# A Catalog of the Diversity and Ubiquity of Metabolic Organelles in Bacteria

**DOI:** 10.1101/2021.01.25.427685

**Authors:** Markus Sutter, Matthew R. Melnicki, Frederik Schulz, Tanja Woyke, Cheryl A. Kerfeld

## Abstract

Bacterial microcompartments (BMCs) are organelles that segregate segments of metabolic pathways, which are incompatible with surrounding metabolism. In contrast to their eukaryotic counterparts, the selectively permeable membrane of BMCs, the shell, is composed of protein. While the sequestered enzymes vary among functionally distinct BMCs, the proteins that form diverse BMC shells are structurally homologous; this enables the bioinformatic identification of the organelles by locating genes encoding shell proteins, which are typically proximal to those for the encapsulated enzymes. With recent advances in genome‐resolved metagenomics and the emphasis on “microbial dark matter”, many new genome sequences from diverse and obscure bacterial clades have become available. We find the number of identifiable BMC loci has increased twenty‐fold since the last comprehensive census of 2014. Moreover, the addition of new types we uncovered doubles the number of distinct BMC types known. These expand the range of catalysis encapsulated in BMCs, underscoring that there is dark biochemistry that is compartmentalized in bacterial organelles yet to be discovered through genome sequencing. Our comprehensive catalog of BMCs provides a framework for their identification, correlation with bacterial niche adaptation, and experimental characterization, and broadens the foundation for the development of BMC‐based nanoarchitectures for biomedical and bioengineering applications.

## Main

Bacterial microcompartments (BMCs) are metabolic organelles that consist entirely of protein; a modular shell surrounds an enzymatically active core, with the shell functioning as a semipermeable membrane for substrates and products (Figure 1a). BMCs were discovered by electron microscopy as polyhedral structures in Cyanobacteria [1]. Carboxysomes enhance CO_2_ fixation [2] in all cyanobacteria and some chemoautotrophs by encapsulating RuBisCO together with carbonic anhydrase (CA) to concentrate the substrate CO_2_ (Figure 1b). Much later, similar structures were observed in heterotrophs, however only when grown in the presence of the substrate of those BMCs, ethanolamine or 1,2‐propanediol [3]. DNA sequencing confirmed that the shell proteins of those BMCs are similar to those of the carboxysomes, a fact that has enabled finding a multitude of BMCs with the advent of genomic sequencing [4, 5]. The majority of BMCs are catabolic and are known as metabolosomes. Many diverse types share a common core chemistry of a signature enzyme, that generates an aldehyde, an ubiquitous pfam00171 aldehyde dehydrogenase (AldDh) to oxidize it as well as a phosphotransacylase (PTAC) to generate an acyl‐phosphate and an alcohol dehydrogenase (AlcDh) for cofactor regeneration (Figure 1c) [6]. Targeting of enzymes into the lumen of many BMCs, including the beta‐carboxysome [7], typically proceeds via encapsulation peptides (EPs), which consist of a ∼15‐20 amino acid amphipathic alpha‐helix that is connected to the N‐ or C‐termini of cargo proteins via a flexible linker [8].

**Figure 1.**
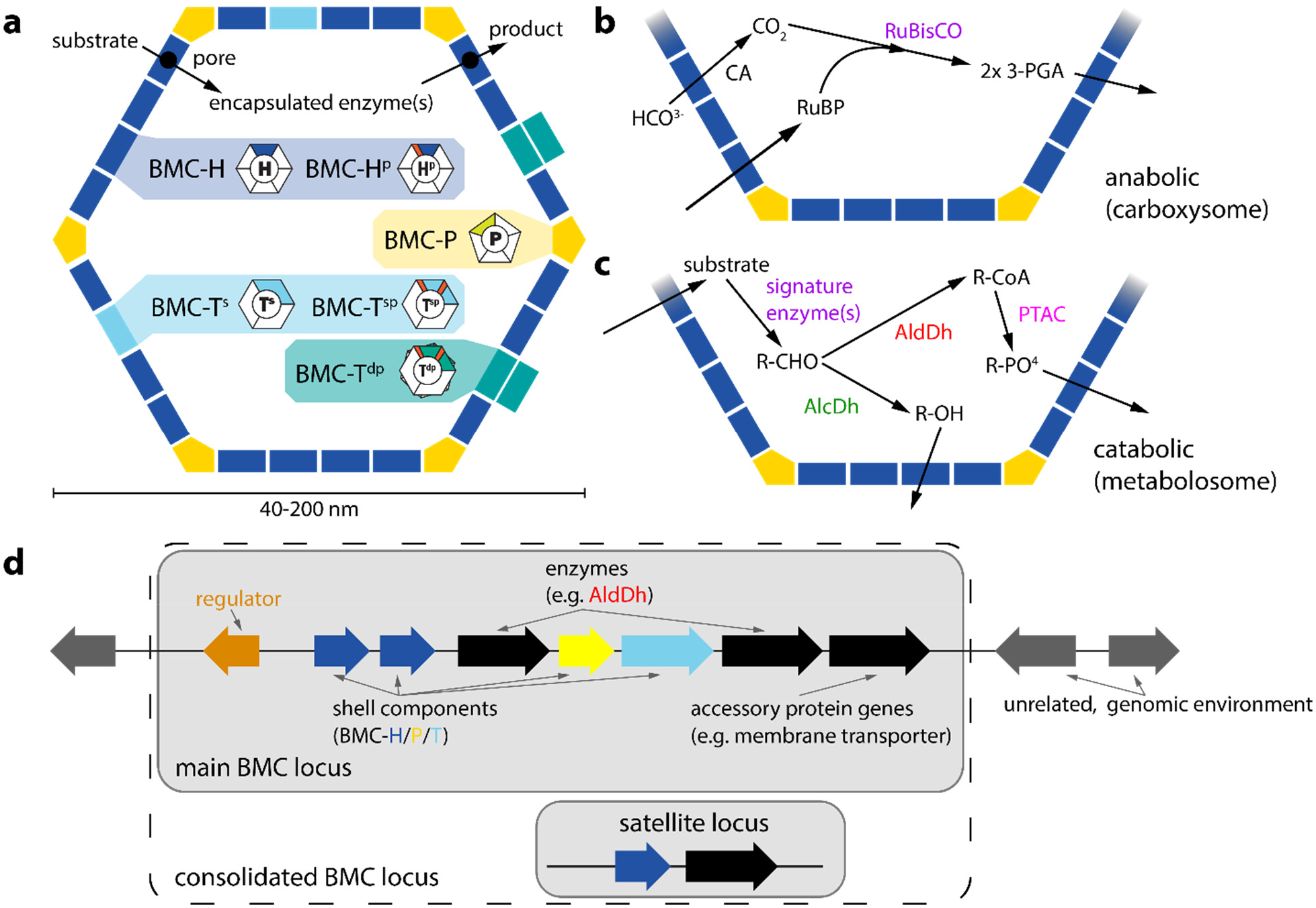
Generalized BMC structure, function and chromosomal organization of component genes. **a**, Overview of a BMC shell and types of shell protein components. **BMC‐P**: pentamer, pfam03319 domain (Supplementary Fig. 1a); **BMC‐H**: hexamer, pfam00936 domain(Supplementary Fig. 1b); **BMC‐H**^**p**^: circularly permuted variant of BMC‐H with two secondary structure elements translocated from the C‐ to the N‐terminus (Supplementary Fig. 1e); **BMC‐T**^**s**^,: standard trimer, fusion of two pfam00936 domains (Supplementary Fig. 1c); **BMC‐T**^**sp**^: permuted BMC‐T^s^, each pfam00936 domain contains a circular permutation as in BMC‐H^p^ (Supplementary Fig. 1f); **BMC‐T**^**dp**^: a permuted BMC‐T variant in which two trimers dimerize across their concave faces to form an interior chamber (Supplementary Fig. 1d). The pores in BMC‐T^dp^ trimers are relatively large (14 Å in diameter), gated by conformational changes in the surrounding sidechains and are predicted to serve as conduits for larger metabolites [45, 46, 47]. **b**, Simplified reactions of anabolic and catabolic BMCs. CA: carbonic anhydrase, RuBP: ribulose 1,5‐ bisphosphate, 3‐PGA: 3‐phosphoglycerate, AlcDh: alcohol dehydrogenase, AldDh: aldehyde dehydrogenase, PTAC: phosphotransacylase, CoA: coenzyme A. **c**, Typical BMC locus consisting of genes for shell proteins (blue, cyan, yellow), enzymes (black, colored according to enzyme type in detailed diagrams of Supplementary Data: BMC types and Supplementary Data: Locus diagrams), regulators (orange) and ancillary proteins such as cell membrane transporters for substrates; The combination of main and satellite BMC locus is termed a consolidated BMC locus.

BMCs have a polyhedral, often icosahedral shape and are about 40‐200 nm in diameter [9]. Structures of model shells confirmed icosahedral symmetry with pentagons occupying 12 vertices and hexagons forming the facets [10, 11, 12, 13] (Figure 1a). The pentagons (BMC‐P) are formed by five subunits of the pfam03319 fold and have the shape of a truncated pyramid [14] (Supplementary Fig. 1a). BMC‐H proteins contain the pfam00936 domain and form almost perfect hexagons and with a concave and convex side [15] (Supplementary Fig. 1b). The facets of shells also contain BMC‐T proteins, which are genetic fusions of two pfam00936 domains that form trimers that in size and shape resemble the hexamers. Variations of the pfam00936 domain that are the result of a circular permutation of the primary structure expand the number of distinct hexagonal building blocks to five (Figure 1a, Supplementary Fig. 1b‐f).

In 2014, a comprehensive bioinformatic survey identified 357 BMC loci representative of 30 types/subtypes across 23 bacterial phyla [4]. This cataloging of BMC loci within genomic databases was feasible, because both pfam03319 and pfam00936 protein folds are unique to BMCs and are often encoded together with their cargo proteins in chromosomal loci. Within the last six years, the number of metagenomic datasets has increased nearly an order of magnitude [16]. In addition to the microbiome data that have become available, the extraction of tens of thousands of genomes from the uncultivated majority of microorganisms has been facilitated through recent advances of genome‐resolved metagenomics [17]. Mining these massive new datasets, we here compiled a database of more than 7000 BMC loci that cluster into 68 BMC types or subtypes, including 29 new functional BMC types or subtypes. BMC loci are widespread, now evident in 45 phyla across the bacterial tree of life. Collectively our results show that the known BMC functional diversity and distribution at the phylum level has essentially doubled in the last six years and foregrounds the widespread occurrence of bacterial organelles in microbial dark matter.

## Results

### BMC shell protein and locus analysis

We hypothesized that BMC diversity has greatly expanded since the previous survey (2014) due to growth in genome sequencing of microbial diversity, and in particular that of uncultivated clades. We compiled and curated an in‐house dataset of all putative BMC loci based on the UniProt Knowledgebase (UniProtKB) with data as recent as March 2020. After retrieving all available BMC shell protein sequences, we collected the sequences for neighboring genes encoded within 12 ORFs from any shell protein gene, which covers all BMC locus related genes in previously known loci and, in retrospect, in all new BMC loci. Contiguous genes were classified as a “main locus” when they contained at least one BMC‐H and one BMC‐P gene, to distinguish from “satellite loci” [4]. Satellite loci were combined with the main locus to form a consolidated locus (Figure 1d).

The previous comprehensive survey [4] relied on pfam co‐occurrence. Many of the new BMC loci would be difficult to categorize by this method, because they contained few recognizably type‐specific proteins other than the shell components (unlike the well‐characterized PDU and EUT loci with around 20 total gene products; Figure 2a). We therefore sought to improve BMC type clustering by subclassifying all pfam00936 and pfam03319 shell proteins using a phylogenomic approach [18]. Trees were built for representative sequences from each of the six shell protein types: BMC‐H, BMC‐P, BMC‐T^s^, BMC‐T^dp^, BMC‐ H^p^ and BMC‐T^sp^ (Figure 1a, Figure 3). Subclades for each shell type were identified visually, usually containing a long internal stem, and were each assigned a unique color name chosen from the xkcd color survey (https://xkcd.com/color/rgb/), using names from related color families for adjacent subclades within each major clade (for high resolution trees with full annotations, see Supplementary Data: Shell protein trees). The sequences comprising each of the colored subclades were then used to calculate a profile Hidden Markov Model (HMM) [19] for each color group and combined with HMMs derived from proteins common to BMC loci that were clustered by protein pfam information (Supplementary Fig. 2a). Decoupling the identity of a shell protein from a specific BMC type allowed us to make unbiased observations of shell proteins that are similar, despite being constituents of functionally distinct BMCs. We find that for many loci, the component shell proteins are drawn from across the tree, revealing functional and evolutionary relationships among distinct BMC types (described below).

**Figure 2.**
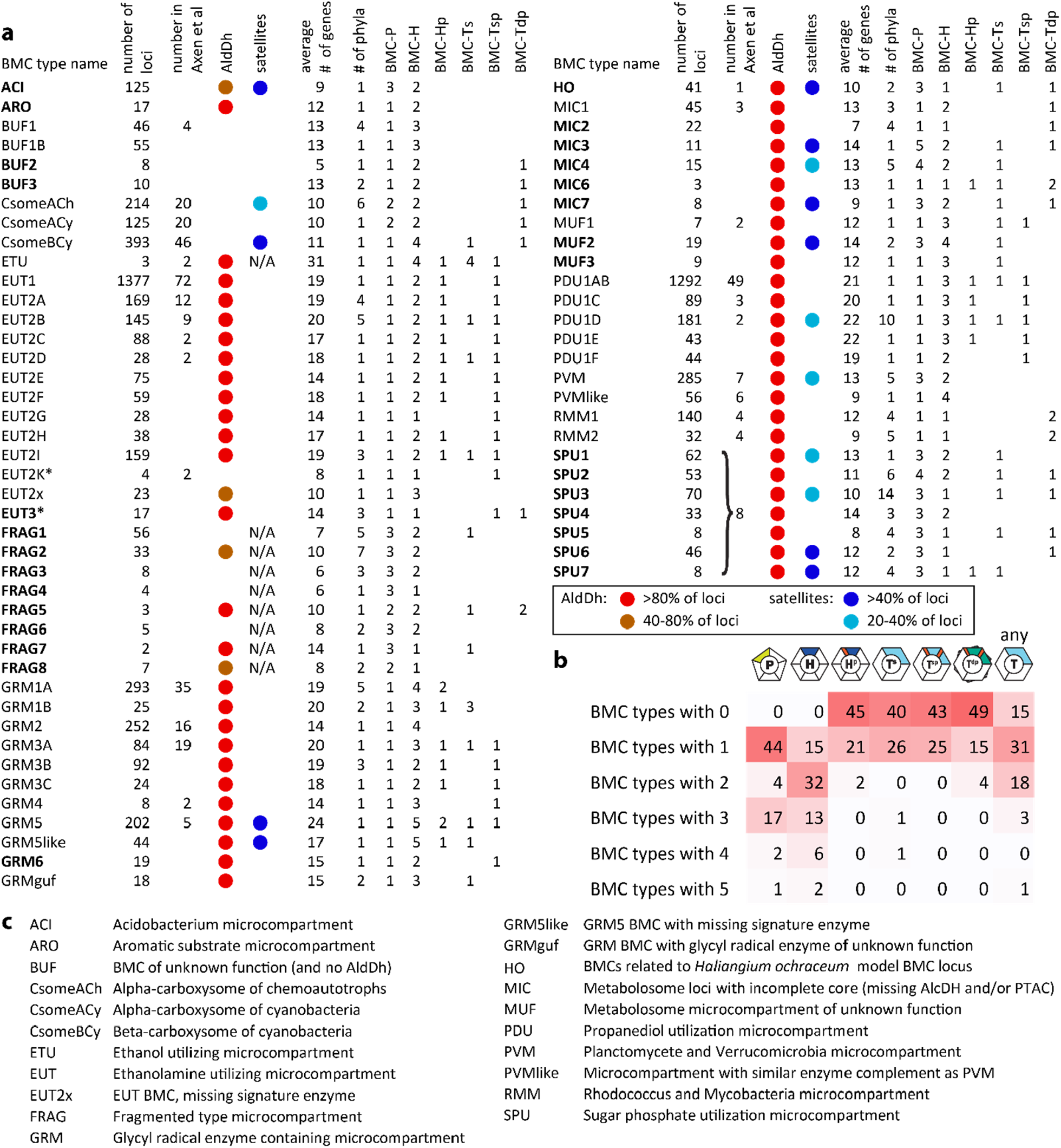
Overview of BMC types and shell protein content found across all bacteria. **a**, Table of number of loci for each BMC type in this study and in Axen et al. 2014 [4], AldDh occurrence, prevalence of satellite loci, average number of genes in the locus, number of observed phyla and number of each type of shell protein. Major new BMC types or subtypes identified in this study in bold. The asterisk denotes the name change of EUT3 (Axen et al. 2014, [4]) to EUT2K because the present analysis indicates EUT2K is not a major new type. **b**, Numbers of each type of shell protein across BMC types. **c**, Explanation of BMC type abbreviations.

**Figure 3.**
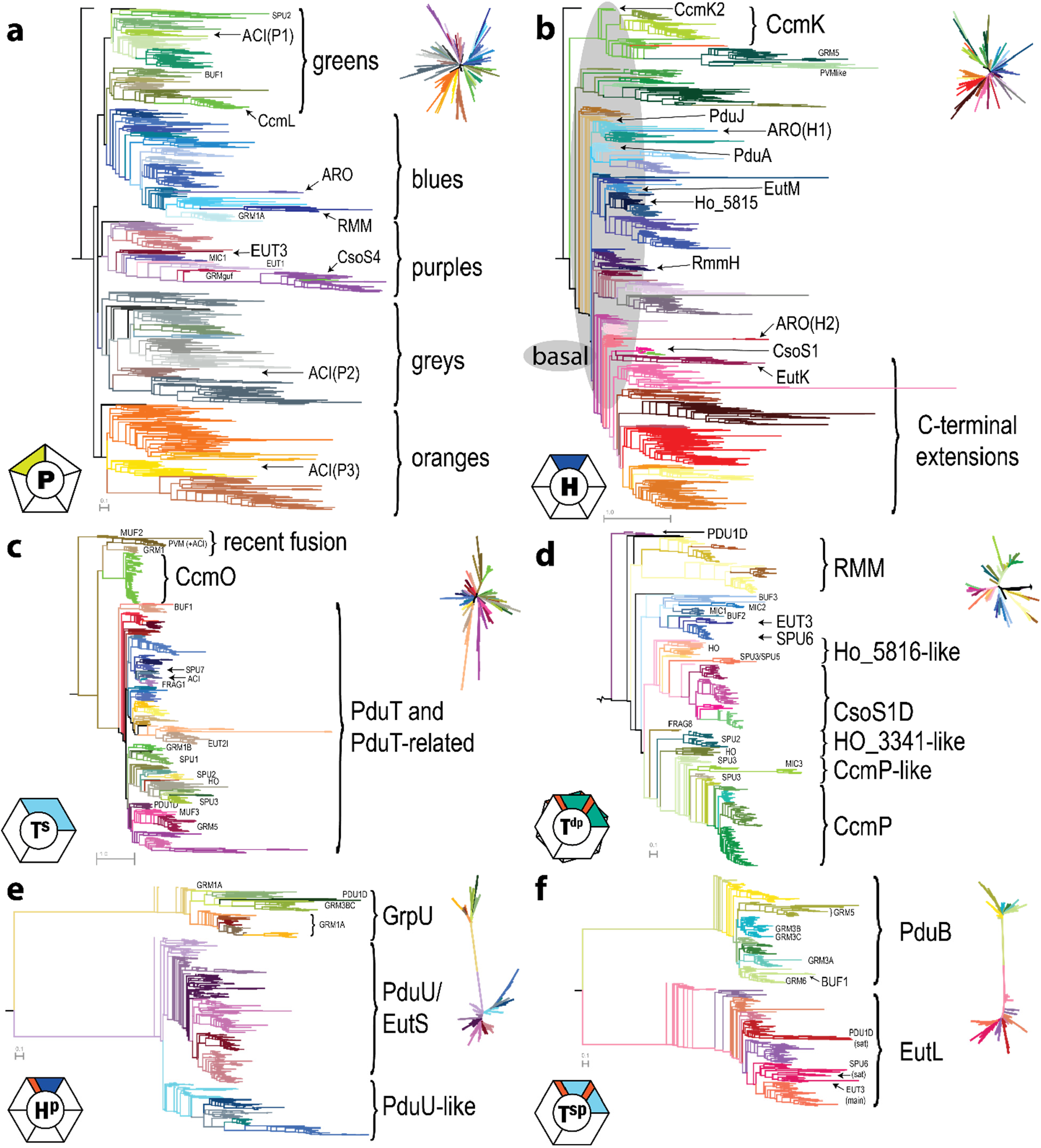
Phylogenetic maximum likelihood trees for the six types of shell protein: **a**, BMC‐P, **b**, BMC‐H with region of basal hexamers encompassed by gray shading, **c**, BMC‐T^s^, **d**, BMC‐T^dp^, **e**, BMC‐H^p^ and **f**, BMC‐T^sp^. Representative sequences were selected by removing redundancy. Full hi‐resolution images of each tree with annotation labels for all terminal nodes are available as Supplementary Data: High resolution shell protein trees. All caps labels refer to the predominantly associated locus with a certain BMC type, mixed case labels denote a specific, previously characterized protein found in that clade. For BMC‐P the major clades are labeled by color. Unrooted versions of the trees are shown in the top right corner of each panel.

The combination of detailed shell protein, enzyme and accessory protein HMMs allowed us to cluster loci into distinct BMC types and subtypes (Supplementary Fig. 2a, see Materials and Methods for details). Subtypes generally share most enzymatic components but differ in gene order and/or shell protein content. For example, the sugar‐phosphate utilization (SPU loci) can be separated into seven distinct subtypes (Supplementary Fig. 2b). For the naming of the loci we adopted and expanded the nomenclature from Axen et al [4] (Figure 2c). New BMCs were named either by a distinguishing feature such as predominant occurrence in a certain taxon of organisms (e.g. ACI for Acidobacteria) or the organism that gave rise to a model system (e.g. HO for *Haliangium ochraceum*), or putative class of BMC substrate (e.g. ARO for aromatic substrate). In absence of any potentially defining feature, BMCs were classified under broad groupings such as Metabolosomes of unknown function (MUF), Metabolosomes with an incomplete core (MIC) or BMC of unknown function (BUF) (Figure 2c). In this analysis we have added 29 new major BMC types or subtypes as well as 10 new subtypes for several established loci (Figure 2a). Representative locus diagrams and a short description of the common components for a total of 68 types are listed in Supplementary Data: BMC types.

### Overview of BMC distribution and characteristics

BMCs occur widely across the bacterial domain (Figure 4), they are now evident in 45 different phyla, compared to 23 phyla from an analysis in 2014 [4]. Some BMCs are confined to a single phylum (e.g. ACI, ARO, BUF2) but most are found across multiple phyla (Figure 4). Our analysis revealed a large BMC functional diversity in certain phyla, such as Proteobacteria, Actinobacteria and Firmicutes (Figure 4), reflecting the importance of metabolic flexibility across disparate niches. To get an estimate of the overall prevalence of BMCs we performed a shell protein HMM search against IMG/M [16] and GEM [17] genomes and found hits in 20% of them.

**Figure 4.**
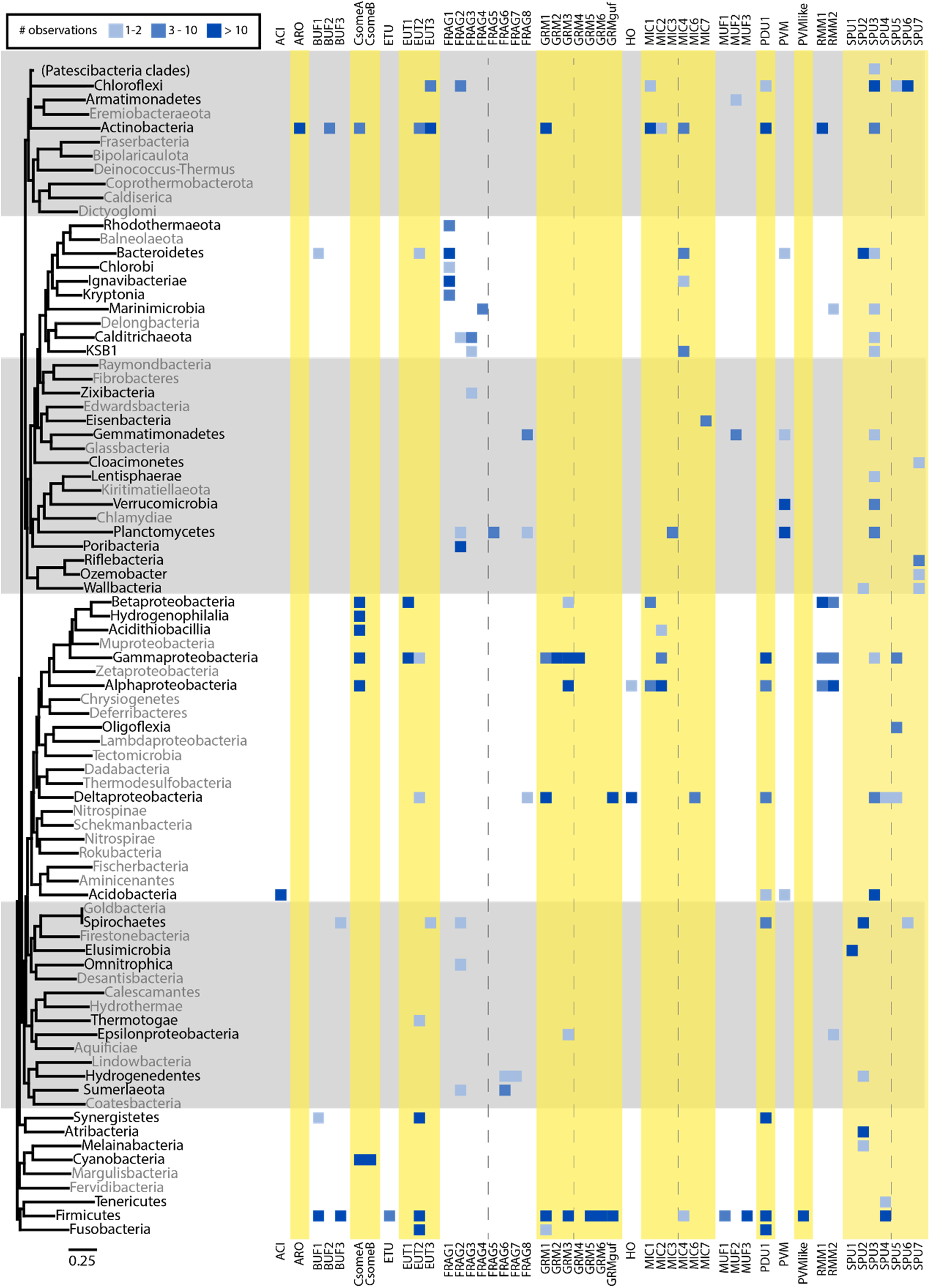
Distribution of BMC types in 45 bacterial phyla. Phylogenetic tree representing all phylum‐level taxonomic groups was generated by aligning a set of 56 different marker proteins. Phyla lacking BMC loci shown in grey and different numbers of BMCs in blue. Alternate yellow/grey blocks and dashed lines are added for visual guidance.

A core enzyme of many functionally distinct metabolosomes is the AldDh (Figure 2a). A phylogenetic tree of representative sequences reveals that the AldDh is specific to BMC functional type (Figure 5). AldDh from BMCs with similar substrates cluster on the tree and, accordingly, can be used in prediction of potential substrates for unknown BMCs by looking at the closest AldDh homologs. AldDh typically have an encapsulation peptide on either the N‐ and C‐ termini; strikingly the two major branches of the tree also are distinct in the location of the EP extensions (Figure 5); type I have EP at their C‐terminus, type II at the N‐terminus. The most parsimonious interpretation of these data is that the acquisition of a sequence extension that serves to facilitate encapsulation is an ancient innovation that arose independently twice in the evolution of BMCs.

**Figure 5.**
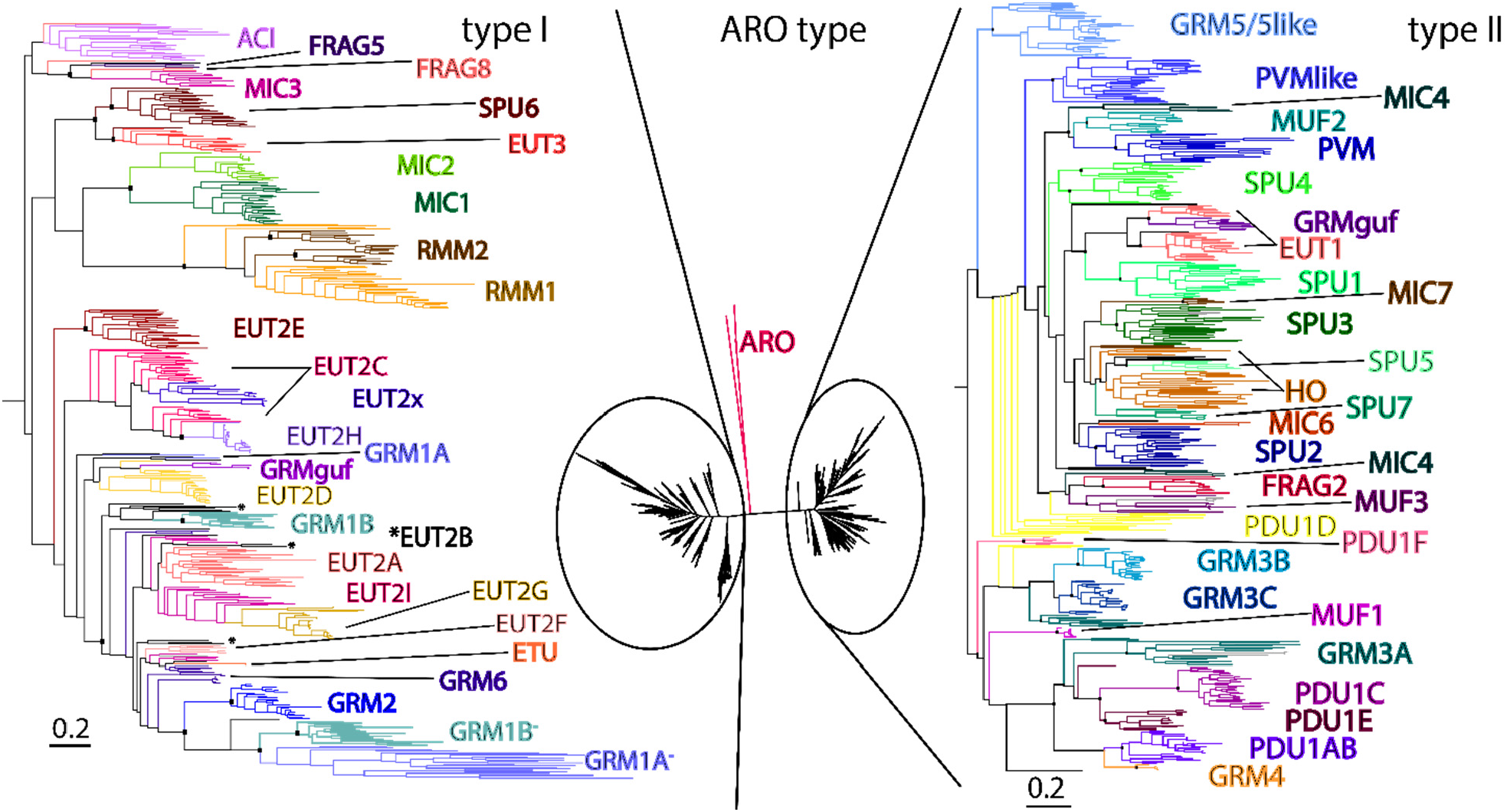
Phylogenetic tree of pfam00171 aldehyde dehydrogenases from BMC loci. Sequence redundancy was reduced by limiting to 30 sequences per BMC type (based on percent identity). Three major groups were identified: type I, which is dominated by EUT2, belongs to IPR013357, and contains a C‐terminal EP (left); the ARO type outlier clade; and type II, which belongs to IPR012408, contains an N‐terminal EP, and contains a larger diversity of BMC types, relative to type I AldDhs. Branch lengths are scaled by the number of substitutions per site. Bootstrap values for important nodes are represented as black squares (above 50%). GRM1A and GRM1B loci contain a putatively inactive second copy (marked with a “‐” superscript). EUT2B AldDh marked with an asterisk are found in different basal locations.

In addition to enzymes, BMC loci also contain genes for functions that support the expression and activity of the organelle like transcriptional regulators and cell membrane transporters for the substrate. Like AldDh, these gene products also can provide clues as to the function of a BMC, by comparison to non‐ BMC related homologs. For example, for one of the new BMC types that we identified here involves an aromatic substrate (ARO), a SWISS‐MODEL search [20] with the protein sequence of the regulator reveals the top characterized hit as a regulator of catechol degradation, which is consistent with the enzymes found in that BMC locus (Supplementary Data: BMC types).

When comparing the shell protein inventories across the BMC types (Figure 2a), both unexpected differences and new patterns emerge. On average, BMC loci contain 1.7 BMC‐P, 2.2 BMC‐H, 0.4 BMC‐Hp, 0.5 BMC‐T^s^, 0.4 BMC‐Tsp and 0.3 BMC‐T^dp^. The distribution reveals that BMC‐P most commonly occur singly or as three paralogs (Figure 2b). Correlating the BMC‐P found in the loci that have three copies with their location on the BMC‐P tree (Figure 3a), we find a “BMC‐P triplet” pattern: one member each from the grey and orange major clades and a third member from one of the other clades (Supplementary Fig. 3). Many BMC loci also contain multiples of BMC‐H (Figure 2a,b) and for a specific BMC type, those do not necessarily cluster together on a phylogenetic tree. A reason for this could be that the different paralogs fulfill a function that is shared across BMC types. The wide variety of different compositions of the BMC shell highlights its modular construction from building blocks that have the same size and shape, but different permselectivities.

### New and expanded variants of BMC loci

One of the most prominent expanded BMC types is the SPU (sugar phosphate utilization), first noted in 2014 through identification of only eight representatives [4]. We now find more than 280 members distributed across 26 bacterial phyla (Figure 4), establishing SPU as one of the most prevalent BMC types. Our clustering has identified seven distinct subtypes (Supplementary Fig. 2b, Supplementary Data: BMC types) that all share two sugar phosphate processing enzymes (pfam01791, pfam02502).The DeoC‐type pfam01791 aldolase converts 2‐deoxy‐D‐ribose to glyceraldehyde‐3‐phosphate and acetaldehyde which can then be processed by the AldDh to acetyl‐CoA. The AldDh of SPU6 is close to EUT3 and the other SPU types are on the same major branch as EUT1, which both process acetaldehyde (Figure 5). The pfam02502 is of the RpiB type [21] that isomerizes ribose‐5‐phosphate. Collectively, these data suggest potential function of this BMC type is metabolizing the products of DNA degradation, presumably from the ubiquitous detritus available in diverse environments.

The HO BMC from *Haliangium ochraceum* has been the primary model system for structural studies of the shell [10, 12, 22]. We have identified similar loci in 40 other genomes but the function of this organelle remains enigmatic; the HO AldDh is most closely related to those of SPU5 and SPU7 (Figure 5), all loci encode the characteristic BMC‐P triplet, and share similar types of BMC‐T^dp^ (Figure 3d). Some genomes containing the HO loci have a pfam01791 sugar processing enzyme in a different genomic location and some of these orthologs contain an EP‐like N‐terminal extension and could therefore be encapsulated in an HO BMC. A similar function as SPU BMCs, the catabolism of nucleic acid, is consistent with the their presence in Myxobacteria that that are known for degrading biomacromolecules [23].

Another functional type that has substantially increased in membership are the PVMs [24] now with 285 representatives (Figure 2), as compared to seven found in 2014 [4]. This reflects the increased attention to Verrucomicrobia species for their role in the global carbon cycle as degraders of complex algal and bacterial cell wall polysaccharides [25, 26]. For example, Lentimonas species devote 4% of their proteome to the degradation of fucoidan, the major cell wall polysaccharide of brown algae into fucose, which is catabolized in the PVM BMC [26]. The PVM‐like BMC that shares the PVM aldolase signature enzyme that processes sugar derivatives also gained 50 new members. We expect more PVM and PVM‐like BMCs to be discovered with increased attention on sequencing bacteria that degrade complex polysaccharides, and we find their shell protein components prevalent in searches of metagenomes from environmental samples (see below).

Our analysis discovered several completely new BMC types. One, ARO, for its predicted aromatic substrate, is found in the Micromonosporales and Pseudonocardiales orders of Actinobacteria. The ARO locus contains two pfam02900 ring‐opening oxygenases and a set of enzymes related to the degradation of aromatic aldehyde compounds (Supplementary Data: BMC types). A possible initial substrate is 2‐ aminophenol based on the assignment of related pfam00171 AldDh as aminomuconate‐semialdehyde dehydrogenases. Unlike most other AldDhs, the ARO AldDh does not contain a detectable encapsulation peptide, and constitutes its own group falling between both major groups on the phylogenetic tree on the tree (Figure 5). Likewise, the ARO shell protein composition is among the simplest observed: consisting of one BMC‐P and two distinct types of BMC‐H. The three ARO shell proteins are all found in late‐branching subclades that are strongly divergent from other shell proteins (Figure 3a,b), suggesting that the predicted catabolism of aromatic compounds and the involvement of an AldDh remote from others found in BMCs has imposed a distinctive function on these shell proteins that is under evolutionary constraint.

Another major new type with 125 members, ACI, is found exclusively in Acidobacteria, primarily in only two of the 26 Subdivisions, Subdivision 4 and Subdivision 6. While Acidobacteria are found across differing ecosystems [27] they are of particular relevance to the soil environment, as they can comprise up to 60% of the soil bacterial community [28]. The locus contains a pfam00596 class II aldolase with a C‐terminal EP, a hydroxyacid dehydrogenase (pfam00389/02826), a triplet of BMC‐P proteins, at least one BMC‐H and several proteins of unknown function (Supplementary Data: BMC types). Its AldDh is consistently found on a satellite locus and is phylogenetically similar only to MIC3, another uncharacterized BMC type (Figure 5). The pfam00596 aldolase is also observed in the PVM and GRM5 types that process L‐fuculose and L‐rhamnulose phosphate so a function related to carbohydrate degradation can be proposed.

We discovered several enigmatic new BMC types that lack a defined locus organization but are able to be grouped based solely on shell protein composition. These FRAG BMCs are composed of shell protein genes encoded by as many as six different genomic locations. One common feature of FRAG BMCs is the presence of a BMC‐P triplet (Figure 3a, Supplementary Fig. 3). Most of these genes are remote from genes for known enzymes, although a few have proximal AldDh (Figure 5, Supplementary Data: BMC types), that map to uncharacterized AldDhs in different parts of the tree (Figure 5). FRAG BMCs are found in diverse organisms, including the Ignavibacteriae and closely related phyla, the Gemmatimonadetes, the Planctomycetes and a large number of candidate phyla (Figure 4). In addition, we have found several more distinct BMC types of unknown function: BUF2, BUF3, GRM6, MIC2‐7, MUF2 and MUF3 (Supplementary Data: BMC types). There is a large number of new metabolosome loci that encode a pfam00171 AldDh yet do not have an obvious signature enzyme that generates that aldehyde (ACI, FRAG1‐7, HO, MIC2‐7, MUF2‐3; Figure 2a,c). Assignment of function for these and other functionally cryptic loci could be approached by combining genetic, biochemical studies and metabolic arrays focused on the AldDH and the shell proteins, as done for the identification of the fucose/rhamnose substrate for PVM BMCs [24].

### BMC type co‐occurrence and horizontal gene transfer

About 80% of BMC‐containing genomes encode only one locus, but a substantial number encode two (20%) or more (2%) loci. The most frequent co‐occurring pair are the PDU and EUT loci (Supplementary Table 1), providing catabolism for ethanolamine and 1,2‐propanediol. In *Salmonella enterica*, this combination has been shown to be regulated mutually exclusively to prevent formation of mixed BMCs [29]. Most of the genomes encoding three or more BMC types are combinations of EUT, PDU and GRM (Supplementary Table 2). The most extreme examples of the potential to form multiple metabolic organelles is the genome of the Firmicute *Maledivibacter halophilus* that contains six loci: two EUT2B and one each of PDU1D, GRM1A, BUF1 and BUF3. This organism is found in anoxic hypersaline sediment [30]. All of the organisms with three or more BMC types are either Proteobacteria or Firmicutes, predominantly from the Enterobacterales or Clostridiales orders. Members of these orders are various human and other animal pathogens, and are likewise abundant in aquatic and soil ecosystems. The ability to form multiple functionally distinct BMCs confers metabolic potential and flexibility; for example, the prevalence of multiple BMCs in the Firmicutes (Clostridiaceae, Ruminococcaceae) likely contributes to their ability to break down of complex polysaccharides [31].

BMC loci are genetic modules, a compact organization of the structural, regulatory and ancillary components necessary for BMC function. As such, they are an ideal target for horizontal gene transfer (HGT). One example of a possibly recent HGT event involves a EUT2I locus in *Oceanotoga teriensis*, the only BMC instance in the Thermotogae phylum. About 13% of the proteome of this organism shares >30% sequence identity with BLAST hits against Firmicutes (img.jgi.doe.gov) so the BMC locus is likely part of large horizontal gene transfer from a Firmicute. The closest relative to the BMC in this organism according to locus scoring is a EUT2I from *Clostridium scatologenes*, indicating a potential origin. In the case of RMM2 there are two phylum outliers, one in Epsilonproteobacteria and one in Candidatus Marinimicrobia (Supplementary Data: Locus diagrams). Both of them have phage integrase proteins (pfam13356 and pfam00589 domains) right next to the BMC locus, indicating a likely transfer via a phage vector.

### Satellite loci

We define satellite loci as loci distal from the main locus and containing either shell proteins only, or a combination of shell proteins and enzymes (Figure 1d). Some satellite loci seem to be obligately distal from main BMC locus, such as the CcmK3/K4 paralogs of the beta‐carboxysome [32, 33]; separate regulation of its expression may serve as a means to tune shell permeability under changing environmental conditions [32]; alteration of shell permeability may be a general function for shell proteins encoded in satellite loci of many types of BMCs (Figure 2). Other satellite loci appear to have arisen as fissions from the main locus, such as the satellite locus for type I HO BMCs that contains two BMC‐P with an aldolase, which are found in the main locus of type II HO BMCs (Supplementary Data: BMC types). For HO, SPU3 and SPU6 BMCs some satellite loci resemble “EUT modules”, consisting of the ethanolamine degradation signature enzymes EutA/B/C and a EutL type BMC‐T^sp^ shell protein (Supplementary Fig. 4). Because those BMC types are not expected to primarily process ethanolamine, this could represent a functional extension of the main BMC to use ethanolamine as an alternate substrate, with the BMC‐T^sp^ acting as a shell protein that facilitates entry and the EutA/B/C enzymes to process it. In the case of SPU6 there is even indication of integration of the satellite locus into the main locus, replacing the SPU type signature enzymes and the resulting locus is similar to that of EUT3 (Supplementary Fig. 4). EUT3 is a new type of ethanolamine utilization BMC with an AldDh that is phylogenetically distinct from the ones from both EUT1 and EUT2 loci (Figure 5) and it contains a BMC‐T^dp^ shell protein, unlike any other known EUT type BMC (Figure 3d). Phylogenetic trees validate the link between the two locus types; they are on the same major branch of the AldDh tree (Figure 5) and shell proteins like the BMC‐T^dp^ are also adjacent, with the members of the fused locus found at the base of the SPU6 part of the branch (Figure 3d), another example of how the expanded survey plausibly recounts BMC evolutionary history.

### Shell protein trees and BMC identification

From all the BMC loci we have collected more than 40,000 shell protein sequences. Almost 19,000 of them are unique, highlighting their diversity despite a common function to form BMC shells. Among them are about 4900 BMC‐P, 8000 BMC‐H, 1550 BMC‐H^p^, 1700 BMC‐T^s^, 1600 BMC‐T^sp^ and 1000 BMC‐T^dp^. A further reduced set of those was used to build phylogenetic trees that illustrate their diversity (Figure 3).

The BMC‐P proteins resolve into five major clades: green, blue, purple, grey and orange (Figure 3a). Representatives of grey and orange clades always co‐occur as two of three members of a BMC‐P triplet. The third member of a given triplet varies across locus types, and is drawn from the green, blue or purple clades (Supplementary Fig. 3). The BMC‐P proteins from these three clades frequently occur as the sole BMC‐P gene in loci that lack BMC‐P paralogs. The only exception is the alpha‐carboxysomal CsoS4A and CsoS4B, BMC‐P proteins that form a long stem of the purple clade; they always co‐occur as pair. The multiplicity of BMC‐P paralogs is unexpected because only 12 pentamers are needed to cap polyhedral structures and hints at an additional functional role for the vertex proteins.

The defining, conserved primary structure of the BMC‐H proteins, the pfam00936 domain is about 80 amino acids in length. However, extensions of up to 200 residues are observed, most frequently at the C‐terminus and those BMC‐H cluster on the phylogenetic tree (Figure 3b). Each BMC locus type contains at least one BMC‐H found close to the base of the tree (Figure 3b). We refer to these as “basal” BMC‐H proteins, because they share sequence motifs including the highly conserved inter‐hexamer interface residue motifs KAAN and (P/A)RPH of the facets [10]. Those likely constitute the bulk of the BMC shell facets while other BMC‐H have more specialized functions.

Before the availability of genomic data, BMC‐T^dp^ proteins were thought to be a rarity because their occurrence was limited to carboxysomal members CcmP and CsoS1D, as the two metabolosome model systems, EUT1 and PDU1, lack these proteins. In our analysis we find BMC‐T^dp^ proteins in a large variety of BMC types (Figure 2a). The BMC‐T^dp^ tree can be divided into four major clades (Figure 3d). The yellow clade contains members from the RMM1 and RMM2 BMC types. An unusual outlier is found at the base of the tree with homologs found in PDU1D loci from Proteobacteria and a single Acidobacterium; no other PDU type BMCs contain BMC‐T^dp^ shell proteins. The blue major clade contains a variety of uncharacterized BMCs (MIC12/BUF23) as well as EUT3 and SPU6 members that are in adjacent clades. The largest number of sequences are found in two clades that can be characterized by the presence of carboxysomal members. The purple clade contains the alpha‐carboxysomal CsoS1D and a number of CsoS1D‐like proteins from mainly HO and SPU3 and SPU5 type loci. The green major clade contains the beta‐ carboxysomal CcmP as well as CcmP‐like proteins from HO, MIC3, SPU2 and SPU3 type loci. The proximity of the BMC‐T^dp^ of SPU to carboxysome ones indicates parallels with regard to the molecules that enter or leave the shell through these proteins. Sugar phosphates seem a likely candidate that both BMC types have in common, however BMC‐T^dp^ from the other two major clades do not all share that type of substrate. It is possible that those BMCs have adopted the gated shell proteins for other purposes, such as large cofactors. Universally conserved across all types are the residues for the gating mechanism, indicating that this is a crucial function of the BMC‐T^dp^.

Shell proteins are diagnostic of the potential to form BMCs, in contrast to the enzymes found in BMCs, which have homologs that are not BMC‐related. Scoring a proteome with the collection of BMC‐type specific shell protein HMMs allows for quick assessment of both the presence of BMCs and initial identification due to their specificity, predicted from the color‐type combinations of shell proteins present (Figure 3 and Supplementary Data: BMC types). We made a preliminary survey for the prevalence of BMCs in metagenomes. The shell protein HMMs derived from our locus collection were used to score data from 26,948 metagenomes in IMG/M [16], finding hits in 15,604 of them, and the total shell protein gene count adds up to more than 1.7M. Using the assumption that the co‐occurrence of the type‐specific BMC‐H and BMC‐P colors are indicative of BMC functional type (or types that use the same or closely related substrates) we can make initial predictions the kinds of encapsulated metabolism to be found in metagenomes/microbiomes (Supplementary Fig. 6). The distribution of BMC types across bacterial clades, such as the prominent occurrence of PVM and SPU, provides a diagnostic marker of the specialized metabolism required in diverse nutritional landscapes of environmental samples. For example, the Anaerolineae class of Chloroflexi that contain SPU6 BMCs are prominent in environments characterized by anaerobic digestion [34] and the Planctomycetes harboring PVM BMCs, despite growing very slowly, are one of the dominant bacterial species in algal blooms [35].

## Discussion

By compiling over 40,000 BMC shell protein genes and surveying their genomic context, we find that the number of identifiable metabolic organelles has, over the last seven years, increased 20‐fold and representatives are found across 45 phyla. The phylogenetic classification of shell proteins combined with the BMC type assignment shows that describing BMC shell proteins based on a specific locus type (e.g. PDU) is of limited usefulness as the growing number of functionally distinct BMC loci encode shell proteins from across the phylogenetic tree. Our phylogenetic classification also enables us to predict the basal hexamer(s) for a given locus (Figure 3b), likely to form the bulk of the BMC shell facets. As such, it likely conducts key metabolites; in conjunction with structural modeling focused on pore size and charge these data can be useful for predicting the first substrate of the encapsulated chemistry [36]. In addition, we predict that basal BMC‐H proteins are broadly interchangeable among functionally distinct BMCs, consistent with results of previous efforts to construct chimeric shells that, in retrospect, involved basal BMC‐H proteins [37, 38]. The color combinations of shell proteins found in loci provides a guide to their compatibility in assembly and can be used for designing shells for metabolic engineering [39, 40, 41].

Our analysis also sheds light on the evolutionary history of BMCs. In addition to the widespread HGT of loci evident from their distribution across phyla (Figure 4), individual shell proteins provide links, such as the BMC‐T^dp^ of EUT3 and SPU6. Likewise, with our observations we can propose steps in the evolution of the encapsulated catalysis; in the canonical PDU1AB and the recently described GRM4 [36, 42], the colors of all shell the same in both loci (Supplementary Data: Locus diagrams). Comparison of the locus diagrams reveals the same gene order, with the PDU signature enzymes interchanged with the GRM4 signature enzyme. Such a transformation, without altering the shell protein composition, is facilitated by sharing the initial substrate, 1,2‐propanediol. This highlights the role of the shell proteins in shaping the encapsulated catalytic potential. Moreover, it suggests that finding the closest homologs from known BMCs, and the combination of shell protein color types, may be useful in predicting function in the absence of any information about the encapsulated enzymes, as in our preliminary survey of metagenomic data (Supplementary Fig. 6).

With the continued emphasis on sequencing using cultivation‐independent methods, which capture candidate phyla that are suggested to be the majority of bacteria [43], we expect that the role of BMCs in predicting the metabolic potential of environmental samples will become even more prominent. For example, in this study, relative to the data available in 2014, we find the large expansion of the occurrence of SPU loci that is now represented by more than 280 loci across 26 bacterial phyla (8 members in 2014; Figure 5). Their saprophytic function, and the availability of detritus in environmental samples, likely accounts for their emergence. We expect that with continued focus on genome sequencing of the members of diverse ecosystems that the occurrence of FRAG loci will eventually be shown to correlate with an environmental factor that reveals the function of FRAG BMCs. These and other newly discovered functional types (Figure 2) expand the types of predicted chemistry performed by BMCs and indicate that there is additional encapsulated dark biochemistry to be found.

Many microbial communities across Earth’s biomes thrive in highly competitive environments, where their success to utilize resources and adapt under stable or changing environmental conditions will determine the fate of their genetic persistence. Our study finds BMCs in about 20% of all sequenced bacterial genomes. Their broad phylogenetic distribution across 45 bacterial phyla and wide environmental distribution spanning diverse ecosystems, underscores the important role they play in allowing bacteria to thrive in otherwise inaccessible environments. In addition to their prominence in candidate phyla and environmental samples, the importance of BMCs in organisms of the human microbiome and their link to dysbiosis is also becoming apparent. For example, of the eleven bacterial species most prominently associated with complicated urinary tract infections [44], eight contain BMC loci, including PDU, EUT, GRM1, GRM2, GRM3 and FRAG; several of these commonly co‐occur within a single genome. In the urinary tract, in addition to the breakdown of ethanolamine and 1,2‐propanediol, the catabolism of choline via GRM1 and GRM2, likely serves as a carbon, nitrogen and energy source that confers a competitive advantage to the uropathogens. More broadly, a species’ potential to form multiple distinct BMCs functionally parallels the prevalence of BMCs in coexisting community members, foregrounds their previously undescribed role in providing metabolic flexibility as a driver for niche expansion. In addition to providing the foundation for understanding the native functions of BMCs in natural ecosystem function and dysbiosis, our catalog also provides insight into the diversity of metabolic compartmentalization and the evolutionary steps that can inform engineering strategies.

## Materials and Methods

### Locus identification, clustering and HMM generation

The complete Uniprot database was searched with shell protein pfams (PF03319 and PF00936), interpro domain identifiers (IPR000249, IPR004992, IPR009193, IPR009307, IPR013501, IPR014076, IPR014077, IPR020808, IPR030983, IPR030984), prosite identifier PS01139, and smart ID SM0877, accessed as recently as March 2020. Adjacent proteins based on the numerical suffix were then obtained by downloading gene entries within 12 loci of the extracted gene identifier. Pfam tags were extracted from the uniprot annotations for initial HMM generation from proteins of the same pfam. HMMs were generated by aligning the sequences with clustalw 2.1 [48], trimming with trimAl 1.2rev59 with parameters ‐gt 0.6 ‐cons 30 ‐w 3 [49] and HMMs built with hmmbuild from the HMMER package version 3.1b2 [50]. HMMs were calculated analogously for distinct shell proteins (see shell protein section below) and we then scored every locus protein against this combined HMM library, allowing us to represent each consolidated BMC locus as a string of identifier elements derived from the best‐scoring HMM for each protein. To cluster the BMC loci and identify BMC types, we calculated pairwise correlation scores across all loci. Pairwise locus‐locus scores were calculated amongst all loci using a python script based on the sum of two values: the total number of HMMs found in both loci; and the length of the longest sequence of consecutive HMM matches, multiplied by a weighting factor of 10. Data was then imported in Cytoscape 3.7.2 [51] to visualize the locus clustering. See Supplementary Fig. 5 for an all‐vs‐all BMC types visualization. Clusters were manually matched to known BMC types and unknown types were assigned new identifiers. In a second round, the proteins that were not assigned a pfam in Uniprot and did not match another protein in the HMM library were collected and the whole set was then clustered with MMseqs2 (mmseqs easy‐cluster [52]). The clusters were then manually inspected and promising candidates were selected based on occurrence within a specific BMC type, distance to shell proteins and direction of translation. The clusters were then analyzed and HMMs were generated from those proteins to identify them in the locus analysis. The overall prevalence of BMCs in genomes was assessed by scoring 64,495 isolate genomes, 1667 single cell genomes (IMG/M [16], April 2020) and 9,331 metagenome assembled genomes (GEM [17], February 2021) with a shell protein HMM library using hmmsearch [50] with a cutoff of 1E‐20.

### Locus visualization

The HMM library was used to score each BMC type separately to generate type‐specific HMMs using only the sequence from one type. This type‐specific HMM was then used to score loci with hmmsearch and locus data was visualized using a python script (Supplementary Data: Locus diagrams). The directionality of the gene coding for the proteins is shown as an arrow. While this data is not present in Uniprot files it can be determined by extracting the EMBL database identifier and parsing a corresponding DNA file downloaded through the European Nucleotide Archive (ENA). To determine the presence of encapsulation peptides (EPs) we collected sequences of known proteins with EPs and used the EP portion to generate HMMs. A separate HMM was generated for each BMC locus type and protein family. The combined HMM library was then used to identify potential encapsulation peptides. Despite the low sequence conservation and short length of EPs this method is quite sensitive, however manual inspection of the results is still necessary.

### Shell protein phylogenies

An initial set of 6408 BMC‐P protein sequences was made non‐redundant to 70% identity using the usearch –cluster_fast algorithm (v.11.0 [53]), resulting in a set of 1183 unique sequences. An initial alignment with muscle [54] was manually edited upon visual inspection with Jalview [55] to prune fragments and problematic sequences likely arising from genome sequencing or gene modeling errors. Sequences were then realigned using MAFFT‐linsi [56] and uninformative columns were removed with BMGE (‐h 0.8 –g 0.05) [57]. ModelTest‐NG [58] was used to determine the best‐scoring substitution model, LG4M, which was then used to construct a maximum likelihood tree using RAxML‐NG (v0.6.0) [59]. A similar approach was used to collapse the redundancy of BMC‐H proteins (95% identity), BMC‐H^p^: (95%), BMC‐T^s^ (90%), BMC‐T^dp^ (90%), BMC‐T^sp^ (90%); higher thresholds for these were necessary because of the higher overall homology. Trees were examined using Archaeopteryx (www.phylosoft.org/archaeopteryx) and significant clades were manually identified based on a general criterion of having a long internal stem. They were then assigned unique color names from the XKCD color survey (https://xkcd.com/color/rgb/) and colored with their corresponding RGB hexcodes, selecting similar colors for nearby clades. The sequences from each color‐based clade were then subdivided by the initial BMC locus type assignments and used to generate subtype‐specific HMMs for scoring the entire BMC dataset. The phylogenies were later re‐examined with regards to final locus type assignments and annotated for functional correspondences to the clades or subclans (Figure 3). Vector‐quality images of the six phylogenies are provided as supplementary material with sequence identifiers and color assignments provided, for legibility upon manual zoom in a PDF reader (Supplementary Data: High resolution shell protein trees).

### Tree of bacterial phyla

A set of 56 universal single‐copy marker proteins [60, 61] was used to build a phylogenetic tree for the domain Bacteria based on a representative dataset that included one genome for each bacterial order present in the IMG/M database ([16]; accessed March 2020). Marker proteins were identified with hmmsearch (version 3.1b2) using a specific HMM for each marker. Genomes lacking a substantial proportion of marker proteins (more than 26) or which had additional copies of more than five single‐copy markers were removed from the dataset. For each marker, proteins were extracted, alignments built with MAFFT‐linsi (v7.294b, [62]) and subsequently trimmed with BMGE (v1.12, [57]) using BLOSUM30. Single protein alignments were then concatenated resulting in an alignment of 10,755 sites. Maximum likelihood phylogenies were inferred with FastTree2 [63] using the options: ‐spr 4 ‐mlacc 2 ‐slownni ‐wag. The phylogenetic tree was pruned to keep only 1 representative genome for each phylum and then visualized using the ete3 package [64].

## Supporting information

Supplementary Data: HMM library

Supplementary Data: High resolution shell protein trees

Supplementary Data: HMM names table

Supplementary Data: Locus diagrams

Supplementary Data: BMC types

## Acknowledgements

This work was supported by the National Institutes of Health, National Institute of Allergy and Infectious Diseases (NIAID) grant 1R01AI114975‐01 and the U.S. Department of Energy, Basic Energy Sciences, Contract DE‐FG02‐91ER20021. We thank Henning Kirst, Jan Zarzycki and Stephanie A. Eichorst for helpful discussions, and Emiley Eloe‐Fadrosh and Neha Varghese for custom HMM searches against the IMG/M database. Parts of this study were performed by the US Department of Energy Joint Genome Institute, a DOE Office of Science User Facility, under Contract No. DE‐AC02–05CH11231 and made use of resources of the National Energy Research Scientific Computing Center, which is supported by the Office of Science of the US Department of Energy under Contract No. DE‐AC02–05CH11231.

## Supplementary Files

Supplementary Figures and Tables

Supplementary Data: BMC types

Supplementary Data: Locus diagrams

Supplementary Data: High resolution shell protein trees

Supplementary Table: HMM names table

Supplementary Data: HMM library

## References

1. Drews, G. and W. Niklowitz, Beiträge zur Cytologie der Blaualgen. II. Zentroplasma und granuläre Einschlüsse von Phormidium uncinatum. Archiv für Mikrobiologie, 1956. 24(2): p. 147‐162.

2. Shively, J.M., F. Ball, D.H. Brown, and R.E. Saunders, Functional organelles in prokaryotes: polyhedral inclusions (carboxysomes) of Thiobacillus neapolitanus. Science, 1973. 182(4112): p. 584‐6.

3. Shively, J.M., C.E. Bradburne, H.C. Aldrich, T.A. Bobik, J.L. Mehlman, S. Jin, and S.H. Baker, Sequence homologs of the carboxysomal polypeptide CsoS1 of the thiobacilli are present in cyanobacteria and enteric bacteria that form carboxysomes ‐ polyhedral bodies. Canadian Journal of Botany‐Revue Canadienne De Botanique, 1998. 76(6): p. 906‐916.

4. Axen, S.D., O. Erbilgin, and C.A. Kerfeld, A taxonomy of bacterial microcompartment loci constructed by a novel scoring method. PLoS Comput Biol, 2014. 10(10): p. e1003898.

5. Jorda, J., D. Lopez, N.M. Wheatley, and T.O. Yeates, Using comparative genomics to uncover new kinds of protein‐based metabolic organelles in bacteria. Protein Science, 2013. 22(2): p. 179‐195.

6. Kerfeld, C.A. and O. Erbilgin, Bacterial microcompartments and the modular construction of microbial metabolism. Trends Microbiol, 2015. 23(1): p. 22‐34.

7. Kinney, J.N., A. Salmeen, F. Cai, and C.A. Kerfeld, Elucidating essential role of conserved carboxysomal protein CcmN reveals common feature of bacterial microcompartment assembly. J Biol Chem, 2012. 287(21): p. 17729‐36.

8. Aussignargues, C., B.C. Paasch, R. Gonzalez‐Esquer, O. Erbilgin, and C.A. Kerfeld, Bacterial microcompartment assembly: The key role of encapsulation peptides. Commun Integr Biol, 2015. 8(3): p. e1039755.

9. Kerfeld, C.A., C. Aussignargues, J. Zarzycki, F. Cai, and M. Sutter, Bacterial microcompartments. Nat Rev Microbiol, 2018. 16(5): p. 277‐290.

10. Sutter, M., B. Greber, C. Aussignargues, and C.A. Kerfeld, Assembly principles and structure of a 6.5‐MDa bacterial microcompartment shell. Science, 2017. 356(6344): p. 1293‐1297.

11. Kalnins, G., E.E. Cesle, J. Jansons, J. Liepins, A. Filimonenko, and K. Tars, Encapsulation mechanisms and structural studies of GRM2 bacterial microcompartment particles. Nat Commun, 2020. 11(1): p. 388.

12. Greber, B.J., M. Sutter, and C.A. Kerfeld, The Plasticity of Molecular Interactions Governs Bacterial Microcompartment Shell Assembly. Structure, 2019. 27(5): p. 749‐763 e4.

13. Sutter, M., T.G. Laughlin, N.B. Sloan, D. Serwas, K.M. Davies, and C.A. Kerfeld, Structure of a Synthetic beta‐Carboxysome Shell. Plant Physiol, 2019. 181(3): p. 1050‐1058.

14. Tanaka, S., C.A. Kerfeld, M.R. Sawaya, F. Cai, S. Heinhorst, G.C. Cannon, and T.O. Yeates, Atomic‐level models of the bacterial carboxysome shell. Science, 2008. 319(5866): p. 1083‐6.

15. Kerfeld, C.A., M.R. Sawaya, S. Tanaka, C.V. Nguyen, M. Phillips, M. Beeby, and T.O. Yeates, Protein structures forming the shell of primitive bacterial organelles. Science, 2005. 309(5736): p. 936‐8.

16. Chen, I.A., K. Chu, K. Palaniappan, A. Ratner, J. Huang, M. Huntemann, P. Hajek, S. Ritter, N. Varghese, R. Seshadri, S. Roux, T. Woyke, E.A. Eloe‐Fadrosh, N.N. Ivanova, and N.C. Kyrpides, The IMG/M data management and analysis system v.6.0: new tools and advanced capabilities. Nucleic Acids Res, 2021. 49(D1): p. D751‐D763.

17. Nayfach, S., S. Roux, R. Seshadri, D. Udwary, N. Varghese, F. Schulz, D. Wu, D. Paez‐Espino, I.M. Chen, M. Huntemann, K. Palaniappan, J. Ladau, S. Mukherjee, T.B.K. Reddy, T. Nielsen, E. Kirton, J.P. Faria, J.N. Edirisinghe, C.S. Henry, S.P. Jungbluth, D. Chivian, P. Dehal, E.M. Wood‐Charlson, A.P. Arkin, S.G. Tringe, A. Visel, I.M.D. Consortium, T. Woyke, N.J. Mouncey, N.N. Ivanova, N.C. Kyrpides, and E.A. Eloe‐Fadrosh, A genomic catalog of Earth’s microbiomes. Nat Biotechnol, 2020.

18. Sjolander, K., Phylogenomic inference of protein molecular function: advances and challenges. Bioinformatics, 2004. 20(2): p. 170‐9.

19. Krogh, A., M. Brown, I.S. Mian, K. Sjolander, and D. Haussler, Hidden Markov models in computational biology. Applications to protein modeling. J Mol Biol, 1994. 235(5): p. 1501‐31.

20. Waterhouse, A., M. Bertoni, S. Bienert, G. Studer, G. Tauriello, R. Gumienny, F.T. Heer, T.A.P. de Beer, C. Rempfer, L. Bordoli, R. Lepore, and T. Schwede, SWISS‐MODEL: homology modelling of protein structures and complexes. Nucleic Acids Res, 2018. 46(W1): p. W296‐W303.

21. Zhang, R.G., C.E. Andersson, T. Skarina, E. Evdokimova, A.M. Edwards, A. Joachimiak, A. Savchenko, and S.L. Mowbray, The 2.2 A resolution structure of RpiB/AlsB from Escherichia coli illustrates a new approach to the ribose‐5‐phosphate isomerase reaction. J Mol Biol, 2003. 332(5): p. 1083‐94.

22. Aussignargues, C., M.E. Pandelia, M. Sutter, J.S. Plegaria, J. Zarzycki, A. Turmo, J. Huang, D.C. Ducat, E.L. Hegg, B.R. Gibney, and C.A. Kerfeld, Structure and Function of a Bacterial Microcompartment Shell Protein Engineered to Bind a [4Fe‐4S] Cluster. Journal of the American Chemical Society, 2016. 138(16): p. 5262‐70.

23. Mohr, K.I., Diversity of Myxobacteria‐We Only See the Tip of the Iceberg. Microorganisms, 2018. 6(3).

24. Erbilgin, O., K.L. McDonald, and C.A. Kerfeld, Characterization of a planctomycetal organelle: a novel bacterial microcompartment for the aerobic degradation of plant saccharides. Appl Environ Microbiol, 2014. 80(7): p. 2193‐205.

25. Sizikov, S., I. Burgsdorf, K.M. Handley, M. Lahyani, M. Haber, and L. Steindler, Characterization of sponge‐associated Verrucomicrobia: microcompartment‐based sugar utilization and enhanced toxin‐antitoxin modules as features of host‐associated Opitutales. Environ Microbiol, 2020. 22(11): p. 4669‐4688.

26. Sichert, A., C.H. Corzett, M.S. Schechter, F. Unfried, S. Markert, D. Becher, A. Fernandez‐Guerra, M. Liebeke, T. Schweder, M.F. Polz, and J.H. Hehemann, Verrucomicrobia use hundreds of enzymes to digest the algal polysaccharide fucoidan. Nature Microbiology, 2020. 5(8): p. 1026‐+.

27. Losey, N.A., B.S. Stevenson, H.J. Busse, J.S.S. Damste, W.I.C. Rijpstra, S. Rudd, and P.A. Lawson, Thermoanaerobaculum aquaticum gen. nov., sp nov., the first cultivated member of Acidobacteria subdivision 23, isolated from a hot spring. International Journal of Systematic and Evolutionary Microbiology, 2013. 63: p. 4149‐4157.

28. Fierer, N., Embracing the unknown: disentangling the complexities of the soil microbiome. Nature Reviews Microbiology, 2017. 15(10): p. 579‐590.

29. Sturms, R., N.A. Streauslin, S. Cheng, and T.A. Bobik, In Salmonella enterica, Ethanolamine Utilization Is Repressed by 1,2‐Propanediol To Prevent Detrimental Mixing of Components of Two Different Bacterial Microcompartments. J Bacteriol, 2015. 197(14): p. 2412‐21.

30. Fendrich, C., H. Hippe, and G. Gottschalk, Clostridium‐Halophilium Sp‐Nov and Clostridium‐Litorale Sp‐Nov, an Obligate Halophilic and a Marine Species Degrading Betaine in the Stickland Reaction. Archives of Microbiology, 1990. 154(2): p. 127‐132.

31. Flint, H.J., K.P. Scott, S.H. Duncan, P. Louis, and E. Forano, Microbial degradation of complex carbohydrates in the gut. Gut Microbes, 2012. 3(4): p. 289‐306.

32. Sommer, M., M. Sutter, S. Gupta, H. Kirst, A. Turmo, S. Lechno‐Yossef, R.L. Burton, C. Saechao, N.B. Sloan, X. Cheng, L.G. Chan, C.J. Petzold, M. Fuentes‐Cabrera, C.Y. Ralston, and C.A. Kerfeld, Heterohexamers Formed by CcmK3 and CcmK4 Increase the Complexity of Beta Carboxysome Shells. Plant Physiology, 2019. 179(1): p. 156.

33. Sommer, M., F. Cai, M. Melnicki, and C.A. Kerfeld, beta‐Carboxysome bioinformatics: identification and evolution of new bacterial microcompartment protein gene classes and core locus constraints. J Exp Bot, 2017. 68(14): p. 3841‐3855.

34. Xia, Y., Y.B. Wang, Y. Wang, F.Y.L. Chin, and T. Zhang, Cellular adhesiveness and cellulolytic capacity in Anaerolineae revealed by omics‐based genome interpretation. Biotechnology for Biofuels, 2016. 9.

35. Cai, F., S.L. Bernstein, S.C. Wilson, and C.A. Kerfeld, Production and Characterization of Synthetic Carboxysome Shells with Incorporated Luminal Proteins. Plant Physiol, 2016. 170(3): p. 1868‐77.

36. Zarzycki, J., O. Erbilgin, and C.A. Kerfeld, Bioinformatic characterization of glycyl radical enzyme‐associated bacterial microcompartments. Appl Environ Microbiol, 2015. 81(24): p. 8315‐29.

37. Cai, F., M. Sutter, S.L. Bernstein, J.N. Kinney, and C.A. Kerfeld, Engineering Bacterial Microcompartment Shells: Chimeric Shell Proteins and Chimeric Carboxysome Shells. ACS Synthetic Biology, 2015. 4(4): p. 444‐453.

38. Slininger Lee, M.F., C.M. Jakobson, and D. Tullman‐Ercek, Evidence for Improved Encapsulated Pathway Behavior in a Bacterial Microcompartment through Shell Protein Engineering. ACS Synthetic Biology, 2017.

39. Frank, S., A.D. Lawrence, M.B. Prentice, and M.J. Warren, Bacterial microcompartments moving into a synthetic biological world. Journal of Biotechnology, 2013. 163(2): p. 273‐279.

40. Kerfeld, C.A. and M. Sutter, Engineered bacterial microcompartments: apps for programming metabolism. Current Opinion in Biotechnology, 2020. 65: p. 225‐232.

41. Kirst, H. and C.A. Kerfeld, Bacterial microcompartments: catalysis‐enhancing metabolic modules for next generation metabolic and biomedical engineering. Bmc Biology, 2019. 17(1).

42. Ferlez, B., M. Sutter, and C.A. Kerfeld, Glycyl Radical Enzyme‐Associated Microcompartments: Redox‐Replete Bacterial Organelles. mBio, 2019. 10(1).

43. Castelle, C.J. and J.F. Banfield, Major New Microbial Groups Expand Diversity and Alter our Understanding of the Tree of Life. Cell, 2018. 172(6): p. 1181‐1197.

44. Flores‐Mireles, A.L., J.N. Walker, M. Caparon, and S.J. Hultgren, Urinary tract infections: epidemiology, mechanisms of infection and treatment options. Nature Reviews Microbiology, 2015. 13(5): p. 269‐284.

45. Klein, M.G., P. Zwart, S.C. Bagby, F. Cai, S.W. Chisholm, S. Heinhorst, G.C. Cannon, and C.A. Kerfeld, Identification and structural analysis of a novel carboxysome shell protein with implications for metabolite transport. Journal of Molecular Biology, 2009. 392(2): p. 319‐33.

46. Cai, F., M. Sutter, J.C. Cameron, D.N. Stanley, J.N. Kinney, and C.A. Kerfeld, The Structure of CcmP, a Tandem Bacterial Microcompartment Domain Protein from the beta‐Carboxysome, Forms a Subcompartment Within a Microcompartment. Journal of Biological Chemistry, 2013. 288(22): p. 16055‐16063.

47. Larsson, A.M., D. Hasse, K. Valegard, and I. Andersson, Crystal structures of beta‐carboxysome shell protein CcmP: ligand binding correlates with the closed or open central pore. J Exp Bot, 2017. 68(14): p. 3857‐3867.

48. Larkin, M.A., G. Blackshields, N.P. Brown, R. Chenna, P.A. McGettigan, H. McWilliam, F. Valentin, I.M. Wallace, A. Wilm, R. Lopez, J.D. Thompson, T.J. Gibson, and D.G. Higgins, Clustal W and clustal X version 2.0. Bioinformatics, 2007. 23(21): p. 2947‐2948.

49. Capella‐Gutierrez, S.J. M. Silla‐Martinez, and T. Gabaldon, trimAl: a tool for automated alignment trimming in large‐scale phylogenetic analyses. Bioinformatics, 2009. 25(15): p. 1972‐1973.

50. Eddy, S.R., Accelerated Profile HMM Searches. Plos Computational Biology, 2011. 7(10).

51. Shannon, P., A. Markiel, O. Ozier, N.S. Baliga, J.T. Wang, D. Ramage, N. Amin, B. Schwikowski, and T. Ideker, Cytoscape: A software environment for integrated models of biomolecular interaction networks. Genome Research, 2003. 13(11): p. 2498‐2504.

52. Steinegger, M. and J. Soding, MMseqs2 enables sensitive protein sequence searching for the analysis of massive data sets. Nature Biotechnology, 2017. 35(11): p. 1026‐1028.

53. Edgar, R.C., Search and clustering orders of magnitude faster than BLAST. Bioinformatics, 2010. 26(19): p. 2460‐2461.

54. Edgar, R.C., MUSCLE: multiple sequence alignment with high accuracy and high throughput. Nucleic Acids Research, 2004. 32(5): p. 1792‐1797.

55. Waterhouse, A.M., J.B. Procter, D.M.A. Martin, M. Clamp, and G.J. Barton, Jalview Version 2‐a multiple sequence alignment editor and analysis workbench. Bioinformatics, 2009. 25(9): p. 1189‐ 1191.

56. Yamada, K.D., K. Tomii, and K. Katoh, Application of the MAFFT sequence alignment program to large data‐reexamination of the usefulness of chained guide trees. Bioinformatics, 2016. 32(21): p. 3246‐3251.

57. Criscuolo, A. and S. Gribaldo, BMGE (Block Mapping and Gathering with Entropy): a new software for selection of phylogenetic informative regions from multiple sequence alignments. BMC Evol Biol, 2010. 10: p. 210.

58. Darriba, D., D. Posada, A.M. Kozlov, A. Stamatakis, B. Morel, and T. Flouri, ModelTest‐NG: A New and Scalable Tool for the Selection of DNA and Protein Evolutionary Models. Molecular Biology and Evolution, 2020. 37(1): p. 291‐294.

59. Kozlov, A.M., D. Darriba, T. Flouri, B. Morel, and A. Stamatakis, RAxML‐NG: a fast, scalable and user‐friendly tool for maximum likelihood phylogenetic inference. Bioinformatics, 2019. 35(21): p. 4453‐4455.

60. Yu, F.B., P.C. Blainey, F. Schulz, T. Woyke, M.A. Horowitz, and S.R. Quake, Microfluidic‐based mini‐metagenomics enables discovery of novel microbial lineages from complex environmental samples. Elife, 2017. 6.

61. Eloe‐Fadrosh, E.A., D. Paez‐Espino, J. Jarett, P.F. Dunfield, B.P. Hedlund, A.E. Dekas, S.E. Grasby, A.L. Brady, H. Dong, B.R. Briggs, W.J. Li, D. Goudeau, R. Malmstrom, A. Pati, J. Pett‐Ridge, E.M. Rubin, T. Woyke, N.C. Kyrpides, and N.N. Ivanova, Global metagenomic survey reveals a new bacterial candidate phylum in geothermal springs. Nat Commun, 2016. 7: p. 10476.

62. Katoh, K. and D.M. Standley, A simple method to control over‐alignment in the MAFFT multiple sequence alignment program. Bioinformatics, 2016. 32(13): p. 1933‐42.

63. Price, M.N., P.S. Dehal, and A.P. Arkin, FastTree 2‐‐approximately maximum‐likelihood trees for large alignments. PLoS One, 2010. 5(3): p. e9490.

64. Huerta‐Cepas, J., F. Serra, and P. Bork, ETE 3: Reconstruction, Analysis, and Visualization of Phylogenomic Data. Mol Biol Evol, 2016. 33(6): p. 1635‐8.

