## Supplementary Data: High resolution shell protein trees for "A Catalog of the Diversity and Ubiquity of Metabolic Organelles in Bacteria"

The six shell protein phylogenies are presented here as zoomable vector-quality images, in order to legibly present sequence identifiers and other relevant annotations. All sequence labels begin with the genomic locus tag, which serves as a unique identifier, followed by a curated taxonomic identifier for the source genome (typically the genus, strain ID, locus-prefix (if necessary), and phylum (provided in parentheses)). The assigned BMC locus type is then indicated ("bmcType="), followed by a qualifier in parentheses to indicate whether the shell protein is part of the "main" locus or is encoded on a satellite locus ("sat"). Sequences which were later omitted from the analysis during subsequent iterations of the BMC locus-locus scoring procedure are indicated ("omitted"); for these shell proteins, the BMC type reported corresponds to the initial assignments and thus may not be accurate, or may contain unresolvable locus type categories (e.g., UA for unassigned, PDU_EUT, EUT2_GRM1, etc). The amino acid length of the full sequence is also provided, followed by the uniprot ID (enclosed in square-brackets). When available, a curated gene "symbol" is provided to help point out functionally-characterized genes and their homologs. A four-character PDB code is also provided for solved 3D structures, or their closest sequence homolog (when PDB code is followed by an apostrophe). Finally, the original color name (corresponding to the RGB hexcodes from the XKCD color survey (https://xkcd.com/color/rgb/), as reported within the Python package, Seaborn), followed by the numerical "subclan" division, when available.

Contents:


| | | |
|---|---|---|
| p. 2 | BMC-P | pentamers |
| p. 3 | BMC-H | standard hexamers |
| p. 4 | BMC-Hp | permuted hexamers (GrpU/EutS/PduU/CutR-related) |
| p. 5 | BMC-Ts | standard tandem-domains (PduT/CcmO/Hoch5812-related) |
| p. 6 | BMC-Tsp | permuted tandem-domains (PduB/EutL-related) |
| p. 7 | BMC-Tdp | dimeric permuted tandem-domains (CcmP/CsoS1D/MSM0271+MSM0275/Hoch3341/Hoch5816-related) |

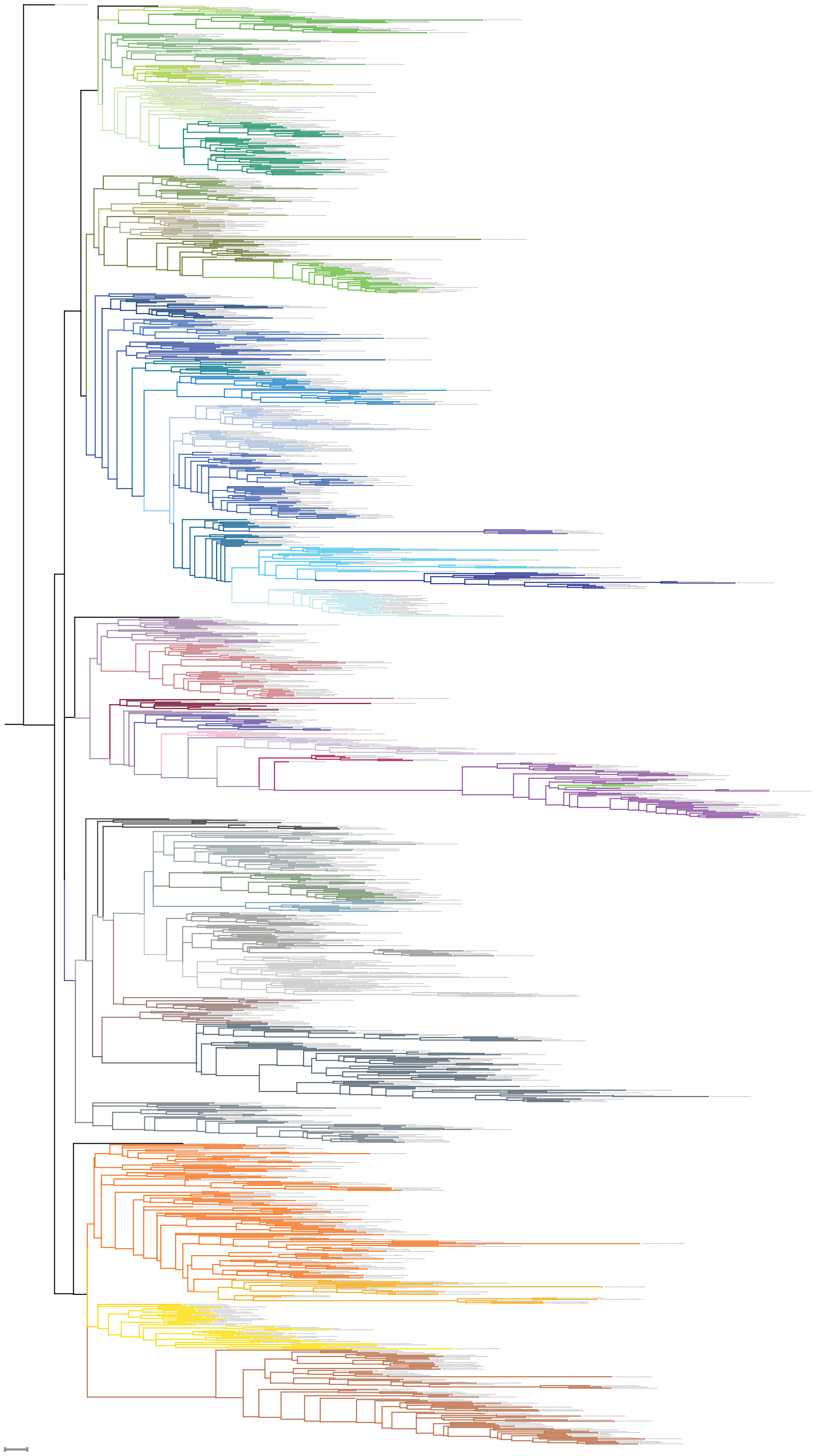

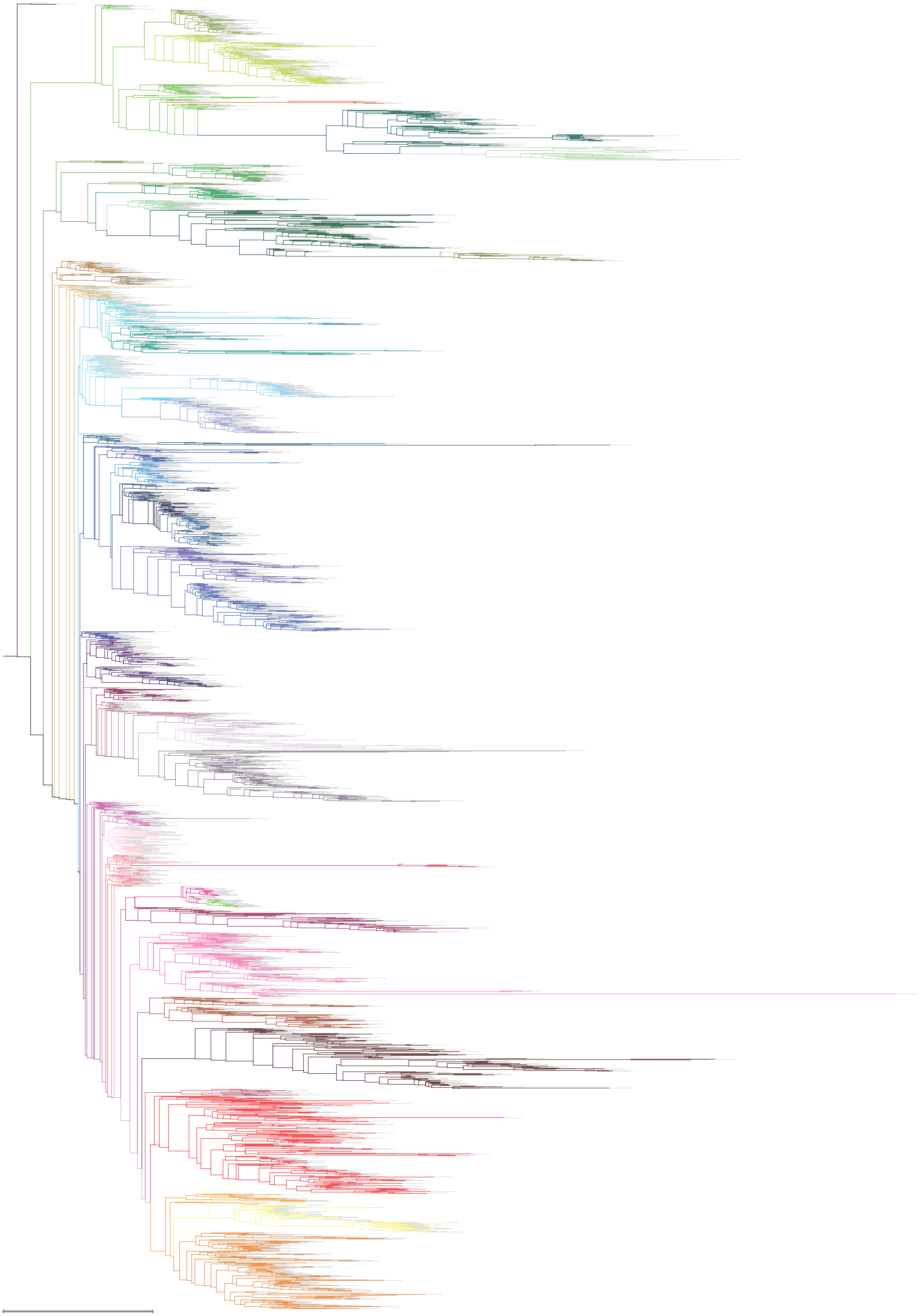

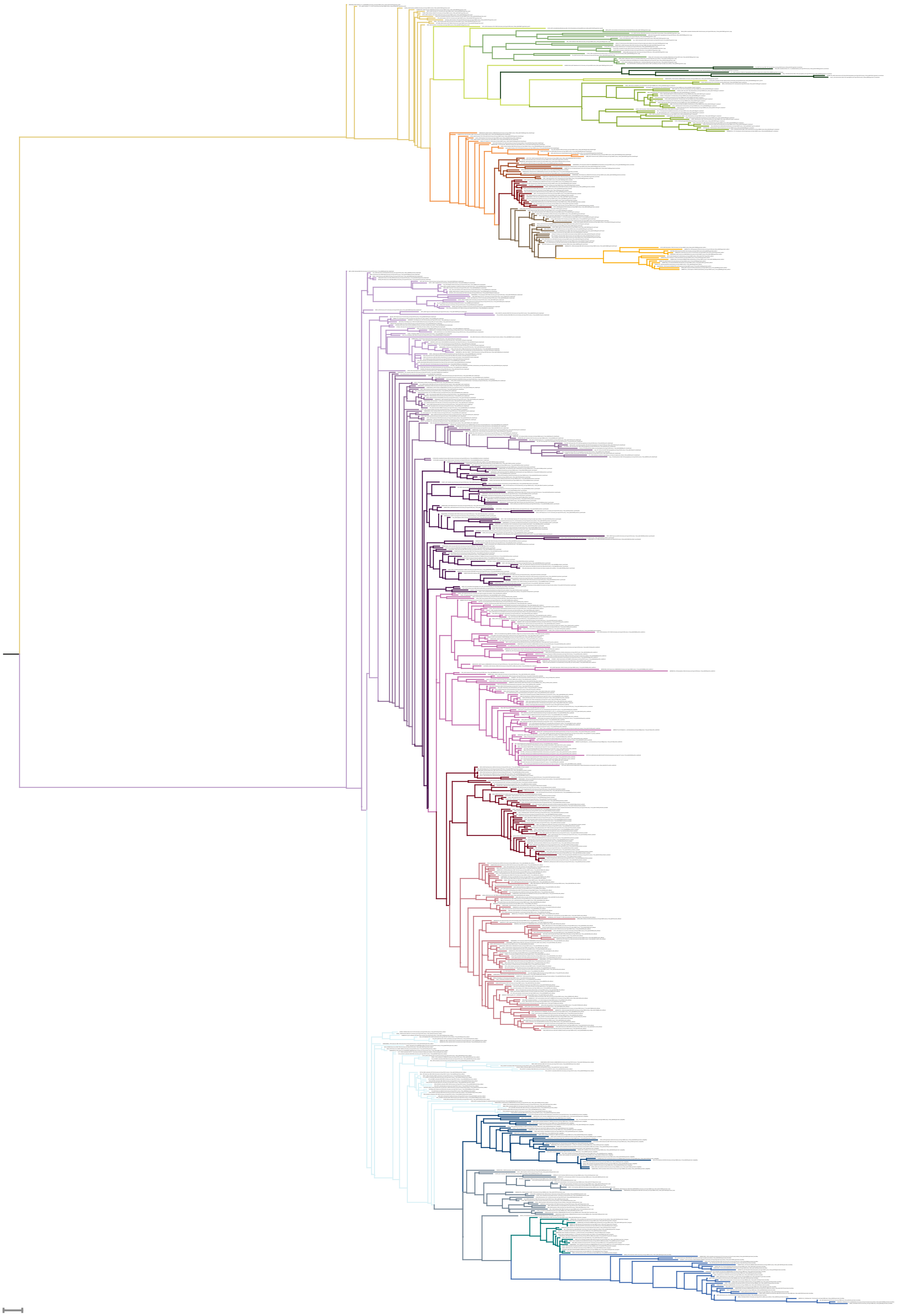

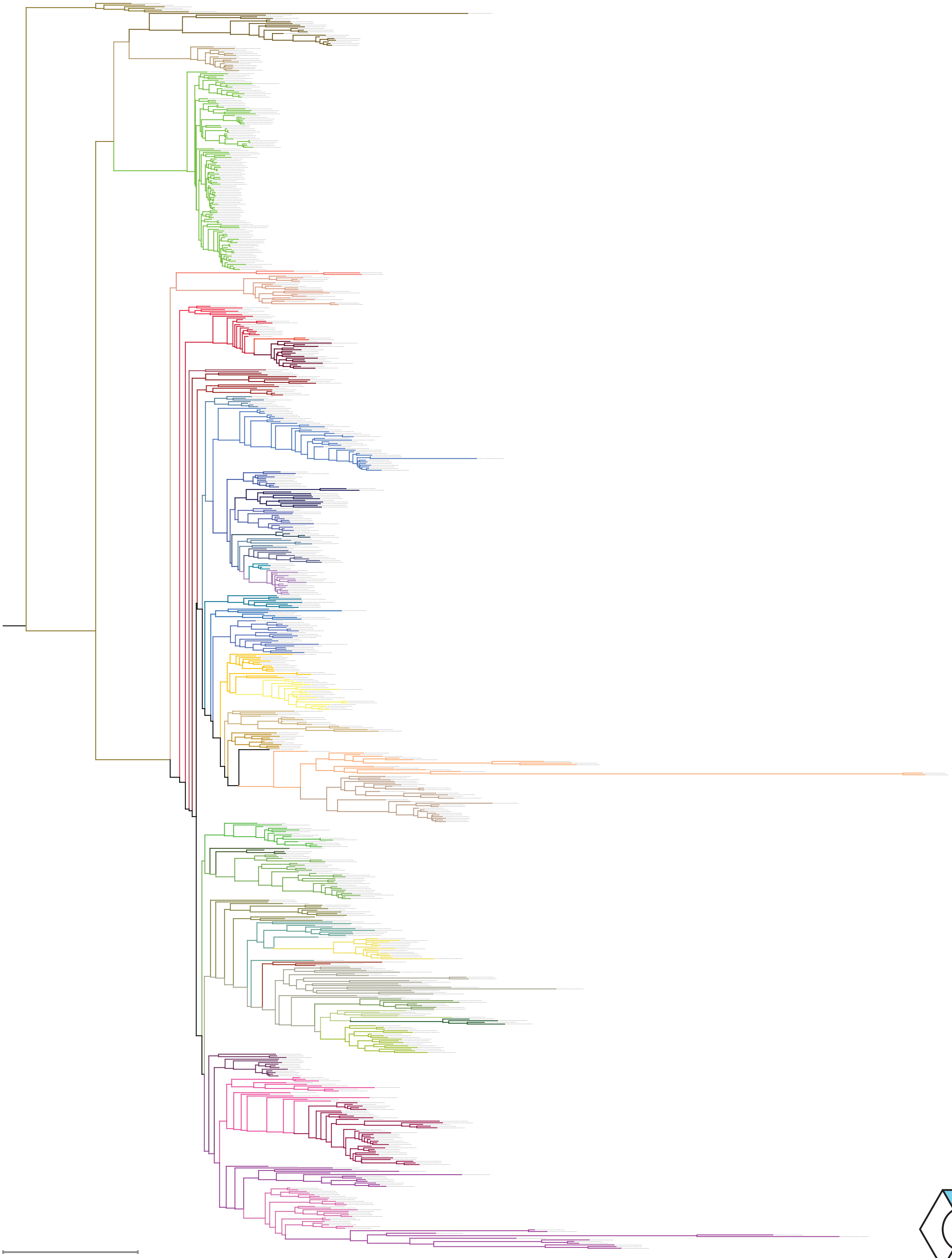

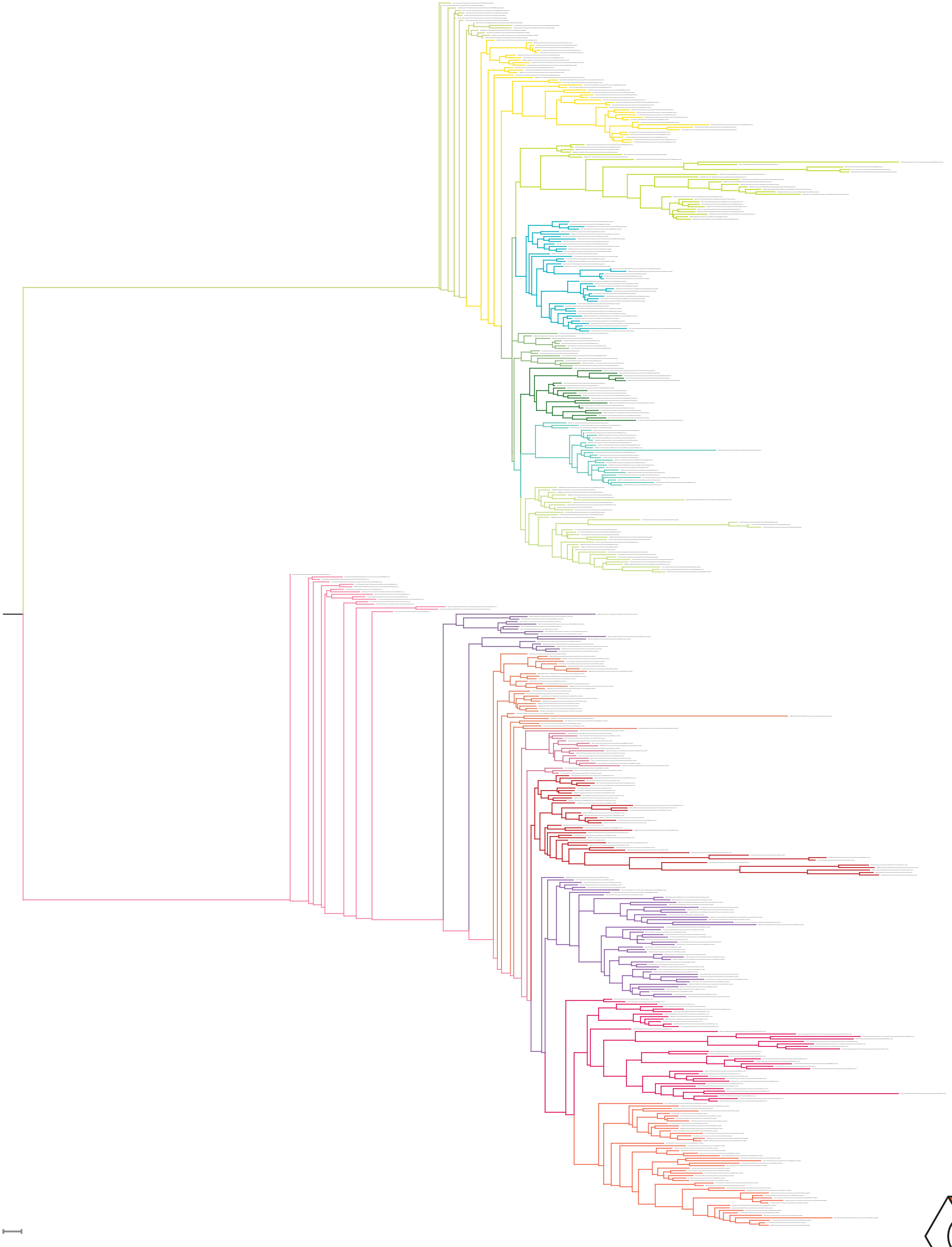

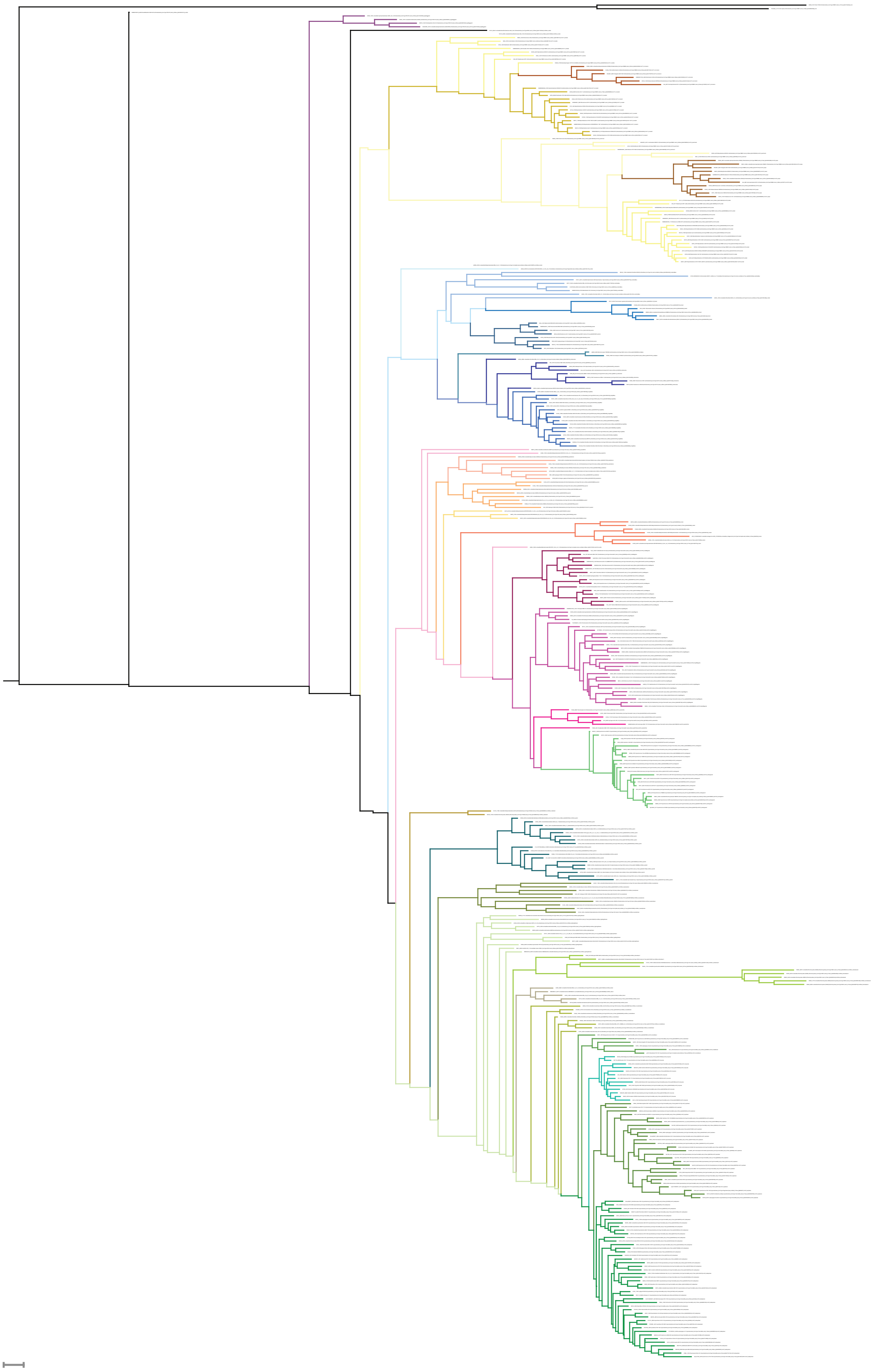
