## Supplementary Data: Locus diagrams for "A Catalog of the Diversity and Ubiquity of Metabolic Organelles in Bacteria"

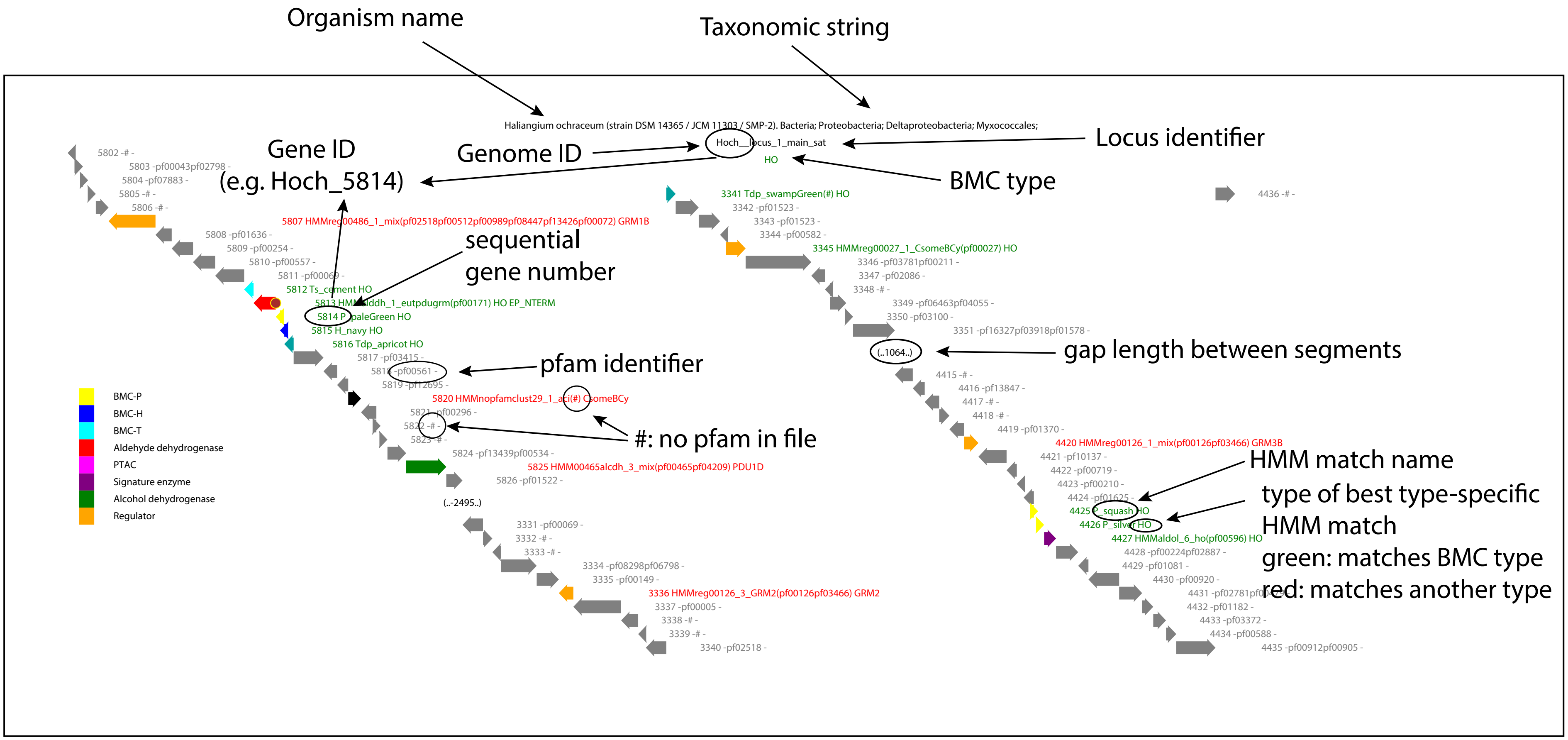

Acidobacteria bacterium RIFCSPHIGHO2_02_FULL_67_57. Bacteria; Acidobacteria.

A3B65__locus_1_main_sat

ACI

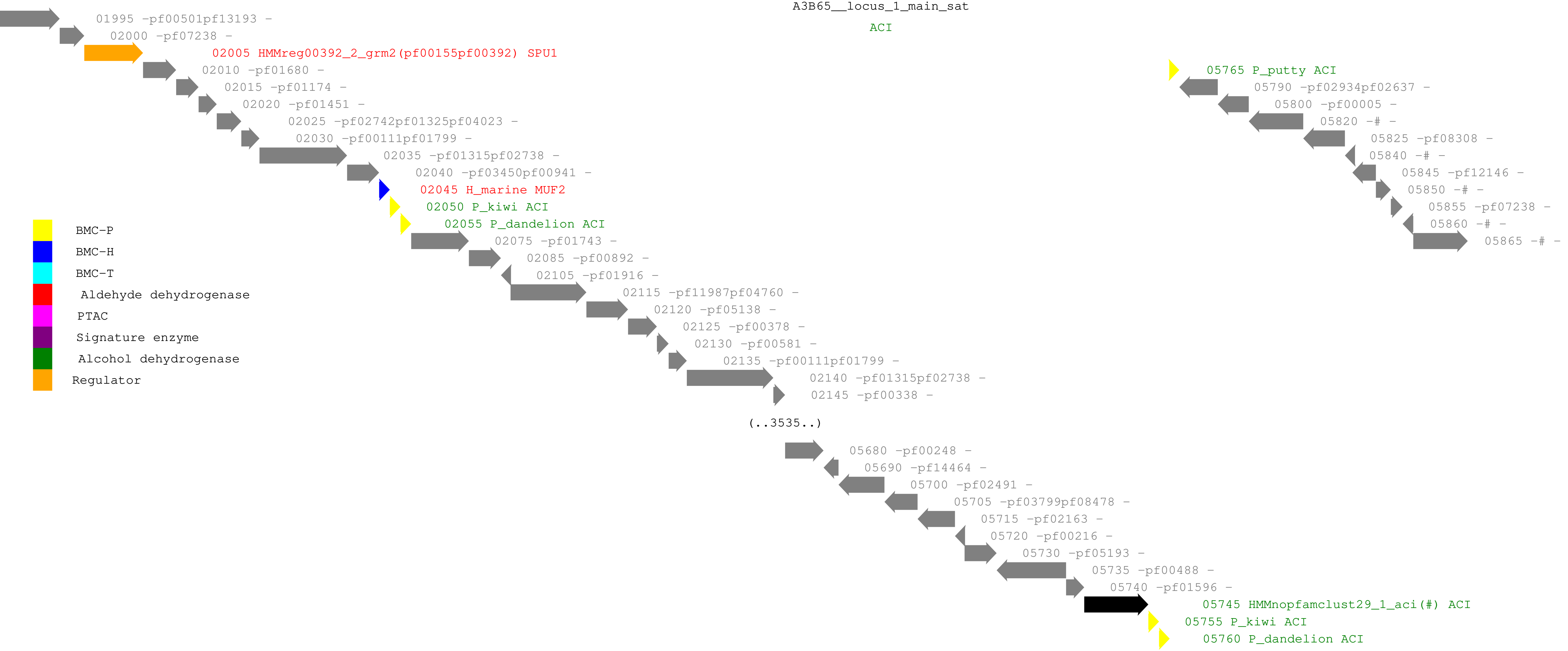

Acidobacteria bacterium RIFCSPHIGHO2_12_FULL_67_30. Bacteria; Acidobacteria.

A3D93__locus_1

ACI

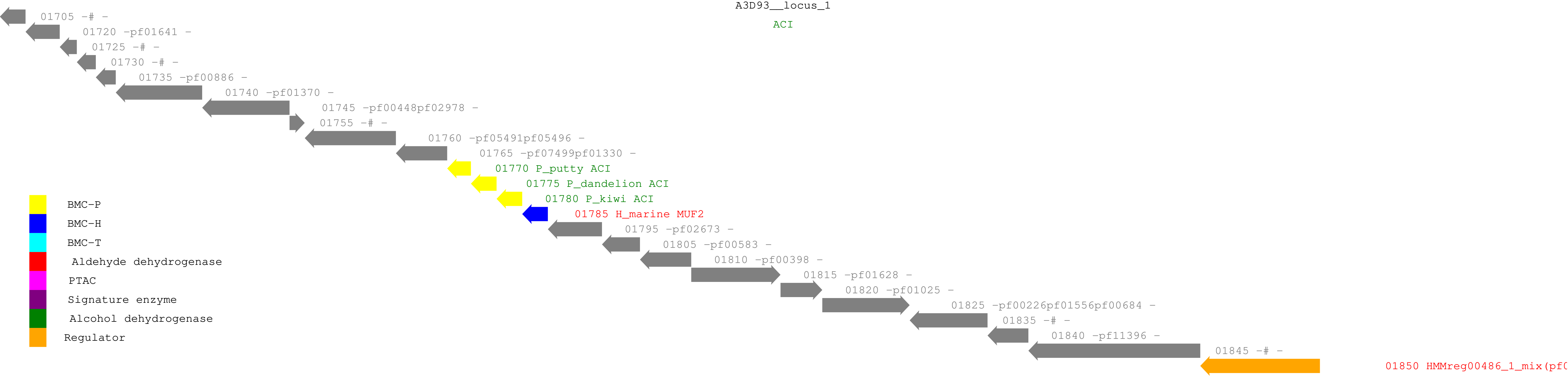

Acidobacteria bacterium RIFCSPLOWO2_12_FULL_66_21. Bacteria; Acidobacteria.

A3G21__locus_1_main_sat

ACI

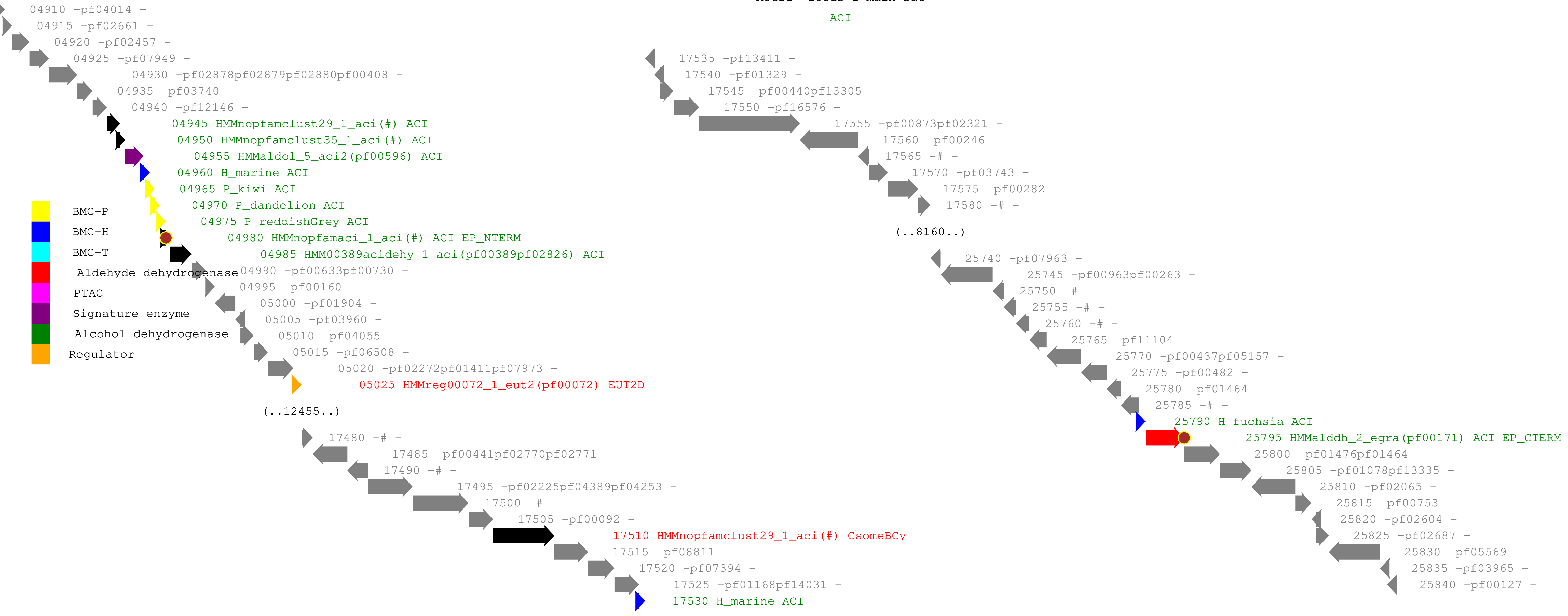

Acidobacteria bacterium RIFCSPLOWO2_02_FULL_67_21. Bacteria; Acidobacteria.

A3H29__locus_1_main_sat

ACI

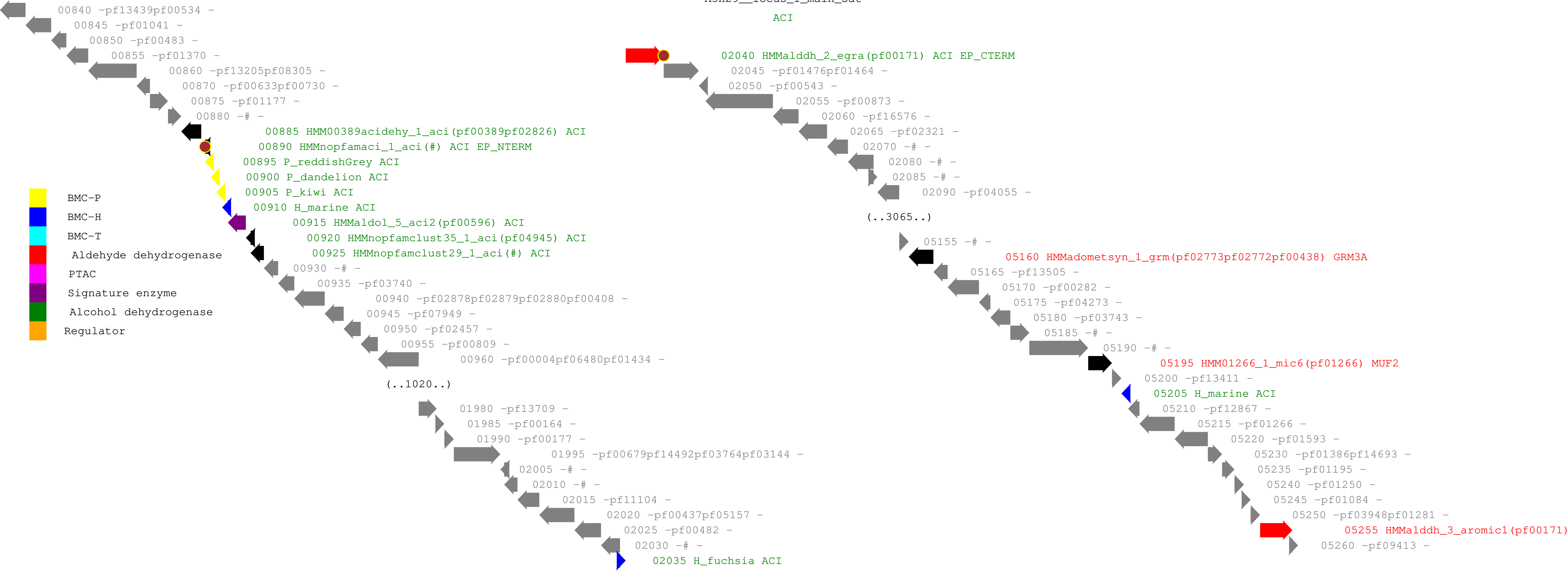

Acidobacteria bacterium RIFCSPLOWO2_02_FULL_64_15. Bacteria; Acidobacteria.

A3H95__locus_1_main_sat

ACI

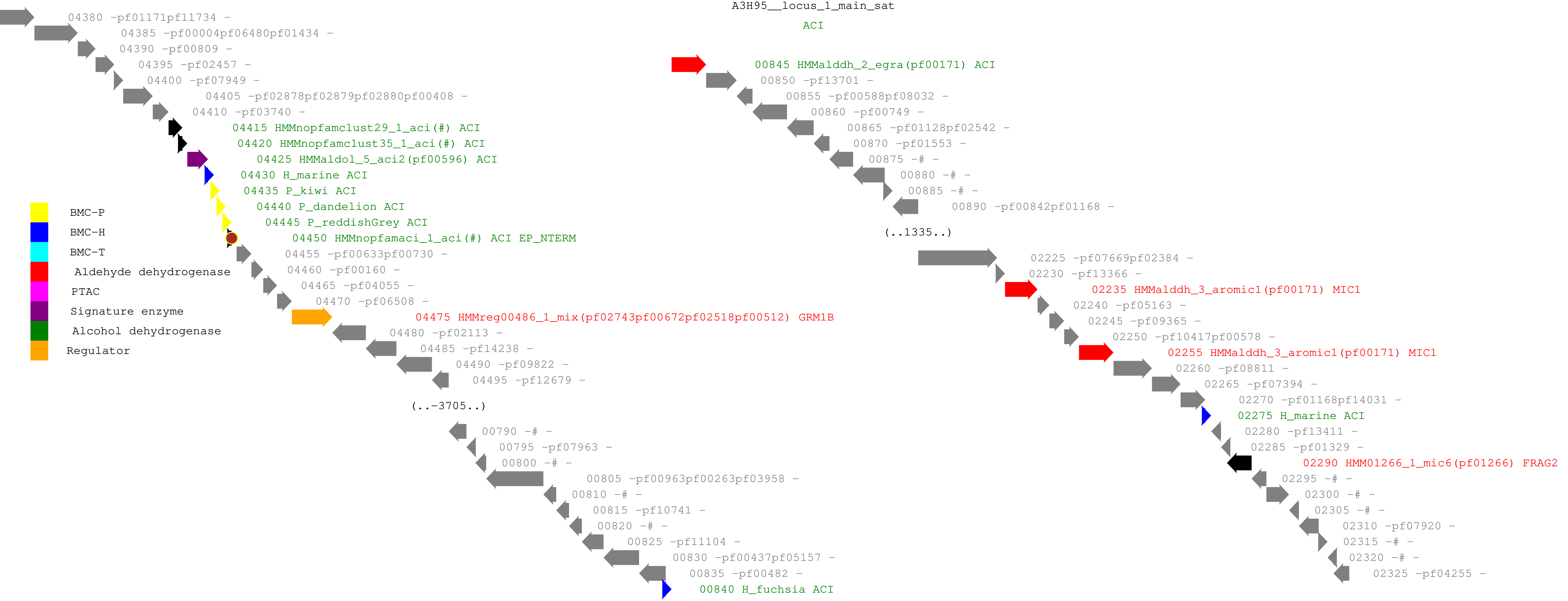

Acidobacteria bacterium RIFCSPLOWO2_02_FULL_65_29. Bacteria; Acidobacteria.

A3H97__locus_1_main_sat

ACI

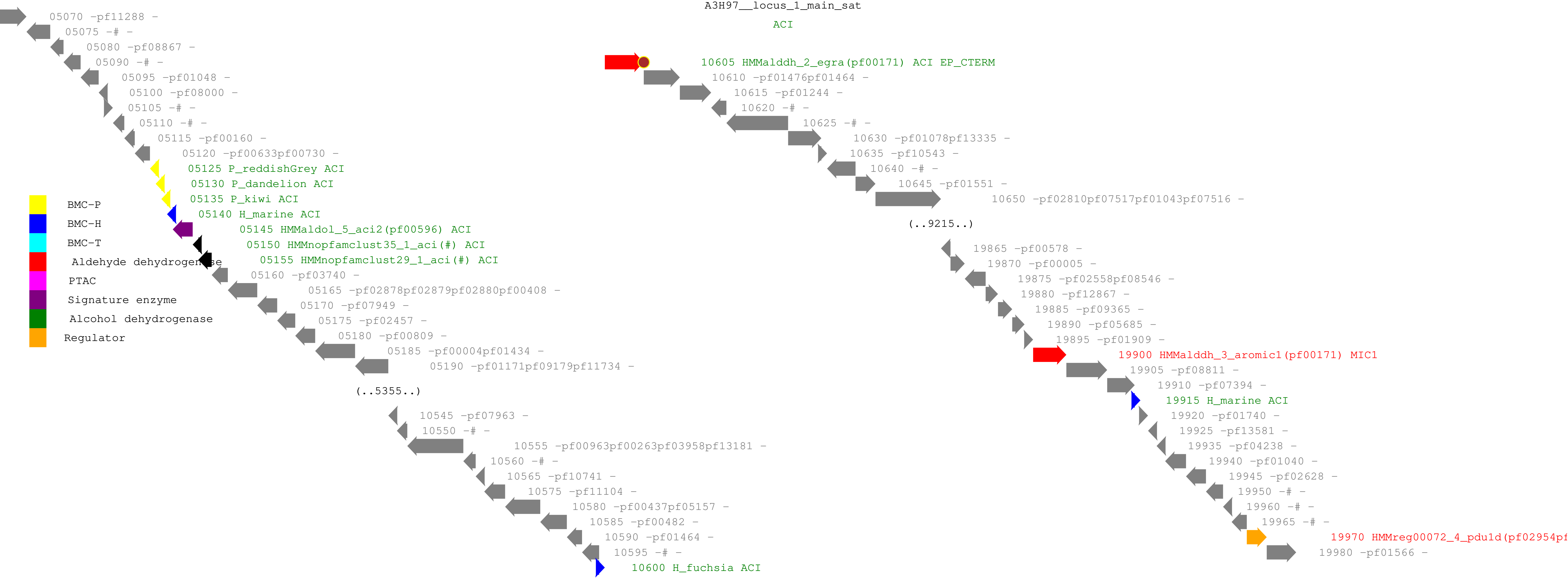

Acidobacteria bacterium RIFCSPLOWO2_02_FULL_68_18. Bacteria; Acidobacteria.

A3I61__locus_1_main_sat

ACI

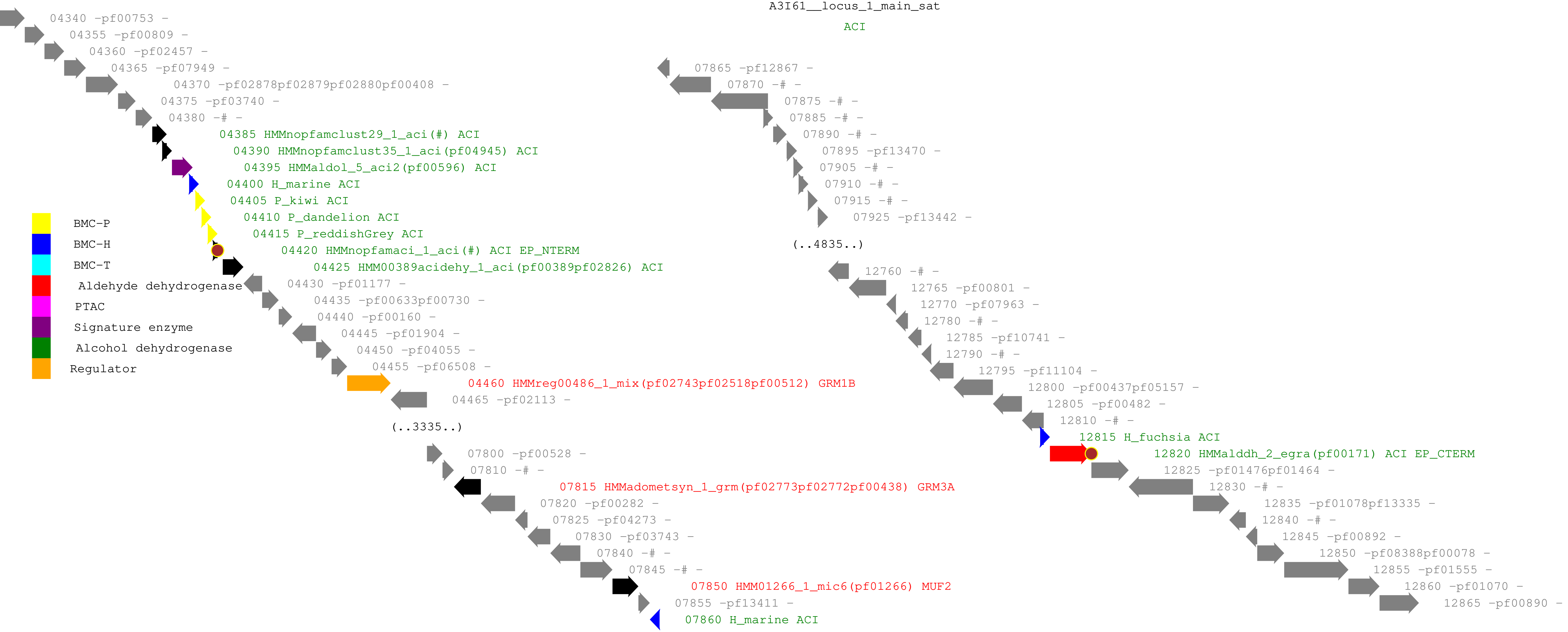

Acidobacteria bacterium RIFCSPLOWO2_12_FULL_67_14b. Bacteria; Acidobacteria.

A3J29__locus_1_main_sat

ACI

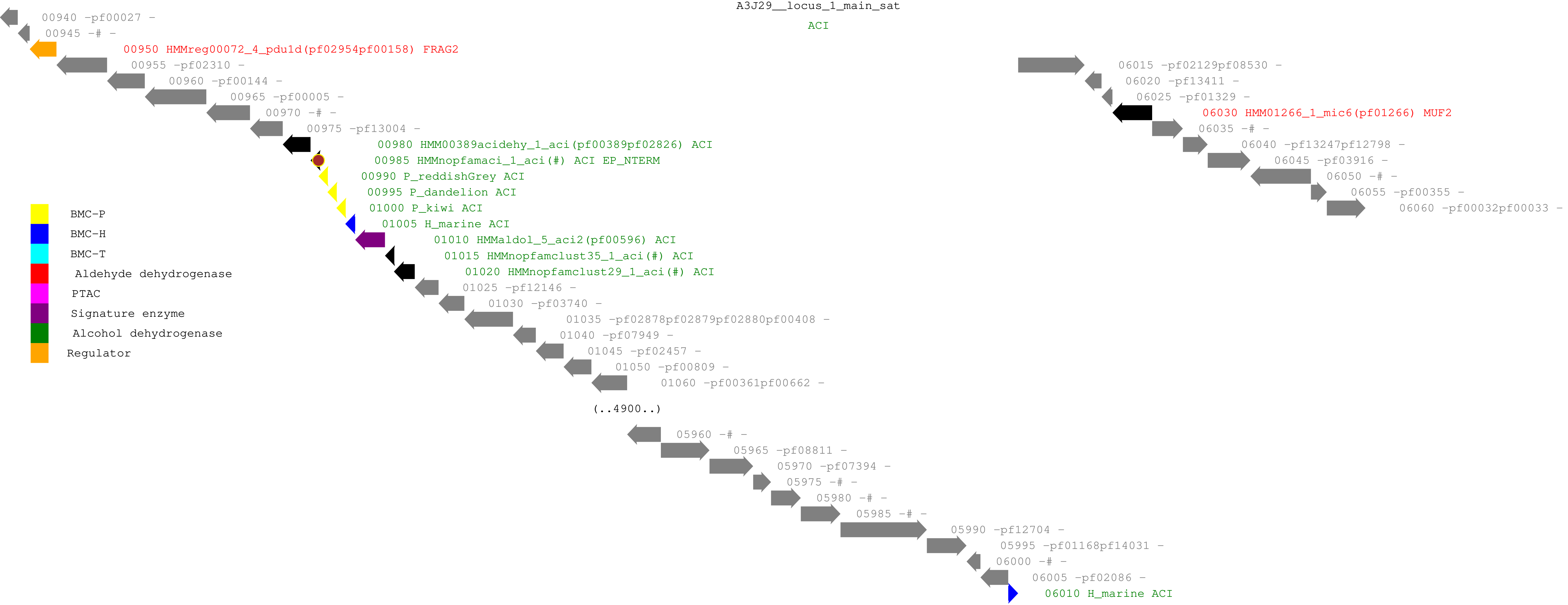

Acidobacteria bacterium SCN 69-37. Bacteria; Acidobacteria.

ABS36__locus_1_main_sat

ACI

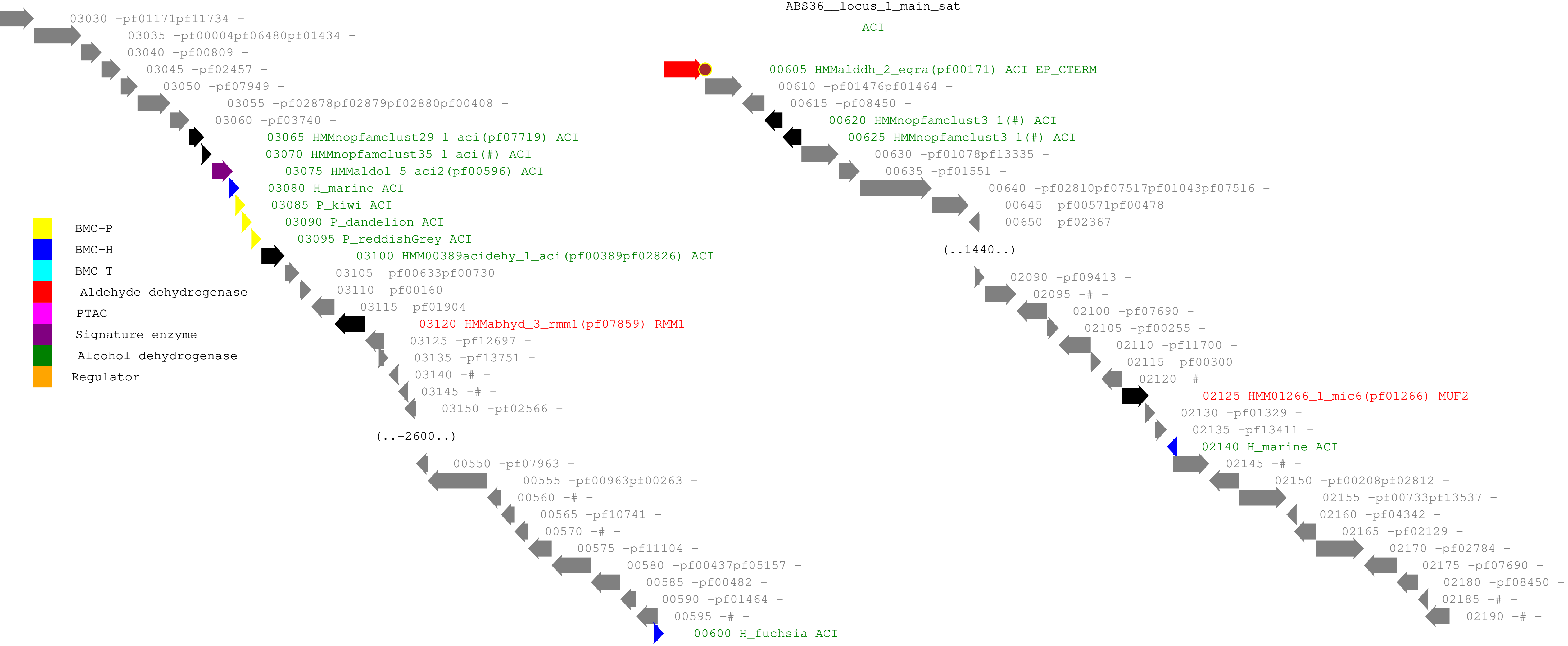

Solibacter usitatus (strain Ellin6076). Bacteria; Acidobacteria; Solibacteres; Solibacterales;

Acid__locus_1_main_sat

ACI

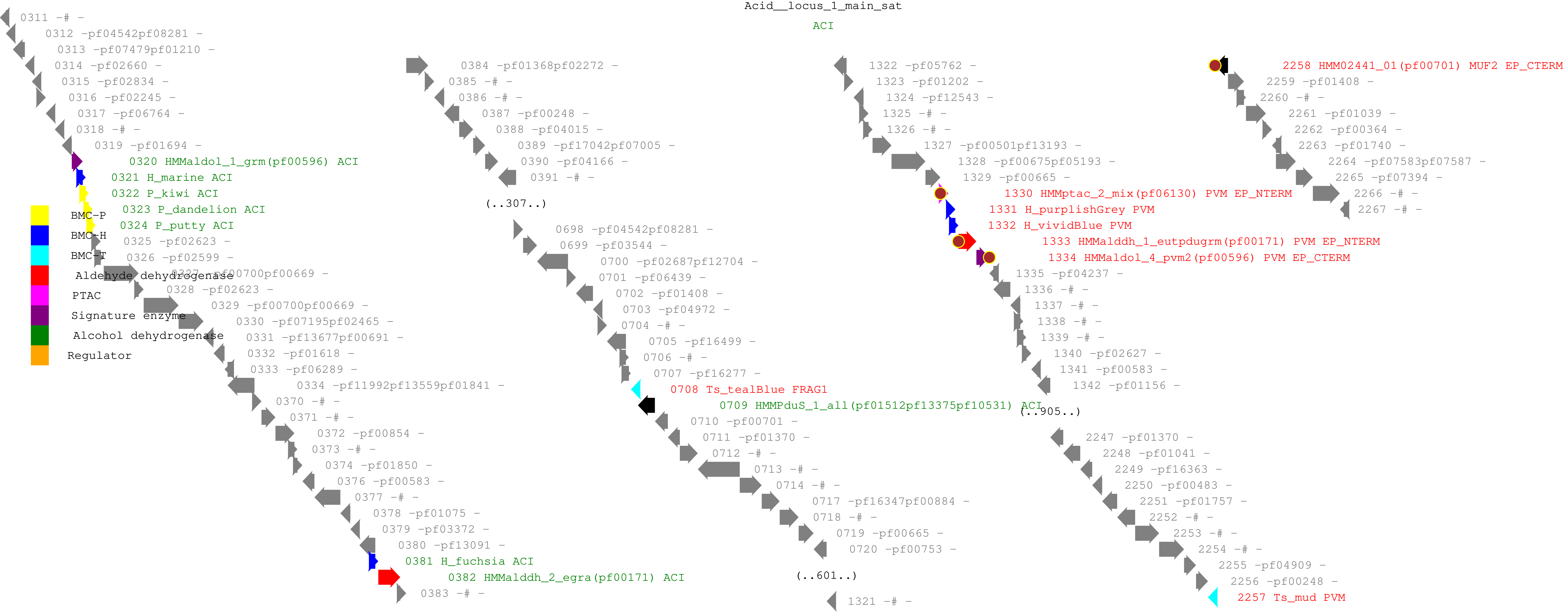

Acidobacteria bacterium 13_1_20CM_2_65_9. Bacteria; Acidobacteria.

AUF76__locus_1_main_sat

ACI

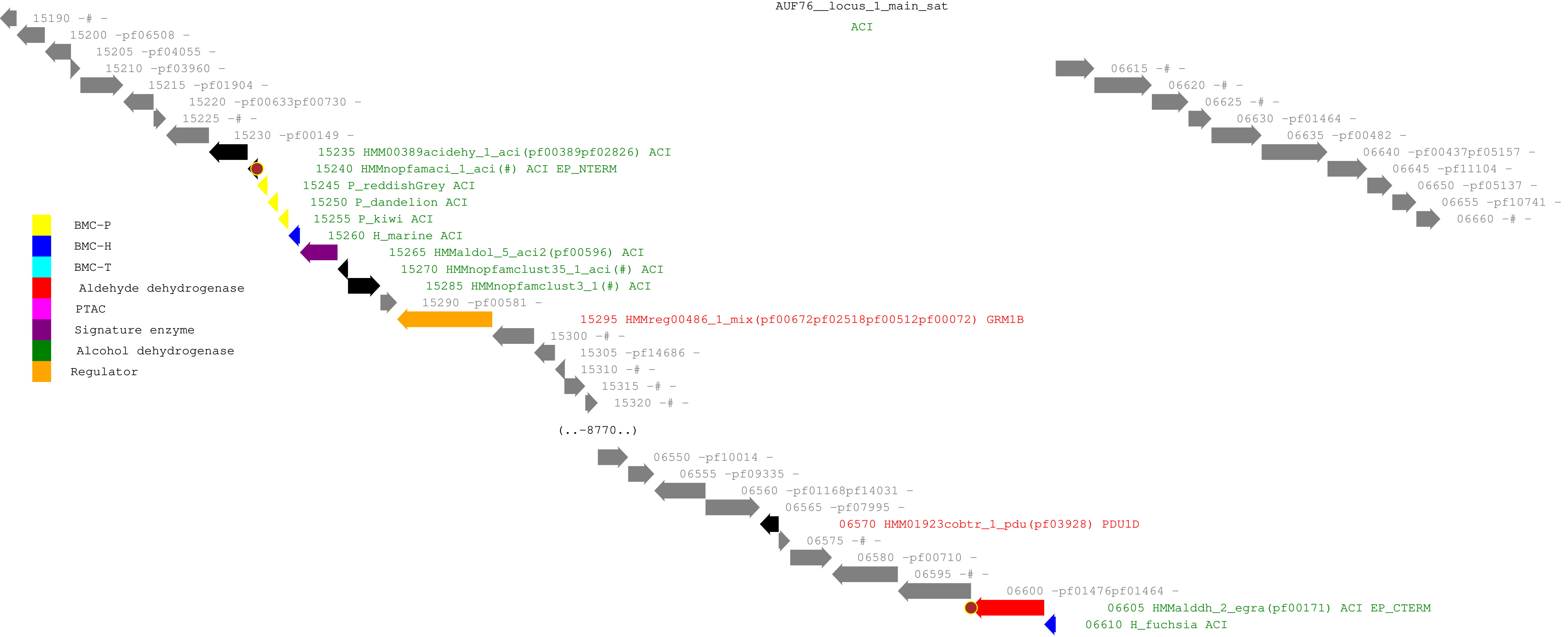

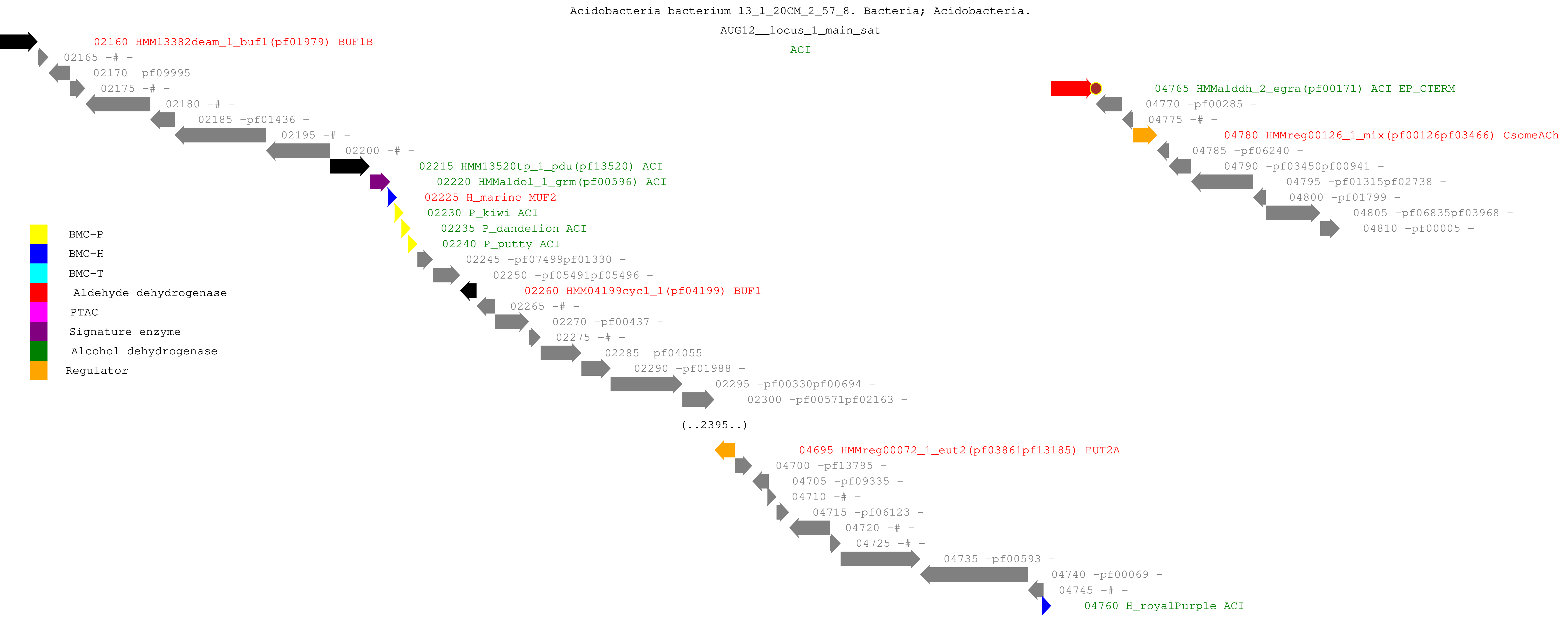

Acidobacteria bacterium 13_1_20CM_2_57_8. Bacteria; Acidobacteria.
AUG12__locus_1_main_sat
ACI
02160 HMM13382deam_1_buf1(pf01979) BUF1B
02165 -# -
02170 -pf09995 -
02175 -# -
02180 -# -
02185 -pf01436 -
02195 -# -
02200 -# -
02215 HMM13520tp_1_pdu(pf13520) ACI
02220 HMMaldol_1_grm(pf00596) ACI
02225 H_marine MUF2
02230 P_kiwi ACI
02235 P_dandelion ACI
02240 P_putty ACI
02245 -pf07499pf01330 -
02250 -pf05491pf05496 -
02260 HMM04199cycl_1(pf04199) BUF1
02265 -# -
02270 -pf00437 -
02275 -# -
02285 -pf04055 -
02290 -pf01988 -
02295 -pf00330pf00694 -
02300 -pf00571pf02163 -
(..2395..)
04695 HMMreg00072_1_eut2(pf03861pf13185) EUT2A
04700 -pf13795 -
04705 -pf09335 -
04710 -# -
04715 -pf06123 -
04720 -# -
04725 -# -
04735 -pf00593 -
04740 -pf00069 -
04745 -# -
04760 H_royalPurple ACI
04765 HMMalddh_2_egra(pf00171) ACI EP_CTERM
04770 -pf00285 -
04775 -# -
04780 HMMreg00126_1_mix(pf00126pf03466) CsomeACh
04785 -pf06240 -
04790 -pf03450pf00941 -
04795 -pf01315pf02738 -
04800 -pf01799 -
04805 -pf06835pf03968 -
04810 -pf00005 -
BMC-P
BMC-H
BMC-T
Aldehyde dehydrogenase
PTAC
Signature enzyme
Alcohol dehydrogenase
Regulator

Acidobacteria bacterium 13_1_20CM_3_53_8. Bacteria; Acidobacteria.

AUG51__locus_1_main_sat

ACI

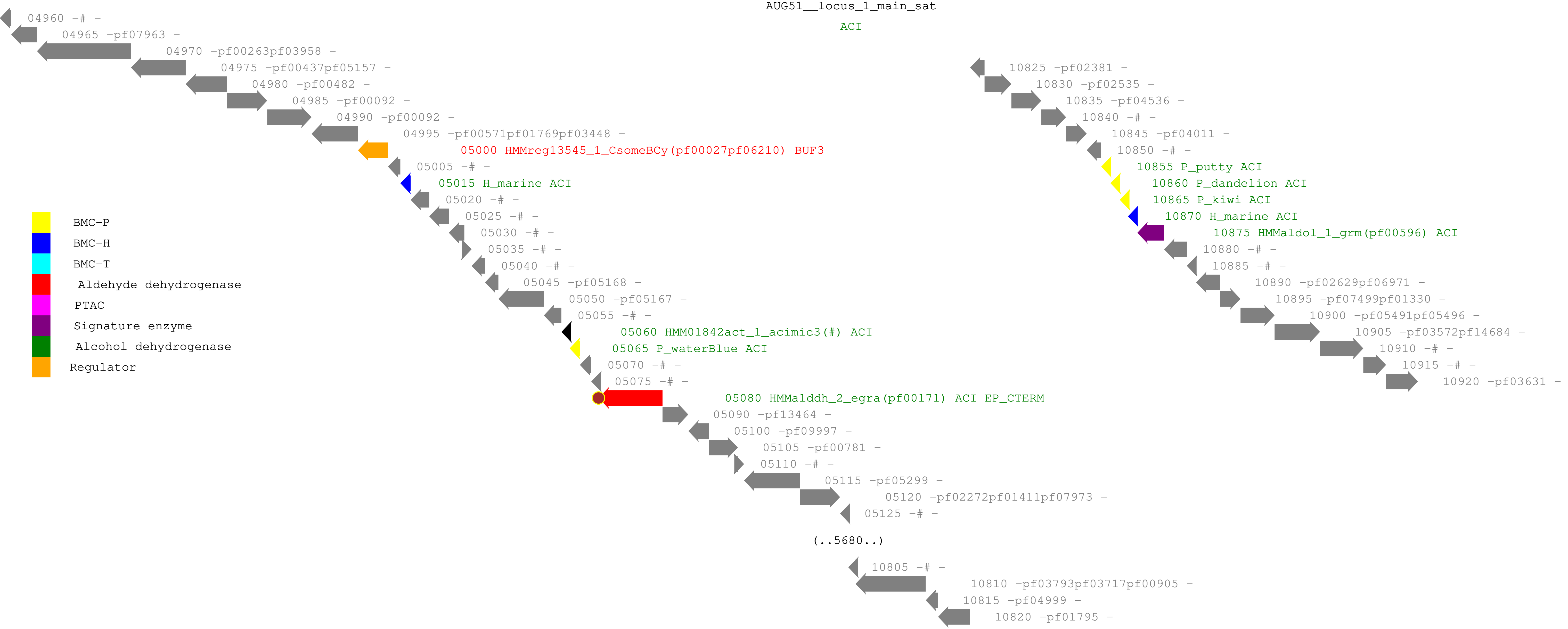

Acidobacteria bacterium 13_1_40CM_56_16. Bacteria; Acidobacteria.

AUH28__locus_1_main_sat

ACI

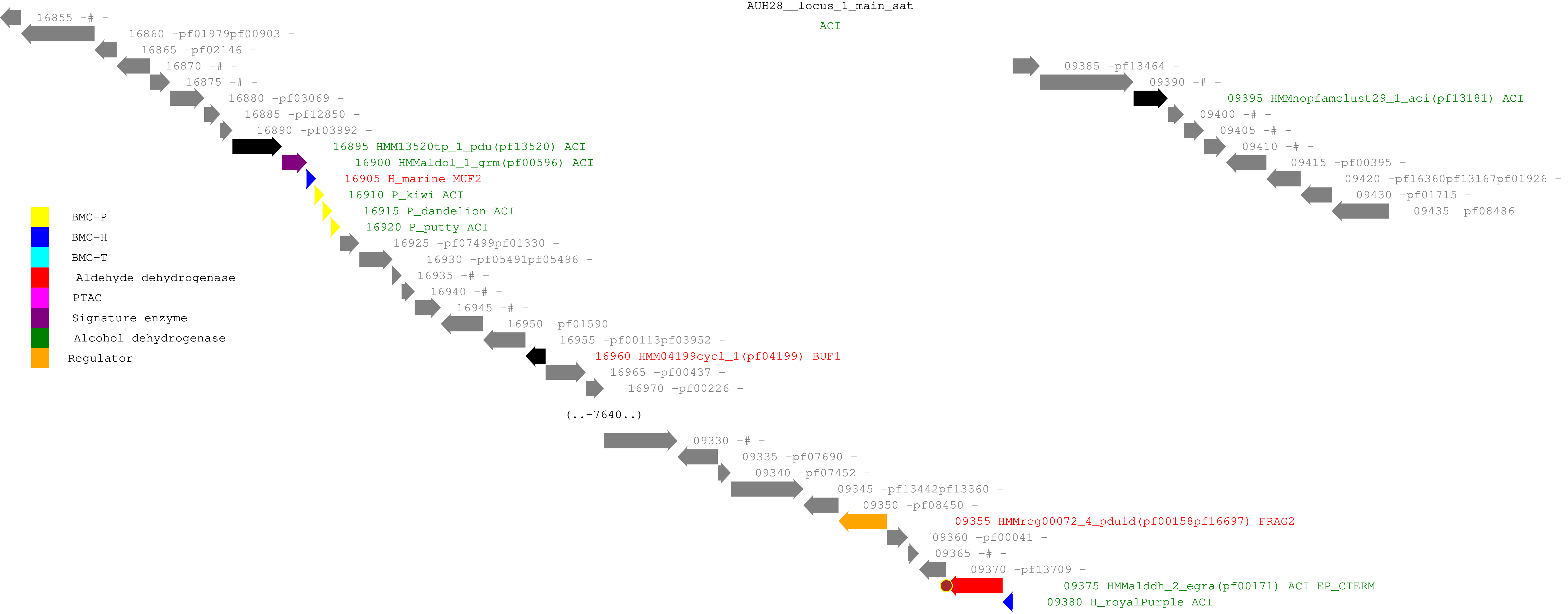

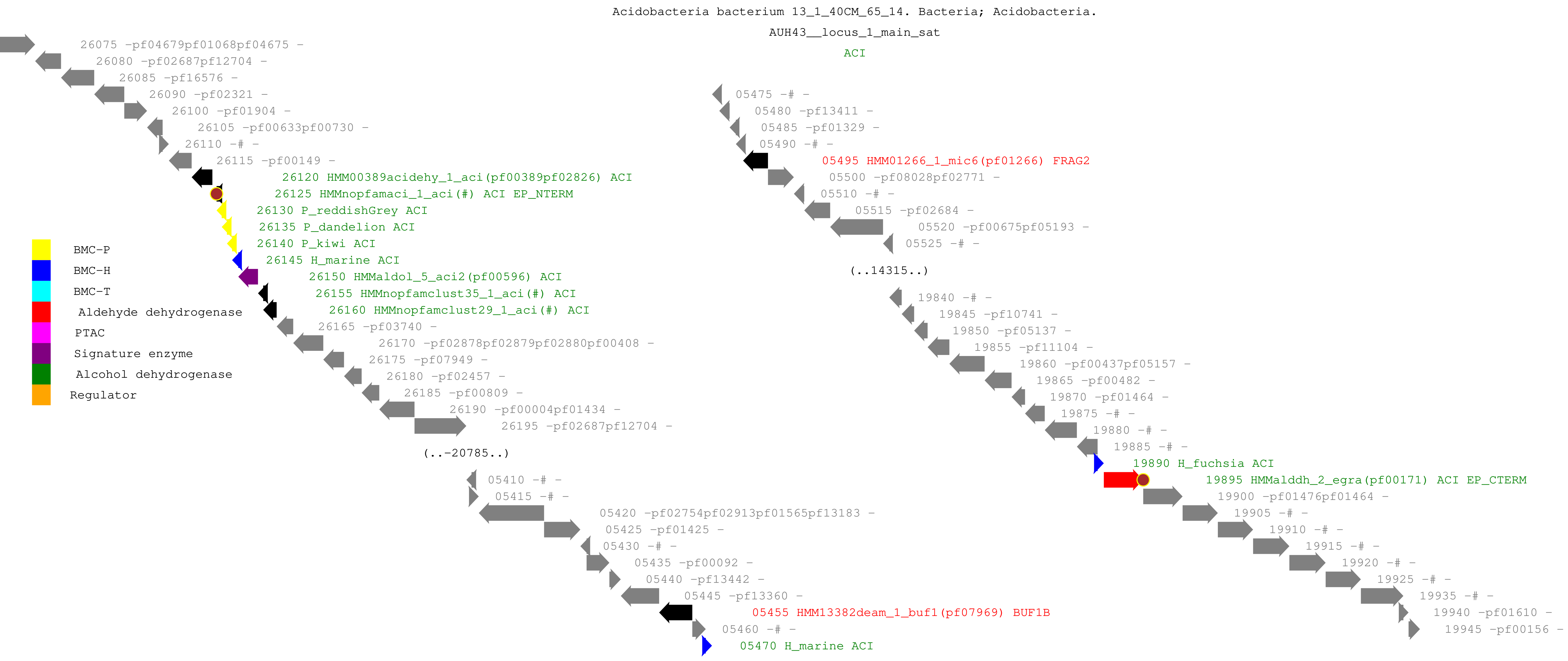

Acidobacteria bacterium 13_1_40CM_65_14. Bacteria; Acidobacteria.
AUH43__locus_1_main_sat
ACI
26075 -pf04679pf01068pf04675 -
26080 -pf02687pf12704 -
26085 -pf16576 -
26090 -pf02321 -
26100 -pf01904 -
26105 -pf00633pf00730 -
26110 -# -
26115 -pf00149 -
26120 HMM00389acidehy_1_aci(pf00389pf02826) ACI
26125 HMMnopfamaci_1_aci(#) ACI EP_NTERM
26130 P_reddishGrey ACI
26135 P_dandelion ACI
26140 P_kiwi ACI
26145 H_marine ACI
26150 HMMaldol_5_aci2(pf00596) ACI
26155 HMMnopfamclust35_1_aci(#) ACI
26160 HMMnopfamclust29_1_aci(#) ACI
26165 -pf03740 -
26170 -pf02878pf02879pf02880pf00408 -
26175 -pf07949 -
26180 -pf02457 -
26185 -pf00809 -
26190 -pf00004pf01434 -
26195 -pf02687pf12704 -
(..-20785..)
05410 -# -
05415 -# -
05420 -pf02754pf02913pf01565pf13183 -
05425 -pf01425 -
05430 -# -
05435 -pf00092 -
05440 -pf13442 -
05445 -pf13360 -
05455 HMM13382deam_1_buf1(pf07969) BUF1B
05460 -# -
05470 H_marine ACI
05475 -# -
05480 -pf13411 -
05485 -pf01329 -
05490 -# -
05495 HMM01266_1_mic6(pf01266) FRAG2
05500 -pf08028pf02771 -
05510 -# -
05515 -pf02684 -
05520 -pf00675pf05193 -
05525 -# -
(..14315..)
19840 -# -
19845 -pf10741 -
19850 -pf05137 -
19855 -pf11104 -
19860 -pf00437pf05157 -
19865 -pf00482 -
19870 -pf01464 -
19875 -# -
19880 -# -
19885 -# -
19890 H_fuchsia ACI
19895 HMMalddh_2_egra(pf00171) ACI EP_CTERM
19900 -pf01476pf01464 -
19905 -# -
19910 -# -
19915 -# -
19920 -# -
19925 -# -
19935 -# -
19940 -pf01610 -
19945 -pf00156 -
BMC-P
BMC-H
BMC-T
Aldehyde dehydrogenase
PTAC
Signature enzyme
Alcohol dehydrogenase
Regulator

Acidobacteria bacterium 13_1_40CM_4_65_8. Bacteria; Acidobacteria.

AUH72__locus_1_main_sat

ACI

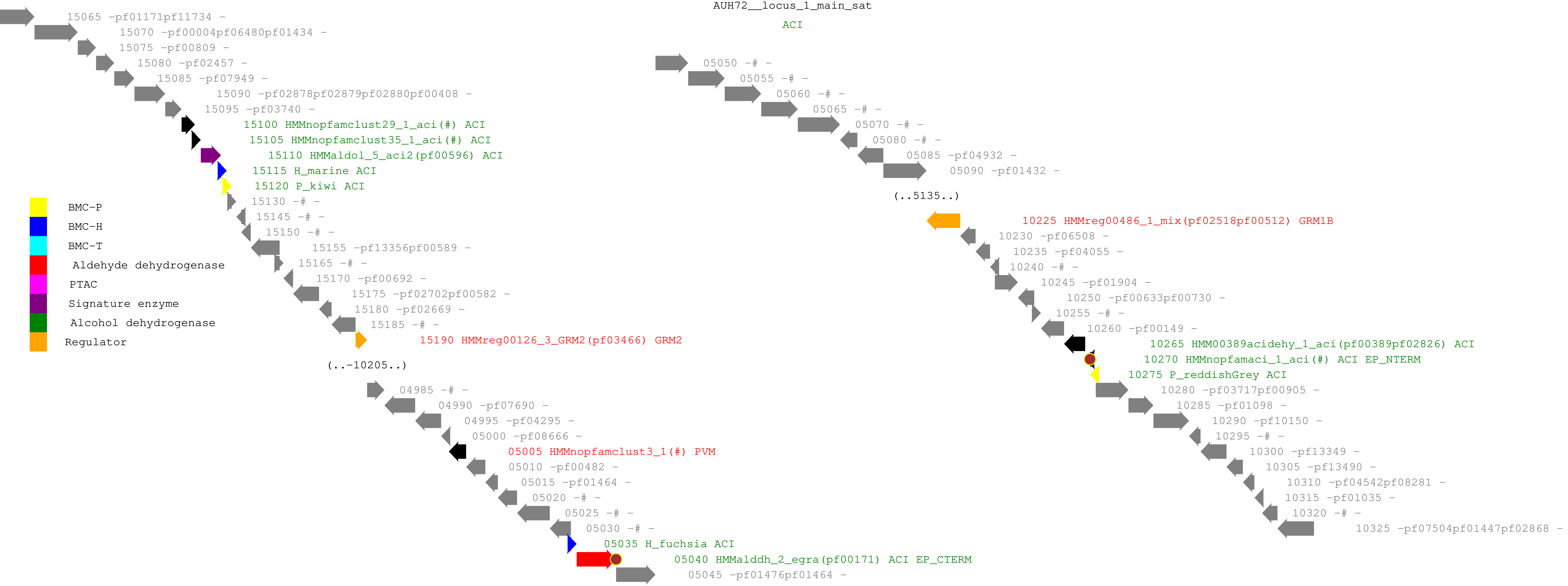

Acidobacteria bacterium 13_1_40CM_2_56_11. Bacteria; Acidobacteria.

AUI45__locus_1

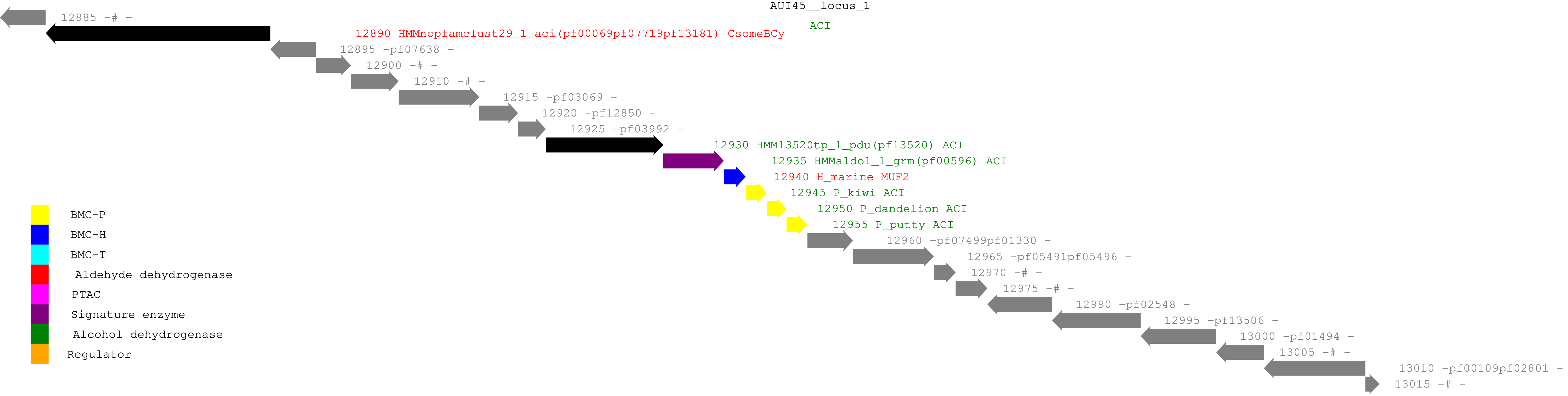

Acidobacteria bacterium 13_1_40CM_2_64_6. Bacteria; Acidobacteria.

AUI64__locus_1_main_sat

ACI

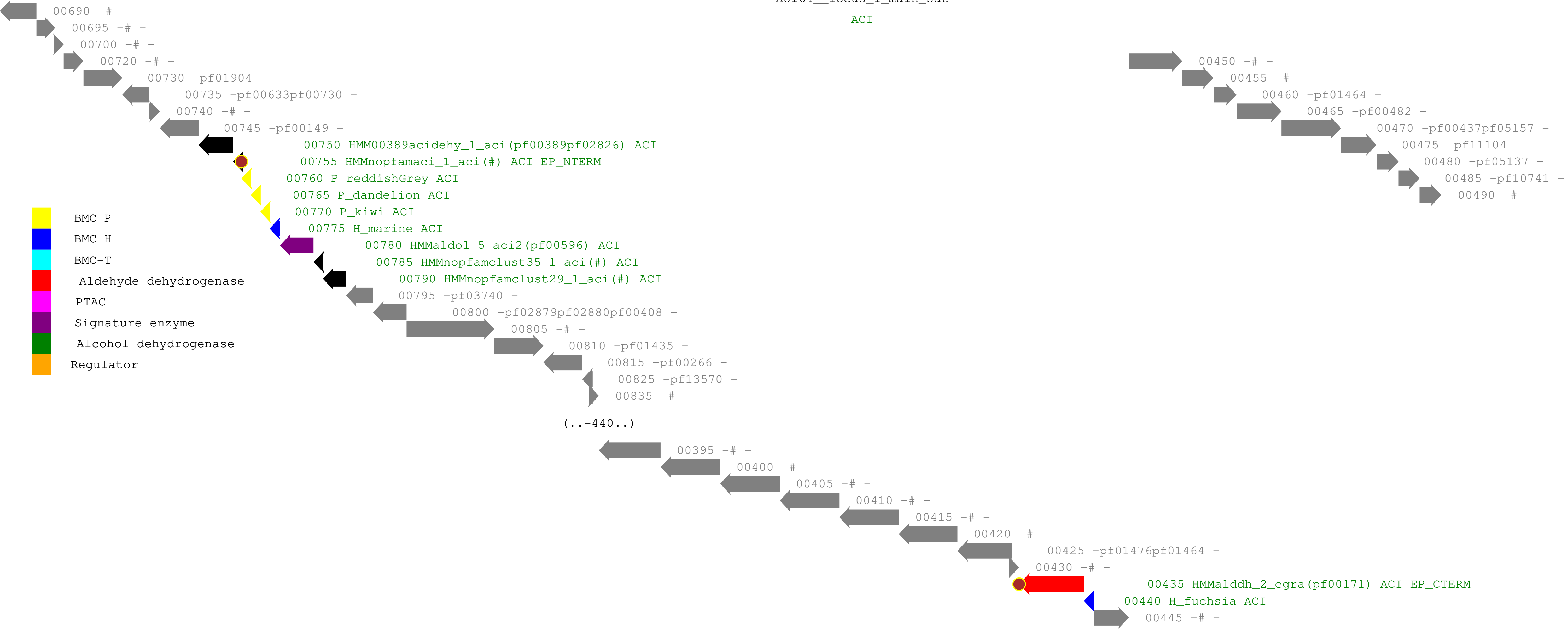

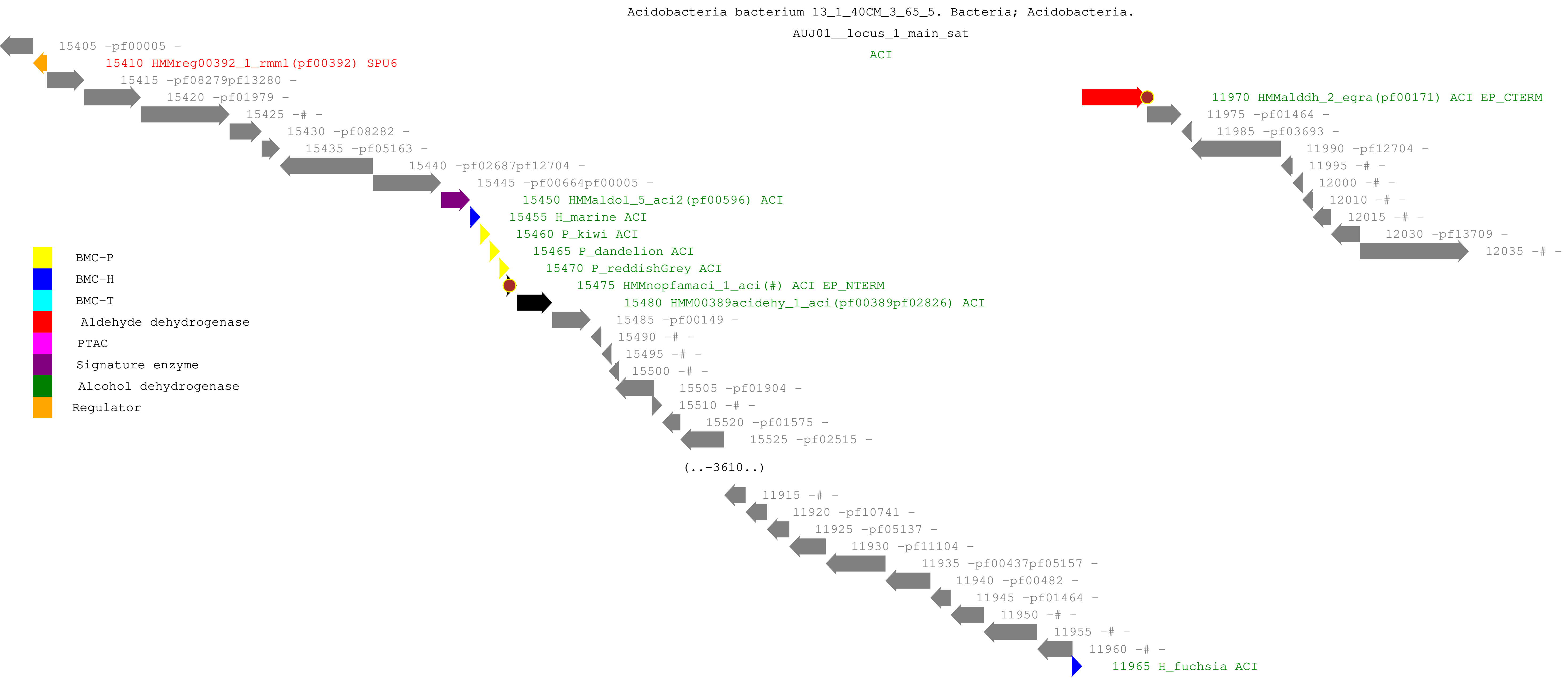
Acidobacteria bacterium 13_1_40CM_3_65_5. Bacteria; Acidobacteria.
AUJ01__locus_1_main_sat
ACI
15405 -pf00005 -
15410 HMMreg00392_1_rmm1(pf00392) SPU6
15415 -pf08279pf13280 -
15420 -pf01979 -
15425 -# -
15430 -pf08282 -
15435 -pf05163 -
15440 -pf02687pf12704 -
15445 -pf00664pf00005 -
15450 HMMaldol_5_aci2(pf00596) ACI
15455 H_marine ACI
15460 P_kiwi ACI
15465 P_dandelion ACI
15470 P_reddishGrey ACI
15475 HMMnopfamaci_1_aci(#) ACI EP_NTERM
15480 HMM00389acidehy_1_aci(pf00389pf02826) ACI
15485 -pf00149 -
15490 -# -
15495 -# -
15500 -# -
15505 -pf01904 -
15510 -# -
15520 -pf01575 -
15525 -pf02515 -
(..-3610..)
11915 -# -
11920 -pf10741 -
11925 -pf05137 -
11930 -pf11104 -
11935 -pf00437pf05157 -
11940 -pf00482 -
11945 -pf01464 -
11950 -# -
11955 -# -
11960 -# -
11965 H_fuchsia ACI
11970 HMMalddh_2_egra(pf00171) ACI EP_CTERM
11975 -pf01464 -
11985 -pf03693 -
11990 -pf12704 -
11995 -# -
12000 -# -
12010 -# -
12015 -# -
12030 -pf13709 -
12035 -# -
BMC-P
BMC-H
BMC-T
Aldehyde dehydrogenase
PTAC
Signature enzyme
Alcohol dehydrogenase
Regulator

Acidobacteria bacterium 37-65-4. Bacteria; Acidobacteria.

B7X11__locus_1_main_sat

ACI

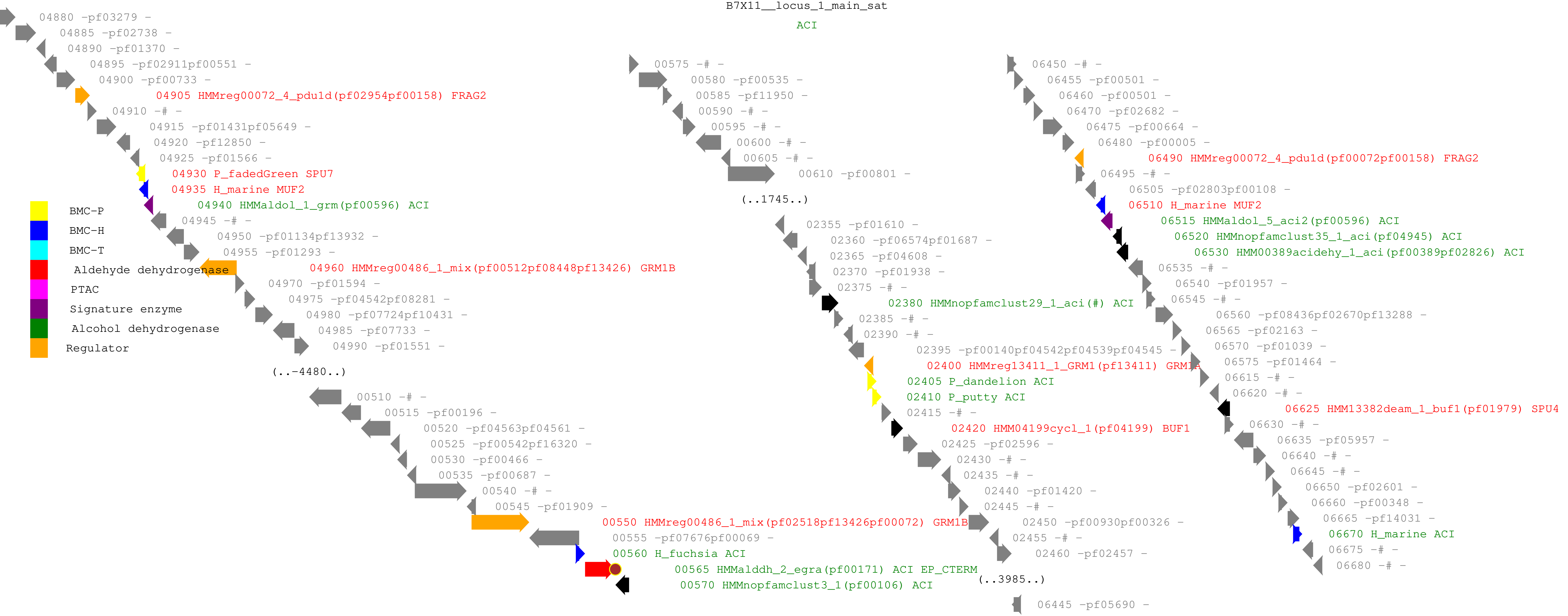

Acidobacteria bacterium. Bacteria; Acidobacteria.

C5B44__locus_1_main_sat

ACI

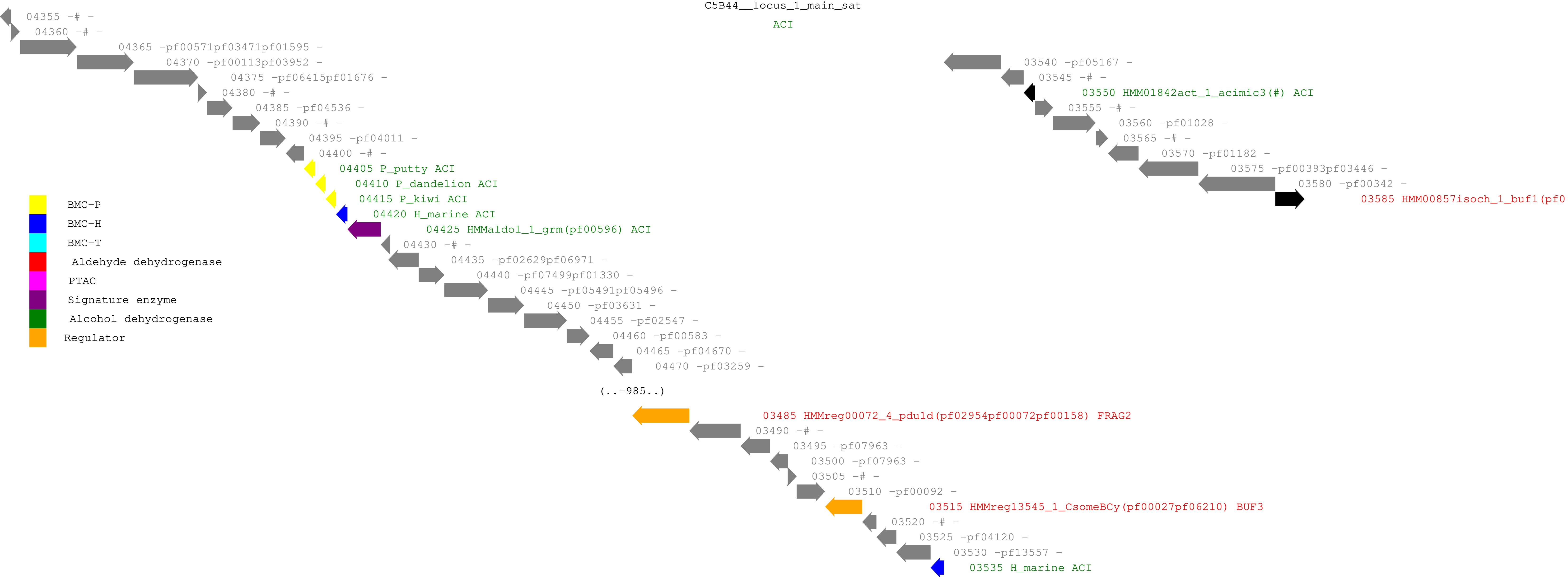

Solibacteres bacterium. Bacteria; Acidobacteria; Solibacteres.

C5B51__locus_1_main_sat

ACI

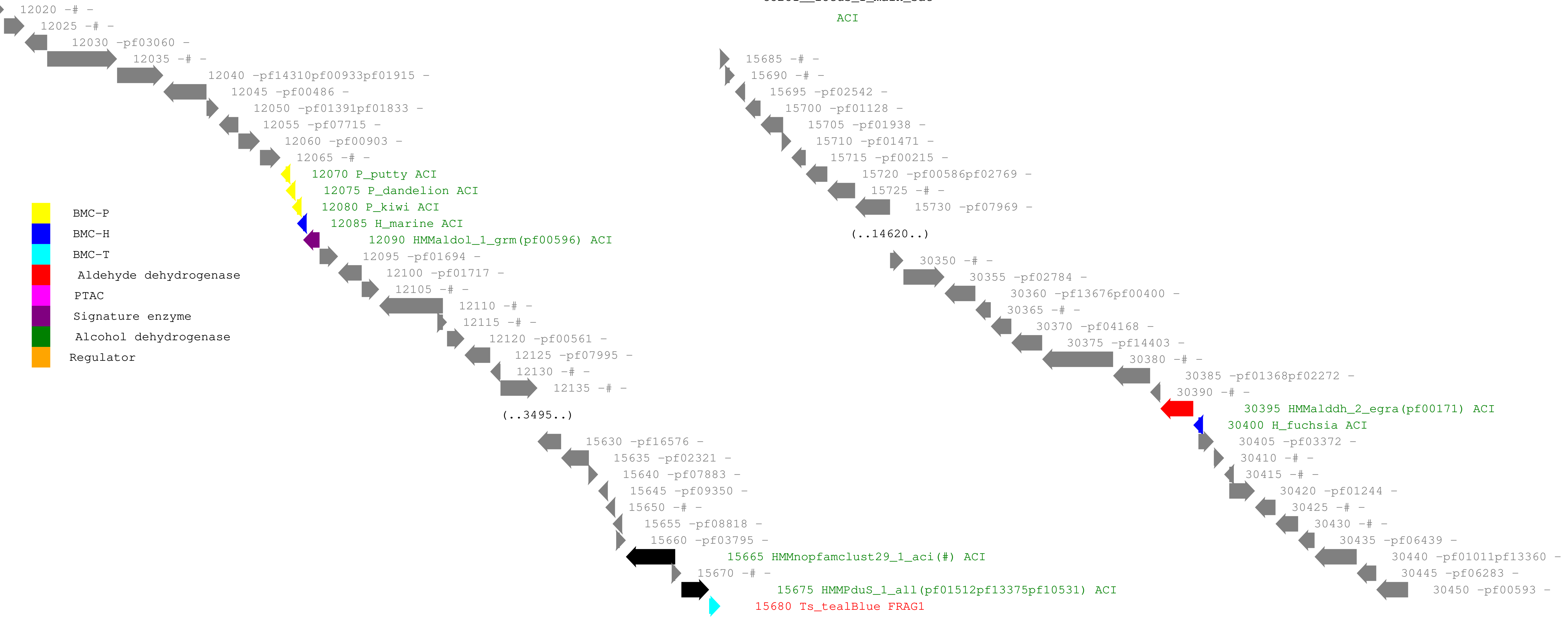

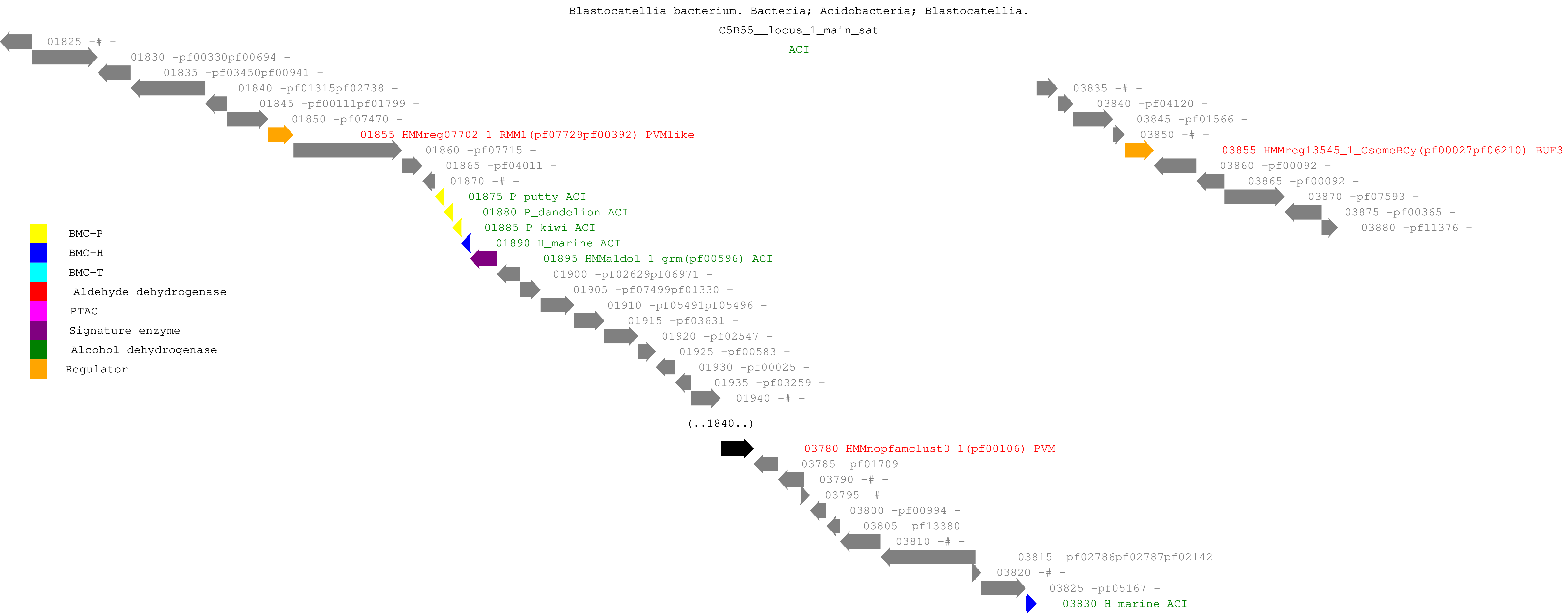
Blastocatellia bacterium. Bacteria; Acidobacteria; Blastocatellia.
C5B55__locus_1_main_sat
ACI
01825 -# -
01830 -pf00330pf00694 -
01835 -pf03450pf00941 -
01840 -pf01315pf02738 -
01845 -pf00111pf01799 -
01850 -pf07470 -
01855 HMMreg07702_1_RMM1(pf07729pf00392) PVMlike
01860 -pf07715 -
01865 -pf04011 -
01870 -# -
01875 P_putty ACI
01880 P_dandelion ACI
01885 P_kiwi ACI
01890 H_marine ACI
01895 HMMaldol_1_grm(pf00596) ACI
01900 -pf02629pf06971 -
01905 -pf07499pf01330 -
01910 -pf05491pf05496 -
01915 -pf03631 -
01920 -pf02547 -
01925 -pf00583 -
01930 -pf00025 -
01935 -pf03259 -
01940 -# -
(..1840..)
03780 HMMnopfamclust3_1(pf00106) PVM
03785 -pf01709 -
03790 -# -
03795 -# -
03800 -pf00994 -
03805 -pf13380 -
03810 -# -
03815 -pf02786pf02787pf02142 -
03820 -# -
03825 -pf05167 -
03830 H_marine ACI
03835 -# -
03840 -pf04120 -
03845 -pf01566 -
03850 -# -
03855 HMMreg13545_1_CsomeBCy(pf00027pf06210) BUF3
03860 -pf00092 -
03865 -pf00092 -
03870 -pf07593 -
03875 -pf00365 -
03880 -pf11376 -
BMC-P
BMC-H
BMC-T
Aldehyde dehydrogenase
PTAC
Signature enzyme
Alcohol dehydrogenase
Regulator

Acidobacterium sp. Bacteria; Acidobacteria; Acidobacteriales; Acidobacteriaceae;

CK533__locus_1_main_sat

ACI

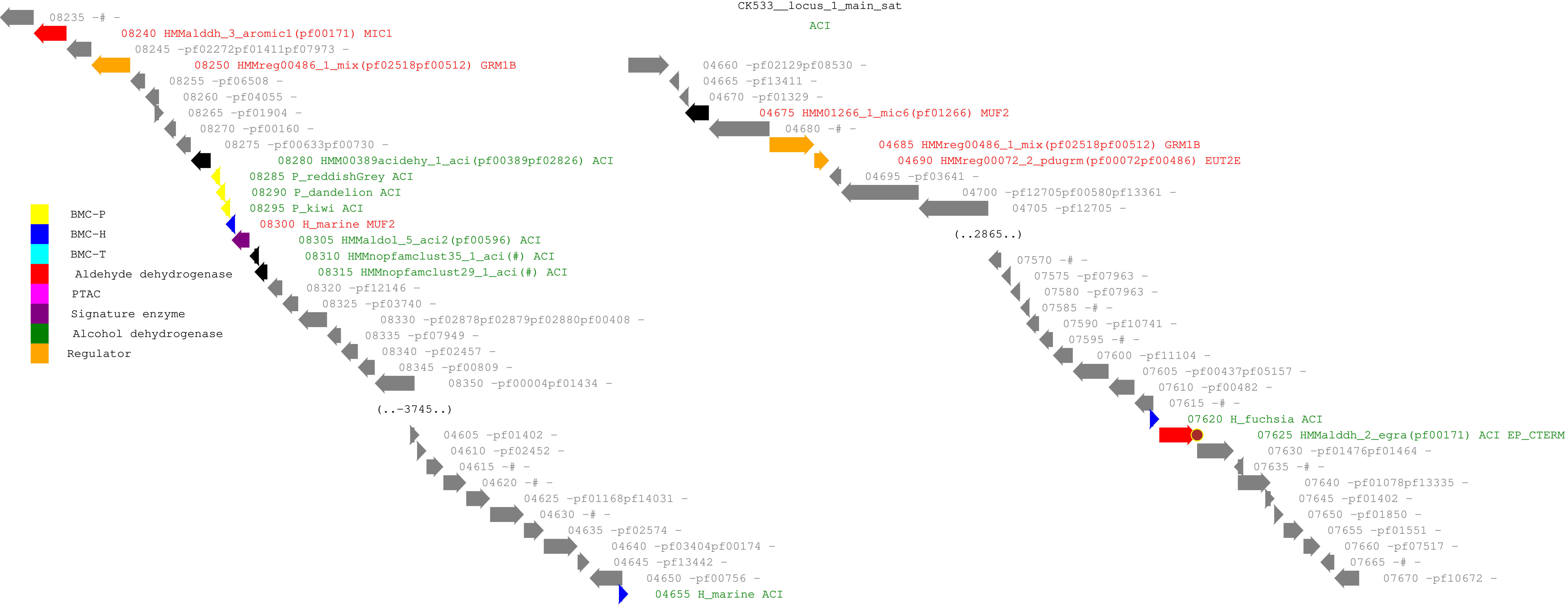

Acidobacteria bacterium. Bacteria; Acidobacteria.

CL471__locus_1_main_sat

ACI

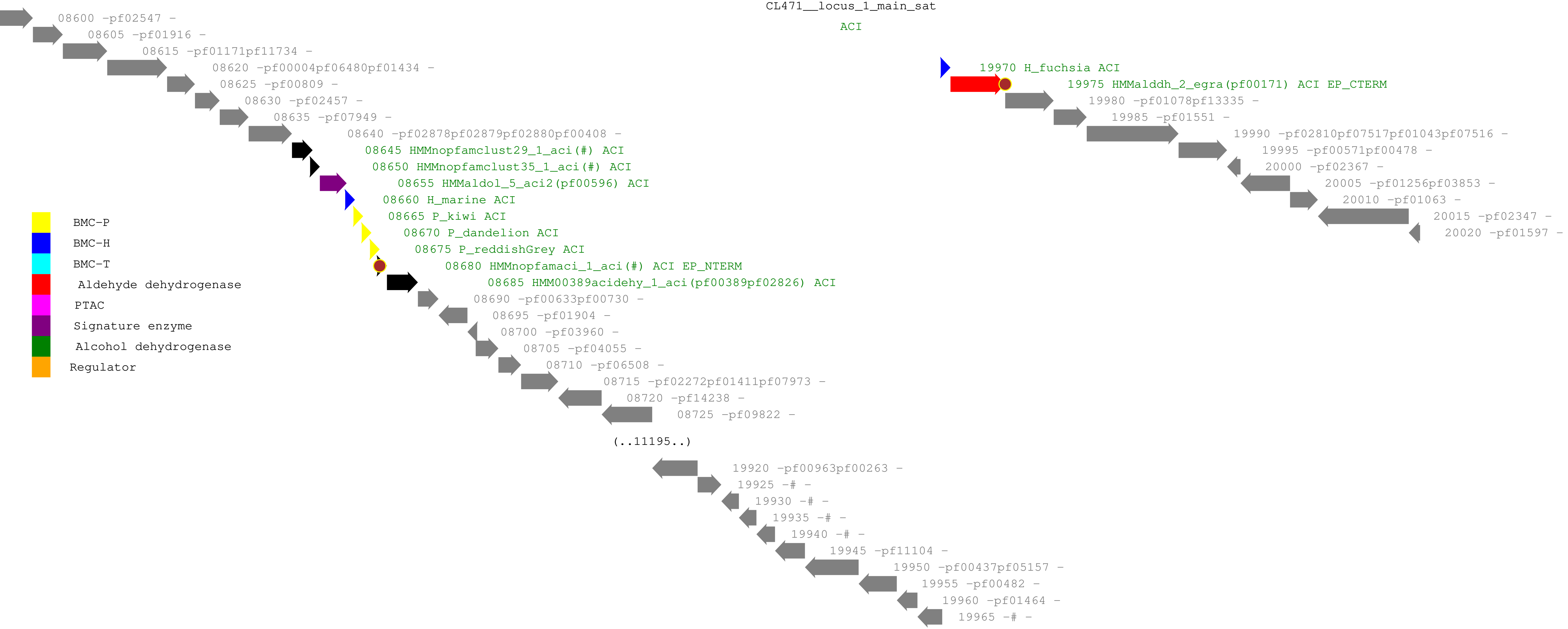


(..11195..)

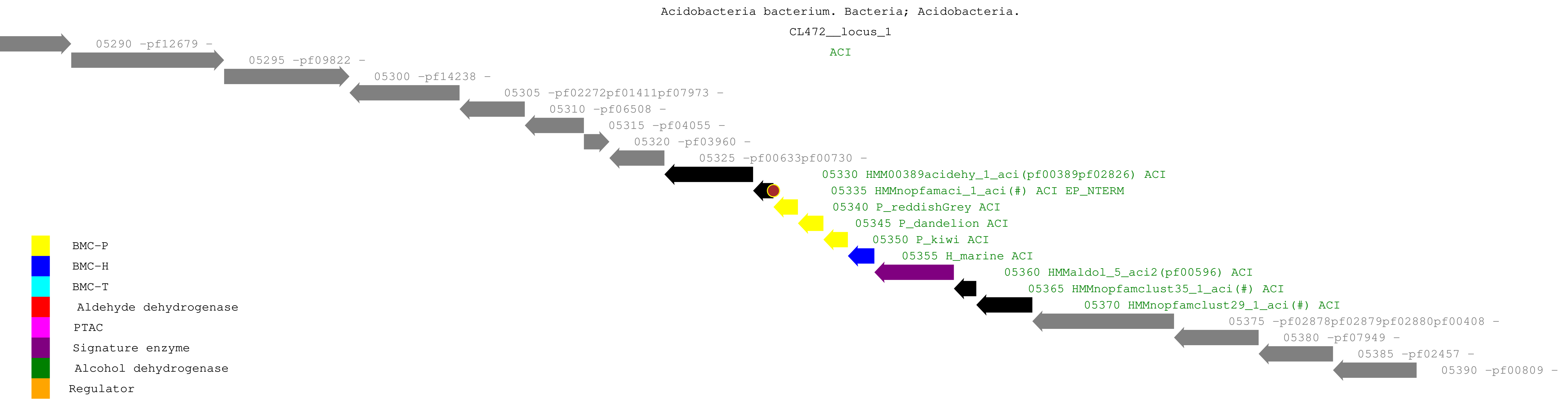
Acidobacteria bacterium. Bacteria; Acidobacteria.
CL472__locus_1
ACI
05290 -pf12679 -
05295 -pf09822 -
05300 -pf14238 -
05305 -pf02272pf01411pf07973 -
05310 -pf06508 -
05315 -pf04055 -
05320 -pf03960 -
05325 -pf00633pf00730 -
05330 HMM00389acidehy_1_aci(pf00389pf02826) ACI
05335 HMMnopfamaci_1_aci(#) ACI EP_NTERM
05340 P_reddishGrey ACI
05345 P_dandelion ACI
05350 P_kiwi ACI
05355 H_marine ACI
05360 HMMaldol_5_aci2(pf00596) ACI
05365 HMMnopfamclust35_1_aci(#) ACI
05370 HMMnopfamclust29_1_aci(#) ACI
05375 -pf02878pf02879pf02880pf00408 -
05380 -pf07949 -
05385 -pf02457 -
05390 -pf00809 -
BMC-P
BMC-H
BMC-T
Aldehyde dehydrogenase
PTAC
Signature enzyme
Alcohol dehydrogenase
Regulator

Acidobacteria bacterium. Bacteria; Acidobacteria.

CL475__locus_1

ACI

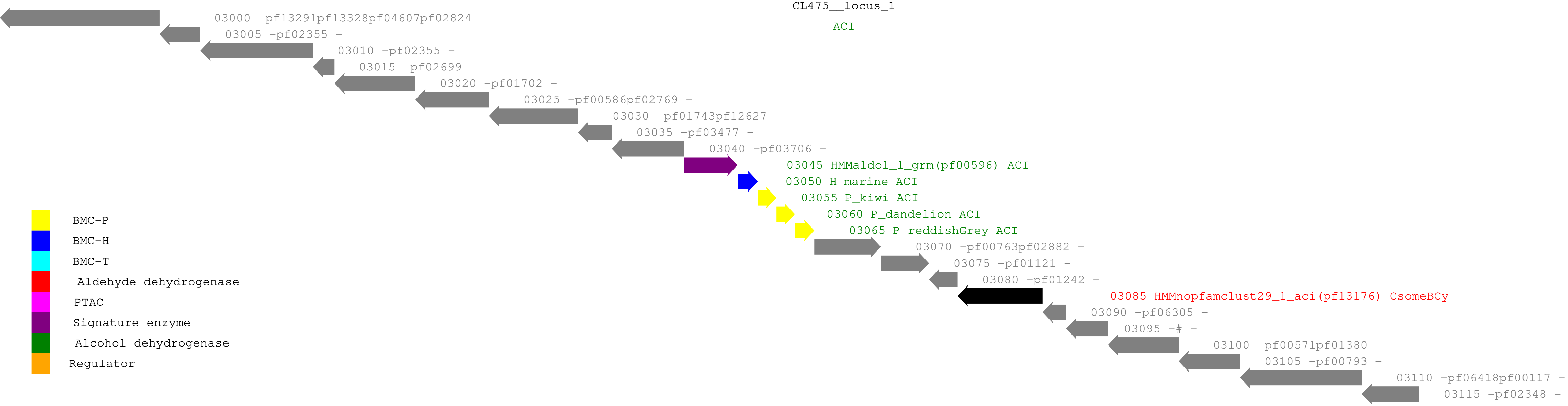

Acidobacteria bacterium. Bacteria; Acidobacteria.

CL476__locus_1_main_sat

ACI

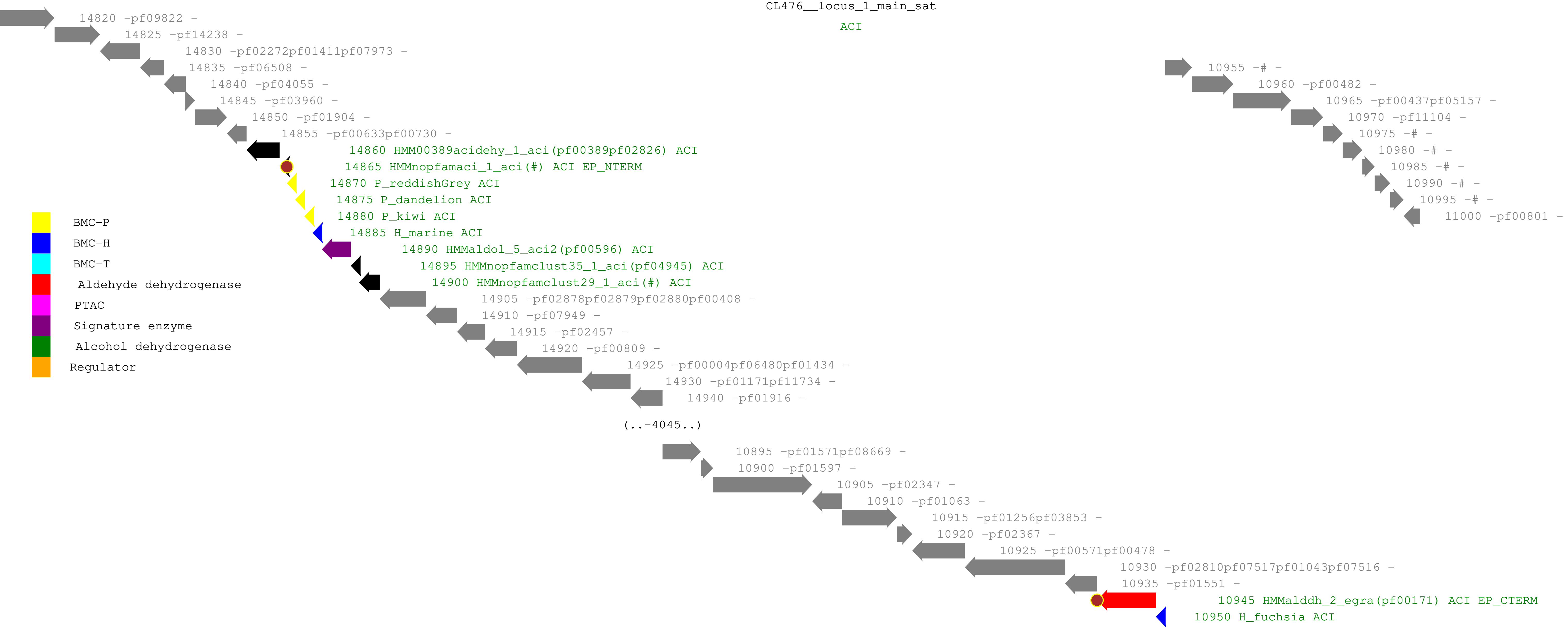

Acidobacteria bacterium. Bacteria; Acidobacteria.

CL477__locus_1_main_sat

ACI

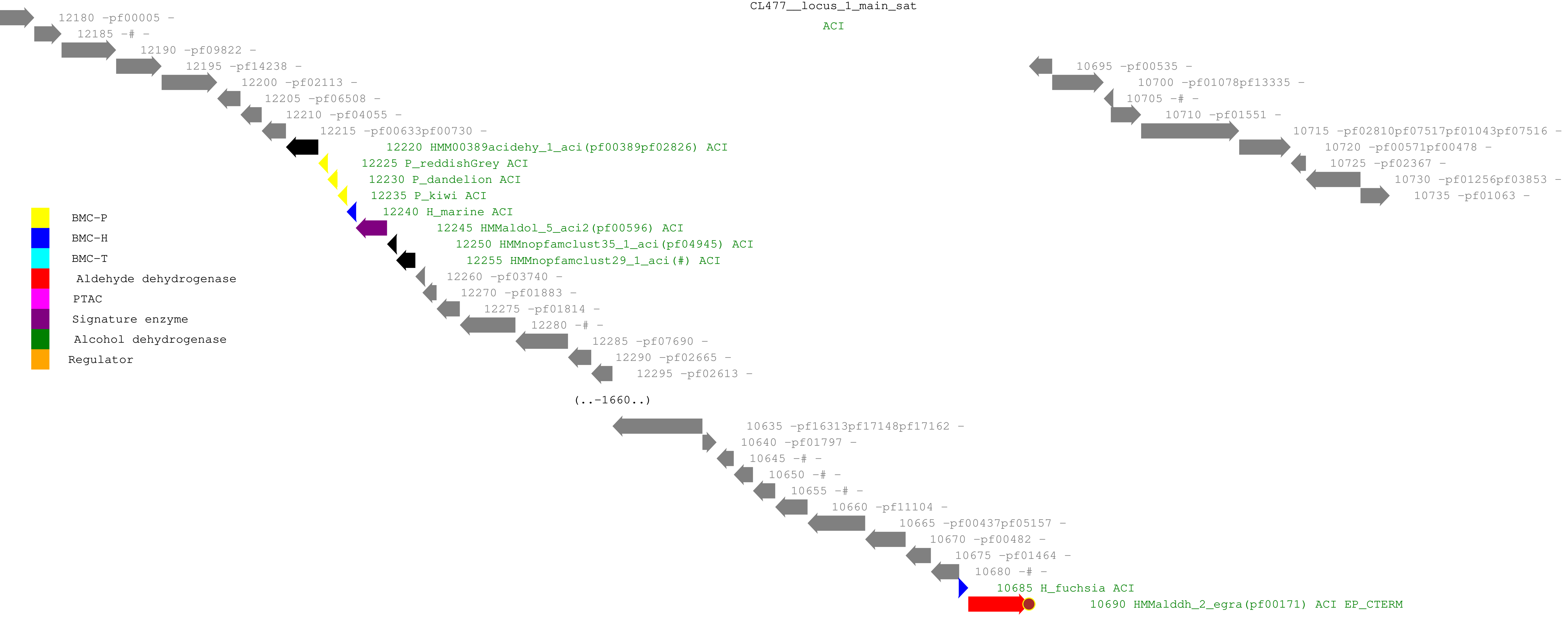

Acidobacteria bacterium. Bacteria; Acidobacteria.

CL478__locus_1_main_sat

ACI

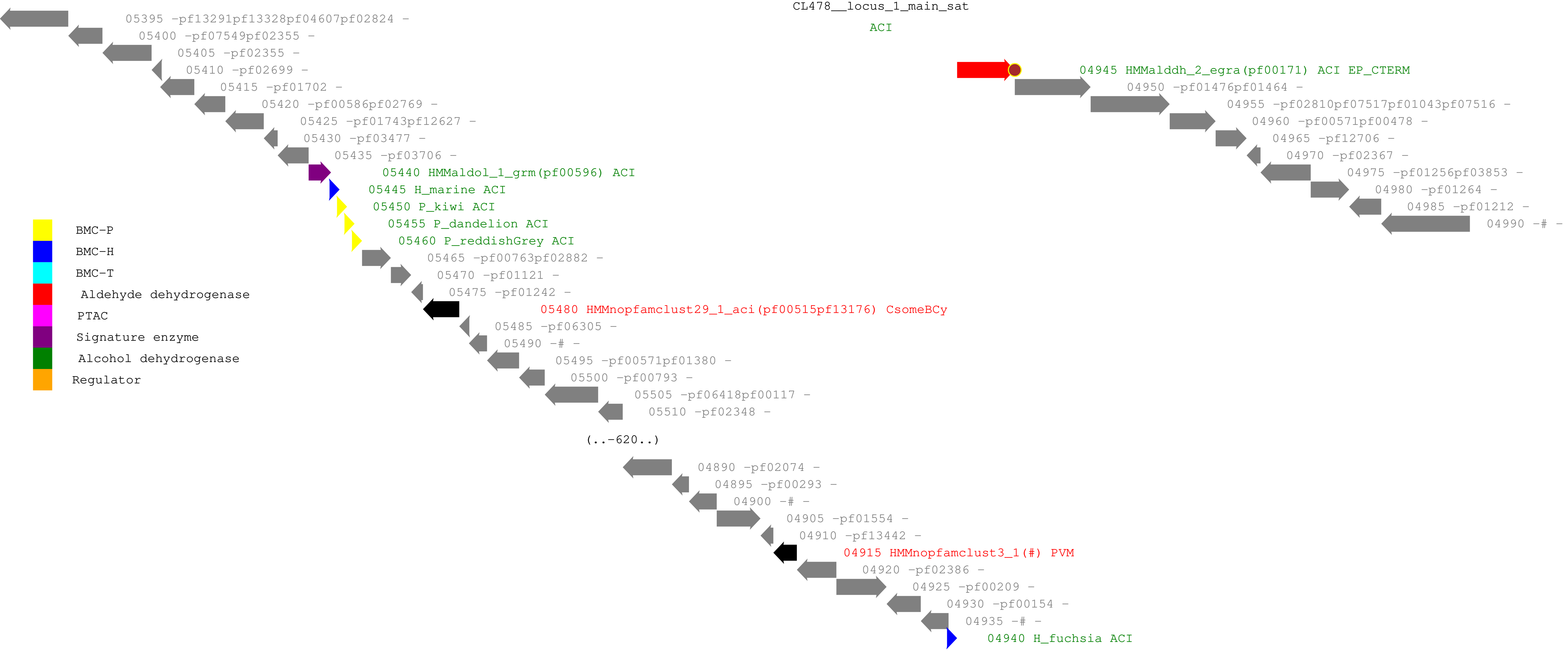

Acidobacteria bacterium. Bacteria; Acidobacteria.

CL481__locus_1_main_sat

ACI

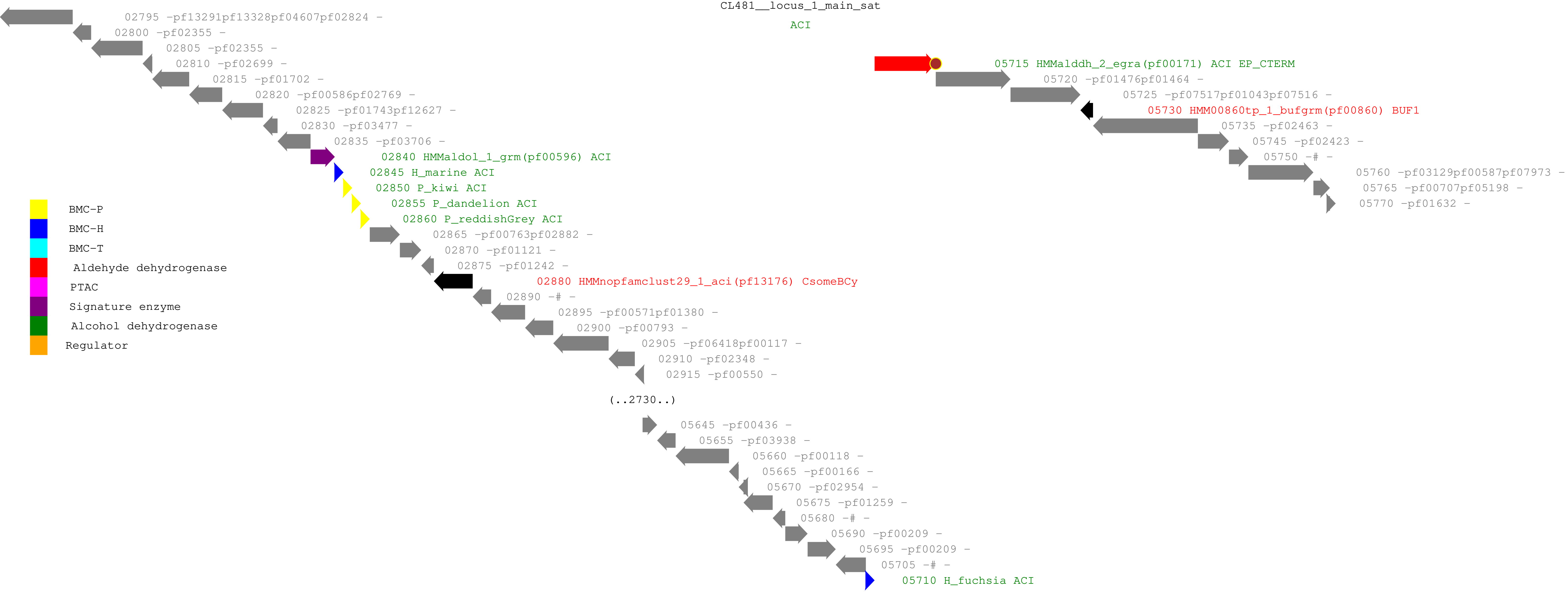

Acidobacteria bacterium. Bacteria; Acidobacteria.

CL482__locus_1

ACI

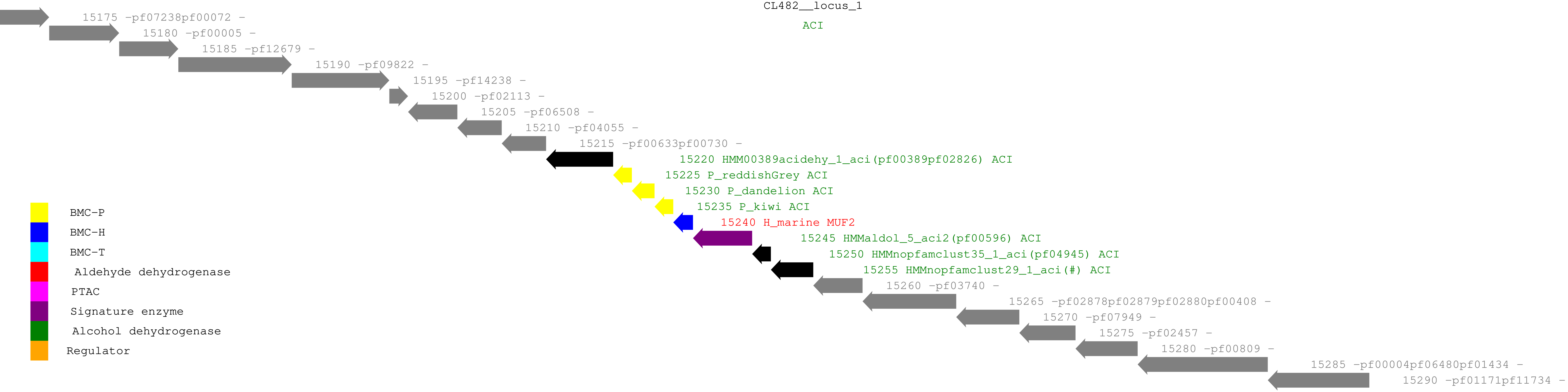

Acidobacteria bacterium. Bacteria; Acidobacteria.

CL483__locus_1_main_sat

ACI

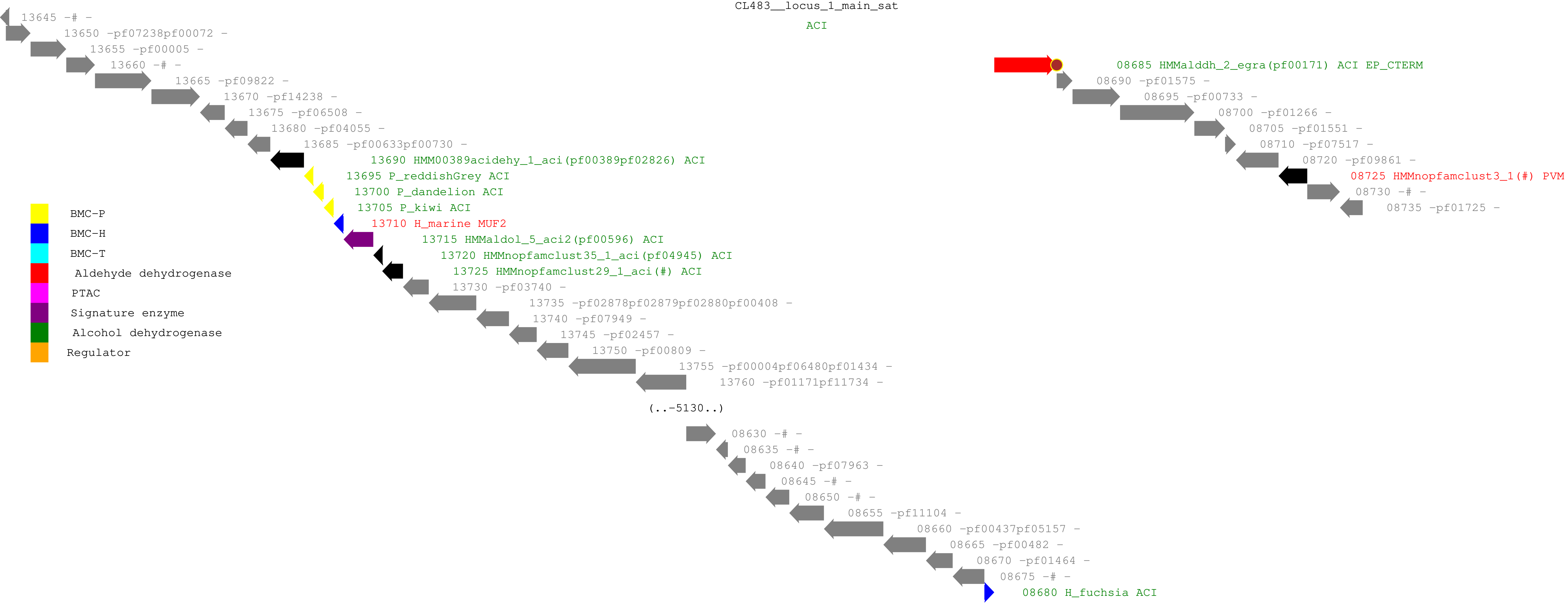

Acidobacteria bacterium. Bacteria; Acidobacteria.

CL484__locus_1_main_sat

ACI

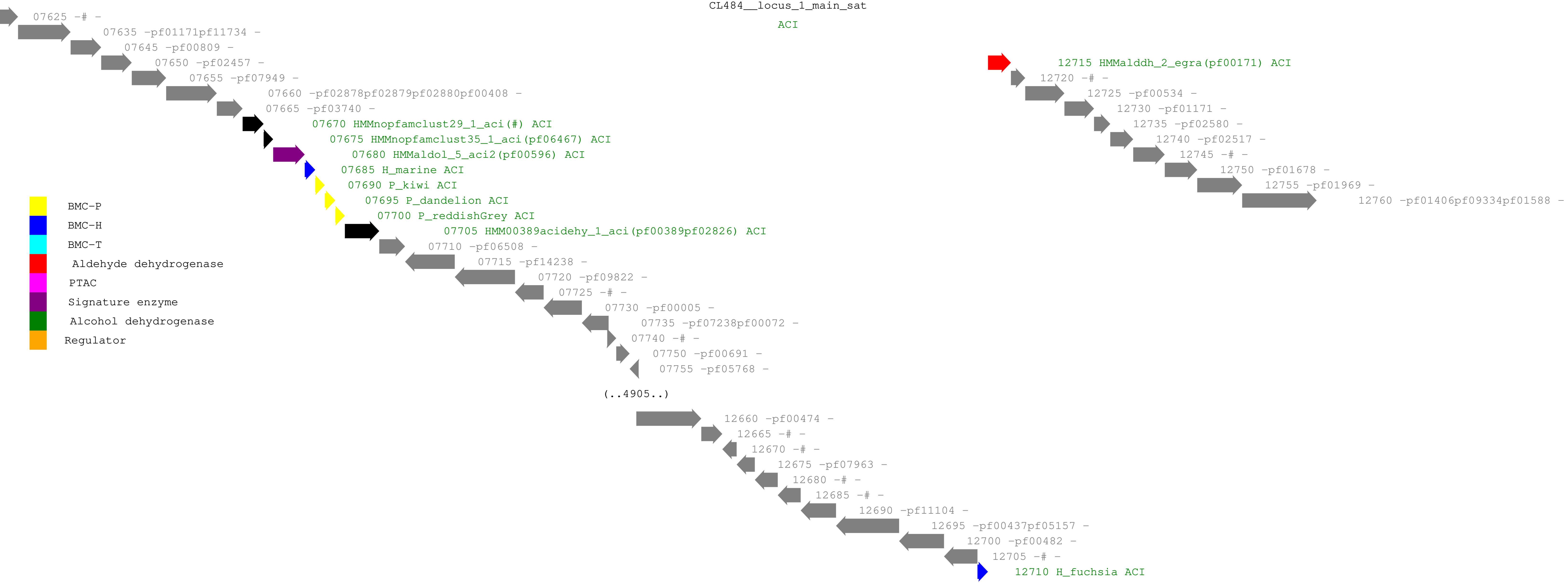

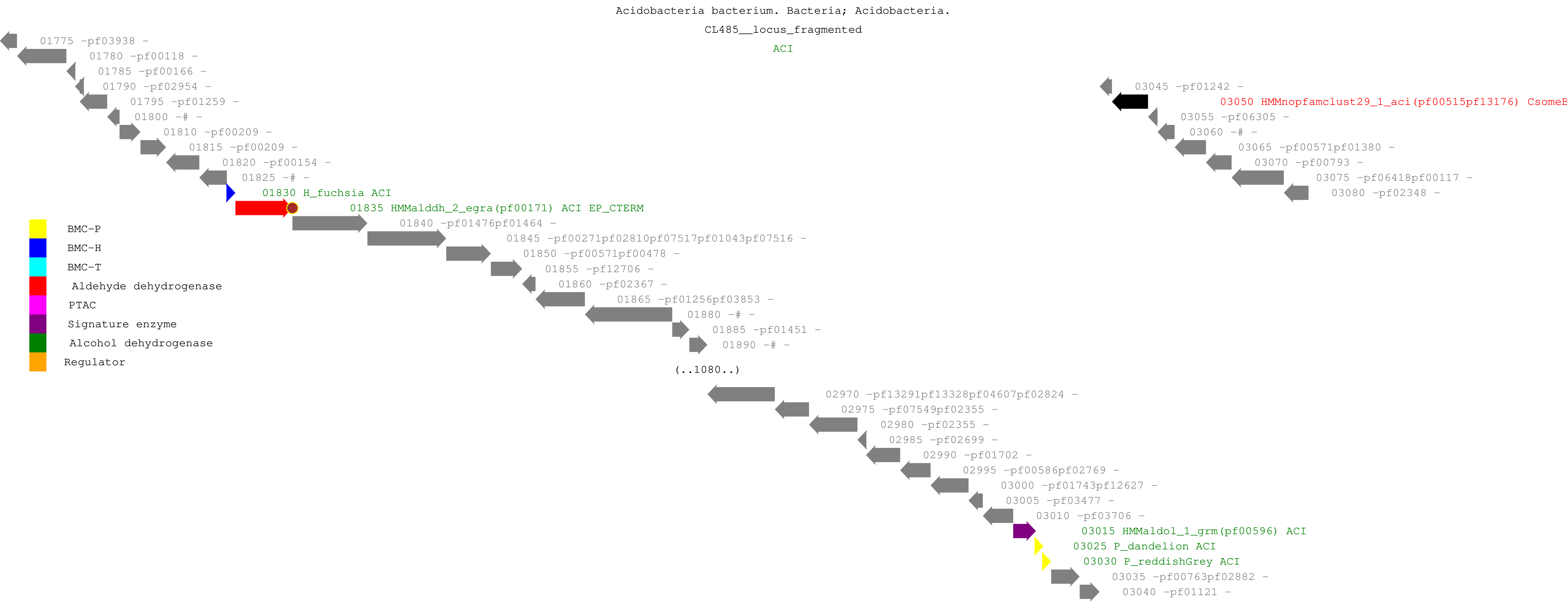

Acidobacteria bacterium. Bacteria; Acidobacteria.
CL485__locus_fragmented
ACI
01775 -pf03938 -
01780 -pf00118 -
01785 -pf00166 -
01790 -pf02954 -
01795 -pf01259 -
01800 -# -
01810 -pf00209 -
01815 -pf00209 -
01820 -pf00154 -
01825 -# -
01830 H_fuchsia ACI
01835 HMMalddh_2_egra(pf00171) ACI EP_CTERM
01840 -pf01476pf01464 -
01845 -pf00271pf02810pf07517pf01043pf07516 -
01850 -pf00571pf00478 -
01855 -pf12706 -
01860 -pf02367 -
01865 -pf01256pf03853 -
01880 -# -
01885 -pf01451 -
01890 -# -
(..1080..)
02970 -pf13291pf13328pf04607pf02824 -
02975 -pf07549pf02355 -
02980 -pf02355 -
02985 -pf02699 -
02990 -pf01702 -
02995 -pf00586pf02769 -
03000 -pf01743pf12627 -
03005 -pf03477 -
03010 -pf03706 -
03015 HMMaldol_1_grm(pf00596) ACI
03025 P_dandelion ACI
03030 P_reddishGrey ACI
03035 -pf00763pf02882 -
03040 -pf01121 -
03045 -pf01242 -
03050 HMMnopfamclust29_1_aci(pf00515pf13176) CsomeE
03055 -pf06305 -
03060 -# -
03065 -pf00571pf01380 -
03070 -pf00793 -
03075 -pf06418pf00117 -
03080 -pf02348 -
BMC-P
BMC-H
BMC-T
Aldehyde dehydrogenase
PTAC
Signature enzyme
Alcohol dehydrogenase
Regulator

Acidobacteria bacterium. Bacteria; Acidobacteria.

CL486__locus_1_main_sat

ACI

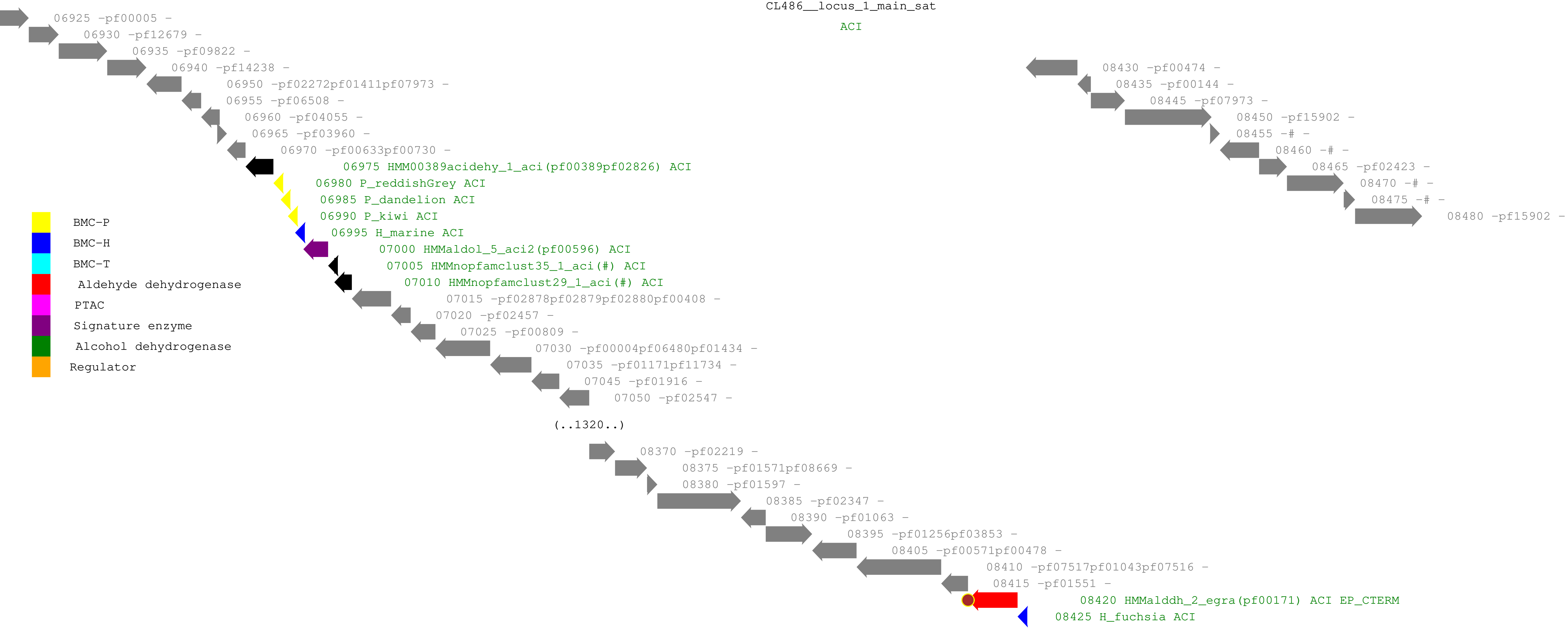

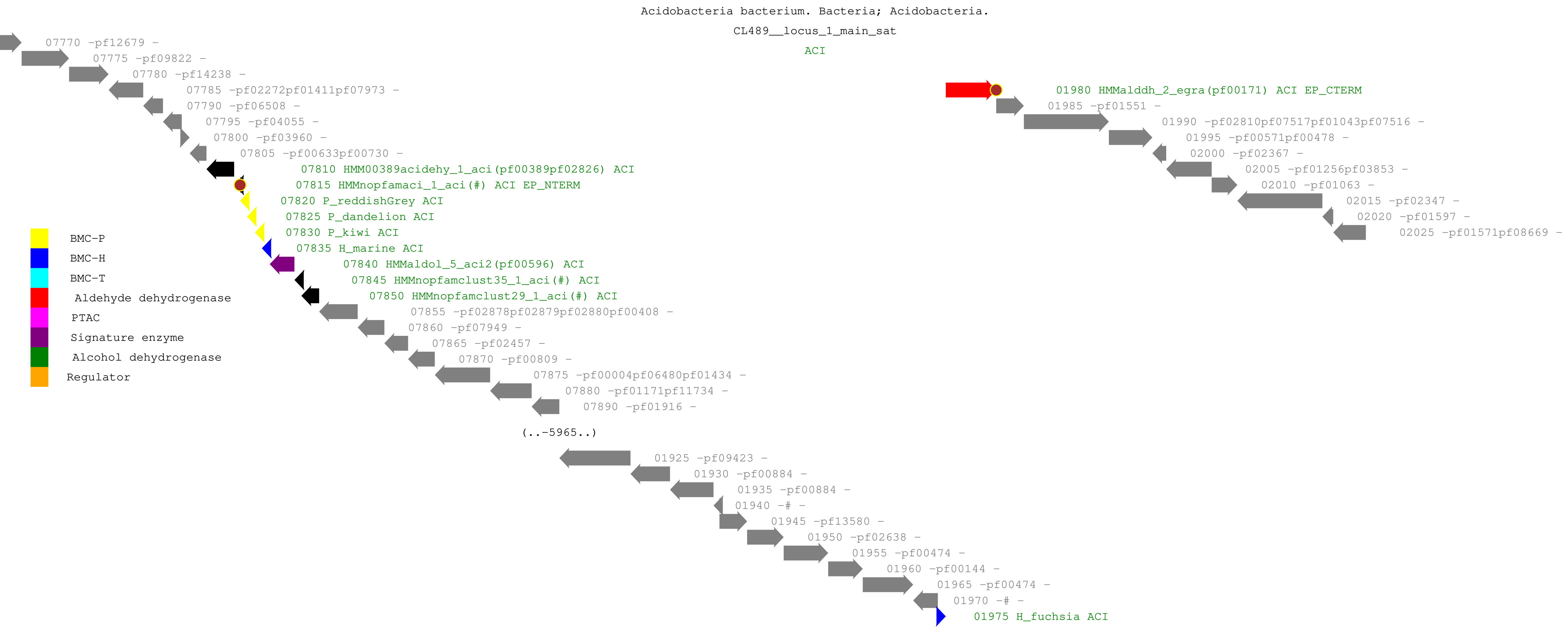
Acidobacteria bacterium. Bacteria; Acidobacteria.
CL489__locus_1_main_sat
ACI
07770 -pf12679 -
07775 -pf09822 -
07780 -pf14238 -
07785 -pf02272pf01411pf07973 -
07790 -pf06508 -
07795 -pf04055 -
07800 -pf03960 -
07805 -pf00633pf00730 -
07810 HMM00389acidehy_1_aci(pf00389pf02826) ACI
07815 HMMnopfamaci_1_aci(#) ACI EP_NTERM
07820 P_reddishGrey ACI
07825 P_dandelion ACI
07830 P_kiwi ACI
07835 H_marine ACI
07840 HMMaldol_5_aci2(pf00596) ACI
07845 HMMnopfamclust35_1_aci(#) ACI
07850 HMMnopfamclust29_1_aci(#) ACI
07855 -pf02878pf02879pf02880pf00408 -
07860 -pf07949 -
07865 -pf02457 -
07870 -pf00809 -
07875 -pf00004pf06480pf01434 -
07880 -pf01171pf11734 -
07890 -pf01916 -
(..-5965..)
01925 -pf09423 -
01930 -pf00884 -
01935 -pf00884 -
01940 -# -
01945 -pf13580 -
01950 -pf02638 -
01955 -pf00474 -
01960 -pf00144 -
01965 -pf00474 -
01970 -# -
01975 H_fuchsia ACI
01980 HMMaldh_2_egra(pf00171) ACI EP_CTERM
01985 -pf01551 -
01990 -pf02810pf07517pf01043pf07516 -
01995 -pf00571pf00478 -
02000 -pf02367 -
02005 -pf01256pf03853 -
02010 -pf01063 -
02015 -pf02347 -
02020 -pf01597 -
02025 -pf01571pf08669 -
BMC-P
BMC-H
BMC-T
Aldehyde dehydrogenase
PTAC
Signature enzyme
Alcohol dehydrogenase
Regulator

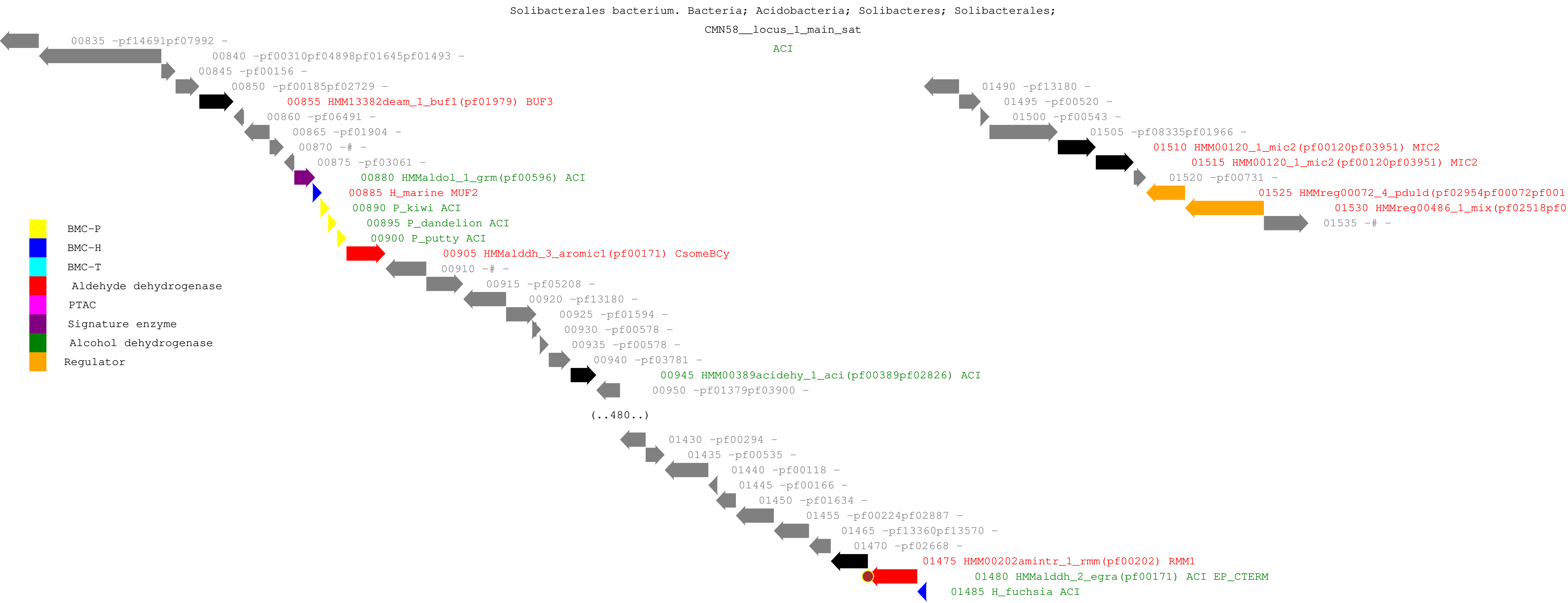
Solibacterales bacterium. Bacteria; Acidobacteria; Solibacteres; Solibacterales;
CMN58__locus_1_main_sat
ACI
00835 -pf14691pf07992 -
00840 -pf00310pf04898pf01645pf01493 -
00845 -pf00156 -
00850 -pf00185pf02729 -
00855 HMM13382deam_1_buf1(pf01979) BUF3
00860 -pf06491 -
00865 -pf01904 -
00870 -# -
00875 -pf03061 -
00880 HMMaldol_1_grm(pf00596) ACI
00885 H_marine MUF2
00890 P_kiwi ACI
00895 P_dandelion ACI
00900 P_putty ACI
00905 HMMalddh_3_aromic1(pf00171) CsomeBCy
00910 -# -
00915 -pf05208 -
00920 -pf13180 -
00925 -pf01594 -
00930 -pf00578 -
00935 -pf00578 -
00940 -pf03781 -
00945 HMM00389acidehy_1_aci(pf00389pf02826) ACI
00950 -pf01379pf03900 -
(..480..)
01430 -pf00294 -
01435 -pf00535 -
01440 -pf00118 -
01445 -pf00166 -
01450 -pf01634 -
01455 -pf00224pf02887 -
01465 -pf13360pf13570 -
01470 -pf02668 -
01475 HMM00202amintr_1_rmm(pf00202) RMM1
01480 HMMalddh_2_egra(pf00171) ACI EP_CTERM
01485 H_fuchsia ACI
01490 -pf13180 -
01495 -pf00520 -
01500 -pf00543 -
01505 -pf08335pf01966 -
01510 HMM00120_1_mic2(pf00120pf03951) MIC2
01515 HMM00120_1_mic2(pf00120pf03951) MIC2
01520 -pf00731 -
01525 HMMreg00072_4_pdu1d(pf02954pf00072pf001
01530 HMMreg00486_1_mix(pf02518pf0
01535 -# -
BMC-P
BMC-H
BMC-T
Aldehyde dehydrogenase
PTAC
Signature enzyme
Alcohol dehydrogenase
Regulator

Woeseiaceae bacterium. Bacteria; Proteobacteria; Gammaproteobacteria; Chromatiales;

CMP01__locus_1_main_sat

ACI

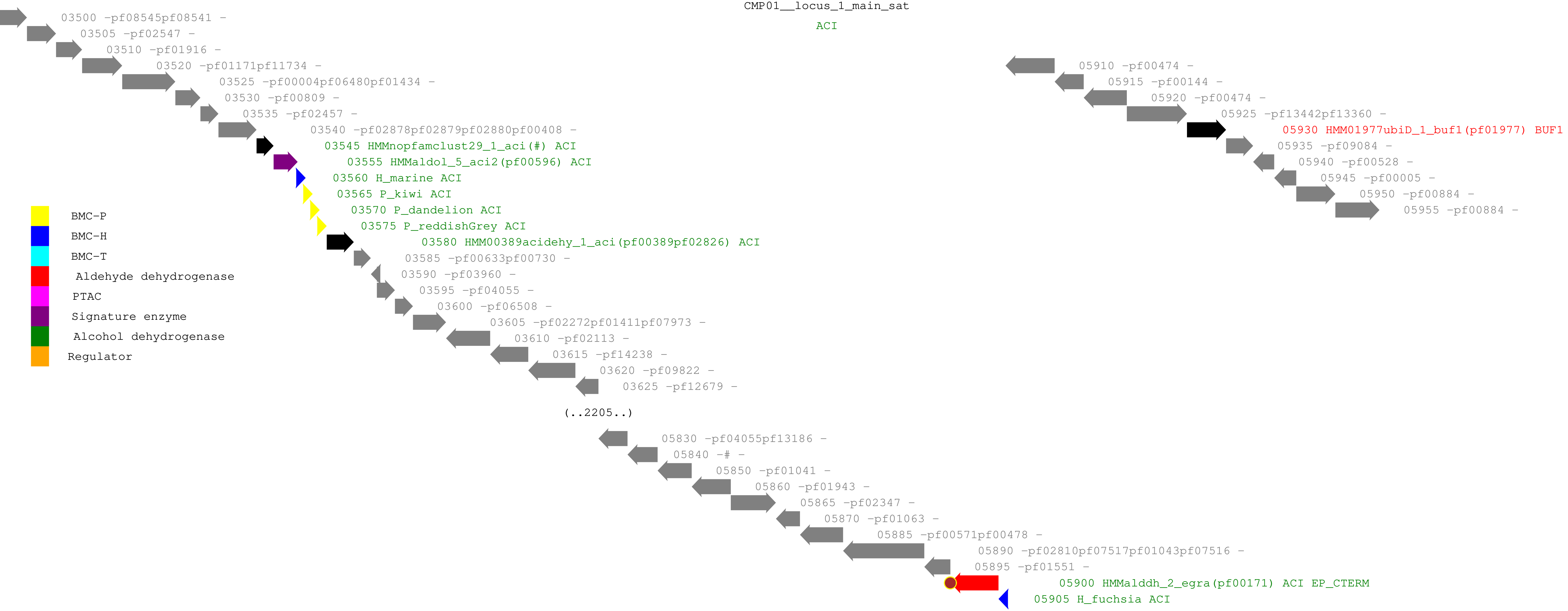

Acidobacteria bacterium CG_4_9_14_3_um_filter_49_7. Bacteria; Acidobacteria.

CO090__locus_1_main_sat

ACI

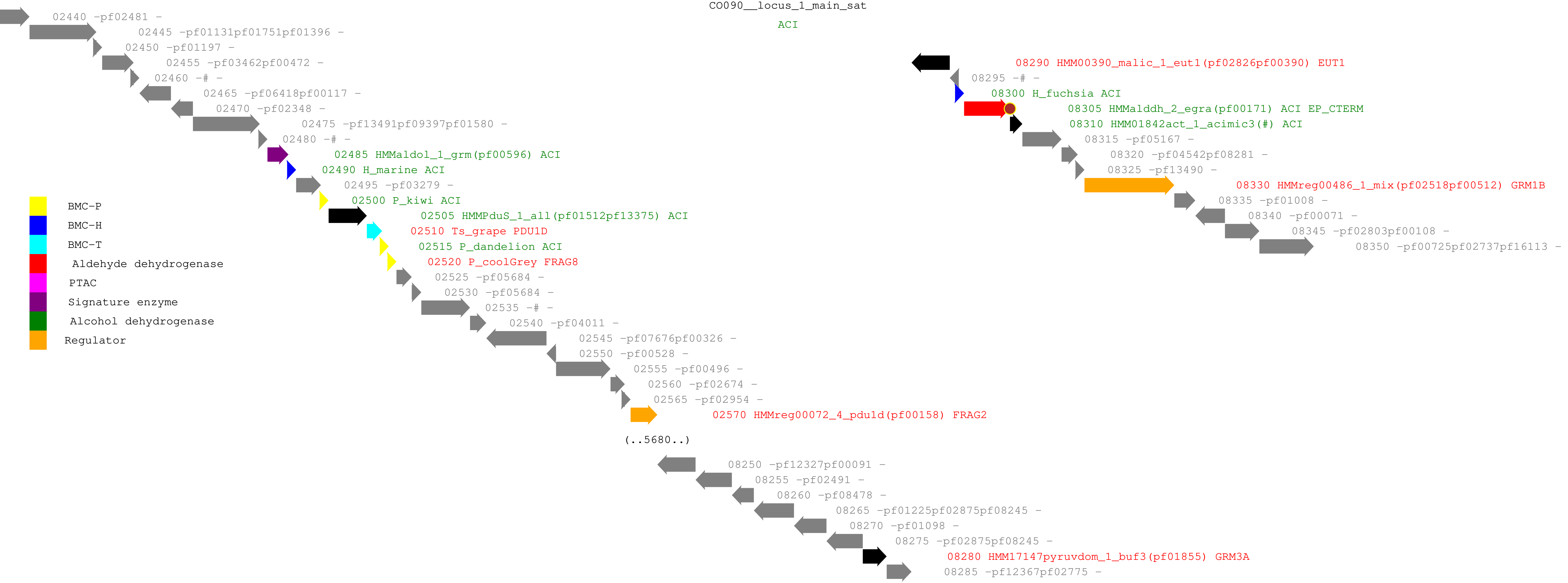

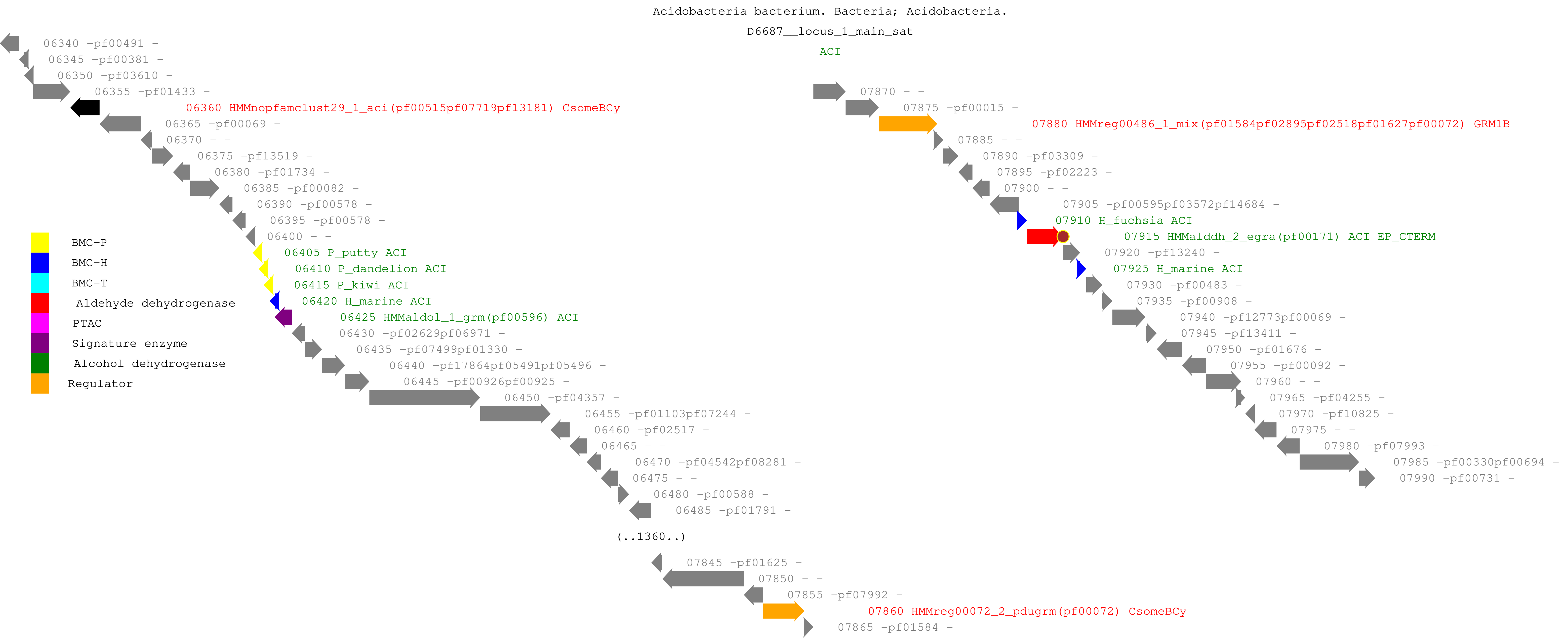
Acidobacteria bacterium. Bacteria; Acidobacteria.
D6687__locus_1_main_sat
ACI
06340 -pf00491 -
06345 -pf00381 -
06350 -pf03610 -
06355 -pf01433 -
06360 HMMnopfamclust29_1_aci(pf00515pf07719pf13181) CsomeBCy
06365 -pf00069 -
06370 - -
06375 -pf13519 -
06380 -pf01734 -
06385 -pf00082 -
06390 -pf00578 -
06395 -pf00578 -
06400 - -
06405 P_putty ACI
06410 P_dandelion ACI
06415 P_kiwi ACI
06420 H_marine ACI
06425 HMMaldol_1_grm(pf00596) ACI
06430 -pf02629pf06971 -
06435 -pf07499pf01330 -
06440 -pf17864pf05491pf05496 -
06445 -pf00926pf00925 -
06450 -pf04357 -
06455 -pf01103pf07244 -
06460 -pf02517 -
06465 - -
06470 -pf04542pf08281 -
06475 - -
06480 -pf00588 -
06485 -pf01791 -
(..1360..)
07845 -pf01625 -
07850 - -
07855 -pf07992 -
07860 HMMreg00072_2_pdugrm(pf00072) CsomeBCy
07865 -pf01584 -
07870 - -
07875 -pf00015 -
07880 HMMreg00486_1_mix(pf01584pf02895pf02518pf01627pf00072) GRM1B
07885 - -
07890 -pf03309 -
07895 -pf02223 -
07900 - -
07905 -pf00595pf03572pf14684 -
07910 H_fuchsia ACI
07915 HMMalddh_2_egra(pf00171) ACI EP_CTERM
07920 -pf13240 -
07925 H_marine ACI
07930 -pf00483 -
07935 -pf00908 -
07940 -pf12773pf00069 -
07945 -pf13411 -
07950 -pf01676 -
07955 -pf00092 -
07960 - -
07965 -pf04255 -
07970 -pf10825 -
07975 - -
07980 -pf07993 -
07985 -pf00330pf00694 -
07990 -pf00731 -
BMC-P
BMC-H
BMC-T
Aldehyde dehydrogenase
PTAC
Signature enzyme
Alcohol dehydrogenase
Regulator

Acidobacteria bacterium. Bacteria; Acidobacteria.

D6723__locus_1_main_sat

ACI

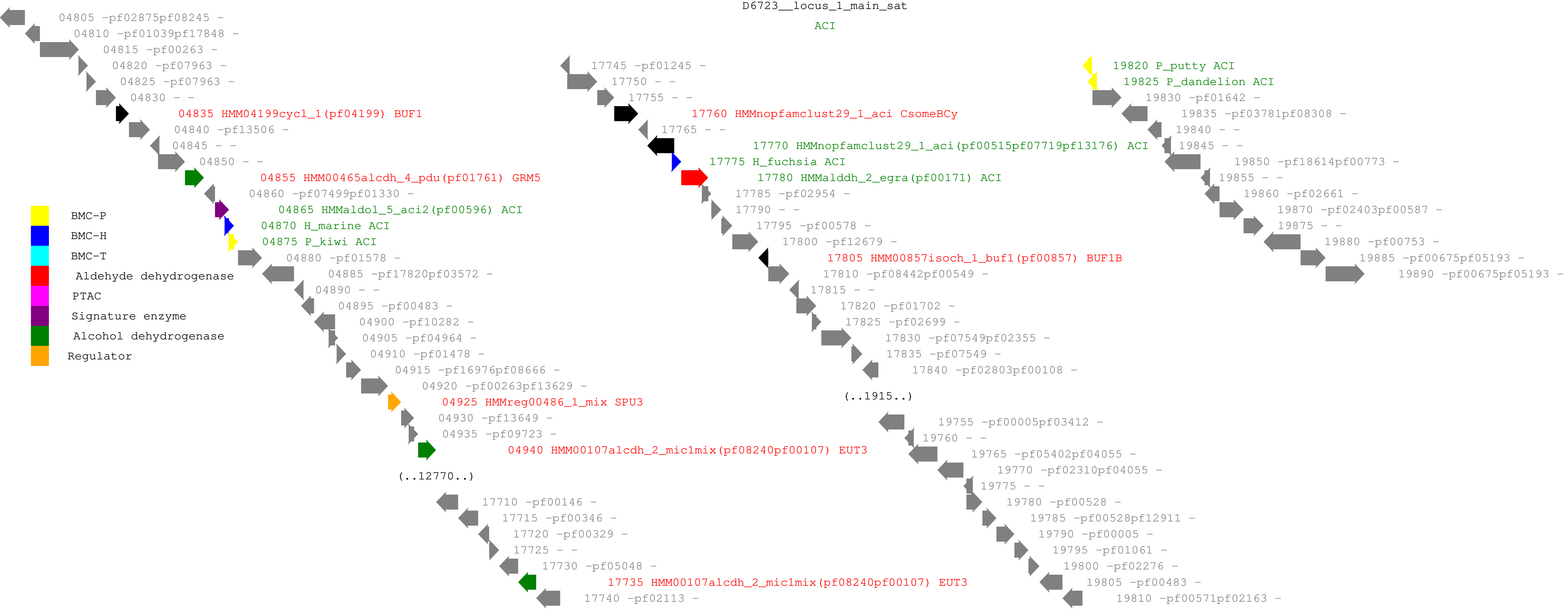

Acidobacteria bacterium. Bacteria; Acidobacteria.

D6735__locus_1_main_sat

ACI

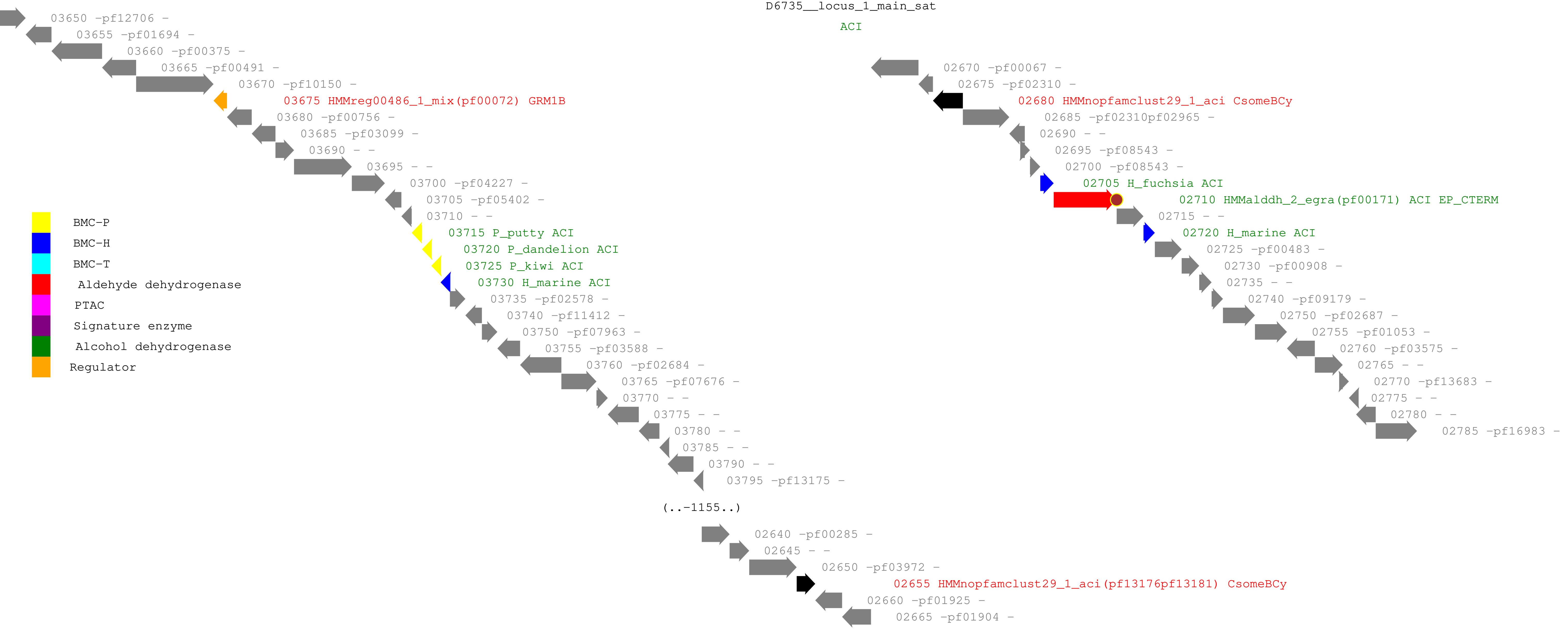

Blastocatellia bacterium. Bacteria; Acidobacteria; Blastocatellia.

DC047__locus_1_main_sat

ACI

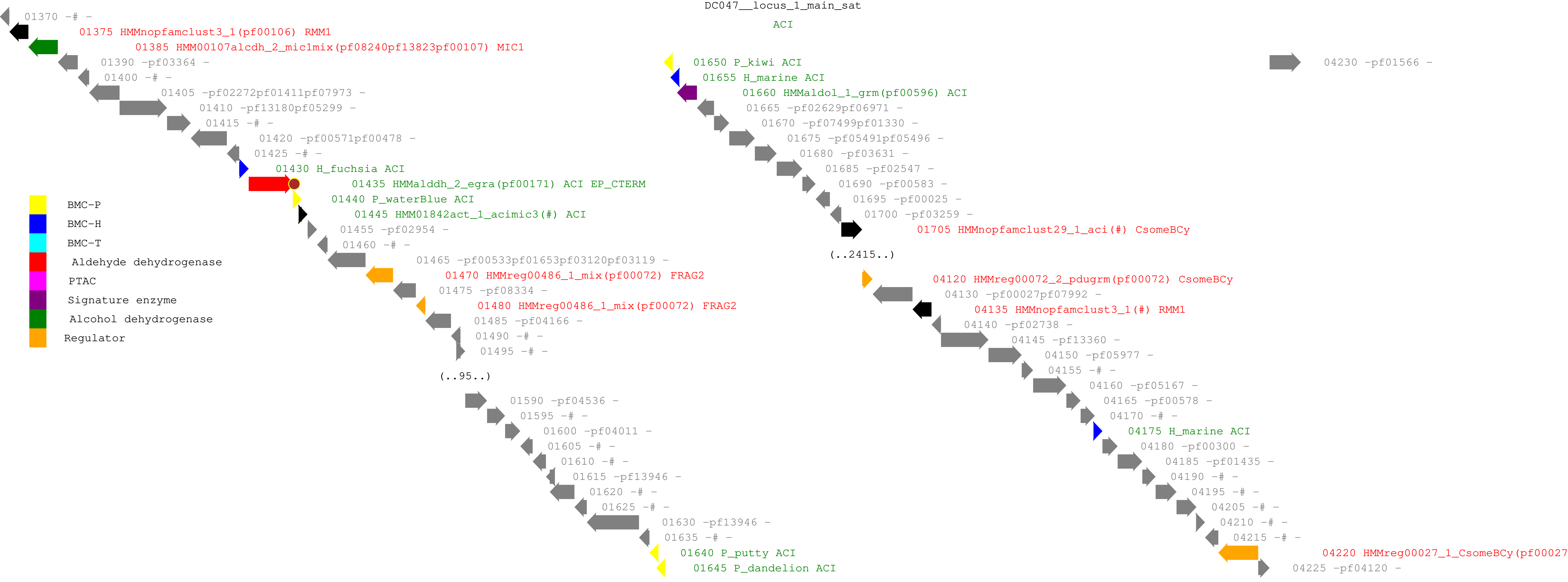

Blastocatellia bacterium. Bacteria; Acidobacteria; Blastocatellia.

DC054__locus_1_main_sat

ACI

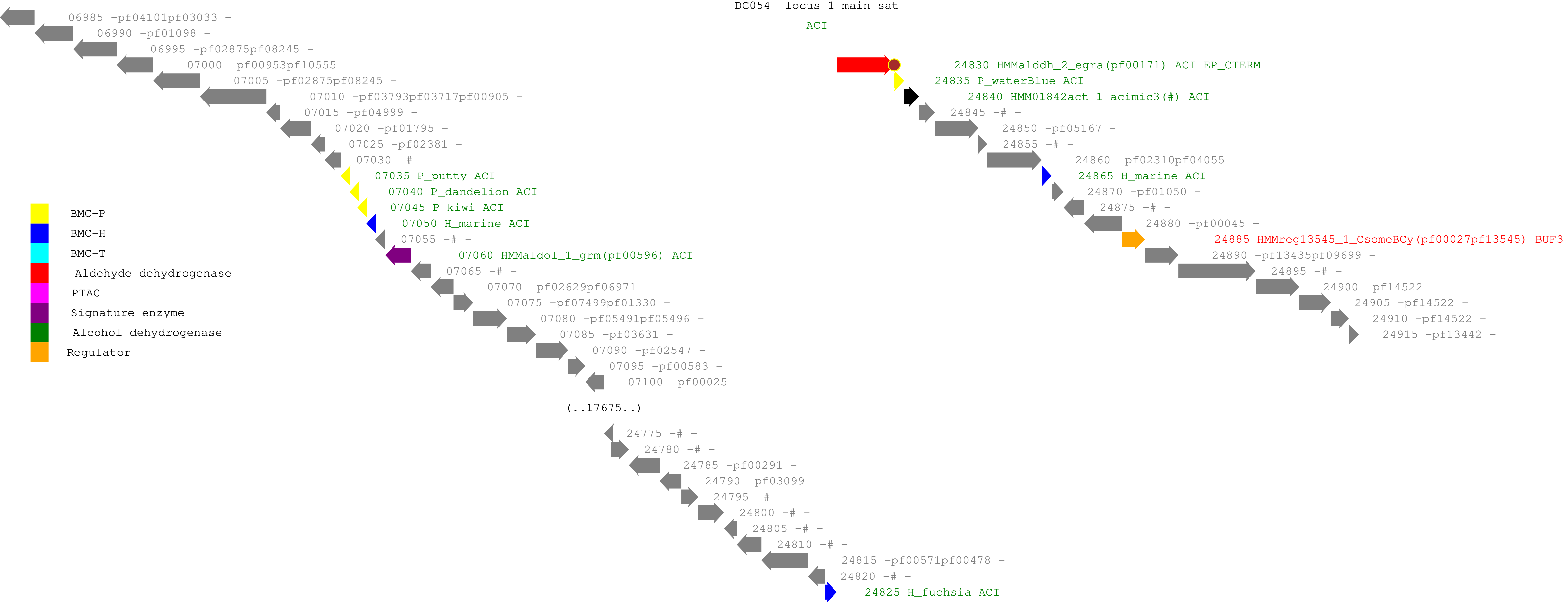

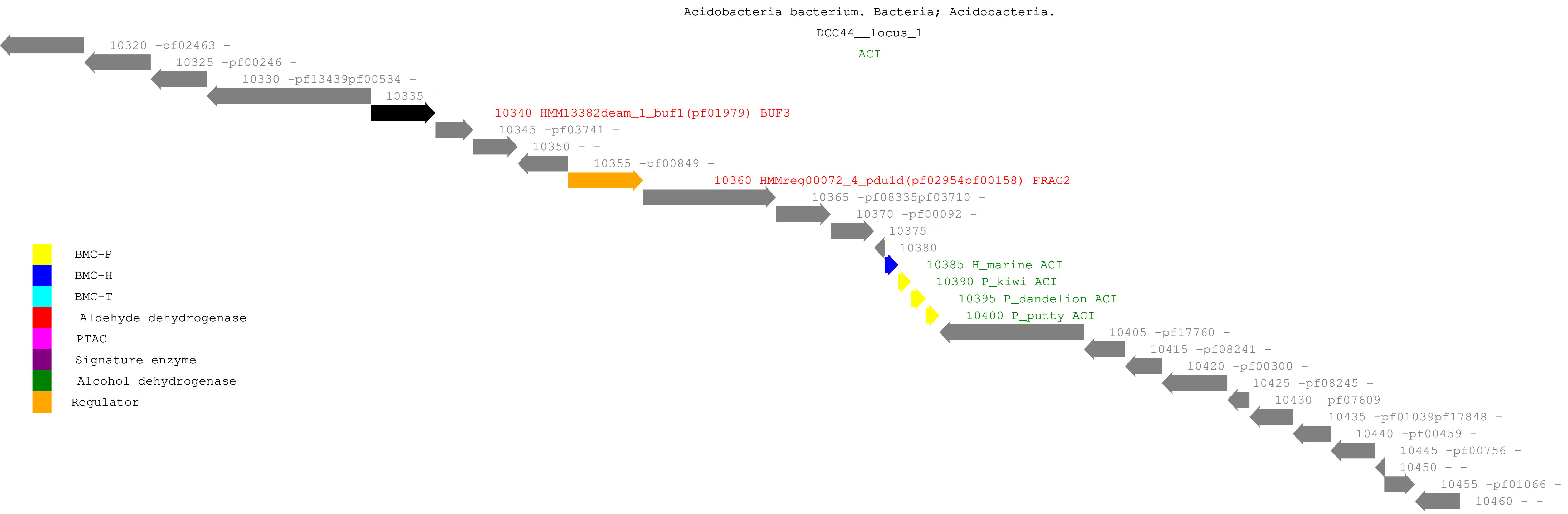

Acidobacteria bacterium. Bacteria; Acidobacteria.
DCC44__locus_1
ACI
10320 -pf02463 -
10325 -pf00246 -
10330 -pf13439pf00534 -
10335 - -
10340 HMM13382deam_1_buf1(pf01979) BUF3
10345 -pf03741 -
10350 - -
10355 -pf00849 -
10360 HMMreg00072_4_pdu1d(pf02954pf00158) FRAG2
10365 -pf08335pf03710 -
10370 -pf00092 -
10375 - -
10380 - -
10385 H_marine ACI
10390 P_kiwi ACI
10395 P_dandelion ACI
10400 P_putty ACI
10405 -pf17760 -
10415 -pf08241 -
10420 -pf00300 -
10425 -pf08245 -
10430 -pf07609 -
10435 -pf01039pf17848 -
10440 -pf00459 -
10445 -pf00756 -
10450 - -
10455 -pf01066 -
10460 - -
BMC-P
BMC-H
BMC-T
Aldehyde dehydrogenase
PTAC
Signature enzyme
Alcohol dehydrogenase
Regulator

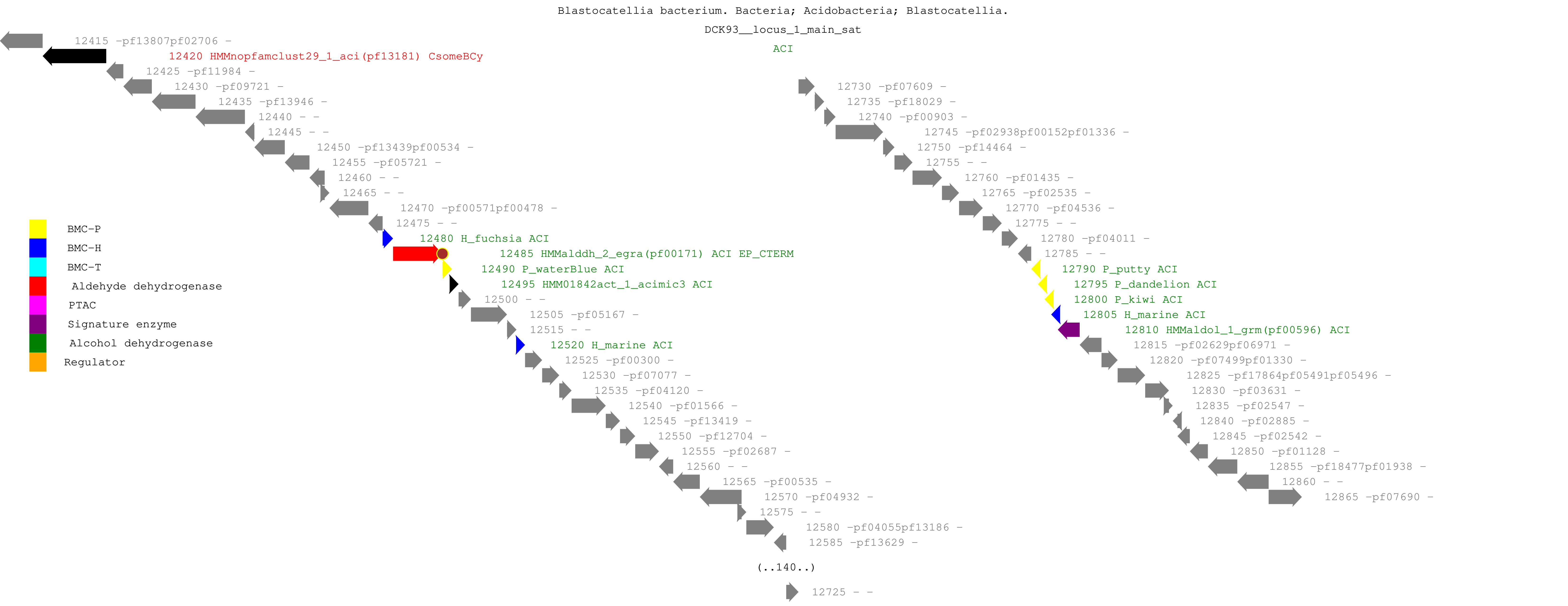

Blastocatellia bacterium. Bacteria; Acidobacteria; Blastocatellia.
DCK93__locus_1_main_sat
ACI
12415 -pf13807pf02706 -
12420 HMMnopfamclust29_1_aci(pf13181) CsomeBCy
12425 -pf11984 -
12430 -pf09721 -
12435 -pf13946 -
12440 - -
12445 - -
12450 -pf13439pf00534 -
12455 -pf05721 -
12460 - -
12465 - -
12470 -pf00571pf00478 -
12475 - -
12480 H_fuchsia ACI
12485 HMMalddh_2_egra(pf00171) ACI EP_CTERM
12490 P_waterBlue ACI
12495 HMM01842act_1_acimic3 ACI
12500 - -
12505 -pf05167 -
12515 - -
12520 H_marine ACI
12525 -pf00300 -
12530 -pf07077 -
12535 -pf04120 -
12540 -pf01566 -
12545 -pf13419 -
12550 -pf12704 -
12555 -pf02687 -
12560 - -
12565 -pf00535 -
12570 -pf04932 -
12575 - -
12580 -pf04055pf13186 -
12585 -pf13629 -
(..140..)
12725 - -
12730 -pf07609 -
12735 -pf18029 -
12740 -pf00903 -
12745 -pf02938pf00152pf01336 -
12750 -pf14464 -
12755 - -
12760 -pf01435 -
12765 -pf02535 -
12770 -pf04536 -
12775 - -
12780 -pf04011 -
12785 - -
12790 P_putty ACI
12795 P_dandelion ACI
12800 P_kiwi ACI
12805 H_marine ACI
12810 HMMaldol_1_grm(pf00596) ACI
12815 -pf02629pf06971 -
12820 -pf07499pf01330 -
12825 -pf17864pf05491pf05496 -
12830 -pf03631 -
12835 -pf02547 -
12840 -pf02885 -
12845 -pf02542 -
12850 -pf01128 -
12855 -pf18477pf01938 -
12860 - -
12865 -pf07690 -
BMC-P
BMC-H
BMC-T
Aldehyde dehydrogenase
PTAC
Signature enzyme
Alcohol dehydrogenase
Regulator

Blastocatellia bacterium. Bacteria; Acidobacteria; Blastocatellia.

DCK99__locus_1_main_sat

ACI

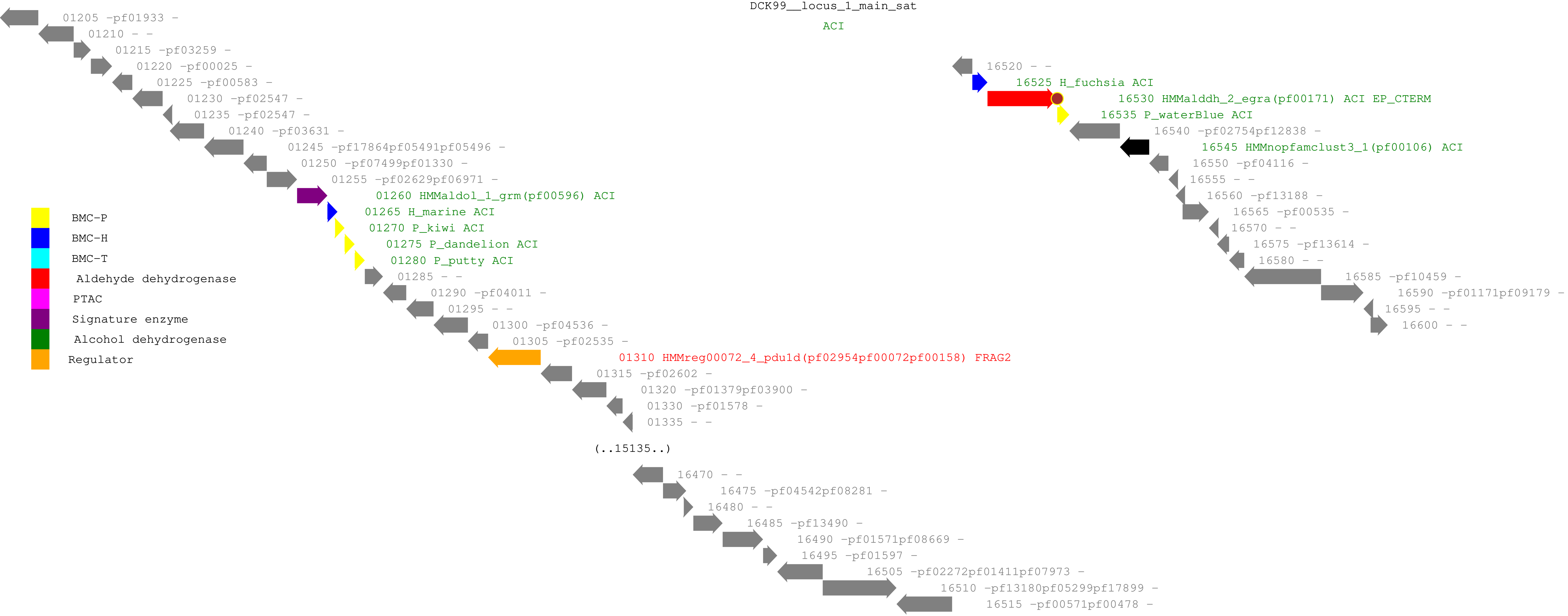

Solibacterales bacterium. Bacteria; Acidobacteria; Solibacteres; Solibacterales;

DCY80__locus_fragmented

ACI

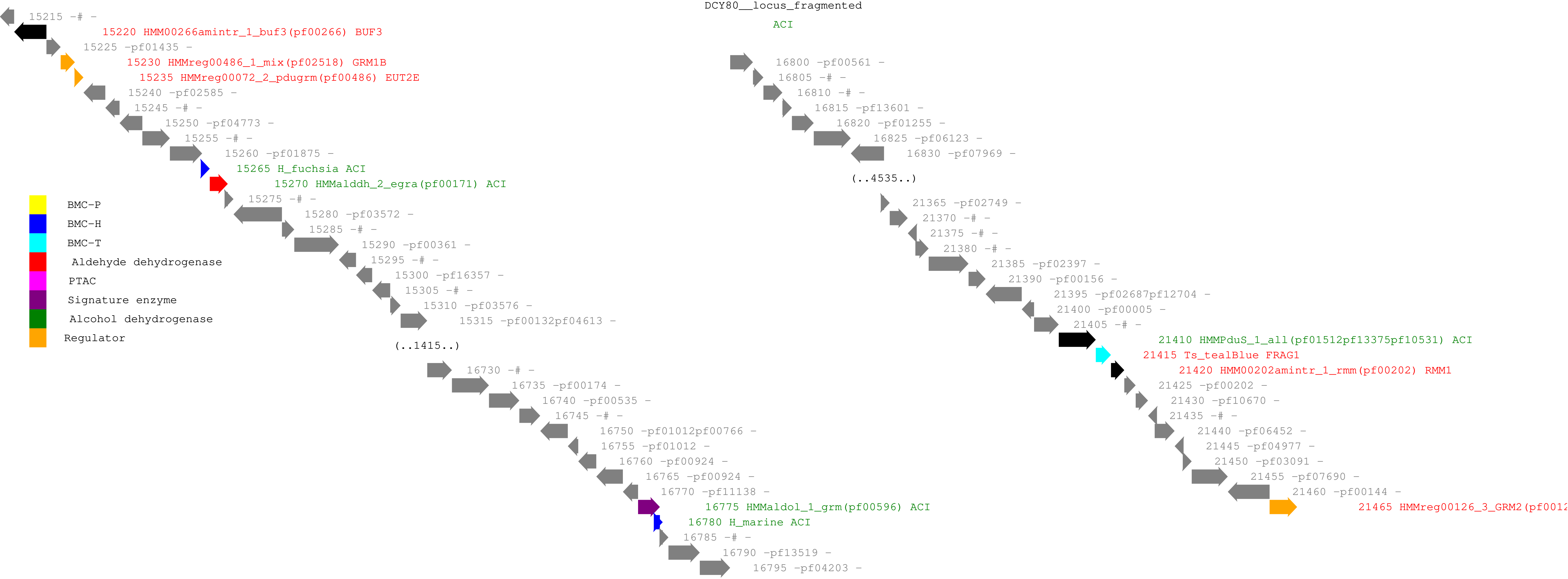

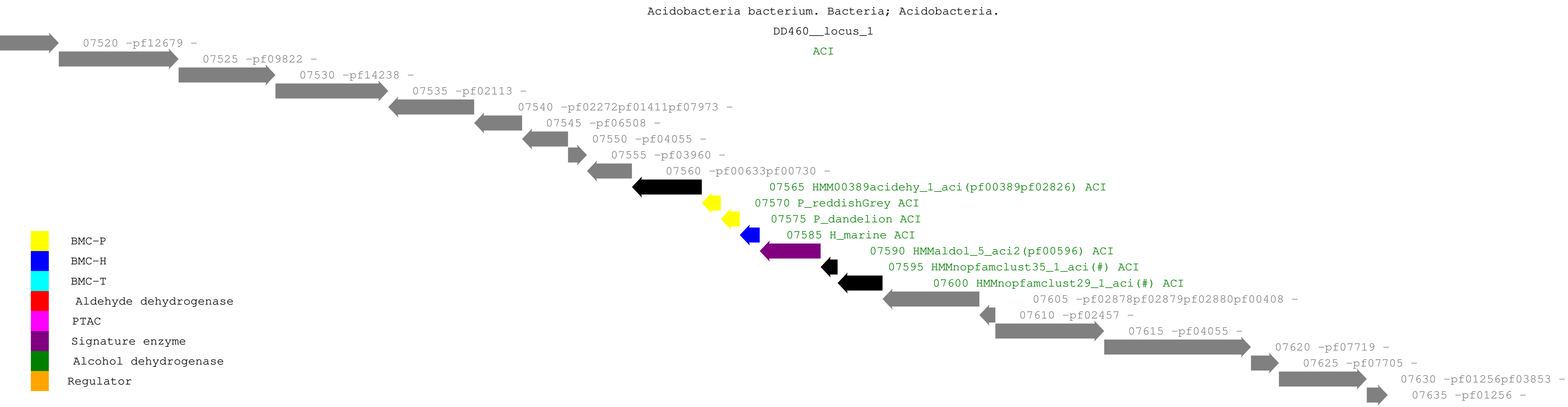
Acidobacteria bacterium. Bacteria; Acidobacteria.
DD460__locus_1
ACI
07520 -pf12679 -
07525 -pf09822 -
07530 -pf14238 -
07535 -pf02113 -
07540 -pf02272pf01411pf07973 -
07545 -pf06508 -
07550 -pf04055 -
07555 -pf03960 -
07560 -pf00633pf00730 -
07565 HMM00389acidehy_1_aci(pf00389pf02826) ACI
07570 P_reddishGrey ACI
07575 P_dandelion ACI
07585 H_marine ACI
07590 HMMaldol_5_aci2(pf00596) ACI
07595 HMMnopfamclust35_1_aci(#) ACI
07600 HMMnopfamclust29_1_aci(#) ACI
07605 -pf02878pf02879pf02880pf00408 -
07610 -pf02457 -
07615 -pf04055 -
07620 -pf07719 -
07625 -pf07705 -
07630 -pf01256pf03853 -
07635 -pf01256 -
BMC-P
BMC-H
BMC-T
Aldehyde dehydrogenase
PTAC
Signature enzyme
Alcohol dehydrogenase
Regulator

Blastocatellia bacterium. Bacteria; Acidobacteria; Blastocatellia.

DDW24__locus_1_main_sat

ACI

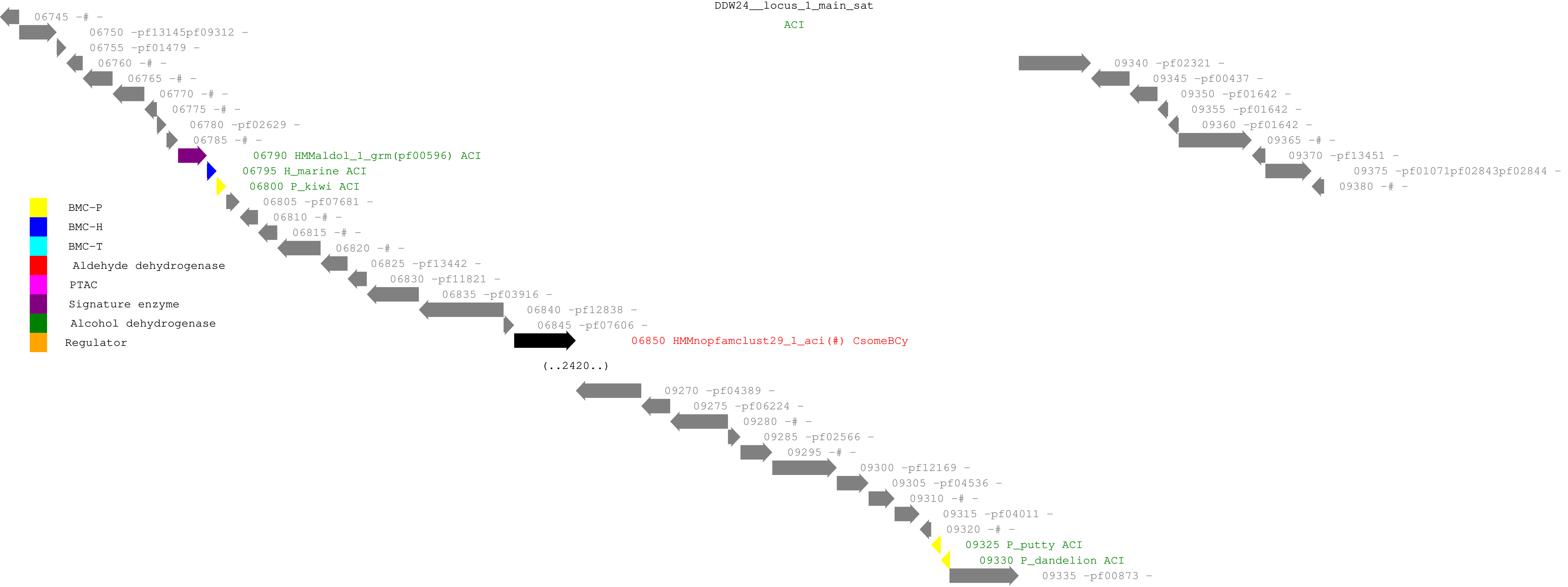

Blastocatellia bacterium. Bacteria; Acidobacteria; Blastocatellia.

DEA22__locus_1_main_sat

ACI

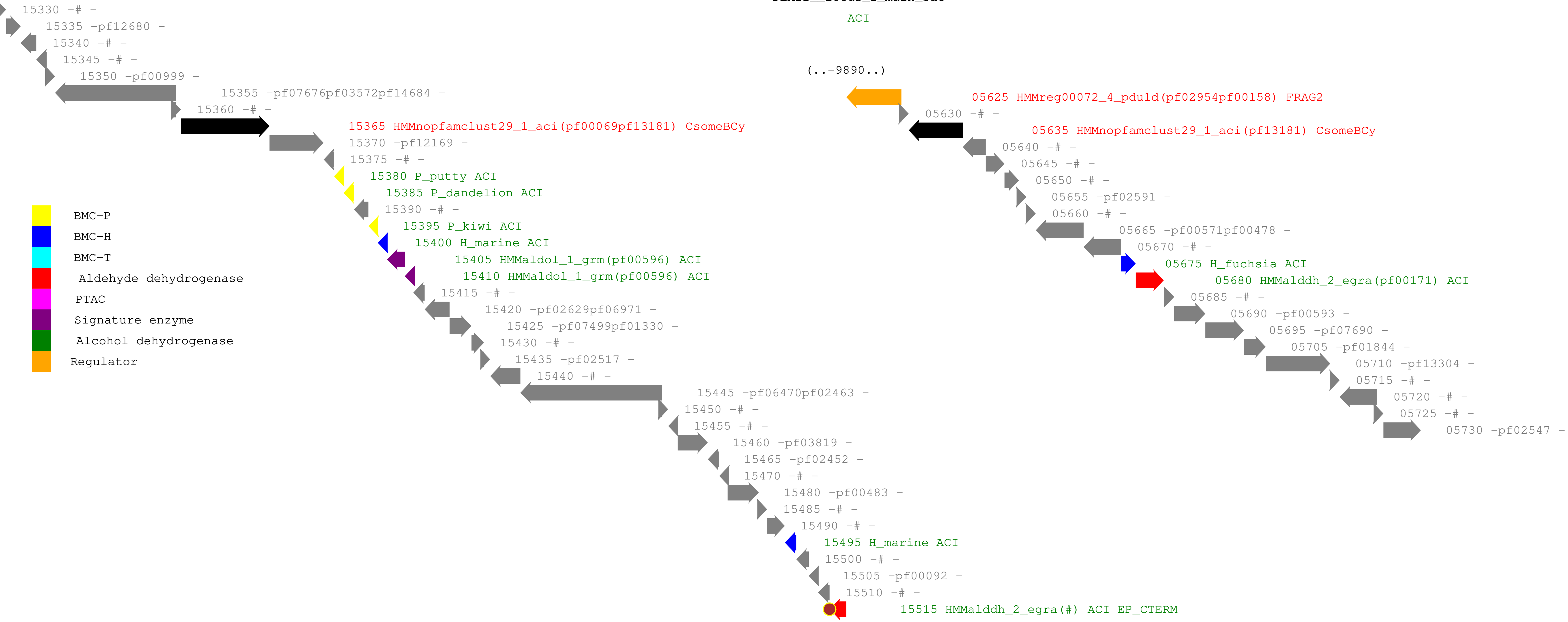

Solibacterales bacterium. Bacteria; Acidobacteria; Solibacteres; Solibacterales;

DEH78__locus_1_main_sat

ACI

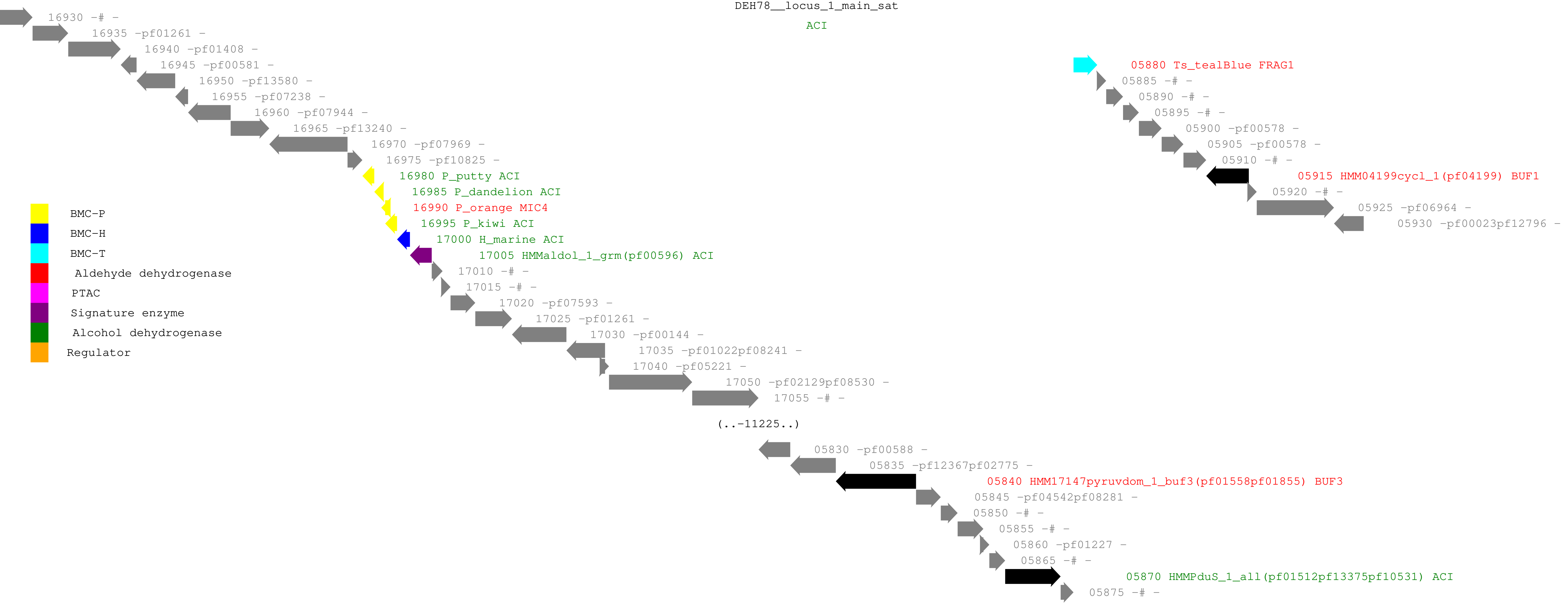

Blastocatellia bacterium. Bacteria; Acidobacteria; Blastocatellia.

DEP46__locus_1_main_sat

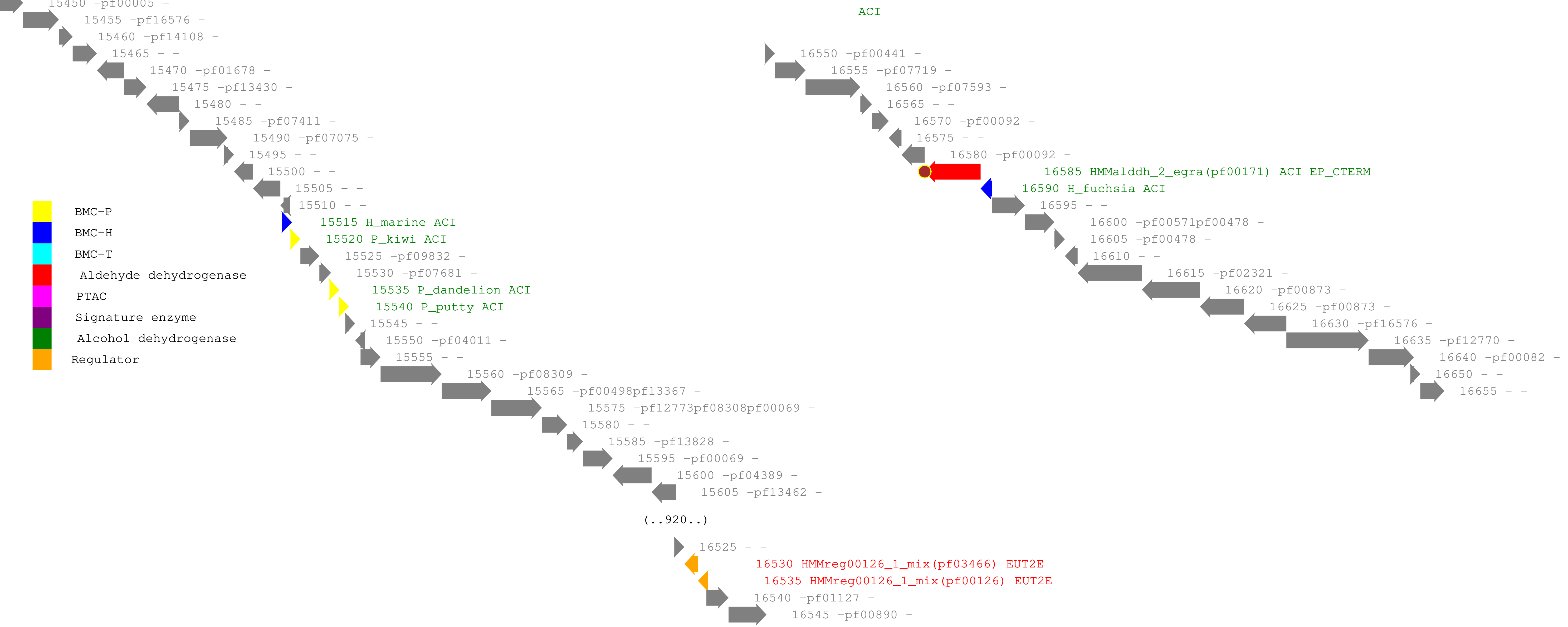

Acidobacteria bacterium. Bacteria; Acidobacteria.

DEQ98__locus_1

ACI

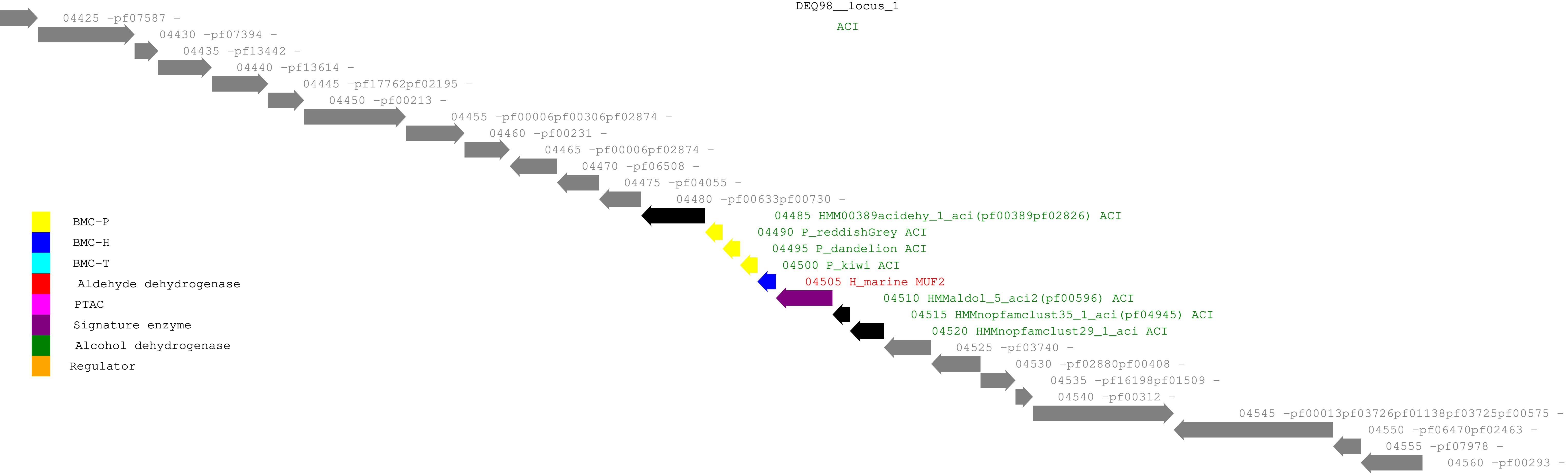

Blastocatellia bacterium. Bacteria; Acidobacteria; Blastocatellia.

DHU55__locus_1_main_sat

ACI

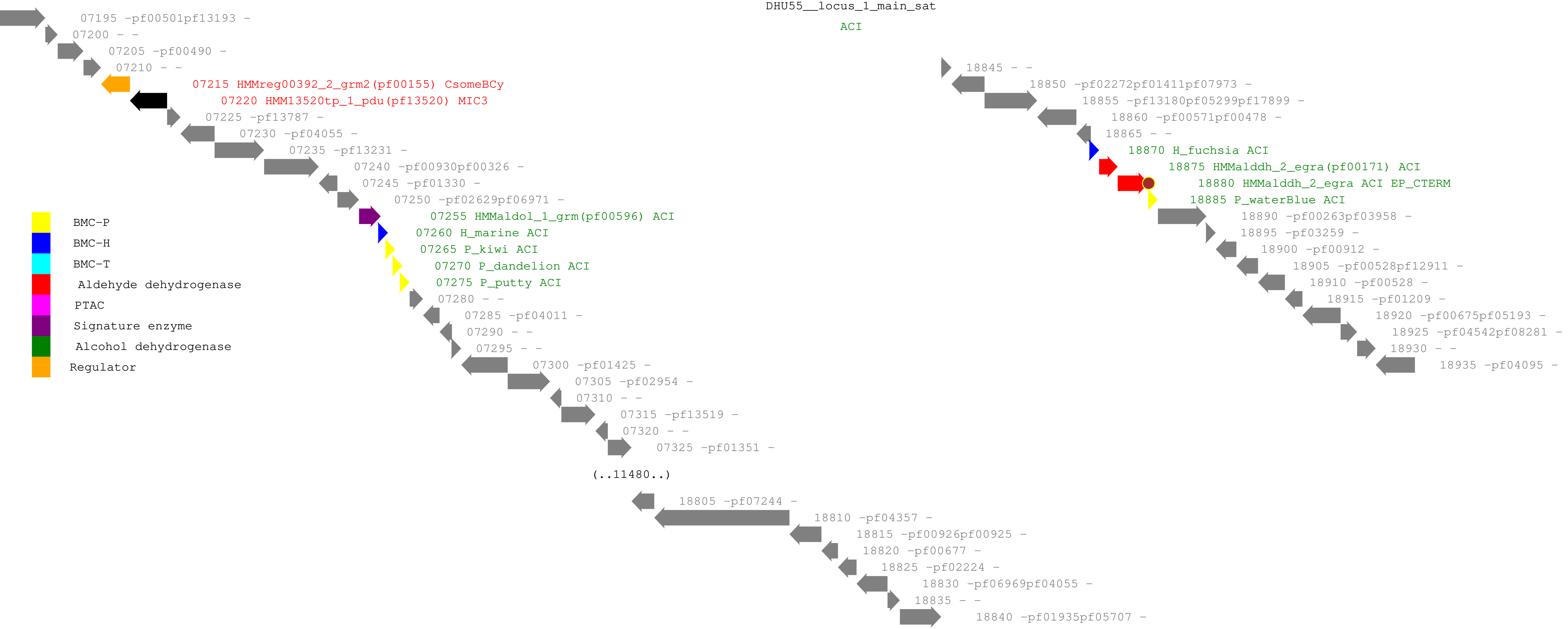

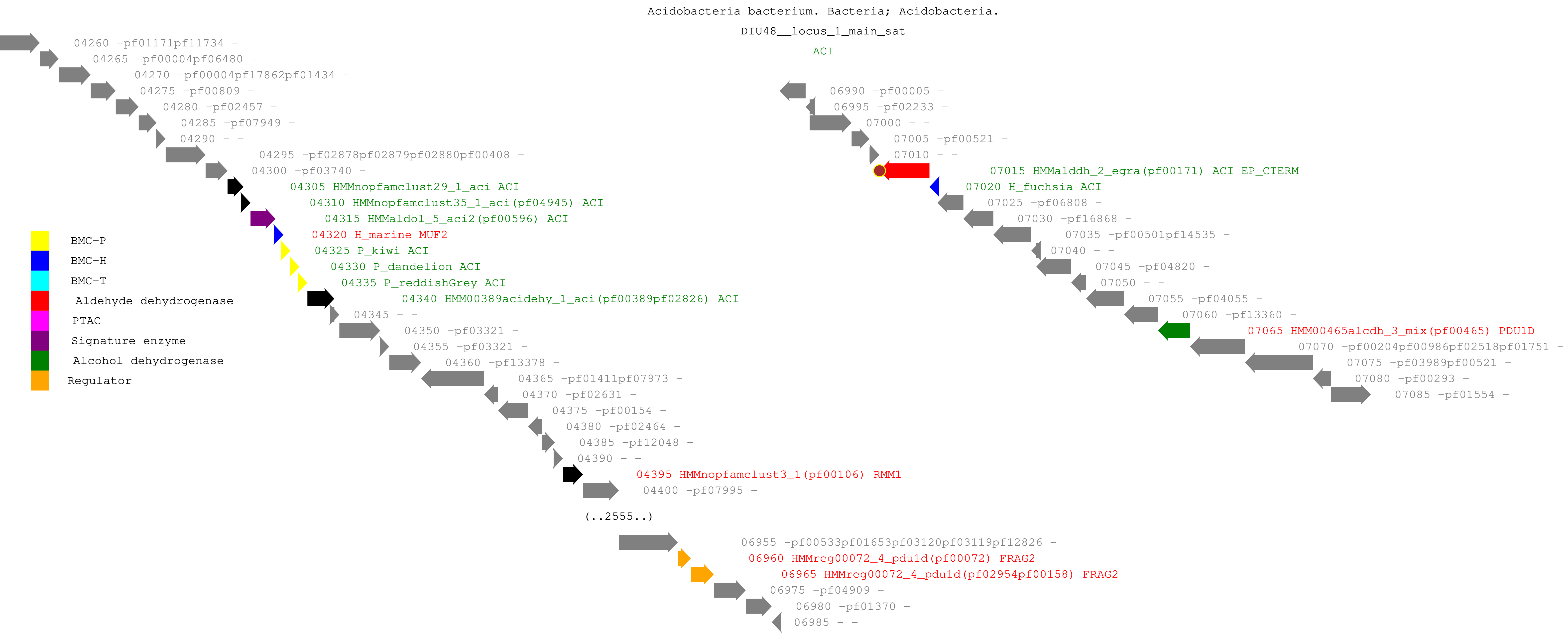

Acidobacteria bacterium. Bacteria; Acidobacteria.
DIU48__locus_1_main_sat
ACI
04260 -pf01171pf11734 -
04265 -pf00004pf06480 -
04270 -pf00004pf17862pf01434 -
04275 -pf00809 -
04280 -pf02457 -
04285 -pf07949 -
04290 - -
04295 -pf02878pf02879pf02880pf00408 -
04300 -pf03740 -
04305 HMMnopfamclust29_1_aci ACI
04310 HMMnopfamclust35_1_aci(pf04945) ACI
04315 HMMaldol_5_aci2(pf00596) ACI
04320 H_marine MUF2
04325 P_kiwi ACI
04330 P_dandelion ACI
04335 P_reddishGrey ACI
04340 HMM00389acidehy_1_aci(pf00389pf02826) ACI
04345 - -
04350 -pf03321 -
04355 -pf03321 -
04360 -pf13378 -
04365 -pf01411pf07973 -
04370 -pf02631 -
04375 -pf00154 -
04380 -pf02464 -
04385 -pf12048 -
04390 - -
04395 HMMnopfamclust3_1(pf00106) RMM1
04400 -pf07995 -
(..2555..)
06955 -pf00533pf01653pf03120pf03119pf12826 -
06960 HMMreg00072_4_pdu1d(pf00072) FRAG2
06965 HMMreg00072_4_pdu1d(pf02954pf00158) FRAG2
06975 -pf04909 -
06980 -pf01370 -
06985 - -
06990 -pf00005 -
06995 -pf02233 -
07000 - -
07005 -pf00521 -
07010 - -
07015 HMMalddh_2_egra(pf00171) ACI EP_CTERM
07020 H_fuchsia ACI
07025 -pf06808 -
07030 -pf16868 -
07035 -pf00501pf14535 -
07040 - -
07045 -pf04820 -
07050 - -
07055 -pf04055 -
07060 -pf13360 -
07065 HMM00465alcdh_3_mix(pf00465) PDU1D
07070 -pf00204pf00986pf02518pf01751 -
07075 -pf03989pf00521 -
07080 -pf00293 -
07085 -pf01554 -
BMC-P
BMC-H
BMC-T
Aldehyde dehydrogenase
PTAC
Signature enzyme
Alcohol dehydrogenase
Regulator

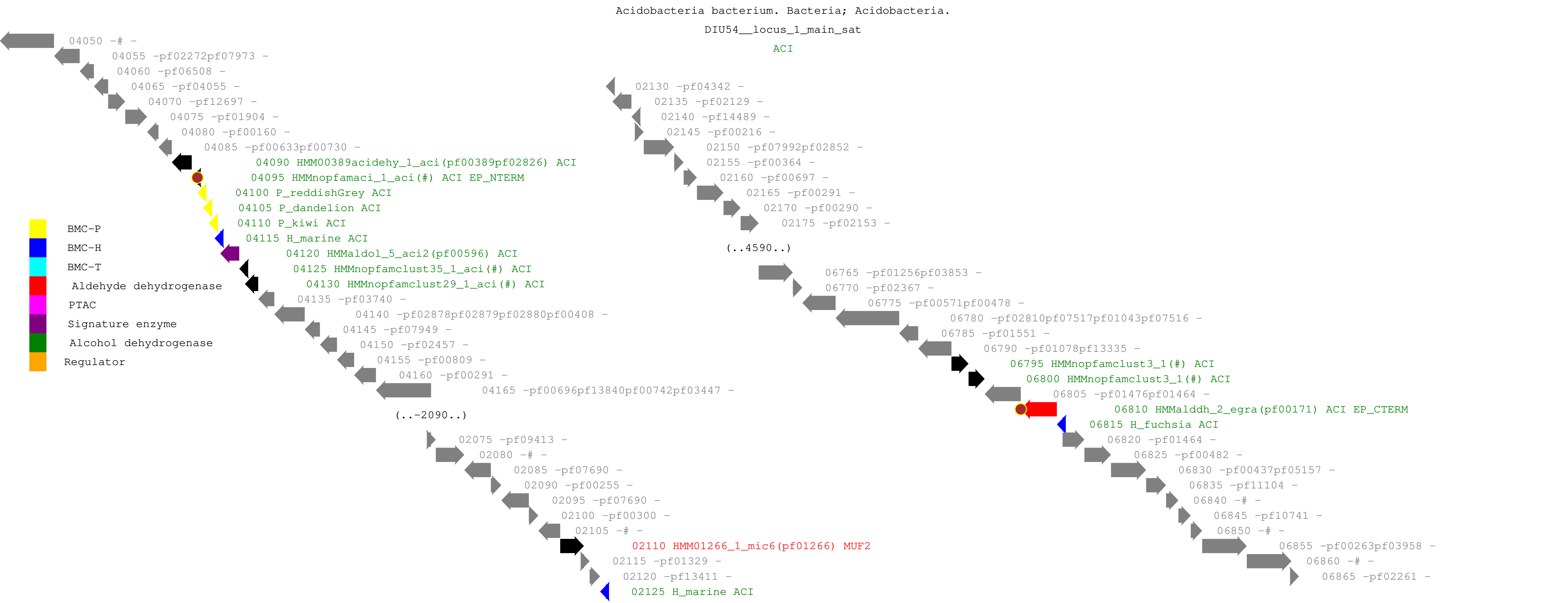
Acidobacteria bacterium. Bacteria; Acidobacteria.
DIU54__locus_1_main_sat
ACI
04050 -# -
04055 -pf02272pf07973 -
04060 -pf06508 -
04065 -pf04055 -
04070 -pf12697 -
04075 -pf01904 -
04080 -pf00160 -
04085 -pf00633pf00730 -
04090 HMM00389acidehy_1_aci(pf00389pf02826) ACI
04095 HMMnopfamaci_1_aci(#) ACI EP_NTERM
04100 P_reddishGrey ACI
04105 P_dandelion ACI
04110 P_kiwi ACI
04115 H_marine ACI
04120 HMMaldol_5_aci2(pf00596) ACI
04125 HMMnopfamclust35_1_aci(#) ACI
04130 HMMnopfamclust29_1_aci(#) ACI
04135 -pf03740 -
04140 -pf02878pf02879pf02880pf00408 -
04145 -pf07949 -
04150 -pf02457 -
04155 -pf00809 -
04160 -pf00291 -
04165 -pf00696pf13840pf00742pf03447 -
(..-2090..)
02075 -pf09413 -
02080 -# -
02085 -pf07690 -
02090 -pf00255 -
02095 -pf07690 -
02100 -pf00300 -
02105 -# -
02110 HMM01266_1_mic6(pf01266) MUF2
02115 -pf01329 -
02120 -pf13411 -
02125 H_marine ACI
02130 -pf04342 -
02135 -pf02129 -
02140 -pf14489 -
02145 -pf00216 -
02150 -pf07992pf02852 -
02155 -pf00364 -
02160 -pf00697 -
02165 -pf00291 -
02170 -pf00290 -
02175 -pf02153 -
(..4590..)
06765 -pf01256pf03853 -
06770 -pf02367 -
06775 -pf00571pf00478 -
06780 -pf02810pf07517pf01043pf07516 -
06785 -pf01551 -
06790 -pf01078pf13335 -
06795 HMMnopfamclust3_1(#) ACI
06800 HMMnopfamclust3_1(#) ACI
06805 -pf01476pf01464 -
06810 HMMalddh_2_egra(pf00171) ACI EP_CTERM
06815 H_fuchsia ACI
06820 -pf01464 -
06825 -pf00482 -
06830 -pf00437pf05157 -
06835 -pf11104 -
06840 -# -
06845 -pf10741 -
06850 -# -
06855 -pf00263pf03958 -
06860 -# -
06865 -pf02261 -
BMC-P
BMC-H
BMC-T
Aldehyde dehydrogenase
PTAC
Signature enzyme
Alcohol dehydrogenase
Regulator

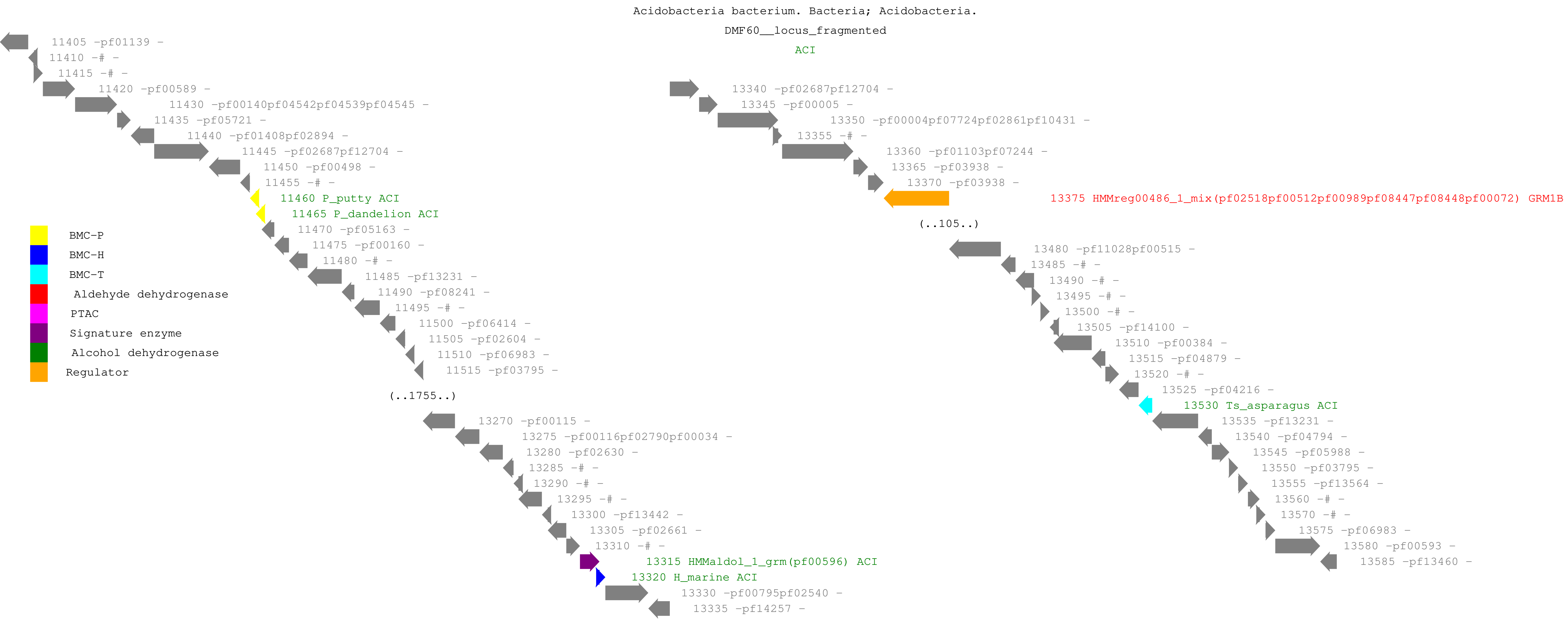

Acidobacteria bacterium. Bacteria; Acidobacteria.
DMF60__locus_fragmented
ACI
11405 -pf01139 -
11410 -# -
11415 -# -
11420 -pf00589 -
11430 -pf00140pf04542pf04539pf04545 -
11435 -pf05721 -
11440 -pf01408pf02894 -
11445 -pf02687pf12704 -
11450 -pf00498 -
11455 -# -
11460 P_putty ACI
11465 P_dandelion ACI
11470 -pf05163 -
11475 -pf00160 -
11480 -# -
11485 -pf13231 -
11490 -pf08241 -
11495 -# -
11500 -pf06414 -
11505 -pf02604 -
11510 -pf06983 -
11515 -pf03795 -
(..1755..)
13270 -pf00115 -
13275 -pf00116pf02790pf00034 -
13280 -pf02630 -
13285 -# -
13290 -# -
13295 -# -
13300 -pf13442 -
13305 -pf02661 -
13310 -# -
13315 HMMaldol_1_grm(pf00596) ACI
13320 H_marine ACI
13330 -pf00795pf02540 -
13335 -pf14257 -
BMC-P
BMC-H
BMC-T
Aldehyde dehydrogenase
PTAC
Signature enzyme
Alcohol dehydrogenase
Regulator
13340 -pf02687pf12704 -
13345 -pf00005 -
13350 -pf00004pf07724pf02861pf10431 -
13355 -# -
13360 -pf01103pf07244 -
13365 -pf03938 -
13370 -pf03938 -
13375 HMMreg00486_1_mix(pf02518pf00512pf00989pf08447pf08448pf00072) GRM1B
(..105..)
13480 -pf11028pf00515 -
13485 -# -
13490 -# -
13495 -# -
13500 -# -
13505 -pf14100 -
13510 -pf00384 -
13515 -pf04879 -
13520 -# -
13525 -pf04216 -
13530 Ts_asparagus ACI
13535 -pf13231 -
13540 -pf04794 -
13545 -pf05988 -
13550 -pf03795 -
13555 -pf13564 -
13560 -# -
13570 -# -
13575 -pf06983 -
13580 -pf00593 -
13585 -pf13460 -

Blastocatellia bacterium AA13. Bacteria; Acidobacteria; Blastocatellia.

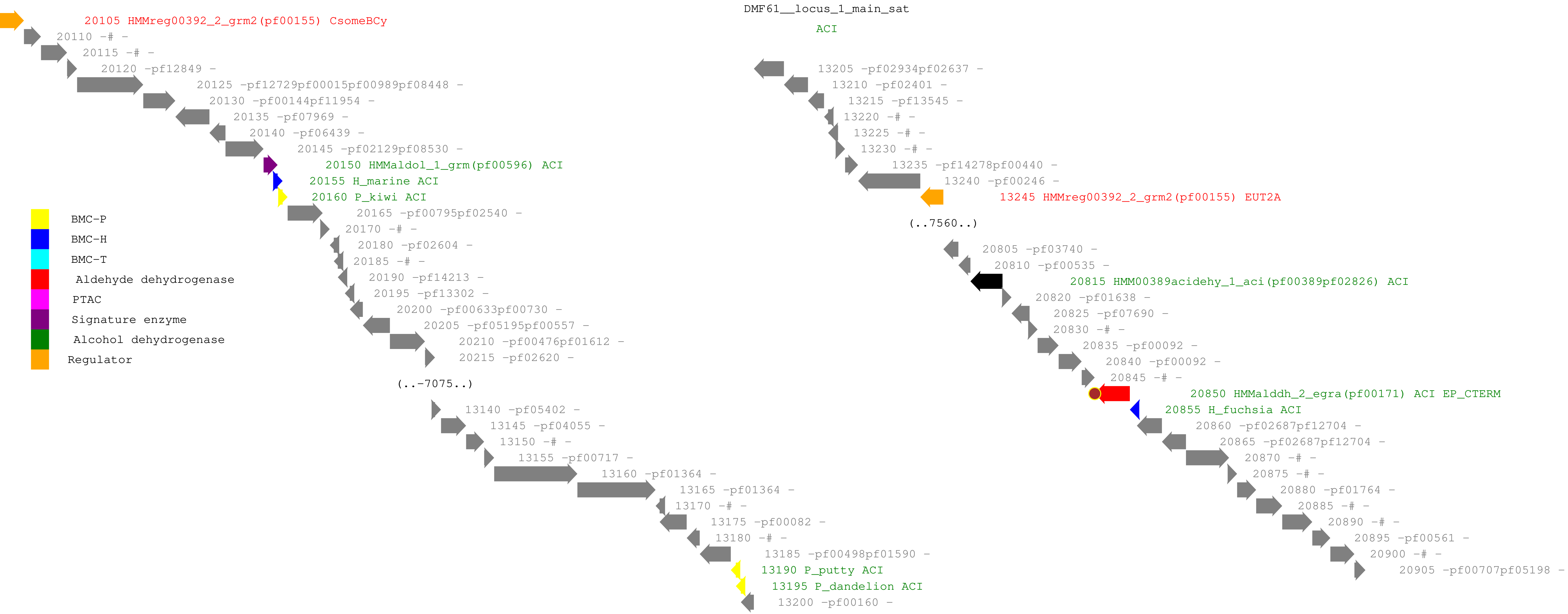

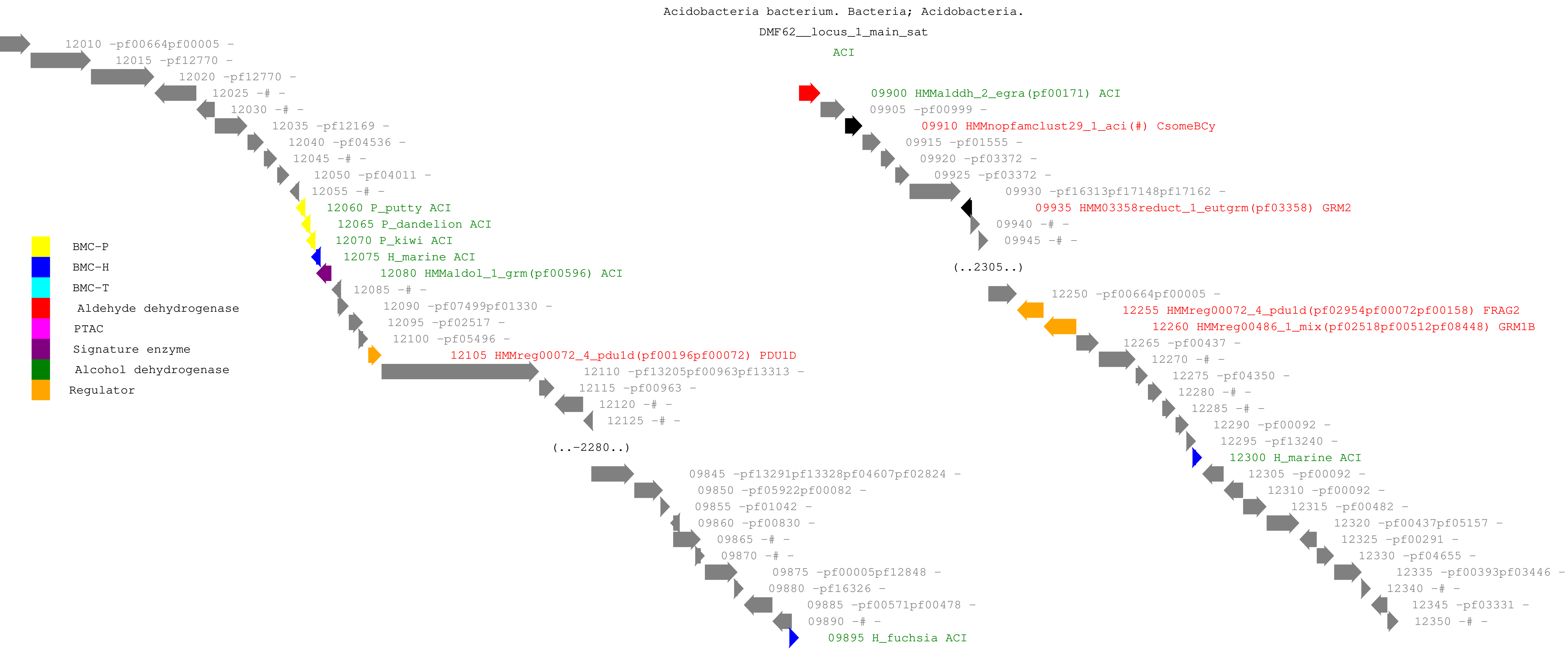

Acidobacteria bacterium. Bacteria; Acidobacteria.
DMF62__locus_1_main_sat
ACI
12010 -pf00664pf00005 -
12015 -pf12770 -
12020 -pf12770 -
12025 -# -
12030 -# -
12035 -pf12169 -
12040 -pf04536 -
12045 -# -
12050 -pf04011 -
12055 -# -
12060 P_putty ACI
12065 P_dandelion ACI
12070 P_kiwi ACI
12075 H_marine ACI
12080 HMMaldol_1_grm(pf00596) ACI
12085 -# -
12090 -pf07499pf01330 -
12095 -pf02517 -
12100 -pf05496 -
12105 HMMreg00072_4_pdu1d(pf00196pf00072) PDU1D
12110 -pf13205pf00963pf13313 -
12115 -pf00963 -
12120 -# -
12125 -# -
(..-2280..)
09845 -pf13291pf13328pf04607pf02824 -
09850 -pf05922pf00082 -
09855 -pf01042 -
09860 -pf00830 -
09865 -# -
09870 -# -
09875 -pf00005pf12848 -
09880 -pf16326 -
09885 -pf00571pf00478 -
09890 -# -
09895 H_fuchsia ACI
09900 HMMaldh_2_egra(pf00171) ACI
09905 -pf00999 -
09910 HMMnopfamclust29_1_aci(#) CsomeBCy
09915 -pf01555 -
09920 -pf03372 -
09925 -pf03372 -
09930 -pf16313pf17148pf17162 -
09935 HMM03358reduct_1_eutgrm(pf03358) GRM2
09940 -# -
09945 -# -
(..2305..)
12250 -pf00664pf00005 -
12255 HMMreg00072_4_pdu1d(pf02954pf00072pf00158) FRAG2
12260 HMMreg00486_1_mix(pf02518pf00512pf08448) GRM1B
12265 -pf00437 -
12270 -# -
12275 -pf04350 -
12280 -# -
12285 -# -
12290 -pf00092 -
12295 -pf13240 -
12300 H_marine ACI
12305 -pf00092 -
12310 -pf00092 -
12315 -pf00482 -
12320 -pf00437pf05157 -
12325 -pf00291 -
12330 -pf04655 -
12335 -pf00393pf03446 -
12340 -# -
12345 -pf03331 -
12350 -# -
BMC-P
BMC-H
BMC-T
Aldehyde dehydrogenase
PTAC
Signature enzyme
Alcohol dehydrogenase
Regulator

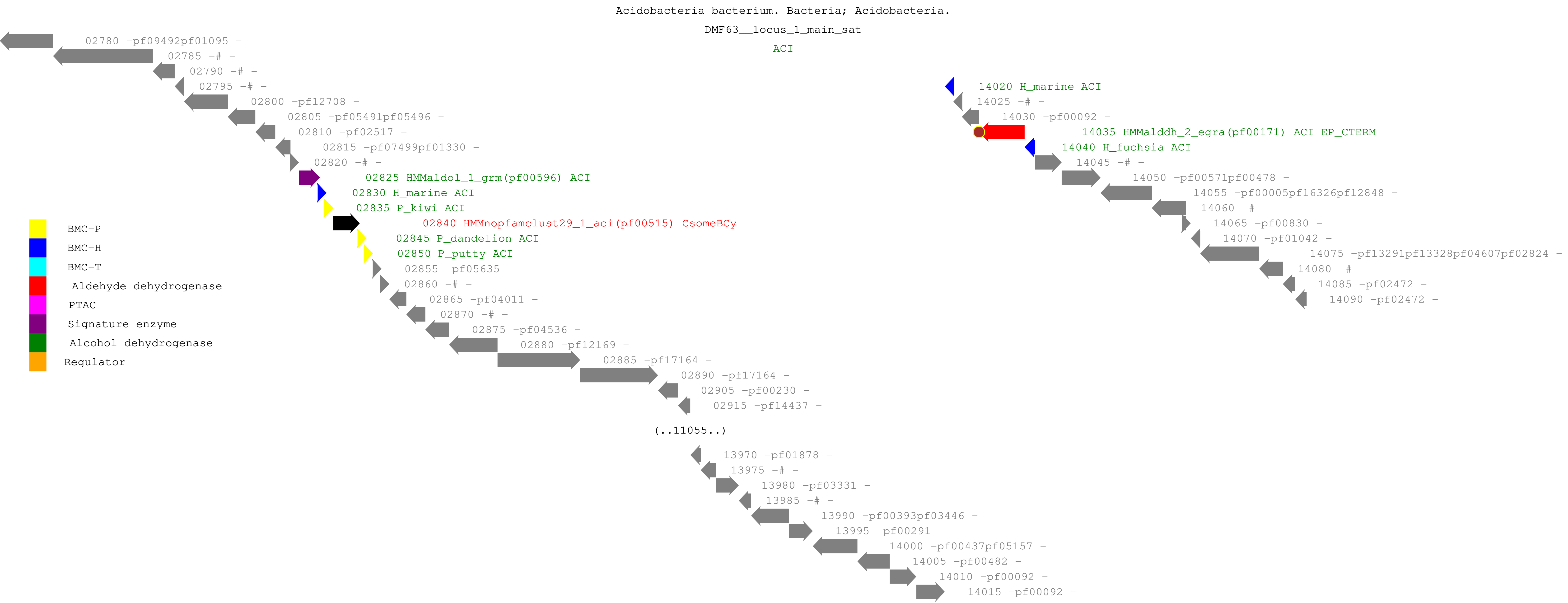

Acidobacteria bacterium. Bacteria; Acidobacteria.
DMF63__locus_1_main_sat
ACI
02780 -pf09492pf01095 -
02785 -# -
02790 -# -
02795 -# -
02800 -pf12708 -
02805 -pf05491pf05496 -
02810 -pf02517 -
02815 -pf07499pf01330 -
02820 -# -
02825 HMMaldol_1_grm(pf00596) ACI
02830 H_marine ACI
02835 P_kiwi ACI
02840 HMMnopfamclust29_1_aci(pf00515) CsomeBCy
02845 P_dandelion ACI
02850 P_putty ACI
02855 -pf05635 -
02860 -# -
02865 -pf04011 -
02870 -# -
02875 -pf04536 -
02880 -pf12169 -
02885 -pf17164 -
02890 -pf17164 -
02905 -pf00230 -
02915 -pf14437 -
(..11055..)
13970 -pf01878 -
13975 -# -
13980 -pf03331 -
13985 -# -
13990 -pf00393pf03446 -
13995 -pf00291 -
14000 -pf00437pf05157 -
14005 -pf00482 -
14010 -pf00092 -
14015 -pf00092 -
14020 H_marine ACI
14025 -# -
14030 -pf00092 -
14035 HMMalddh_2_egra(pf00171) ACI EP_CTERM
14040 H_fuchsia ACI
14045 -# -
14050 -pf00571pf00478 -
14055 -pf00005pf16326pf12848 -
14060 -# -
14065 -pf00830 -
14070 -pf01042 -
14075 -pf13291pf13328pf04607pf02824 -
14080 -# -
14085 -pf02472 -
14090 -pf02472 -
BMC-P
BMC-H
BMC-T
Aldehyde dehydrogenase
PTAC
Signature enzyme
Alcohol dehydrogenase
Regulator

Acidobacteria bacterium. Bacteria; Acidobacteria.

DMF64__locus_1_main_sat

ACI

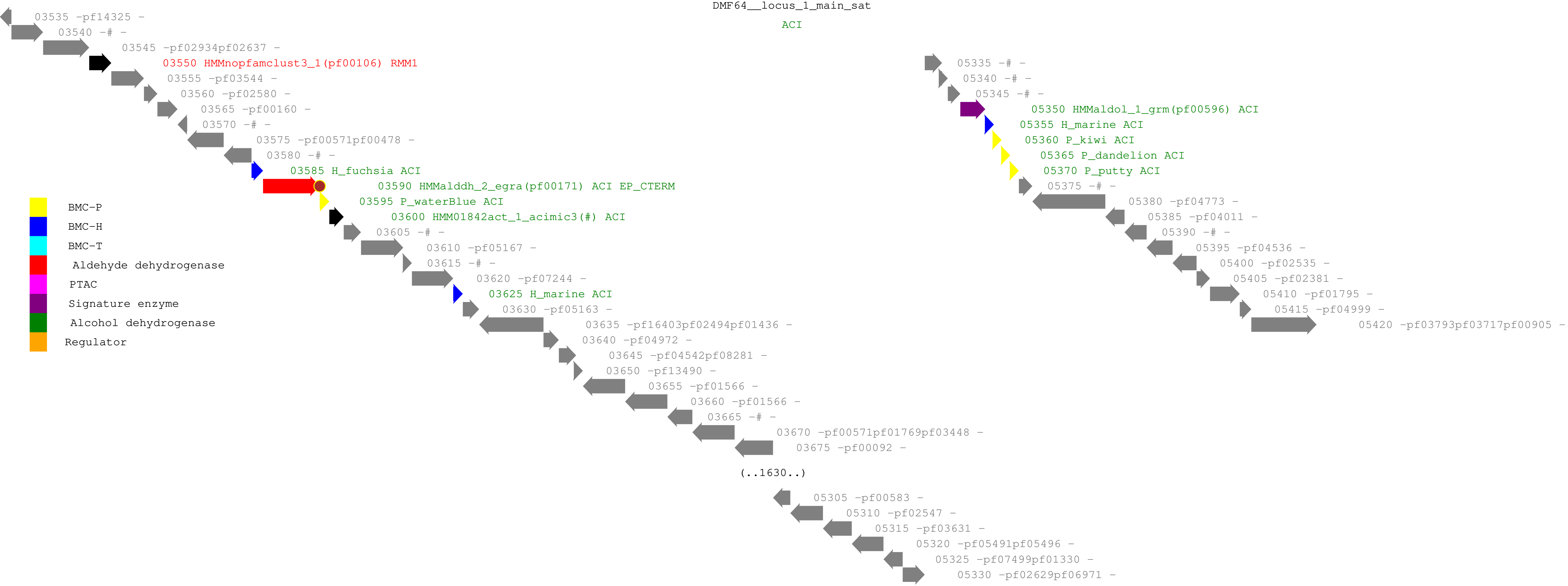

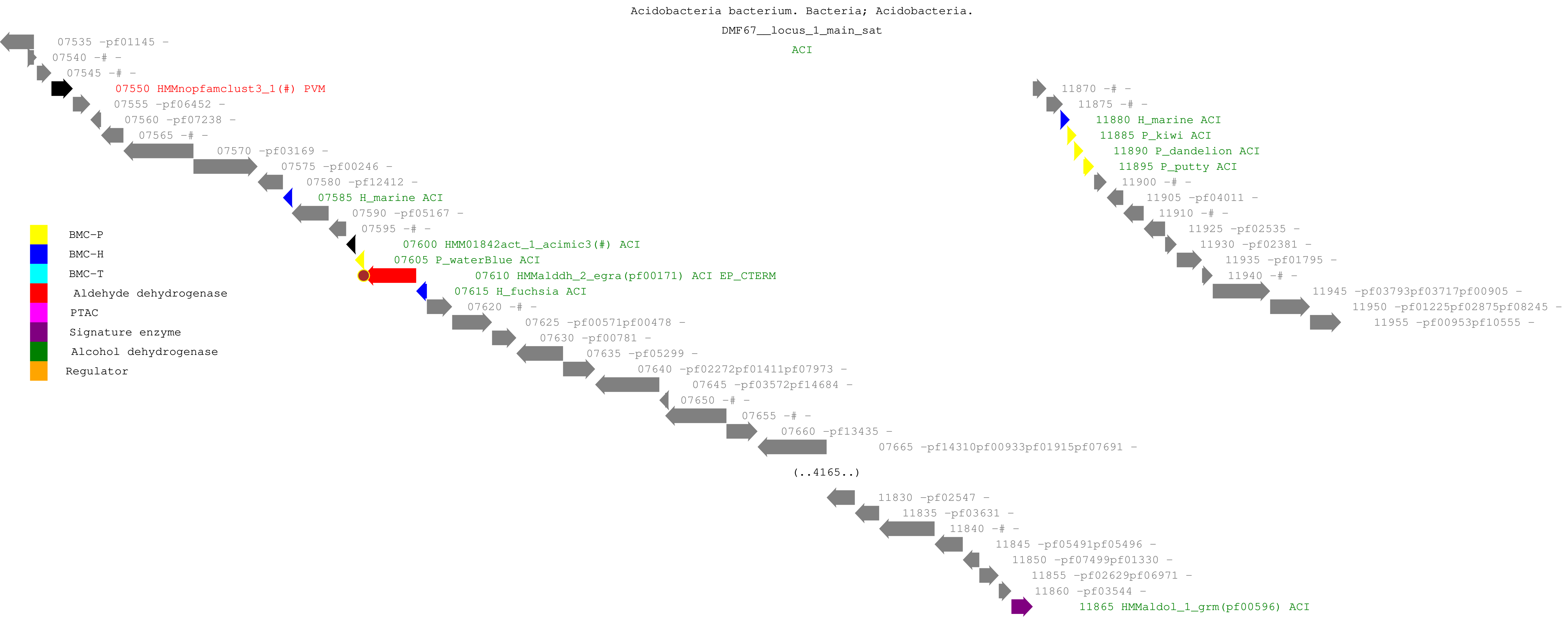

Acidobacteria bacterium. Bacteria; Acidobacteria.
DMF67__locus_1_main_sat
ACI
07535 -pf01145 -
07540 -# -
07545 -# -
07550 HMMnopfamclust3_1(#) PVM
07555 -pf06452 -
07560 -pf07238 -
07565 -# -
07570 -pf03169 -
07575 -pf00246 -
07580 -pf12412 -
07585 H_marine ACI
07590 -pf05167 -
07595 -# -
07600 HMM01842act_1_acimic3(#) ACI
07605 P_waterBlue ACI
07610 HMMalddh_2_egra(pf00171) ACI EP_CTERM
07615 H_fuchsia ACI
07620 -# -
07625 -pf00571pf00478 -
07630 -pf00781 -
07635 -pf05299 -
07640 -pf02272pf01411pf07973 -
07645 -pf03572pf14684 -
07650 -# -
07655 -# -
07660 -pf13435 -
07665 -pf14310pf00933pf01915pf07691 -
(..4165..)
11830 -pf02547 -
11835 -pf03631 -
11840 -# -
11845 -pf05491pf05496 -
11850 -pf07499pf01330 -
11855 -pf02629pf06971 -
11860 -pf03544 -
11865 HMMaldol_1_grm(pf00596) ACI
11870 -# -
11875 -# -
11880 H_marine ACI
11885 P_kiwi ACI
11890 P_dandelion ACI
11895 P_putty ACI
11900 -# -
11905 -pf04011 -
11910 -# -
11925 -pf02535 -
11930 -pf02381 -
11935 -pf01795 -
11940 -# -
11945 -pf03793pf03717pf00905 -
11950 -pf01225pf02875pf08245 -
11955 -pf00953pf10555 -
BMC-P
BMC-H
BMC-T
Aldehyde dehydrogenase
PTAC
Signature enzyme
Alcohol dehydrogenase
Regulator

Acidobacteria bacterium. Bacteria; Acidobacteria.

DMF68__locus_1_main_sat

ACI

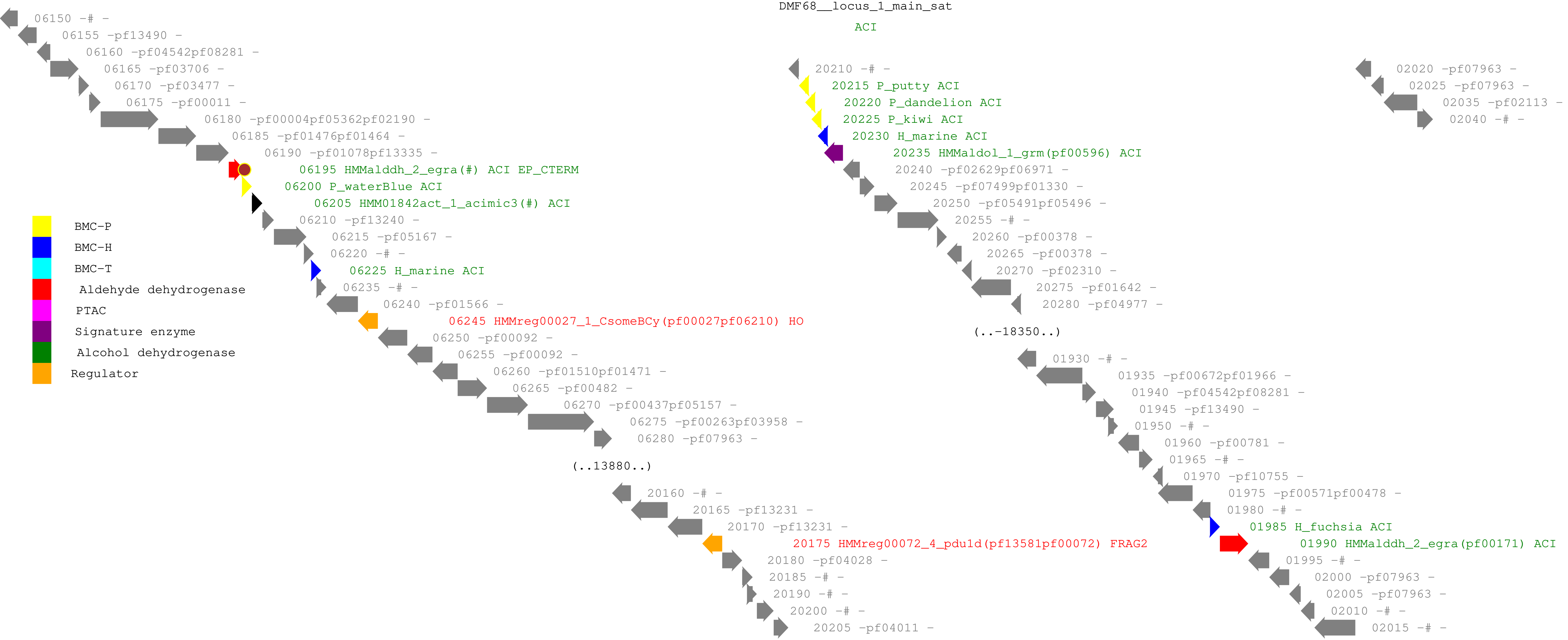

Acidobacteria bacterium. Bacteria; Acidobacteria.

DMF69__locus_1_main_sat

ACI

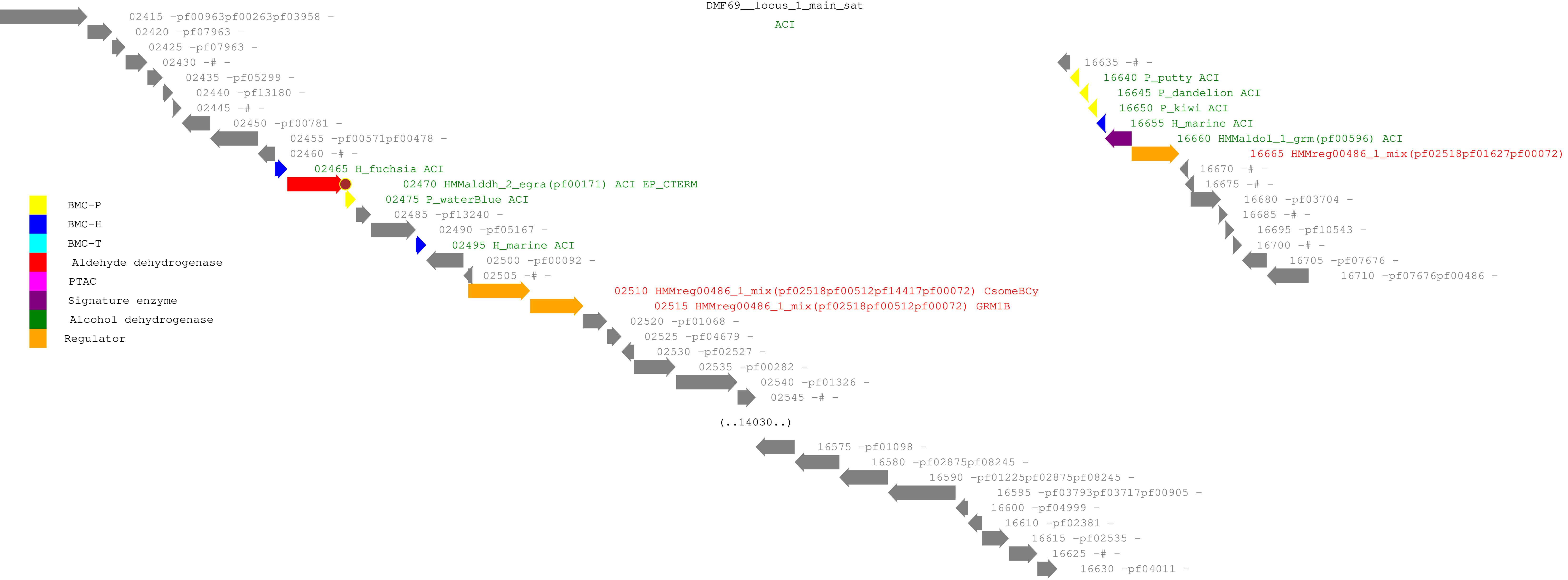

Acidobacteria bacterium. Bacteria; Acidobacteria.

DMF70__locus_1_main_sat

ACI

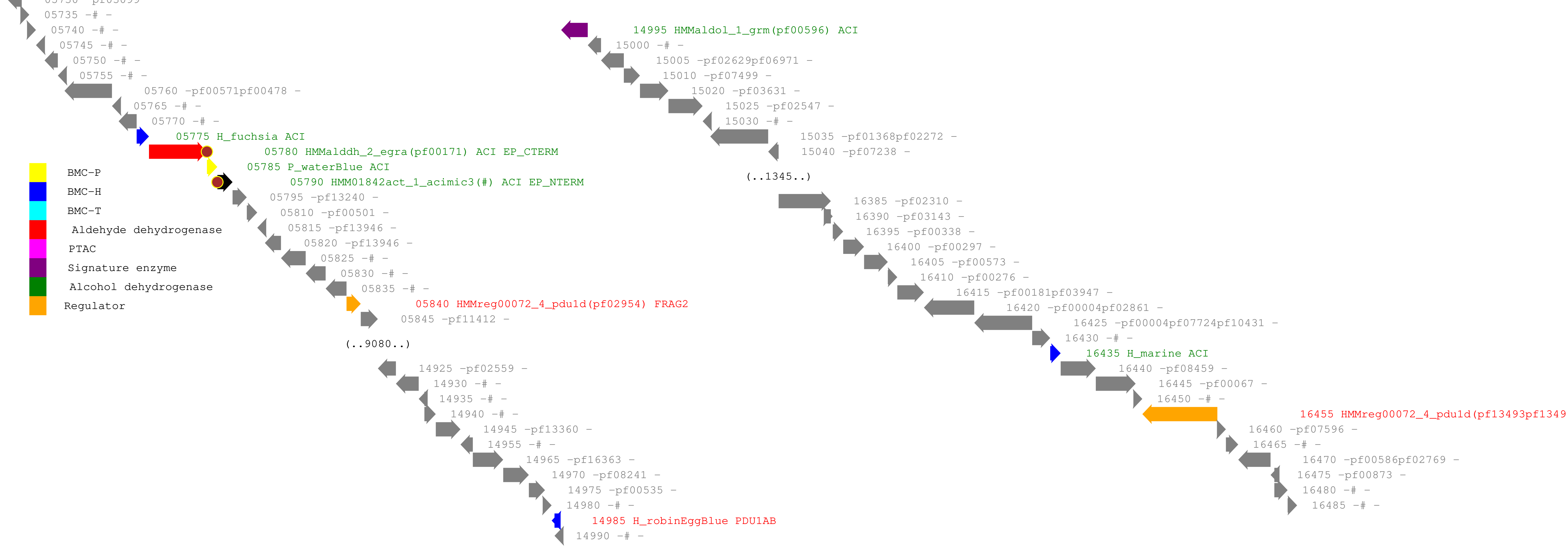

Acidobacteria bacterium. Bacteria; Acidobacteria.

DMF71__locus_1_main_sat

ACI

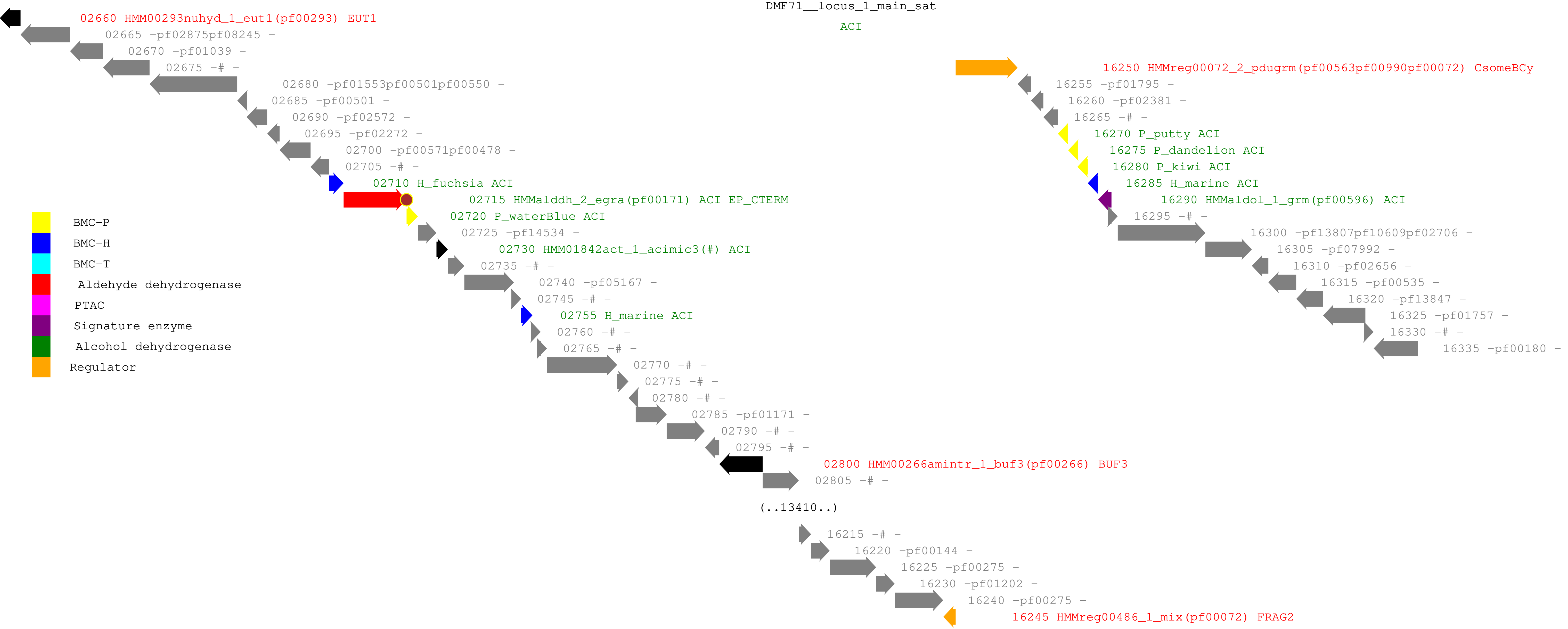

Acidobacteria bacterium. Bacteria; Acidobacteria.

DMF72__locus_1_main_sat

ACI

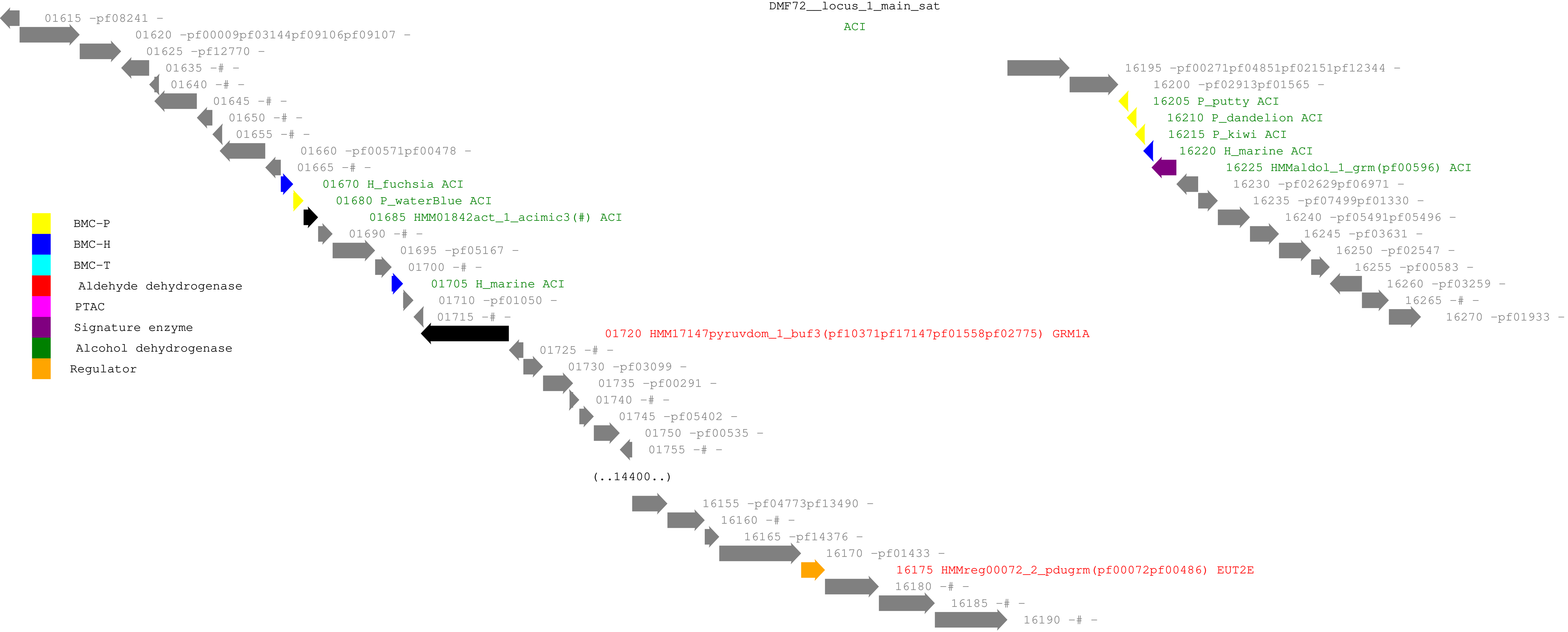

Acidobacteria bacterium. Bacteria; Acidobacteria.

DMF73__locus_1

ACI

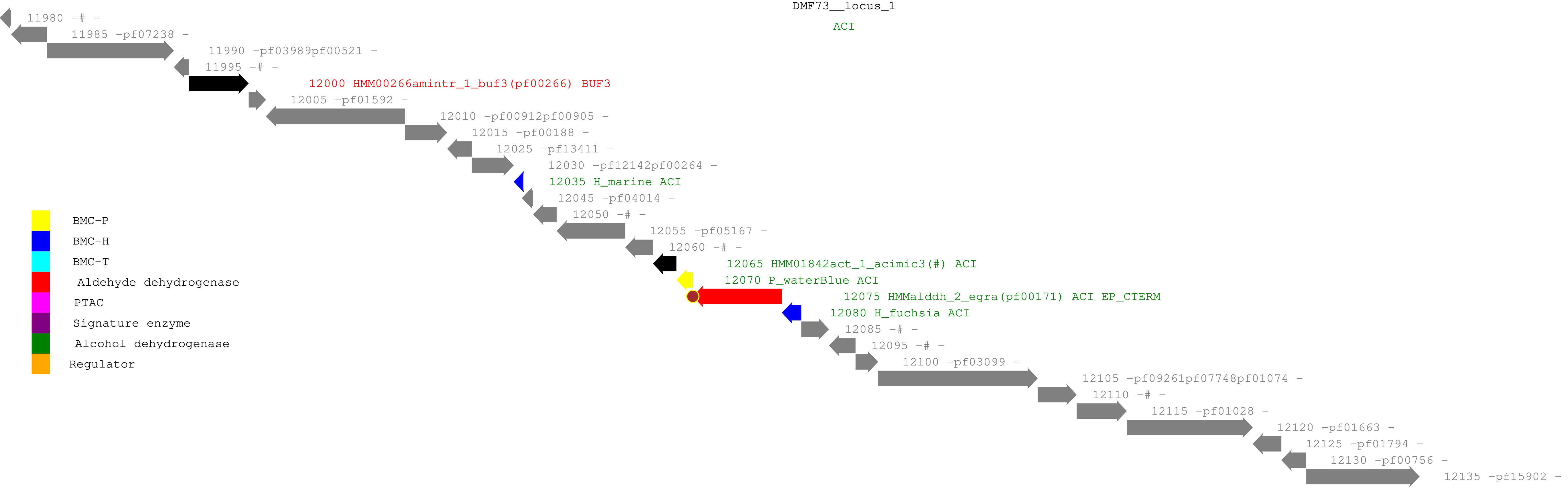

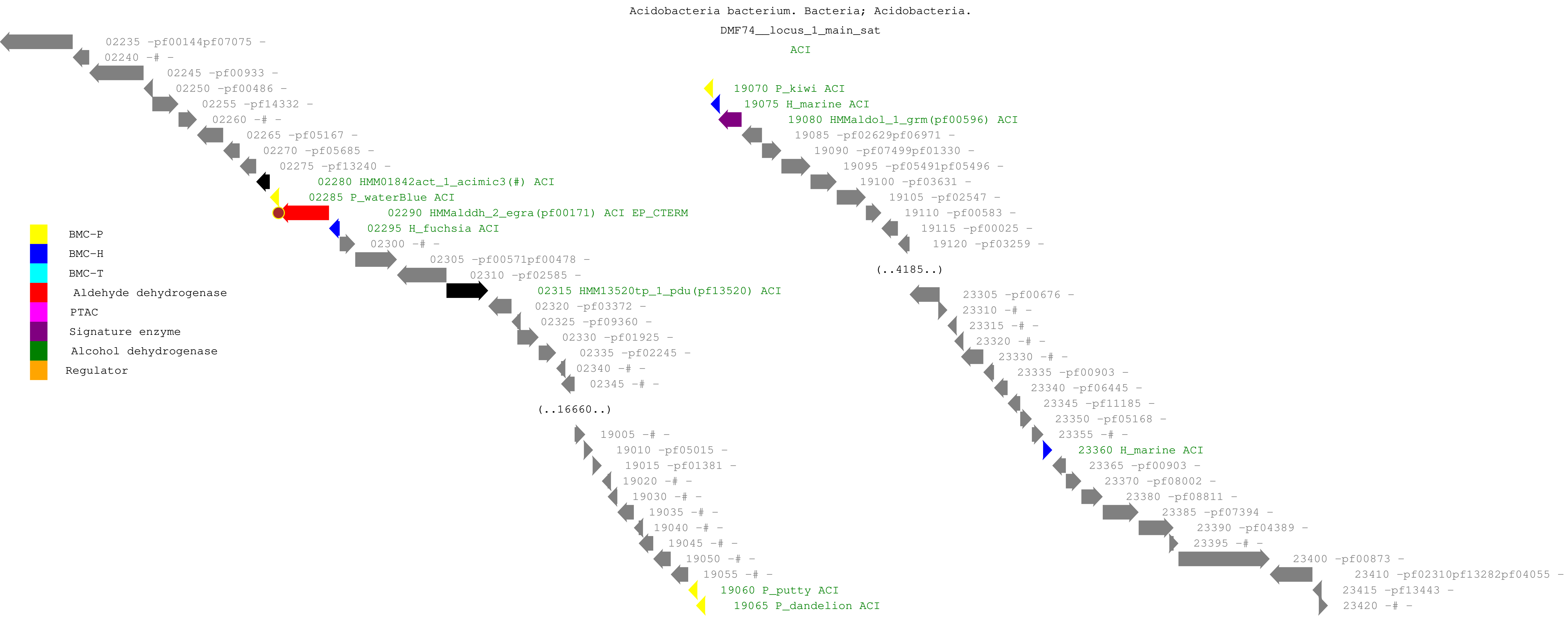

Acidobacteria bacterium. Bacteria; Acidobacteria.
DMF74__locus_1_main_sat
ACI
02235 -pf00144pf07075 -
02240 -# -
02245 -pf00933 -
02250 -pf00486 -
02255 -pf14332 -
02260 -# -
02265 -pf05167 -
02270 -pf05685 -
02275 -pf13240 -
02280 HMM01842act_1_acimic3(#) ACI
02285 P_waterBlue ACI
02290 HMMalddh_2_egra(pf00171) ACI EP_CTERM
02295 H_fuchsia ACI
02300 -# -
02305 -pf00571pf00478 -
02310 -pf02585 -
02315 HMM13520tp_1_pdu(pf13520) ACI
02320 -pf03372 -
02325 -pf09360 -
02330 -pf01925 -
02335 -pf02245 -
02340 -# -
02345 -# -
(..16660..)
19005 -# -
19010 -pf05015 -
19015 -pf01381 -
19020 -# -
19030 -# -
19035 -# -
19040 -# -
19045 -# -
19050 -# -
19055 -# -
19060 P_putty ACI
19065 P_dandelion ACI
19070 P_kiwi ACI
19075 H_marine ACI
19080 HMMaldol_1_grm(pf00596) ACI
19085 -pf02629pf06971 -
19090 -pf07499pf01330 -
19095 -pf05491pf05496 -
19100 -pf03631 -
19105 -pf02547 -
19110 -pf00583 -
19115 -pf00025 -
19120 -pf03259 -
(..4185..)
23305 -pf00676 -
23310 -# -
23315 -# -
23320 -# -
23330 -# -
23335 -pf00903 -
23340 -pf06445 -
23345 -pf11185 -
23350 -pf05168 -
23355 -# -
23360 H_marine ACI
23365 -pf00903 -
23370 -pf08002 -
23380 -pf08811 -
23385 -pf07394 -
23390 -pf04389 -
23395 -# -
23400 -pf00873 -
23410 -pf02310pf13282pf04055 -
23415 -pf13443 -
23420 -# -
BMC-P
BMC-H
BMC-T
Aldehyde dehydrogenase
PTAC
Signature enzyme
Alcohol dehydrogenase
Regulator

Acidobacteria bacterium. Bacteria; Acidobacteria.

DMF75__locus_1

ACI

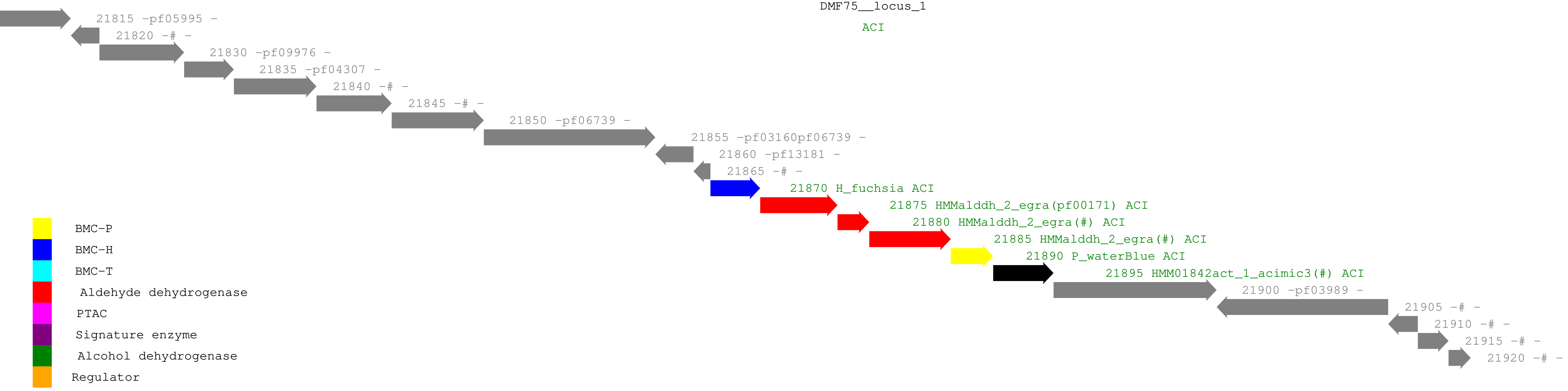

Acidobacteria bacterium. Bacteria; Acidobacteria.

DMF76__locus_1_main_sat

ACI

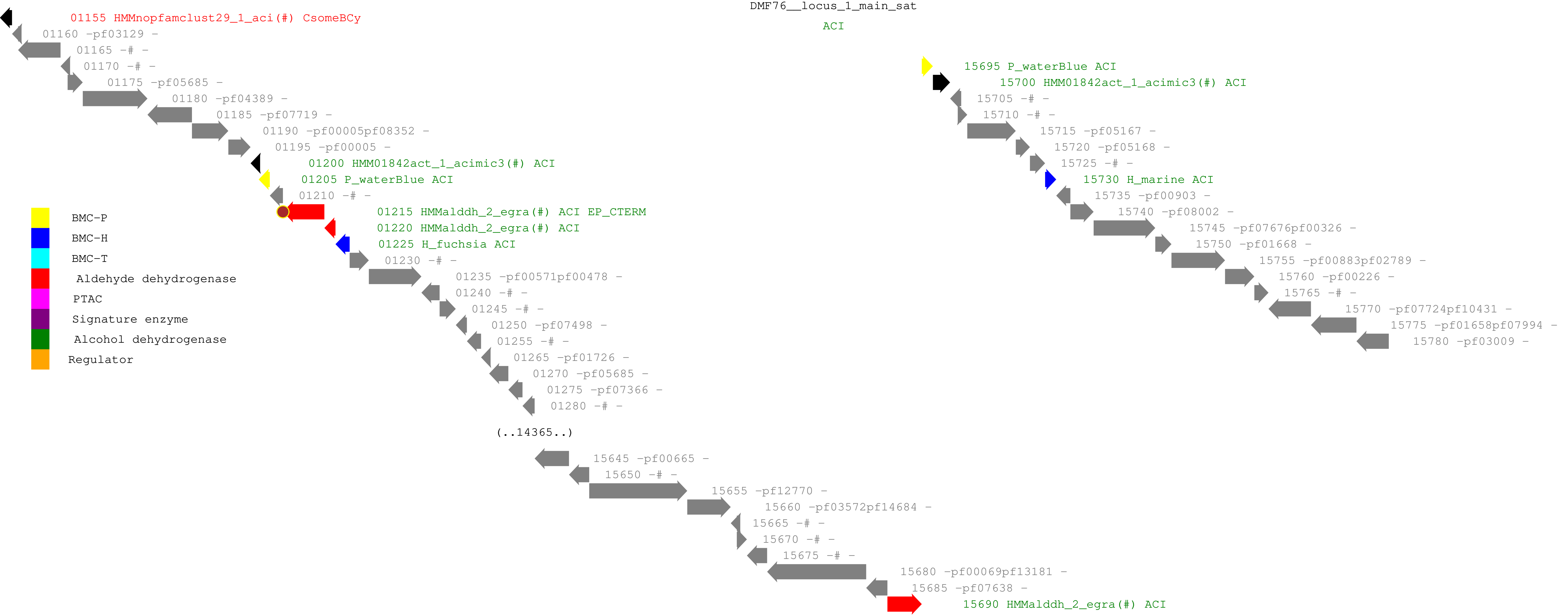

Acidobacteria bacterium. Bacteria; Acidobacteria.

DMF76__locus_3

ACI

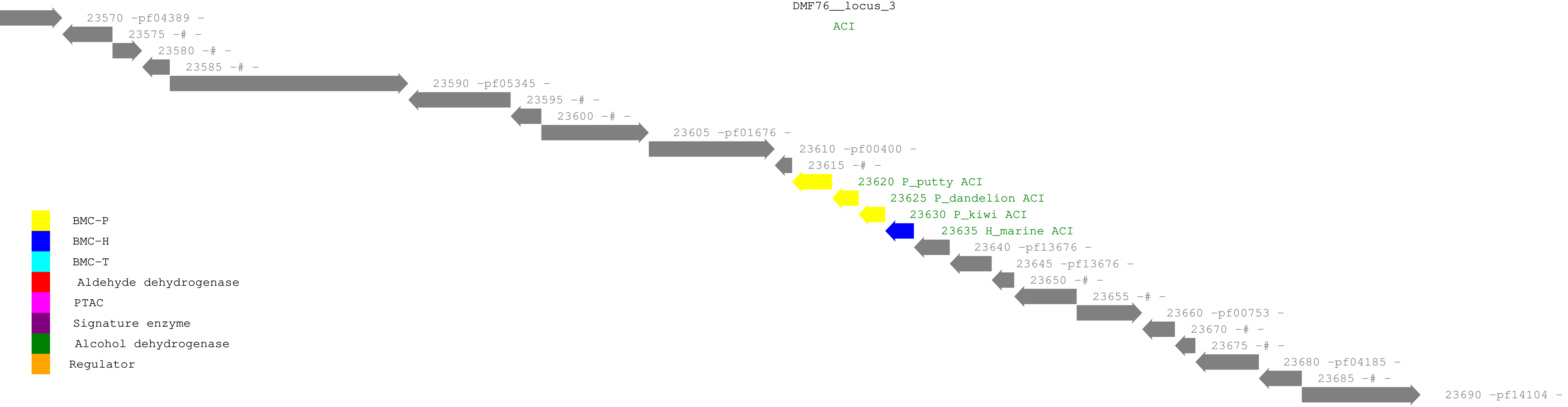

Acidobacteria bacterium. Bacteria; Acidobacteria.

DMF77__locus_1_main_sat

ACI

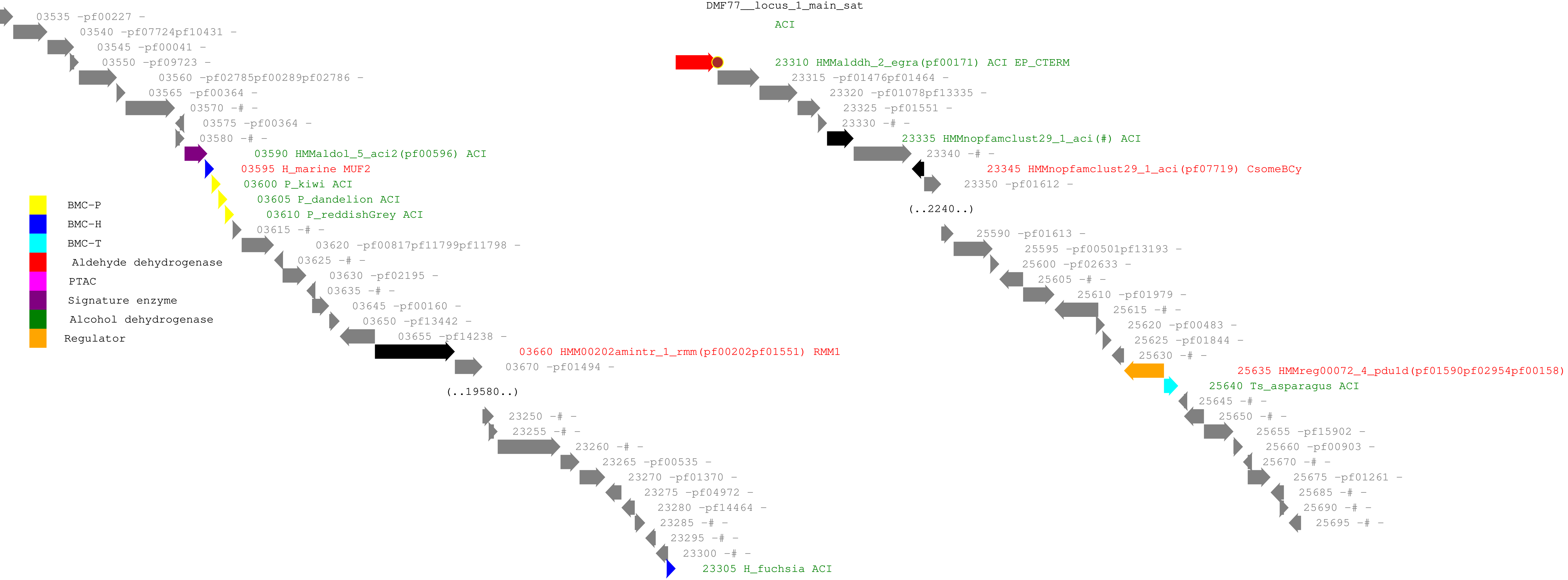

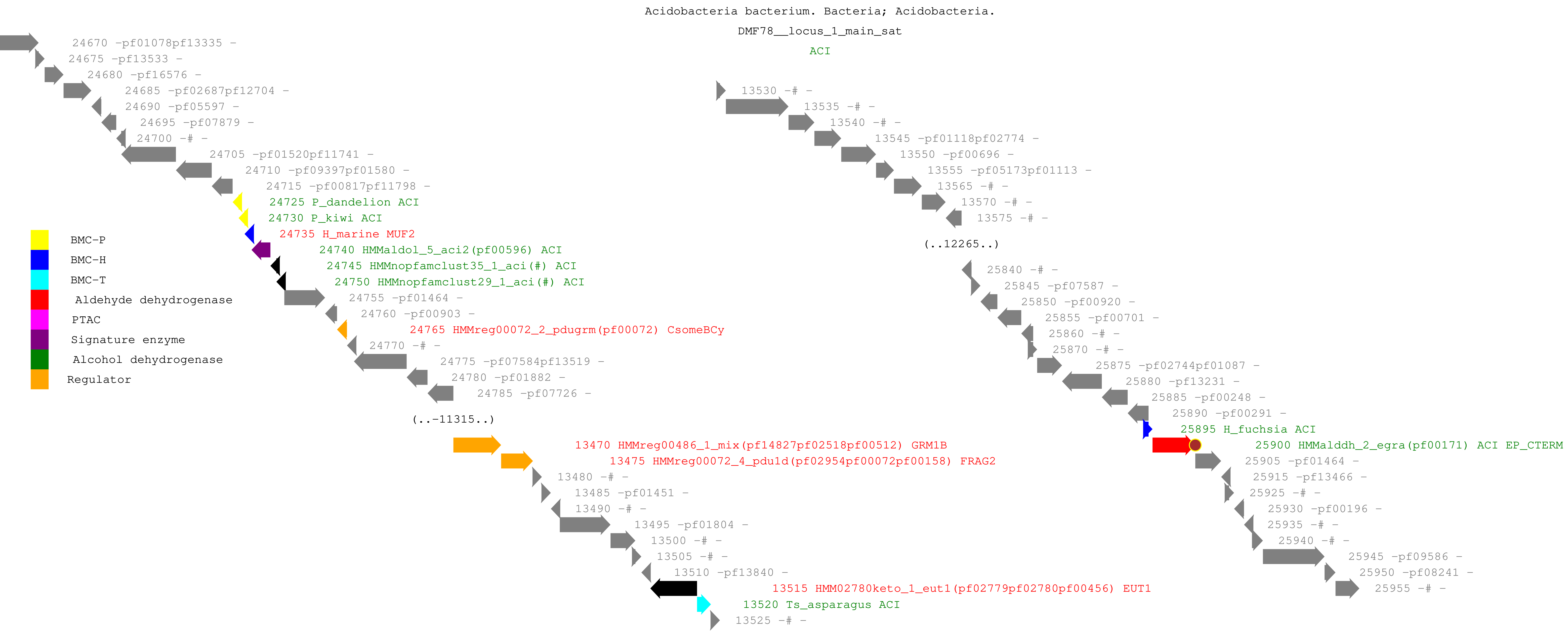
Acidobacteria bacterium. Bacteria; Acidobacteria.
DMF78__locus_1_main_sat
ACI
24670 -pf01078pf13335 -
24675 -pf13533 -
24680 -pf16576 -
24685 -pf02687pf12704 -
24690 -pf05597 -
24695 -pf07879 -
24700 -# -
24705 -pf01520pf11741 -
24710 -pf09397pf01580 -
24715 -pf00817pf11798 -
24725 P_dandelion ACI
24730 P_kiwi ACI
24735 H_marine MUF2
24740 HMMaldol_5_aci2(pf00596) ACI
24745 HMMnopfamclust35_1_aci(#) ACI
24750 HMMnopfamclust29_1_aci(#) ACI
24755 -pf01464 -
24760 -pf00903 -
24765 HMMreg00072_2_pdugrm(pf00072) CsomeBCy
24770 -# -
24775 -pf07584pf13519 -
24780 -pf01882 -
24785 -pf07726 -
(..-11315..)
13470 HMMreg00486_1_mix(pf14827pf02518pf00512) GRM1B
13475 HMMreg00072_4_pdu1d(pf02954pf00072pf00158) FRAG2
13480 -# -
13485 -pf01451 -
13490 -# -
13495 -pf01804 -
13500 -# -
13505 -# -
13510 -pf13840 -
13515 HMM02780keto_1_eut1(pf02779pf02780pf00456) EUT1
13520 Ts_asparagus ACI
13525 -# -
13530 -# -
13535 -# -
13540 -# -
13545 -pf01118pf02774 -
13550 -pf00696 -
13555 -pf05173pf01113 -
13565 -# -
13570 -# -
13575 -# -
(..12265..)
25840 -# -
25845 -pf07587 -
25850 -pf00920 -
25855 -pf00701 -
25860 -# -
25870 -# -
25875 -pf02744pf01087 -
25880 -pf13231 -
25885 -pf00248 -
25890 -pf00291 -
25895 H_fuchsia ACI
25900 HMMalddh_2_egra(pf00171) ACI EP_CTERM
25905 -pf01464 -
25915 -pf13466 -
25925 -# -
25930 -pf00196 -
25935 -# -
25940 -# -
25945 -pf09586 -
25950 -pf08241 -
25955 -# -
BMC-P
BMC-H
BMC-T
Aldehyde dehydrogenase
PTAC
Signature enzyme
Alcohol dehydrogenase
Regulator

Acidobacteria bacterium. Bacteria; Acidobacteria.

DMF79__locus_1_main_sat

ACI

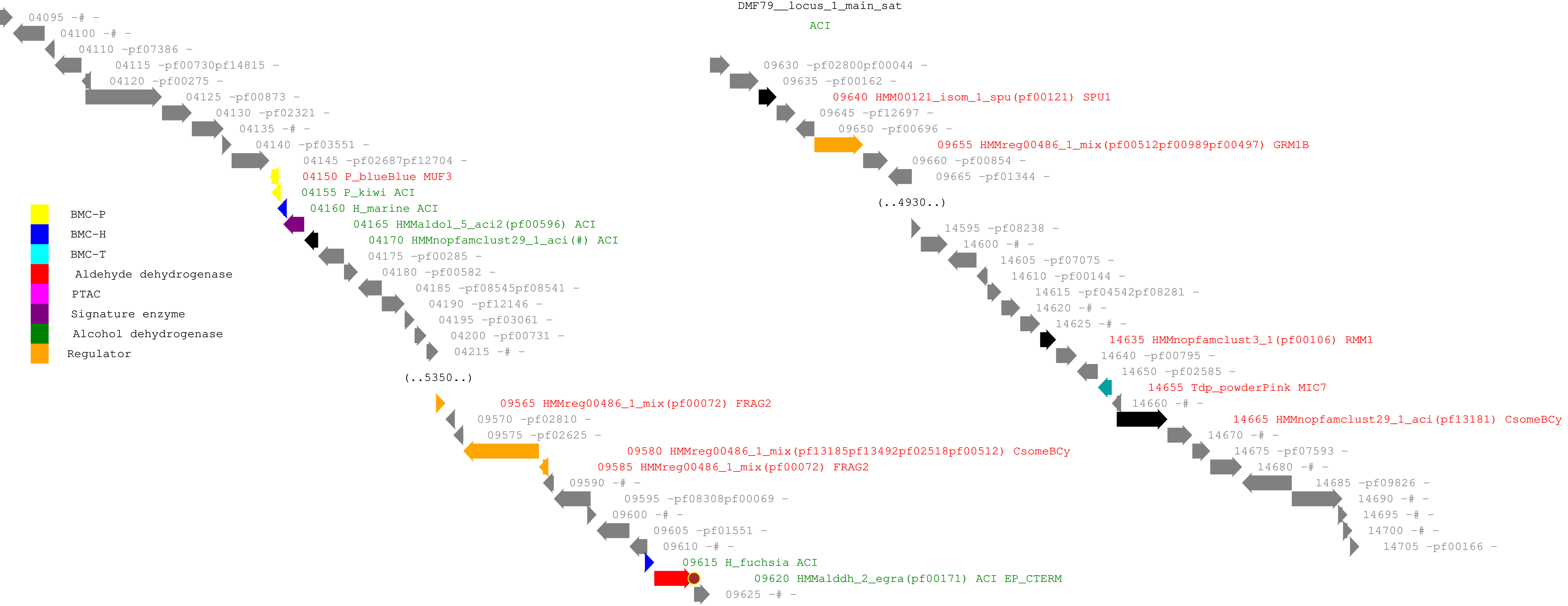

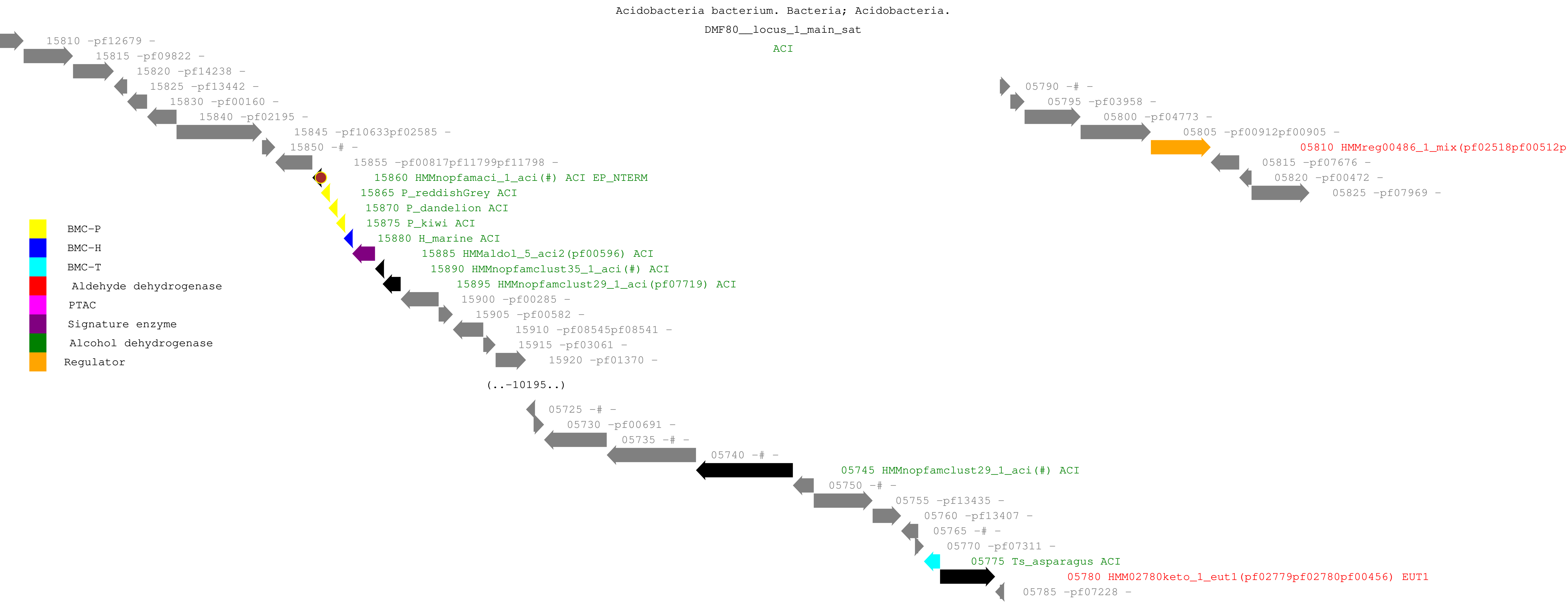
Acidobacteria bacterium. Bacteria; Acidobacteria.
DMF80__locus_1_main_sat
ACI
15810 -pf12679 -
15815 -pf09822 -
15820 -pf14238 -
15825 -pf13442 -
15830 -pf00160 -
15840 -pf02195 -
15845 -pf10633pf02585 -
15850 -# -
15855 -pf00817pf11799pf11798 -
15860 HMMnopfamaci_1_aci(#) ACI EP_NTERM
15865 P_reddishGrey ACI
15870 P_dandelion ACI
15875 P_kiwi ACI
15880 H_marine ACI
15885 HMMaldol_5_aci2(pf00596) ACI
15890 HMMnopfamclust35_1_aci(#) ACI
15895 HMMnopfamclust29_1_aci(pf07719) ACI
15900 -pf00285 -
15905 -pf00582 -
15910 -pf08545pf08541 -
15915 -pf03061 -
15920 -pf01370 -
(..-10195..)
05725 -# -
05730 -pf00691 -
05735 -# -
05740 -# -
05745 HMMnopfamclust29_1_aci(#) ACI
05750 -# -
05755 -pf13435 -
05760 -pf13407 -
05765 -# -
05770 -pf07311 -
05775 Ts_asparagus ACI
05780 HMM02780keto_1_eut1(pf02779pf02780pf00456) EUT1
05785 -pf07228 -
05790 -# -
05795 -pf03958 -
05800 -pf04773 -
05805 -pf00912pf00905 -
05810 HMMreg00486_1_mix(pf02518pf00512p
05815 -pf07676 -
05820 -pf00472 -
05825 -pf07969 -
BMC-P
BMC-H
BMC-T
Aldehyde dehydrogenase
PTAC
Signature enzyme
Alcohol dehydrogenase
Regulator

Acidobacteria bacterium. Bacteria; Acidobacteria.

DMF81__locus_fragmented

ACI

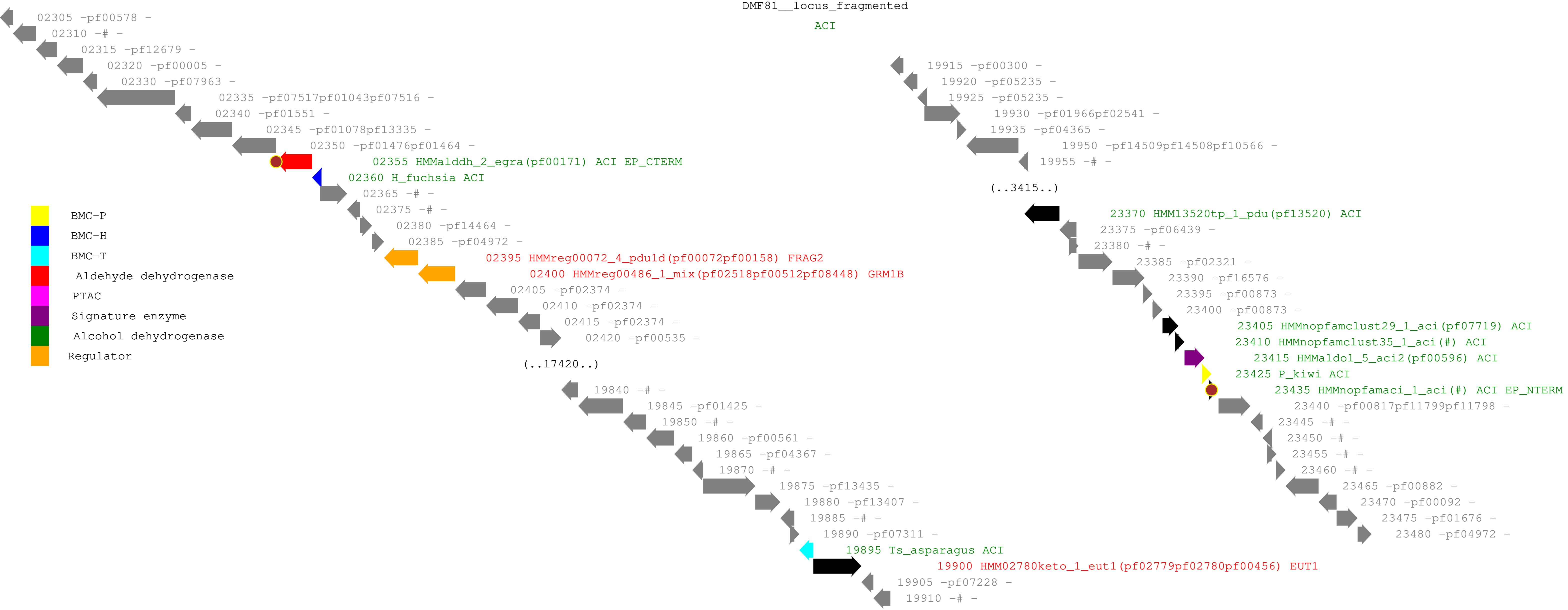

Acidobacteria bacterium. Bacteria; Acidobacteria.

DMF82__locus_1_main_sat

ACI

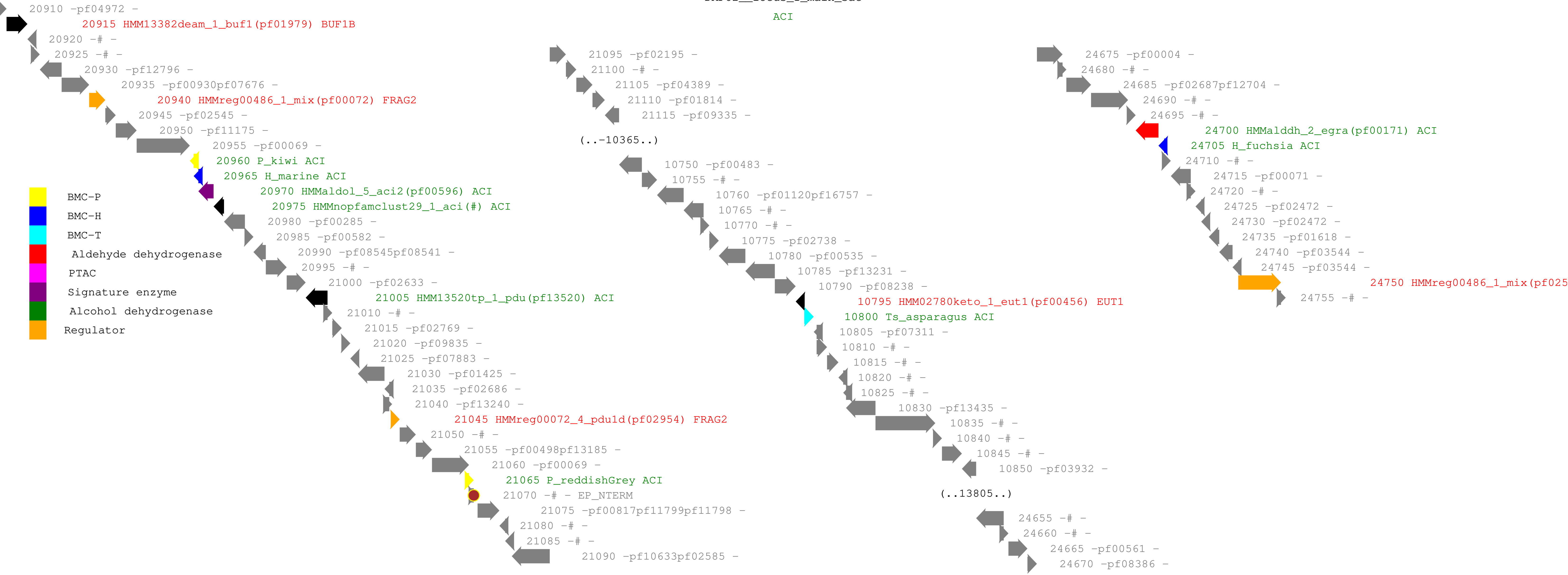

Acidobacteria bacterium. Bacteria; Acidobacteria.

DMF83__locus_1_main_sat

ACI

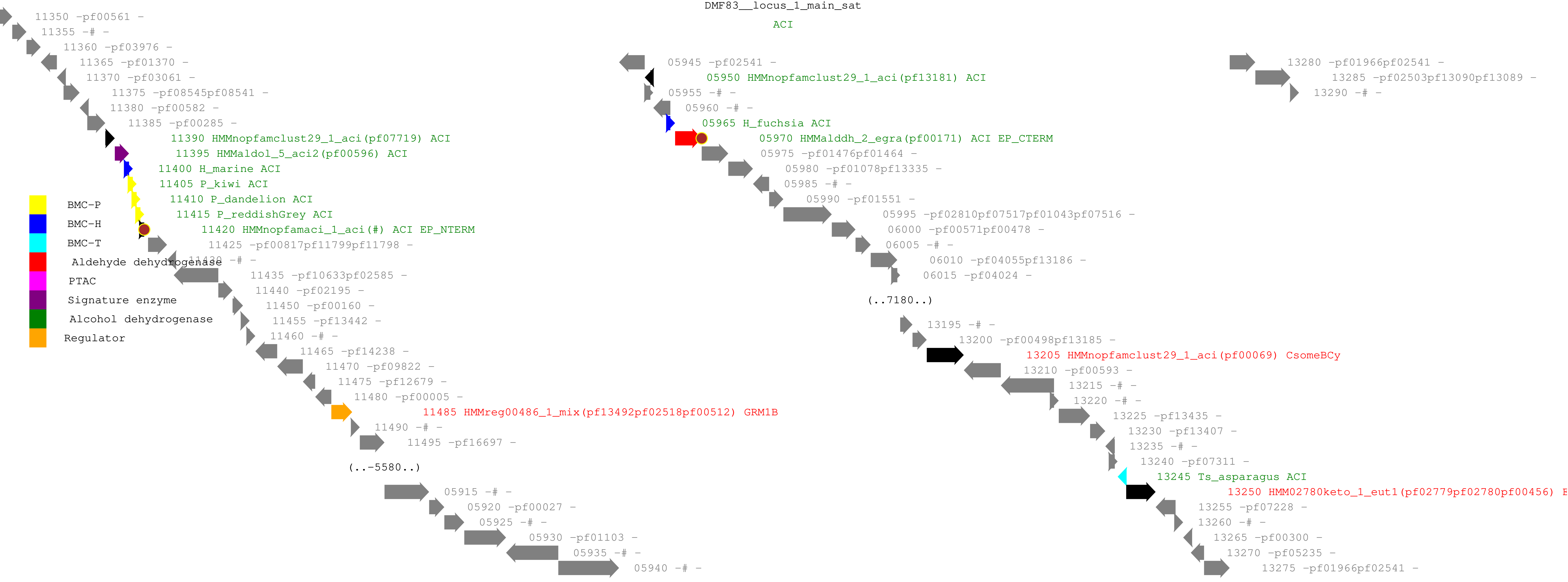

Acidobacteria bacterium. Bacteria; Acidobacteria.

DMF84__locus_1_main_sat

ACI

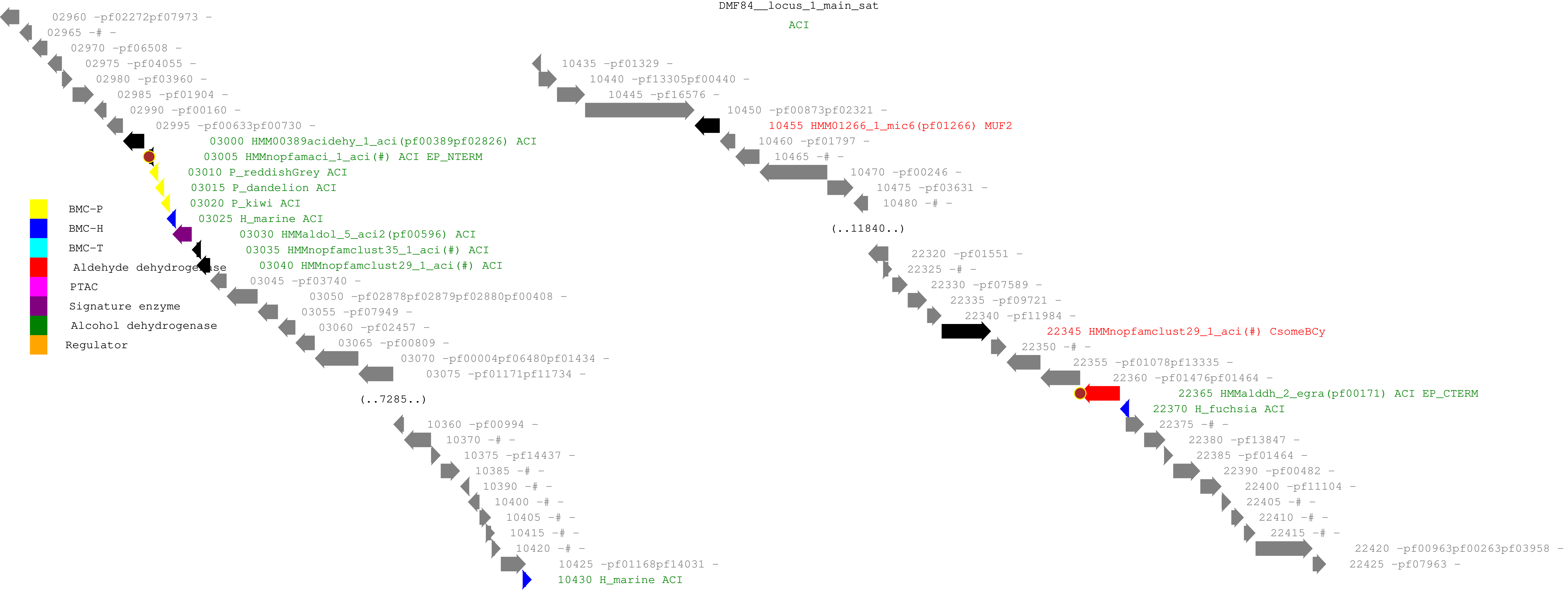

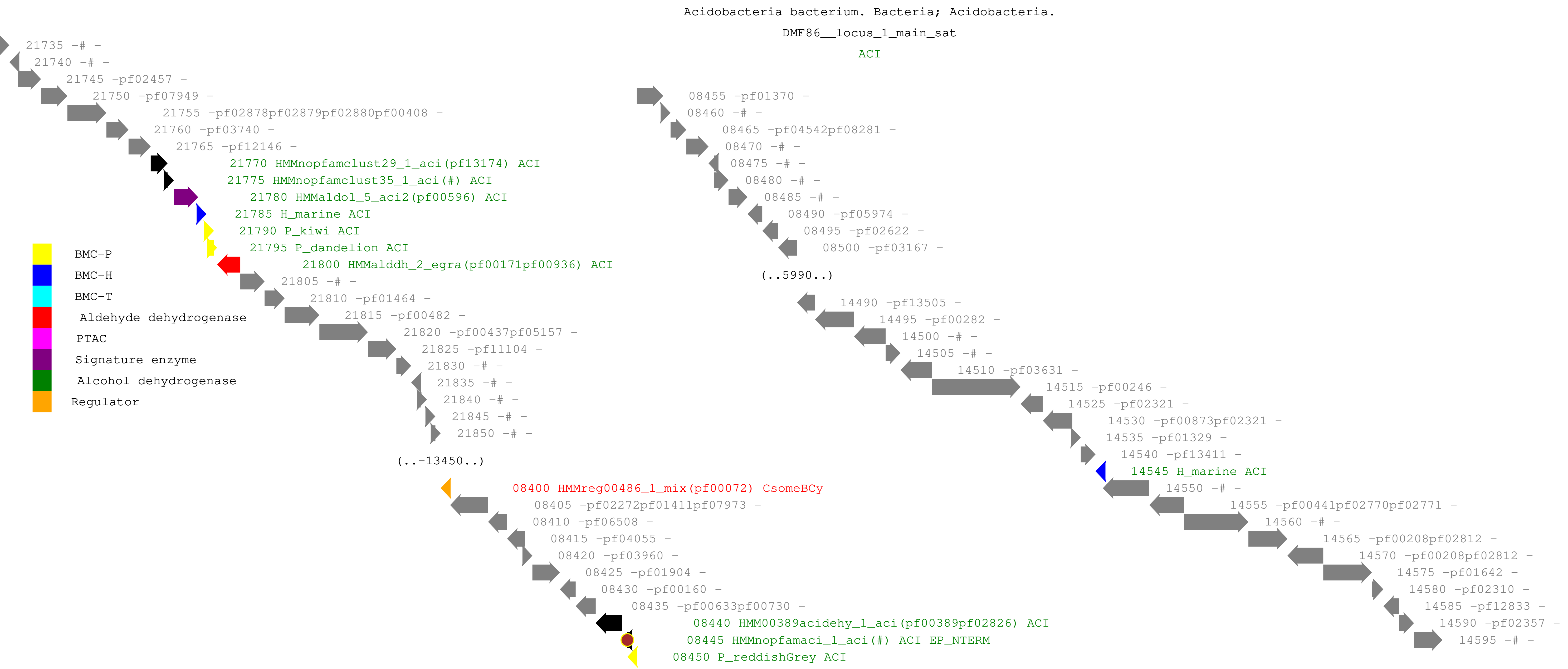

Acidobacteria bacterium. Bacteria; Acidobacteria.
DMF86__locus_1_main_sat
ACI
21735 -# -
21740 -# -
21745 -pf02457 -
21750 -pf07949 -
21755 -pf02878pf02879pf02880pf00408 -
21760 -pf03740 -
21765 -pf12146 -
21770 HMMnopfamclust29_1_aci(pf13174) ACI
21775 HMMnopfamclust35_1_aci(#) ACI
21780 HMMaldol_5_aci2(pf00596) ACI
21785 H_marine ACI
21790 P_kiwi ACI
21795 P_dandelion ACI
21800 HMMalddh_2_egra(pf00171pf00936) ACI
21805 -# -
21810 -pf01464 -
21815 -pf00482 -
21820 -pf00437pf05157 -
21825 -pf11104 -
21830 -# -
21835 -# -
21840 -# -
21845 -# -
21850 -# -
(..-13450..)
08400 HMMreg00486_1_mix(pf00072) CsomeBCy
08405 -pf02272pf01411pf07973 -
08410 -pf06508 -
08415 -pf04055 -
08420 -pf03960 -
08425 -pf01904 -
08430 -pf00160 -
08435 -pf00633pf00730 -
08440 HMM00389acidehy_1_aci(pf00389pf02826) ACI
08445 HMMnopfamaci_1_aci(#) ACI EP_NTERM
08450 P_reddishGrey ACI
08455 -pf01370 -
08460 -# -
08465 -pf04542pf08281 -
08470 -# -
08475 -# -
08480 -# -
08485 -# -
08490 -pf05974 -
08495 -pf02622 -
08500 -pf03167 -
(..5990..)
14490 -pf13505 -
14495 -pf00282 -
14500 -# -
14505 -# -
14510 -pf03631 -
14515 -pf00246 -
14525 -pf02321 -
14530 -pf00873pf02321 -
14535 -pf01329 -
14540 -pf13411 -
14545 H_marine ACI
14550 -# -
14555 -pf00441pf02770pf02771 -
14560 -# -
14565 -pf00208pf02812 -
14570 -pf00208pf02812 -
14575 -pf01642 -
14580 -pf02310 -
14585 -pf12833 -
14590 -pf02357 -
14595 -# -
BMC-P
BMC-H
BMC-T
Aldehyde dehydrogenase
PTAC
Signature enzyme
Alcohol dehydrogenase
Regulator

Acidobacteria bacterium. Bacteria; Acidobacteria.

DMF87__locus_1_main_sat

ACI

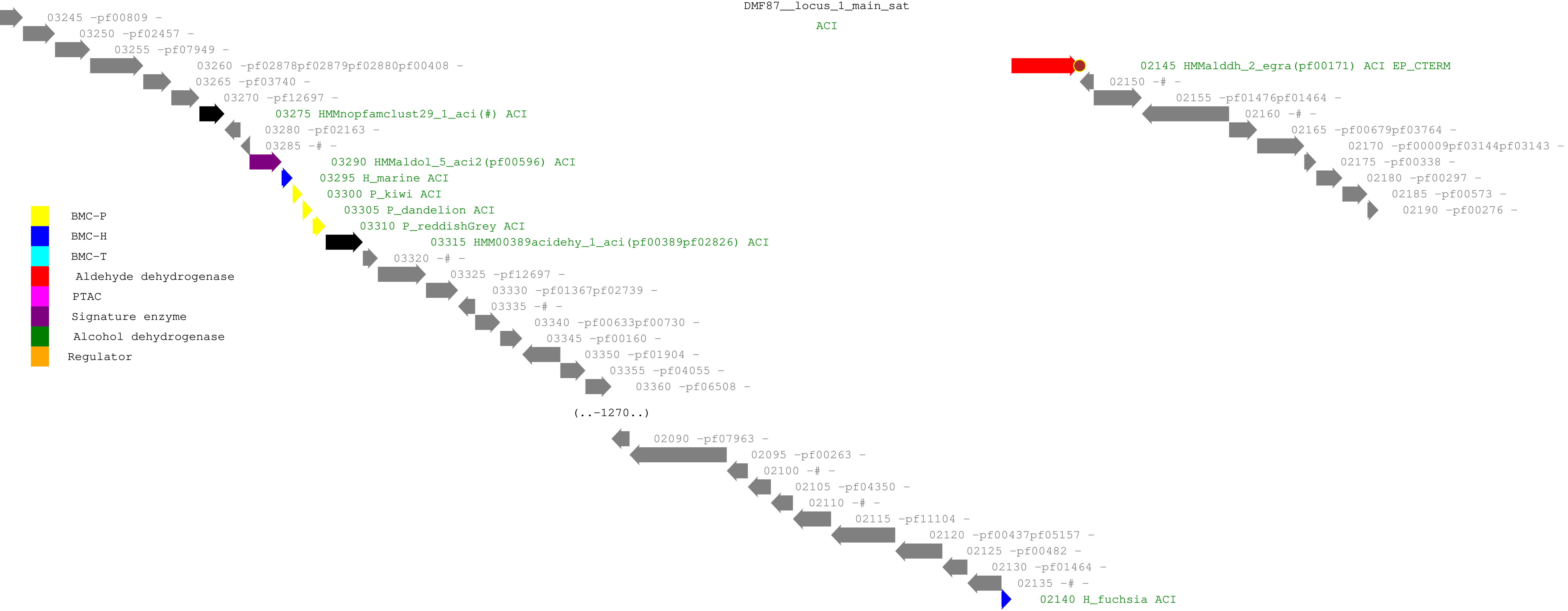

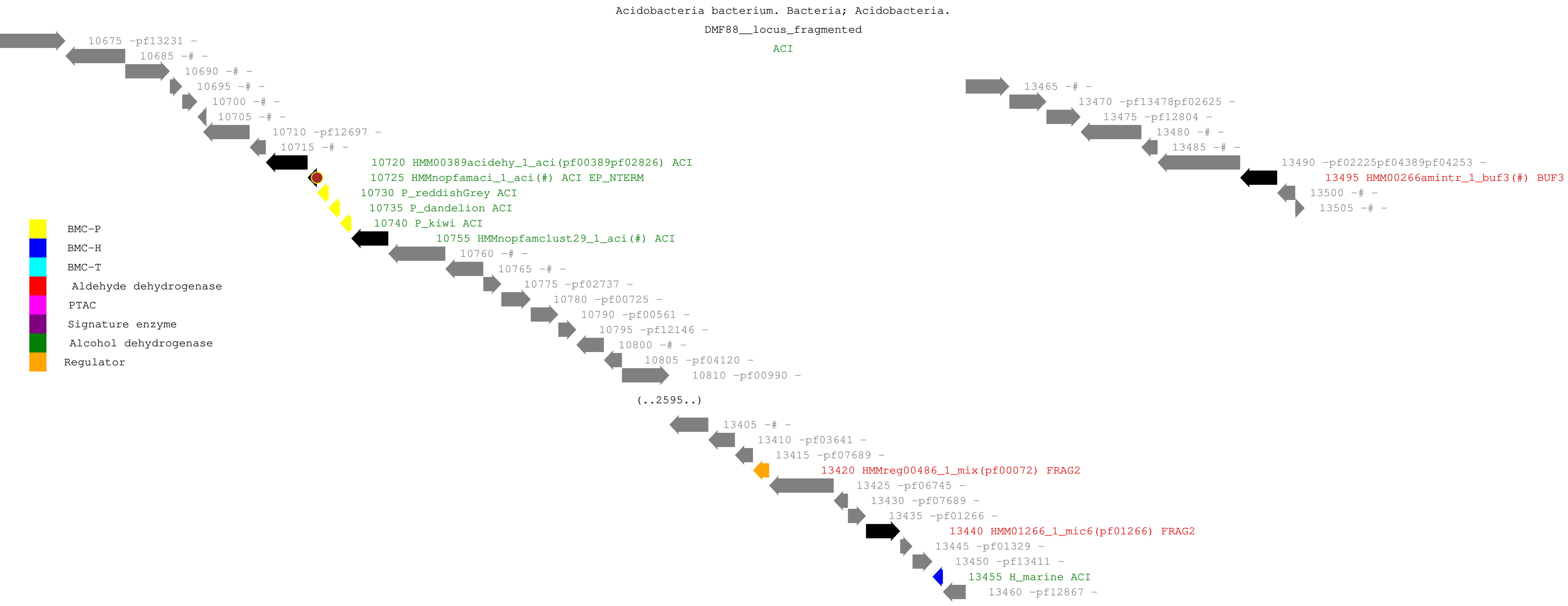
Acidobacteria bacterium. Bacteria; Acidobacteria.
DMF88__locus_fragmented
ACI
10675 -pf13231 -
10685 -# -
10690 -# -
10695 -# -
10700 -# -
10705 -# -
10710 -pf12697 -
10715 -# -
10720 HMM00389acidehy_1_aci(pf00389pf02826) ACI
10725 HMMnopfamaci_1_aci(#) ACI EP_NTERM
10730 P_reddishGrey ACI
10735 P_dandelion ACI
10740 P_kiwi ACI
10755 HMMnopfamclust29_1_aci(#) ACI
10760 -# -
10765 -# -
10775 -pf02737 -
10780 -pf00725 -
10790 -pf00561 -
10795 -pf12146 -
10800 -# -
10805 -pf04120 -
10810 -pf00990 -
(..2595..)
13405 -# -
13410 -pf03641 -
13415 -pf07689 -
13420 HMMreg00486_1_mix(pf00072) FRAG2
13425 -pf06745 -
13430 -pf07689 -
13435 -pf01266 -
13440 HMM01266_1_mic6(pf01266) FRAG2
13445 -pf01329 -
13450 -pf13411 -
13455 H_marine ACI
13460 -pf12867 -
13465 -# -
13470 -pf13478pf02625 -
13475 -pf12804 -
13480 -# -
13485 -# -
13490 -pf02225pf04389pf04253 -
13495 HMM00266amintr_1_buf3(#) BUF3
13500 -# -
13505 -# -
BMC-P
BMC-H
BMC-T
Aldehyde dehydrogenase
PTAC
Signature enzyme
Alcohol dehydrogenase
Regulator

Acidobacteria bacterium. Bacteria; Acidobacteria.

DMF90__locus_1_main_sat

ACI

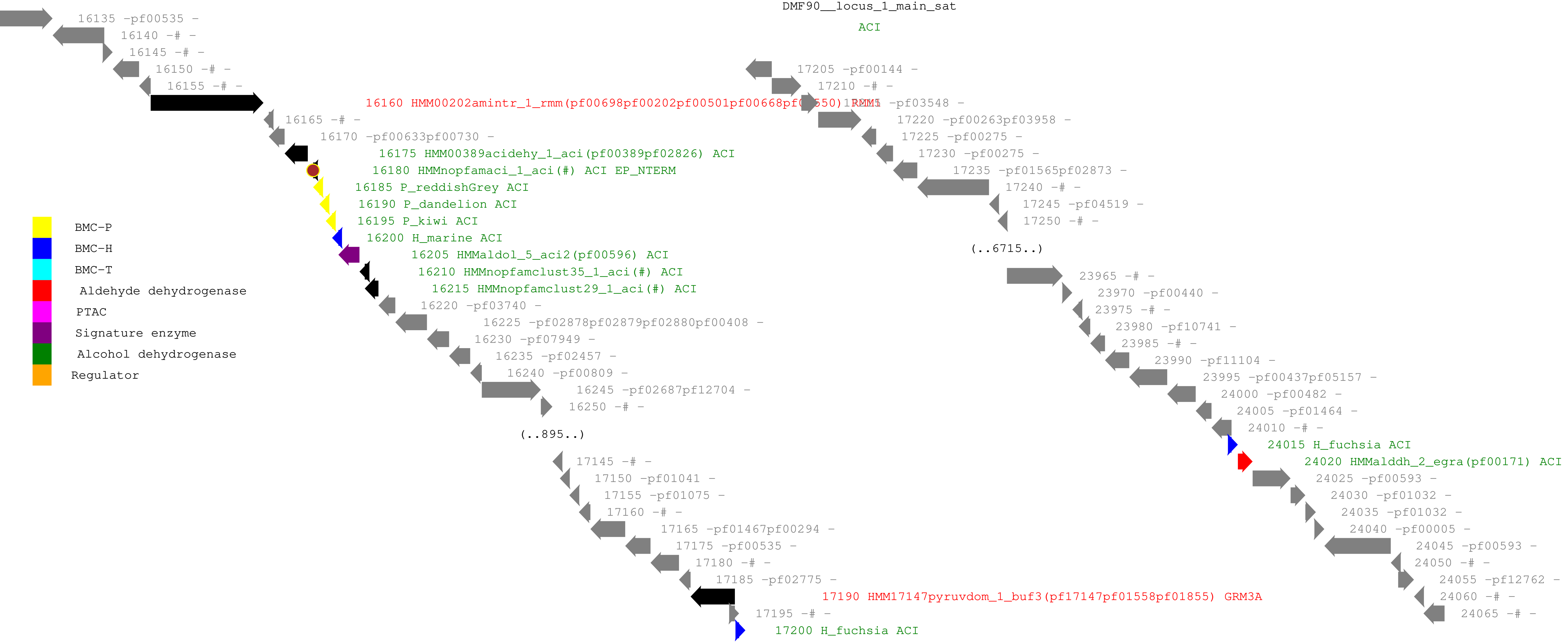

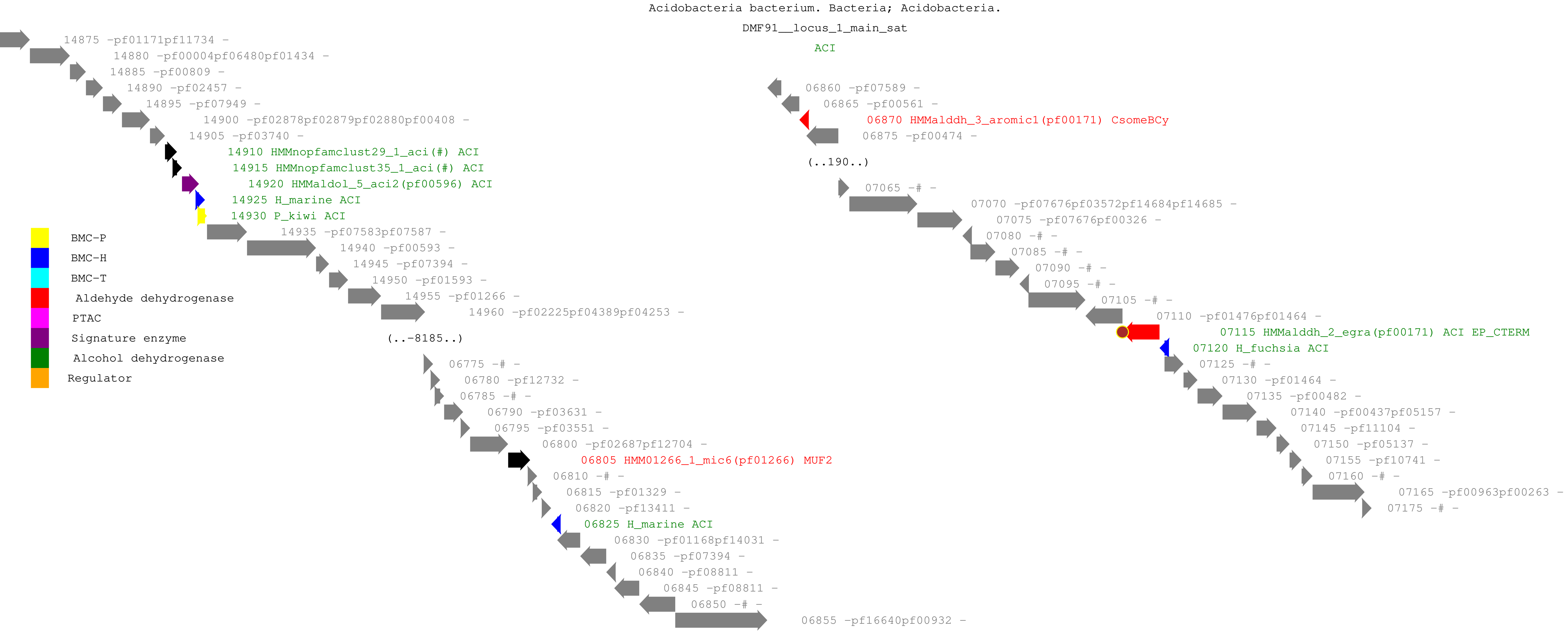

Acidobacteria bacterium. Bacteria; Acidobacteria.
DMF91__locus_1_main_sat
ACI
14875 -pf01171pf11734 -
14880 -pf00004pf06480pf01434 -
14885 -pf00809 -
14890 -pf02457 -
14895 -pf07949 -
14900 -pf02878pf02879pf02880pf00408 -
14905 -pf03740 -
14910 HMMnopfamclust29_1_aci(#) ACI
14915 HMMnopfamclust35_1_aci(#) ACI
14920 HMMaldol_5_aci2(pf00596) ACI
14925 H_marine ACI
14930 P_kiwi ACI
14935 -pf07583pf07587 -
14940 -pf00593 -
14945 -pf07394 -
14950 -pf01593 -
14955 -pf01266 -
14960 -pf02225pf04389pf04253 -
(..-8185..)
06775 -# -
06780 -pf12732 -
06785 -# -
06790 -pf03631 -
06795 -pf03551 -
06800 -pf02687pf12704 -
06805 HMM01266_1_mic6(pf01266) MUF2
06810 -# -
06815 -pf01329 -
06820 -pf13411 -
06825 H_marine ACI
06830 -pf01168pf14031 -
06835 -pf07394 -
06840 -pf08811 -
06845 -pf08811 -
06850 -# -
06855 -pf16640pf00932 -
06860 -pf07589 -
06865 -pf00561 -
06870 HMMalddh_3_aromic1(pf00171) CsomeBCy
06875 -pf00474 -
(..190..)
07065 -# -
07070 -pf07676pf03572pf14684pf14685 -
07075 -pf07676pf00326 -
07080 -# -
07085 -# -
07090 -# -
07095 -# -
07105 -# -
07110 -pf01476pf01464 -
07115 HMMalddh_2_egra(pf00171) ACI EP_CTERM
07120 H_fuchsia ACI
07125 -# -
07130 -pf01464 -
07135 -pf00482 -
07140 -pf00437pf05157 -
07145 -pf11104 -
07150 -pf05137 -
07155 -pf10741 -
07160 -# -
07165 -pf00963pf00263 -
07175 -# -
BMC-P
BMC-H
BMC-T
Aldehyde dehydrogenase
PTAC
Signature enzyme
Alcohol dehydrogenase
Regulator

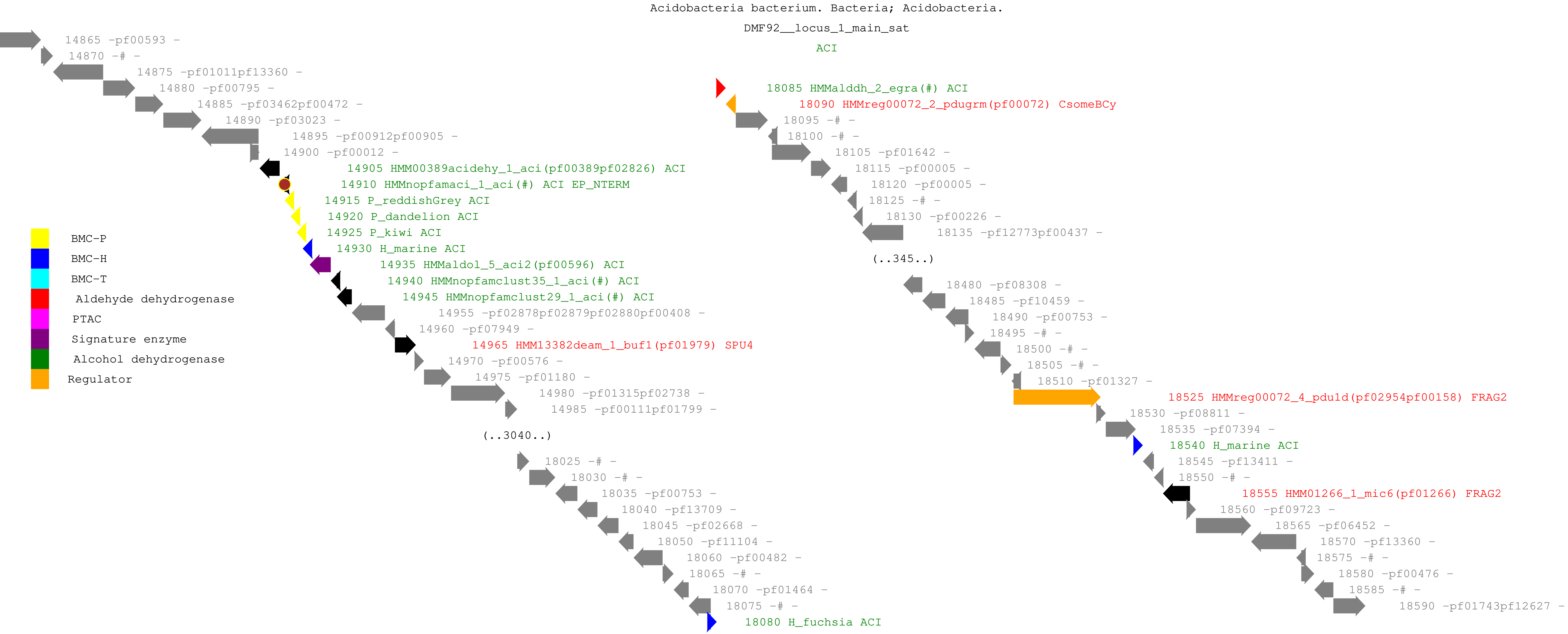
Acidobacteria bacterium. Bacteria; Acidobacteria.
DMF92__locus_1_main_sat
ACI
14865 -pf00593 -
14870 -# -
14875 -pf01011pf13360 -
14880 -pf00795 -
14885 -pf03462pf00472 -
14890 -pf03023 -
14895 -pf00912pf00905 -
14900 -pf00012 -
14905 HMM00389acidehy_1_aci(pf00389pf02826) ACI
14910 HMMnopfamaci_1_aci(#) ACI EP_NTERM
14915 P_reddishGrey ACI
14920 P_dandelion ACI
14925 P_kiwi ACI
14930 H_marine ACI
14935 HMMaldol_5_aci2(pf00596) ACI
14940 HMMnopfamclust35_1_aci(#) ACI
14945 HMMnopfamclust29_1_aci(#) ACI
14955 -pf02878pf02879pf02880pf00408 -
14960 -pf07949 -
14965 HMM13382deam_1_buf1(pf01979) SPU4
14970 -pf00576 -
14975 -pf01180 -
14980 -pf01315pf02738 -
14985 -pf00111pf01799 -
(..3040..)
18025 -# -
18030 -# -
18035 -pf00753 -
18040 -pf13709 -
18045 -pf02668 -
18050 -pf11104 -
18060 -pf00482 -
18065 -# -
18070 -pf01464 -
18075 -# -
18080 H_fuchsia ACI
18085 HMMaldh_2_egra(#) ACI
18090 HMMreg00072_2_pdugrm(pf00072) CsomeBCy
18095 -# -
18100 -# -
18105 -pf01642 -
18115 -pf00005 -
18120 -pf00005 -
18125 -# -
18130 -pf00226 -
18135 -pf12773pf00437 -
(..345..)
18480 -pf08308 -
18485 -pf10459 -
18490 -pf00753 -
18495 -# -
18500 -# -
18505 -# -
18510 -pf01327 -
18525 HMMreg00072_4_pdu1d(pf02954pf00158) FRAG2
18530 -pf08811 -
18535 -pf07394 -
18540 H_marine ACI
18545 -pf13411 -
18550 -# -
18555 HMM01266_1_mic6(pf01266) FRAG2
18560 -pf09723 -
18565 -pf06452 -
18570 -pf13360 -
18575 -# -
18580 -pf00476 -
18585 -# -
18590 -pf01743pf12627 -
BMC-P
BMC-H
BMC-T
Aldehyde dehydrogenase
PTAC
Signature enzyme
Alcohol dehydrogenase
Regulator

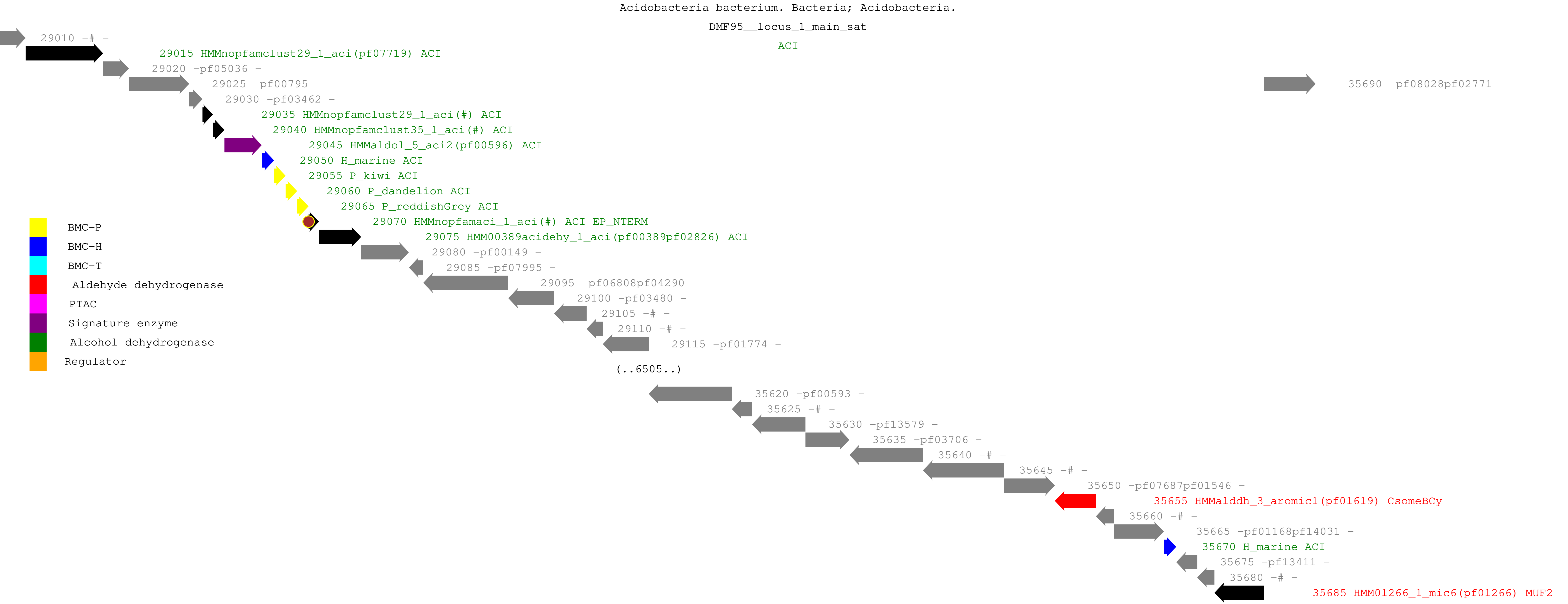
Acidobacteria bacterium. Bacteria; Acidobacteria.
DMF95__locus_1_main_sat
ACI
29010 -# -
29015 HMMnopfamclust29_1_aci(pf07719) ACI
29020 -pf05036 -
29025 -pf00795 -
29030 -pf03462 -
29035 HMMnopfamclust29_1_aci(#) ACI
29040 HMMnopfamclust35_1_aci(#) ACI
29045 HMMaldol_5_aci2(pf00596) ACI
29050 H_marine ACI
29055 P_kiwi ACI
29060 P_dandelion ACI
29065 P_reddishGrey ACI
29070 HMMnopfamaci_1_aci(#) ACI EP_NTERM
29075 HMM00389acidehy_1_aci(pf00389pf02826) ACI
29080 -pf00149 -
29085 -pf07995 -
29095 -pf06808pf04290 -
29100 -pf03480 -
29105 -# -
29110 -# -
29115 -pf01774 -
(..6505..)
35620 -pf00593 -
35625 -# -
35630 -pf13579 -
35635 -pf03706 -
35640 -# -
35645 -# -
35650 -pf07687pf01546 -
35655 HMMaldh_3_aromic1(pf01619) CsomeBCy
35660 -# -
35665 -pf01168pf14031 -
35670 H_marine ACI
35675 -pf13411 -
35680 -# -
35685 HMM01266_1_mic6(pf01266) MUF2
35690 -pf08028pf02771 -
BMC-P
BMC-H
BMC-T
Aldehyde dehydrogenase
PTAC
Signature enzyme
Alcohol dehydrogenase
Regulator

Acidobacteria bacterium. Bacteria; Acidobacteria.

DMF96__locus_1_main_sat

ACI

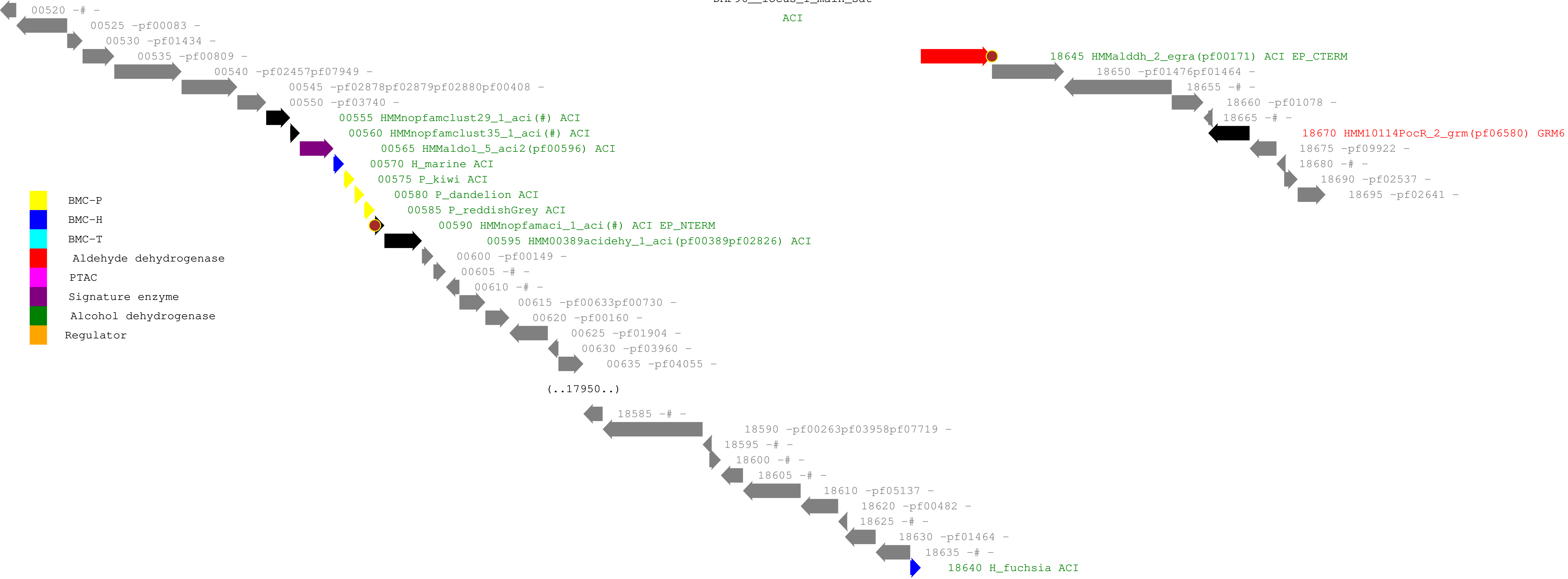

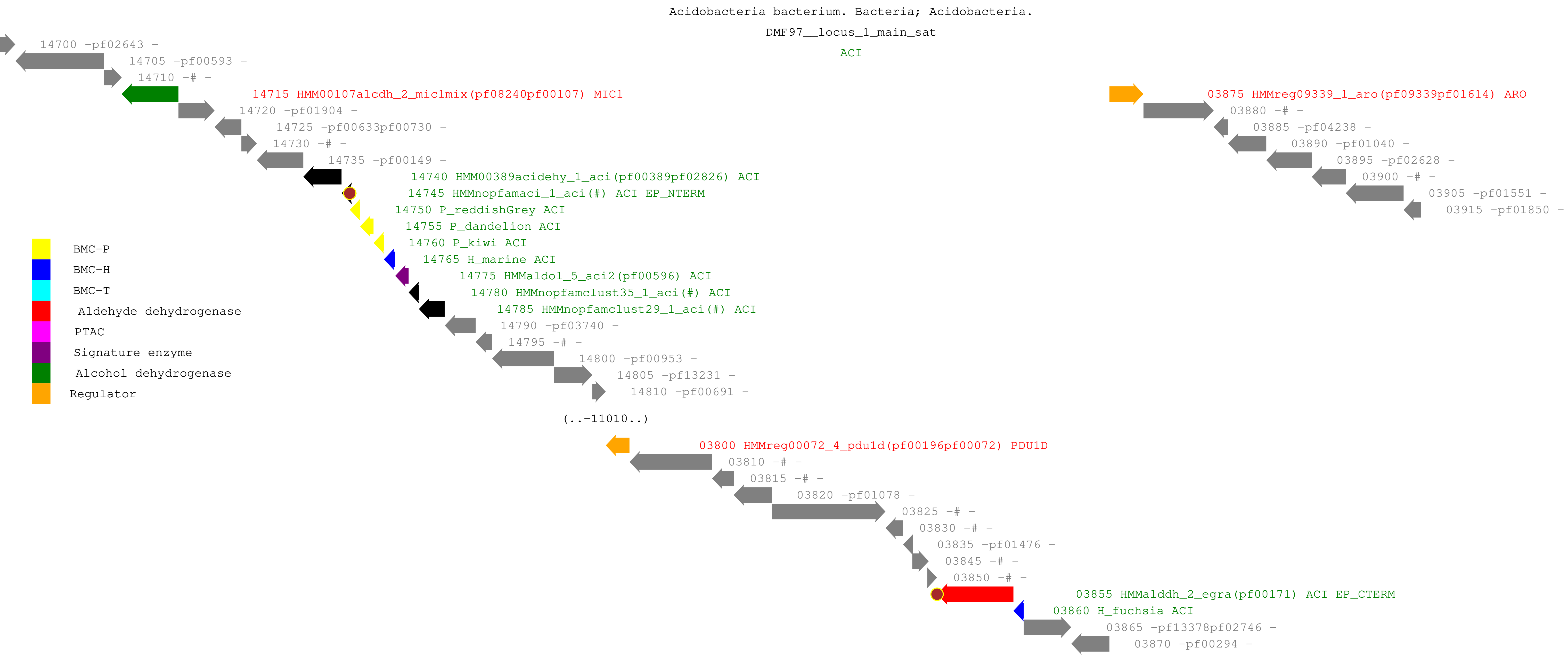

Acidobacteria bacterium. Bacteria; Acidobacteria.
DMF97__locus_1_main_sat
ACI
14700 -pf02643 -
14705 -pf00593 -
14710 -# -
14715 HMM00107alcdh_2_mic1mix(pf08240pf00107) MIC1
14720 -pf01904 -
14725 -pf00633pf00730 -
14730 -# -
14735 -pf00149 -
14740 HMM00389acidehy_1_aci(pf00389pf02826) ACI
14745 HMMnopfamaci_1_aci(#) ACI EP_NTERM
14750 P_reddishGrey ACI
14755 P_dandelion ACI
14760 P_kiwi ACI
14765 H_marine ACI
14775 HMMaldol_5_aci2(pf00596) ACI
14780 HMMnopfamclust35_1_aci(#) ACI
14785 HMMnopfamclust29_1_aci(#) ACI
14790 -pf03740 -
14795 -# -
14800 -pf00953 -
14805 -pf13231 -
14810 -pf00691 -
(..-11010..)
03800 HMMreg00072_4_pdu1d(pf00196pf00072) PDU1D
03810 -# -
03815 -# -
03820 -pf01078 -
03825 -# -
03830 -# -
03835 -pf01476 -
03845 -# -
03850 -# -
03855 HMMalddh_2_egra(pf00171) ACI EP_CTERM
03860 H_fuchsia ACI
03865 -pf13378pf02746 -
03870 -pf00294 -
03875 HMMreg09339_1_aro(pf09339pf01614) ARO
03880 -# -
03885 -pf04238 -
03890 -pf01040 -
03895 -pf02628 -
03900 -# -
03905 -pf01551 -
03915 -pf01850 -
BMC-P
BMC-H
BMC-T
Aldehyde dehydrogenase
PTAC
Signature enzyme
Alcohol dehydrogenase
Regulator

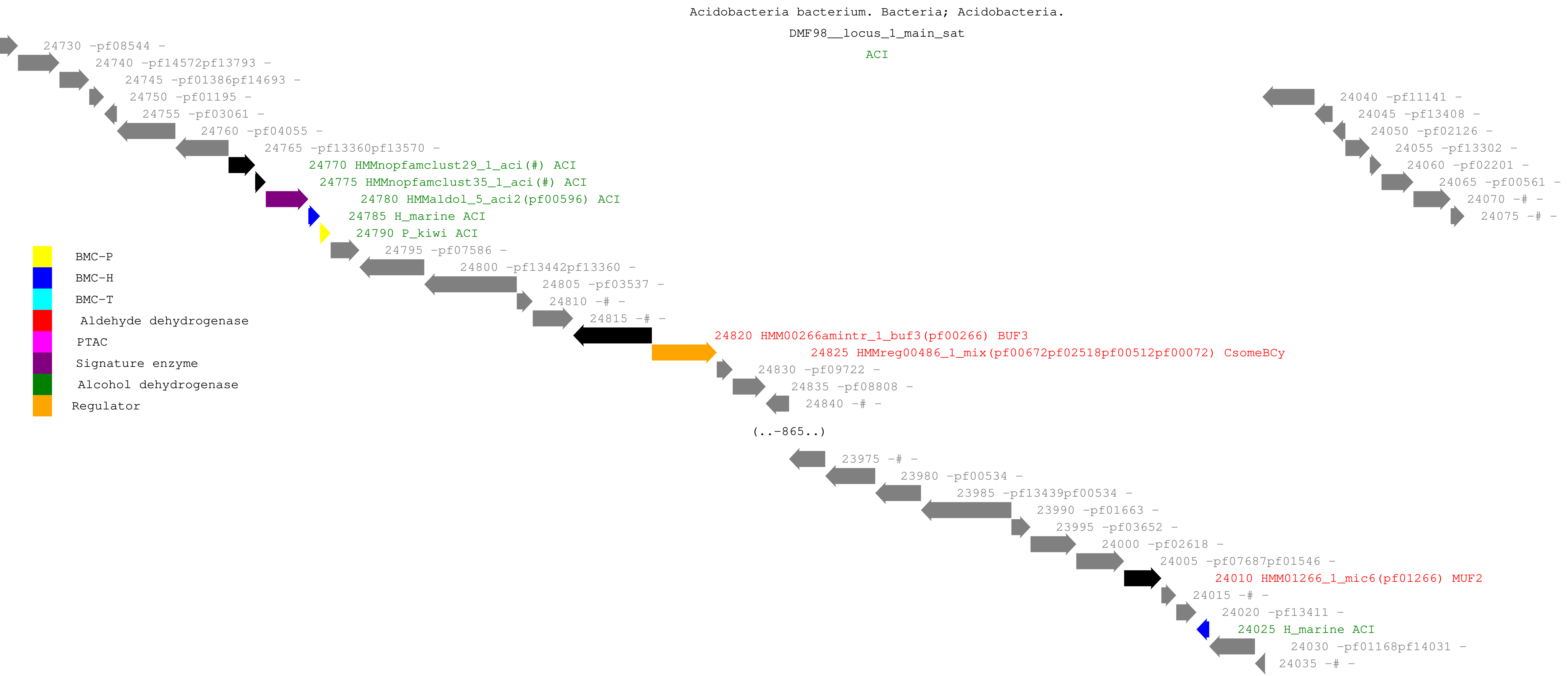

Acidobacteria bacterium. Bacteria; Acidobacteria.
DMF98__locus_1_main_sat
ACI
24730 -pf08544 -
24740 -pf14572pf13793 -
24745 -pf01386pf14693 -
24750 -pf01195 -
24755 -pf03061 -
24760 -pf04055 -
24765 -pf13360pf13570 -
24770 HMMnopfamclust29_1_aci(#) ACI
24775 HMMnopfamclust35_1_aci(#) ACI
24780 HMMaldol_5_aci2(pf00596) ACI
24785 H_marine ACI
24790 P_kiwi ACI
24795 -pf07586 -
24800 -pf13442pf13360 -
24805 -pf03537 -
24810 -# -
24815 -# -
24820 HMM00266amintr_1_buf3(pf00266) BUF3
24825 HMMreg00486_1_mix(pf00672pf02518pf00512pf00072) CsomeBCy
24830 -pf09722 -
24835 -pf08808 -
24840 -# -
(..-865..)
23975 -# -
23980 -pf00534 -
23985 -pf13439pf00534 -
23990 -pf01663 -
23995 -pf03652 -
24000 -pf02618 -
24005 -pf07687pf01546 -
24010 HMM01266_1_mic6(pf01266) MUF2
24015 -# -
24020 -pf13411 -
24025 H_marine ACI
24030 -pf01168pf14031 -
24035 -# -
24040 -pf11141 -
24045 -pf13408 -
24050 -pf02126 -
24055 -pf13302 -
24060 -pf02201 -
24065 -pf00561 -
24070 -# -
24075 -# -
BMC-P
BMC-H
BMC-T
Aldehyde dehydrogenase
PTAC
Signature enzyme
Alcohol dehydrogenase
Regulator

Acidobacteria bacterium. Bacteria; Acidobacteria.

DMG00__locus_1_main_sat

ACI

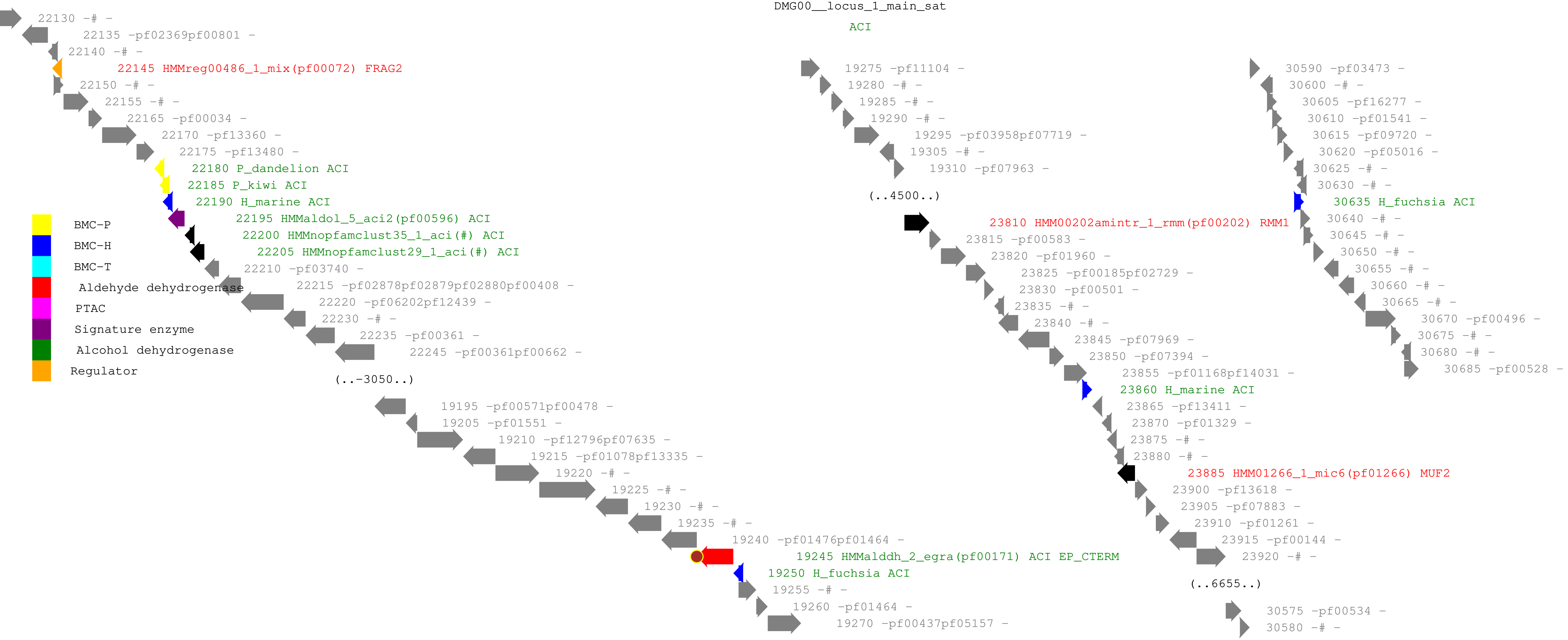

Acidobacteria bacterium. Bacteria; Acidobacteria.

DMG01__locus_1_main_sat

ACI

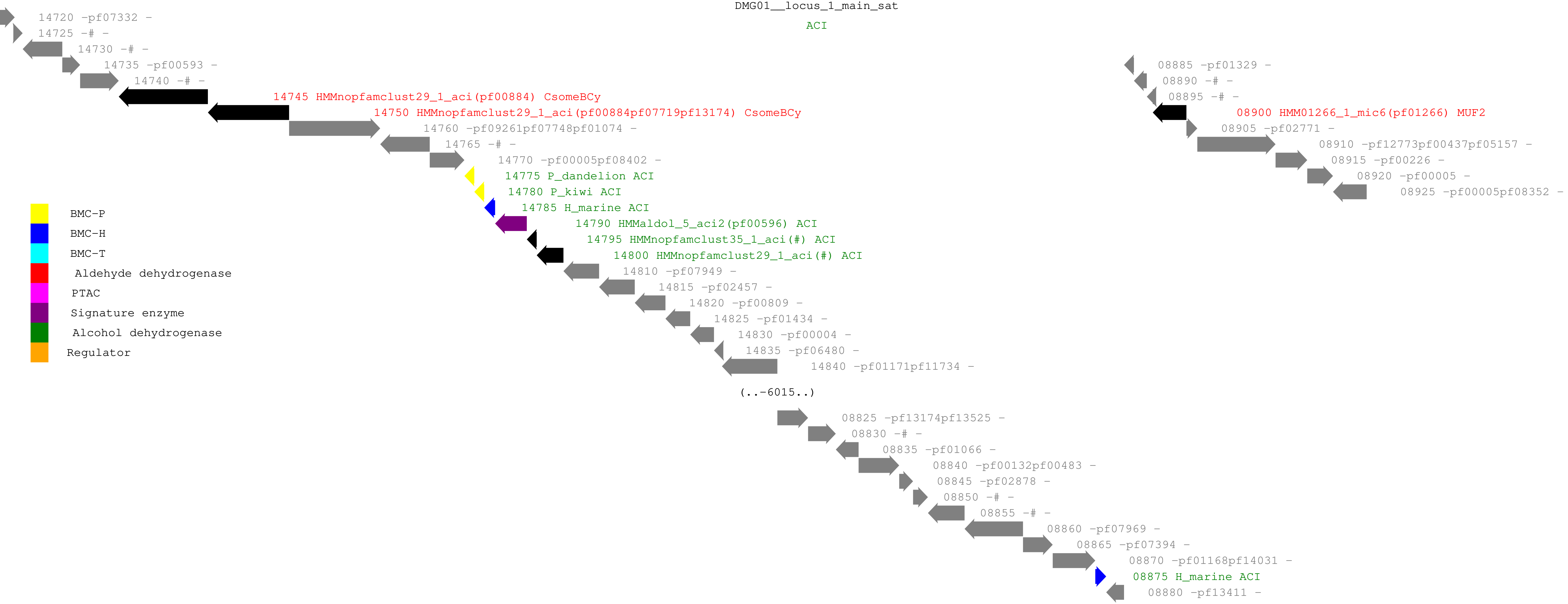

Acidobacteria bacterium. Bacteria; Acidobacteria.

DMG02__locus_1_main_sat

ACI

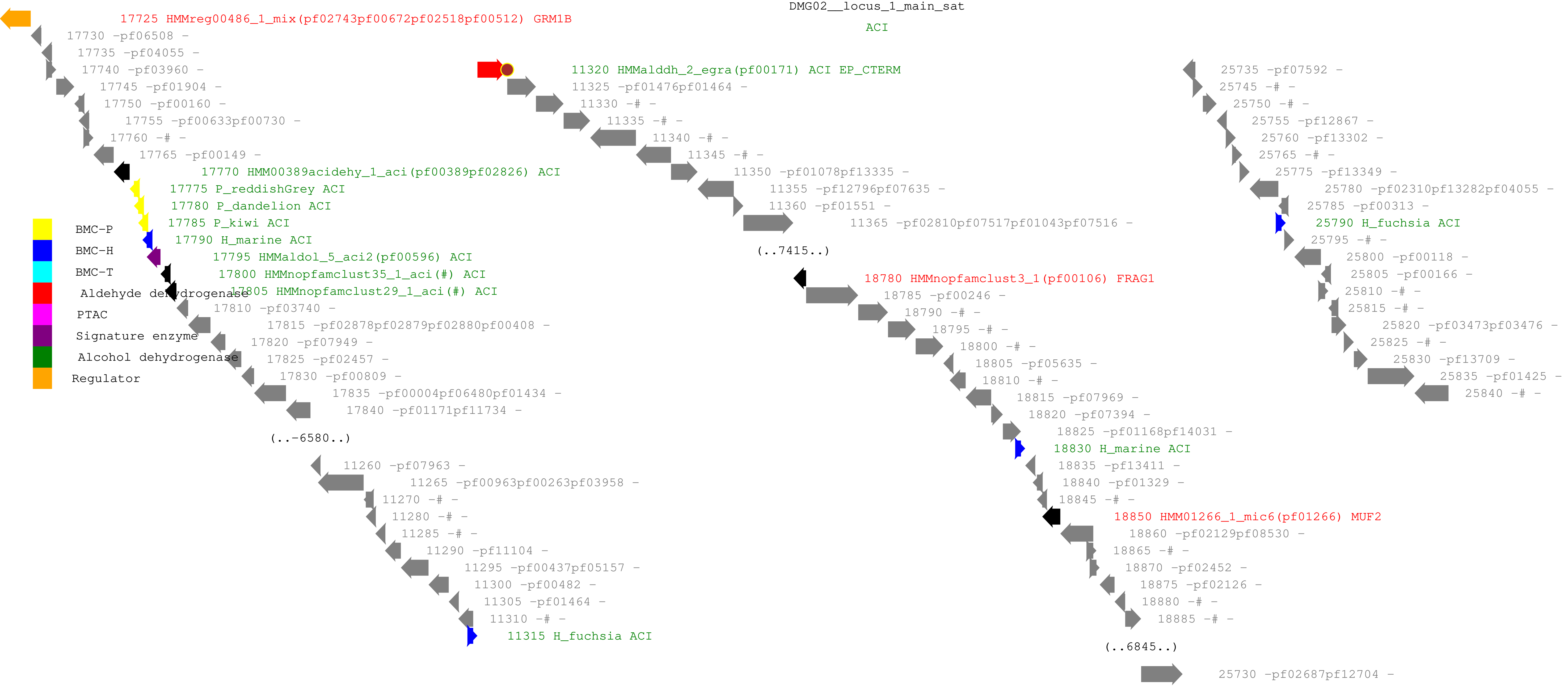

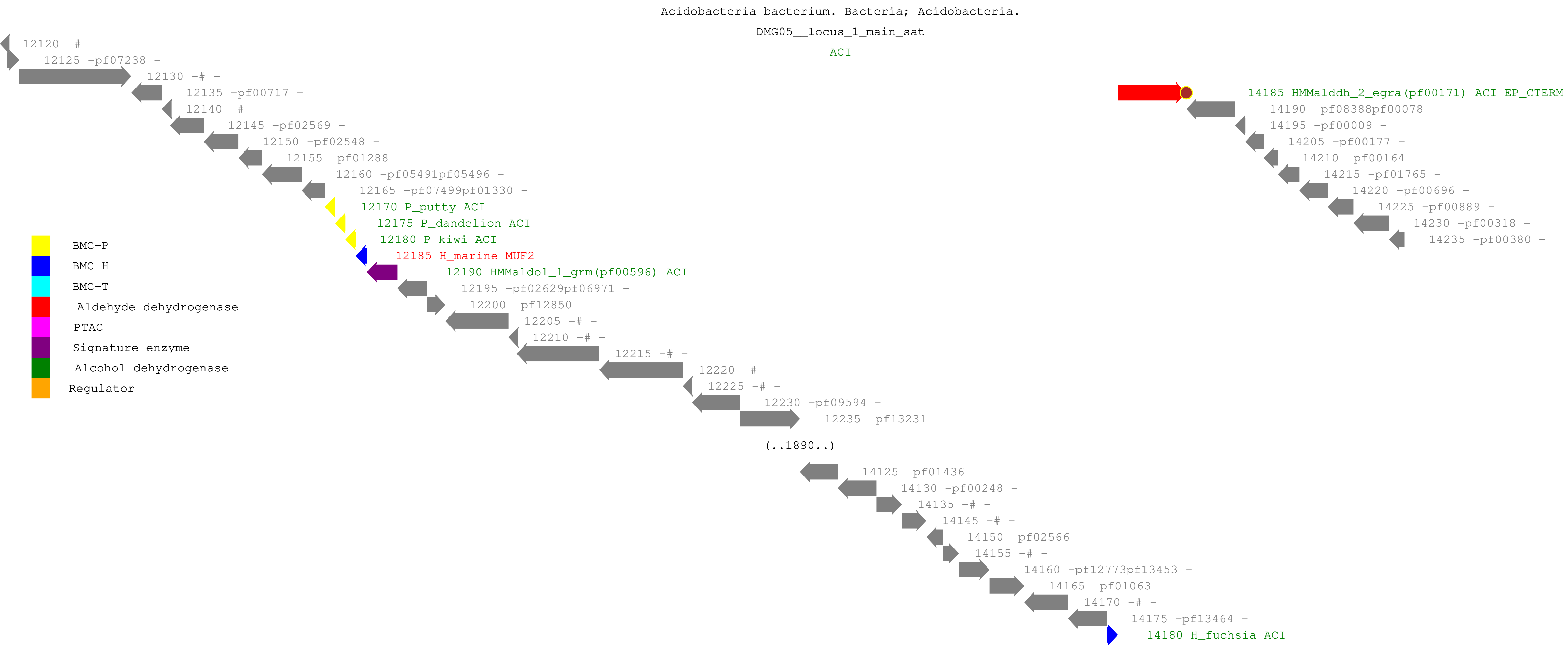

Acidobacteria bacterium. Bacteria; Acidobacteria.
DMG05__locus_1_main_sat
ACI
12120 -# -
12125 -pf07238 -
12130 -# -
12135 -pf00717 -
12140 -# -
12145 -pf02569 -
12150 -pf02548 -
12155 -pf01288 -
12160 -pf05491pf05496 -
12165 -pf07499pf01330 -
12170 P_putty ACI
12175 P_dandelion ACI
12180 P_kiwi ACI
12185 H_marine MUF2
12190 HMMaldol_1_grm(pf00596) ACI
12195 -pf02629pf06971 -
12200 -pf12850 -
12205 -# -
12210 -# -
12215 -# -
12220 -# -
12225 -# -
12230 -pf09594 -
12235 -pf13231 -
(..1890..)
14125 -pf01436 -
14130 -pf00248 -
14135 -# -
14145 -# -
14150 -pf02566 -
14155 -# -
14160 -pf12773pf13453 -
14165 -pf01063 -
14170 -# -
14175 -pf13464 -
14180 H_fuchsia ACI
14185 HMMalddh_2_egra(pf00171) ACI EP_CTERM
14190 -pf08388pf00078 -
14195 -pf00009 -
14205 -pf00177 -
14210 -pf00164 -
14215 -pf01765 -
14220 -pf00696 -
14225 -pf00889 -
14230 -pf00318 -
14235 -pf00380 -
BMC-P
BMC-H
BMC-T
Aldehyde dehydrogenase
PTAC
Signature enzyme
Alcohol dehydrogenase
Regulator

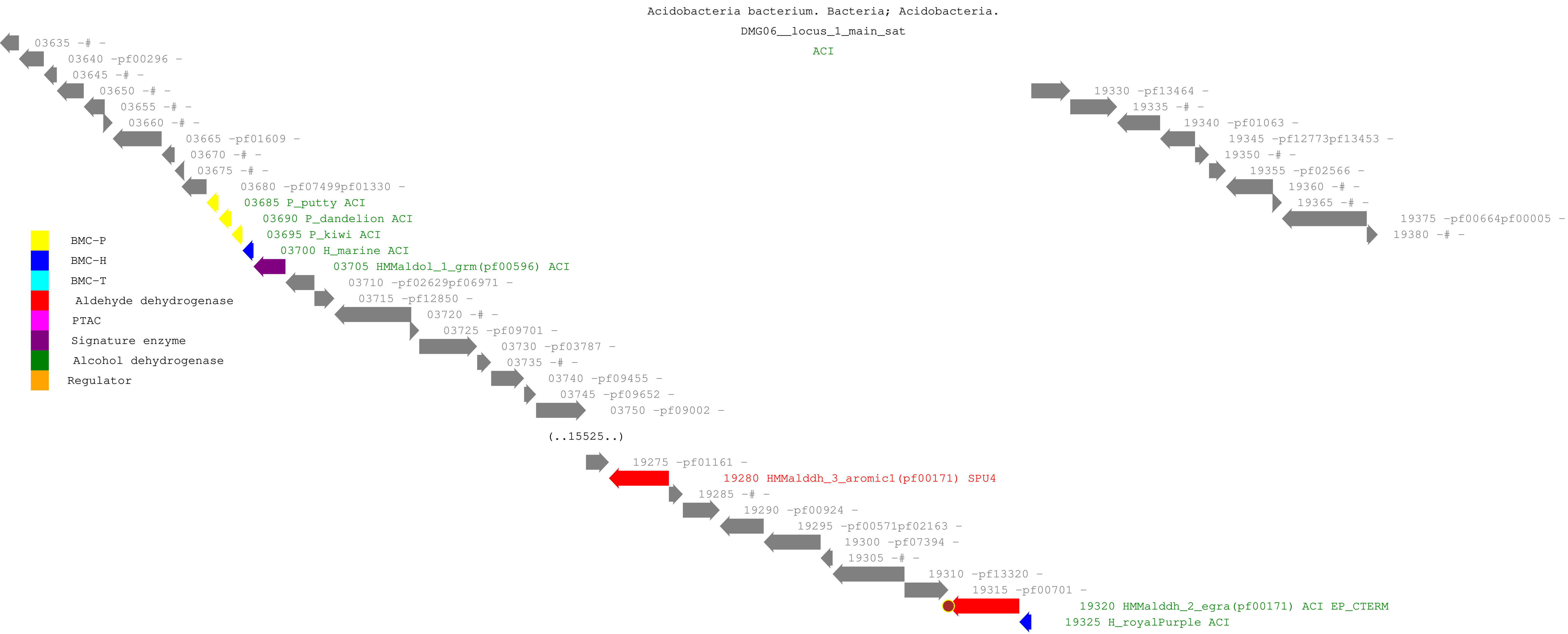
Acidobacteria bacterium. Bacteria; Acidobacteria.
DMG06__locus_1_main_sat
ACI
03635 -# -
03640 -pf00296 -
03645 -# -
03650 -# -
03655 -# -
03660 -# -
03665 -pf01609 -
03670 -# -
03675 -# -
03680 -pf07499pf01330 -
03685 P_putty ACI
03690 P_dandelion ACI
03695 P_kiwi ACI
03700 H_marine ACI
03705 HMMaldol_1_grm(pf00596) ACI
03710 -pf02629pf06971 -
03715 -pf12850 -
03720 -# -
03725 -pf09701 -
03730 -pf03787 -
03735 -# -
03740 -pf09455 -
03745 -pf09652 -
03750 -pf09002 -
(..15525..)
19275 -pf01161 -
19280 HMMalddh_3_aromic1(pf00171) SPU4
19285 -# -
19290 -pf00924 -
19295 -pf00571pf02163 -
19300 -pf07394 -
19305 -# -
19310 -pf13320 -
19315 -pf00701 -
19320 HMMalddh_2_egra(pf00171) ACI EP_CTERM
19325 H_royalPurple ACI
19330 -pf13464 -
19335 -# -
19340 -pf01063 -
19345 -pf12773pf13453 -
19350 -# -
19355 -pf02566 -
19360 -# -
19365 -# -
19375 -pf00664pf00005 -
19380 -# -
BMC-P
BMC-H
BMC-T
Aldehyde dehydrogenase
PTAC
Signature enzyme
Alcohol dehydrogenase
Regulator

Acidobacteria bacterium. Bacteria; Acidobacteria.

DMG07__locus_fragmented

ACI

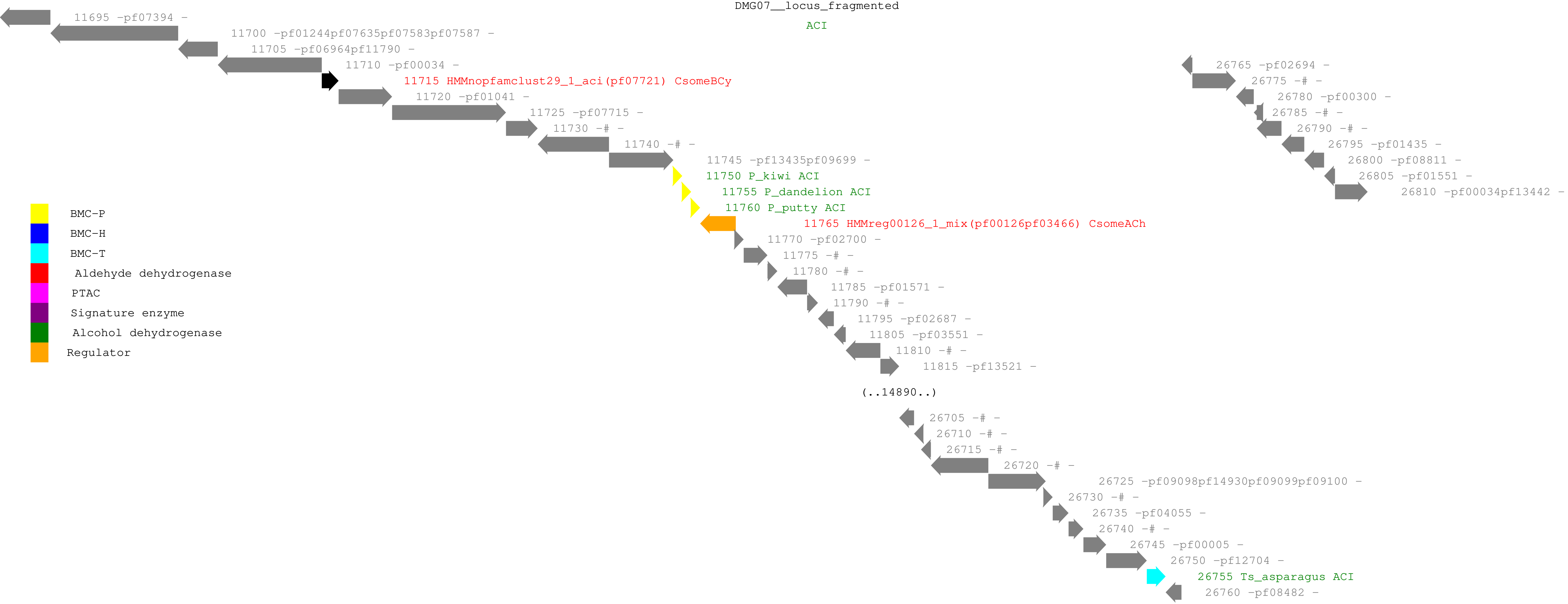

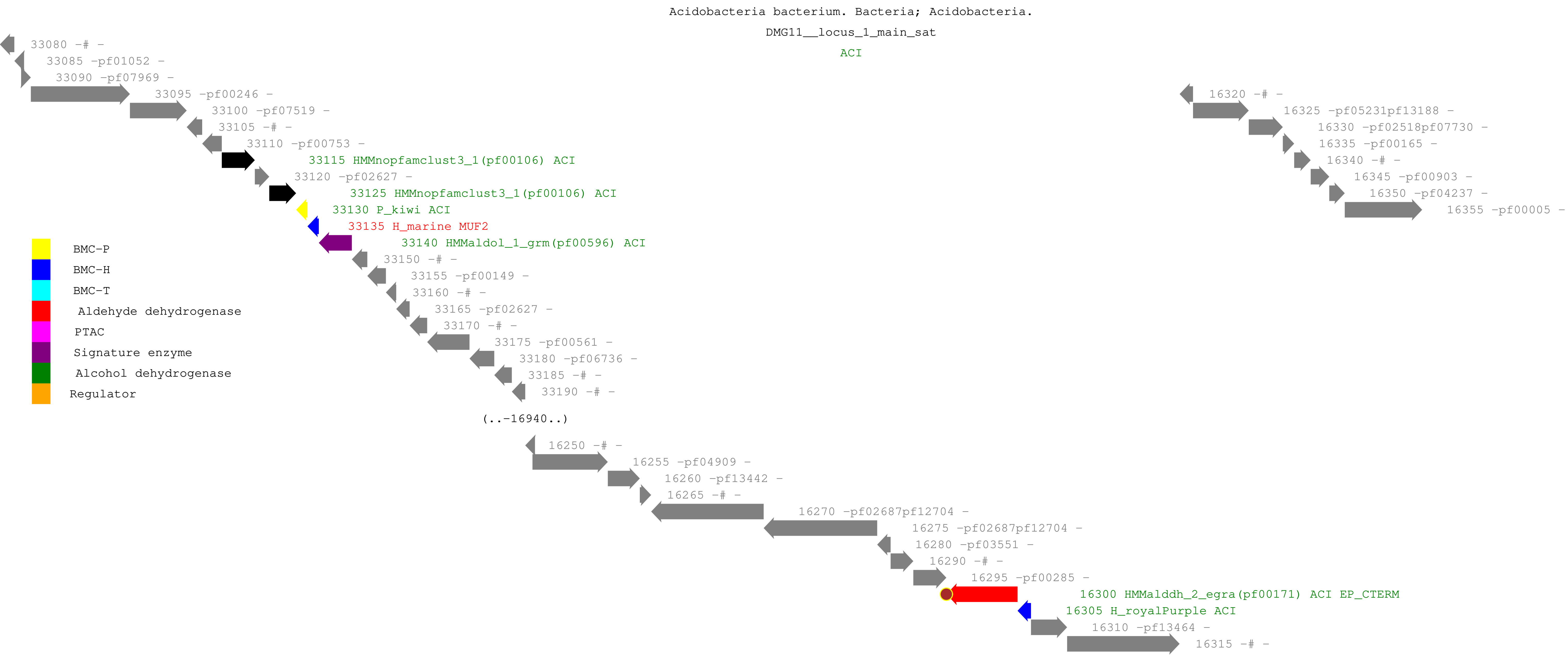

Acidobacteria bacterium. Bacteria; Acidobacteria.
DMG11__locus_1_main_sat
ACI
33080 -# -
33085 -pf01052 -
33090 -pf07969 -
33095 -pf00246 -
33100 -pf07519 -
33105 -# -
33110 -pf00753 -
33115 HMMnopfamclust3_1(pf00106) ACI
33120 -pf02627 -
33125 HMMnopfamclust3_1(pf00106) ACI
33130 P_kiwi ACI
33135 H_marine MUF2
33140 HMMaldol_1_grm(pf00596) ACI
33150 -# -
33155 -pf00149 -
33160 -# -
33165 -pf02627 -
33170 -# -
33175 -pf00561 -
33180 -pf06736 -
33185 -# -
33190 -# -
(..-16940..)
16250 -# -
16255 -pf04909 -
16260 -pf13442 -
16265 -# -
16270 -pf02687pf12704 -
16275 -pf02687pf12704 -
16280 -pf03551 -
16290 -# -
16295 -pf00285 -
16300 HMMalddh_2_egra(pf00171) ACI EP_CTERM
16305 H_royalPurple ACI
16310 -pf13464 -
16315 -# -
16320 -# -
16325 -pf05231pf13188 -
16330 -pf02518pf07730 -
16335 -pf00165 -
16340 -# -
16345 -pf00903 -
16350 -pf04237 -
16355 -pf00005 -
BMC-P
BMC-H
BMC-T
Aldehyde dehydrogenase
PTAC
Signature enzyme
Alcohol dehydrogenase
Regulator

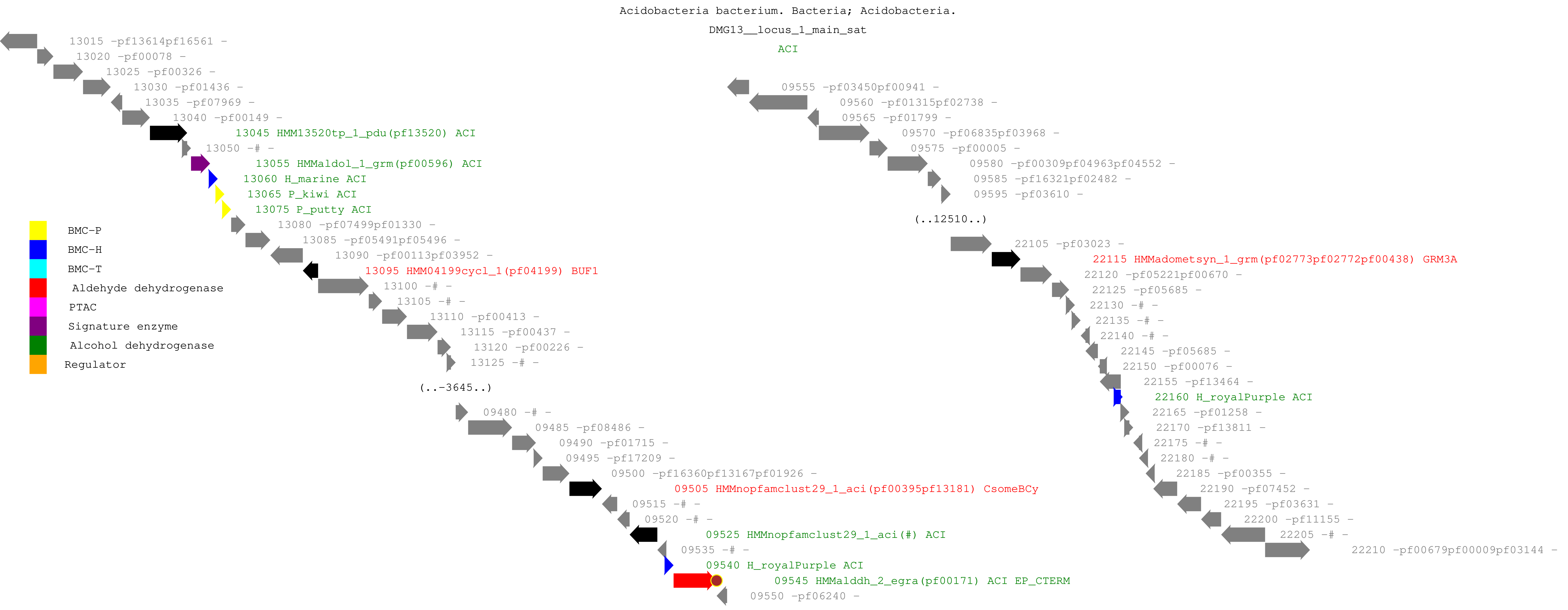

Acidobacteria bacterium. Bacteria; Acidobacteria.
DMG13__locus_1_main_sat
ACI
13015 -pf13614pf16561 -
13020 -pf00078 -
13025 -pf00326 -
13030 -pf01436 -
13035 -pf07969 -
13040 -pf00149 -
13045 HMM13520tp_1_pdu(pf13520) ACI
13050 -# -
13055 HMMaldol_1_grm(pf00596) ACI
13060 H_marine ACI
13065 P_kiwi ACI
13075 P_putty ACI
13080 -pf07499pf01330 -
13085 -pf05491pf05496 -
13090 -pf00113pf03952 -
13095 HMM04199cycl_1(pf04199) BUF1
13100 -# -
13105 -# -
13110 -pf00413 -
13115 -pf00437 -
13120 -pf00226 -
13125 -# -
(..-3645..)
09480 -# -
09485 -pf08486 -
09490 -pf01715 -
09495 -pf17209 -
09500 -pf16360pf13167pf01926 -
09505 HMMnopfamclust29_1_aci(pf00395pf13181) CsomeBCy
09515 -# -
09520 -# -
09525 HMMnopfamclust29_1_aci(#) ACI
09535 -# -
09540 H_royalPurple ACI
09545 HMMalddh_2_egra(pf00171) ACI EP_CTERM
09550 -pf06240 -
09555 -pf03450pf00941 -
09560 -pf01315pf02738 -
09565 -pf01799 -
09570 -pf06835pf03968 -
09575 -pf00005 -
09580 -pf00309pf04963pf04552 -
09585 -pf16321pf02482 -
09595 -pf03610 -
(..12510..)
22105 -pf03023 -
22115 HMMadometsyn_1_grm(pf02773pf02772pf00438) GRM3A
22120 -pf05221pf00670 -
22125 -pf05685 -
22130 -# -
22135 -# -
22140 -# -
22145 -pf05685 -
22150 -pf00076 -
22155 -pf13464 -
22160 H_royalPurple ACI
22165 -pf01258 -
22170 -pf13811 -
22175 -# -
22180 -# -
22185 -pf00355 -
22190 -pf07452 -
22195 -pf03631 -
22200 -pf11155 -
22205 -# -
22210 -pf00679pf00009pf03144 -
BMC-P
BMC-H
BMC-T
Aldehyde dehydrogenase
PTAC
Signature enzyme
Alcohol dehydrogenase
Regulator

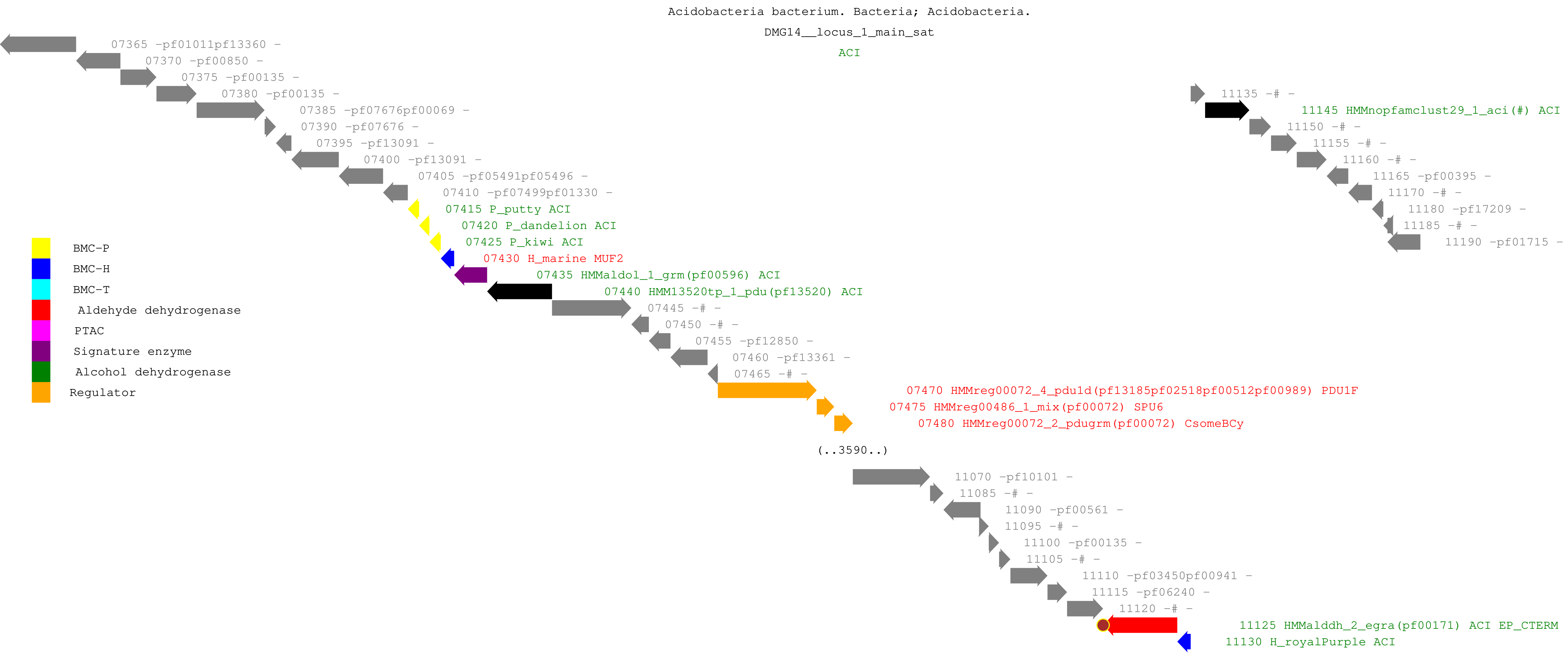
Acidobacteria bacterium. Bacteria; Acidobacteria.
DMG14__locus_1_main_sat
ACI
07365 -pf01011pf13360 -
07370 -pf00850 -
07375 -pf00135 -
07380 -pf00135 -
07385 -pf07676pf00069 -
07390 -pf07676 -
07395 -pf13091 -
07400 -pf13091 -
07405 -pf05491pf05496 -
07410 -pf07499pf01330 -
07415 P_putty ACI
07420 P_dandelion ACI
07425 P_kiwi ACI
07430 H_marine MUF2
07435 HMMaldol_1_grm(pf00596) ACI
07440 HMM13520tp_1_pdu(pf13520) ACI
07445 -# -
07450 -# -
07455 -pf12850 -
07460 -pf13361 -
07465 -# -
07470 HMMreg00072_4_pdu1d(pf13185pf02518pf00512pf00989) PDU1F
07475 HMMreg00486_1_mix(pf00072) SPU6
07480 HMMreg00072_2_pdugrm(pf00072) CsomeBCy
(..3590..)
11070 -pf10101 -
11085 -# -
11090 -pf00561 -
11095 -# -
11100 -pf00135 -
11105 -# -
11110 -pf03450pf00941 -
11115 -pf06240 -
11120 -# -
11125 HMMaldh_2_egra(pf00171) ACI EP_CTERM
11130 H_royalPurple ACI
11135 -# -
11145 HMMnopfamclust29_1_aci(#) ACI
11150 -# -
11155 -# -
11160 -# -
11165 -pf00395 -
11170 -# -
11180 -pf17209 -
11185 -# -
11190 -pf01715 -
BMC-P
BMC-H
BMC-T
Aldehyde dehydrogenase
PTAC
Signature enzyme
Alcohol dehydrogenase
Regulator

Acidobacteria bacterium. Bacteria; Acidobacteria.

DMG15__locus_1_main_sat

ACI

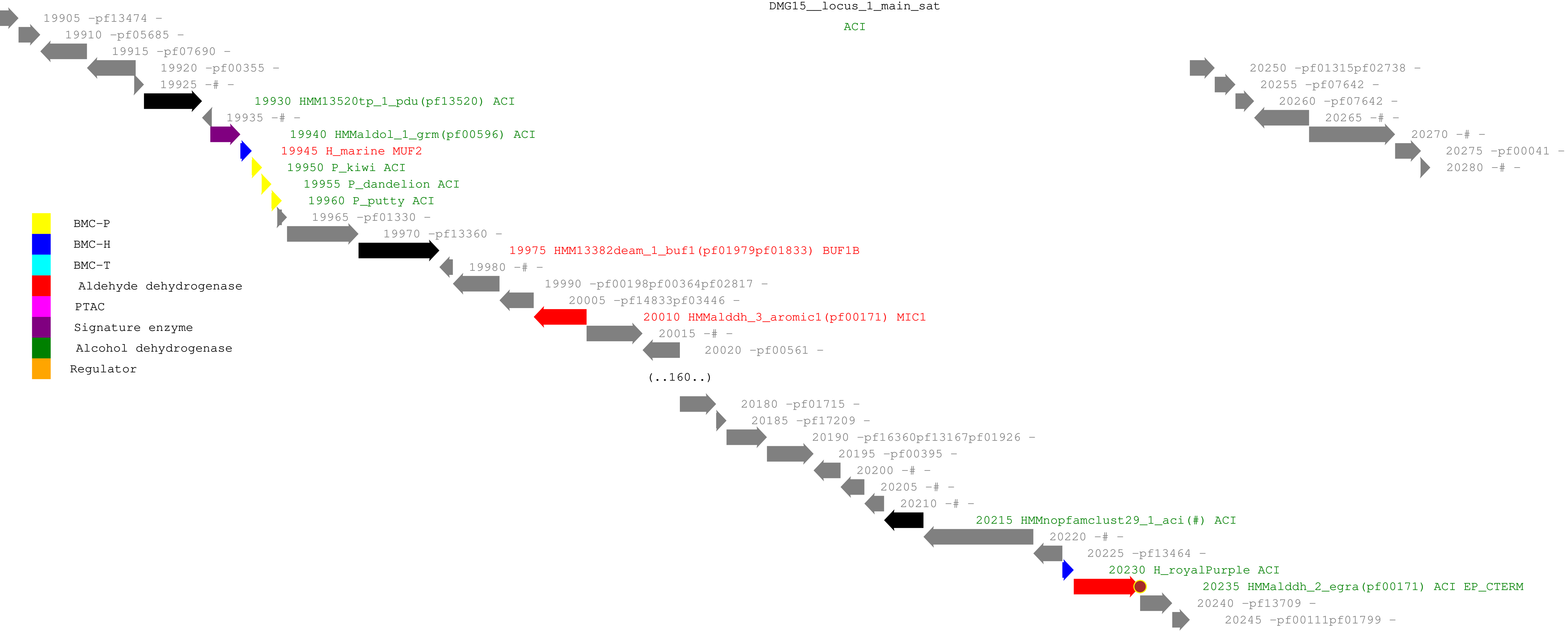

Acidobacteria bacterium. Bacteria; Acidobacteria.

DMG16__locus_1_main_sat

ACI

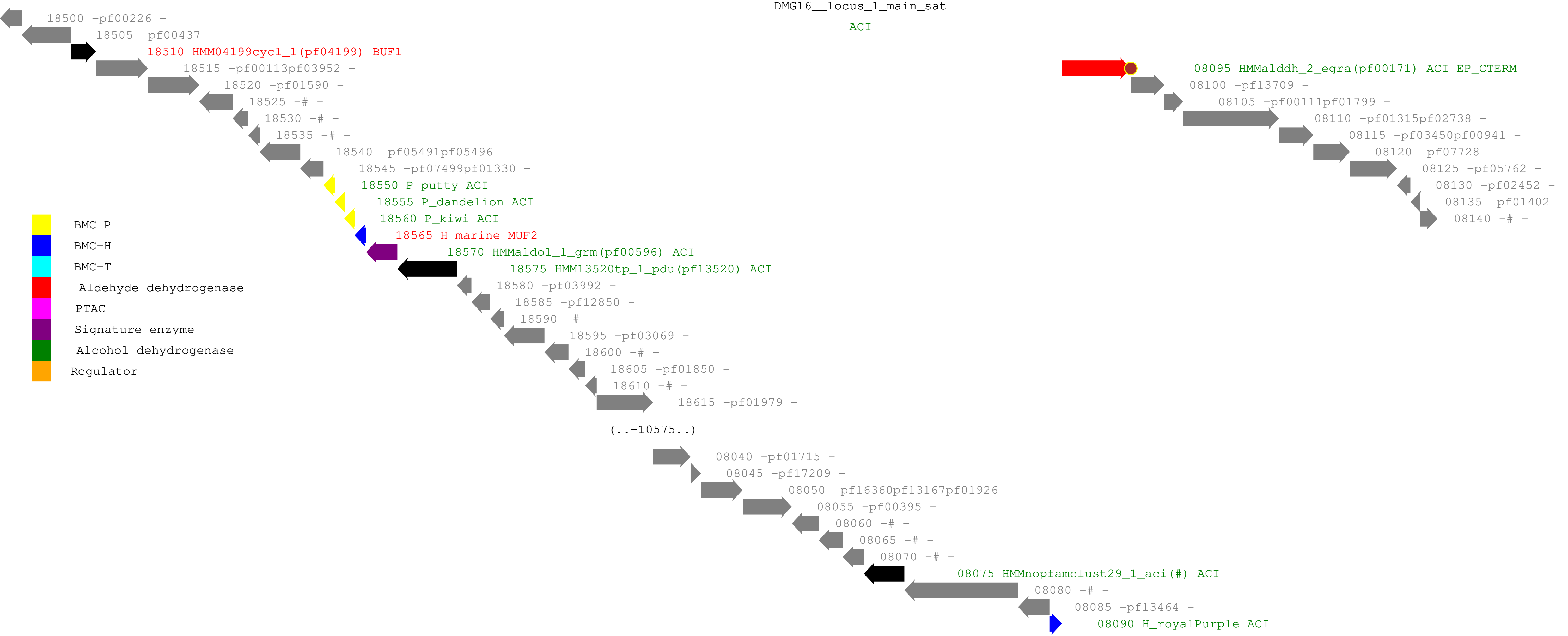

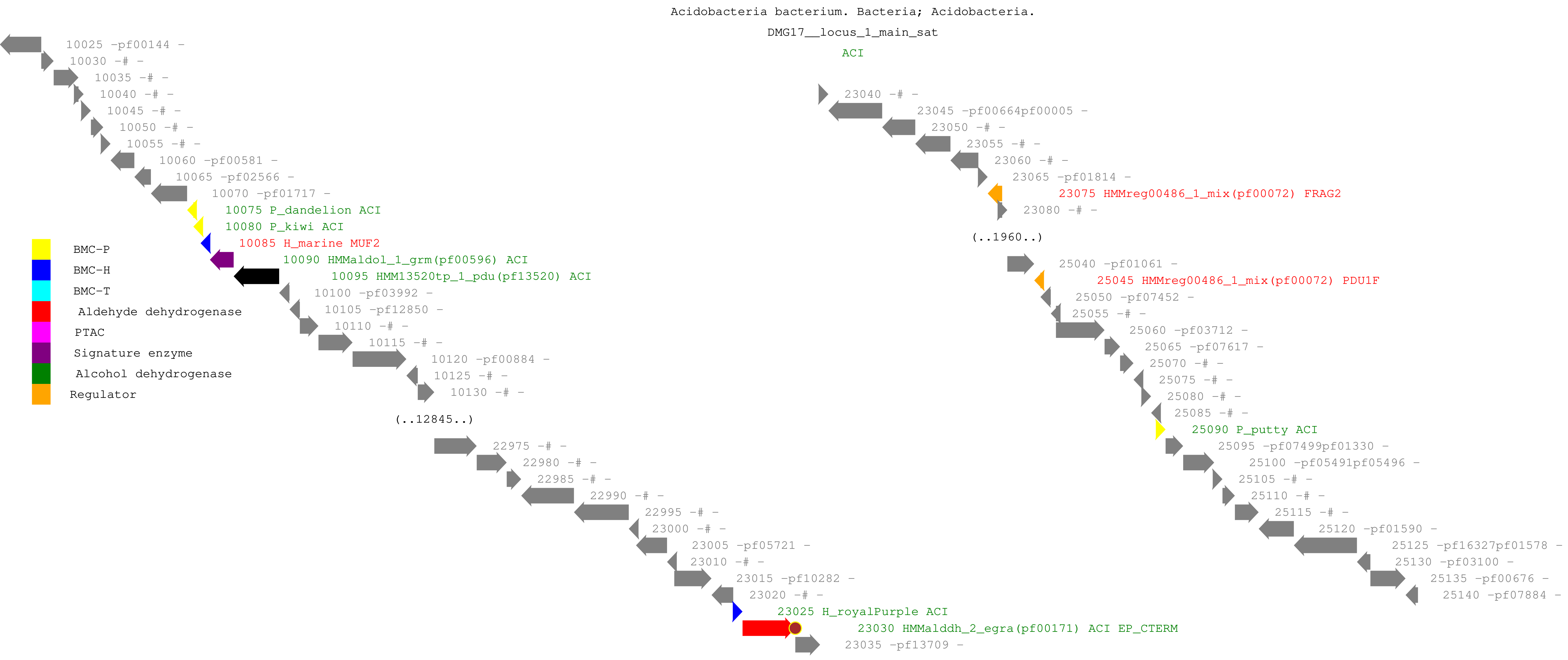
Acidobacteria bacterium. Bacteria; Acidobacteria.
DMG17__locus_1_main_sat
ACI
10025 -pf00144 -
10030 -# -
10035 -# -
10040 -# -
10045 -# -
10050 -# -
10055 -# -
10060 -pf00581 -
10065 -pf02566 -
10070 -pf01717 -
10075 P_dandelion ACI
10080 P_kiwi ACI
10085 H_marine MUF2
10090 HMMaldol_1_grm(pf00596) ACI
10095 HMM13520tp_1_pdu(pf13520) ACI
10100 -pf03992 -
10105 -pf12850 -
10110 -# -
10115 -# -
10120 -pf00884 -
10125 -# -
10130 -# -
(..12845..)
22975 -# -
22980 -# -
22985 -# -
22990 -# -
22995 -# -
23000 -# -
23005 -pf05721 -
23010 -# -
23015 -pf10282 -
23020 -# -
23025 H_royalPurple ACI
23030 HMMalddh_2_egra(pf00171) ACI EP_CTERM
23035 -pf13709 -
23040 -# -
23045 -pf00664pf00005 -
23050 -# -
23055 -# -
23060 -# -
23065 -pf01814 -
23075 HMMreg00486_1_mix(pf00072) FRAG2
23080 -# -
(..1960..)
25040 -pf01061 -
25045 HMMreg00486_1_mix(pf00072) PDU1F
25050 -pf07452 -
25055 -# -
25060 -pf03712 -
25065 -pf07617 -
25070 -# -
25075 -# -
25080 -# -
25085 -# -
25090 P_putty ACI
25095 -pf07499pf01330 -
25100 -pf05491pf05496 -
25105 -# -
25110 -# -
25115 -# -
25120 -pf01590 -
25125 -pf16327pf01578 -
25130 -pf03100 -
25135 -pf00676 -
25140 -pf07884 -
BMC-P
BMC-H
BMC-T
Aldehyde dehydrogenase
PTAC
Signature enzyme
Alcohol dehydrogenase
Regulator

Acidobacteria bacterium. Bacteria; Acidobacteria.

DMG19__locus_1_main_sat

ACI

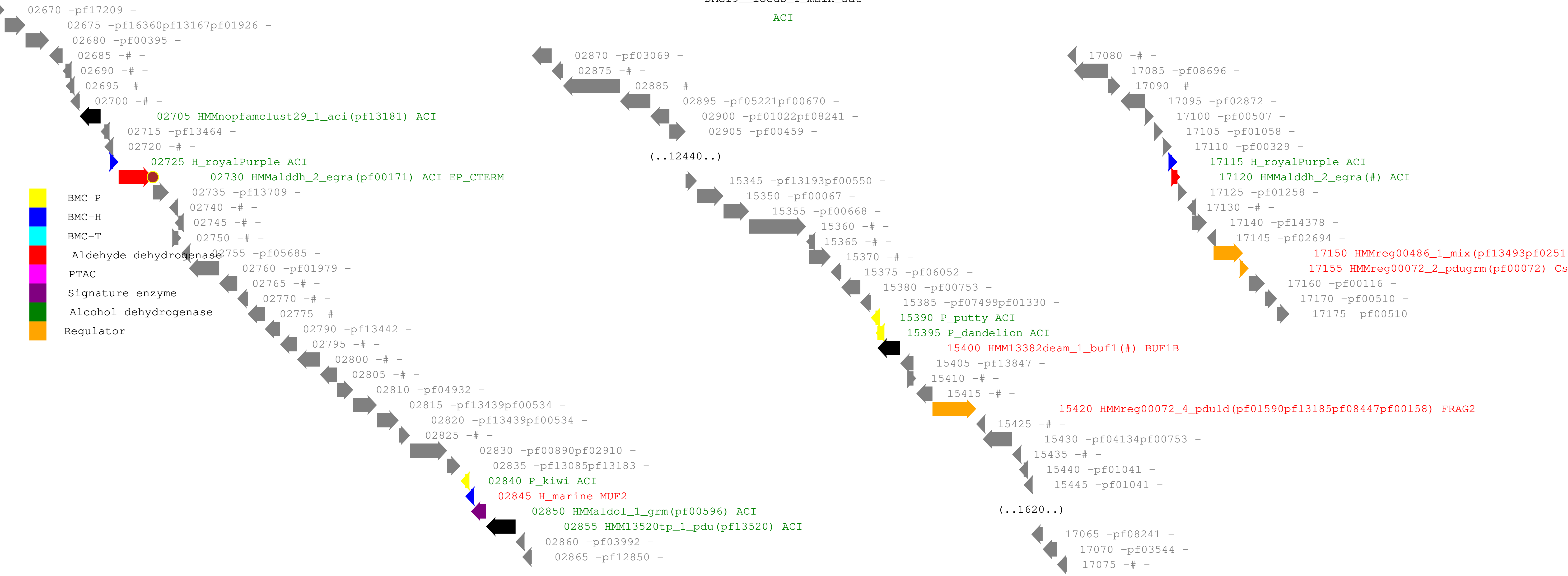

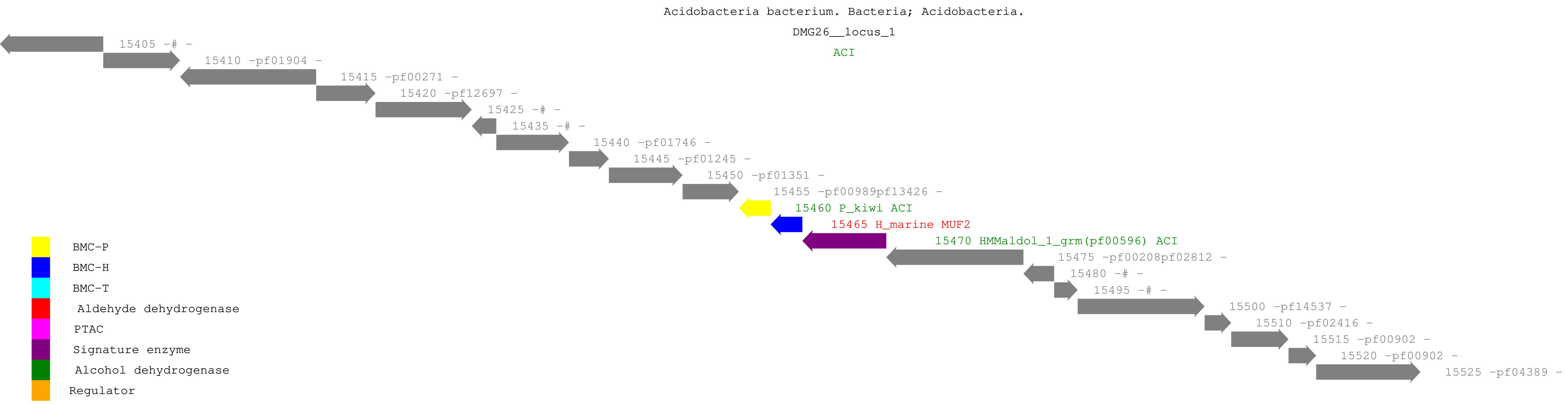
Acidobacteria bacterium. Bacteria; Acidobacteria.
DMG26__locus_1
ACI
15405 -# -
15410 -pf01904 -
15415 -pf00271 -
15420 -pf12697 -
15425 -# -
15435 -# -
15440 -pf01746 -
15445 -pf01245 -
15450 -pf01351 -
15455 -pf00989pf13426 -
15460 P_kiwi ACI
15465 H_marine MUF2
15470 HMMaldol_1_grm(pf00596) ACI
15475 -pf00208pf02812 -
15480 -# -
15495 -# -
15500 -pf14537 -
15510 -pf02416 -
15515 -pf00902 -
15520 -pf00902 -
15525 -pf04389 -
BMC-P
BMC-H
BMC-T
Aldehyde dehydrogenase
PTAC
Signature enzyme
Alcohol dehydrogenase
Regulator

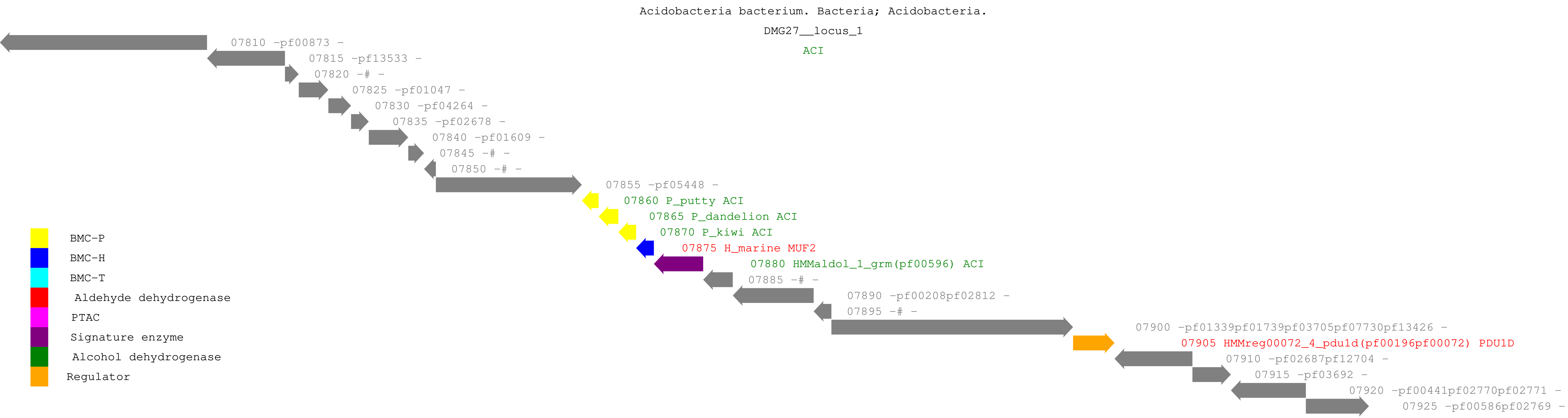

Acidobacteria bacterium. Bacteria; Acidobacteria.
DMG27__locus_1
ACI
07810 -pf00873 -
07815 -pf13533 -
07820 -# -
07825 -pf01047 -
07830 -pf04264 -
07835 -pf02678 -
07840 -pf01609 -
07845 -# -
07850 -# -
07855 -pf05448 -
07860 P_putty ACI
07865 P_dandelion ACI
07870 P_kiwi ACI
07875 H_marine MUF2
07880 HMMaldol_1_grm(pf00596) ACI
07885 -# -
07890 -pf00208pf02812 -
07895 -# -
07900 -pf01339pf01739pf03705pf07730pf13426 -
07905 HMMreg00072_4_pdu1d(pf00196pf00072) PDU1D
07910 -pf02687pf12704 -
07915 -pf03692 -
07920 -pf00441pf02770pf02771 -
07925 -pf00586pf02769 -
BMC-P
BMC-H
BMC-T
Aldehyde dehydrogenase
PTAC
Signature enzyme
Alcohol dehydrogenase
Regulator

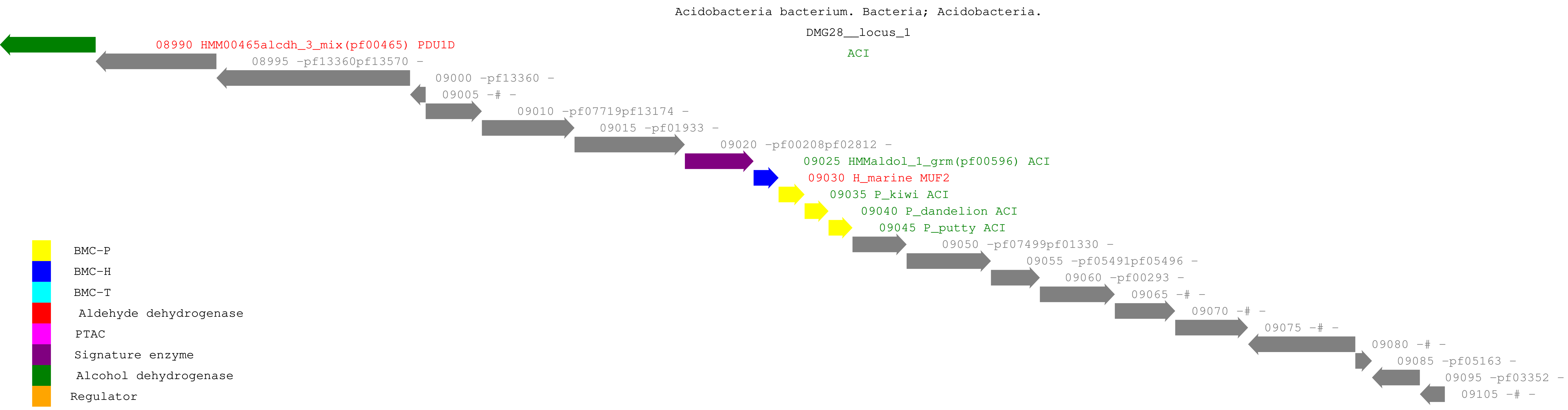

Acidobacteria bacterium. Bacteria; Acidobacteria.
DMG28__locus_1
ACI
08990 HMM00465alcdh_3_mix(pf00465) PDU1D
08995 -pf13360pf13570 -
09000 -pf13360 -
09005 -# -
09010 -pf07719pf13174 -
09015 -pf01933 -
09020 -pf00208pf02812 -
09025 HMMaldol_1_grm(pf00596) ACI
09030 H_marine MUF2
09035 P_kiwi ACI
09040 P_dandelion ACI
09045 P_putty ACI
09050 -pf07499pf01330 -
09055 -pf05491pf05496 -
09060 -pf00293 -
09065 -# -
09070 -# -
09075 -# -
09080 -# -
09085 -pf05163 -
09095 -pf03352 -
09105 -# -
BMC-P
BMC-H
BMC-T
Aldehyde dehydrogenase
PTAC
Signature enzyme
Alcohol dehydrogenase
Regulator

Acidobacteria bacterium. Bacteria; Acidobacteria.

DMG29__locus_1_main_sat

ACI

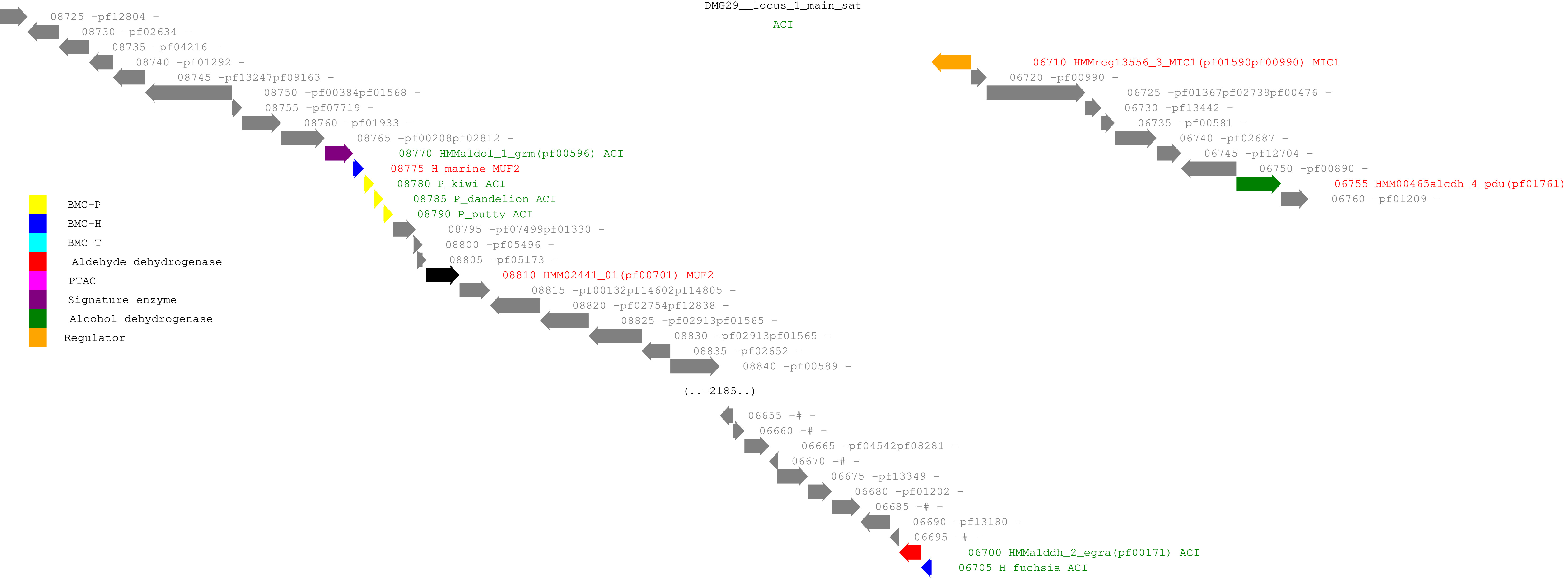

Acidobacteria bacterium. Bacteria; Acidobacteria.

DMG57__locus_1_main_sat

ACI

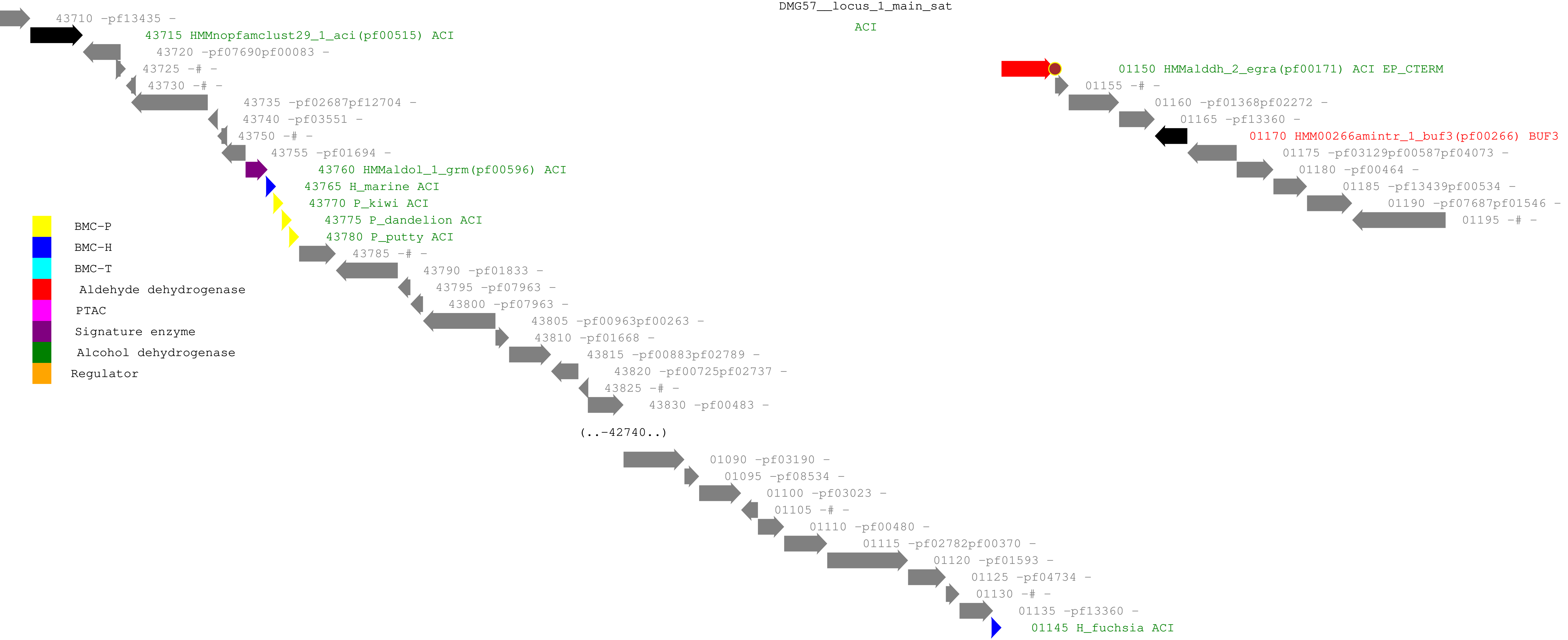

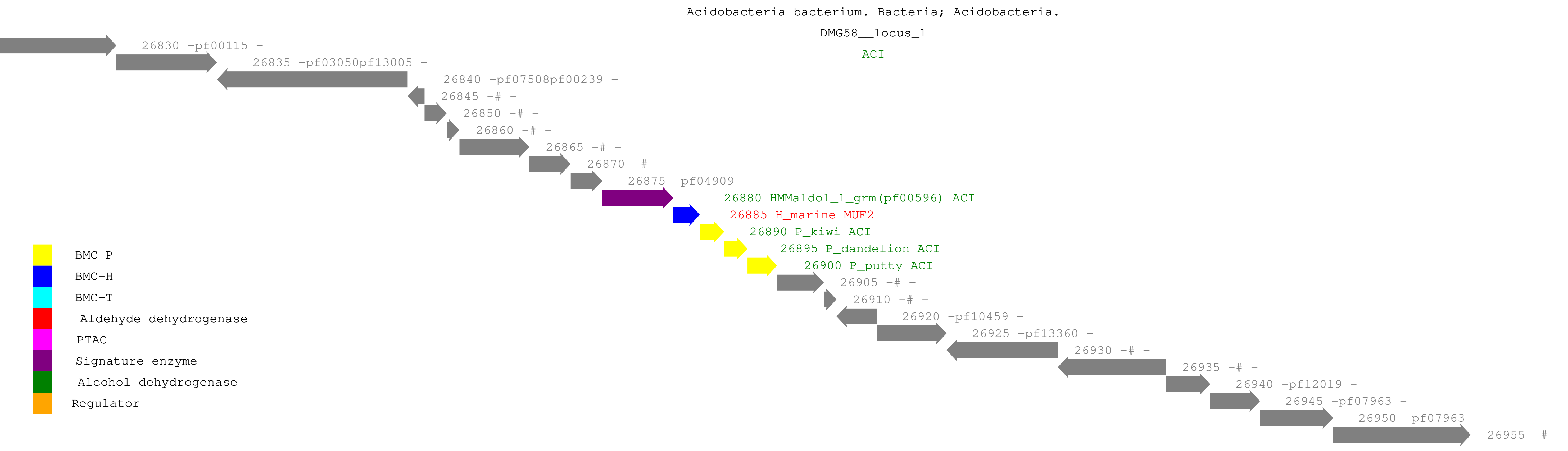
Acidobacteria bacterium. Bacteria; Acidobacteria.
DMG58__locus_1
ACI
26830 -pf00115 -
26835 -pf03050pf13005 -
26840 -pf07508pf00239 -
26845 -# -
26850 -# -
26860 -# -
26865 -# -
26870 -# -
26875 -pf04909 -
26880 HMMaldol_1_grm(pf00596) ACI
26885 H_marine MUF2
26890 P_kiwi ACI
26895 P_dandelion ACI
26900 P_putty ACI
26905 -# -
26910 -# -
26920 -pf10459 -
26925 -pf13360 -
26930 -# -
26935 -# -
26940 -pf12019 -
26945 -pf07963 -
26950 -pf07963 -
26955 -# -
BMC-P
BMC-H
BMC-T
Aldehyde dehydrogenase
PTAC
Signature enzyme
Alcohol dehydrogenase
Regulator

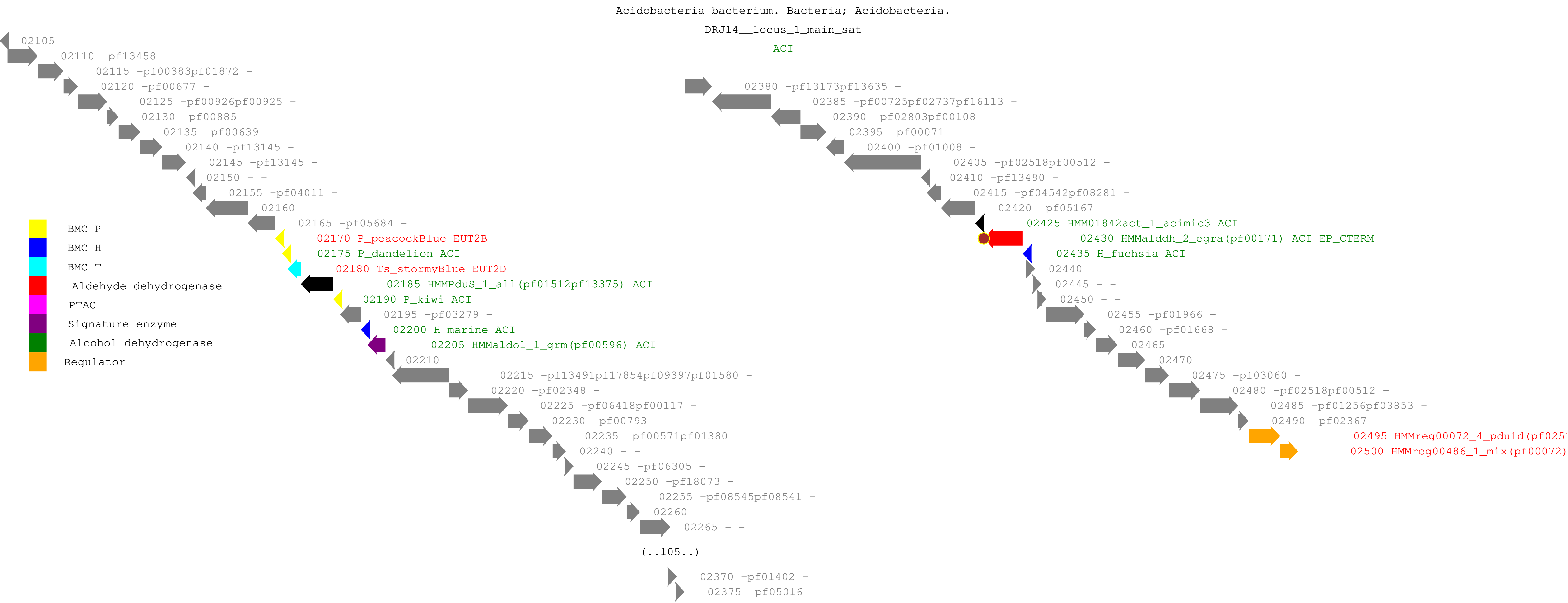

Acidobacteria bacterium. Bacteria; Acidobacteria.
DRJ14__locus_1_main_sat
ACI
02105 - -
02110 -pf13458 -
02115 -pf00383pf01872 -
02120 -pf00677 -
02125 -pf00926pf00925 -
02130 -pf00885 -
02135 -pf00639 -
02140 -pf13145 -
02145 -pf13145 -
02150 - -
02155 -pf04011 -
02160 - -
02165 -pf05684 -
02170 P_peacockBlue EUT2B
02175 P_dandelion ACI
02180 Ts_stormyBlue EUT2D
02185 HMMPduS_1_all(pf01512pf13375) ACI
02190 P_kiwi ACI
02195 -pf03279 -
02200 H_marine ACI
02205 HMMaldol_1_grm(pf00596) ACI
02210 - -
02215 -pf13491pf17854pf09397pf01580 -
02220 -pf02348 -
02225 -pf06418pf00117 -
02230 -pf00793 -
02235 -pf00571pf01380 -
02240 - -
02245 -pf06305 -
02250 -pf18073 -
02255 -pf08545pf08541 -
02260 - -
02265 - -
(..105..)
02370 -pf01402 -
02375 -pf05016 -
02380 -pf13173pf13635 -
02385 -pf00725pf02737pf16113 -
02390 -pf02803pf00108 -
02395 -pf00071 -
02400 -pf01008 -
02405 -pf02518pf00512 -
02410 -pf13490 -
02415 -pf04542pf08281 -
02420 -pf05167 -
02425 HMM01842act_1_acimic3 ACI
02430 HMMalddh_2_egra(pf00171) ACI EP_CTERM
02435 H_fuchsia ACI
02440 - -
02445 - -
02450 - -
02455 -pf01966 -
02460 -pf01668 -
02465 - -
02470 - -
02475 -pf03060 -
02480 -pf02518pf00512 -
02485 -pf01256pf03853 -
02490 -pf02367 -
02495 HMMreg00072_4_pdu1d(pf025
02500 HMMreg00486_1_mix(pf00072)
BMC-P
BMC-H
BMC-T
Aldehyde dehydrogenase
PTAC
Signature enzyme
Alcohol dehydrogenase
Regulator

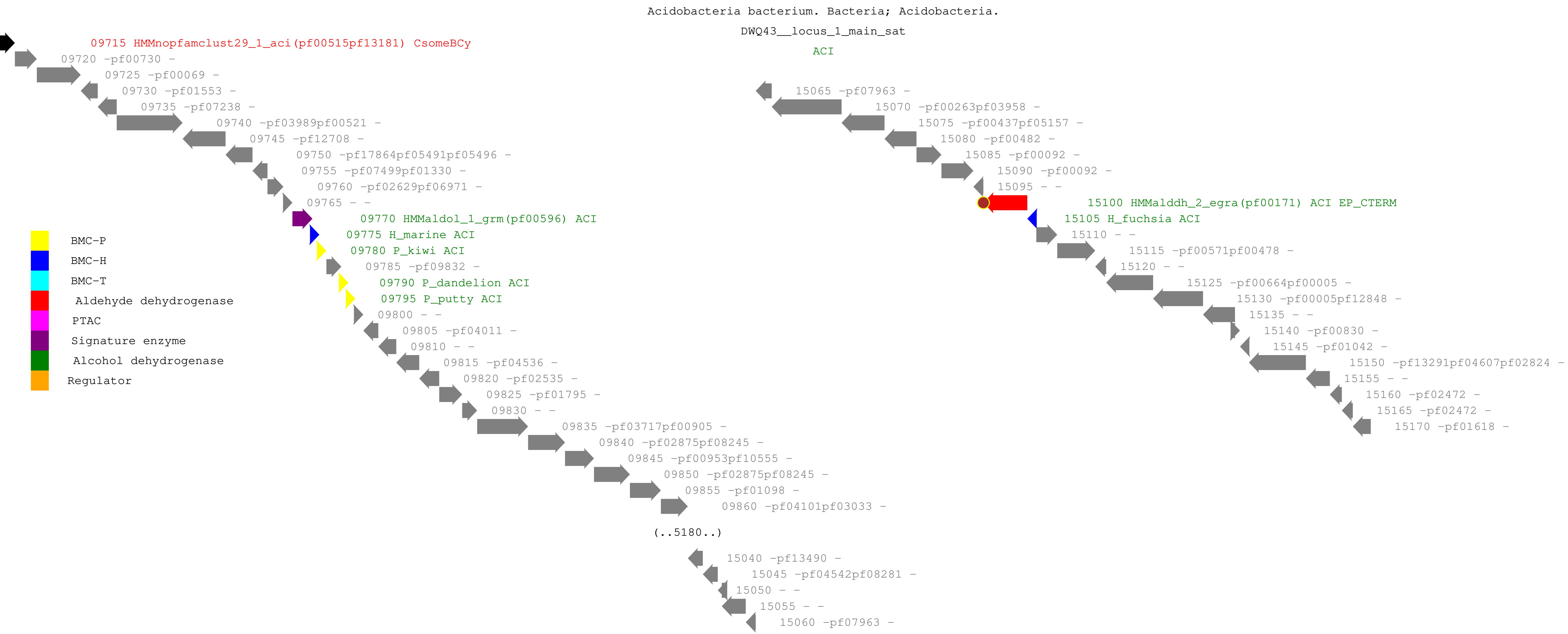
Acidobacteria bacterium. Bacteria; Acidobacteria.
DWQ43__locus_1_main_sat
ACI
09715 HMMnopfamclust29_1_aci(pf00515pf13181) CsomeBCy
09720 -pf00730 -
09725 -pf00069 -
09730 -pf01553 -
09735 -pf07238 -
09740 -pf03989pf00521 -
09745 -pf12708 -
09750 -pf17864pf05491pf05496 -
09755 -pf07499pf01330 -
09760 -pf02629pf06971 -
09765 - -
09770 HMMaldol_1_grm(pf00596) ACI
09775 H_marine ACI
09780 P_kiwi ACI
09785 -pf09832 -
09790 P_dandelion ACI
09795 P_putty ACI
09800 - -
09805 -pf04011 -
09810 - -
09815 -pf04536 -
09820 -pf02535 -
09825 -pf01795 -
09830 - -
09835 -pf03717pf00905 -
09840 -pf02875pf08245 -
09845 -pf00953pf10555 -
09850 -pf02875pf08245 -
09855 -pf01098 -
09860 -pf04101pf03033 -
(..5180..)
15040 -pf13490 -
15045 -pf04542pf08281 -
15050 - -
15055 - -
15060 -pf07963 -
15065 -pf07963 -
15070 -pf00263pf03958 -
15075 -pf00437pf05157 -
15080 -pf00482 -
15085 -pf00092 -
15090 -pf00092 -
15095 - -
15100 HMMalddh_2_egra(pf00171) ACI EP_CTERM
15105 H_fuchsia ACI
15110 - -
15115 -pf00571pf00478 -
15120 - -
15125 -pf00664pf00005 -
15130 -pf00005pf12848 -
15135 - -
15140 -pf00830 -
15145 -pf01042 -
15150 -pf13291pf04607pf02824 -
15155 - -
15160 -pf02472 -
15165 -pf02472 -
15170 -pf01618 -
BMC-P
BMC-H
BMC-T
Aldehyde dehydrogenase
PTAC
Signature enzyme
Alcohol dehydrogenase
Regulator

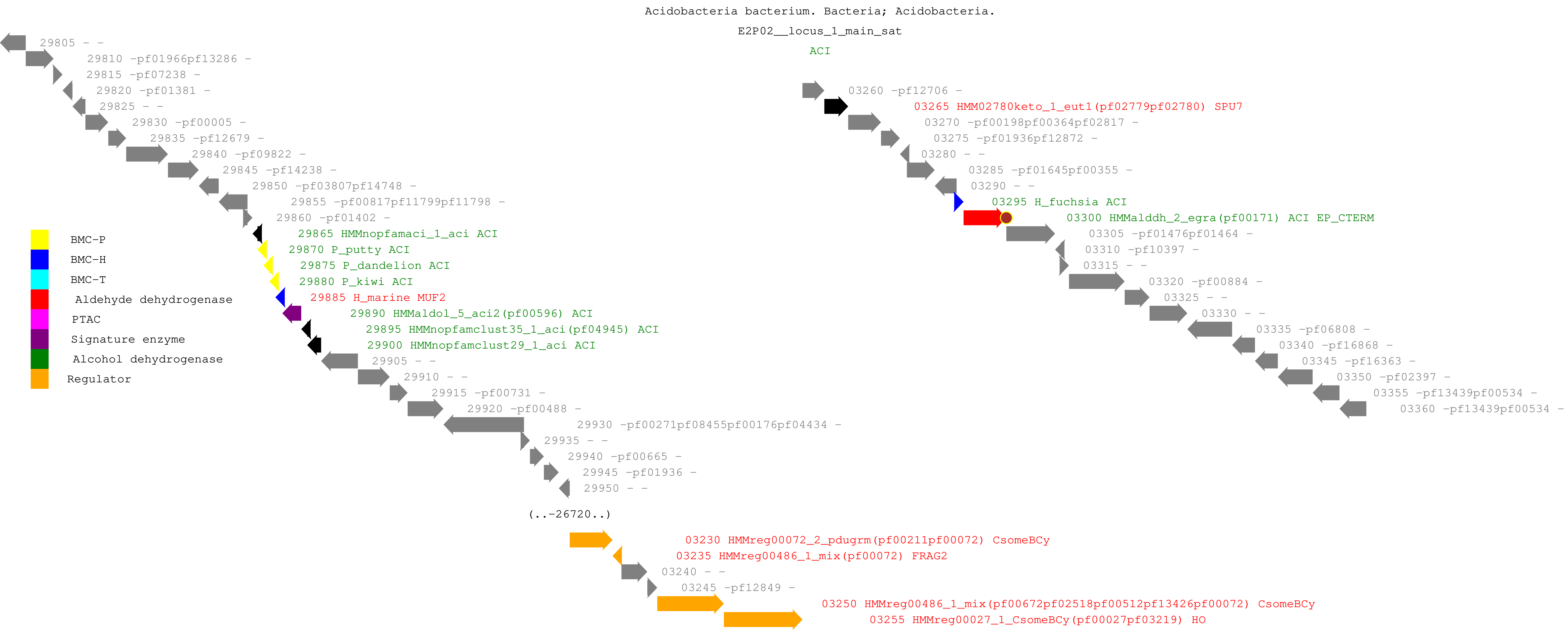

Acidobacteria bacterium. Bacteria; Acidobacteria.
E2P02__locus_1_main_sat
ACI
29805 - -
29810 -pf01966pf13286 -
29815 -pf07238 -
29820 -pf01381 -
29825 - -
29830 -pf00005 -
29835 -pf12679 -
29840 -pf09822 -
29845 -pf14238 -
29850 -pf03807pf14748 -
29855 -pf00817pf11799pf11798 -
29860 -pf01402 -
29865 HMMnopfamaci_1_aci ACI
29870 P_putty ACI
29875 P_dandelion ACI
29880 P_kiwi ACI
29885 H_marine MUF2
29890 HMMaldol_5_aci2(pf00596) ACI
29895 HMMnopfamclust35_1_aci(pf04945) ACI
29900 HMMnopfamclust29_1_aci ACI
29905 - -
29910 - -
29915 -pf00731 -
29920 -pf00488 -
29930 -pf00271pf08455pf00176pf04434 -
29935 - -
29940 -pf00665 -
29945 -pf01936 -
29950 - -
(..-26720..)
03230 HMMreg00072_2_pdugrm(pf00211pf00072) CsomeBCy
03235 HMMreg00486_1_mix(pf00072) FRAG2
03240 - -
03245 -pf12849 -
03250 HMMreg00486_1_mix(pf00672pf02518pf00512pf13426pf00072) CsomeBCy
03255 HMMreg00027_1_CsomeBCy(pf00027pf03219) HO
03260 -pf12706 -
03265 HMM02780keto_1_eut1(pf02779pf02780) SPU7
03270 -pf00198pf00364pf02817 -
03275 -pf01936pf12872 -
03280 - -
03285 -pf01645pf00355 -
03290 - -
03295 H_fuchsia ACI
03300 HMMalddh_2_egra(pf00171) ACI EP_CTERM
03305 -pf01476pf01464 -
03310 -pf10397 -
03315 - -
03320 -pf00884 -
03325 - -
03330 - -
03335 -pf06808 -
03340 -pf16868 -
03345 -pf16363 -
03350 -pf02397 -
03355 -pf13439pf00534 -
03360 -pf13439pf00534 -
BMC-P
BMC-H
BMC-T
Aldehyde dehydrogenase
PTAC
Signature enzyme
Alcohol dehydrogenase
Regulator

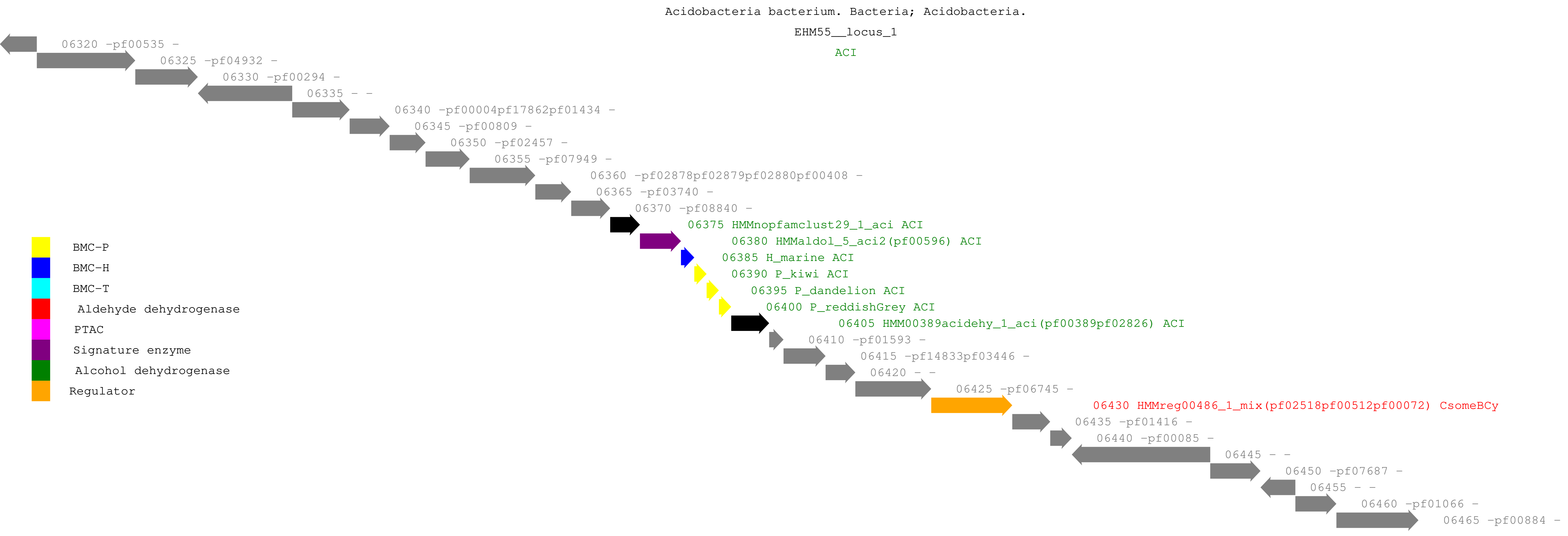

Acidobacteria bacterium. Bacteria; Acidobacteria.
EHM55__locus_1
ACI
06320 -pf00535 -
06325 -pf04932 -
06330 -pf00294 -
06335 - -
06340 -pf00004pf17862pf01434 -
06345 -pf00809 -
06350 -pf02457 -
06355 -pf07949 -
06360 -pf02878pf02879pf02880pf00408 -
06365 -pf03740 -
06370 -pf08840 -
06375 HMMnopfamclust29_1_aci ACI
06380 HMMaldol_5_aci2(pf00596) ACI
06385 H_marine ACI
06390 P_kiwi ACI
06395 P_dandelion ACI
06400 P_reddishGrey ACI
06405 HMM00389acidehy_1_aci(pf00389pf02826) ACI
06410 -pf01593 -
06415 -pf14833pf03446 -
06420 - -
06425 -pf06745 -
06430 HMMreg00486_1_mix(pf02518pf00512pf00072) CsomeBCy
06435 -pf01416 -
06440 -pf00085 -
06445 - -
06450 -pf07687 -
06455 - -
06460 -pf01066 -
06465 -pf00884 -
BMC-P
BMC-H
BMC-T
Aldehyde dehydrogenase
PTAC
Signature enzyme
Alcohol dehydrogenase
Regulator

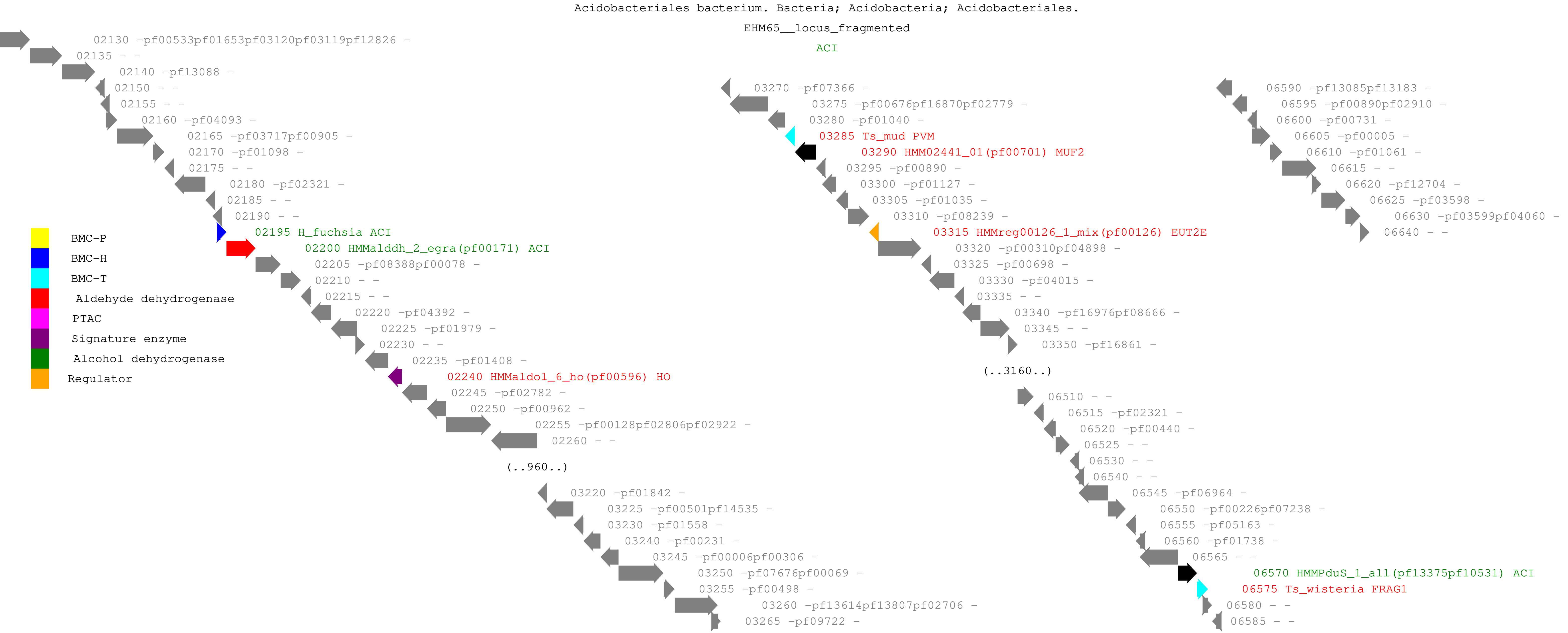
Acidobacteriales bacterium. Bacteria; Acidobacteria; Acidobacteriales.
EHM65__locus_fragmented
ACI
02130 -pf00533pf01653pf03120pf03119pf12826 -
02135 - -
02140 -pf13088 -
02150 - -
02155 - -
02160 -pf04093 -
02165 -pf03717pf00905 -
02170 -pf01098 -
02175 - -
02180 -pf02321 -
02185 - -
02190 - -
02195 H_fuchsia ACI
02200 HMMalddh_2_egra(pf00171) ACI
02205 -pf08388pf00078 -
02210 - -
02215 - -
02220 -pf04392 -
02225 -pf01979 -
02230 - -
02235 -pf01408 -
02240 HMMaldol_6_ho(pf00596) HO
02245 -pf02782 -
02250 -pf00962 -
02255 -pf00128pf02806pf02922 -
02260 - -
(..960..)
03220 -pf01842 -
03225 -pf00501pf14535 -
03230 -pf01558 -
03240 -pf00231 -
03245 -pf00006pf00306 -
03250 -pf07676pf00069 -
03255 -pf00498 -
03260 -pf13614pf13807pf02706 -
03265 -pf09722 -
03270 -pf07366 -
03275 -pf00676pf16870pf02779 -
03280 -pf01040 -
03285 Ts_mud PVM
03290 HMM02441_01(pf00701) MUF2
03295 -pf00890 -
03300 -pf01127 -
03305 -pf01035 -
03310 -pf08239 -
03315 HMMreg00126_1_mix(pf00126) EUT2E
03320 -pf00310pf04898 -
03325 -pf00698 -
03330 -pf04015 -
03335 - -
03340 -pf16976pf08666 -
03345 - -
03350 -pf16861 -
(..3160..)
06510 - -
06515 -pf02321 -
06520 -pf00440 -
06525 - -
06530 - -
06540 - -
06545 -pf06964 -
06550 -pf00226pf07238 -
06555 -pf05163 -
06560 -pf01738 -
06565 - -
06570 HMMPduS_1_all(pf13375pf10531) ACI
06575 Ts_wisteria FRAG1
06580 - -
06585 - -
06590 -pf13085pf13183 -
06595 -pf00890pf02910 -
06600 -pf00731 -
06605 -pf00005 -
06610 -pf01061 -
06615 - -
06620 -pf12704 -
06625 -pf03598 -
06630 -pf03599pf04060 -
06640 - -
BMC-P
BMC-H
BMC-T
Aldehyde dehydrogenase
PTAC
Signature enzyme
Alcohol dehydrogenase
Regulator

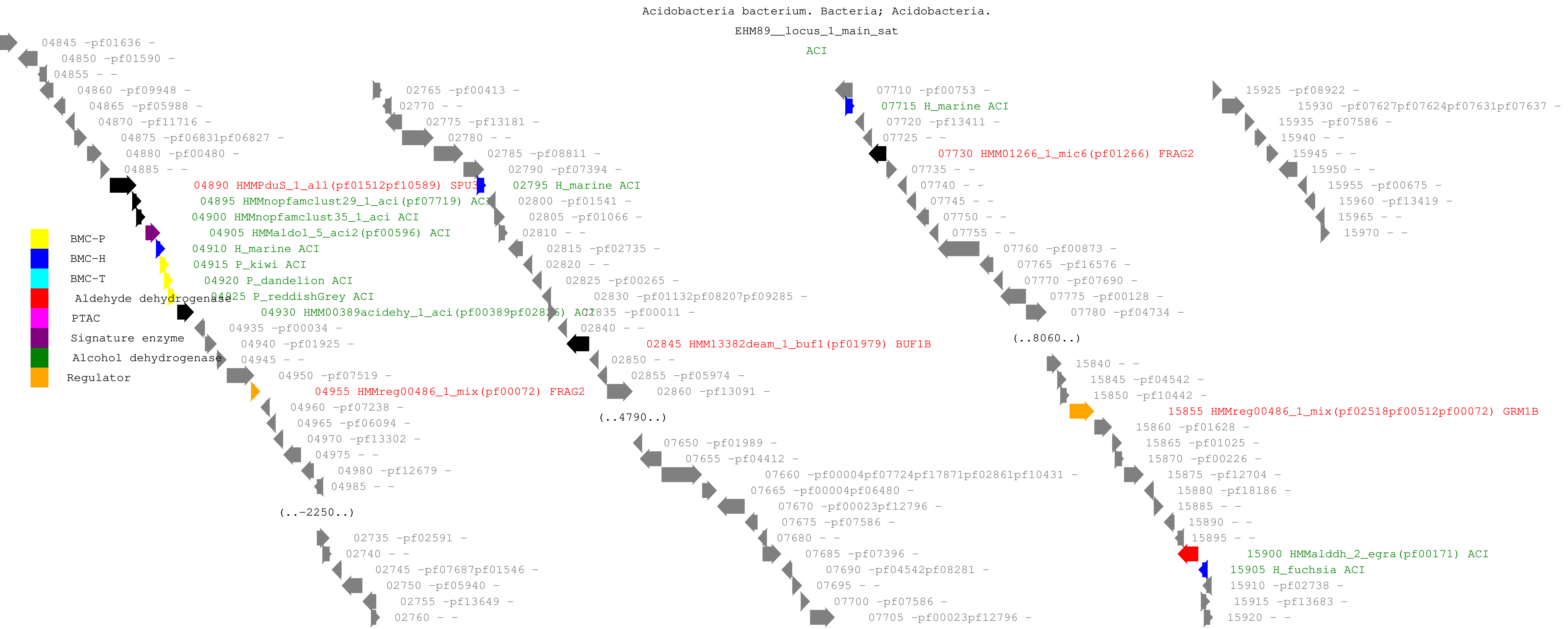

Acidobacteria bacterium. Bacteria; Acidobacteria.
EHM89__locus_1_main_sat
ACI
BMC-P
BMC-H
BMC-T
Aldehyde dehydrogenase
PTAC
Signature enzyme
Alcohol dehydrogenase
Regulator
04845 -pf01636 -
04850 -pf01590 -
04855 - -
04860 -pf09948 -
04865 -pf05988 -
04870 -pf11716 -
04875 -pf06831pf06827 -
04880 -pf00480 -
04885 - -
04890 HMMPduS_1_all(pf01512pf10589) SPU3
04895 HMMnopfamclust29_1_aci(pf07719) ACI
04900 HMMnopfamclust35_1_aci ACI
04905 HMMaldol_5_aci2(pf00596) ACI
04910 H_marine ACI
04915 P_kiwi ACI
04920 P_dandelion ACI
04925 P_reddishGrey ACI
04930 HMM00389acidehy_1_aci(pf00389pf02826) ACI
04935 -pf00034 -
04940 -pf01925 -
04945 - -
04950 -pf07519 -
04955 HMMreg00486_1_mix(pf00072) FRAG2
04960 -pf07238 -
04965 -pf06094 -
04970 -pf13302 -
04975 - -
04980 -pf12679 -
04985 - -
(..-2250..)
02735 -pf02591 -
02740 - -
02745 -pf07687pf01546 -
02750 -pf05940 -
02755 -pf13649 -
02760 - -
02765 -pf00413 -
02770 - -
02775 -pf13181 -
02780 - -
02785 -pf08811 -
02790 -pf07394 -
02795 H_marine ACI
02800 -pf01541 -
02805 -pf01066 -
02810 - -
02815 -pf02735 -
02820 - -
02825 -pf00265 -
02830 -pf01132pf08207pf09285 -
02835 -pf00011 -
02840 - -
02845 HMM13382deam_1_buf1(pf01979) BUF1B
02850 - -
02855 -pf05974 -
02860 -pf13091 -
(..4790..)
07650 -pf01989 -
07655 -pf04412 -
07660 -pf00004pf07724pf17871pf02861pf10431 -
07665 -pf00004pf06480 -
07670 -pf00023pf12796 -
07675 -pf07586 -
07680 - -
07685 -pf07396 -
07690 -pf04542pf08281 -
07695 - -
07700 -pf07586 -
07705 -pf00023pf12796 -
07710 -pf00753 -
07715 H_marine ACI
07720 -pf13411 -
07725 - -
07730 HMM01266_1_mic6(pf01266) FRAG2
07735 - -
07740 - -
07745 - -
07750 - -
07755 - -
07760 -pf00873 -
07765 -pf16576 -
07770 -pf07690 -
07775 -pf00128 -
07780 -pf04734 -
(..8060..)
15840 - -
15845 -pf04542 -
15850 -pf10442 -
15855 HMMreg00486_1_mix(pf02518pf00512pf00072) GRM1B
15860 -pf01628 -
15865 -pf01025 -
15870 -pf00226 -
15875 -pf12704 -
15880 -pf18186 -
15885 - -
15890 - -
15895 - -
15900 HMMaldh_2_egra(pf00171) ACI
15905 H_fuchsia ACI
15910 -pf02738 -
15915 -pf13683 -
15920 - -
15925 -pf08922 -
15930 -pf07627pf07624pf07631pf07637 -
15935 -pf07586 -
15940 - -
15945 - -
15950 - -
15955 -pf00675 -
15960 -pf13419 -
15965 - -
15970 - -

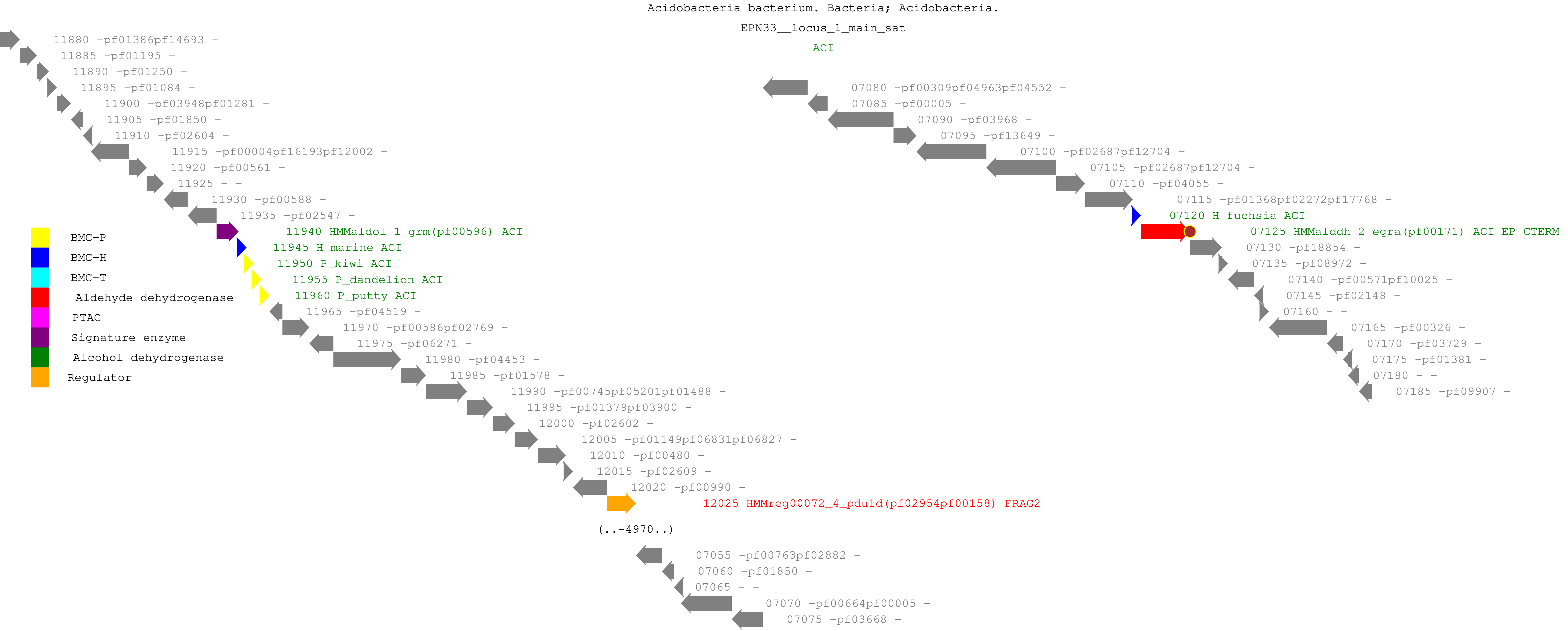
Acidobacteria bacterium. Bacteria; Acidobacteria.
EPN33__locus_1_main_sat
ACI
11880 -pf01386pf14693 -
11885 -pf01195 -
11890 -pf01250 -
11895 -pf01084 -
11900 -pf03948pf01281 -
11905 -pf01850 -
11910 -pf02604 -
11915 -pf00004pf16193pf12002 -
11920 -pf00561 -
11925 - -
11930 -pf00588 -
11935 -pf02547 -
11940 HMMaldol_1_grm(pf00596) ACI
11945 H_marine ACI
11950 P_kiwi ACI
11955 P_dandelion ACI
11960 P_putty ACI
11965 -pf04519 -
11970 -pf00586pf02769 -
11975 -pf06271 -
11980 -pf04453 -
11985 -pf01578 -
11990 -pf00745pf05201pf01488 -
11995 -pf01379pf03900 -
12000 -pf02602 -
12005 -pf01149pf06831pf06827 -
12010 -pf00480 -
12015 -pf02609 -
12020 -pf00990 -
12025 HMMreg00072_4_pdu1d(pf02954pf00158) FRAG2
(..-4970..)
07055 -pf00763pf02882 -
07060 -pf01850 -
07065 - -
07070 -pf00664pf00005 -
07075 -pf03668 -
07080 -pf00309pf04963pf04552 -
07085 -pf00005 -
07090 -pf03968 -
07095 -pf13649 -
07100 -pf02687pf12704 -
07105 -pf02687pf12704 -
07110 -pf04055 -
07115 -pf01368pf02272pf17768 -
07120 H_fuchsia ACI
07125 HMMalddh_2_egra(pf00171) ACI EP_CTERM
07130 -pf18854 -
07135 -pf08972 -
07140 -pf00571pf10025 -
07145 -pf02148 -
07160 - -
07165 -pf00326 -
07170 -pf03729 -
07175 -pf01381 -
07180 - -
07185 -pf09907 -
BMC-P
BMC-H
BMC-T
Aldehyde dehydrogenase
PTAC
Signature enzyme
Alcohol dehydrogenase
Regulator

Acidobacteria bacterium. Bacteria; Acidobacteria.

EPO35__locus_1_main_sat

ACI

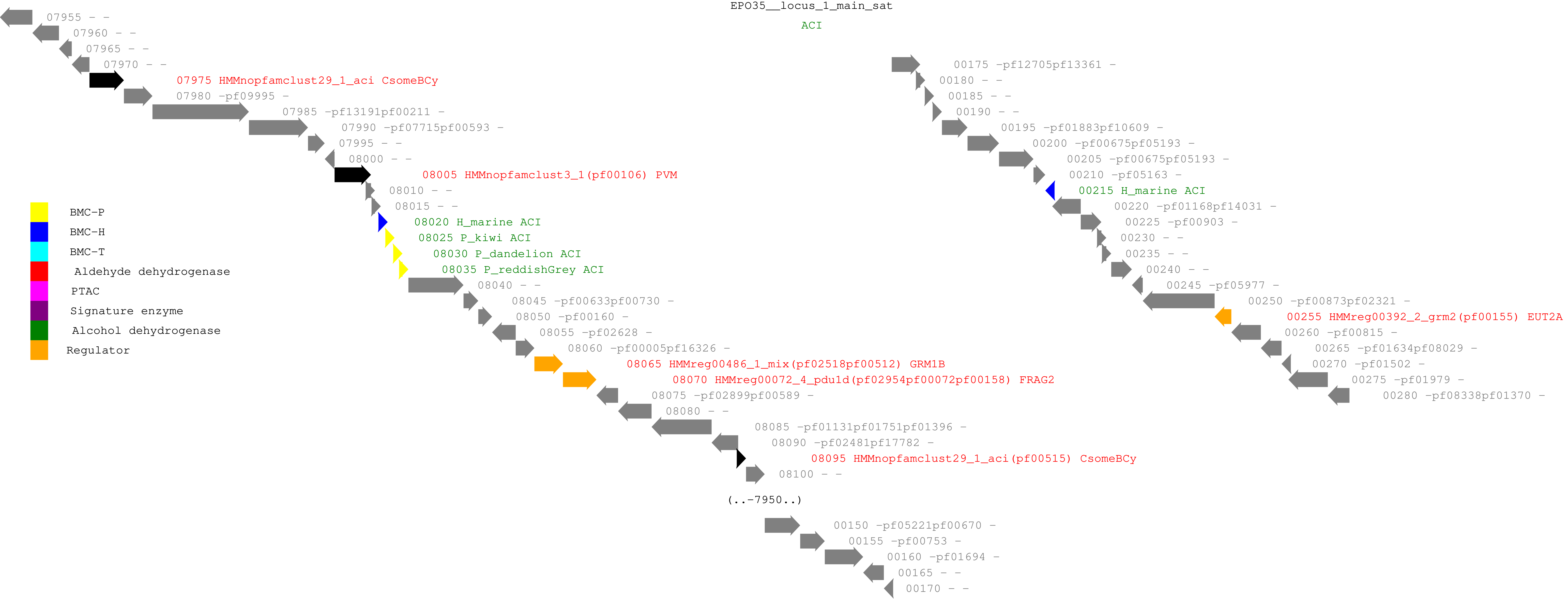

Acidobacteria bacterium DSM 100886. Bacteria; Acidobacteria; Acidobacteria subdivision 6.

LuPra__locus_1_main_sat

ACI

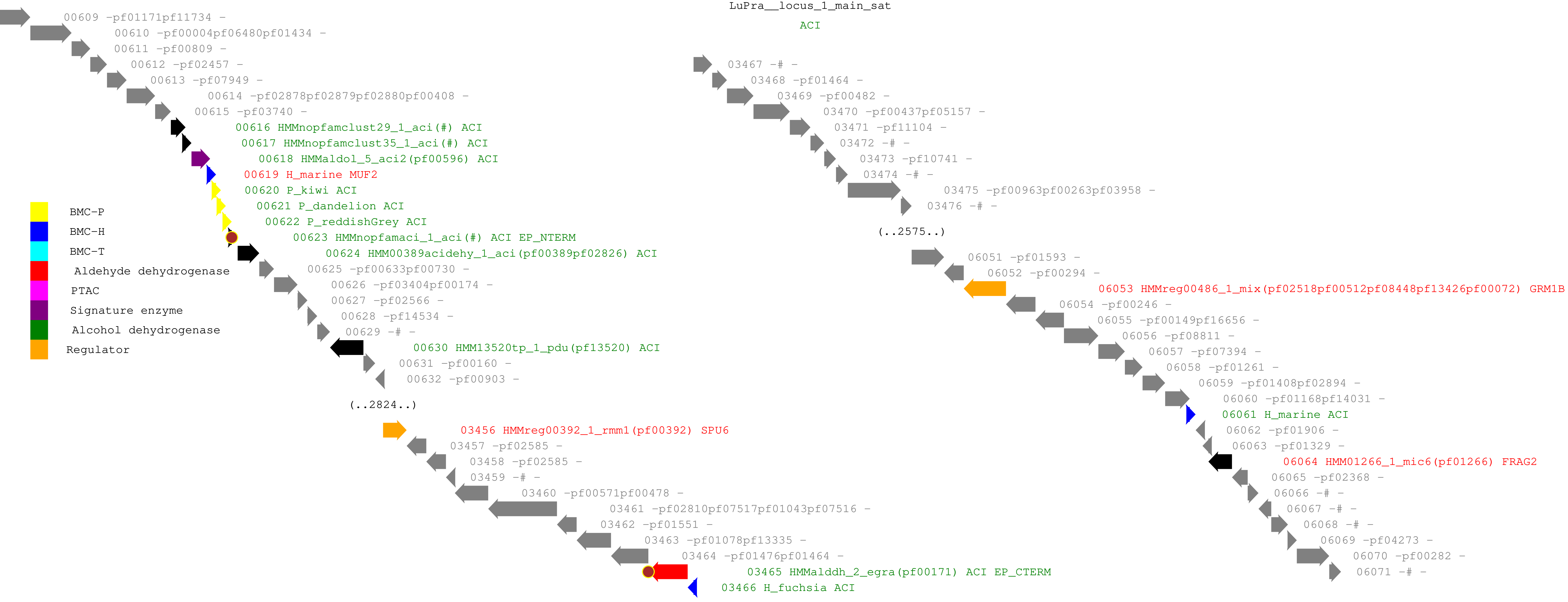

Pyrinomonas methylaliphatogenes. Bacteria; Acidobacteria; Blastocatellia; Pyrinomonas.

PYK22__locus_1_main_sat

ACI

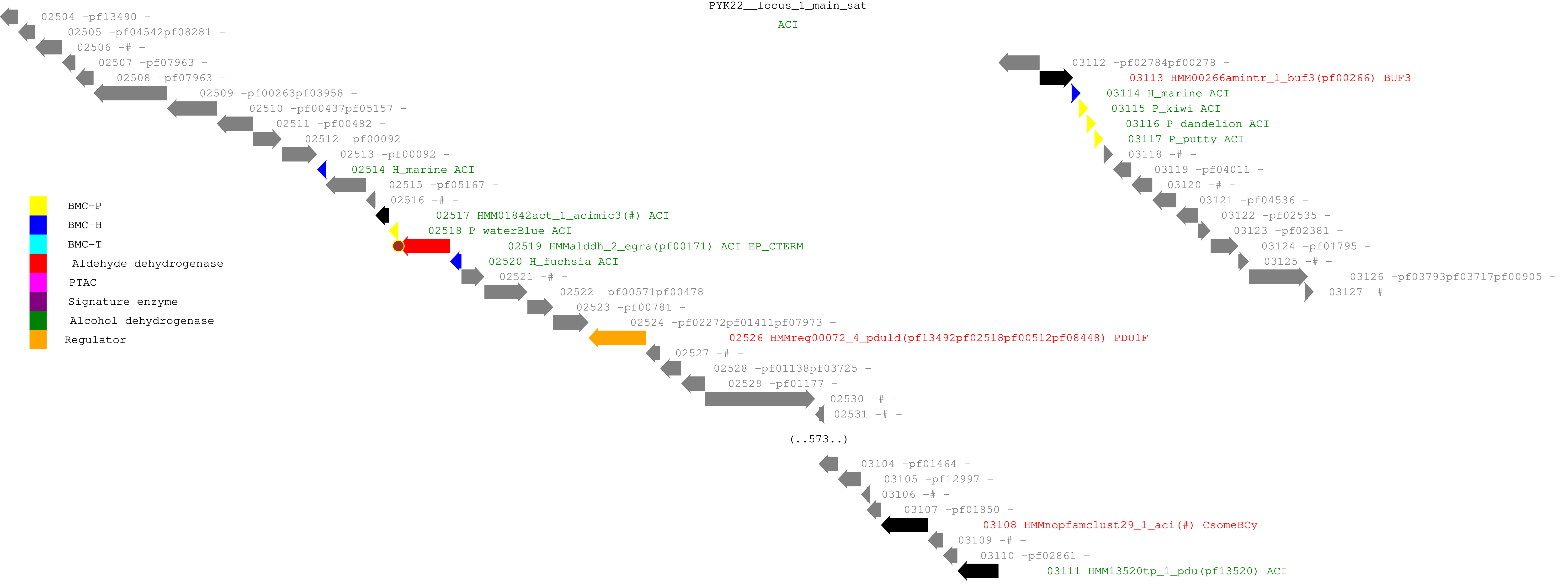

Candidatus Sulfopaludibacter sp. SbA3. Bacteria; Acidobacteria; Solibacteres; Solibacterales;

SBA3__locus_1_main_sat

ACI

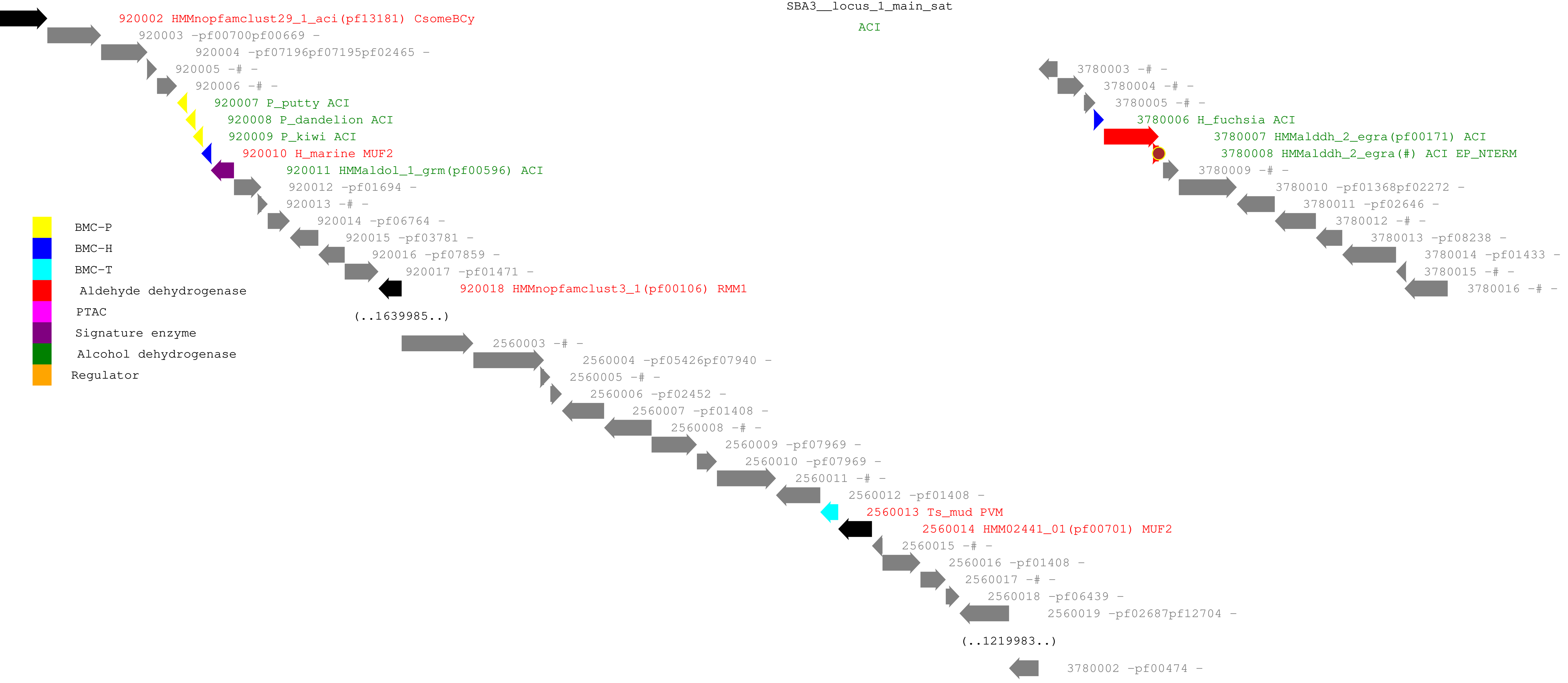

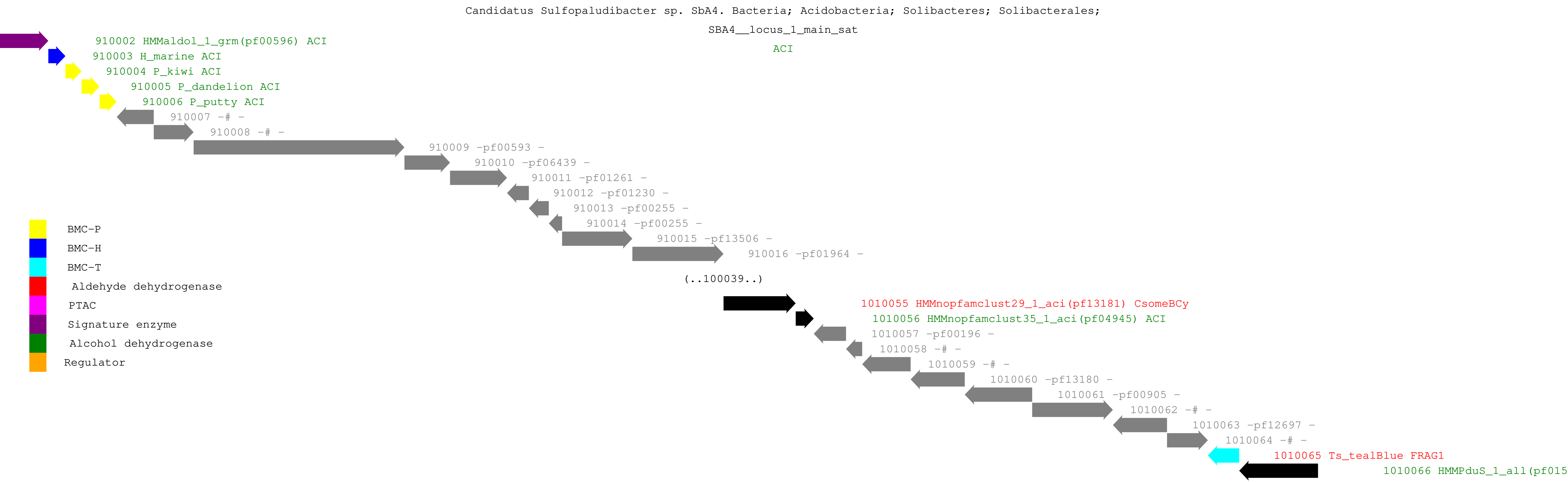
Candidatus Sulfopaludibacter sp. SbA4. Bacteria; Acidobacteria; Solibacteres; Solibacterales;
SBA4__locus_1_main_sat
ACI
910002 HMMaldol_1_grm(pf00596) ACI
910003 H_marine ACI
910004 P_kiwi ACI
910005 P_dandelion ACI
910006 P_putty ACI
910007 -# -
910008 -# -
910009 -pf00593 -
910010 -pf06439 -
910011 -pf01261 -
910012 -pf01230 -
910013 -pf00255 -
910014 -pf00255 -
910015 -pf13506 -
910016 -pf01964 -
(..100039..)
1010055 HMMnopfamclust29_1_aci(pf13181) CsomeBCy
1010056 HMMnopfamclust35_1_aci(pf04945) ACI
1010057 -pf00196 -
1010058 -# -
1010059 -# -
1010060 -pf13180 -
1010061 -pf00905 -
1010062 -# -
1010063 -pf12697 -
1010064 -# -
1010065 Ts_tealBlue FRAG1
1010066 HMMPduS_1_all(pf015
BMC-P
BMC-H
BMC-T
Aldehyde dehydrogenase
PTAC
Signature enzyme
Alcohol dehydrogenase
Regulator

Candidatus Sulfopaludibacter sp. SbA6. Bacteria; Acidobacteria; Solibacteres; Solibacterales;

SBA6__locus_1

ACI

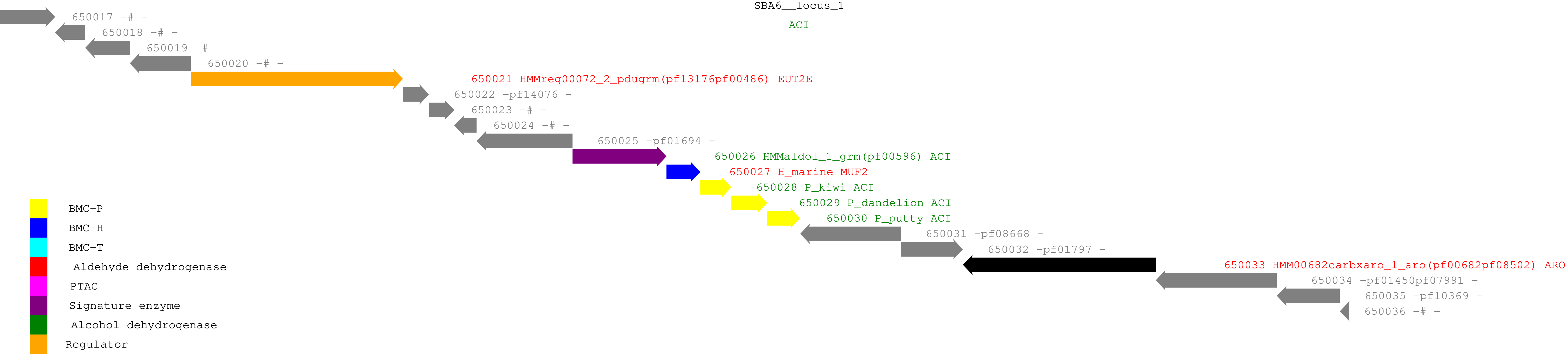

Acidobacteria bacterium OLB17. Bacteria; Acidobacteria.

UZ17_ACD_locus_1_main_sat

ACI

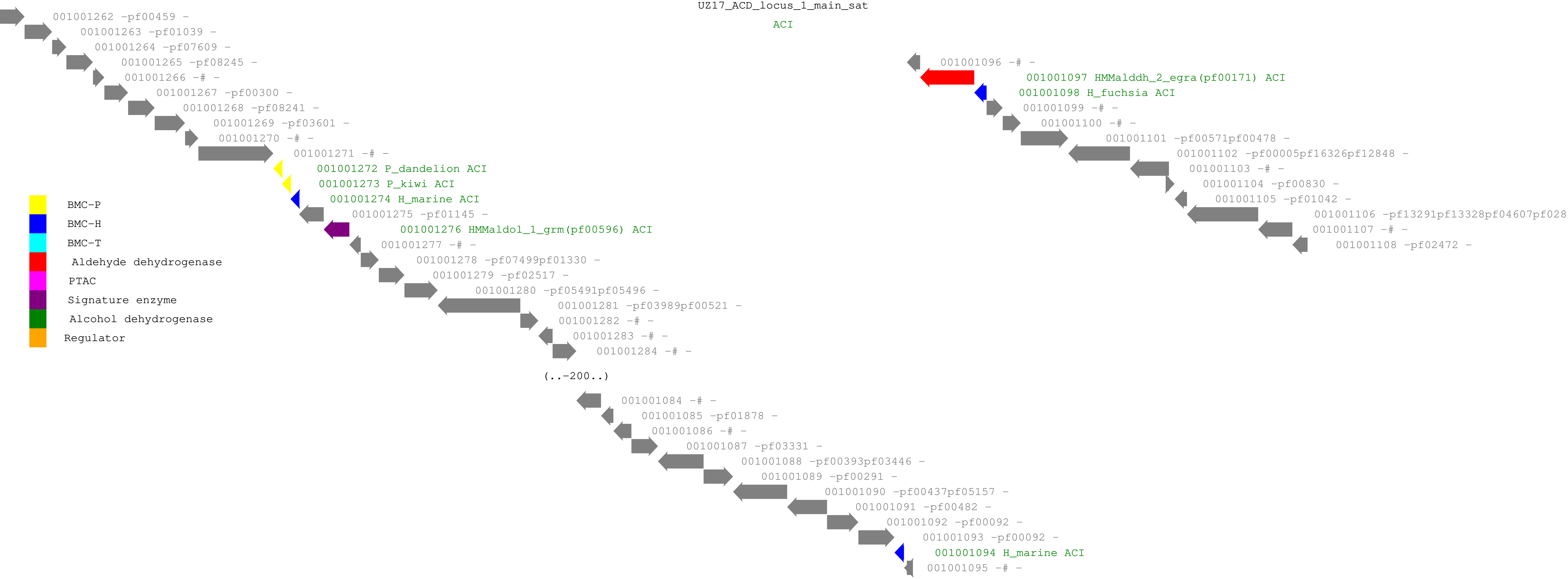

Pseudonocardia sp. SCN 72-86. Bacteria; Actinobacteria; Pseudonocardiales; Pseudonocardiaceae;

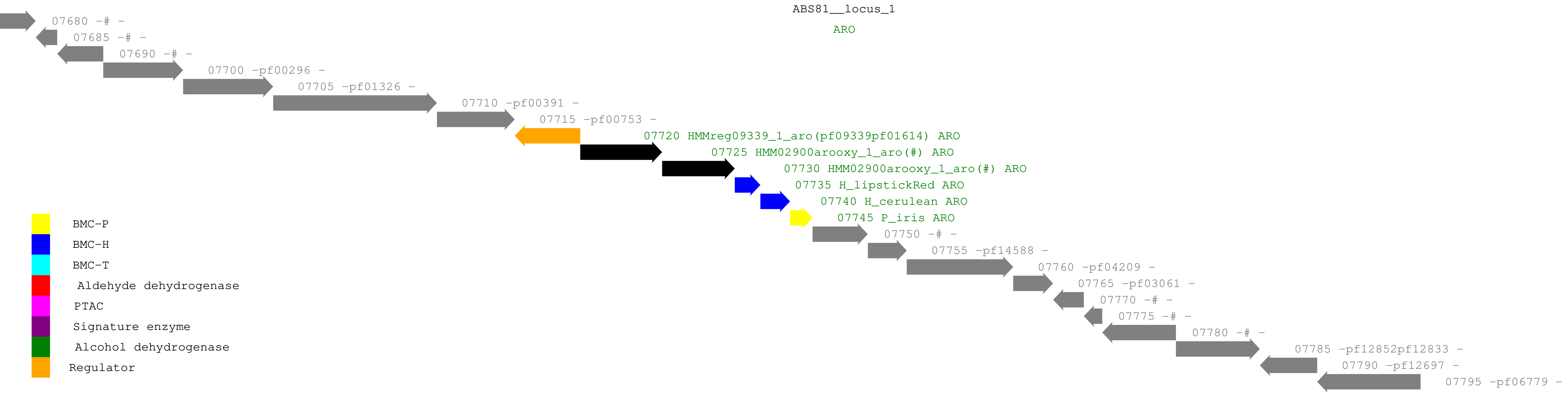

Micromonospora rosaria. Bacteria; Actinobacteria; Micromonosporales; Micromonosporaceae;

AWW66__locus_1

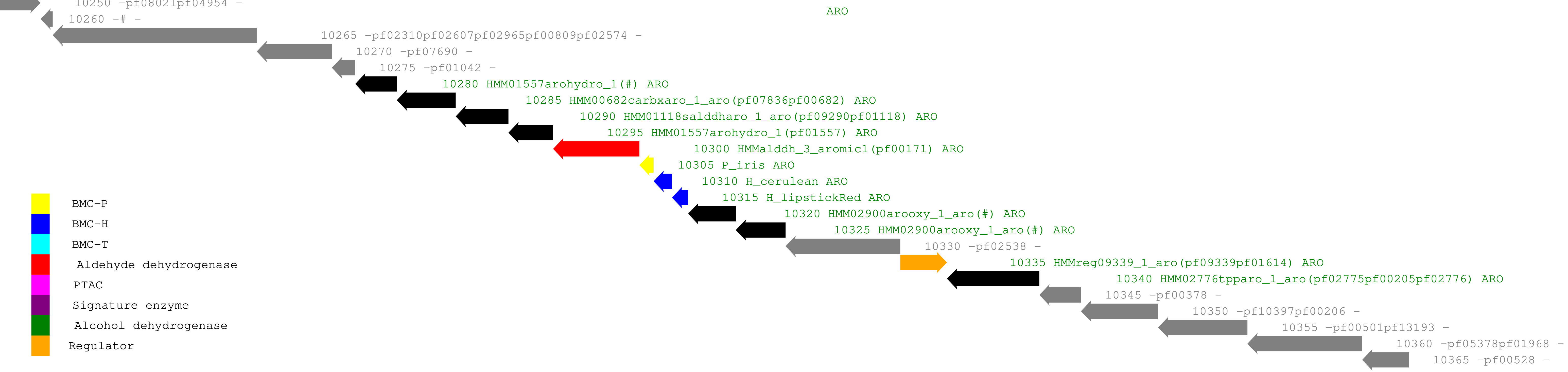

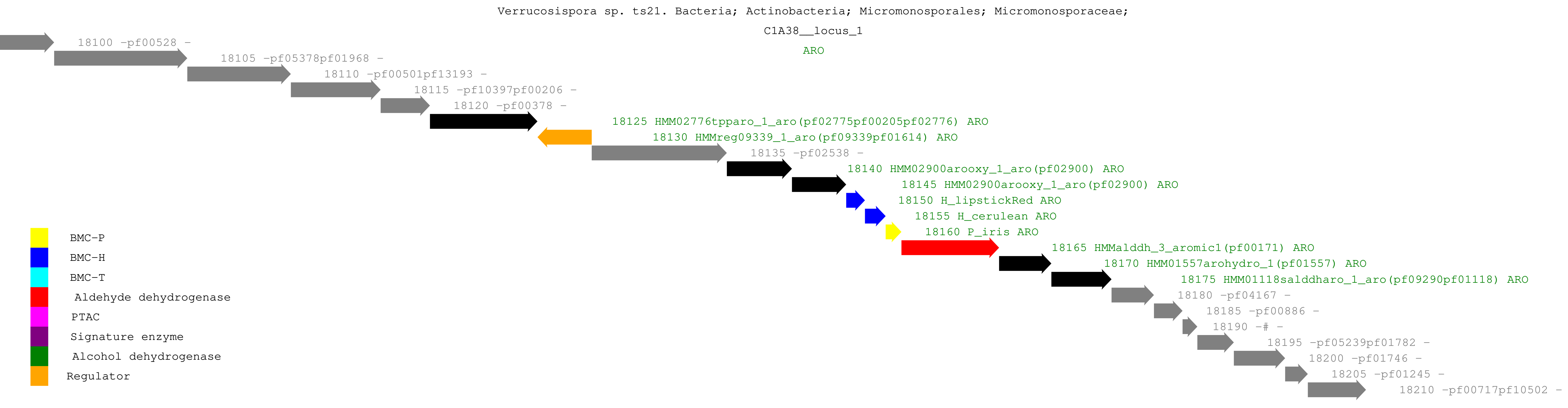
Verrucosispora sp. ts21. Bacteria; Actinobacteria; Micromonosporales; Micromonosporaceae;
C1A38__locus_1
ARO
18100 -pf00528 -
18105 -pf05378pf01968 -
18110 -pf00501pf13193 -
18115 -pf10397pf00206 -
18120 -pf00378 -
18125 HMM02776tpparo_1_aro(pf02775pf00205pf02776) ARO
18130 HMMreg09339_1_aro(pf09339pf01614) ARO
18135 -pf02538 -
18140 HMM02900arooxy_1_aro(pf02900) ARO
18145 HMM02900arooxy_1_aro(pf02900) ARO
18150 H_lipstickRed ARO
18155 H_cerulean ARO
18160 P_iris ARO
18165 HMMalddh_3_aromic1(pf00171) ARO
18170 HMM01557arohydro_1(pf01557) ARO
18175 HMM01118salddharo_1_aro(pf09290pf01118) ARO
18180 -pf04167 -
18185 -pf00886 -
18190 -# -
18195 -pf05239pf01782 -
18200 -pf01746 -
18205 -pf01245 -
18210 -pf00717pf10502 -
BMC-P
BMC-H
BMC-T
Aldehyde dehydrogenase
PTAC
Signature enzyme
Alcohol dehydrogenase
Regulator

Actinomycetospora cinnamomea. Bacteria; Actinobacteria; Pseudonocardiales; Pseudonocardiaceae;

C8D89__locus_2

ARO

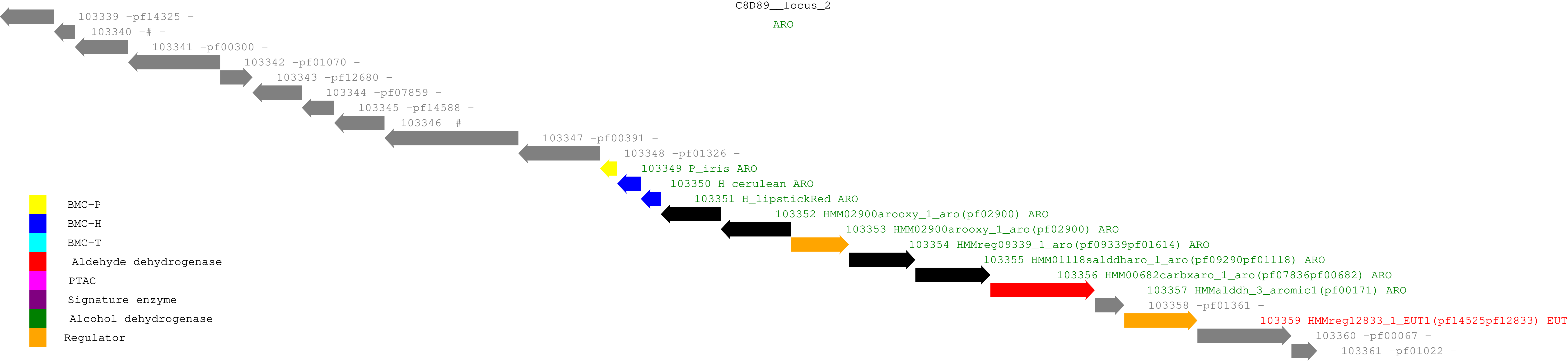

Micromonospora sp. CNZ309. Bacteria; Actinobacteria; Micromonosporales; Micromonosporaceae;

CLT72__locus_1

ARO

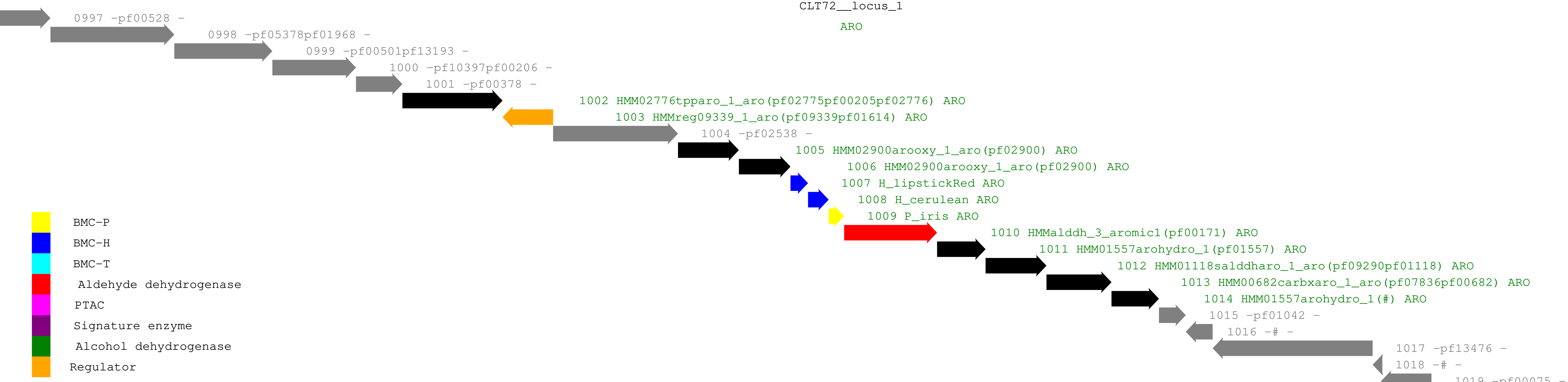

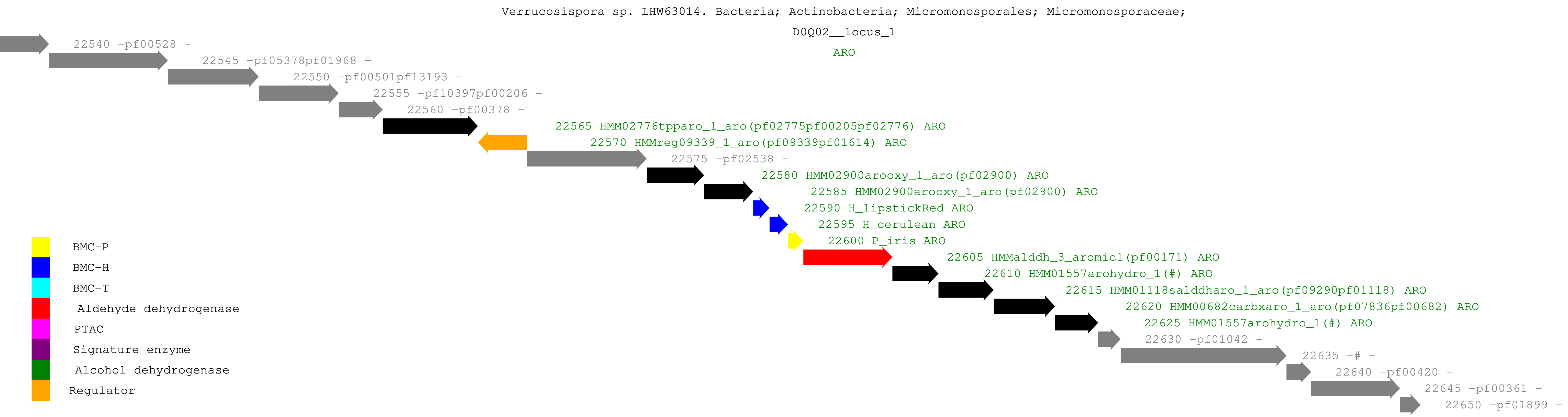

Verrucosispora sp. LHW63014. Bacteria; Actinobacteria; Micromonosporales; Micromonosporaceae;
D0Q02__locus_1
ARO
22540 -pf00528 -
22545 -pf05378pf01968 -
22550 -pf00501pf13193 -
22555 -pf10397pf00206 -
22560 -pf00378 -
22565 HMM02776tpparo_1_aro(pf02775pf00205pf02776) ARO
22570 HMMreg09339_1_aro(pf09339pf01614) ARO
22575 -pf02538 -
22580 HMM02900arooxy_1_aro(pf02900) ARO
22585 HMM02900arooxy_1_aro(pf02900) ARO
22590 H_lipstickRed ARO
22595 H_cerulean ARO
22600 P_iris ARO
22605 HMMalddh_3_aromic1(pf00171) ARO
22610 HMM01557arohydro_1(#) ARO
22615 HMM01118salddharo_1_aro(pf09290pf01118) ARO
22620 HMM00682carbxaro_1_aro(pf07836pf00682) ARO
22625 HMM01557arohydro_1(#) ARO
22630 -pf01042 -
22635 -# -
22640 -pf00420 -
22645 -pf00361 -
22650 -pf01899 -
BMC-P
BMC-H
BMC-T
Aldehyde dehydrogenase
PTAC
Signature enzyme
Alcohol dehydrogenase
Regulator

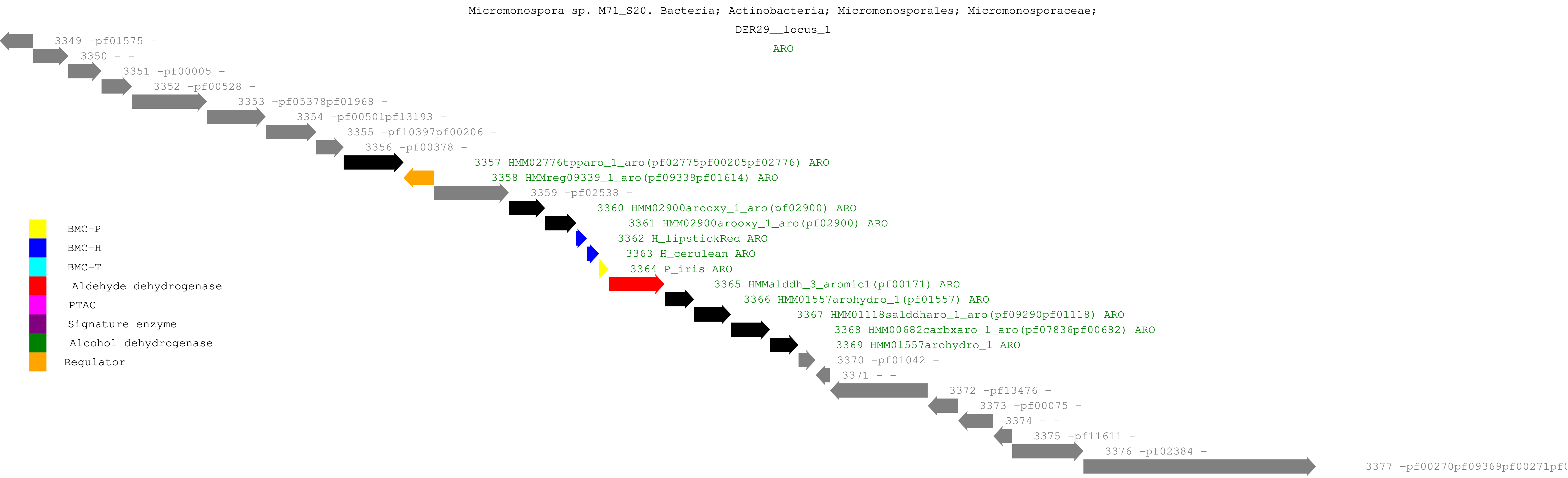
Micromonospora sp. M71_S20. Bacteria; Actinobacteria; Micromonosporales; Micromonosporaceae;
DER29__locus_1
ARO
3349 -pf01575 -
3350 - -
3351 -pf00005 -
3352 -pf00528 -
3353 -pf05378pf01968 -
3354 -pf00501pf13193 -
3355 -pf10397pf00206 -
3356 -pf00378 -
3357 HMM02776tpparo_1_aro(pf02775pf00205pf02776) ARO
3358 HMMreg09339_1_aro(pf09339pf01614) ARO
3359 -pf02538 -
3360 HMM02900arooxy_1_aro(pf02900) ARO
3361 HMM02900arooxy_1_aro(pf02900) ARO
3362 H_lipstickRed ARO
3363 H_cerulean ARO
3364 P_iris ARO
3365 HMMalddh_3_aromic1(pf00171) ARO
3366 HMM01557arohydro_1(pf01557) ARO
3367 HMM01118salddharo_1_aro(pf09290pf01118) ARO
3368 HMM00682carbxaro_1_aro(pf07836pf00682) ARO
3369 HMM01557arohydro_1 ARO
3370 -pf01042 -
3371 - -
3372 -pf13476 -
3373 -pf00075 -
3374 - -
3375 -pf11611 -
3376 -pf02384 -
3377 -pf00270pf09369pf00271pf
BMC-P
BMC-H
BMC-T
Aldehyde dehydrogenase
PTAC
Signature enzyme
Alcohol dehydrogenase
Regulator

Verrucosispora sp. FIM060022. Bacteria; Actinobacteria; Micromonosporales; Micromonosporaceae;

EG812__locus_1

ARO

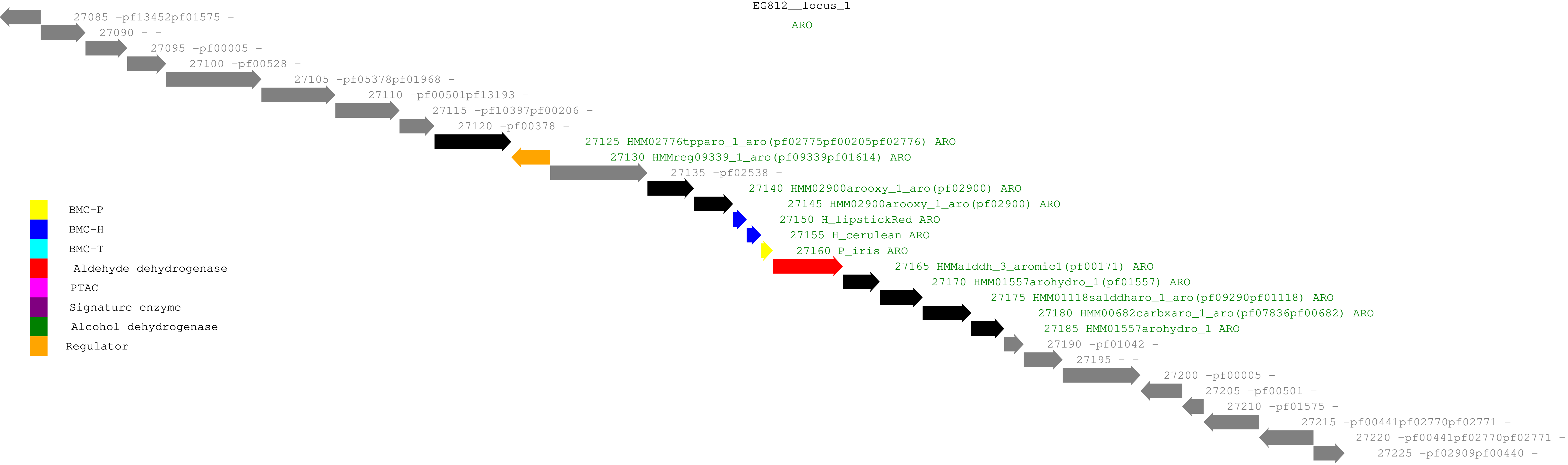

Actinomycetospora succinea. Bacteria; Actinobacteria; Pseudonocardiales; Pseudonocardiaceae;

EV188__locus_1

ARO

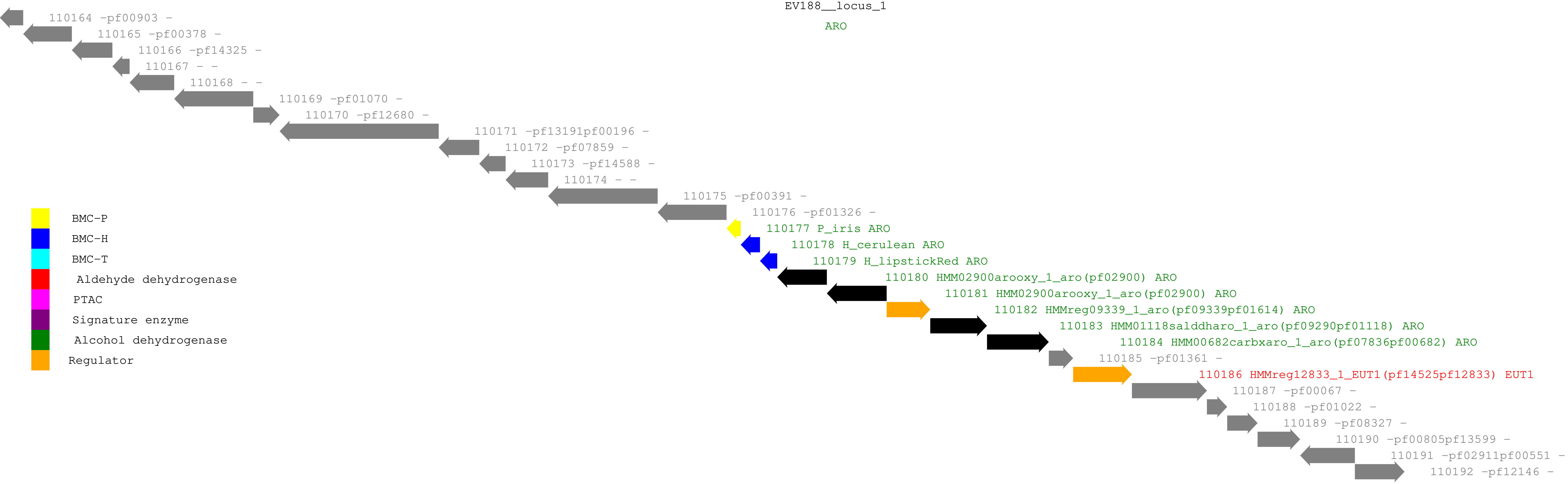

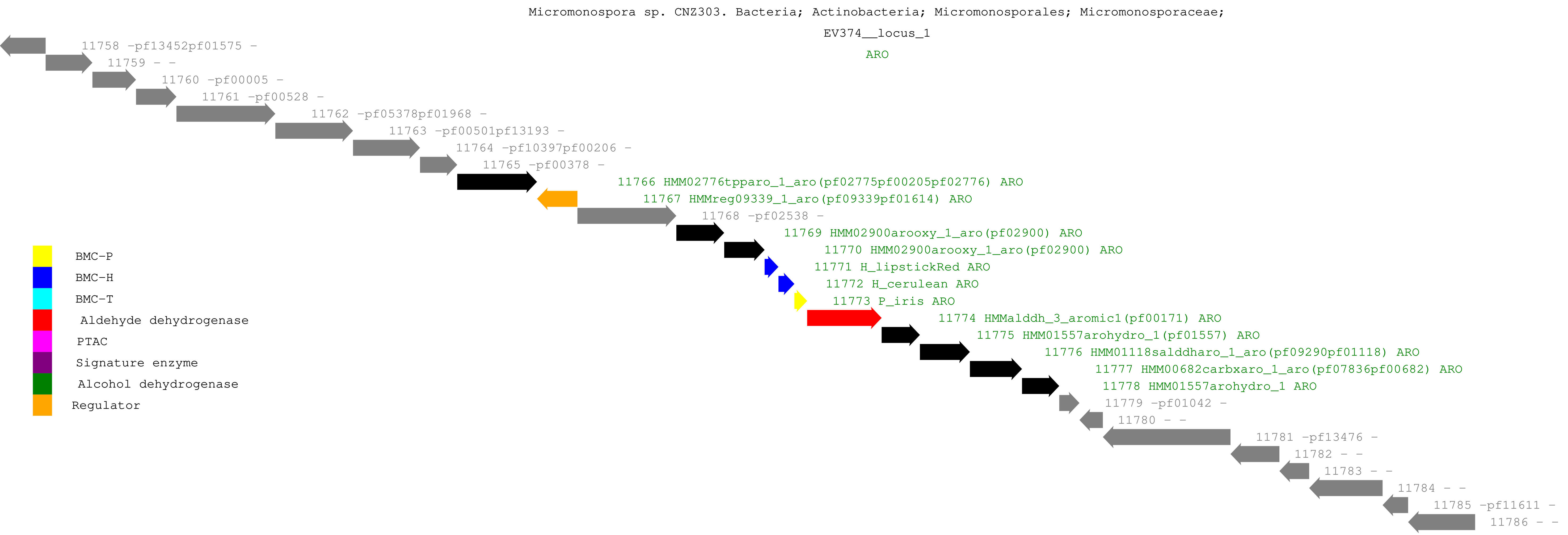
Micromonospora sp. CNZ303. Bacteria; Actinobacteria; Micromonosporales; Micromonosporaceae;
EV374__locus_1
ARO
11758 -pf13452pf01575 -
11759 - -
11760 -pf00005 -
11761 -pf00528 -
11762 -pf05378pf01968 -
11763 -pf00501pf13193 -
11764 -pf10397pf00206 -
11765 -pf00378 -
11766 HMM02776tpparo_1_aro(pf02775pf00205pf02776) ARO
11767 HMMreg09339_1_aro(pf09339pf01614) ARO
11768 -pf02538 -
11769 HMM02900arooxy_1_aro(pf02900) ARO
11770 HMM02900arooxy_1_aro(pf02900) ARO
11771 H_lipstickRed ARO
11772 H_cerulean ARO
11773 P_iris ARO
11774 HMMaldh_3_aromic1(pf00171) ARO
11775 HMM01557arohydro_1(pf01557) ARO
11776 HMM01118salddharo_1_aro(pf09290pf01118) ARO
11777 HMM00682carbxaro_1_aro(pf07836pf00682) ARO
11778 HMM01557arohydro_1 ARO
11779 -pf01042 -
11780 - -
11781 -pf13476 -
11782 - -
11783 - -
11784 - -
11785 -pf11611 -
11786 - -
BMC-P
BMC-H
BMC-T
Aldehyde dehydrogenase
PTAC
Signature enzyme
Alcohol dehydrogenase
Regulator

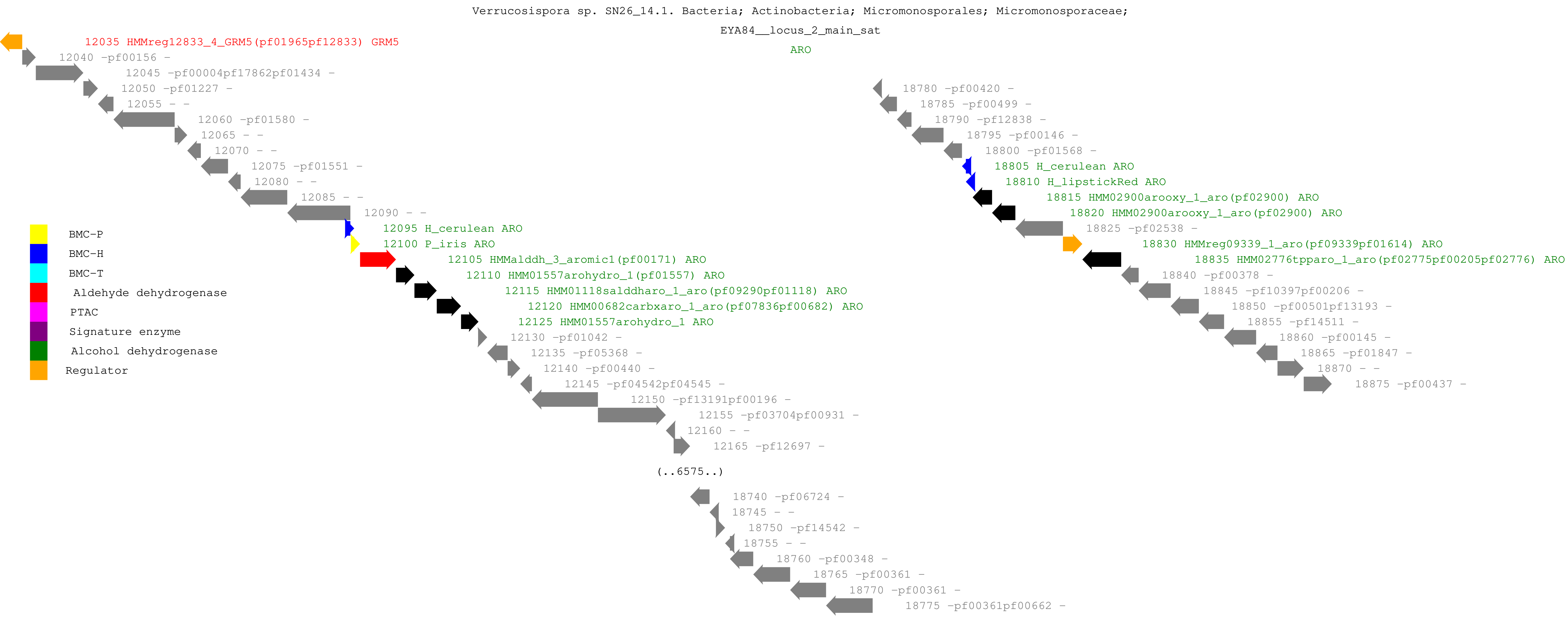
Verrucosispora sp. SN26_14.1. Bacteria; Actinobacteria; Micromonosporales; Micromonosporaceae;
EYA84__locus_2_main_sat
ARO
12035 HMMreg12833_4_GRM5(pf01965pf12833) GRM5
12040 -pf00156 -
12045 -pf00004pf17862pf01434 -
12050 -pf01227 -
12055 - -
12060 -pf01580 -
12065 - -
12070 - -
12075 -pf01551 -
12080 - -
12085 - -
12090 - -
12095 H_cerulean ARO
12100 P_iris ARO
12105 HMMalddh_3_aromic1(pf00171) ARO
12110 HMM01557arohydro_1(pf01557) ARO
12115 HMM01118salddharo_1_aro(pf09290pf01118) ARO
12120 HMM00682carbxaro_1_aro(pf07836pf00682) ARO
12125 HMM01557arohydro_1 ARO
12130 -pf01042 -
12135 -pf05368 -
12140 -pf00440 -
12145 -pf04542pf04545 -
12150 -pf13191pf00196 -
12155 -pf03704pf00931 -
12160 - -
12165 -pf12697 -
(..6575..)
18740 -pf06724 -
18745 - -
18750 -pf14542 -
18755 - -
18760 -pf00348 -
18765 -pf00361 -
18770 -pf00361 -
18775 -pf00361pf00662 -
18780 -pf00420 -
18785 -pf00499 -
18790 -pf12838 -
18795 -pf00146 -
18800 -pf01568 -
18805 H_cerulean ARO
18810 H_lipstickRed ARO
18815 HMM02900arooxy_1_aro(pf02900) ARO
18820 HMM02900arooxy_1_aro(pf02900) ARO
18825 -pf02538 -
18830 HMMreg09339_1_aro(pf09339pf01614) ARO
18835 HMM02776tpparo_1_aro(pf02775pf00205pf02776) ARO
18840 -pf00378 -
18845 -pf10397pf00206 -
18850 -pf00501pf13193 -
18855 -pf14511 -
18860 -pf00145 -
18865 -pf01847 -
18870 - -
18875 -pf00437 -
BMC-P
BMC-H
BMC-T
Aldehyde dehydrogenase
PTAC
Signature enzyme
Alcohol dehydrogenase
Regulator

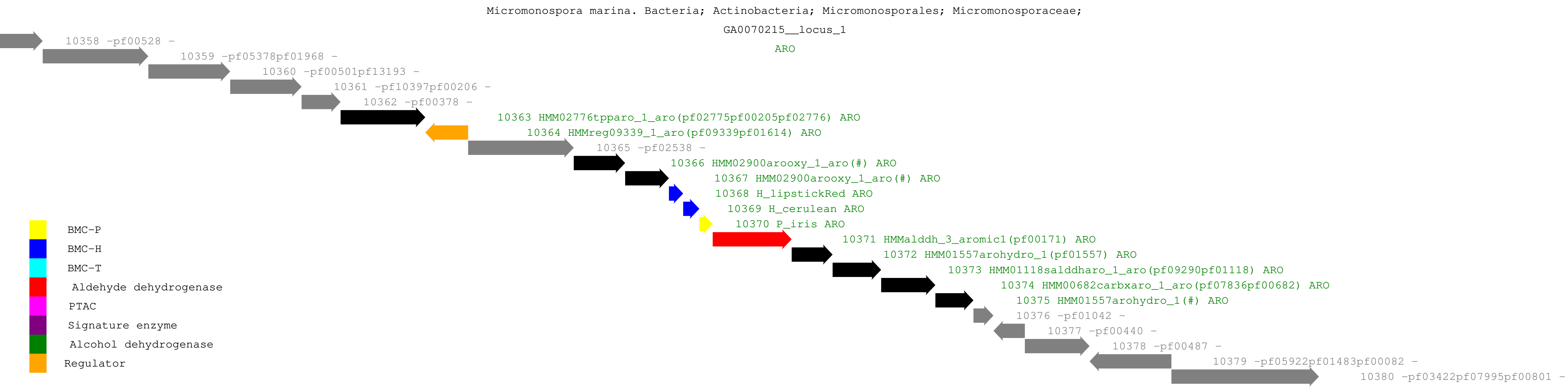
Micromonospora marina. Bacteria; Actinobacteria; Micromonosporales; Micromonosporaceae;
GA0070215__locus_1
ARO
10358 -pf00528 -
10359 -pf05378pf01968 -
10360 -pf00501pf13193 -
10361 -pf10397pf00206 -
10362 -pf00378 -
10363 HMM02776tpparo_1_aro(pf02775pf00205pf02776) ARO
10364 HMMreg09339_1_aro(pf09339pf01614) ARO
10365 -pf02538 -
10366 HMM02900arooxy_1_aro(#) ARO
10367 HMM02900arooxy_1_aro(#) ARO
10368 H_lipstickRed ARO
10369 H_cerulean ARO
10370 P_iris ARO
10371 HMMalddh_3_aromic1(pf00171) ARO
10372 HMM01557arohydro_1(pf01557) ARO
10373 HMM01118salddharo_1_aro(pf09290pf01118) ARO
10374 HMM00682carbxaro_1_aro(pf07836pf00682) ARO
10375 HMM01557arohydro_1(#) ARO
10376 -pf01042 -
10377 -pf00440 -
10378 -pf00487 -
10379 -pf05922pf01483pf00082 -
10380 -pf03422pf07995pf00801 -
BMC-P
BMC-H
BMC-T
Aldehyde dehydrogenase
PTAC
Signature enzyme
Alcohol dehydrogenase
Regulator

Micromonospora echinofusca. Bacteria; Actinobacteria; Micromonosporales; Micromonosporaceae;

GA0070610__locus_1

ARO

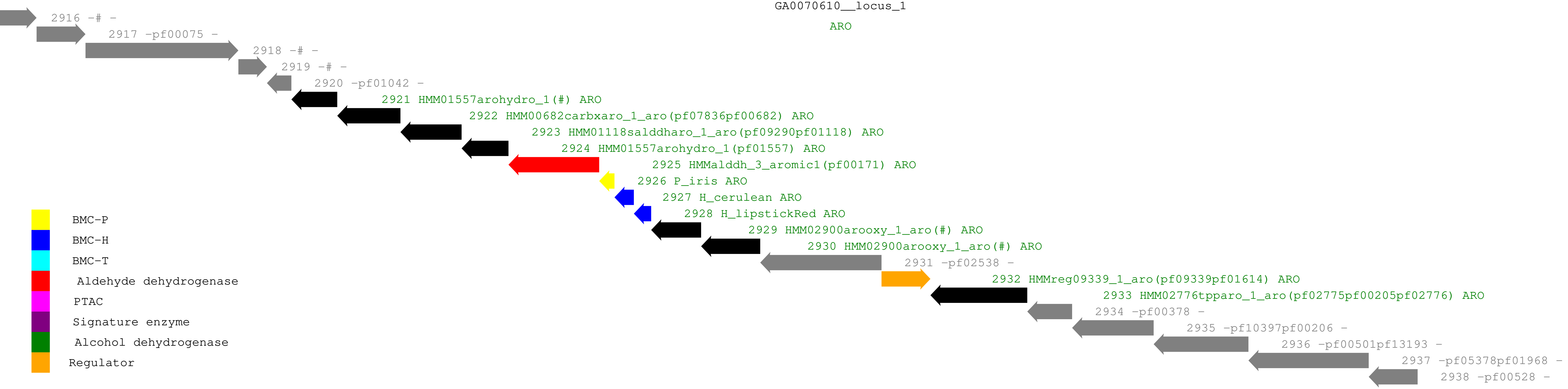

Pseudonocardia asaccharolytica DSM 44247 = NBRC 16224. Bacteria; Actinobacteria; Pseudonocardiales; Pseudonocardiaceae;

PA7__locus_2

ARO

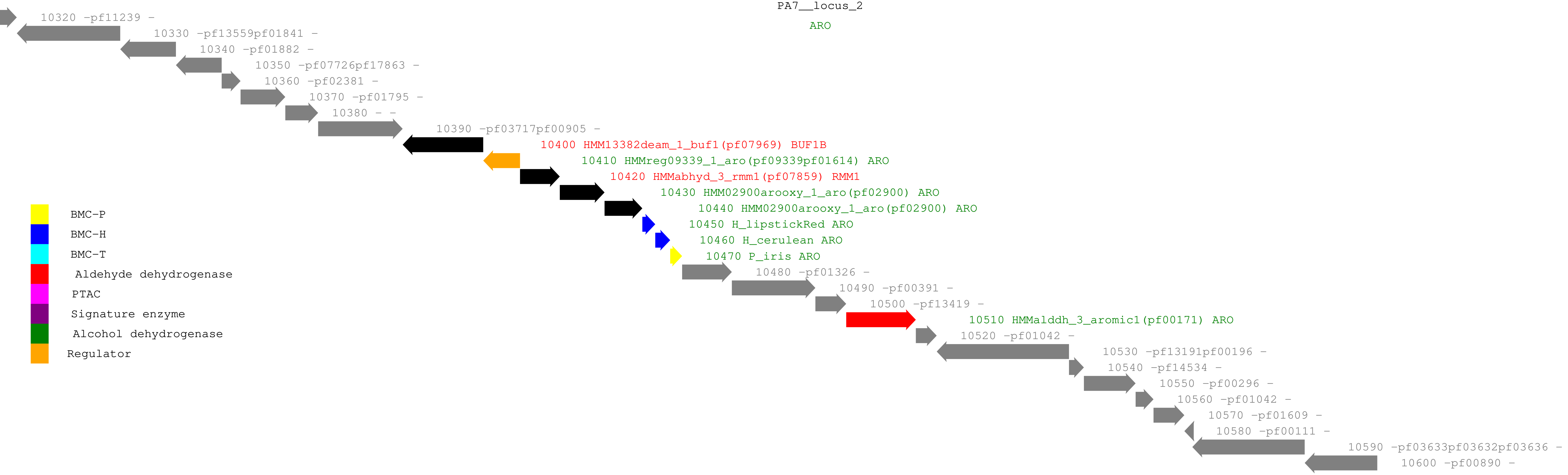

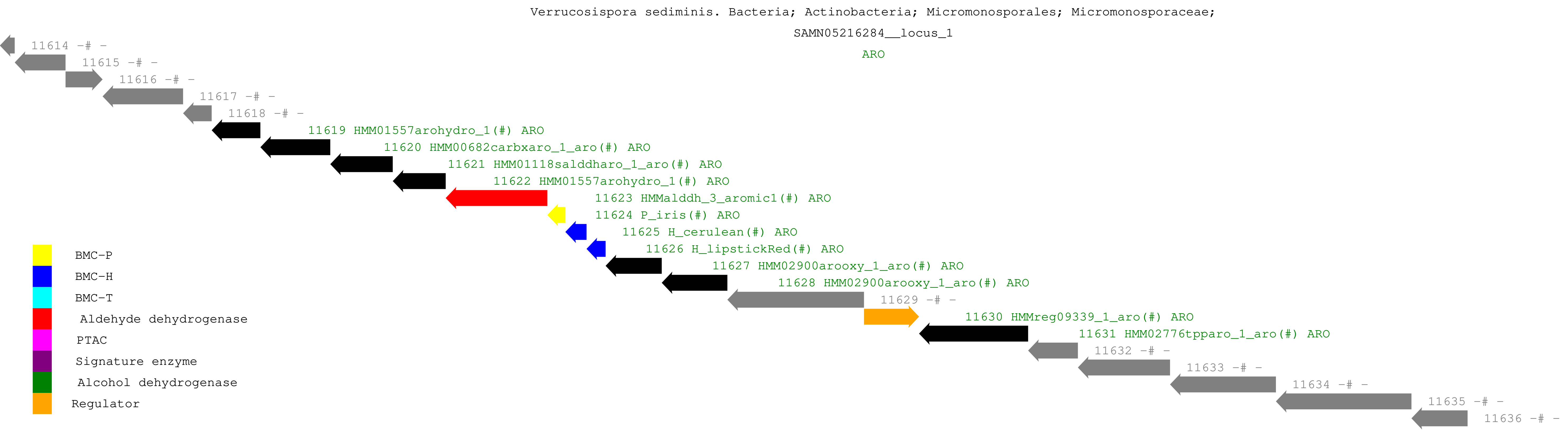
Verrucosispora sediminis. Bacteria; Actinobacteria; Micromonosporales; Micromonosporaceae;
SAMN05216284__locus_1
ARO
11614 -# -
11615 -# -
11616 -# -
11617 -# -
11618 -# -
11619 HMM01557arohydro_1(#) ARO
11620 HMM00682carbxaro_1_aro(#) ARO
11621 HMM01118salddharo_1_aro(#) ARO
11622 HMM01557arohydro_1(#) ARO
11623 HMMalddh_3_aromic1(#) ARO
11624 P_iris(#) ARO
11625 H_cerulean(#) ARO
11626 H_lipstickRed(#) ARO
11627 HMM02900arooxy_1_aro(#) ARO
11628 HMM02900arooxy_1_aro(#) ARO
11629 -# -
11630 HMMreg09339_1_aro(#) ARO
11631 HMM02776tpparo_1_aro(#) ARO
11632 -# -
11633 -# -
11634 -# -
11635 -# -
11636 -# -
BMC-P
BMC-H
BMC-T
Aldehyde dehydrogenase
PTAC
Signature enzyme
Alcohol dehydrogenase
Regulator

Kibdelosporangium aridum. Bacteria; Actinobacteria; Pseudonocardiales; Pseudonocardiaceae;

SAMN05661093__locus_1

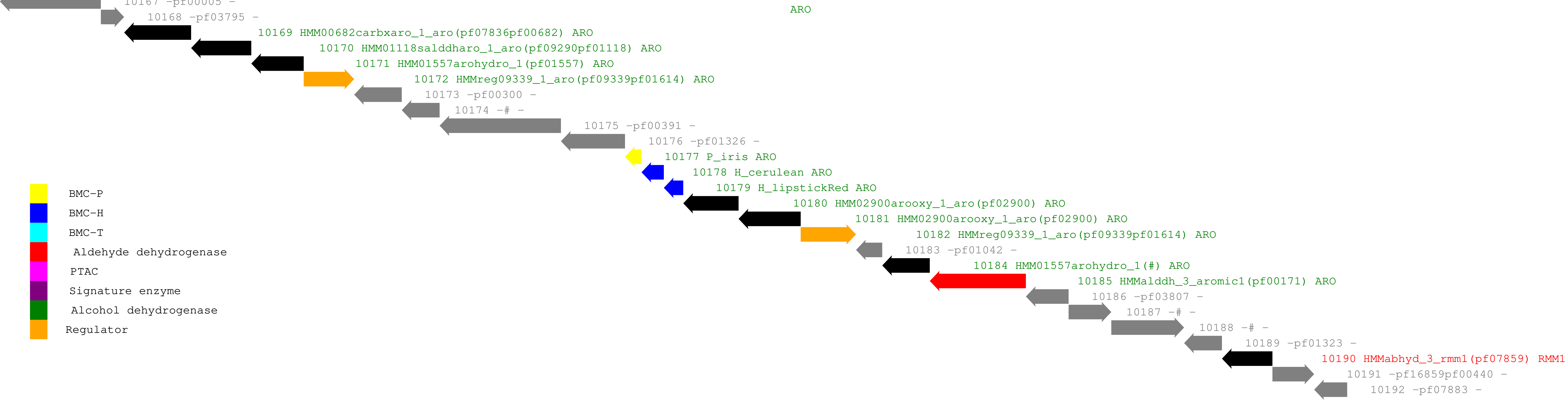

Verrucosispora maris (strain AB-18-032). Bacteria; Actinobacteria; Micromonosporales; Micromonosporaceae;

VAB18032__locus_1

ARO

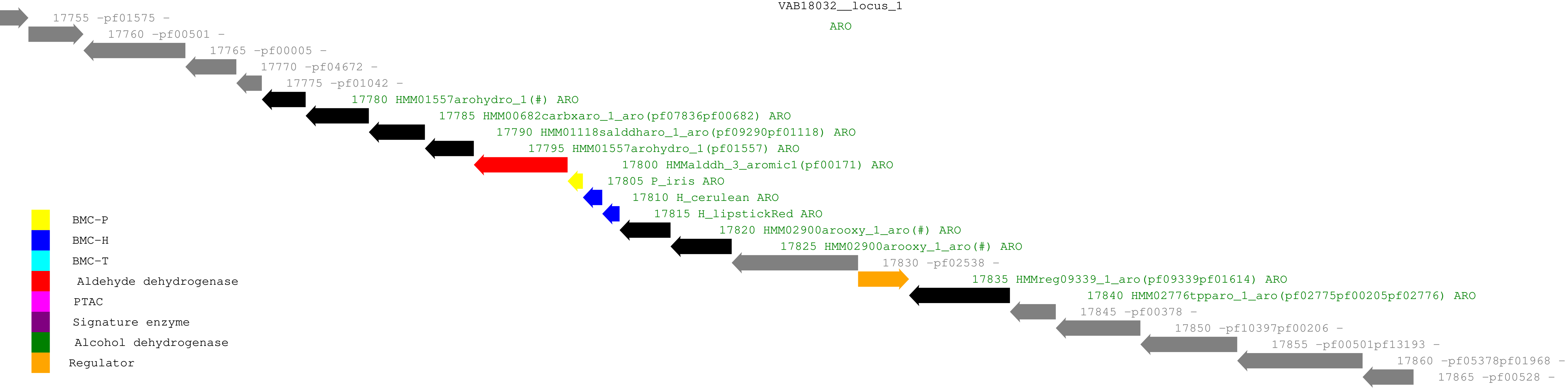

Caloramator mitchellensis. Bacteria; Firmicutes; Clostridia; Clostridiales; Clostridiaceae;

ABG79__locus_1

BUF1

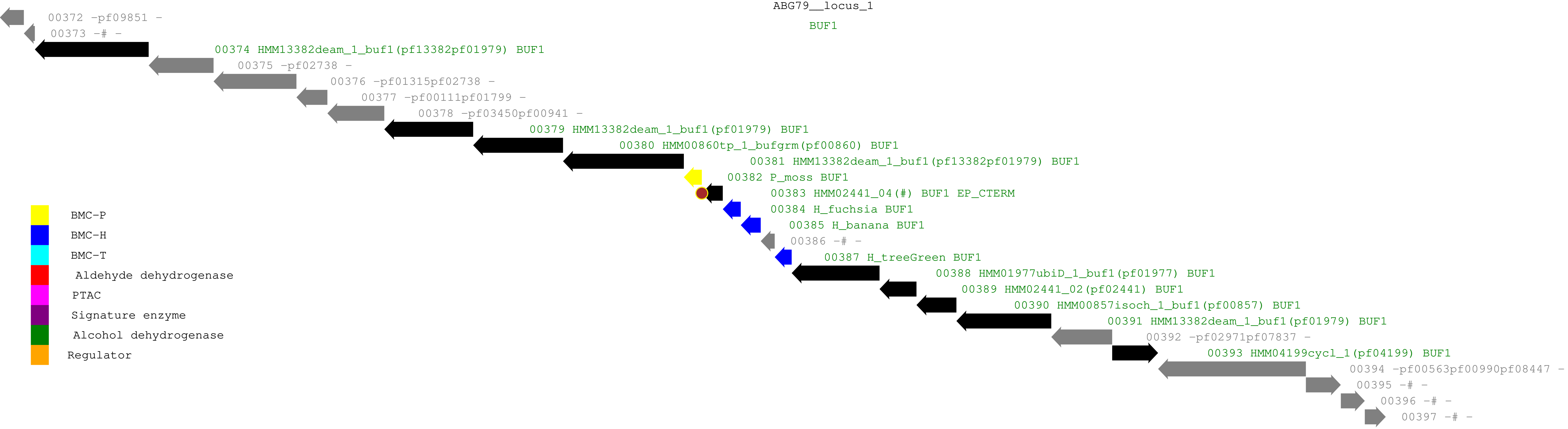

Alkaliphilus metalliredigens (strain QYMF). Bacteria; Firmicutes; Clostridia; Clostridiales; Clostridiaceae;

Amet__locus_3

BUF1

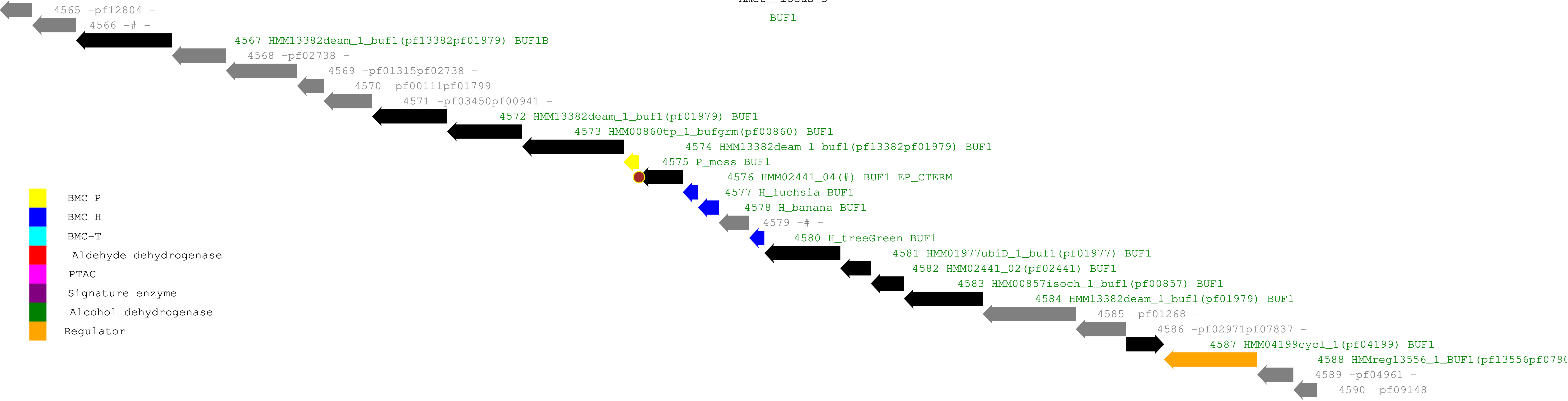

Thermotalea metallivorans. Bacteria; Firmicutes; Clostridia; Clostridiales; Clostridiaceae;

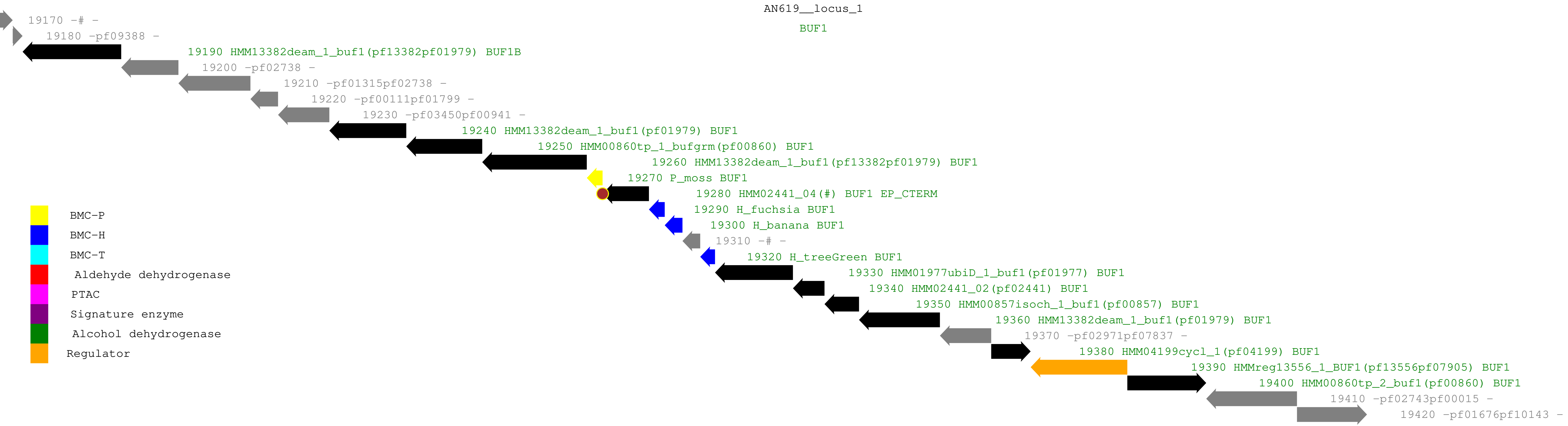

Paraclostridium bifermentans (Clostridium bifermentans). Bacteria; Firmicutes; Clostridia; Clostridiales;

B2H97__locus_2

BUF1

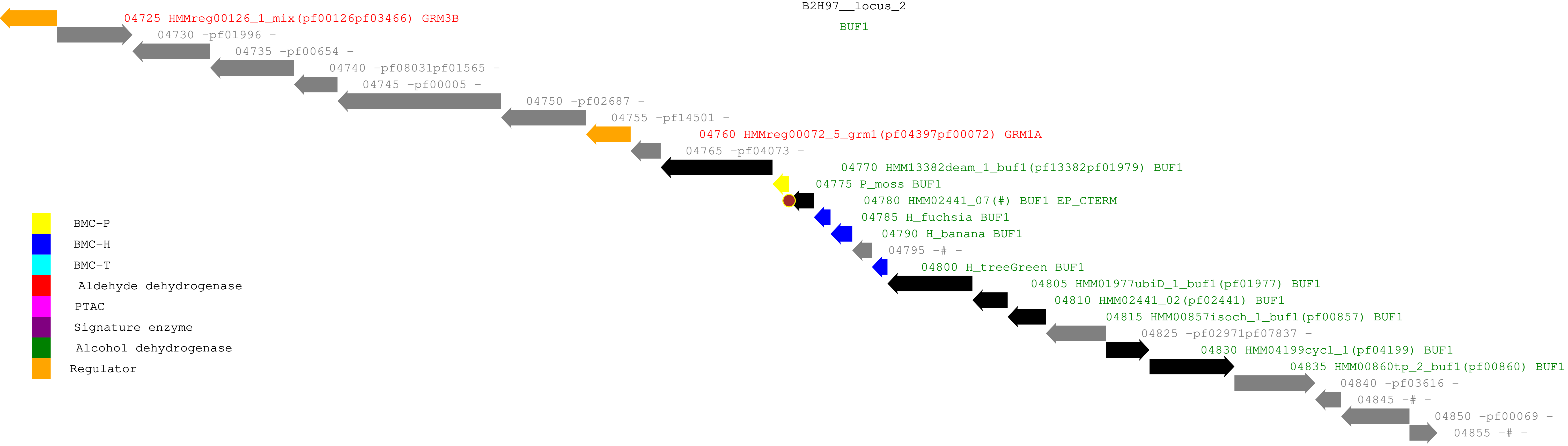

Bacteroidetes bacterium 4572_77. Bacteria; Bacteroidetes.

B7C24__locus_2

BUF1

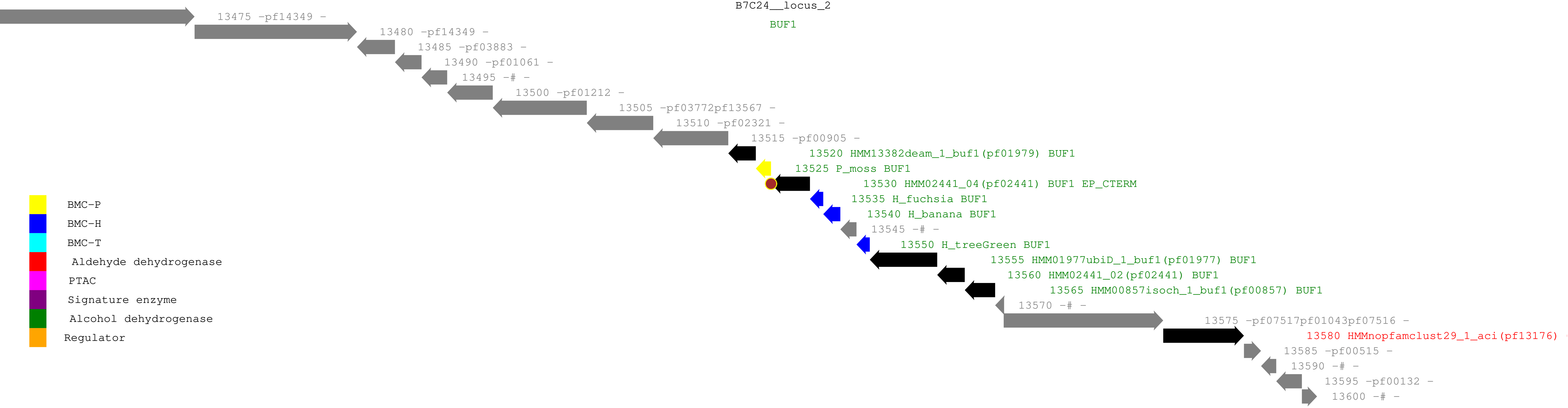

Thermohalobacter berrensis. Bacteria; Firmicutes; Clostridia; Clostridiales; Clostridiaceae;

BET03__locus_2

BUF1

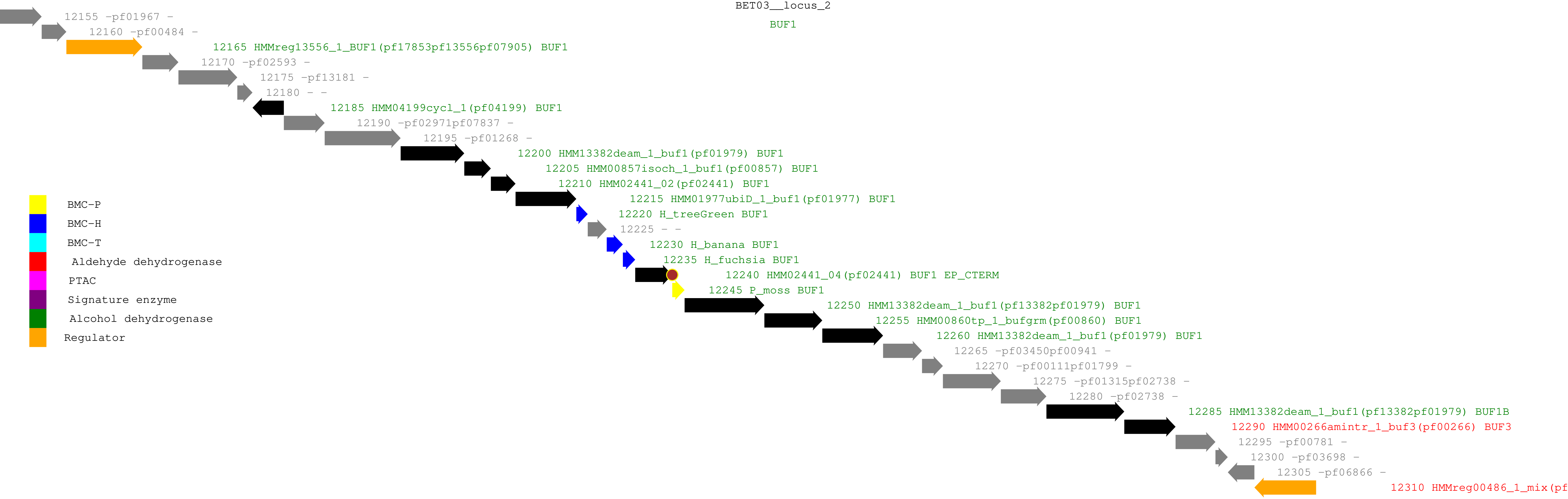

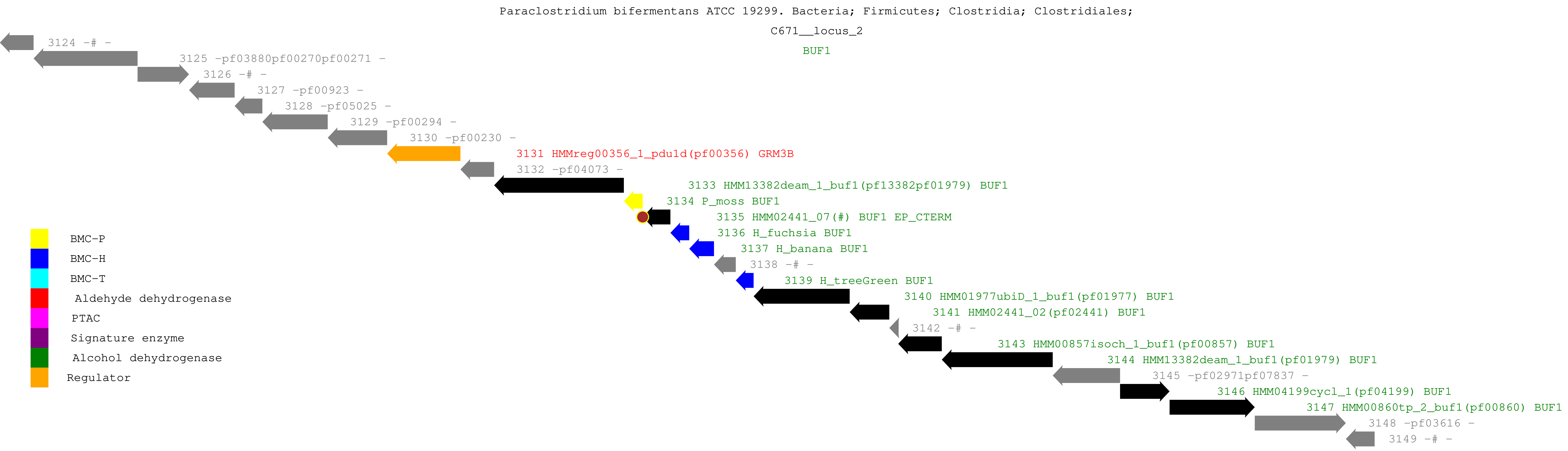
Paraclostridium bifermentans ATCC 19299. Bacteria; Firmicutes; Clostridia; Clostridiales;
C671__locus_2
BUF1
3124 -# -
3125 -pf03880pf00270pf00271 -
3126 -# -
3127 -pf00923 -
3128 -pf05025 -
3129 -pf00294 -
3130 -pf00230 -
3131 HMMreg00356_1_pdu1d(pf00356) GRM3B
3132 -pf04073 -
3133 HMM13382deam_1_buf1(pf13382pf01979) BUF1
3134 P_moss BUF1
3135 HMM02441_07(#) BUF1 EP_CTERM
3136 H_fuchsia BUF1
3137 H_banana BUF1
3138 -# -
3139 H_treeGreen BUF1
3140 HMM01977ubiD_1_buf1(pf01977) BUF1
3141 HMM02441_02(pf02441) BUF1
3142 -# -
3143 HMM00857isoch_1_buf1(pf00857) BUF1
3144 HMM13382deam_1_buf1(pf01979) BUF1
3145 -pf02971pf07837 -
3146 HMM04199cycl_1(pf04199) BUF1
3147 HMM00860tp_2_buf1(pf00860) BUF1
3148 -pf03616 -
3149 -# -
BMC-P
BMC-H
BMC-T
Aldehyde dehydrogenase
PTAC
Signature enzyme
Alcohol dehydrogenase
Regulator

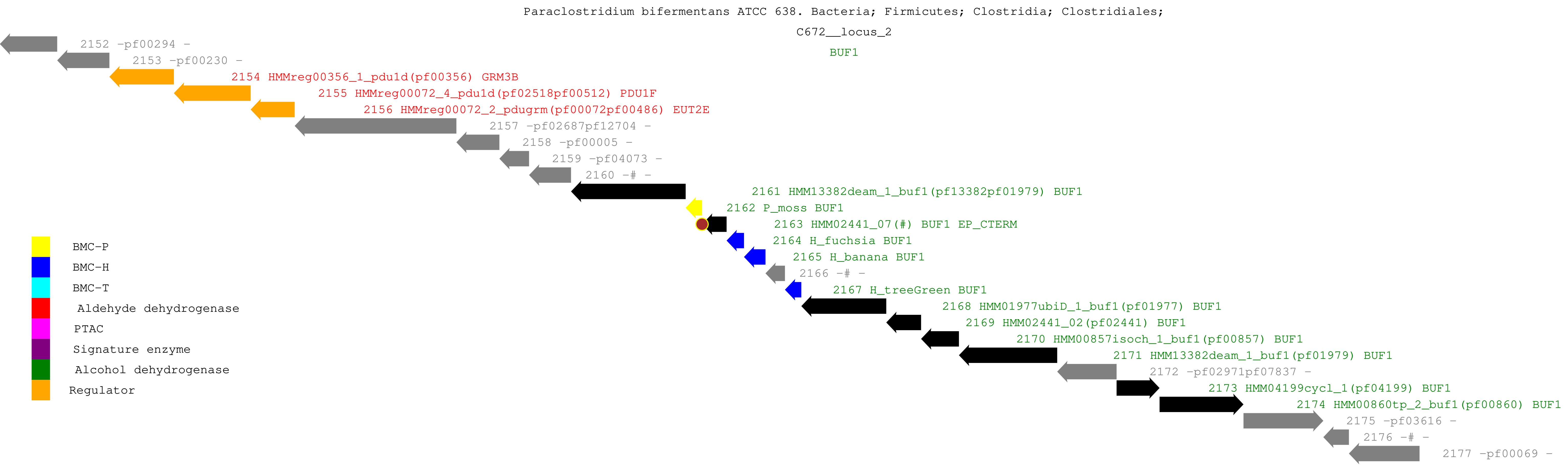
Paraclostridium bifermentans ATCC 638. Bacteria; Firmicutes; Clostridia; Clostridiales;
C672__locus_2
BUF1
2152 -pf00294 -
2153 -pf00230 -
2154 HMMreg00356_1_pdu1d(pf00356) GRM3B
2155 HMMreg00072_4_pdu1d(pf02518pf00512) PDU1F
2156 HMMreg00072_2_pdugrm(pf00072pf00486) EUT2E
2157 -pf02687pf12704 -
2158 -pf00005 -
2159 -pf04073 -
2160 -# -
2161 HMM13382deam_1_buf1(pf13382pf01979) BUF1
2162 P_moss BUF1
2163 HMM02441_07(#) BUF1 EP_CTERM
2164 H_fuchsia BUF1
2165 H_banana BUF1
2166 -# -
2167 H_treeGreen BUF1
2168 HMM01977ubiD_1_buf1(pf01977) BUF1
2169 HMM02441_02(pf02441) BUF1
2170 HMM00857isoch_1_buf1(pf00857) BUF1
2171 HMM13382deam_1_buf1(pf01979) BUF1
2172 -pf02971pf07837 -
2173 HMM04199cycl_1(pf04199) BUF1
2174 HMM00860tp_2_buf1(pf00860) BUF1
2175 -pf03616 -
2176 -# -
2177 -pf00069 -
BMC-P
BMC-H
BMC-T
Aldehyde dehydrogenase
PTAC
Signature enzyme
Alcohol dehydrogenase
Regulator

Caloramator australicus RC3. Bacteria; Firmicutes; Clostridia; Clostridiales; Clostridiaceae;

CAAU__locus_1

BUF1

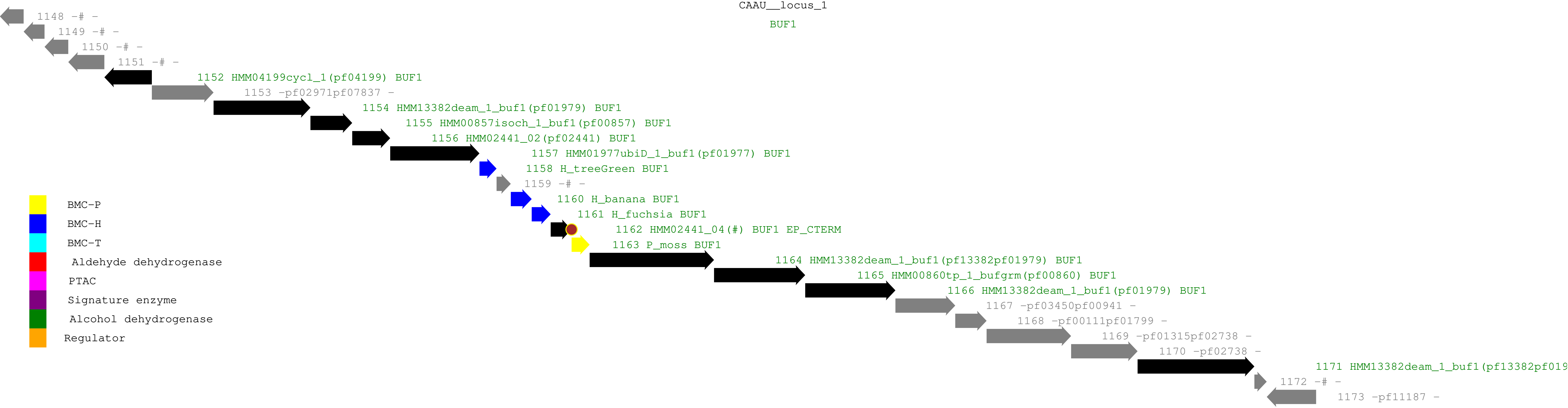

Anaeromicrobium sediminis. Bacteria; Firmicutes; Clostridia; Clostridiales; Clostridiaceae;

CCE28__locus_1

BUF1

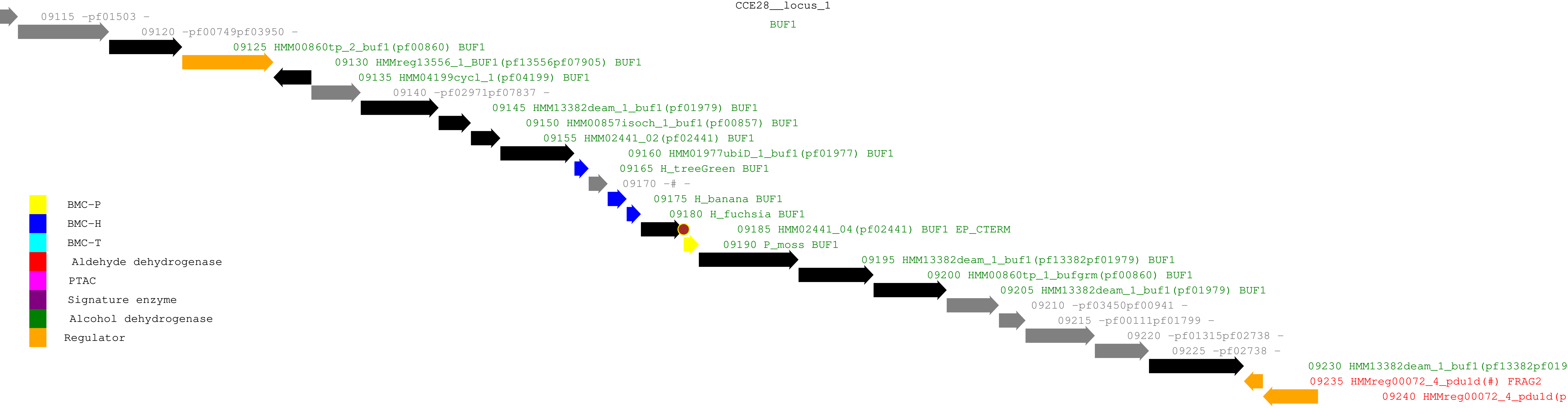

Carboxydothermus hydrogenoformans (strain ATCC BAA-161 / DSM 6008 / Bacteria; Firmicutes; Clostridia; Thermoanaerobacterales;

CHY__locus_1

BUF1

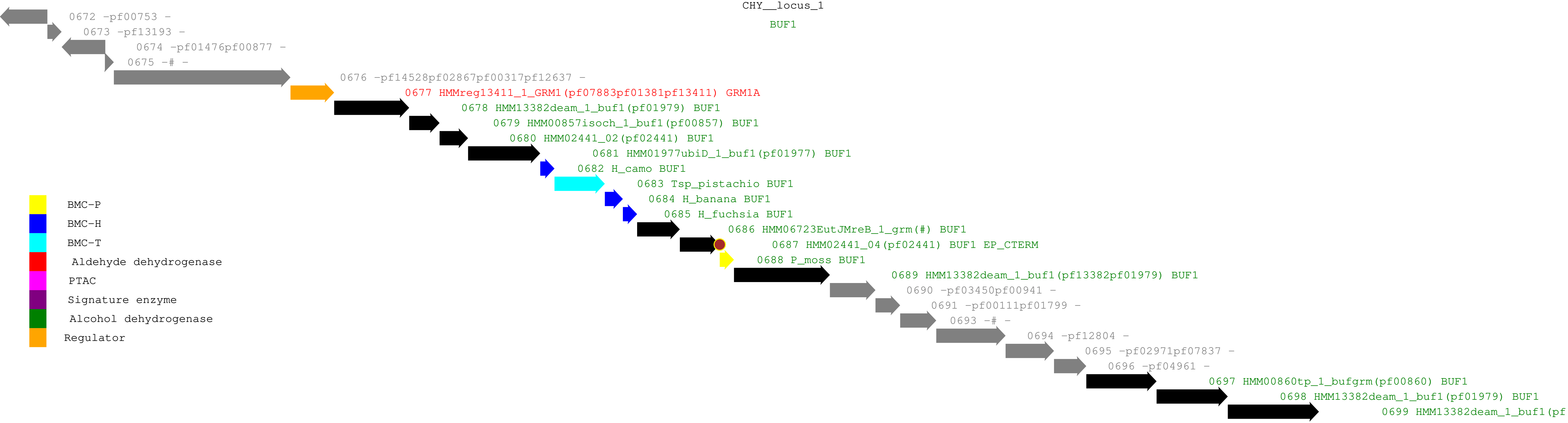

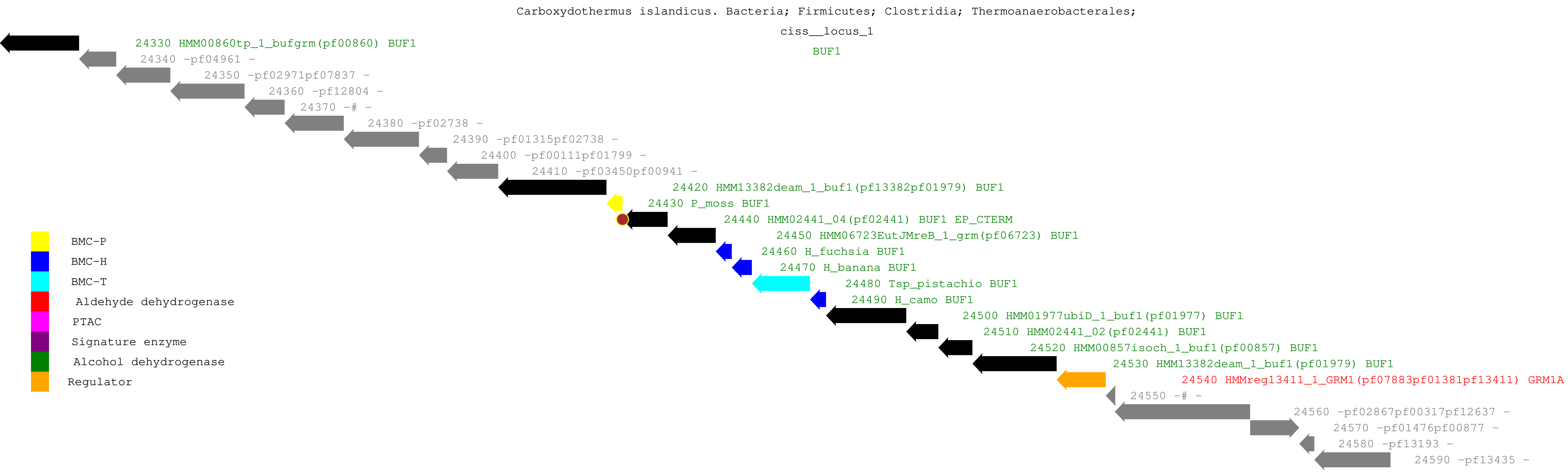
Carboxydothermus islandicus. Bacteria; Firmicutes; Clostridia; Thermoanaerobacterales;
ciss__locus_1
BUF1
24330 HMM00860tp_1_bufgrm(pf00860) BUF1
24340 -pf04961 -
24350 -pf02971pf07837 -
24360 -pf12804 -
24370 -# -
24380 -pf02738 -
24390 -pf01315pf02738 -
24400 -pf00111pf01799 -
24410 -pf03450pf00941 -
24420 HMM13382deam_1_buf1(pf13382pf01979) BUF1
24430 P_moss BUF1
24440 HMM02441_04(pf02441) BUF1 EP_CTERM
24450 HMM06723EutJMreB_1_grm(pf06723) BUF1
24460 H_fuchsia BUF1
24470 H_banana BUF1
24480 Tsp_pistachio BUF1
24490 H_camo BUF1
24500 HMM01977ubiD_1_buf1(pf01977) BUF1
24510 HMM02441_02(pf02441) BUF1
24520 HMM00857isoch_1_buf1(pf00857) BUF1
24530 HMM13382deam_1_buf1(pf01979) BUF1
24540 HMMreg13411_1_GRM1(pf07883pf01381pf13411) GRM1A
24550 -# -
24560 -pf02867pf00317pf12637 -
24570 -pf01476pf00877 -
24580 -pf13193 -
24590 -pf13435 -
BMC-P
BMC-H
BMC-T
Aldehyde dehydrogenase
PTAC
Signature enzyme
Alcohol dehydrogenase
Regulator

Tissierella sp. P1. Bacteria; Firmicutes; Tissierellia; Tissierellales; Tissierellaceae;

CIW83__locus_2

BUF1

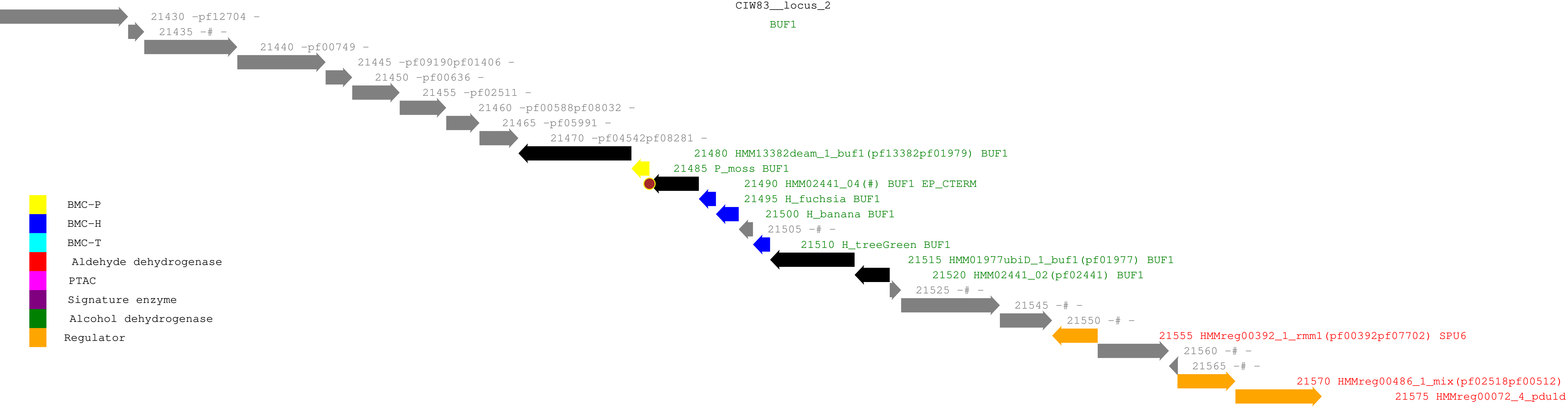

Clostridium cylindrosporum DSM 605. Bacteria; Firmicutes; Clostridia; Clostridiales; Clostridiaceae;

CLCY_4c_locus_1

BUF1

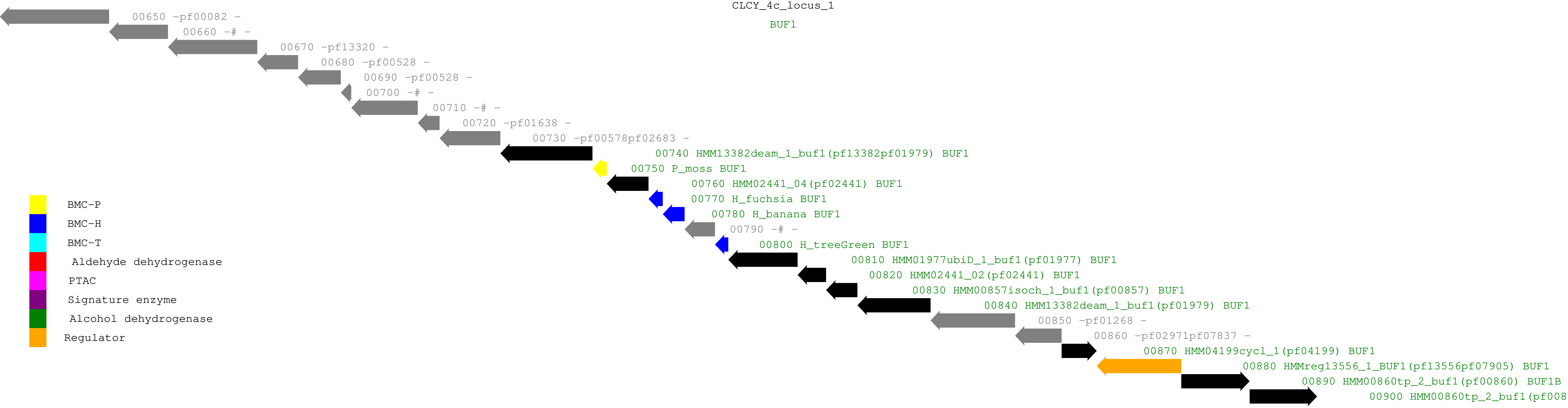

Alkaliphilus oremlandii (strain OhILAs) (Clostridium oremlandii Bacteria; Firmicutes; Clostridia; Clostridiales; Clostridiaceae;

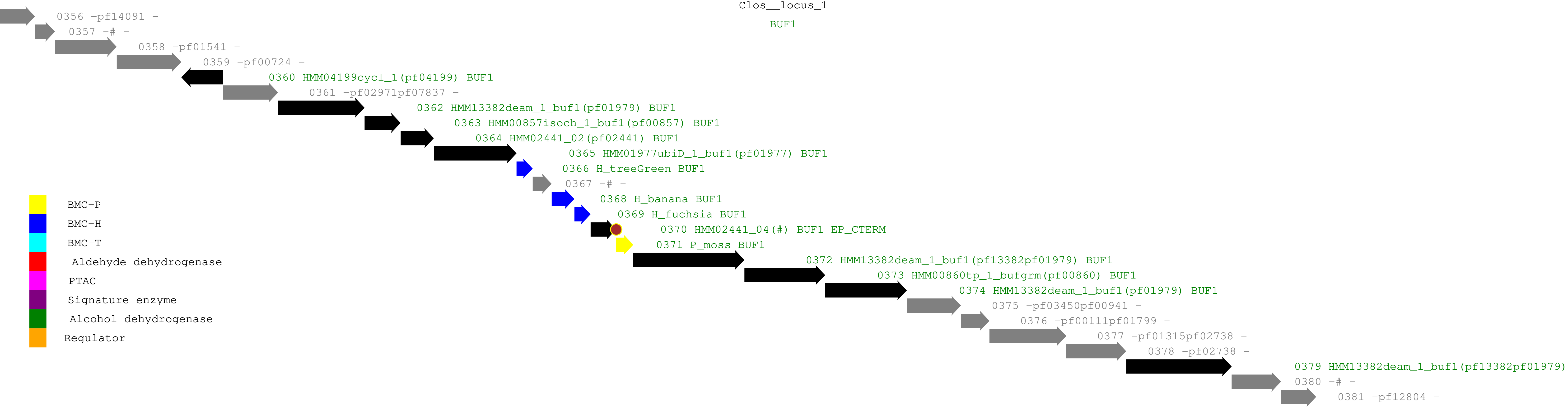

Clostridium purinilyticum. Bacteria; Firmicutes; Clostridia; Clostridiales; Gottschalkia.

CLPU_29c_locus_1

BUF1

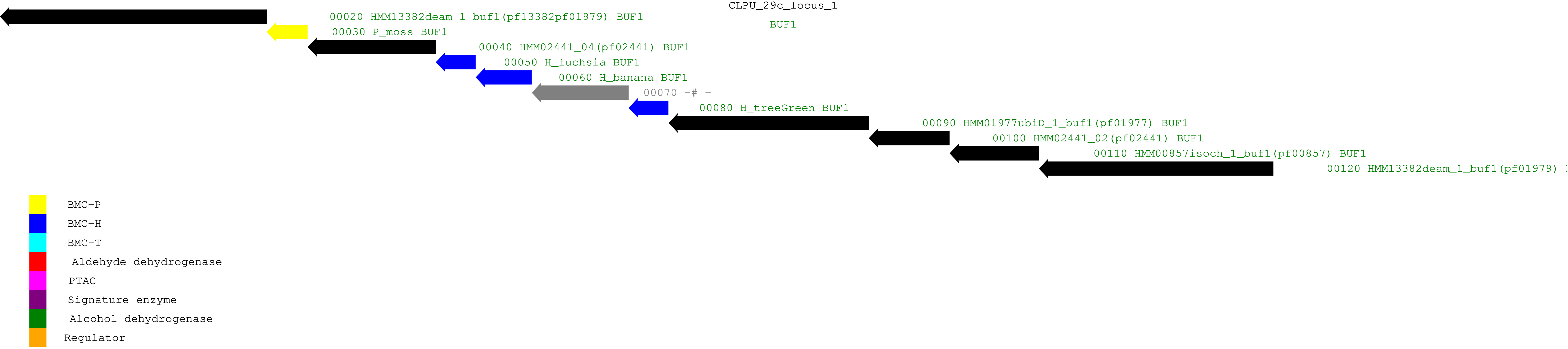

Alkaliphilus sp. Bacteria; Firmicutes; Clostridia; Clostridiales; Clostridiaceae;

COA82__locus_1

BUF1

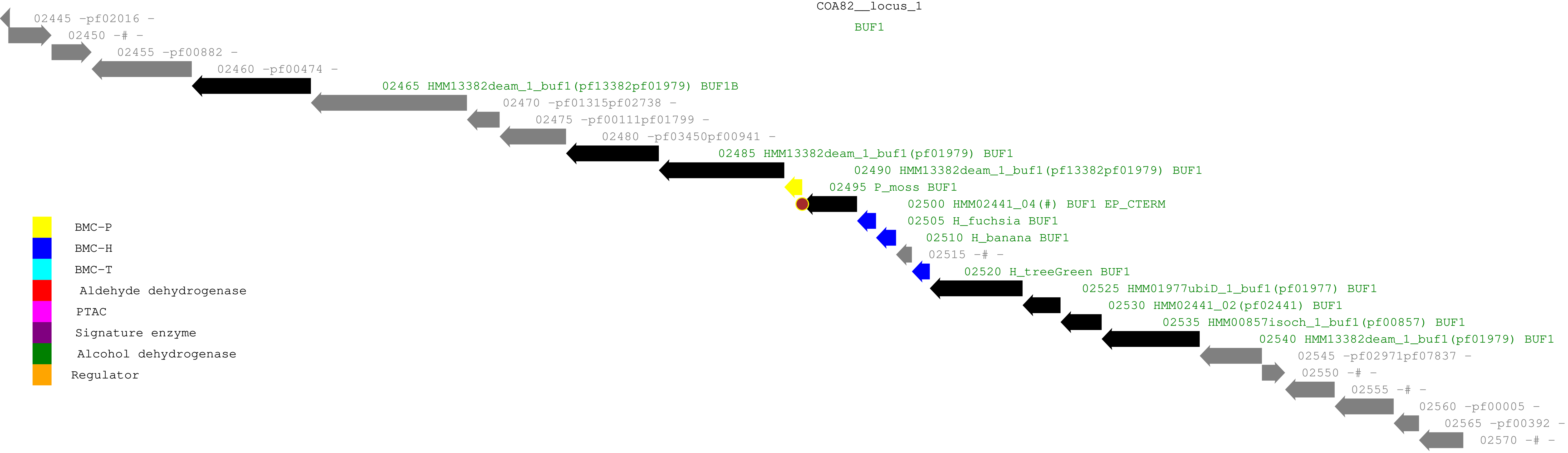

Carboxydothermus pertinax. Bacteria; Firmicutes; Clostridia; Thermoanaerobacterales;

cpu__locus_1

BUF1

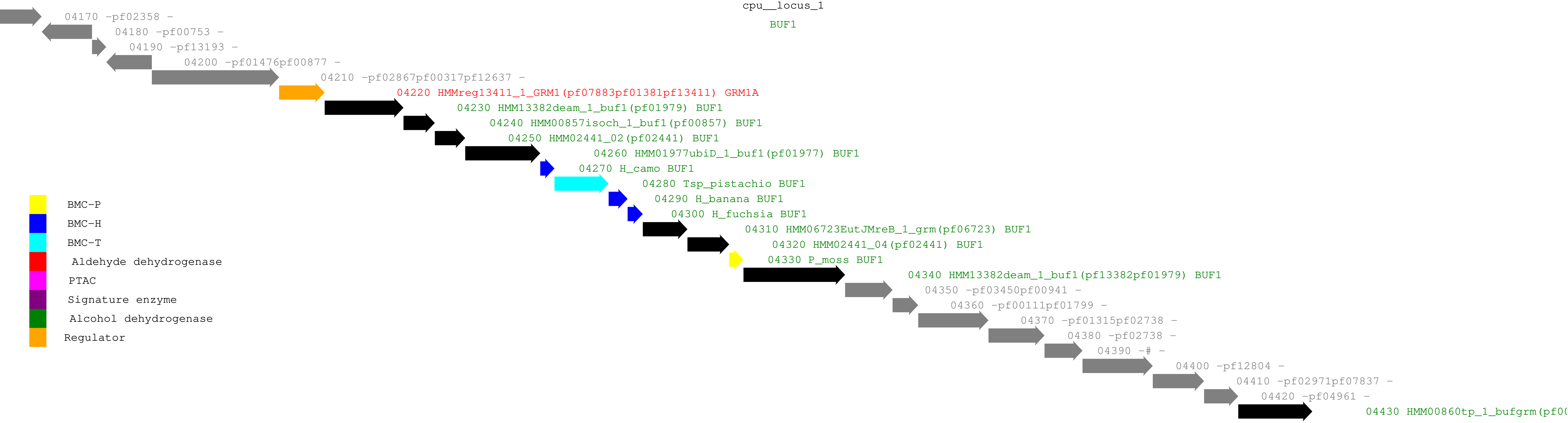

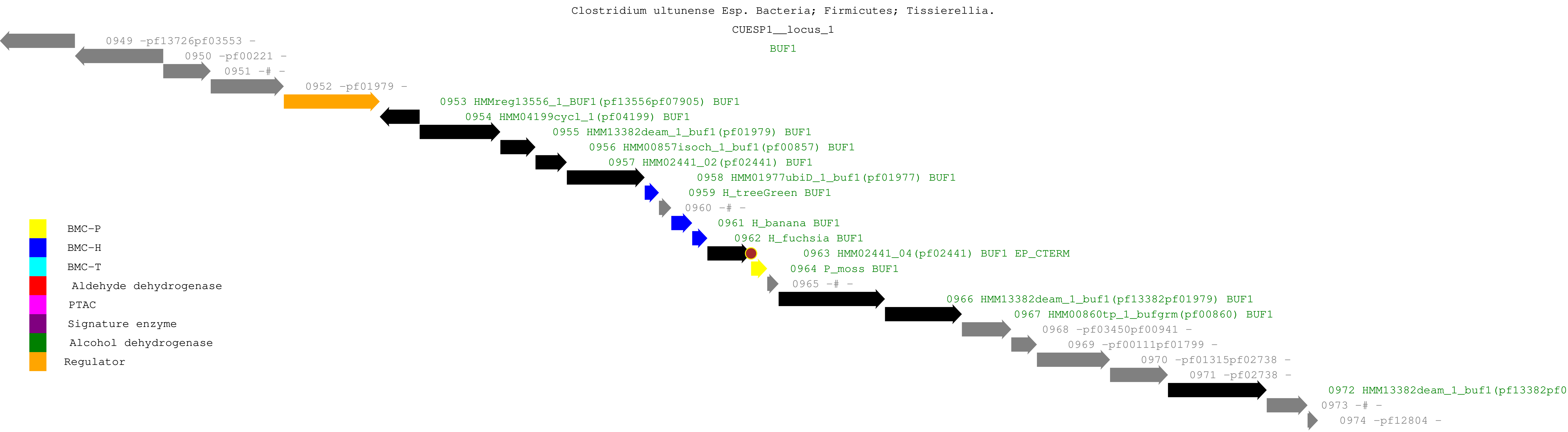
Clostridium ultunense Esp. Bacteria; Firmicutes; Tissierellia.
CUESP1__locus_1
BUF1
0949 -pf13726pf03553 -
0950 -pf00221 -
0951 -# -
0952 -pf01979 -
0953 HMMreg13556_1_BUF1(pf13556pf07905) BUF1
0954 HMM04199cycl_1(pf04199) BUF1
0955 HMM13382deam_1_buf1(pf01979) BUF1
0956 HMM00857isoch_1_buf1(pf00857) BUF1
0957 HMM02441_02(pf02441) BUF1
0958 HMM01977ubiD_1_buf1(pf01977) BUF1
0959 H_treeGreen BUF1
0960 -# -
0961 H_banana BUF1
0962 H_fuchsia BUF1
0963 HMM02441_04(pf02441) BUF1 EP_CTERM
0964 P_moss BUF1
0965 -# -
0966 HMM13382deam_1_buf1(pf13382pf01979) BUF1
0967 HMM00860tp_1_bufgrm(pf00860) BUF1
0968 -pf03450pf00941 -
0969 -pf00111pf01799 -
0970 -pf01315pf02738 -
0971 -pf02738 -
0972 HMM13382deam_1_buf1(pf13382pf0
0973 -# -
0974 -pf12804 -
BMC-P
BMC-H
BMC-T
Aldehyde dehydrogenase
PTAC
Signature enzyme
Alcohol dehydrogenase
Regulator

Clostridium acidurici (strain ATCC 7906 / CIP 104303 / DSM 604 / NCIMB Bacteria; Firmicutes; Clostridia; Clostridiales; Gottschalkia.

Curi_c_locus_1_main_sat

BUF1

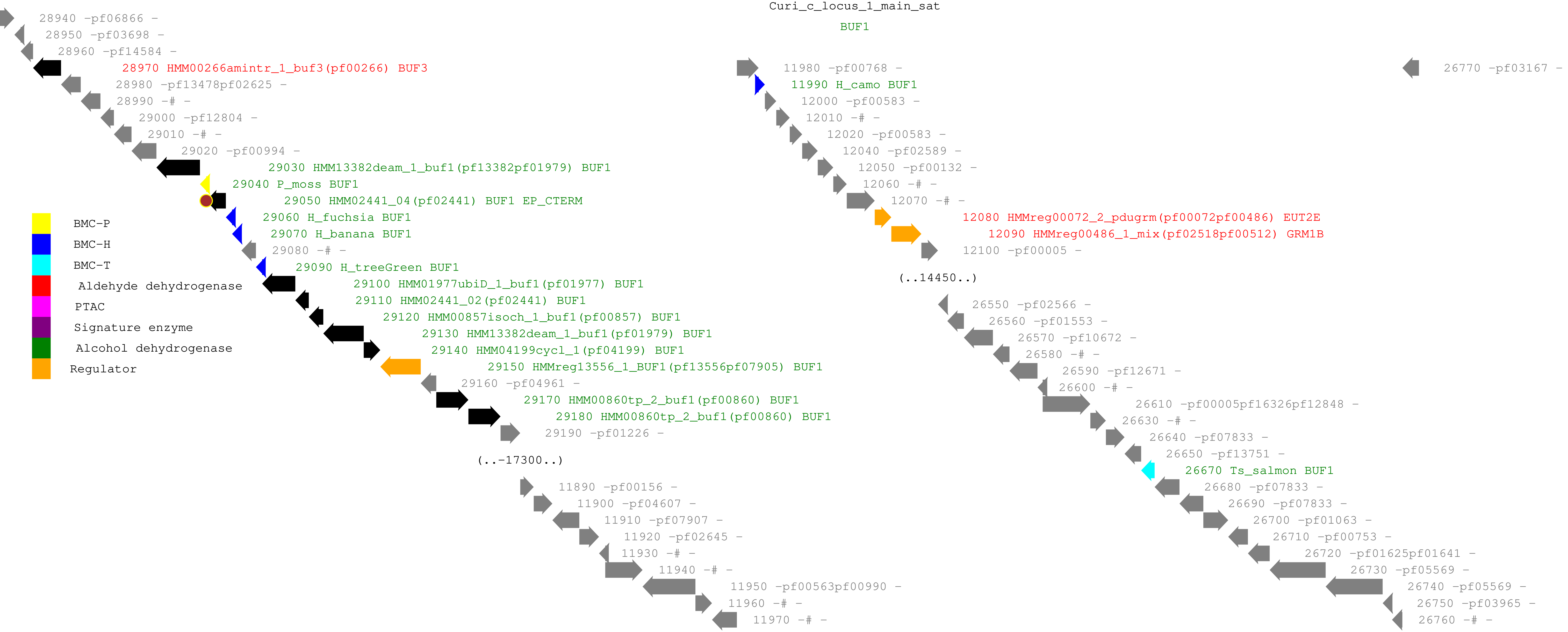

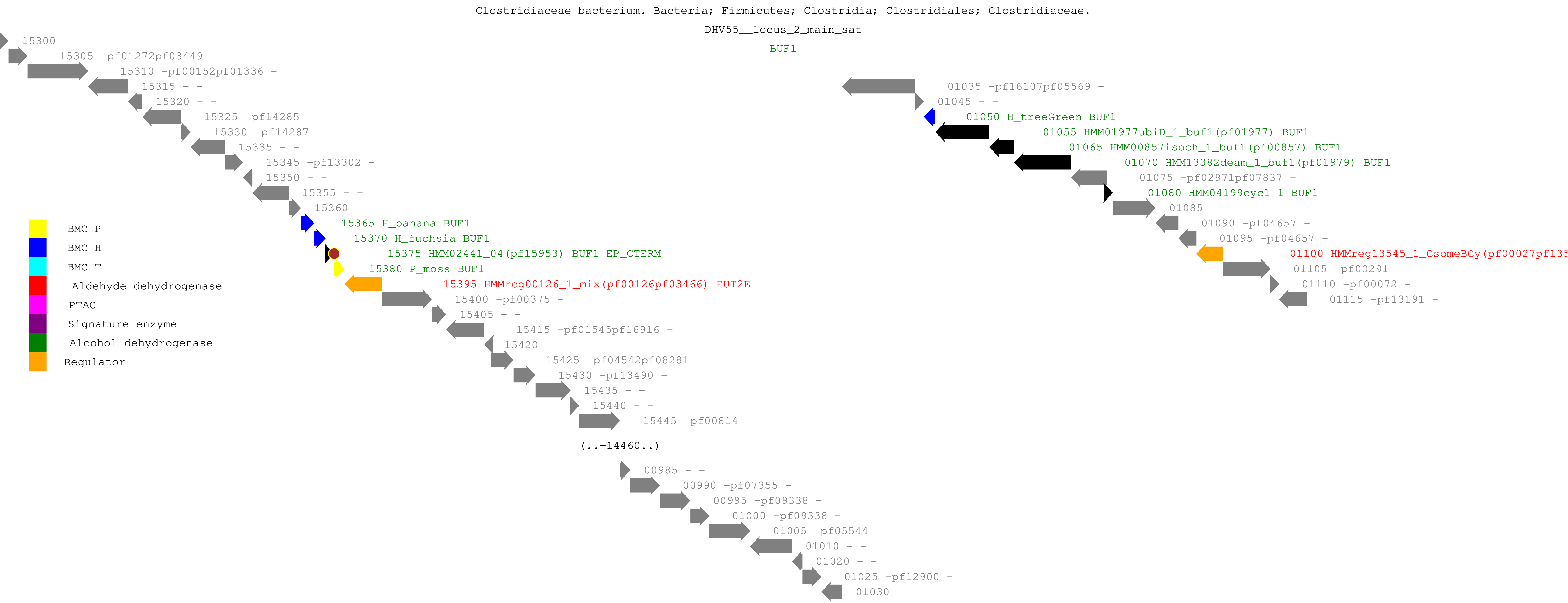
Clostridiaceae bacterium. Bacteria; Firmicutes; Clostridia; Clostridiales; Clostridiaceae.
DHV55__locus_2_main_sat
BUF1
15300 - -
15305 -pf01272pf03449 -
15310 -pf00152pf01336 -
15315 - -
15320 - -
15325 -pf14285 -
15330 -pf14287 -
15335 - -
15345 -pf13302 -
15350 - -
15355 - -
15360 - -
15365 H_banana BUF1
15370 H_fuchsia BUF1
15375 HMM02441_04(pf15953) BUF1 EP_CTERM
15380 P_moss BUF1
15395 HMMreg00126_1_mix(pf00126pf03466) EUT2E
15400 -pf00375 -
15405 - -
15415 -pf01545pf16916 -
15420 - -
15425 -pf04542pf08281 -
15430 -pf13490 -
15435 - -
15440 - -
15445 -pf00814 -
(..-14460..)
00985 - -
00990 -pf07355 -
00995 -pf09338 -
01000 -pf09338 -
01005 -pf05544 -
01010 - -
01020 - -
01025 -pf12900 -
01030 - -
01035 -pf16107pf05569 -
01045 - -
01050 H_treeGreen BUF1
01055 HMM01977ubiD_1_buf1(pf01977) BUF1
01065 HMM00857isoch_1_buf1(pf00857) BUF1
01070 HMM13382deam_1_buf1(pf01979) BUF1
01075 -pf02971pf07837 -
01080 HMM04199cycl_1 BUF1
01085 - -
01090 -pf04657 -
01095 -pf04657 -
01100 HMMreg13545_1_CsomeBCy(pf00027pf135
01105 -pf00291 -
01110 -pf00072 -
01115 -pf13191 -
BMC-P
BMC-H
BMC-T
Aldehyde dehydrogenase
PTAC
Signature enzyme
Alcohol dehydrogenase
Regulator

Acinetobacter sp. RIT592. Bacteria; Proteobacteria; Gammaproteobacteria; Pseudomonadales;

DVA85__locus_2

BUF1

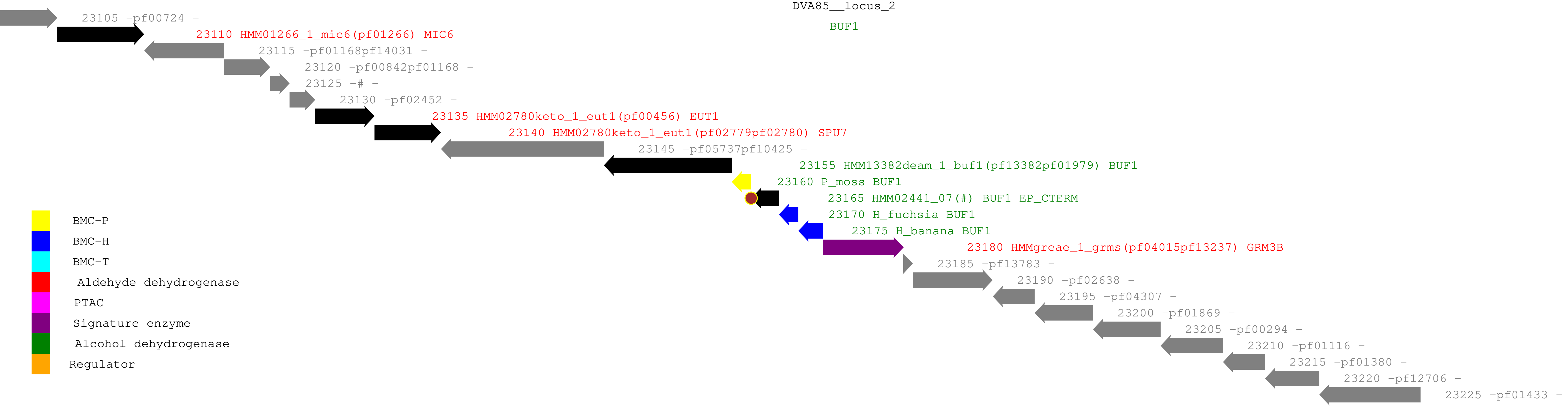

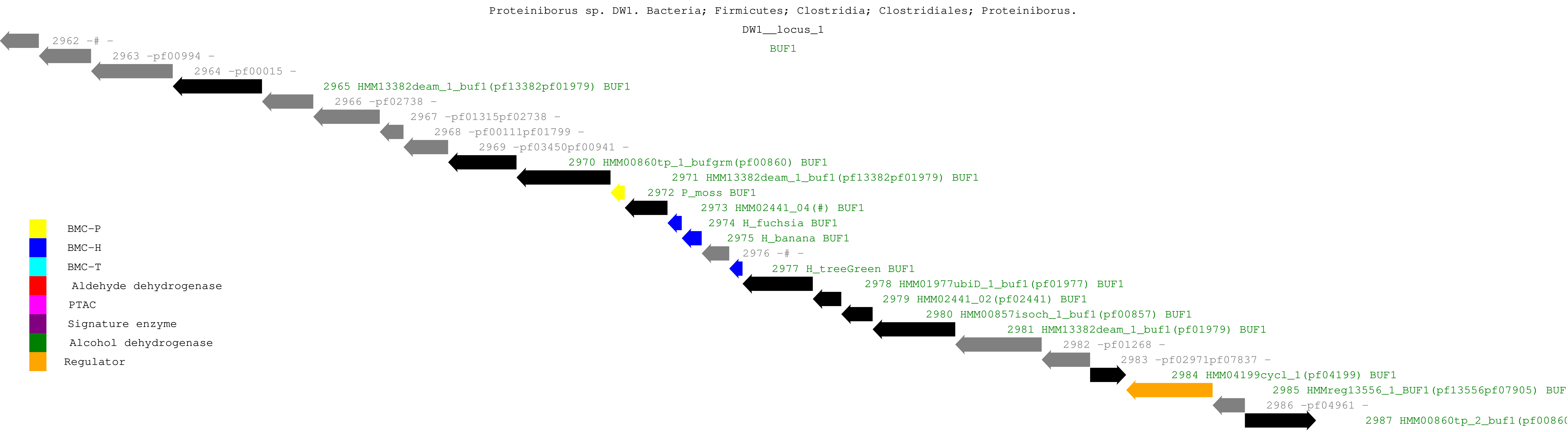
Proteiniborus sp. DW1. Bacteria; Firmicutes; Clostridia; Clostridiales; Proteiniborus.
DW1__locus_1
BUF1
2962 -# -
2963 -pf00994 -
2964 -pf00015 -
2965 HMM13382deam_1_buf1(pf13382pf01979) BUF1
2966 -pf02738 -
2967 -pf01315pf02738 -
2968 -pf00111pf01799 -
2969 -pf03450pf00941 -
2970 HMM00860tp_1_bufgrm(pf00860) BUF1
2971 HMM13382deam_1_buf1(pf13382pf01979) BUF1
2972 P_moss BUF1
2973 HMM02441_04(#) BUF1
2974 H_fuchsia BUF1
2975 H_banana BUF1
2976 -# -
2977 H_treeGreen BUF1
2978 HMM01977ubiD_1_buf1(pf01977) BUF1
2979 HMM02441_02(pf02441) BUF1
2980 HMM00857isoch_1_buf1(pf00857) BUF1
2981 HMM13382deam_1_buf1(pf01979) BUF1
2982 -pf01268 -
2983 -pf02971pf07837 -
2984 HMM04199cycl_1(pf04199) BUF1
2985 HMMreg13556_1_BUF1(pf13556pf07905) BUF
2986 -pf04961 -
2987 HMM00860tp_2_buf1(pf0086
BMC-P
BMC-H
BMC-T
Aldehyde dehydrogenase
PTAC
Signature enzyme
Alcohol dehydrogenase
Regulator

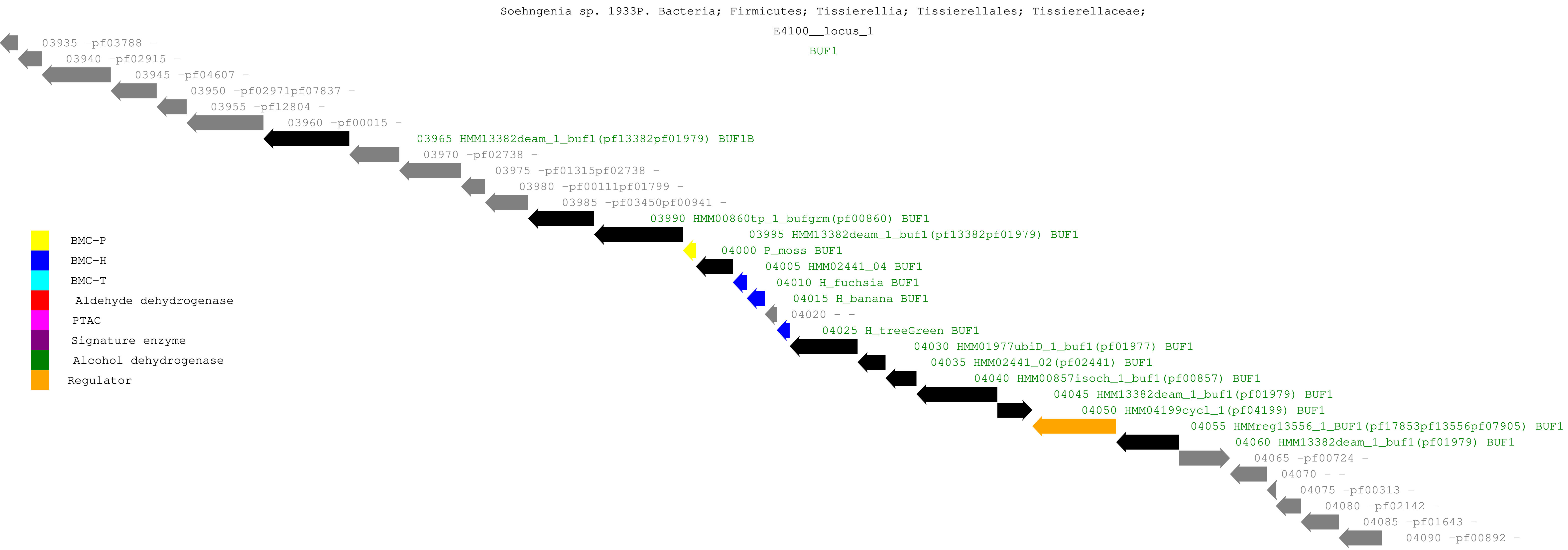

Soehngenia sp. 1933P. Bacteria; Firmicutes; Tissierellia; Tissierellales; Tissierellaceae;
E4100__locus_1
BUF1
03935 -pf03788 -
03940 -pf02915 -
03945 -pf04607 -
03950 -pf02971pf07837 -
03955 -pf12804 -
03960 -pf00015 -
03965 HMM13382deam_1_buf1(pf13382pf01979) BUF1B
03970 -pf02738 -
03975 -pf01315pf02738 -
03980 -pf00111pf01799 -
03985 -pf03450pf00941 -
03990 HMM00860tp_1_bufgrm(pf00860) BUF1
03995 HMM13382deam_1_buf1(pf13382pf01979) BUF1
04000 P_moss BUF1
04005 HMM02441_04 BUF1
04010 H_fuchsia BUF1
04015 H_banana BUF1
04020 - -
04025 H_treeGreen BUF1
04030 HMM01977ubiD_1_buf1(pf01977) BUF1
04035 HMM02441_02(pf02441) BUF1
04040 HMM00857isoch_1_buf1(pf00857) BUF1
04045 HMM13382deam_1_buf1(pf01979) BUF1
04050 HMM04199cycl_1(pf04199) BUF1
04055 HMMreg13556_1_BUF1(pf17853pf13556pf07905) BUF1
04060 HMM13382deam_1_buf1(pf01979) BUF1
04065 -pf00724 -
04070 - -
04075 -pf00313 -
04080 -pf02142 -
04085 -pf01643 -
04090 -pf00892 -
BMC-P
BMC-H
BMC-T
Aldehyde dehydrogenase
PTAC
Signature enzyme
Alcohol dehydrogenase
Regulator

Tissierella creatinini. Bacteria; Firmicutes; Tissierellia; Tissierellales; Tissierellaceae;

E9840__locus_1

BUF1

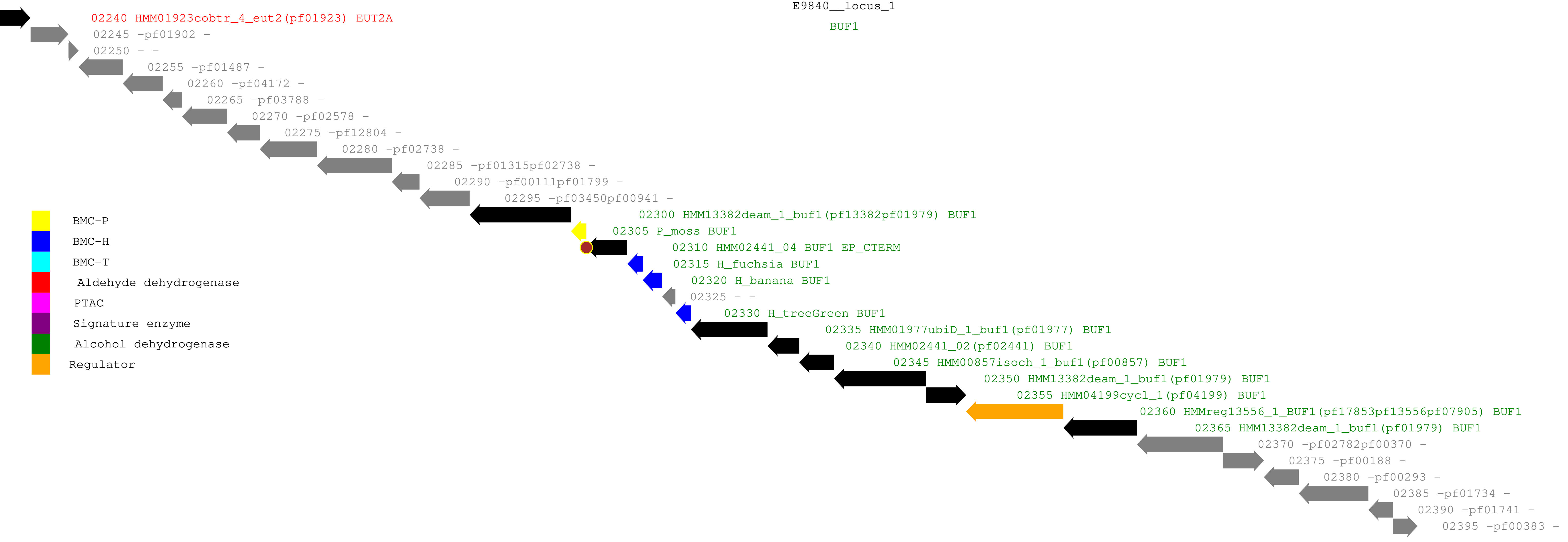

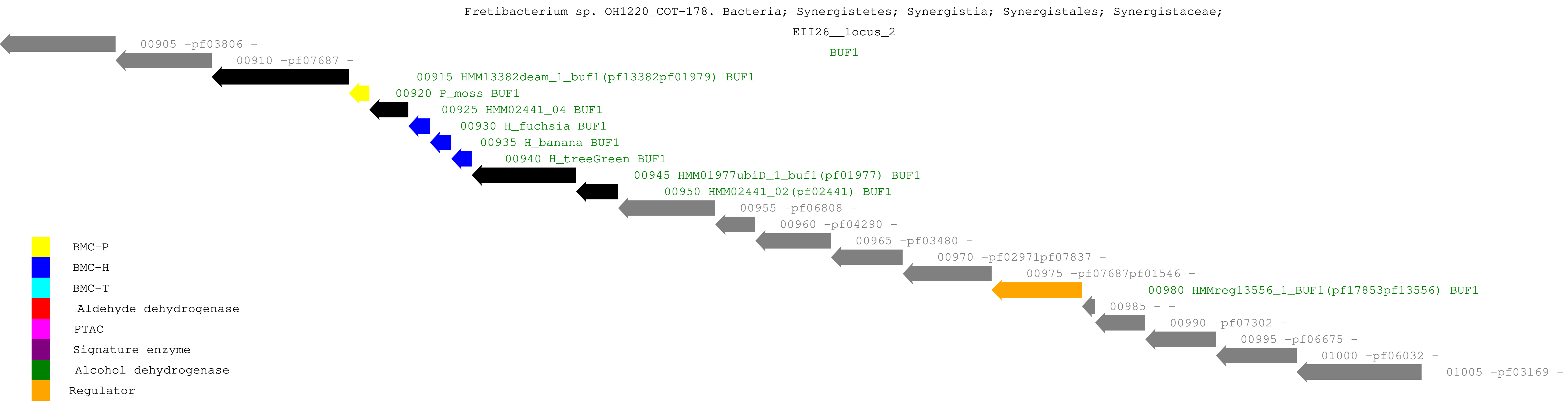
Fretibacterium sp. OH1220_COT-178. Bacteria; Synergistetes; Synergistia; Synergistales; Synergistaceae;
EII26__locus_2
BUF1
00905 -pf03806 -
00910 -pf07687 -
00915 HMM13382deam_1_buf1(pf13382pf01979) BUF1
00920 P_moss BUF1
00925 HMM02441_04 BUF1
00930 H_fuchsia BUF1
00935 H_banana BUF1
00940 H_treeGreen BUF1
00945 HMM01977ubiD_1_buf1(pf01977) BUF1
00950 HMM02441_02(pf02441) BUF1
00955 -pf06808 -
00960 -pf04290 -
00965 -pf03480 -
00970 -pf02971pf07837 -
00975 -pf07687pf01546 -
00980 HMMreg13556_1_BUF1(pf17853pf13556) BUF1
00985 - -
00990 -pf07302 -
00995 -pf06675 -
01000 -pf06032 -
01005 -pf03169 -
BMC-P
BMC-H
BMC-T
Aldehyde dehydrogenase
PTAC
Signature enzyme
Alcohol dehydrogenase
Regulator

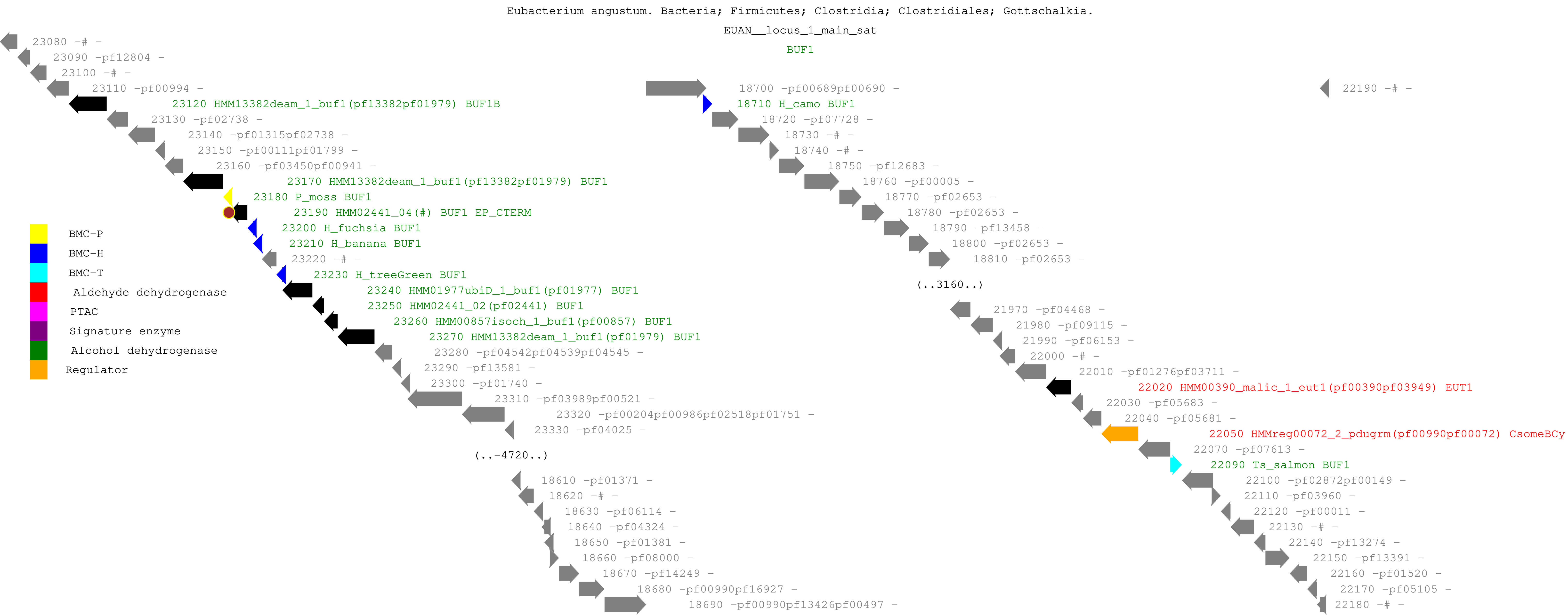
Eubacterium angustum. Bacteria; Firmicutes; Clostridia; Clostridiales; Gottschalkia.
EUAN__locus_1_main_sat
BUF1
23080 -# -
23090 -pf12804 -
23100 -# -
23110 -pf00994 -
23120 HMM13382deam_1_buf1(pf13382pf01979) BUF1B
23130 -pf02738 -
23140 -pf01315pf02738 -
23150 -pf00111pf01799 -
23160 -pf03450pf00941 -
23170 HMM13382deam_1_buf1(pf13382pf01979) BUF1
23180 P_moss BUF1
23190 HMM02441_04(#) BUF1 EP_CTERM
23200 H_fuchsia BUF1
23210 H_banana BUF1
23220 -# -
23230 H_treeGreen BUF1
23240 HMM01977ubiD_1_buf1(pf01977) BUF1
23250 HMM02441_02(pf02441) BUF1
23260 HMM00857isoch_1_buf1(pf00857) BUF1
23270 HMM13382deam_1_buf1(pf01979) BUF1
23280 -pf04542pf04539pf04545 -
23290 -pf13581 -
23300 -pf01740 -
23310 -pf03989pf00521 -
23320 -pf00204pf00986pf02518pf01751 -
23330 -pf04025 -
(..-4720..)
18610 -pf01371 -
18620 -# -
18630 -pf06114 -
18640 -pf04324 -
18650 -pf01381 -
18660 -pf08000 -
18670 -pf14249 -
18680 -pf00990pf16927 -
18690 -pf00990pf13426pf00497 -
BMC-P
BMC-H
BMC-T
Aldehyde dehydrogenase
PTAC
Signature enzyme
Alcohol dehydrogenase
Regulator
18700 -pf00689pf00690 -
18710 H_camo BUF1
18720 -pf07728 -
18730 -# -
18740 -# -
18750 -pf12683 -
18760 -pf00005 -
18770 -pf02653 -
18780 -pf02653 -
18790 -pf13458 -
18800 -pf02653 -
18810 -pf02653 -
(..3160..)
21970 -pf04468 -
21980 -pf09115 -
21990 -pf06153 -
22000 -# -
22010 -pf01276pf03711 -
22020 HMM00390_malic_1_eut1(pf00390pf03949) EUT1
22030 -pf05683 -
22040 -pf05681 -
22050 HMMreg00072_2_pdugrm(pf00990pf00072) CsomeBCy
22070 -pf07613 -
22090 Ts_salmon BUF1
22100 -pf02872pf00149 -
22110 -pf03960 -
22120 -pf00011 -
22130 -# -
22140 -pf13274 -
22150 -pf13391 -
22160 -pf01520 -
22170 -pf05105 -
22180 -# -
22190 -# -

Marinisporobacter balticus. Bacteria; Firmicutes; Clostridia; Clostridiales; Clostridiaceae;

EV214__locus_2

BUF1

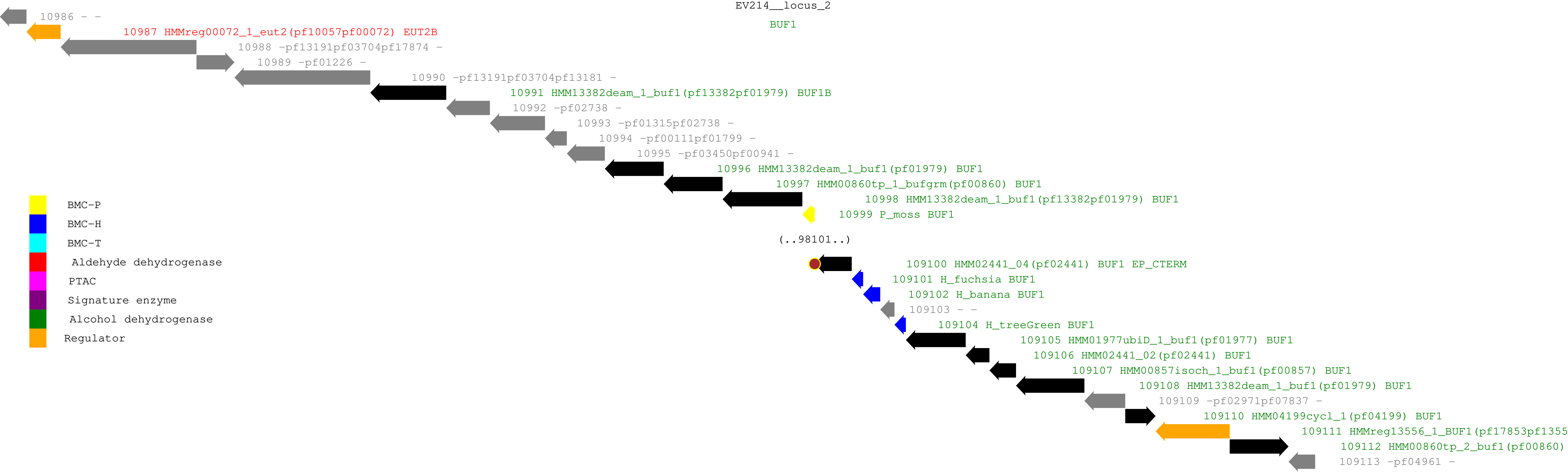

Crassaminicella sp. SY095. Bacteria; Firmicutes; Clostridia; Clostridiales; Clostridiaceae;

FQB35__locus_3

BUF1

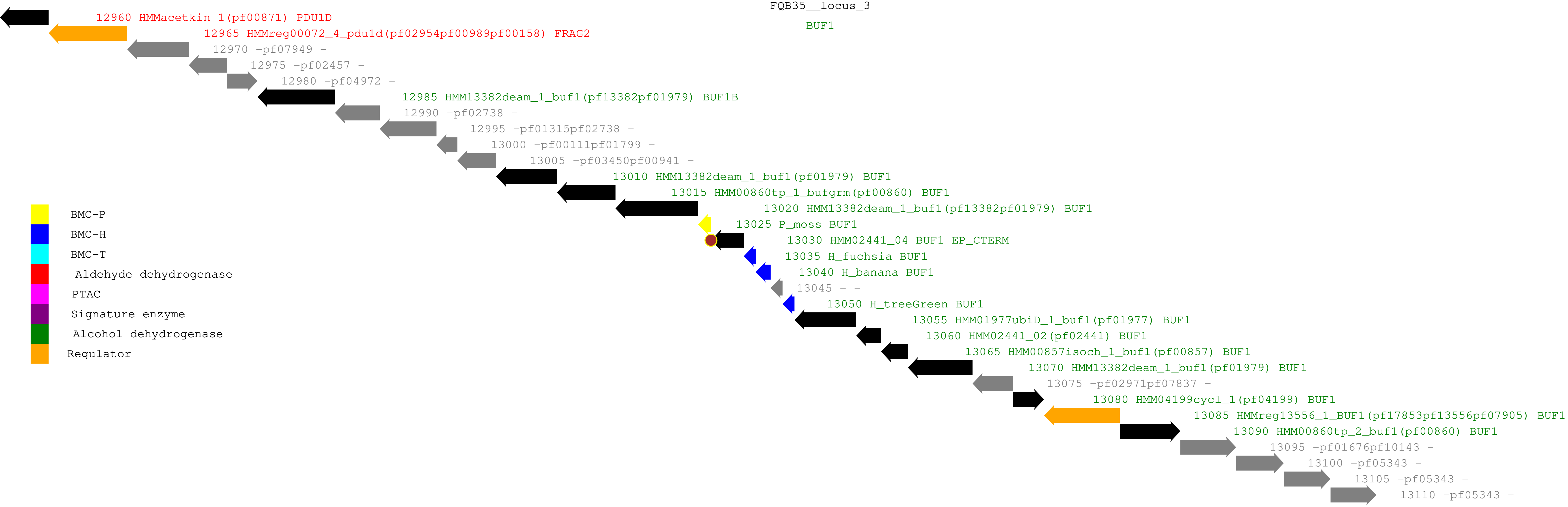

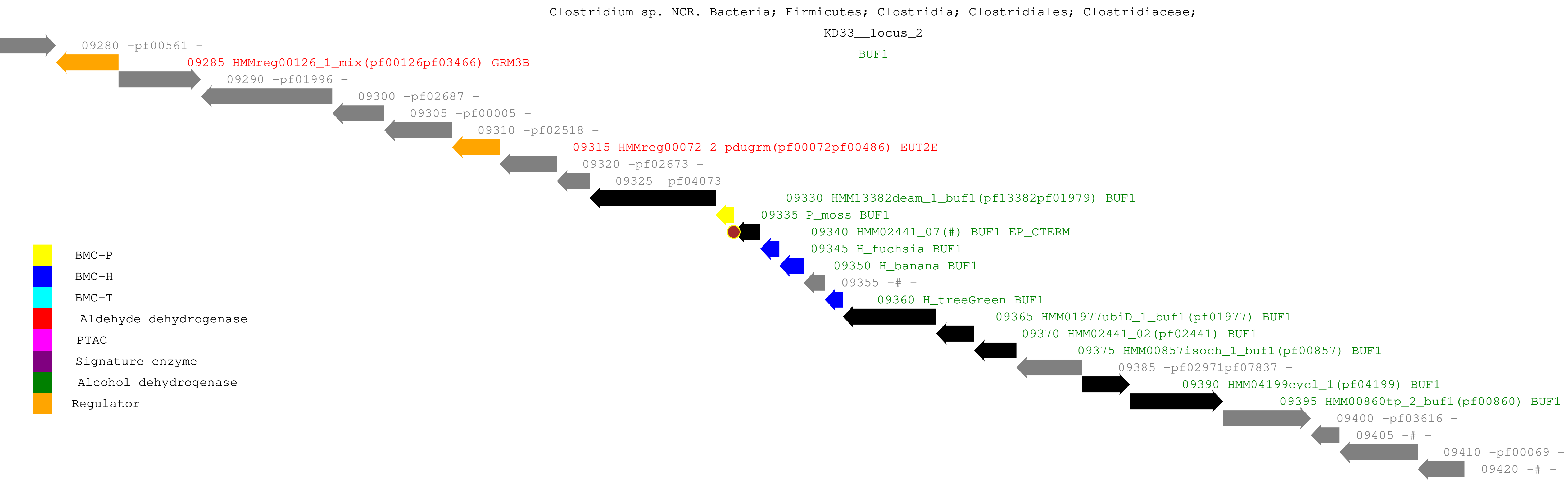
Clostridium sp. NCR. Bacteria; Firmicutes; Clostridia; Clostridiales; Clostridiaceae;
KD33__locus_2
BUF1
09280 -pf00561 -
09285 HMMreg00126_1_mix(pf00126pf03466) GRM3B
09290 -pf01996 -
09300 -pf02687 -
09305 -pf00005 -
09310 -pf02518 -
09315 HMMreg00072_2_pdugrm(pf00072pf00486) EUT2E
09320 -pf02673 -
09325 -pf04073 -
09330 HMM13382deam_1_buf1(pf13382pf01979) BUF1
09335 P_moss BUF1
09340 HMM02441_07(#) BUF1 EP_CTERM
09345 H_fuchsia BUF1
09350 H_banana BUF1
09355 -# -
09360 H_treeGreen BUF1
09365 HMM01977ubiD_1_buf1(pf01977) BUF1
09370 HMM02441_02(pf02441) BUF1
09375 HMM00857isoch_1_buf1(pf00857) BUF1
09385 -pf02971pf07837 -
09390 HMM04199cycl_1(pf04199) BUF1
09395 HMM00860tp_2_buf1(pf00860) BUF1
09400 -pf03616 -
09405 -# -
09410 -pf00069 -
09420 -# -
BMC-P
BMC-H
BMC-T
Aldehyde dehydrogenase
PTAC
Signature enzyme
Alcohol dehydrogenase
Regulator

Tissierella praeacuta. Bacteria; Firmicutes; Tissierellia; Tissierellales; Tissierellaceae;

NCTC11158__locus_1

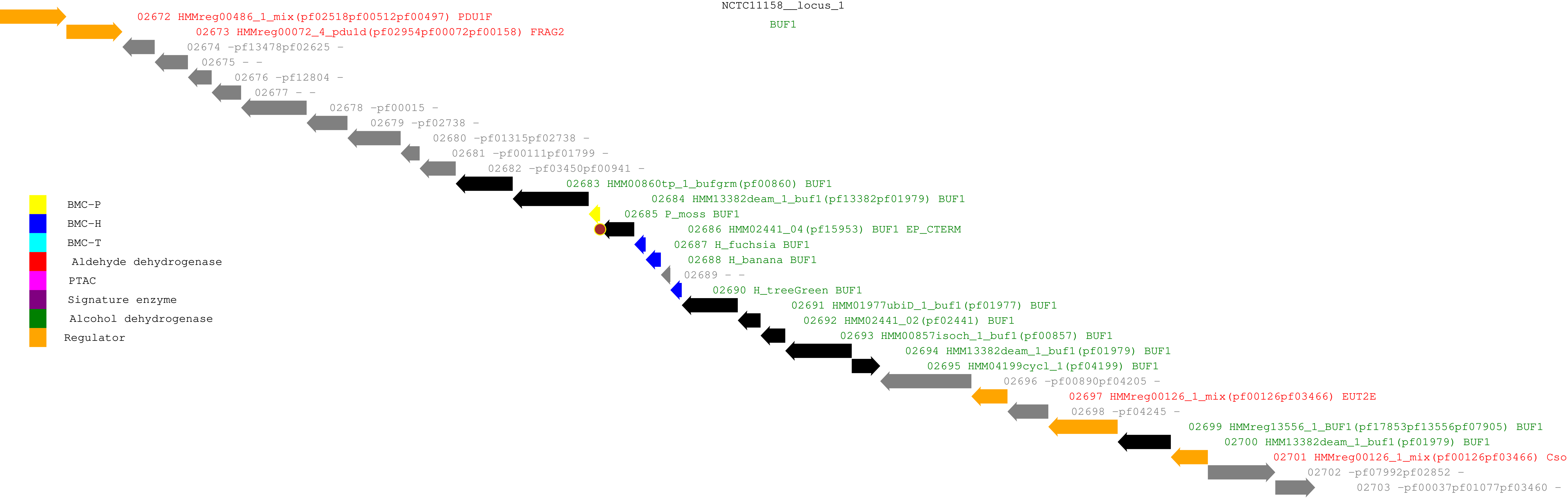

Maledivibacter halophilus. Bacteria; Firmicutes; Clostridia; Clostridiales; Clostridiaceae;

SAMN02194393__locus_3

BUF1

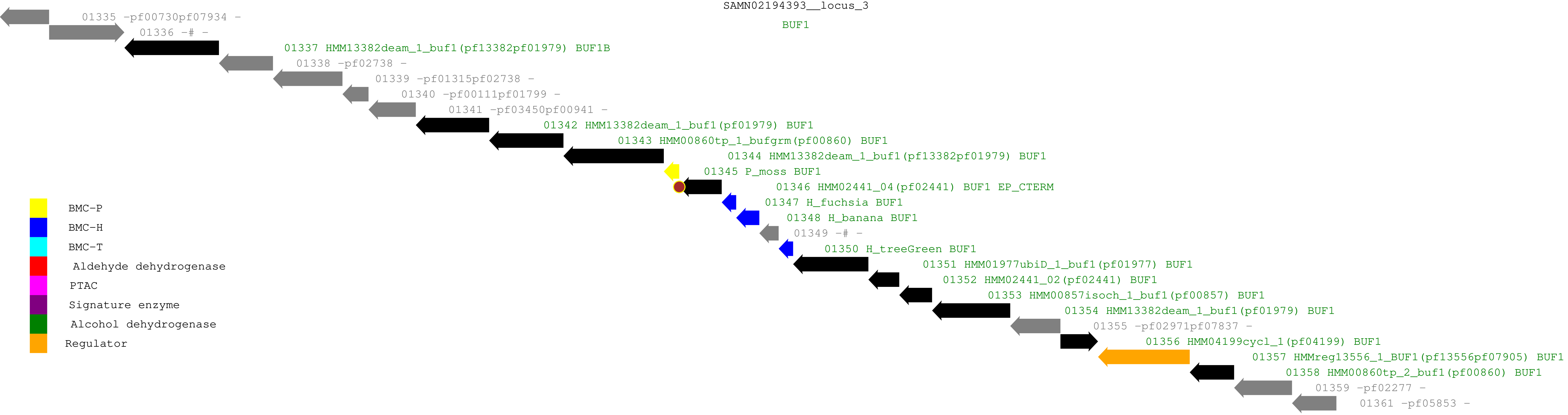

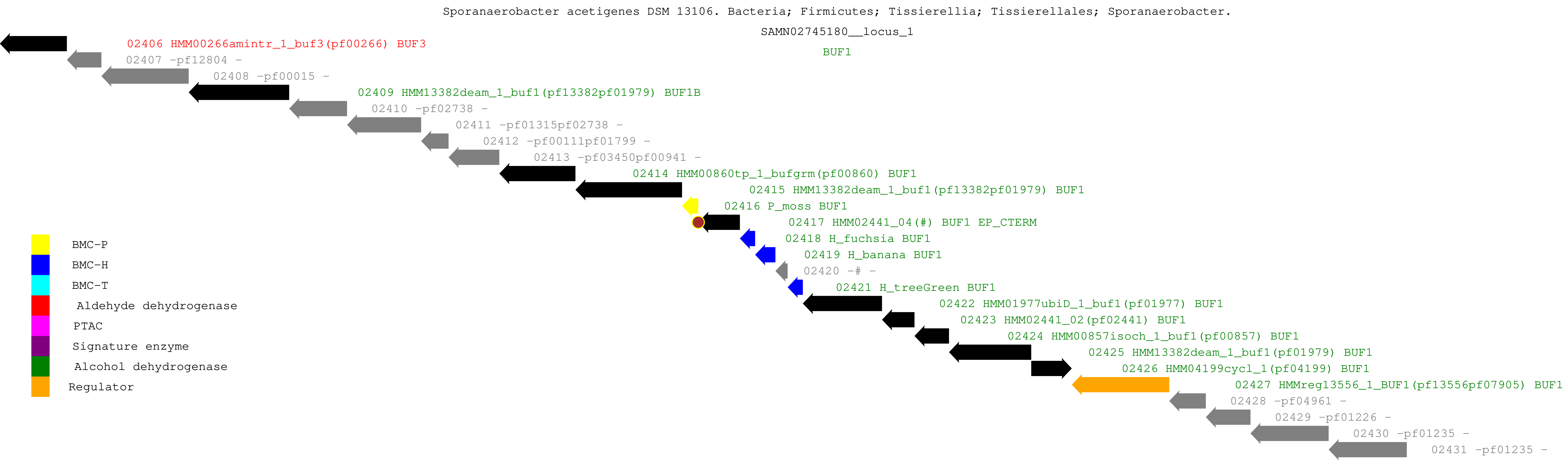
Sporanaerobacter acetigenes DSM 13106. Bacteria; Firmicutes; Tissierellia; Tissierellales; Sporanaerobacter.
SAMN02745180__locus_1
BUF1
02406 HMM00266amintr_1_buf3(pf00266) BUF3
02407 -pf12804 -
02408 -pf00015 -
02409 HMM13382deam_1_buf1(pf13382pf01979) BUF1B
02410 -pf02738 -
02411 -pf01315pf02738 -
02412 -pf00111pf01799 -
02413 -pf03450pf00941 -
02414 HMM00860tp_1_bufgrm(pf00860) BUF1
02415 HMM13382deam_1_buf1(pf13382pf01979) BUF1
02416 P_moss BUF1
02417 HMM02441_04(#) BUF1 EP_CTERM
02418 H_fuchsia BUF1
02419 H_banana BUF1
02420 -# -
02421 H_treeGreen BUF1
02422 HMM01977ubiD_1_buf1(pf01977) BUF1
02423 HMM02441_02(pf02441) BUF1
02424 HMM00857isoch_1_buf1(pf00857) BUF1
02425 HMM13382deam_1_buf1(pf01979) BUF1
02426 HMM04199cycl_1(pf04199) BUF1
02427 HMMreg13556_1_BUF1(pf13556pf07905) BUF1
02428 -pf04961 -
02429 -pf01226 -
02430 -pf01235 -
02431 -pf01235 -
BMC-P
BMC-H
BMC-T
Aldehyde dehydrogenase
PTAC
Signature enzyme
Alcohol dehydrogenase
Regulator

Tissierella praeacuta DSM 18095. Bacteria; Firmicutes; Tissierellia; Tissierellales; Tissierellaceae;

SAMN02745784__locus_1

BUF1

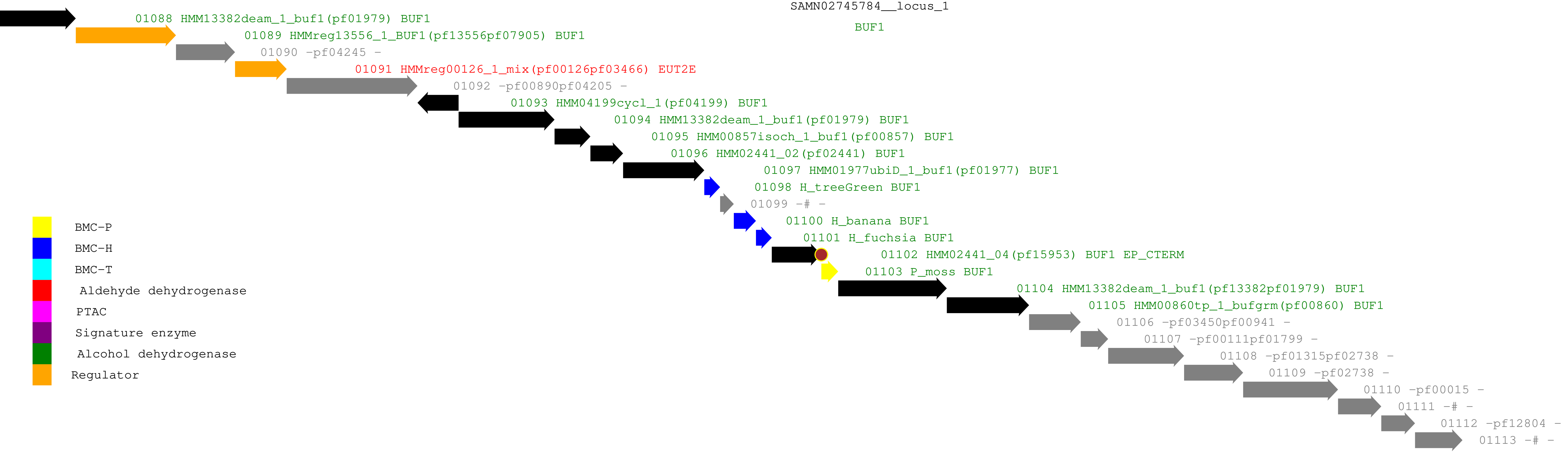

Acidaminobacter hydrogenoformans DSM 2784. Bacteria; Firmicutes; Clostridia; Clostridiales;

SAMN03080599__locus_1_main_sat

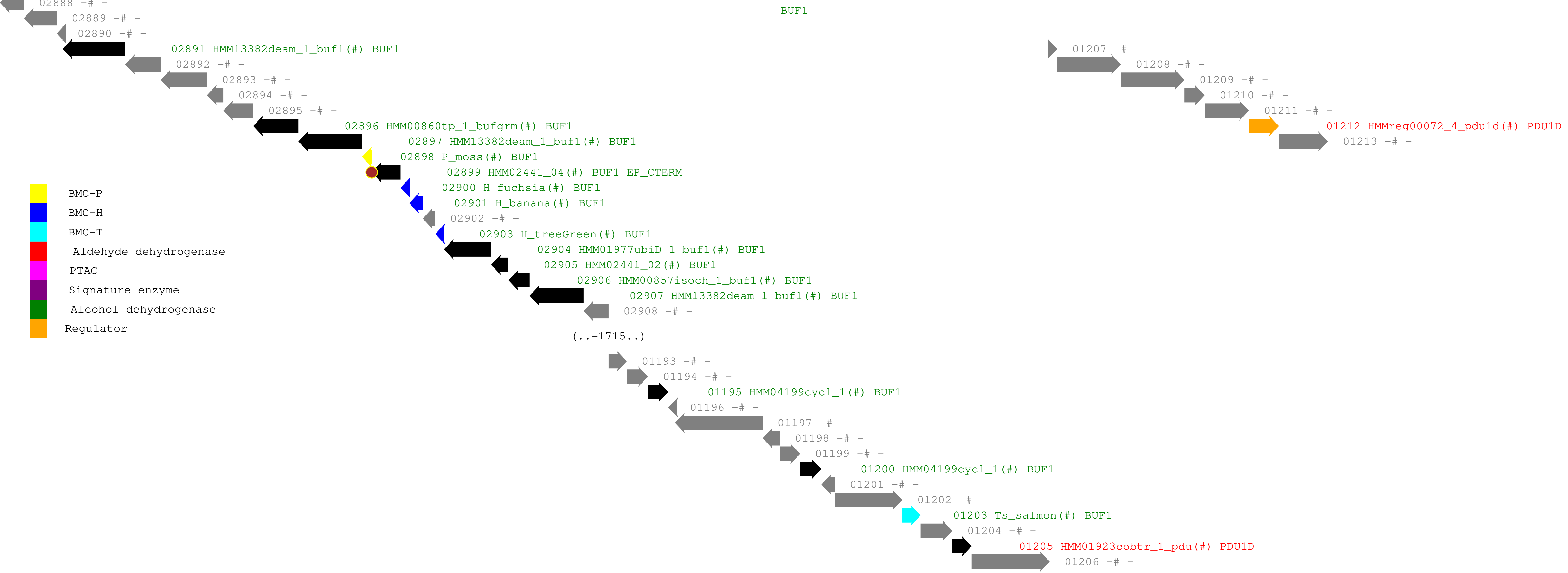

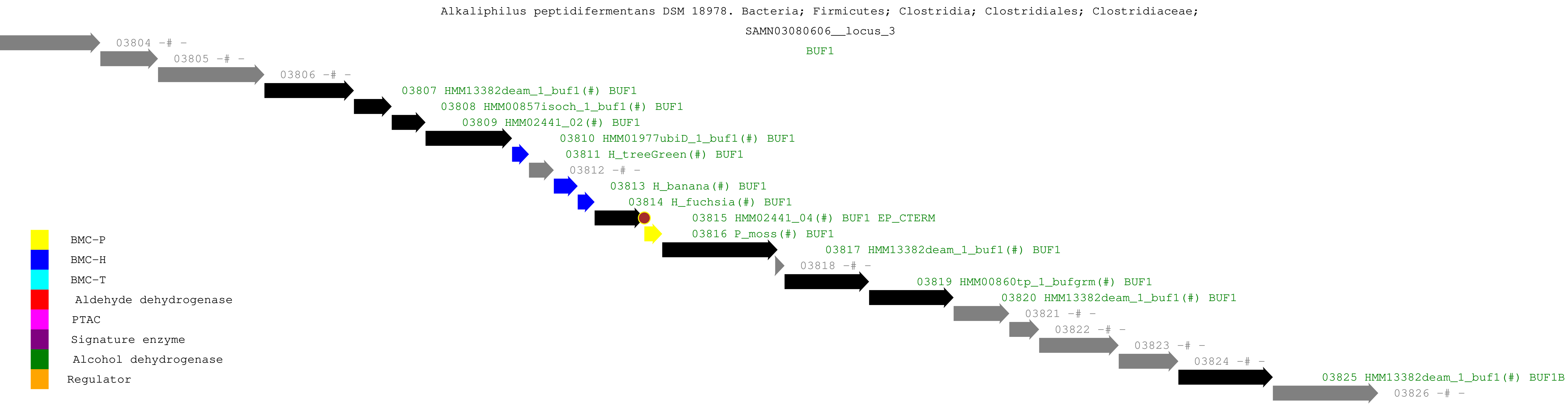

Alkaliphilus peptidifermentans DSM 18978. Bacteria; Firmicutes; Clostridia; Clostridiales; Clostridiaceae;
SAMN03080606__locus_3
BUF1
03804 -# -
03805 -# -
03806 -# -
03807 HMM13382deam_1_buf1(#) BUF1
03808 HMM00857isoch_1_buf1(#) BUF1
03809 HMM02441_02(#) BUF1
03810 HMM01977ubiD_1_buf1(#) BUF1
03811 H_treeGreen(#) BUF1
03812 -# -
03813 H_banana(#) BUF1
03814 H_fuchsia(#) BUF1
03815 HMM02441_04(#) BUF1 EP_CTERM
03816 P_moss(#) BUF1
03817 HMM13382deam_1_buf1(#) BUF1
03818 -# -
03819 HMM00860tp_1_bufgrm(#) BUF1
03820 HMM13382deam_1_buf1(#) BUF1
03821 -# -
03822 -# -
03823 -# -
03824 -# -
03825 HMM13382deam_1_buf1(#) BUF1B
03826 -# -
BMC-P
BMC-H
BMC-T
Aldehyde dehydrogenase
PTAC
Signature enzyme
Alcohol dehydrogenase
Regulator

Romboutsia lituseburensis DSM 797. Bacteria; Firmicutes; Clostridia; Clostridiales;

SAMN04515677__locus_2

BUF1

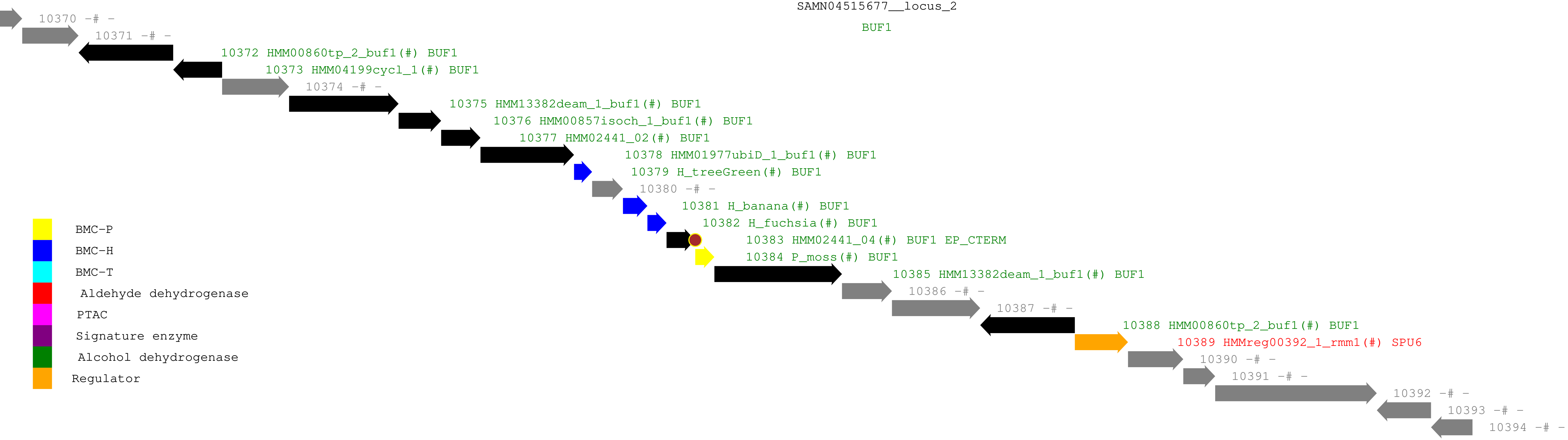

Tindallia californiensis. Bacteria; Firmicutes; Clostridia; Clostridiales; Clostridiaceae;

SAMN05192546__locus_2

BUF1

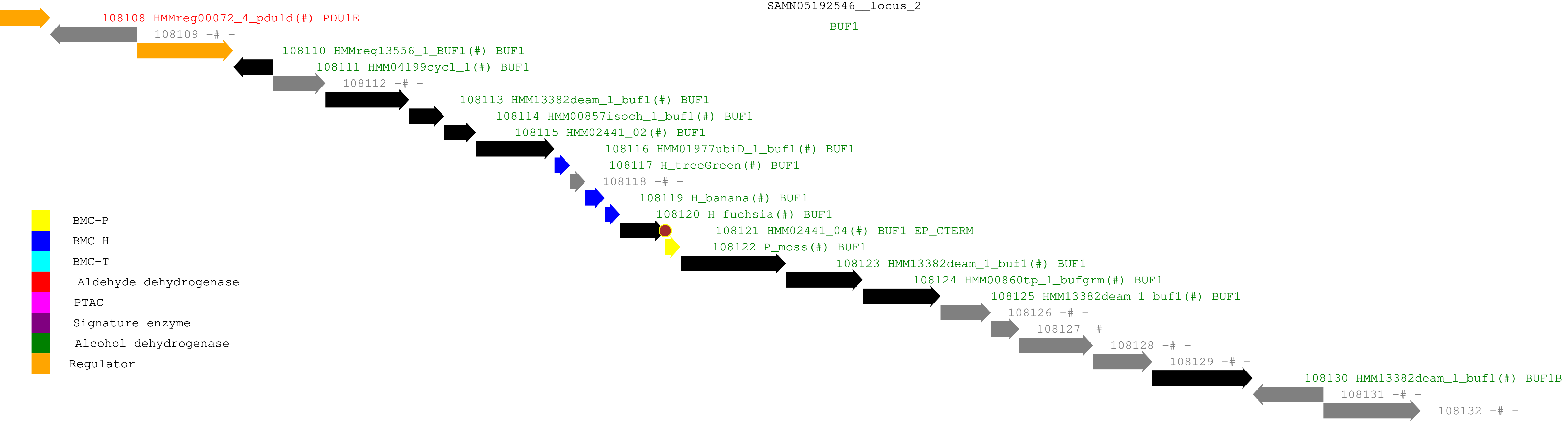

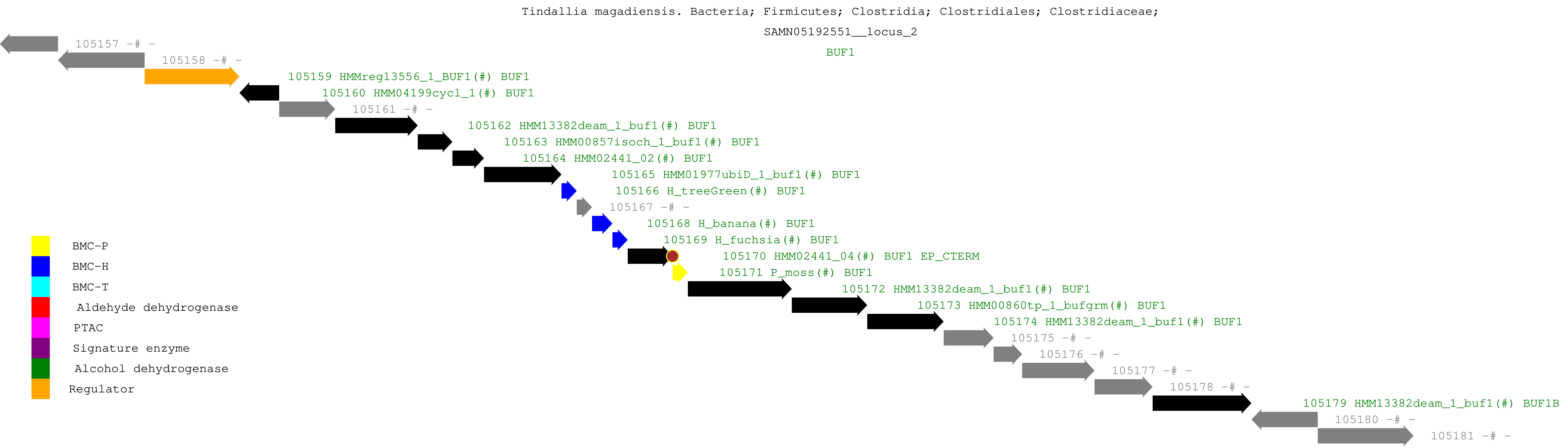

Tindallia magadiensis. Bacteria; Firmicutes; Clostridia; Clostridiales; Clostridiaceae;
SAMN05192551__locus_2
BUF1
105157 -# -
105158 -# -
105159 HMMreg13556_1_BUF1(#) BUF1
105160 HMM04199cycl_1(#) BUF1
105161 -# -
105162 HMM13382deam_1_buf1(#) BUF1
105163 HMM00857isoch_1_buf1(#) BUF1
105164 HMM02441_02(#) BUF1
105165 HMM01977ubiD_1_buf1(#) BUF1
105166 H_treeGreen(#) BUF1
105167 -# -
105168 H_banana(#) BUF1
105169 H_fuchsia(#) BUF1
105170 HMM02441_04(#) BUF1 EP_CTERM
105171 P_moss(#) BUF1
105172 HMM13382deam_1_buf1(#) BUF1
105173 HMM00860tp_1_bufgrm(#) BUF1
105174 HMM13382deam_1_buf1(#) BUF1
105175 -# -
105176 -# -
105177 -# -
105178 -# -
105179 HMM13382deam_1_buf1(#) BUF1B
105180 -# -
105181 -# -
BMC-P
BMC-H
BMC-T
Aldehyde dehydrogenase
PTAC
Signature enzyme
Alcohol dehydrogenase
Regulator

Anaerovirgula multivorans. Bacteria; Firmicutes; Clostridia; Clostridiales; Anaerovirgula.

SAMN05446037__locus_1

BUF1

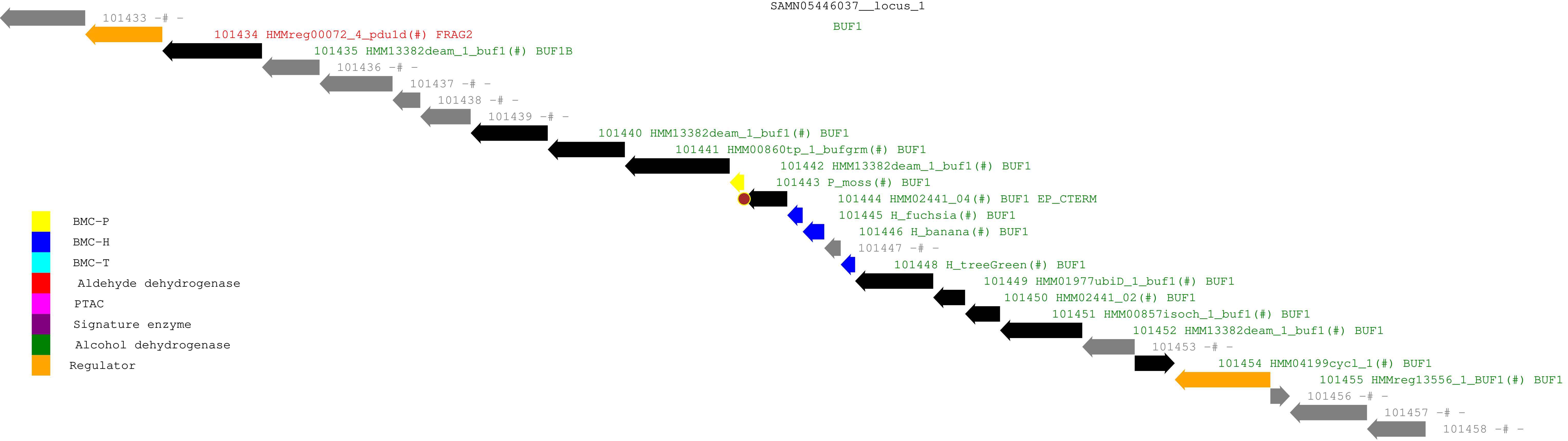

Natronincola peptidivorans. Bacteria; Firmicutes; Clostridia; Clostridiales; Clostridiaceae;

SAMN05660297__locus_4

BUF1

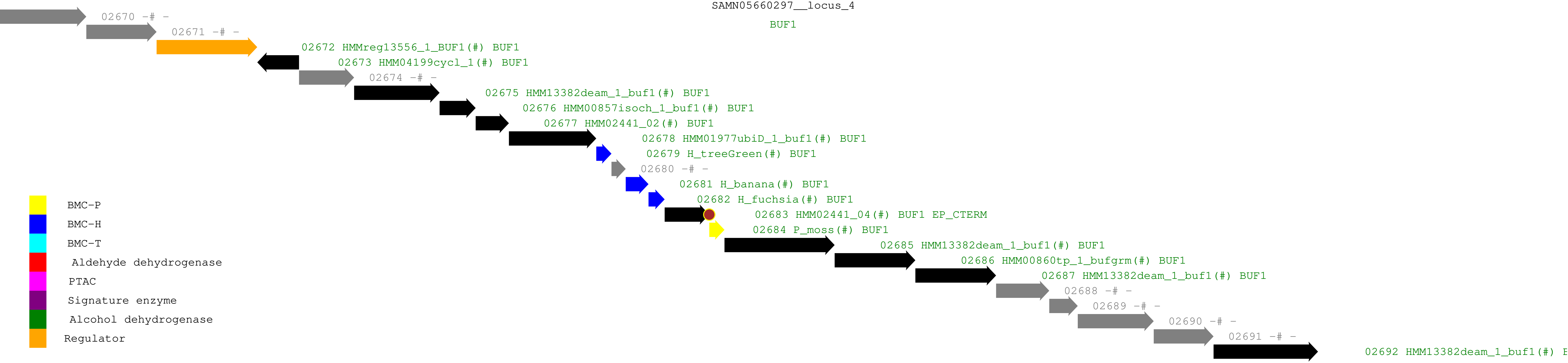

Proteiniborus ethanoligenes. Bacteria; Firmicutes; Clostridia; Clostridiales; Proteiniborus.

SAMN05660462__locus_1

BUF1

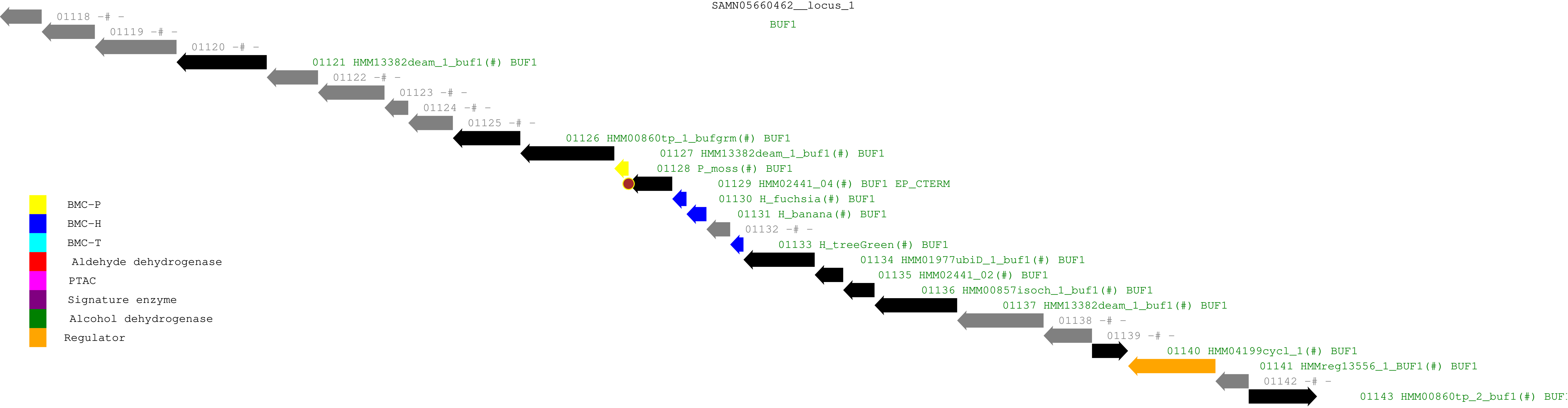

Caloramator fervidus. Bacteria; Firmicutes; Clostridia; Clostridiales; Clostridiaceae;

SAMN05660865__locus_1

BUF1

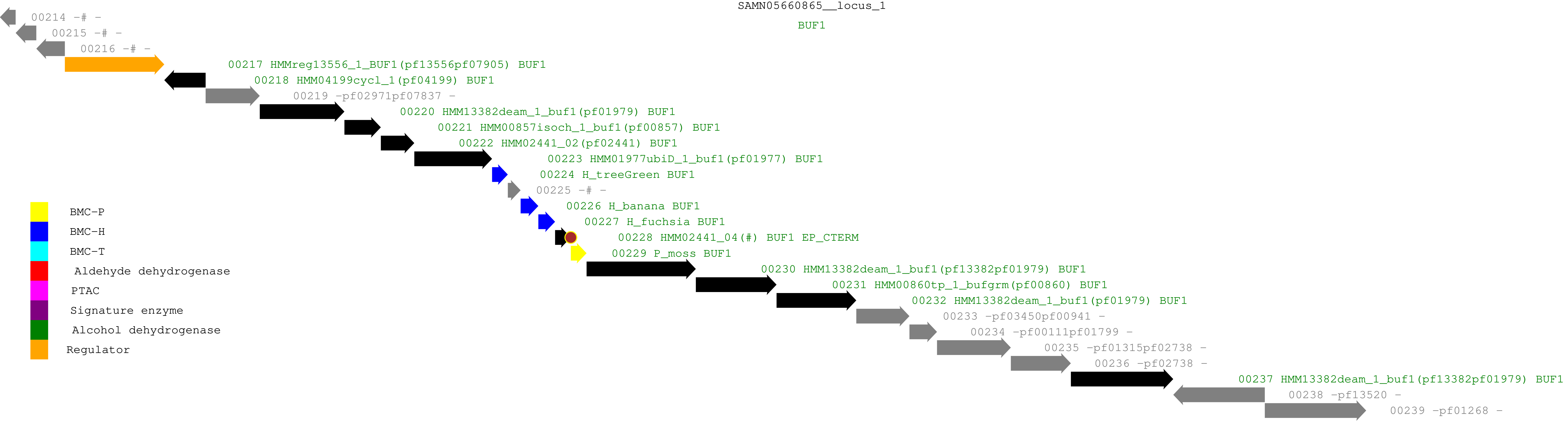

Tepidimicrobium xylanilyticum. Bacteria; Firmicutes; Tissierellia; Tepidimicrobium.

SAMN05660923__locus_1

BUF1

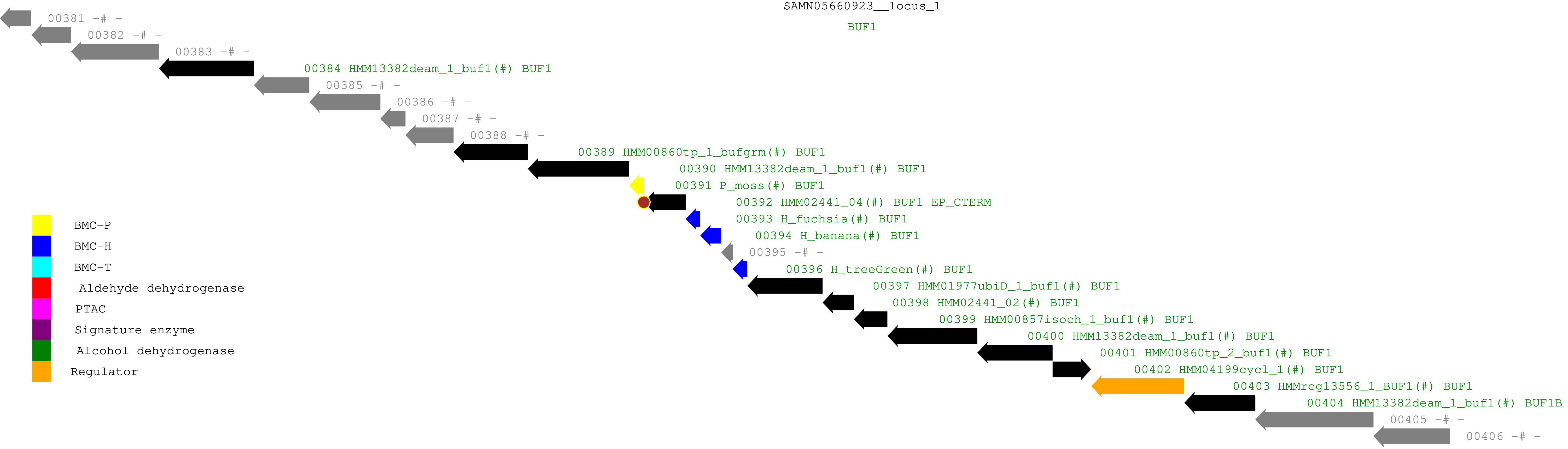

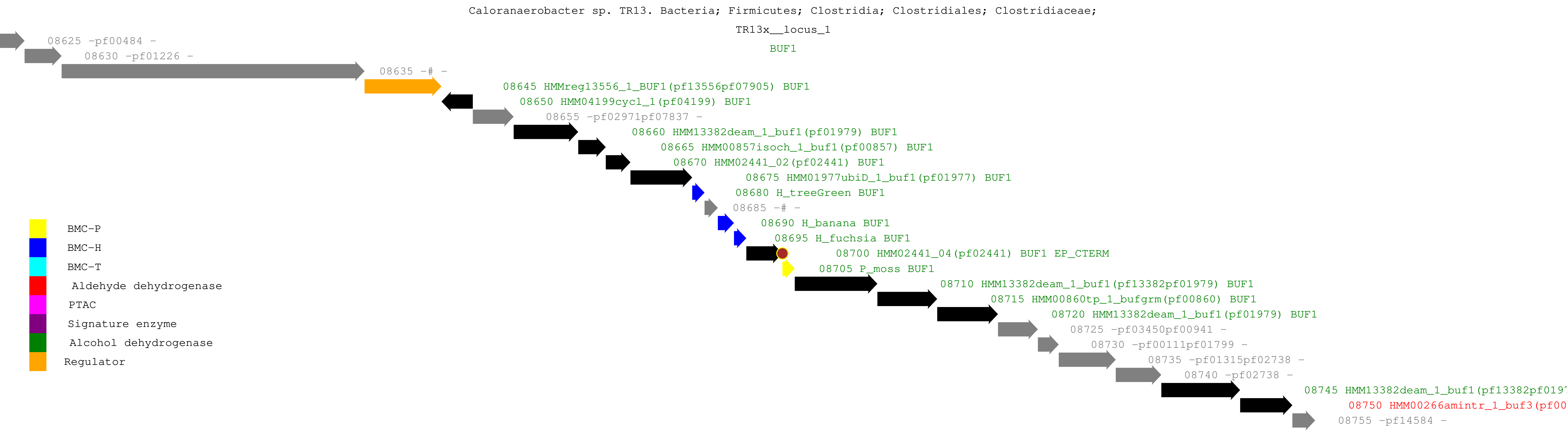

Caloranaerobacter sp. TR13. Bacteria; Firmicutes; Clostridia; Clostridiales; Clostridiaceae;
TR13x__locus_1
BUF1
08625 -pf00484 -
08630 -pf01226 -
08635 -# -
08645 HMMreg13556_1_BUF1(pf13556pf07905) BUF1
08650 HMM04199cycl_1(pf04199) BUF1
08655 -pf02971pf07837 -
08660 HMM13382deam_1_buf1(pf01979) BUF1
08665 HMM00857isoch_1_buf1(pf00857) BUF1
08670 HMM02441_02(pf02441) BUF1
08675 HMM01977ubiD_1_buf1(pf01977) BUF1
08680 H_treeGreen BUF1
08685 -# -
08690 H_banana BUF1
08695 H_fuchsia BUF1
08700 HMM02441_04(pf02441) BUF1 EP_CTERM
08705 P_moss BUF1
08710 HMM13382deam_1_buf1(pf13382pf01979) BUF1
08715 HMM00860tp_1_bufgrm(pf00860) BUF1
08720 HMM13382deam_1_buf1(pf01979) BUF1
08725 -pf03450pf00941 -
08730 -pf00111pf01799 -
08735 -pf01315pf02738 -
08740 -pf02738 -
08755 -pf14584 -
BMC-P
BMC-H
BMC-T
Aldehyde dehydrogenase
PTAC
Signature enzyme
Alcohol dehydrogenase
Regulator

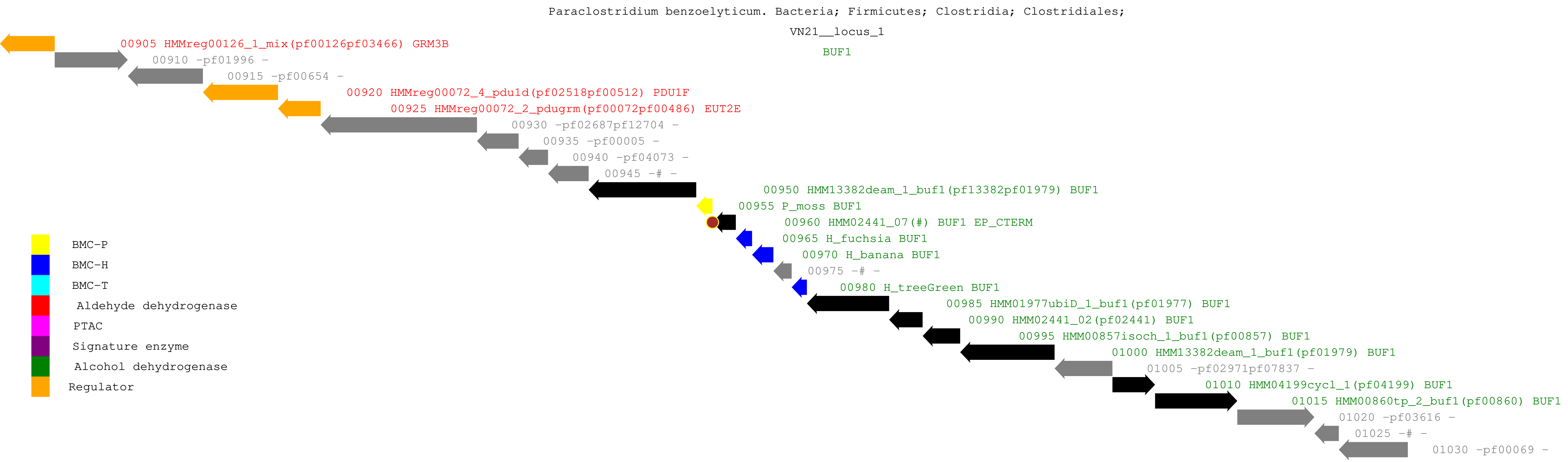
Paraclostridium benzoelyticum. Bacteria; Firmicutes; Clostridia; Clostridiales;
VN21__locus_1
BUF1
00905 HMMreg00126_1_mix(pf00126pf03466) GRM3B
00910 -pf01996 -
00915 -pf00654 -
00920 HMMreg00072_4_pdu1d(pf02518pf00512) PDU1F
00925 HMMreg00072_2_pdugrm(pf00072pf00486) EUT2E
00930 -pf02687pf12704 -
00935 -pf00005 -
00940 -pf04073 -
00945 -# -
00950 HMM13382deam_1_buf1(pf13382pf01979) BUF1
00955 P_moss BUF1
00960 HMM02441_07(#) BUF1 EP_CTERM
00965 H_fuchsia BUF1
00970 H_banana BUF1
00975 -# -
00980 H_treeGreen BUF1
00985 HMM01977ubiD_1_buf1(pf01977) BUF1
00990 HMM02441_02(pf02441) BUF1
00995 HMM00857isoch_1_buf1(pf00857) BUF1
01000 HMM13382deam_1_buf1(pf01979) BUF1
01005 -pf02971pf07837 -
01010 HMM04199cycl_1(pf04199) BUF1
01015 HMM00860tp_2_buf1(pf00860) BUF1
01020 -pf03616 -
01025 -# -
01030 -pf00069 -
BMC-P
BMC-H
BMC-T
Aldehyde dehydrogenase
PTAC
Signature enzyme
Alcohol dehydrogenase
Regulator

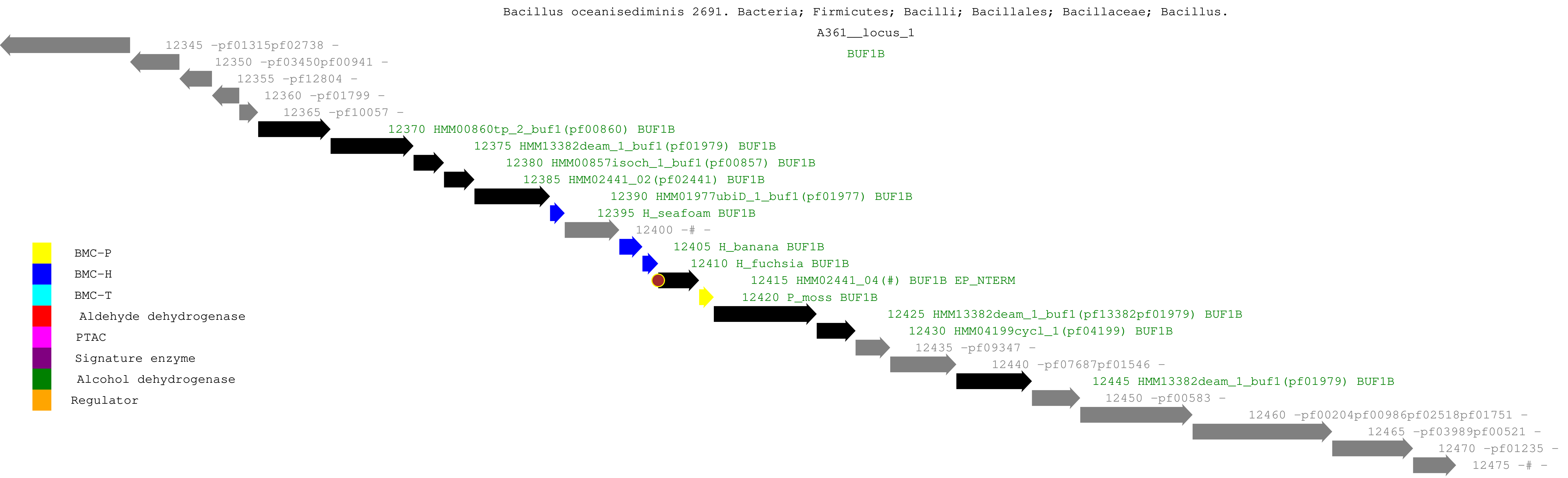

Bacillus oceanisediminis 2691. Bacteria; Firmicutes; Bacilli; Bacillales; Bacillaceae; Bacillus.
A361__locus_1
BUF1B
12345 -pf01315pf02738 -
12350 -pf03450pf00941 -
12355 -pf12804 -
12360 -pf01799 -
12365 -pf10057 -
12370 HMM00860tp_2_buf1(pf00860) BUF1B
12375 HMM13382deam_1_buf1(pf01979) BUF1B
12380 HMM00857isoch_1_buf1(pf00857) BUF1B
12385 HMM02441_02(pf02441) BUF1B
12390 HMM01977ubiD_1_buf1(pf01977) BUF1B
12395 H_seafoam BUF1B
12400 -# -
12405 H_banana BUF1B
12410 H_fuchsia BUF1B
12415 HMM02441_04(#) BUF1B EP_NTERM
12420 P_moss BUF1B
12425 HMM13382deam_1_buf1(pf13382pf01979) BUF1B
12430 HMM04199cycl_1(pf04199) BUF1B
12435 -pf09347 -
12440 -pf07687pf01546 -
12445 HMM13382deam_1_buf1(pf01979) BUF1B
12450 -pf00583 -
12460 -pf00204pf00986pf02518pf01751 -
12465 -pf03989pf00521 -
12470 -pf01235 -
12475 -# -
BMC-P
BMC-H
BMC-T
Aldehyde dehydrogenase
PTAC
Signature enzyme
Alcohol dehydrogenase
Regulator

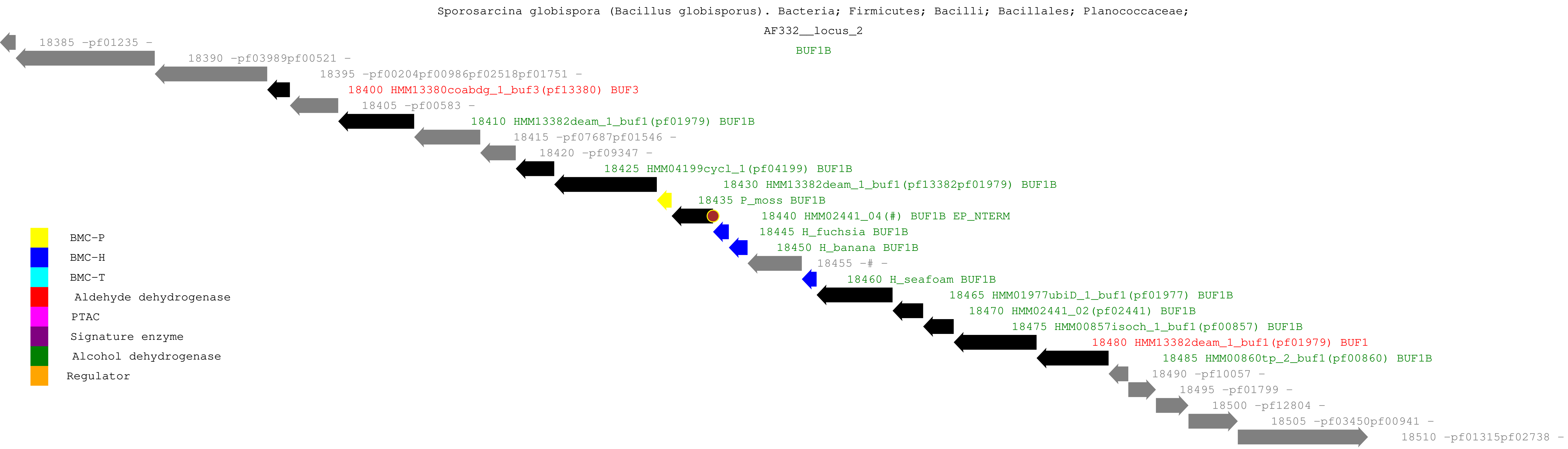

Sporosarcina globispora (Bacillus globisporus). Bacteria; Firmicutes; Bacilli; Bacillales; Planococcaceae;
AF332__locus_2
BUF1B
18385 -pf01235 -
18390 -pf03989pf00521 -
18395 -pf00204pf00986pf02518pf01751 -
18400 HMM13380coabdg_1_buf3(pf13380) BUF3
18405 -pf00583 -
18410 HMM13382deam_1_buf1(pf01979) BUF1B
18415 -pf07687pf01546 -
18420 -pf09347 -
18425 HMM04199cycl_1(pf04199) BUF1B
18430 HMM13382deam_1_buf1(pf13382pf01979) BUF1B
18435 P_moss BUF1B
18440 HMM02441_04(#) BUF1B EP_NTERM
18445 H_fuchsia BUF1B
18450 H_banana BUF1B
18455 -# -
18460 H_seafoam BUF1B
18465 HMM01977ubiD_1_buf1(pf01977) BUF1B
18470 HMM02441_02(pf02441) BUF1B
18475 HMM00857isoch_1_buf1(pf00857) BUF1B
18480 HMM13382deam_1_buf1(pf01979) BUF1
18485 HMM00860tp_2_buf1(pf00860) BUF1B
18490 -pf10057 -
18495 -pf01799 -
18500 -pf12804 -
18505 -pf03450pf00941 -
18510 -pf01315pf02738 -
BMC-P
BMC-H
BMC-T
Aldehyde dehydrogenase
PTAC
Signature enzyme
Alcohol dehydrogenase
Regulator

Aneurinibacillus migulanus (Bacillus migulanus). Bacteria; Firmicutes; Bacilli; Bacillales; Paenibacillaceae;

AF333__locus_2

BUF1B

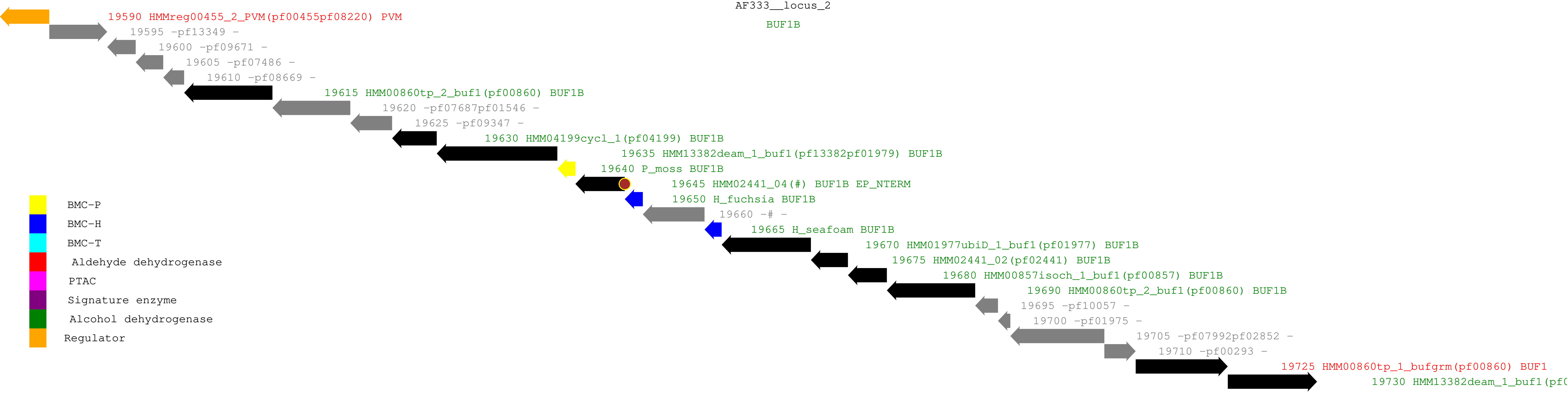

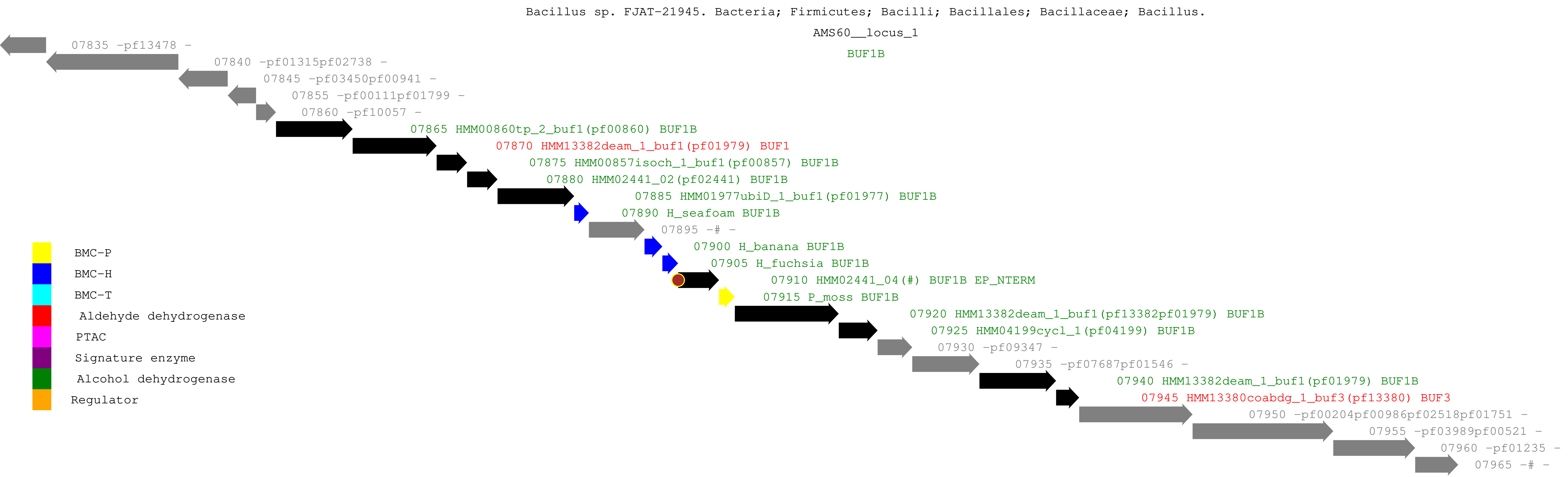

Bacillus sp. FJAT-21945. Bacteria; Firmicutes; Bacilli; Bacillales; Bacillaceae; Bacillus.
AMS60__locus_1
BUF1B
07835 -pf13478 -
07840 -pf01315pf02738 -
07845 -pf03450pf00941 -
07855 -pf00111pf01799 -
07860 -pf10057 -
07865 HMM00860tp_2_buf1(pf00860) BUF1B
07870 HMM13382deam_1_buf1(pf01979) BUF1
07875 HMM00857isoch_1_buf1(pf00857) BUF1B
07880 HMM02441_02(pf02441) BUF1B
07885 HMM01977ubiD_1_buf1(pf01977) BUF1B
07890 H_seafoam BUF1B
07895 -# -
07900 H_banana BUF1B
07905 H_fuchsia BUF1B
07910 HMM02441_04(#) BUF1B EP_NTERM
07915 P_moss BUF1B
07920 HMM13382deam_1_buf1(pf13382pf01979) BUF1B
07925 HMM04199cycl_1(pf04199) BUF1B
07930 -pf09347 -
07935 -pf07687pf01546 -
07940 HMM13382deam_1_buf1(pf01979) BUF1B
07945 HMM13380coabdg_1_buf3(pf13380) BUF3
07950 -pf00204pf00986pf02518pf01751 -
07955 -pf03989pf00521 -
07960 -pf01235 -
07965 -# -
BMC-P
BMC-H
BMC-T
Aldehyde dehydrogenase
PTAC
Signature enzyme
Alcohol dehydrogenase
Regulator

Bacillus solani. Bacteria; Firmicutes; Bacilli; Bacillales; Bacillaceae; Bacillus.

AN957__locus_1

BUF1B

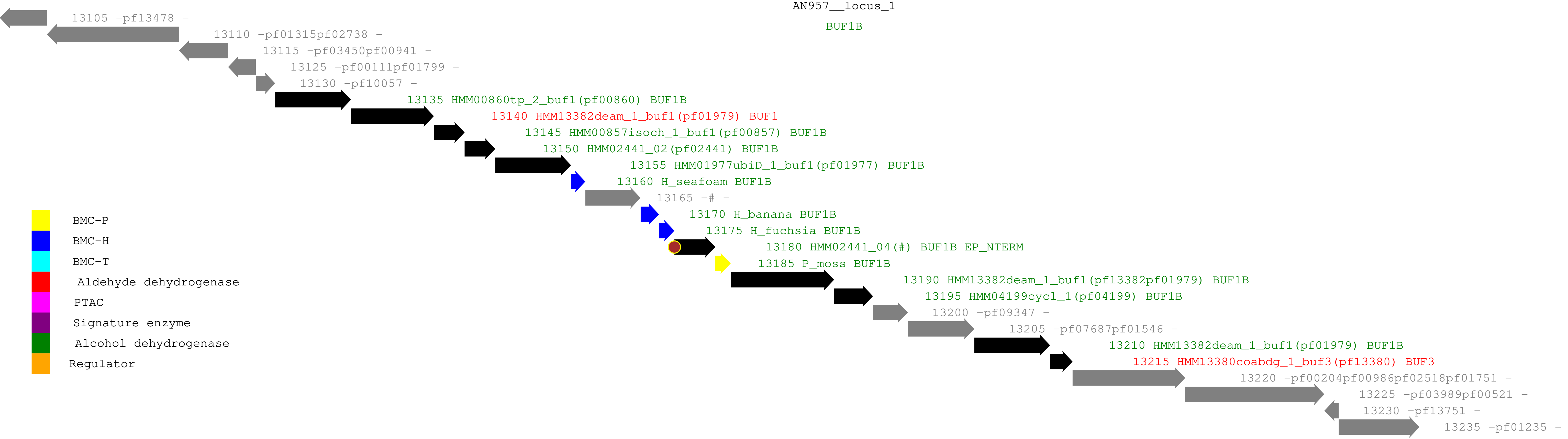

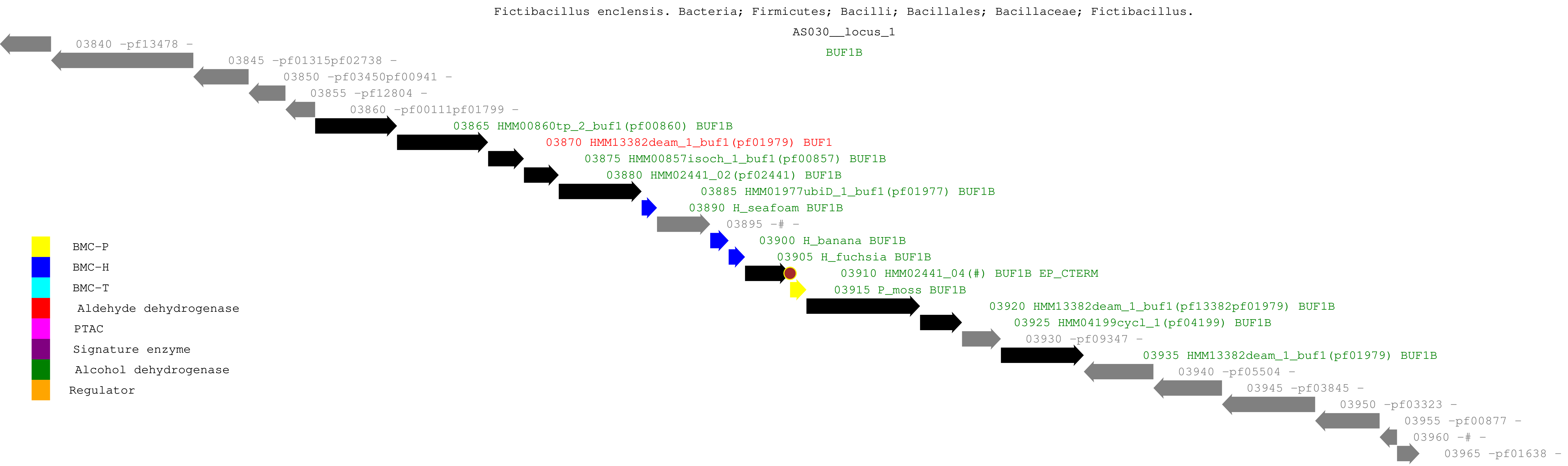

Fictibacillus enclensis. Bacteria; Firmicutes; Bacilli; Bacillales; Bacillaceae; Fictibacillus.
AS030__locus_1
BUF1B
03840 -pf13478 -
03845 -pf01315pf02738 -
03850 -pf03450pf00941 -
03855 -pf12804 -
03860 -pf00111pf01799 -
03865 HMM00860tp_2_buf1(pf00860) BUF1B
03870 HMM13382deam_1_buf1(pf01979) BUF1
03875 HMM00857isoch_1_buf1(pf00857) BUF1B
03880 HMM02441_02(pf02441) BUF1B
03885 HMM01977ubiD_1_buf1(pf01977) BUF1B
03890 H_seafoam BUF1B
03895 -# -
03900 H_banana BUF1B
03905 H_fuchsia BUF1B
03910 HMM02441_04(#) BUF1B EP_CTERM
03915 P_moss BUF1B
03920 HMM13382deam_1_buf1(pf13382pf01979) BUF1B
03925 HMM04199cycl_1(pf04199) BUF1B
03930 -pf09347 -
03935 HMM13382deam_1_buf1(pf01979) BUF1B
03940 -pf05504 -
03945 -pf03845 -
03950 -pf03323 -
03955 -pf00877 -
03960 -# -
03965 -pf01638 -
BMC-P
BMC-H
BMC-T
Aldehyde dehydrogenase
PTAC
Signature enzyme
Alcohol dehydrogenase
Regulator

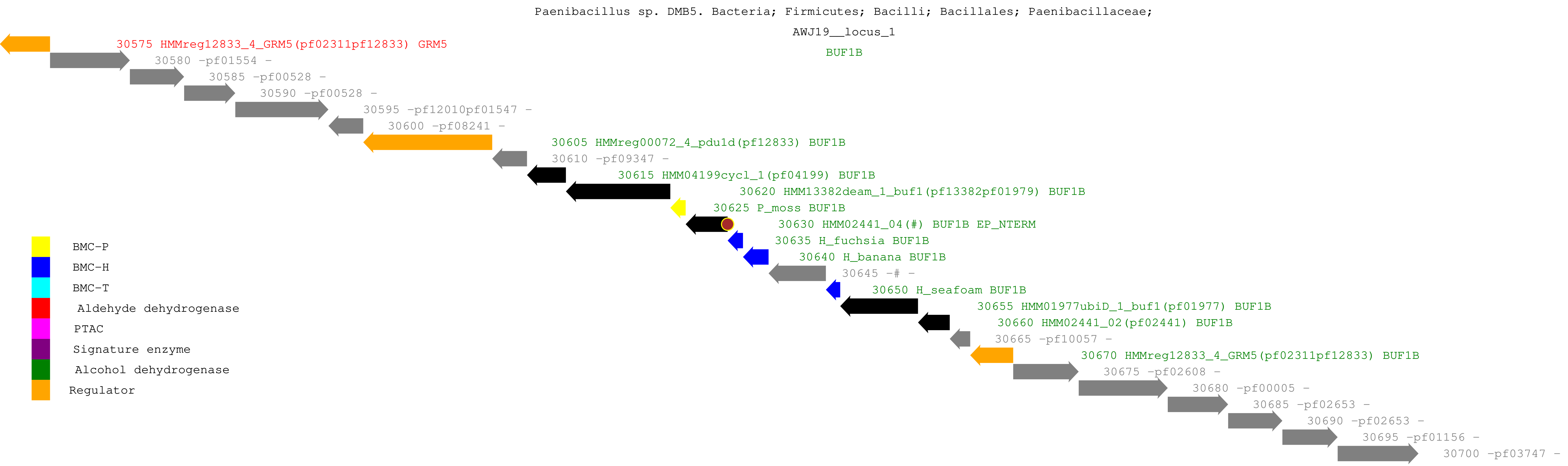

Paenibacillus sp. DMB5. Bacteria; Firmicutes; Bacilli; Bacillales; Paenibacillaceae;
AWJ19__locus_1
BUF1B
30575 HMMreg12833_4_GRM5(pf02311pf12833) GRM5
30580 -pf01554 -
30585 -pf00528 -
30590 -pf00528 -
30595 -pf12010pf01547 -
30600 -pf08241 -
30605 HMMreg00072_4_pdu1d(pf12833) BUF1B
30610 -pf09347 -
30615 HMM04199cycl_1(pf04199) BUF1B
30620 HMM13382deam_1_buf1(pf13382pf01979) BUF1B
30625 P_moss BUF1B
30630 HMM02441_04(#) BUF1B EP_NTERM
30635 H_fuchsia BUF1B
30640 H_banana BUF1B
30645 -# -
30650 H_seafoam BUF1B
30655 HMM01977ubiD_1_buf1(pf01977) BUF1B
30660 HMM02441_02(pf02441) BUF1B
30665 -pf10057 -
30670 HMMreg12833_4_GRM5(pf02311pf12833) BUF1B
30675 -pf02608 -
30680 -pf00005 -
30685 -pf02653 -
30690 -pf02653 -
30695 -pf01156 -
30700 -pf03747 -
BMC-P
BMC-H
BMC-T
Aldehyde dehydrogenase
PTAC
Signature enzyme
Alcohol dehydrogenase
Regulator

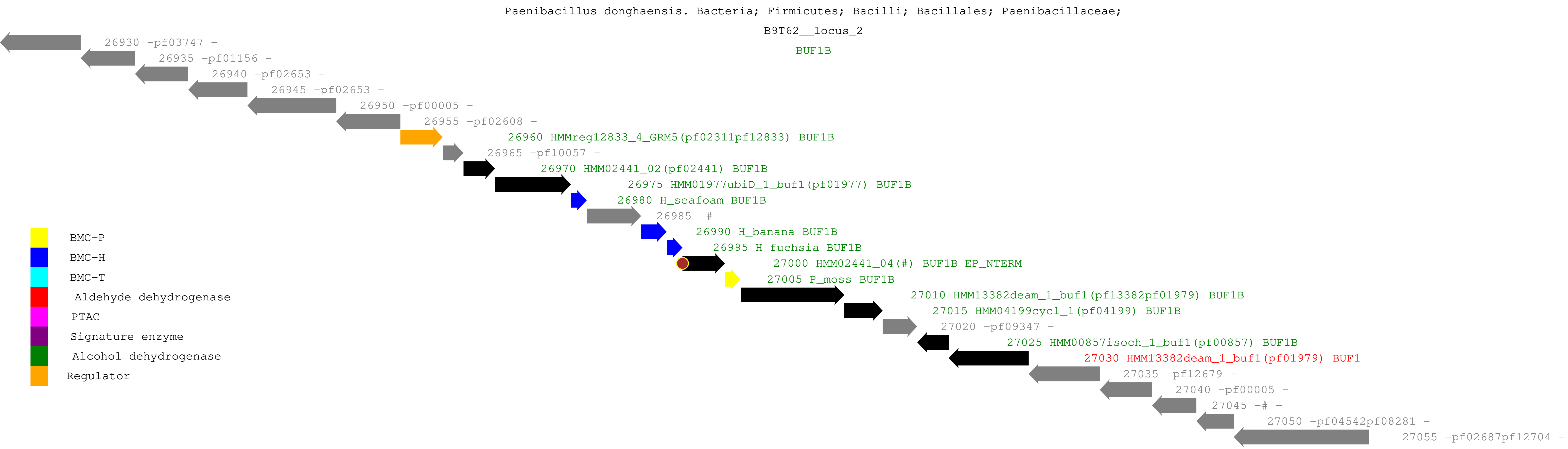
Paenibacillus donghaensis. Bacteria; Firmicutes; Bacilli; Bacillales; Paenibacillaceae;
B9T62__locus_2
BUF1B
26930 -pf03747 -
26935 -pf01156 -
26940 -pf02653 -
26945 -pf02653 -
26950 -pf00005 -
26955 -pf02608 -
26960 HMMreg12833_4_GRM5(pf02311pf12833) BUF1B
26965 -pf10057 -
26970 HMM02441_02(pf02441) BUF1B
26975 HMM01977ubiD_1_buf1(pf01977) BUF1B
26980 H_seafoam BUF1B
26985 -# -
26990 H_banana BUF1B
26995 H_fuchsia BUF1B
27000 HMM02441_04(#) BUF1B EP_NTERM
27005 P_moss BUF1B
27010 HMM13382deam_1_buf1(pf13382pf01979) BUF1B
27015 HMM04199cycl_1(pf04199) BUF1B
27020 -pf09347 -
27025 HMM00857isoch_1_buf1(pf00857) BUF1B
27030 HMM13382deam_1_buf1(pf01979) BUF1
27035 -pf12679 -
27040 -pf00005 -
27045 -# -
27050 -pf04542pf08281 -
27055 -pf02687pf12704 -
BMC-P
BMC-H
BMC-T
Aldehyde dehydrogenase
PTAC
Signature enzyme
Alcohol dehydrogenase
Regulator

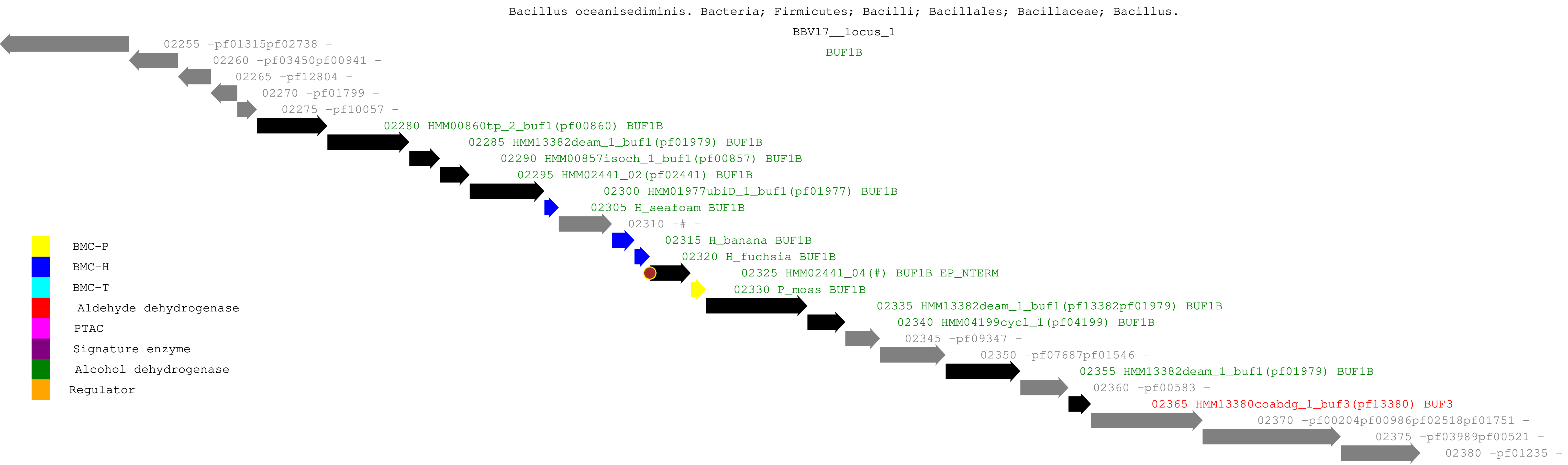

Bacillus oceanisediminis. Bacteria; Firmicutes; Bacilli; Bacillales; Bacillaceae; Bacillus.
BBV17__locus_1
BUF1B
02255 -pf01315pf02738 -
02260 -pf03450pf00941 -
02265 -pf12804 -
02270 -pf01799 -
02275 -pf10057 -
02280 HMM00860tp_2_buf1(pf00860) BUF1B
02285 HMM13382deam_1_buf1(pf01979) BUF1B
02290 HMM00857isoch_1_buf1(pf00857) BUF1B
02295 HMM02441_02(pf02441) BUF1B
02300 HMM01977ubiD_1_buf1(pf01977) BUF1B
02305 H_seafoam BUF1B
02310 -# -
02315 H_banana BUF1B
02320 H_fuchsia BUF1B
02325 HMM02441_04(#) BUF1B EP_NTERM
02330 P_moss BUF1B
02335 HMM13382deam_1_buf1(pf13382pf01979) BUF1B
02340 HMM04199cycl_1(pf04199) BUF1B
02345 -pf09347 -
02350 -pf07687pf01546 -
02355 HMM13382deam_1_buf1(pf01979) BUF1B
02360 -pf00583 -
02365 HMM13380coabdg_1_buf3(pf13380) BUF3
02370 -pf00204pf00986pf02518pf01751 -
02375 -pf03989pf00521 -
02380 -pf01235 -
BMC-P
BMC-H
BMC-T
Aldehyde dehydrogenase
PTAC
Signature enzyme
Alcohol dehydrogenase
Regulator

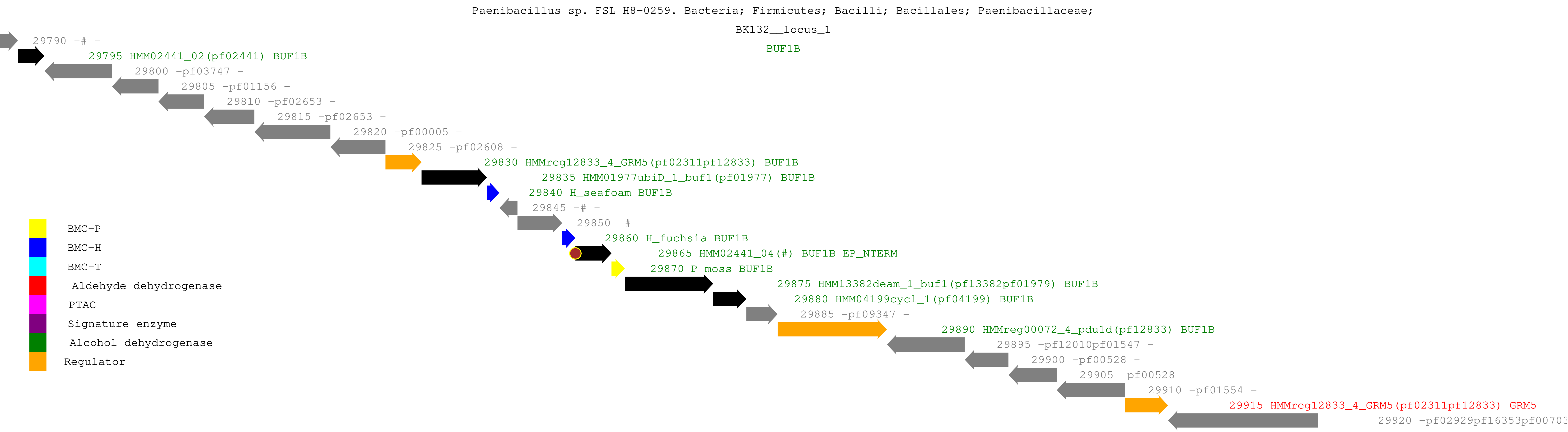
Paenibacillus sp. FSL H8-0259. Bacteria; Firmicutes; Bacilli; Bacillales; Paenibacillaceae;
BK132__locus_1
BUF1B
29790 -# -
29795 HMM02441_02(pf02441) BUF1B
29800 -pf03747 -
29805 -pf01156 -
29810 -pf02653 -
29815 -pf02653 -
29820 -pf00005 -
29825 -pf02608 -
29830 HMMreg12833_4_GRM5(pf02311pf12833) BUF1B
29835 HMM01977ubiD_1_buf1(pf01977) BUF1B
29840 H_seafoam BUF1B
29845 -# -
29850 -# -
29860 H_fuchsia BUF1B
29865 HMM02441_04(#) BUF1B EP_NTERM
29870 P_moss BUF1B
29875 HMM13382deam_1_buf1(pf13382pf01979) BUF1B
29880 HMM04199cycl_1(pf04199) BUF1B
29885 -pf09347 -
29890 HMMreg00072_4_pdu1d(pf12833) BUF1B
29895 -pf12010pf01547 -
29900 -pf00528 -
29905 -pf00528 -
29910 -pf01554 -
29915 HMMreg12833_4_GRM5(pf02311pf12833) GRM5
29920 -pf02929pf16353pf00703
BMC-P
BMC-H
BMC-T
Aldehyde dehydrogenase
PTAC
Signature enzyme
Alcohol dehydrogenase
Regulator

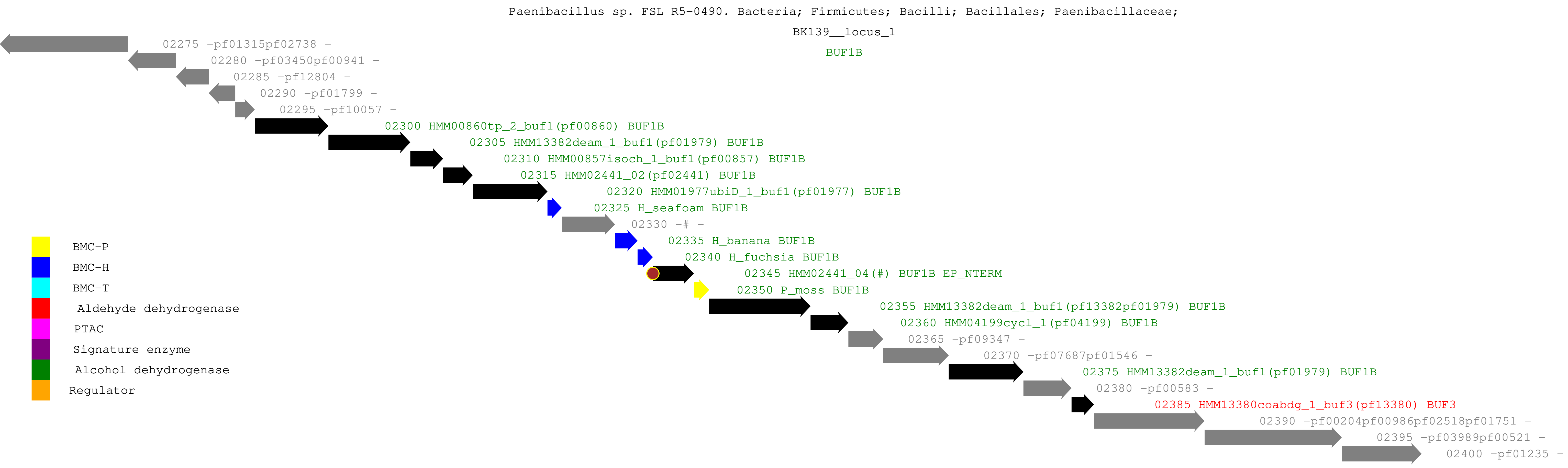
Paenibacillus sp. FSL R5-0490. Bacteria; Firmicutes; Bacilli; Bacillales; Paenibacillaceae;
BK139__locus_1
BUF1B
02275 -pf01315pf02738 -
02280 -pf03450pf00941 -
02285 -pf12804 -
02290 -pf01799 -
02295 -pf10057 -
02300 HMM00860tp_2_buf1(pf00860) BUF1B
02305 HMM13382deam_1_buf1(pf01979) BUF1B
02310 HMM00857isoch_1_buf1(pf00857) BUF1B
02315 HMM02441_02(pf02441) BUF1B
02320 HMM01977ubiD_1_buf1(pf01977) BUF1B
02325 H_seafoam BUF1B
02330 -# -
02335 H_banana BUF1B
02340 H_fuchsia BUF1B
02345 HMM02441_04(#) BUF1B EP_NTERM
02350 P_moss BUF1B
02355 HMM13382deam_1_buf1(pf13382pf01979) BUF1B
02360 HMM04199cycl_1(pf04199) BUF1B
02365 -pf09347 -
02370 -pf07687pf01546 -
02375 HMM13382deam_1_buf1(pf01979) BUF1B
02380 -pf00583 -
02385 HMM13380coabdg_1_buf3(pf13380) BUF3
02390 -pf00204pf00986pf02518pf01751 -
02395 -pf03989pf00521 -
02400 -pf01235 -
BMC-P
BMC-H
BMC-T
Aldehyde dehydrogenase
PTAC
Signature enzyme
Alcohol dehydrogenase
Regulator

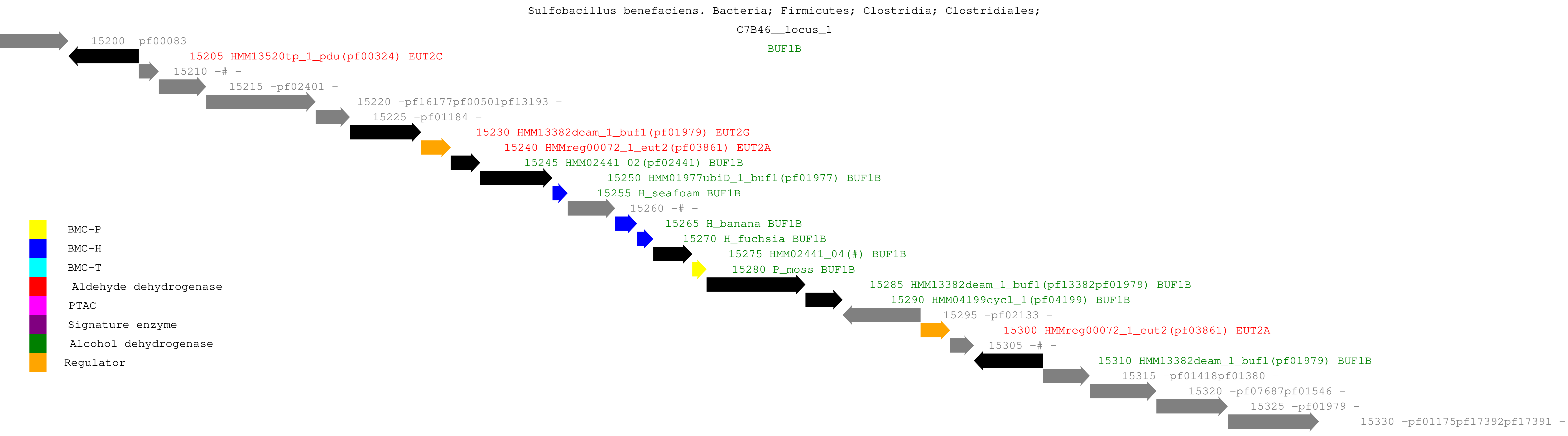

Sulfobacillus benefaciens. Bacteria; Firmicutes; Clostridia; Clostridiales;
C7B46__locus_1
BUF1B
15200 -pf00083 -
15205 HMM13520tp_1_pdu(pf00324) EUT2C
15210 -# -
15215 -pf02401 -
15220 -pf16177pf00501pf13193 -
15225 -pf01184 -
15230 HMM13382deam_1_buf1(pf01979) EUT2G
15240 HMMreg00072_1_eut2(pf03861) EUT2A
15245 HMM02441_02(pf02441) BUF1B
15250 HMM01977ubiD_1_buf1(pf01977) BUF1B
15255 H_seafoam BUF1B
15260 -# -
15265 H_banana BUF1B
15270 H_fuchsia BUF1B
15275 HMM02441_04(#) BUF1B
15280 P_moss BUF1B
15285 HMM13382deam_1_buf1(pf13382pf01979) BUF1B
15290 HMM04199cycl_1(pf04199) BUF1B
15295 -pf02133 -
15300 HMMreg00072_1_eut2(pf03861) EUT2A
15305 -# -
15310 HMM13382deam_1_buf1(pf01979) BUF1B
15315 -pf01418pf01380 -
15320 -pf07687pf01546 -
15325 -pf01979 -
15330 -pf01175pf17392pf17391 -
BMC-P
BMC-H
BMC-T
Aldehyde dehydrogenase
PTAC
Signature enzyme
Alcohol dehydrogenase
Regulator

Bacillus sp. 7894-2. Bacteria; Firmicutes; Bacilli; Bacillales; Bacillaceae; Bacillus.

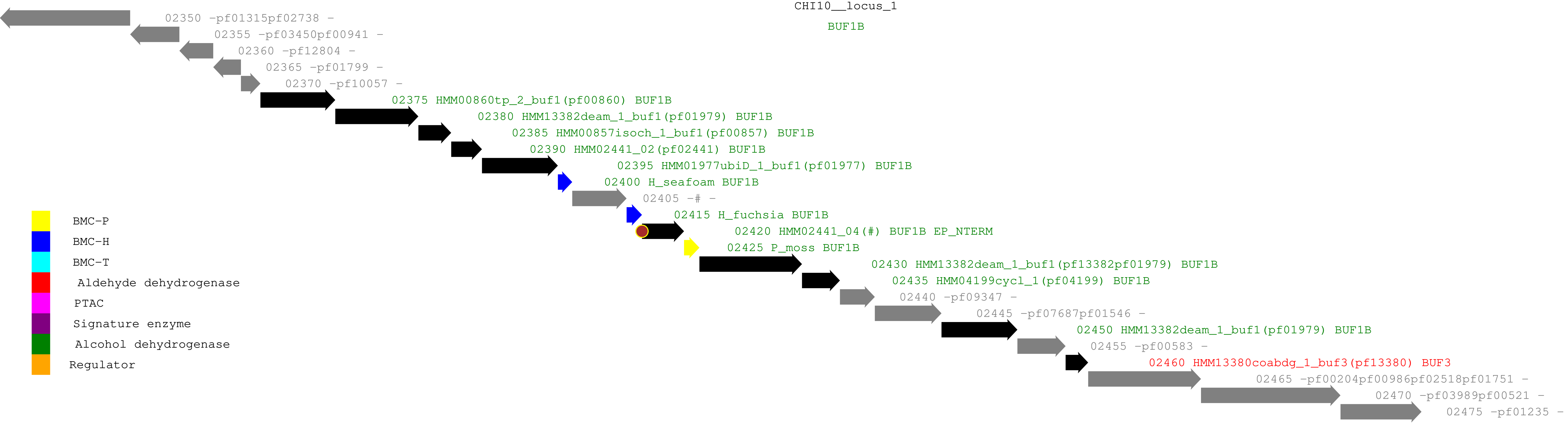

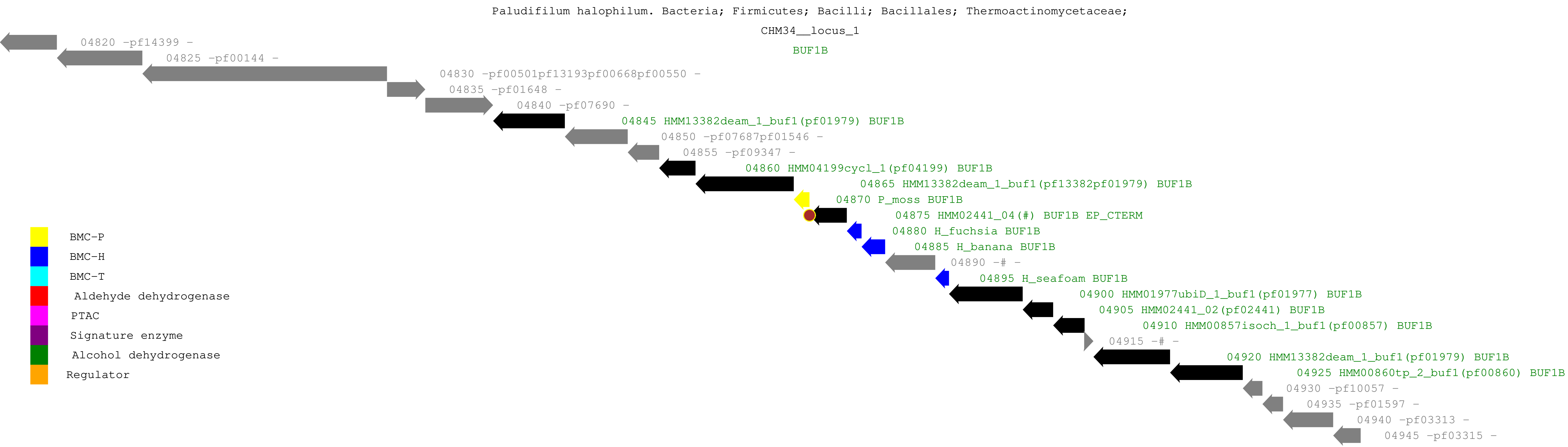
Paludifilum halophilum. Bacteria; Firmicutes; Bacilli; Bacillales; Thermoactinomycetaceae;
CHM34__locus_1
BUF1B
04820 -pf14399 -
04825 -pf00144 -
04830 -pf00501pf13193pf00668pf00550 -
04835 -pf01648 -
04840 -pf07690 -
04845 HMM13382deam_1_buf1(pf01979) BUF1B
04850 -pf07687pf01546 -
04855 -pf09347 -
04860 HMM04199cycl_1(pf04199) BUF1B
04865 HMM13382deam_1_buf1(pf13382pf01979) BUF1B
04870 P_moss BUF1B
04875 HMM02441_04(#) BUF1B EP_CTERM
04880 H_fuchsia BUF1B
04885 H_banana BUF1B
04890 -# -
04895 H_seafoam BUF1B
04900 HMM01977ubiD_1_buf1(pf01977) BUF1B
04905 HMM02441_02(pf02441) BUF1B
04910 HMM00857isoch_1_buf1(pf00857) BUF1B
04915 -# -
04920 HMM13382deam_1_buf1(pf01979) BUF1B
04925 HMM00860tp_2_buf1(pf00860) BUF1B
04930 -pf10057 -
04935 -pf01597 -
04940 -pf03313 -
04945 -pf03315 -
BMC-P
BMC-H
BMC-T
Aldehyde dehydrogenase
PTAC
Signature enzyme
Alcohol dehydrogenase
Regulator

Bacillus mesonae. Bacteria; Firmicutes; Bacilli; Bacillales; Bacillaceae; Bacillus.

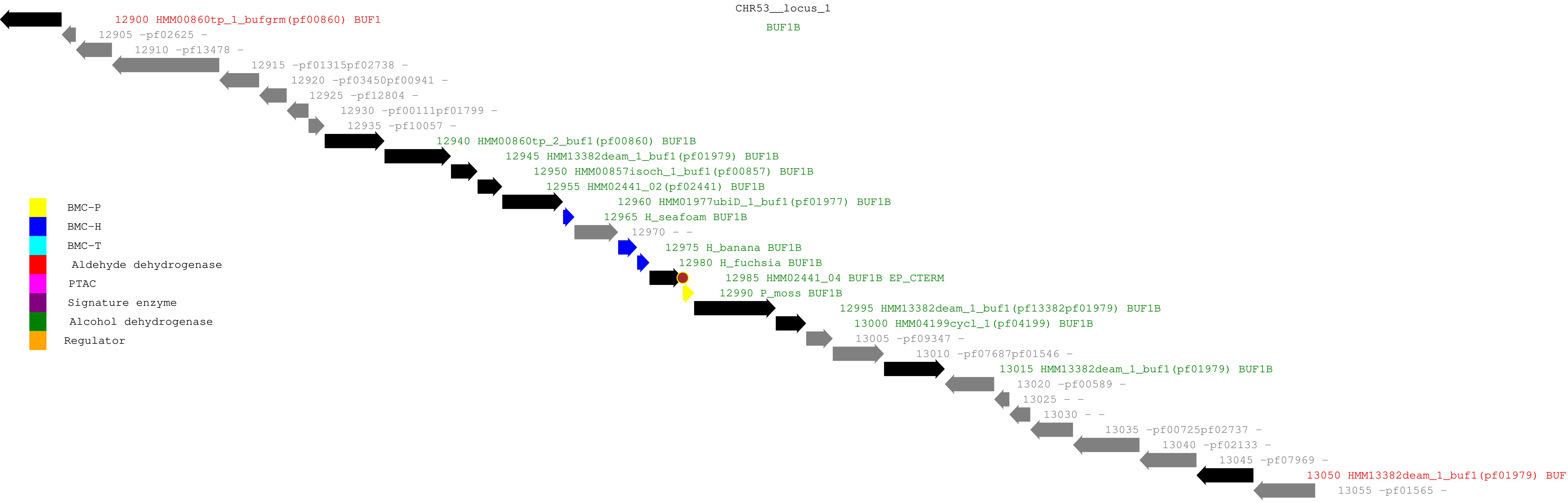

Virgibacillus sp. IO3-P2-C2. Bacteria; Firmicutes; Bacilli; Bacillales; Bacillaceae; Virgibacillus.

CIL03__locus_1

BUF1B

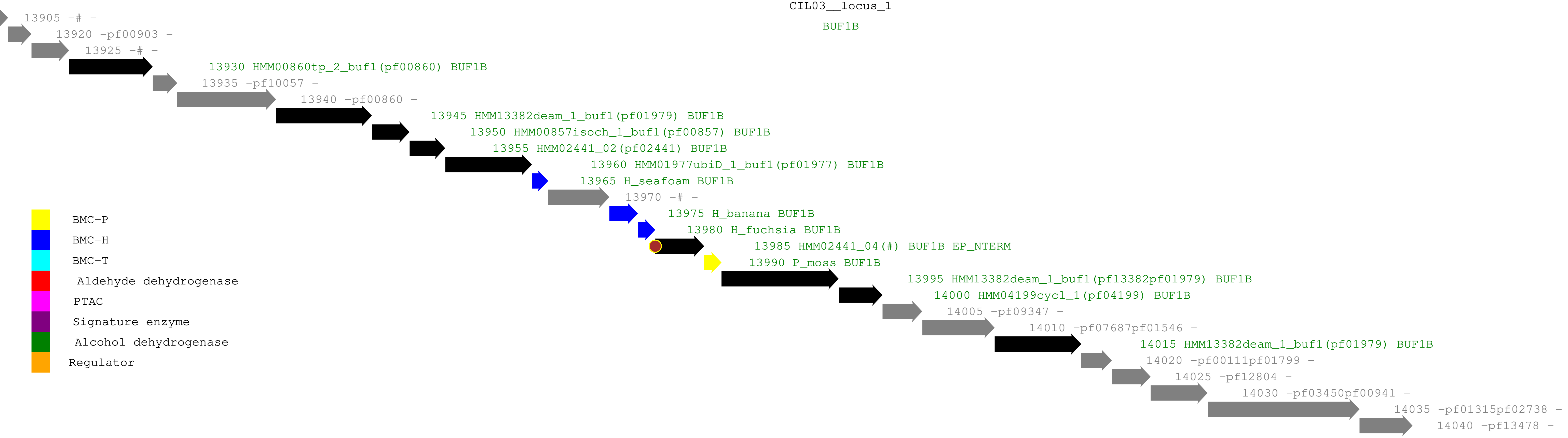

Virgibacillus sp. IO3-P3-H5. Bacteria; Firmicutes; Bacilli; Bacillales; Bacillaceae; Virgibacillus.

CIL05__locus_1

BUF1B

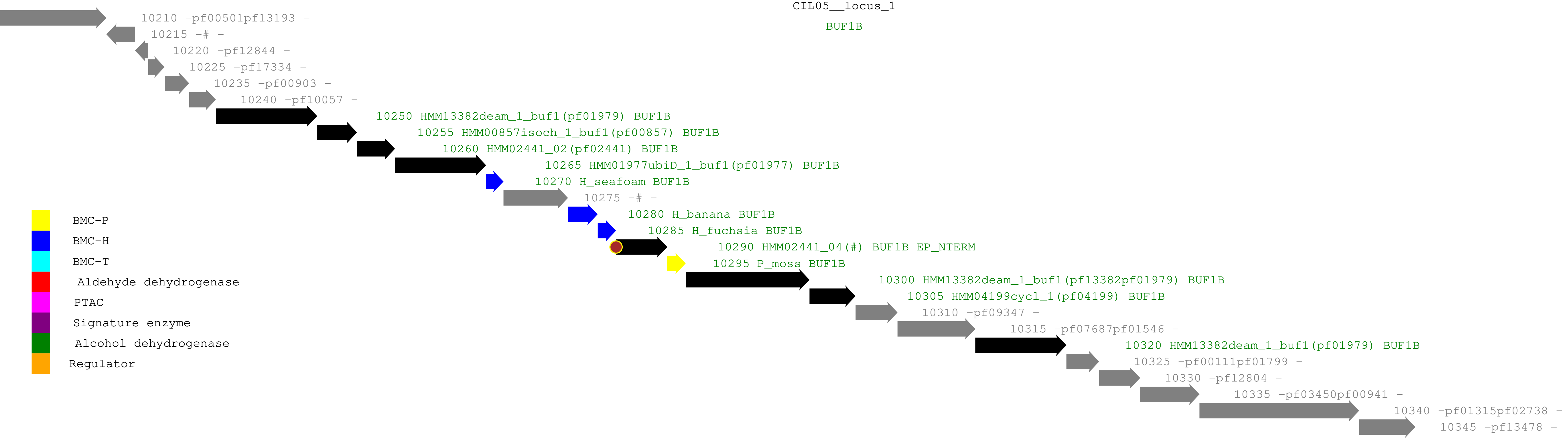

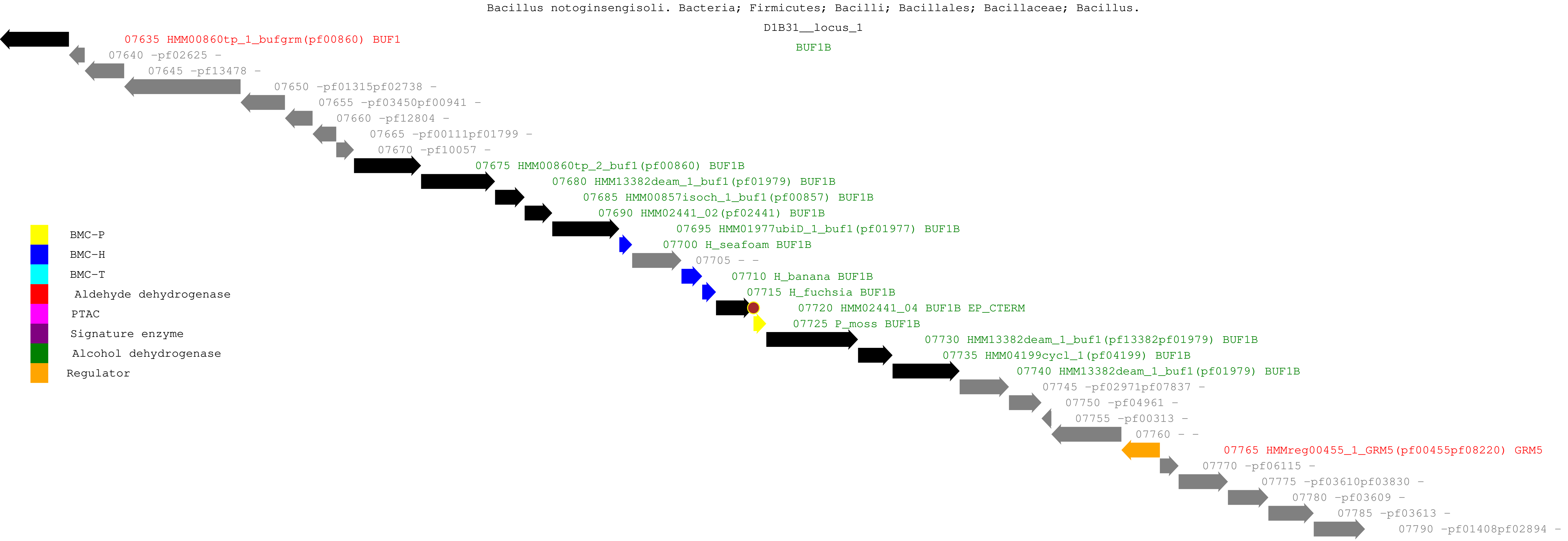

Bacillus notoginsengisoli. Bacteria; Firmicutes; Bacilli; Bacillales; Bacillaceae; Bacillus.
D1B31__locus_1
BUF1B
07635 HMM00860tp_1_bufgrm(pf00860) BUF1
07640 -pf02625 -
07645 -pf13478 -
07650 -pf01315pf02738 -
07655 -pf03450pf00941 -
07660 -pf12804 -
07665 -pf00111pf01799 -
07670 -pf10057 -
07675 HMM00860tp_2_buf1(pf00860) BUF1B
07680 HMM13382deam_1_buf1(pf01979) BUF1B
07685 HMM00857isoch_1_buf1(pf00857) BUF1B
07690 HMM02441_02(pf02441) BUF1B
07695 HMM01977ubiD_1_buf1(pf01977) BUF1B
07700 H_seafoam BUF1B
07705 - -
07710 H_banana BUF1B
07715 H_fuchsia BUF1B
07720 HMM02441_04 BUF1B EP_CTERM
07725 P_moss BUF1B
07730 HMM13382deam_1_buf1(pf13382pf01979) BUF1B
07735 HMM04199cycl_1(pf04199) BUF1B
07740 HMM13382deam_1_buf1(pf01979) BUF1B
07745 -pf02971pf07837 -
07750 -pf04961 -
07755 -pf00313 -
07760 - -
07765 HMMreg00455_1_GRM5(pf00455pf08220) GRM5
07770 -pf06115 -
07775 -pf03610pf03830 -
07780 -pf03609 -
07785 -pf03613 -
07790 -pf01408pf02894 -
BMC-P
BMC-H
BMC-T
Aldehyde dehydrogenase
PTAC
Signature enzyme
Alcohol dehydrogenase
Regulator

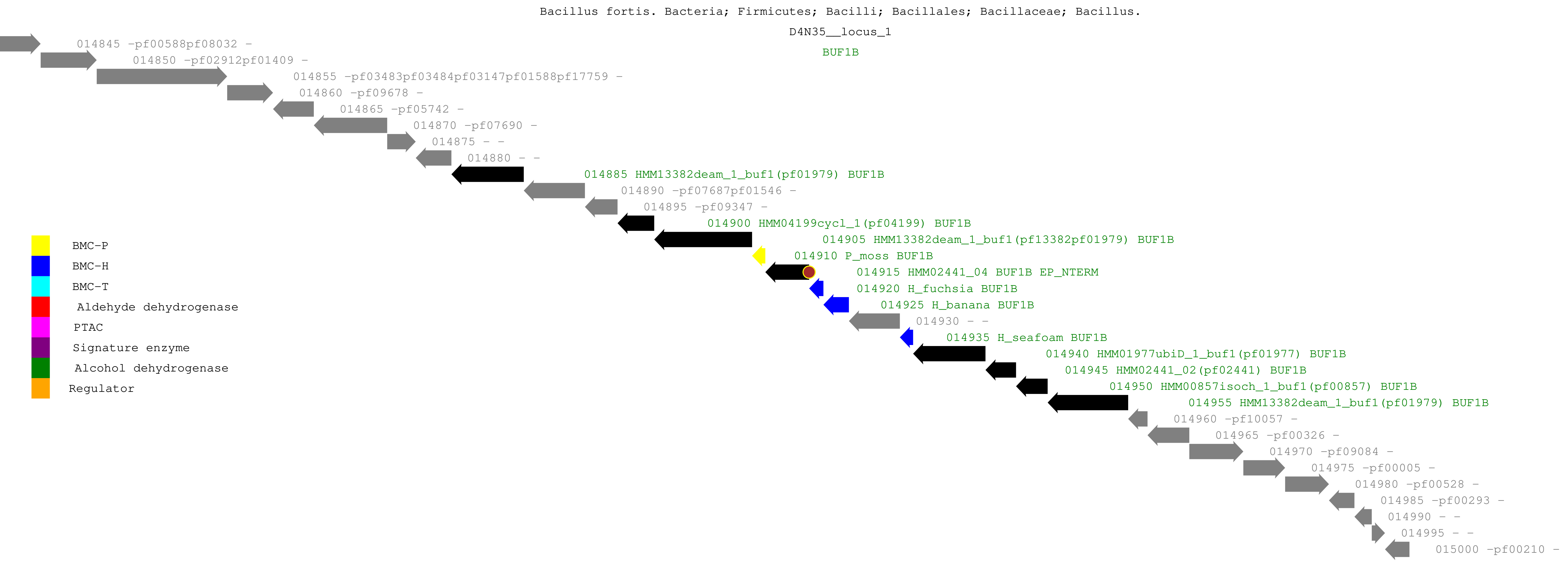

Bacillus fortis. Bacteria; Firmicutes; Bacilli; Bacillales; Bacillaceae; Bacillus.
D4N35__locus_1
BUF1B
014845 -pf00588pf08032 -
014850 -pf02912pf01409 -
014855 -pf03483pf03484pf03147pf01588pf17759 -
014860 -pf09678 -
014865 -pf05742 -
014870 -pf07690 -
014875 - -
014880 - -
014885 HMM13382deam_1_buf1(pf01979) BUF1B
014890 -pf07687pf01546 -
014895 -pf09347 -
014900 HMM04199cycl_1(pf04199) BUF1B
014905 HMM13382deam_1_buf1(pf13382pf01979) BUF1B
014910 P_moss BUF1B
014915 HMM02441_04 BUF1B EP_NTERM
014920 H_fuchsia BUF1B
014925 H_banana BUF1B
014930 - -
014935 H_seafoam BUF1B
014940 HMM01977ubiD_1_buf1(pf01977) BUF1B
014945 HMM02441_02(pf02441) BUF1B
014950 HMM00857isoch_1_buf1(pf00857) BUF1B
014955 HMM13382deam_1_buf1(pf01979) BUF1B
014960 -pf10057 -
014965 -pf00326 -
014970 -pf09084 -
014975 -pf00005 -
014980 -pf00528 -
014985 -pf00293 -
014990 - -
014995 - -
015000 -pf00210 -
BMC-P
BMC-H
BMC-T
Aldehyde dehydrogenase
PTAC
Signature enzyme
Alcohol dehydrogenase
Regulator

Bacillus sp. 3-2-2. Bacteria; Firmicutes; Bacilli; Bacillales; Bacillaceae; Bacillus.

D4T97__locus_1

BUF1B

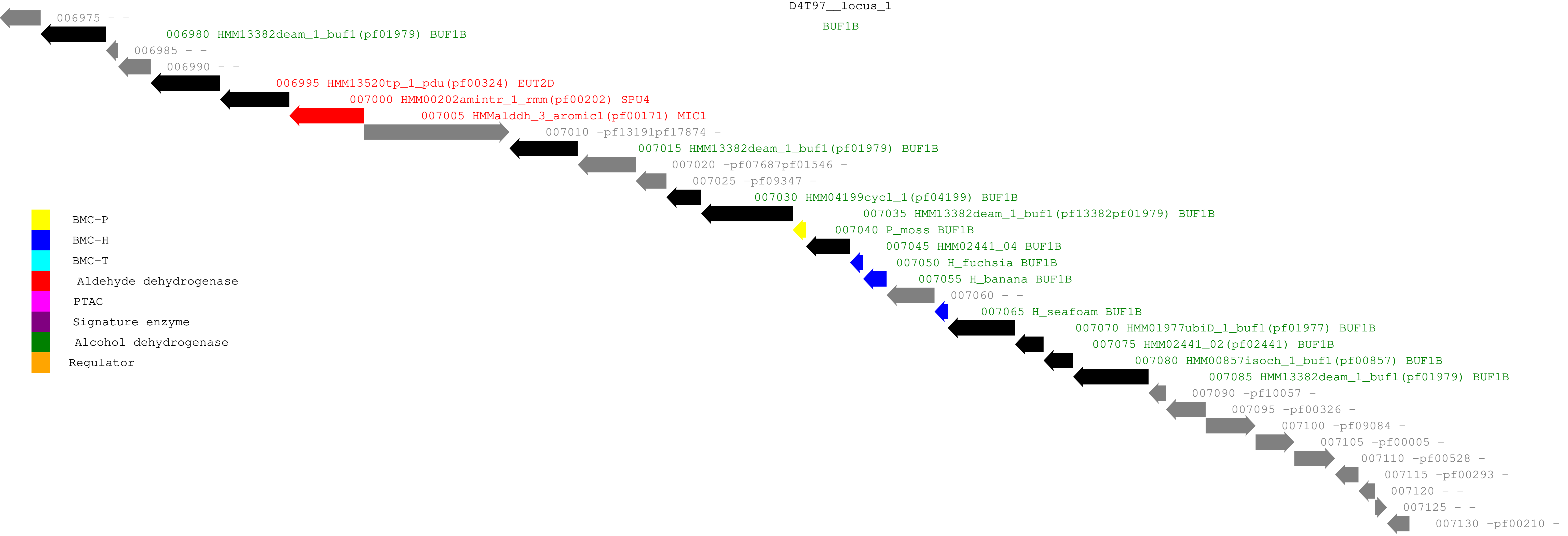

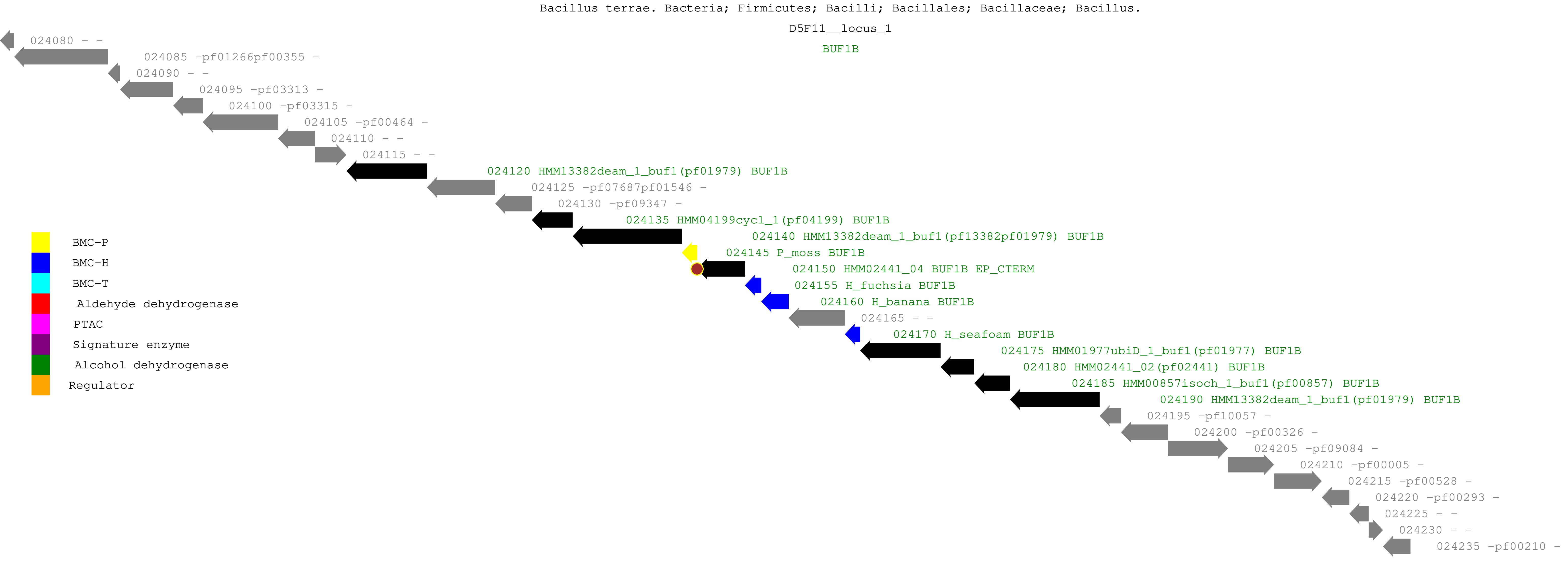
Bacillus terrae. Bacteria; Firmicutes; Bacilli; Bacillales; Bacillaceae; Bacillus.
D5F11__locus_1
BUF1B
024080 - -
024085 -pf01266pf00355 -
024090 - -
024095 -pf03313 -
024100 -pf03315 -
024105 -pf00464 -
024110 - -
024115 - -
024120 HMM13382deam_1_buf1(pf01979) BUF1B
024125 -pf07687pf01546 -
024130 -pf09347 -
024135 HMM04199cycl_1(pf04199) BUF1B
024140 HMM13382deam_1_buf1(pf13382pf01979) BUF1B
024145 P_moss BUF1B
024150 HMM02441_04 BUF1B EP_CTERM
024155 H_fuchsia BUF1B
024160 H_banana BUF1B
024165 - -
024170 H_seafoam BUF1B
024175 HMM01977ubiD_1_buf1(pf01977) BUF1B
024180 HMM02441_02(pf02441) BUF1B
024185 HMM00857isoch_1_buf1(pf00857) BUF1B
024190 HMM13382deam_1_buf1(pf01979) BUF1B
024195 -pf10057 -
024200 -pf00326 -
024205 -pf09084 -
024210 -pf00005 -
024215 -pf00528 -
024220 -pf00293 -
024225 - -
024230 - -
024235 -pf00210 -
BMC-P
BMC-H
BMC-T
Aldehyde dehydrogenase
PTAC
Signature enzyme
Alcohol dehydrogenase
Regulator

Bacillus bacterium. Bacteria; Firmicutes; Bacilli; Bacillales; Bacillaceae; Bacillus.

DEO65__locus_1

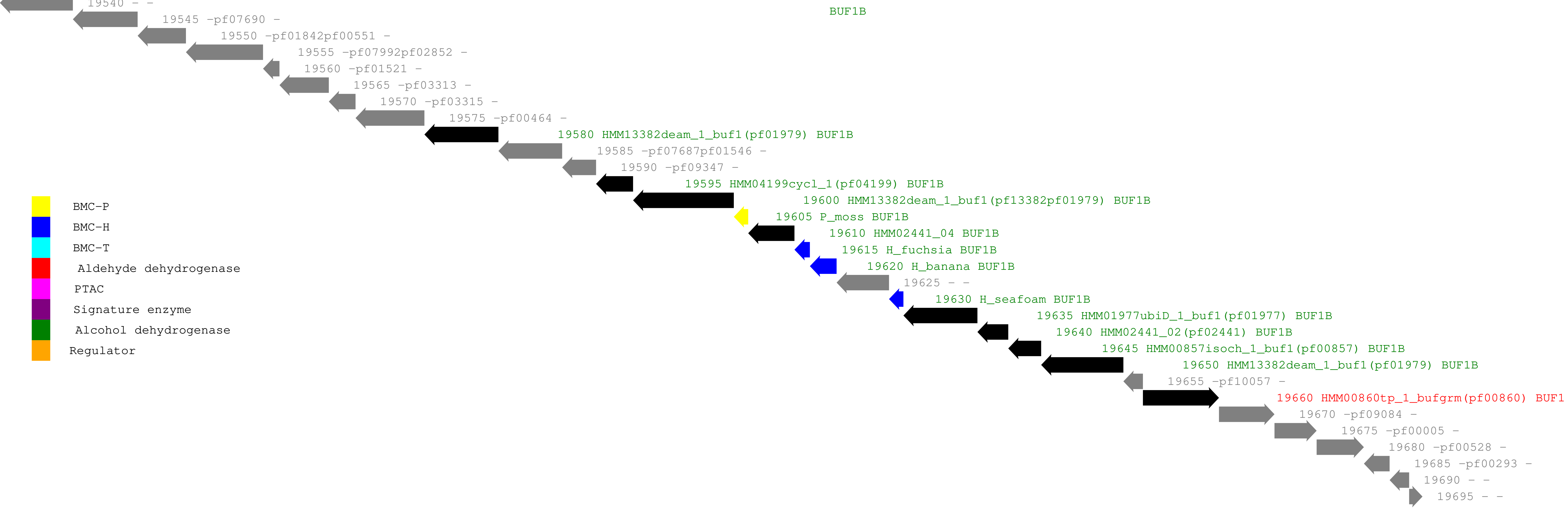

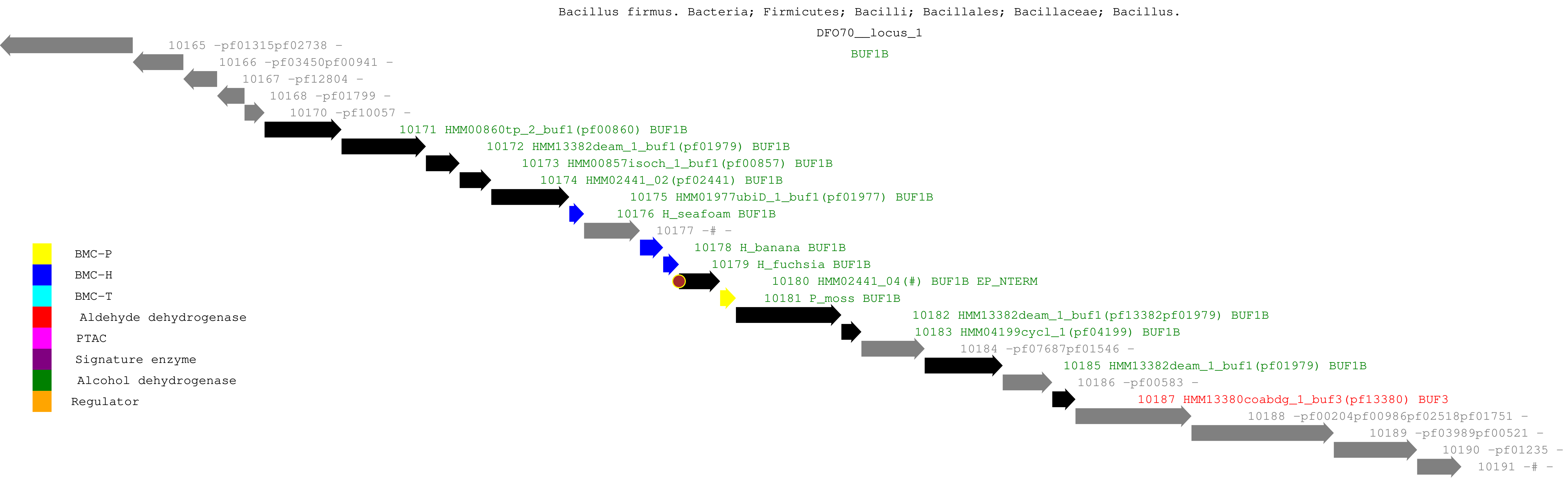
Bacillus firmus. Bacteria; Firmicutes; Bacilli; Bacillales; Bacillaceae; Bacillus.
DFO70__locus_1
BUF1B
10165 -pf01315pf02738 -
10166 -pf03450pf00941 -
10167 -pf12804 -
10168 -pf01799 -
10170 -pf10057 -
10171 HMM00860tp_2_buf1(pf00860) BUF1B
10172 HMM13382deam_1_buf1(pf01979) BUF1B
10173 HMM00857isoch_1_buf1(pf00857) BUF1B
10174 HMM02441_02(pf02441) BUF1B
10175 HMM01977ubiD_1_buf1(pf01977) BUF1B
10176 H_seafoam BUF1B
10177 -# -
10178 H_banana BUF1B
10179 H_fuchsia BUF1B
10180 HMM02441_04(#) BUF1B EP_NTERM
10181 P_moss BUF1B
10182 HMM13382deam_1_buf1(pf13382pf01979) BUF1B
10183 HMM04199cycl_1(pf04199) BUF1B
10184 -pf07687pf01546 -
10185 HMM13382deam_1_buf1(pf01979) BUF1B
10186 -pf00583 -
10187 HMM13380coabdg_1_buf3(pf13380) BUF3
10188 -pf00204pf00986pf02518pf01751 -
10189 -pf03989pf00521 -
10190 -pf01235 -
10191 -# -
BMC-P
BMC-H
BMC-T
Aldehyde dehydrogenase
PTAC
Signature enzyme
Alcohol dehydrogenase
Regulator

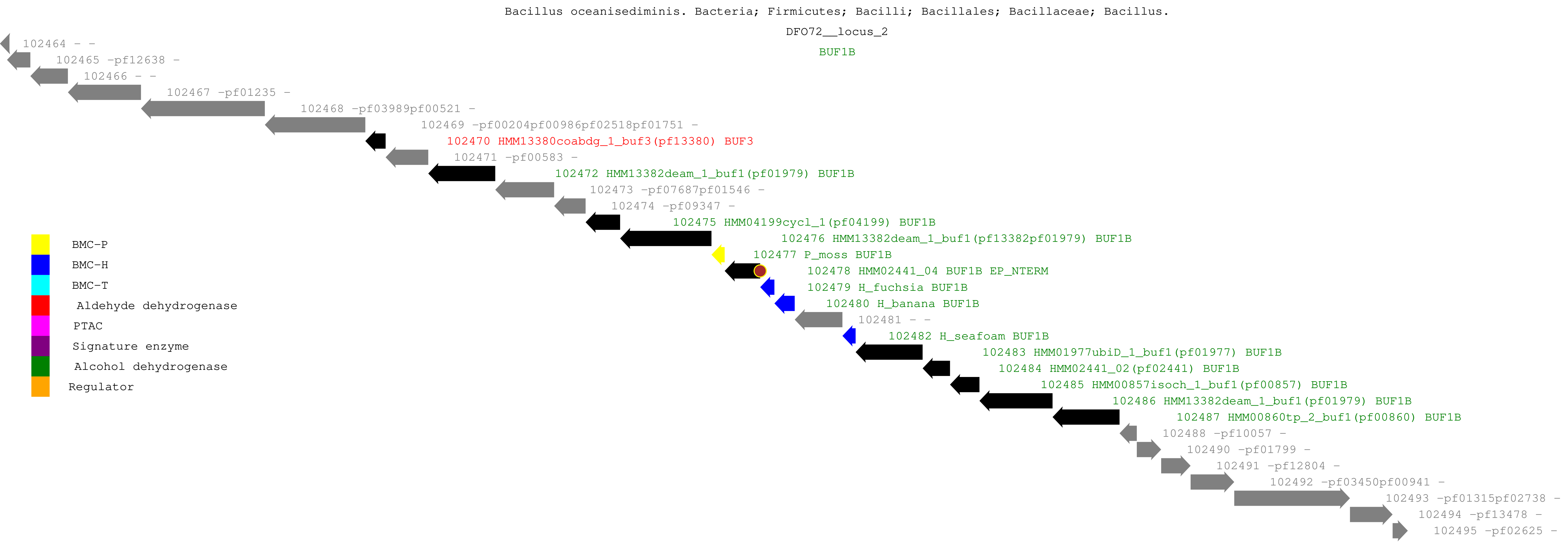
Bacillus oceanisediminis. Bacteria; Firmicutes; Bacilli; Bacillales; Bacillaceae; Bacillus.
DFO72__locus_2
BUF1B
102464 - -
102465 -pf12638 -
102466 - -
102467 -pf01235 -
102468 -pf03989pf00521 -
102469 -pf00204pf00986pf02518pf01751 -
102470 HMM13380coabdg_1_buf3(pf13380) BUF3
102471 -pf00583 -
102472 HMM13382deam_1_buf1(pf01979) BUF1B
102473 -pf07687pf01546 -
102474 -pf09347 -
102475 HMM04199cycl_1(pf04199) BUF1B
102476 HMM13382deam_1_buf1(pf13382pf01979) BUF1B
102477 P_moss BUF1B
102478 HMM02441_04 BUF1B EP_NTERM
102479 H_fuchsia BUF1B
102480 H_banana BUF1B
102481 - -
102482 H_seafoam BUF1B
102483 HMM01977ubiD_1_buf1(pf01977) BUF1B
102484 HMM02441_02(pf02441) BUF1B
102485 HMM00857isoch_1_buf1(pf00857) BUF1B
102486 HMM13382deam_1_buf1(pf01979) BUF1B
102487 HMM00860tp_2_buf1(pf00860) BUF1B
102488 -pf10057 -
102490 -pf01799 -
102491 -pf12804 -
102492 -pf03450pf00941 -
102493 -pf01315pf02738 -
102494 -pf13478 -
102495 -pf02625 -
BMC-P
BMC-H
BMC-T
Aldehyde dehydrogenase
PTAC
Signature enzyme
Alcohol dehydrogenase
Regulator

Bacillus oceanisediminis. Bacteria; Firmicutes; Bacilli; Bacillales; Bacillaceae; Bacillus.

DFO73__locus_2_main_sat

BUF1B

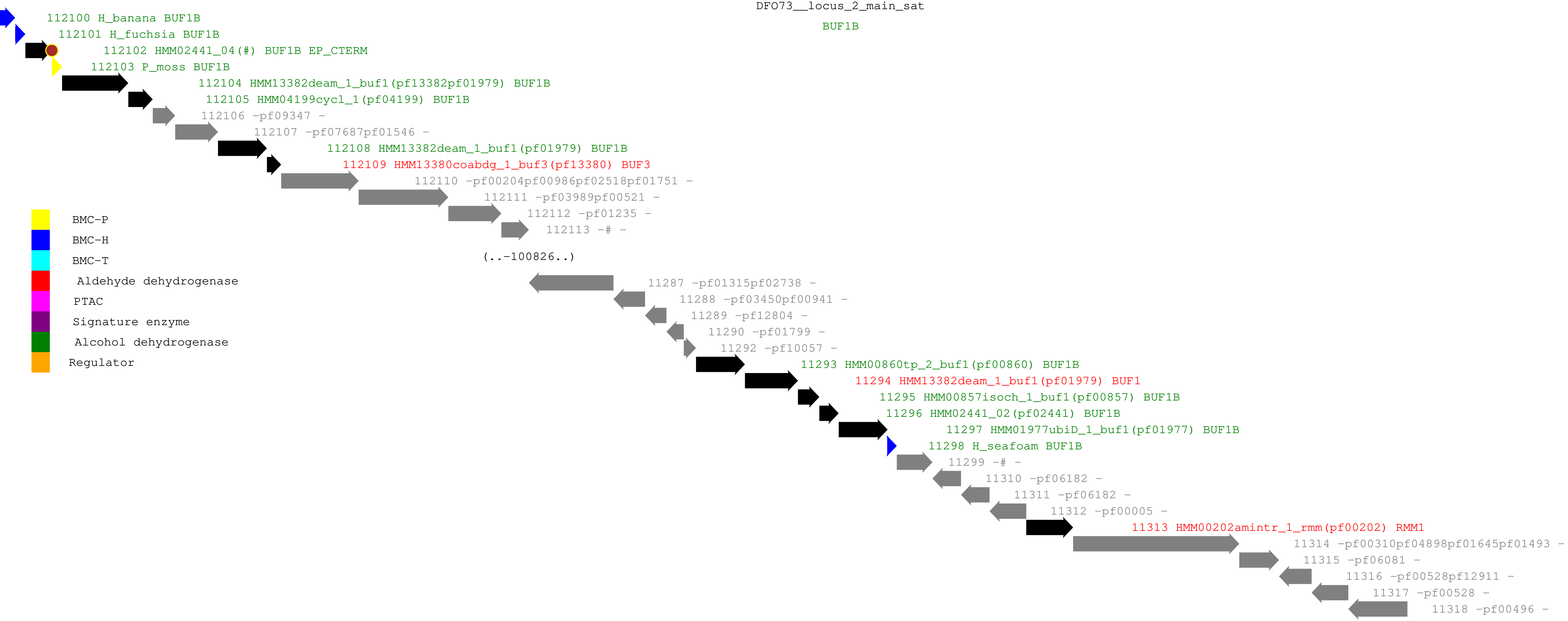

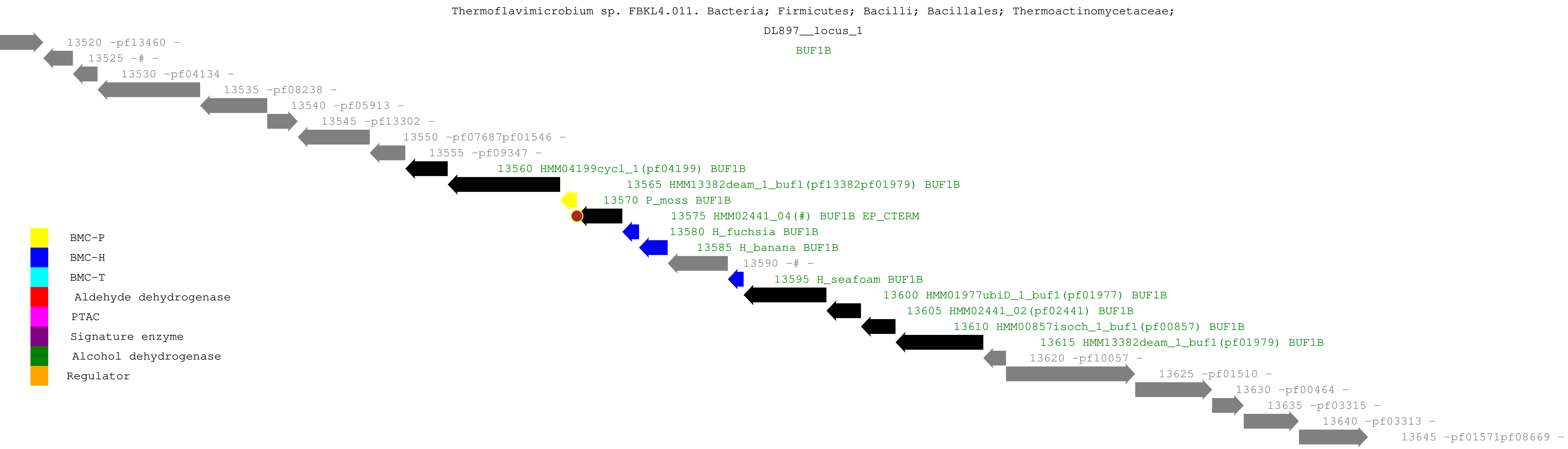
Thermoflavimicrobium sp. FBKL4.011. Bacteria; Firmicutes; Bacilli; Bacillales; Thermoactinomycetaceae;
DL897__locus_1
BUF1B
13520 -pf13460 -
13525 -# -
13530 -pf04134 -
13535 -pf08238 -
13540 -pf05913 -
13545 -pf13302 -
13550 -pf07687pf01546 -
13555 -pf09347 -
13560 HMM04199cycl_1(pf04199) BUF1B
13565 HMM13382deam_1_buf1(pf13382pf01979) BUF1B
13570 P_moss BUF1B
13575 HMM02441_04(#) BUF1B EP_CTERM
13580 H_fuchsia BUF1B
13585 H_banana BUF1B
13590 -# -
13595 H_seafoam BUF1B
13600 HMM01977ubiD_1_buf1(pf01977) BUF1B
13605 HMM02441_02(pf02441) BUF1B
13610 HMM00857isoch_1_buf1(pf00857) BUF1B
13615 HMM13382deam_1_buf1(pf01979) BUF1B
13620 -pf10057 -
13625 -pf01510 -
13630 -pf00464 -
13635 -pf03315 -
13640 -pf03313 -
13645 -pf01571pf08669 -
BMC-P
BMC-H
BMC-T
Aldehyde dehydrogenase
PTAC
Signature enzyme
Alcohol dehydrogenase
Regulator

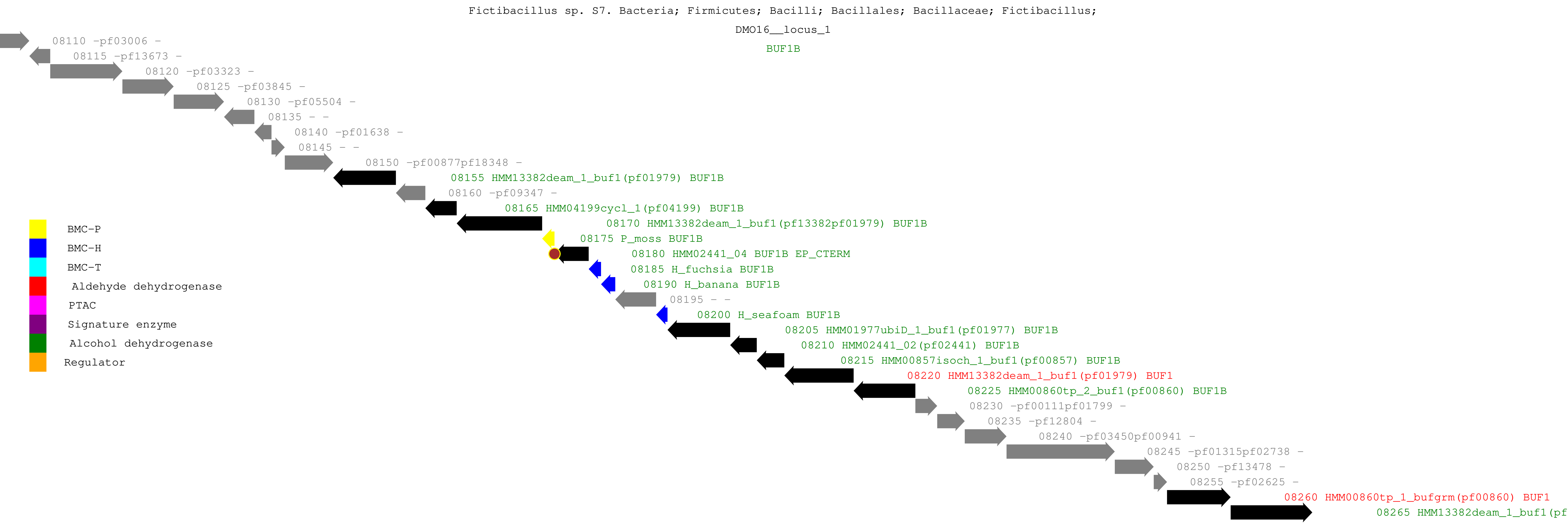
Fictibacillus sp. S7. Bacteria; Firmicutes; Bacilli; Bacillales; Bacillaceae; Fictibacillus;
DMO16__locus_1
BUF1B
08110 -pf03006 -
08115 -pf13673 -
08120 -pf03323 -
08125 -pf03845 -
08130 -pf05504 -
08135 - -
08140 -pf01638 -
08145 - -
08150 -pf00877pf18348 -
08155 HMM13382deam_1_buf1(pf01979) BUF1B
08160 -pf09347 -
08165 HMM04199cycl_1(pf04199) BUF1B
08170 HMM13382deam_1_buf1(pf13382pf01979) BUF1B
08175 P_moss BUF1B
08180 HMM02441_04 BUF1B EP_CTERM
08185 H_fuchsia BUF1B
08190 H_banana BUF1B
08195 - -
08200 H_seafoam BUF1B
08205 HMM01977ubiD_1_buf1(pf01977) BUF1B
08210 HMM02441_02(pf02441) BUF1B
08215 HMM00857isoch_1_buf1(pf00857) BUF1B
08220 HMM13382deam_1_buf1(pf01979) BUF1
08225 HMM00860tp_2_buf1(pf00860) BUF1B
08230 -pf00111pf01799 -
08235 -pf12804 -
08240 -pf03450pf00941 -
08245 -pf01315pf02738 -
08250 -pf13478 -
08255 -pf02625 -
08260 HMM00860tp_1_bufgrm(pf00860) BUF1
08265 HMM13382deam_1_buf1(pf
BMC-P
BMC-H
BMC-T
Aldehyde dehydrogenase
PTAC
Signature enzyme
Alcohol dehydrogenase
Regulator

Bacillus praedii. Bacteria; Firmicutes; Bacilli; Bacillales; Bacillaceae; Bacillus.

E0Y62__locus_2

BUF1B

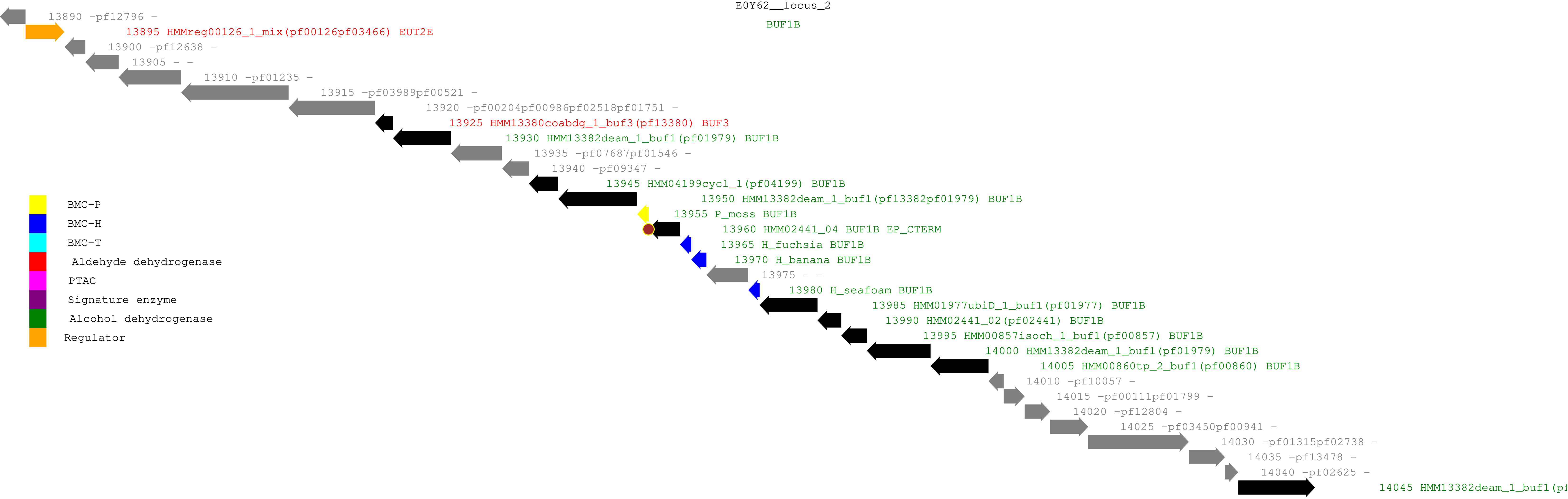

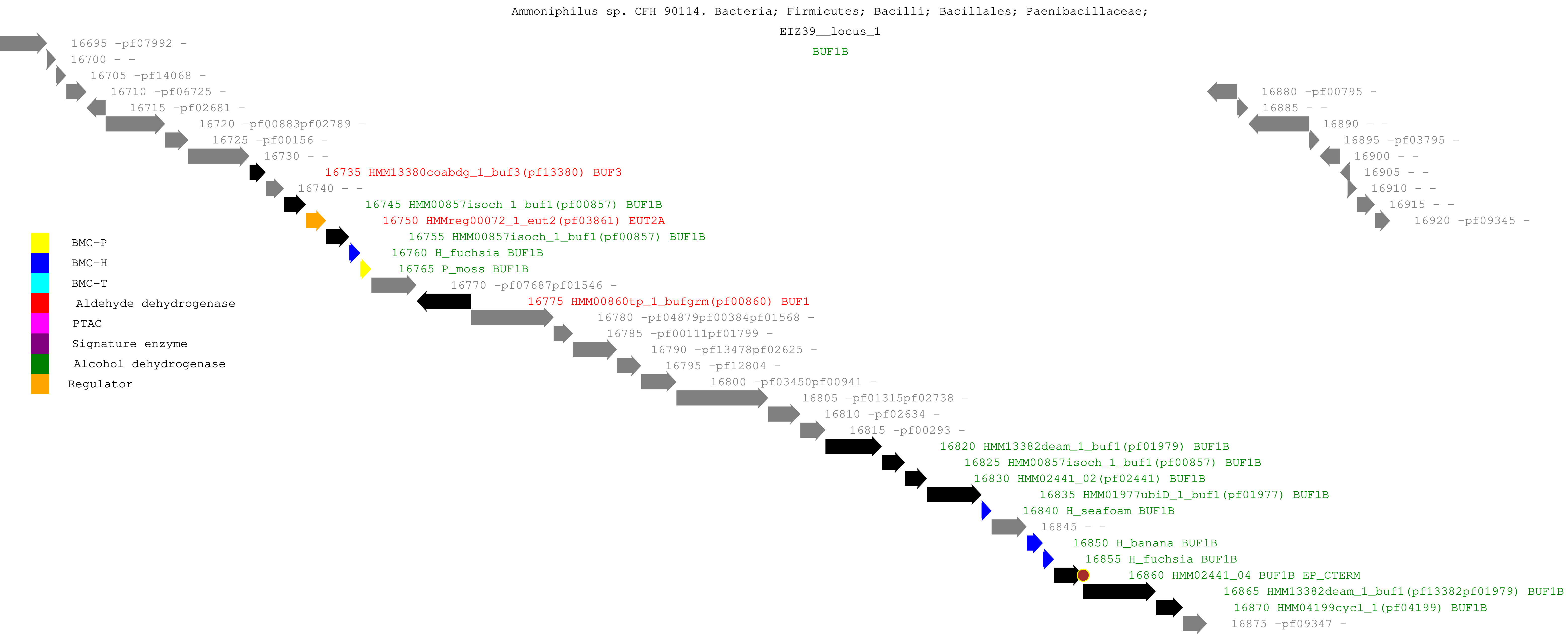

Ammoniphilus sp. CFH 90114. Bacteria; Firmicutes; Bacilli; Bacillales; Paenibacillaceae;
EIZ39__locus_1
BUF1B
16695 -pf07992 -
16700 - -
16705 -pf14068 -
16710 -pf06725 -
16715 -pf02681 -
16720 -pf00883pf02789 -
16725 -pf00156 -
16730 - -
16735 HMM13380coabdg_1_buf3(pf13380) BUF3
16740 - -
16745 HMM00857isoch_1_buf1(pf00857) BUF1B
16750 HMMreg00072_1_eut2(pf03861) EUT2A
16755 HMM00857isoch_1_buf1(pf00857) BUF1B
16760 H_fuchsia BUF1B
16765 P_moss BUF1B
16770 -pf07687pf01546 -
16775 HMM00860tp_1_bufgrm(pf00860) BUF1
16780 -pf04879pf00384pf01568 -
16785 -pf00111pf01799 -
16790 -pf13478pf02625 -
16795 -pf12804 -
16800 -pf03450pf00941 -
16805 -pf01315pf02738 -
16810 -pf02634 -
16815 -pf00293 -
16820 HMM13382deam_1_buf1(pf01979) BUF1B
16825 HMM00857isoch_1_buf1(pf00857) BUF1B
16830 HMM02441_02(pf02441) BUF1B
16835 HMM01977ubiD_1_buf1(pf01977) BUF1B
16840 H_seafoam BUF1B
16845 - -
16850 H_banana BUF1B
16855 H_fuchsia BUF1B
16860 HMM02441_04 BUF1B EP_CTERM
16865 HMM13382deam_1_buf1(pf13382pf01979) BUF1B
16870 HMM04199cycl_1(pf04199) BUF1B
16875 -pf09347 -
16880 -pf00795 -
16885 - -
16890 - -
16895 -pf03795 -
16900 - -
16905 - -
16910 - -
16915 - -
16920 -pf09345 -
BMC-P
BMC-H
BMC-T
Aldehyde dehydrogenase
PTAC
Signature enzyme
Alcohol dehydrogenase
Regulator

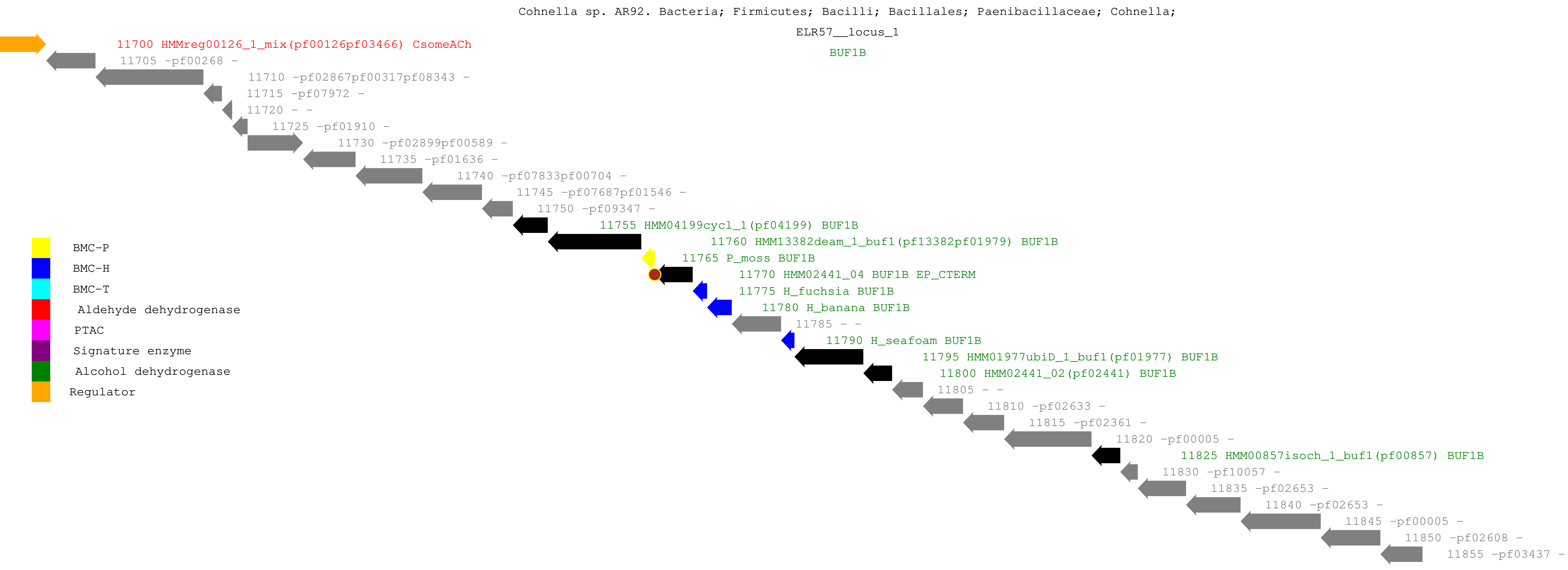
Cohnella sp. AR92. Bacteria; Firmicutes; Bacilli; Bacillales; Paenibacillaceae; Cohnella;
ELR57__locus_1
BUF1B
11700 HMMreg00126_1_mix(pf00126pf03466) CsomeACh
11705 -pf00268 -
11710 -pf02867pf00317pf08343 -
11715 -pf07972 -
11720 - -
11725 -pf01910 -
11730 -pf02899pf00589 -
11735 -pf01636 -
11740 -pf07833pf00704 -
11745 -pf07687pf01546 -
11750 -pf09347 -
11755 HMM04199cycl_1(pf04199) BUF1B
11760 HMM13382deam_1_buf1(pf13382pf01979) BUF1B
11765 P_moss BUF1B
11770 HMM02441_04 BUF1B EP_CTERM
11775 H_fuchsia BUF1B
11780 H_banana BUF1B
11785 - -
11790 H_seafoam BUF1B
11795 HMM01977ubiD_1_buf1(pf01977) BUF1B
11800 HMM02441_02(pf02441) BUF1B
11805 - -
11810 -pf02633 -
11815 -pf02361 -
11820 -pf00005 -
11825 HMM00857isoch_1_buf1(pf00857) BUF1B
11830 -pf10057 -
11835 -pf02653 -
11840 -pf02653 -
11845 -pf00005 -
11850 -pf02608 -
11855 -pf03437 -
BMC-P
BMC-H
BMC-T
Aldehyde dehydrogenase
PTAC
Signature enzyme
Alcohol dehydrogenase
Regulator

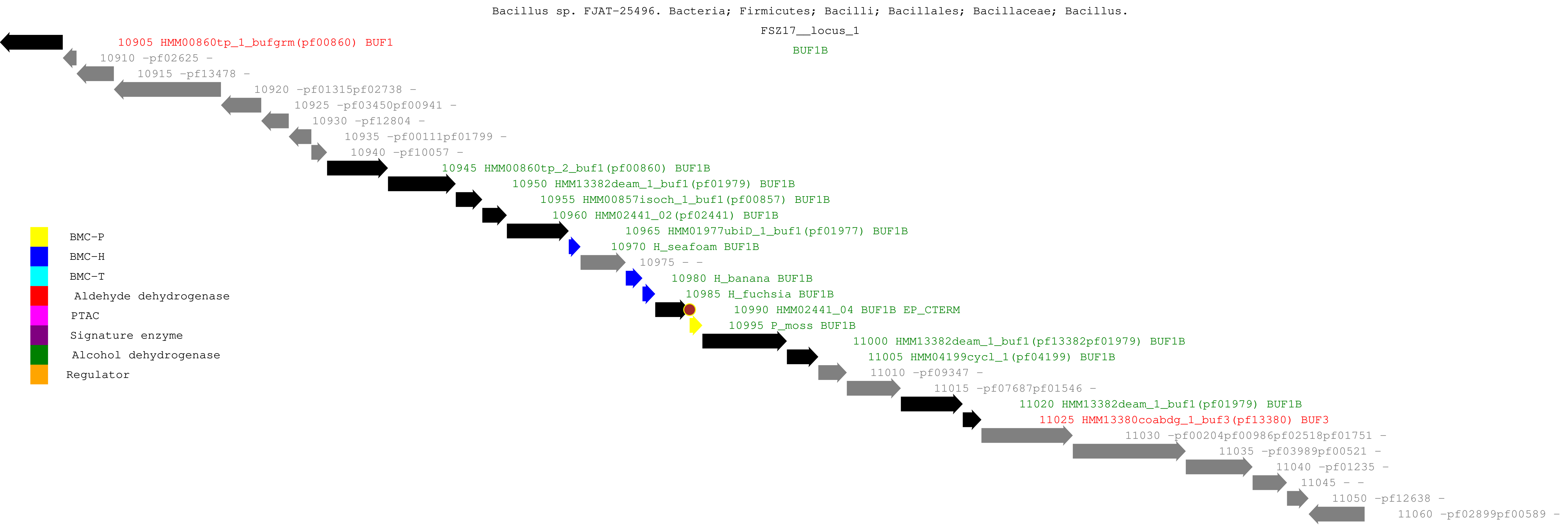

Bacillus sp. FJAT-25496. Bacteria; Firmicutes; Bacilli; Bacillales; Bacillaceae; Bacillus.
FSZ17__locus_1
BUF1B
10905 HMM00860tp_1_bufgrm(pf00860) BUF1
10910 -pf02625 -
10915 -pf13478 -
10920 -pf01315pf02738 -
10925 -pf03450pf00941 -
10930 -pf12804 -
10935 -pf00111pf01799 -
10940 -pf10057 -
10945 HMM00860tp_2_buf1(pf00860) BUF1B
10950 HMM13382deam_1_buf1(pf01979) BUF1B
10955 HMM00857isoch_1_buf1(pf00857) BUF1B
10960 HMM02441_02(pf02441) BUF1B
10965 HMM01977ubiD_1_buf1(pf01977) BUF1B
10970 H_seafoam BUF1B
10975 - -
10980 H_banana BUF1B
10985 H_fuchsia BUF1B
10990 HMM02441_04 BUF1B EP_CTERM
10995 P_moss BUF1B
11000 HMM13382deam_1_buf1(pf13382pf01979) BUF1B
11005 HMM04199cycl_1(pf04199) BUF1B
11010 -pf09347 -
11015 -pf07687pf01546 -
11020 HMM13382deam_1_buf1(pf01979) BUF1B
11025 HMM13380coabdg_1_buf3(pf13380) BUF3
11030 -pf00204pf00986pf02518pf01751 -
11035 -pf03989pf00521 -
11040 -pf01235 -
11045 - -
11050 -pf12638 -
11060 -pf02899pf00589 -
BMC-P
BMC-H
BMC-T
Aldehyde dehydrogenase
PTAC
Signature enzyme
Alcohol dehydrogenase
Regulator

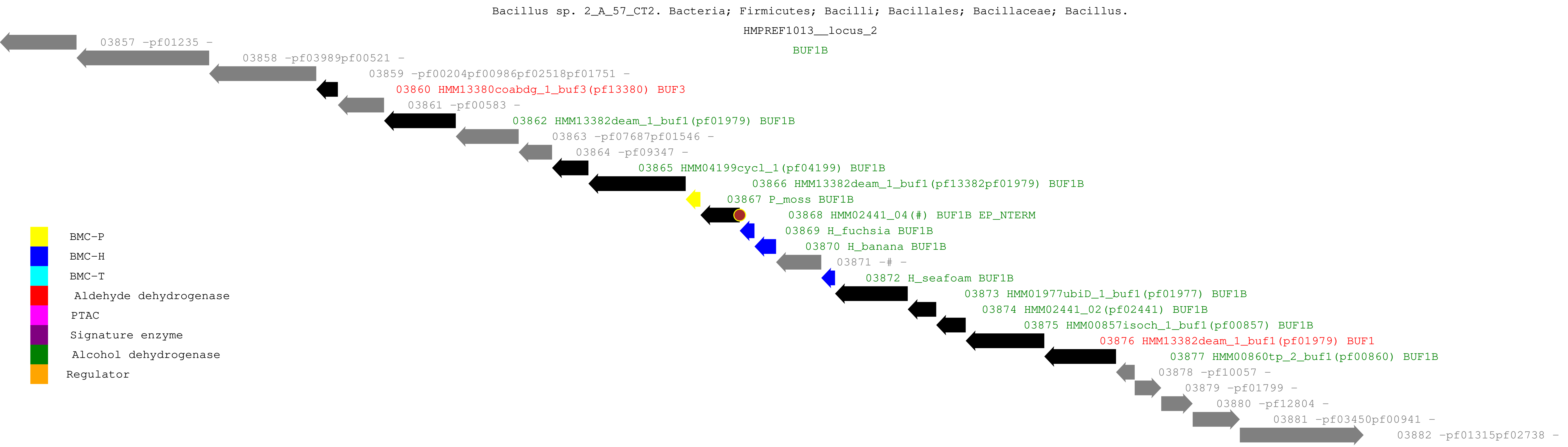
Bacillus sp. 2_A_57_CT2. Bacteria; Firmicutes; Bacilli; Bacillales; Bacillaceae; Bacillus.
HMPREF1013__locus_2
BUF1B
03857 -pf01235 -
03858 -pf03989pf00521 -
03859 -pf00204pf00986pf02518pf01751 -
03860 HMM13380coabdg_1_buf3(pf13380) BUF3
03861 -pf00583 -
03862 HMM13382deam_1_buf1(pf01979) BUF1B
03863 -pf07687pf01546 -
03864 -pf09347 -
03865 HMM04199cycl_1(pf04199) BUF1B
03866 HMM13382deam_1_buf1(pf13382pf01979) BUF1B
03867 P_moss BUF1B
03868 HMM02441_04(#) BUF1B EP_NTERM
03869 H_fuchsia BUF1B
03870 H_banana BUF1B
03871 -# -
03872 H_seafoam BUF1B
03873 HMM01977ubiD_1_buf1(pf01977) BUF1B
03874 HMM02441_02(pf02441) BUF1B
03875 HMM00857isoch_1_buf1(pf00857) BUF1B
03876 HMM13382deam_1_buf1(pf01979) BUF1
03877 HMM00860tp_2_buf1(pf00860) BUF1B
03878 -pf10057 -
03879 -pf01799 -
03880 -pf12804 -
03881 -pf03450pf00941 -
03882 -pf01315pf02738 -
BMC-P
BMC-H
BMC-T
Aldehyde dehydrogenase
PTAC
Signature enzyme
Alcohol dehydrogenase
Regulator

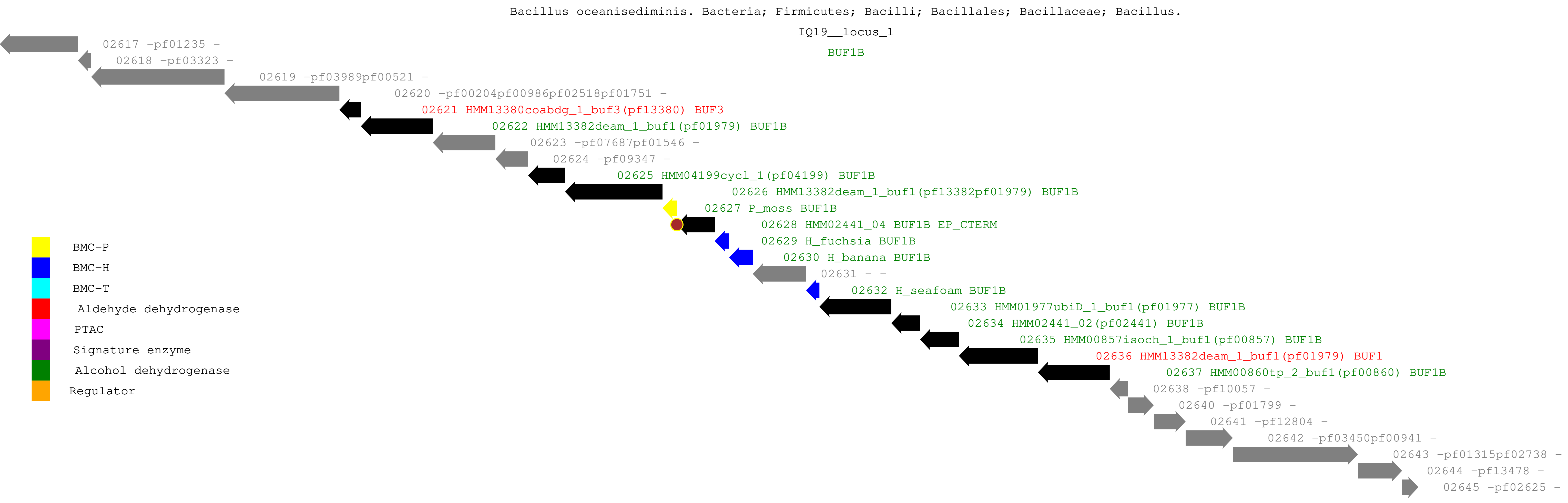
Bacillus oceanisediminis. Bacteria; Firmicutes; Bacilli; Bacillales; Bacillaceae; Bacillus.
IQ19__locus_1
BUF1B
02617 -pf01235 -
02618 -pf03323 -
02619 -pf03989pf00521 -
02620 -pf00204pf00986pf02518pf01751 -
02621 HMM13380coabdg_1_buf3(pf13380) BUF3
02622 HMM13382deam_1_buf1(pf01979) BUF1B
02623 -pf07687pf01546 -
02624 -pf09347 -
02625 HMM04199cycl_1(pf04199) BUF1B
02626 HMM13382deam_1_buf1(pf13382pf01979) BUF1B
02627 P_moss BUF1B
02628 HMM02441_04 BUF1B EP_CTERM
02629 H_fuchsia BUF1B
02630 H_banana BUF1B
02631 - -
02632 H_seafoam BUF1B
02633 HMM01977ubiD_1_buf1(pf01977) BUF1B
02634 HMM02441_02(pf02441) BUF1B
02635 HMM00857isoch_1_buf1(pf00857) BUF1B
02636 HMM13382deam_1_buf1(pf01979) BUF1
02637 HMM00860tp_2_buf1(pf00860) BUF1B
02638 -pf10057 -
02640 -pf01799 -
02641 -pf12804 -
02642 -pf03450pf00941 -
02643 -pf01315pf02738 -
02644 -pf13478 -
02645 -pf02625 -
BMC-P
BMC-H
BMC-T
Aldehyde dehydrogenase
PTAC
Signature enzyme
Alcohol dehydrogenase
Regulator

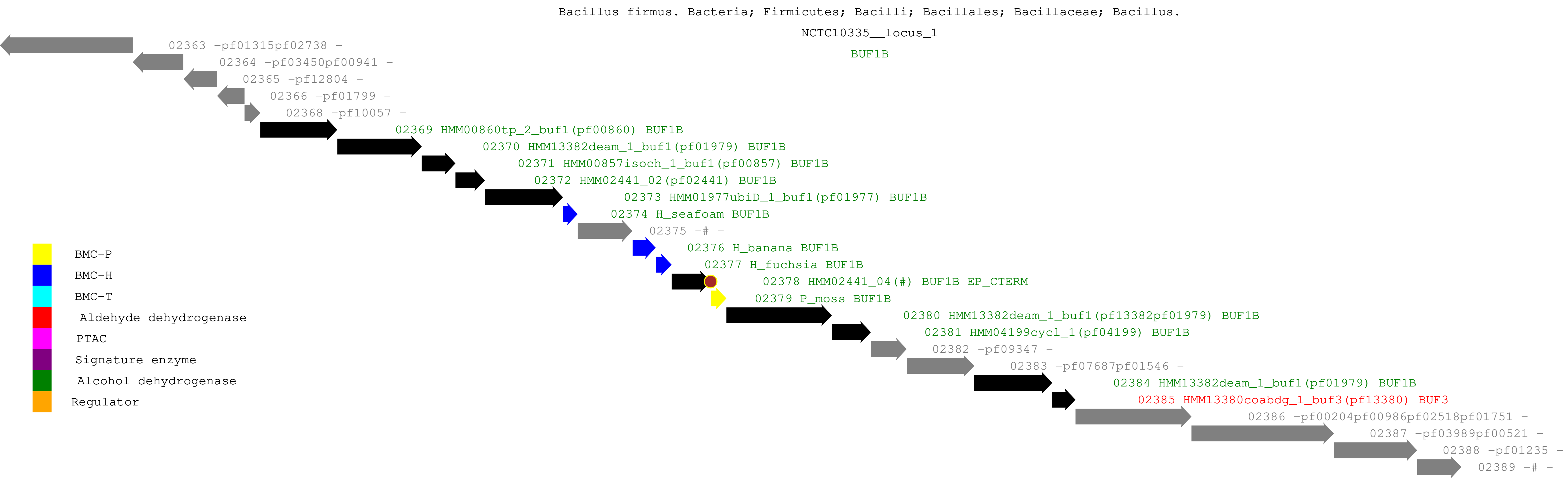

Bacillus firmus. Bacteria; Firmicutes; Bacilli; Bacillales; Bacillaceae; Bacillus.
NCTC10335__locus_1
BUF1B
02363 -pf01315pf02738 -
02364 -pf03450pf00941 -
02365 -pf12804 -
02366 -pf01799 -
02368 -pf10057 -
02369 HMM00860tp_2_buf1(pf00860) BUF1B
02370 HMM13382deam_1_buf1(pf01979) BUF1B
02371 HMM00857isoch_1_buf1(pf00857) BUF1B
02372 HMM02441_02(pf02441) BUF1B
02373 HMM01977ubiD_1_buf1(pf01977) BUF1B
02374 H_seafoam BUF1B
02375 -# -
02376 H_banana BUF1B
02377 H_fuchsia BUF1B
02378 HMM02441_04(#) BUF1B EP_CTERM
02379 P_moss BUF1B
02380 HMM13382deam_1_buf1(pf13382pf01979) BUF1B
02381 HMM04199cycl_1(pf04199) BUF1B
02382 -pf09347 -
02383 -pf07687pf01546 -
02384 HMM13382deam_1_buf1(pf01979) BUF1B
02385 HMM13380coabdg_1_buf3(pf13380) BUF3
02386 -pf00204pf00986pf02518pf01751 -
02387 -pf03989pf00521 -
02388 -pf01235 -
02389 -# -
BMC-P
BMC-H
BMC-T
Aldehyde dehydrogenase
PTAC
Signature enzyme
Alcohol dehydrogenase
Regulator

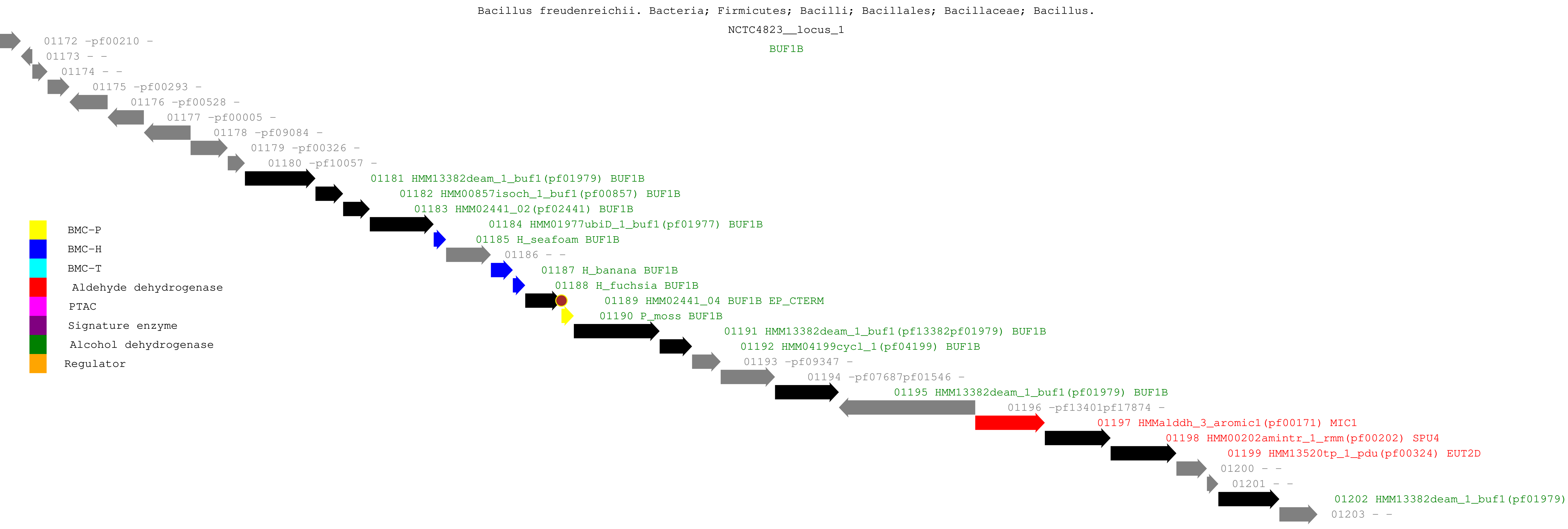

Bacillus freudenreichii. Bacteria; Firmicutes; Bacilli; Bacillales; Bacillaceae; Bacillus.
NCTC4823__locus_1
BUF1B
01172 -pf00210 -
01173 - -
01174 - -
01175 -pf00293 -
01176 -pf00528 -
01177 -pf00005 -
01178 -pf09084 -
01179 -pf00326 -
01180 -pf10057 -
01181 HMM13382deam_1_buf1(pf01979) BUF1B
01182 HMM00857isoch_1_buf1(pf00857) BUF1B
01183 HMM02441_02(pf02441) BUF1B
01184 HMM01977ubiD_1_buf1(pf01977) BUF1B
01185 H_seafoam BUF1B
01186 - -
01187 H_banana BUF1B
01188 H_fuchsia BUF1B
01189 HMM02441_04 BUF1B EP_CTERM
01190 P_moss BUF1B
01191 HMM13382deam_1_buf1(pf13382pf01979) BUF1B
01192 HMM04199cycl_1(pf04199) BUF1B
01193 -pf09347 -
01194 -pf07687pf01546 -
01195 HMM13382deam_1_buf1(pf01979) BUF1B
01196 -pf13401pf17874 -
01197 HMMalddh_3_aromic1(pf00171) MIC1
01198 HMM00202amintr_1_rmm(pf00202) SPU4
01199 HMM13520tp_1_pdu(pf00324) EUT2D
01200 - -
01201 - -
01202 HMM13382deam_1_buf1(pf01979)
01203 - -
BMC-P
BMC-H
BMC-T
Aldehyde dehydrogenase
PTAC
Signature enzyme
Alcohol dehydrogenase
Regulator

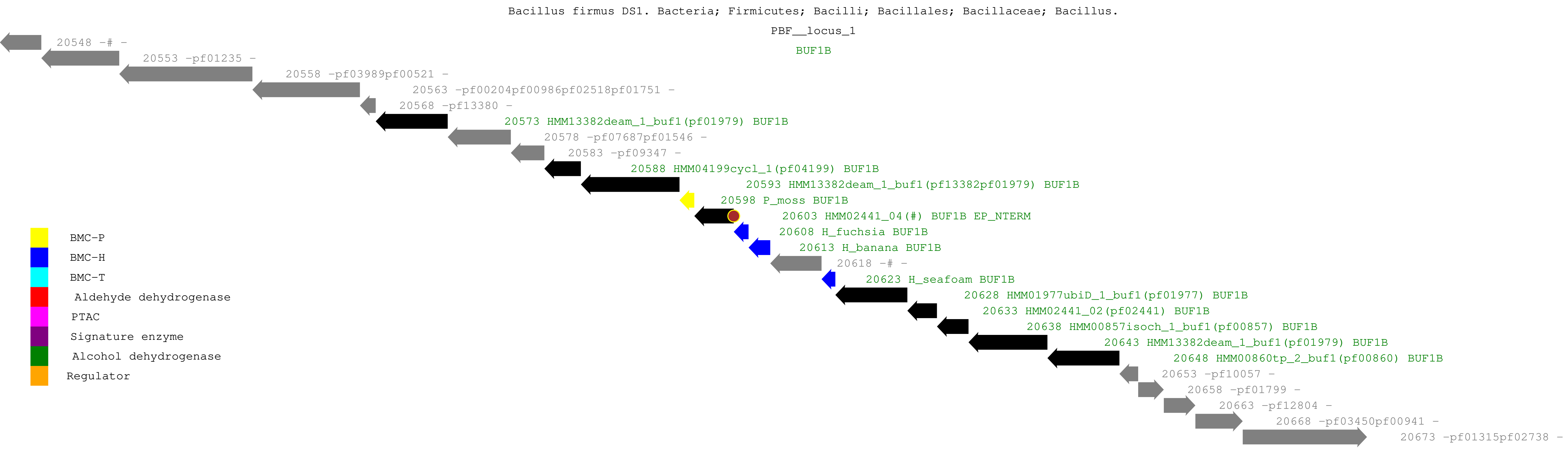
Bacillus firmus DS1. Bacteria; Firmicutes; Bacilli; Bacillales; Bacillaceae; Bacillus.
PBF__locus_1
BUF1B
20548 -# -
20553 -pf01235 -
20558 -pf03989pf00521 -
20563 -pf00204pf00986pf02518pf01751 -
20568 -pf13380 -
20573 HMM13382deam_1_buf1(pf01979) BUF1B
20578 -pf07687pf01546 -
20583 -pf09347 -
20588 HMM04199cycl_1(pf04199) BUF1B
20593 HMM13382deam_1_buf1(pf13382pf01979) BUF1B
20598 P_moss BUF1B
20603 HMM02441_04(#) BUF1B EP_NTERM
20608 H_fuchsia BUF1B
20613 H_banana BUF1B
20618 -# -
20623 H_seafoam BUF1B
20628 HMM01977ubiD_1_buf1(pf01977) BUF1B
20633 HMM02441_02(pf02441) BUF1B
20638 HMM00857isoch_1_buf1(pf00857) BUF1B
20643 HMM13382deam_1_buf1(pf01979) BUF1B
20648 HMM00860tp_2_buf1(pf00860) BUF1B
20653 -pf10057 -
20658 -pf01799 -
20663 -pf12804 -
20668 -pf03450pf00941 -
20673 -pf01315pf02738 -
BMC-P
BMC-H
BMC-T
Aldehyde dehydrogenase
PTAC
Signature enzyme
Alcohol dehydrogenase
Regulator

Paenibacillus borealis. Bacteria; Firmicutes; Bacilli; Bacillales; Paenibacillaceae;

PBOR__locus_1

BUF1B

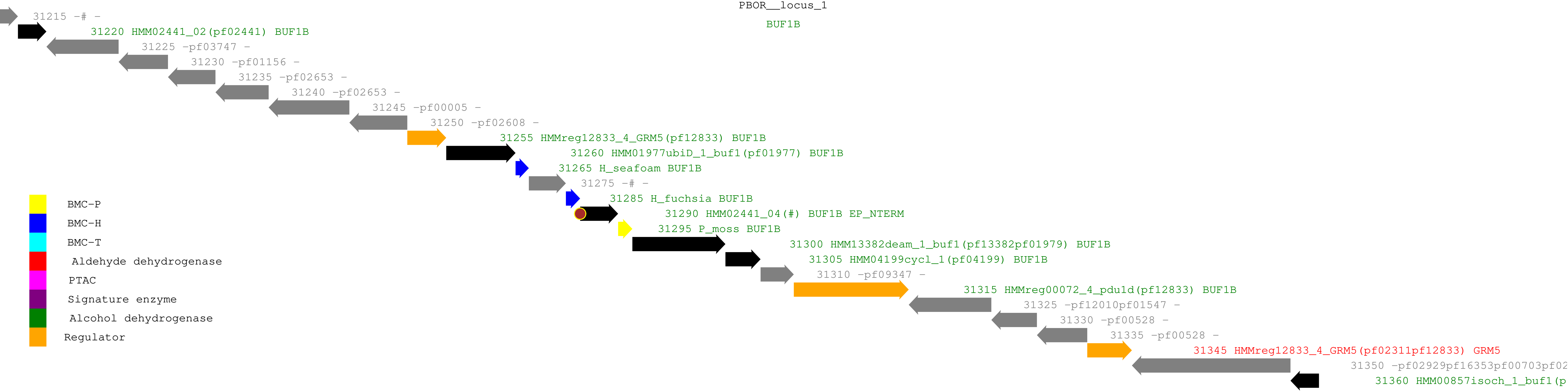

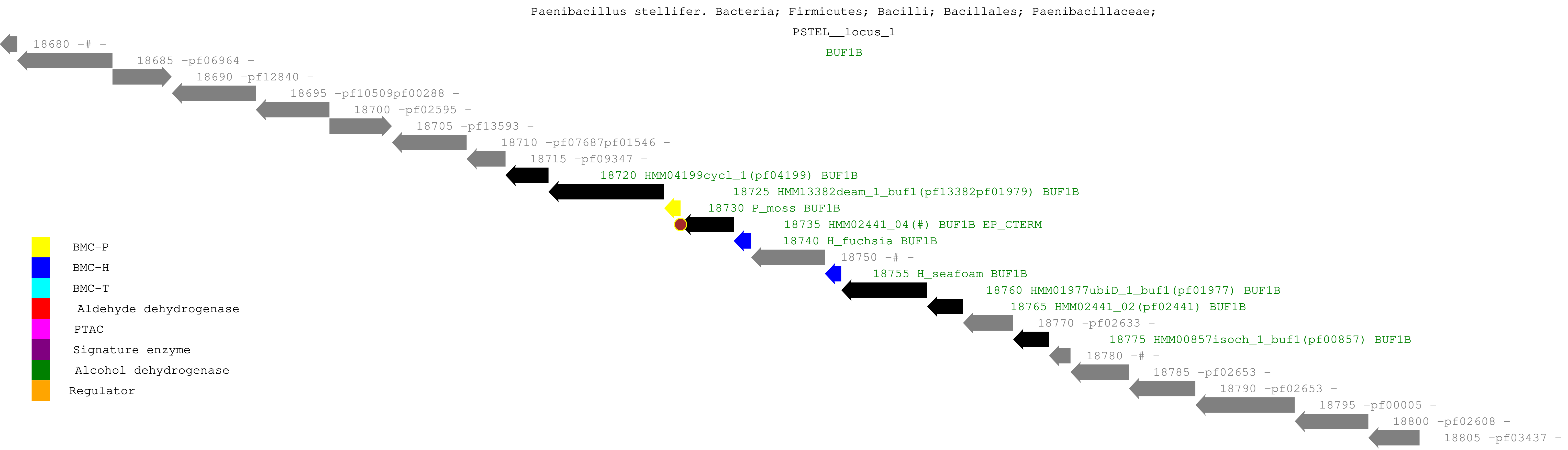
Paenibacillus stellifer. Bacteria; Firmicutes; Bacilli; Bacillales; Paenibacillaceae;
PSTEL__locus_1
BUF1B
18680 -# -
18685 -pf06964 -
18690 -pf12840 -
18695 -pf10509pf00288 -
18700 -pf02595 -
18705 -pf13593 -
18710 -pf07687pf01546 -
18715 -pf09347 -
18720 HMM04199cycl_1(pf04199) BUF1B
18725 HMM13382deam_1_buf1(pf13382pf01979) BUF1B
18730 P_moss BUF1B
18735 HMM02441_04(#) BUF1B EP_CTERM
18740 H_fuchsia BUF1B
18750 -# -
18755 H_seafoam BUF1B
18760 HMM01977ubiD_1_buf1(pf01977) BUF1B
18765 HMM02441_02(pf02441) BUF1B
18770 -pf02633 -
18775 HMM00857isoch_1_buf1(pf00857) BUF1B
18780 -# -
18785 -pf02653 -
18790 -pf02653 -
18795 -pf00005 -
18800 -pf02608 -
18805 -pf03437 -
BMC-P
BMC-H
BMC-T
Aldehyde dehydrogenase
PTAC
Signature enzyme
Alcohol dehydrogenase
Regulator

Paenibacillus sp. IHB B 3415. Bacteria; Firmicutes; Bacilli; Bacillales; Paenibacillaceae;

QW71__locus_1

BUF1B

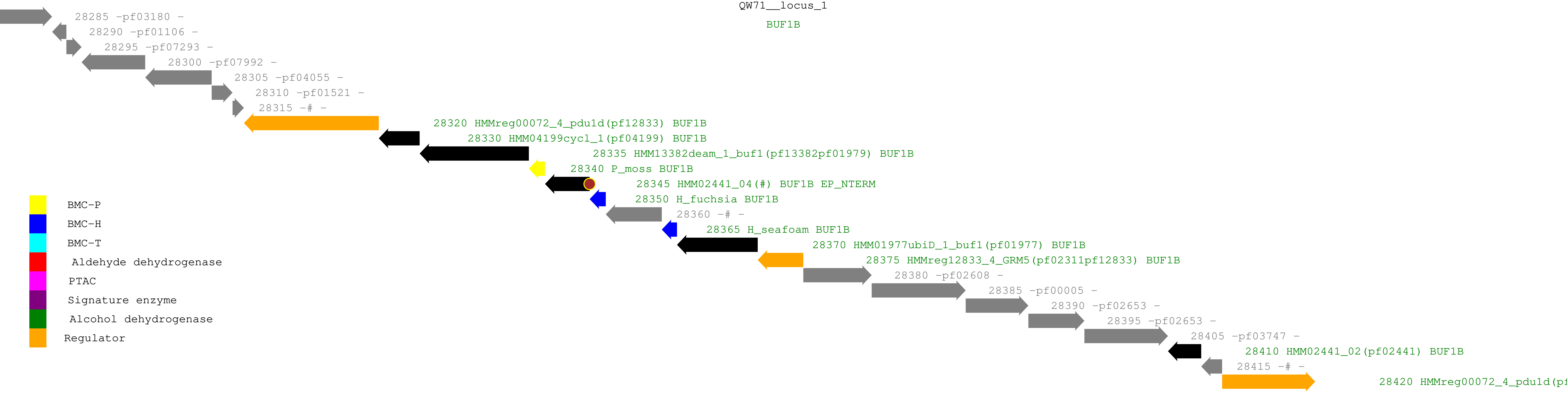

Paenibacillus sp. FSL R5-0912. Bacteria; Firmicutes; Bacilli; Bacillales; Paenibacillaceae;

R50912__locus_1

BUF1B

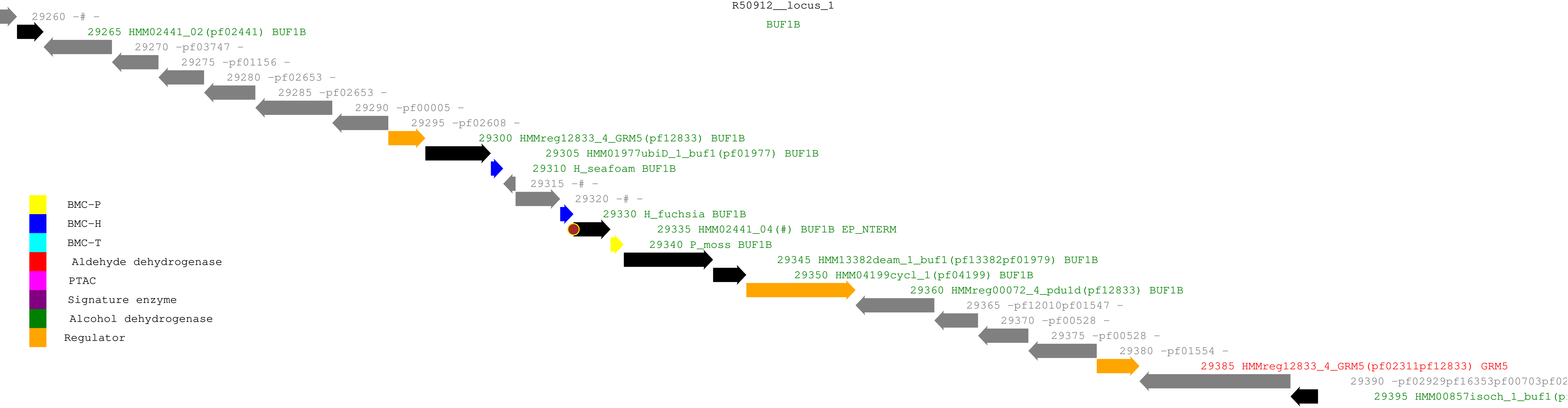

Paenibacillus sp. FSL R7-0331. Bacteria; Firmicutes; Bacilli; Bacillales; Paenibacillaceae;

R70331__locus_1

BUF1B

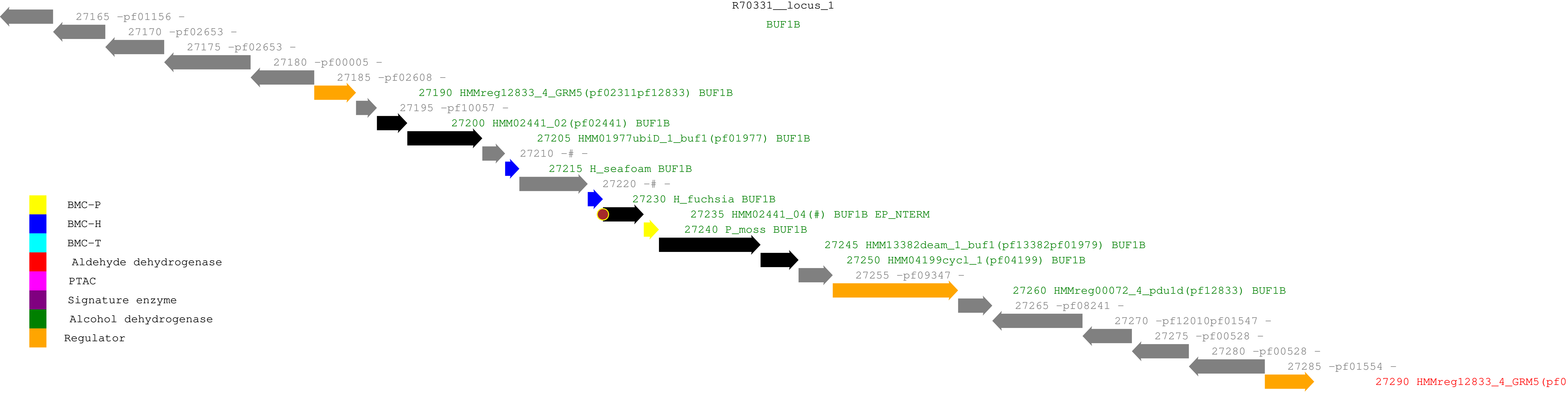

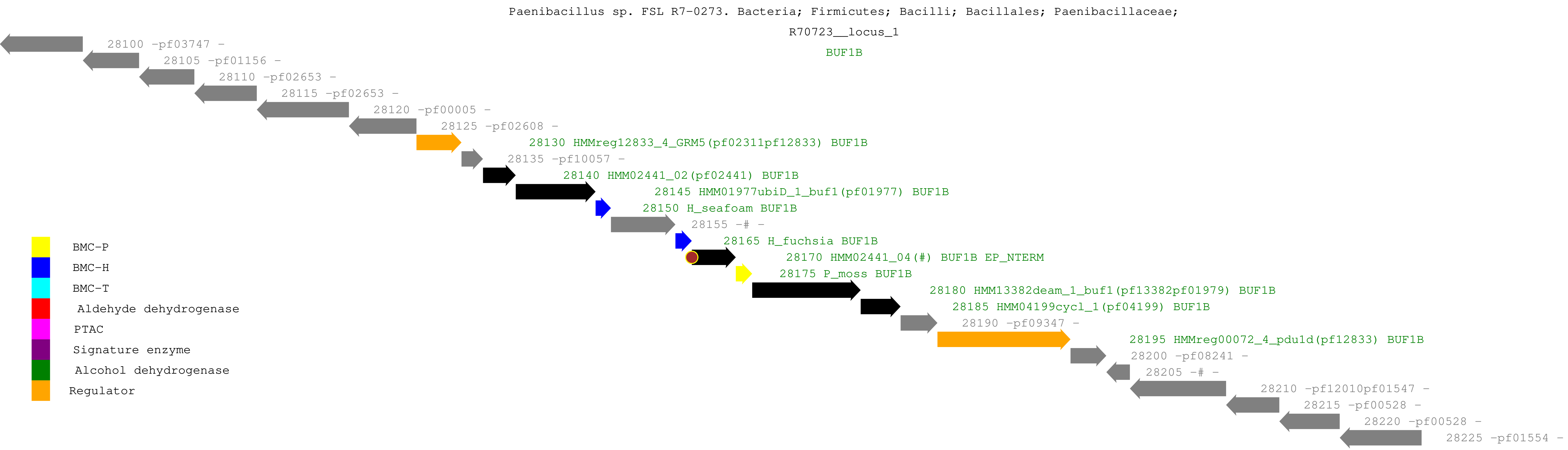
Paenibacillus sp. FSL R7-0273. Bacteria; Firmicutes; Bacilli; Bacillales; Paenibacillaceae;
R70723__locus_1
BUF1B
28100 -pf03747 -
28105 -pf01156 -
28110 -pf02653 -
28115 -pf02653 -
28120 -pf00005 -
28125 -pf02608 -
28130 HMMreg12833_4_GRM5(pf02311pf12833) BUF1B
28135 -pf10057 -
28140 HMM02441_02(pf02441) BUF1B
28145 HMM01977ubiD_1_buf1(pf01977) BUF1B
28150 H_seafoam BUF1B
28155 -# -
28165 H_fuchsia BUF1B
28170 HMM02441_04(#) BUF1B EP_NTERM
28175 P_moss BUF1B
28180 HMM13382deam_1_buf1(pf13382pf01979) BUF1B
28185 HMM04199cycl_1(pf04199) BUF1B
28190 -pf09347 -
28195 HMMreg00072_4_pdu1d(pf12833) BUF1B
28200 -pf08241 -
28205 -# -
28210 -pf12010pf01547 -
28215 -pf00528 -
28220 -pf00528 -
28225 -pf01554 -
BMC-P
BMC-H
BMC-T
Aldehyde dehydrogenase
PTAC
Signature enzyme
Alcohol dehydrogenase
Regulator

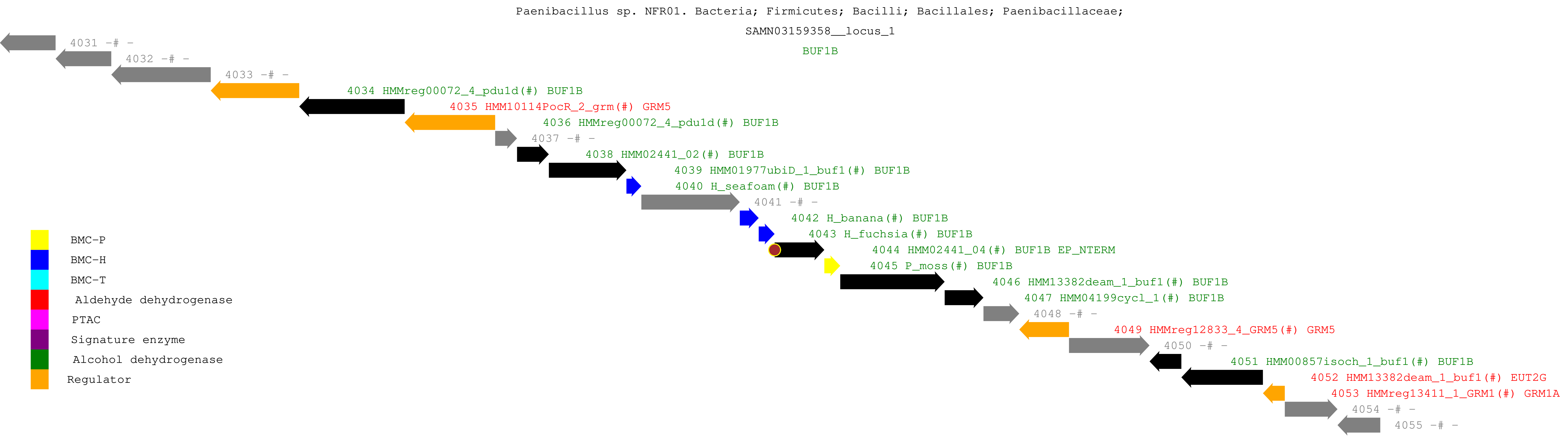

Paenibacillus sp. NFR01. Bacteria; Firmicutes; Bacilli; Bacillales; Paenibacillaceae;
SAMN03159358__locus_1
BUF1B
4031 -# -
4032 -# -
4033 -# -
4034 HMMreg00072_4_pdu1d(#) BUF1B
4035 HMM10114PocR_2_grm(#) GRM5
4036 HMMreg00072_4_pdu1d(#) BUF1B
4037 -# -
4038 HMM02441_02(#) BUF1B
4039 HMM01977ubiD_1_buf1(#) BUF1B
4040 H_seafoam(#) BUF1B
4041 -# -
4042 H_banana(#) BUF1B
4043 H_fuchsia(#) BUF1B
4044 HMM02441_04(#) BUF1B EP_NTERM
4045 P_moss(#) BUF1B
4046 HMM13382deam_1_buf1(#) BUF1B
4047 HMM04199cycl_1(#) BUF1B
4048 -# -
4049 HMMreg12833_4_GRM5(#) GRM5
4050 -# -
4051 HMM00857isoch_1_buf1(#) BUF1B
4052 HMM13382deam_1_buf1(#) EUT2G
4053 HMMreg13411_1_GRM1(#) GRM1A
4054 -# -
4055 -# -
BMC-P
BMC-H
BMC-T
Aldehyde dehydrogenase
PTAC
Signature enzyme
Alcohol dehydrogenase
Regulator

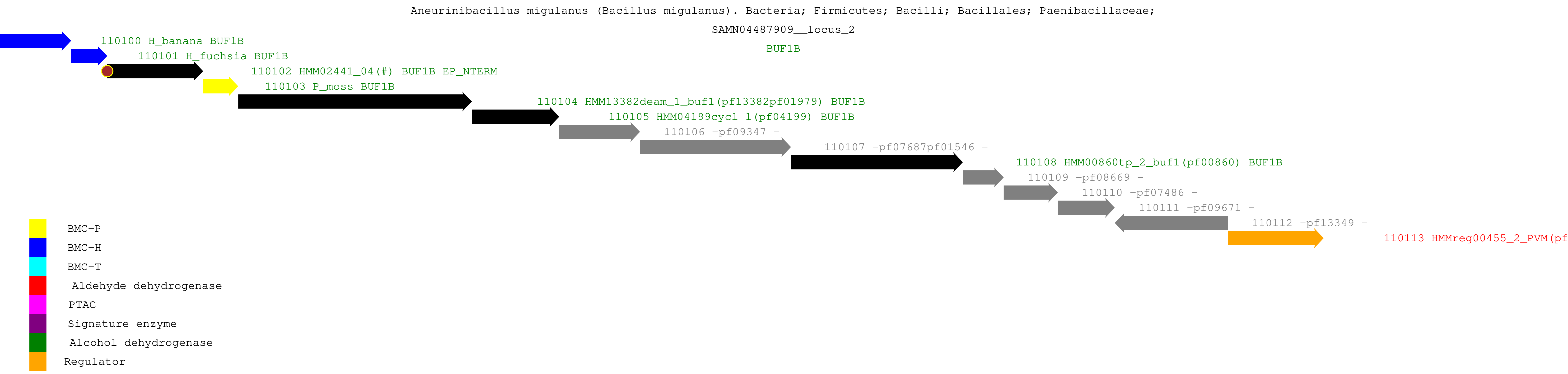

Aneurinibacillus migulanus (Bacillus migulanus). Bacteria; Firmicutes; Bacilli; Bacillales; Paenibacillaceae;
SAMN04487909__locus_2
BUF1B
110100 H_banana BUF1B
110101 H_fuchsia BUF1B
110102 HMM02441_04(#) BUF1B EP_NTERM
110103 P_moss BUF1B
110104 HMM13382deam_1_buf1(pf13382pf01979) BUF1B
110105 HMM04199cycl_1(pf04199) BUF1B
110106 -pf09347 -
110107 -pf07687pf01546 -
110108 HMM00860tp_2_buf1(pf00860) BUF1B
110109 -pf08669 -
110110 -pf07486 -
110111 -pf09671 -
110112 -pf13349 -
110113 HMMreg00455_2_PVM(pf
BMC-P
BMC-H
BMC-T
Aldehyde dehydrogenase
PTAC
Signature enzyme
Alcohol dehydrogenase
Regulator

Natribacillus halophilus. Bacteria; Firmicutes; Bacilli; Bacillales; Bacillaceae; Natribacillus.

SAMN04488123__locus_1

BUF1B

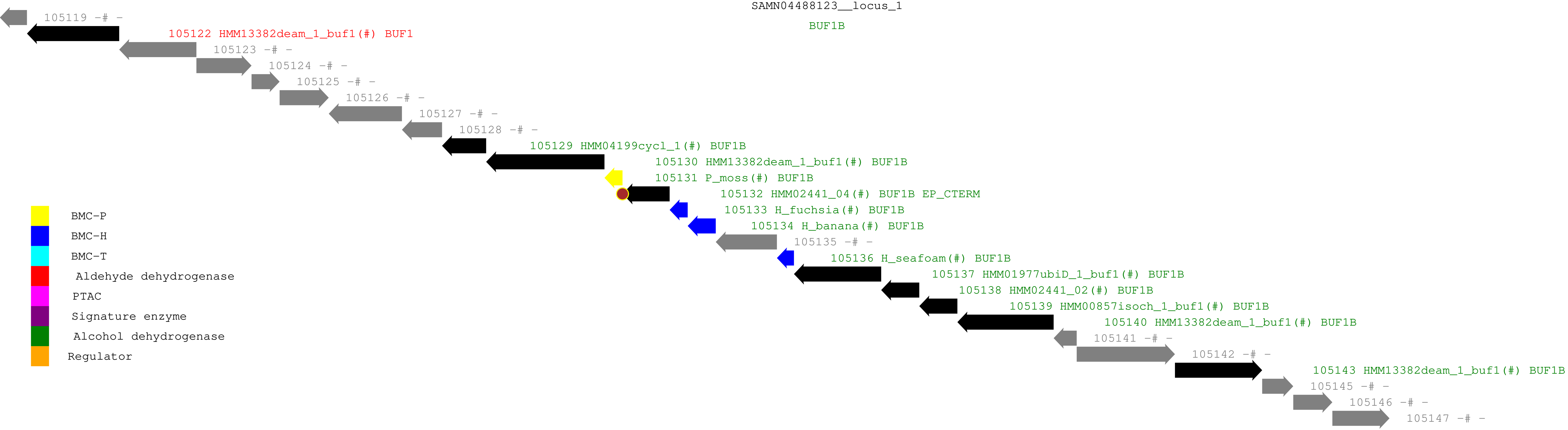

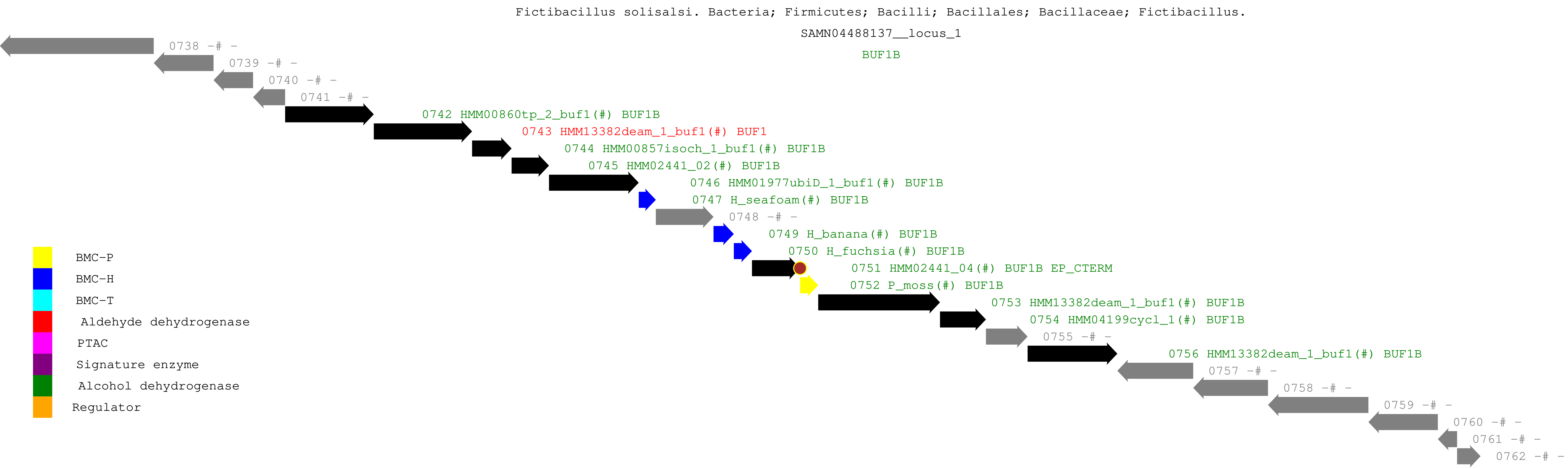

Fictibacillus solisalsi. Bacteria; Firmicutes; Bacilli; Bacillales; Bacillaceae; Fictibacillus.
SAMN04488137__locus_1
BUF1B
0738 -# -
0739 -# -
0740 -# -
0741 -# -
0742 HMM00860tp_2_buf1(#) BUF1B
0743 HMM13382deam_1_buf1(#) BUF1
0744 HMM00857isoch_1_buf1(#) BUF1B
0745 HMM02441_02(#) BUF1B
0746 HMM01977ubiD_1_buf1(#) BUF1B
0747 H_seafoam(#) BUF1B
0748 -# -
0749 H_banana(#) BUF1B
0750 H_fuchsia(#) BUF1B
0751 HMM02441_04(#) BUF1B EP_CTERM
0752 P_moss(#) BUF1B
0753 HMM13382deam_1_buf1(#) BUF1B
0754 HMM04199cycl_1(#) BUF1B
0755 -# -
0756 HMM13382deam_1_buf1(#) BUF1B
0757 -# -
0758 -# -
0759 -# -
0760 -# -
0761 -# -
0762 -# -
BMC-P
BMC-H
BMC-T
Aldehyde dehydrogenase
PTAC
Signature enzyme
Alcohol dehydrogenase
Regulator

Fontibacillus panacisegetis. Bacteria; Firmicutes; Bacilli; Bacillales; Paenibacillaceae;

SAMN04488542__locus_1

BUF1B

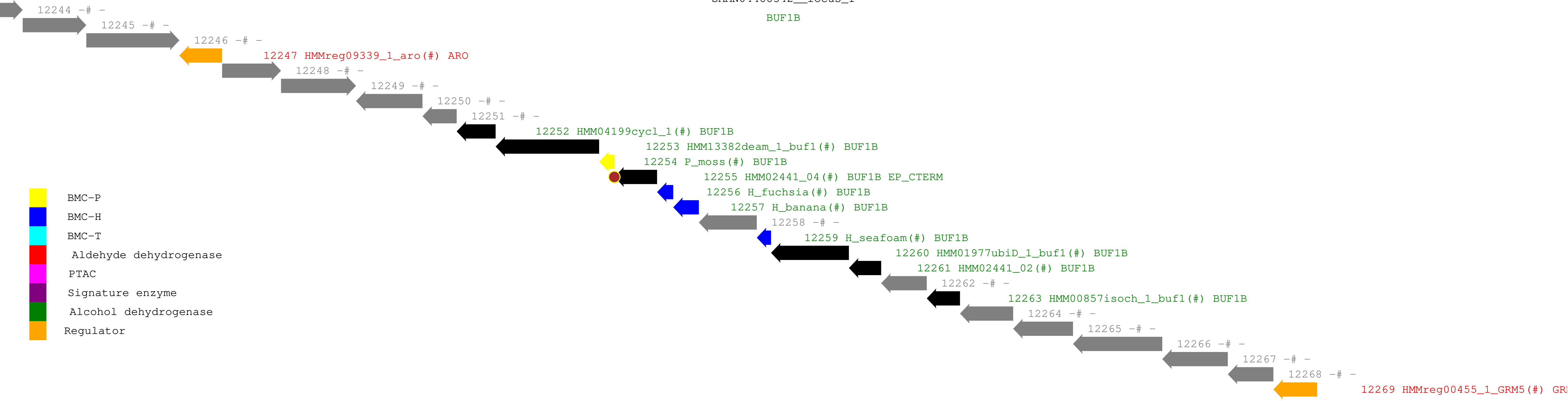

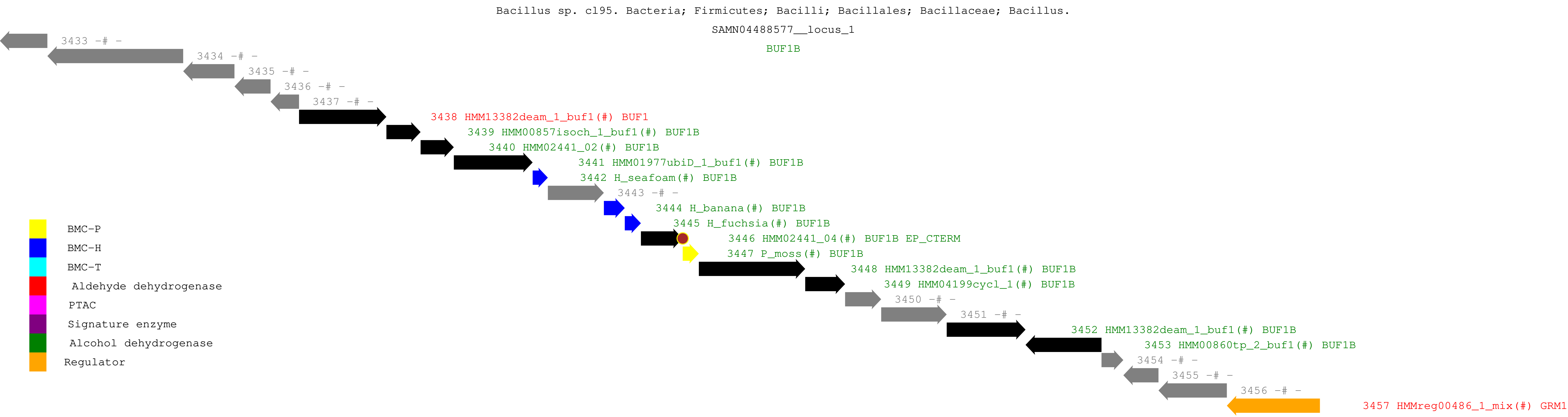

Bacillus sp. cl95. Bacteria; Firmicutes; Bacilli; Bacillales; Bacillaceae; Bacillus.
SAMN04488577__locus_1
BUF1B
3433 -# -
3434 -# -
3435 -# -
3436 -# -
3437 -# -
3438 HMM13382deam_1_buf1(#) BUF1
3439 HMM00857isoch_1_buf1(#) BUF1B
3440 HMM02441_02(#) BUF1B
3441 HMM01977ubiD_1_buf1(#) BUF1B
3442 H_seafoam(#) BUF1B
3443 -# -
3444 H_banana(#) BUF1B
3445 H_fuchsia(#) BUF1B
3446 HMM02441_04(#) BUF1B EP_CTERM
3447 P_moss(#) BUF1B
3448 HMM13382deam_1_buf1(#) BUF1B
3449 HMM04199cycl_1(#) BUF1B
3450 -# -
3451 -# -
3452 HMM13382deam_1_buf1(#) BUF1B
3453 HMM00860tp_2_buf1(#) BUF1B
3454 -# -
3455 -# -
3456 -# -
3457 HMMreg00486_1_mix(#) GRM1
BMC-P
BMC-H
BMC-T
Aldehyde dehydrogenase
PTAC
Signature enzyme
Alcohol dehydrogenase
Regulator

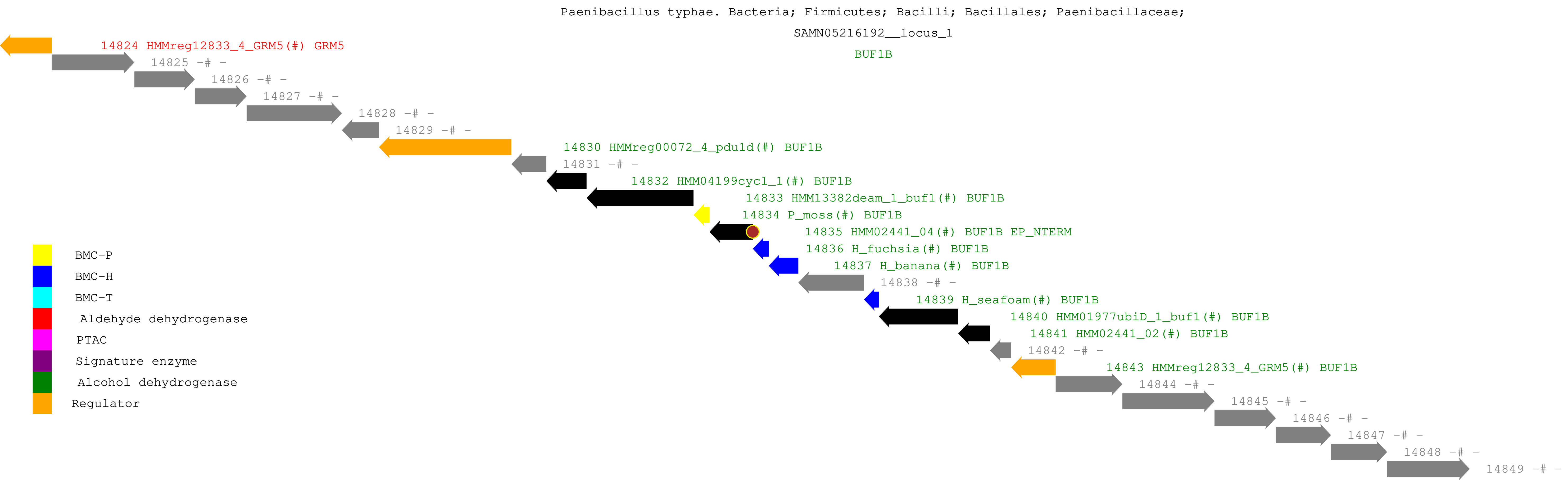
Paenibacillus typhae. Bacteria; Firmicutes; Bacilli; Bacillales; Paenibacillaceae;
SAMN05216192__locus_1
BUF1B
14824 HMMreg12833_4_GRM5(#) GRM5
14825 -# -
14826 -# -
14827 -# -
14828 -# -
14829 -# -
14830 HMMreg00072_4_pdu1d(#) BUF1B
14831 -# -
14832 HMM04199cycl_1(#) BUF1B
14833 HMM13382deam_1_buf1(#) BUF1B
14834 P_moss(#) BUF1B
14835 HMM02441_04(#) BUF1B EP_NTERM
14836 H_fuchsia(#) BUF1B
14837 H_banana(#) BUF1B
14838 -# -
14839 H_seafoam(#) BUF1B
14840 HMM01977ubiD_1_buf1(#) BUF1B
14841 HMM02441_02(#) BUF1B
14842 -# -
14843 HMMreg12833_4_GRM5(#) BUF1B
14844 -# -
14845 -# -
14846 -# -
14847 -# -
14848 -# -
14849 -# -
BMC-P
BMC-H
BMC-T
Aldehyde dehydrogenase
PTAC
Signature enzyme
Alcohol dehydrogenase
Regulator

Ornithinibacillus halophilus. Bacteria; Firmicutes; Bacilli; Bacillales; Bacillaceae;

SAMN05216225__locus_1

BUF1B

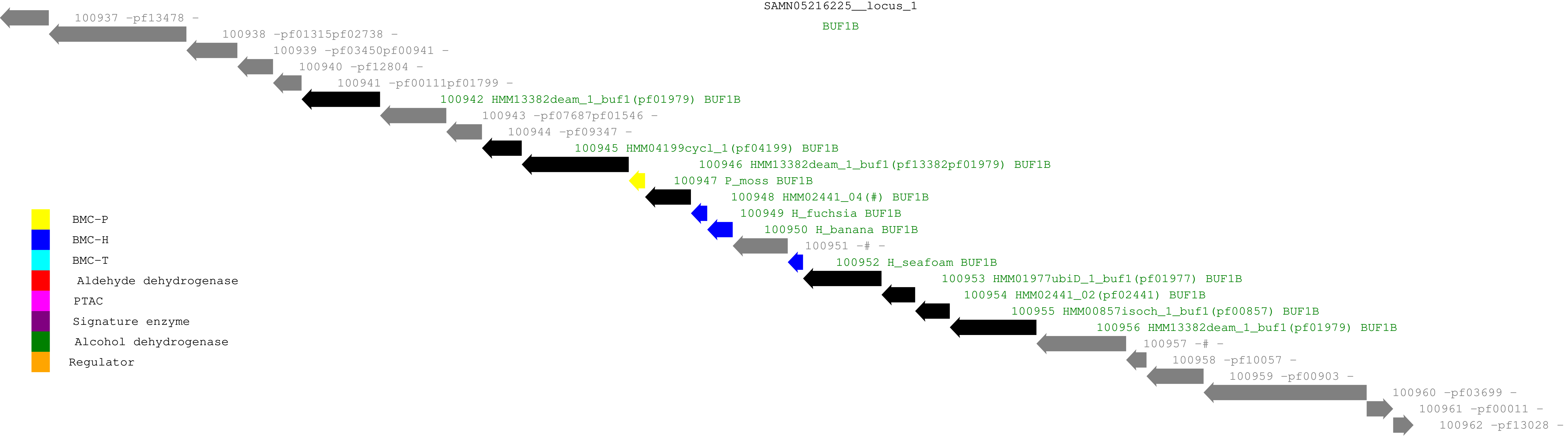

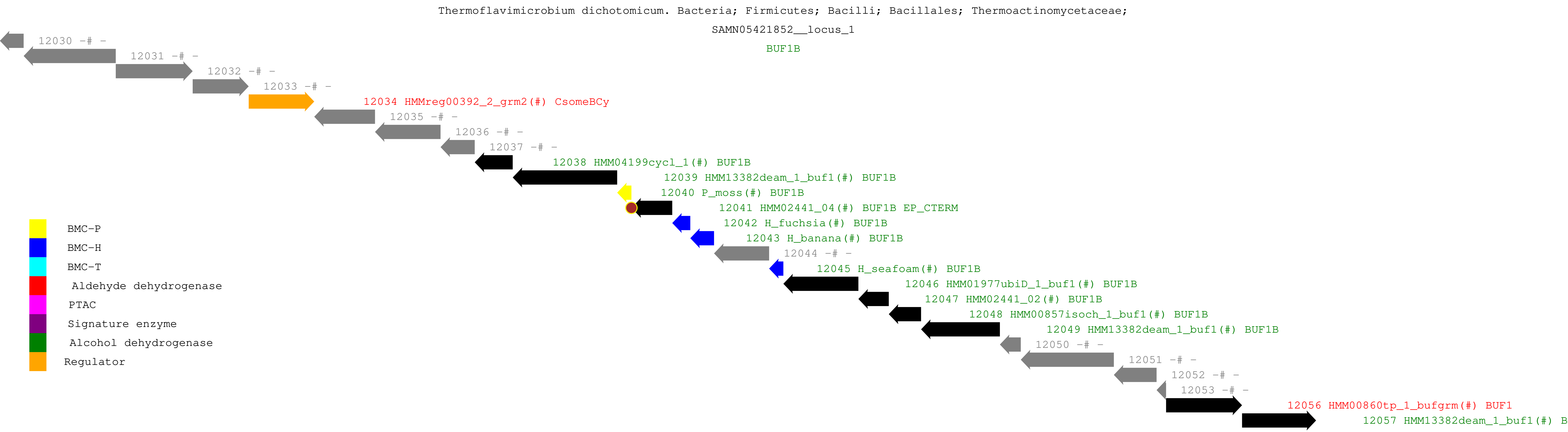
Thermoflavimicrobium dichotomicum. Bacteria; Firmicutes; Bacilli; Bacillales; Thermoactinomycetaceae;
SAMN05421852__locus_1
BUF1B
12030 -# -
12031 -# -
12032 -# -
12033 -# -
12034 HMMreg00392_2_grm2(#) CsomeBCy
12035 -# -
12036 -# -
12037 -# -
12038 HMM04199cycl_1(#) BUF1B
12039 HMM13382deam_1_buf1(#) BUF1B
12040 P_moss(#) BUF1B
12041 HMM02441_04(#) BUF1B EP_CTERM
12042 H_fuchsia(#) BUF1B
12043 H_banana(#) BUF1B
12044 -# -
12045 H_seafoam(#) BUF1B
12046 HMM01977ubiD_1_buf1(#) BUF1B
12047 HMM02441_02(#) BUF1B
12048 HMM00857isoch_1_buf1(#) BUF1B
12049 HMM13382deam_1_buf1(#) BUF1B
12050 -# -
12051 -# -
12052 -# -
12053 -# -
12056 HMM00860tp_1_bufgrm(#) BUF1
12057 HMM13382deam_1_buf1(#) B
BMC-P
BMC-H
BMC-T
Aldehyde dehydrogenase
PTAC
Signature enzyme
Alcohol dehydrogenase
Regulator

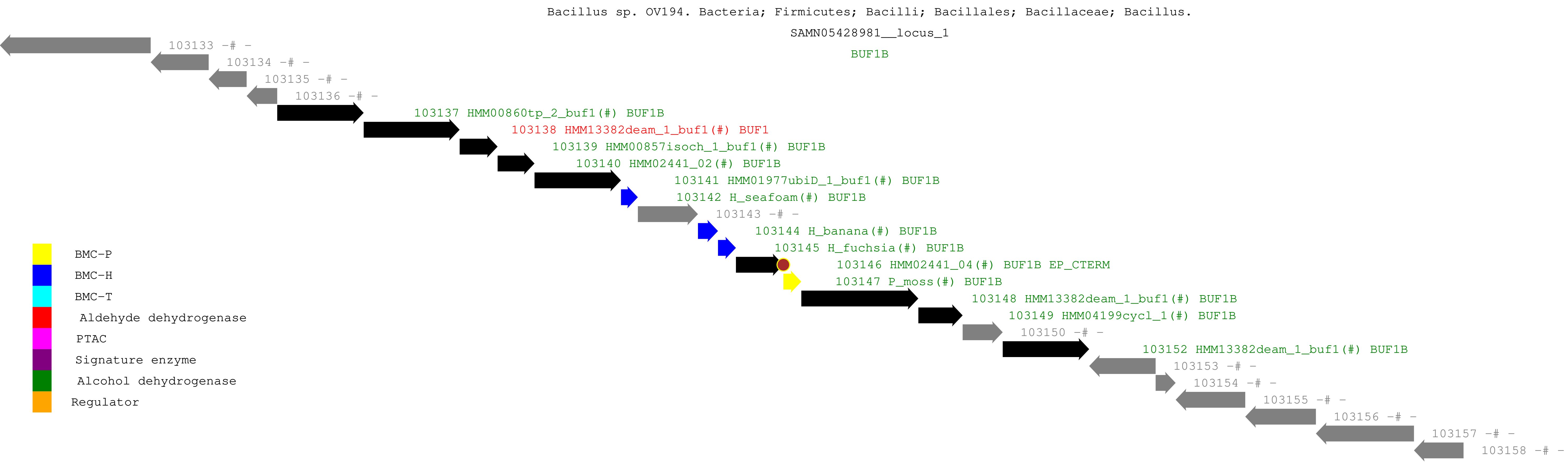
Bacillus sp. OV194. Bacteria; Firmicutes; Bacilli; Bacillales; Bacillaceae; Bacillus.
SAMN05428981__locus_1
BUF1B
103133 -# -
103134 -# -
103135 -# -
103136 -# -
103137 HMM00860tp_2_buf1(#) BUF1B
103138 HMM13382deam_1_buf1(#) BUF1
103139 HMM00857isoch_1_buf1(#) BUF1B
103140 HMM02441_02(#) BUF1B
103141 HMM01977ubiD_1_buf1(#) BUF1B
103142 H_seafoam(#) BUF1B
103143 -# -
103144 H_banana(#) BUF1B
103145 H_fuchsia(#) BUF1B
103146 HMM02441_04(#) BUF1B EP_CTERM
103147 P_moss(#) BUF1B
103148 HMM13382deam_1_buf1(#) BUF1B
103149 HMM04199cycl_1(#) BUF1B
103150 -# -
103152 HMM13382deam_1_buf1(#) BUF1B
103153 -# -
103154 -# -
103155 -# -
103156 -# -
103157 -# -
103158 -# -
BMC-P
BMC-H
BMC-T
Aldehyde dehydrogenase
PTAC
Signature enzyme
Alcohol dehydrogenase
Regulator

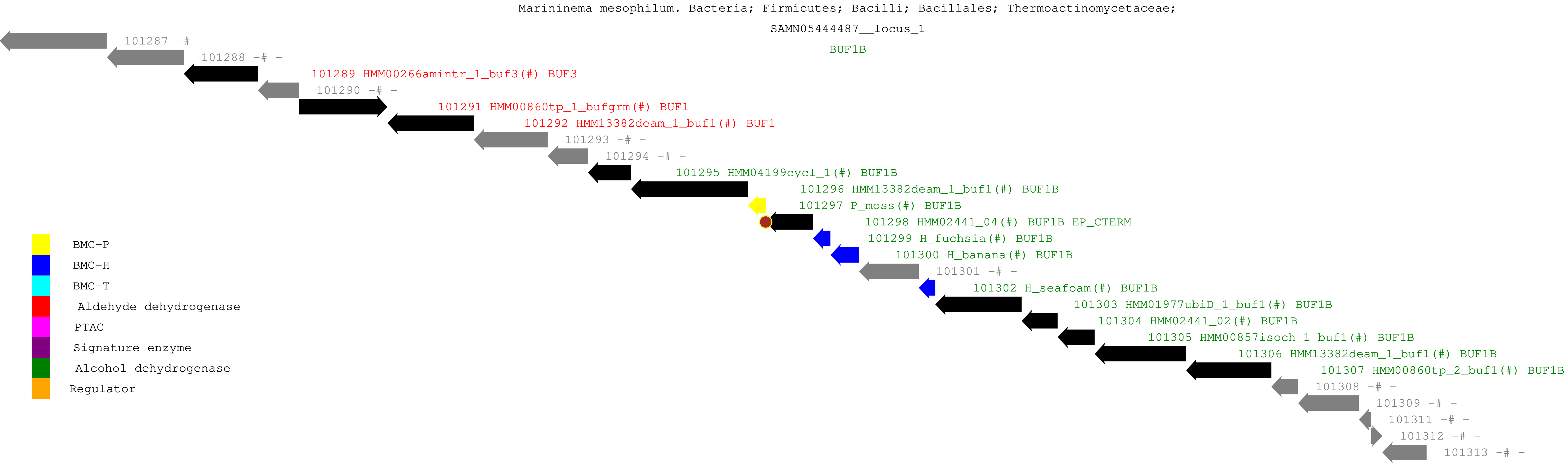

Marininema mesophilum. Bacteria; Firmicutes; Bacilli; Bacillales; Thermoactinomycetaceae;
SAMN05444487__locus_1
BUF1B
101287 -# -
101288 -# -
101289 HMM00266amintr_1_buf3(#) BUF3
101290 -# -
101291 HMM00860tp_1_bufgrm(#) BUF1
101292 HMM13382deam_1_buf1(#) BUF1
101293 -# -
101294 -# -
101295 HMM04199cycl_1(#) BUF1B
101296 HMM13382deam_1_buf1(#) BUF1B
101297 P_moss(#) BUF1B
101298 HMM02441_04(#) BUF1B EP_CTERM
101299 H_fuchsia(#) BUF1B
101300 H_banana(#) BUF1B
101301 -# -
101302 H_seafoam(#) BUF1B
101303 HMM01977ubiD_1_buf1(#) BUF1B
101304 HMM02441_02(#) BUF1B
101305 HMM00857isoch_1_buf1(#) BUF1B
101306 HMM13382deam_1_buf1(#) BUF1B
101307 HMM00860tp_2_buf1(#) BUF1B
101308 -# -
101309 -# -
101311 -# -
101312 -# -
101313 -# -
BMC-P
BMC-H
BMC-T
Aldehyde dehydrogenase
PTAC
Signature enzyme
Alcohol dehydrogenase
Regulator

Marininema halotolerans. Bacteria; Firmicutes; Bacilli; Bacillales; Thermoactinomycetaceae;

SAMN05444972__locus_1

BUF1B

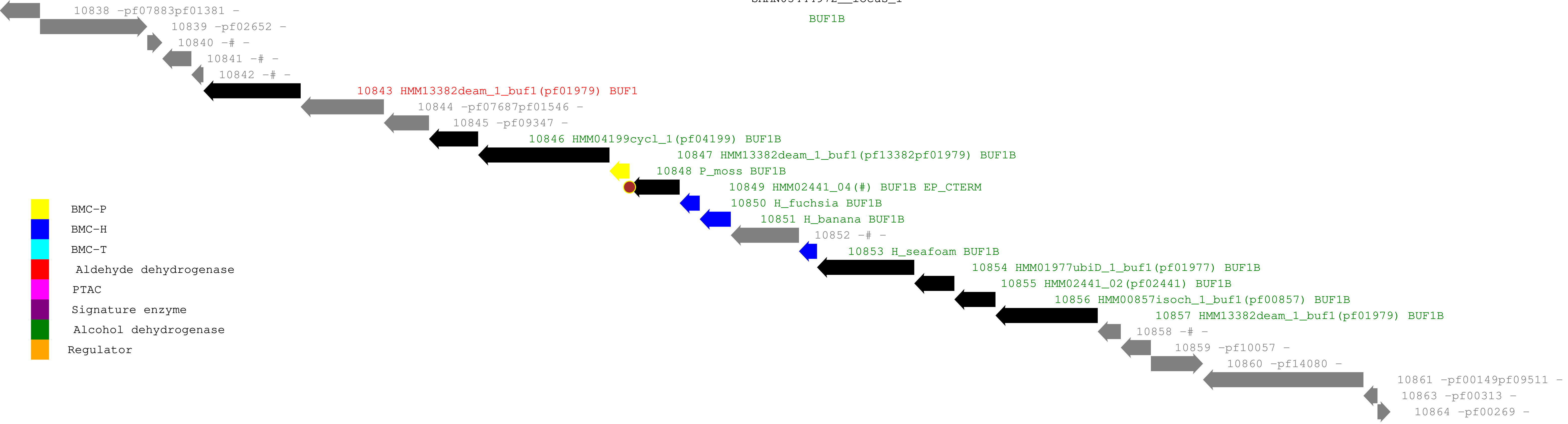

Bacillus firmus. Bacteria; Firmicutes; Bacilli; Bacillales; Bacillaceae; Bacillus.

VL14__locus_1

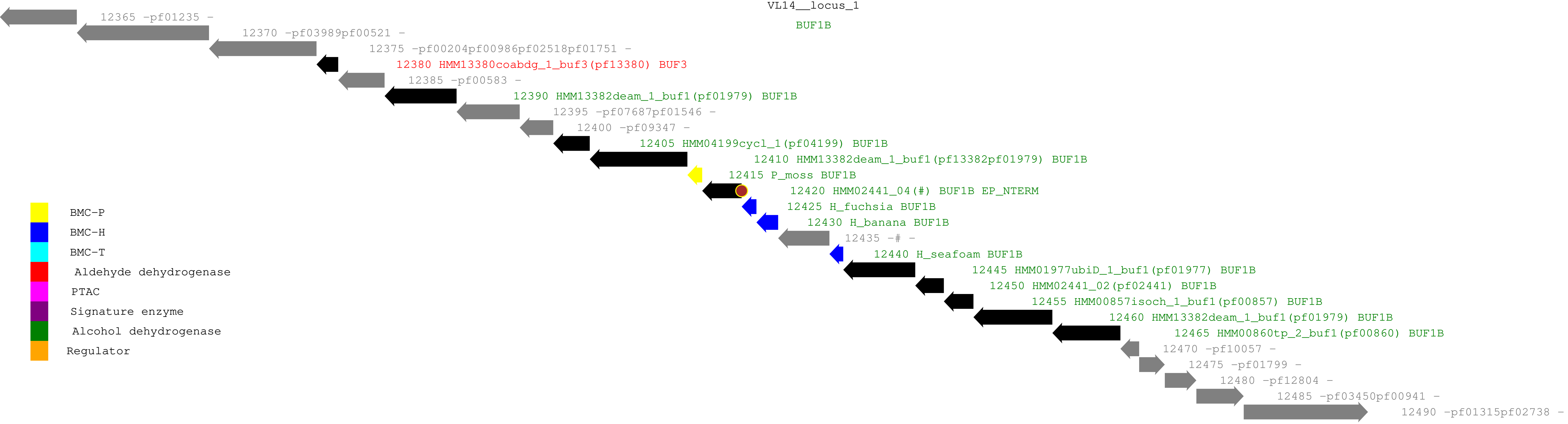

Micromonospora sp. TSRI0369. Bacteria; Actinobacteria; Micromonosporales; Micromonosporaceae;

AMK25__locus_1

BUF2

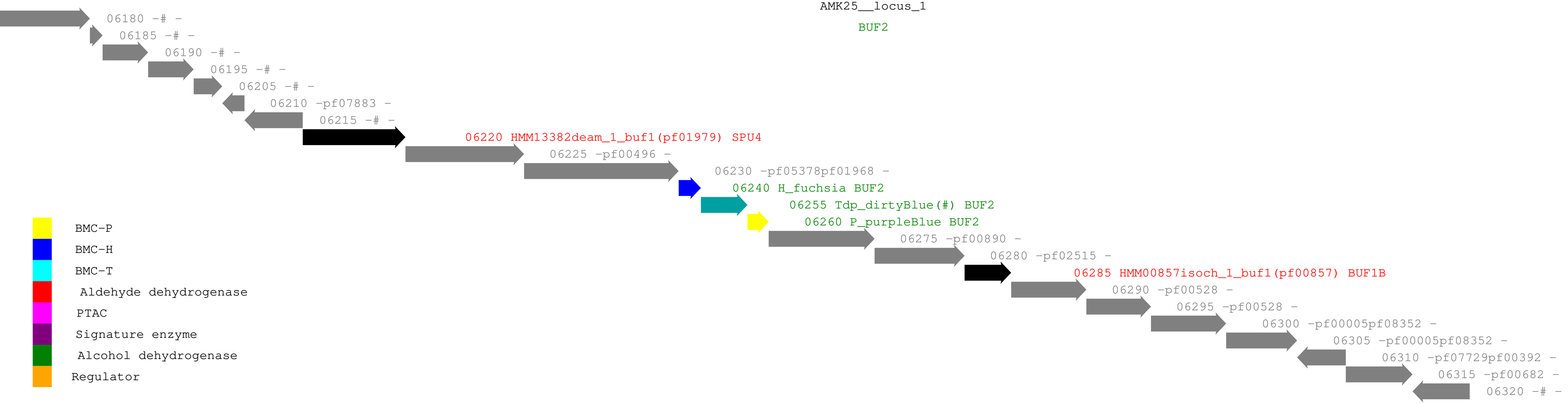

Micromonospora tulbaghiae. Bacteria; Actinobacteria; Micromonosporales; Micromonosporaceae;

CSH63__locus_1

BUF2

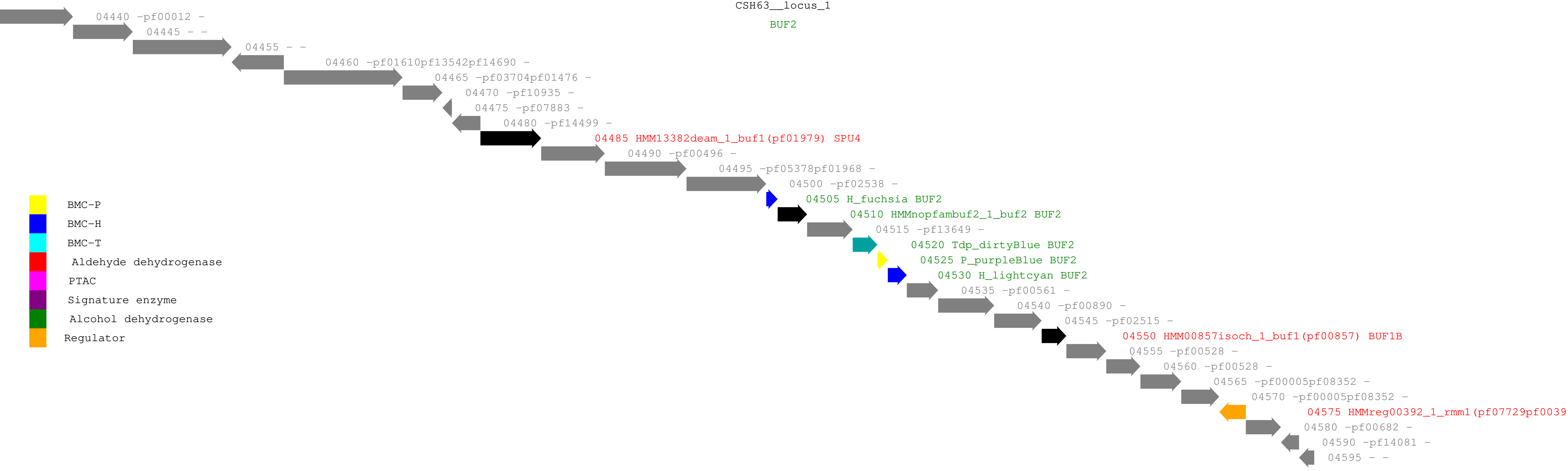

Verrucosispora sp. LHW63014. Bacteria; Actinobacteria; Micromonosporales; Micromonosporaceae;

D0Q02__locus_2

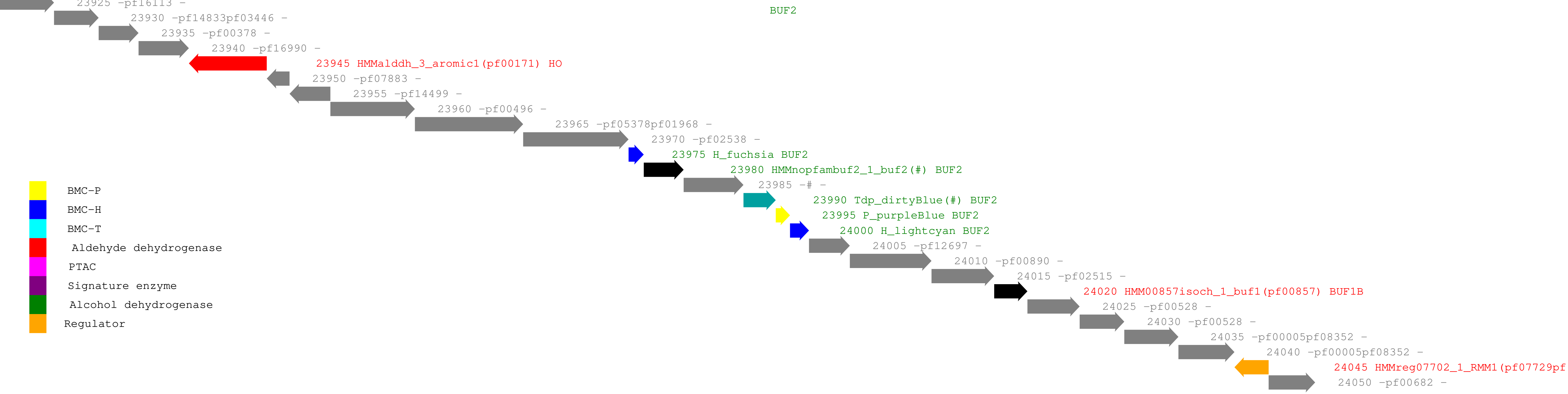

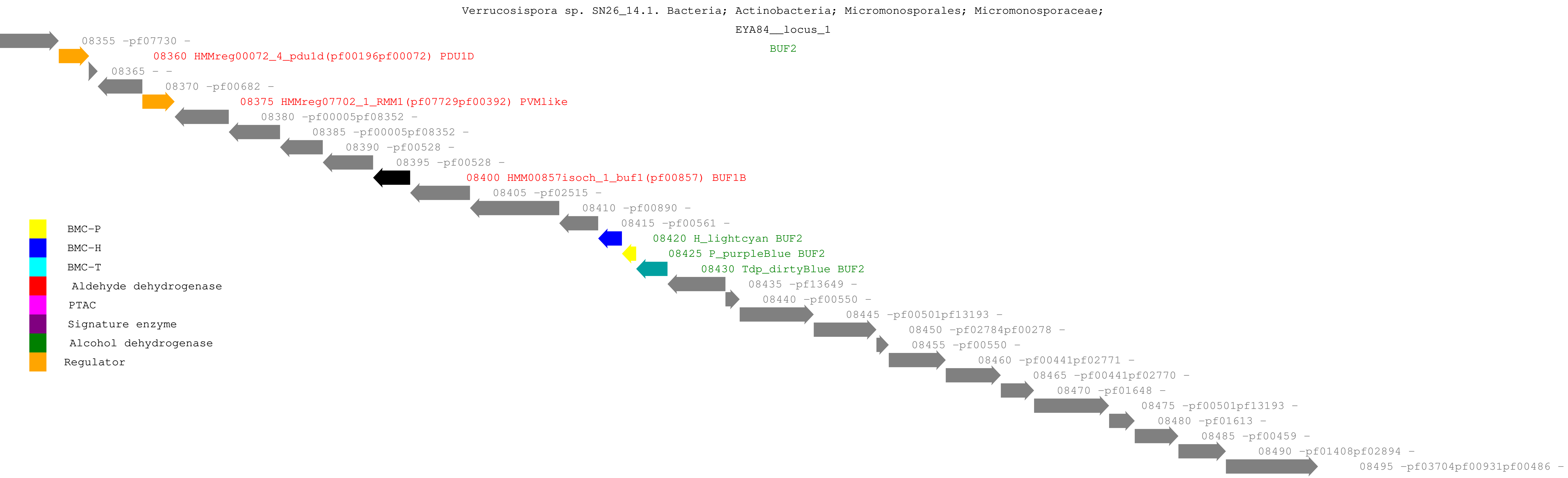
Verrucosispora sp. SN26_14.1. Bacteria; Actinobacteria; Micromonosporales; Micromonosporaceae;
EYA84__locus_1
BUF2
08355 -pf07730 -
08360 HMMreg00072_4_pdu1d(pf00196pf00072) PDU1D
08365 - -
08370 -pf00682 -
08375 HMMreg07702_1_RMM1(pf07729pf00392) PVMlike
08380 -pf00005pf08352 -
08385 -pf00005pf08352 -
08390 -pf00528 -
08395 -pf00528 -
08400 HMM00857isoch_1_buf1(pf00857) BUF1B
08405 -pf02515 -
08410 -pf00890 -
08415 -pf00561 -
08420 H_lightcyan BUF2
08425 P_purpleBlue BUF2
08430 Tdp_dirtyBlue BUF2
08435 -pf13649 -
08440 -pf00550 -
08445 -pf00501pf13193 -
08450 -pf02784pf00278 -
08455 -pf00550 -
08460 -pf00441pf02771 -
08465 -pf00441pf02770 -
08470 -pf01648 -
08475 -pf00501pf13193 -
08480 -pf01613 -
08485 -pf00459 -
08490 -pf01408pf02894 -
08495 -pf03704pf00931pf00486 -
BMC-P
BMC-H
BMC-T
Aldehyde dehydrogenase
PTAC
Signature enzyme
Alcohol dehydrogenase
Regulator

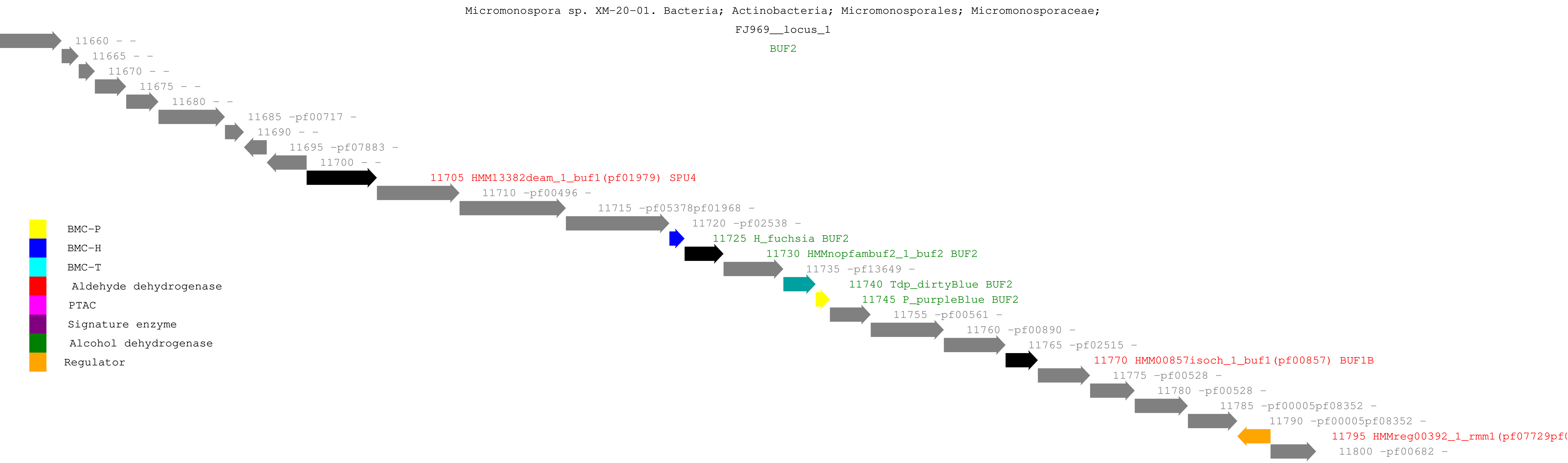
Micromonospora sp. XM-20-01. Bacteria; Actinobacteria; Micromonosporales; Micromonosporaceae;
FJ969__locus_1
BUF2
11660 - -
11665 - -
11670 - -
11675 - -
11680 - -
11685 -pf00717 -
11690 - -
11695 -pf07883 -
11700 - -
11705 HMM13382deam_1_buf1(pf01979) SPU4
11710 -pf00496 -
11715 -pf05378pf01968 -
11720 -pf02538 -
11725 H_fuchsia BUF2
11730 HMMnopfambuf2_1_buf2 BUF2
11735 -pf13649 -
11740 Tdp_dirtyBlue BUF2
11745 P_purpleBlue BUF2
11755 -pf00561 -
11760 -pf00890 -
11765 -pf02515 -
11770 HMM00857isoch_1_buf1(pf00857) BUF1B
11775 -pf00528 -
11780 -pf00528 -
11785 -pf00005pf08352 -
11790 -pf00005pf08352 -
11795 HMMreg00392_1_rmm1(pf07729pf0
11800 -pf00682 -
BMC-P
BMC-H
BMC-T
Aldehyde dehydrogenase
PTAC
Signature enzyme
Alcohol dehydrogenase
Regulator

Micromonospora sp. HM134. Bacteria; Actinobacteria; Micromonosporales; Micromonosporaceae;

FJK98__locus_1

BUF2

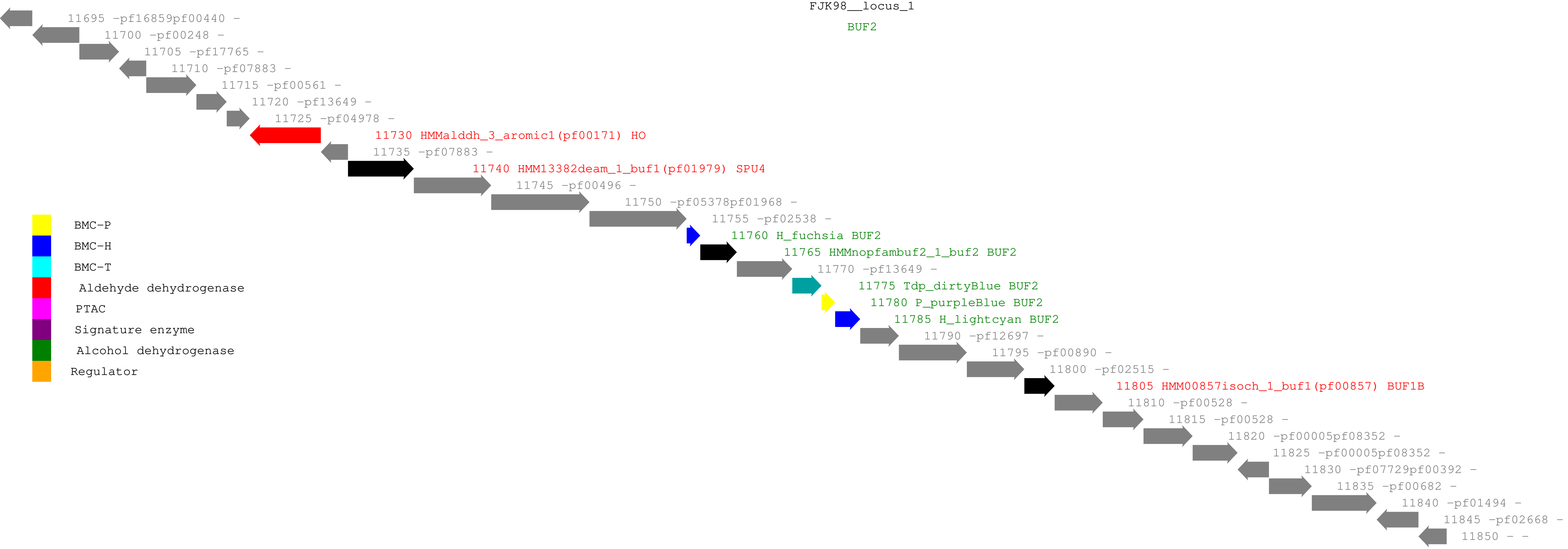

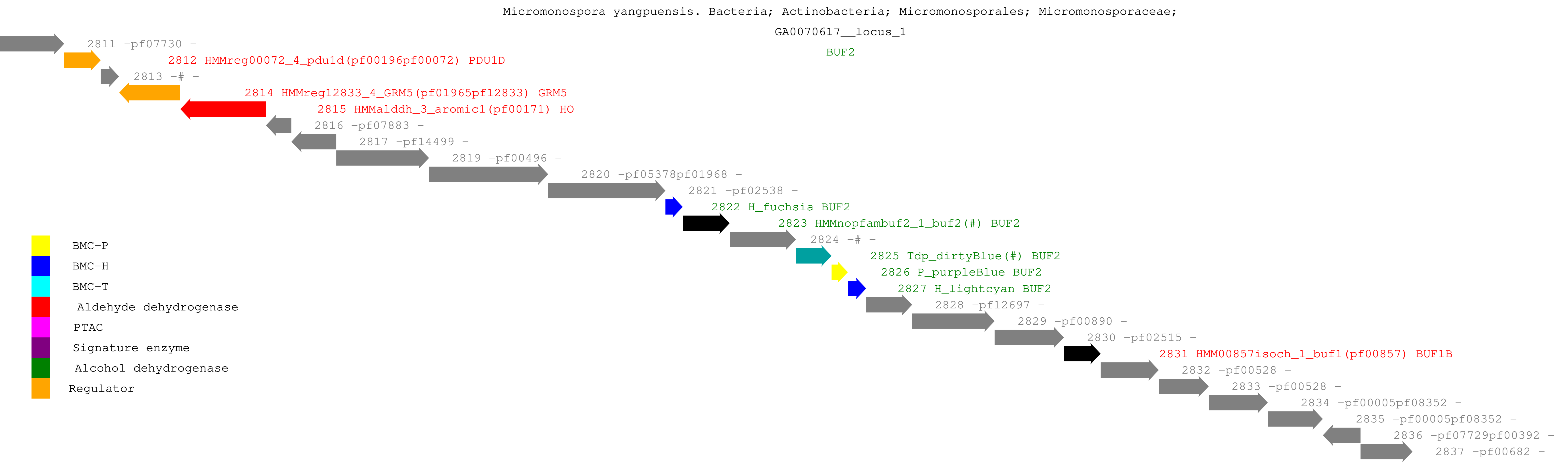

Micromonospora yangpuensis. Bacteria; Actinobacteria; Micromonosporales; Micromonosporaceae;
GA0070617__locus_1
BUF2
2811 -pf07730 -
2812 HMMreg00072_4_pdu1d(pf00196pf00072) PDU1D
2813 -# -
2814 HMMreg12833_4_GRM5(pf01965pf12833) GRM5
2815 HMMalddh_3_aromic1(pf00171) HO
2816 -pf07883 -
2817 -pf14499 -
2819 -pf00496 -
2820 -pf05378pf01968 -
2821 -pf02538 -
2822 H_fuchsia BUF2
2823 HMMnopfambuf2_1_buf2(#) BUF2
2824 -# -
2825 Tdp_dirtyBlue(#) BUF2
2826 P_purpleBlue BUF2
2827 H_lightcyan BUF2
2828 -pf12697 -
2829 -pf00890 -
2830 -pf02515 -
2831 HMM00857isoch_1_buf1(pf00857) BUF1B
2832 -pf00528 -
2833 -pf00528 -
2834 -pf00005pf08352 -
2835 -pf00005pf08352 -
2836 -pf07729pf00392 -
2837 -pf00682 -
BMC-P
BMC-H
BMC-T
Aldehyde dehydrogenase
PTAC
Signature enzyme
Alcohol dehydrogenase
Regulator

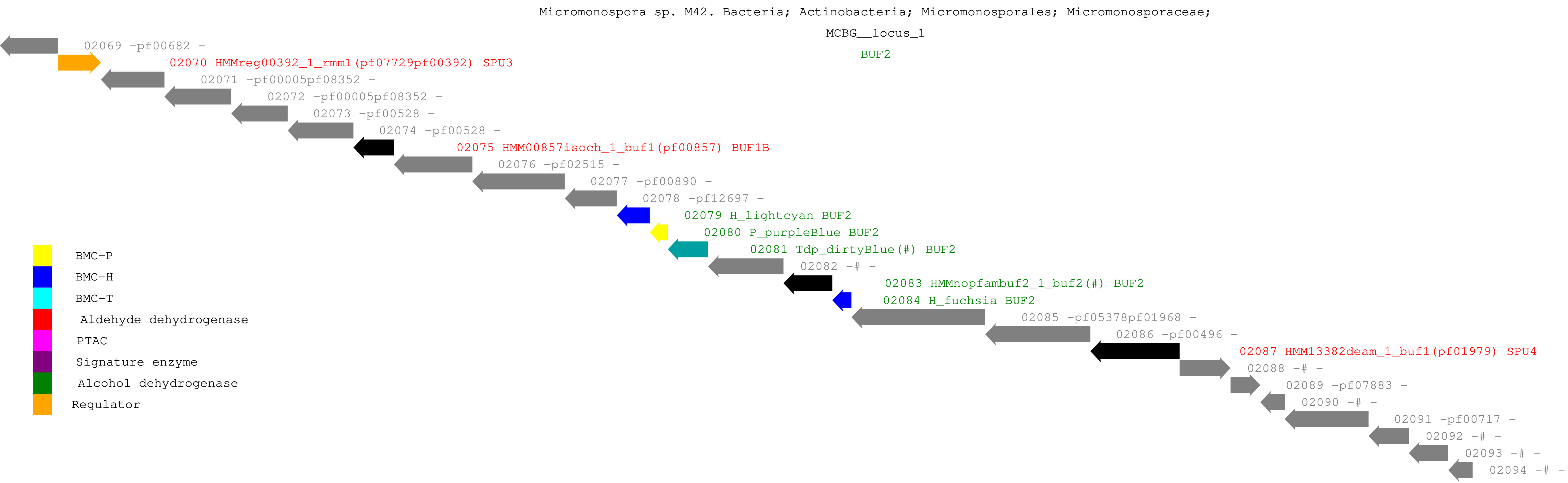
Micromonospora sp. M42. Bacteria; Actinobacteria; Micromonosporales; Micromonosporaceae;
MCBG__locus_1
BUF2
02069 -pf00682 -
02070 HMMreg00392_1_rmm1(pf07729pf00392) SPU3
02071 -pf00005pf08352 -
02072 -pf00005pf08352 -
02073 -pf00528 -
02074 -pf00528 -
02075 HMM00857isoch_1_buf1(pf00857) BUF1B
02076 -pf02515 -
02077 -pf00890 -
02078 -pf12697 -
02079 H_lightcyan BUF2
02080 P_purpleBlue BUF2
02081 Tdp_dirtyBlue(#) BUF2
02082 -# -
02083 HMMnopfambuf2_1_buf2(#) BUF2
02084 H_fuchsia BUF2
02085 -pf05378pf01968 -
02086 -pf00496 -
02087 HMM13382deam_1_buf1(pf01979) SPU4
02088 -# -
02089 -pf07883 -
02090 -# -
02091 -pf00717 -
02092 -# -
02093 -# -
02094 -# -
BMC-P
BMC-H
BMC-T
Aldehyde dehydrogenase
PTAC
Signature enzyme
Alcohol dehydrogenase
Regulator

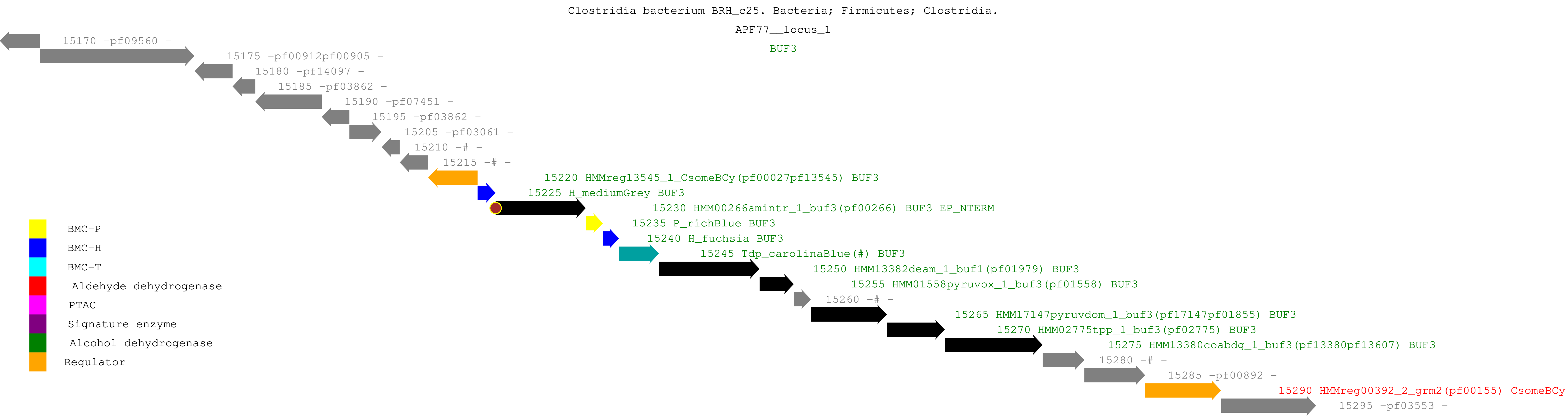
Clostridia bacterium BRH_c25. Bacteria; Firmicutes; Clostridia.
APF77__locus_1
BUF3
15170 -pf09560 -
15175 -pf00912pf00905 -
15180 -pf14097 -
15185 -pf03862 -
15190 -pf07451 -
15195 -pf03862 -
15205 -pf03061 -
15210 -# -
15215 -# -
15220 HMMreg13545_1_CsomeBCy(pf00027pf13545) BUF3
15225 H_mediumGrey BUF3
15230 HMM00266amintr_1_buf3(pf00266) BUF3 EP_NTERM
15235 P_richBlue BUF3
15240 H_fuchsia BUF3
15245 Tdp_carolinaBlue(#) BUF3
15250 HMM13382deam_1_buf1(pf01979) BUF3
15255 HMM01558pyruvox_1_buf3(pf01558) BUF3
15260 -# -
15265 HMM17147pyruvdom_1_buf3(pf17147pf01855) BUF3
15270 HMM02775tpp_1_buf3(pf02775) BUF3
15275 HMM13380coabdg_1_buf3(pf13380pf13607) BUF3
15280 -# -
15285 -pf00892 -
15290 HMMreg00392_2_grm2(pf00155) CsomeBCy
15295 -pf03553 -
BMC-P
BMC-H
BMC-T
Aldehyde dehydrogenase
PTAC
Signature enzyme
Alcohol dehydrogenase
Regulator

Firmicutes bacterium. Bacteria; Firmicutes.

C4551__locus_1

BUF3

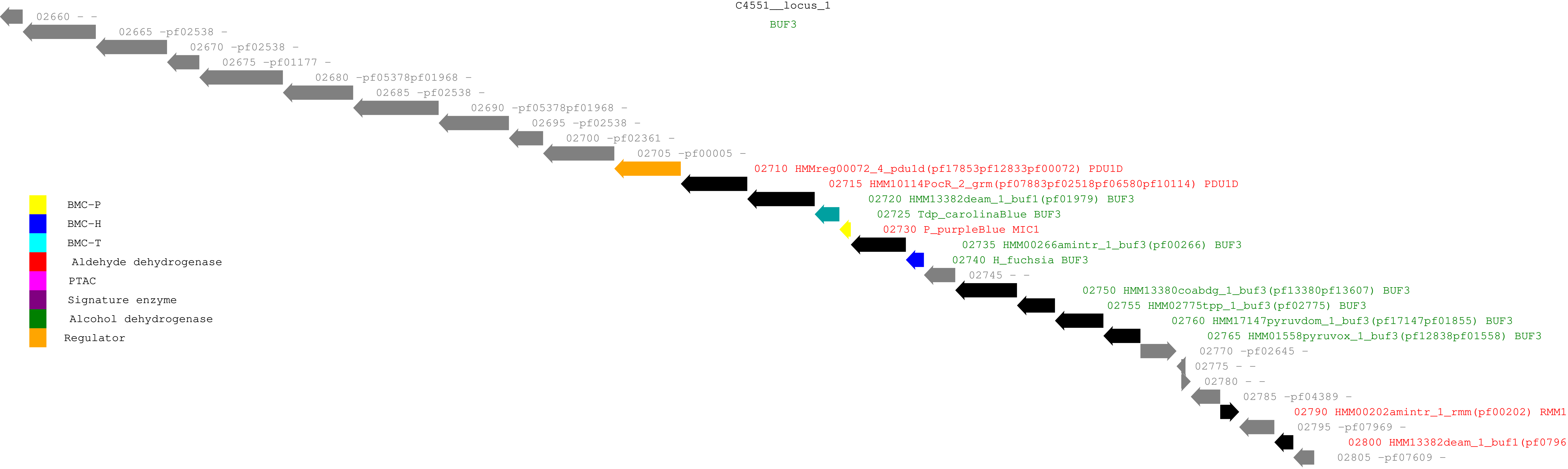

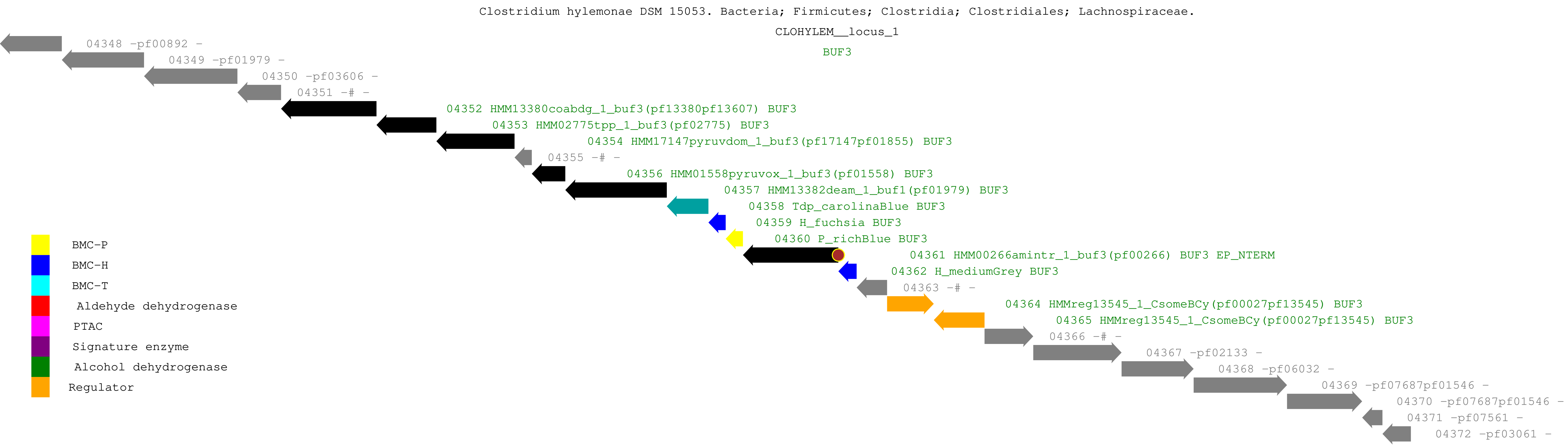
Clostridium hylemonae DSM 15053. Bacteria; Firmicutes; Clostridia; Clostridiales; Lachnospiraceae.
CLOHYLEM__locus_1
BUF3
04348 -pf00892 -
04349 -pf01979 -
04350 -pf03606 -
04351 -# -
04352 HMM13380coabdg_1_buf3(pf13380pf13607) BUF3
04353 HMM02775tpp_1_buf3(pf02775) BUF3
04354 HMM17147pyruvdom_1_buf3(pf17147pf01855) BUF3
04355 -# -
04356 HMM01558pyruvox_1_buf3(pf01558) BUF3
04357 HMM13382deam_1_buf1(pf01979) BUF3
04358 Tdp_carolinaBlue BUF3
04359 H_fuchsia BUF3
04360 P_richBlue BUF3
04361 HMM00266amintr_1_buf3(pf00266) BUF3 EP_NTERM
04362 H_mediumGrey BUF3
04363 -# -
04364 HMMreg13545_1_CsomeBCy(pf00027pf13545) BUF3
04365 HMMreg13545_1_CsomeBCy(pf00027pf13545) BUF3
04366 -# -
04367 -pf02133 -
04368 -pf06032 -
04369 -pf07687pf01546 -
04370 -pf07687pf01546 -
04371 -pf07561 -
04372 -pf03061 -
BMC-P
BMC-H
BMC-T
Aldehyde dehydrogenase
PTAC
Signature enzyme
Alcohol dehydrogenase
Regulator

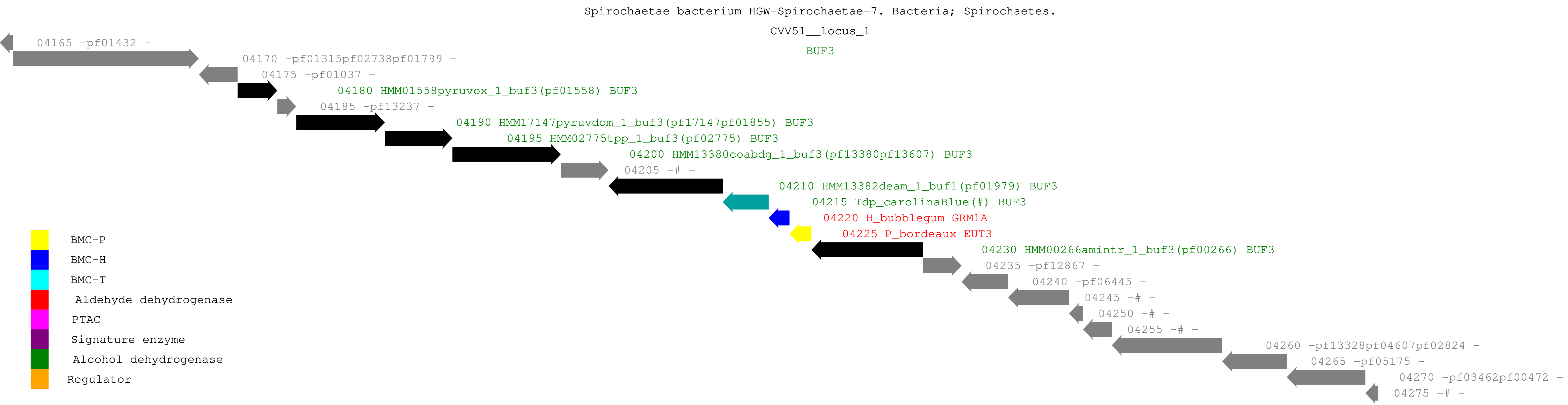

Spirochaetae bacterium HGW-Spirochaetae-7. Bacteria; Spirochaetes.
CVV51__locus_1
BUF3
04165 -pf01432 -
04170 -pf01315pf02738pf01799 -
04175 -pf01037 -
04180 HMM01558pyruvox_1_buf3(pf01558) BUF3
04185 -pf13237 -
04190 HMM17147pyruvdom_1_buf3(pf17147pf01855) BUF3
04195 HMM02775tpp_1_buf3(pf02775) BUF3
04200 HMM13380coabdg_1_buf3(pf13380pf13607) BUF3
04205 -# -
04210 HMM13382deam_1_buf1(pf01979) BUF3
04215 Tdp_carolinaBlue(#) BUF3
04220 H_bubblegum GRM1A
04225 P_bordeaux EUT3
04230 HMM00266amintr_1_buf3(pf00266) BUF3
04235 -pf12867 -
04240 -pf06445 -
04245 -# -
04250 -# -
04255 -# -
04260 -pf13328pf04607pf02824 -
04265 -pf05175 -
04270 -pf03462pf00472 -
04275 -# -
BMC-P
BMC-H
BMC-T
Aldehyde dehydrogenase
PTAC
Signature enzyme
Alcohol dehydrogenase
Regulator

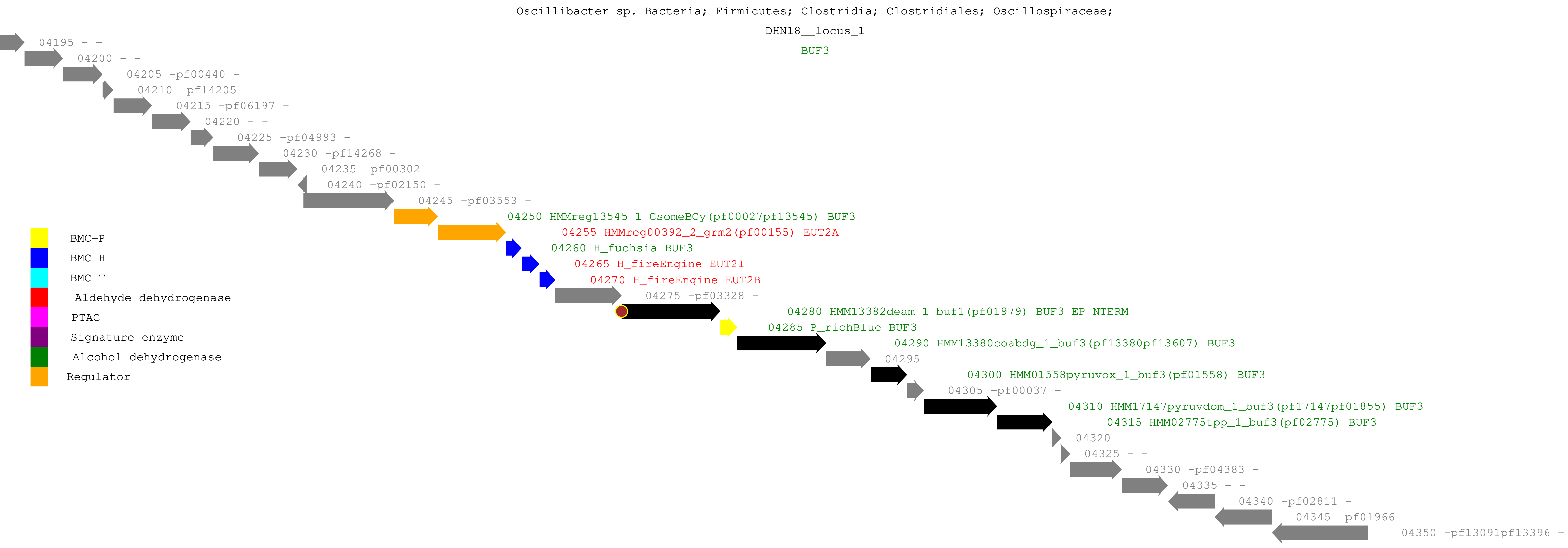
Oscillibacter sp. Bacteria; Firmicutes; Clostridia; Clostridiales; Oscillospiraceae;
DHN18__locus_1
BUF3
04195 - -
04200 - -
04205 -pf00440 -
04210 -pf14205 -
04215 -pf06197 -
04220 - -
04225 -pf04993 -
04230 -pf14268 -
04235 -pf00302 -
04240 -pf02150 -
04245 -pf03553 -
04250 HMMreg13545_1_CsomeBCy(pf00027pf13545) BUF3
04255 HMMreg00392_2_grm2(pf00155) EUT2A
04260 H_fuchsia BUF3
04265 H_fireEngine EUT2I
04270 H_fireEngine EUT2B
04275 -pf03328 -
04280 HMM13382deam_1_buf1(pf01979) BUF3 EP_NTERM
04285 P_richBlue BUF3
04290 HMM13380coabdg_1_buf3(pf13380pf13607) BUF3
04295 - -
04300 HMM01558pyruvox_1_buf3(pf01558) BUF3
04305 -pf00037 -
04310 HMM17147pyruvdom_1_buf3(pf17147pf01855) BUF3
04315 HMM02775tpp_1_buf3(pf02775) BUF3
04320 - -
04325 - -
04330 -pf04383 -
04335 - -
04340 -pf02811 -
04345 -pf01966 -
04350 -pf13091pf13396 -
BMC-P
BMC-H
BMC-T
Aldehyde dehydrogenase
PTAC
Signature enzyme
Alcohol dehydrogenase
Regulator

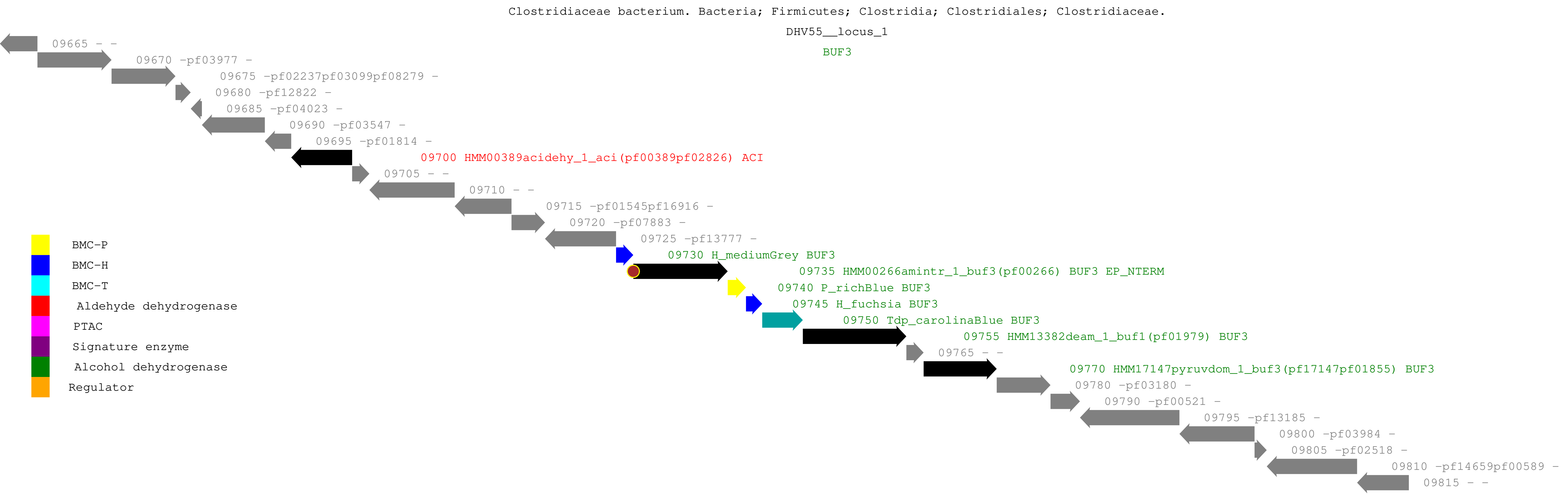

Clostridiaceae bacterium. Bacteria; Firmicutes; Clostridia; Clostridiales; Clostridiaceae.
DHV55__locus_1
BUF3
09665 - -
09670 -pf03977 -
09675 -pf02237pf03099pf08279 -
09680 -pf12822 -
09685 -pf04023 -
09690 -pf03547 -
09695 -pf01814 -
09700 HMM00389acidehy_1_aci(pf00389pf02826) ACI
09705 - -
09710 - -
09715 -pf01545pf16916 -
09720 -pf07883 -
09725 -pf13777 -
09730 H_mediumGrey BUF3
09735 HMM00266amintr_1_buf3(pf00266) BUF3 EP_NTERM
09740 P_richBlue BUF3
09745 H_fuchsia BUF3
09750 Tdp_carolinaBlue BUF3
09755 HMM13382deam_1_buf1(pf01979) BUF3
09765 - -
09770 HMM17147pyruvdom_1_buf3(pf17147pf01855) BUF3
09780 -pf03180 -
09790 -pf00521 -
09795 -pf13185 -
09800 -pf03984 -
09805 -pf02518 -
09810 -pf14659pf00589 -
09815 - -
BMC-P
BMC-H
BMC-T
Aldehyde dehydrogenase
PTAC
Signature enzyme
Alcohol dehydrogenase
Regulator

Clostridium sp. AF15-17LB. Bacteria; Firmicutes; Clostridia; Clostridiales; Clostridiaceae;

DWW31__locus_2

BUF3

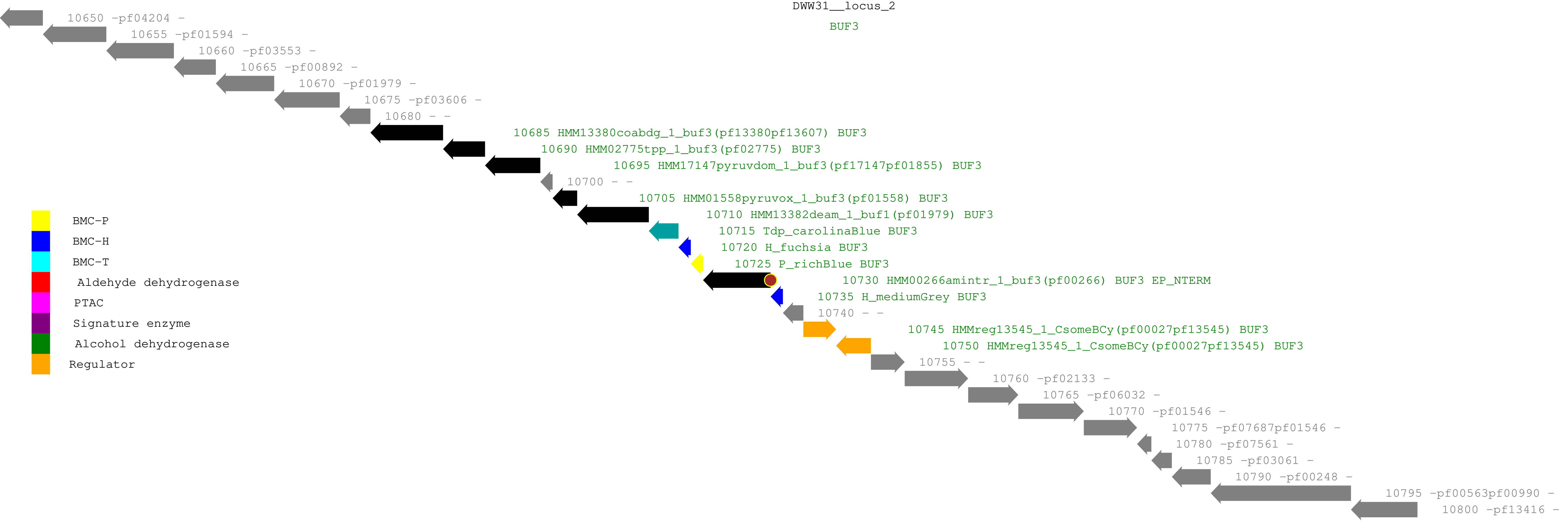

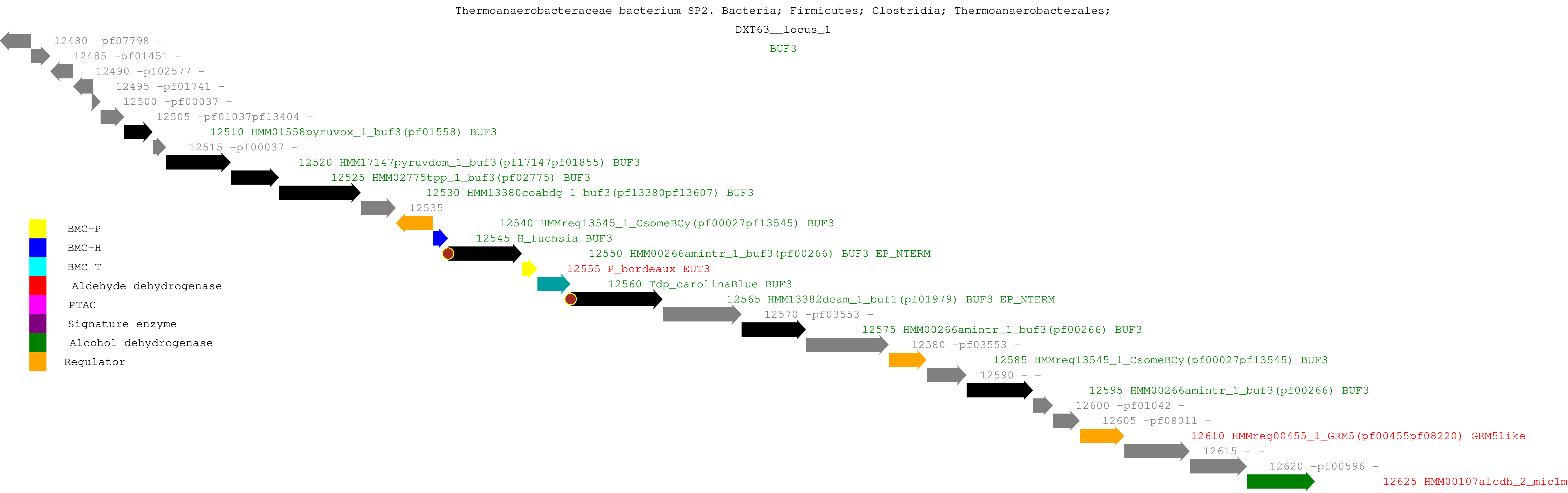
Thermoanaerobacteraceae bacterium SP2. Bacteria; Firmicutes; Clostridia; Thermoanaerobacterales;
DXT63__locus_1
BUF3
12480 -pf07798 -
12485 -pf01451 -
12490 -pf02577 -
12495 -pf01741 -
12500 -pf00037 -
12505 -pf01037pf13404 -
12510 HMM01558pyruvox_1_buf3(pf01558) BUF3
12515 -pf00037 -
12520 HMM17147pyruvdom_1_buf3(pf17147pf01855) BUF3
12525 HMM02775tpp_1_buf3(pf02775) BUF3
12530 HMM13380coabdg_1_buf3(pf13380pf13607) BUF3
12535 - -
12540 HMMreg13545_1_CsomeBCy(pf00027pf13545) BUF3
12545 H_fuchsia BUF3
12550 HMM00266amintr_1_buf3(pf00266) BUF3 EP_NTERM
12555 P_bordeaux EUT3
12560 Tdp_carolinaBlue BUF3
12565 HMM13382deam_1_buf1(pf01979) BUF3 EP_NTERM
12570 -pf03553 -
12575 HMM00266amintr_1_buf3(pf00266) BUF3
12580 -pf03553 -
12585 HMMreg13545_1_CsomeBCy(pf00027pf13545) BUF3
12590 - -
12595 HMM00266amintr_1_buf3(pf00266) BUF3
12600 -pf01042 -
12605 -pf08011 -
12610 HMMreg00455_1_GRM5(pf00455pf08220) GRM5like
12615 - -
12620 -pf00596 -
12625 HMM00107alcdh_2_mic1m
BMC-P
BMC-H
BMC-T
Aldehyde dehydrogenase
PTAC
Signature enzyme
Alcohol dehydrogenase
Regulator

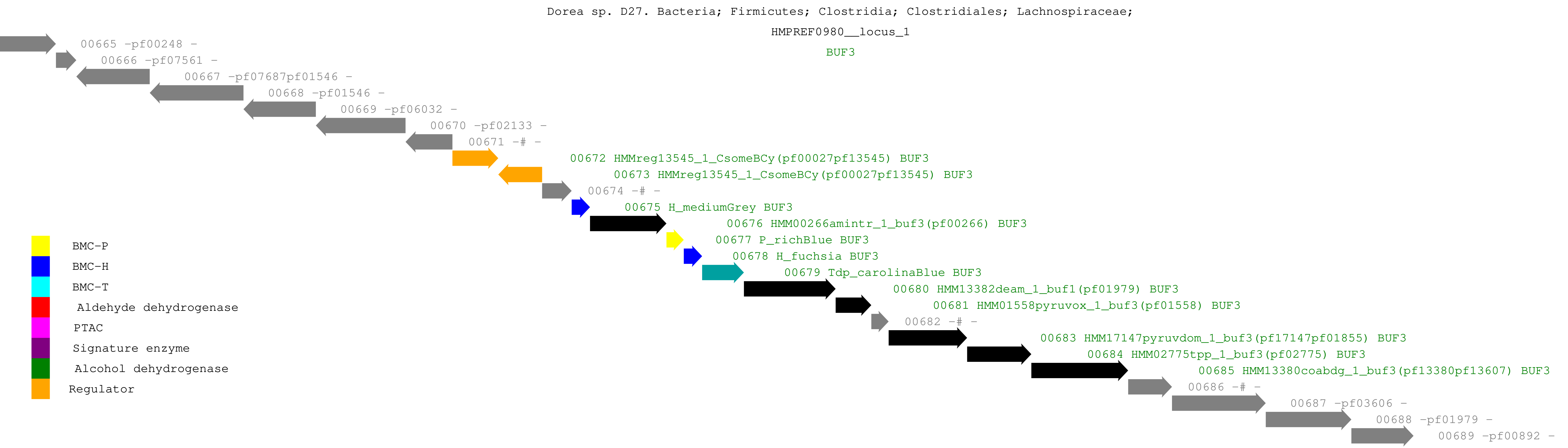

Dorea sp. D27. Bacteria; Firmicutes; Clostridia; Clostridiales; Lachnospiraceae;
HMPREF0980__locus_1
BUF3
00665 -pf00248 -
00666 -pf07561 -
00667 -pf07687pf01546 -
00668 -pf01546 -
00669 -pf06032 -
00670 -pf02133 -
00671 -# -
00672 HMMreg13545_1_CsomeBCy(pf00027pf13545) BUF3
00673 HMMreg13545_1_CsomeBCy(pf00027pf13545) BUF3
00674 -# -
00675 H_mediumGrey BUF3
00676 HMM00266amintr_1_buf3(pf00266) BUF3
00677 P_richBlue BUF3
00678 H_fuchsia BUF3
00679 Tdp_carolinaBlue BUF3
00680 HMM13382deam_1_buf1(pf01979) BUF3
00681 HMM01558pyruvox_1_buf3(pf01558) BUF3
00682 -# -
00683 HMM17147pyruvdom_1_buf3(pf17147pf01855) BUF3
00684 HMM02775tpp_1_buf3(pf02775) BUF3
00685 HMM13380coabdg_1_buf3(pf13380pf13607) BUF3
00686 -# -
00687 -pf03606 -
00688 -pf01979 -
00689 -pf00892 -
BMC-P
BMC-H
BMC-T
Aldehyde dehydrogenase
PTAC
Signature enzyme
Alcohol dehydrogenase
Regulator

Maledivibacter halophilus. Bacteria; Firmicutes; Clostridia; Clostridiales; Clostridiaceae;

SAMN02194393__locus_1

BUF3

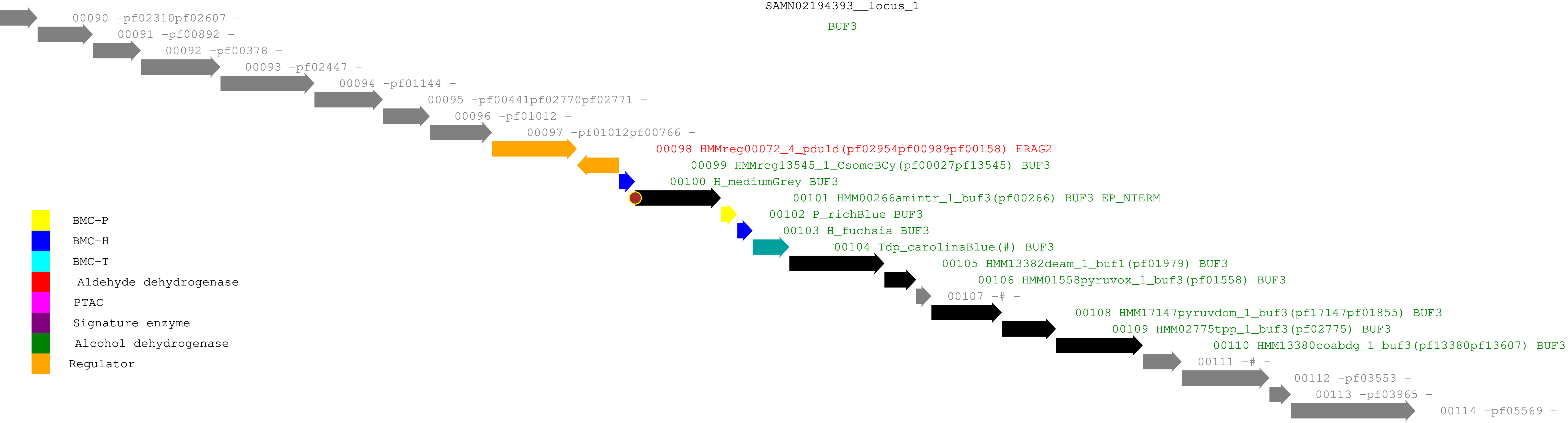

Hydrogenophilales bacterium RIFOXYD1_FULL_62_11. Bacteria; Proteobacteria; Betaproteobacteria; Hydrogenophilales.

A2580__locus_1_main_sat

CsomeACh

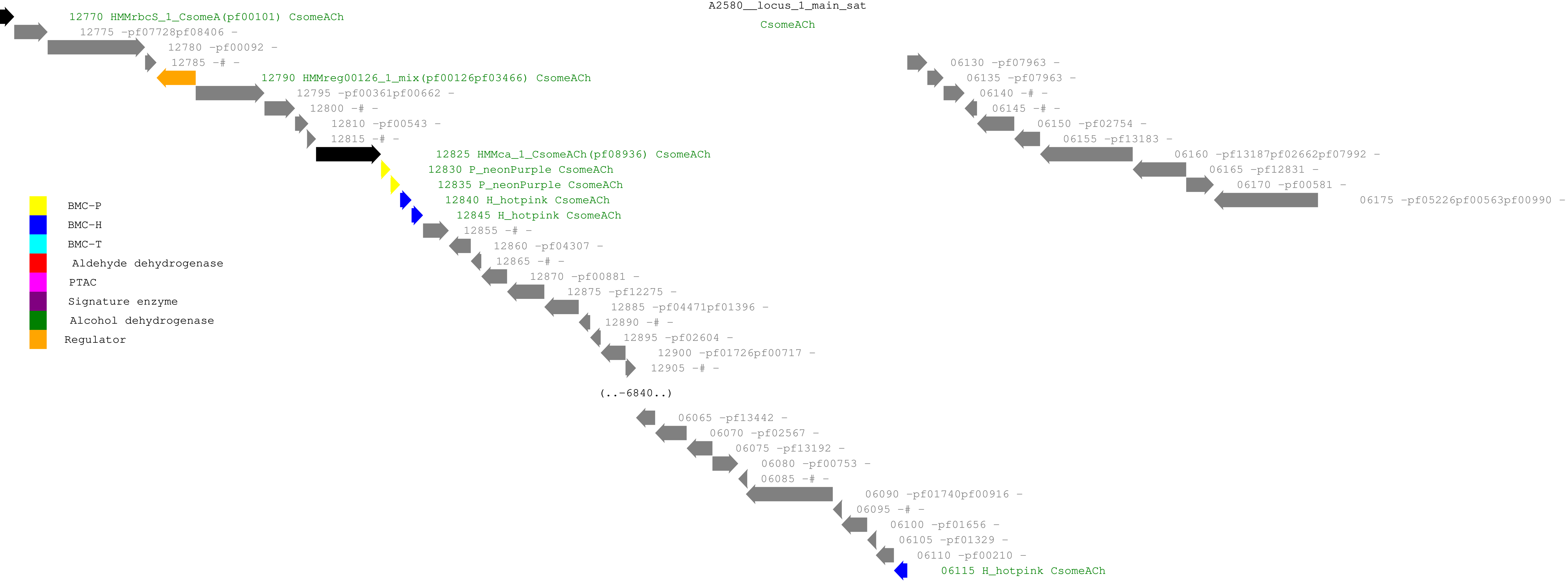

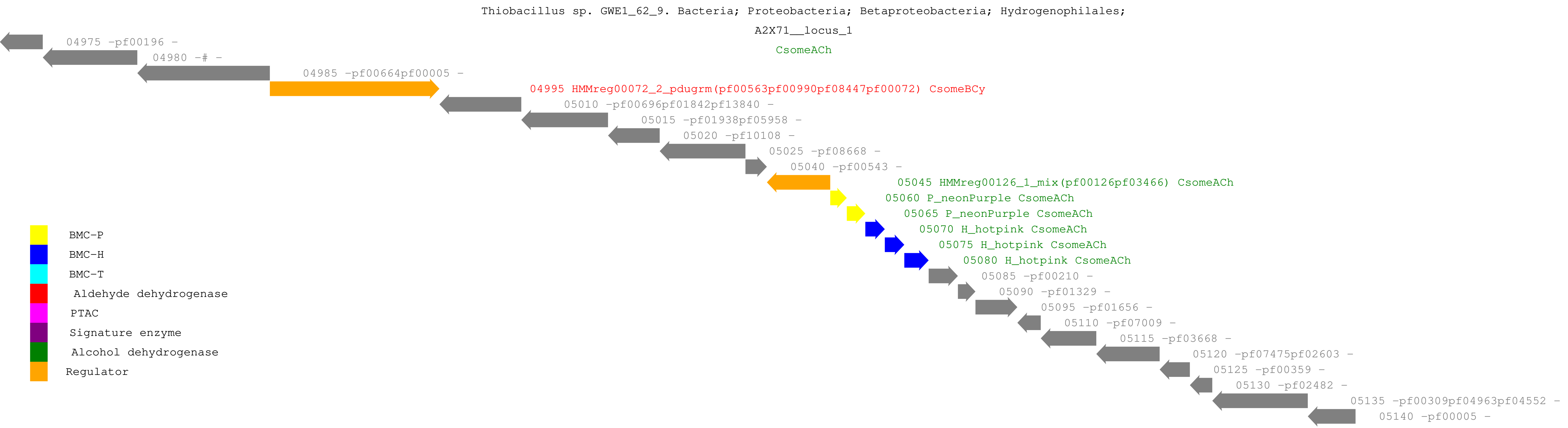

Thiobacillus sp. GWE1_62_9. Bacteria; Proteobacteria; Betaproteobacteria; Hydrogenophilales;
A2X71__locus_1
CsomeACh
04975 -pf00196 -
04980 -# -
04985 -pf00664pf00005 -
04995 HMMreg00072_2_pdugrm(pf00563pf00990pf08447pf00072) CsomeBCy
05010 -pf00696pf01842pf13840 -
05015 -pf01938pf05958 -
05020 -pf10108 -
05025 -pf08668 -
05040 -pf00543 -
05045 HMMreg00126_1_mix(pf00126pf03466) CsomeACh
05060 P_neonPurple CsomeACh
05065 P_neonPurple CsomeACh
05070 H_hotpink CsomeACh
05075 H_hotpink CsomeACh
05080 H_hotpink CsomeACh
05085 -pf00210 -
05090 -pf01329 -
05095 -pf01656 -
05110 -pf07009 -
05115 -pf03668 -
05120 -pf07475pf02603 -
05125 -pf00359 -
05130 -pf02482 -
05135 -pf00309pf04963pf04552 -
05140 -pf00005 -
BMC-P
BMC-H
BMC-T
Aldehyde dehydrogenase
PTAC
Signature enzyme
Alcohol dehydrogenase
Regulator

Acidithiobacillus ferrooxidans (Thiobacillus ferrooxidans). Bacteria; Proteobacteria; Acidithiobacillia; Acidithiobacillales;

A4H96__locus_fragmented

CsomeACh

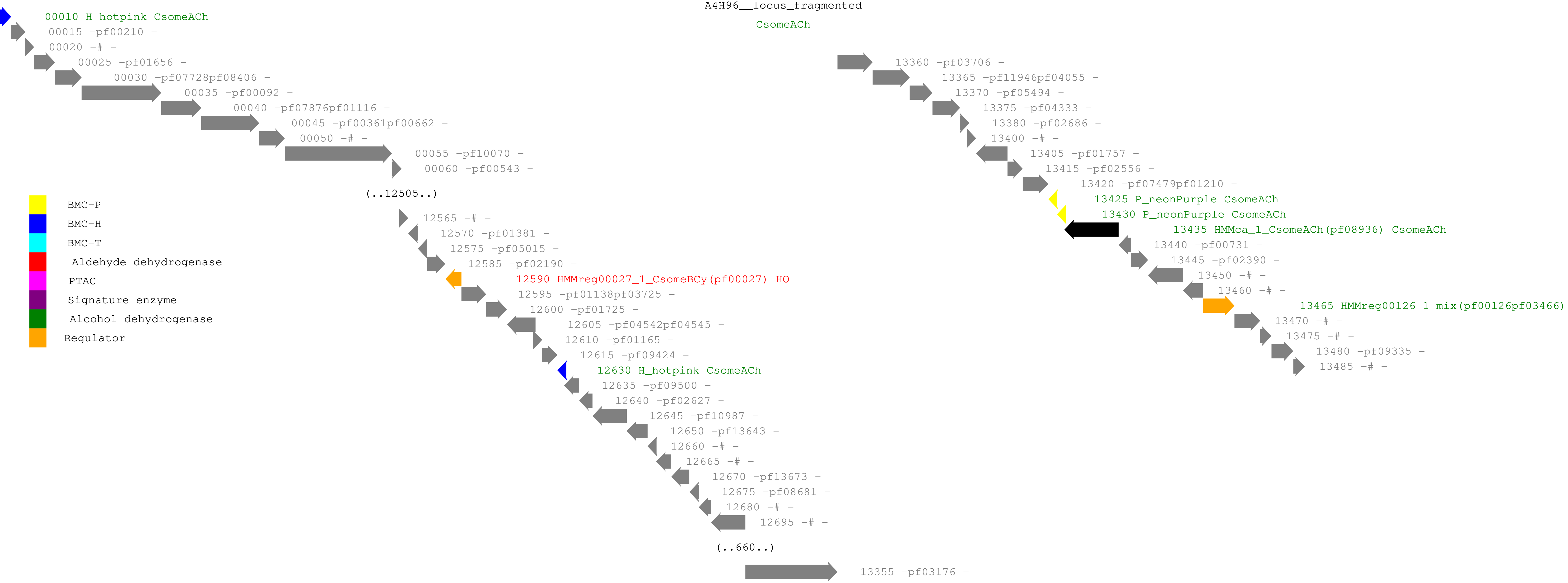

Acidithiobacillus ferrivorans. Bacteria; Proteobacteria; Acidithiobacillia; Acidithiobacillales;

A4U49__locus_1

CsomeACh

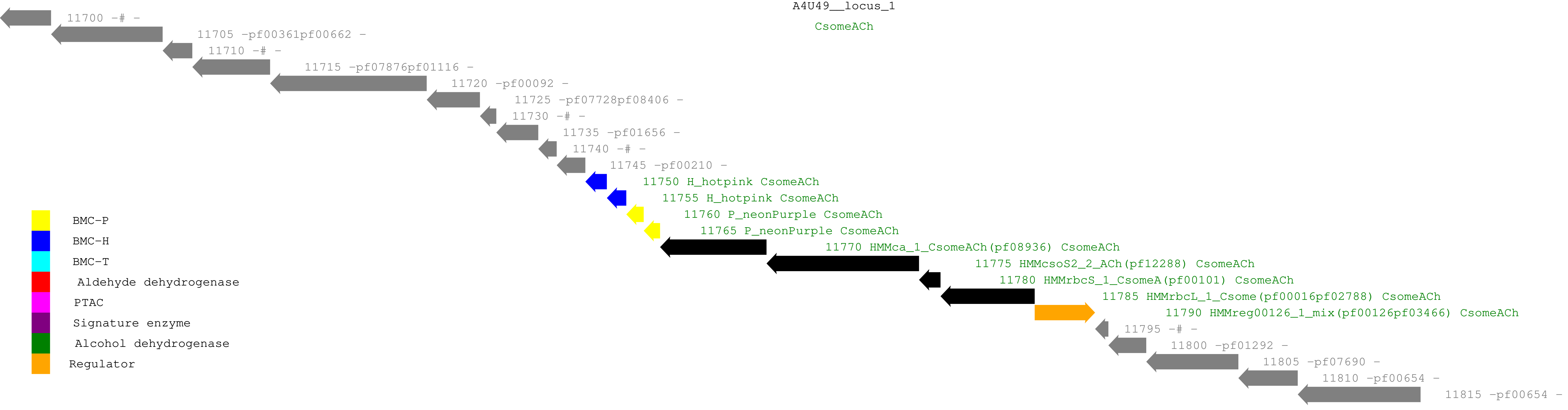

Acidithiobacillus thiooxidans (Thiobacillus thiooxidans). Bacteria; Proteobacteria; Acidithiobacillia; Acidithiobacillales;

A6M23__locus_1_main_sat

CsomeACh

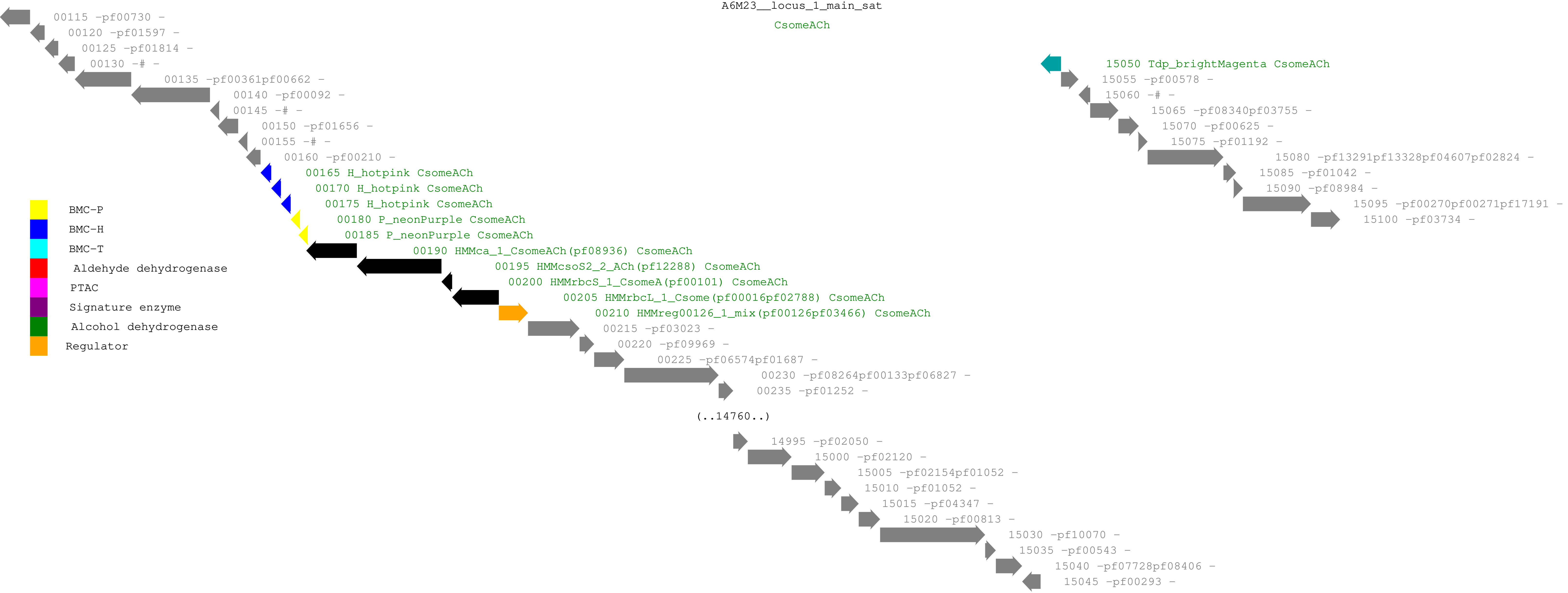

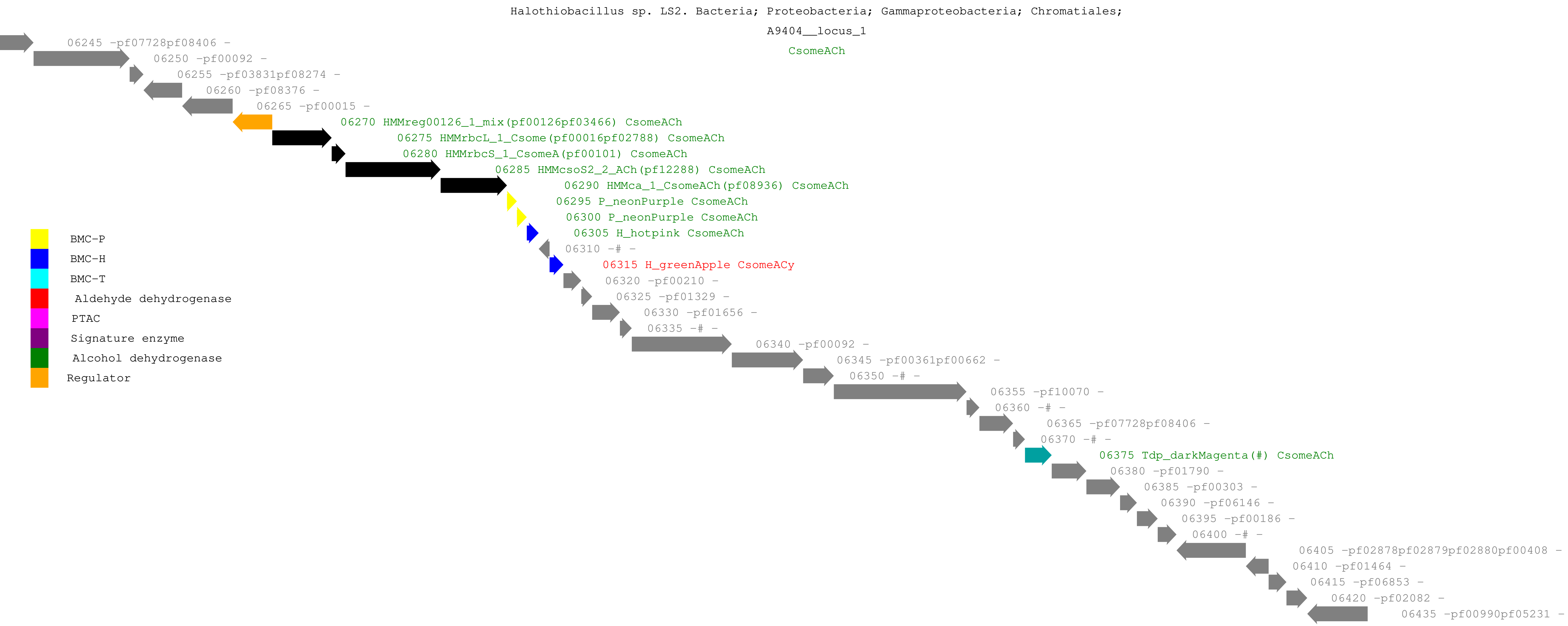
Halothiobacillus sp. LS2. Bacteria; Proteobacteria; Gammaproteobacteria; Chromatiales;
A9404__locus_1
CsomeACh
06245 -pf07728pf08406 -
06250 -pf00092 -
06255 -pf03831pf08274 -
06260 -pf08376 -
06265 -pf00015 -
06270 HMMreg00126_1_mix(pf00126pf03466) CsomeACh
06275 HMMrbcL_1_Csome(pf00016pf02788) CsomeACh
06280 HMMrbcS_1_CsomeA(pf00101) CsomeACh
06285 HMMcsoS2_2_ACh(pf12288) CsomeACh
06290 HMMca_1_CsomeACh(pf08936) CsomeACh
06295 P_neonPurple CsomeACh
06300 P_neonPurple CsomeACh
06305 H_hotpink CsomeACh
06310 -# -
06315 H_greenApple CsomeACy
06320 -pf00210 -
06325 -pf01329 -
06330 -pf01656 -
06335 -# -
06340 -pf00092 -
06345 -pf00361pf00662 -
06350 -# -
06355 -pf10070 -
06360 -# -
06365 -pf07728pf08406 -
06370 -# -
06375 Tdp_darkMagenta(#) CsomeACh
06380 -pf01790 -
06385 -pf00303 -
06390 -pf06146 -
06395 -pf00186 -
06400 -# -
06405 -pf02878pf02879pf02880pf00408 -
06410 -pf01464 -
06415 -pf06853 -
06420 -pf02082 -
06435 -pf00990pf05231 -
BMC-P
BMC-H
BMC-T
Aldehyde dehydrogenase
PTAC
Signature enzyme
Alcohol dehydrogenase
Regulator

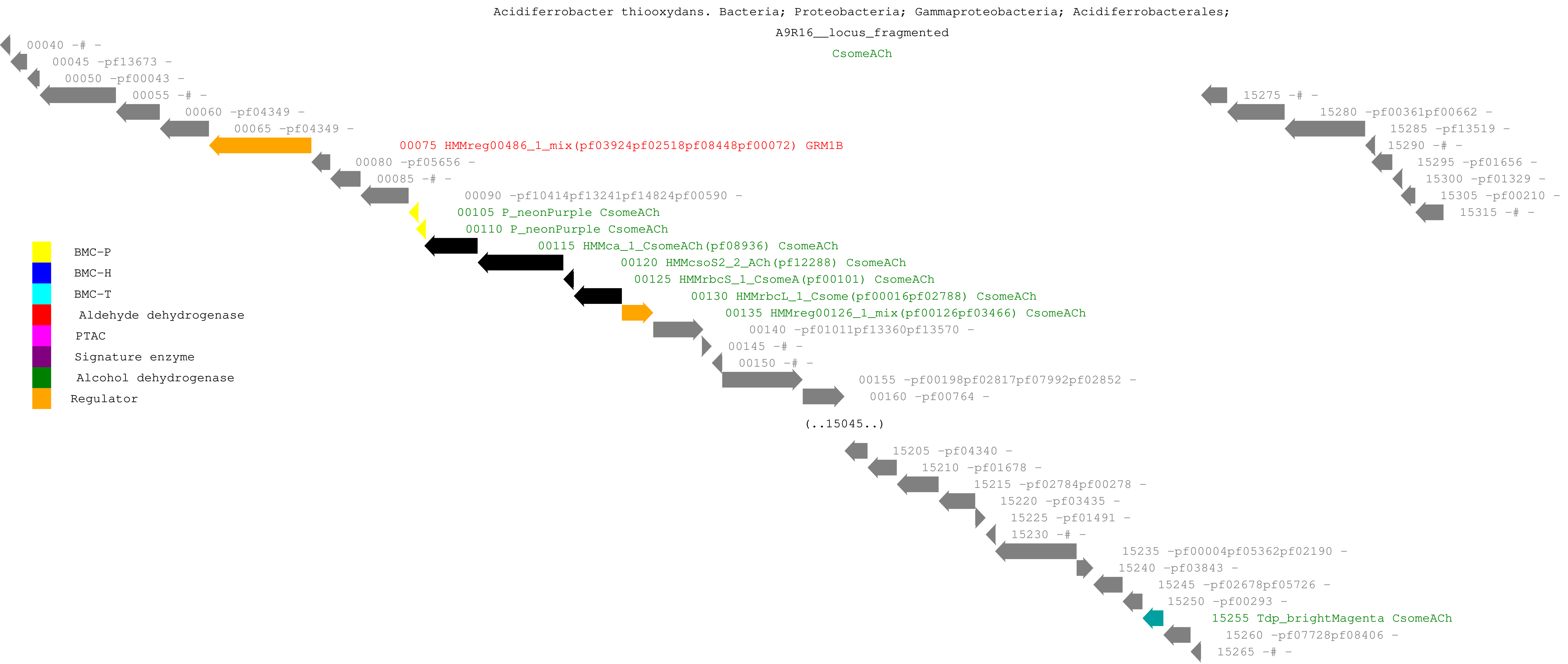

Acidiferrobacter thiooxydans. Bacteria; Proteobacteria; Gammaproteobacteria; Acidiferrobacterales;
A9R16__locus_fragmented
CsomeACh
00040 -# -
00045 -pf13673 -
00050 -pf00043 -
00055 -# -
00060 -pf04349 -
00065 -pf04349 -
00075 HMMreg00486_1_mix(pf03924pf02518pf08448pf00072) GRM1B
00080 -pf05656 -
00085 -# -
00090 -pf10414pf13241pf14824pf00590 -
00105 P_neonPurple CsomeACh
00110 P_neonPurple CsomeACh
00115 HMMca_1_CsomeACh(pf08936) CsomeACh
00120 HMMcsoS2_2_ACh(pf12288) CsomeACh
00125 HMMrbcS_1_CsomeA(pf00101) CsomeACh
00130 HMMrbcL_1_Csome(pf00016pf02788) CsomeACh
00135 HMMreg00126_1_mix(pf00126pf03466) CsomeACh
00140 -pf01011pf13360pf13570 -
00145 -# -
00150 -# -
00155 -pf00198pf02817pf07992pf02852 -
00160 -pf00764 -
(..15045..)
15205 -pf04340 -
15210 -pf01678 -
15215 -pf02784pf00278 -
15220 -pf03435 -
15225 -pf01491 -
15230 -# -
15235 -pf00004pf05362pf02190 -
15240 -pf03843 -
15245 -pf02678pf05726 -
15250 -pf00293 -
15255 Tdp_brightMagenta CsomeACh
15260 -pf07728pf08406 -
15265 -# -
15275 -# -
15280 -pf00361pf00662 -
15285 -pf13519 -
15290 -# -
15295 -pf01656 -
15300 -pf01329 -
15305 -pf00210 -
15315 -# -
BMC-P
BMC-H
BMC-T
Aldehyde dehydrogenase
PTAC
Signature enzyme
Alcohol dehydrogenase
Regulator

Nitrosomonadales bacterium SCN 54-20. Bacteria; Proteobacteria; Betaproteobacteria; Nitrosomonadales.

ABS69__locus_1

CsomeACh

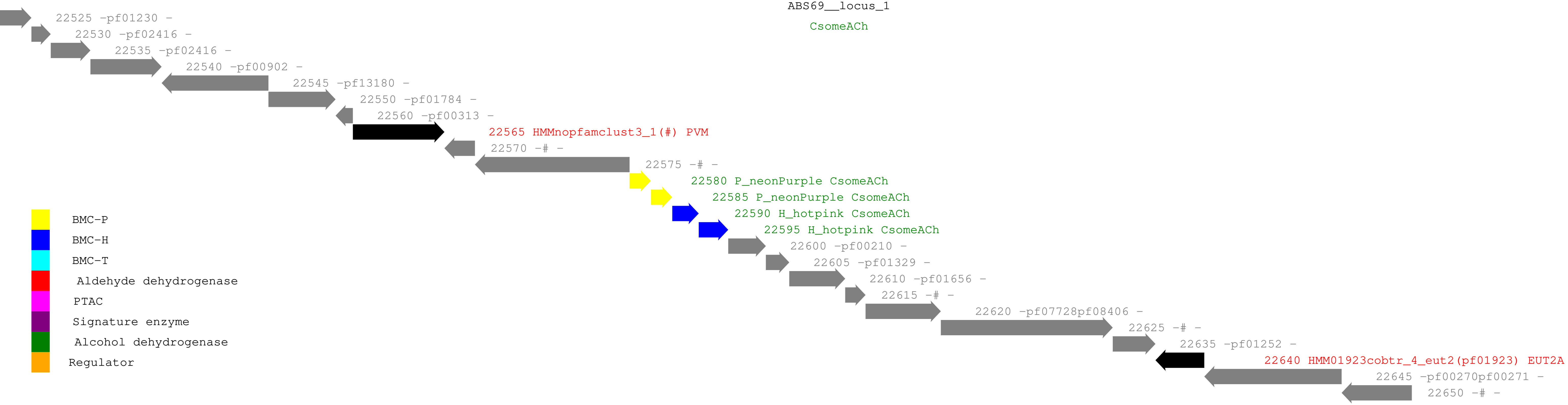

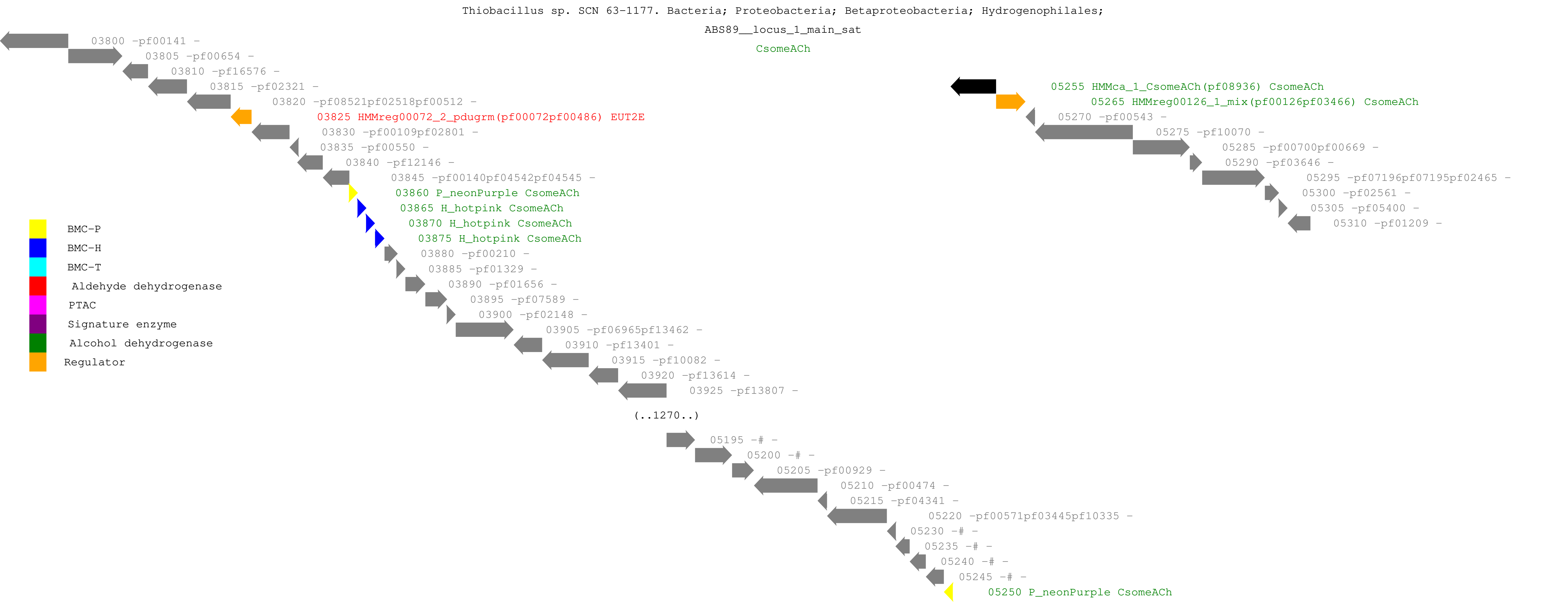
Thiobacillus sp. SCN 63-1177. Bacteria; Proteobacteria; Betaproteobacteria; Hydrogenophilales;
ABS89__locus_1_main_sat
CsomeACh
03800 -pf00141 -
03805 -pf00654 -
03810 -pf16576 -
03815 -pf02321 -
03820 -pf08521pf02518pf00512 -
03825 HMMreg00072_2_pdugrm(pf00072pf00486) EUT2E
03830 -pf00109pf02801 -
03835 -pf00550 -
03840 -pf12146 -
03845 -pf00140pf04542pf04545 -
03860 P_neonPurple CsomeACh
03865 H_hotpink CsomeACh
03870 H_hotpink CsomeACh
03875 H_hotpink CsomeACh
03880 -pf00210 -
03885 -pf01329 -
03890 -pf01656 -
03895 -pf07589 -
03900 -pf02148 -
03905 -pf06965pf13462 -
03910 -pf13401 -
03915 -pf10082 -
03920 -pf13614 -
03925 -pf13807 -
(..1270..)
05195 -# -
05200 -# -
05205 -pf00929 -
05210 -pf00474 -
05215 -pf04341 -
05220 -pf00571pf03445pf10335 -
05230 -# -
05235 -# -
05240 -# -
05245 -# -
05250 P_neonPurple CsomeACh
05255 HMMca_1_CsomeACh(pf08936) CsomeACh
05265 HMMreg00126_1_mix(pf00126pf03466) CsomeACh
05270 -pf00543 -
05275 -pf10070 -
05285 -pf00700pf00669 -
05290 -pf03646 -
05295 -pf07196pf07195pf02465 -
05300 -pf02561 -
05305 -pf05400 -
05310 -pf01209 -
BMC-P
BMC-H
BMC-T
Aldehyde dehydrogenase
PTAC
Signature enzyme
Alcohol dehydrogenase
Regulator

Thiobacillus sp. SCN 63-374. Bacteria; Proteobacteria; Betaproteobacteria; Hydrogenophilales;

ABS92__locus_1_main_sat

CsomeACh

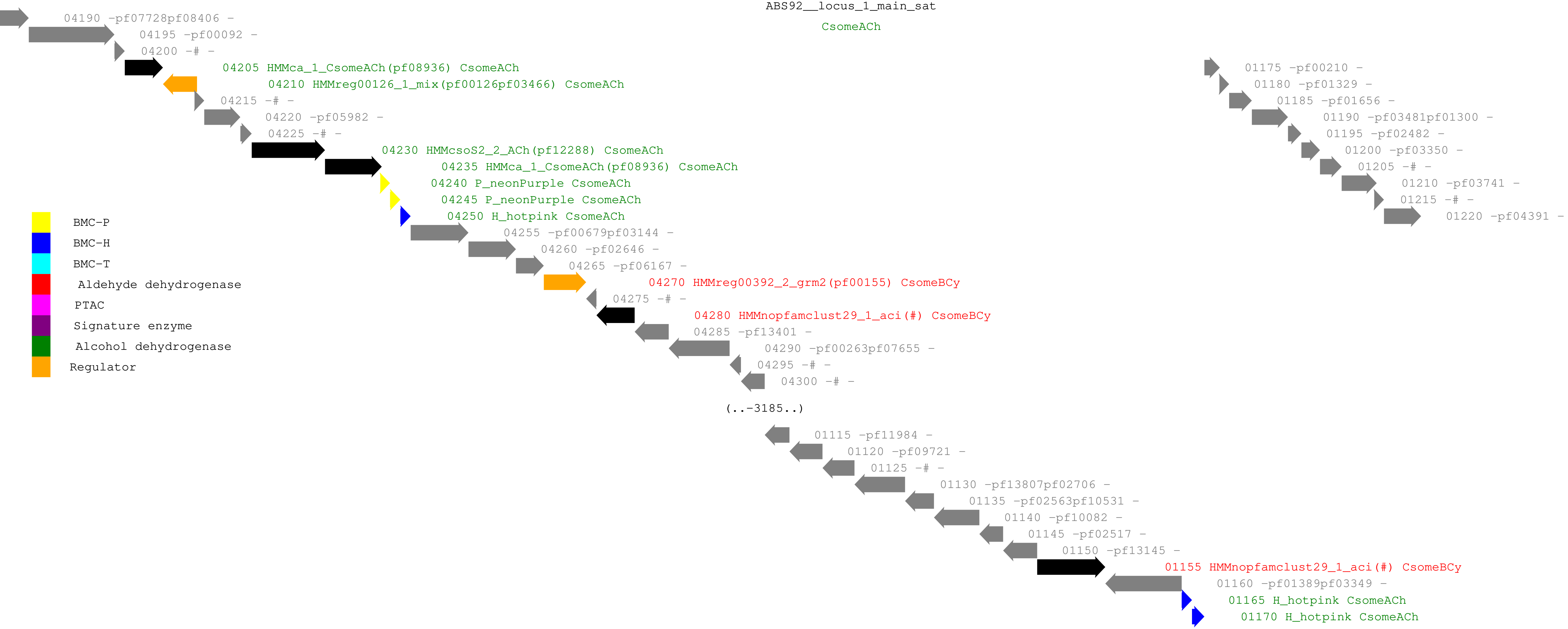

Pseudonocardia sp. SCN 73-27. Bacteria; Actinobacteria; Pseudonocardiales; Pseudonocardiaceae;

ABT15__locus_1

CsomeACh

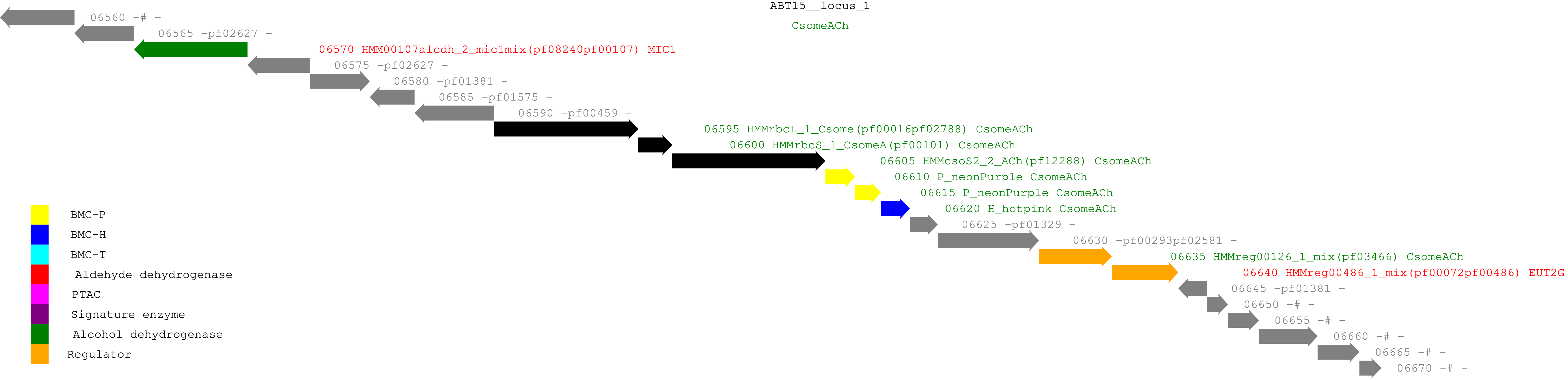

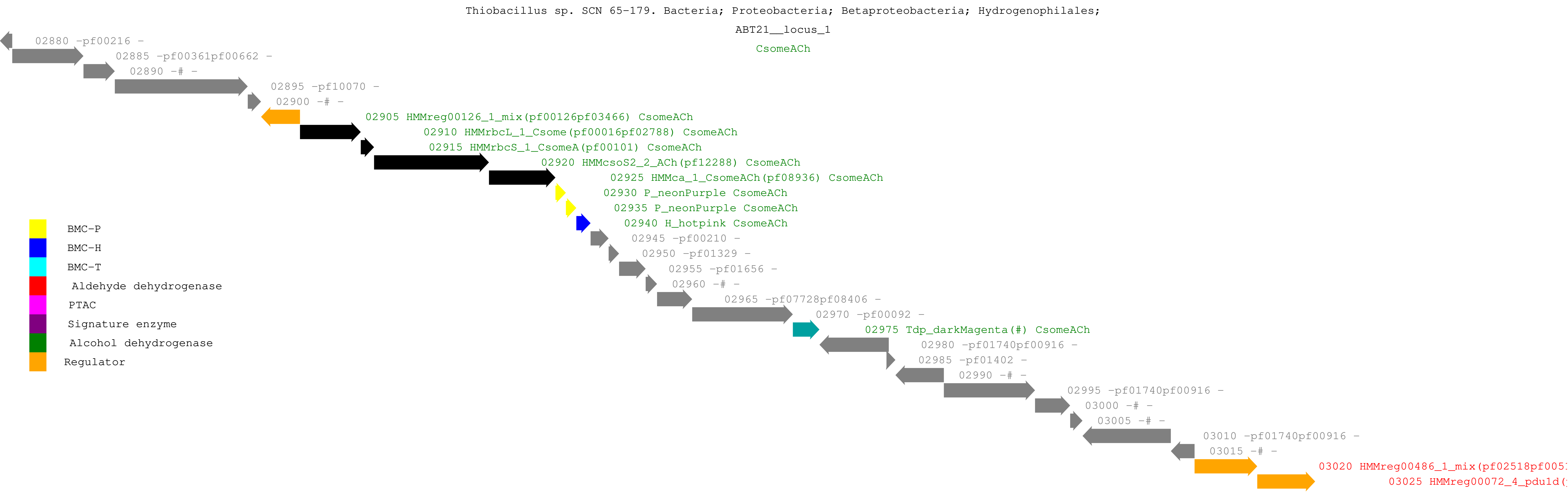

Thiobacillus sp. SCN 65-179. Bacteria; Proteobacteria; Betaproteobacteria; Hydrogenophilales;
ABT21__locus_1
CsomeACh
02880 -pf00216 -
02885 -pf00361pf00662 -
02890 -# -
02895 -pf10070 -
02900 -# -
02905 HMMreg00126_1_mix(pf00126pf03466) CsomeACh
02910 HMMrbcL_1_Csome(pf00016pf02788) CsomeACh
02915 HMMrbcS_1_CsomeA(pf00101) CsomeACh
02920 HMMcsoS2_2_ACh(pf12288) CsomeACh
02925 HMMca_1_CsomeACh(pf08936) CsomeACh
02930 P_neonPurple CsomeACh
02935 P_neonPurple CsomeACh
02940 H_hotpink CsomeACh
02945 -pf00210 -
02950 -pf01329 -
02955 -pf01656 -
02960 -# -
02965 -pf07728pf08406 -
02970 -pf00092 -
02975 Tdp_darkMagenta(#) CsomeACh
02980 -pf01740pf00916 -
02985 -pf01402 -
02990 -# -
02995 -pf01740pf00916 -
03000 -# -
03005 -# -
03010 -pf01740pf00916 -
03015 -# -
03020 HMMreg00486_1_mix(pf02518pf005
03025 HMMreg00072_4_pdu1d(
BMC-P
BMC-H
BMC-T
Aldehyde dehydrogenase
PTAC
Signature enzyme
Alcohol dehydrogenase
Regulator

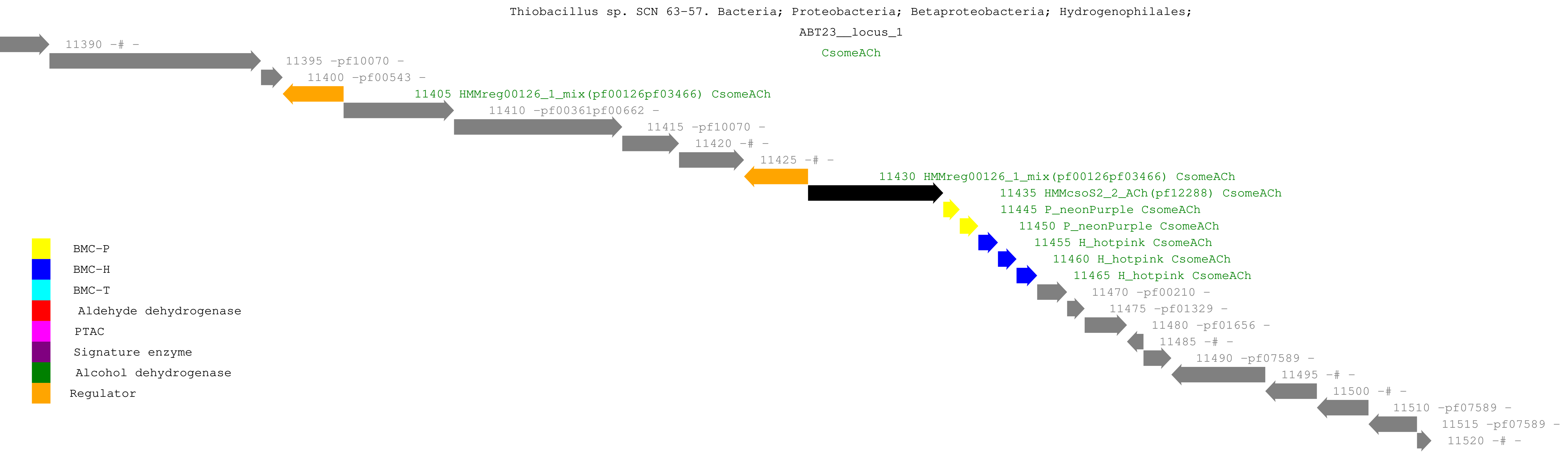

Thiobacillus sp. SCN 63-57. Bacteria; Proteobacteria; Betaproteobacteria; Hydrogenophilales;
ABT23__locus_1
CsomeACh
11390 -# -
11395 -pf10070 -
11400 -pf00543 -
11405 HMMreg00126_1_mix(pf00126pf03466) CsomeACh
11410 -pf00361pf00662 -
11415 -pf10070 -
11420 -# -
11425 -# -
11430 HMMreg00126_1_mix(pf00126pf03466) CsomeACh
11435 HMMcsoS2_2_ACh(pf12288) CsomeACh
11445 P_neonPurple CsomeACh
11450 P_neonPurple CsomeACh
11455 H_hotpink CsomeACh
11460 H_hotpink CsomeACh
11465 H_hotpink CsomeACh
11470 -pf00210 -
11475 -pf01329 -
11480 -pf01656 -
11485 -# -
11490 -pf07589 -
11495 -# -
11500 -# -
11510 -pf07589 -
11515 -pf07589 -
11520 -# -
BMC-P
BMC-H
BMC-T
Aldehyde dehydrogenase
PTAC
Signature enzyme
Alcohol dehydrogenase
Regulator

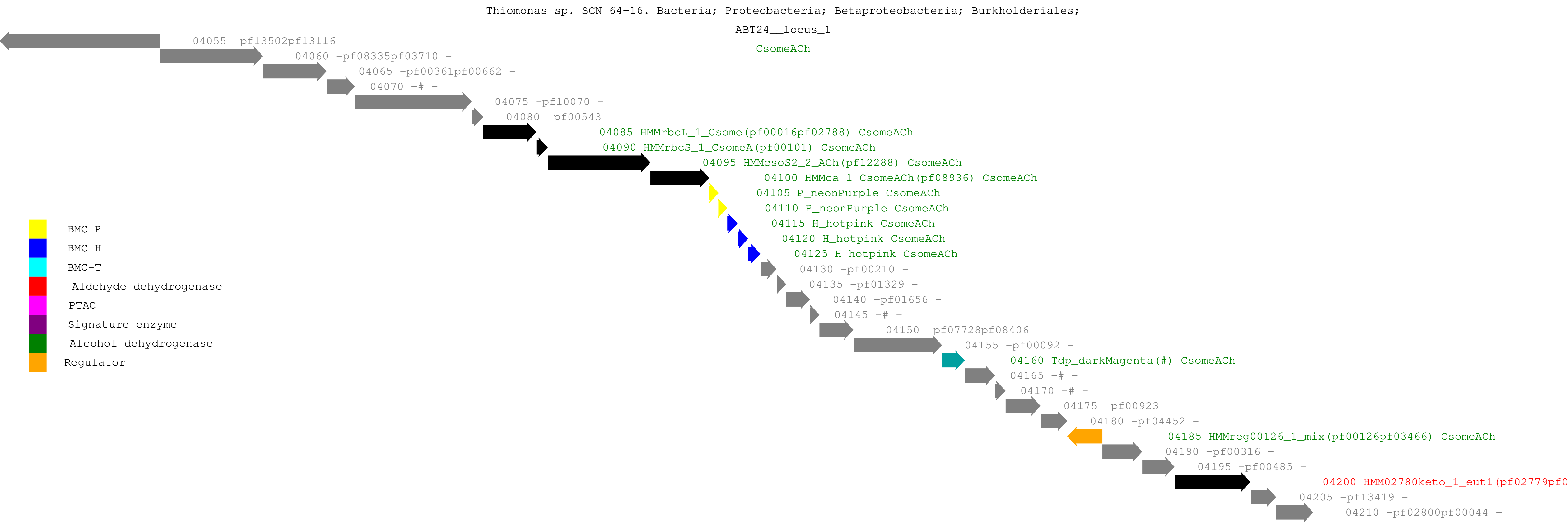

Thiomonas sp. SCN 64-16. Bacteria; Proteobacteria; Betaproteobacteria; Burkholderiales;
ABT24__locus_1
CsomeACh
04055 -pf13502pf13116 -
04060 -pf08335pf03710 -
04065 -pf00361pf00662 -
04070 -# -
04075 -pf10070 -
04080 -pf00543 -
04085 HMMrbcL_1_Csome(pf00016pf02788) CsomeACh
04090 HMMrbcS_1_CsomeA(pf00101) CsomeACh
04095 HMMcsoS2_2_ACh(pf12288) CsomeACh
04100 HMMca_1_CsomeACh(pf08936) CsomeACh
04105 P_neonPurple CsomeACh
04110 P_neonPurple CsomeACh
04115 H_hotpink CsomeACh
04120 H_hotpink CsomeACh
04125 H_hotpink CsomeACh
04130 -pf00210 -
04135 -pf01329 -
04140 -pf01656 -
04145 -# -
04150 -pf07728pf08406 -
04155 -pf00092 -
04160 Tdp_darkMagenta(#) CsomeACh
04165 -# -
04170 -# -
04175 -pf00923 -
04180 -pf04452 -
04185 HMMreg00126_1_mix(pf00126pf03466) CsomeACh
04190 -pf00316 -
04195 -pf00485 -
04200 HMM02780keto_1_eut1(pf02779pf0
04205 -pf13419 -
04210 -pf02800pf00044 -
BMC-P
BMC-H
BMC-T
Aldehyde dehydrogenase
PTAC
Signature enzyme
Alcohol dehydrogenase
Regulator

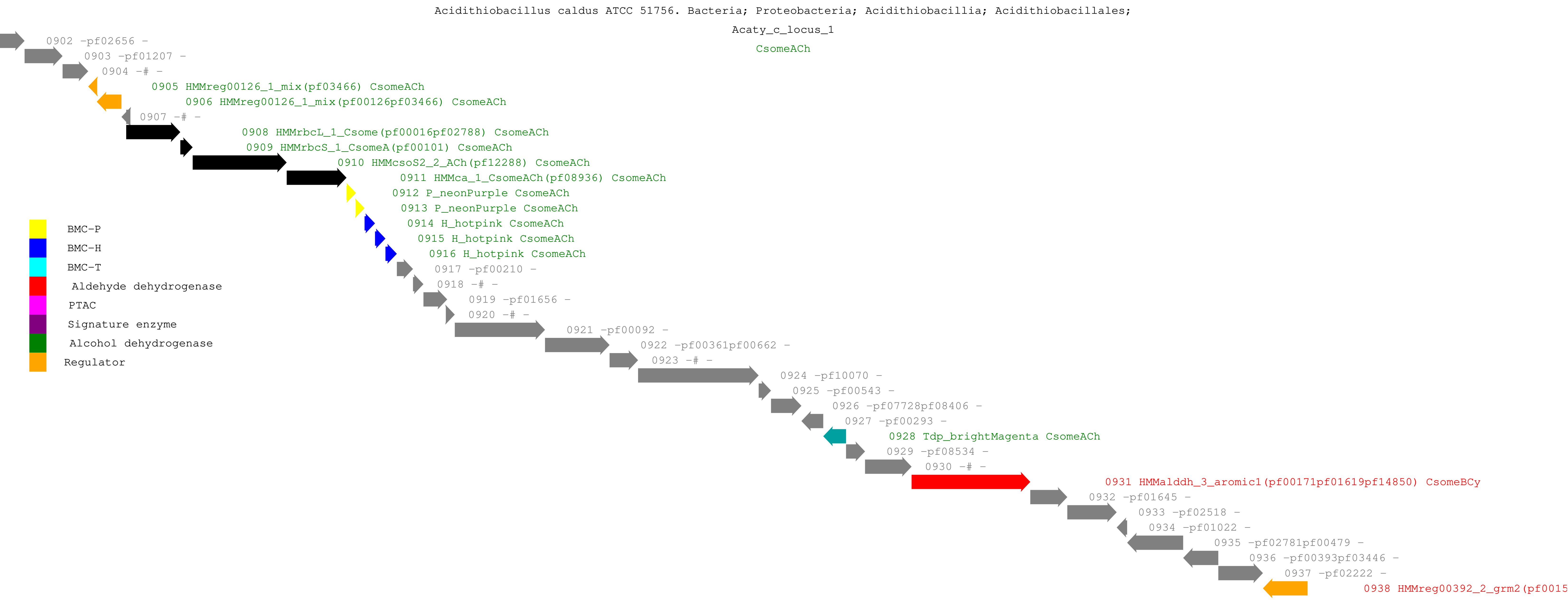
Acidithiobacillus caldus ATCC 51756. Bacteria; Proteobacteria; Acidithiobacillia; Acidithiobacillales;
Acaty_c_locus_1
CsomeACh
0902 -pf02656 -
0903 -pf01207 -
0904 -# -
0905 HMMreg00126_1_mix(pf03466) CsomeACh
0906 HMMreg00126_1_mix(pf00126pf03466) CsomeACh
0907 -# -
0908 HMMrbcL_1_Csome(pf00016pf02788) CsomeACh
0909 HMMrbcS_1_CsomeA(pf00101) CsomeACh
0910 HMMcsoS2_2_ACh(pf12288) CsomeACh
0911 HMMca_1_CsomeACh(pf08936) CsomeACh
0912 P_neonPurple CsomeACh
0913 P_neonPurple CsomeACh
0914 H_hotpink CsomeACh
0915 H_hotpink CsomeACh
0916 H_hotpink CsomeACh
0917 -pf00210 -
0918 -# -
0919 -pf01656 -
0920 -# -
0921 -pf00092 -
0922 -pf00361pf00662 -
0923 -# -
0924 -pf10070 -
0925 -pf00543 -
0926 -pf07728pf08406 -
0927 -pf00293 -
0928 Tdp_brightMagenta CsomeACh
0929 -pf08534 -
0930 -# -
0931 HMMaldh_3_aromic1(pf00171pf01619pf14850) CsomeBCy
0932 -pf01645 -
0933 -pf02518 -
0934 -pf01022 -
0935 -pf02781pf00479 -
0936 -pf00393pf03446 -
0937 -pf02222 -
0938 HMMreg00392_2_grm2(pf0015
BMC-P
BMC-H
BMC-T
Aldehyde dehydrogenase
PTAC
Signature enzyme
Alcohol dehydrogenase
Regulator

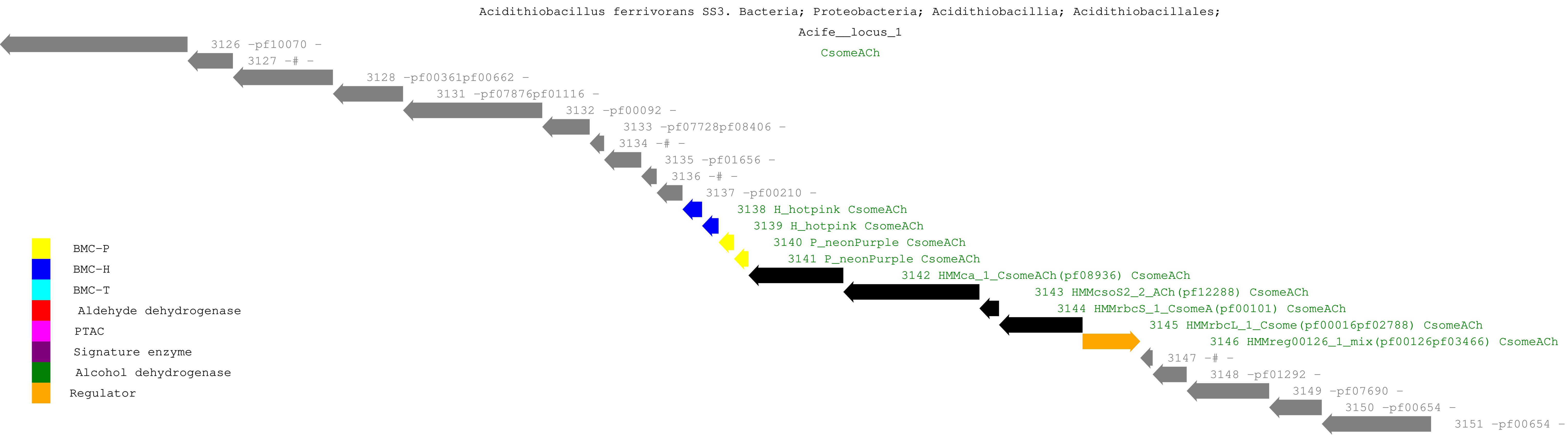

Acidithiobacillus ferrivorans SS3. Bacteria; Proteobacteria; Acidithiobacillia; Acidithiobacillales;
Acife__locus_1
CsomeACh
3126 -pf10070 -
3127 -# -
3128 -pf00361pf00662 -
3131 -pf07876pf01116 -
3132 -pf00092 -
3133 -pf07728pf08406 -
3134 -# -
3135 -pf01656 -
3136 -# -
3137 -pf00210 -
3138 H_hotpink CsomeACh
3139 H_hotpink CsomeACh
3140 P_neonPurple CsomeACh
3141 P_neonPurple CsomeACh
3142 HMMca_1_CsomeACh(pf08936) CsomeACh
3143 HMMcsoS2_2_ACh(pf12288) CsomeACh
3144 HMMrbcS_1_CsomeA(pf00101) CsomeACh
3145 HMMrbcL_1_Csome(pf00016pf02788) CsomeACh
3146 HMMreg00126_1_mix(pf00126pf03466) CsomeACh
3147 -# -
3148 -pf01292 -
3149 -pf07690 -
3150 -pf00654 -
3151 -pf00654 -
BMC-P
BMC-H
BMC-T
Aldehyde dehydrogenase
PTAC
Signature enzyme
Alcohol dehydrogenase
Regulator

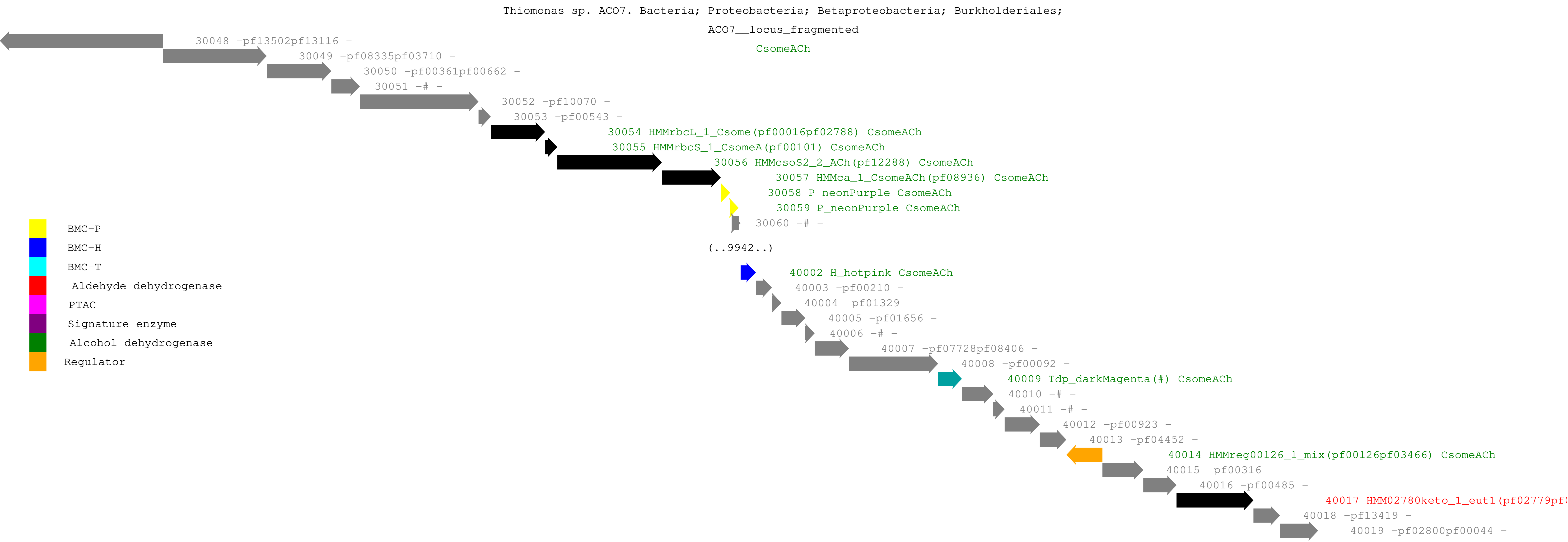
Thiomonas sp. ACO7. Bacteria; Proteobacteria; Betaproteobacteria; Burkholderiales;
ACO7__locus_fragmented
CsomeACh
30048 -pf13502pf13116 -
30049 -pf08335pf03710 -
30050 -pf00361pf00662 -
30051 -# -
30052 -pf10070 -
30053 -pf00543 -
30054 HMMrbcL_1_Csome(pf00016pf02788) CsomeACh
30055 HMMrbcS_1_CsomeA(pf00101) CsomeACh
30056 HMMcsoS2_2_ACh(pf12288) CsomeACh
30057 HMMca_1_CsomeACh(pf08936) CsomeACh
30058 P_neonPurple CsomeACh
30059 P_neonPurple CsomeACh
30060 -# -
(..9942..)
40002 H_hotpink CsomeACh
40003 -pf00210 -
40004 -pf01329 -
40005 -pf01656 -
40006 -# -
40007 -pf07728pf08406 -
40008 -pf00092 -
40009 Tdp_darkMagenta(#) CsomeACh
40010 -# -
40011 -# -
40012 -pf00923 -
40013 -pf04452 -
40014 HMMreg00126_1_mix(pf00126pf03466) CsomeACh
40015 -pf00316 -
40016 -pf00485 -
40017 HMM02780keto_1_eut1(pf02779pf
40018 -pf13419 -
40019 -pf02800pf00044 -
BMC-P
BMC-H
BMC-T
Aldehyde dehydrogenase
PTAC
Signature enzyme
Alcohol dehydrogenase
Regulator

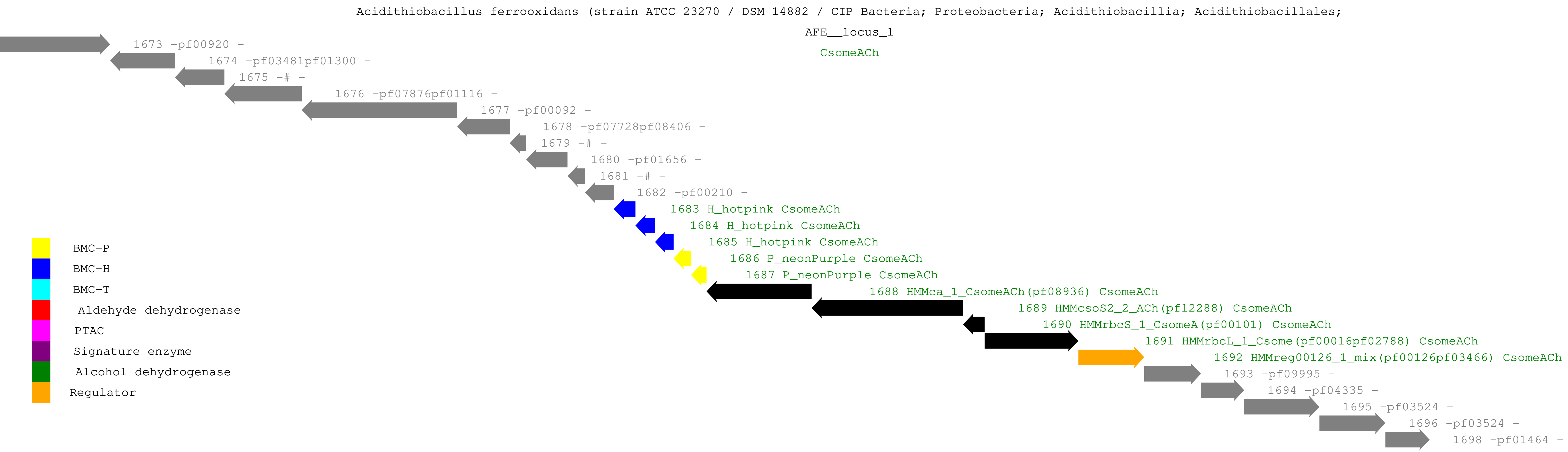
Acidithiobacillus ferrooxidans (strain ATCC 23270 / DSM 14882 / CIP Bacteria; Proteobacteria; Acidithiobacillia; Acidithiobacillales;
AFE__locus_1
CsomeACh
1673 -pf00920 -
1674 -pf03481pf01300 -
1675 -# -
1676 -pf07876pf01116 -
1677 -pf00092 -
1678 -pf07728pf08406 -
1679 -# -
1680 -pf01656 -
1681 -# -
1682 -pf00210 -
1683 H_hotpink CsomeACh
1684 H_hotpink CsomeACh
1685 H_hotpink CsomeACh
1686 P_neonPurple CsomeACh
1687 P_neonPurple CsomeACh
1688 HMMca_1_CsomeACh(pf08936) CsomeACh
1689 HMMcsoS2_2_ACh(pf12288) CsomeACh
1690 HMMrbcS_1_CsomeA(pf00101) CsomeACh
1691 HMMrbcL_1_Csome(pf00016pf02788) CsomeACh
1692 HMMreg00126_1_mix(pf00126pf03466) CsomeACh
1693 -pf09995 -
1694 -pf04335 -
1695 -pf03524 -
1696 -pf03524 -
1698 -pf01464 -
BMC-P
BMC-H
BMC-T
Aldehyde dehydrogenase
PTAC
Signature enzyme
Alcohol dehydrogenase
Regulator

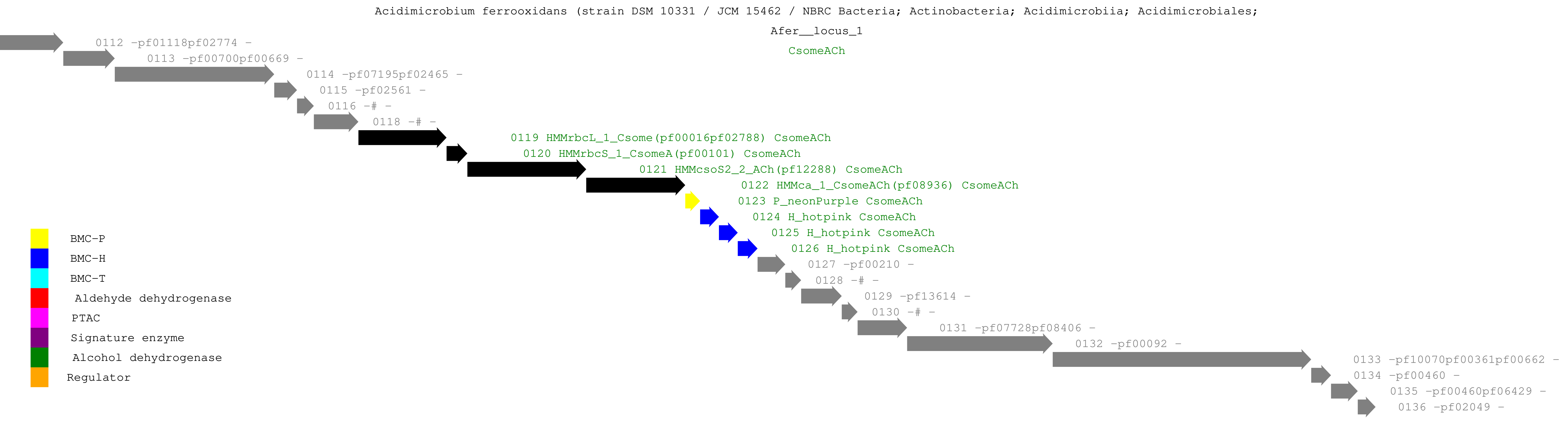

Acidimicrobium ferrooxidans (strain DSM 10331 / JCM 15462 / NBRC Bacteria; Actinobacteria; Acidimicrobiia; Acidimicrobiales;
Afer__locus_1
CsomeACh
0112 -pf01118pf02774 -
0113 -pf00700pf00669 -
0114 -pf07195pf02465 -
0115 -pf02561 -
0116 -# -
0118 -# -
0119 HMMrbcL_1_Csome(pf00016pf02788) CsomeACh
0120 HMMrbcS_1_CsomeA(pf00101) CsomeACh
0121 HMMcsoS2_2_ACh(pf12288) CsomeACh
0122 HMMca_1_CsomeACh(pf08936) CsomeACh
0123 P_neonPurple CsomeACh
0124 H_hotpink CsomeACh
0125 H_hotpink CsomeACh
0126 H_hotpink CsomeACh
0127 -pf00210 -
0128 -# -
0129 -pf13614 -
0130 -# -
0131 -pf07728pf08406 -
0132 -pf00092 -
0133 -pf10070pf00361pf00662 -
0134 -pf00460 -
0135 -pf00460pf06429 -
0136 -pf02049 -
BMC-P
BMC-H
BMC-T
Aldehyde dehydrogenase
PTAC
Signature enzyme
Alcohol dehydrogenase
Regulator

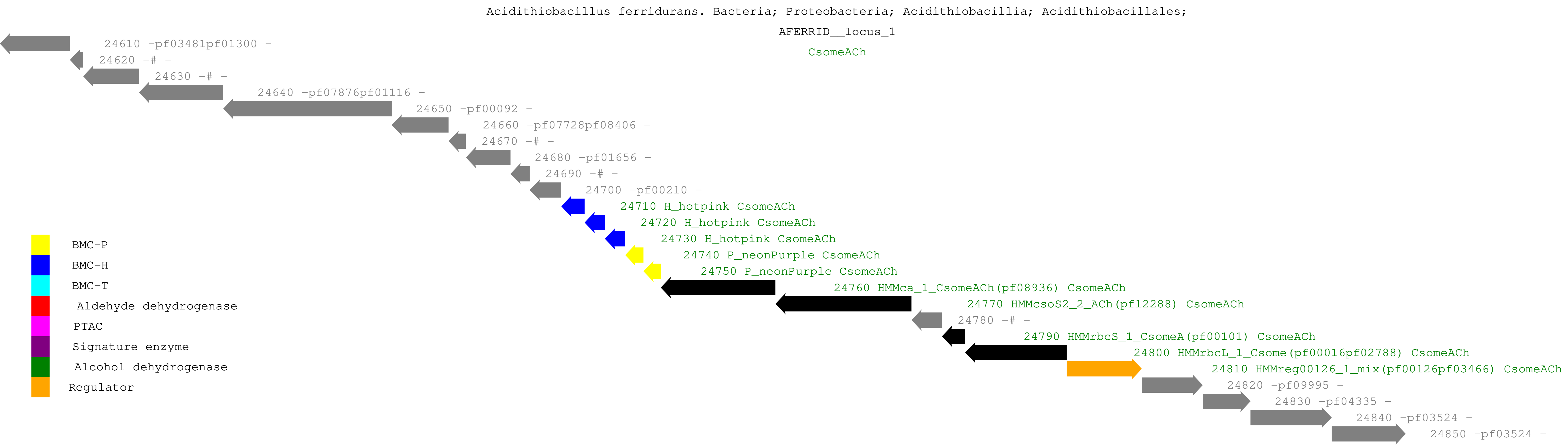
Acidithiobacillus ferridurans. Bacteria; Proteobacteria; Acidithiobacillia; Acidithiobacillales;
AFERRID__locus_1
CsomeACh
24610 -pf03481pf01300 -
24620 -# -
24630 -# -
24640 -pf07876pf01116 -
24650 -pf00092 -
24660 -pf07728pf08406 -
24670 -# -
24680 -pf01656 -
24690 -# -
24700 -pf00210 -
24710 H_hotpink CsomeACh
24720 H_hotpink CsomeACh
24730 H_hotpink CsomeACh
24740 P_neonPurple CsomeACh
24750 P_neonPurple CsomeACh
24760 HMMca_1_CsomeACh(pf08936) CsomeACh
24770 HMMcsoS2_2_ACh(pf12288) CsomeACh
24780 -# -
24790 HMMrbcS_1_CsomeA(pf00101) CsomeACh
24800 HMMrbcL_1_Csome(pf00016pf02788) CsomeACh
24810 HMMreg00126_1_mix(pf00126pf03466) CsomeACh
24820 -pf09995 -
24830 -pf04335 -
24840 -pf03524 -
24850 -pf03524 -
BMC-P
BMC-H
BMC-T
Aldehyde dehydrogenase
PTAC
Signature enzyme
Alcohol dehydrogenase
Regulator

Acidithiobacillus ferrivorans. Bacteria; Proteobacteria; Acidithiobacillia; Acidithiobacillales;

AFERRI__locus_1

CsomeACh

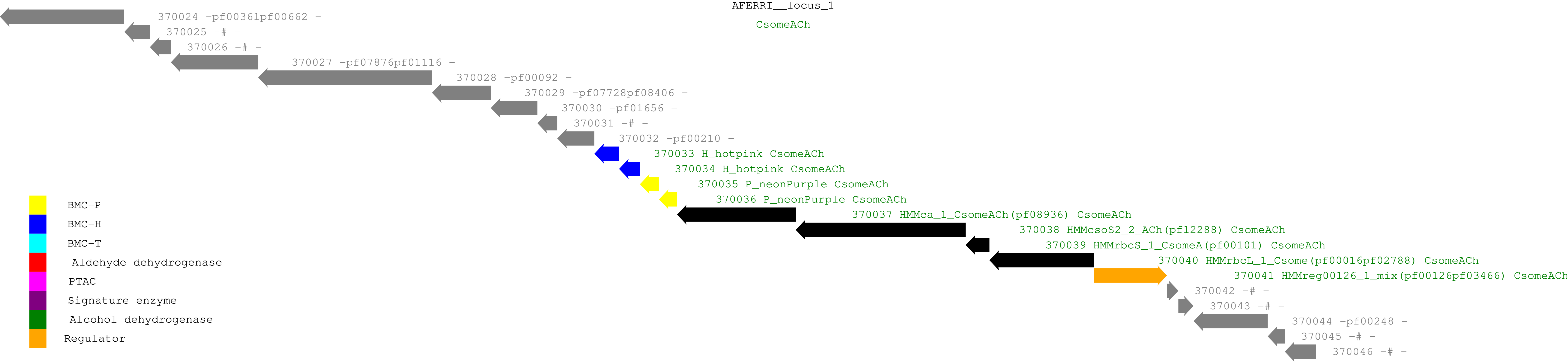

Allochromatium vinosum (strain ATCC 17899 / DSM 180 / NBRC 103801 / Bacteria; Proteobacteria; Gammaproteobacteria; Chromatiales;

Alvin__locus_1_main_sat

CsomeACh

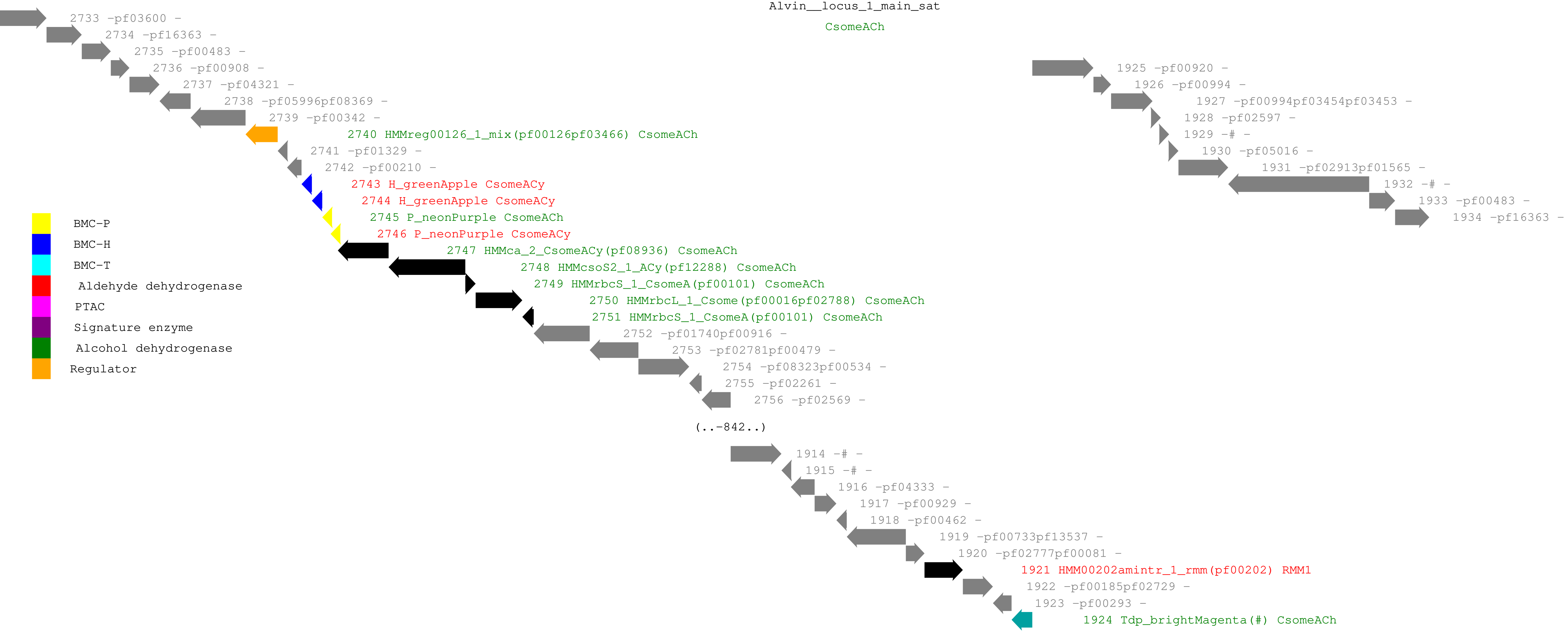

Gammaproteobacteria bacterium SG8_15. Bacteria; Proteobacteria; Gammaproteobacteria.

AMJ55__locus_1_main_sat

CsomeACh

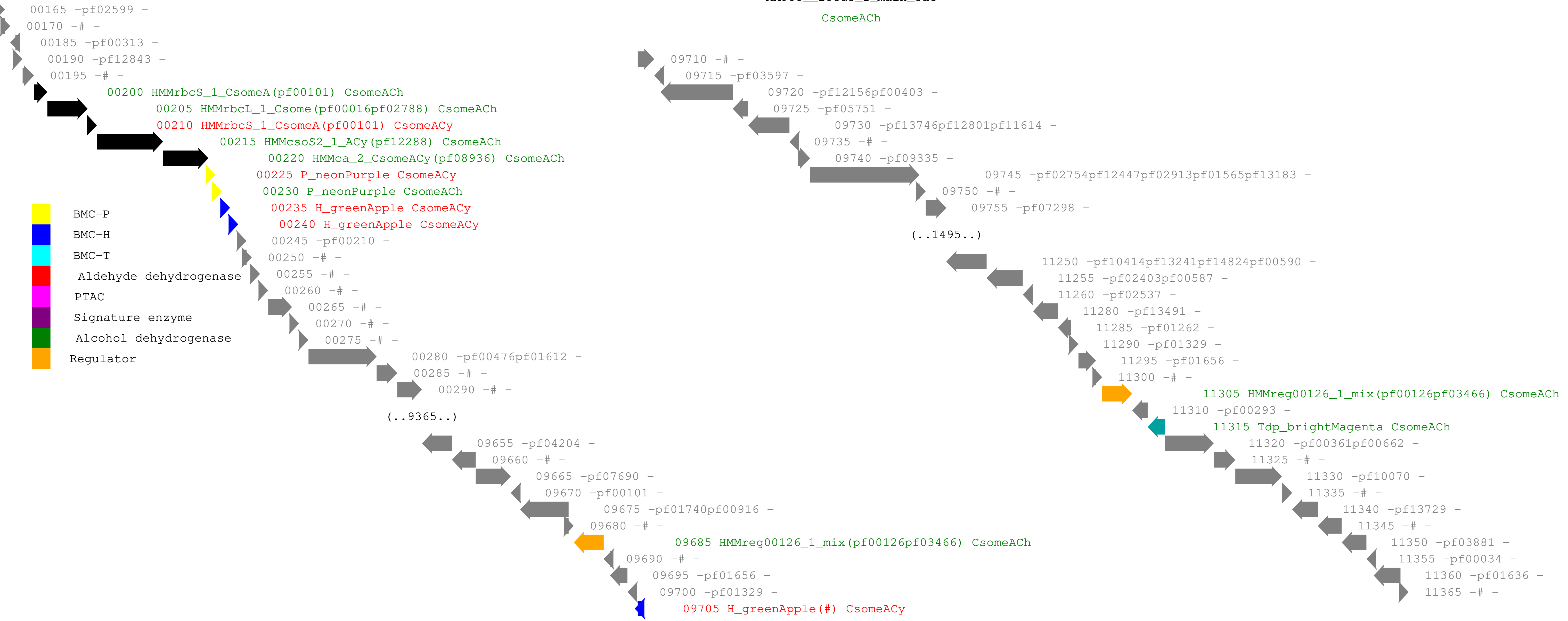

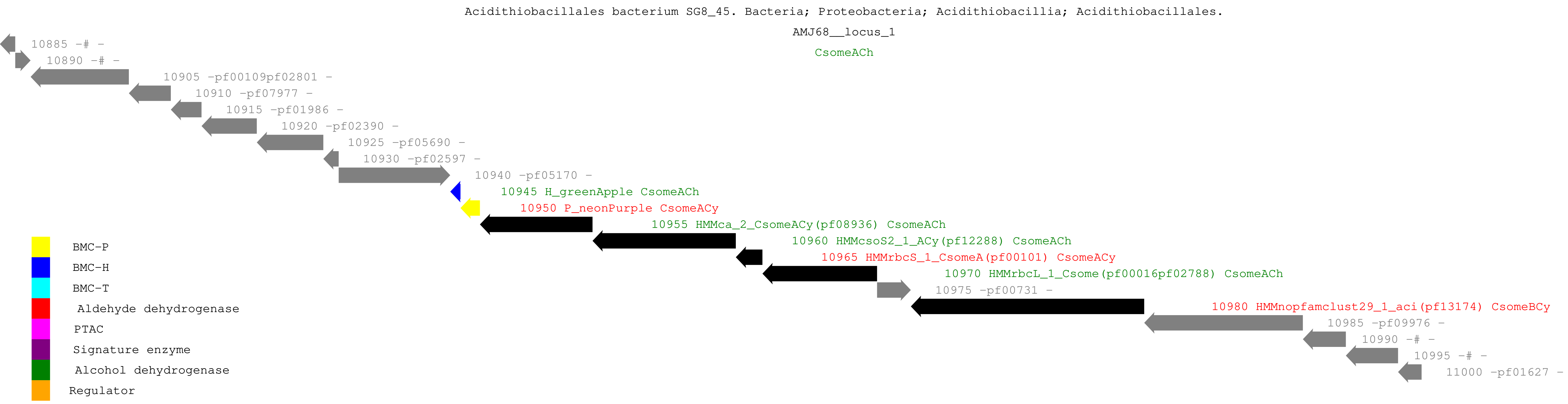

Acidithiobacillales bacterium SG8_45. Bacteria; Proteobacteria; Acidithiobacillia; Acidithiobacillales.
AMJ68__locus_1
CsomeACh
10885 -# -
10890 -# -
10905 -pf00109pf02801 -
10910 -pf07977 -
10915 -pf01986 -
10920 -pf02390 -
10925 -pf05690 -
10930 -pf02597 -
10940 -pf05170 -
10945 H_greenApple CsomeACh
10950 P_neonPurple CsomeACy
10955 HMMca_2_CsomeACy(pf08936) CsomeACh
10960 HMMcsoS2_1_ACy(pf12288) CsomeACh
10965 HMMrbcS_1_CsomeA(pf00101) CsomeACy
10970 HMMrbcL_1_Csome(pf00016pf02788) CsomeACh
10975 -pf00731 -
10980 HMMnopfamclust29_1_aci(pf13174) CsomeBCy
10985 -pf09976 -
10990 -# -
10995 -# -
11000 -pf01627 -
BMC-P
BMC-H
BMC-T
Aldehyde dehydrogenase
PTAC
Signature enzyme
Alcohol dehydrogenase
Regulator

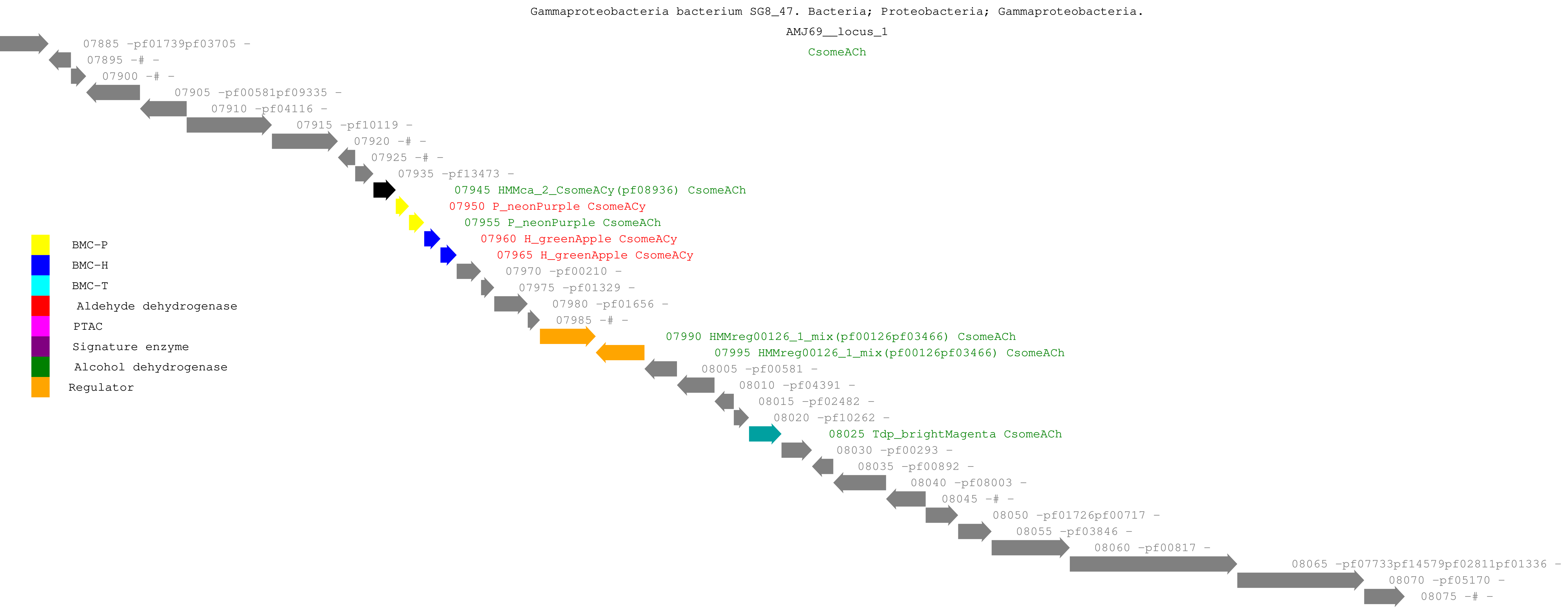
Gammaproteobacteria bacterium SG8_47. Bacteria; Proteobacteria; Gammaproteobacteria.
AMJ69__locus_1
CsomeACh
07885 -pf01739pf03705 -
07895 -# -
07900 -# -
07905 -pf00581pf09335 -
07910 -pf04116 -
07915 -pf10119 -
07920 -# -
07925 -# -
07935 -pf13473 -
07945 HMMca_2_CsomeACy(pf08936) CsomeACh
07950 P_neonPurple CsomeACy
07955 P_neonPurple CsomeACh
07960 H_greenApple CsomeACy
07965 H_greenApple CsomeACy
07970 -pf00210 -
07975 -pf01329 -
07980 -pf01656 -
07985 -# -
07990 HMMreg00126_1_mix(pf00126pf03466) CsomeACh
07995 HMMreg00126_1_mix(pf00126pf03466) CsomeACh
08005 -pf00581 -
08010 -pf04391 -
08015 -pf02482 -
08020 -pf10262 -
08025 Tdp_brightMagenta CsomeACh
08030 -pf00293 -
08035 -pf00892 -
08040 -pf08003 -
08045 -# -
08050 -pf01726pf00717 -
08055 -pf03846 -
08060 -pf00817 -
08065 -pf07733pf14579pf02811pf01336 -
08070 -pf05170 -
08075 -# -
BMC-P
BMC-H
BMC-T
Aldehyde dehydrogenase
PTAC
Signature enzyme
Alcohol dehydrogenase
Regulator

Thiotrichales bacterium SG8_50. Bacteria; Proteobacteria; Gammaproteobacteria; Thiotrichales;

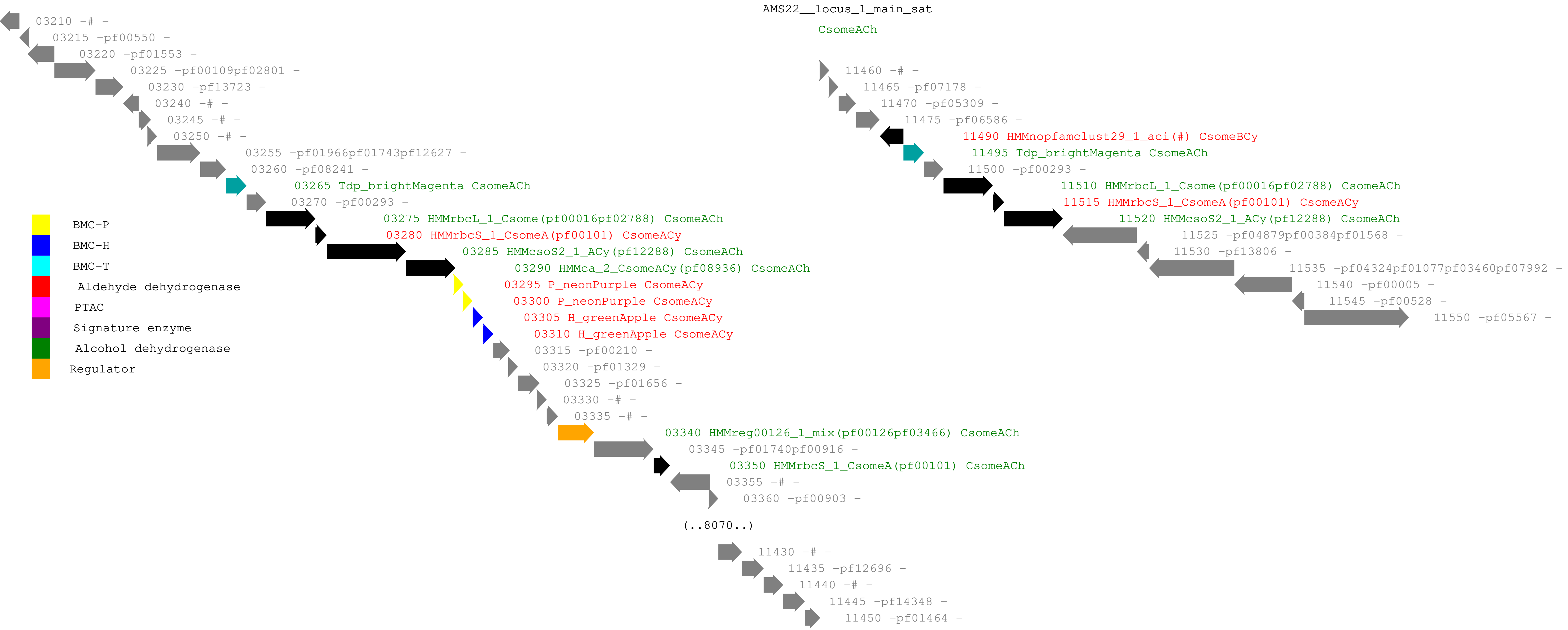

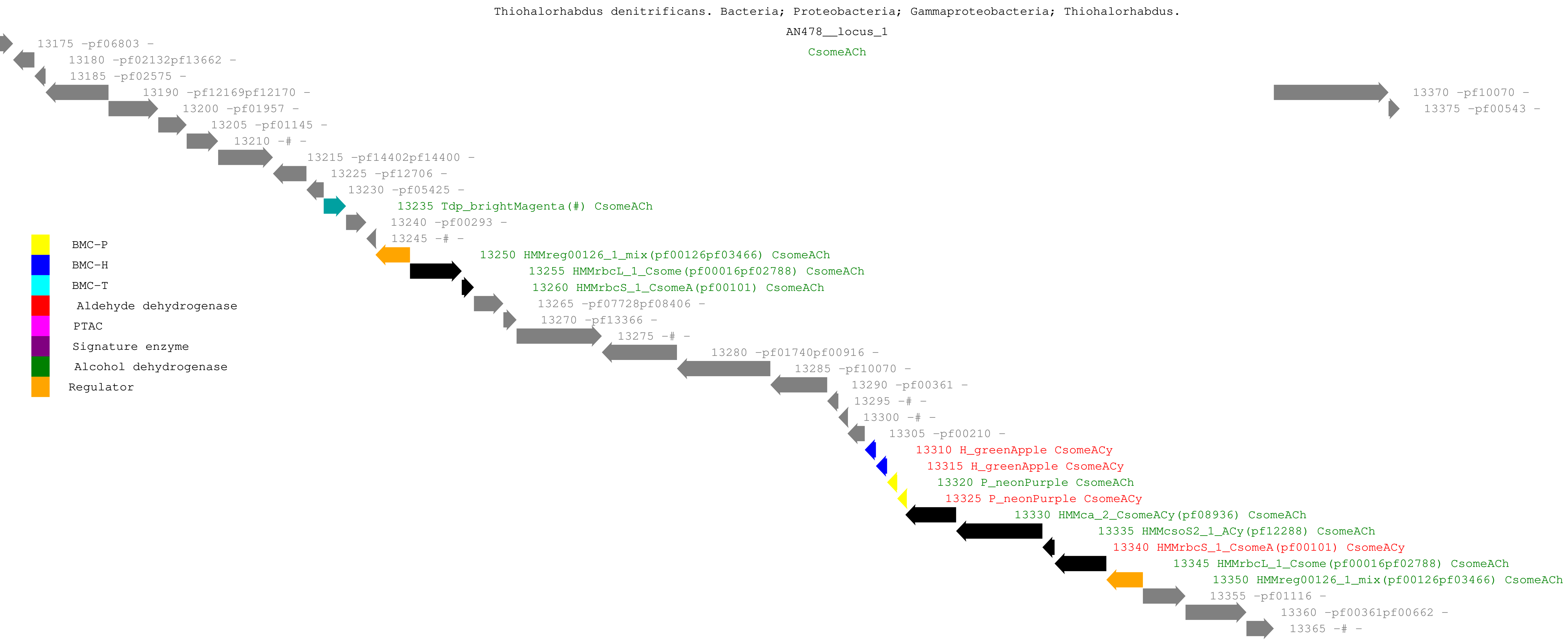

Thiohalorhabdus denitrificans. Bacteria; Proteobacteria; Gammaproteobacteria; Thiohalorhabdus.
AN478__locus_1
CsomeACh
13175 -pf06803 -
13180 -pf02132pf13662 -
13185 -pf02575 -
13190 -pf12169pf12170 -
13200 -pf01957 -
13205 -pf01145 -
13210 -# -
13215 -pf14402pf14400 -
13225 -pf12706 -
13230 -pf05425 -
13235 Tdp_brightMagenta(#) CsomeACh
13240 -pf00293 -
13245 -# -
13250 HMMreg00126_1_mix(pf00126pf03466) CsomeACh
13255 HMMrbcL_1_Csome(pf00016pf02788) CsomeACh
13260 HMMrbcS_1_CsomeA(pf00101) CsomeACh
13265 -pf07728pf08406 -
13270 -pf13366 -
13275 -# -
13280 -pf01740pf00916 -
13285 -pf10070 -
13290 -pf00361 -
13295 -# -
13300 -# -
13305 -pf00210 -
13310 H_greenApple CsomeACy
13315 H_greenApple CsomeACy
13320 P_neonPurple CsomeACh
13325 P_neonPurple CsomeACy
13330 HMMca_2_CsomeACy(pf08936) CsomeACh
13335 HMMcsoS2_1_ACy(pf12288) CsomeACh
13340 HMMrbcS_1_CsomeA(pf00101) CsomeACy
13345 HMMrbcL_1_Csome(pf00016pf02788) CsomeACh
13350 HMMreg00126_1_mix(pf00126pf03466) CsomeACh
13355 -pf01116 -
13360 -pf00361pf00662 -
13365 -# -
13370 -pf10070 -
13375 -pf00543 -
BMC-P
BMC-H
BMC-T
Aldehyde dehydrogenase
PTAC
Signature enzyme
Alcohol dehydrogenase
Regulator

Acidithiobacillus caldus (strain SM-1). Bacteria; Proteobacteria; Acidithiobacillia; Acidithiobacillales;

Atc__locus_1

CsomeACh

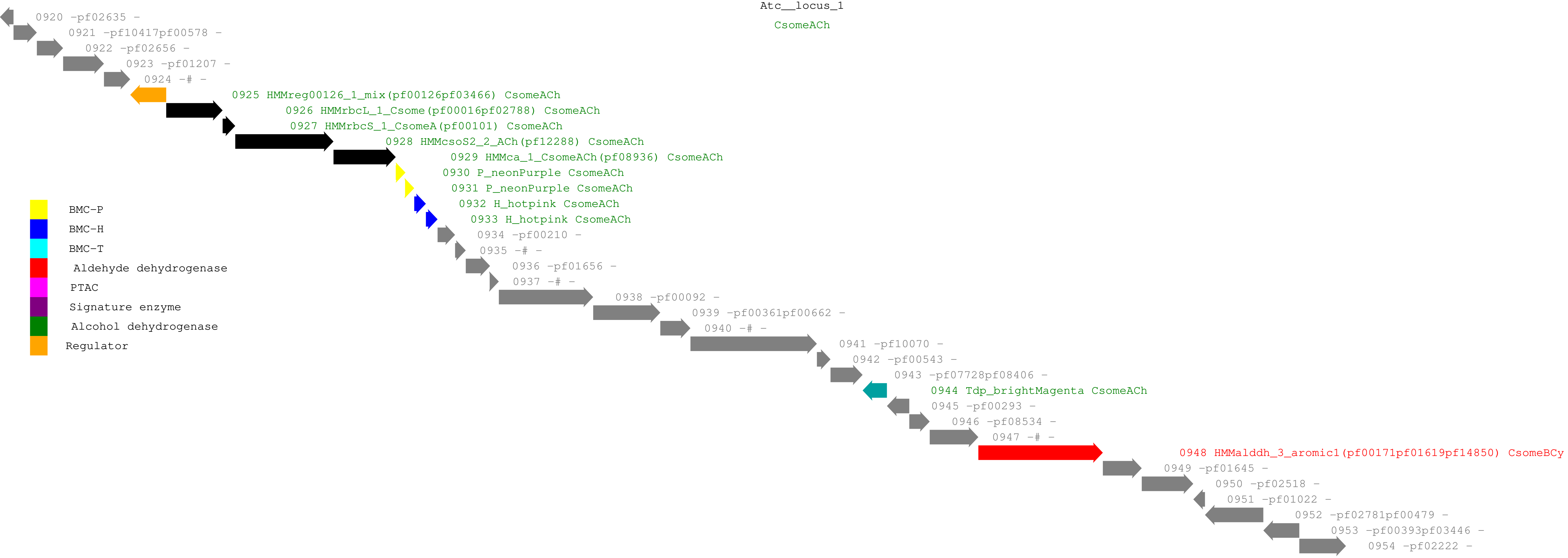

Loktanella sp. 22II-4b. Bacteria; Proteobacteria; Alphaproteobacteria; Rhodobacterales;

ATO5__locus_1

CsomeACh

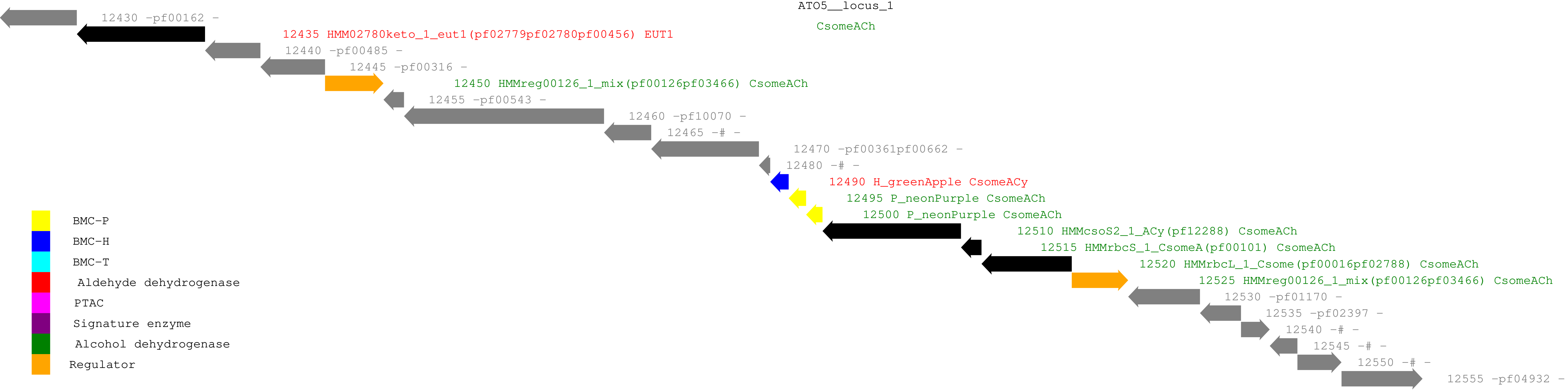

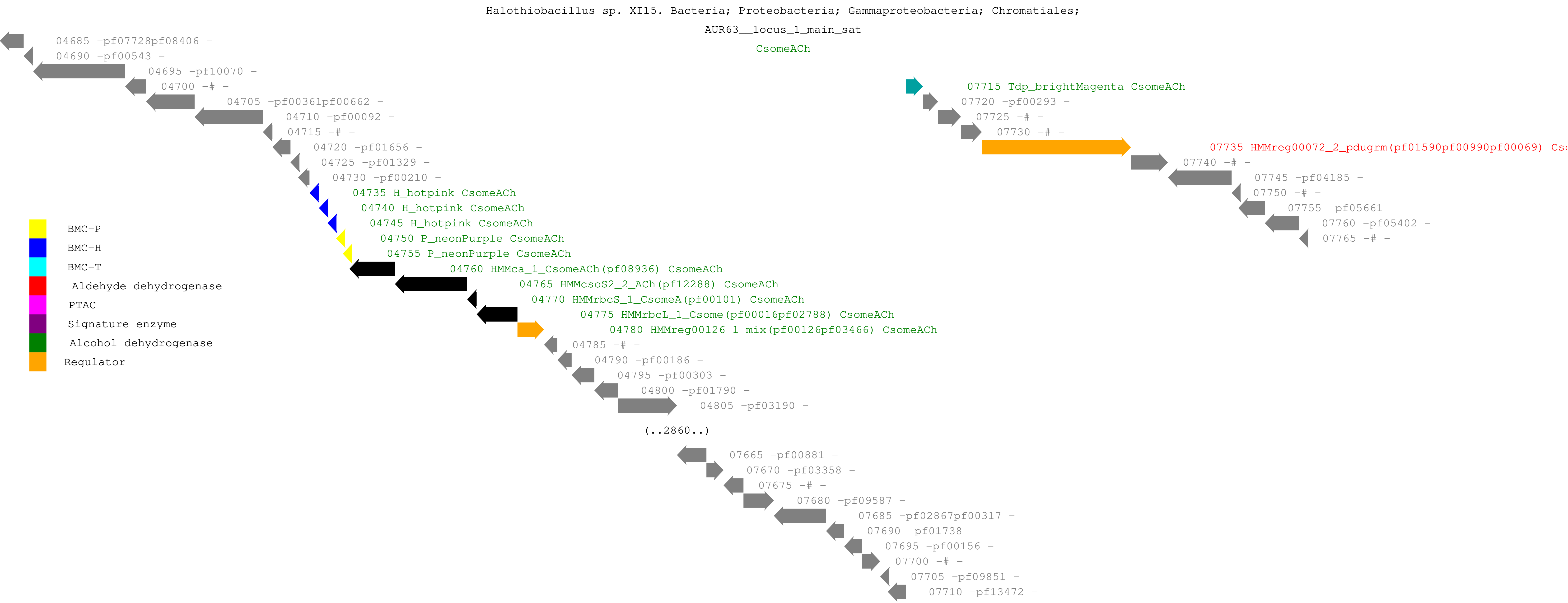
Halothiobacillus sp. XI15. Bacteria; Proteobacteria; Gammaproteobacteria; Chromatiales;
AUR63__locus_1_main_sat
CsomeACh
04685 -pf07728pf08406 -
04690 -pf00543 -
04695 -pf10070 -
04700 -# -
04705 -pf00361pf00662 -
04710 -pf00092 -
04715 -# -
04720 -pf01656 -
04725 -pf01329 -
04730 -pf00210 -
04735 H_hotpink CsomeACh
04740 H_hotpink CsomeACh
04745 H_hotpink CsomeACh
04750 P_neonPurple CsomeACh
04755 P_neonPurple CsomeACh
04760 HMMca_1_CsomeACh(pf08936) CsomeACh
04765 HMMcsoS2_2_ACh(pf12288) CsomeACh
04770 HMMrbcS_1_CsomeA(pf00101) CsomeACh
04775 HMMrbcL_1_Csome(pf00016pf02788) CsomeACh
04780 HMMreg00126_1_mix(pf00126pf03466) CsomeACh
04785 -# -
04790 -pf00186 -
04795 -pf00303 -
04800 -pf01790 -
04805 -pf03190 -
(..2860..)
07665 -pf00881 -
07670 -pf03358 -
07675 -# -
07680 -pf09587 -
07685 -pf02867pf00317 -
07690 -pf01738 -
07695 -pf00156 -
07700 -# -
07705 -pf09851 -
07710 -pf13472 -
07715 Tdp_brightMagenta CsomeACh
07720 -pf00293 -
07725 -# -
07730 -# -
07735 HMMreg00072_2_pdugrm(pf01590pf00990pf00069) Cs
07740 -# -
07745 -pf04185 -
07750 -# -
07755 -pf05661 -
07760 -pf05402 -
07765 -# -
BMC-P
BMC-H
BMC-T
Aldehyde dehydrogenase
PTAC
Signature enzyme
Alcohol dehydrogenase
Regulator

Thiomicrospira sp. WB1. Bacteria; Proteobacteria; Gammaproteobacteria; Thiotrichales;

AVO41__locus_1

CsomeACh

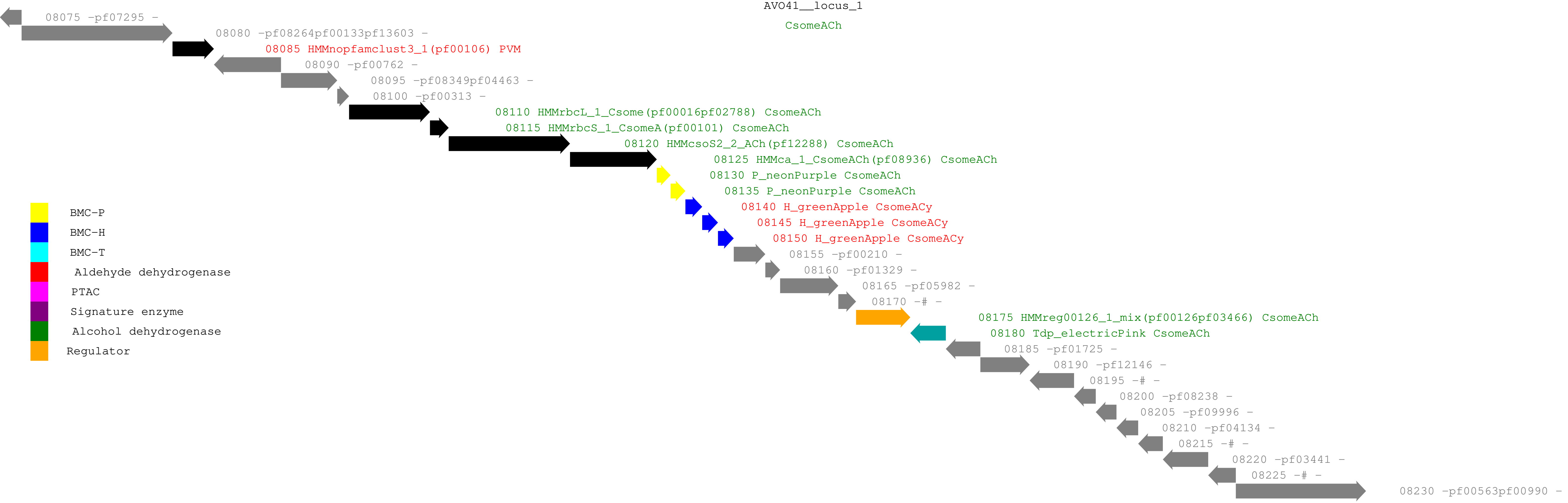

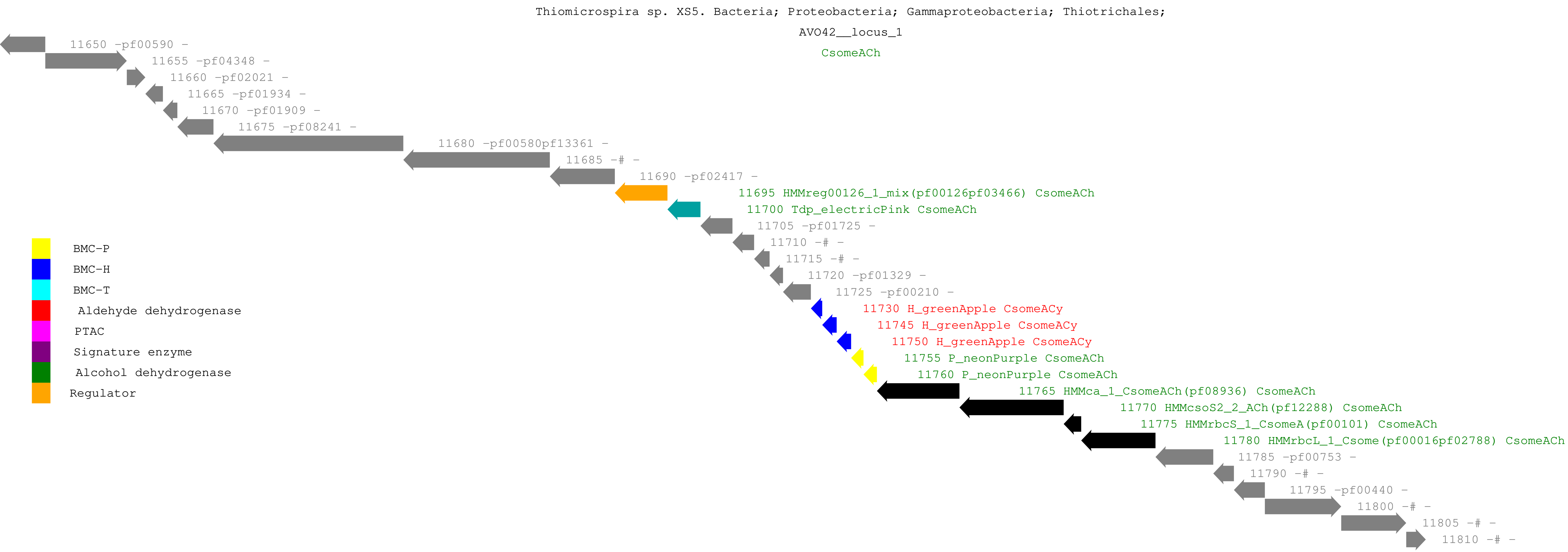
Thiomicrospira sp. XS5. Bacteria; Proteobacteria; Gammaproteobacteria; Thiotrichales;
AVO42__locus_1
CsomeACh
11650 -pf00590 -
11655 -pf04348 -
11660 -pf02021 -
11665 -pf01934 -
11670 -pf01909 -
11675 -pf08241 -
11680 -pf00580pf13361 -
11685 -# -
11690 -pf02417 -
11695 HMMreg00126_1_mix(pf00126pf03466) CsomeACh
11700 Tdp_electricPink CsomeACh
11705 -pf01725 -
11710 -# -
11715 -# -
11720 -pf01329 -
11725 -pf00210 -
11730 H_greenApple CsomeACy
11745 H_greenApple CsomeACy
11750 H_greenApple CsomeACy
11755 P_neonPurple CsomeACh
11760 P_neonPurple CsomeACh
11765 HMMca_1_CsomeACh(pf08936) CsomeACh
11770 HMMcsoS2_2_ACh(pf12288) CsomeACh
11775 HMMrbcS_1_CsomeA(pf00101) CsomeACh
11780 HMMrbcL_1_Csome(pf00016pf02788) CsomeACh
11785 -pf00753 -
11790 -# -
11795 -pf00440 -
11800 -# -
11805 -# -
11810 -# -
BMC-P
BMC-H
BMC-T
Aldehyde dehydrogenase
PTAC
Signature enzyme
Alcohol dehydrogenase
Regulator

Candidatus Gallionella acididurans. Bacteria; Proteobacteria; Betaproteobacteria; Gallionellales;

AWT59__locus_1_main_sat

CsomeACh

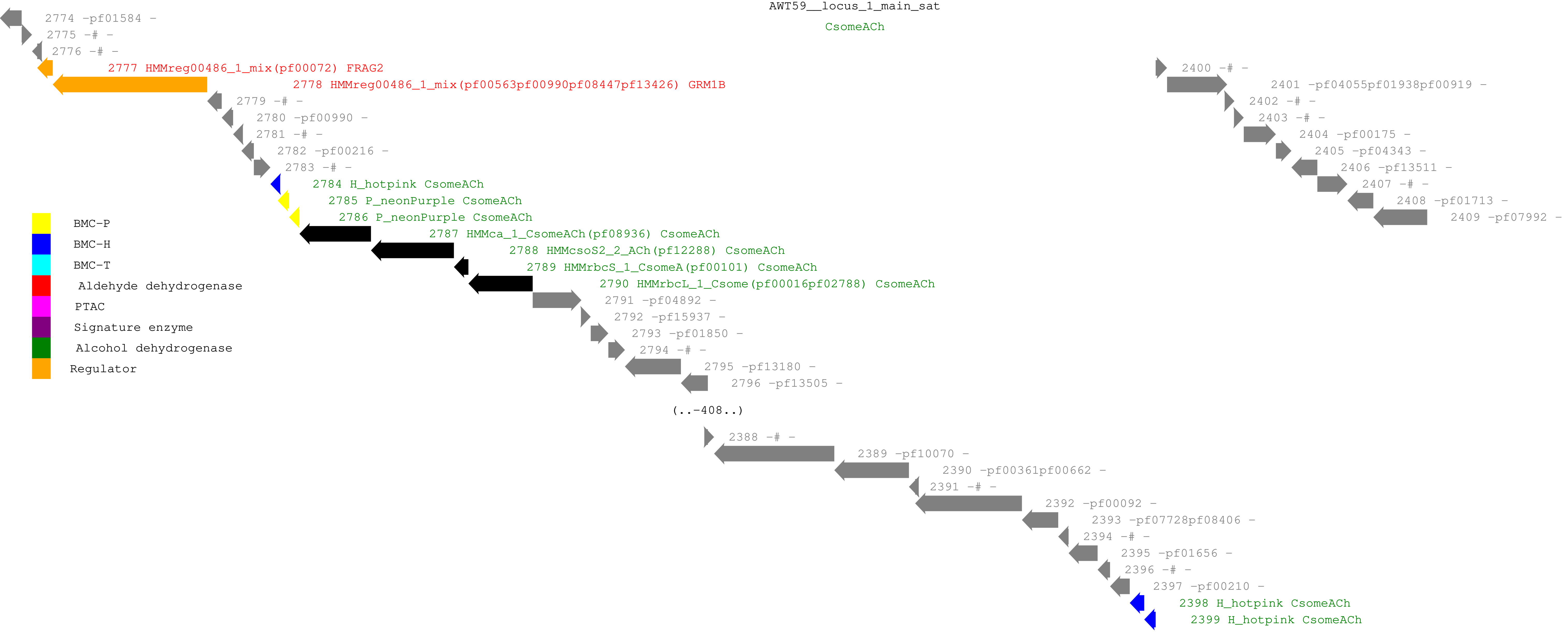

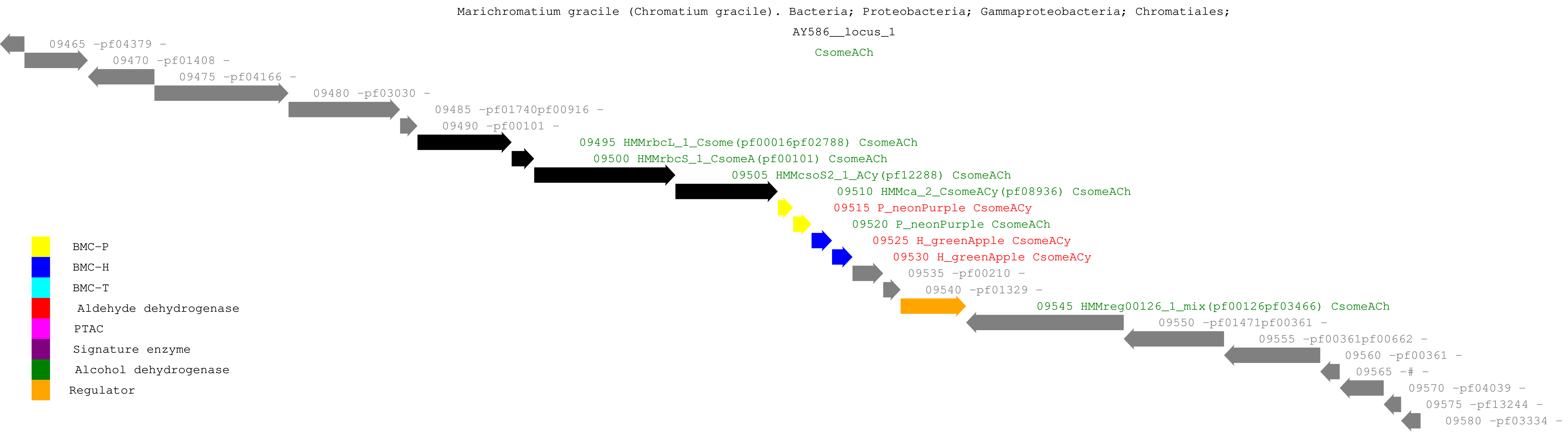
Marichromatium gracile (Chromatium gracile). Bacteria; Proteobacteria; Gammaproteobacteria; Chromatiales;
AY586__locus_1
CsomeACh
09465 -pf04379 -
09470 -pf01408 -
09475 -pf04166 -
09480 -pf03030 -
09485 -pf01740pf00916 -
09490 -pf00101 -
09495 HMMrbcL_1_Csome(pf00016pf02788) CsomeACh
09500 HMMrbcS_1_CsomeA(pf00101) CsomeACh
09505 HMMcsoS2_1_ACy(pf12288) CsomeACh
09510 HMMca_2_CsomeACy(pf08936) CsomeACh
09515 P_neonPurple CsomeACy
09520 P_neonPurple CsomeACh
09525 H_greenApple CsomeACy
09530 H_greenApple CsomeACy
09535 -pf00210 -
09540 -pf01329 -
09545 HMMreg00126_1_mix(pf00126pf03466) CsomeACh
09550 -pf01471pf00361 -
09555 -pf00361pf00662 -
09560 -pf00361 -
09565 -# -
09570 -pf04039 -
09575 -pf13244 -
09580 -pf03334 -
BMC-P
BMC-H
BMC-T
Aldehyde dehydrogenase
PTAC
Signature enzyme
Alcohol dehydrogenase
Regulator

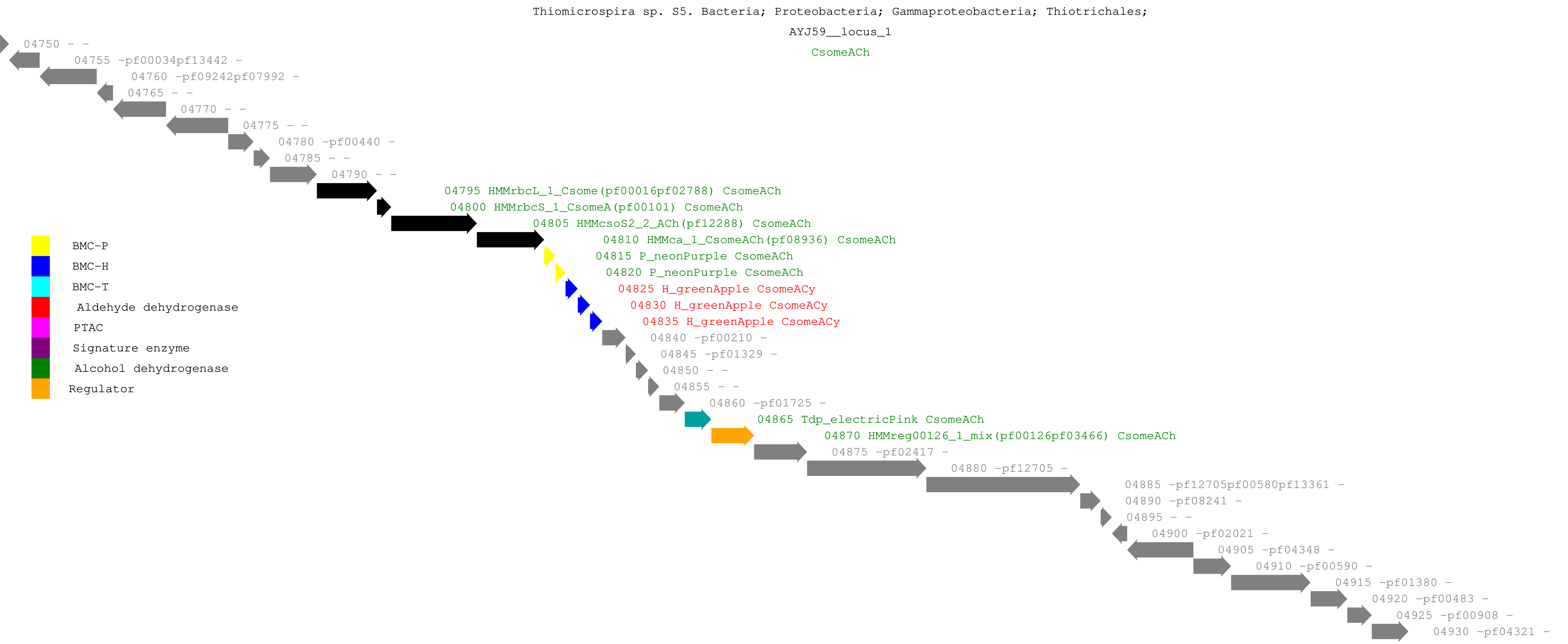
Thiomicrospira sp. S5. Bacteria; Proteobacteria; Gammaproteobacteria; Thiotrichales;
AYJ59__locus_1
CsomeACh
04750 - -
04755 -pf00034pf13442 -
04760 -pf09242pf07992 -
04765 - -
04770 - -
04775 - -
04780 -pf00440 -
04785 - -
04790 - -
04795 HMMrbcL_1_Csome(pf00016pf02788) CsomeACh
04800 HMMrbcS_1_CsomeA(pf00101) CsomeACh
04805 HMMcsoS2_2_ACh(pf12288) CsomeACh
04810 HMMca_1_CsomeACh(pf08936) CsomeACh
04815 P_neonPurple CsomeACh
04820 P_neonPurple CsomeACh
04825 H_greenApple CsomeACy
04830 H_greenApple CsomeACy
04835 H_greenApple CsomeACy
04840 -pf00210 -
04845 -pf01329 -
04850 - -
04855 - -
04860 -pf01725 -
04865 Tdp_electricPink CsomeACh
04870 HMMreg00126_1_mix(pf00126pf03466) CsomeACh
04875 -pf02417 -
04880 -pf12705 -
04885 -pf12705pf00580pf13361 -
04890 -pf08241 -
04895 - -
04900 -pf02021 -
04905 -pf04348 -
04910 -pf00590 -
04915 -pf01380 -
04920 -pf00483 -
04925 -pf00908 -
04930 -pf04321 -
BMC-P
BMC-H
BMC-T
Aldehyde dehydrogenase
PTAC
Signature enzyme
Alcohol dehydrogenase
Regulator

Thioalkalivibrio versutus. Bacteria; Proteobacteria; Gammaproteobacteria; Chromatiales;

B0684__locus_1_main_sat

CsomeACh

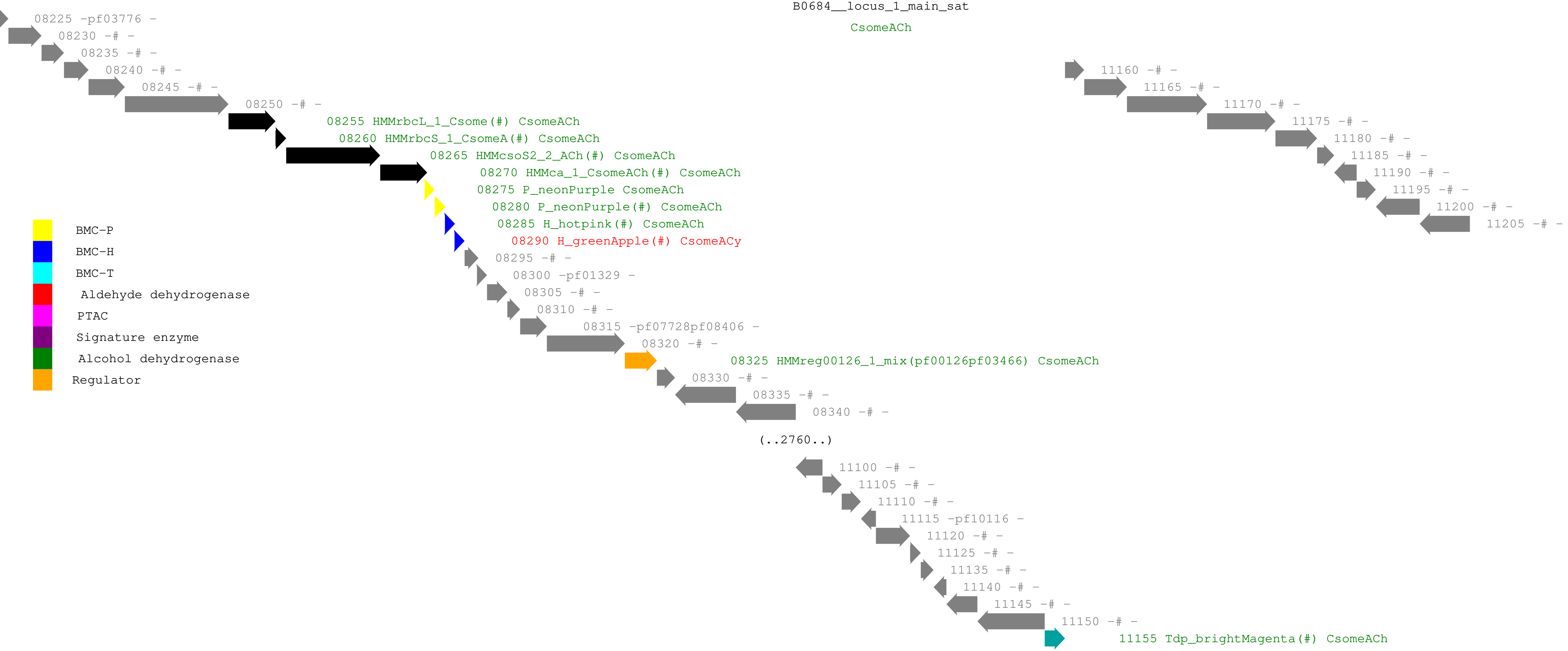

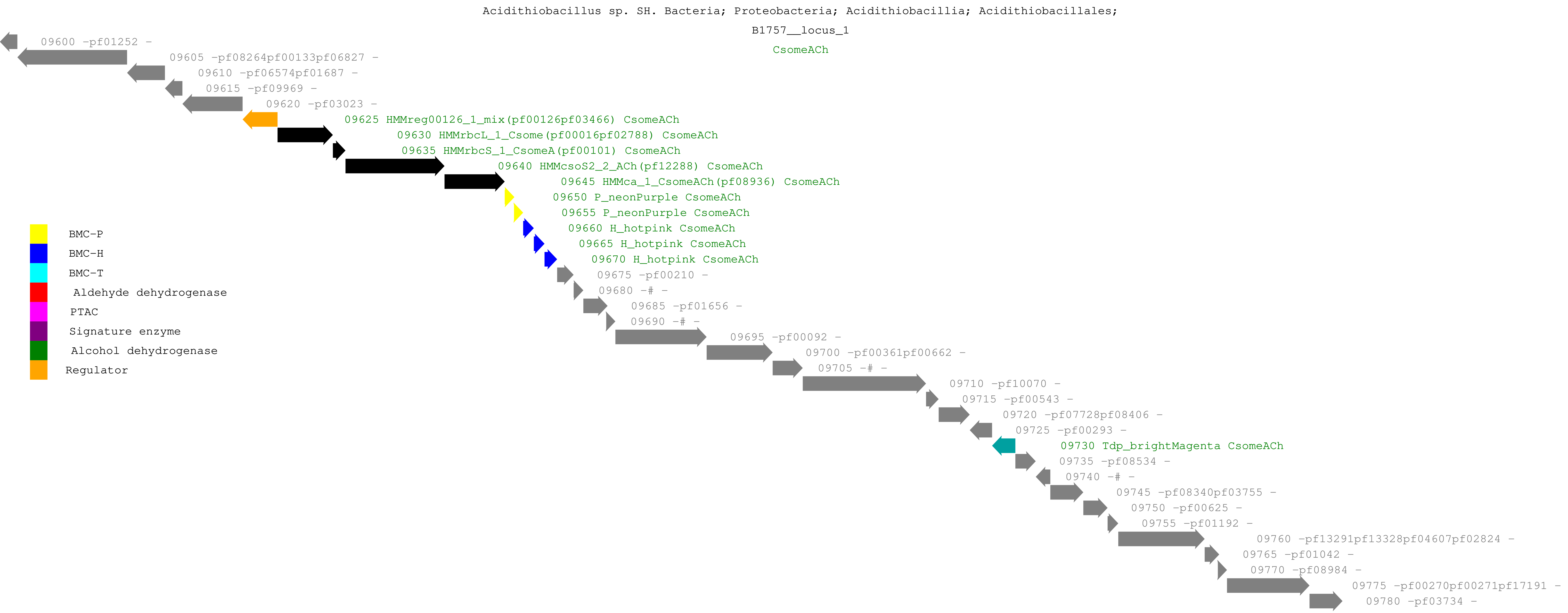
Acidithiobacillus sp. SH. Bacteria; Proteobacteria; Acidithiobacillia; Acidithiobacillales;
B1757__locus_1
CsomeACh
09600 -pf01252 -
09605 -pf08264pf00133pf06827 -
09610 -pf06574pf01687 -
09615 -pf09969 -
09620 -pf03023 -
09625 HMMreg00126_1_mix(pf00126pf03466) CsomeACh
09630 HMMrbcL_1_Csome(pf00016pf02788) CsomeACh
09635 HMMrbcS_1_CsomeA(pf00101) CsomeACh
09640 HMMcsoS2_2_ACh(pf12288) CsomeACh
09645 HMMca_1_CsomeACh(pf08936) CsomeACh
09650 P_neonPurple CsomeACh
09655 P_neonPurple CsomeACh
09660 H_hotpink CsomeACh
09665 H_hotpink CsomeACh
09670 H_hotpink CsomeACh
09675 -pf00210 -
09680 -# -
09685 -pf01656 -
09690 -# -
09695 -pf00092 -
09700 -pf00361pf00662 -
09705 -# -
09710 -pf10070 -
09715 -pf00543 -
09720 -pf07728pf08406 -
09725 -pf00293 -
09730 Tdp_brightMagenta CsomeACh
09735 -pf08534 -
09740 -# -
09745 -pf08340pf03755 -
09750 -pf00625 -
09755 -pf01192 -
09760 -pf13291pf13328pf04607pf02824 -
09765 -pf01042 -
09770 -pf08984 -
09775 -pf00270pf00271pf17191 -
09780 -pf03734 -
BMC-P
BMC-H
BMC-T
Aldehyde dehydrogenase
PTAC
Signature enzyme
Alcohol dehydrogenase
Regulator

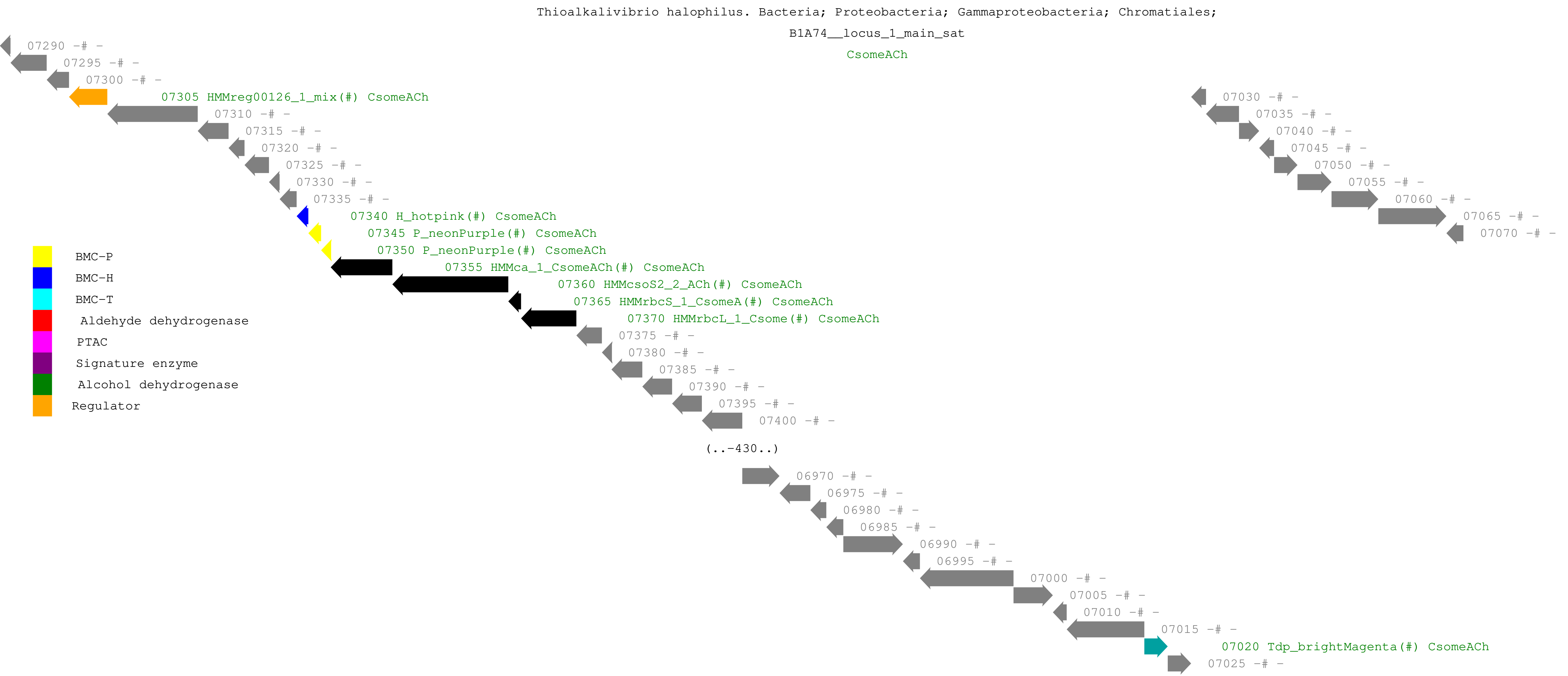
Thioalkalivibrio halophilus. Bacteria; Proteobacteria; Gammaproteobacteria; Chromatiales;
B1A74__locus_1_main_sat
CsomeACh
07290 -# -
07295 -# -
07300 -# -
07305 HMMreg00126_1_mix(#) CsomeACh
07310 -# -
07315 -# -
07320 -# -
07325 -# -
07330 -# -
07335 -# -
07340 H_hotpink(#) CsomeACh
07345 P_neonPurple(#) CsomeACh
07350 P_neonPurple(#) CsomeACh
07355 HMMca_1_CsomeACh(#) CsomeACh
07360 HMMcsoS2_2_ACh(#) CsomeACh
07365 HMMrbcS_1_CsomeA(#) CsomeACh
07370 HMMrbcL_1_Csome(#) CsomeACh
07375 -# -
07380 -# -
07385 -# -
07390 -# -
07395 -# -
07400 -# -
(..-430..)
06970 -# -
06975 -# -
06980 -# -
06985 -# -
06990 -# -
06995 -# -
07000 -# -
07005 -# -
07010 -# -
07015 -# -
07020 Tdp_brightMagenta(#) CsomeACh
07025 -# -
07030 -# -
07035 -# -
07040 -# -
07045 -# -
07050 -# -
07055 -# -
07060 -# -
07065 -# -
07070 -# -
BMC-P
BMC-H
BMC-T
Aldehyde dehydrogenase
PTAC
Signature enzyme
Alcohol dehydrogenase
Regulator

mine drainage metagenome. unclassified sequences; metagenomes; ecological metagenomes.

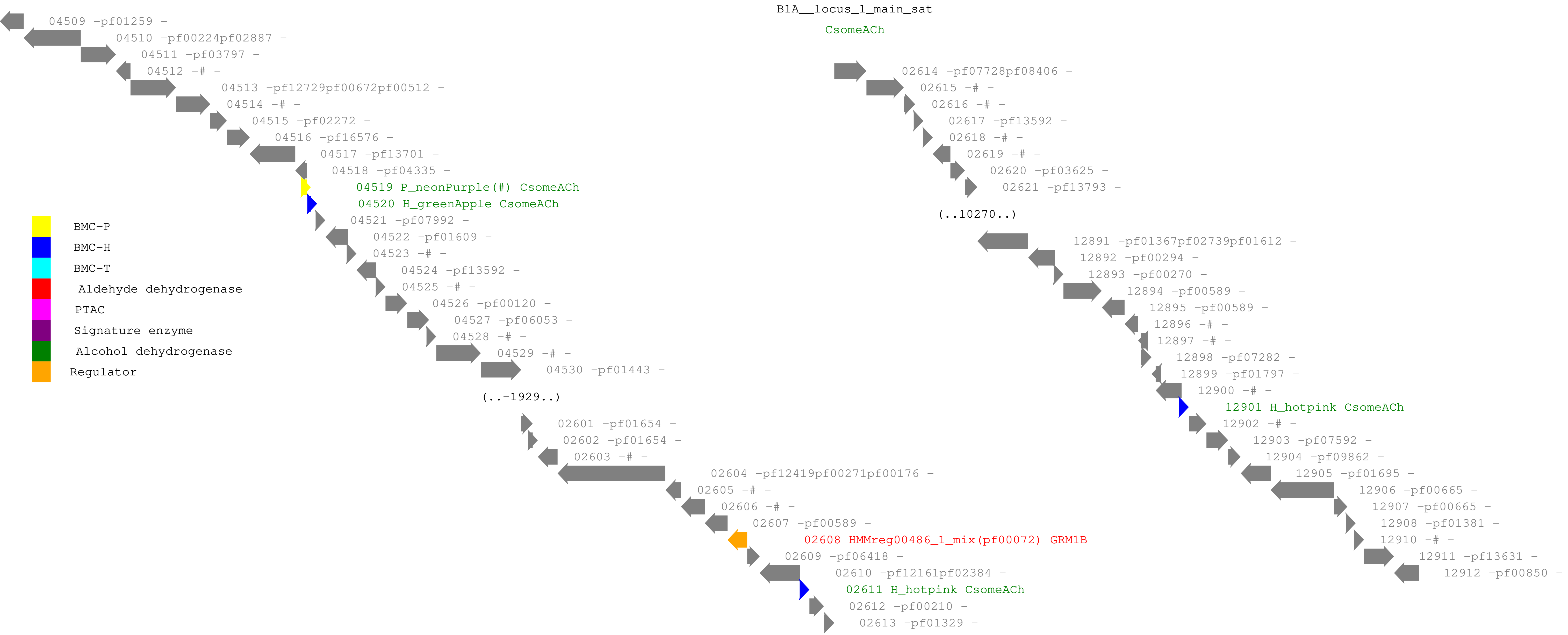

Thioalkalivibrio denitrificans. Bacteria; Proteobacteria; Gammaproteobacteria; Chromatiales;

B1C78__locus_1

CsomeACh

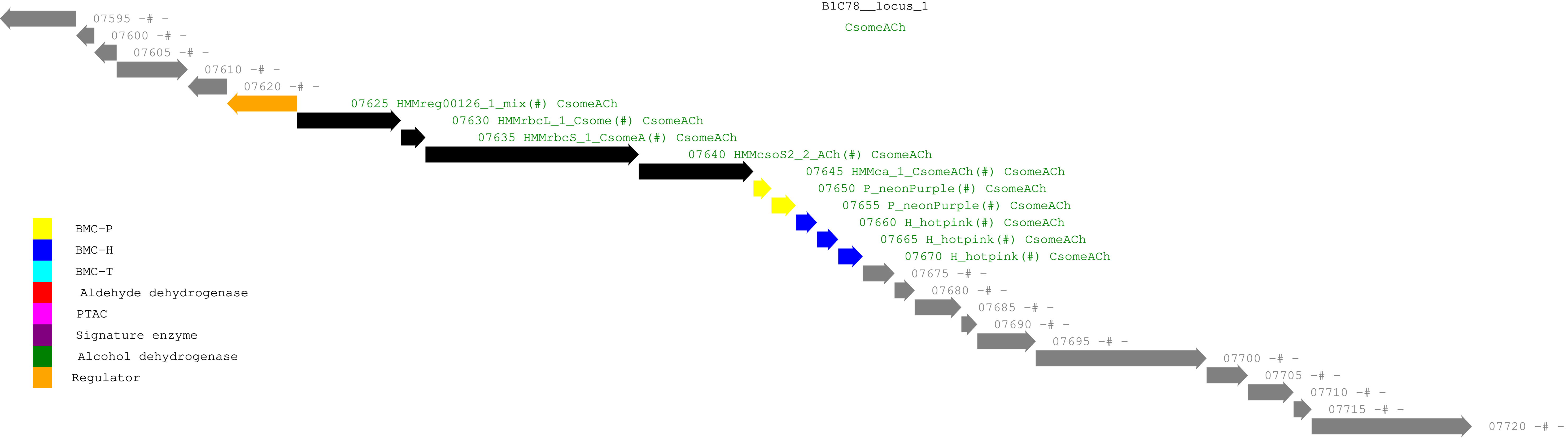

Nitrobacter vulgaris. Bacteria; Proteobacteria; Alphaproteobacteria; Rhizobiales;

B2M20__locus_1

CsomeACh

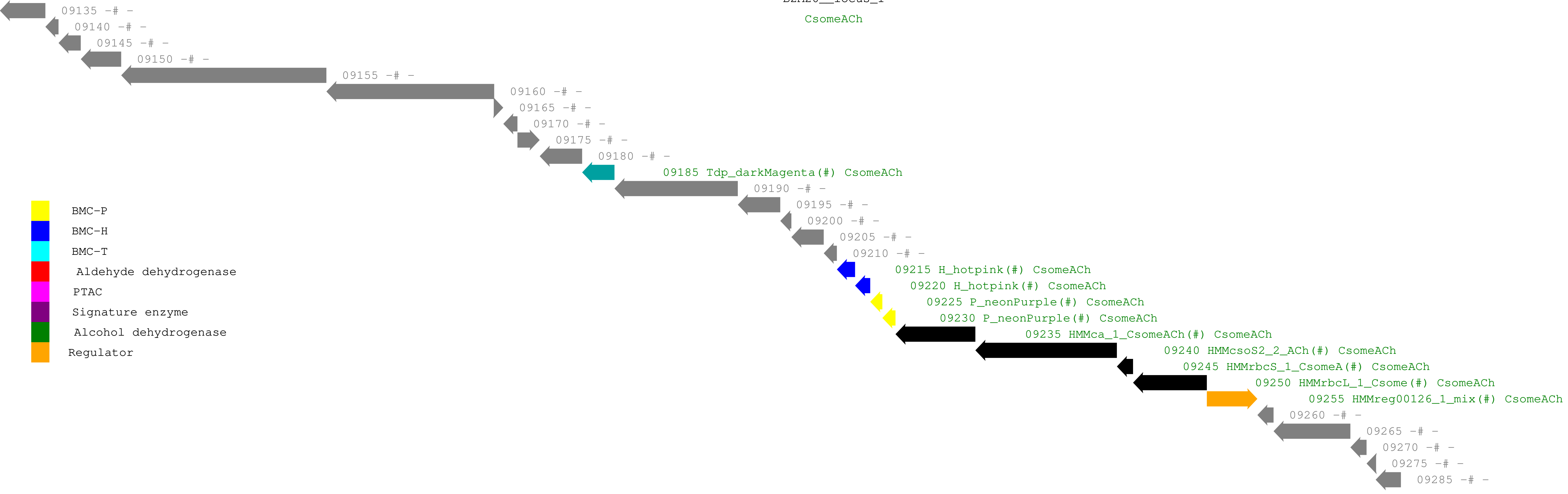

Burkholderiales bacterium 21-58-4. Bacteria; Proteobacteria; Betaproteobacteria; Burkholderiales.

B7X10__locus_1

CsomeACh

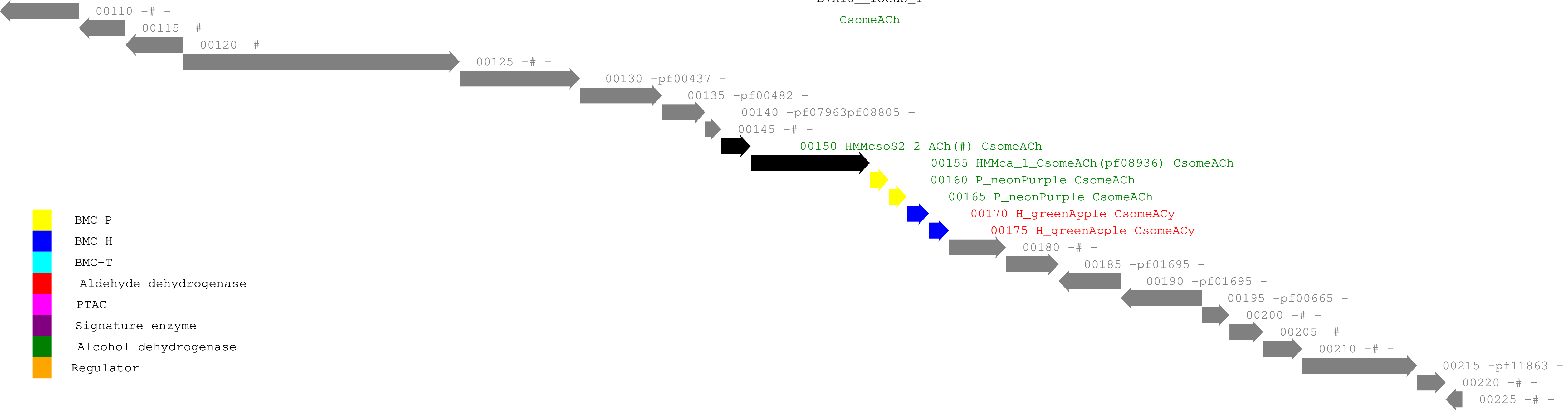

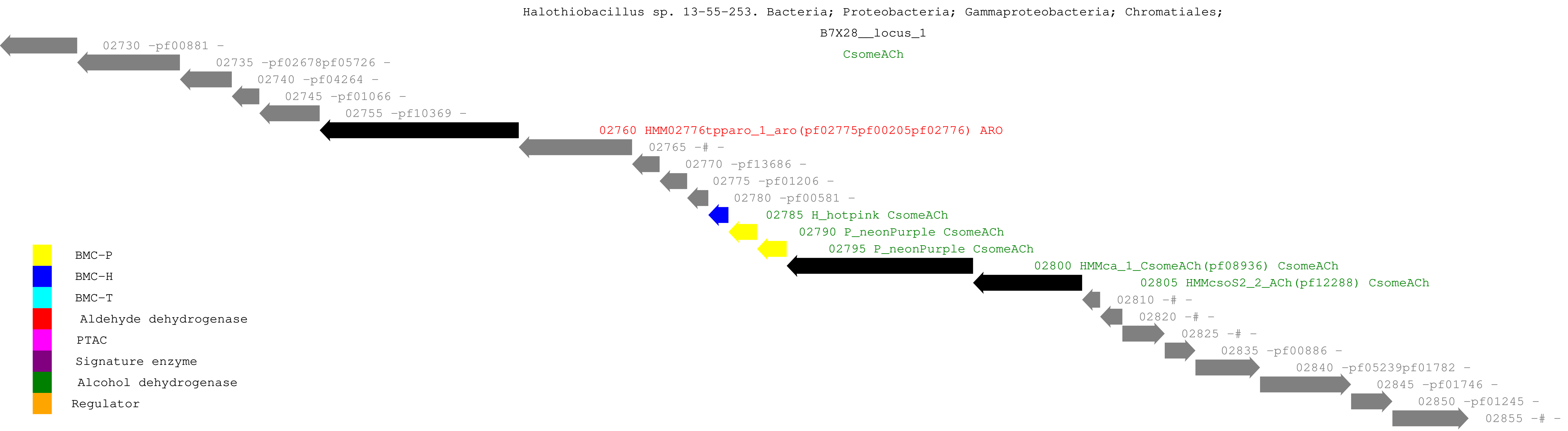
Halothiobacillus sp. 13-55-253. Bacteria; Proteobacteria; Gammaproteobacteria; Chromatiales;
B7X28__locus_1
CsomeACh
02730 -pf00881 -
02735 -pf02678pf05726 -
02740 -pf04264 -
02745 -pf01066 -
02755 -pf10369 -
02760 HMM02776tpparo_1_aro(pf02775pf00205pf02776) ARO
02765 -# -
02770 -pf13686 -
02775 -pf01206 -
02780 -pf00581 -
02785 H_hotpink CsomeACh
02790 P_neonPurple CsomeACh
02795 P_neonPurple CsomeACh
02800 HMMca_1_CsomeACh(pf08936) CsomeACh
02805 HMMcsoS2_2_ACh(pf12288) CsomeACh
02810 -# -
02820 -# -
02825 -# -
02835 -pf00886 -
02840 -pf05239pf01782 -
02845 -pf01746 -
02850 -pf01245 -
02855 -# -
BMC-P
BMC-H
BMC-T
Aldehyde dehydrogenase
PTAC
Signature enzyme
Alcohol dehydrogenase
Regulator

Halothiobacillus sp. 13-55-115. Bacteria; Proteobacteria; Gammaproteobacteria; Chromatiales;

B7X29__locus_1_main_sat

CsomeACh

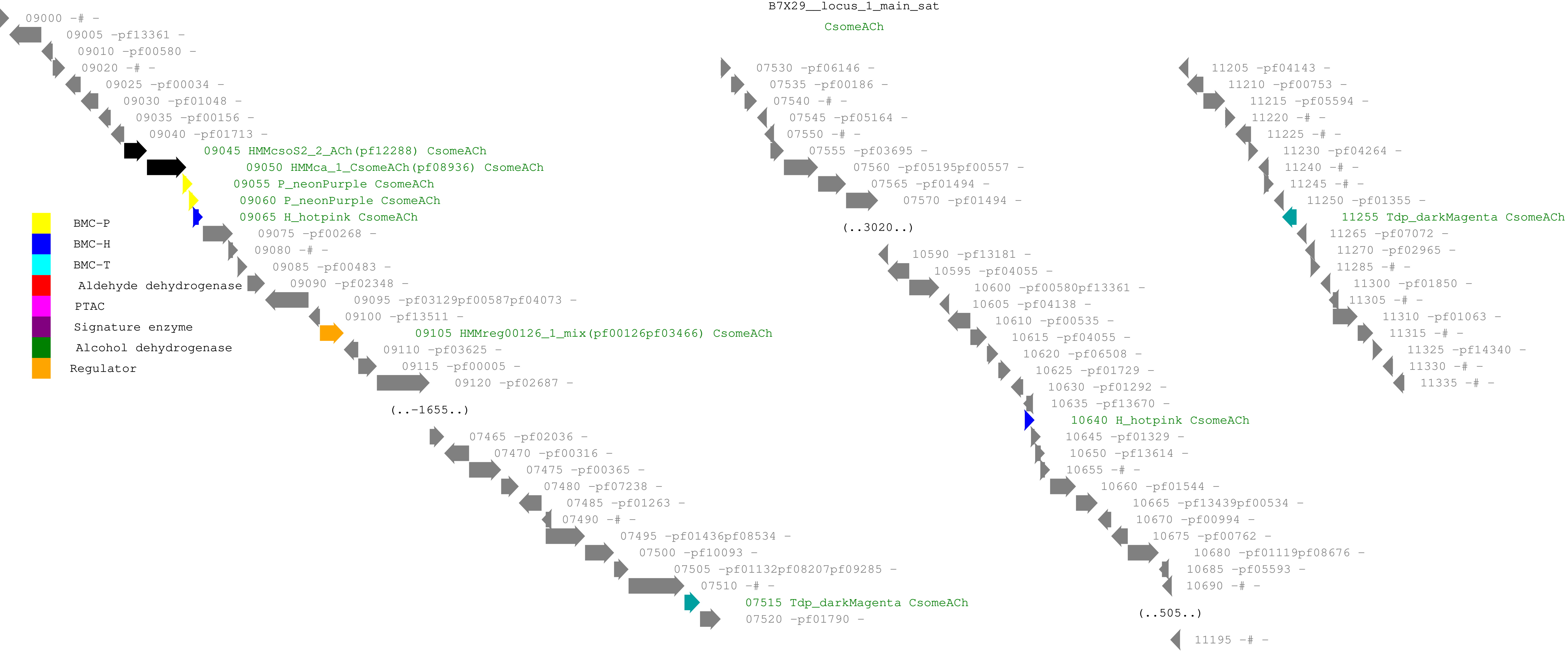

Thiomonas sp. 13-64-67. Bacteria; Proteobacteria; Betaproteobacteria; Burkholderiales;

B7X30__locus_1

CsomeACh

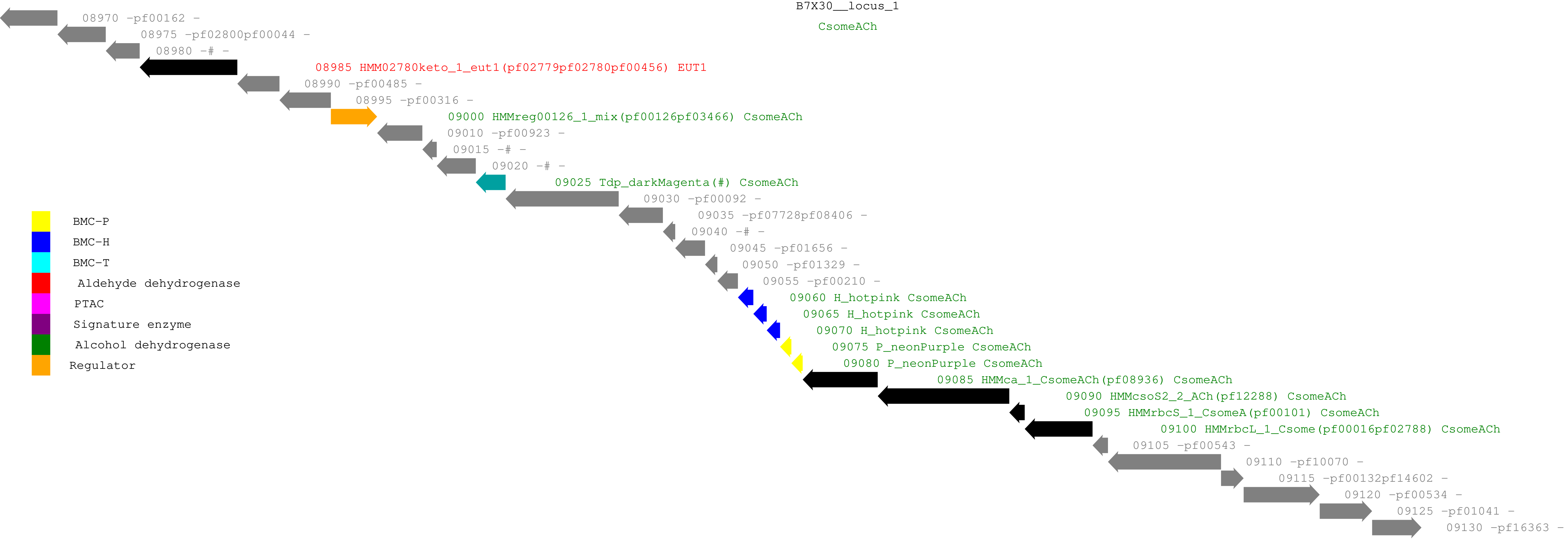

Thiomonas sp. 13-66-29. Bacteria; Proteobacteria; Betaproteobacteria; Burkholderiales;

B7X31__locus_1_main_sat

CsomeACh

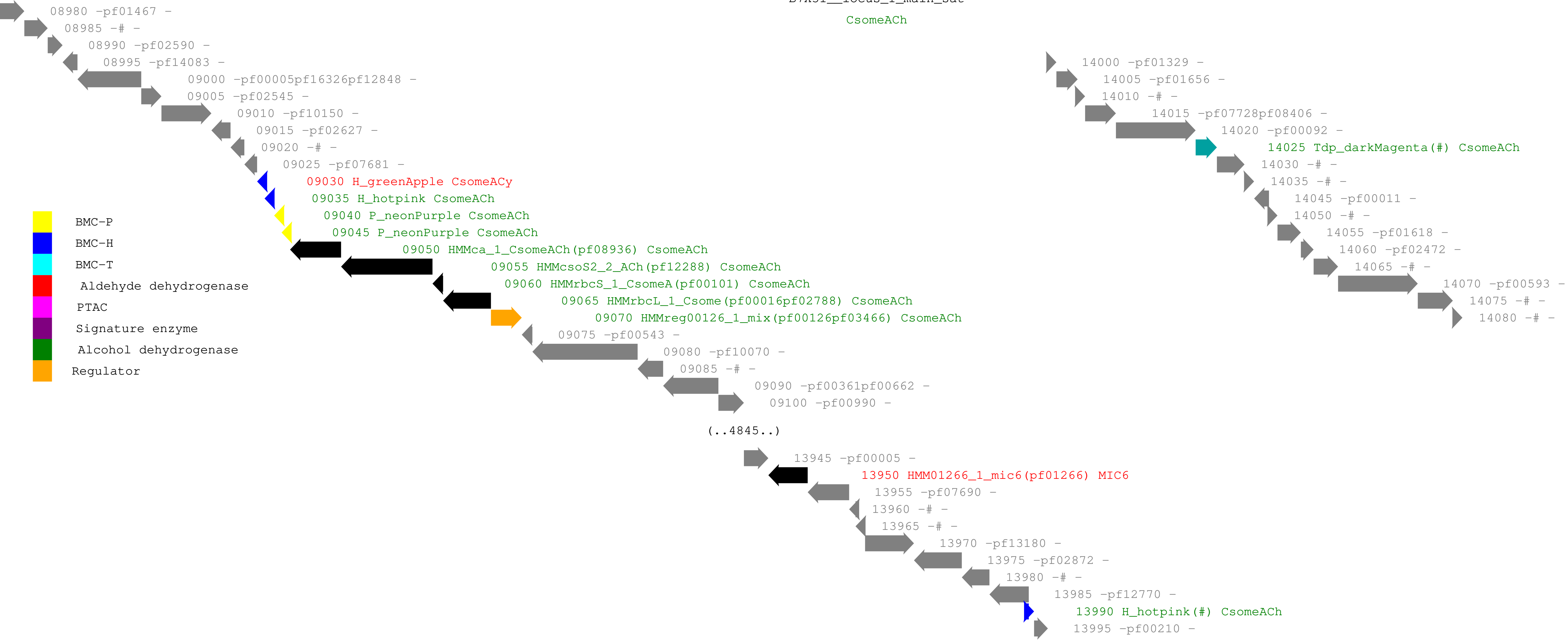

Thiomonas sp. 14-64-326. Bacteria; Proteobacteria; Betaproteobacteria; Burkholderiales;

B7X36__locus_1

CsomeACh

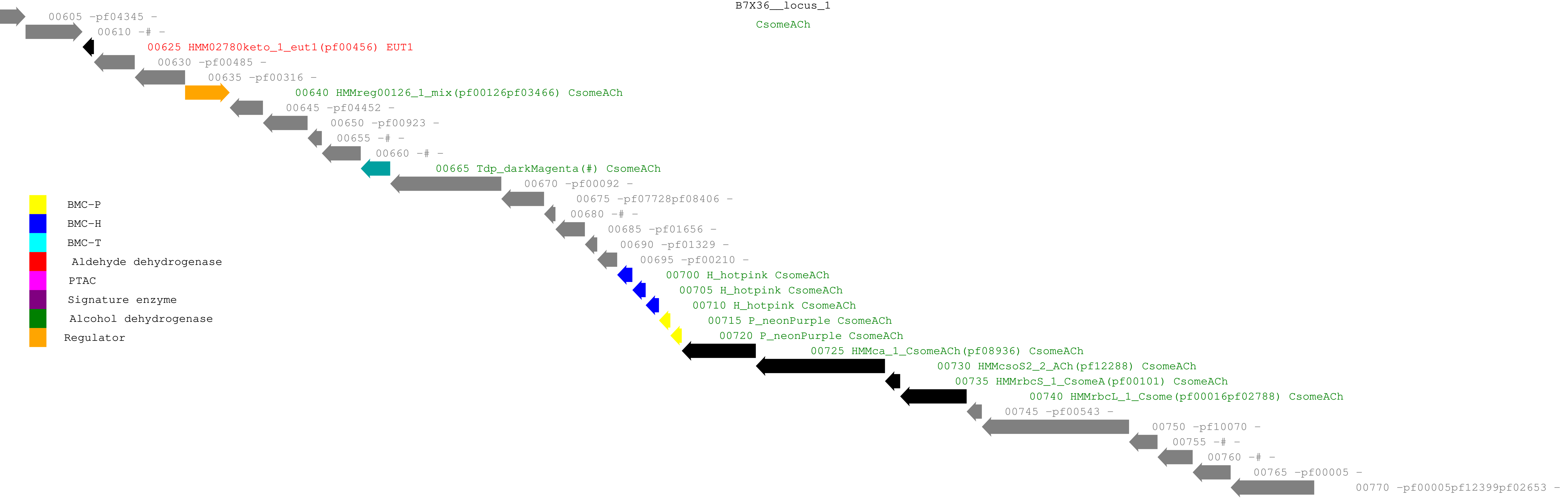

Halothiobacillus sp. 14-55-98. Bacteria; Proteobacteria; Gammaproteobacteria; Chromatiales;

B7X37__locus_1

CsomeACh

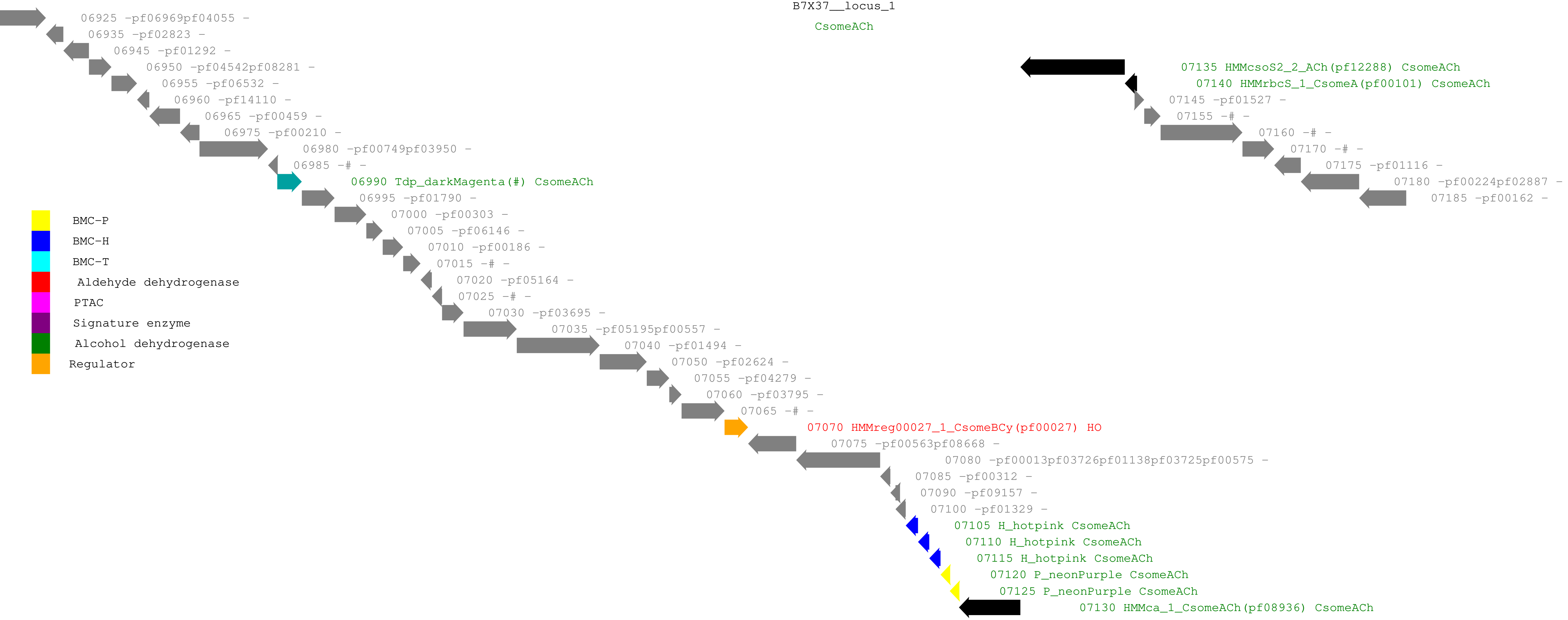

Thiomonas sp. 15-63-373. Bacteria; Proteobacteria; Betaproteobacteria; Burkholderiales;

B7X43__locus_1

CsomeACh

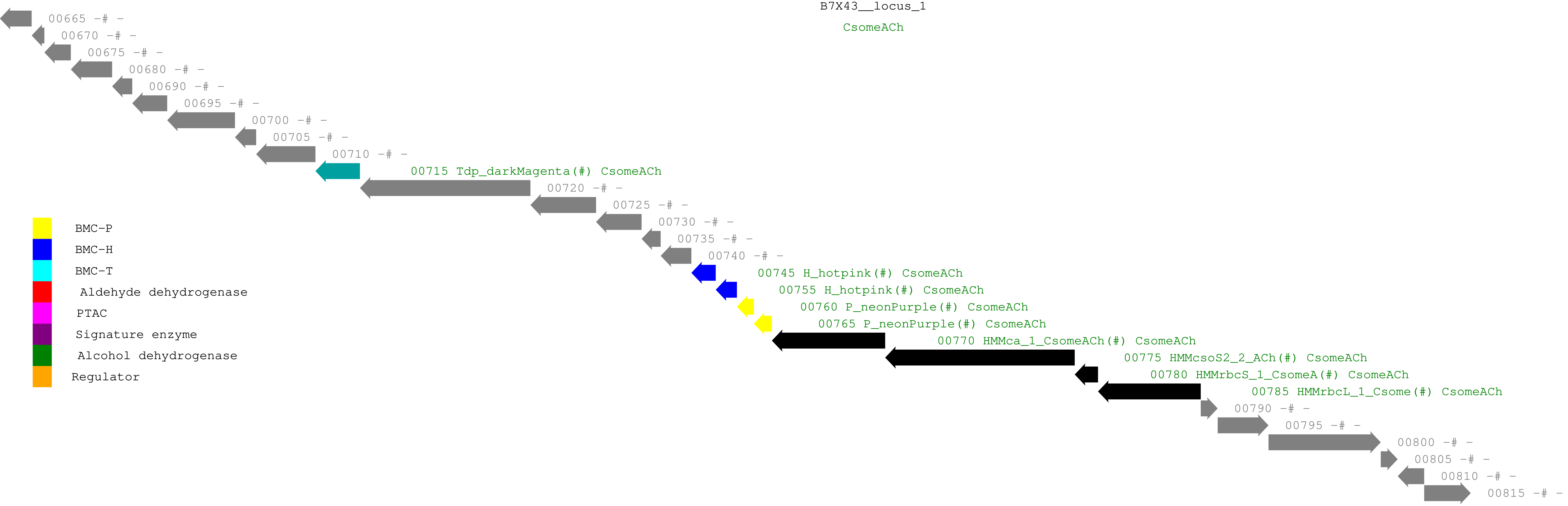

Halothiobacillus sp. 15-55-196. Bacteria; Proteobacteria; Gammaproteobacteria; Chromatiales;

B7X44__locus_1

CsomeACh

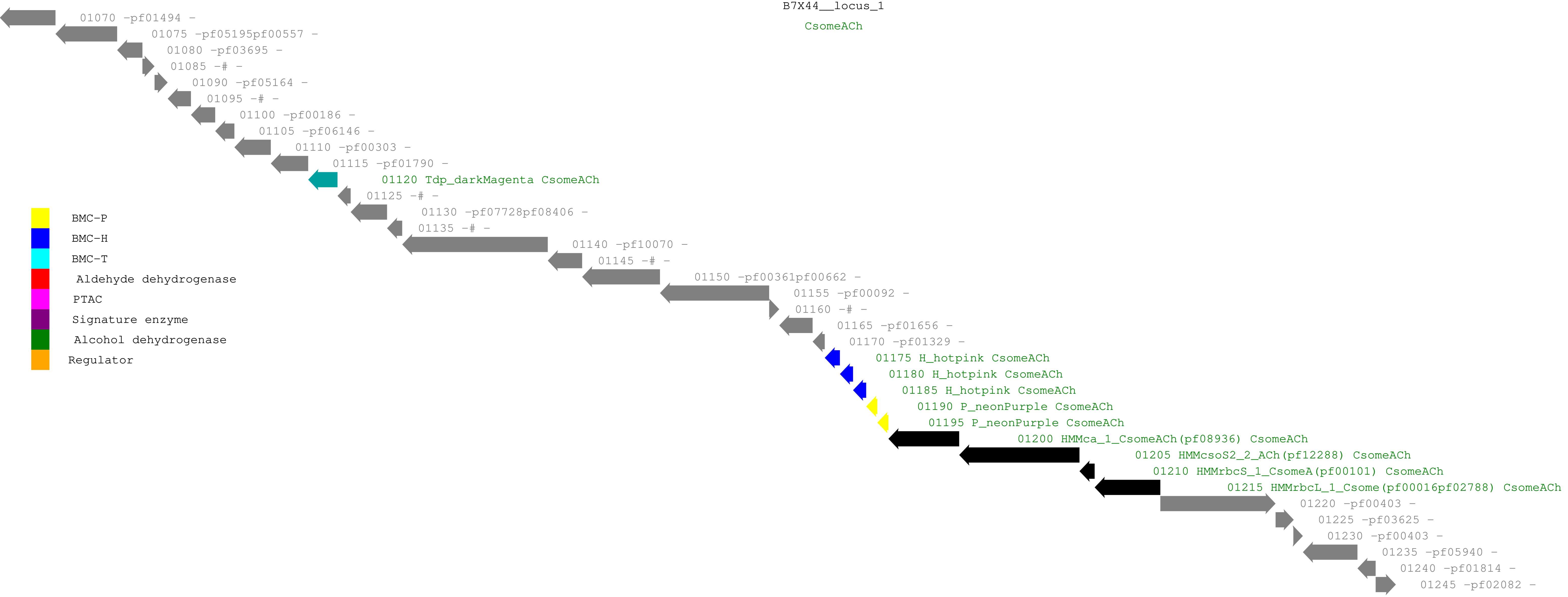

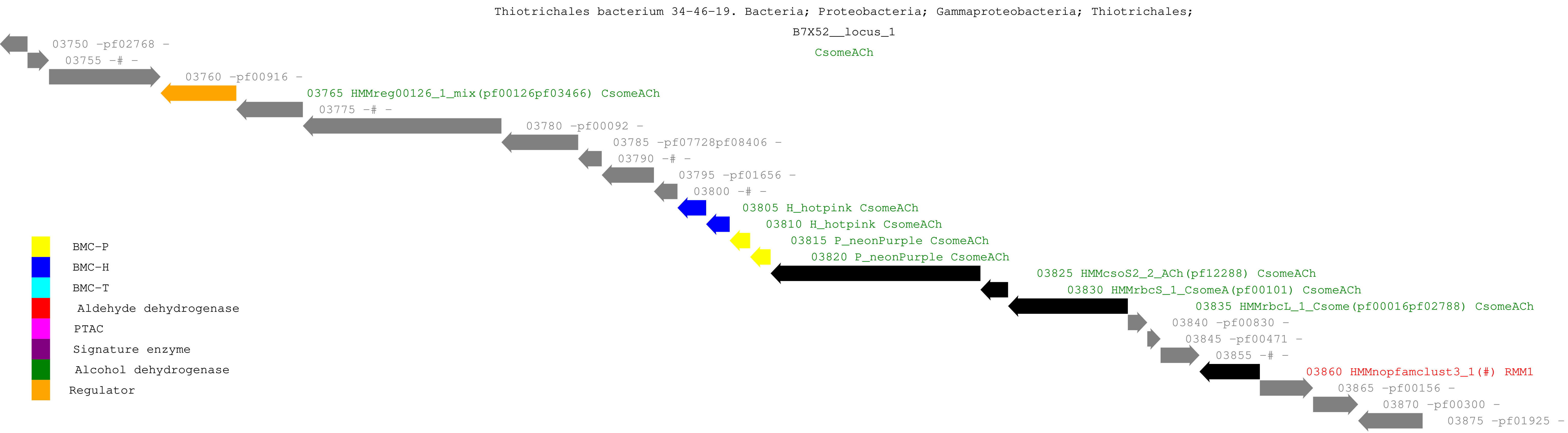
Thiotrichales bacterium 34-46-19. Bacteria; Proteobacteria; Gammaproteobacteria; Thiotrichales;
B7X52__locus_1
CsomeACh
03750 -pf02768 -
03755 -# -
03760 -pf00916 -
03765 HMMreg00126_1_mix(pf00126pf03466) CsomeACh
03775 -# -
03780 -pf00092 -
03785 -pf07728pf08406 -
03790 -# -
03795 -pf01656 -
03800 -# -
03805 H_hotpink CsomeACh
03810 H_hotpink CsomeACh
03815 P_neonPurple CsomeACh
03820 P_neonPurple CsomeACh
03825 HMMcsoS2_2_ACh(pf12288) CsomeACh
03830 HMMrbcS_1_CsomeA(pf00101) CsomeACh
03835 HMMrbcL_1_Csome(pf00016pf02788) CsomeACh
03840 -pf00830 -
03845 -pf00471 -
03855 -# -
03860 HMMnopfamclust3_1(#) RMM1
03865 -pf00156 -
03870 -pf00300 -
03875 -pf01925 -
BMC-P
BMC-H
BMC-T
Aldehyde dehydrogenase
PTAC
Signature enzyme
Alcohol dehydrogenase
Regulator

Hydrogenophilales bacterium 17-61-76. Bacteria; Proteobacteria; Hydrogenophilalia; Hydrogenophilales.

B7X81__locus_1

CsomeACh

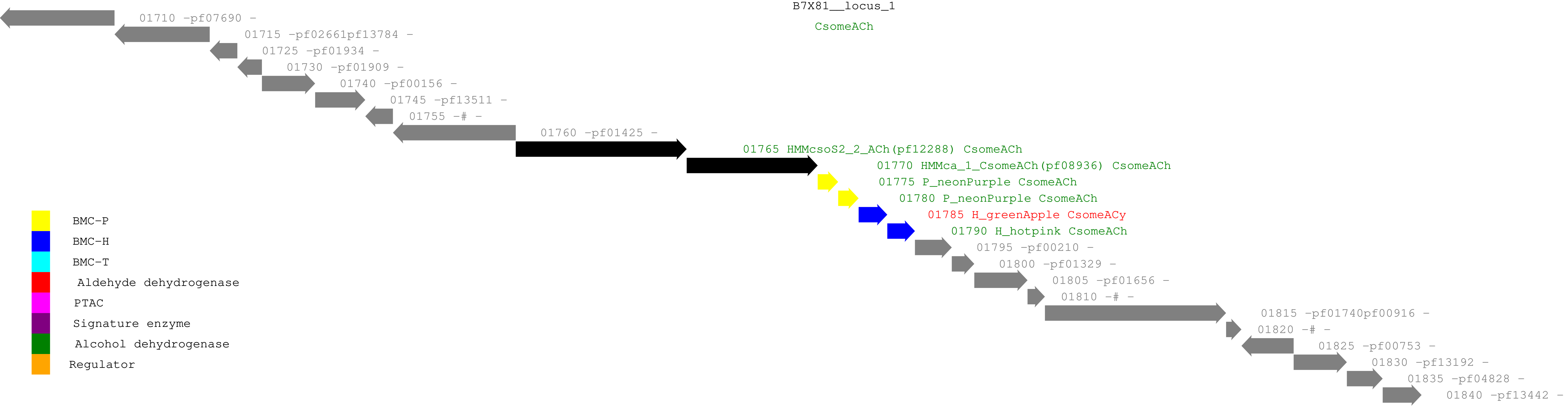

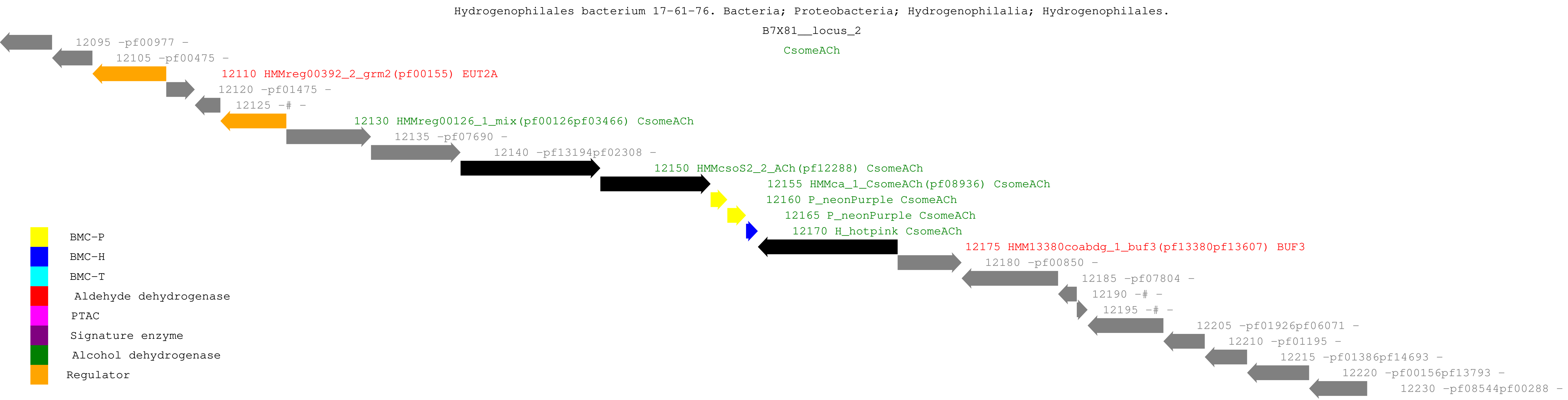
Hydrogenophilales bacterium 17-61-76. Bacteria; Proteobacteria; Hydrogenophilalia; Hydrogenophilales.
B7X81__locus_2
CsomeACh
12095 -pf00977 -
12105 -pf00475 -
12110 HMMreg00392_2_grm2(pf00155) EUT2A
12120 -pf01475 -
12125 -# -
12130 HMMreg00126_1_mix(pf00126pf03466) CsomeACh
12135 -pf07690 -
12140 -pf13194pf02308 -
12150 HMMcsoS2_2_ACh(pf12288) CsomeACh
12155 HMMca_1_CsomeACh(pf08936) CsomeACh
12160 P_neonPurple CsomeACh
12165 P_neonPurple CsomeACh
12170 H_hotpink CsomeACh
12175 HMM13380coabdg_1_buf3(pf13380pf13607) BUF3
12180 -pf00850 -
12185 -pf07804 -
12190 -# -
12195 -# -
12205 -pf01926pf06071 -
12210 -pf01195 -
12215 -pf01386pf14693 -
12220 -pf00156pf13793 -
12230 -pf08544pf00288 -
BMC-P
BMC-H
BMC-T
Aldehyde dehydrogenase
PTAC
Signature enzyme
Alcohol dehydrogenase
Regulator

Hydrogenophilales bacterium 17-64-34. Bacteria; Proteobacteria; Hydrogenophilalia; Hydrogenophilales.

B7X87__locus_1

CsomeACh

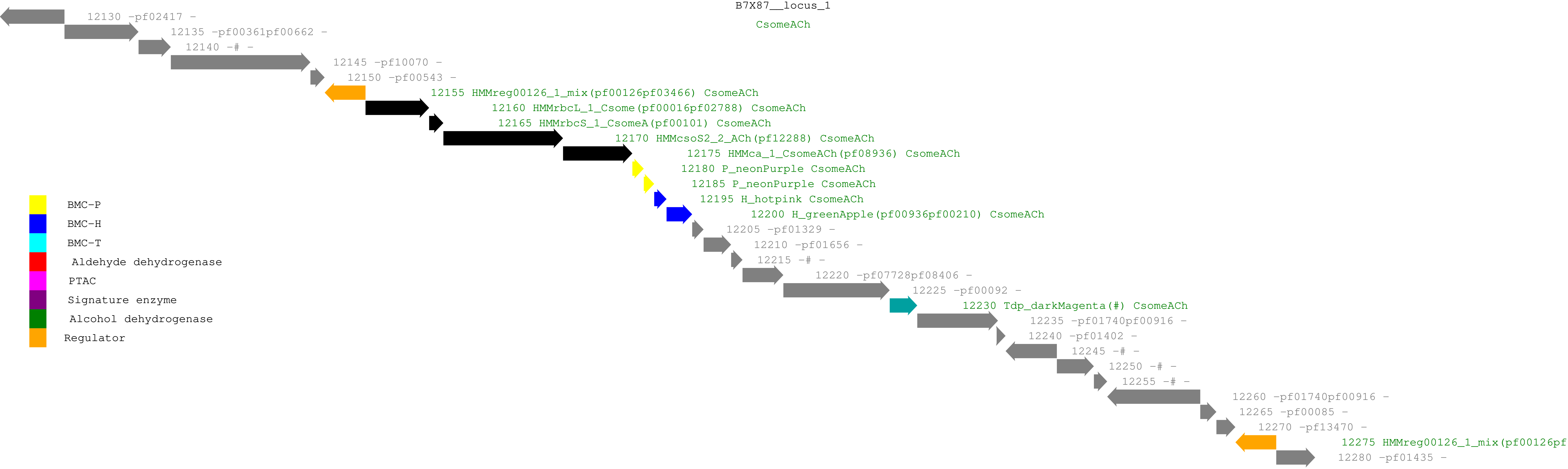

Hydrogenophilales bacterium 17-64-11. Bacteria; Proteobacteria; Hydrogenophilalia; Hydrogenophilales.

B7X91__locus_1

CsomeACh

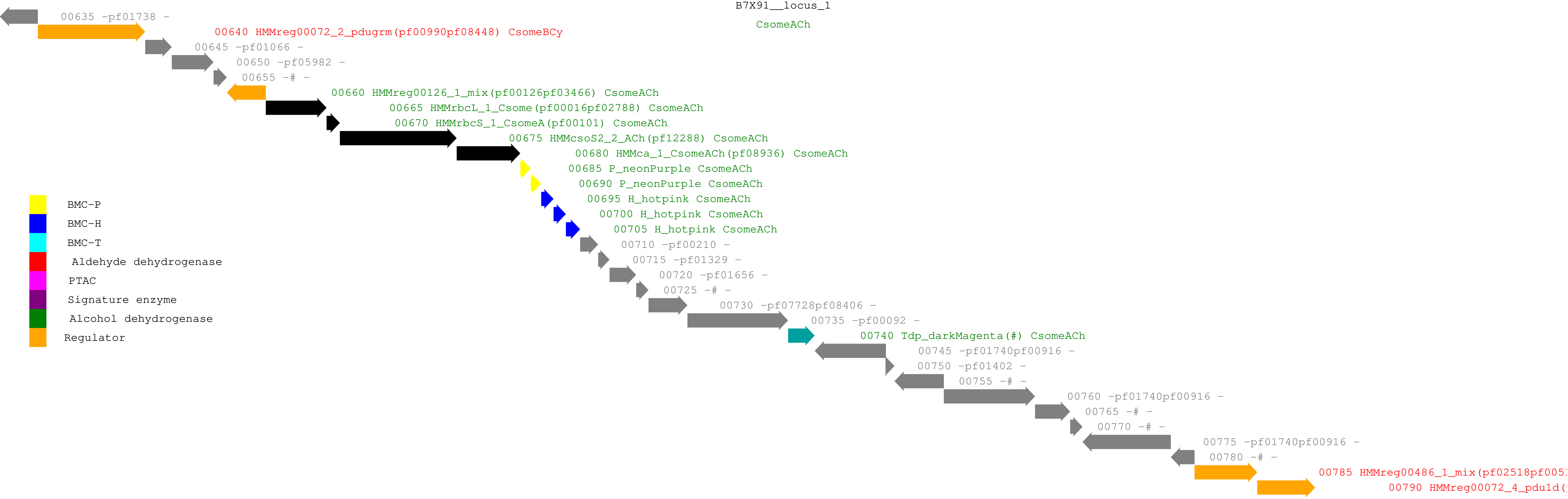

Hydrogenophilales bacterium 17-61-9. Bacteria; Proteobacteria; Hydrogenophilalia; Hydrogenophilales.

B7X93__locus_1

CsomeACh

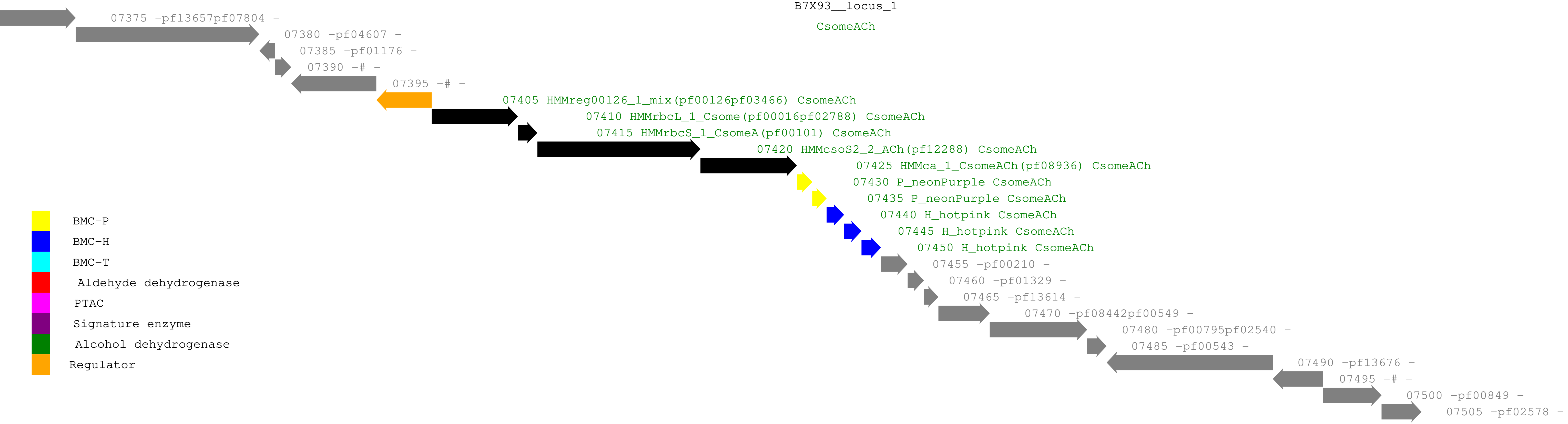

Halothiobacillus sp. 24-54-40. Bacteria; Proteobacteria; Gammaproteobacteria; Chromatiales;

B7Y07__locus_1

CsomeACh

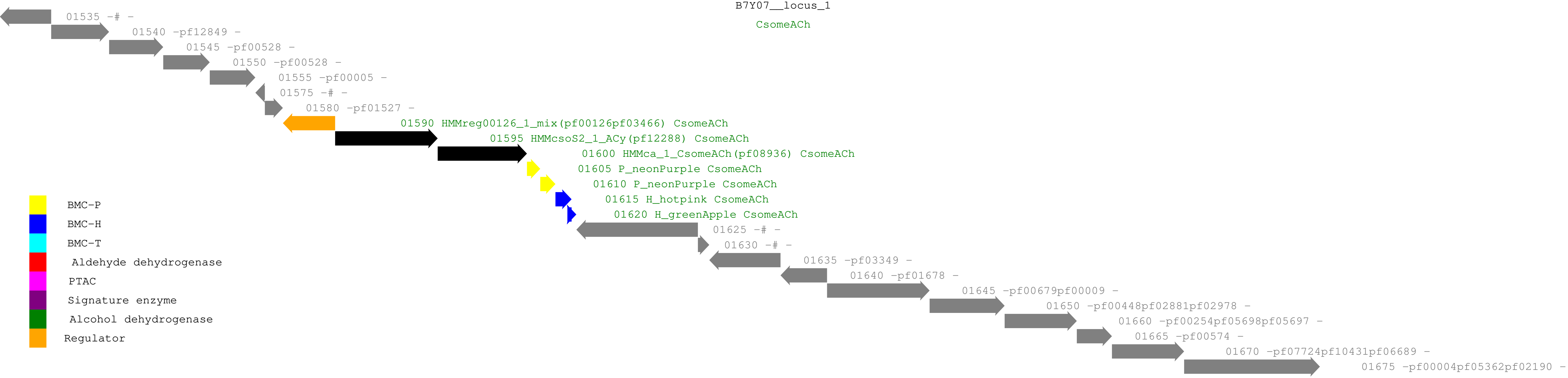

Hydrogenophilales bacterium 16-61-112. Bacteria; Proteobacteria; Hydrogenophilalia; Hydrogenophilales.

B7Y21__locus_1

CsomeACh

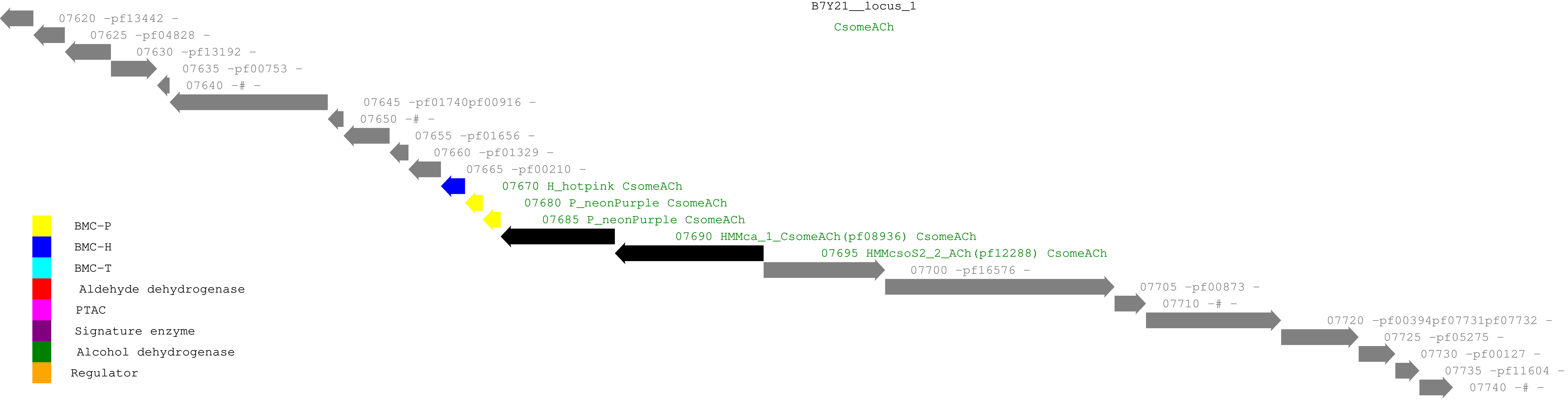

Thiotrichales bacterium 16-46-22. Bacteria; Proteobacteria; Gammaproteobacteria; Thiotrichales;

B7Y29__locus_1

CsomeACh

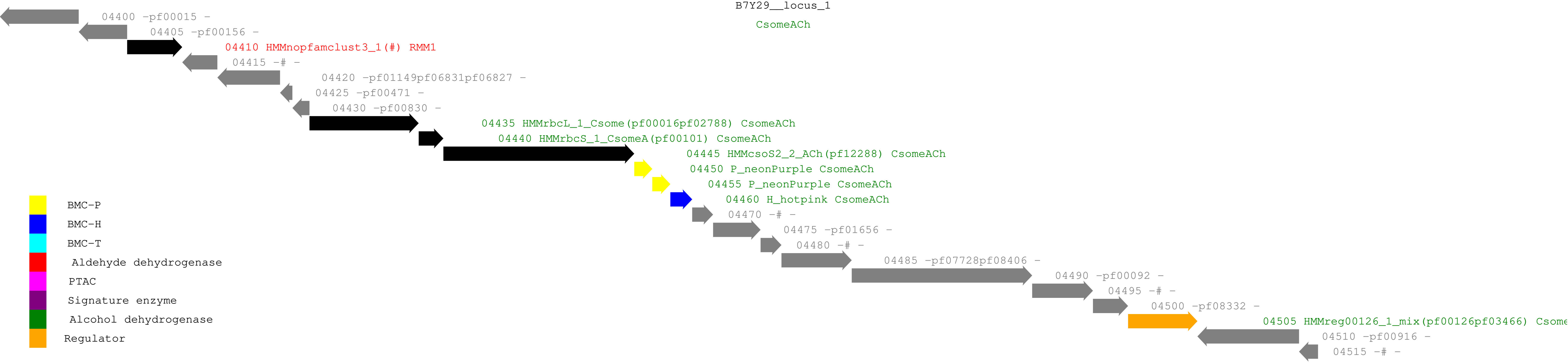

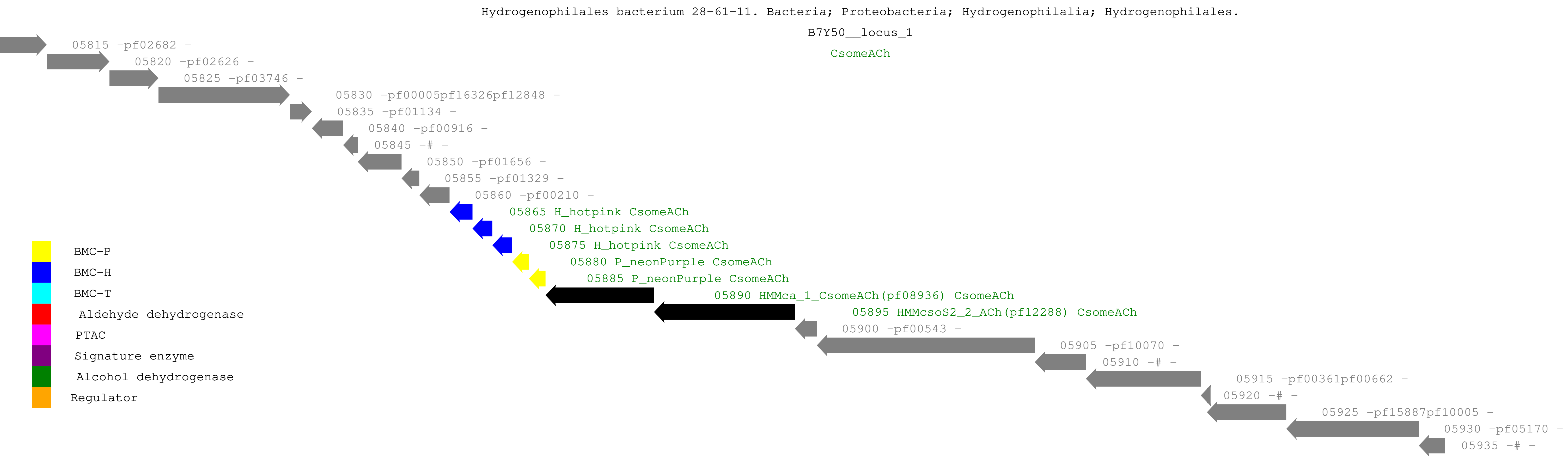
Hydrogenophilales bacterium 28-61-11. Bacteria; Proteobacteria; Hydrogenophilalia; Hydrogenophilales.
B7Y50__locus_1
CsomeACh
05815 -pf02682 -
05820 -pf02626 -
05825 -pf03746 -
05830 -pf00005pf16326pf12848 -
05835 -pf01134 -
05840 -pf00916 -
05845 -# -
05850 -pf01656 -
05855 -pf01329 -
05860 -pf00210 -
05865 H_hotpink CsomeACh
05870 H_hotpink CsomeACh
05875 H_hotpink CsomeACh
05880 P_neonPurple CsomeACh
05885 P_neonPurple CsomeACh
05890 HMMca_1_CsomeACh(pf08936) CsomeACh
05895 HMMcsoS2_2_ACh(pf12288) CsomeACh
05900 -pf00543 -
05905 -pf10070 -
05910 -# -
05915 -pf00361pf00662 -
05920 -# -
05925 -pf15887pf10005 -
05930 -pf05170 -
05935 -# -
BMC-P
BMC-H
BMC-T
Aldehyde dehydrogenase
PTAC
Signature enzyme
Alcohol dehydrogenase
Regulator

Thiotrichales bacterium 35-46-9. Bacteria; Proteobacteria; Gammaproteobacteria; Thiotrichales;

B7Y68__locus_1

CsomeACh

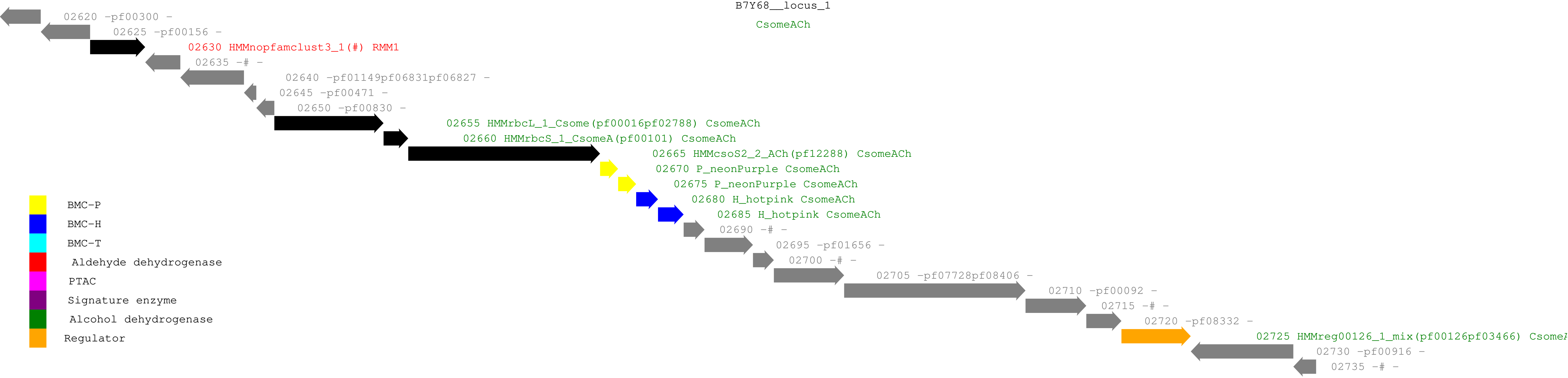

Hydrogenophilales bacterium 32-62-9. Bacteria; Proteobacteria; Hydrogenophilalia; Hydrogenophilales.

B7Z03__locus_1_main_sat

CsomeACh

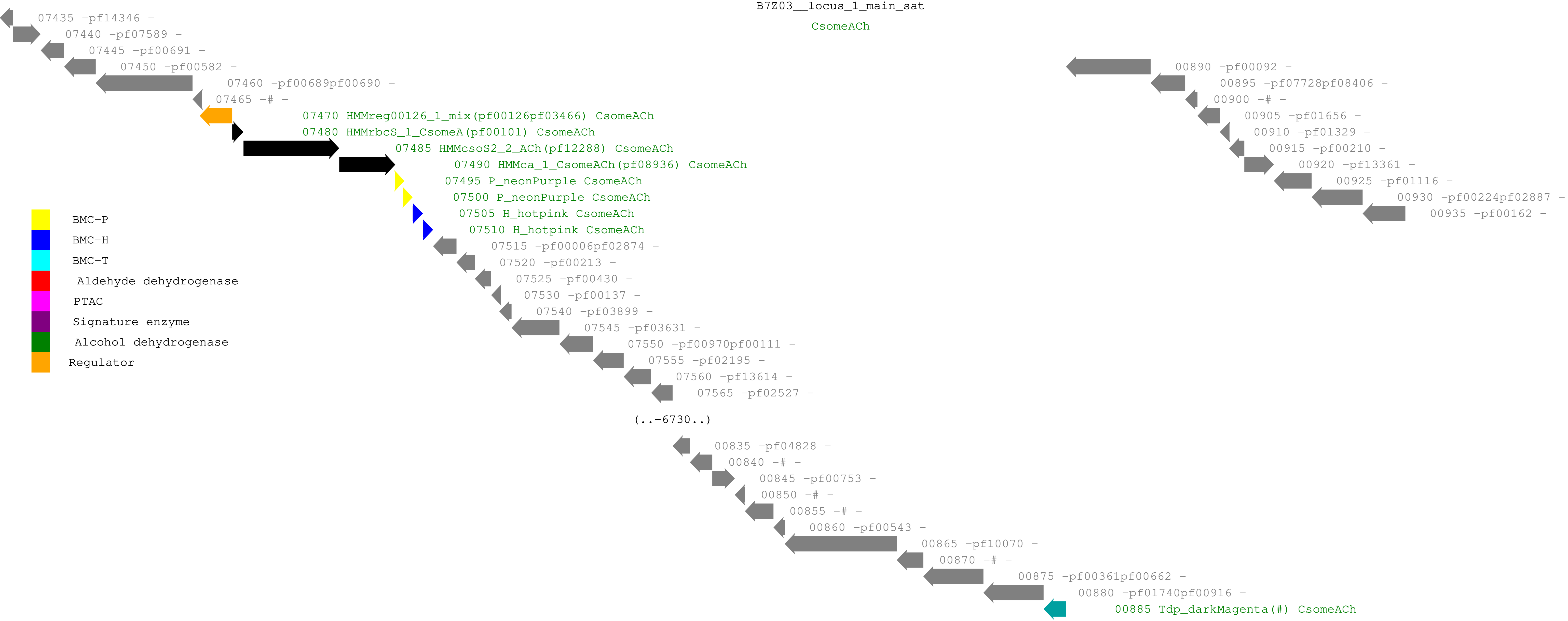

Thiotrichales bacterium 32-46-8. Bacteria; Proteobacteria; Gammaproteobacteria; Thiotrichales;

B7Z05__locus_1

CsomeACh

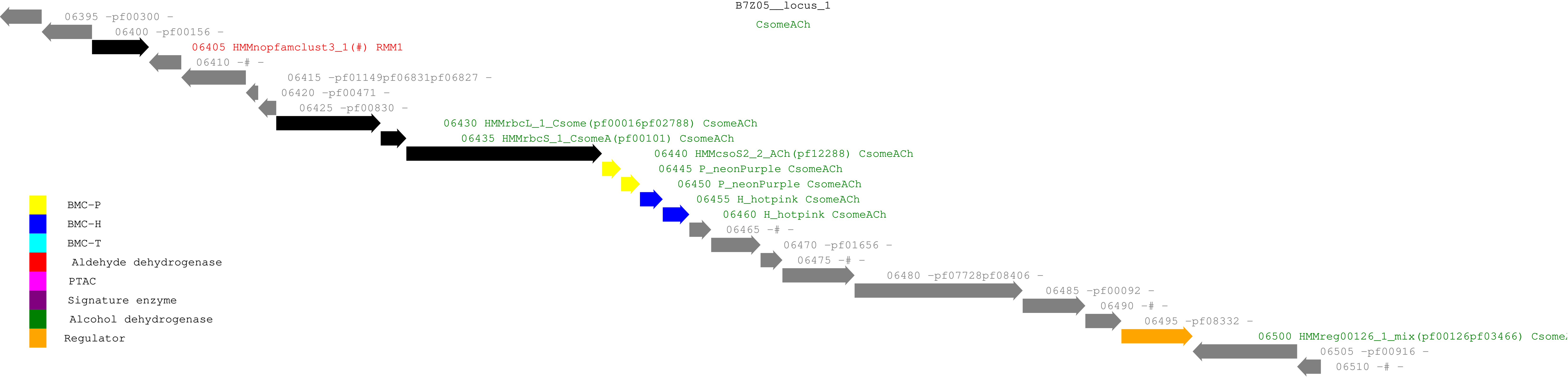

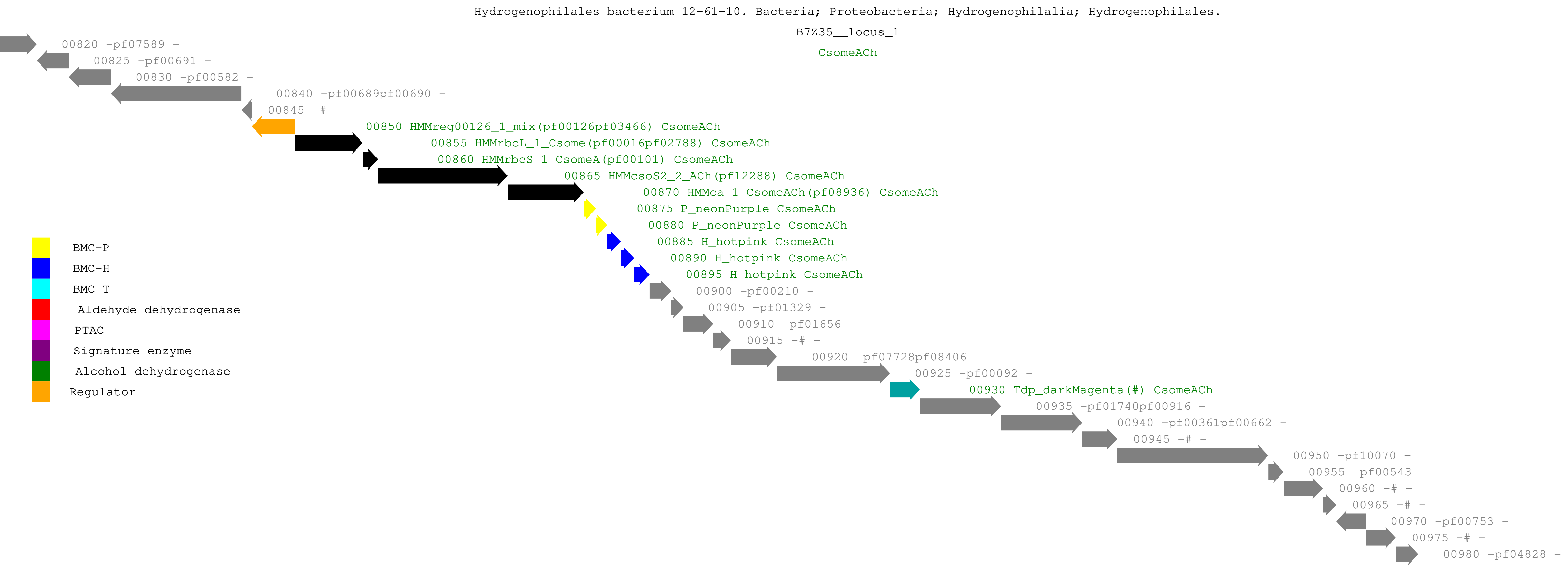

Hydrogenophilales bacterium 12-61-10. Bacteria; Proteobacteria; Hydrogenophilalia; Hydrogenophilales.
B7Z35__locus_1
CsomeACh
00820 -pf07589 -
00825 -pf00691 -
00830 -pf00582 -
00840 -pf00689pf00690 -
00845 -# -
00850 HMMreg00126_1_mix(pf00126pf03466) CsomeACh
00855 HMMrbcL_1_Csome(pf00016pf02788) CsomeACh
00860 HMMrbcS_1_CsomeA(pf00101) CsomeACh
00865 HMMcsoS2_2_ACh(pf12288) CsomeACh
00870 HMMca_1_CsomeACh(pf08936) CsomeACh
00875 P_neonPurple CsomeACh
00880 P_neonPurple CsomeACh
00885 H_hotpink CsomeACh
00890 H_hotpink CsomeACh
00895 H_hotpink CsomeACh
00900 -pf00210 -
00905 -pf01329 -
00910 -pf01656 -
00915 -# -
00920 -pf07728pf08406 -
00925 -pf00092 -
00930 Tdp_darkMagenta(#) CsomeACh
00935 -pf01740pf00916 -
00940 -pf00361pf00662 -
00945 -# -
00950 -pf10070 -
00955 -pf00543 -
00960 -# -
00965 -# -
00970 -pf00753 -
00975 -# -
00980 -pf04828 -
BMC-P
BMC-H
BMC-T
Aldehyde dehydrogenase
PTAC
Signature enzyme
Alcohol dehydrogenase
Regulator

Ferrovum sp. 37-45-19. Bacteria; Proteobacteria; Betaproteobacteria; Ferrovales; Ferrovaceae;

B7Z60__locus_1_main_sat

CsomeACh

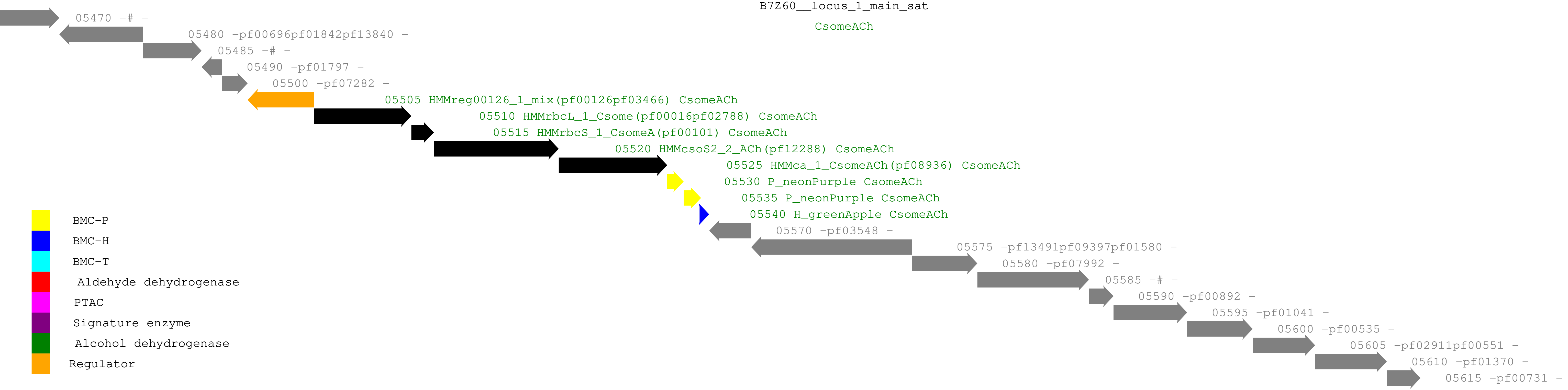

Chromatiales bacterium 21-64-14. Bacteria; Proteobacteria; Gammaproteobacteria; Chromatiales;

B7Z66__locus_1

CsomeACh

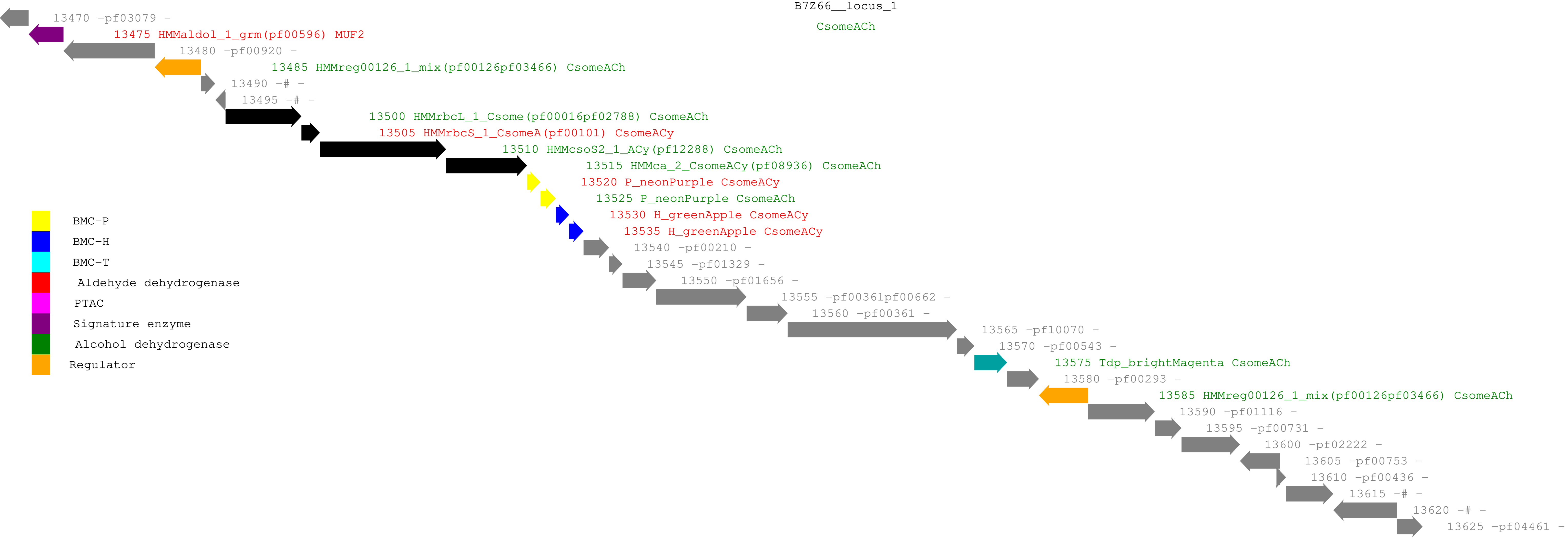

Acidithiobacillus ferrivorans. Bacteria; Proteobacteria; Acidithiobacillia; Acidithiobacillales;

B7Z70__locus_1_main_sat

CsomeACh

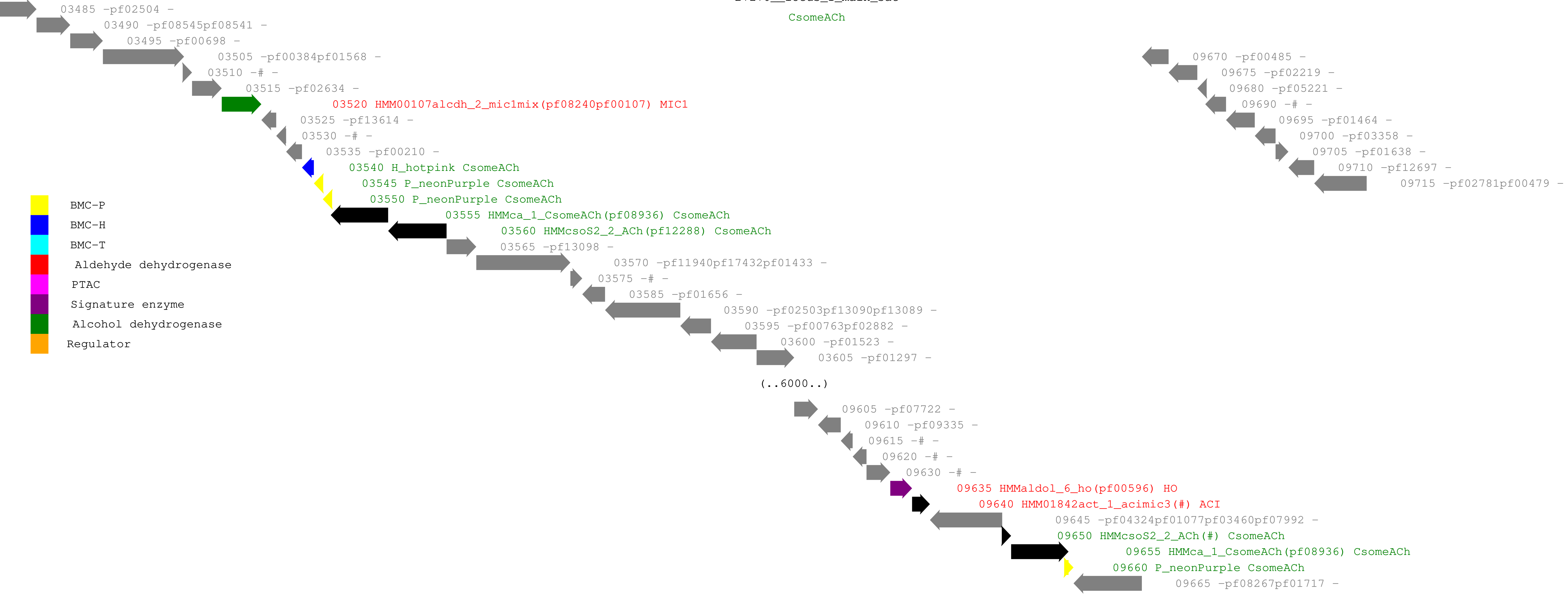

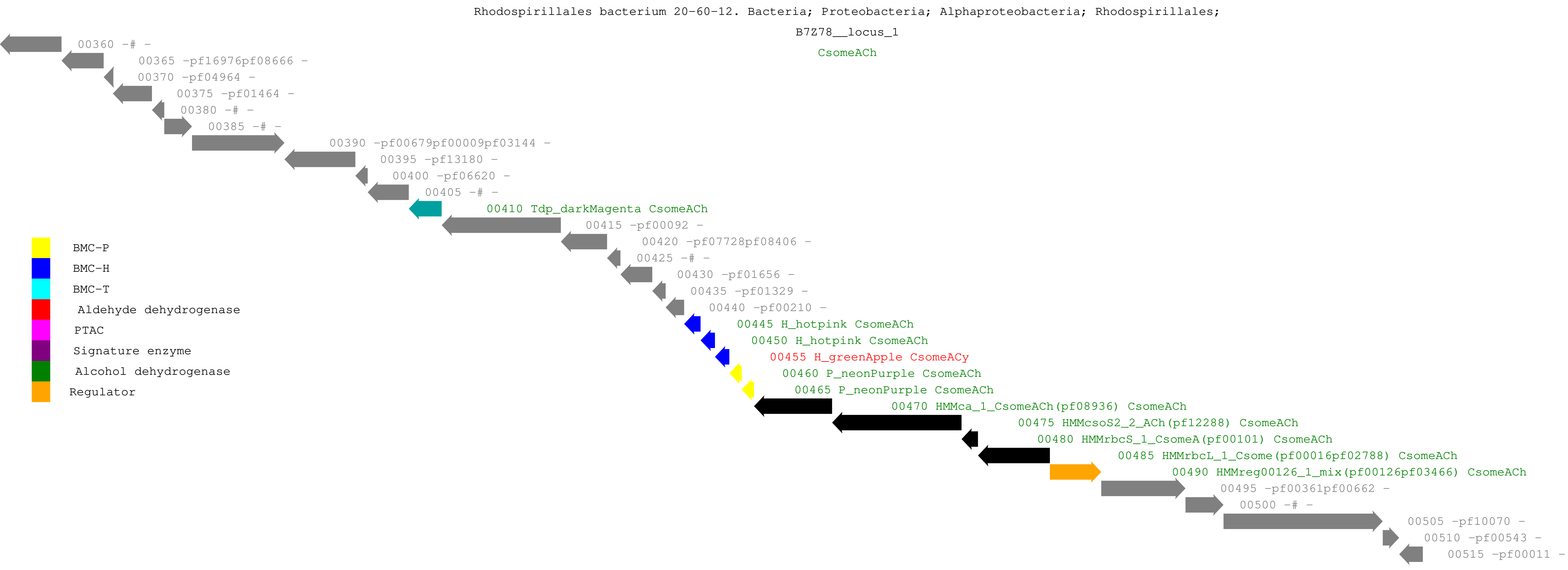
Rhodospirillales bacterium 20-60-12. Bacteria; Proteobacteria; Alphaproteobacteria; Rhodospirillales;
B7Z78__locus_1
CsomeACh
00360 -# -
00365 -pf16976pf08666 -
00370 -pf04964 -
00375 -pf01464 -
00380 -# -
00385 -# -
00390 -pf00679pf00009pf03144 -
00395 -pf13180 -
00400 -pf06620 -
00405 -# -
00410 Tdp_darkMagenta CsomeACh
00415 -pf00092 -
00420 -pf07728pf08406 -
00425 -# -
00430 -pf01656 -
00435 -pf01329 -
00440 -pf00210 -
00445 H_hotpink CsomeACh
00450 H_hotpink CsomeACh
00455 H_greenApple CsomeACy
00460 P_neonPurple CsomeACh
00465 P_neonPurple CsomeACh
00470 HMMca_1_CsomeACh(pf08936) CsomeACh
00475 HMMcsoS2_2_ACh(pf12288) CsomeACh
00480 HMMrbcS_1_CsomeA(pf00101) CsomeACh
00485 HMMrbcL_1_Csome(pf00016pf02788) CsomeACh
00490 HMMreg00126_1_mix(pf00126pf03466) CsomeACh
00495 -pf00361pf00662 -
00500 -# -
00505 -pf10070 -
00510 -pf00543 -
00515 -pf00011 -
BMC-P
BMC-H
BMC-T
Aldehyde dehydrogenase
PTAC
Signature enzyme
Alcohol dehydrogenase
Regulator

Thiomonas sp. 20-64-5. Bacteria; Proteobacteria; Betaproteobacteria; Burkholderiales;

B7Z83__locus_fragmented

CsomeACh

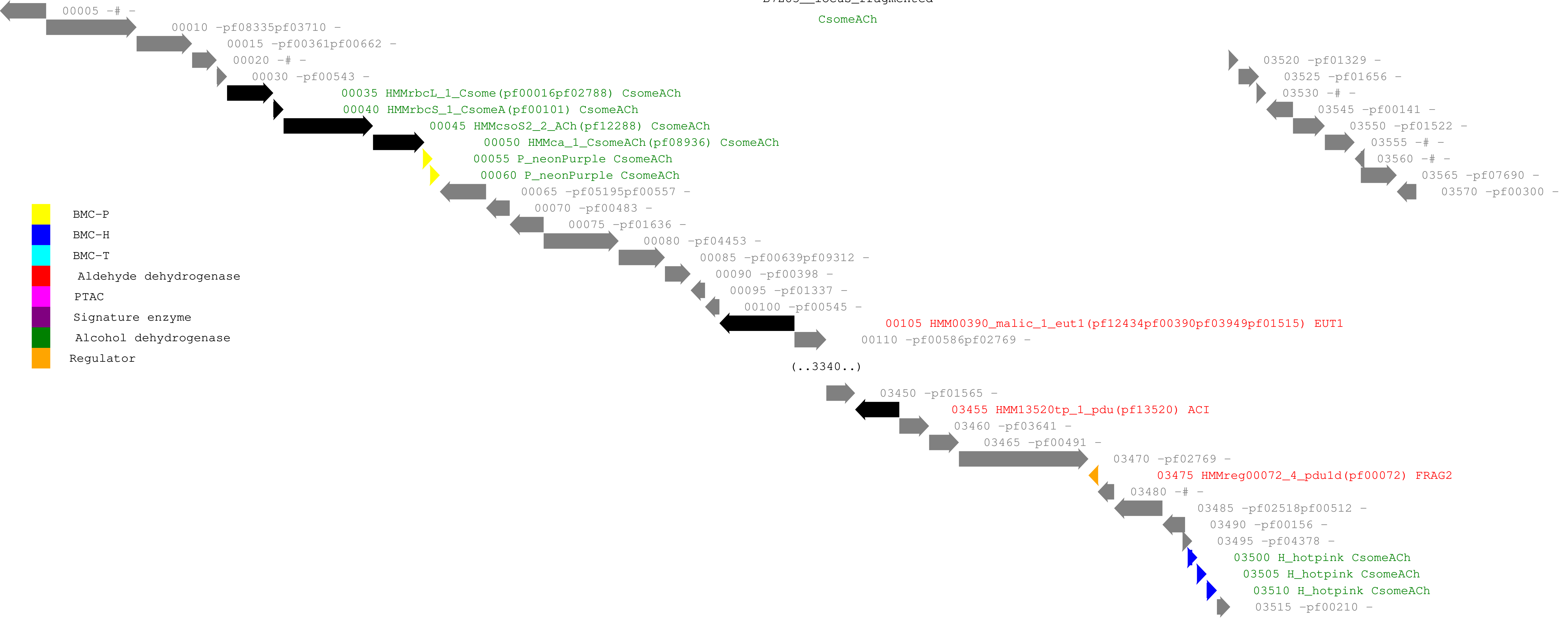

Acidithiobacillus caldus. Bacteria; Proteobacteria; Acidithiobacillia; Acidithiobacillales;

BAE27__locus_fragmented

CsomeACh

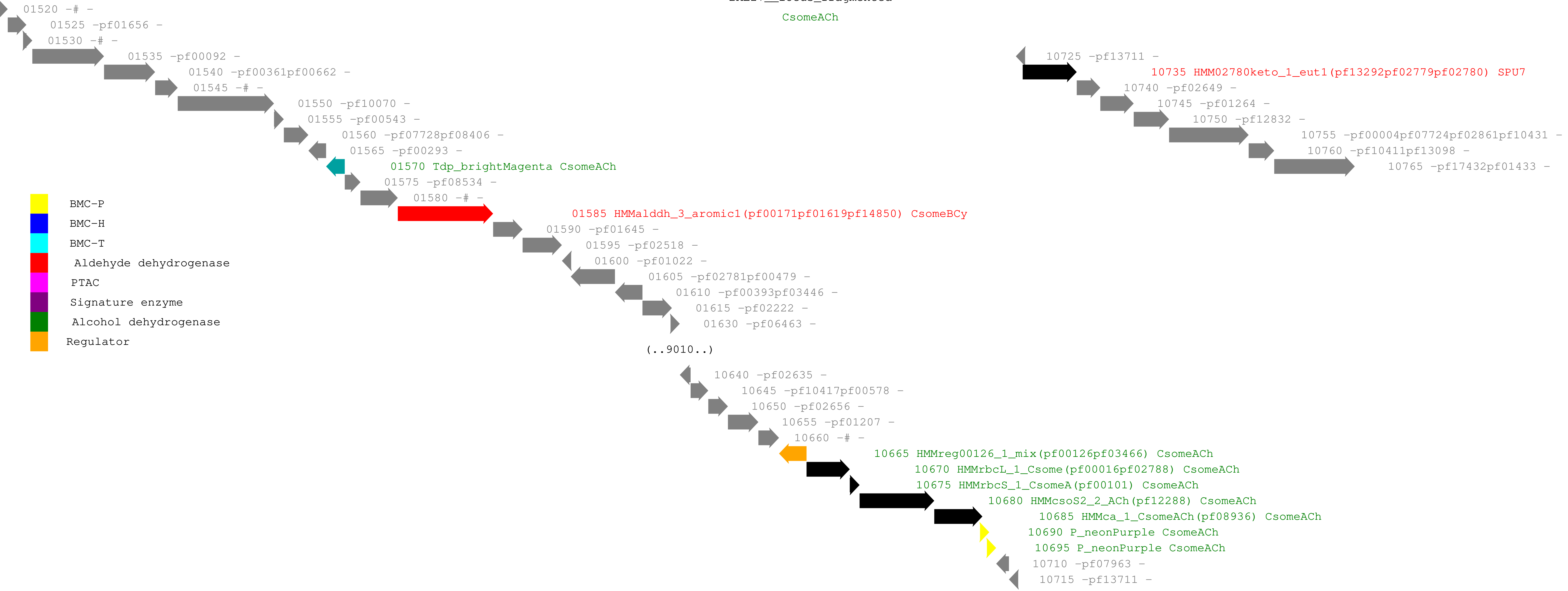

Bradyrhizobium sp. (strain BTAi1 / ATCC BAA-1182). Bacteria; Proteobacteria; Alphaproteobacteria; Rhizobiales;

BBta__locus_1

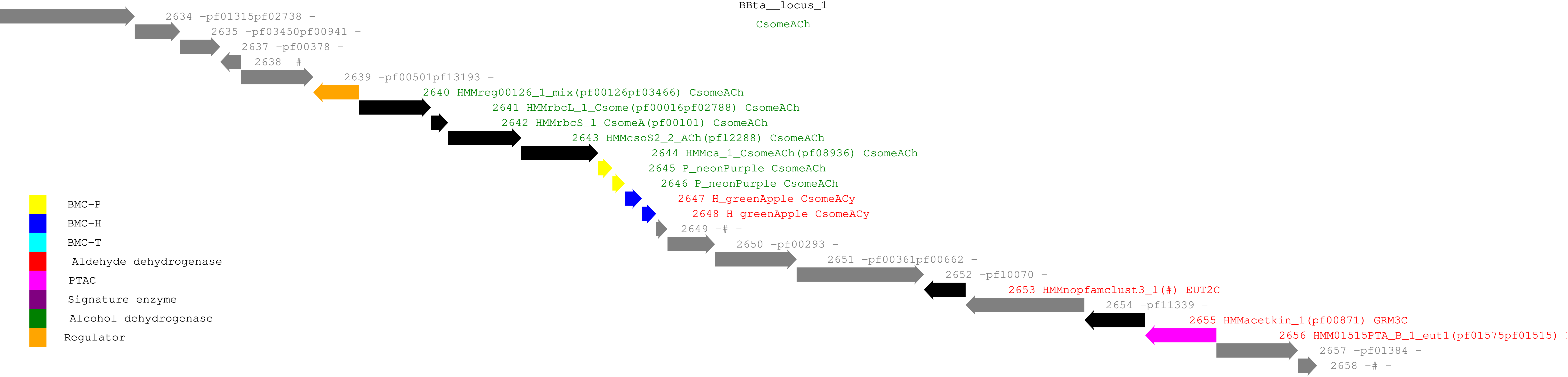

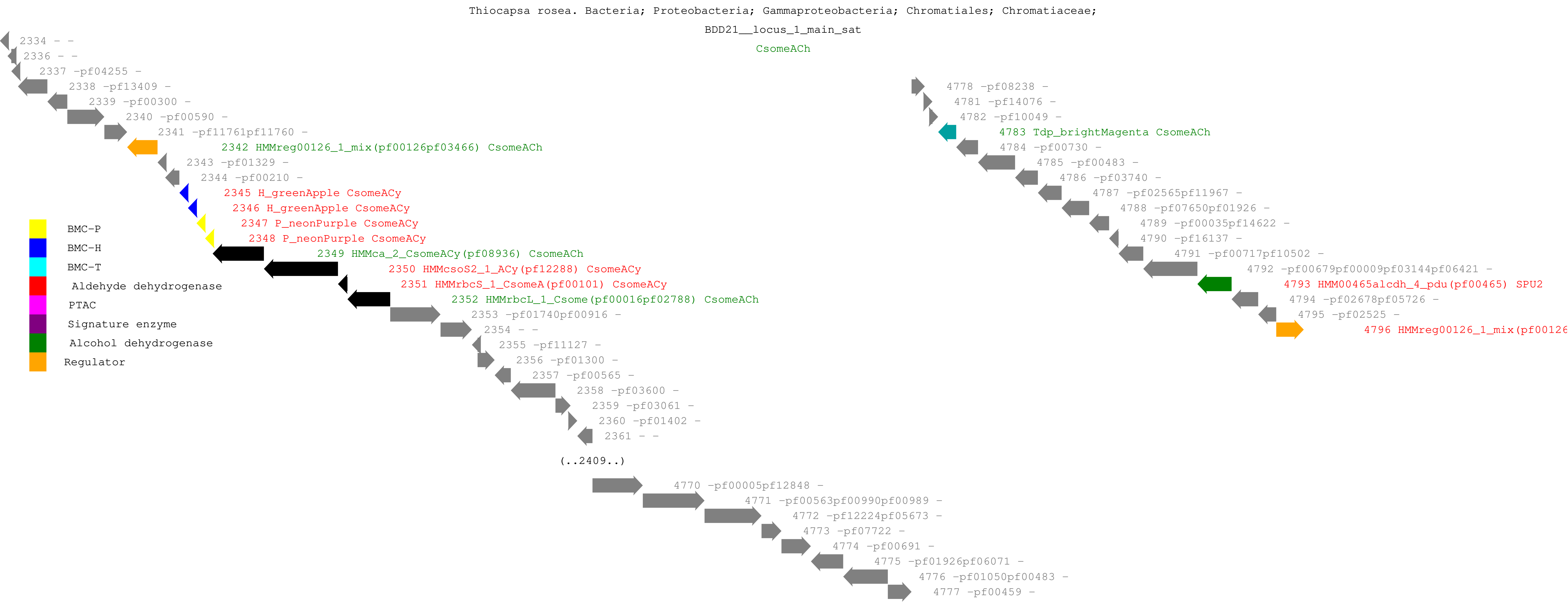

Thiocapsa rosea. Bacteria; Proteobacteria; Gammaproteobacteria; Chromatiales; Chromatiaceae;
BDD21__locus_1_main_sat
CsomeACh
2334 - -
2336 - -
2337 -pf04255 -
2338 -pf13409 -
2339 -pf00300 -
2340 -pf00590 -
2341 -pf11761pf11760 -
2342 HMMreg00126_1_mix(pf00126pf03466) CsomeACh
2343 -pf01329 -
2344 -pf00210 -
2345 H_greenApple CsomeACy
2346 H_greenApple CsomeACy
2347 P_neonPurple CsomeACy
2348 P_neonPurple CsomeACy
2349 HMMca_2_CsomeACy(pf08936) CsomeACh
2350 HMMcsoS2_1_ACy(pf12288) CsomeACy
2351 HMMrbcS_1_CsomeA(pf00101) CsomeACy
2352 HMMrbcL_1_Csome(pf00016pf02788) CsomeACh
2353 -pf01740pf00916 -
2354 - -
2355 -pf11127 -
2356 -pf01300 -
2357 -pf00565 -
2358 -pf03600 -
2359 -pf03061 -
2360 -pf01402 -
2361 - -
(..2409..)
4770 -pf00005pf12848 -
4771 -pf00563pf00990pf00989 -
4772 -pf12224pf05673 -
4773 -pf07722 -
4774 -pf00691 -
4775 -pf01926pf06071 -
4776 -pf01050pf00483 -
4777 -pf00459 -
4778 -pf08238 -
4781 -pf14076 -
4782 -pf10049 -
4783 Tdp_brightMagenta CsomeACh
4784 -pf00730 -
4785 -pf00483 -
4786 -pf03740 -
4787 -pf02565pf11967 -
4788 -pf07650pf01926 -
4789 -pf00035pf14622 -
4790 -pf16137 -
4791 -pf00717pf10502 -
4792 -pf00679pf00009pf03144pf06421 -
4793 HMM00465alcdh_4_pdu(pf00465) SPU2
4794 -pf02678pf05726 -
4795 -pf02525 -
4796 HMMreg00126_1_mix(pf00126
BMC-P
BMC-H
BMC-T
Aldehyde dehydrogenase
PTAC
Signature enzyme
Alcohol dehydrogenase
Regulator

Nitrobacter sp. 62-13. Bacteria; Proteobacteria; Alphaproteobacteria; Rhizobiales;

BGN91__locus_fragmented

CsomeACh

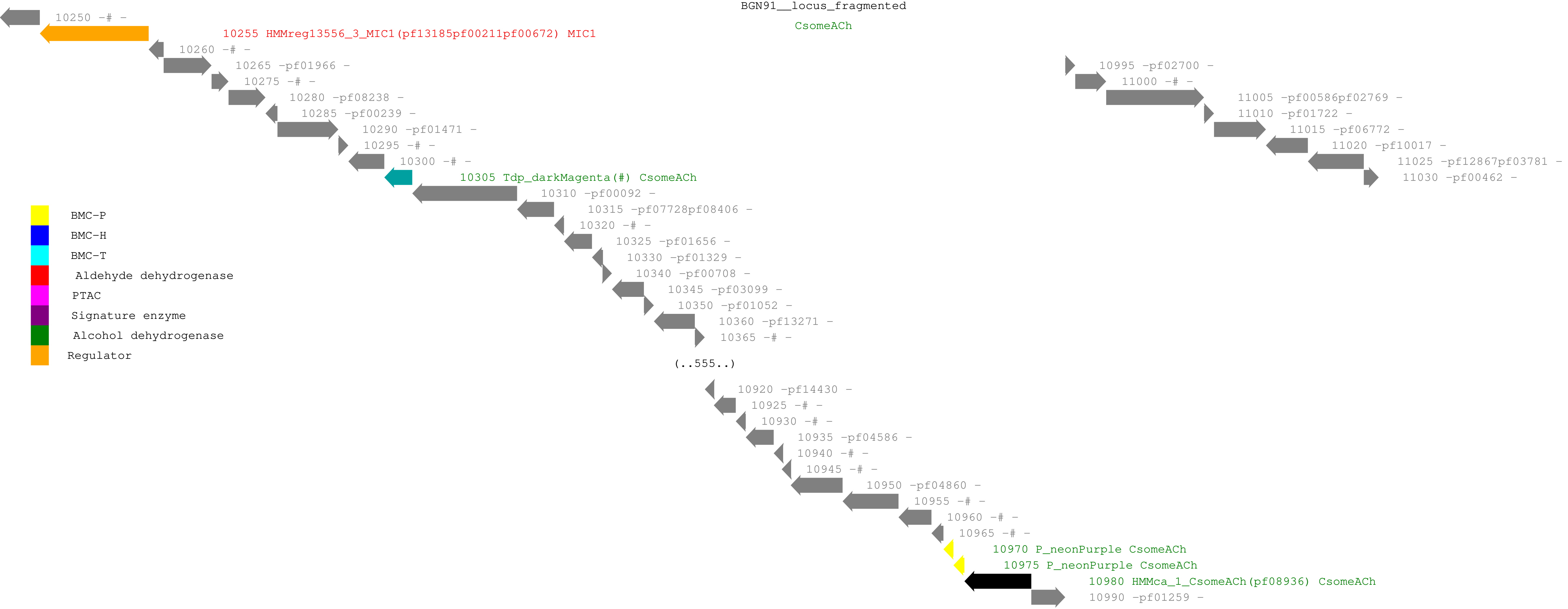

Nitrobacter sp. 62-23. Bacteria; Proteobacteria; Alphaproteobacteria; Rhizobiales;

BGO16__locus_1_main_sat

CsomeACh

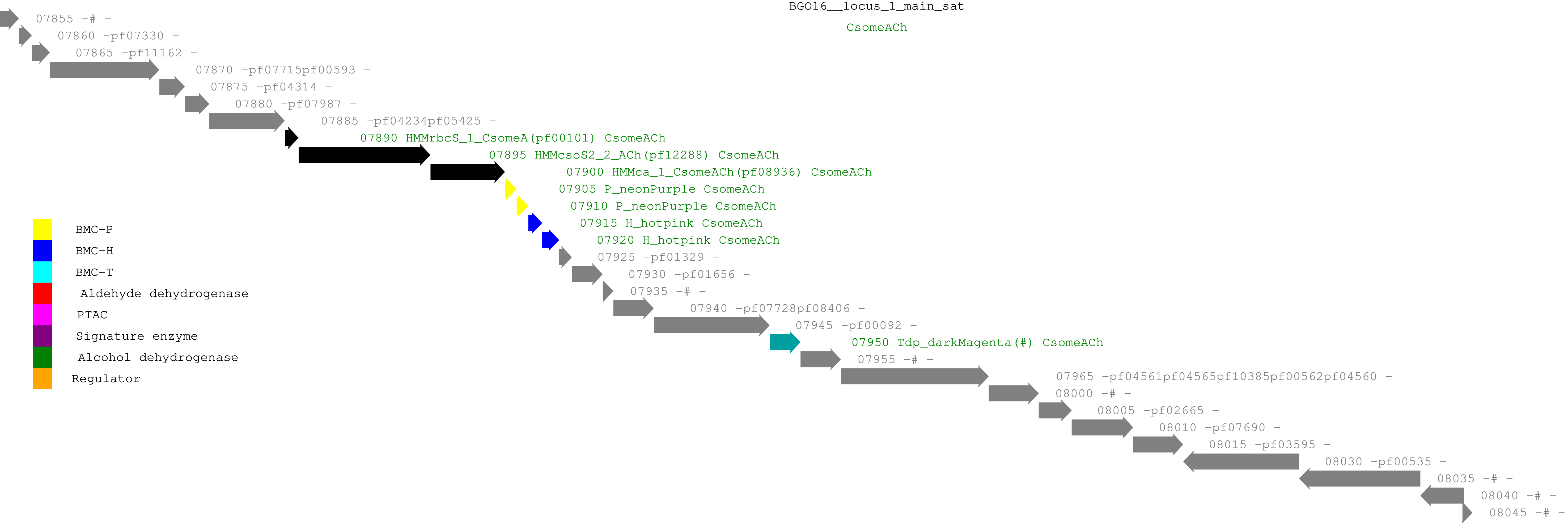

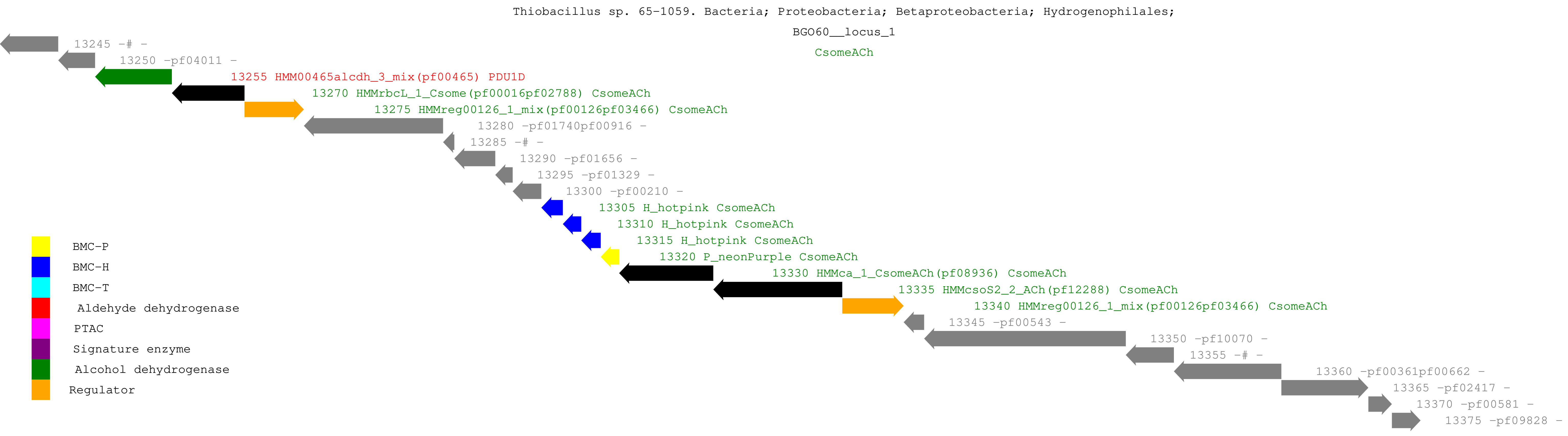
Thiobacillus sp. 65-1059. Bacteria; Proteobacteria; Betaproteobacteria; Hydrogenophilales;
BGO60__locus_1
CsomeACh
13245 -# -
13250 -pf04011 -
13255 HMM00465alcdh_3_mix(pf00465) PDU1D
13270 HMMrbcL_1_Csome(pf00016pf02788) CsomeACh
13275 HMMreg00126_1_mix(pf00126pf03466) CsomeACh
13280 -pf01740pf00916 -
13285 -# -
13290 -pf01656 -
13295 -pf01329 -
13300 -pf00210 -
13305 H_hotpink CsomeACh
13310 H_hotpink CsomeACh
13315 H_hotpink CsomeACh
13320 P_neonPurple CsomeACh
13330 HMMca_1_CsomeACh(pf08936) CsomeACh
13335 HMMcsoS2_2_ACh(pf12288) CsomeACh
13340 HMMreg00126_1_mix(pf00126pf03466) CsomeACh
13345 -pf00543 -
13350 -pf10070 -
13355 -# -
13360 -pf00361pf00662 -
13365 -pf02417 -
13370 -pf00581 -
13375 -pf09828 -
BMC-P
BMC-H
BMC-T
Aldehyde dehydrogenase
PTAC
Signature enzyme
Alcohol dehydrogenase
Regulator

Thiobacillus sp. 65-1402. Bacteria; Proteobacteria; Betaproteobacteria; Hydrogenophilales;

BGO62__locus_1

CsomeACh

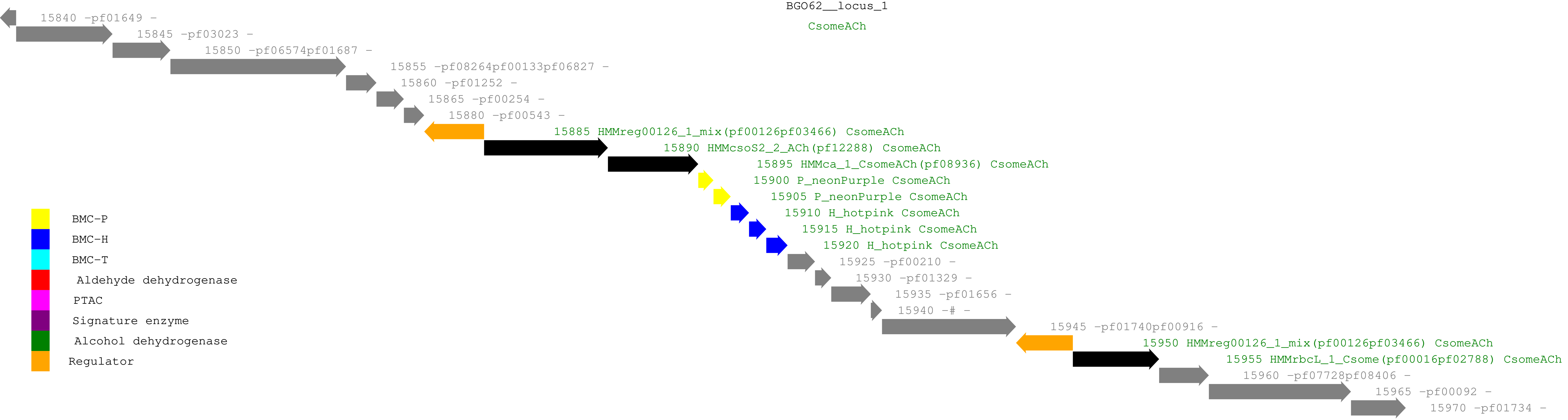

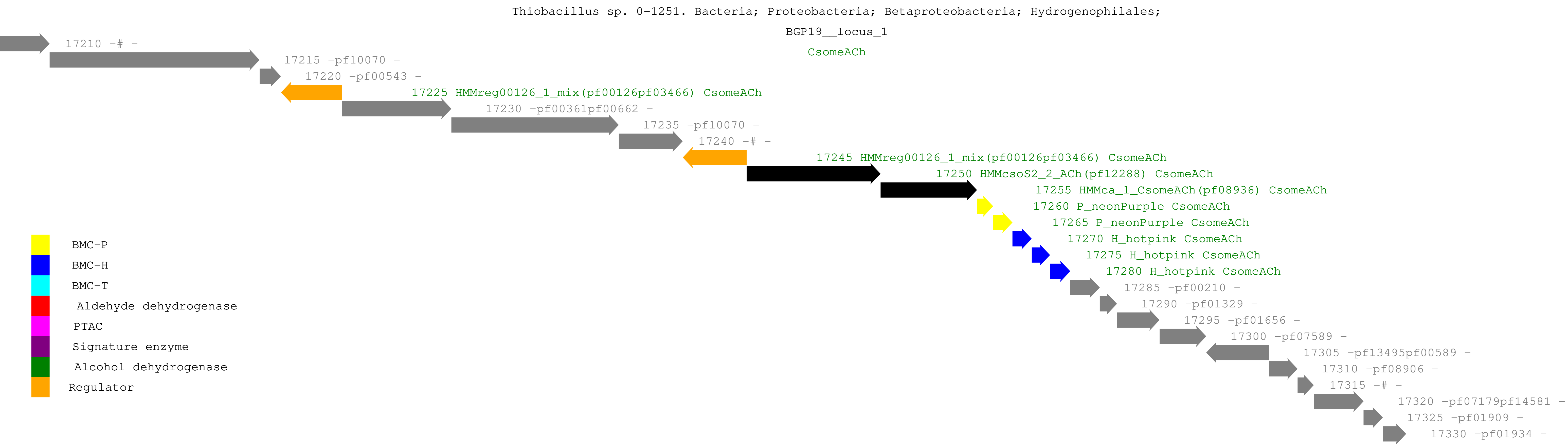
Thiobacillus sp. 0-1251. Bacteria; Proteobacteria; Betaproteobacteria; Hydrogenophilales;
BGP19__locus_1
CsomeACh
17210 -# -
17215 -pf10070 -
17220 -pf00543 -
17225 HMMreg00126_1_mix(pf00126pf03466) CsomeACh
17230 -pf00361pf00662 -
17235 -pf10070 -
17240 -# -
17245 HMMreg00126_1_mix(pf00126pf03466) CsomeACh
17250 HMMcsoS2_2_ACh(pf12288) CsomeACh
17255 HMMca_1_CsomeACh(pf08936) CsomeACh
17260 P_neonPurple CsomeACh
17265 P_neonPurple CsomeACh
17270 H_hotpink CsomeACh
17275 H_hotpink CsomeACh
17280 H_hotpink CsomeACh
17285 -pf00210 -
17290 -pf01329 -
17295 -pf01656 -
17300 -pf07589 -
17305 -pf13495pf00589 -
17310 -pf08906 -
17315 -# -
17320 -pf07179pf14581 -
17325 -pf01909 -
17330 -pf01934 -
BMC-P
BMC-H
BMC-T
Aldehyde dehydrogenase
PTAC
Signature enzyme
Alcohol dehydrogenase
Regulator

Acidihalobacter prosperus. Bacteria; Proteobacteria; Gammaproteobacteria; Chromatiales;

BI364__locus_1

CsomeACh

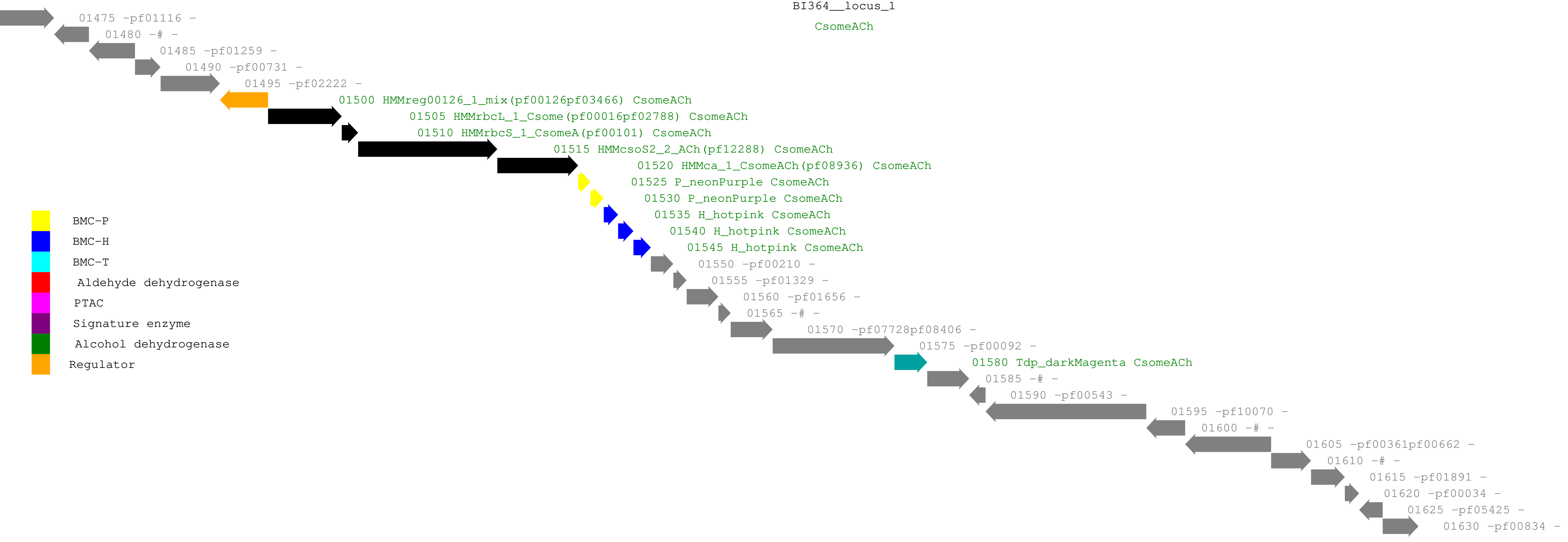

Acidihalobacter prosperus. Bacteria; Proteobacteria; Gammaproteobacteria; Chromatiales;

BJI67__locus_1

CsomeACh

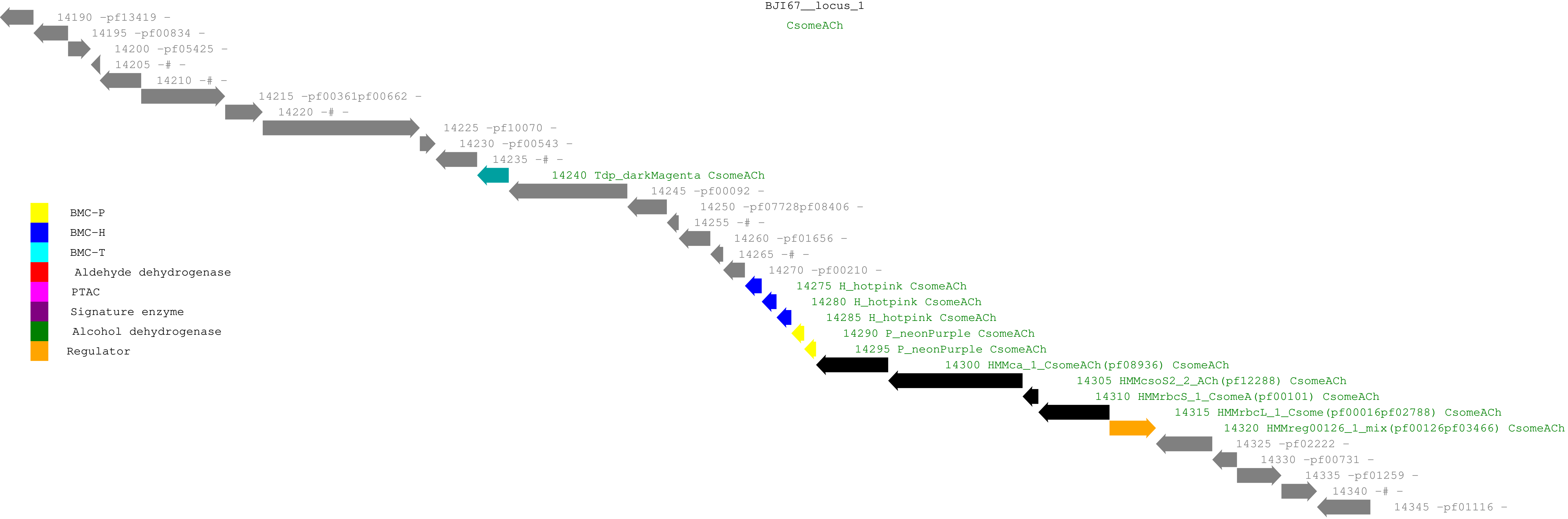

Bradyrhizobium sp. (strain ORS 278). Bacteria; Proteobacteria; Alphaproteobacteria; Rhizobiales;

BRADO_locus_1

CsomeACh

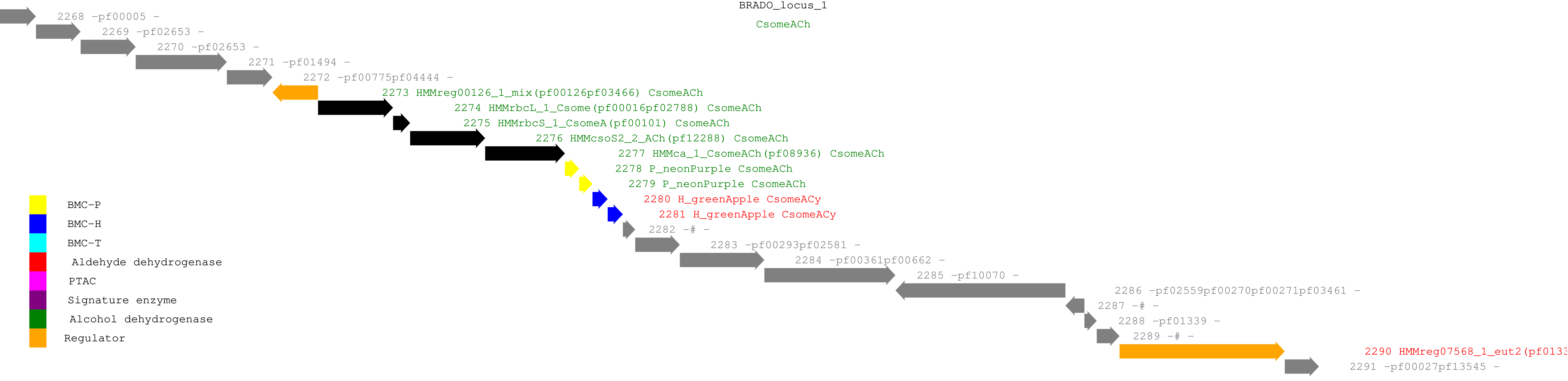

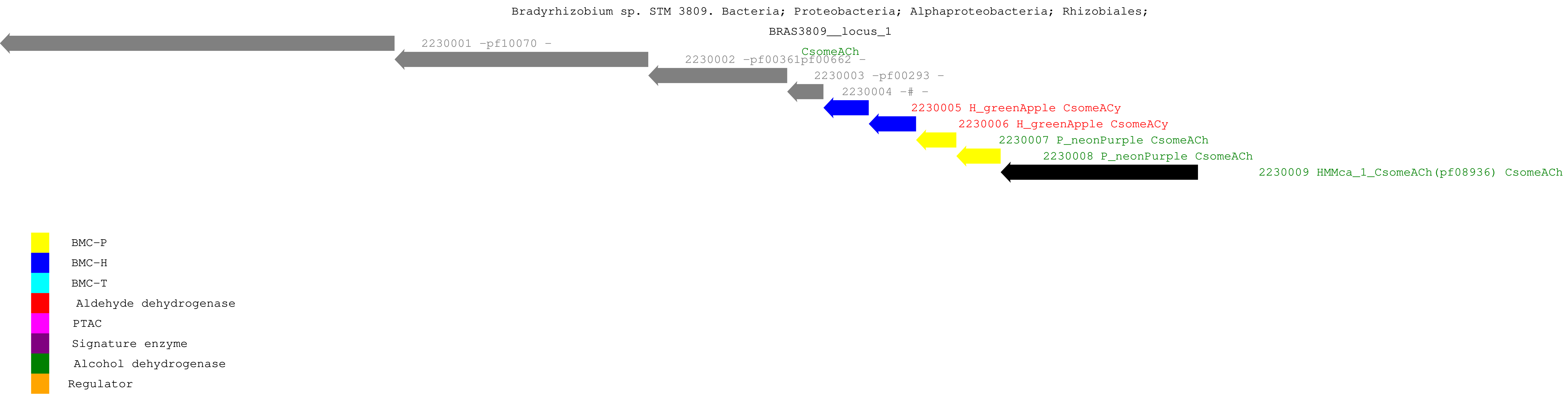

Bradyrhizobium sp. STM 3809. Bacteria; Proteobacteria; Alphaproteobacteria; Rhizobiales;
BRAS3809__locus_1
CsomeACh
2230001 -pf10070 -
2230002 -pf00361pf00662 -
2230003 -pf00293 -
2230004 -# -
2230005 H_greenApple CsomeACy
2230006 H_greenApple CsomeACy
2230007 P_neonPurple CsomeACh
2230008 P_neonPurple CsomeACh
2230009 HMMca_1_CsomeACh(pf08936) CsomeACh
BMC-P
BMC-H
BMC-T
Aldehyde dehydrogenase
PTAC
Signature enzyme
Alcohol dehydrogenase
Regulator

Xuhuaishuia manganoxidans. Bacteria; Proteobacteria; Alphaproteobacteria; Rhodobacterales;

BV394__locus_1

CsomeACh

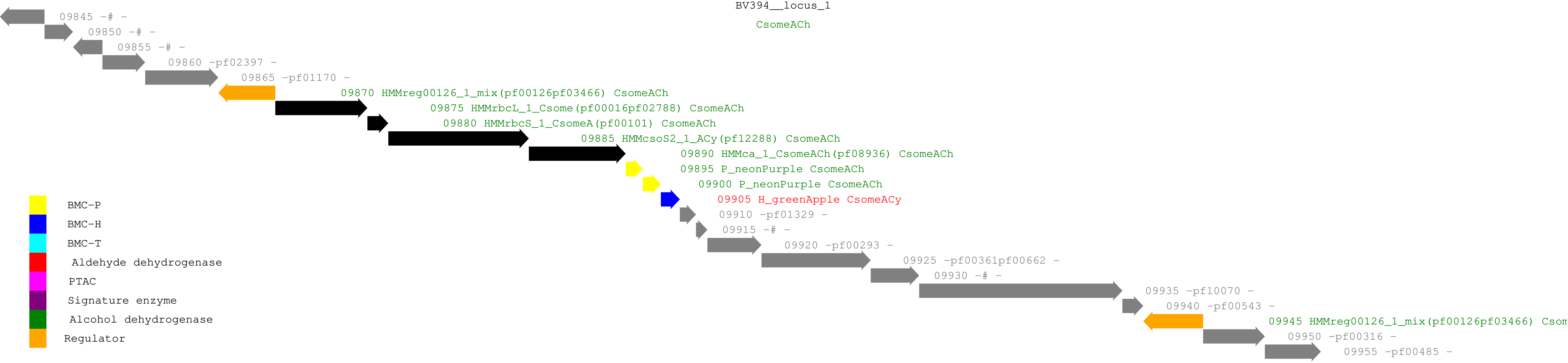

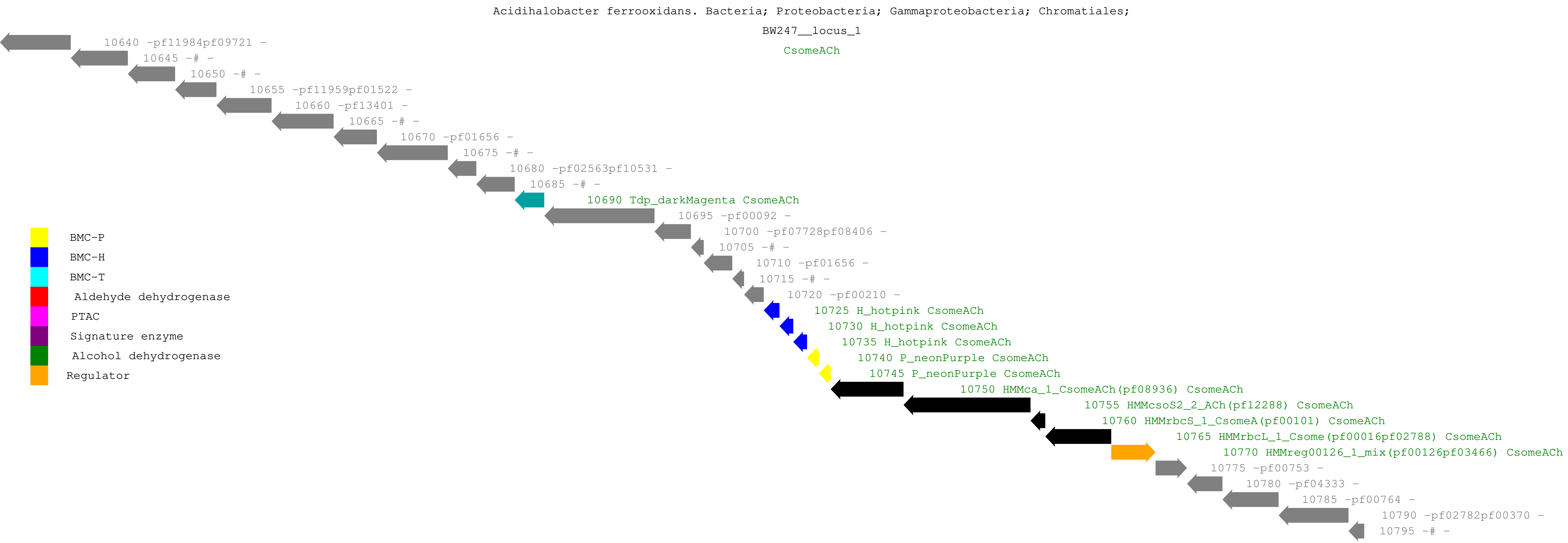
Acidihalobacter ferrooxidans. Bacteria; Proteobacteria; Gammaproteobacteria; Chromatiales;
BW247__locus_1
CsomeACh
10640 -pf11984pf09721 -
10645 -# -
10650 -# -
10655 -pf11959pf01522 -
10660 -pf13401 -
10665 -# -
10670 -pf01656 -
10675 -# -
10680 -pf02563pf10531 -
10685 -# -
10690 Tdp_darkMagenta CsomeACh
10695 -pf00092 -
10700 -pf07728pf08406 -
10705 -# -
10710 -pf01656 -
10715 -# -
10720 -pf00210 -
10725 H_hotpink CsomeACh
10730 H_hotpink CsomeACh
10735 H_hotpink CsomeACh
10740 P_neonPurple CsomeACh
10745 P_neonPurple CsomeACh
10750 HMMca_1_CsomeACh(pf08936) CsomeACh
10755 HMMcsoS2_2_ACh(pf12288) CsomeACh
10760 HMMrbcS_1_CsomeA(pf00101) CsomeACh
10765 HMMrbcL_1_Csome(pf00016pf02788) CsomeACh
10770 HMMreg00126_1_mix(pf00126pf03466) CsomeACh
10775 -pf00753 -
10780 -pf04333 -
10785 -pf00764 -
10790 -pf02782pf00370 -
10795 -# -
BMC-P
BMC-H
BMC-T
Aldehyde dehydrogenase
PTAC
Signature enzyme
Alcohol dehydrogenase
Regulator

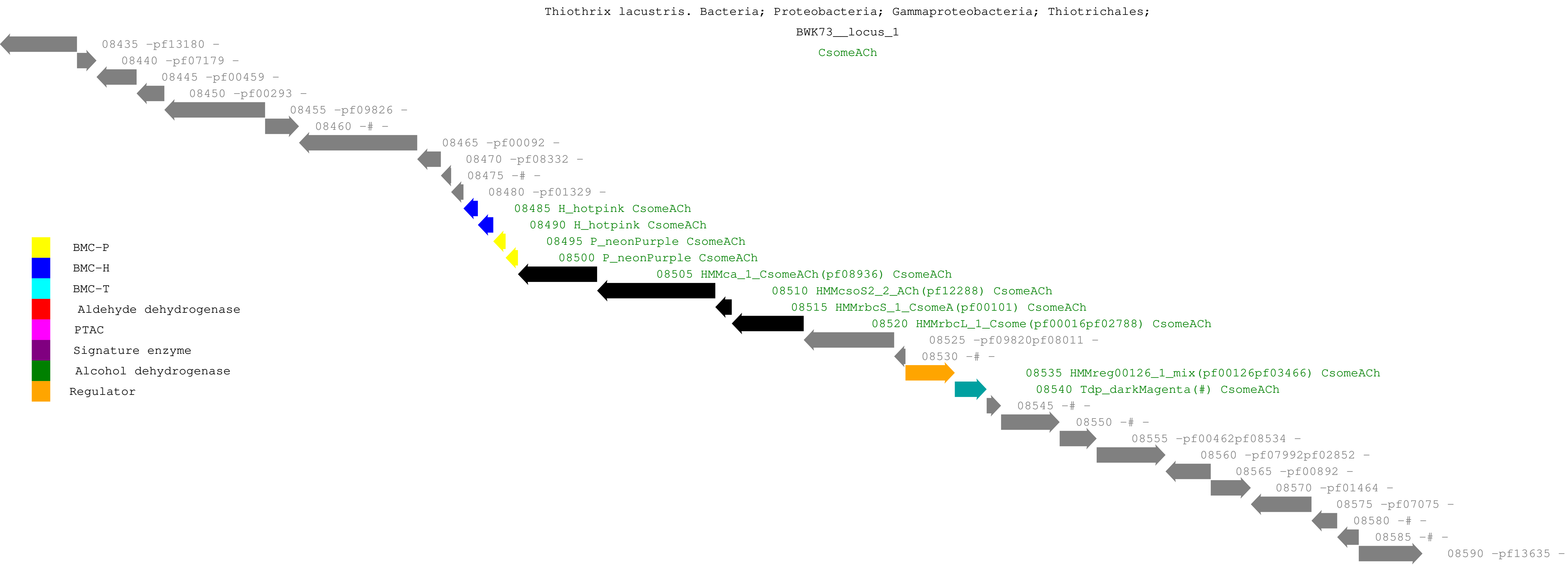
Thiothrix lacustris. Bacteria; Proteobacteria; Gammaproteobacteria; Thiotrichales;
BWK73__locus_1
CsomeACh
08435 -pf13180 -
08440 -pf07179 -
08445 -pf00459 -
08450 -pf00293 -
08455 -pf09826 -
08460 -# -
08465 -pf00092 -
08470 -pf08332 -
08475 -# -
08480 -pf01329 -
08485 H_hotpink CsomeACh
08490 H_hotpink CsomeACh
08495 P_neonPurple CsomeACh
08500 P_neonPurple CsomeACh
08505 HMMca_1_CsomeACh(pf08936) CsomeACh
08510 HMMcsoS2_2_ACh(pf12288) CsomeACh
08515 HMMrbcS_1_CsomeA(pf00101) CsomeACh
08520 HMMrbcL_1_Csome(pf00016pf02788) CsomeACh
08525 -pf09820pf08011 -
08530 -# -
08535 HMMreg00126_1_mix(pf00126pf03466) CsomeACh
08540 Tdp_darkMagenta(#) CsomeACh
08545 -# -
08550 -# -
08555 -pf00462pf08534 -
08560 -pf07992pf02852 -
08565 -pf00892 -
08570 -pf01464 -
08575 -pf07075 -
08580 -# -
08585 -# -
08590 -pf13635 -
BMC-P
BMC-H
BMC-T
Aldehyde dehydrogenase
PTAC
Signature enzyme
Alcohol dehydrogenase
Regulator

Acidithiobacillus ferrooxidans (Thiobacillus ferrooxidans). Bacteria; Proteobacteria; Acidithiobacillia; Acidithiobacillales;

C3R74__locus_1

CsomeACh

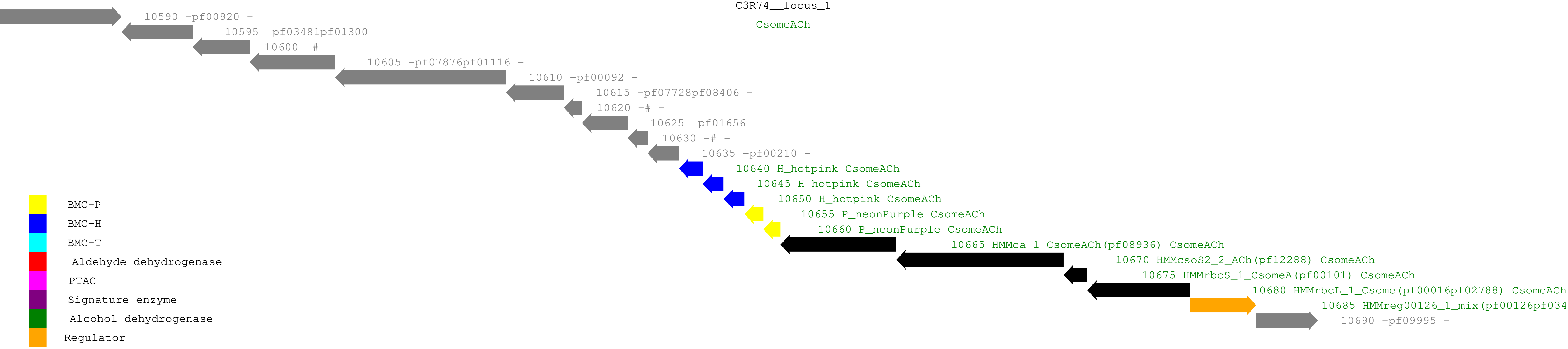

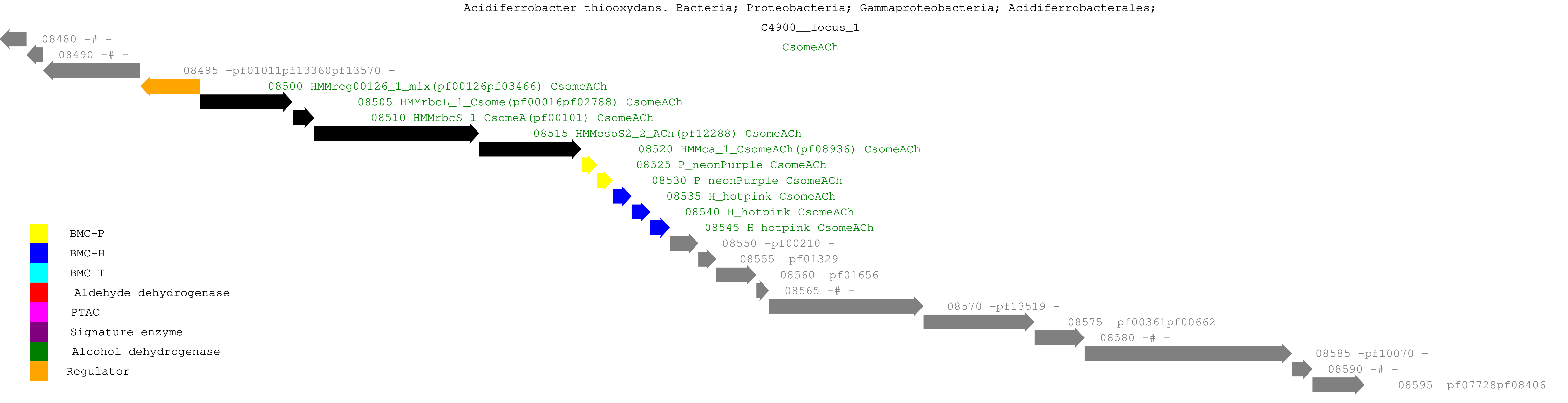

Acidiferrobacter thiooxydans. Bacteria; Proteobacteria; Gammaproteobacteria; Acidiferrobacterales;
C4900__locus_1
CsomeACh
08480 -# -
08490 -# -
08495 -pf01011pf13360pf13570 -
08500 HMMreg00126_1_mix(pf00126pf03466) CsomeACh
08505 HMMrbcL_1_Csome(pf00016pf02788) CsomeACh
08510 HMMrbcS_1_CsomeA(pf00101) CsomeACh
08515 HMMcsoS2_2_ACh(pf12288) CsomeACh
08520 HMMca_1_CsomeACh(pf08936) CsomeACh
08525 P_neonPurple CsomeACh
08530 P_neonPurple CsomeACh
08535 H_hotpink CsomeACh
08540 H_hotpink CsomeACh
08545 H_hotpink CsomeACh
08550 -pf00210 -
08555 -pf01329 -
08560 -pf01656 -
08565 -# -
08570 -pf13519 -
08575 -pf00361pf00662 -
08580 -# -
08585 -pf10070 -
08590 -# -
08595 -pf07728pf08406 -
BMC-P
BMC-H
BMC-T
Aldehyde dehydrogenase
PTAC
Signature enzyme
Alcohol dehydrogenase
Regulator

Acidiferrobacter sp. SPIII_3. Bacteria; Proteobacteria; Gammaproteobacteria; Acidiferrobacterales;

C4901__locus_1

CsomeACh

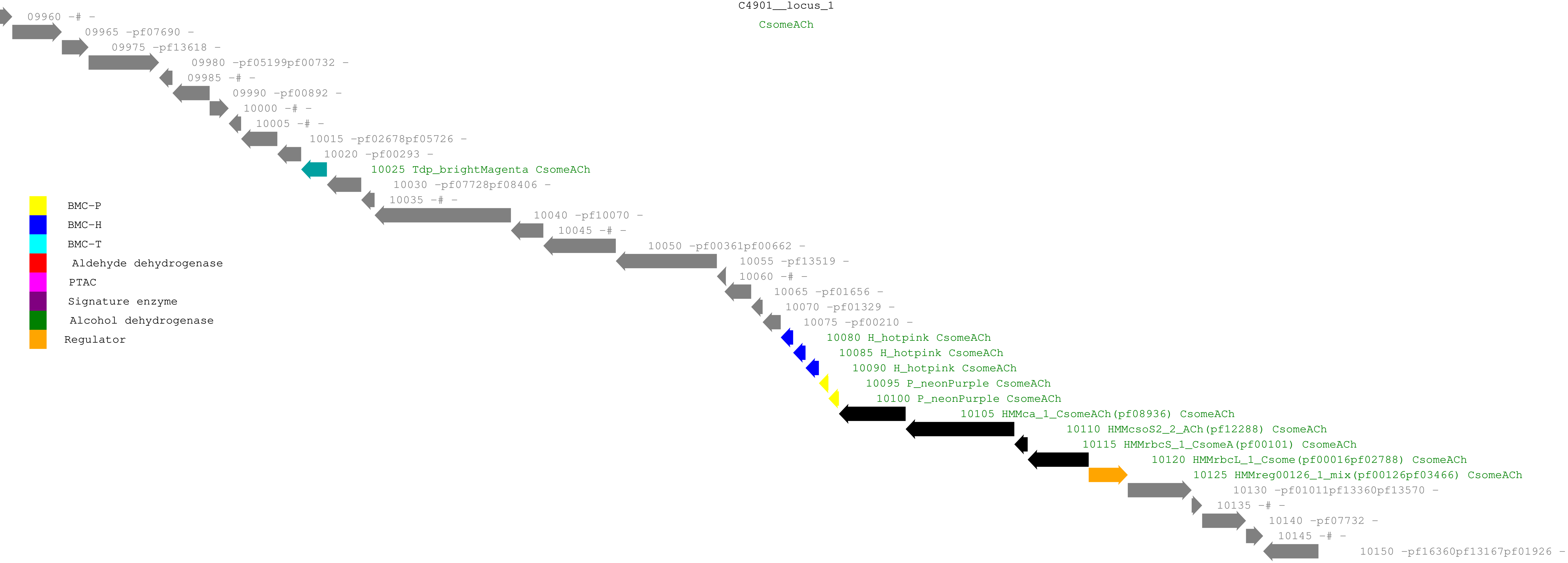

Nitrosospira sp. Nsp5. Bacteria; Proteobacteria; Betaproteobacteria; Nitrosomonadales;

C8R32__locus_1

CsomeACh

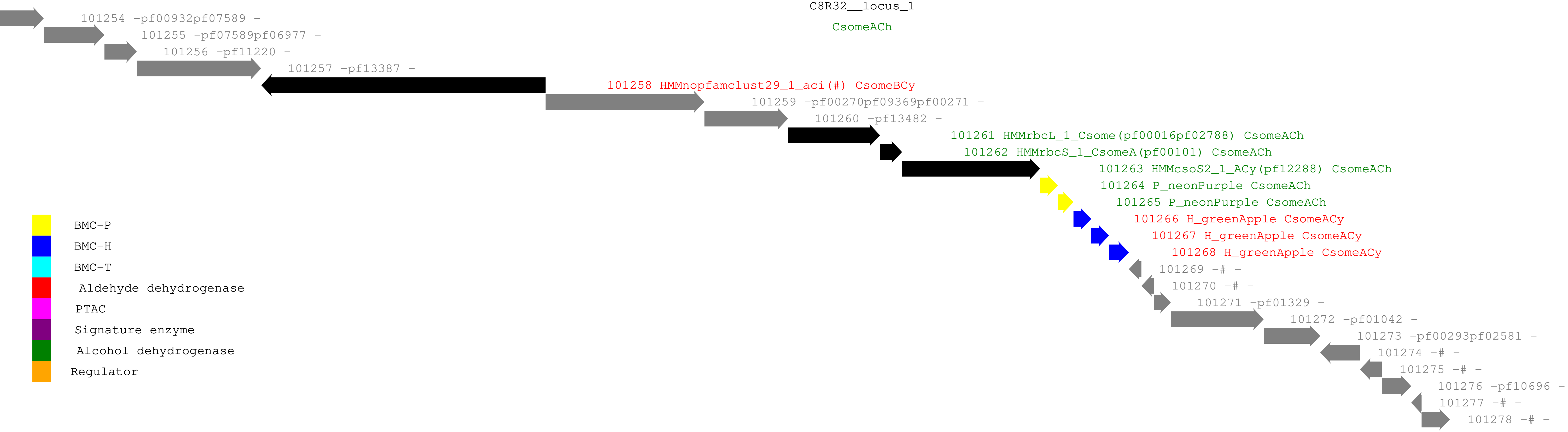

Nitrosomonas sp. Nm143. Bacteria; Proteobacteria; Betaproteobacteria; Nitrosomonadales;

C8R33__locus_1

CsomeACh

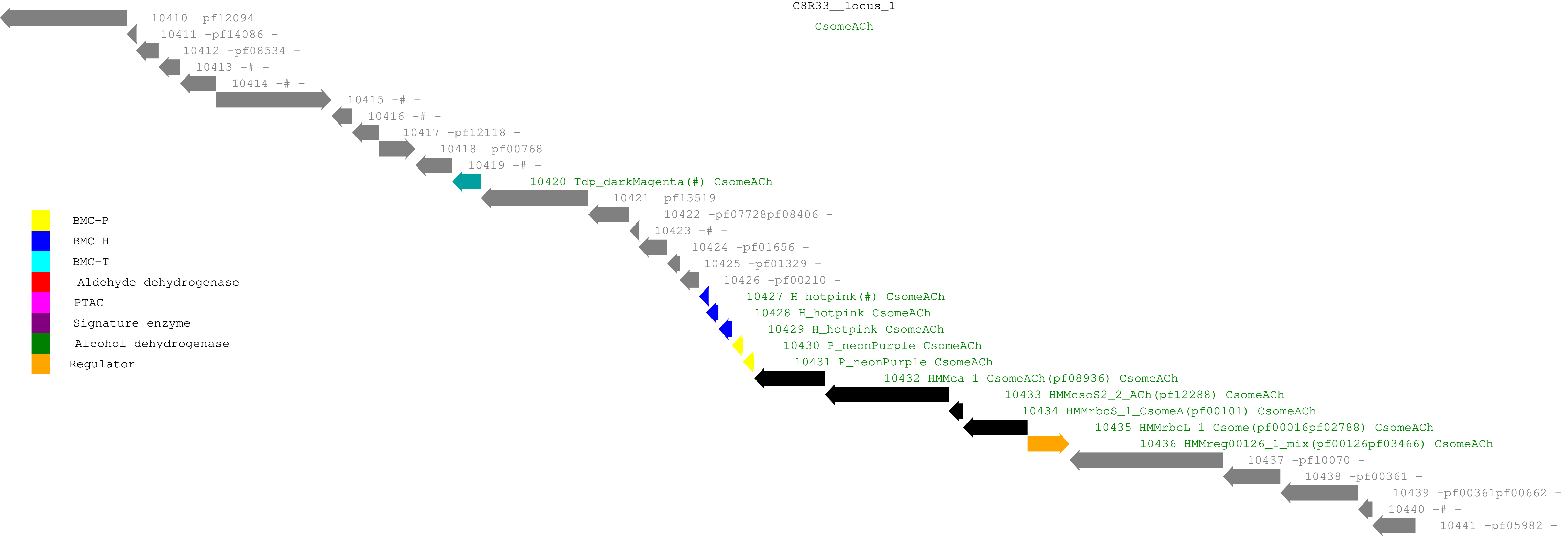

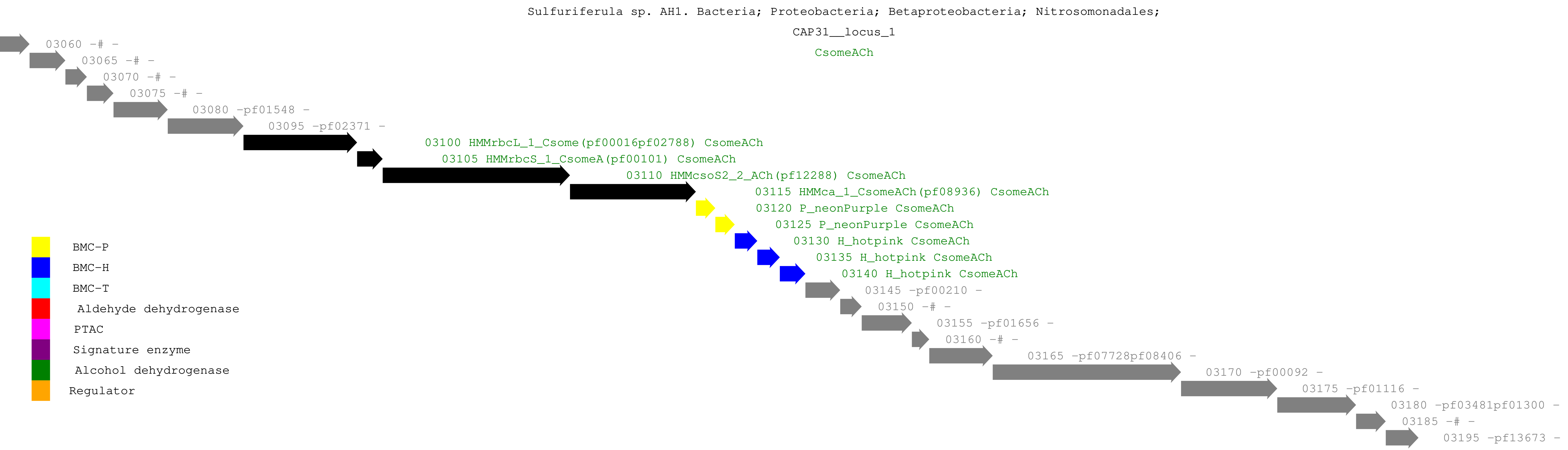
Sulfuriferula sp. AH1. Bacteria; Proteobacteria; Betaproteobacteria; Nitrosomonadales;
CAP31__locus_1
CsomeACh
03060 -# -
03065 -# -
03070 -# -
03075 -# -
03080 -pf01548 -
03095 -pf02371 -
03100 HMMrbcL_1_Csome(pf00016pf02788) CsomeACh
03105 HMMrbcS_1_CsomeA(pf00101) CsomeACh
03110 HMMcsoS2_2_ACh(pf12288) CsomeACh
03115 HMMca_1_CsomeACh(pf08936) CsomeACh
03120 P_neonPurple CsomeACh
03125 P_neonPurple CsomeACh
03130 H_hotpink CsomeACh
03135 H_hotpink CsomeACh
03140 H_hotpink CsomeACh
03145 -pf00210 -
03150 -# -
03155 -pf01656 -
03160 -# -
03165 -pf07728pf08406 -
03170 -pf00092 -
03175 -pf01116 -
03180 -pf03481pf01300 -
03185 -# -
03195 -pf13673 -
BMC-P
BMC-H
BMC-T
Aldehyde dehydrogenase
PTAC
Signature enzyme
Alcohol dehydrogenase
Regulator

mine drainage metagenome. unclassified sequences; metagenomes; ecological metagenomes.

CARN2__locus_1

CsomeACh

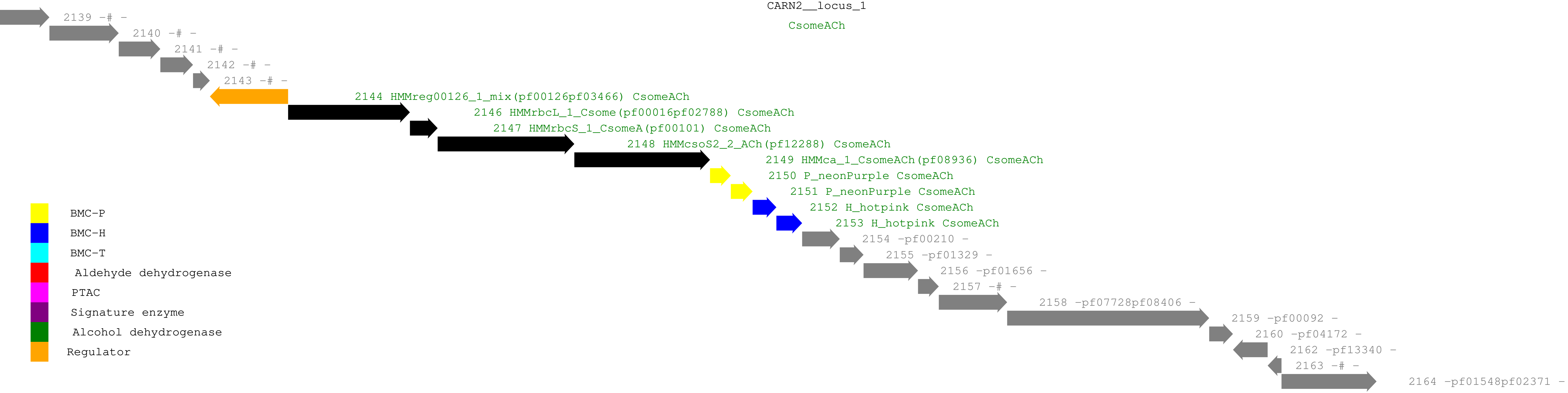

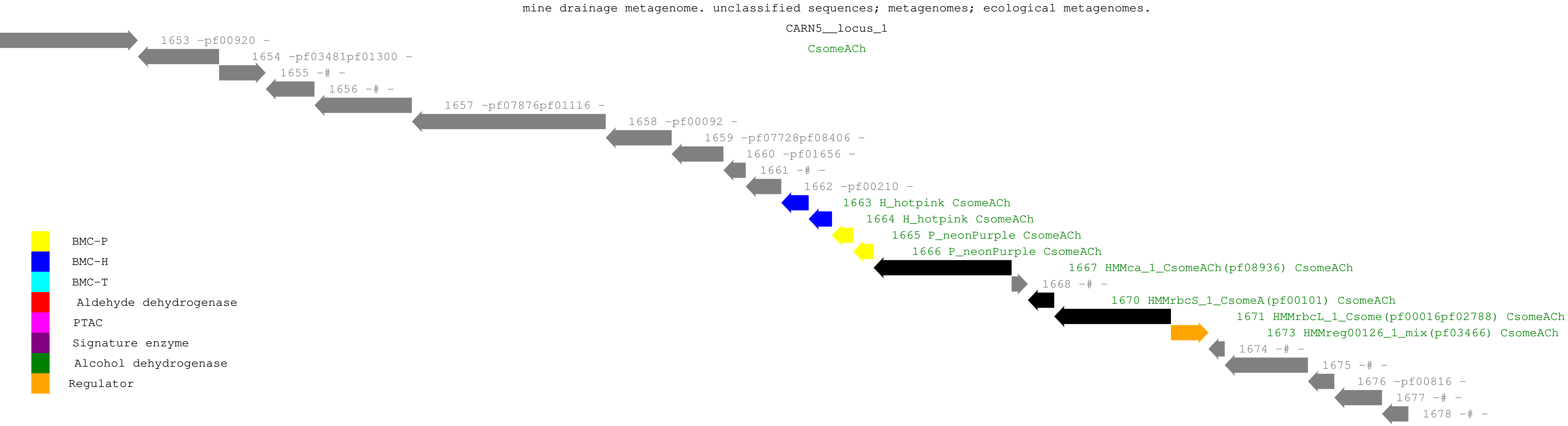
mine drainage metagenome. unclassified sequences; metagenomes; ecological metagenomes.
CARN5__locus_1
CsomeACh
1653 -pf00920 -
1654 -pf03481pf01300 -
1655 -# -
1656 -# -
1657 -pf07876pf01116 -
1658 -pf00092 -
1659 -pf07728pf08406 -
1660 -pf01656 -
1661 -# -
1662 -pf00210 -
1663 H_hotpink CsomeACh
1664 H_hotpink CsomeACh
1665 P_neonPurple CsomeACh
1666 P_neonPurple CsomeACh
1667 HMMca_1_CsomeACh(pf08936) CsomeACh
1668 -# -
1670 HMMrbcS_1_CsomeA(pf00101) CsomeACh
1671 HMMrbcL_1_Csome(pf00016pf02788) CsomeACh
1673 HMMreg00126_1_mix(pf03466) CsomeACh
1674 -# -
1675 -# -
1676 -pf00816 -
1677 -# -
1678 -# -
BMC-P
BMC-H
BMC-T
Aldehyde dehydrogenase
PTAC
Signature enzyme
Alcohol dehydrogenase
Regulator

mine drainage metagenome. unclassified sequences; metagenomes; ecological metagenomes.

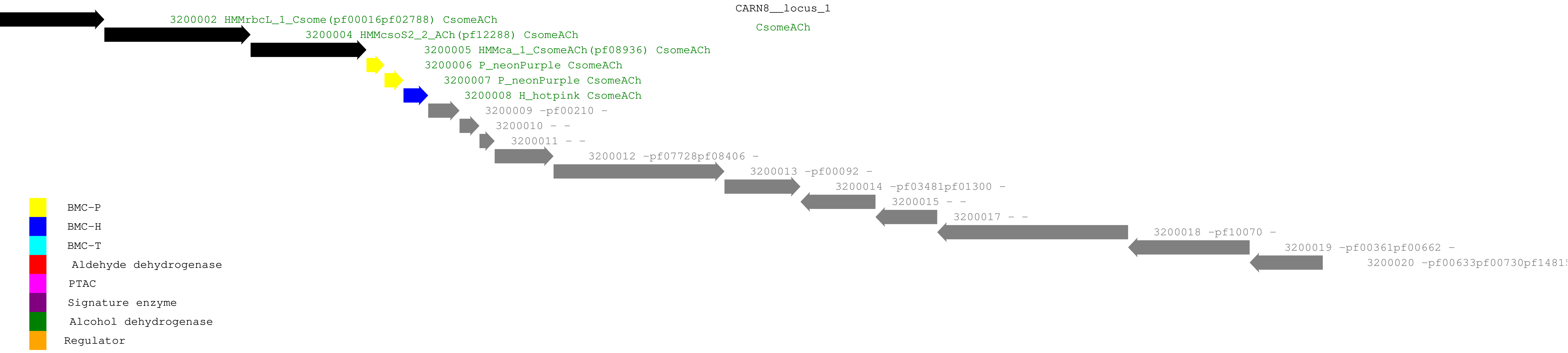

Oceanospirillales bacterium TMED59. Bacteria; Proteobacteria; Gammaproteobacteria; Oceanospirillales;

CBC19__locus_1

CsomeACh

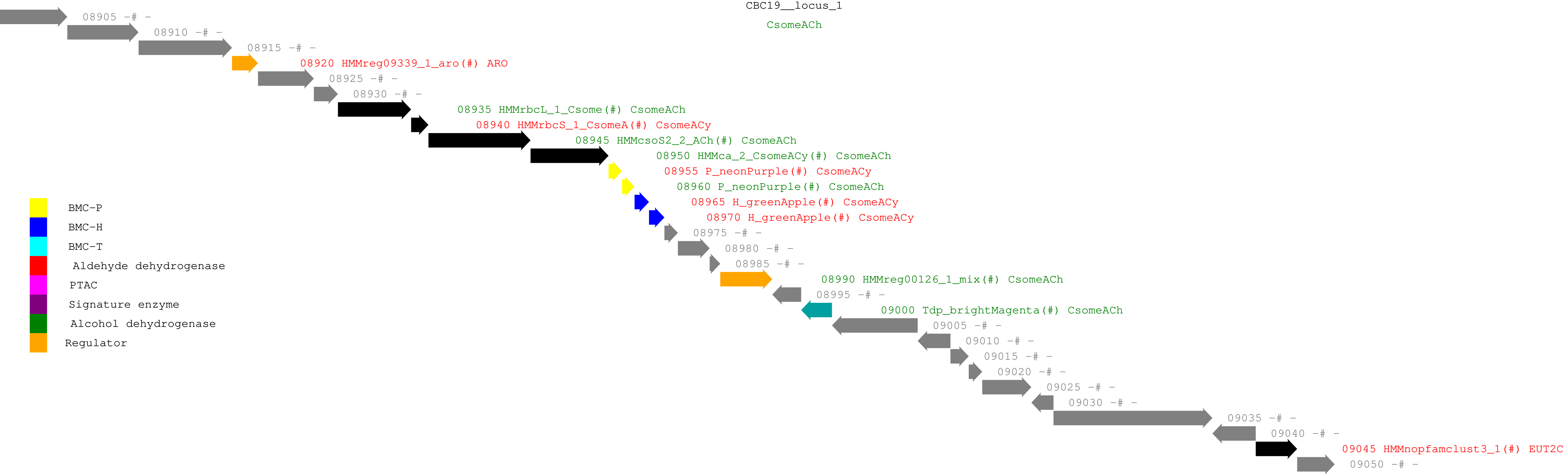

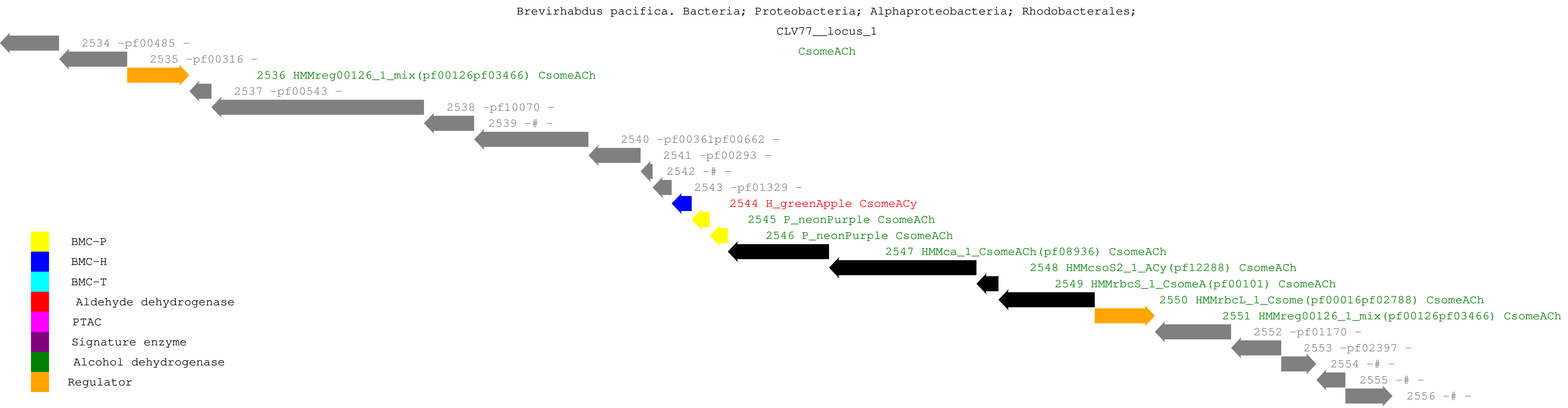
Brevirhabdus pacifica. Bacteria; Proteobacteria; Alphaproteobacteria; Rhodobacterales;
CLV77__locus_1
CsomeACh
2534 -pf00485 -
2535 -pf00316 -
2536 HMMreg00126_1_mix(pf00126pf03466) CsomeACh
2537 -pf00543 -
2538 -pf10070 -
2539 -# -
2540 -pf00361pf00662 -
2541 -pf00293 -
2542 -# -
2543 -pf01329 -
2544 H_greenApple CsomeACy
2545 P_neonPurple CsomeACh
2546 P_neonPurple CsomeACh
2547 HMMca_1_CsomeACh(pf08936) CsomeACh
2548 HMMcsoS2_1_ACy(pf12288) CsomeACh
2549 HMMrbcS_1_CsomeA(pf00101) CsomeACh
2550 HMMrbcL_1_Csome(pf00016pf02788) CsomeACh
2551 HMMreg00126_1_mix(pf00126pf03466) CsomeACh
2552 -pf01170 -
2553 -pf02397 -
2554 -# -
2555 -# -
2556 -# -
BMC-P
BMC-H
BMC-T
Aldehyde dehydrogenase
PTAC
Signature enzyme
Alcohol dehydrogenase
Regulator

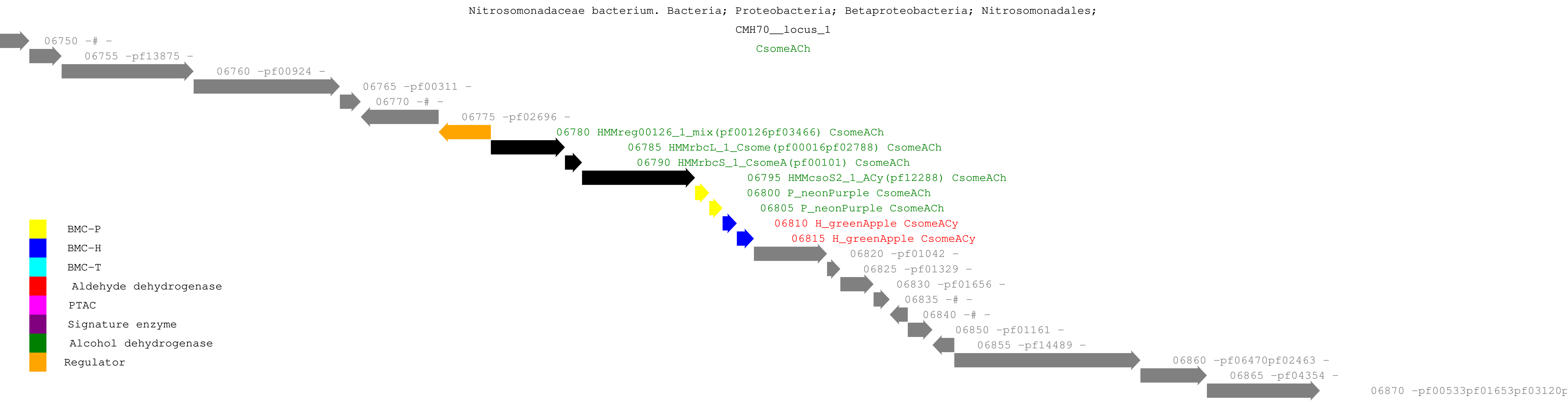

Nitrosomonadaceae bacterium. Bacteria; Proteobacteria; Betaproteobacteria; Nitrosomonadales;
CMH70__locus_1
CsomeACh
06750 -# -
06755 -pf13875 -
06760 -pf00924 -
06765 -pf00311 -
06770 -# -
06775 -pf02696 -
06780 HMMreg00126_1_mix(pf00126pf03466) CsomeACh
06785 HMMrbcL_1_Csome(pf00016pf02788) CsomeACh
06790 HMMrbcS_1_CsomeA(pf00101) CsomeACh
06795 HMMcsoS2_1_ACy(pf12288) CsomeACh
06800 P_neonPurple CsomeACh
06805 P_neonPurple CsomeACh
06810 H_greenApple CsomeACy
06815 H_greenApple CsomeACy
06820 -pf01042 -
06825 -pf01329 -
06830 -pf01656 -
06835 -# -
06840 -# -
06850 -pf01161 -
06855 -pf14489 -
06860 -pf06470pf02463 -
06865 -pf04354 -
06870 -pf00533pf01653pf03120p
BMC-P
BMC-H
BMC-T
Aldehyde dehydrogenase
PTAC
Signature enzyme
Alcohol dehydrogenase
Regulator

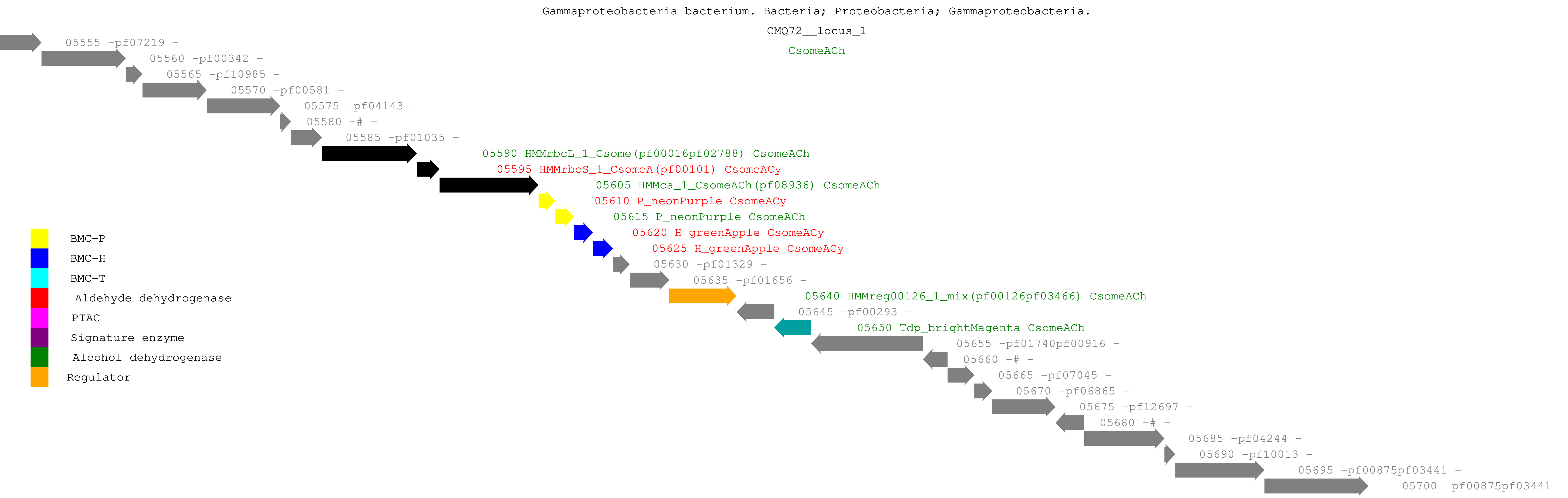

Gammaproteobacteria bacterium. Bacteria; Proteobacteria; Gammaproteobacteria.
CMQ72__locus_1
CsomeACh
05555 -pf07219 -
05560 -pf00342 -
05565 -pf10985 -
05570 -pf00581 -
05575 -pf04143 -
05580 -# -
05585 -pf01035 -
05590 HMMrbcL_1_Csome(pf00016pf02788) CsomeACh
05595 HMMrbcS_1_CsomeA(pf00101) CsomeACy
05605 HMMca_1_CsomeACh(pf08936) CsomeACh
05610 P_neonPurple CsomeACy
05615 P_neonPurple CsomeACh
05620 H_greenApple CsomeACy
05625 H_greenApple CsomeACy
05630 -pf01329 -
05635 -pf01656 -
05640 HMMreg00126_1_mix(pf00126pf03466) CsomeACh
05645 -pf00293 -
05650 Tdp_brightMagenta CsomeACh
05655 -pf01740pf00916 -
05660 -# -
05665 -pf07045 -
05670 -pf06865 -
05675 -pf12697 -
05680 -# -
05685 -pf04244 -
05690 -pf10013 -
05695 -pf00875pf03441 -
05700 -pf00875pf03441 -
BMC-P
BMC-H
BMC-T
Aldehyde dehydrogenase
PTAC
Signature enzyme
Alcohol dehydrogenase
Regulator

Gammaproteobacteria bacterium. Bacteria; Proteobacteria; Gammaproteobacteria.

CMQ84__locus_1

CsomeACh

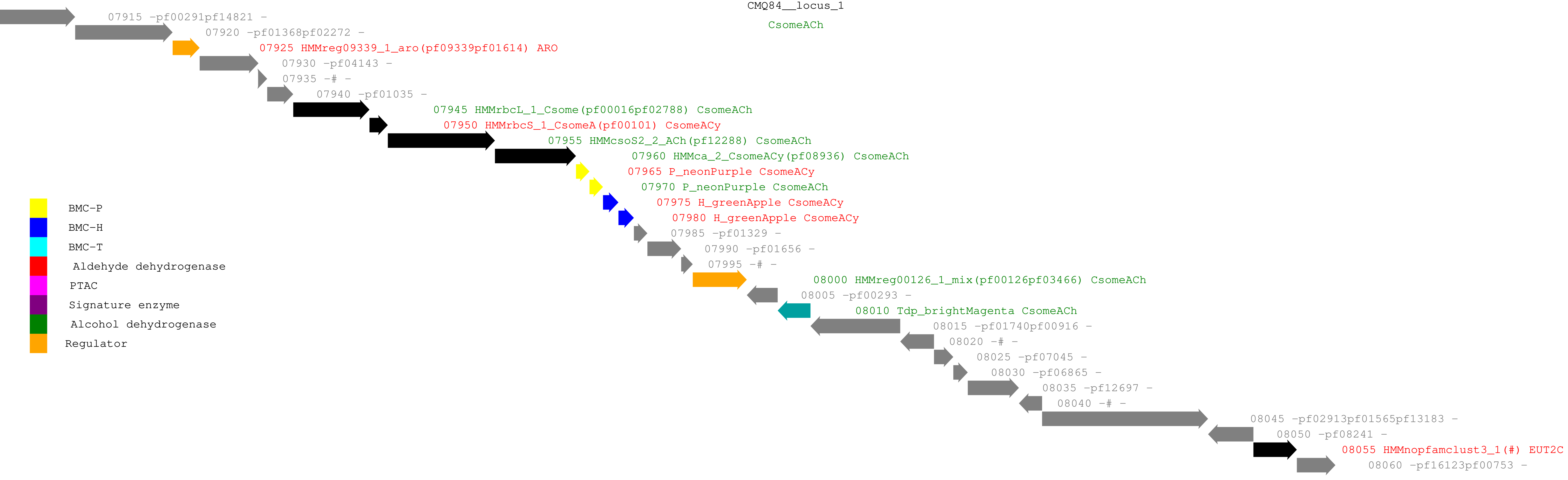

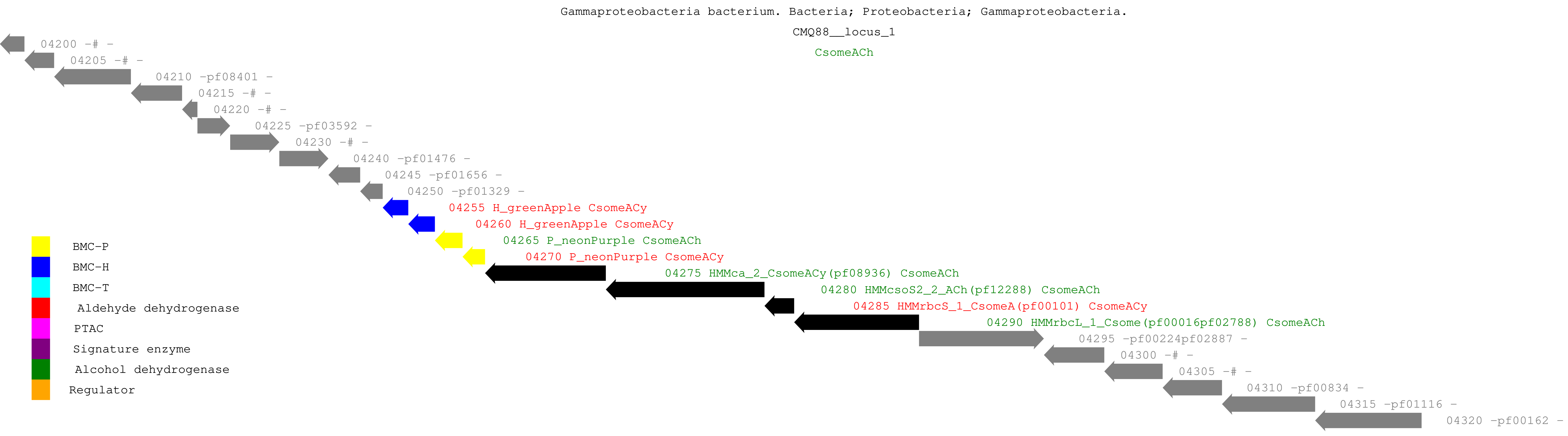

Gammaproteobacteria bacterium. Bacteria; Proteobacteria; Gammaproteobacteria.
CMQ88__locus_1
CsomeACh
04200 -# -
04205 -# -
04210 -pf08401 -
04215 -# -
04220 -# -
04225 -pf03592 -
04230 -# -
04240 -pf01476 -
04245 -pf01656 -
04250 -pf01329 -
04255 H_greenApple CsomeACy
04260 H_greenApple CsomeACy
04265 P_neonPurple CsomeACh
04270 P_neonPurple CsomeACy
04275 HMMca_2_CsomeACy(pf08936) CsomeACh
04280 HMMcsoS2_2_ACh(pf12288) CsomeACh
04285 HMMrbcS_1_CsomeA(pf00101) CsomeACy
04290 HMMrbcL_1_Csome(pf00016pf02788) CsomeACh
04295 -pf00224pf02887 -
04300 -# -
04305 -# -
04310 -pf00834 -
04315 -pf01116 -
04320 -pf00162 -
BMC-P
BMC-H
BMC-T
Aldehyde dehydrogenase
PTAC
Signature enzyme
Alcohol dehydrogenase
Regulator

Thiotrichaceae bacterium. Bacteria; Proteobacteria; Gammaproteobacteria; Thiotrichales;

COA54__locus_1

CsomeACh

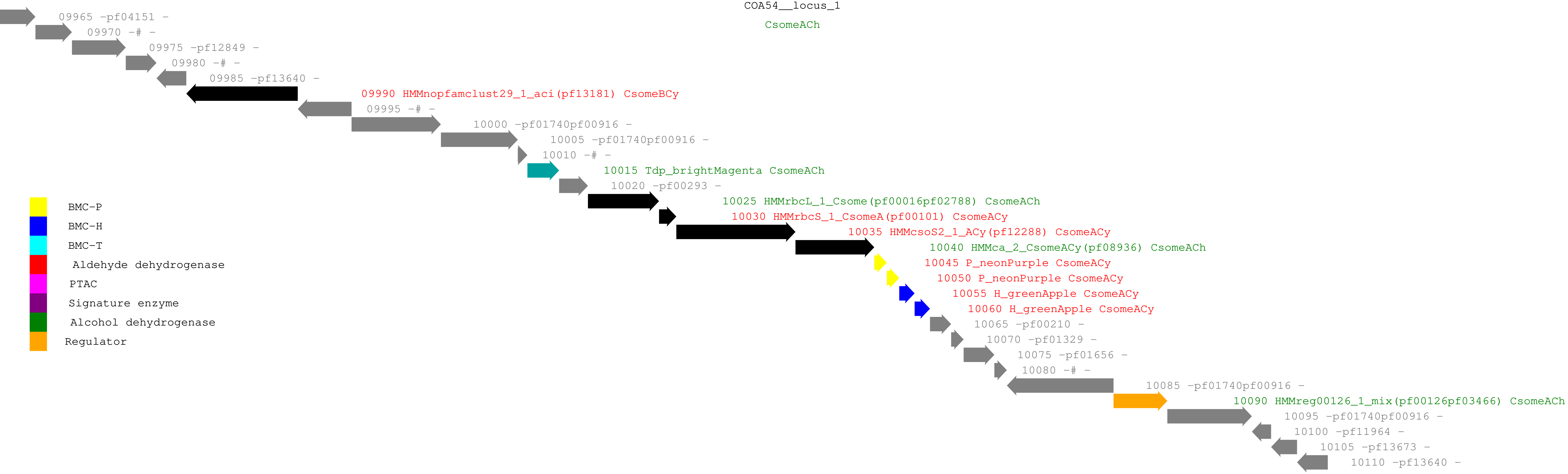

Thiotrichaceae bacterium. Bacteria; Proteobacteria; Gammaproteobacteria; Thiotrichales;

COB33__locus_1

CsomeACh

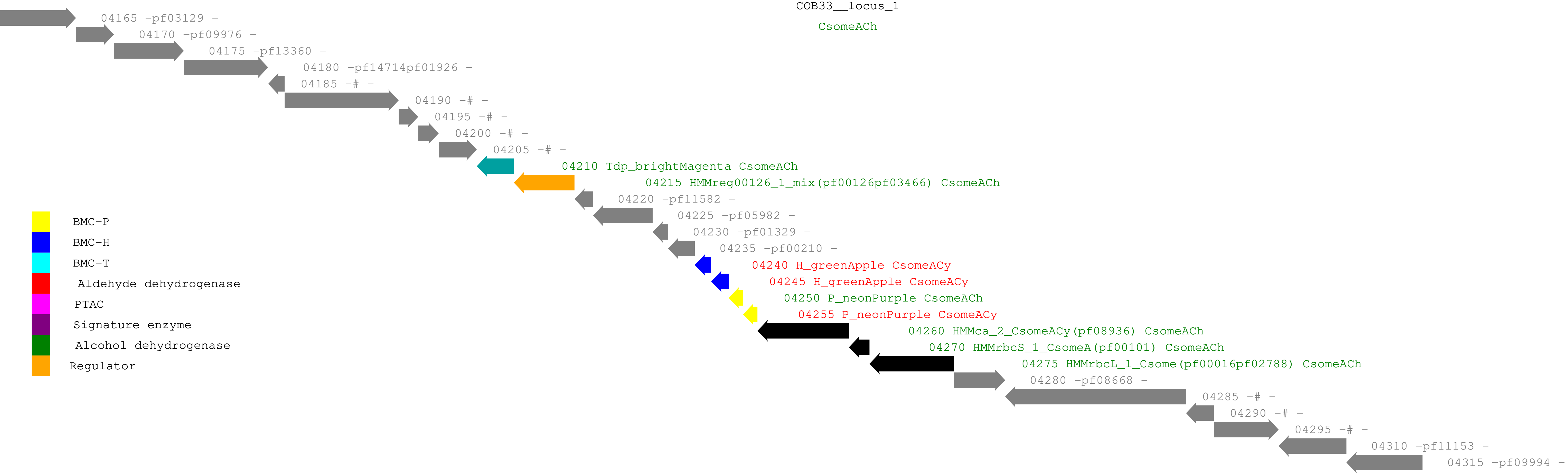

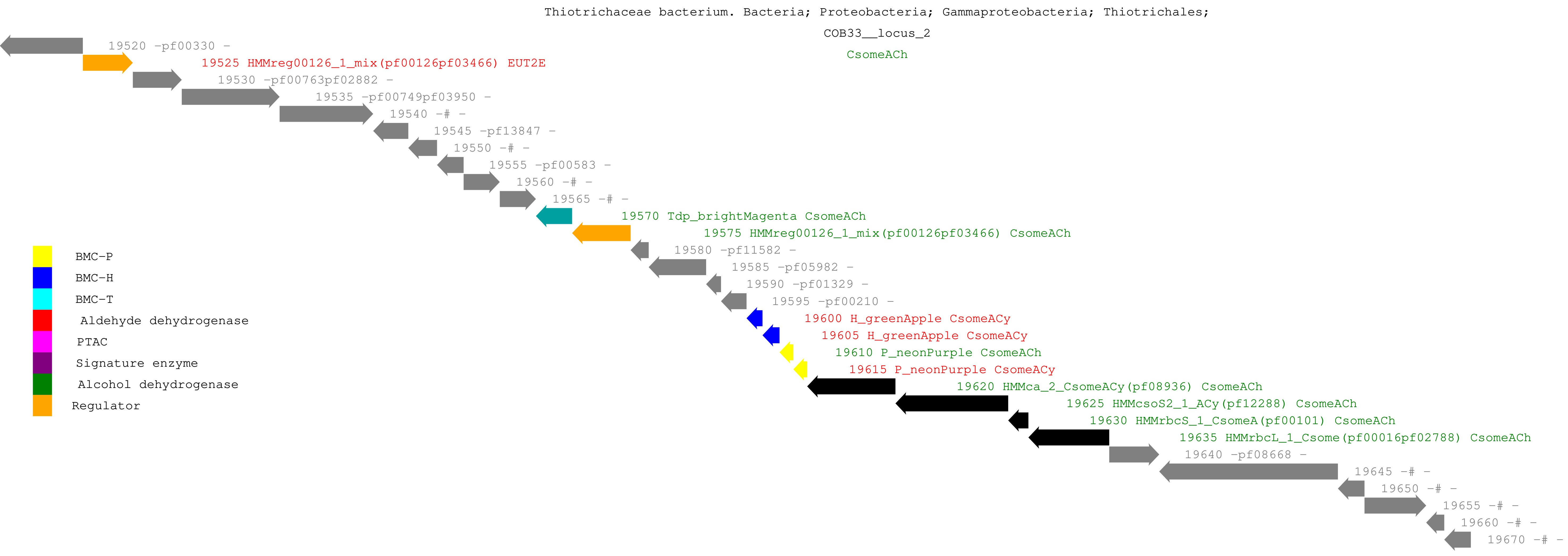
Thiotrichaceae bacterium. Bacteria; Proteobacteria; Gammaproteobacteria; Thiotrichales;
COB33__locus_2
CsomeACh
19520 -pf00330 -
19525 HMMreg00126_1_mix(pf00126pf03466) EUT2E
19530 -pf00763pf02882 -
19535 -pf00749pf03950 -
19540 -# -
19545 -pf13847 -
19550 -# -
19555 -pf00583 -
19560 -# -
19565 -# -
19570 Tdp_brightMagenta CsomeACh
19575 HMMreg00126_1_mix(pf00126pf03466) CsomeACh
19580 -pf11582 -
19585 -pf05982 -
19590 -pf01329 -
19595 -pf00210 -
19600 H_greenApple CsomeACy
19605 H_greenApple CsomeACy
19610 P_neonPurple CsomeACh
19615 P_neonPurple CsomeACy
19620 HMMca_2_CsomeACy(pf08936) CsomeACh
19625 HMMcsoS2_1_ACy(pf12288) CsomeACh
19630 HMMrbcS_1_CsomeA(pf00101) CsomeACh
19635 HMMrbcL_1_Csome(pf00016pf02788) CsomeACh
19640 -pf08668 -
19645 -# -
19650 -# -
19655 -# -
19660 -# -
19670 -# -
BMC-P
BMC-H
BMC-T
Aldehyde dehydrogenase
PTAC
Signature enzyme
Alcohol dehydrogenase
Regulator

Gammaproteobacteria bacterium. Bacteria; Proteobacteria; Gammaproteobacteria.

COC09__locus_1

CsomeACh

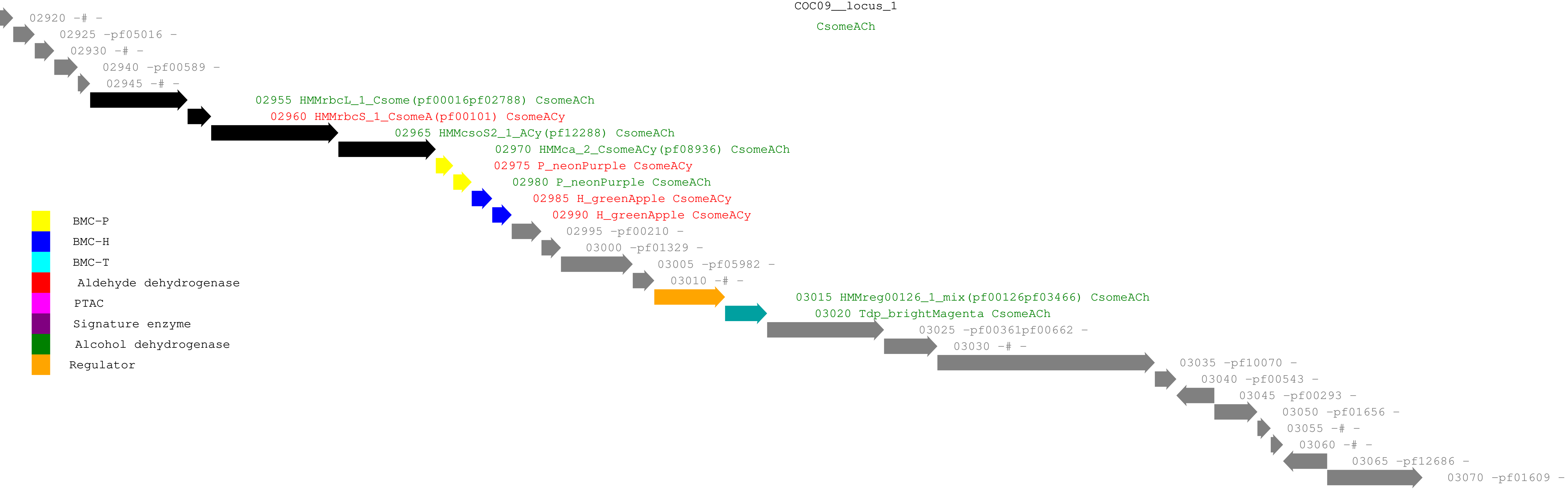

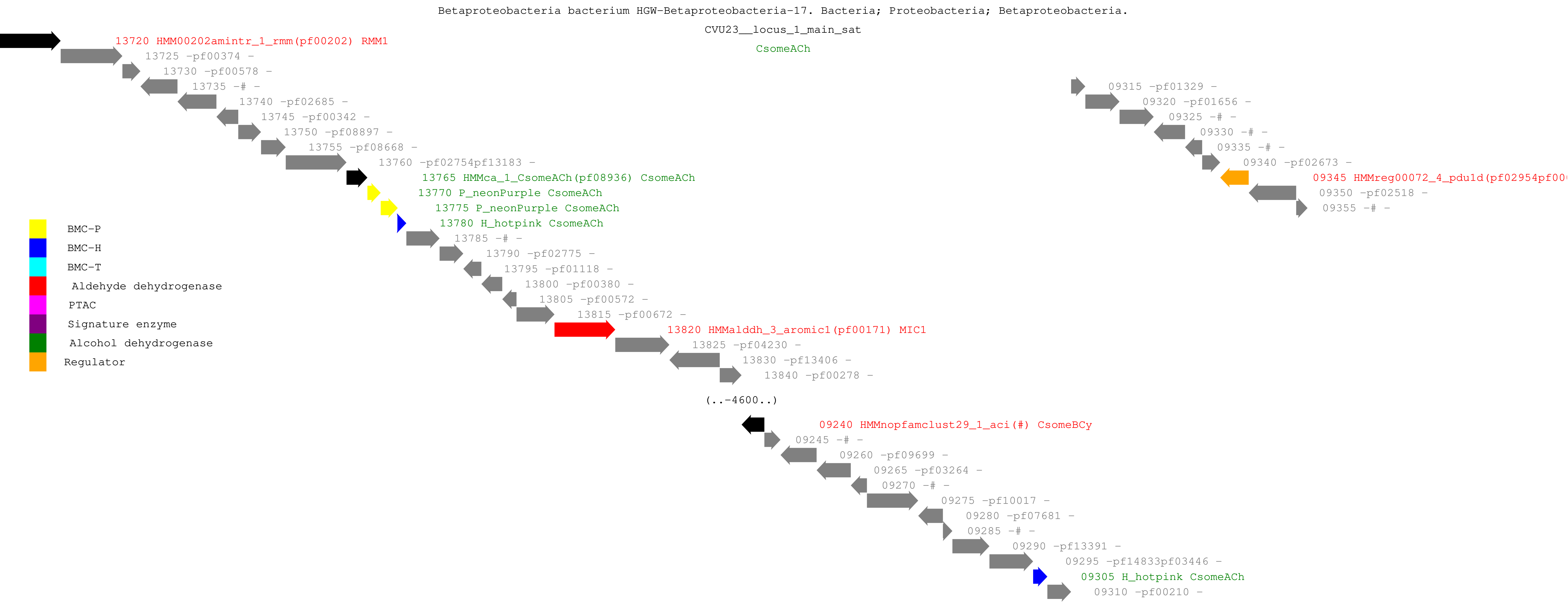

Betaproteobacteria bacterium HGW-Betaproteobacteria-17. Bacteria; Proteobacteria; Betaproteobacteria.
CVU23__locus_1_main_sat
CsomeACh
13720 HMM00202amintr_1_rmm(pf00202) RMM1
13725 -pf00374 -
13730 -pf00578 -
13735 -# -
13740 -pf02685 -
13745 -pf00342 -
13750 -pf08897 -
13755 -pf08668 -
13760 -pf02754pf13183 -
13765 HMMca_1_CsomeACh(pf08936) CsomeACh
13770 P_neonPurple CsomeACh
13775 P_neonPurple CsomeACh
13780 H_hotpink CsomeACh
13785 -# -
13790 -pf02775 -
13795 -pf01118 -
13800 -pf00380 -
13805 -pf00572 -
13815 -pf00672 -
13820 HMMaldh_3_aromic1(pf00171) MIC1
13825 -pf04230 -
13830 -pf13406 -
13840 -pf00278 -
(..-4600..)
09240 HMMnopfamclust29_1_aci(#) CsomeBCy
09245 -# -
09260 -pf09699 -
09265 -pf03264 -
09270 -# -
09275 -pf10017 -
09280 -pf07681 -
09285 -# -
09290 -pf13391 -
09295 -pf14833pf03446 -
09305 H_hotpink CsomeACh
09310 -pf00210 -
09315 -pf01329 -
09320 -pf01656 -
09325 -# -
09330 -# -
09335 -# -
09340 -pf02673 -
09345 HMMreg00072_4_pdu1d(pf02954pf00
09350 -pf02518 -
09355 -# -
BMC-P
BMC-H
BMC-T
Aldehyde dehydrogenase
PTAC
Signature enzyme
Alcohol dehydrogenase
Regulator

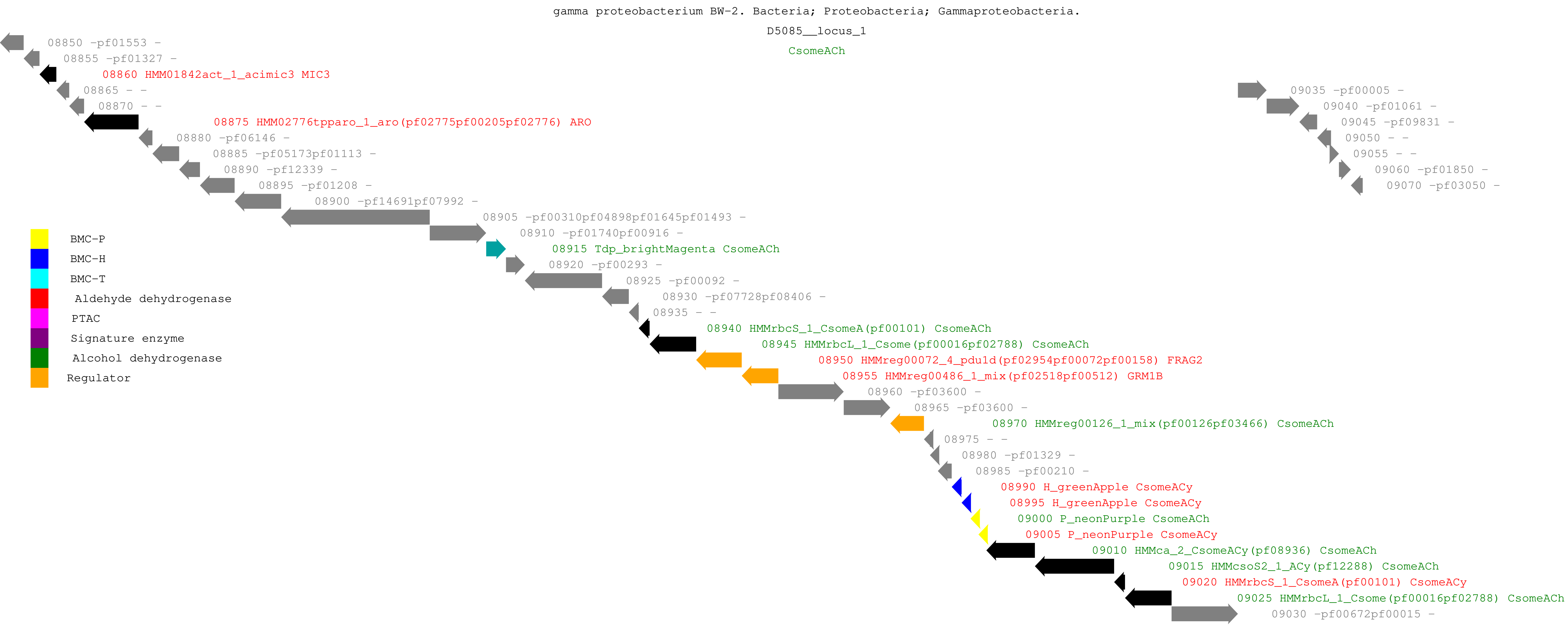
gamma proteobacterium BW-2. Bacteria; Proteobacteria; Gammaproteobacteria.
D5085__locus_1
CsomeACh
08850 -pf01553 -
08855 -pf01327 -
08860 HMM01842act_1_acimic3 MIC3
08865 - -
08870 - -
08875 HMM02776tpparo_1_aro(pf02775pf00205pf02776) ARO
08880 -pf06146 -
08885 -pf05173pf01113 -
08890 -pf12339 -
08895 -pf01208 -
08900 -pf14691pf07992 -
08905 -pf00310pf04898pf01645pf01493 -
08910 -pf01740pf00916 -
08915 Tdp_brightMagenta CsomeACh
08920 -pf00293 -
08925 -pf00092 -
08930 -pf07728pf08406 -
08935 - -
08940 HMMrbcS_1_CsomeA(pf00101) CsomeACh
08945 HMMrbcL_1_Csome(pf00016pf02788) CsomeACh
08950 HMMreg00072_4_pduld(pf02954pf00072pf00158) FRAG2
08955 HMMreg00486_1_mix(pf02518pf00512) GRM1B
08960 -pf03600 -
08965 -pf03600 -
08970 HMMreg00126_1_mix(pf00126pf03466) CsomeACh
08975 - -
08980 -pf01329 -
08985 -pf00210 -
08990 H_greenApple CsomeACy
08995 H_greenApple CsomeACy
09000 P_neonPurple CsomeACh
09005 P_neonPurple CsomeACy
09010 HMMca_2_CsomeACy(pf08936) CsomeACh
09015 HMMcsoS2_1_ACy(pf12288) CsomeACh
09020 HMMrbcS_1_CsomeA(pf00101) CsomeACy
09025 HMMrbcL_1_Csome(pf00016pf02788) CsomeACh
09030 -pf00672pf00015 -
09035 -pf00005 -
09040 -pf01061 -
09045 -pf09831 -
09050 - -
09055 - -
09060 -pf01850 -
09070 -pf03050 -
BMC-P
BMC-H
BMC-T
Aldehyde dehydrogenase
PTAC
Signature enzyme
Alcohol dehydrogenase
Regulator

Guyparkeria sp. SCN-R1. Bacteria; Proteobacteria; Gammaproteobacteria; Chromatiales;

D5687__locus_1_main_sat

CsomeACh

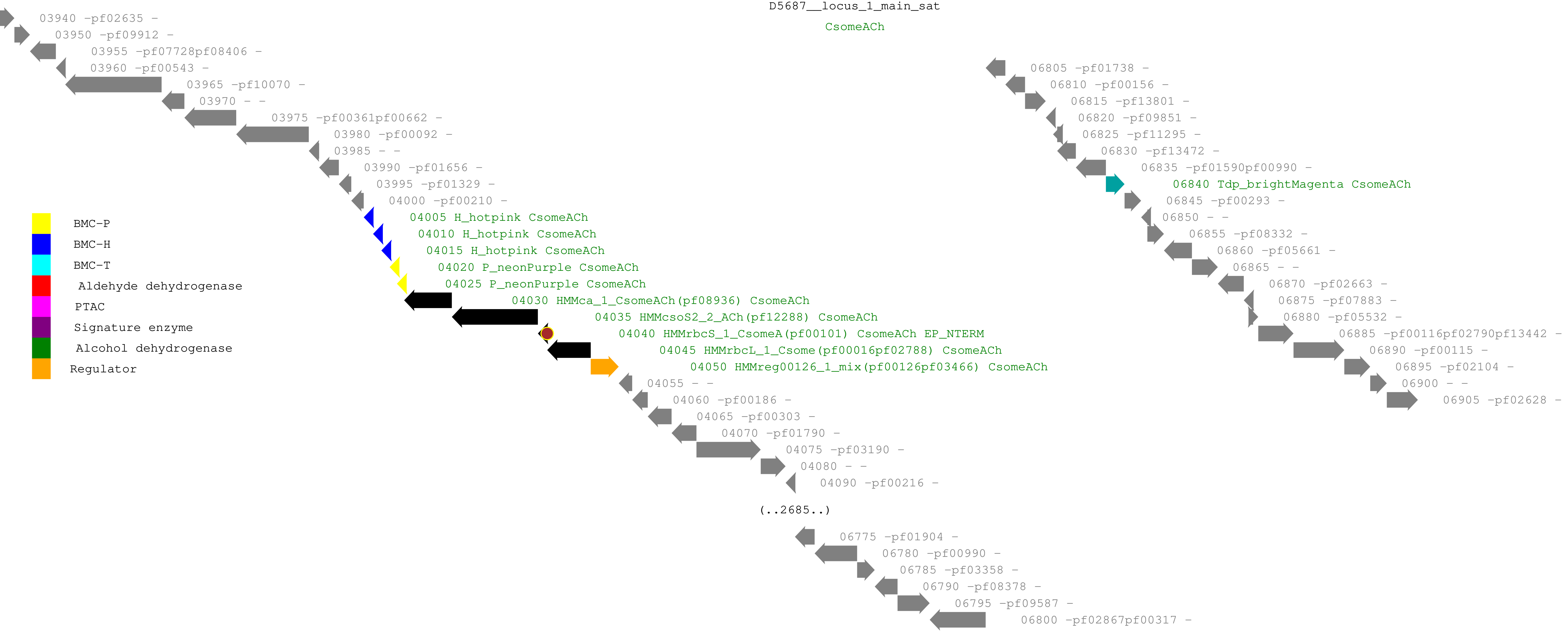

Thiohalobacter thiocyanaticus. Bacteria; Proteobacteria; Gammaproteobacteria; Chromatiales;

D6C00__locus_1

CsomeACh

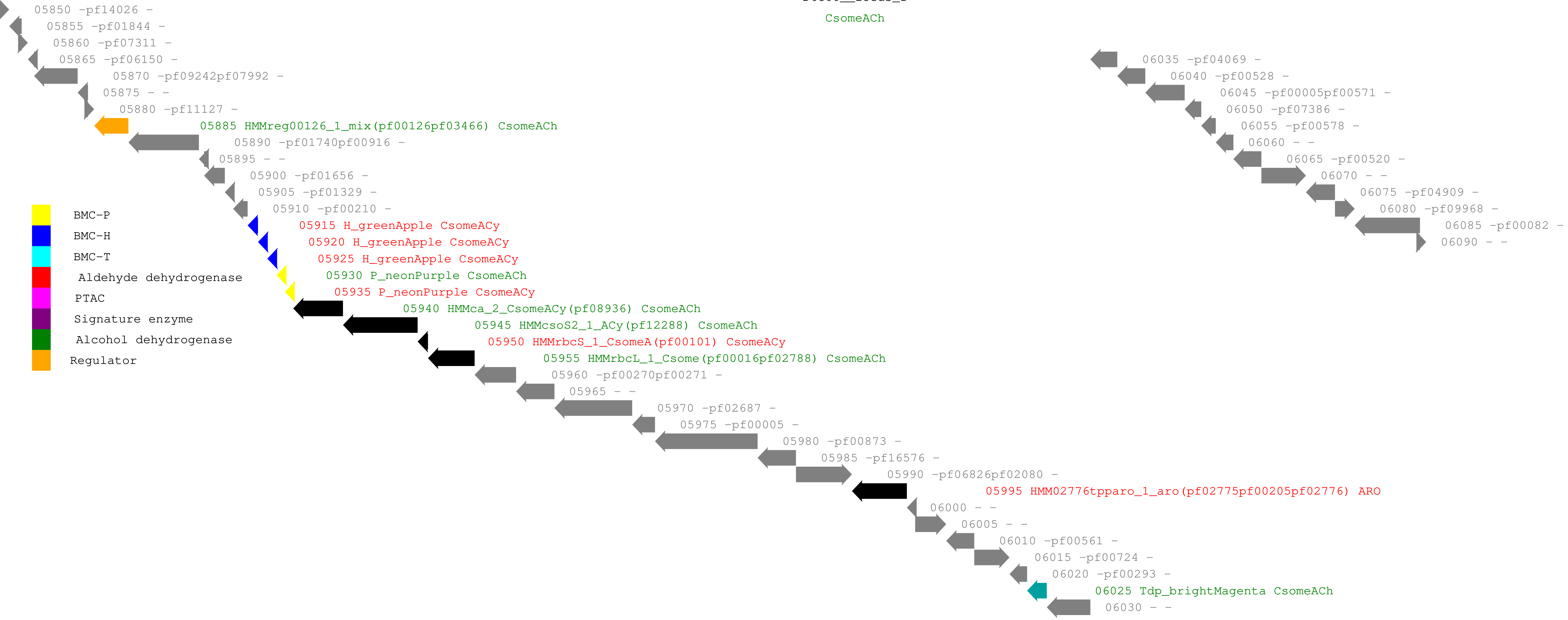

Thiorhodococcus sp. AK35. Bacteria; Proteobacteria; Gammaproteobacteria; Chromatiales;

D779__locus_1

CsomeACh

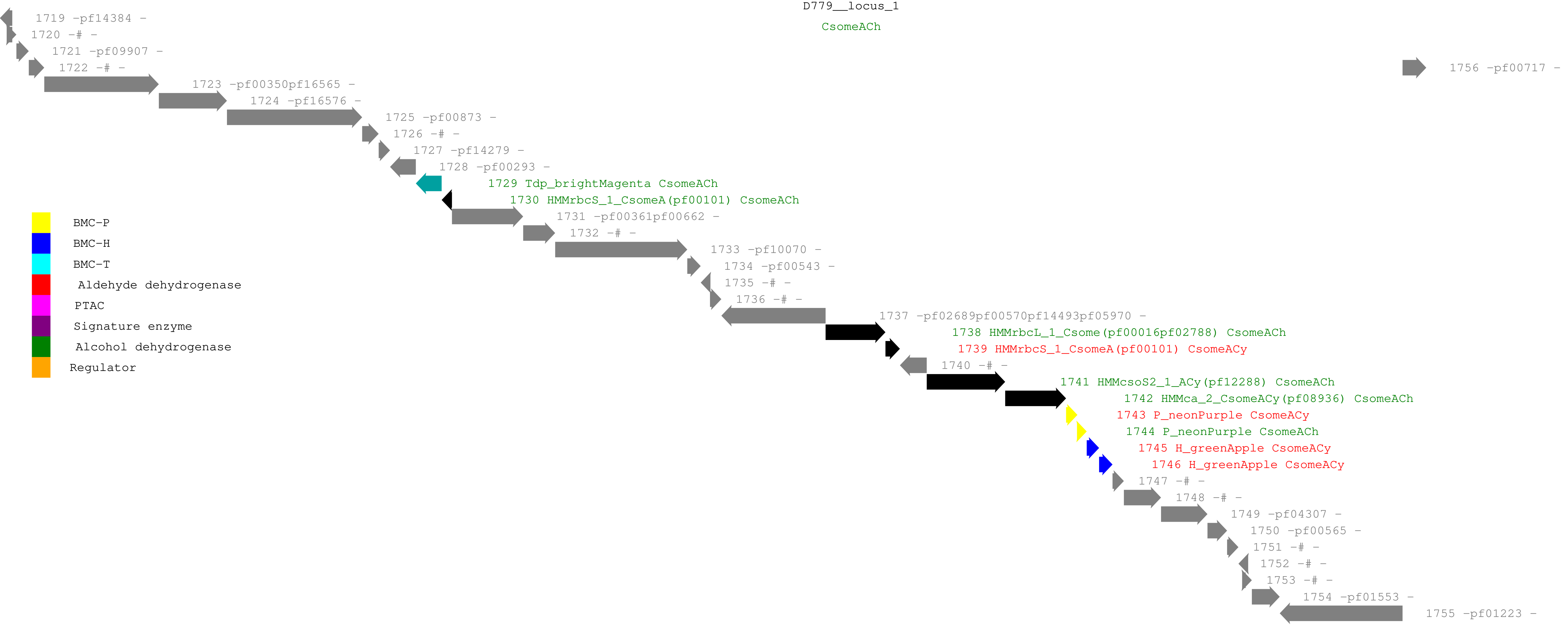

Thioalkalivibrio sp. ALE21. Bacteria; Proteobacteria; Gammaproteobacteria; Chromatiales;

D893__locus_1_main_sat

CsomeACh

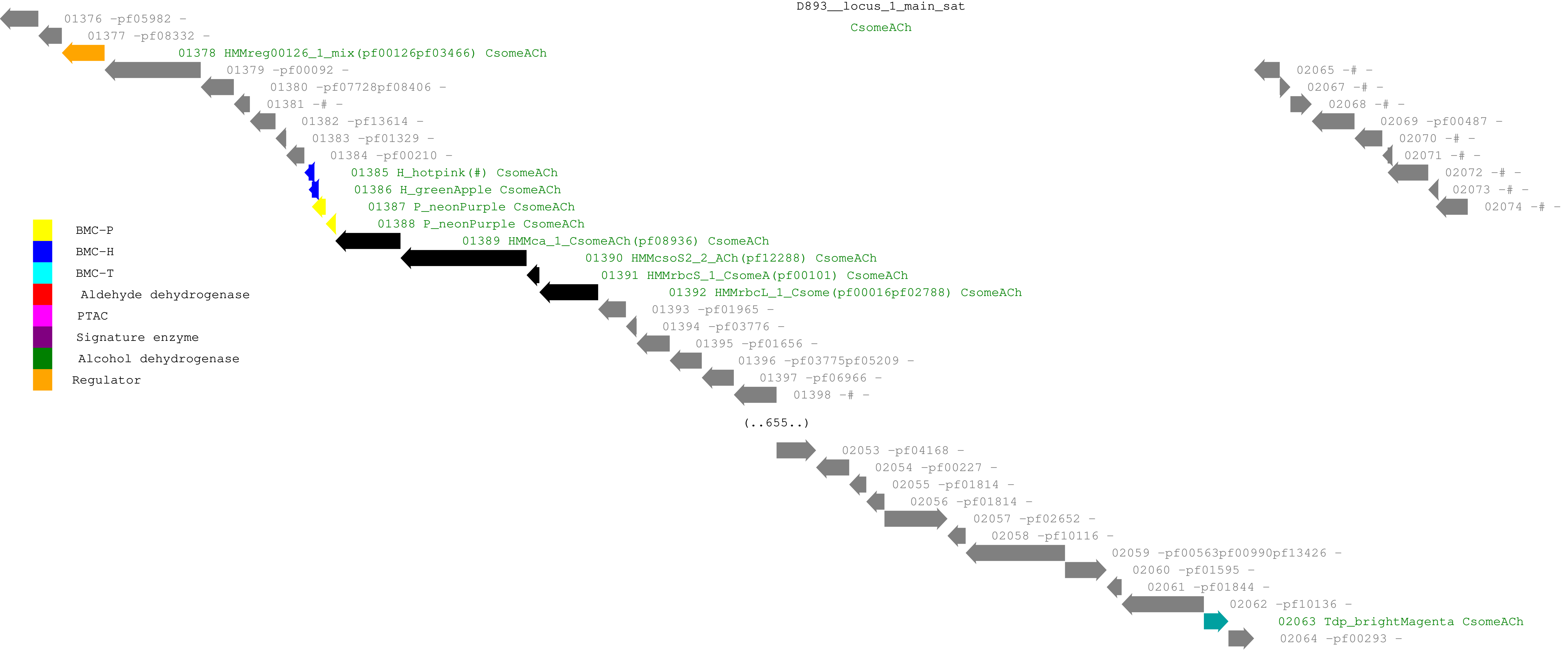

Gammaproteobacteria bacterium. Bacteria; Proteobacteria; Gammaproteobacteria.

DCE57__locus_1

CsomeACh

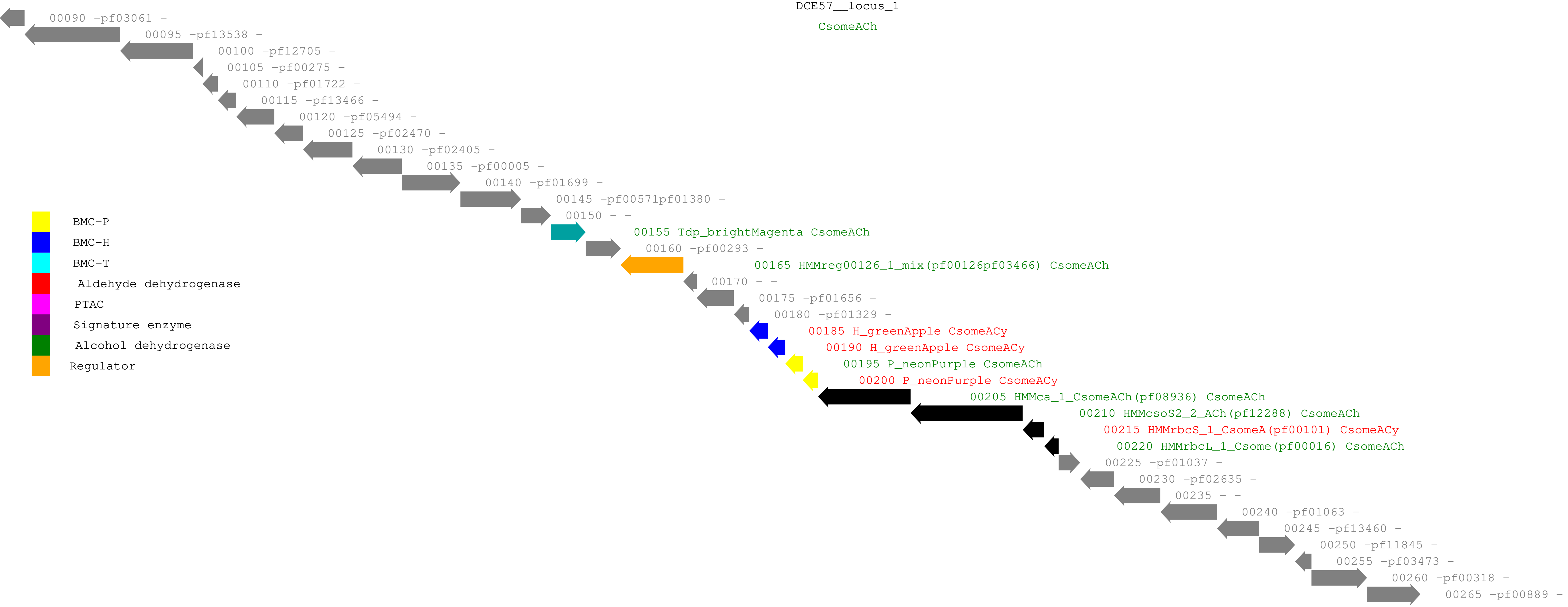

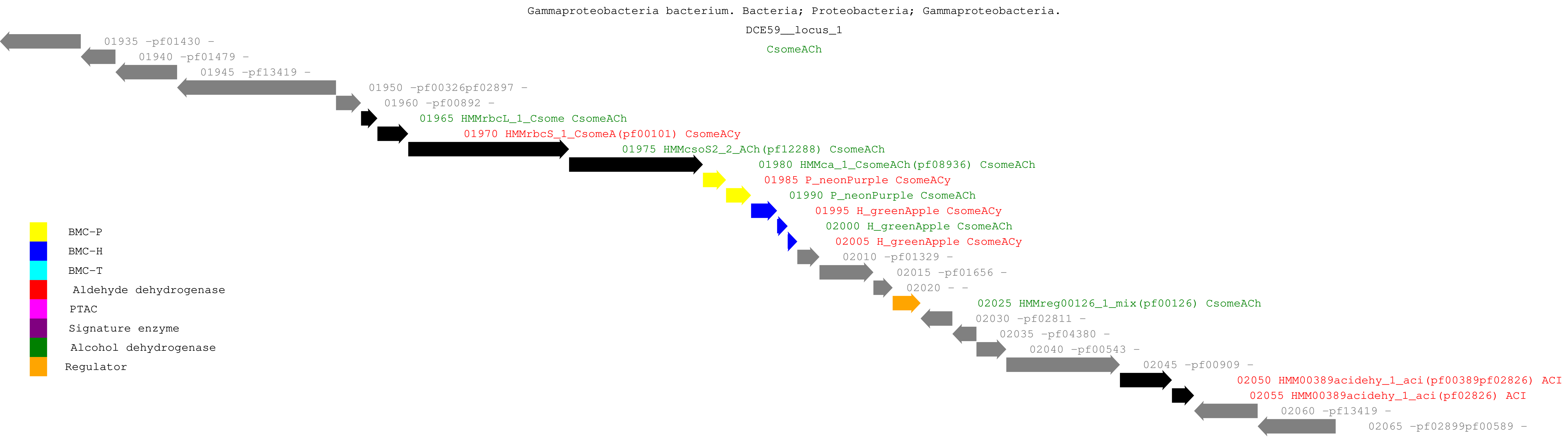

Gammaproteobacteria bacterium. Bacteria; Proteobacteria; Gammaproteobacteria.
DCE59__locus_1
CsomeACh
01935 -pf01430 -
01940 -pf01479 -
01945 -pf13419 -
01950 -pf00326pf02897 -
01960 -pf00892 -
01965 HMMrbcL_1_Csome CsomeACh
01970 HMMrbcS_1_CsomeA(pf00101) CsomeACy
01975 HMMcsoS2_2_ACh(pf12288) CsomeACh
01980 HMMca_1_CsomeACh(pf08936) CsomeACh
01985 P_neonPurple CsomeACy
01990 P_neonPurple CsomeACh
01995 H_greenApple CsomeACy
02000 H_greenApple CsomeACh
02005 H_greenApple CsomeACy
02010 -pf01329 -
02015 -pf01656 -
02020 - -
02025 HMMreg00126_1_mix(pf00126) CsomeACh
02030 -pf02811 -
02035 -pf04380 -
02040 -pf00543 -
02045 -pf00909 -
02050 HMM00389acidehy_1_aci(pf00389pf02826) ACI
02055 HMM00389acidehy_1_aci(pf02826) ACI
02060 -pf13419 -
02065 -pf02899pf00589 -
BMC-P
BMC-H
BMC-T
Aldehyde dehydrogenase
PTAC
Signature enzyme
Alcohol dehydrogenase
Regulator

Thiomicrospira sp. Bacteria; Proteobacteria; Gammaproteobacteria; Thiotrichales;

DD716__locus_fragmented

CsomeACh

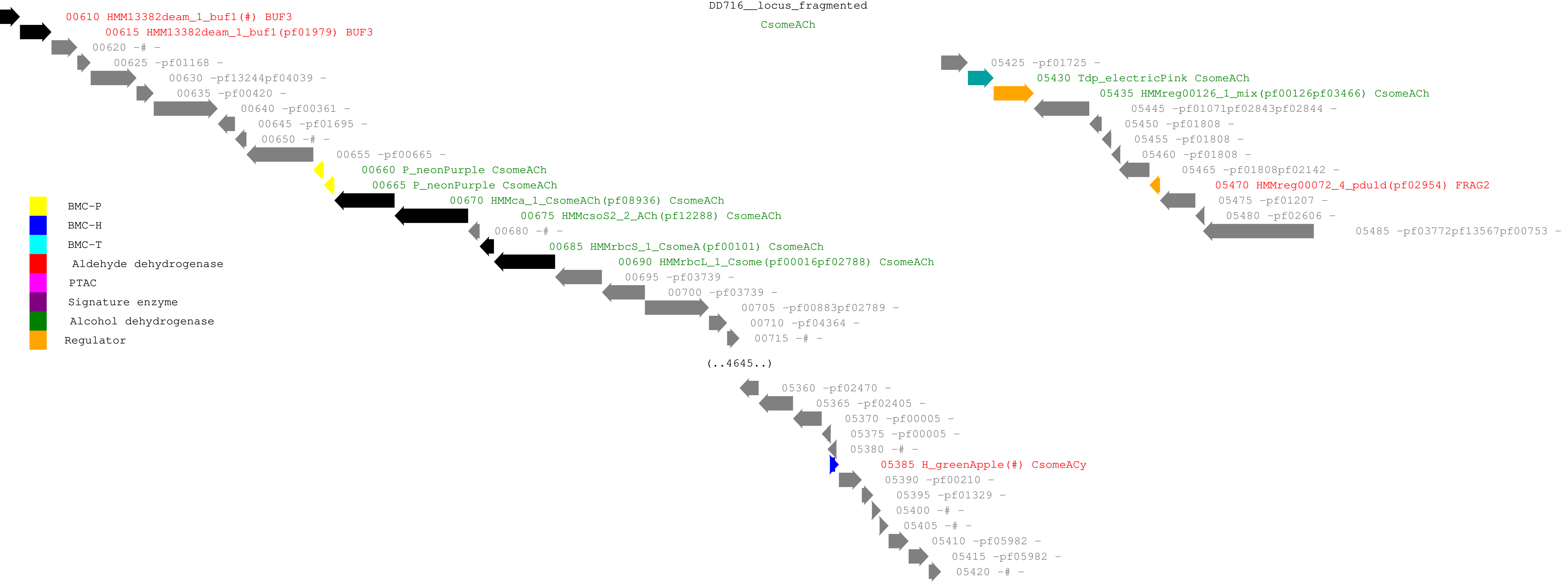

Gammaproteobacteria bacterium. Bacteria; Proteobacteria; Gammaproteobacteria.

DD979__locus_1

CsomeACh

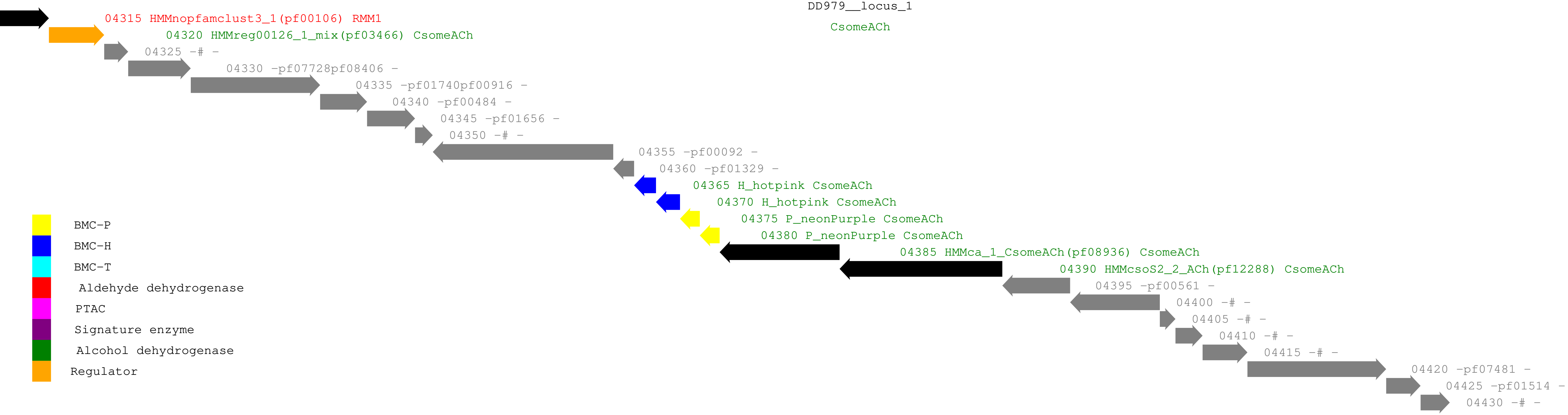

Chromatiaceae bacterium. Bacteria; Proteobacteria; Gammaproteobacteria; Chromatiales;

DDY14__locus_1_main_sat

CsomeACh

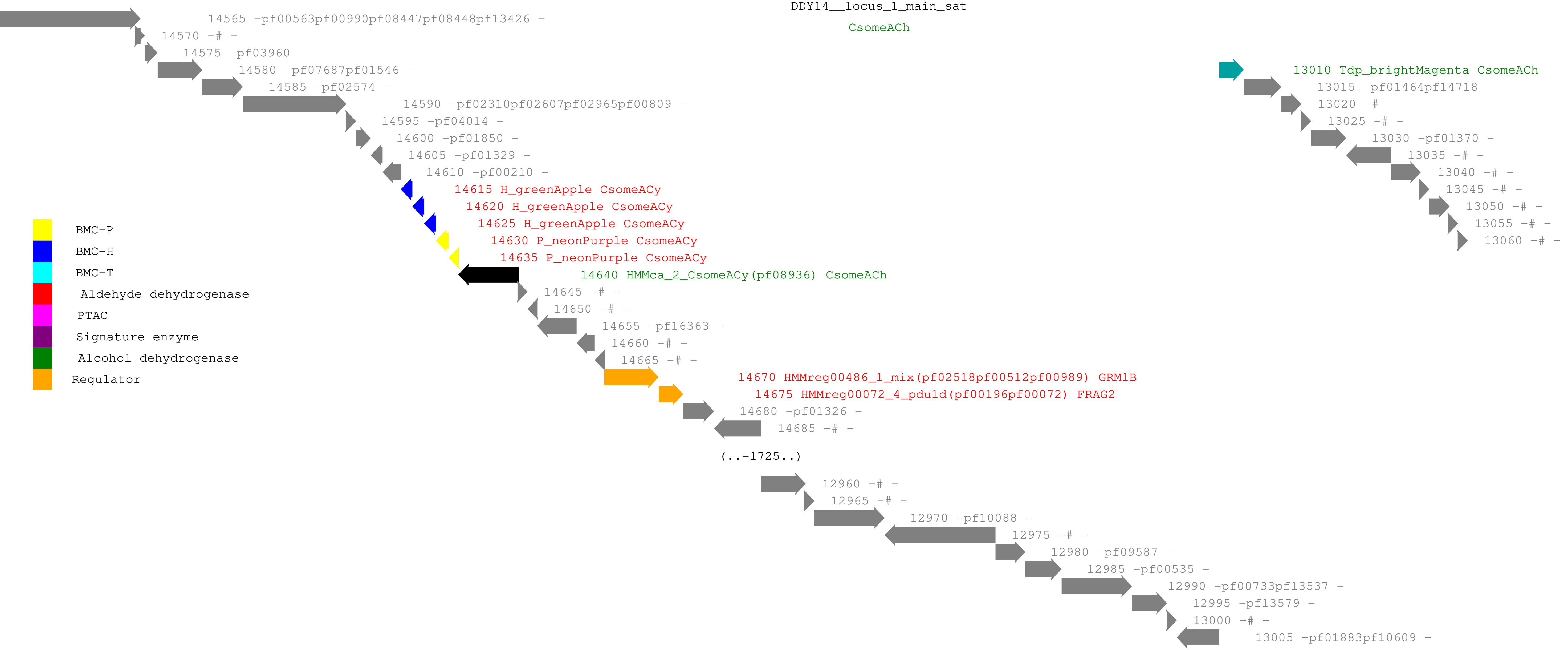

Thiobacillus sp. Bacteria; Proteobacteria; Betaproteobacteria; Nitrosomonadales;

DEB56__locus_1

CsomeACh

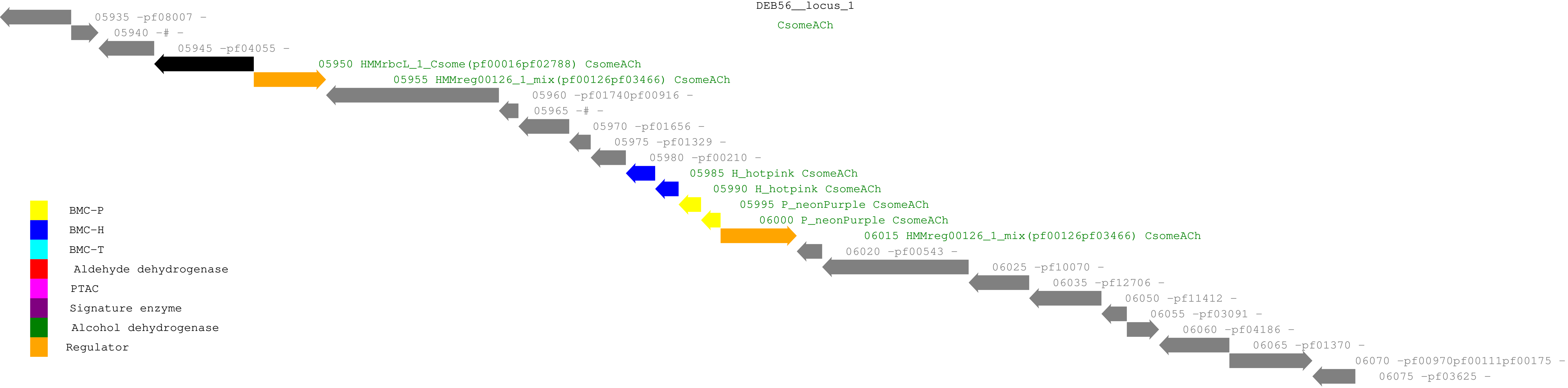

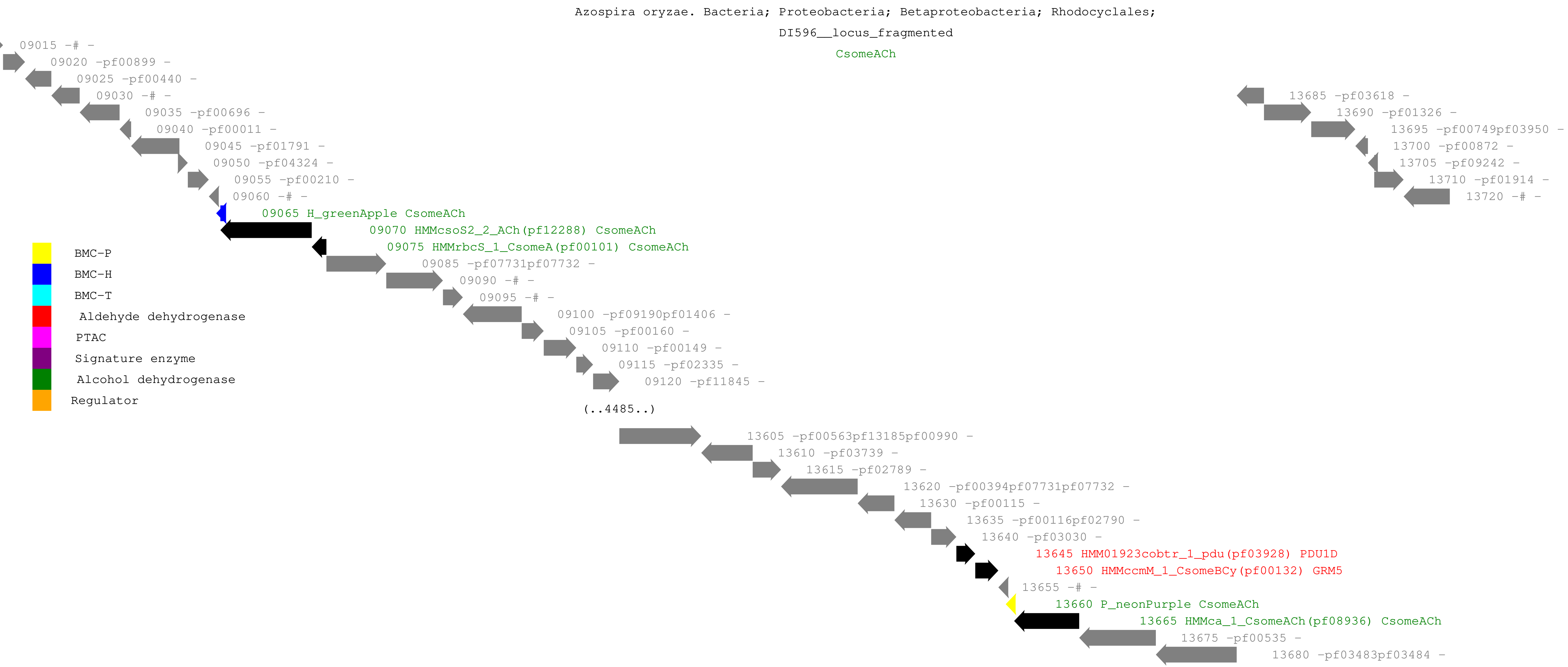
Azospira oryzae. Bacteria; Proteobacteria; Betaproteobacteria; Rhodocyclales;
DI596__locus_fragmented
CsomeACh
09015 -# -
09020 -pf00899 -
09025 -pf00440 -
09030 -# -
09035 -pf00696 -
09040 -pf00011 -
09045 -pf01791 -
09050 -pf04324 -
09055 -pf00210 -
09060 -# -
09065 H_greenApple CsomeACh
09070 HMMcsoS2_2_ACh(pf12288) CsomeACh
09075 HMMrbcS_1_CsomeA(pf00101) CsomeACh
09085 -pf07731pf07732 -
09090 -# -
09095 -# -
09100 -pf09190pf01406 -
09105 -pf00160 -
09110 -pf00149 -
09115 -pf02335 -
09120 -pf11845 -
(..4485..)
13605 -pf00563pf13185pf00990 -
13610 -pf03739 -
13615 -pf02789 -
13620 -pf00394pf07731pf07732 -
13630 -pf00115 -
13635 -pf00116pf02790 -
13640 -pf03030 -
13645 HMM01923cobtr_1_pdu(pf03928) PDU1D
13650 HMMccmM_1_CsomeBCy(pf00132) GRM5
13655 -# -
13660 P_neonPurple CsomeACh
13665 HMMca_1_CsomeACh(pf08936) CsomeACh
13675 -pf00535 -
13680 -pf03483pf03484 -
13685 -pf03618 -
13690 -pf01326 -
13695 -pf00749pf03950 -
13700 -pf00872 -
13705 -pf09242 -
13710 -pf01914 -
13720 -# -
BMC-P
BMC-H
BMC-T
Aldehyde dehydrogenase
PTAC
Signature enzyme
Alcohol dehydrogenase
Regulator

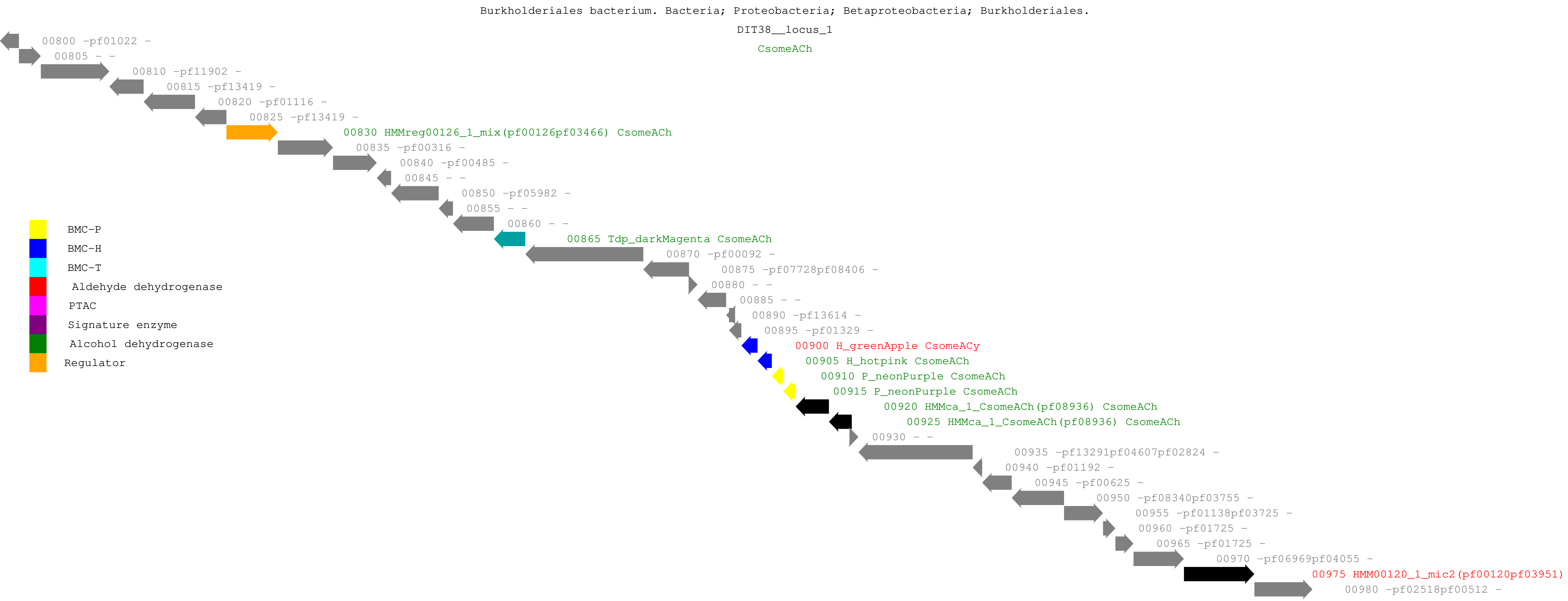
Burkholderiales bacterium. Bacteria; Proteobacteria; Betaproteobacteria; Burkholderiales.
DIT38__locus_1
CsomeACh
00800 -pf01022 -
00805 - -
00810 -pf11902 -
00815 -pf13419 -
00820 -pf01116 -
00825 -pf13419 -
00830 HMMreg00126_1_mix(pf00126pf03466) CsomeACh
00835 -pf00316 -
00840 -pf00485 -
00845 - -
00850 -pf05982 -
00855 - -
00860 - -
00865 Tdp_darkMagenta CsomeACh
00870 -pf00092 -
00875 -pf07728pf08406 -
00880 - -
00885 - -
00890 -pf13614 -
00895 -pf01329 -
00900 H_greenApple CsomeACy
00905 H_hotpink CsomeACh
00910 P_neonPurple CsomeACh
00915 P_neonPurple CsomeACh
00920 HMMca_1_CsomeACh(pf08936) CsomeACh
00925 HMMca_1_CsomeACh(pf08936) CsomeACh
00930 - -
00935 -pf13291pf04607pf02824 -
00940 -pf01192 -
00945 -pf00625 -
00950 -pf08340pf03755 -
00955 -pf01138pf03725 -
00960 -pf01725 -
00965 -pf01725 -
00970 -pf06969pf04055 -
00975 HMM00120_1_mic2(pf00120pf03951)
00980 -pf02518pf00512 -
BMC-P
BMC-H
BMC-T
Aldehyde dehydrogenase
PTAC
Signature enzyme
Alcohol dehydrogenase
Regulator

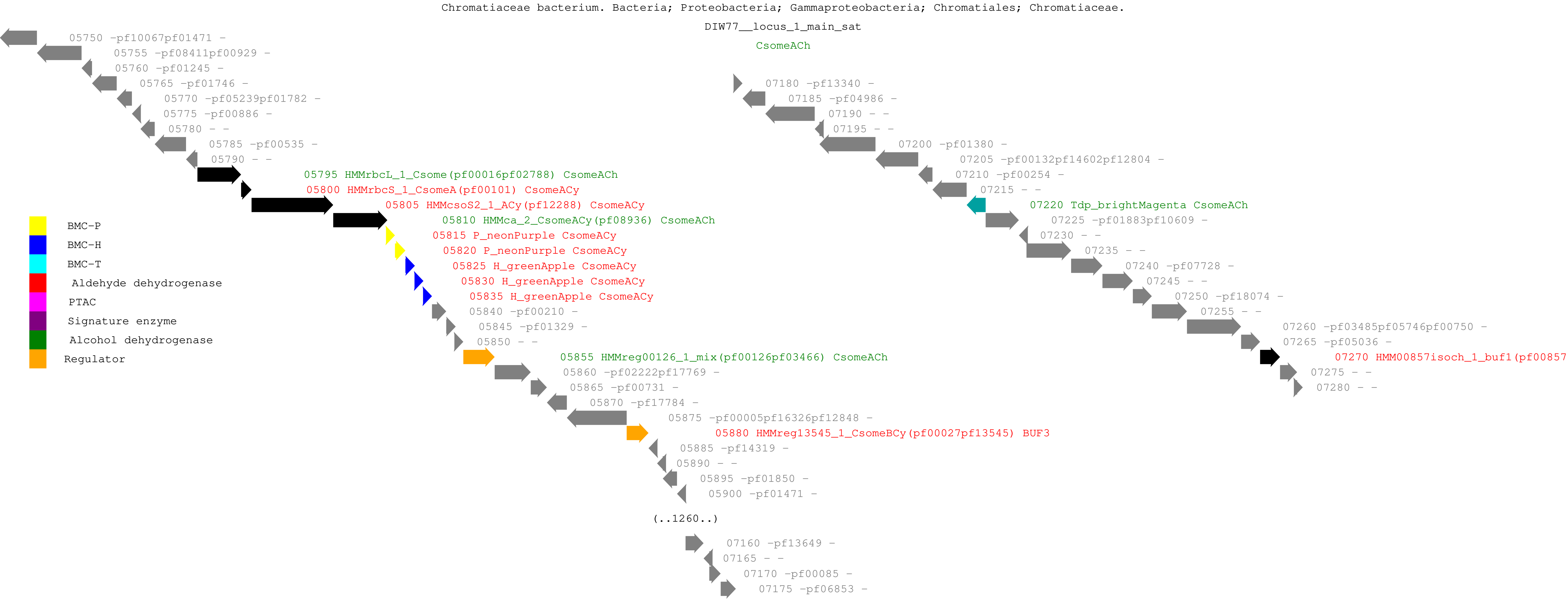

Chromatiaceae bacterium. Bacteria; Proteobacteria; Gammaproteobacteria; Chromatiales; Chromatiaceae.
DIW77__locus_1_main_sat
CsomeACh
05750 -pf10067pf01471 -
05755 -pf08411pf00929 -
05760 -pf01245 -
05765 -pf01746 -
05770 -pf05239pf01782 -
05775 -pf00886 -
05780 - -
05785 -pf00535 -
05790 - -
05795 HMMrbcL_1_Csome(pf00016pf02788) CsomeACh
05800 HMMrbcS_1_CsomeA(pf00101) CsomeACy
05805 HMMcsoS2_1_ACy(pf12288) CsomeACy
05810 HMMca_2_CsomeACy(pf08936) CsomeACh
05815 P_neonPurple CsomeACy
05820 P_neonPurple CsomeACy
05825 H_greenApple CsomeACy
05830 H_greenApple CsomeACy
05835 H_greenApple CsomeACy
05840 -pf00210 -
05845 -pf01329 -
05850 - -
05855 HMMreg00126_1_mix(pf00126pf03466) CsomeACh
05860 -pf02222pf17769 -
05865 -pf00731 -
05870 -pf17784 -
05875 -pf00005pf16326pf12848 -
05880 HMMreg13545_1_CsomeBCy(pf00027pf13545) BUF3
05885 -pf14319 -
05890 - -
05895 -pf01850 -
05900 -pf01471 -
(..1260..)
07160 -pf13649 -
07165 - -
07170 -pf00085 -
07175 -pf06853 -
07180 -pf13340 -
07185 -pf04986 -
07190 - -
07195 - -
07200 -pf01380 -
07205 -pf00132pf14602pf12804 -
07210 -pf00254 -
07215 - -
07220 Tdp_brightMagenta CsomeACh
07225 -pf01883pf10609 -
07230 - -
07235 - -
07240 -pf07728 -
07245 - -
07250 -pf18074 -
07255 - -
07260 -pf03485pf05746pf00750 -
07265 -pf05036 -
07270 HMM00857isoch_1_buf1(pf00857
07275 - -
07280 - -
BMC-P
BMC-H
BMC-T
Aldehyde dehydrogenase
PTAC
Signature enzyme
Alcohol dehydrogenase
Regulator

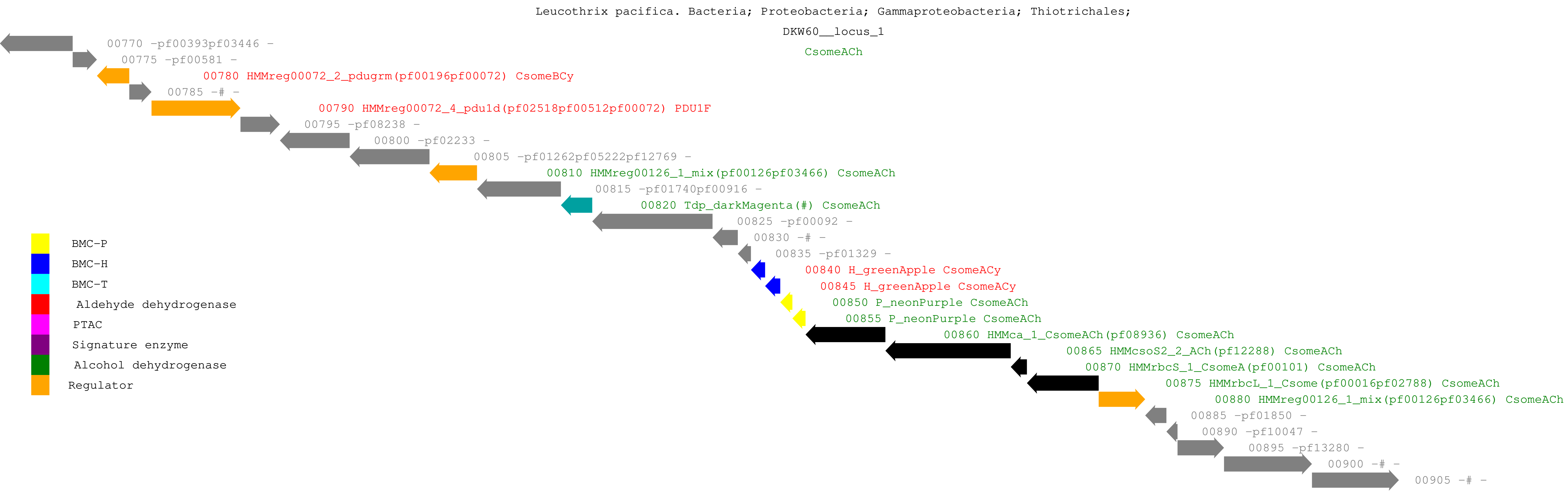
Leucothrix pacifica. Bacteria; Proteobacteria; Gammaproteobacteria; Thiotrichales;
DKW60__locus_1
CsomeACh
00770 -pf00393pf03446 -
00775 -pf00581 -
00780 HMMreg00072_2_pdugrm(pf00196pf00072) CsomeBCy
00785 -# -
00790 HMMreg00072_4_pdu1d(pf02518pf00512pf00072) PDU1F
00795 -pf08238 -
00800 -pf02233 -
00805 -pf01262pf05222pf12769 -
00810 HMMreg00126_1_mix(pf00126pf03466) CsomeACh
00815 -pf01740pf00916 -
00820 Tdp_darkMagenta(#) CsomeACh
00825 -pf00092 -
00830 -# -
00835 -pf01329 -
00840 H_greenApple CsomeACy
00845 H_greenApple CsomeACy
00850 P_neonPurple CsomeACh
00855 P_neonPurple CsomeACh
00860 HMMca_1_CsomeACh(pf08936) CsomeACh
00865 HMMcsoS2_2_ACh(pf12288) CsomeACh
00870 HMMrbcS_1_CsomeA(pf00101) CsomeACh
00875 HMMrbcL_1_Csome(pf00016pf02788) CsomeACh
00880 HMMreg00126_1_mix(pf00126pf03466) CsomeACh
00885 -pf01850 -
00890 -pf10047 -
00895 -pf13280 -
00900 -# -
00905 -# -
BMC-P
BMC-H
BMC-T
Aldehyde dehydrogenase
PTAC
Signature enzyme
Alcohol dehydrogenase
Regulator

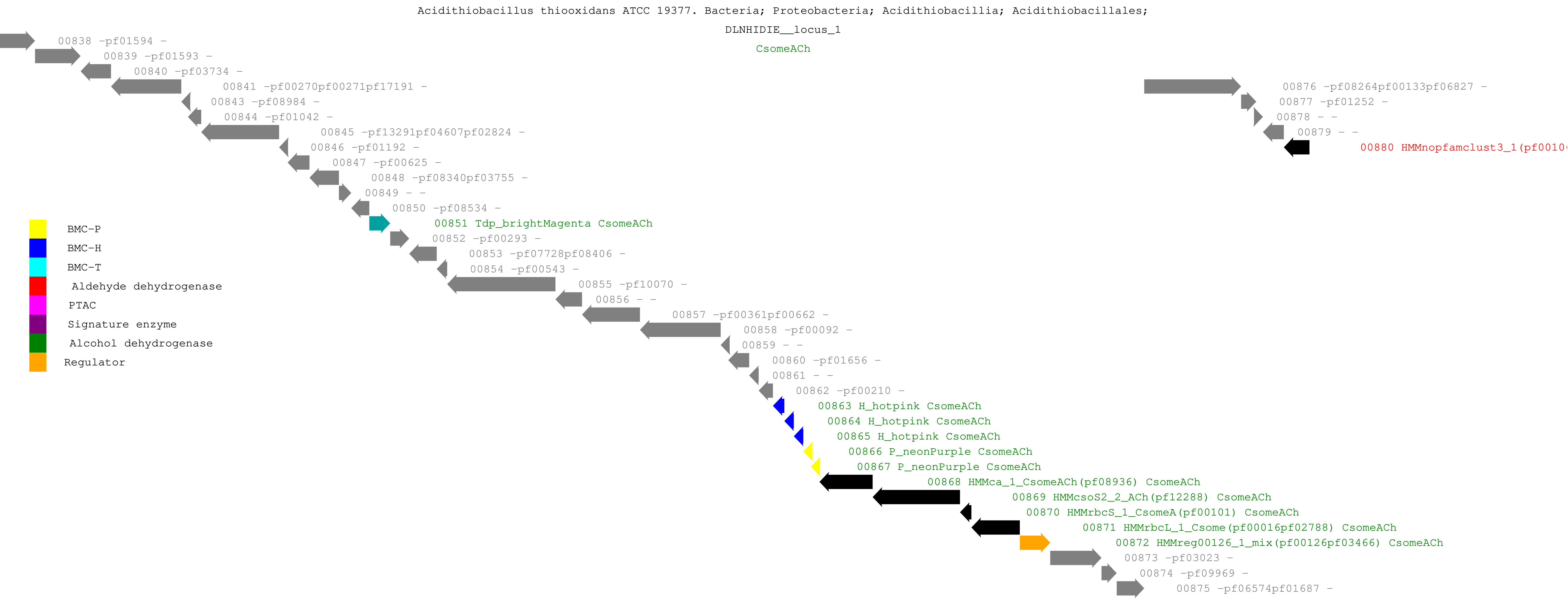

Acidithiobacillus thiooxidans ATCC 19377. Bacteria; Proteobacteria; Acidithiobacillia; Acidithiobacillales;
DLNHIDIE__locus_1
CsomeACh
00838 -pf01594 -
00839 -pf01593 -
00840 -pf03734 -
00841 -pf00270pf00271pf17191 -
00843 -pf08984 -
00844 -pf01042 -
00845 -pf13291pf04607pf02824 -
00846 -pf01192 -
00847 -pf00625 -
00848 -pf08340pf03755 -
00849 - -
00850 -pf08534 -
00851 Tdp_brightMagenta CsomeACh
00852 -pf00293 -
00853 -pf07728pf08406 -
00854 -pf00543 -
00855 -pf10070 -
00856 - -
00857 -pf00361pf00662 -
00858 -pf00092 -
00859 - -
00860 -pf01656 -
00861 - -
00862 -pf00210 -
00863 H_hotpink CsomeACh
00864 H_hotpink CsomeACh
00865 H_hotpink CsomeACh
00866 P_neonPurple CsomeACh
00867 P_neonPurple CsomeACh
00868 HMMca_1_CsomeACh(pf08936) CsomeACh
00869 HMMcsoS2_2_ACh(pf12288) CsomeACh
00870 HMMrbcS_1_CsomeA(pf00101) CsomeACh
00871 HMMrbcL_1_Csome(pf00016pf02788) CsomeACh
00872 HMMreg00126_1_mix(pf00126pf03466) CsomeACh
00873 -pf03023 -
00874 -pf09969 -
00875 -pf06574pf01687 -
00876 -pf08264pf00133pf06827 -
00877 -pf01252 -
00878 - -
00879 - -
00880 HMMnopfamclust3_1(pf0010
BMC-P
BMC-H
BMC-T
Aldehyde dehydrogenase
PTAC
Signature enzyme
Alcohol dehydrogenase
Regulator

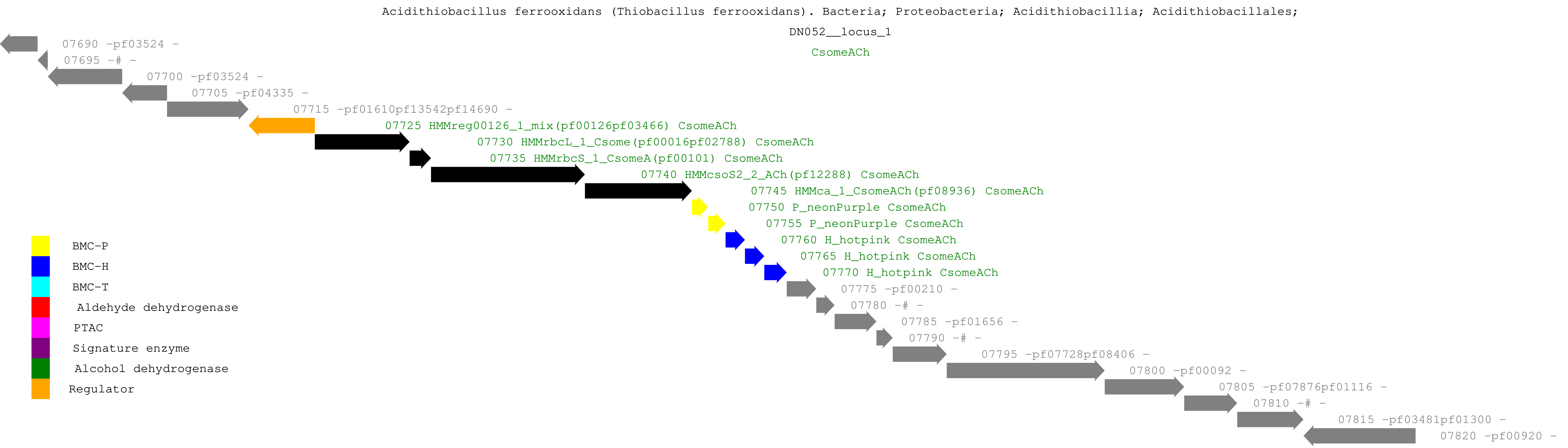
Acidithiobacillus ferrooxidans (Thiobacillus ferrooxidans). Bacteria; Proteobacteria; Acidithiobacillia; Acidithiobacillales;
DN052__locus_1
CsomeACh
07690 -pf03524 -
07695 -# -
07700 -pf03524 -
07705 -pf04335 -
07715 -pf01610pf13542pf14690 -
07725 HMMreg00126_1_mix(pf00126pf03466) CsomeACh
07730 HMMrbcL_1_Csome(pf00016pf02788) CsomeACh
07735 HMMrbcS_1_CsomeA(pf00101) CsomeACh
07740 HMMcsoS2_2_ACh(pf12288) CsomeACh
07745 HMMca_1_CsomeACh(pf08936) CsomeACh
07750 P_neonPurple CsomeACh
07755 P_neonPurple CsomeACh
07760 H_hotpink CsomeACh
07765 H_hotpink CsomeACh
07770 H_hotpink CsomeACh
07775 -pf00210 -
07780 -# -
07785 -pf01656 -
07790 -# -
07795 -pf07728pf08406 -
07800 -pf00092 -
07805 -pf07876pf01116 -
07810 -# -
07815 -pf03481pf01300 -
07820 -pf00920 -
BMC-P
BMC-H
BMC-T
Aldehyde dehydrogenase
PTAC
Signature enzyme
Alcohol dehydrogenase
Regulator

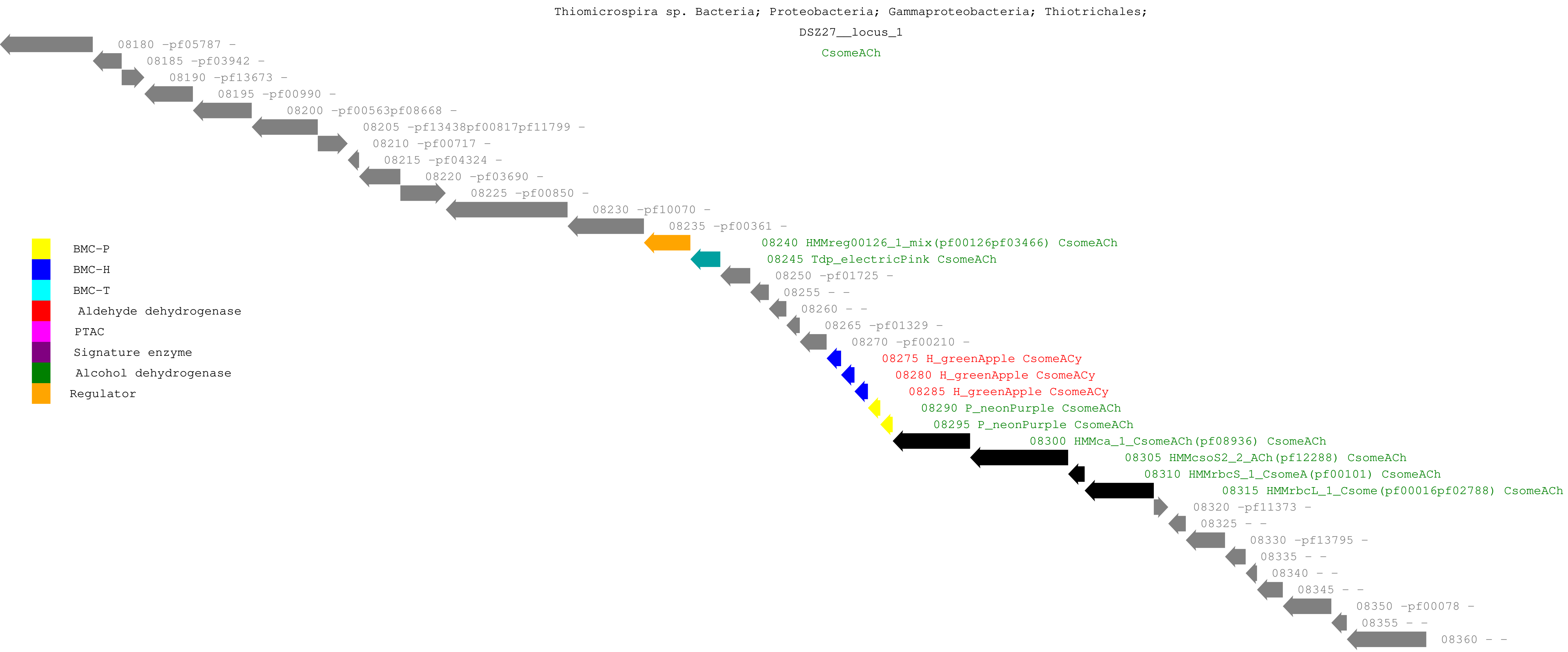
Thiomicrospira sp. Bacteria; Proteobacteria; Gammaproteobacteria; Thiotrichales;
DSZ27__locus_1
CsomeACh
08180 -pf05787 -
08185 -pf03942 -
08190 -pf13673 -
08195 -pf00990 -
08200 -pf00563pf08668 -
08205 -pf13438pf00817pf11799 -
08210 -pf00717 -
08215 -pf04324 -
08220 -pf03690 -
08225 -pf00850 -
08230 -pf10070 -
08235 -pf00361 -
08240 HMMreg00126_1_mix(pf00126pf03466) CsomeACh
08245 Tdp_electricPink CsomeACh
08250 -pf01725 -
08255 - -
08260 - -
08265 -pf01329 -
08270 -pf00210 -
08275 H_greenApple CsomeACy
08280 H_greenApple CsomeACy
08285 H_greenApple CsomeACy
08290 P_neonPurple CsomeACh
08295 P_neonPurple CsomeACh
08300 HMMca_1_CsomeACh(pf08936) CsomeACh
08305 HMMcsoS2_2_ACh(pf12288) CsomeACh
08310 HMMrbcS_1_CsomeA(pf00101) CsomeACh
08315 HMMrbcL_1_Csome(pf00016pf02788) CsomeACh
08320 -pf11373 -
08325 - -
08330 -pf13795 -
08335 - -
08340 - -
08345 - -
08350 -pf00078 -
08355 - -
08360 - -
BMC-P
BMC-H
BMC-T
Aldehyde dehydrogenase
PTAC
Signature enzyme
Alcohol dehydrogenase
Regulator

Burkholderiales bacterium GJ-E10. Bacteria; Proteobacteria; Betaproteobacteria; Burkholderiales.

E10__locus_1

CsomeACh

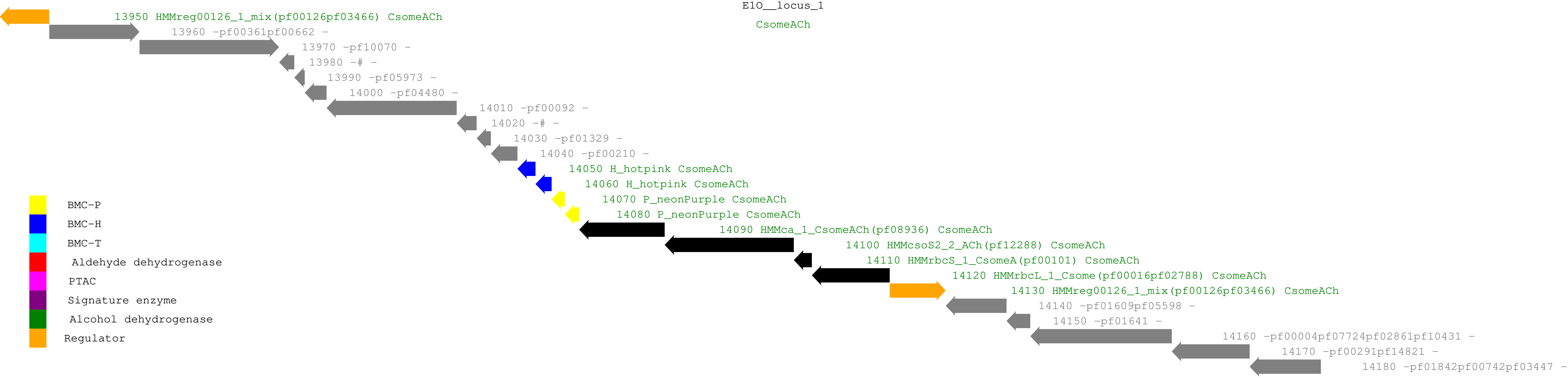

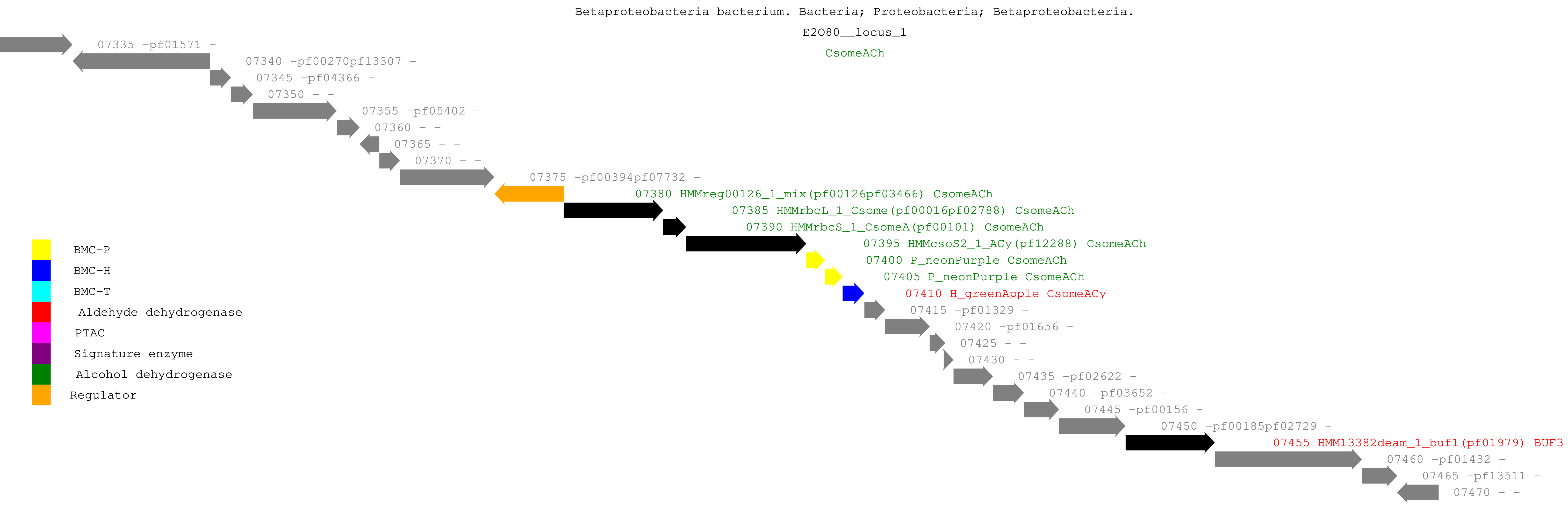

Betaproteobacteria bacterium. Bacteria; Proteobacteria; Betaproteobacteria.
E2O80__locus_1
CsomeACh
07335 -pf01571 -
07340 -pf00270pf13307 -
07345 -pf04366 -
07350 - -
07355 -pf05402 -
07360 - -
07365 - -
07370 - -
07375 -pf00394pf07732 -
07380 HMMreg00126_1_mix(pf00126pf03466) CsomeACh
07385 HMMrbcL_1_Csome(pf00016pf02788) CsomeACh
07390 HMMrbcS_1_CsomeA(pf00101) CsomeACh
07395 HMMcsoS2_1_ACy(pf12288) CsomeACh
07400 P_neonPurple CsomeACh
07405 P_neonPurple CsomeACh
07410 H_greenApple CsomeACy
07415 -pf01329 -
07420 -pf01656 -
07425 - -
07430 - -
07435 -pf02622 -
07440 -pf03652 -
07445 -pf00156 -
07450 -pf00185pf02729 -
07455 HMM13382deam_1_buf1(pf01979) BUF3
07460 -pf01432 -
07465 -pf13511 -
07470 - -
BMC-P
BMC-H
BMC-T
Aldehyde dehydrogenase
PTAC
Signature enzyme
Alcohol dehydrogenase
Regulator

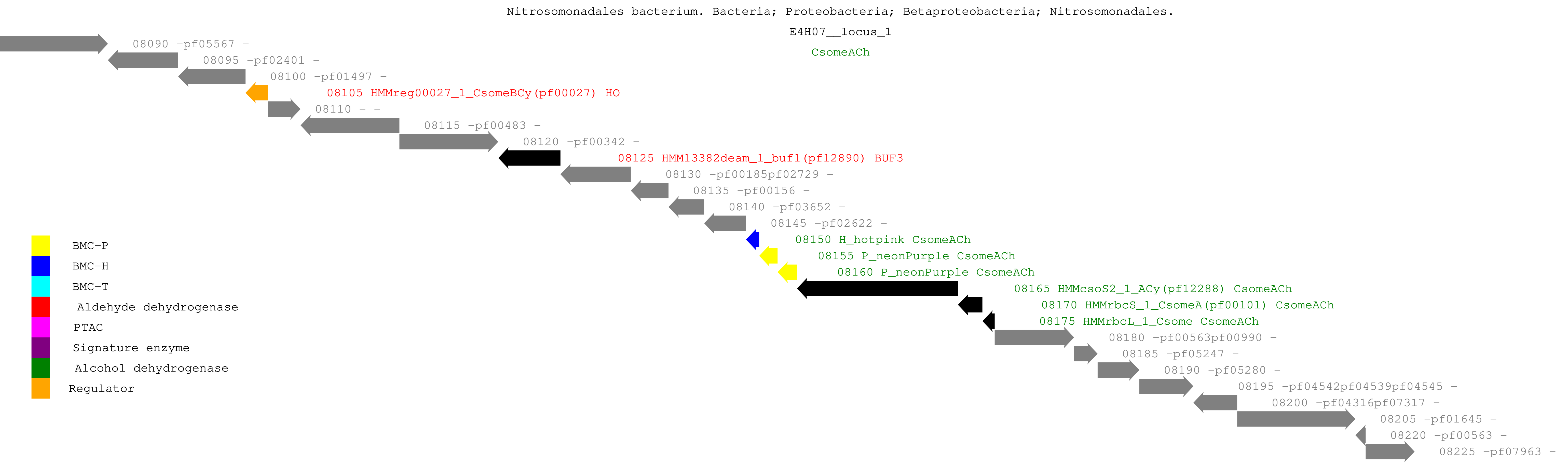

Nitrosomonadales bacterium. Bacteria; Proteobacteria; Betaproteobacteria; Nitrosomonadales.
E4H07__locus_1
CsomeACh
08090 -pf05567 -
08095 -pf02401 -
08100 -pf01497 -
08105 HMMreg00027_1_CsomeBCy(pf00027) HO
08110 - -
08115 -pf00483 -
08120 -pf00342 -
08125 HMM13382deam_1_buf1(pf12890) BUF3
08130 -pf00185pf02729 -
08135 -pf00156 -
08140 -pf03652 -
08145 -pf02622 -
08150 H_hotpink CsomeACh
08155 P_neonPurple CsomeACh
08160 P_neonPurple CsomeACh
08165 HMMcsoS2_1_ACy(pf12288) CsomeACh
08170 HMMrbcS_1_CsomeA(pf00101) CsomeACh
08175 HMMrbcL_1_Csome CsomeACh
08180 -pf00563pf00990 -
08185 -pf05247 -
08190 -pf05280 -
08195 -pf04542pf04539pf04545 -
08200 -pf04316pf07317 -
08205 -pf01645 -
08220 -pf00563 -
08225 -pf07963 -
BMC-P
BMC-H
BMC-T
Aldehyde dehydrogenase
PTAC
Signature enzyme
Alcohol dehydrogenase
Regulator

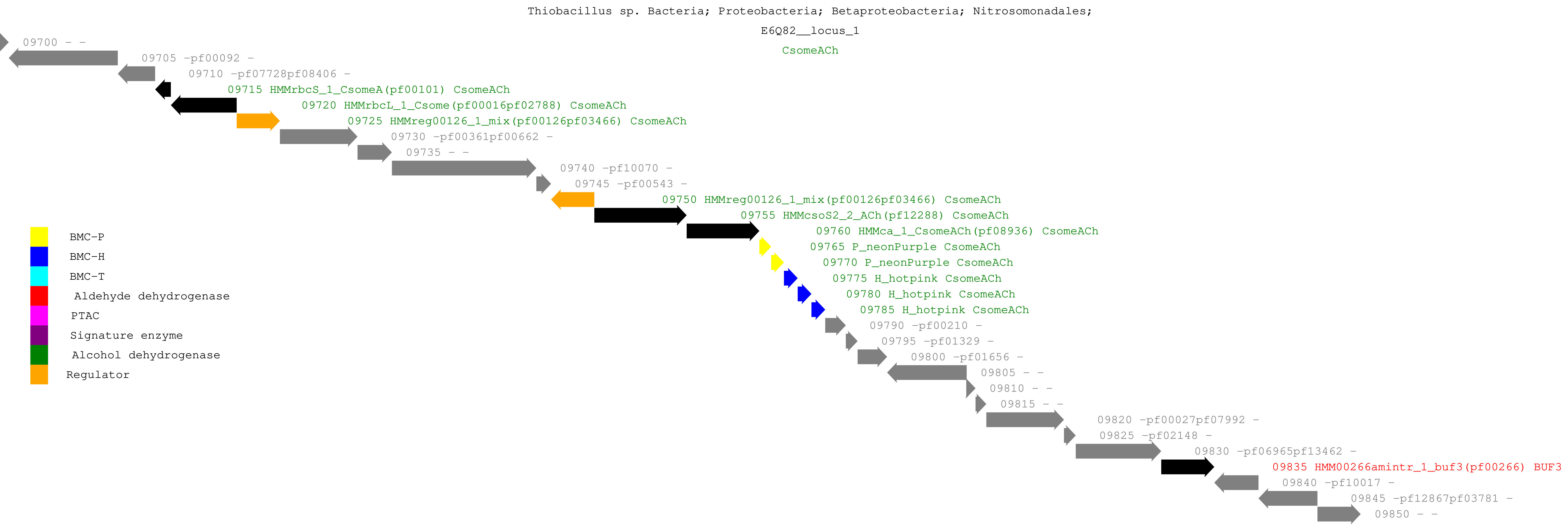
Thiobacillus sp. Bacteria; Proteobacteria; Betaproteobacteria; Nitrosomonadales;
E6Q82__locus_1
CsomeACh
09700 - -
09705 -pf00092 -
09710 -pf07728pf08406 -
09715 HMMrbcS_1_CsomeA(pf00101) CsomeACh
09720 HMMrbcL_1_Csome(pf00016pf02788) CsomeACh
09725 HMMreg00126_1_mix(pf00126pf03466) CsomeACh
09730 -pf00361pf00662 -
09735 - -
09740 -pf10070 -
09745 -pf00543 -
09750 HMMreg00126_1_mix(pf00126pf03466) CsomeACh
09755 HMMcsoS2_2_ACh(pf12288) CsomeACh
09760 HMMca_1_CsomeACh(pf08936) CsomeACh
09765 P_neonPurple CsomeACh
09770 P_neonPurple CsomeACh
09775 H_hotpink CsomeACh
09780 H_hotpink CsomeACh
09785 H_hotpink CsomeACh
09790 -pf00210 -
09795 -pf01329 -
09800 -pf01656 -
09805 - -
09810 - -
09815 - -
09820 -pf00027pf07992 -
09825 -pf02148 -
09830 -pf06965pf13462 -
09835 HMM00266amintr_1_buf3(pf00266) BUF3
09840 -pf10017 -
09845 -pf12867pf03781 -
09850 - -
BMC-P
BMC-H
BMC-T
Aldehyde dehydrogenase
PTAC
Signature enzyme
Alcohol dehydrogenase
Regulator

Acidithiobacillus sulfuriphilus. Bacteria; Proteobacteria; Acidithiobacillia; Acidithiobacillales;

EC580__locus_1

CsomeACh

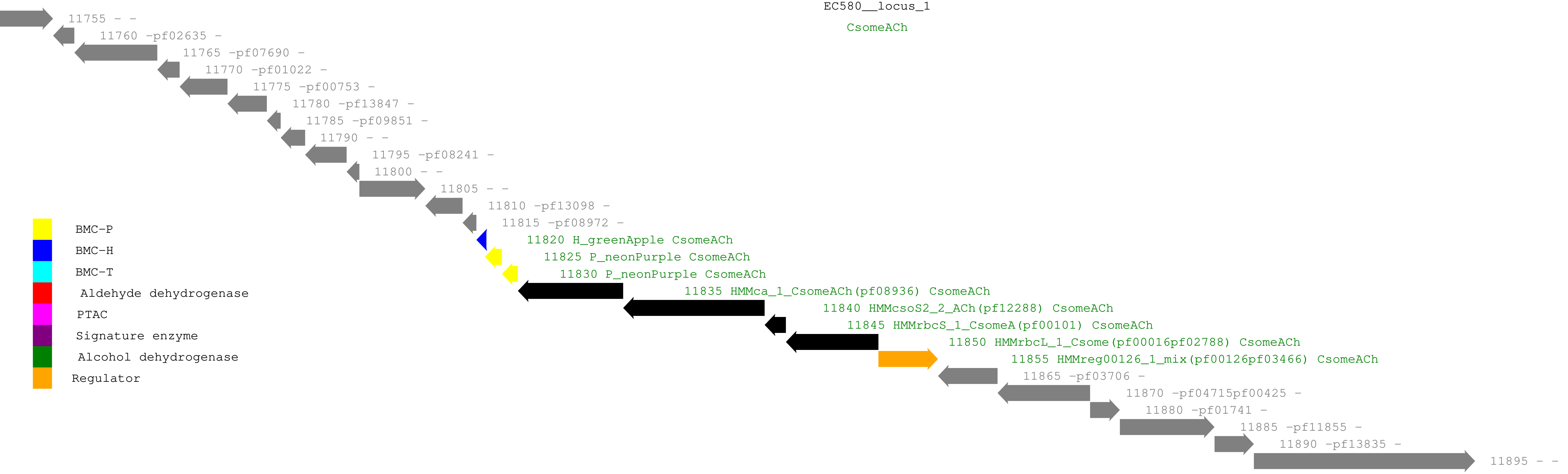

Ectothiorhodospira sp. BSL-9. Bacteria; Proteobacteria; Gammaproteobacteria; Chromatiales;

ECTOBSL9__locus_1_main_sat

CsomeACh

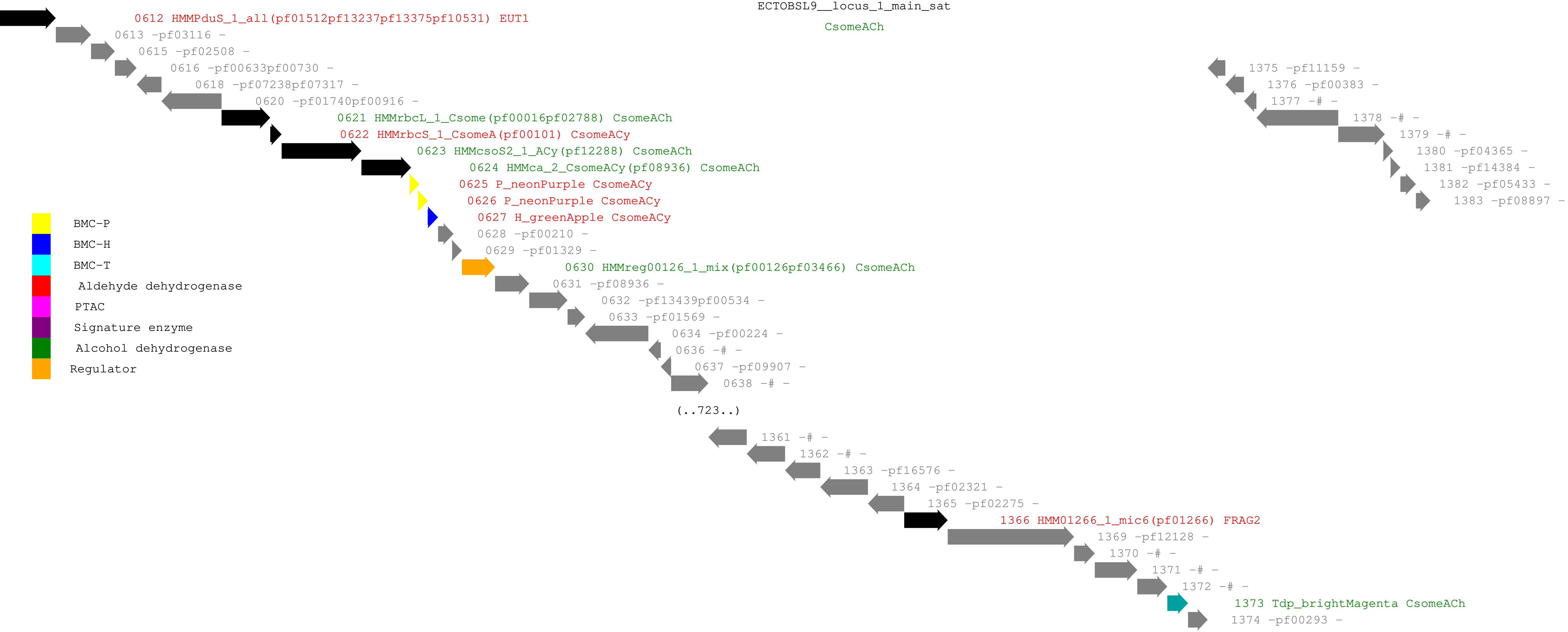

Thiohalophilus thiocyanatoxydans. Bacteria; Proteobacteria; Gammaproteobacteria; Chromatiales;

EDC23__locus_1

CsomeACh

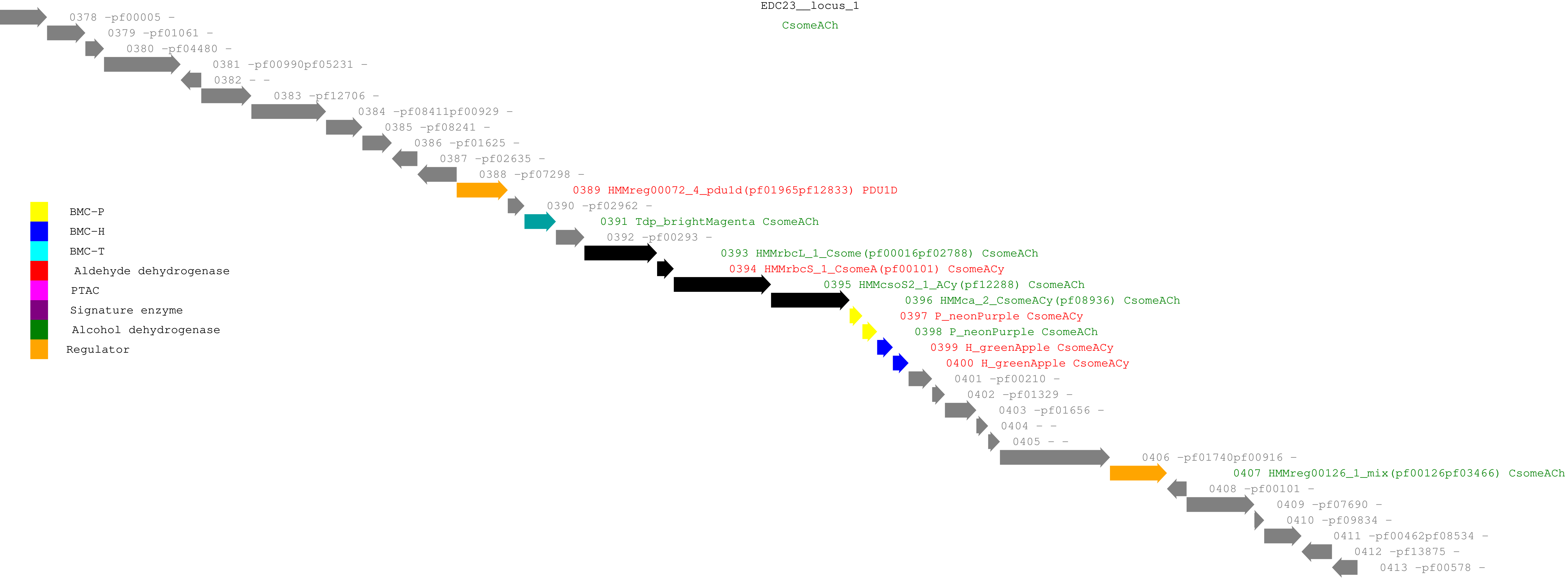

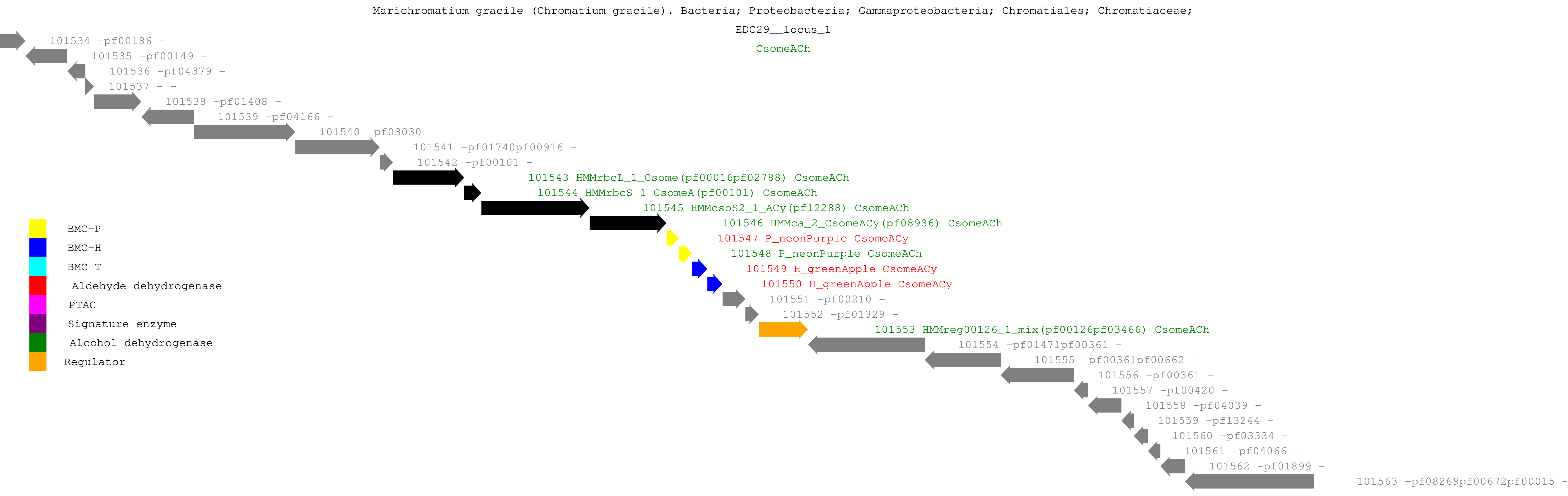
Marichromatium gracile (Chromatium gracile). Bacteria; Proteobacteria; Gammaproteobacteria; Chromatiales; Chromatiaceae;
EDC29__locus_1
CsomeACh
101534 -pf00186 -
101535 -pf00149 -
101536 -pf04379 -
101537 - -
101538 -pf01408 -
101539 -pf04166 -
101540 -pf03030 -
101541 -pf01740pf00916 -
101542 -pf00101 -
101543 HMMrbcL_1_Csome(pf00016pf02788) CsomeACh
101544 HMMrbcS_1_CsomeA(pf00101) CsomeACh
101545 HMMcsoS2_1_ACy(pf12288) CsomeACh
101546 HMMca_2_CsomeACy(pf08936) CsomeACh
101547 P_neonPurple CsomeACy
101548 P_neonPurple CsomeACh
101549 H_greenApple CsomeACy
101550 H_greenApple CsomeACy
101551 -pf00210 -
101552 -pf01329 -
101553 HMMreg00126_1_mix(pf00126pf03466) CsomeACh
101554 -pf01471pf00361 -
101555 -pf00361pf00662 -
101556 -pf00361 -
101557 -pf00420 -
101558 -pf04039 -
101559 -pf13244 -
101560 -pf03334 -
101561 -pf04066 -
101562 -pf01899 -
101563 -pf08269pf00672pf00015 -
BMC-P
BMC-H
BMC-T
Aldehyde dehydrogenase
PTAC
Signature enzyme
Alcohol dehydrogenase
Regulator

Tepidimonas ignava. Bacteria; Proteobacteria; Betaproteobacteria; Burkholderiales; Tepidimonas.

EDC36__locus_1

CsomeACh

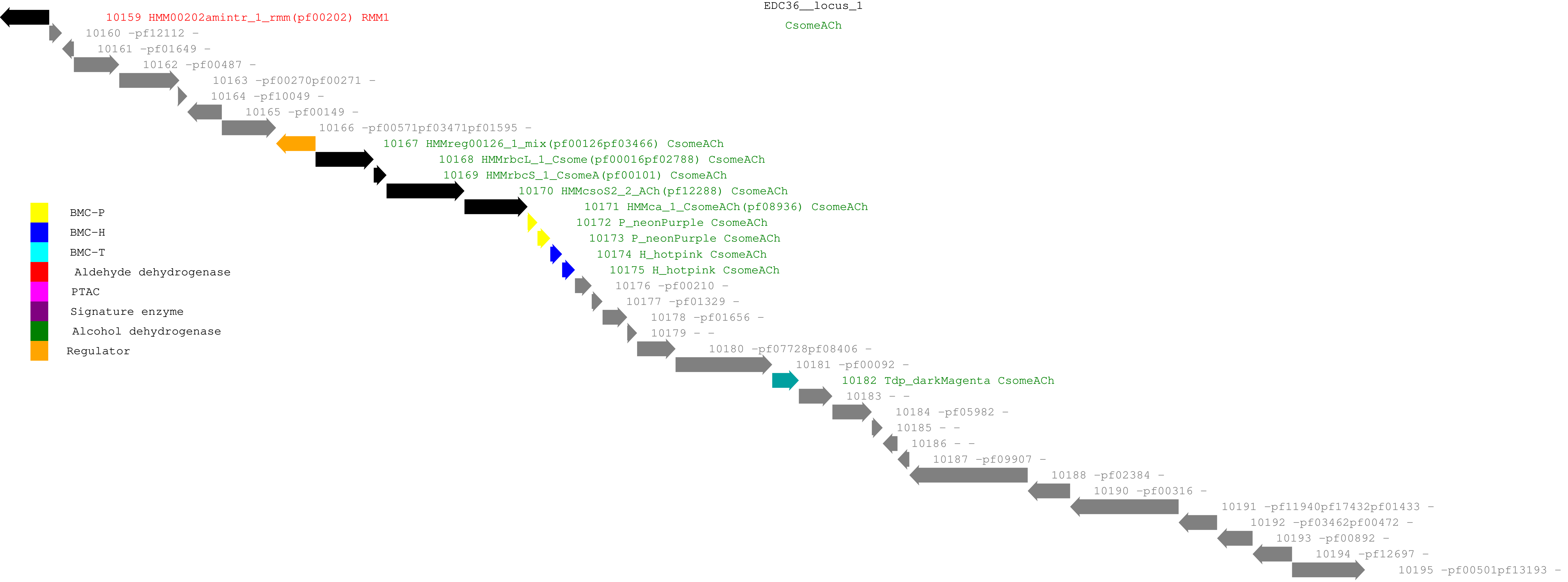

Sulfuritortus calidifontis. Bacteria; Proteobacteria; Betaproteobacteria; Nitrosomonadales;

EDC61__locus_1

CsomeACh

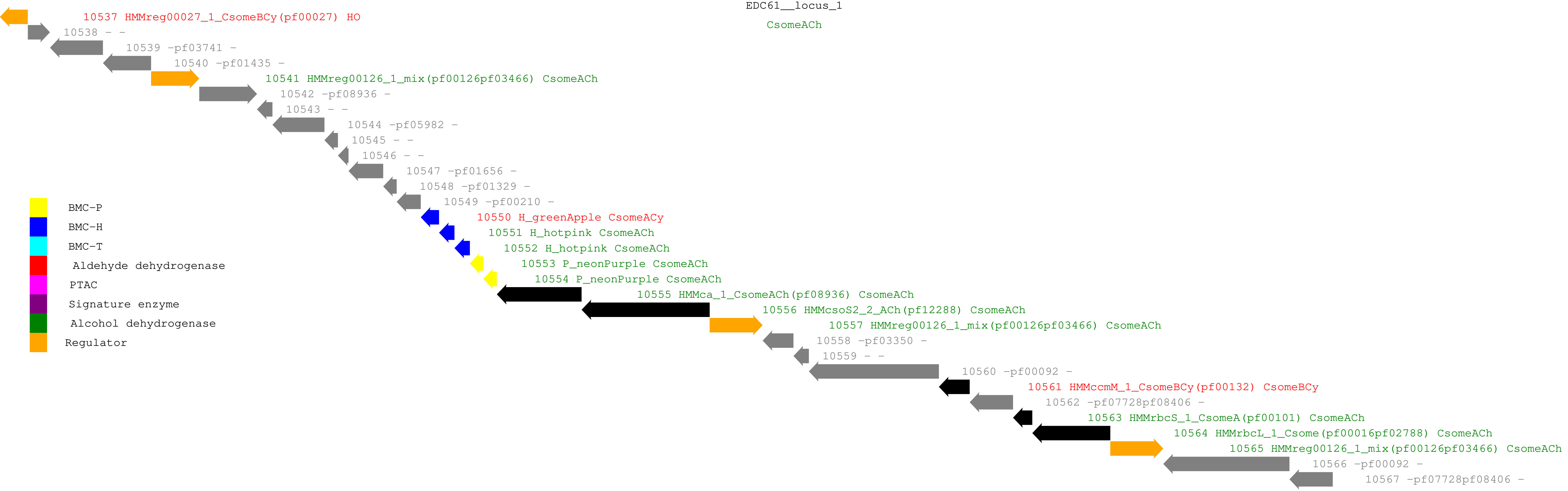

Sulfurirhabdus autotrophica. Bacteria; Proteobacteria; Betaproteobacteria; Nitrosomonadales;

EDC63__locus_1

CsomeACh

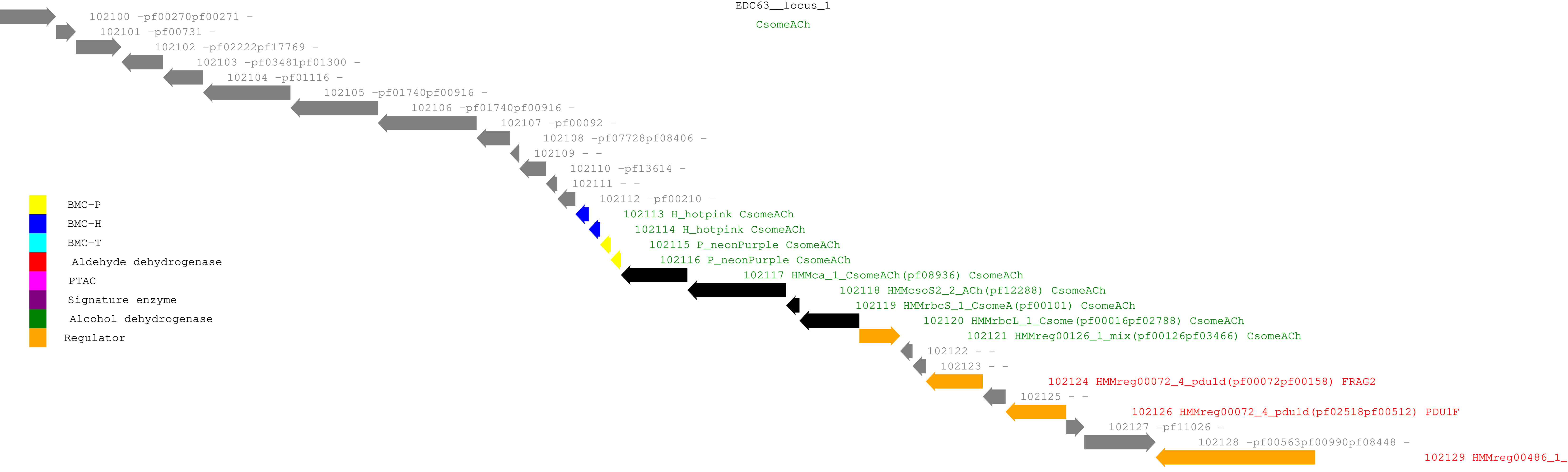

Hydrogenovibrio marinus. Bacteria; Proteobacteria; Gammaproteobacteria; Thiotrichales;

EI16__locus_1

CsomeACh

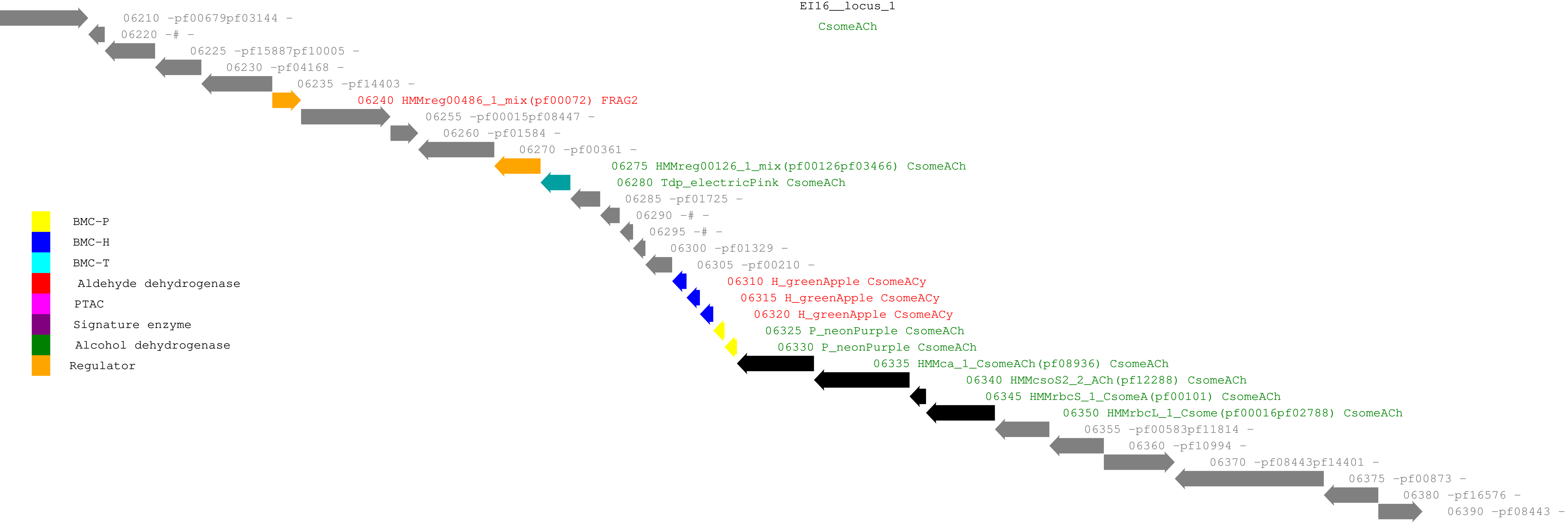

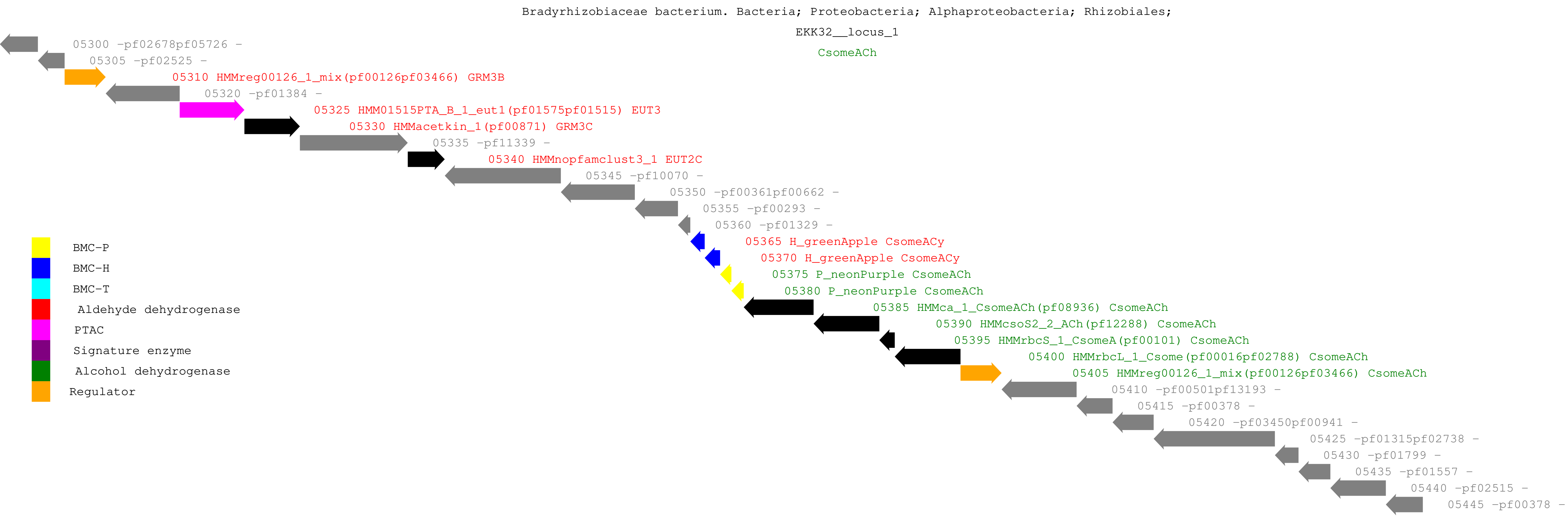
Bradyrhizobiaceae bacterium. Bacteria; Proteobacteria; Alphaproteobacteria; Rhizobiales;
EKK32__locus_1
CsomeACh
05300 -pf02678pf05726 -
05305 -pf02525 -
05310 HMMreg00126_1_mix(pf00126pf03466) GRM3B
05320 -pf01384 -
05325 HMM01515PTA_B_1_eut1(pf01575pf01515) EUT3
05330 HMMacetkin_1(pf00871) GRM3C
05335 -pf11339 -
05340 HMMnopfamclust3_1 EUT2C
05345 -pf10070 -
05350 -pf00361pf00662 -
05355 -pf00293 -
05360 -pf01329 -
05365 H_greenApple CsomeACy
05370 H_greenApple CsomeACy
05375 P_neonPurple CsomeACh
05380 P_neonPurple CsomeACh
05385 HMMca_1_CsomeACh(pf08936) CsomeACh
05390 HMMcsoS2_2_ACh(pf12288) CsomeACh
05395 HMMrbcS_1_CsomeA(pf00101) CsomeACh
05400 HMMrbcL_1_Csome(pf00016pf02788) CsomeACh
05405 HMMreg00126_1_mix(pf00126pf03466) CsomeACh
05410 -pf00501pf13193 -
05415 -pf00378 -
05420 -pf03450pf00941 -
05425 -pf01315pf02738 -
05430 -pf01799 -
05435 -pf01557 -
05440 -pf02515 -
05445 -pf00378 -
BMC-P
BMC-H
BMC-T
Aldehyde dehydrogenase
PTAC
Signature enzyme
Alcohol dehydrogenase
Regulator

Pseudonocardiaceae bacterium. Bacteria; Actinobacteria; Pseudonocardiales; Pseudonocardiaceae.

EKK42__locus_2

CsomeACh

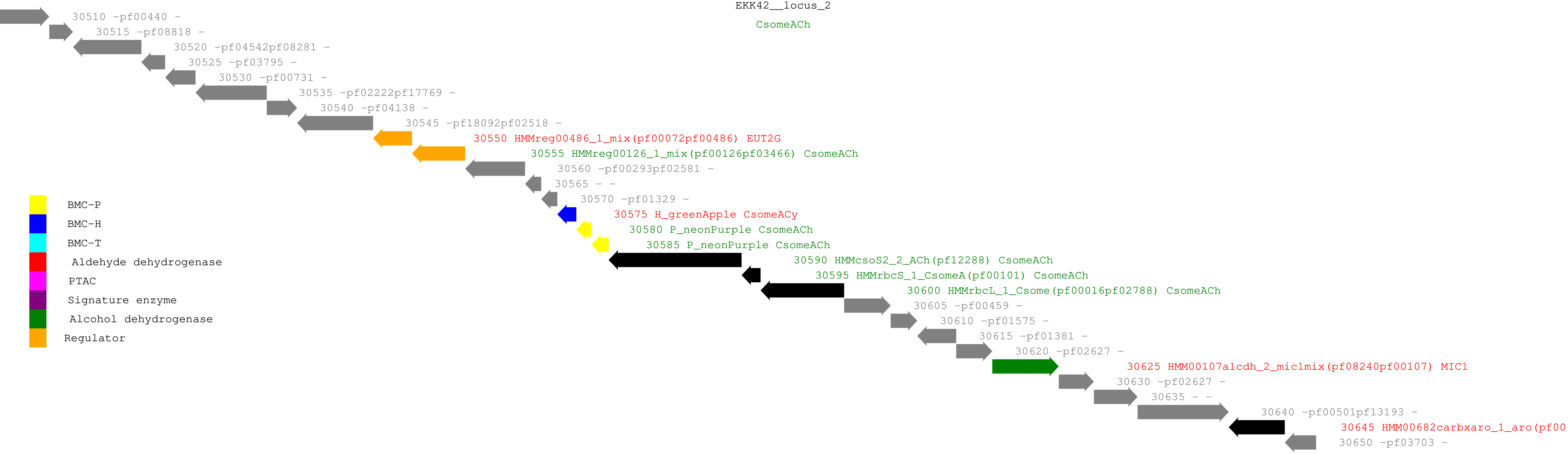

Gallionellaceae bacterium. Bacteria; Proteobacteria; Betaproteobacteria; Nitrosomonadales;

EPO42__locus_1_main_sat

CsomeACh

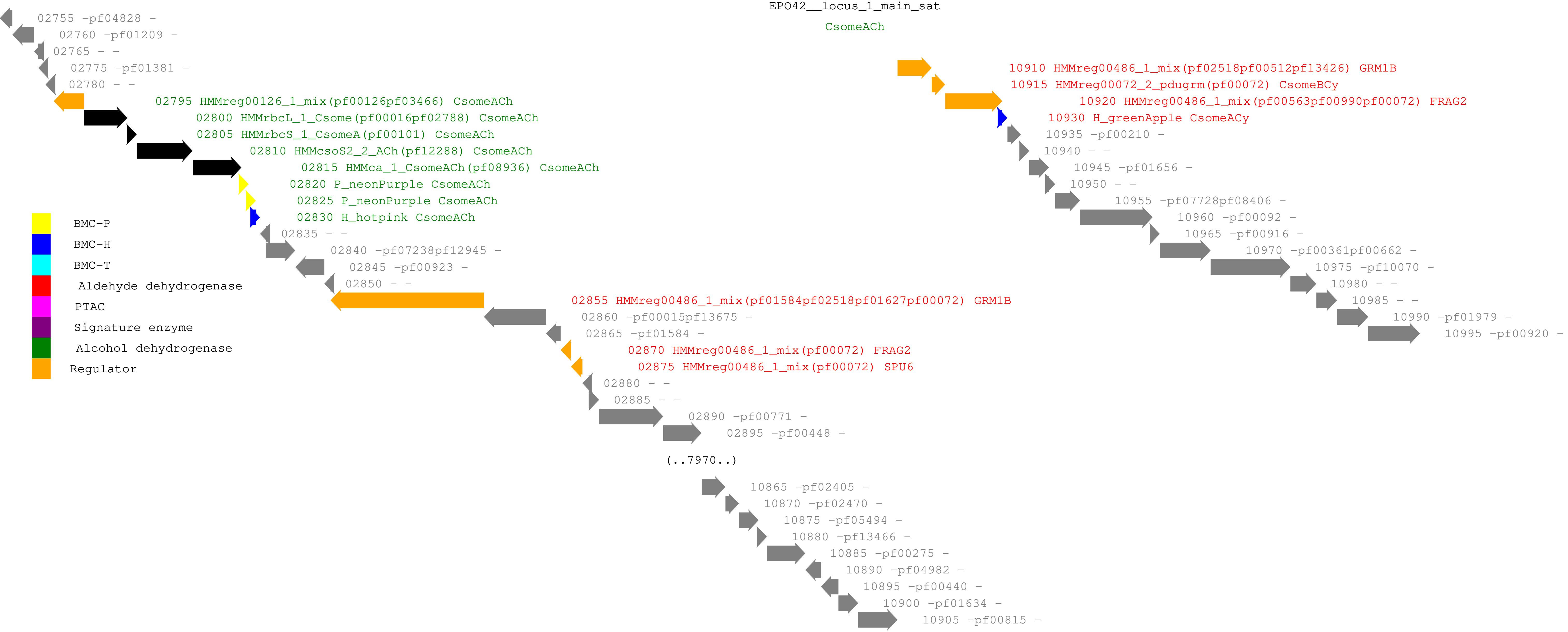

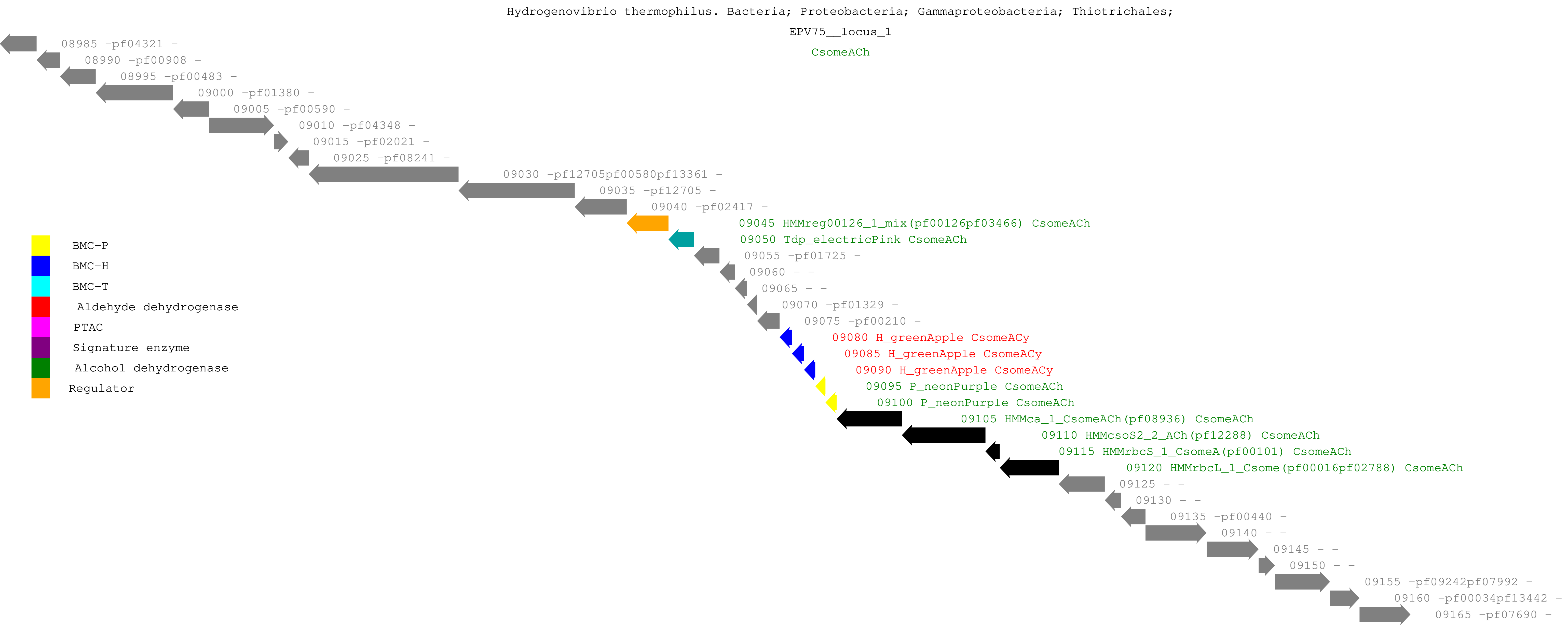

Hydrogenovibrio thermophilus. Bacteria; Proteobacteria; Gammaproteobacteria; Thiotrichales;
EPV75__locus_1
CsomeACh
08985 -pf04321 -
08990 -pf00908 -
08995 -pf00483 -
09000 -pf01380 -
09005 -pf00590 -
09010 -pf04348 -
09015 -pf02021 -
09025 -pf08241 -
09030 -pf12705pf00580pf13361 -
09035 -pf12705 -
09040 -pf02417 -
09045 HMMreg00126_1_mix(pf00126pf03466) CsomeACh
09050 Tdp_electricPink CsomeACh
09055 -pf01725 -
09060 - -
09065 - -
09070 -pf01329 -
09075 -pf00210 -
09080 H_greenApple CsomeACy
09085 H_greenApple CsomeACy
09090 H_greenApple CsomeACy
09095 P_neonPurple CsomeACh
09100 P_neonPurple CsomeACh
09105 HMMca_1_CsomeACh(pf08936) CsomeACh
09110 HMMcsoS2_2_ACh(pf12288) CsomeACh
09115 HMMrbcS_1_CsomeA(pf00101) CsomeACh
09120 HMMrbcL_1_Csome(pf00016pf02788) CsomeACh
09125 - -
09130 - -
09135 -pf00440 -
09140 - -
09145 - -
09150 - -
09155 -pf09242pf07992 -
09160 -pf00034pf13442 -
09165 -pf07690 -
BMC-P
BMC-H
BMC-T
Aldehyde dehydrogenase
PTAC
Signature enzyme
Alcohol dehydrogenase
Regulator

Litoricolaceae bacterium. Bacteria; Proteobacteria; Gammaproteobacteria; Oceanospirillales;

EVA60__locus_1

CsomeACh

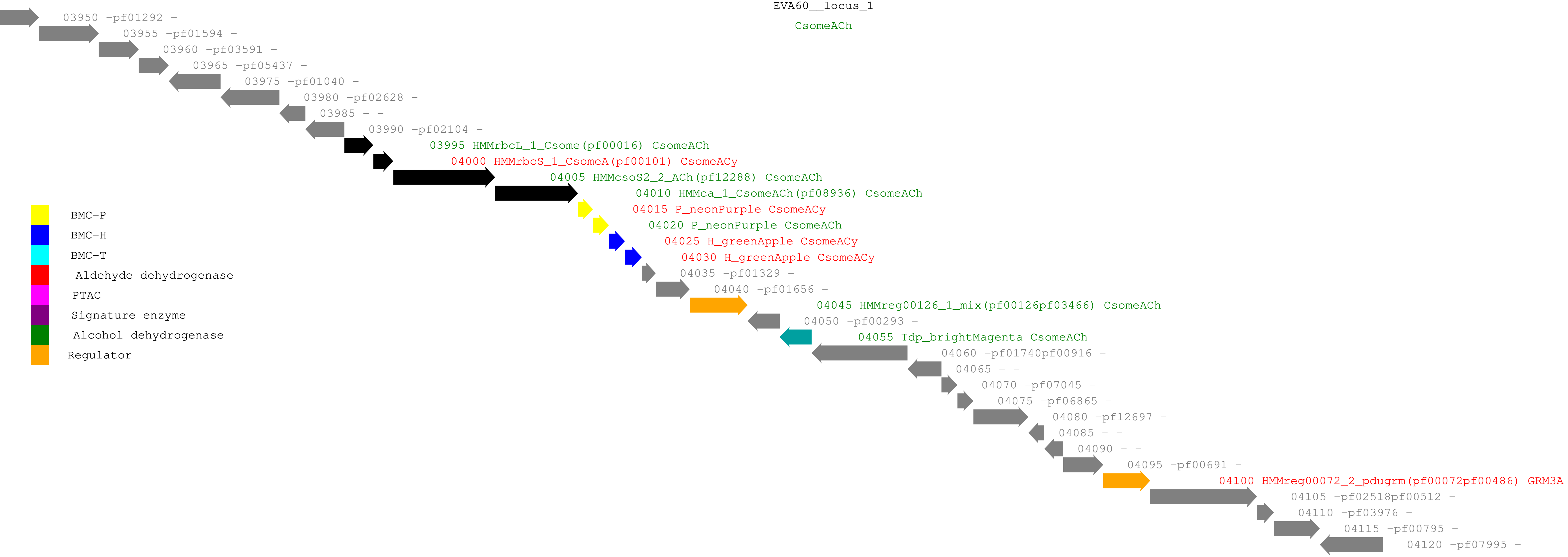

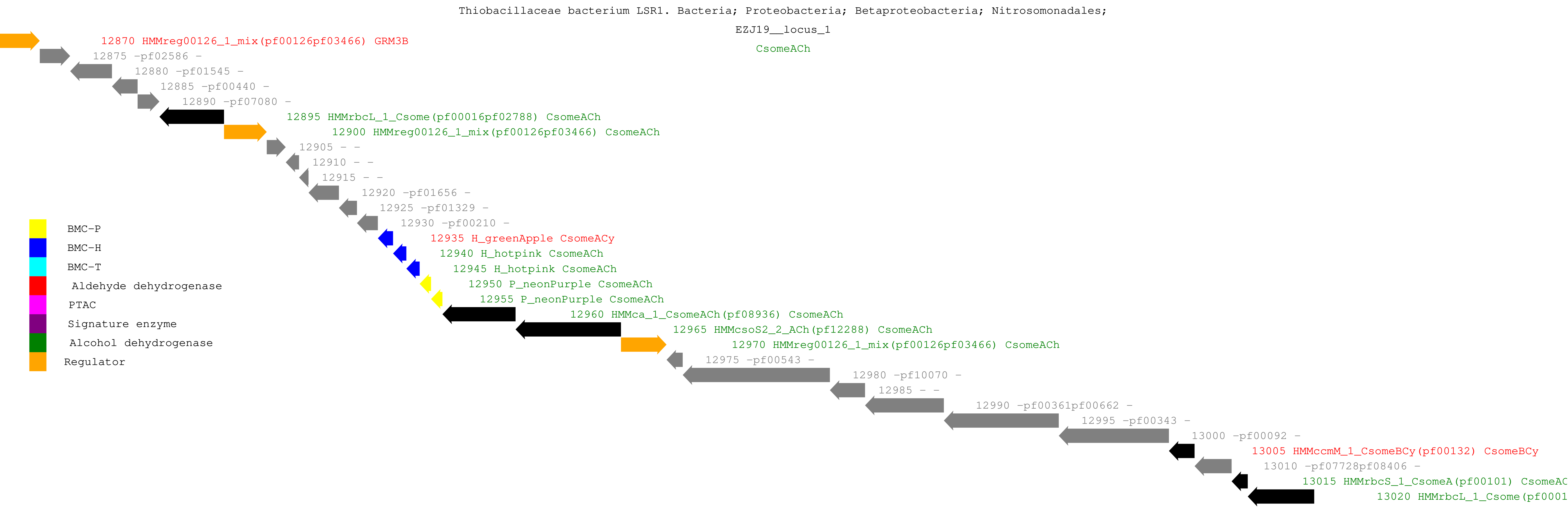

Thiobacillaceae bacterium LSR1. Bacteria; Proteobacteria; Betaproteobacteria; Nitrosomonadales;
EZJ19__locus_1
CsomeACh
12870 HMMreg00126_1_mix(pf00126pf03466) GRM3B
12875 -pf02586 -
12880 -pf01545 -
12885 -pf00440 -
12890 -pf07080 -
12895 HMMrbcL_1_Csome(pf00016pf02788) CsomeACh
12900 HMMreg00126_1_mix(pf00126pf03466) CsomeACh
12905 - -
12910 - -
12915 - -
12920 -pf01656 -
12925 -pf01329 -
12930 -pf00210 -
12935 H_greenApple CsomeACy
12940 H_hotpink CsomeACh
12945 H_hotpink CsomeACh
12950 P_neonPurple CsomeACh
12955 P_neonPurple CsomeACh
12960 HMMca_1_CsomeACh(pf08936) CsomeACh
12965 HMMcsoS2_2_ACh(pf12288) CsomeACh
12970 HMMreg00126_1_mix(pf00126pf03466) CsomeACh
12975 -pf00543 -
12980 -pf10070 -
12985 - -
12990 -pf00361pf00662 -
12995 -pf00343 -
13000 -pf00092 -
13005 HMMccmM_1_CsomeBCy(pf00132) CsomeBCy
13010 -pf07728pf08406 -
13015 HMMrbcS_1_CsomeA(pf00101) CsomeAC
13020 HMMrbcL_1_Csome(pf0001
BMC-P
BMC-H
BMC-T
Aldehyde dehydrogenase
PTAC
Signature enzyme
Alcohol dehydrogenase
Regulator

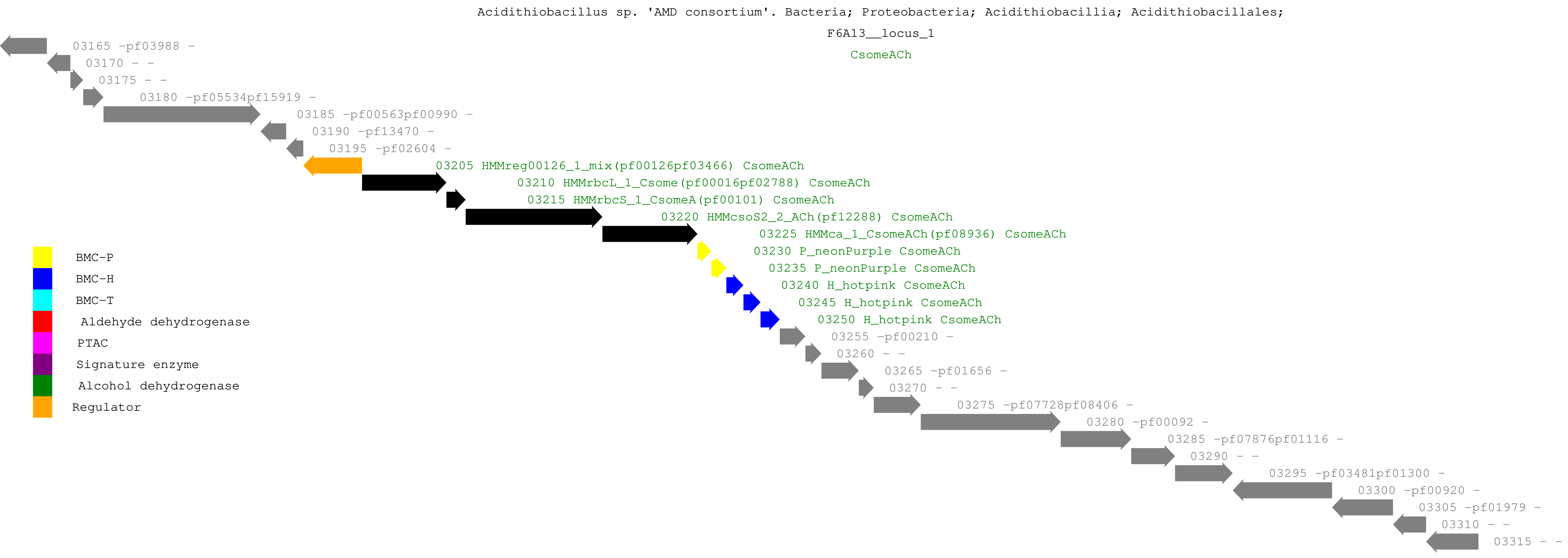
Acidithiobacillus sp. 'AMD consortium'. Bacteria; Proteobacteria; Acidithiobacillia; Acidithiobacillales;
F6A13__locus_1
CsomeACh
03165 -pf03988 -
03170 - -
03175 - -
03180 -pf05534pf15919 -
03185 -pf00563pf00990 -
03190 -pf13470 -
03195 -pf02604 -
03205 HMMreg00126_1_mix(pf00126pf03466) CsomeACh
03210 HMMrbcL_1_Csome(pf00016pf02788) CsomeACh
03215 HMMrbcS_1_CsomeA(pf00101) CsomeACh
03220 HMMcsoS2_2_ACh(pf12288) CsomeACh
03225 HMMca_1_CsomeACh(pf08936) CsomeACh
03230 P_neonPurple CsomeACh
03235 P_neonPurple CsomeACh
03240 H_hotpink CsomeACh
03245 H_hotpink CsomeACh
03250 H_hotpink CsomeACh
03255 -pf00210 -
03260 - -
03265 -pf01656 -
03270 - -
03275 -pf07728pf08406 -
03280 -pf00092 -
03285 -pf07876pf01116 -
03290 - -
03295 -pf03481pf01300 -
03300 -pf00920 -
03305 -pf01979 -
03310 - -
03315 - -
BMC-P
BMC-H
BMC-T
Aldehyde dehydrogenase
PTAC
Signature enzyme
Alcohol dehydrogenase
Regulator

Acidithiobacillus sp. 'AMD consortium'. Bacteria; Proteobacteria; Acidithiobacillia; Acidithiobacillales;

F6A13__locus_2

CsomeACh

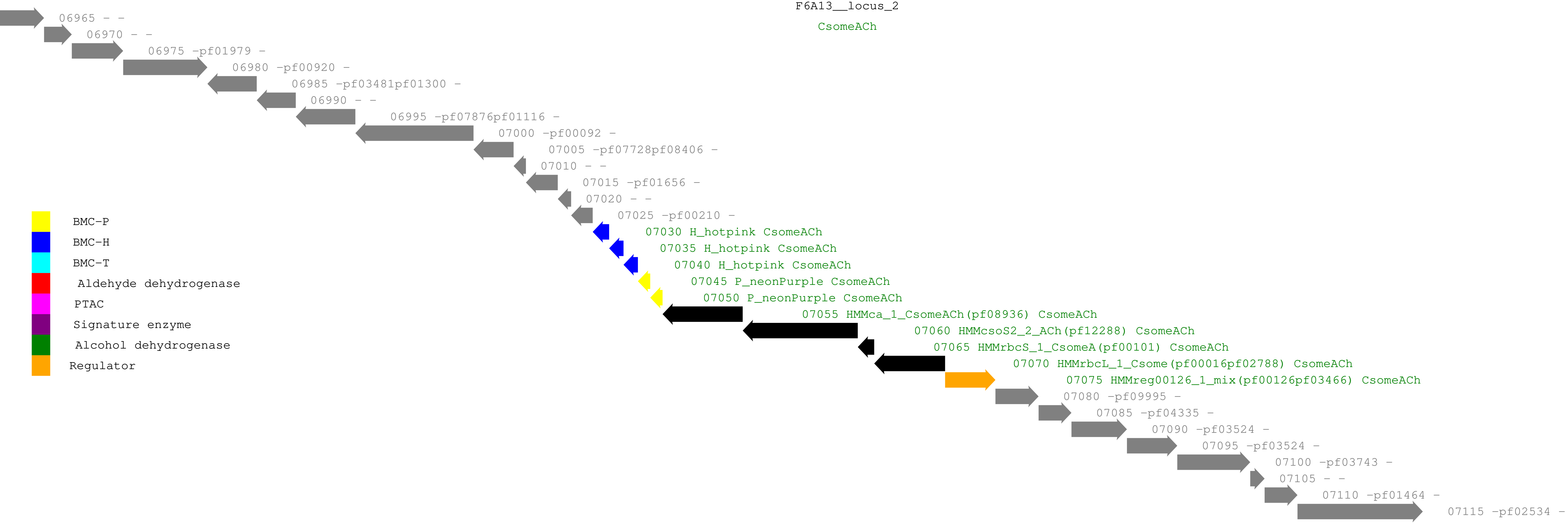

Halothiobacillus sp. SB14A. Bacteria; Proteobacteria; Gammaproteobacteria; Chromatiales;

FAZ79__locus_1_main_sat

CsomeACh

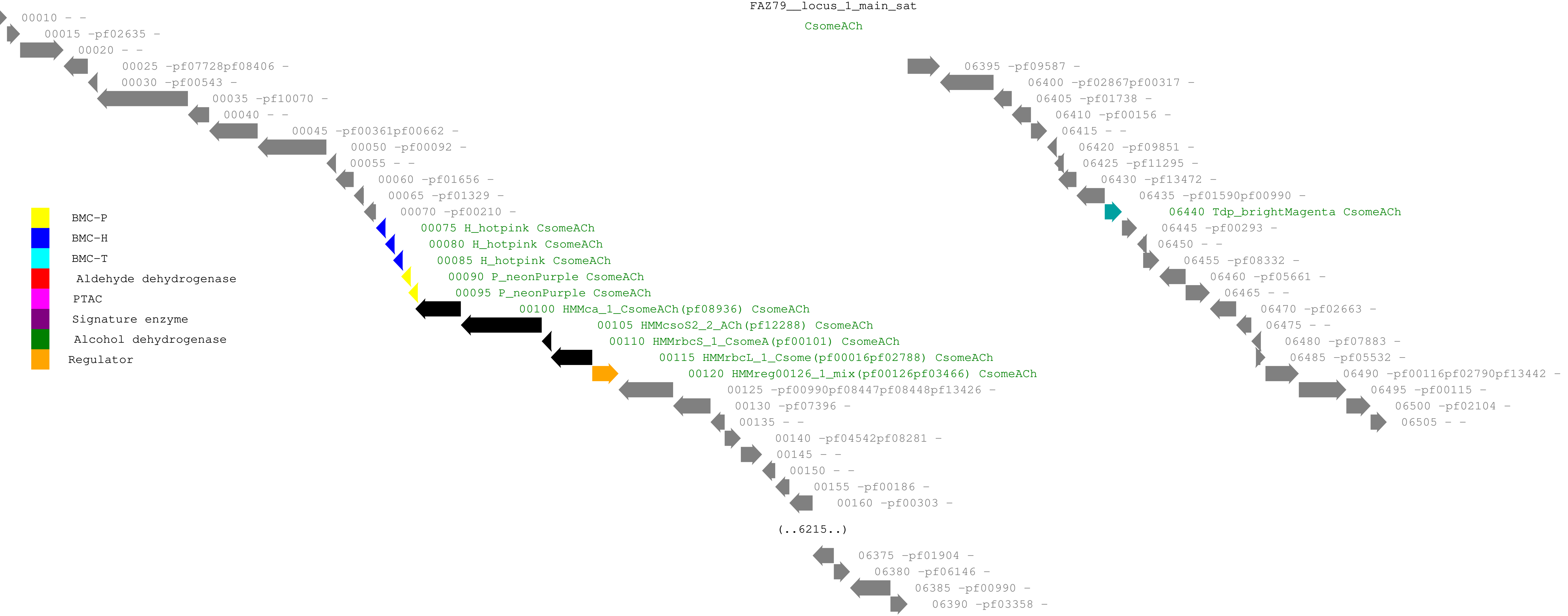

Halothiobacillaceae bacterium. Bacteria; Proteobacteria; Gammaproteobacteria; Chromatiales;

FD130__locus_1_main_sat

CsomeACh

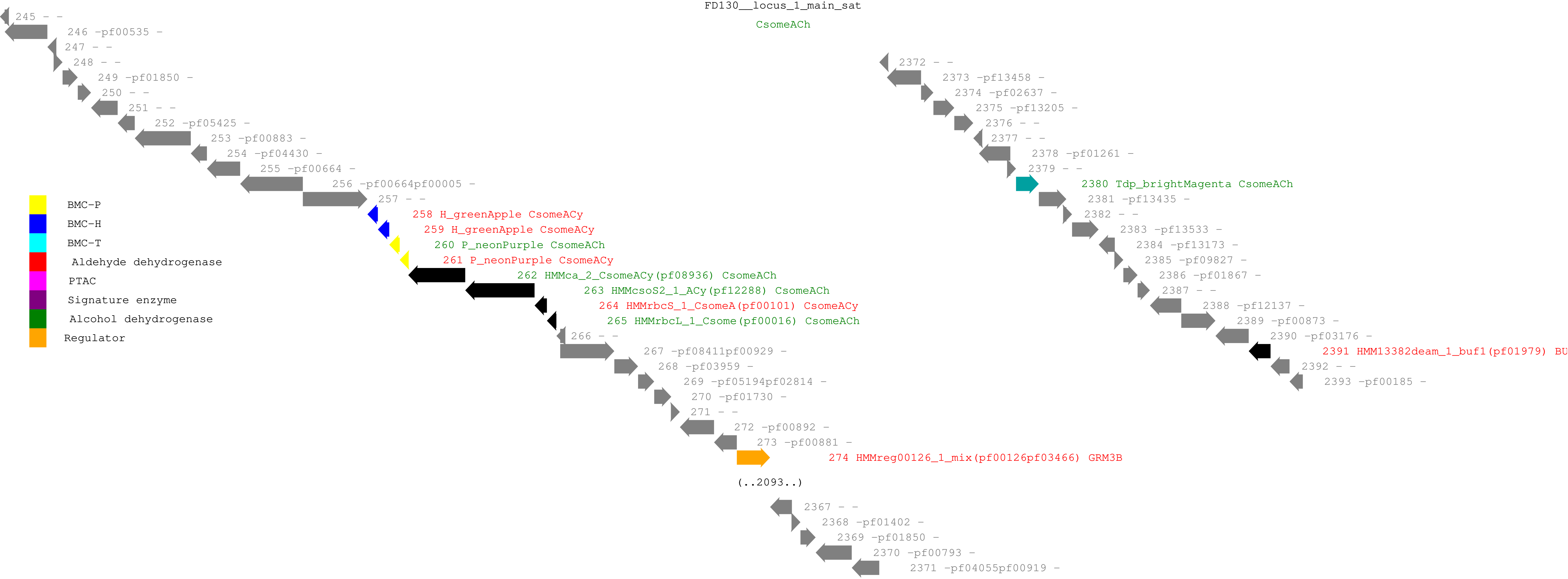

Thiomicrorhabdus sp. G1. Bacteria; Proteobacteria; Gammaproteobacteria; Thiotrichales;

FE785__locus_1_main_sat

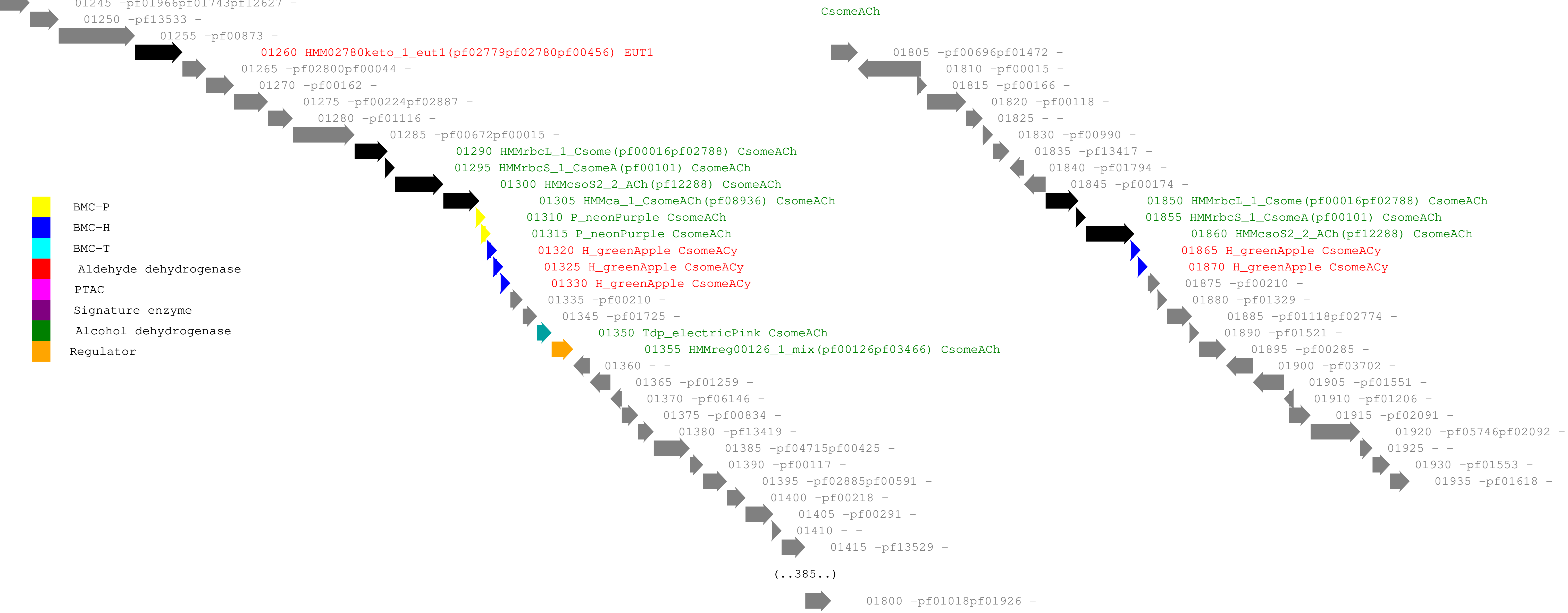

Ferrovum sp. Z-31. Bacteria; Proteobacteria; Betaproteobacteria; Ferrovales; Ferrovaceae;

FEMY__locus_1

CsomeACh

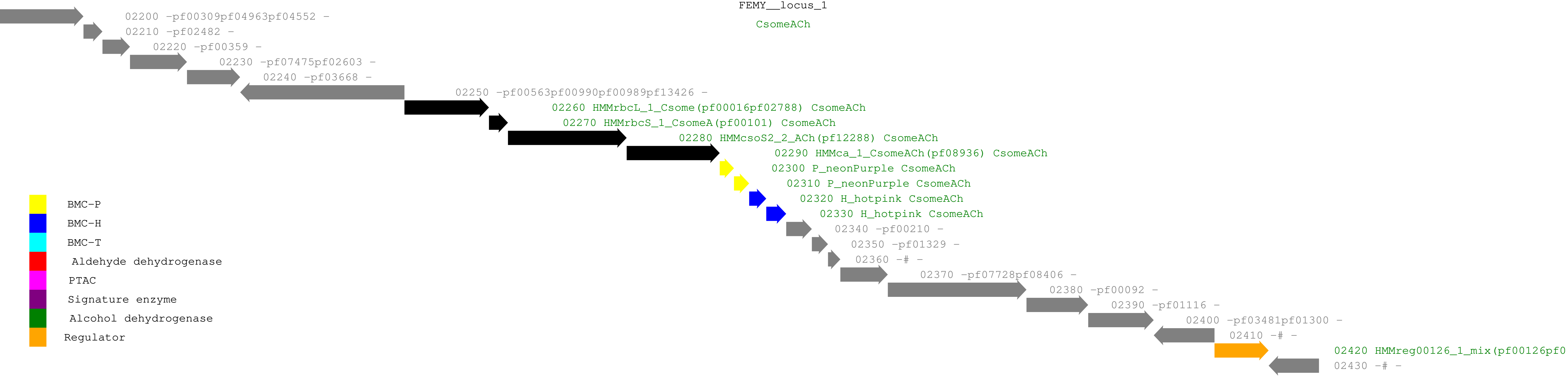

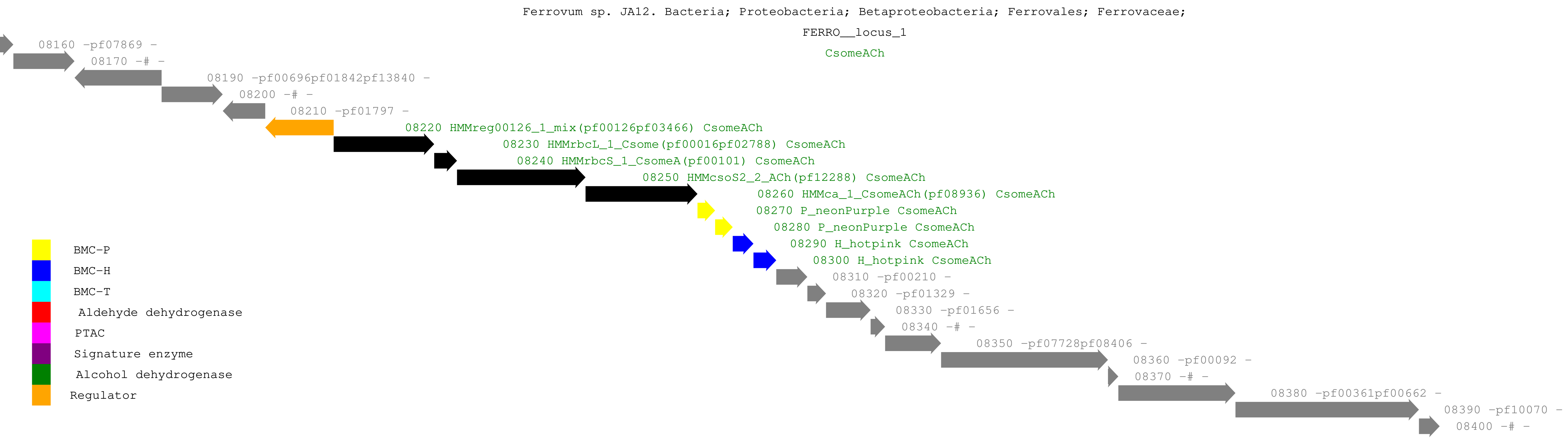

Ferrovum sp. JA12. Bacteria; Proteobacteria; Betaproteobacteria; Ferrovales; Ferrovaceae;
FERRO__locus_1
CsomeACh
08160 -pf07869 -
08170 -# -
08190 -pf00696pf01842pf13840 -
08200 -# -
08210 -pf01797 -
08220 HMMreg00126_1_mix(pf00126pf03466) CsomeACh
08230 HMMrbcL_1_Csome(pf00016pf02788) CsomeACh
08240 HMMrbcS_1_CsomeA(pf00101) CsomeACh
08250 HMMcsoS2_2_ACh(pf12288) CsomeACh
08260 HMMca_1_CsomeACh(pf08936) CsomeACh
08270 P_neonPurple CsomeACh
08280 P_neonPurple CsomeACh
08290 H_hotpink CsomeACh
08300 H_hotpink CsomeACh
08310 -pf00210 -
08320 -pf01329 -
08330 -pf01656 -
08340 -# -
08350 -pf07728pf08406 -
08360 -pf00092 -
08370 -# -
08380 -pf00361pf00662 -
08390 -pf10070 -
08400 -# -
BMC-P
BMC-H
BMC-T
Aldehyde dehydrogenase
PTAC
Signature enzyme
Alcohol dehydrogenase
Regulator

Thiohalobacter thiocyanaticus. Bacteria; Proteobacteria; Gammaproteobacteria; Thiohalobacter.

FOKN1__locus_1

CsomeACh

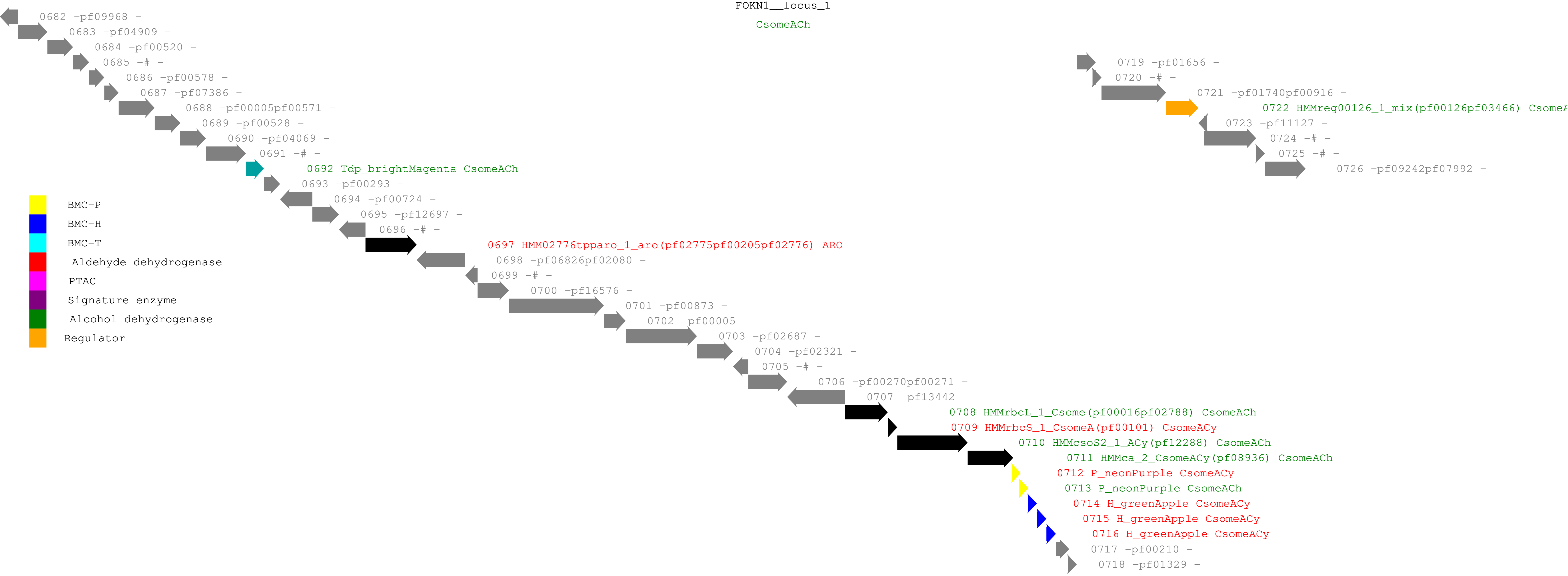

Hydrogenophilales bacterium SM250. Bacteria; Proteobacteria; Hydrogenophilalia; Hydrogenophilales.

FR698__locus_1

CsomeACh

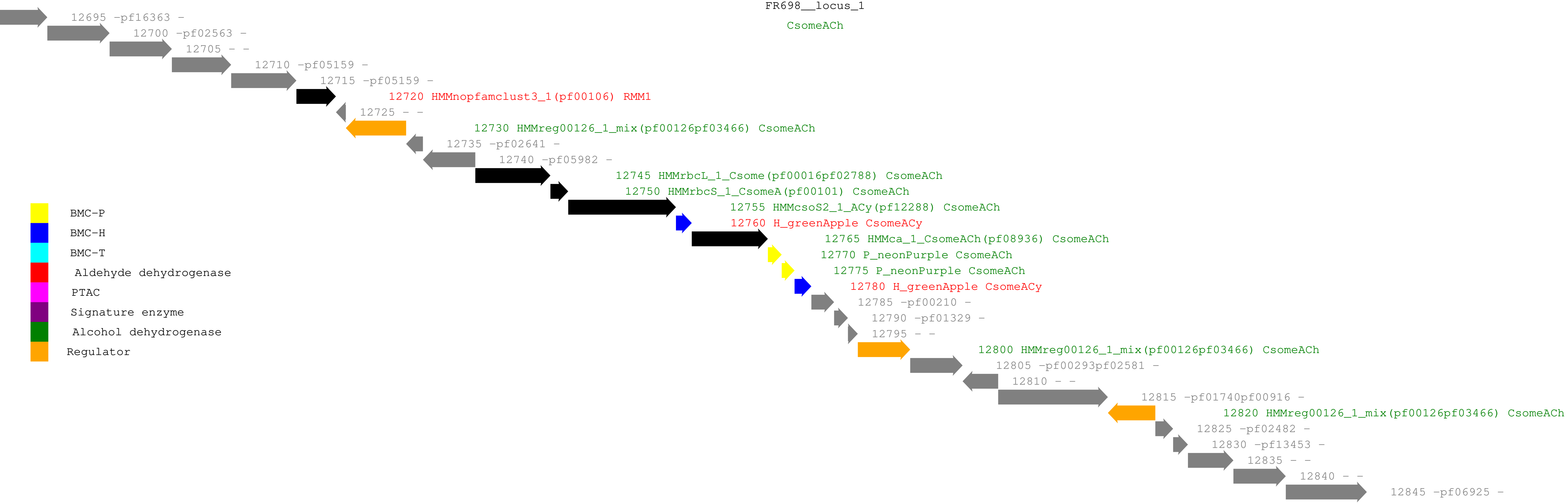

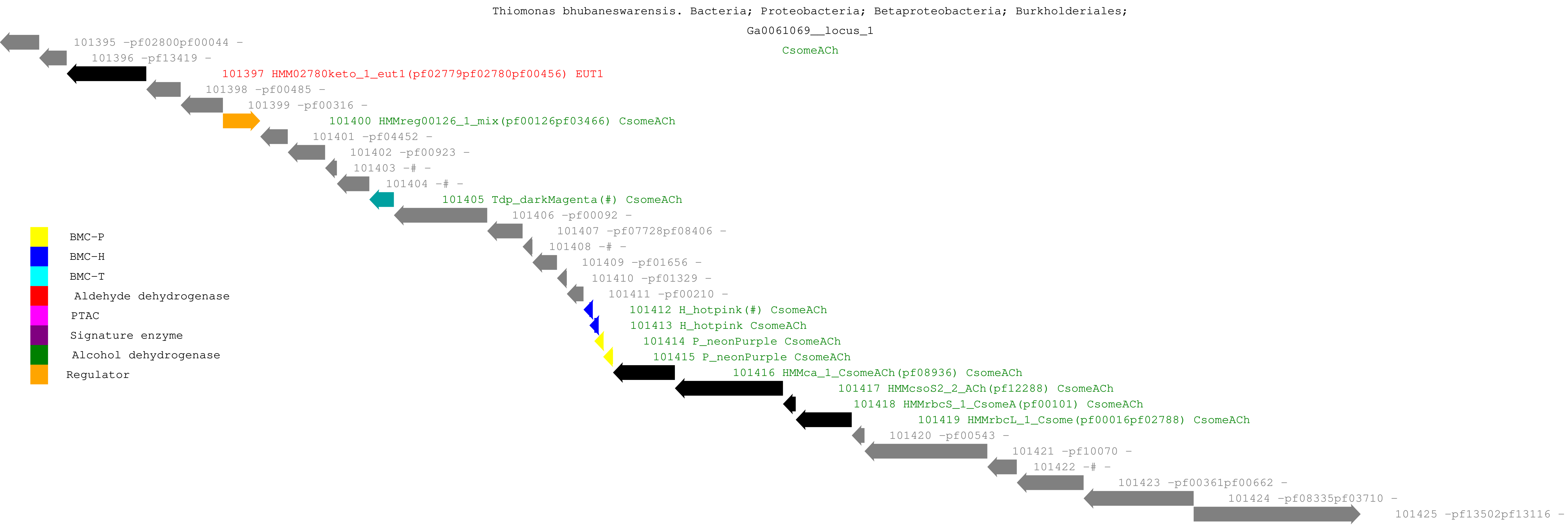
Thiomonas bhubaneswarensis. Bacteria; Proteobacteria; Betaproteobacteria; Burkholderiales;
Ga0061069__locus_1
CsomeACh
101395 -pf02800pf00044 -
101396 -pf13419 -
101397 HMM02780keto_1_eut1(pf02779pf02780pf00456) EUT1
101398 -pf00485 -
101399 -pf00316 -
101400 HMMreg00126_1_mix(pf00126pf03466) CsomeACh
101401 -pf04452 -
101402 -pf00923 -
101403 -# -
101404 -# -
101405 Tdp_darkMagenta(#) CsomeACh
101406 -pf00092 -
101407 -pf07728pf08406 -
101408 -# -
101409 -pf01656 -
101410 -pf01329 -
101411 -pf00210 -
101412 H_hotpink(#) CsomeACh
101413 H_hotpink CsomeACh
101414 P_neonPurple CsomeACh
101415 P_neonPurple CsomeACh
101416 HMMca_1_CsomeACh(pf08936) CsomeACh
101417 HMMcsoS2_2_ACh(pf12288) CsomeACh
101418 HMMrbcS_1_CsomeA(pf00101) CsomeACh
101419 HMMrbcL_1_Csome(pf00016pf02788) CsomeACh
101420 -pf00543 -
101421 -pf10070 -
101422 -# -
101423 -pf00361pf00662 -
101424 -pf08335pf03710 -
101425 -pf13502pf13116 -
BMC-P
BMC-H
BMC-T
Aldehyde dehydrogenase
PTAC
Signature enzyme
Alcohol dehydrogenase
Regulator

mine drainage metagenome. unclassified sequences; metagenomes; ecological metagenomes.

GALL__locus_2

CsomeACh

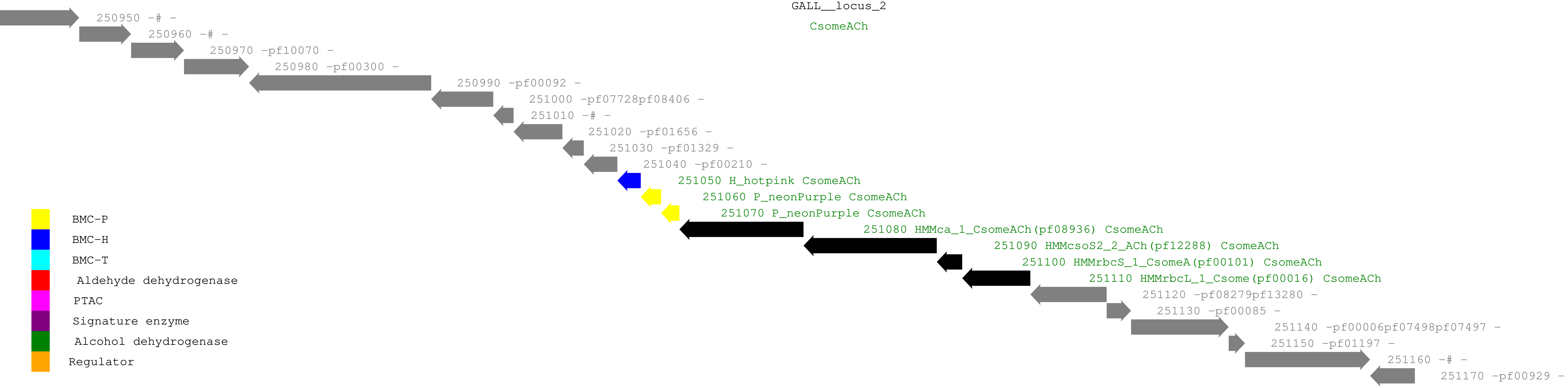

Acidithiobacillus sp. GGI-221. Bacteria; Proteobacteria; Acidithiobacillia; Acidithiobacillales;

GGI1__locus_fragmented

CsomeACh

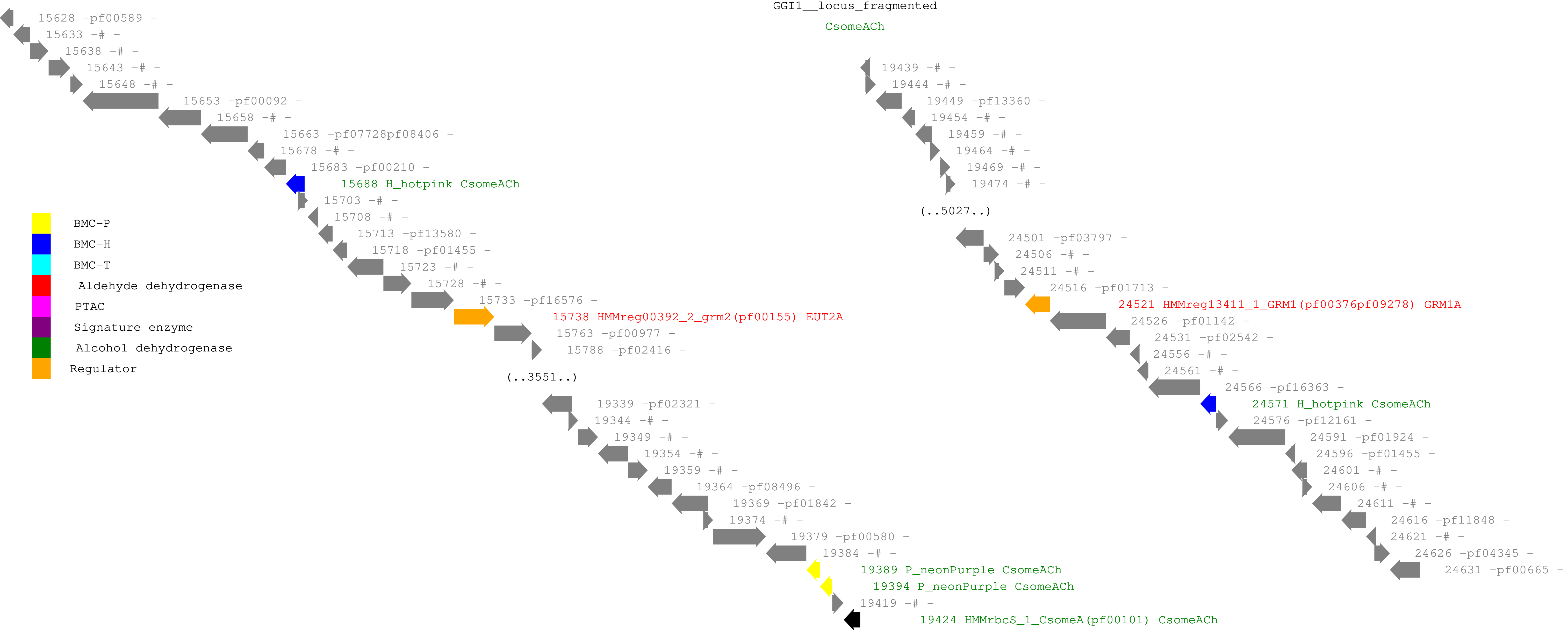

Hydrogenovibrio crunogenus. Bacteria; Proteobacteria; Gammaproteobacteria; Thiotrichales;

GHNINEIG__locus_1

CsomeACh

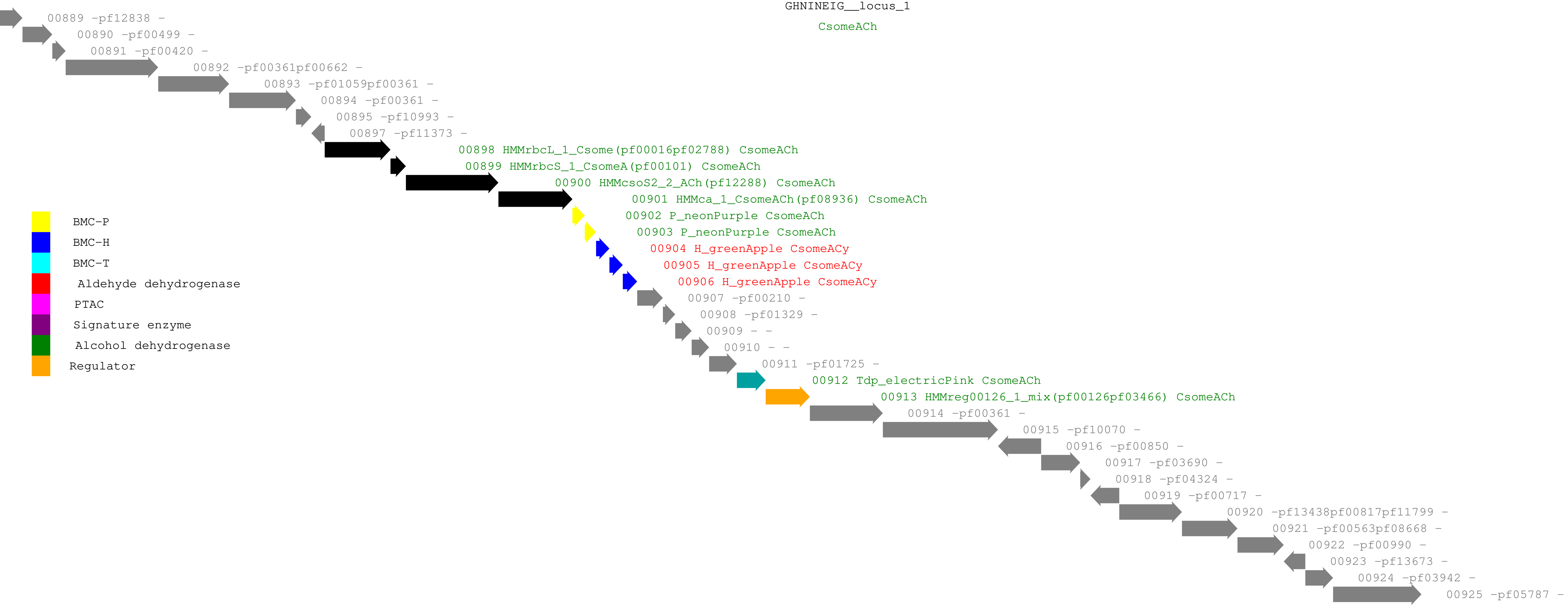

Halothiobacillus neapolitanus (strain ATCC 23641 / c2) (Thiobacillus Bacteria; Proteobacteria; Gammaproteobacteria; Chromatiales;

Hneap__locus_1

CsomeACh

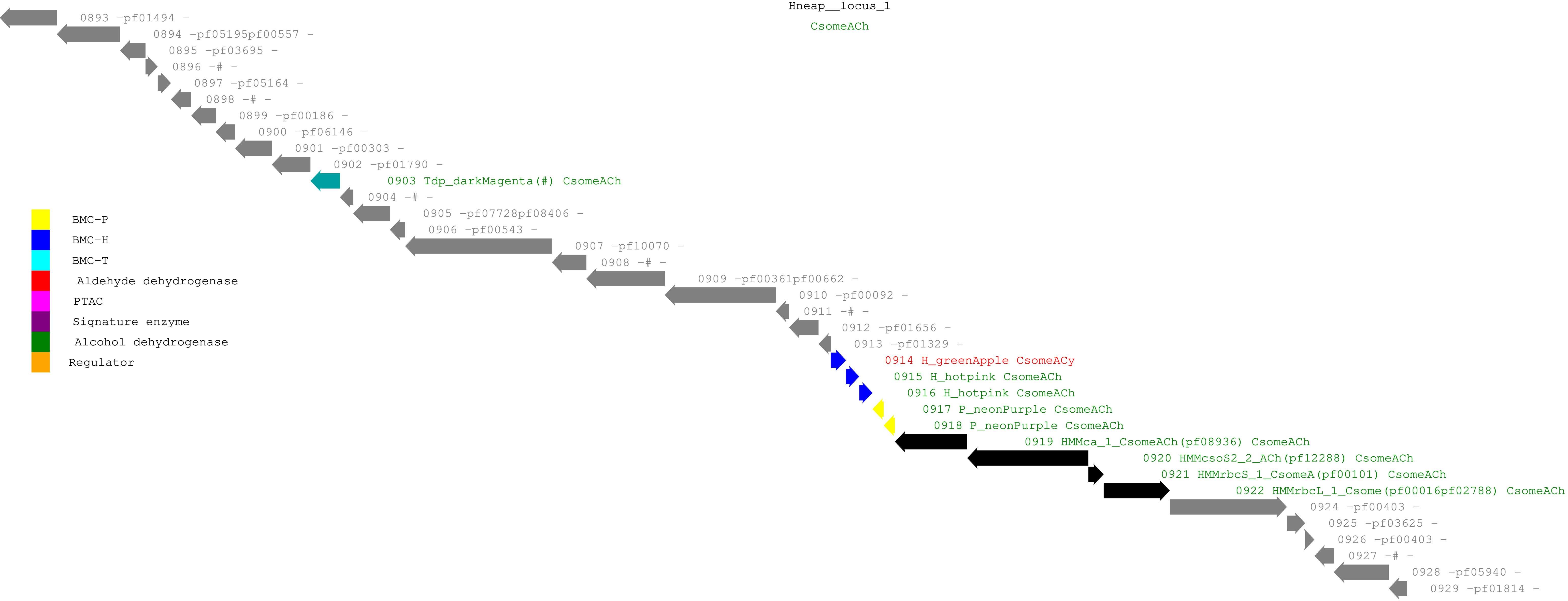

Nitrosomonadaceae bacterium. Bacteria; Proteobacteria; Betaproteobacteria; Nitrosomonadales;

LBMAG32__locus_1

CsomeACh

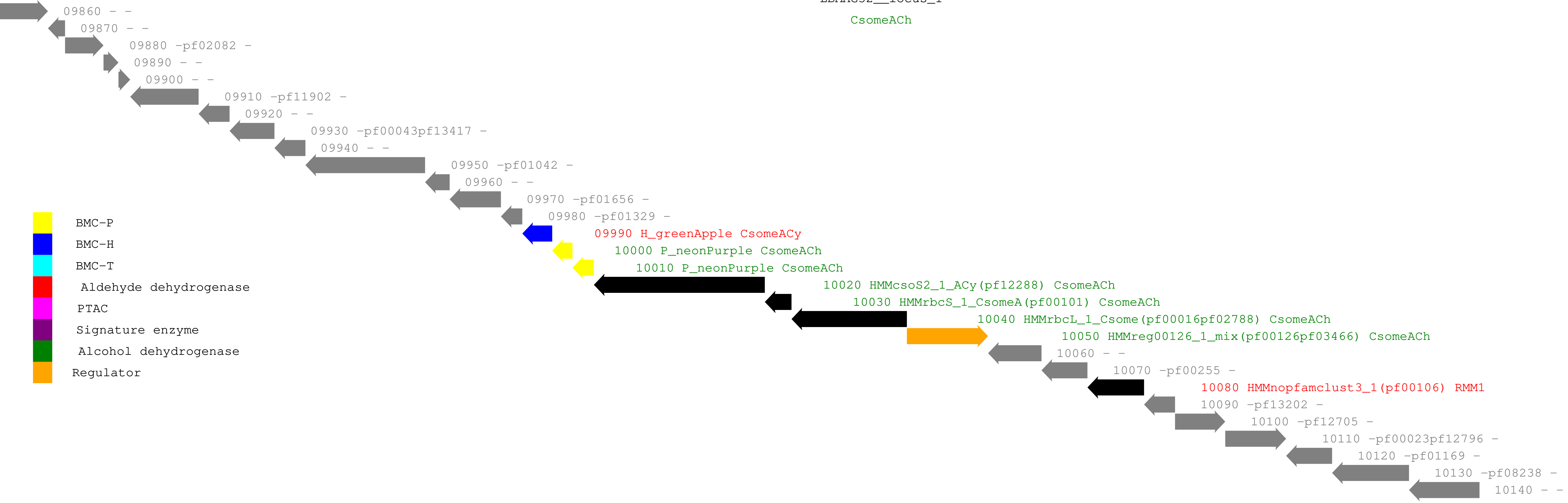

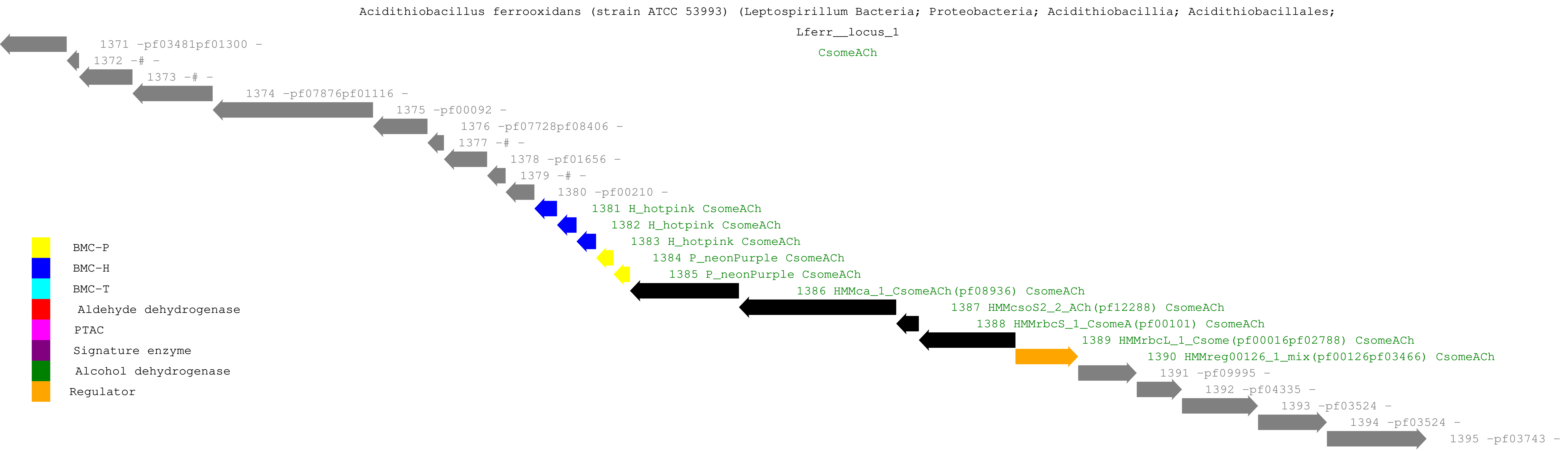
Acidithiobacillus ferrooxidans (strain ATCC 53993) (Leptospirillum Bacteria; Proteobacteria; Acidithiobacillia; Acidithiobacillales;
Lferr__locus_1
CsomeACh
1371 -pf03481pf01300 -
1372 -# -
1373 -# -
1374 -pf07876pf01116 -
1375 -pf00092 -
1376 -pf07728pf08406 -
1377 -# -
1378 -pf01656 -
1379 -# -
1380 -pf00210 -
1381 H_hotpink CsomeACh
1382 H_hotpink CsomeACh
1383 H_hotpink CsomeACh
1384 P_neonPurple CsomeACh
1385 P_neonPurple CsomeACh
1386 HMMca_1_CsomeACh(pf08936) CsomeACh
1387 HMMcsoS2_2_ACh(pf12288) CsomeACh
1388 HMMrbcS_1_CsomeA(pf00101) CsomeACh
1389 HMMrbcL_1_Csome(pf00016pf02788) CsomeACh
1390 HMMreg00126_1_mix(pf00126pf03466) CsomeACh
1391 -pf09995 -
1392 -pf04335 -
1393 -pf03524 -
1394 -pf03524 -
1395 -pf03743 -
BMC-P
BMC-H
BMC-T
Aldehyde dehydrogenase
PTAC
Signature enzyme
Alcohol dehydrogenase
Regulator

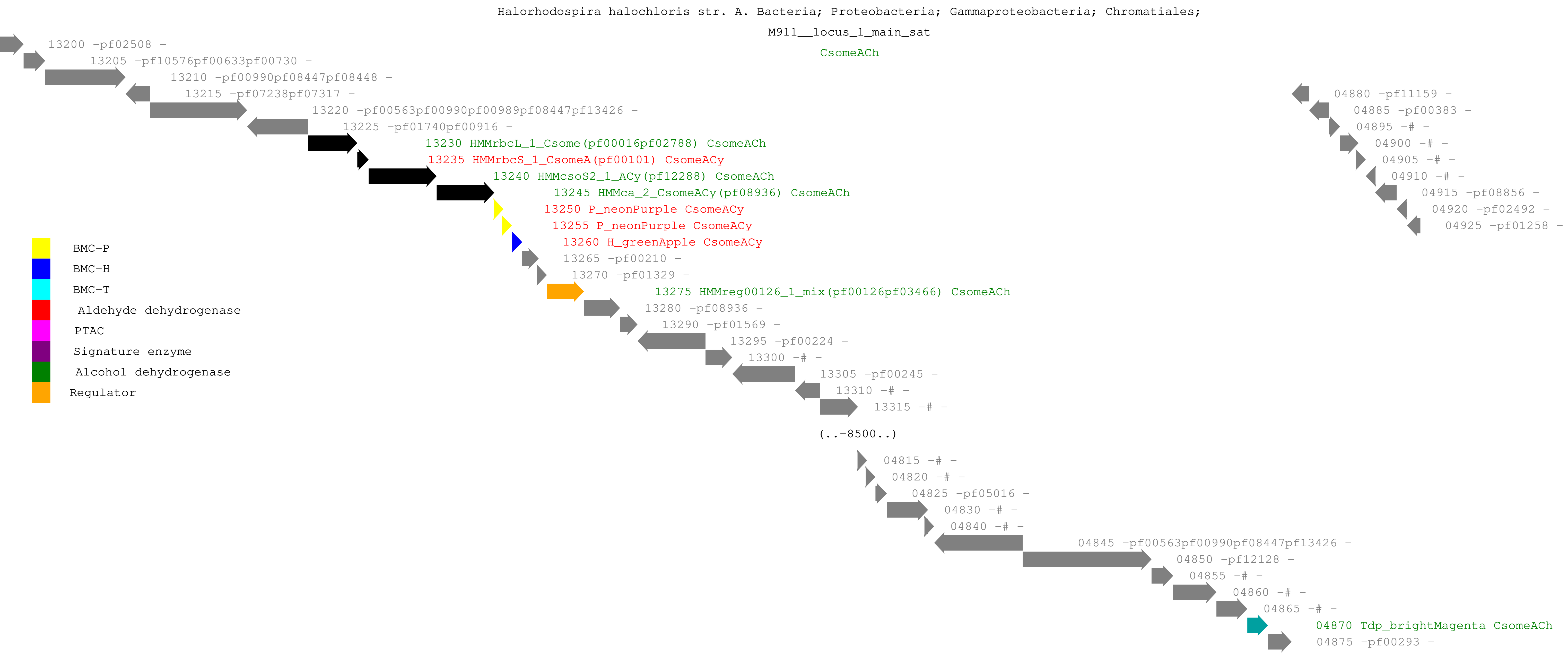
Halorhodospira halochloris str. A. Bacteria; Proteobacteria; Gammaproteobacteria; Chromatiales;
M911__locus_1_main_sat
CsomeACh
13200 -pf02508 -
13205 -pf10576pf00633pf00730 -
13210 -pf00990pf08447pf08448 -
13215 -pf07238pf07317 -
13220 -pf00563pf00990pf00989pf08447pf13426 -
13225 -pf01740pf00916 -
13230 HMMrbcL_1_Csome(pf00016pf02788) CsomeACh
13235 HMMrbcS_1_CsomeA(pf00101) CsomeACy
13240 HMMcsoS2_1_ACy(pf12288) CsomeACh
13245 HMMca_2_CsomeACy(pf08936) CsomeACh
13250 P_neonPurple CsomeACy
13255 P_neonPurple CsomeACy
13260 H_greenApple CsomeACy
13265 -pf00210 -
13270 -pf01329 -
13275 HMMreg00126_1_mix(pf00126pf03466) CsomeACh
13280 -pf08936 -
13290 -pf01569 -
13295 -pf00224 -
13300 -# -
13305 -pf00245 -
13310 -# -
13315 -# -
(..-8500..)
04815 -# -
04820 -# -
04825 -pf05016 -
04830 -# -
04840 -# -
04845 -pf00563pf00990pf08447pf13426 -
04850 -pf12128 -
04855 -# -
04860 -# -
04865 -# -
04870 Tdp_brightMagenta CsomeACh
04875 -pf00293 -
04880 -pf11159 -
04885 -pf00383 -
04895 -# -
04900 -# -
04905 -# -
04910 -# -
04915 -pf08856 -
04920 -pf02492 -
04925 -pf01258 -
BMC-P
BMC-H
BMC-T
Aldehyde dehydrogenase
PTAC
Signature enzyme
Alcohol dehydrogenase
Regulator

Marichromatium purpuratum 984. Bacteria; Proteobacteria; Gammaproteobacteria; Chromatiales;

MARPU__locus_1

CsomeACh

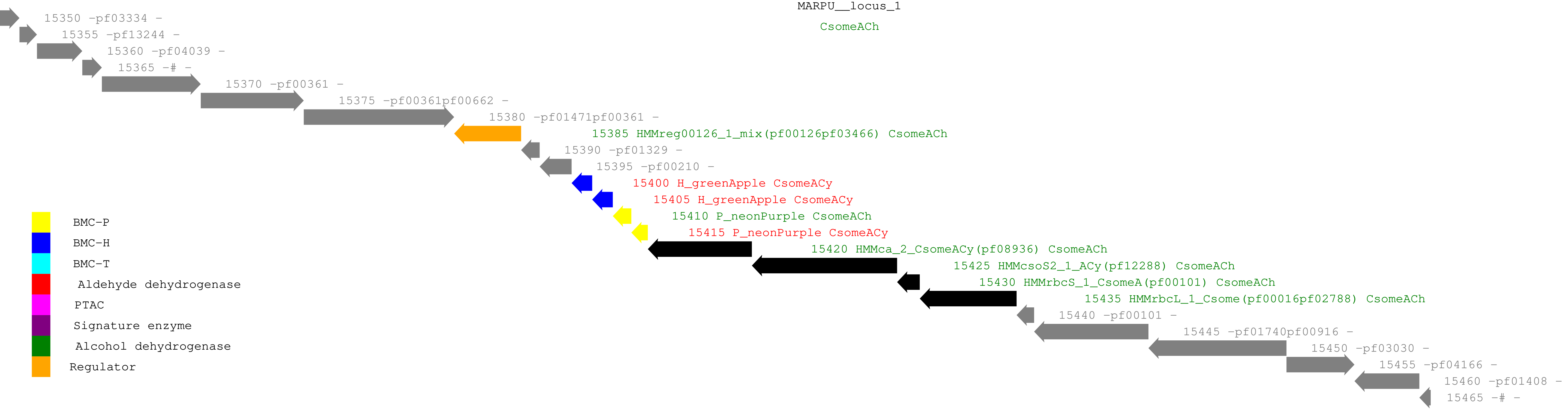

hydrothermal vent metagenome. unclassified sequences; metagenomes; ecological metagenomes.

MNBD_GAMMA04-_locus_1_main_sat

CsomeACh

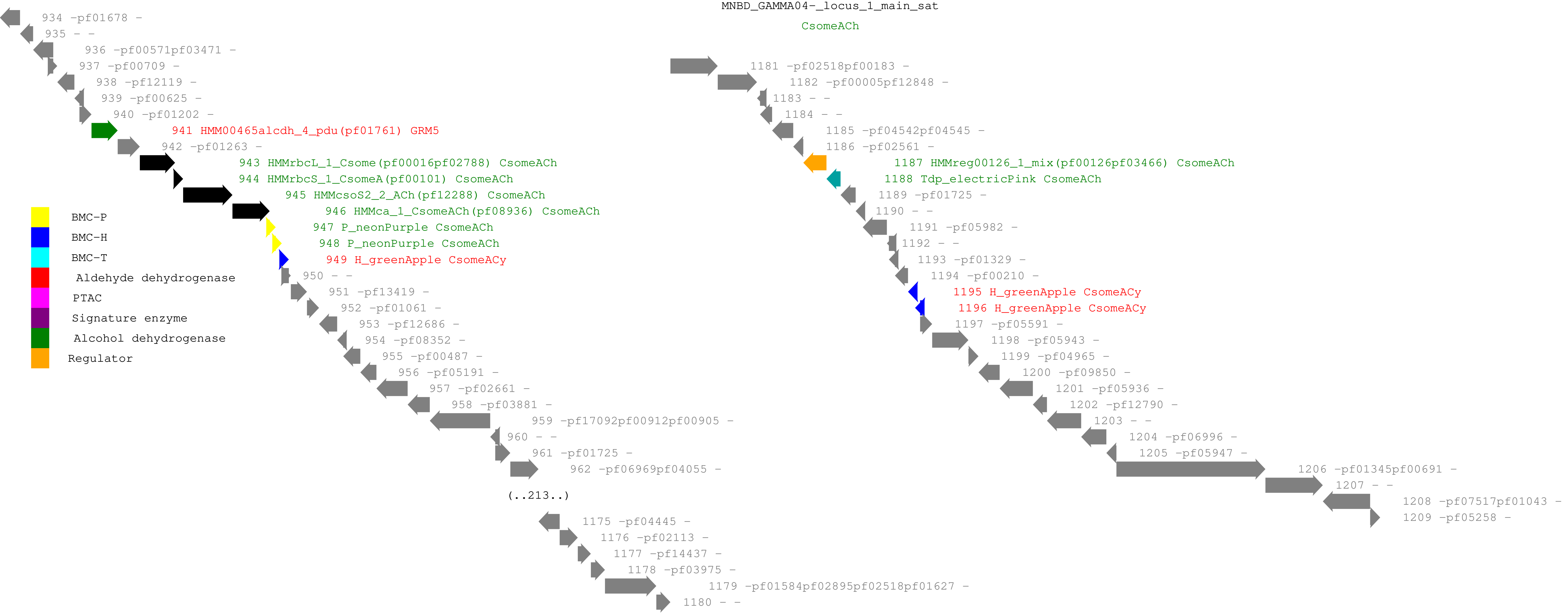

hydrothermal vent metagenome. unclassified sequences; metagenomes; ecological metagenomes.

MNBD_GAMMA16-_locus_1

CsomeACh

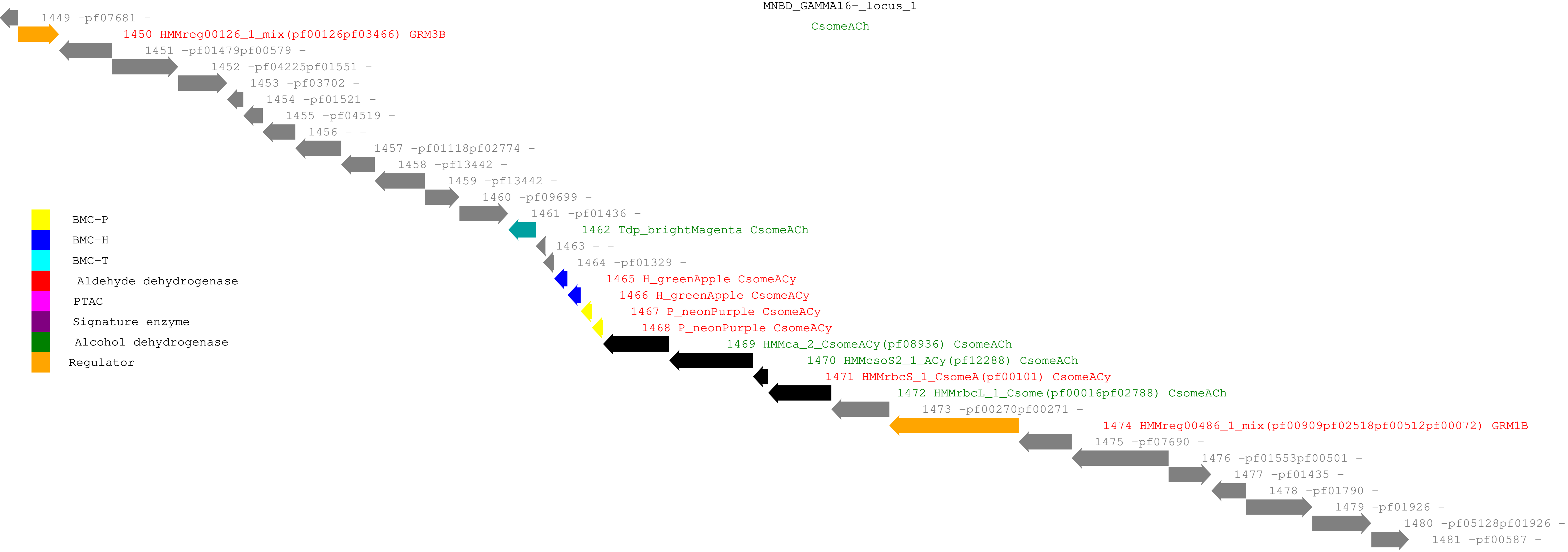

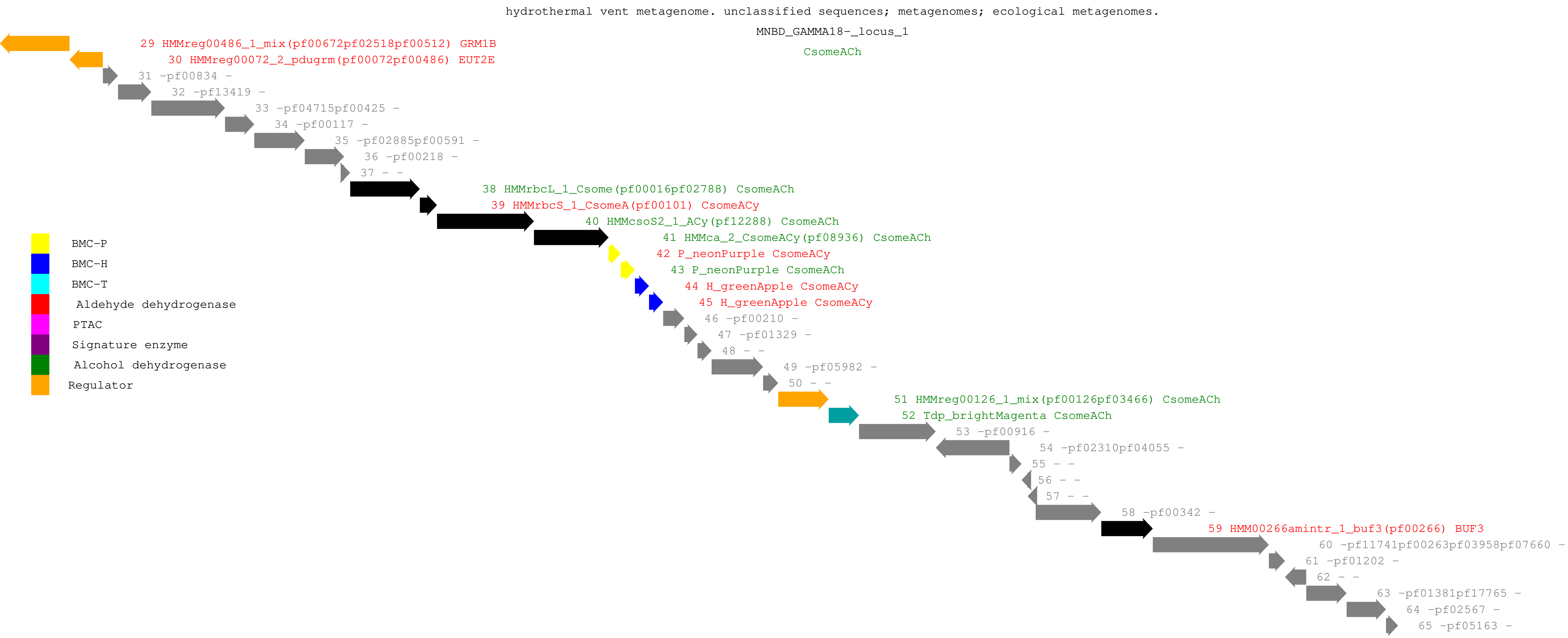

hydrothermal vent metagenome. unclassified sequences; metagenomes; ecological metagenomes.
MNBD_GAMMA18-_locus_1
CsomeACh
29 HMMreg00486_1_mix(pf00672pf02518pf00512) GRM1B
30 HMMreg00072_2_pdugrm(pf00072pf00486) EUT2E
31 -pf00834 -
32 -pf13419 -
33 -pf04715pf00425 -
34 -pf00117 -
35 -pf02885pf00591 -
36 -pf00218 -
37 - -
38 HMMrbcL_1_Csome(pf00016pf02788) CsomeACh
39 HMMrbcS_1_CsomeA(pf00101) CsomeACy
40 HMMcsoS2_1_ACy(pf12288) CsomeACh
41 HMMca_2_CsomeACy(pf08936) CsomeACh
42 P_neonPurple CsomeACy
43 P_neonPurple CsomeACh
44 H_greenApple CsomeACy
45 H_greenApple CsomeACy
46 -pf00210 -
47 -pf01329 -
48 - -
49 -pf05982 -
50 - -
51 HMMreg00126_1_mix(pf00126pf03466) CsomeACh
52 Tdp_brightMagenta CsomeACh
53 -pf00916 -
54 -pf02310pf04055 -
55 - -
56 - -
57 - -
58 -pf00342 -
59 HMM00266amintr_1_buf3(pf00266) BUF3
60 -pf11741pf00263pf03958pf07660 -
61 -pf01202 -
62 - -
63 -pf01381pf17765 -
64 -pf02567 -
65 -pf05163 -
BMC-P
BMC-H
BMC-T
Aldehyde dehydrogenase
PTAC
Signature enzyme
Alcohol dehydrogenase
Regulator

hydrothermal vent metagenome. unclassified sequences; metagenomes; ecological metagenomes.

MNBD_GAMMA20-_locus_1

CsomeACh

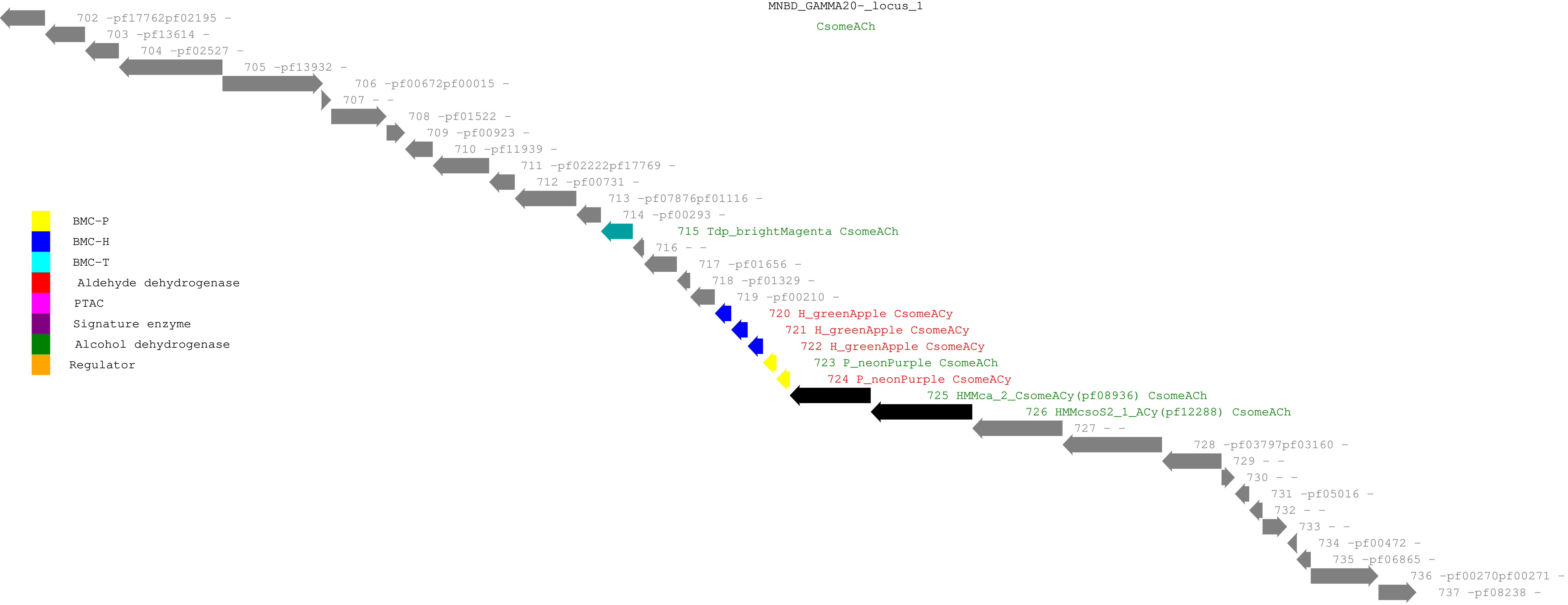

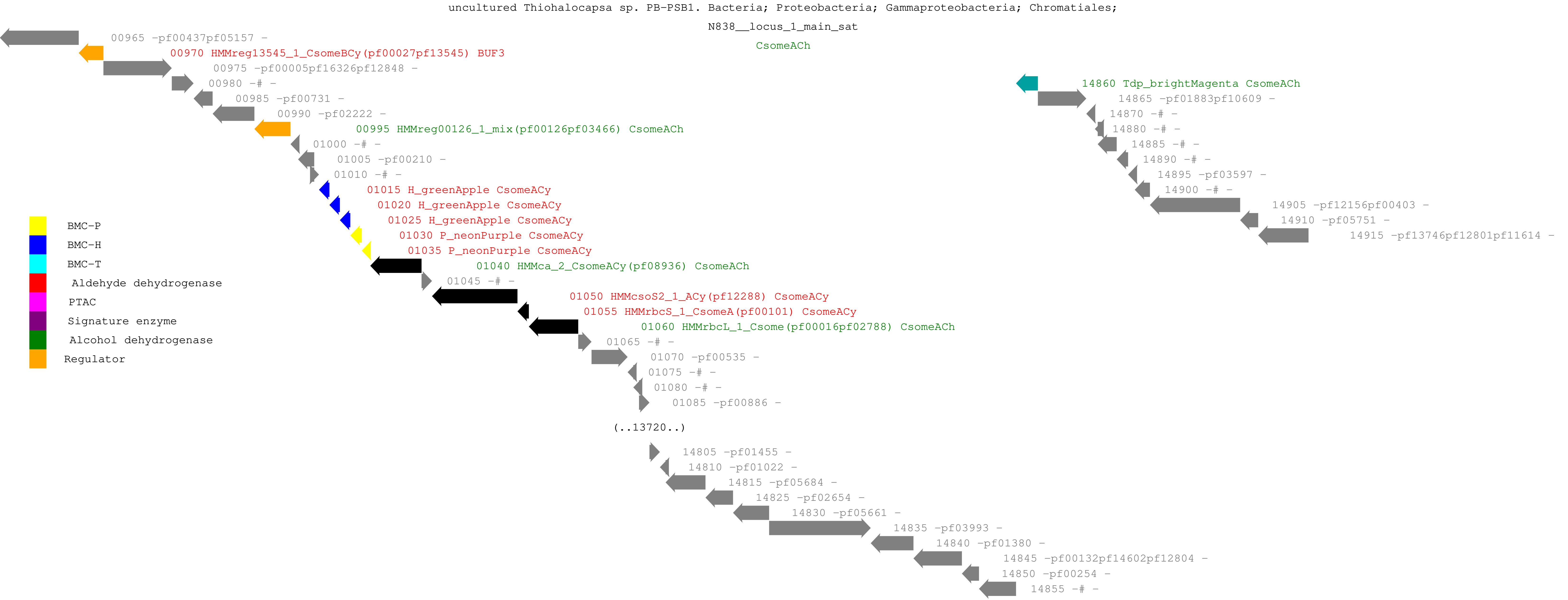

uncultured Thiohalocapsa sp. PB-PSB1. Bacteria; Proteobacteria; Gammaproteobacteria; Chromatiales;
N838__locus_1_main_sat
CsomeACh
00965 -pf00437pf05157 -
00970 HMMreg13545_1_CsomeBCy(pf00027pf13545) BUF3
00975 -pf00005pf16326pf12848 -
00980 -# -
00985 -pf00731 -
00990 -pf02222 -
00995 HMMreg00126_1_mix(pf00126pf03466) CsomeACh
01000 -# -
01005 -pf00210 -
01010 -# -
01015 H_greenApple CsomeACy
01020 H_greenApple CsomeACy
01025 H_greenApple CsomeACy
01030 P_neonPurple CsomeACy
01035 P_neonPurple CsomeACy
01040 HMMca_2_CsomeACy(pf08936) CsomeACh
01045 -# -
01050 HMMcsoS2_1_ACy(pf12288) CsomeACy
01055 HMMrbcS_1_CsomeA(pf00101) CsomeACy
01060 HMMrbcL_1_Csome(pf00016pf02788) CsomeACh
01065 -# -
01070 -pf00535 -
01075 -# -
01080 -# -
01085 -pf00886 -
(..13720..)
14805 -pf01455 -
14810 -pf01022 -
14815 -pf05684 -
14825 -pf02654 -
14830 -pf05661 -
14835 -pf03993 -
14840 -pf01380 -
14845 -pf00132pf14602pf12804 -
14850 -pf00254 -
14855 -# -
14860 Tdp_brightMagenta CsomeACh
14865 -pf01883pf10609 -
14870 -# -
14880 -# -
14885 -# -
14890 -# -
14895 -pf03597 -
14900 -# -
14905 -pf12156pf00403 -
14910 -pf05751 -
14915 -pf13746pf12801pf11614 -
BMC-P
BMC-H
BMC-T
Aldehyde dehydrogenase
PTAC
Signature enzyme
Alcohol dehydrogenase
Regulator

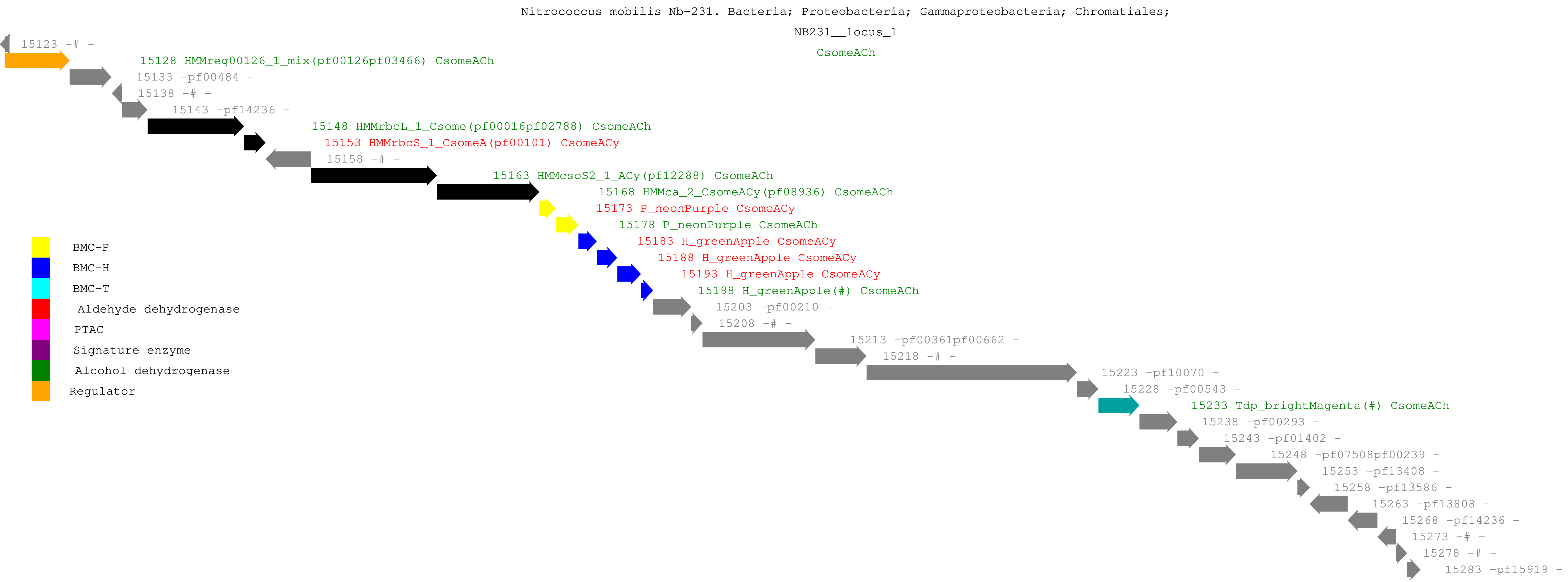
Nitrococcus mobilis Nb-231. Bacteria; Proteobacteria; Gammaproteobacteria; Chromatiales;
NB231__locus_1
CsomeACh
15123 -# -
15128 HMMreg00126_1_mix(pf00126pf03466) CsomeACh
15133 -pf00484 -
15138 -# -
15143 -pf14236 -
15148 HMMrbcL_1_Csome(pf00016pf02788) CsomeACh
15153 HMMrbcS_1_CsomeA(pf00101) CsomeACy
15158 -# -
15163 HMMcsoS2_1_ACy(pf12288) CsomeACh
15168 HMMca_2_CsomeACy(pf08936) CsomeACh
15173 P_neonPurple CsomeACy
15178 P_neonPurple CsomeACh
15183 H_greenApple CsomeACy
15188 H_greenApple CsomeACy
15193 H_greenApple CsomeACy
15198 H_greenApple(#) CsomeACh
15203 -pf00210 -
15208 -# -
15213 -pf00361pf00662 -
15218 -# -
15223 -pf10070 -
15228 -pf00543 -
15233 Tdp_brightMagenta(#) CsomeACh
15238 -pf00293 -
15243 -pf01402 -
15248 -pf07508pf00239 -
15253 -pf13408 -
15258 -pf13586 -
15263 -pf13808 -
15268 -pf14236 -
15273 -# -
15278 -# -
15283 -pf15919 -
BMC-P
BMC-H
BMC-T
Aldehyde dehydrogenase
PTAC
Signature enzyme
Alcohol dehydrogenase
Regulator

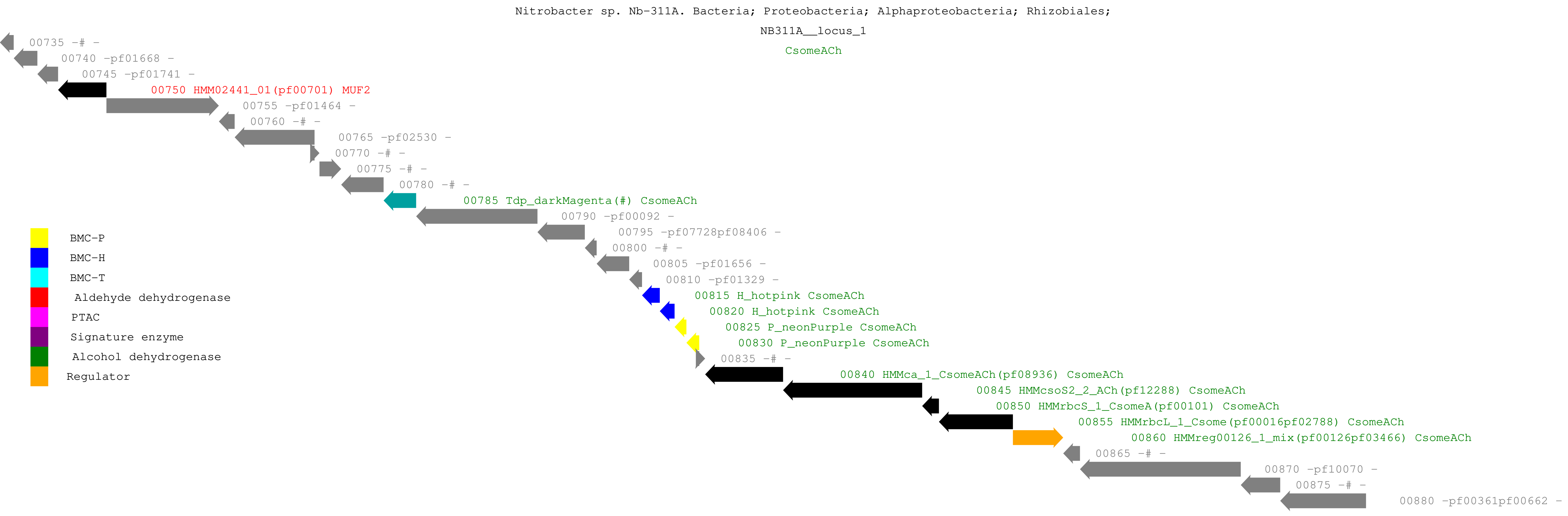
Nitrobacter sp. Nb-311A. Bacteria; Proteobacteria; Alphaproteobacteria; Rhizobiales;
NB311A__locus_1
CsomeACh
00735 -# -
00740 -pf01668 -
00745 -pf01741 -
00750 HMM02441_01(pf00701) MUF2
00755 -pf01464 -
00760 -# -
00765 -pf02530 -
00770 -# -
00775 -# -
00780 -# -
00785 Tdp_darkMagenta(#) CsomeACh
00790 -pf00092 -
00795 -pf07728pf08406 -
00800 -# -
00805 -pf01656 -
00810 -pf01329 -
00815 H_hotpink CsomeACh
00820 H_hotpink CsomeACh
00825 P_neonPurple CsomeACh
00830 P_neonPurple CsomeACh
00835 -# -
00840 HMMca_1_CsomeACh(pf08936) CsomeACh
00845 HMMcsoS2_2_ACh(pf12288) CsomeACh
00850 HMMrbcS_1_CsomeA(pf00101) CsomeACh
00855 HMMrbcL_1_Csome(pf00016pf02788) CsomeACh
00860 HMMreg00126_1_mix(pf00126pf03466) CsomeACh
00865 -# -
00870 -pf10070 -
00875 -# -
00880 -pf00361pf00662 -
BMC-P
BMC-H
BMC-T
Aldehyde dehydrogenase
PTAC
Signature enzyme
Alcohol dehydrogenase
Regulator

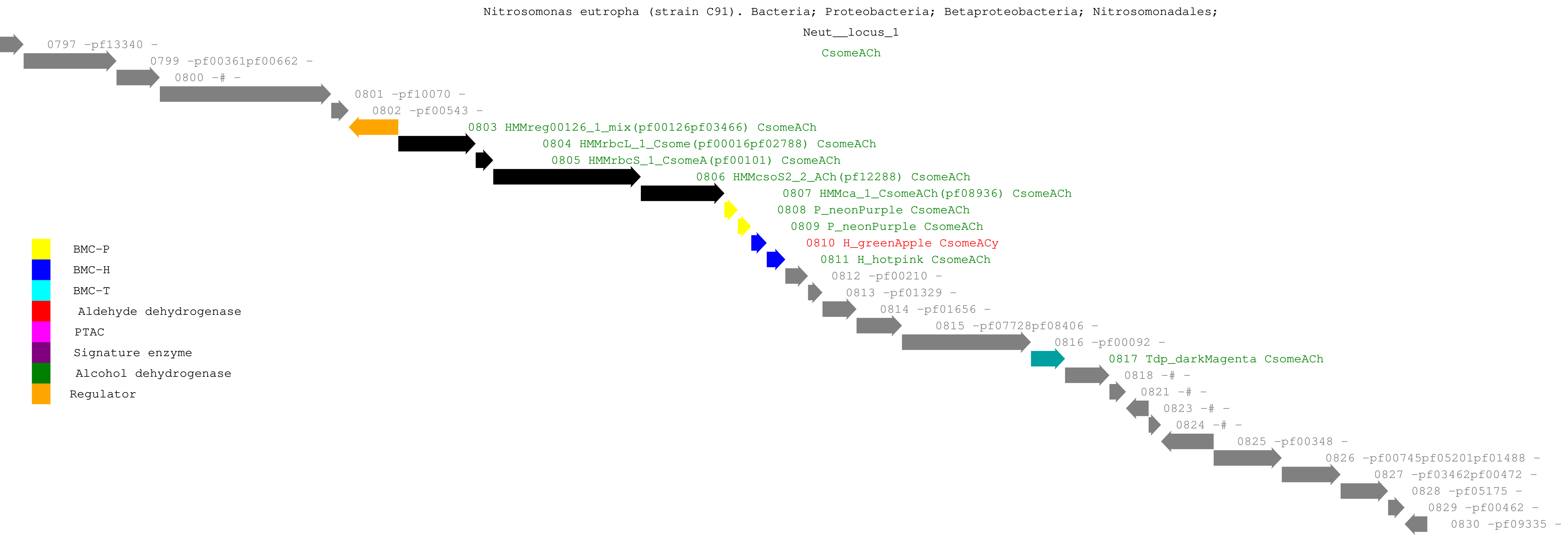
Nitrosomonas eutropha (strain C91). Bacteria; Proteobacteria; Betaproteobacteria; Nitrosomonadales;
Neut__locus_1
CsomeACh
0797 -pf13340 -
0799 -pf00361pf00662 -
0800 -# -
0801 -pf10070 -
0802 -pf00543 -
0803 HMMreg00126_1_mix(pf00126pf03466) CsomeACh
0804 HMMrbcL_1_Csome(pf00016pf02788) CsomeACh
0805 HMMrbcS_1_CsomeA(pf00101) CsomeACh
0806 HMMcsoS2_2_ACh(pf12288) CsomeACh
0807 HMMca_1_CsomeACh(pf08936) CsomeACh
0808 P_neonPurple CsomeACh
0809 P_neonPurple CsomeACh
0810 H_greenApple CsomeACy
0811 H_hotpink CsomeACh
0812 -pf00210 -
0813 -pf01329 -
0814 -pf01656 -
0815 -pf07728pf08406 -
0816 -pf00092 -
0817 Tdp_darkMagenta CsomeACh
0818 -# -
0821 -# -
0823 -# -
0824 -# -
0825 -pf00348 -
0826 -pf00745pf05201pf01488 -
0827 -pf03462pf00472 -
0828 -pf05175 -
0829 -pf00462 -
0830 -pf09335 -
BMC-P
BMC-H
BMC-T
Aldehyde dehydrogenase
PTAC
Signature enzyme
Alcohol dehydrogenase
Regulator

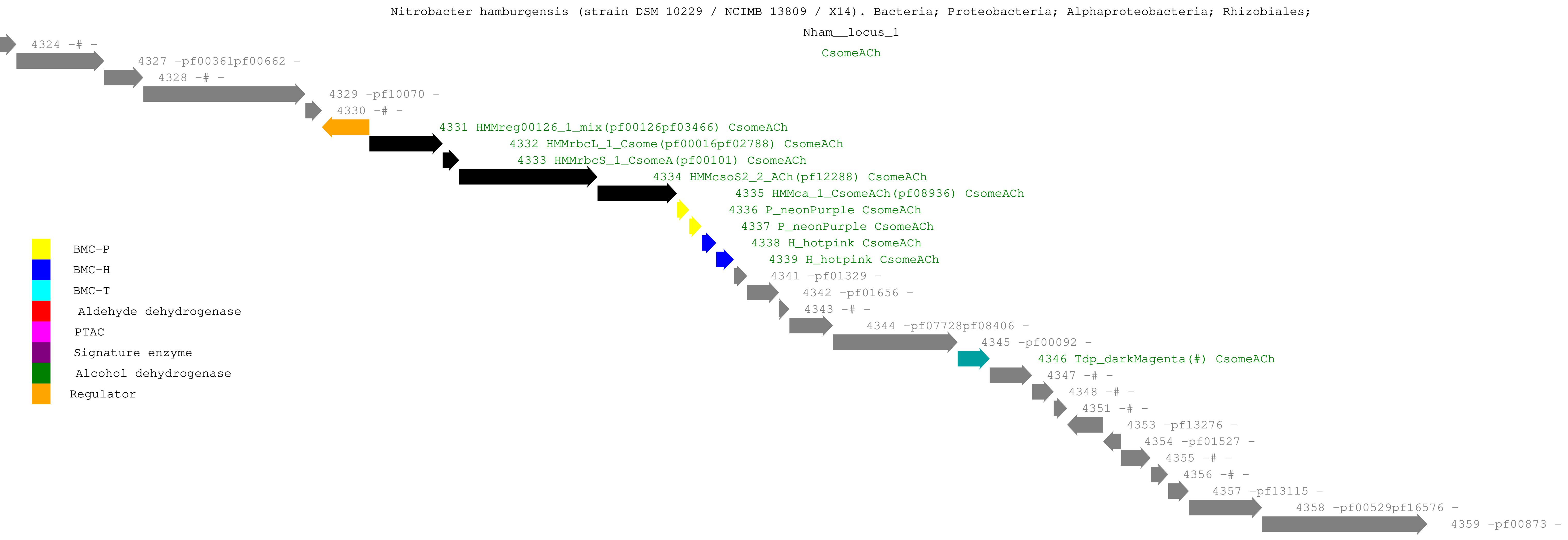
Nitrobacter hamburgensis (strain DSM 10229 / NCIMB 13809 / X14). Bacteria; Proteobacteria; Alphaproteobacteria; Rhizobiales;
Nham__locus_1
CsomeACh
4324 -# -
4327 -pf00361pf00662 -
4328 -# -
4329 -pf10070 -
4330 -# -
4331 HMMreg00126_1_mix(pf00126pf03466) CsomeACh
4332 HMMrbcL_1_Csome(pf00016pf02788) CsomeACh
4333 HMMrbcS_1_CsomeA(pf00101) CsomeACh
4334 HMMcsoS2_2_ACh(pf12288) CsomeACh
4335 HMMca_1_CsomeACh(pf08936) CsomeACh
4336 P_neonPurple CsomeACh
4337 P_neonPurple CsomeACh
4338 H_hotpink CsomeACh
4339 H_hotpink CsomeACh
4341 -pf01329 -
4342 -pf01656 -
4343 -# -
4344 -pf07728pf08406 -
4345 -pf00092 -
4346 Tdp_darkMagenta(#) CsomeACh
4347 -# -
4348 -# -
4351 -# -
4353 -pf13276 -
4354 -pf01527 -
4355 -# -
4356 -# -
4357 -pf13115 -
4358 -pf00529pf16576 -
4359 -pf00873 -
BMC-P
BMC-H
BMC-T
Aldehyde dehydrogenase
PTAC
Signature enzyme
Alcohol dehydrogenase
Regulator

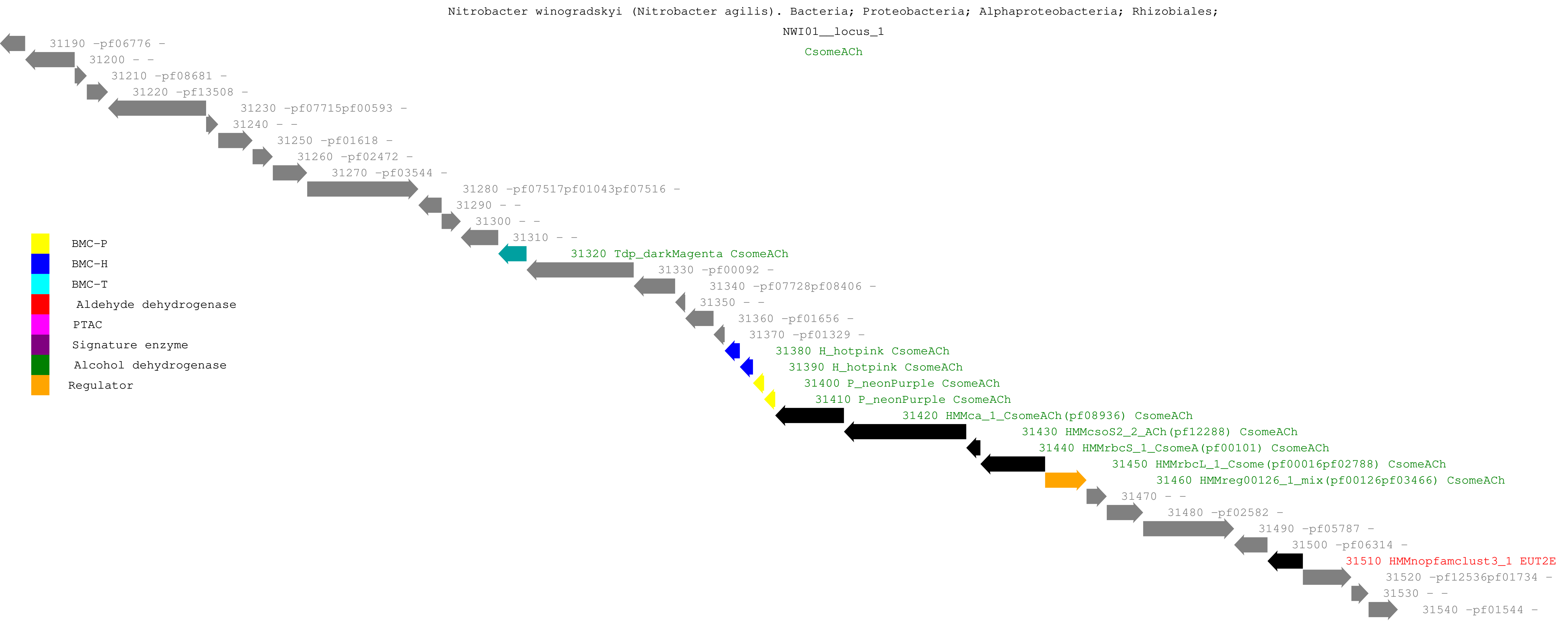

Nitrobacter winogradskyi (Nitrobacter agilis). Bacteria; Proteobacteria; Alphaproteobacteria; Rhizobiales;
NWI01__locus_1
CsomeACh
31190 -pf06776 -
31200 - -
31210 -pf08681 -
31220 -pf13508 -
31230 -pf07715pf00593 -
31240 - -
31250 -pf01618 -
31260 -pf02472 -
31270 -pf03544 -
31280 -pf07517pf01043pf07516 -
31290 - -
31300 - -
31310 - -
31320 Tdp_darkMagenta CsomeACh
31330 -pf00092 -
31340 -pf07728pf08406 -
31350 - -
31360 -pf01656 -
31370 -pf01329 -
31380 H_hotpink CsomeACh
31390 H_hotpink CsomeACh
31400 P_neonPurple CsomeACh
31410 P_neonPurple CsomeACh
31420 HMMca_1_CsomeACh(pf08936) CsomeACh
31430 HMMcsoS2_2_ACh(pf12288) CsomeACh
31440 HMMrbcS_1_CsomeA(pf00101) CsomeACh
31450 HMMrbcL_1_Csome(pf00016pf02788) CsomeACh
31460 HMMreg00126_1_mix(pf00126pf03466) CsomeACh
31470 - -
31480 -pf02582 -
31490 -pf05787 -
31500 -pf06314 -
31510 HMMnopfamclust3_1 EUT2E
31520 -pf12536pf01734 -
31530 - -
31540 -pf01544 -
BMC-P
BMC-H
BMC-T
Aldehyde dehydrogenase
PTAC
Signature enzyme
Alcohol dehydrogenase
Regulator

Nitrobacter winogradskyi (strain ATCC 25391 / DSM 10237 / CIP 104748 / Bacteria; Proteobacteria; Alphaproteobacteria; Rhizobiales;

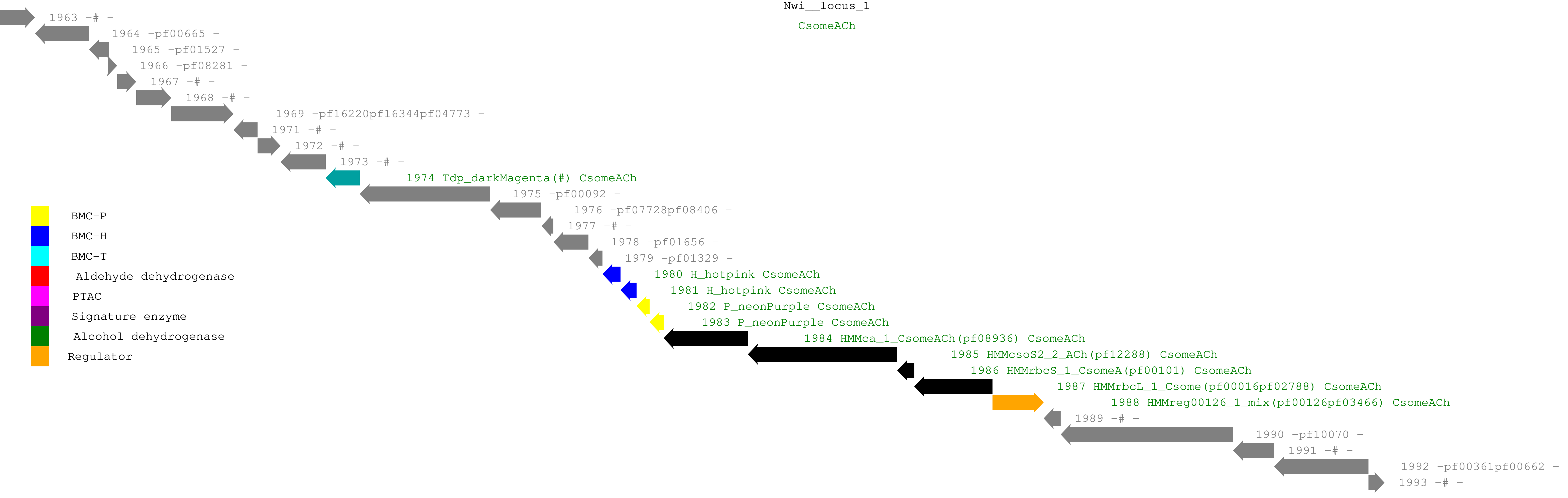

Ferrithrix thermotolerans DSM 19514. Bacteria; Actinobacteria; Acidimicrobiia; Acidimicrobiales;

SAMN02745225__locus_1_main_sat

CsomeACh

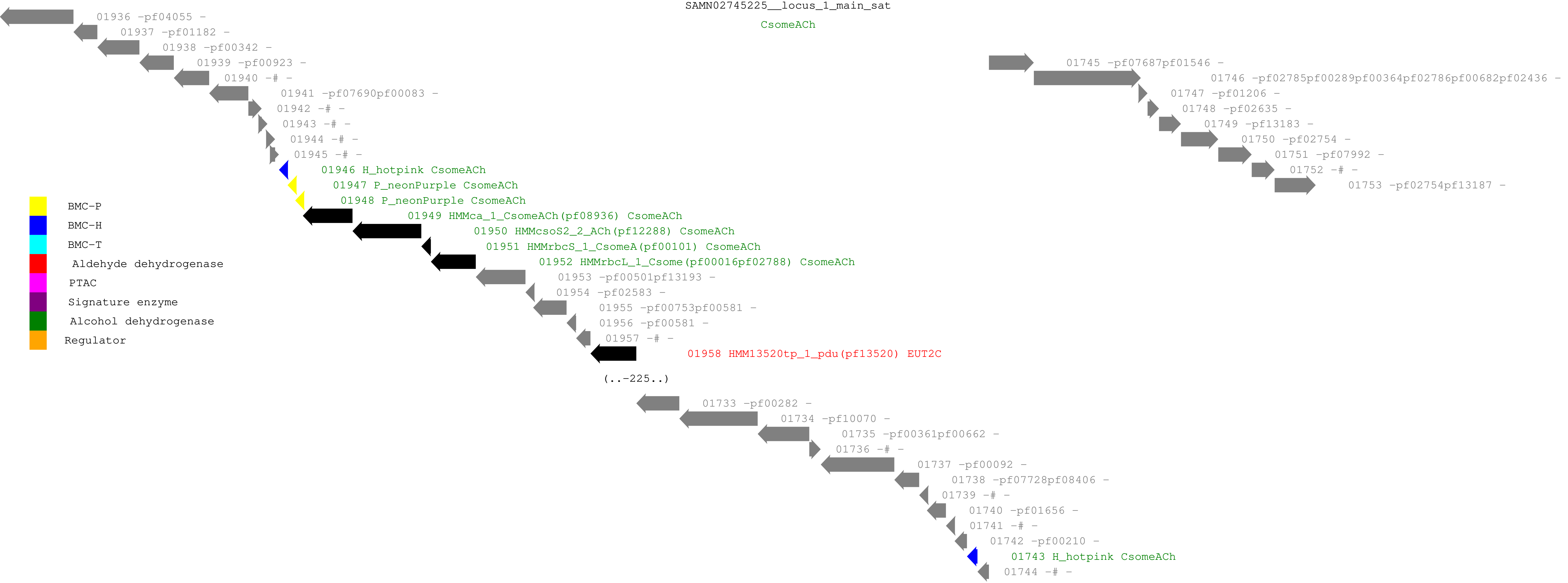

Thiomicrospira sp. ALE5. Bacteria; Proteobacteria; Gammaproteobacteria; Thiotrichales;

SAMN03092900__locus_1

CsomeACh

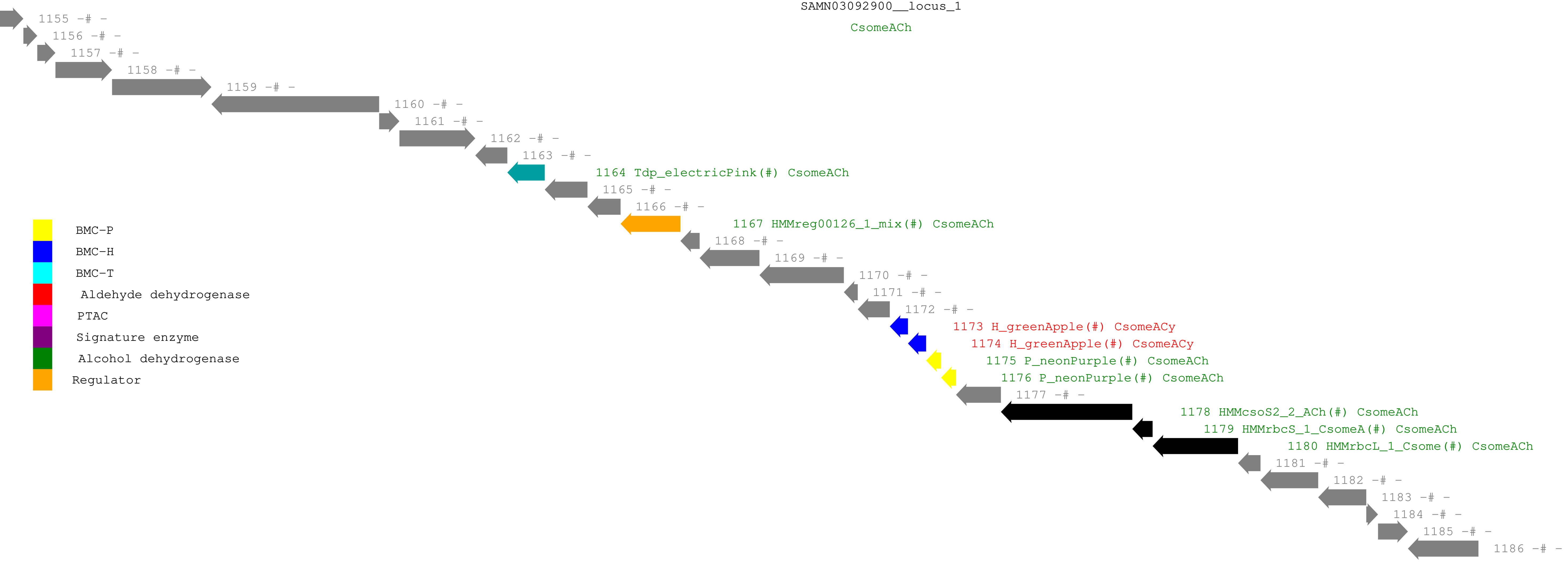

Roseivivax sediminis. Bacteria; Proteobacteria; Alphaproteobacteria; Rhodobacterales;

SAMN04515678__locus_1

CsomeACh

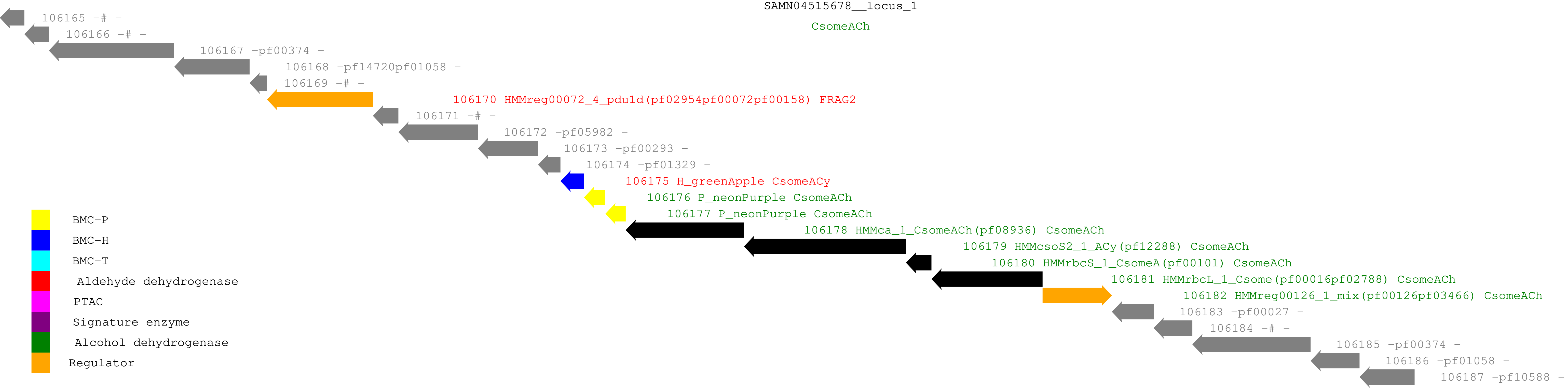

Nitrosomonas europha. Bacteria; Proteobacteria; Betaproteobacteria; Nitrosomonadales;

SAMN05216339__locus_fragmented

CsomeACh

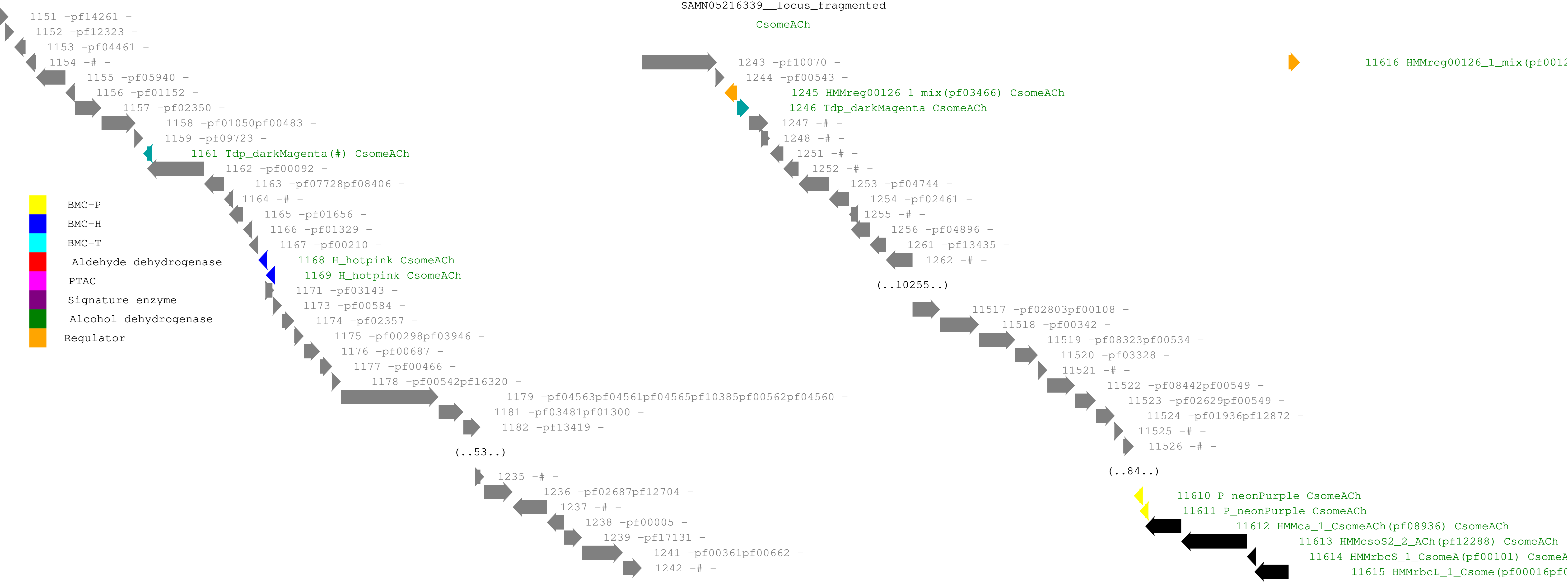

Nitrosomonas eutropha. Bacteria; Proteobacteria; Betaproteobacteria; Nitrosomonadales;

SAMN05216379__locus_1

CsomeACh

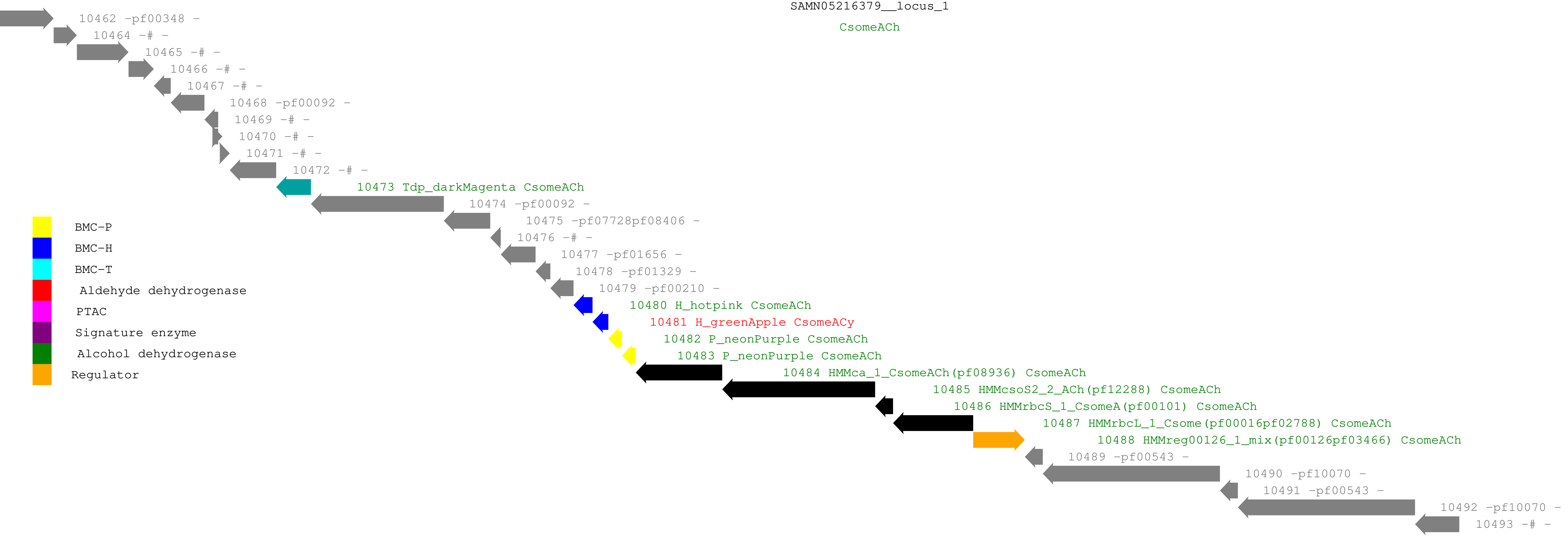

Nitrosospira multiformis. Bacteria; Proteobacteria; Betaproteobacteria; Nitrosomonadales;

SAMN05216402__locus_1

CsomeACh

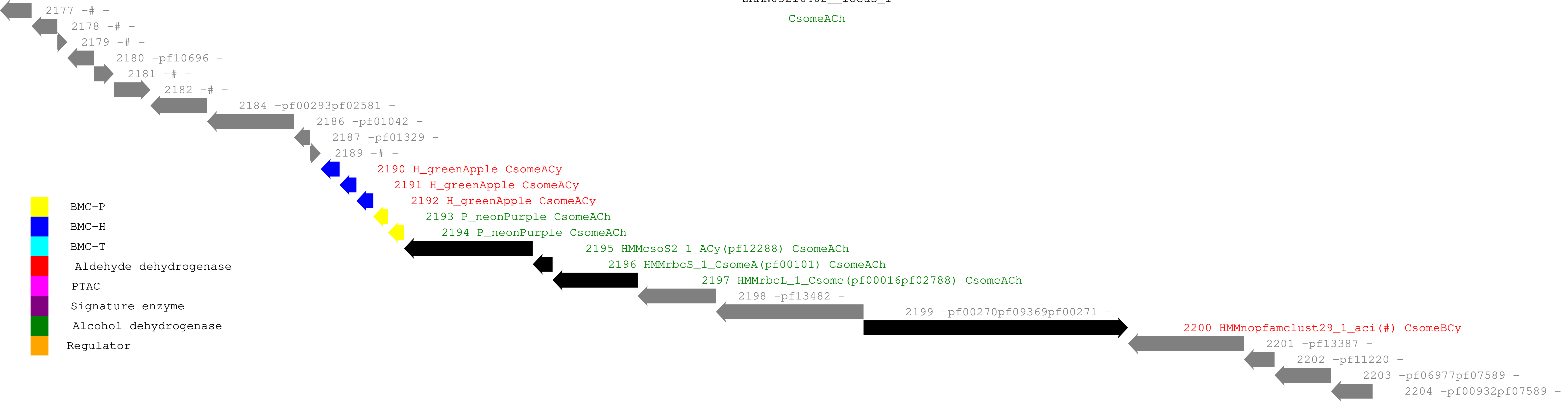

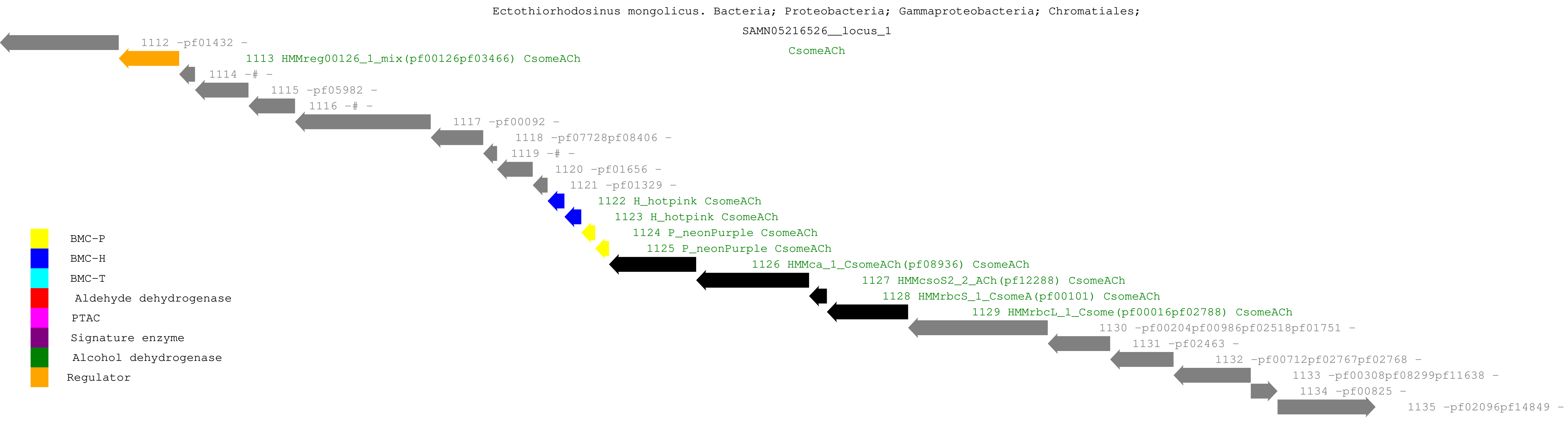

Ectothiorhodosinus mongolicus. Bacteria; Proteobacteria; Gammaproteobacteria; Chromatiales;
SAMN05216526__locus_1
CsomeACh
1112 -pf01432 -
1113 HMMreg00126_1_mix(pf00126pf03466) CsomeACh
1114 -# -
1115 -pf05982 -
1116 -# -
1117 -pf00092 -
1118 -pf07728pf08406 -
1119 -# -
1120 -pf01656 -
1121 -pf01329 -
1122 H_hotpink CsomeACh
1123 H_hotpink CsomeACh
1124 P_neonPurple CsomeACh
1125 P_neonPurple CsomeACh
1126 HMMca_1_CsomeACh(pf08936) CsomeACh
1127 HMMcsoS2_2_ACh(pf12288) CsomeACh
1128 HMMrbcS_1_CsomeA(pf00101) CsomeACh
1129 HMMrbcL_1_Csome(pf00016pf02788) CsomeACh
1130 -pf00204pf00986pf02518pf01751 -
1131 -pf02463 -
1132 -pf00712pf02767pf02768 -
1133 -pf00308pf08299pf11638 -
1134 -pf00825 -
1135 -pf02096pf14849 -
BMC-P
BMC-H
BMC-T
Aldehyde dehydrogenase
PTAC
Signature enzyme
Alcohol dehydrogenase
Regulator

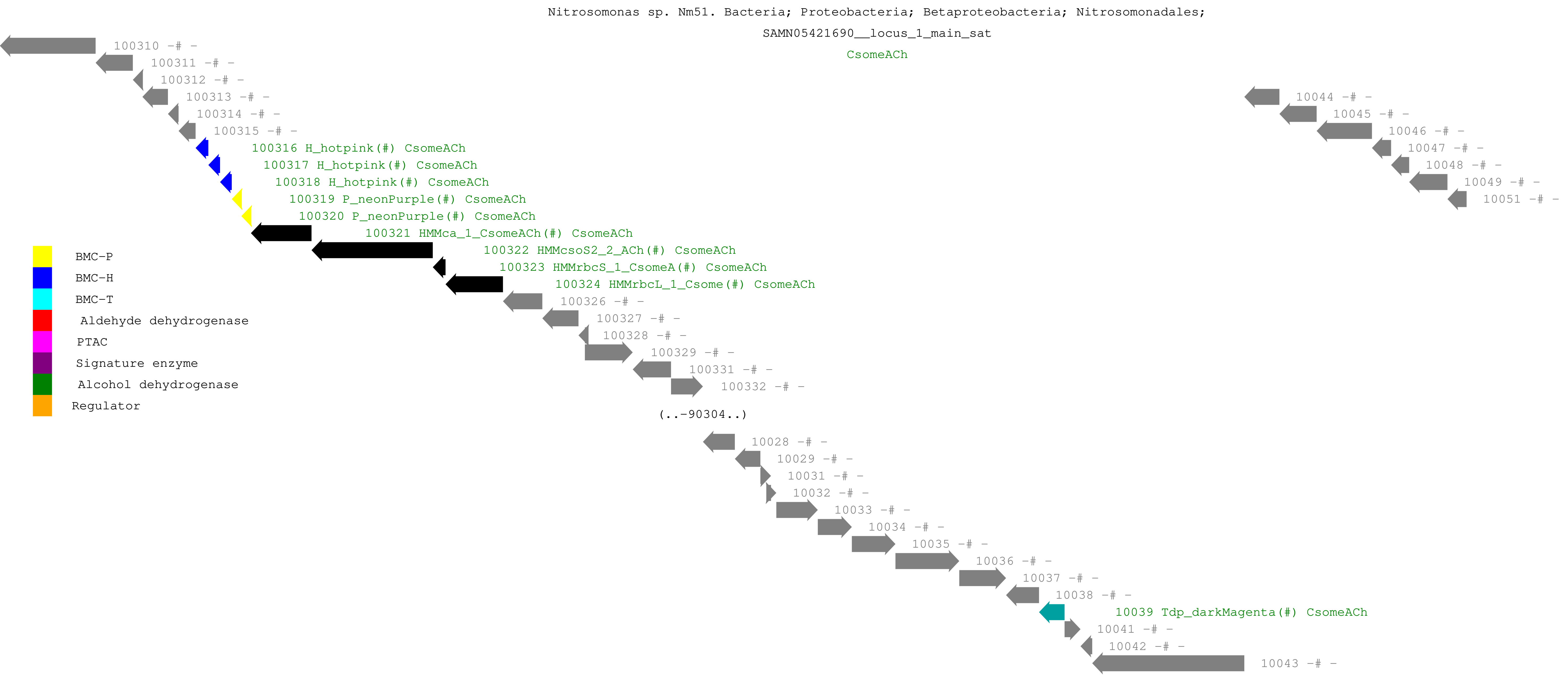

Nitrosomonas sp. Nm51. Bacteria; Proteobacteria; Betaproteobacteria; Nitrosomonadales;
SAMN05421690__locus_1_main_sat
CsomeACh
100310 -# -
100311 -# -
100312 -# -
100313 -# -
100314 -# -
100315 -# -
100316 H_hotpink(#) CsomeACh
100317 H_hotpink(#) CsomeACh
100318 H_hotpink(#) CsomeACh
100319 P_neonPurple(#) CsomeACh
100320 P_neonPurple(#) CsomeACh
100321 HMMca_1_CsomeACh(#) CsomeACh
100322 HMMcsoS2_2_ACh(#) CsomeACh
100323 HMMrbcS_1_CsomeA(#) CsomeACh
100324 HMMrbcL_1_Csome(#) CsomeACh
100326 -# -
100327 -# -
100328 -# -
100329 -# -
100331 -# -
100332 -# -
(..-90304..)
10028 -# -
10029 -# -
10031 -# -
10032 -# -
10033 -# -
10034 -# -
10035 -# -
10036 -# -
10037 -# -
10038 -# -
10039 Tdp_darkMagenta(#) CsomeACh
10041 -# -
10042 -# -
10043 -# -
10044 -# -
10045 -# -
10046 -# -
10047 -# -
10048 -# -
10049 -# -
10051 -# -
BMC-P
BMC-H
BMC-T
Aldehyde dehydrogenase
PTAC
Signature enzyme
Alcohol dehydrogenase
Regulator

Ectothiorhodospira magna. Bacteria; Proteobacteria; Gammaproteobacteria; Chromatiales;

SAMN05421693__locus_1_main_sat

CsomeACh

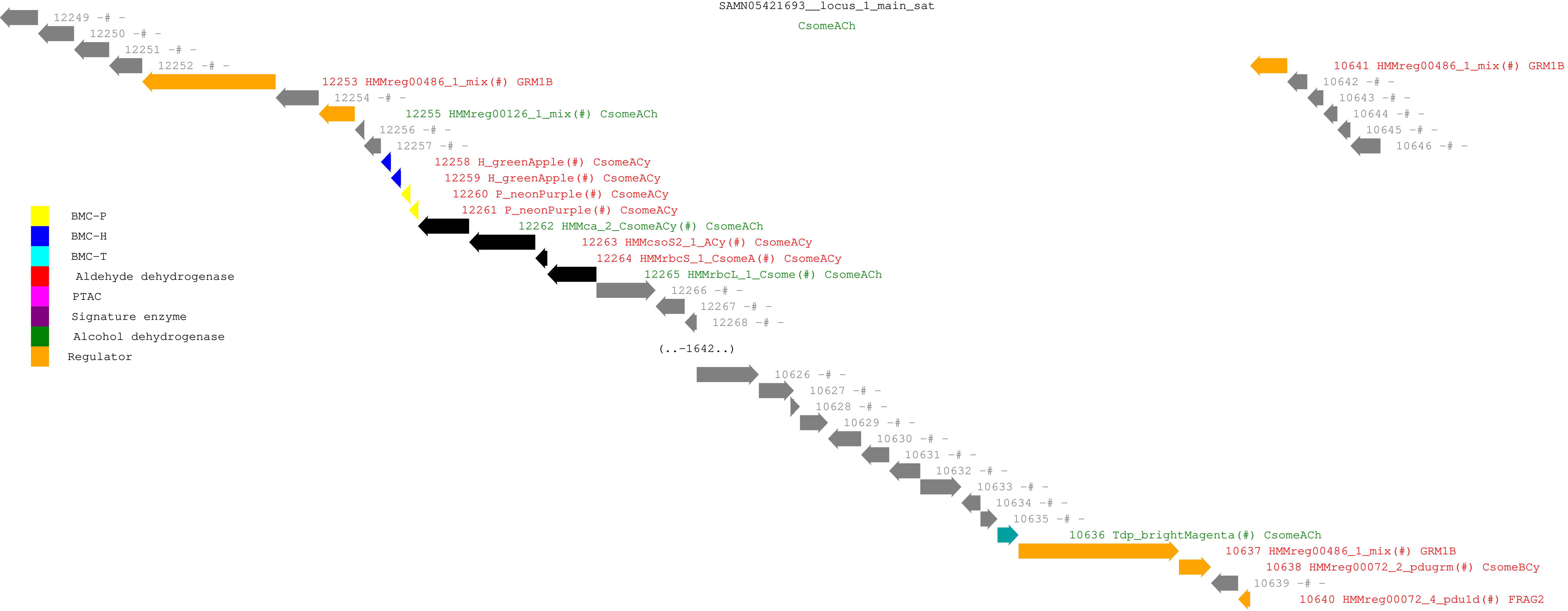

Thiocapsa roseopersicina. Bacteria; Proteobacteria; Gammaproteobacteria; Chromatiales;

SAMN05421783__locus_1_main_sat

CsomeACh

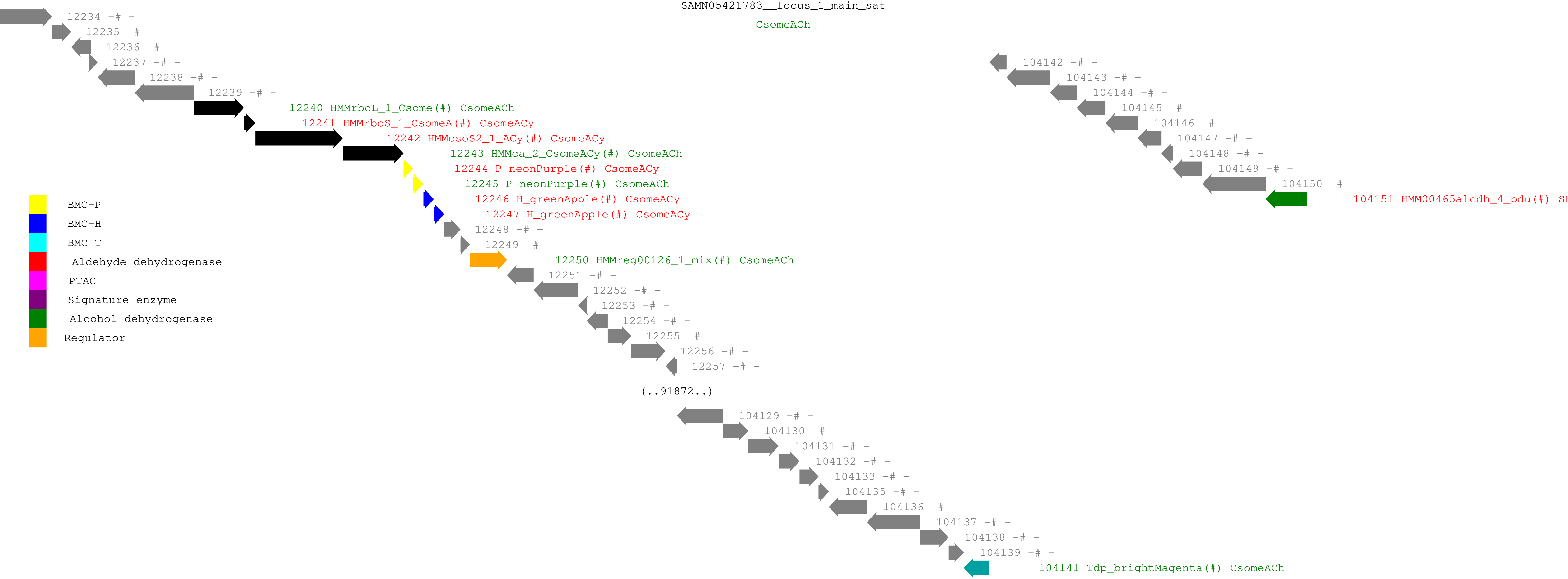

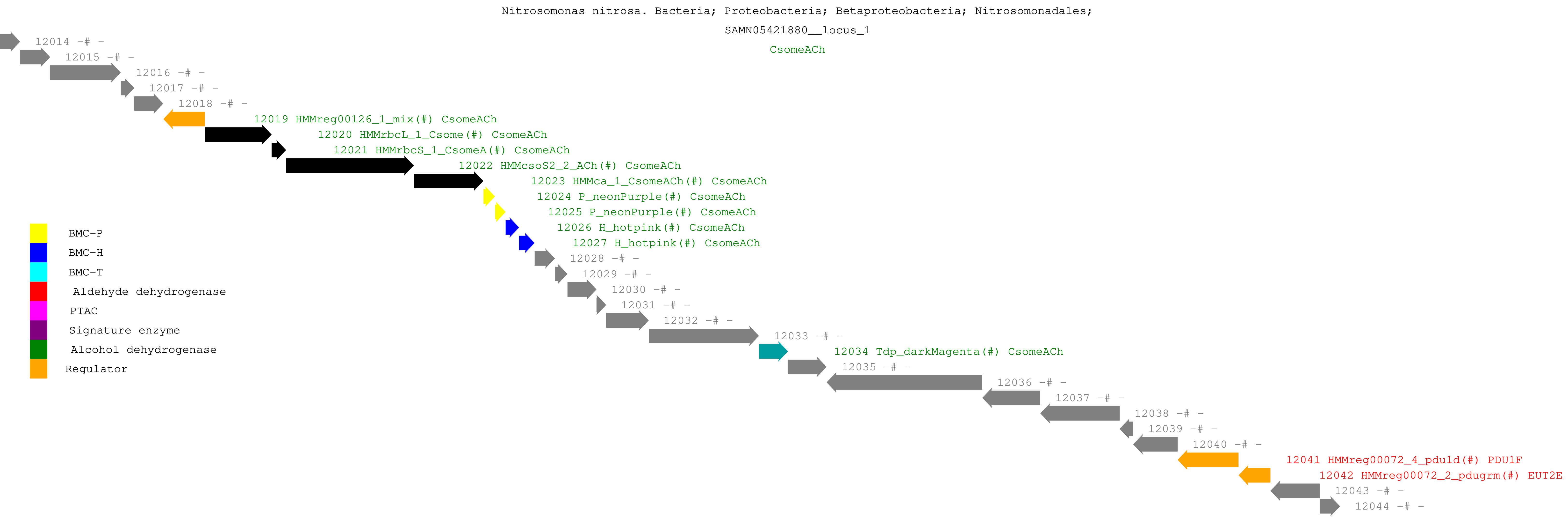

Nitrosomonas nitrosa. Bacteria; Proteobacteria; Betaproteobacteria; Nitrosomonadales;
SAMN05421880__locus_1
CsomeACh
12014 -# -
12015 -# -
12016 -# -
12017 -# -
12018 -# -
12019 HMMreg00126_1_mix(#) CsomeACh
12020 HMMrbcL_1_Csome(#) CsomeACh
12021 HMMrbcS_1_CsomeA(#) CsomeACh
12022 HMMcsoS2_2_ACh(#) CsomeACh
12023 HMMca_1_CsomeACh(#) CsomeACh
12024 P_neonPurple(#) CsomeACh
12025 P_neonPurple(#) CsomeACh
12026 H_hotpink(#) CsomeACh
12027 H_hotpink(#) CsomeACh
12028 -# -
12029 -# -
12030 -# -
12031 -# -
12032 -# -
12033 -# -
12034 Tdp_darkMagenta(#) CsomeACh
12035 -# -
12036 -# -
12037 -# -
12038 -# -
12039 -# -
12040 -# -
12041 HMMreg00072_4_pdu1d(#) PDU1F
12042 HMMreg00072_2_pdugrm(#) EUT2E
12043 -# -
12044 -# -
BMC-P
BMC-H
BMC-T
Aldehyde dehydrogenase
PTAC
Signature enzyme
Alcohol dehydrogenase
Regulator

Nitrosomonas halophila. Bacteria; Proteobacteria; Betaproteobacteria; Nitrosomonadales;

SAMN05421881__locus_1_main_sat

CsomeACh

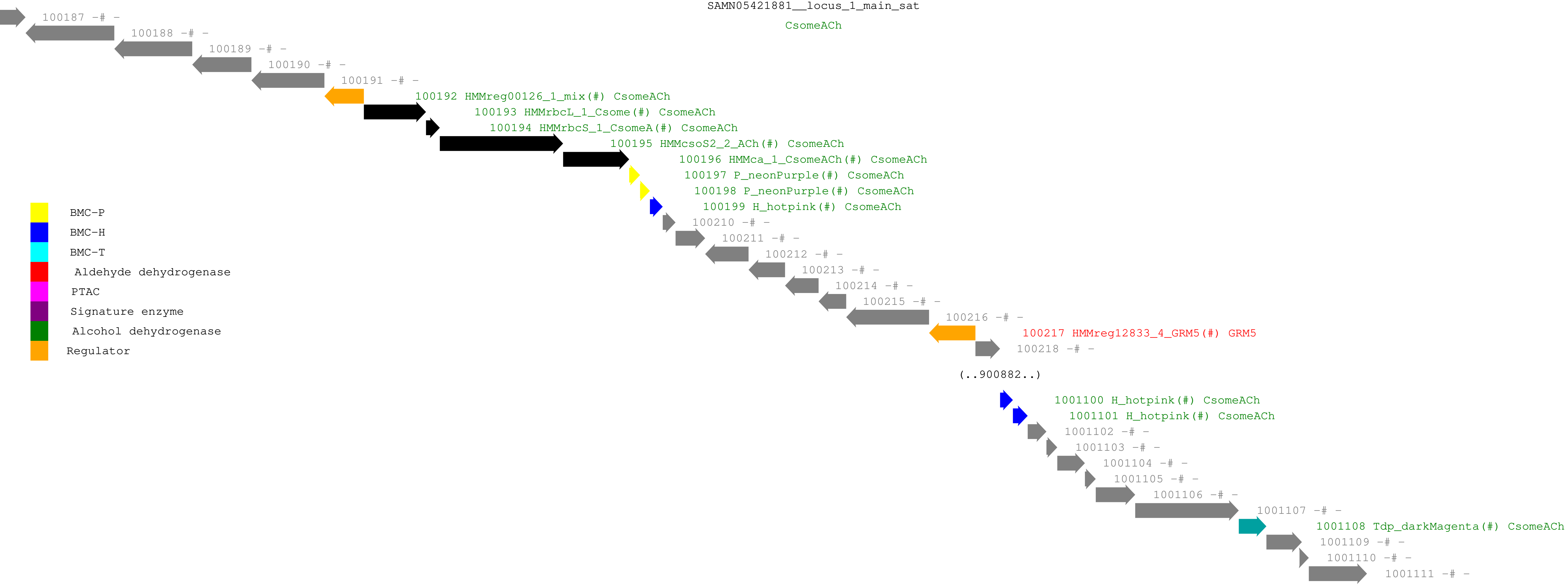

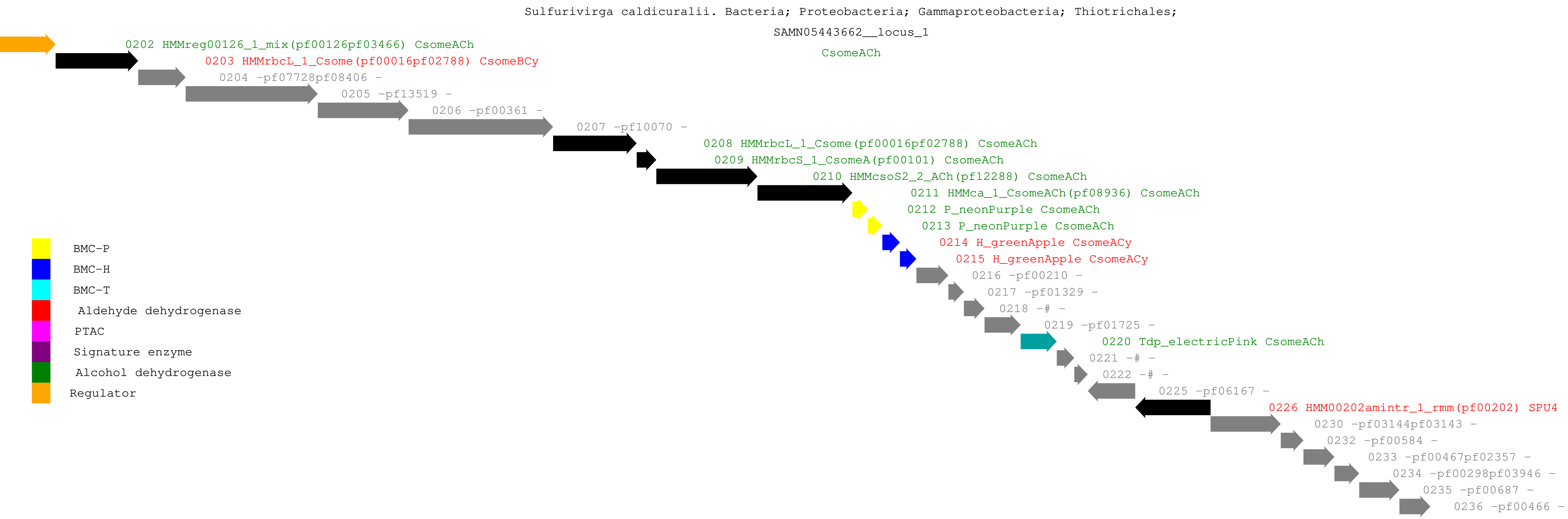
Sulfurivirga caldicuralii. Bacteria; Proteobacteria; Gammaproteobacteria; Thiotrichales;
SAMN05443662__locus_1
CsomeACh
0202 HMMreg00126_1_mix(pf00126pf03466) CsomeACh
0203 HMMrbcL_1_Csome(pf00016pf02788) CsomeBCy
0204 -pf07728pf08406 -
0205 -pf13519 -
0206 -pf00361 -
0207 -pf10070 -
0208 HMMrbcL_1_Csome(pf00016pf02788) CsomeACh
0209 HMMrbcS_1_CsomeA(pf00101) CsomeACh
0210 HMMcsoS2_2_ACh(pf12288) CsomeACh
0211 HMMca_1_CsomeACh(pf08936) CsomeACh
0212 P_neonPurple CsomeACh
0213 P_neonPurple CsomeACh
0214 H_greenApple CsomeACy
0215 H_greenApple CsomeACy
0216 -pf00210 -
0217 -pf01329 -
0218 -# -
0219 -pf01725 -
0220 Tdp_electricPink CsomeACh
0221 -# -
0222 -# -
0225 -pf06167 -
0226 HMM00202amintr_1_rmm(pf00202) SPU4
0230 -pf03144pf03143 -
0232 -pf00584 -
0233 -pf00467pf02357 -
0234 -pf00298pf03946 -
0235 -pf00687 -
0236 -pf00466 -
BMC-P
BMC-H
BMC-T
Aldehyde dehydrogenase
PTAC
Signature enzyme
Alcohol dehydrogenase
Regulator

Ectothiorhodospira marina. Bacteria; Proteobacteria; Gammaproteobacteria; Chromatiales;

SAMN05444515__locus_1_main_sat

CsomeACh

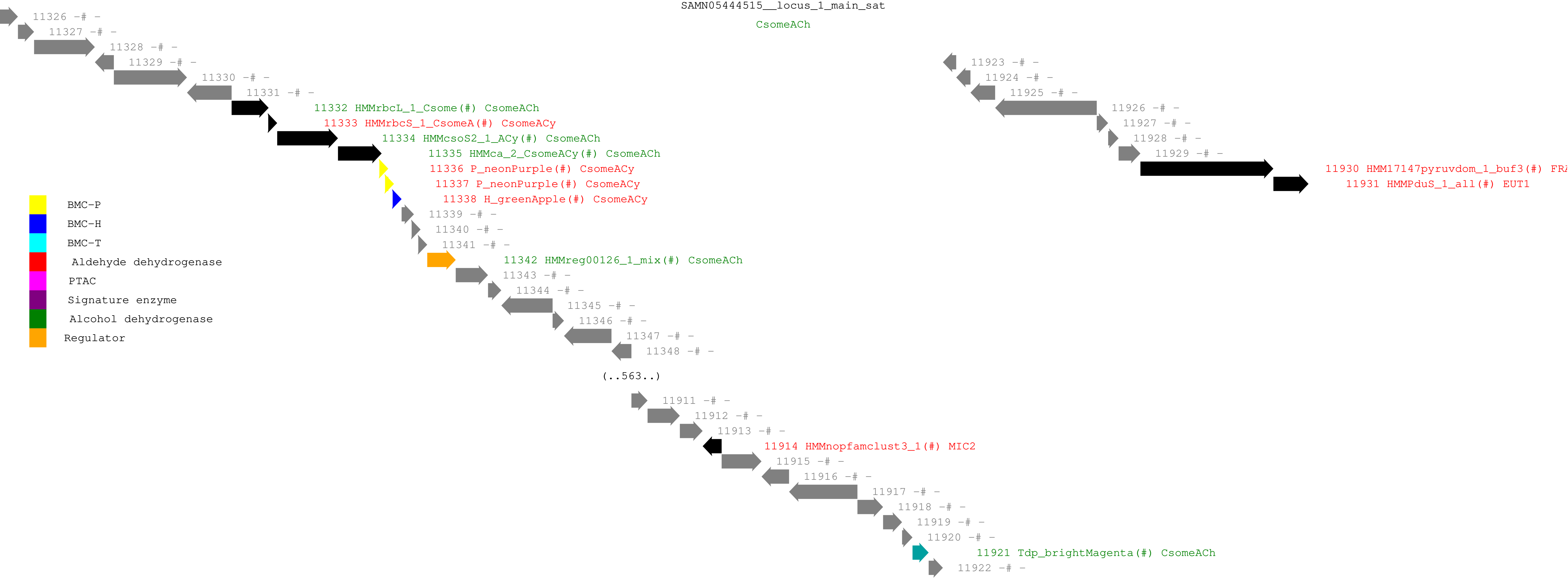

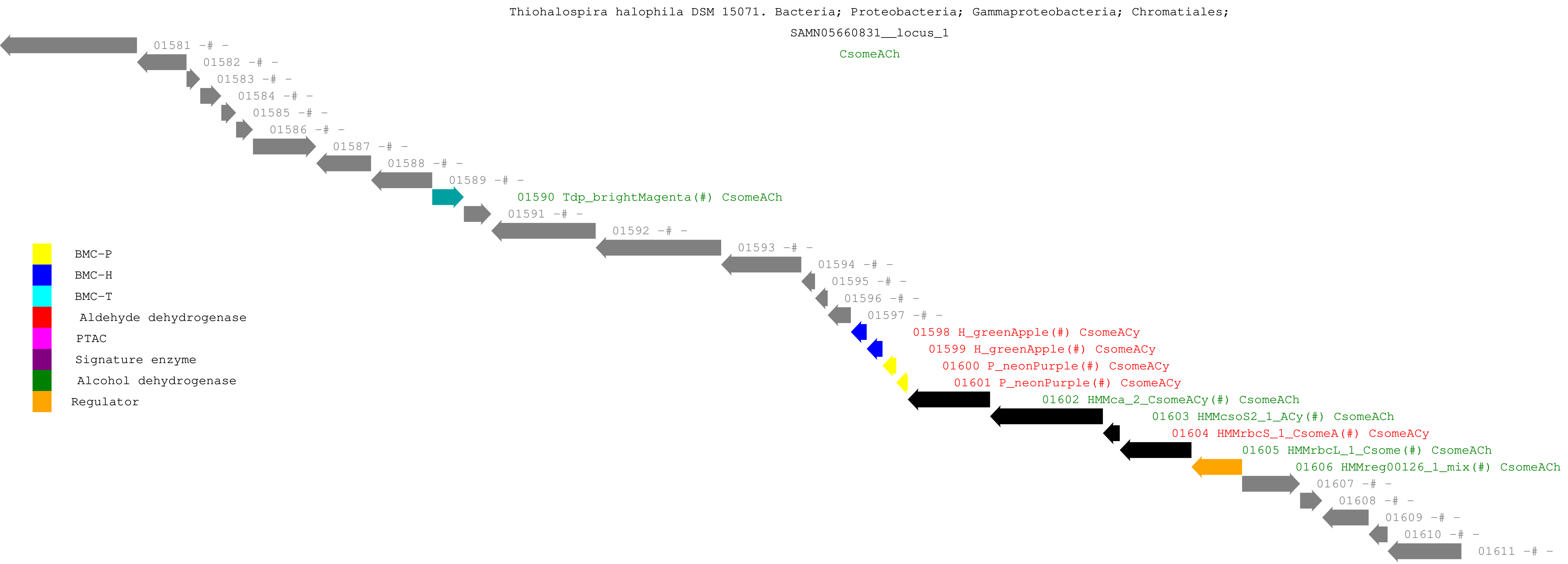
Thiohalospira halophila DSM 15071. Bacteria; Proteobacteria; Gammaproteobacteria; Chromatiales;
SAMN05660831__locus_1
CsomeACh
01581 -# -
01582 -# -
01583 -# -
01584 -# -
01585 -# -
01586 -# -
01587 -# -
01588 -# -
01589 -# -
01590 Tdp_brightMagenta(#) CsomeACh
01591 -# -
01592 -# -
01593 -# -
01594 -# -
01595 -# -
01596 -# -
01597 -# -
01598 H_greenApple(#) CsomeACy
01599 H_greenApple(#) CsomeACy
01600 P_neonPurple(#) CsomeACy
01601 P_neonPurple(#) CsomeACy
01602 HMMca_2_CsomeACy(#) CsomeACh
01603 HMMcsoS2_1_ACy(#) CsomeACh
01604 HMMrbcS_1_CsomeA(#) CsomeACy
01605 HMMrbcL_1_Csome(#) CsomeACh
01606 HMMreg00126_1_mix(#) CsomeACh
01607 -# -
01608 -# -
01609 -# -
01610 -# -
01611 -# -
BMC-P
BMC-H
BMC-T
Aldehyde dehydrogenase
PTAC
Signature enzyme
Alcohol dehydrogenase
Regulator

Thiothrix caldifontis. Bacteria; Proteobacteria; Gammaproteobacteria; Thiotrichales;

SAMN05660964__locus_1

CsomeACh

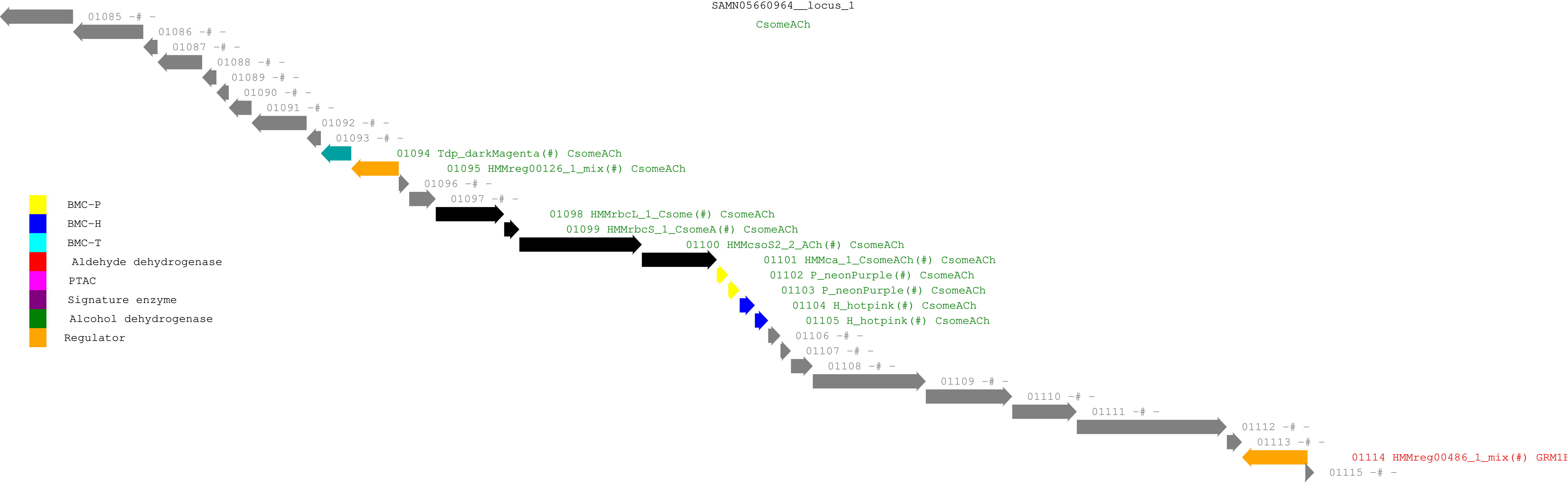

Sulfuriferula multivorans. Bacteria; Proteobacteria; Betaproteobacteria; Nitrosomonadales;

SFMTTN__locus_1_main_sat

CsomeACh

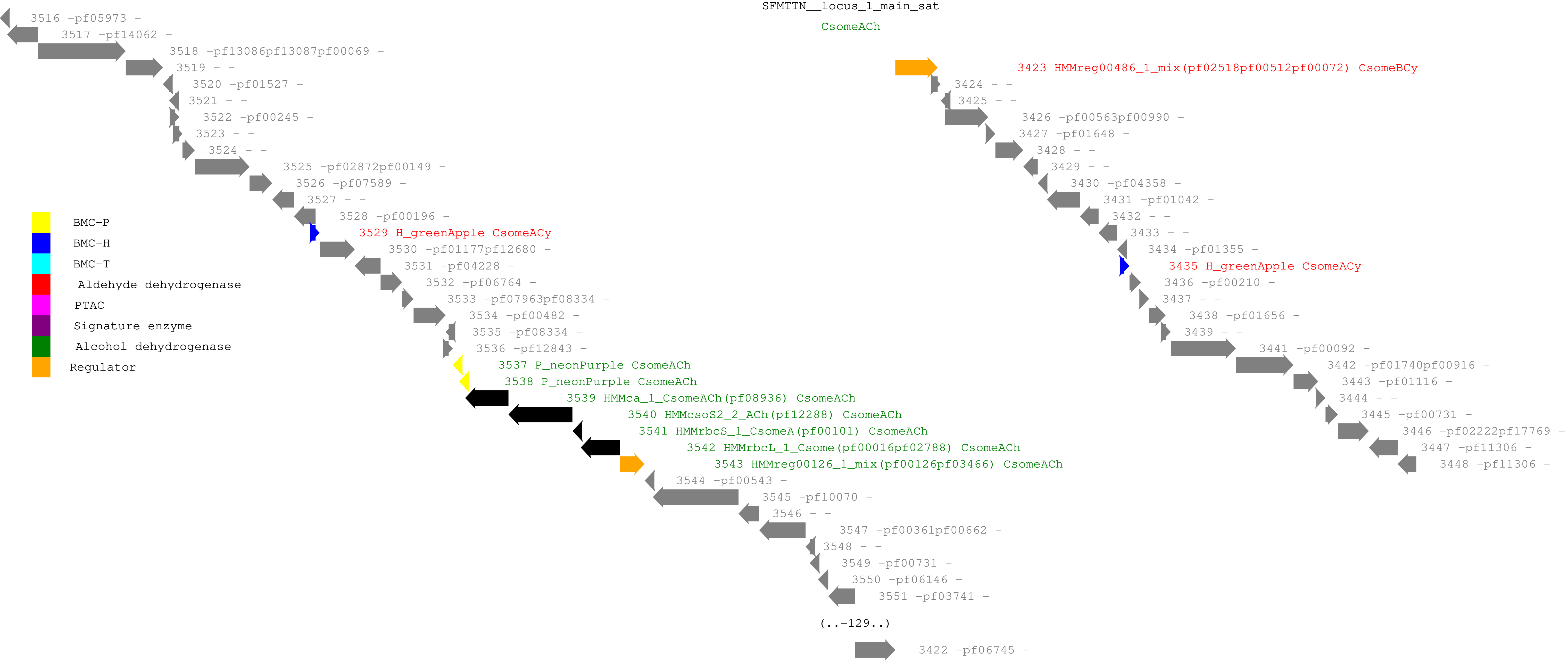

Comamonadaceae bacterium B1. Bacteria; Proteobacteria; Betaproteobacteria; Burkholderiales;

SMCB__locus_1

CsomeACh

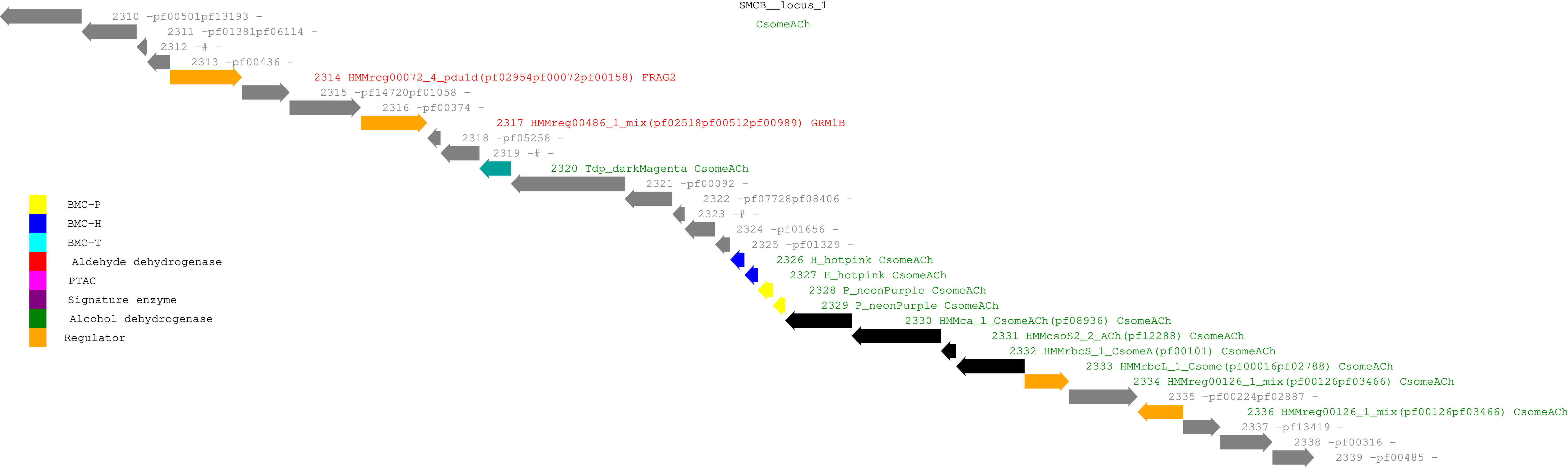

Comamonadaceae bacterium A1. Bacteria; Proteobacteria; Betaproteobacteria; Burkholderiales;

SRAA__locus_1

CsomeACh

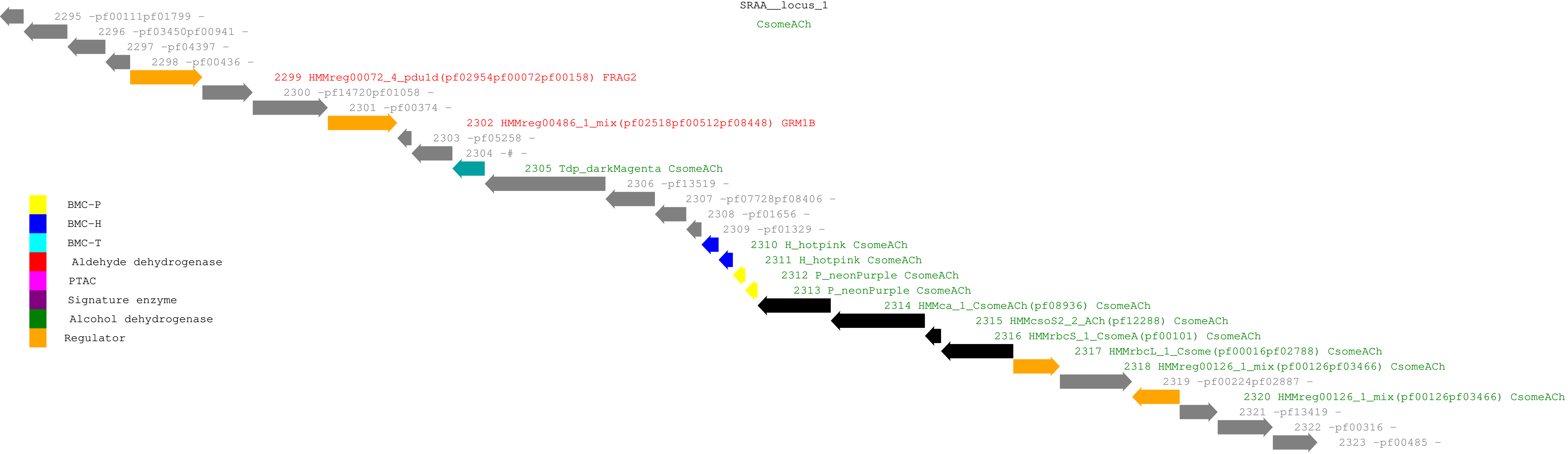

Tepidimonas alkaliphilus. Bacteria; Proteobacteria; Betaproteobacteria; Burkholderiales; Tepidimonas.

Talka__locus_fragmented

CsomeACh

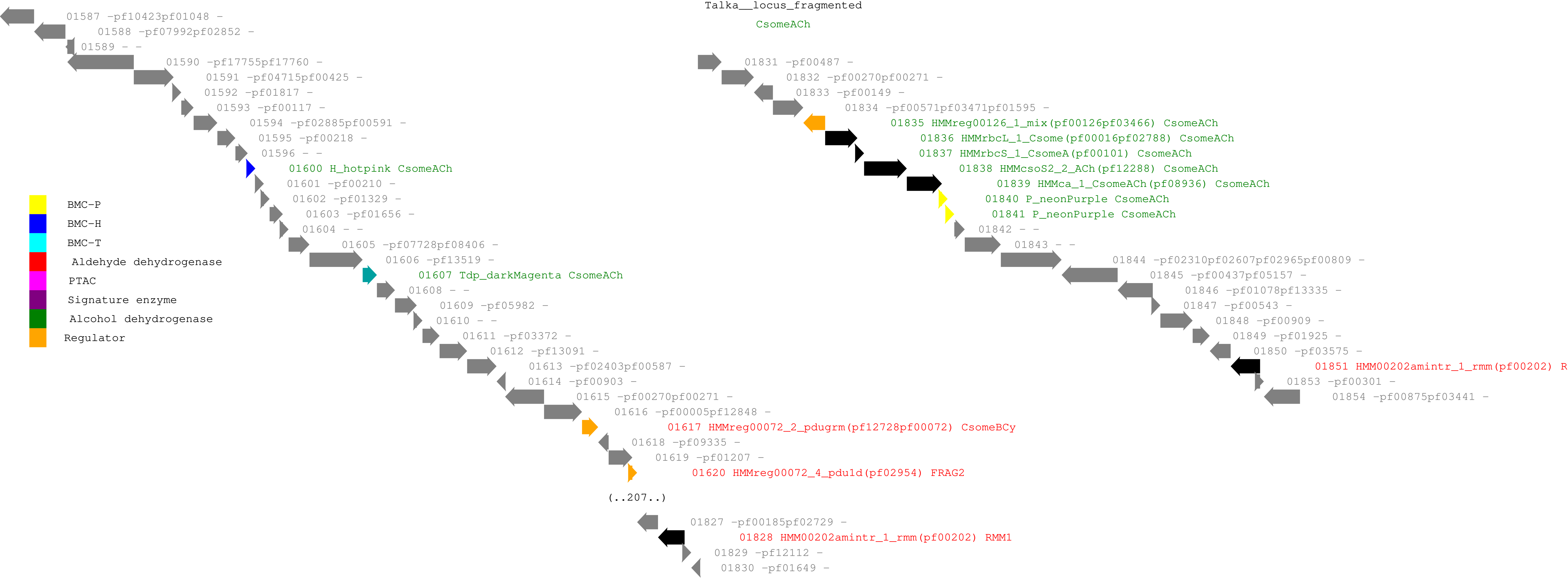

Tepidimonas aquatica. Bacteria; Proteobacteria; Betaproteobacteria; Burkholderiales; Tepidimonas.

Taqua__locus_1

CsomeACh

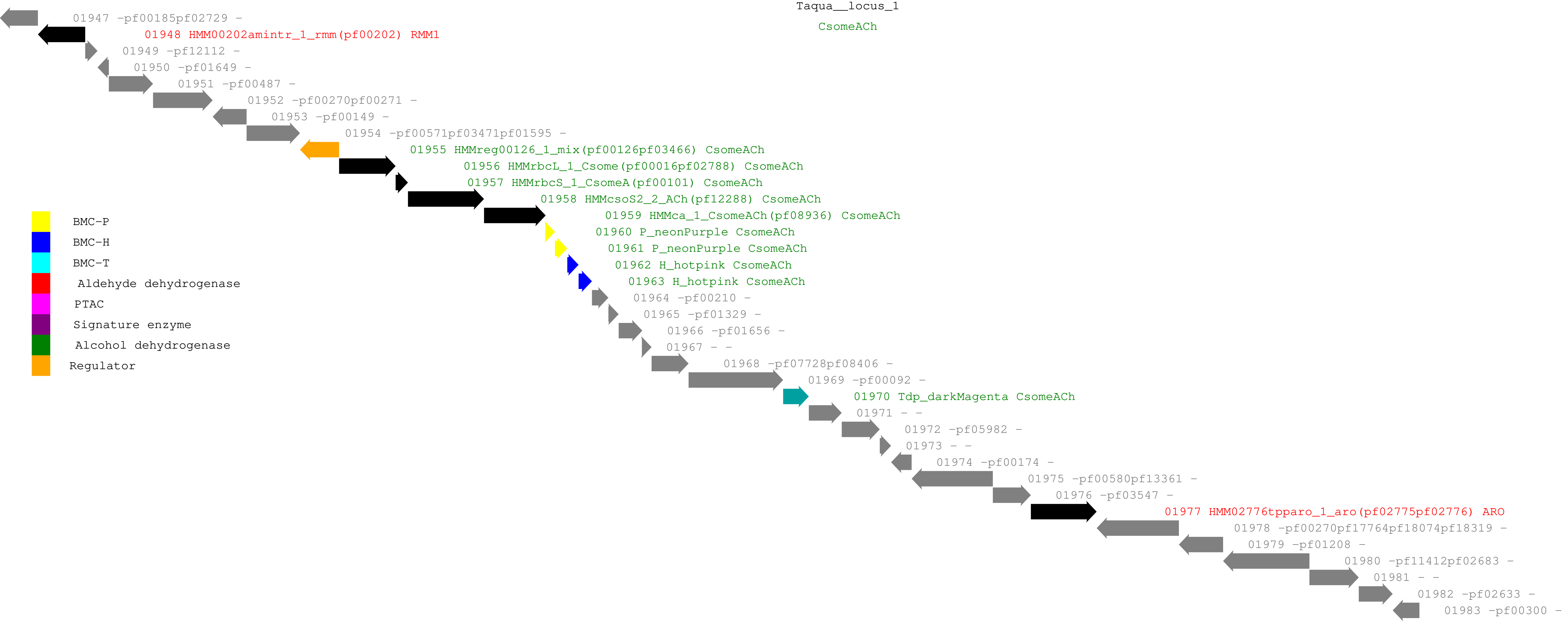

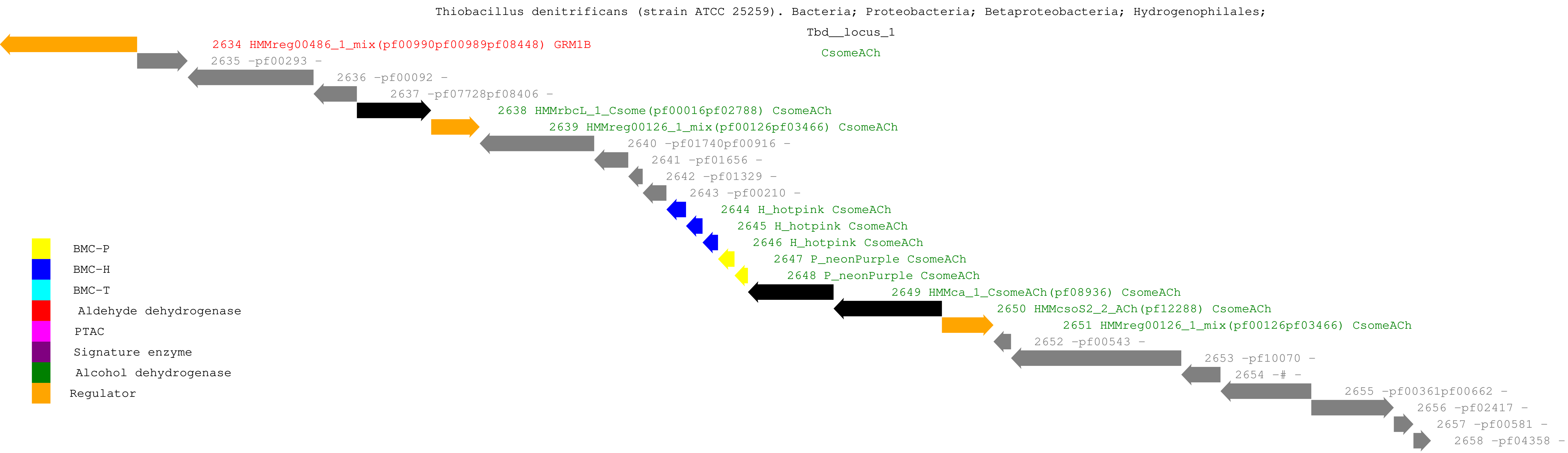
Thiobacillus denitrificans (strain ATCC 25259). Bacteria; Proteobacteria; Betaproteobacteria; Hydrogenophilales;
Tbd__locus_1
CsomeACh
2634 HMMreg00486_1_mix(pf00990pf00989pf08448) GRM1B
2635 -pf00293 -
2636 -pf00092 -
2637 -pf07728pf08406 -
2638 HMMrbcL_1_Csome(pf00016pf02788) CsomeACh
2639 HMMreg00126_1_mix(pf00126pf03466) CsomeACh
2640 -pf01740pf00916 -
2641 -pf01656 -
2642 -pf01329 -
2643 -pf00210 -
2644 H_hotpink CsomeACh
2645 H_hotpink CsomeACh
2646 H_hotpink CsomeACh
2647 P_neonPurple CsomeACh
2648 P_neonPurple CsomeACh
2649 HMMca_1_CsomeACh(pf08936) CsomeACh
2650 HMMcsoS2_2_ACh(pf12288) CsomeACh
2651 HMMreg00126_1_mix(pf00126pf03466) CsomeACh
2652 -pf00543 -
2653 -pf10070 -
2654 -# -
2655 -pf00361pf00662 -
2656 -pf02417 -
2657 -pf00581 -
2658 -pf04358 -
BMC-P
BMC-H
BMC-T
Aldehyde dehydrogenase
PTAC
Signature enzyme
Alcohol dehydrogenase
Regulator

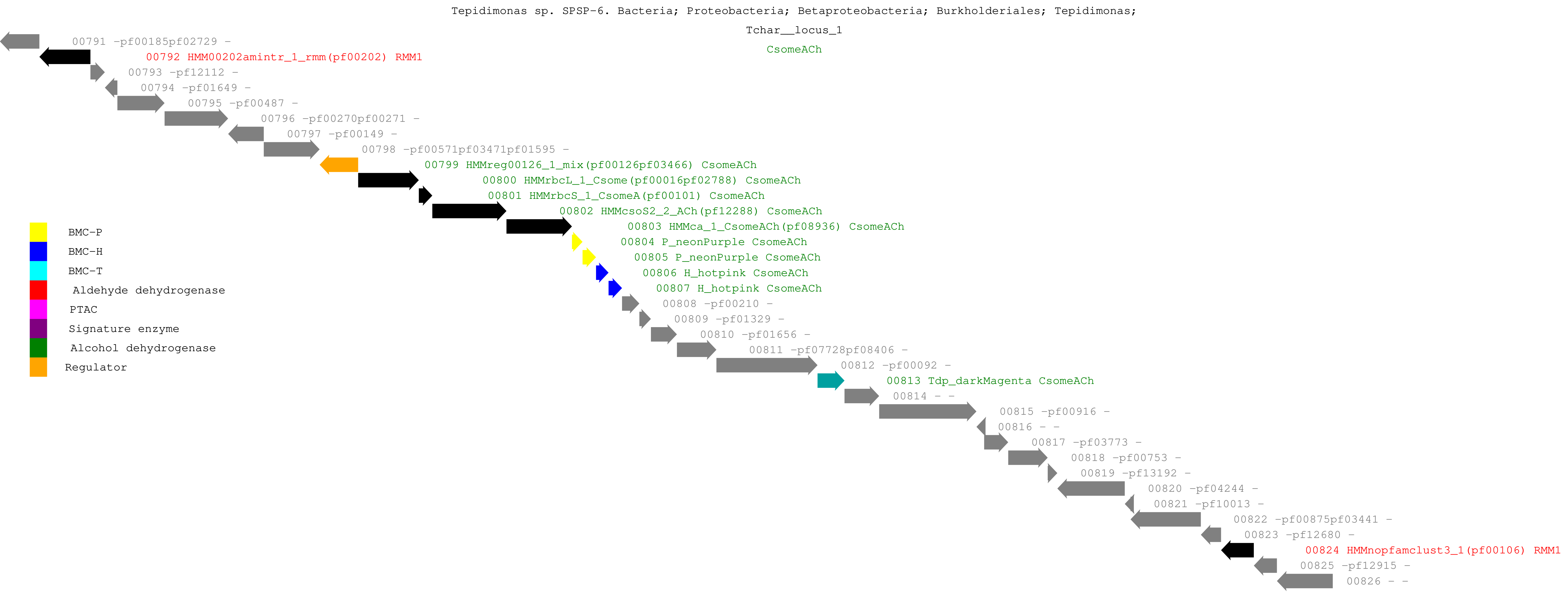
Tepidimonas sp. SPSP-6. Bacteria; Proteobacteria; Betaproteobacteria; Burkholderiales; Tepidimonas;
Tchar__locus_1
CsomeACh
00791 -pf00185pf02729 -
00792 HMM00202amintr_1_rmm(pf00202) RMM1
00793 -pf12112 -
00794 -pf01649 -
00795 -pf00487 -
00796 -pf00270pf00271 -
00797 -pf00149 -
00798 -pf00571pf03471pf01595 -
00799 HMMreg00126_1_mix(pf00126pf03466) CsomeACh
00800 HMMrbcL_1_Csome(pf00016pf02788) CsomeACh
00801 HMMrbcS_1_CsomeA(pf00101) CsomeACh
00802 HMMcsoS2_2_ACh(pf12288) CsomeACh
00803 HMMca_1_CsomeACh(pf08936) CsomeACh
00804 P_neonPurple CsomeACh
00805 P_neonPurple CsomeACh
00806 H_hotpink CsomeACh
00807 H_hotpink CsomeACh
00808 -pf00210 -
00809 -pf01329 -
00810 -pf01656 -
00811 -pf07728pf08406 -
00812 -pf00092 -
00813 Tdp_darkMagenta CsomeACh
00814 - -
00815 -pf00916 -
00816 - -
00817 -pf03773 -
00818 -pf00753 -
00819 -pf13192 -
00820 -pf04244 -
00821 -pf10013 -
00822 -pf00875pf03441 -
00823 -pf12680 -
00824 HMMnopfamclust3_1(pf00106) RMM1
00825 -pf12915 -
00826 - -
BMC-P
BMC-H
BMC-T
Aldehyde dehydrogenase
PTAC
Signature enzyme
Alcohol dehydrogenase
Regulator

Thiomicrospira crunogena (strain XCL-2). Bacteria; Proteobacteria; Gammaproteobacteria; Thiotrichales;

Tcr__locus_1

CsomeACh

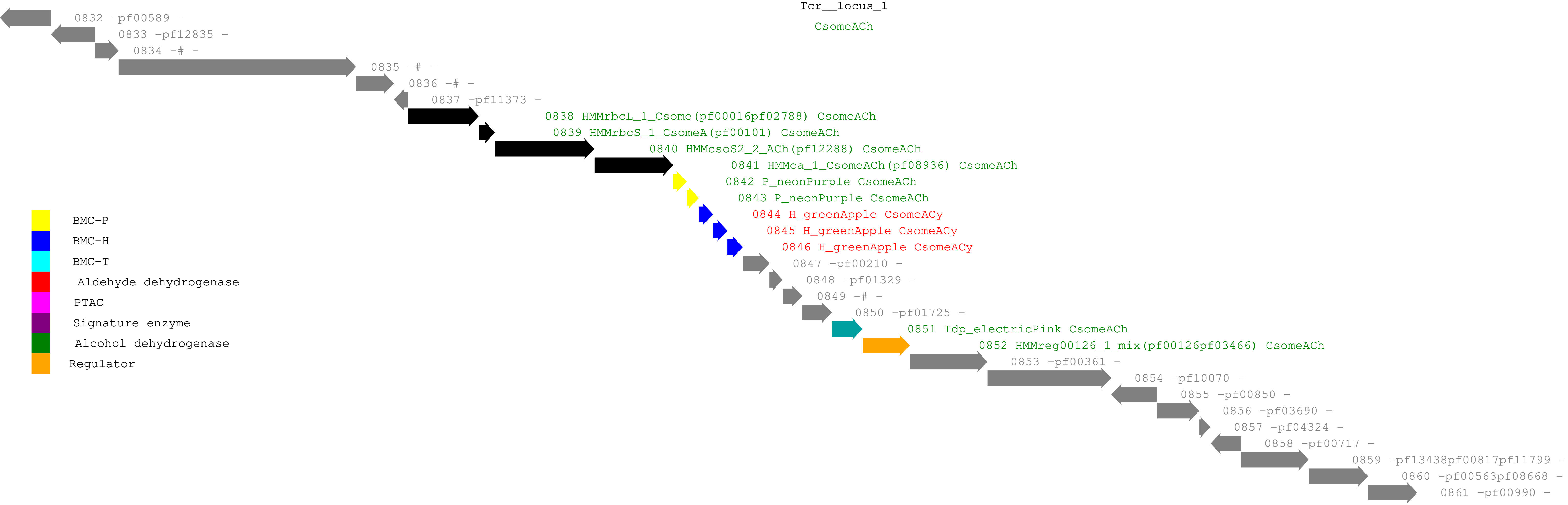

Candidatus Tenderia electrophaga. Bacteria; Proteobacteria; Gammaproteobacteria; Chromatiales;

Tel__locus_1

CsomeACh

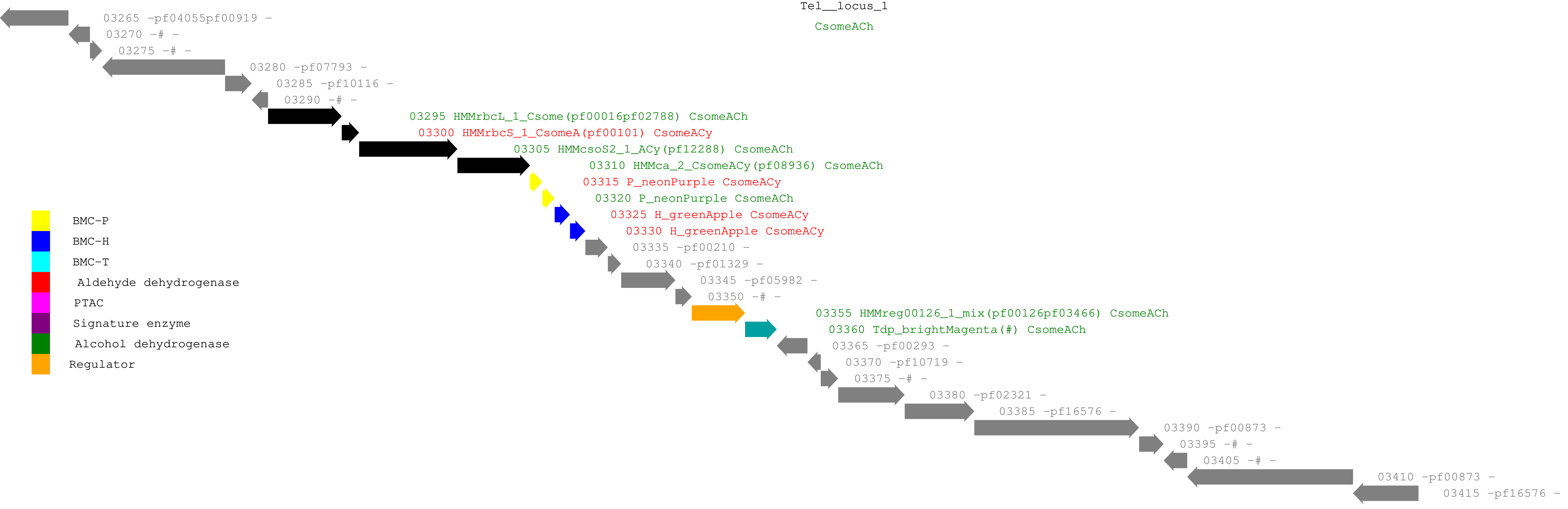

Thioalkalivibrio sulfidiphilus (strain HL-EbGR7). Bacteria; Proteobacteria; Gammaproteobacteria; Chromatiales;

Tgr7__locus_1_main_sat

CsomeACh

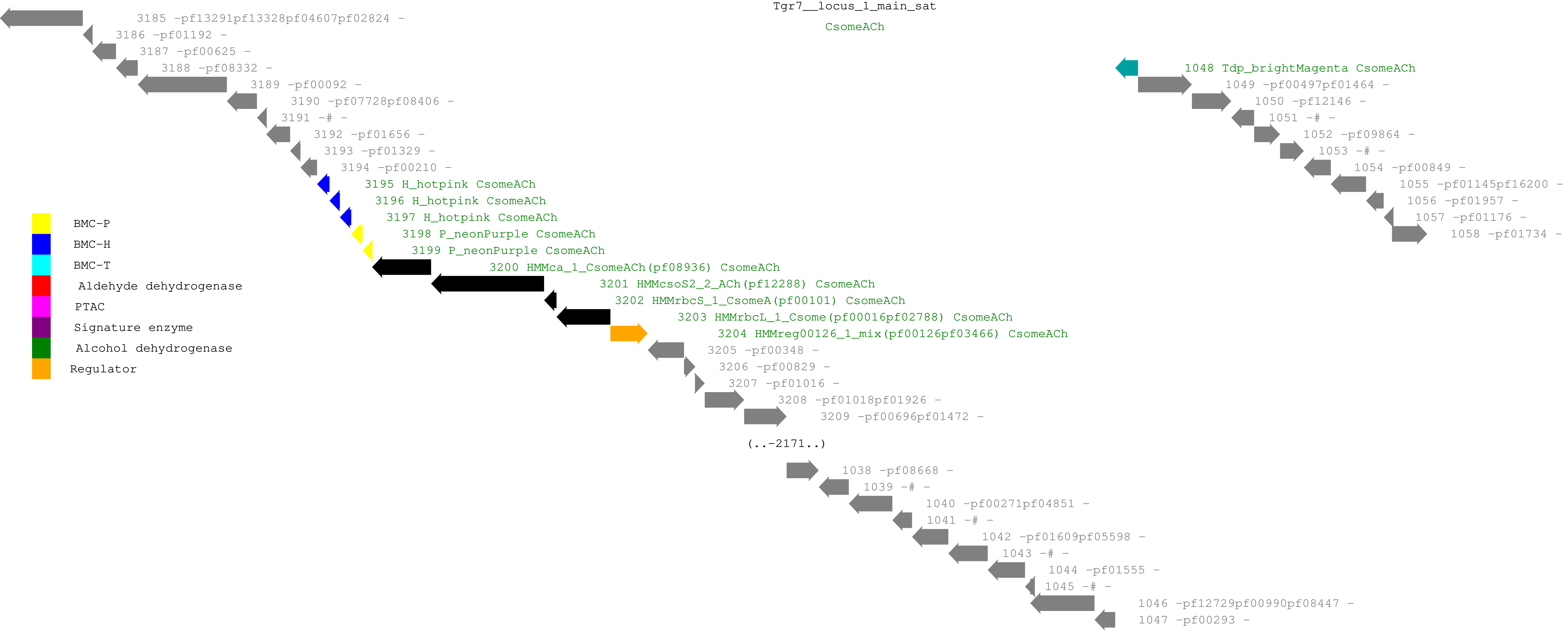

Thiorhodovibrio sp. 970. Bacteria; Proteobacteria; Gammaproteobacteria; Chromatiales;

Thi970DRAFT__locus_1_main_sat

CsomeACh

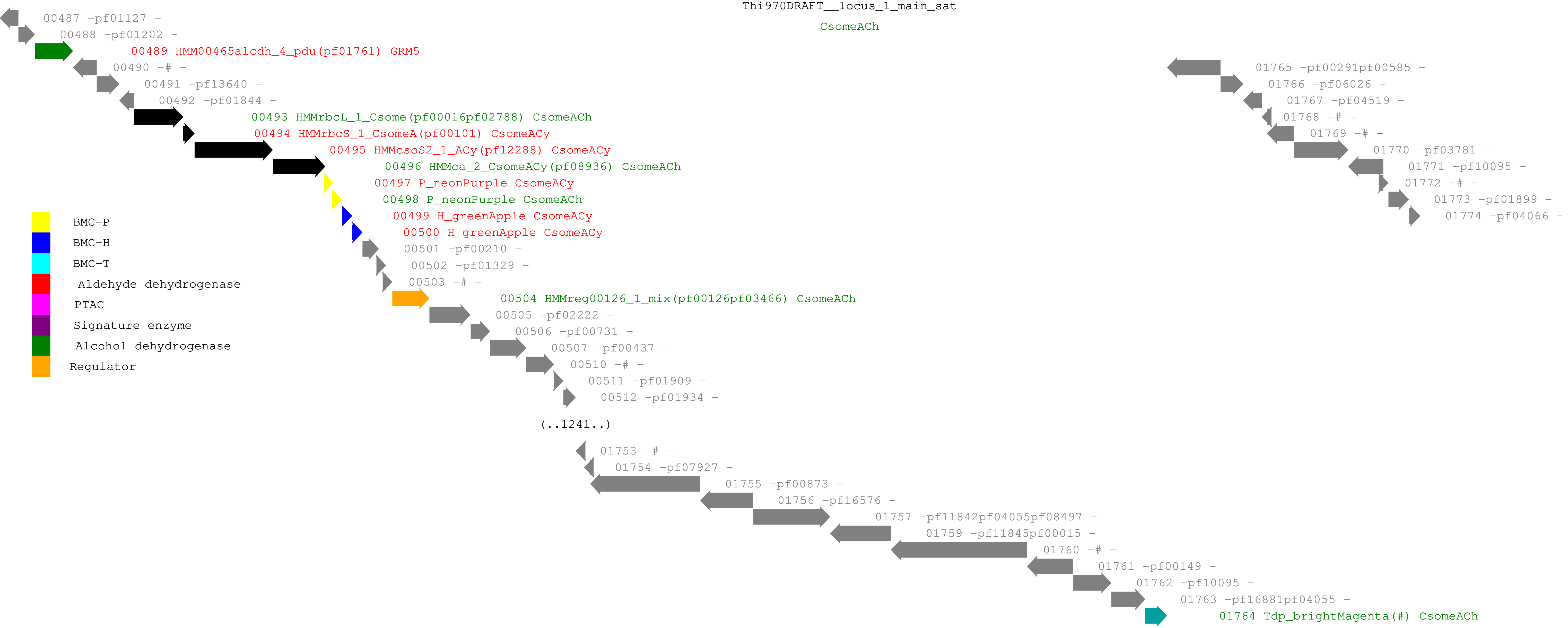

Thioalkalimicrobium aerophilum AL3. Bacteria; Proteobacteria; Gammaproteobacteria; Thiotrichales;

THIAE__locus_1

CsomeACh

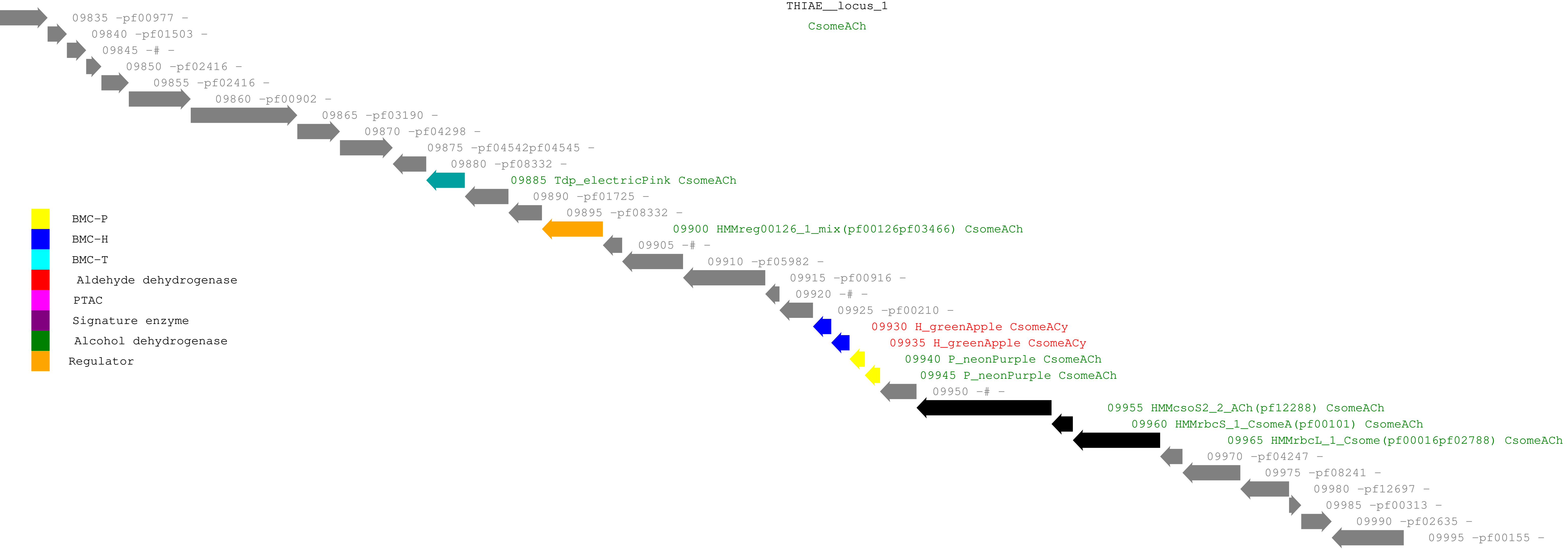

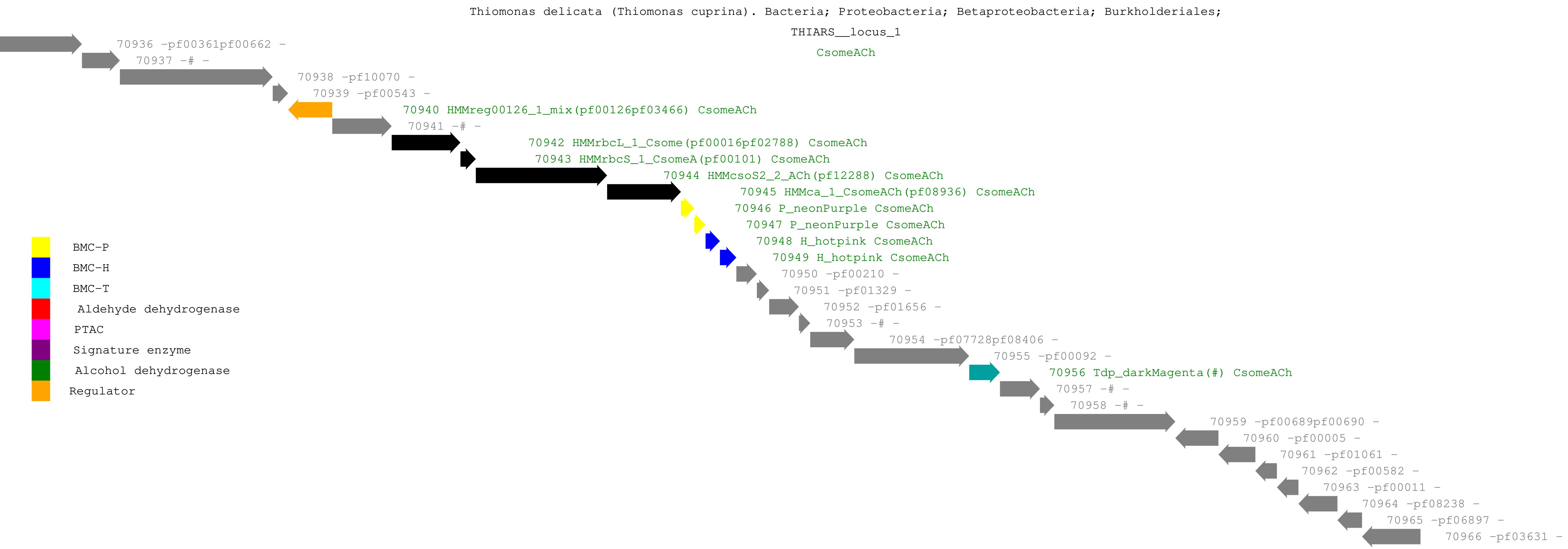
Thiomonas delicata (Thiomonas cuprina). Bacteria; Proteobacteria; Betaproteobacteria; Burkholderiales;
THIARS__locus_1
CsomeACh
70936 -pf00361pf00662 -
70937 -# -
70938 -pf10070 -
70939 -pf00543 -
70940 HMMreg00126_1_mix(pf00126pf03466) CsomeACh
70941 -# -
70942 HMMrbcL_1_Csome(pf00016pf02788) CsomeACh
70943 HMMrbcS_1_CsomeA(pf00101) CsomeACh
70944 HMMcsoS2_2_ACh(pf12288) CsomeACh
70945 HMMca_1_CsomeACh(pf08936) CsomeACh
70946 P_neonPurple CsomeACh
70947 P_neonPurple CsomeACh
70948 H_hotpink CsomeACh
70949 H_hotpink CsomeACh
70950 -pf00210 -
70951 -pf01329 -
70952 -pf01656 -
70953 -# -
70954 -pf07728pf08406 -
70955 -pf00092 -
70956 Tdp_darkMagenta(#) CsomeACh
70957 -# -
70958 -# -
70959 -pf00689pf00690 -
70960 -pf00005 -
70961 -pf01061 -
70962 -pf00582 -
70963 -pf00011 -
70964 -pf08238 -
70965 -pf06897 -
70966 -pf03631 -
BMC-P
BMC-H
BMC-T
Aldehyde dehydrogenase
PTAC
Signature enzyme
Alcohol dehydrogenase
Regulator

Thioalkalimicrobium cyclicum (strain DSM 14477 / JCM 11371 / ALM1). Bacteria; Proteobacteria; Gammaproteobacteria; Thiotrichales;

Thicy__locus_1

CsomeACh

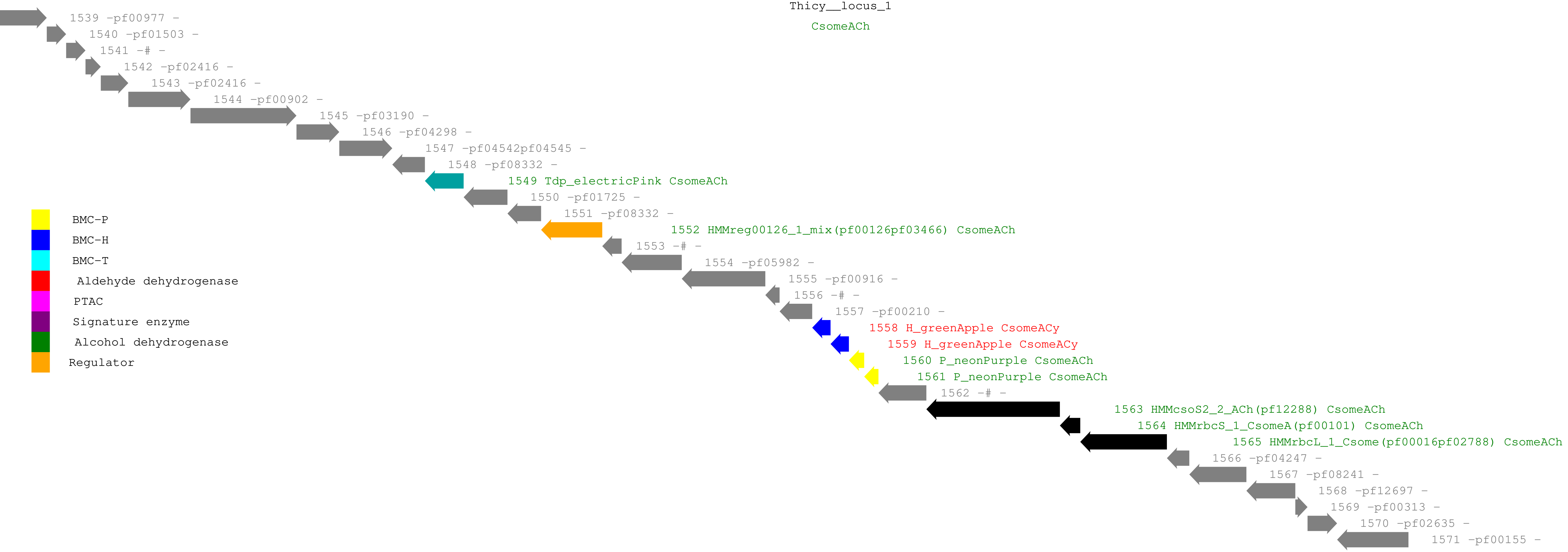

Thiorhodococcus drewsii AZ1. Bacteria; Proteobacteria; Gammaproteobacteria; Chromatiales;

ThidrDRAFT__locus_1

CsomeACh

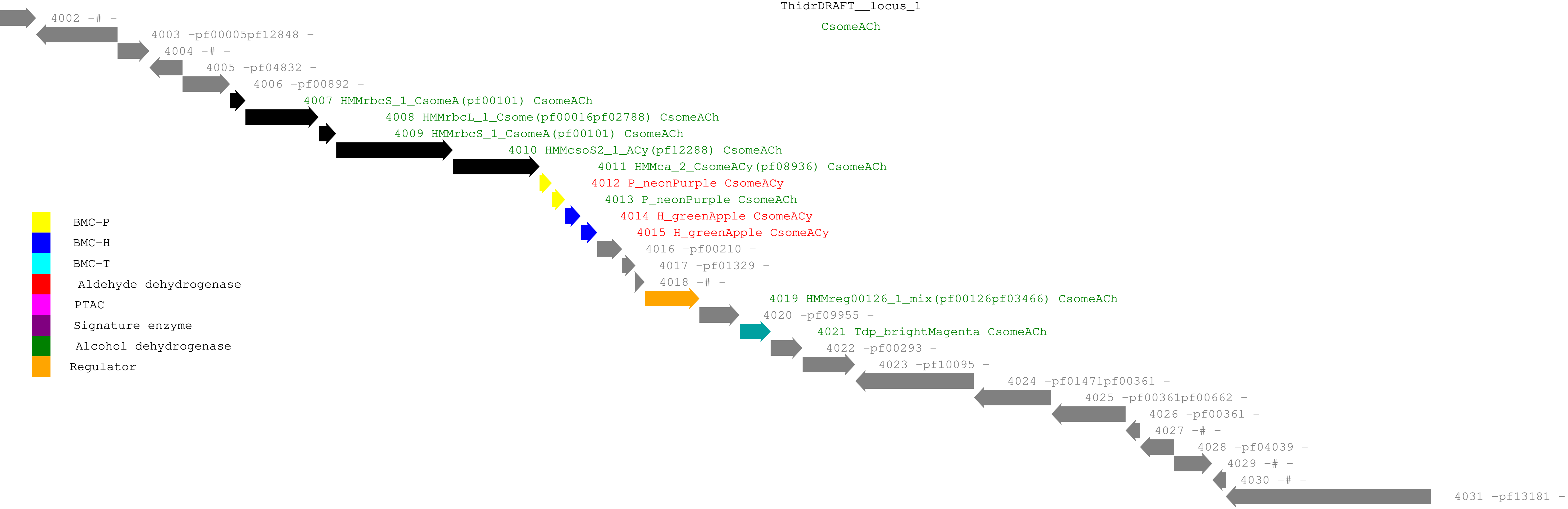

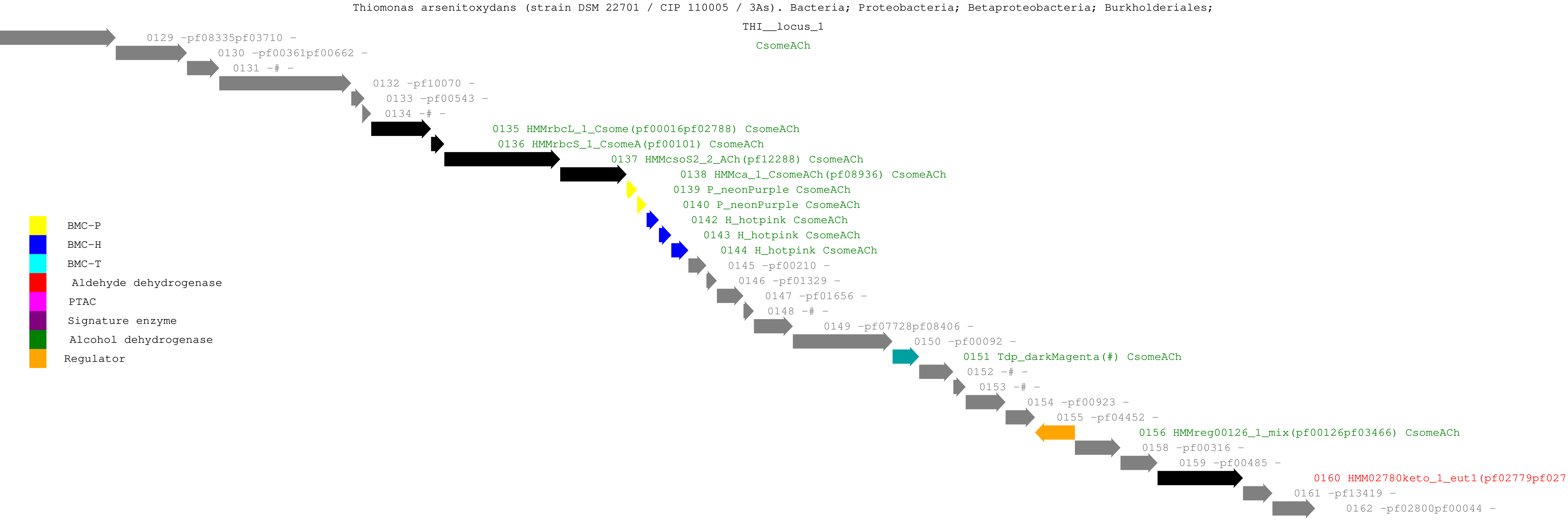
Thiomonas arsenitoxydans (strain DSM 22701 / CIP 110005 / 3As). Bacteria; Proteobacteria; Betaproteobacteria; Burkholderiales;
THI__locus_1
CsomeACh
0129 -pf08335pf03710 -
0130 -pf00361pf00662 -
0131 -# -
0132 -pf10070 -
0133 -pf00543 -
0134 -# -
0135 HMMrbcL_1_Csome(pf00016pf02788) CsomeACh
0136 HMMrbcS_1_CsomeA(pf00101) CsomeACh
0137 HMMcsoS2_2_ACh(pf12288) CsomeACh
0138 HMMca_1_CsomeACh(pf08936) CsomeACh
0139 P_neonPurple CsomeACh
0140 P_neonPurple CsomeACh
0142 H_hotpink CsomeACh
0143 H_hotpink CsomeACh
0144 H_hotpink CsomeACh
0145 -pf00210 -
0146 -pf01329 -
0147 -pf01656 -
0148 -# -
0149 -pf07728pf08406 -
0150 -pf00092 -
0151 Tdp_darkMagenta(#) CsomeACh
0152 -# -
0153 -# -
0154 -pf00923 -
0155 -pf04452 -
0156 HMMreg00126_1_mix(pf00126pf03466) CsomeACh
0158 -pf00316 -
0159 -pf00485 -
0160 HMM02780keto_1_eut1(pf02779pf027
0161 -pf13419 -
0162 -pf02800pf00044 -
BMC-P
BMC-H
BMC-T
Aldehyde dehydrogenase
PTAC
Signature enzyme
Alcohol dehydrogenase
Regulator

Thiocapsa marina 5811. Bacteria; Proteobacteria; Gammaproteobacteria; Chromatiales;

ThimaDRAFT__locus_1_main_sat

CsomeACh

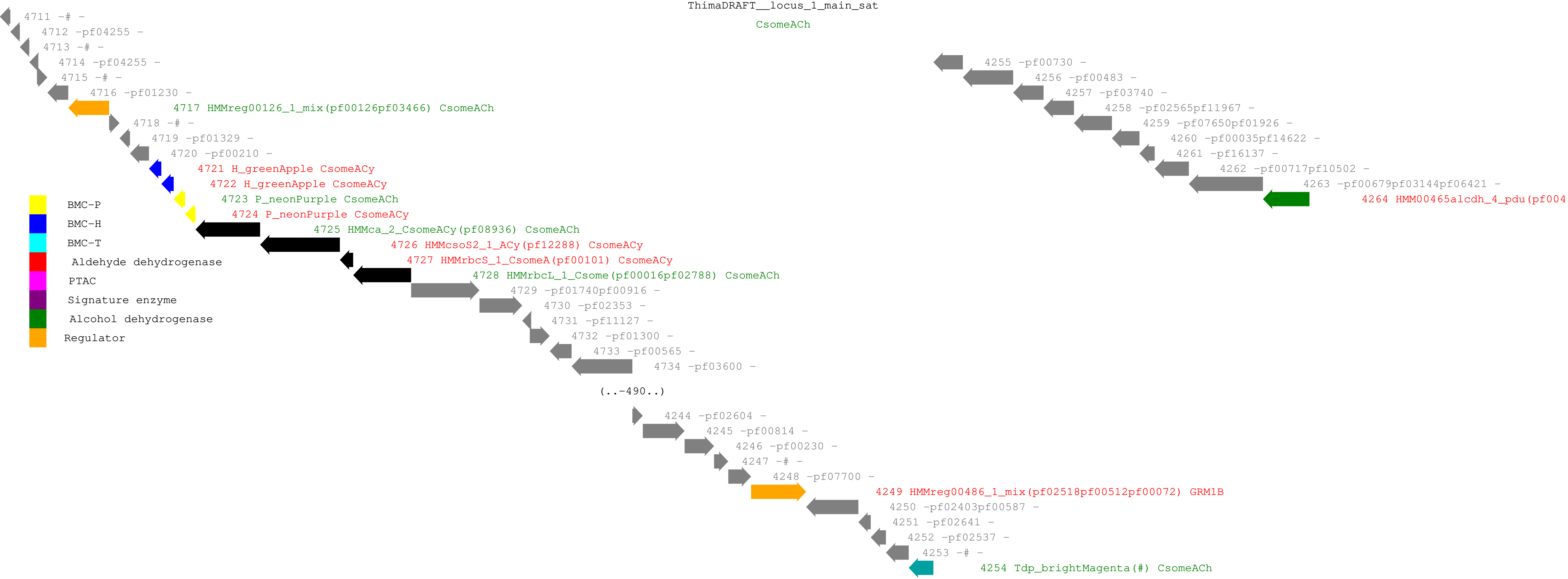

Thioflavicoccus mobilis 8321. Bacteria; Proteobacteria; Gammaproteobacteria; Chromatiales;

Thimo__locus_1

CsomeACh

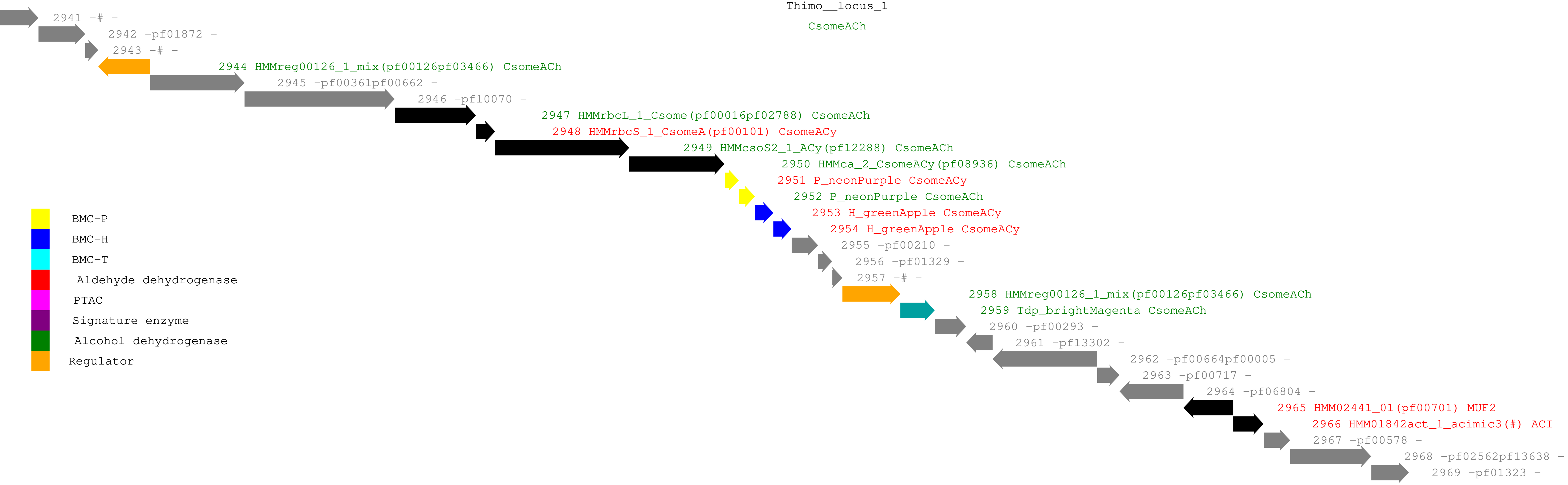

Thiothrix nivea (strain ATCC 35100 / DSM 5205 / JP2). Bacteria; Proteobacteria; Gammaproteobacteria; Thiotrichales;

Thini__locus_1

CsomeACh

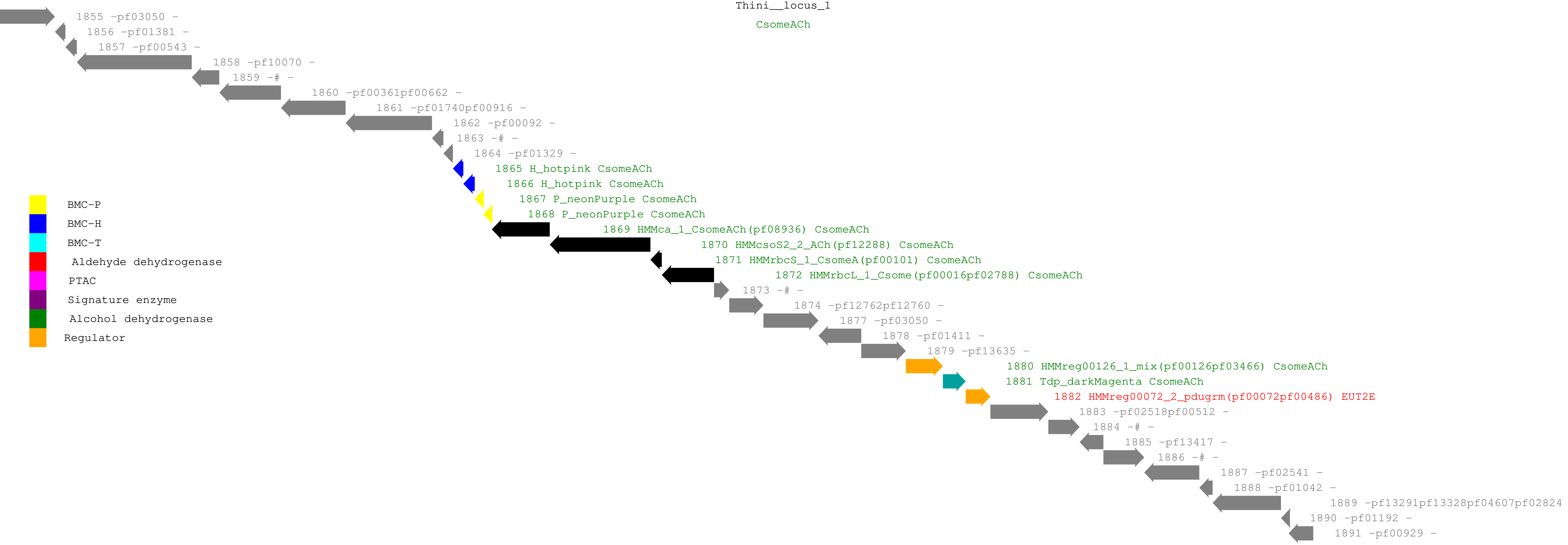

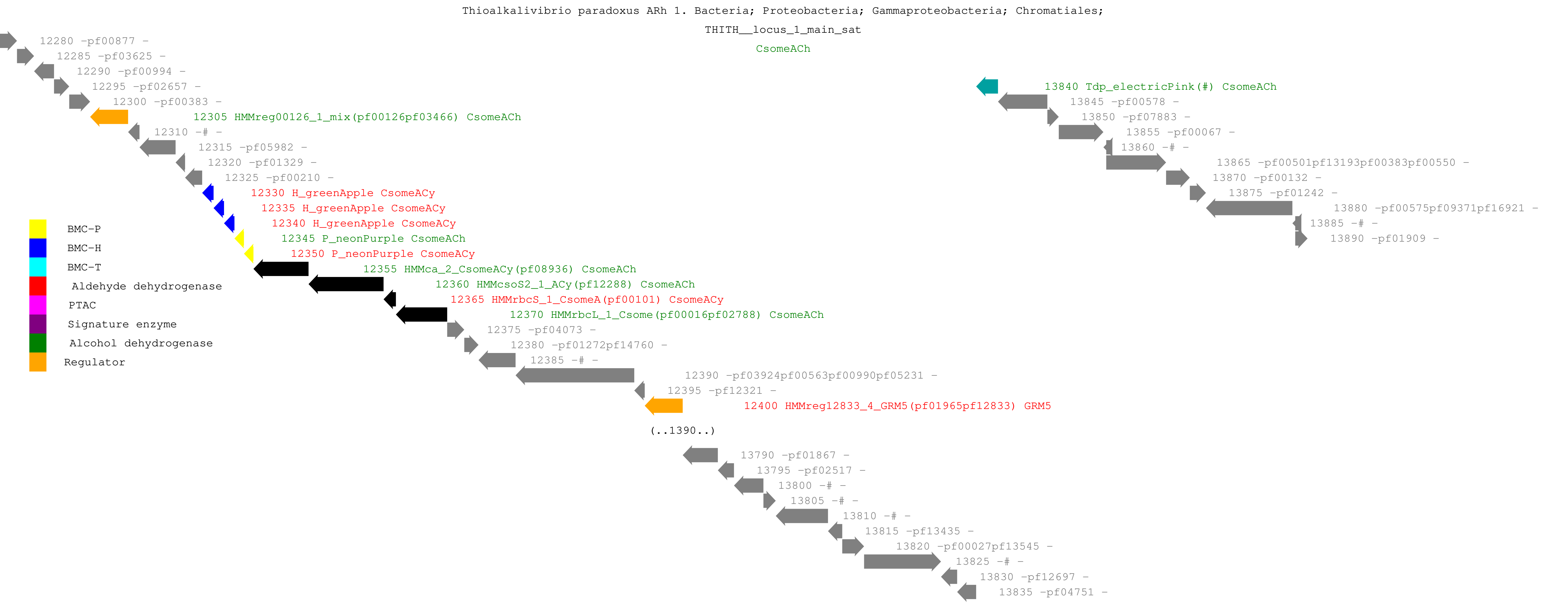
Thioalkalivibrio paradoxus ARh 1. Bacteria; Proteobacteria; Gammaproteobacteria; Chromatiales;
THITH__locus_1_main_sat
CsomeACh
12280 -pf00877 -
12285 -pf03625 -
12290 -pf00994 -
12295 -pf02657 -
12300 -pf00383 -
12305 HMMreg00126_1_mix(pf00126pf03466) CsomeACh
12310 -# -
12315 -pf05982 -
12320 -pf01329 -
12325 -pf00210 -
12330 H_greenApple CsomeACy
12335 H_greenApple CsomeACy
12340 H_greenApple CsomeACy
12345 P_neonPurple CsomeACh
12350 P_neonPurple CsomeACy
12355 HMMca_2_CsomeACy(pf08936) CsomeACh
12360 HMMcsoS2_1_ACy(pf12288) CsomeACh
12365 HMMrbcS_1_CsomeA(pf00101) CsomeACy
12370 HMMrbcL_1_Csome(pf00016pf02788) CsomeACh
12375 -pf04073 -
12380 -pf01272pf14760 -
12385 -# -
12390 -pf03924pf00563pf00990pf05231 -
12395 -pf12321 -
12400 HMMreg12833_4_GRM5(pf01965pf12833) GRM5
(..1390..)
13790 -pf01867 -
13795 -pf02517 -
13800 -# -
13805 -# -
13810 -# -
13815 -pf13435 -
13820 -pf00027pf13545 -
13825 -# -
13830 -pf12697 -
13835 -pf04751 -
13840 Tdp_electricPink(#) CsomeACh
13845 -pf00578 -
13850 -pf07883 -
13855 -pf00067 -
13860 -# -
13865 -pf00501pf13193pf00383pf00550 -
13870 -pf00132 -
13875 -pf01242 -
13880 -pf00575pf09371pf16921 -
13885 -# -
13890 -pf01909 -
BMC-P
BMC-H
BMC-T
Aldehyde dehydrogenase
PTAC
Signature enzyme
Alcohol dehydrogenase
Regulator

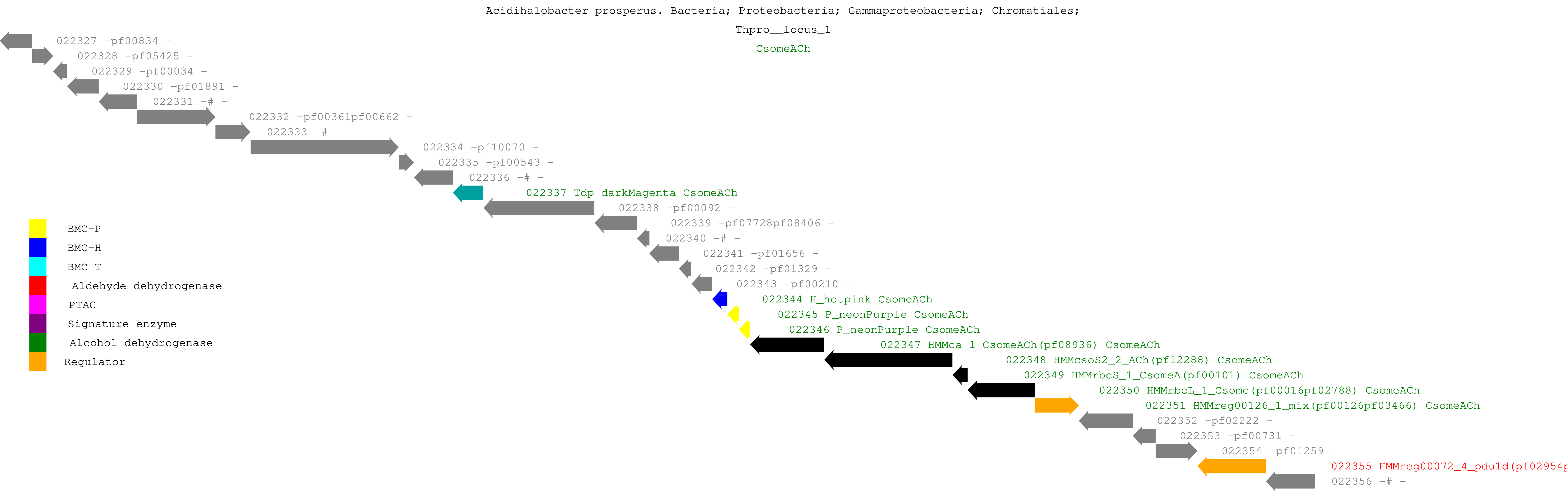
Acidihalobacter prosperus. Bacteria; Proteobacteria; Gammaproteobacteria; Chromatiales;
Thpro__locus_1
CsomeACh
022327 -pf00834 -
022328 -pf05425 -
022329 -pf00034 -
022330 -pf01891 -
022331 -# -
022332 -pf00361pf00662 -
022333 -# -
022334 -pf10070 -
022335 -pf00543 -
022336 -# -
022337 Tdp_darkMagenta CsomeACh
022338 -pf00092 -
022339 -pf07728pf08406 -
022340 -# -
022341 -pf01656 -
022342 -pf01329 -
022343 -pf00210 -
022344 H_hotpink CsomeACh
022345 P_neonPurple CsomeACh
022346 P_neonPurple CsomeACh
022347 HMMca_1_CsomeACh(pf08936) CsomeACh
022348 HMMcsoS2_2_ACh(pf12288) CsomeACh
022349 HMMrbcS_1_CsomeA(pf00101) CsomeACh
022350 HMMrbcL_1_Csome(pf00016pf02788) CsomeACh
022351 HMMreg00126_1_mix(pf00126pf03466) CsomeACh
022352 -pf02222 -
022353 -pf00731 -
022354 -pf01259 -
022355 HMMreg00072_4_pdu1d(pf02954p
022356 -# -
BMC-P
BMC-H
BMC-T
Aldehyde dehydrogenase
PTAC
Signature enzyme
Alcohol dehydrogenase
Regulator

Candidatus Thiodictyon syntrophicum. Bacteria; Proteobacteria; Gammaproteobacteria; Chromatiales;

THSYN__locus_1

CsomeACh

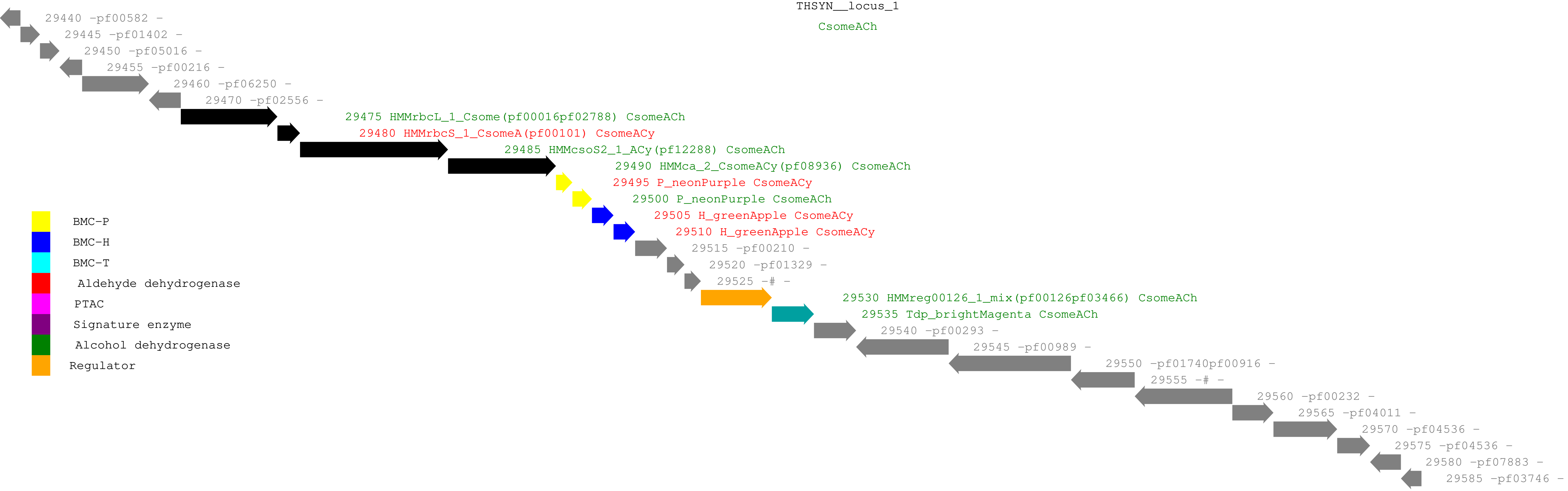

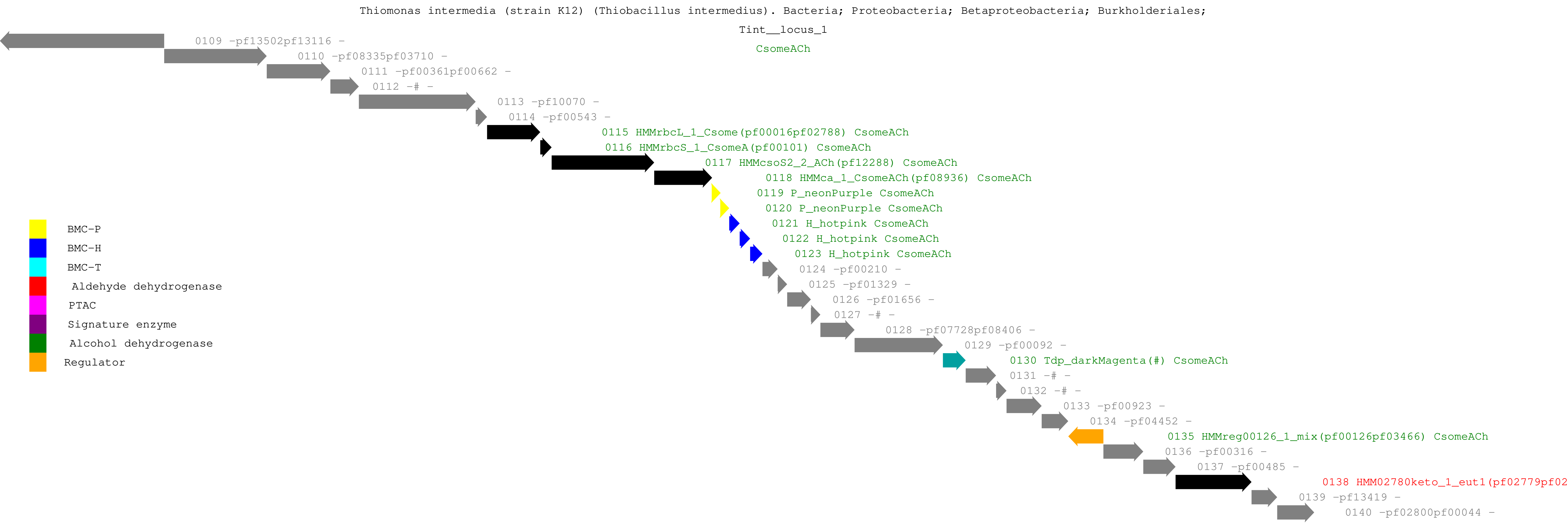

Thiomonas intermedia (strain K12) (Thiobacillus intermedius). Bacteria; Proteobacteria; Betaproteobacteria; Burkholderiales;
Tint__locus_1
CsomeACh
0109 -pf13502pf13116 -
0110 -pf08335pf03710 -
0111 -pf00361pf00662 -
0112 -# -
0113 -pf10070 -
0114 -pf00543 -
0115 HMMrbcL_1_Csome(pf00016pf02788) CsomeACh
0116 HMMrbcS_1_CsomeA(pf00101) CsomeACh
0117 HMMcsoS2_2_ACh(pf12288) CsomeACh
0118 HMMca_1_CsomeACh(pf08936) CsomeACh
0119 P_neonPurple CsomeACh
0120 P_neonPurple CsomeACh
0121 H_hotpink CsomeACh
0122 H_hotpink CsomeACh
0123 H_hotpink CsomeACh
0124 -pf00210 -
0125 -pf01329 -
0126 -pf01656 -
0127 -# -
0128 -pf07728pf08406 -
0129 -pf00092 -
0130 Tdp_darkMagenta(#) CsomeACh
0131 -# -
0132 -# -
0133 -pf00923 -
0134 -pf04452 -
0135 HMMreg00126_1_mix(pf00126pf03466) CsomeACh
0136 -pf00316 -
0137 -pf00485 -
0138 HMM02780keto_1_eut1(pf02779pf02
0139 -pf13419 -
0140 -pf02800pf00044 -
BMC-P
BMC-H
BMC-T
Aldehyde dehydrogenase
PTAC
Signature enzyme
Alcohol dehydrogenase
Regulator

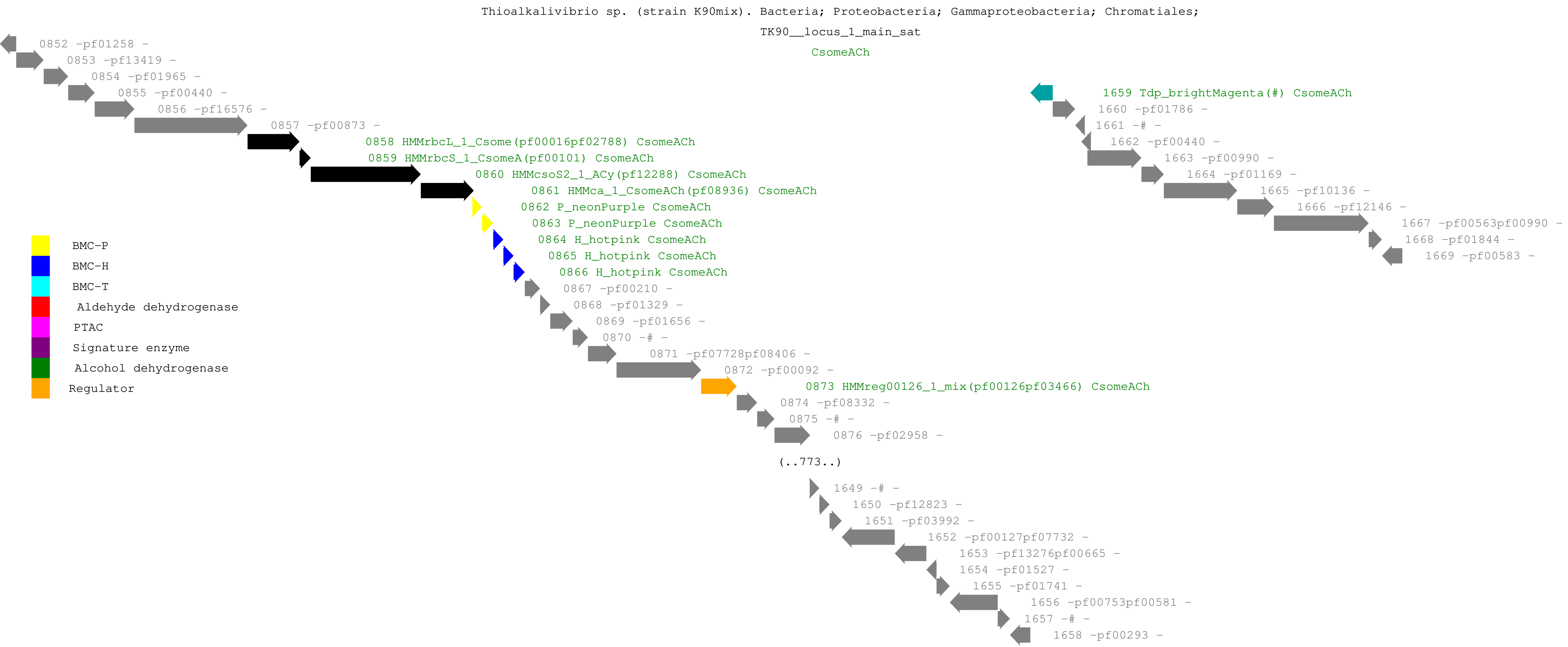
Thioalkalivibrio sp. (strain K90mix). Bacteria; Proteobacteria; Gammaproteobacteria; Chromatiales;
TK90__locus_1_main_sat
CsomeACh
0852 -pf01258 -
0853 -pf13419 -
0854 -pf01965 -
0855 -pf00440 -
0856 -pf16576 -
0857 -pf00873 -
0858 HMMrbcL_1_Csome(pf00016pf02788) CsomeACh
0859 HMMrbcS_1_CsomeA(pf00101) CsomeACh
0860 HMMcsoS2_1_ACy(pf12288) CsomeACh
0861 HMMca_1_CsomeACh(pf08936) CsomeACh
0862 P_neonPurple CsomeACh
0863 P_neonPurple CsomeACh
0864 H_hotpink CsomeACh
0865 H_hotpink CsomeACh
0866 H_hotpink CsomeACh
0867 -pf00210 -
0868 -pf01329 -
0869 -pf01656 -
0870 -# -
0871 -pf07728pf08406 -
0872 -pf00092 -
0873 HMMreg00126_1_mix(pf00126pf03466) CsomeACh
0874 -pf08332 -
0875 -# -
0876 -pf02958 -
(..773..)
1649 -# -
1650 -pf12823 -
1651 -pf03992 -
1652 -pf00127pf07732 -
1653 -pf13276pf00665 -
1654 -pf01527 -
1655 -pf01741 -
1656 -pf00753pf00581 -
1657 -# -
1658 -pf00293 -
1659 Tdp_brightMagenta(#) CsomeACh
1660 -pf01786 -
1661 -# -
1662 -pf00440 -
1663 -pf00990 -
1664 -pf01169 -
1665 -pf10136 -
1666 -pf12146 -
1667 -pf00563pf00990 -
1668 -pf01844 -
1669 -pf00583 -
BMC-P
BMC-H
BMC-T
Aldehyde dehydrogenase
PTAC
Signature enzyme
Alcohol dehydrogenase
Regulator

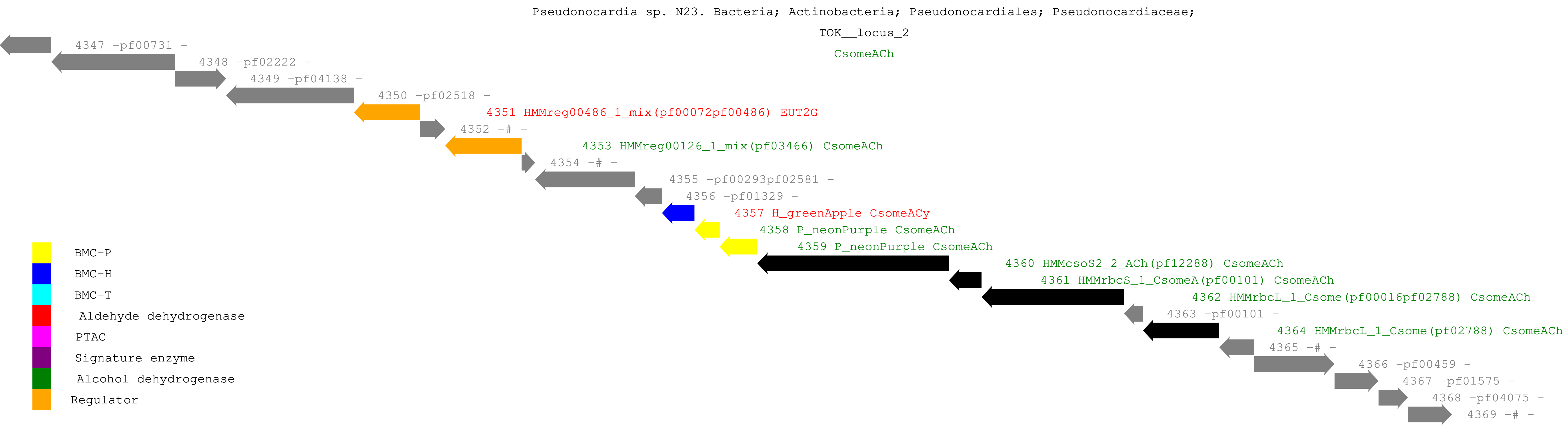
Pseudonocardia sp. N23. Bacteria; Actinobacteria; Pseudonocardiales; Pseudonocardiaceae;
TOK__locus_2
CsomeACh
4347 -pf00731 -
4348 -pf02222 -
4349 -pf04138 -
4350 -pf02518 -
4351 HMMreg00486_1_mix(pf00072pf00486) EUT2G
4352 -# -
4353 HMMreg00126_1_mix(pf03466) CsomeACh
4354 -# -
4355 -pf00293pf02581 -
4356 -pf01329 -
4357 H_greenApple CsomeACy
4358 P_neonPurple CsomeACh
4359 P_neonPurple CsomeACh
4360 HMMcsoS2_2_ACh(pf12288) CsomeACh
4361 HMMrbcS_1_CsomeA(pf00101) CsomeACh
4362 HMMrbcL_1_Csome(pf00016pf02788) CsomeACh
4363 -pf00101 -
4364 HMMrbcL_1_Csome(pf02788) CsomeACh
4365 -# -
4366 -pf00459 -
4367 -pf01575 -
4368 -pf04075 -
4369 -# -
BMC-P
BMC-H
BMC-T
Aldehyde dehydrogenase
PTAC
Signature enzyme
Alcohol dehydrogenase
Regulator

Tepidimonas taiwanensis. Bacteria; Proteobacteria; Betaproteobacteria; Burkholderiales; Tepidimonas.

Ttaiw__locus_1

CsomeACh

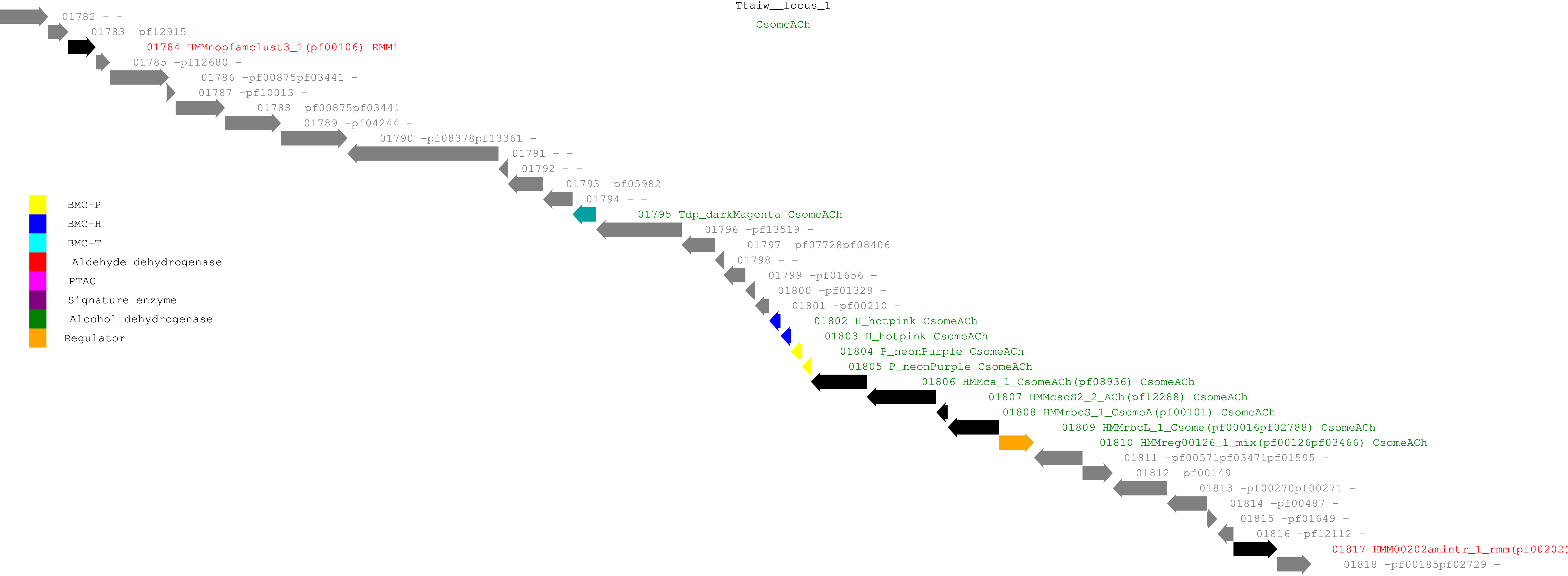

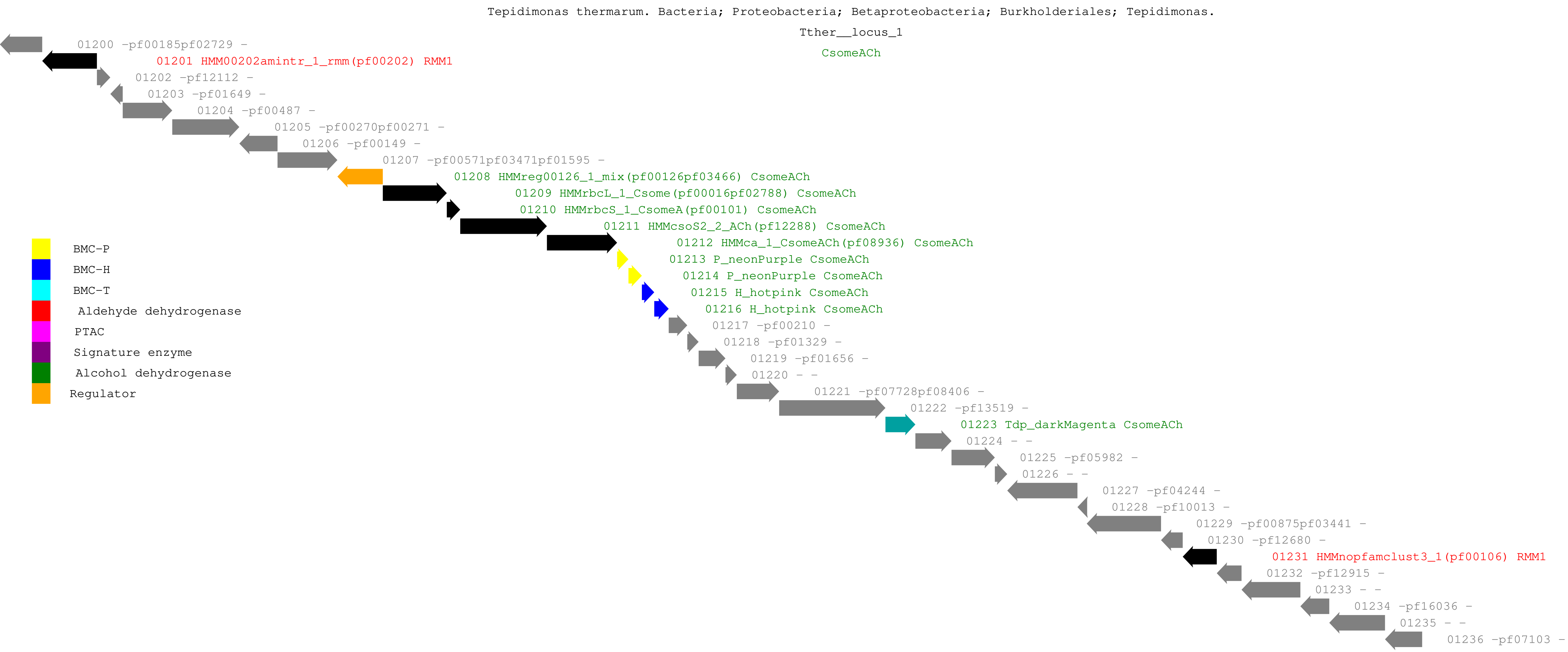
Tepidimonas thermarum. Bacteria; Proteobacteria; Betaproteobacteria; Burkholderiales; Tepidimonas.
Tther__locus_1
CsomeACh
01200 -pf00185pf02729 -
01201 HMM00202amintr_1_rmm(pf00202) RMM1
01202 -pf12112 -
01203 -pf01649 -
01204 -pf00487 -
01205 -pf00270pf00271 -
01206 -pf00149 -
01207 -pf00571pf03471pf01595 -
01208 HMMreg00126_1_mix(pf00126pf03466) CsomeACh
01209 HMMrbcL_1_Csome(pf00016pf02788) CsomeACh
01210 HMMrbcS_1_CsomeA(pf00101) CsomeACh
01211 HMMcsoS2_2_ACh(pf12288) CsomeACh
01212 HMMca_1_CsomeACh(pf08936) CsomeACh
01213 P_neonPurple CsomeACh
01214 P_neonPurple CsomeACh
01215 H_hotpink CsomeACh
01216 H_hotpink CsomeACh
01217 -pf00210 -
01218 -pf01329 -
01219 -pf01656 -
01220 - -
01221 -pf07728pf08406 -
01222 -pf13519 -
01223 Tdp_darkMagenta CsomeACh
01224 - -
01225 -pf05982 -
01226 - -
01227 -pf04244 -
01228 -pf10013 -
01229 -pf00875pf03441 -
01230 -pf12680 -
01231 HMMnopfamclust3_1(pf00106) RMM1
01232 -pf12915 -
01233 - -
01234 -pf16036 -
01235 - -
01236 -pf07103 -
BMC-P
BMC-H
BMC-T
Aldehyde dehydrogenase
PTAC
Signature enzyme
Alcohol dehydrogenase
Regulator

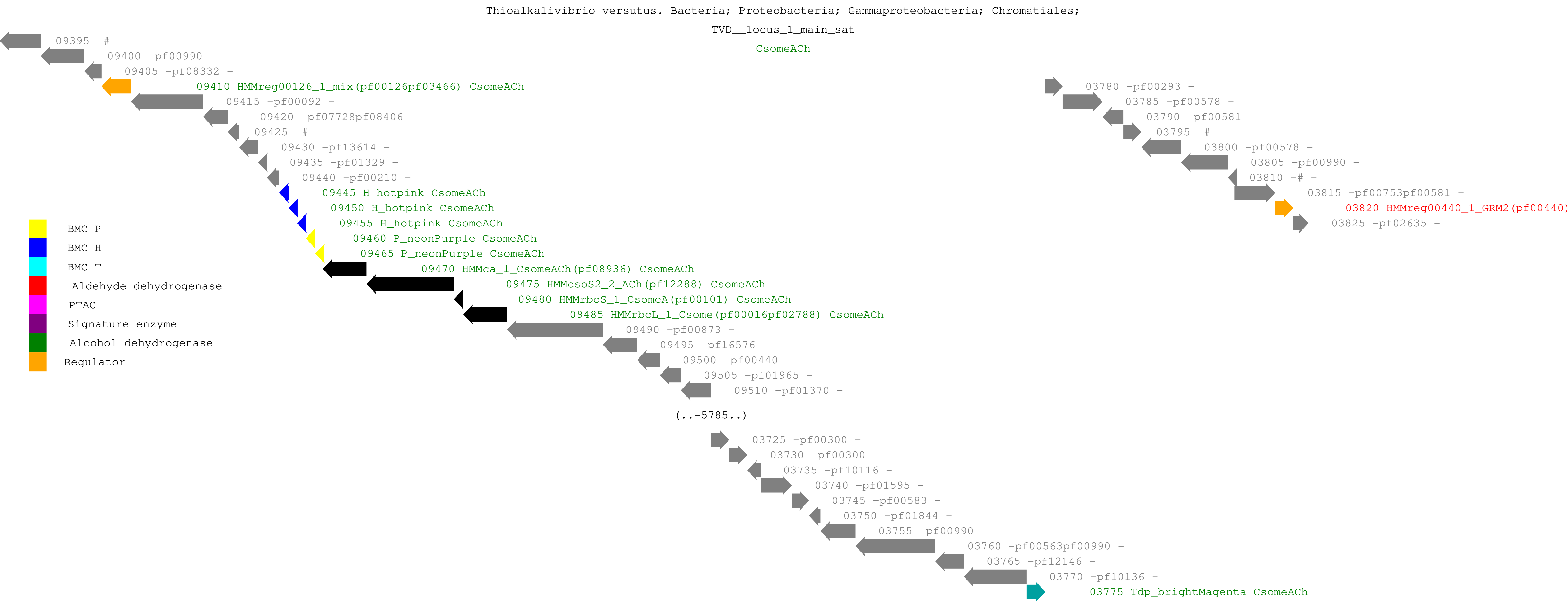
Thioalkalivibrio versutus. Bacteria; Proteobacteria; Gammaproteobacteria; Chromatiales;
TVD__locus_1_main_sat
CsomeACh
09395 -# -
09400 -pf00990 -
09405 -pf08332 -
09410 HMMreg00126_1_mix(pf00126pf03466) CsomeACh
09415 -pf00092 -
09420 -pf07728pf08406 -
09425 -# -
09430 -pf13614 -
09435 -pf01329 -
09440 -pf00210 -
09445 H_hotpink CsomeACh
09450 H_hotpink CsomeACh
09455 H_hotpink CsomeACh
09460 P_neonPurple CsomeACh
09465 P_neonPurple CsomeACh
09470 HMMca_1_CsomeACh(pf08936) CsomeACh
09475 HMMcsoS2_2_ACh(pf12288) CsomeACh
09480 HMMrbcS_1_CsomeA(pf00101) CsomeACh
09485 HMMrbcL_1_Csome(pf00016pf02788) CsomeACh
09490 -pf00873 -
09495 -pf16576 -
09500 -pf00440 -
09505 -pf01965 -
09510 -pf01370 -
(..-5785..)
03725 -pf00300 -
03730 -pf00300 -
03735 -pf10116 -
03740 -pf01595 -
03745 -pf00583 -
03750 -pf01844 -
03755 -pf00990 -
03760 -pf00563pf00990 -
03765 -pf12146 -
03770 -pf10136 -
03775 Tdp_brightMagenta CsomeACh
03780 -pf00293 -
03785 -pf00578 -
03790 -pf00581 -
03795 -# -
03800 -pf00578 -
03805 -pf00990 -
03810 -# -
03815 -pf00753pf00581 -
03820 HMMreg00440_1_GRM2(pf00440)
03825 -pf02635 -
BMC-P
BMC-H
BMC-T
Aldehyde dehydrogenase
PTAC
Signature enzyme
Alcohol dehydrogenase
Regulator

Thioalkalivibrio nitratireducens (strain DSM 14787 / UNIQEM 213 / Bacteria; Proteobacteria; Gammaproteobacteria; Chromatiales;

TVNIR__locus_1_main_sat

CsomeACh

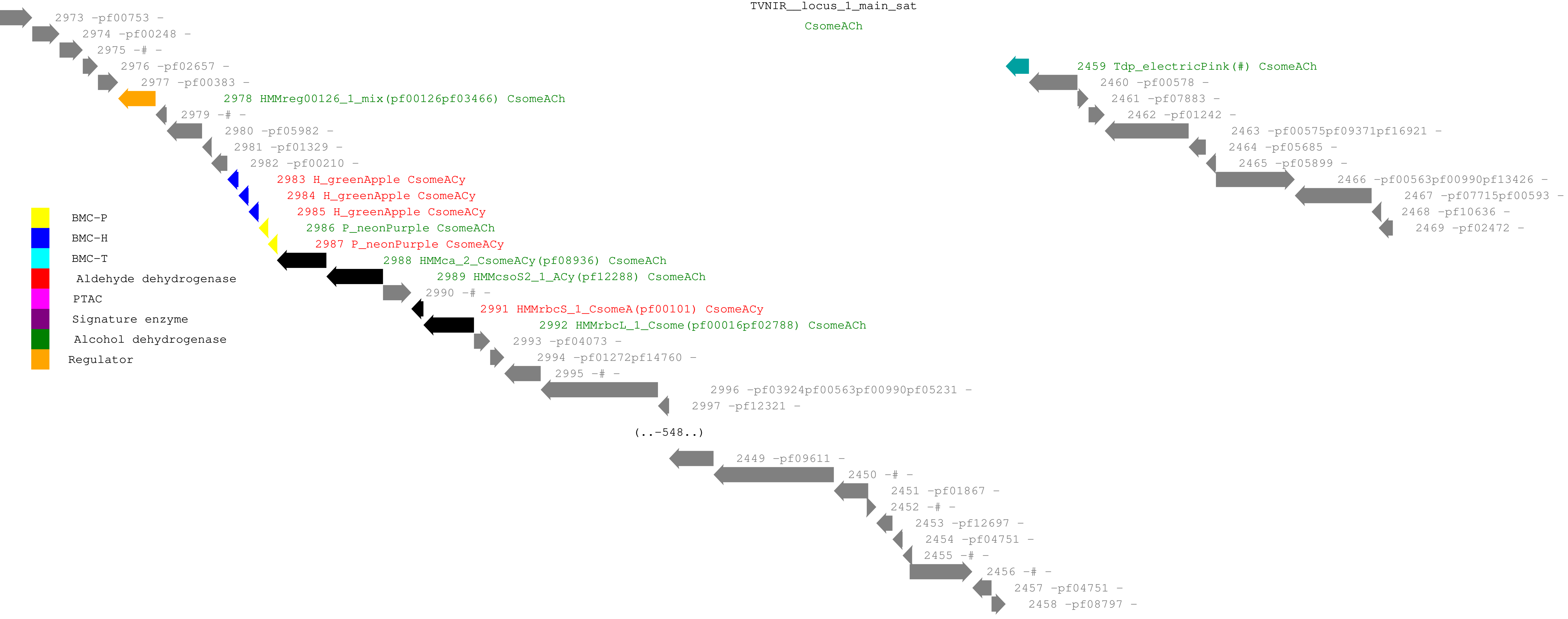

Prochlorococcus marinus (strain AS9601). Bacteria; Cyanobacteria; Synechococcales; Prochloraceae;

A9601__locus_1

CsomeACy

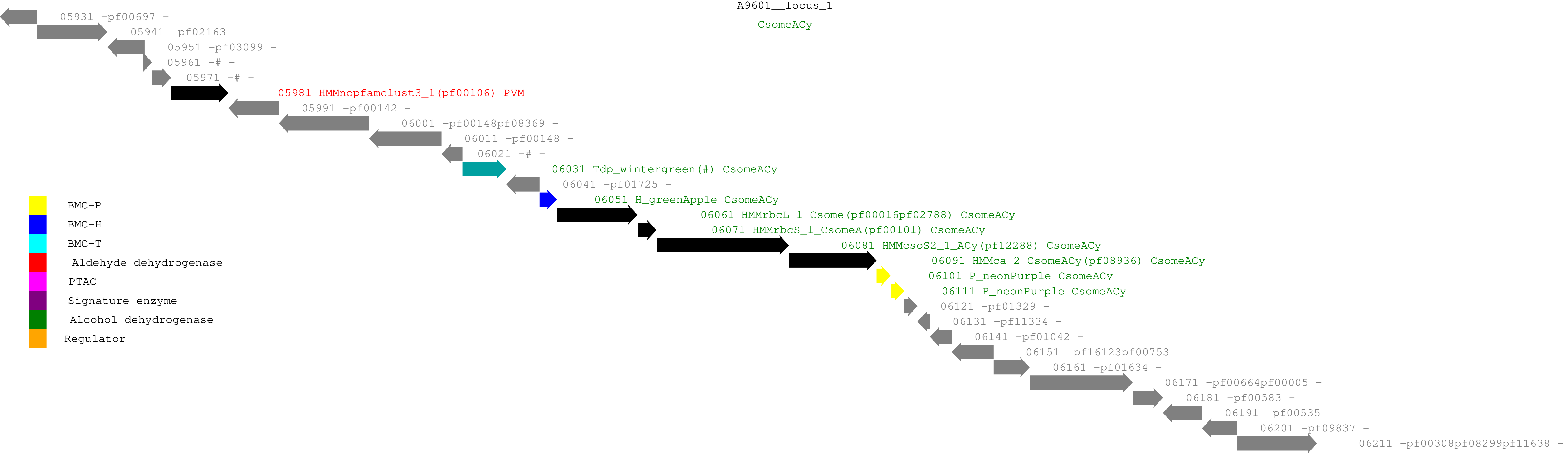

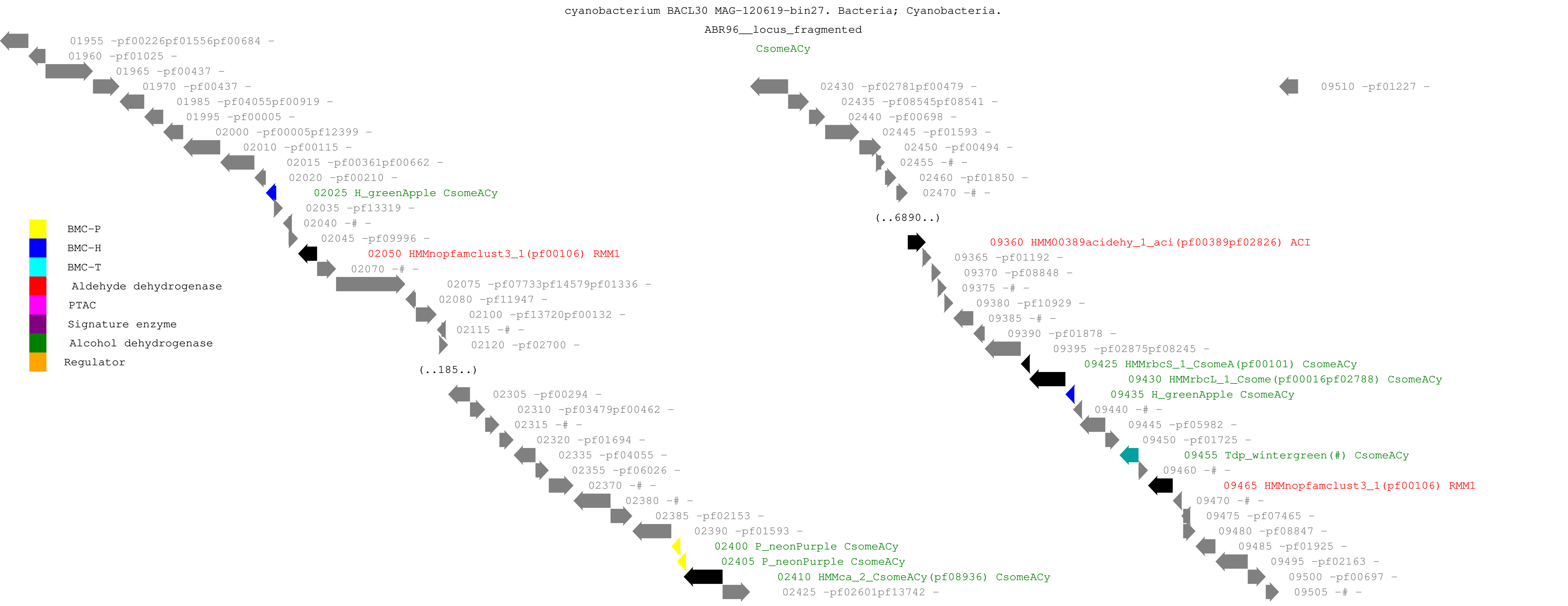

cyanobacterium BACL30 MAG-120619-bin27. Bacteria; Cyanobacteria.
ABR96__locus_fragmented
CsomeACy
01955 -pf00226pf01556pf00684 -
01960 -pf01025 -
01965 -pf00437 -
01970 -pf00437 -
01985 -pf04055pf00919 -
01995 -pf00005 -
02000 -pf00005pf12399 -
02010 -pf00115 -
02015 -pf00361pf00662 -
02020 -pf00210 -
02025 H_greenApple CsomeACy
02035 -pf13319 -
02040 -# -
02045 -pf09996 -
02050 HMMnopfamclust3_1(pf00106) RMM1
02070 -# -
02075 -pf07733pf14579pf01336 -
02080 -pf11947 -
02100 -pf13720pf00132 -
02115 -# -
02120 -pf02700 -
(..185..)
02305 -pf00294 -
02310 -pf03479pf00462 -
02315 -# -
02320 -pf01694 -
02335 -pf04055 -
02355 -pf06026 -
02370 -# -
02380 -# -
02385 -pf02153 -
02390 -pf01593 -
02400 P_neonPurple CsomeACy
02405 P_neonPurple CsomeACy
02410 HMMca_2_CsomeACy(pf08936) CsomeACy
02425 -pf02601pf13742 -
02430 -pf02781pf00479 -
02435 -pf08545pf08541 -
02440 -pf00698 -
02445 -pf01593 -
02450 -pf00494 -
02455 -# -
02460 -pf01850 -
02470 -# -
(..6890..)
09360 HMM00389acidehy_1_aci(pf00389pf02826) ACI
09365 -pf01192 -
09370 -pf08848 -
09375 -# -
09380 -pf10929 -
09385 -# -
09390 -pf01878 -
09395 -pf02875pf08245 -
09425 HMMrbcS_1_CsomeA(pf00101) CsomeACy
09430 HMMrbcL_1_Csome(pf00016pf02788) CsomeACy
09435 H_greenApple CsomeACy
09440 -# -
09445 -pf05982 -
09450 -pf01725 -
09455 Tdp_wintergreen(#) CsomeACy
09460 -# -
09465 HMMnopfamclust3_1(pf00106) RMM1
09470 -# -
09475 -pf07465 -
09480 -pf08847 -
09485 -pf01925 -
09495 -pf02163 -
09500 -pf00697 -
09505 -# -
09510 -pf01227 -
BMC-P
BMC-H
BMC-T
Aldehyde dehydrogenase
PTAC
Signature enzyme
Alcohol dehydrogenase
Regulator

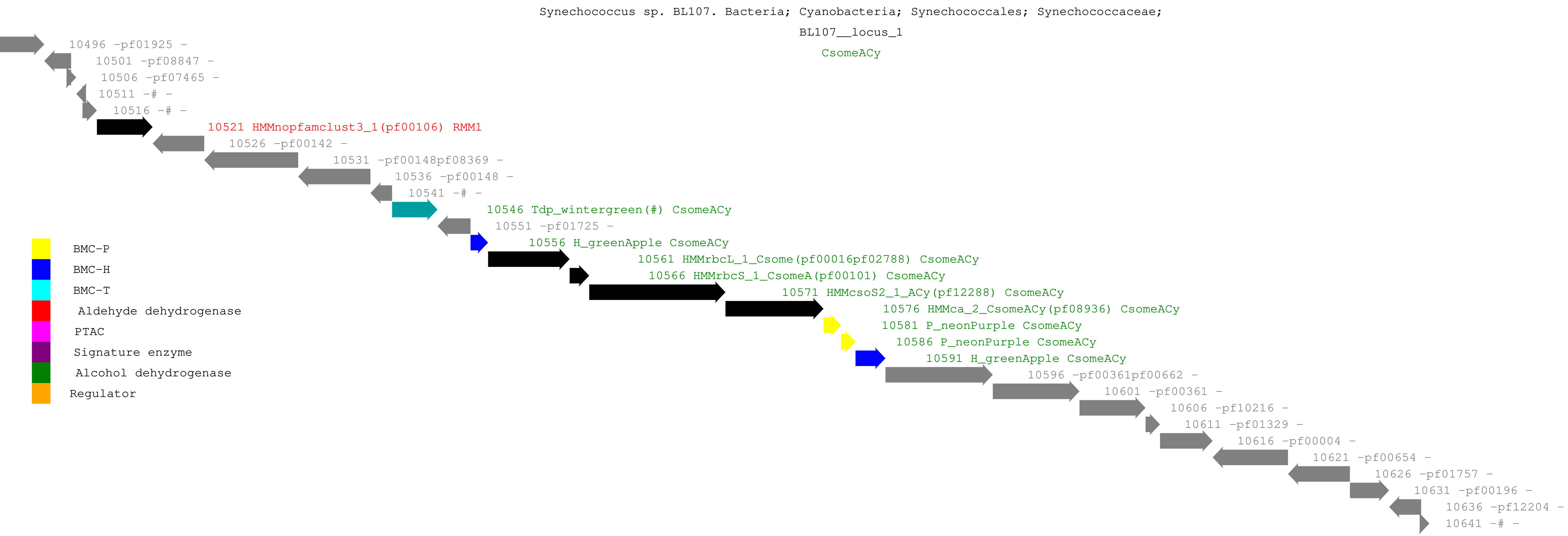

Synechococcus sp. BL107. Bacteria; Cyanobacteria; Synechococcales; Synechococcaceae;
BL107__locus_1
CsomeACy
10496 -pf01925 -
10501 -pf08847 -
10506 -pf07465 -
10511 -# -
10516 -# -
10521 HMMnopfamclust3_1(pf00106) RMM1
10526 -pf00142 -
10531 -pf00148pf08369 -
10536 -pf00148 -
10541 -# -
10546 Tdp_wintergreen(#) CsomeACy
10551 -pf01725 -
10556 H_greenApple CsomeACy
10561 HMMrbcL_1_Csome(pf00016pf02788) CsomeACy
10566 HMMrbcS_1_CsomeA(pf00101) CsomeACy
10571 HMMcsoS2_1_ACy(pf12288) CsomeACy
10576 HMMca_2_CsomeACy(pf08936) CsomeACy
10581 P_neonPurple CsomeACy
10586 P_neonPurple CsomeACy
10591 H_greenApple CsomeACy
10596 -pf00361pf00662 -
10601 -pf00361 -
10606 -pf10216 -
10611 -pf01329 -
10616 -pf00004 -
10621 -pf00654 -
10626 -pf01757 -
10631 -pf00196 -
10636 -pf12204 -
10641 -# -
BMC-P
BMC-H
BMC-T
Aldehyde dehydrogenase
PTAC
Signature enzyme
Alcohol dehydrogenase
Regulator

Synechococcus sp. SynAce01. Bacteria; Cyanobacteria; Synechococcales; Synechococcaceae;

BM449__locus_1

CsomeACy

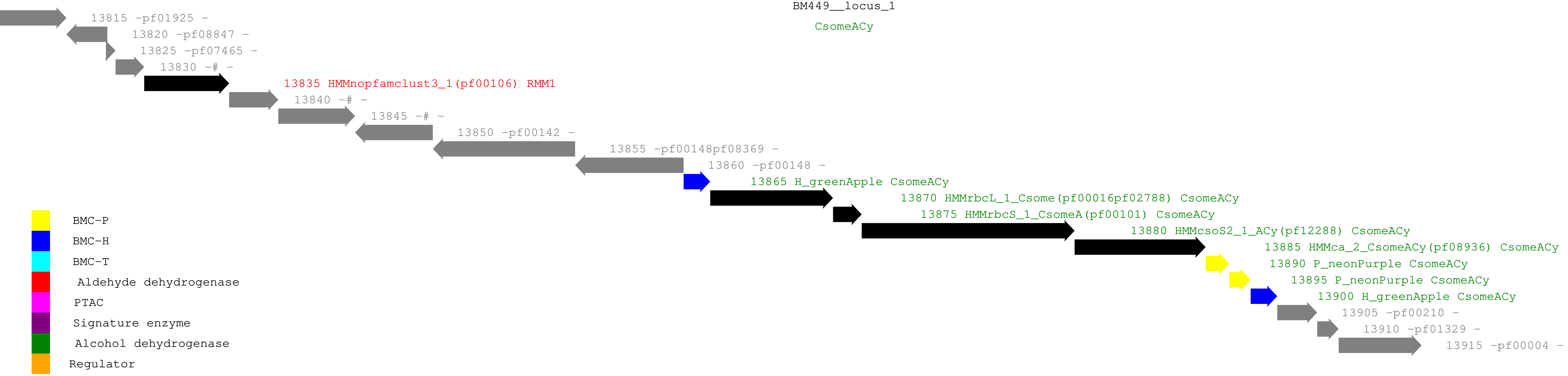

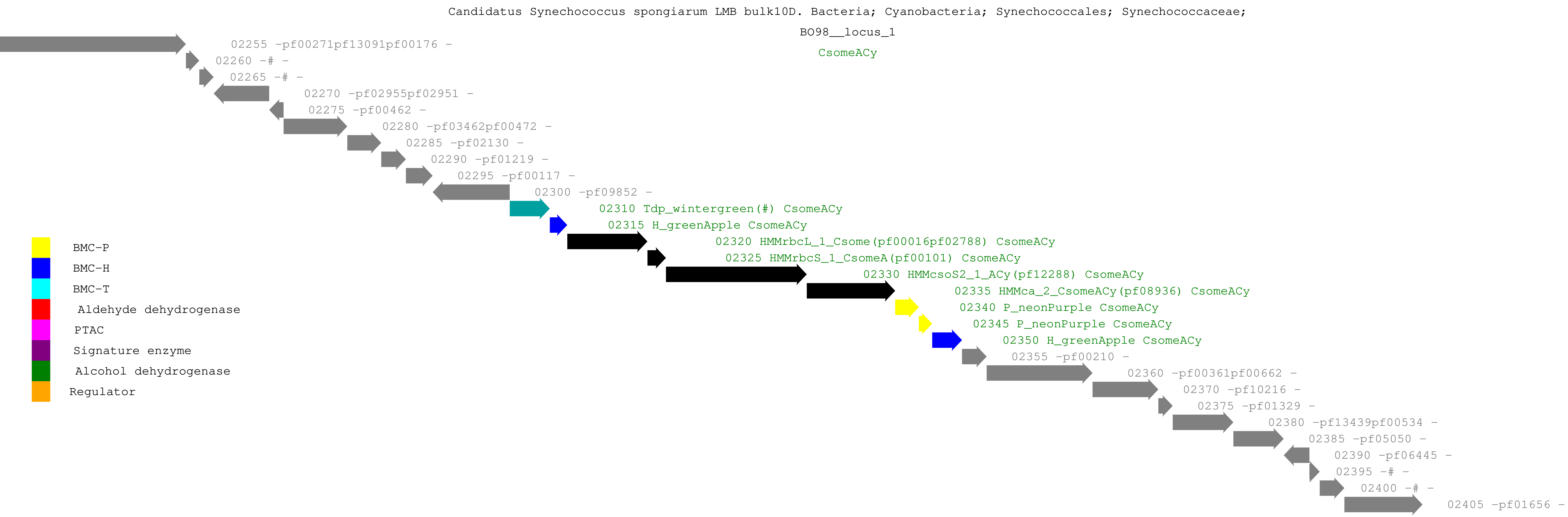

Candidatus Synechococcus spongiarum LMB bulk10D. Bacteria; Cyanobacteria; Synechococcales; Synechococcaceae;
BO98__locus_1
CsomeACy
02255 -pf00271pf13091pf00176 -
02260 -# -
02265 -# -
02270 -pf02955pf02951 -
02275 -pf00462 -
02280 -pf03462pf00472 -
02285 -pf02130 -
02290 -pf01219 -
02295 -pf00117 -
02300 -pf09852 -
02310 Tdp_wintergreen(#) CsomeACy
02315 H_greenApple CsomeACy
02320 HMMrbcL_1_Csome(pf00016pf02788) CsomeACy
02325 HMMrbcS_1_CsomeA(pf00101) CsomeACy
02330 HMMcsoS2_1_ACy(pf12288) CsomeACy
02335 HMMca_2_CsomeACy(pf08936) CsomeACy
02340 P_neonPurple CsomeACy
02345 P_neonPurple CsomeACy
02350 H_greenApple CsomeACy
02355 -pf00210 -
02360 -pf00361pf00662 -
02370 -pf10216 -
02375 -pf01329 -
02380 -pf13439pf00534 -
02385 -pf05050 -
02390 -pf06445 -
02395 -# -
02400 -# -
02405 -pf01656 -
BMC-P
BMC-H
BMC-T
Aldehyde dehydrogenase
PTAC
Signature enzyme
Alcohol dehydrogenase
Regulator

Prochlorococcus sp. RS01. A0A1Q1EUC2_9PROC Unreviewed; 464 AA.

BS620__locus_1

CsomeACy

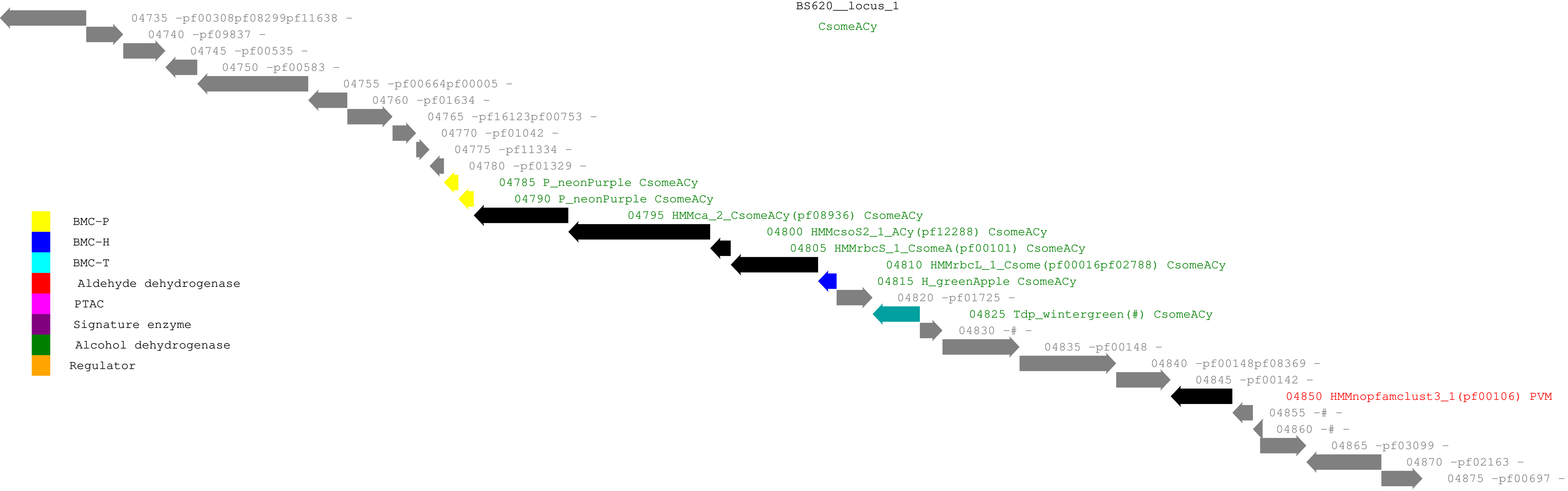

Prochlorococcus sp. RS50. A0A1Q1ERN1_9PROC Unreviewed; 221 AA.

BSR22__locus_1

CsomeACy

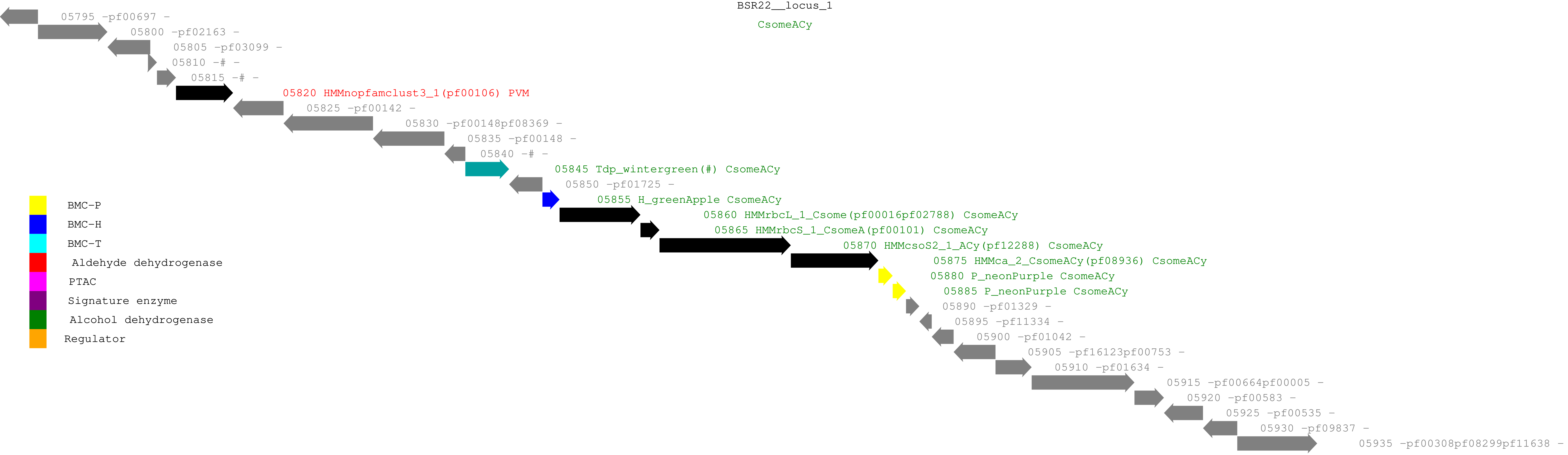

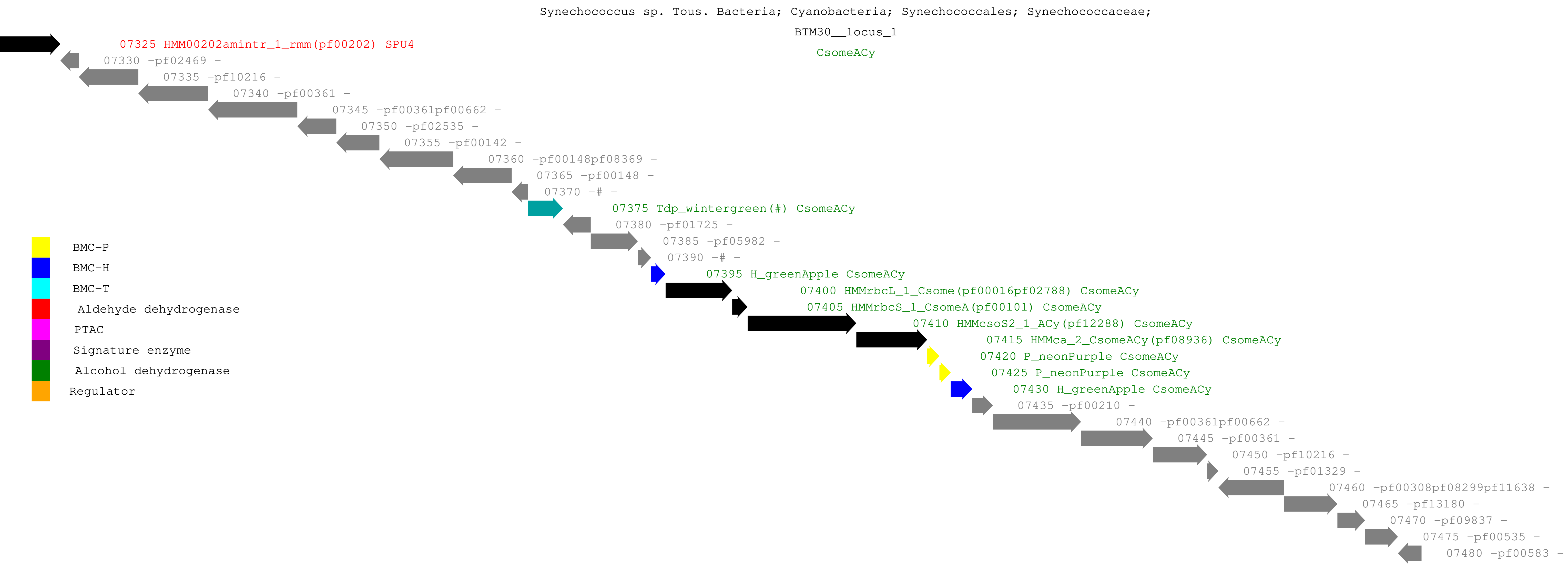
Synechococcus sp. Tous. Bacteria; Cyanobacteria; Synechococcales; Synechococcaceae;
BTM30__locus_1
CsomeACy
07325 HMM00202amintr_1_rmm(pf00202) SPU4
07330 -pf02469 -
07335 -pf10216 -
07340 -pf00361 -
07345 -pf00361pf00662 -
07350 -pf02535 -
07355 -pf00142 -
07360 -pf00148pf08369 -
07365 -pf00148 -
07370 -# -
07375 Tdp_wintergreen(#) CsomeACy
07380 -pf01725 -
07385 -pf05982 -
07390 -# -
07395 H_greenApple CsomeACy
07400 HMMrbcL_1_Csome(pf00016pf02788) CsomeACy
07405 HMMrbcS_1_CsomeA(pf00101) CsomeACy
07410 HMMcsoS2_1_ACy(pf12288) CsomeACy
07415 HMMca_2_CsomeACy(pf08936) CsomeACy
07420 P_neonPurple CsomeACy
07425 P_neonPurple CsomeACy
07430 H_greenApple CsomeACy
07435 -pf00210 -
07440 -pf00361pf00662 -
07445 -pf00361 -
07450 -pf10216 -
07455 -pf01329 -
07460 -pf00308pf08299pf11638 -
07465 -pf13180 -
07470 -pf09837 -
07475 -pf00535 -
07480 -pf00583 -
BMC-P
BMC-H
BMC-T
Aldehyde dehydrogenase
PTAC
Signature enzyme
Alcohol dehydrogenase
Regulator

Candidatus Synechococcus spongiarum LMB bulk15N. Bacteria; Cyanobacteria; Synechococcales; Synechococcaceae;

BV53__locus_1

CsomeACy

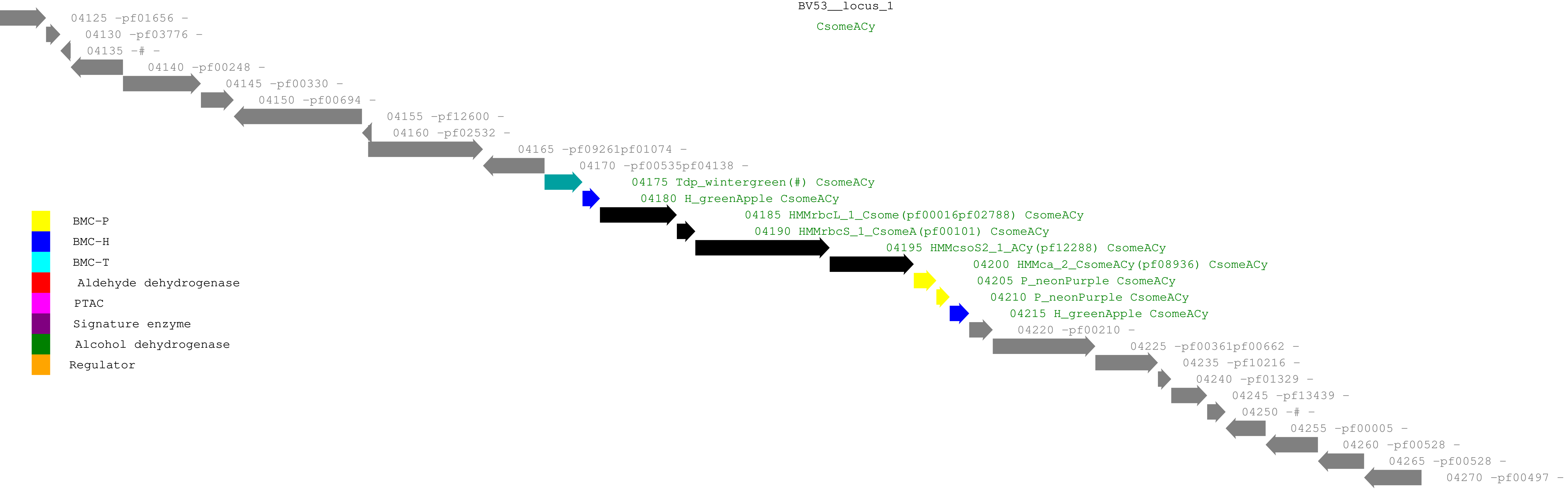

Candidatus Synechococcus spongiarum LMB bulk15M. Bacteria; Cyanobacteria; Synechococcales; Synechococcaceae;

BV61__locus_1_main_sat

CsomeACy

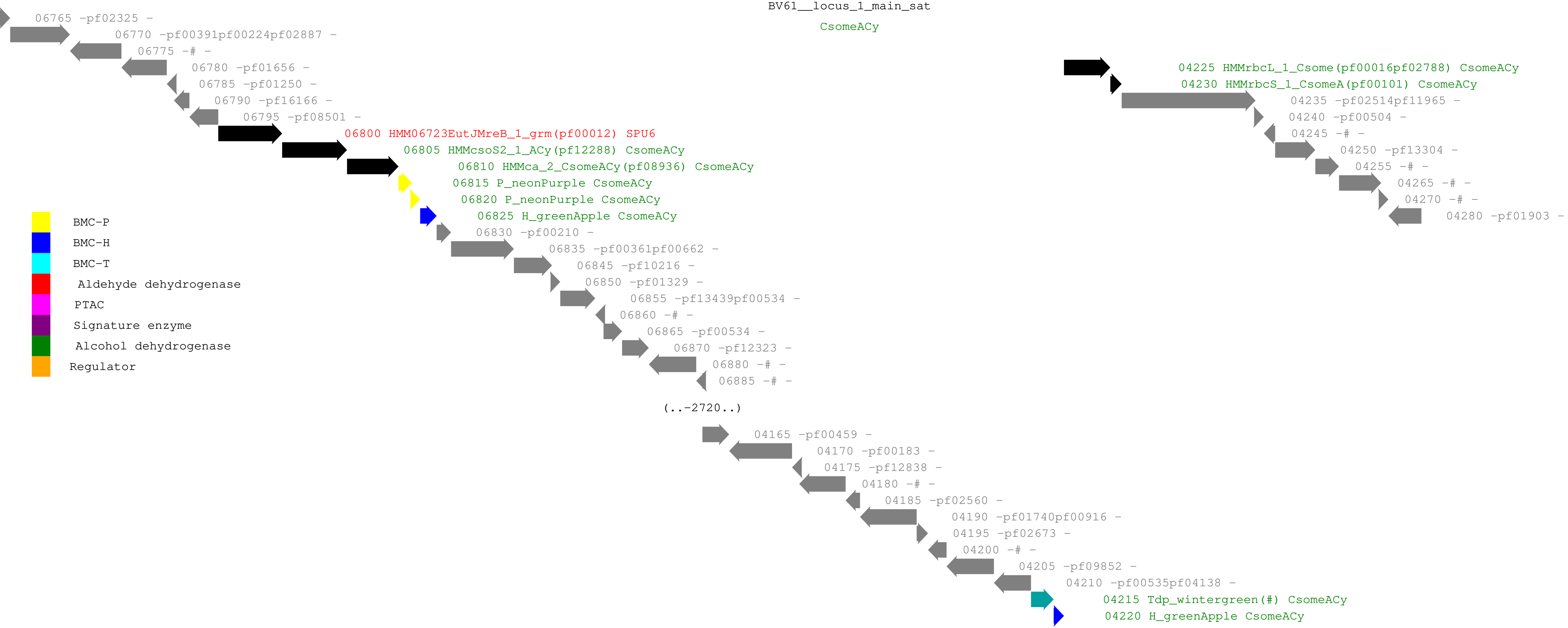

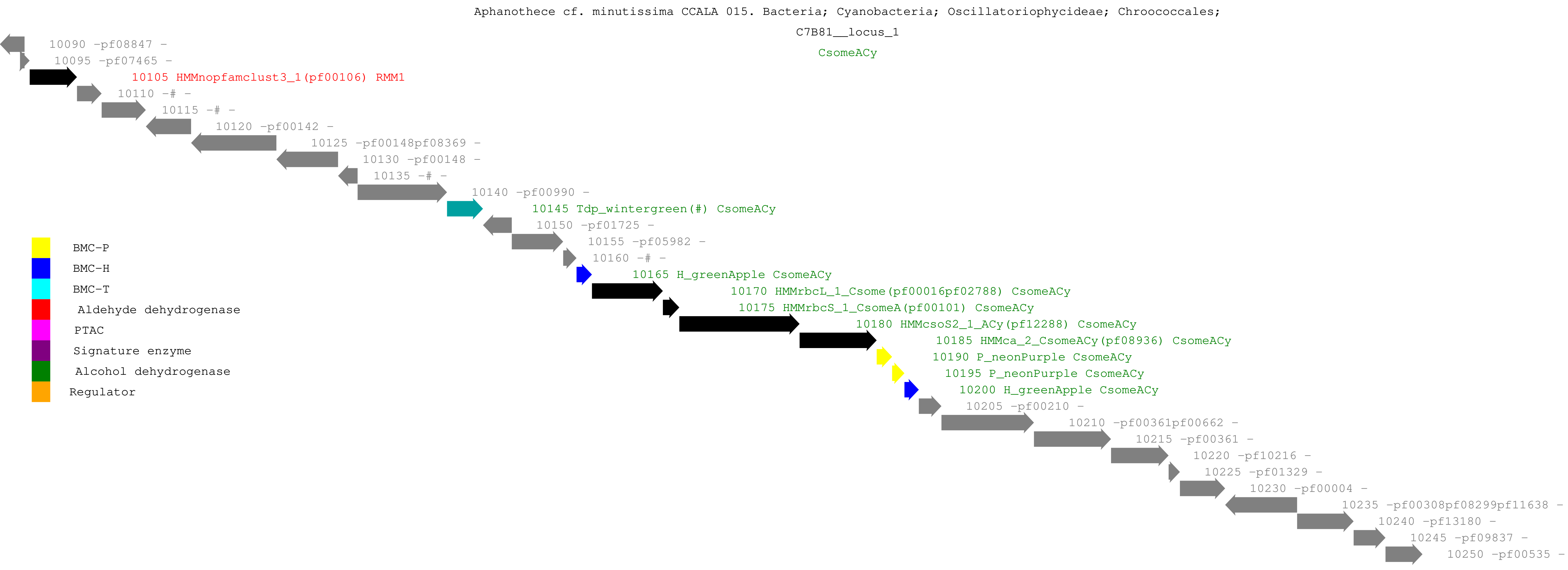
Aphanothece cf. minutissima CCALA 015. Bacteria; Cyanobacteria; Oscillatoriophycideae; Chroococcales;
C7B81__locus_1
CsomeACy
10090 -pf08847 -
10095 -pf07465 -
10105 HMMnopfamclust3_1(pf00106) RMM1
10110 -# -
10115 -# -
10120 -pf00142 -
10125 -pf00148pf08369 -
10130 -pf00148 -
10135 -# -
10140 -pf00990 -
10145 Tdp_wintergreen(#) CsomeACy
10150 -pf01725 -
10155 -pf05982 -
10160 -# -
10165 H_greenApple CsomeACy
10170 HMMrbcL_1_Csome(pf00016pf02788) CsomeACy
10175 HMMrbcS_1_CsomeA(pf00101) CsomeACy
10180 HMMcsoS2_1_ACy(pf12288) CsomeACy
10185 HMMca_2_CsomeACy(pf08936) CsomeACy
10190 P_neonPurple CsomeACy
10195 P_neonPurple CsomeACy
10200 H_greenApple CsomeACy
10205 -pf00210 -
10210 -pf00361pf00662 -
10215 -pf00361 -
10220 -pf10216 -
10225 -pf01329 -
10230 -pf00004 -
10235 -pf00308pf08299pf11638 -
10240 -pf13180 -
10245 -pf09837 -
10250 -pf00535 -
BMC-P
BMC-H
BMC-T
Aldehyde dehydrogenase
PTAC
Signature enzyme
Alcohol dehydrogenase
Regulator

Candidatus Synechococcus lacustris. Bacteria; Cyanobacteria; Synechococcales; Synechococcaceae;

C7K08__locus_1

CsomeACy

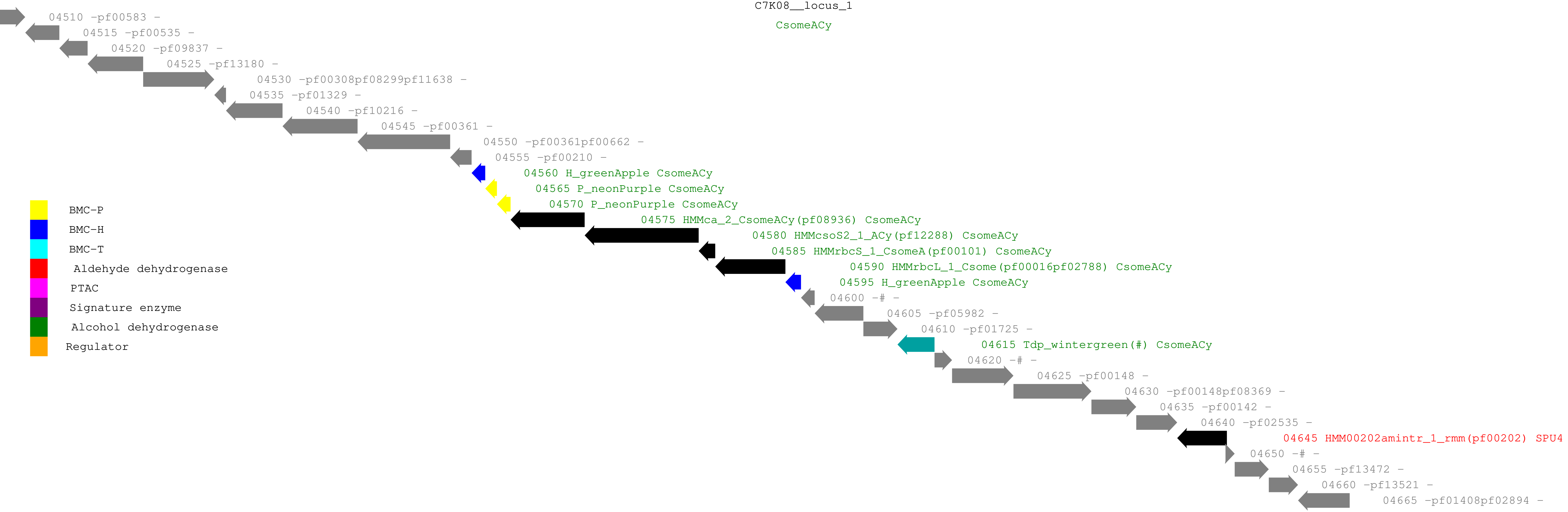

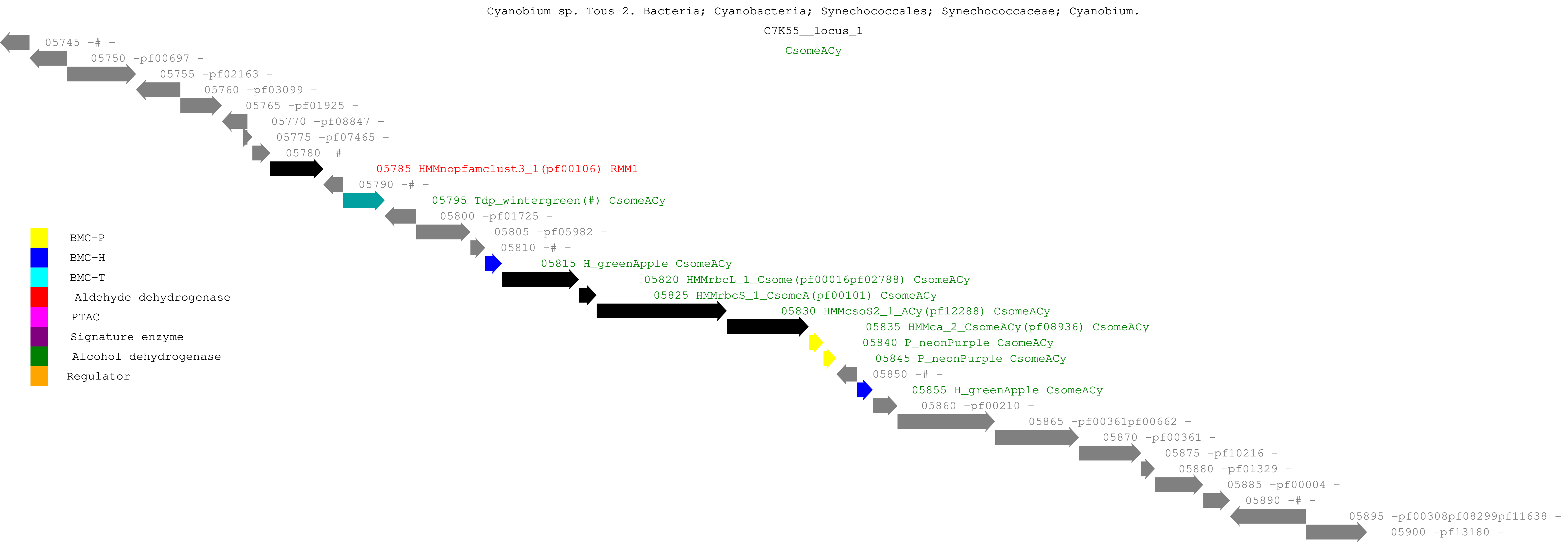
Cyanobium sp. Tous-2. Bacteria; Cyanobacteria; Synechococcales; Synechococcaceae; Cyanobium.
C7K55__locus_1
CsomeACy
05745 -# -
05750 -pf00697 -
05755 -pf02163 -
05760 -pf03099 -
05765 -pf01925 -
05770 -pf08847 -
05775 -pf07465 -
05780 -# -
05785 HMMnopfamclust3_1(pf00106) RMM1
05790 -# -
05795 Tdp_wintergreen(#) CsomeACy
05800 -pf01725 -
05805 -pf05982 -
05810 -# -
05815 H_greenApple CsomeACy
05820 HMMrbcL_1_Csome(pf00016pf02788) CsomeACy
05825 HMMrbcS_1_CsomeA(pf00101) CsomeACy
05830 HMMcsoS2_1_ACy(pf12288) CsomeACy
05835 HMMca_2_CsomeACy(pf08936) CsomeACy
05840 P_neonPurple CsomeACy
05845 P_neonPurple CsomeACy
05850 -# -
05855 H_greenApple CsomeACy
05860 -pf00210 -
05865 -pf00361pf00662 -
05870 -pf00361 -
05875 -pf10216 -
05880 -pf01329 -
05885 -pf00004 -
05890 -# -
05895 -pf00308pf08299pf11638 -
05900 -pf13180 -
BMC-P
BMC-H
BMC-T
Aldehyde dehydrogenase
PTAC
Signature enzyme
Alcohol dehydrogenase
Regulator

Synechococcus sp. CB0101. Bacteria; Cyanobacteria; Synechococcales; Synechococcaceae; Synechococcus;

CB0101__locus_1

CsomeACy

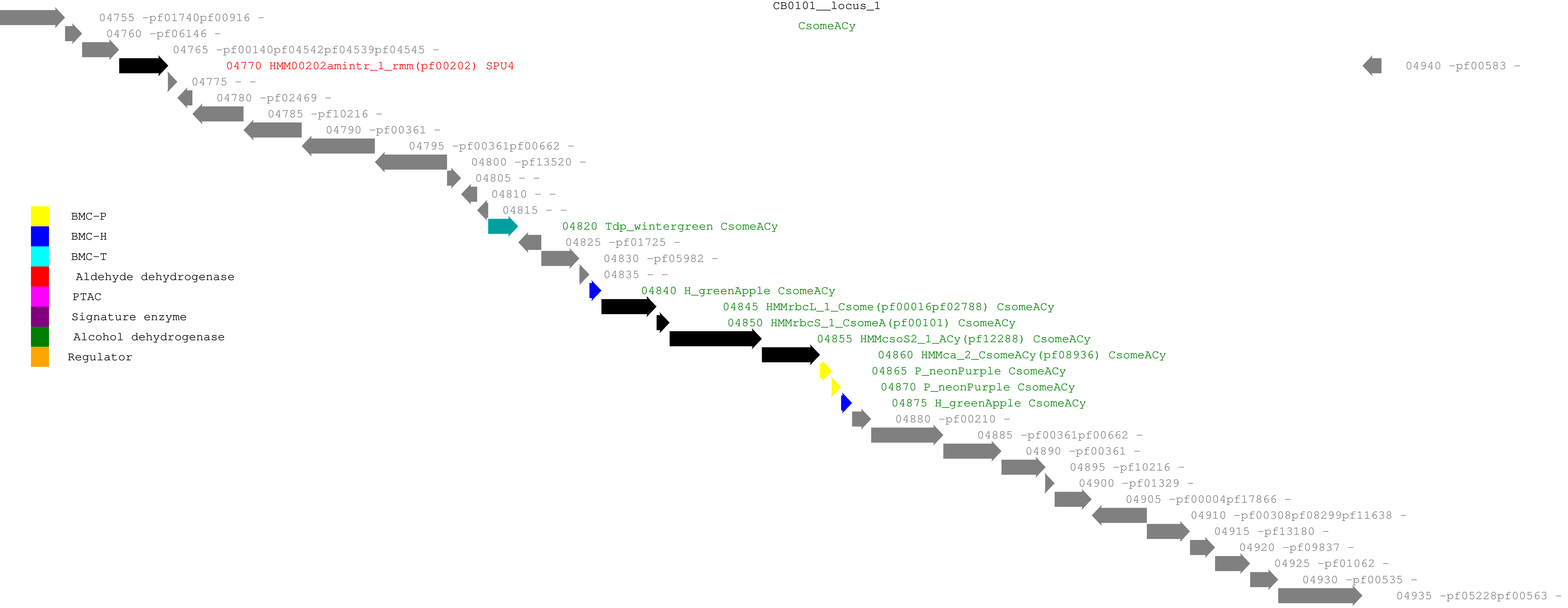

Phycisphaera sp. TMED9. Bacteria; Planctomycetes; Phycisphaerae; Phycisphaerales;

CBB69__locus_1

CsomeACy

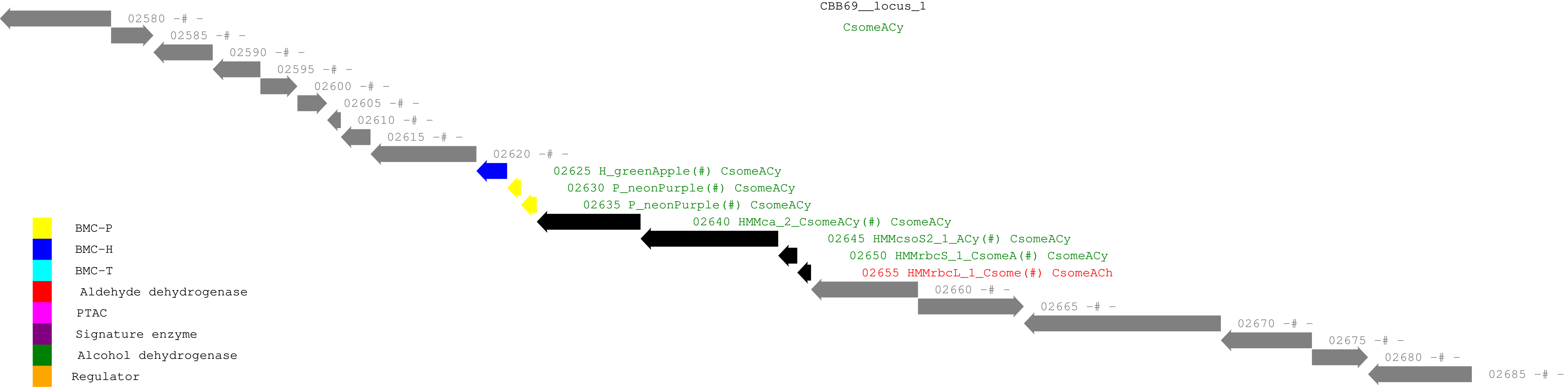

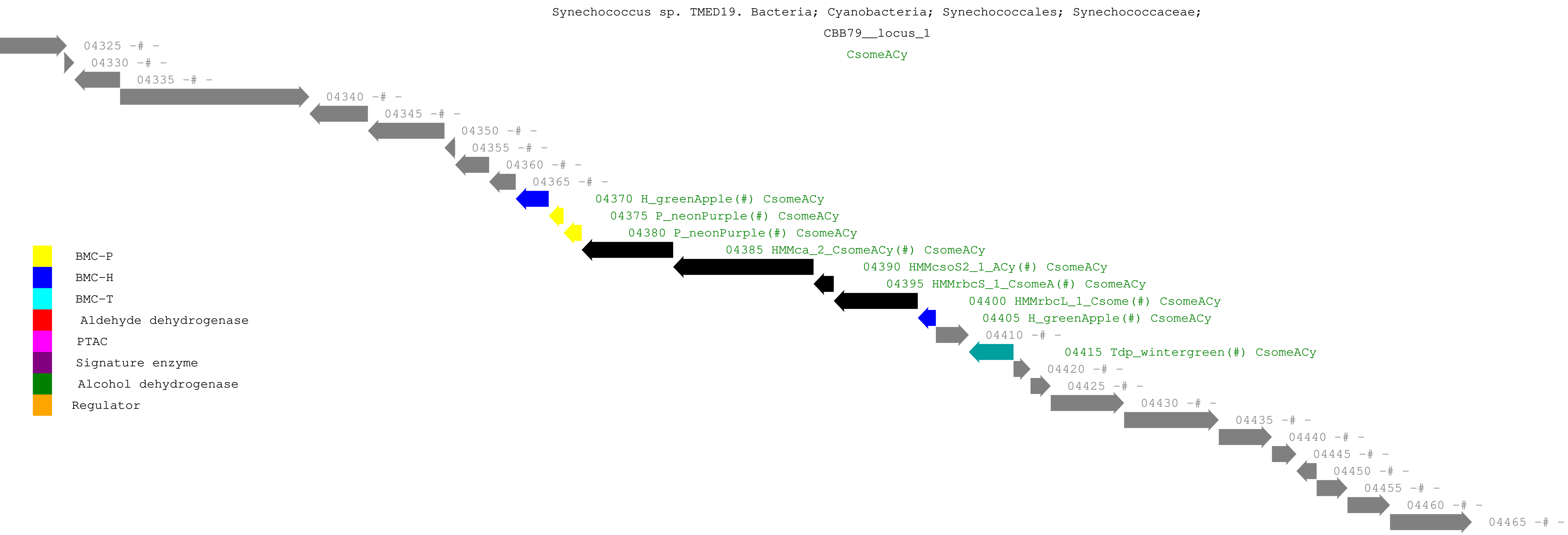
Synechococcus sp. TMED19. Bacteria; Cyanobacteria; Synechococcales; Synechococcaceae;
CBB79__locus_1
CsomeACy
04325 -# -
04330 -# -
04335 -# -
04340 -# -
04345 -# -
04350 -# -
04355 -# -
04360 -# -
04365 -# -
04370 H_greenApple(#) CsomeACy
04375 P_neonPurple(#) CsomeACy
04380 P_neonPurple(#) CsomeACy
04385 HMMca_2_CsomeACy(#) CsomeACy
04390 HMMcsoS2_1_ACy(#) CsomeACy
04395 HMMrbcS_1_CsomeA(#) CsomeACy
04400 HMMrbcL_1_Csome(#) CsomeACy
04405 H_greenApple(#) CsomeACy
04410 -# -
04415 Tdp_wintergreen(#) CsomeACy
04420 -# -
04425 -# -
04430 -# -
04435 -# -
04440 -# -
04445 -# -
04450 -# -
04455 -# -
04460 -# -
04465 -# -
BMC-P
BMC-H
BMC-T
Aldehyde dehydrogenase
PTAC
Signature enzyme
Alcohol dehydrogenase
Regulator

Synechococcus sp. TMED20. Bacteria; Cyanobacteria; Synechococcales; Synechococcaceae; Synechococcus;

CBB80__locus_1

CsomeACy

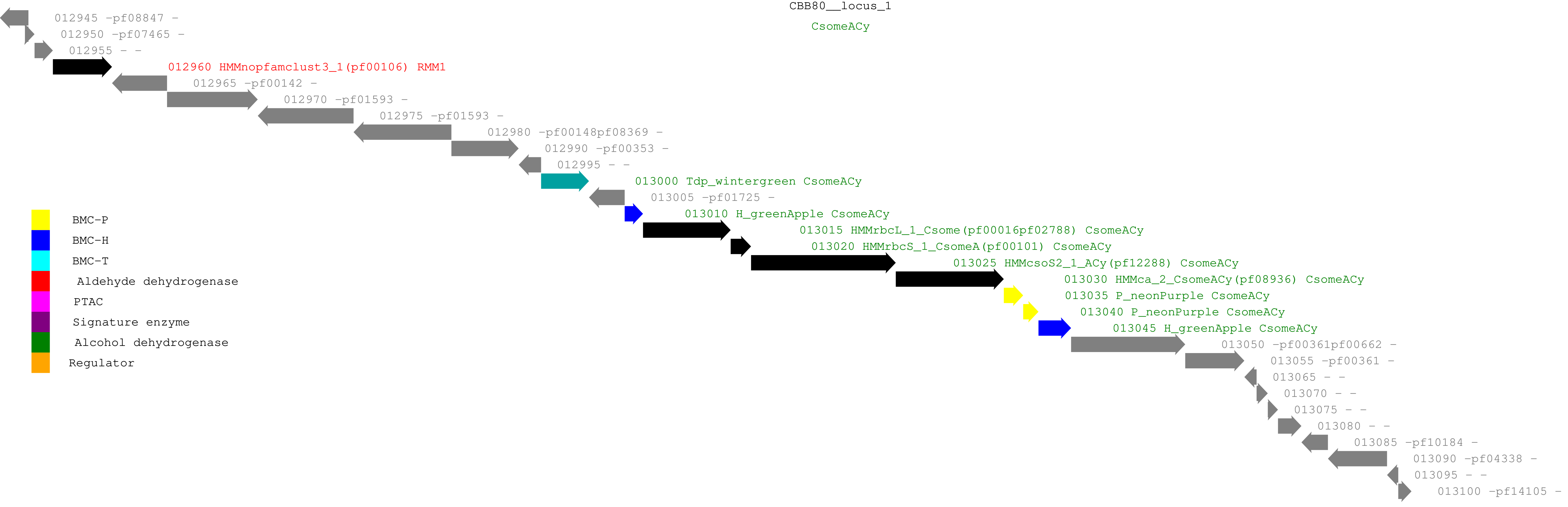

Synechococcus sp. TMED66. Bacteria; Cyanobacteria; Synechococcales; Synechococcaceae;

CBC26__locus_1_main_sat

CsomeACy

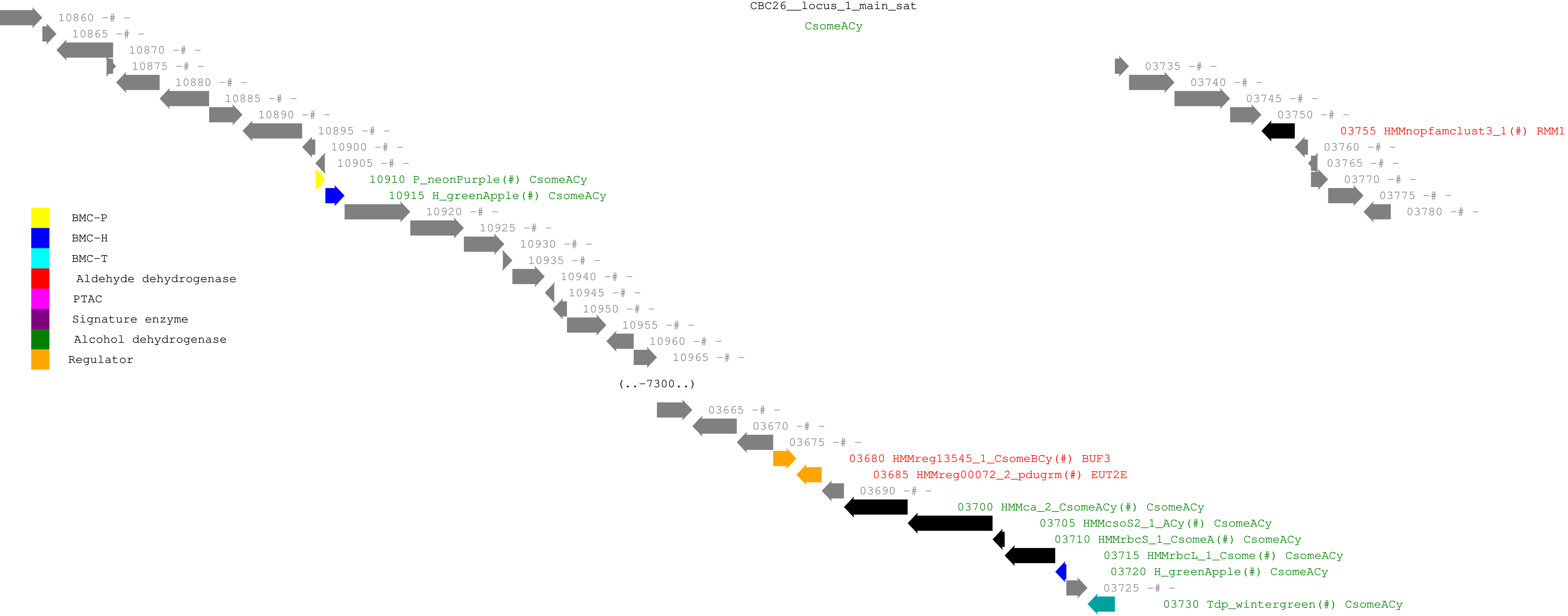

Synechococcus sp. TMED90. Bacteria; Cyanobacteria; Synechococcales; Synechococcaceae;

CBC50__locus_1_main_sat

CsomeACy

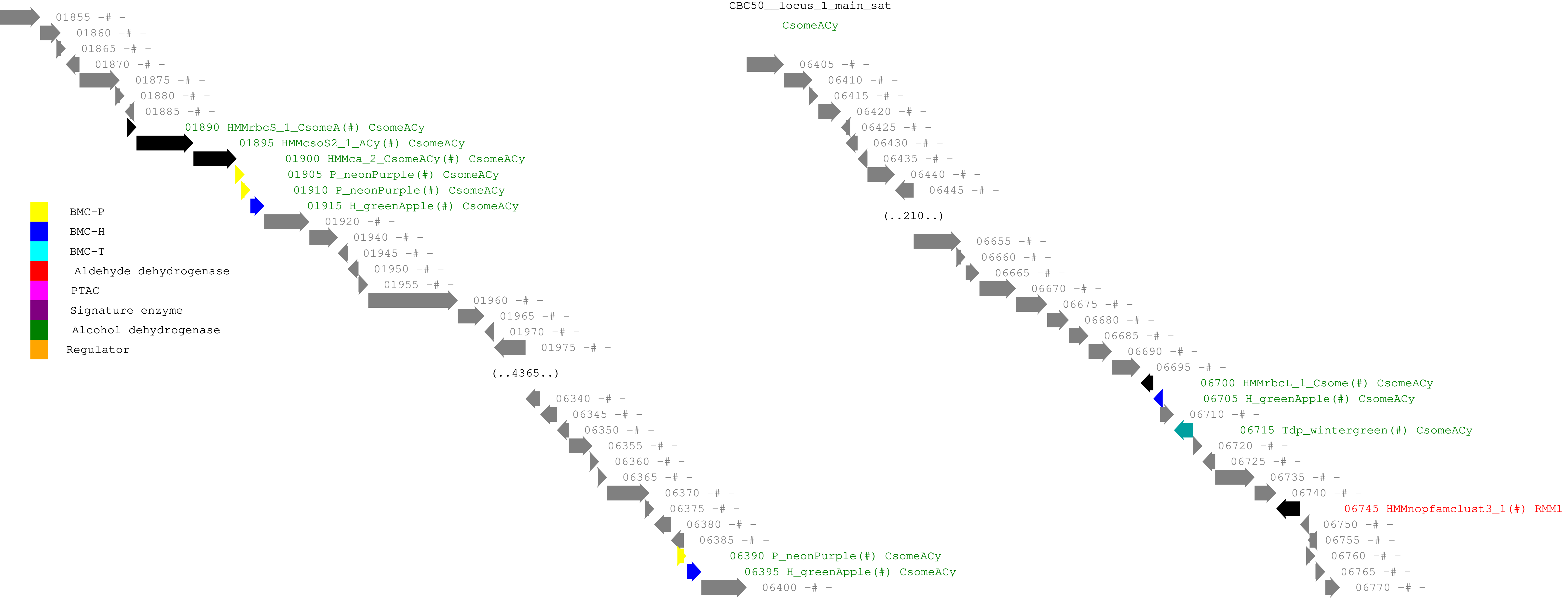

Synechococcus sp. TMED155. Bacteria; Cyanobacteria; Synechococcales; Synechococcaceae; Synechococcus;

CBD15__locus_1

CsomeACy

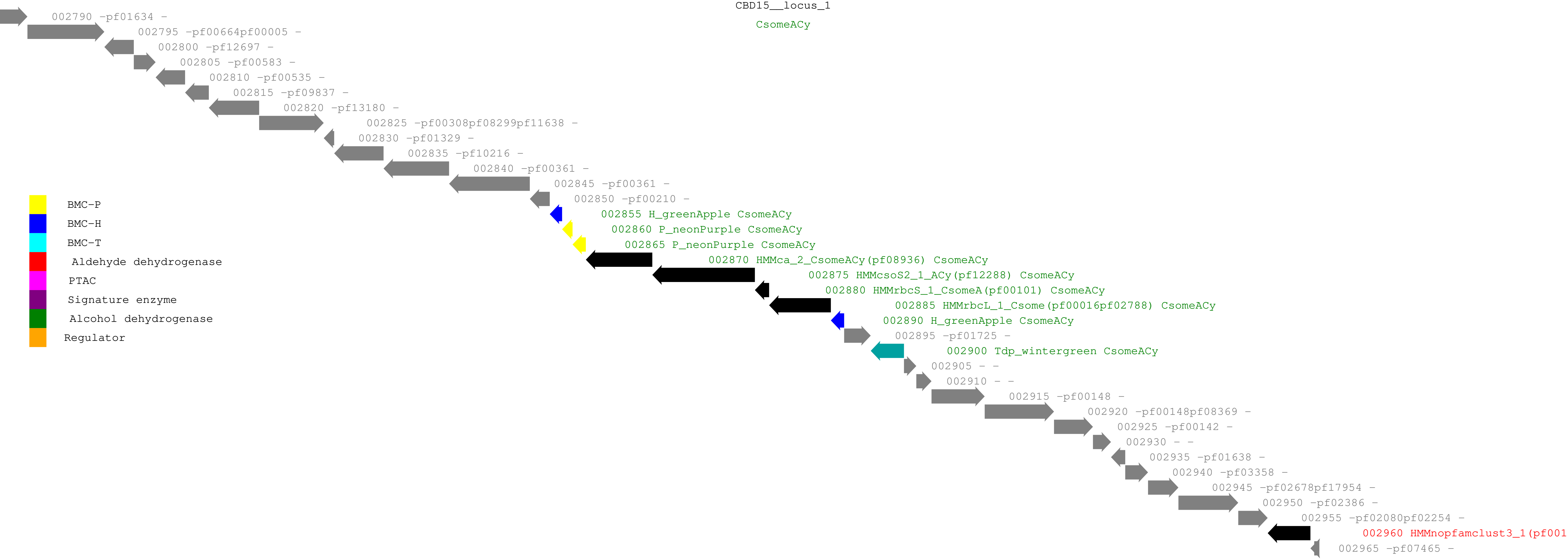

Cyanobacteria bacterium TMED177. Bacteria; Cyanobacteria.

CBD37__locus_1

CsomeACy

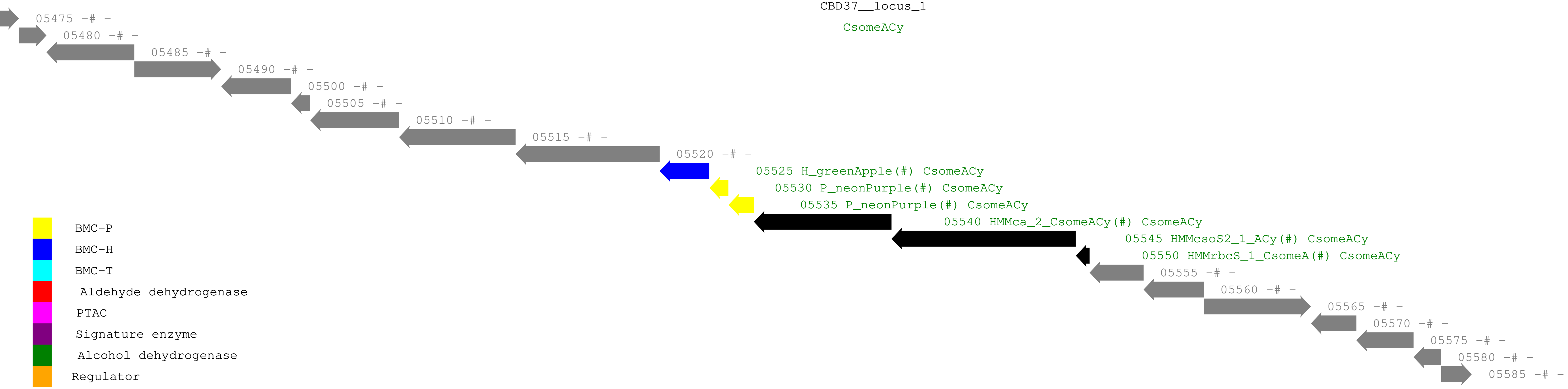

Cyanobacteria bacterium TMED188. Bacteria; Cyanobacteria.

CBD48__locus_1

CsomeACy

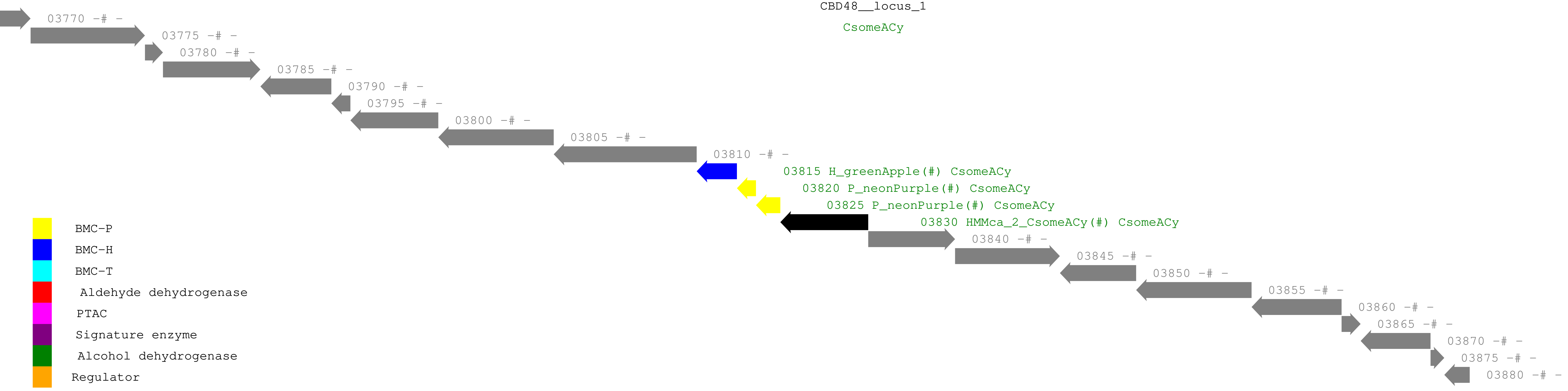

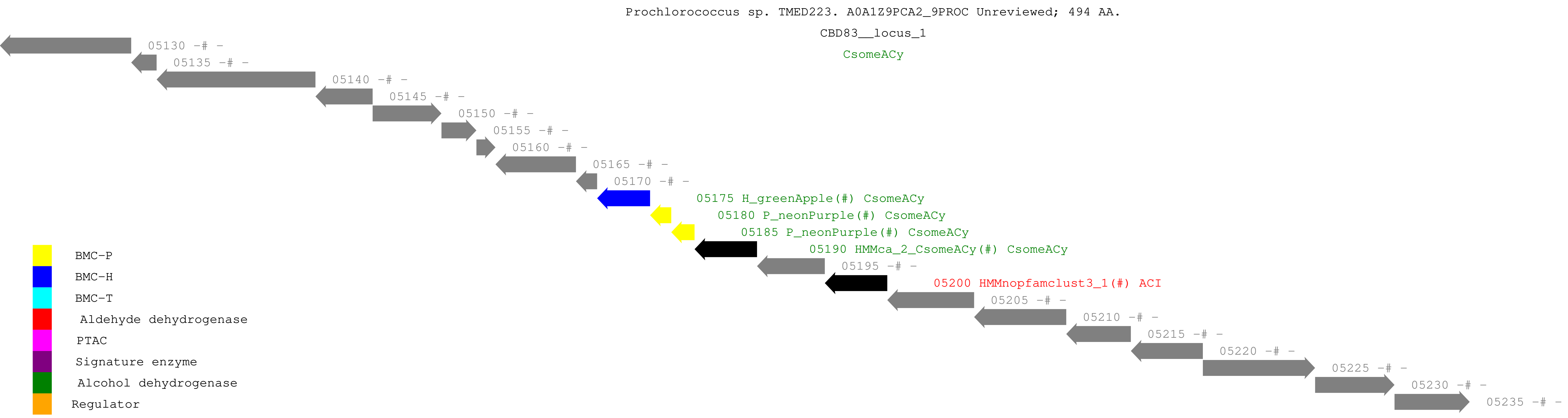
Prochlorococcus sp. TMED223. A0A1Z9PCA2_9PROC Unreviewed; 494 AA.
CBD83__locus_1
CsomeACy
05130 -# -
05135 -# -
05140 -# -
05145 -# -
05150 -# -
05155 -# -
05160 -# -
05165 -# -
05170 -# -
05175 H_greenApple(#) CsomeACy
05180 P_neonPurple(#) CsomeACy
05185 P_neonPurple(#) CsomeACy
05190 HMMca_2_CsomeACy(#) CsomeACy
05195 -# -
05200 HMMnopfamclust3_1(#) ACI
05205 -# -
05210 -# -
05215 -# -
05220 -# -
05225 -# -
05230 -# -
05235 -# -
BMC-P
BMC-H
BMC-T
Aldehyde dehydrogenase
PTAC
Signature enzyme
Alcohol dehydrogenase
Regulator

Cyanobacteria bacterium TMED229. Bacteria; Cyanobacteria.

CBD89__locus_1_main_sat

CsomeACy

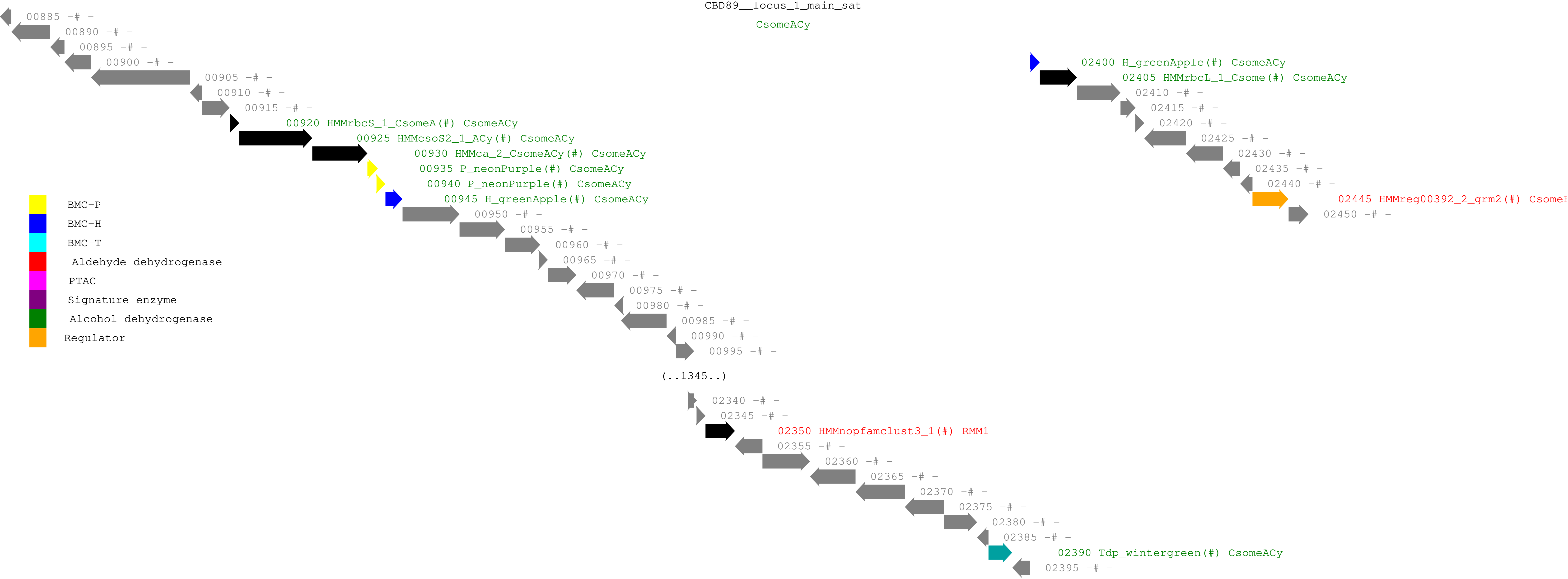

Cyanobium sp. NIES-981. Bacteria; Cyanobacteria; Synechococcales; Synechococcaceae; Cyanobium.

CBM981__locus_1

CsomeACy

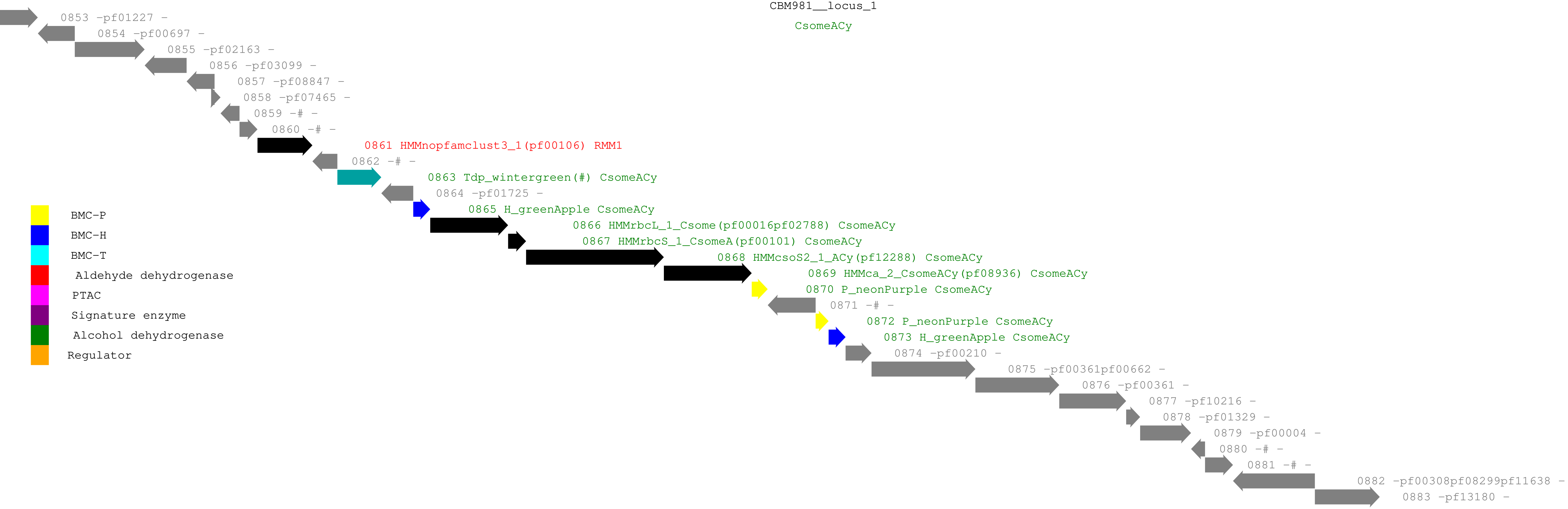

Synechococcus sp. Baikal-G1. Bacteria; Cyanobacteria; Synechococcales; Synechococcaceae;

CK536__locus_1

CsomeACy

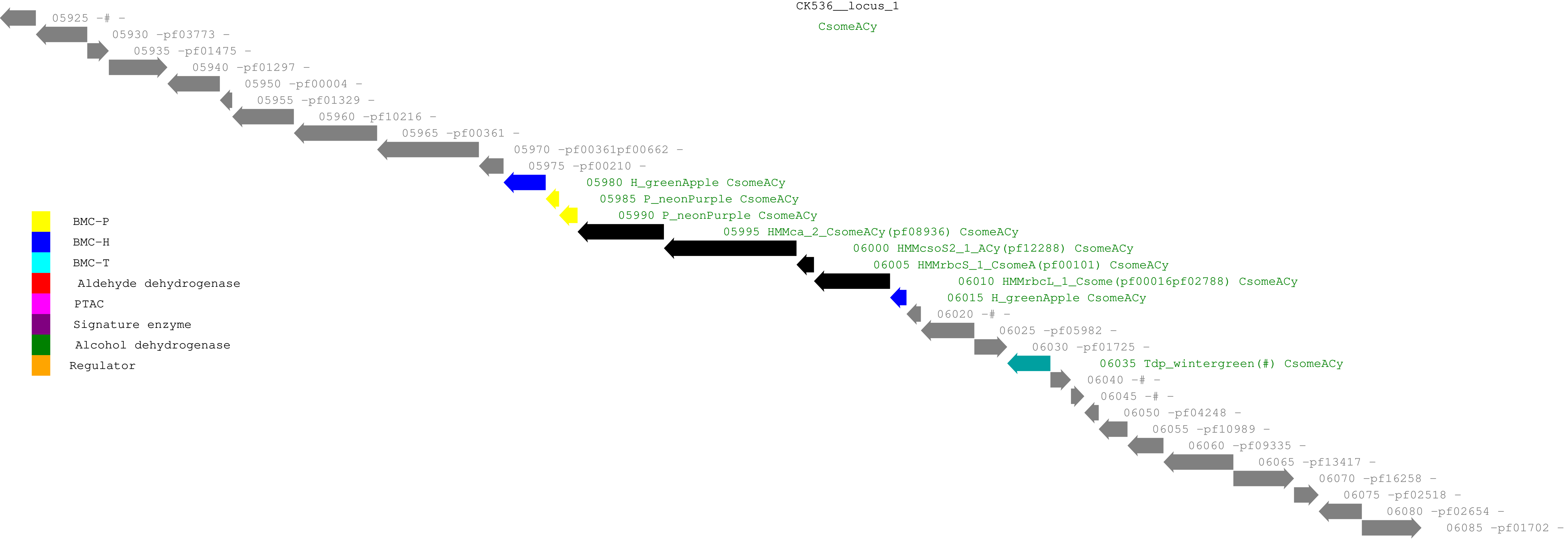

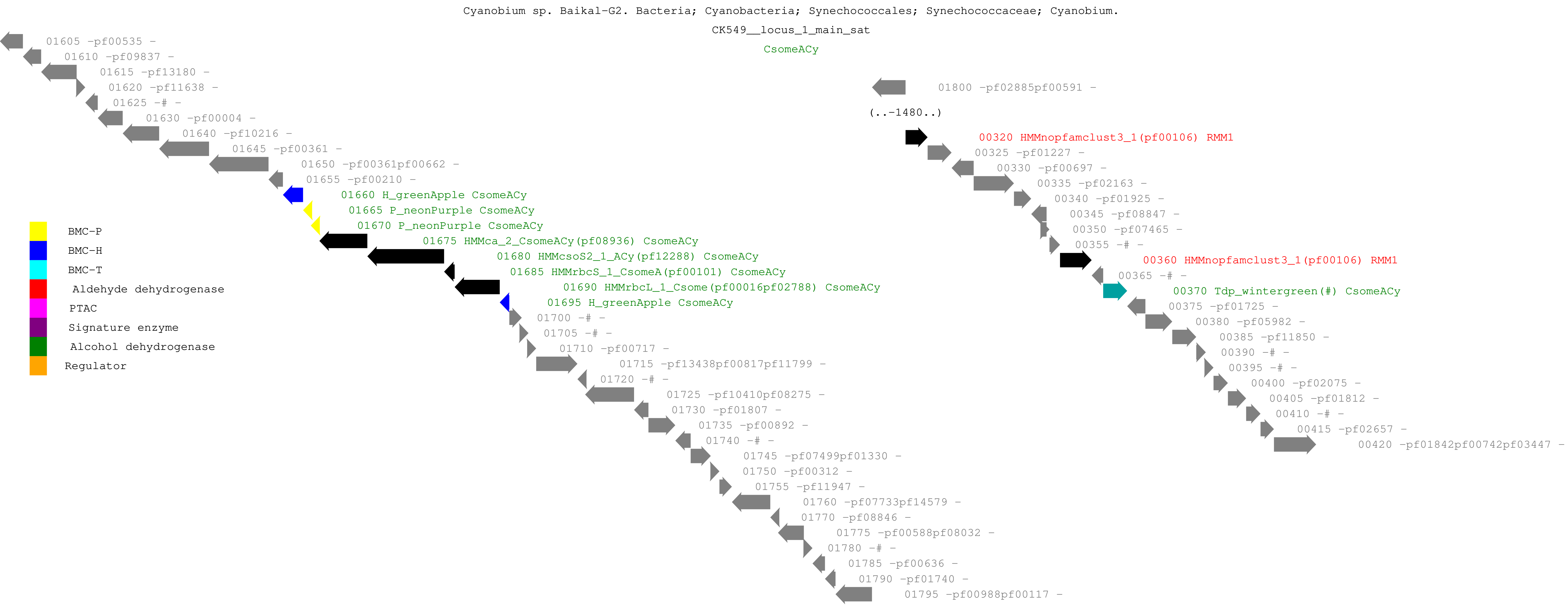
Cyanobium sp. Baikal-G2. Bacteria; Cyanobacteria; Synechococcales; Synechococcaceae; Cyanobium.
CK549__locus_1_main_sat
CsomeACy
01605 -pf00535 -
01610 -pf09837 -
01615 -pf13180 -
01620 -pf11638 -
01625 -# -
01630 -pf00004 -
01640 -pf10216 -
01645 -pf00361 -
01650 -pf00361pf00662 -
01655 -pf00210 -
01660 H_greenApple CsomeACy
01665 P_neonPurple CsomeACy
01670 P_neonPurple CsomeACy
01675 HMMca_2_CsomeACy(pf08936) CsomeACy
01680 HMMcsoS2_1_ACy(pf12288) CsomeACy
01685 HMMrbcS_1_CsomeA(pf00101) CsomeACy
01690 HMMrbcL_1_Csome(pf00016pf02788) CsomeACy
01695 H_greenApple CsomeACy
01700 -# -
01705 -# -
01710 -pf00717 -
01715 -pf13438pf00817pf11799 -
01720 -# -
01725 -pf10410pf08275 -
01730 -pf01807 -
01735 -pf00892 -
01740 -# -
01745 -pf07499pf01330 -
01750 -pf00312 -
01755 -pf11947 -
01760 -pf07733pf14579 -
01770 -pf08846 -
01775 -pf00588pf08032 -
01780 -# -
01785 -pf00636 -
01790 -pf01740 -
01795 -pf00988pf00117 -
01800 -pf02885pf00591 -
(..-1480..)
00320 HMMnopfamclust3_1(pf00106) RMM1
00325 -pf01227 -
00330 -pf00697 -
00335 -pf02163 -
00340 -pf01925 -
00345 -pf08847 -
00350 -pf07465 -
00355 -# -
00360 HMMnopfamclust3_1(pf00106) RMM1
00365 -# -
00370 Tdp_wintergreen(#) CsomeACy
00375 -pf01725 -
00380 -pf05982 -
00385 -pf11850 -
00390 -# -
00395 -# -
00400 -pf02075 -
00405 -pf01812 -
00410 -# -
00415 -pf02657 -
00420 -pf01842pf00742pf03447 -
BMC-P
BMC-H
BMC-T
Aldehyde dehydrogenase
PTAC
Signature enzyme
Alcohol dehydrogenase
Regulator

Cyanobium sp. NAT70. Bacteria; Cyanobacteria; Synechococcales; Synechococcaceae; Cyanobium.

CL862__locus_1_main_sat

CsomeACy

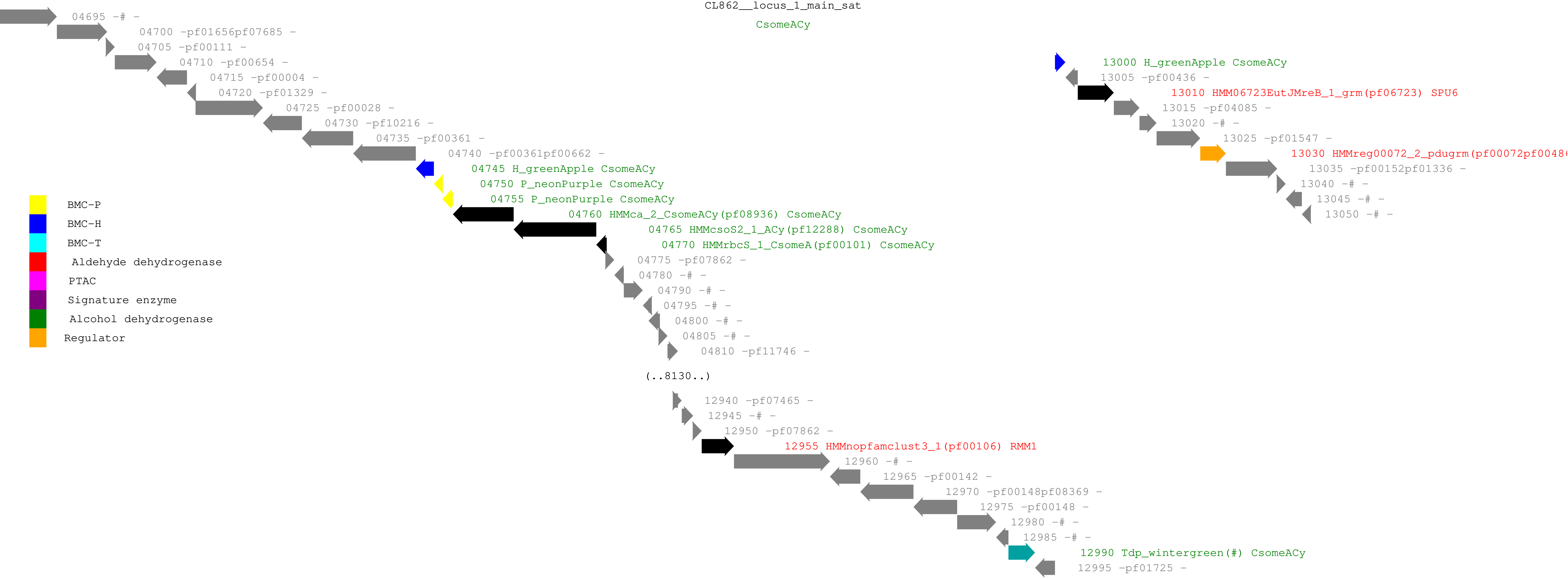

Prochlorococcus sp. MED105. A0A2D7TIP7_9PROC Unreviewed; 112 AA.

CMK47__locus_1

CsomeACy

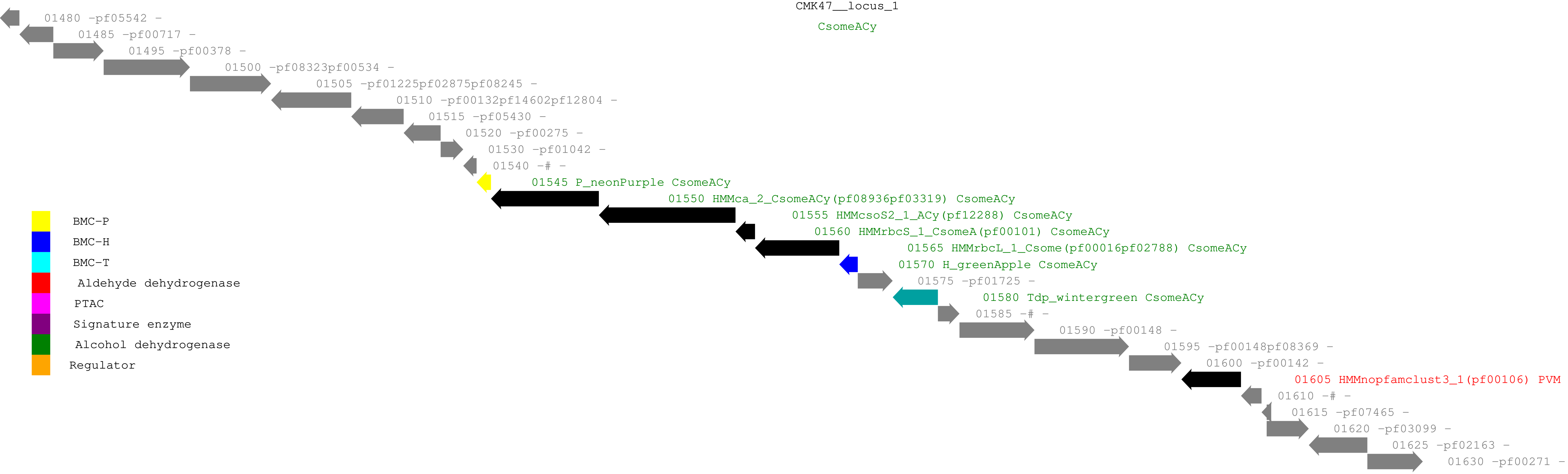

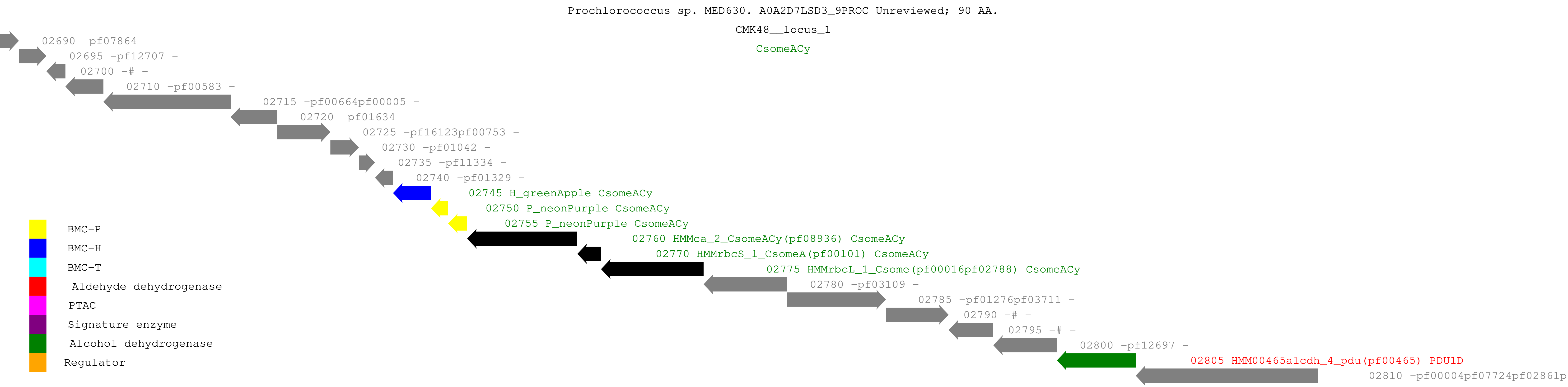
Prochlorococcus sp. MED630. A0A2D7LSD3_9PROC Unreviewed; 90 AA.
CMK48__locus_1
CsomeACy
02690 -pf07864 -
02695 -pf12707 -
02700 -# -
02710 -pf00583 -
02715 -pf00664pf00005 -
02720 -pf01634 -
02725 -pf16123pf00753 -
02730 -pf01042 -
02735 -pf11334 -
02740 -pf01329 -
02745 H_greenApple CsomeACy
02750 P_neonPurple CsomeACy
02755 P_neonPurple CsomeACy
02760 HMMca_2_CsomeACy(pf08936) CsomeACy
02770 HMMrbcS_1_CsomeA(pf00101) CsomeACy
02775 HMMrbcL_1_Csome(pf00016pf02788) CsomeACy
02780 -pf03109 -
02785 -pf01276pf03711 -
02790 -# -
02795 -# -
02800 -pf12697 -
02805 HMM00465alcdh_4_pdu(pf00465) PDU1D
02810 -pf00004pf07724pf02861p
BMC-P
BMC-H
BMC-T
Aldehyde dehydrogenase
PTAC
Signature enzyme
Alcohol dehydrogenase
Regulator

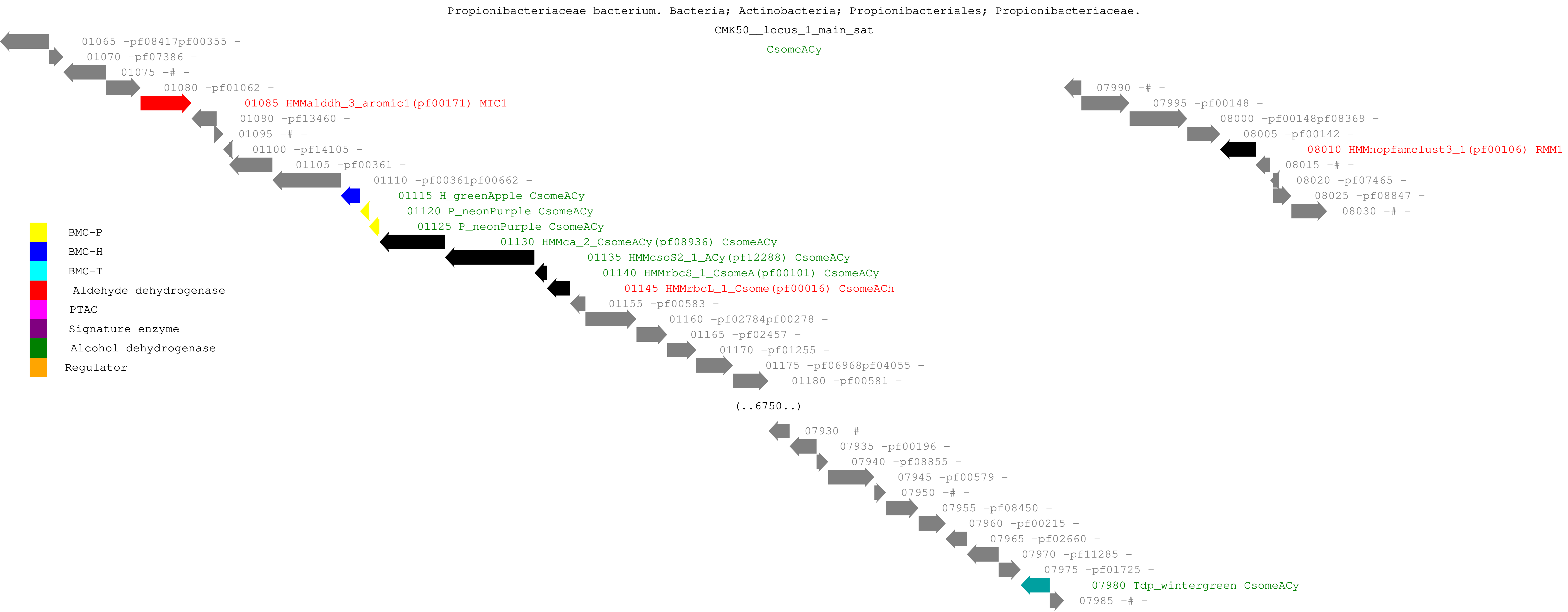
Propionibacteriaceae bacterium. Bacteria; Actinobacteria; Propionibacteriales; Propionibacteriaceae.
CMK50__locus_1_main_sat
CsomeACy
01065 -pf08417pf00355 -
01070 -pf07386 -
01075 -# -
01080 -pf01062 -
01085 HMMalddh_3_aromic1(pf00171) MIC1
01090 -pf13460 -
01095 -# -
01100 -pf14105 -
01105 -pf00361 -
01110 -pf00361pf00662 -
01115 H_greenApple CsomeACy
01120 P_neonPurple CsomeACy
01125 P_neonPurple CsomeACy
01130 HMMca_2_CsomeACy(pf08936) CsomeACy
01135 HMMcsoS2_1_ACy(pf12288) CsomeACy
01140 HMMrbcS_1_CsomeA(pf00101) CsomeACy
01145 HMMrbcL_1_Csome(pf00016) CsomeACh
01155 -pf00583 -
01160 -pf02784pf00278 -
01165 -pf02457 -
01170 -pf01255 -
01175 -pf06968pf04055 -
01180 -pf00581 -
(..6750..)
07930 -# -
07935 -pf00196 -
07940 -pf08855 -
07945 -pf00579 -
07950 -# -
07955 -pf08450 -
07960 -pf00215 -
07965 -pf02660 -
07970 -pf11285 -
07975 -pf01725 -
07980 Tdp_wintergreen CsomeACy
07985 -# -
07990 -# -
07995 -pf00148 -
08000 -pf00148pf08369 -
08005 -pf00142 -
08010 HMMnopfamclust3_1(pf00106) RMM1
08015 -# -
08020 -pf07465 -
08025 -pf08847 -
08030 -# -
BMC-P
BMC-H
BMC-T
Aldehyde dehydrogenase
PTAC
Signature enzyme
Alcohol dehydrogenase
Regulator

Synechococcus sp. ARS1019. Bacteria; Cyanobacteria; Synechococcales; Synechococcaceae;

CMN91__locus_1_main_sat

CsomeACy

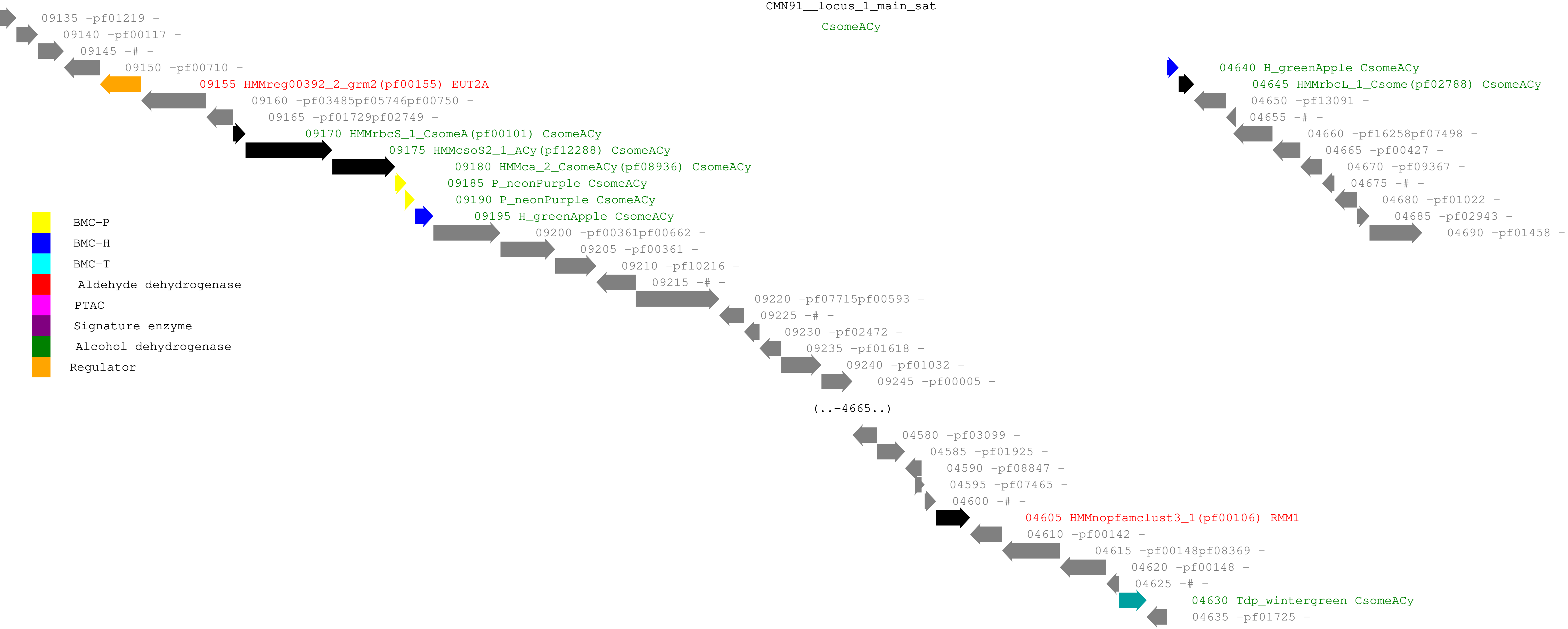

Synechococcus sp. CPC35. Bacteria; Cyanobacteria; Synechococcales; Synechococcaceae;

CMN93__locus_1_main_sat

CsomeACy

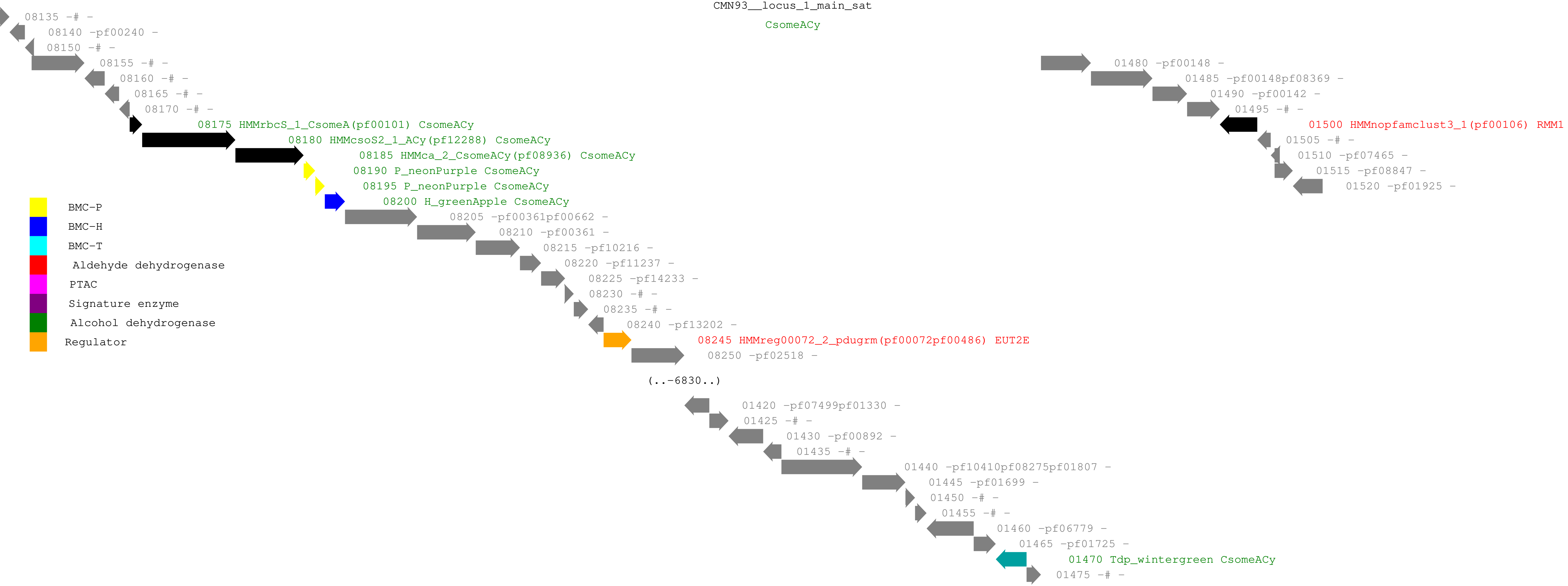

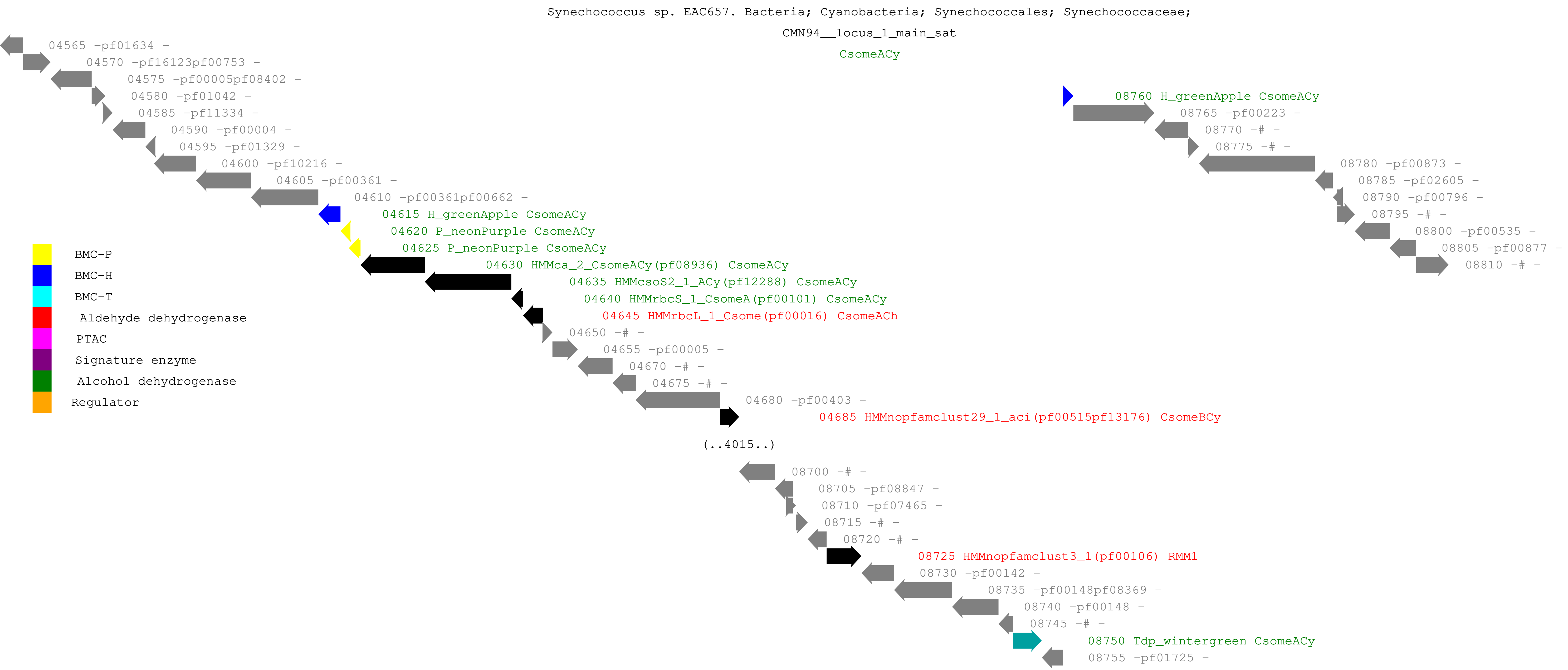

Synechococcus sp. EAC657. Bacteria; Cyanobacteria; Synechococcales; Synechococcaceae;
CMN94__locus_1_main_sat
CsomeACy
04565 -pf01634 -
04570 -pf16123pf00753 -
04575 -pf00005pf08402 -
04580 -pf01042 -
04585 -pf11334 -
04590 -pf00004 -
04595 -pf01329 -
04600 -pf10216 -
04605 -pf00361 -
04610 -pf00361pf00662 -
04615 H_greenApple CsomeACy
04620 P_neonPurple CsomeACy
04625 P_neonPurple CsomeACy
04630 HMMca_2_CsomeACy(pf08936) CsomeACy
04635 HMMcsoS2_1_ACy(pf12288) CsomeACy
04640 HMMrbcS_1_CsomeA(pf00101) CsomeACy
04645 HMMrbcL_1_Csome(pf00016) CsomeACh
04650 -# -
04655 -pf00005 -
04670 -# -
04675 -# -
04680 -pf00403 -
04685 HMMnopfamclust29_1_aci(pf00515pf13176) CsomeBCy
(..4015..)
08700 -# -
08705 -pf08847 -
08710 -pf07465 -
08715 -# -
08720 -# -
08725 HMMnopfamclust3_1(pf00106) RMM1
08730 -pf00142 -
08735 -pf00148pf08369 -
08740 -pf00148 -
08745 -# -
08750 Tdp_wintergreen CsomeACy
08755 -pf01725 -
08760 H_greenApple CsomeACy
08765 -pf00223 -
08770 -# -
08775 -# -
08780 -pf00873 -
08785 -pf02605 -
08790 -pf00796 -
08795 -# -
08800 -pf00535 -
08805 -pf00877 -
08810 -# -
BMC-P
BMC-H
BMC-T
Aldehyde dehydrogenase
PTAC
Signature enzyme
Alcohol dehydrogenase
Regulator

Synechococcus sp. MED650. Bacteria; Cyanobacteria; Synechococcales; Synechococcaceae;

CMN95__locus_1_main_sat

CsomeACy

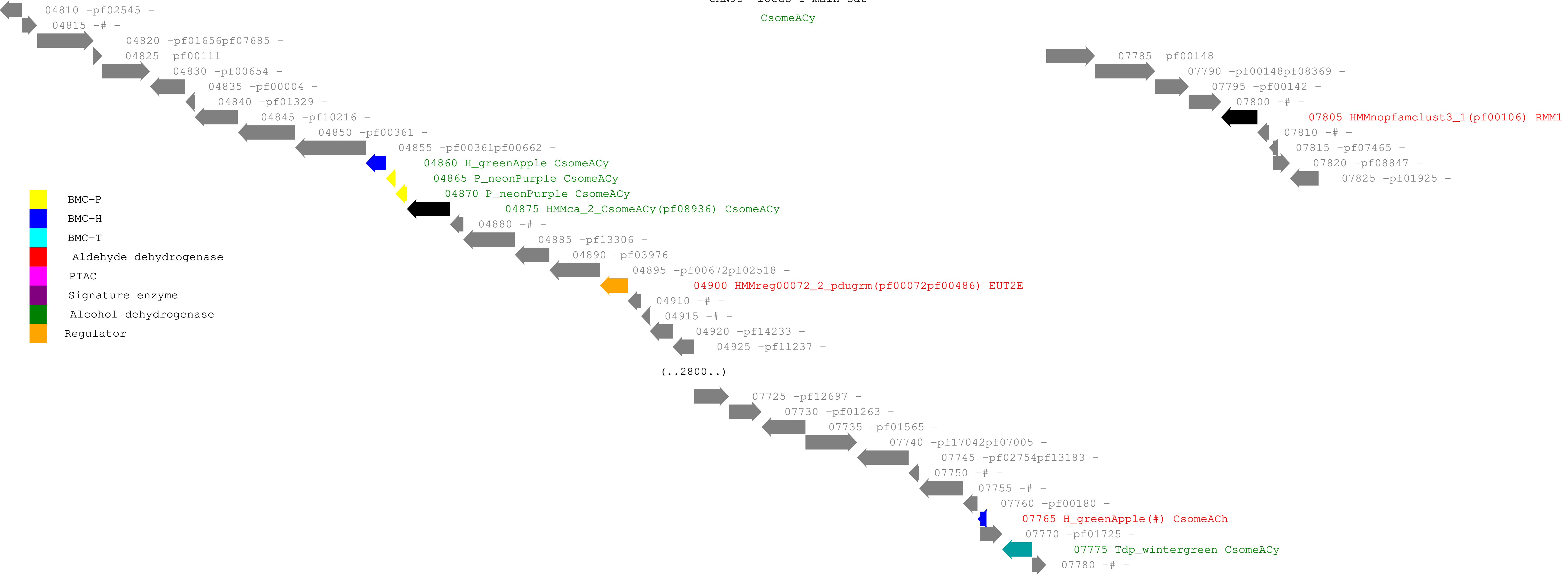

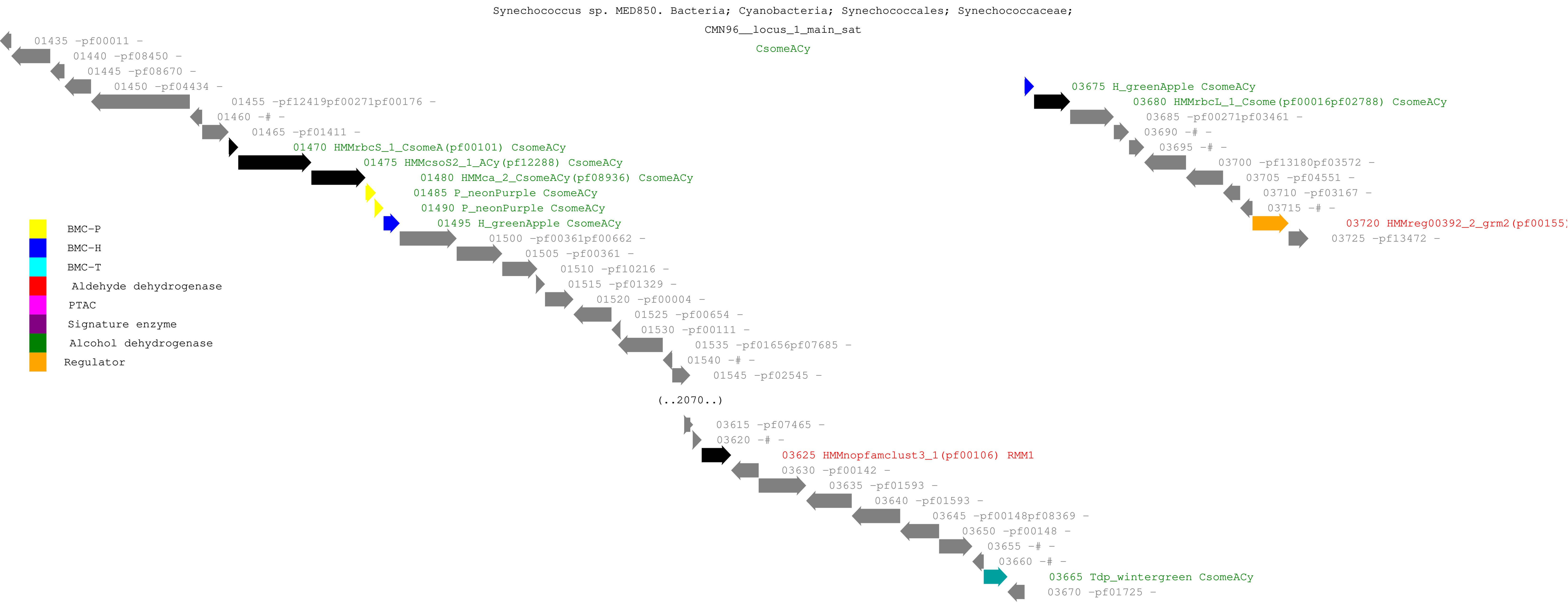
Synechococcus sp. MED850. Bacteria; Cyanobacteria; Synechococcales; Synechococcaceae;
CMN96__locus_1_main_sat
CsomeACy
01435 -pf00011 -
01440 -pf08450 -
01445 -pf08670 -
01450 -pf04434 -
01455 -pf12419pf00271pf00176 -
01460 -# -
01465 -pf01411 -
01470 HMMrbcS_1_CsomeA(pf00101) CsomeACy
01475 HMMcsoS2_1_ACy(pf12288) CsomeACy
01480 HMMca_2_CsomeACy(pf08936) CsomeACy
01485 P_neonPurple CsomeACy
01490 P_neonPurple CsomeACy
01495 H_greenApple CsomeACy
01500 -pf00361pf00662 -
01505 -pf00361 -
01510 -pf10216 -
01515 -pf01329 -
01520 -pf00004 -
01525 -pf00654 -
01530 -pf00111 -
01535 -pf01656pf07685 -
01540 -# -
01545 -pf02545 -
(..2070..)
03615 -pf07465 -
03620 -# -
03625 HMMnopfamclust3_1(pf00106) RMM1
03630 -pf00142 -
03635 -pf01593 -
03640 -pf01593 -
03645 -pf00148pf08369 -
03650 -pf00148 -
03655 -# -
03660 -# -
03665 Tdp_wintergreen CsomeACy
03670 -pf01725 -
03675 H_greenApple CsomeACy
03680 HMMrbcL_1_Csome(pf00016pf02788) CsomeACy
03685 -pf00271pf03461 -
03690 -# -
03695 -# -
03700 -pf13180pf03572 -
03705 -pf04551 -
03710 -pf03167 -
03715 -# -
03720 HMMreg00392_2_grm2(pf00155
03725 -pf13472 -
BMC-P
BMC-H
BMC-T
Aldehyde dehydrogenase
PTAC
Signature enzyme
Alcohol dehydrogenase
Regulator

Synechococcus sp. NAT40. Bacteria; Cyanobacteria; Synechococcales; Synechococcaceae;

CMN97__locus_1

CsomeACy

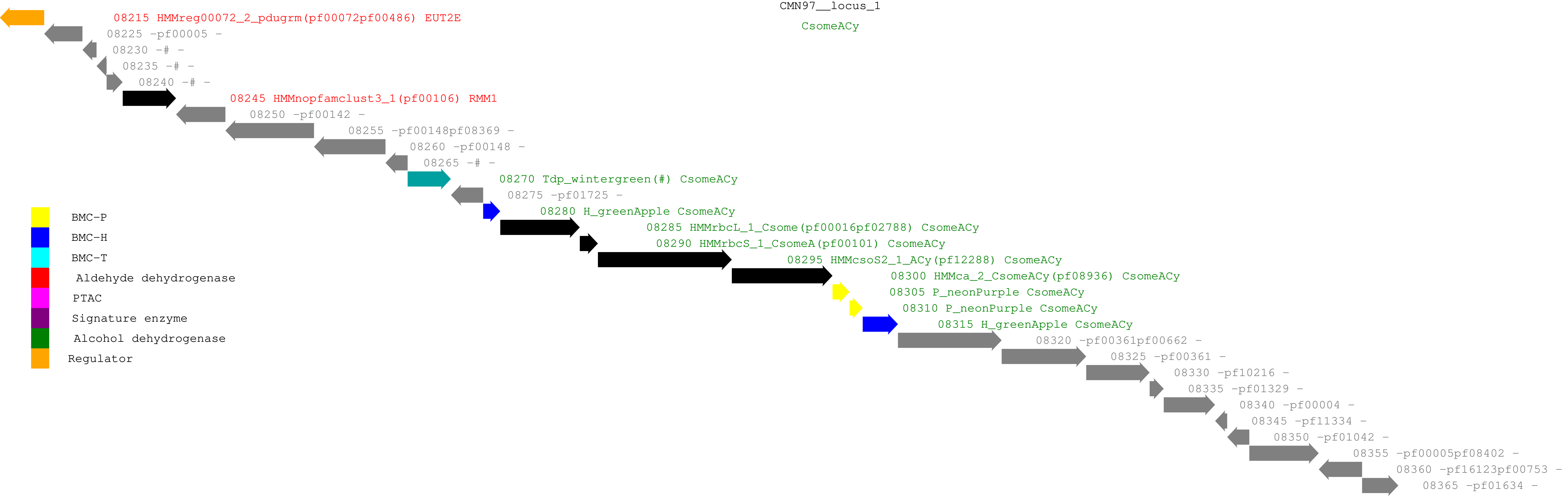

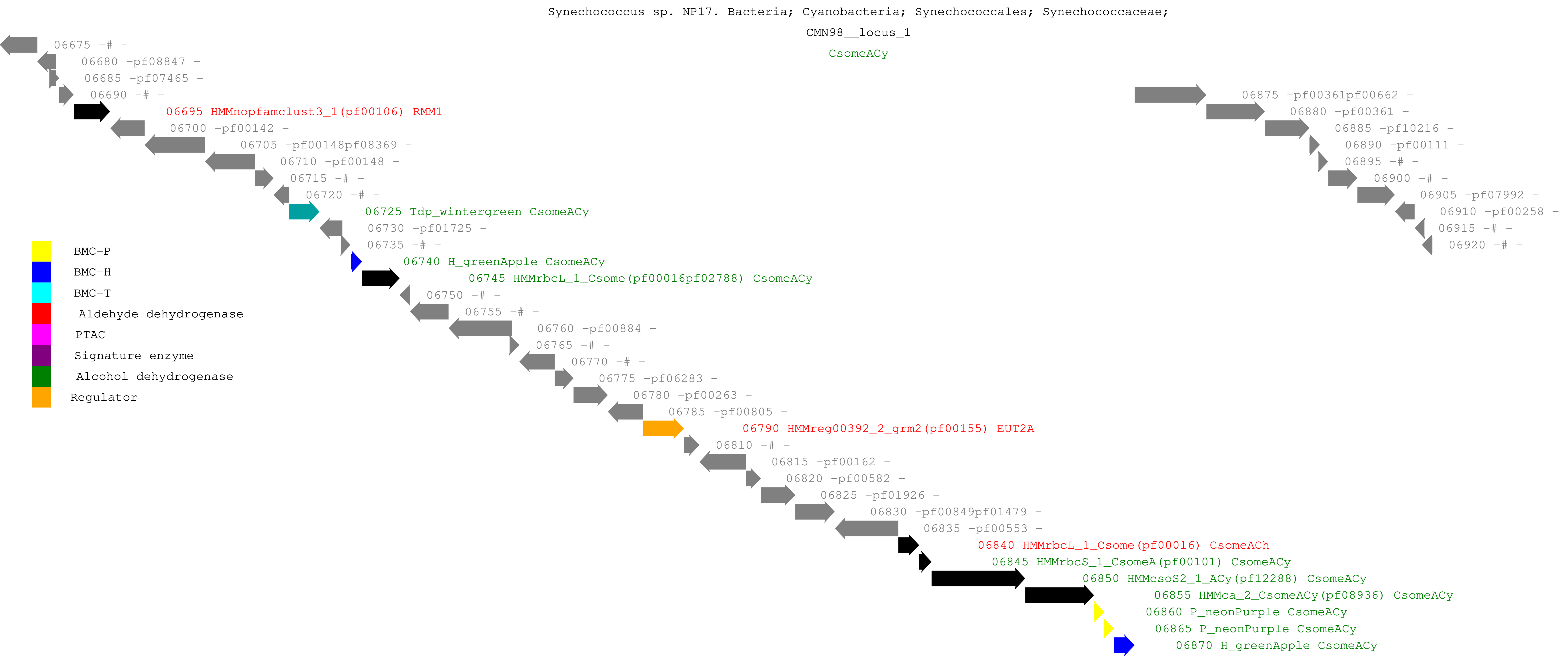

Synechococcus sp. NP17. Bacteria; Cyanobacteria; Synechococcales; Synechococcaceae;
CMN98__locus_1
CsomeACy
06675 -# -
06680 -pf08847 -
06685 -pf07465 -
06690 -# -
06695 HMMnopfamclust3_1(pf00106) RMM1
06700 -pf00142 -
06705 -pf00148pf08369 -
06710 -pf00148 -
06715 -# -
06720 -# -
06725 Tdp_wintergreen CsomeACy
06730 -pf01725 -
06735 -# -
06740 H_greenApple CsomeACy
06745 HMMrbcL_1_Csome(pf00016pf02788) CsomeACy
06750 -# -
06755 -# -
06760 -pf00884 -
06765 -# -
06770 -# -
06775 -pf06283 -
06780 -pf00263 -
06785 -pf00805 -
06790 HMMreg00392_2_grm2(pf00155) EUT2A
06810 -# -
06815 -pf00162 -
06820 -pf00582 -
06825 -pf01926 -
06830 -pf00849pf01479 -
06835 -pf00553 -
06840 HMMrbcL_1_Csome(pf00016) CsomeACh
06845 HMMrbcS_1_CsomeA(pf00101) CsomeACy
06850 HMMcsoS2_1_ACy(pf12288) CsomeACy
06855 HMMca_2_CsomeACy(pf08936) CsomeACy
06860 P_neonPurple CsomeACy
06865 P_neonPurple CsomeACy
06870 H_greenApple CsomeACy
06875 -pf00361pf00662 -
06880 -pf00361 -
06885 -pf10216 -
06890 -pf00111 -
06895 -# -
06900 -# -
06905 -pf07992 -
06910 -pf00258 -
06915 -# -
06920 -# -
BMC-P
BMC-H
BMC-T
Aldehyde dehydrogenase
PTAC
Signature enzyme
Alcohol dehydrogenase
Regulator

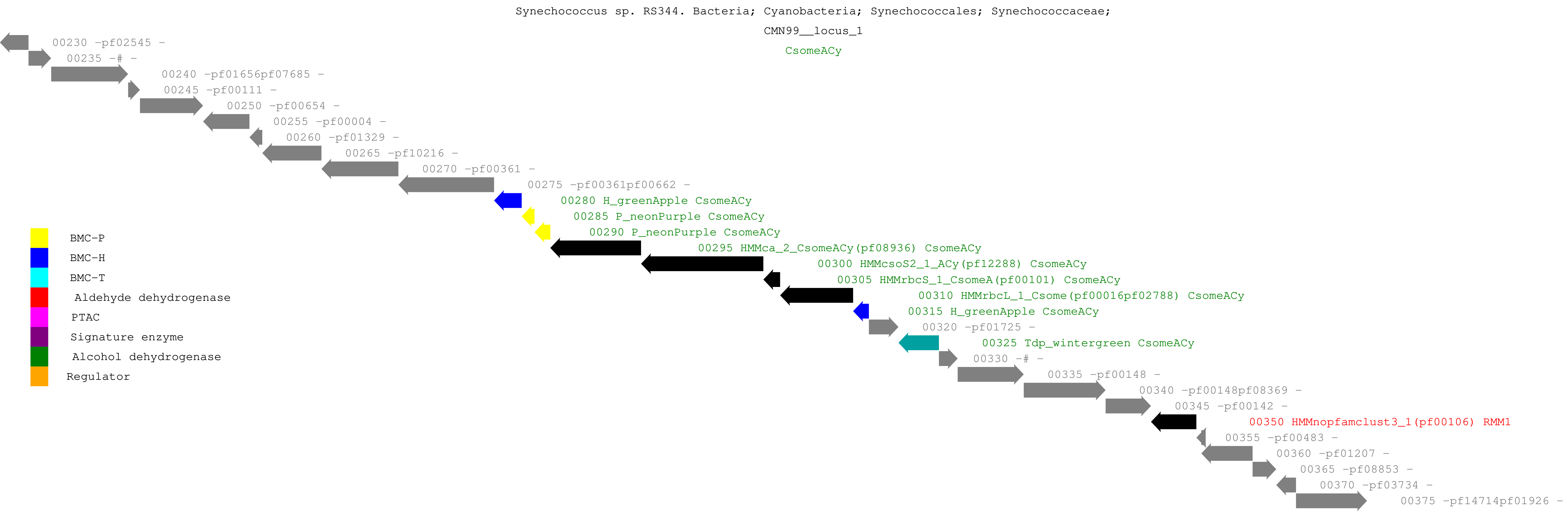
Synechococcus sp. RS344. Bacteria; Cyanobacteria; Synechococcales; Synechococcaceae;
CMN99__locus_1
CsomeACy
00230 -pf02545 -
00235 -# -
00240 -pf01656pf07685 -
00245 -pf00111 -
00250 -pf00654 -
00255 -pf00004 -
00260 -pf01329 -
00265 -pf10216 -
00270 -pf00361 -
00275 -pf00361pf00662 -
00280 H_greenApple CsomeACy
00285 P_neonPurple CsomeACy
00290 P_neonPurple CsomeACy
00295 HMMca_2_CsomeACy(pf08936) CsomeACy
00300 HMMcsoS2_1_ACy(pf12288) CsomeACy
00305 HMMrbcS_1_CsomeA(pf00101) CsomeACy
00310 HMMrbcL_1_Csome(pf00016pf02788) CsomeACy
00315 H_greenApple CsomeACy
00320 -pf01725 -
00325 Tdp_wintergreen CsomeACy
00330 -# -
00335 -pf00148 -
00340 -pf00148pf08369 -
00345 -pf00142 -
00350 HMMnopfamclust3_1(pf00106) RMM1
00355 -pf00483 -
00360 -pf01207 -
00365 -pf08853 -
00370 -pf03734 -
00375 -pf14714pf01926 -
BMC-P
BMC-H
BMC-T
Aldehyde dehydrogenase
PTAC
Signature enzyme
Alcohol dehydrogenase
Regulator

Cyanobium sp. PCC 7001. Bacteria; Cyanobacteria; Synechococcales; Synechococcaceae; Cyanobium.

CPCC7001__locus_fragmented

CsomeACy

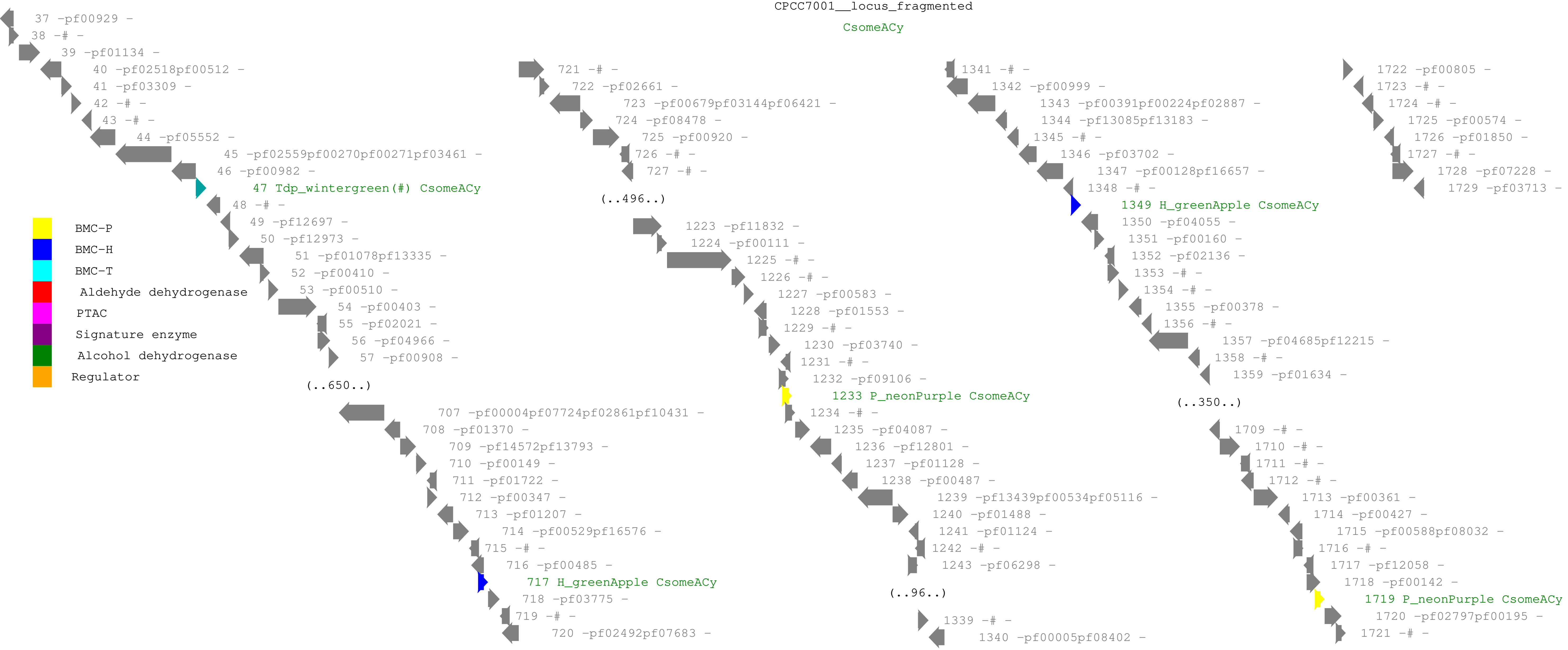

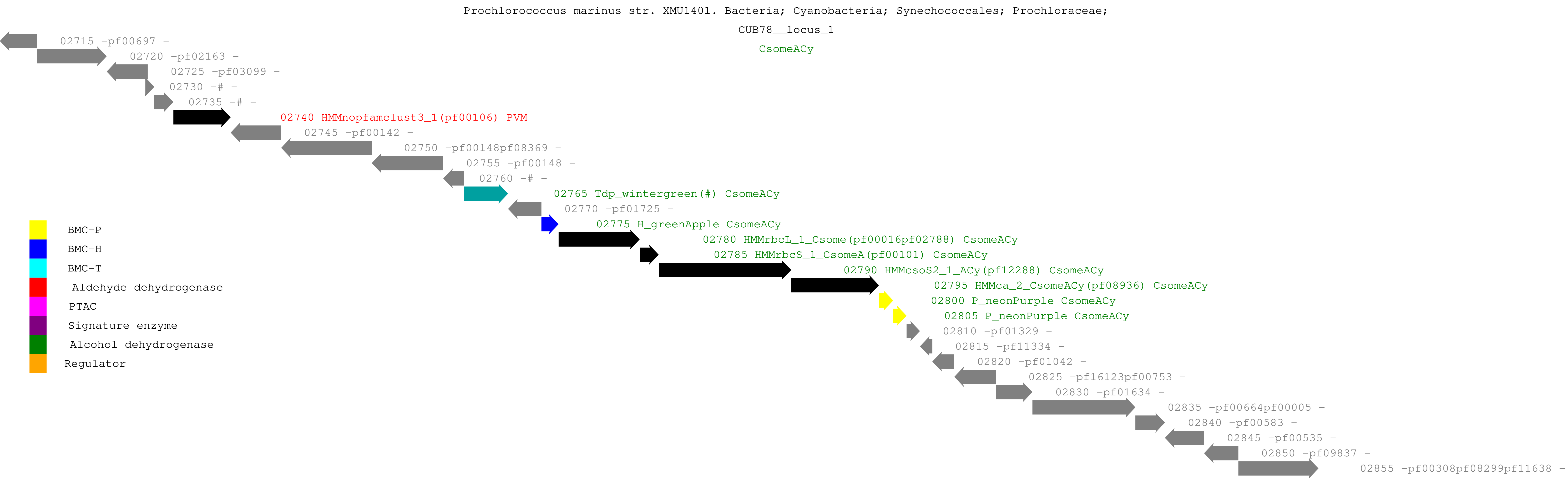
Prochlorococcus marinus str. XMU1401. Bacteria; Cyanobacteria; Synechococcales; Prochloraceae;
CUB78__locus_1
CsomeACy
02715 -pf00697 -
02720 -pf02163 -
02725 -pf03099 -
02730 -# -
02735 -# -
02740 HMMnopfamclust3_1(pf00106) PVM
02745 -pf00142 -
02750 -pf00148pf08369 -
02755 -pf00148 -
02760 -# -
02765 Tdp_wintergreen(#) CsomeACy
02770 -pf01725 -
02775 H_greenApple CsomeACy
02780 HMMrbcL_1_Csome(pf00016pf02788) CsomeACy
02785 HMMrbcS_1_CsomeA(pf00101) CsomeACy
02790 HMMcsoS2_1_ACy(pf12288) CsomeACy
02795 HMMca_2_CsomeACy(pf08936) CsomeACy
02800 P_neonPurple CsomeACy
02805 P_neonPurple CsomeACy
02810 -pf01329 -
02815 -pf11334 -
02820 -pf01042 -
02825 -pf16123pf00753 -
02830 -pf01634 -
02835 -pf00664pf00005 -
02840 -pf00583 -
02845 -pf00535 -
02850 -pf09837 -
02855 -pf00308pf08299pf11638 -
BMC-P
BMC-H
BMC-T
Aldehyde dehydrogenase
PTAC
Signature enzyme
Alcohol dehydrogenase
Regulator

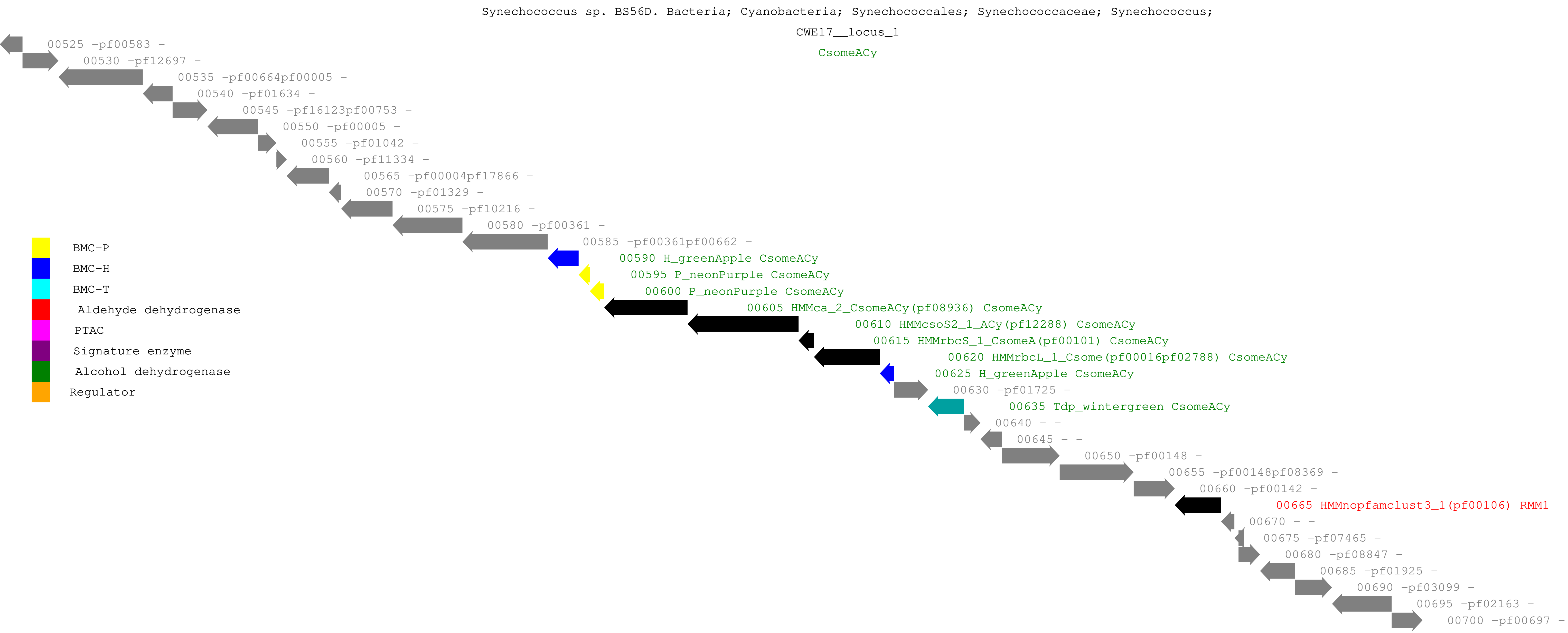
Synechococcus sp. BS56D. Bacteria; Cyanobacteria; Synechococcales; Synechococcaceae; Synechococcus;
CWE17__locus_1
CsomeACy
00525 -pf00583 -
00530 -pf12697 -
00535 -pf00664pf00005 -
00540 -pf01634 -
00545 -pf16123pf00753 -
00550 -pf00005 -
00555 -pf01042 -
00560 -pf11334 -
00565 -pf00004pf17866 -
00570 -pf01329 -
00575 -pf10216 -
00580 -pf00361 -
00585 -pf00361pf00662 -
00590 H_greenApple CsomeACy
00595 P_neonPurple CsomeACy
00600 P_neonPurple CsomeACy
00605 HMMca_2_CsomeACy(pf08936) CsomeACy
00610 HMMcsoS2_1_ACy(pf12288) CsomeACy
00615 HMMrbcS_1_CsomeA(pf00101) CsomeACy
00620 HMMrbcL_1_Csome(pf00016pf02788) CsomeACy
00625 H_greenApple CsomeACy
00630 -pf01725 -
00635 Tdp_wintergreen CsomeACy
00640 - -
00645 - -
00650 -pf00148 -
00655 -pf00148pf08369 -
00660 -pf00142 -
00665 HMMnopfamclust3_1(pf00106) RMM1
00670 - -
00675 -pf07465 -
00680 -pf08847 -
00685 -pf01925 -
00690 -pf03099 -
00695 -pf02163 -
00700 -pf00697 -
BMC-P
BMC-H
BMC-T
Aldehyde dehydrogenase
PTAC
Signature enzyme
Alcohol dehydrogenase
Regulator

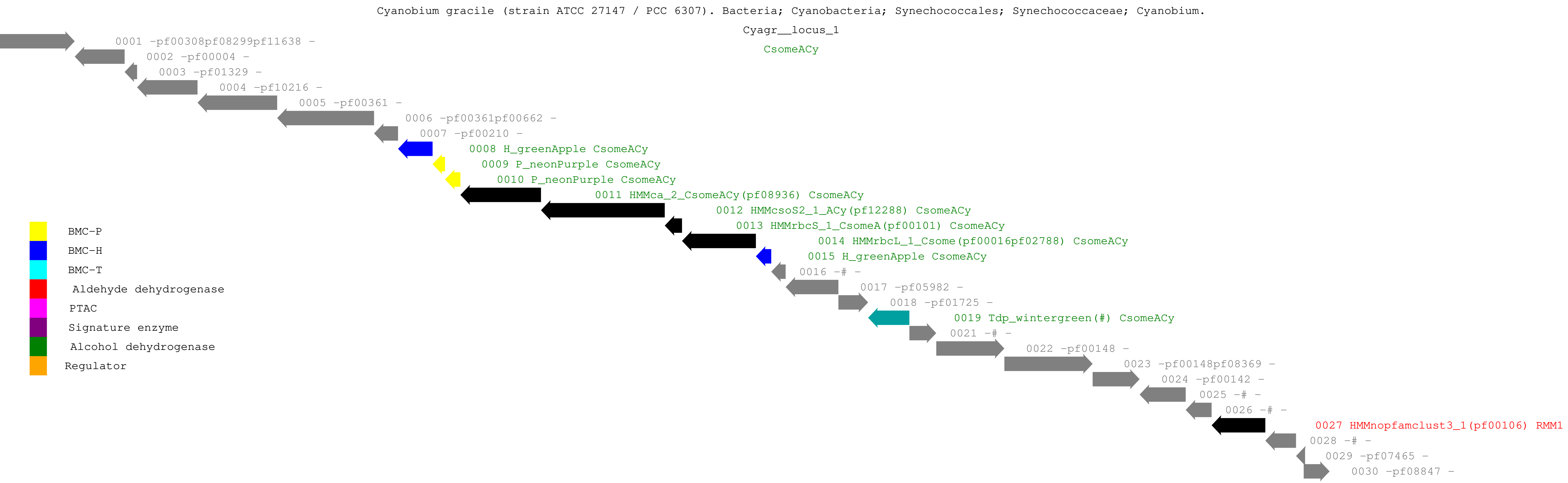
Cyanobium gracile (strain ATCC 27147 / PCC 6307). Bacteria; Cyanobacteria; Synechococcales; Synechococcaceae; Cyanobium.
Cyagr__locus_1
CsomeACy
0001 -pf00308pf08299pf11638 -
0002 -pf00004 -
0003 -pf01329 -
0004 -pf10216 -
0005 -pf00361 -
0006 -pf00361pf00662 -
0007 -pf00210 -
0008 H_greenApple CsomeACy
0009 P_neonPurple CsomeACy
0010 P_neonPurple CsomeACy
0011 HMMca_2_CsomeACy(pf08936) CsomeACy
0012 HMMcsoS2_1_ACy(pf12288) CsomeACy
0013 HMMrbcS_1_CsomeA(pf00101) CsomeACy
0014 HMMrbcL_1_Csome(pf00016pf02788) CsomeACy
0015 H_greenApple CsomeACy
0016 -# -
0017 -pf05982 -
0018 -pf01725 -
0019 Tdp_wintergreen(#) CsomeACy
0021 -# -
0022 -pf00148 -
0023 -pf00148pf08369 -
0024 -pf00142 -
0025 -# -
0026 -# -
0027 HMMnopfamclust3_1(pf00106) RMM1
0028 -# -
0029 -pf07465 -
0030 -pf08847 -
BMC-P
BMC-H
BMC-T
Aldehyde dehydrogenase
PTAC
Signature enzyme
Alcohol dehydrogenase
Regulator

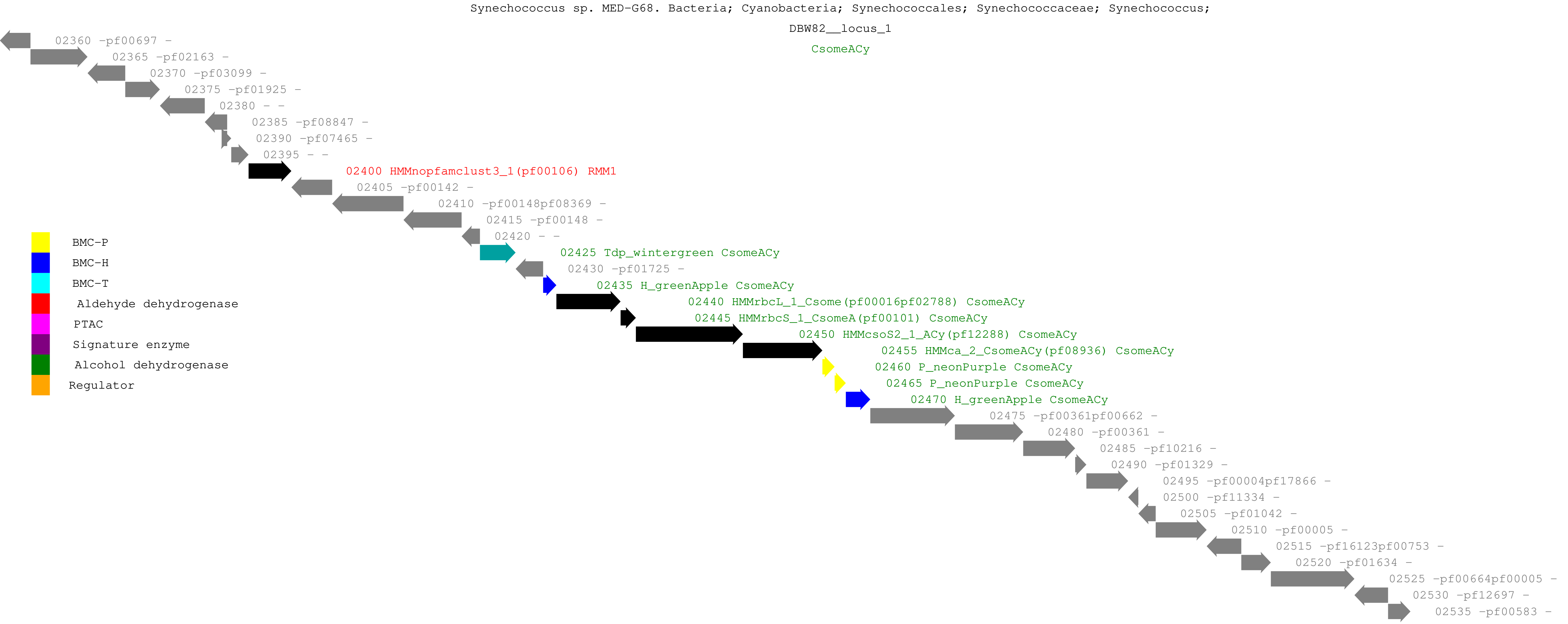
Synechococcus sp. MED-G68. Bacteria; Cyanobacteria; Synechococcales; Synechococcaceae; Synechococcus;
DBW82__locus_1
CsomeACy
02360 -pf00697 -
02365 -pf02163 -
02370 -pf03099 -
02375 -pf01925 -
02380 - -
02385 -pf08847 -
02390 -pf07465 -
02395 - -
02400 HMMnopfamclust3_1(pf00106) RMM1
02405 -pf00142 -
02410 -pf00148pf08369 -
02415 -pf00148 -
02420 - -
02425 Tdp_wintergreen CsomeACy
02430 -pf01725 -
02435 H_greenApple CsomeACy
02440 HMMrbcL_1_Csome(pf00016pf02788) CsomeACy
02445 HMMrbcS_1_CsomeA(pf00101) CsomeACy
02450 HMMcsoS2_1_ACy(pf12288) CsomeACy
02455 HMMca_2_CsomeACy(pf08936) CsomeACy
02460 P_neonPurple CsomeACy
02465 P_neonPurple CsomeACy
02470 H_greenApple CsomeACy
02475 -pf00361pf00662 -
02480 -pf00361 -
02485 -pf10216 -
02490 -pf01329 -
02495 -pf00004pf17866 -
02500 -pf11334 -
02505 -pf01042 -
02510 -pf00005 -
02515 -pf16123pf00753 -
02520 -pf01634 -
02525 -pf00664pf00005 -
02530 -pf12697 -
02535 -pf00583 -
BMC-P
BMC-H
BMC-T
Aldehyde dehydrogenase
PTAC
Signature enzyme
Alcohol dehydrogenase
Regulator

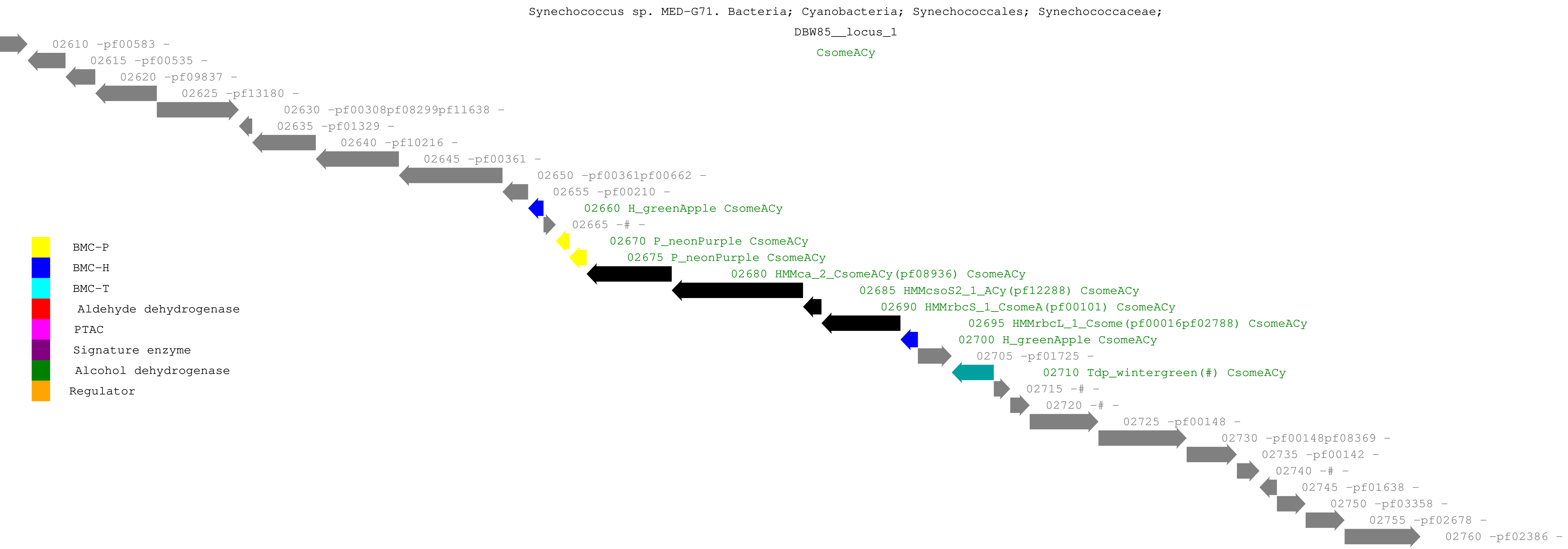
Synechococcus sp. MED-G71. Bacteria; Cyanobacteria; Synechococcales; Synechococcaceae;
DBW85__locus_1
CsomeACy
02610 -pf00583 -
02615 -pf00535 -
02620 -pf09837 -
02625 -pf13180 -
02630 -pf00308pf08299pf11638 -
02635 -pf01329 -
02640 -pf10216 -
02645 -pf00361 -
02650 -pf00361pf00662 -
02655 -pf00210 -
02660 H_greenApple CsomeACy
02665 -# -
02670 P_neonPurple CsomeACy
02675 P_neonPurple CsomeACy
02680 HMMca_2_CsomeACy(pf08936) CsomeACy
02685 HMMcsoS2_1_ACy(pf12288) CsomeACy
02690 HMMrbcS_1_CsomeA(pf00101) CsomeACy
02695 HMMrbcL_1_Csome(pf00016pf02788) CsomeACy
02700 H_greenApple CsomeACy
02705 -pf01725 -
02710 Tdp_wintergreen(#) CsomeACy
02715 -# -
02720 -# -
02725 -pf00148 -
02730 -pf00148pf08369 -
02735 -pf00142 -
02740 -# -
02745 -pf01638 -
02750 -pf03358 -
02755 -pf02678 -
02760 -pf02386 -
BMC-P
BMC-H
BMC-T
Aldehyde dehydrogenase
PTAC
Signature enzyme
Alcohol dehydrogenase
Regulator

Cyanobium sp. Bacteria; Cyanobacteria; Synechococcales; Synechococcaceae; Cyanobium.

DCF18__locus_1_main_sat

CsomeACy

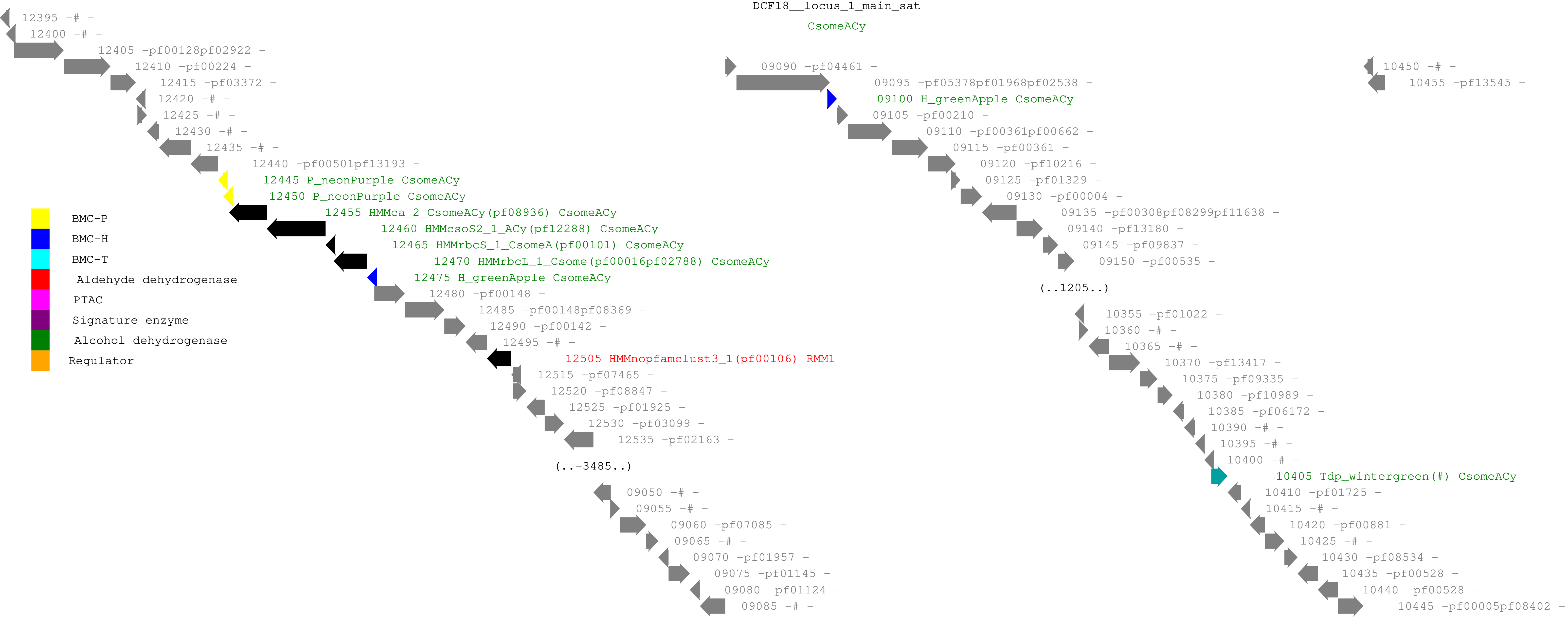

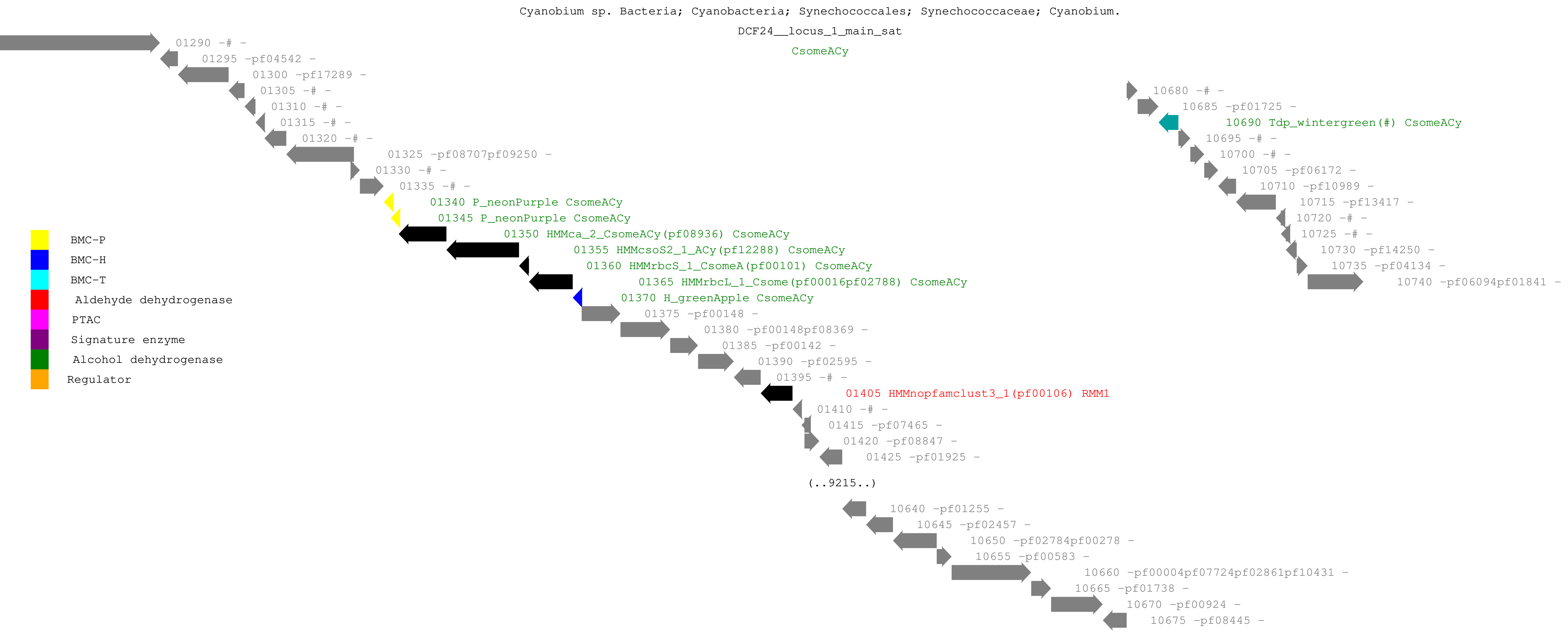

Cyanobium sp. Bacteria; Cyanobacteria; Synechococcales; Synechococcaceae; Cyanobium.
DCF24__locus_1_main_sat
CsomeACy
01290 -# -
01295 -pf04542 -
01300 -pf17289 -
01305 -# -
01310 -# -
01315 -# -
01320 -# -
01325 -pf08707pf09250 -
01330 -# -
01335 -# -
01340 P_neonPurple CsomeACy
01345 P_neonPurple CsomeACy
01350 HMMca_2_CsomeACy(pf08936) CsomeACy
01355 HMMcsoS2_1_ACy(pf12288) CsomeACy
01360 HMMrbcS_1_CsomeA(pf00101) CsomeACy
01365 HMMrbcL_1_Csome(pf00016pf02788) CsomeACy
01370 H_greenApple CsomeACy
01375 -pf00148 -
01380 -pf00148pf08369 -
01385 -pf00142 -
01390 -pf02595 -
01395 -# -
01405 HMMnopfamclust3_1(pf00106) RMM1
01410 -# -
01415 -pf07465 -
01420 -pf08847 -
01425 -pf01925 -
(..9215..)
10640 -pf01255 -
10645 -pf02457 -
10650 -pf02784pf00278 -
10655 -pf00583 -
10660 -pf00004pf07724pf02861pf10431 -
10665 -pf01738 -
10670 -pf00924 -
10675 -pf08445 -
10680 -# -
10685 -pf01725 -
10690 Tdp_wintergreen(#) CsomeACy
10695 -# -
10700 -# -
10705 -pf06172 -
10710 -pf10989 -
10715 -pf13417 -
10720 -# -
10725 -# -
10730 -pf14250 -
10735 -pf04134 -
10740 -pf06094pf01841 -
BMC-P
BMC-H
BMC-T
Aldehyde dehydrogenase
PTAC
Signature enzyme
Alcohol dehydrogenase
Regulator

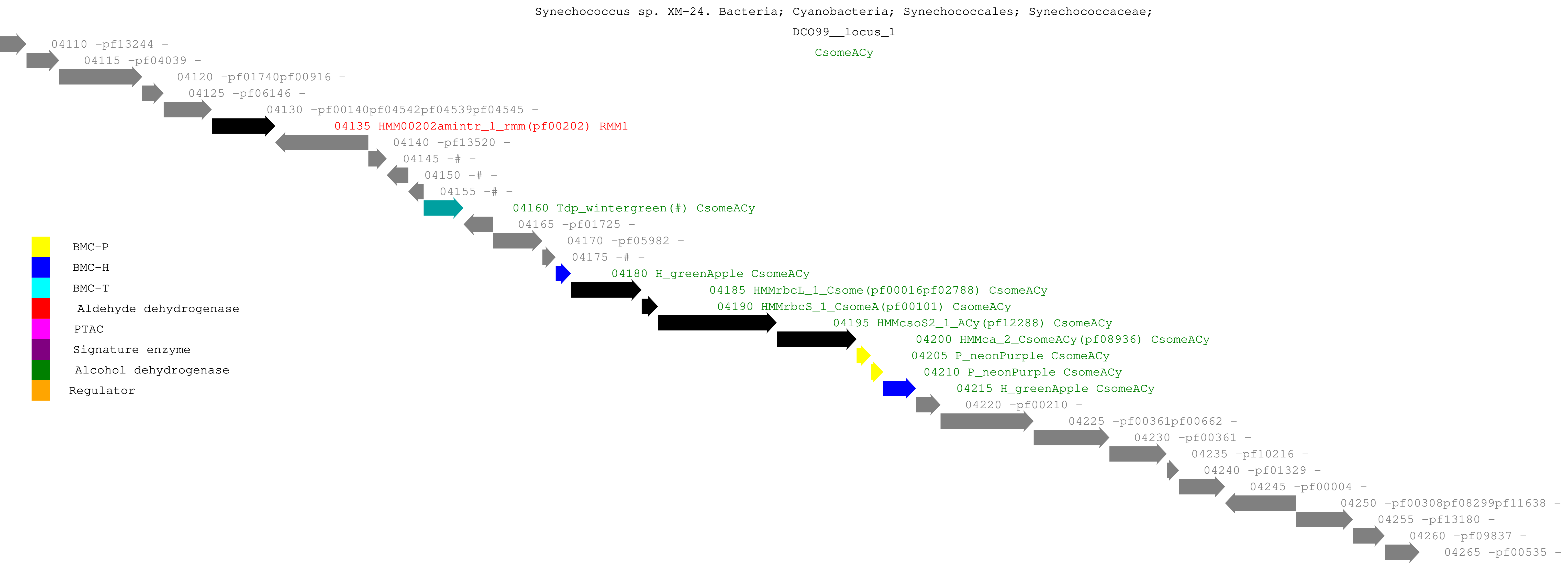
Synechococcus sp. XM-24. Bacteria; Cyanobacteria; Synechococcales; Synechococcaceae;
DCO99__locus_1
CsomeACy
04110 -pf13244 -
04115 -pf04039 -
04120 -pf01740pf00916 -
04125 -pf06146 -
04130 -pf00140pf04542pf04539pf04545 -
04135 HMM00202amintr_1_rmm(pf00202) RMM1
04140 -pf13520 -
04145 -# -
04150 -# -
04155 -# -
04160 Tdp_wintergreen(#) CsomeACy
04165 -pf01725 -
04170 -pf05982 -
04175 -# -
04180 H_greenApple CsomeACy
04185 HMMrbcL_1_Csome(pf00016pf02788) CsomeACy
04190 HMMrbcS_1_CsomeA(pf00101) CsomeACy
04195 HMMcsoS2_1_ACy(pf12288) CsomeACy
04200 HMMca_2_CsomeACy(pf08936) CsomeACy
04205 P_neonPurple CsomeACy
04210 P_neonPurple CsomeACy
04215 H_greenApple CsomeACy
04220 -pf00210 -
04225 -pf00361pf00662 -
04230 -pf00361 -
04235 -pf10216 -
04240 -pf01329 -
04245 -pf00004 -
04250 -pf00308pf08299pf11638 -
04255 -pf13180 -
04260 -pf09837 -
04265 -pf00535 -
BMC-P
BMC-H
BMC-T
Aldehyde dehydrogenase
PTAC
Signature enzyme
Alcohol dehydrogenase
Regulator

Synechococcus sp. UBA8638. Bacteria; Cyanobacteria; Synechococcales; Synechococcaceae;

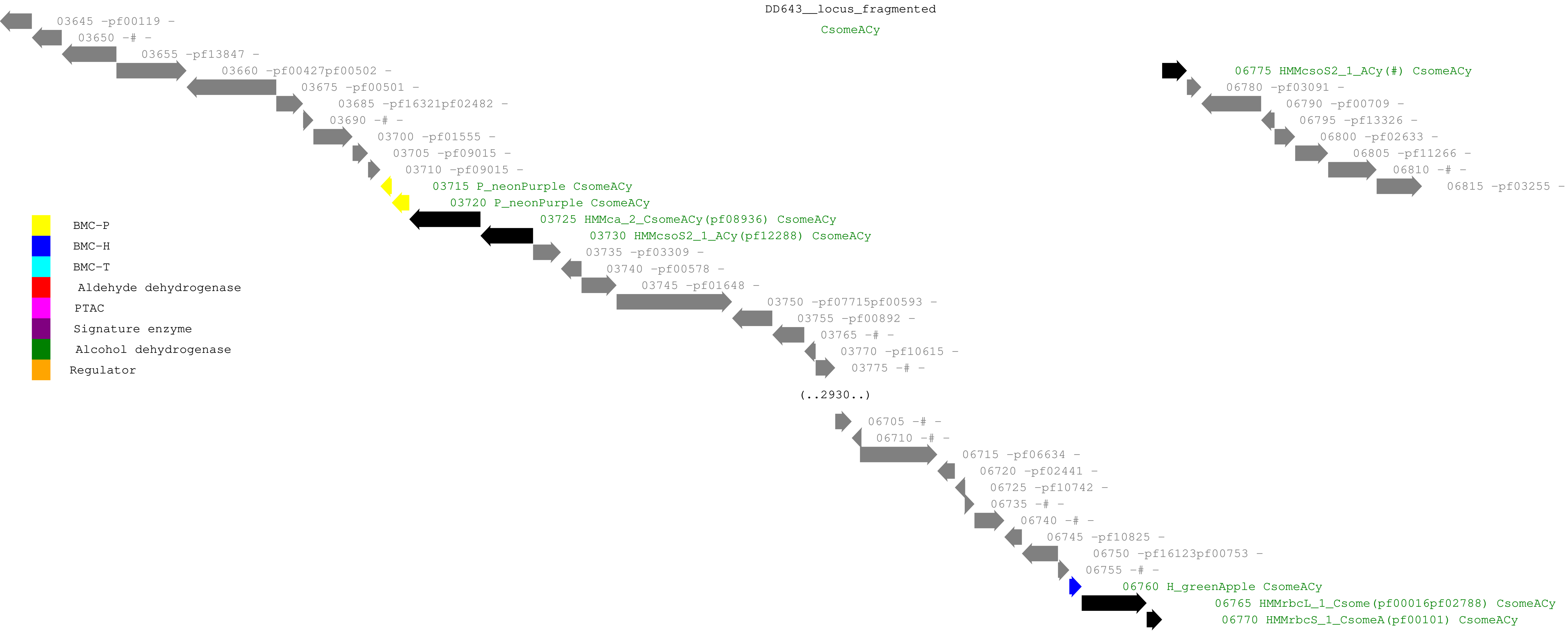

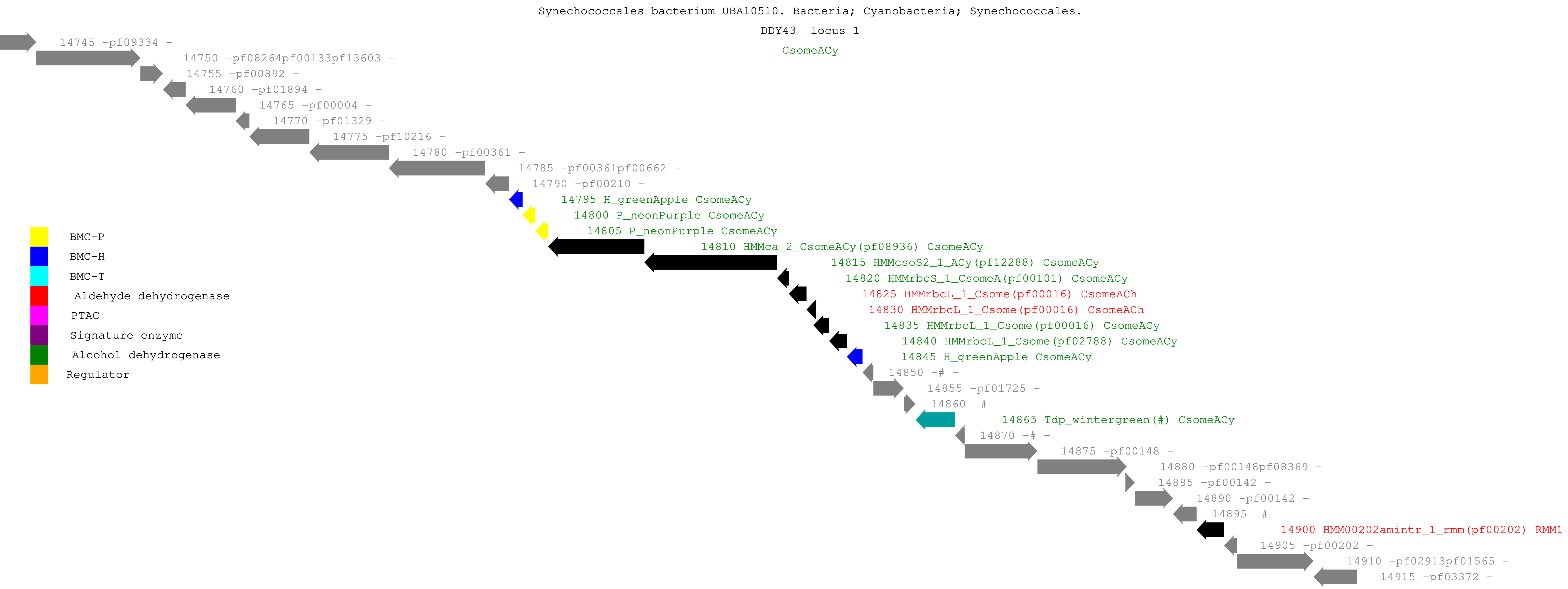
Synechococcales bacterium UBA10510. Bacteria; Cyanobacteria; Synechococcales.
DDY43__locus_1
CsomeACy
14745 -pf09334 -
14750 -pf08264pf00133pf13603 -
14755 -pf00892 -
14760 -pf01894 -
14765 -pf00004 -
14770 -pf01329 -
14775 -pf10216 -
14780 -pf00361 -
14785 -pf00361pf00662 -
14790 -pf00210 -
14795 H_greenApple CsomeACy
14800 P_neonPurple CsomeACy
14805 P_neonPurple CsomeACy
14810 HMMca_2_CsomeACy(pf08936) CsomeACy
14815 HMMcsoS2_1_ACy(pf12288) CsomeACy
14820 HMMrbcS_1_CsomeA(pf00101) CsomeACy
14825 HMMrbcL_1_Csome(pf00016) CsomeACh
14830 HMMrbcL_1_Csome(pf00016) CsomeACh
14835 HMMrbcL_1_Csome(pf00016) CsomeACy
14840 HMMrbcL_1_Csome(pf02788) CsomeACy
14845 H_greenApple CsomeACy
14850 -# -
14855 -pf01725 -
14860 -# -
14865 Tdp_wintergreen(#) CsomeACy
14870 -# -
14875 -pf00148 -
14880 -pf00148pf08369 -
14885 -pf00142 -
14890 -pf00142 -
14895 -# -
14900 HMM00202amintr_1_rmm(pf00202) RMM1
14905 -pf00202 -
14910 -pf02913pf01565 -
14915 -pf03372 -
BMC-P
BMC-H
BMC-T
Aldehyde dehydrogenase
PTAC
Signature enzyme
Alcohol dehydrogenase
Regulator

Synechococcales bacterium UBA8138. Bacteria; Cyanobacteria; Synechococcales.

DEB19__locus_1

CsomeACy

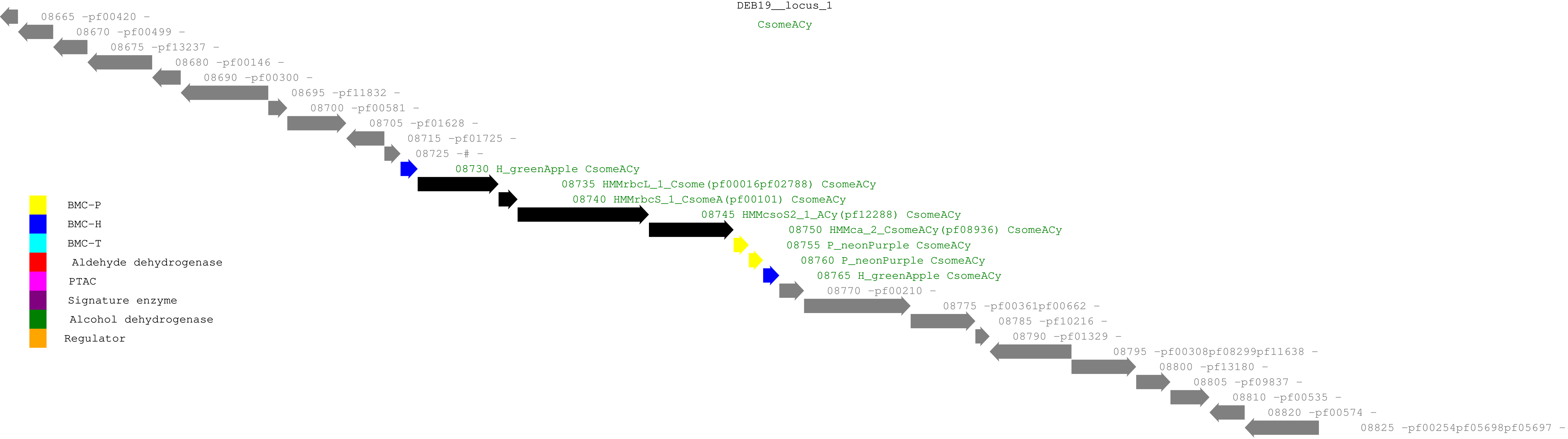

Synechococcales bacterium UBA8647. Bacteria; Cyanobacteria; Synechococcales.

DEO90__locus_1_main_sat

CsomeACy

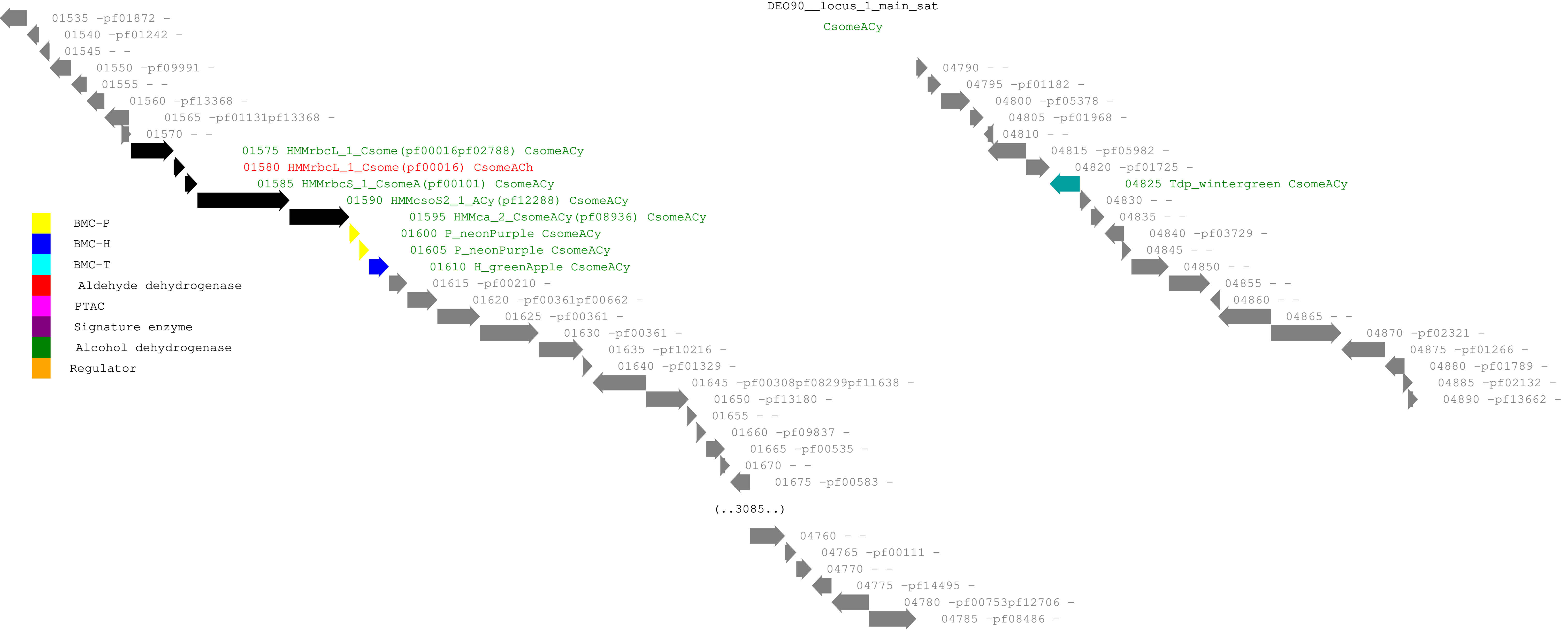

Synechococcales bacterium UBA12195. Bacteria; Cyanobacteria; Synechococcales.

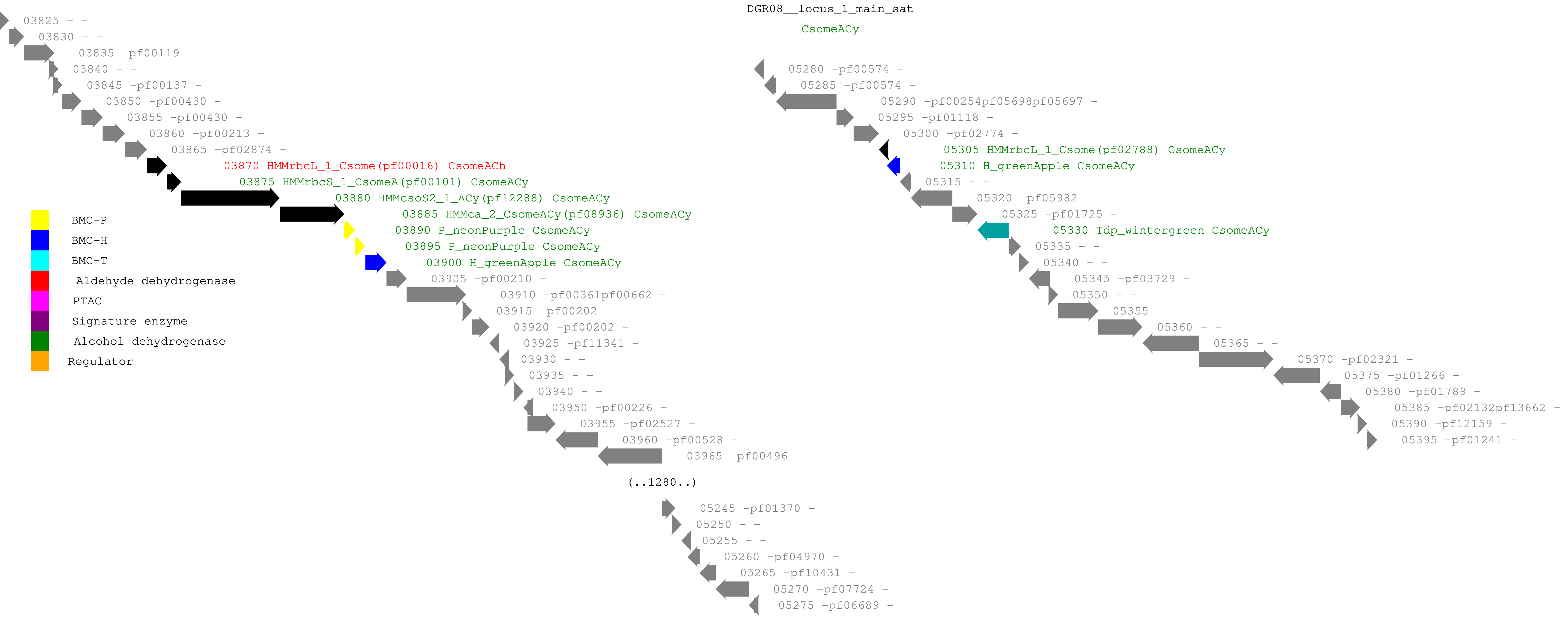

Synechococcus sp. UBA8071. Bacteria; Cyanobacteria; Synechococcales; Synechococcaceae; Synechococcus;

DIT12__locus_1

CsomeACy

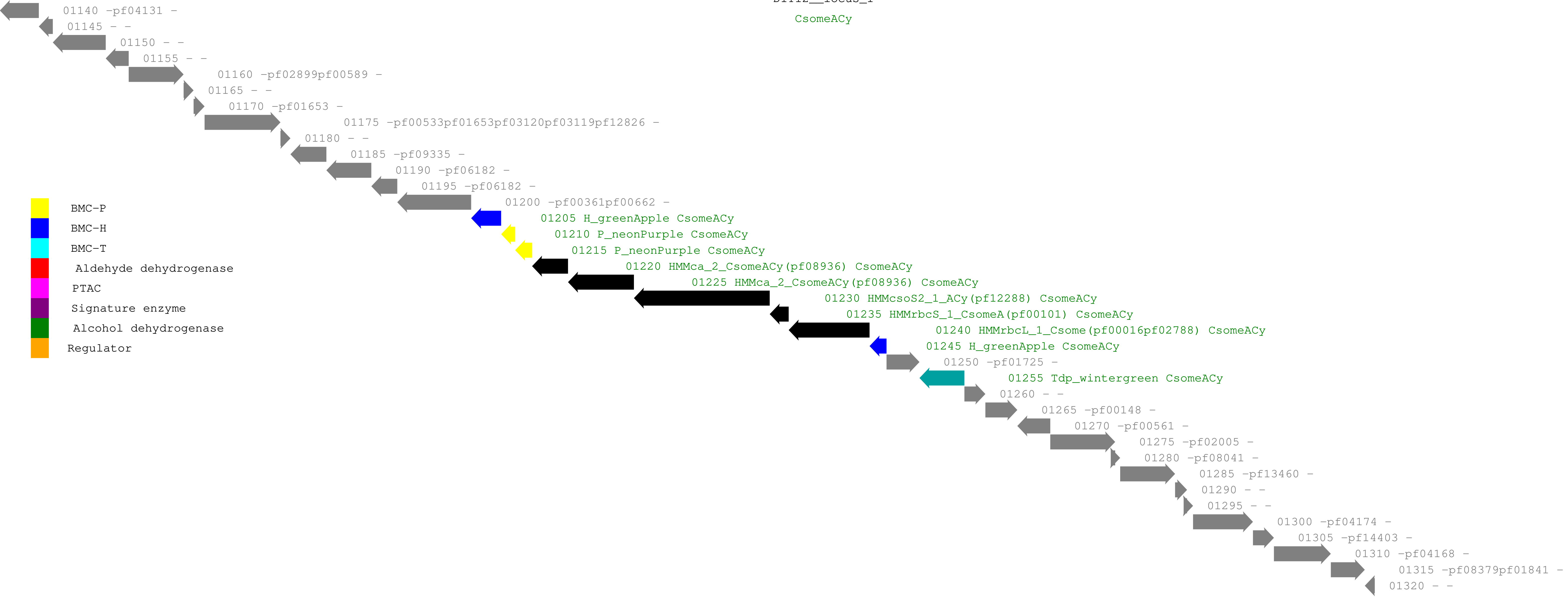

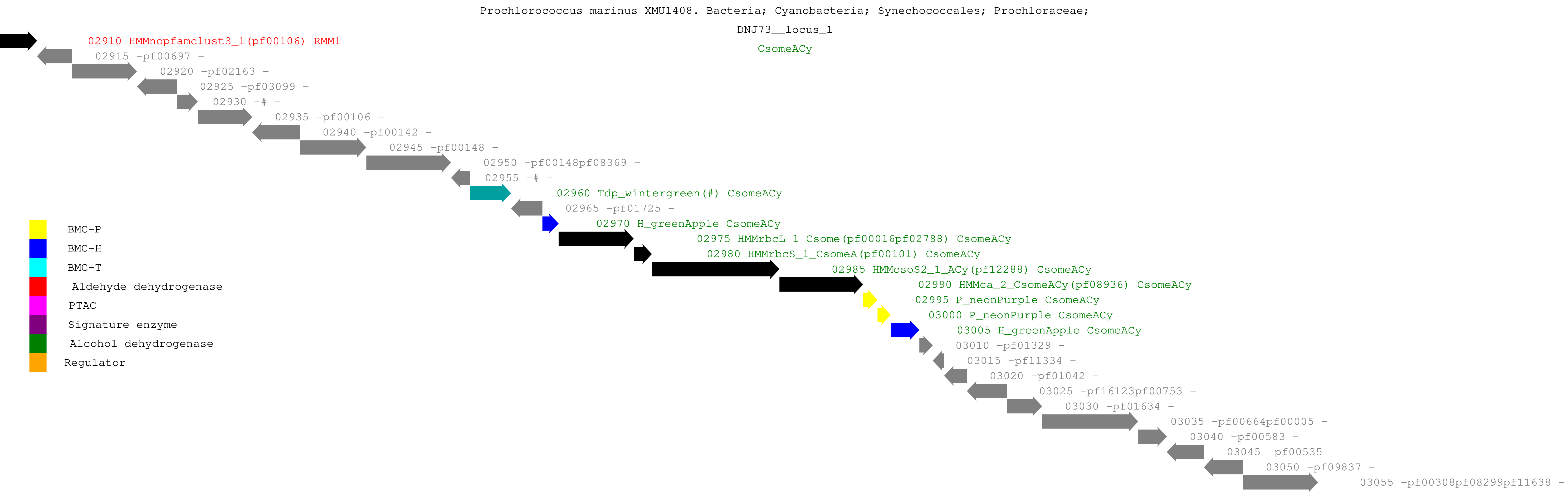
Prochlorococcus marinus XMU1408. Bacteria; Cyanobacteria; Synechococcales; Prochloraceae;
DNJ73__locus_1
CsomeACy
02910 HMMnopfamclust3_1(pf00106) RMM1
02915 -pf00697 -
02920 -pf02163 -
02925 -pf03099 -
02930 -# -
02935 -pf00106 -
02940 -pf00142 -
02945 -pf00148 -
02950 -pf00148pf08369 -
02955 -# -
02960 Tdp_wintergreen(#) CsomeACy
02965 -pf01725 -
02970 H_greenApple CsomeACy
02975 HMMrbcL_1_Csome(pf00016pf02788) CsomeACy
02980 HMMrbcS_1_CsomeA(pf00101) CsomeACy
02985 HMMcsoS2_1_ACy(pf12288) CsomeACy
02990 HMMca_2_CsomeACy(pf08936) CsomeACy
02995 P_neonPurple CsomeACy
03000 P_neonPurple CsomeACy
03005 H_greenApple CsomeACy
03010 -pf01329 -
03015 -pf11334 -
03020 -pf01042 -
03025 -pf16123pf00753 -
03030 -pf01634 -
03035 -pf00664pf00005 -
03040 -pf00583 -
03045 -pf00535 -
03050 -pf09837 -
03055 -pf00308pf08299pf11638 -
BMC-P
BMC-H
BMC-T
Aldehyde dehydrogenase
PTAC
Signature enzyme
Alcohol dehydrogenase
Regulator

Cyanobium sp. CACIAM 14. Bacteria; Cyanobacteria; Synechococcales; Synechococcaceae; Cyanobium.

ER33__locus_1_main_sat

CsomeACy

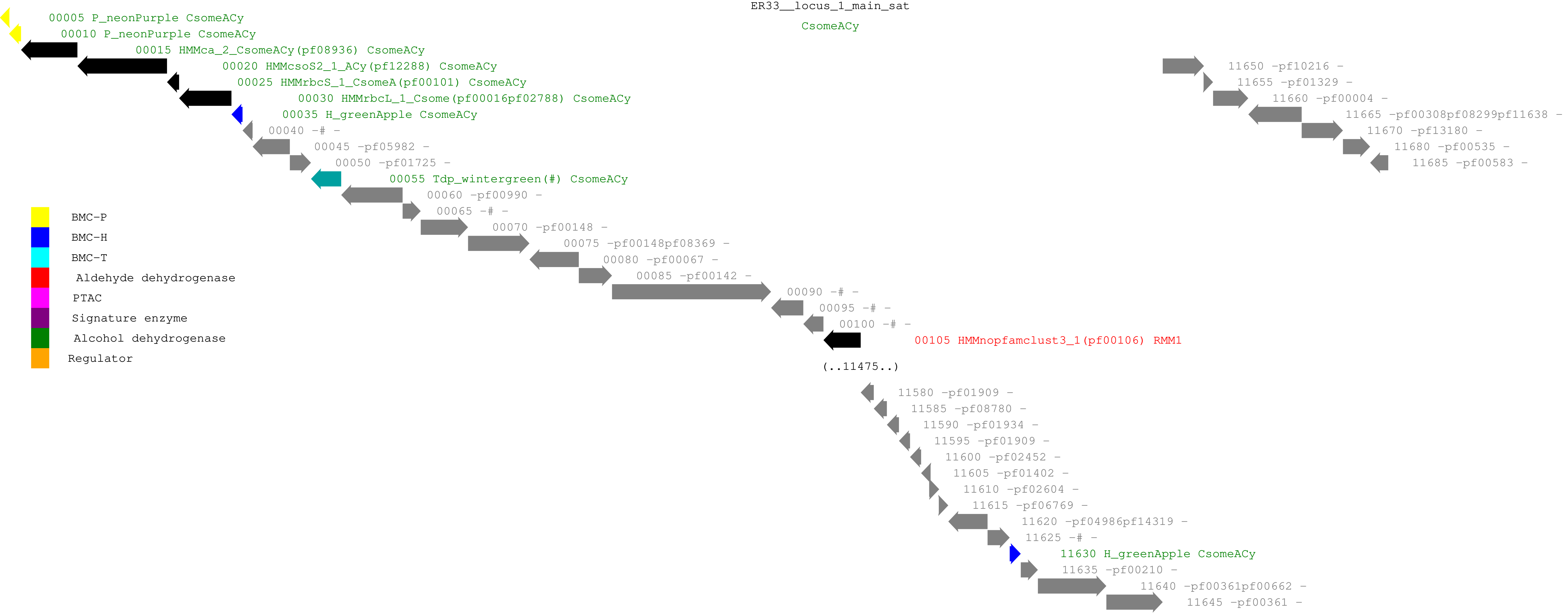

Aphanocapsa feldmannii 277cV. Bacteria; Cyanobacteria; Synechococcales; Merismopediaceae; Aphanocapsa.

ERJ67__locus_1

CsomeACy

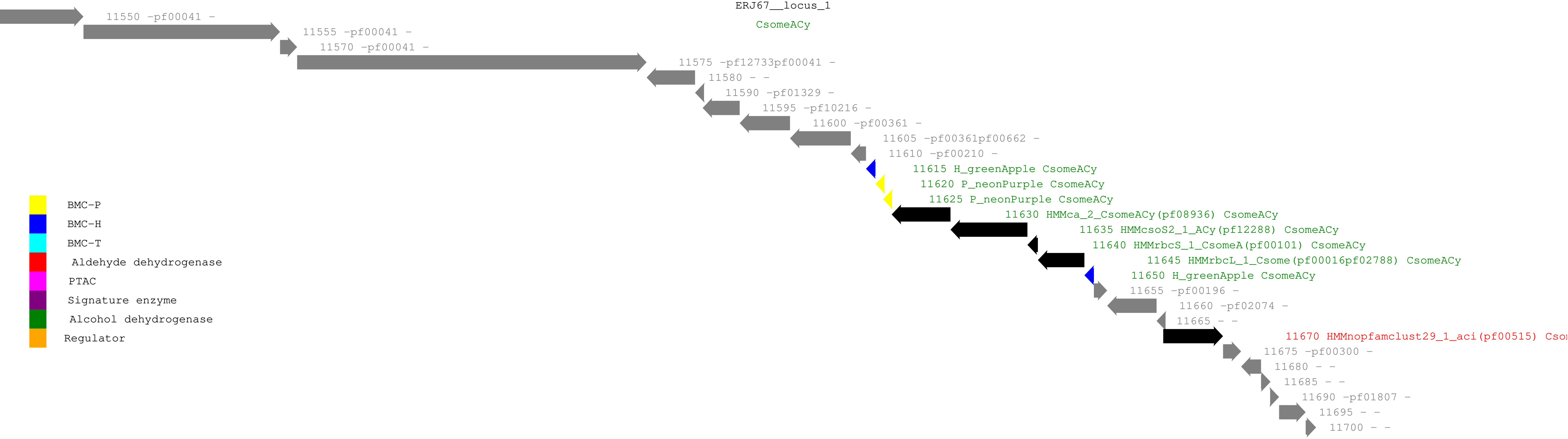

Aphanocapsa feldmannii 277cI. Bacteria; Cyanobacteria; Synechococcales; Merismopediaceae; Aphanocapsa.

ERJ68__locus_1_main_sat

CsomeACy

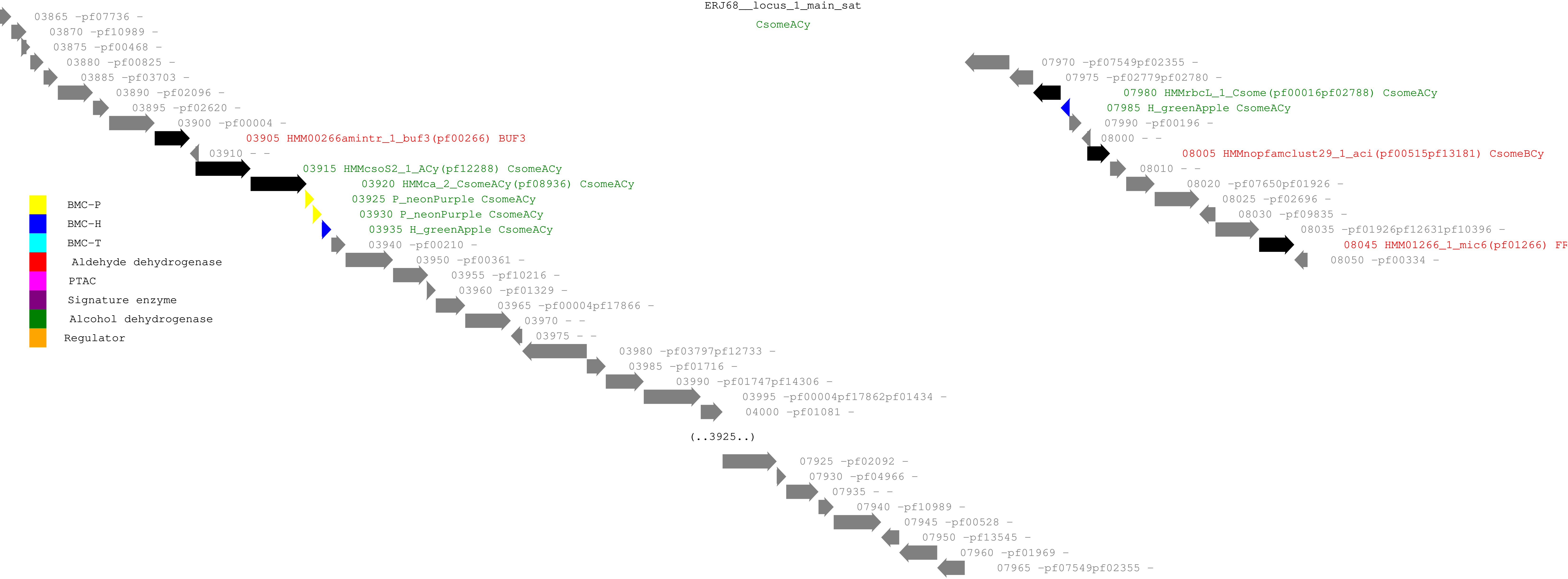

Prochlorococcus marinus str. GP2. Bacteria; Cyanobacteria; Synechococcales; Prochloraceae;

EU91__locus_1

CsomeACy

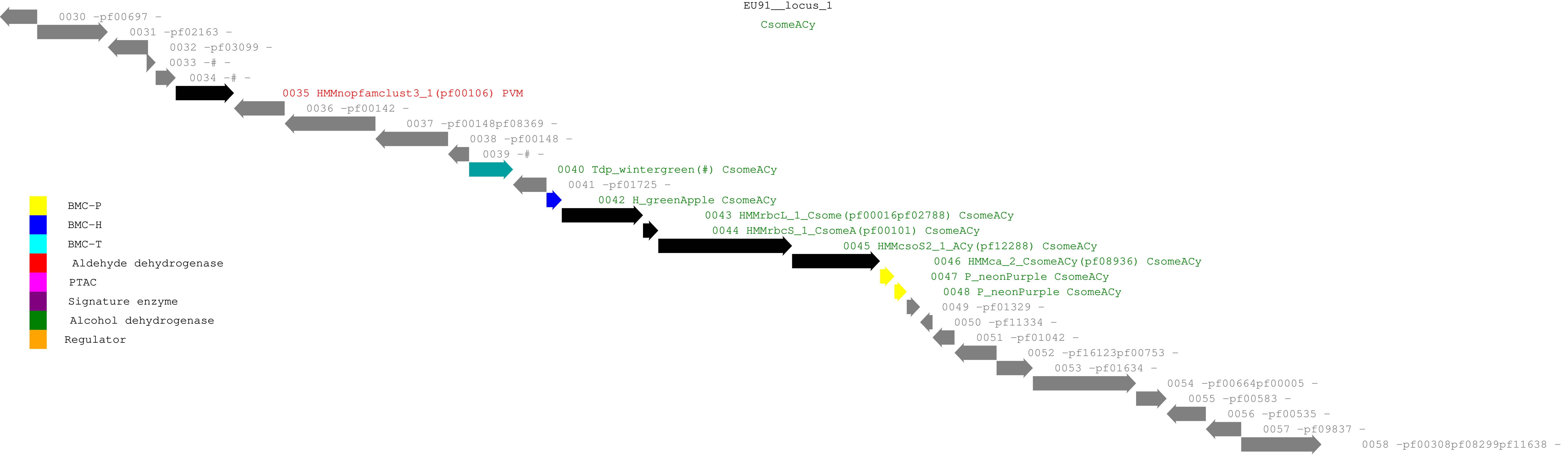

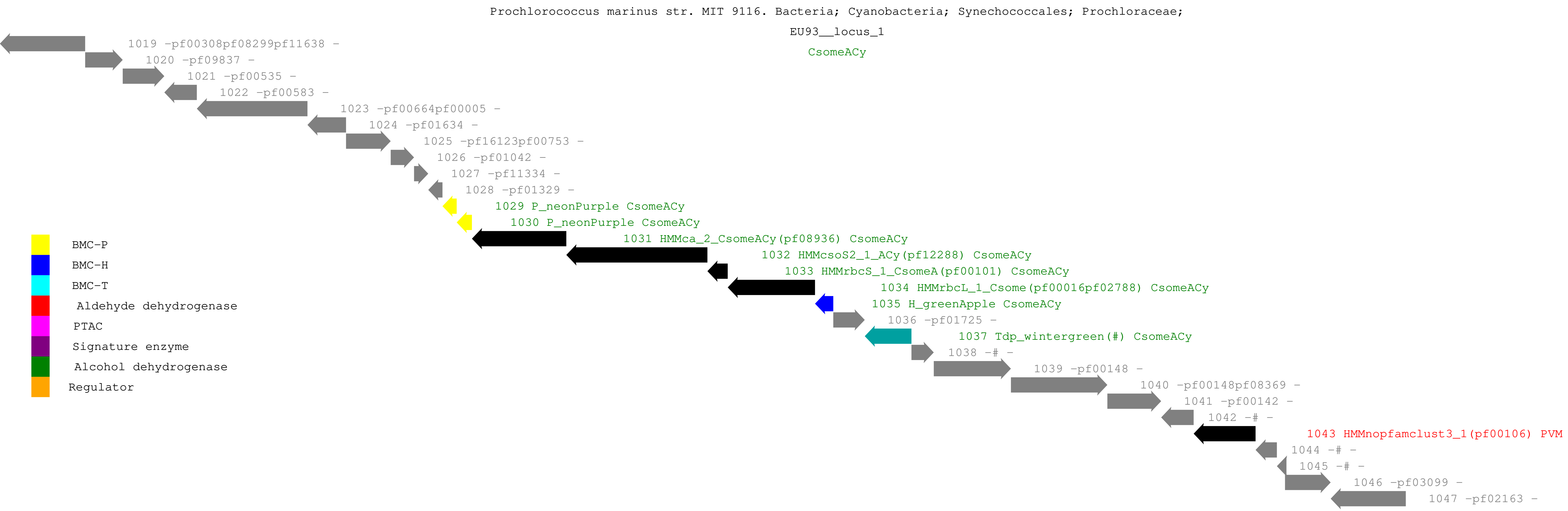

Prochlorococcus marinus str. MIT 9116. Bacteria; Cyanobacteria; Synechococcales; Prochloraceae;
EU93__locus_1
CsomeACy
1019 -pf00308pf08299pf11638 -
1020 -pf09837 -
1021 -pf00535 -
1022 -pf00583 -
1023 -pf00664pf00005 -
1024 -pf01634 -
1025 -pf16123pf00753 -
1026 -pf01042 -
1027 -pf11334 -
1028 -pf01329 -
1029 P_neonPurple CsomeACy
1030 P_neonPurple CsomeACy
1031 HMMca_2_CsomeACy(pf08936) CsomeACy
1032 HMMcsoS2_1_ACy(pf12288) CsomeACy
1033 HMMrbcS_1_CsomeA(pf00101) CsomeACy
1034 HMMrbcL_1_Csome(pf00016pf02788) CsomeACy
1035 H_greenApple CsomeACy
1036 -pf01725 -
1037 Tdp_wintergreen(#) CsomeACy
1038 -# -
1039 -pf00148 -
1040 -pf00148pf08369 -
1041 -pf00142 -
1042 -# -
1043 HMMnopfamclust3_1(pf00106) PVM
1044 -# -
1045 -# -
1046 -pf03099 -
1047 -pf02163 -
BMC-P
BMC-H
BMC-T
Aldehyde dehydrogenase
PTAC
Signature enzyme
Alcohol dehydrogenase
Regulator

Prochlorococcus marinus str. MIT 9201. Bacteria; Cyanobacteria; Synechococcales; Prochloraceae;

EU95__locus_1

CsomeACy

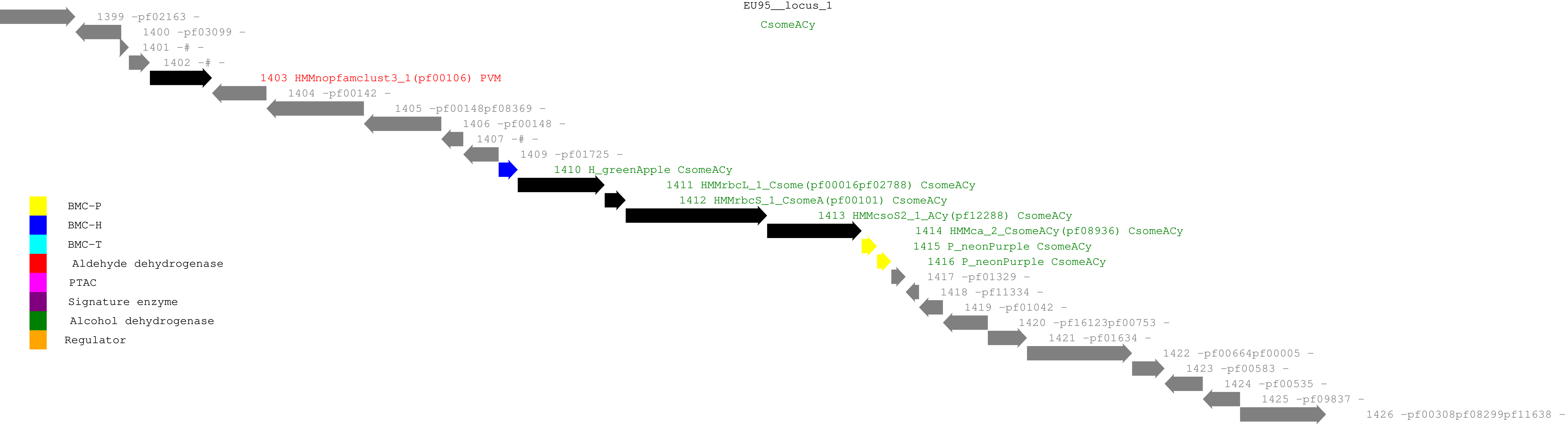

Prochlorococcus marinus str. MIT 9302. Bacteria; Cyanobacteria; Synechococcales; Prochloraceae;

EU96__locus_1

CsomeACy

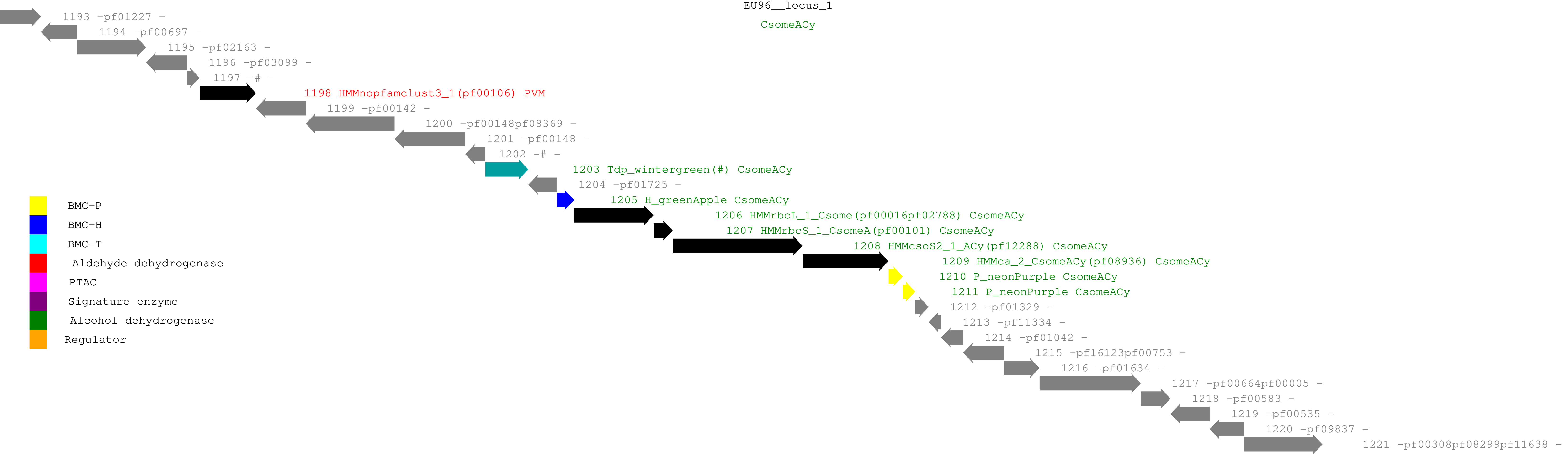

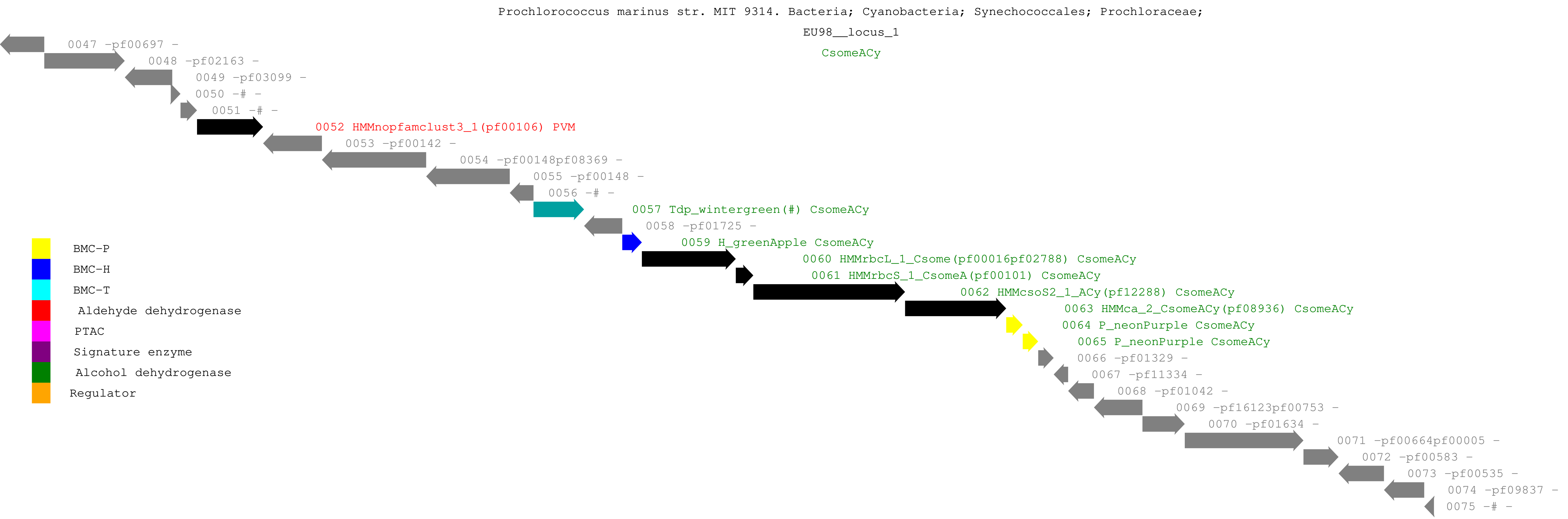
Prochlorococcus marinus str. MIT 9314. Bacteria; Cyanobacteria; Synechococcales; Prochloraceae;
EU98__locus_1
CsomeACy
0047 -pf00697 -
0048 -pf02163 -
0049 -pf03099 -
0050 -# -
0051 -# -
0052 HMMnopfamclust3_1(pf00106) PVM
0053 -pf00142 -
0054 -pf00148pf08369 -
0055 -pf00148 -
0056 -# -
0057 Tdp_wintergreen(#) CsomeACy
0058 -pf01725 -
0059 H_greenApple CsomeACy
0060 HMMrbcL_1_Csome(pf00016pf02788) CsomeACy
0061 HMMrbcS_1_CsomeA(pf00101) CsomeACy
0062 HMMcsoS2_1_ACy(pf12288) CsomeACy
0063 HMMca_2_CsomeACy(pf08936) CsomeACy
0064 P_neonPurple CsomeACy
0065 P_neonPurple CsomeACy
0066 -pf01329 -
0067 -pf11334 -
0068 -pf01042 -
0069 -pf16123pf00753 -
0070 -pf01634 -
0071 -pf00664pf00005 -
0072 -pf00583 -
0073 -pf00535 -
0074 -pf09837 -
0075 -# -
BMC-P
BMC-H
BMC-T
Aldehyde dehydrogenase
PTAC
Signature enzyme
Alcohol dehydrogenase
Regulator

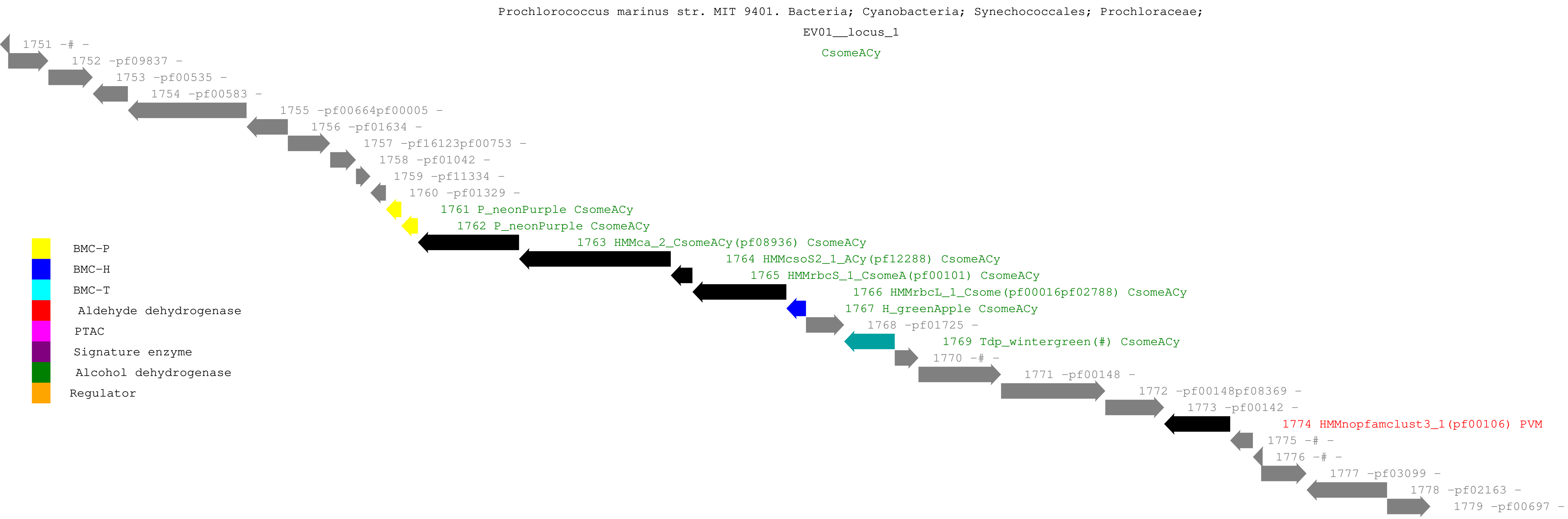
Prochlorococcus marinus str. MIT 9401. Bacteria; Cyanobacteria; Synechococcales; Prochloraceae;
EV01__locus_1
CsomeACy
1751 -# -
1752 -pf09837 -
1753 -pf00535 -
1754 -pf00583 -
1755 -pf00664pf00005 -
1756 -pf01634 -
1757 -pf16123pf00753 -
1758 -pf01042 -
1759 -pf11334 -
1760 -pf01329 -
1761 P_neonPurple CsomeACy
1762 P_neonPurple CsomeACy
1763 HMMca_2_CsomeACy(pf08936) CsomeACy
1764 HMMcsoS2_1_ACy(pf12288) CsomeACy
1765 HMMrbcS_1_CsomeA(pf00101) CsomeACy
1766 HMMrbcL_1_Csome(pf00016pf02788) CsomeACy
1767 H_greenApple CsomeACy
1768 -pf01725 -
1769 Tdp_wintergreen(#) CsomeACy
1770 -# -
1771 -pf00148 -
1772 -pf00148pf08369 -
1773 -pf00142 -
1774 HMMnopfamclust3_1(pf00106) PVM
1775 -# -
1776 -# -
1777 -pf03099 -
1778 -pf02163 -
1779 -pf00697 -
BMC-P
BMC-H
BMC-T
Aldehyde dehydrogenase
PTAC
Signature enzyme
Alcohol dehydrogenase
Regulator

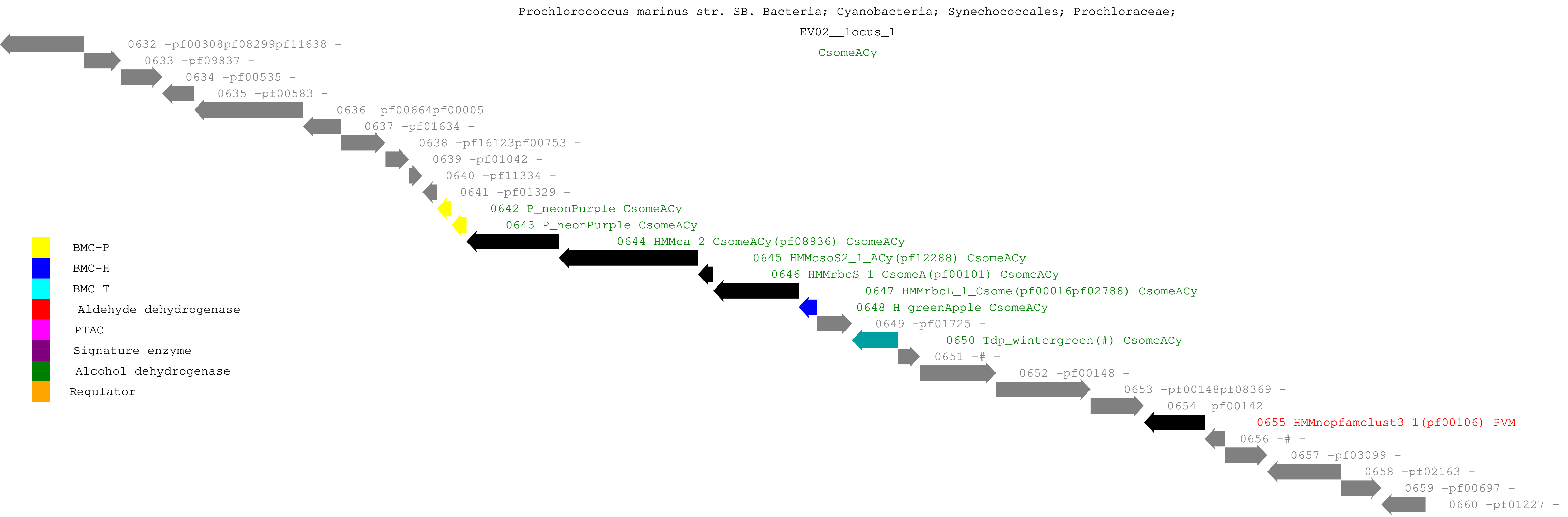
Prochlorococcus marinus str. SB. Bacteria; Cyanobacteria; Synechococcales; Prochloraceae;
EV02__locus_1
CsomeACy
0632 -pf00308pf08299pf11638 -
0633 -pf09837 -
0634 -pf00535 -
0635 -pf00583 -
0636 -pf00664pf00005 -
0637 -pf01634 -
0638 -pf16123pf00753 -
0639 -pf01042 -
0640 -pf11334 -
0641 -pf01329 -
0642 P_neonPurple CsomeACy
0643 P_neonPurple CsomeACy
0644 HMMca_2_CsomeACy(pf08936) CsomeACy
0645 HMMcsoS2_1_ACy(pf12288) CsomeACy
0646 HMMrbcS_1_CsomeA(pf00101) CsomeACy
0647 HMMrbcL_1_Csome(pf00016pf02788) CsomeACy
0648 H_greenApple CsomeACy
0649 -pf01725 -
0650 Tdp_wintergreen(#) CsomeACy
0651 -# -
0652 -pf00148 -
0653 -pf00148pf08369 -
0654 -pf00142 -
0655 HMMnopfamclust3_1(pf00106) PVM
0656 -# -
0657 -pf03099 -
0658 -pf02163 -
0659 -pf00697 -
0660 -pf01227 -
BMC-P
BMC-H
BMC-T
Aldehyde dehydrogenase
PTAC
Signature enzyme
Alcohol dehydrogenase
Regulator

Prochlorococcus marinus str. PAC1. Bacteria; Cyanobacteria; Synechococcales; Prochloraceae;

EV03__locus_1

CsomeACy

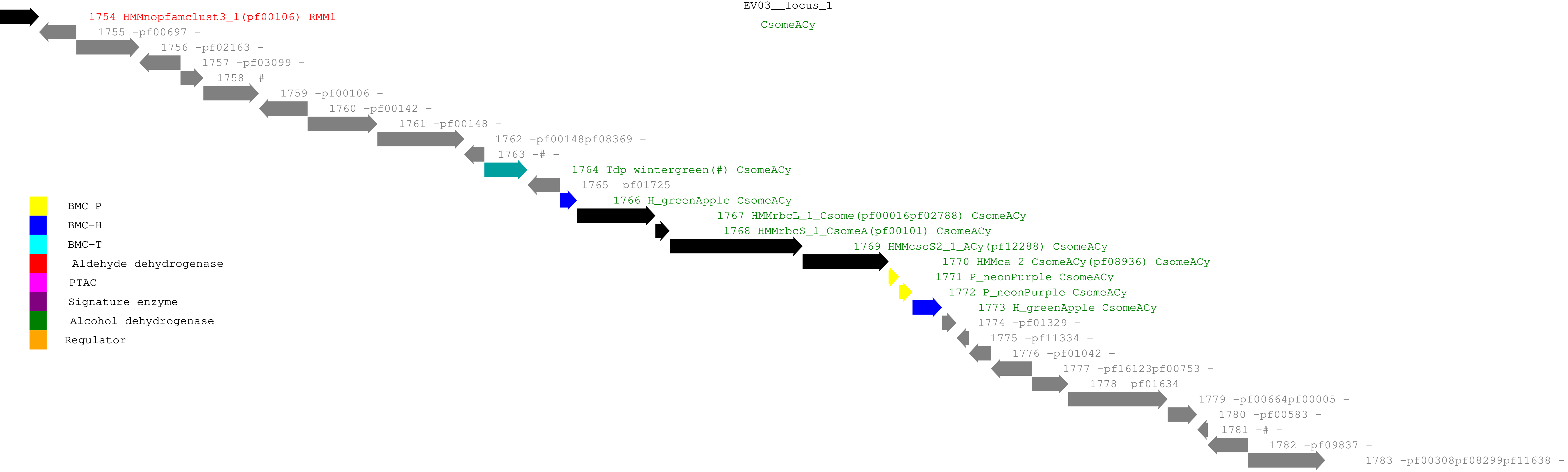

Prochlorococcus sp. MIT 0601. A0A0A2BFG3_9PROC Unreviewed; 466 AA.

EV05__locus_1

CsomeACy

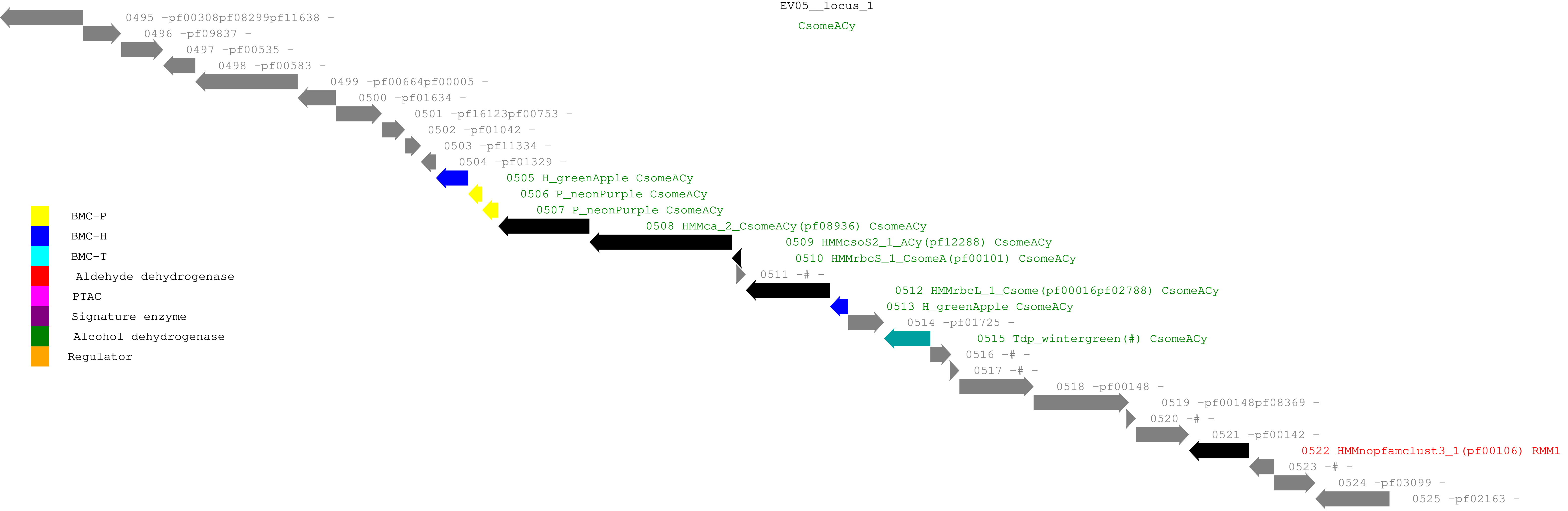

Prochlorococcus sp. MIT 0602. A0A0A2BQP8_9PROC Unreviewed; 222 AA.

EV06__locus_1

CsomeACy

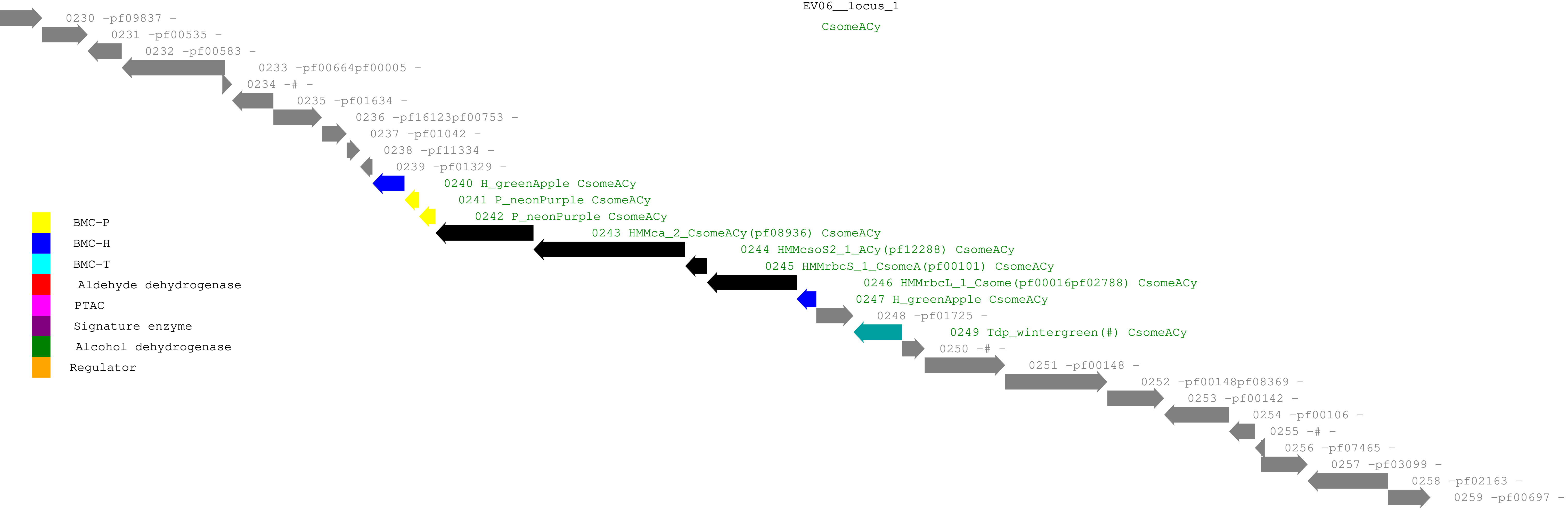

Prochlorococcus sp. MIT 0701. A0A0A2CMC6_9PROC Unreviewed; 227 AA.

EV12__locus_1

CsomeACy

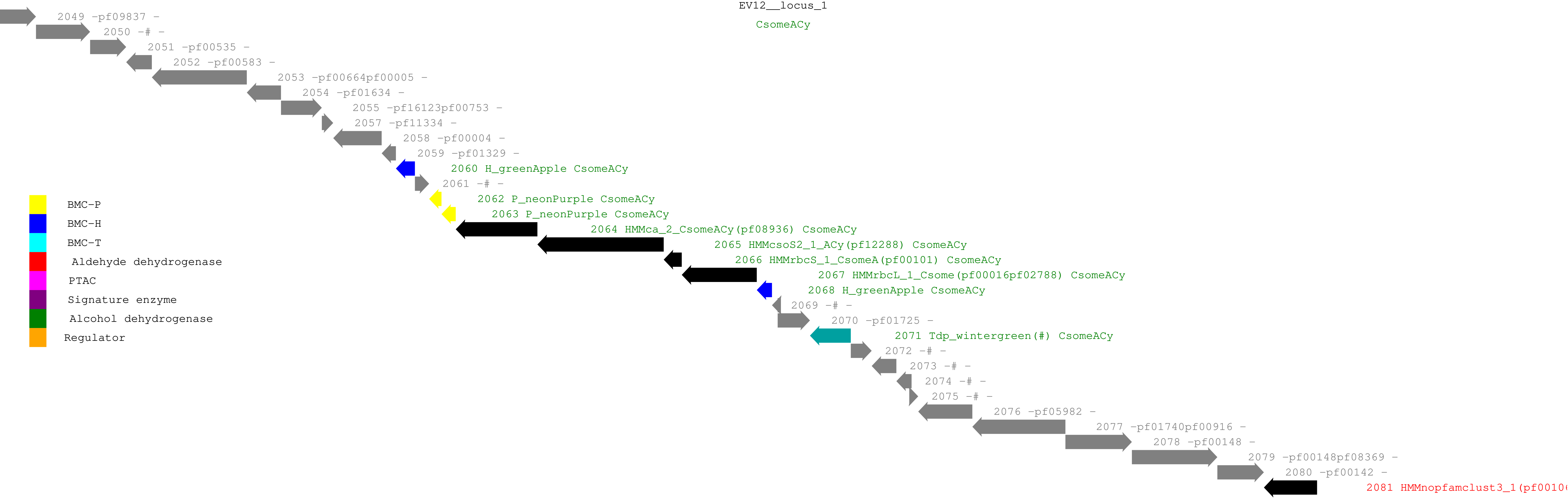

Prochlorococcus sp. MED-G132. A0A520PZ09_9PROC Unreviewed; 547 AA.

EVA79__locus_1

CsomeACy

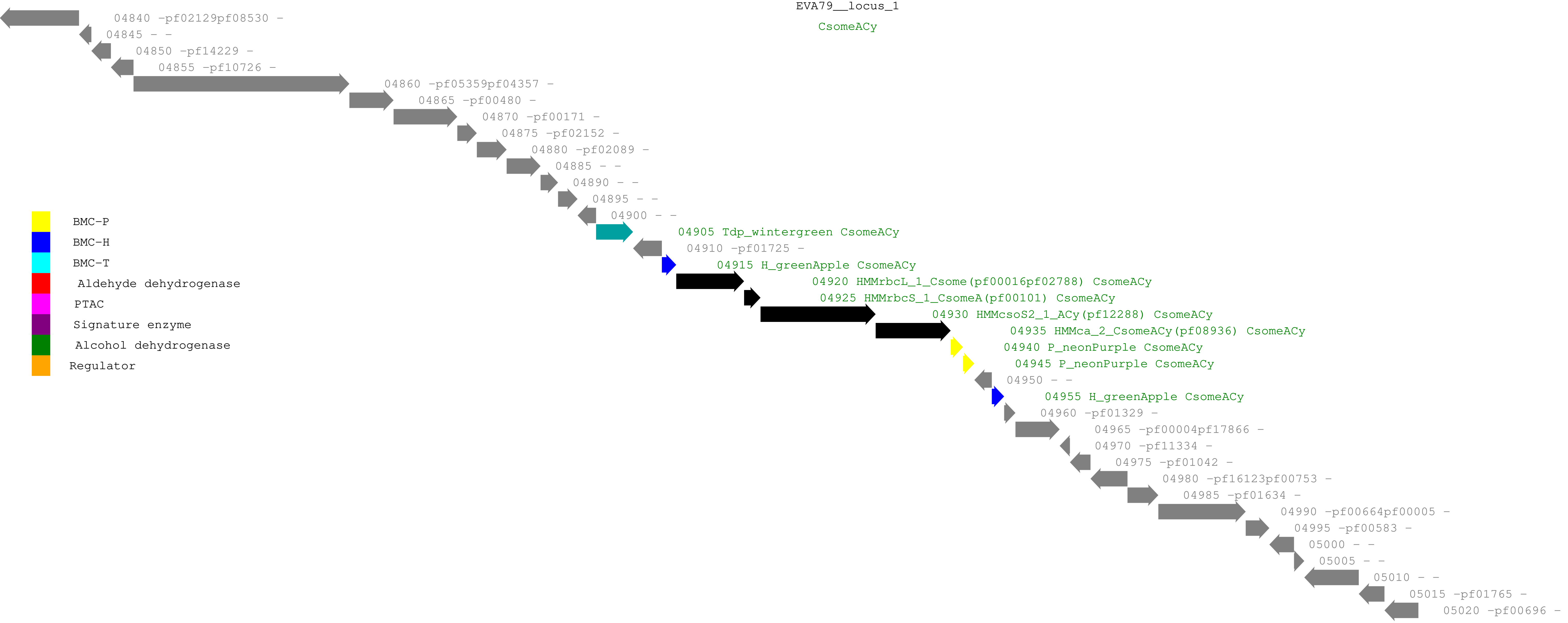

Synechococcus sp. MED-G134. Bacteria; Cyanobacteria; Synechococcales; Synechococcaceae; Synechococcus;

EVB07__locus_1_main_sat

CsomeACy

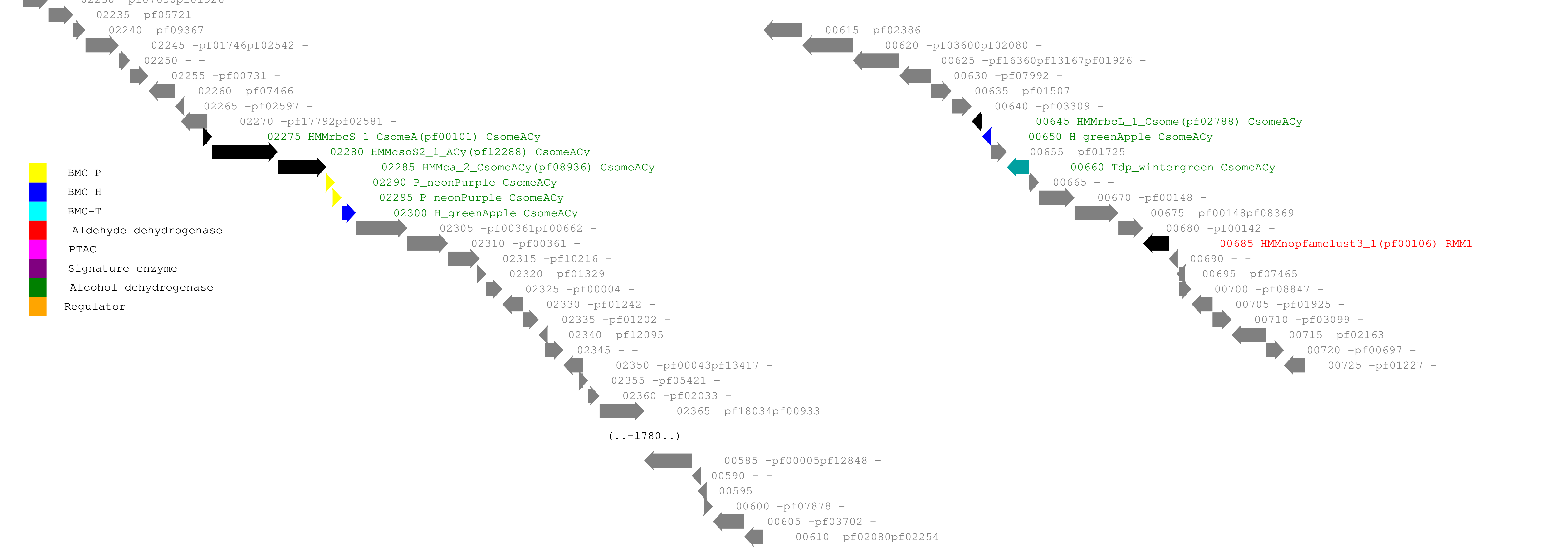

Synechococcus sp. RSCCF101. Bacteria; Cyanobacteria; Synechococcales; Synechococcaceae; Synechococcus;

EVJ50__locus_1

CsomeACy

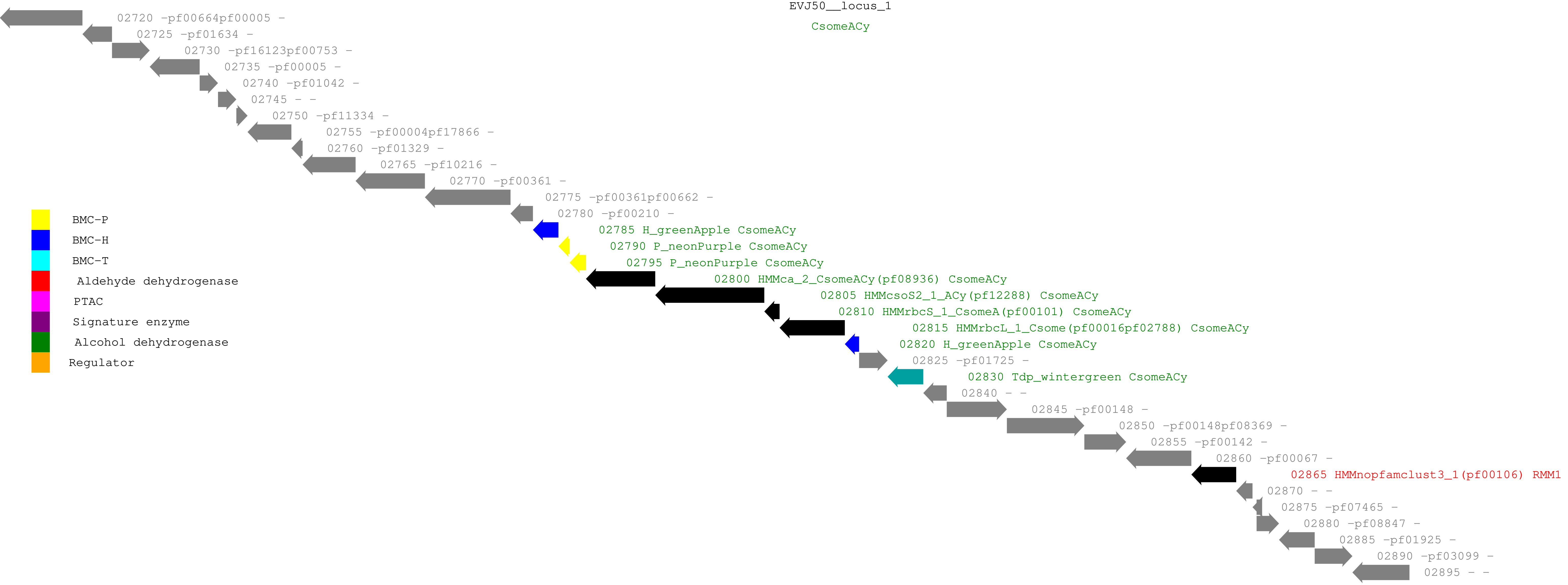

Prochlorococcus sp. MIT 0604. A0A089P2Z2_9PROC Unreviewed; 227 AA.

EW14__locus_1

CsomeACy

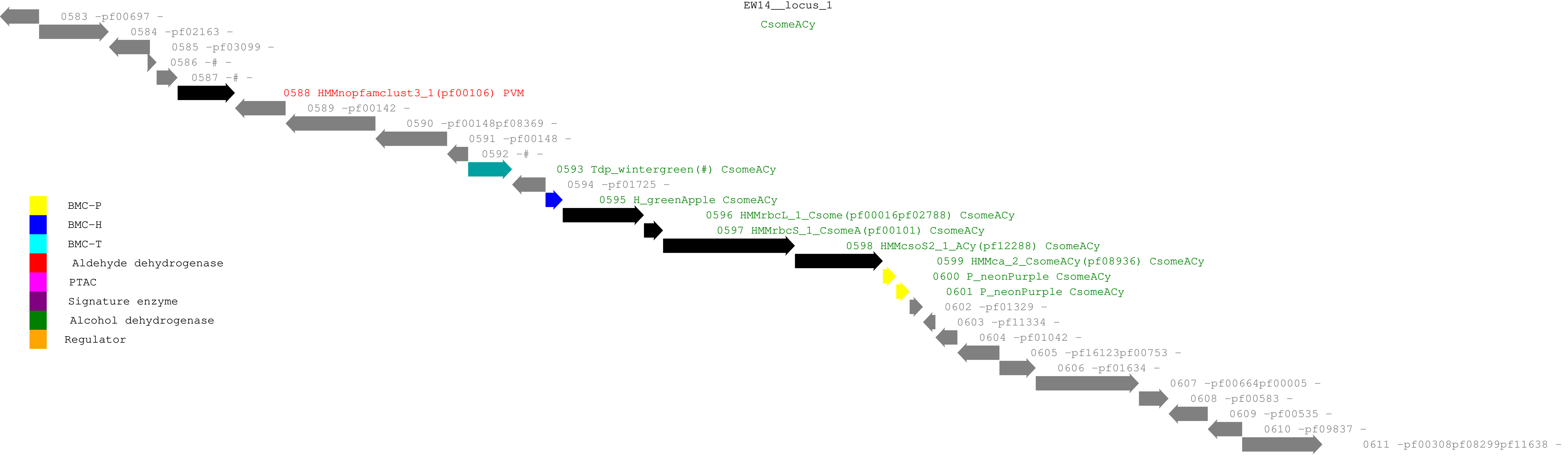

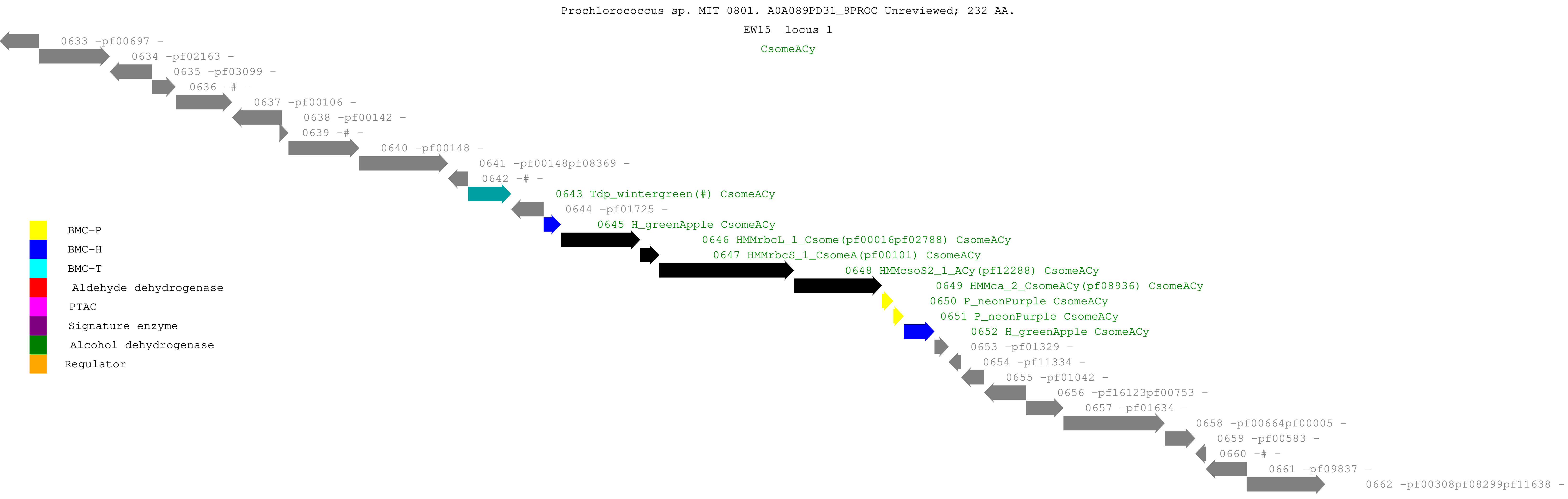

Prochlorococcus sp. MIT 0801. A0A089PD31_9PROC Unreviewed; 232 AA.
EW15__locus_1
CsomeACy
0633 -pf00697 -
0634 -pf02163 -
0635 -pf03099 -
0636 -# -
0637 -pf00106 -
0638 -pf00142 -
0639 -# -
0640 -pf00148 -
0641 -pf00148pf08369 -
0642 -# -
0643 Tdp_wintergreen(#) CsomeACy
0644 -pf01725 -
0645 H_greenApple CsomeACy
0646 HMMrbcL_1_Csome(pf00016pf02788) CsomeACy
0647 HMMrbcS_1_CsomeA(pf00101) CsomeACy
0648 HMMcsoS2_1_ACy(pf12288) CsomeACy
0649 HMMca_2_CsomeACy(pf08936) CsomeACy
0650 P_neonPurple CsomeACy
0651 P_neonPurple CsomeACy
0652 H_greenApple CsomeACy
0653 -pf01329 -
0654 -pf11334 -
0655 -pf01042 -
0656 -pf16123pf00753 -
0657 -pf01634 -
0658 -pf00664pf00005 -
0659 -pf00583 -
0660 -# -
0661 -pf09837 -
0662 -pf00308pf08299pf11638 -
BMC-P
BMC-H
BMC-T
Aldehyde dehydrogenase
PTAC
Signature enzyme
Alcohol dehydrogenase
Regulator

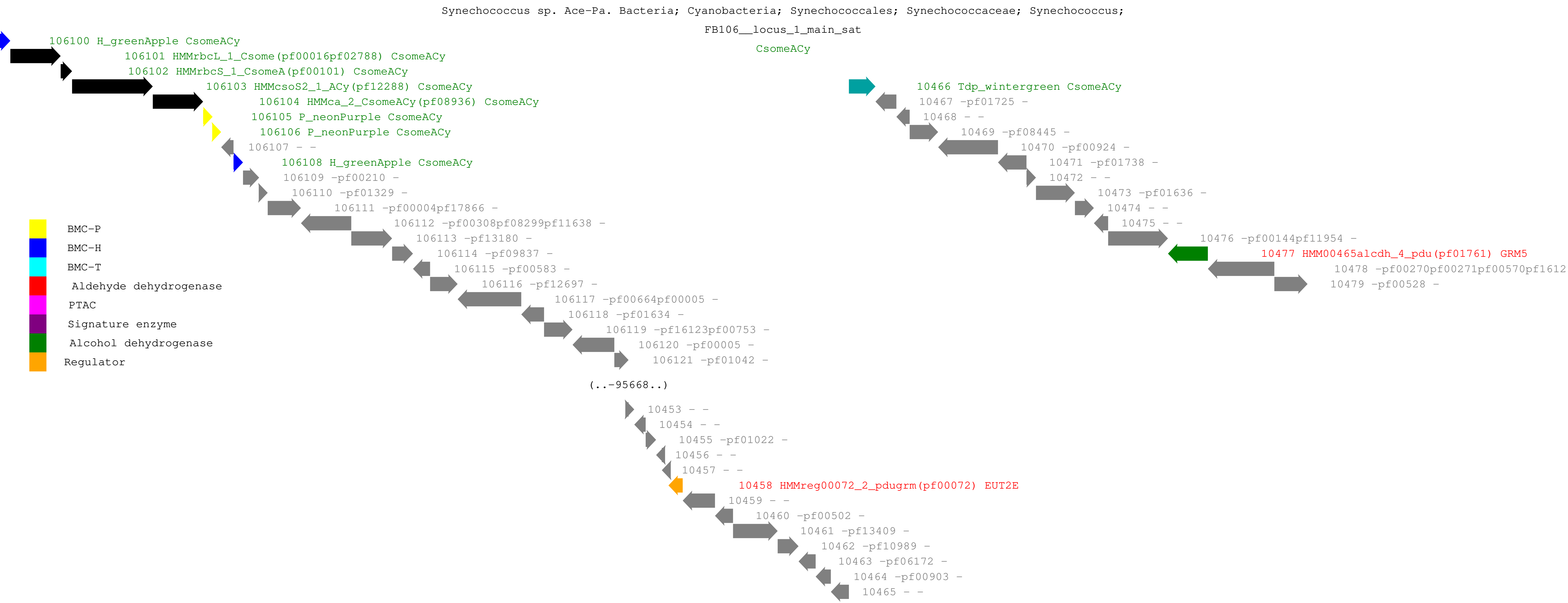
Synechococcus sp. Ace-Pa. Bacteria; Cyanobacteria; Synechococcales; Synechococcaceae; Synechococcus;
FB106__locus_1_main_sat
CsomeACy
106100 H_greenApple CsomeACy
106101 HMMrbcL_1_Csome(pf00016pf02788) CsomeACy
106102 HMMrbcS_1_CsomeA(pf00101) CsomeACy
106103 HMMcsoS2_1_ACy(pf12288) CsomeACy
106104 HMMca_2_CsomeACy(pf08936) CsomeACy
106105 P_neonPurple CsomeACy
106106 P_neonPurple CsomeACy
106107 - -
106108 H_greenApple CsomeACy
106109 -pf00210 -
106110 -pf01329 -
106111 -pf00004pf17866 -
106112 -pf00308pf08299pf11638 -
106113 -pf13180 -
106114 -pf09837 -
106115 -pf00583 -
106116 -pf12697 -
106117 -pf00664pf00005 -
106118 -pf01634 -
106119 -pf16123pf00753 -
106120 -pf00005 -
106121 -pf01042 -
(..-95668..)
10453 - -
10454 - -
10455 -pf01022 -
10456 - -
10457 - -
10458 HMMreg00072_2_pdugrm(pf00072) EUT2E
10459 - -
10460 -pf00502 -
10461 -pf13409 -
10462 -pf10989 -
10463 -pf06172 -
10464 -pf00903 -
10465 - -
10466 Tdp_wintergreen CsomeACy
10467 -pf01725 -
10468 - -
10469 -pf08445 -
10470 -pf00924 -
10471 -pf01738 -
10472 - -
10473 -pf01636 -
10474 - -
10475 - -
10476 -pf00144pf11954 -
10477 HMM00465alcdh_4_pdu(pf01761) GRM5
10478 -pf00270pf00271pf00570pf1612
10479 -pf00528 -
BMC-P
BMC-H
BMC-T
Aldehyde dehydrogenase
PTAC
Signature enzyme
Alcohol dehydrogenase
Regulator

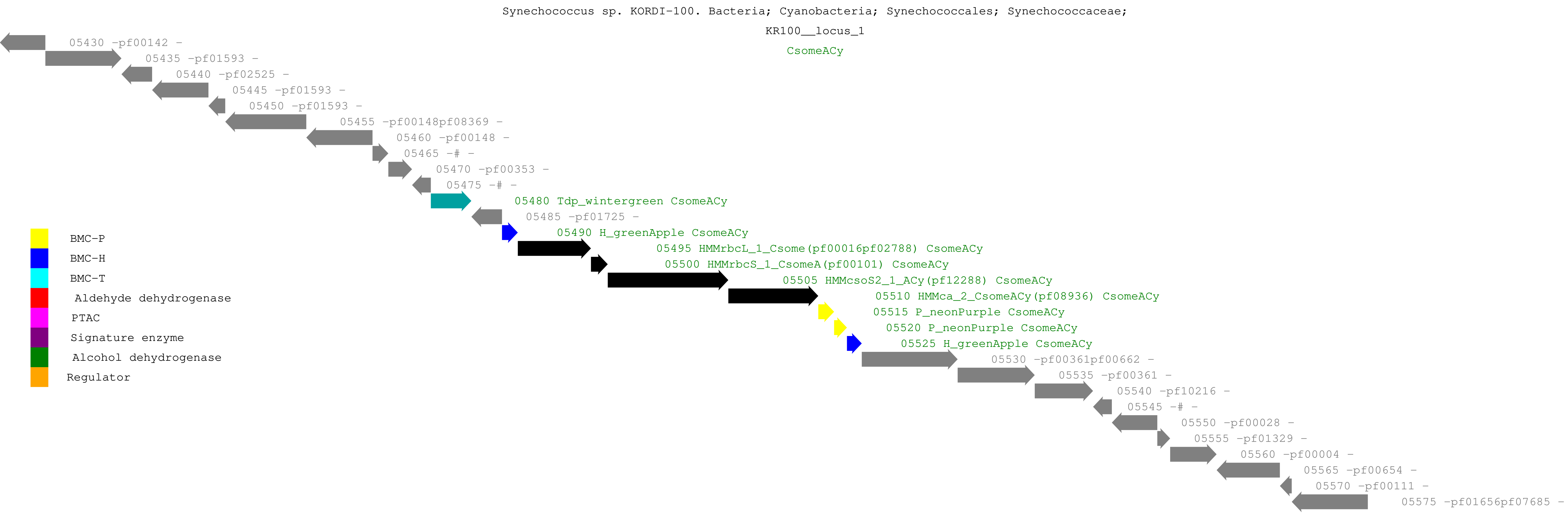
Synechococcus sp. KORDI-100. Bacteria; Cyanobacteria; Synechococcales; Synechococcaceae;
KR100__locus_1
CsomeACy
05430 -pf00142 -
05435 -pf01593 -
05440 -pf02525 -
05445 -pf01593 -
05450 -pf01593 -
05455 -pf00148pf08369 -
05460 -pf00148 -
05465 -# -
05470 -pf00353 -
05475 -# -
05480 Tdp_wintergreen CsomeACy
05485 -pf01725 -
05490 H_greenApple CsomeACy
05495 HMMrbcL_1_Csome(pf00016pf02788) CsomeACy
05500 HMMrbcS_1_CsomeA(pf00101) CsomeACy
05505 HMMcsoS2_1_ACy(pf12288) CsomeACy
05510 HMMca_2_CsomeACy(pf08936) CsomeACy
05515 P_neonPurple CsomeACy
05520 P_neonPurple CsomeACy
05525 H_greenApple CsomeACy
05530 -pf00361pf00662 -
05535 -pf00361 -
05540 -pf10216 -
05545 -# -
05550 -pf00028 -
05555 -pf01329 -
05560 -pf00004 -
05565 -pf00654 -
05570 -pf00111 -
05575 -pf01656pf07685 -
BMC-P
BMC-H
BMC-T
Aldehyde dehydrogenase
PTAC
Signature enzyme
Alcohol dehydrogenase
Regulator

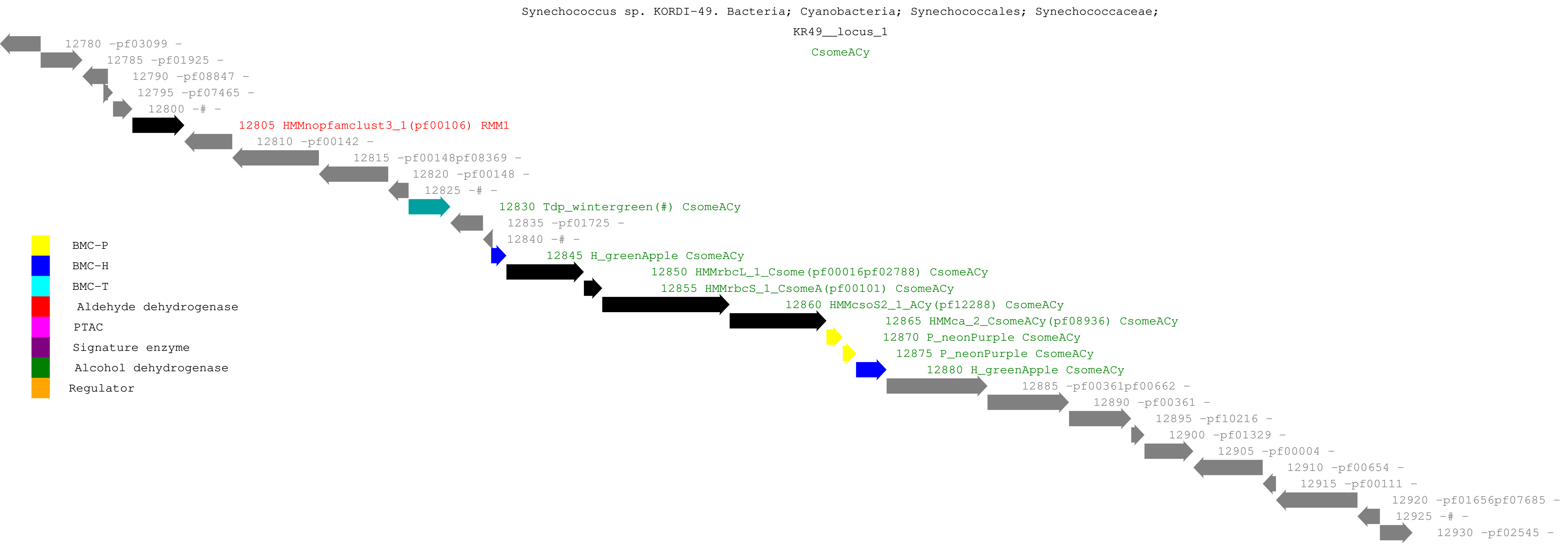
Synechococcus sp. KORDI-49. Bacteria; Cyanobacteria; Synechococcales; Synechococcaceae;
KR49__locus_1
CsomeACy
12780 -pf03099 -
12785 -pf01925 -
12790 -pf08847 -
12795 -pf07465 -
12800 -# -
12805 HMMnopfamclust3_1(pf00106) RMM1
12810 -pf00142 -
12815 -pf00148pf08369 -
12820 -pf00148 -
12825 -# -
12830 Tdp_wintergreen(#) CsomeACy
12835 -pf01725 -
12840 -# -
12845 H_greenApple CsomeACy
12850 HMMrbcL_1_Csome(pf00016pf02788) CsomeACy
12855 HMMrbcS_1_CsomeA(pf00101) CsomeACy
12860 HMMcsoS2_1_ACy(pf12288) CsomeACy
12865 HMMca_2_CsomeACy(pf08936) CsomeACy
12870 P_neonPurple CsomeACy
12875 P_neonPurple CsomeACy
12880 H_greenApple CsomeACy
12885 -pf00361pf00662 -
12890 -pf00361 -
12895 -pf10216 -
12900 -pf01329 -
12905 -pf00004 -
12910 -pf00654 -
12915 -pf00111 -
12920 -pf01656pf07685 -
12925 -# -
12930 -pf02545 -
BMC-P
BMC-H
BMC-T
Aldehyde dehydrogenase
PTAC
Signature enzyme
Alcohol dehydrogenase
Regulator

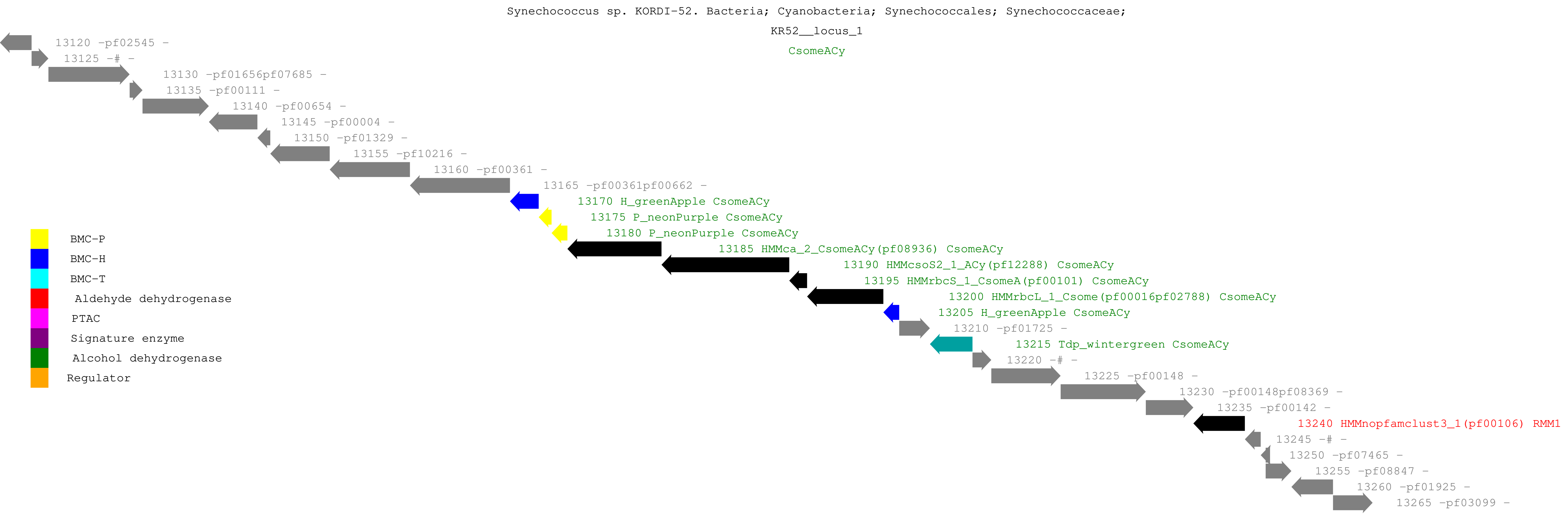
Synechococcus sp. KORDI-52. Bacteria; Cyanobacteria; Synechococcales; Synechococcaceae;
KR52__locus_1
CsomeACy
13120 -pf02545 -
13125 -# -
13130 -pf01656pf07685 -
13135 -pf00111 -
13140 -pf00654 -
13145 -pf00004 -
13150 -pf01329 -
13155 -pf10216 -
13160 -pf00361 -
13165 -pf00361pf00662 -
13170 H_greenApple CsomeACy
13175 P_neonPurple CsomeACy
13180 P_neonPurple CsomeACy
13185 HMMca_2_CsomeACy(pf08936) CsomeACy
13190 HMMcsoS2_1_ACy(pf12288) CsomeACy
13195 HMMrbcS_1_CsomeA(pf00101) CsomeACy
13200 HMMrbcL_1_Csome(pf00016pf02788) CsomeACy
13205 H_greenApple CsomeACy
13210 -pf01725 -
13215 Tdp_wintergreen CsomeACy
13220 -# -
13225 -pf00148 -
13230 -pf00148pf08369 -
13235 -pf00142 -
13240 HMMnopfamclust3_1(pf00106) RMM1
13245 -# -
13250 -pf07465 -
13255 -pf08847 -
13260 -pf01925 -
13265 -pf03099 -
BMC-P
BMC-H
BMC-T
Aldehyde dehydrogenase
PTAC
Signature enzyme
Alcohol dehydrogenase
Regulator

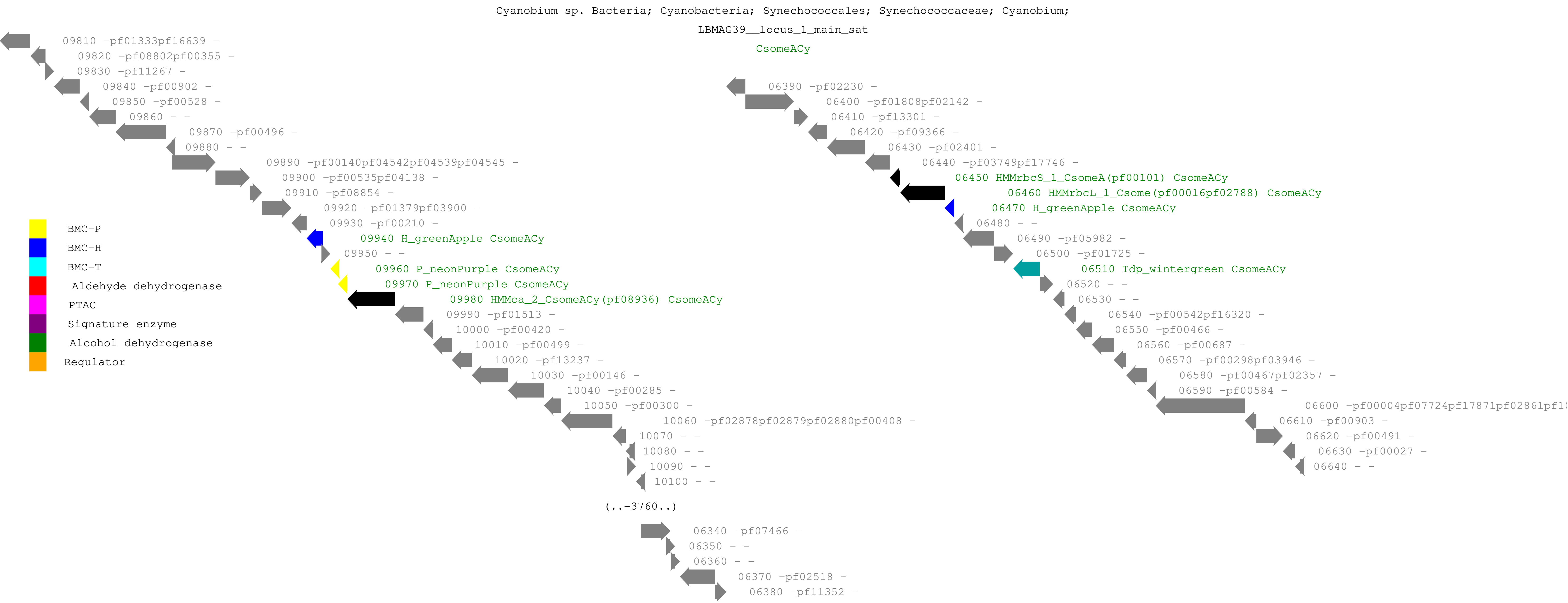
Cyanobium sp. Bacteria; Cyanobacteria; Synechococcales; Synechococcaceae; Cyanobium;
LBMAG39__locus_1_main_sat
CsomeACy
09810 -pf01333pf16639 -
09820 -pf08802pf00355 -
09830 -pf11267 -
09840 -pf00902 -
09850 -pf00528 -
09860 - -
09870 -pf00496 -
09880 - -
09890 -pf00140pf04542pf04539pf04545 -
09900 -pf00535pf04138 -
09910 -pf08854 -
09920 -pf01379pf03900 -
09930 -pf00210 -
09940 H_greenApple CsomeACy
09950 - -
09960 P_neonPurple CsomeACy
09970 P_neonPurple CsomeACy
09980 HMMca_2_CsomeACy(pf08936) CsomeACy
09990 -pf01513 -
10000 -pf00420 -
10010 -pf00499 -
10020 -pf13237 -
10030 -pf00146 -
10040 -pf00285 -
10050 -pf00300 -
10060 -pf02878pf02879pf02880pf00408 -
10070 - -
10080 - -
10090 - -
10100 - -
(..-3760..)
06340 -pf07466 -
06350 - -
06360 - -
06370 -pf02518 -
06380 -pf11352 -
06390 -pf02230 -
06400 -pf01808pf02142 -
06410 -pf13301 -
06420 -pf09366 -
06430 -pf02401 -
06440 -pf03749pf17746 -
06450 HMMrbcS_1_CsomeA(pf00101) CsomeACy
06460 HMMrbcL_1_Csome(pf00016pf02788) CsomeACy
06470 H_greenApple CsomeACy
06480 - -
06490 -pf05982 -
06500 -pf01725 -
06510 Tdp_wintergreen CsomeACy
06520 - -
06530 - -
06540 -pf00542pf16320 -
06550 -pf00466 -
06560 -pf00687 -
06570 -pf00298pf03946 -
06580 -pf00467pf02357 -
06590 -pf00584 -
06600 -pf00004pf07724pf17871pf02861pf1
06610 -pf00903 -
06620 -pf00491 -
06630 -pf00027 -
06640 - -
BMC-P
BMC-H
BMC-T
Aldehyde dehydrogenase
PTAC
Signature enzyme
Alcohol dehydrogenase
Regulator

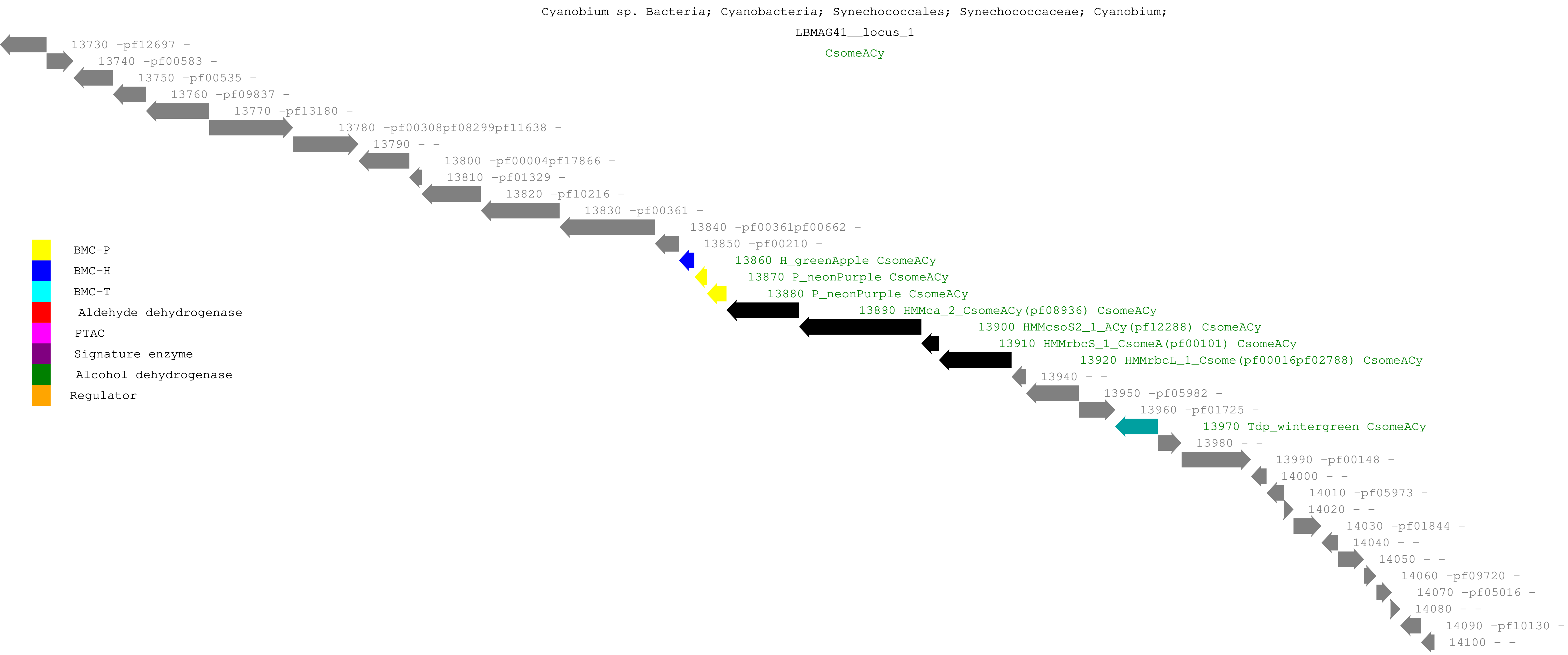
Cyanobium sp. Bacteria; Cyanobacteria; Synechococcales; Synechococcaceae; Cyanobium;
LBMAG41__locus_1
CsomeACy
13730 -pf12697 -
13740 -pf00583 -
13750 -pf00535 -
13760 -pf09837 -
13770 -pf13180 -
13780 -pf00308pf08299pf11638 -
13790 - -
13800 -pf00004pf17866 -
13810 -pf01329 -
13820 -pf10216 -
13830 -pf00361 -
13840 -pf00361pf00662 -
13850 -pf00210 -
13860 H_greenApple CsomeACy
13870 P_neonPurple CsomeACy
13880 P_neonPurple CsomeACy
13890 HMMca_2_CsomeACy(pf08936) CsomeACy
13900 HMMcsoS2_1_ACy(pf12288) CsomeACy
13910 HMMrbcS_1_CsomeA(pf00101) CsomeACy
13920 HMMrbcL_1_Csome(pf00016pf02788) CsomeACy
13940 - -
13950 -pf05982 -
13960 -pf01725 -
13970 Tdp_wintergreen CsomeACy
13980 - -
13990 -pf00148 -
14000 - -
14010 -pf05973 -
14020 - -
14030 -pf01844 -
14040 - -
14050 - -
14060 -pf09720 -
14070 -pf05016 -
14080 - -
14090 -pf10130 -
14100 - -
BMC-P
BMC-H
BMC-T
Aldehyde dehydrogenase
PTAC
Signature enzyme
Alcohol dehydrogenase
Regulator

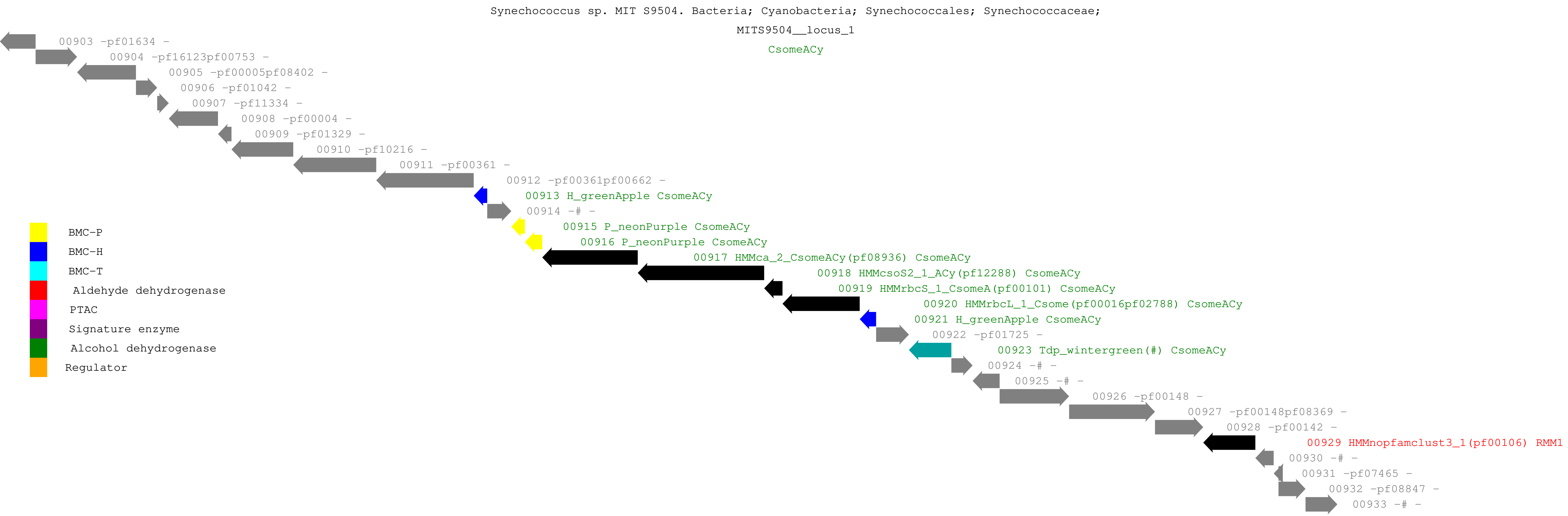
Synechococcus sp. MIT S9504. Bacteria; Cyanobacteria; Synechococcales; Synechococcaceae;
MITS9504__locus_1
CsomeACy
00903 -pf01634 -
00904 -pf16123pf00753 -
00905 -pf00005pf08402 -
00906 -pf01042 -
00907 -pf11334 -
00908 -pf00004 -
00909 -pf01329 -
00910 -pf10216 -
00911 -pf00361 -
00912 -pf00361pf00662 -
00913 H_greenApple CsomeACy
00914 -# -
00915 P_neonPurple CsomeACy
00916 P_neonPurple CsomeACy
00917 HMMca_2_CsomeACy(pf08936) CsomeACy
00918 HMMcsoS2_1_ACy(pf12288) CsomeACy
00919 HMMrbcS_1_CsomeA(pf00101) CsomeACy
00920 HMMrbcL_1_Csome(pf00016pf02788) CsomeACy
00921 H_greenApple CsomeACy
00922 -pf01725 -
00923 Tdp_wintergreen(#) CsomeACy
00924 -# -
00925 -# -
00926 -pf00148 -
00927 -pf00148pf08369 -
00928 -pf00142 -
00929 HMMnopfamclust3_1(pf00106) RMM1
00930 -# -
00931 -pf07465 -
00932 -pf08847 -
00933 -# -
BMC-P
BMC-H
BMC-T
Aldehyde dehydrogenase
PTAC
Signature enzyme
Alcohol dehydrogenase
Regulator

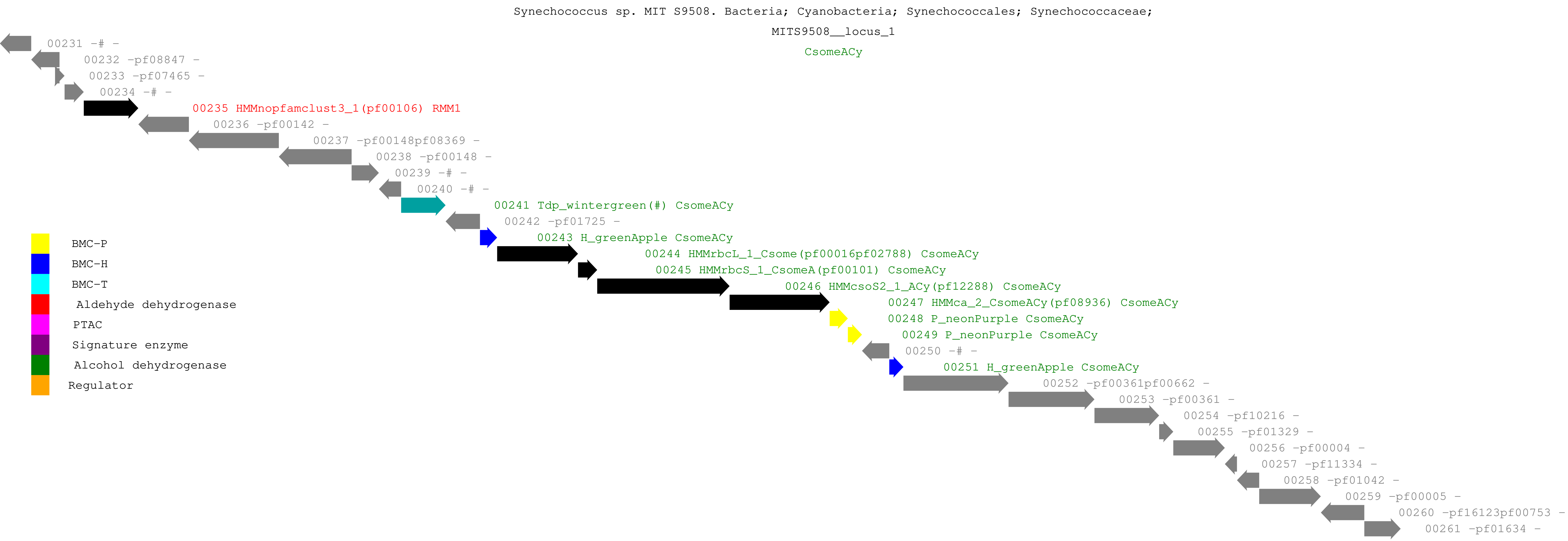

Synechococcus sp. MIT S9508. Bacteria; Cyanobacteria; Synechococcales; Synechococcaceae;
MITS9508__locus_1
CsomeACy
00231 -# -
00232 -pf08847 -
00233 -pf07465 -
00234 -# -
00235 HMMnopfamclust3_1(pf00106) RMM1
00236 -pf00142 -
00237 -pf00148pf08369 -
00238 -pf00148 -
00239 -# -
00240 -# -
00241 Tdp_wintergreen(#) CsomeACy
00242 -pf01725 -
00243 H_greenApple CsomeACy
00244 HMMrbcL_1_Csome(pf00016pf02788) CsomeACy
00245 HMMrbcS_1_CsomeA(pf00101) CsomeACy
00246 HMMcsoS2_1_ACy(pf12288) CsomeACy
00247 HMMca_2_CsomeACy(pf08936) CsomeACy
00248 P_neonPurple CsomeACy
00249 P_neonPurple CsomeACy
00250 -# -
00251 H_greenApple CsomeACy
00252 -pf00361pf00662 -
00253 -pf00361 -
00254 -pf10216 -
00255 -pf01329 -
00256 -pf00004 -
00257 -pf11334 -
00258 -pf01042 -
00259 -pf00005 -
00260 -pf16123pf00753 -
00261 -pf01634 -
BMC-P
BMC-H
BMC-T
Aldehyde dehydrogenase
PTAC
Signature enzyme
Alcohol dehydrogenase
Regulator

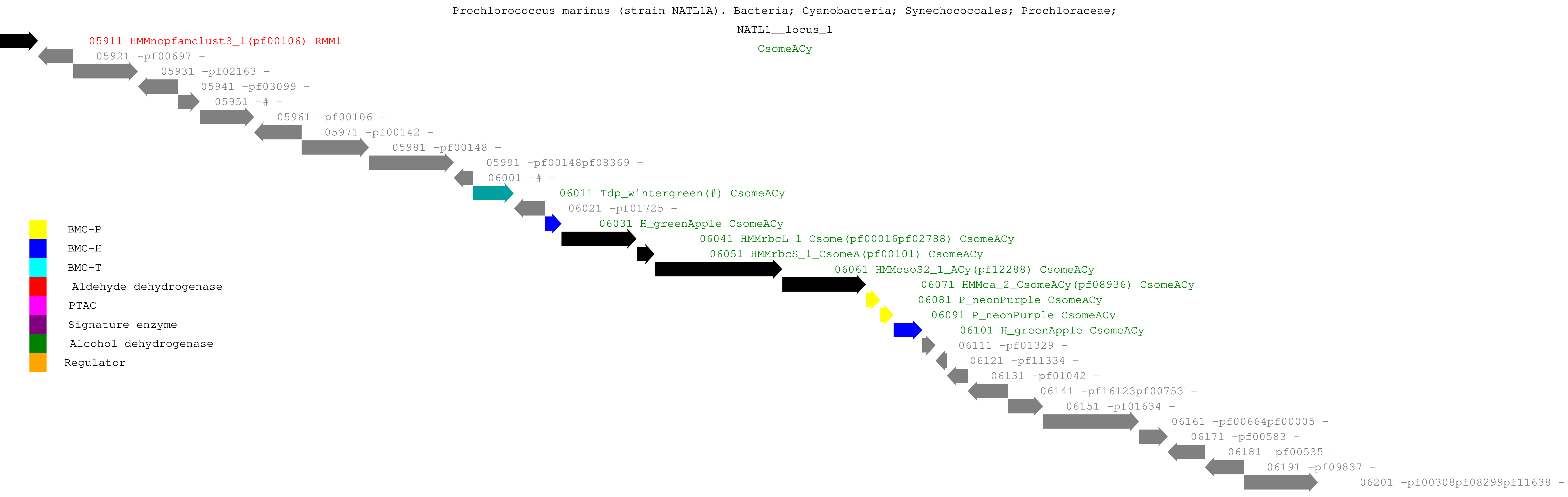

Prochlorococcus marinus (strain NATL1A). Bacteria; Cyanobacteria; Synechococcales; Prochloraceae;
NATL1__locus_1
CsomeACy
05911 HMMnopfamclust3_1(pf00106) RMM1
05921 -pf00697 -
05931 -pf02163 -
05941 -pf03099 -
05951 -# -
05961 -pf00106 -
05971 -pf00142 -
05981 -pf00148 -
05991 -pf00148pf08369 -
06001 -# -
06011 Tdp_wintergreen(#) CsomeACy
06021 -pf01725 -
06031 H_greenApple CsomeACy
06041 HMMrbcL_1_Csome(pf00016pf02788) CsomeACy
06051 HMMrbcS_1_CsomeA(pf00101) CsomeACy
06061 HMMcsoS2_1_ACy(pf12288) CsomeACy
06071 HMMca_2_CsomeACy(pf08936) CsomeACy
06081 P_neonPurple CsomeACy
06091 P_neonPurple CsomeACy
06101 H_greenApple CsomeACy
06111 -pf01329 -
06121 -pf11334 -
06131 -pf01042 -
06141 -pf16123pf00753 -
06151 -pf01634 -
06161 -pf00664pf00005 -
06171 -pf00583 -
06181 -pf00535 -
06191 -pf09837 -
06201 -pf00308pf08299pf11638 -
BMC-P
BMC-H
BMC-T
Aldehyde dehydrogenase
PTAC
Signature enzyme
Alcohol dehydrogenase
Regulator

cyanobiont of Ornithocercus magnificus. Bacteria; Cyanobacteria; Synechococcales.

OMCYN__locus_1

CsomeACy

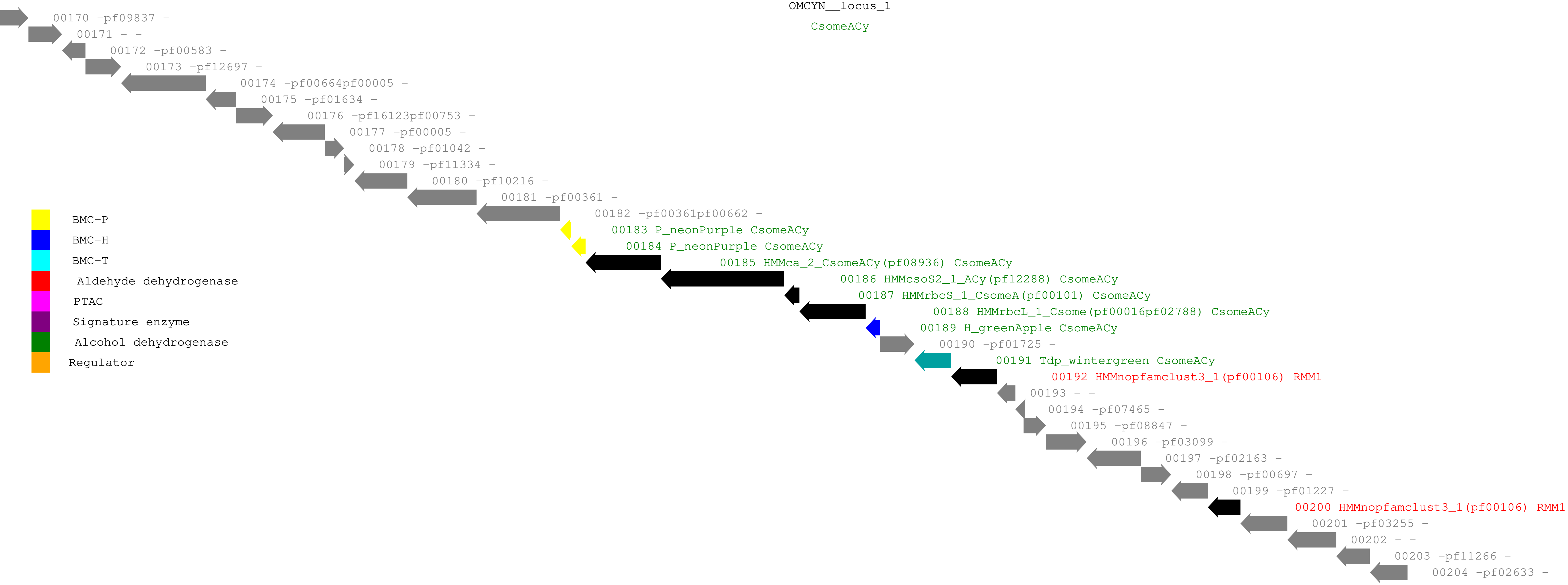

Prochlorococcus marinus str. MIT 9202. Bacteria; Cyanobacteria; Synechococcales; Prochloraceae;

P9202__locus_fragmented

CsomeACy

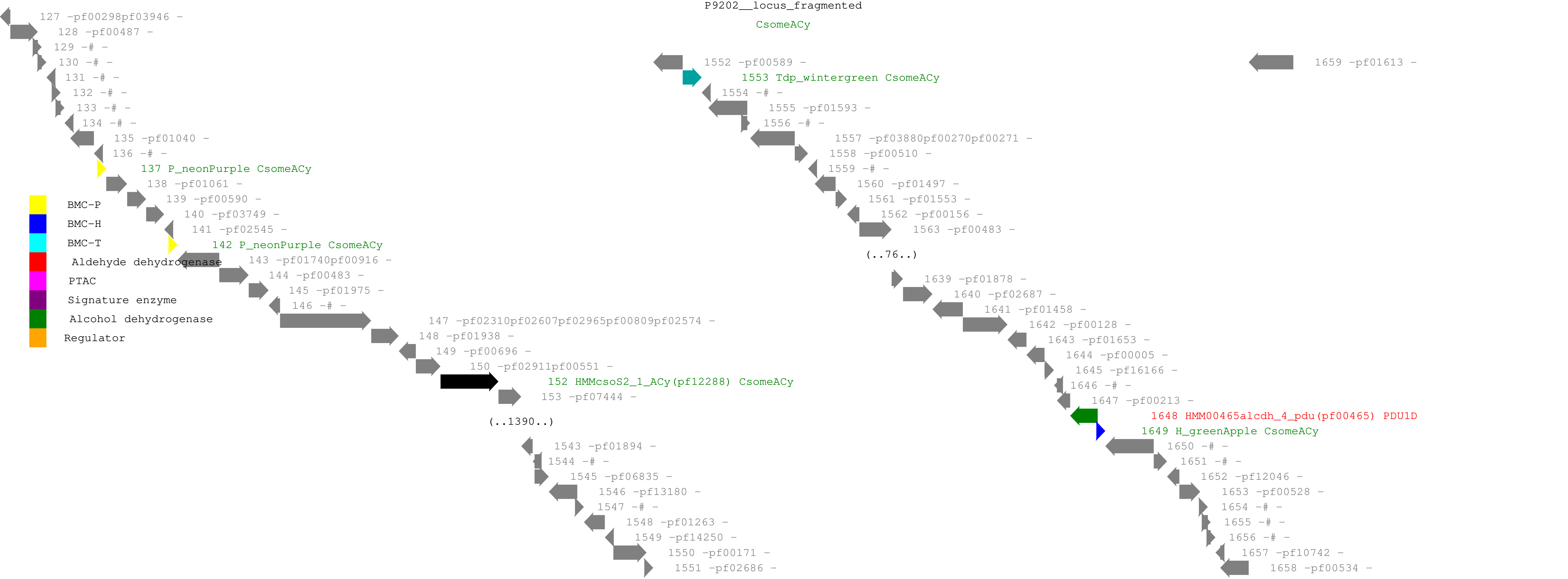

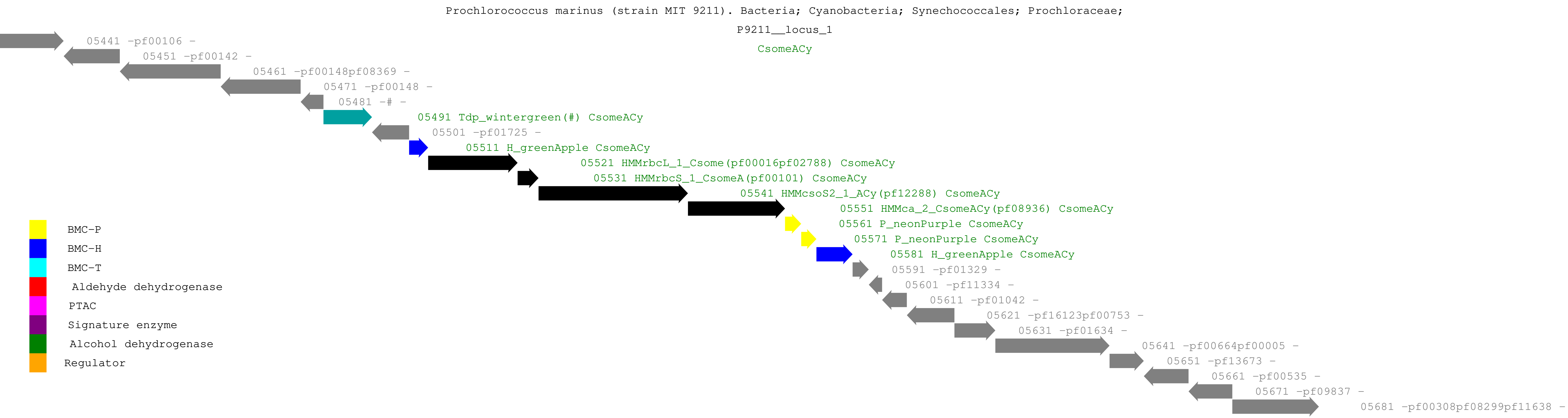
Prochlorococcus marinus (strain MIT 9211). Bacteria; Cyanobacteria; Synechococcales; Prochloraceae;
P9211__locus_1
CsomeACy
05441 -pf00106 -
05451 -pf00142 -
05461 -pf00148pf08369 -
05471 -pf00148 -
05481 -# -
05491 Tdp_wintergreen(#) CsomeACy
05501 -pf01725 -
05511 H_greenApple CsomeACy
05521 HMMrbcL_1_Csome(pf00016pf02788) CsomeACy
05531 HMMrbcS_1_CsomeA(pf00101) CsomeACy
05541 HMMcsoS2_1_ACy(pf12288) CsomeACy
05551 HMMca_2_CsomeACy(pf08936) CsomeACy
05561 P_neonPurple CsomeACy
05571 P_neonPurple CsomeACy
05581 H_greenApple CsomeACy
05591 -pf01329 -
05601 -pf11334 -
05611 -pf01042 -
05621 -pf16123pf00753 -
05631 -pf01634 -
05641 -pf00664pf00005 -
05651 -pf13673 -
05661 -pf00535 -
05671 -pf09837 -
05681 -pf00308pf08299pf11638 -
BMC-P
BMC-H
BMC-T
Aldehyde dehydrogenase
PTAC
Signature enzyme
Alcohol dehydrogenase
Regulator

Prochlorococcus marinus (strain MIT 9215). Bacteria; Cyanobacteria; Synechococcales; Prochloraceae;

P9215__locus_1

CsomeACy

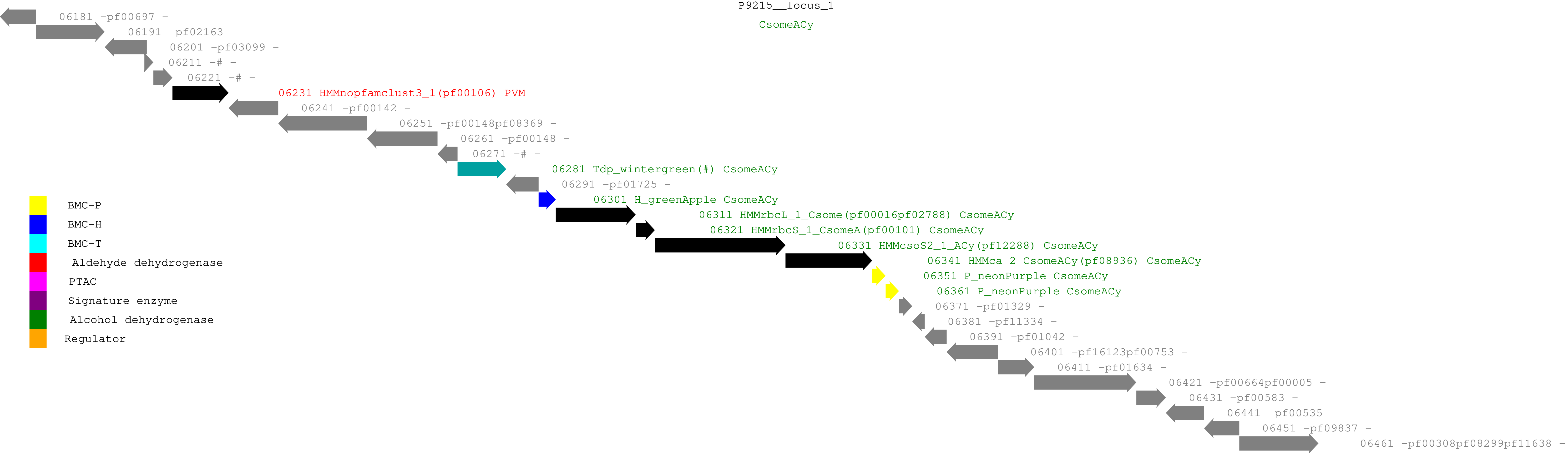

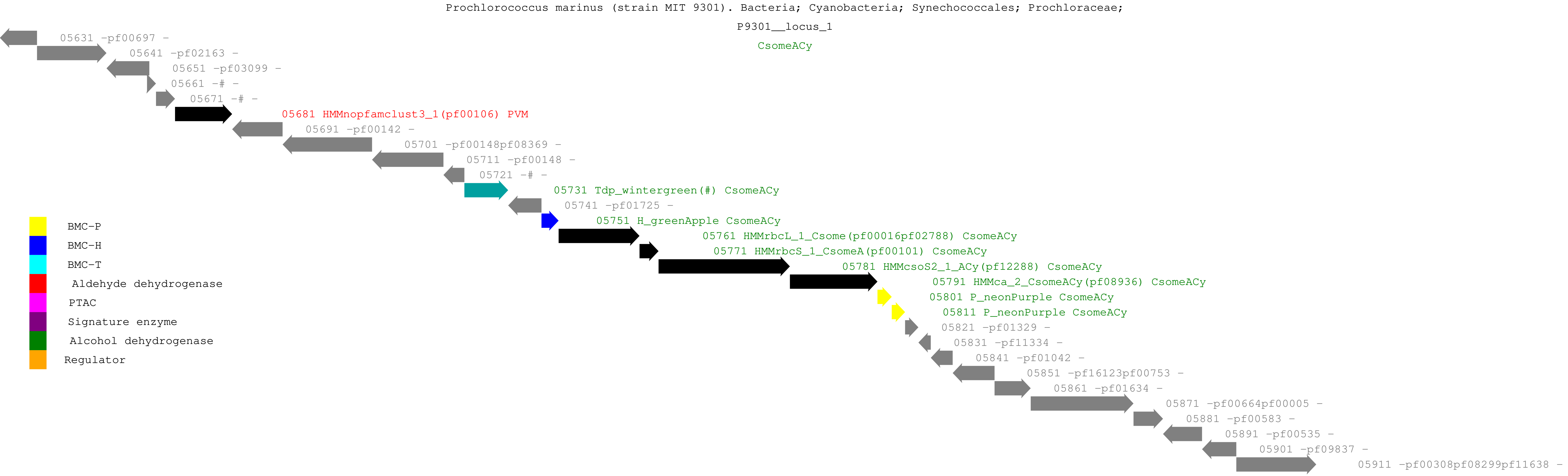
Prochlorococcus marinus (strain MIT 9301). Bacteria; Cyanobacteria; Synechococcales; Prochloraceae;
P9301__locus_1
CsomeACy
05631 -pf00697 -
05641 -pf02163 -
05651 -pf03099 -
05661 -# -
05671 -# -
05681 HMMnopfamclust3_1(pf00106) PVM
05691 -pf00142 -
05701 -pf00148pf08369 -
05711 -pf00148 -
05721 -# -
05731 Tdp_wintergreen(#) CsomeACy
05741 -pf01725 -
05751 H_greenApple CsomeACy
05761 HMMrbcL_1_Csome(pf00016pf02788) CsomeACy
05771 HMMrbcS_1_CsomeA(pf00101) CsomeACy
05781 HMMcsoS2_1_ACy(pf12288) CsomeACy
05791 HMMca_2_CsomeACy(pf08936) CsomeACy
05801 P_neonPurple CsomeACy
05811 P_neonPurple CsomeACy
05821 -pf01329 -
05831 -pf11334 -
05841 -pf01042 -
05851 -pf16123pf00753 -
05861 -pf01634 -
05871 -pf00664pf00005 -
05881 -pf00583 -
05891 -pf00535 -
05901 -pf09837 -
05911 -pf00308pf08299pf11638 -
BMC-P
BMC-H
BMC-T
Aldehyde dehydrogenase
PTAC
Signature enzyme
Alcohol dehydrogenase
Regulator

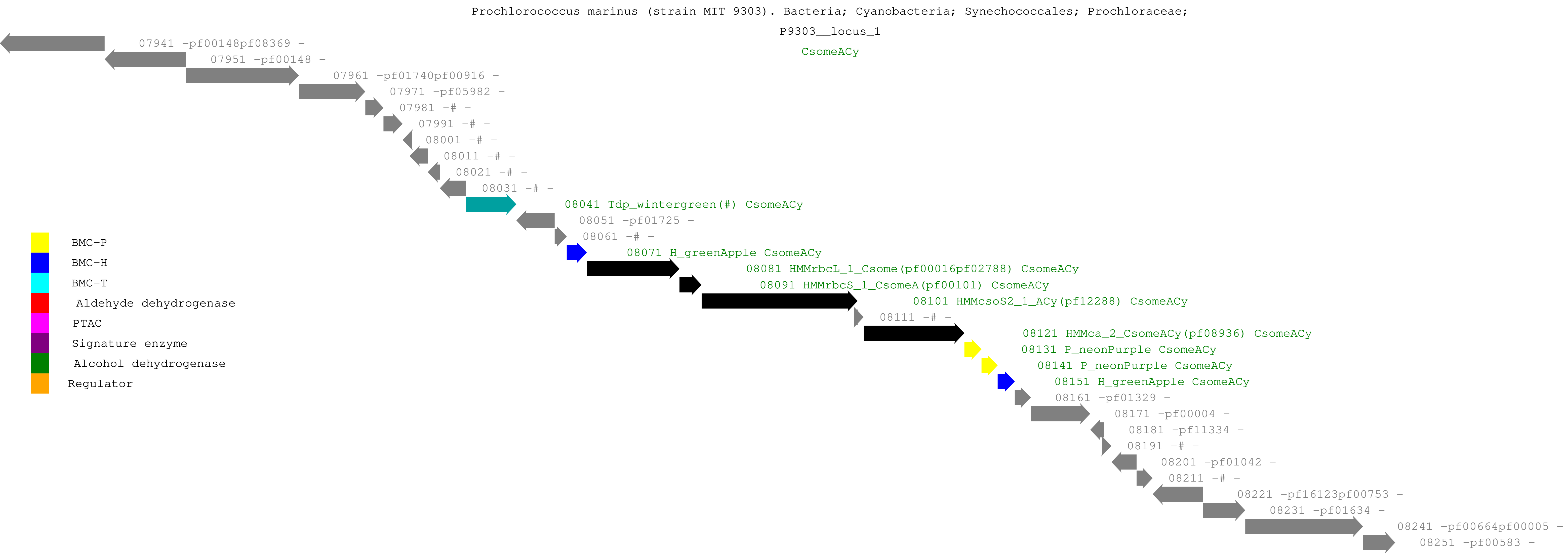

Prochlorococcus marinus (strain MIT 9303). Bacteria; Cyanobacteria; Synechococcales; Prochloraceae;
P9303__locus_1
CsomeACy
07941 -pf00148pf08369 -
07951 -pf00148 -
07961 -pf01740pf00916 -
07971 -pf05982 -
07981 -# -
07991 -# -
08001 -# -
08011 -# -
08021 -# -
08031 -# -
08041 Tdp_wintergreen(#) CsomeACy
08051 -pf01725 -
08061 -# -
08071 H_greenApple CsomeACy
08081 HMMrbcL_1_Csome(pf00016pf02788) CsomeACy
08091 HMMrbcS_1_CsomeA(pf00101) CsomeACy
08101 HMMcsoS2_1_ACy(pf12288) CsomeACy
08111 -# -
08121 HMMca_2_CsomeACy(pf08936) CsomeACy
08131 P_neonPurple CsomeACy
08141 P_neonPurple CsomeACy
08151 H_greenApple CsomeACy
08161 -pf01329 -
08171 -pf00004 -
08181 -pf11334 -
08191 -# -
08201 -pf01042 -
08211 -# -
08221 -pf16123pf00753 -
08231 -pf01634 -
08241 -pf00664pf00005 -
08251 -pf00583 -
BMC-P
BMC-H
BMC-T
Aldehyde dehydrogenase
PTAC
Signature enzyme
Alcohol dehydrogenase
Regulator

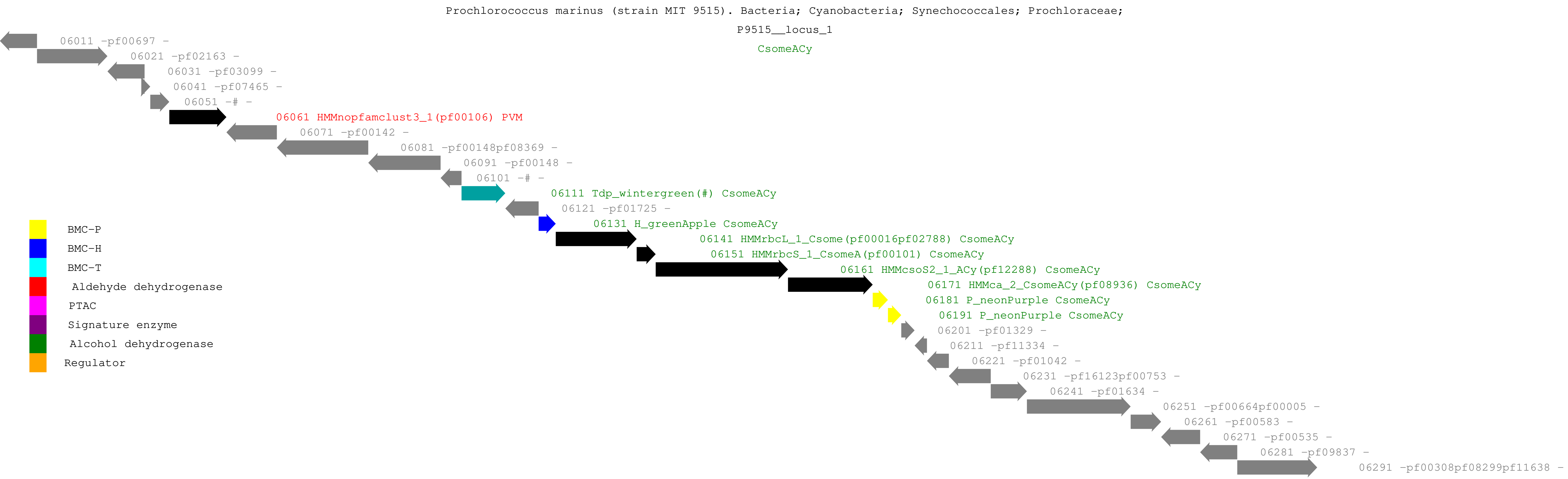
Prochlorococcus marinus (strain MIT 9515). Bacteria; Cyanobacteria; Synechococcales; Prochloraceae;
P9515__locus_1
CsomeACy
06011 -pf00697 -
06021 -pf02163 -
06031 -pf03099 -
06041 -pf07465 -
06051 -# -
06061 HMMnopfamclust3_1(pf00106) PVM
06071 -pf00142 -
06081 -pf00148pf08369 -
06091 -pf00148 -
06101 -# -
06111 Tdp_wintergreen(#) CsomeACy
06121 -pf01725 -
06131 H_greenApple CsomeACy
06141 HMMrbcL_1_Csome(pf00016pf02788) CsomeACy
06151 HMMrbcS_1_CsomeA(pf00101) CsomeACy
06161 HMMcsoS2_1_ACy(pf12288) CsomeACy
06171 HMMca_2_CsomeACy(pf08936) CsomeACy
06181 P_neonPurple CsomeACy
06191 P_neonPurple CsomeACy
06201 -pf01329 -
06211 -pf11334 -
06221 -pf01042 -
06231 -pf16123pf00753 -
06241 -pf01634 -
06251 -pf00664pf00005 -
06261 -pf00583 -
06271 -pf00535 -
06281 -pf09837 -
06291 -pf00308pf08299pf11638 -
BMC-P
BMC-H
BMC-T
Aldehyde dehydrogenase
PTAC
Signature enzyme
Alcohol dehydrogenase
Regulator

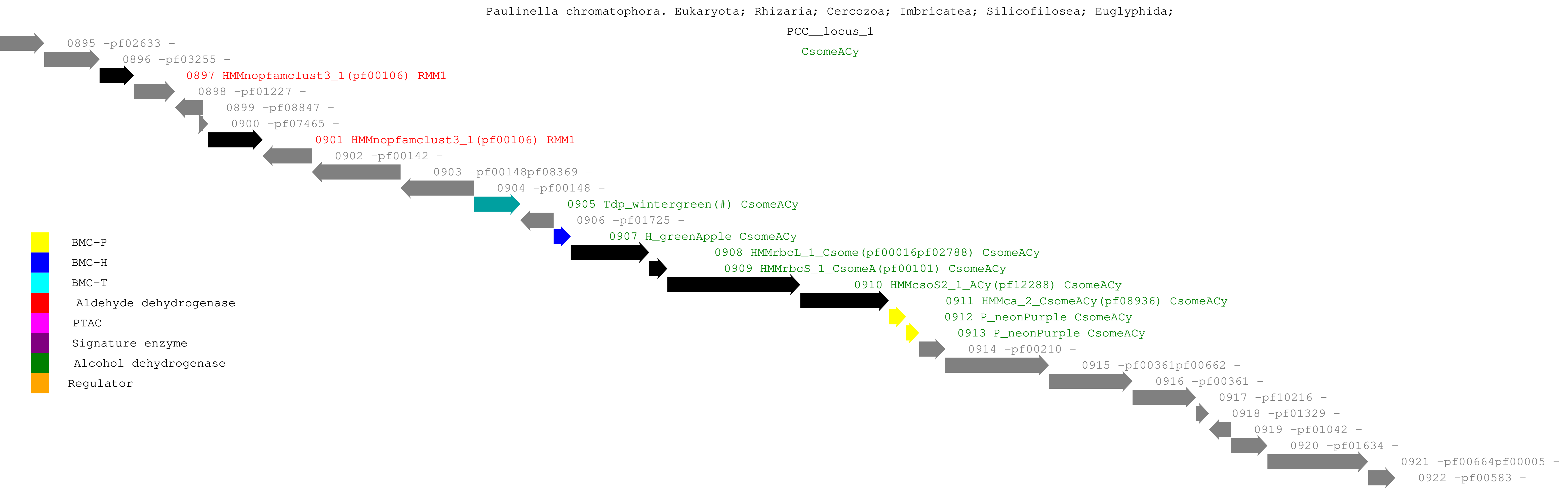
Paulinella chromatophora. Eukaryota; Rhizaria; Cercozoa; Imbricatea; Silicofilosea; Euglyphida;
PCC__locus_1
CsomeACy
0895 -pf02633 -
0896 -pf03255 -
0897 HMMnopfamclust3_1(pf00106) RMM1
0898 -pf01227 -
0899 -pf08847 -
0900 -pf07465 -
0901 HMMnopfamclust3_1(pf00106) RMM1
0902 -pf00142 -
0903 -pf00148pf08369 -
0904 -pf00148 -
0905 Tdp_wintergreen(#) CsomeACy
0906 -pf01725 -
0907 H_greenApple CsomeACy
0908 HMMrbcL_1_Csome(pf00016pf02788) CsomeACy
0909 HMMrbcS_1_CsomeA(pf00101) CsomeACy
0910 HMMcsoS2_1_ACy(pf12288) CsomeACy
0911 HMMca_2_CsomeACy(pf08936) CsomeACy
0912 P_neonPurple CsomeACy
0913 P_neonPurple CsomeACy
0914 -pf00210 -
0915 -pf00361pf00662 -
0916 -pf00361 -
0917 -pf10216 -
0918 -pf01329 -
0919 -pf01042 -
0920 -pf01634 -
0921 -pf00664pf00005 -
0922 -pf00583 -
BMC-P
BMC-H
BMC-T
Aldehyde dehydrogenase
PTAC
Signature enzyme
Alcohol dehydrogenase
Regulator

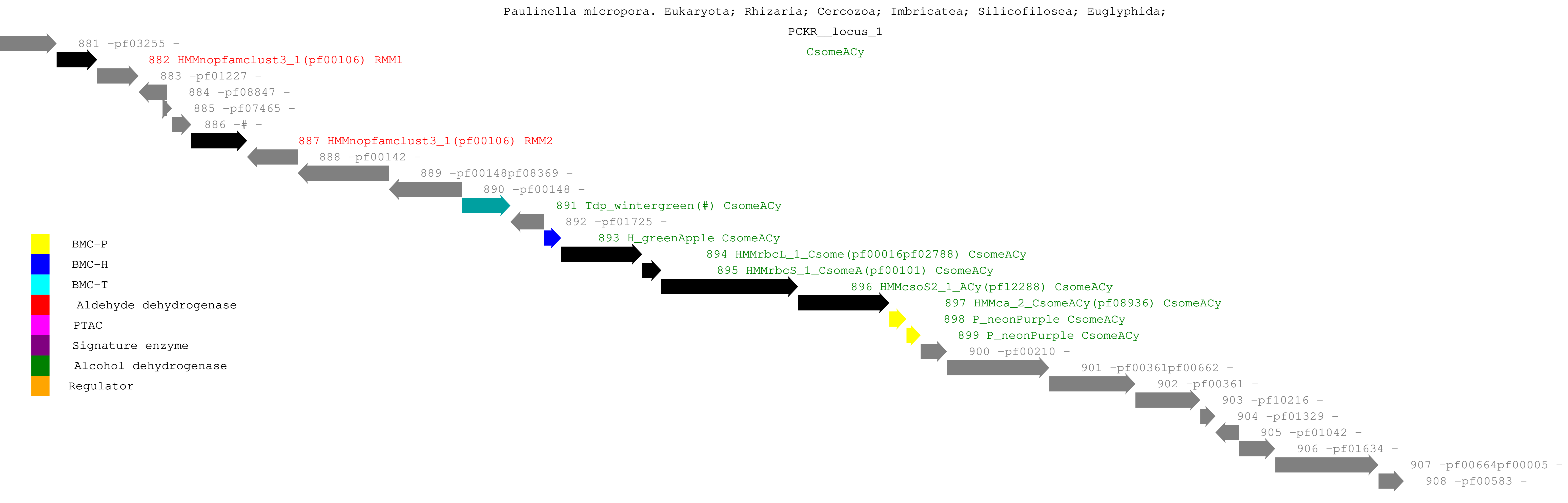

Paulinella micropora. Eukaryota; Rhizaria; Cercozoa; Imbricatea; Silicofilosea; Euglyphida;
PCKR__locus_1
CsomeACy
881 -pf03255 -
882 HMMnopfamclust3_1(pf00106) RMM1
883 -pf01227 -
884 -pf08847 -
885 -pf07465 -
886 -# -
887 HMMnopfamclust3_1(pf00106) RMM2
888 -pf00142 -
889 -pf00148pf08369 -
890 -pf00148 -
891 Tdp_wintergreen(#) CsomeACy
892 -pf01725 -
893 H_greenApple CsomeACy
894 HMMrbcL_1_Csome(pf00016pf02788) CsomeACy
895 HMMrbcS_1_CsomeA(pf00101) CsomeACy
896 HMMcsoS2_1_ACy(pf12288) CsomeACy
897 HMMca_2_CsomeACy(pf08936) CsomeACy
898 P_neonPurple CsomeACy
899 P_neonPurple CsomeACy
900 -pf00210 -
901 -pf00361pf00662 -
902 -pf00361 -
903 -pf10216 -
904 -pf01329 -
905 -pf01042 -
906 -pf01634 -
907 -pf00664pf00005 -
908 -pf00583 -
BMC-P
BMC-H
BMC-T
Aldehyde dehydrogenase
PTAC
Signature enzyme
Alcohol dehydrogenase
Regulator

Prochlorococcus sp. MIT 1303. A0A163G6N1_9PROC Unreviewed; 105 AA.

PMIT1303__locus_1

CsomeACy

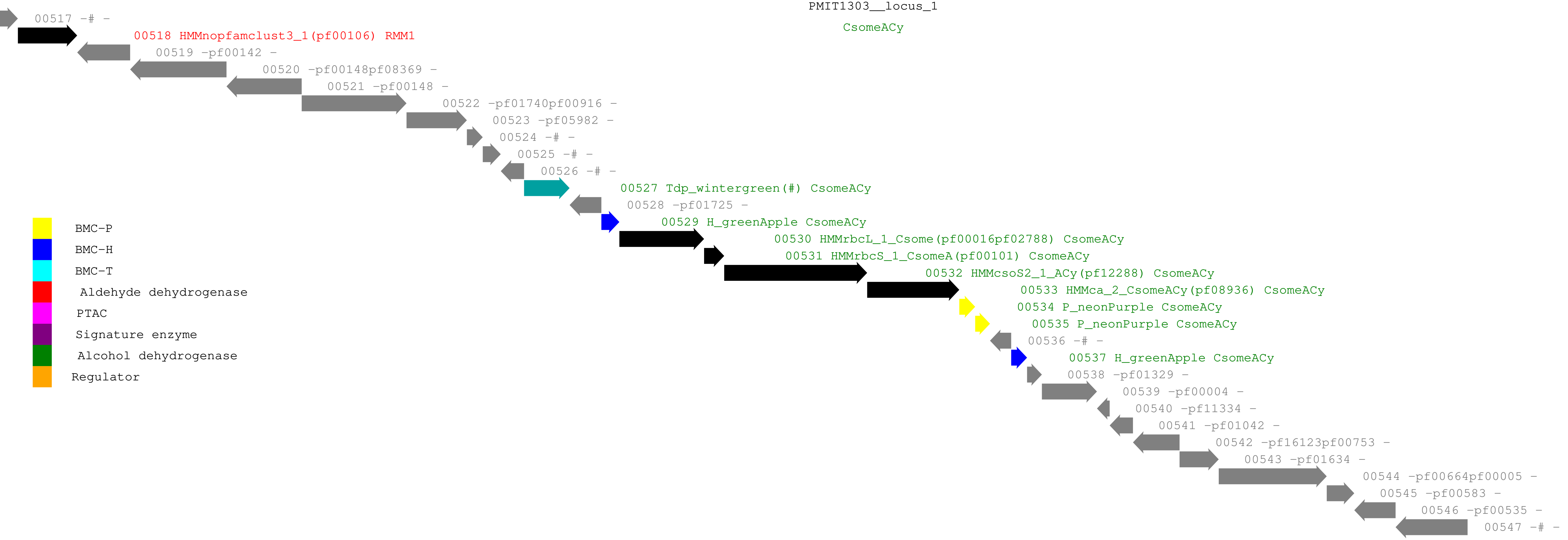

Prochlorococcus sp. MIT 1306. A0A163C295_9PROC Unreviewed; 477 AA.

PMIT1306__locus_1

CsomeACy

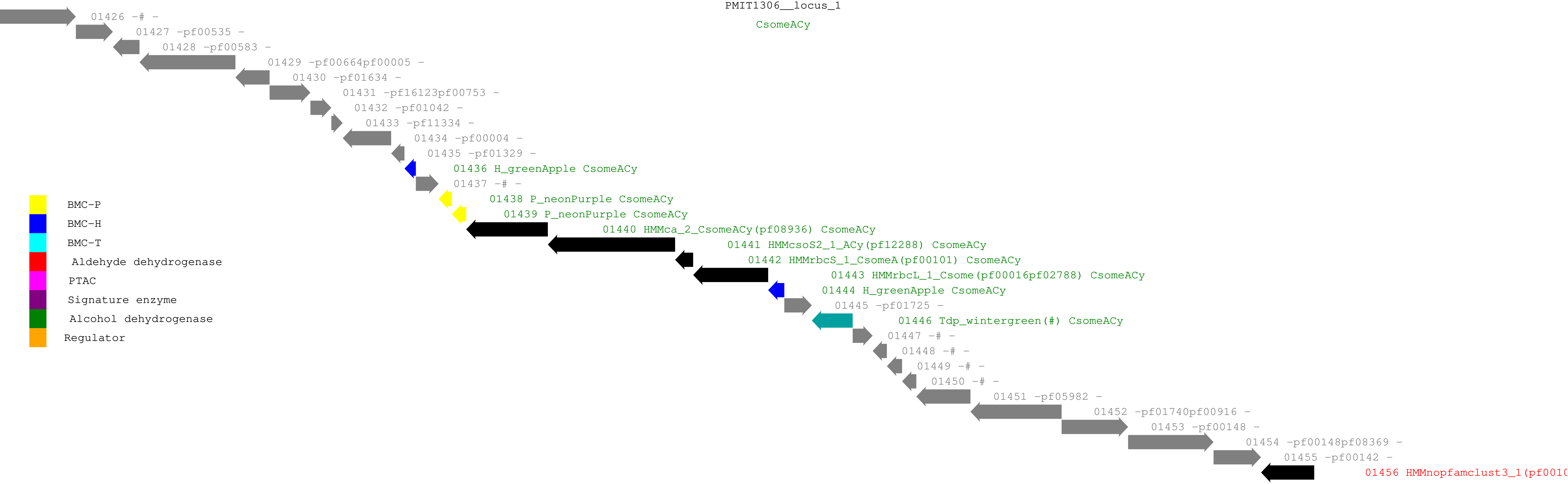

Prochlorococcus marinus str. MIT 1318. Bacteria; Cyanobacteria; Synechococcales; Prochloraceae;

PMIT1318__locus_1

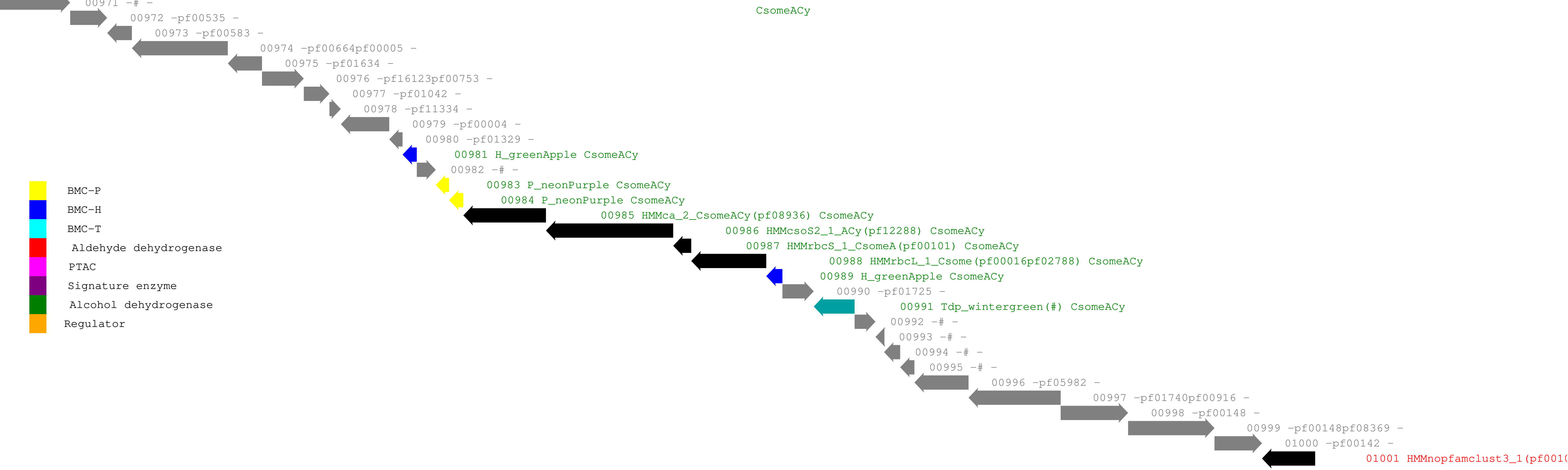

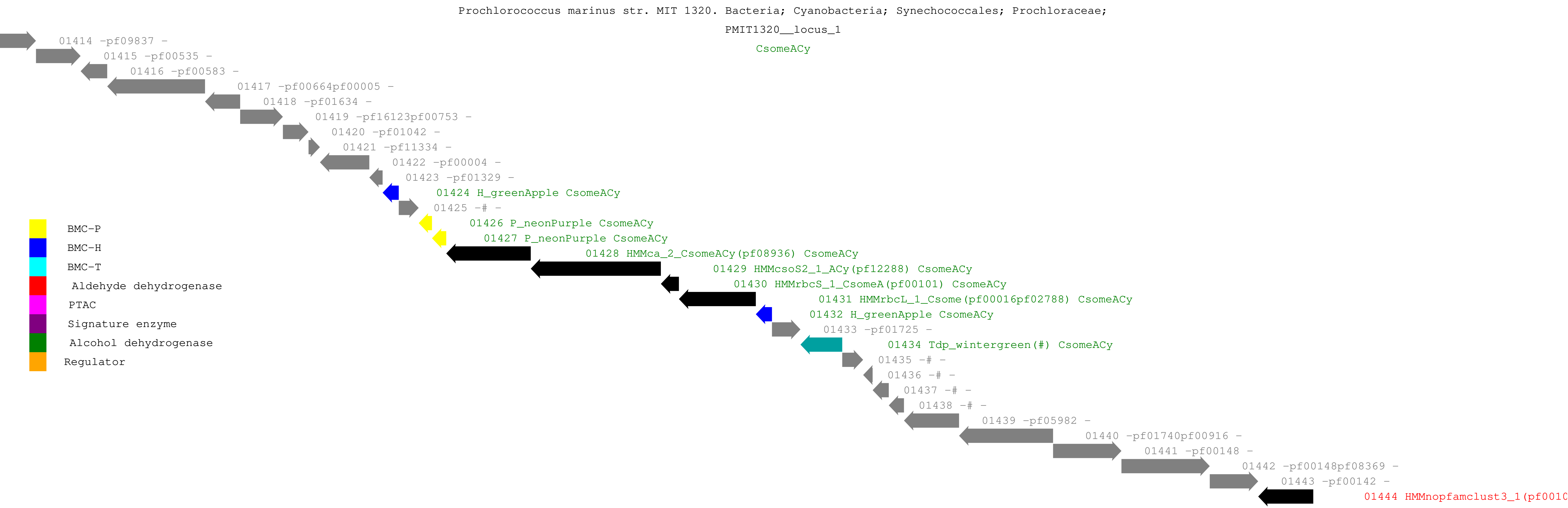
Prochlorococcus marinus str. MIT 1320. Bacteria; Cyanobacteria; Synechococcales; Prochloraceae;
PMIT1320__locus_1
CsomeACy
01414 -pf09837 -
01415 -pf00535 -
01416 -pf00583 -
01417 -pf00664pf00005 -
01418 -pf01634 -
01419 -pf16123pf00753 -
01420 -pf01042 -
01421 -pf11334 -
01422 -pf00004 -
01423 -pf01329 -
01424 H_greenApple CsomeACy
01425 -# -
01426 P_neonPurple CsomeACy
01427 P_neonPurple CsomeACy
01428 HMMca_2_CsomeACy(pf08936) CsomeACy
01429 HMMcsoS2_1_ACy(pf12288) CsomeACy
01430 HMMrbcS_1_CsomeA(pf00101) CsomeACy
01431 HMMrbcL_1_Csome(pf00016pf02788) CsomeACy
01432 H_greenApple CsomeACy
01433 -pf01725 -
01434 Tdp_wintergreen(#) CsomeACy
01435 -# -
01436 -# -
01437 -# -
01438 -# -
01439 -pf05982 -
01440 -pf01740pf00916 -
01441 -pf00148 -
01442 -pf00148pf08369 -
01443 -pf00142 -
01444 HMMnopfamclust3_1(pf0010
BMC-P
BMC-H
BMC-T
Aldehyde dehydrogenase
PTAC
Signature enzyme
Alcohol dehydrogenase
Regulator

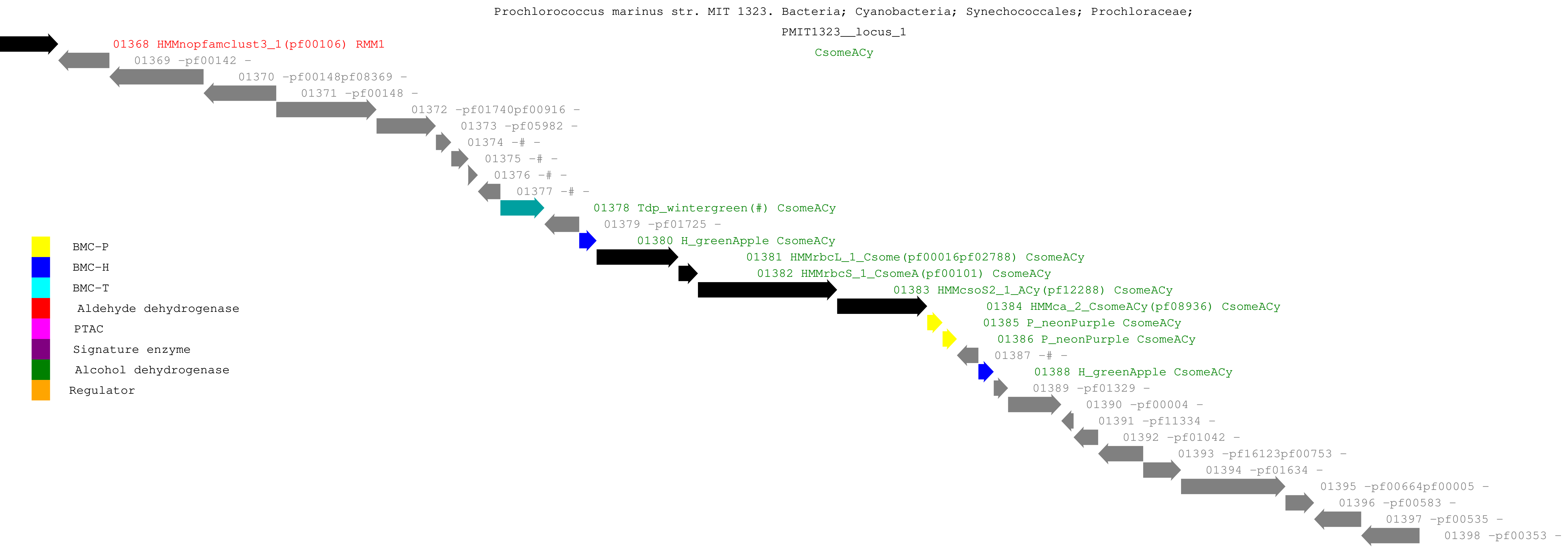
Prochlorococcus marinus str. MIT 1323. Bacteria; Cyanobacteria; Synechococcales; Prochloraceae;
PMIT1323__locus_1
CsomeACy
01368 HMMnopfamclust3_1(pf00106) RMM1
01369 -pf00142 -
01370 -pf00148pf08369 -
01371 -pf00148 -
01372 -pf01740pf00916 -
01373 -pf05982 -
01374 -# -
01375 -# -
01376 -# -
01377 -# -
01378 Tdp_wintergreen(#) CsomeACy
01379 -pf01725 -
01380 H_greenApple CsomeACy
01381 HMMrbcL_1_Csome(pf00016pf02788) CsomeACy
01382 HMMrbcS_1_CsomeA(pf00101) CsomeACy
01383 HMMcsoS2_1_ACy(pf12288) CsomeACy
01384 HMMca_2_CsomeACy(pf08936) CsomeACy
01385 P_neonPurple CsomeACy
01386 P_neonPurple CsomeACy
01387 -# -
01388 H_greenApple CsomeACy
01389 -pf01329 -
01390 -pf00004 -
01391 -pf11334 -
01392 -pf01042 -
01393 -pf16123pf00753 -
01394 -pf01634 -
01395 -pf00664pf00005 -
01396 -pf00583 -
01397 -pf00535 -
01398 -pf00353 -
BMC-P
BMC-H
BMC-T
Aldehyde dehydrogenase
PTAC
Signature enzyme
Alcohol dehydrogenase
Regulator

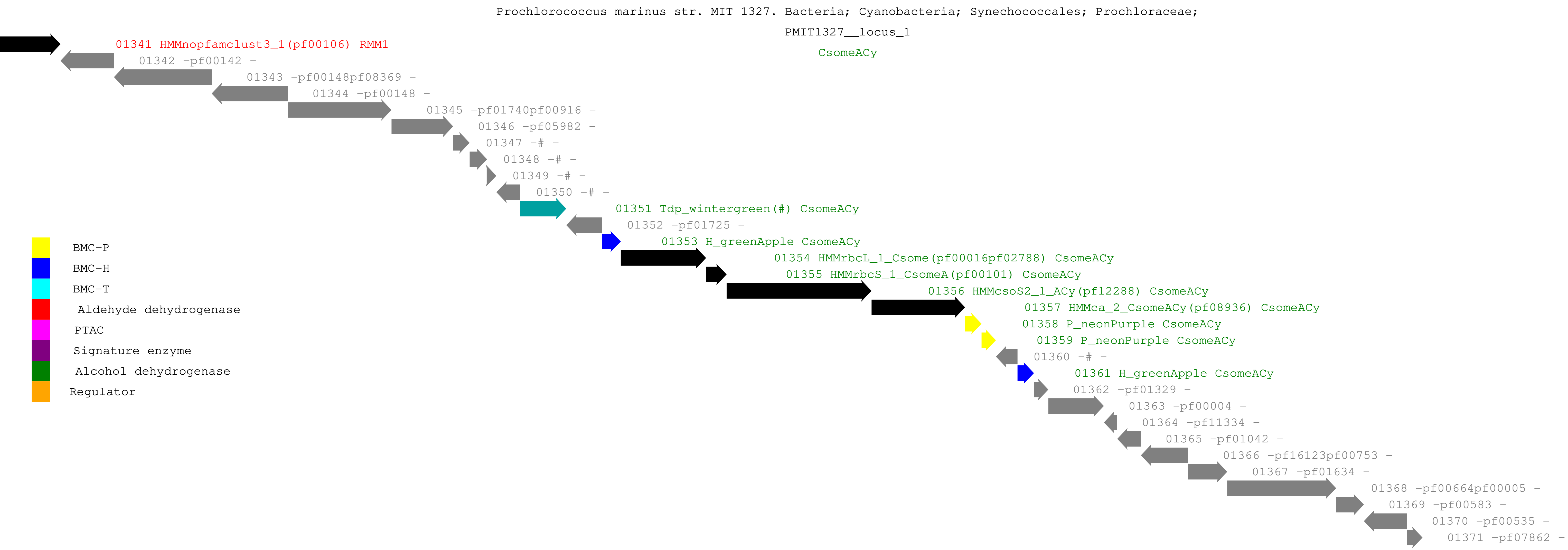
Prochlorococcus marinus str. MIT 1327. Bacteria; Cyanobacteria; Synechococcales; Prochloraceae;
PMIT1327__locus_1
CsomeACy
01341 HMMnopfamclust3_1(pf00106) RMM1
01342 -pf00142 -
01343 -pf00148pf08369 -
01344 -pf00148 -
01345 -pf01740pf00916 -
01346 -pf05982 -
01347 -# -
01348 -# -
01349 -# -
01350 -# -
01351 Tdp_wintergreen(#) CsomeACy
01352 -pf01725 -
01353 H_greenApple CsomeACy
01354 HMMrbcL_1_Csome(pf00016pf02788) CsomeACy
01355 HMMrbcS_1_CsomeA(pf00101) CsomeACy
01356 HMMcsoS2_1_ACy(pf12288) CsomeACy
01357 HMMca_2_CsomeACy(pf08936) CsomeACy
01358 P_neonPurple CsomeACy
01359 P_neonPurple CsomeACy
01360 -# -
01361 H_greenApple CsomeACy
01362 -pf01329 -
01363 -pf00004 -
01364 -pf11334 -
01365 -pf01042 -
01366 -pf16123pf00753 -
01367 -pf01634 -
01368 -pf00664pf00005 -
01369 -pf00583 -
01370 -pf00535 -
01371 -pf07862 -
BMC-P
BMC-H
BMC-T
Aldehyde dehydrogenase
PTAC
Signature enzyme
Alcohol dehydrogenase
Regulator

Prochlorococcus marinus str. MIT 1342. Bacteria; Cyanobacteria; Synechococcales; Prochloraceae;

PMIT1342__locus_1

CsomeACy

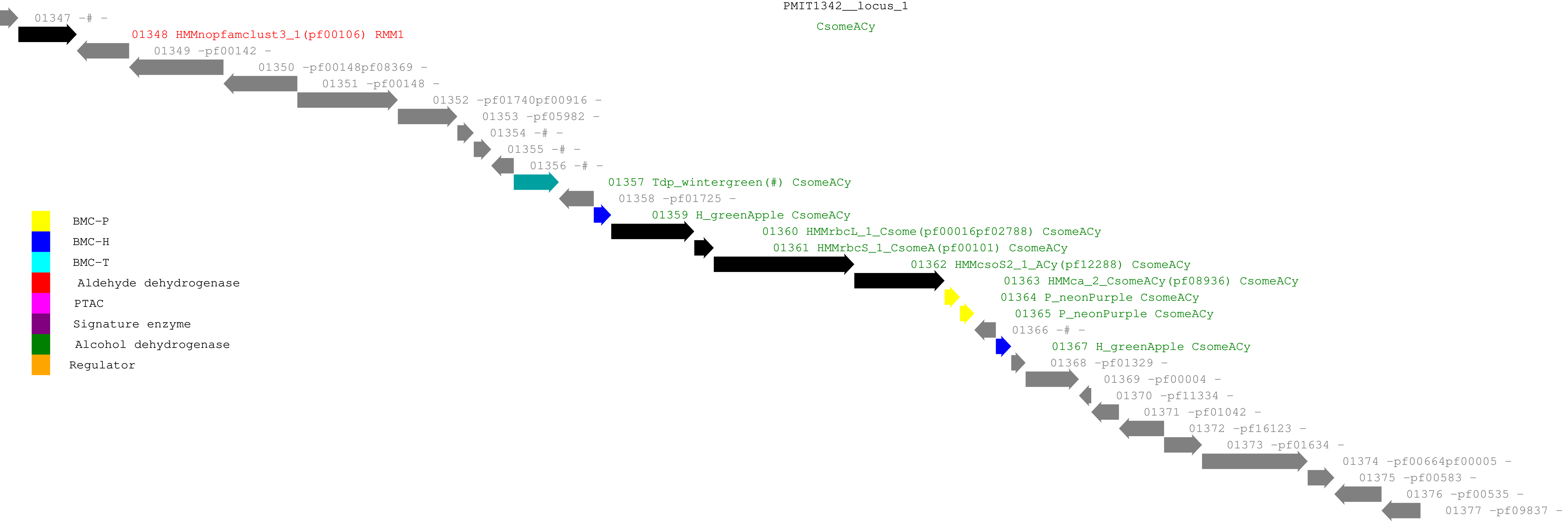

Prochlorococcus marinus subsp. pastoris (strain CCMP1986 / NIES-2087 / Bacteria; Cyanobacteria; Synechococcales; Prochloraceae;

PMM_locus_1

CsomeACy

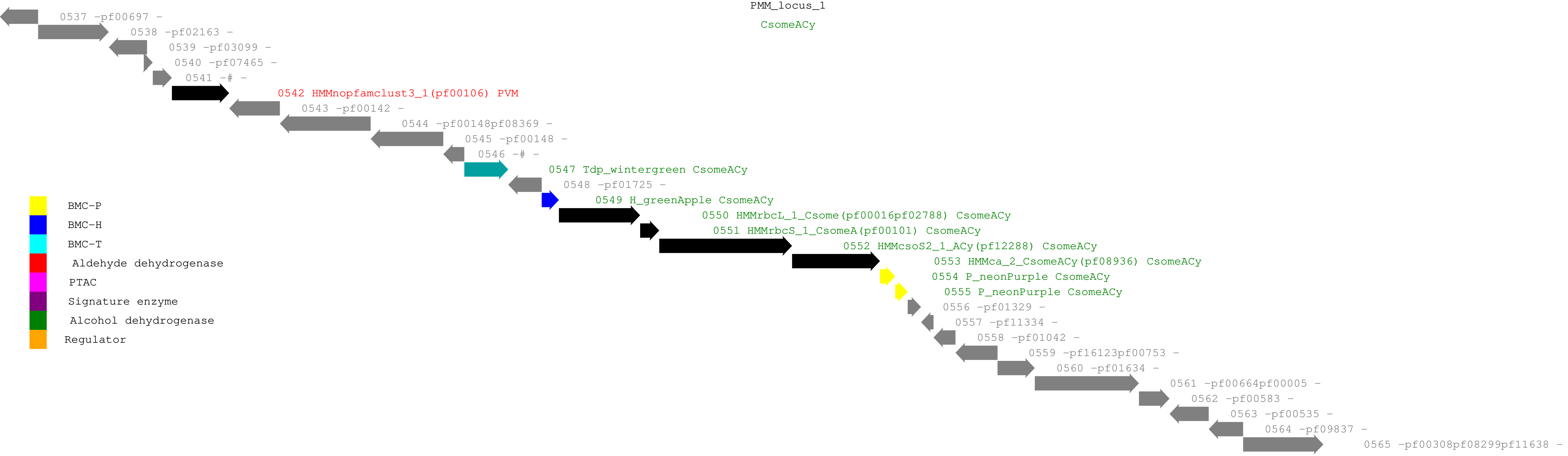

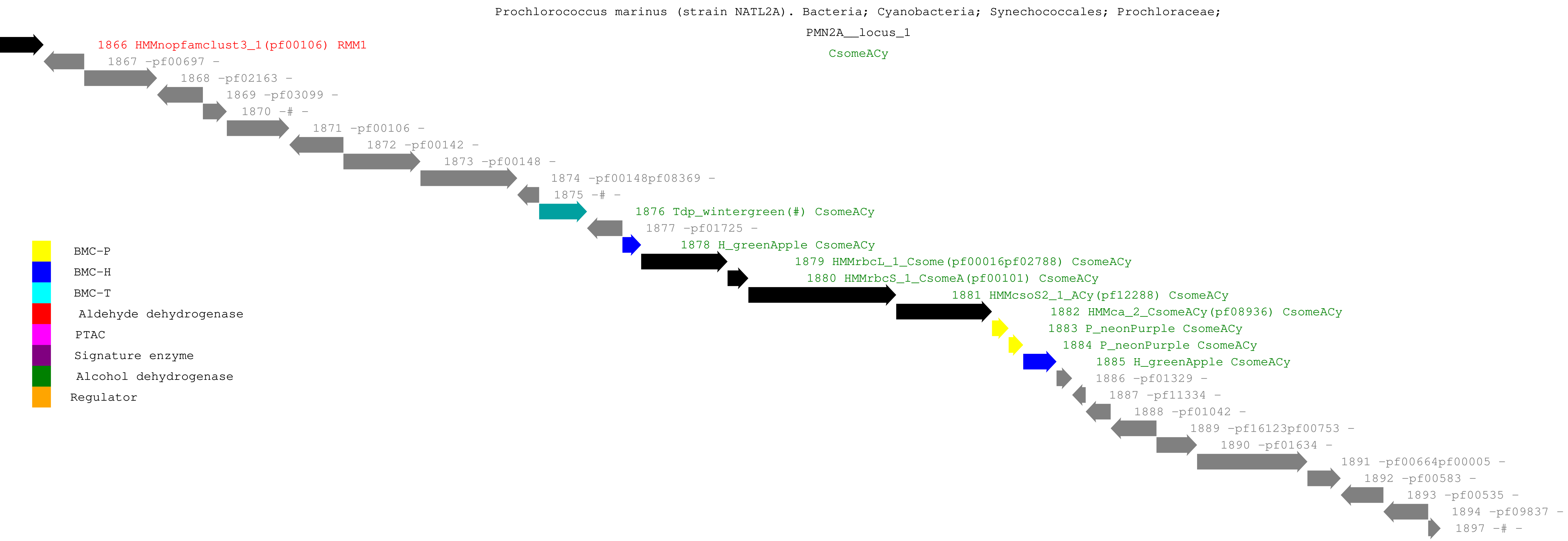
Prochlorococcus marinus (strain NATL2A). Bacteria; Cyanobacteria; Synechococcales; Prochloraceae;
PMN2A__locus_1
CsomeACy
1866 HMMnopfamclust3_1(pf00106) RMM1
1867 -pf00697 -
1868 -pf02163 -
1869 -pf03099 -
1870 -# -
1871 -pf00106 -
1872 -pf00142 -
1873 -pf00148 -
1874 -pf00148pf08369 -
1875 -# -
1876 Tdp_wintergreen(#) CsomeACy
1877 -pf01725 -
1878 H_greenApple CsomeACy
1879 HMMrbcL_1_Csome(pf00016pf02788) CsomeACy
1880 HMMrbcS_1_CsomeA(pf00101) CsomeACy
1881 HMMcsoS2_1_ACy(pf12288) CsomeACy
1882 HMMca_2_CsomeACy(pf08936) CsomeACy
1883 P_neonPurple CsomeACy
1884 P_neonPurple CsomeACy
1885 H_greenApple CsomeACy
1886 -pf01329 -
1887 -pf11334 -
1888 -pf01042 -
1889 -pf16123pf00753 -
1890 -pf01634 -
1891 -pf00664pf00005 -
1892 -pf00583 -
1893 -pf00535 -
1894 -pf09837 -
1897 -# -
BMC-P
BMC-H
BMC-T
Aldehyde dehydrogenase
PTAC
Signature enzyme
Alcohol dehydrogenase
Regulator

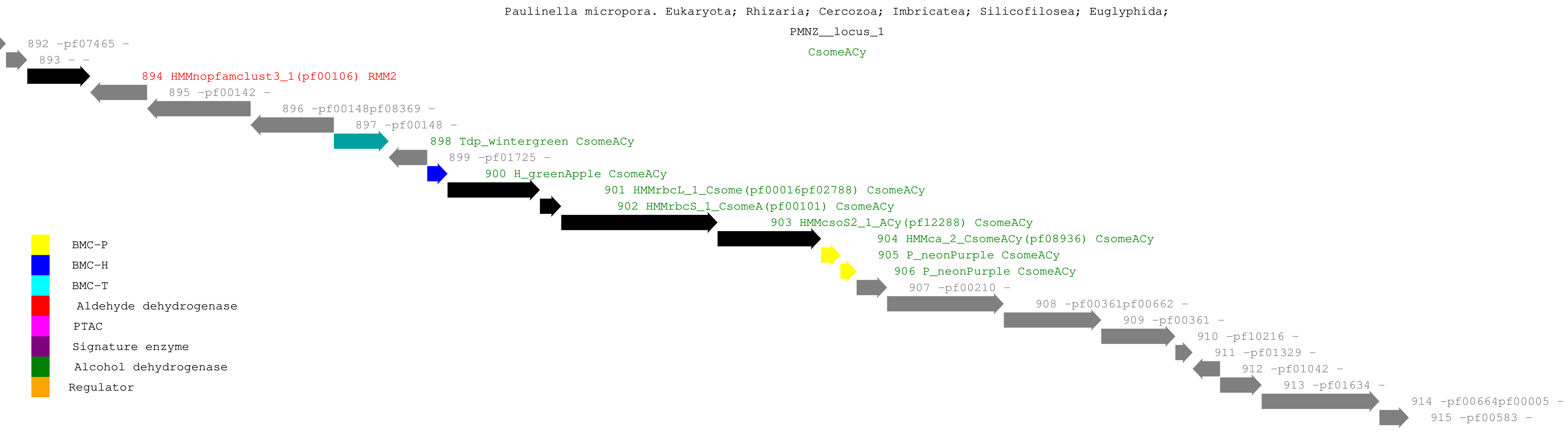
Paulinella micropora. Eukaryota; Rhizaria; Cercozoa; Imbricatea; Silicofilosea; Euglyphida;
PMNZ__locus_1
CsomeACy
892 -pf07465 -
893 - -
894 HMMnopfamclust3_1(pf00106) RMM2
895 -pf00142 -
896 -pf00148pf08369 -
897 -pf00148 -
898 Tdp_wintergreen CsomeACy
899 -pf01725 -
900 H_greenApple CsomeACy
901 HMMrbcL_1_Csome(pf00016pf02788) CsomeACy
902 HMMrbcS_1_CsomeA(pf00101) CsomeACy
903 HMMcsoS2_1_ACy(pf12288) CsomeACy
904 HMMca_2_CsomeACy(pf08936) CsomeACy
905 P_neonPurple CsomeACy
906 P_neonPurple CsomeACy
907 -pf00210 -
908 -pf00361pf00662 -
909 -pf00361 -
910 -pf10216 -
911 -pf01329 -
912 -pf01042 -
913 -pf01634 -
914 -pf00664pf00005 -
915 -pf00583 -
BMC-P
BMC-H
BMC-T
Aldehyde dehydrogenase
PTAC
Signature enzyme
Alcohol dehydrogenase
Regulator

Prochlorococcus marinus (strain MIT 9312). Bacteria; Cyanobacteria; Synechococcales; Prochloraceae;

PMT9312__locus_1

CsomeACy

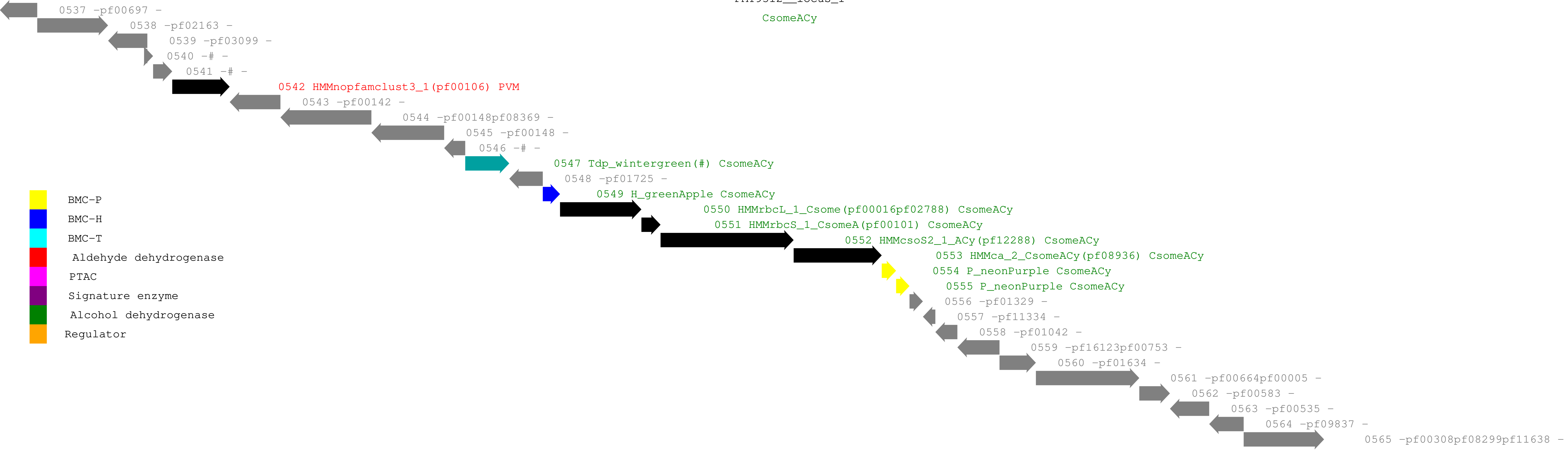

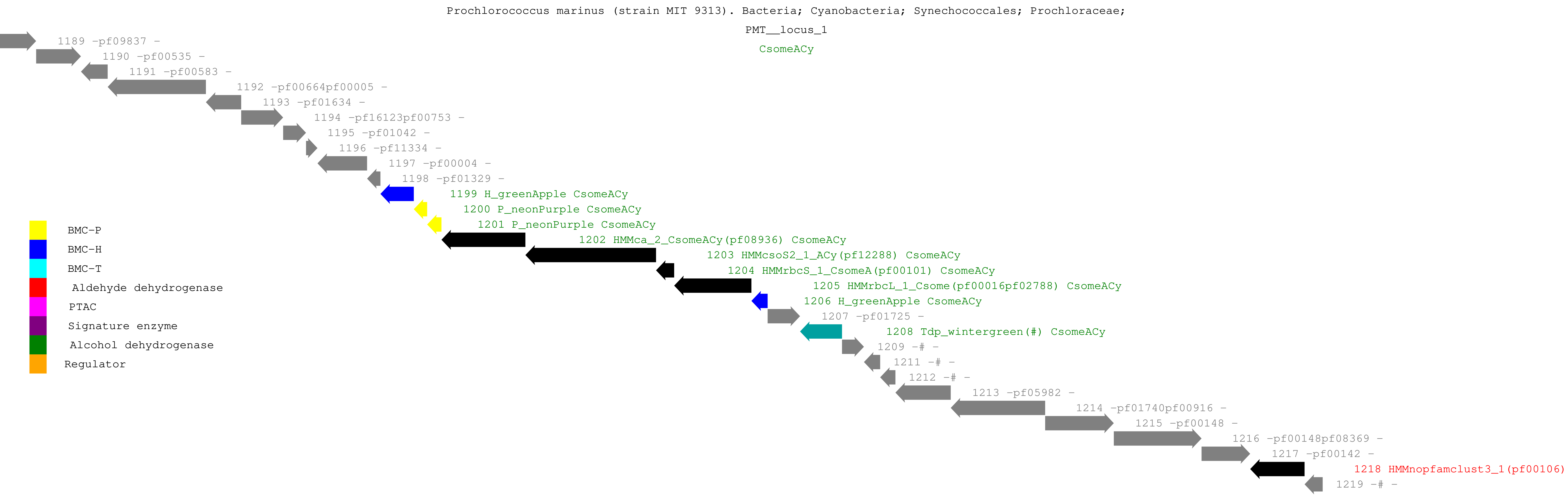

Prochlorococcus marinus (strain MIT 9313). Bacteria; Cyanobacteria; Synechococcales; Prochloraceae;
PMT__locus_1
CsomeACy
1189 -pf09837 -
1190 -pf00535 -
1191 -pf00583 -
1192 -pf00664pf00005 -
1193 -pf01634 -
1194 -pf16123pf00753 -
1195 -pf01042 -
1196 -pf11334 -
1197 -pf00004 -
1198 -pf01329 -
1199 H_greenApple CsomeACy
1200 P_neonPurple CsomeACy
1201 P_neonPurple CsomeACy
1202 HMMca_2_CsomeACy(pf08936) CsomeACy
1203 HMMcsoS2_1_ACy(pf12288) CsomeACy
1204 HMMrbcS_1_CsomeA(pf00101) CsomeACy
1205 HMMrbcL_1_Csome(pf00016pf02788) CsomeACy
1206 H_greenApple CsomeACy
1207 -pf01725 -
1208 Tdp_wintergreen(#) CsomeACy
1209 -# -
1211 -# -
1212 -# -
1213 -pf05982 -
1214 -pf01740pf00916 -
1215 -pf00148 -
1216 -pf00148pf08369 -
1217 -pf00142 -
1218 HMMnopfamclust3_1(pf00106)
1219 -# -
BMC-P
BMC-H
BMC-T
Aldehyde dehydrogenase
PTAC
Signature enzyme
Alcohol dehydrogenase
Regulator

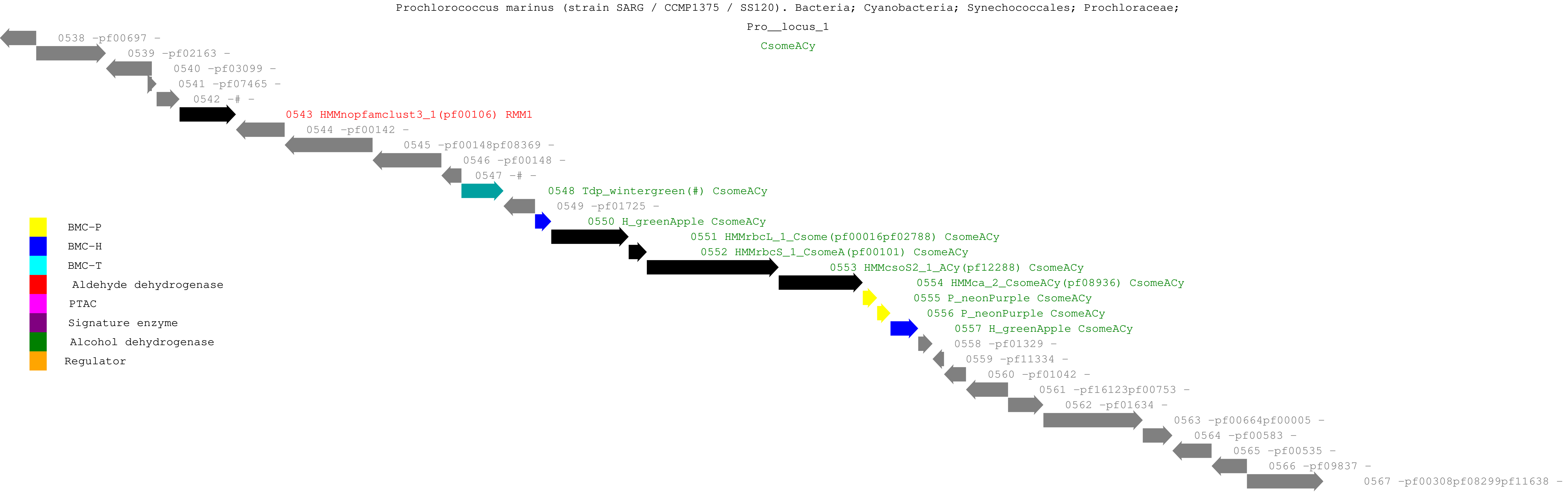
Prochlorococcus marinus (strain SARG / CCMP1375 / SS120). Bacteria; Cyanobacteria; Synechococcales; Prochloraceae;
Pro__locus_1
CsomeACy
0538 -pf00697 -
0539 -pf02163 -
0540 -pf03099 -
0541 -pf07465 -
0542 -# -
0543 HMMnopfamclust3_1(pf00106) RMM1
0544 -pf00142 -
0545 -pf00148pf08369 -
0546 -pf00148 -
0547 -# -
0548 Tdp_wintergreen(#) CsomeACy
0549 -pf01725 -
0550 H_greenApple CsomeACy
0551 HMMrbcL_1_Csome(pf00016pf02788) CsomeACy
0552 HMMrbcS_1_CsomeA(pf00101) CsomeACy
0553 HMMcsoS2_1_ACy(pf12288) CsomeACy
0554 HMMca_2_CsomeACy(pf08936) CsomeACy
0555 P_neonPurple CsomeACy
0556 P_neonPurple CsomeACy
0557 H_greenApple CsomeACy
0558 -pf01329 -
0559 -pf11334 -
0560 -pf01042 -
0561 -pf16123pf00753 -
0562 -pf01634 -
0563 -pf00664pf00005 -
0564 -pf00583 -
0565 -pf00535 -
0566 -pf09837 -
0567 -pf00308pf08299pf11638 -
BMC-P
BMC-H
BMC-T
Aldehyde dehydrogenase
PTAC
Signature enzyme
Alcohol dehydrogenase
Regulator

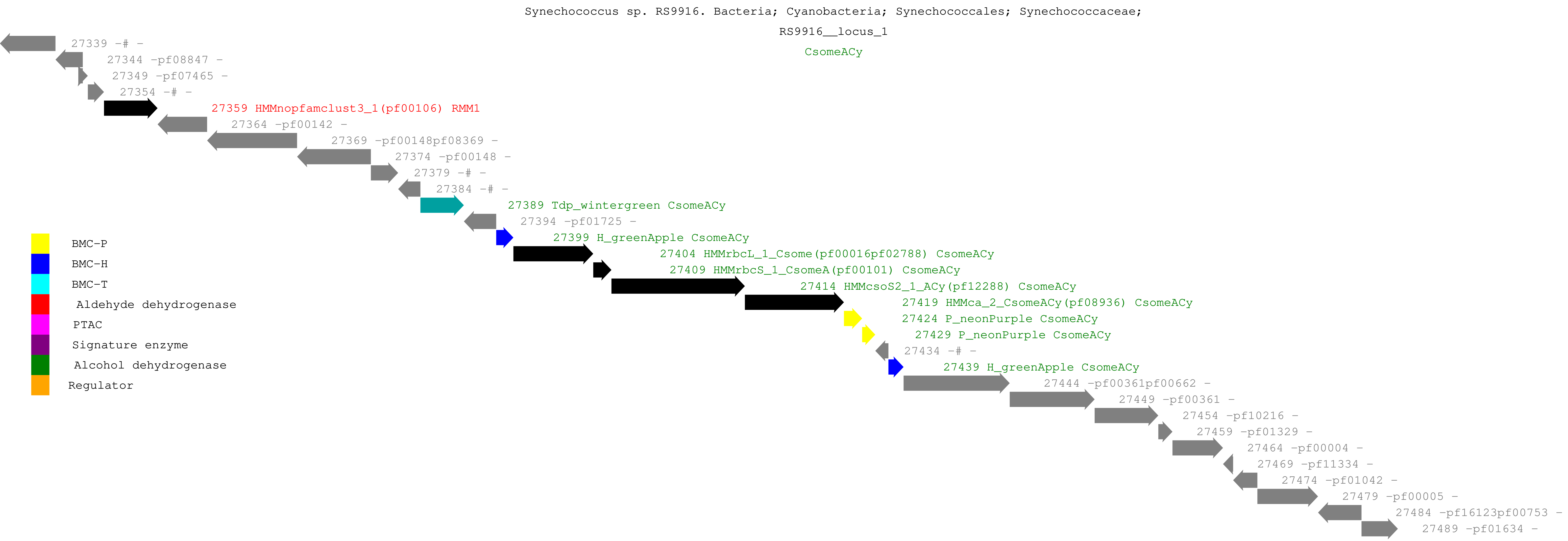
Synechococcus sp. RS9916. Bacteria; Cyanobacteria; Synechococcales; Synechococcaceae;
RS9916__locus_1
CsomeACy
27339 -# -
27344 -pf08847 -
27349 -pf07465 -
27354 -# -
27359 HMMnopfamclust3_1(pf00106) RMM1
27364 -pf00142 -
27369 -pf00148pf08369 -
27374 -pf00148 -
27379 -# -
27384 -# -
27389 Tdp_wintergreen CsomeACy
27394 -pf01725 -
27399 H_greenApple CsomeACy
27404 HMMrbcL_1_Csome(pf00016pf02788) CsomeACy
27409 HMMrbcS_1_CsomeA(pf00101) CsomeACy
27414 HMMcsoS2_1_ACy(pf12288) CsomeACy
27419 HMMca_2_CsomeACy(pf08936) CsomeACy
27424 P_neonPurple CsomeACy
27429 P_neonPurple CsomeACy
27434 -# -
27439 H_greenApple CsomeACy
27444 -pf00361pf00662 -
27449 -pf00361 -
27454 -pf10216 -
27459 -pf01329 -
27464 -pf00004 -
27469 -pf11334 -
27474 -pf01042 -
27479 -pf00005 -
27484 -pf16123pf00753 -
27489 -pf01634 -
BMC-P
BMC-H
BMC-T
Aldehyde dehydrogenase
PTAC
Signature enzyme
Alcohol dehydrogenase
Regulator

Synechococcus sp. RS9917. Bacteria; Cyanobacteria; Synechococcales; Synechococcaceae;

RS9917__locus_1

CsomeACy

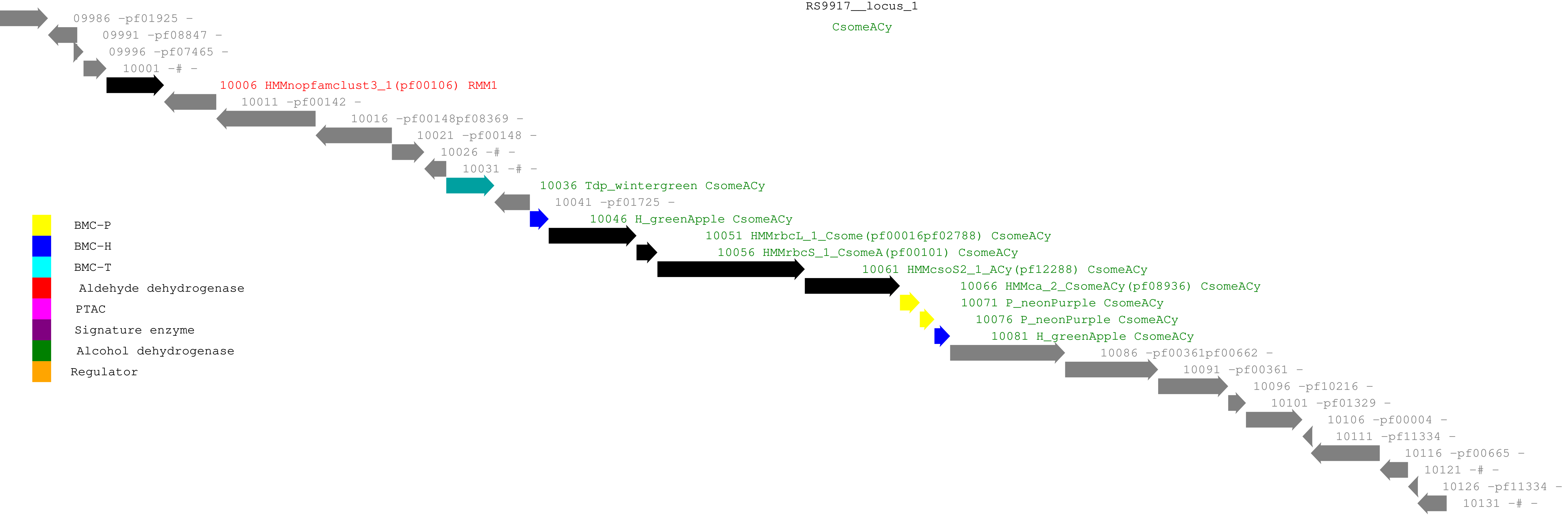

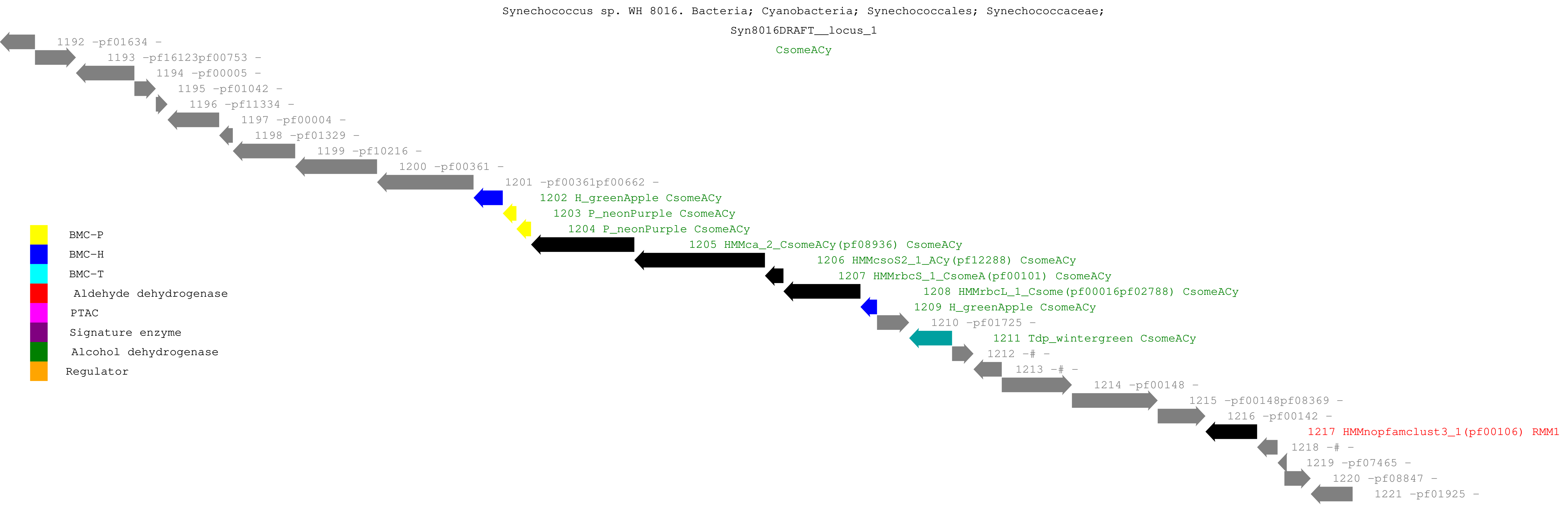
Synechococcus sp. WH 8016. Bacteria; Cyanobacteria; Synechococcales; Synechococcaceae;
Syn8016DRAFT__locus_1
CsomeACy
1192 -pf01634 -
1193 -pf16123pf00753 -
1194 -pf00005 -
1195 -pf01042 -
1196 -pf11334 -
1197 -pf00004 -
1198 -pf01329 -
1199 -pf10216 -
1200 -pf00361 -
1201 -pf00361pf00662 -
1202 H_greenApple CsomeACy
1203 P_neonPurple CsomeACy
1204 P_neonPurple CsomeACy
1205 HMMca_2_CsomeACy(pf08936) CsomeACy
1206 HMMcsoS2_1_ACy(pf12288) CsomeACy
1207 HMMrbcS_1_CsomeA(pf00101) CsomeACy
1208 HMMrbcL_1_Csome(pf00016pf02788) CsomeACy
1209 H_greenApple CsomeACy
1210 -pf01725 -
1211 Tdp_wintergreen CsomeACy
1212 -# -
1213 -# -
1214 -pf00148 -
1215 -pf00148pf08369 -
1216 -pf00142 -
1217 HMMnopfamclust3_1(pf00106) RMM1
1218 -# -
1219 -pf07465 -
1220 -pf08847 -
1221 -pf01925 -
BMC-P
BMC-H
BMC-T
Aldehyde dehydrogenase
PTAC
Signature enzyme
Alcohol dehydrogenase
Regulator

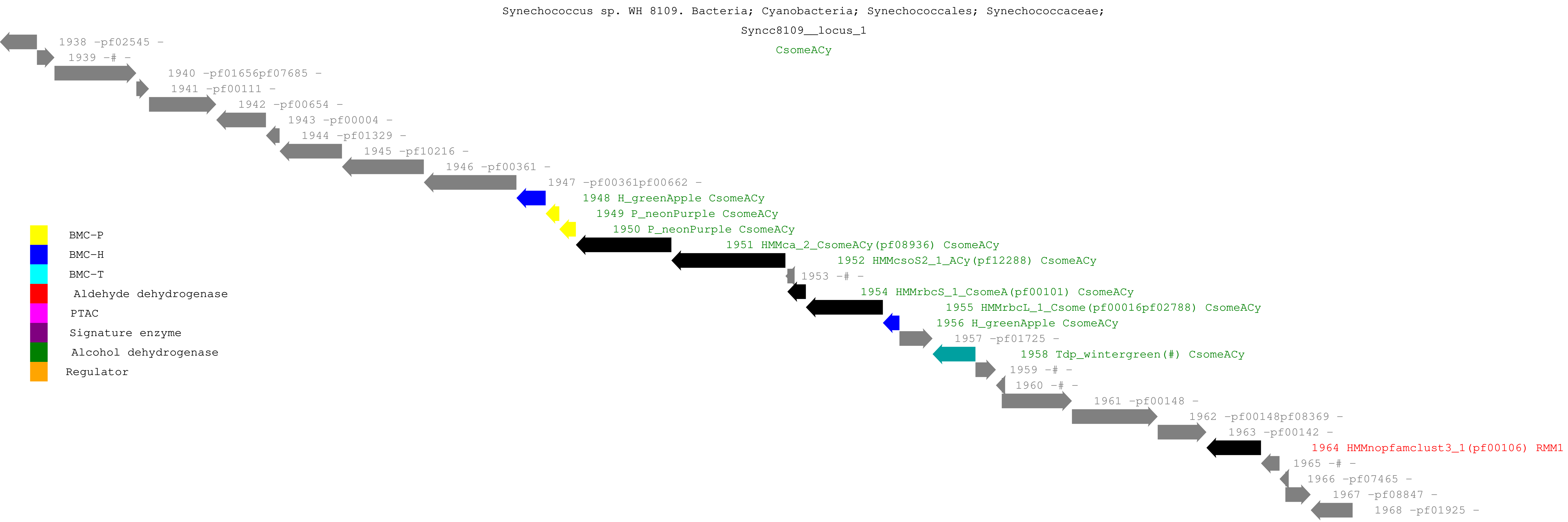

Synechococcus sp. WH 8109. Bacteria; Cyanobacteria; Synechococcales; Synechococcaceae;
Syncc8109__locus_1
CsomeACy
1938 -pf02545 -
1939 -# -
1940 -pf01656pf07685 -
1941 -pf00111 -
1942 -pf00654 -
1943 -pf00004 -
1944 -pf01329 -
1945 -pf10216 -
1946 -pf00361 -
1947 -pf00361pf00662 -
1948 H_greenApple CsomeACy
1949 P_neonPurple CsomeACy
1950 P_neonPurple CsomeACy
1951 HMMca_2_CsomeACy(pf08936) CsomeACy
1952 HMMcsoS2_1_ACy(pf12288) CsomeACy
1953 -# -
1954 HMMrbcS_1_CsomeA(pf00101) CsomeACy
1955 HMMrbcL_1_Csome(pf00016pf02788) CsomeACy
1956 H_greenApple CsomeACy
1957 -pf01725 -
1958 Tdp_wintergreen(#) CsomeACy
1959 -# -
1960 -# -
1961 -pf00148 -
1962 -pf00148pf08369 -
1963 -pf00142 -
1964 HMMnopfamclust3_1(pf00106) RMM1
1965 -# -
1966 -pf07465 -
1967 -pf08847 -
1968 -pf01925 -
BMC-P
BMC-H
BMC-T
Aldehyde dehydrogenase
PTAC
Signature enzyme
Alcohol dehydrogenase
Regulator

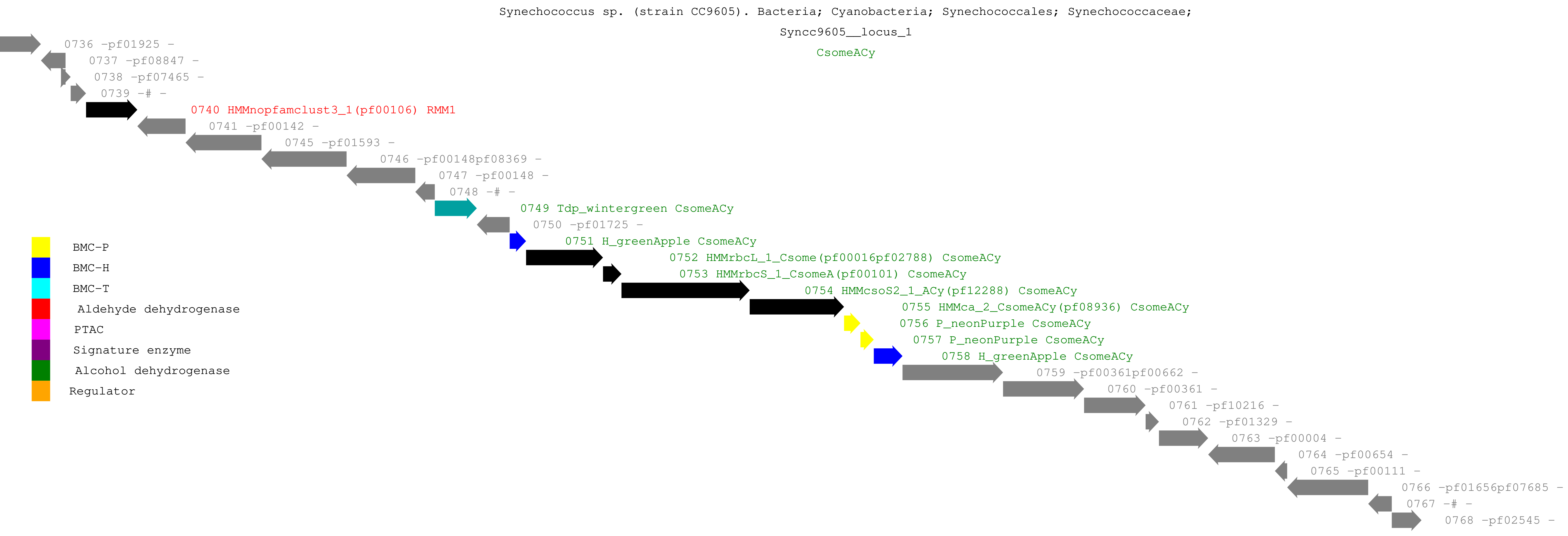
Synechococcus sp. (strain CC9605). Bacteria; Cyanobacteria; Synechococcales; Synechococcaceae;
Syncc9605__locus_1
CsomeACy
0736 -pf01925 -
0737 -pf08847 -
0738 -pf07465 -
0739 -# -
0740 HMMnopfamclust3_1(pf00106) RMM1
0741 -pf00142 -
0745 -pf01593 -
0746 -pf00148pf08369 -
0747 -pf00148 -
0748 -# -
0749 Tdp_wintergreen CsomeACy
0750 -pf01725 -
0751 H_greenApple CsomeACy
0752 HMMrbcL_1_Csome(pf00016pf02788) CsomeACy
0753 HMMrbcS_1_CsomeA(pf00101) CsomeACy
0754 HMMcsoS2_1_ACy(pf12288) CsomeACy
0755 HMMca_2_CsomeACy(pf08936) CsomeACy
0756 P_neonPurple CsomeACy
0757 P_neonPurple CsomeACy
0758 H_greenApple CsomeACy
0759 -pf00361pf00662 -
0760 -pf00361 -
0761 -pf10216 -
0762 -pf01329 -
0763 -pf00004 -
0764 -pf00654 -
0765 -pf00111 -
0766 -pf01656pf07685 -
0767 -# -
0768 -pf02545 -
BMC-P
BMC-H
BMC-T
Aldehyde dehydrogenase
PTAC
Signature enzyme
Alcohol dehydrogenase
Regulator

Synechococcus sp. (strain CC9902). Bacteria; Cyanobacteria; Synechococcales; Synechococcaceae;

Syncc9902__locus_1

CsomeACy

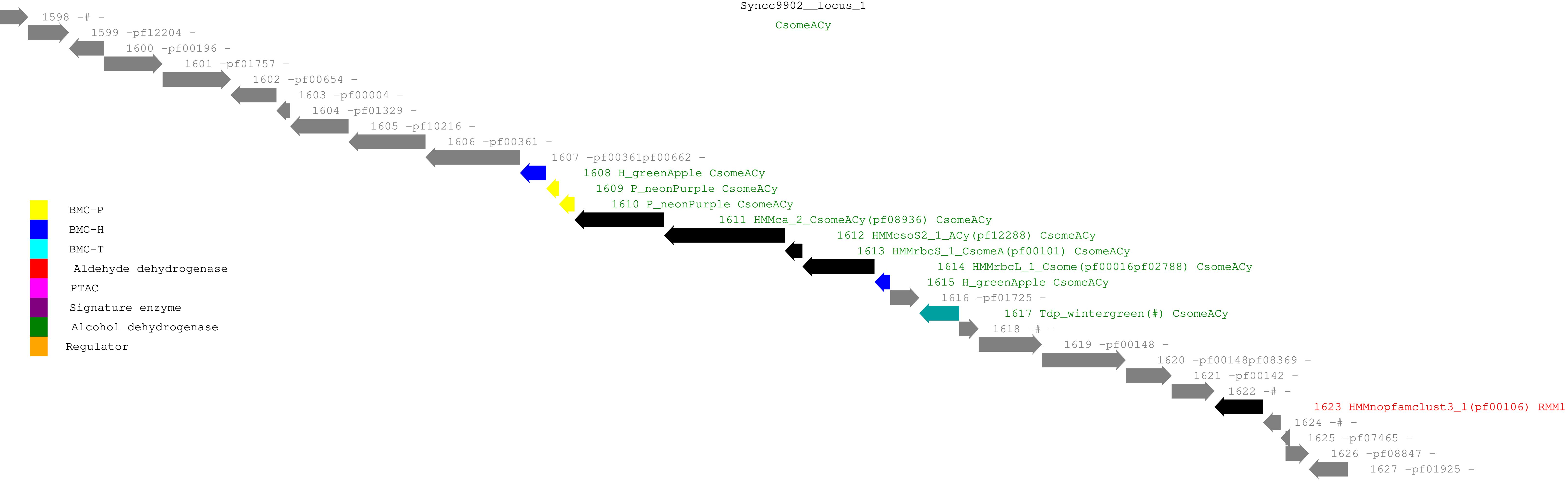

Synechococcus sp. (strain CC9311). Bacteria; Cyanobacteria; Synechococcales; Synechococcaceae;

sync__locus_1

CsomeACy

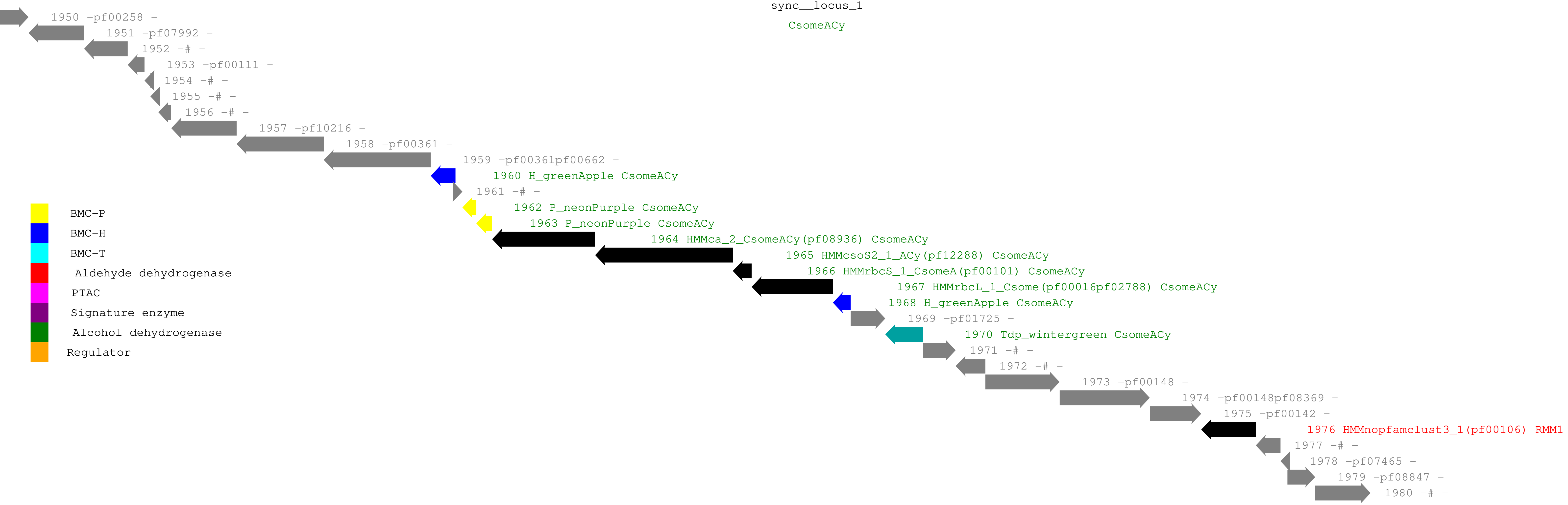

Synechococcus sp. GFB01. Bacteria; Cyanobacteria; Synechococcales; Synechococcaceae;

SYNGFB01__locus_1_main_sat

CsomeACy

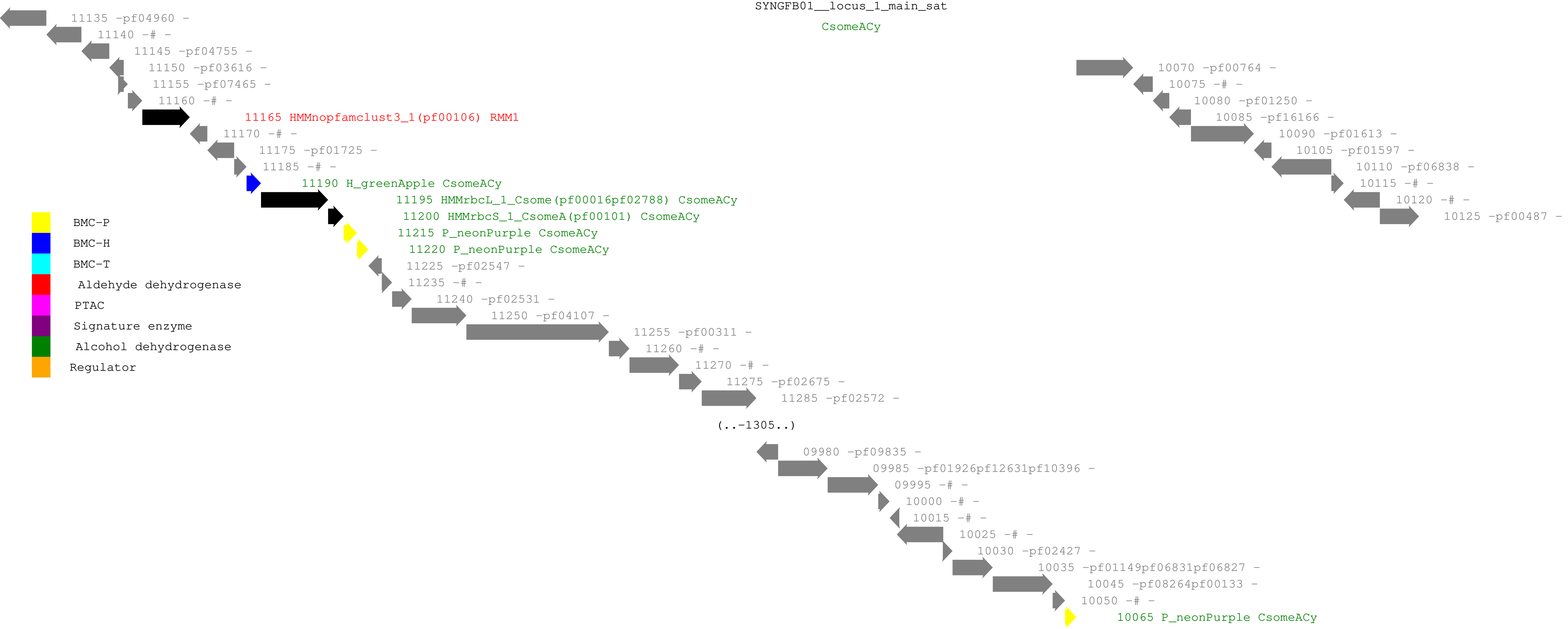

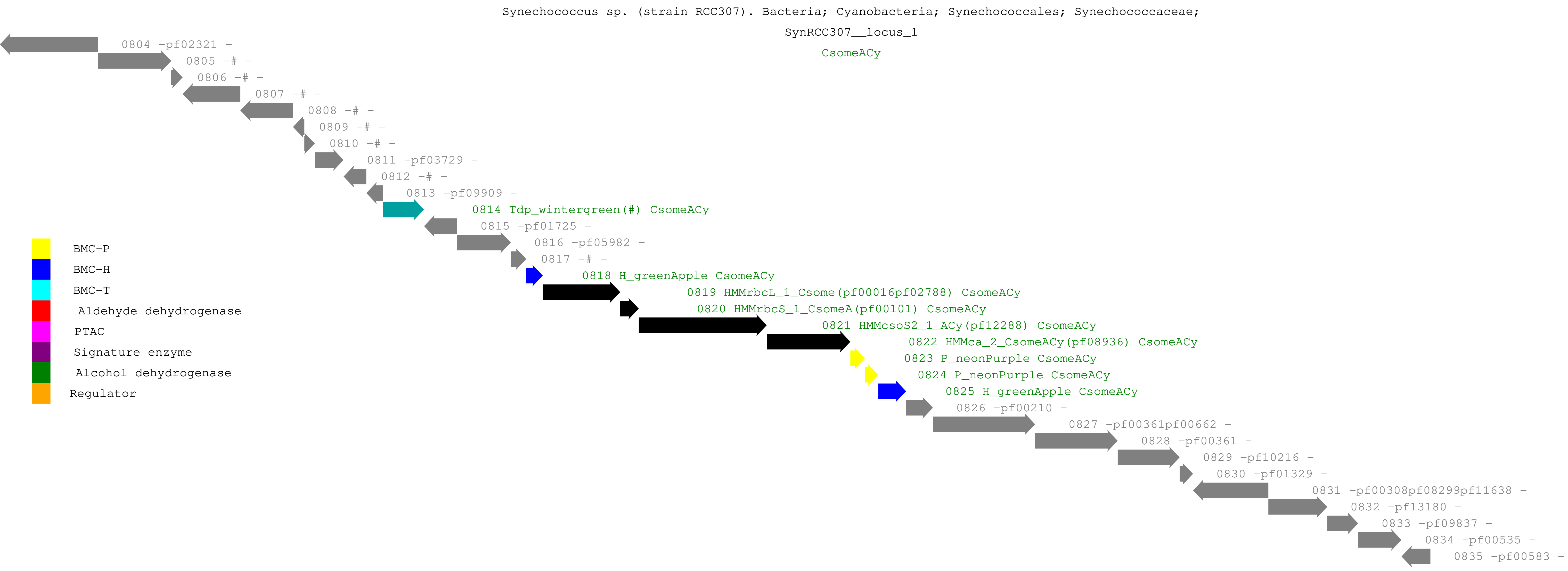
Synechococcus sp. (strain RCC307). Bacteria; Cyanobacteria; Synechococcales; Synechococcaceae;
SynRCC307__locus_1
CsomeACy
0804 -pf02321 -
0805 -# -
0806 -# -
0807 -# -
0808 -# -
0809 -# -
0810 -# -
0811 -pf03729 -
0812 -# -
0813 -pf09909 -
0814 Tdp_wintergreen(#) CsomeACy
0815 -pf01725 -
0816 -pf05982 -
0817 -# -
0818 H_greenApple CsomeACy
0819 HMMrbcL_1_Csome(pf00016pf02788) CsomeACy
0820 HMMrbcS_1_CsomeA(pf00101) CsomeACy
0821 HMMcsoS2_1_ACy(pf12288) CsomeACy
0822 HMMca_2_CsomeACy(pf08936) CsomeACy
0823 P_neonPurple CsomeACy
0824 P_neonPurple CsomeACy
0825 H_greenApple CsomeACy
0826 -pf00210 -
0827 -pf00361pf00662 -
0828 -pf00361 -
0829 -pf10216 -
0830 -pf01329 -
0831 -pf00308pf08299pf11638 -
0832 -pf13180 -
0833 -pf09837 -
0834 -pf00535 -
0835 -pf00583 -
BMC-P
BMC-H
BMC-T
Aldehyde dehydrogenase
PTAC
Signature enzyme
Alcohol dehydrogenase
Regulator

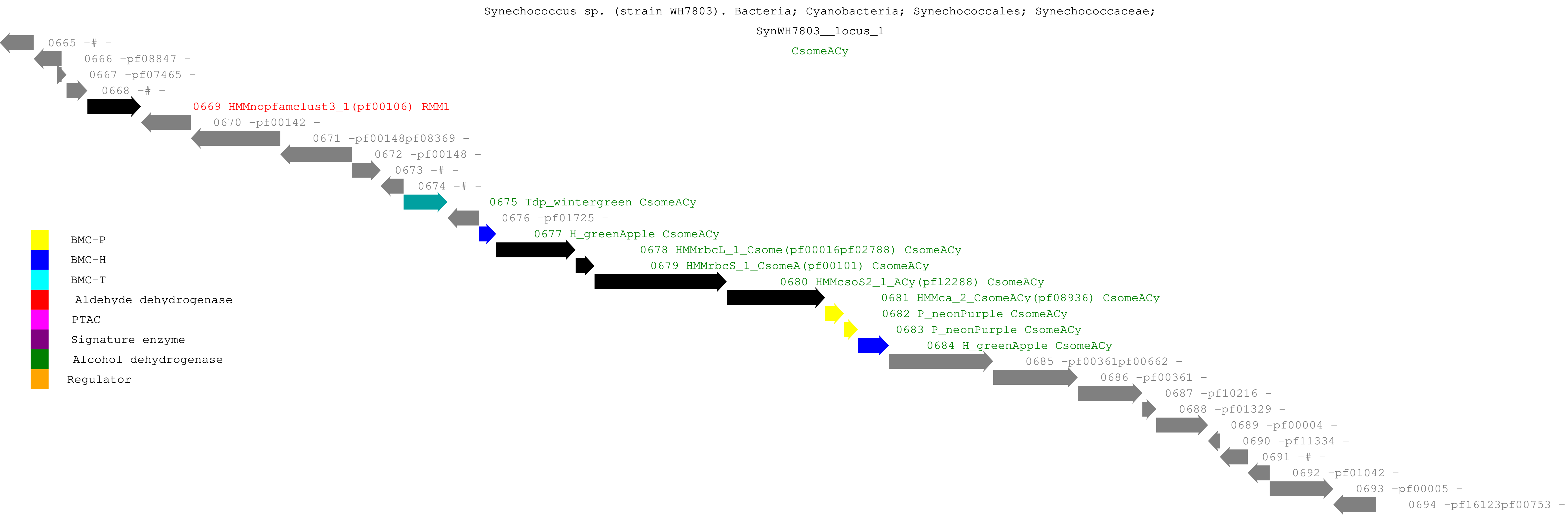
Synechococcus sp. (strain WH7803). Bacteria; Cyanobacteria; Synechococcales; Synechococcaceae;
SynWH7803__locus_1
CsomeACy
0665 -# -
0666 -pf08847 -
0667 -pf07465 -
0668 -# -
0669 HMMnopfamclust3_1(pf00106) RMM1
0670 -pf00142 -
0671 -pf00148pf08369 -
0672 -pf00148 -
0673 -# -
0674 -# -
0675 Tdp_wintergreen CsomeACy
0676 -pf01725 -
0677 H_greenApple CsomeACy
0678 HMMrbcL_1_Csome(pf00016pf02788) CsomeACy
0679 HMMrbcS_1_CsomeA(pf00101) CsomeACy
0680 HMMcsoS2_1_ACy(pf12288) CsomeACy
0681 HMMca_2_CsomeACy(pf08936) CsomeACy
0682 P_neonPurple CsomeACy
0683 P_neonPurple CsomeACy
0684 H_greenApple CsomeACy
0685 -pf00361pf00662 -
0686 -pf00361 -
0687 -pf10216 -
0688 -pf01329 -
0689 -pf00004 -
0690 -pf11334 -
0691 -# -
0692 -pf01042 -
0693 -pf00005 -
0694 -pf16123pf00753 -
BMC-P
BMC-H
BMC-T
Aldehyde dehydrogenase
PTAC
Signature enzyme
Alcohol dehydrogenase
Regulator

Synechococcus sp. WH 8101. Bacteria; Cyanobacteria; Synechococcales; Synechococcaceae; Synechococcus;

SynWH8101__locus_1

CsomeACy

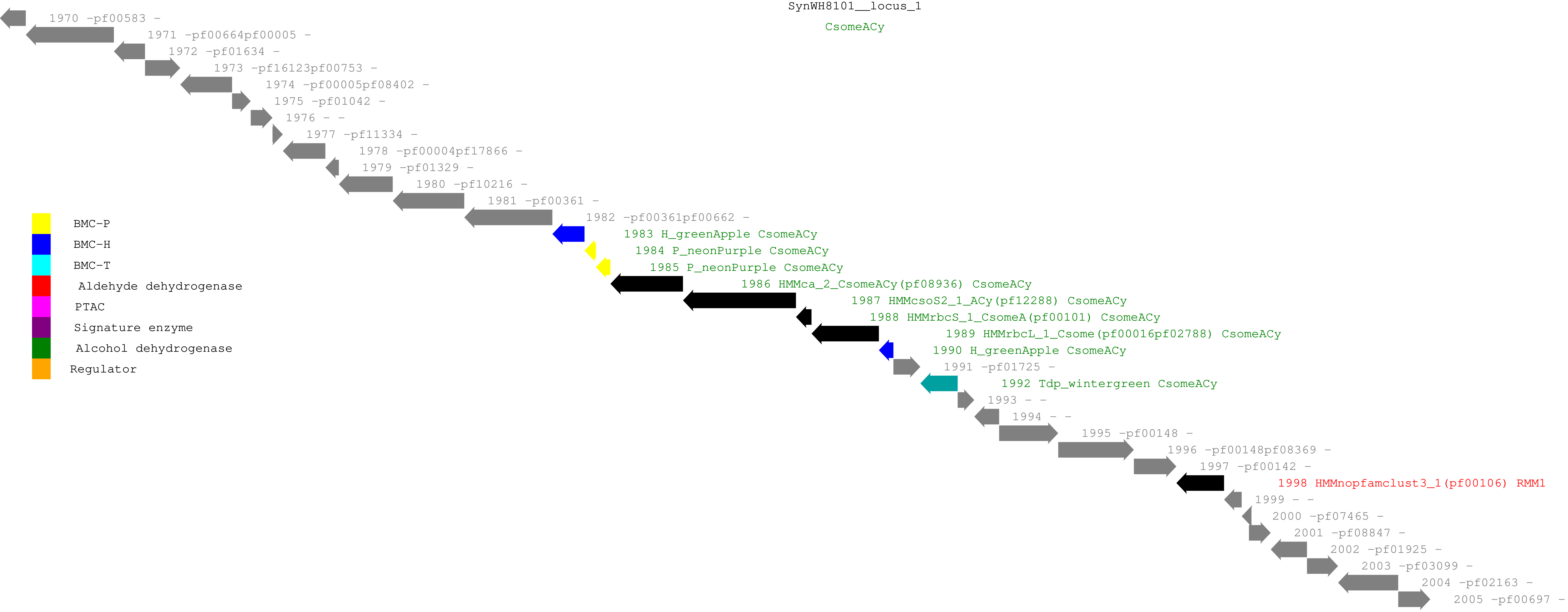

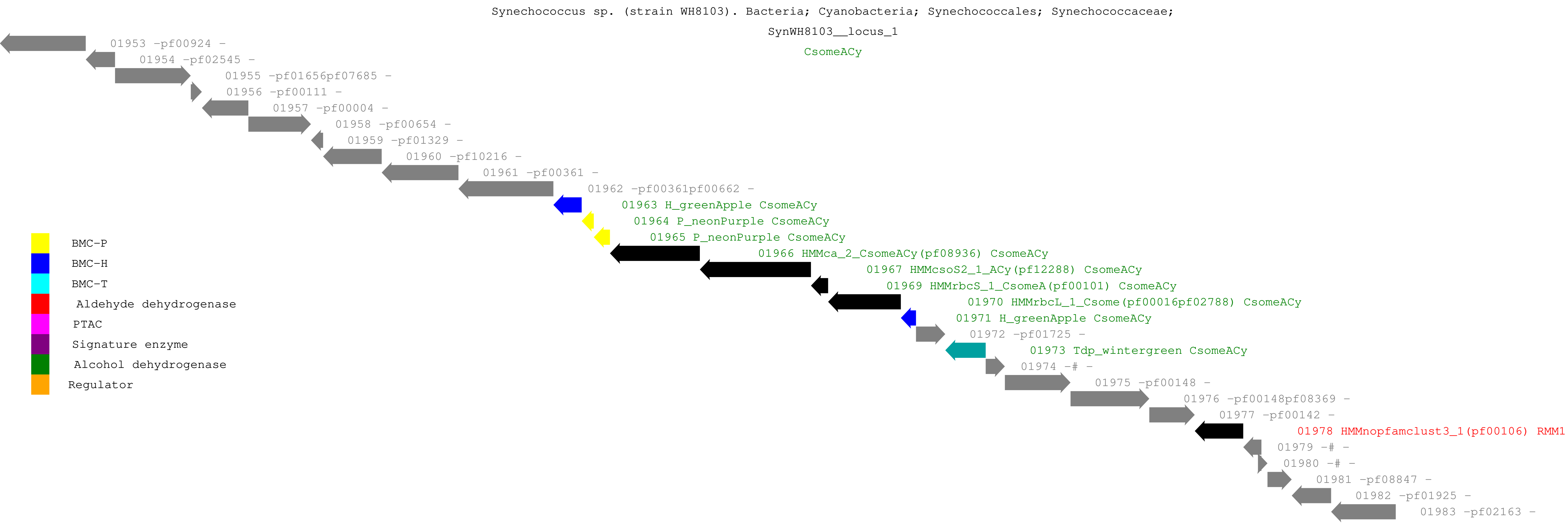
Synechococcus sp. (strain WH8103). Bacteria; Cyanobacteria; Synechococcales; Synechococcaceae;
SynWH8103__locus_1
CsomeACy
01953 -pf00924 -
01954 -pf02545 -
01955 -pf01656pf07685 -
01956 -pf00111 -
01957 -pf00004 -
01958 -pf00654 -
01959 -pf01329 -
01960 -pf10216 -
01961 -pf00361 -
01962 -pf00361pf00662 -
01963 H_greenApple CsomeACy
01964 P_neonPurple CsomeACy
01965 P_neonPurple CsomeACy
01966 HMMca_2_CsomeACy(pf08936) CsomeACy
01967 HMMcsoS2_1_ACy(pf12288) CsomeACy
01969 HMMrbcS_1_CsomeA(pf00101) CsomeACy
01970 HMMrbcL_1_Csome(pf00016pf02788) CsomeACy
01971 H_greenApple CsomeACy
01972 -pf01725 -
01973 Tdp_wintergreen CsomeACy
01974 -# -
01975 -pf00148 -
01976 -pf00148pf08369 -
01977 -pf00142 -
01978 HMMnopfamclust3_1(pf00106) RMM1
01979 -# -
01980 -# -
01981 -pf08847 -
01982 -pf01925 -
01983 -pf02163 -
BMC-P
BMC-H
BMC-T
Aldehyde dehydrogenase
PTAC
Signature enzyme
Alcohol dehydrogenase
Regulator

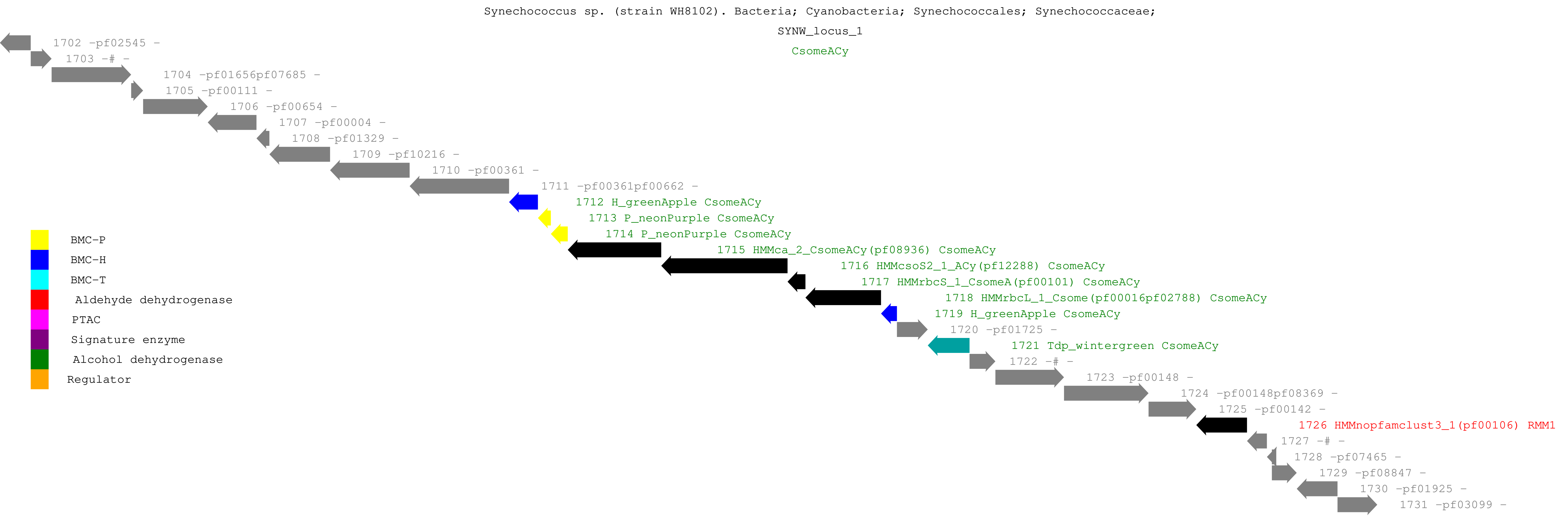

Synechococcus sp. (strain WH8102). Bacteria; Cyanobacteria; Synechococcales; Synechococcaceae;
SYNW_locus_1
CsomeACy
1702 -pf02545 -
1703 -# -
1704 -pf01656pf07685 -
1705 -pf00111 -
1706 -pf00654 -
1707 -pf00004 -
1708 -pf01329 -
1709 -pf10216 -
1710 -pf00361 -
1711 -pf00361pf00662 -
1712 H_greenApple CsomeACy
1713 P_neonPurple CsomeACy
1714 P_neonPurple CsomeACy
1715 HMMca_2_CsomeACy(pf08936) CsomeACy
1716 HMMcsoS2_1_ACy(pf12288) CsomeACy
1717 HMMrbcS_1_CsomeA(pf00101) CsomeACy
1718 HMMrbcL_1_Csome(pf00016pf02788) CsomeACy
1719 H_greenApple CsomeACy
1720 -pf01725 -
1721 Tdp_wintergreen CsomeACy
1722 -# -
1723 -pf00148 -
1724 -pf00148pf08369 -
1725 -pf00142 -
1726 HMMnopfamclust3_1(pf00106) RMM1
1727 -# -
1728 -pf07465 -
1729 -pf08847 -
1730 -pf01925 -
1731 -pf03099 -
BMC-P
BMC-H
BMC-T
Aldehyde dehydrogenase
PTAC
Signature enzyme
Alcohol dehydrogenase
Regulator

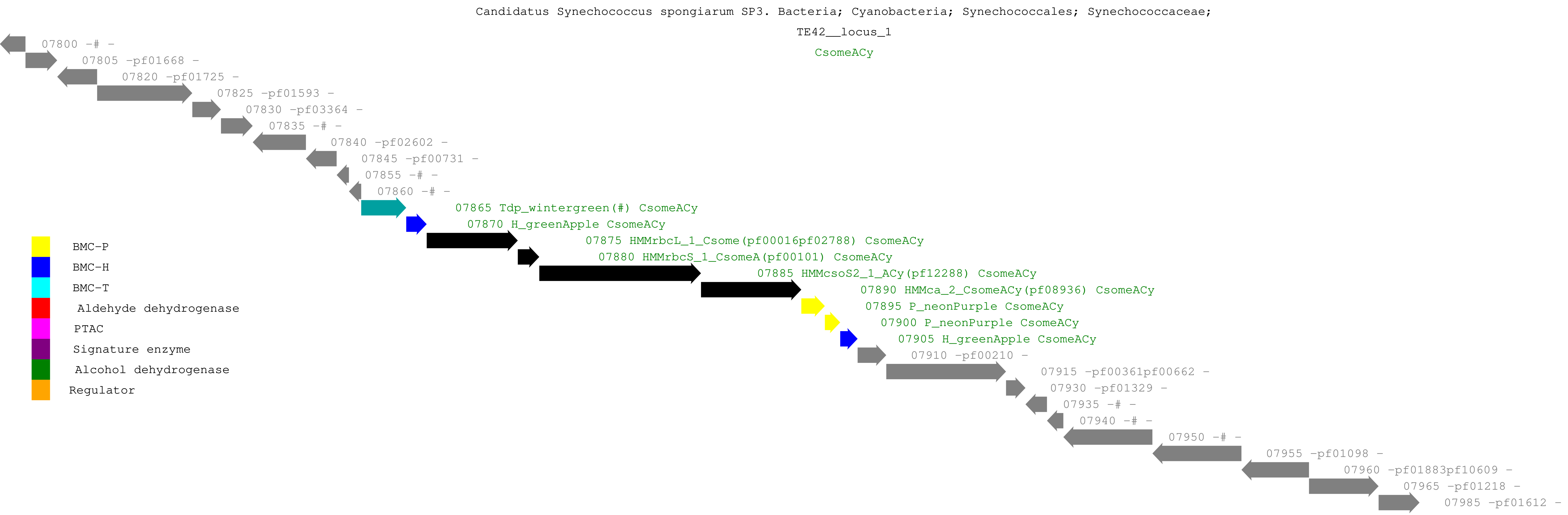

Candidatus Synechococcus spongiarum SP3. Bacteria; Cyanobacteria; Synechococcales; Synechococcaceae;
TE42__locus_1
CsomeACy
07800 -# -
07805 -pf01668 -
07820 -pf01725 -
07825 -pf01593 -
07830 -pf03364 -
07835 -# -
07840 -pf02602 -
07845 -pf00731 -
07855 -# -
07860 -# -
07865 Tdp_wintergreen(#) CsomeACy
07870 H_greenApple CsomeACy
07875 HMMrbcL_1_Csome(pf00016pf02788) CsomeACy
07880 HMMrbcS_1_CsomeA(pf00101) CsomeACy
07885 HMMcsoS2_1_ACy(pf12288) CsomeACy
07890 HMMca_2_CsomeACy(pf08936) CsomeACy
07895 P_neonPurple CsomeACy
07900 P_neonPurple CsomeACy
07905 H_greenApple CsomeACy
07910 -pf00210 -
07915 -pf00361pf00662 -
07930 -pf01329 -
07935 -# -
07940 -# -
07950 -# -
07955 -pf01098 -
07960 -pf01883pf10609 -
07965 -pf01218 -
07985 -pf01612 -
BMC-P
BMC-H
BMC-T
Aldehyde dehydrogenase
PTAC
Signature enzyme
Alcohol dehydrogenase
Regulator

Candidatus Synechococcus spongiarum 142. Bacteria; Cyanobacteria; Synechococcales; Synechococcaceae;

TH68__locus_1

CsomeACy

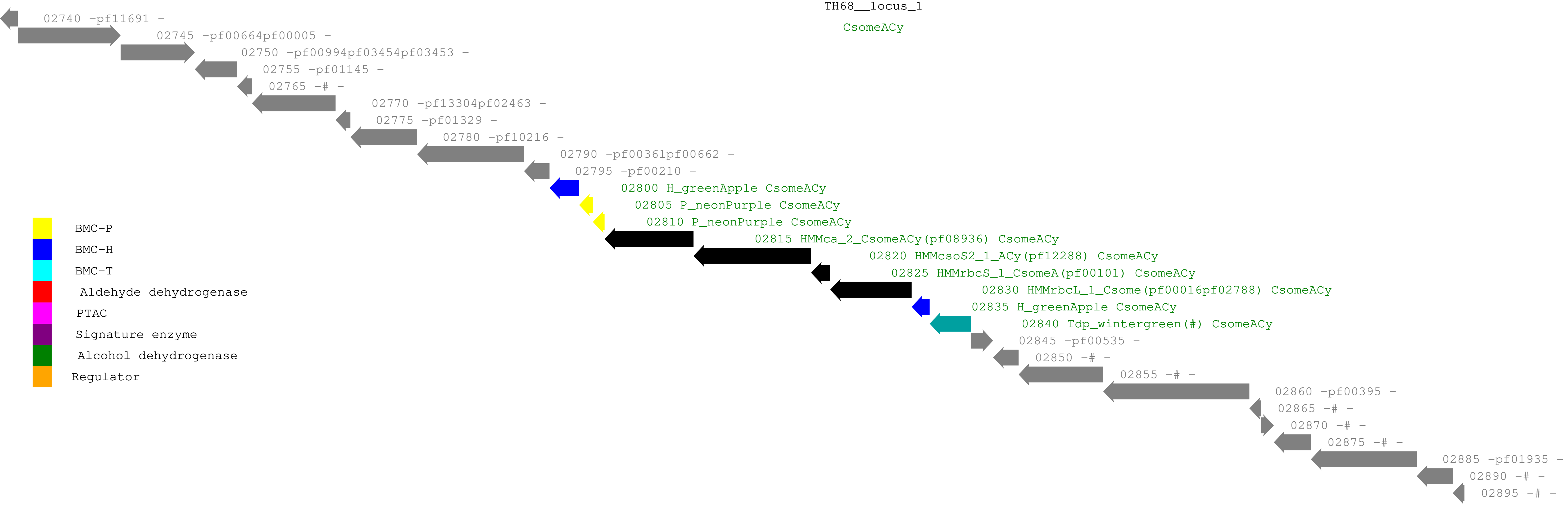

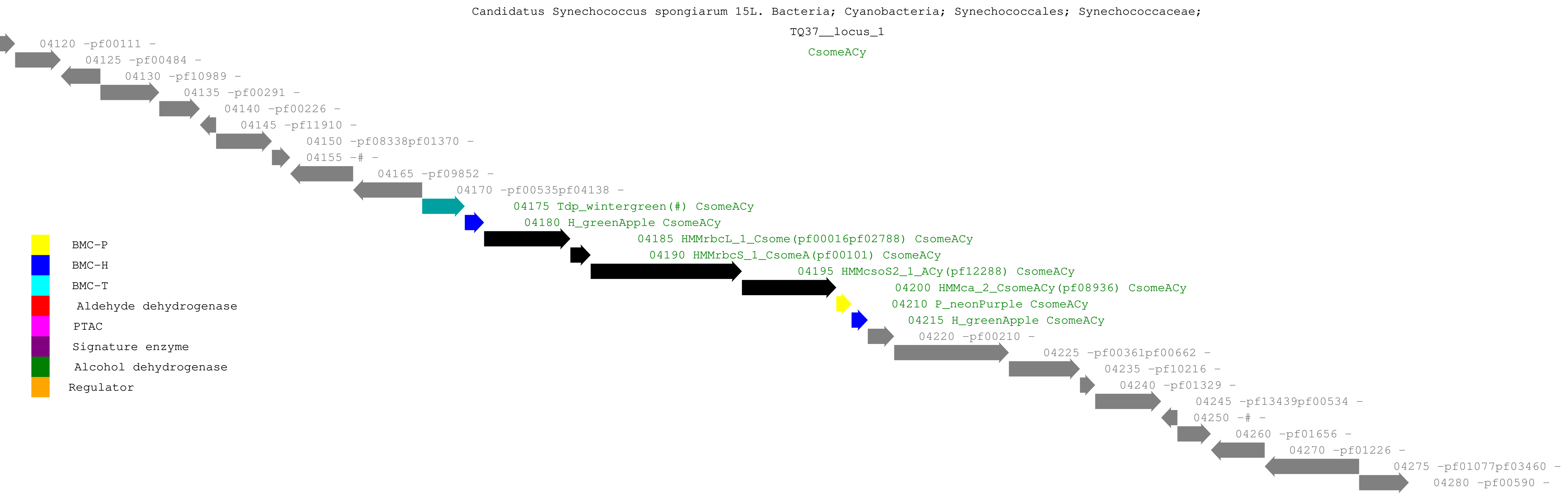

Candidatus Synechococcus spongiarum 15L. Bacteria; Cyanobacteria; Synechococcales; Synechococcaceae;
TQ37__locus_1
CsomeACy
04120 -pf00111 -
04125 -pf00484 -
04130 -pf10989 -
04135 -pf00291 -
04140 -pf00226 -
04145 -pf11910 -
04150 -pf08338pf01370 -
04155 -# -
04165 -pf09852 -
04170 -pf00535pf04138 -
04175 Tdp_wintergreen(#) CsomeACy
04180 H_greenApple CsomeACy
04185 HMMrbcL_1_Csome(pf00016pf02788) CsomeACy
04190 HMMrbcS_1_CsomeA(pf00101) CsomeACy
04195 HMMcsoS2_1_ACy(pf12288) CsomeACy
04200 HMMca_2_CsomeACy(pf08936) CsomeACy
04210 P_neonPurple CsomeACy
04215 H_greenApple CsomeACy
04220 -pf00210 -
04225 -pf00361pf00662 -
04235 -pf10216 -
04240 -pf01329 -
04245 -pf13439pf00534 -
04250 -# -
04260 -pf01656 -
04270 -pf01226 -
04275 -pf01077pf03460 -
04280 -pf00590 -
BMC-P
BMC-H
BMC-T
Aldehyde dehydrogenase
PTAC
Signature enzyme
Alcohol dehydrogenase
Regulator

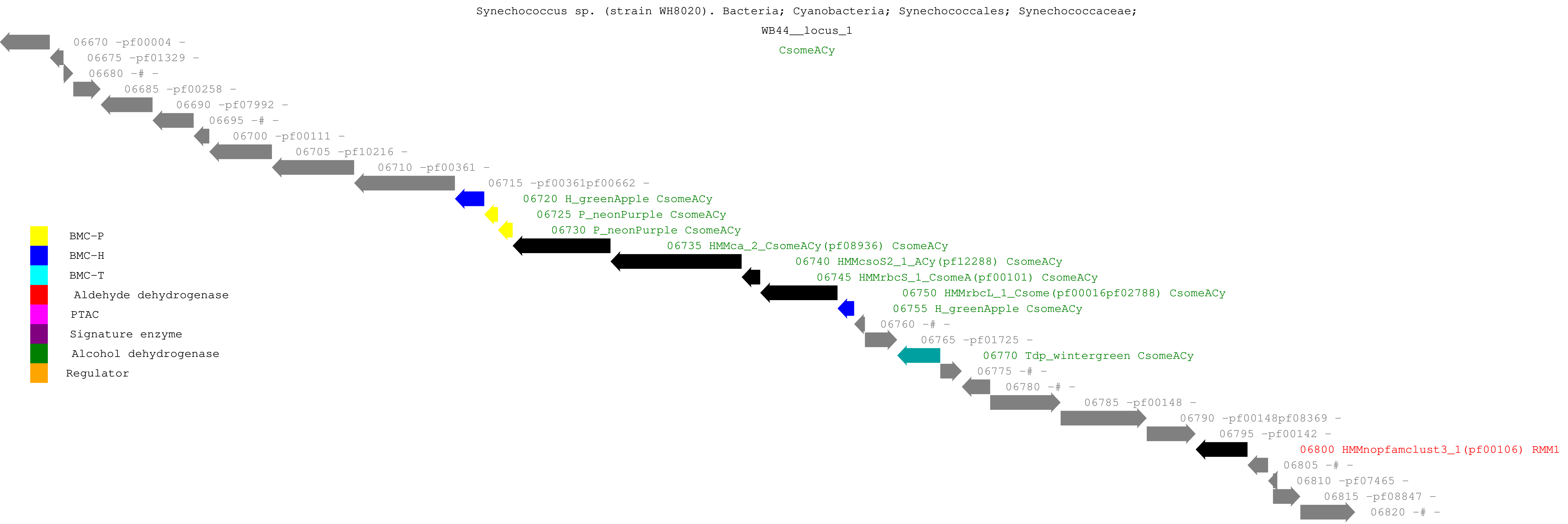
Synechococcus sp. (strain WH8020). Bacteria; Cyanobacteria; Synechococcales; Synechococcaceae;
WB44__locus_1
CsomeACy
06670 -pf00004 -
06675 -pf01329 -
06680 -# -
06685 -pf00258 -
06690 -pf07992 -
06695 -# -
06700 -pf00111 -
06705 -pf10216 -
06710 -pf00361 -
06715 -pf00361pf00662 -
06720 H_greenApple CsomeACy
06725 P_neonPurple CsomeACy
06730 P_neonPurple CsomeACy
06735 HMMca_2_CsomeACy(pf08936) CsomeACy
06740 HMMcsoS2_1_ACy(pf12288) CsomeACy
06745 HMMrbcS_1_CsomeA(pf00101) CsomeACy
06750 HMMrbcL_1_Csome(pf00016pf02788) CsomeACy
06755 H_greenApple CsomeACy
06760 -# -
06765 -pf01725 -
06770 Tdp_wintergreen CsomeACy
06775 -# -
06780 -# -
06785 -pf00148 -
06790 -pf00148pf08369 -
06795 -pf00142 -
06800 HMMnopfamclust3_1(pf00106) RMM1
06805 -# -
06810 -pf07465 -
06815 -pf08847 -
06820 -# -
BMC-P
BMC-H
BMC-T
Aldehyde dehydrogenase
PTAC
Signature enzyme
Alcohol dehydrogenase
Regulator

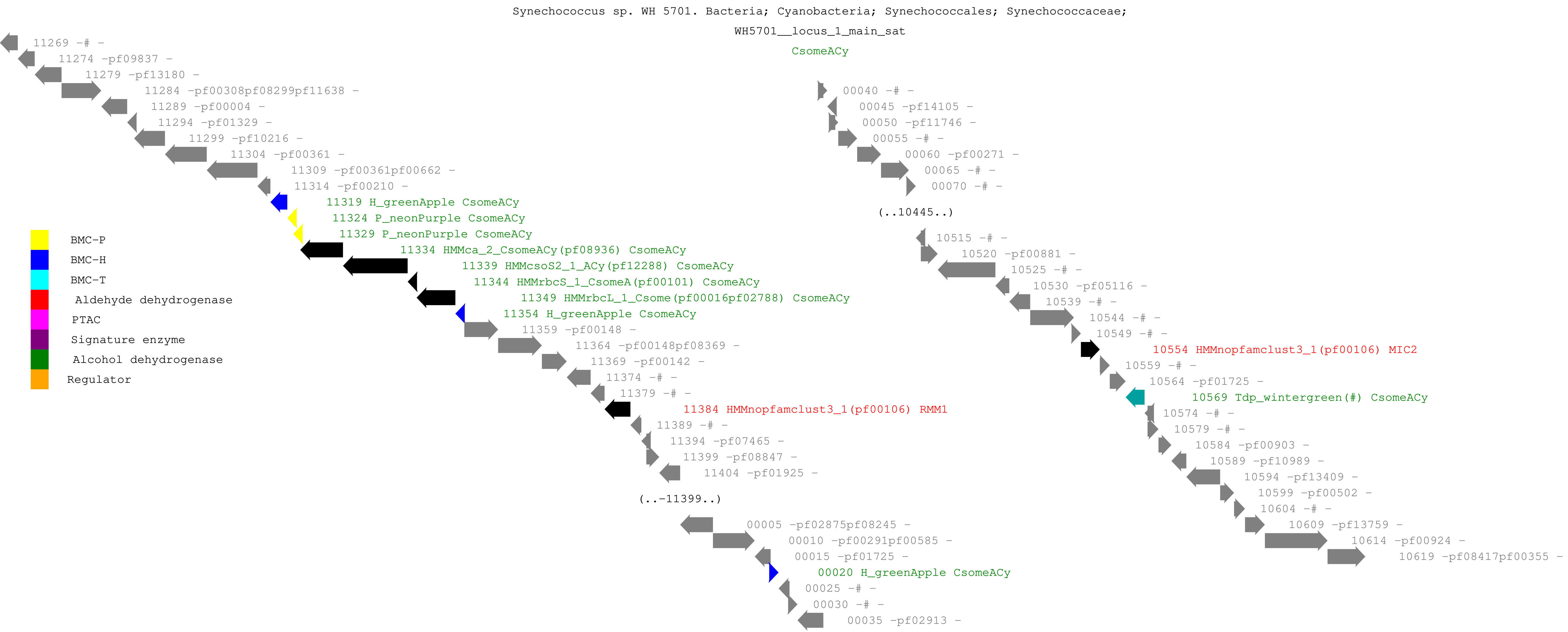

Synechococcus sp. WH 5701. Bacteria; Cyanobacteria; Synechococcales; Synechococcaceae;
WH5701__locus_1_main_sat
CsomeACy
11269 -# -
11274 -pf09837 -
11279 -pf13180 -
11284 -pf00308pf08299pf11638 -
11289 -pf00004 -
11294 -pf01329 -
11299 -pf10216 -
11304 -pf00361 -
11309 -pf00361pf00662 -
11314 -pf00210 -
11319 H_greenApple CsomeACy
11324 P_neonPurple CsomeACy
11329 P_neonPurple CsomeACy
11334 HMMca_2_CsomeACy(pf08936) CsomeACy
11339 HMMcsoS2_1_ACy(pf12288) CsomeACy
11344 HMMrbcS_1_CsomeA(pf00101) CsomeACy
11349 HMMrbcL_1_Csome(pf00016pf02788) CsomeACy
11354 H_greenApple CsomeACy
11359 -pf00148 -
11364 -pf00148pf08369 -
11369 -pf00142 -
11374 -# -
11379 -# -
11384 HMMnopfamclust3_1(pf00106) RMM1
11389 -# -
11394 -pf07465 -
11399 -pf08847 -
11404 -pf01925 -
(..-11399..)
00005 -pf02875pf08245 -
00010 -pf00291pf00585 -
00015 -pf01725 -
00020 H_greenApple CsomeACy
00025 -# -
00030 -# -
00035 -pf02913 -
00040 -# -
00045 -pf14105 -
00050 -pf11746 -
00055 -# -
00060 -pf00271 -
00065 -# -
00070 -# -
(..10445..)
10515 -# -
10520 -pf00881 -
10525 -# -
10530 -pf05116 -
10539 -# -
10544 -# -
10549 -# -
10554 HMMnopfamclust3_1(pf00106) MIC2
10559 -# -
10564 -pf01725 -
10569 Tdp_wintergreen(#) CsomeACy
10574 -# -
10579 -# -
10584 -pf00903 -
10589 -pf10989 -
10594 -pf13409 -
10599 -pf00502 -
10604 -# -
10609 -pf13759 -
10614 -pf00924 -
10619 -pf08417pf00355 -
BMC-P
BMC-H
BMC-T
Aldehyde dehydrogenase
PTAC
Signature enzyme
Alcohol dehydrogenase
Regulator

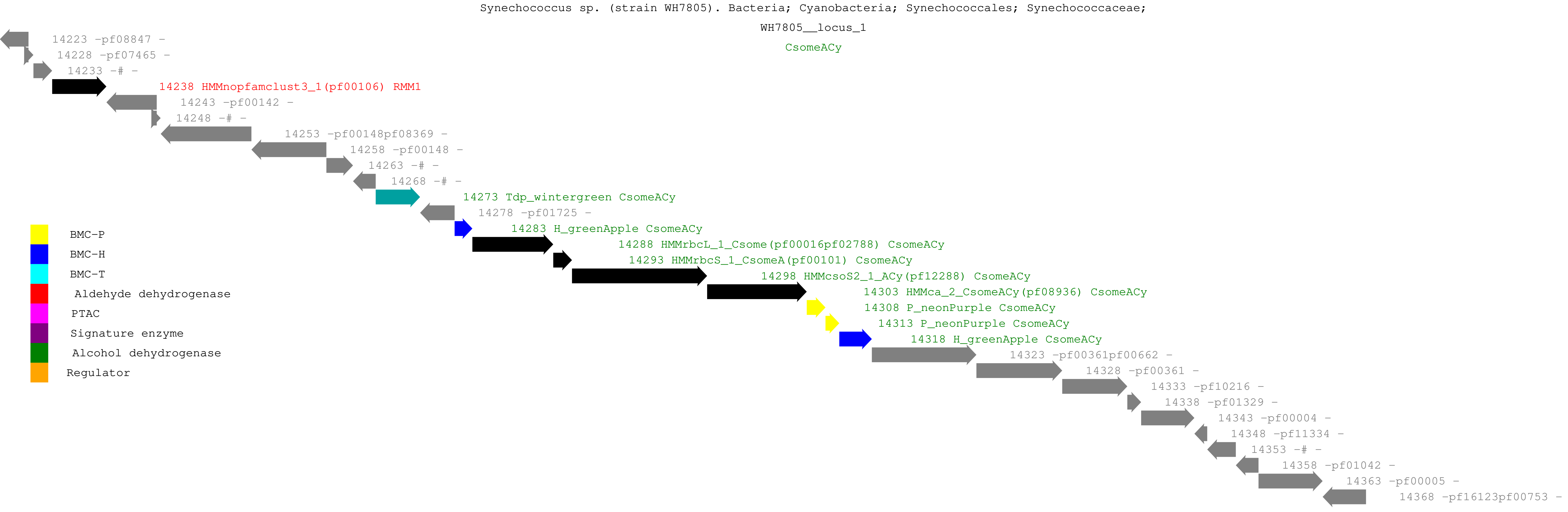
Synechococcus sp. (strain WH7805). Bacteria; Cyanobacteria; Synechococcales; Synechococcaceae;
WH7805__locus_1
CsomeACy
14223 -pf08847 -
14228 -pf07465 -
14233 -# -
14238 HMMnopfamclust3_1(pf00106) RMM1
14243 -pf00142 -
14248 -# -
14253 -pf00148pf08369 -
14258 -pf00148 -
14263 -# -
14268 -# -
14273 Tdp_wintergreen CsomeACy
14278 -pf01725 -
14283 H_greenApple CsomeACy
14288 HMMrbcL_1_Csome(pf00016pf02788) CsomeACy
14293 HMMrbcS_1_CsomeA(pf00101) CsomeACy
14298 HMMcsoS2_1_ACy(pf12288) CsomeACy
14303 HMMca_2_CsomeACy(pf08936) CsomeACy
14308 P_neonPurple CsomeACy
14313 P_neonPurple CsomeACy
14318 H_greenApple CsomeACy
14323 -pf00361pf00662 -
14328 -pf00361 -
14333 -pf10216 -
14338 -pf01329 -
14343 -pf00004 -
14348 -pf11334 -
14353 -# -
14358 -pf01042 -
14363 -pf00005 -
14368 -pf16123pf00753 -
BMC-P
BMC-H
BMC-T
Aldehyde dehydrogenase
PTAC
Signature enzyme
Alcohol dehydrogenase
Regulator

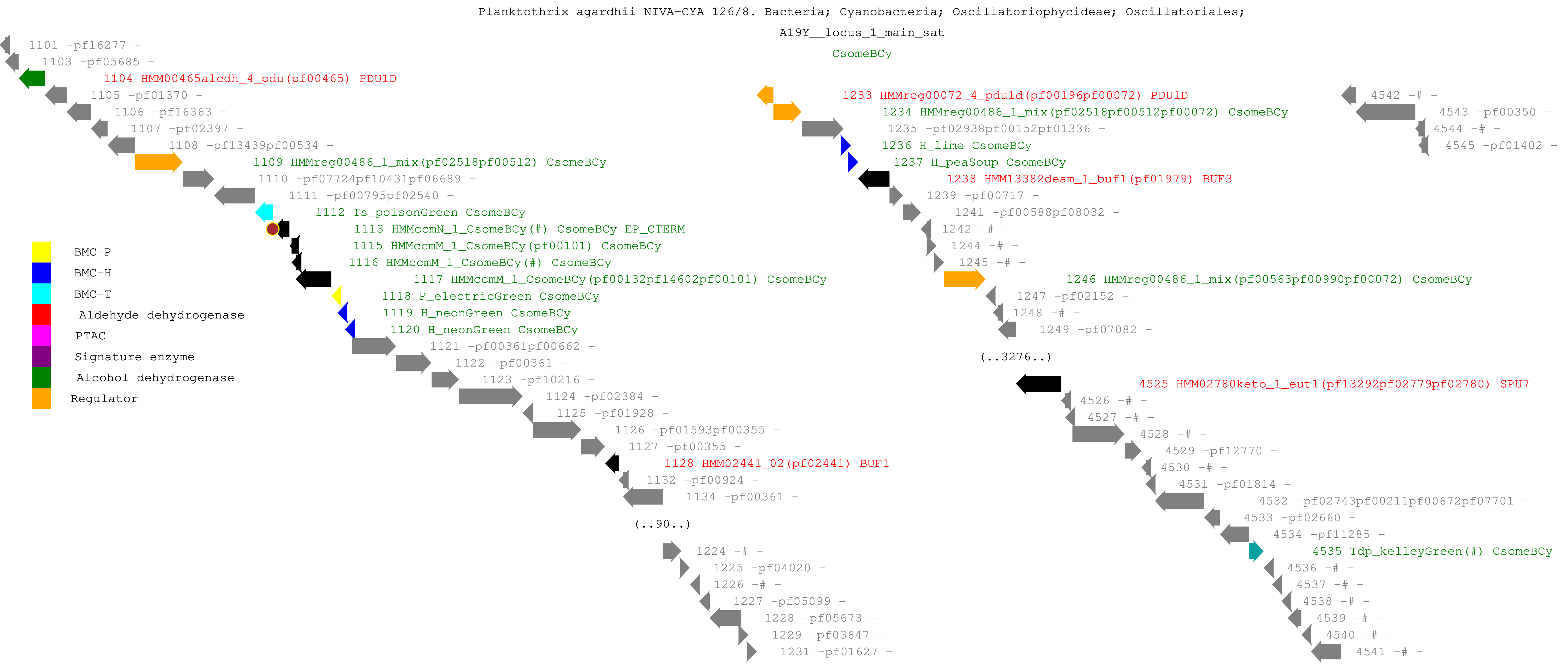
Planktothrix agardhii NIVA-CYA 126/8. Bacteria; Cyanobacteria; Oscillatoriophycideae; Oscillatoriales;
A19Y__locus_1_main_sat
CsomeBCy
1101 -pf16277 -
1103 -pf05685 -
1104 HMM00465alcdh_4_pdu(pf00465) PDU1D
1105 -pf01370 -
1106 -pf16363 -
1107 -pf02397 -
1108 -pf13439pf00534 -
1109 HMMreg00486_1_mix(pf02518pf00512) CsomeBCy
1110 -pf07724pf10431pf06689 -
1111 -pf00795pf02540 -
1112 Ts_poisonGreen CsomeBCy
1113 HMMccmN_1_CsomeBCy(#) CsomeBCy EP_CTERM
1115 HMMccmM_1_CsomeBCy(pf00101) CsomeBCy
1116 HMMccmM_1_CsomeBCy(#) CsomeBCy
1117 HMMccmM_1_CsomeBCy(pf00132pf14602pf00101) CsomeBCy
1118 P_electricGreen CsomeBCy
1119 H_neonGreen CsomeBCy
1120 H_neonGreen CsomeBCy
1121 -pf00361pf00662 -
1122 -pf00361 -
1123 -pf10216 -
1124 -pf02384 -
1125 -pf01928 -
1126 -pf01593pf00355 -
1127 -pf00355 -
1128 HMM02441_02(pf02441) BUF1
1132 -pf00924 -
1134 -pf00361 -
(..90..)
1224 -# -
1225 -pf04020 -
1226 -# -
1227 -pf05099 -
1228 -pf05673 -
1229 -pf03647 -
1231 -pf01627 -
BMC-P
BMC-H
BMC-T
Aldehyde dehydrogenase
PTAC
Signature enzyme
Alcohol dehydrogenase
Regulator
1233 HMMreg00072_4_pdu1d(pf00196pf00072) PDU1D
1234 HMMreg00486_1_mix(pf02518pf00512pf00072) CsomeBCy
1235 -pf02938pf00152pf01336 -
1236 H_lime CsomeBCy
1237 H_peaSoup CsomeBCy
1238 HMM13382deam_1_buf1(pf01979) BUF3
1239 -pf00717 -
1241 -pf00588pf08032 -
1242 -# -
1244 -# -
1245 -# -
1246 HMMreg00486_1_mix(pf00563pf00990pf00072) CsomeBCy
1247 -pf02152 -
1248 -# -
1249 -pf07082 -
(..3276..)
4525 HMM02780keto_1_eut1(pf13292pf02779pf02780) SPU7
4526 -# -
4527 -# -
4528 -# -
4529 -pf12770 -
4530 -# -
4531 -pf01814 -
4532 -pf02743pf00211pf00672pf07701 -
4533 -pf02660 -
4534 -pf11285 -
4535 Tdp_kelleyGreen(#) CsomeBCy
4536 -# -
4537 -# -
4538 -# -
4539 -# -
4540 -# -
4541 -# -
4542 -# -
4543 -pf00350 -
4544 -# -
4545 -pf01402 -

Nodularia spumigena CENA596. Bacteria; Cyanobacteria; Nostocales; Aphanizomenonaceae; Nodularia.

A2T98__locus_1_main_sat

CsomeBCy

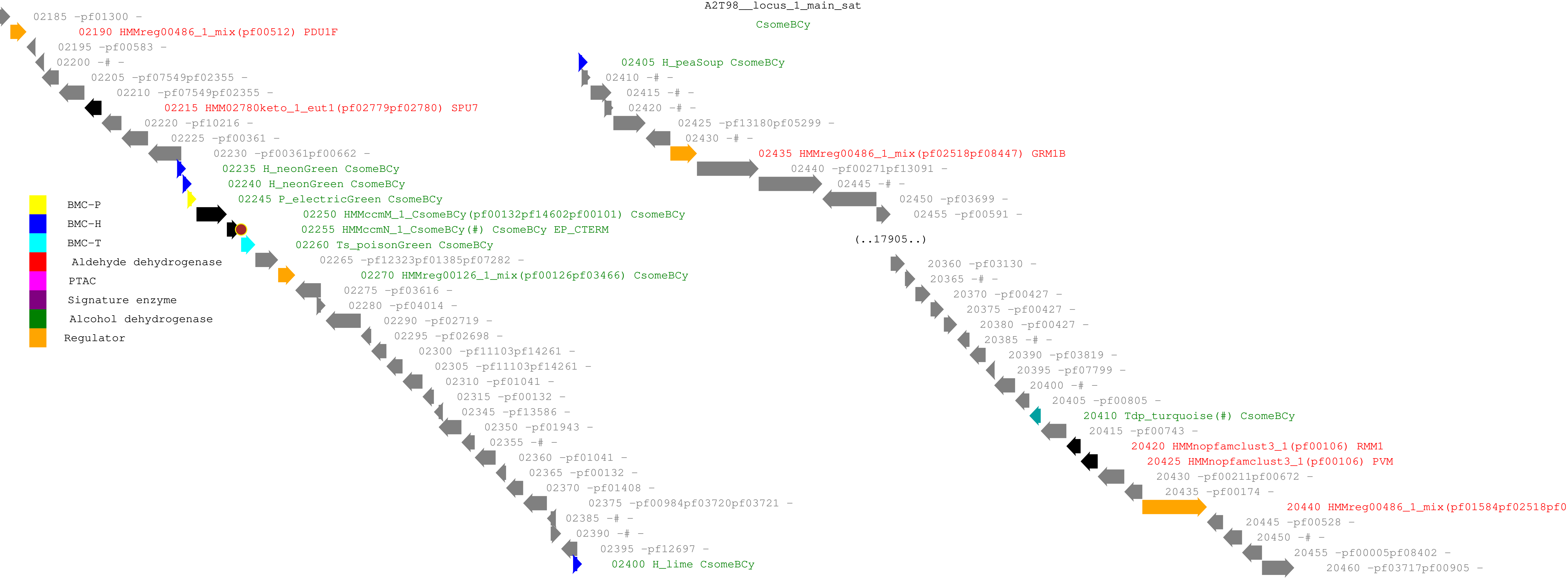

Nostoc sp. KVJ20. Bacteria; Cyanobacteria; Nostocales; Nostocaceae; Nostoc.

A4S05__locus_1_main_sat

CsomeBCy

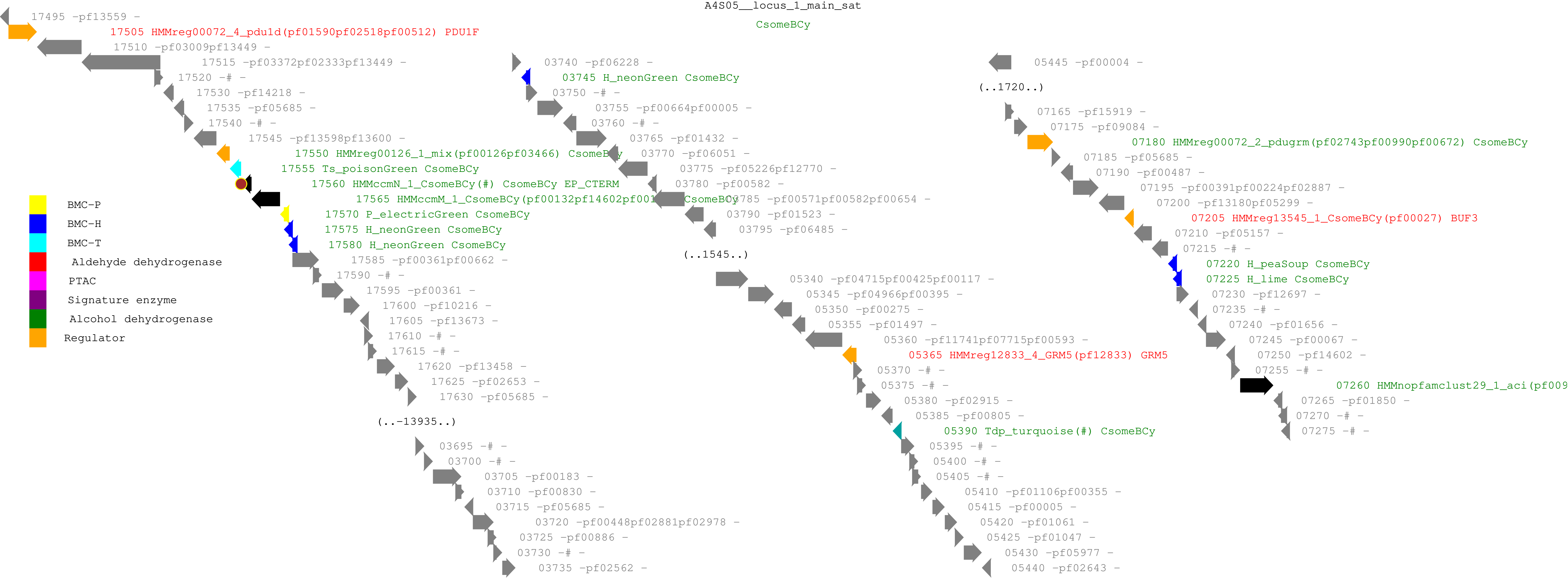

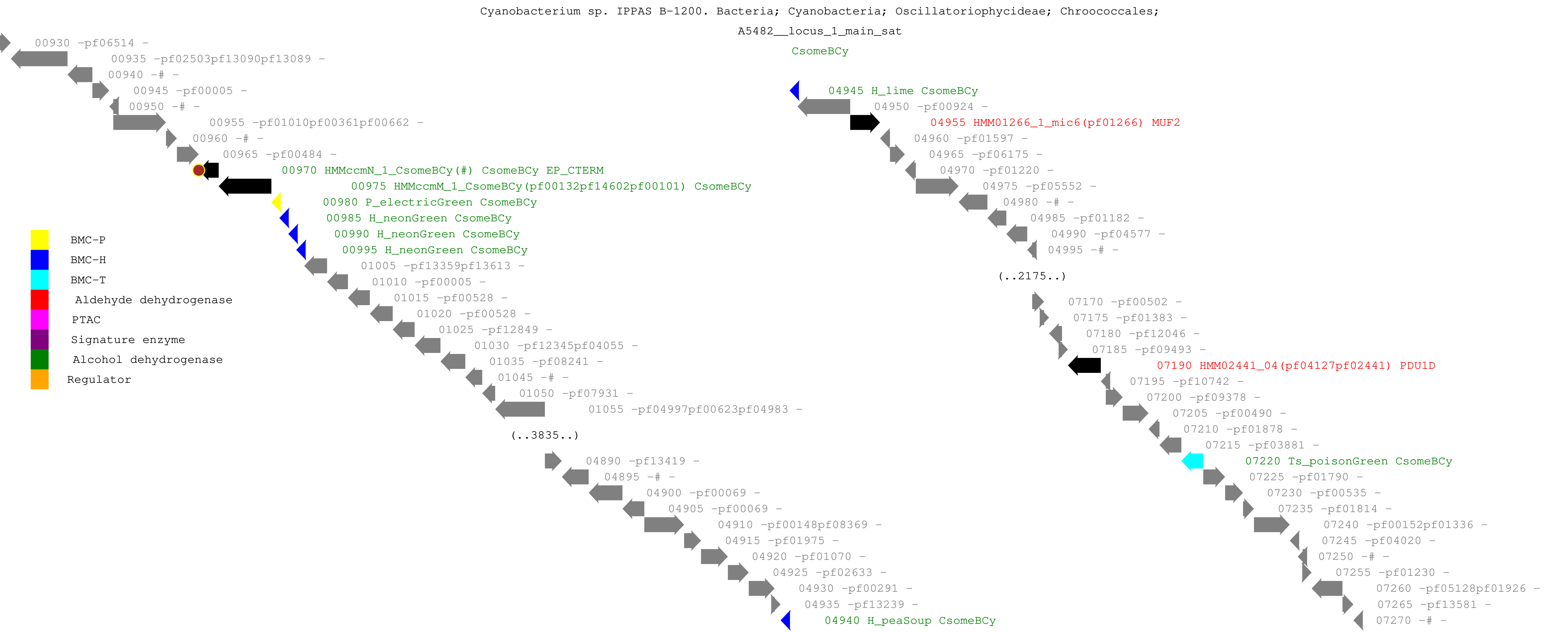

Cyanobacterium sp. IPPAS B-1200. Bacteria; Cyanobacteria; Oscillatoriophycideae; Chroococcales;
A5482__locus_1_main_sat
CsomeBCy
00930 -pf06514 -
00935 -pf02503pf13090pf13089 -
00940 -# -
00945 -pf00005 -
00950 -# -
00955 -pf01010pf00361pf00662 -
00960 -# -
00965 -pf00484 -
00970 HMMccmN_1_CsomeBCy(#) CsomeBCy EP_CTERM
00975 HMMccmM_1_CsomeBCy(pf00132pf14602pf00101) CsomeBCy
00980 P_electricGreen CsomeBCy
00985 H_neonGreen CsomeBCy
00990 H_neonGreen CsomeBCy
00995 H_neonGreen CsomeBCy
01005 -pf13359pf13613 -
01010 -pf00005 -
01015 -pf00528 -
01020 -pf00528 -
01025 -pf12849 -
01030 -pf12345pf04055 -
01035 -pf08241 -
01045 -# -
01050 -pf07931 -
01055 -pf04997pf00623pf04983 -
(..3835..)
04890 -pf13419 -
04895 -# -
04900 -pf00069 -
04905 -pf00069 -
04910 -pf00148pf08369 -
04915 -pf01975 -
04920 -pf01070 -
04925 -pf02633 -
04930 -pf00291 -
04935 -pf13239 -
04940 H_peaSoup CsomeBCy
04945 H_lime CsomeBCy
04950 -pf00924 -
04955 HMM01266_1_mic6(pf01266) MUF2
04960 -pf01597 -
04965 -pf06175 -
04970 -pf01220 -
04975 -pf05552 -
04980 -# -
04985 -pf01182 -
04990 -pf04577 -
04995 -# -
(..2175..)
07170 -pf00502 -
07175 -pf01383 -
07180 -pf12046 -
07185 -pf09493 -
07190 HMM02441_04(pf04127pf02441) PDU1D
07195 -pf10742 -
07200 -pf09378 -
07205 -pf00490 -
07210 -pf01878 -
07215 -pf03881 -
07220 Ts_poisonGreen CsomeBCy
07225 -pf01790 -
07230 -pf00535 -
07235 -pf01814 -
07240 -pf00152pf01336 -
07245 -pf04020 -
07250 -# -
07255 -pf01230 -
07260 -pf05128pf01926 -
07265 -pf13581 -
07270 -# -
BMC-P
BMC-H
BMC-T
Aldehyde dehydrogenase
PTAC
Signature enzyme
Alcohol dehydrogenase
Regulator

Nostoc punctiforme NIES-2108. Bacteria; Cyanobacteria; Nostocales; Nostocaceae; Nostoc.

A6769__locus_1_main_sat

CsomeBCy

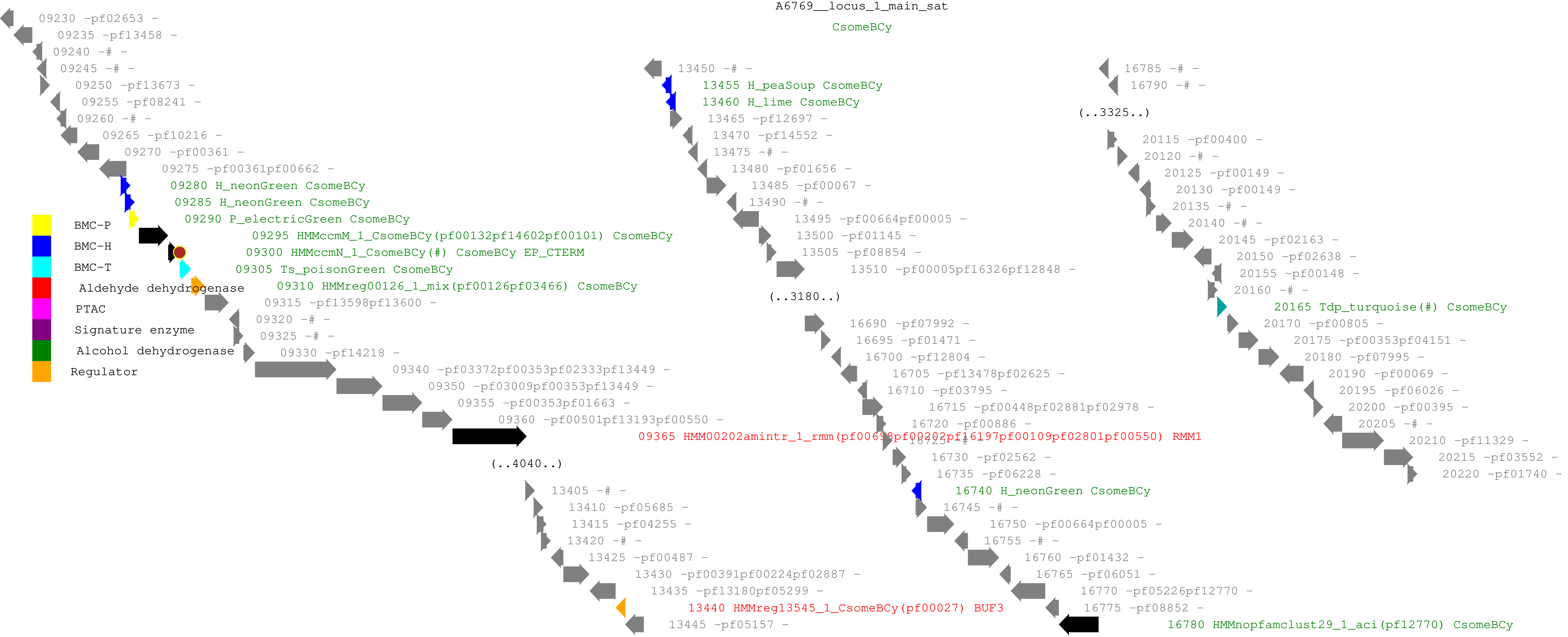

Nostoc minutum NIES-26. Bacteria; Cyanobacteria; Nostocales; Nostocaceae; Nostoc.

A6770__locus_1_main_sat

CsomeBCy

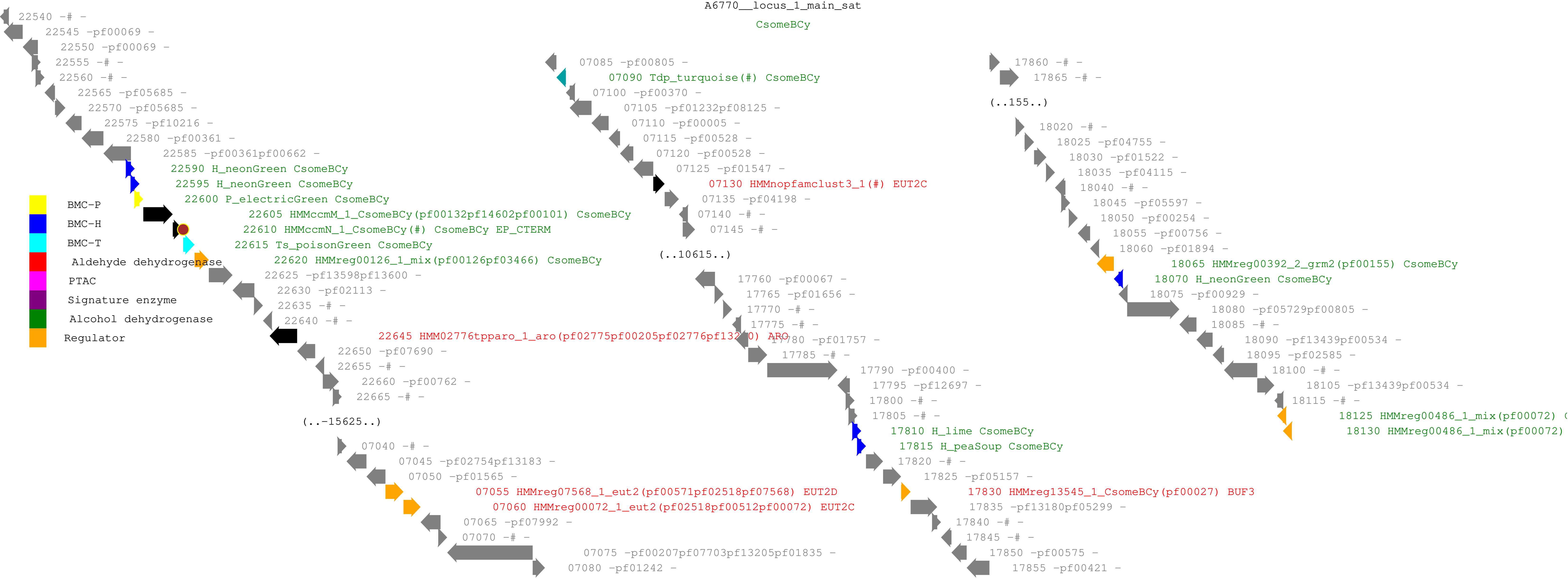

Nostoc sp. ATCC 43529. Bacteria; Cyanobacteria; Nostocales; Nostocaceae; Nostoc.

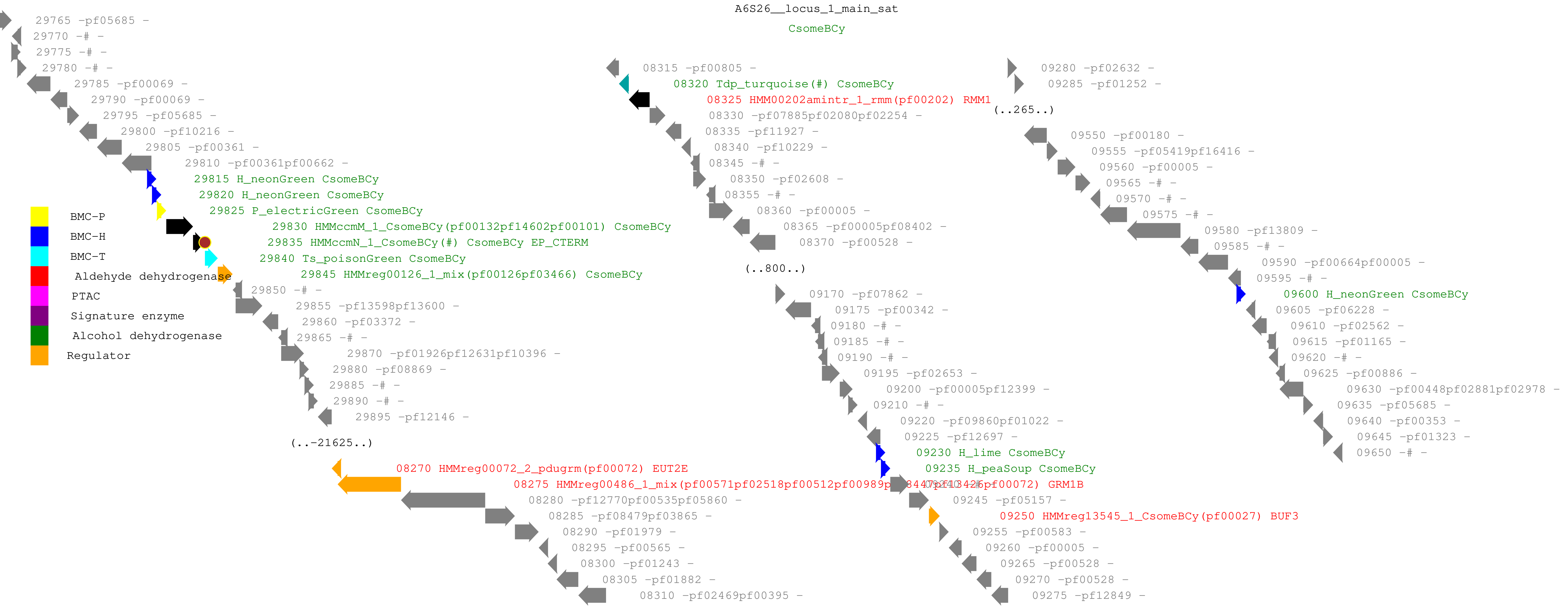

Nostoc sp. ATCC 53789. Bacteria; Cyanobacteria; Nostocales; Nostocaceae; Nostoc.

A6V25__locus_1_main_sat

CsomeBCy

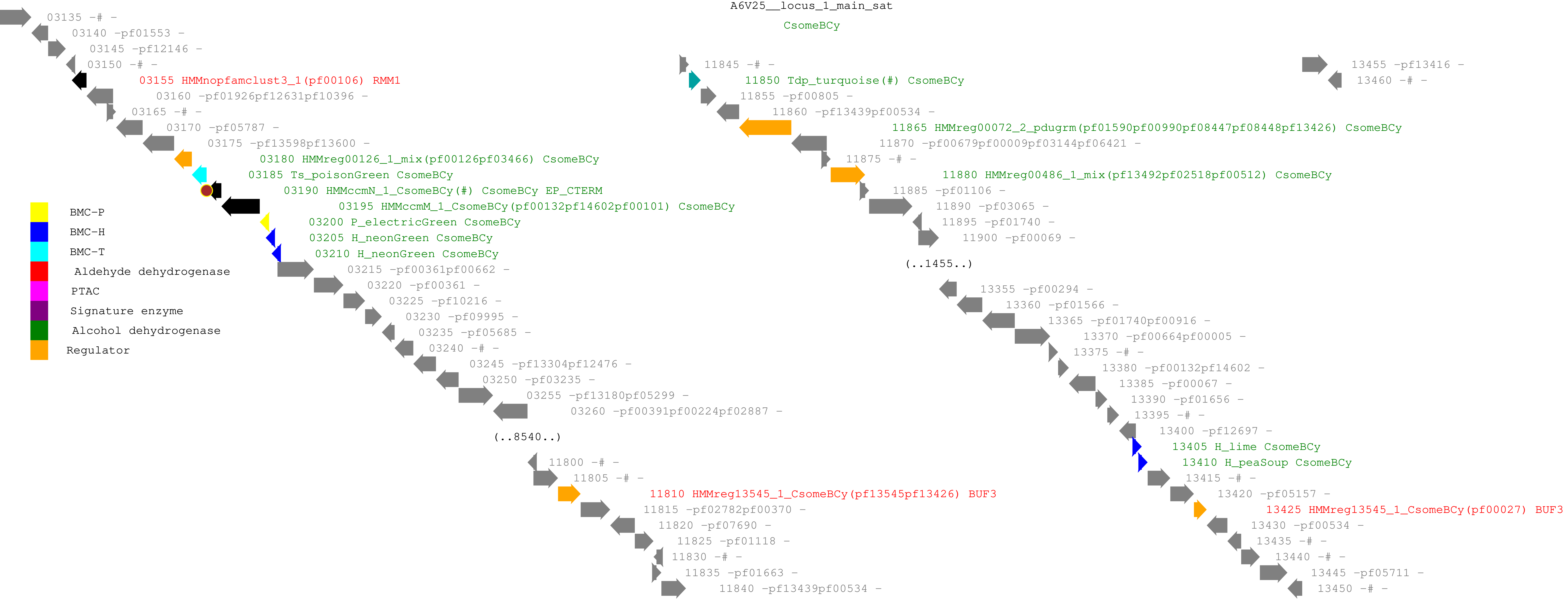

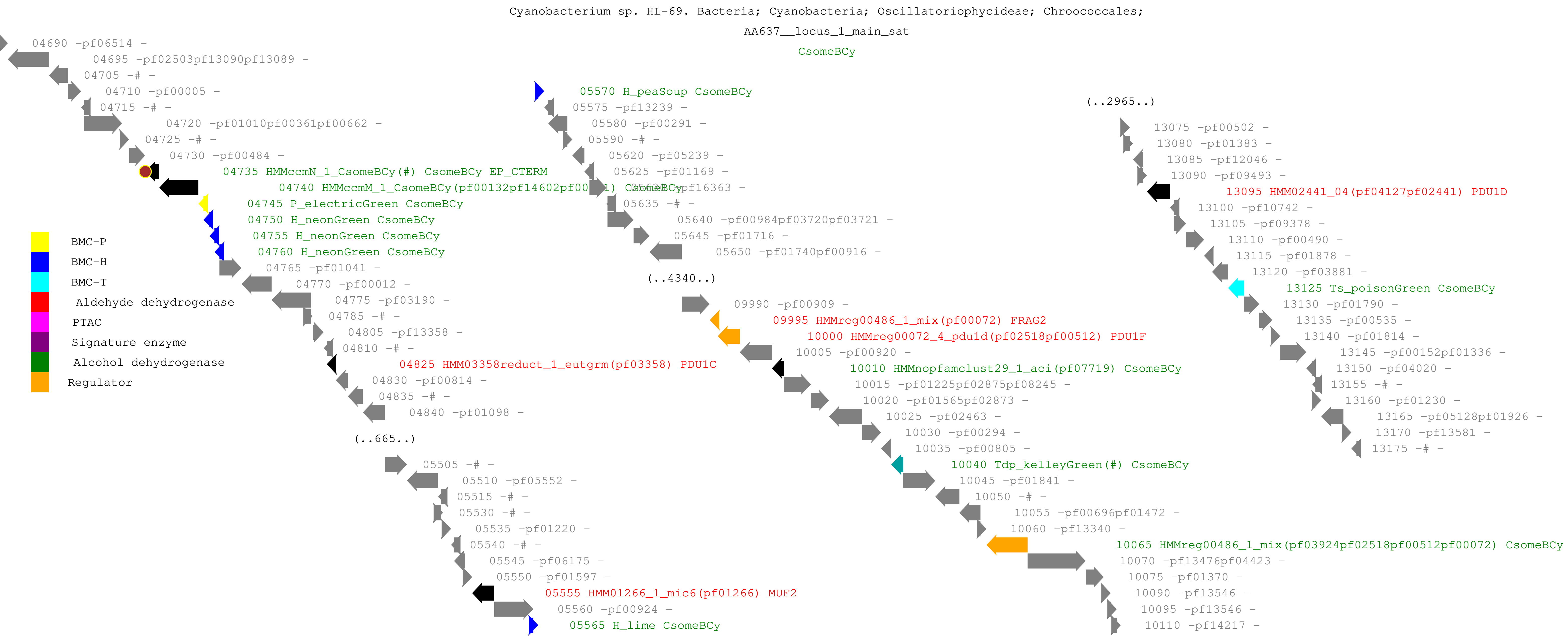
Cyanobacterium sp. HL-69. Bacteria; Cyanobacteria; Oscillatoriophycideae; Chroococcales;
AA637__locus_1_main_sat
CsomeBCy
04690 -pf06514 -
04695 -pf02503pf13090pf13089 -
04705 -# -
04710 -pf00005 -
04715 -# -
04720 -pf01010pf00361pf00662 -
04725 -# -
04730 -pf00484 -
04735 HMMccmN_1_CsomeBCy(#) CsomeBCy EP_CTERM
04740 HMMccmM_1_CsomeBCy(pf00132pf14602pf00[illegible]1) CsomeBCy
04745 P_electricGreen CsomeBCy
04750 H_neonGreen CsomeBCy
04755 H_neonGreen CsomeBCy
04760 H_neonGreen CsomeBCy
04765 -pf01041 -
04770 -pf00012 -
04775 -pf03190 -
04785 -# -
04805 -pf13358 -
04810 -# -
04825 HMM03358reduct_1_eutgrm(pf03358) PDU1C
04830 -pf00814 -
04835 -# -
04840 -pf01098 -
(..665..)
05505 -# -
05510 -pf05552 -
05515 -# -
05530 -# -
05535 -pf01220 -
05540 -# -
05545 -pf06175 -
05550 -pf01597 -
05555 HMM01266_1_mic6(pf01266) MUF2
05560 -pf00924 -
05565 H_lime CsomeBCy
05570 H_peaSoup CsomeBCy
05575 -pf13239 -
05580 -pf00291 -
05590 -# -
05620 -pf05239 -
05625 -pf01169 -
[illegible]pf16363 -
05635 -# -
05640 -pf00984pf03720pf03721 -
05645 -pf01716 -
05650 -pf01740pf00916 -
(..4340..)
09990 -pf00909 -
09995 HMMreg00486_1_mix(pf00072) FRAG2
10000 HMMreg00072_4_pdu1d(pf02518pf00512) PDU1F
10005 -pf00920 -
10010 HMMnopfamclust29_1_aci(pf07719) CsomeBCy
10015 -pf01225pf02875pf08245 -
10020 -pf01565pf02873 -
10025 -pf02463 -
10030 -pf00294 -
10035 -pf00805 -
10040 Tdp_kelleyGreen(#) CsomeBCy
10045 -pf01841 -
10050 -# -
10055 -pf00696pf01472 -
10060 -pf13340 -
10065 HMMreg00486_1_mix(pf03924pf02518pf00512pf00072) CsomeBCy
10070 -pf13476pf04423 -
10075 -pf01370 -
10090 -pf13546 -
10095 -pf13546 -
10110 -pf14217 -
(..2965..)
13075 -pf00502 -
13080 -pf01383 -
13085 -pf12046 -
13090 -pf09493 -
13095 HMM02441_04(pf04127pf02441) PDU1D
13100 -pf10742 -
13105 -pf09378 -
13110 -pf00490 -
13115 -pf01878 -
13120 -pf03881 -
13125 Ts_poisonGreen CsomeBCy
13130 -pf01790 -
13135 -pf00535 -
13140 -pf01814 -
13145 -pf00152pf01336 -
13150 -pf04020 -
13155 -# -
13160 -pf01230 -
13165 -pf05128pf01926 -
13170 -pf13581 -
13175 -# -
BMC-P
BMC-H
BMC-T
Aldehyde dehydrogenase
PTAC
Signature enzyme
Alcohol dehydrogenase
Regulator

Anabaena sp. WA102. Bacteria; Cyanobacteria; Nostocales; Nostocaceae; Anabaena.

AA650__locus_1_main_sat

CsomeBCy

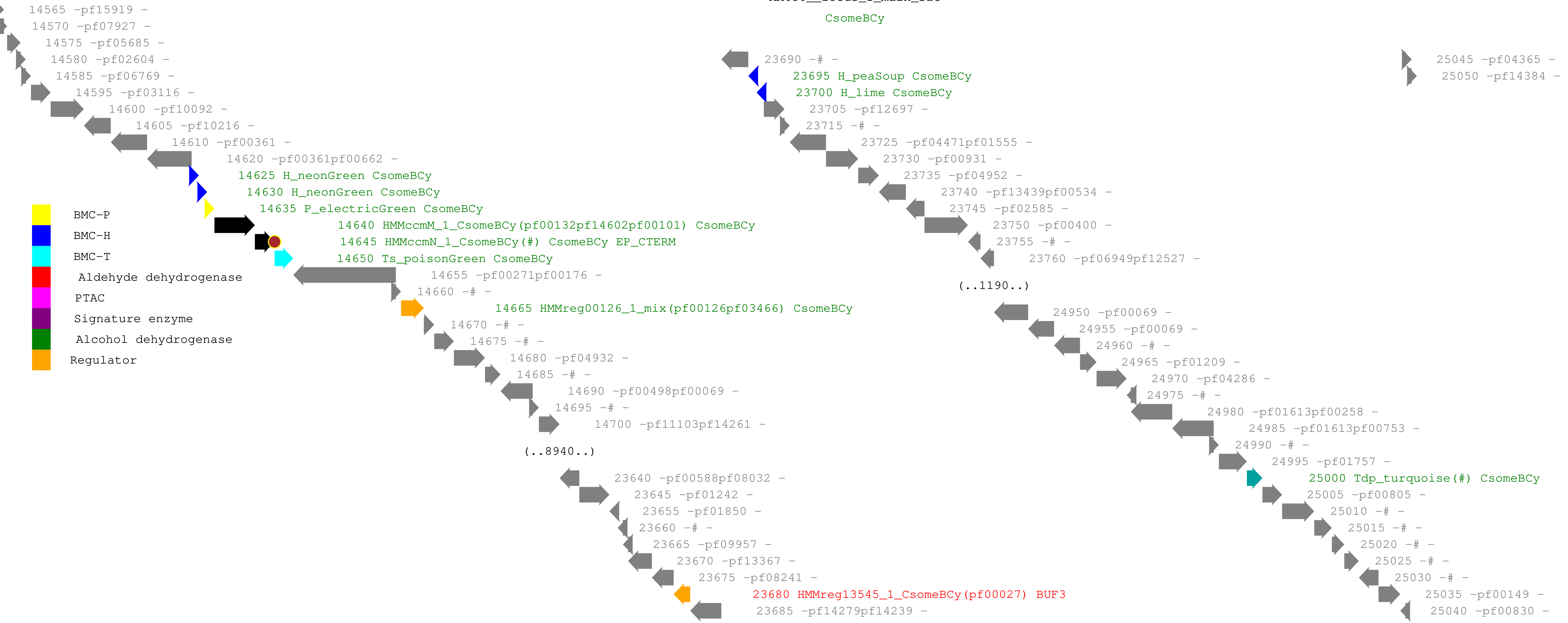

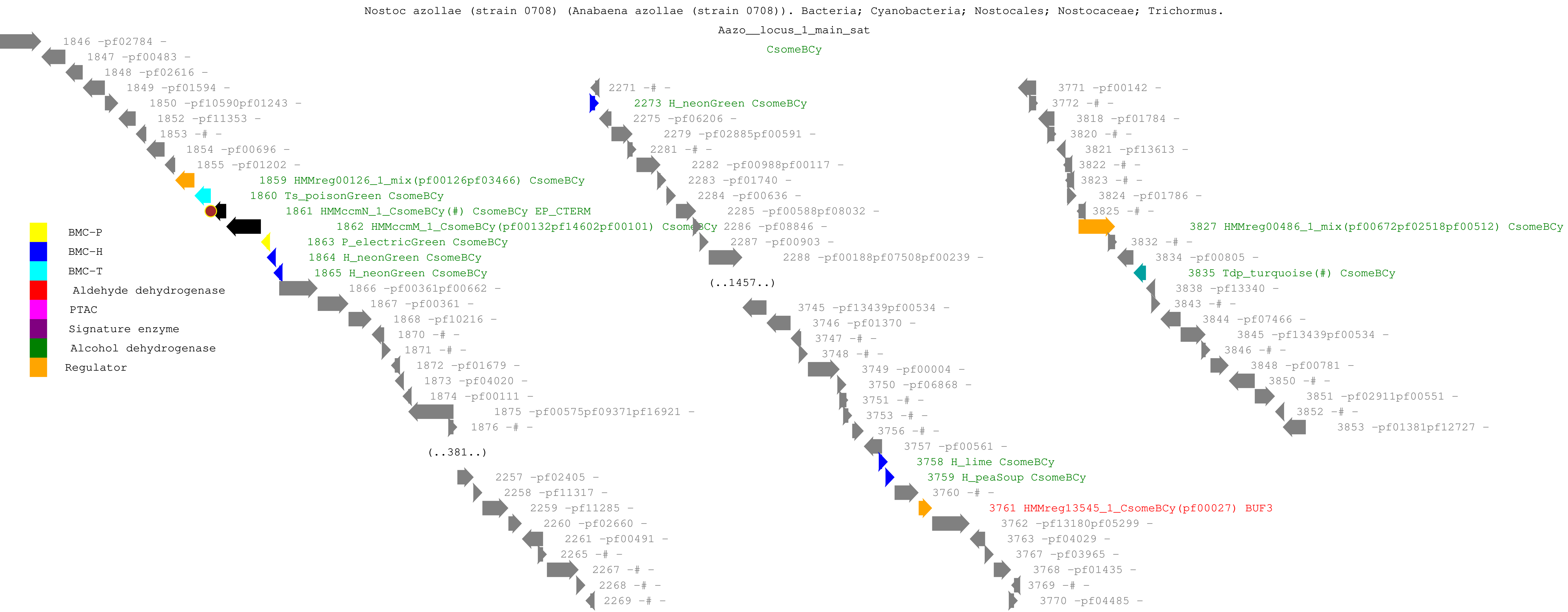

Nostoc azollae (strain 0708) (Anabaena azollae (strain 0708)). Bacteria; Cyanobacteria; Nostocales; Nostocaceae; Trichormus.
Aazo__locus_1_main_sat
CsomeBCy
1846 -pf02784 -
1847 -pf00483 -
1848 -pf02616 -
1849 -pf01594 -
1850 -pf10590pf01243 -
1852 -pf11353 -
1853 -# -
1854 -pf00696 -
1855 -pf01202 -
1859 HMMreg00126_1_mix(pf00126pf03466) CsomeBCy
1860 Ts_poisonGreen CsomeBCy
1861 HMMccmN_1_CsomeBCy(#) CsomeBCy EP_CTERM
1862 HMMccmM_1_CsomeBCy(pf00132pf14602pf00101) CsomeBCy
1863 P_electricGreen CsomeBCy
1864 H_neonGreen CsomeBCy
1865 H_neonGreen CsomeBCy
1866 -pf00361pf00662 -
1867 -pf00361 -
1868 -pf10216 -
1870 -# -
1871 -# -
1872 -pf01679 -
1873 -pf04020 -
1874 -pf00111 -
1875 -pf00575pf09371pf16921 -
1876 -# -
(..381..)
2257 -pf02405 -
2258 -pf11317 -
2259 -pf11285 -
2260 -pf02660 -
2261 -pf00491 -
2265 -# -
2267 -# -
2268 -# -
2269 -# -
BMC-P
BMC-H
BMC-T
Aldehyde dehydrogenase
PTAC
Signature enzyme
Alcohol dehydrogenase
Regulator
2271 -# -
2273 H_neonGreen CsomeBCy
2275 -pf06206 -
2279 -pf02885pf00591 -
2281 -# -
2282 -pf00988pf00117 -
2283 -pf01740 -
2284 -pf00636 -
2285 -pf00588pf08032 -
2286 -pf08846 -
2287 -pf00903 -
2288 -pf00188pf07508pf00239 -
(..1457..)
3745 -pf13439pf00534 -
3746 -pf01370 -
3747 -# -
3748 -# -
3749 -pf00004 -
3750 -pf06868 -
3751 -# -
3753 -# -
3756 -# -
3757 -pf00561 -
3758 H_lime CsomeBCy
3759 H_peaSoup CsomeBCy
3760 -# -
3761 HMMreg13545_1_CsomeBCy(pf00027) BUF3
3762 -pf13180pf05299 -
3763 -pf04029 -
3767 -pf03965 -
3768 -pf01435 -
3769 -# -
3770 -pf04485 -
3771 -pf00142 -
3772 -# -
3818 -pf01784 -
3820 -# -
3821 -pf13613 -
3822 -# -
3823 -# -
3824 -pf01786 -
3825 -# -
3827 HMMreg00486_1_mix(pf00672pf02518pf00512) CsomeBCy
3832 -# -
3834 -pf00805 -
3835 Tdp_turquoise(#) CsomeBCy
3838 -pf13340 -
3843 -# -
3844 -pf07466 -
3845 -pf13439pf00534 -
3846 -# -
3848 -pf00781 -
3850 -# -
3851 -pf02911pf00551 -
3852 -# -
3853 -pf01381pf12727 -

Pseudanabaena sp. ABRG5-3. Bacteria; Cyanobacteria; Synechococcales; Pseudanabaenaceae;

ABRG53__locus_1_main_sat

CsomeBCy

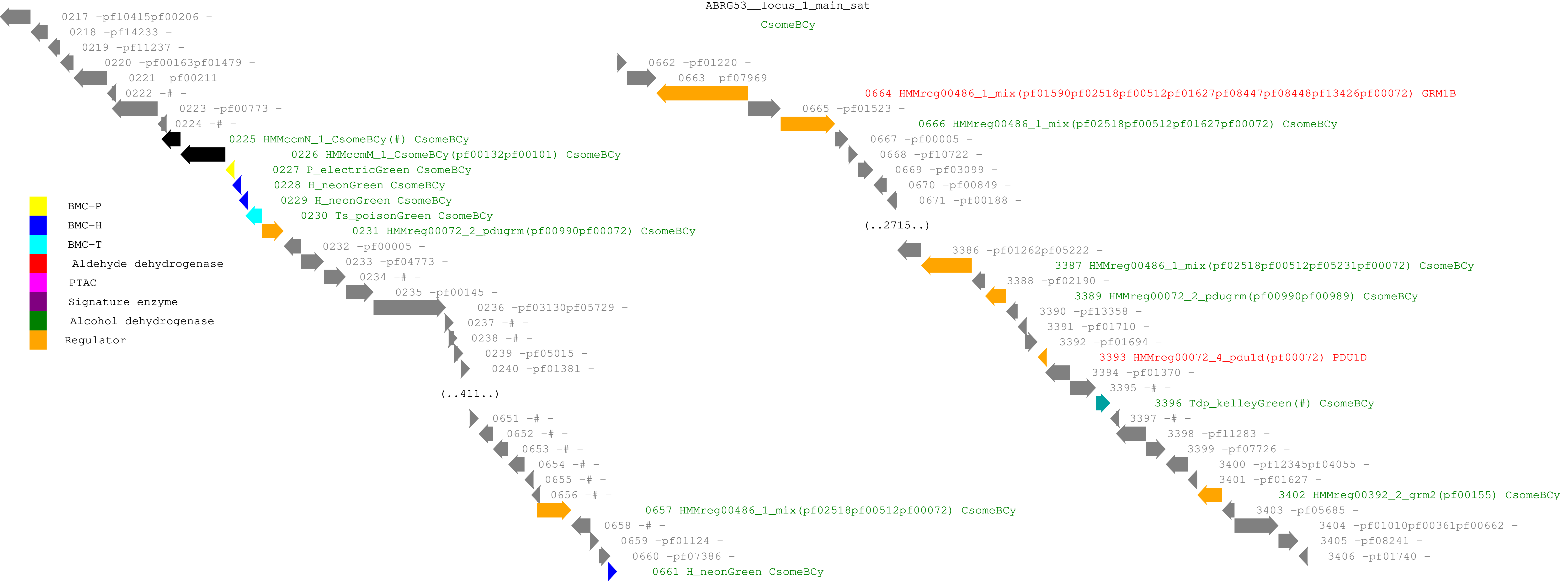

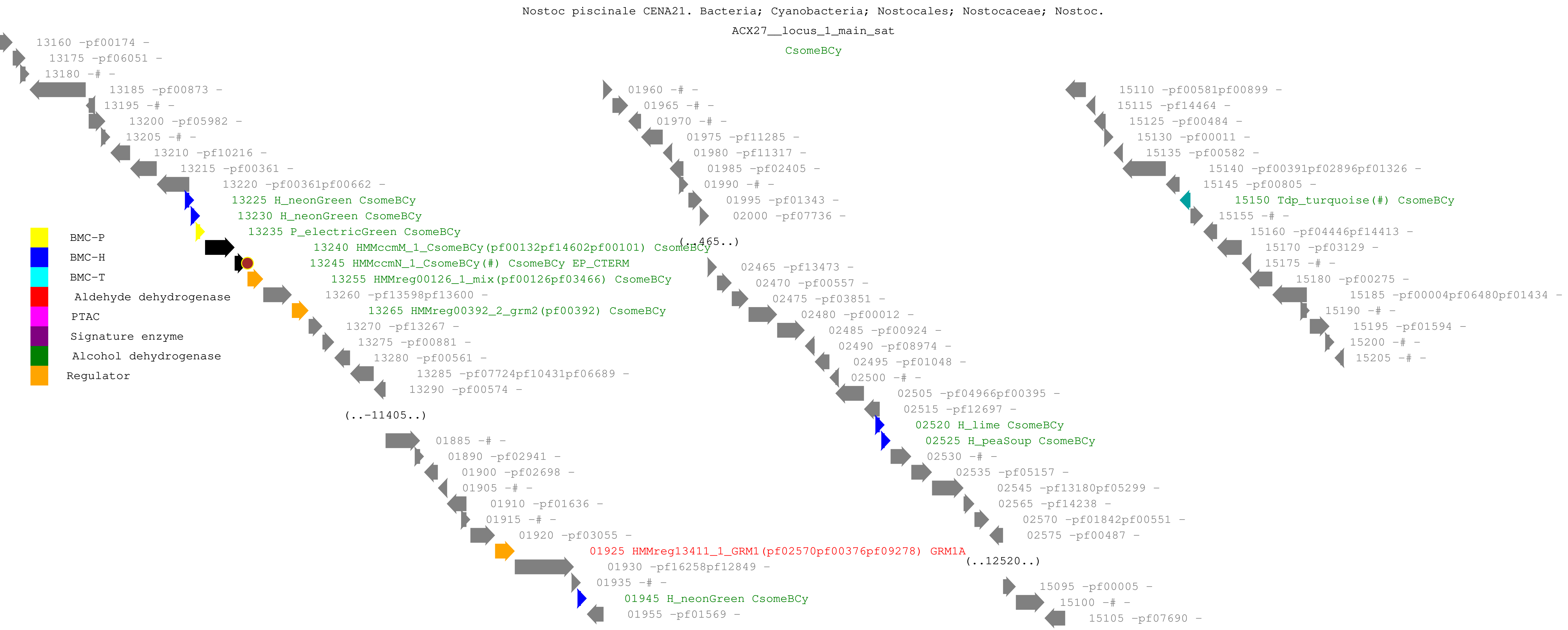
Nostoc piscinale CENA21. Bacteria; Cyanobacteria; Nostocales; Nostocaceae; Nostoc.
ACX27__locus_1_main_sat
CsomeBCy
13160 -pf00174 -
13175 -pf06051 -
13180 -# -
13185 -pf00873 -
13195 -# -
13200 -pf05982 -
13205 -# -
13210 -pf10216 -
13215 -pf00361 -
13220 -pf00361pf00662 -
13225 H_neonGreen CsomeBCy
13230 H_neonGreen CsomeBCy
13235 P_electricGreen CsomeBCy
13240 HMMccmM_1_CsomeBCy(pf00132pf14602pf00101) CsomeBCy
13245 HMMccmN_1_CsomeBCy(#) CsomeBCy EP_CTERM
13255 HMMreg00126_1_mix(pf00126pf03466) CsomeBCy
13260 -pf13598pf13600 -
13265 HMMreg00392_2_grm2(pf00392) CsomeBCy
13270 -pf13267 -
13275 -pf00881 -
13280 -pf00561 -
13285 -pf07724pf10431pf06689 -
13290 -pf00574 -
(..-11405..)
01885 -# -
01890 -pf02941 -
01900 -pf02698 -
01905 -# -
01910 -pf01636 -
01915 -# -
01920 -pf03055 -
01925 HMMreg13411_1_GRM1(pf02570pf00376pf09278) GRM1A
01930 -pf16258pf12849 -
01935 -# -
01945 H_neonGreen CsomeBCy
01955 -pf01569 -
BMC-P
BMC-H
BMC-T
Aldehyde dehydrogenase
PTAC
Signature enzyme
Alcohol dehydrogenase
Regulator
01960 -# -
01965 -# -
01970 -# -
01975 -pf11285 -
01980 -pf11317 -
01985 -pf02405 -
01990 -# -
01995 -pf01343 -
02000 -pf07736 -
(..465..)
02465 -pf13473 -
02470 -pf00557 -
02475 -pf03851 -
02480 -pf00012 -
02485 -pf00924 -
02490 -pf08974 -
02495 -pf01048 -
02500 -# -
02505 -pf04966pf00395 -
02515 -pf12697 -
02520 H_lime CsomeBCy
02525 H_peaSoup CsomeBCy
02530 -# -
02535 -pf05157 -
02545 -pf13180pf05299 -
02565 -pf14238 -
02570 -pf01842pf00551 -
02575 -pf00487 -
(..12520..)
15095 -pf00005 -
15100 -# -
15105 -pf07690 -
15110 -pf00581pf00899 -
15115 -pf14464 -
15125 -pf00484 -
15130 -pf00011 -
15135 -pf00582 -
15140 -pf00391pf02896pf01326 -
15145 -pf00805 -
15150 Tdp_turquoise(#) CsomeBCy
15155 -# -
15160 -pf04446pf14413 -
15170 -pf03129 -
15175 -# -
15180 -pf00275 -
15185 -pf00004pf06480pf01434 -
15190 -# -
15195 -pf01594 -
15200 -# -
15205 -# -

Hydrocoleum sp. CS-953. Bacteria; Cyanobacteria; Oscillatoriophycideae; Oscillatoriales;

AFK68__locus_1_main_sat

CsomeBCy

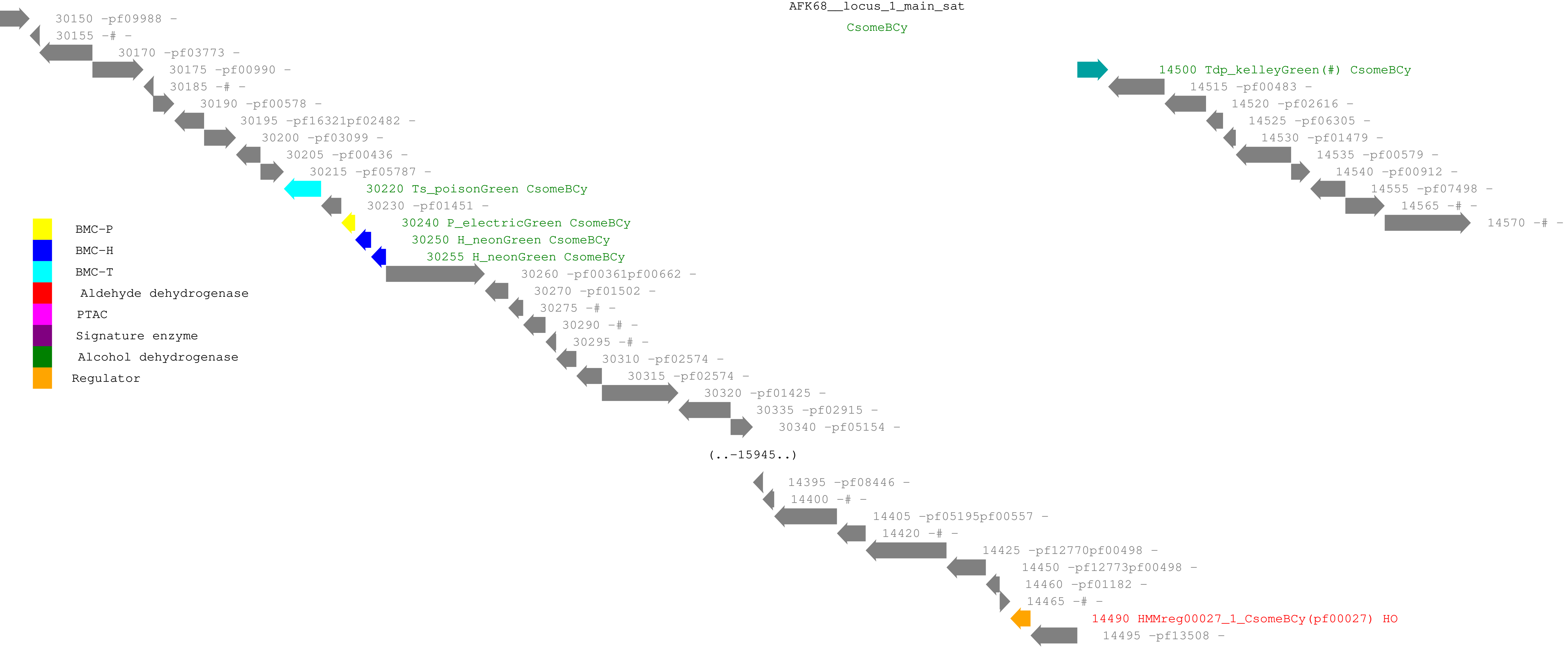

Acaryochloris marina (strain MBIC 11017). Bacteria; Cyanobacteria; Synechococcales; Acaryochloridaceae;
AM1__locus_1_main_sat
CsomeBCy

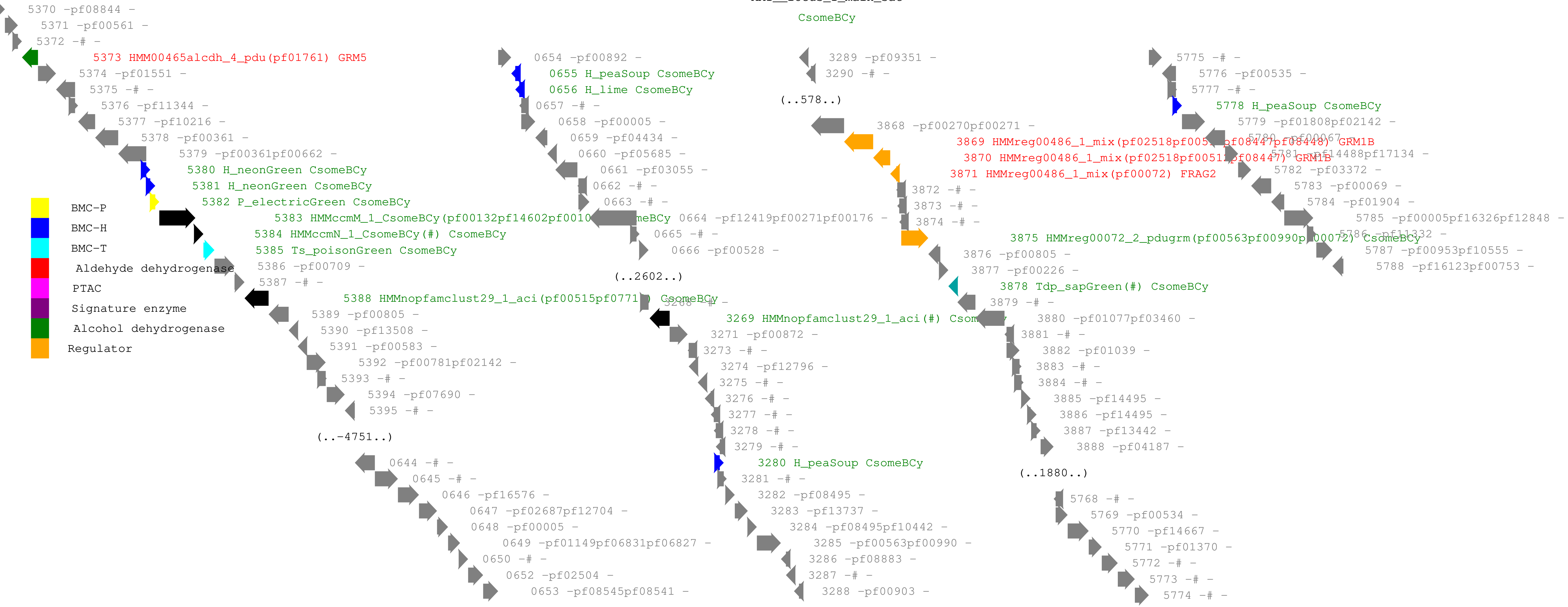

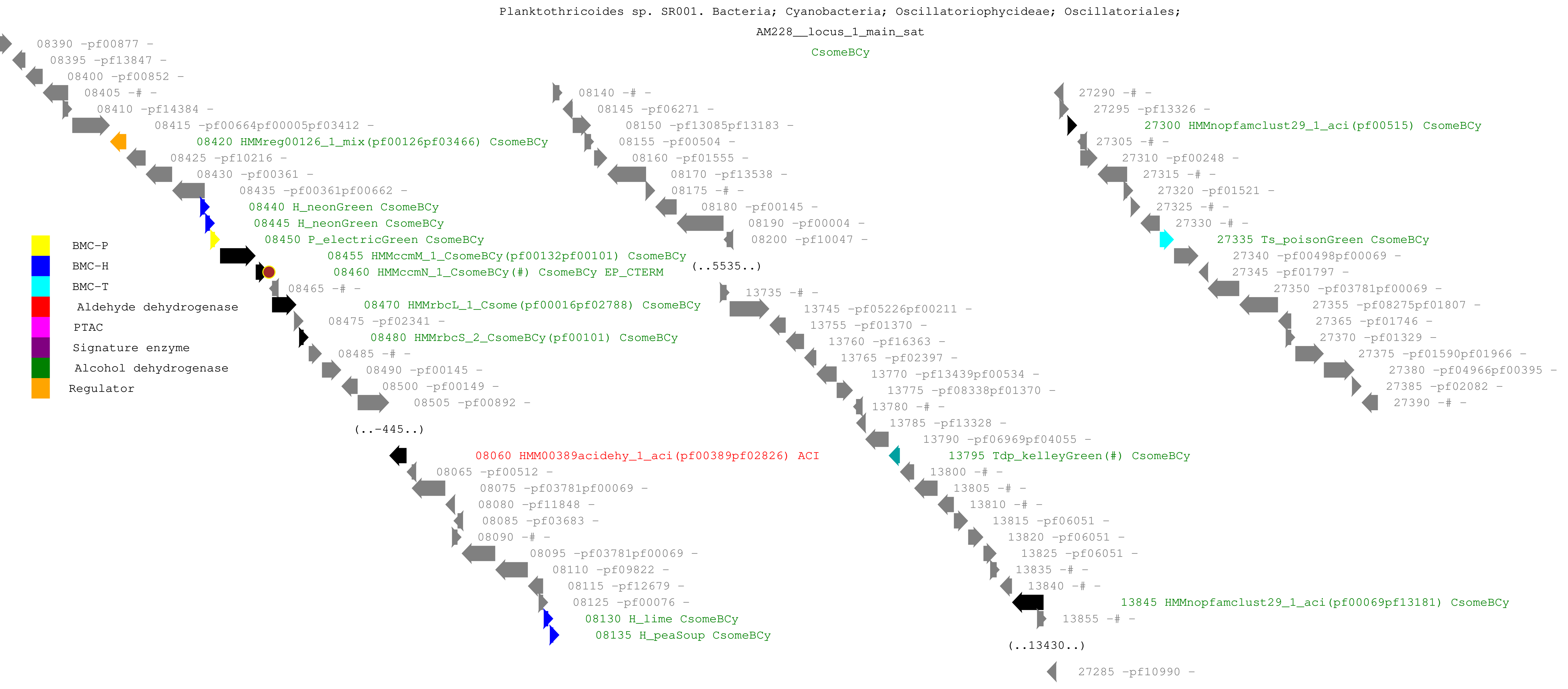

Planktothricoides sp. SR001. Bacteria; Cyanobacteria; Oscillatoriophycideae; Oscillatoriales;
AM228__locus_1_main_sat
CsomeBCy
08390 -pf00877 -
08395 -pf13847 -
08400 -pf00852 -
08405 -# -
08410 -pf14384 -
08415 -pf00664pf00005pf03412 -
08420 HMMreg00126_1_mix(pf00126pf03466) CsomeBCy
08425 -pf10216 -
08430 -pf00361 -
08435 -pf00361pf00662 -
08440 H_neonGreen CsomeBCy
08445 H_neonGreen CsomeBCy
08450 P_electricGreen CsomeBCy
08455 HMMccmM_1_CsomeBCy(pf00132pf00101) CsomeBCy
08460 HMMccmN_1_CsomeBCy(#) CsomeBCy EP_CTERM
08465 -# -
08470 HMMrbcL_1_Csome(pf00016pf02788) CsomeBCy
08475 -pf02341 -
08480 HMMrbcS_2_CsomeBCy(pf00101) CsomeBCy
08485 -# -
08490 -pf00145 -
08500 -pf00149 -
08505 -pf00892 -
(..-445..)
08060 HMM00389acidehy_1_aci(pf00389pf02826) ACI
08065 -pf00512 -
08075 -pf03781pf00069 -
08080 -pf11848 -
08085 -pf03683 -
08090 -# -
08095 -pf03781pf00069 -
08110 -pf09822 -
08115 -pf12679 -
08125 -pf00076 -
08130 H_lime CsomeBCy
08135 H_peaSoup CsomeBCy
08140 -# -
08145 -pf06271 -
08150 -pf13085pf13183 -
08155 -pf00504 -
08160 -pf01555 -
08170 -pf13538 -
08175 -# -
08180 -pf00145 -
08190 -pf00004 -
08200 -pf10047 -
(..5535..)
13735 -# -
13745 -pf05226pf00211 -
13755 -pf01370 -
13760 -pf16363 -
13765 -pf02397 -
13770 -pf13439pf00534 -
13775 -pf08338pf01370 -
13780 -# -
13785 -pf13328 -
13790 -pf06969pf04055 -
13795 Tdp_kelleyGreen(#) CsomeBCy
13800 -# -
13805 -# -
13810 -# -
13815 -pf06051 -
13820 -pf06051 -
13825 -pf06051 -
13835 -# -
13840 -# -
13845 HMMnopfamclust29_1_aci(pf00069pf13181) CsomeBCy
13855 -# -
(..13430..)
27285 -pf10990 -
27290 -# -
27295 -pf13326 -
27300 HMMnopfamclust29_1_aci(pf00515) CsomeBCy
27305 -# -
27310 -pf00248 -
27315 -# -
27320 -pf01521 -
27325 -# -
27330 -# -
27335 Ts_poisonGreen CsomeBCy
27340 -pf00498pf00069 -
27345 -pf01797 -
27350 -pf03781pf00069 -
27355 -pf08275pf01807 -
27365 -pf01746 -
27370 -pf01329 -
27375 -pf01590pf01966 -
27380 -pf04966pf00395 -
27385 -pf02082 -
27390 -# -
BMC-P
BMC-H
BMC-T
Aldehyde dehydrogenase
PTAC
Signature enzyme
Alcohol dehydrogenase
Regulator

Arthrospira maxima CS-328. Bacteria; Cyanobacteria; Oscillatoriophycideae; Oscillatoriales;

AmaxDRAFT__locus_1_main_sat

CsomeBCy

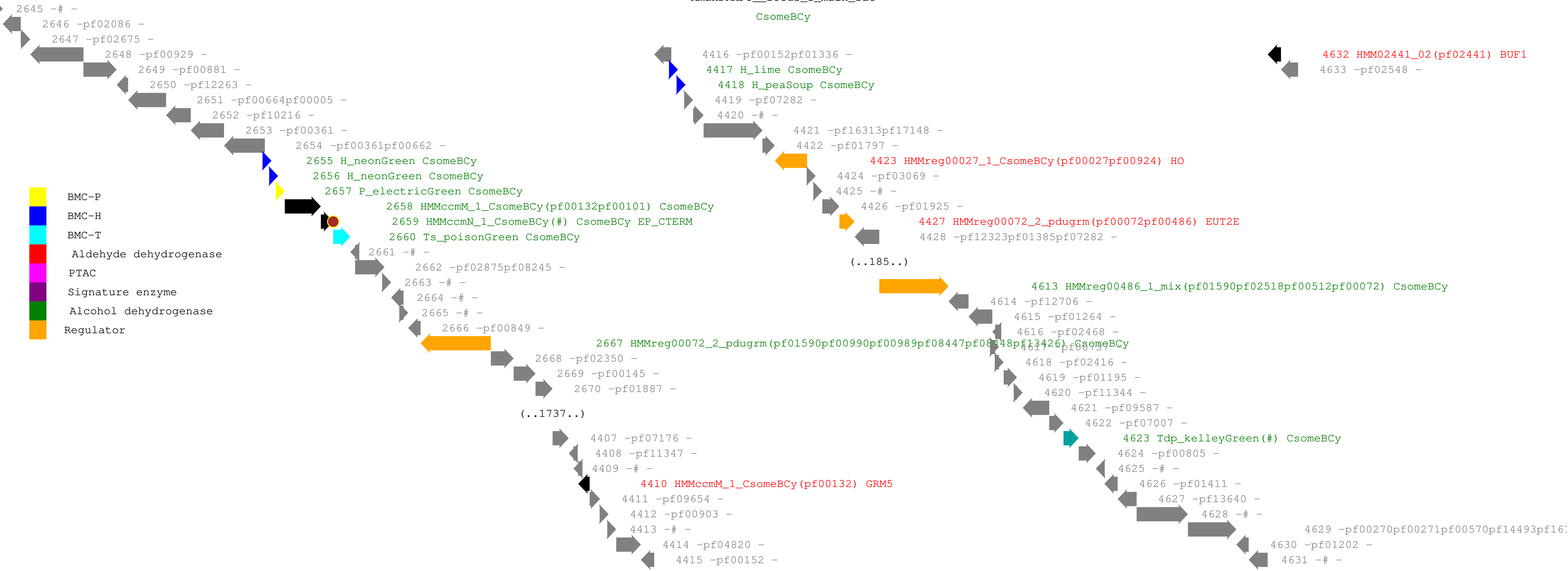

Hapalosiphon sp. MRB220. Bacteria; Cyanobacteria; Nostocales; Hapalosiphonaceae; Hapalosiphon.

AMR41__locus_1_main_sat

CsomeBCy

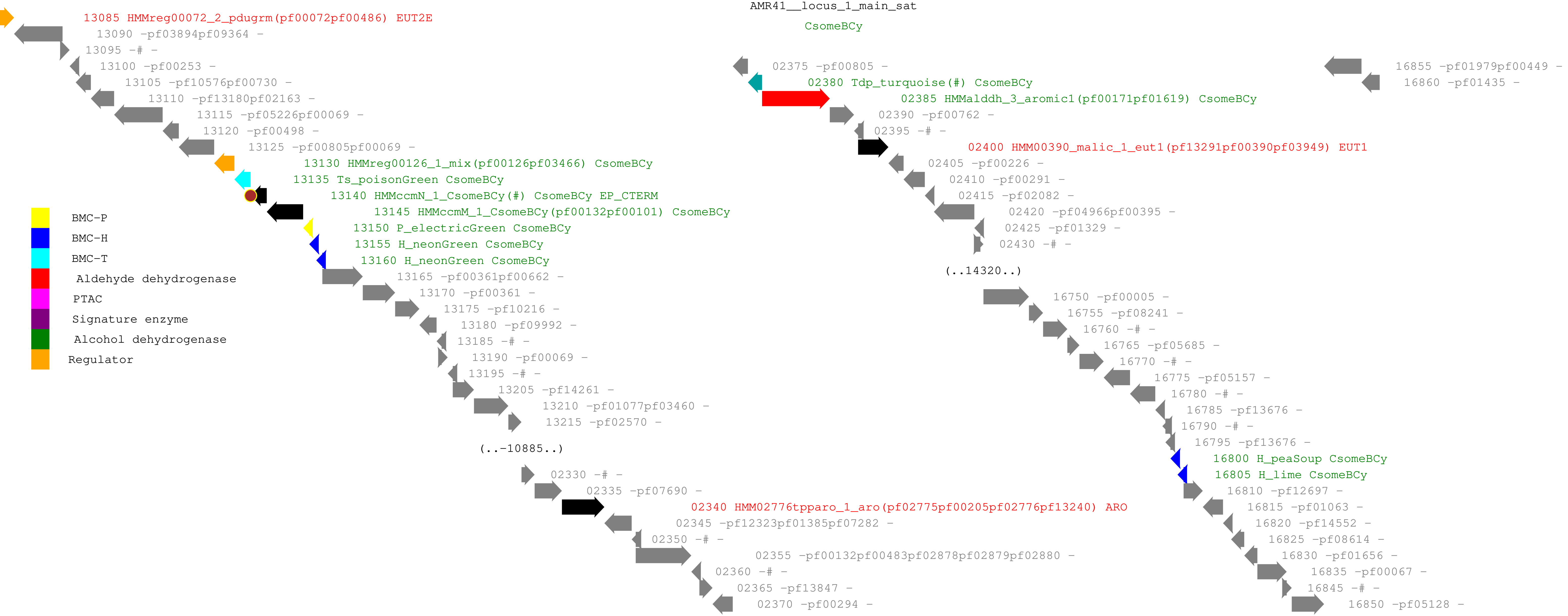

Limnothrix sp. PR1529. Bacteria; Cyanobacteria; Synechococcales; Pseudanabaenaceae;

AMR42__locus_1_main_sat

CsomeBCy

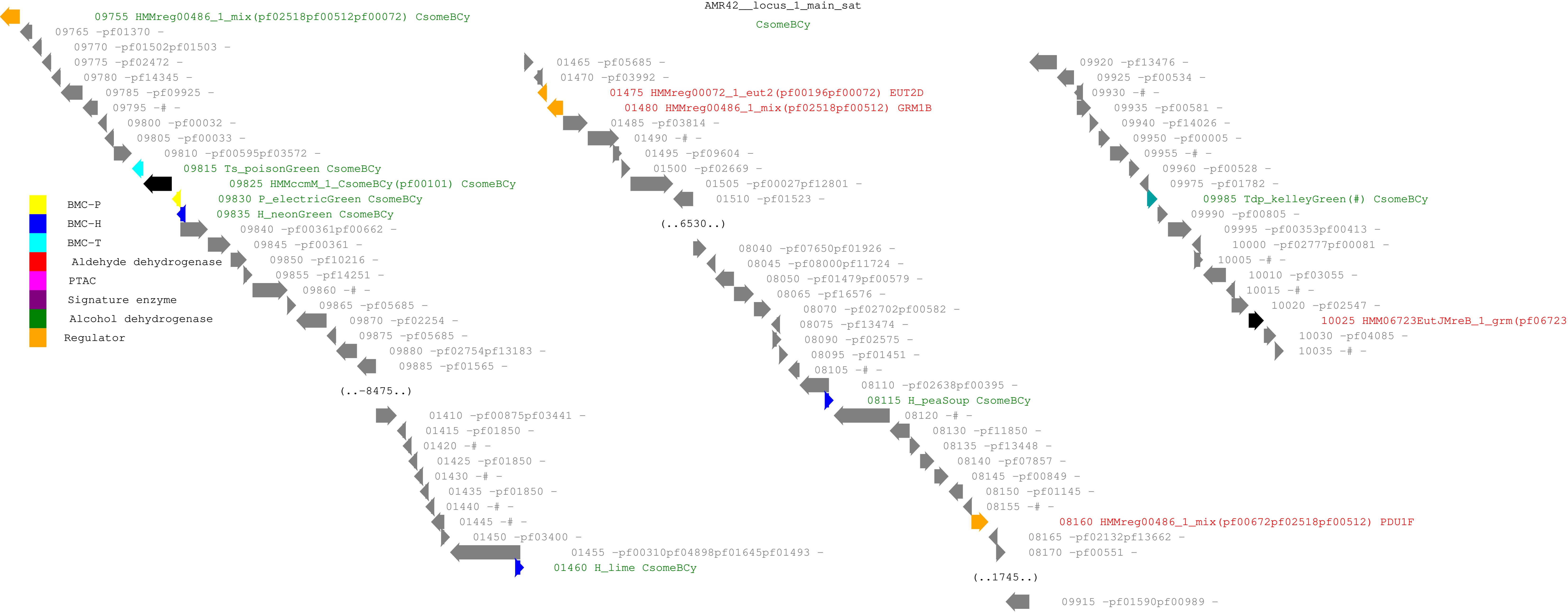

Aphanizomenon flos-aquae LD13. Bacteria; Cyanobacteria; Nostocales; Aphanizomenonaceae;

AN481__locus_1_main_sat

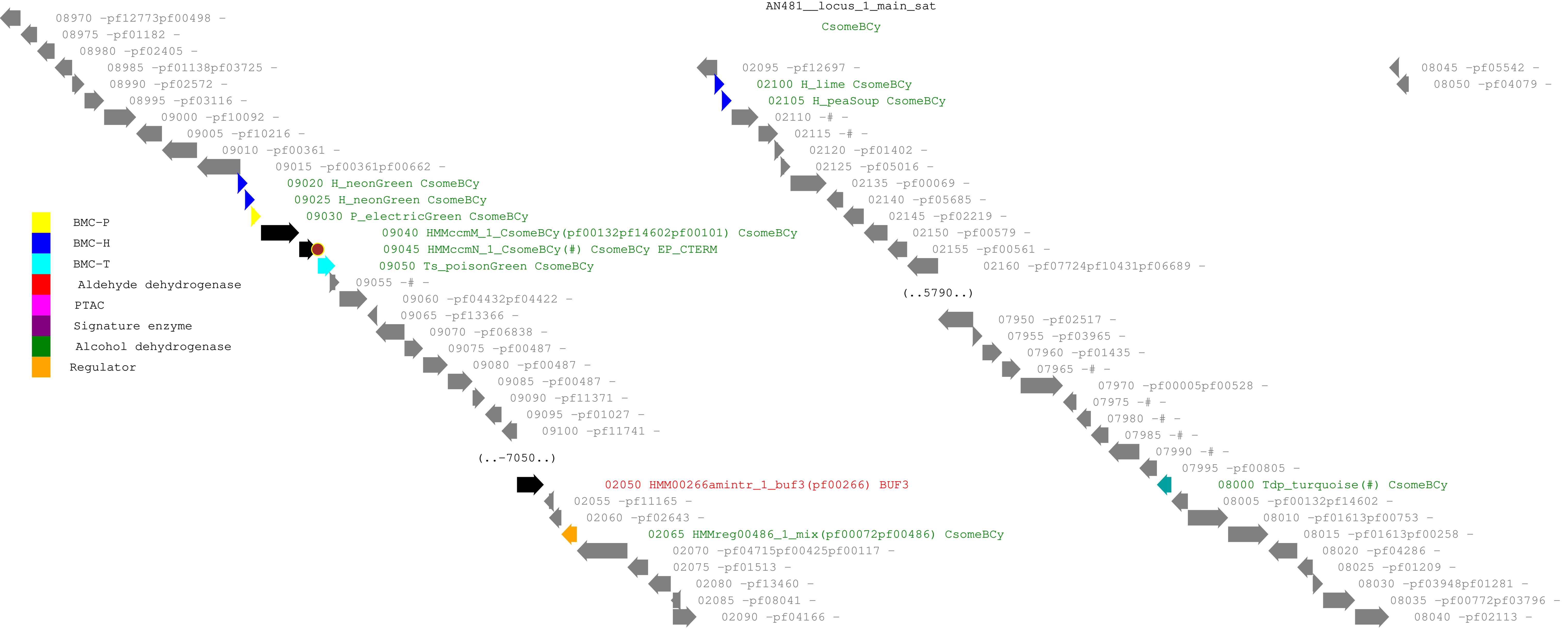

Anabaena sp. LE011-02. Bacteria; Cyanobacteria; Nostocales; Nostocaceae; Anabaena.

AN482__locus_1_main_sat

CsomeBCy

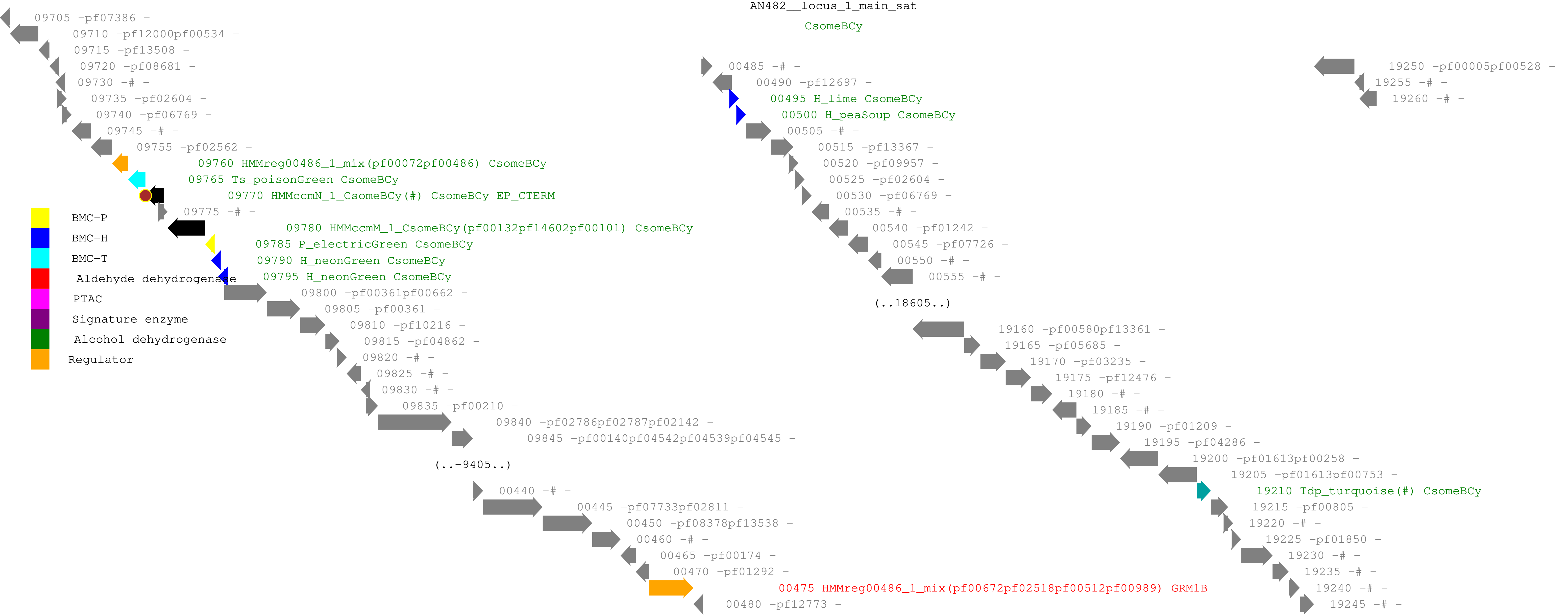

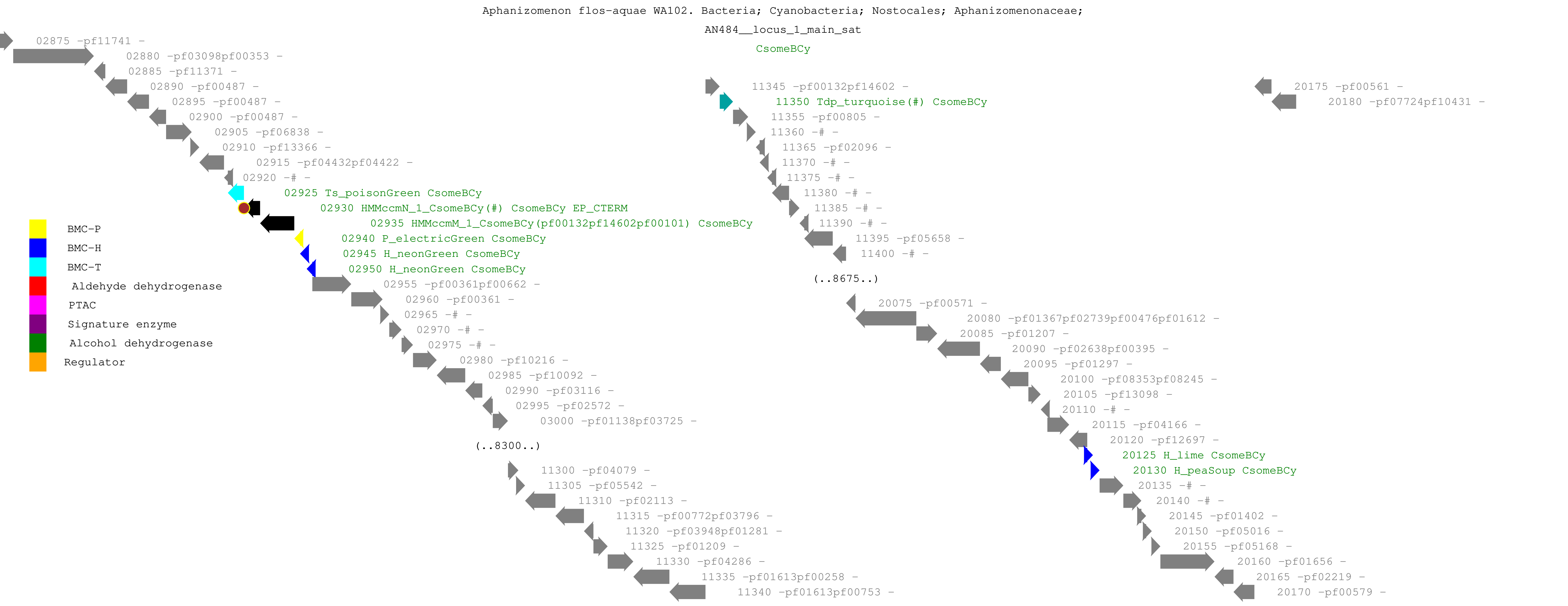

Aphanizomenon flos-aquae WA102. Bacteria; Cyanobacteria; Nostocales; Aphanizomenonaceae;
AN484__locus_1_main_sat
CsomeBCy
02875 -pf11741 -
02880 -pf03098pf00353 -
02885 -pf11371 -
02890 -pf00487 -
02895 -pf00487 -
02900 -pf00487 -
02905 -pf06838 -
02910 -pf13366 -
02915 -pf04432pf04422 -
02920 -# -
02925 Ts_poisonGreen CsomeBCy
02930 HMMccmN_1_CsomeBCy(#) CsomeBCy EP_CTERM
02935 HMMccmM_1_CsomeBCy(pf00132pf14602pf00101) CsomeBCy
02940 P_electricGreen CsomeBCy
02945 H_neonGreen CsomeBCy
02950 H_neonGreen CsomeBCy
02955 -pf00361pf00662 -
02960 -pf00361 -
02965 -# -
02970 -# -
02975 -# -
02980 -pf10216 -
02985 -pf10092 -
02990 -pf03116 -
02995 -pf02572 -
03000 -pf01138pf03725 -
(..8300..)
11300 -pf04079 -
11305 -pf05542 -
11310 -pf02113 -
11315 -pf00772pf03796 -
11320 -pf03948pf01281 -
11325 -pf01209 -
11330 -pf04286 -
11335 -pf01613pf00258 -
11340 -pf01613pf00753 -
11345 -pf00132pf14602 -
11350 Tdp_turquoise(#) CsomeBCy
11355 -pf00805 -
11360 -# -
11365 -pf02096 -
11370 -# -
11375 -# -
11380 -# -
11385 -# -
11390 -# -
11395 -pf05658 -
11400 -# -
(..8675..)
20075 -pf00571 -
20080 -pf01367pf02739pf00476pf01612 -
20085 -pf01207 -
20090 -pf02638pf00395 -
20095 -pf01297 -
20100 -pf08353pf08245 -
20105 -pf13098 -
20110 -# -
20115 -pf04166 -
20120 -pf12697 -
20125 H_lime CsomeBCy
20130 H_peaSoup CsomeBCy
20135 -# -
20140 -# -
20145 -pf01402 -
20150 -pf05016 -
20155 -pf05168 -
20160 -pf01656 -
20165 -pf02219 -
20170 -pf00579 -
20175 -pf00561 -
20180 -pf07724pf10431 -
BMC-P
BMC-H
BMC-T
Aldehyde dehydrogenase
PTAC
Signature enzyme
Alcohol dehydrogenase
Regulator

Anabaena sp. MDT14b. Bacteria; Cyanobacteria; Nostocales; Nostocaceae; Anabaena.

AN485__locus_1_main_sat

CsomeBCy

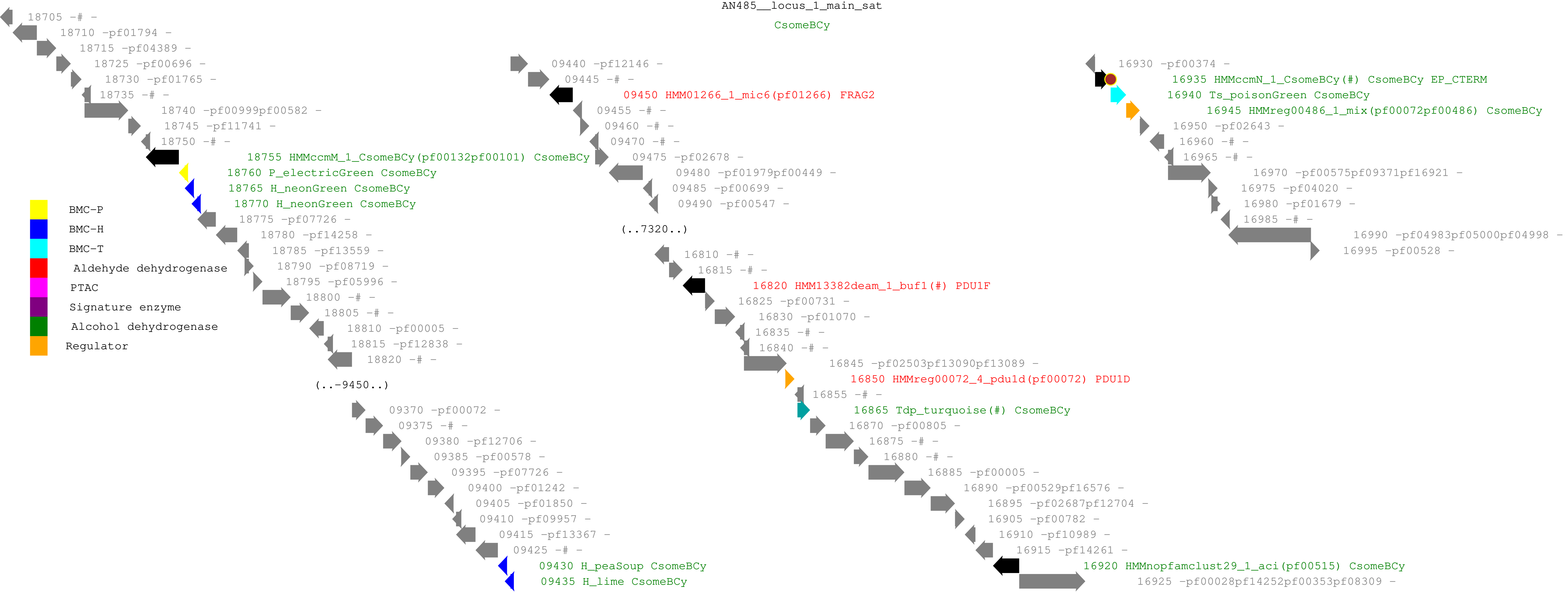

Anabaena sp. CRKS33. Bacteria; Cyanobacteria; Nostocales; Nostocaceae; Anabaena.

AN487__locus_1_main_sat

CsomeBCy

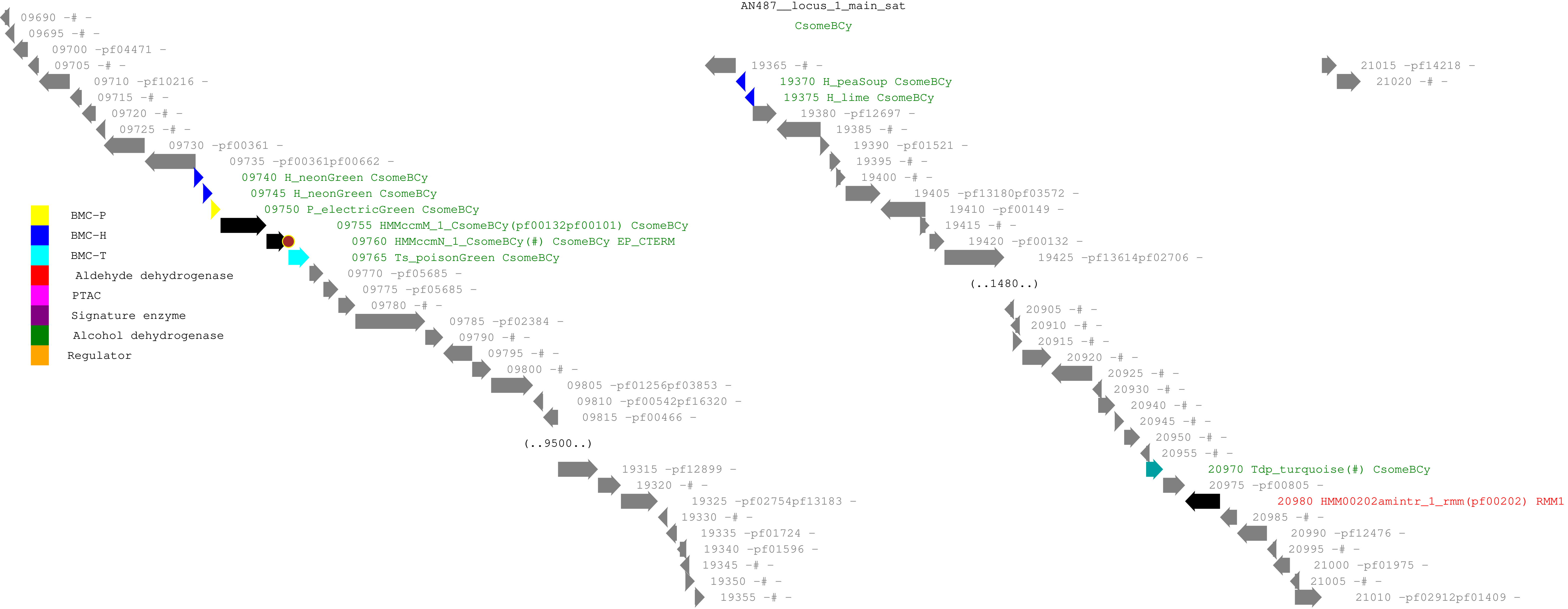

Anabaena sp. WA113. Bacteria; Cyanobacteria; Nostocales; Nostocaceae; Anabaena.

AN488__locus_1_main_sat

CsomeBCy

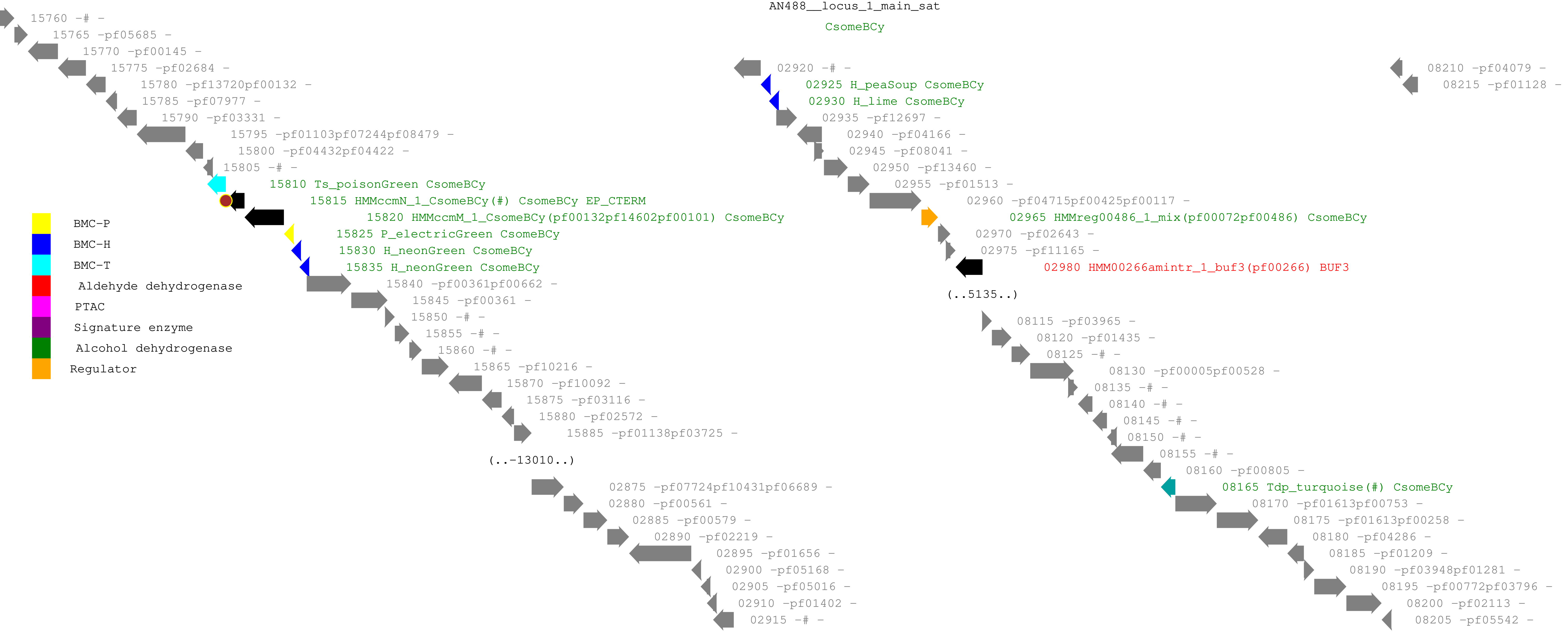

Anabaena sp. 39858. Bacteria; Cyanobacteria; Nostocales; Nostocaceae; Anabaena.

AN489__locus_1_main_sat

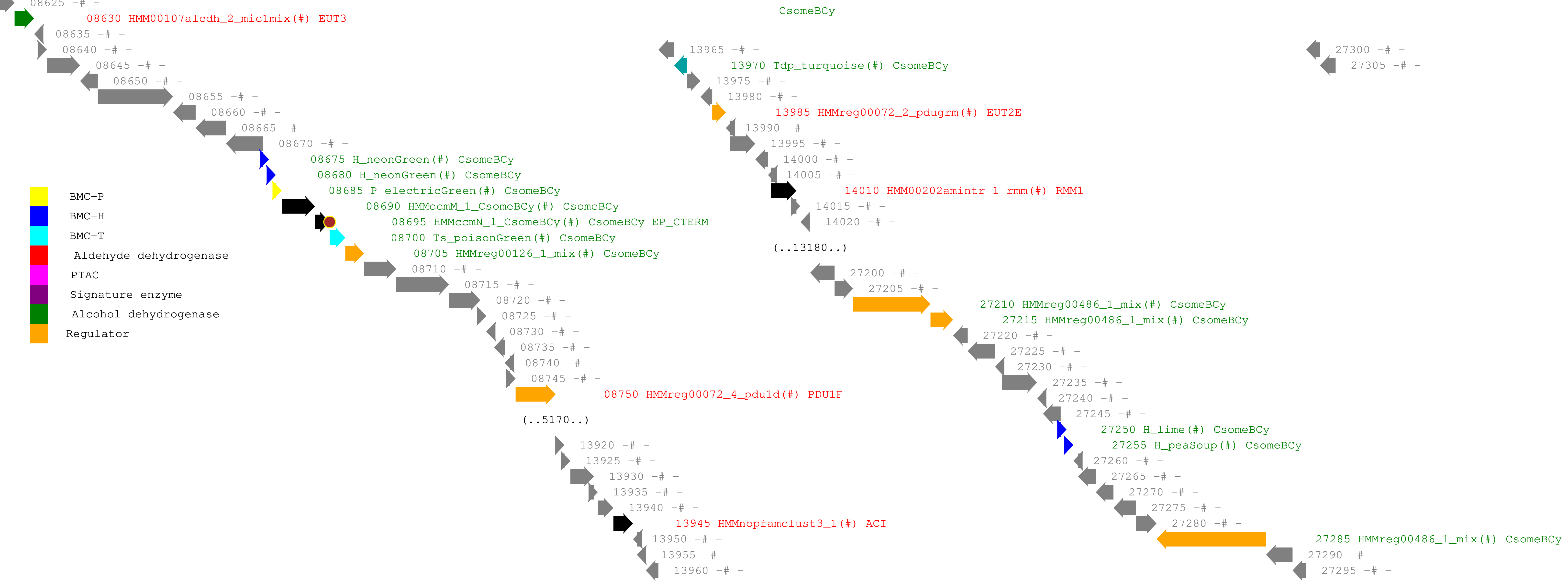

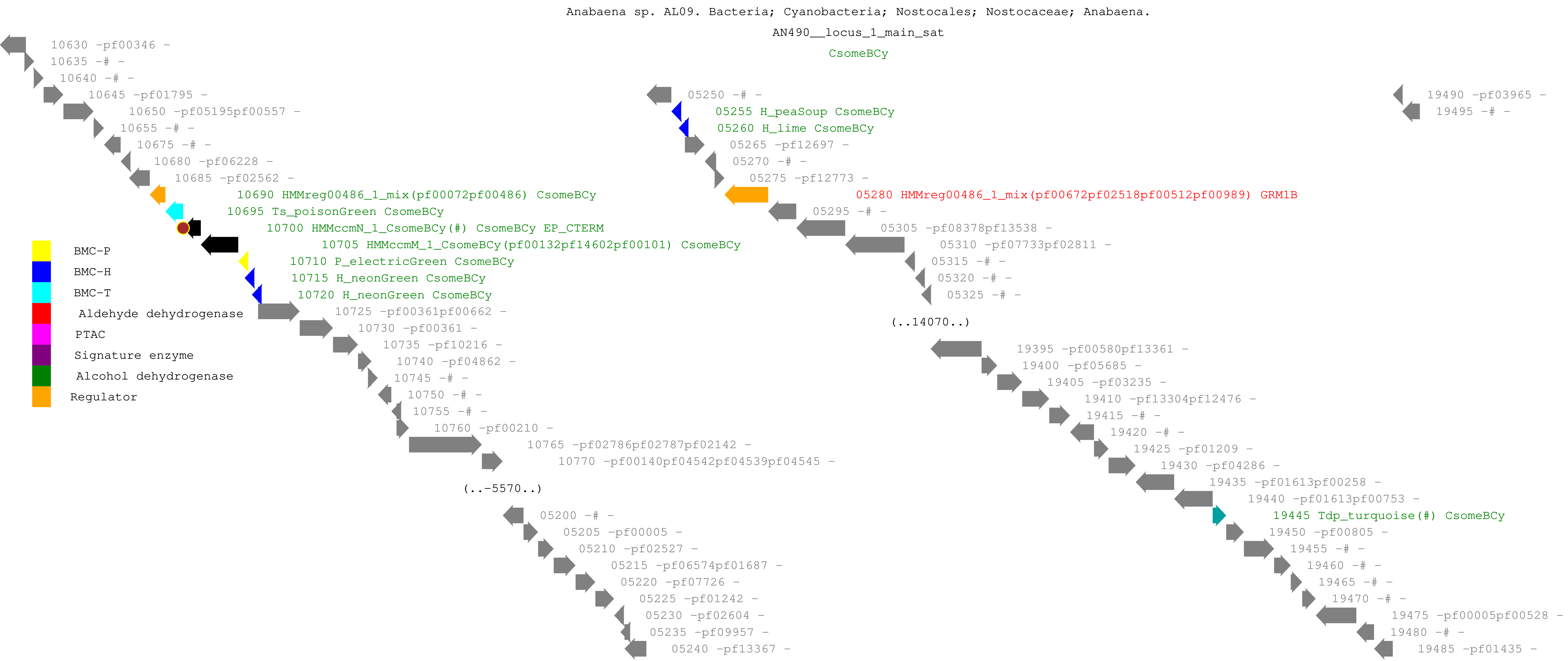

Anabaena sp. AL09. Bacteria; Cyanobacteria; Nostocales; Nostocaceae; Anabaena.
AN490__locus_1_main_sat
CsomeBCy
10630 -pf00346 -
10635 -# -
10640 -# -
10645 -pf01795 -
10650 -pf05195pf00557 -
10655 -# -
10675 -# -
10680 -pf06228 -
10685 -pf02562 -
10690 HMMreg00486_1_mix(pf00072pf00486) CsomeBCy
10695 Ts_poisonGreen CsomeBCy
10700 HMMccmN_1_CsomeBCy(#) CsomeBCy EP_CTERM
10705 HMMccmM_1_CsomeBCy(pf00132pf14602pf00101) CsomeBCy
10710 P_electricGreen CsomeBCy
10715 H_neonGreen CsomeBCy
10720 H_neonGreen CsomeBCy
10725 -pf00361pf00662 -
10730 -pf00361 -
10735 -pf10216 -
10740 -pf04862 -
10745 -# -
10750 -# -
10755 -# -
10760 -pf00210 -
10765 -pf02786pf02787pf02142 -
10770 -pf00140pf04542pf04539pf04545 -
(..-5570..)
05200 -# -
05205 -pf00005 -
05210 -pf02527 -
05215 -pf06574pf01687 -
05220 -pf07726 -
05225 -pf01242 -
05230 -pf02604 -
05235 -pf09957 -
05240 -pf13367 -
05250 -# -
05255 H_peaSoup CsomeBCy
05260 H_lime CsomeBCy
05265 -pf12697 -
05270 -# -
05275 -pf12773 -
05280 HMMreg00486_1_mix(pf00672pf02518pf00512pf00989) GRM1B
05295 -# -
05305 -pf08378pf13538 -
05310 -pf07733pf02811 -
05315 -# -
05320 -# -
05325 -# -
(..14070..)
19395 -pf00580pf13361 -
19400 -pf05685 -
19405 -pf03235 -
19410 -pf13304pf12476 -
19415 -# -
19420 -# -
19425 -pf01209 -
19430 -pf04286 -
19435 -pf01613pf00258 -
19440 -pf01613pf00753 -
19445 Tdp_turquoise(#) CsomeBCy
19450 -pf00805 -
19455 -# -
19460 -# -
19465 -# -
19470 -# -
19475 -pf00005pf00528 -
19480 -# -
19485 -pf01435 -
19490 -pf03965 -
19495 -# -
BMC-P
BMC-H
BMC-T
Aldehyde dehydrogenase
PTAC
Signature enzyme
Alcohol dehydrogenase
Regulator

Anabaena sp. 90. Bacteria; Cyanobacteria; Nostocales; Nostocaceae; Anabaena.

ANA_C_locus_1_main_sat

CsomeBCy

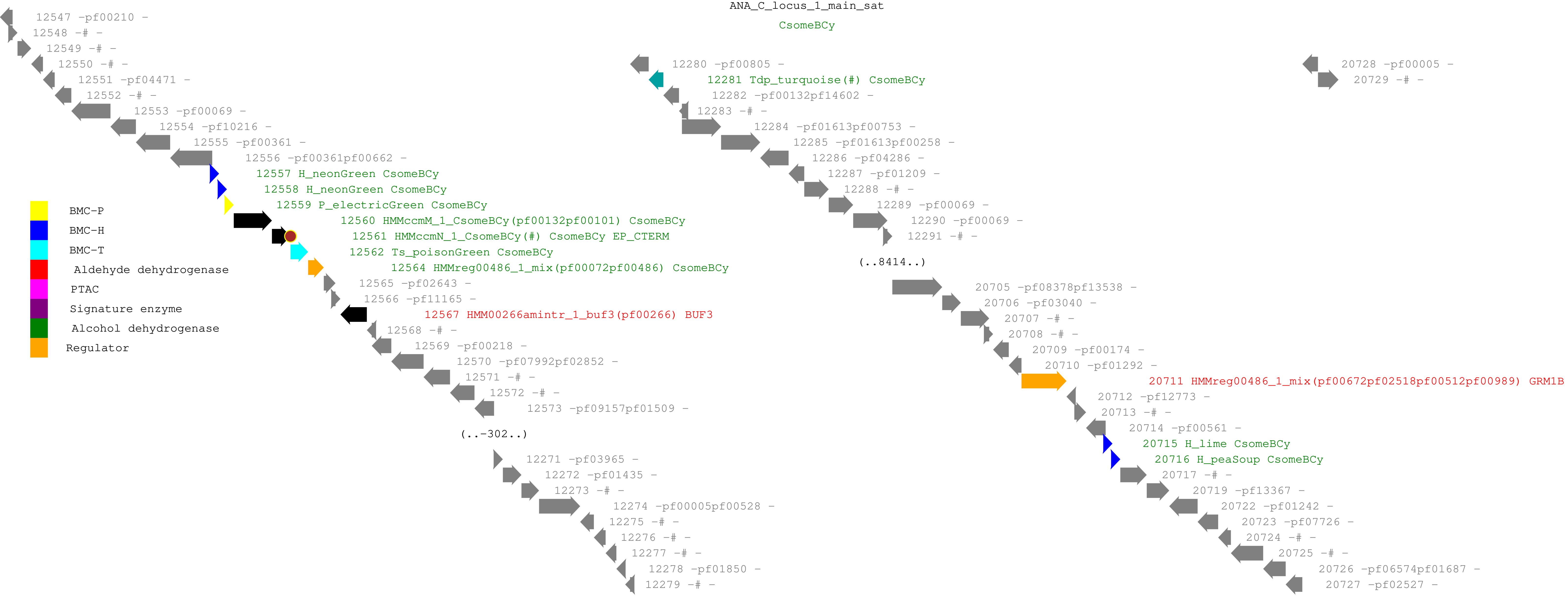

Anabaena cylindrica (strain ATCC 27899 / PCC 7122). Bacteria; Cyanobacteria; Nostocales; Nostocaceae; Anabaena.

Anacy__locus_1_main_sat

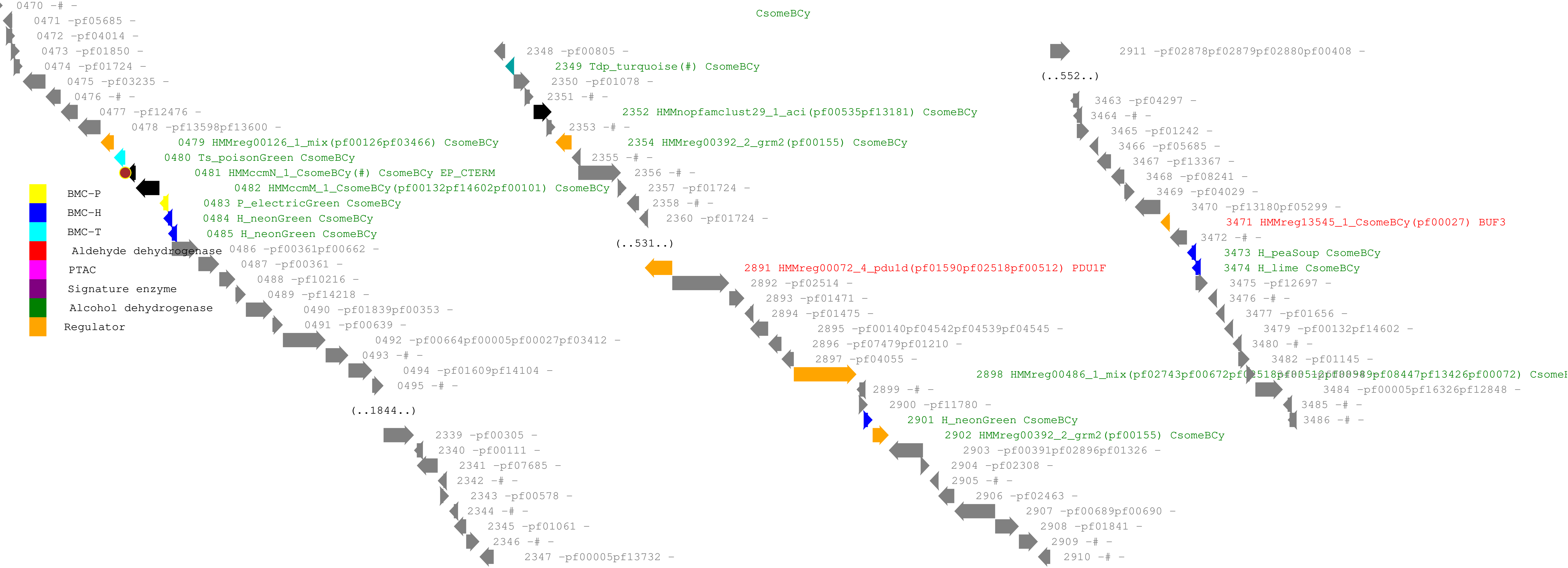

Chrysosporum ovalisporum. Bacteria; Cyanobacteria; Nostocales; Aphanizomenonaceae; Chrysosporum.

apha__locus_1_main_sat

CsomeBCy

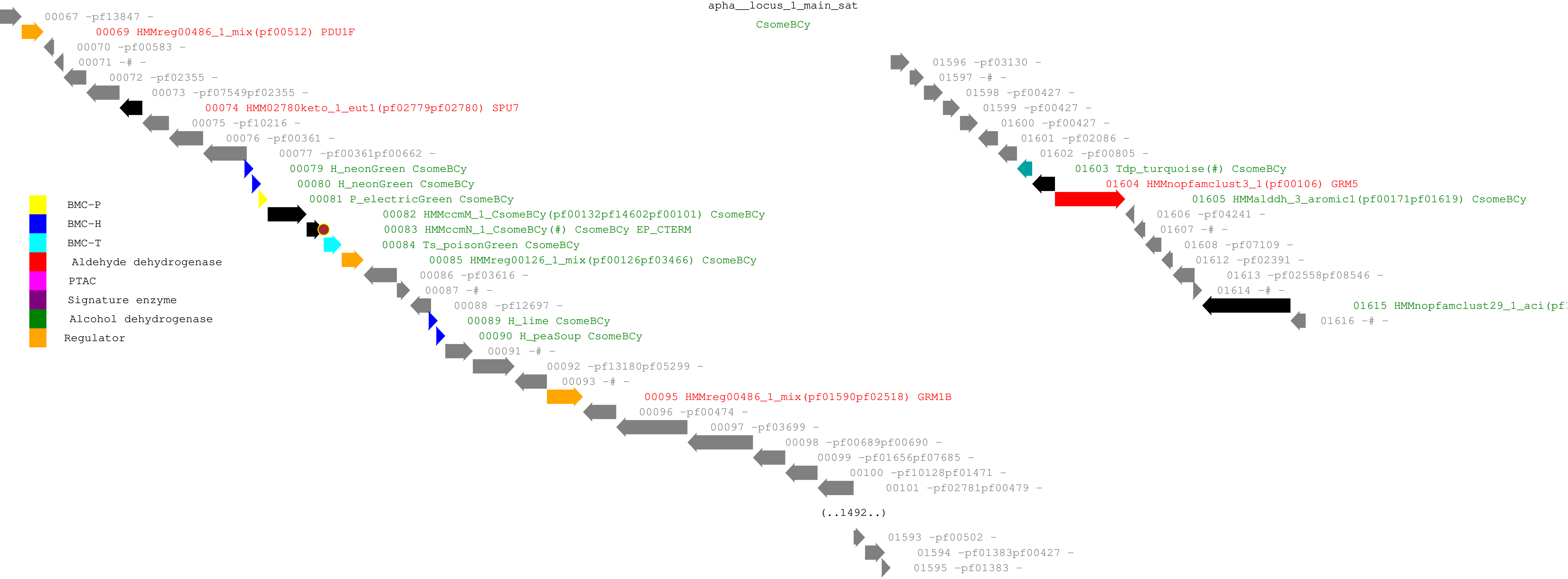

Arthrospira sp. PCC 8005. Bacteria; Cyanobacteria; Oscillatoriophycideae; Oscillatoriales;

ARTHRO__locus_1_main_sat

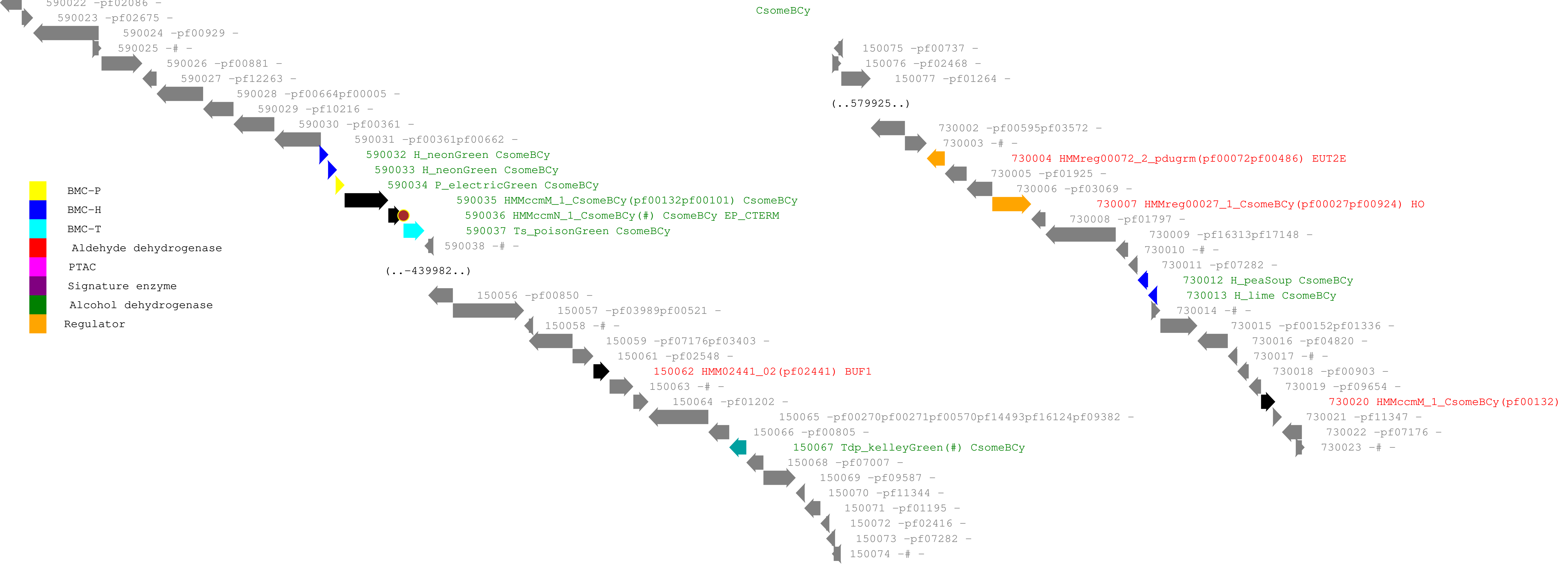

Aphanothece sacrum FPU1. Bacteria; Cyanobacteria; Oscillatoriophycideae; Chroococcales;

AsFPU1__locus_1_main_sat

CsomeBCy

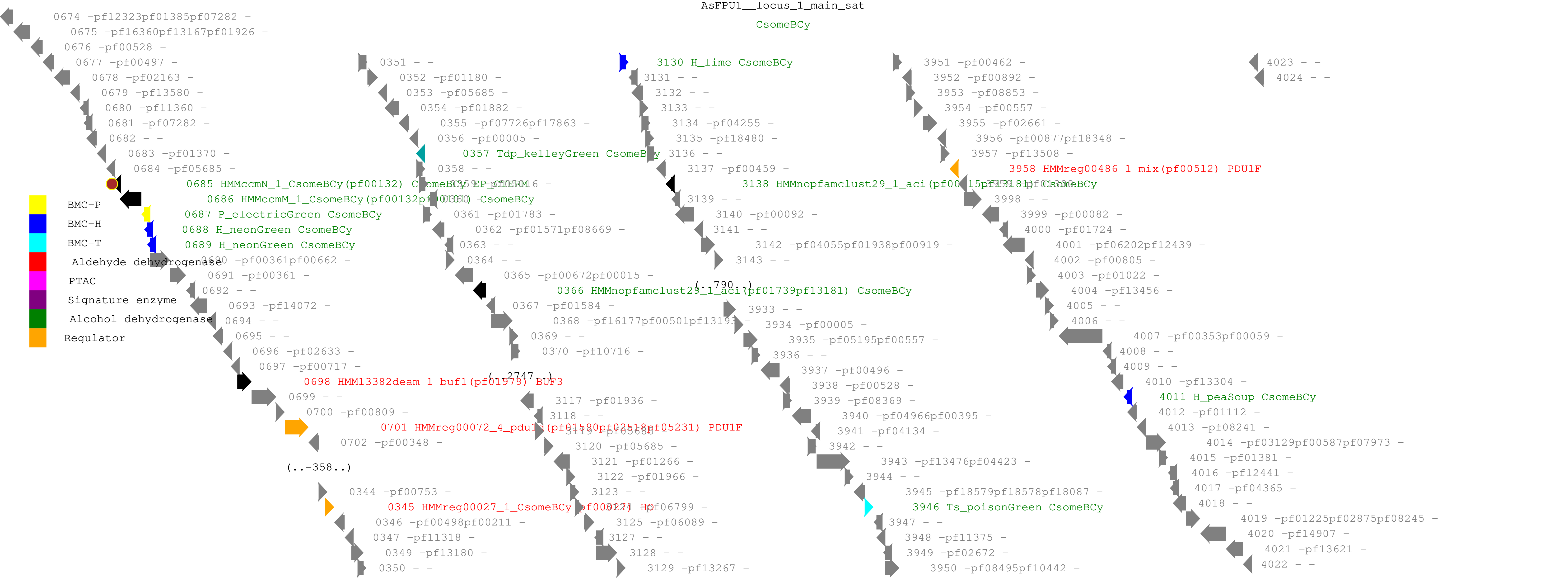

Cylindrospermopsis sp. CR12. Bacteria; Cyanobacteria; Nostocales; Aphanizomenonaceae;

ASL19__locus_1_main_sat

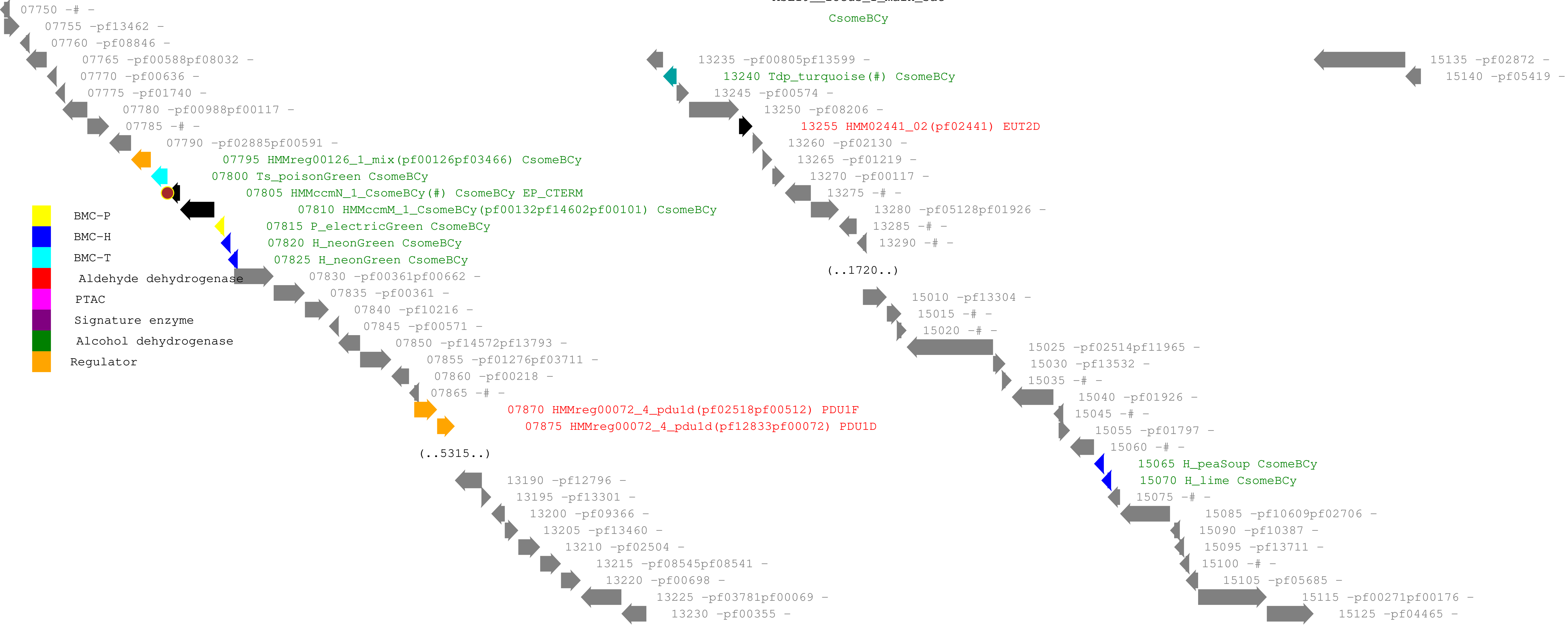

Oscillatoriales cyanobacterium CG2_30_40_61. Bacteria; Cyanobacteria; Oscillatoriophycideae; Oscillatoriales;

AUK43__locus_1_main_sat

CsomeBCy

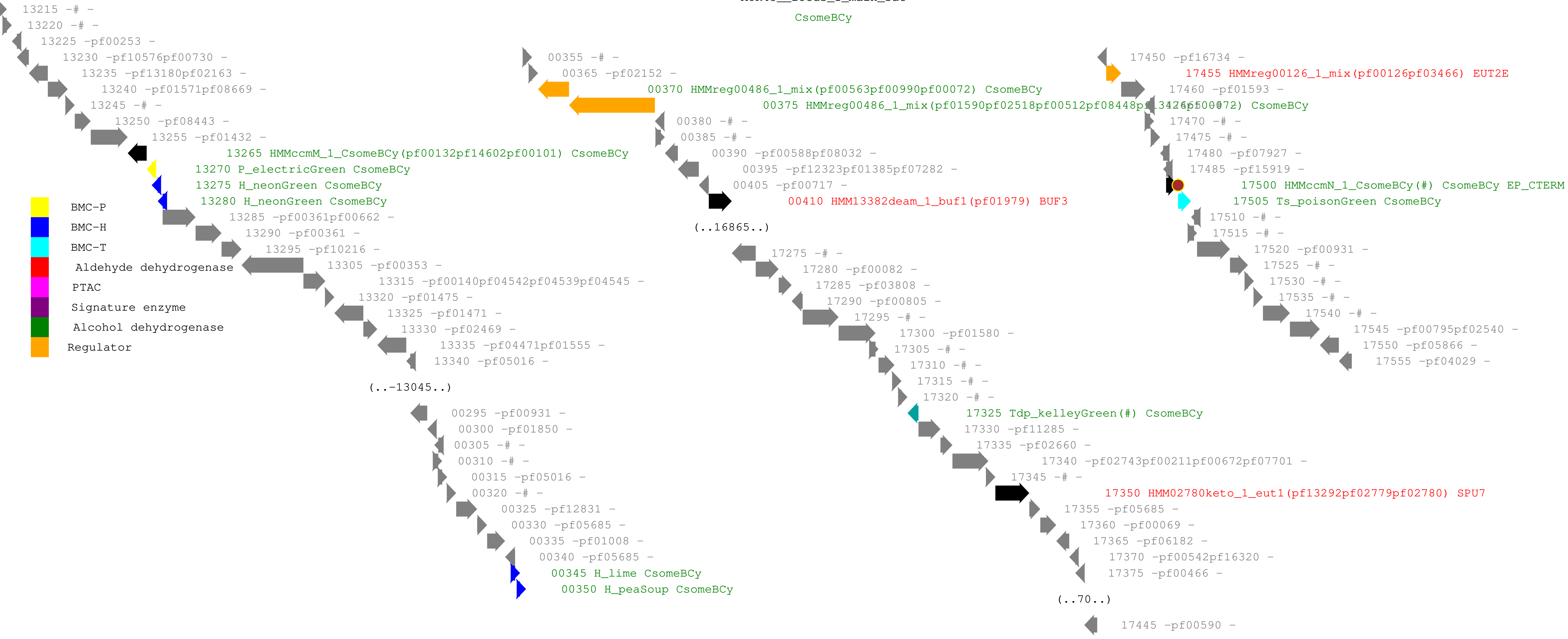

Oscillatoriales cyanobacterium CG2_30_44_21. Bacteria; Cyanobacteria; Oscillatoriophycideae; Oscillatoriales;

AUK48__locus_1_main_sat

CsomeBCy

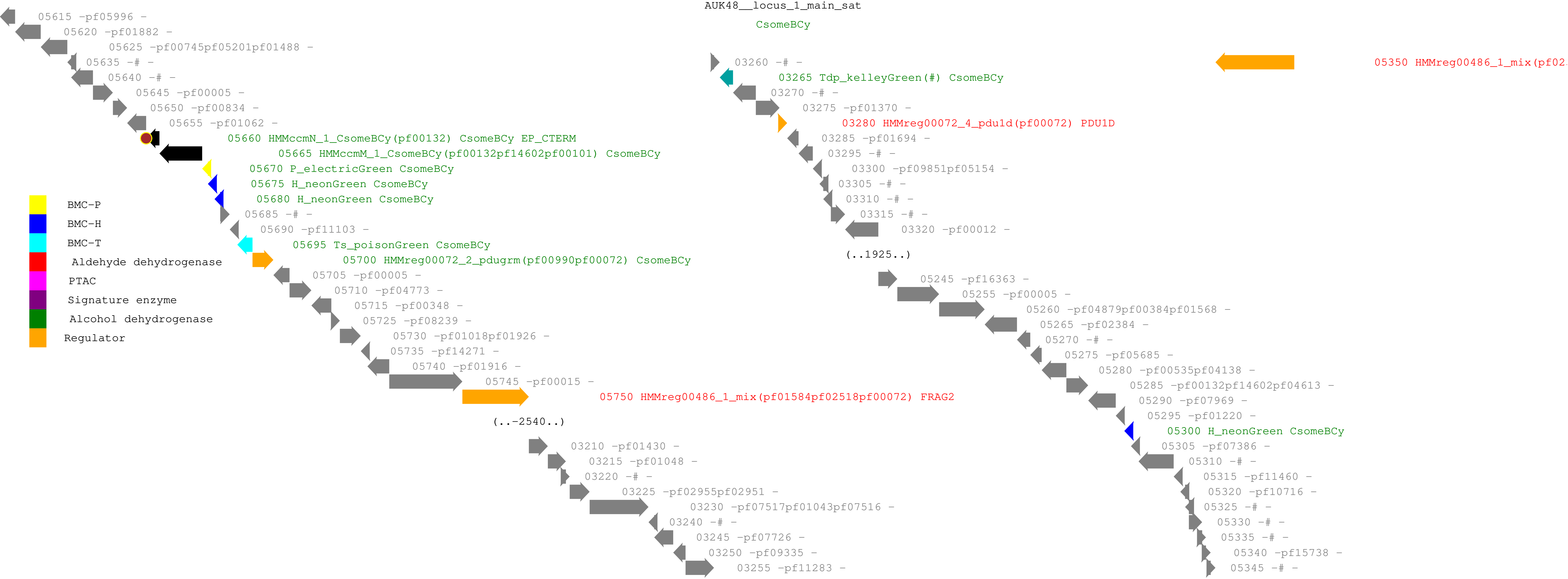

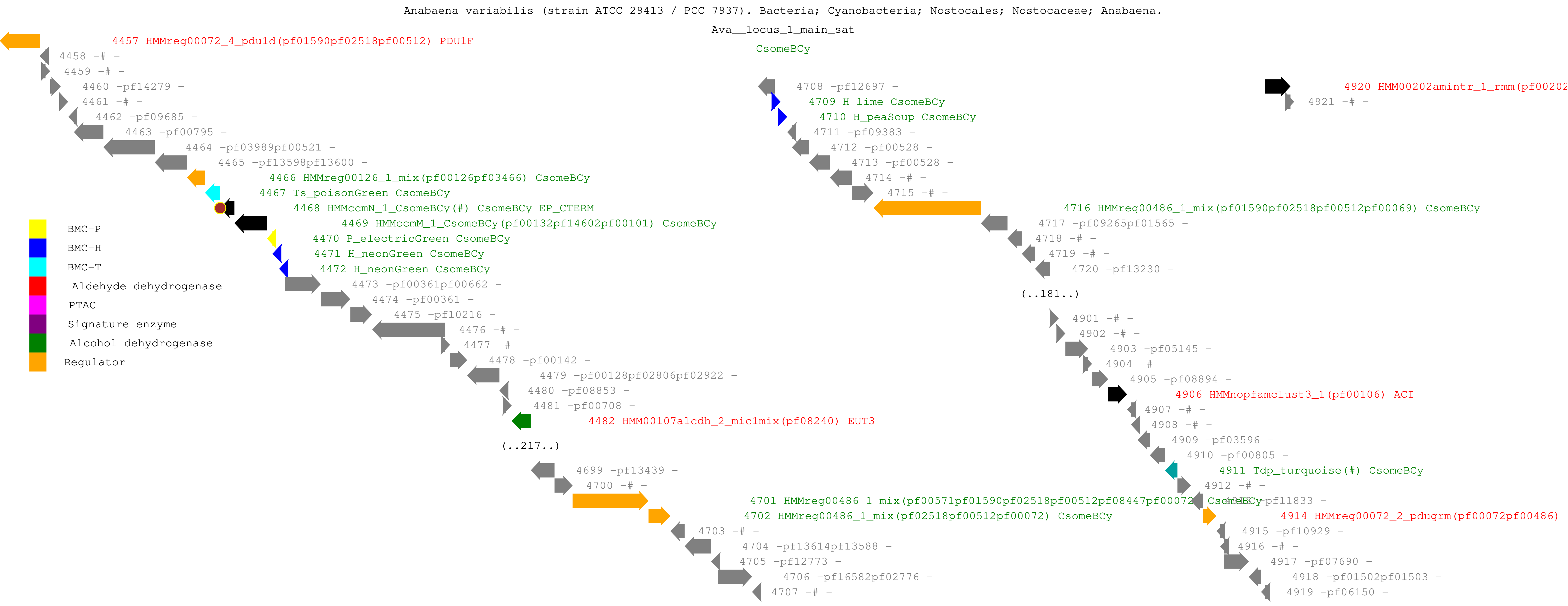
Anabaena variabilis (strain ATCC 29413 / PCC 7937). Bacteria; Cyanobacteria; Nostocales; Nostocaceae; Anabaena.
Ava__locus_1_main_sat
CsomeBCy
4457 HMMreg00072_4_pdu1d(pf01590pf02518pf00512) PDU1F
4458 -# -
4459 -# -
4460 -pf14279 -
4461 -# -
4462 -pf09685 -
4463 -pf00795 -
4464 -pf03989pf00521 -
4465 -pf13598pf13600 -
4466 HMMreg00126_1_mix(pf00126pf03466) CsomeBCy
4467 Ts_poisonGreen CsomeBCy
4468 HMMccmN_1_CsomeBCy(#) CsomeBCy EP_CTERM
4469 HMMccmM_1_CsomeBCy(pf00132pf14602pf00101) CsomeBCy
4470 P_electricGreen CsomeBCy
4471 H_neonGreen CsomeBCy
4472 H_neonGreen CsomeBCy
4473 -pf00361pf00662 -
4474 -pf00361 -
4475 -pf10216 -
4476 -# -
4477 -# -
4478 -pf00142 -
4479 -pf00128pf02806pf02922 -
4480 -pf08853 -
4481 -pf00708 -
4482 HMM00107alcdh_2_mic1mix(pf08240) EUT3
(..217..)
4699 -pf13439 -
4700 -# -
4701 HMMreg00486_1_mix(pf00571pf01590pf02518pf00512pf08447pf00072) CsomeBCy
4702 HMMreg00486_1_mix(pf02518pf00512pf00072) CsomeBCy
4703 -# -
4704 -pf13614pf13588 -
4705 -pf12773 -
4706 -pf16582pf02776 -
4707 -# -
4708 -pf12697 -
4709 H_lime CsomeBCy
4710 H_peaSoup CsomeBCy
4711 -pf09383 -
4712 -pf00528 -
4713 -pf00528 -
4714 -# -
4715 -# -
4716 HMMreg00486_1_mix(pf01590pf02518pf00512pf00069) CsomeBCy
4717 -pf09265pf01565 -
4718 -# -
4719 -# -
4720 -pf13230 -
(..181..)
4901 -# -
4902 -# -
4903 -pf05145 -
4904 -# -
4905 -pf08894 -
4906 HMMnopfamclust3_1(pf00106) ACI
4907 -# -
4908 -# -
4909 -pf03596 -
4910 -pf00805 -
4911 Tdp_turquoise(#) CsomeBCy
4912 -# -
4913 -pf11833 -
4914 HMMreg00072_2_pdugrm(pf00072pf00486)
4915 -pf10929 -
4916 -# -
4917 -pf07690 -
4918 -pf01502pf01503 -
4919 -pf06150 -
4920 HMM00202amintr_1_rmm(pf00202
4921 -# -
BMC-P
BMC-H
BMC-T
Aldehyde dehydrogenase
PTAC
Signature enzyme
Alcohol dehydrogenase
Regulator

Synechococcus sp. PCC 7003. Bacteria; Cyanobacteria; Synechococcales; Synechococcaceae;

AWQ21__locus_1_main_sat

CsomeBCy

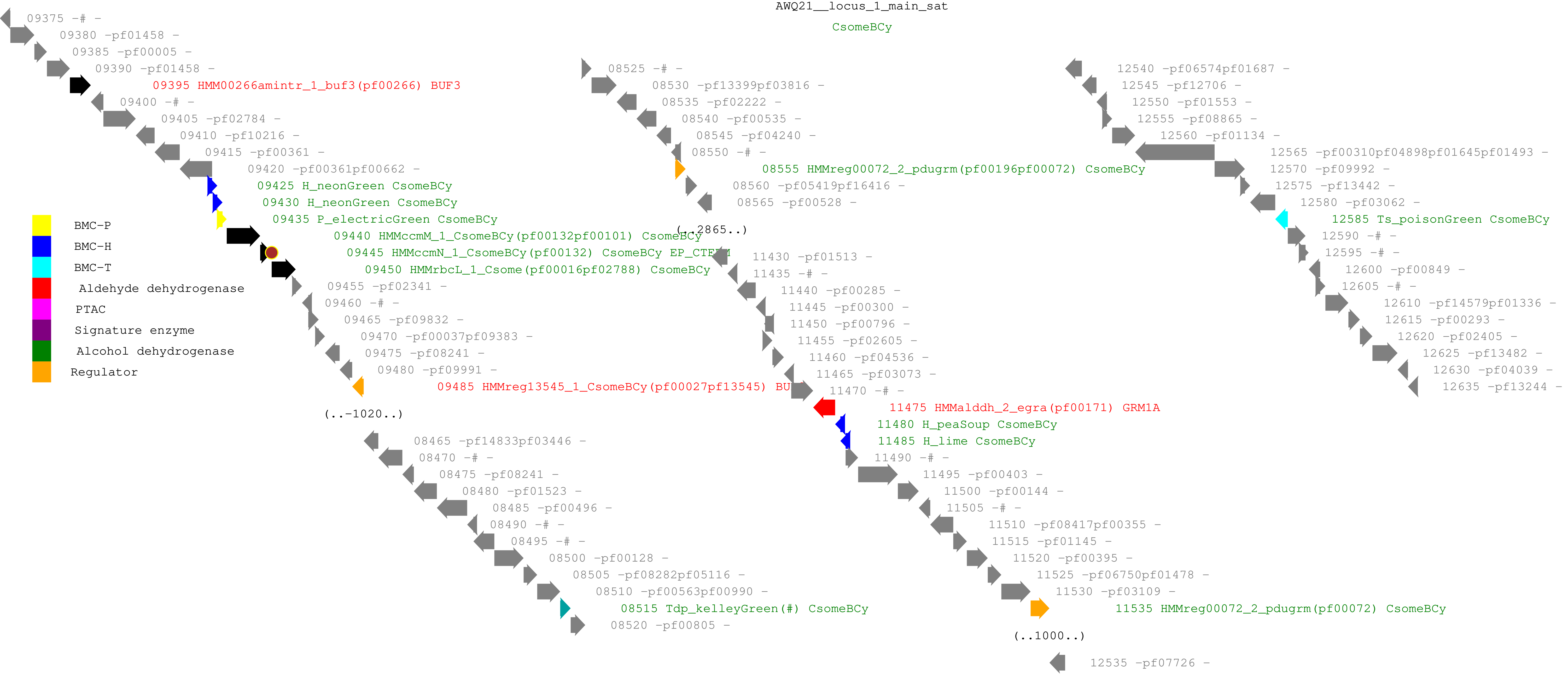

Synechococcus sp. PCC 7117. Bacteria; Cyanobacteria; Synechococcales; Synechococcaceae;

AWQ22__locus_1_main_sat

CsomeBCy

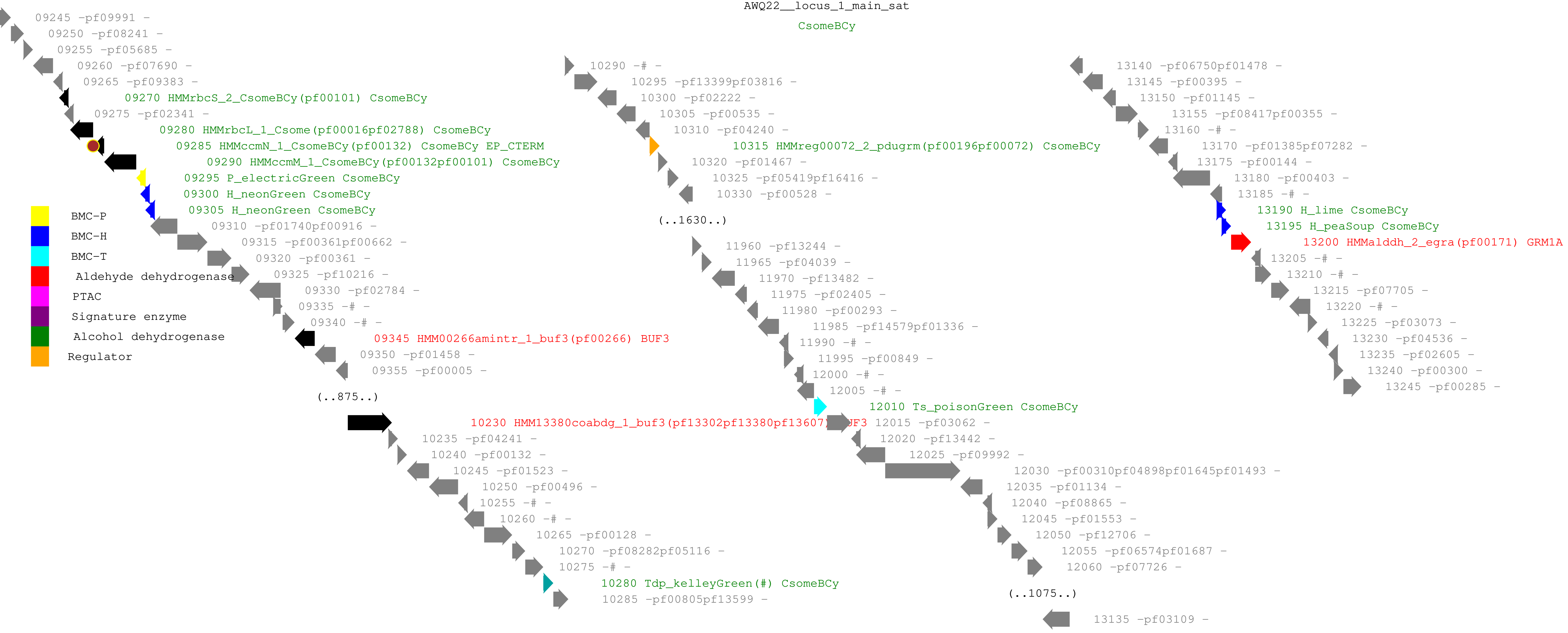

Phormidium willei BDU 130791. Bacteria; Cyanobacteria; Oscillatoriophycideae; Oscillatoriales;

AY600__locus_1_main_sat

CsomeBCy

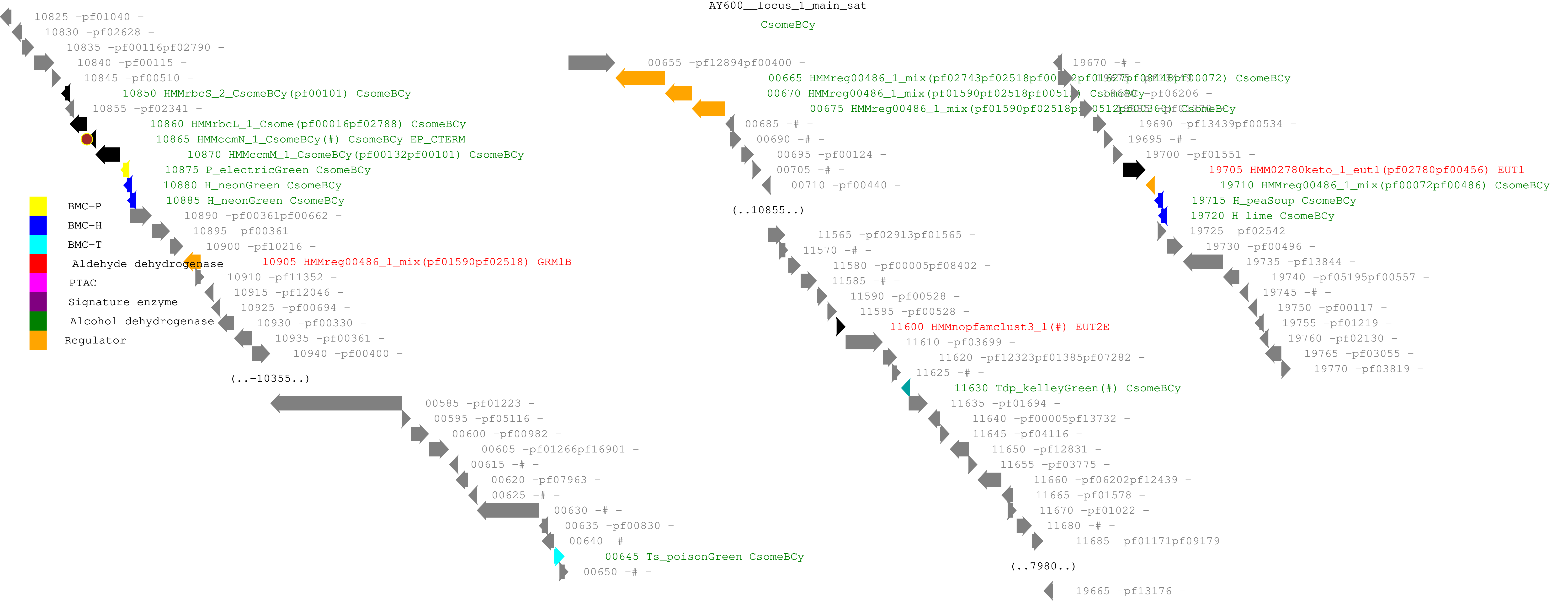

Alkalinema sp. CACIAM 70d. Bacteria; Cyanobacteria; Synechococcales; Pseudanabaenaceae;

B0A82__locus_1_main_sat

CsomeBCy

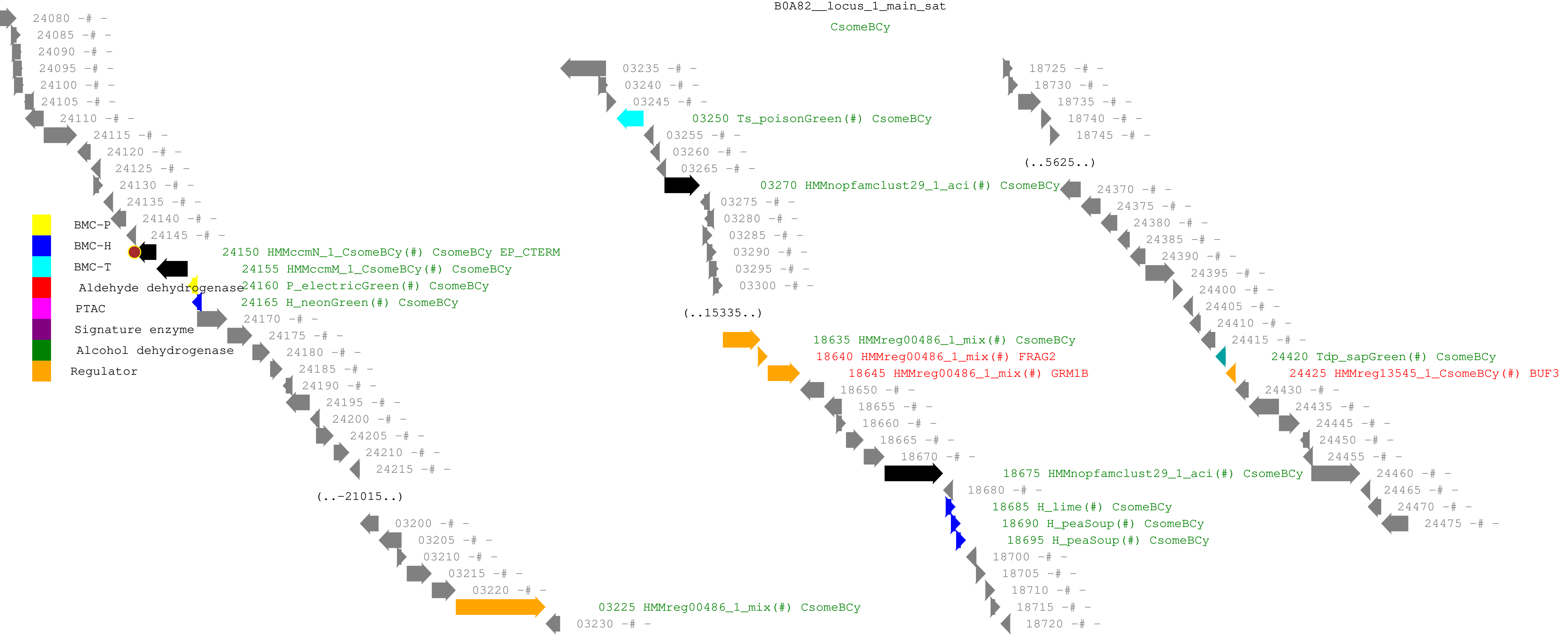

Chroococcidiopsis sp. TS-821. Bacteria; Cyanobacteria; Chroococcidiopsidales;

B1A85__locus_1_main_sat

CsomeBCy

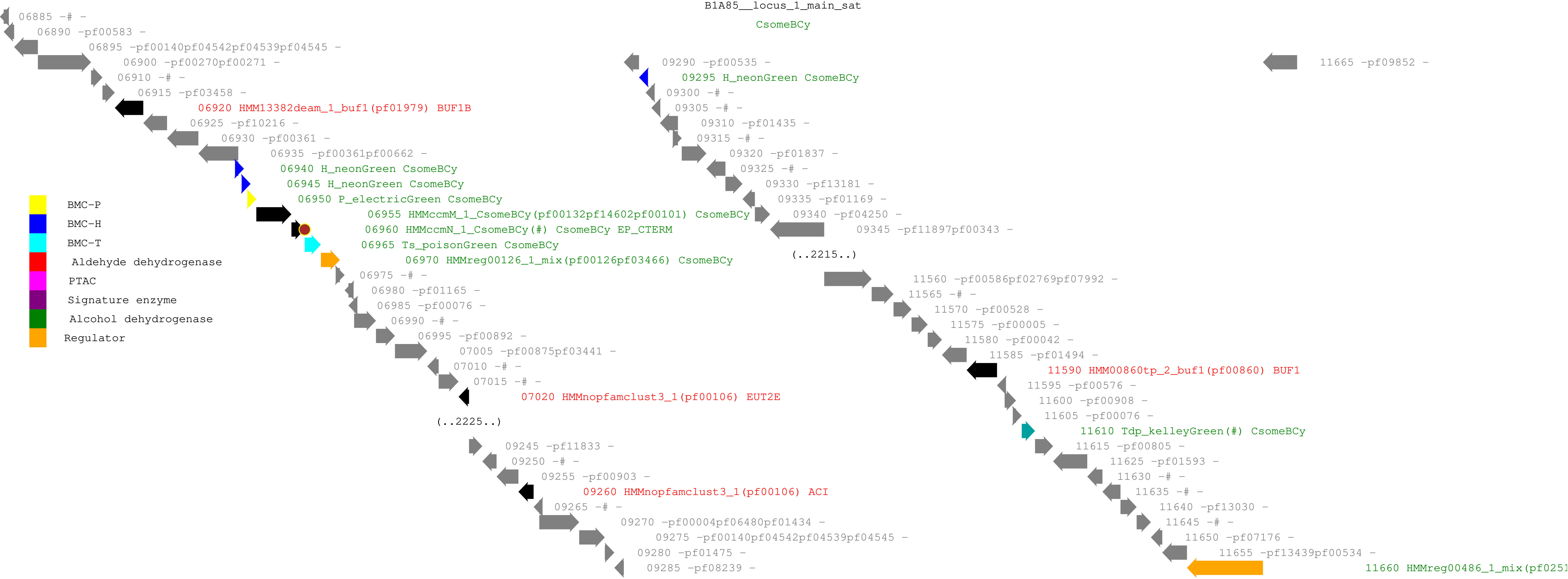

Microcystis aeruginosa KW. Bacteria; Cyanobacteria; Oscillatoriophycideae; Chroococcales;

B1L04__locus_1_main_sat

CsomeBCy

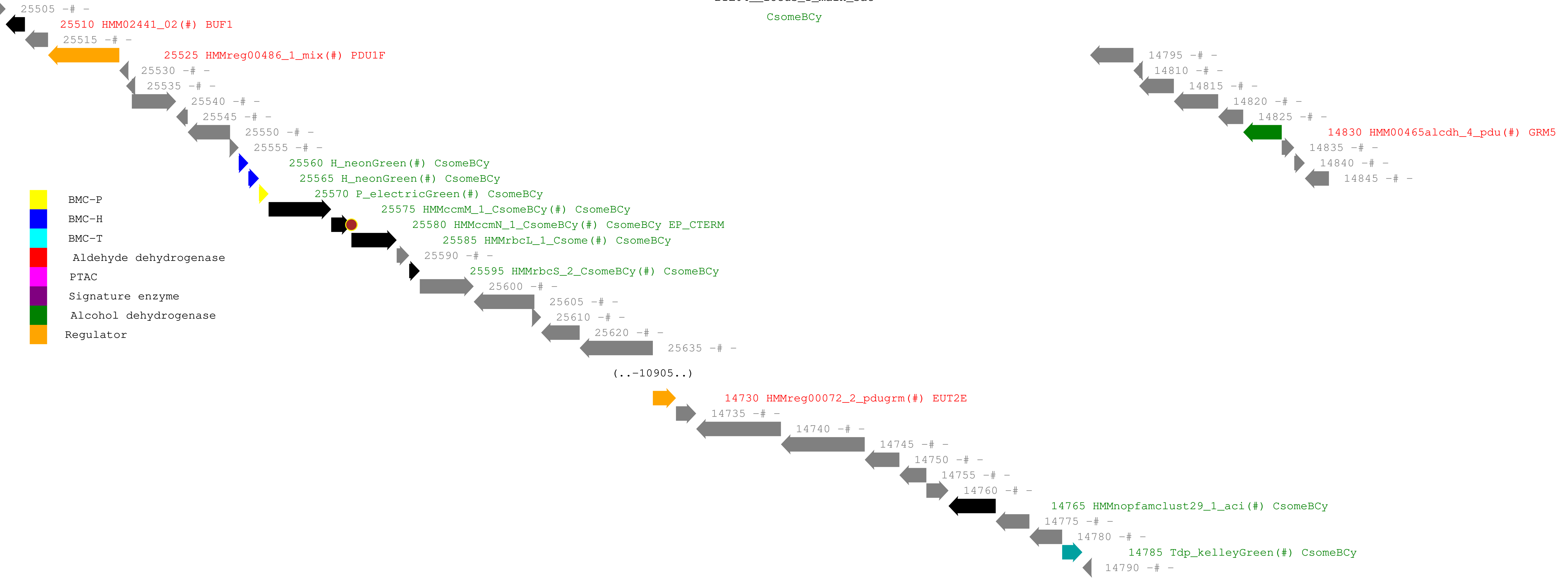

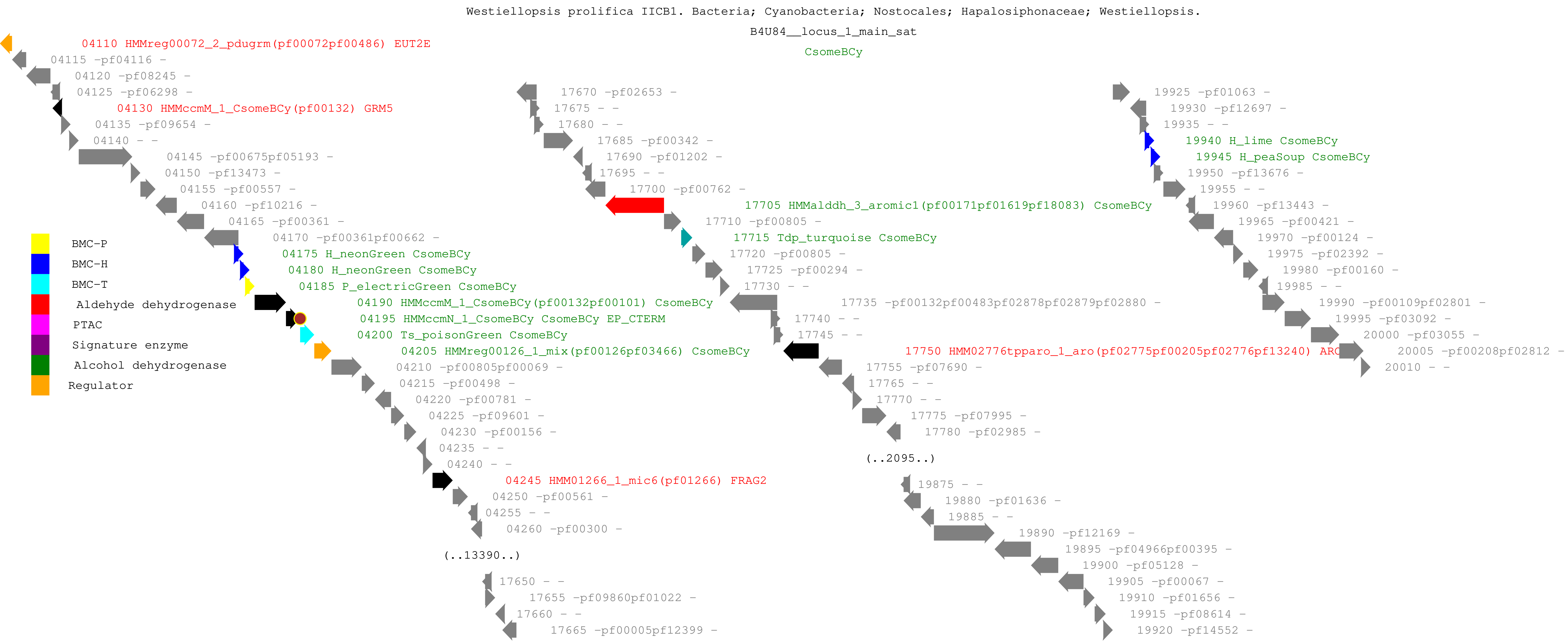

Westiellopsis prolifica IICB1. Bacteria; Cyanobacteria; Nostocales; Hapalosiphonaceae; Westiellopsis.
B4U84__locus_1_main_sat
CsomeBCy
04110 HMMreg00072_2_pdugrm(pf00072pf00486) EUT2E
04115 -pf04116 -
04120 -pf08245 -
04125 -pf06298 -
04130 HMMccmM_1_CsomeBCy(pf00132) GRM5
04135 -pf09654 -
04140 - -
04145 -pf00675pf05193 -
04150 -pf13473 -
04155 -pf00557 -
04160 -pf10216 -
04165 -pf00361 -
04170 -pf00361pf00662 -
04175 H_neonGreen CsomeBCy
04180 H_neonGreen CsomeBCy
04185 P_electricGreen CsomeBCy
04190 HMMccmM_1_CsomeBCy(pf00132pf00101) CsomeBCy
04195 HMMccmN_1_CsomeBCy CsomeBCy EP_CTERM
04200 Ts_poisonGreen CsomeBCy
04205 HMMreg00126_1_mix(pf00126pf03466) CsomeBCy
04210 -pf00805pf00069 -
04215 -pf00498 -
04220 -pf00781 -
04225 -pf09601 -
04230 -pf00156 -
04235 - -
04240 - -
04245 HMM01266_1_mic6(pf01266) FRAG2
04250 -pf00561 -
04255 - -
04260 -pf00300 -
(..13390..)
17650 - -
17655 -pf09860pf01022 -
17660 - -
17665 -pf00005pf12399 -
BMC-P
BMC-H
BMC-T
Aldehyde dehydrogenase
PTAC
Signature enzyme
Alcohol dehydrogenase
Regulator
17670 -pf02653 -
17675 - -
17680 - -
17685 -pf00342 -
17690 -pf01202 -
17695 - -
17700 -pf00762 -
17705 HMMalddh_3_aromic1(pf00171pf01619pf18083) CsomeBCy
17710 -pf00805 -
17715 Tdp_turquoise CsomeBCy
17720 -pf00805 -
17725 -pf00294 -
17730 - -
17735 -pf00132pf00483pf02878pf02879pf02880 -
17740 - -
17745 - -
17750 HMM02776tpparo_1_aro(pf02775pf00205pf02776pf13240) AR
17755 -pf07690 -
17765 - -
17770 - -
17775 -pf07995 -
17780 -pf02985 -
(..2095..)
19875 - -
19880 -pf01636 -
19885 - -
19890 -pf12169 -
19895 -pf04966pf00395 -
19900 -pf05128 -
19905 -pf00067 -
19910 -pf01656 -
19915 -pf08614 -
19920 -pf14552 -
19925 -pf01063 -
19930 -pf12697 -
19935 - -
19940 H_lime CsomeBCy
19945 H_peaSoup CsomeBCy
19950 -pf13676 -
19955 - -
19960 -pf13443 -
19965 -pf00421 -
19970 -pf00124 -
19975 -pf02392 -
19980 -pf00160 -
19985 - -
19990 -pf00109pf02801 -
19995 -pf03092 -
20000 -pf03055 -
20005 -pf00208pf02812 -
20010 - -

Microcystis sp. MC19. Bacteria; Cyanobacteria; Oscillatoriophycideae; Chroococcales;

B5D77__locus_1_main_sat

CsomeBCy

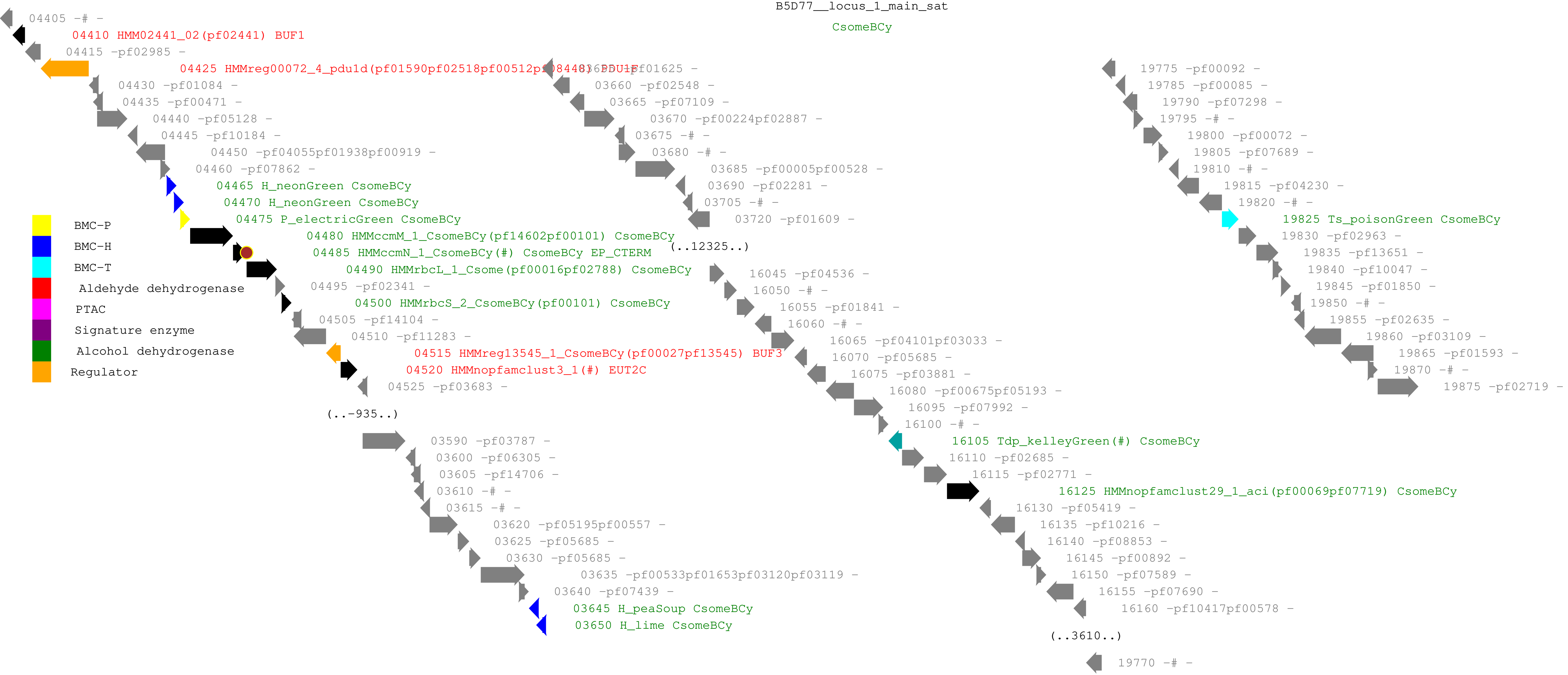

cyanobacterium TDX16. Bacteria; Cyanobacteria.

B7486__locus_2

CsomeBCy

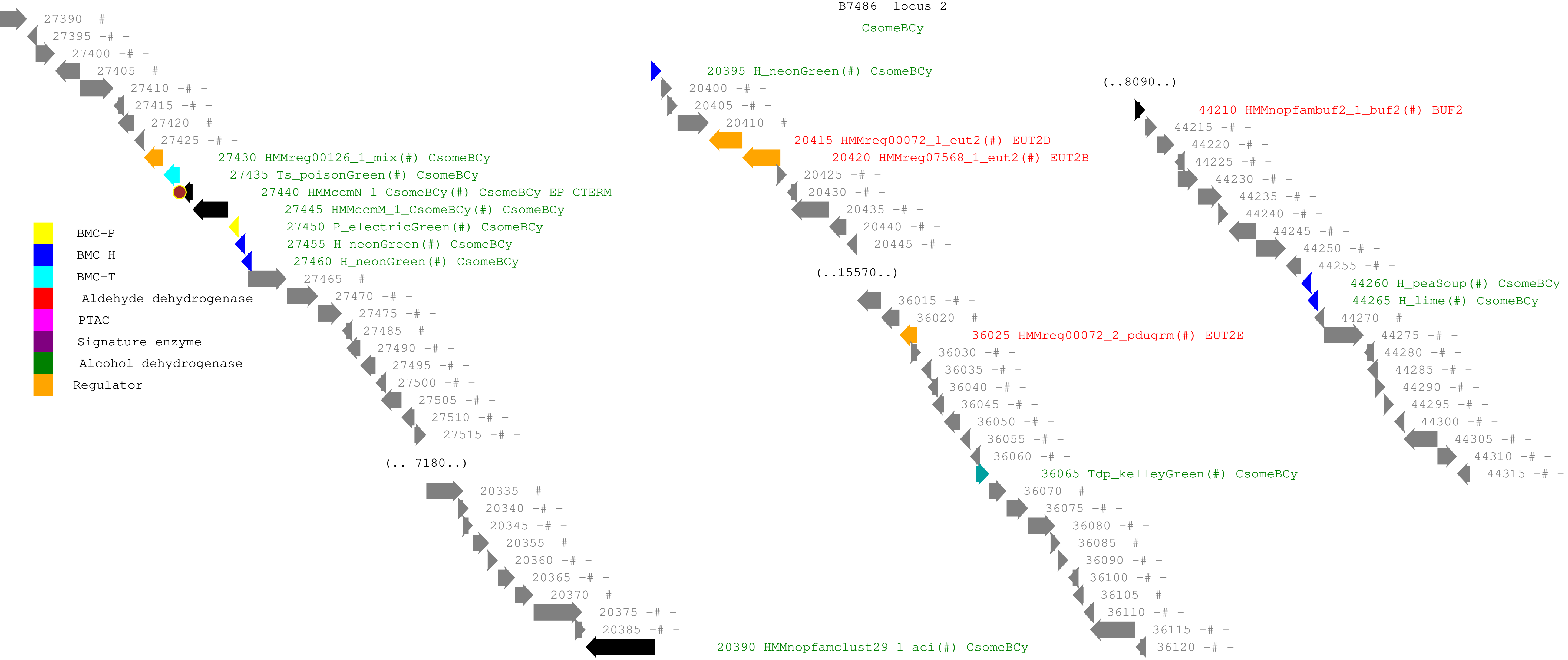

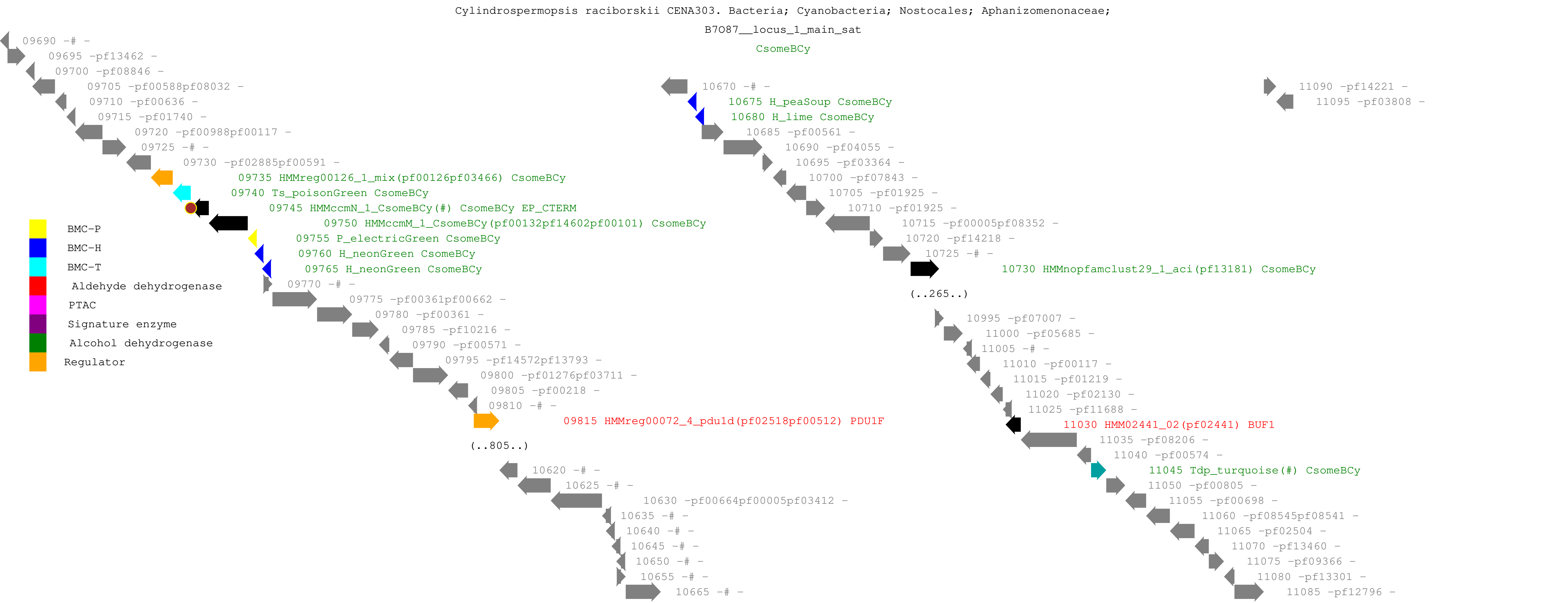
Cylindrospermopsis raciborskii CENA303. Bacteria; Cyanobacteria; Nostocales; Aphanizomenonaceae;
B7087__locus_1_main_sat
CsomeBCy
09690 -# -
09695 -pf13462 -
09700 -pf08846 -
09705 -pf00588pf08032 -
09710 -pf00636 -
09715 -pf01740 -
09720 -pf00988pf00117 -
09725 -# -
09730 -pf02885pf00591 -
09735 HMMreg00126_1_mix(pf00126pf03466) CsomeBCy
09740 Ts_poisonGreen CsomeBCy
09745 HMMccmN_1_CsomeBCy(#) CsomeBCy EP_CTERM
09750 HMMccmM_1_CsomeBCy(pf00132pf14602pf00101) CsomeBCy
09755 P_electricGreen CsomeBCy
09760 H_neonGreen CsomeBCy
09765 H_neonGreen CsomeBCy
09770 -# -
09775 -pf00361pf00662 -
09780 -pf00361 -
09785 -pf10216 -
09790 -pf00571 -
09795 -pf14572pf13793 -
09800 -pf01276pf03711 -
09805 -pf00218 -
09810 -# -
09815 HMMreg00072_4_pdu1d(pf02518pf00512) PDU1F
(..805..)
10620 -# -
10625 -# -
10630 -pf00664pf00005pf03412 -
10635 -# -
10640 -# -
10645 -# -
10650 -# -
10655 -# -
10665 -# -
10670 -# -
10675 H_peaSoup CsomeBCy
10680 H_lime CsomeBCy
10685 -pf00561 -
10690 -pf04055 -
10695 -pf03364 -
10700 -pf07843 -
10705 -pf01925 -
10710 -pf01925 -
10715 -pf00005pf08352 -
10720 -pf14218 -
10725 -# -
10730 HMMnopfamclust29_1_aci(pf13181) CsomeBCy
(..265..)
10995 -pf07007 -
11000 -pf05685 -
11005 -# -
11010 -pf00117 -
11015 -pf01219 -
11020 -pf02130 -
11025 -pf11688 -
11030 HMM02441_02(pf02441) BUF1
11035 -pf08206 -
11040 -pf00574 -
11045 Tdp_turquoise(#) CsomeBCy
11050 -pf00805 -
11055 -pf00698 -
11060 -pf08545pf08541 -
11065 -pf02504 -
11070 -pf13460 -
11075 -pf09366 -
11080 -pf13301 -
11085 -pf12796 -
11090 -pf14221 -
11095 -pf03808 -
BMC-P
BMC-H
BMC-T
Aldehyde dehydrogenase
PTAC
Signature enzyme
Alcohol dehydrogenase
Regulator

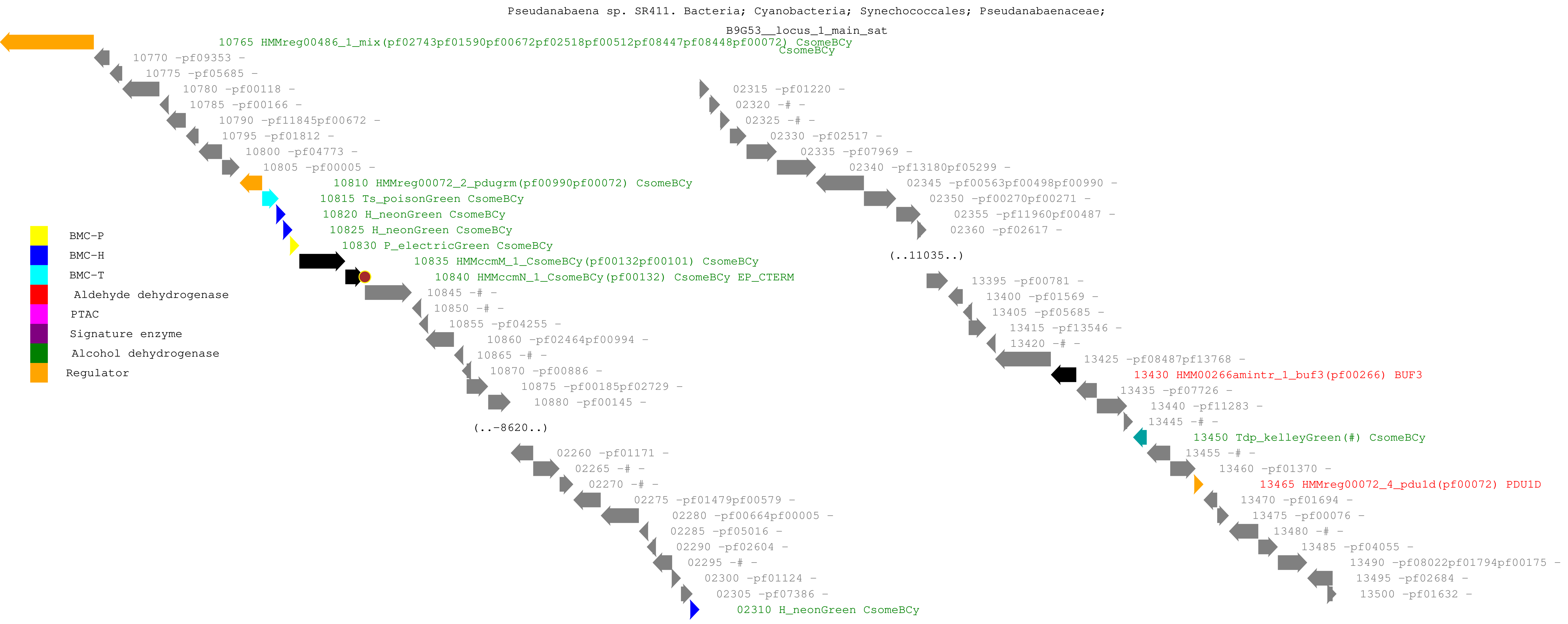
Pseudanabaena sp. SR411. Bacteria; Cyanobacteria; Synechococcales; Pseudanabaenaceae;
B9G53__locus_1_main_sat
CsomeBCy
10765 HMMreg00486_1_mix(pf02743pf01590pf00672pf02518pf00512pf08447pf08448pf00072) CsomeBCy
10770 -pf09353 -
10775 -pf05685 -
10780 -pf00118 -
10785 -pf00166 -
10790 -pf11845pf00672 -
10795 -pf01812 -
10800 -pf04773 -
10805 -pf00005 -
10810 HMMreg00072_2_pdugrm(pf00990pf00072) CsomeBCy
10815 Ts_poisonGreen CsomeBCy
10820 H_neonGreen CsomeBCy
10825 H_neonGreen CsomeBCy
10830 P_electricGreen CsomeBCy
10835 HMMccmM_1_CsomeBCy(pf00132pf00101) CsomeBCy
10840 HMMccmN_1_CsomeBCy(pf00132) CsomeBCy EP_CTERM
10845 -# -
10850 -# -
10855 -pf04255 -
10860 -pf02464pf00994 -
10865 -# -
10870 -pf00886 -
10875 -pf00185pf02729 -
10880 -pf00145 -
(..-8620..)
02260 -pf01171 -
02265 -# -
02270 -# -
02275 -pf01479pf00579 -
02280 -pf00664pf00005 -
02285 -pf05016 -
02290 -pf02604 -
02295 -# -
02300 -pf01124 -
02305 -pf07386 -
02310 H_neonGreen CsomeBCy
02315 -pf01220 -
02320 -# -
02325 -# -
02330 -pf02517 -
02335 -pf07969 -
02340 -pf13180pf05299 -
02345 -pf00563pf00498pf00990 -
02350 -pf00270pf00271 -
02355 -pf11960pf00487 -
02360 -pf02617 -
(..11035..)
13395 -pf00781 -
13400 -pf01569 -
13405 -pf05685 -
13415 -pf13546 -
13420 -# -
13425 -pf08487pf13768 -
13430 HMM00266amintr_1_buf3(pf00266) BUF3
13435 -pf07726 -
13440 -pf11283 -
13445 -# -
13450 Tdp_kelleyGreen(#) CsomeBCy
13455 -# -
13460 -pf01370 -
13465 HMMreg00072_4_pdu1d(pf00072) PDU1D
13470 -pf01694 -
13475 -pf00076 -
13480 -# -
13485 -pf04055 -
13490 -pf08022pf01794pf00175 -
13495 -pf02684 -
13500 -pf01632 -
BMC-P
BMC-H
BMC-T
Aldehyde dehydrogenase
PTAC
Signature enzyme
Alcohol dehydrogenase
Regulator

Arthrospira sp. O9.13F. Bacteria; Cyanobacteria; Oscillatoriophycideae; Oscillatoriales;

B9S53__locus_1_main_sat

CsomeBCy

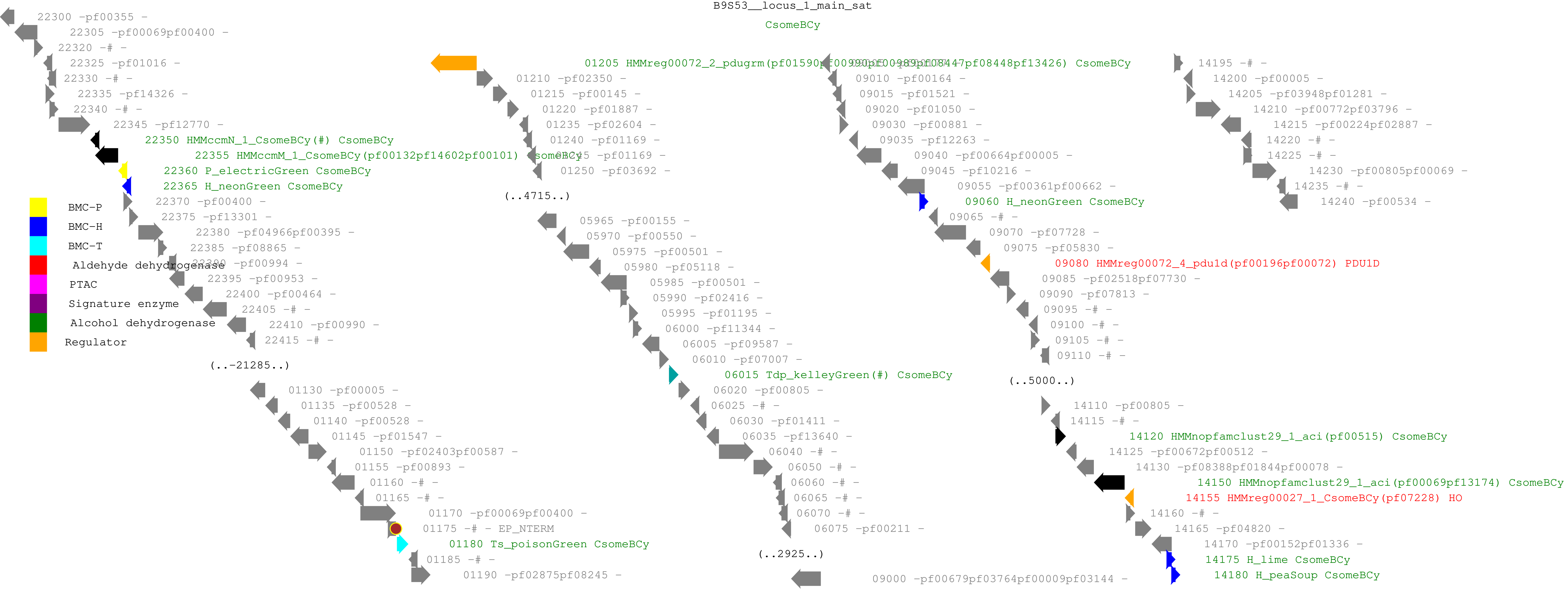

Mastigocoleus testarum BC008. Bacteria; Cyanobacteria; Nostocales; Hapalosiphonaceae; Mastigocoleus.

BC008__locus_1_main_sat

CsomeBCy

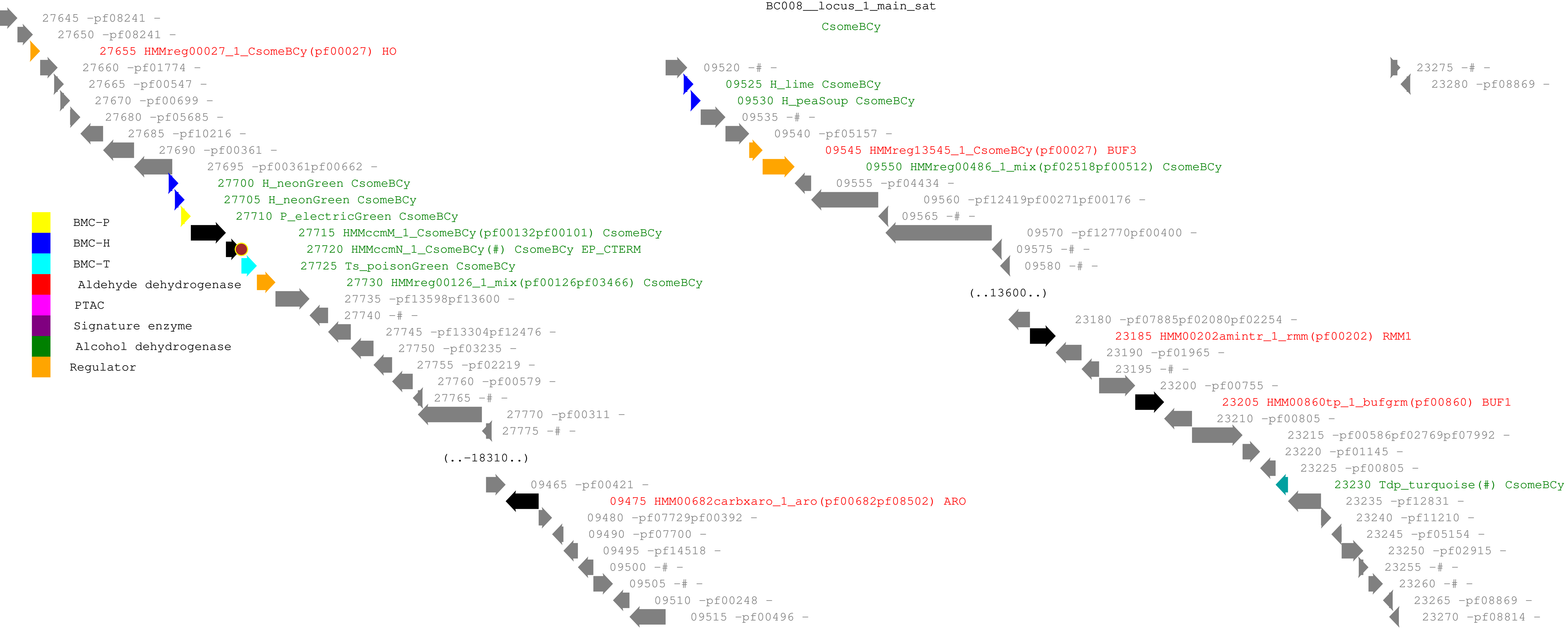

Nostoc sp. MBR 210. Bacteria; Cyanobacteria; Nostocales; Nostocaceae; Nostoc.

BCD64__locus_1_main_sat

CsomeBCy

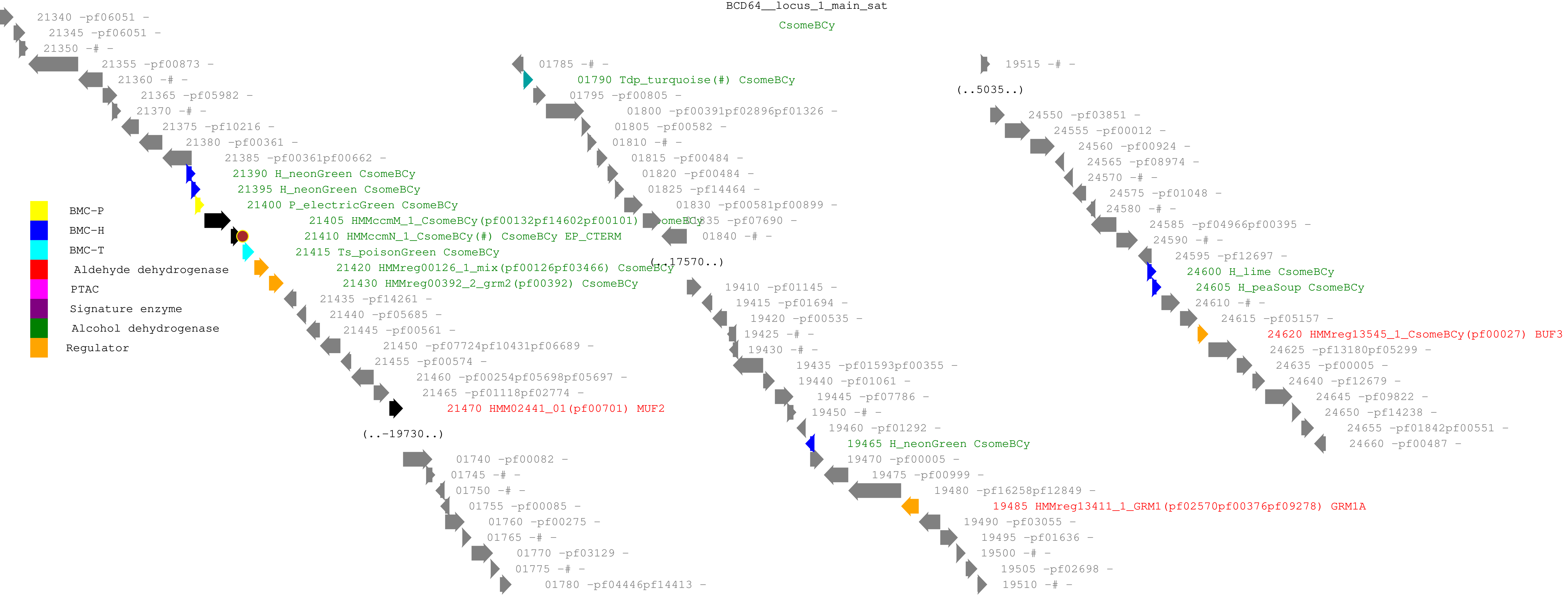

Oscillatoriales cyanobacterium USR001. Bacteria; Cyanobacteria; Oscillatoriophycideae; Oscillatoriales;

BCD67__locus_1_main_sat

CsomeBCy

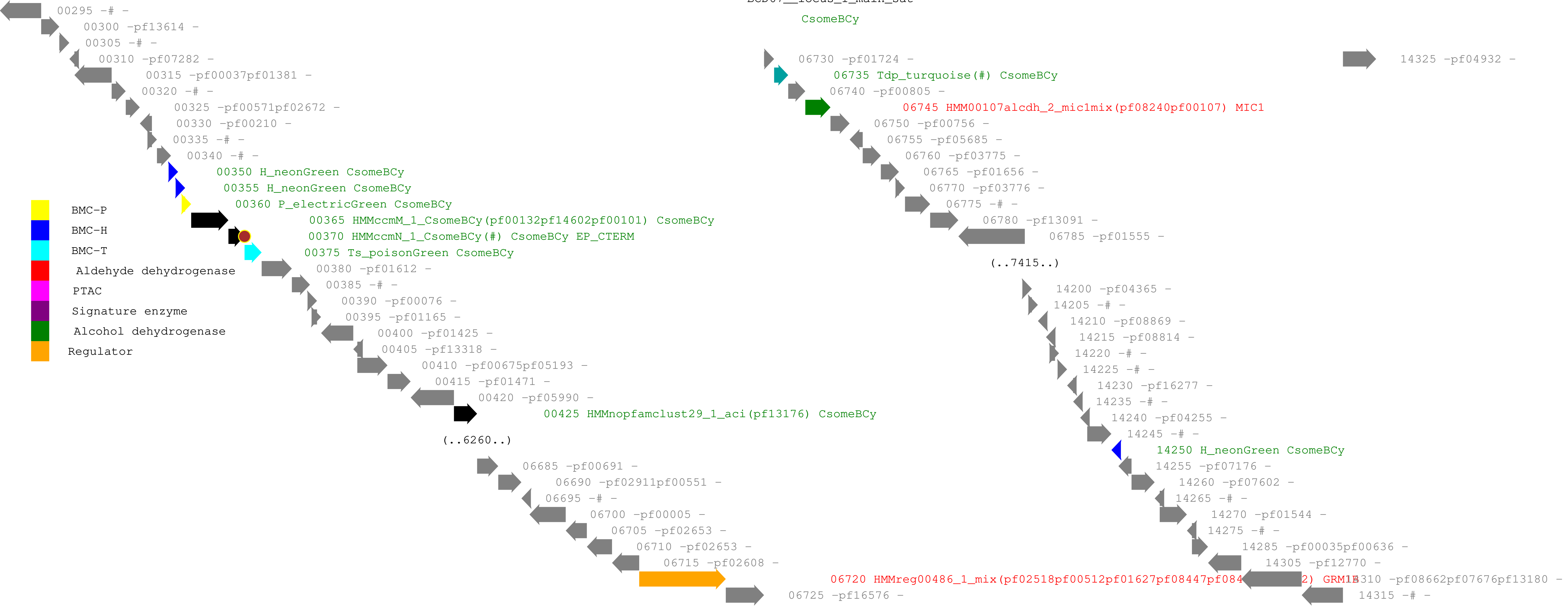

Limnothrix sp. P13C2. Bacteria; Cyanobacteria; Synechococcales; Pseudanabaenaceae;

BCR12__locus_1_main_sat

CsomeBCy

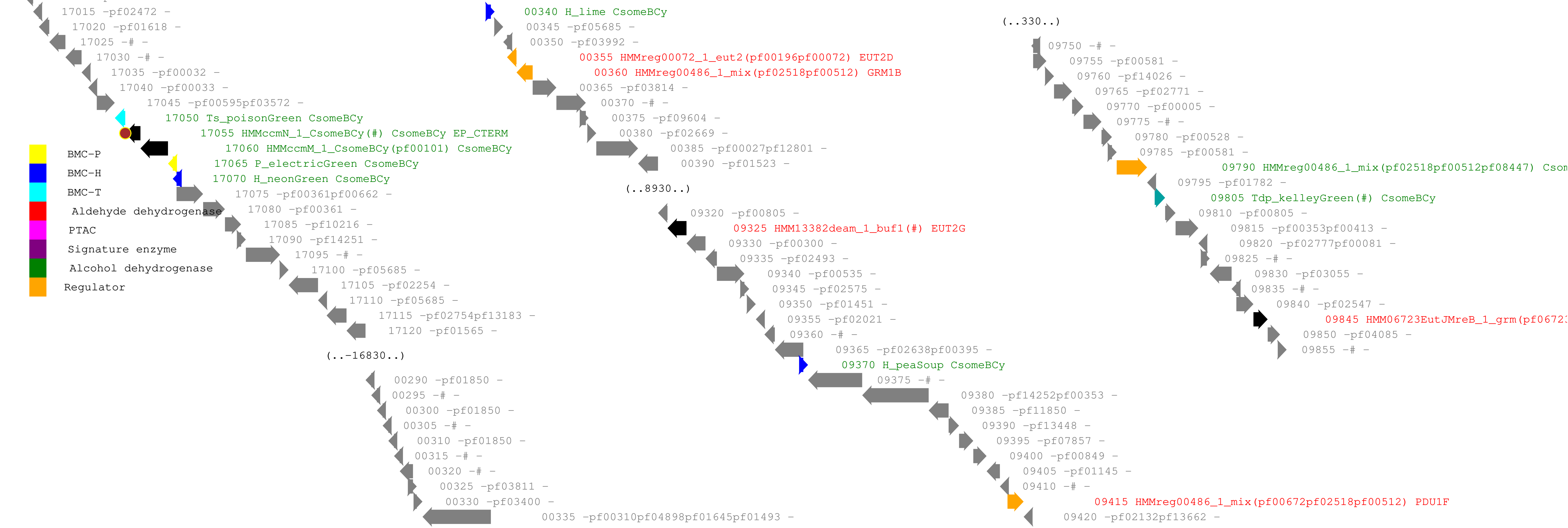

Cylindrospermopsis raciborskii MVCC14. Bacteria; Cyanobacteria; Nostocales; Aphanizomenonaceae;

BCV64__locus_1_main_sat

CsomeBCy

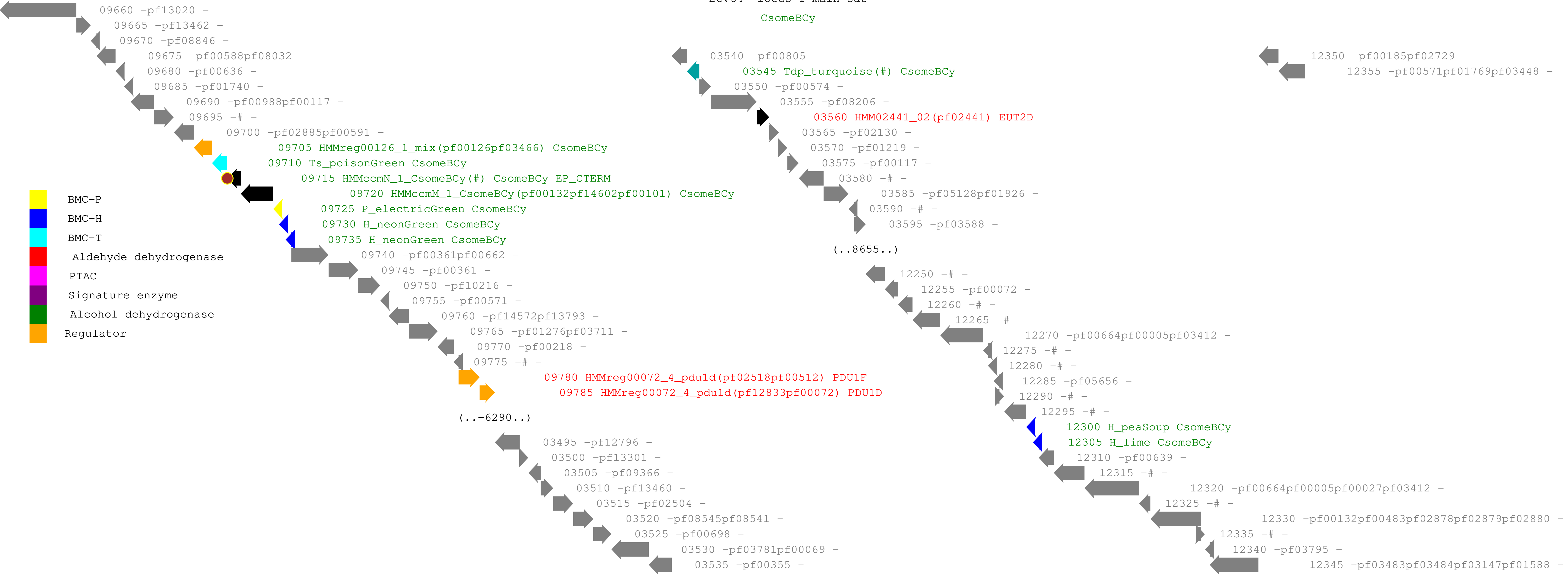

Halothece sp. (strain PCC 7418) (Synechococcus sp. (strain PCC 7418)). Bacteria; Cyanobacteria; Oscillatoriophycideae; Chroococcales;

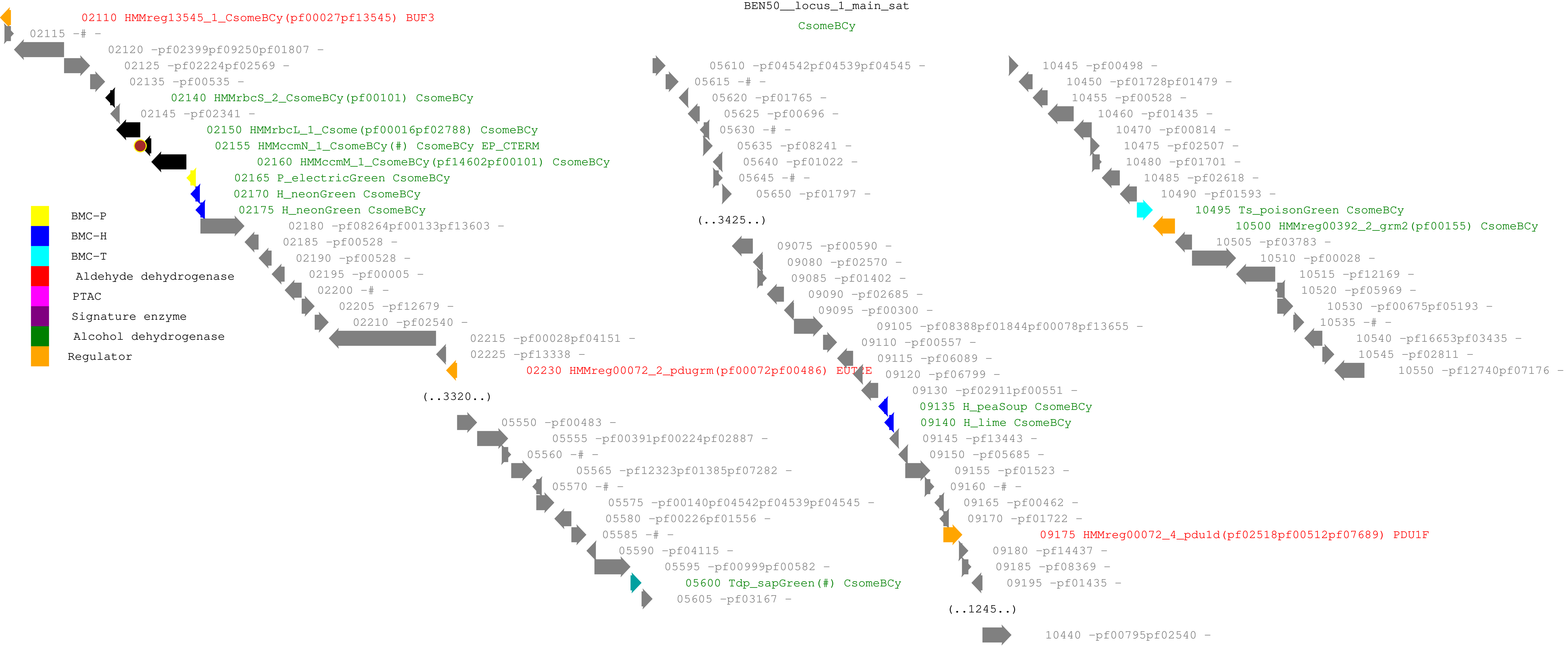

Bacillus subtilis BEST7613. Bacteria; Firmicutes; Bacilli; Bacillales; Bacillaceae; Bacillus.

BEST7613__locus_1_main_sat

CsomeBCy

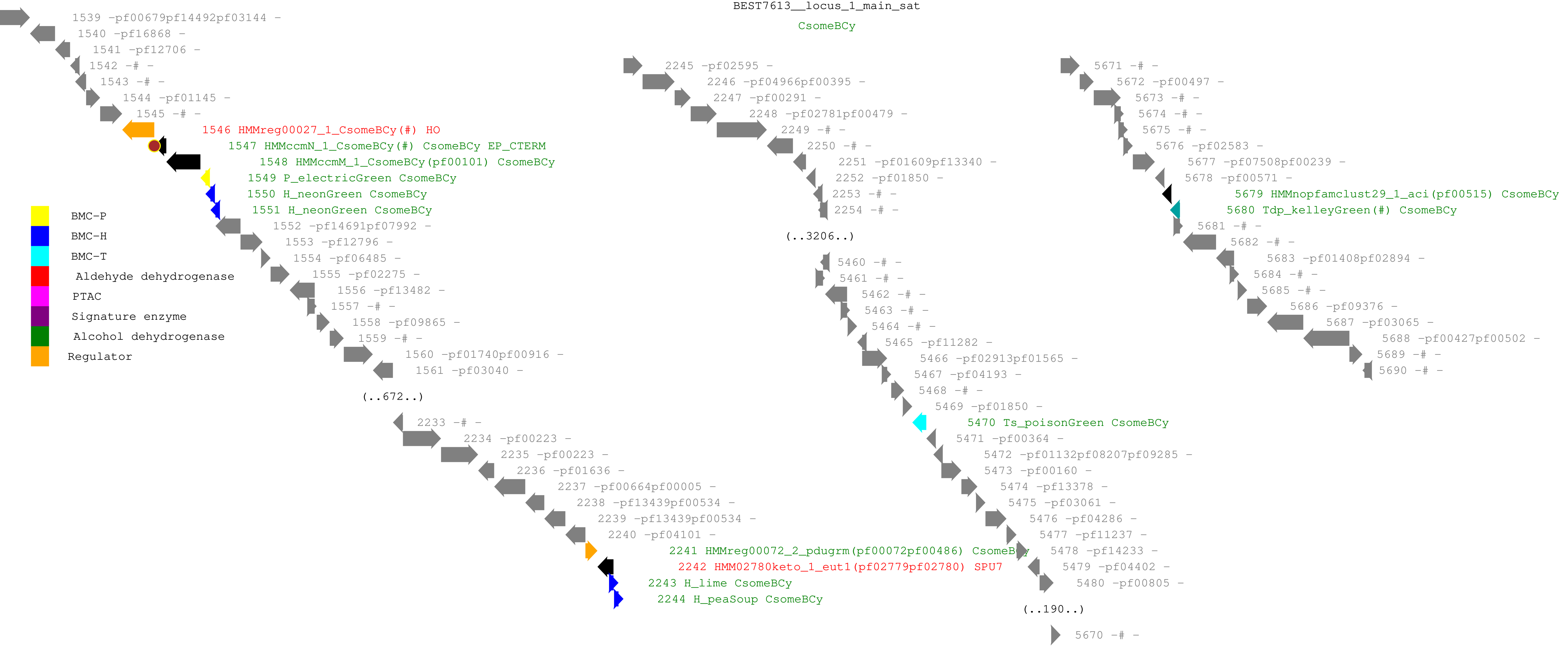

Microcystis aeruginosa NIES-98. Bacteria; Cyanobacteria; Oscillatoriophycideae; Chroococcales;

BFG60__locus_1_main_sat

CsomeBCy

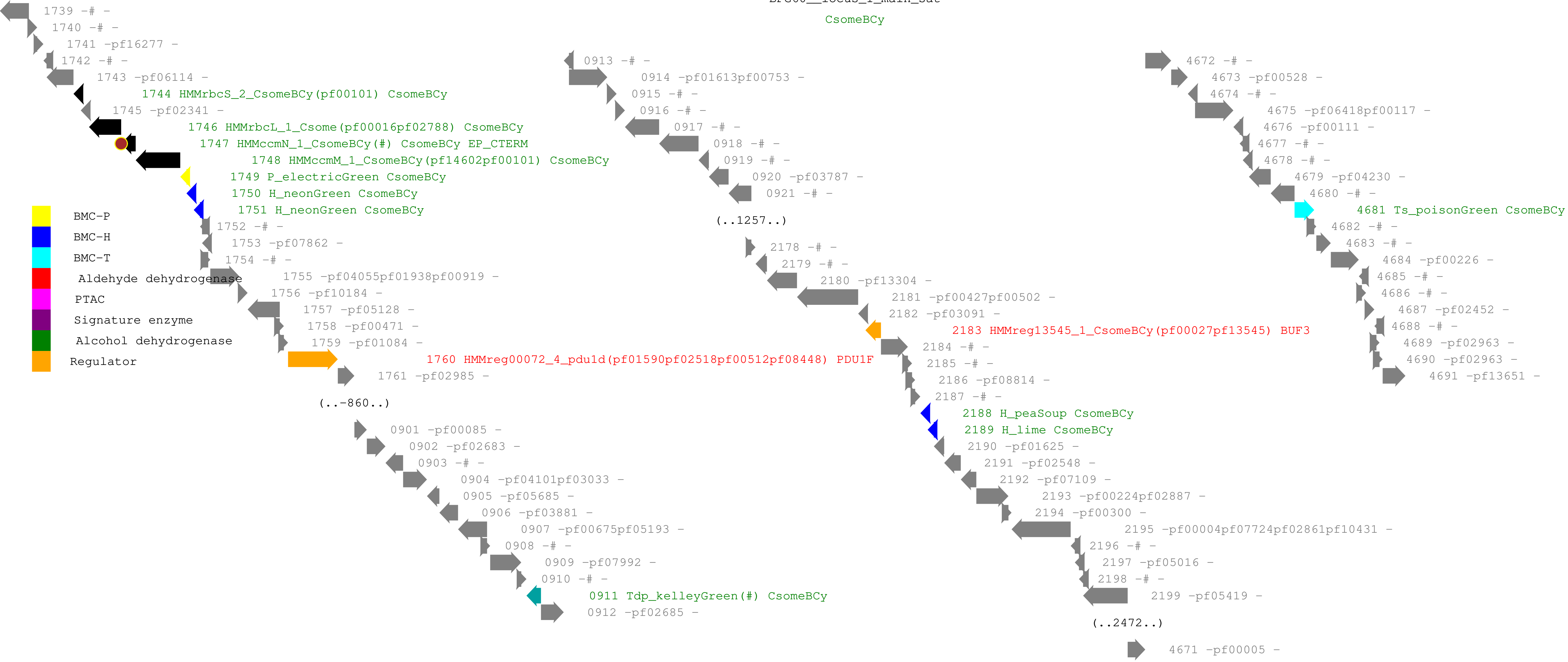

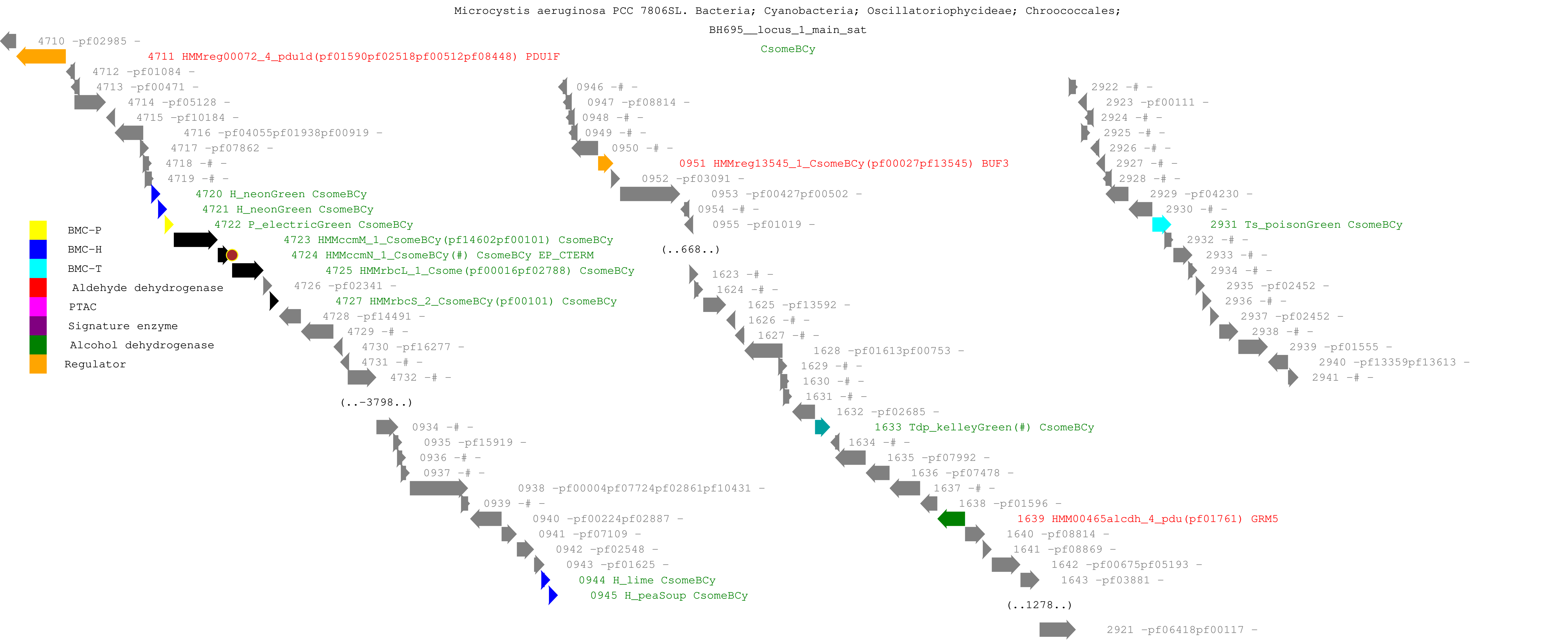
Microcystis aeruginosa PCC 7806SL. Bacteria; Cyanobacteria; Oscillatoriophycideae; Chroococcales;
BH695__locus_1_main_sat
CsomeBCy
4710 -pf02985 -
4711 HMMreg00072_4_pdu1d(pf01590pf02518pf00512pf08448) PDU1F
4712 -pf01084 -
4713 -pf00471 -
4714 -pf05128 -
4715 -pf10184 -
4716 -pf04055pf01938pf00919 -
4717 -pf07862 -
4718 -# -
4719 -# -
4720 H_neonGreen CsomeBCy
4721 H_neonGreen CsomeBCy
4722 P_electricGreen CsomeBCy
4723 HMMccmM_1_CsomeBCy(pf14602pf00101) CsomeBCy
4724 HMMccmN_1_CsomeBCy(#) CsomeBCy EP_CTERM
4725 HMMrbcL_1_Csome(pf00016pf02788) CsomeBCy
4726 -pf02341 -
4727 HMMrbcS_2_CsomeBCy(pf00101) CsomeBCy
4728 -pf14491 -
4729 -# -
4730 -pf16277 -
4731 -# -
4732 -# -
(..-3798..)
0934 -# -
0935 -pf15919 -
0936 -# -
0937 -# -
0938 -pf00004pf07724pf02861pf10431 -
0939 -# -
0940 -pf00224pf02887 -
0941 -pf07109 -
0942 -pf02548 -
0943 -pf01625 -
0944 H_lime CsomeBCy
0945 H_peaSoup CsomeBCy
BMC-P
BMC-H
BMC-T
Aldehyde dehydrogenase
PTAC
Signature enzyme
Alcohol dehydrogenase
Regulator
0946 -# -
0947 -pf08814 -
0948 -# -
0949 -# -
0950 -# -
0951 HMMreg13545_1_CsomeBCy(pf00027pf13545) BUF3
0952 -pf03091 -
0953 -pf00427pf00502 -
0954 -# -
0955 -pf01019 -
(..668..)
1623 -# -
1624 -# -
1625 -pf13592 -
1626 -# -
1627 -# -
1628 -pf01613pf00753 -
1629 -# -
1630 -# -
1631 -# -
1632 -pf02685 -
1633 Tdp_kelleyGreen(#) CsomeBCy
1634 -# -
1635 -pf07992 -
1636 -pf07478 -
1637 -# -
1638 -pf01596 -
1639 HMM00465alcdh_4_pdu(pf01761) GRM5
1640 -pf08814 -
1641 -pf08869 -
1642 -pf00675pf05193 -
1643 -pf03881 -
(..1278..)
2921 -pf06418pf00117 -
2922 -# -
2923 -pf00111 -
2924 -# -
2925 -# -
2926 -# -
2927 -# -
2928 -# -
2929 -pf04230 -
2930 -# -
2931 Ts_poisonGreen CsomeBCy
2932 -# -
2933 -# -
2934 -# -
2935 -pf02452 -
2936 -# -
2937 -pf02452 -
2938 -# -
2939 -pf01555 -
2940 -pf13359pf13613 -
2941 -# -

Desertifilum sp. IPPAS B-1220. Bacteria; Cyanobacteria; Oscillatoriophycideae; Oscillatoriales;

BH720__locus_1_main_sat

CsomeBCy

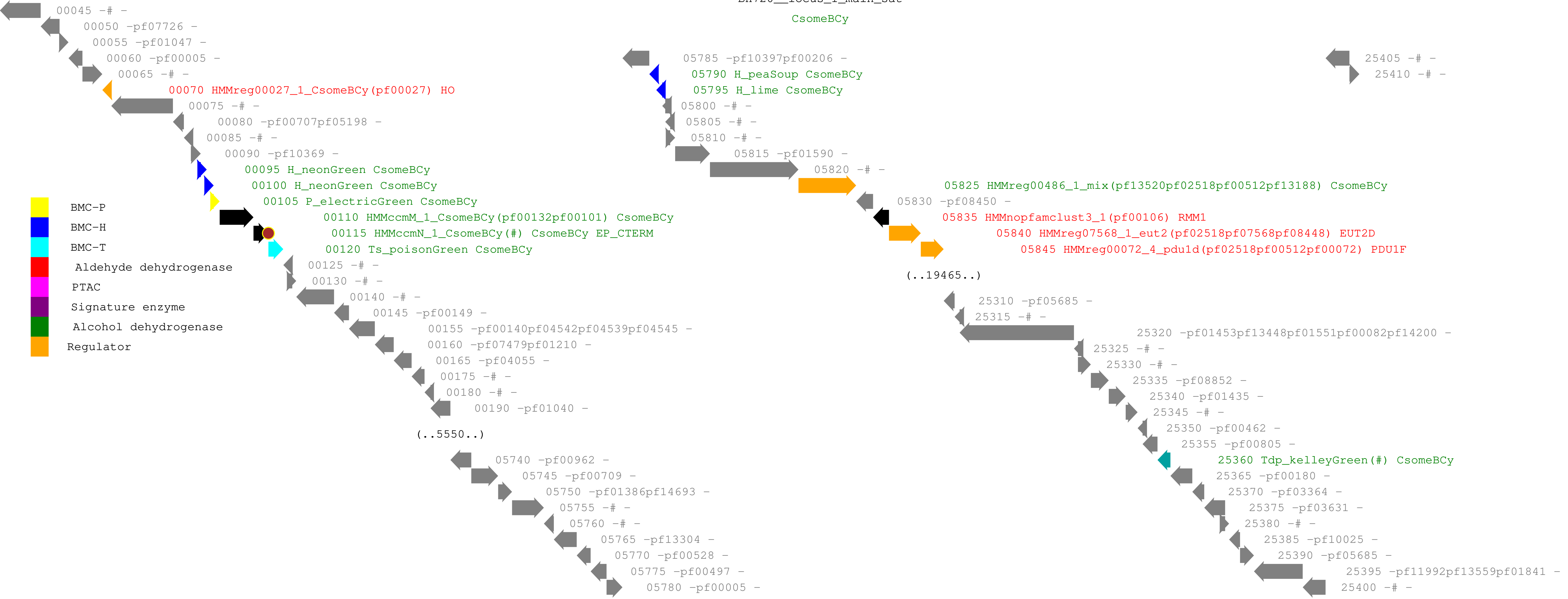

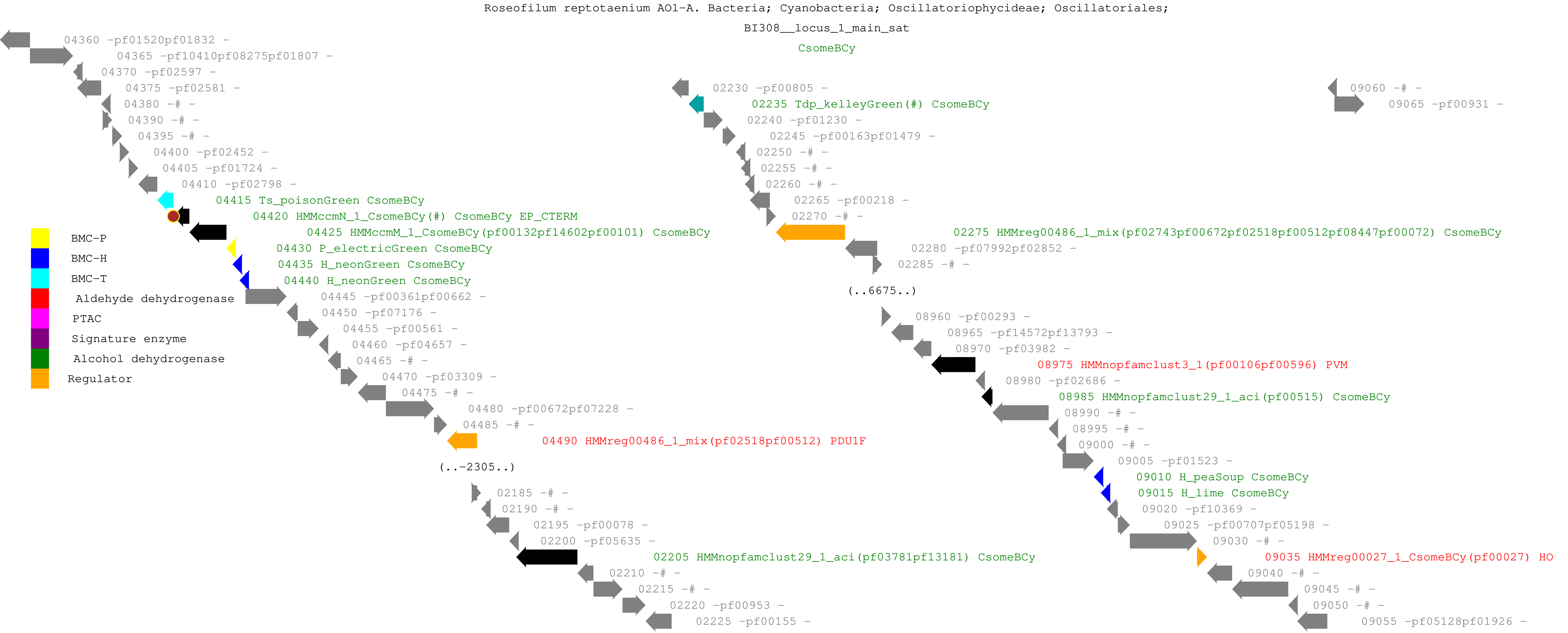
Roseofilum reptotaenium AO1-A. Bacteria; Cyanobacteria; Oscillatoriophycideae; Oscillatoriales;
BI308__locus_1_main_sat
CsomeBCy
04360 -pf01520pf01832 -
04365 -pf10410pf08275pf01807 -
04370 -pf02597 -
04375 -pf02581 -
04380 -# -
04390 -# -
04395 -# -
04400 -pf02452 -
04405 -pf01724 -
04410 -pf02798 -
04415 Ts_poisonGreen CsomeBCy
04420 HMMccmN_1_CsomeBCy(#) CsomeBCy EP_CTERM
04425 HMMccmM_1_CsomeBCy(pf00132pf14602pf00101) CsomeBCy
04430 P_electricGreen CsomeBCy
04435 H_neonGreen CsomeBCy
04440 H_neonGreen CsomeBCy
04445 -pf00361pf00662 -
04450 -pf07176 -
04455 -pf00561 -
04460 -pf04657 -
04465 -# -
04470 -pf03309 -
04475 -# -
04480 -pf00672pf07228 -
04485 -# -
04490 HMMreg00486_1_mix(pf02518pf00512) PDU1F
(..-2305..)
02185 -# -
02190 -# -
02195 -pf00078 -
02200 -pf05635 -
02205 HMMnopfamclust29_1_aci(pf03781pf13181) CsomeBCy
02210 -# -
02215 -# -
02220 -pf00953 -
02225 -pf00155 -
02230 -pf00805 -
02235 Tdp_kelleyGreen(#) CsomeBCy
02240 -pf01230 -
02245 -pf00163pf01479 -
02250 -# -
02255 -# -
02260 -# -
02265 -pf00218 -
02270 -# -
02275 HMMreg00486_1_mix(pf02743pf00672pf02518pf00512pf08447pf00072) CsomeBCy
02280 -pf07992pf02852 -
02285 -# -
(..6675..)
08960 -pf00293 -
08965 -pf14572pf13793 -
08970 -pf03982 -
08975 HMMnopfamclust3_1(pf00106pf00596) PVM
08980 -pf02686 -
08985 HMMnopfamclust29_1_aci(pf00515) CsomeBCy
08990 -# -
08995 -# -
09000 -# -
09005 -pf01523 -
09010 H_peaSoup CsomeBCy
09015 H_lime CsomeBCy
09020 -pf10369 -
09025 -pf00707pf05198 -
09030 -# -
09035 HMMreg00027_1_CsomeBCy(pf00027) HO
09040 -# -
09045 -# -
09050 -# -
09055 -pf05128pf01926 -
09060 -# -
09065 -pf00931 -
BMC-P
BMC-H
BMC-T
Aldehyde dehydrogenase
PTAC
Signature enzyme
Alcohol dehydrogenase
Regulator

Limnothrix sp. CACIAM 69d. Bacteria; Cyanobacteria; Synechococcales; Pseudanabaenaceae; Limnothrix;

BJG00__locus_1_main_sat

CsomeBCy

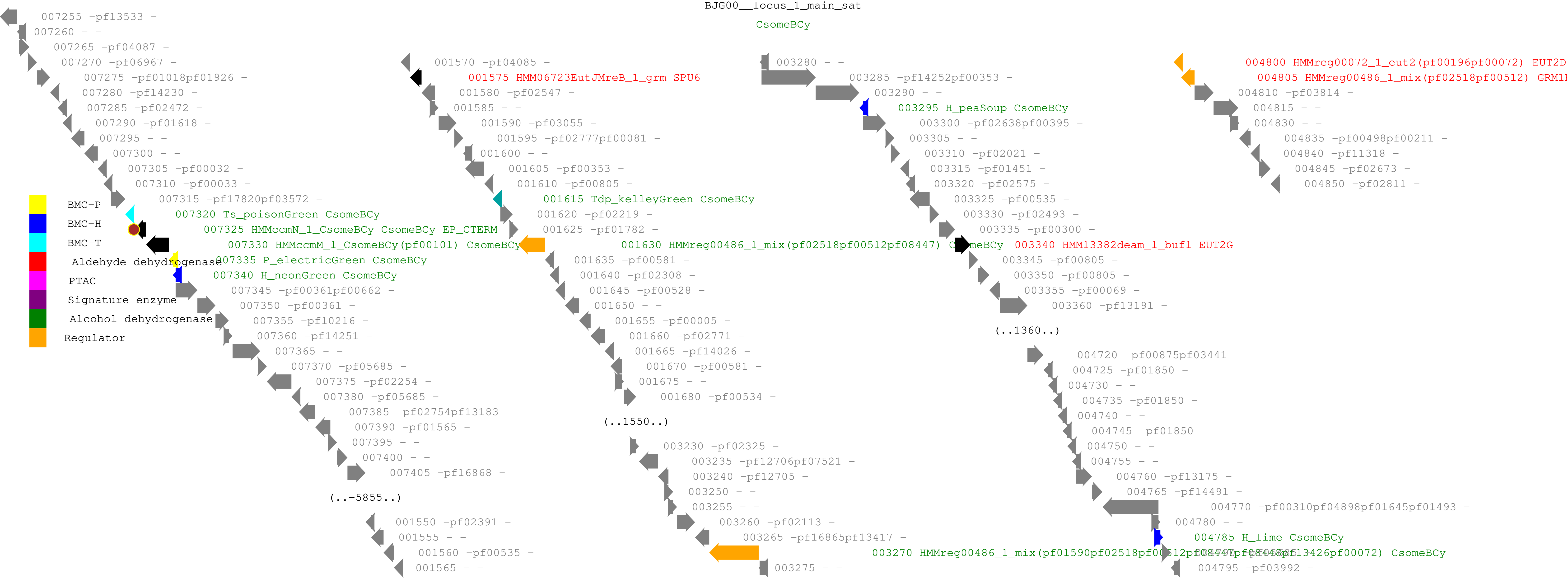

Moorea producens PAL-8-15-08-1. Bacteria; Cyanobacteria; Oscillatoriophycideae; Oscillatoriales;

BJP34__locus_1_main_sat

CsomeBCy

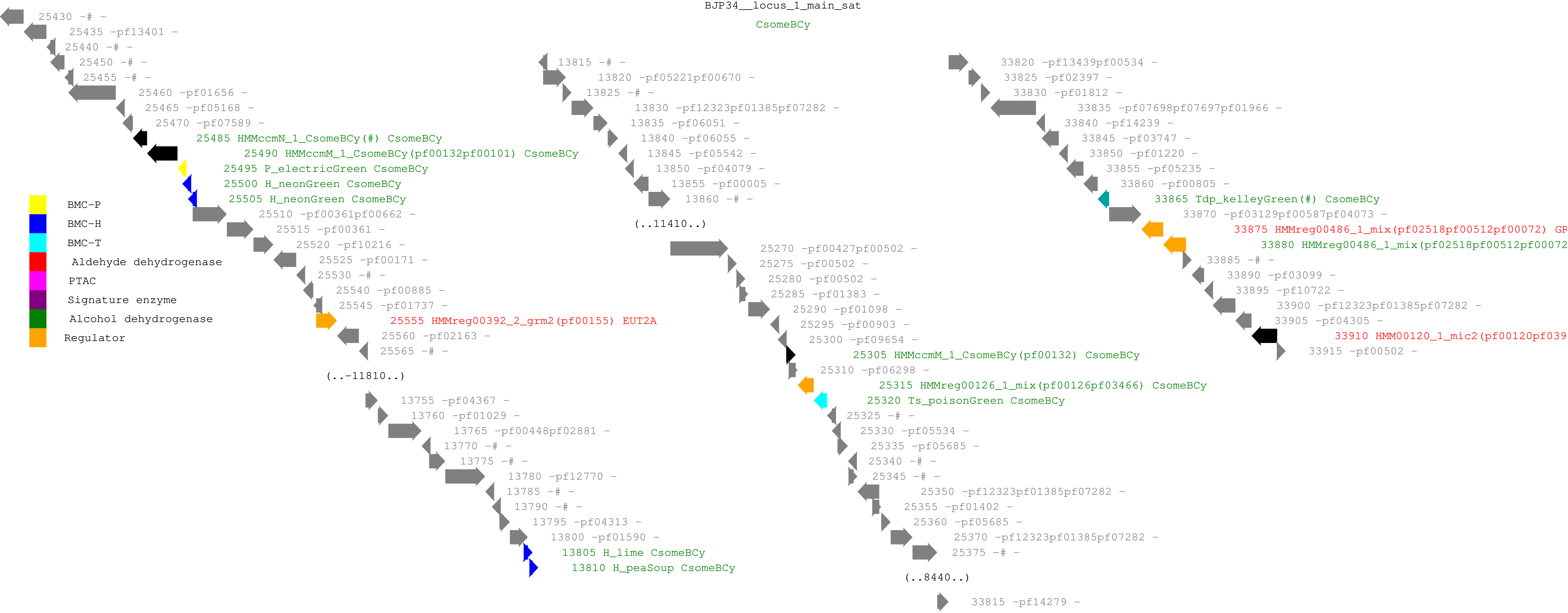

Moorea producens JHB. Bacteria; Cyanobacteria; Oscillatoriophycideae; Oscillatoriales;

BJP36__locus_1_main_sat

CsomeBCy

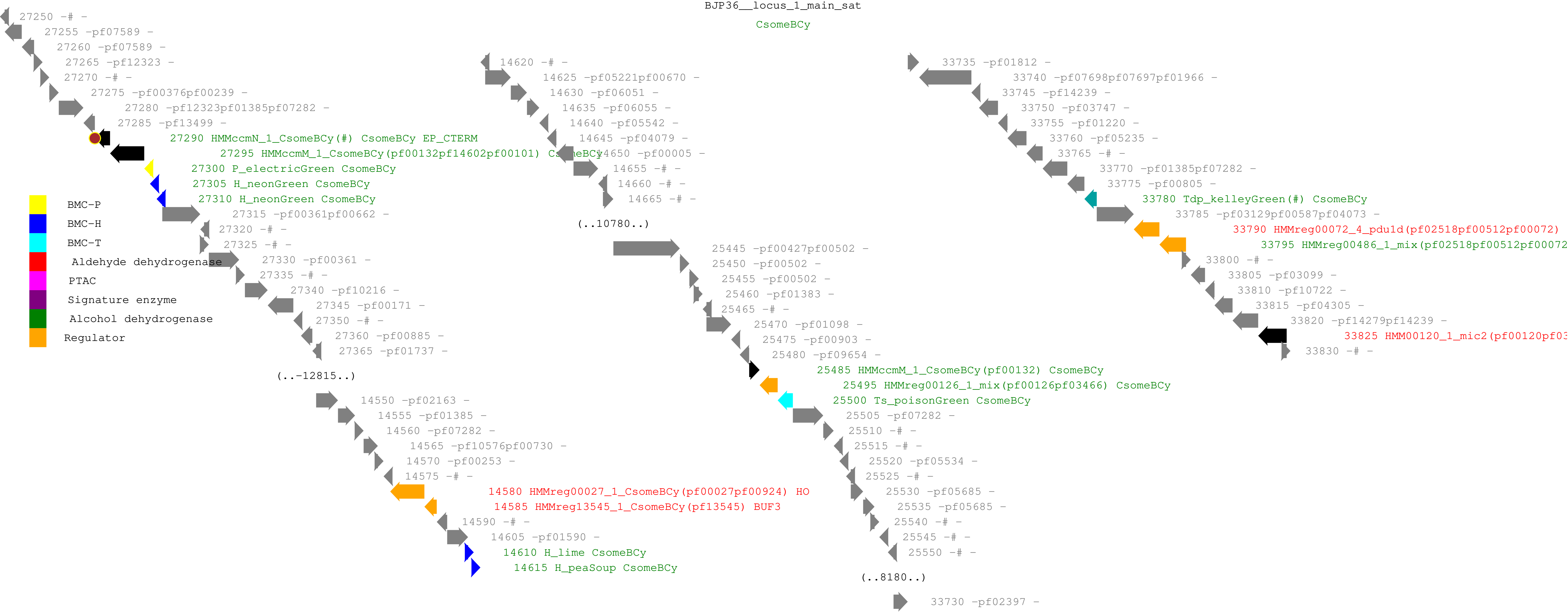

Moorea bouillonii PNG. Bacteria; Cyanobacteria; Oscillatoriophycideae; Oscillatoriales;

BJP37__locus_1_main_sat

CsomeBCy

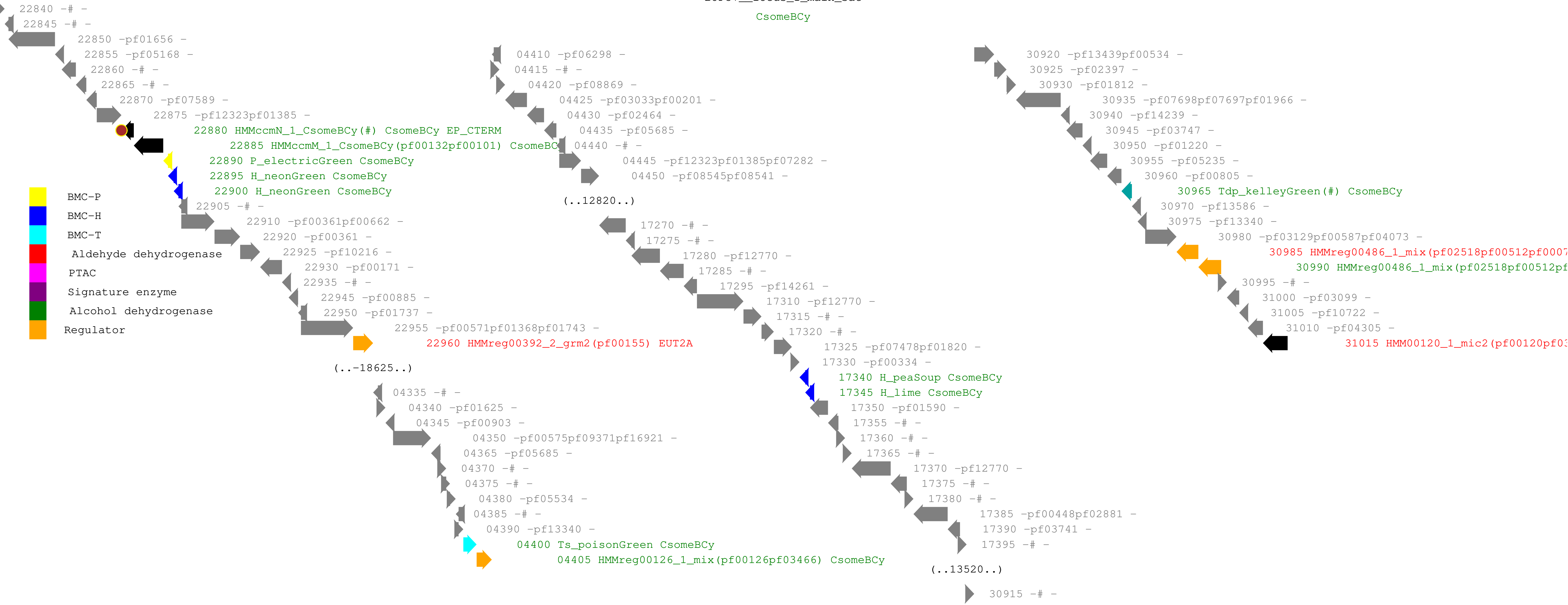

Mastigocladus laminosus UU774. Bacteria; Cyanobacteria; Nostocales; Hapalosiphonaceae; Mastigocladus.

BLD44__locus_1_main_sat

CsomeBCy

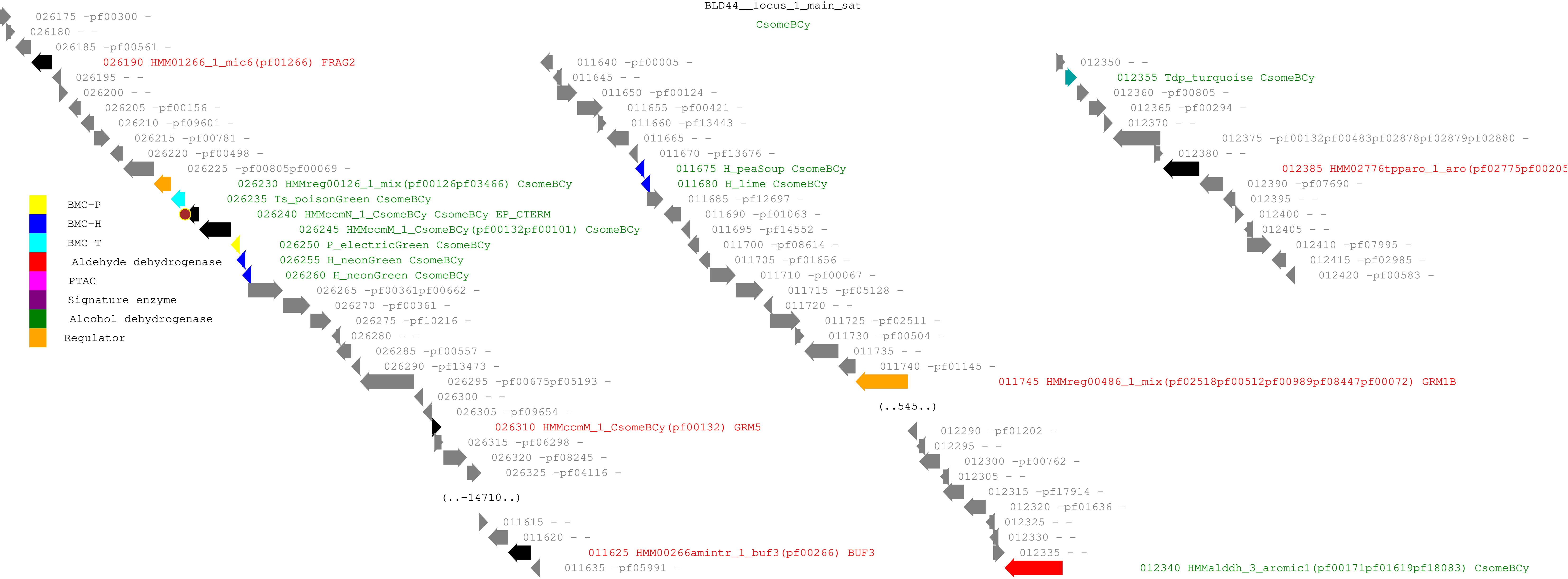

Dolichospermum sp. UHCC 0315A. Bacteria; Cyanobacteria; Nostocales; Aphanizomenonaceae; Dolichospermum;

BMF77__locus_1_main_sat

CsomeBCy

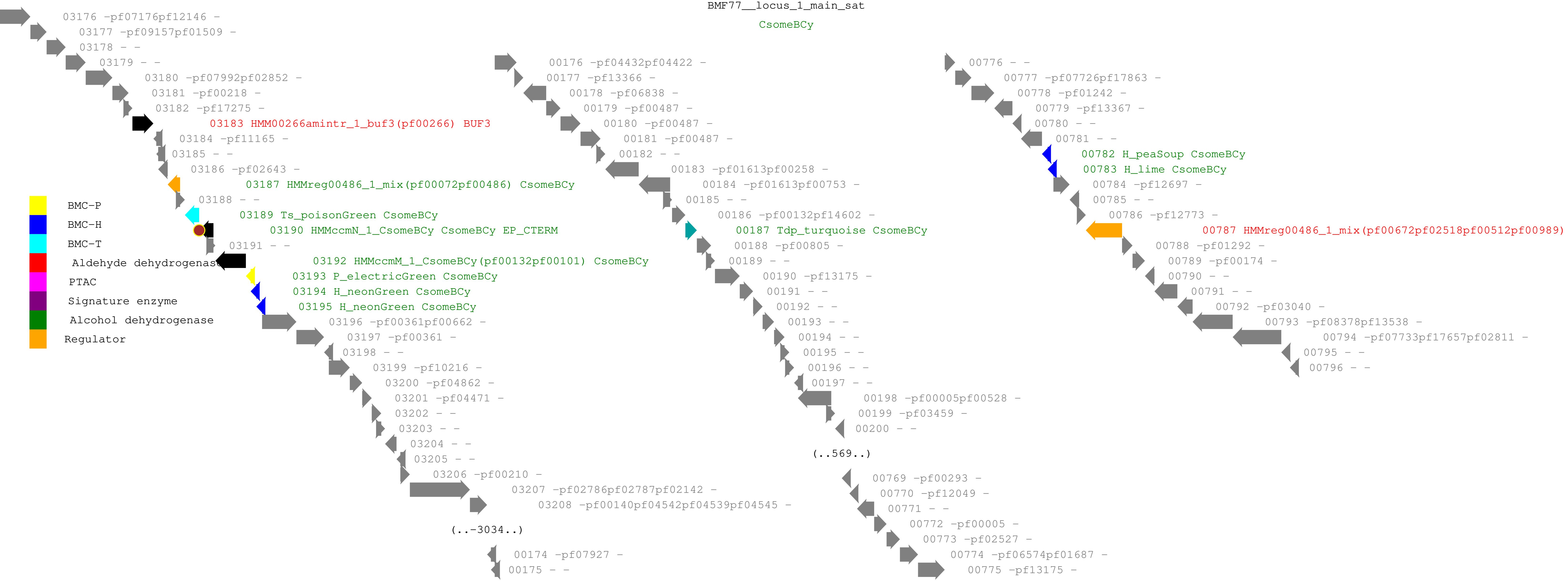

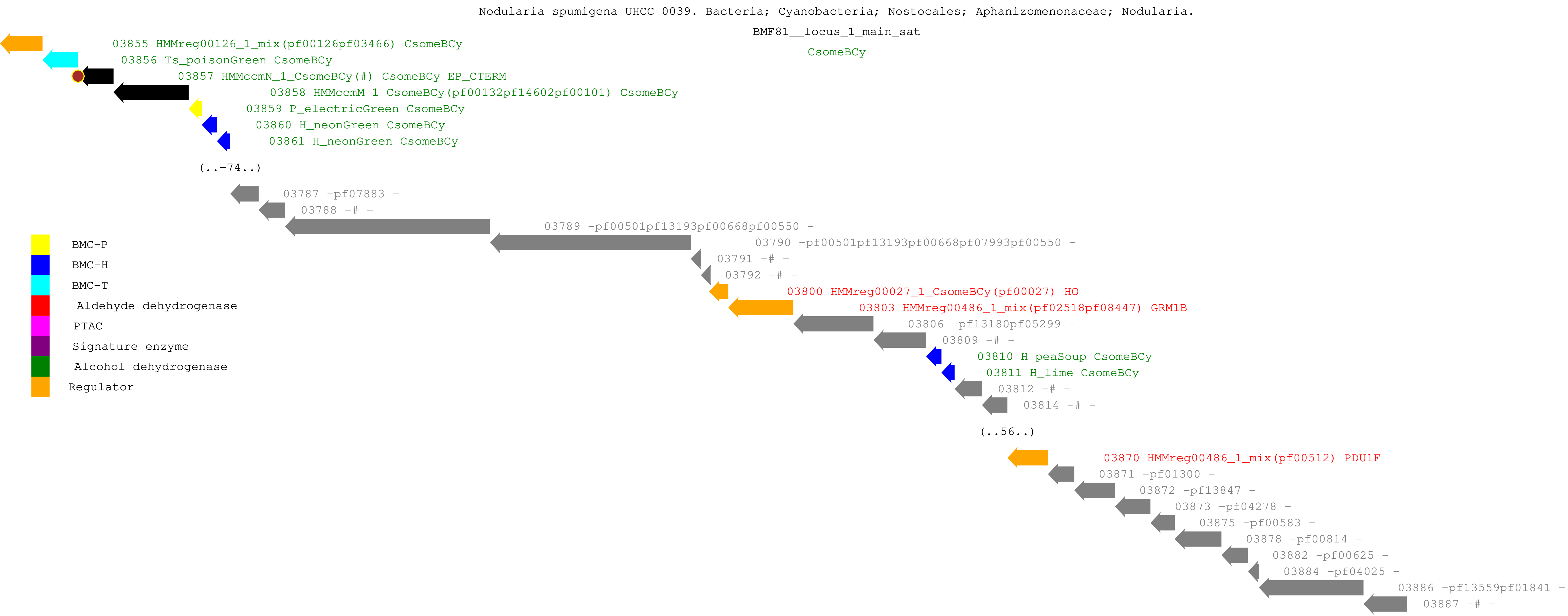
Nodularia spumigena UHCC 0039. Bacteria; Cyanobacteria; Nostocales; Aphanizomenonaceae; Nodularia.
BMF81__locus_1_main_sat
CsomeBCy
03855 HMMreg00126_1_mix(pf00126pf03466) CsomeBCy
03856 Ts_poisonGreen CsomeBCy
03857 HMMccmN_1_CsomeBCy(#) CsomeBCy EP_CTERM
03858 HMMccmM_1_CsomeBCy(pf00132pf14602pf00101) CsomeBCy
03859 P_electricGreen CsomeBCy
03860 H_neonGreen CsomeBCy
03861 H_neonGreen CsomeBCy
(..-74..)
03787 -pf07883 -
03788 -# -
03789 -pf00501pf13193pf00668pf00550 -
03790 -pf00501pf13193pf00668pf07993pf00550 -
03791 -# -
03792 -# -
03800 HMMreg00027_1_CsomeBCy(pf00027) HO
03803 HMMreg00486_1_mix(pf02518pf08447) GRM1B
03806 -pf13180pf05299 -
03809 -# -
03810 H_peaSoup CsomeBCy
03811 H_lime CsomeBCy
03812 -# -
03814 -# -
(..56..)
03870 HMMreg00486_1_mix(pf00512) PDU1F
03871 -pf01300 -
03872 -pf13847 -
03873 -pf04278 -
03875 -pf00583 -
03878 -pf00814 -
03882 -pf00625 -
03884 -pf04025 -
03886 -pf13559pf01841 -
03887 -# -
BMC-P
BMC-H
BMC-T
Aldehyde dehydrogenase
PTAC
Signature enzyme
Alcohol dehydrogenase
Regulator

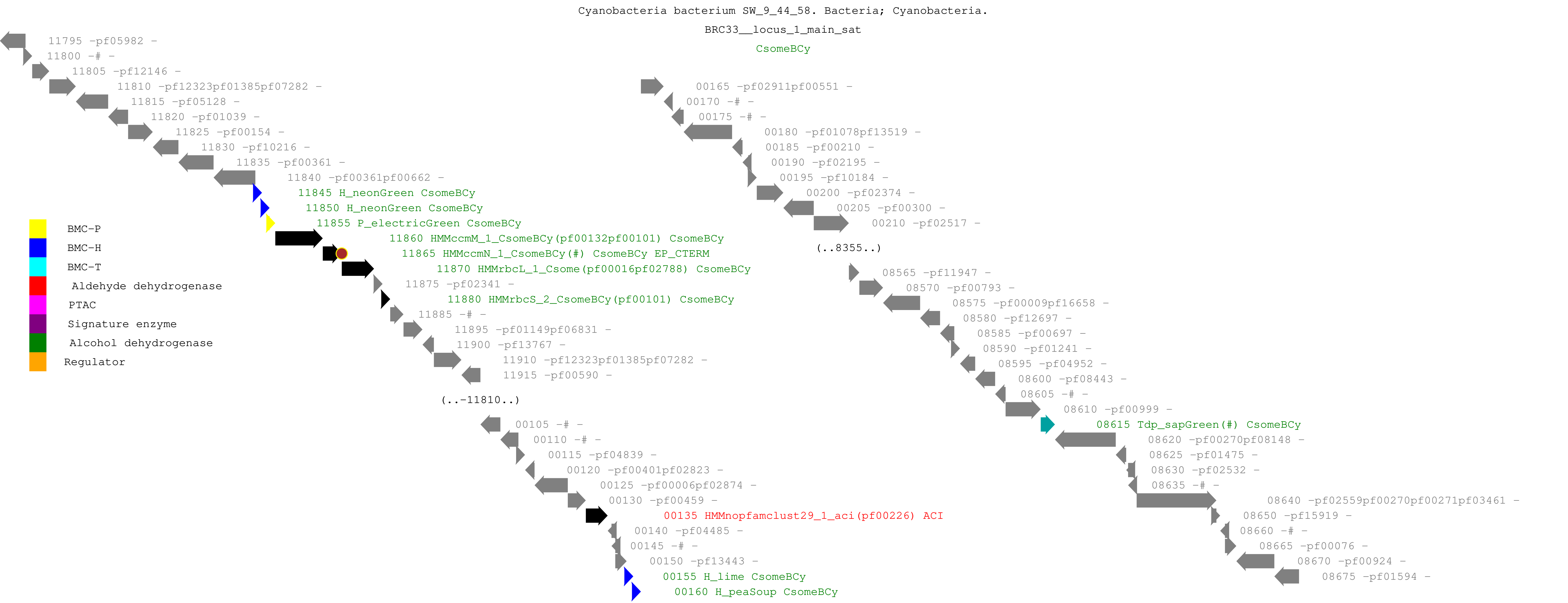

Cyanobacteria bacterium SW_9_44_58. Bacteria; Cyanobacteria.
BRC33__locus_1_main_sat
CsomeBCy
11795 -pf05982 -
11800 -# -
11805 -pf12146 -
11810 -pf12323pf01385pf07282 -
11815 -pf05128 -
11820 -pf01039 -
11825 -pf00154 -
11830 -pf10216 -
11835 -pf00361 -
11840 -pf00361pf00662 -
11845 H_neonGreen CsomeBCy
11850 H_neonGreen CsomeBCy
11855 P_electricGreen CsomeBCy
11860 HMMccmM_1_CsomeBCy(pf00132pf00101) CsomeBCy
11865 HMMccmN_1_CsomeBCy(#) CsomeBCy EP_CTERM
11870 HMMrbcL_1_Csome(pf00016pf02788) CsomeBCy
11875 -pf02341 -
11880 HMMrbcS_2_CsomeBCy(pf00101) CsomeBCy
11885 -# -
11895 -pf01149pf06831 -
11900 -pf13767 -
11910 -pf12323pf01385pf07282 -
11915 -pf00590 -
(..-11810..)
00105 -# -
00110 -# -
00115 -pf04839 -
00120 -pf00401pf02823 -
00125 -pf00006pf02874 -
00130 -pf00459 -
00135 HMMnopfamclust29_1_aci(pf00226) ACI
00140 -pf04485 -
00145 -# -
00150 -pf13443 -
00155 H_lime CsomeBCy
00160 H_peaSoup CsomeBCy
00165 -pf02911pf00551 -
00170 -# -
00175 -# -
00180 -pf01078pf13519 -
00185 -pf00210 -
00190 -pf02195 -
00195 -pf10184 -
00200 -pf02374 -
00205 -pf00300 -
00210 -pf02517 -
(..8355..)
08565 -pf11947 -
08570 -pf00793 -
08575 -pf00009pf16658 -
08580 -pf12697 -
08585 -pf00697 -
08590 -pf01241 -
08595 -pf04952 -
08600 -pf08443 -
08605 -# -
08610 -pf00999 -
08615 Tdp_sapGreen(#) CsomeBCy
08620 -pf00270pf08148 -
08625 -pf01475 -
08630 -pf02532 -
08635 -# -
08640 -pf02559pf00270pf00271pf03461 -
08650 -pf15919 -
08660 -# -
08665 -pf00076 -
08670 -pf00924 -
08675 -pf01594 -
BMC-P
BMC-H
BMC-T
Aldehyde dehydrogenase
PTAC
Signature enzyme
Alcohol dehydrogenase
Regulator

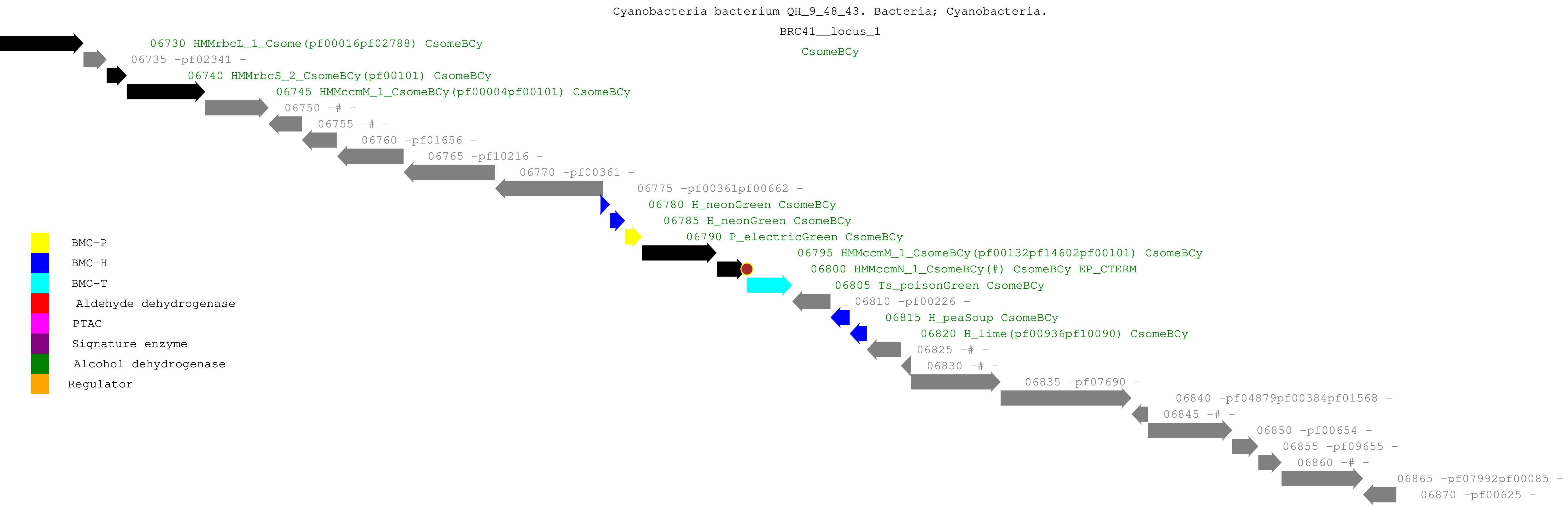
Cyanobacteria bacterium QH_9_48_43. Bacteria; Cyanobacteria.
BRC41__locus_1
CsomeBCy
06730 HMMrbcL_1_Csome(pf00016pf02788) CsomeBCy
06735 -pf02341 -
06740 HMMrbcS_2_CsomeBCy(pf00101) CsomeBCy
06745 HMMccmM_1_CsomeBCy(pf00004pf00101) CsomeBCy
06750 -# -
06755 -# -
06760 -pf01656 -
06765 -pf10216 -
06770 -pf00361 -
06775 -pf00361pf00662 -
06780 H_neonGreen CsomeBCy
06785 H_neonGreen CsomeBCy
06790 P_electricGreen CsomeBCy
06795 HMMccmM_1_CsomeBCy(pf00132pf14602pf00101) CsomeBCy
06800 HMMccmN_1_CsomeBCy(#) CsomeBCy EP_CTERM
06805 Ts_poisonGreen CsomeBCy
06810 -pf00226 -
06815 H_peaSoup CsomeBCy
06820 H_lime(pf00936pf10090) CsomeBCy
06825 -# -
06830 -# -
06835 -pf07690 -
06840 -pf04879pf00384pf01568 -
06845 -# -
06850 -pf00654 -
06855 -pf09655 -
06860 -# -
06865 -pf07992pf00085 -
06870 -pf00625 -
BMC-P
BMC-H
BMC-T
Aldehyde dehydrogenase
PTAC
Signature enzyme
Alcohol dehydrogenase
Regulator

Synechococcus lividus PCC 6715. Bacteria; Cyanobacteria; Synechococcales; Synechococcaceae;

BRW62__locus_1_main_sat

CsomeBCy

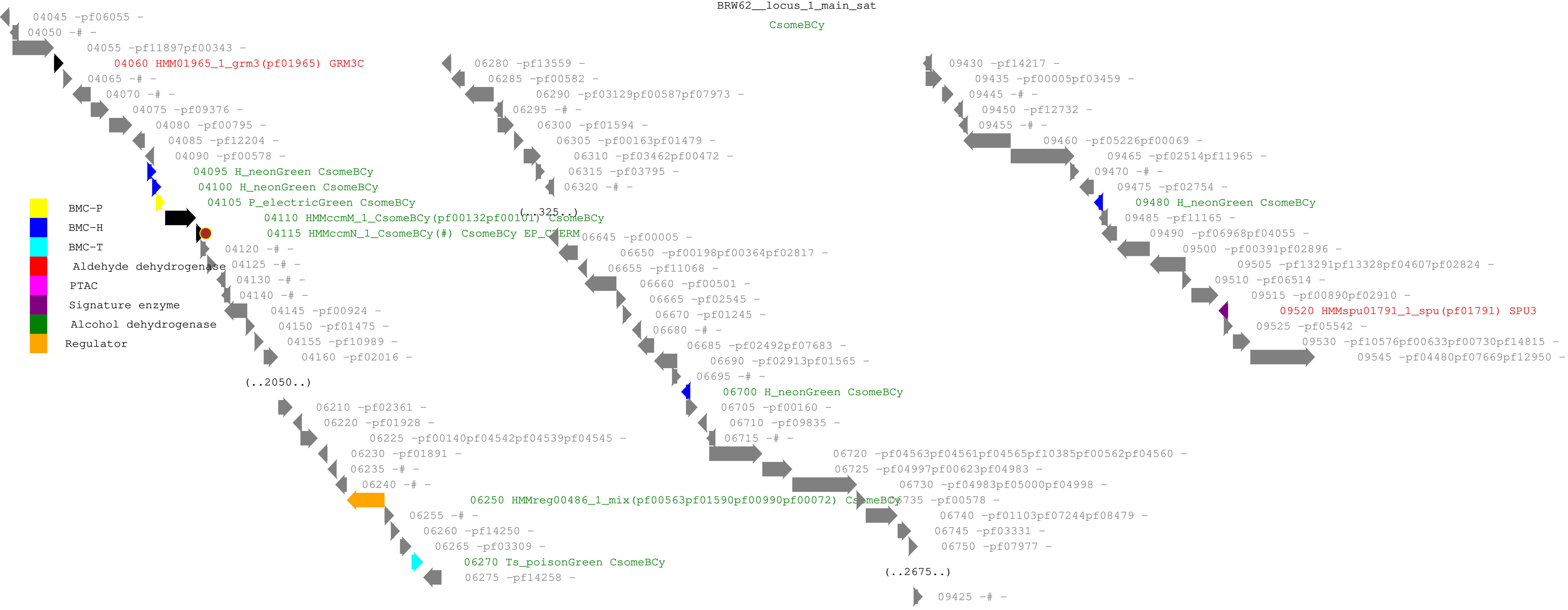

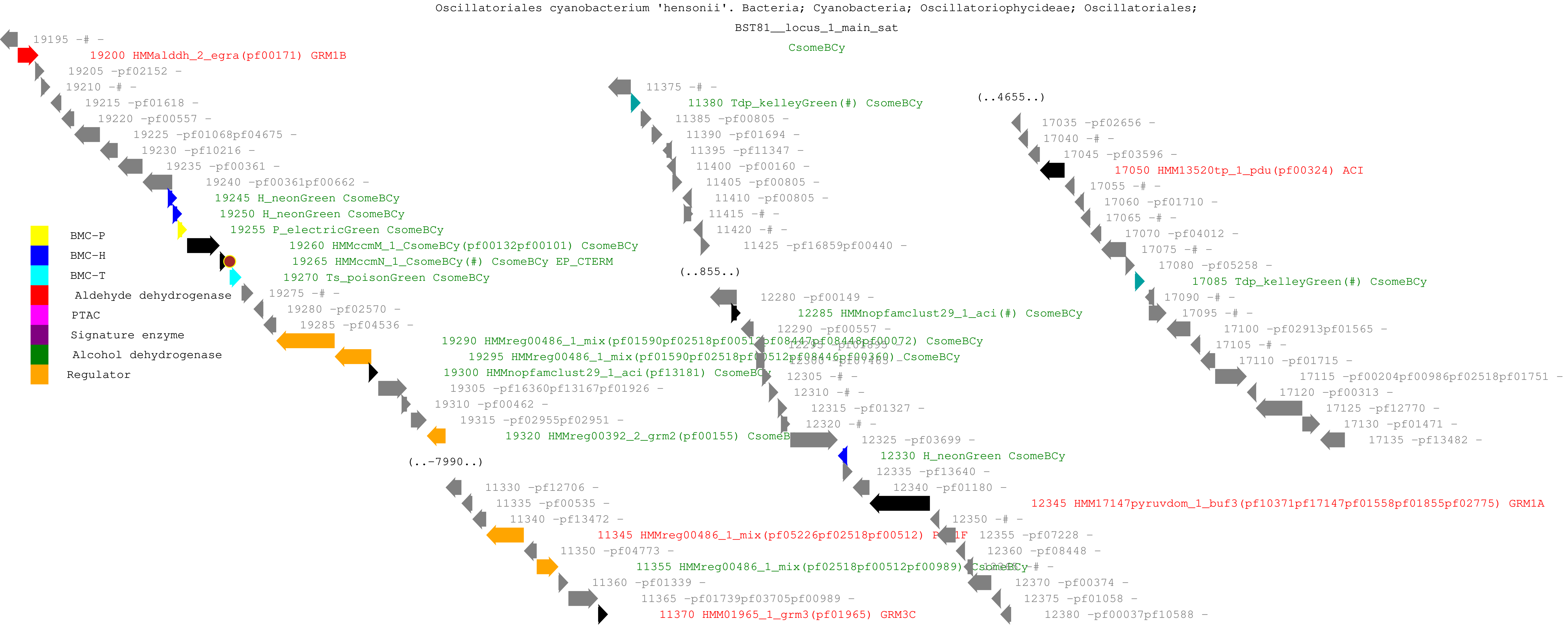
Oscillatoriales cyanobacterium 'hensonii'. Bacteria; Cyanobacteria; Oscillatoriophycideae; Oscillatoriales;
BST81__locus_1_main_sat
CsomeBCy
19195 -# -
19200 HMMalddh_2_egra(pf00171) GRM1B
19205 -pf02152 -
19210 -# -
19215 -pf01618 -
19220 -pf00557 -
19225 -pf01068pf04675 -
19230 -pf10216 -
19235 -pf00361 -
19240 -pf00361pf00662 -
19245 H_neonGreen CsomeBCy
19250 H_neonGreen CsomeBCy
19255 P_electricGreen CsomeBCy
19260 HMMccmM_1_CsomeBCy(pf00132pf00101) CsomeBCy
19265 HMMccmN_1_CsomeBCy(#) CsomeBCy EP_CTERM
19270 Ts_poisonGreen CsomeBCy
19275 -# -
19280 -pf02570 -
19285 -pf04536 -
19290 HMMreg00486_1_mix(pf01590pf02518pf00512pf08447pf08448pf00072) CsomeBCy
19295 HMMreg00486_1_mix(pf01590pf02518pf00512pf08446pf00360) CsomeBCy
19300 HMMnopfamclust29_1_aci(pf13181) CsomeBCy
19305 -pf16360pf13167pf01926 -
19310 -pf00462 -
19315 -pf02955pf02951 -
19320 HMMreg00392_2_grm2(pf00155) CsomeB
(..-7990..)
11330 -pf12706 -
11335 -pf00535 -
11340 -pf13472 -
11345 HMMreg00486_1_mix(pf05226pf02518pf00512)
11350 -pf04773 -
11355 HMMreg00486_1_mix(pf02518pf00512pf00989) CsomeBCy
11360 -pf01339 -
11365 -pf01739pf03705pf00989 -
11370 HMM01965_1_grm3(pf01965) GRM3C
11375 -# -
11380 Tdp_kelleyGreen(#) CsomeBCy
11385 -pf00805 -
11390 -pf01694 -
11395 -pf11347 -
11400 -pf00160 -
11405 -pf00805 -
11410 -pf00805 -
11415 -# -
11420 -# -
11425 -pf16859pf00440 -
(..855..)
12280 -pf00149 -
12285 HMMnopfamclust29_1_aci(#) CsomeBCy
12290 -pf00557 -
12305 -# -
12310 -# -
12315 -pf01327 -
12320 -# -
12325 -pf03699 -
12330 H_neonGreen CsomeBCy
12335 -pf13640 -
12340 -pf01180 -
12345 HMM17147pyruvdom_1_buf3(pf10371pf17147pf01558pf01855pf02775) GRM1A
12350 -# -
12355 -pf07228 -
12360 -pf08448 -
12370 -pf00374 -
12375 -pf01058 -
12380 -pf00037pf10588 -
(..4655..)
17035 -pf02656 -
17040 -# -
17045 -pf03596 -
17050 HMM13520tp_1_pdu(pf00324) ACI
17055 -# -
17060 -pf01710 -
17065 -# -
17070 -pf04012 -
17075 -# -
17080 -pf05258 -
17085 Tdp_kelleyGreen(#) CsomeBCy
17090 -# -
17095 -# -
17100 -pf02913pf01565 -
17105 -# -
17110 -pf01715 -
17115 -pf00204pf00986pf02518pf01751 -
17120 -pf00313 -
17125 -pf12770 -
17130 -pf01471 -
17135 -pf13482 -
BMC-P
BMC-H
BMC-T
Aldehyde dehydrogenase
PTAC
Signature enzyme
Alcohol dehydrogenase
Regulator

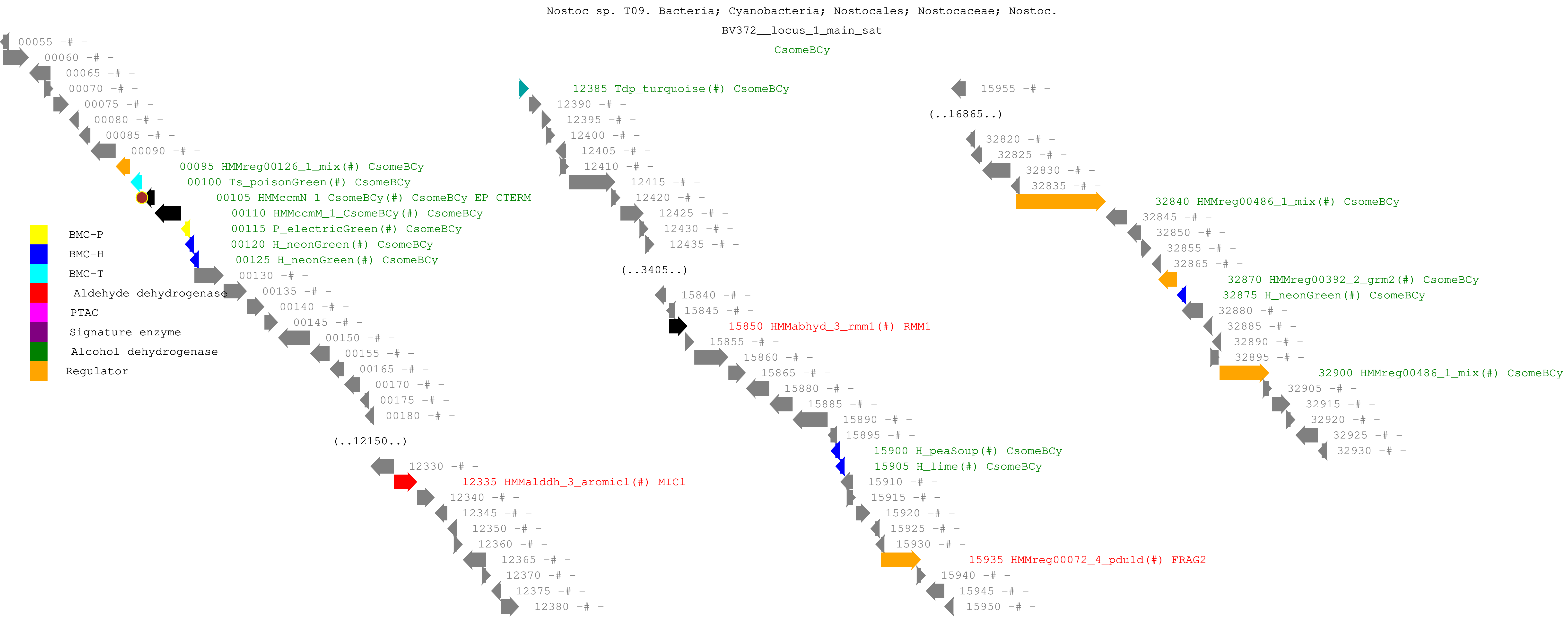

Nostoc sp. T09. Bacteria; Cyanobacteria; Nostocales; Nostocaceae; Nostoc.
BV372__locus_1_main_sat
CsomeBCy
00055 -# -
00060 -# -
00065 -# -
00070 -# -
00075 -# -
00080 -# -
00085 -# -
00090 -# -
00095 HMMreg00126_1_mix(#) CsomeBCy
00100 Ts_poisonGreen(#) CsomeBCy
00105 HMMccmN_1_CsomeBCy(#) CsomeBCy EP_CTERM
00110 HMMccmM_1_CsomeBCy(#) CsomeBCy
00115 P_electricGreen(#) CsomeBCy
00120 H_neonGreen(#) CsomeBCy
00125 H_neonGreen(#) CsomeBCy
00130 -# -
00135 -# -
00140 -# -
00145 -# -
00150 -# -
00155 -# -
00165 -# -
00170 -# -
00175 -# -
00180 -# -
(..12150..)
12330 -# -
12335 HMMaldh_3_aromic1(#) MIC1
12340 -# -
12345 -# -
12350 -# -
12360 -# -
12365 -# -
12370 -# -
12375 -# -
12380 -# -
12385 Tdp_turquoise(#) CsomeBCy
12390 -# -
12395 -# -
12400 -# -
12405 -# -
12410 -# -
12415 -# -
12420 -# -
12425 -# -
12430 -# -
12435 -# -
(..3405..)
15840 -# -
15845 -# -
15850 HMMabhyd_3_rmm1(#) RMM1
15855 -# -
15860 -# -
15865 -# -
15880 -# -
15885 -# -
15890 -# -
15895 -# -
15900 H_peaSoup(#) CsomeBCy
15905 H_lime(#) CsomeBCy
15910 -# -
15915 -# -
15920 -# -
15925 -# -
15930 -# -
15935 HMMreg00072_4_pdu1d(#) FRAG2
15940 -# -
15945 -# -
15950 -# -
15955 -# -
(..16865..)
32820 -# -
32825 -# -
32830 -# -
32835 -# -
32840 HMMreg00486_1_mix(#) CsomeBCy
32845 -# -
32850 -# -
32855 -# -
32865 -# -
32870 HMMreg00392_2_grm2(#) CsomeBCy
32875 H_neonGreen(#) CsomeBCy
32880 -# -
32885 -# -
32890 -# -
32895 -# -
32900 HMMreg00486_1_mix(#) CsomeBCy
32905 -# -
32915 -# -
32920 -# -
32925 -# -
32930 -# -
BMC-P
BMC-H
BMC-T
Aldehyde dehydrogenase
PTAC
Signature enzyme
Alcohol dehydrogenase
Regulator

Nostoc sp. 106C. Bacteria; Cyanobacteria; Nostocales; Nostocaceae; Nostoc.

BV375__locus_1_main_sat

CsomeBCy

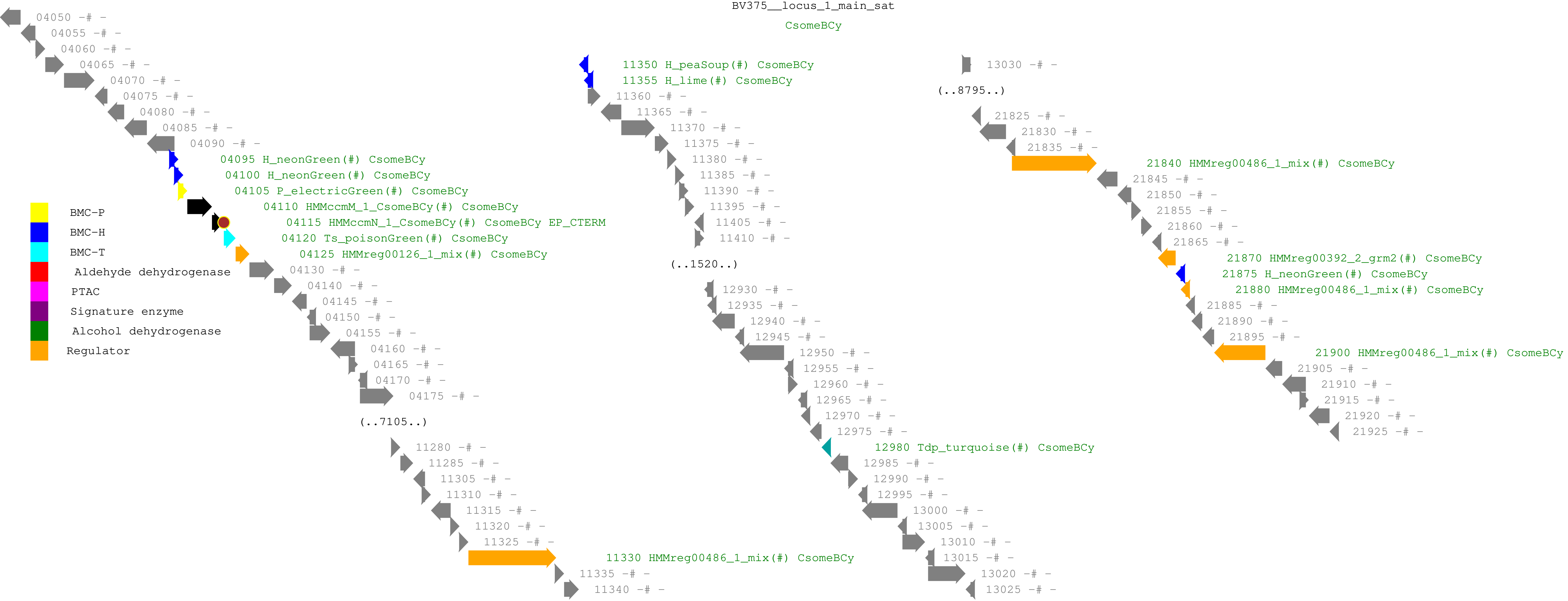

Nostoc sp. RF31Y. Bacteria; Cyanobacteria; Nostocales; Nostocaceae; Nostoc.

BV378__locus_1_main_sat

CsomeBCy

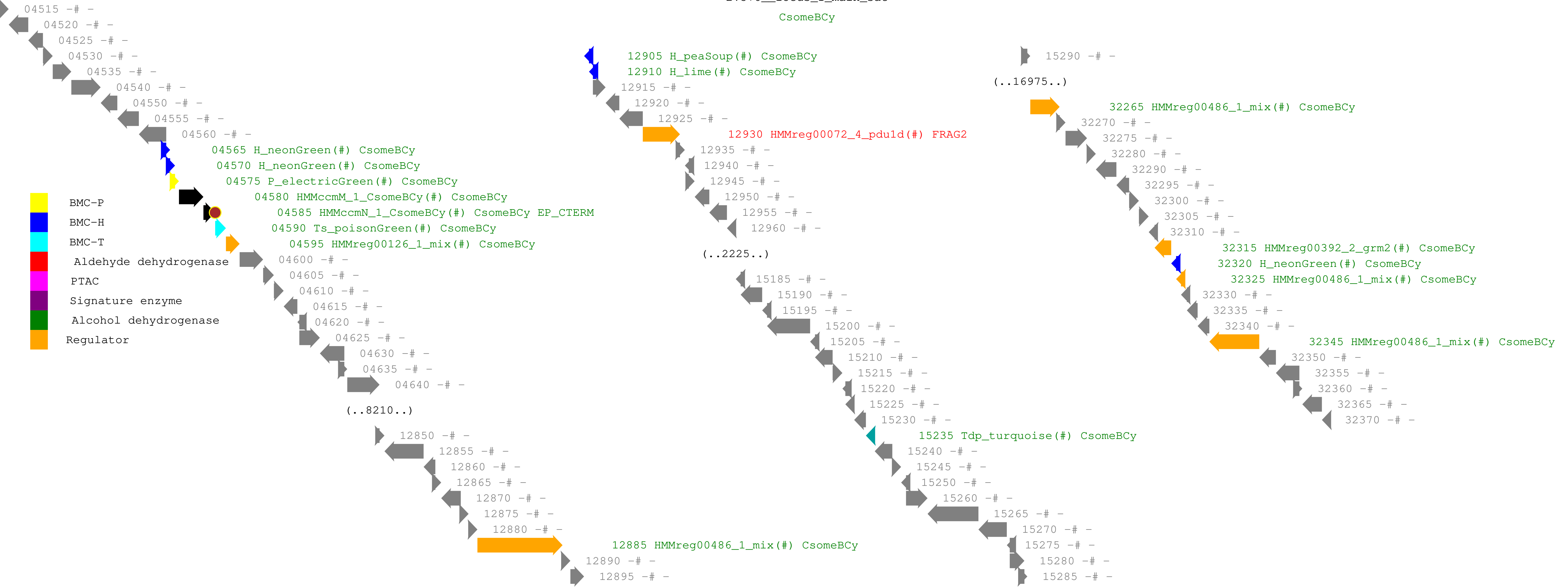

Nostocales cyanobacterium HT-58-2. Bacteria; Cyanobacteria; Nostocales.

BZZ01__locus_1_main_sat

CsomeBCy

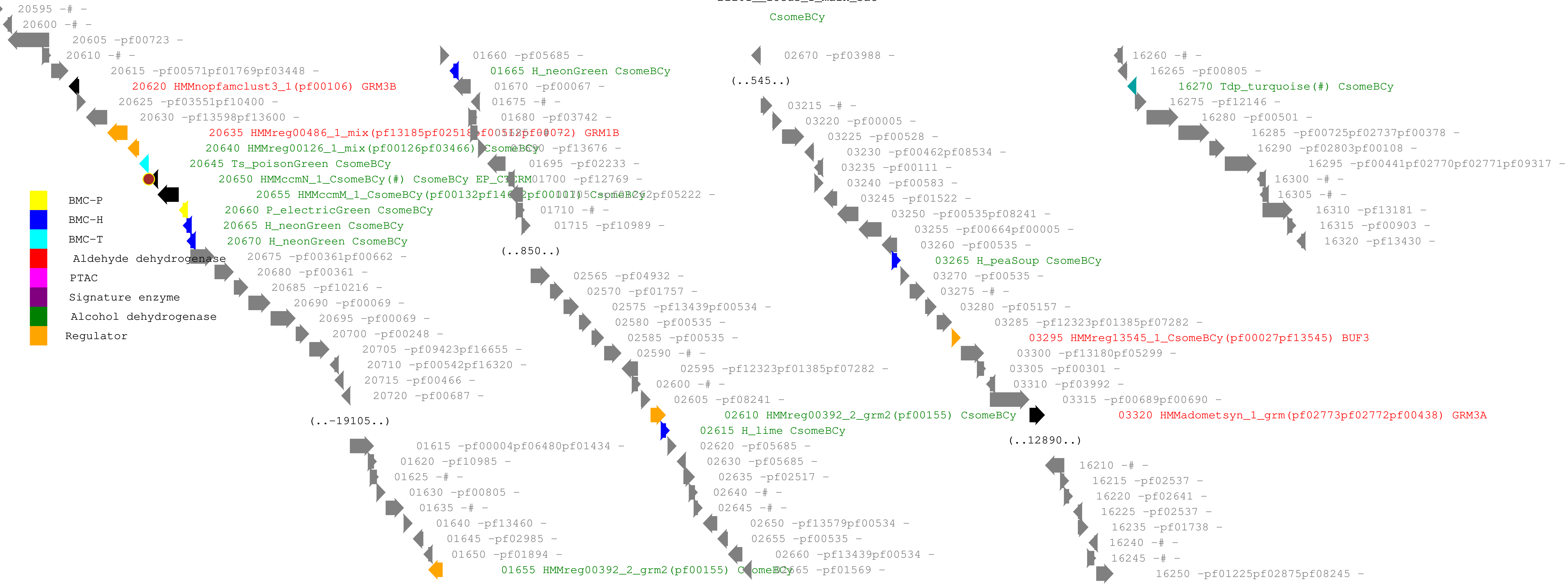

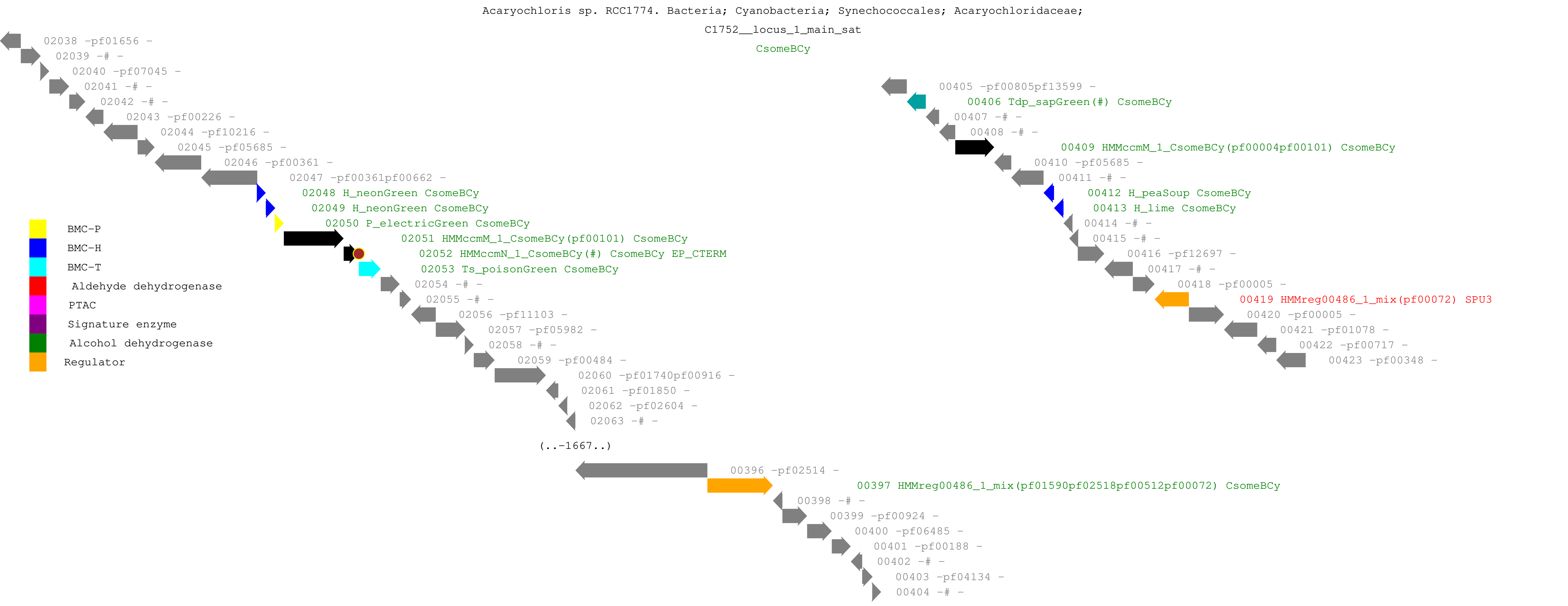
Acaryochloris sp. RCC1774. Bacteria; Cyanobacteria; Synechococcales; Acaryochloridaceae;
C1752__locus_1_main_sat
CsomeBCy
02038 -pf01656 -
02039 -# -
02040 -pf07045 -
02041 -# -
02042 -# -
02043 -pf00226 -
02044 -pf10216 -
02045 -pf05685 -
02046 -pf00361 -
02047 -pf00361pf00662 -
02048 H_neonGreen CsomeBCy
02049 H_neonGreen CsomeBCy
02050 P_electricGreen CsomeBCy
02051 HMMccmM_1_CsomeBCy(pf00101) CsomeBCy
02052 HMMccmN_1_CsomeBCy(#) CsomeBCy EP_CTERM
02053 Ts_poisonGreen CsomeBCy
02054 -# -
02055 -# -
02056 -pf11103 -
02057 -pf05982 -
02058 -# -
02059 -pf00484 -
02060 -pf01740pf00916 -
02061 -pf01850 -
02062 -pf02604 -
02063 -# -
(..-1667..)
00396 -pf02514 -
00397 HMMreg00486_1_mix(pf01590pf02518pf00512pf00072) CsomeBCy
00398 -# -
00399 -pf00924 -
00400 -pf06485 -
00401 -pf00188 -
00402 -# -
00403 -pf04134 -
00404 -# -
00405 -pf00805pf13599 -
00406 Tdp_sapGreen(#) CsomeBCy
00407 -# -
00408 -# -
00409 HMMccmM_1_CsomeBCy(pf00004pf00101) CsomeBCy
00410 -pf05685 -
00411 -# -
00412 H_peaSoup CsomeBCy
00413 H_lime CsomeBCy
00414 -# -
00415 -# -
00416 -pf12697 -
00417 -# -
00418 -pf00005 -
00419 HMMreg00486_1_mix(pf00072) SPU3
00420 -pf00005 -
00421 -pf01078 -
00422 -pf00717 -
00423 -pf00348 -
BMC-P
BMC-H
BMC-T
Aldehyde dehydrogenase
PTAC
Signature enzyme
Alcohol dehydrogenase
Regulator

Hapalosiphonaceae cyanobacterium JJU2. Bacteria; Cyanobacteria; Nostocales; Hapalosiphonaceae.

C6Y22__locus_1_main_sat

CsomeBCy

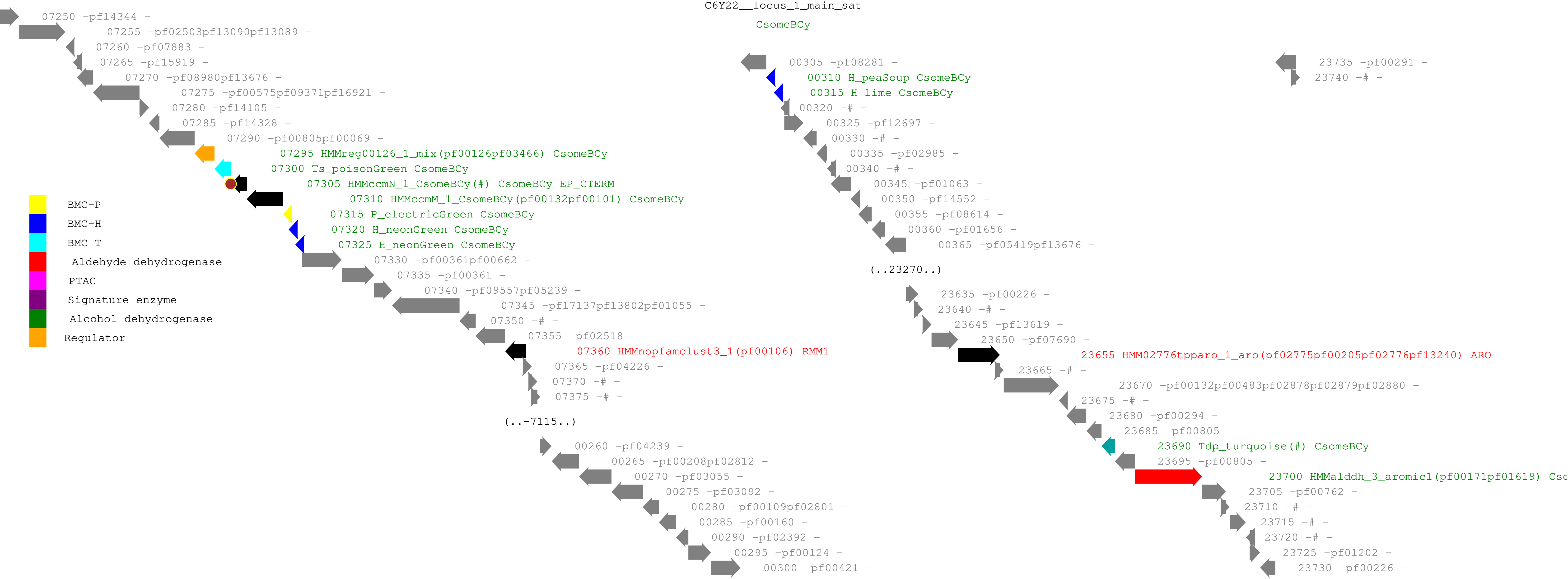

filamentous cyanobacterium CCP5. Bacteria; Cyanobacteria.

C7271__locus_1_main_sat

CsomeBCy

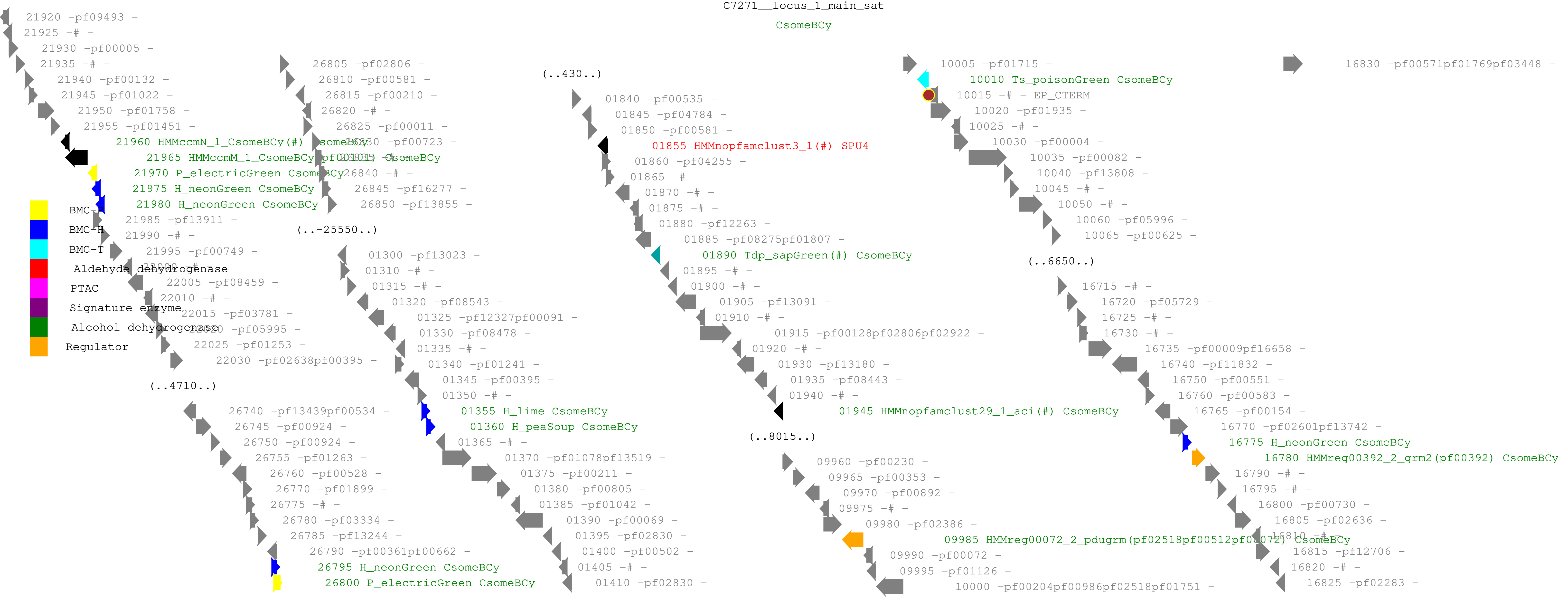

filamentous cyanobacterium CCT1. Bacteria; Cyanobacteria.

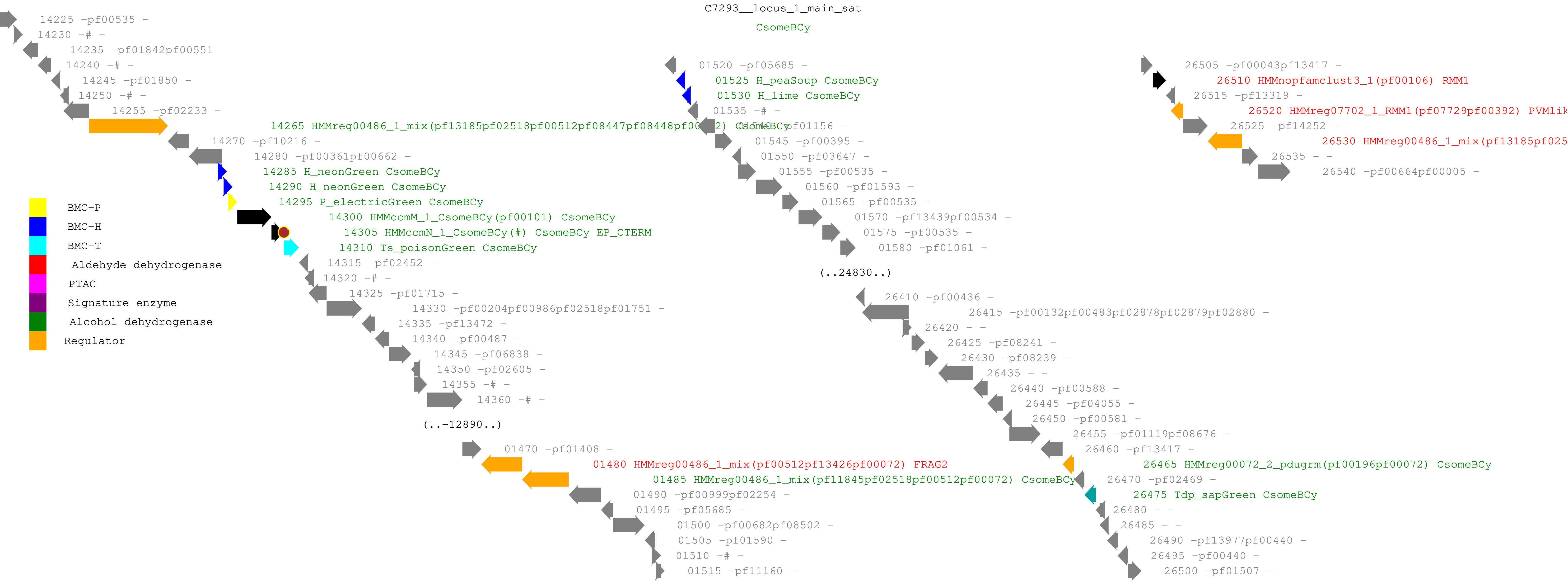

Microcystis aeruginosa DIANCHI905. Bacteria; Cyanobacteria; Oscillatoriophycideae; Chroococcales;

C789__locus_1_main_sat

CsomeBCy

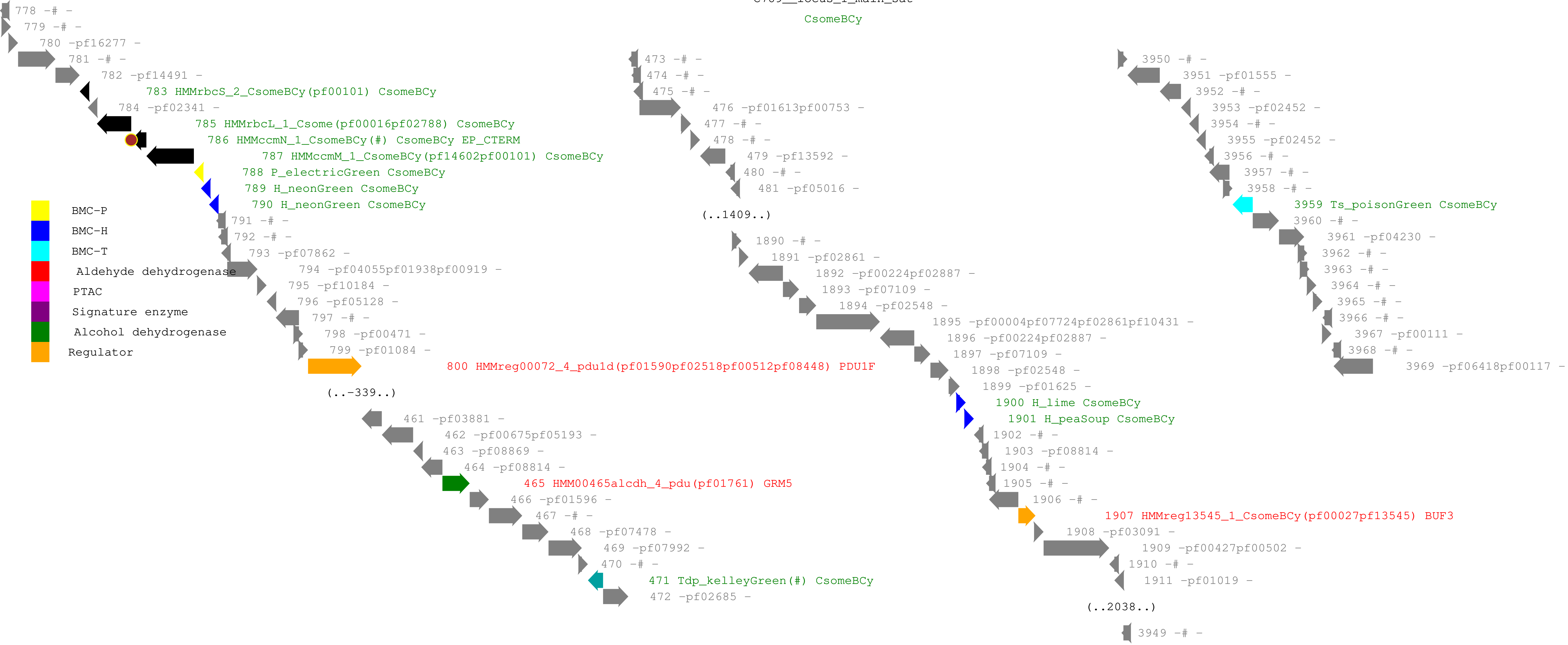

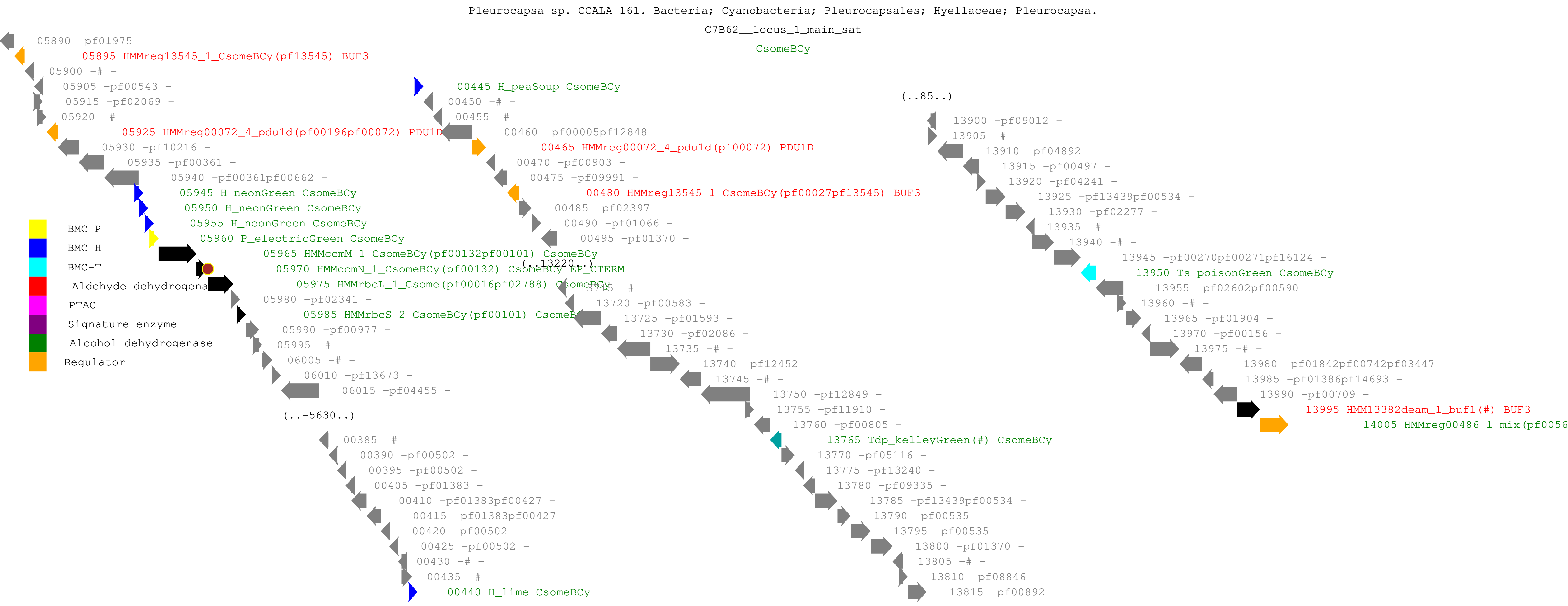

Pleurocapsa sp. CCALA 161. Bacteria; Cyanobacteria; Pleurocapsales; Hyellaceae; Pleurocapsa.
C7B62__locus_1_main_sat
CsomeBCy
05890 -pf01975 -
05895 HMMreg13545_1_CsomeBCy(pf13545) BUF3
05900 -# -
05905 -pf00543 -
05915 -pf02069 -
05920 -# -
05925 HMMreg00072_4_pdu1d(pf00196pf00072) PDU1D
05930 -pf10216 -
05935 -pf00361 -
05940 -pf00361pf00662 -
05945 H_neonGreen CsomeBCy
05950 H_neonGreen CsomeBCy
05955 H_neonGreen CsomeBCy
05960 P_electricGreen CsomeBCy
05965 HMMccmM_1_CsomeBCy(pf00132pf00101) CsomeBCy
05970 HMMccmN_1_CsomeBCy(pf00132) CsomeBCy EP_CTERM
05975 HMMrbcL_1_Csome(pf00016pf02788) CsomeBCy
05980 -pf02341 -
05985 HMMrbcS_2_CsomeBCy(pf00101) CsomeB
05990 -pf00977 -
05995 -# -
06005 -# -
06010 -pf13673 -
06015 -pf04455 -
(..-5630..)
00385 -# -
00390 -pf00502 -
00395 -pf00502 -
00405 -pf01383 -
00410 -pf01383pf00427 -
00415 -pf01383pf00427 -
00420 -pf00502 -
00425 -pf00502 -
00430 -# -
00435 -# -
00440 H_lime CsomeBCy
00445 H_peaSoup CsomeBCy
00450 -# -
00455 -# -
00460 -pf00005pf12848 -
00465 HMMreg00072_4_pdu1d(pf00072) PDU1D
00470 -pf00903 -
00475 -pf09991 -
00480 HMMreg13545_1_CsomeBCy(pf00027pf13545) BUF3
00485 -pf02397 -
00490 -pf01066 -
00495 -pf01370 -
(..13220..)
13715 -# -
13720 -pf00583 -
13725 -pf01593 -
13730 -pf02086 -
13735 -# -
13740 -pf12452 -
13745 -# -
13750 -pf12849 -
13755 -pf11910 -
13760 -pf00805 -
13765 Tdp_kelleyGreen(#) CsomeBCy
13770 -pf05116 -
13775 -pf13240 -
13780 -pf09335 -
13785 -pf13439pf00534 -
13790 -pf00535 -
13795 -pf00535 -
13800 -pf01370 -
13805 -# -
13810 -pf08846 -
13815 -pf00892 -
(..85..)
13900 -pf09012 -
13905 -# -
13910 -pf04892 -
13915 -pf00497 -
13920 -pf04241 -
13925 -pf13439pf00534 -
13930 -pf02277 -
13935 -# -
13940 -# -
13945 -pf00270pf00271pf16124 -
13950 Ts_poisonGreen CsomeBCy
13955 -pf02602pf00590 -
13960 -# -
13965 -pf01904 -
13970 -pf00156 -
13975 -# -
13980 -pf01842pf00742pf03447 -
13985 -pf01386pf14693 -
13990 -pf00709 -
13995 HMM13382deam_1_buf1(#) BUF3
14005 HMMreg00486_1_mix(pf0056
BMC-P
BMC-H
BMC-T
Aldehyde dehydrogena
PTAC
Signature enzyme
Alcohol dehydrogenase
Regulator

Merismopedia glauca CCAP 1448/3. Bacteria; Cyanobacteria; Synechococcales; Merismopediaceae;

C7B64__locus_1_main_sat

CsomeBCy

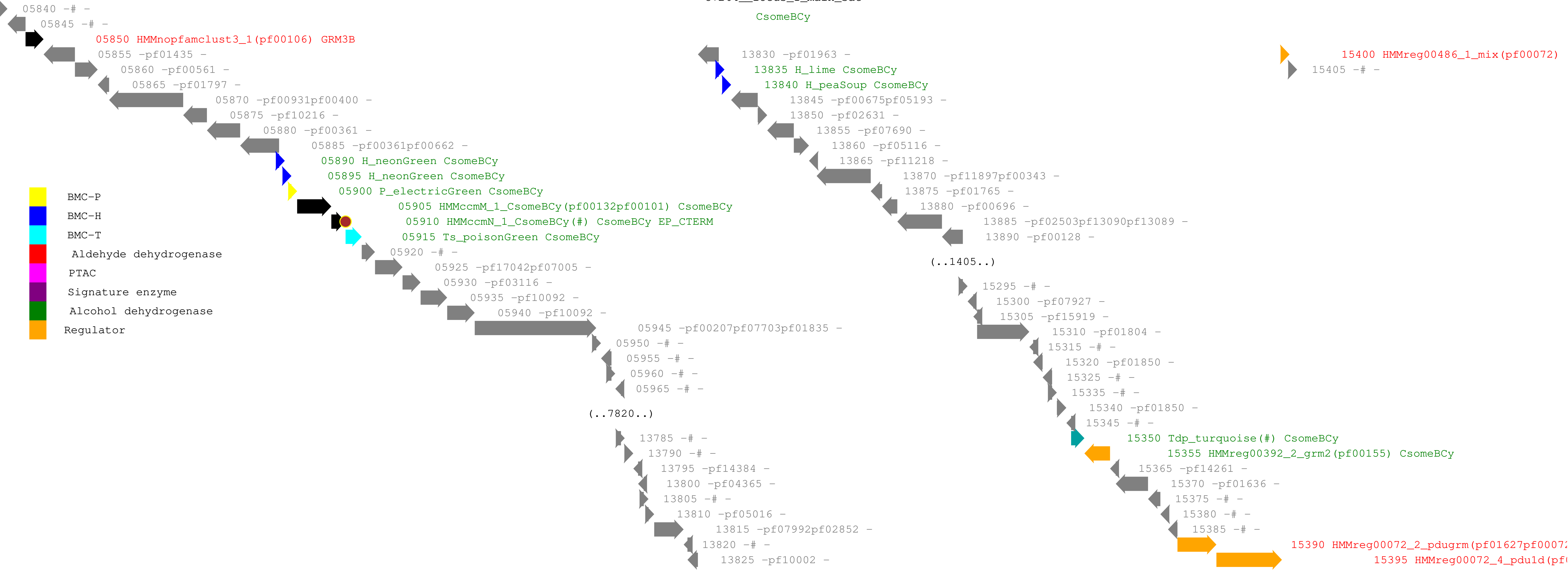

Phormidesmis priestleyi ULC007. Bacteria; Cyanobacteria; Synechococcales; Leptolyngbyaceae;

C7B65__locus_1_main_sat

CsomeBCy

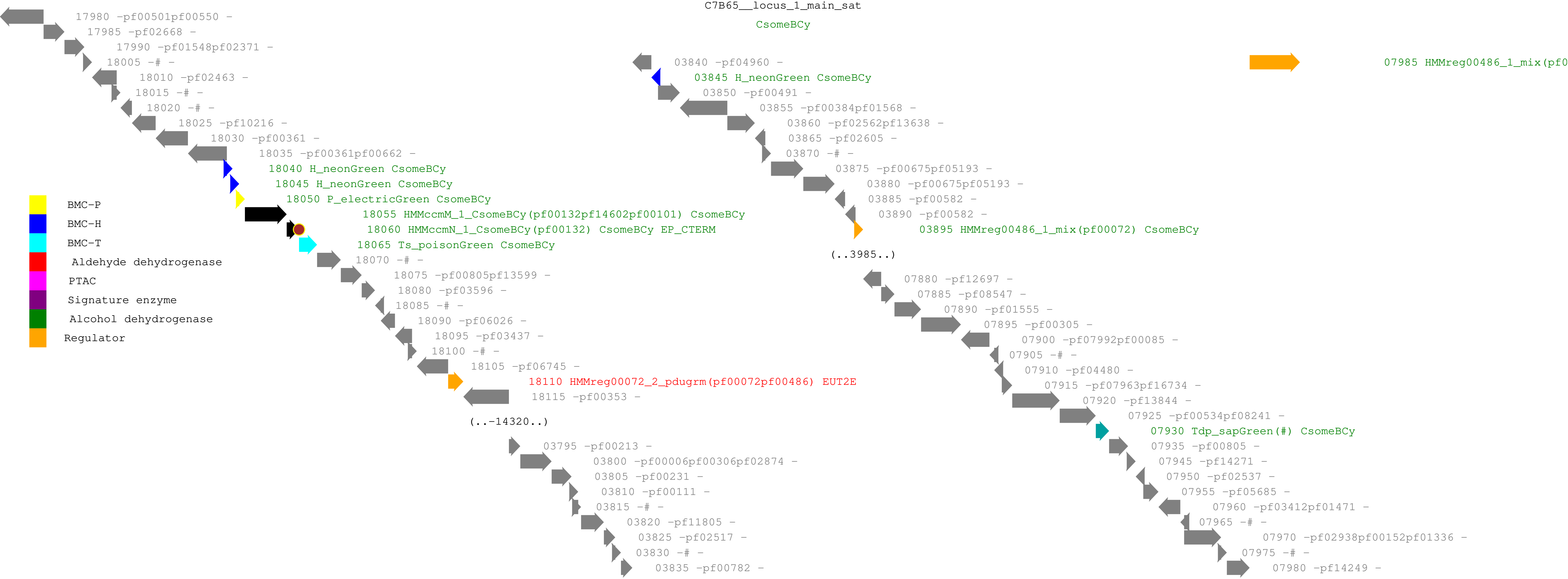

filamentous cyanobacterium Phorm 6. Bacteria; Cyanobacteria.

C7B67__locus_1_main_sat

CsomeBCy

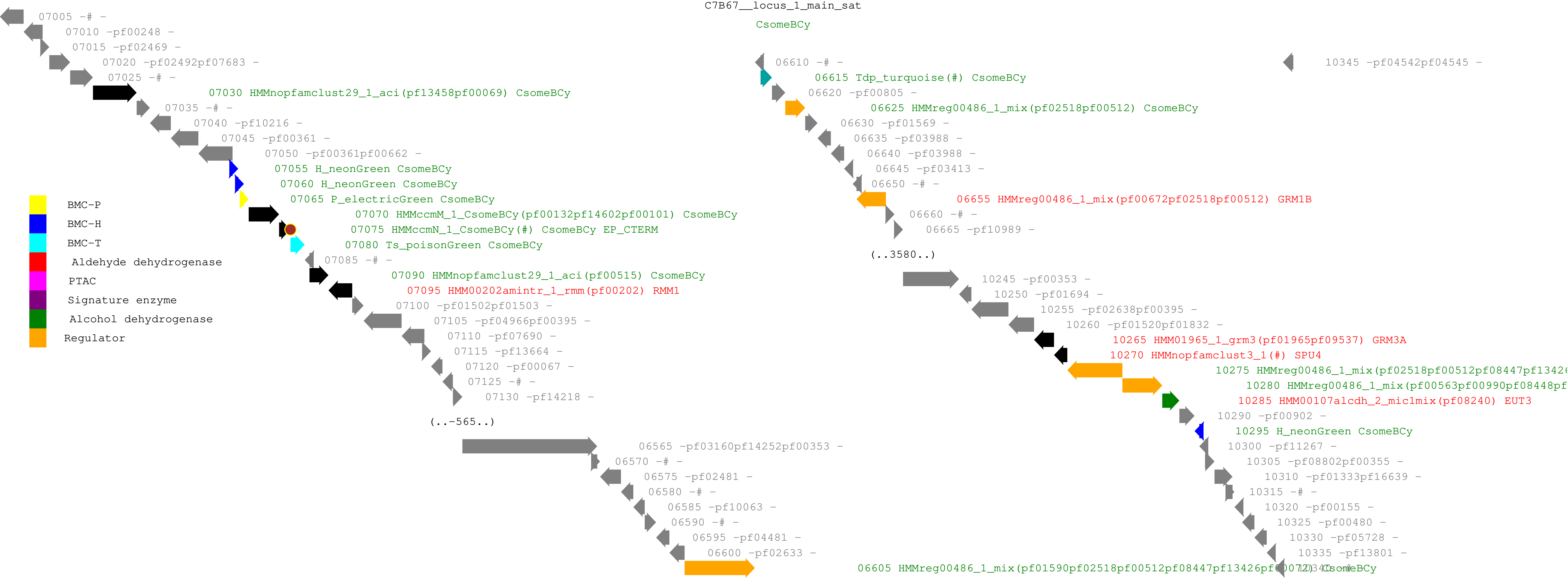

filamentous cyanobacterium Phorm 46. Bacteria; Cyanobacteria.

C7B69__locus_1_main_sat

CsomeBCy

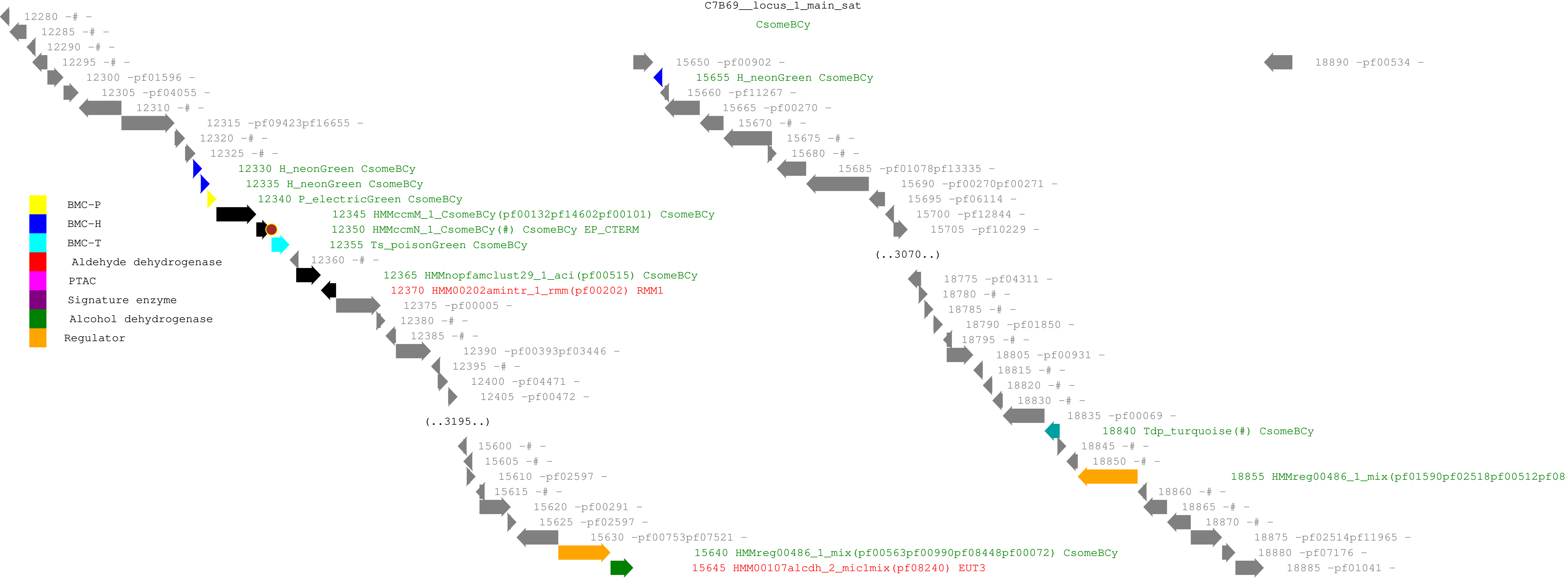

Chlorogloea sp. CCALA 695. Bacteria; Cyanobacteria; Oscillatoriophycideae; Chroococcales;

C7B70__locus_1_main_sat

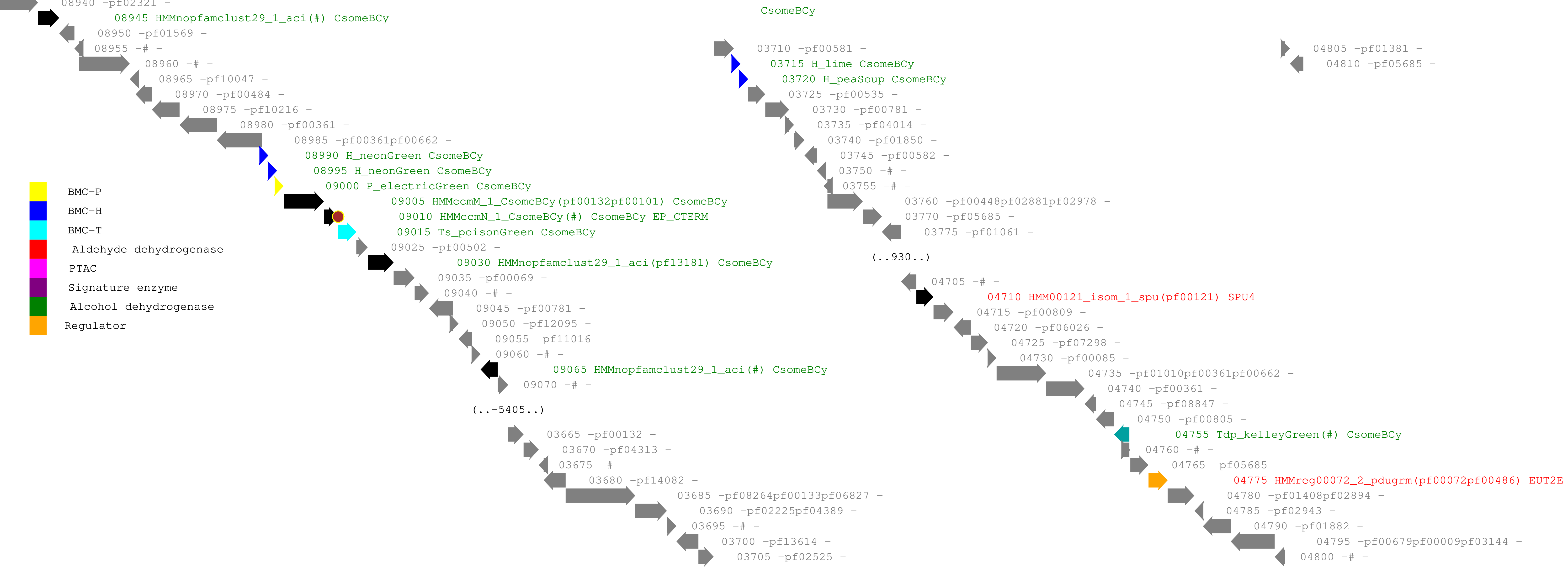

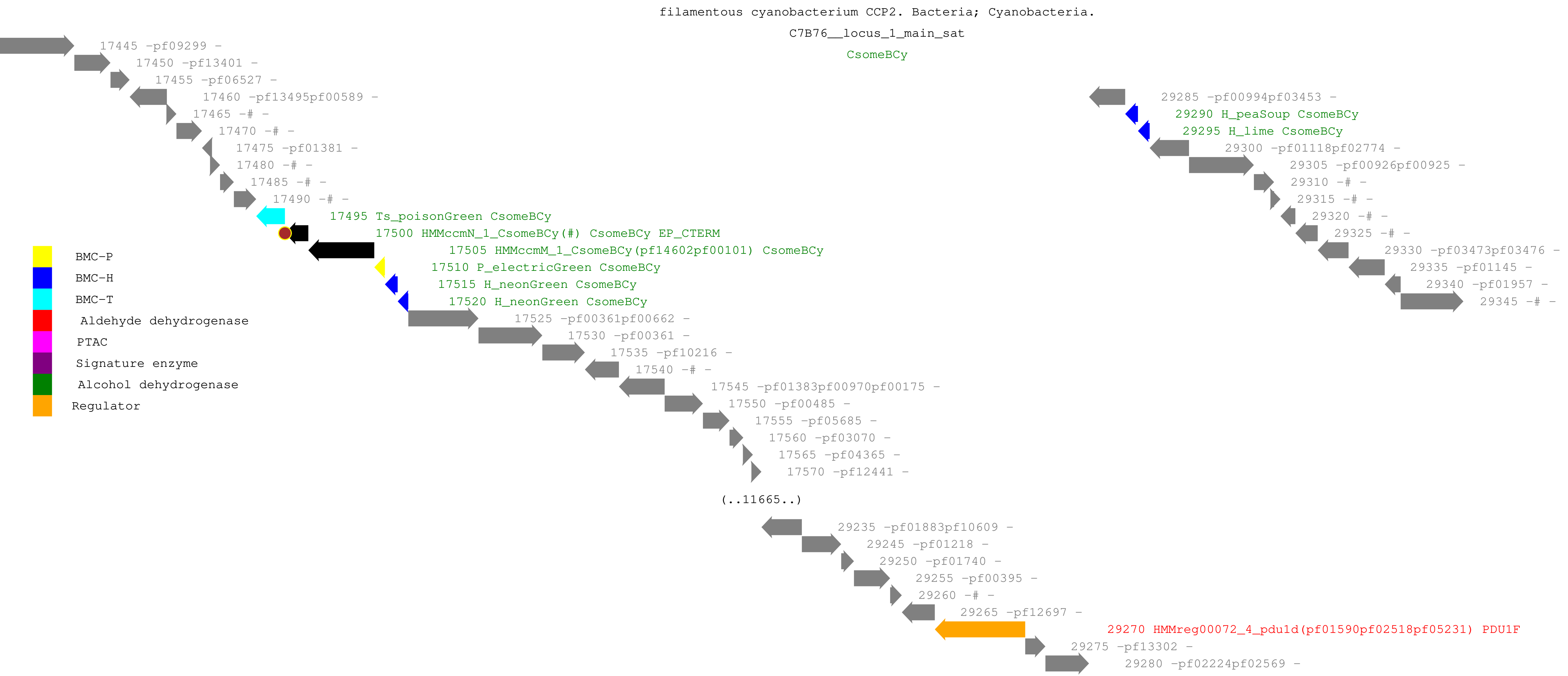
filamentous cyanobacterium CCP2. Bacteria; Cyanobacteria.
C7B76__locus_1_main_sat
CsomeBCy
17445 -pf09299 -
17450 -pf13401 -
17455 -pf06527 -
17460 -pf13495pf00589 -
17465 -# -
17470 -# -
17475 -pf01381 -
17480 -# -
17485 -# -
17490 -# -
17495 Ts_poisonGreen CsomeBCy
17500 HMMccmN_1_CsomeBCy(#) CsomeBCy EP_CTERM
17505 HMMccmM_1_CsomeBCy(pf14602pf00101) CsomeBCy
17510 P_electricGreen CsomeBCy
17515 H_neonGreen CsomeBCy
17520 H_neonGreen CsomeBCy
17525 -pf00361pf00662 -
17530 -pf00361 -
17535 -pf10216 -
17540 -# -
17545 -pf01383pf00970pf00175 -
17550 -pf00485 -
17555 -pf05685 -
17560 -pf03070 -
17565 -pf04365 -
17570 -pf12441 -
(..11665..)
29235 -pf01883pf10609 -
29245 -pf01218 -
29250 -pf01740 -
29255 -pf00395 -
29260 -# -
29265 -pf12697 -
29270 HMMreg00072_4_pdu1d(pf01590pf02518pf05231) PDU1F
29275 -pf13302 -
29280 -pf02224pf02569 -
29285 -pf00994pf03453 -
29290 H_peaSoup CsomeBCy
29295 H_lime CsomeBCy
29300 -pf01118pf02774 -
29305 -pf00926pf00925 -
29310 -# -
29315 -# -
29320 -# -
29325 -# -
29330 -pf03473pf03476 -
29335 -pf01145 -
29340 -pf01957 -
29345 -# -
BMC-P
BMC-H
BMC-T
Aldehyde dehydrogenase
PTAC
Signature enzyme
Alcohol dehydrogenase
Regulator

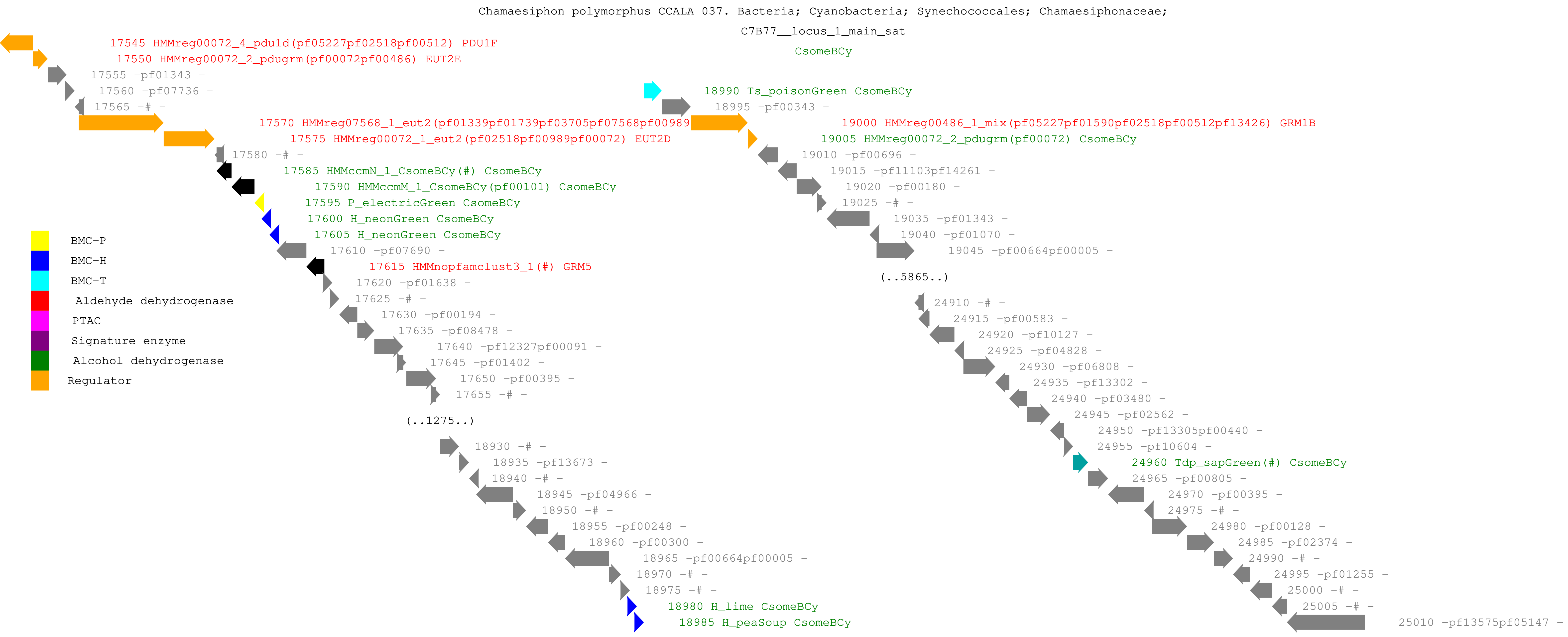
Chamaesiphon polymorphus CCALA 037. Bacteria; Cyanobacteria; Synechococcales; Chamaesiphonaceae;
C7B77__locus_1_main_sat
CsomeBCy
17545 HMMreg00072_4_pdu1d(pf05227pf02518pf00512) PDU1F
17550 HMMreg00072_2_pdugrm(pf00072pf00486) EUT2E
17555 -pf01343 -
17560 -pf07736 -
17565 -# -
17570 HMMreg07568_1_eut2(pf01339pf01739pf03705pf07568pf00989
17575 HMMreg00072_1_eut2(pf02518pf00989pf00072) EUT2D
17580 -# -
17585 HMMccmN_1_CsomeBCy(#) CsomeBCy
17590 HMMccmM_1_CsomeBCy(pf00101) CsomeBCy
17595 P_electricGreen CsomeBCy
17600 H_neonGreen CsomeBCy
17605 H_neonGreen CsomeBCy
17610 -pf07690 -
17615 HMMnopfamclust3_1(#) GRM5
17620 -pf01638 -
17625 -# -
17630 -pf00194 -
17635 -pf08478 -
17640 -pf12327pf00091 -
17645 -pf01402 -
17650 -pf00395 -
17655 -# -
(..1275..)
18930 -# -
18935 -pf13673 -
18940 -# -
18945 -pf04966 -
18950 -# -
18955 -pf00248 -
18960 -pf00300 -
18965 -pf00664pf00005 -
18970 -# -
18975 -# -
18980 H_lime CsomeBCy
18985 H_peaSoup CsomeBCy
18990 Ts_poisonGreen CsomeBCy
18995 -pf00343 -
19000 HMMreg00486_1_mix(pf05227pf01590pf02518pf00512pf13426) GRM1B
19005 HMMreg00072_2_pdugrm(pf00072) CsomeBCy
19010 -pf00696 -
19015 -pf11103pf14261 -
19020 -pf00180 -
19025 -# -
19035 -pf01343 -
19040 -pf01070 -
19045 -pf00664pf00005 -
(..5865..)
24910 -# -
24915 -pf00583 -
24920 -pf10127 -
24925 -pf04828 -
24930 -pf06808 -
24935 -pf13302 -
24940 -pf03480 -
24945 -pf02562 -
24950 -pf13305pf00440 -
24955 -pf10604 -
24960 Tdp_sapGreen(#) CsomeBCy
24965 -pf00805 -
24970 -pf00395 -
24975 -# -
24980 -pf00128 -
24985 -pf02374 -
24990 -# -
24995 -pf01255 -
25000 -# -
25005 -# -
25010 -pf13575pf05147 -
BMC-P
BMC-H
BMC-T
Aldehyde dehydrogenase
PTAC
Signature enzyme
Alcohol dehydrogenase
Regulator

Chroococcidiopsis cubana CCALA 043. Bacteria; Cyanobacteria; Chroococcidiopsidales;

C7B79__locus_1_main_sat

CsomeBCy

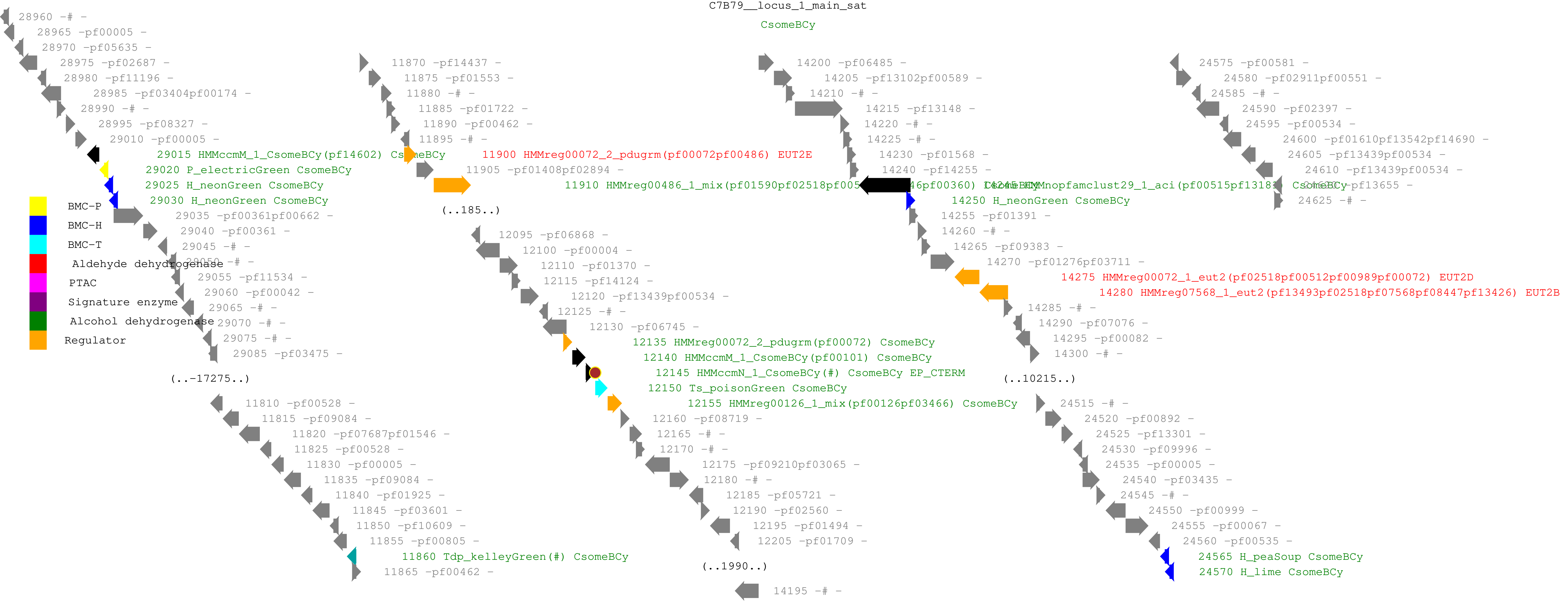

Cyanosarcina cf. burmensis CCALA 770. Bacteria; Cyanobacteria; Oscillatoriophycideae; Chroococcales;

C7B80__locus_1_main_sat

CsomeBCy

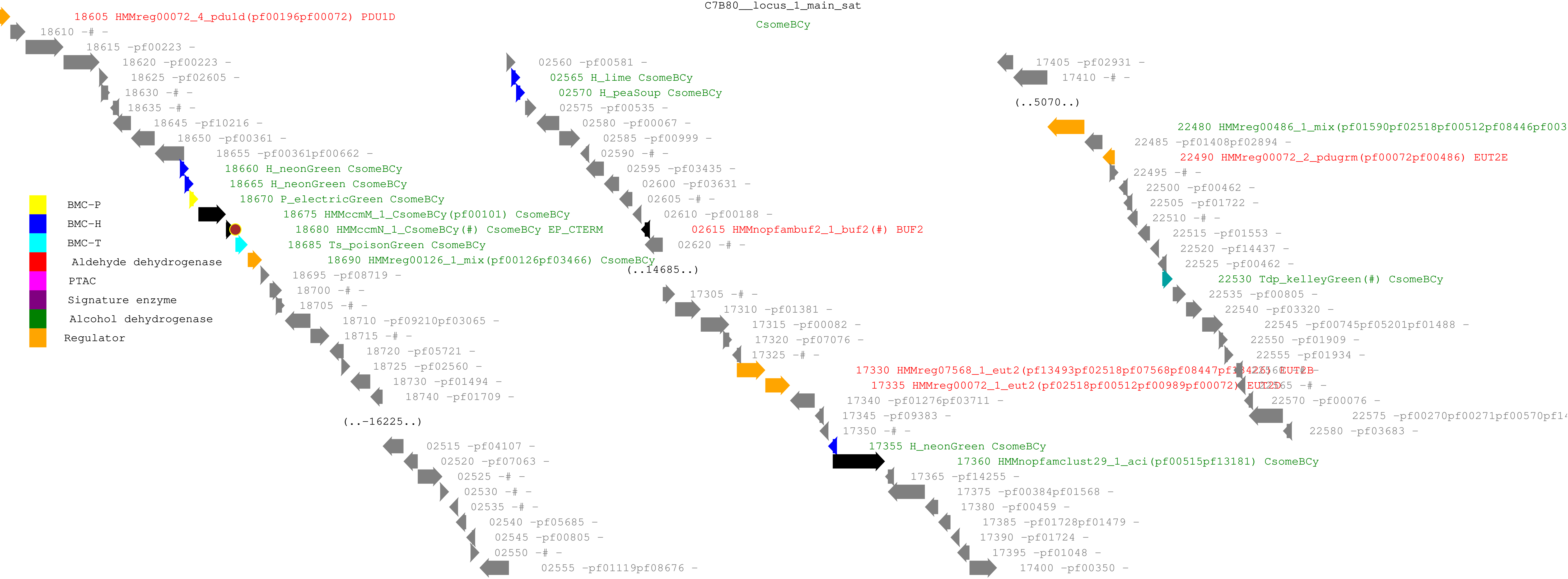

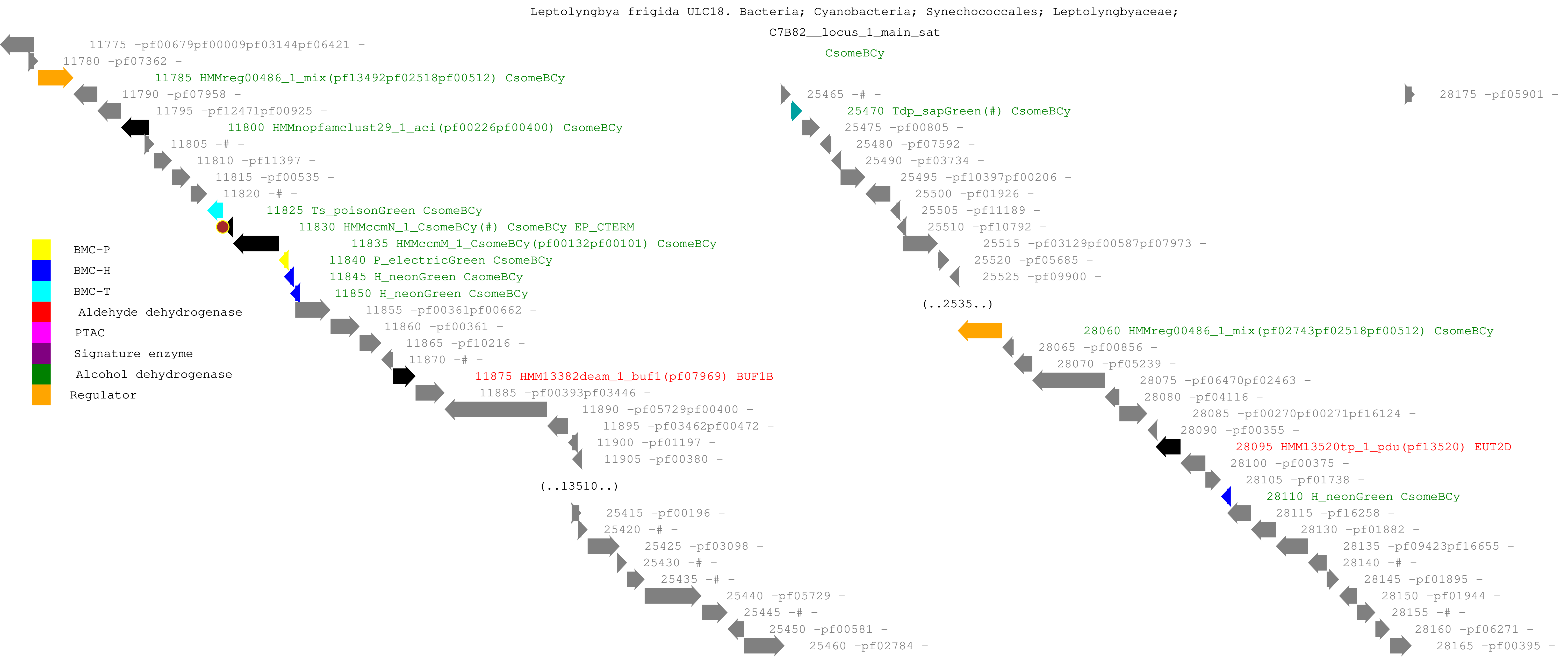
Leptolyngbya frigida ULC18. Bacteria; Cyanobacteria; Synechococcales; Leptolyngbyaceae;
C7B82__locus_1_main_sat
CsomeBCy
11775 -pf00679pf00009pf03144pf06421 -
11780 -pf07362 -
11785 HMMreg00486_1_mix(pf13492pf02518pf00512) CsomeBCy
11790 -pf07958 -
11795 -pf12471pf00925 -
11800 HMMnopfamclust29_1_aci(pf00226pf00400) CsomeBCy
11805 -# -
11810 -pf11397 -
11815 -pf00535 -
11820 -# -
11825 Ts_poisonGreen CsomeBCy
11830 HMMccmN_1_CsomeBCy(#) CsomeBCy EP_CTERM
11835 HMMccmM_1_CsomeBCy(pf00132pf00101) CsomeBCy
11840 P_electricGreen CsomeBCy
11845 H_neonGreen CsomeBCy
11850 H_neonGreen CsomeBCy
11855 -pf00361pf00662 -
11860 -pf00361 -
11865 -pf10216 -
11870 -# -
11875 HMM13382deam_1_buf1(pf07969) BUF1B
11885 -pf00393pf03446 -
11890 -pf05729pf00400 -
11895 -pf03462pf00472 -
11900 -pf01197 -
11905 -pf00380 -
(..13510..)
25415 -pf00196 -
25420 -# -
25425 -pf03098 -
25430 -# -
25435 -# -
25440 -pf05729 -
25445 -# -
25450 -pf00581 -
25460 -pf02784 -
25465 -# -
25470 Tdp_sapGreen(#) CsomeBCy
25475 -pf00805 -
25480 -pf07592 -
25490 -pf03734 -
25495 -pf10397pf00206 -
25500 -pf01926 -
25505 -pf11189 -
25510 -pf10792 -
25515 -pf03129pf00587pf07973 -
25520 -pf05685 -
25525 -pf09900 -
(..2535..)
28060 HMMreg00486_1_mix(pf02743pf02518pf00512) CsomeBCy
28065 -pf00856 -
28070 -pf05239 -
28075 -pf06470pf02463 -
28080 -pf04116 -
28085 -pf00270pf00271pf16124 -
28090 -pf00355 -
28095 HMM13520tp_1_pdu(pf13520) EUT2D
28100 -pf00375 -
28105 -pf01738 -
28110 H_neonGreen CsomeBCy
28115 -pf16258 -
28130 -pf01882 -
28135 -pf09423pf16655 -
28140 -# -
28145 -pf01895 -
28150 -pf01944 -
28155 -# -
28160 -pf06271 -
28165 -pf00395 -
28175 -pf05901 -
BMC-P
BMC-H
BMC-T
Aldehyde dehydrogenase
PTAC
Signature enzyme
Alcohol dehydrogenase
Regulator

Aphanothece hegewaldii CCALA 016. Bacteria; Cyanobacteria; Oscillatoriophycideae; Chroococcales;

C7H19__locus_1_main_sat

CsomeBCy

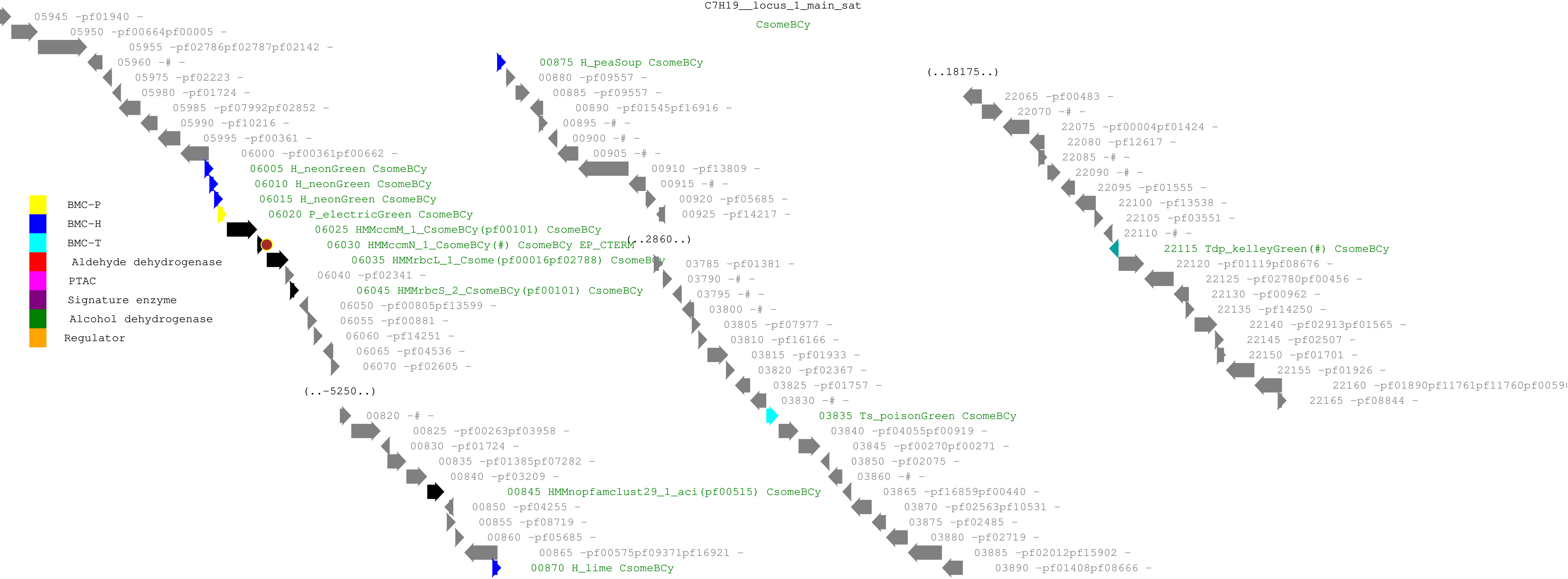

Chroococcidiopsis sp. CCALA 051. Bacteria; Cyanobacteria; Chroococcidiopsidales;

C7Y66__locus_1_main_sat

CsomeBCy

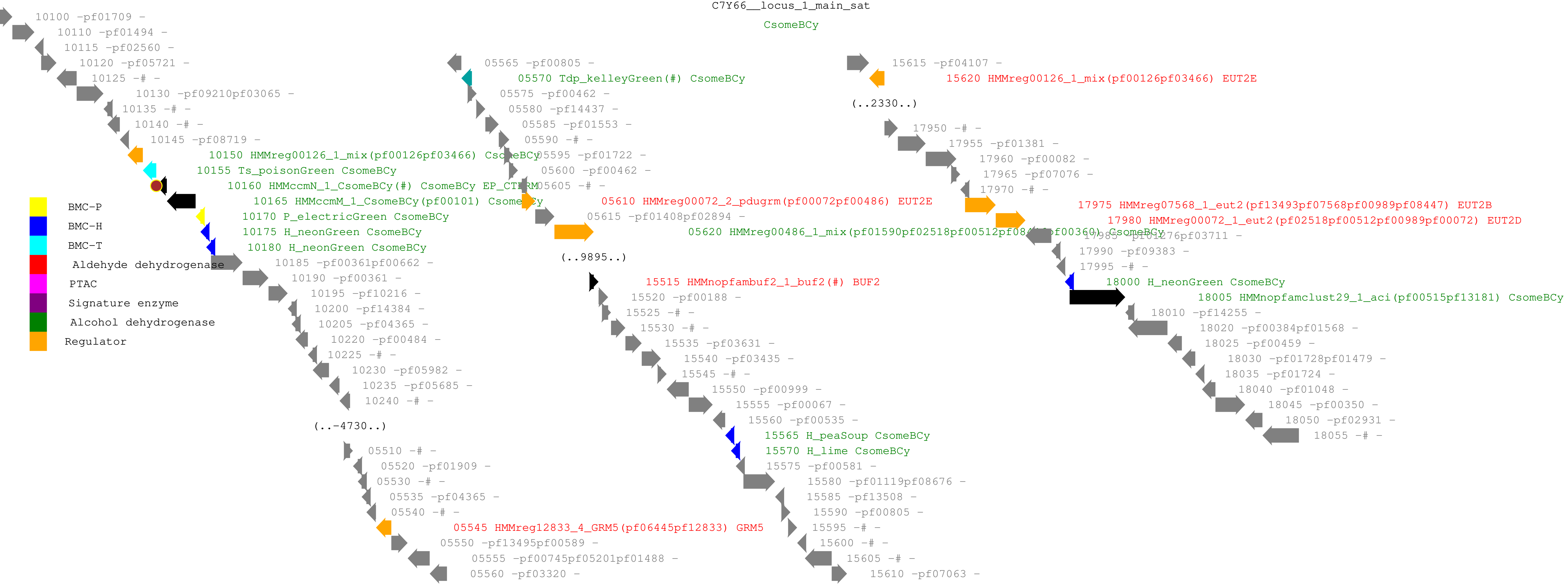

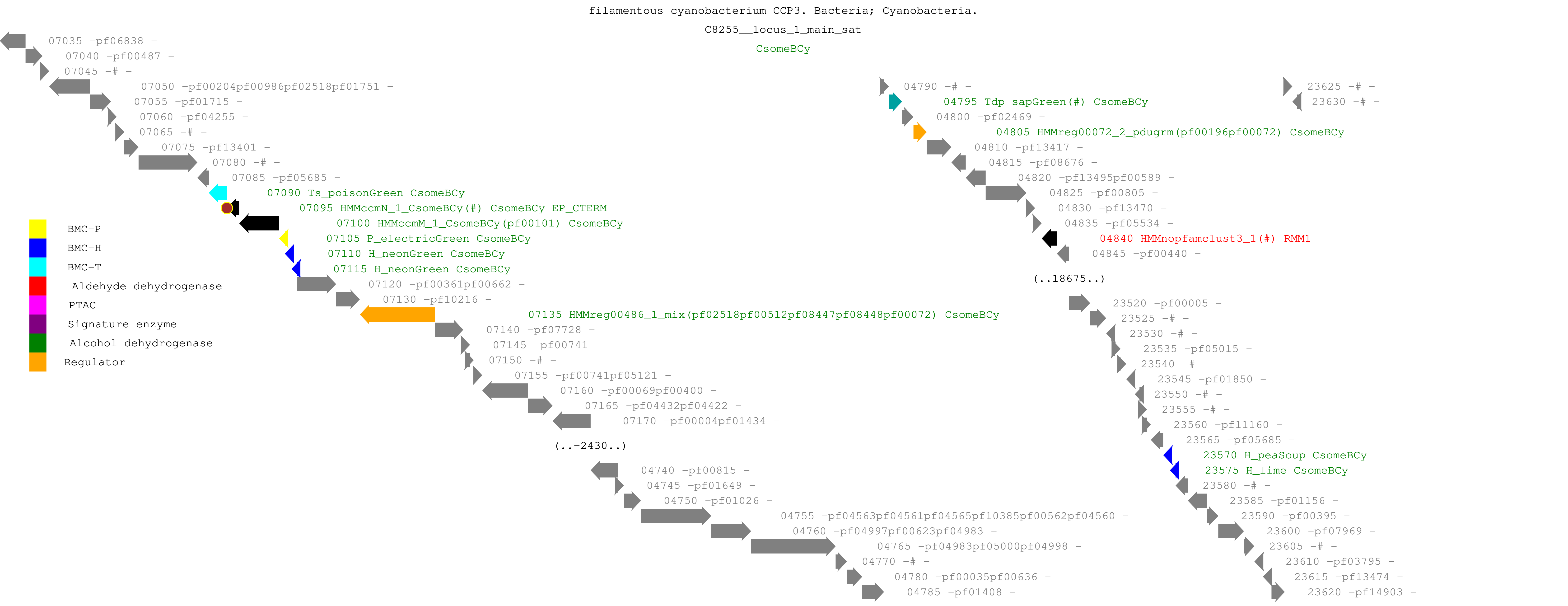
filamentous cyanobacterium CCP3. Bacteria; Cyanobacteria.
C8255__locus_1_main_sat
CsomeBCy
07035 -pf06838 -
07040 -pf00487 -
07045 -# -
07050 -pf00204pf00986pf02518pf01751 -
07055 -pf01715 -
07060 -pf04255 -
07065 -# -
07075 -pf13401 -
07080 -# -
07085 -pf05685 -
07090 Ts_poisonGreen CsomeBCy
07095 HMMccmN_1_CsomeBCy(#) CsomeBCy EP_CTERM
07100 HMMccmM_1_CsomeBCy(pf00101) CsomeBCy
07105 P_electricGreen CsomeBCy
07110 H_neonGreen CsomeBCy
07115 H_neonGreen CsomeBCy
07120 -pf00361pf00662 -
07130 -pf10216 -
07135 HMMreg00486_1_mix(pf02518pf00512pf08447pf08448pf00072) CsomeBCy
07140 -pf07728 -
07145 -pf00741 -
07150 -# -
07155 -pf00741pf05121 -
07160 -pf00069pf00400 -
07165 -pf04432pf04422 -
07170 -pf00004pf01434 -
(..-2430..)
04740 -pf00815 -
04745 -pf01649 -
04750 -pf01026 -
04755 -pf04563pf04561pf04565pf10385pf00562pf04560 -
04760 -pf04997pf00623pf04983 -
04765 -pf04983pf05000pf04998 -
04770 -# -
04780 -pf00035pf00636 -
04785 -pf01408 -
04790 -# -
04795 Tdp_sapGreen(#) CsomeBCy
04800 -pf02469 -
04805 HMMreg00072_2_pdugrm(pf00196pf00072) CsomeBCy
04810 -pf13417 -
04815 -pf08676 -
04820 -pf13495pf00589 -
04825 -pf00805 -
04830 -pf13470 -
04835 -pf05534 -
04840 HMMnopfamclust3_1(#) RMM1
04845 -pf00440 -
(..18675..)
23520 -pf00005 -
23525 -# -
23530 -# -
23535 -pf05015 -
23540 -# -
23545 -pf01850 -
23550 -# -
23555 -# -
23560 -pf11160 -
23565 -pf05685 -
23570 H_peaSoup CsomeBCy
23575 H_lime CsomeBCy
23580 -# -
23585 -pf01156 -
23590 -pf00395 -
23600 -pf07969 -
23605 -# -
23610 -pf03795 -
23615 -pf13474 -
23620 -pf14903 -
23625 -# -
23630 -# -
BMC-P
BMC-H
BMC-T
Aldehyde dehydrogenase
PTAC
Signature enzyme
Alcohol dehydrogenase
Regulator

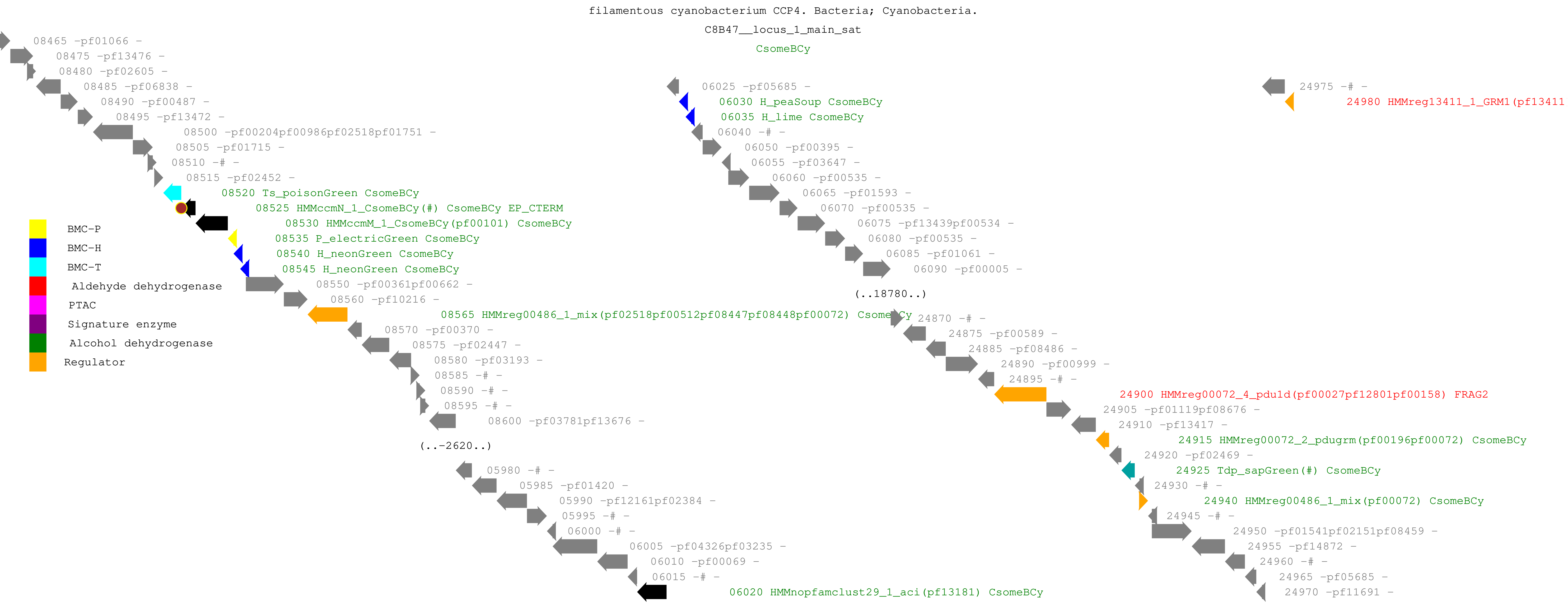
filamentous cyanobacterium CCP4. Bacteria; Cyanobacteria.
C8B47__locus_1_main_sat
CsomeBCy
08465 -pf01066 -
08475 -pf13476 -
08480 -pf02605 -
08485 -pf06838 -
08490 -pf00487 -
08495 -pf13472 -
08500 -pf00204pf00986pf02518pf01751 -
08505 -pf01715 -
08510 -# -
08515 -pf02452 -
08520 Ts_poisonGreen CsomeBCy
08525 HMMccmN_1_CsomeBCy(#) CsomeBCy EP_CTERM
08530 HMMccmM_1_CsomeBCy(pf00101) CsomeBCy
08535 P_electricGreen CsomeBCy
08540 H_neonGreen CsomeBCy
08545 H_neonGreen CsomeBCy
08550 -pf00361pf00662 -
08560 -pf10216 -
08565 HMMreg00486_1_mix(pf02518pf00512pf08447pf08448pf00072) CsomeBCy
08570 -pf00370 -
08575 -pf02447 -
08580 -pf03193 -
08585 -# -
08590 -# -
08595 -# -
08600 -pf03781pf13676 -
(..-2620..)
05980 -# -
05985 -pf01420 -
05990 -pf12161pf02384 -
05995 -# -
06000 -# -
06005 -pf04326pf03235 -
06010 -pf00069 -
06015 -# -
06020 HMMnopfamclust29_1_aci(pf13181) CsomeBCy
06025 -pf05685 -
06030 H_peaSoup CsomeBCy
06035 H_lime CsomeBCy
06040 -# -
06050 -pf00395 -
06055 -pf03647 -
06060 -pf00535 -
06065 -pf01593 -
06070 -pf00535 -
06075 -pf13439pf00534 -
06080 -pf00535 -
06085 -pf01061 -
06090 -pf00005 -
(..18780..)
24870 -# -
24875 -pf00589 -
24885 -pf08486 -
24890 -pf00999 -
24895 -# -
24900 HMMreg00072_4_pdu1d(pf00027pf12801pf00158) FRAG2
24905 -pf01119pf08676 -
24910 -pf13417 -
24915 HMMreg00072_2_pdugrm(pf00196pf00072) CsomeBCy
24920 -pf02469 -
24925 Tdp_sapGreen(#) CsomeBCy
24930 -# -
24940 HMMreg00486_1_mix(pf00072) CsomeBCy
24945 -# -
24950 -pf01541pf02151pf08459 -
24955 -pf14872 -
24960 -# -
24965 -pf05685 -
24970 -pf11691 -
24975 -# -
24980 HMMreg13411_1_GRM1(pf13411
BMC-P
BMC-H
BMC-T
Aldehyde dehydrogenase
PTAC
Signature enzyme
Alcohol dehydrogenase
Regulator

Calothrix sp. PCC 6303. Bacteria; Cyanobacteria; Nostocales; Rivulariaceae; Calothrix.

Cal6303__locus_1_main_sat

CsomeBCy

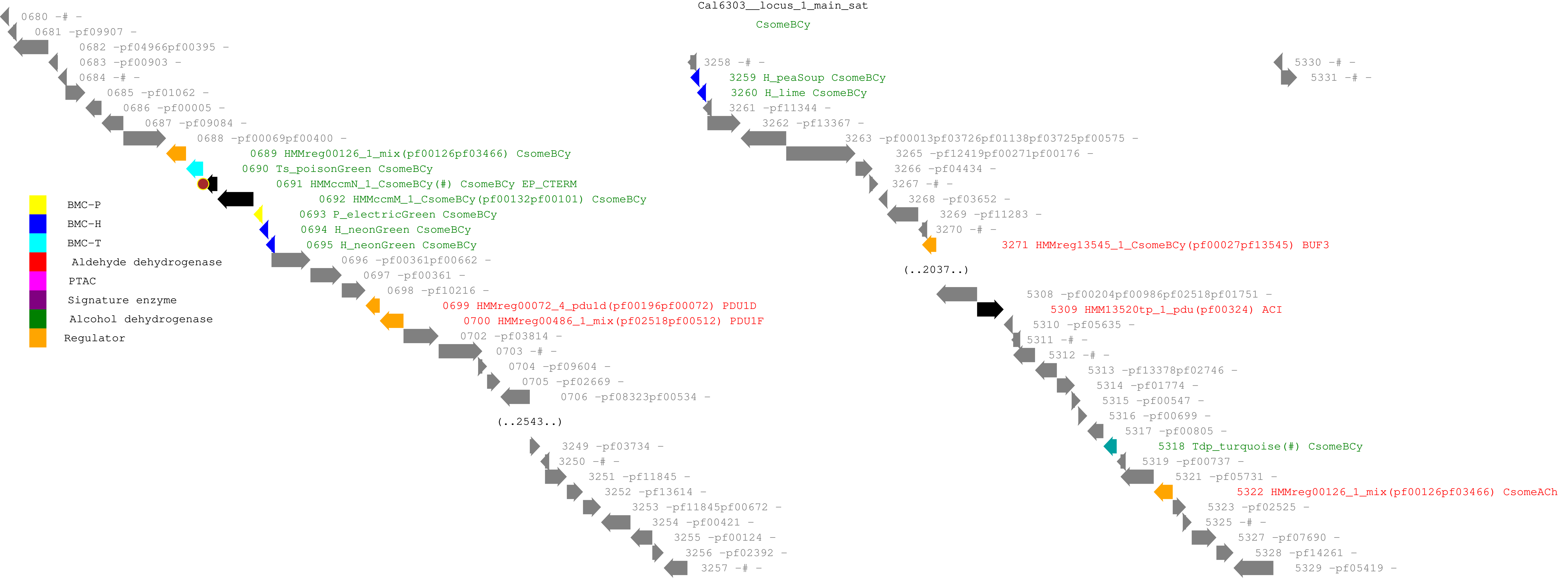

Calothrix sp. PCC 7507. Bacteria; Cyanobacteria; Nostocales; Rivulariaceae; Calothrix.

Cal7507__locus_1_main_sat

CsomeBCy

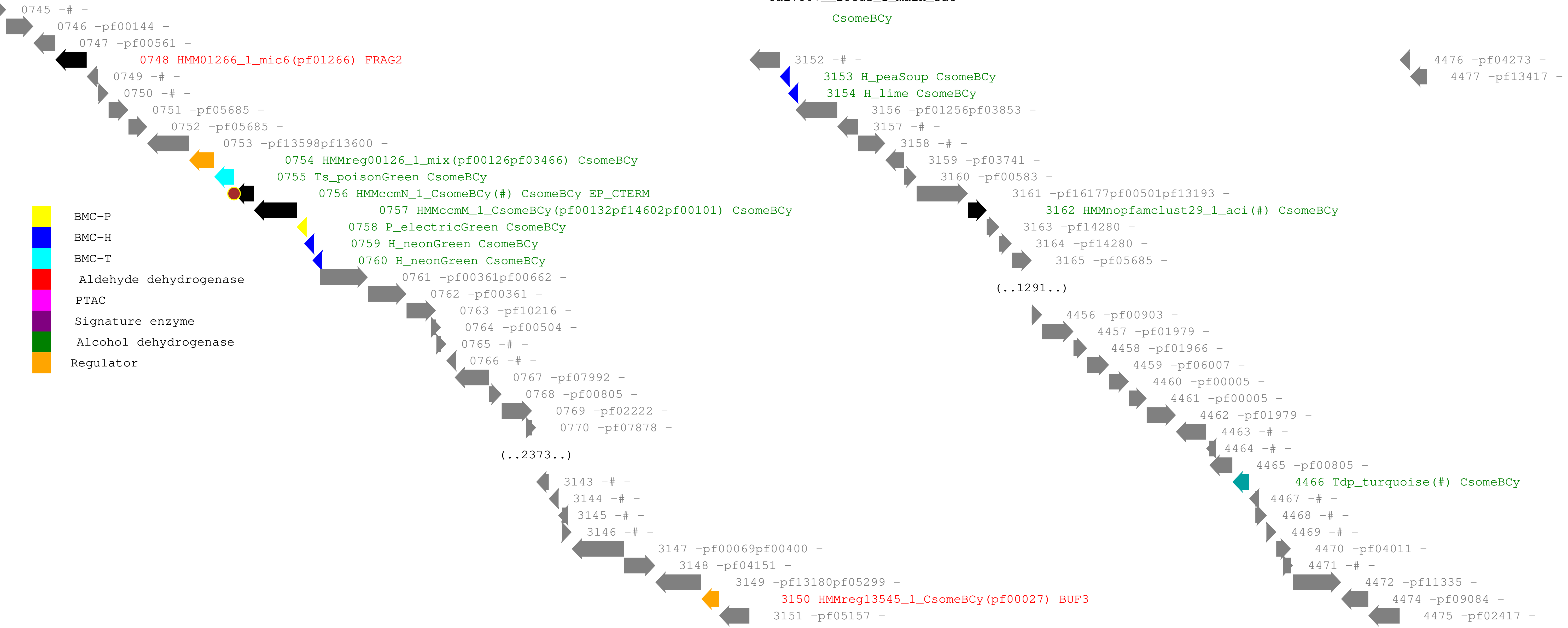

Fischerella thermalis WC341. Bacteria; Cyanobacteria; Nostocales; Hapalosiphonaceae; Fischerella.

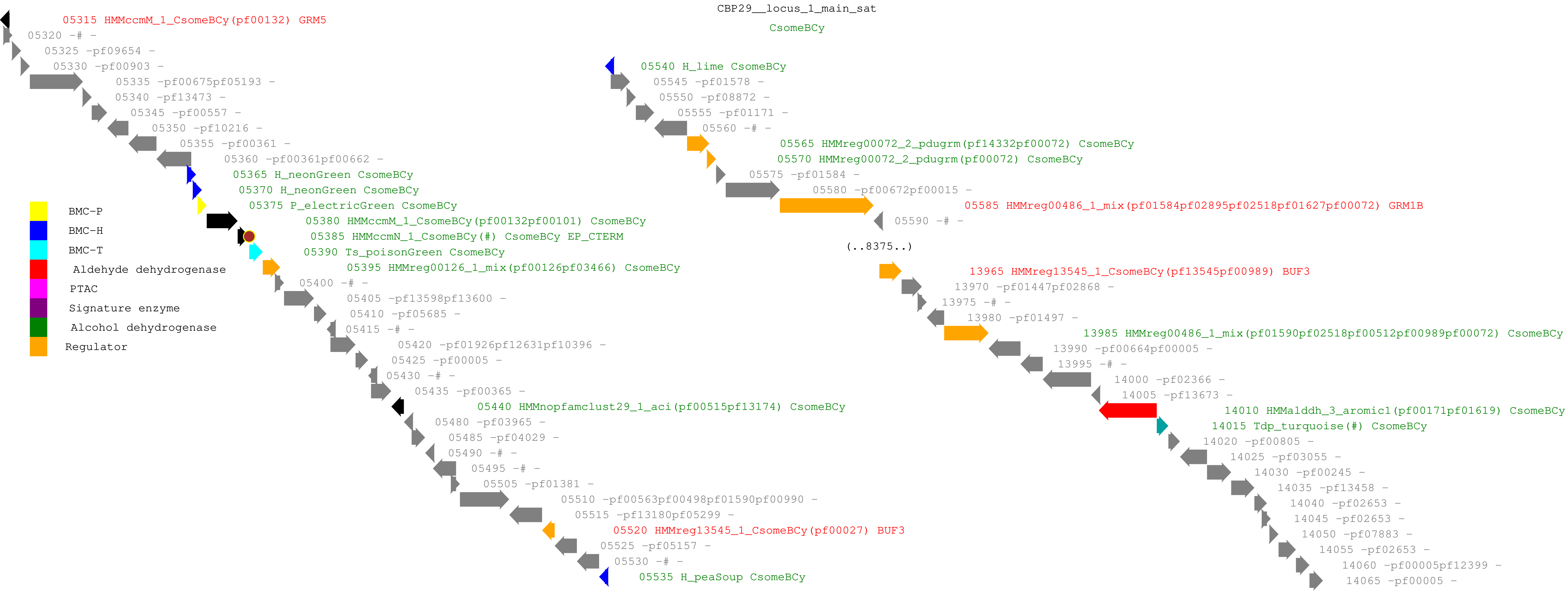

Cyanothece sp. (strain ATCC 51142). Bacteria; Cyanobacteria; Oscillatoriophycideae; Oscillatoriales;

cce__locus_1_main_sat

CsomeBCy

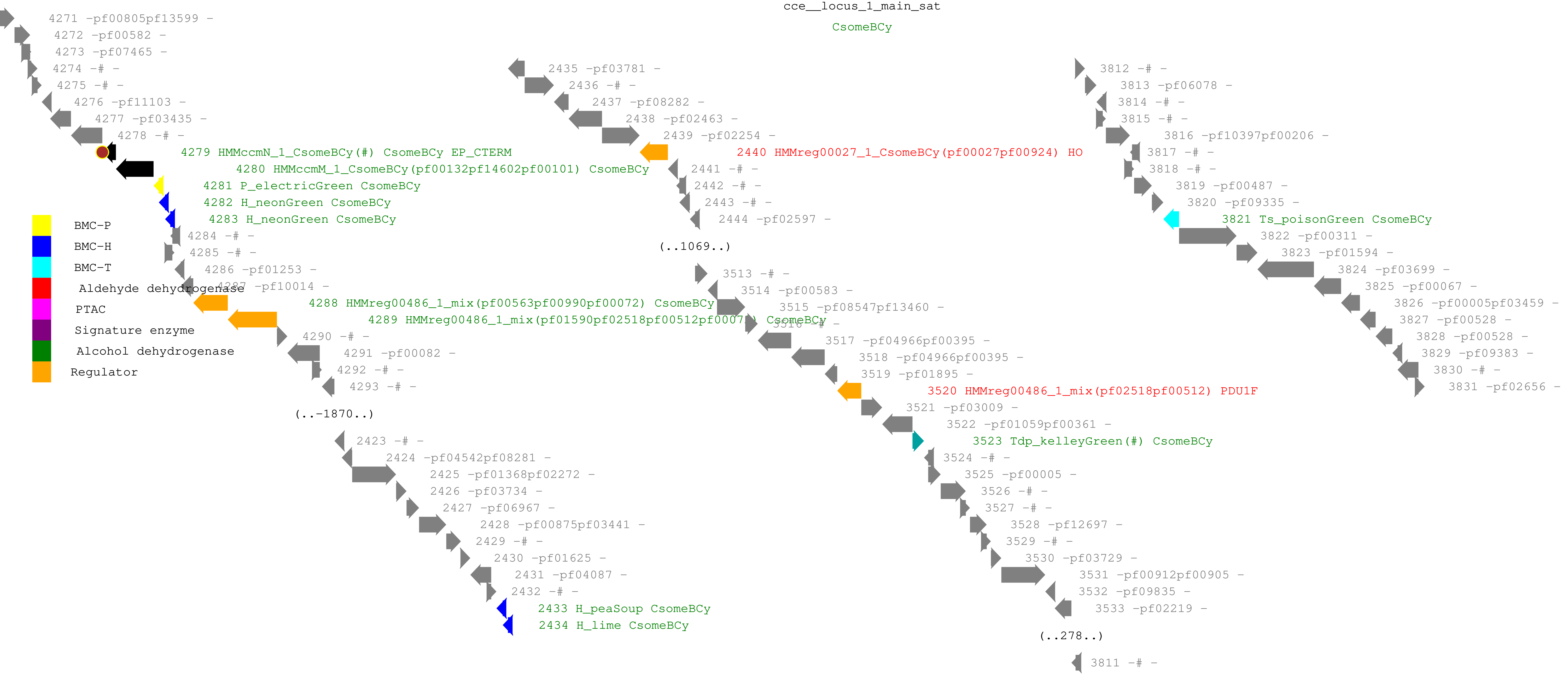

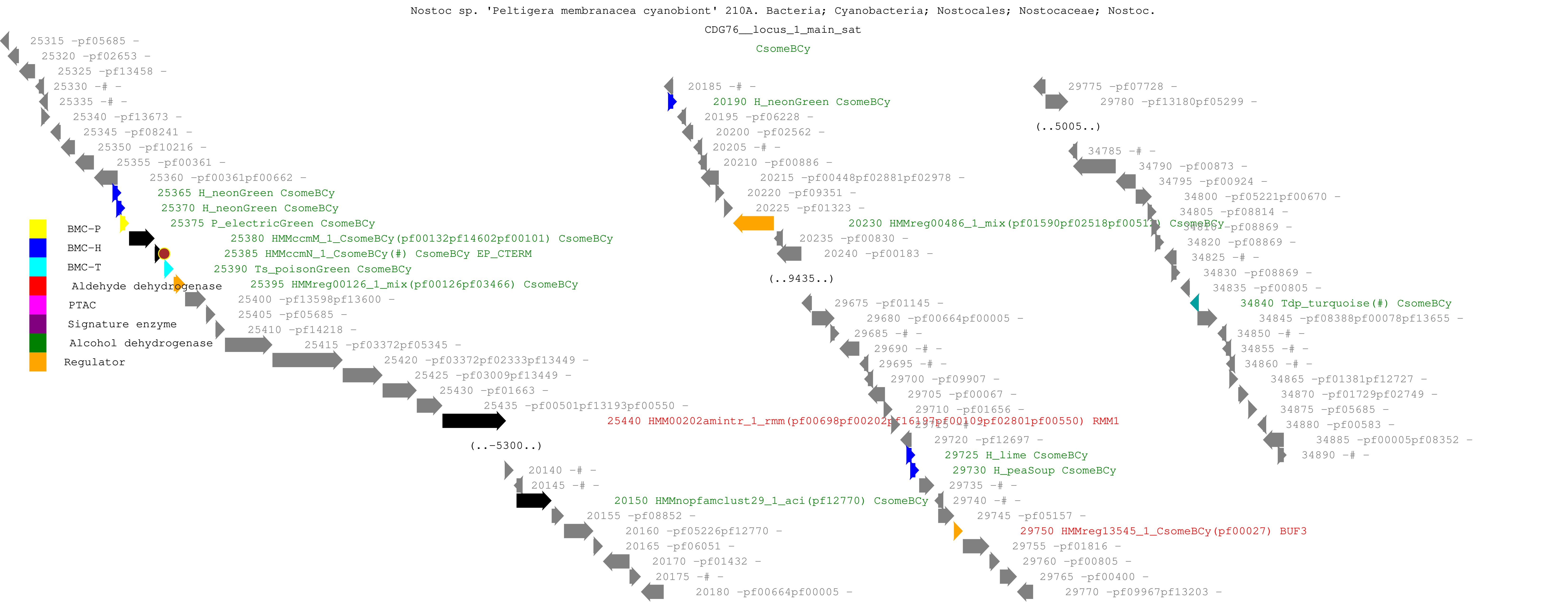
Nostoc sp. 'Peltigera membranacea cyanobiont' 210A. Bacteria; Cyanobacteria; Nostocales; Nostocaceae; Nostoc.
CDG76__locus_1_main_sat
CsomeBCy
BMC-P
BMC-H
BMC-T
Aldehyde dehydrogenase
PTAC
Signature enzyme
Alcohol dehydrogenase
Regulator
25315 -pf05685 -
25320 -pf02653 -
25325 -pf13458 -
25330 -# -
25335 -# -
25340 -pf13673 -
25345 -pf08241 -
25350 -pf10216 -
25355 -pf00361 -
25360 -pf00361pf00662 -
25365 H_neonGreen CsomeBCy
25370 H_neonGreen CsomeBCy
25375 P_electricGreen CsomeBCy
25380 HMMccmM_1_CsomeBCy(pf00132pf14602pf00101) CsomeBCy
25385 HMMccmN_1_CsomeBCy(#) CsomeBCy EP_CTERM
25390 Ts_poisonGreen CsomeBCy
25395 HMMreg00126_1_mix(pf00126pf03466) CsomeBCy
25400 -pf13598pf13600 -
25405 -pf05685 -
25410 -pf14218 -
25415 -pf03372pf05345 -
25420 -pf03372pf02333pf13449 -
25425 -pf03009pf13449 -
25430 -pf01663 -
25435 -pf00501pf13193pf00550 -
25440 HMM00202amintr_1_rmm(pf00698pf00202pf16197pf00109pf02801pf00550) RMM1
(..-5300..)
20140 -# -
20145 -# -
20150 HMMnopfamclust29_1_aci(pf12770) CsomeBCy
20155 -pf08852 -
20160 -pf05226pf12770 -
20165 -pf06051 -
20170 -pf01432 -
20175 -# -
20180 -pf00664pf00005 -
20185 -# -
20190 H_neonGreen CsomeBCy
20195 -pf06228 -
20200 -pf02562 -
20205 -# -
20210 -pf00886 -
20215 -pf00448pf02881pf02978 -
20220 -pf09351 -
20225 -pf01323 -
20230 HMMreg00486_1_mix(pf01590pf02518pf00512) CsomeBCy
20235 -pf00830 -
20240 -pf00183 -
(..9435..)
29675 -pf01145 -
29680 -pf00664pf00005 -
29685 -# -
29690 -# -
29695 -# -
29700 -pf09907 -
29705 -pf00067 -
29710 -pf01656 -
29715 -# -
29720 -pf12697 -
29725 H_lime CsomeBCy
29730 H_peaSoup CsomeBCy
29735 -# -
29740 -# -
29745 -pf05157 -
29750 HMMreg13545_1_CsomeBCy(pf00027) BUF3
29755 -pf01816 -
29760 -pf00805 -
29765 -pf00400 -
29770 -pf09967pf13203 -
29775 -pf07728 -
29780 -pf13180pf05299 -
(..5005..)
34785 -# -
34790 -pf00873 -
34795 -pf00924 -
34800 -pf05221pf00670 -
34805 -pf08814 -
34810 -pf08869 -
34820 -pf08869 -
34825 -# -
34830 -pf08869 -
34835 -pf00805 -
34840 Tdp_turquoise(#) CsomeBCy
34845 -pf08388pf00078pf13655 -
34850 -# -
34855 -# -
34860 -# -
34865 -pf01381pf12727 -
34870 -pf01729pf02749 -
34875 -pf05685 -
34880 -pf00583 -
34885 -pf00005pf08352 -
34890 -# -

Nostoc sp. 'Peltigera membranacea cyanobiont' 213. Bacteria; Cyanobacteria; Nostocales; Nostocaceae; Nostoc.

CDG77__locus_1_main_sat

CsomeBCy

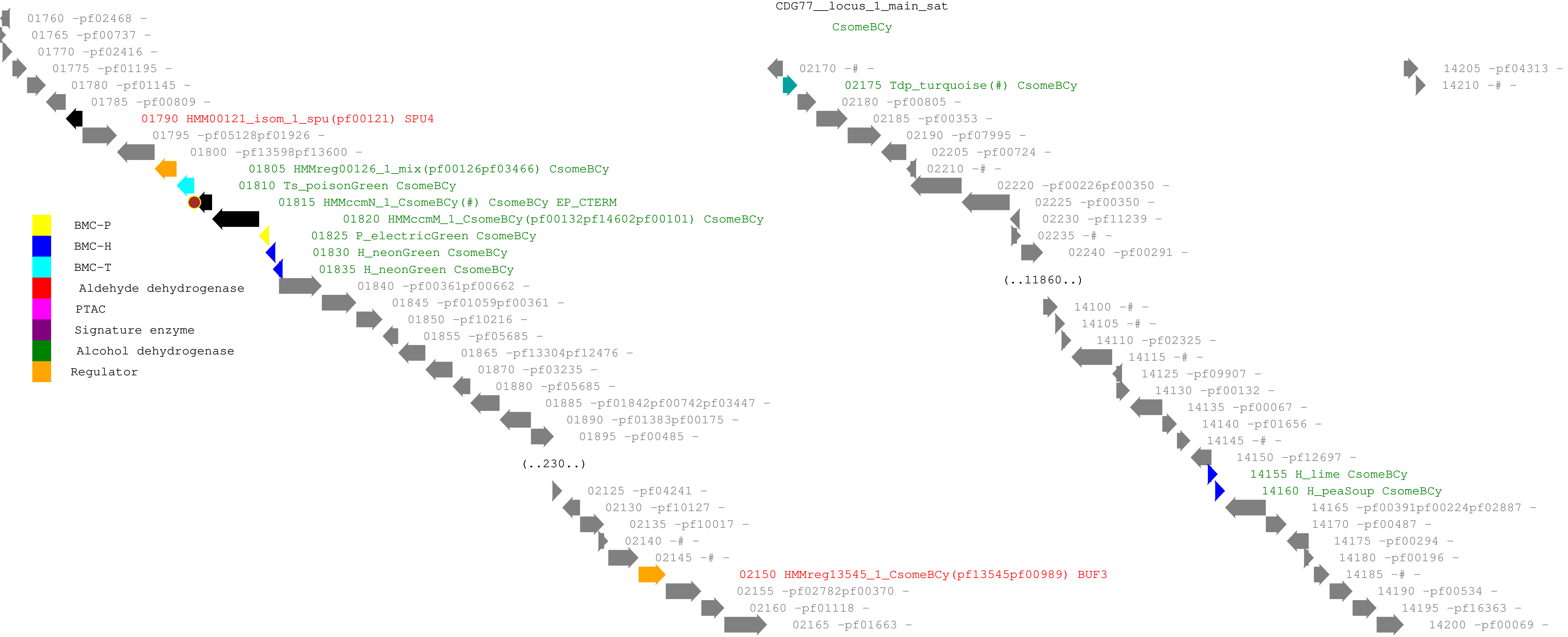

Nostoc sp. 'Peltigera membranacea cyanobiont' 232. Bacteria; Cyanobacteria; Nostocales; Nostocaceae; Nostoc.

CDG79__locus_1_main_sat

CsomeBCy

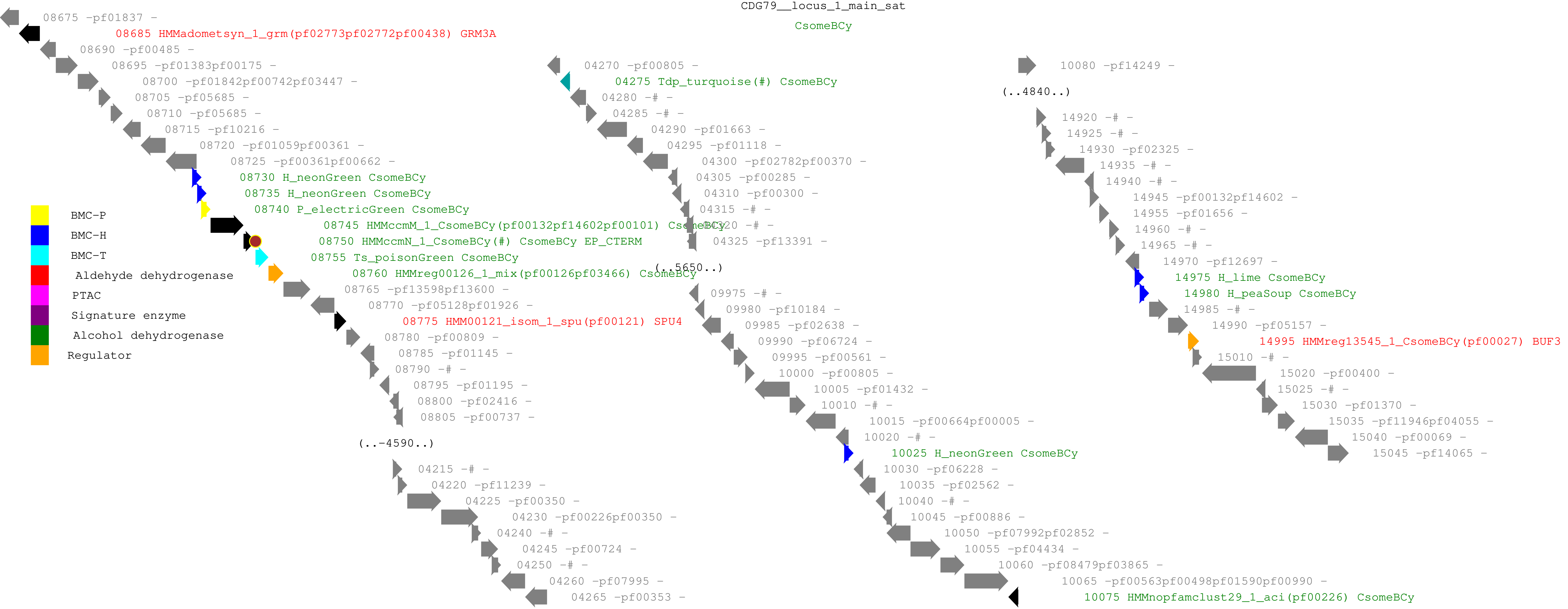

Fischerella thermalis CCMEE 5201. Bacteria; Cyanobacteria; Nostocales; Hapalosiphonaceae; Fischerella.

CEN39__locus_1_main_sat

CsomeBCy

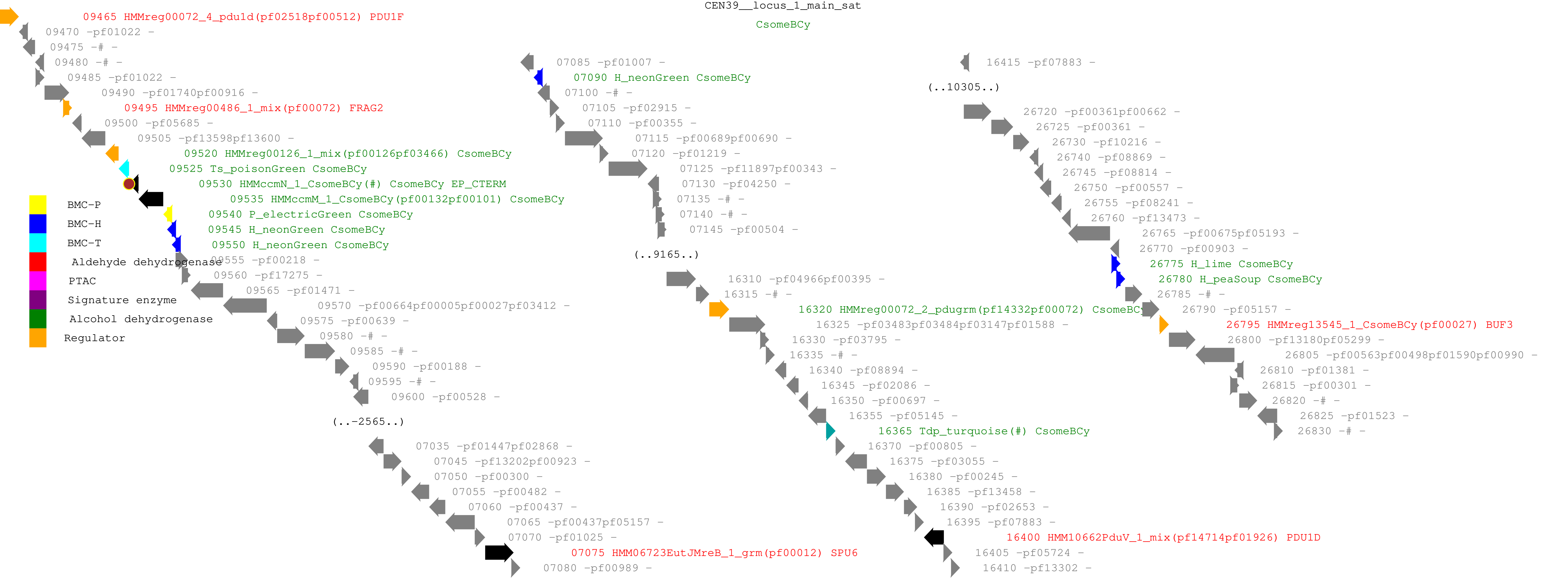

Fischerella thermalis CCMEE 5205. Bacteria; Cyanobacteria; Nostocales; Hapalosiphonaceae; Fischerella.

CEN40__locus_1_main_sat

CsomeBCy

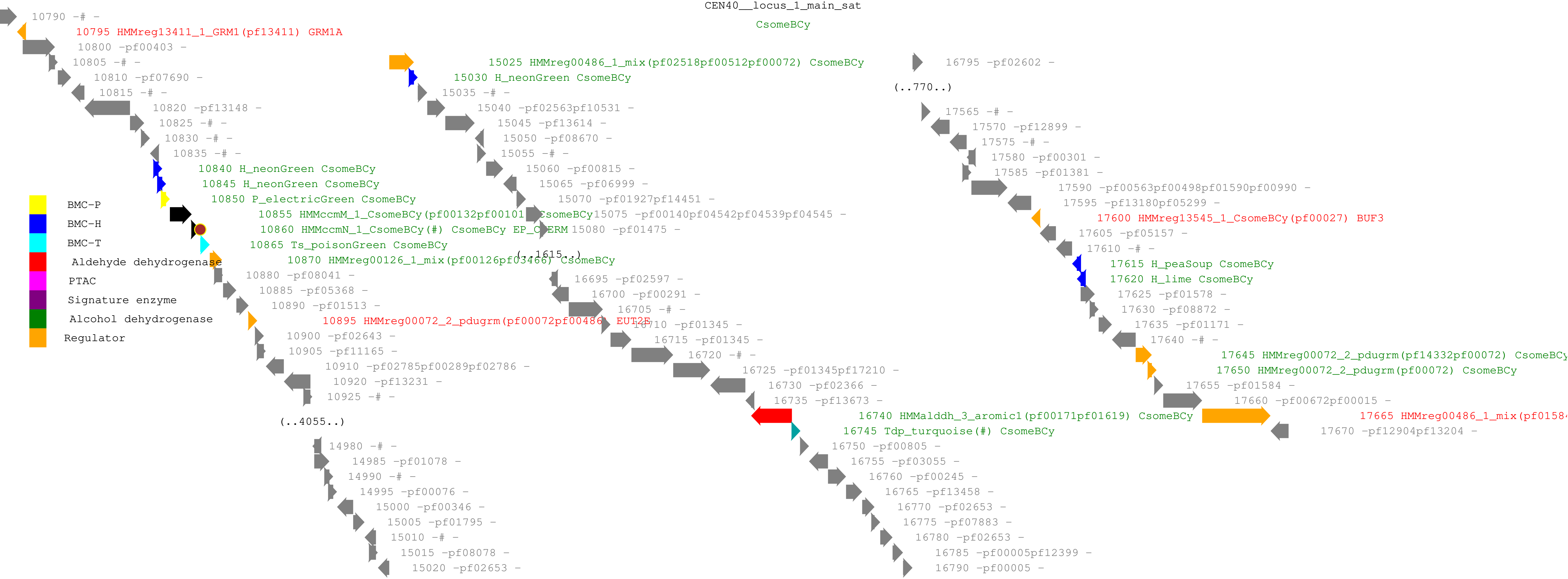

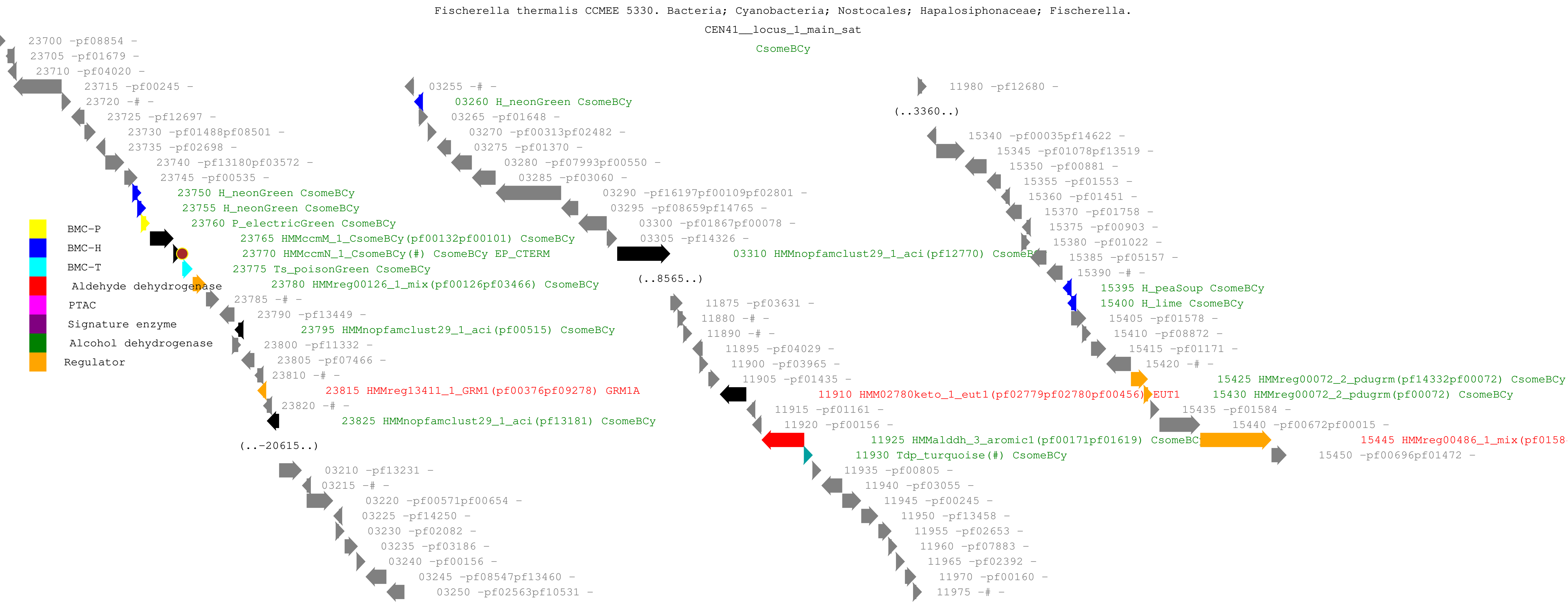

Fischerella thermalis CCMEE 5330. Bacteria; Cyanobacteria; Nostocales; Hapalosiphonaceae; Fischerella.
CEN41__locus_1_main_sat
CsomeBCy
23700 -pf08854 -
23705 -pf01679 -
23710 -pf04020 -
23715 -pf00245 -
23720 -# -
23725 -pf12697 -
23730 -pf01488pf08501 -
23735 -pf02698 -
23740 -pf13180pf03572 -
23745 -pf00535 -
23750 H_neonGreen CsomeBCy
23755 H_neonGreen CsomeBCy
23760 P_electricGreen CsomeBCy
23765 HMMccmM_1_CsomeBCy(pf00132pf00101) CsomeBCy
23770 HMMccmN_1_CsomeBCy(#) CsomeBCy EP_CTERM
23775 Ts_poisonGreen CsomeBCy
23780 HMMreg00126_1_mix(pf00126pf03466) CsomeBCy
23785 -# -
23790 -pf13449 -
23795 HMMnopfamclust29_1_aci(pf00515) CsomeBCy
23800 -pf11332 -
23805 -pf07466 -
23810 -# -
23815 HMMreg13411_1_GRM1(pf00376pf09278) GRM1A
23820 -# -
23825 HMMnopfamclust29_1_aci(pf13181) CsomeBCy
(..-20615..)
03210 -pf13231 -
03215 -# -
03220 -pf00571pf00654 -
03225 -pf14250 -
03230 -pf02082 -
03235 -pf03186 -
03240 -pf00156 -
03245 -pf08547pf13460 -
03250 -pf02563pf10531 -
03255 -# -
03260 H_neonGreen CsomeBCy
03265 -pf01648 -
03270 -pf00313pf02482 -
03275 -pf01370 -
03280 -pf07993pf00550 -
03285 -pf03060 -
03290 -pf16197pf00109pf02801 -
03295 -pf08659pf14765 -
03300 -pf01867pf00078 -
03305 -pf14326 -
03310 HMMnopfamclust29_1_aci(pf12770) CsomeBC
(..8565..)
11875 -pf03631 -
11880 -# -
11890 -# -
11895 -pf04029 -
11900 -pf03965 -
11905 -pf01435 -
11910 HMM02780keto_1_eut1(pf02779pf02780pf00456) EUT1
11915 -pf01161 -
11920 -pf00156 -
11925 HMMalddh_3_aromic1(pf00171pf01619) CsomeBC
11930 Tdp_turquoise(#) CsomeBCy
11935 -pf00805 -
11940 -pf03055 -
11945 -pf00245 -
11950 -pf13458 -
11955 -pf02653 -
11960 -pf07883 -
11965 -pf02392 -
11970 -pf00160 -
11975 -# -
11980 -pf12680 -
(..3360..)
15340 -pf00035pf14622 -
15345 -pf01078pf13519 -
15350 -pf00881 -
15355 -pf01553 -
15360 -pf01451 -
15370 -pf01758 -
15375 -pf00903 -
15380 -pf01022 -
15385 -pf05157 -
15390 -# -
15395 H_peaSoup CsomeBCy
15400 H_lime CsomeBCy
15405 -pf01578 -
15410 -pf08872 -
15415 -pf01171 -
15420 -# -
15425 HMMreg00072_2_pdugrm(pf14332pf00072) CsomeBCy
15430 HMMreg00072_2_pdugrm(pf00072) CsomeBCy
15435 -pf01584 -
15440 -pf00672pf00015 -
15445 HMMreg00486_1_mix(pf0158
15450 -pf00696pf01472 -
BMC-P
BMC-H
BMC-T
Aldehyde dehydrogenase
PTAC
Signature enzyme
Alcohol dehydrogenase
Regulator

Fischerella muscicola CCMEE 5323. Bacteria; Cyanobacteria; Nostocales; Hapalosiphonaceae; Fischerella.

CEN44__locus_1_main_sat

CsomeBCy

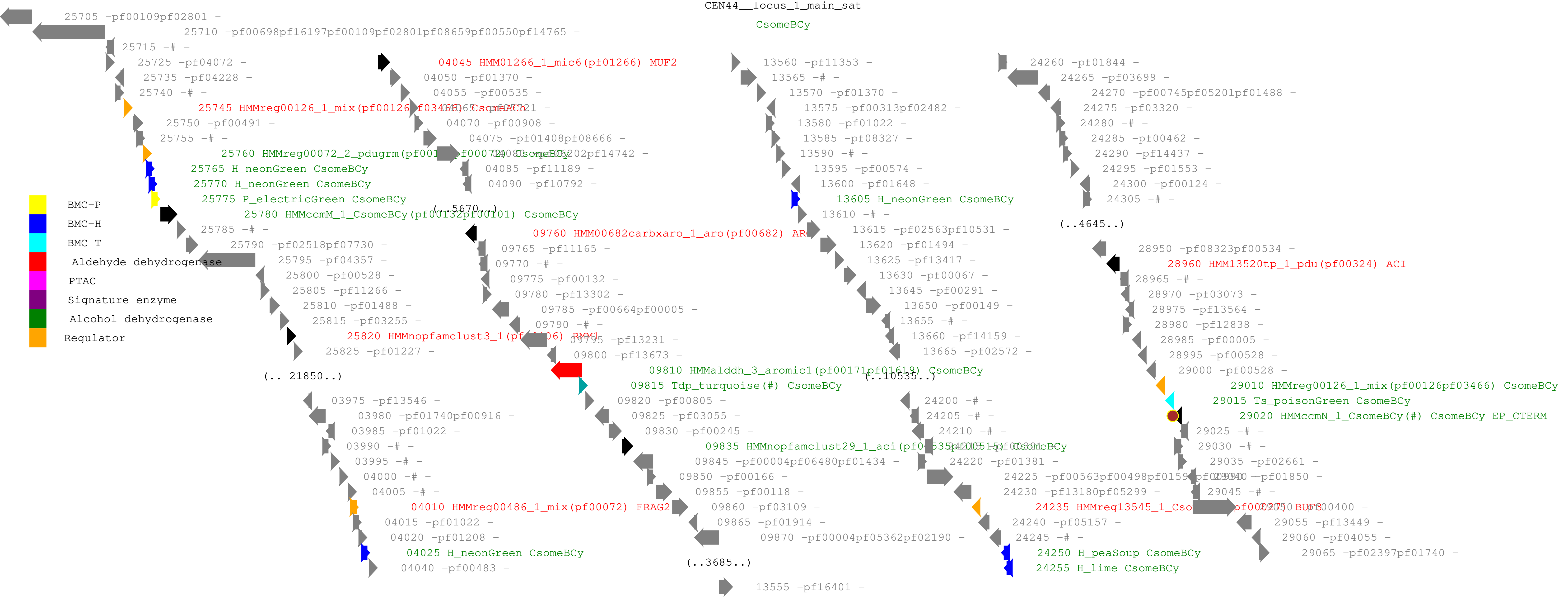

Fischerella thermalis CCMEE 5318. Bacteria; Cyanobacteria; Nostocales; Hapalosiphonaceae; Fischerella.

CEN46__locus_1_main_sat

CsomeBCy

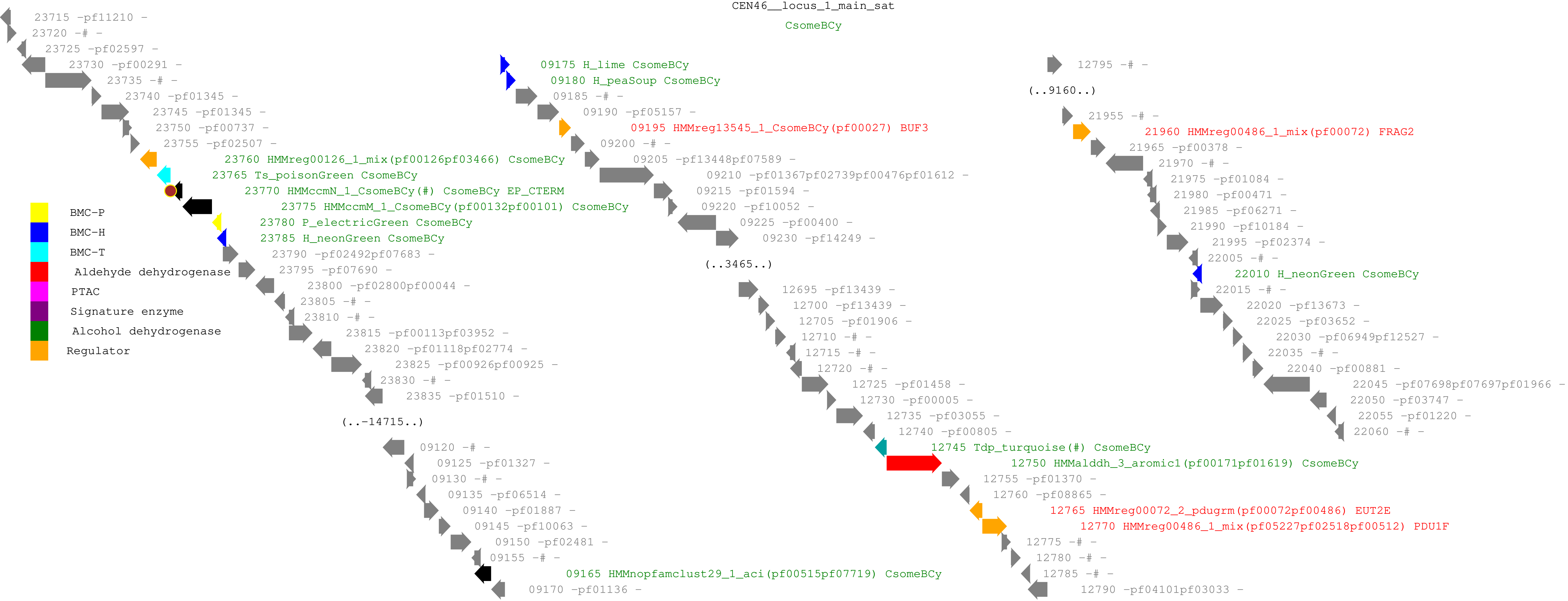

Fischerella thermalis CCMEE 5319. Bacteria; Cyanobacteria; Nostocales; Hapalosiphonaceae; Fischerella.

CEN47__locus_1_main_sat

CsomeBCy

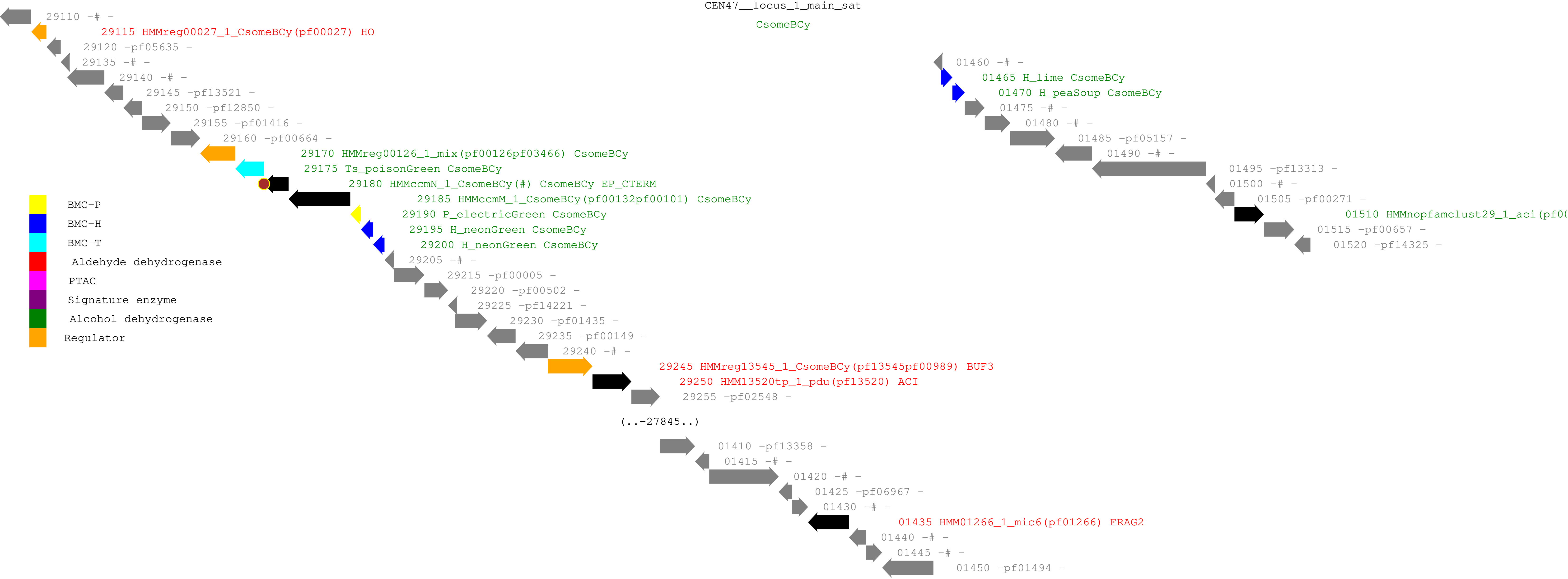

Fischerella thermalis CCMEE 5273. Bacteria; Cyanobacteria; Nostocales; Hapalosiphonaceae; Fischerella.

CEN49__locus_1_main_sat

CsomeBCy

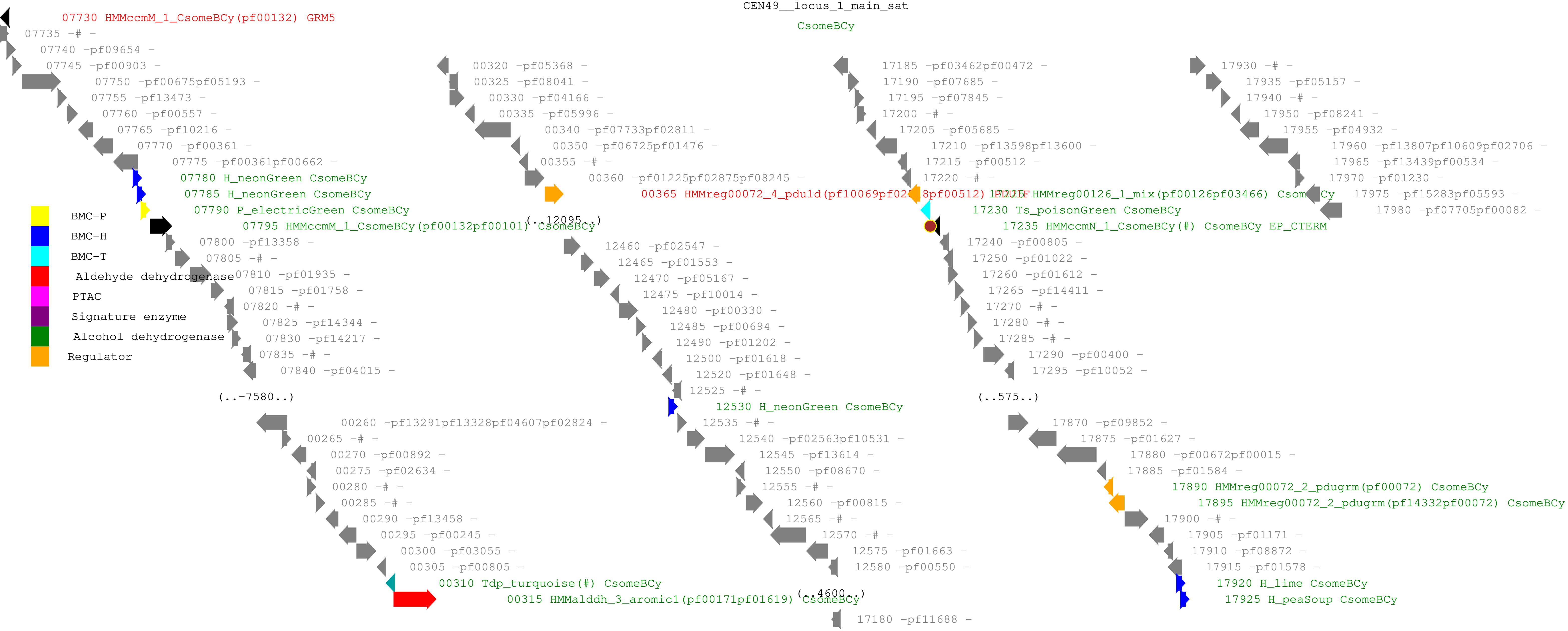

Fischerella thermalis CCMEE 5268. Bacteria; Cyanobacteria; Nostocales; Hapalosiphonaceae; Fischerella.

CEN50__locus_1_main_sat

CsomeBCy

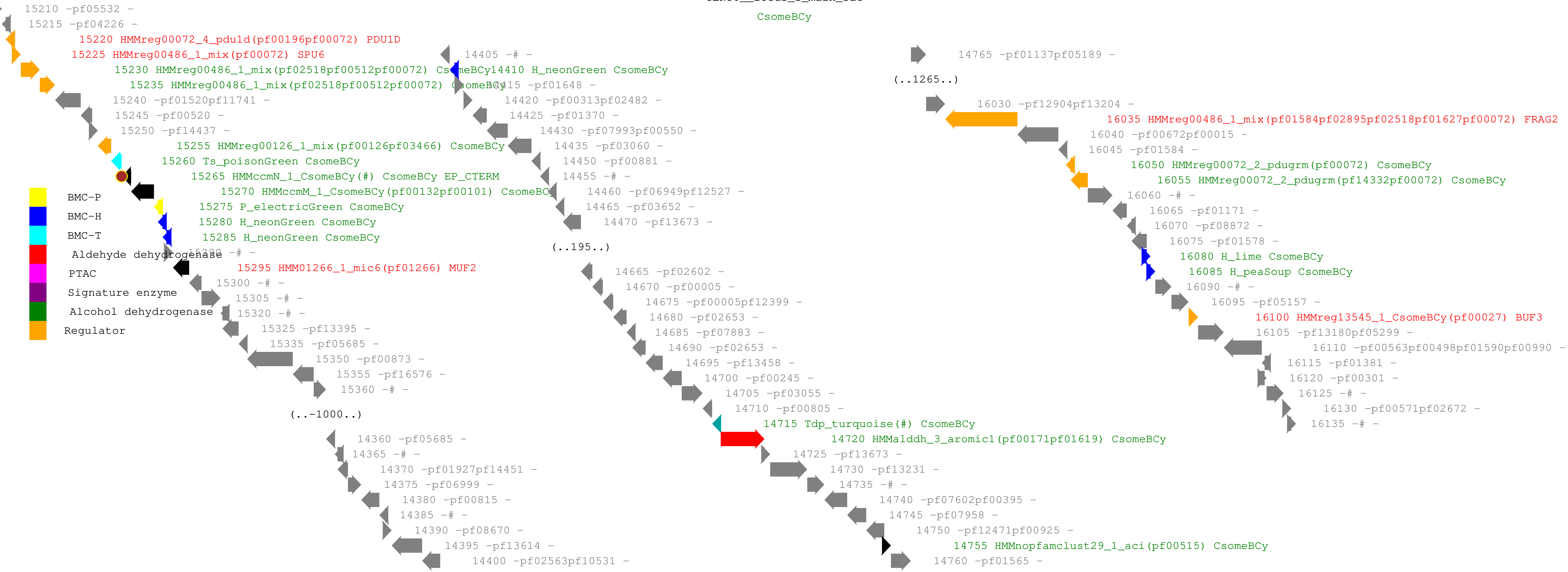

Cylindrospermopsis raciborskii S05. Bacteria; Cyanobacteria; Nostocales; Aphanizomenonaceae;

CEP08__locus_1_main_sat

CsomeBCy

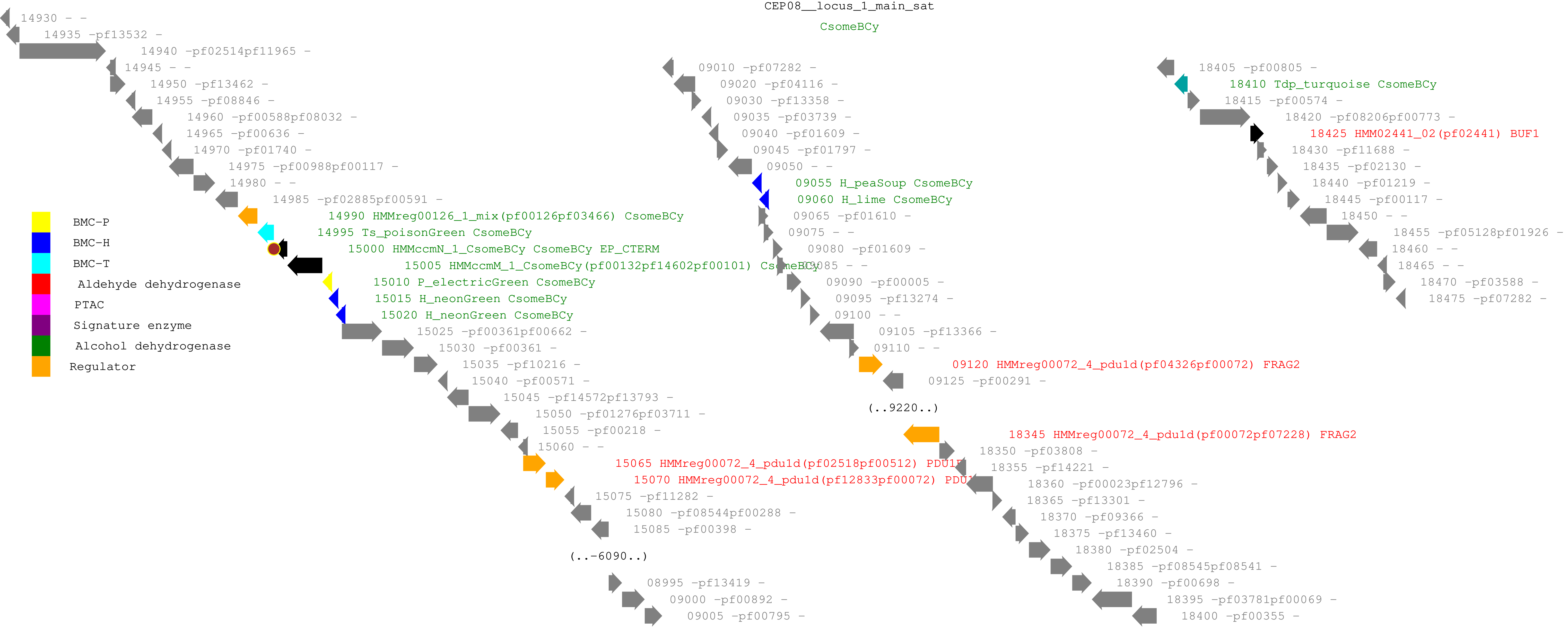

Chamaesiphon minutus PCC 6605. Bacteria; Cyanobacteria; Synechococcales; Chamaesiphonaceae;

Cha6605__locus_1_main_sat

CsomeBCy

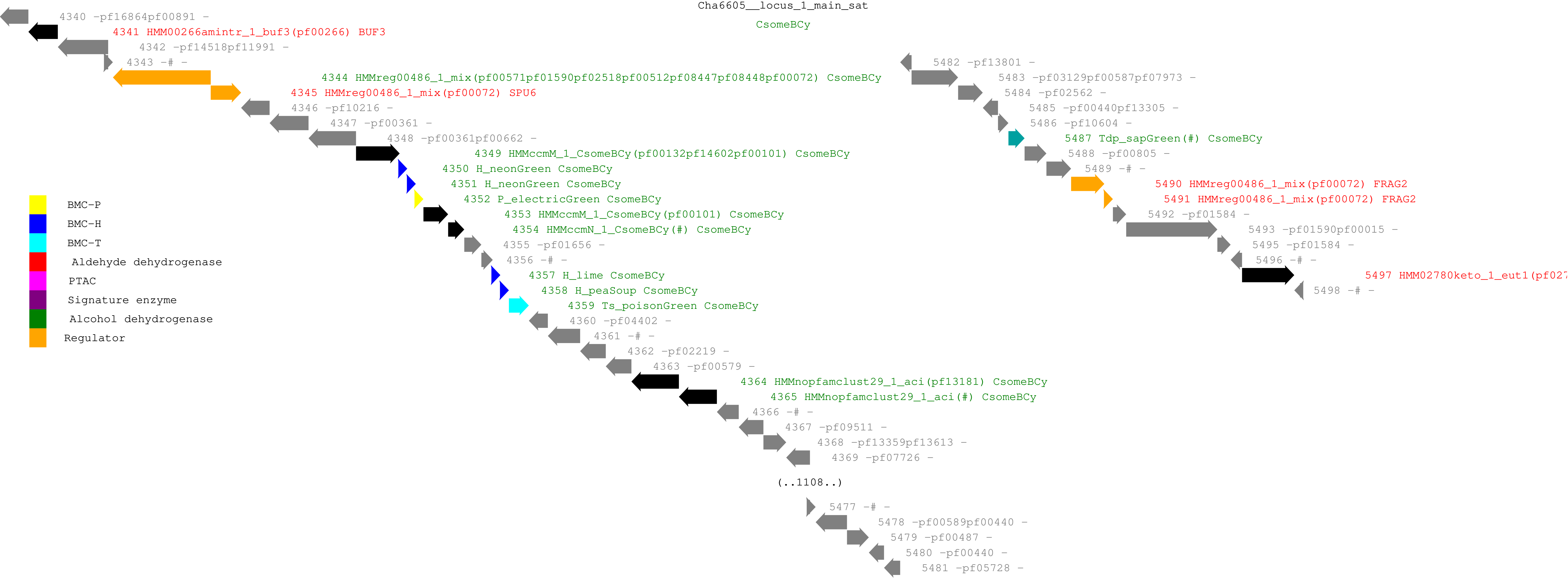

Chroococcidiopsis thermalis PCC 7203. Bacteria; Cyanobacteria; Chroococcidiopsidales;

Chro__locus_1_main_sat

CsomeBCy

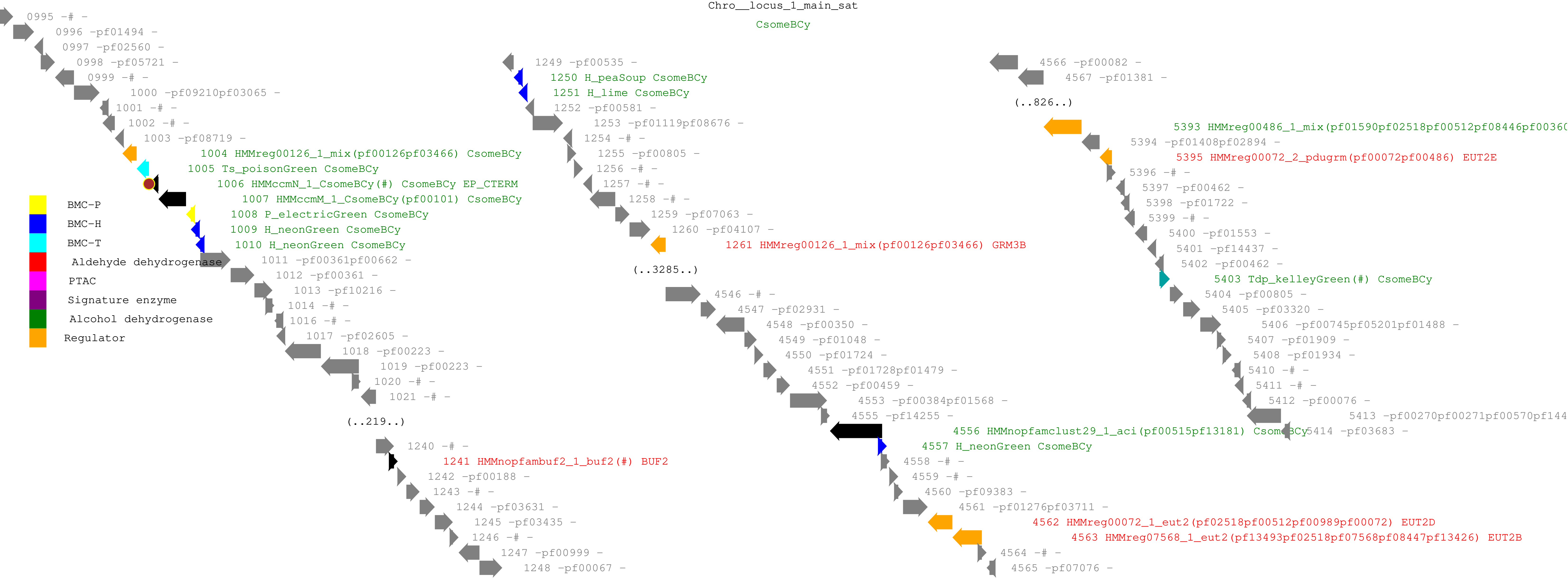

Calothrix elsteri CCALA 953. Bacteria; Cyanobacteria; Nostocales; Rivulariaceae; Calothrix.

CK510__locus_1_main_sat

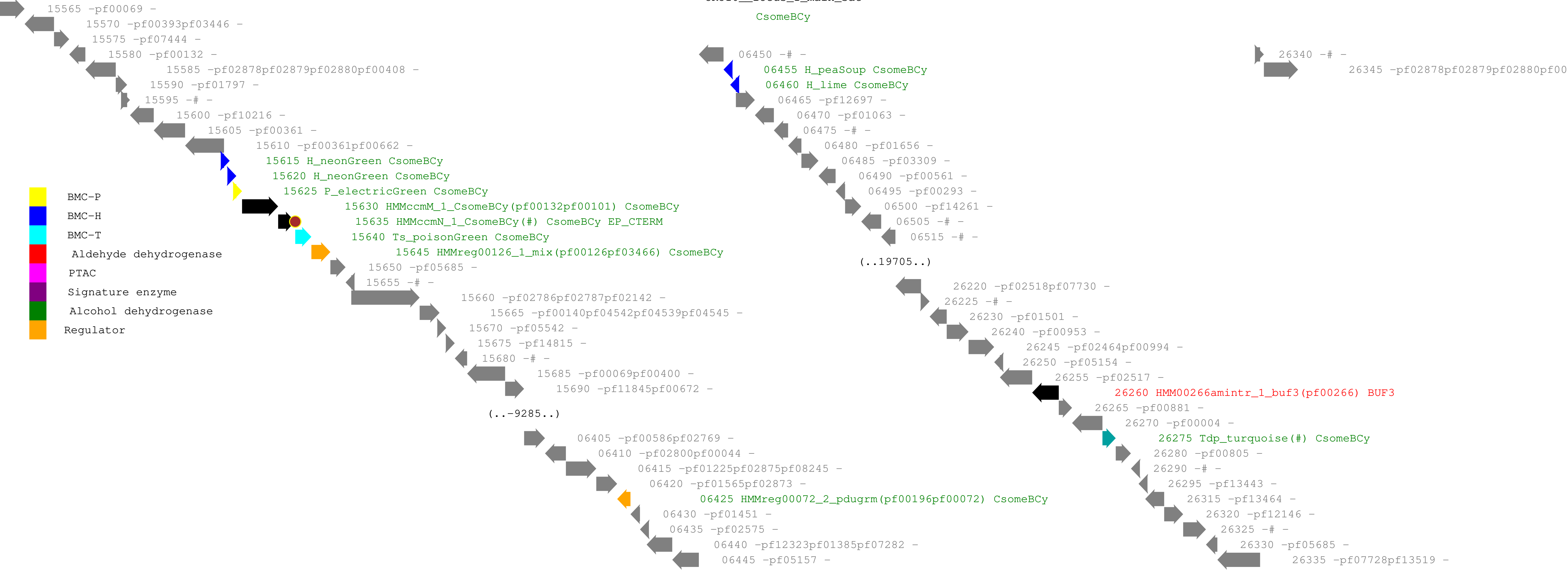

Nostoc sp. 'Peltigera malacea cyanobiont' DB3992. Bacteria; Cyanobacteria; Nostocales; Nostocaceae; Nostoc.

CK516__locus_1_main_sat

CsomeBCy

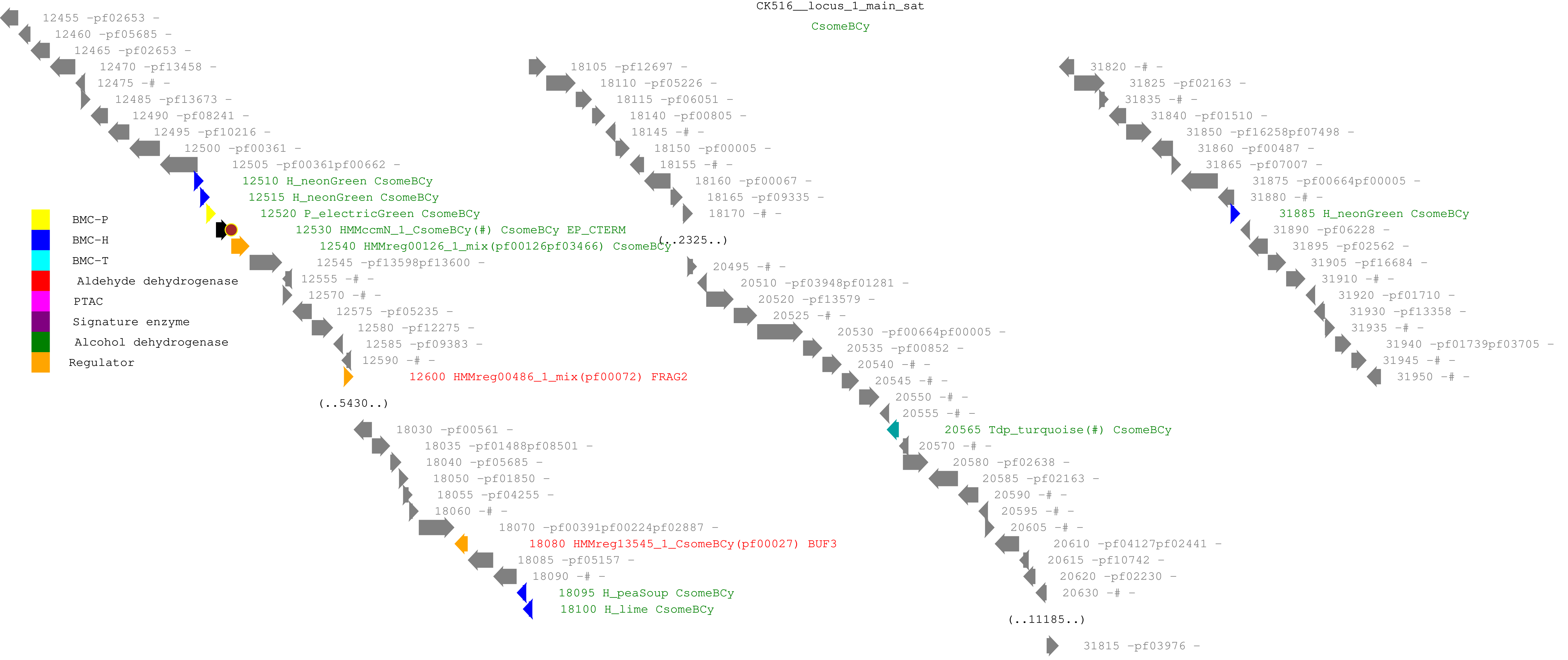

Nostoc sp. CENA543. Bacteria; Cyanobacteria; Nostocales; Nostocaceae; Nostoc.

CLI64__locus_1_main_sat

CsomeBCy

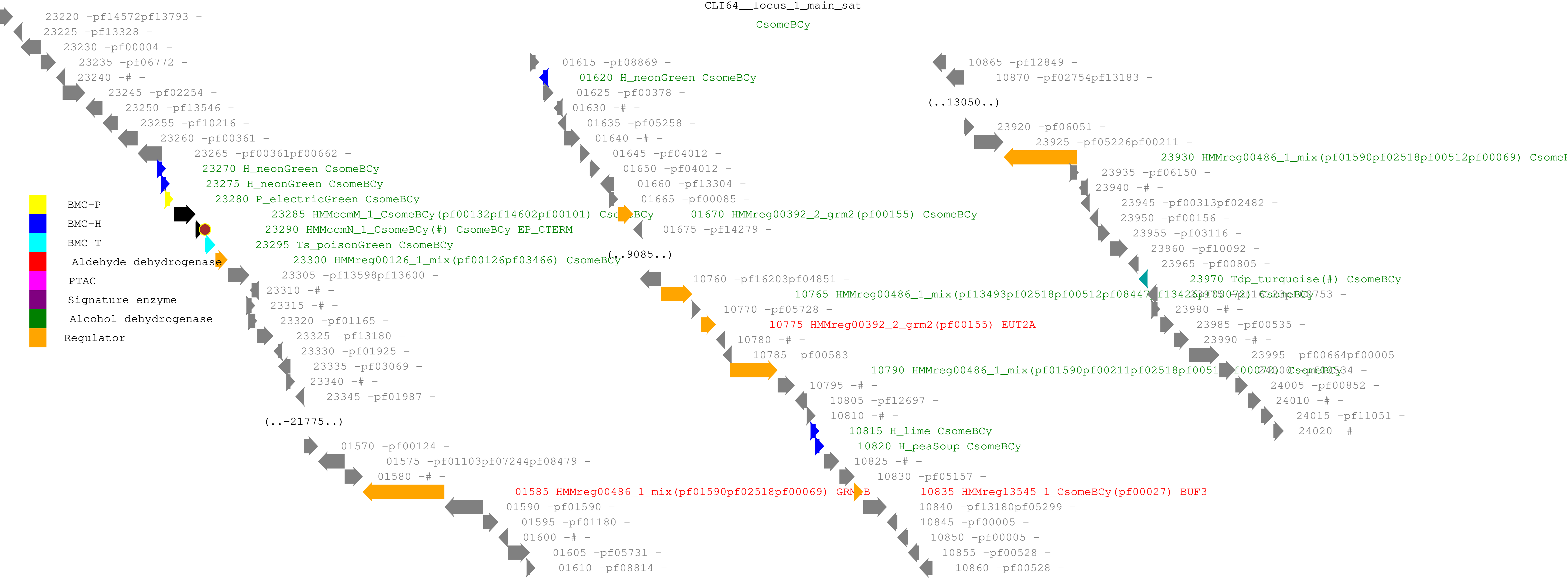

Nostoc flagelliforme CCNUN1. Bacteria; Cyanobacteria; Nostocales; Nostocaceae; Nostoc.

COO91__locus_1_main_sat

CsomeBCy

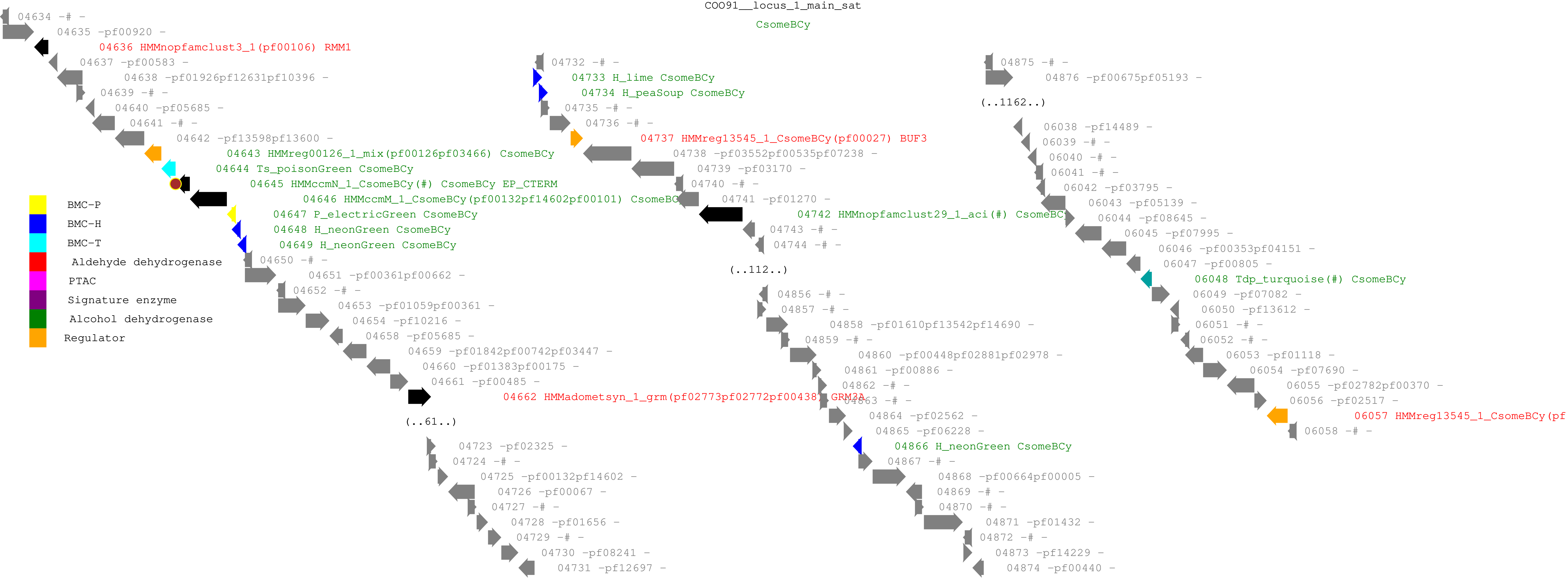

Tychonema bourrellyi FEM_GT703. Bacteria; Cyanobacteria; Oscillatoriophycideae; Oscillatoriales;

CP500__locus_1_main_sat

CsomeBCy

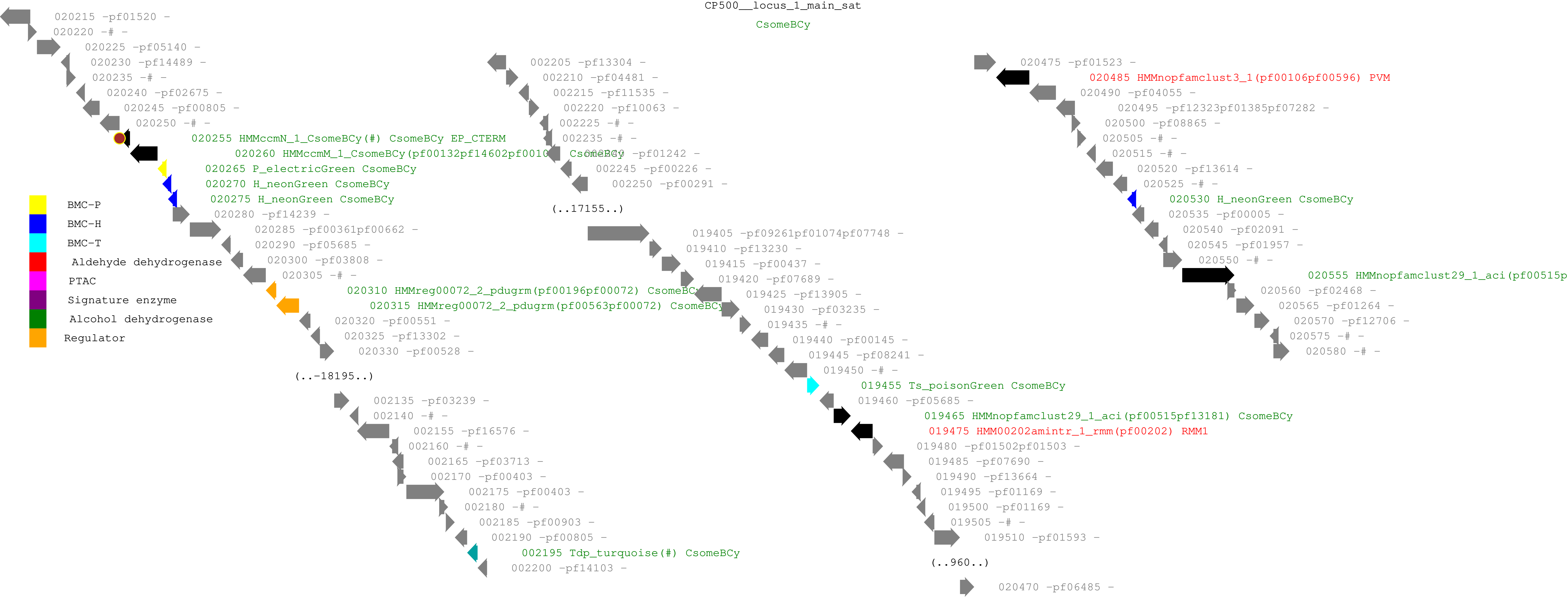

Cylindrospermopsis raciborskii CS-505. Bacteria; Cyanobacteria; Nostocales; Aphanizomenonaceae;

CRC__locus_1_main_sat

CsomeBCy

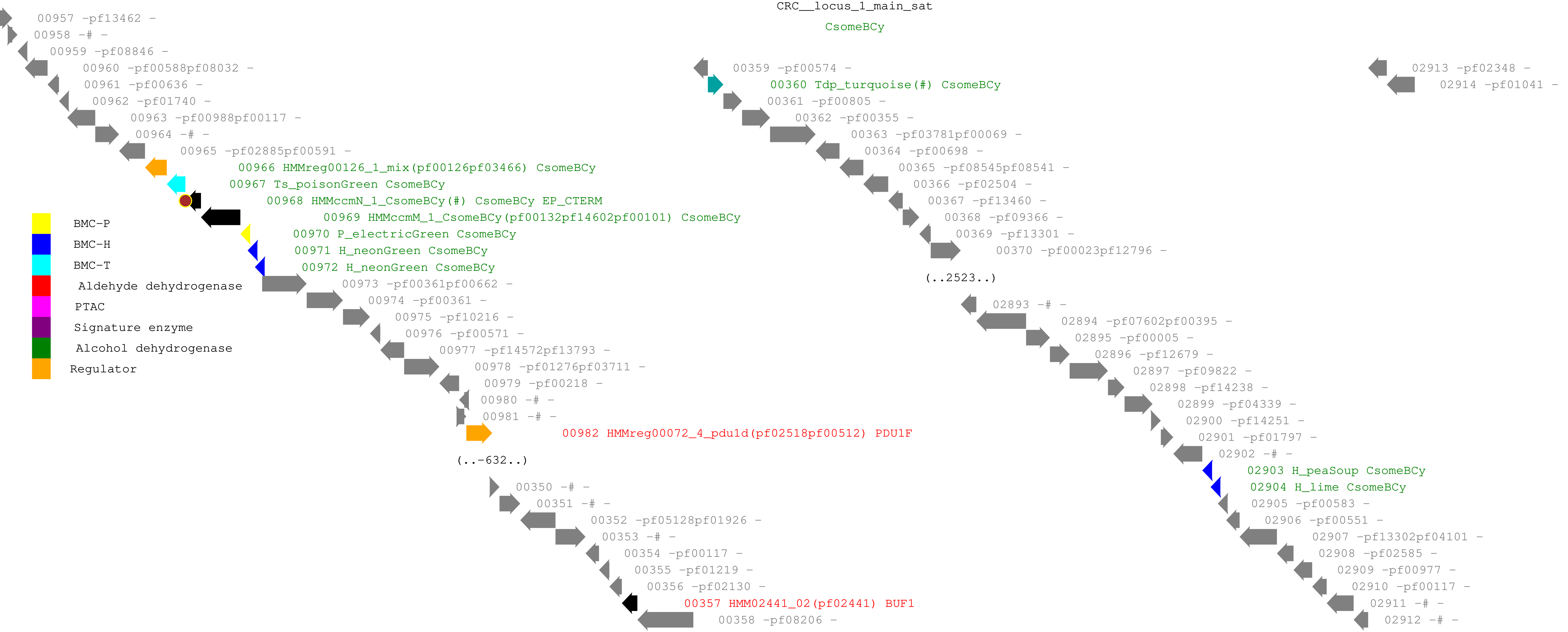

Raphidiopsis brookii D9. Bacteria; Cyanobacteria; Nostocales; Aphanizomenonaceae; Raphidiopsis.

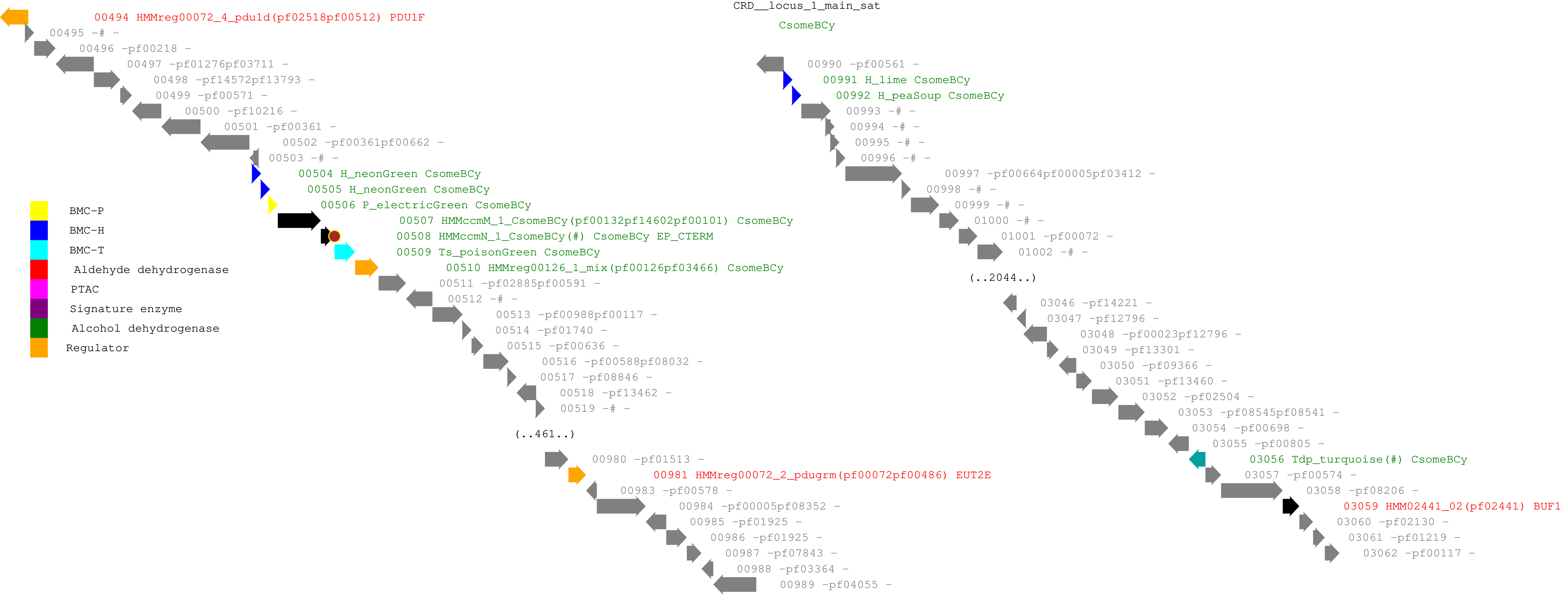

Crinalium epipsammum PCC 9333. Bacteria; Cyanobacteria; Oscillatoriophycideae; Oscillatoriales;

Cri9333__locus_1_main_sat

CsomeBCy

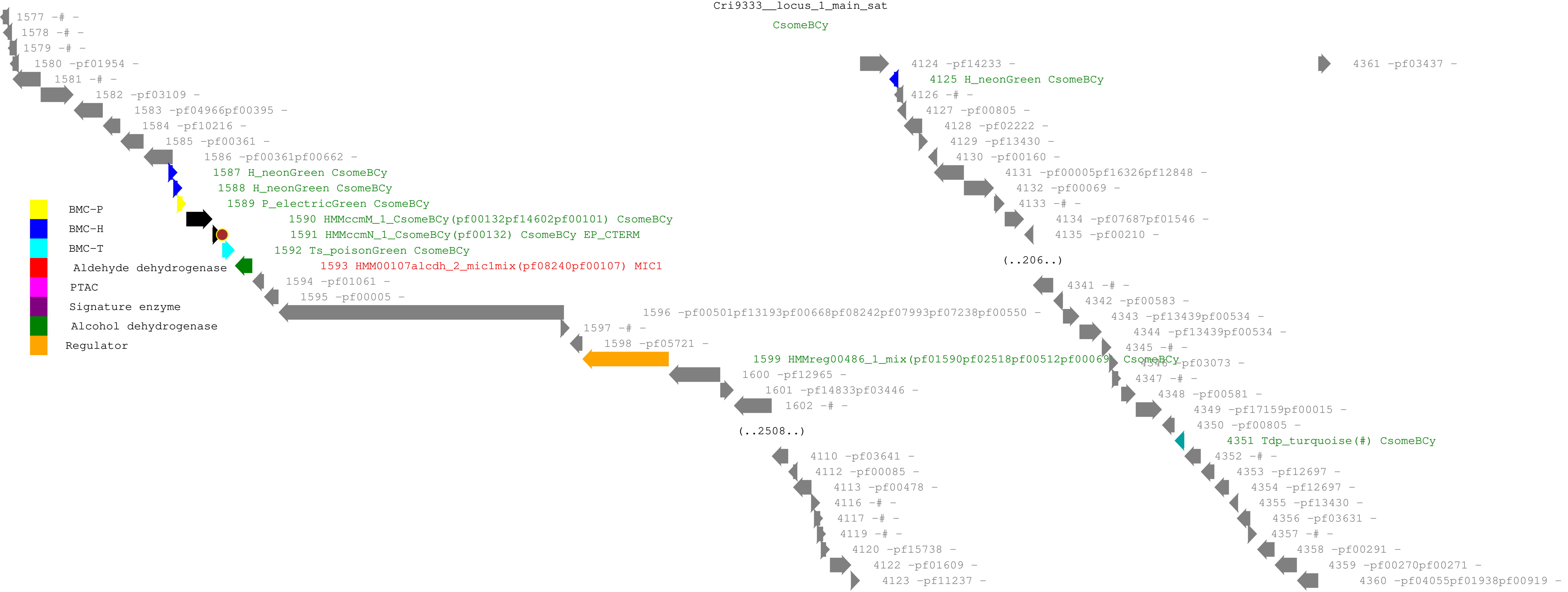

Chroococcales cyanobacterium IPPAS B-1203. Bacteria; Cyanobacteria; Oscillatoriophycideae; Chroococcales.

CSQ79__locus_1_main_sat

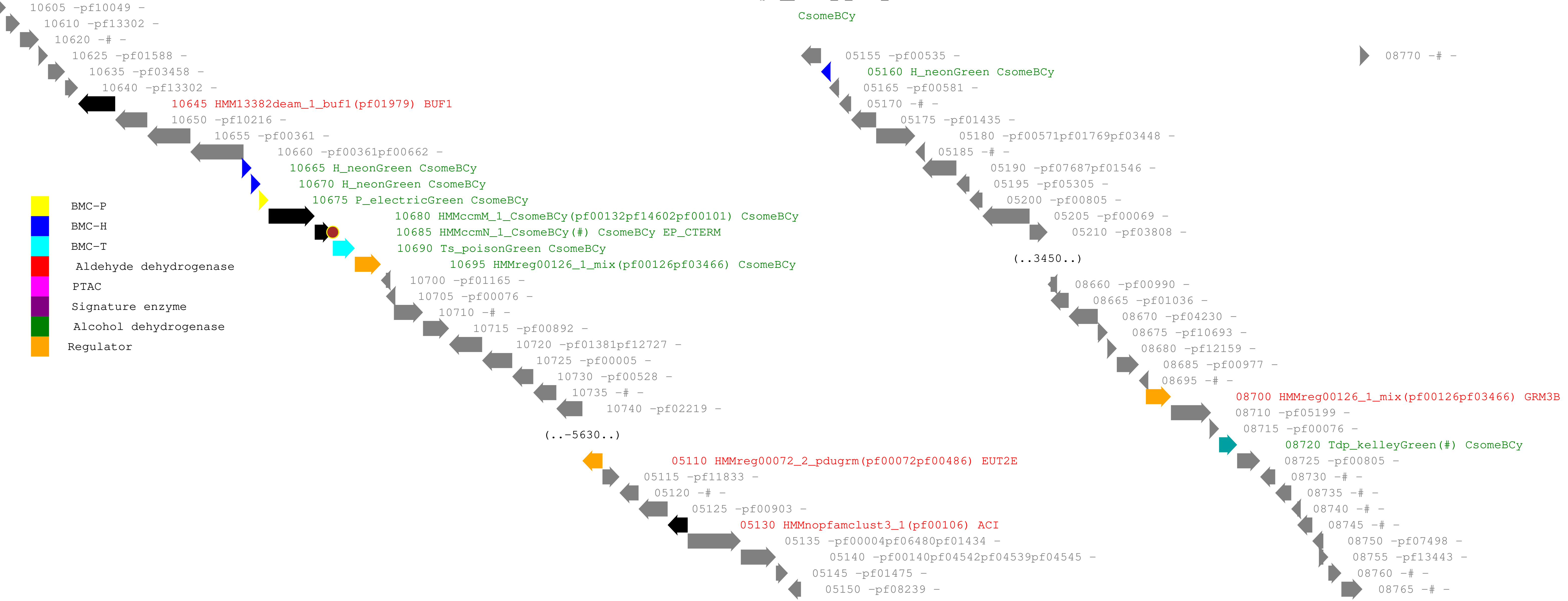

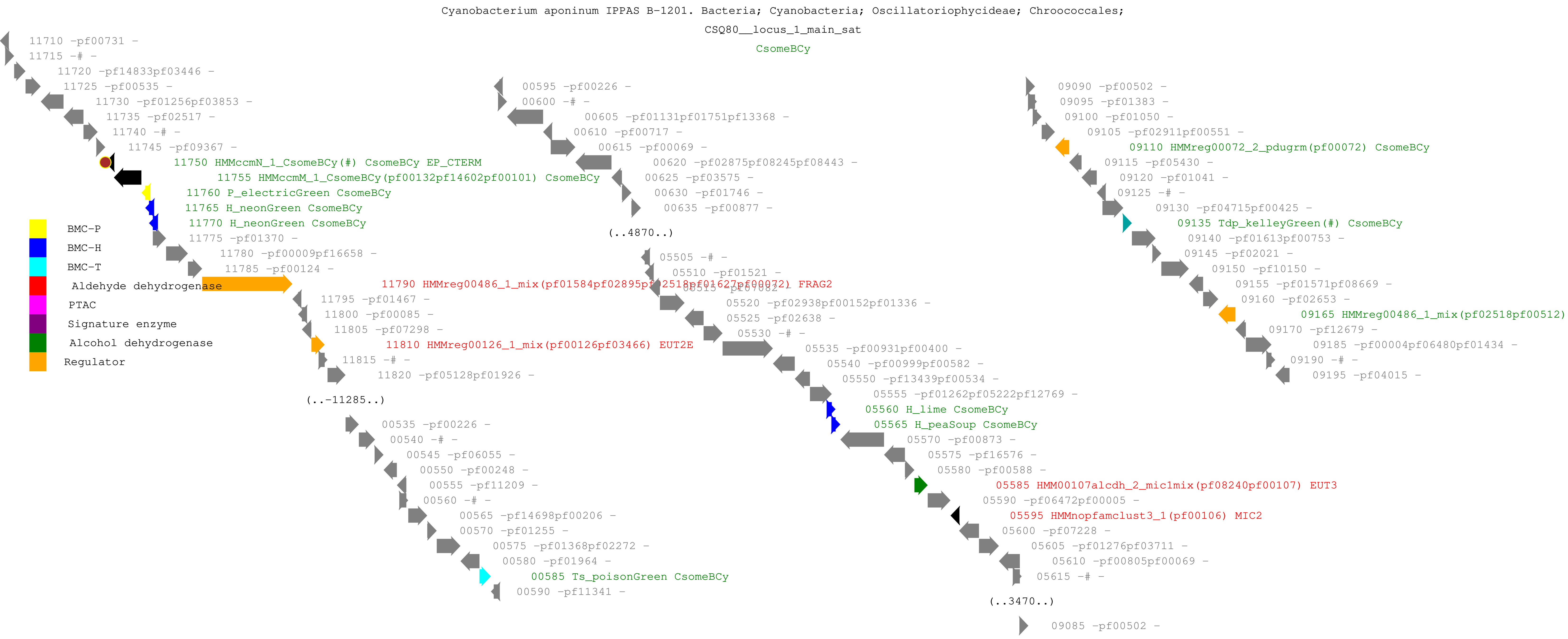
Cyanobacterium aponinum IPPAS B-1201. Bacteria; Cyanobacteria; Oscillatoriophycideae; Chroococcales;
CSQ80__locus_1_main_sat
CsomeBCy
11710 -pf00731 -
11715 -# -
11720 -pf14833pf03446 -
11725 -pf00535 -
11730 -pf01256pf03853 -
11735 -pf02517 -
11740 -# -
11745 -pf09367 -
11750 HMMccmN_1_CsomeBCy(#) CsomeBCy EP_CTERM
11755 HMMccmM_1_CsomeBCy(pf00132pf14602pf00101) CsomeBCy
11760 P_electricGreen CsomeBCy
11765 H_neonGreen CsomeBCy
11770 H_neonGreen CsomeBCy
11775 -pf01370 -
11780 -pf00009pf16658 -
11785 -pf00124 -
11790 HMMreg00486_1_mix(pf01584pf02895pf02518pf01627pf00072) FRAG2
11795 -pf01467 -
11800 -pf00085 -
11805 -pf07298 -
11810 HMMreg00126_1_mix(pf00126pf03466) EUT2E
11815 -# -
11820 -pf05128pf01926 -
(..-11285..)
00535 -pf00226 -
00540 -# -
00545 -pf06055 -
00550 -pf00248 -
00555 -pf11209 -
00560 -# -
00565 -pf14698pf00206 -
00570 -pf01255 -
00575 -pf01368pf02272 -
00580 -pf01964 -
00585 Ts_poisonGreen CsomeBCy
00590 -pf11341 -
BMC-P
BMC-H
BMC-T
Aldehyde dehydrogenase
PTAC
Signature enzyme
Alcohol dehydrogenase
Regulator
00595 -pf00226 -
00600 -# -
00605 -pf01131pf01751pf13368 -
00610 -pf00717 -
00615 -pf00069 -
00620 -pf02875pf08245pf08443 -
00625 -pf03575 -
00630 -pf01746 -
00635 -pf00877 -
(..4870..)
05505 -# -
05510 -pf01521 -
05515 -pf07082 -
05520 -pf02938pf00152pf01336 -
05525 -pf02638 -
05530 -# -
05535 -pf00931pf00400 -
05540 -pf00999pf00582 -
05550 -pf13439pf00534 -
05555 -pf01262pf05222pf12769 -
05560 H_lime CsomeBCy
05565 H_peaSoup CsomeBCy
05570 -pf00873 -
05575 -pf16576 -
05580 -pf00588 -
05585 HMM00107alcdh_2_mic1mix(pf08240pf00107) EUT3
05590 -pf06472pf00005 -
05595 HMMnopfamclust3_1(pf00106) MIC2
05600 -pf07228 -
05605 -pf01276pf03711 -
05610 -pf00805pf00069 -
05615 -# -
(..3470..)
09085 -pf00502 -
09090 -pf00502 -
09095 -pf01383 -
09100 -pf01050 -
09105 -pf02911pf00551 -
09110 HMMreg00072_2_pdugrm(pf00072) CsomeBCy
09115 -pf05430 -
09120 -pf01041 -
09125 -# -
09130 -pf04715pf00425 -
09135 Tdp_kelleyGreen(#) CsomeBCy
09140 -pf01613pf00753 -
09145 -pf02021 -
09150 -pf10150 -
09155 -pf01571pf08669 -
09160 -pf02653 -
09165 HMMreg00486_1_mix(pf02518pf00512)
09170 -pf12679 -
09185 -pf00004pf06480pf01434 -
09190 -# -
09195 -pf04015 -

Cuspidothrix issatschenkoi CHARLIE-1. Bacteria; Cyanobacteria; Nostocales; Aphanizomenonaceae; Cuspidothrix.

CUN59__locus_1_main_sat

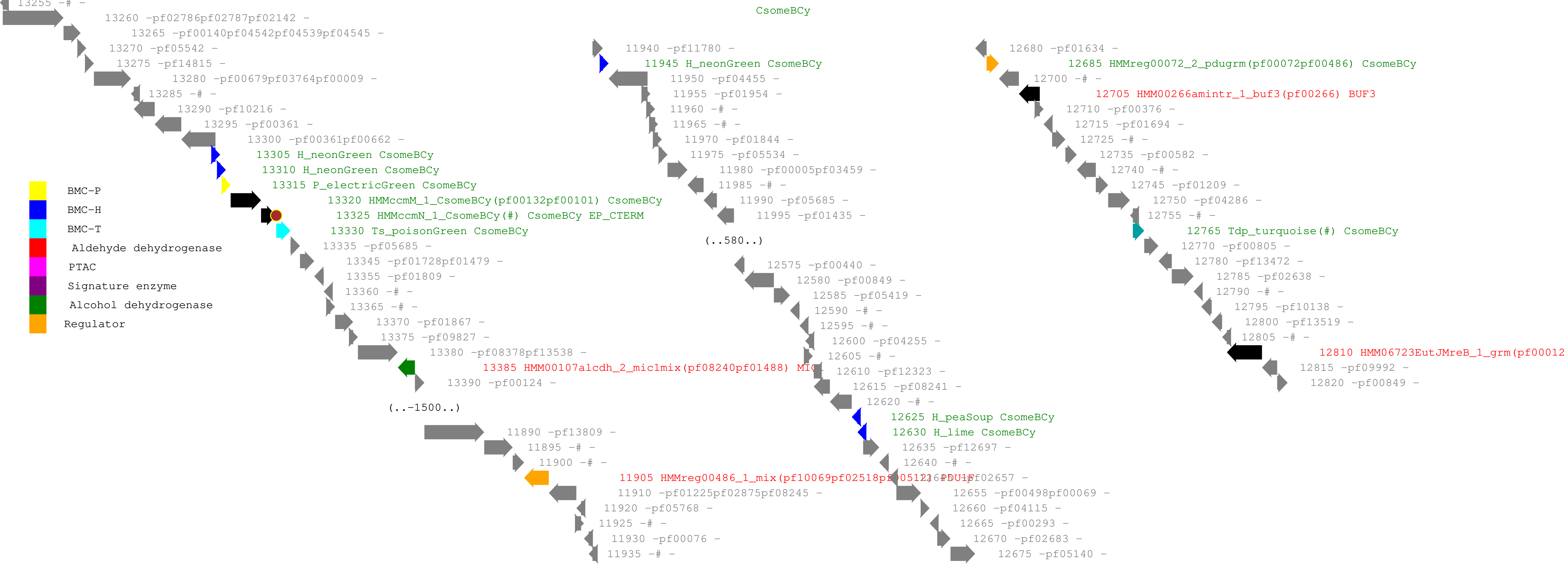

Cyanobacteria bacterium M5B4. Bacteria; Cyanobacteria.

CV045__locus_fragmented

CsomeBCy

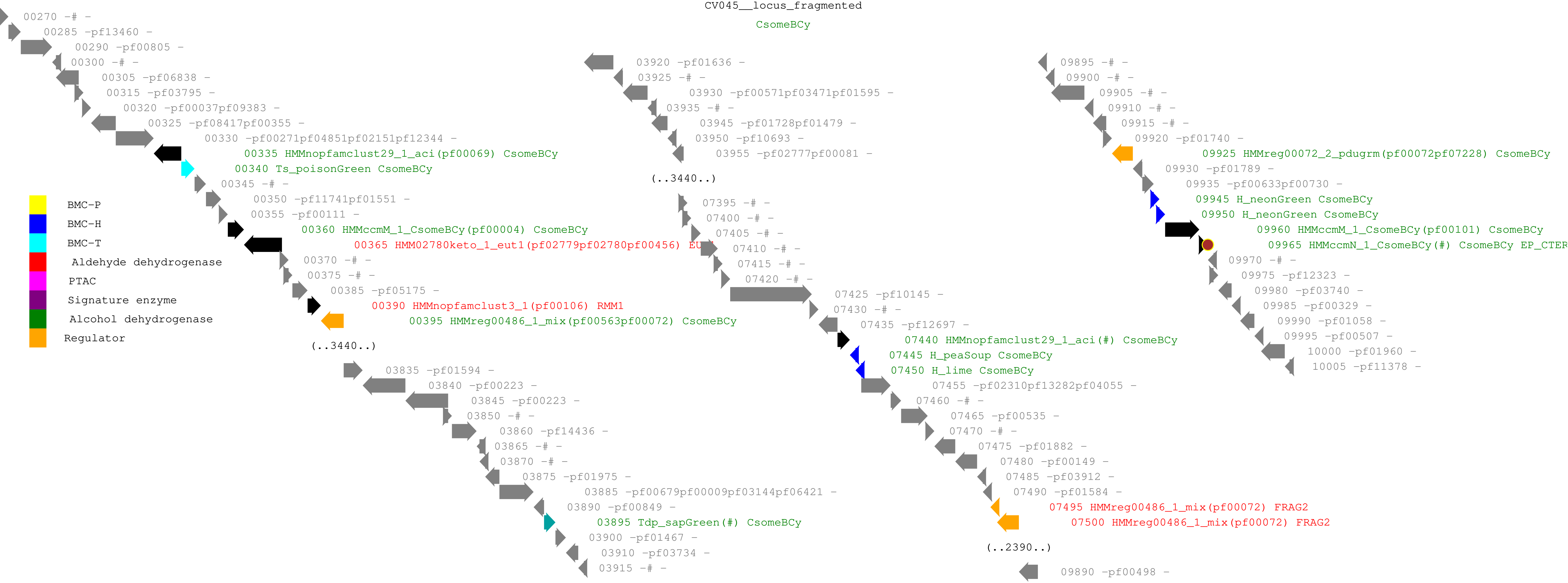

Crocosphaera watsonii WH 8501. Bacteria; Cyanobacteria; Oscillatoriophycideae; Chroococcales;

CwatDRAFT__locus_1_main_sat

CsomeBCy

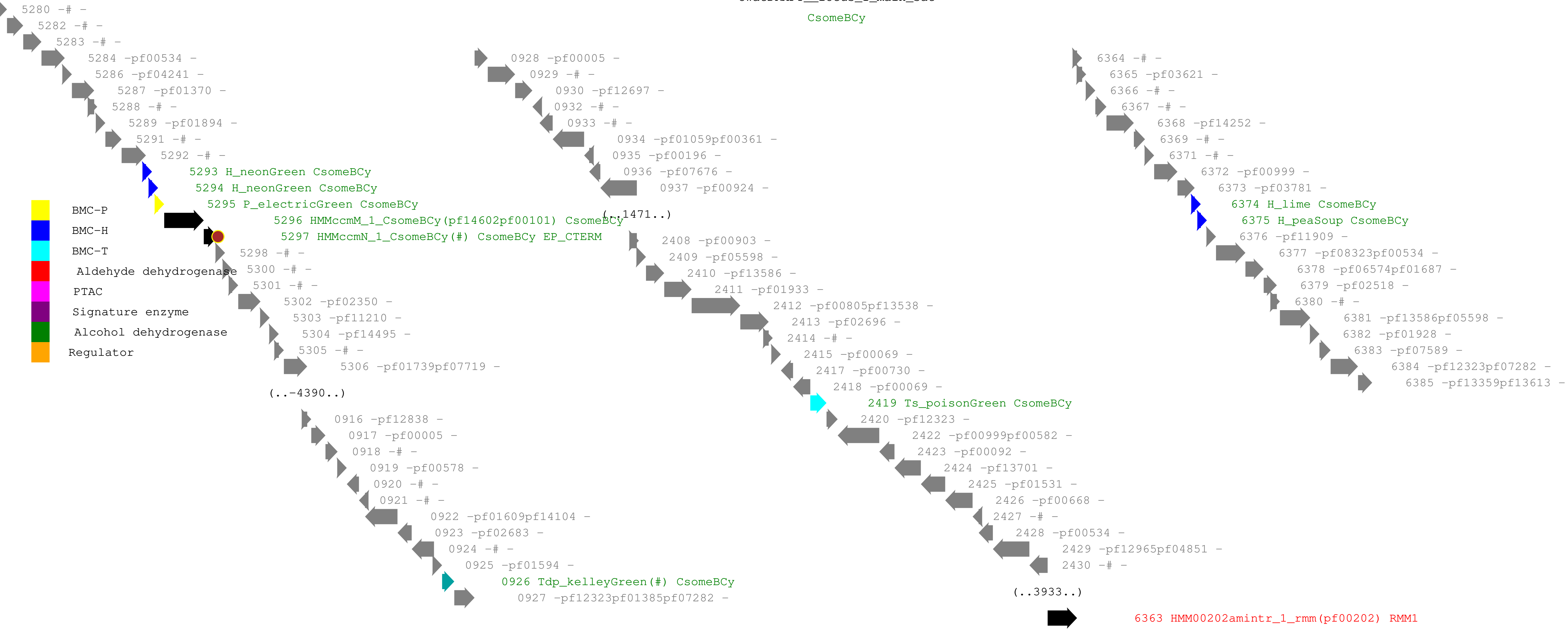

Crocosphaera watsonii WH 0003. Bacteria; Cyanobacteria; Oscillatoriophycideae; Chroococcales;

CWATWH0003__locus_1_main_sat

CsomeBCy

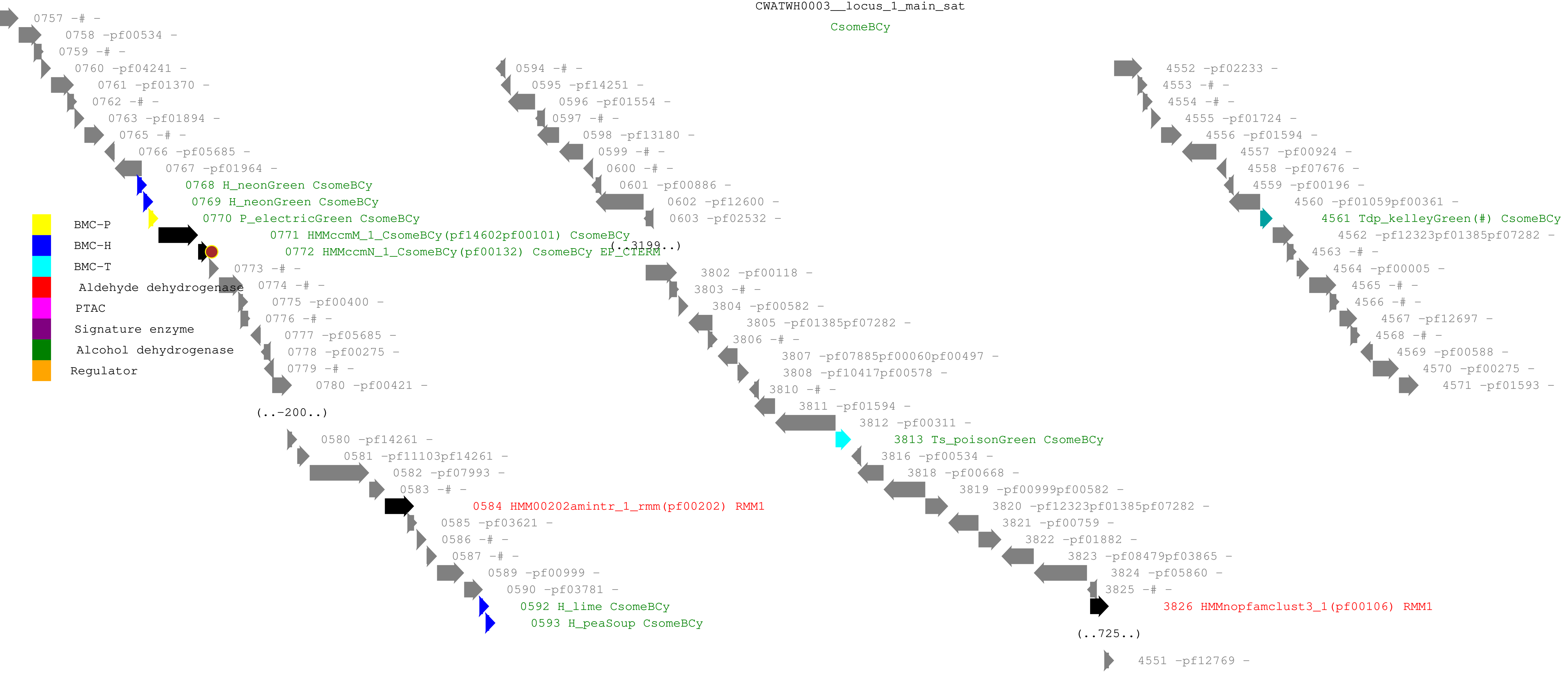

Crocosphaera watsonii WH 0005. Bacteria; Cyanobacteria; Oscillatoriophycideae; Chroococcales;

CWATWH0005__locus_fragmented

CsomeBCy

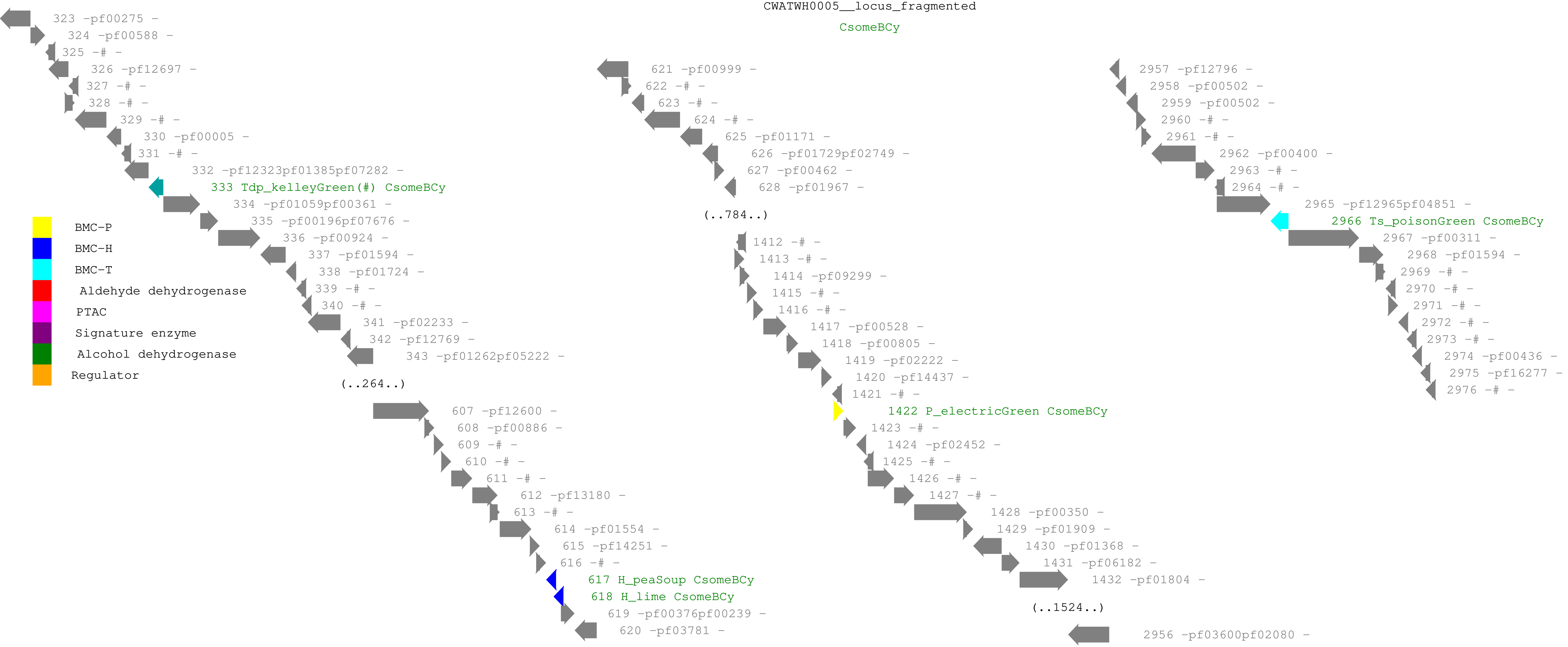

Crocosphaera watsonii WH 0401. Bacteria; Cyanobacteria; Oscillatoriophycideae; Chroococcales;

CWATWH0401__locus_1_main_sat

CsomeBCy

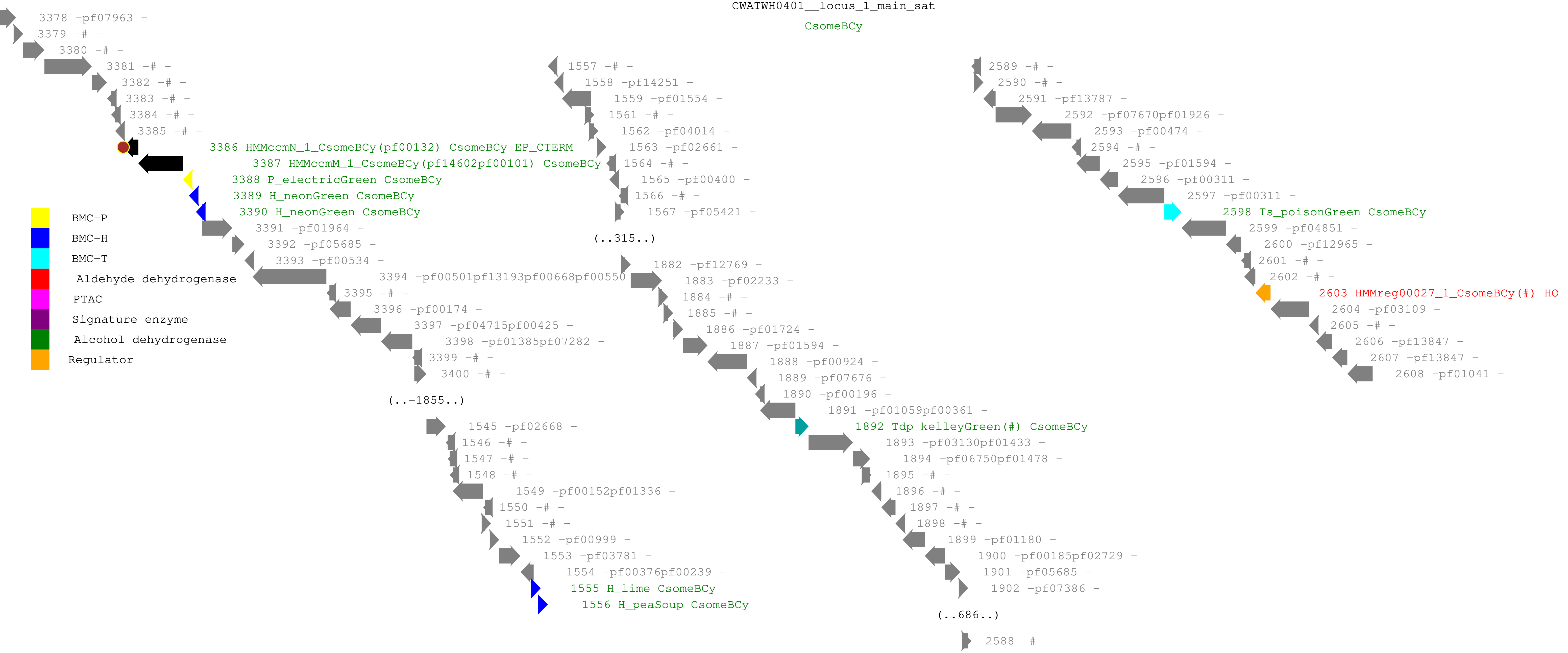

Crocosphaera watsonii WH 0402. Bacteria; Cyanobacteria; Oscillatoriophycideae; Chroococcales;

CWATWH0402__locus_1_main_sat

CsomeBCy

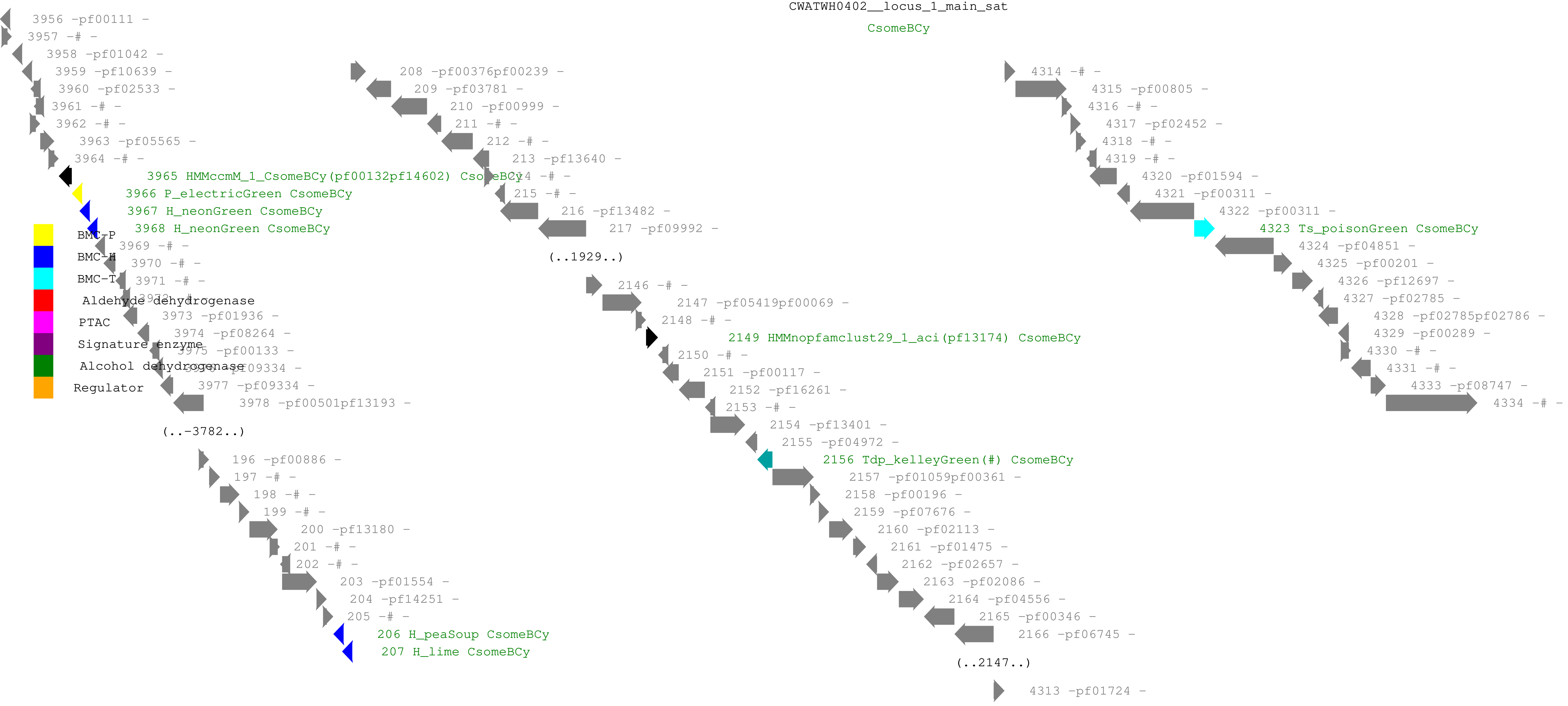

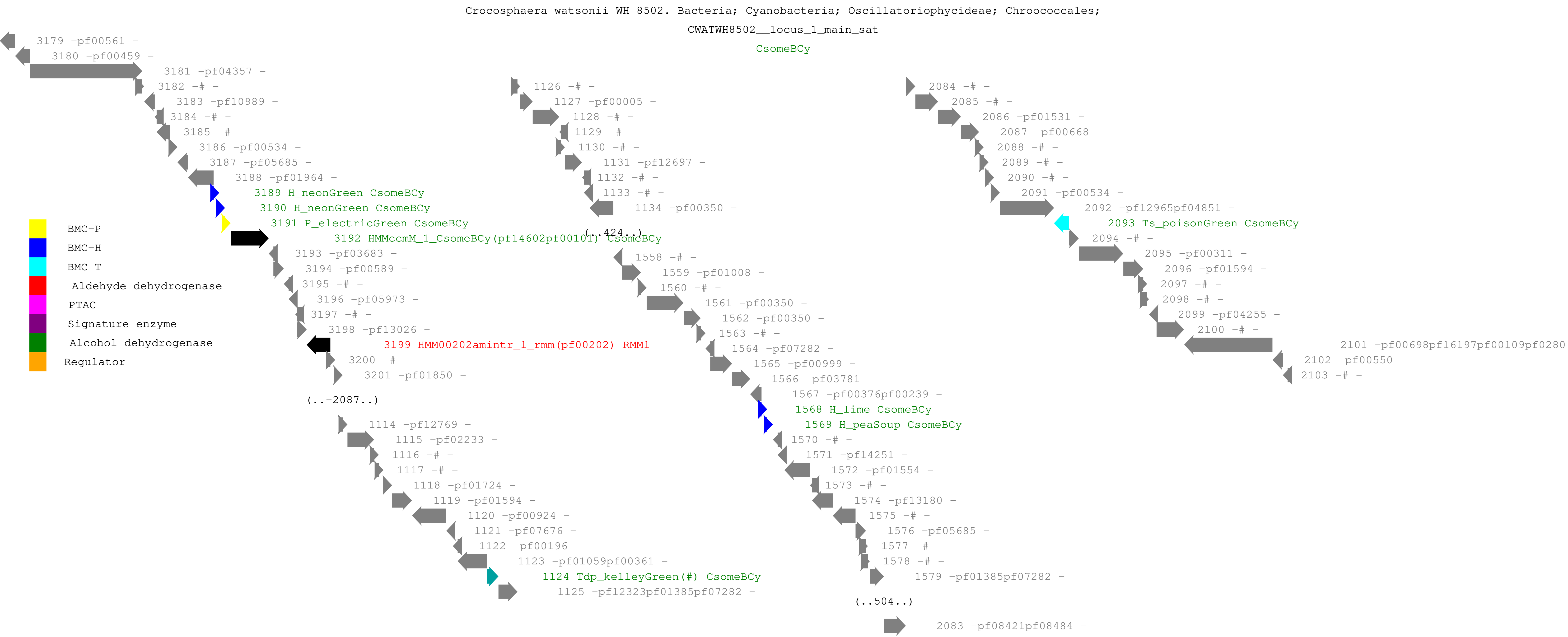
Crocosphaera watsonii WH 8502. Bacteria; Cyanobacteria; Oscillatoriophycideae; Chroococcales;
CWATWH8502__locus_1_main_sat
CsomeBCy
3179 -pf00561 -
3180 -pf00459 -
3181 -pf04357 -
3182 -# -
3183 -pf10989 -
3184 -# -
3185 -# -
3186 -pf00534 -
3187 -pf05685 -
3188 -pf01964 -
3189 H_neonGreen CsomeBCy
3190 H_neonGreen CsomeBCy
3191 P_electricGreen CsomeBCy
3192 HMMccmM_1_CsomeBCy(pf14602pf00101) CsomeBCy
3193 -pf03683 -
3194 -pf00589 -
3195 -# -
3196 -pf05973 -
3197 -# -
3198 -pf13026 -
3199 HMM00202amintr_1_rmm(pf00202) RMM1
3200 -# -
3201 -pf01850 -
(..-2087..)
1114 -pf12769 -
1115 -pf02233 -
1116 -# -
1117 -# -
1118 -pf01724 -
1119 -pf01594 -
1120 -pf00924 -
1121 -pf07676 -
1122 -pf00196 -
1123 -pf01059pf00361 -
1124 Tdp_kelleyGreen(#) CsomeBCy
1125 -pf12323pf01385pf07282 -
1126 -# -
1127 -pf00005 -
1128 -# -
1129 -# -
1130 -# -
1131 -pf12697 -
1132 -# -
1133 -# -
1134 -pf00350 -
(..424..)
1558 -# -
1559 -pf01008 -
1560 -# -
1561 -pf00350 -
1562 -pf00350 -
1563 -# -
1564 -pf07282 -
1565 -pf00999 -
1566 -pf03781 -
1567 -pf00376pf00239 -
1568 H_lime CsomeBCy
1569 H_peaSoup CsomeBCy
1570 -# -
1571 -pf14251 -
1572 -pf01554 -
1573 -# -
1574 -pf13180 -
1575 -# -
1576 -pf05685 -
1577 -# -
1578 -# -
1579 -pf01385pf07282 -
(..504..)
2083 -pf08421pf08484 -
2084 -# -
2085 -# -
2086 -pf01531 -
2087 -pf00668 -
2088 -# -
2089 -# -
2090 -# -
2091 -pf00534 -
2092 -pf12965pf04851 -
2093 Ts_poisonGreen CsomeBCy
2094 -# -
2095 -pf00311 -
2096 -pf01594 -
2097 -# -
2098 -# -
2099 -pf04255 -
2100 -# -
2101 -pf00698pf16197pf00109pf0280
2102 -pf00550 -
2103 -# -
BMC-P
BMC-H
BMC-T
Aldehyde dehydrogenase
PTAC
Signature enzyme
Alcohol dehydrogenase
Regulator

Cyanothece sp. CCY0110. Bacteria; Cyanobacteria; Oscillatoriophycideae; Oscillatoriales;

CY0110__locus_1_main_sat

CsomeBCy

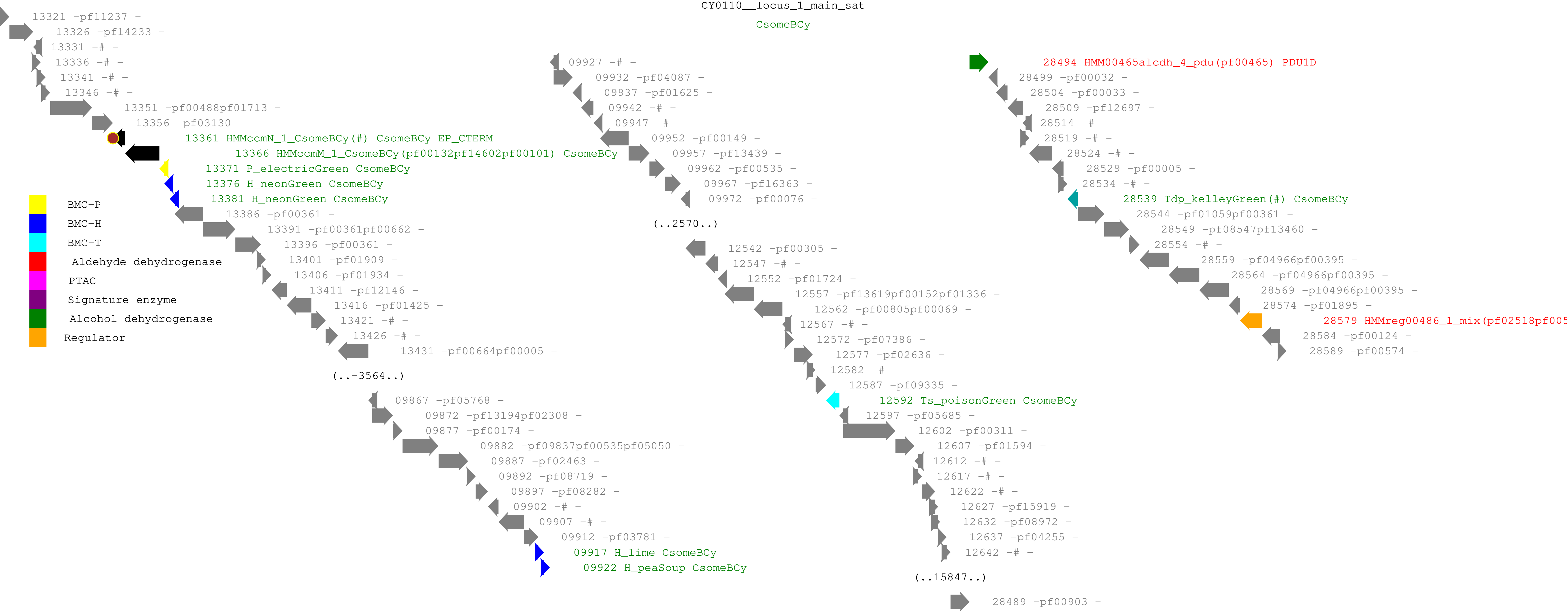

Synechococcus sp. (strain JA-3-3Ab) (Cyanobacteria bacterium Bacteria; Cyanobacteria; Synechococcales; Synechococcaceae;

CYA__locus_1_main_sat

CsomeBCy

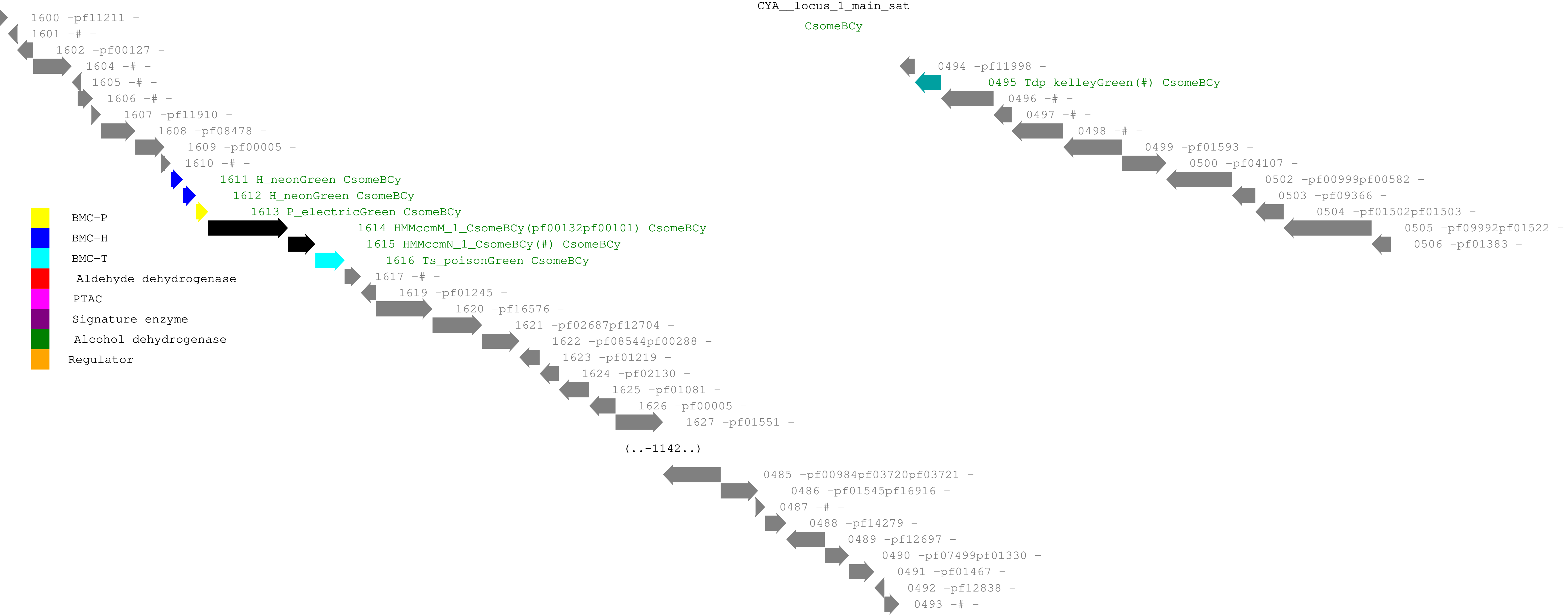

Cyanobacterium aponinum (strain PCC 10605). Bacteria; Cyanobacteria; Oscillatoriophycideae; Chroococcales;

Cyan10605__locus_1_main_sat

CsomeBCy

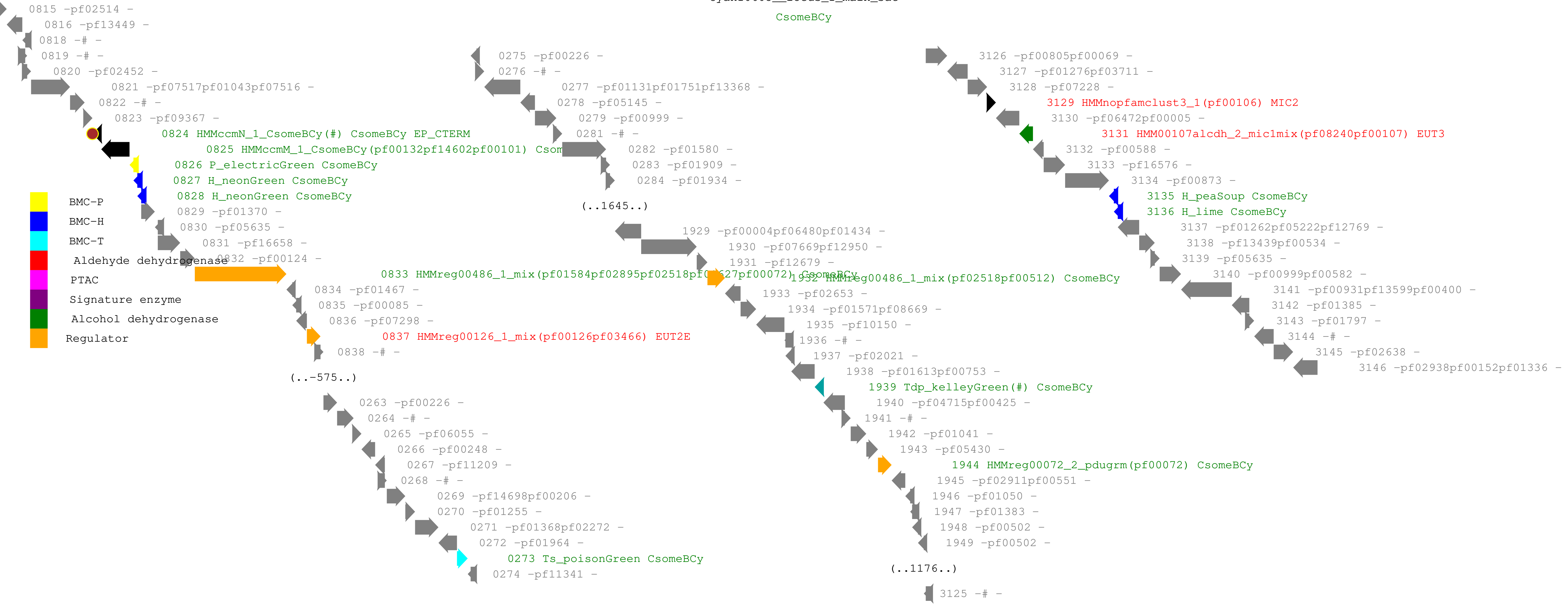

Cyanothece sp. (strain PCC 7425 / ATCC 29141). Bacteria; Cyanobacteria; Oscillatoriophycideae; Oscillatoriales;

Cyan7425__locus_1_main_sat

CsomeBCy

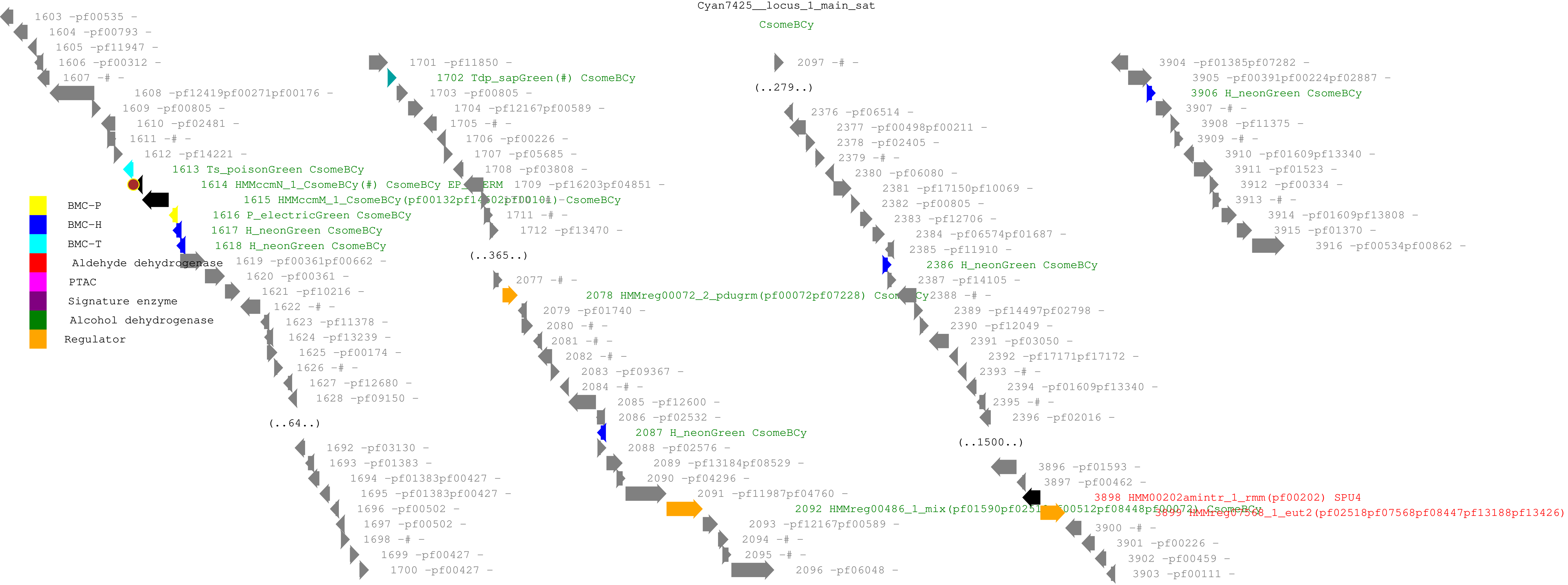

Cyanothece sp. (strain PCC 7822). Bacteria; Cyanobacteria; Oscillatoriophycideae; Oscillatoriales;

Cyan7822__locus_1_main_sat

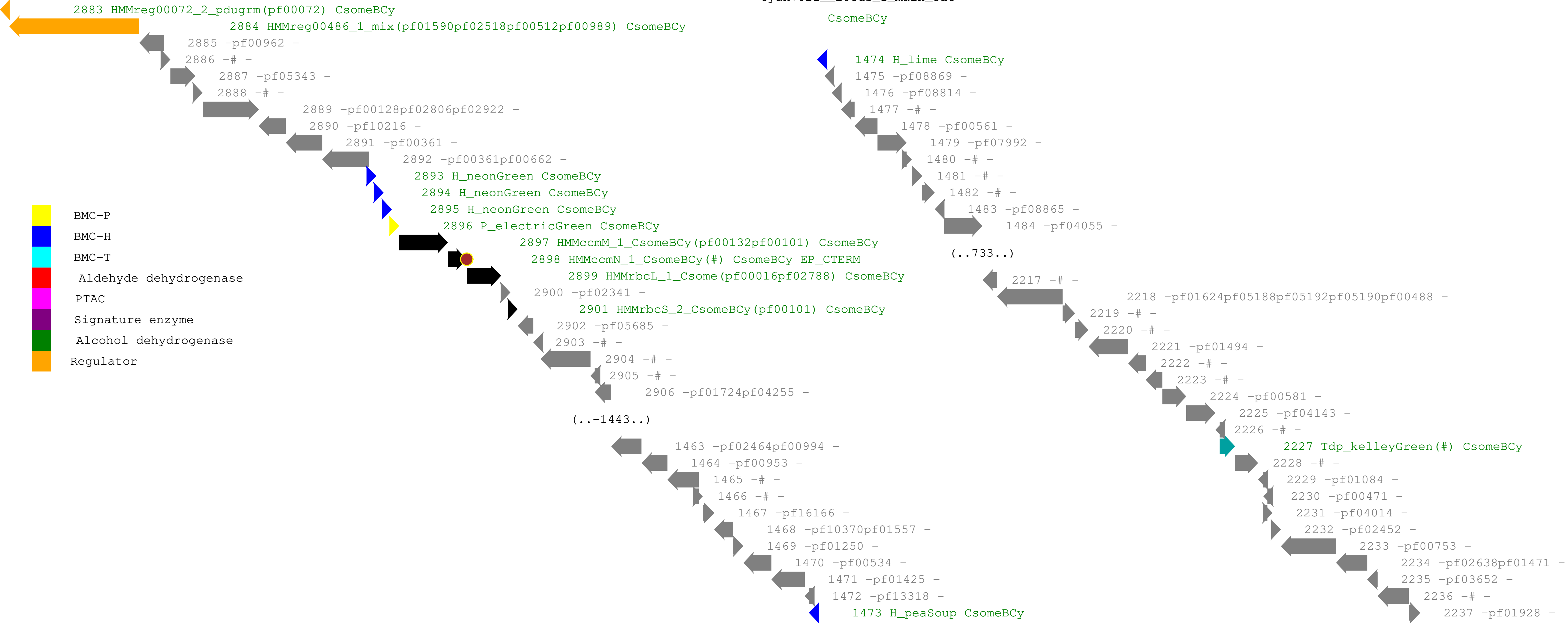

Cyanobacterium stanieri (strain ATCC 29140 / PCC 7202). Bacteria; Cyanobacteria; Oscillatoriophycideae; Chroococcales;

Cyast__locus_1_main_sat

CsomeBCy

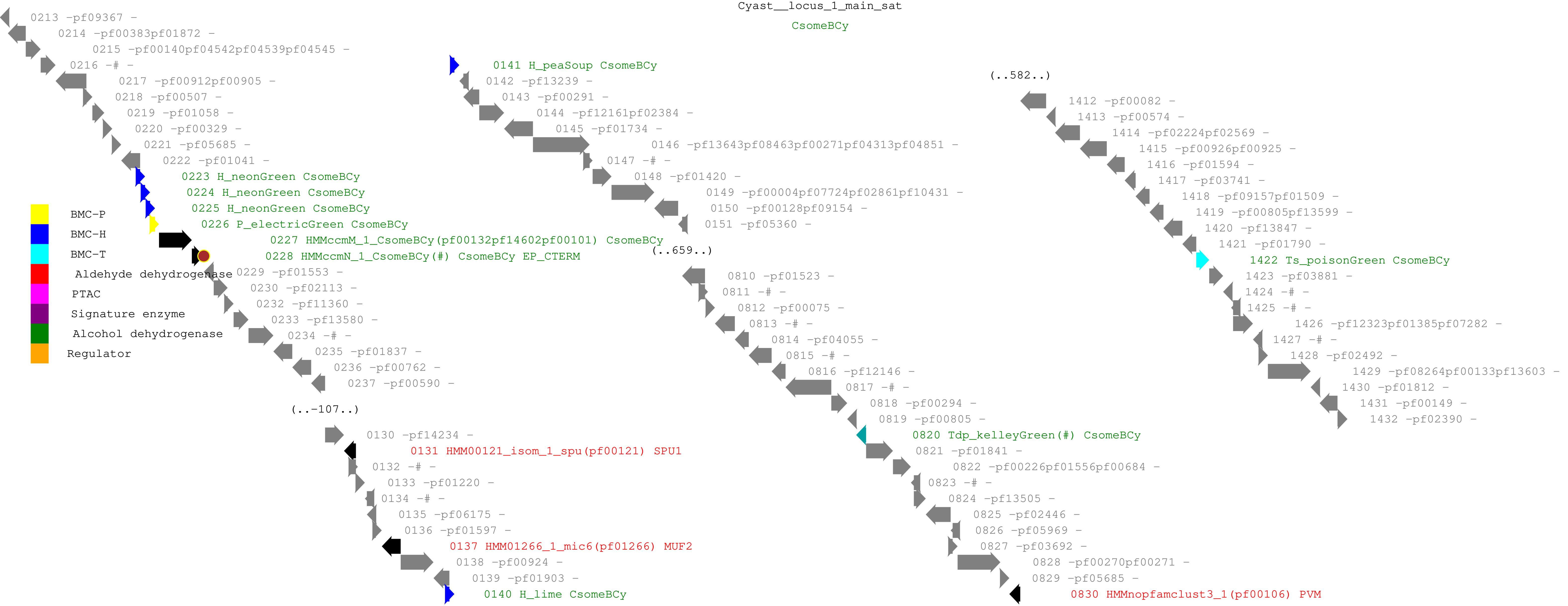

Synechococcus sp. (strain JA-2-3B'a(2-13)) (Cyanobacteria bacterium Bacteria; Cyanobacteria; Synechococcales; Synechococcaceae;

CYB__locus_1_main_sat

CsomeBCy

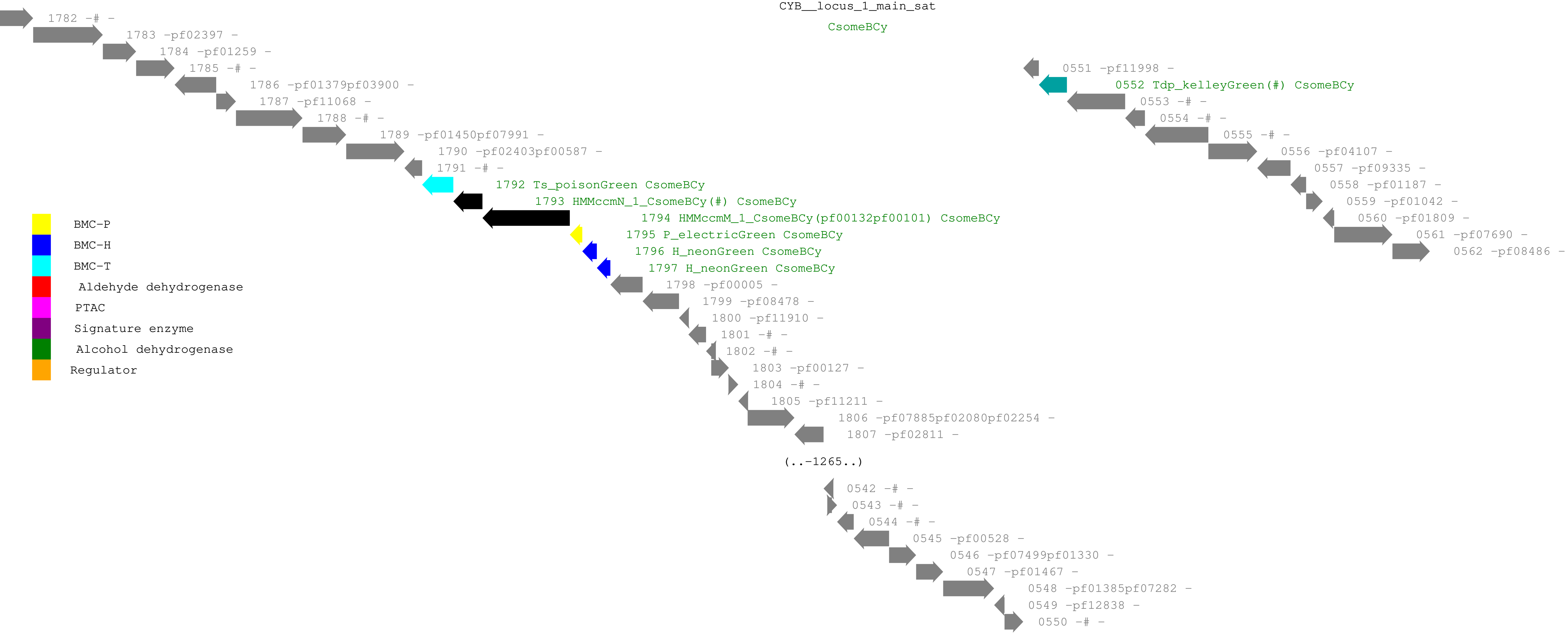

Cylindrospermum stagnale PCC 7417. Bacteria; Cyanobacteria; Nostocales; Nostocaceae; Cylindrospermum.

Cylst__locus_1_main_sat

CsomeBCy

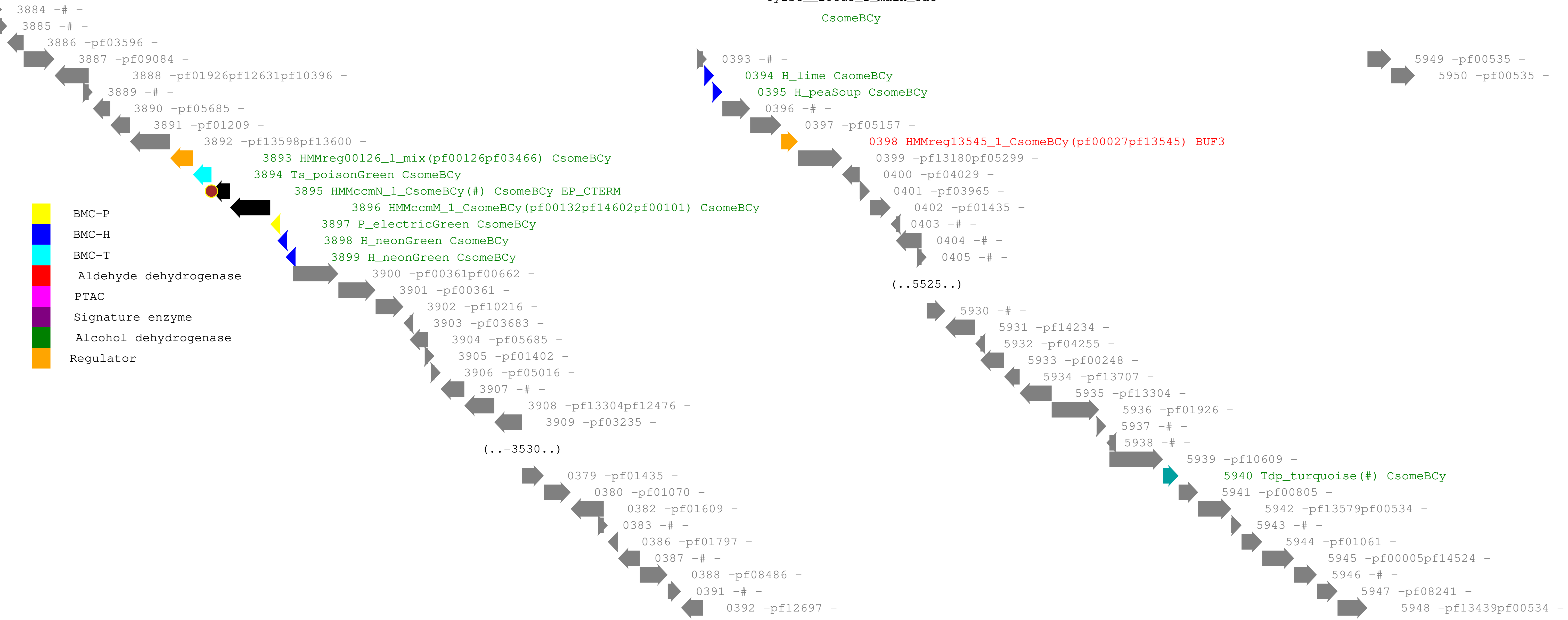

Synechocystis sp. (strain PCC 6714) (Aphanocapsa sp. (strain PCC Bacteria; Cyanobacteria; Synechococcales; Merismopediaceae;

D082__locus_1_main_sat

CsomeBCy

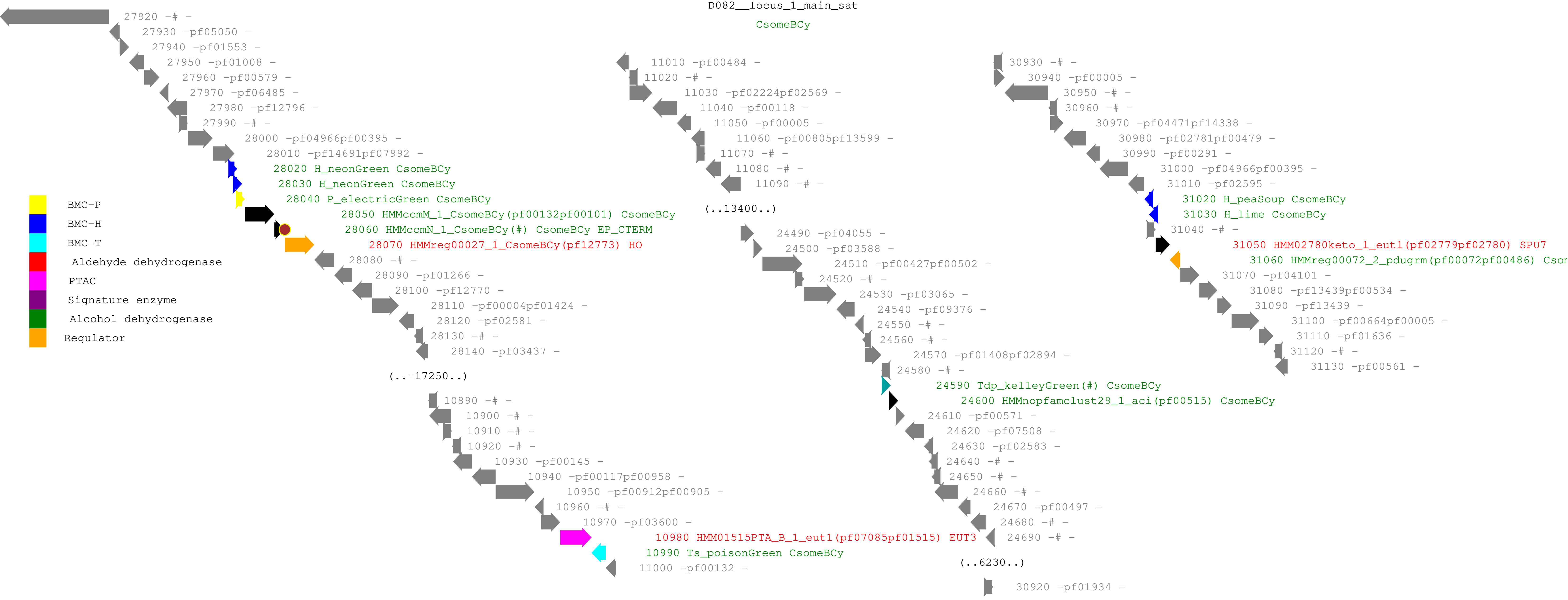

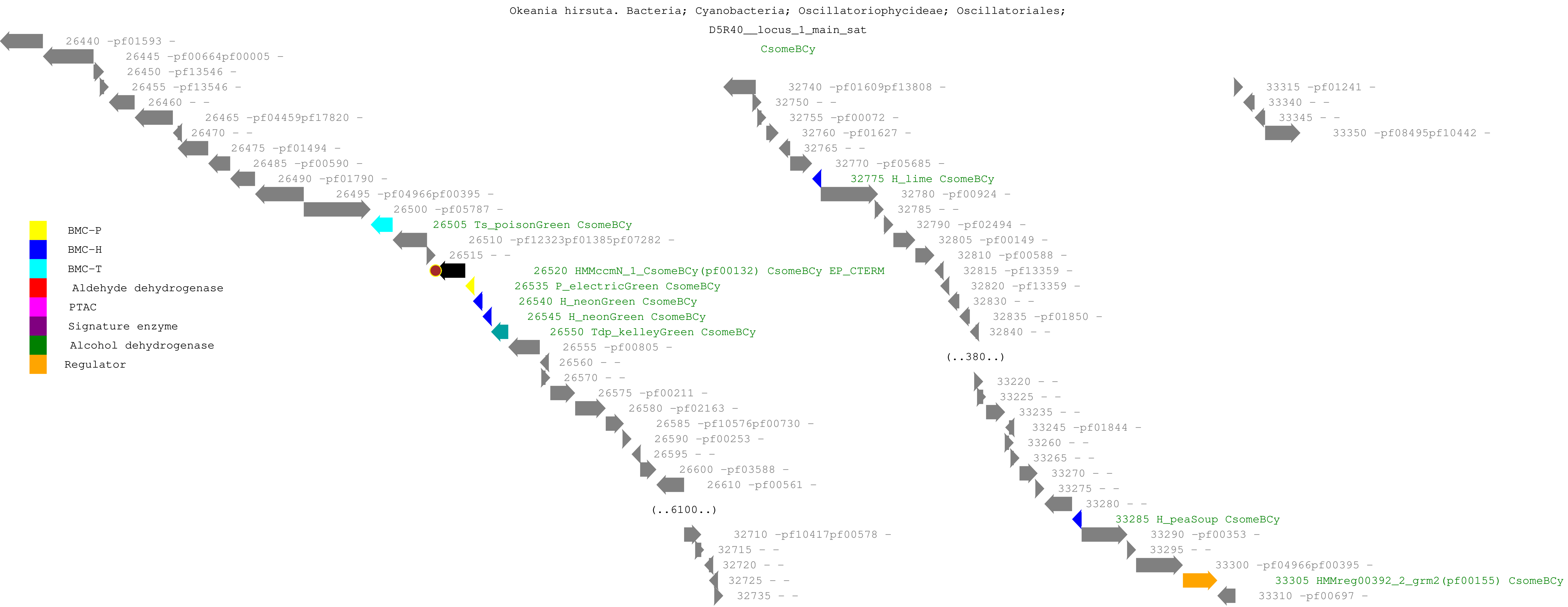

Okeania hirsuta. Bacteria; Cyanobacteria; Oscillatoriophycideae; Oscillatoriales;
D5R40__locus_1_main_sat
CsomeBCy
26440 -pf01593 -
26445 -pf00664pf00005 -
26450 -pf13546 -
26455 -pf13546 -
26460 - -
26465 -pf04459pf17820 -
26470 - -
26475 -pf01494 -
26485 -pf00590 -
26490 -pf01790 -
26495 -pf04966pf00395 -
26500 -pf05787 -
26505 Ts_poisonGreen CsomeBCy
26510 -pf12323pf01385pf07282 -
26515 - -
26520 HMMccmN_1_CsomeBCy(pf00132) CsomeBCy EP_CTERM
26535 P_electricGreen CsomeBCy
26540 H_neonGreen CsomeBCy
26545 H_neonGreen CsomeBCy
26550 Tdp_kelleyGreen CsomeBCy
26555 -pf00805 -
26560 - -
26570 - -
26575 -pf00211 -
26580 -pf02163 -
26585 -pf10576pf00730 -
26590 -pf00253 -
26595 - -
26600 -pf03588 -
26610 -pf00561 -
(..6100..)
32710 -pf10417pf00578 -
32715 - -
32720 - -
32725 - -
32735 - -
32740 -pf01609pf13808 -
32750 - -
32755 -pf00072 -
32760 -pf01627 -
32765 - -
32770 -pf05685 -
32775 H_lime CsomeBCy
32780 -pf00924 -
32785 - -
32790 -pf02494 -
32805 -pf00149 -
32810 -pf00588 -
32815 -pf13359 -
32820 -pf13359 -
32830 - -
32835 -pf01850 -
32840 - -
(..380..)
33220 - -
33225 - -
33235 - -
33245 -pf01844 -
33260 - -
33265 - -
33270 - -
33275 - -
33280 - -
33285 H_peaSoup CsomeBCy
33290 -pf00353 -
33295 - -
33300 -pf04966pf00395 -
33305 HMMreg00392_2_grm2(pf00155) CsomeBCy
33310 -pf00697 -
33315 -pf01241 -
33340 - -
33345 - -
33350 -pf08495pf10442 -
BMC-P
BMC-H
BMC-T
Aldehyde dehydrogenase
PTAC
Signature enzyme
Alcohol dehydrogenase
Regulator

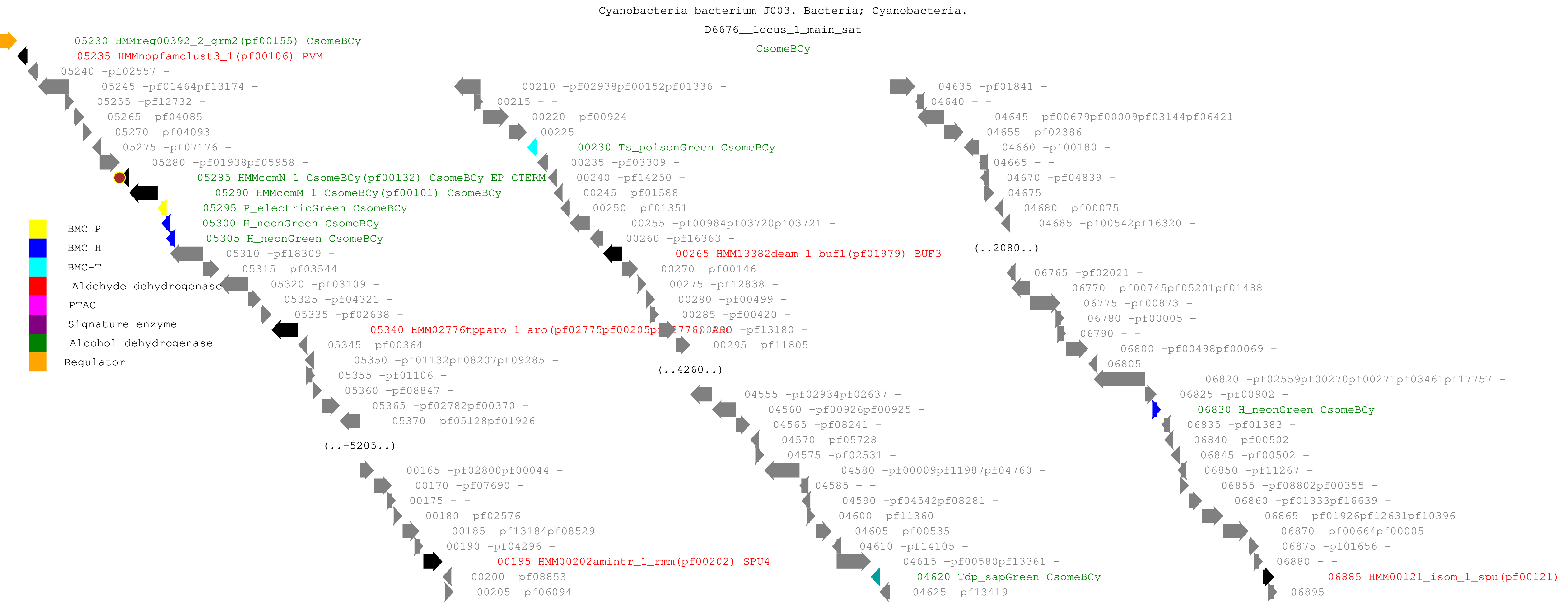

Cyanobacteria bacterium J003. Bacteria; Cyanobacteria.
D6676__locus_1_main_sat
CsomeBCy
05230 HMMreg00392_2_grm2(pf00155) CsomeBCy
05235 HMMnopfamclust3_1(pf00106) PVM
05240 -pf02557 -
05245 -pf01464pf13174 -
05255 -pf12732 -
05265 -pf04085 -
05270 -pf04093 -
05275 -pf07176 -
05280 -pf01938pf05958 -
05285 HMMccmN_1_CsomeBCy(pf00132) CsomeBCy EP_CTERM
05290 HMMccmM_1_CsomeBCy(pf00101) CsomeBCy
05295 P_electricGreen CsomeBCy
05300 H_neonGreen CsomeBCy
05305 H_neonGreen CsomeBCy
05310 -pf18309 -
05315 -pf03544 -
05320 -pf03109 -
05325 -pf04321 -
05335 -pf02638 -
05340 HMM02776tpparo_1_aro(pf02775pf00205pf02776) ARO
05345 -pf00364 -
05350 -pf01132pf08207pf09285 -
05355 -pf01106 -
05360 -pf08847 -
05365 -pf02782pf00370 -
05370 -pf05128pf01926 -
(..-5205..)
00165 -pf02800pf00044 -
00170 -pf07690 -
00175 - -
00180 -pf02576 -
00185 -pf13184pf08529 -
00190 -pf04296 -
00195 HMM00202amintr_1_rmm(pf00202) SPU4
00200 -pf08853 -
00205 -pf06094 -
00210 -pf02938pf00152pf01336 -
00215 - -
00220 -pf00924 -
00225 - -
00230 Ts_poisonGreen CsomeBCy
00235 -pf03309 -
00240 -pf14250 -
00245 -pf01588 -
00250 -pf01351 -
00255 -pf00984pf03720pf03721 -
00260 -pf16363 -
00265 HMM13382deam_1_buf1(pf01979) BUF3
00270 -pf00146 -
00275 -pf12838 -
00280 -pf00499 -
00285 -pf00420 -
00290 -pf13180 -
00295 -pf11805 -
(..4260..)
04555 -pf02934pf02637 -
04560 -pf00926pf00925 -
04565 -pf08241 -
04570 -pf05728 -
04575 -pf02531 -
04580 -pf00009pf11987pf04760 -
04585 - -
04590 -pf04542pf08281 -
04600 -pf11360 -
04605 -pf00535 -
04610 -pf14105 -
04615 -pf00580pf13361 -
04620 Tdp_sapGreen CsomeBCy
04625 -pf13419 -
04635 -pf01841 -
04640 - -
04645 -pf00679pf00009pf03144pf06421 -
04655 -pf02386 -
04660 -pf00180 -
04665 - -
04670 -pf04839 -
04675 - -
04680 -pf00075 -
04685 -pf00542pf16320 -
(..2080..)
06765 -pf02021 -
06770 -pf00745pf05201pf01488 -
06775 -pf00873 -
06780 -pf00005 -
06790 - -
06800 -pf00498pf00069 -
06805 - -
06820 -pf02559pf00270pf00271pf03461pf17757 -
06825 -pf00902 -
06830 H_neonGreen CsomeBCy
06835 -pf01383 -
06840 -pf00502 -
06845 -pf00502 -
06850 -pf11267 -
06855 -pf08802pf00355 -
06860 -pf01333pf16639 -
06865 -pf01926pf12631pf10396 -
06870 -pf00664pf00005 -
06875 -pf01656 -
06880 - -
06885 HMM00121_isom_1_spu(pf00121)
06895 - -
BMC-P
BMC-H
BMC-T
Aldehyde dehydrogenase
PTAC
Signature enzyme
Alcohol dehydrogenase
Regulator

Cyanobacteria bacterium J007. Bacteria; Cyanobacteria.

D6680__locus_1_main_sat

CsomeBCy

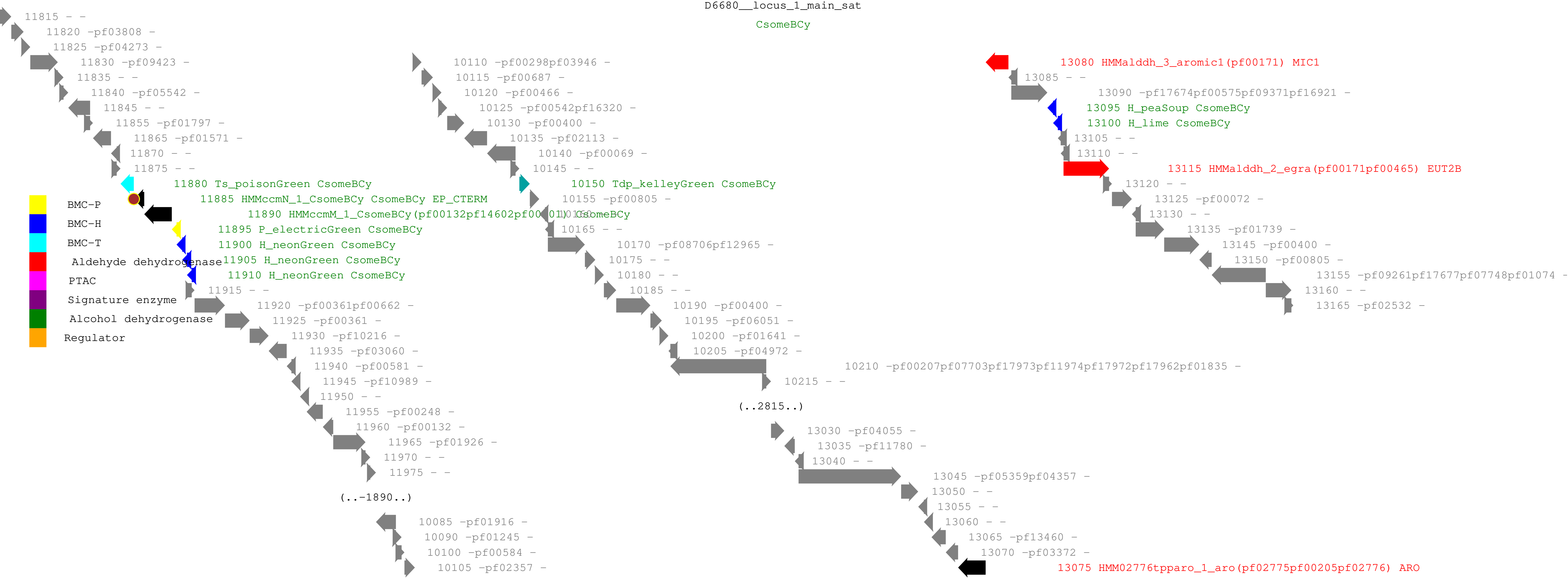

Cyanobacteria bacterium J055. Bacteria; Cyanobacteria.

D6728__locus_1_main_sat

CsomeBCy

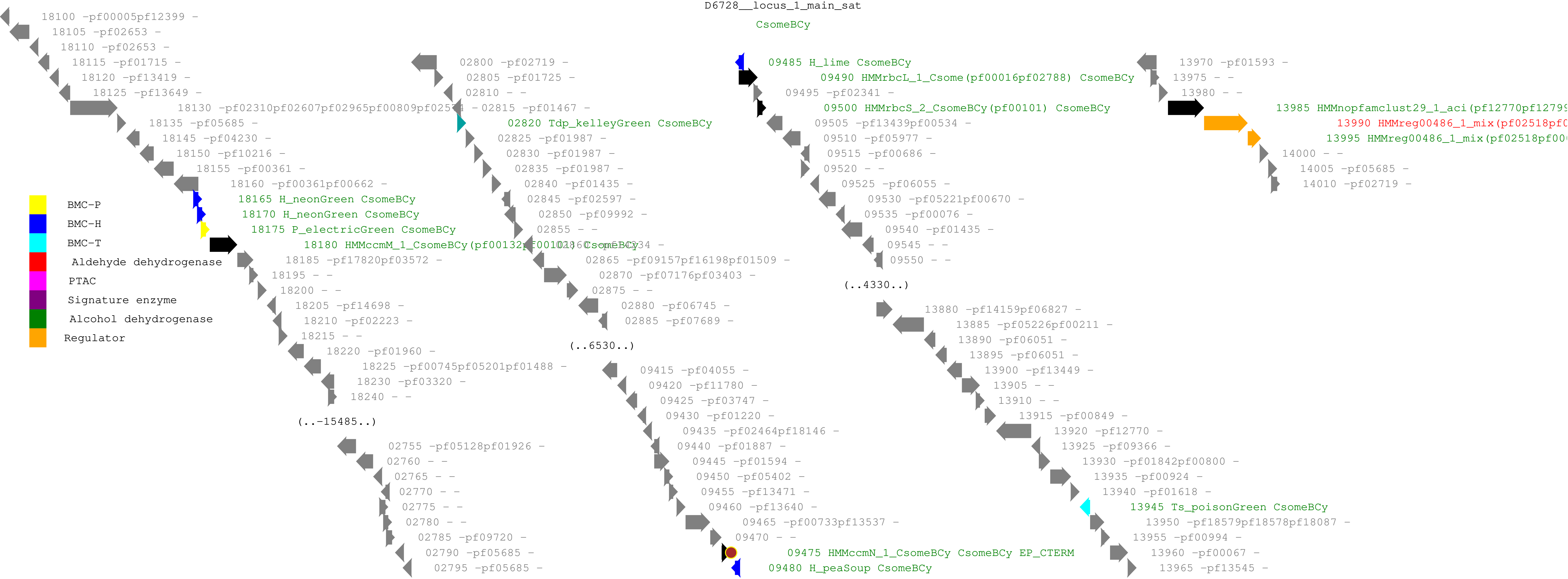

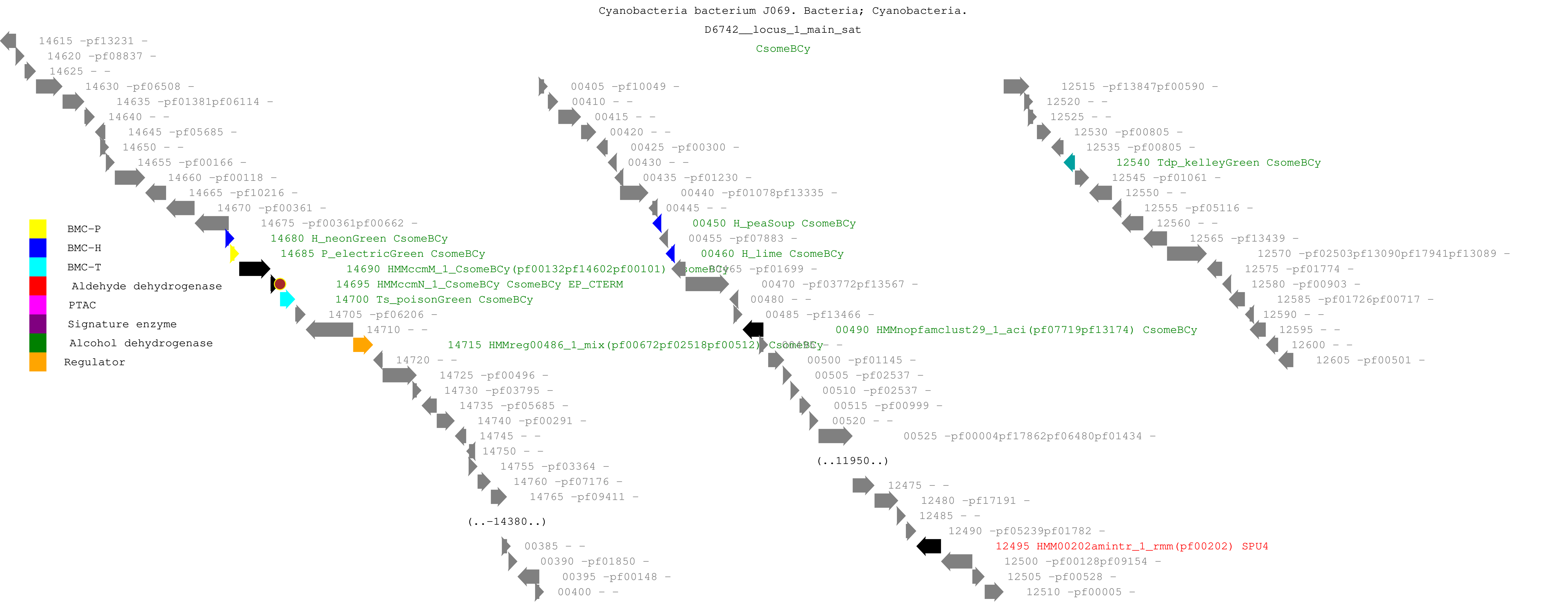

Cyanobacteria bacterium J069. Bacteria; Cyanobacteria.
D6742__locus_1_main_sat
CsomeBCy
BMC-P
BMC-H
BMC-T
Aldehyde dehydrogenase
PTAC
Signature enzyme
Alcohol dehydrogenase
Regulator
14615 -pf13231 -
14620 -pf08837 -
14625 - -
14630 -pf06508 -
14635 -pf01381pf06114 -
14640 - -
14645 -pf05685 -
14650 - -
14655 -pf00166 -
14660 -pf00118 -
14665 -pf10216 -
14670 -pf00361 -
14675 -pf00361pf00662 -
14680 H_neonGreen CsomeBCy
14685 P_electricGreen CsomeBCy
14690 HMMccmM_1_CsomeBCy(pf00132pf14602pf00101)
14695 HMMccmN_1_CsomeBCy CsomeBCy EP_CTERM
14700 Ts_poisonGreen CsomeBCy
14705 -pf06206 -
14710 - -
14715 HMMreg00486_1_mix(pf00672pf02518pf00512) CsomeBCy
14720 - -
14725 -pf00496 -
14730 -pf03795 -
14735 -pf05685 -
14740 -pf00291 -
14745 - -
14750 - -
14755 -pf03364 -
14760 -pf07176 -
14765 -pf09411 -
(..-14380..)
00385 - -
00390 -pf01850 -
00395 -pf00148 -
00400 - -
00405 -pf10049 -
00410 - -
00415 - -
00420 - -
00425 -pf00300 -
00430 - -
00435 -pf01230 -
00440 -pf01078pf13335 -
00445 - -
00450 H_peaSoup CsomeBCy
00455 -pf07883 -
00460 H_lime CsomeBCy
00465 -pf01699 -
00470 -pf03772pf13567 -
00480 - -
00485 -pf13466 -
00490 HMMnopfamclust29_1_aci(pf07719pf13174) CsomeBCy
00495 - -
00500 -pf01145 -
00505 -pf02537 -
00510 -pf02537 -
00515 -pf00999 -
00520 - -
00525 -pf00004pf17862pf06480pf01434 -
(..11950..)
12475 - -
12480 -pf17191 -
12485 - -
12490 -pf05239pf01782 -
12495 HMM00202amintr_1_rmm(pf00202) SPU4
12500 -pf00128pf09154 -
12505 -pf00528 -
12510 -pf00005 -
12515 -pf13847pf00590 -
12520 - -
12525 - -
12530 -pf00805 -
12535 -pf00805 -
12540 Tdp_kelleyGreen CsomeBCy
12545 -pf01061 -
12550 - -
12555 -pf05116 -
12560 - -
12565 -pf13439 -
12570 -pf02503pf13090pf17941pf13089 -
12575 -pf01774 -
12580 -pf00903 -
12585 -pf01726pf00717 -
12590 - -
12595 - -
12600 - -
12605 -pf00501 -

Cyanobacteria bacterium J083. Bacteria; Cyanobacteria.

D6756__locus_1_main_sat

CsomeBCy

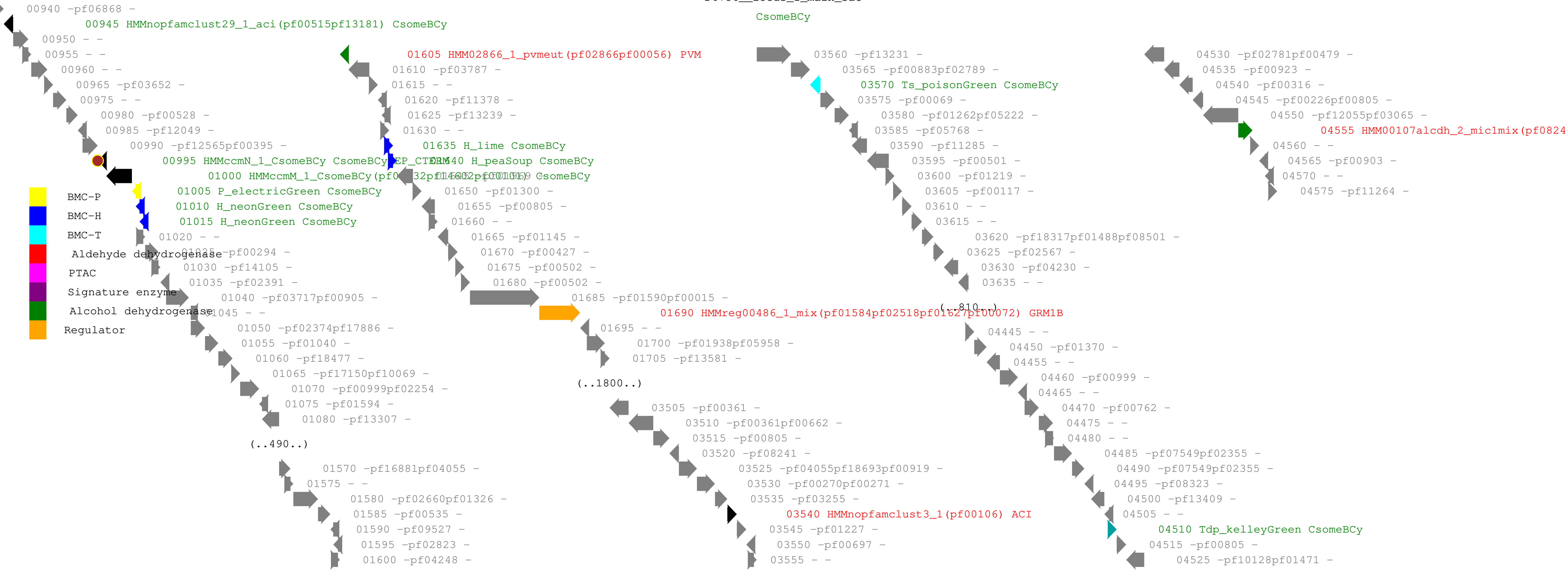

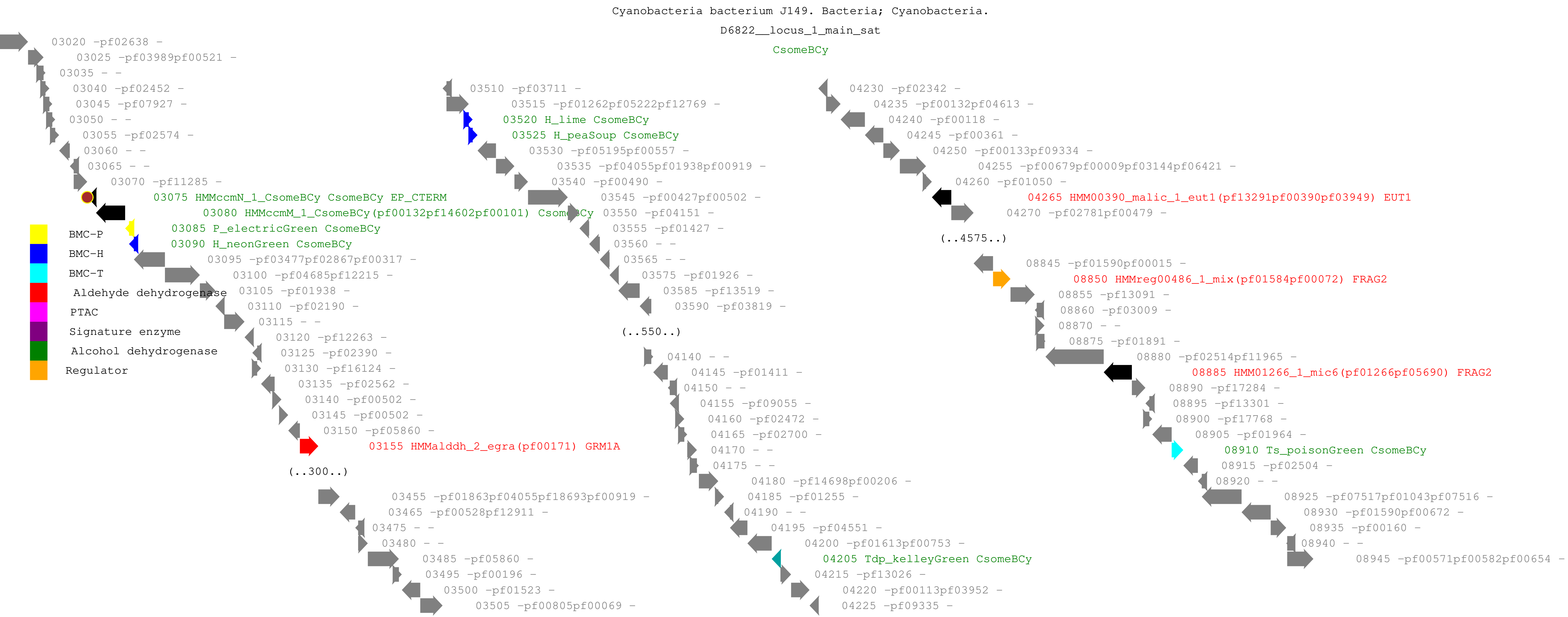

Cyanobacteria bacterium J149. Bacteria; Cyanobacteria.
D6822__locus_1_main_sat
CsomeBCy
03020 -pf02638 -
03025 -pf03989pf00521 -
03035 - -
03040 -pf02452 -
03045 -pf07927 -
03050 - -
03055 -pf02574 -
03060 - -
03065 - -
03070 -pf11285 -
03075 HMMccmN_1_CsomeBCy CsomeBCy EP_CTERM
03080 HMMccmM_1_CsomeBCy(pf00132pf14602pf00101) CsomeBCy
03085 P_electricGreen CsomeBCy
03090 H_neonGreen CsomeBCy
03095 -pf03477pf02867pf00317 -
03100 -pf04685pf12215 -
03105 -pf01938 -
03110 -pf02190 -
03115 - -
03120 -pf12263 -
03125 -pf02390 -
03130 -pf16124 -
03135 -pf02562 -
03140 -pf00502 -
03145 -pf00502 -
03150 -pf05860 -
03155 HMMaldh_2_egra(pf00171) GRM1A
(..300..)
03455 -pf01863pf04055pf18693pf00919 -
03465 -pf00528pf12911 -
03475 - -
03480 - -
03485 -pf05860 -
03495 -pf00196 -
03500 -pf01523 -
03505 -pf00805pf00069 -
BMC-P
BMC-H
BMC-T
Aldehyde dehydrogenase
PTAC
Signature enzyme
Alcohol dehydrogenase
Regulator
03510 -pf03711 -
03515 -pf01262pf05222pf12769 -
03520 H_lime CsomeBCy
03525 H_peaSoup CsomeBCy
03530 -pf05195pf00557 -
03535 -pf04055pf01938pf00919 -
03540 -pf00490 -
03545 -pf00427pf00502 -
03550 -pf04151 -
03555 -pf01427 -
03560 - -
03565 - -
03575 -pf01926 -
03585 -pf13519 -
03590 -pf03819 -
(..550..)
04140 - -
04145 -pf01411 -
04150 - -
04155 -pf09055 -
04160 -pf02472 -
04165 -pf02700 -
04170 - -
04175 - -
04180 -pf14698pf00206 -
04185 -pf01255 -
04190 - -
04195 -pf04551 -
04200 -pf01613pf00753 -
04205 Tdp_kelleyGreen CsomeBCy
04215 -pf13026 -
04220 -pf00113pf03952 -
04225 -pf09335 -
04230 -pf02342 -
04235 -pf00132pf04613 -
04240 -pf00118 -
04245 -pf00361 -
04250 -pf00133pf09334 -
04255 -pf00679pf00009pf03144pf06421 -
04260 -pf01050 -
04265 HMM00390_malic_1_eut1(pf13291pf00390pf03949) EUT1
04270 -pf02781pf00479 -
(..4575..)
08845 -pf01590pf00015 -
08850 HMMreg00486_1_mix(pf01584pf00072) FRAG2
08855 -pf13091 -
08860 -pf03009 -
08870 - -
08875 -pf01891 -
08880 -pf02514pf11965 -
08885 HMM01266_1_mic6(pf01266pf05690) FRAG2
08890 -pf17284 -
08895 -pf13301 -
08900 -pf17768 -
08905 -pf01964 -
08910 Ts_poisonGreen CsomeBCy
08915 -pf02504 -
08920 - -
08925 -pf07517pf01043pf07516 -
08930 -pf01590pf00672 -
08935 -pf00160 -
08940 - -
08945 -pf00571pf00582pf00654 -

Tolypothrix bouteillei VB521301. Bacteria; Cyanobacteria; Nostocales; Tolypothrichaceae; Tolypothrix.

DA73__locus_1_main_sat

CsomeBCy

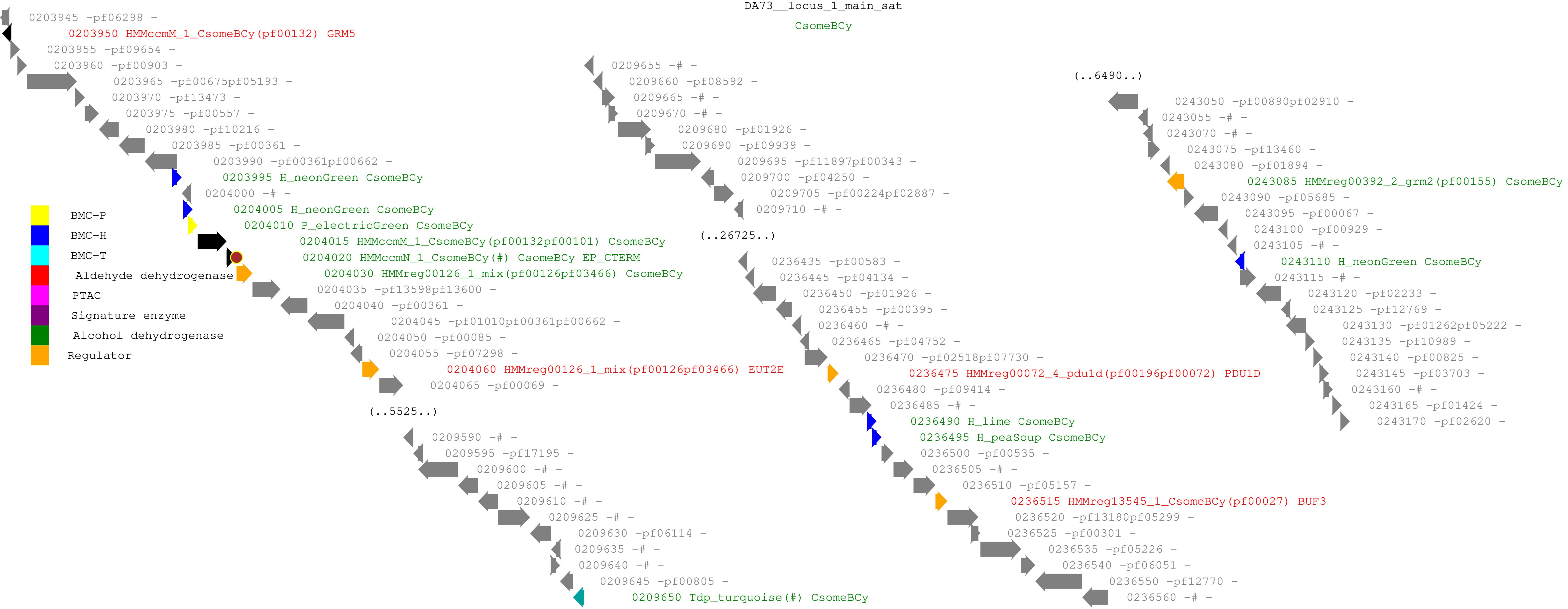

Dactylococcopsis salina PCC 8305. Bacteria; Cyanobacteria; Synechococcales; Synechococcaceae;

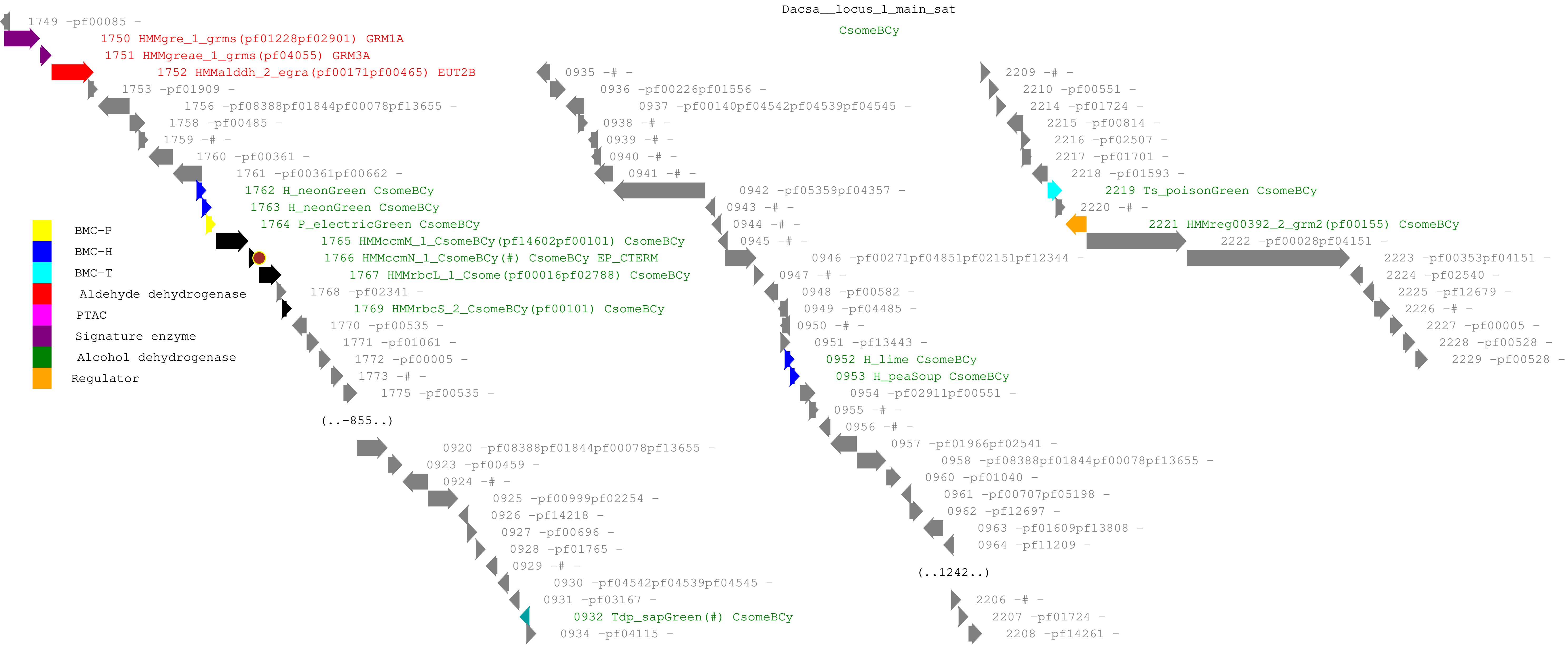

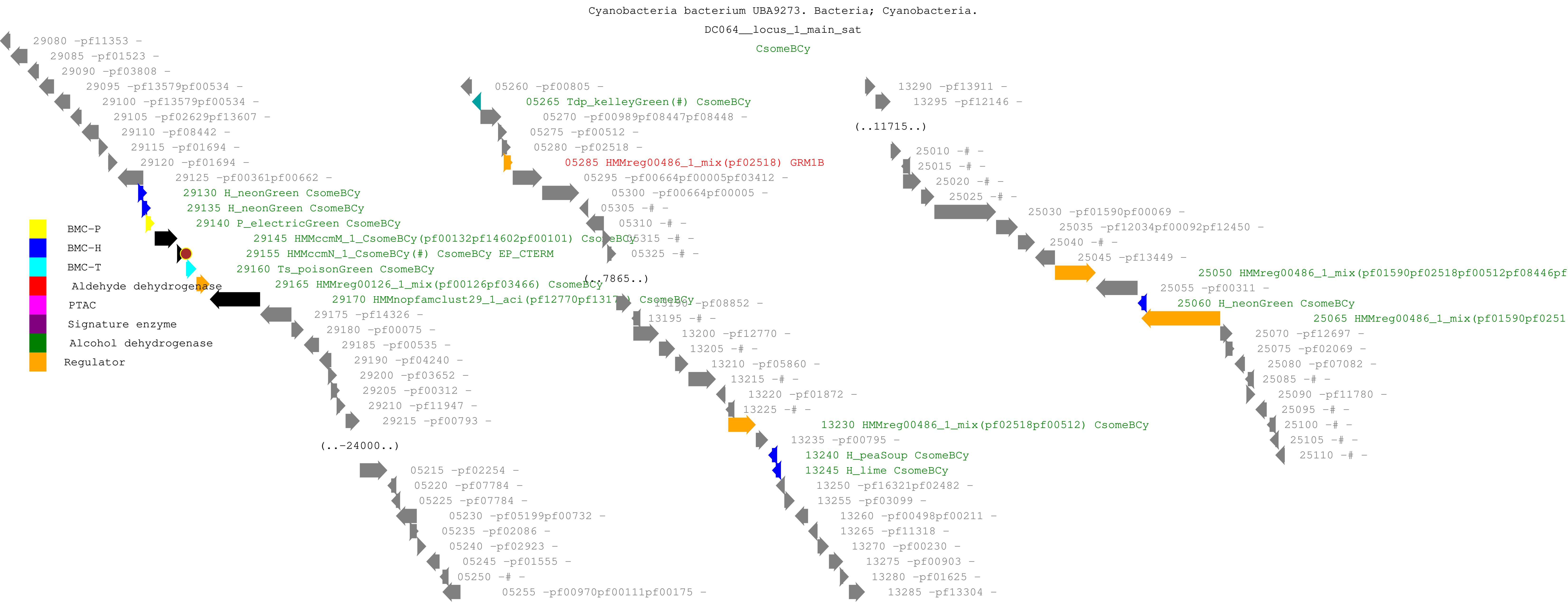

Cyanobacteria bacterium UBA9273. Bacteria; Cyanobacteria.
DC064__locus_1_main_sat
CsomeBCy
BMC-P
BMC-H
BMC-T
Aldehyde dehydrogenase
PTAC
Signature enzyme
Alcohol dehydrogenase
Regulator
29080 -pf11353 -
29085 -pf01523 -
29090 -pf03808 -
29095 -pf13579pf00534 -
29100 -pf13579pf00534 -
29105 -pf02629pf13607 -
29110 -pf08442 -
29115 -pf01694 -
29120 -pf01694 -
29125 -pf00361pf00662 -
29130 H_neonGreen CsomeBCy
29135 H_neonGreen CsomeBCy
29140 P_electricGreen CsomeBCy
29145 HMMccmM_1_CsomeBCy(pf00132pf14602pf00101) CsomeBCy
29155 HMMccmN_1_CsomeBCy(#) CsomeBCy EP_CTERM
29160 Ts_poisonGreen CsomeBCy
29165 HMMreg00126_1_mix(pf00126pf03466) CsomeBCy
29170 HMMnopfamclust29_1_aci(pf12770pf1317 CsomeBCy
29175 -pf14326 -
29180 -pf00075 -
29185 -pf00535 -
29190 -pf04240 -
29200 -pf03652 -
29205 -pf00312 -
29210 -pf11947 -
29215 -pf00793 -
(..-24000..)
05215 -pf02254 -
05220 -pf07784 -
05225 -pf07784 -
05230 -pf05199pf00732 -
05235 -pf02086 -
05240 -pf02923 -
05245 -pf01555 -
05250 -# -
05255 -pf00970pf00111pf00175 -
05260 -pf00805 -
05265 Tdp_kelleyGreen(#) CsomeBCy
05270 -pf00989pf08447pf08448 -
05275 -pf00512 -
05280 -pf02518 -
05285 HMMreg00486_1_mix(pf02518) GRM1B
05295 -pf00664pf00005pf03412 -
05300 -pf00664pf00005 -
05305 -# -
05310 -# -
05315 -# -
05325 -# -
(..7865..)
13190 -pf08852 -
13195 -# -
13200 -pf12770 -
13205 -# -
13210 -pf05860 -
13215 -# -
13220 -pf01872 -
13225 -# -
13230 HMMreg00486_1_mix(pf02518pf00512) CsomeBCy
13235 -pf00795 -
13240 H_peaSoup CsomeBCy
13245 H_lime CsomeBCy
13250 -pf16321pf02482 -
13255 -pf03099 -
13260 -pf00498pf00211 -
13265 -pf11318 -
13270 -pf00230 -
13275 -pf00903 -
13280 -pf01625 -
13285 -pf13304 -
13290 -pf13911 -
13295 -pf12146 -
(..11715..)
25010 -# -
25015 -# -
25020 -# -
25025 -# -
25030 -pf01590pf00069 -
25035 -pf12034pf00092pf12450 -
25040 -# -
25045 -pf13449 -
25050 HMMreg00486_1_mix(pf01590pf02518pf00512pf08446pf
25055 -pf00311 -
25060 H_neonGreen CsomeBCy
25065 HMMreg00486_1_mix(pf01590pf0251
25070 -pf12697 -
25075 -pf02069 -
25080 -pf07082 -
25085 -# -
25090 -pf11780 -
25095 -# -
25100 -# -
25105 -# -
25110 -# -

Cyanobacteria bacterium UBA8553. Bacteria; Cyanobacteria.

DCE56__locus_1_main_sat

CsomeBCy

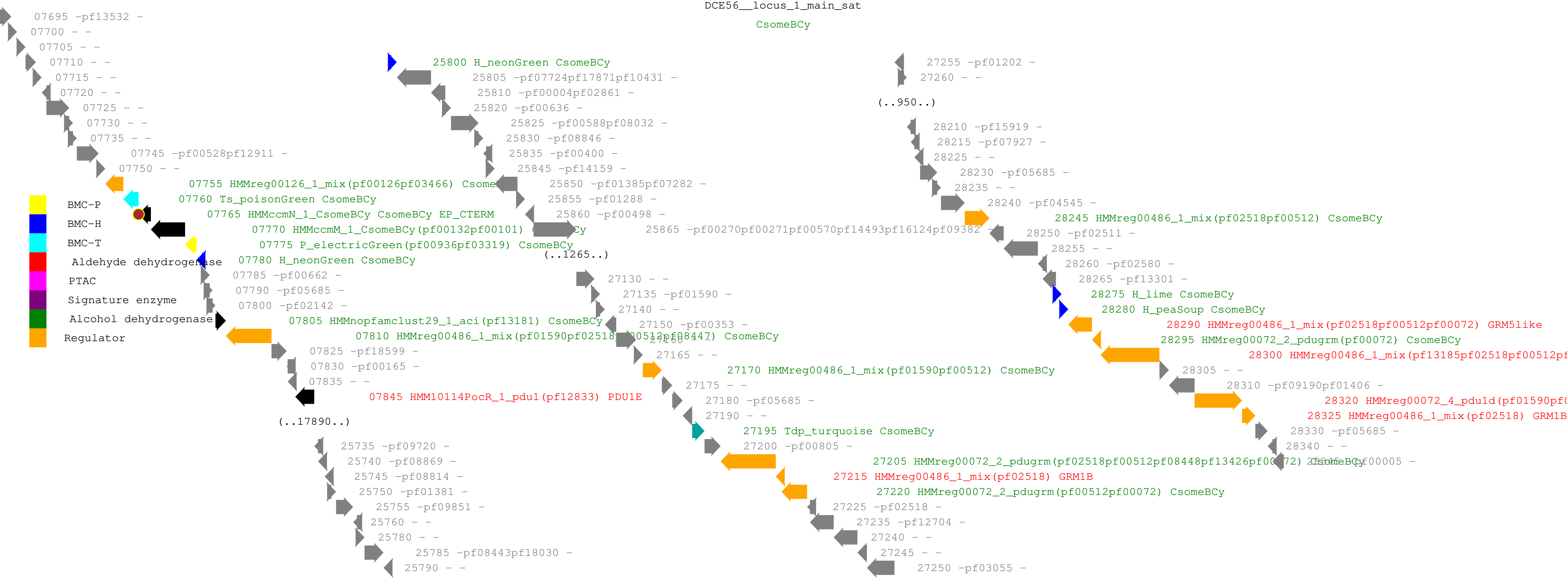

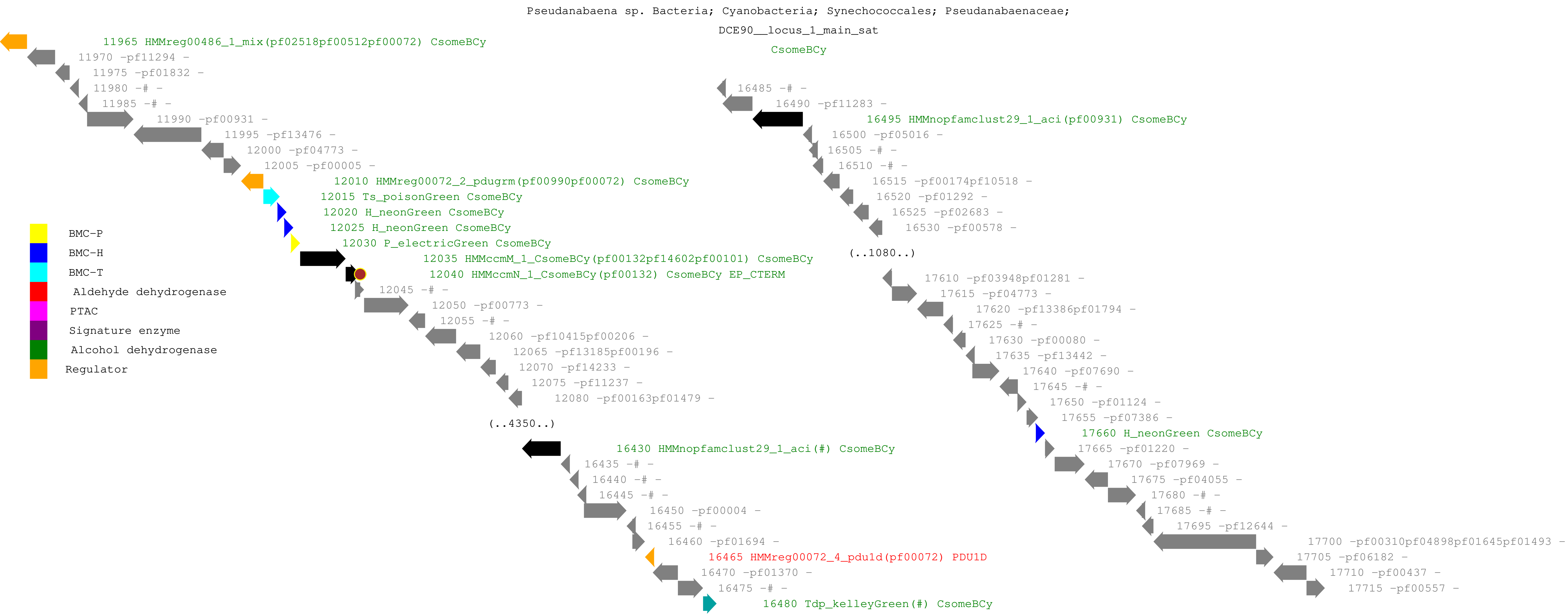
Pseudanabaena sp. Bacteria; Cyanobacteria; Synechococcales; Pseudanabaenaceae;
DCE90__locus_1_main_sat
CsomeBCy
11965 HMMreg00486_1_mix(pf02518pf00512pf00072) CsomeBCy
11970 -pf11294 -
11975 -pf01832 -
11980 -# -
11985 -# -
11990 -pf00931 -
11995 -pf13476 -
12000 -pf04773 -
12005 -pf00005 -
12010 HMMreg00072_2_pdugrm(pf00990pf00072) CsomeBCy
12015 Ts_poisonGreen CsomeBCy
12020 H_neonGreen CsomeBCy
12025 H_neonGreen CsomeBCy
12030 P_electricGreen CsomeBCy
12035 HMMccmM_1_CsomeBCy(pf00132pf14602pf00101) CsomeBCy
12040 HMMccmN_1_CsomeBCy(pf00132) CsomeBCy EP_CTERM
12045 -# -
12050 -pf00773 -
12055 -# -
12060 -pf10415pf00206 -
12065 -pf13185pf00196 -
12070 -pf14233 -
12075 -pf11237 -
12080 -pf00163pf01479 -
(..4350..)
16430 HMMnopfamclust29_1_aci(#) CsomeBCy
16435 -# -
16440 -# -
16445 -# -
16450 -pf00004 -
16455 -# -
16460 -pf01694 -
16465 HMMreg00072_4_pdu1d(pf00072) PDU1D
16470 -pf01370 -
16475 -# -
16480 Tdp_kelleyGreen(#) CsomeBCy
BMC-P
BMC-H
BMC-T
Aldehyde dehydrogenase
PTAC
Signature enzyme
Alcohol dehydrogenase
Regulator
16485 -# -
16490 -pf11283 -
16495 HMMnopfamclust29_1_aci(pf00931) CsomeBCy
16500 -pf05016 -
16505 -# -
16510 -# -
16515 -pf00174pf10518 -
16520 -pf01292 -
16525 -pf02683 -
16530 -pf00578 -
(..1080..)
17610 -pf03948pf01281 -
17615 -pf04773 -
17620 -pf13386pf01794 -
17625 -# -
17630 -pf00080 -
17635 -pf13442 -
17640 -pf07690 -
17645 -# -
17650 -pf01124 -
17655 -pf07386 -
17660 H_neonGreen CsomeBCy
17665 -pf01220 -
17670 -pf07969 -
17675 -pf04055 -
17680 -# -
17685 -# -
17695 -pf12644 -
17700 -pf00310pf04898pf01645pf01493 -
17705 -pf06182 -
17710 -pf00437 -
17715 -pf00557 -

Snowella sp. Bacteria; Cyanobacteria; Synechococcales; Coelosphaeriaceae; Snowella.

DCF12__locus_1_main_sat

CsomeBCy

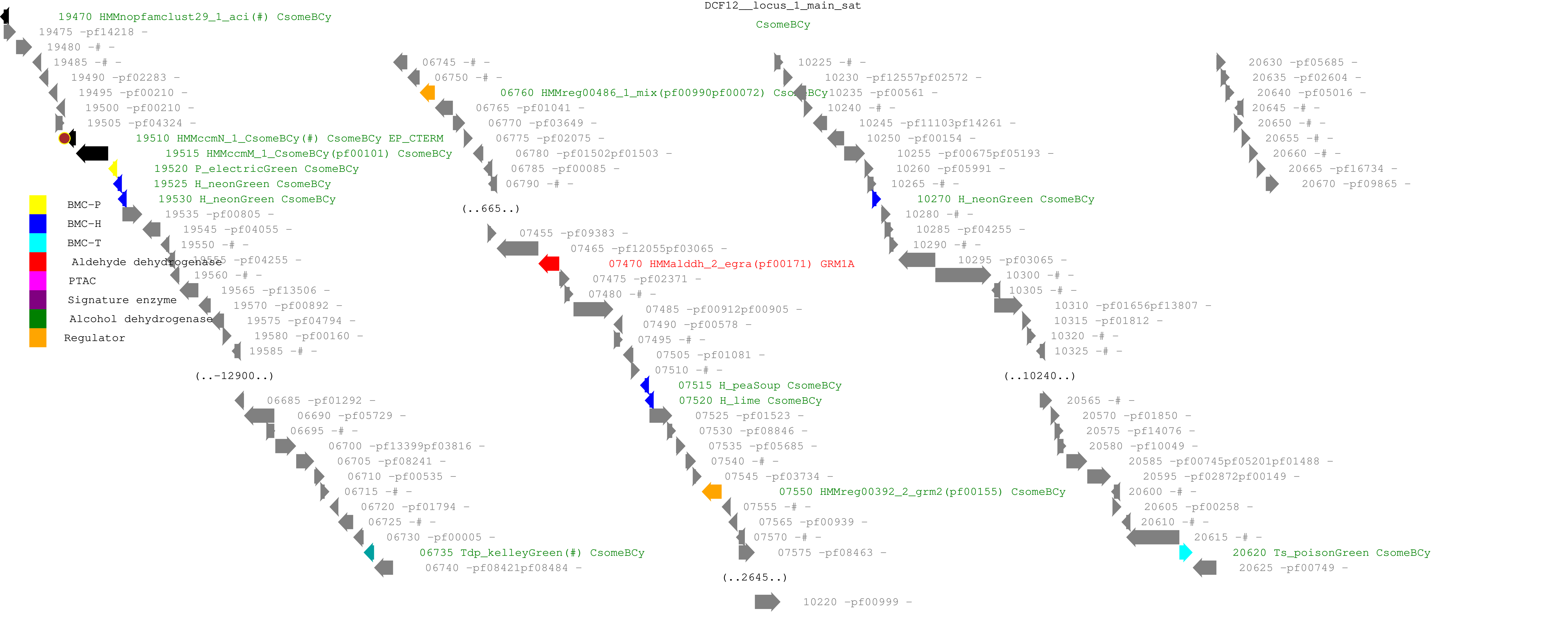

Phormidesmis priestleyi. Bacteria; Cyanobacteria; Synechococcales; Leptolyngbyaceae;

DCF15__locus_1_main_sat

CsomeBCy

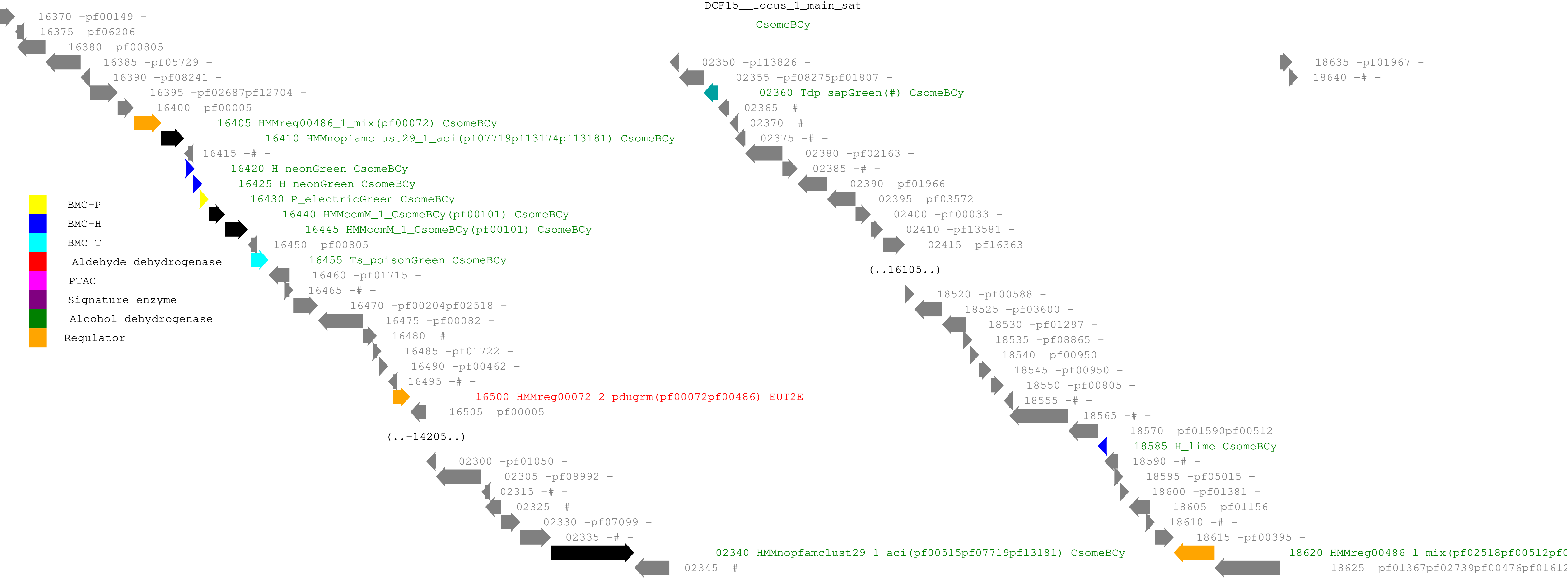

Leptolyngbya antarctica. Bacteria; Cyanobacteria; Synechococcales; Leptolyngbyaceae;

DCF17__locus_1_main_sat

CsomeBCy

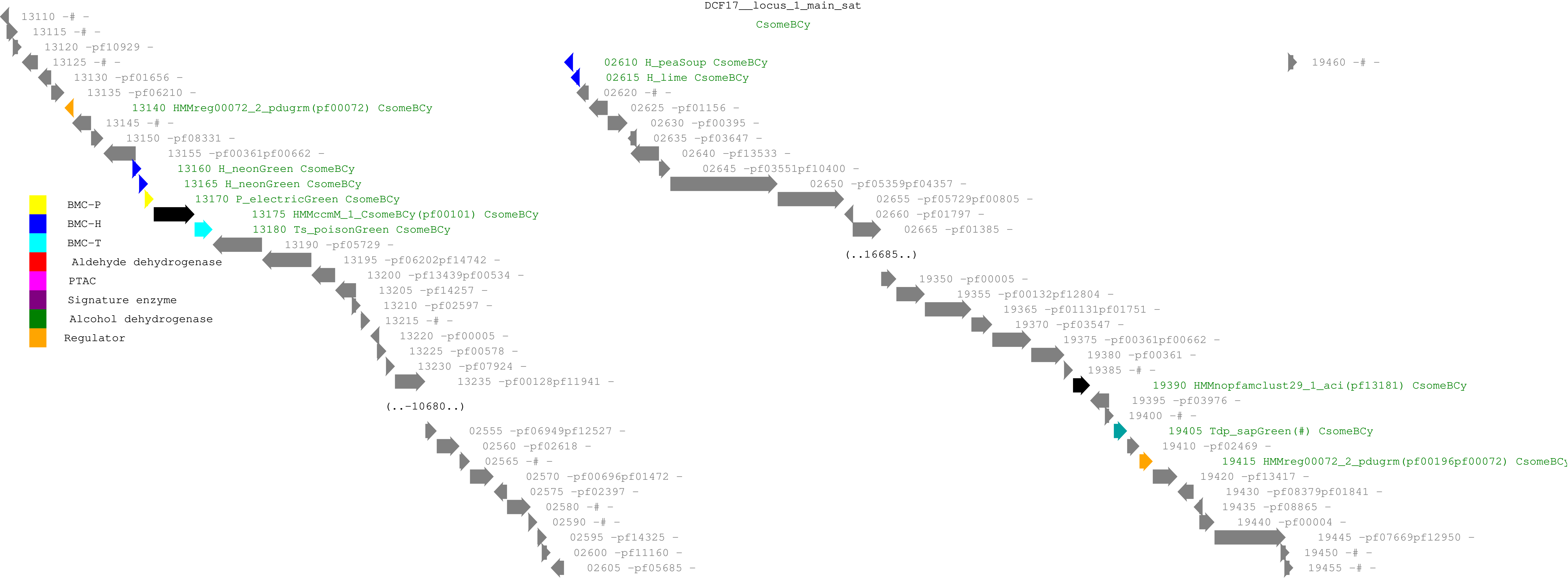

Pseudanabaena frigida. Bacteria; Cyanobacteria; Synechococcales; Pseudanabaenaceae;

DCF19__locus_fragmented

CsomeBCy

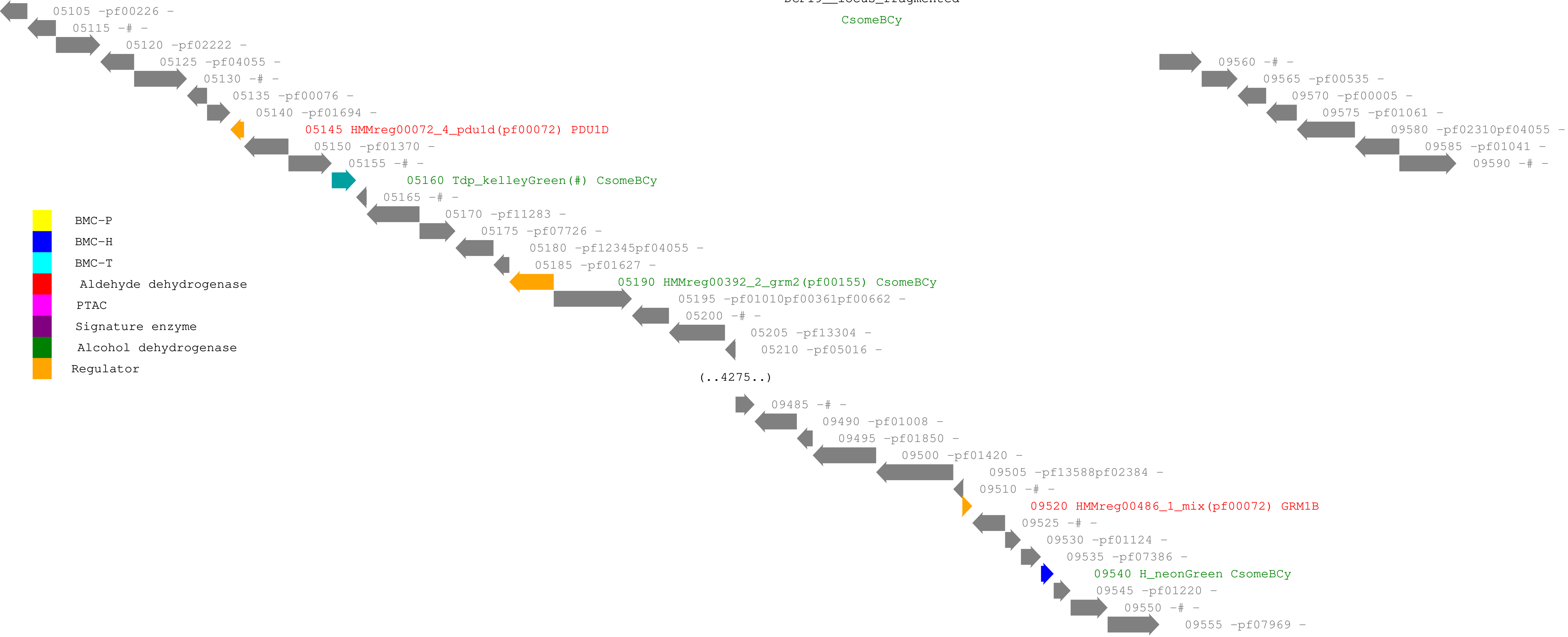

Pseudanabaena sp. Bacteria; Cyanobacteria; Synechococcales; Pseudanabaenaceae;

DCF20__locus_1_main_sat

CsomeBCy

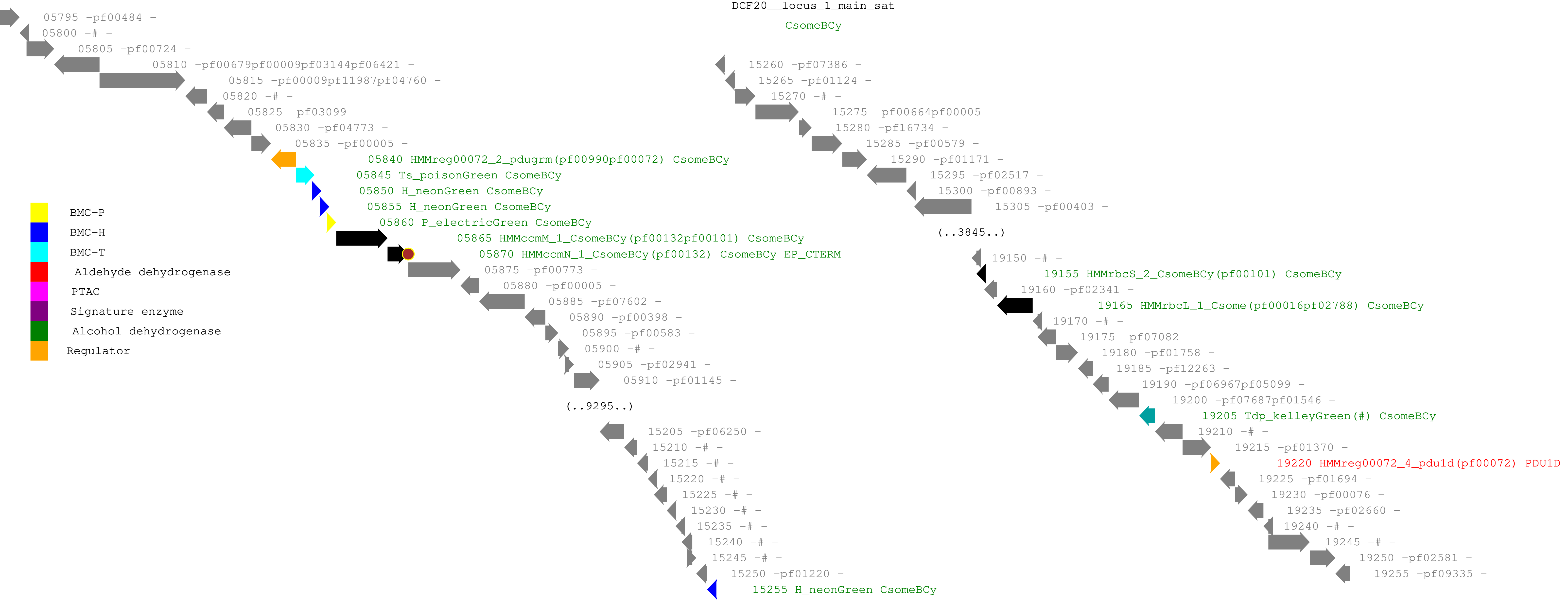

Leptolyngbya sp. Bacteria; Cyanobacteria; Synechococcales; Leptolyngbyaceae;

DCF21__locus_1_main_sat

CsomeBCy

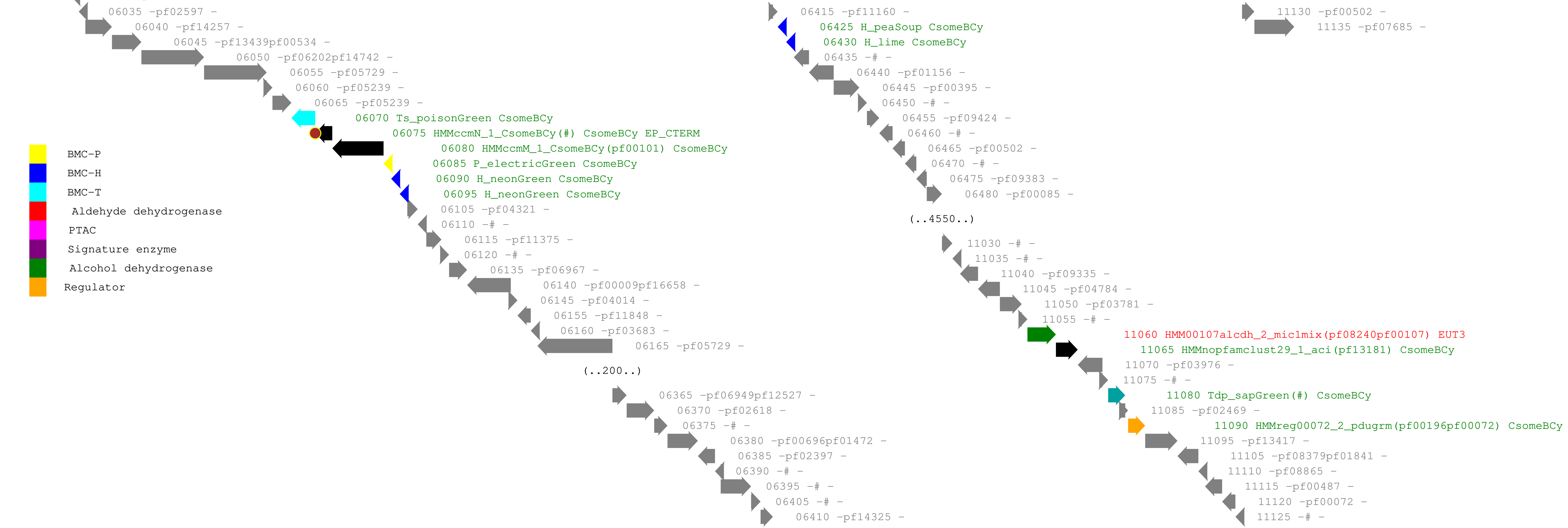

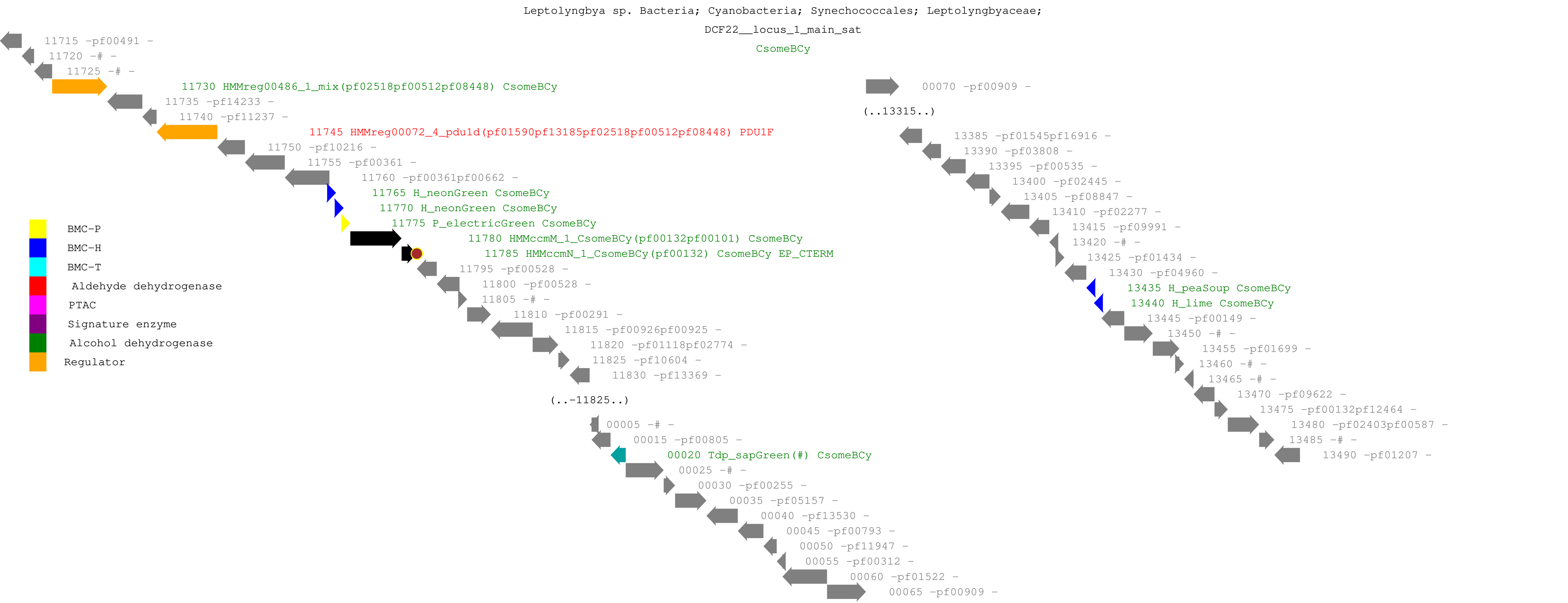

Leptolyngbya sp. Bacteria; Cyanobacteria; Synechococcales; Leptolyngbyaceae;
DCF22__locus_1_main_sat
CsomeBCy
11715 -pf00491 -
11720 -# -
11725 -# -
11730 HMMreg00486_1_mix(pf02518pf00512pf08448) CsomeBCy
11735 -pf14233 -
11740 -pf11237 -
11745 HMMreg00072_4_pdu1d(pf01590pf13185pf02518pf00512pf08448) PDU1F
11750 -pf10216 -
11755 -pf00361 -
11760 -pf00361pf00662 -
11765 H_neonGreen CsomeBCy
11770 H_neonGreen CsomeBCy
11775 P_electricGreen CsomeBCy
11780 HMMccmM_1_CsomeBCy(pf00132pf00101) CsomeBCy
11785 HMMccmN_1_CsomeBCy(pf00132) CsomeBCy EP_CTERM
11795 -pf00528 -
11800 -pf00528 -
11805 -# -
11810 -pf00291 -
11815 -pf00926pf00925 -
11820 -pf01118pf02774 -
11825 -pf10604 -
11830 -pf13369 -
(..-11825..)
00005 -# -
00015 -pf00805 -
00020 Tdp_sapGreen(#) CsomeBCy
00025 -# -
00030 -pf00255 -
00035 -pf05157 -
00040 -pf13530 -
00045 -pf00793 -
00050 -pf11947 -
00055 -pf00312 -
00060 -pf01522 -
00065 -pf00909 -
00070 -pf00909 -
(..13315..)
13385 -pf01545pf16916 -
13390 -pf03808 -
13395 -pf00535 -
13400 -pf02445 -
13405 -pf08847 -
13410 -pf02277 -
13415 -pf09991 -
13420 -# -
13425 -pf01434 -
13430 -pf04960 -
13435 H_peaSoup CsomeBCy
13440 H_lime CsomeBCy
13445 -pf00149 -
13450 -# -
13455 -pf01699 -
13460 -# -
13465 -# -
13470 -pf09622 -
13475 -pf00132pf12464 -
13480 -pf02403pf00587 -
13485 -# -
13490 -pf01207 -
BMC-P
BMC-H
BMC-T
Aldehyde dehydrogenase
PTAC
Signature enzyme
Alcohol dehydrogenase
Regulator

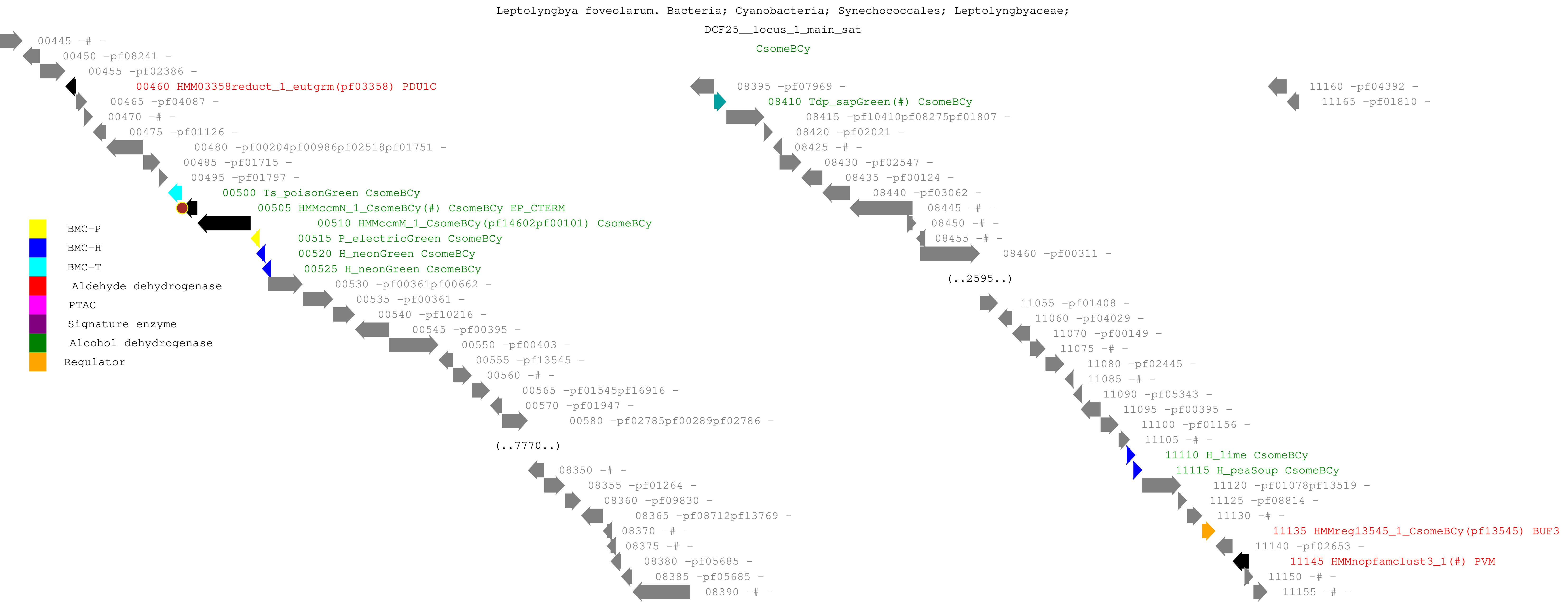
Leptolyngbya foveolarum. Bacteria; Cyanobacteria; Synechococcales; Leptolyngbyaceae;
DCF25__locus_1_main_sat
CsomeBCy
00445 -# -
00450 -pf08241 -
00455 -pf02386 -
00460 HMM03358reduct_1_eutgrm(pf03358) PDU1C
00465 -pf04087 -
00470 -# -
00475 -pf01126 -
00480 -pf00204pf00986pf02518pf01751 -
00485 -pf01715 -
00495 -pf01797 -
00500 Ts_poisonGreen CsomeBCy
00505 HMMccmN_1_CsomeBCy(#) CsomeBCy EP_CTERM
00510 HMMccmM_1_CsomeBCy(pf14602pf00101) CsomeBCy
00515 P_electricGreen CsomeBCy
00520 H_neonGreen CsomeBCy
00525 H_neonGreen CsomeBCy
00530 -pf00361pf00662 -
00535 -pf00361 -
00540 -pf10216 -
00545 -pf00395 -
00550 -pf00403 -
00555 -pf13545 -
00560 -# -
00565 -pf01545pf16916 -
00570 -pf01947 -
00580 -pf02785pf00289pf02786 -
(..7770..)
08350 -# -
08355 -pf01264 -
08360 -pf09830 -
08365 -pf08712pf13769 -
08370 -# -
08375 -# -
08380 -pf05685 -
08385 -pf05685 -
08390 -# -
08395 -pf07969 -
08410 Tdp_sapGreen(#) CsomeBCy
08415 -pf10410pf08275pf01807 -
08420 -pf02021 -
08425 -# -
08430 -pf02547 -
08435 -pf00124 -
08440 -pf03062 -
08445 -# -
08450 -# -
08455 -# -
08460 -pf00311 -
(..2595..)
11055 -pf01408 -
11060 -pf04029 -
11070 -pf00149 -
11075 -# -
11080 -pf02445 -
11085 -# -
11090 -pf05343 -
11095 -pf00395 -
11100 -pf01156 -
11105 -# -
11110 H_lime CsomeBCy
11115 H_peaSoup CsomeBCy
11120 -pf01078pf13519 -
11125 -pf08814 -
11130 -# -
11135 HMMreg13545_1_CsomeBCy(pf13545) BUF3
11140 -pf02653 -
11145 HMMnopfamclust3_1(#) PVM
11150 -# -
11155 -# -
11160 -pf04392 -
11165 -pf01810 -
BMC-P
BMC-H
BMC-T
Aldehyde dehydrogenase
PTAC
Signature enzyme
Alcohol dehydrogenase
Regulator

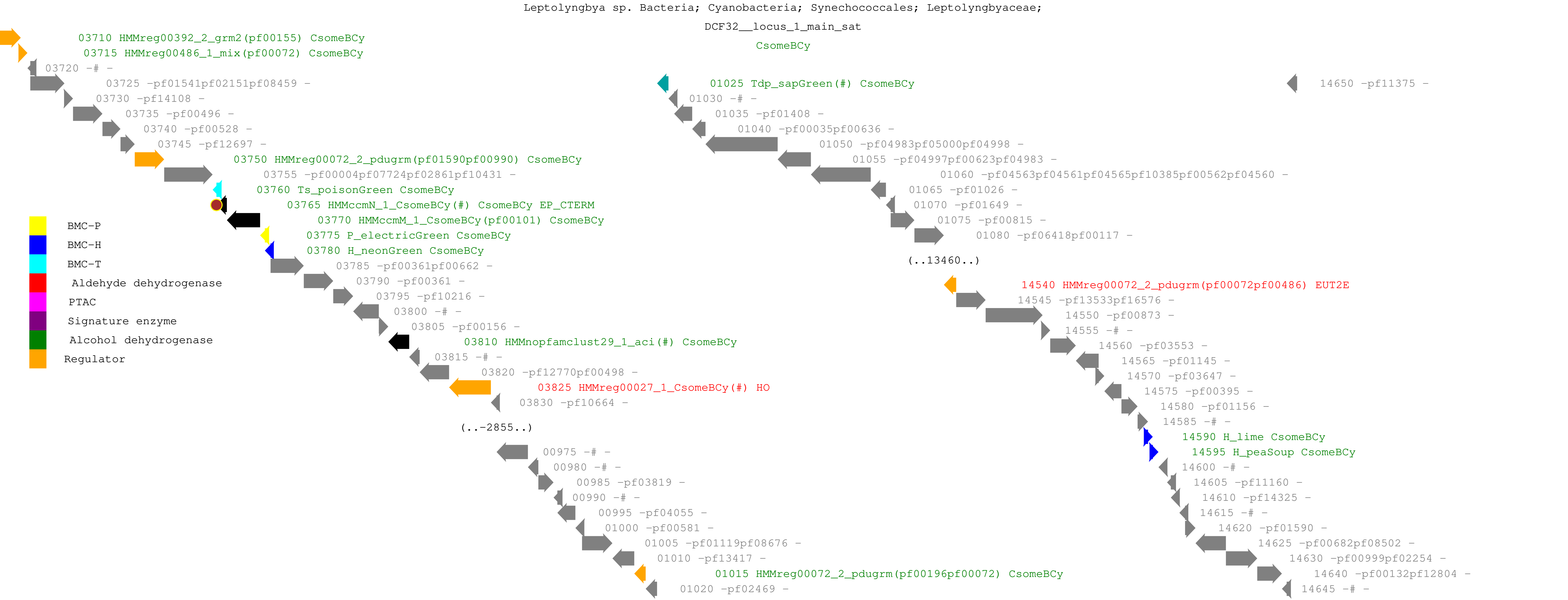
Leptolyngbya sp. Bacteria; Cyanobacteria; Synechococcales; Leptolyngbyaceae;
DCF32__locus_1_main_sat
CsomeBCy
03710 HMMreg00392_2_grm2(pf00155) CsomeBCy
03715 HMMreg00486_1_mix(pf00072) CsomeBCy
03720 -# -
03725 -pf01541pf02151pf08459 -
03730 -pf14108 -
03735 -pf00496 -
03740 -pf00528 -
03745 -pf12697 -
03750 HMMreg00072_2_pdugrm(pf01590pf00990) CsomeBCy
03755 -pf00004pf07724pf02861pf10431 -
03760 Ts_poisonGreen CsomeBCy
03765 HMMccmN_1_CsomeBCy(#) CsomeBCy EP_CTERM
03770 HMMccmM_1_CsomeBCy(pf00101) CsomeBCy
03775 P_electricGreen CsomeBCy
03780 H_neonGreen CsomeBCy
03785 -pf00361pf00662 -
03790 -pf00361 -
03795 -pf10216 -
03800 -# -
03805 -pf00156 -
03810 HMMnopfamclust29_1_aci(#) CsomeBCy
03815 -# -
03820 -pf12770pf00498 -
03825 HMMreg00027_1_CsomeBCy(#) HO
03830 -pf10664 -
(..-2855..)
00975 -# -
00980 -# -
00985 -pf03819 -
00990 -# -
00995 -pf04055 -
01000 -pf00581 -
01005 -pf01119pf08676 -
01010 -pf13417 -
01015 HMMreg00072_2_pdugrm(pf00196pf00072) CsomeBCy
01020 -pf02469 -
01025 Tdp_sapGreen(#) CsomeBCy
01030 -# -
01035 -pf01408 -
01040 -pf00035pf00636 -
01050 -pf04983pf05000pf04998 -
01055 -pf04997pf00623pf04983 -
01060 -pf04563pf04561pf04565pf10385pf00562pf04560 -
01065 -pf01026 -
01070 -pf01649 -
01075 -pf00815 -
01080 -pf06418pf00117 -
(..13460..)
14540 HMMreg00072_2_pdugrm(pf00072pf00486) EUT2E
14545 -pf13533pf16576 -
14550 -pf00873 -
14555 -# -
14560 -pf03553 -
14565 -pf01145 -
14570 -pf03647 -
14575 -pf00395 -
14580 -pf01156 -
14585 -# -
14590 H_lime CsomeBCy
14595 H_peaSoup CsomeBCy
14600 -# -
14605 -pf11160 -
14610 -pf14325 -
14615 -# -
14620 -pf01590 -
14625 -pf00682pf08502 -
14630 -pf00999pf02254 -
14640 -pf00132pf12804 -
14645 -# -
14650 -pf11375 -
BMC-P
BMC-H
BMC-T
Aldehyde dehydrogenase
PTAC
Signature enzyme
Alcohol dehydrogenase
Regulator

Cyanothece sp. UBA12306. Bacteria; Cyanobacteria; Oscillatoriophycideae; Oscillatoriales;

DCF68__locus_1_main_sat

CsomeBCy

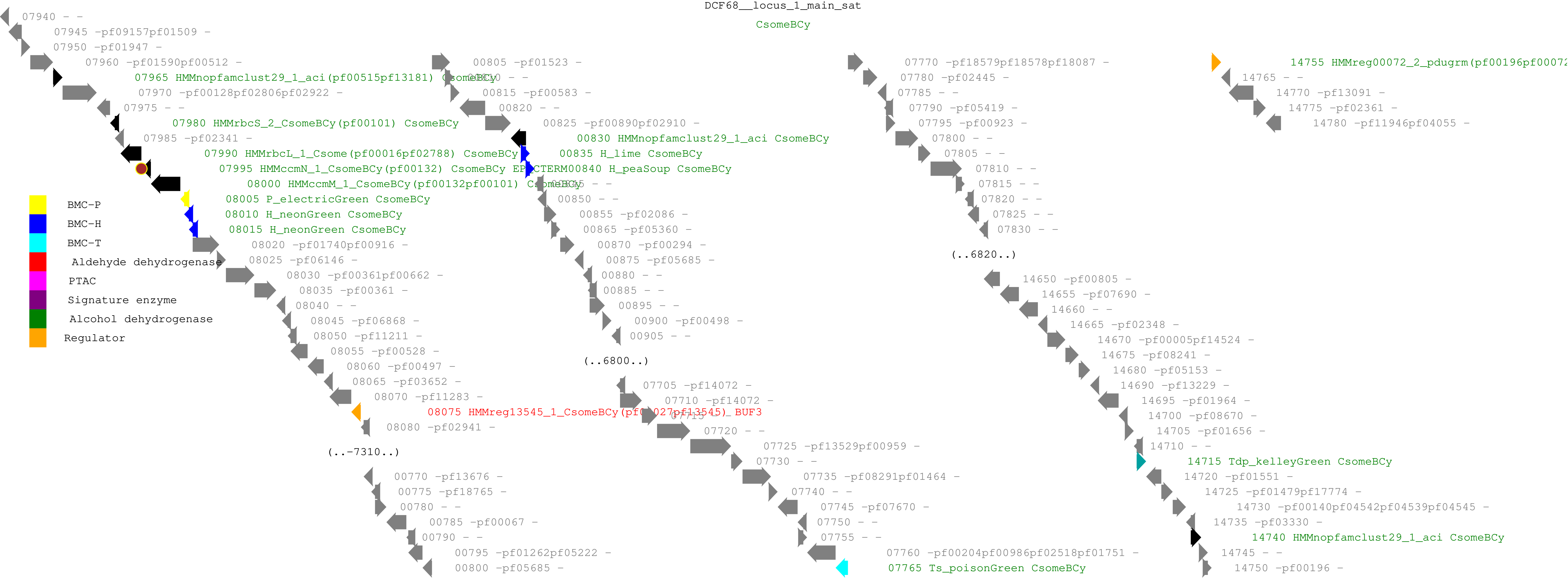

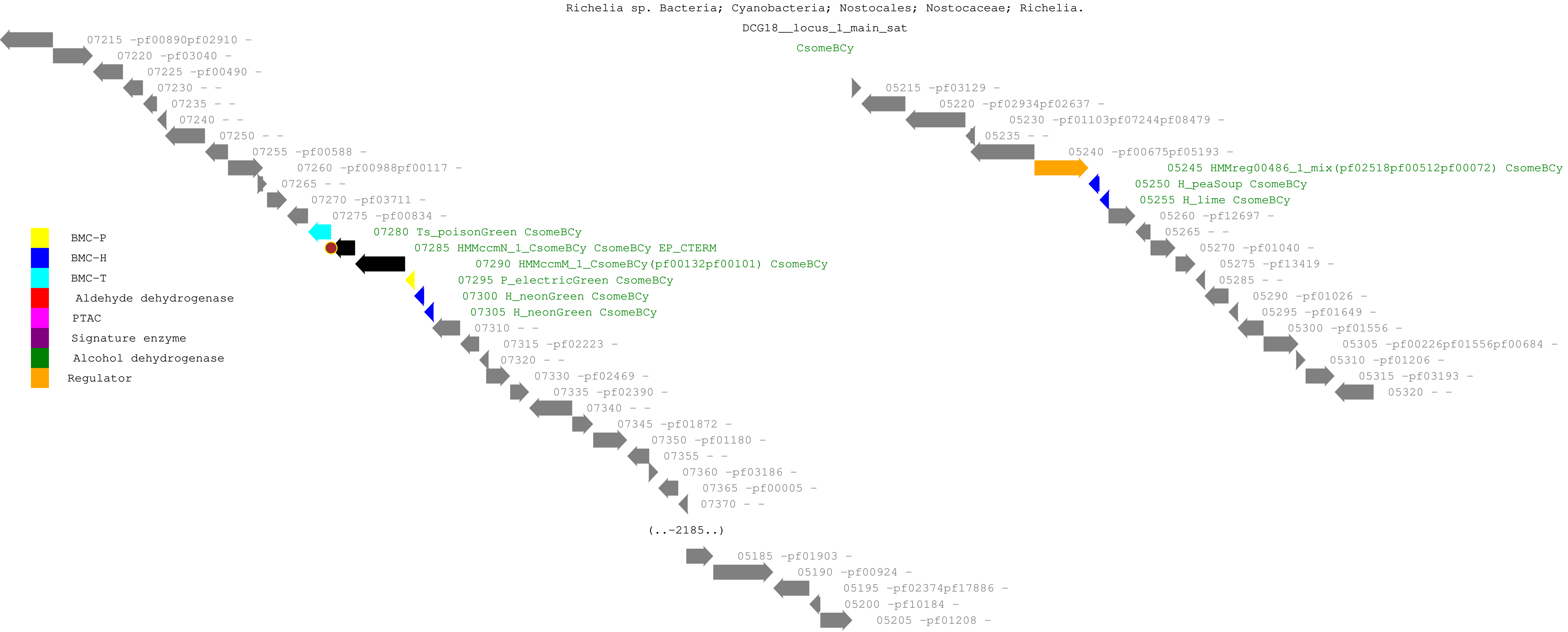
Richelia sp. Bacteria; Cyanobacteria; Nostocales; Nostocaceae; Richelia.
DCG18__locus_1_main_sat
CsomeBCy
07215 -pf00890pf02910 -
07220 -pf03040 -
07225 -pf00490 -
07230 - -
07235 - -
07240 - -
07250 - -
07255 -pf00588 -
07260 -pf00988pf00117 -
07265 - -
07270 -pf03711 -
07275 -pf00834 -
07280 Ts_poisonGreen CsomeBCy
07285 HMMccmN_1_CsomeBCy CsomeBCy EP_CTERM
07290 HMMccmM_1_CsomeBCy(pf00132pf00101) CsomeBCy
07295 P_electricGreen CsomeBCy
07300 H_neonGreen CsomeBCy
07305 H_neonGreen CsomeBCy
07310 - -
07315 -pf02223 -
07320 - -
07330 -pf02469 -
07335 -pf02390 -
07340 - -
07345 -pf01872 -
07350 -pf01180 -
07355 - -
07360 -pf03186 -
07365 -pf00005 -
07370 - -
(..-2185..)
05185 -pf01903 -
05190 -pf00924 -
05195 -pf02374pf17886 -
05200 -pf10184 -
05205 -pf01208 -
05215 -pf03129 -
05220 -pf02934pf02637 -
05230 -pf01103pf07244pf08479 -
05235 - -
05240 -pf00675pf05193 -
05245 HMMreg00486_1_mix(pf02518pf00512pf00072) CsomeBCy
05250 H_peaSoup CsomeBCy
05255 H_lime CsomeBCy
05260 -pf12697 -
05265 - -
05270 -pf01040 -
05275 -pf13419 -
05285 - -
05290 -pf01026 -
05295 -pf01649 -
05300 -pf01556 -
05305 -pf00226pf01556pf00684 -
05310 -pf01206 -
05315 -pf03193 -
05320 - -
BMC-P
BMC-H
BMC-T
Aldehyde dehydrogenase
PTAC
Signature enzyme
Alcohol dehydrogenase
Regulator

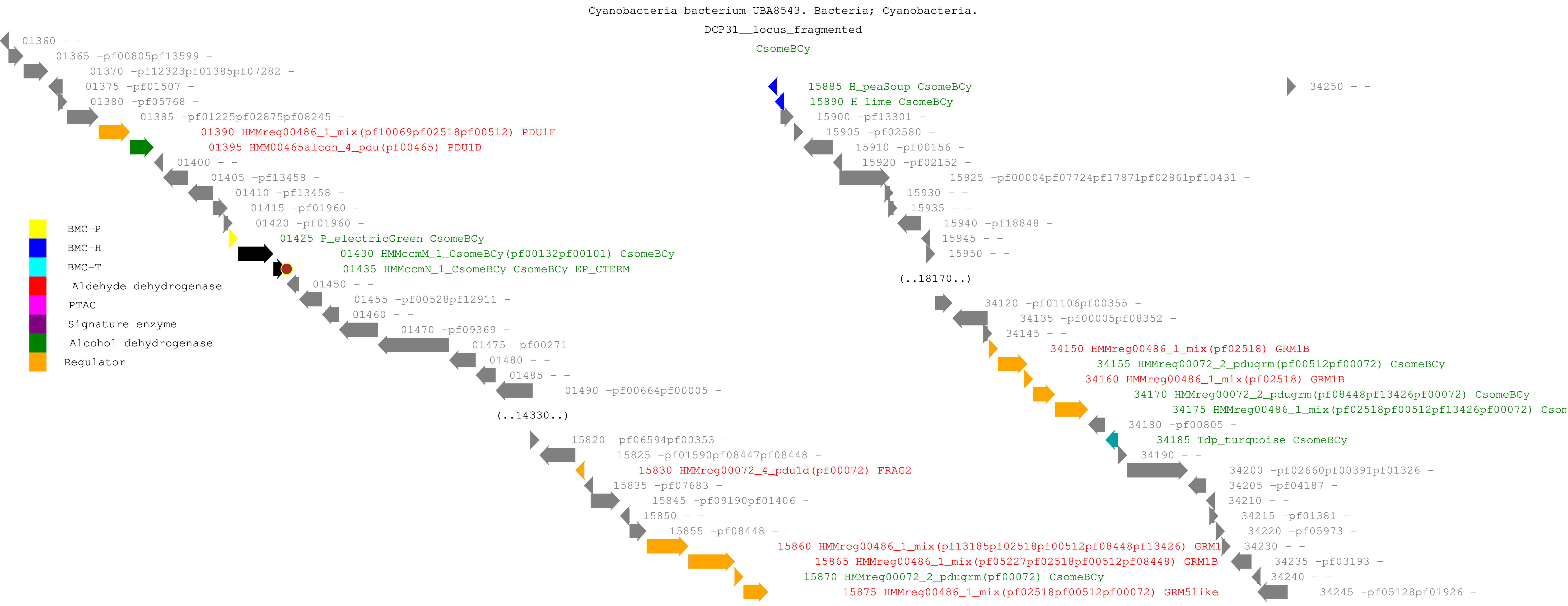
Cyanobacteria bacterium UBA8543. Bacteria; Cyanobacteria.
DCP31__locus_fragmented
CsomeBCy
01360 - -
01365 -pf00805pf13599 -
01370 -pf12323pf01385pf07282 -
01375 -pf01507 -
01380 -pf05768 -
01385 -pf01225pf02875pf08245 -
01390 HMMreg00486_1_mix(pf10069pf02518pf00512) PDU1F
01395 HMM00465alcdh_4_pdu(pf00465) PDU1D
01400 - -
01405 -pf13458 -
01410 -pf13458 -
01415 -pf01960 -
01420 -pf01960 -
01425 P_electricGreen CsomeBCy
01430 HMMccmM_1_CsomeBCy(pf00132pf00101) CsomeBCy
01435 HMMccmN_1_CsomeBCy CsomeBCy EP_CTERM
01450 - -
01455 -pf00528pf12911 -
01460 - -
01470 -pf09369 -
01475 -pf00271 -
01480 - -
01485 - -
01490 -pf00664pf00005 -
(..14330..)
15820 -pf06594pf00353 -
15825 -pf01590pf08447pf08448 -
15830 HMMreg00072_4_pdu1d(pf00072) FRAG2
15835 -pf07683 -
15845 -pf09190pf01406 -
15850 - -
15855 -pf08448 -
15860 HMMreg00486_1_mix(pf13185pf02518pf00512pf08448pf13426) GRM1
15865 HMMreg00486_1_mix(pf05227pf02518pf00512pf08448) GRM1B
15870 HMMreg00072_2_pdugrm(pf00072) CsomeBCy
15875 HMMreg00486_1_mix(pf02518pf00512pf00072) GRM5like
BMC-P
BMC-H
BMC-T
Aldehyde dehydrogenase
PTAC
Signature enzyme
Alcohol dehydrogenase
Regulator
15885 H_peaSoup CsomeBCy
15890 H_lime CsomeBCy
15900 -pf13301 -
15905 -pf02580 -
15910 -pf00156 -
15920 -pf02152 -
15925 -pf00004pf07724pf17871pf02861pf10431 -
15930 - -
15935 - -
15940 -pf18848 -
15945 - -
15950 - -
(..18170..)
34120 -pf01106pf00355 -
34135 -pf00005pf08352 -
34145 - -
34150 HMMreg00486_1_mix(pf02518) GRM1B
34155 HMMreg00072_2_pdugrm(pf00512pf00072) CsomeBCy
34160 HMMreg00486_1_mix(pf02518) GRM1B
34170 HMMreg00072_2_pdugrm(pf08448pf13426pf00072) CsomeBCy
34175 HMMreg00486_1_mix(pf02518pf00512pf13426pf00072) Csom
34180 -pf00805 -
34185 Tdp_turquoise CsomeBCy
34190 - -
34200 -pf02660pf00391pf01326 -
34205 -pf04187 -
34210 - -
34215 -pf01381 -
34220 -pf05973 -
34230 - -
34235 -pf03193 -
34240 - -
34245 -pf05128pf01926 -
34250 - -

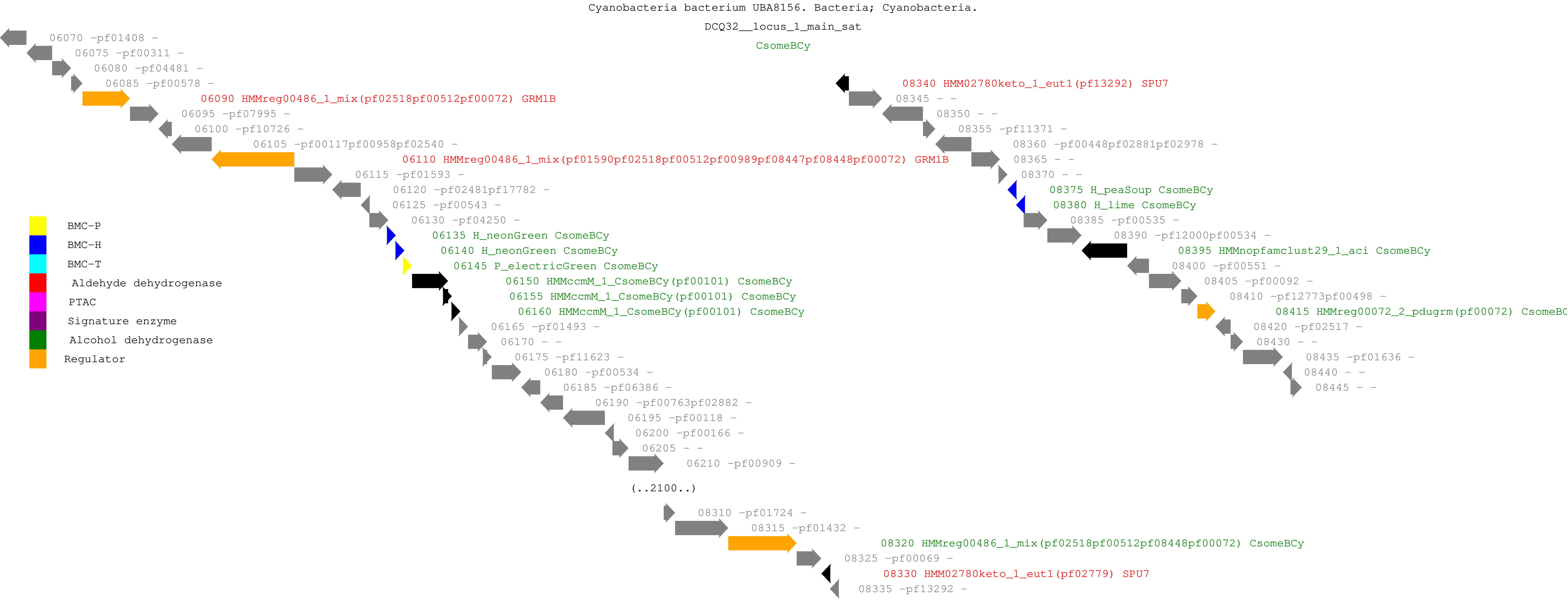

Cyanobacteria bacterium UBA8156. Bacteria; Cyanobacteria.
DCQ32__locus_1_main_sat
CsomeBCy
06070 -pf01408 -
06075 -pf00311 -
06080 -pf04481 -
06085 -pf00578 -
06090 HMMreg00486_1_mix(pf02518pf00512pf00072) GRM1B
06095 -pf07995 -
06100 -pf10726 -
06105 -pf00117pf00958pf02540 -
06110 HMMreg00486_1_mix(pf01590pf02518pf00512pf00989pf08447pf08448pf00072) GRM1B
06115 -pf01593 -
06120 -pf02481pf17782 -
06125 -pf00543 -
06130 -pf04250 -
06135 H_neonGreen CsomeBCy
06140 H_neonGreen CsomeBCy
06145 P_electricGreen CsomeBCy
06150 HMMccmM_1_CsomeBCy(pf00101) CsomeBCy
06155 HMMccmM_1_CsomeBCy(pf00101) CsomeBCy
06160 HMMccmM_1_CsomeBCy(pf00101) CsomeBCy
06165 -pf01493 -
06170 - -
06175 -pf11623 -
06180 -pf00534 -
06185 -pf06386 -
06190 -pf00763pf02882 -
06195 -pf00118 -
06200 -pf00166 -
06205 - -
06210 -pf00909 -
(..2100..)
08310 -pf01724 -
08315 -pf01432 -
08320 HMMreg00486_1_mix(pf02518pf00512pf08448pf00072) CsomeBCy
08325 -pf00069 -
08330 HMM02780keto_1_eut1(pf02779) SPU7
08335 -pf13292 -
08340 HMM02780keto_1_eut1(pf13292) SPU7
08345 - -
08350 - -
08355 -pf11371 -
08360 -pf00448pf02881pf02978 -
08365 - -
08370 - -
08375 H_peaSoup CsomeBCy
08380 H_lime CsomeBCy
08385 -pf00535 -
08390 -pf12000pf00534 -
08395 HMMnopfamclust29_1_aci CsomeBCy
08400 -pf00551 -
08405 -pf00092 -
08410 -pf12773pf00498 -
08415 HMMreg00072_2_pdugrm(pf00072) CsomeBC
08420 -pf02517 -
08430 - -
08435 -pf01636 -
08440 - -
08445 - -
BMC-P
BMC-H
BMC-T
Aldehyde dehydrogenase
PTAC
Signature enzyme
Alcohol dehydrogenase
Regulator

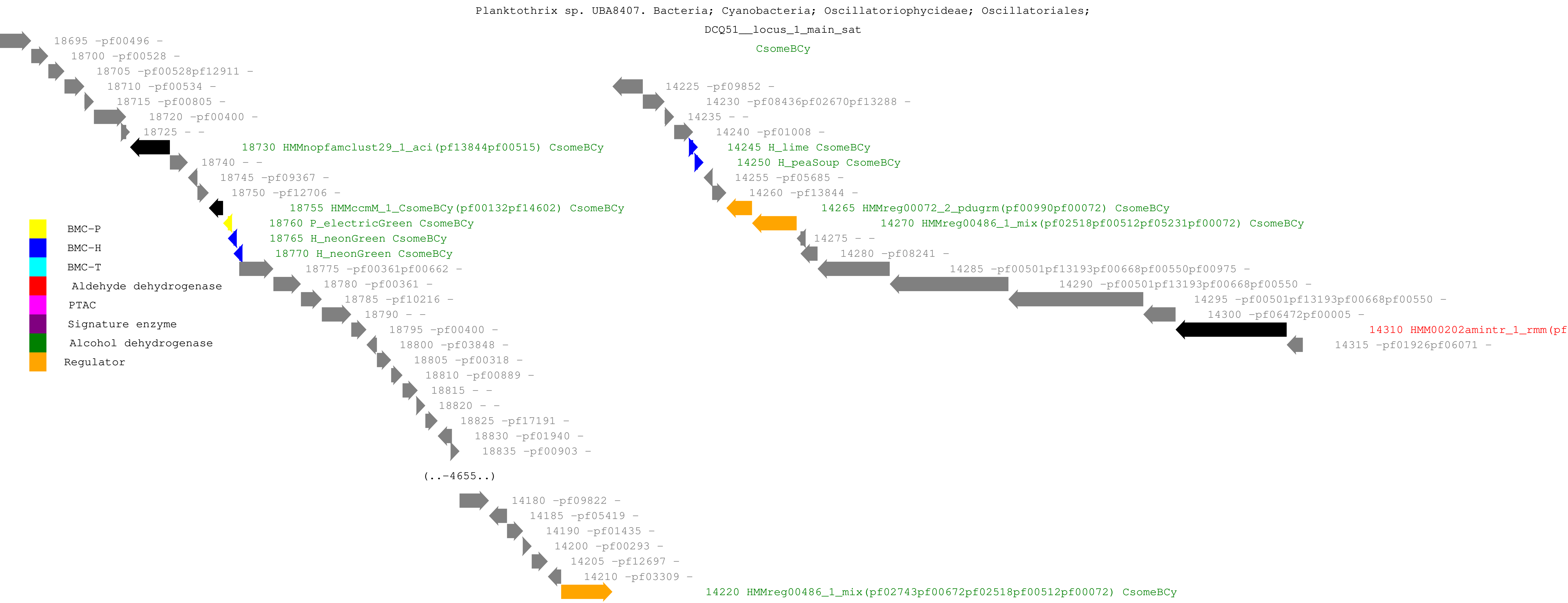
Planktothrix sp. UBA8407. Bacteria; Cyanobacteria; Oscillatoriophycideae; Oscillatoriales;
DCQ51__locus_1_main_sat
CsomeBCy
18695 -pf00496 -
18700 -pf00528 -
18705 -pf00528pf12911 -
18710 -pf00534 -
18715 -pf00805 -
18720 -pf00400 -
18725 - -
18730 HMMnopfamclust29_1_aci(pf13844pf00515) CsomeBCy
18740 - -
18745 -pf09367 -
18750 -pf12706 -
18755 HMMccmM_1_CsomeBCy(pf00132pf14602) CsomeBCy
18760 P_electricGreen CsomeBCy
18765 H_neonGreen CsomeBCy
18770 H_neonGreen CsomeBCy
18775 -pf00361pf00662 -
18780 -pf00361 -
18785 -pf10216 -
18790 - -
18795 -pf00400 -
18800 -pf03848 -
18805 -pf00318 -
18810 -pf00889 -
18815 - -
18820 - -
18825 -pf17191 -
18830 -pf01940 -
18835 -pf00903 -
(..-4655..)
14180 -pf09822 -
14185 -pf05419 -
14190 -pf01435 -
14200 -pf00293 -
14205 -pf12697 -
14210 -pf03309 -
14220 HMMreg00486_1_mix(pf02743pf00672pf02518pf00512pf00072) CsomeBCy
14225 -pf09852 -
14230 -pf08436pf02670pf13288 -
14235 - -
14240 -pf01008 -
14245 H_lime CsomeBCy
14250 H_peaSoup CsomeBCy
14255 -pf05685 -
14260 -pf13844 -
14265 HMMreg00072_2_pdugrm(pf00990pf00072) CsomeBCy
14270 HMMreg00486_1_mix(pf02518pf00512pf05231pf00072) CsomeBCy
14275 - -
14280 -pf08241 -
14285 -pf00501pf13193pf00668pf00550pf00975 -
14290 -pf00501pf13193pf00668pf00550 -
14295 -pf00501pf13193pf00668pf00550 -
14300 -pf06472pf00005 -
14310 HMM00202amintr_1_rmm(pf
14315 -pf01926pf06071 -
BMC-P
BMC-H
BMC-T
Aldehyde dehydrogenase
PTAC
Signature enzyme
Alcohol dehydrogenase
Regulator

Planktothrix sp. UBA8402. Bacteria; Cyanobacteria; Oscillatoriophycideae; Oscillatoriales;

DCQ63__locus_fragmented

CsomeBCy

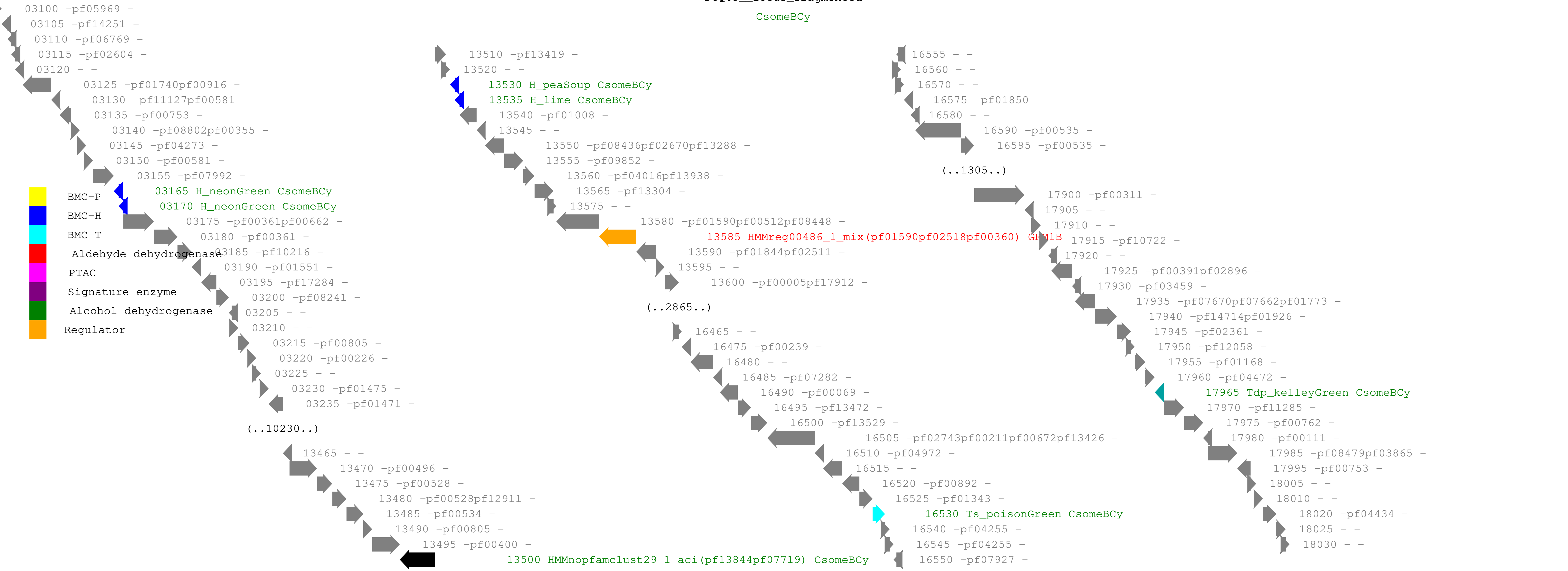

Microcoleaceae bacterium UBA11344. Bacteria; Cyanobacteria; Oscillatoriophycideae; Oscillatoriales;

DCS91__locus_1_main_sat

CsomeBCy

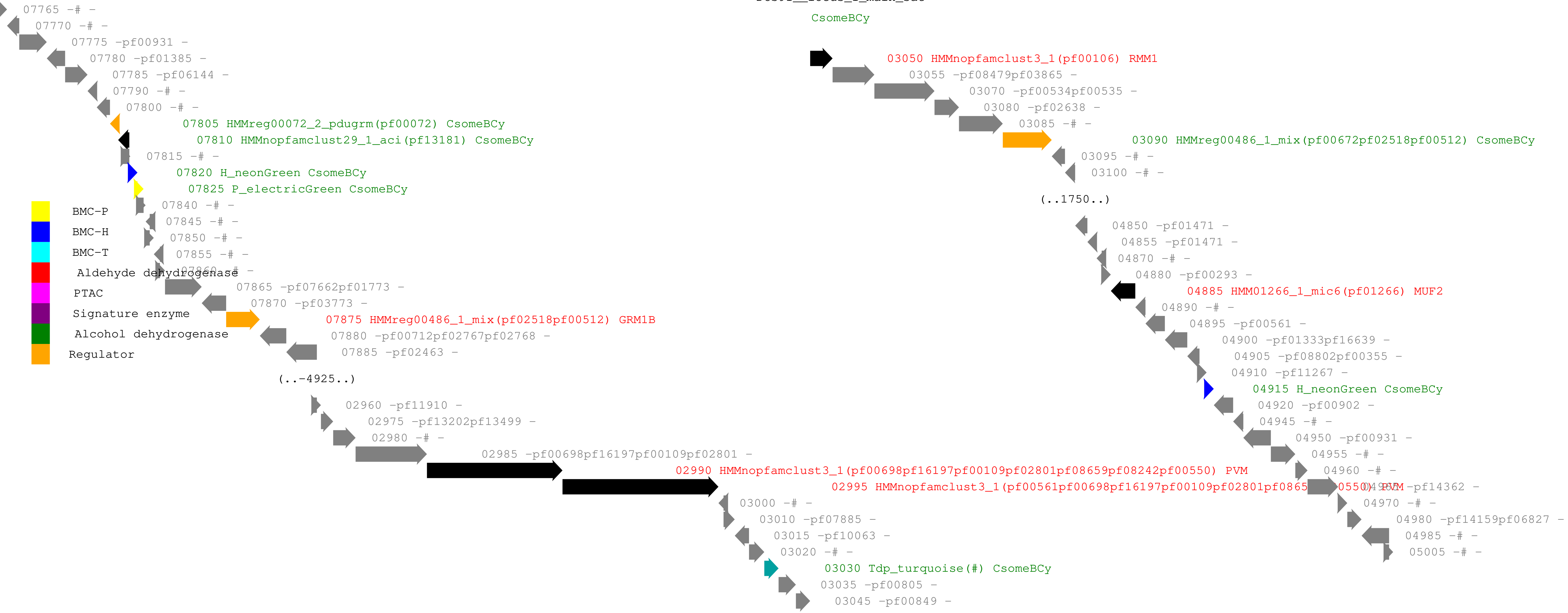

Cyanobacteria bacterium UBA11372. Bacteria; Cyanobacteria.

DCY88__locus_fragmented

CsomeBCy

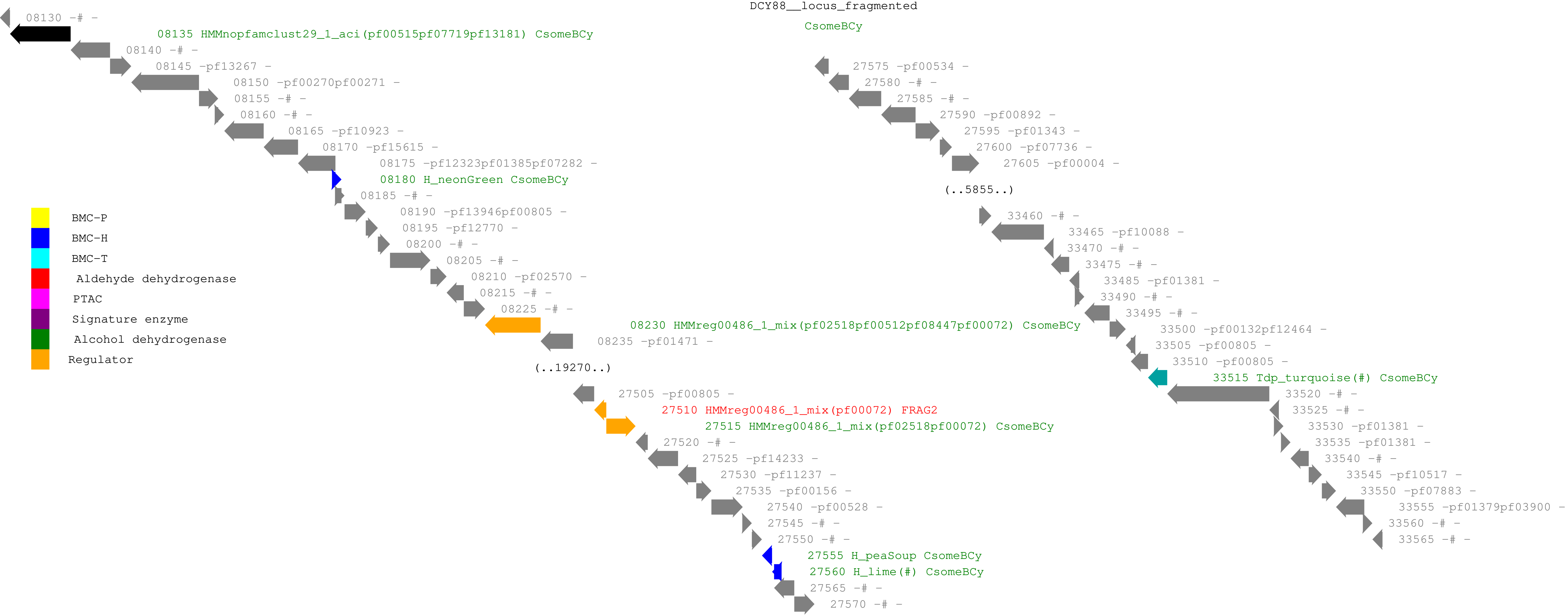

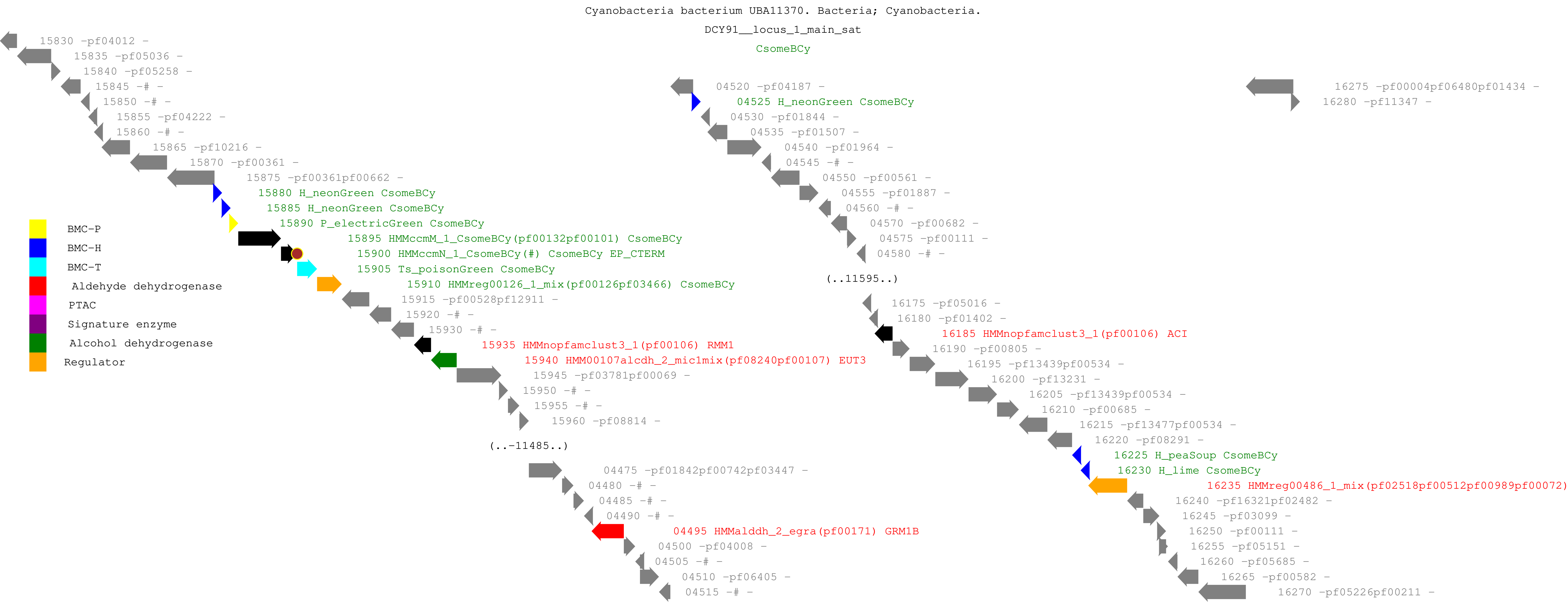
Cyanobacteria bacterium UBA11370. Bacteria; Cyanobacteria.
DCY91__locus_1_main_sat
CsomeBCy
15830 -pf04012 -
15835 -pf05036 -
15840 -pf05258 -
15845 -# -
15850 -# -
15855 -pf04222 -
15860 -# -
15865 -pf10216 -
15870 -pf00361 -
15875 -pf00361pf00662 -
15880 H_neonGreen CsomeBCy
15885 H_neonGreen CsomeBCy
15890 P_electricGreen CsomeBCy
15895 HMMccmM_1_CsomeBCy(pf00132pf00101) CsomeBCy
15900 HMMccmN_1_CsomeBCy(#) CsomeBCy EP_CTERM
15905 Ts_poisonGreen CsomeBCy
15910 HMMreg00126_1_mix(pf00126pf03466) CsomeBCy
15915 -pf00528pf12911 -
15920 -# -
15930 -# -
15935 HMMnopfamclust3_1(pf00106) RMM1
15940 HMM00107alcdh_2_mic1mix(pf08240pf00107) EUT3
15945 -pf03781pf00069 -
15950 -# -
15955 -# -
15960 -pf08814 -
(..-11485..)
04475 -pf01842pf00742pf03447 -
04480 -# -
04485 -# -
04490 -# -
04495 HMMaldh_2_egra(pf00171) GRM1B
04500 -pf04008 -
04505 -# -
04510 -pf06405 -
04515 -# -
04520 -pf04187 -
04525 H_neonGreen CsomeBCy
04530 -pf01844 -
04535 -pf01507 -
04540 -pf01964 -
04545 -# -
04550 -pf00561 -
04555 -pf01887 -
04560 -# -
04570 -pf00682 -
04575 -pf00111 -
04580 -# -
(..11595..)
16175 -pf05016 -
16180 -pf01402 -
16185 HMMnopfamclust3_1(pf00106) ACI
16190 -pf00805 -
16195 -pf13439pf00534 -
16200 -pf13231 -
16205 -pf13439pf00534 -
16210 -pf00685 -
16215 -pf13477pf00534 -
16220 -pf08291 -
16225 H_peaSoup CsomeBCy
16230 H_lime CsomeBCy
16235 HMMreg00486_1_mix(pf02518pf00512pf00989pf00072)
16240 -pf16321pf02482 -
16245 -pf03099 -
16250 -pf00111 -
16255 -pf05151 -
16260 -pf05685 -
16265 -pf00582 -
16270 -pf05226pf00211 -
16275 -pf00004pf06480pf01434 -
16280 -pf11347 -
BMC-P
BMC-H
BMC-T
Aldehyde dehydrogenase
PTAC
Signature enzyme
Alcohol dehydrogenase
Regulator

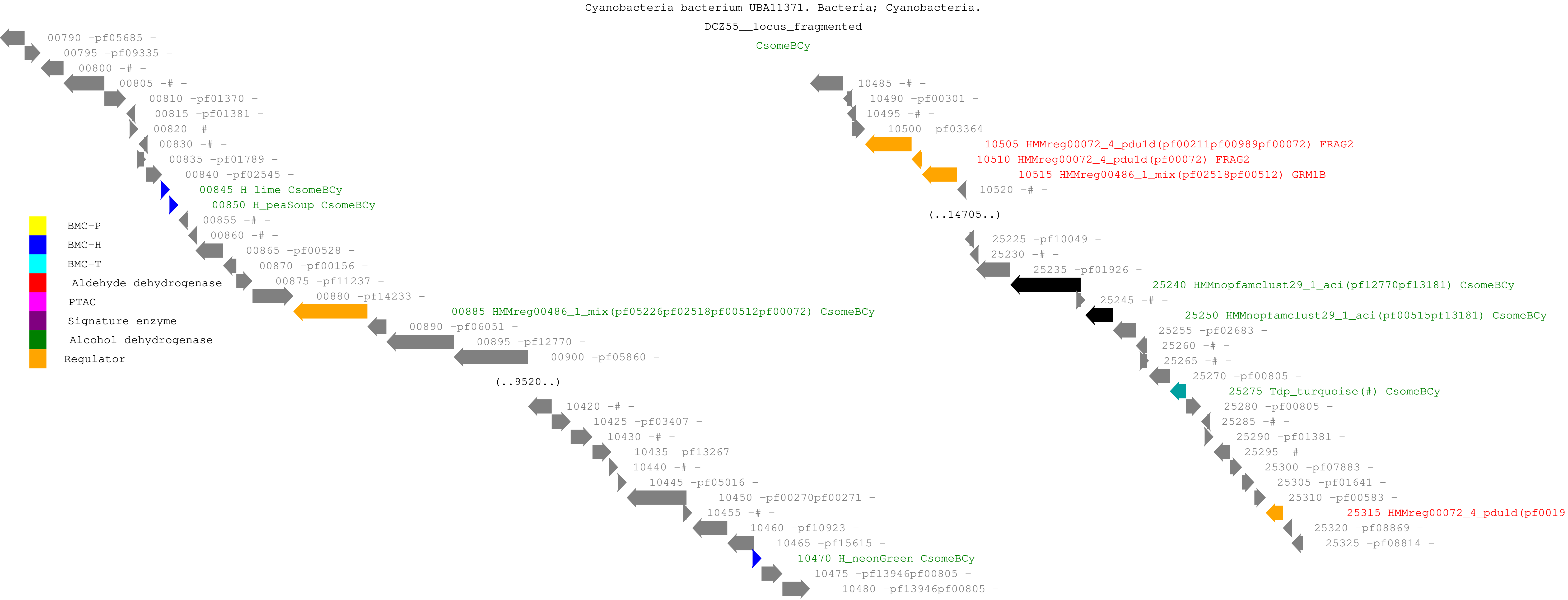

Cyanobacteria bacterium UBA11371. Bacteria; Cyanobacteria.
DCZ55__locus_fragmented
CsomeBCy
00790 -pf05685 -
00795 -pf09335 -
00800 -# -
00805 -# -
00810 -pf01370 -
00815 -pf01381 -
00820 -# -
00830 -# -
00835 -pf01789 -
00840 -pf02545 -
00845 H_lime CsomeBCy
00850 H_peaSoup CsomeBCy
00855 -# -
00860 -# -
00865 -pf00528 -
00870 -pf00156 -
00875 -pf11237 -
00880 -pf14233 -
00885 HMMreg00486_1_mix(pf05226pf02518pf00512pf00072) CsomeBCy
00890 -pf06051 -
00895 -pf12770 -
00900 -pf05860 -
(..9520..)
10420 -# -
10425 -pf03407 -
10430 -# -
10435 -pf13267 -
10440 -# -
10445 -pf05016 -
10450 -pf00270pf00271 -
10455 -# -
10460 -pf10923 -
10465 -pf15615 -
10470 H_neonGreen CsomeBCy
10475 -pf13946pf00805 -
10480 -pf13946pf00805 -
10485 -# -
10490 -pf00301 -
10495 -# -
10500 -pf03364 -
10505 HMMreg00072_4_pdu1d(pf00211pf00989pf00072) FRAG2
10510 HMMreg00072_4_pdu1d(pf00072) FRAG2
10515 HMMreg00486_1_mix(pf02518pf00512) GRM1B
10520 -# -
(..14705..)
25225 -pf10049 -
25230 -# -
25235 -pf01926 -
25240 HMMnopfamclust29_1_aci(pf12770pf13181) CsomeBCy
25245 -# -
25250 HMMnopfamclust29_1_aci(pf00515pf13181) CsomeBCy
25255 -pf02683 -
25260 -# -
25265 -# -
25270 -pf00805 -
25275 Tdp_turquoise(#) CsomeBCy
25280 -pf00805 -
25285 -# -
25290 -pf01381 -
25295 -# -
25300 -pf07883 -
25305 -pf01641 -
25310 -pf00583 -
25315 HMMreg00072_4_pdu1d(pf0019
25320 -pf08869 -
25325 -pf08814 -
BMC-P
BMC-H
BMC-T
Aldehyde dehydrogenase
PTAC
Signature enzyme
Alcohol dehydrogenase
Regulator

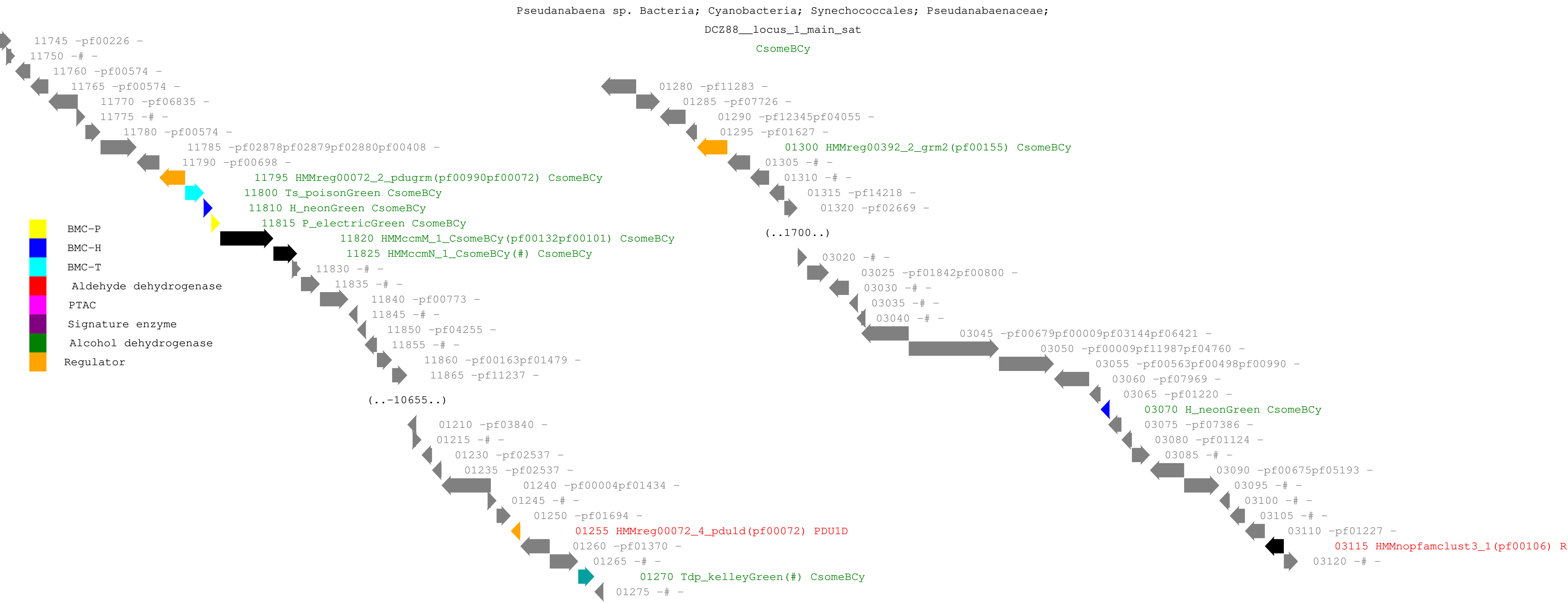
Pseudanabaena sp. Bacteria; Cyanobacteria; Synechococcales; Pseudanabaenaceae;
DCZ88__locus_1_main_sat
CsomeBCy
11745 -pf00226 -
11750 -# -
11760 -pf00574 -
11765 -pf00574 -
11770 -pf06835 -
11775 -# -
11780 -pf00574 -
11785 -pf02878pf02879pf02880pf00408 -
11790 -pf00698 -
11795 HMMreg00072_2_pdugrm(pf00990pf00072) CsomeBCy
11800 Ts_poisonGreen CsomeBCy
11810 H_neonGreen CsomeBCy
11815 P_electricGreen CsomeBCy
11820 HMMccmM_1_CsomeBCy(pf00132pf00101) CsomeBCy
11825 HMMccmN_1_CsomeBCy(#) CsomeBCy
11830 -# -
11835 -# -
11840 -pf00773 -
11845 -# -
11850 -pf04255 -
11855 -# -
11860 -pf00163pf01479 -
11865 -pf11237 -
(..-10655..)
01210 -pf03840 -
01215 -# -
01230 -pf02537 -
01235 -pf02537 -
01240 -pf00004pf01434 -
01245 -# -
01250 -pf01694 -
01255 HMMreg00072_4_pdu1d(pf00072) PDU1D
01260 -pf01370 -
01265 -# -
01270 Tdp_kelleyGreen(#) CsomeBCy
01275 -# -
BMC-P
BMC-H
BMC-T
Aldehyde dehydrogenase
PTAC
Signature enzyme
Alcohol dehydrogenase
Regulator
01280 -pf11283 -
01285 -pf07726 -
01290 -pf12345pf04055 -
01295 -pf01627 -
01300 HMMreg00392_2_grm2(pf00155) CsomeBCy
01305 -# -
01310 -# -
01315 -pf14218 -
01320 -pf02669 -
(..1700..)
03020 -# -
03025 -pf01842pf00800 -
03030 -# -
03035 -# -
03040 -# -
03045 -pf00679pf00009pf03144pf06421 -
03050 -pf00009pf11987pf04760 -
03055 -pf00563pf00498pf00990 -
03060 -pf07969 -
03065 -pf01220 -
03070 H_neonGreen CsomeBCy
03075 -pf07386 -
03080 -pf01124 -
03085 -# -
03090 -pf00675pf05193 -
03095 -# -
03100 -# -
03105 -# -
03110 -pf01227 -
03115 HMMnopfamclust3_1(pf00106) R
03120 -# -

Cyanobacteria bacterium UBA11166. Bacteria; Cyanobacteria.

DD000__locus_1_main_sat

CsomeBCy

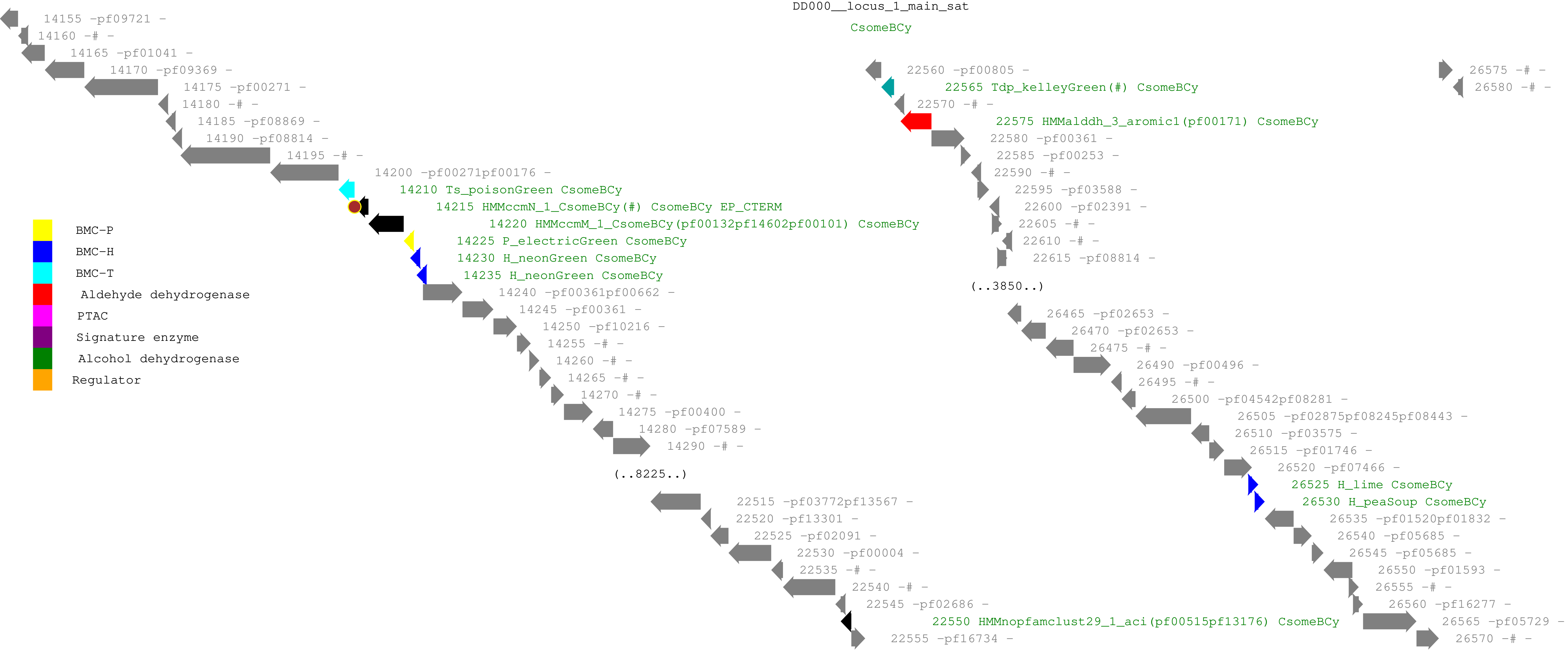

Microcoleaceae bacterium UBA10368. Bacteria; Cyanobacteria; Oscillatoriophycideae; Oscillatoriales;

DD001__locus_fragmented

CsomeBCy

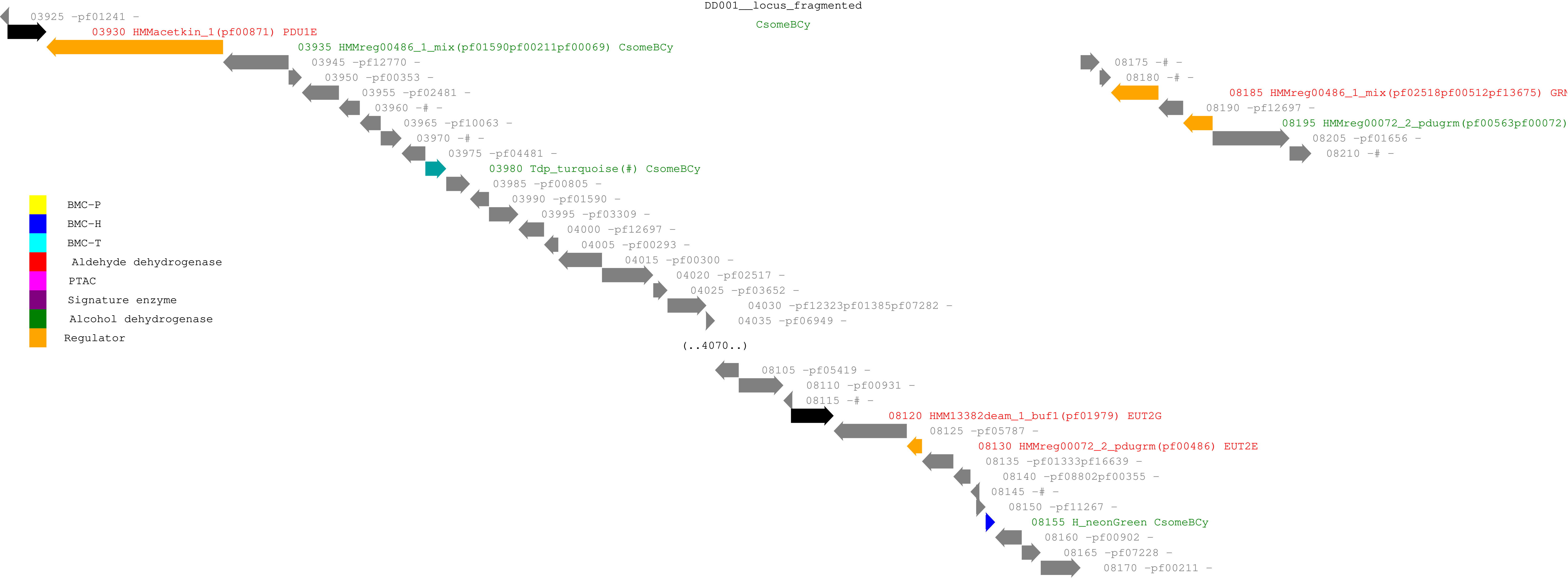

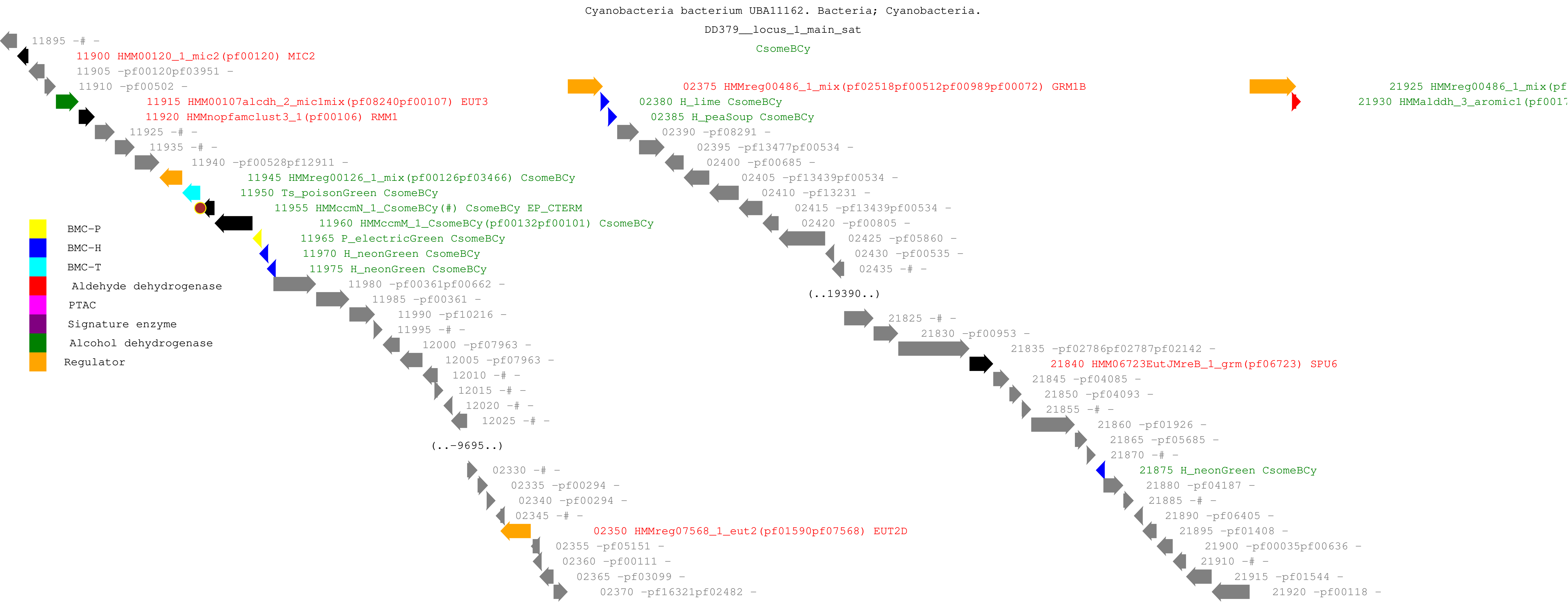
Cyanobacteria bacterium UBA11162. Bacteria; Cyanobacteria.
DD379__locus_1_main_sat
CsomeBCy
11895 -# -
11900 HMM00120_1_mic2(pf00120) MIC2
11905 -pf00120pf03951 -
11910 -pf00502 -
11915 HMM00107alcdh_2_mic1mix(pf08240pf00107) EUT3
11920 HMMnopfamclust3_1(pf00106) RMM1
11925 -# -
11935 -# -
11940 -pf00528pf12911 -
11945 HMMreg00126_1_mix(pf00126pf03466) CsomeBCy
11950 Ts_poisonGreen CsomeBCy
11955 HMMccmN_1_CsomeBCy(#) CsomeBCy EP_CTERM
11960 HMMccmM_1_CsomeBCy(pf00132pf00101) CsomeBCy
11965 P_electricGreen CsomeBCy
11970 H_neonGreen CsomeBCy
11975 H_neonGreen CsomeBCy
11980 -pf00361pf00662 -
11985 -pf00361 -
11990 -pf10216 -
11995 -# -
12000 -pf07963 -
12005 -pf07963 -
12010 -# -
12015 -# -
12020 -# -
12025 -# -
(..-9695..)
02330 -# -
02335 -pf00294 -
02340 -pf00294 -
02345 -# -
02350 HMMreg07568_1_eut2(pf01590pf07568) EUT2D
02355 -pf05151 -
02360 -pf00111 -
02365 -pf03099 -
02370 -pf16321pf02482 -
02375 HMMreg00486_1_mix(pf02518pf00512pf00989pf00072) GRM1B
02380 H_lime CsomeBCy
02385 H_peaSoup CsomeBCy
02390 -pf08291 -
02395 -pf13477pf00534 -
02400 -pf00685 -
02405 -pf13439pf00534 -
02410 -pf13231 -
02415 -pf13439pf00534 -
02420 -pf00805 -
02425 -pf05860 -
02430 -pf00535 -
02435 -# -
(..19390..)
21825 -# -
21830 -pf00953 -
21835 -pf02786pf02787pf02142 -
21840 HMM06723EutJMreB_1_grm(pf06723) SPU6
21845 -pf04085 -
21850 -pf04093 -
21855 -# -
21860 -pf01926 -
21865 -pf05685 -
21870 -# -
21875 H_neonGreen CsomeBCy
21880 -pf04187 -
21885 -# -
21890 -pf06405 -
21895 -pf01408 -
21900 -pf00035pf00636 -
21910 -# -
21915 -pf01544 -
21920 -pf00118 -
21925 HMMreg00486_1_mix(pf
21930 HMMaldh_3_aromic1(pf0017
BMC-P
BMC-H
BMC-T
Aldehyde dehydrogenase
PTAC
Signature enzyme
Alcohol dehydrogenase
Regulator

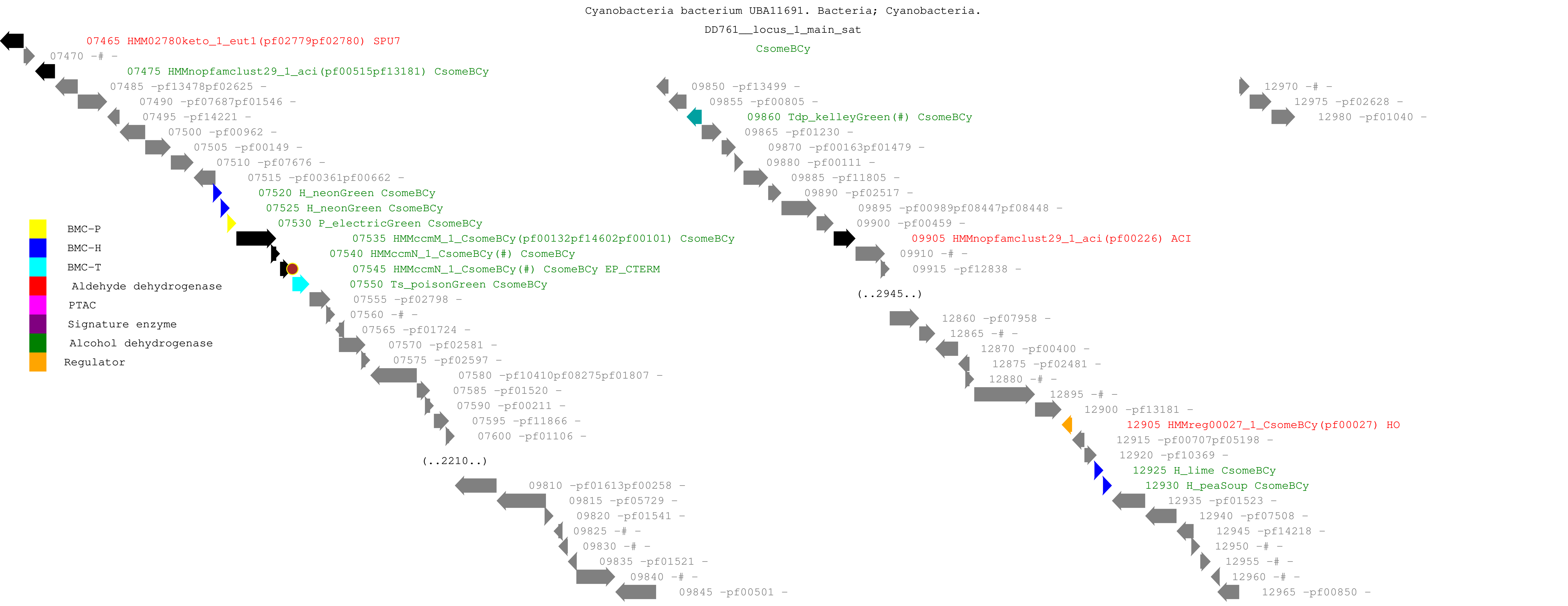

Cyanobacteria bacterium UBA11691. Bacteria; Cyanobacteria.
DD761__locus_1_main_sat
CsomeBCy
07465 HMM02780keto_1_eut1(pf02779pf02780) SPU7
07470 -# -
07475 HMMnopfamclust29_1_aci(pf00515pf13181) CsomeBCy
07485 -pf13478pf02625 -
07490 -pf07687pf01546 -
07495 -pf14221 -
07500 -pf00962 -
07505 -pf00149 -
07510 -pf07676 -
07515 -pf00361pf00662 -
07520 H_neonGreen CsomeBCy
07525 H_neonGreen CsomeBCy
07530 P_electricGreen CsomeBCy
07535 HMMccmM_1_CsomeBCy(pf00132pf14602pf00101) CsomeBCy
07540 HMMccmN_1_CsomeBCy(#) CsomeBCy
07545 HMMccmN_1_CsomeBCy(#) CsomeBCy EP_CTERM
07550 Ts_poisonGreen CsomeBCy
07555 -pf02798 -
07560 -# -
07565 -pf01724 -
07570 -pf02581 -
07575 -pf02597 -
07580 -pf10410pf08275pf01807 -
07585 -pf01520 -
07590 -pf00211 -
07595 -pf11866 -
07600 -pf01106 -
(..2210..)
09810 -pf01613pf00258 -
09815 -pf05729 -
09820 -pf01541 -
09825 -# -
09830 -# -
09835 -pf01521 -
09840 -# -
09845 -pf00501 -
BMC-P
BMC-H
BMC-T
Aldehyde dehydrogenase
PTAC
Signature enzyme
Alcohol dehydrogenase
Regulator
09850 -pf13499 -
09855 -pf00805 -
09860 Tdp_kelleyGreen(#) CsomeBCy
09865 -pf01230 -
09870 -pf00163pf01479 -
09880 -pf00111 -
09885 -pf11805 -
09890 -pf02517 -
09895 -pf00989pf08447pf08448 -
09900 -pf00459 -
09905 HMMnopfamclust29_1_aci(pf00226) ACI
09910 -# -
09915 -pf12838 -
(..2945..)
12860 -pf07958 -
12865 -# -
12870 -pf00400 -
12875 -pf02481 -
12880 -# -
12895 -# -
12900 -pf13181 -
12905 HMMreg00027_1_CsomeBCy(pf00027) HO
12915 -pf00707pf05198 -
12920 -pf10369 -
12925 H_lime CsomeBCy
12930 H_peaSoup CsomeBCy
12935 -pf01523 -
12940 -pf07508 -
12945 -pf14218 -
12950 -# -
12955 -# -
12960 -# -
12965 -pf00850 -
12970 -# -
12975 -pf02628 -
12980 -pf01040 -

Cyanobacteria bacterium UBA11368. Bacteria; Cyanobacteria.

DD990__locus_1_main_sat

CsomeBCy

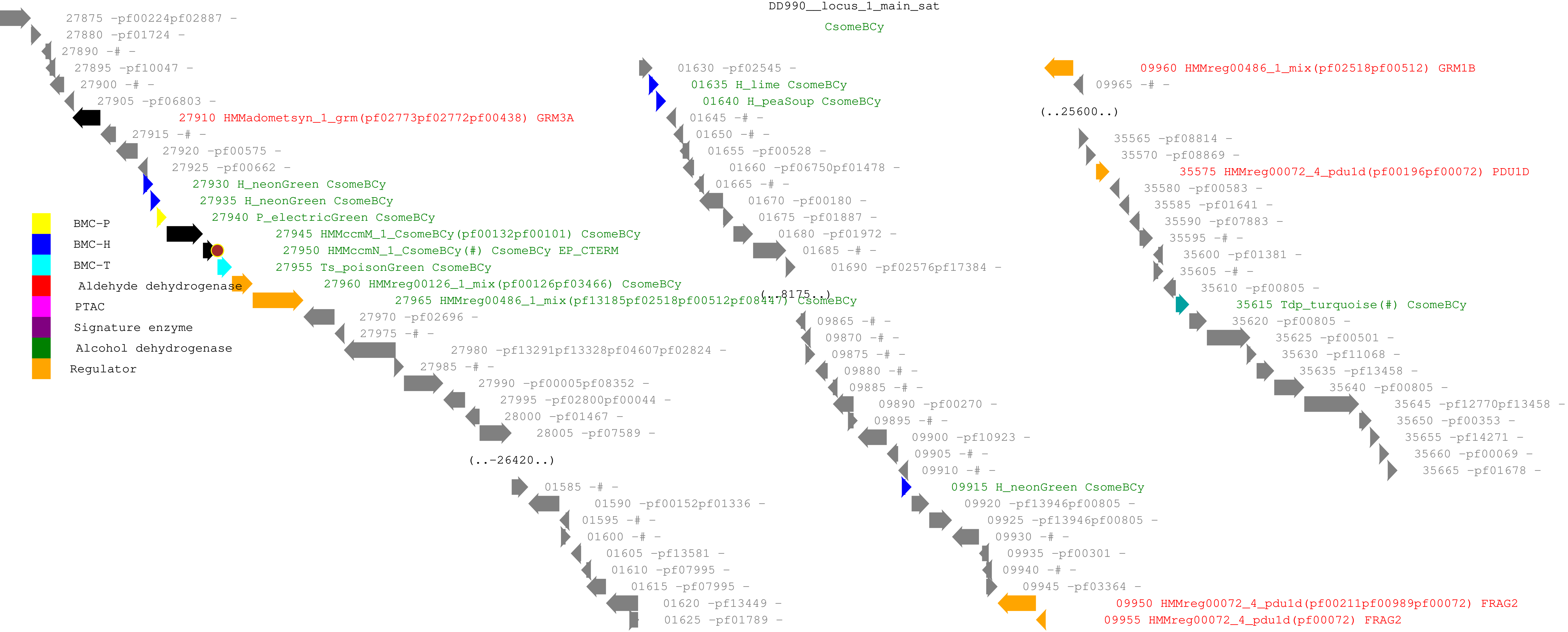

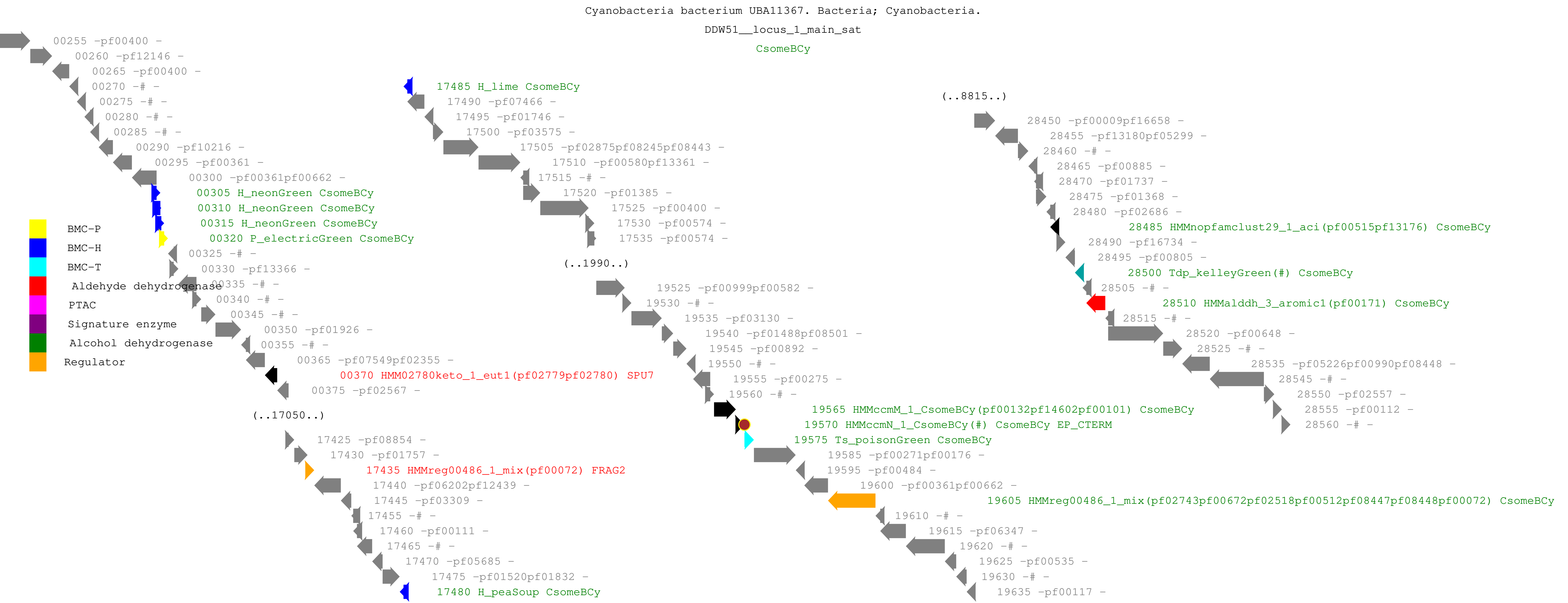

Cyanobacteria bacterium UBA11367. Bacteria; Cyanobacteria.
DDW51__locus_1_main_sat
CsomeBCy
00255 -pf00400 -
00260 -pf12146 -
00265 -pf00400 -
00270 -# -
00275 -# -
00280 -# -
00285 -# -
00290 -pf10216 -
00295 -pf00361 -
00300 -pf00361pf00662 -
00305 H_neonGreen CsomeBCy
00310 H_neonGreen CsomeBCy
00315 H_neonGreen CsomeBCy
00320 P_electricGreen CsomeBCy
00325 -# -
00330 -pf13366 -
00335 -# -
00340 -# -
00345 -# -
00350 -pf01926 -
00355 -# -
00365 -pf07549pf02355 -
00370 HMM02780keto_1_eut1(pf02779pf02780) SPU7
00375 -pf02567 -
(..17050..)
17425 -pf08854 -
17430 -pf01757 -
17435 HMMreg00486_1_mix(pf00072) FRAG2
17440 -pf06202pf12439 -
17445 -pf03309 -
17455 -# -
17460 -pf00111 -
17465 -# -
17470 -pf05685 -
17475 -pf01520pf01832 -
17480 H_peaSoup CsomeBCy
17485 H_lime CsomeBCy
17490 -pf07466 -
17495 -pf01746 -
17500 -pf03575 -
17505 -pf02875pf08245pf08443 -
17510 -pf00580pf13361 -
17515 -# -
17520 -pf01385 -
17525 -pf00400 -
17530 -pf00574 -
17535 -pf00574 -
(..1990..)
19525 -pf00999pf00582 -
19530 -# -
19535 -pf03130 -
19540 -pf01488pf08501 -
19545 -pf00892 -
19550 -# -
19555 -pf00275 -
19560 -# -
19565 HMMccmM_1_CsomeBCy(pf00132pf14602pf00101) CsomeBCy
19570 HMMccmN_1_CsomeBCy(#) CsomeBCy EP_CTERM
19575 Ts_poisonGreen CsomeBCy
19585 -pf00271pf00176 -
19595 -pf00484 -
19600 -pf00361pf00662 -
19605 HMMreg00486_1_mix(pf02743pf00672pf02518pf00512pf08447pf08448pf00072) CsomeBCy
19610 -# -
19615 -pf06347 -
19620 -# -
19625 -pf00535 -
19630 -# -
19635 -pf00117 -
(..8815..)
28450 -pf00009pf16658 -
28455 -pf13180pf05299 -
28460 -# -
28465 -pf00885 -
28470 -pf01737 -
28475 -pf01368 -
28480 -pf02686 -
28485 HMMnopfamclust29_1_aci(pf00515pf13176) CsomeBCy
28490 -pf16734 -
28495 -pf00805 -
28500 Tdp_kelleyGreen(#) CsomeBCy
28505 -# -
28510 HMMaldd h_3_aromic1(pf00171) CsomeBCy
28515 -# -
28520 -pf00648 -
28525 -# -
28535 -pf05226pf00990pf08448 -
28545 -# -
28550 -pf02557 -
28555 -pf00112 -
28560 -# -
BMC-P
BMC-H
BMC-T
Aldehyde dehydrogenase
PTAC
Signature enzyme
Alcohol dehydrogenase
Regulator

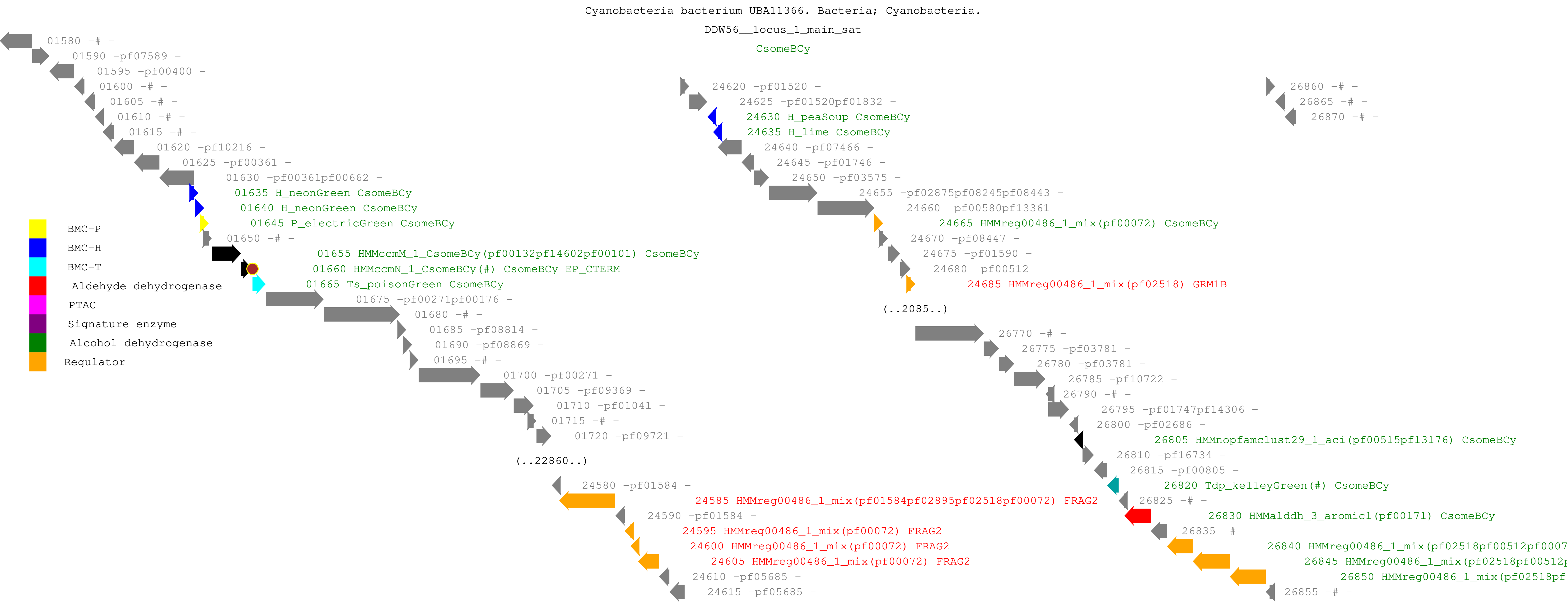
Cyanobacteria bacterium UBA11366. Bacteria; Cyanobacteria.
DDW56__locus_1_main_sat
CsomeBCy
BMC-P
BMC-H
BMC-T
Aldehyde dehydrogenase
PTAC
Signature enzyme
Alcohol dehydrogenase
Regulator
01580 -# -
01590 -pf07589 -
01595 -pf00400 -
01600 -# -
01605 -# -
01610 -# -
01615 -# -
01620 -pf10216 -
01625 -pf00361 -
01630 -pf00361pf00662 -
01635 H_neonGreen CsomeBCy
01640 H_neonGreen CsomeBCy
01645 P_electricGreen CsomeBCy
01650 -# -
01655 HMMccmM_1_CsomeBCy(pf00132pf14602pf00101) CsomeBCy
01660 HMMccmN_1_CsomeBCy(#) CsomeBCy EP_CTERM
01665 Ts_poisonGreen CsomeBCy
01675 -pf00271pf00176 -
01680 -# -
01685 -pf08814 -
01690 -pf08869 -
01695 -# -
01700 -pf00271 -
01705 -pf09369 -
01710 -pf01041 -
01715 -# -
01720 -pf09721 -
(..22860..)
24580 -pf01584 -
24585 HMMreg00486_1_mix(pf01584pf02895pf02518pf00072) FRAG2
24590 -pf01584 -
24595 HMMreg00486_1_mix(pf00072) FRAG2
24600 HMMreg00486_1_mix(pf00072) FRAG2
24605 HMMreg00486_1_mix(pf00072) FRAG2
24610 -pf05685 -
24615 -pf05685 -
24620 -pf01520 -
24625 -pf01520pf01832 -
24630 H_peaSoup CsomeBCy
24635 H_lime CsomeBCy
24640 -pf07466 -
24645 -pf01746 -
24650 -pf03575 -
24655 -pf02875pf08245pf08443 -
24660 -pf00580pf13361 -
24665 HMMreg00486_1_mix(pf00072) CsomeBCy
24670 -pf08447 -
24675 -pf01590 -
24680 -pf00512 -
24685 HMMreg00486_1_mix(pf02518) GRM1B
(..2085..)
26770 -# -
26775 -pf03781 -
26780 -pf03781 -
26785 -pf10722 -
26790 -# -
26795 -pf01747pf14306 -
26800 -pf02686 -
26805 HMMnopfamclust29_1_aci(pf00515pf13176) CsomeBCy
26810 -pf16734 -
26815 -pf00805 -
26820 Tdp_kelleyGreen(#) CsomeBCy
26825 -# -
26830 HMMalddh_3_aromic1(pf00171) CsomeBCy
26835 -# -
26840 HMMreg00486_1_mix(pf02518pf00512pf0007
26845 HMMreg00486_1_mix(pf02518pf00512p
26850 HMMreg00486_1_mix(pf02518pf
26855 -# -
26860 -# -
26865 -# -
26870 -# -

Cyanobacteria bacterium UBA11369. Bacteria; Cyanobacteria.

DDW76__locus_1_main_sat

CsomeBCy

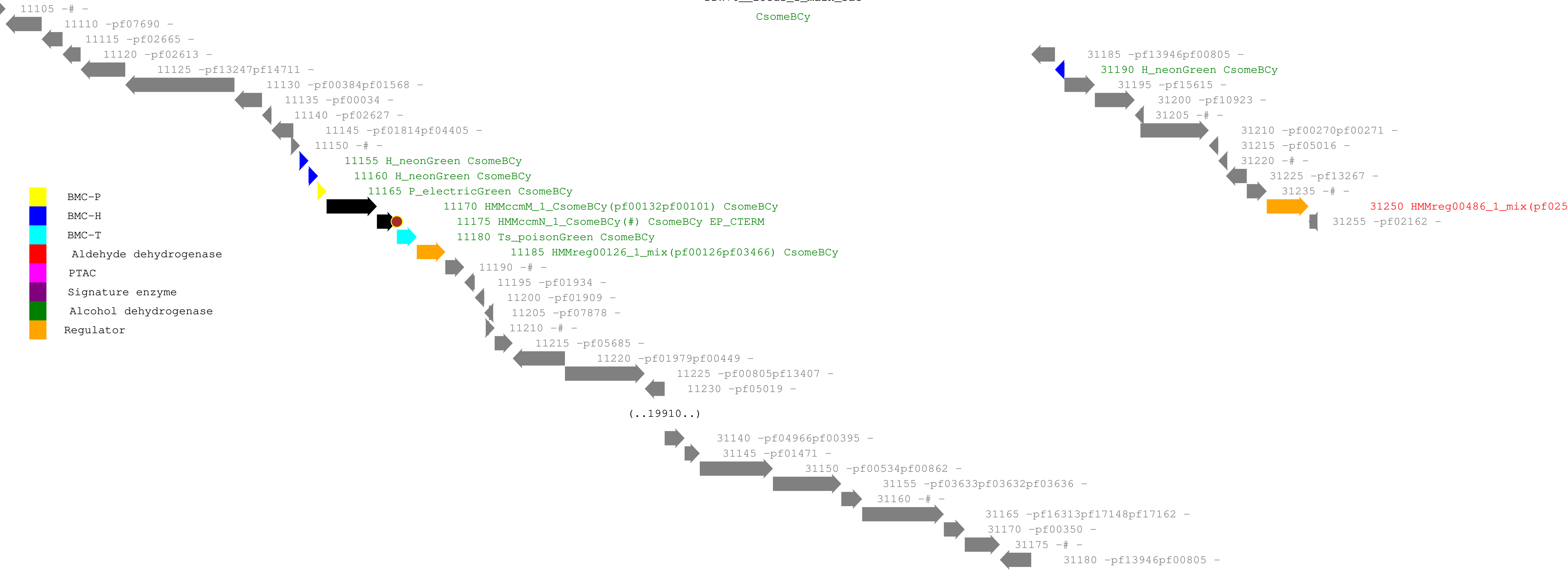

Planktothrix sp. UBA10369. Bacteria; Cyanobacteria; Oscillatoriophycideae; Oscillatoriales;

DDZ60__locus_1_main_sat

CsomeBCy

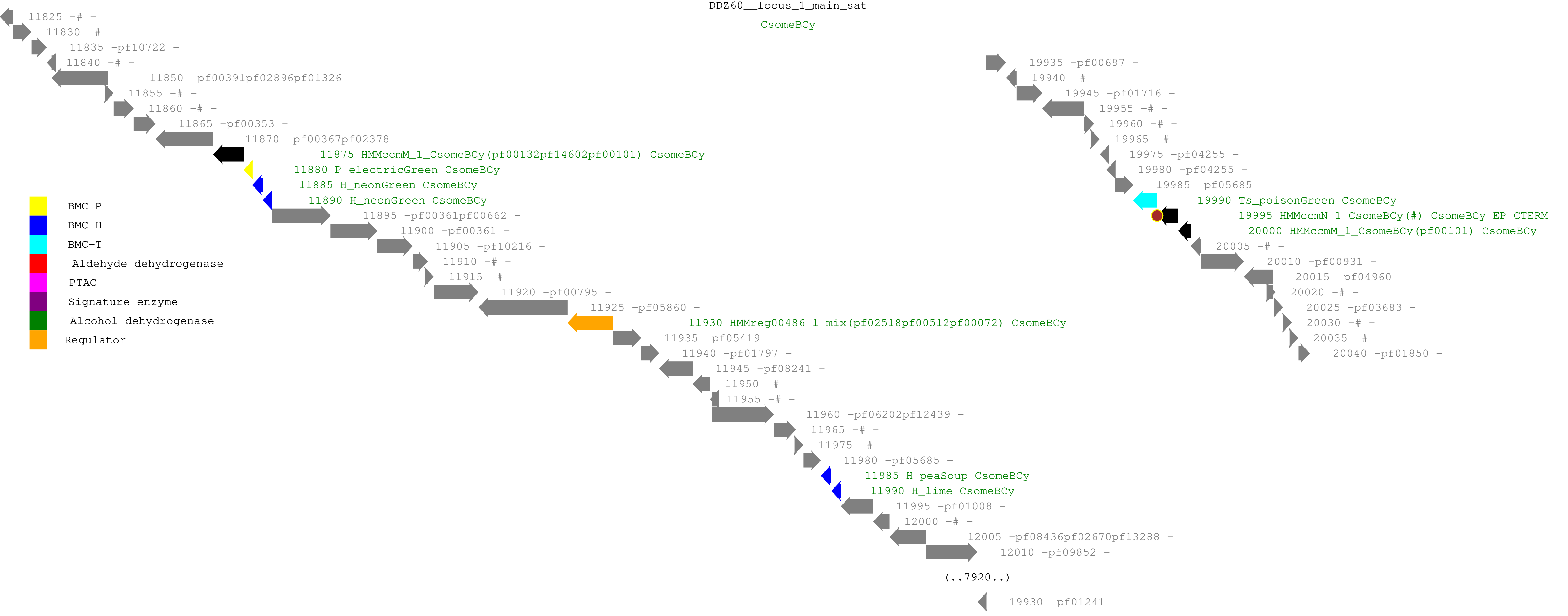

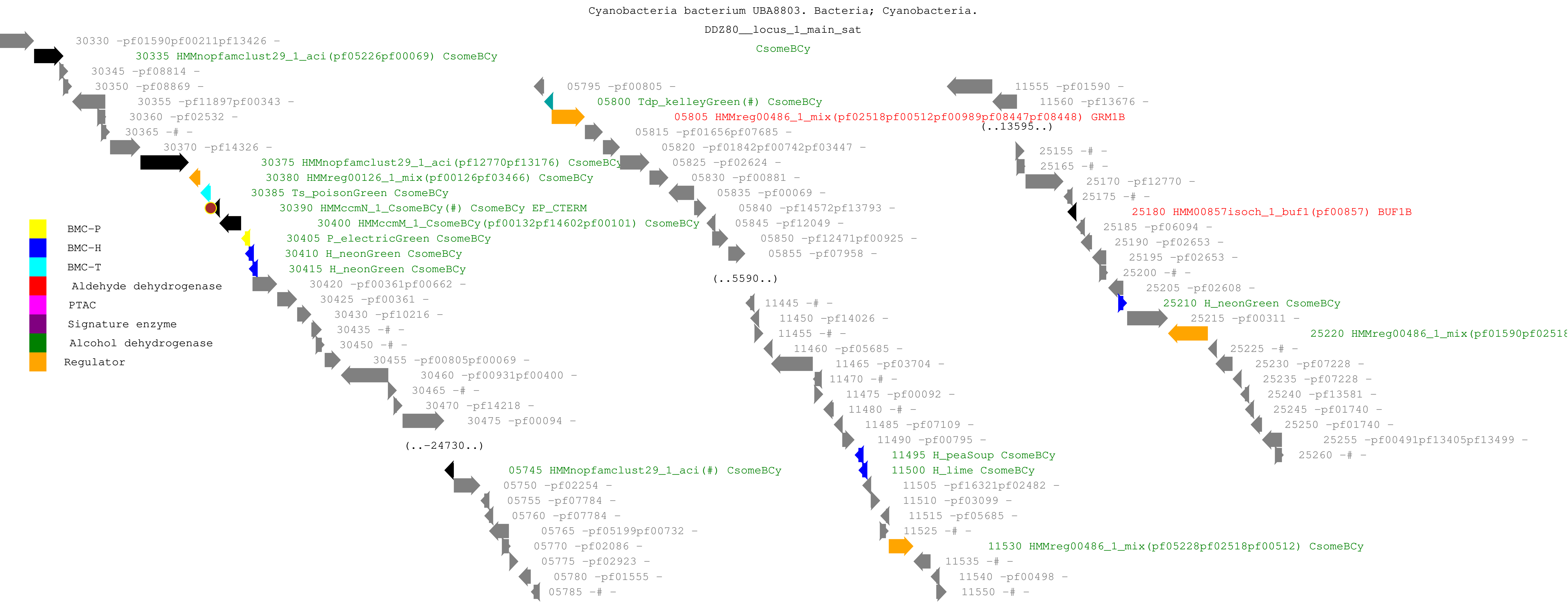
Cyanobacteria bacterium UBA8803. Bacteria; Cyanobacteria.
DDZ80__locus_1_main_sat
CsomeBCy
30330 -pf01590pf00211pf13426 -
30335 HMMnopfamclust29_1_aci(pf05226pf00069) CsomeBCy
30345 -pf08814 -
30350 -pf08869 -
30355 -pf11897pf00343 -
30360 -pf02532 -
30365 -# -
30370 -pf14326 -
30375 HMMnopfamclust29_1_aci(pf12770pf13176) CsomeBCy
30380 HMMreg00126_1_mix(pf00126pf03466) CsomeBCy
30385 Ts_poisonGreen CsomeBCy
30390 HMMccmN_1_CsomeBCy(#) CsomeBCy EP_CTERM
30400 HMMccmM_1_CsomeBCy(pf00132pf14602pf00101) CsomeBCy
30405 P_electricGreen CsomeBCy
30410 H_neonGreen CsomeBCy
30415 H_neonGreen CsomeBCy
30420 -pf00361pf00662 -
30425 -pf00361 -
30430 -pf10216 -
30435 -# -
30450 -# -
30455 -pf00805pf00069 -
30460 -pf00931pf00400 -
30465 -# -
30470 -pf14218 -
30475 -pf00094 -
(..-24730..)
05745 HMMnopfamclust29_1_aci(#) CsomeBCy
05750 -pf02254 -
05755 -pf07784 -
05760 -pf07784 -
05765 -pf05199pf00732 -
05770 -pf02086 -
05775 -pf02923 -
05780 -pf01555 -
05785 -# -
05795 -pf00805 -
05800 Tdp_kelleyGreen(#) CsomeBCy
05805 HMMreg00486_1_mix(pf02518pf00512pf00989pf08447pf08448) GRM1B
05815 -pf01656pf07685 -
05820 -pf01842pf00742pf03447 -
05825 -pf02624 -
05830 -pf00881 -
05835 -pf00069 -
05840 -pf14572pf13793 -
05845 -pf12049 -
05850 -pf12471pf00925 -
05855 -pf07958 -
(..5590..)
11445 -# -
11450 -pf14026 -
11455 -# -
11460 -pf05685 -
11465 -pf03704 -
11470 -# -
11475 -pf00092 -
11480 -# -
11485 -pf07109 -
11490 -pf00795 -
11495 H_peaSoup CsomeBCy
11500 H_lime CsomeBCy
11505 -pf16321pf02482 -
11510 -pf03099 -
11515 -pf05685 -
11525 -# -
11530 HMMreg00486_1_mix(pf05228pf02518pf00512) CsomeBCy
11535 -# -
11540 -pf00498 -
11550 -# -
11555 -pf01590 -
11560 -pf13676 -
(..13595..)
25155 -# -
25165 -# -
25170 -pf12770 -
25175 -# -
25180 HMM00857isoch_1_buf1(pf00857) BUF1B
25185 -pf06094 -
25190 -pf02653 -
25195 -pf02653 -
25200 -# -
25205 -pf02608 -
25210 H_neonGreen CsomeBCy
25215 -pf00311 -
25220 HMMreg00486_1_mix(pf01590pf02518
25225 -# -
25230 -pf07228 -
25235 -pf07228 -
25240 -pf13581 -
25245 -pf01740 -
25250 -pf01740 -
25255 -pf00491pf13405pf13499 -
25260 -# -
BMC-P
BMC-H
BMC-T
Aldehyde dehydrogenase
PTAC
Signature enzyme
Alcohol dehydrogenase
Regulator

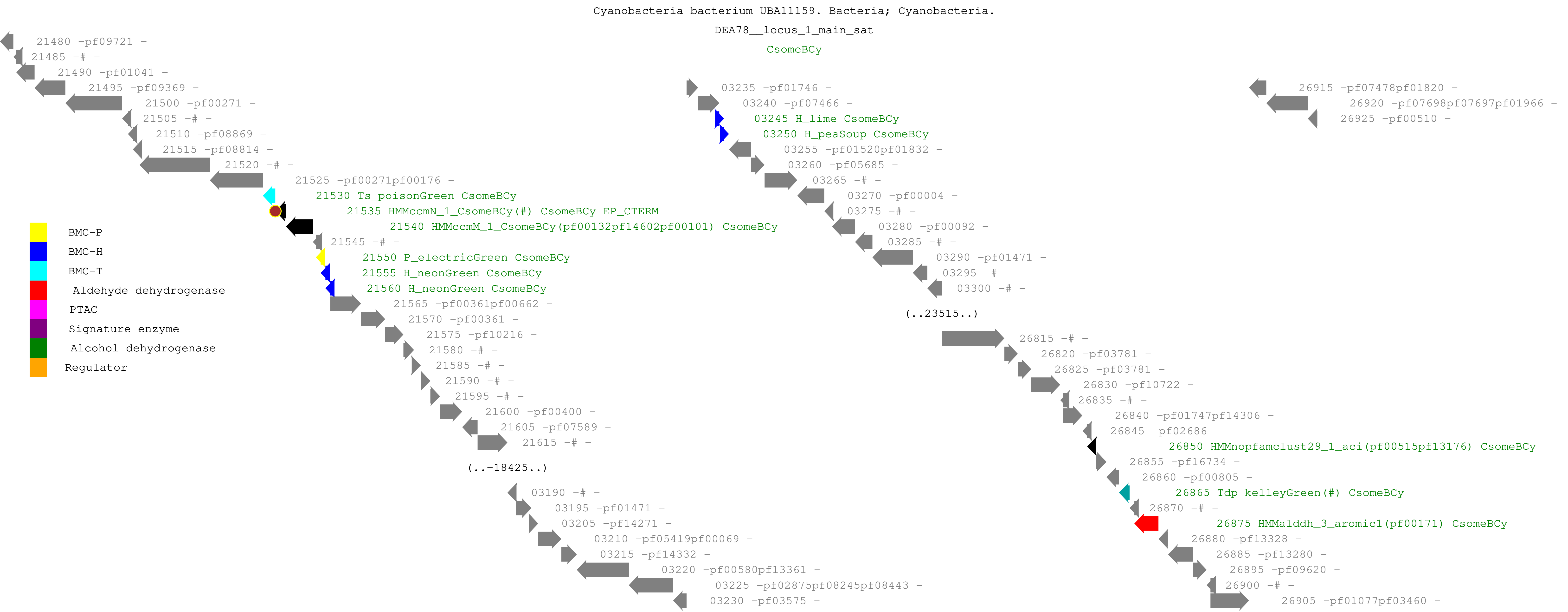

Cyanobacteria bacterium UBA11159. Bacteria; Cyanobacteria.
DEA78__locus_1_main_sat
CsomeBCy
21480 -pf09721 -
21485 -# -
21490 -pf01041 -
21495 -pf09369 -
21500 -pf00271 -
21505 -# -
21510 -pf08869 -
21515 -pf08814 -
21520 -# -
21525 -pf00271pf00176 -
21530 Ts_poisonGreen CsomeBCy
21535 HMMccmN_1_CsomeBCy(#) CsomeBCy EP_CTERM
21540 HMMccmM_1_CsomeBCy(pf00132pf14602pf00101) CsomeBCy
21545 -# -
21550 P_electricGreen CsomeBCy
21555 H_neonGreen CsomeBCy
21560 H_neonGreen CsomeBCy
21565 -pf00361pf00662 -
21570 -pf00361 -
21575 -pf10216 -
21580 -# -
21585 -# -
21590 -# -
21595 -# -
21600 -pf00400 -
21605 -pf07589 -
21615 -# -
(..-18425..)
03190 -# -
03195 -pf01471 -
03205 -pf14271 -
03210 -pf05419pf00069 -
03215 -pf14332 -
03220 -pf00580pf13361 -
03225 -pf02875pf08245pf08443 -
03230 -pf03575 -
BMC-P
BMC-H
BMC-T
Aldehyde dehydrogenase
PTAC
Signature enzyme
Alcohol dehydrogenase
Regulator
03235 -pf01746 -
03240 -pf07466 -
03245 H_lime CsomeBCy
03250 H_peaSoup CsomeBCy
03255 -pf01520pf01832 -
03260 -pf05685 -
03265 -# -
03270 -pf00004 -
03275 -# -
03280 -pf00092 -
03285 -# -
03290 -pf01471 -
03295 -# -
03300 -# -
(..23515..)
26815 -# -
26820 -pf03781 -
26825 -pf03781 -
26830 -pf10722 -
26835 -# -
26840 -pf01747pf14306 -
26845 -pf02686 -
26850 HMMnopfamclust29_1_aci(pf00515pf13176) CsomeBCy
26855 -pf16734 -
26860 -pf00805 -
26865 Tdp_kelleyGreen(#) CsomeBCy
26870 -# -
26875 HMMaldh_3_aromic1(pf00171) CsomeBCy
26880 -pf13328 -
26885 -pf13280 -
26895 -pf09620 -
26900 -# -
26905 -pf01077pf03460 -
26915 -pf07478pf01820 -
26920 -pf07698pf07697pf01966 -
26925 -pf00510 -

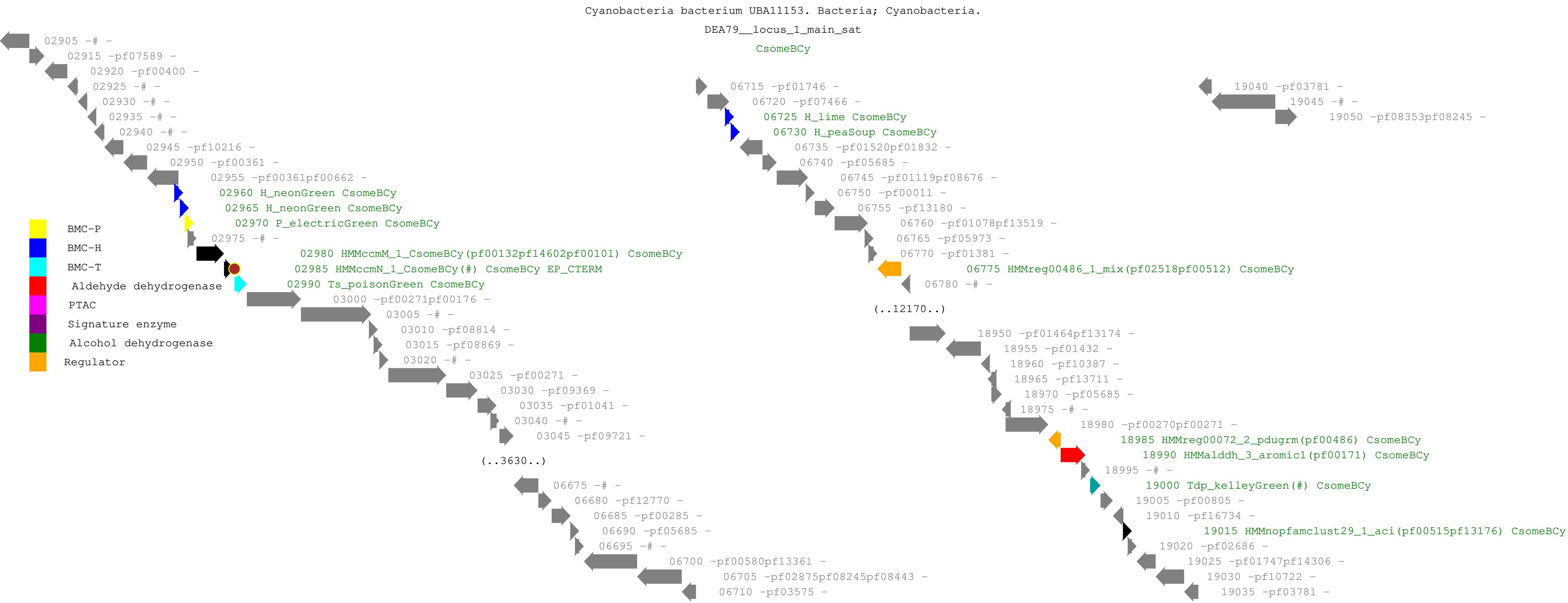

Cyanobacteria bacterium UBA11153. Bacteria; Cyanobacteria.
DEA79__locus_1_main_sat
CsomeBCy
02905 -# -
02915 -pf07589 -
02920 -pf00400 -
02925 -# -
02930 -# -
02935 -# -
02940 -# -
02945 -pf10216 -
02950 -pf00361 -
02955 -pf00361pf00662 -
02960 H_neonGreen CsomeBCy
02965 H_neonGreen CsomeBCy
02970 P_electricGreen CsomeBCy
02975 -# -
02980 HMMccmM_1_CsomeBCy(pf00132pf14602pf00101) CsomeBCy
02985 HMMccmN_1_CsomeBCy(#) CsomeBCy EP_CTERM
02990 Ts_poisonGreen CsomeBCy
03000 -pf00271pf00176 -
03005 -# -
03010 -pf08814 -
03015 -pf08869 -
03020 -# -
03025 -pf00271 -
03030 -pf09369 -
03035 -pf01041 -
03040 -# -
03045 -pf09721 -
(..3630..)
06675 -# -
06680 -pf12770 -
06685 -pf00285 -
06690 -pf05685 -
06695 -# -
06700 -pf00580pf13361 -
06705 -pf02875pf08245pf08443 -
06710 -pf03575 -
06715 -pf01746 -
06720 -pf07466 -
06725 H_lime CsomeBCy
06730 H_peaSoup CsomeBCy
06735 -pf01520pf01832 -
06740 -pf05685 -
06745 -pf01119pf08676 -
06750 -pf00011 -
06755 -pf13180 -
06760 -pf01078pf13519 -
06765 -pf05973 -
06770 -pf01381 -
06775 HMMreg00486_1_mix(pf02518pf00512) CsomeBCy
06780 -# -
(..12170..)
18950 -pf01464pf13174 -
18955 -pf01432 -
18960 -pf10387 -
18965 -pf13711 -
18970 -pf05685 -
18975 -# -
18980 -pf00270pf00271 -
18985 HMMreg00072_2_pdugrm(pf00486) CsomeBCy
18990 HMMalddh_3_aromic1(pf00171) CsomeBCy
18995 -# -
19000 Tdp_kelleyGreen(#) CsomeBCy
19005 -pf00805 -
19010 -pf16734 -
19015 HMMnopfamclust29_1_aci(pf00515pf13176) CsomeBCy
19020 -pf02686 -
19025 -pf01747pf14306 -
19030 -pf10722 -
19035 -pf03781 -
19040 -pf03781 -
19045 -# -
19050 -pf08353pf08245 -
BMC-P
BMC-H
BMC-T
Aldehyde dehydrogenase
PTAC
Signature enzyme
Alcohol dehydrogenase
Regulator

Oscillatoriales bacterium UBA8482. Bacteria; Cyanobacteria; Oscillatoriophycideae; Oscillatoriales;

DEF27__locus_fragmented

CsomeBCy

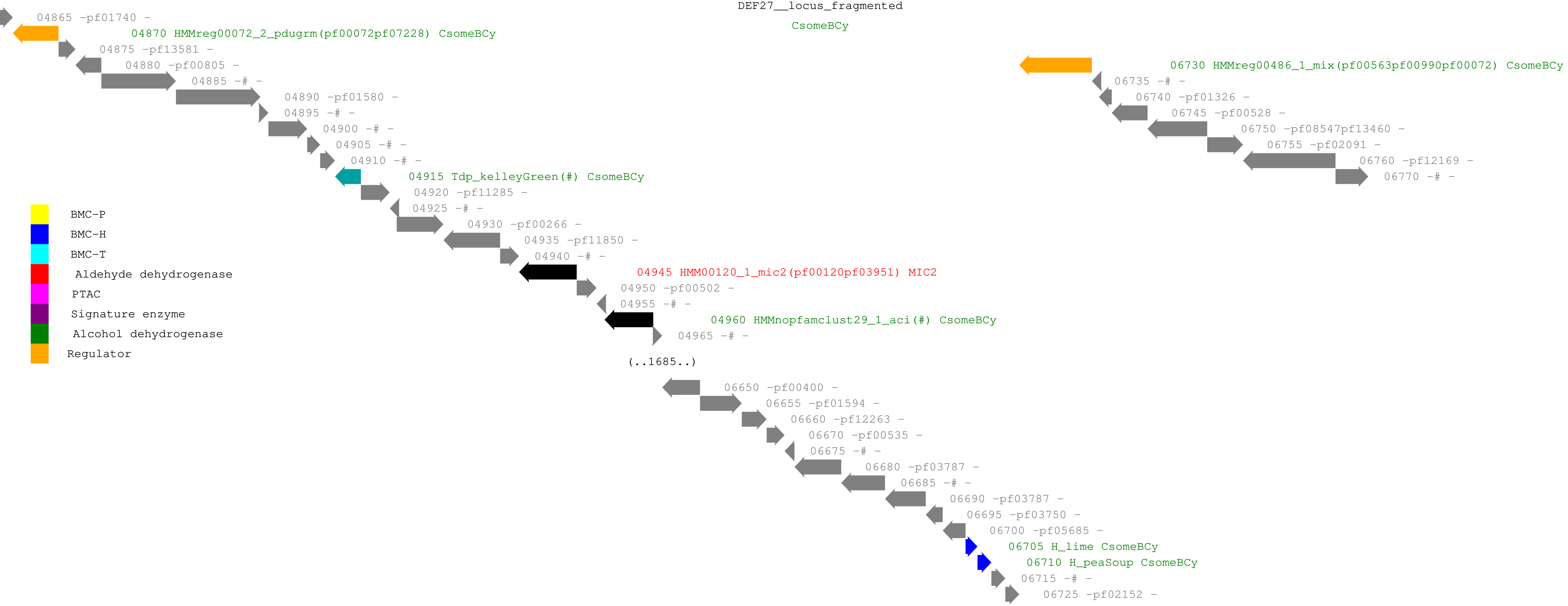

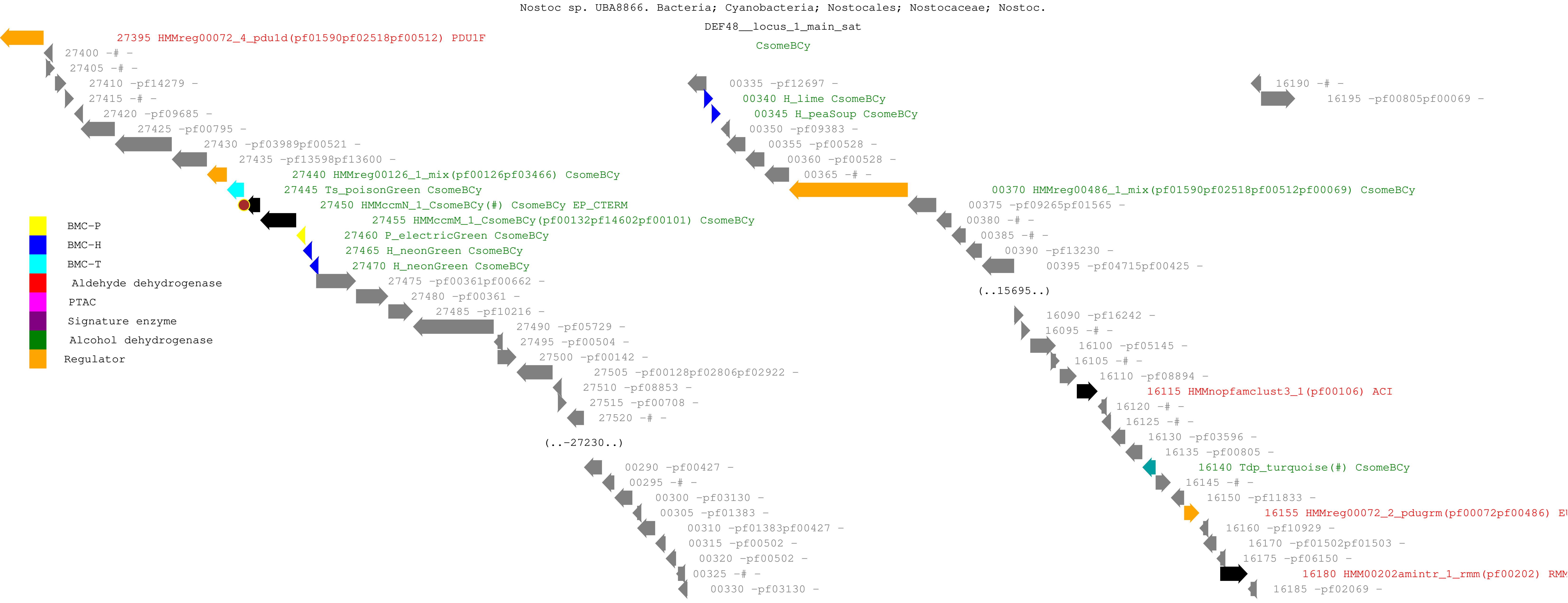

Nostoc sp. UBA8866. Bacteria; Cyanobacteria; Nostocales; Nostocaceae; Nostoc.
DEF48__locus_1_main_sat
CsomeBCy
27395 HMMreg00072_4_pdu1d(pf01590pf02518pf00512) PDU1F
27400 -# -
27405 -# -
27410 -pf14279 -
27415 -# -
27420 -pf09685 -
27425 -pf00795 -
27430 -pf03989pf00521 -
27435 -pf13598pf13600 -
27440 HMMreg00126_1_mix(pf00126pf03466) CsomeBCy
27445 Ts_poisonGreen CsomeBCy
27450 HMMccmN_1_CsomeBCy(#) CsomeBCy EP_CTERM
27455 HMMccmM_1_CsomeBCy(pf00132pf14602pf00101) CsomeBCy
27460 P_electricGreen CsomeBCy
27465 H_neonGreen CsomeBCy
27470 H_neonGreen CsomeBCy
27475 -pf00361pf00662 -
27480 -pf00361 -
27485 -pf10216 -
27490 -pf05729 -
27495 -pf00504 -
27500 -pf00142 -
27505 -pf00128pf02806pf02922 -
27510 -pf08853 -
27515 -pf00708 -
27520 -# -
(..-27230..)
00290 -pf00427 -
00295 -# -
00300 -pf03130 -
00305 -pf01383 -
00310 -pf01383pf00427 -
00315 -pf00502 -
00320 -pf00502 -
00325 -# -
00330 -pf03130 -
BMC-P
BMC-H
BMC-T
Aldehyde dehydrogenase
PTAC
Signature enzyme
Alcohol dehydrogenase
Regulator
00335 -pf12697 -
00340 H_lime CsomeBCy
00345 H_peaSoup CsomeBCy
00350 -pf09383 -
00355 -pf00528 -
00360 -pf00528 -
00365 -# -
00370 HMMreg00486_1_mix(pf01590pf02518pf00512pf00069) CsomeBCy
00375 -pf09265pf01565 -
00380 -# -
00385 -# -
00390 -pf13230 -
00395 -pf04715pf00425 -
(..15695..)
16090 -pf16242 -
16095 -# -
16100 -pf05145 -
16105 -# -
16110 -pf08894 -
16115 HMMnopfamclust3_1(pf00106) ACI
16120 -# -
16125 -# -
16130 -pf03596 -
16135 -pf00805 -
16140 Tdp_turquoise(#) CsomeBCy
16145 -# -
16150 -pf11833 -
16155 HMMreg00072_2_pdugrm(pf00072pf00486) EU
16160 -pf10929 -
16170 -pf01502pf01503 -
16175 -pf06150 -
16180 HMM00202amintr_1_rmm(pf00202) RMM
16185 -pf02069 -
16190 -# -
16195 -pf00805pf00069 -

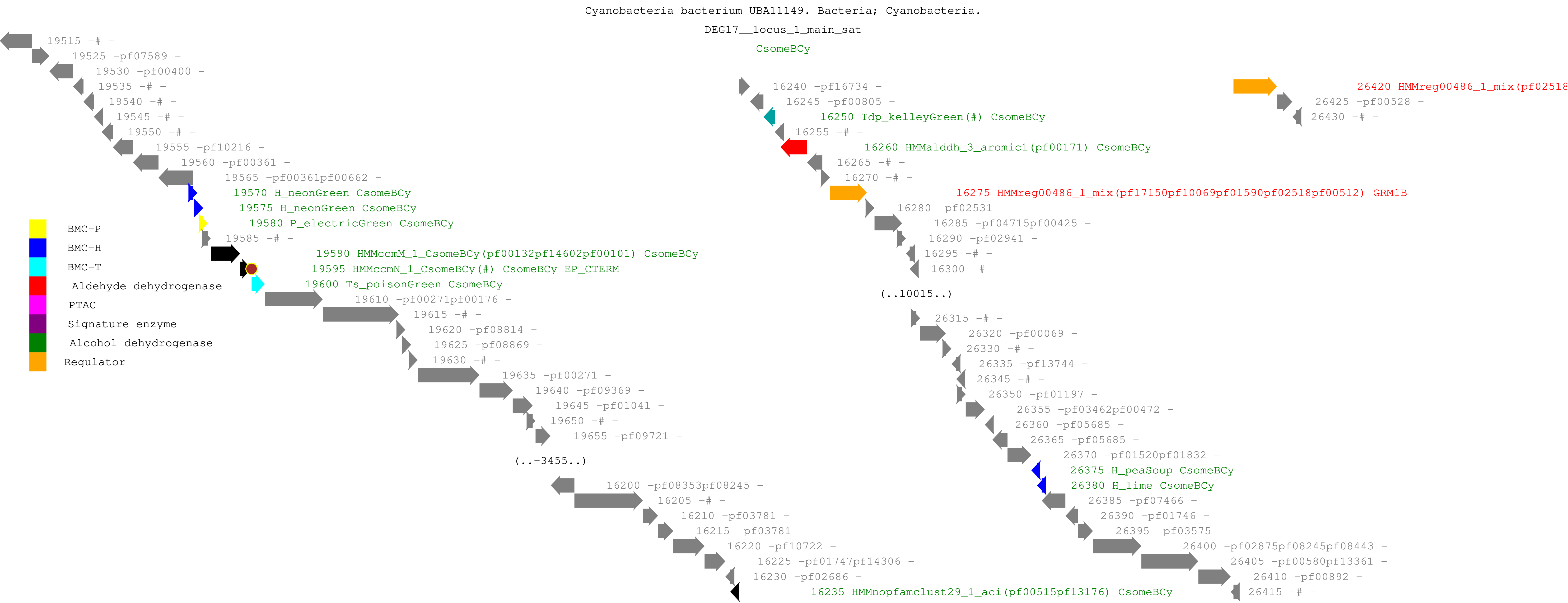
Cyanobacteria bacterium UBA11149. Bacteria; Cyanobacteria.
DEG17__locus_1_main_sat
CsomeBCy
19515 -# -
19525 -pf07589 -
19530 -pf00400 -
19535 -# -
19540 -# -
19545 -# -
19550 -# -
19555 -pf10216 -
19560 -pf00361 -
19565 -pf00361pf00662 -
19570 H_neonGreen CsomeBCy
19575 H_neonGreen CsomeBCy
19580 P_electricGreen CsomeBCy
19585 -# -
19590 HMMccmM_1_CsomeBCy(pf00132pf14602pf00101) CsomeBCy
19595 HMMccmN_1_CsomeBCy(#) CsomeBCy EP_CTERM
19600 Ts_poisonGreen CsomeBCy
19610 -pf00271pf00176 -
19615 -# -
19620 -pf08814 -
19625 -pf08869 -
19630 -# -
19635 -pf00271 -
19640 -pf09369 -
19645 -pf01041 -
19650 -# -
19655 -pf09721 -
(..-3455..)
16200 -pf08353pf08245 -
16205 -# -
16210 -pf03781 -
16215 -pf03781 -
16220 -pf10722 -
16225 -pf01747pf14306 -
16230 -pf02686 -
16235 HMMnopfamclust29_1_aci(pf00515pf13176) CsomeBCy
16240 -pf16734 -
16245 -pf00805 -
16250 Tdp_kelleyGreen(#) CsomeBCy
16255 -# -
16260 HMMaldh_3_aromic1(pf00171) CsomeBCy
16265 -# -
16270 -# -
16275 HMMreg00486_1_mix(pf17150pf10069pf01590pf02518pf00512) GRM1B
16280 -pf02531 -
16285 -pf04715pf00425 -
16290 -pf02941 -
16295 -# -
16300 -# -
(..10015..)
26315 -# -
26320 -pf00069 -
26330 -# -
26335 -pf13744 -
26345 -# -
26350 -pf01197 -
26355 -pf03462pf00472 -
26360 -pf05685 -
26365 -pf05685 -
26370 -pf01520pf01832 -
26375 H_peaSoup CsomeBCy
26380 H_lime CsomeBCy
26385 -pf07466 -
26390 -pf01746 -
26395 -pf03575 -
26400 -pf02875pf08245pf08443 -
26405 -pf00580pf13361 -
26410 -pf00892 -
26415 -# -
26420 HMMreg00486_1_mix(pf02518
26425 -pf00528 -
26430 -# -
BMC-P
BMC-H
BMC-T
Aldehyde dehydrogenase
PTAC
Signature enzyme
Alcohol dehydrogenase
Regulator

Cyanobacteria bacterium UBA11148. Bacteria; Cyanobacteria.

DEG47__locus_1_main_sat

CsomeBCy

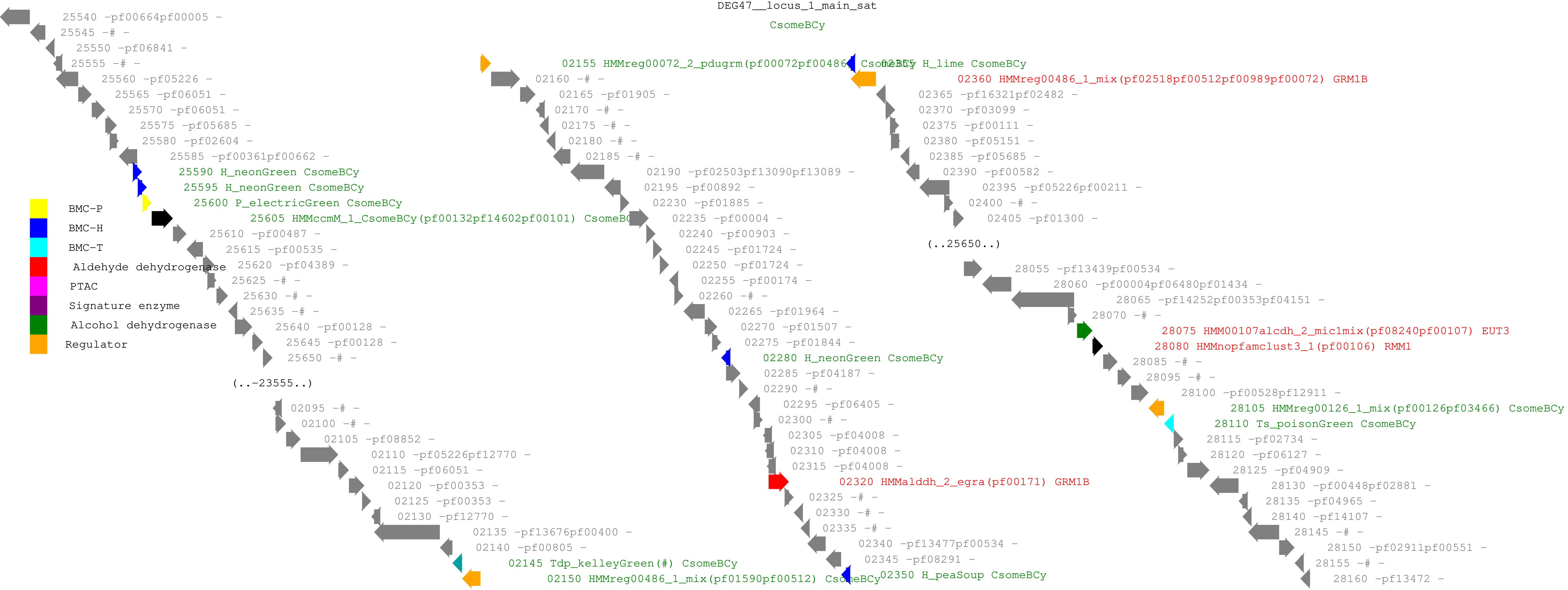

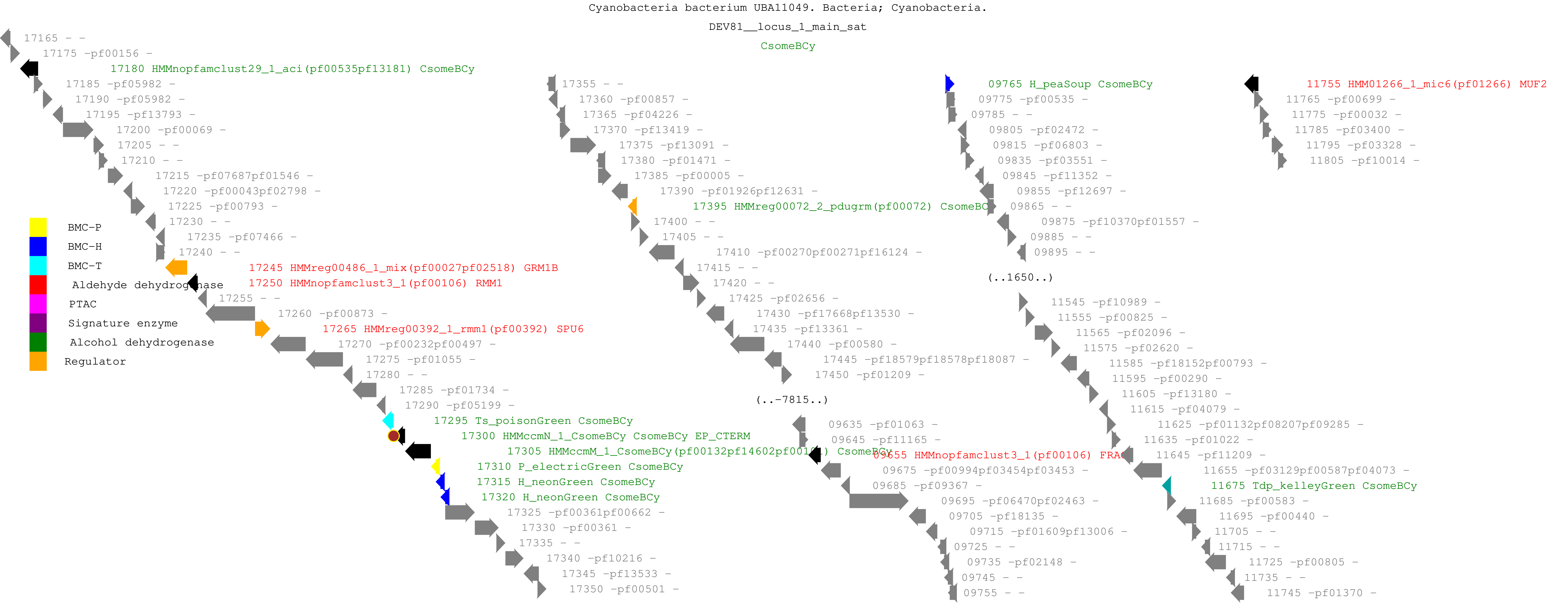
Cyanobacteria bacterium UBA11049. Bacteria; Cyanobacteria.
DEV81__locus_1_main_sat
CsomeBCy
17165 - -
17175 -pf00156 -
17180 HMMnopfamclust29_1_aci(pf00535pf13181) CsomeBCy
17185 -pf05982 -
17190 -pf05982 -
17195 -pf13793 -
17200 -pf00069 -
17205 - -
17210 - -
17215 -pf07687pf01546 -
17220 -pf00043pf02798 -
17225 -pf00793 -
17230 - -
17235 -pf07466 -
17240 - -
17245 HMMreg00486_1_mix(pf00027pf02518) GRM1B
17250 HMMnopfamclust3_1(pf00106) RMM1
17255 - -
17260 -pf00873 -
17265 HMMreg00392_1_rmm1(pf00392) SPU6
17270 -pf00232pf00497 -
17275 -pf01055 -
17280 - -
17285 -pf01734 -
17290 -pf05199 -
17295 Ts_poisonGreen CsomeBCy
17300 HMMccmN_1_CsomeBCy CsomeBCy EP_CTERM
17305 HMMccmM_1_CsomeBCy(pf00132pf14602pf00101) CsomeBCy
17310 P_electricGreen CsomeBCy
17315 H_neonGreen CsomeBCy
17320 H_neonGreen CsomeBCy
17325 -pf00361pf00662 -
17330 -pf00361 -
17335 - -
17340 -pf10216 -
17345 -pf13533 -
17350 -pf00501 -
BMC-P
BMC-H
BMC-T
Aldehyde dehydrogenase
PTAC
Signature enzyme
Alcohol dehydrogenase
Regulator
17355 - -
17360 -pf00857 -
17365 -pf04226 -
17370 -pf13419 -
17375 -pf13091 -
17380 -pf01471 -
17385 -pf00005 -
17390 -pf01926pf12631 -
17395 HMMreg00072_2_pdugrm(pf00072) CsomeBCy
17400 - -
17405 - -
17410 -pf00270pf00271pf16124 -
17415 - -
17420 - -
17425 -pf02656 -
17430 -pf17668pf13530 -
17435 -pf13361 -
17440 -pf00580 -
17445 -pf18579pf18578pf18087 -
17450 -pf01209 -
(..-7815..)
09635 -pf01063 -
09645 -pf11165 -
09655 HMMnopfamclust3_1(pf00106) FRA
09675 -pf00994pf03454pf03453 -
09685 -pf09367 -
09695 -pf06470pf02463 -
09705 -pf18135 -
09715 -pf01609pf13006 -
09725 - -
09735 -pf02148 -
09745 - -
09755 - -
09765 H_peaSoup CsomeBCy
09775 -pf00535 -
09785 - -
09805 -pf02472 -
09815 -pf06803 -
09835 -pf03551 -
09845 -pf11352 -
09855 -pf12697 -
09865 - -
09875 -pf10370pf01557 -
09885 - -
09895 - -
(..1650..)
11545 -pf10989 -
11555 -pf00825 -
11565 -pf02096 -
11575 -pf02620 -
11585 -pf18152pf00793 -
11595 -pf00290 -
11605 -pf13180 -
11615 -pf04079 -
11625 -pf01132pf08207pf09285 -
11635 -pf01022 -
11645 -pf11209 -
11655 -pf03129pf00587pf04073 -
11675 Tdp_kelleyGreen CsomeBCy
11685 -pf00583 -
11695 -pf00440 -
11705 - -
11715 - -
11725 -pf00805 -
11735 - -
11745 -pf01370 -
11755 HMM01266_1_mic6(pf01266) MUF2
11765 -pf00699 -
11775 -pf00032 -
11785 -pf03400 -
11795 -pf03328 -
11805 -pf10014 -

Microcoleaceae bacterium UBA9251. Bacteria; Cyanobacteria; Oscillatoriophycideae; Oscillatoriales;

DGO89__locus_fragmented

CsomeBCy

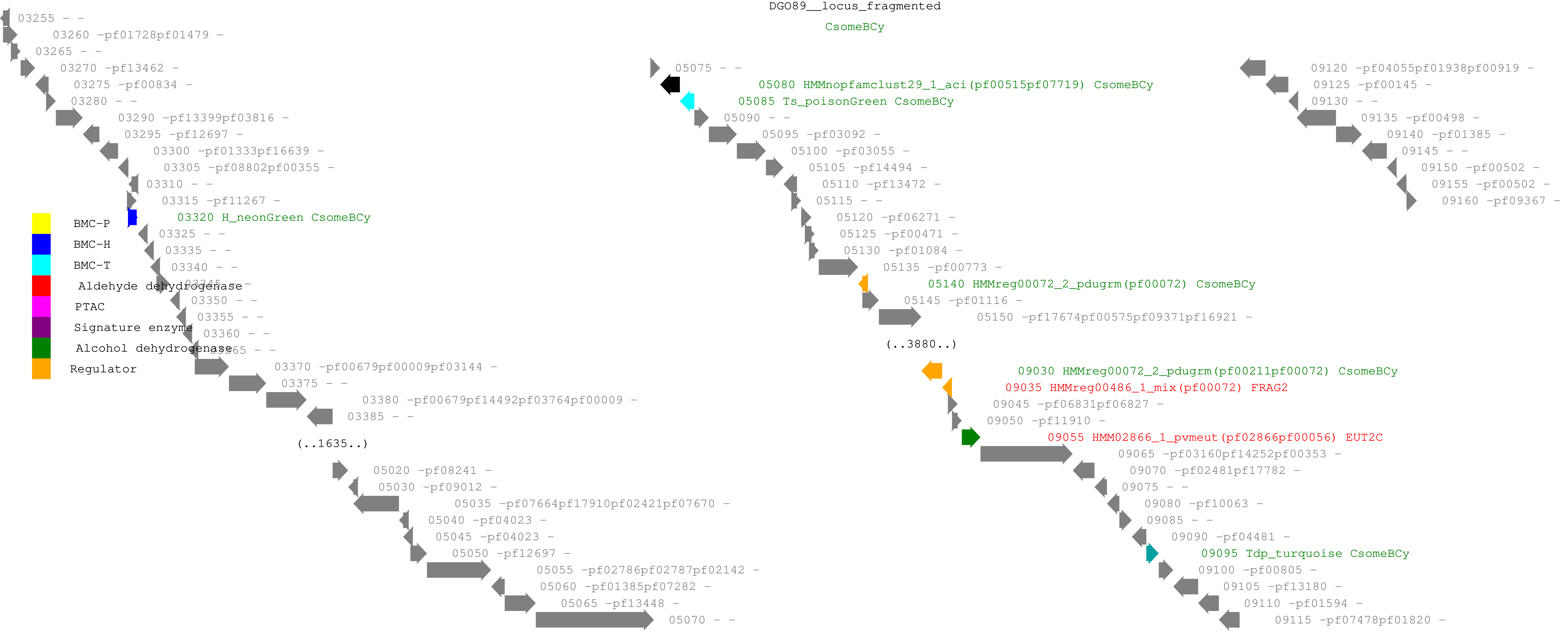

Anabaena sp. UBA12330. Bacteria; Cyanobacteria; Nostocales; Nostocaceae; Anabaena;

DIU28__locus_1_main_sat

CsomeBCy

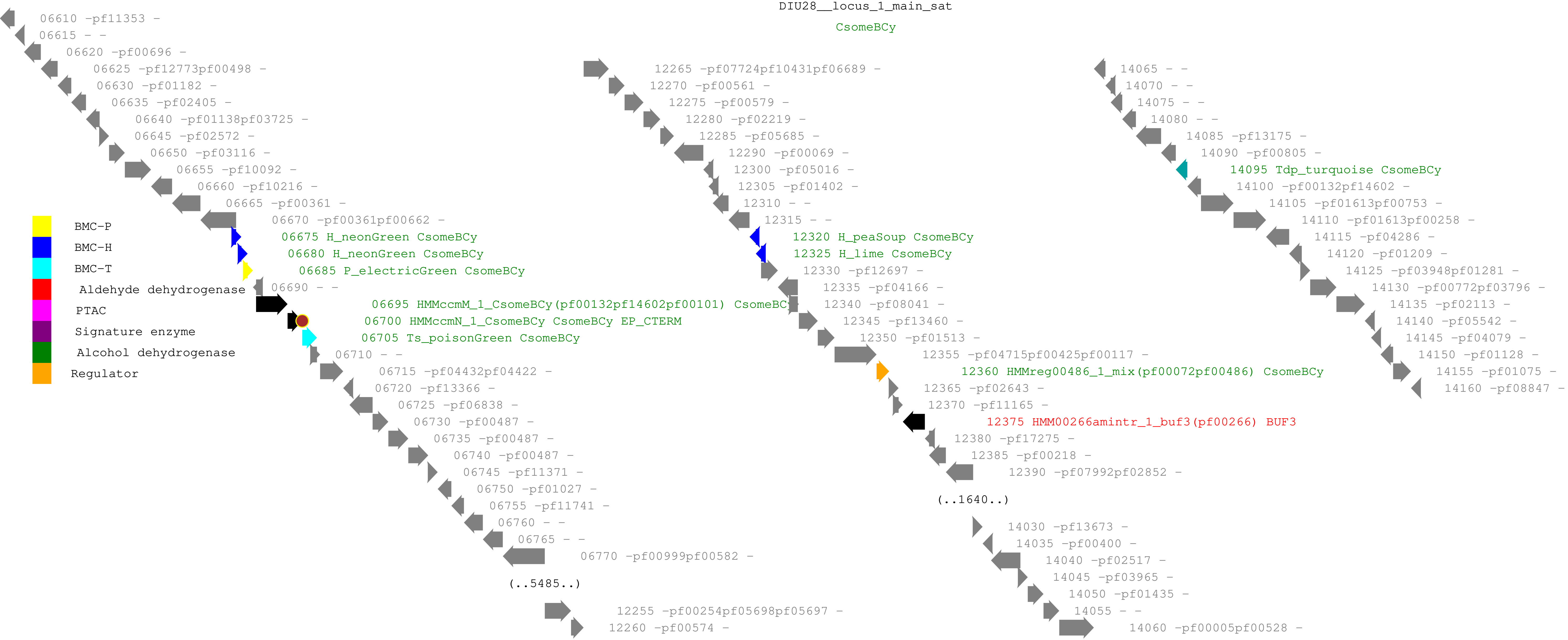

Neosynechococcus sphagnicola sy1. Bacteria; Cyanobacteria; Synechococcales; Leptolyngbyaceae;

DO97__locus_1_main_sat

CsomeBCy

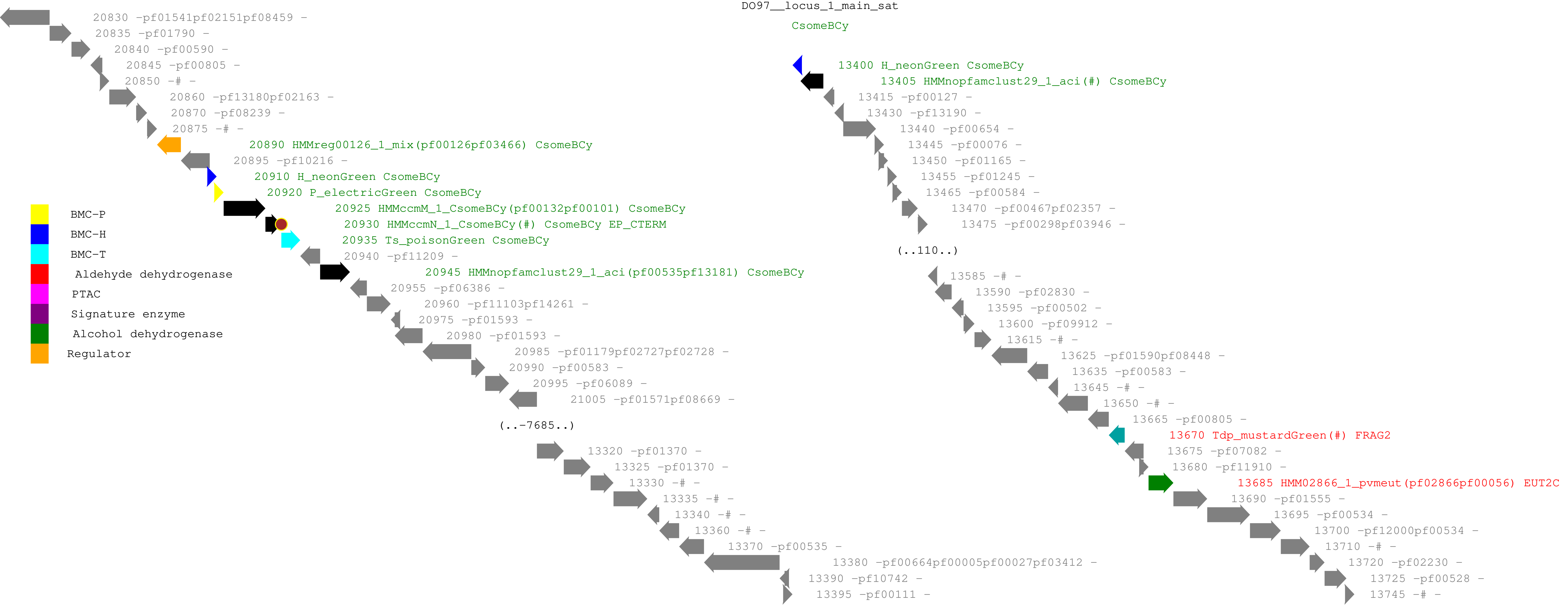

Synechococcus elongatus PCC 11801. Bacteria; Cyanobacteria; Synechococcales; Synechococcaceae; Synechococcus.

DOP62__locus_1_main_sat

CsomeBCy

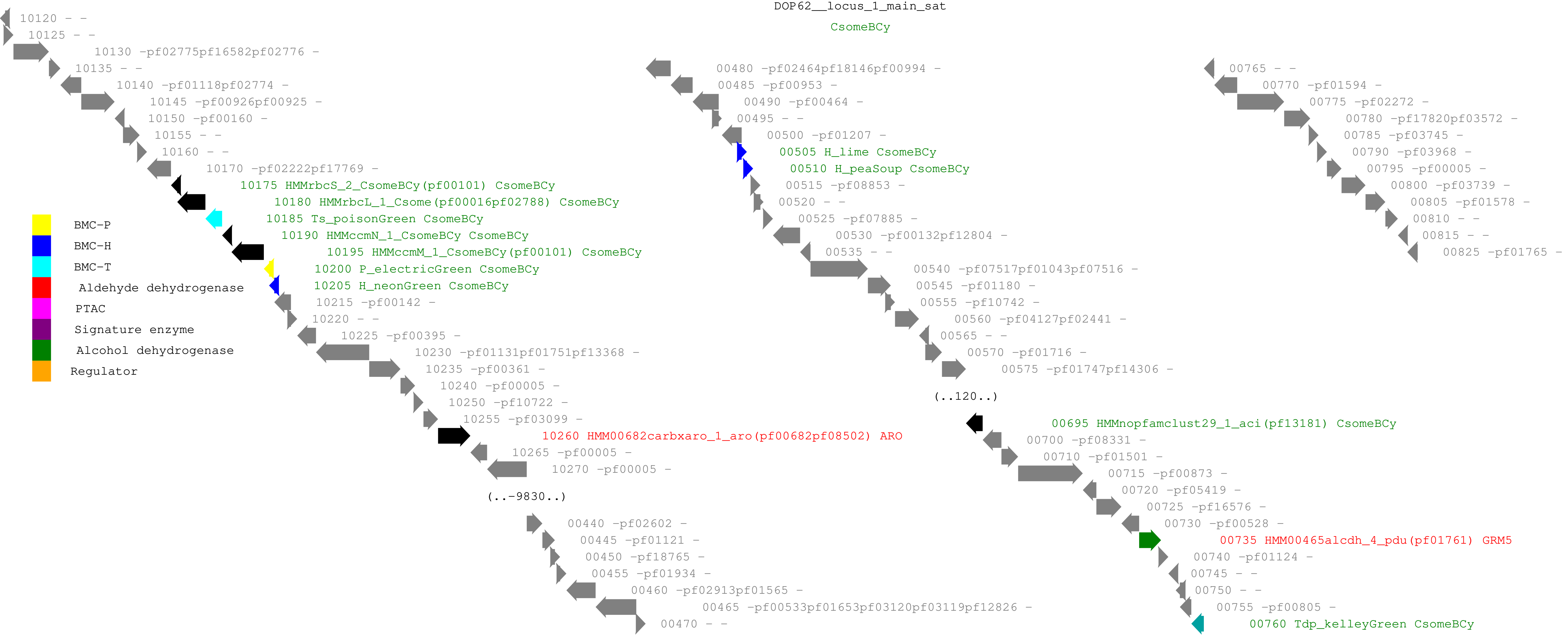

Calothrix desertica PCC 7102. Bacteria; Cyanobacteria; Nostocales; Calotrichaceae; Calothrix.

DSM106972__locus_1_main_sat

CsomeBCy

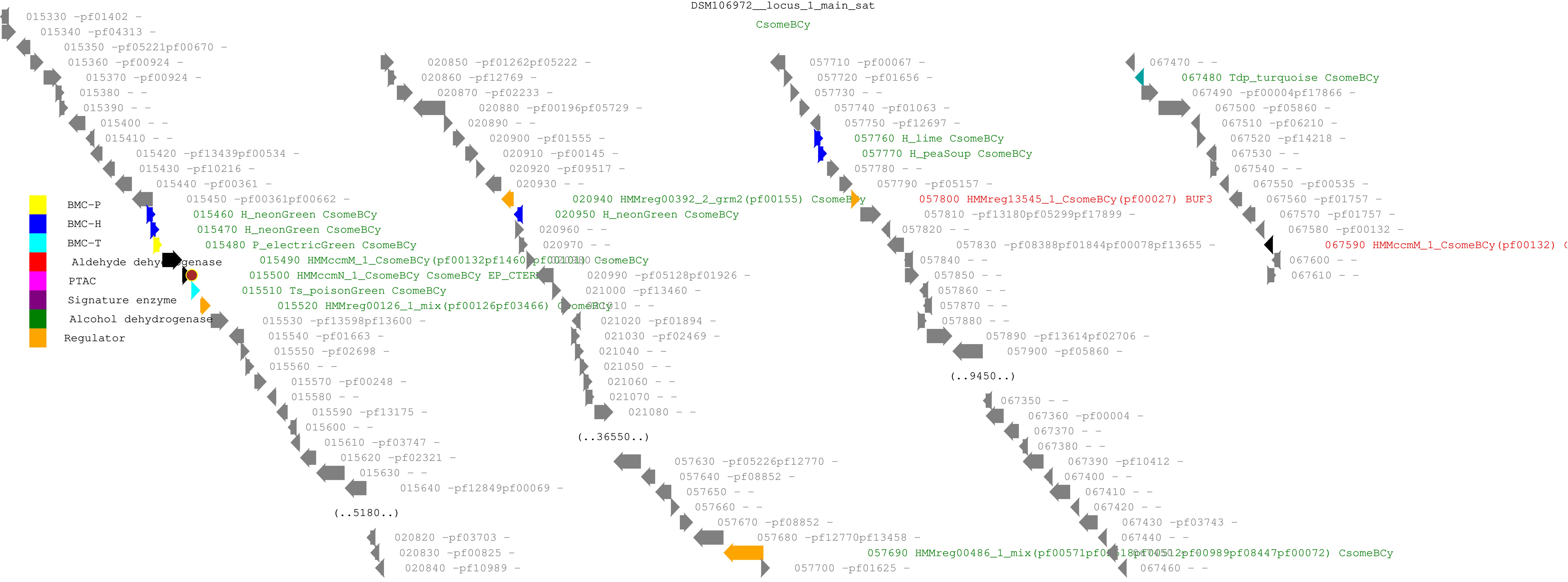

Trichormus variabilis SAG 1403-4b. Bacteria; Cyanobacteria; Nostocales; Nostocaceae; Trichormus.

DSM107003__locus_1_main_sat

CsomeBCy

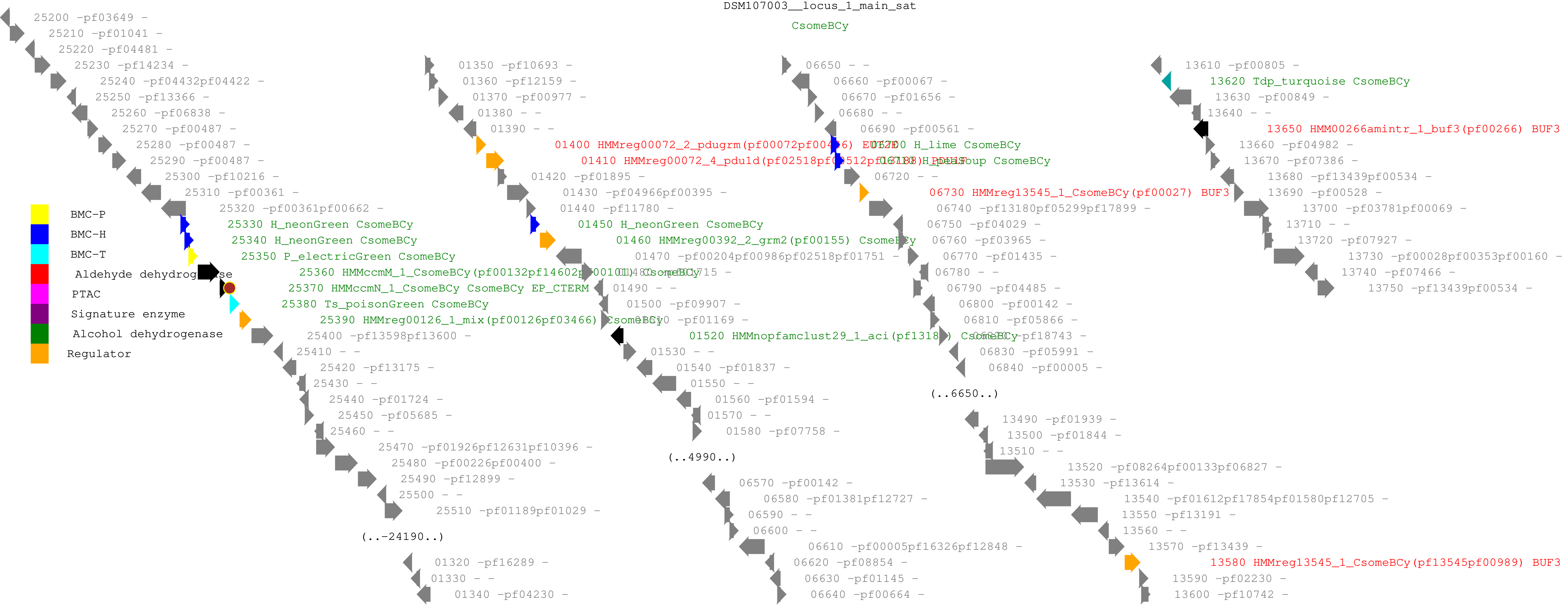

Chroococcidiopsis cubana SAG 39.79. Bacteria; Cyanobacteria; Chroococcidiopsidales; Chroococcidiopsidaceae;

DSM107010__locus_1_main_sat

CsomeBCy

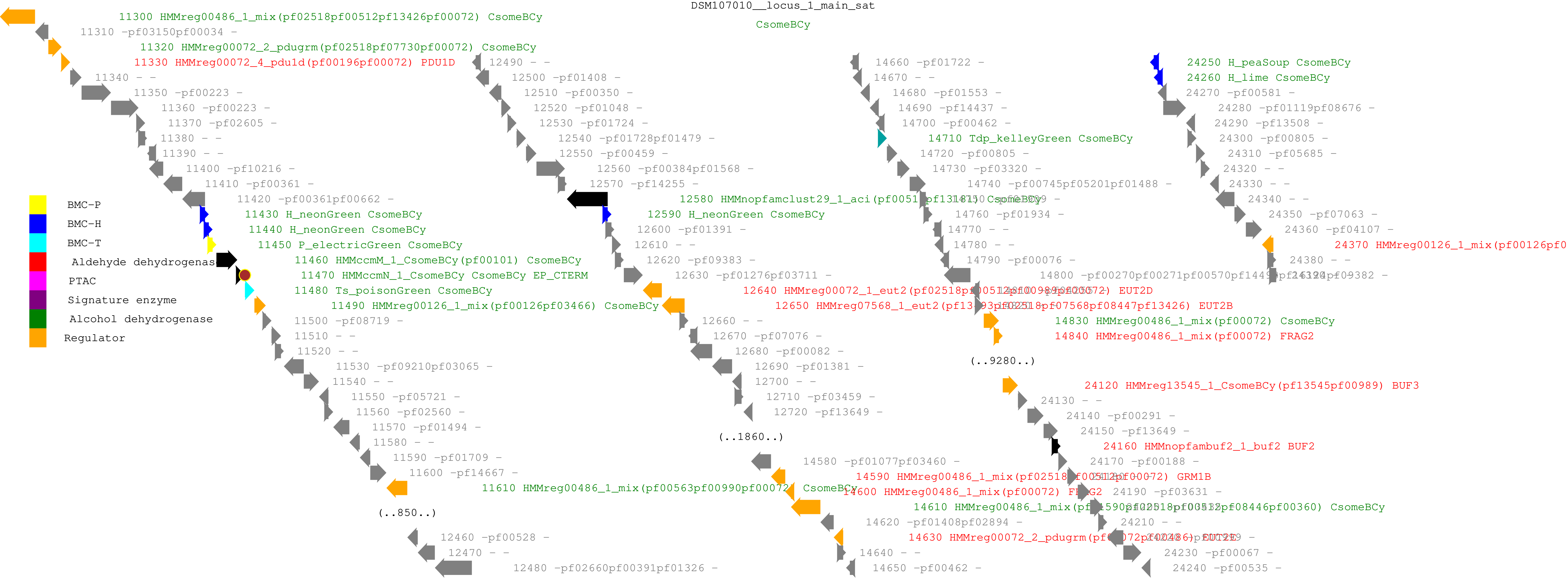

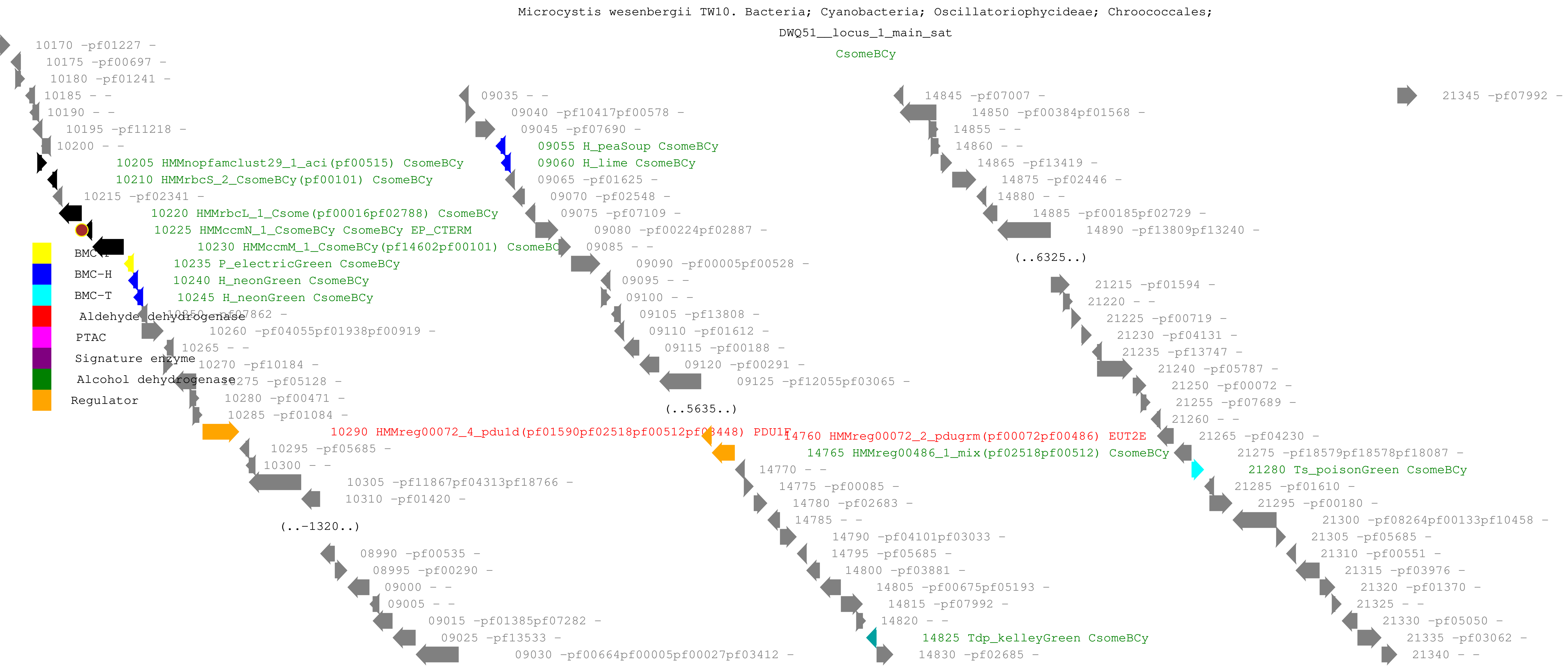

Microcystis wesenbergii TW10. Bacteria; Cyanobacteria; Oscillatoriophycideae; Chroococcales;
DWQ51__locus_1_main_sat
CsomeBCy
10170 -pf01227 -
10175 -pf00697 -
10180 -pf01241 -
10185 - -
10190 - -
10195 -pf11218 -
10200 - -
10205 HMMnopfamclust29_1_aci(pf00515) CsomeBCy
10210 HMMrbcS_2_CsomeBCy(pf00101) CsomeBCy
10215 -pf02341 -
10220 HMMrbcL_1_Csome(pf00016pf02788) CsomeBCy
10225 HMMccmN_1_CsomeBCy CsomeBCy EP_CTERM
10230 HMMccmM_1_CsomeBCy(pf14602pf00101) CsomeBCy
10235 P_electricGreen CsomeBCy
10240 H_neonGreen CsomeBCy
10245 H_neonGreen CsomeBCy
10250 -pf07862 -
10260 -pf04055pf01938pf00919 -
10265 - -
10270 -pf10184 -
10275 -pf05128 -
10280 -pf00471 -
10285 -pf01084 -
10290 HMMreg00072_4_pdu1d(pf01590pf02518pf00512pf03448) PDU1F
10295 -pf05685 -
10300 - -
10305 -pf11867pf04313pf18766 -
10310 -pf01420 -
(..-1320..)
08990 -pf00535 -
08995 -pf00290 -
09000 - -
09005 - -
09015 -pf01385pf07282 -
09025 -pf13533 -
09030 -pf00664pf00005pf00027pf03412 -
09035 - -
09040 -pf10417pf00578 -
09045 -pf07690 -
09055 H_peaSoup CsomeBCy
09060 H_lime CsomeBCy
09065 -pf01625 -
09070 -pf02548 -
09075 -pf07109 -
09080 -pf00224pf02887 -
09085 - -
09090 -pf00005pf00528 -
09095 - -
09100 - -
09105 -pf13808 -
09110 -pf01612 -
09115 -pf00188 -
09120 -pf00291 -
09125 -pf12055pf03065 -
(..5635..)
14760 HMMreg00072_2_pdugrm(pf00072pf00486) EUT2E
14765 HMMreg00486_1_mix(pf02518pf00512) CsomeBCy
14770 - -
14775 -pf00085 -
14780 -pf02683 -
14785 - -
14790 -pf04101pf03033 -
14795 -pf05685 -
14800 -pf03881 -
14805 -pf00675pf05193 -
14815 -pf07992 -
14820 - -
14825 Tdp_kelleyGreen CsomeBCy
14830 -pf02685 -
14845 -pf07007 -
14850 -pf00384pf01568 -
14855 - -
14860 - -
14865 -pf13419 -
14875 -pf02446 -
14880 - -
14885 -pf00185pf02729 -
14890 -pf13809pf13240 -
(..6325..)
21215 -pf01594 -
21220 - -
21225 -pf00719 -
21230 -pf04131 -
21235 -pf13747 -
21240 -pf05787 -
21250 -pf00072 -
21255 -pf07689 -
21260 - -
21265 -pf04230 -
21275 -pf18579pf18578pf18087 -
21280 Ts_poisonGreen CsomeBCy
21285 -pf01610 -
21295 -pf00180 -
21300 -pf08264pf00133pf10458 -
21305 -pf05685 -
21310 -pf00551 -
21315 -pf03976 -
21320 -pf01370 -
21325 - -
21330 -pf05050 -
21335 -pf03062 -
21340 - -
21345 -pf07992 -
BMC-P
BMC-H
BMC-T
Aldehyde dehydrogenase
PTAC
Signature enzyme
Alcohol dehydrogenase
Regulator

Microcystis flos-aquae DF17. Bacteria; Cyanobacteria; Oscillatoriophycideae; Chroococcales;

DWQ53__locus_1_main_sat

CsomeBCy

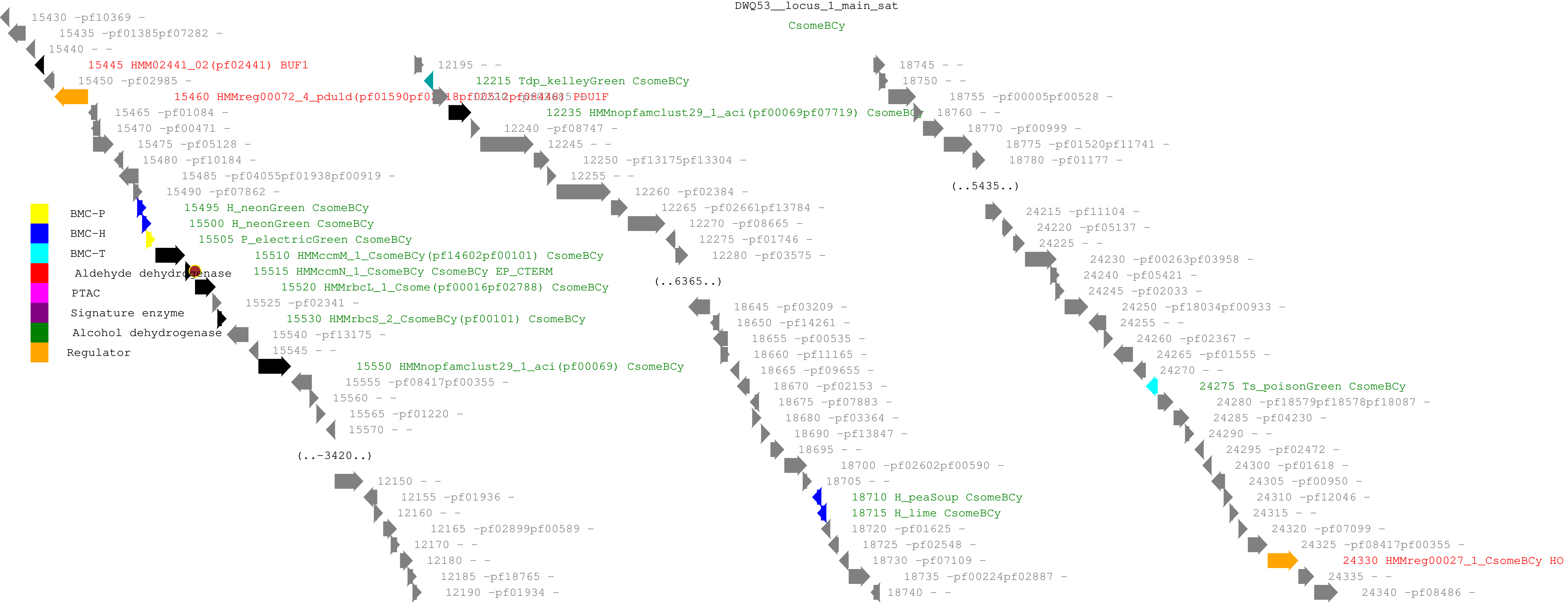

Microcystis flos-aquae TF09. Bacteria; Cyanobacteria; Oscillatoriophycideae; Chroococcales;

DWQ54__locus_1_main_sat

CsomeBCy

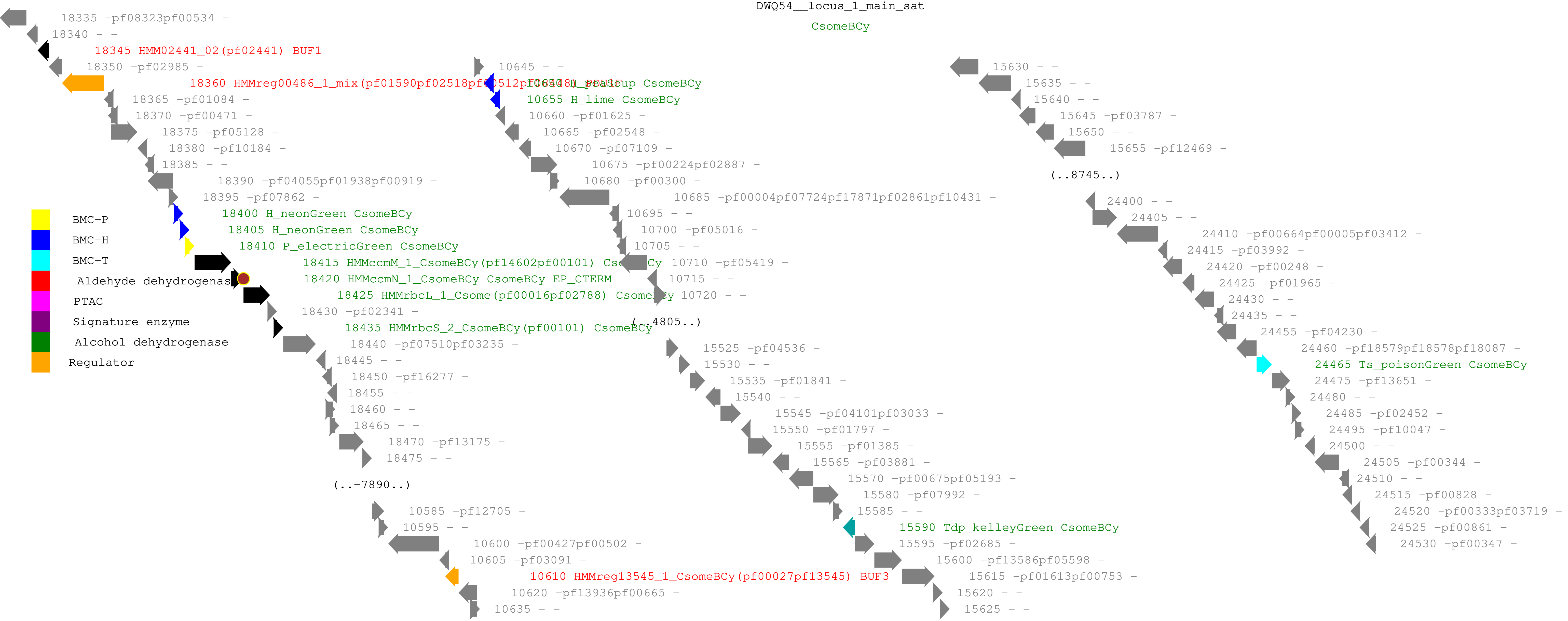

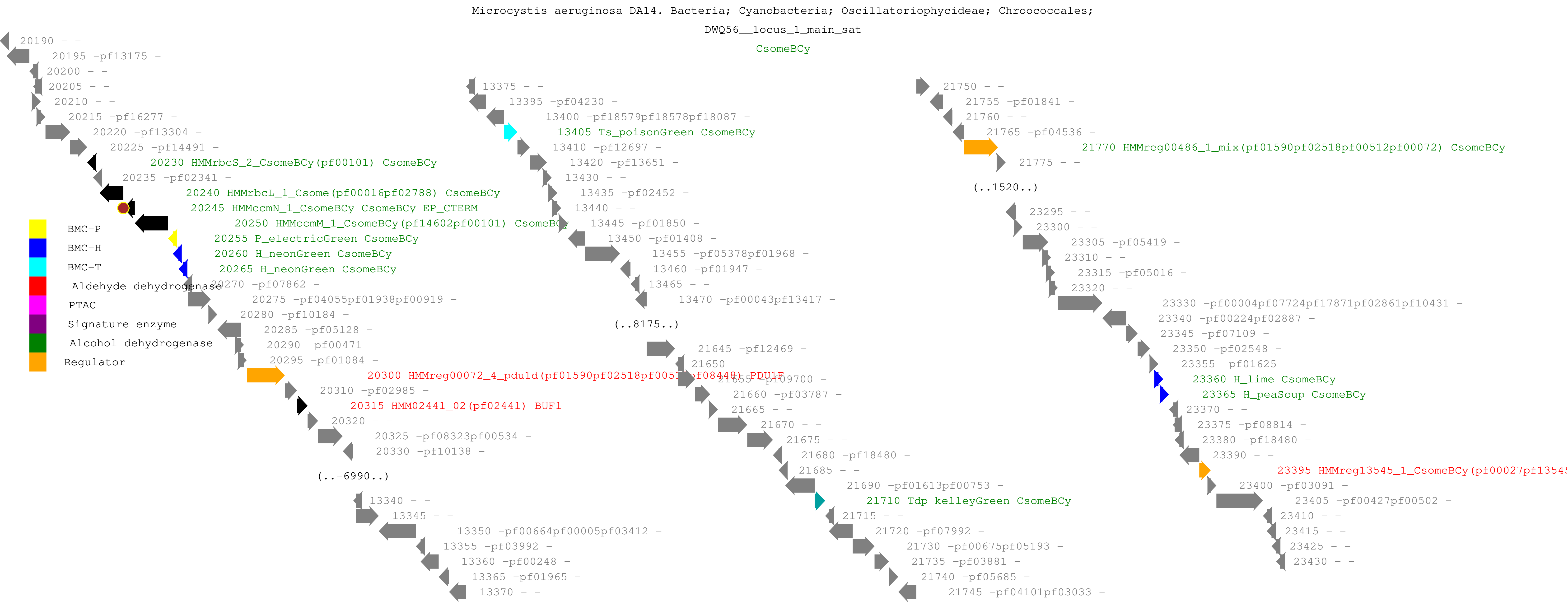

Microcystis aeruginosa DA14. Bacteria; Cyanobacteria; Oscillatoriophycideae; Chroococcales;
DWQ56__locus_1_main_sat
CsomeBCy
BMC-P
BMC-H
BMC-T
Aldehyde dehydrogenase
PTAC
Signature enzyme
Alcohol dehydrogenase
Regulator
20190 - -
20195 -pf13175 -
20200 - -
20205 - -
20210 - -
20215 -pf16277 -
20220 -pf13304 -
20225 -pf14491 -
20230 HMMrbcS_2_CsomeBCy(pf00101) CsomeBCy
20235 -pf02341 -
20240 HMMrbcL_1_Csome(pf00016pf02788) CsomeBCy
20245 HMMccmN_1_CsomeBCy CsomeBCy EP_CTERM
20250 HMMccmM_1_CsomeBCy(pf14602pf00101) CsomeBCy
20255 P_electricGreen CsomeBCy
20260 H_neonGreen CsomeBCy
20265 H_neonGreen CsomeBCy
20270 -pf07862 -
20275 -pf04055pf01938pf00919 -
20280 -pf10184 -
20285 -pf05128 -
20290 -pf00471 -
20295 -pf01084 -
20300 HMMreg00072_4_pdu1d(pf01590pf02518pf0051
20310 -pf02985 -
20315 HMM02441_02(pf02441) BUF1
20320 - -
20325 -pf08323pf00534 -
20330 -pf10138 -
(..-6990..)
13340 - -
13345 - -
13350 -pf00664pf00005pf03412 -
13355 -pf03992 -
13360 -pf00248 -
13365 -pf01965 -
13370 - -
13375 - -
13395 -pf04230 -
13400 -pf18579pf18578pf18087 -
13405 Ts_poisonGreen CsomeBCy
13410 -pf12697 -
13420 -pf13651 -
13430 - -
13435 -pf02452 -
13440 - -
13445 -pf01850 -
13450 -pf01408 -
13455 -pf05378pf01968 -
13460 -pf01947 -
13465 - -
13470 -pf00043pf13417 -
(..8175..)
21645 -pf12469 -
21650 - -
21655 -pf09700 -
21660 -pf03787 -
21665 - -
21670 - -
21675 - -
21680 -pf18480 -
21685 - -
21690 -pf01613pf00753 -
21710 Tdp_kelleyGreen CsomeBCy
21715 - -
21720 -pf07992 -
21730 -pf00675pf05193 -
21735 -pf03881 -
21740 -pf05685 -
21745 -pf04101pf03033 -
21750 - -
21755 -pf01841 -
21760 - -
21765 -pf04536 -
21770 HMMreg00486_1_mix(pf01590pf02518pf00512pf00072) CsomeBCy
21775 - -
(..1520..)
23295 - -
23300 - -
23305 -pf05419 -
23310 - -
23315 -pf05016 -
23320 - -
23330 -pf00004pf07724pf17871pf02861pf10431 -
23340 -pf00224pf02887 -
23345 -pf07109 -
23350 -pf02548 -
23355 -pf01625 -
23360 H_lime CsomeBCy
23365 H_peaSoup CsomeBCy
23370 - -
23375 -pf08814 -
23380 -pf18480 -
23390 - -
23395 HMMreg13545_1_CsomeBCy(pf00027pf1354
23400 -pf03091 -
23405 -pf00427pf00502 -
23410 - -
23415 - -
23425 - -
23430 - -

Microcystis aeruginosa TA09. Bacteria; Cyanobacteria; Oscillatoriophycideae; Chroococcales;

DWQ58__locus_1_main_sat

CsomeBCy

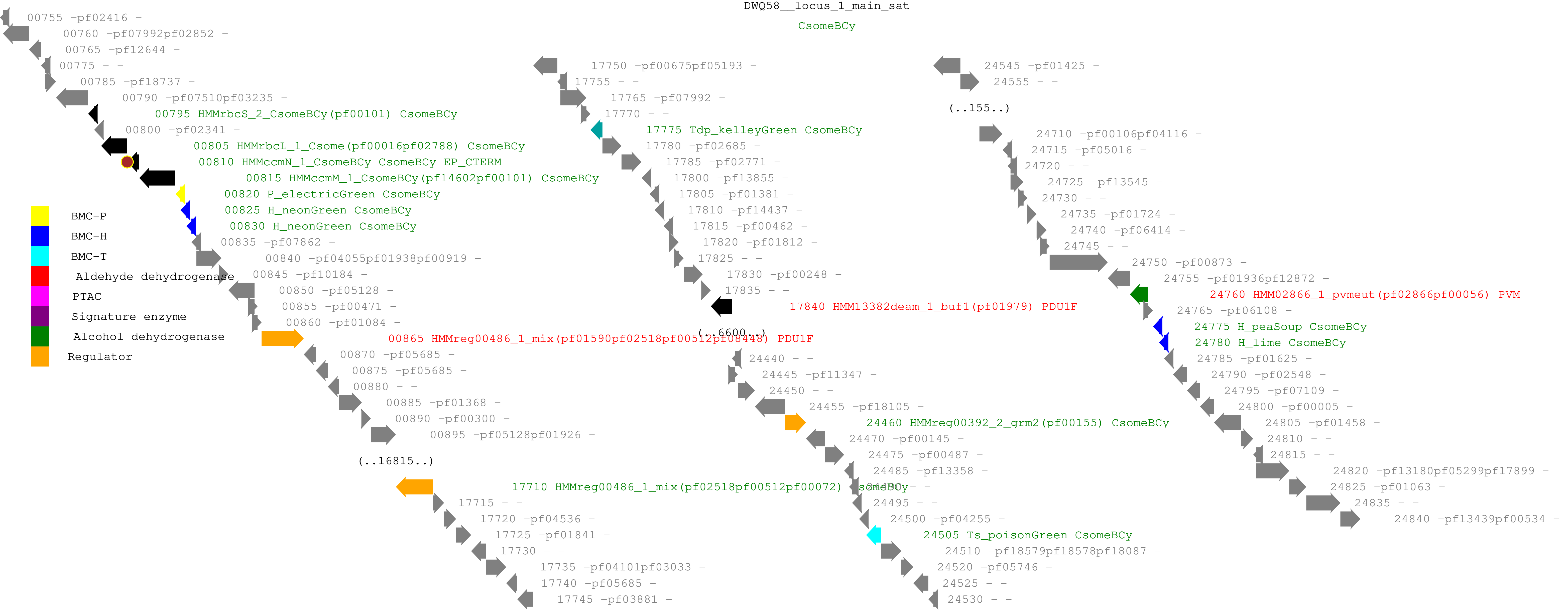

Halomicronema excentricum str. Lakshadweep. Bacteria; Cyanobacteria; Synechococcales; Prochlorotrichaceae;

DYY88__locus_1_main_sat

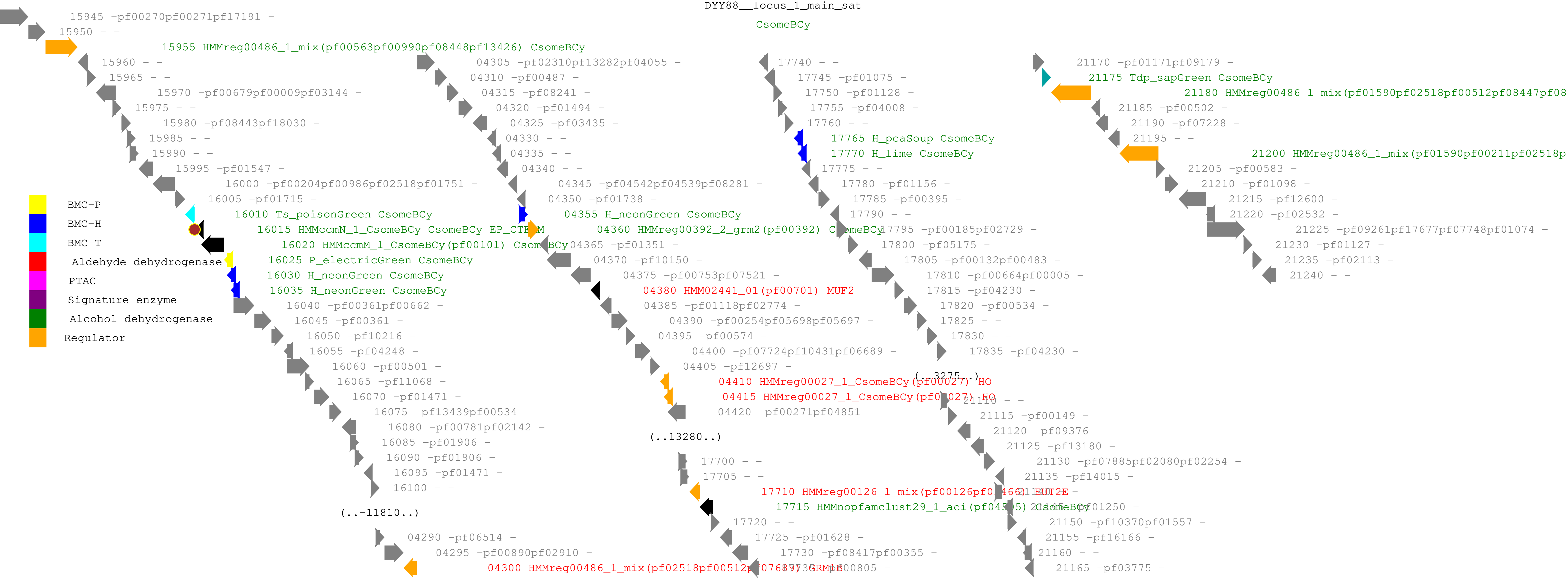

Scytonema millei VB511283. Bacteria; Cyanobacteria; Nostocales; Scytonemataceae; Scytonema.

DYY89__locus_1_main_sat

CsomeBCy

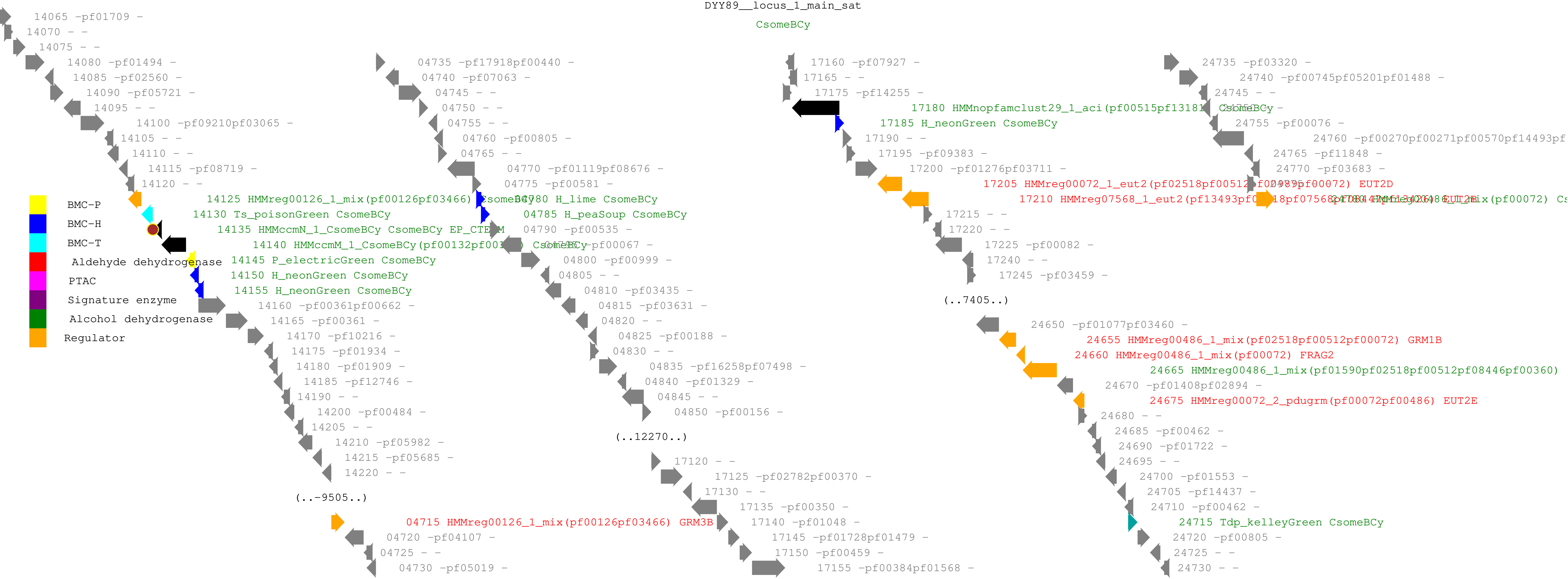

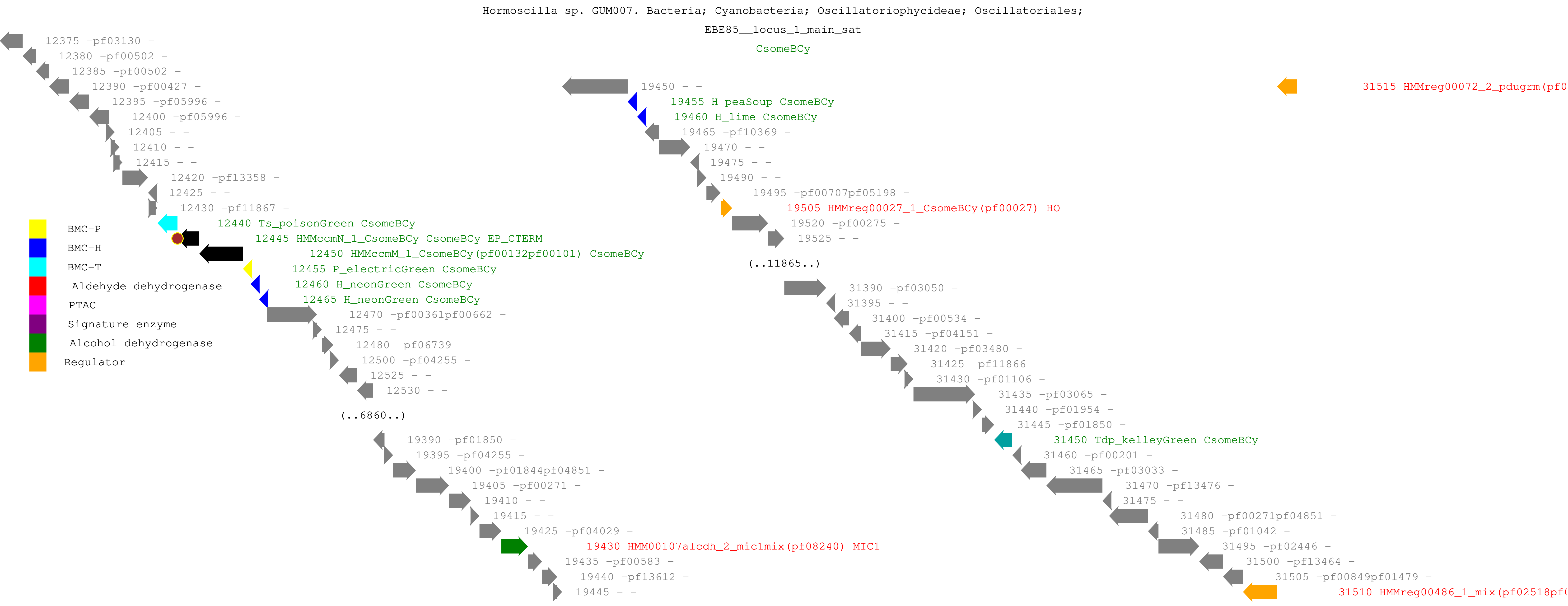
Hormoscilla sp. GUM007. Bacteria; Cyanobacteria; Oscillatoriophycideae; Oscillatoriales;
EBE85__locus_1_main_sat
CsomeBCy
BMC-P
BMC-H
BMC-T
Aldehyde dehydrogenase
PTAC
Signature enzyme
Alcohol dehydrogenase
Regulator
12375 -pf03130 -
12380 -pf00502 -
12385 -pf00502 -
12390 -pf00427 -
12395 -pf05996 -
12400 -pf05996 -
12405 - -
12410 - -
12415 - -
12420 -pf13358 -
12425 - -
12430 -pf11867 -
12440 Ts_poisonGreen CsomeBCy
12445 HMMccmN_1_CsomeBCy CsomeBCy EP_CTERM
12450 HMMccmM_1_CsomeBCy(pf00132pf00101) CsomeBCy
12455 P_electricGreen CsomeBCy
12460 H_neonGreen CsomeBCy
12465 H_neonGreen CsomeBCy
12470 -pf00361pf00662 -
12475 - -
12480 -pf06739 -
12500 -pf04255 -
12525 - -
12530 - -
(..6860..)
19390 -pf01850 -
19395 -pf04255 -
19400 -pf01844pf04851 -
19405 -pf00271 -
19410 - -
19415 - -
19425 -pf04029 -
19430 HMM00107alcdh_2_mic1mix(pf08240) MIC1
19435 -pf00583 -
19440 -pf13612 -
19445 - -
19450 - -
19455 H_peaSoup CsomeBCy
19460 H_lime CsomeBCy
19465 -pf10369 -
19470 - -
19475 - -
19490 - -
19495 -pf00707pf05198 -
19505 HMMreg00027_1_CsomeBCy(pf00027) HO
19520 -pf00275 -
19525 - -
(..11865..)
31390 -pf03050 -
31395 - -
31400 -pf00534 -
31415 -pf04151 -
31420 -pf03480 -
31425 -pf11866 -
31430 -pf01106 -
31435 -pf03065 -
31440 -pf01954 -
31445 -pf01850 -
31450 Tdp_kelleyGreen CsomeBCy
31460 -pf00201 -
31465 -pf03033 -
31470 -pf13476 -
31475 - -
31480 -pf00271pf04851 -
31485 -pf01042 -
31495 -pf02446 -
31500 -pf13464 -
31505 -pf00849pf01479 -
31510 HMMreg00486_1_mix(pf02518pf0
31515 HMMreg00072_2_pdugrm(pf0

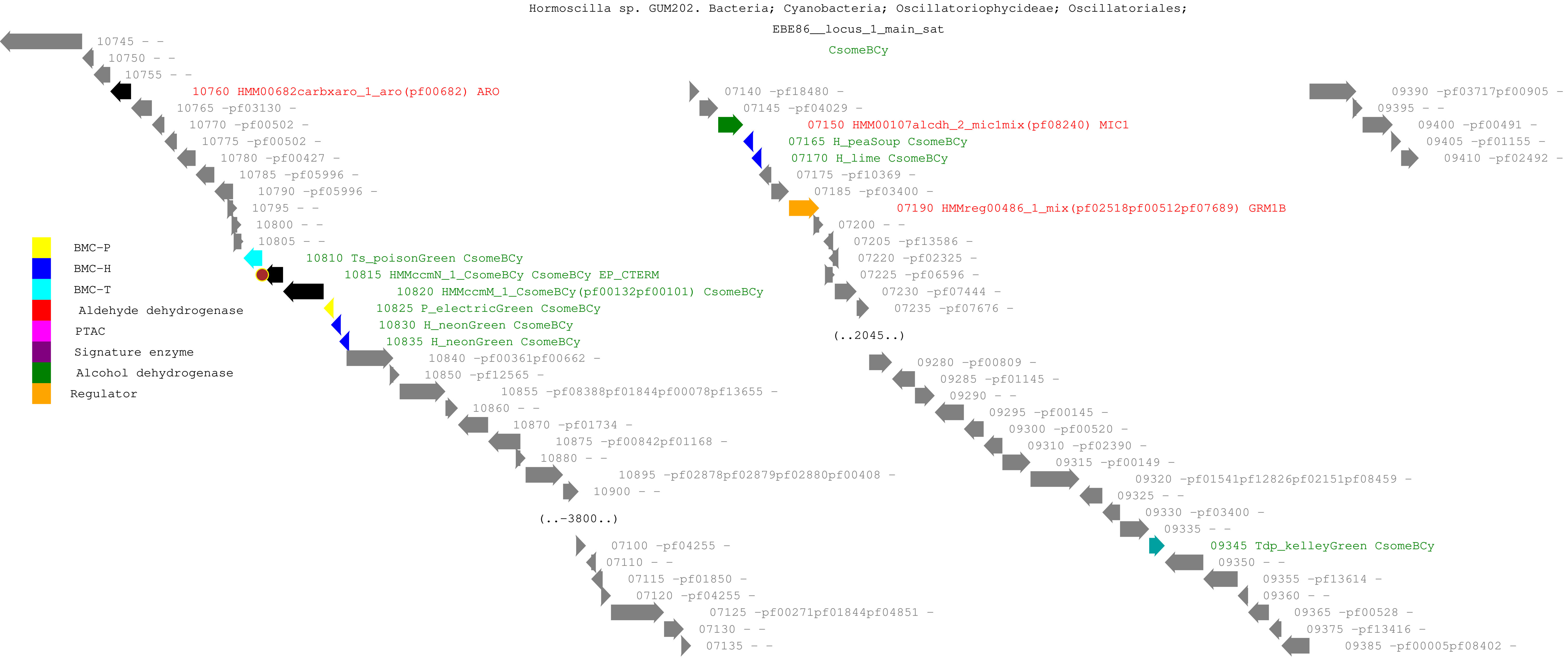
Hormoscilla sp. GUM202. Bacteria; Cyanobacteria; Oscillatoriophycideae; Oscillatoriales;
EBE86__locus_1_main_sat
CsomeBCy
10745 - -
10750 - -
10755 - -
10760 HMM00682carbxaro_1_aro(pf00682) ARO
10765 -pf03130 -
10770 -pf00502 -
10775 -pf00502 -
10780 -pf00427 -
10785 -pf05996 -
10790 -pf05996 -
10795 - -
10800 - -
10805 - -
10810 Ts_poisonGreen CsomeBCy
10815 HMMccmN_1_CsomeBCy CsomeBCy EP_CTERM
10820 HMMccmM_1_CsomeBCy(pf00132pf00101) CsomeBCy
10825 P_electricGreen CsomeBCy
10830 H_neonGreen CsomeBCy
10835 H_neonGreen CsomeBCy
10840 -pf00361pf00662 -
10850 -pf12565 -
10855 -pf08388pf01844pf00078pf13655 -
10860 - -
10870 -pf01734 -
10875 -pf00842pf01168 -
10880 - -
10895 -pf02878pf02879pf02880pf00408 -
10900 - -
(..-3800..)
07100 -pf04255 -
07110 - -
07115 -pf01850 -
07120 -pf04255 -
07125 -pf00271pf01844pf04851 -
07130 - -
07135 - -
07140 -pf18480 -
07145 -pf04029 -
07150 HMM00107alcdh_2_mic1mix(pf08240) MIC1
07165 H_peaSoup CsomeBCy
07170 H_lime CsomeBCy
07175 -pf10369 -
07185 -pf03400 -
07190 HMMreg00486_1_mix(pf02518pf00512pf07689) GRM1B
07200 - -
07205 -pf13586 -
07220 -pf02325 -
07225 -pf06596 -
07230 -pf07444 -
07235 -pf07676 -
(..2045..)
09280 -pf00809 -
09285 -pf01145 -
09290 - -
09295 -pf00145 -
09300 -pf00520 -
09310 -pf02390 -
09315 -pf00149 -
09320 -pf01541pf12826pf02151pf08459 -
09325 - -
09330 -pf03400 -
09335 - -
09345 Tdp_kelleyGreen CsomeBCy
09350 - -
09355 -pf13614 -
09360 - -
09365 -pf00528 -
09375 -pf13416 -
09385 -pf00005pf08402 -
09390 -pf03717pf00905 -
09395 - -
09400 -pf00491 -
09405 -pf01155 -
09410 -pf02492 -
BMC-P
BMC-H
BMC-T
Aldehyde dehydrogenase
PTAC
Signature enzyme
Alcohol dehydrogenase
Regulator

Microcystis aeruginosa FACHB-524. Bacteria; Cyanobacteria; Oscillatoriophycideae; Chroococcales;

ED562__locus_1_main_sat

CsomeBCy

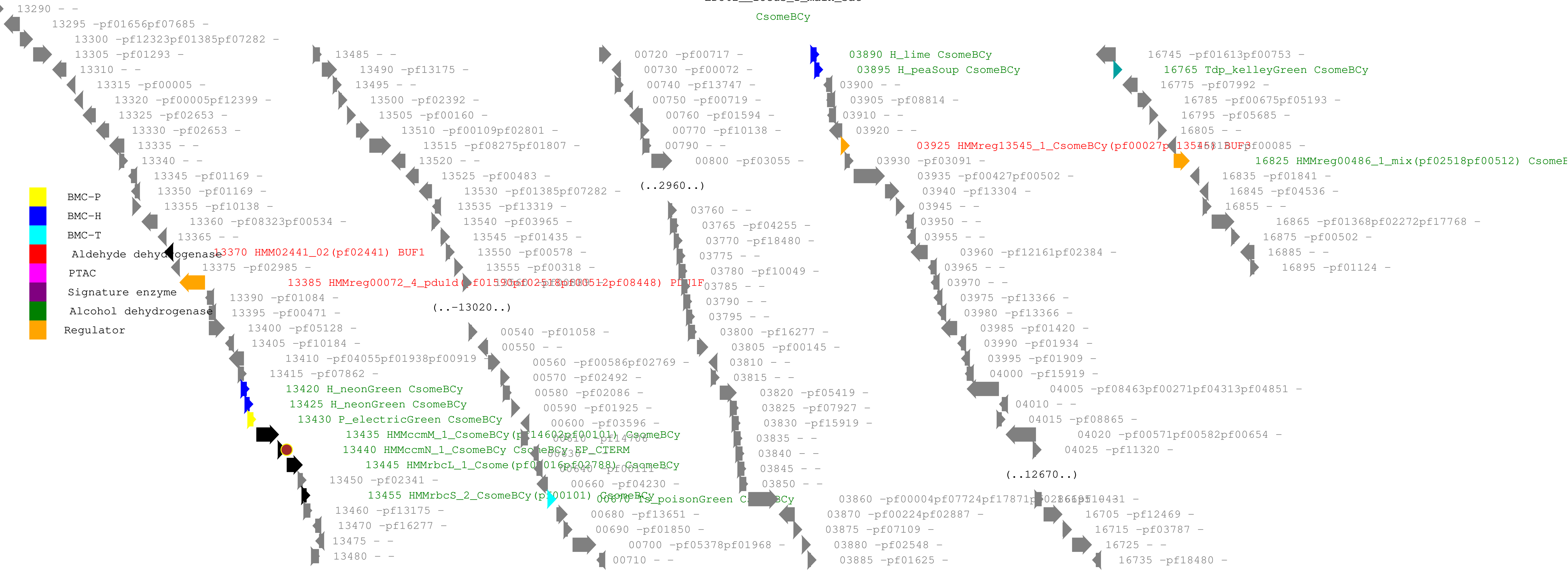

Leptolyngbya sp. IPPAS B-1204. Bacteria; Cyanobacteria; Synechococcales; Leptolyngbyaceae; Leptolyngbya.

EDM05__locus_1_main_sat

CsomeBCy

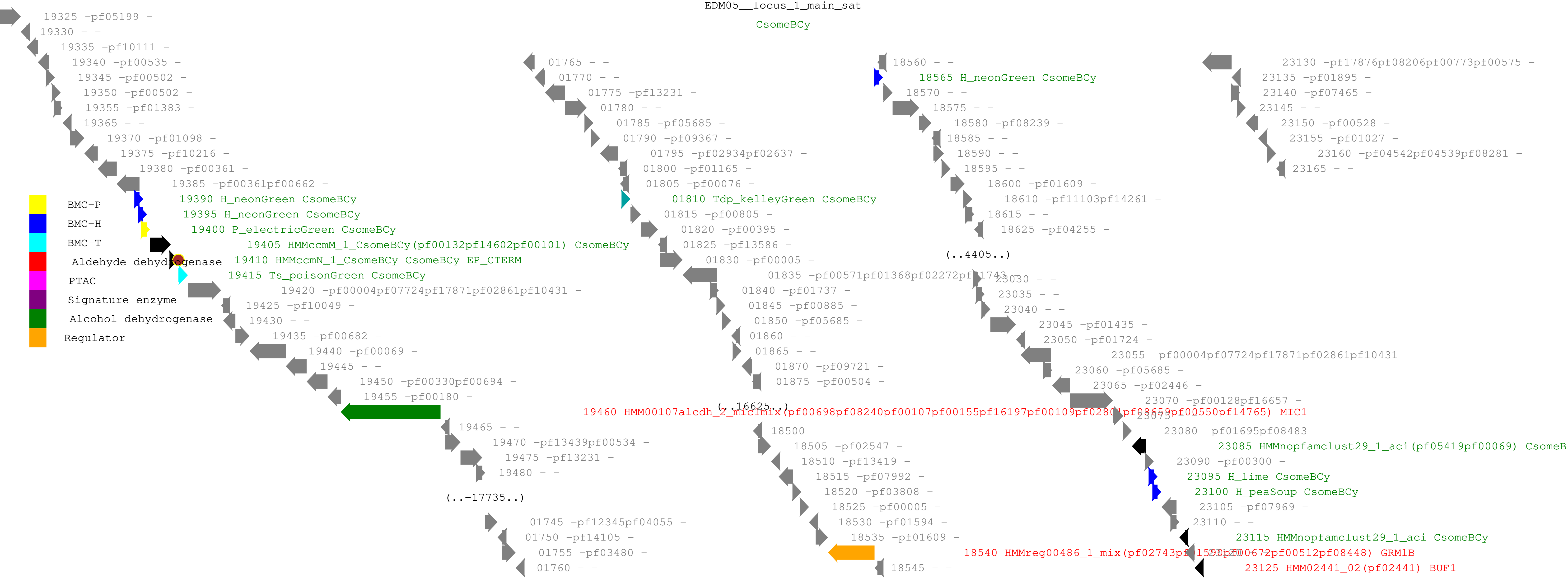

Chroococcales cyanobacterium metabat2.561. Bacteria; Cyanobacteria; Oscillatoriophycideae; Chroococcales.

EHM73__locus_fragmented

CsomeBCy

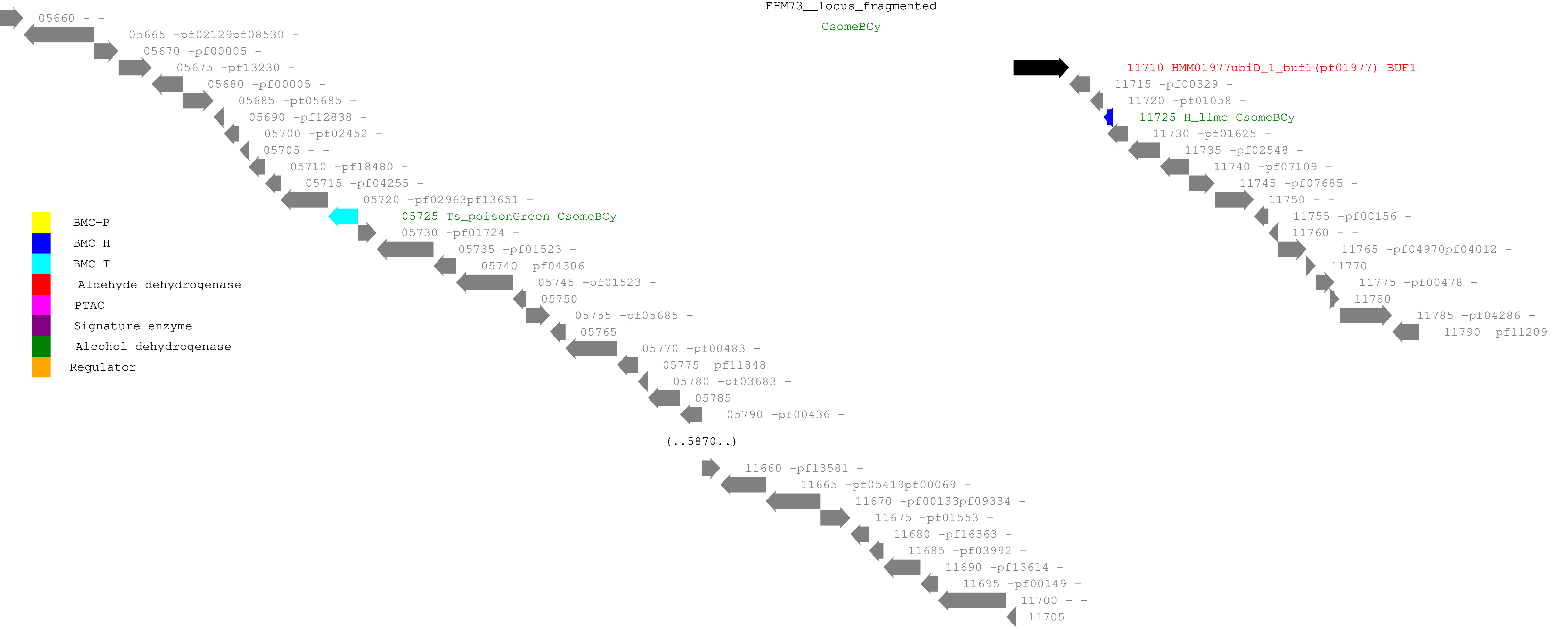

Microcystis flos-aquae Mf_WU_F_19750830_S460. Bacteria; Cyanobacteria; Oscillatoriophycideae; Chroococcales;

EWV40__locus_1_main_sat

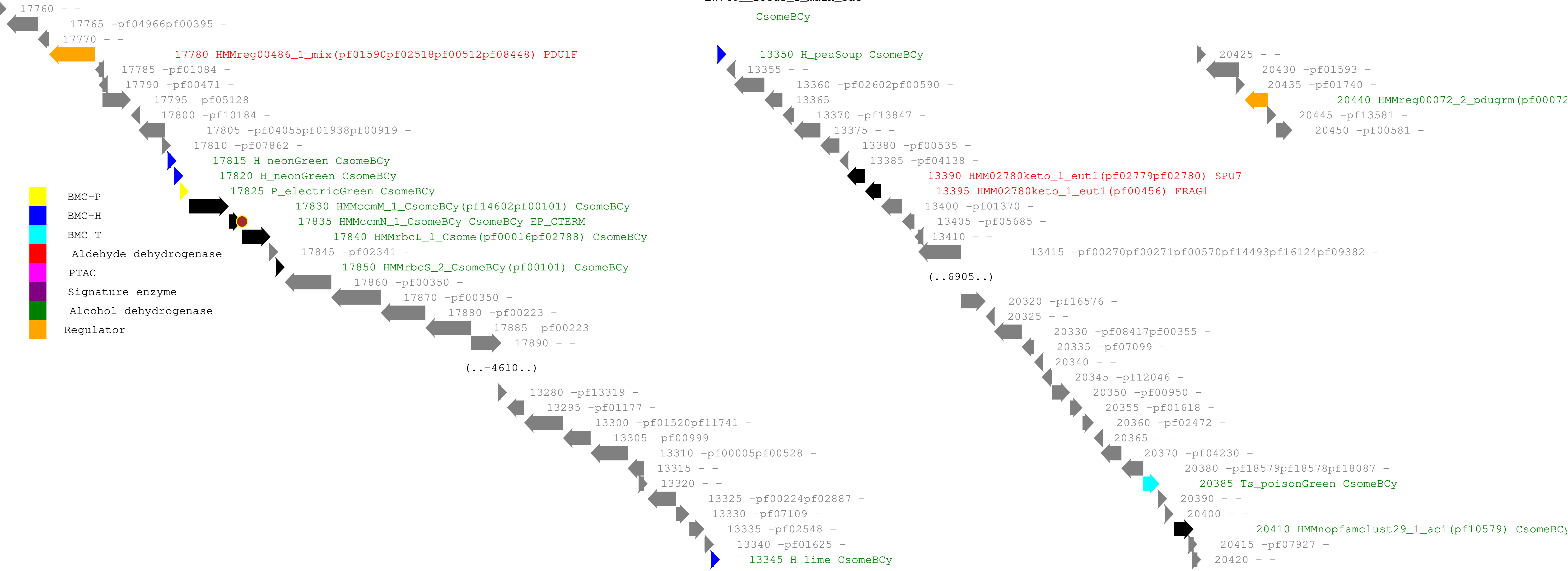

Microcystis flos-aquae Mf_QC_C_20070823_S10D. Bacteria; Cyanobacteria; Oscillatoriophycideae; Chroococcales;

EWV45__locus_1_main_sat

CsomeBCy

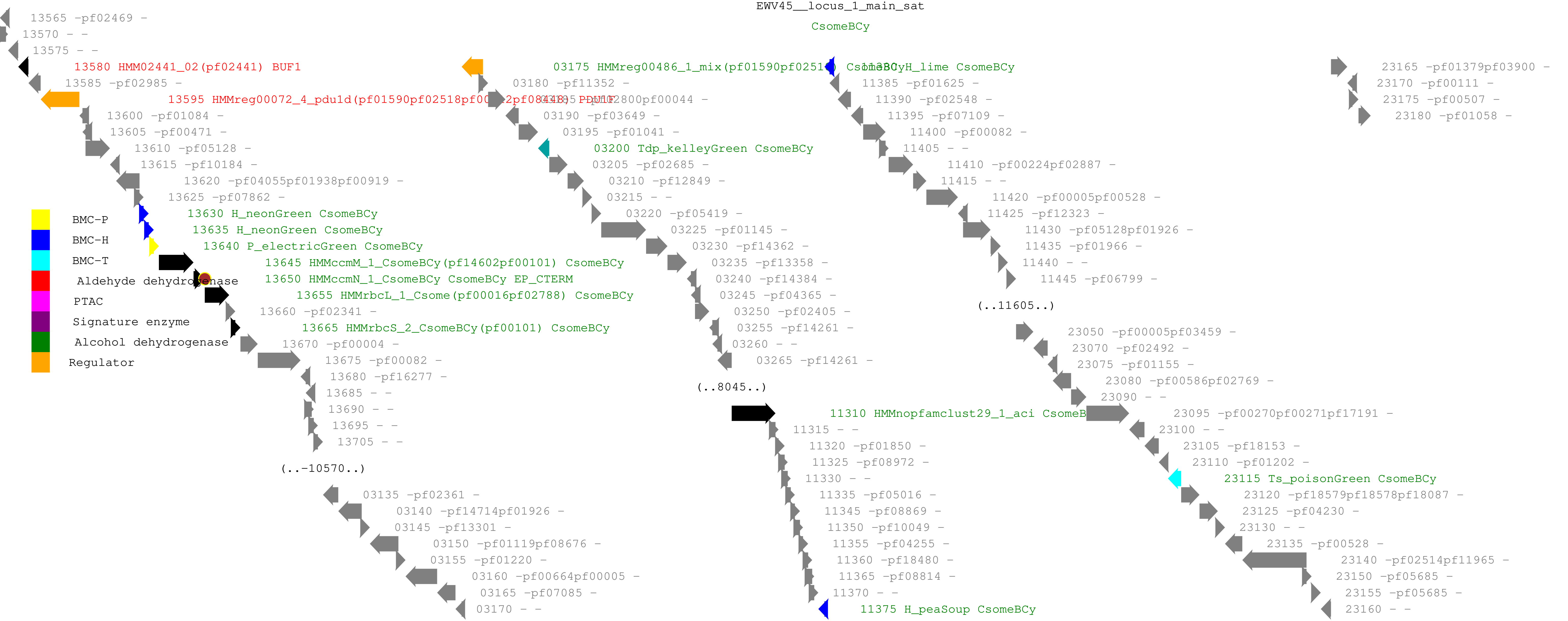

Microcystis panniformis Mp_MB_F_20051200_S9. Bacteria; Cyanobacteria; Oscillatoriophycideae; Chroococcales;

EWV53__locus_1_main_sat

CsomeBCy

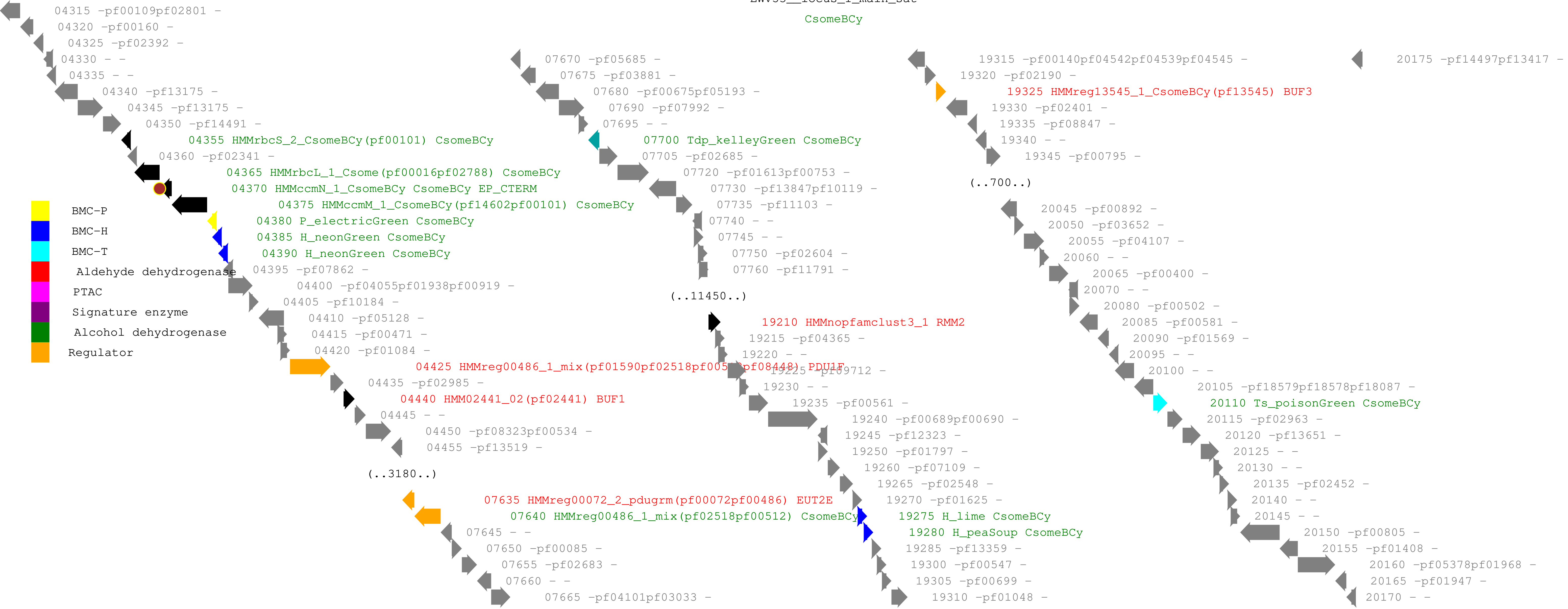

Microcystis novacekii Mn_MB_F_20050700_S1D. Bacteria; Cyanobacteria; Oscillatoriophycideae; Chroococcales;

EWV54__locus_1_main_sat

CsomeBCy

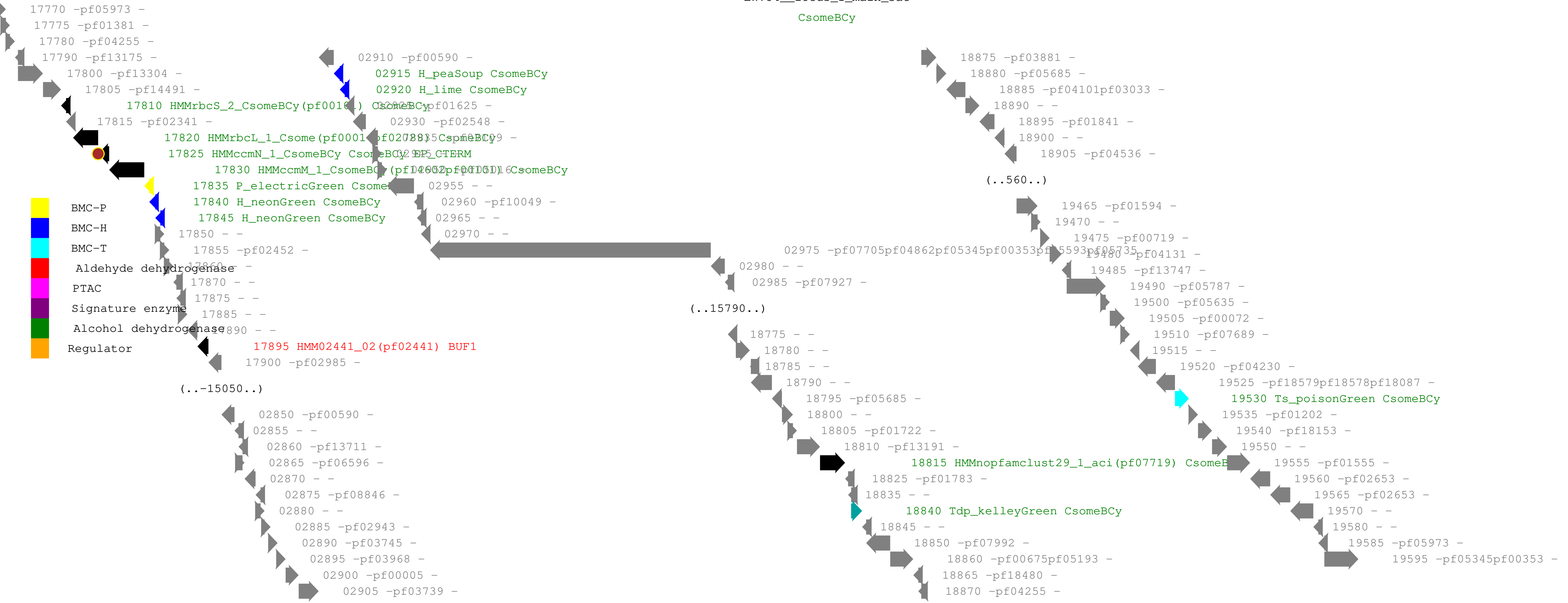

Microcystis aeruginosa Ma_QC_Ch_20071001_S25D. Bacteria; Cyanobacteria; Oscillatoriophycideae; Chroococcales;

EWV57__locus_1_main_sat

CsomeBCy

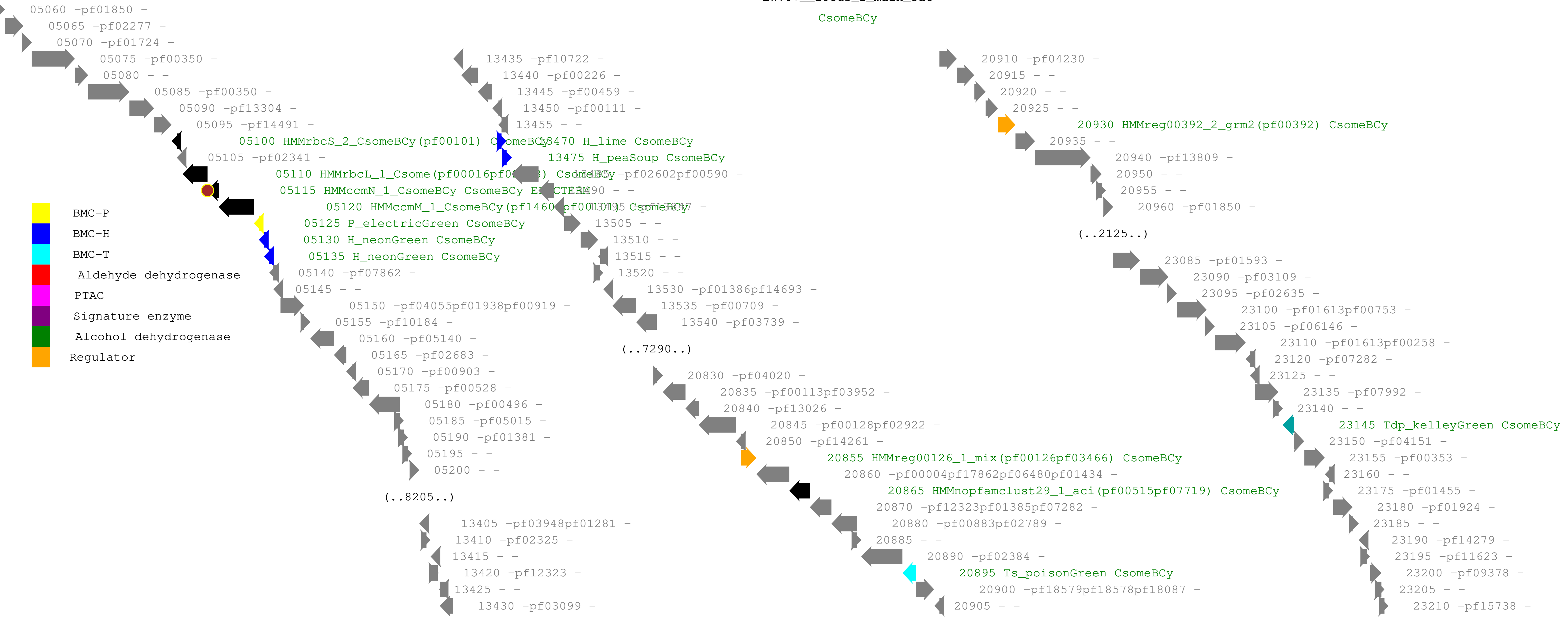

Microcystis sp. Msp_OC_L_20101000_S702. Bacteria; Cyanobacteria; Oscillatoriophycideae; Chroococcales;
EWV60__locus_1_main_sat
CsomeBCy

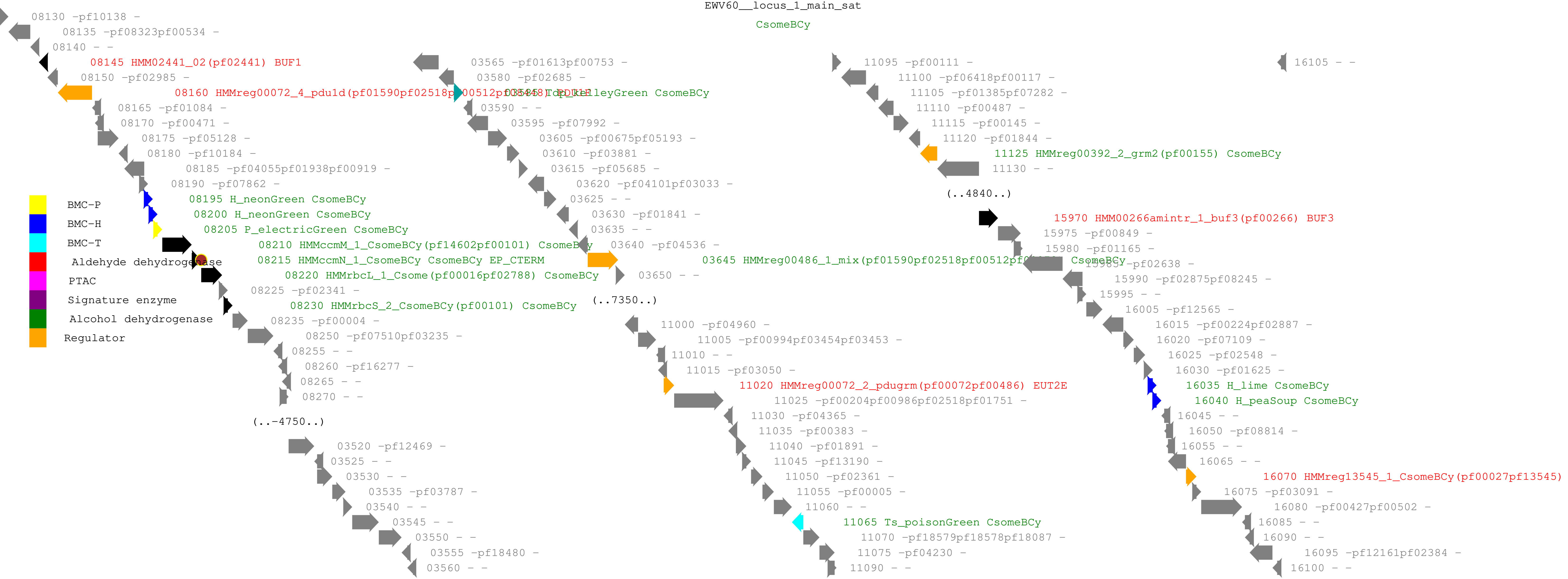

Microcystis aeruginosa Ma_OC_LR_19540900_S633. Bacteria; Cyanobacteria; Oscillatoriophycideae; Chroococcales;

EWV62__locus_1_main_sat

CsomeBCy

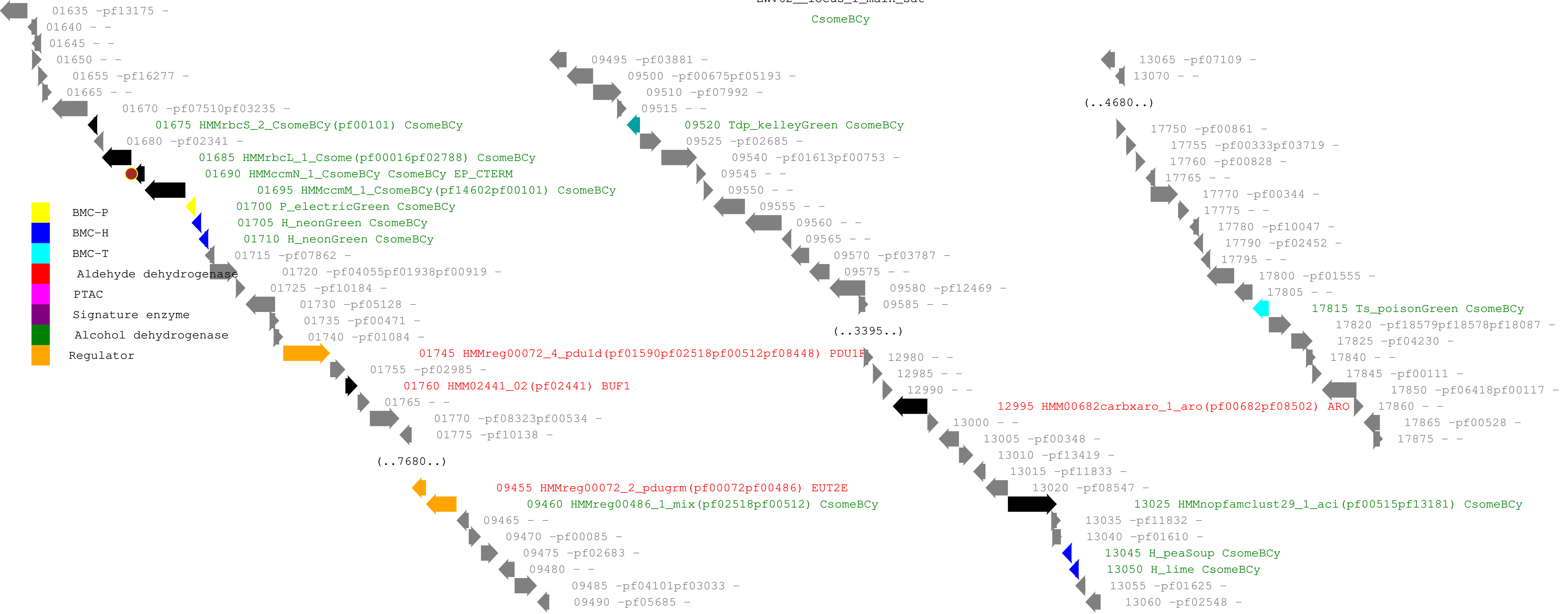

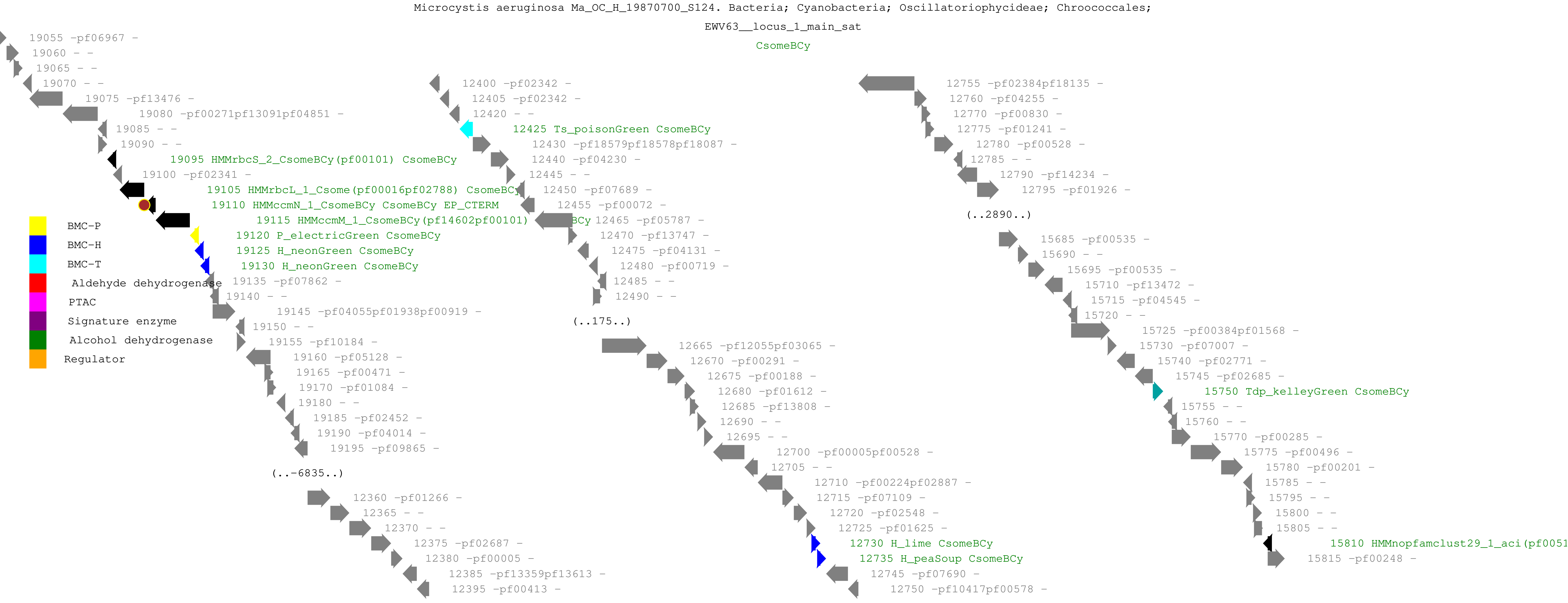
Microcystis aeruginosa Ma_OC_H_19870700_S124. Bacteria; Cyanobacteria; Oscillatoriophycideae; Chroococcales;
EWV63__locus_1_main_sat
CsomeBCy
19055 -pf06967 -
19060 - -
19065 - -
19070 - -
19075 -pf13476 -
19080 -pf00271pf13091pf04851 -
19085 - -
19090 - -
19095 HMMrbcS_2_CsomeBCy(pf00101) CsomeBCy
19100 -pf02341 -
19105 HMMrbcL_1_Csome(pf00016pf02788) CsomeBCy
19110 HMMccmN_1_CsomeBCy CsomeBCy EP_CTERM
19115 HMMccmM_1_CsomeBCy(pf14602pf00101)
19120 P_electricGreen CsomeBCy
19125 H_neonGreen CsomeBCy
19130 H_neonGreen CsomeBCy
19135 -pf07862 -
19140 - -
19145 -pf04055pf01938pf00919 -
19150 - -
19155 -pf10184 -
19160 -pf05128 -
19165 -pf00471 -
19170 -pf01084 -
19180 - -
19185 -pf02452 -
19190 -pf04014 -
19195 -pf09865 -
(..-6835..)
12360 -pf01266 -
12365 - -
12370 - -
12375 -pf02687 -
12380 -pf00005 -
12385 -pf13359pf13613 -
12395 -pf00413 -
BMC-P
BMC-H
BMC-T
Aldehyde dehydrogenase
PTAC
Signature enzyme
Alcohol dehydrogenase
Regulator
12400 -pf02342 -
12405 -pf02342 -
12420 - -
12425 Ts_poisonGreen CsomeBCy
12430 -pf18579pf18578pf18087 -
12440 -pf04230 -
12445 - -
12450 -pf07689 -
12455 -pf00072 -
BCy 12465 -pf05787 -
12470 -pf13747 -
12475 -pf04131 -
12480 -pf00719 -
12485 - -
12490 - -
(..175..)
12665 -pf12055pf03065 -
12670 -pf00291 -
12675 -pf00188 -
12680 -pf01612 -
12685 -pf13808 -
12690 - -
12695 - -
12700 -pf00005pf00528 -
12705 - -
12710 -pf00224pf02887 -
12715 -pf07109 -
12720 -pf02548 -
12725 -pf01625 -
12730 H_lime CsomeBCy
12735 H_peaSoup CsomeBCy
12745 -pf07690 -
12750 -pf10417pf00578 -
12755 -pf02384pf18135 -
12760 -pf04255 -
12770 -pf00830 -
12775 -pf01241 -
12780 -pf00528 -
12785 - -
12790 -pf14234 -
12795 -pf01926 -
(..2890..)
15685 -pf00535 -
15690 - -
15695 -pf00535 -
15710 -pf13472 -
15715 -pf04545 -
15720 - -
15725 -pf00384pf01568 -
15730 -pf07007 -
15740 -pf02771 -
15745 -pf02685 -
15750 Tdp_kelleyGreen CsomeBCy
15755 - -
15760 - -
15770 -pf00285 -
15775 -pf00496 -
15780 -pf00201 -
15785 - -
15795 - -
15800 - -
15805 - -
15810 HMMnopfamclust29_1_aci(pf0051
15815 -pf00248 -

Microcystis sp. M_OC_Ca_00000000_C217Col. Bacteria; Cyanobacteria; Oscillatoriophycideae; Chroococcales;

EWV66__locus_1_main_sat

CsomeBCy

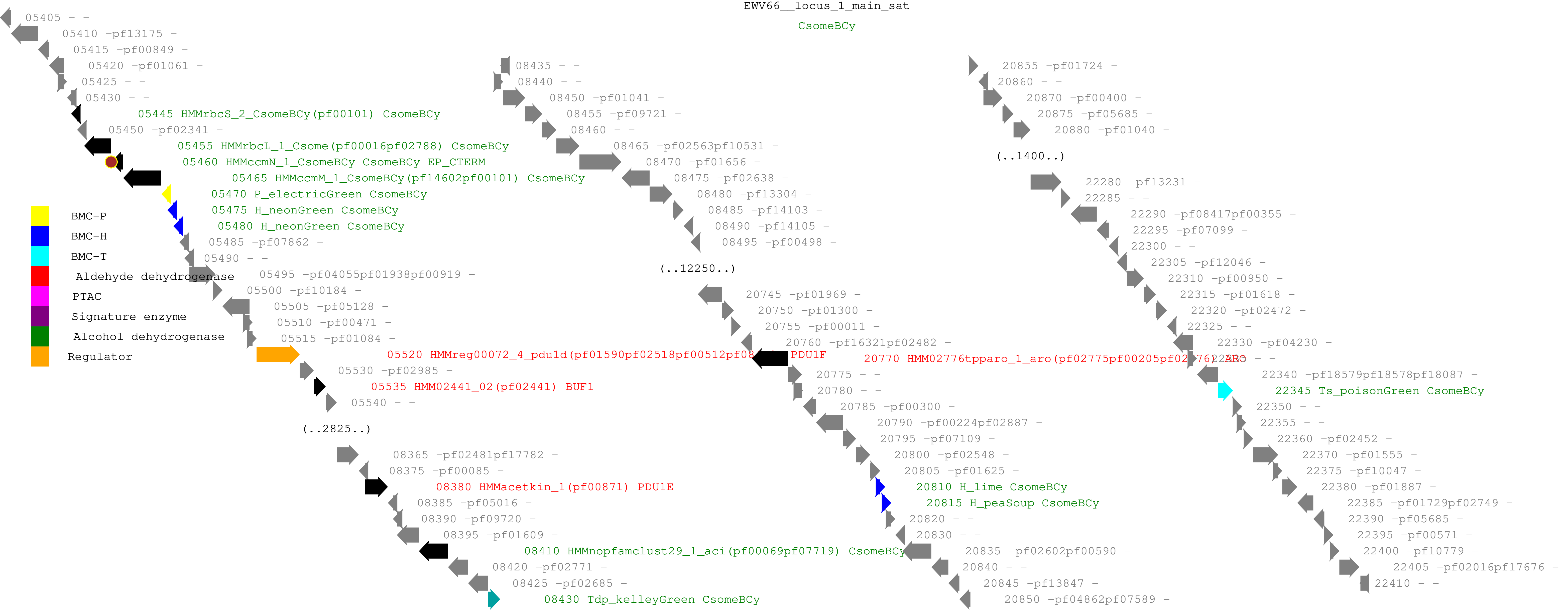

Microcystis sp. M_QC_C_20170808_M9Col. Bacteria; Cyanobacteria; Oscillatoriophycideae; Chroococcales;

EWV68__locus_1_main_sat

CsomeBCy

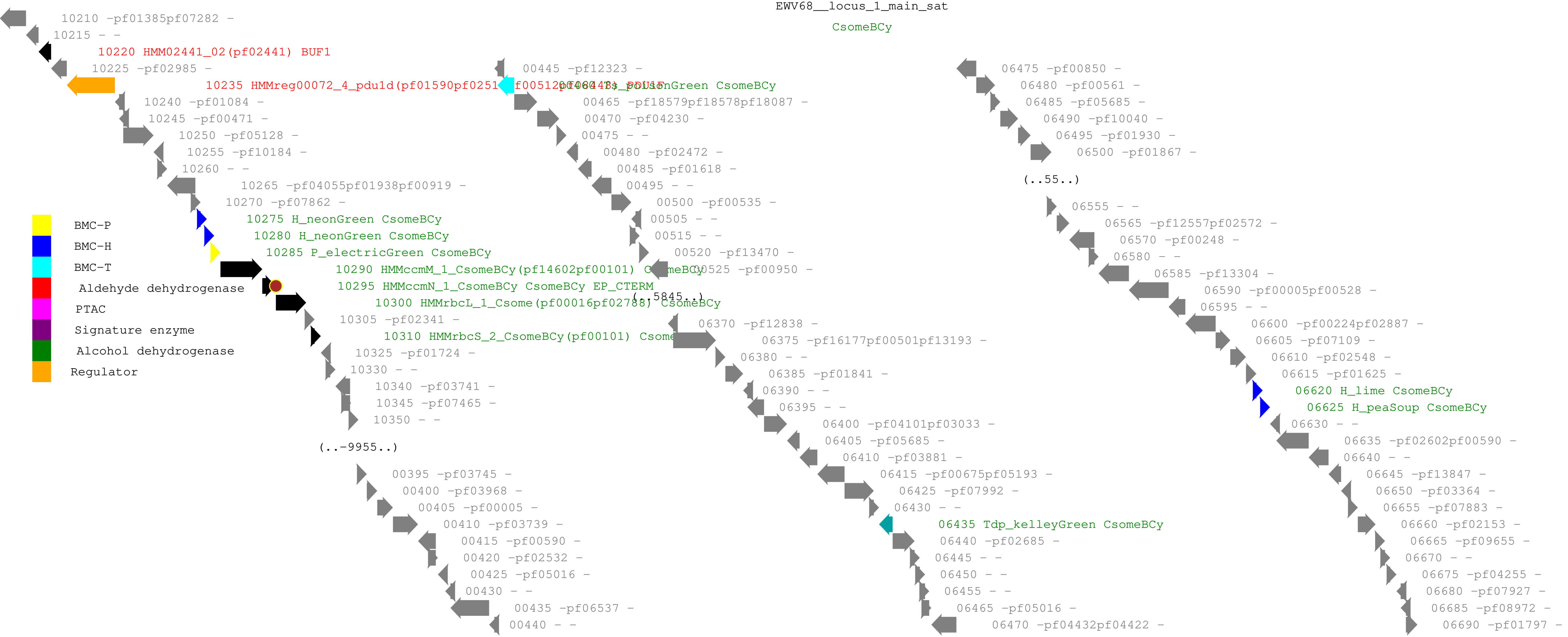

Microcystis wesenbergii Mw_QC_S_20081001_S30D. Bacteria; Cyanobacteria; Oscillatoriophycideae; Chroococcales;

EWV75__locus_1_main_sat

CsomeBCy

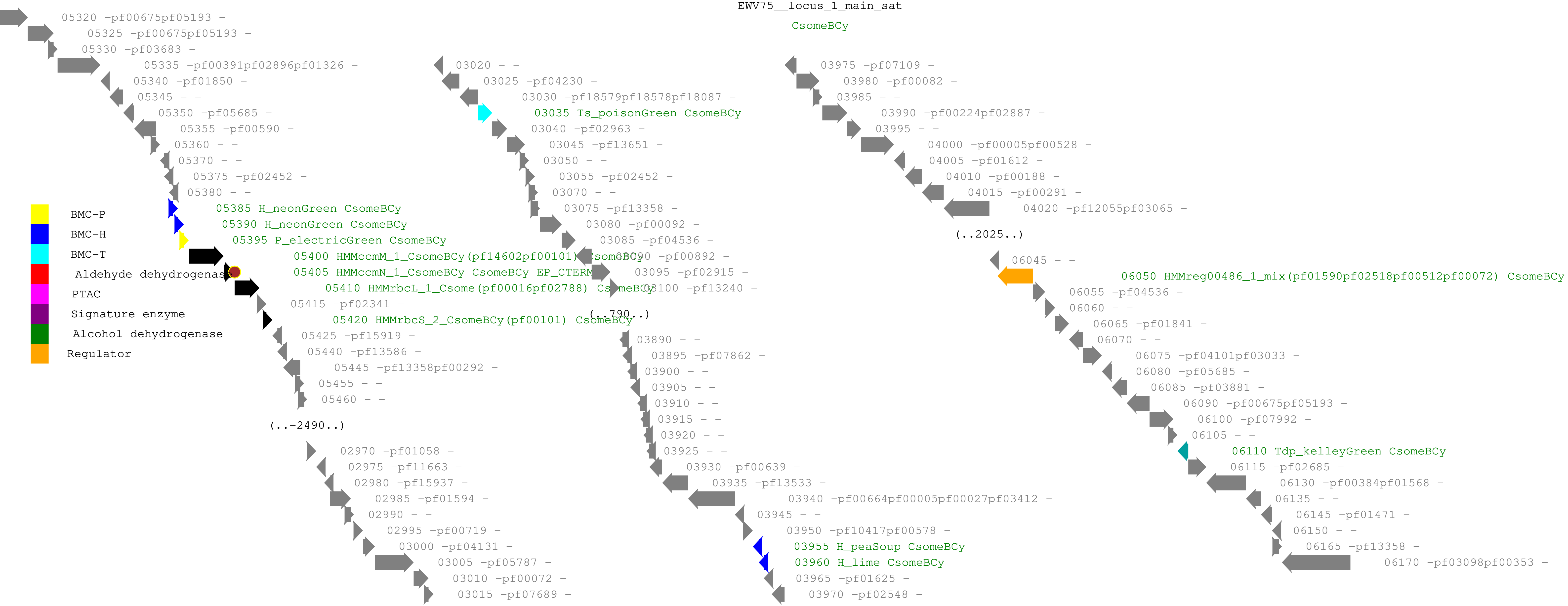

Microcystis viridis Mv_BB_P_19951000_S68D. Bacteria; Cyanobacteria; Oscillatoriophycideae; Chroococcales;

EWV77__locus_1_main_sat

CsomeBCy

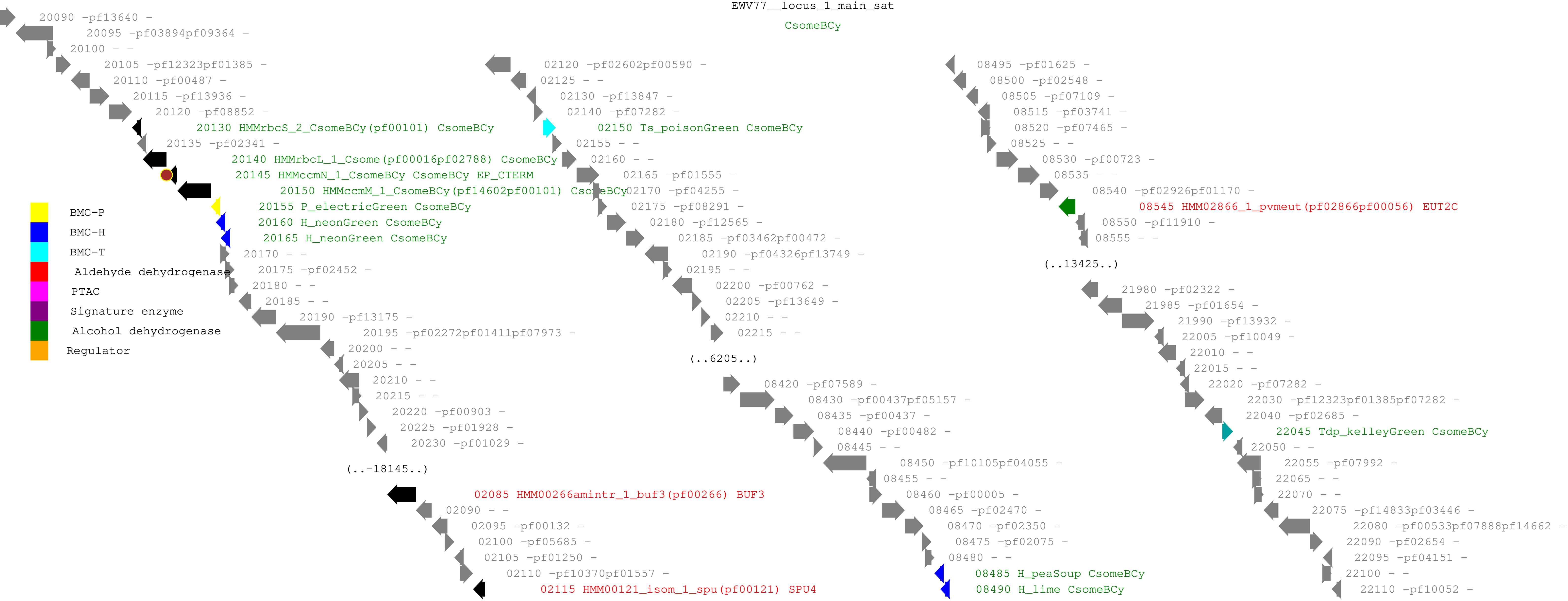

Microcystis aeruginosa Ma_MB_F_20061100_S20D. Bacteria; Cyanobacteria; Oscillatoriophycideae; Chroococcales;

EWV78__locus_1_main_sat

CsomeBCy

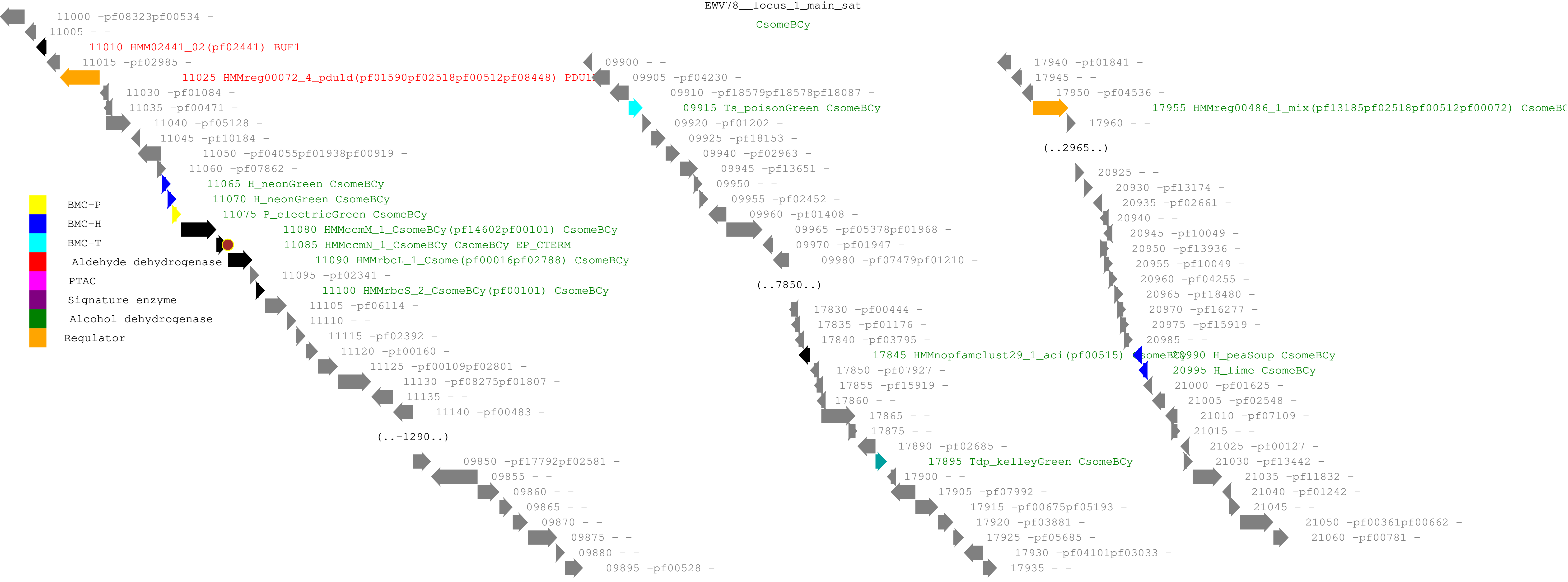

Microcystis aeruginosa Ma_QC_B_20070730_S2. Bacteria; Cyanobacteria; Oscillatoriophycideae; Chroococcales;

EWV80__locus_1_main_sat

CsomeBCy

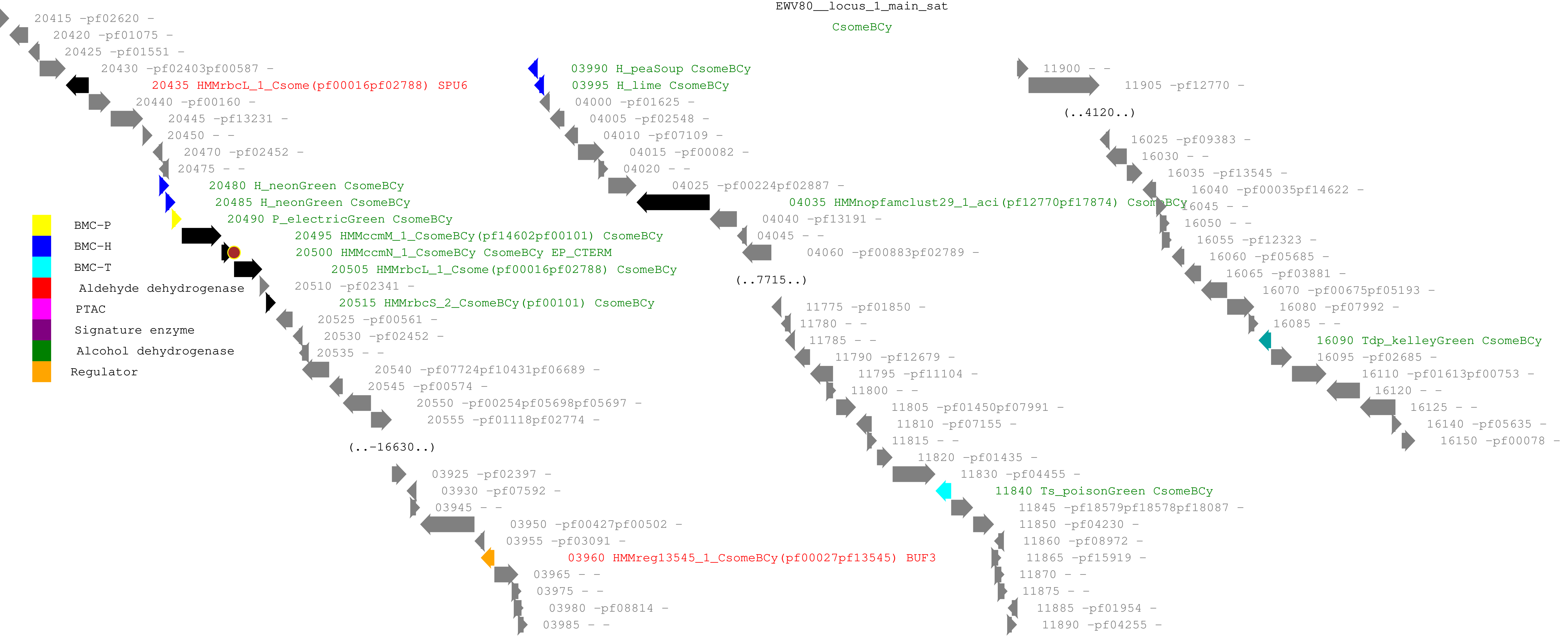

Microcystis aeruginosa Ma_SC_T_19800800_S464. Bacteria; Cyanobacteria; Oscillatoriophycideae; Chroococcales;

EWV81__locus_1_main_sat

CsomeBCy

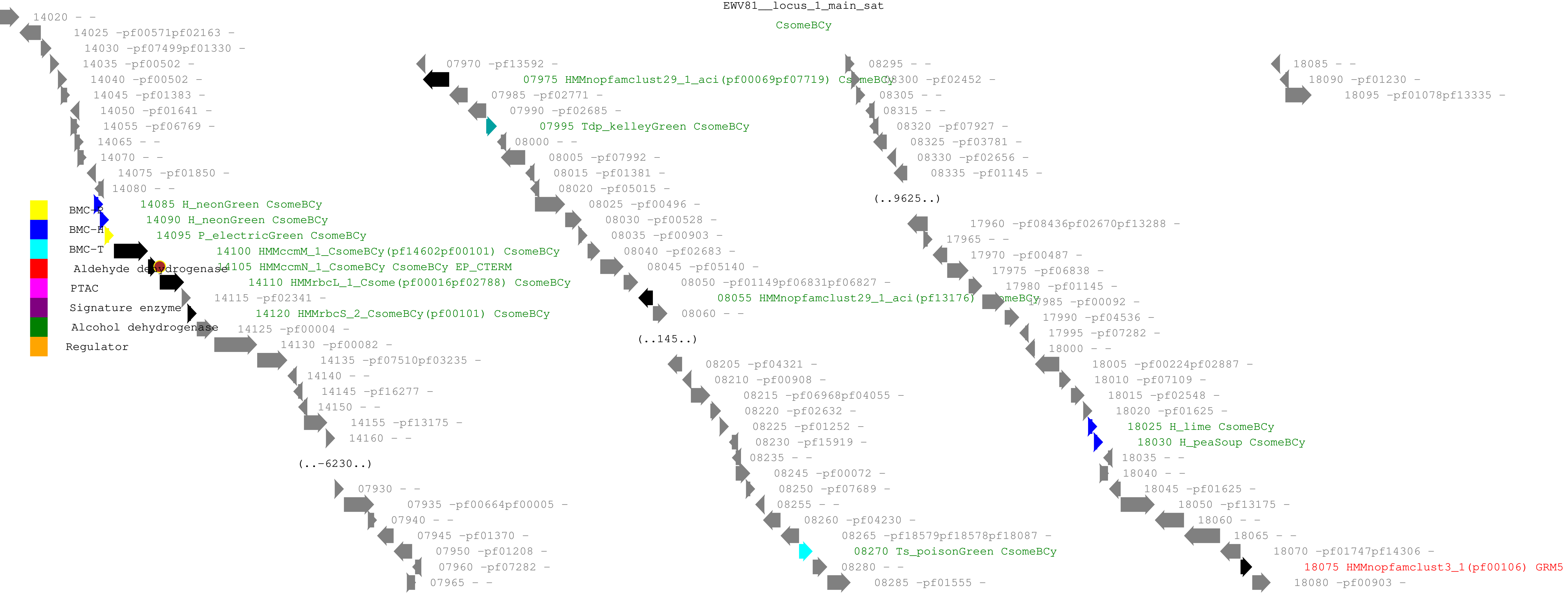

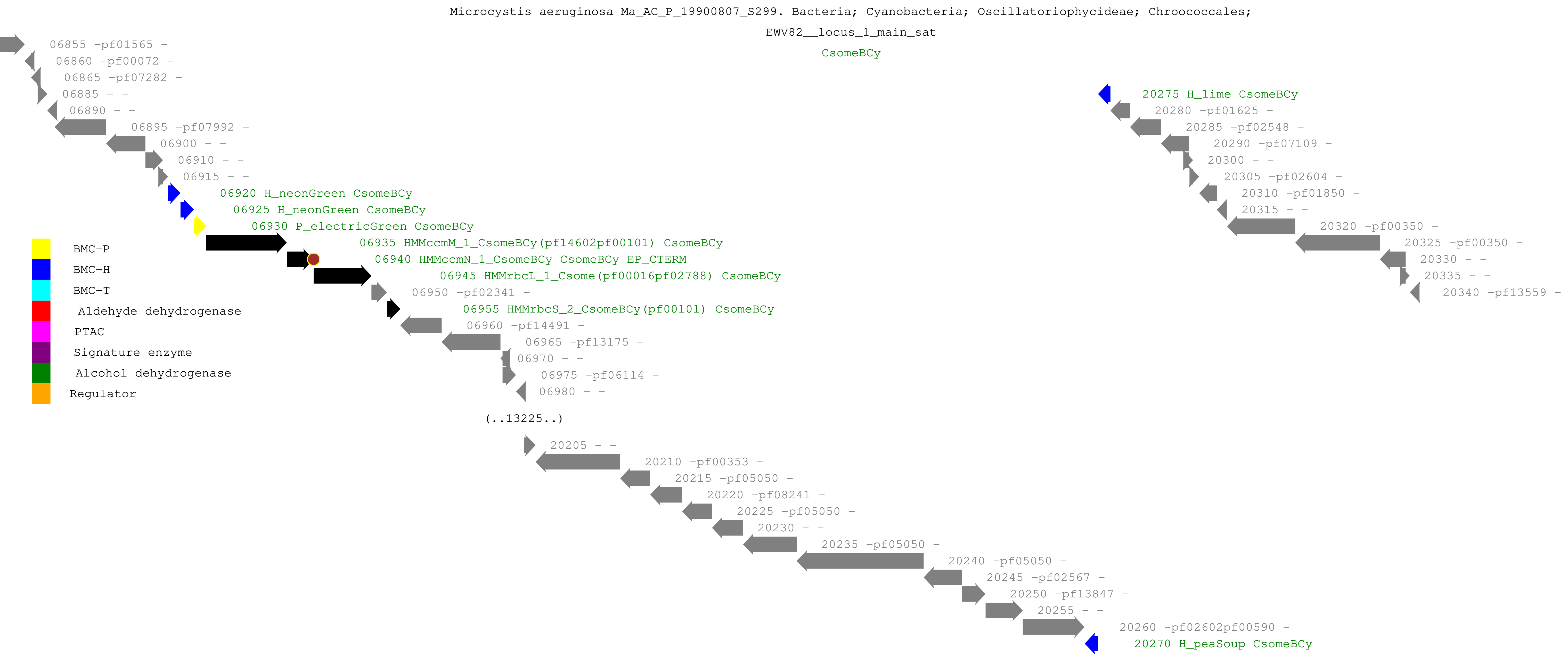

Microcystis aeruginosa Ma_AC_P_19900807_S299. Bacteria; Cyanobacteria; Oscillatoriophycideae; Chroococcales;
EWV82__locus_1_main_sat
CsomeBCy
06855 -pf01565 -
06860 -pf00072 -
06865 -pf07282 -
06885 - -
06890 - -
06895 -pf07992 -
06900 - -
06910 - -
06915 - -
06920 H_neonGreen CsomeBCy
06925 H_neonGreen CsomeBCy
06930 P_electricGreen CsomeBCy
06935 HMMccmM_1_CsomeBCy(pf14602pf00101) CsomeBCy
06940 HMMccmN_1_CsomeBCy CsomeBCy EP_CTERM
06945 HMMrbcL_1_Csome(pf00016pf02788) CsomeBCy
06950 -pf02341 -
06955 HMMrbcS_2_CsomeBCy(pf00101) CsomeBCy
06960 -pf14491 -
06965 -pf13175 -
06970 - -
06975 -pf06114 -
06980 - -
(..13225..)
20205 - -
20210 -pf00353 -
20215 -pf05050 -
20220 -pf08241 -
20225 -pf05050 -
20230 - -
20235 -pf05050 -
20240 -pf05050 -
20245 -pf02567 -
20250 -pf13847 -
20255 - -
20260 -pf02602pf00590 -
20270 H_peaSoup CsomeBCy
20275 H_lime CsomeBCy
20280 -pf01625 -
20285 -pf02548 -
20290 -pf07109 -
20300 - -
20305 -pf02604 -
20310 -pf01850 -
20315 - -
20320 -pf00350 -
20325 -pf00350 -
20330 - -
20335 - -
20340 -pf13559 -
BMC-P
BMC-H
BMC-T
Aldehyde dehydrogenase
PTAC
Signature enzyme
Alcohol dehydrogenase
Regulator

Microcystis sp. M_QC_C_20170808_M3Col. Bacteria; Cyanobacteria; Oscillatoriophycideae; Chroococcales;

EWV84__locus_1_main_sat

CsomeBCy

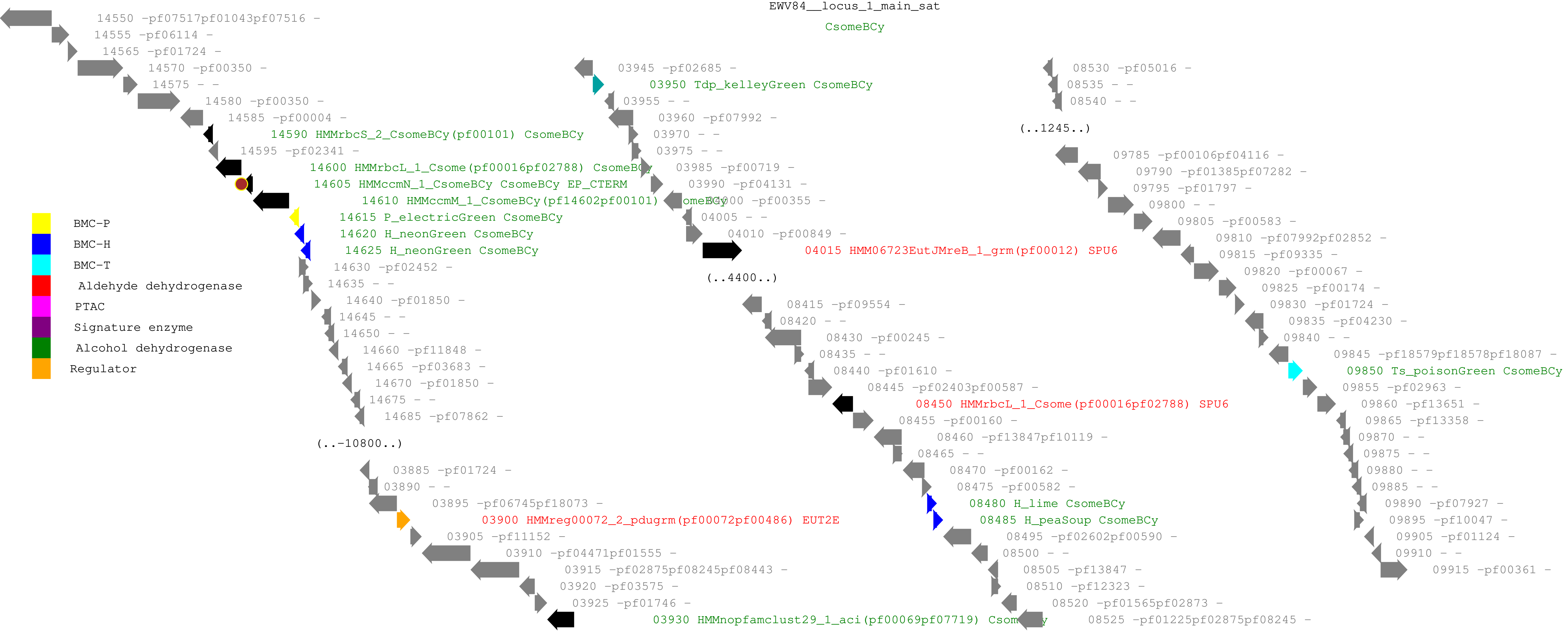

Microcystis aeruginosa Ma_QC_C_20070703_M131. Bacteria; Cyanobacteria; Oscillatoriophycideae; Chroococcales;

EWV85__locus_1_main_sat

CsomeBCy

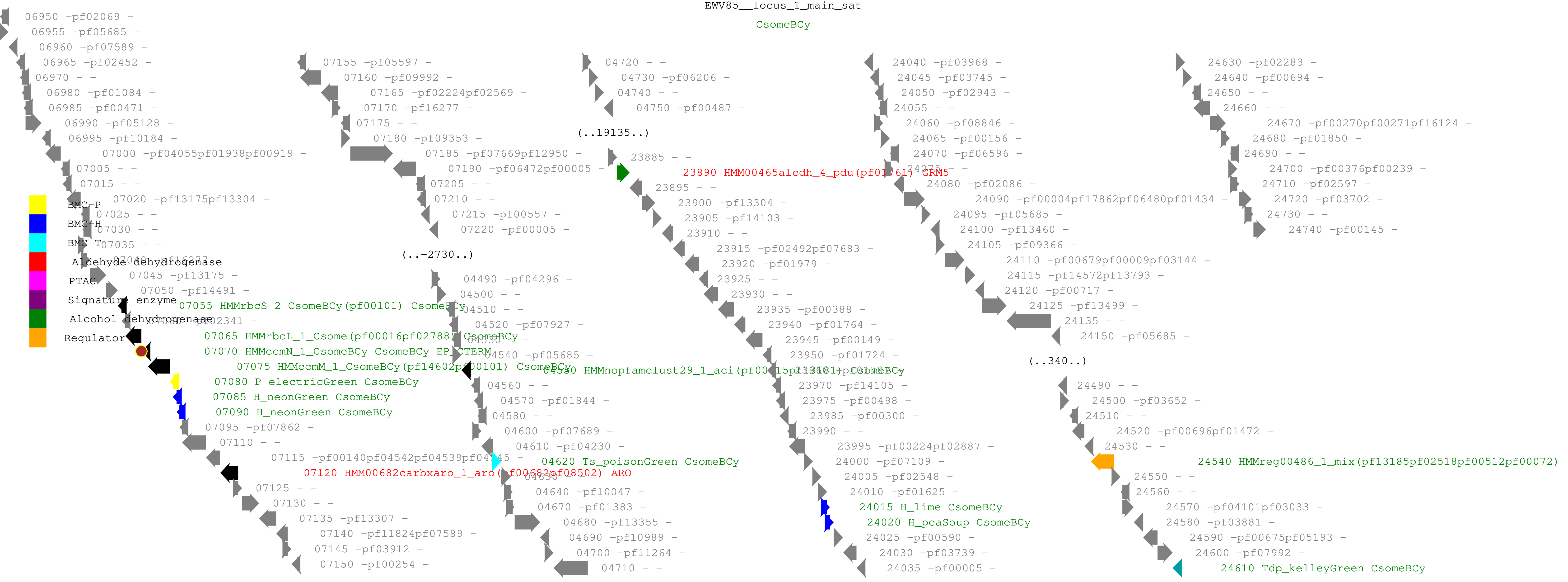

Microcystis wesenbergii Mw_MB_S_20031200_S109D. Bacteria; Cyanobacteria; Oscillatoriophycideae; Chroococcales;

EWV88__locus_1_main_sat

CsomeBCy

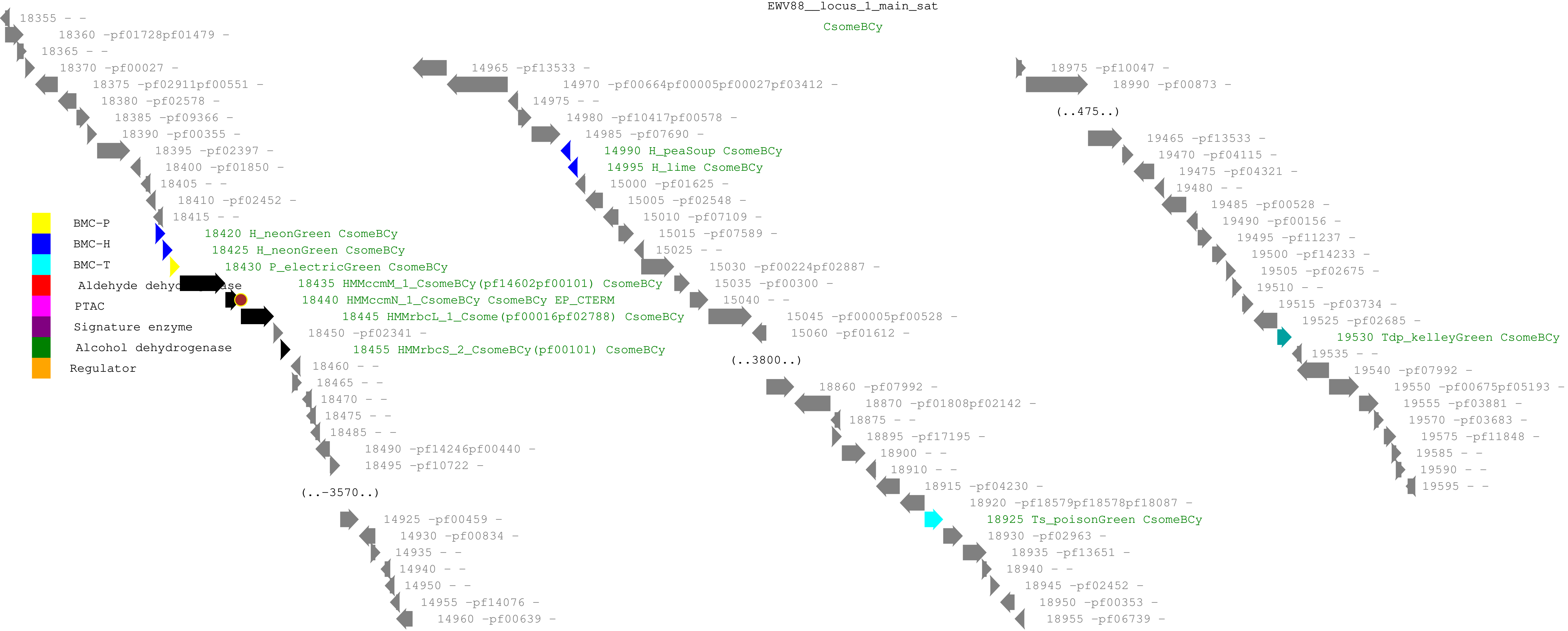

Microcystis aeruginosa Ma_QC_Ca_00000000_S207. Bacteria; Cyanobacteria; Oscillatoriophycideae; Chroococcales;

EWV91__locus_1_main_sat

CsomeBCy

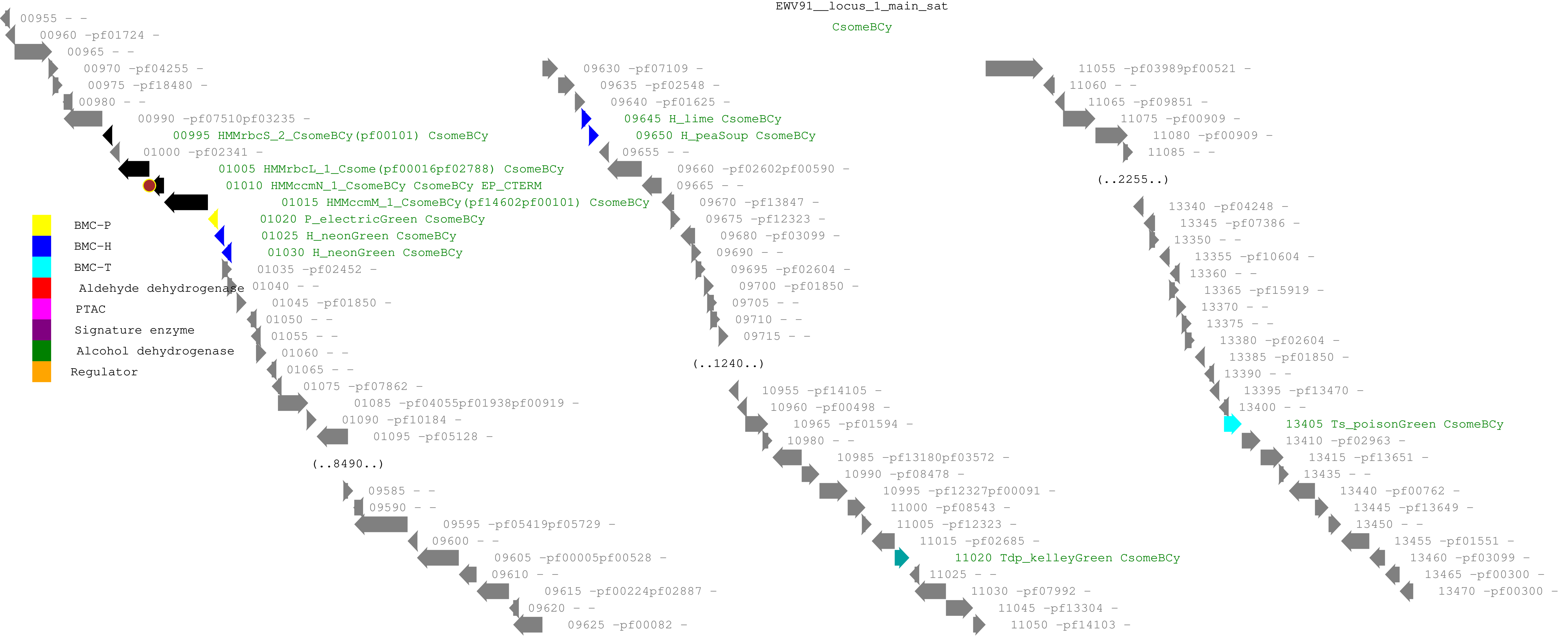

Microcystis aeruginosa Ma_MB_S_20031200_S102. Bacteria; Cyanobacteria; Oscillatoriophycideae; Chroococcales;

EWV92__locus_1_main_sat

CsomeBCy

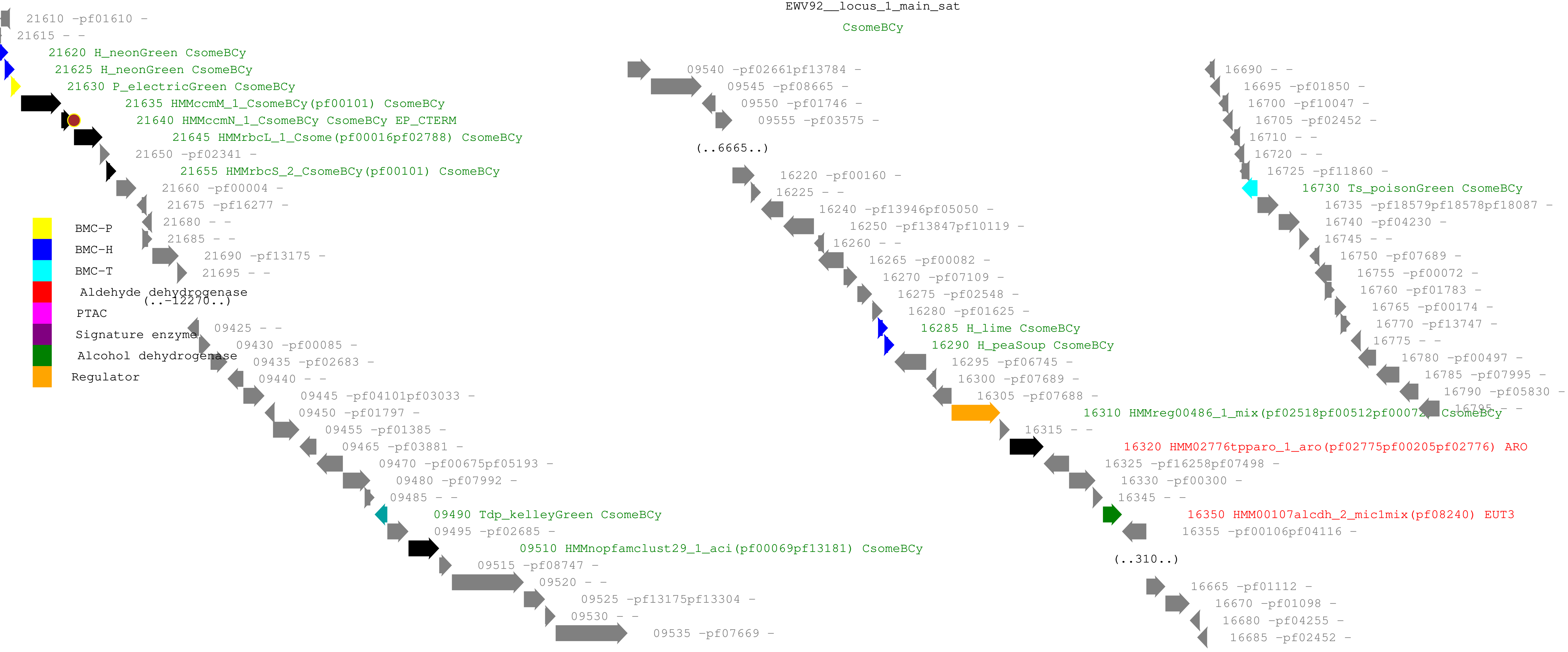

Phormidium sp. SL48-SHIP. Bacteria; Cyanobacteria; Oscillatoriophycideae; Oscillatoriales;

EYR95__locus_1_main_sat

CsomeBCy

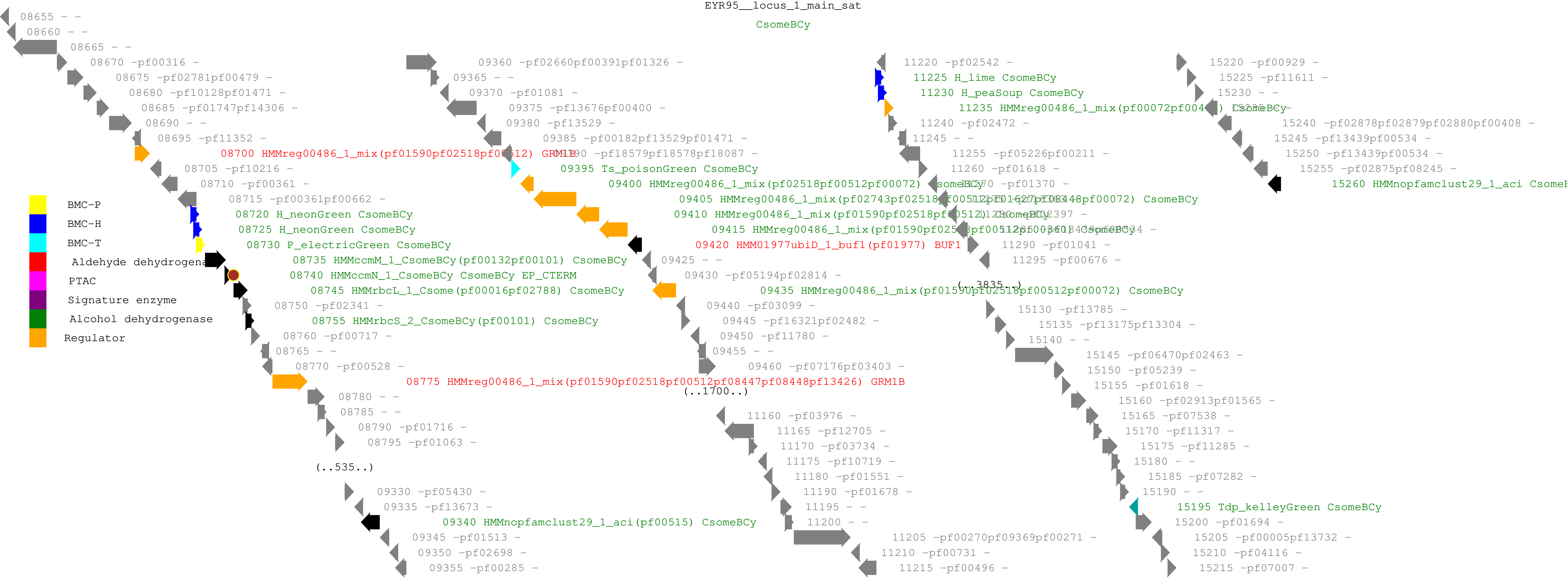

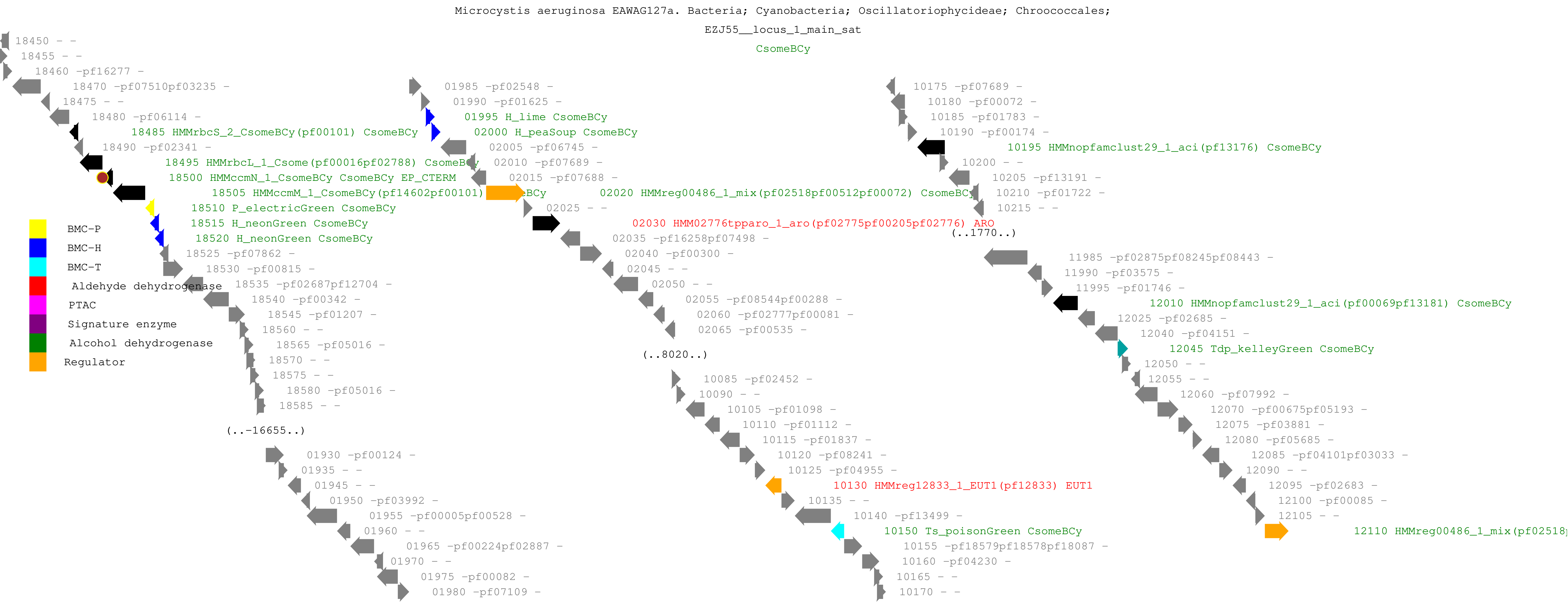

Microcystis aeruginosa EAWAG127a. Bacteria; Cyanobacteria; Oscillatoriophycideae; Chroococcales;
EZJ55__locus_1_main_sat
CsomeBCy
18450 - -
18455 - -
18460 -pf16277 -
18470 -pf07510pf03235 -
18475 - -
18480 -pf06114 -
18485 HMMrbcS_2_CsomeBCy(pf00101) CsomeBCy
18490 -pf02341 -
18495 HMMrbcL_1_Csome(pf00016pf02788) CsomeBCy
18500 HMMccmN_1_CsomeBCy CsomeBCy EP_CTERM
18505 HMMccmM_1_CsomeBCy(pf14602pf00101) CsomeBCy
18510 P_electricGreen CsomeBCy
18515 H_neonGreen CsomeBCy
18520 H_neonGreen CsomeBCy
18525 -pf07862 -
18530 -pf00815 -
18535 -pf02687pf12704 -
18540 -pf00342 -
18545 -pf01207 -
18560 - -
18565 -pf05016 -
18570 - -
18575 - -
18580 -pf05016 -
18585 - -
(..-16655..)
01930 -pf00124 -
01935 - -
01945 - -
01950 -pf03992 -
01955 -pf00005pf00528 -
01960 - -
01965 -pf00224pf02887 -
01970 - -
01975 -pf00082 -
01980 -pf07109 -
BMC-P
BMC-H
BMC-T
Aldehyde dehydrogenase
PTAC
Signature enzyme
Alcohol dehydrogenase
Regulator
01985 -pf02548 -
01990 -pf01625 -
01995 H_lime CsomeBCy
02000 H_peaSoup CsomeBCy
02005 -pf06745 -
02010 -pf07689 -
02015 -pf07688 -
02020 HMMreg00486_1_mix(pf02518pf00512pf00072) CsomeBCy
02025 - -
02030 HMM02776tpparo_1_aro(pf02775pf00205pf02776) ARO
02035 -pf16258pf07498 -
02040 -pf00300 -
02045 - -
02050 - -
02055 -pf08544pf00288 -
02060 -pf02777pf00081 -
02065 -pf00535 -
(..8020..)
10085 -pf02452 -
10090 - -
10105 -pf01098 -
10110 -pf01112 -
10115 -pf01837 -
10120 -pf08241 -
10125 -pf04955 -
10130 HMMreg12833_1_EUT1(pf12833) EUT1
10135 - -
10140 -pf13499 -
10150 Ts_poisonGreen CsomeBCy
10155 -pf18579pf18578pf18087 -
10160 -pf04230 -
10165 - -
10170 - -
10175 -pf07689 -
10180 -pf00072 -
10185 -pf01783 -
10190 -pf00174 -
10195 HMMnopfamclust29_1_aci(pf13176) CsomeBCy
10200 - -
10205 -pf13191 -
10210 -pf01722 -
10215 - -
(..1770..)
11985 -pf02875pf08245pf08443 -
11990 -pf03575 -
11995 -pf01746 -
12010 HMMnopfamclust29_1_aci(pf00069pf13181) CsomeBCy
12025 -pf02685 -
12040 -pf04151 -
12045 Tdp_kelleyGreen CsomeBCy
12050 - -
12055 - -
12060 -pf07992 -
12070 -pf00675pf05193 -
12075 -pf03881 -
12080 -pf05685 -
12085 -pf04101pf03033 -
12090 - -
12095 -pf02683 -
12100 -pf00085 -
12105 - -
12110 HMMreg00486_1_mix(pf02518

Nostoc calcicola FACHB-389. Bacteria; Cyanobacteria; Nostocales; Nostocaceae; Nostoc.

FACHB389__locus_1_main_sat

CsomeBCy

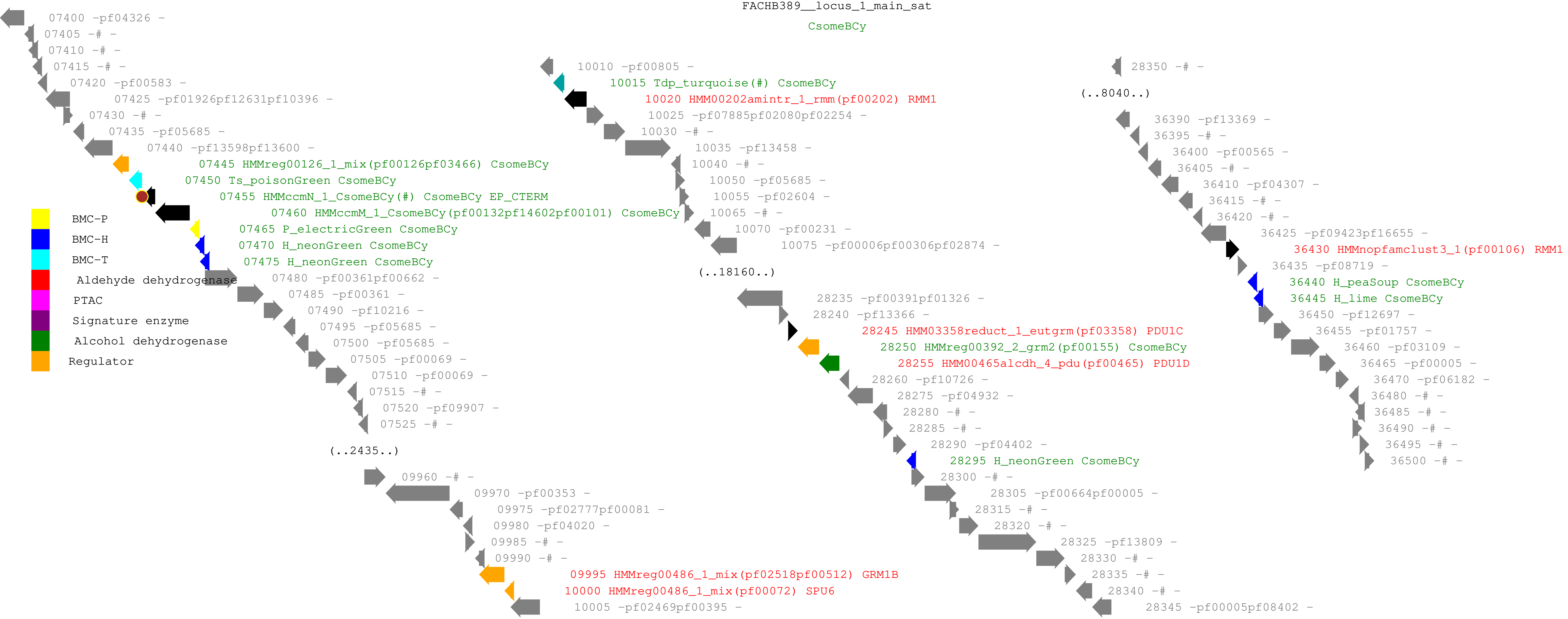

Tolypothrix sp. PCC 7601. Bacteria; Cyanobacteria; Nostocales; Tolypothrichaceae; Tolypothrix.

FDUTEX481__locus_1_main_sat

CsomeBCy

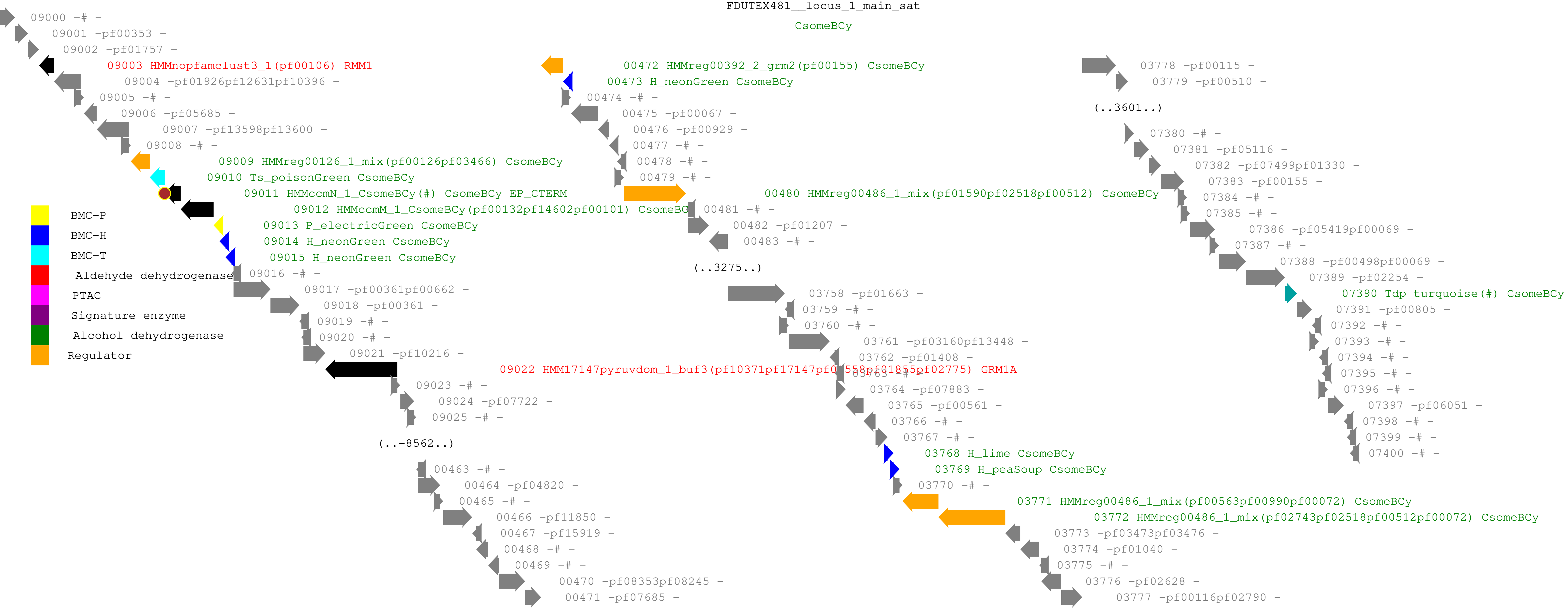

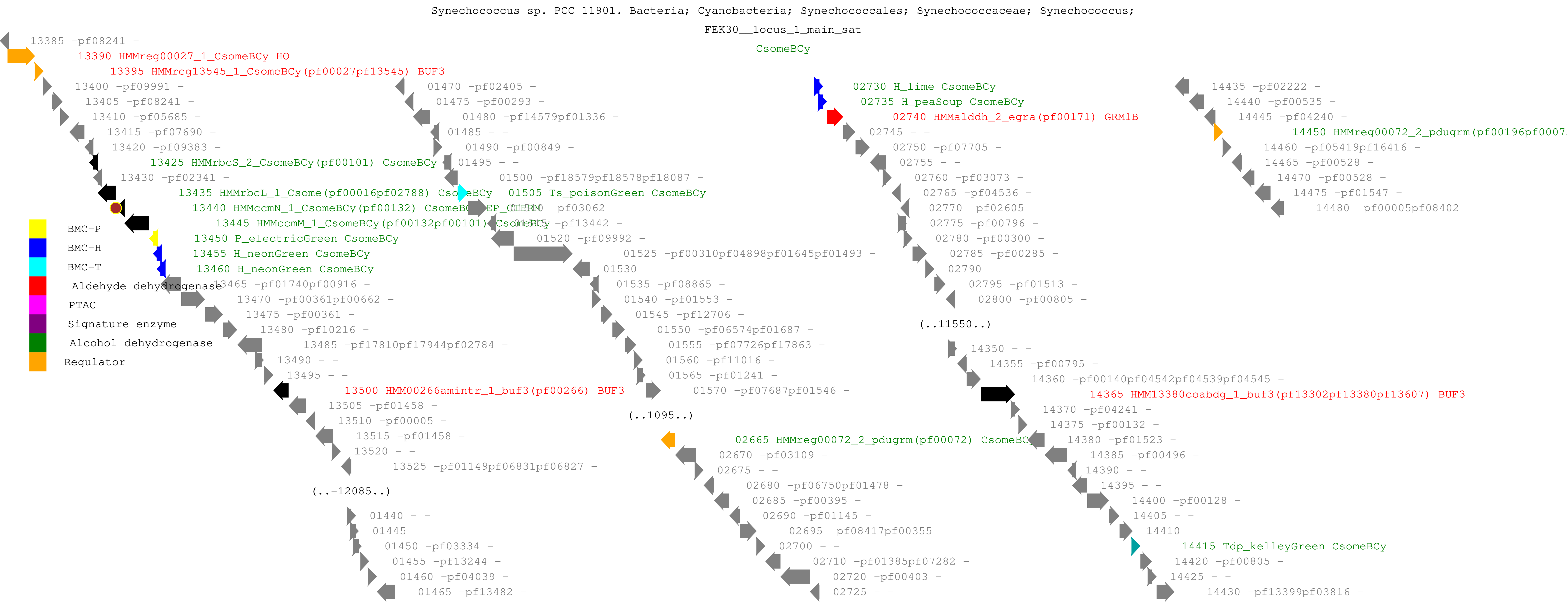

Synechococcus sp. PCC 11901. Bacteria; Cyanobacteria; Synechococcales; Synechococcaceae; Synechococcus;
FEK30__locus_1_main_sat
CsomeBCy
13385 -pf08241 -
13390 HMMreg00027_1_CsomeBCy HO
13395 HMMreg13545_1_CsomeBCy(pf00027pf13545) BUF3
13400 -pf09991 -
13405 -pf08241 -
13410 -pf05685 -
13415 -pf07690 -
13420 -pf09383 -
13425 HMMrbcS_2_CsomeBCy(pf00101) CsomeBCy
13430 -pf02341 -
13435 HMMrbcL_1_Csome(pf00016pf02788) CsomeBCy
13440 HMMccmN_1_CsomeBCy(pf00132) CsomeBCy
13445 HMMccmM_1_CsomeBCy(pf00132pf00101) CsomeBCy
13450 P_electricGreen CsomeBCy
13455 H_neonGreen CsomeBCy
13460 H_neonGreen CsomeBCy
13465 -pf01740pf00916 -
13470 -pf00361pf00662 -
13475 -pf00361 -
13480 -pf10216 -
13485 -pf17810pf17944pf02784 -
13490 - -
13495 - -
13500 HMM00266amintr_1_buf3(pf00266) BUF3
13505 -pf01458 -
13510 -pf00005 -
13515 -pf01458 -
13520 - -
13525 -pf01149pf06831pf06827 -
(..-12085..)
01440 - -
01445 - -
01450 -pf03334 -
01455 -pf13244 -
01460 -pf04039 -
01465 -pf13482 -
BMC-P
BMC-H
BMC-T
Aldehyde dehydrogenase
PTAC
Signature enzyme
Alcohol dehydrogenase
Regulator
01470 -pf02405 -
01475 -pf00293 -
01480 -pf14579pf01336 -
01485 - -
01490 -pf00849 -
01495 - -
01500 -pf18579pf18578pf18087 -
01505 Ts_poisonGreen CsomeBCy
01510 -pf03062 -
EP_CTERM
01515 -pf13442 -
01520 -pf09992 -
01525 -pf00310pf04898pf01645pf01493 -
01530 - -
01535 -pf08865 -
01540 -pf01553 -
01545 -pf12706 -
01550 -pf06574pf01687 -
01555 -pf07726pf17863 -
01560 -pf11016 -
01565 -pf01241 -
01570 -pf07687pf01546 -
(..1095..)
02665 HMMreg00072_2_pdugrm(pf00072) CsomeBCy
02670 -pf03109 -
02675 - -
02680 -pf06750pf01478 -
02685 -pf00395 -
02690 -pf01145 -
02695 -pf08417pf00355 -
02700 - -
02710 -pf01385pf07282 -
02720 -pf00403 -
02725 - -
02730 H_lime CsomeBCy
02735 H_peaSoup CsomeBCy
02740 HMMalddh_2_egra(pf00171) GRM1B
02745 - -
02750 -pf07705 -
02755 - -
02760 -pf03073 -
02765 -pf04536 -
02770 -pf02605 -
02775 -pf00796 -
02780 -pf00300 -
02785 -pf00285 -
02790 - -
02795 -pf01513 -
02800 -pf00805 -
(..11550..)
14350 - -
14355 -pf00795 -
14360 -pf00140pf04542pf04539pf04545 -
14365 HMM13380coabdg_1_buf3(pf13302pf13380pf13607) BUF3
14370 -pf04241 -
14375 -pf00132 -
14380 -pf01523 -
14385 -pf00496 -
14390 - -
14395 - -
14400 -pf00128 -
14405 - -
14410 - -
14415 Tdp_kelleyGreen CsomeBCy
14420 -pf00805 -
14425 - -
14430 -pf13399pf03816 -
14435 -pf02222 -
14440 -pf00535 -
14445 -pf04240 -
14450 HMMreg00072_2_pdugrm(pf00196pf0007
14460 -pf05419pf16416 -
14465 -pf00528 -
14470 -pf00528 -
14475 -pf01547 -
14480 -pf00005pf08402 -

Thermosynechococcus sp. CL-1. Bacteria; Cyanobacteria; Synechococcales; Synechococcaceae;

FFX45__locus_1_main_sat

CsomeBCy

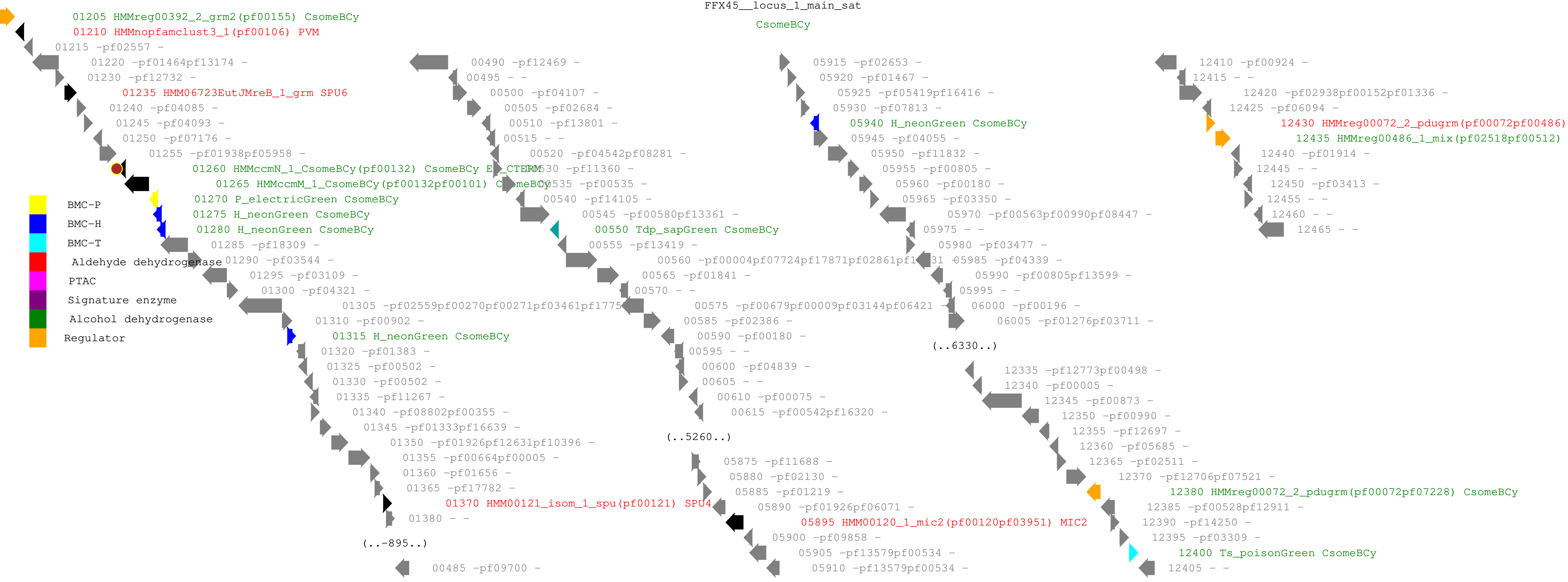

Fischerella sp. NIES-3754. Bacteria; Cyanobacteria; Nostocales; Hapalosiphonaceae; Fischerella.

FIS3754__locus_1_main_sat

CsomeBCy

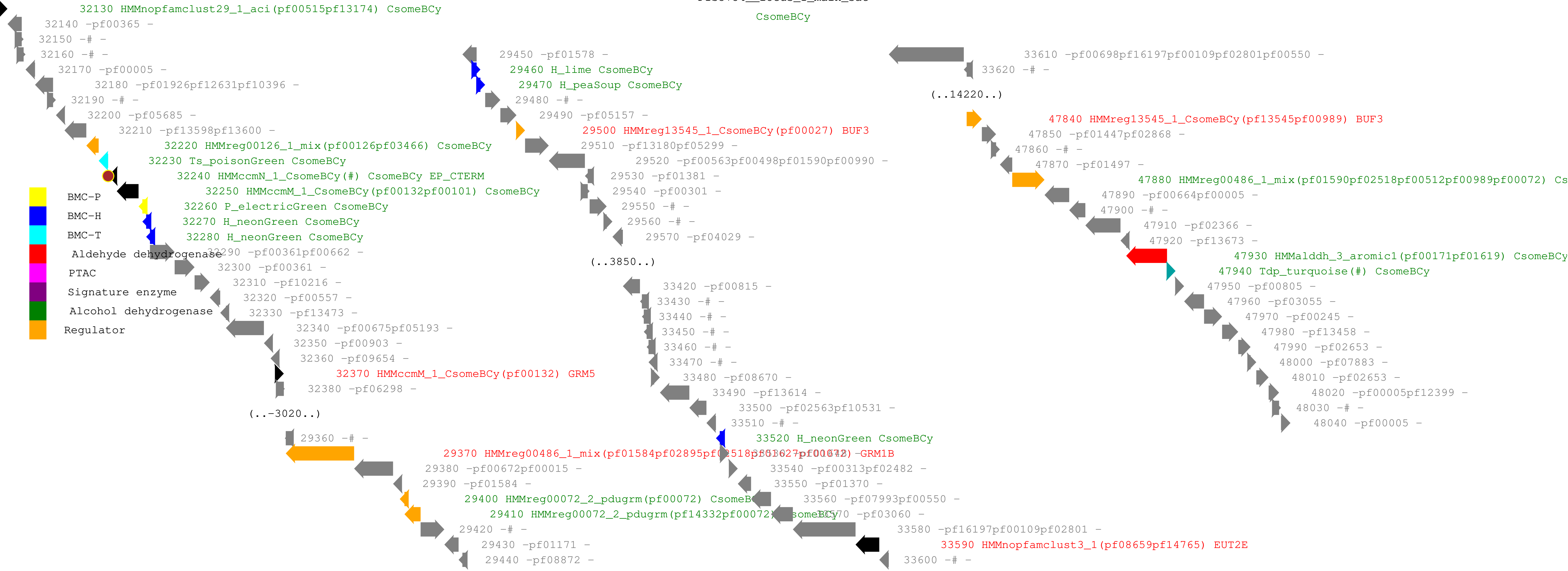

Fischerella sp. JSC-11. Bacteria; Cyanobacteria; Nostocales; Hapalosiphonaceae; Fischerella.

FJSC11DRAFT__locus_1_main_sat

CsomeBCy

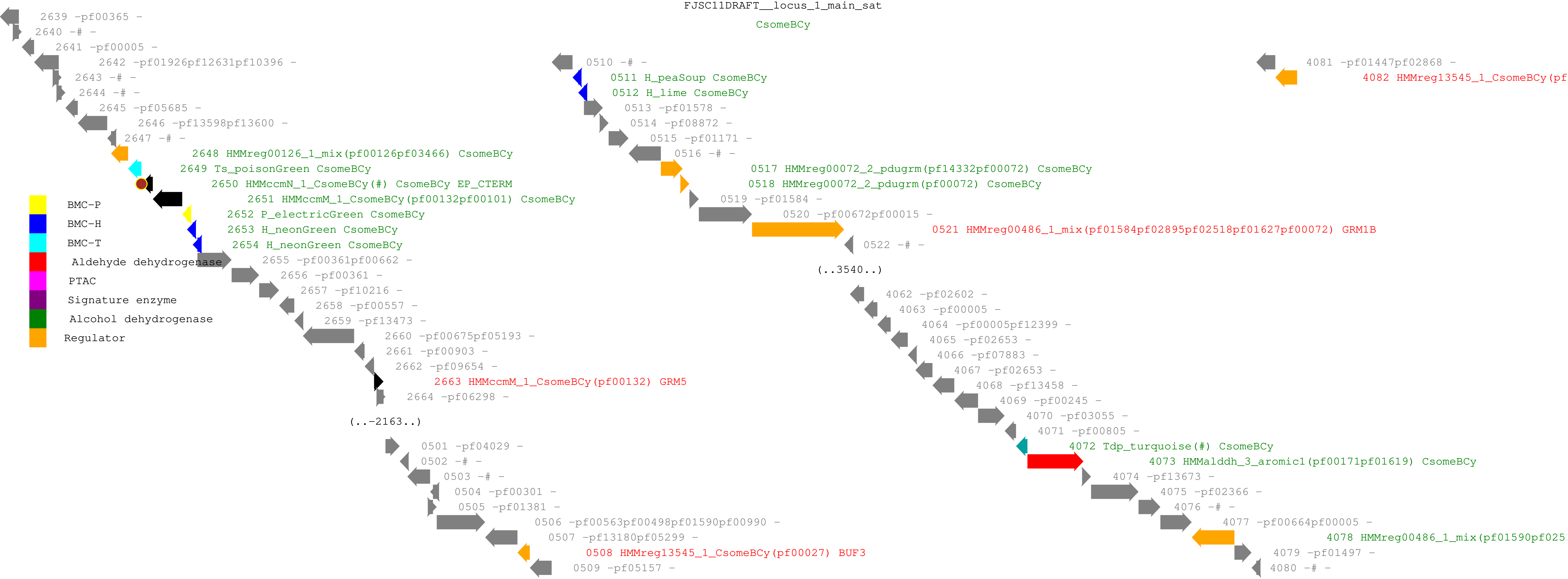

Euhalothece natronophila Z-M001. Bacteria; Cyanobacteria; Oscillatoriophycideae; Chroococcales;

FRE64__locus_1_main_sat

CsomeBCy

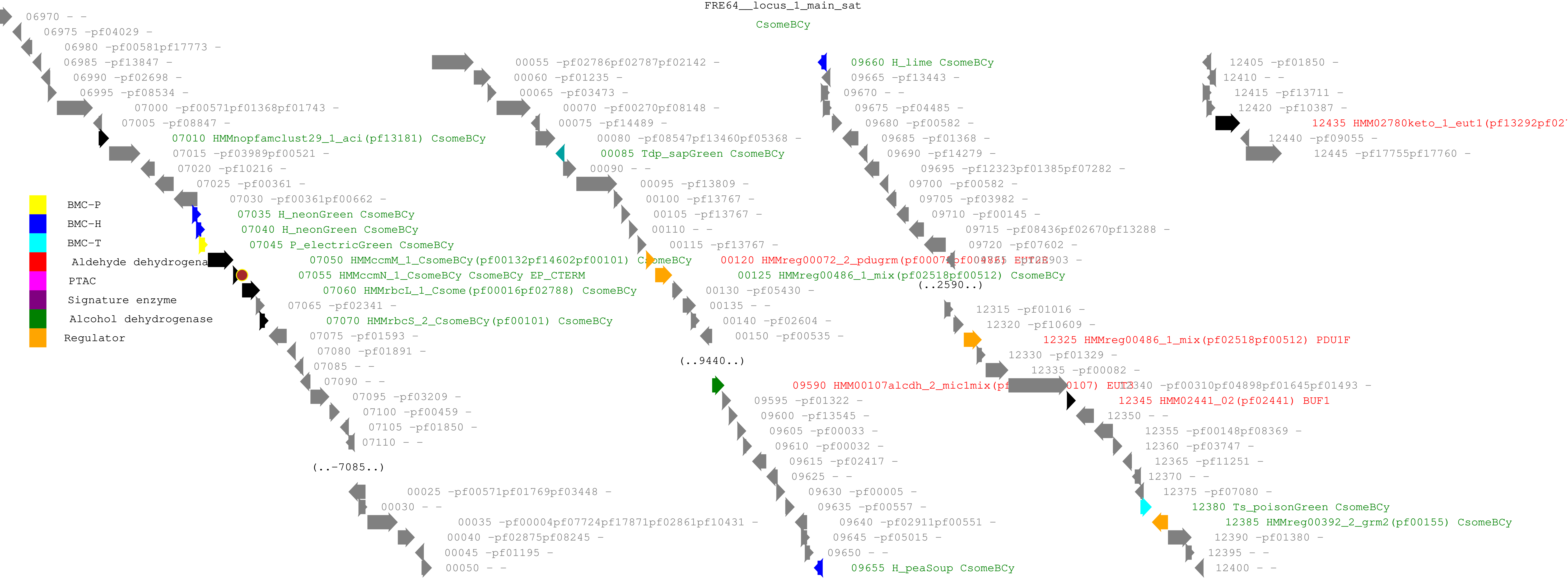

Microcystis aeruginosa KLA2. Bacteria; Cyanobacteria; Oscillatoriophycideae; Chroococcales;

FXO09__locus_1_main_sat

CsomeBCy

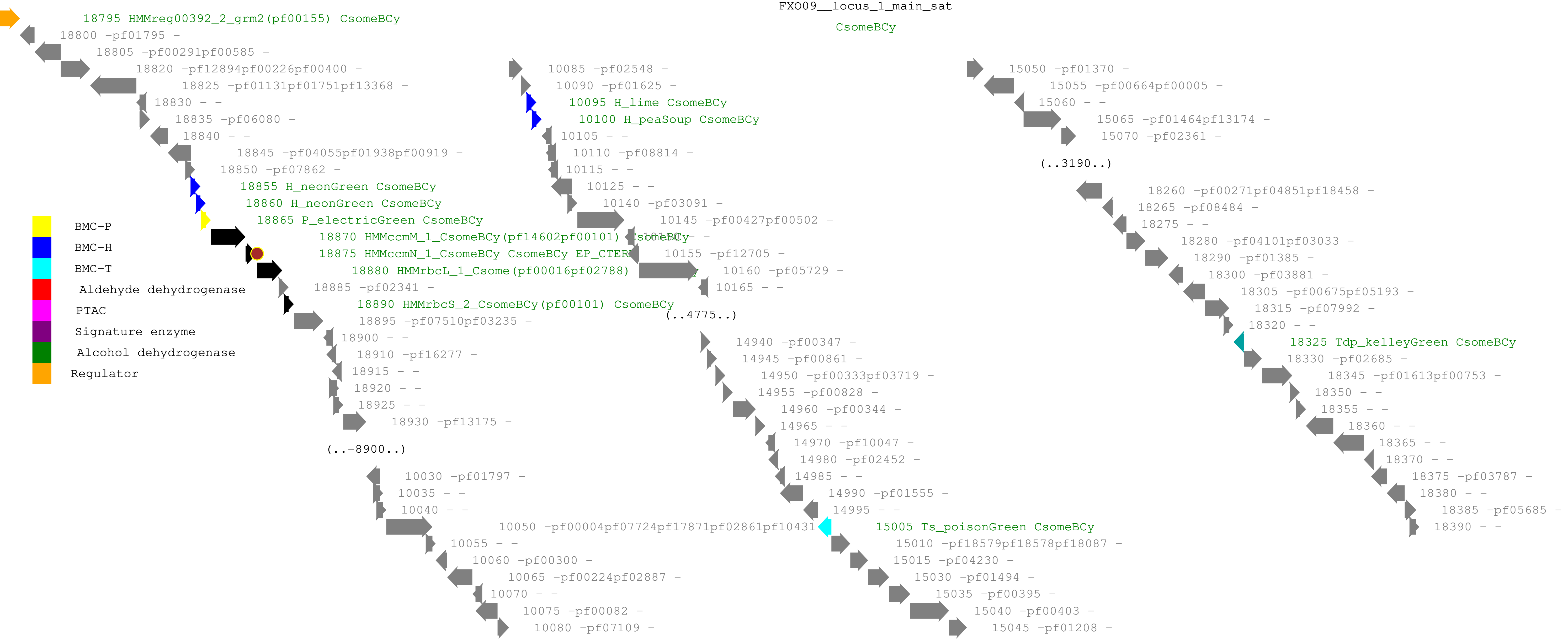

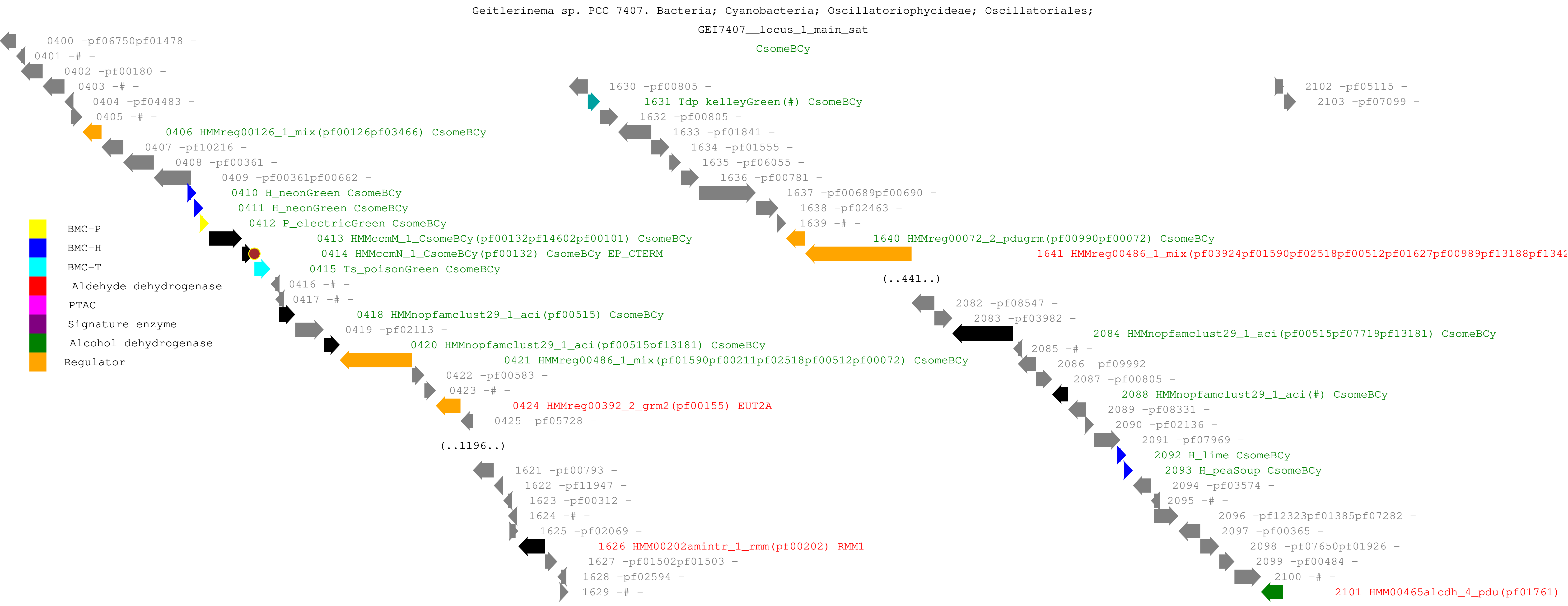

Geitlerinema sp. PCC 7407. Bacteria; Cyanobacteria; Oscillatoriophycideae; Oscillatoriales;
GEI7407__locus_1_main_sat
CsomeBCy
0400 -pf06750pf01478 -
0401 -# -
0402 -pf00180 -
0403 -# -
0404 -pf04483 -
0405 -# -
0406 HMMreg00126_1_mix(pf00126pf03466) CsomeBCy
0407 -pf10216 -
0408 -pf00361 -
0409 -pf00361pf00662 -
0410 H_neonGreen CsomeBCy
0411 H_neonGreen CsomeBCy
0412 P_electricGreen CsomeBCy
0413 HMMccmM_1_CsomeBCy(pf00132pf14602pf00101) CsomeBCy
0414 HMMccmN_1_CsomeBCy(pf00132) CsomeBCy EP_CTERM
0415 Ts_poisonGreen CsomeBCy
0416 -# -
0417 -# -
0418 HMMnopfamclust29_1_aci(pf00515) CsomeBCy
0419 -pf02113 -
0420 HMMnopfamclust29_1_aci(pf00515pf13181) CsomeBCy
0421 HMMreg00486_1_mix(pf01590pf00211pf02518pf00512pf00072) CsomeBCy
0422 -pf00583 -
0423 -# -
0424 HMMreg00392_2_grm2(pf00155) EUT2A
0425 -pf05728 -
(..1196..)
1621 -pf00793 -
1622 -pf11947 -
1623 -pf00312 -
1624 -# -
1625 -pf02069 -
1626 HMM00202amintr_1_rmm(pf00202) RMM1
1627 -pf01502pf01503 -
1628 -pf02594 -
1629 -# -
1630 -pf00805 -
1631 Tdp_kelleyGreen(#) CsomeBCy
1632 -pf00805 -
1633 -pf01841 -
1634 -pf01555 -
1635 -pf06055 -
1636 -pf00781 -
1637 -pf00689pf00690 -
1638 -pf02463 -
1639 -# -
1640 HMMreg00072_2_pdugrm(pf00990pf00072) CsomeBCy
1641 HMMreg00486_1_mix(pf03924pf01590pf02518pf00512pf01627pf00989pf13188pf1342
(..441..)
2082 -pf08547 -
2083 -pf03982 -
2084 HMMnopfamclust29_1_aci(pf00515pf07719pf13181) CsomeBCy
2085 -# -
2086 -pf09992 -
2087 -pf00805 -
2088 HMMnopfamclust29_1_aci(#) CsomeBCy
2089 -pf08331 -
2090 -pf02136 -
2091 -pf07969 -
2092 H_lime CsomeBCy
2093 H_peaSoup CsomeBCy
2094 -pf03574 -
2095 -# -
2096 -pf12323pf01385pf07282 -
2097 -pf00365 -
2098 -pf07650pf01926 -
2099 -pf00484 -
2100 -# -
2101 HMM00465alcdh_4_pdu(pf01761)
2102 -pf05115 -
2103 -pf07099 -
BMC-P
BMC-H
BMC-T
Aldehyde dehydrogenase
PTAC
Signature enzyme
Alcohol dehydrogenase
Regulator

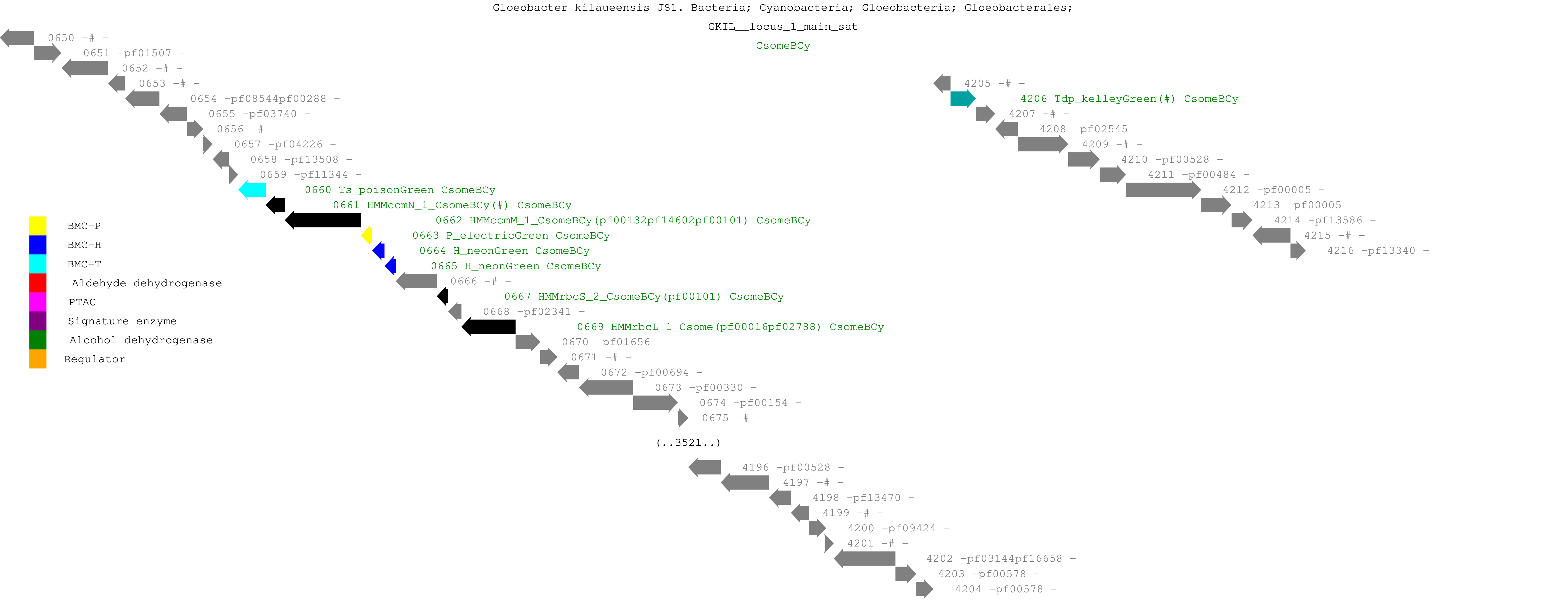
Gloeobacter kilaueensis JS1. Bacteria; Cyanobacteria; Gloeobacteria; Gloeobacterales;
GKIL__locus_1_main_sat
CsomeBCy
0650 -# -
0651 -pf01507 -
0652 -# -
0653 -# -
0654 -pf08544pf00288 -
0655 -pf03740 -
0656 -# -
0657 -pf04226 -
0658 -pf13508 -
0659 -pf11344 -
0660 Ts_poisonGreen CsomeBCy
0661 HMMccmN_1_CsomeBCy(#) CsomeBCy
0662 HMMccmM_1_CsomeBCy(pf00132pf14602pf00101) CsomeBCy
0663 P_electricGreen CsomeBCy
0664 H_neonGreen CsomeBCy
0665 H_neonGreen CsomeBCy
0666 -# -
0667 HMMrbcS_2_CsomeBCy(pf00101) CsomeBCy
0668 -pf02341 -
0669 HMMrbcL_1_Csome(pf00016pf02788) CsomeBCy
0670 -pf01656 -
0671 -# -
0672 -pf00694 -
0673 -pf00330 -
0674 -pf00154 -
0675 -# -
(..3521..)
4196 -pf00528 -
4197 -# -
4198 -pf13470 -
4199 -# -
4200 -pf09424 -
4201 -# -
4202 -pf03144pf16658 -
4203 -pf00578 -
4204 -pf00578 -
4205 -# -
4206 Tdp_kelleyGreen(#) CsomeBCy
4207 -# -
4208 -pf02545 -
4209 -# -
4210 -pf00528 -
4211 -pf00484 -
4212 -pf00005 -
4213 -pf00005 -
4214 -pf13586 -
4215 -# -
4216 -pf13340 -
BMC-P
BMC-H
BMC-T
Aldehyde dehydrogenase
PTAC
Signature enzyme
Alcohol dehydrogenase
Regulator

Gloeomargarita lithophora Alchichica-D10. Bacteria; Cyanobacteria; Gloeomargaritales; Gloeomargaritaceae;

GlitD10__locus_1_main_sat

CsomeBCy

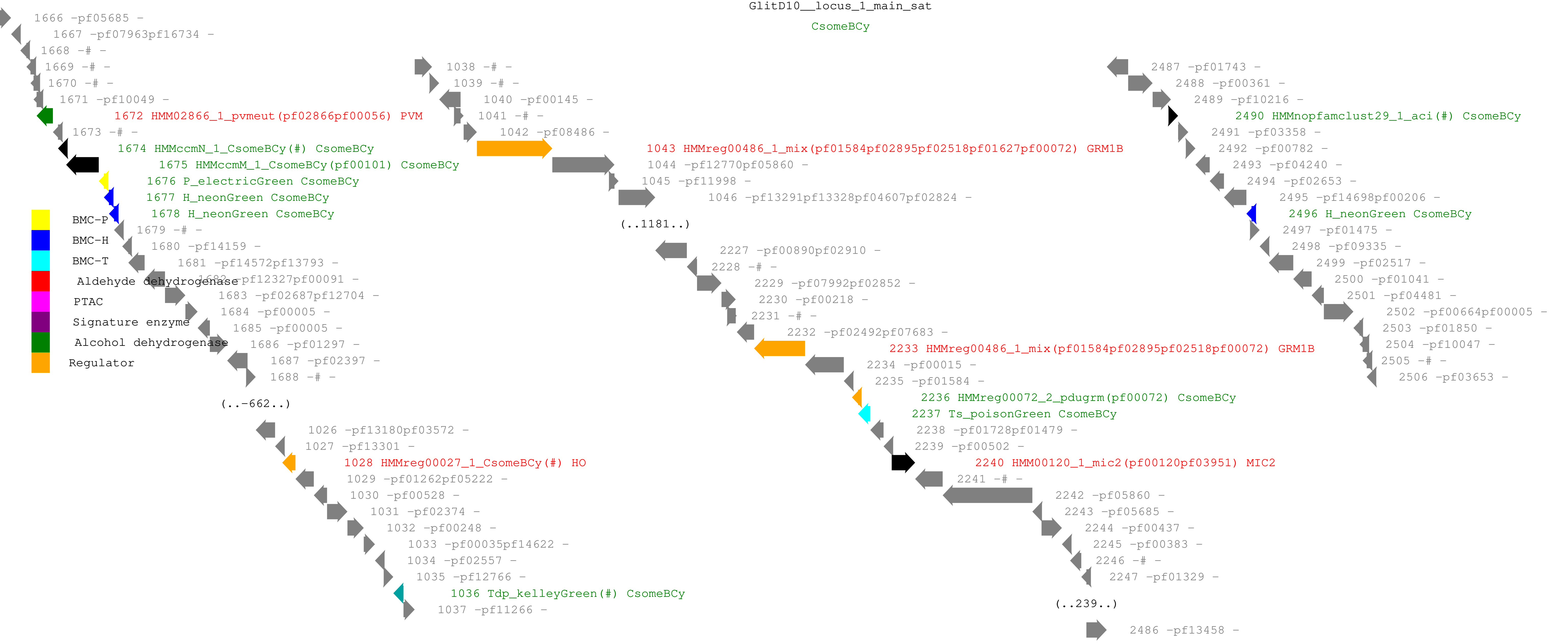

Gloeocapsa sp. PCC 73106. Bacteria; Cyanobacteria; Oscillatoriophycideae; Chroococcales;

GLO73106DRAFT__locus_1_main_sat

CsomeBCy

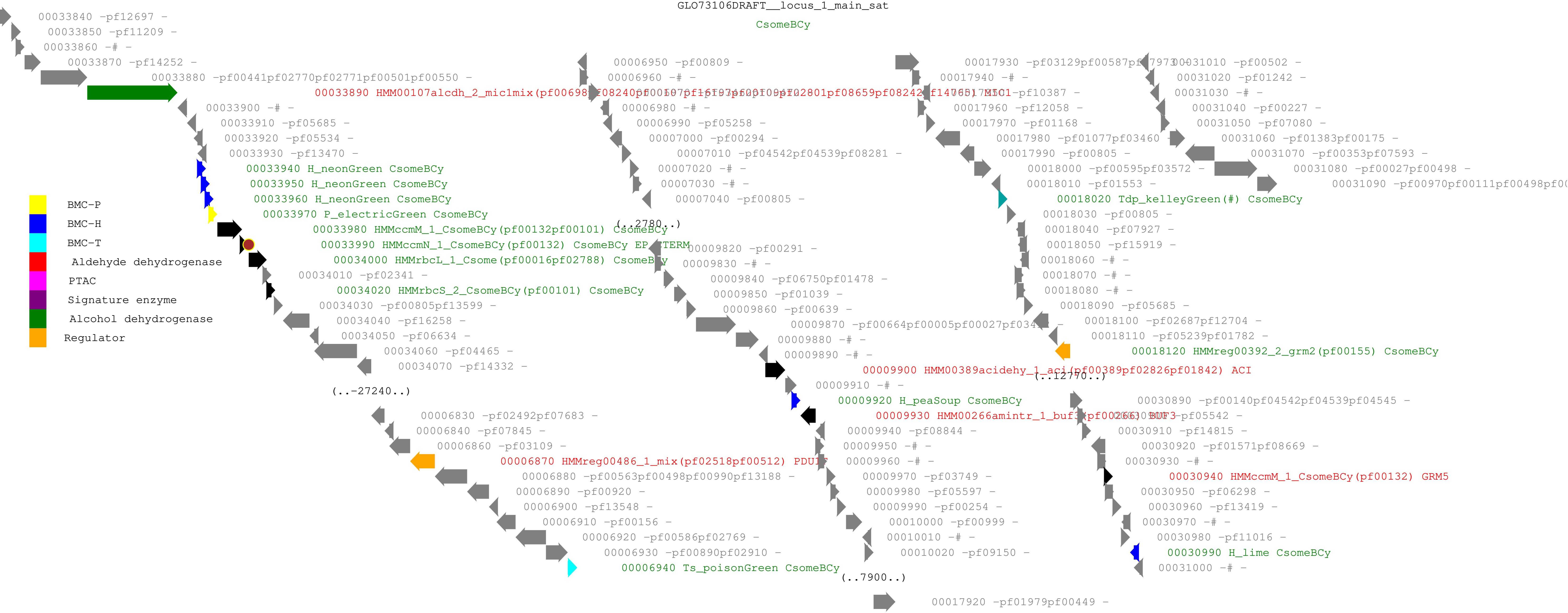

Gloeocapsa sp. PCC 7428. Bacteria; Cyanobacteria; Oscillatoriophycideae; Chroococcales;

Glo7428__locus_1_main_sat

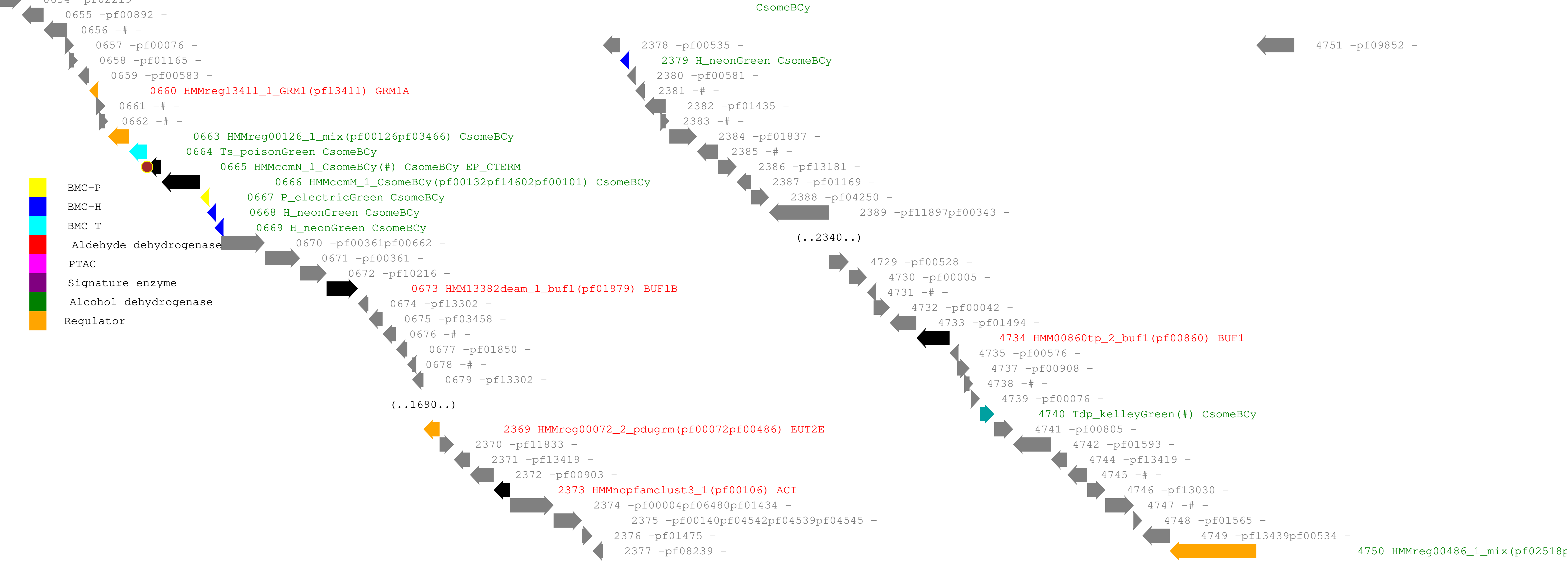

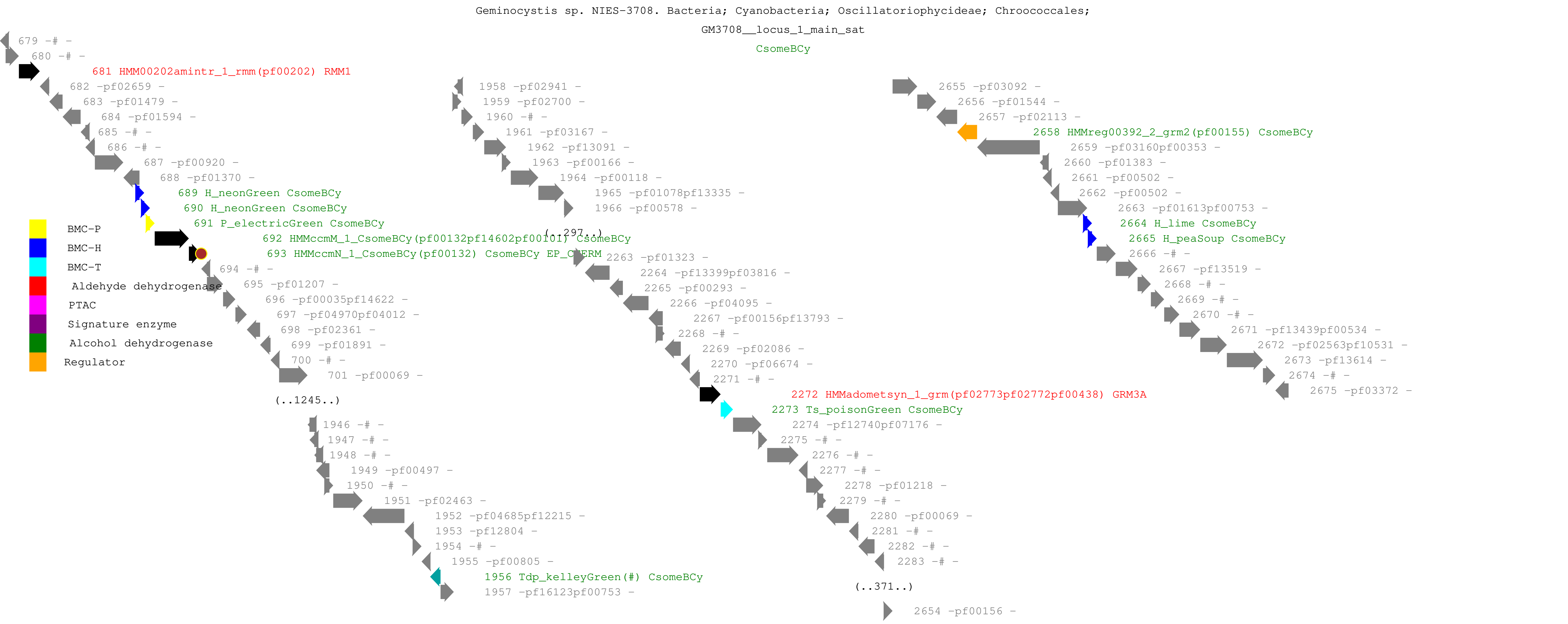
Geminocystis sp. NIES-3708. Bacteria; Cyanobacteria; Oscillatoriophycideae; Chroococcales;
GM3708__locus_1_main_sat
CsomeBCy
679 -# -
680 -# -
681 HMM00202amintr_1_rmm(pf00202) RMM1
682 -pf02659 -
683 -pf01479 -
684 -pf01594 -
685 -# -
686 -# -
687 -pf00920 -
688 -pf01370 -
689 H_neonGreen CsomeBCy
690 H_neonGreen CsomeBCy
691 P_electricGreen CsomeBCy
692 HMMccmM_1_CsomeBCy(pf00132pf14602pf00101) CsomeBCy
693 HMMccmN_1_CsomeBCy(pf00132) CsomeBCy EP_C_TERM
694 -# -
695 -pf01207 -
696 -pf00035pf14622 -
697 -pf04970pf04012 -
698 -pf02361 -
699 -pf01891 -
700 -# -
701 -pf00069 -
(..1245..)
1946 -# -
1947 -# -
1948 -# -
1949 -pf00497 -
1950 -# -
1951 -pf02463 -
1952 -pf04685pf12215 -
1953 -pf12804 -
1954 -# -
1955 -pf00805 -
1956 Tdp_kelleyGreen(#) CsomeBCy
1957 -pf16123pf00753 -
1958 -pf02941 -
1959 -pf02700 -
1960 -# -
1961 -pf03167 -
1962 -pf13091 -
1963 -pf00166 -
1964 -pf00118 -
1965 -pf01078pf13335 -
1966 -pf00578 -
(..297..)
2263 -pf01323 -
2264 -pf13399pf03816 -
2265 -pf00293 -
2266 -pf04095 -
2267 -pf00156pf13793 -
2268 -# -
2269 -pf02086 -
2270 -pf06674 -
2271 -# -
2272 HMMadometsyn_1_grm(pf02773pf02772pf00438) GRM3A
2273 Ts_poisonGreen CsomeBCy
2274 -pf12740pf07176 -
2275 -# -
2276 -# -
2277 -# -
2278 -pf01218 -
2279 -# -
2280 -pf00069 -
2281 -# -
2282 -# -
2283 -# -
(..371..)
2654 -pf00156 -
2655 -pf03092 -
2656 -pf01544 -
2657 -pf02113 -
2658 HMMreg00392_2_grm2(pf00155) CsomeBCy
2659 -pf03160pf00353 -
2660 -pf01383 -
2661 -pf00502 -
2662 -pf00502 -
2663 -pf01613pf00753 -
2664 H_lime CsomeBCy
2665 H_peaSoup CsomeBCy
2666 -# -
2667 -pf13519 -
2668 -# -
2669 -# -
2670 -# -
2671 -pf13439pf00534 -
2672 -pf02563pf10531 -
2673 -pf13614 -
2674 -# -
2675 -pf03372 -
BMC-P
BMC-H
BMC-T
Aldehyde dehydrogenase
PTAC
Signature enzyme
Alcohol dehydrogenase
Regulator

Geminocystis sp. NIES-3709. Bacteria; Cyanobacteria; Oscillatoriophycideae; Chroococcales;

GM3709__locus_1_main_sat

CsomeBCy

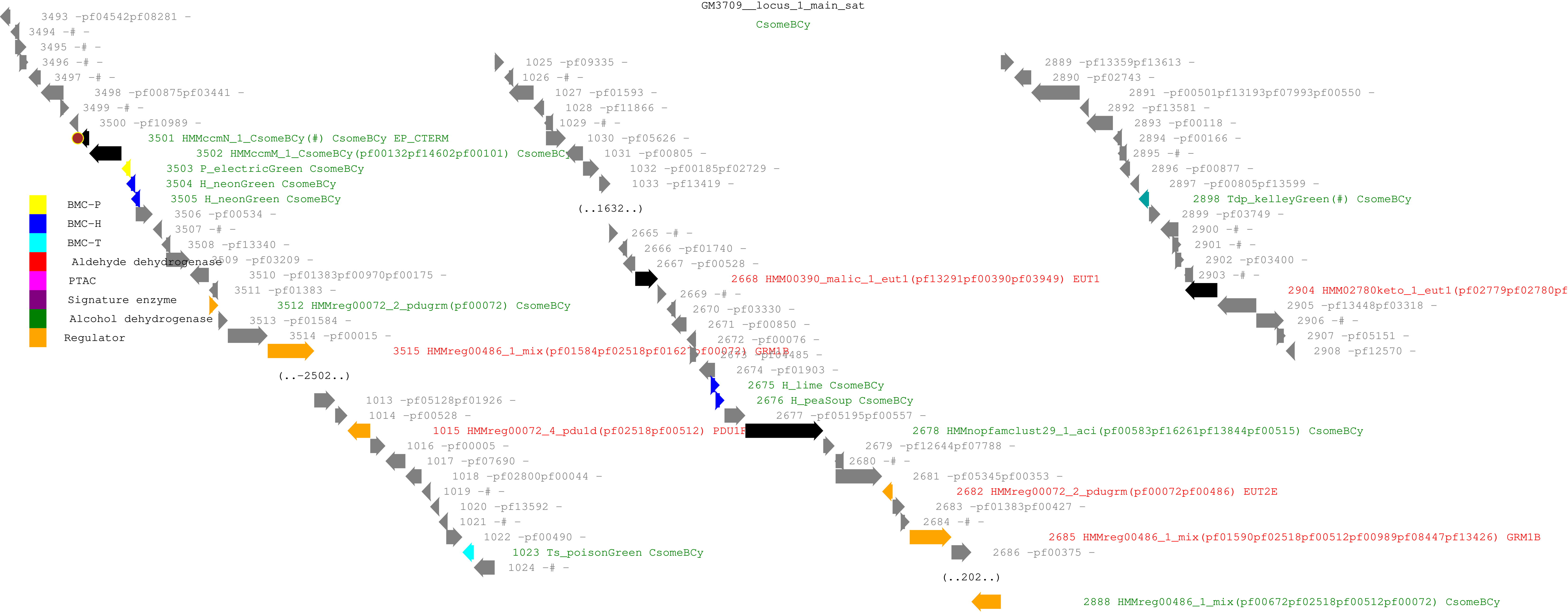

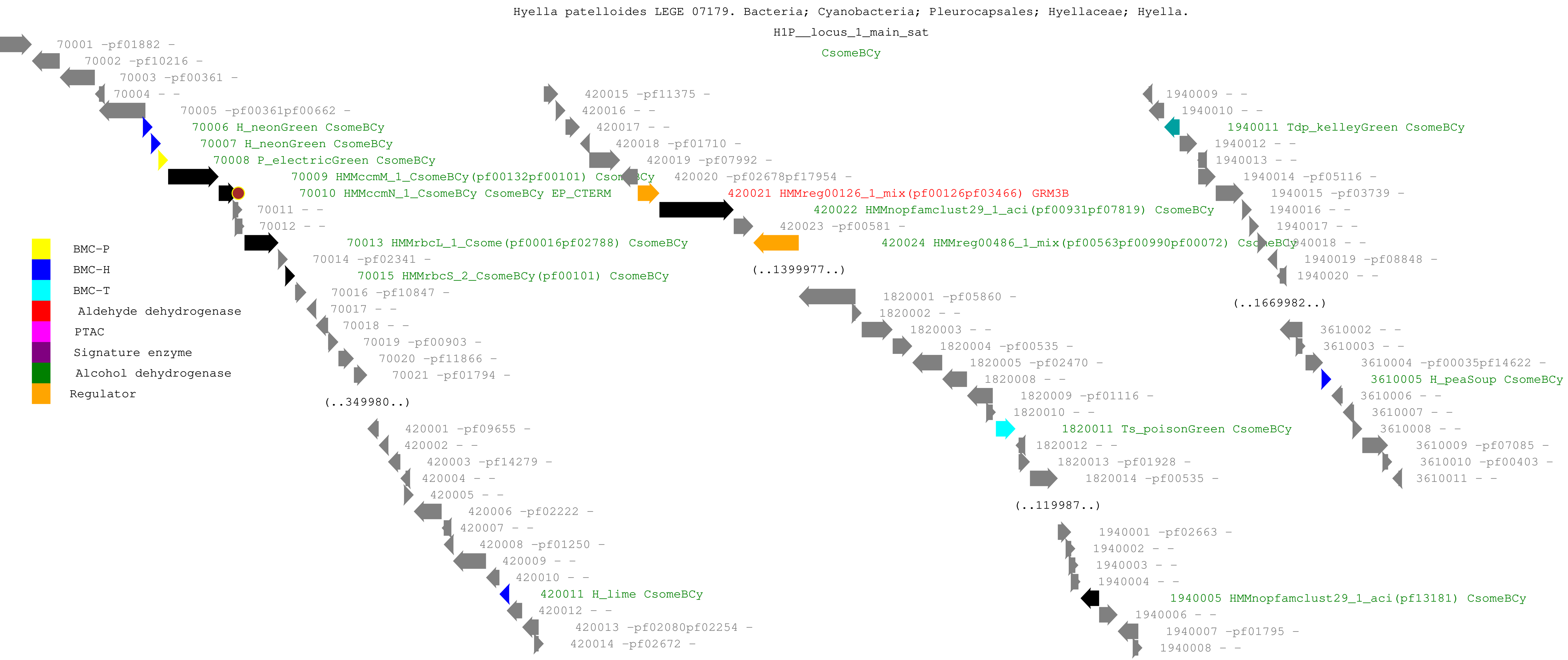
Hyella patelloides LEGE 07179. Bacteria; Cyanobacteria; Pleurocapsales; Hyellaceae; Hyella.
H1P__locus_1_main_sat
CsomeBCy
70001 -pf01882 -
70002 -pf10216 -
70003 -pf00361 -
70004 - -
70005 -pf00361pf00662 -
70006 H_neonGreen CsomeBCy
70007 H_neonGreen CsomeBCy
70008 P_electricGreen CsomeBCy
70009 HMMccmM_1_CsomeBCy(pf00132pf00101) CsomeBCy
70010 HMMccmN_1_CsomeBCy CsomeBCy EP_CTERM
70011 - -
70012 - -
70013 HMMrbcL_1_Csome(pf00016pf02788) CsomeBCy
70014 -pf02341 -
70015 HMMrbcS_2_CsomeBCy(pf00101) CsomeBCy
70016 -pf10847 -
70017 - -
70018 - -
70019 -pf00903 -
70020 -pf11866 -
70021 -pf01794 -
(..349980..)
420001 -pf09655 -
420002 - -
420003 -pf14279 -
420004 - -
420005 - -
420006 -pf02222 -
420007 - -
420008 -pf01250 -
420009 - -
420010 - -
420011 H_lime CsomeBCy
420012 - -
420013 -pf02080pf02254 -
420014 -pf02672 -
420015 -pf11375 -
420016 - -
420017 - -
420018 -pf01710 -
420019 -pf07992 -
420020 -pf02678pf17954 -
420021 HMMreg00126_1_mix(pf00126pf03466) GRM3B
420022 HMMnopfamclust29_1_aci(pf00931pf07819) CsomeBCy
420023 -pf00581 -
420024 HMMreg00486_1_mix(pf00563pf00990pf00072) CsomeBCy
(..1399977..)
1820001 -pf05860 -
1820002 - -
1820003 - -
1820004 -pf00535 -
1820005 -pf02470 -
1820008 - -
1820009 -pf01116 -
1820010 - -
1820011 Ts_poisonGreen CsomeBCy
1820012 - -
1820013 -pf01928 -
1820014 -pf00535 -
(..119987..)
1940001 -pf02663 -
1940002 - -
1940003 - -
1940004 - -
1940005 HMMnopfamclust29_1_aci(pf13181) CsomeBCy
1940006 - -
1940007 -pf01795 -
1940008 - -
1940009 - -
1940010 - -
1940011 Tdp_kelleyGreen CsomeBCy
1940012 - -
1940013 - -
1940014 -pf05116 -
1940015 -pf03739 -
1940016 - -
1940017 - -
1940018 - -
1940019 -pf08848 -
1940020 - -
(..1669982..)
3610002 - -
3610003 - -
3610004 -pf00035pf14622 -
3610005 H_peaSoup CsomeBCy
3610006 - -
3610007 - -
3610008 - -
3610009 -pf07085 -
3610010 -pf00403 -
3610011 - -
BMC-P
BMC-H
BMC-T
Aldehyde dehydrogenase
PTAC
Signature enzyme
Alcohol dehydrogenase
Regulator

Phormidesmis priestleyi Ana. Bacteria; Cyanobacteria; Synechococcales; Leptolyngbyaceae;

HLUCCA11__locus_1_main_sat

CsomeBCy

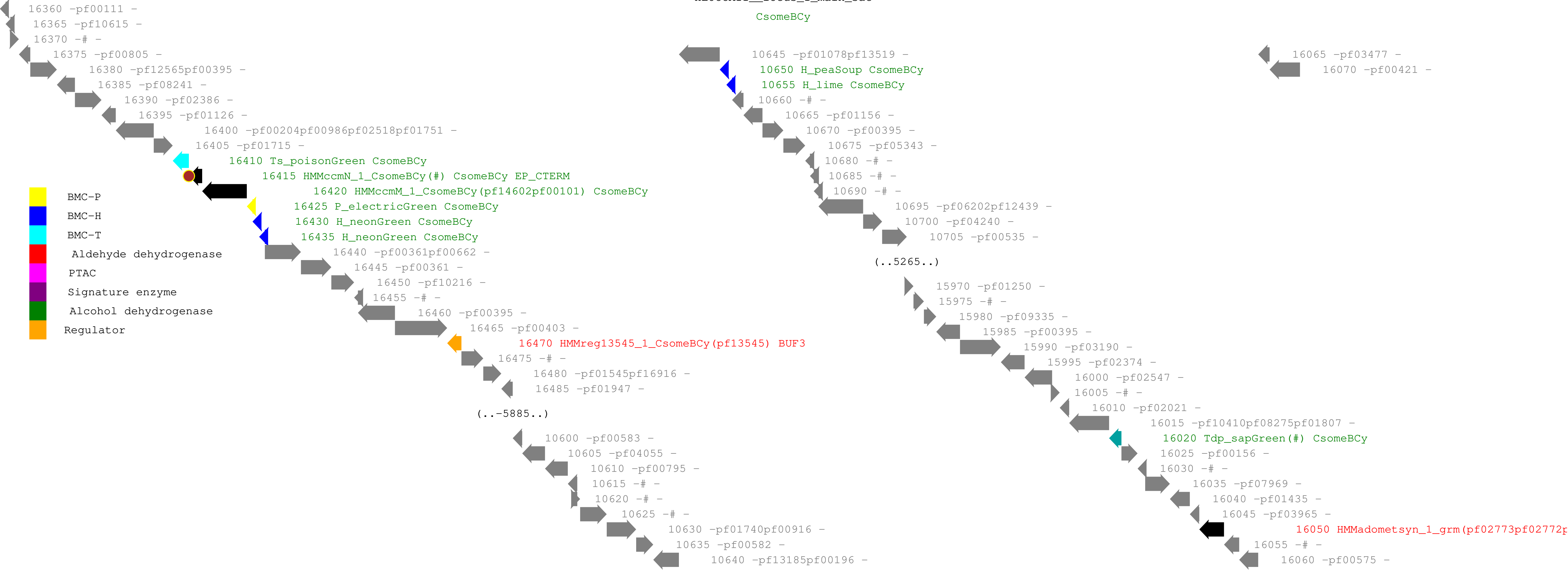

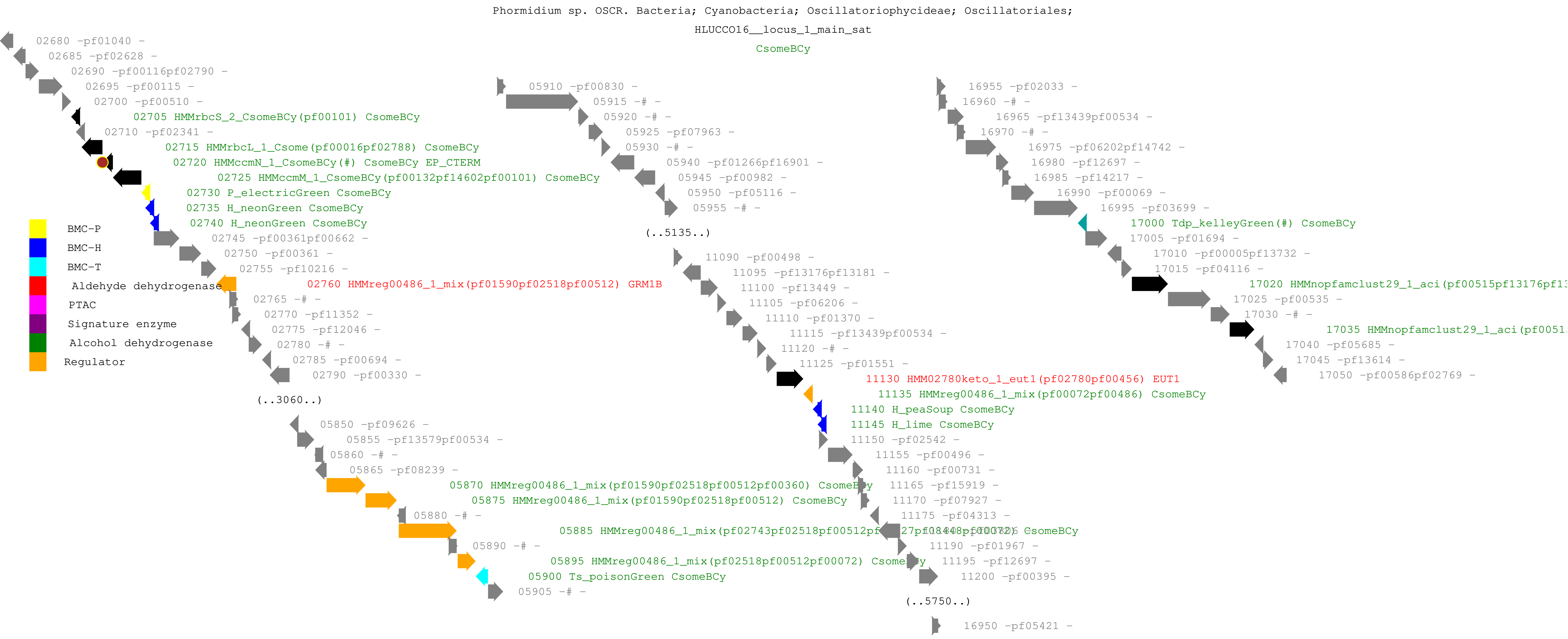

Phormidium sp. OSCR. Bacteria; Cyanobacteria; Oscillatoriophycideae; Oscillatoriales;
HLUCCO16__locus_1_main_sat
CsomeBCy
02680 -pf01040 -
02685 -pf02628 -
02690 -pf00116pf02790 -
02695 -pf00115 -
02700 -pf00510 -
02705 HMMrbcS_2_CsomeBCy(pf00101) CsomeBCy
02710 -pf02341 -
02715 HMMrbcL_1_Csome(pf00016pf02788) CsomeBCy
02720 HMMccmN_1_CsomeBCy(#) CsomeBCy EP_CTERM
02725 HMMccmM_1_CsomeBCy(pf00132pf14602pf00101) CsomeBCy
02730 P_electricGreen CsomeBCy
02735 H_neonGreen CsomeBCy
02740 H_neonGreen CsomeBCy
02745 -pf00361pf00662 -
02750 -pf00361 -
02755 -pf10216 -
02760 HMMreg00486_1_mix(pf01590pf02518pf00512) GRM1B
02765 -# -
02770 -pf11352 -
02775 -pf12046 -
02780 -# -
02785 -pf00694 -
02790 -pf00330 -
(..3060..)
05850 -pf09626 -
05855 -pf13579pf00534 -
05860 -# -
05865 -pf08239 -
05870 HMMreg00486_1_mix(pf01590pf02518pf00512pf00360) CsomeBCy
05875 HMMreg00486_1_mix(pf01590pf02518pf00512) CsomeBCy
05880 -# -
05885 HMMreg00486_1_mix(pf02743pf02518pf00512pf
05890 -# -
05895 HMMreg00486_1_mix(pf02518pf00512pf00072) CsomeBCy
05900 Ts_poisonGreen CsomeBCy
05905 -# -
05910 -pf00830 -
05915 -# -
05920 -# -
05925 -pf07963 -
05930 -# -
05940 -pf01266pf16901 -
05945 -pf00982 -
05950 -pf05116 -
05955 -# -
(..5135..)
11090 -pf00498 -
11095 -pf13176pf13181 -
11100 -pf13449 -
11105 -pf06206 -
11110 -pf01370 -
11115 -pf13439pf00534 -
11120 -# -
11125 -pf01551 -
11130 HMM02780keto_1_eut1(pf02780pf00456) EUT1
11135 HMMreg00486_1_mix(pf00072pf00486) CsomeBCy
11140 H_peaSoup CsomeBCy
11145 H_lime CsomeBCy
11150 -pf02542 -
11155 -pf00496 -
11160 -pf00731 -
11165 -pf15919 -
11170 -pf07927 -
11175 -pf04313 -
11190 -pf01967 -
11195 -pf12697 -
11200 -pf00395 -
(..5750..)
16950 -pf05421 -
16955 -pf02033 -
16960 -# -
16965 -pf13439pf00534 -
16970 -# -
16975 -pf06202pf14742 -
16980 -pf12697 -
16985 -pf14217 -
16990 -pf00069 -
16995 -pf03699 -
17000 Tdp_kelleyGreen(#) CsomeBCy
17005 -pf01694 -
17010 -pf00005pf13732 -
17015 -pf04116 -
17020 HMMnopfamclust29_1_aci(pf00515pf13176pf1
17025 -pf00535 -
17030 -# -
17035 HMMnopfamclust29_1_aci(pf0051
17040 -pf05685 -
17045 -pf13614 -
17050 -pf00586pf02769 -
BMC-P
BMC-H
BMC-T
Aldehyde dehydrogenase
PTAC
Signature enzyme
Alcohol dehydrogenase
Regulator

Calothrix sp. 336/3. Bacteria; Cyanobacteria; Nostocales; Rivulariaceae; Calothrix.

IJ00__locus_1_main_sat

CsomeBCy

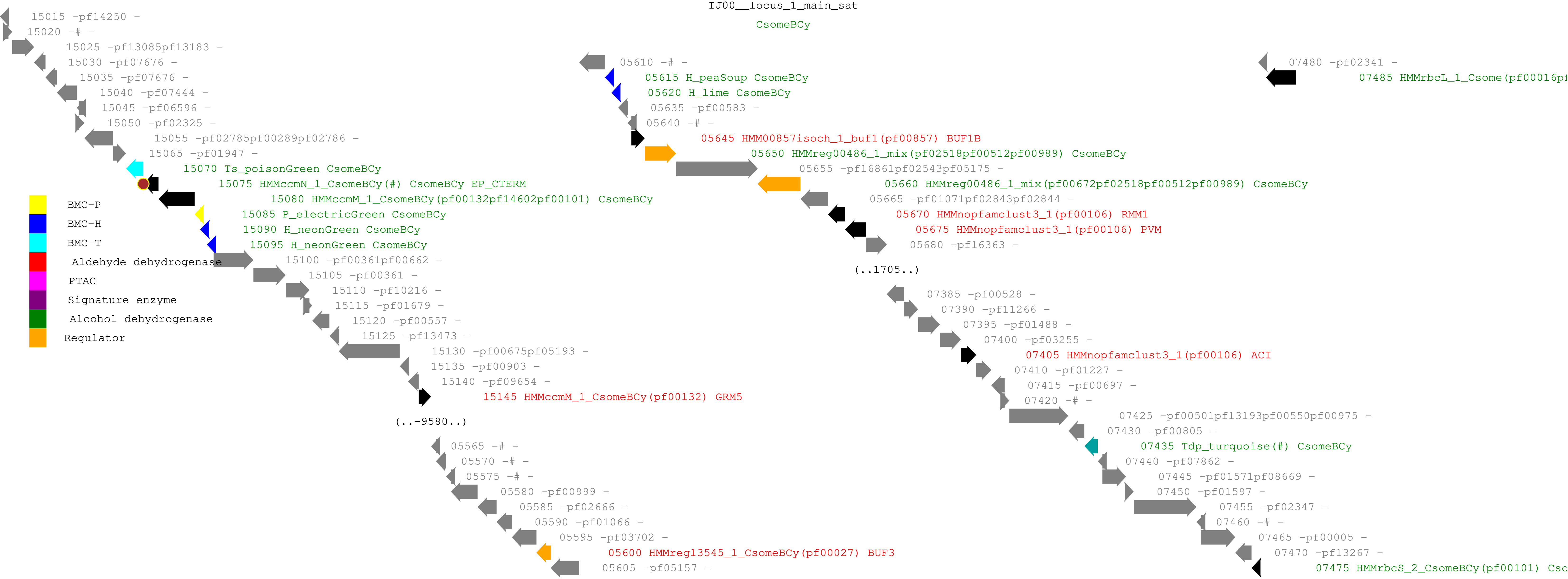

Microcystis aeruginosa PCC 7806. Bacteria; Cyanobacteria; Oscillatoriophycideae; Chroococcales;

IPF__locus_fragmented

CsomeBCy

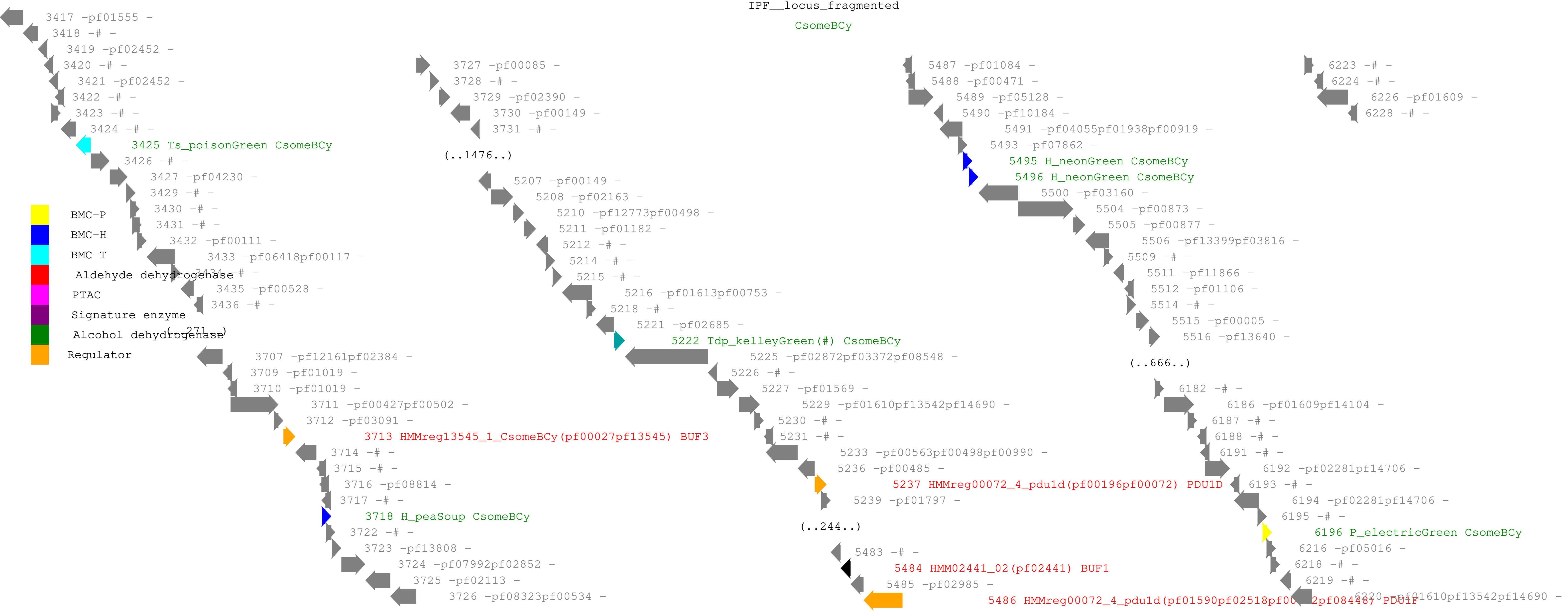

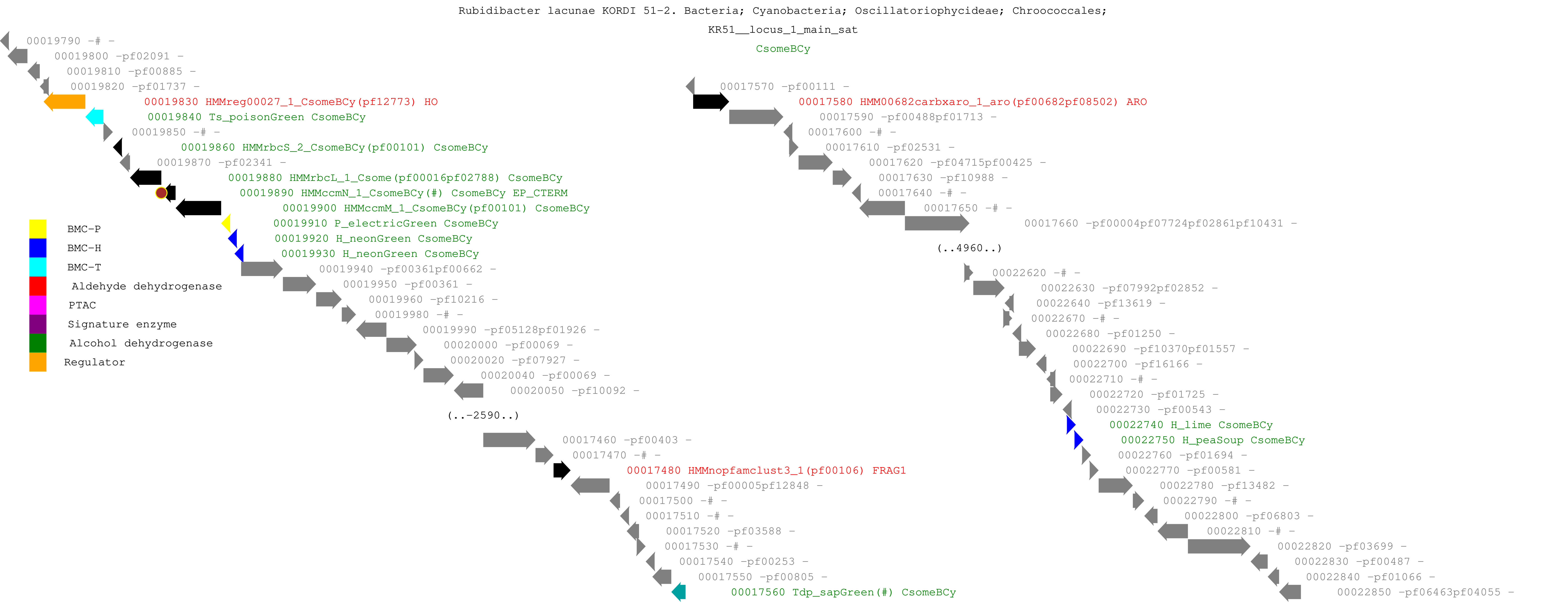
Rubidibacter lacunae KORDI 51-2. Bacteria; Cyanobacteria; Oscillatoriophycideae; Chroococcales;
KR51__locus_1_main_sat
CsomeBCy
00019790 -# -
00019800 -pf02091 -
00019810 -pf00885 -
00019820 -pf01737 -
00019830 HMMreg00027_1_CsomeBCy(pf12773) HO
00019840 Ts_poisonGreen CsomeBCy
00019850 -# -
00019860 HMMrbcS_2_CsomeBCy(pf00101) CsomeBCy
00019870 -pf02341 -
00019880 HMMrbcL_1_Csome(pf00016pf02788) CsomeBCy
00019890 HMMccmN_1_CsomeBCy(#) CsomeBCy EP_CTERM
00019900 HMMccmM_1_CsomeBCy(pf00101) CsomeBCy
00019910 P_electricGreen CsomeBCy
00019920 H_neonGreen CsomeBCy
00019930 H_neonGreen CsomeBCy
00019940 -pf00361pf00662 -
00019950 -pf00361 -
00019960 -pf10216 -
00019980 -# -
00019990 -pf05128pf01926 -
00020000 -pf00069 -
00020020 -pf07927 -
00020040 -pf00069 -
00020050 -pf10092 -
(..-2590..)
00017460 -pf00403 -
00017470 -# -
00017480 HMMnopfamclust3_1(pf00106) FRAG1
00017490 -pf00005pf12848 -
00017500 -# -
00017510 -# -
00017520 -pf03588 -
00017530 -# -
00017540 -pf00253 -
00017550 -pf00805 -
00017560 Tdp_sapGreen(#) CsomeBCy
00017570 -pf00111 -
00017580 HMM00682carbxaro_1_aro(pf00682pf08502) ARO
00017590 -pf00488pf01713 -
00017600 -# -
00017610 -pf02531 -
00017620 -pf04715pf00425 -
00017630 -pf10988 -
00017640 -# -
00017650 -# -
00017660 -pf00004pf07724pf02861pf10431 -
(..4960..)
00022620 -# -
00022630 -pf07992pf02852 -
00022640 -pf13619 -
00022670 -# -
00022680 -pf01250 -
00022690 -pf10370pf01557 -
00022700 -pf16166 -
00022710 -# -
00022720 -pf01725 -
00022730 -pf00543 -
00022740 H_lime CsomeBCy
00022750 H_peaSoup CsomeBCy
00022760 -pf01694 -
00022770 -pf00581 -
00022780 -pf13482 -
00022790 -# -
00022800 -pf06803 -
00022810 -# -
00022820 -pf03699 -
00022830 -pf00487 -
00022840 -pf01066 -
00022850 -pf06463pf04055 -
BMC-P
BMC-H
BMC-T
Aldehyde dehydrogenase
PTAC
Signature enzyme
Alcohol dehydrogenase
Regulator

Lyngbya sp. (strain PCC 8106) (Lyngbya aestuarii (strain CCY9616)). Bacteria; Cyanobacteria; Oscillatoriophycideae; Oscillatoriales;

L8106__locus_1_main_sat

CsomeBCy

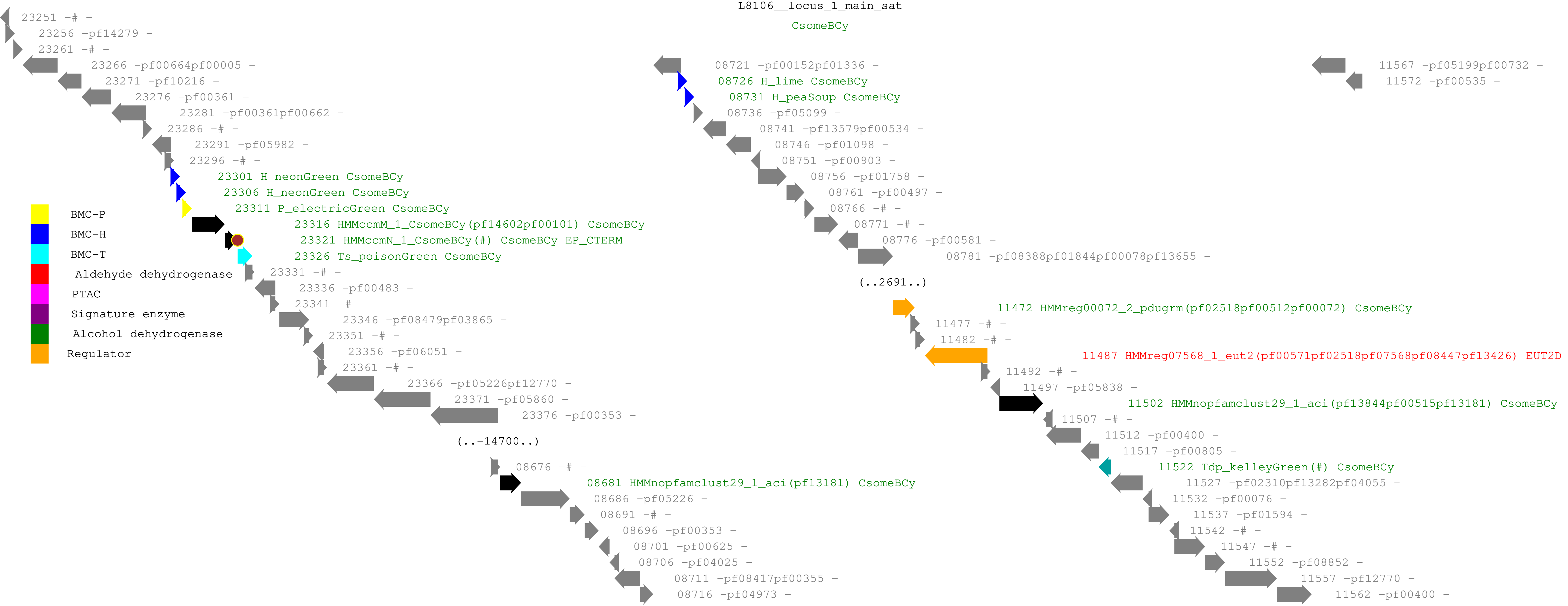

Leptolyngbya sp. NIES-3755. Bacteria; Cyanobacteria; Synechococcales; Leptolyngbyaceae;

LEP3755__locus_1_main_sat

CsomeBCy

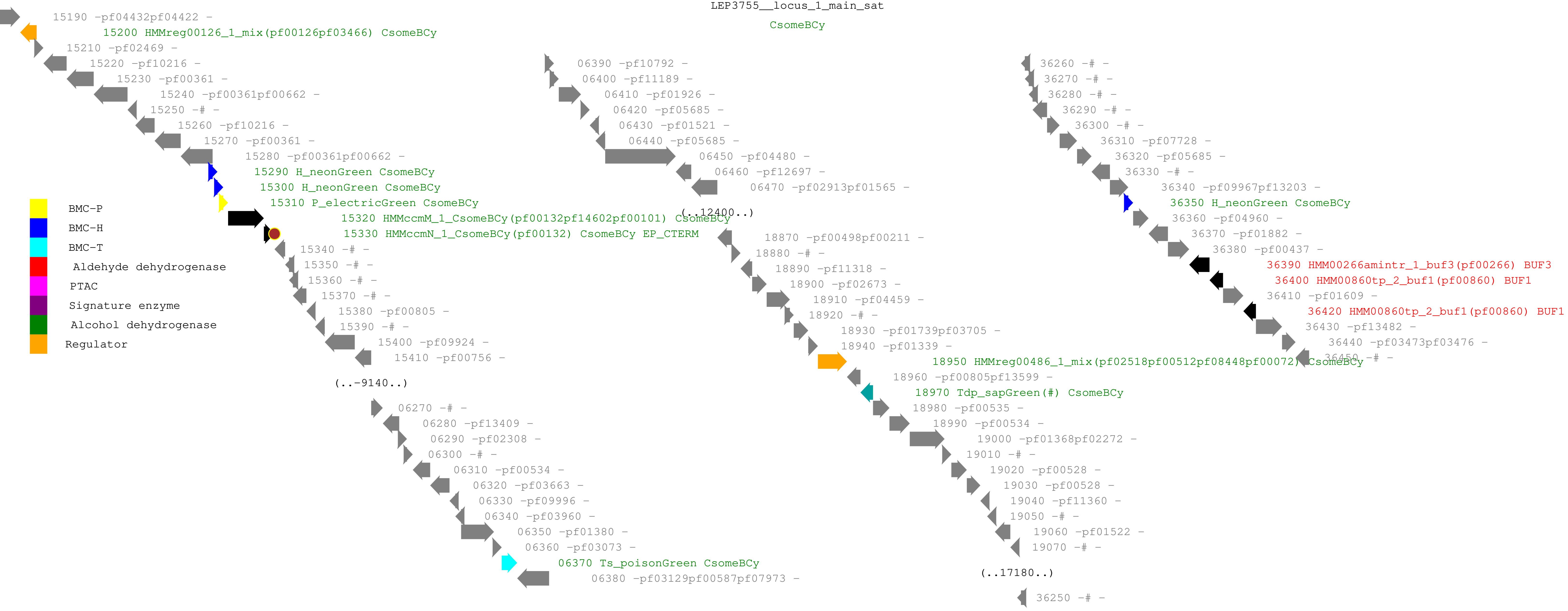

Leptolyngbya sp. PCC 7375. Bacteria; Cyanobacteria; Synechococcales; Leptolyngbyaceae;

Lepto7375DRAFT__locus_1_main_sat

CsomeBCy

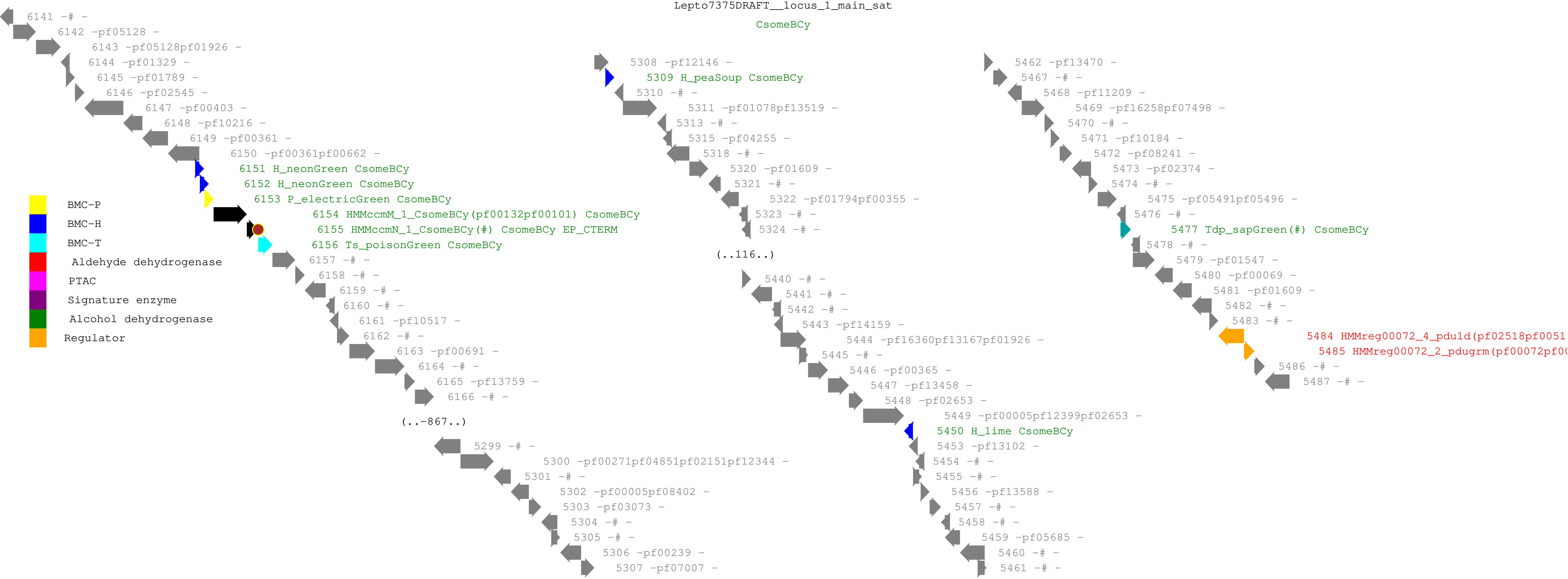

Leptolyngbya sp. PCC 7376. Bacteria; Cyanobacteria; Synechococcales; Leptolyngbyaceae;

Lepto7376__locus_1_main_sat

CsomeBCy

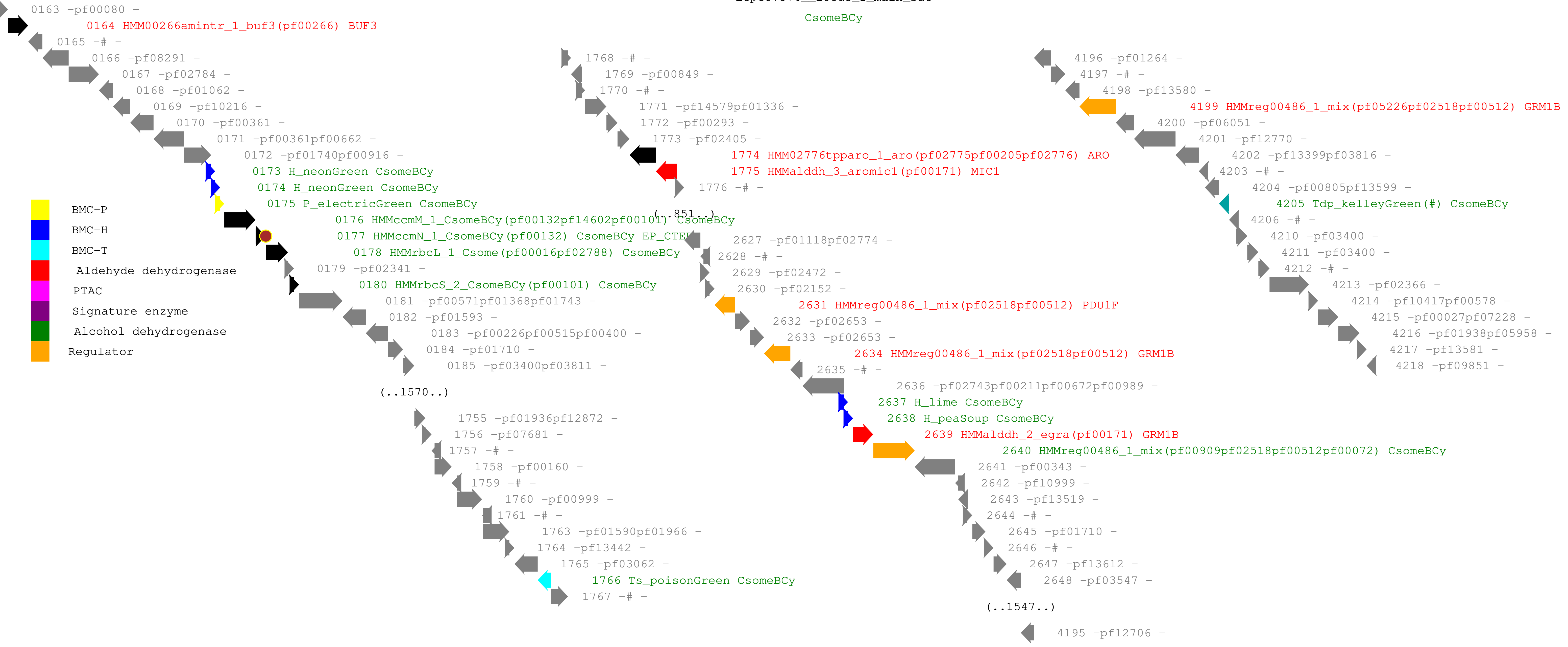

Moorea producens 3L. Bacteria; Cyanobacteria; Oscillatoriophycideae; Oscillatoriales;

LYNGBM3L__locus_1_main_sat

CsomeBCy

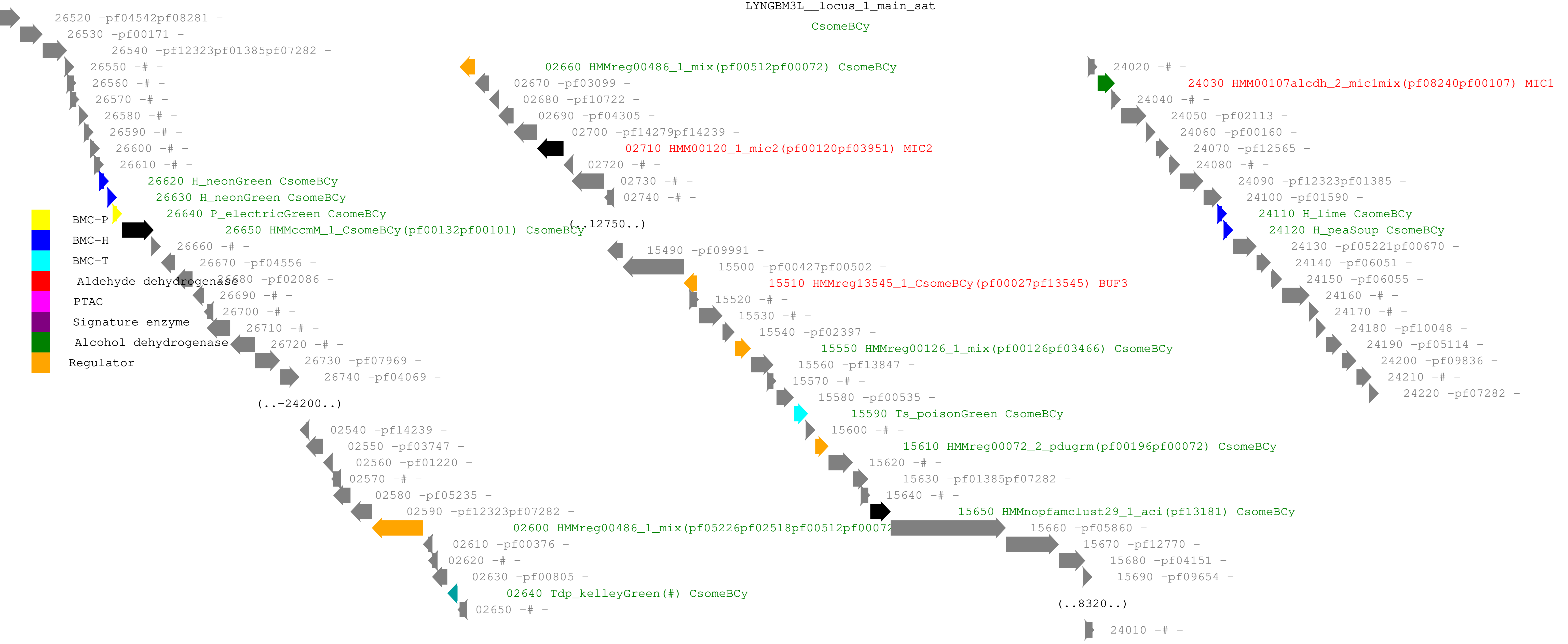

Lyngbya aestuarii BL J. Bacteria; Cyanobacteria; Oscillatoriophycideae; Oscillatoriales;

M595__locus_1_main_sat

CsomeBCy

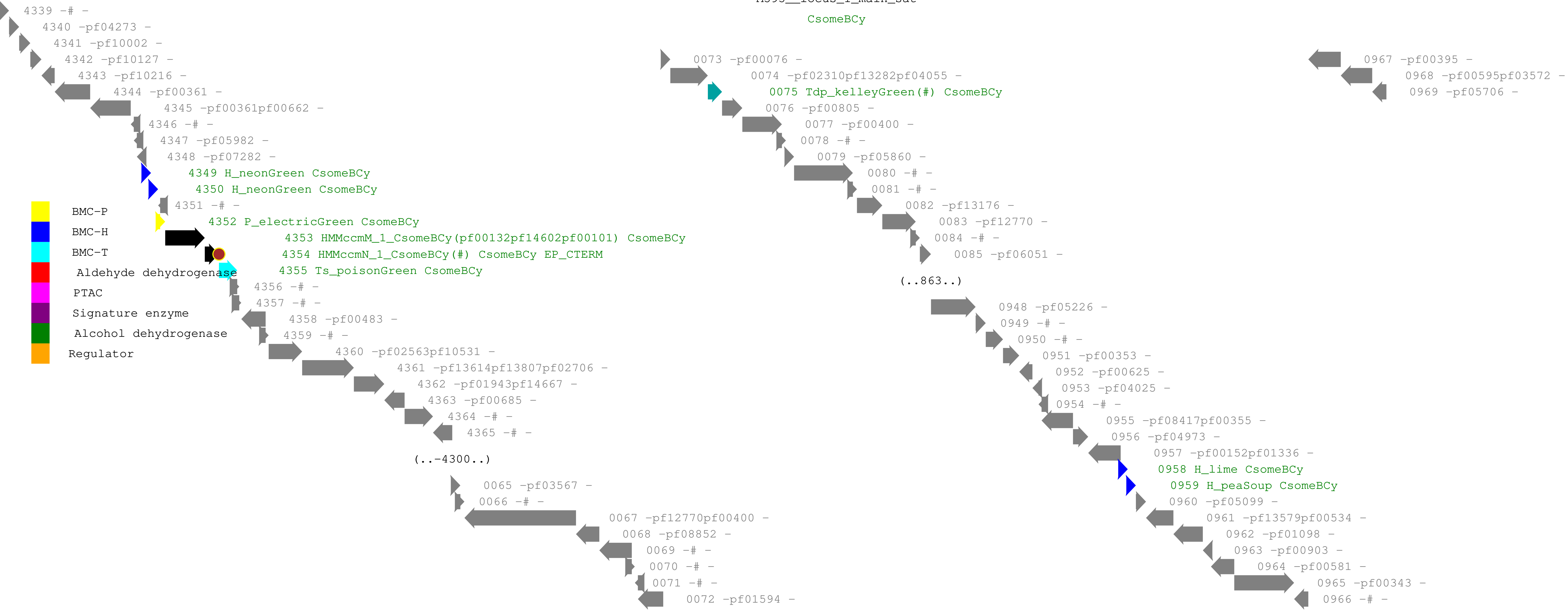

Microcystis aeruginosa 11-30S32. Bacteria; Cyanobacteria; Oscillatoriophycideae; Chroococcales;

MAE30S32__locus_1_main_sat

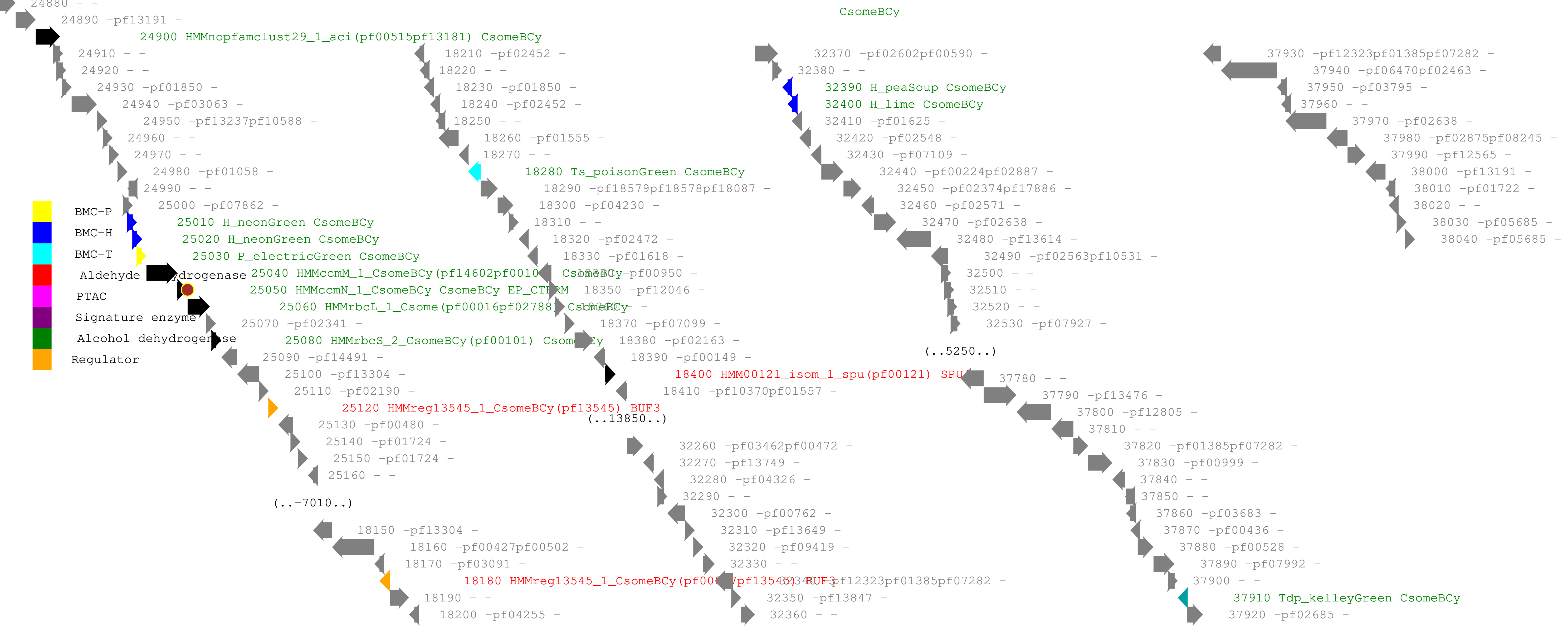

Microcystis aeruginosa (strain NIES-843). Bacteria; Cyanobacteria; Oscillatoriophycideae; Chroococcales;

MAE__locus_1_main_sat

CsomeBCy

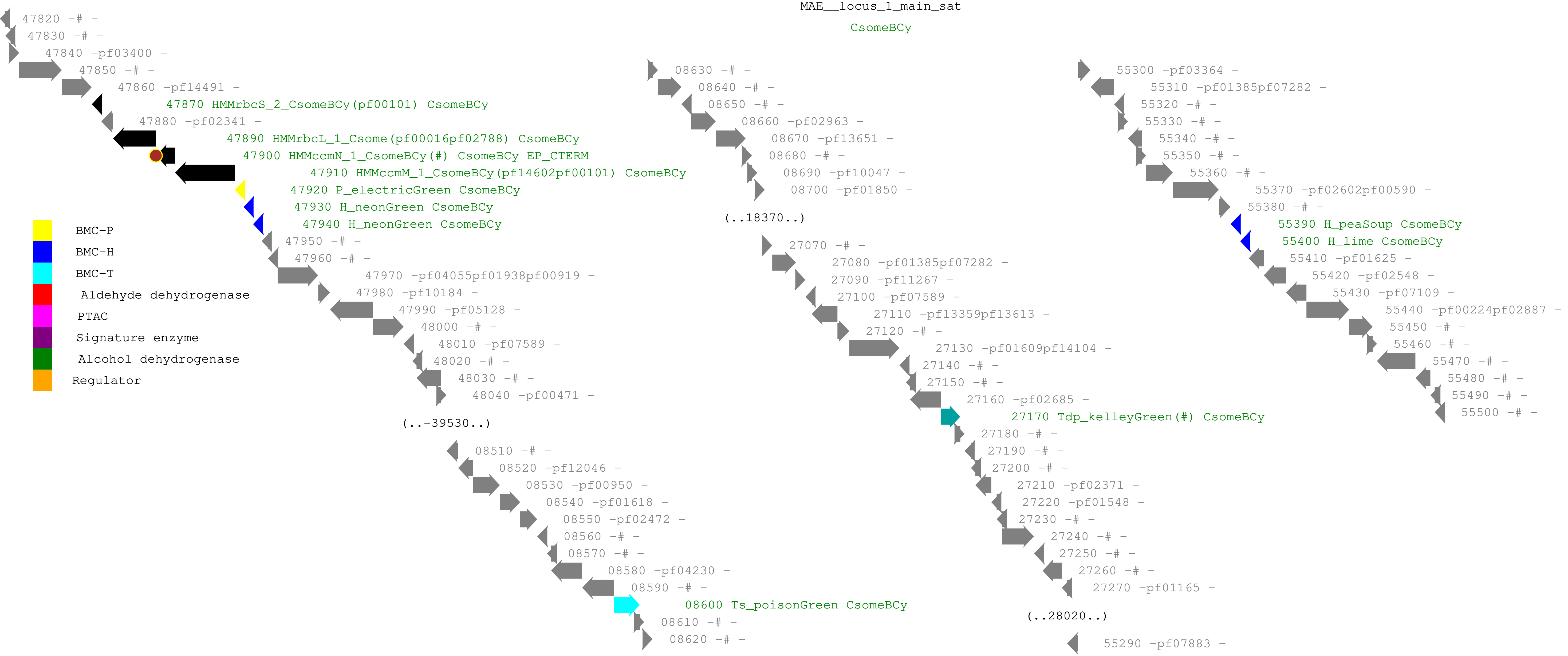

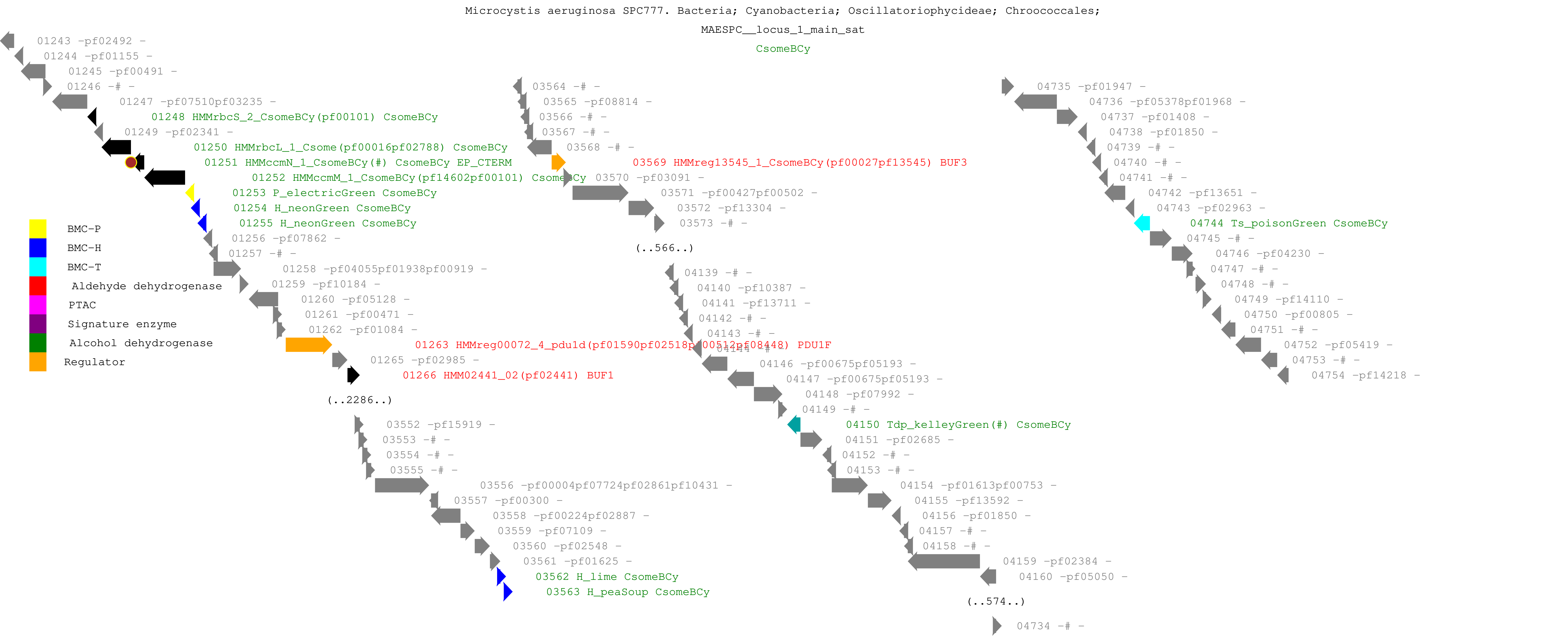

Microcystis aeruginosa SPC777. Bacteria; Cyanobacteria; Oscillatoriophycideae; Chroococcales;
MAESPC__locus_1_main_sat
CsomeBCy
01243 -pf02492 -
01244 -pf01155 -
01245 -pf00491 -
01246 -# -
01247 -pf07510pf03235 -
01248 HMMrbcS_2_CsomeBCy(pf00101) CsomeBCy
01249 -pf02341 -
01250 HMMrbcL_1_Csome(pf00016pf02788) CsomeBCy
01251 HMMccmN_1_CsomeBCy(#) CsomeBCy EP_CTERM
01252 HMMccmM_1_CsomeBCy(pf14602pf00101) CsomeBCy
01253 P_electricGreen CsomeBCy
01254 H_neonGreen CsomeBCy
01255 H_neonGreen CsomeBCy
01256 -pf07862 -
01257 -# -
01258 -pf04055pf01938pf00919 -
01259 -pf10184 -
01260 -pf05128 -
01261 -pf00471 -
01262 -pf01084 -
01263 HMMreg00072_4_pdu1d(pf01590pf02518pf00512pf08448) PDU1F
01265 -pf02985 -
01266 HMM02441_02(pf02441) BUF1
(..2286..)
03552 -pf15919 -
03553 -# -
03554 -# -
03555 -# -
03556 -pf00004pf07724pf02861pf10431 -
03557 -pf00300 -
03558 -pf00224pf02887 -
03559 -pf07109 -
03560 -pf02548 -
03561 -pf01625 -
03562 H_lime CsomeBCy
03563 H_peaSoup CsomeBCy
03564 -# -
03565 -pf08814 -
03566 -# -
03567 -# -
03568 -# -
03569 HMMreg13545_1_CsomeBCy(pf00027pf13545) BUF3
03570 -pf03091 -
03571 -pf00427pf00502 -
03572 -pf13304 -
03573 -# -
(..566..)
04139 -# -
04140 -pf10387 -
04141 -pf13711 -
04142 -# -
04143 -# -
04144 -# -
04146 -pf00675pf05193 -
04147 -pf00675pf05193 -
04148 -pf07992 -
04149 -# -
04150 Tdp_kelleyGreen(#) CsomeBCy
04151 -pf02685 -
04152 -# -
04153 -# -
04154 -pf01613pf00753 -
04155 -pf13592 -
04156 -pf01850 -
04157 -# -
04158 -# -
04159 -pf02384 -
04160 -pf05050 -
(..574..)
04734 -# -
04735 -pf01947 -
04736 -pf05378pf01968 -
04737 -pf01408 -
04738 -pf01850 -
04739 -# -
04740 -# -
04741 -# -
04742 -pf13651 -
04743 -pf02963 -
04744 Ts_poisonGreen CsomeBCy
04745 -# -
04746 -pf04230 -
04747 -# -
04748 -# -
04749 -pf14110 -
04750 -pf00805 -
04751 -# -
04752 -pf05419 -
04753 -# -
04754 -pf14218 -
BMC-P
BMC-H
BMC-T
Aldehyde dehydrogenase
PTAC
Signature enzyme
Alcohol dehydrogenase
Regulator

Coleofasciculus chthonoplastes PCC 7420. Bacteria; Cyanobacteria; Oscillatoriophycideae; Oscillatoriales;

MC7420__locus_1_main_sat

CsomeBCy

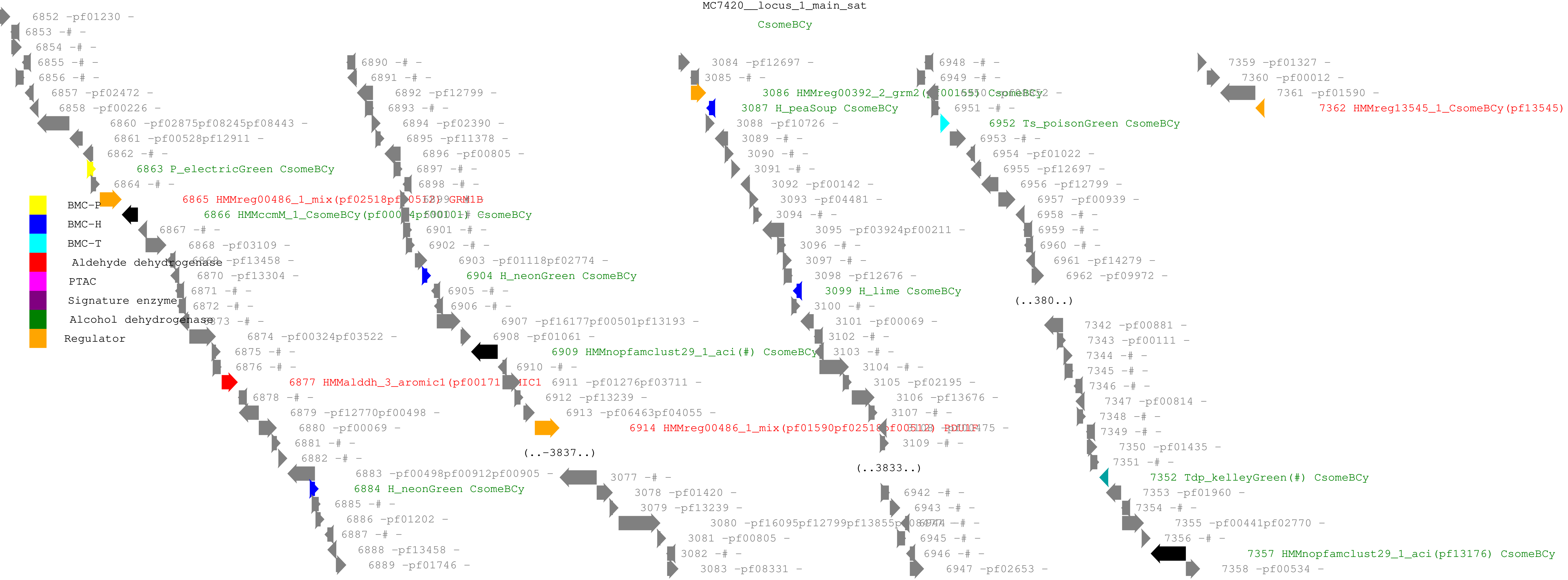

Microcystis aeruginosa NIES-4285. Bacteria; Cyanobacteria; Oscillatoriophycideae; Chroococcales;

MiAbB__locus_1_main_sat

CsomeBCy

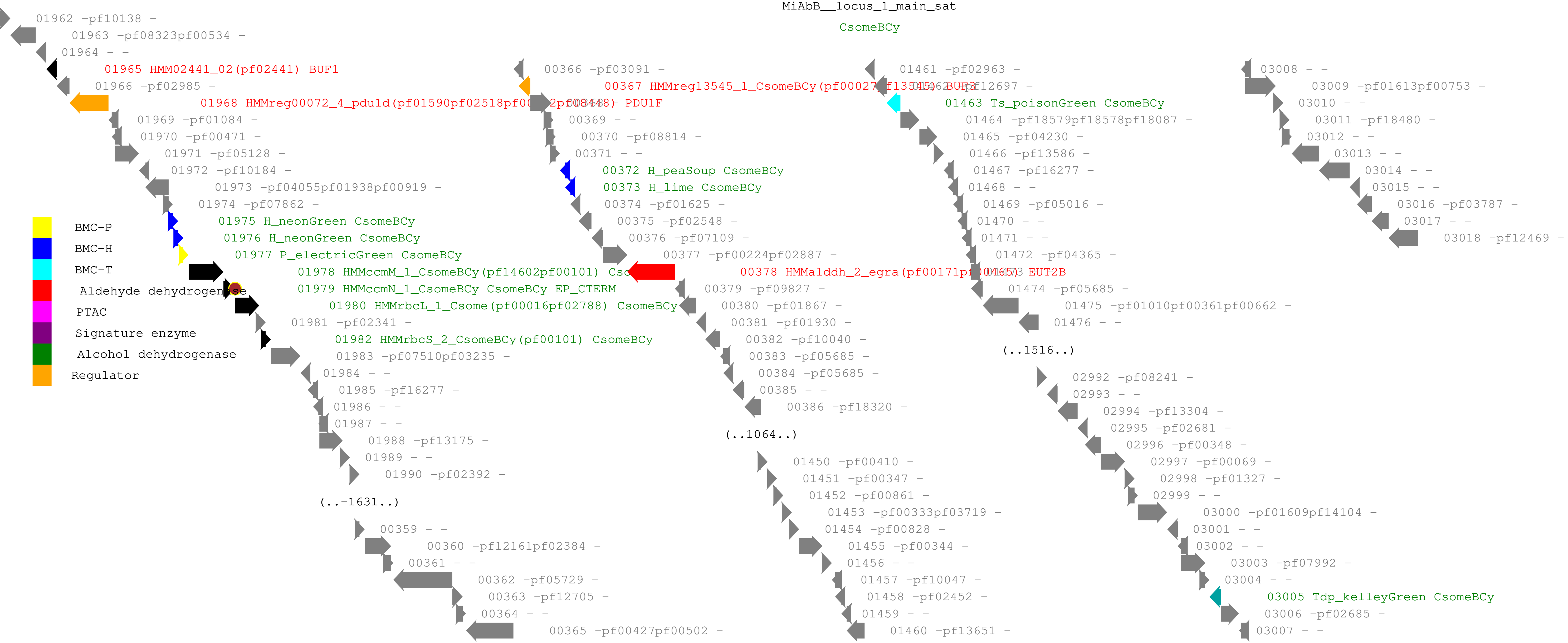

Microcystis aeruginosa NIES-4325. Bacteria; Cyanobacteria; Oscillatoriophycideae; Chroococcales;

MiAbW__locus_1_main_sat

CsomeBCy

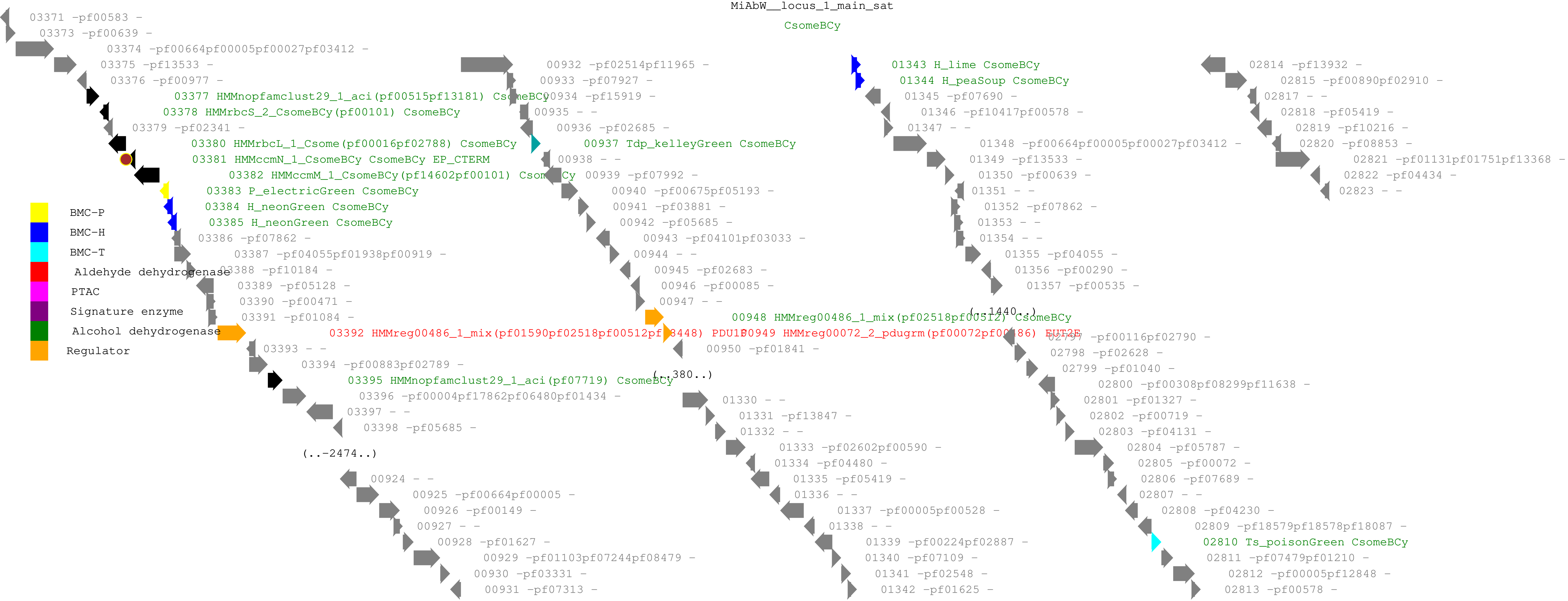

Microcoleus sp. PCC 7113. Bacteria; Cyanobacteria; Oscillatoriophycideae; Oscillatoriales;

Mic7113__locus_1_main_sat

CsomeBCy

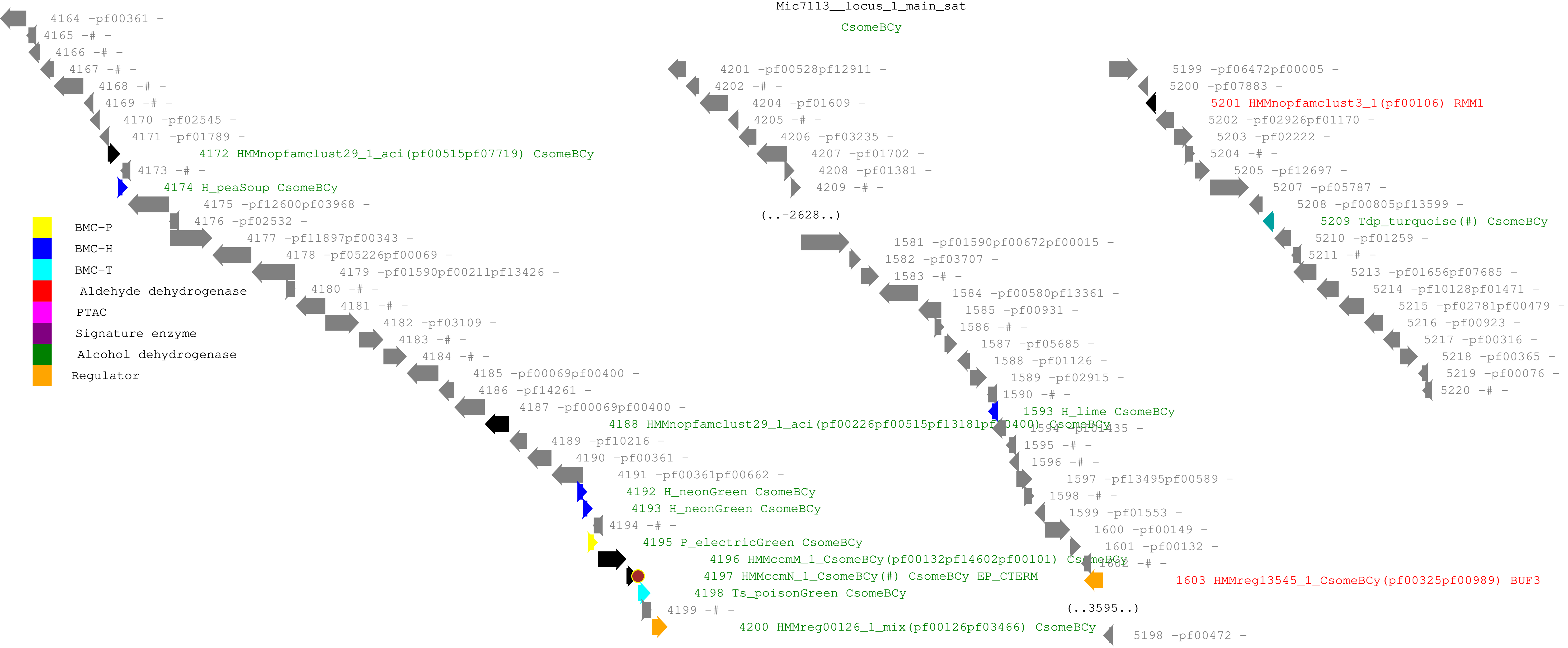

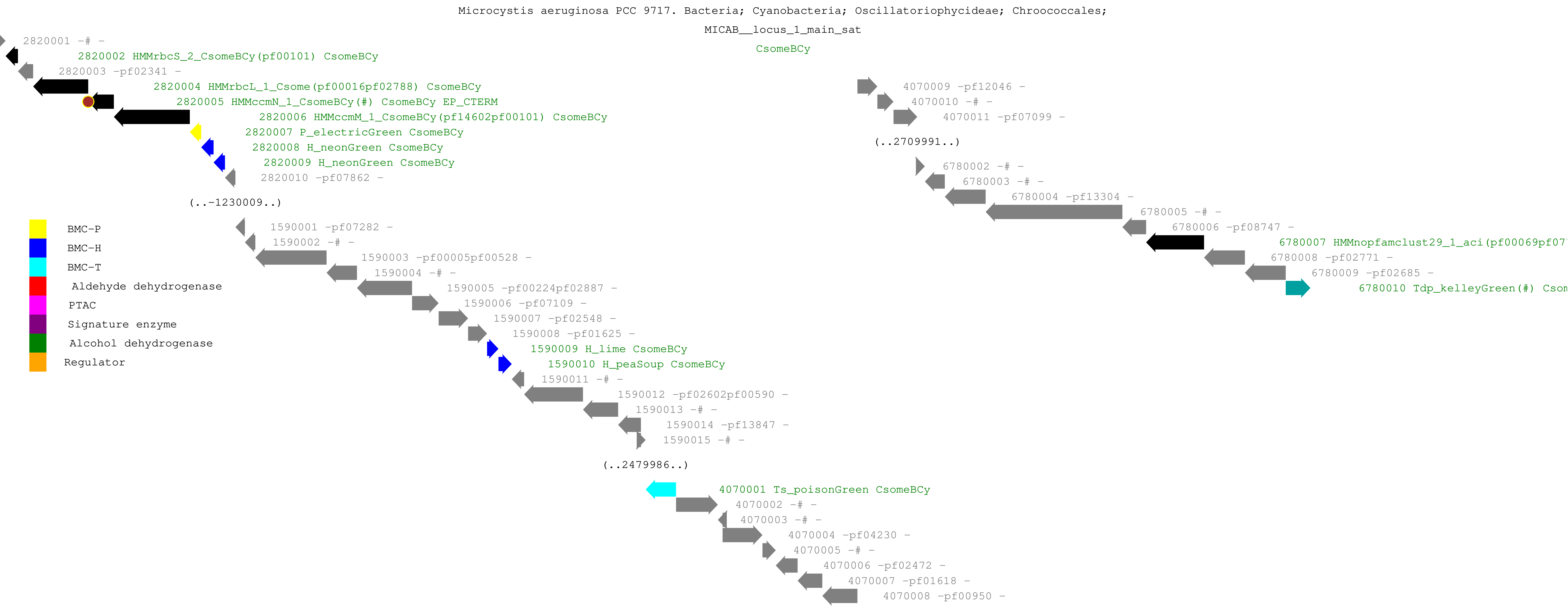
Microcystis aeruginosa PCC 9717. Bacteria; Cyanobacteria; Oscillatoriophycideae; Chroococcales;
MICAB__locus_1_main_sat
CsomeBCy
2820001 -# -
2820002 HMMrbcS_2_CsomeBCy(pf00101) CsomeBCy
2820003 -pf02341 -
2820004 HMMrbcL_1_Csome(pf00016pf02788) CsomeBCy
2820005 HMMccmN_1_CsomeBCy(#) CsomeBCy EP_CTERM
2820006 HMMccmM_1_CsomeBCy(pf14602pf00101) CsomeBCy
2820007 P_electricGreen CsomeBCy
2820008 H_neonGreen CsomeBCy
2820009 H_neonGreen CsomeBCy
2820010 -pf07862 -
(..-1230009..)
BMC-P
BMC-H
BMC-T
Aldehyde dehydrogenase
PTAC
Signature enzyme
Alcohol dehydrogenase
Regulator
1590001 -pf07282 -
1590002 -# -
1590003 -pf00005pf00528 -
1590004 -# -
1590005 -pf00224pf02887 -
1590006 -pf07109 -
1590007 -pf02548 -
1590008 -pf01625 -
1590009 H_lime CsomeBCy
1590010 H_peaSoup CsomeBCy
1590011 -# -
1590012 -pf02602pf00590 -
1590013 -# -
1590014 -pf13847 -
1590015 -# -
(..2479986..)
4070001 Ts_poisonGreen CsomeBCy
4070002 -# -
4070003 -# -
4070004 -pf04230 -
4070005 -# -
4070006 -pf02472 -
4070007 -pf01618 -
4070008 -pf00950 -
4070009 -pf12046 -
4070010 -# -
4070011 -pf07099 -
(..2709991..)
6780002 -# -
6780003 -# -
6780004 -pf13304 -
6780005 -# -
6780006 -pf08747 -
6780007 HMMnopfamclust29_1_aci(pf00069pf07
6780008 -pf02771 -
6780009 -pf02685 -
6780010 Tdp_kelleyGreen(#) Csom

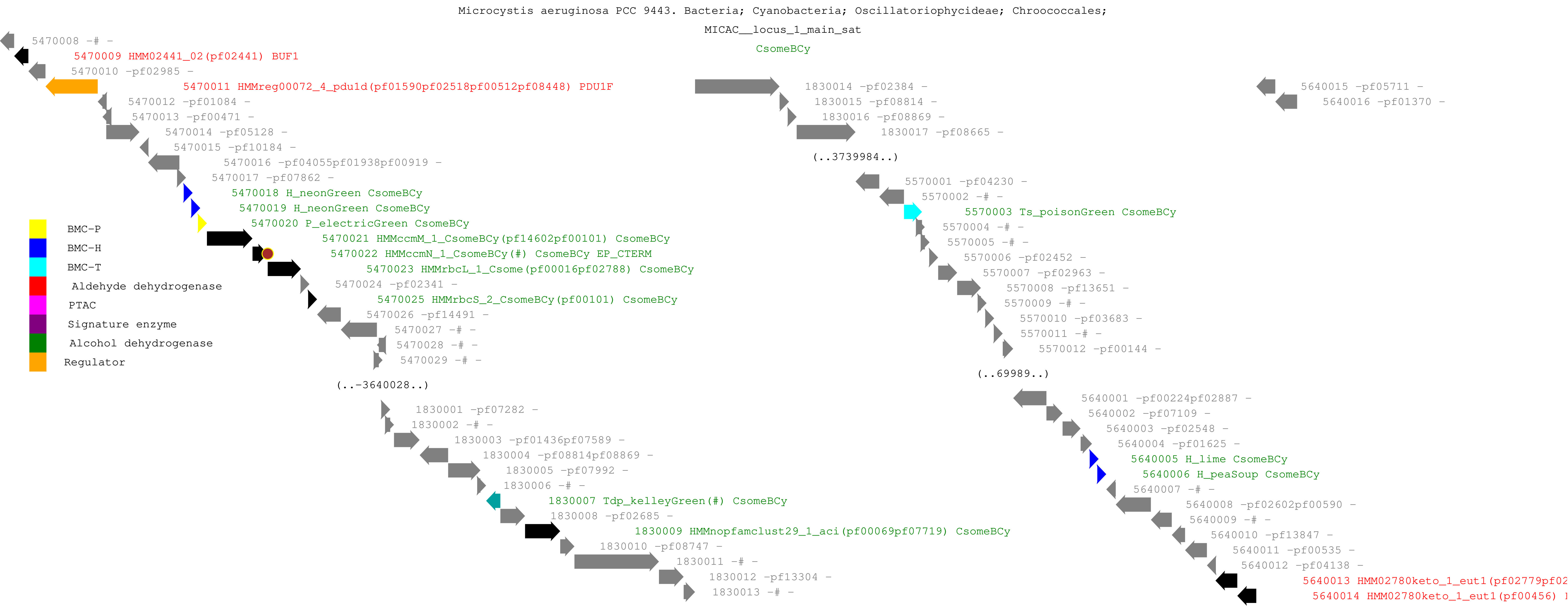
Microcystis aeruginosa PCC 9443. Bacteria; Cyanobacteria; Oscillatoriophycideae; Chroococcales;
MICAC__locus_1_main_sat
CsomeBCy
5470008 -# -
5470009 HMM02441_02(pf02441) BUF1
5470010 -pf02985 -
5470011 HMMreg00072_4_pdu1d(pf01590pf02518pf00512pf08448) PDU1F
5470012 -pf01084 -
5470013 -pf00471 -
5470014 -pf05128 -
5470015 -pf10184 -
5470016 -pf04055pf01938pf00919 -
5470017 -pf07862 -
5470018 H_neonGreen CsomeBCy
5470019 H_neonGreen CsomeBCy
5470020 P_electricGreen CsomeBCy
5470021 HMMccmM_1_CsomeBCy(pf14602pf00101) CsomeBCy
5470022 HMMccmN_1_CsomeBCy(#) CsomeBCy EP_CTERM
5470023 HMMrbcL_1_Csome(pf00016pf02788) CsomeBCy
5470024 -pf02341 -
5470025 HMMrbcS_2_CsomeBCy(pf00101) CsomeBCy
5470026 -pf14491 -
5470027 -# -
5470028 -# -
5470029 -# -
(..-3640028..)
1830001 -pf07282 -
1830002 -# -
1830003 -pf01436pf07589 -
1830004 -pf08814pf08869 -
1830005 -pf07992 -
1830006 -# -
1830007 Tdp_kelleyGreen(#) CsomeBCy
1830008 -pf02685 -
1830009 HMMnopfamclust29_1_aci(pf00069pf07719) CsomeBCy
1830010 -pf08747 -
1830011 -# -
1830012 -pf13304 -
1830013 -# -
BMC-P
BMC-H
BMC-T
Aldehyde dehydrogenase
PTAC
Signature enzyme
Alcohol dehydrogenase
Regulator
1830014 -pf02384 -
1830015 -pf08814 -
1830016 -pf08869 -
1830017 -pf08665 -
(..3739984..)
5570001 -pf04230 -
5570002 -# -
5570003 Ts_poisonGreen CsomeBCy
5570004 -# -
5570005 -# -
5570006 -pf02452 -
5570007 -pf02963 -
5570008 -pf13651 -
5570009 -# -
5570010 -pf03683 -
5570011 -# -
5570012 -pf00144 -
(..69989..)
5640001 -pf00224pf02887 -
5640002 -pf07109 -
5640003 -pf02548 -
5640004 -pf01625 -
5640005 H_lime CsomeBCy
5640006 H_peaSoup CsomeBCy
5640007 -# -
5640008 -pf02602pf00590 -
5640009 -# -
5640010 -pf13847 -
5640011 -pf00535 -
5640012 -pf04138 -
5640013 HMM02780keto_1_eut1(pf02779pf02
5640014 HMM02780keto_1_eut1(pf00456)
5640015 -pf05711 -
5640016 -pf01370 -

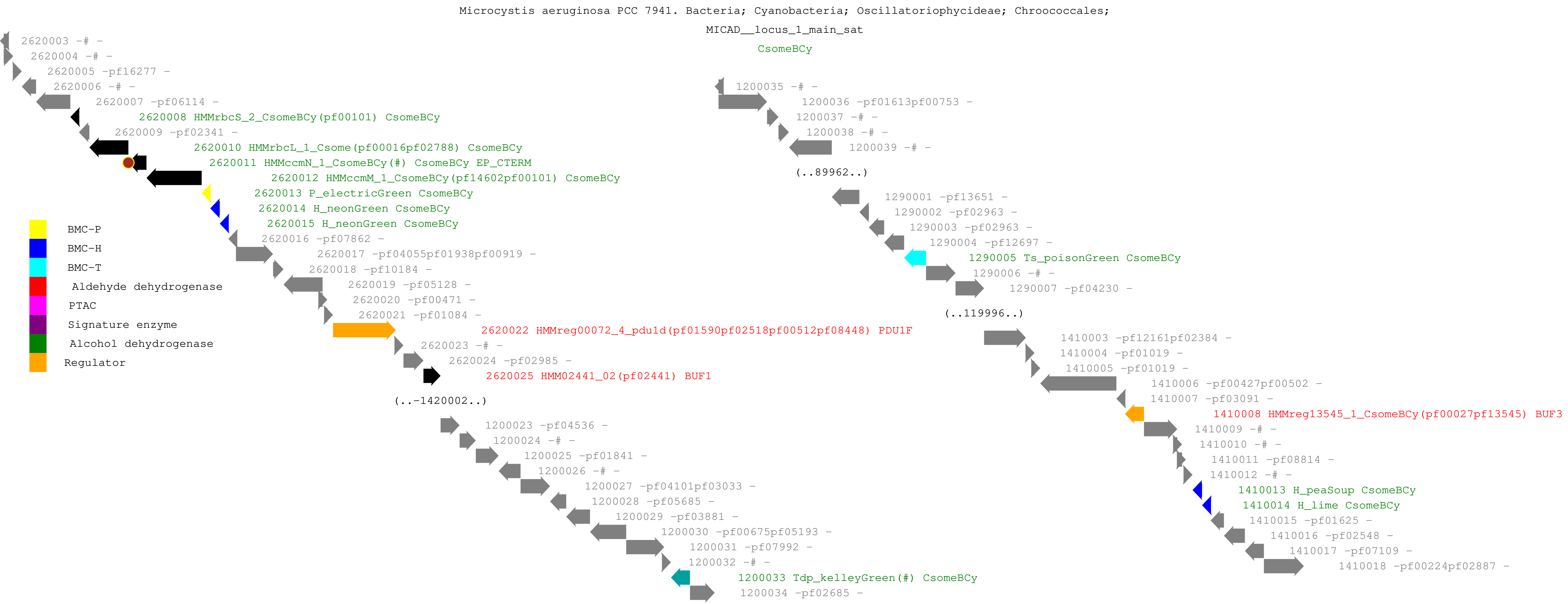

Microcystis aeruginosa PCC 7941. Bacteria; Cyanobacteria; Oscillatoriophycideae; Chroococcales;
MICAD__locus_1_main_sat
CsomeBCy
2620003 -# -
2620004 -# -
2620005 -pf16277 -
2620006 -# -
2620007 -pf06114 -
2620008 HMMrbcS_2_CsomeBCy(pf00101) CsomeBCy
2620009 -pf02341 -
2620010 HMMrbcL_1_Csome(pf00016pf02788) CsomeBCy
2620011 HMMccmN_1_CsomeBCy(#) CsomeBCy EP_CTERM
2620012 HMMccmM_1_CsomeBCy(pf14602pf00101) CsomeBCy
2620013 P_electricGreen CsomeBCy
2620014 H_neonGreen CsomeBCy
2620015 H_neonGreen CsomeBCy
2620016 -pf07862 -
2620017 -pf04055pf01938pf00919 -
2620018 -pf10184 -
2620019 -pf05128 -
2620020 -pf00471 -
2620021 -pf01084 -
2620022 HMMreg00072_4_pdu1d(pf01590pf02518pf00512pf08448) PDU1F
2620023 -# -
2620024 -pf02985 -
2620025 HMM02441_02(pf02441) BUF1
(..-1420002..)
1200023 -pf04536 -
1200024 -# -
1200025 -pf01841 -
1200026 -# -
1200027 -pf04101pf03033 -
1200028 -pf05685 -
1200029 -pf03881 -
1200030 -pf00675pf05193 -
1200031 -pf07992 -
1200032 -# -
1200033 Tdp_kelleyGreen(#) CsomeBCy
1200034 -pf02685 -
1200035 -# -
1200036 -pf01613pf00753 -
1200037 -# -
1200038 -# -
1200039 -# -
(..89962..)
1290001 -pf13651 -
1290002 -pf02963 -
1290003 -pf02963 -
1290004 -pf12697 -
1290005 Ts_poisonGreen CsomeBCy
1290006 -# -
1290007 -pf04230 -
(..119996..)
1410003 -pf12161pf02384 -
1410004 -pf01019 -
1410005 -pf01019 -
1410006 -pf00427pf00502 -
1410007 -pf03091 -
1410008 HMMreg13545_1_CsomeBCy(pf00027pf13545) BUF3
1410009 -# -
1410010 -# -
1410011 -pf08814 -
1410012 -# -
1410013 H_peaSoup CsomeBCy
1410014 H_lime CsomeBCy
1410015 -pf01625 -
1410016 -pf02548 -
1410017 -pf07109 -
1410018 -pf00224pf02887 -
BMC-P
BMC-H
BMC-T
Aldehyde dehydrogenase
PTAC
Signature enzyme
Alcohol dehydrogenase
Regulator

Microcystis aeruginosa PCC 9806. Bacteria; Cyanobacteria; Oscillatoriophycideae; Chroococcales;

MICAE__locus_1_main_sat

CsomeBCy

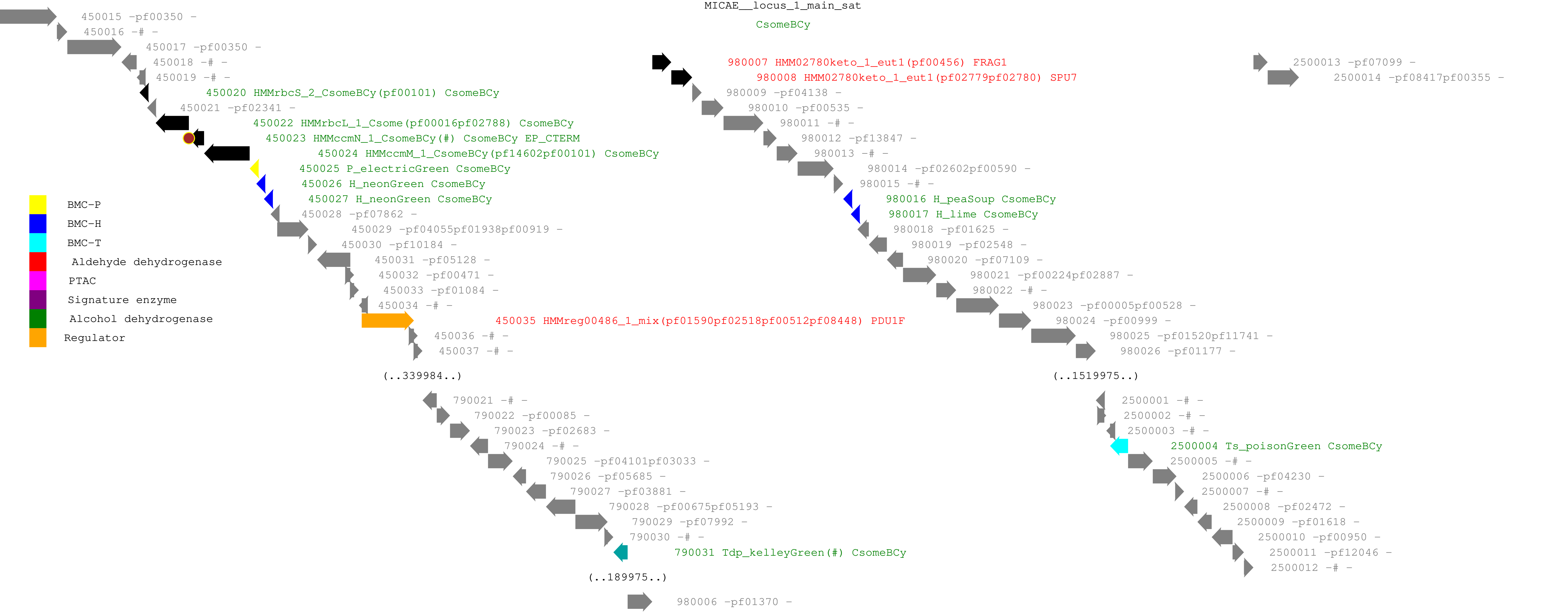

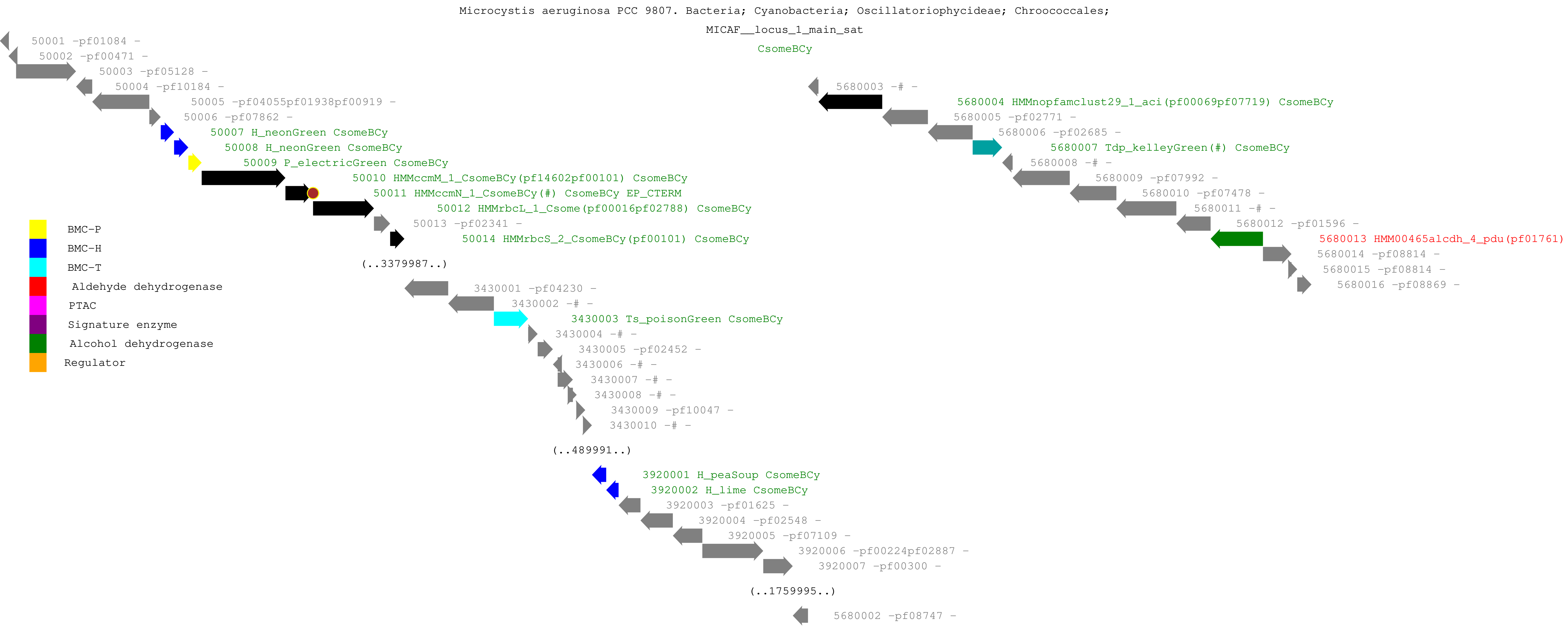
Microcystis aeruginosa PCC 9807. Bacteria; Cyanobacteria; Oscillatoriophycideae; Chroococcales;
MICAF__locus_1_main_sat
CsomeBCy
50001 -pf01084 -
50002 -pf00471 -
50003 -pf05128 -
50004 -pf10184 -
50005 -pf04055pf01938pf00919 -
50006 -pf07862 -
50007 H_neonGreen CsomeBCy
50008 H_neonGreen CsomeBCy
50009 P_electricGreen CsomeBCy
50010 HMMccmM_1_CsomeBCy(pf14602pf00101) CsomeBCy
50011 HMMccmN_1_CsomeBCy(#) CsomeBCy EP_CTERM
50012 HMMrbcL_1_Csome(pf00016pf02788) CsomeBCy
50013 -pf02341 -
50014 HMMrbcS_2_CsomeBCy(pf00101) CsomeBCy
(..3379987..)
3430001 -pf04230 -
3430002 -# -
3430003 Ts_poisonGreen CsomeBCy
3430004 -# -
3430005 -pf02452 -
3430006 -# -
3430007 -# -
3430008 -# -
3430009 -pf10047 -
3430010 -# -
(..489991..)
3920001 H_peaSoup CsomeBCy
3920002 H_lime CsomeBCy
3920003 -pf01625 -
3920004 -pf02548 -
3920005 -pf07109 -
3920006 -pf00224pf02887 -
3920007 -pf00300 -
(..1759995..)
5680002 -pf08747 -
5680003 -# -
5680004 HMMnopfamclust29_1_aci(pf00069pf07719) CsomeBCy
5680005 -pf02771 -
5680006 -pf02685 -
5680007 Tdp_kelleyGreen(#) CsomeBCy
5680008 -# -
5680009 -pf07992 -
5680010 -pf07478 -
5680011 -# -
5680012 -pf01596 -
5680013 HMM00465alcdh_4_pdu(pf01761)
5680014 -pf08814 -
5680015 -pf08814 -
5680016 -pf08869 -
BMC-P
BMC-H
BMC-T
Aldehyde dehydrogenase
PTAC
Signature enzyme
Alcohol dehydrogenase
Regulator

Microcystis aeruginosa PCC 9808. Bacteria; Cyanobacteria; Oscillatoriophycideae; Chroococcales;

MICAG__locus_1_main_sat

CsomeBCy

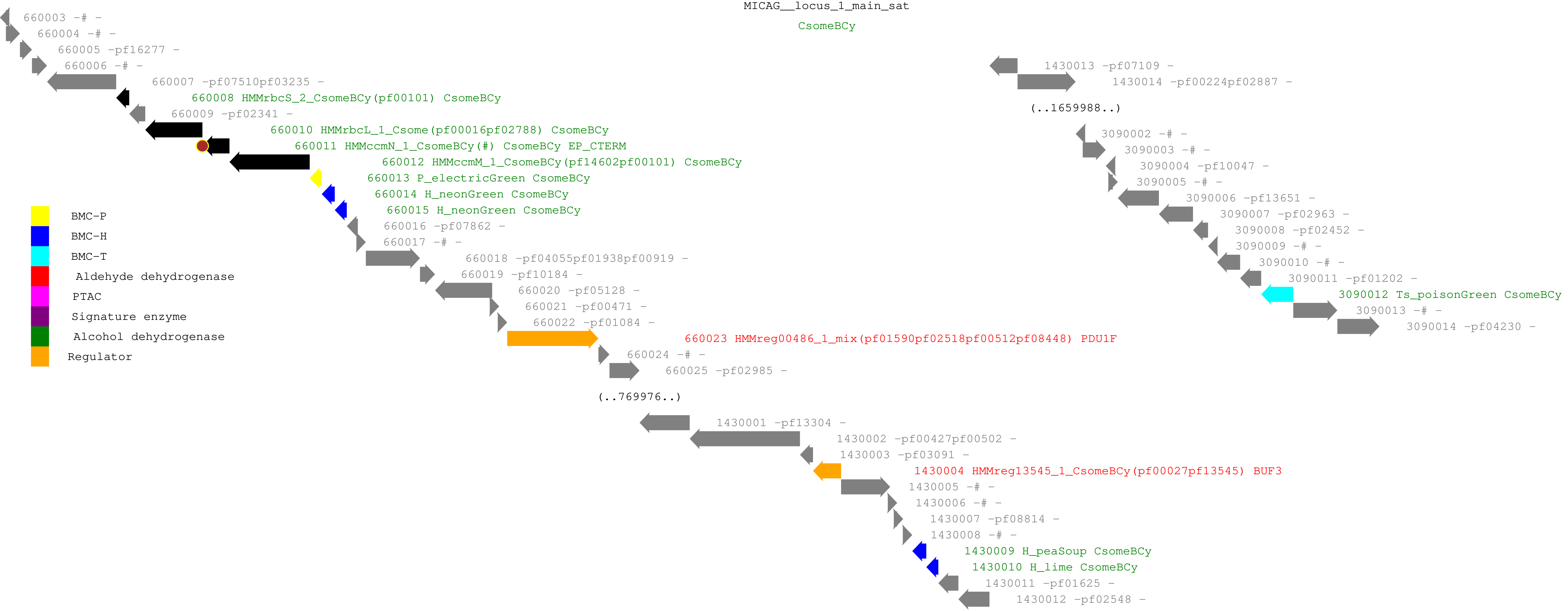

Microcystis aeruginosa PCC 9809. Bacteria; Cyanobacteria; Oscillatoriophycideae; Chroococcales;

MICAH__locus_1_main_sat

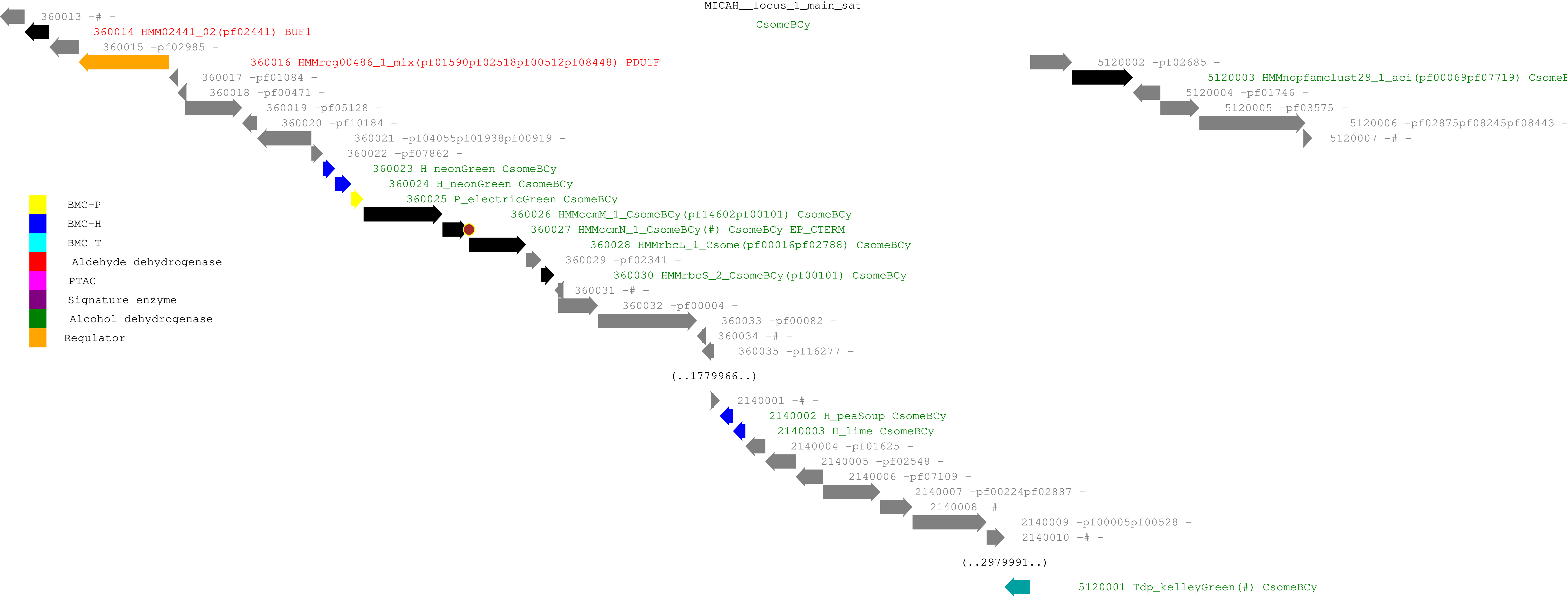

Microcystis sp. T1-4. Bacteria; Cyanobacteria; Oscillatoriophycideae; Chroococcales;

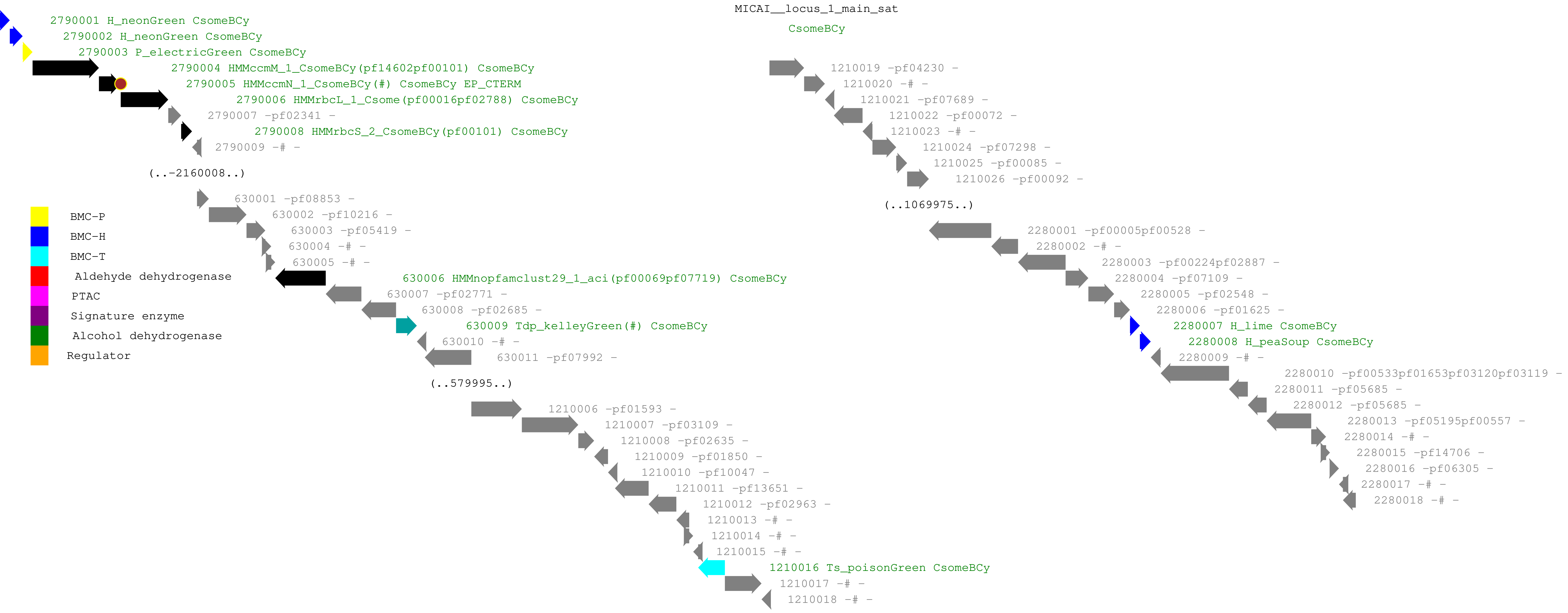

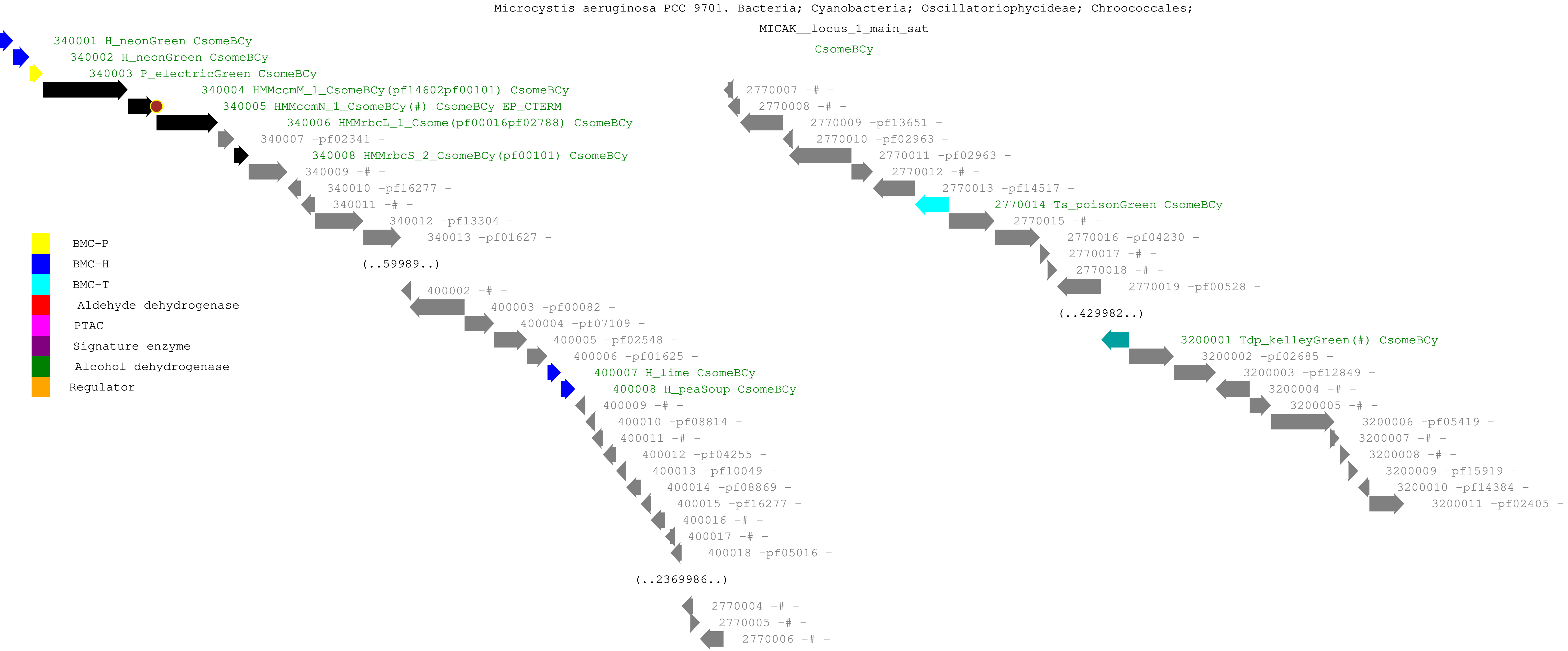
Microcystis aeruginosa PCC 9701. Bacteria; Cyanobacteria; Oscillatoriophycideae; Chroococcales;
MICAK__locus_1_main_sat
CsomeBCy
340001 H_neonGreen CsomeBCy
340002 H_neonGreen CsomeBCy
340003 P_electricGreen CsomeBCy
340004 HMMccmM_1_CsomeBCy(pf14602pf00101) CsomeBCy
340005 HMMccmN_1_CsomeBCy(#) CsomeBCy EP_CTERM
340006 HMMrbcL_1_Csome(pf00016pf02788) CsomeBCy
340007 -pf02341 -
340008 HMMrbcS_2_CsomeBCy(pf00101) CsomeBCy
340009 -# -
340010 -pf16277 -
340011 -# -
340012 -pf13304 -
340013 -pf01627 -
(..59989..)
400002 -# -
400003 -pf00082 -
400004 -pf07109 -
400005 -pf02548 -
400006 -pf01625 -
400007 H_lime CsomeBCy
400008 H_peaSoup CsomeBCy
400009 -# -
400010 -pf08814 -
400011 -# -
400012 -pf04255 -
400013 -pf10049 -
400014 -pf08869 -
400015 -pf16277 -
400016 -# -
400017 -# -
400018 -pf05016 -
(..2369986..)
2770004 -# -
2770005 -# -
2770006 -# -
2770007 -# -
2770008 -# -
2770009 -pf13651 -
2770010 -pf02963 -
2770011 -pf02963 -
2770012 -# -
2770013 -pf14517 -
2770014 Ts_poisonGreen CsomeBCy
2770015 -# -
2770016 -pf04230 -
2770017 -# -
2770018 -# -
2770019 -pf00528 -
(..429982..)
3200001 Tdp_kelleyGreen(#) CsomeBCy
3200002 -pf02685 -
3200003 -pf12849 -
3200004 -# -
3200005 -# -
3200006 -pf05419 -
3200007 -# -
3200008 -# -
3200009 -pf15919 -
3200010 -pf14384 -
3200011 -pf02405 -
BMC-P
BMC-H
BMC-T
Aldehyde dehydrogenase
PTAC
Signature enzyme
Alcohol dehydrogenase
Regulator

Microcystis aeruginosa PCC 9432. Bacteria; Cyanobacteria; Oscillatoriophycideae; Chroococcales;

MICCA__locus_1_main_sat

CsomeBCy

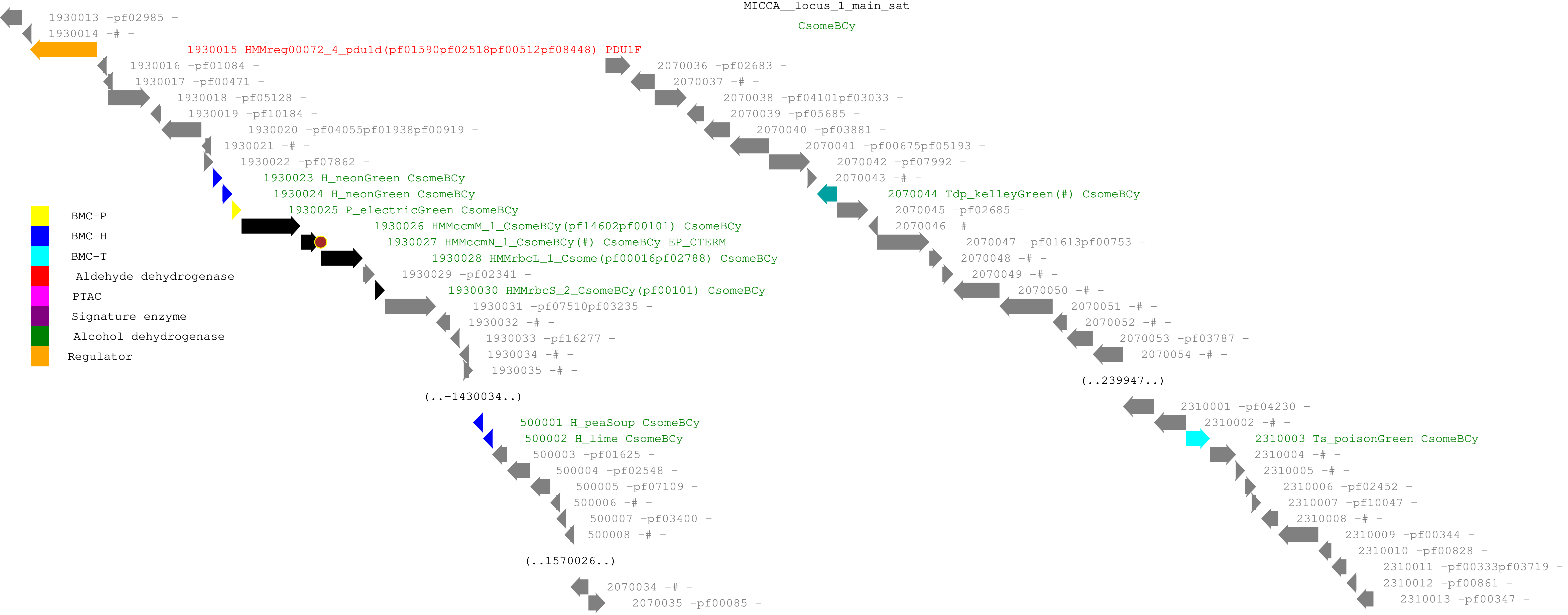

Microcoleus vaginatus FGP-2. Bacteria; Cyanobacteria; Oscillatoriophycideae; Oscillatoriales;

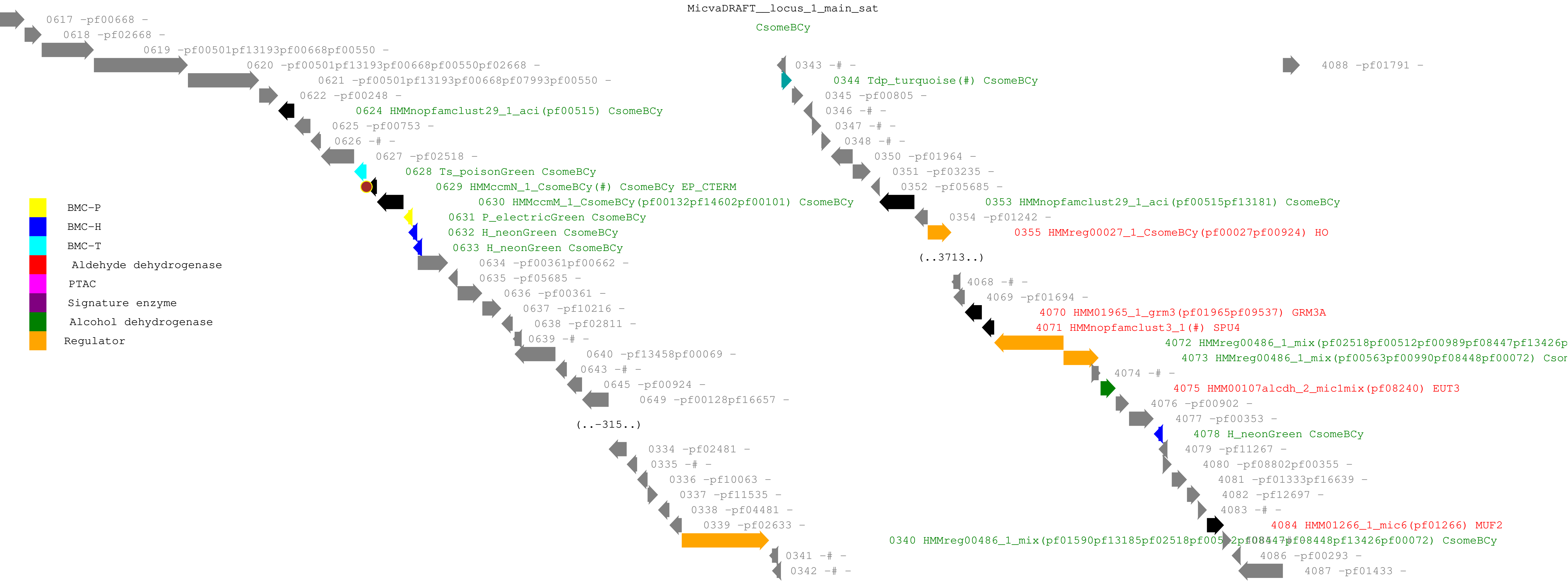

Microcystis aeruginosa NIES-2520. Bacteria; Cyanobacteria; Oscillatoriophycideae; Chroococcales;

MiTe__locus_1_main_sat

CsomeBCy

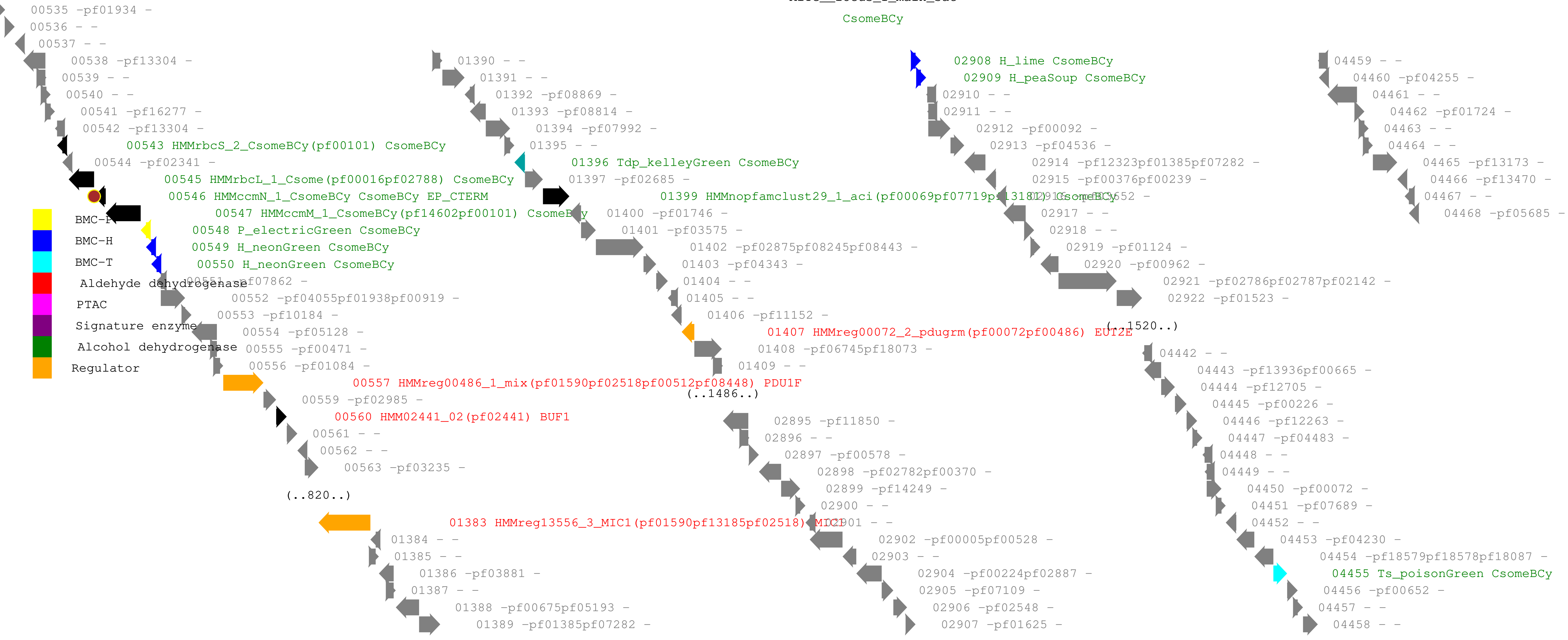

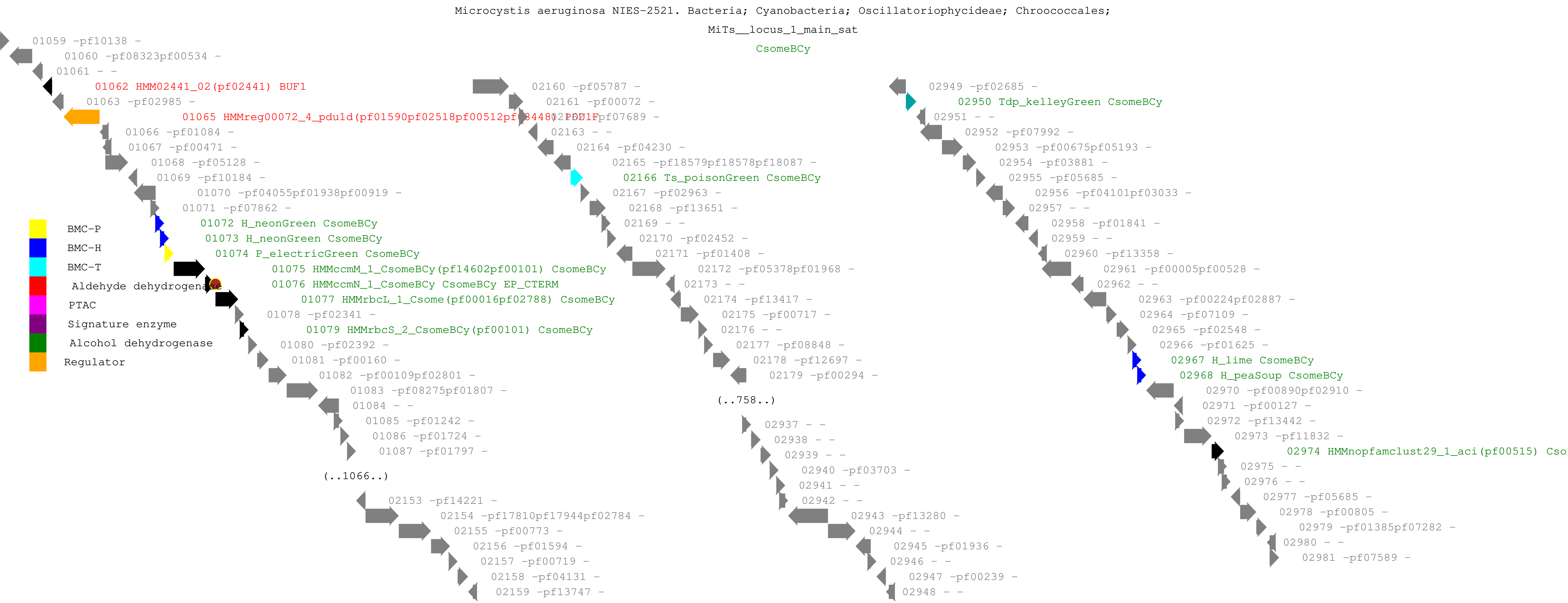

Microcystis aeruginosa NIES-2521. Bacteria; Cyanobacteria; Oscillatoriophycideae; Chroococcales;
MiTs__locus_1_main_sat
CsomeBCy
01059 -pf10138 -
01060 -pf08323pf00534 -
01061 - -
01062 HMM02441_02(pf02441) BUF1
01063 -pf02985 -
01065 HMMreg00072_4_pdu1d(pf01590pf02518pf00512pf
PDU1F
01066 -pf01084 -
01067 -pf00471 -
01068 -pf05128 -
01069 -pf10184 -
01070 -pf04055pf01938pf00919 -
01071 -pf07862 -
01072 H_neonGreen CsomeBCy
01073 H_neonGreen CsomeBCy
01074 P_electricGreen CsomeBCy
01075 HMMccmM_1_CsomeBCy(pf14602pf00101) CsomeBCy
01076 HMMccmN_1_CsomeBCy CsomeBCy EP_CTERM
01077 HMMrbcL_1_Csome(pf00016pf02788) CsomeBCy
01078 -pf02341 -
01079 HMMrbcS_2_CsomeBCy(pf00101) CsomeBCy
01080 -pf02392 -
01081 -pf00160 -
01082 -pf00109pf02801 -
01083 -pf08275pf01807 -
01084 - -
01085 -pf01242 -
01086 -pf01724 -
01087 -pf01797 -
(..1066..)
02153 -pf14221 -
02154 -pf17810pf17944pf02784 -
02155 -pf00773 -
02156 -pf01594 -
02157 -pf00719 -
02158 -pf04131 -
02159 -pf13747 -
02160 -pf05787 -
02161 -pf00072 -
02162 -pf07689 -
02163 - -
02164 -pf04230 -
02165 -pf18579pf18578pf18087 -
02166 Ts_poisonGreen CsomeBCy
02167 -pf02963 -
02168 -pf13651 -
02169 - -
02170 -pf02452 -
02171 -pf01408 -
02172 -pf05378pf01968 -
02173 - -
02174 -pf13417 -
02175 -pf00717 -
02176 - -
02177 -pf08848 -
02178 -pf12697 -
02179 -pf00294 -
(..758..)
02937 - -
02938 - -
02939 - -
02940 -pf03703 -
02941 - -
02942 - -
02943 -pf13280 -
02944 - -
02945 -pf01936 -
02946 - -
02947 -pf00239 -
02948 - -
02949 -pf02685 -
02950 Tdp_kelleyGreen CsomeBCy
02951 - -
02952 -pf07992 -
02953 -pf00675pf05193 -
02954 -pf03881 -
02955 -pf05685 -
02956 -pf04101pf03033 -
02957 - -
02958 -pf01841 -
02959 - -
02960 -pf13358 -
02961 -pf00005pf00528 -
02962 - -
02963 -pf00224pf02887 -
02964 -pf07109 -
02965 -pf02548 -
02966 -pf01625 -
02967 H_lime CsomeBCy
02968 H_peaSoup CsomeBCy
02970 -pf00890pf02910 -
02971 -pf00127 -
02972 -pf13442 -
02973 -pf11832 -
02974 HMMnopfamclust29_1_aci(pf00515) Cso
02975 - -
02976 - -
02977 -pf05685 -
02978 -pf00805 -
02979 -pf01385pf07282 -
02980 - -
02981 -pf07589 -
BMC-P
BMC-H
BMC-T
Aldehyde dehydrogenase
PTAC
Signature enzyme
Alcohol dehydrogenase
Regulator

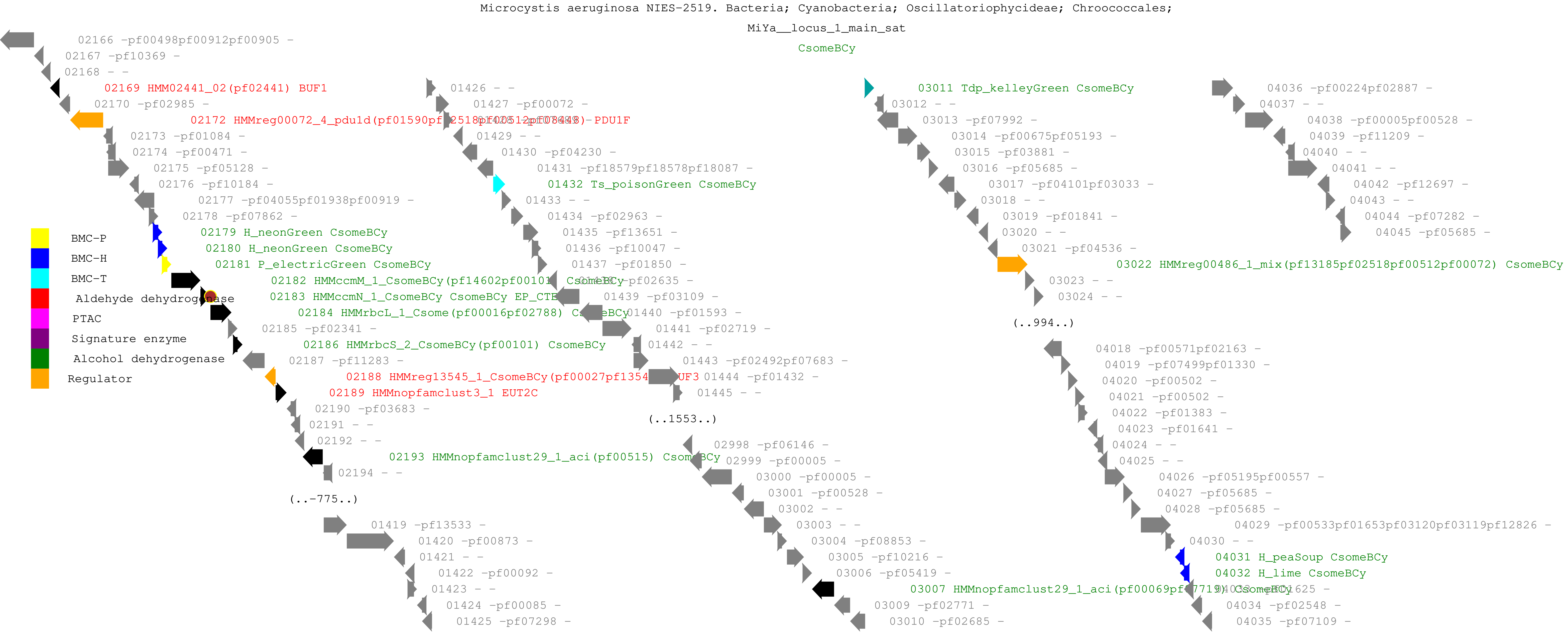

Microcystis aeruginosa NIES-2519. Bacteria; Cyanobacteria; Oscillatoriophycideae; Chroococcales;
MiYa__locus_1_main_sat
CsomeBCy
BMC-P
BMC-H
BMC-T
Aldehyde dehydrogenase
PTAC
Signature enzyme
Alcohol dehydrogenase
Regulator
02166 -pf00498pf00912pf00905 -
02167 -pf10369 -
02168 - -
02169 HMM02441_02(pf02441) BUF1
02170 -pf02985 -
02172 HMMreg00072_4_pdu1d(pf01590pf02518pf00512pf07848) PDU1F
02173 -pf01084 -
02174 -pf00471 -
02175 -pf05128 -
02176 -pf10184 -
02177 -pf04055pf01938pf00919 -
02178 -pf07862 -
02179 H_neonGreen CsomeBCy
02180 H_neonGreen CsomeBCy
02181 P_electricGreen CsomeBCy
02182 HMMccmM_1_CsomeBCy(pf14602pf00101) CsomeBCy
02183 HMMccmN_1_CsomeBCy CsomeBCy EP_CTE
02184 HMMrbcL_1_Csome(pf00016pf02788) CsomeBCy
02185 -pf02341 -
02186 HMMrbcS_2_CsomeBCy(pf00101) CsomeBCy
02187 -pf11283 -
02188 HMMreg13545_1_CsomeBCy(pf00027pf1354 UF3
02189 HMMnopfamclust3_1 EUT2C
02190 -pf03683 -
02191 - -
02192 - -
02193 HMMnopfamclust29_1_aci(pf00515) CsomeBCy
02194 - -
(..-775..)
01419 -pf13533 -
01420 -pf00873 -
01421 - -
01422 -pf00092 -
01423 - -
01424 -pf00085 -
01425 -pf07298 -
01426 - -
01427 -pf00072 -
01429 - -
01430 -pf04230 -
01431 -pf18579pf18578pf18087 -
01432 Ts_poisonGreen CsomeBCy
01433 - -
01434 -pf02963 -
01435 -pf13651 -
01436 -pf10047 -
01437 -pf01850 -
01438 -pf02635 -
01439 -pf03109 -
01440 -pf01593 -
01441 -pf02719 -
01442 - -
01443 -pf02492pf07683 -
01444 -pf01432 -
01445 - -
(..1553..)
02998 -pf06146 -
02999 -pf00005 -
03000 -pf00005 -
03001 -pf00528 -
03002 - -
03003 - -
03004 -pf08853 -
03005 -pf10216 -
03006 -pf05419 -
03007 HMMnopfamclust29_1_aci(pf00069pf07719) CsomeBCy
03009 -pf02771 -
03010 -pf02685 -
03011 Tdp_kelleyGreen CsomeBCy
03012 - -
03013 -pf07992 -
03014 -pf00675pf05193 -
03015 -pf03881 -
03016 -pf05685 -
03017 -pf04101pf03033 -
03018 - -
03019 -pf01841 -
03020 - -
03021 -pf04536 -
03022 HMMreg00486_1_mix(pf13185pf02518pf00512pf00072) CsomeBCy
03023 - -
03024 - -
(..994..)
04018 -pf00571pf02163 -
04019 -pf07499pf01330 -
04020 -pf00502 -
04021 -pf00502 -
04022 -pf01383 -
04023 -pf01641 -
04024 - -
04025 - -
04026 -pf05195pf00557 -
04027 -pf05685 -
04028 -pf05685 -
04029 -pf00533pf01653pf03120pf03119pf12826 -
04030 - -
04031 H_peaSoup CsomeBCy
04032 H_lime CsomeBCy
04033 -pf01625 -
04034 -pf02548 -
04035 -pf07109 -
04036 -pf00224pf02887 -
04037 - -
04038 -pf00005pf00528 -
04039 -pf11209 -
04040 - -
04041 - -
04042 -pf12697 -
04043 - -
04044 -pf07282 -
04045 -pf05685 -

Microcystis aeruginosa Sj. Bacteria; Cyanobacteria; Oscillatoriophycideae; Chroococcales;

MSj__locus_1_main_sat

CsomeBCy

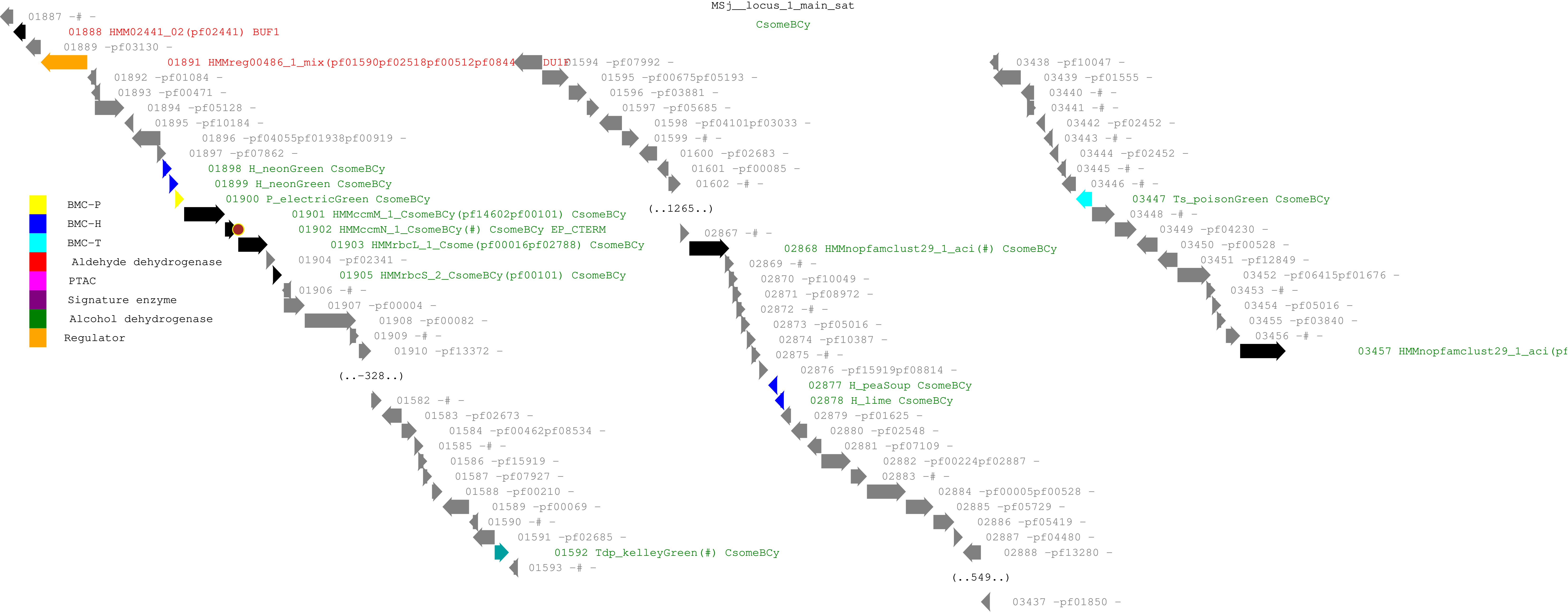

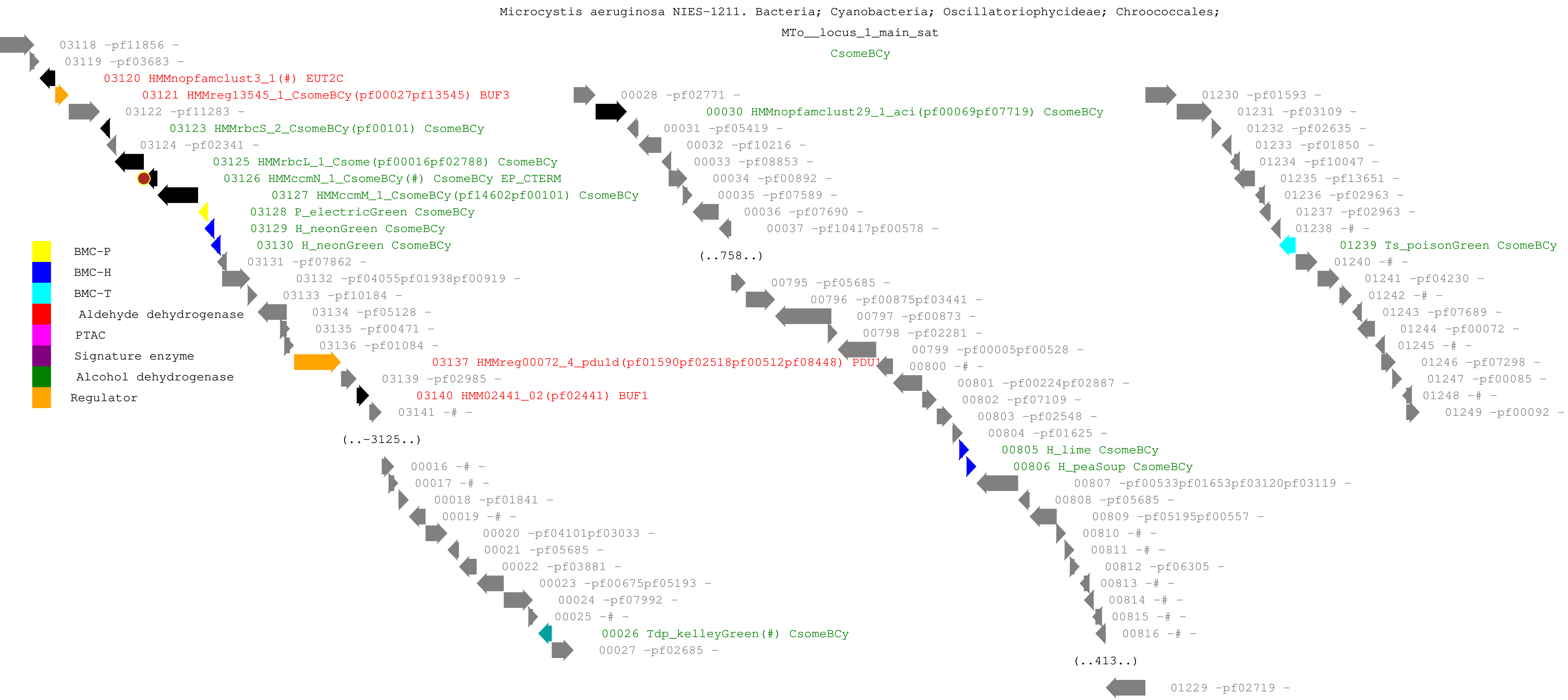
Microcystis aeruginosa NIES-1211. Bacteria; Cyanobacteria; Oscillatoriophycideae; Chroococcales;
MTo__locus_1_main_sat
CsomeBCy
03118 -pf11856 -
03119 -pf03683 -
03120 HMMnopfamclust3_1(#) EUT2C
03121 HMMreg13545_1_CsomeBCy(pf00027pf13545) BUF3
03122 -pf11283 -
03123 HMMrbcS_2_CsomeBCy(pf00101) CsomeBCy
03124 -pf02341 -
03125 HMMrbcL_1_Csome(pf00016pf02788) CsomeBCy
03126 HMMccmN_1_CsomeBCy(#) CsomeBCy EP_CTERM
03127 HMMccmM_1_CsomeBCy(pf14602pf00101) CsomeBCy
03128 P_electricGreen CsomeBCy
03129 H_neonGreen CsomeBCy
03130 H_neonGreen CsomeBCy
03131 -pf07862 -
03132 -pf04055pf01938pf00919 -
03133 -pf10184 -
03134 -pf05128 -
03135 -pf00471 -
03136 -pf01084 -
03137 HMMreg00072_4_pduld(pf01590pf02518pf00512pf08448) PDU1
03139 -pf02985 -
03140 HMM02441_02(pf02441) BUF1
03141 -# -
(..-3125..)
00016 -# -
00017 -# -
00018 -pf01841 -
00019 -# -
00020 -pf04101pf03033 -
00021 -pf05685 -
00022 -pf03881 -
00023 -pf00675pf05193 -
00024 -pf07992 -
00025 -# -
00026 Tdp_kelleyGreen(#) CsomeBCy
00027 -pf02685 -
BMC-P
BMC-H
BMC-T
Aldehyde dehydrogenase
PTAC
Signature enzyme
Alcohol dehydrogenase
Regulator
00028 -pf02771 -
00030 HMMnopfamclust29_1_aci(pf00069pf07719) CsomeBCy
00031 -pf05419 -
00032 -pf10216 -
00033 -pf08853 -
00034 -pf00892 -
00035 -pf07589 -
00036 -pf07690 -
00037 -pf10417pf00578 -
(..758..)
00795 -pf05685 -
00796 -pf00875pf03441 -
00797 -pf00873 -
00798 -pf02281 -
00799 -pf00005pf00528 -
00800 -# -
00801 -pf00224pf02887 -
00802 -pf07109 -
00803 -pf02548 -
00804 -pf01625 -
00805 H_lime CsomeBCy
00806 H_peaSoup CsomeBCy
00807 -pf00533pf01653pf03120pf03119 -
00808 -pf05685 -
00809 -pf05195pf00557 -
00810 -# -
00811 -# -
00812 -pf06305 -
00813 -# -
00814 -# -
00815 -# -
00816 -# -
(..413..)
01229 -pf02719 -
01230 -pf01593 -
01231 -pf03109 -
01232 -pf02635 -
01233 -pf01850 -
01234 -pf10047 -
01235 -pf13651 -
01236 -pf02963 -
01237 -pf02963 -
01238 -# -
01239 Ts_poisonGreen CsomeBCy
01240 -# -
01241 -pf04230 -
01242 -# -
01243 -pf07689 -
01244 -pf00072 -
01245 -# -
01246 -pf07298 -
01247 -pf00085 -
01248 -# -
01249 -pf00092 -

Microcystis viridis NIES-102. Bacteria; Cyanobacteria; Oscillatoriophycideae; Chroococcales;

myaer102__locus_1_main_sat

CsomeBCy

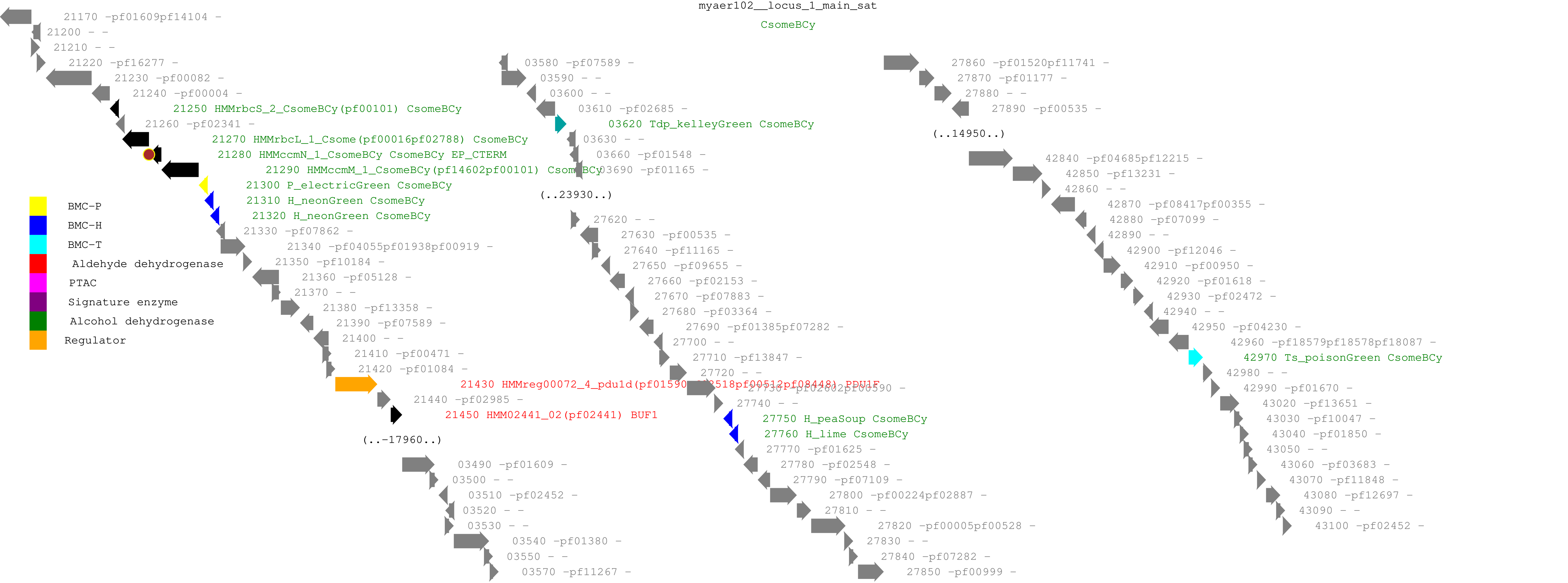

Microcystis aeruginosa NIES-87. Bacteria; Cyanobacteria; Oscillatoriophycideae; Chroococcales;

myaer87__locus_1_main_sat

CsomeBCy

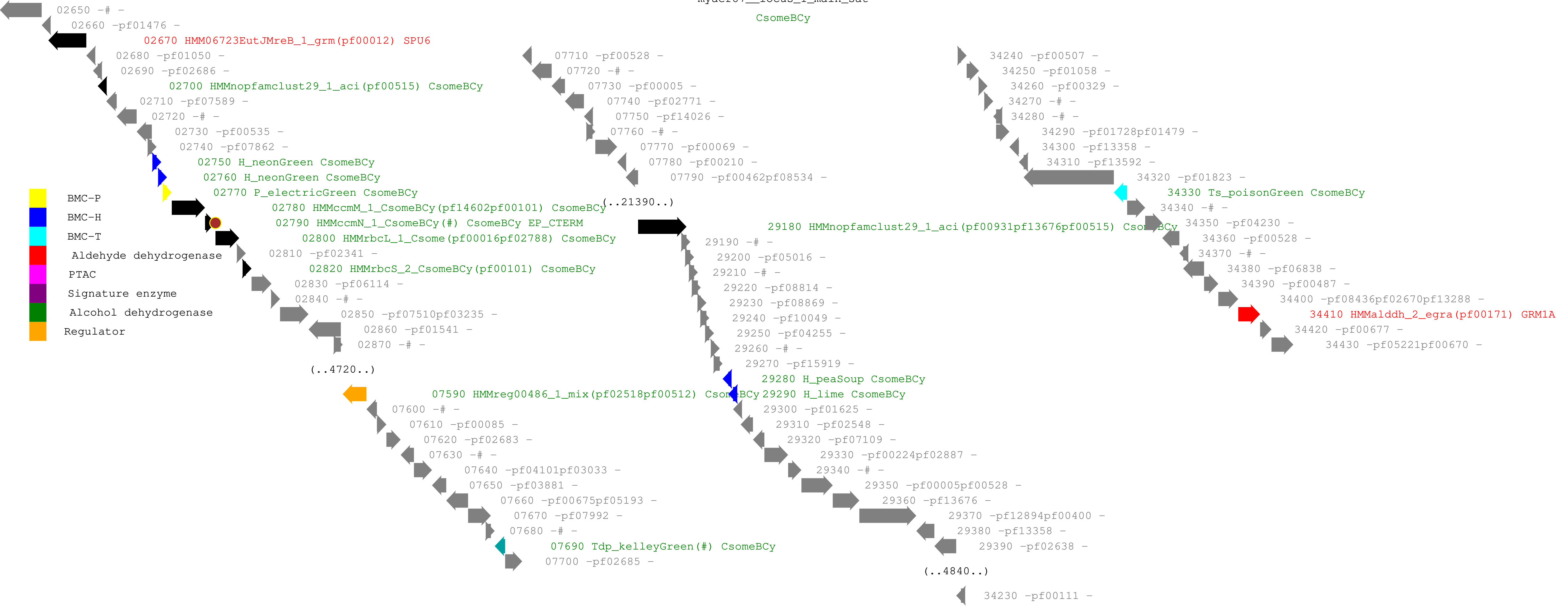

Microcystis aeruginosa NIES-2549. Bacteria; Cyanobacteria; Oscillatoriophycideae; Chroococcales;

MYAER__locus_1_main_sat

CsomeBCy

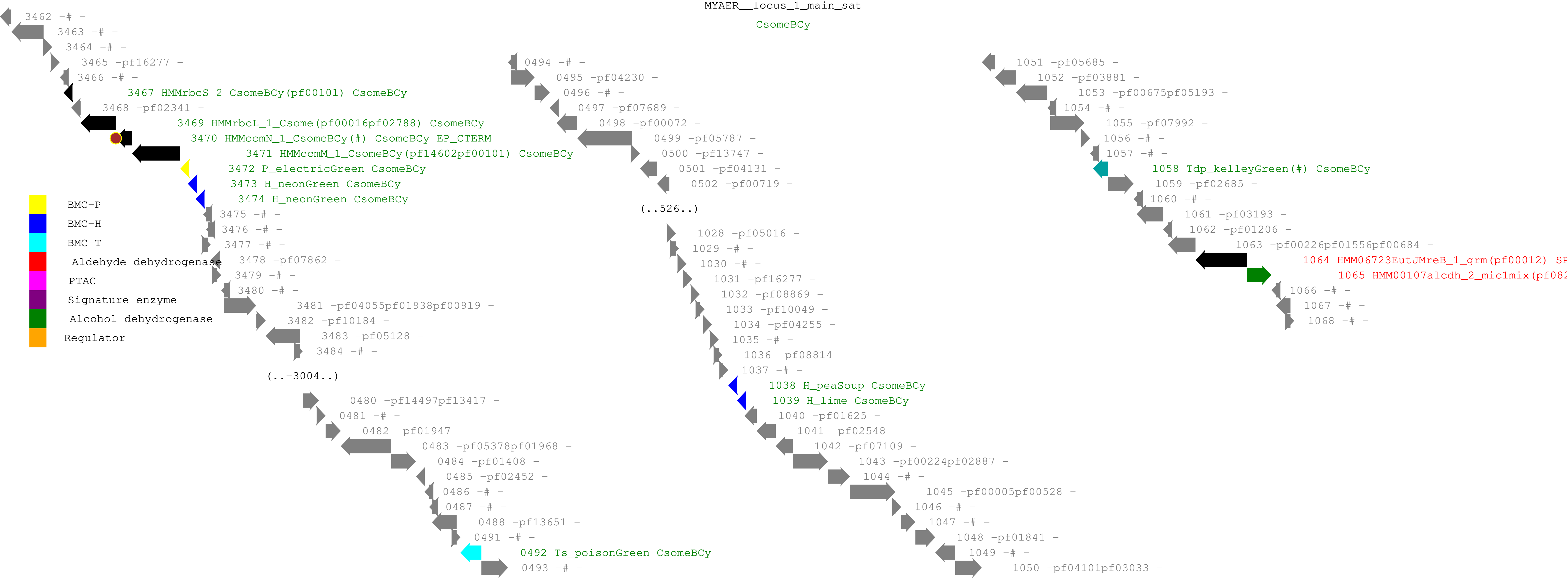

Microcystis sp. 0824. Bacteria; Cyanobacteria; Oscillatoriophycideae; Chroococcales;

N0824__locus_1_main_sat

CsomeBCy

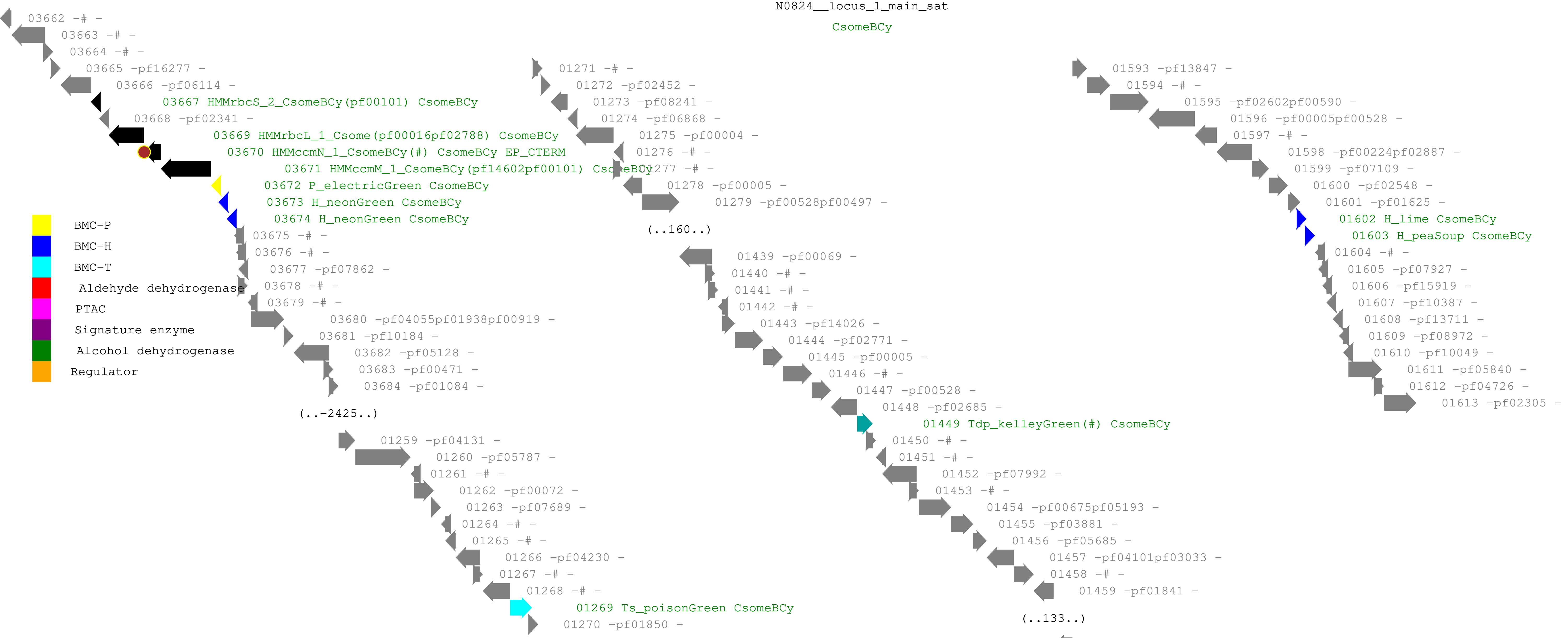

Microcystis aeruginosa NIES-44. Bacteria; Cyanobacteria; Oscillatoriophycideae; Chroococcales;

N44__locus_1_main_sat

CsomeBCy

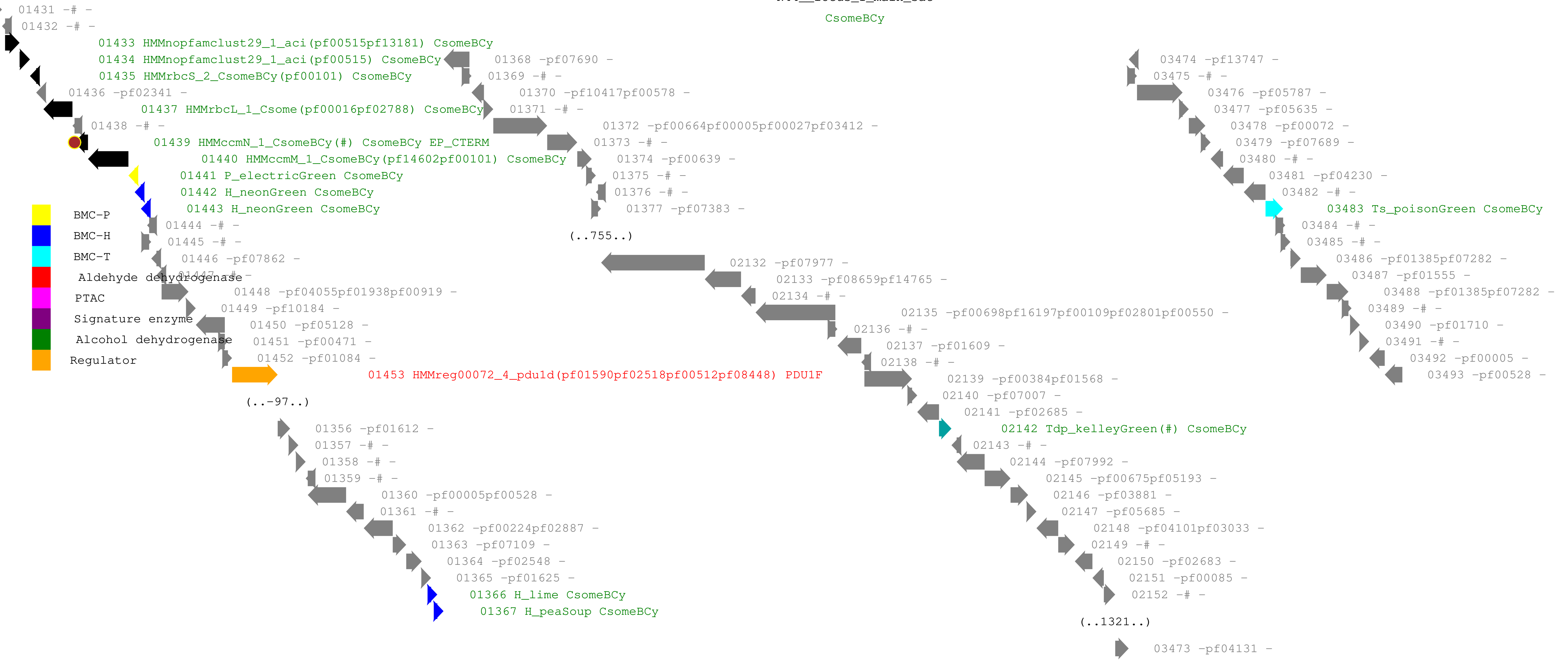

Leptolyngbya sp. Heron Island J. Bacteria; Cyanobacteria; Synechococcales; Leptolyngbyaceae;

N836__locus_1_main_sat

CsomeBCy

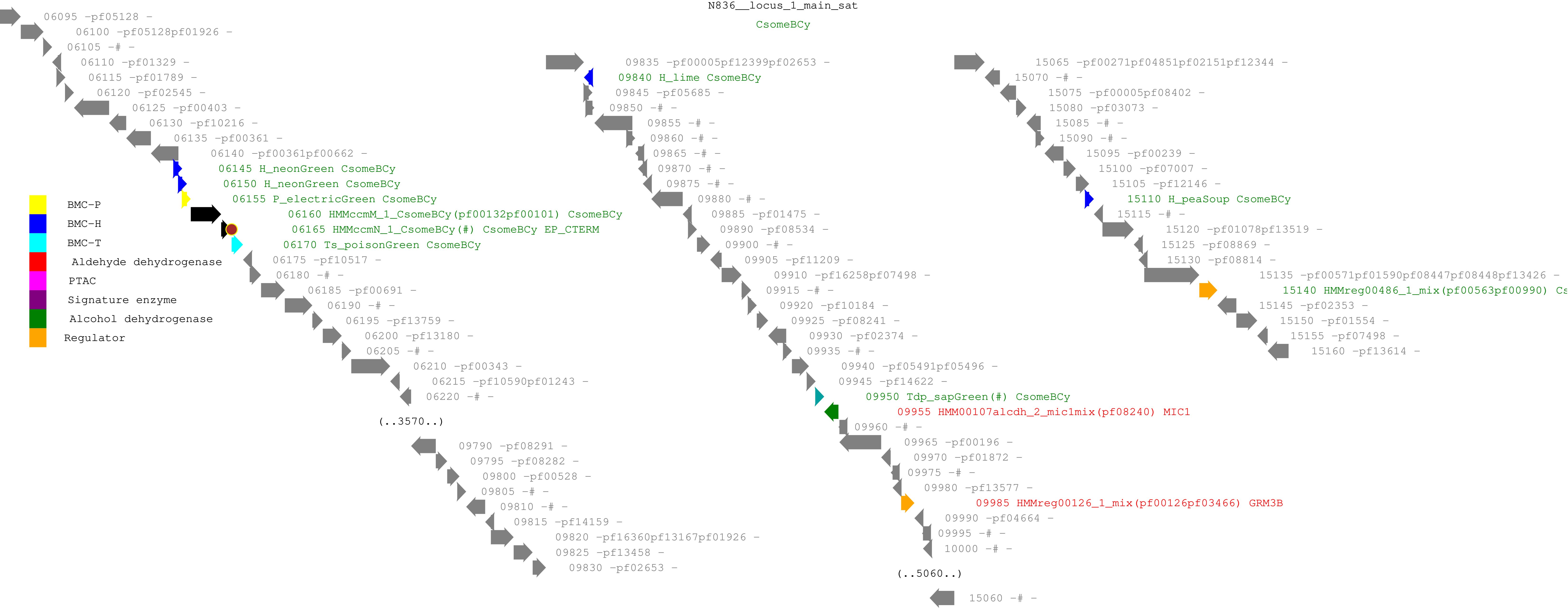

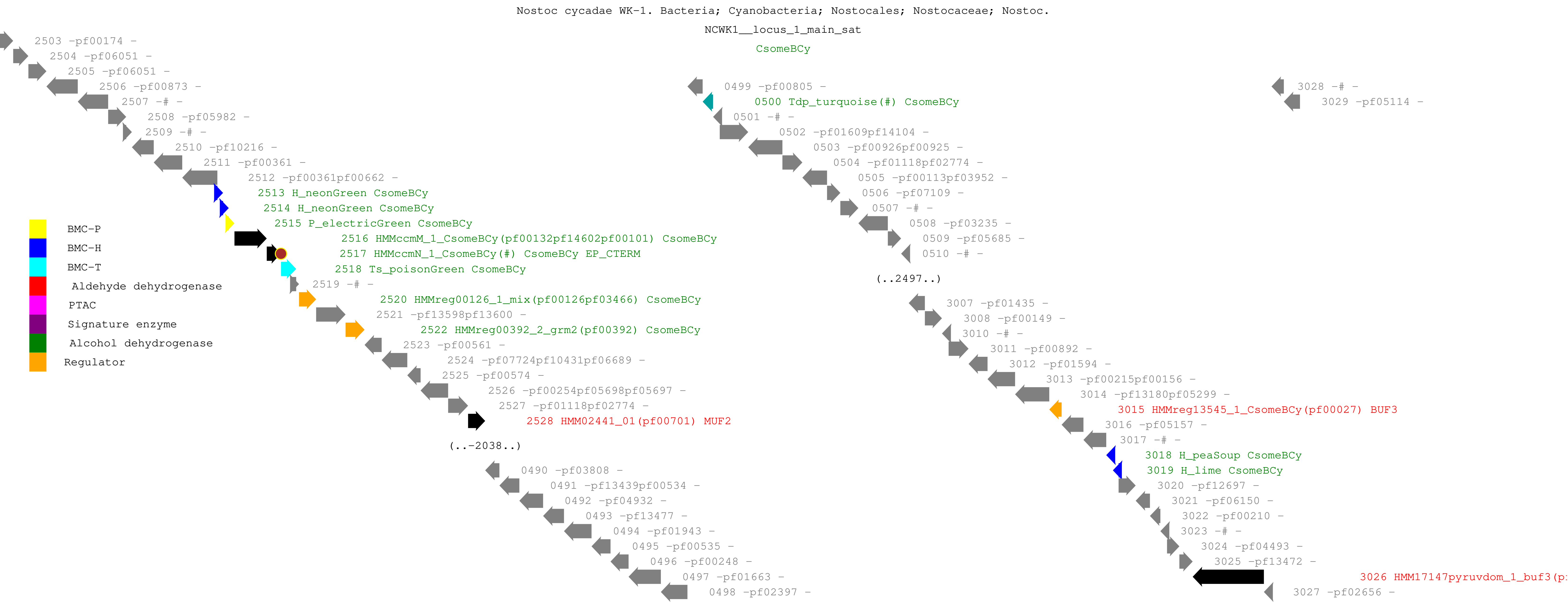
Nostoc cycadae WK-1. Bacteria; Cyanobacteria; Nostocales; Nostocaceae; Nostoc.
NCWK1__locus_1_main_sat
CsomeBCy
2503 -pf00174 -
2504 -pf06051 -
2505 -pf06051 -
2506 -pf00873 -
2507 -# -
2508 -pf05982 -
2509 -# -
2510 -pf10216 -
2511 -pf00361 -
2512 -pf00361pf00662 -
2513 H_neonGreen CsomeBCy
2514 H_neonGreen CsomeBCy
2515 P_electricGreen CsomeBCy
2516 HMMccmM_1_CsomeBCy(pf00132pf14602pf00101) CsomeBCy
2517 HMMccmN_1_CsomeBCy(#) CsomeBCy EP_CTERM
2518 Ts_poisonGreen CsomeBCy
2519 -# -
2520 HMMreg00126_1_mix(pf00126pf03466) CsomeBCy
2521 -pf13598pf13600 -
2522 HMMreg00392_2_grm2(pf00392) CsomeBCy
2523 -pf00561 -
2524 -pf07724pf10431pf06689 -
2525 -pf00574 -
2526 -pf00254pf05698pf05697 -
2527 -pf01118pf02774 -
2528 HMM02441_01(pf00701) MUF2
(..-2038..)
0490 -pf03808 -
0491 -pf13439pf00534 -
0492 -pf04932 -
0493 -pf13477 -
0494 -pf01943 -
0495 -pf00535 -
0496 -pf00248 -
0497 -pf01663 -
0498 -pf02397 -
BMC-P
BMC-H
BMC-T
Aldehyde dehydrogenase
PTAC
Signature enzyme
Alcohol dehydrogenase
Regulator
0499 -pf00805 -
0500 Tdp_turquoise(#) CsomeBCy
0501 -# -
0502 -pf01609pf14104 -
0503 -pf00926pf00925 -
0504 -pf01118pf02774 -
0505 -pf00113pf03952 -
0506 -pf07109 -
0507 -# -
0508 -pf03235 -
0509 -pf05685 -
0510 -# -
(..2497..)
3007 -pf01435 -
3008 -pf00149 -
3010 -# -
3011 -pf00892 -
3012 -pf01594 -
3013 -pf00215pf00156 -
3014 -pf13180pf05299 -
3015 HMMreg13545_1_CsomeBCy(pf00027) BUF3
3016 -pf05157 -
3017 -# -
3018 H_peaSoup CsomeBCy
3019 H_lime CsomeBCy
3020 -pf12697 -
3021 -pf06150 -
3022 -pf00210 -
3023 -# -
3024 -pf04493 -
3025 -pf13472 -
3026 HMM17147pyruvdom_1_buf3(p
3027 -pf02656 -
3028 -# -
3029 -pf05114 -

Chroogloeocystis siderophila 5.2 s.c.1. Bacteria; Cyanobacteria; Oscillatoriophycideae; Chroococcales;

NIES1031__locus_1_main_sat

CsomeBCy

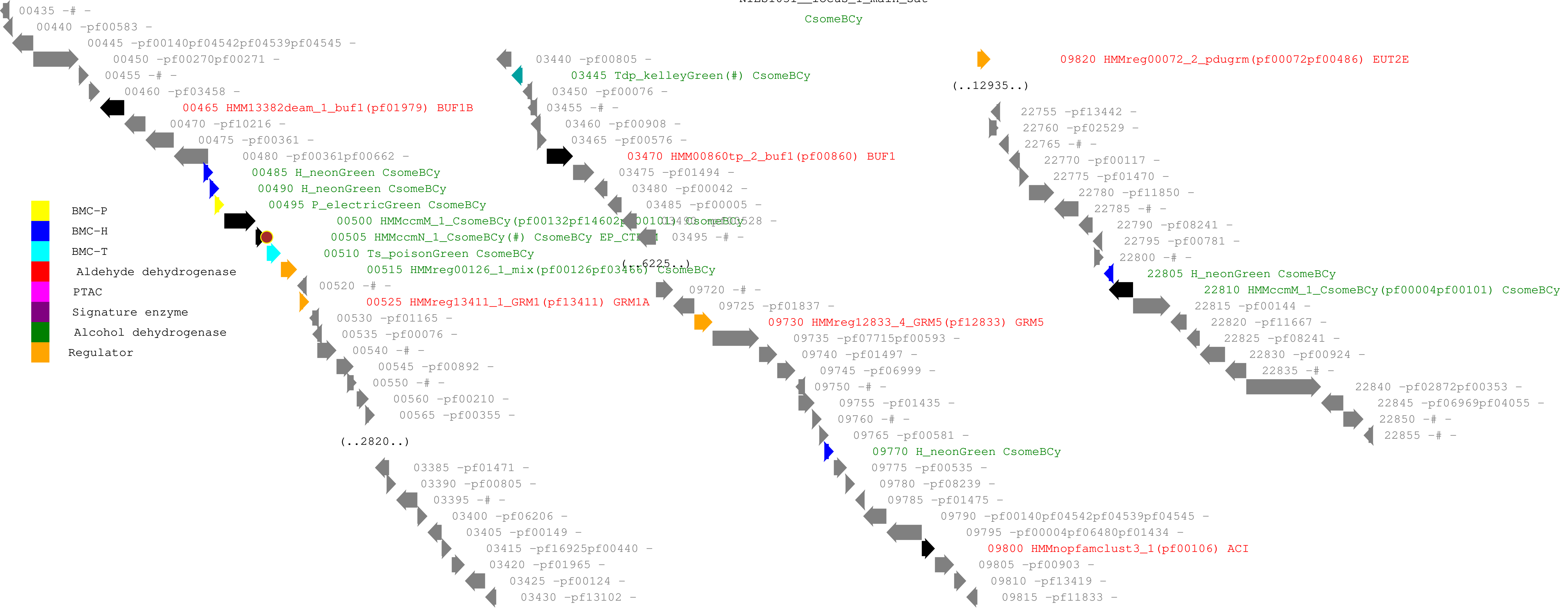

Anabaena cylindrica (strain ATCC 27899 / PCC 7122). Bacteria; Cyanobacteria; Nostocales; Nostocaceae; Anabaena.

NIES19__locus_1_main_sat

CsomeBCy

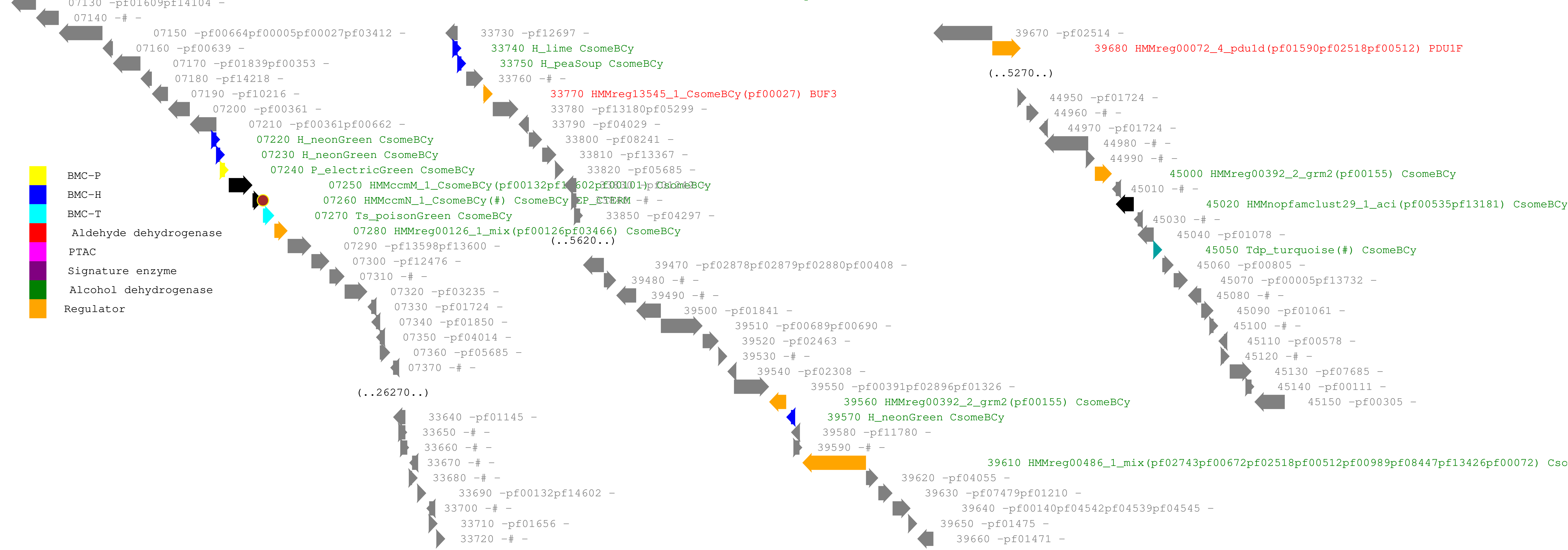

Planktothrix agardhii NIES-204. Bacteria; Cyanobacteria; Oscillatoriophycideae; Oscillatoriales;

NIES204__locus_1_main_sat

CsomeBCy

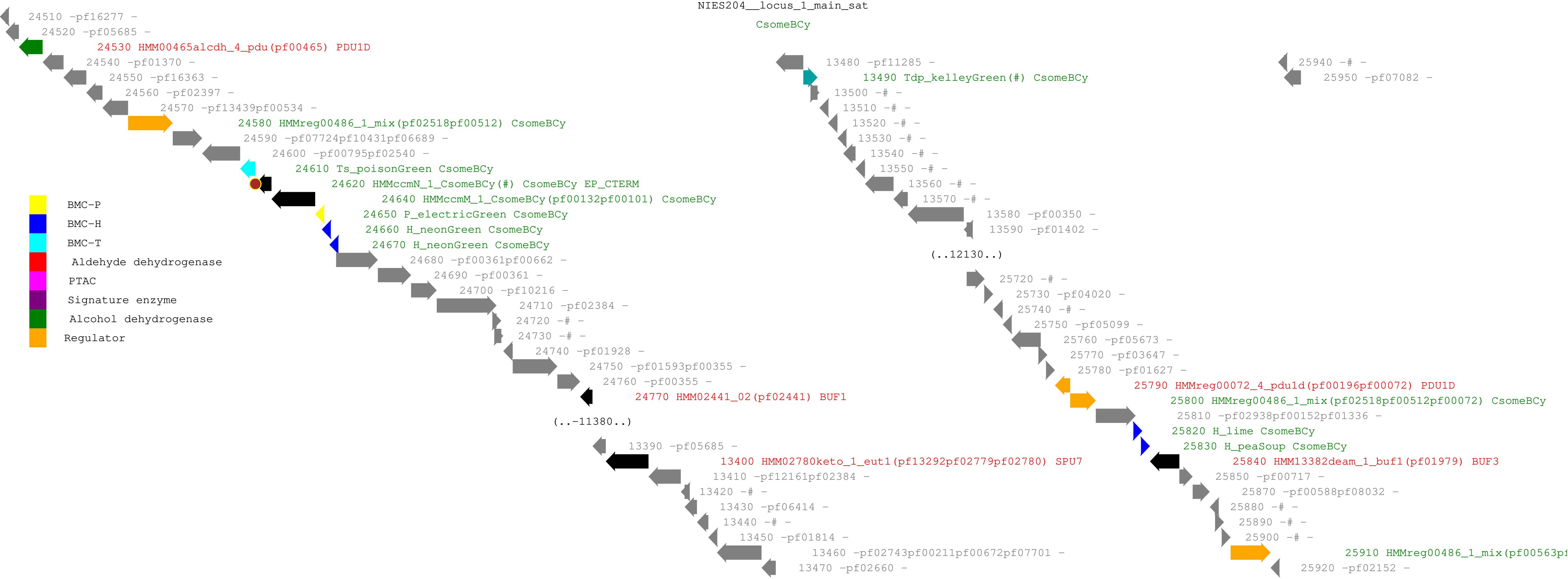

Limnothrix rosea IAM M-220. Bacteria; Cyanobacteria; Synechococcales; Pseudanabaenaceae;

NIES208__locus_1_main_sat

CsomeBCy

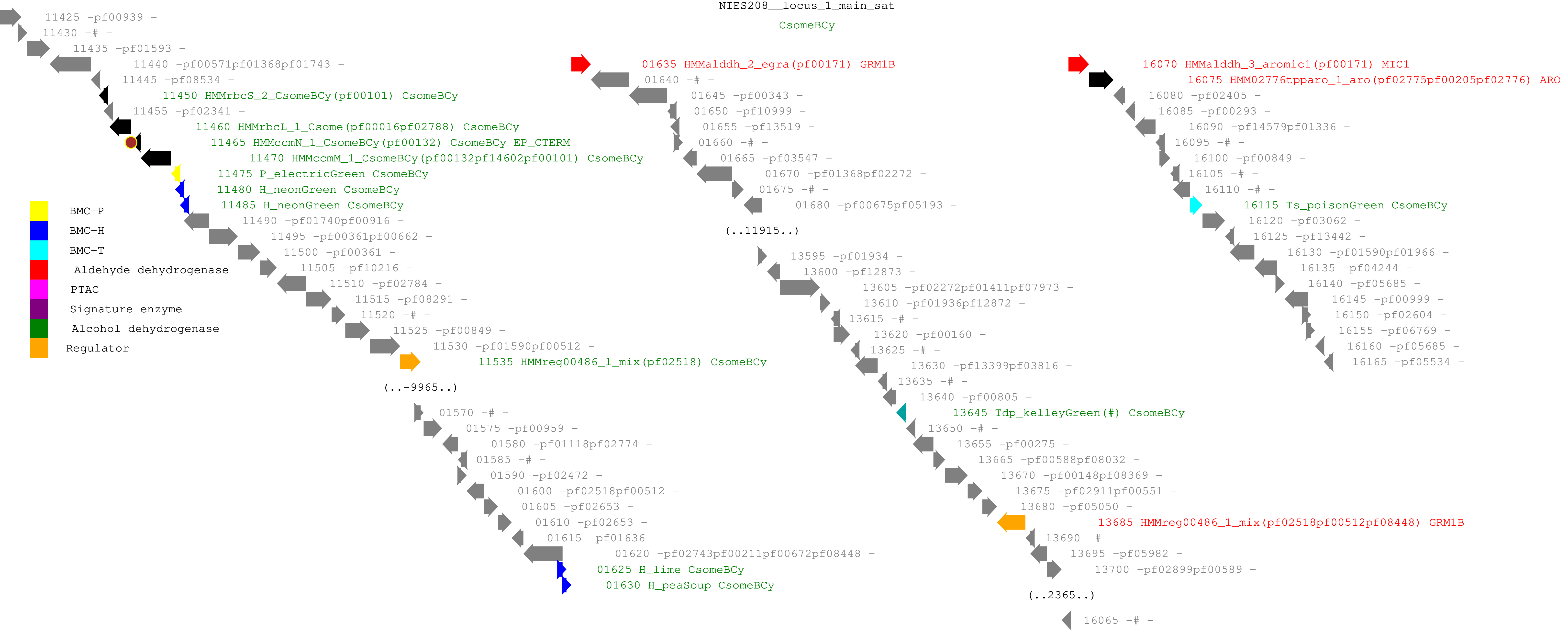

Calothrix sp. NIES-2098. Bacteria; Cyanobacteria; Nostocales; Rivulariaceae; Calothrix.

NIES2098__locus_1_main_sat

CsomeBCy

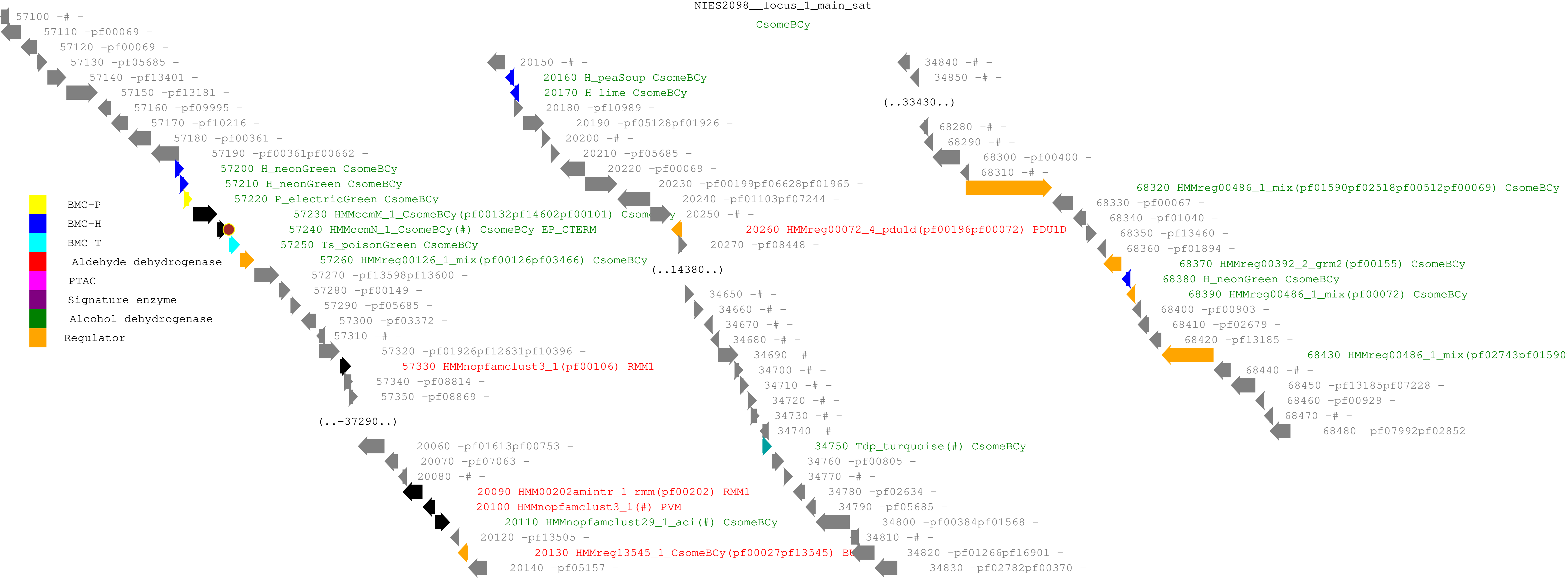

Calothrix sp. NIES-2100. Bacteria; Cyanobacteria; Nostocales; Rivulariaceae; Calothrix.

NIES2100__locus_1_main_sat

CsomeBCy

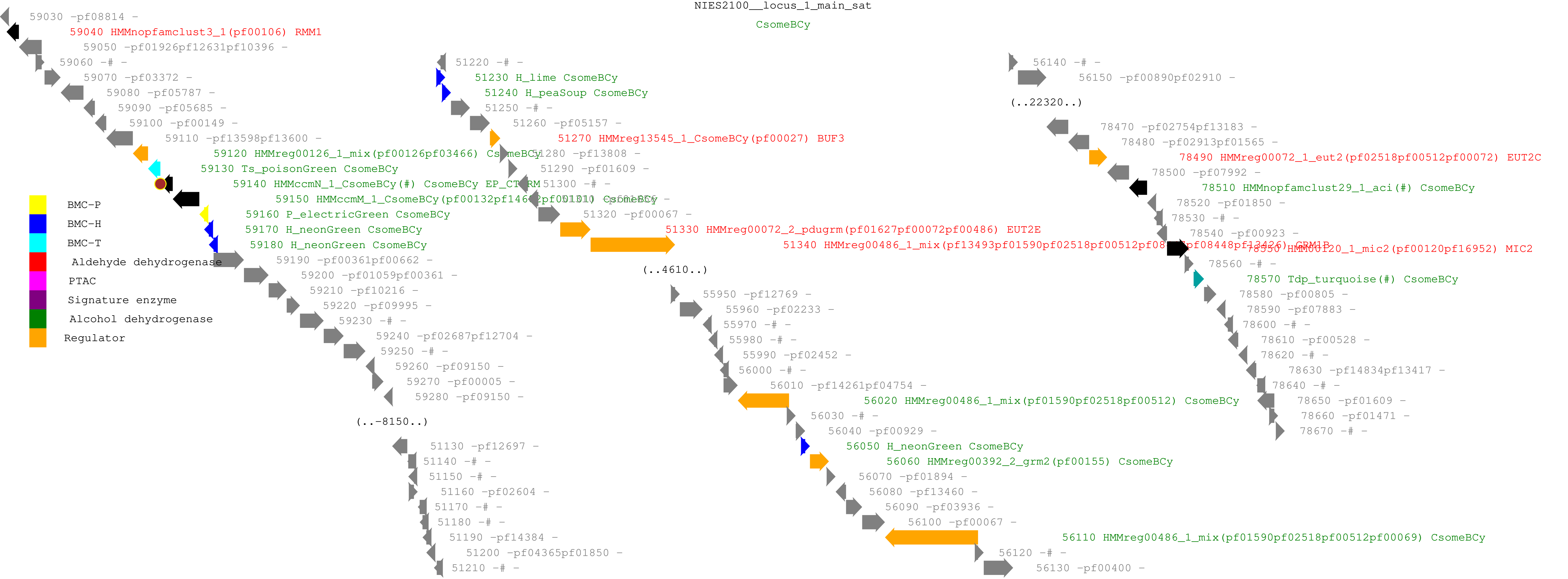

Calothrix sp. HK-06. Bacteria; Cyanobacteria; Nostocales; Rivulariaceae; Calothrix.

NIES2101__locus_1_main_sat

CsomeBCy

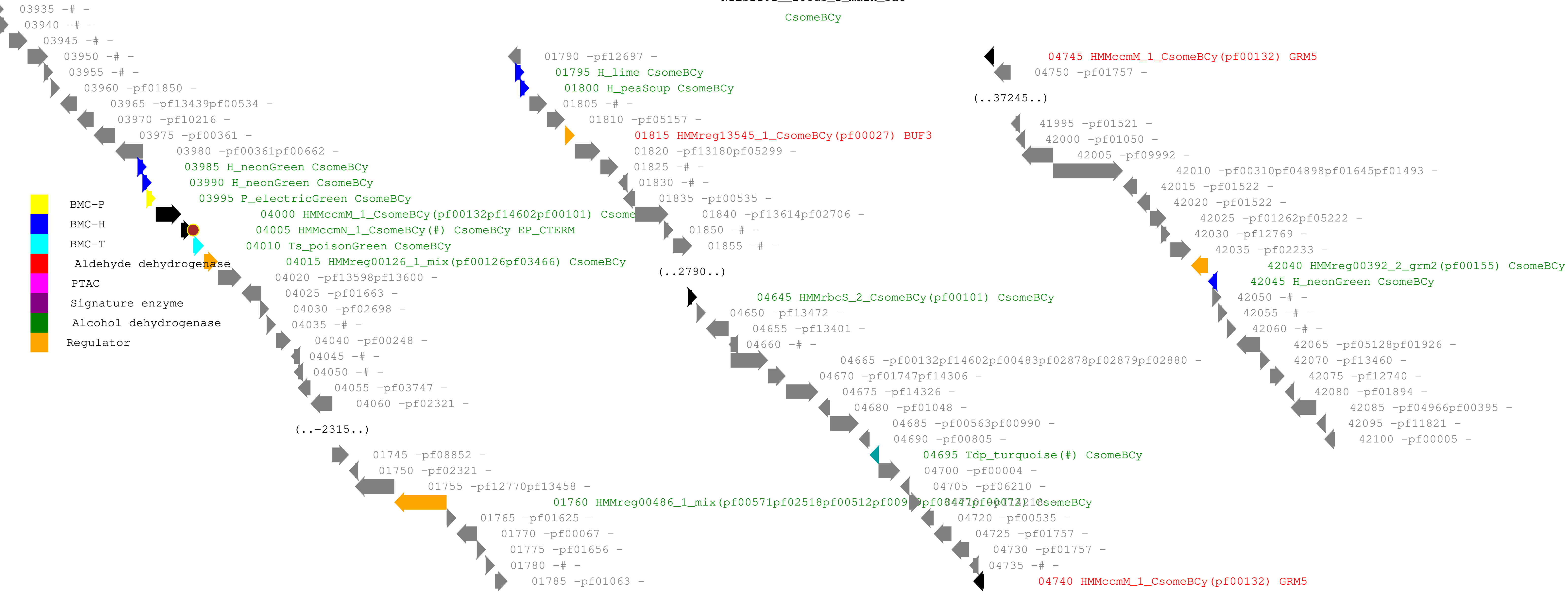

Leptolyngbya sp. NIES-2104. Bacteria; Cyanobacteria; Synechococcales; Leptolyngbyaceae;
NIES2104__locus_1_main_sat
CsomeBCy

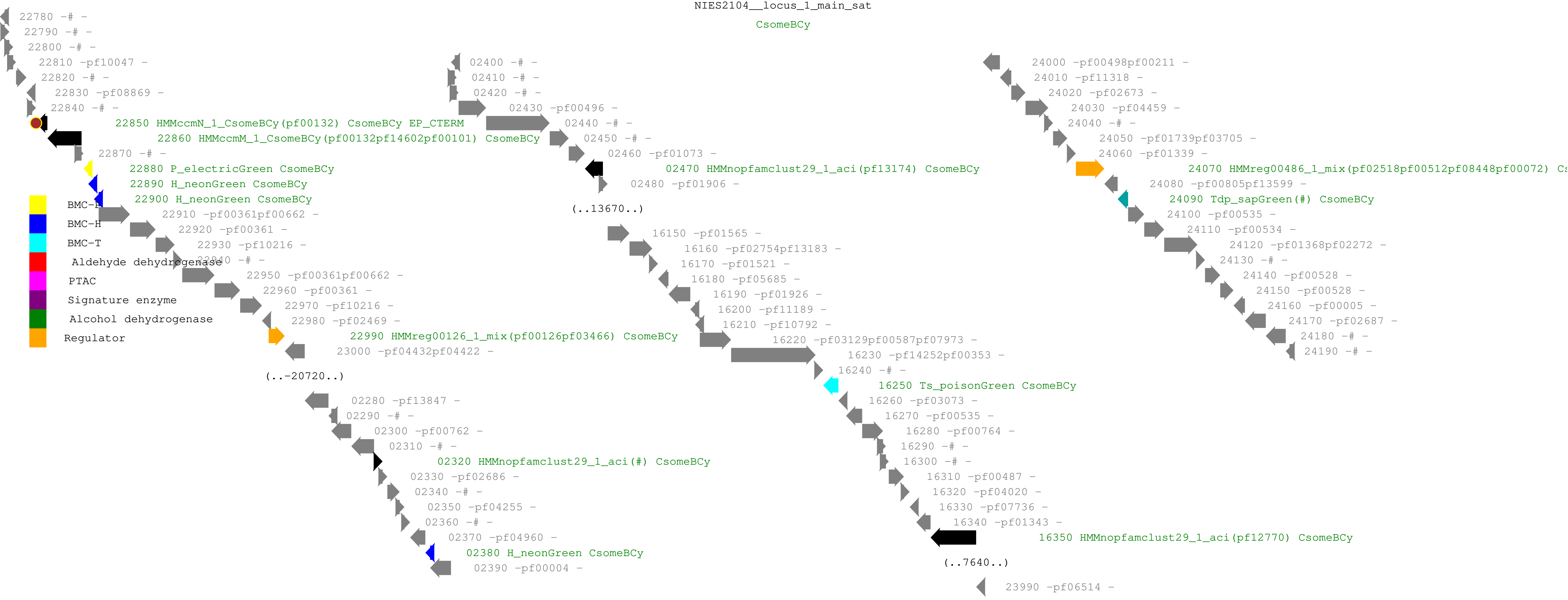

Nostoc carneum NIES-2107. Bacteria; Cyanobacteria; Nostocales; Nostocaceae; Nostoc.

NIES2107__locus_1_main_sat

CsomeBCy

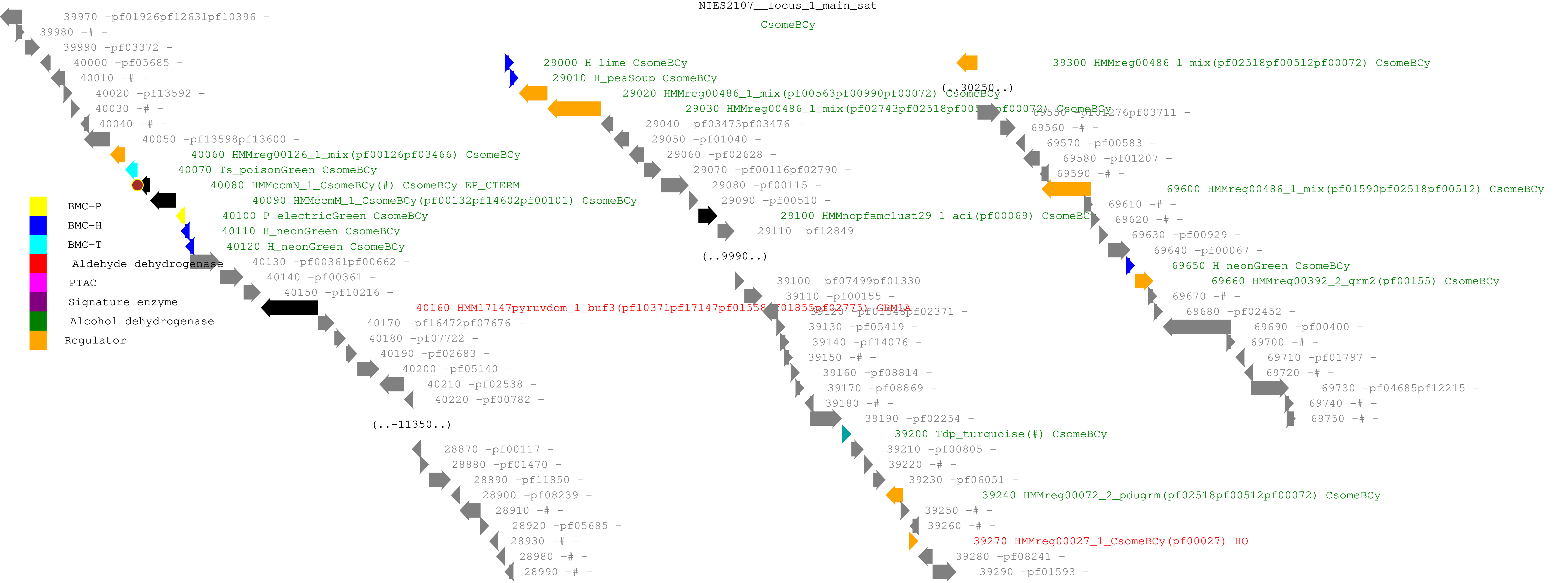

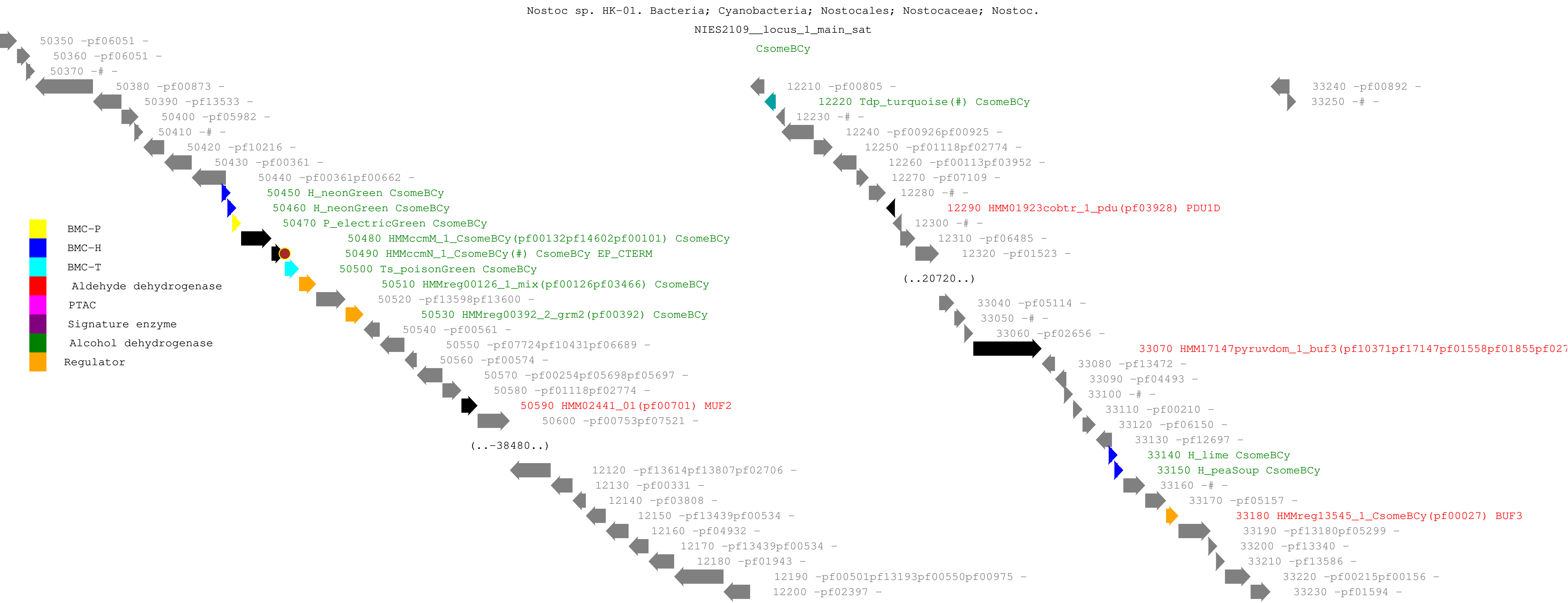
Nostoc sp. HK-01. Bacteria; Cyanobacteria; Nostocales; Nostocaceae; Nostoc.
NIES2109__locus_1_main_sat
CsomeBCy
50350 -pf06051 -
50360 -pf06051 -
50370 -# -
50380 -pf00873 -
50390 -pf13533 -
50400 -pf05982 -
50410 -# -
50420 -pf10216 -
50430 -pf00361 -
50440 -pf00361pf00662 -
50450 H_neonGreen CsomeBCy
50460 H_neonGreen CsomeBCy
50470 P_electricGreen CsomeBCy
50480 HMMccmM_1_CsomeBCy(pf00132pf14602pf00101) CsomeBCy
50490 HMMccmN_1_CsomeBCy(#) CsomeBCy EP_CTERM
50500 Ts_poisonGreen CsomeBCy
50510 HMMreg00126_1_mix(pf00126pf03466) CsomeBCy
50520 -pf13598pf13600 -
50530 HMMreg00392_2_grm2(pf00392) CsomeBCy
50540 -pf00561 -
50550 -pf07724pf10431pf06689 -
50560 -pf00574 -
50570 -pf00254pf05698pf05697 -
50580 -pf01118pf02774 -
50590 HMM02441_01(pf00701) MUF2
50600 -pf00753pf07521 -
(..-38480..)
12120 -pf13614pf13807pf02706 -
12130 -pf00331 -
12140 -pf03808 -
12150 -pf13439pf00534 -
12160 -pf04932 -
12170 -pf13439pf00534 -
12180 -pf01943 -
12190 -pf00501pf13193pf00550pf00975 -
12200 -pf02397 -
BMC-P
BMC-H
BMC-T
Aldehyde dehydrogenase
PTAC
Signature enzyme
Alcohol dehydrogenase
Regulator
12210 -pf00805 -
12220 Tdp_turquoise(#) CsomeBCy
12230 -# -
12240 -pf00926pf00925 -
12250 -pf01118pf02774 -
12260 -pf00113pf03952 -
12270 -pf07109 -
12280 -# -
12290 HMM01923cobtr_1_pdu(pf03928) PDU1D
12300 -# -
12310 -pf06485 -
12320 -pf01523 -
(..20720..)
33040 -pf05114 -
33050 -# -
33060 -pf02656 -
33070 HMM17147pyruvdom_1_buf3(pf10371pf17147pf01558pf01855pf027
33080 -pf13472 -
33090 -pf04493 -
33100 -# -
33110 -pf00210 -
33120 -pf06150 -
33130 -pf12697 -
33140 H_lime CsomeBCy
33150 H_peaSoup CsomeBCy
33160 -# -
33170 -pf05157 -
33180 HMMreg13545_1_CsomeBCy(pf00027) BUF3
33190 -pf13180pf05299 -
33200 -pf13340 -
33210 -pf13586 -
33220 -pf00215pf00156 -
33230 -pf01594 -
33240 -pf00892 -
33250 -# -

Nostoc sp. NIES-2111. Bacteria; Cyanobacteria; Nostocales; Nostocaceae; Nostoc.

NIES2111__locus_1_main_sat

CsomeBCy

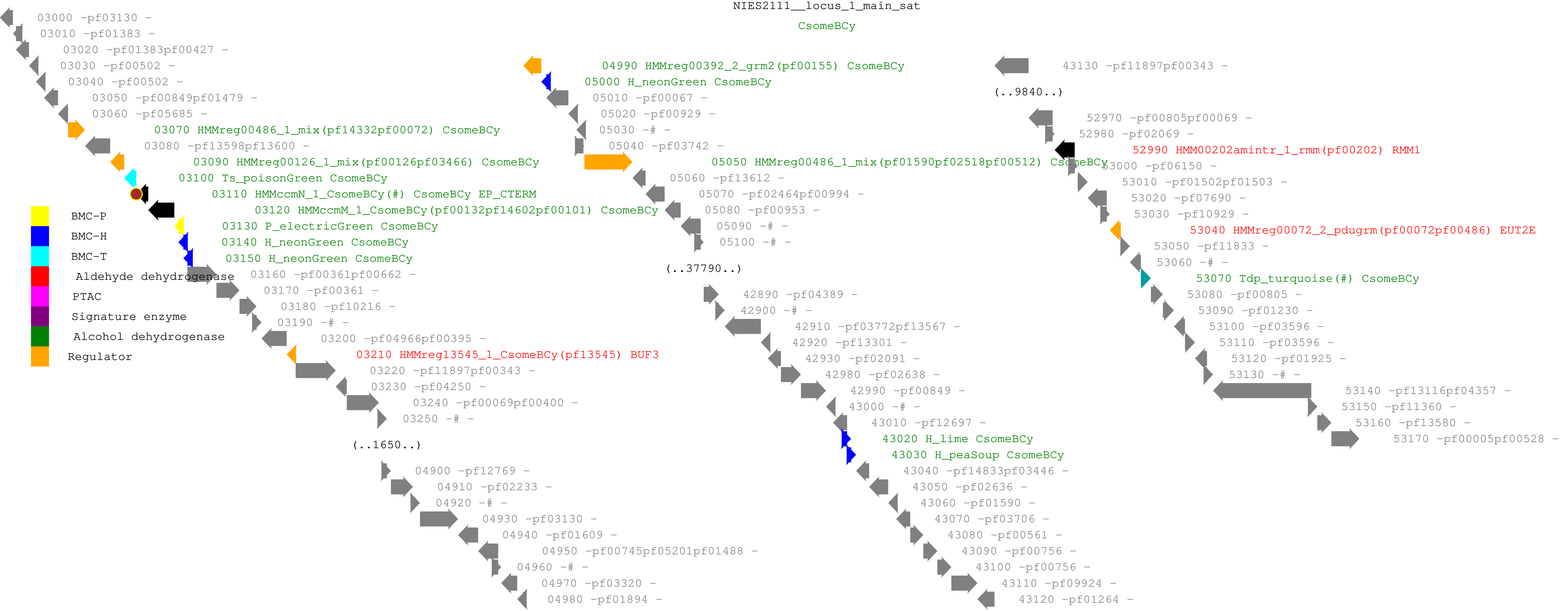

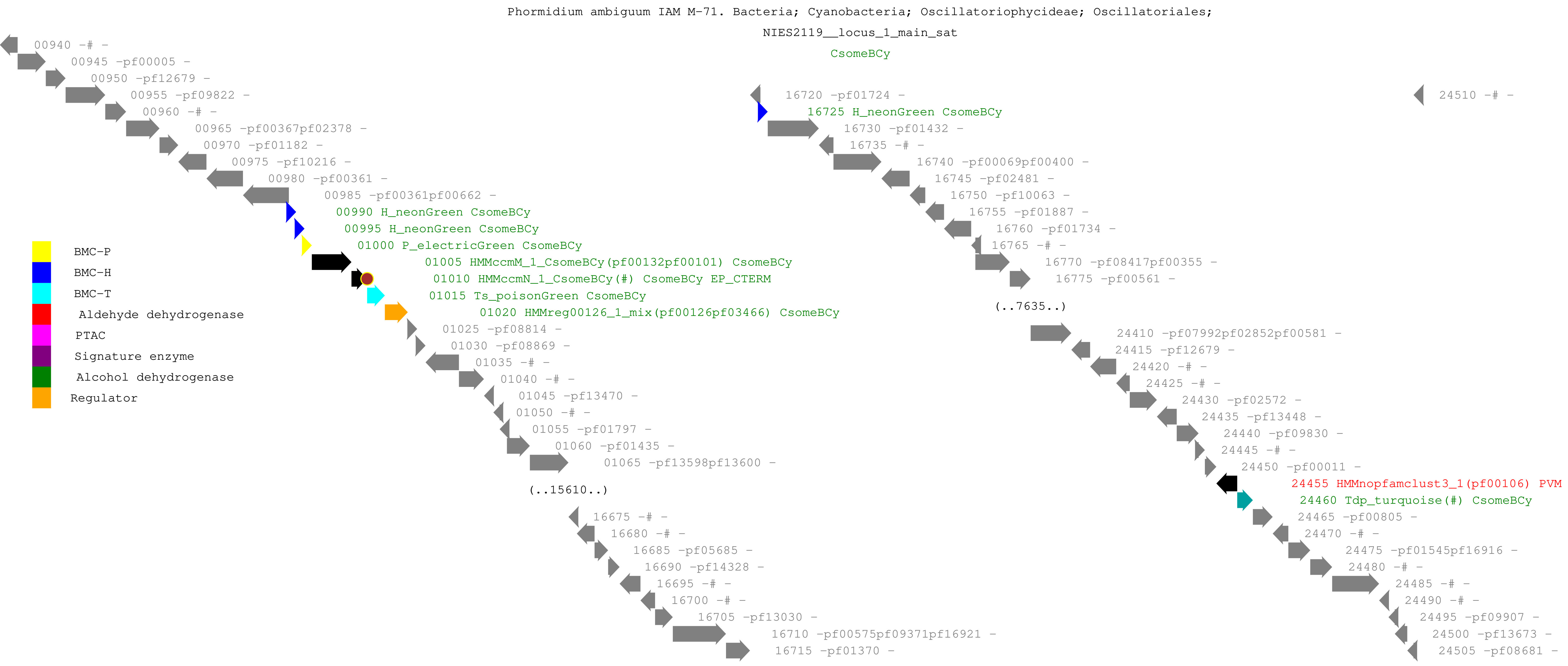

Phormidium ambiguum IAM M-71. Bacteria; Cyanobacteria; Oscillatoriophycideae; Oscillatoriales;
NIES2119__locus_1_main_sat
CsomeBCy
00940 -# -
00945 -pf00005 -
00950 -pf12679 -
00955 -pf09822 -
00960 -# -
00965 -pf00367pf02378 -
00970 -pf01182 -
00975 -pf10216 -
00980 -pf00361 -
00985 -pf00361pf00662 -
00990 H_neonGreen CsomeBCy
00995 H_neonGreen CsomeBCy
01000 P_electricGreen CsomeBCy
01005 HMMccmM_1_CsomeBCy(pf00132pf00101) CsomeBCy
01010 HMMccmN_1_CsomeBCy(#) CsomeBCy EP_CTERM
01015 Ts_poisonGreen CsomeBCy
01020 HMMreg00126_1_mix(pf00126pf03466) CsomeBCy
01025 -pf08814 -
01030 -pf08869 -
01035 -# -
01040 -# -
01045 -pf13470 -
01050 -# -
01055 -pf01797 -
01060 -pf01435 -
01065 -pf13598pf13600 -
(..15610..)
16675 -# -
16680 -# -
16685 -pf05685 -
16690 -pf14328 -
16695 -# -
16700 -# -
16705 -pf13030 -
16710 -pf00575pf09371pf16921 -
16715 -pf01370 -
16720 -pf01724 -
16725 H_neonGreen CsomeBCy
16730 -pf01432 -
16735 -# -
16740 -pf00069pf00400 -
16745 -pf02481 -
16750 -pf10063 -
16755 -pf01887 -
16760 -pf01734 -
16765 -# -
16770 -pf08417pf00355 -
16775 -pf00561 -
(..7635..)
24410 -pf07992pf02852pf00581 -
24415 -pf12679 -
24420 -# -
24425 -# -
24430 -pf02572 -
24435 -pf13448 -
24440 -pf09830 -
24445 -# -
24450 -pf00011 -
24455 HMMnopfamclust3_1(pf00106) PVM
24460 Tdp_turquoise(#) CsomeBCy
24465 -pf00805 -
24470 -# -
24475 -pf01545pf16916 -
24480 -# -
24485 -# -
24490 -# -
24495 -pf09907 -
24500 -pf13673 -
24505 -pf08681 -
24510 -# -
BMC-P
BMC-H
BMC-T
Aldehyde dehydrogenase
PTAC
Signature enzyme
Alcohol dehydrogenase
Regulator

Scytonema sp. HK-05. Bacteria; Cyanobacteria; Nostocales; Scytonemataceae; Scytonema.

NIES2130__locus_1_main_sat

CsomeBCy

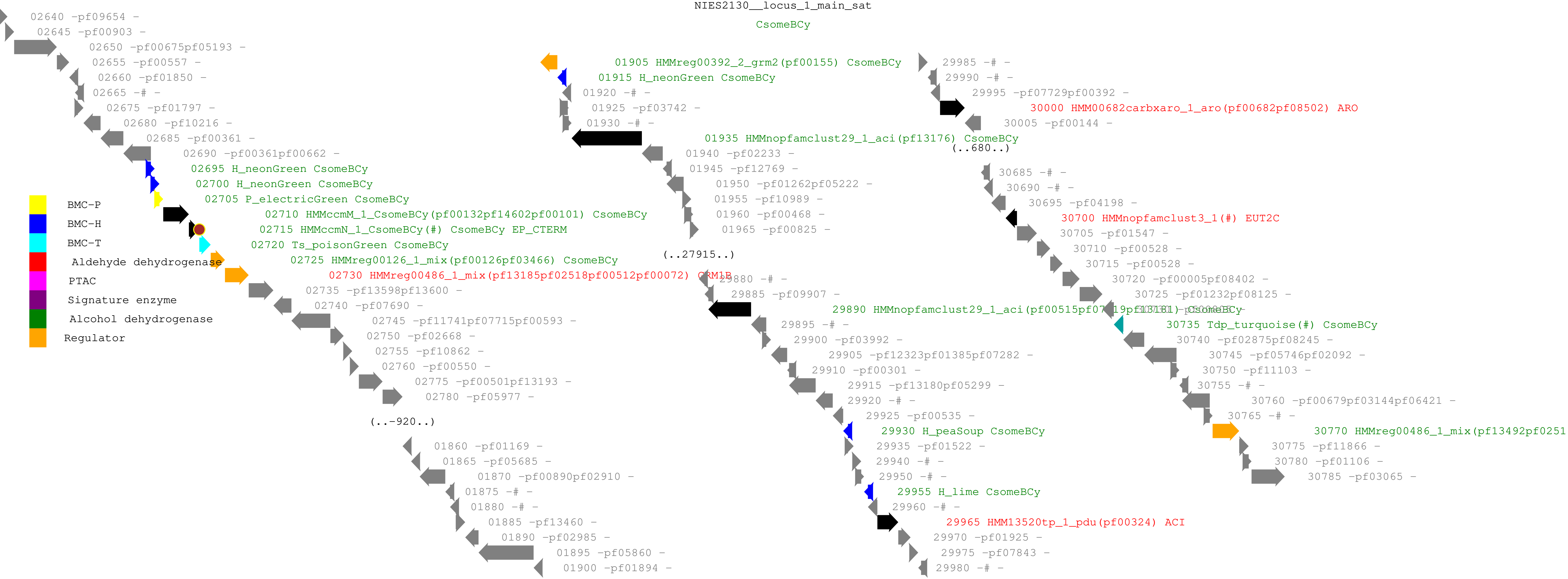

Leptolyngbya boryana NIES-2135. Bacteria; Cyanobacteria; Synechococcales; Leptolyngbyaceae;

NIES2135__locus_1_main_sat

CsomeBCy

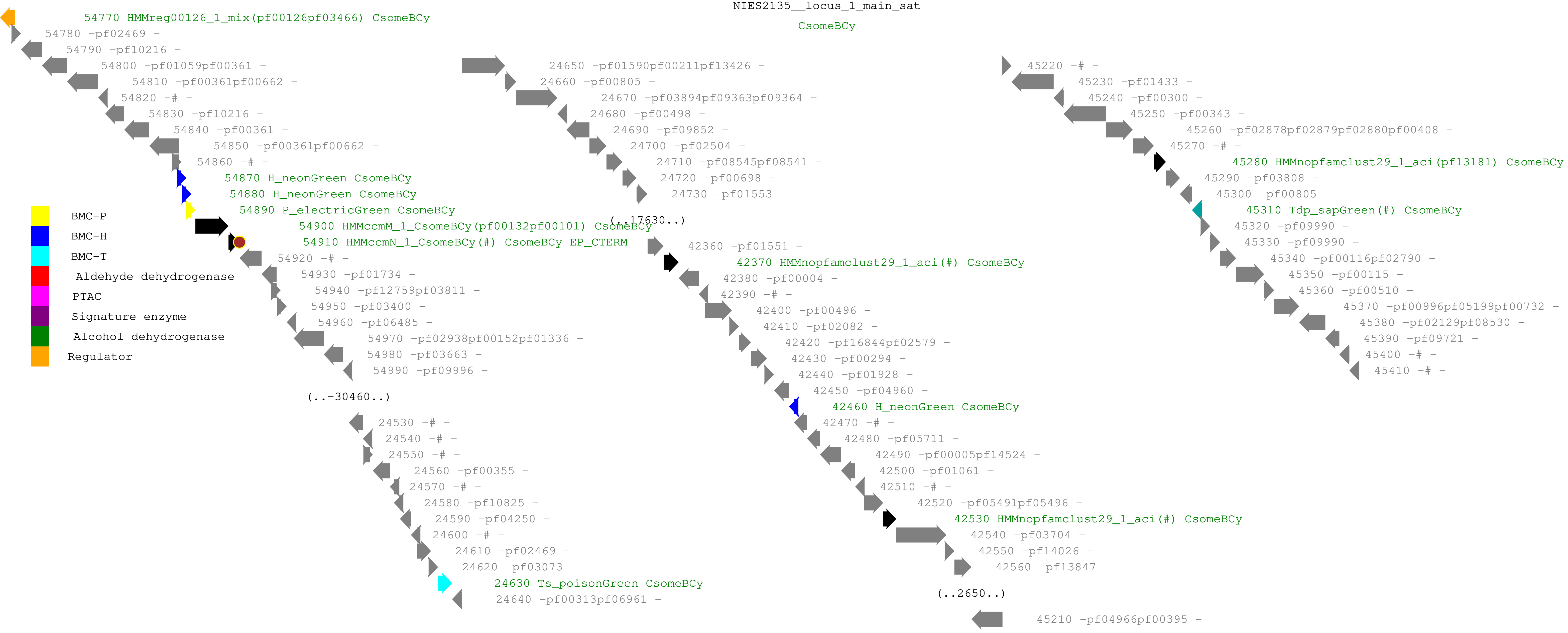

Anabaenopsis circularis NIES-21. Bacteria; Cyanobacteria; Nostocales; Aphanizomenonaceae; Anabaenopsis.

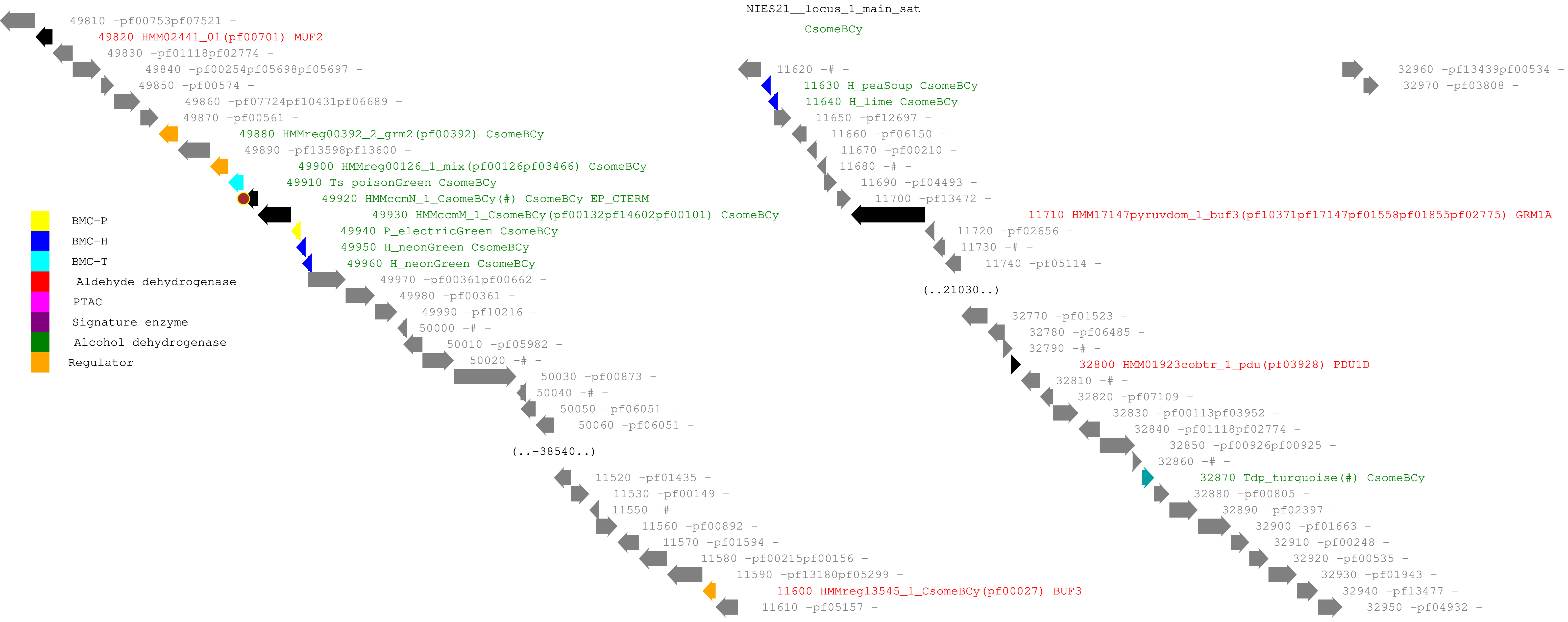

Calothrix brevissima NIES-22. Bacteria; Cyanobacteria; Nostocales; Rivulariaceae; Calothrix.

NIES22__locus_1_main_sat

CsomeBCy

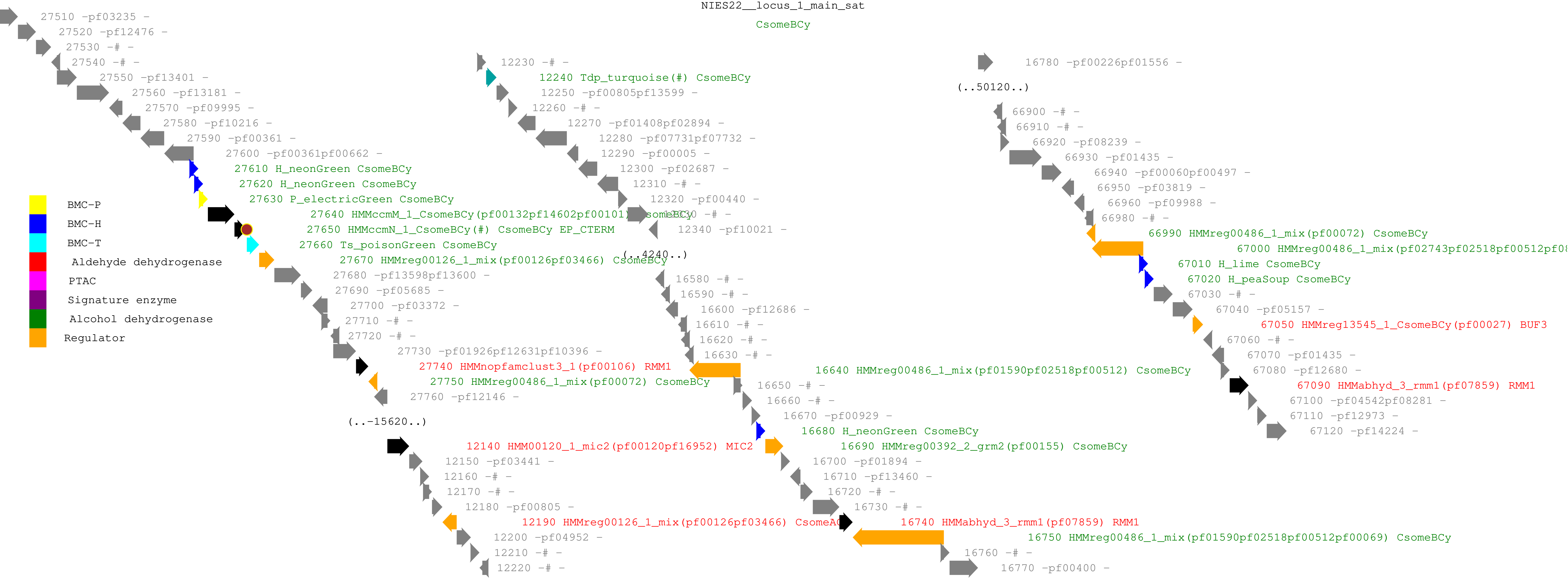

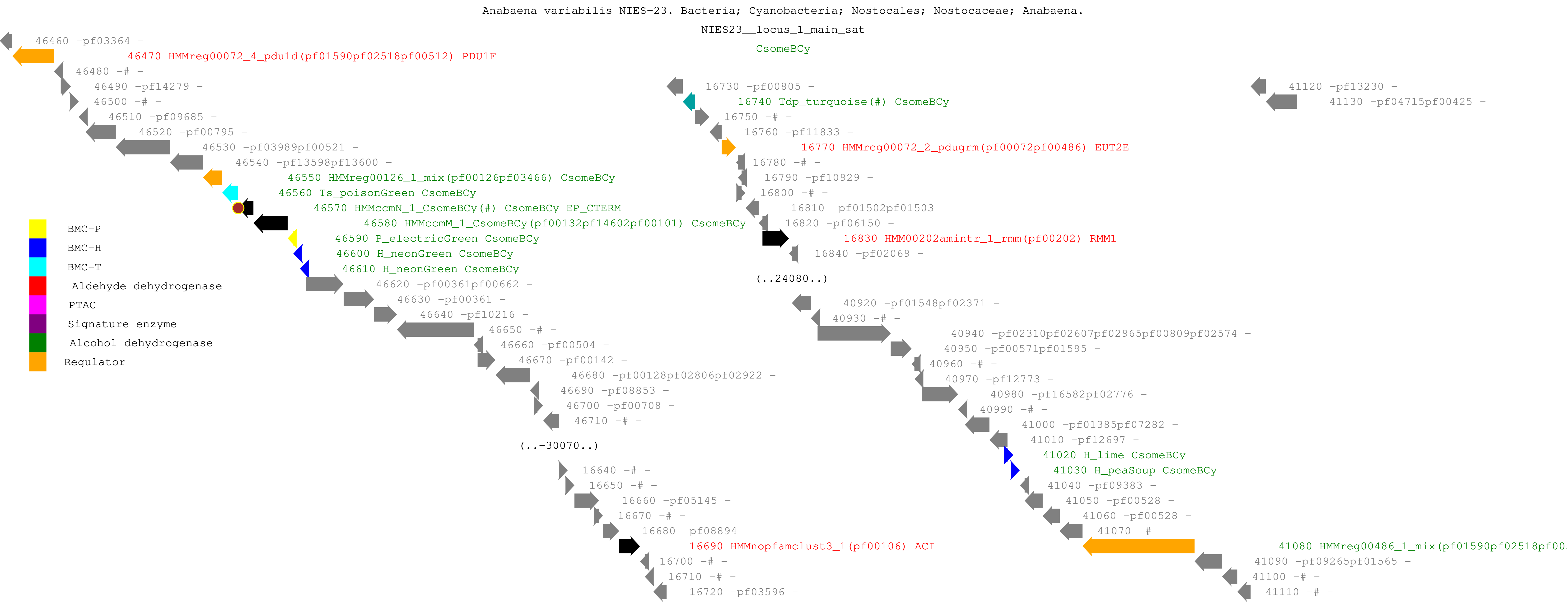
Anabaena variabilis NIES-23. Bacteria; Cyanobacteria; Nostocales; Nostocaceae; Anabaena.
NIES23__locus_1_main_sat
CsomeBCy
46460 -pf03364 -
46470 HMMreg00072_4_pdu1d(pf01590pf02518pf00512) PDU1F
46480 -# -
46490 -pf14279 -
46500 -# -
46510 -pf09685 -
46520 -pf00795 -
46530 -pf03989pf00521 -
46540 -pf13598pf13600 -
46550 HMMreg00126_1_mix(pf00126pf03466) CsomeBCy
46560 Ts_poisonGreen CsomeBCy
46570 HMMccmN_1_CsomeBCy(#) CsomeBCy EP_CTERM
46580 HMMccmM_1_CsomeBCy(pf00132pf14602pf00101) CsomeBCy
46590 P_electricGreen CsomeBCy
46600 H_neonGreen CsomeBCy
46610 H_neonGreen CsomeBCy
46620 -pf00361pf00662 -
46630 -pf00361 -
46640 -pf10216 -
46650 -# -
46660 -pf00504 -
46670 -pf00142 -
46680 -pf00128pf02806pf02922 -
46690 -pf08853 -
46700 -pf00708 -
46710 -# -
(..-30070..)
16640 -# -
16650 -# -
16660 -pf05145 -
16670 -# -
16680 -pf08894 -
16690 HMMnopfamclust3_1(pf00106) ACI
16700 -# -
16710 -# -
16720 -pf03596 -
16730 -pf00805 -
16740 Tdp_turquoise(#) CsomeBCy
16750 -# -
16760 -pf11833 -
16770 HMMreg00072_2_pdugrm(pf00072pf00486) EUT2E
16780 -# -
16790 -pf10929 -
16800 -# -
16810 -pf01502pf01503 -
16820 -pf06150 -
16830 HMM00202amintr_1_rmm(pf00202) RMM1
16840 -pf02069 -
(..24080..)
40920 -pf01548pf02371 -
40930 -# -
40940 -pf02310pf02607pf02965pf00809pf02574 -
40950 -pf00571pf01595 -
40960 -# -
40970 -pf12773 -
40980 -pf16582pf02776 -
40990 -# -
41000 -pf01385pf07282 -
41010 -pf12697 -
41020 H_lime CsomeBCy
41030 H_peaSoup CsomeBCy
41040 -pf09383 -
41050 -pf00528 -
41060 -pf00528 -
41070 -# -
41080 HMMreg00486_1_mix(pf01590pf02518pf00
41090 -pf09265pf01565 -
41100 -# -
41110 -# -
41120 -pf13230 -
41130 -pf04715pf00425 -
BMC-P
BMC-H
BMC-T
Aldehyde dehydrogenase
PTAC
Signature enzyme
Alcohol dehydrogenase
Regulator

Nostoc linckia NIES-25. Bacteria; Cyanobacteria; Nostocales; Nostocaceae; Nostoc.

NIES25__locus_1_main_sat

CsomeBCy

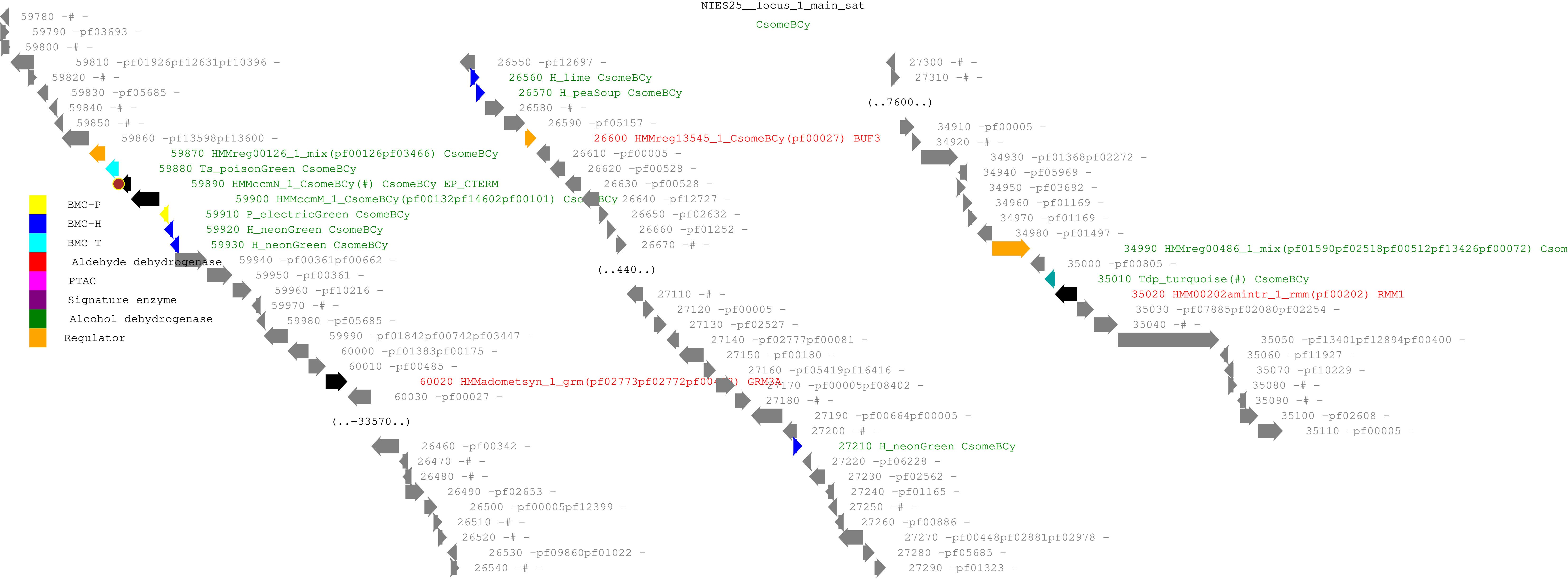

Calothrix parasitica NIES-267. Bacteria; Cyanobacteria; Nostocales; Rivulariaceae; Calothrix.

NIES267__locus_1_main_sat

CsomeBCy

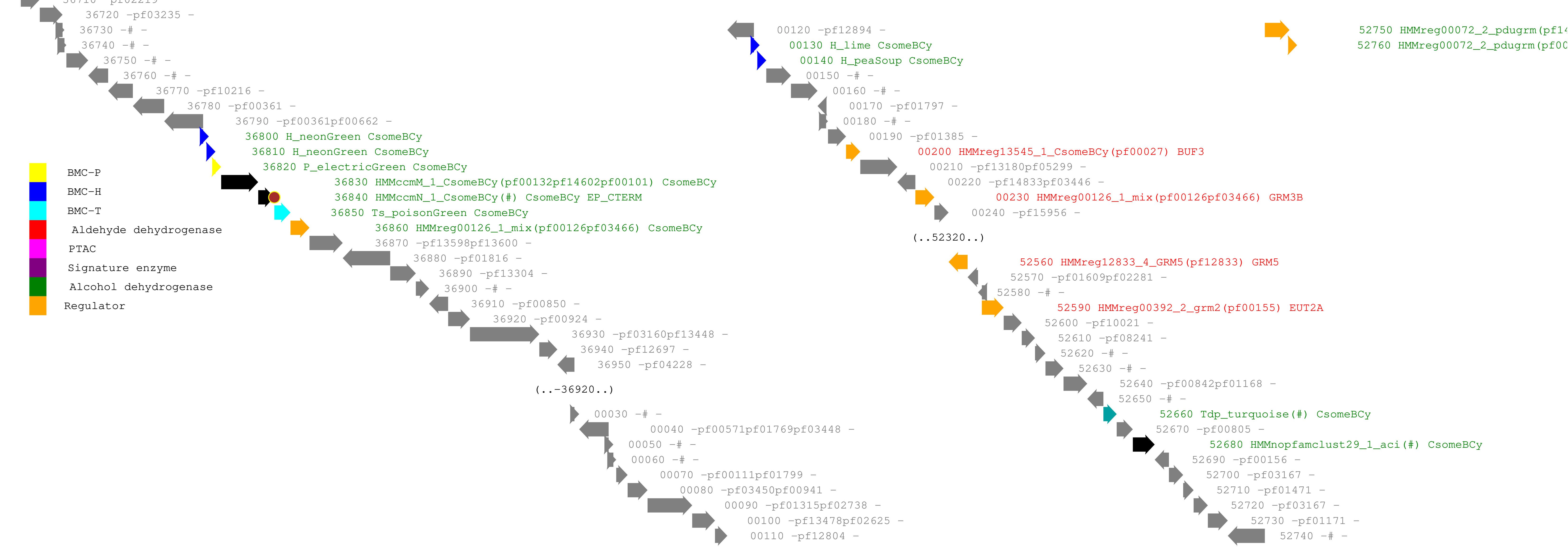

Microcystis aeruginosa NIES-298. Bacteria; Cyanobacteria; Oscillatoriophycideae; Chroococcales;

NIES298__locus_1_main_sat

CsomeBCy

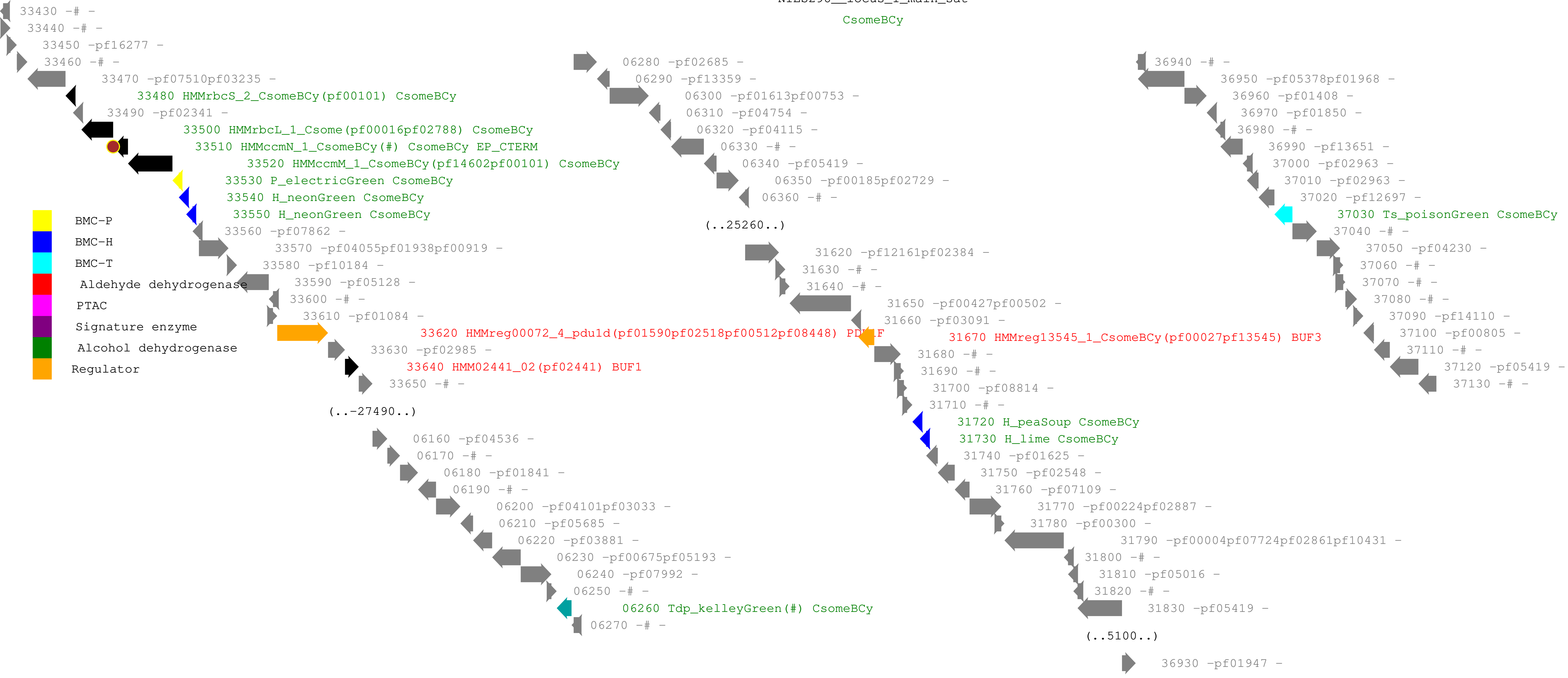

Phormidium tenue NIES-30. Bacteria; Cyanobacteria; Oscillatoriophycideae; Oscillatoriales;

NIES30__locus_1_main_sat

CsomeBCy

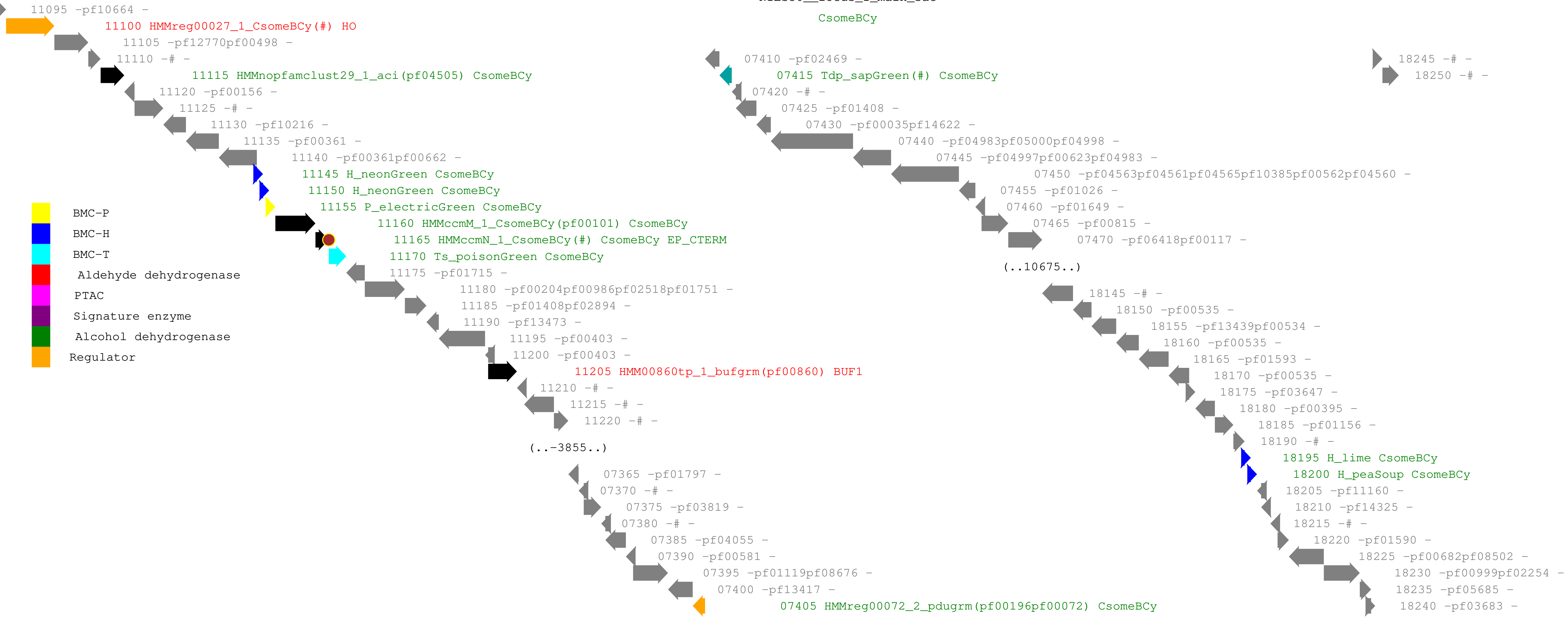

Fremyella diplosiphon NIES-3275. Bacteria; Cyanobacteria; Nostocales; Rivulariaceae; Microchaete.

NIES3275__locus_1_main_sat

CsomeBCy

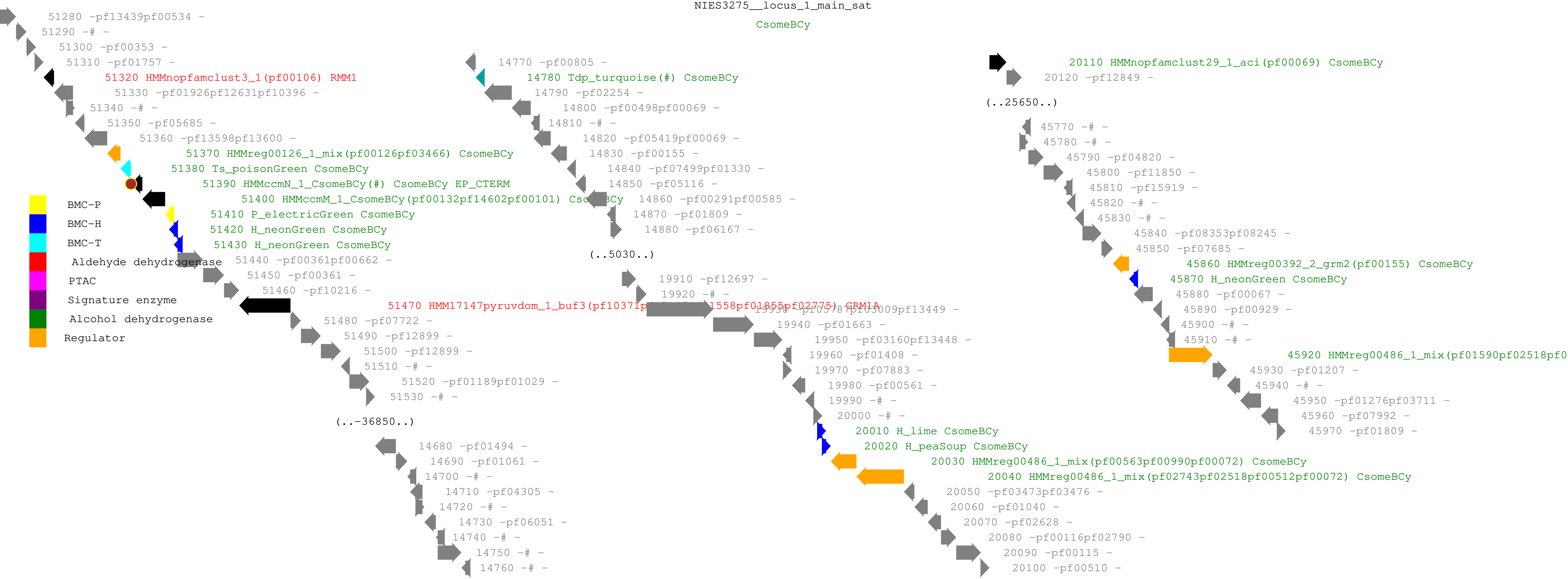

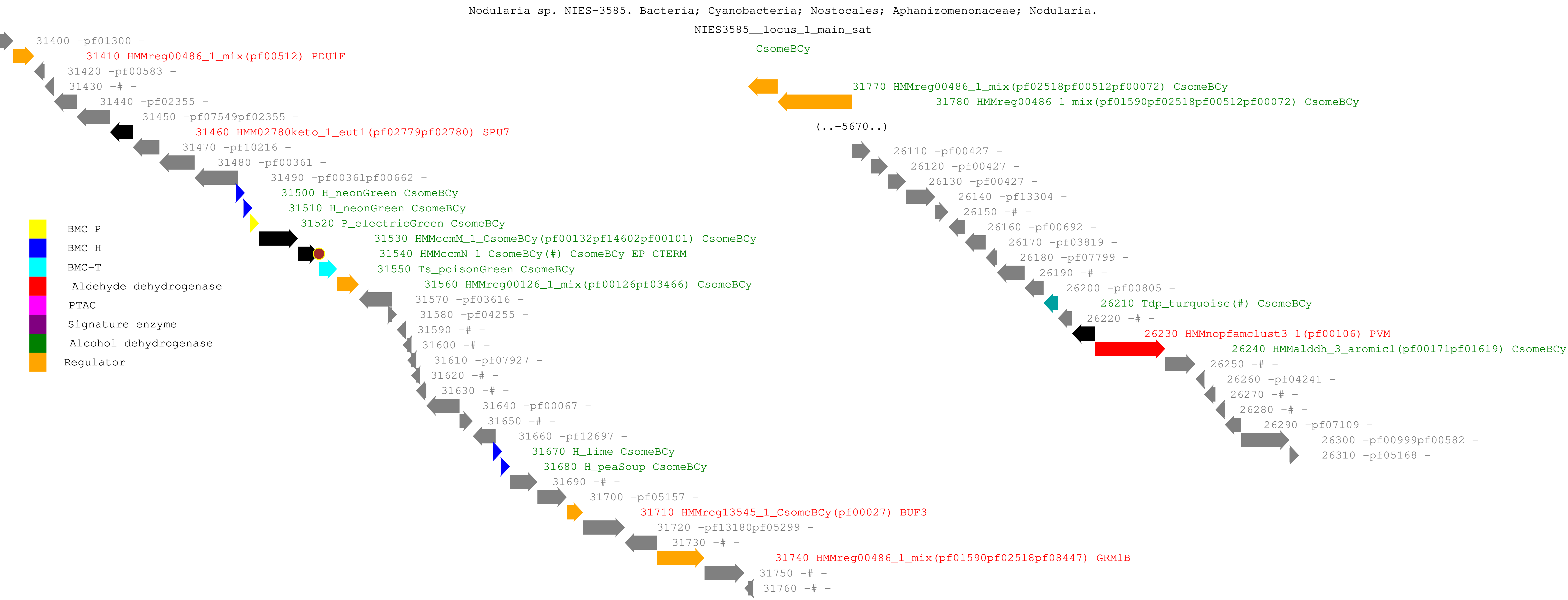
Nodularia sp. NIES-3585. Bacteria; Cyanobacteria; Nostocales; Aphanizomenonaceae; Nodularia.
NIES3585__locus_1_main_sat
CsomeBCy
31400 -pf01300 -
31410 HMMreg00486_1_mix(pf00512) PDU1F
31420 -pf00583 -
31430 -# -
31440 -pf02355 -
31450 -pf07549pf02355 -
31460 HMM02780keto_1_eut1(pf02779pf02780) SPU7
31470 -pf10216 -
31480 -pf00361 -
31490 -pf00361pf00662 -
31500 H_neonGreen CsomeBCy
31510 H_neonGreen CsomeBCy
31520 P_electricGreen CsomeBCy
31530 HMMccmM_1_CsomeBCy(pf00132pf14602pf00101) CsomeBCy
31540 HMMccmN_1_CsomeBCy(#) CsomeBCy EP_CTERM
31550 Ts_poisonGreen CsomeBCy
31560 HMMreg00126_1_mix(pf00126pf03466) CsomeBCy
31570 -pf03616 -
31580 -pf04255 -
31590 -# -
31600 -# -
31610 -pf07927 -
31620 -# -
31630 -# -
31640 -pf00067 -
31650 -# -
31660 -pf12697 -
31670 H_lime CsomeBCy
31680 H_peaSoup CsomeBCy
31690 -# -
31700 -pf05157 -
31710 HMMreg13545_1_CsomeBCy(pf00027) BUF3
31720 -pf13180pf05299 -
31730 -# -
31740 HMMreg00486_1_mix(pf01590pf02518pf08447) GRM1B
31750 -# -
31760 -# -
31770 HMMreg00486_1_mix(pf02518pf00512pf00072) CsomeBCy
31780 HMMreg00486_1_mix(pf01590pf02518pf00512pf00072) CsomeBCy
(..-5670..)
26110 -pf00427 -
26120 -pf00427 -
26130 -pf00427 -
26140 -pf13304 -
26150 -# -
26160 -pf00692 -
26170 -pf03819 -
26180 -pf07799 -
26190 -# -
26200 -pf00805 -
26210 Tdp_turquoise(#) CsomeBCy
26220 -# -
26230 HMMnopfamclust3_1(pf00106) PVM
26240 HMMaldh_3_aromic1(pf00171pf01619) CsomeBCy
26250 -# -
26260 -pf04241 -
26270 -# -
26280 -# -
26290 -pf07109 -
26300 -pf00999pf00582 -
26310 -pf05168 -
BMC-P
BMC-H
BMC-T
Aldehyde dehydrogenase
PTAC
Signature enzyme
Alcohol dehydrogenase
Regulator

Tolypothrix tenuis PCC 7101. Bacteria; Cyanobacteria; Nostocales; Tolypothrichaceae; Tolypothrix.

NIES37__locus_1_main_sat

CsomeBCy

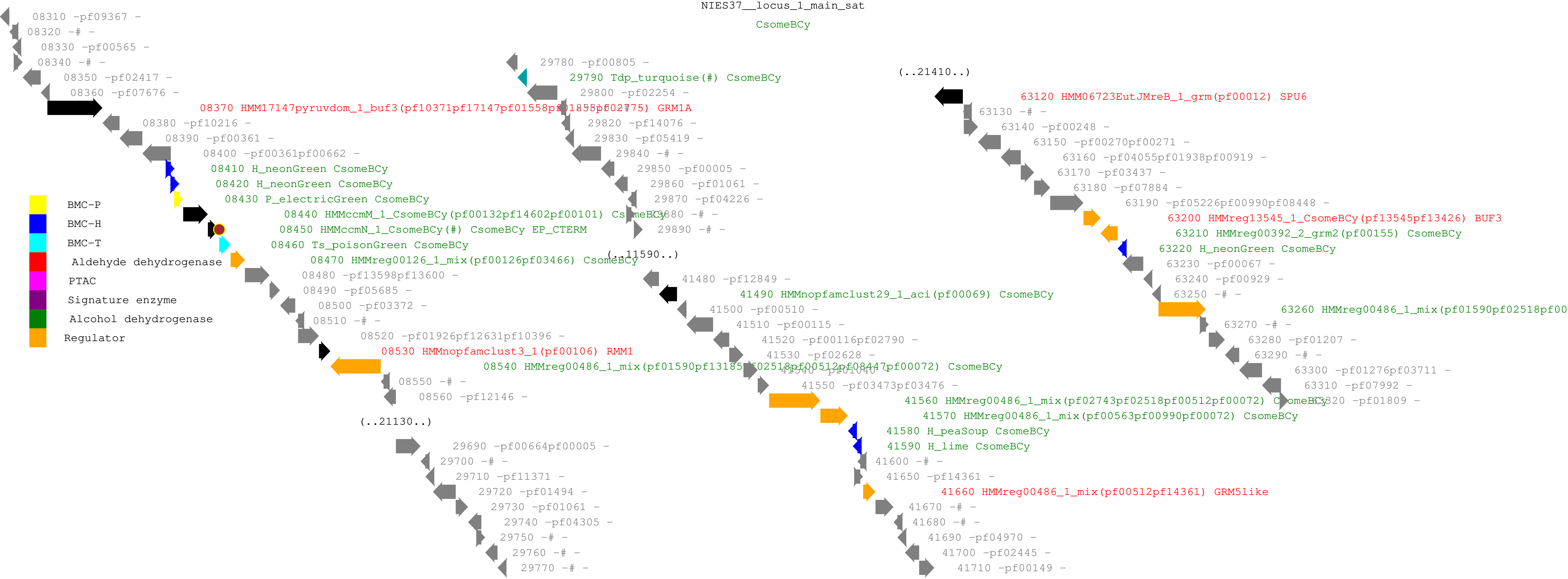

Calothrix sp. NIES-3974. Bacteria; Cyanobacteria; Nostocales; Rivulariaceae; Calothrix.

NIES3974__locus_1_main_sat

CsomeBCy

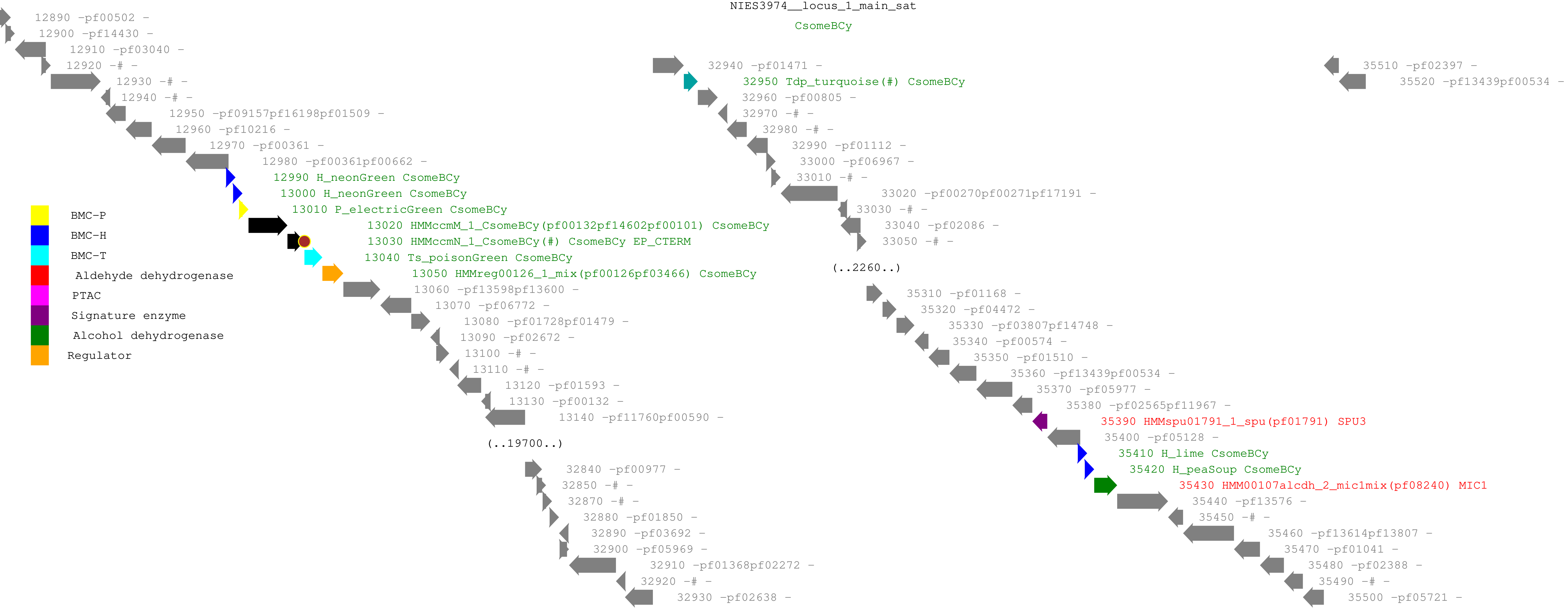

Arthrospira platensis (strain NIES-39 / IAM M-135) (Spirulina Bacteria; Cyanobacteria; Oscillatoriophycideae; Oscillatoriales;

NIES39_K_locus_1

CsomeBCy

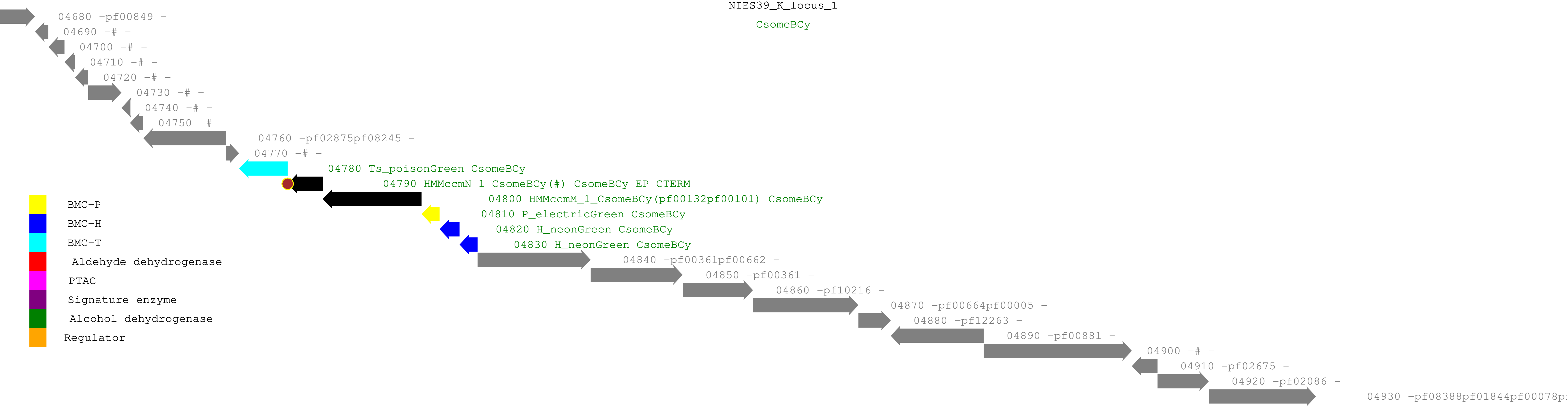

Nostoc commune HK-02. Bacteria; Cyanobacteria; Nostocales; Nostocaceae; Nostoc.

NIES4070__locus_1_main_sat

CsomeBCy

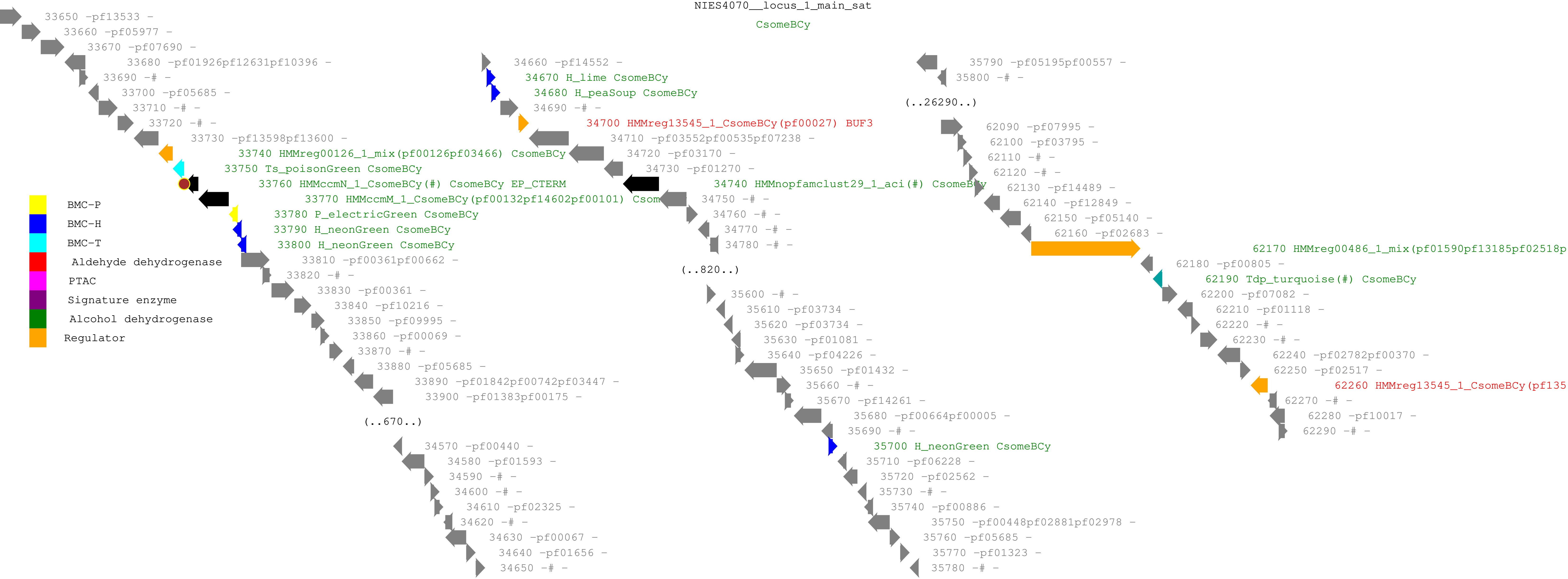

Calothrix sp. NIES-4071. Bacteria; Cyanobacteria; Nostocales; Rivulariaceae; Calothrix.

NIES4071__locus_1_main_sat

CsomeBCy

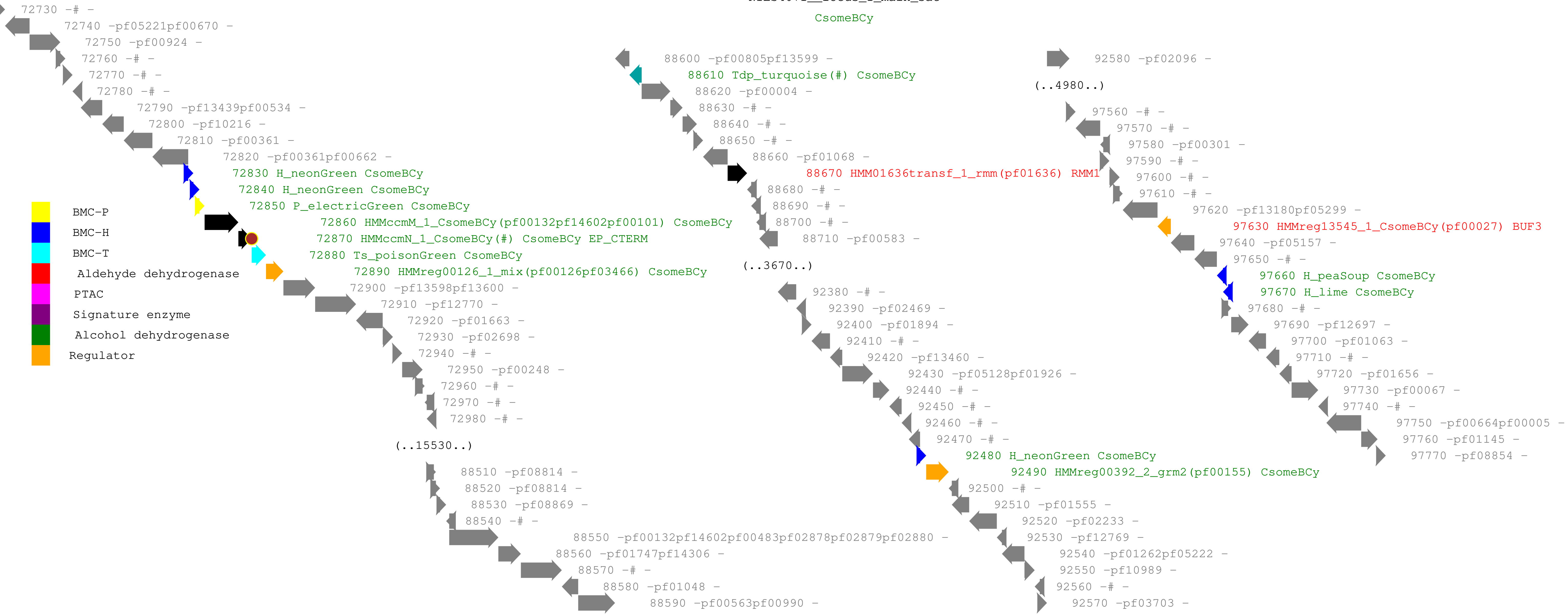

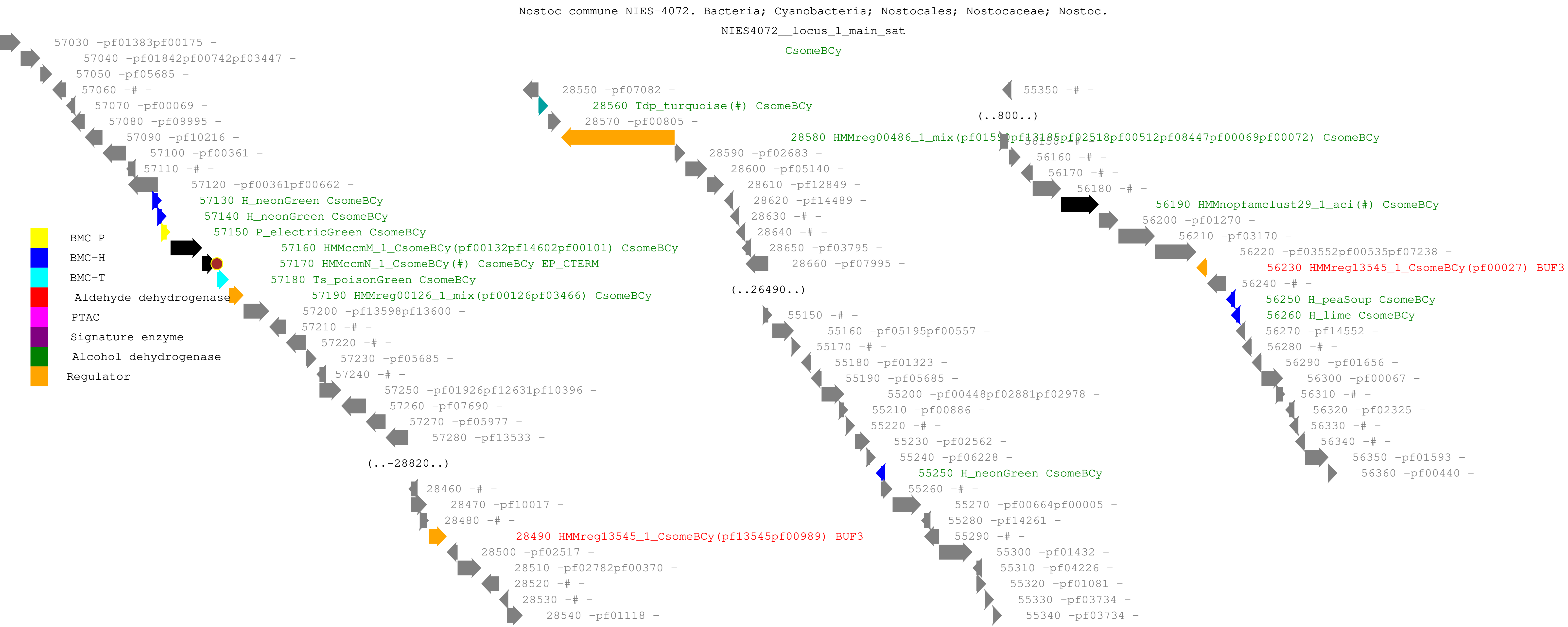
Nostoc commune NIES-4072. Bacteria; Cyanobacteria; Nostocales; Nostocaceae; Nostoc.
NIES4072__locus_1_main_sat
CsomeBCy
57030 -pf01383pf00175 -
57040 -pf01842pf00742pf03447 -
57050 -pf05685 -
57060 -# -
57070 -pf00069 -
57080 -pf09995 -
57090 -pf10216 -
57100 -pf00361 -
57110 -# -
57120 -pf00361pf00662 -
57130 H_neonGreen CsomeBCy
57140 H_neonGreen CsomeBCy
57150 P_electricGreen CsomeBCy
57160 HMMccmM_1_CsomeBCy(pf00132pf14602pf00101) CsomeBCy
57170 HMMccmN_1_CsomeBCy(#) CsomeBCy EP_CTERM
57180 Ts_poisonGreen CsomeBCy
57190 HMMreg00126_1_mix(pf00126pf03466) CsomeBCy
57200 -pf13598pf13600 -
57210 -# -
57220 -# -
57230 -pf05685 -
57240 -# -
57250 -pf01926pf12631pf10396 -
57260 -pf07690 -
57270 -pf05977 -
57280 -pf13533 -
(..-28820..)
28460 -# -
28470 -pf10017 -
28480 -# -
28490 HMMreg13545_1_CsomeBCy(pf13545pf00989) BUF3
28500 -pf02517 -
28510 -pf02782pf00370 -
28520 -# -
28530 -# -
28540 -pf01118 -
28550 -pf07082 -
28560 Tdp_turquoise(#) CsomeBCy
28570 -pf00805 -
28580 HMMreg00486_1_mix(pf01590pf13185pf02518pf00512pf08447pf00069pf00072) CsomeBCy
28590 -pf02683 -
28600 -pf05140 -
28610 -pf12849 -
28620 -pf14489 -
28630 -# -
28640 -# -
28650 -pf03795 -
28660 -pf07995 -
(..26490..)
55150 -# -
55160 -pf05195pf00557 -
55170 -# -
55180 -pf01323 -
55190 -pf05685 -
55200 -pf00448pf02881pf02978 -
55210 -pf00886 -
55220 -# -
55230 -pf02562 -
55240 -pf06228 -
55250 H_neonGreen CsomeBCy
55260 -# -
55270 -pf00664pf00005 -
55280 -pf14261 -
55290 -# -
55300 -pf01432 -
55310 -pf04226 -
55320 -pf01081 -
55330 -pf03734 -
55340 -pf03734 -
55350 -# -
(..800..)
56150 -# -
56160 -# -
56170 -# -
56180 -# -
56190 HMMnopfamclust29_1_aci(#) CsomeBCy
56200 -pf01270 -
56210 -pf03170 -
56220 -pf03552pf00535pf07238 -
56230 HMMreg13545_1_CsomeBCy(pf00027) BUF3
56240 -# -
56250 H_peaSoup CsomeBCy
56260 H_lime CsomeBCy
56270 -pf14552 -
56280 -# -
56290 -pf01656 -
56300 -pf00067 -
56310 -# -
56320 -pf02325 -
56330 -# -
56340 -# -
56350 -pf01593 -
56360 -pf00440 -
BMC-P
BMC-H
BMC-T
Aldehyde dehydrogenase
PTAC
Signature enzyme
Alcohol dehydrogenase
Regulator

Scytonema sp. NIES-4073. Bacteria; Cyanobacteria; Nostocales; Scytonemataceae; Scytonema.

NIES4073__locus_1_main_sat

CsomeBCy

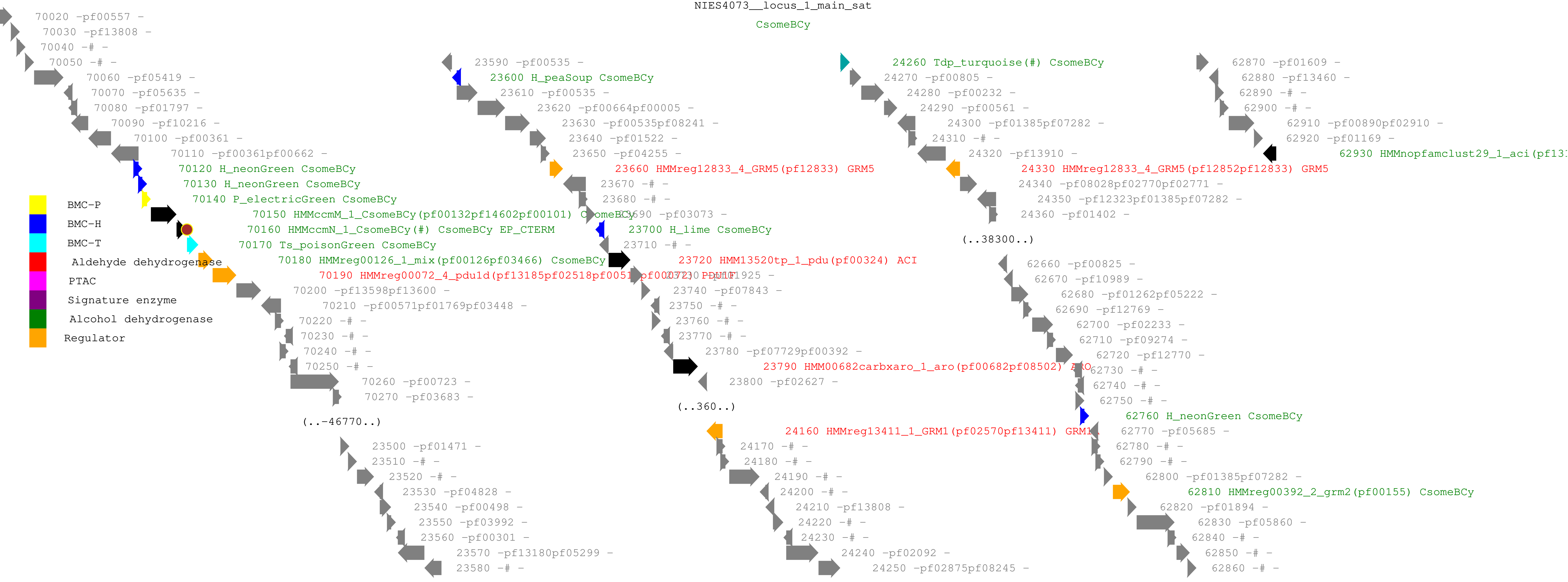

Cylindrospermum sp. NIES-4074. Bacteria; Cyanobacteria; Nostocales; Nostocaceae; Cylindrospermum.

NIES4074__locus_1_main_sat

CsomeBCy

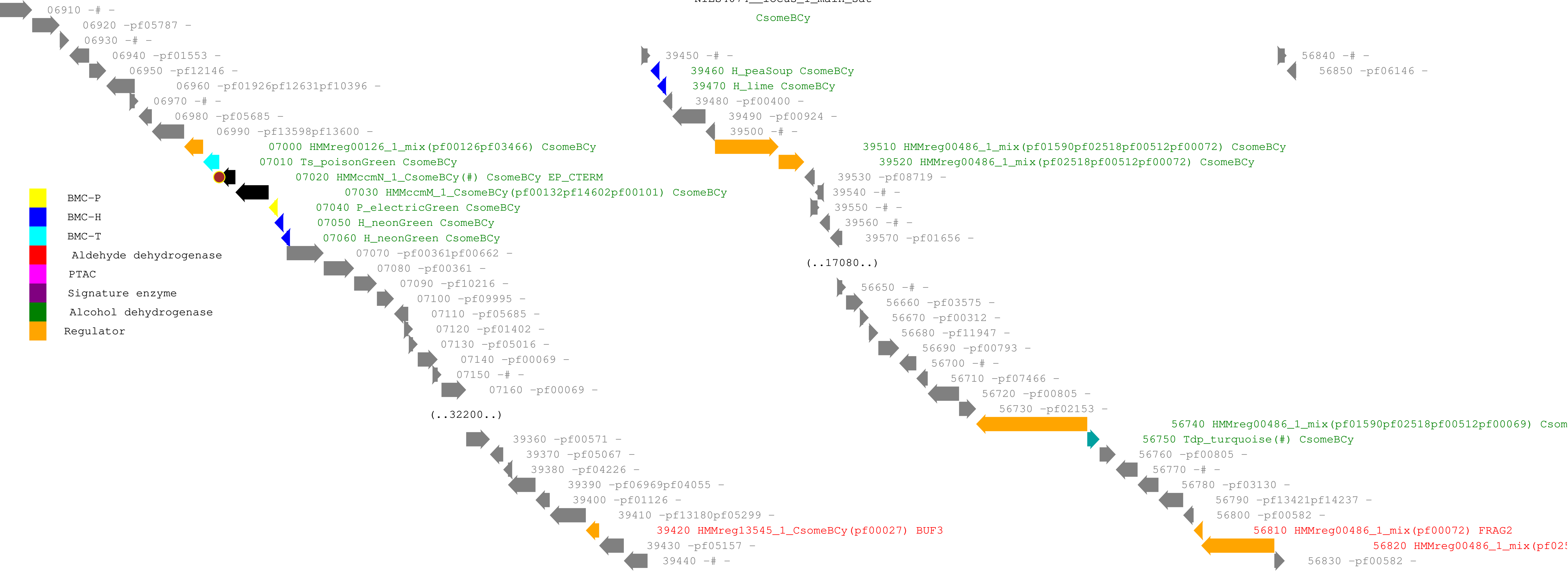

Tolypothrix sp. NIES-4075. Bacteria; Cyanobacteria; Nostocales; Tolypothrichaceae; Tolypothrix.

NIES4075__locus_1_main_sat

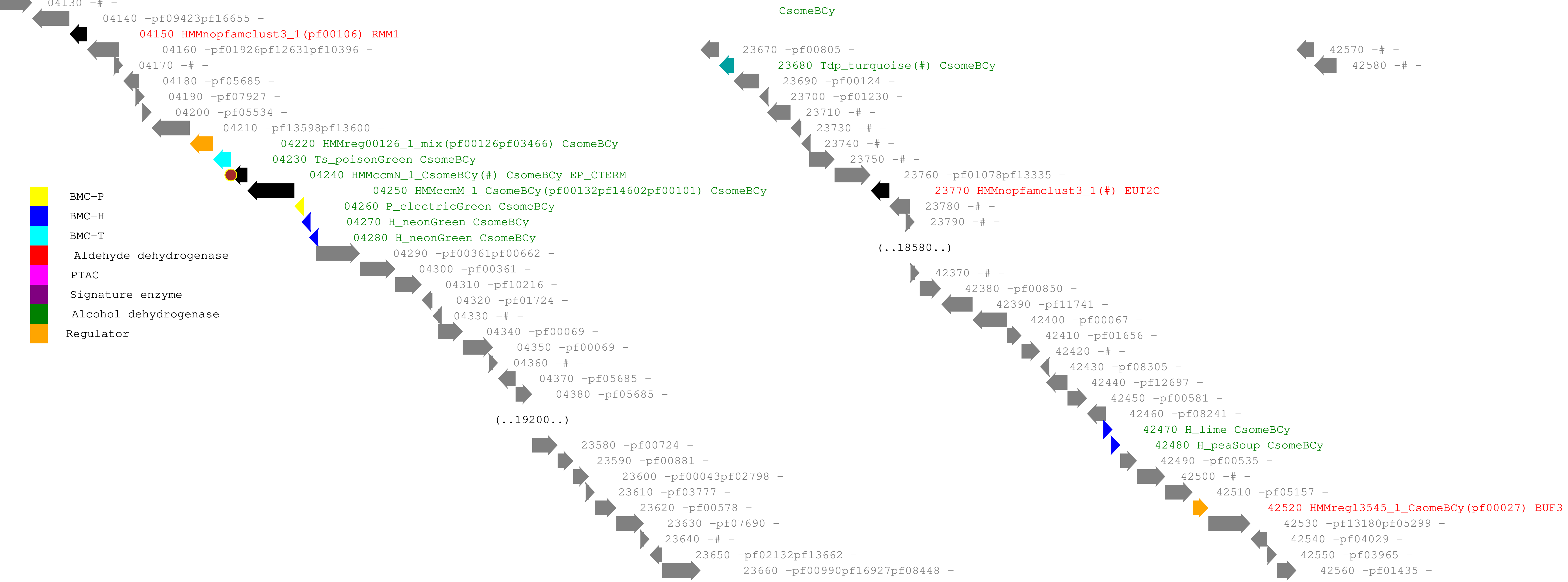

Calothrix sp. NIES-4101. Bacteria; Cyanobacteria; Nostocales; Rivulariaceae; Calothrix.

NIES4101__locus_1_main_sat

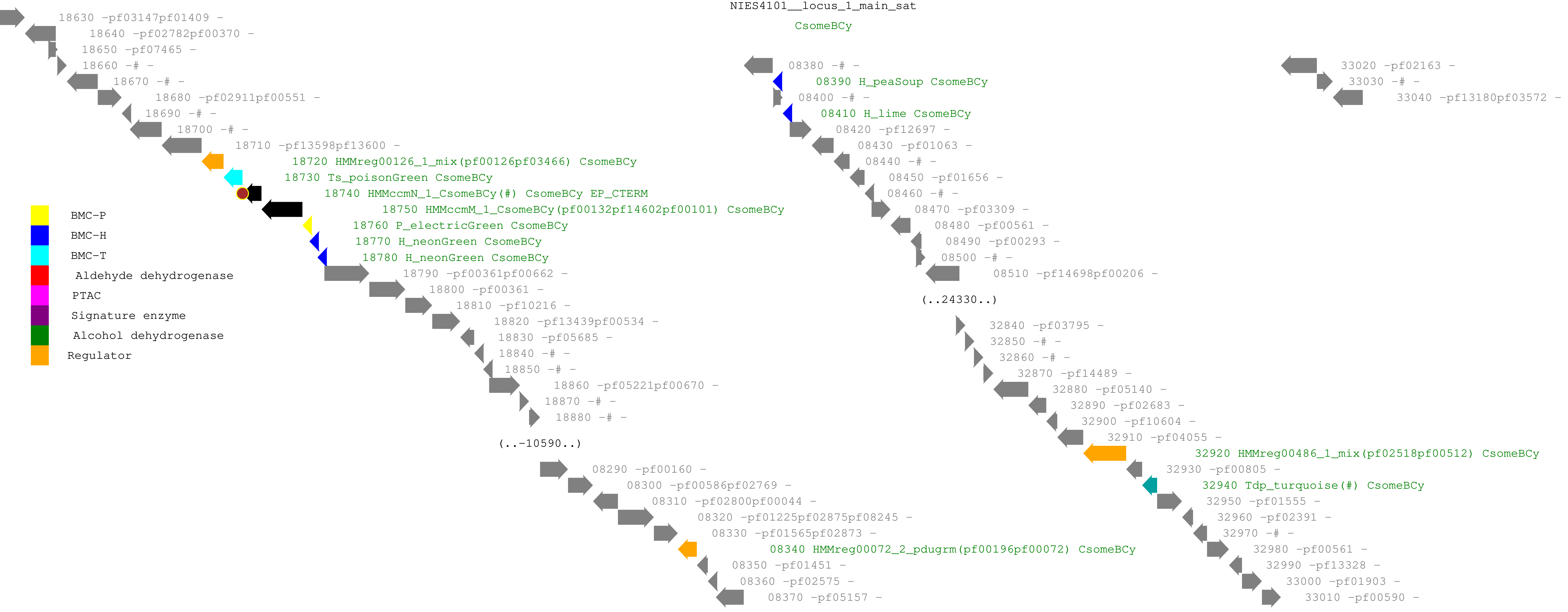

Chondrocystis sp. NIES-4102. Bacteria; Cyanobacteria; Oscillatoriophycideae; Chroococcales;

NIES4102__locus_1_main_sat

CsomeBCy

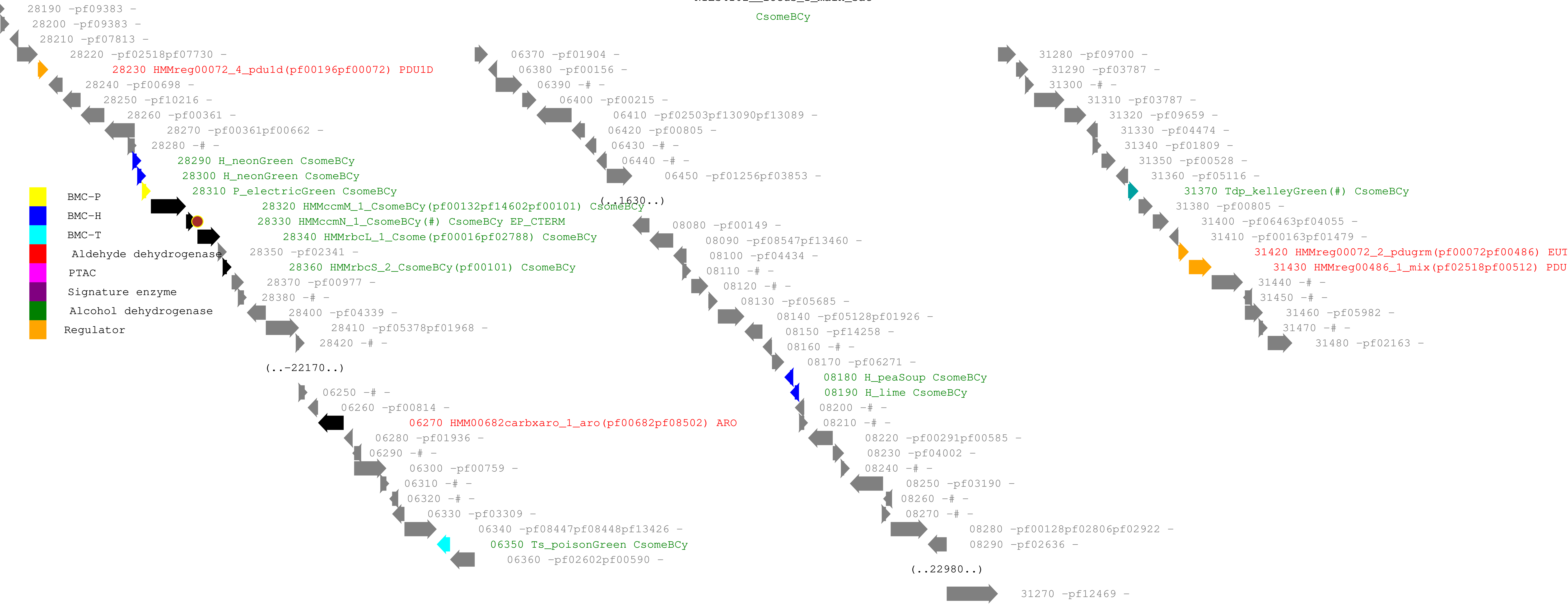

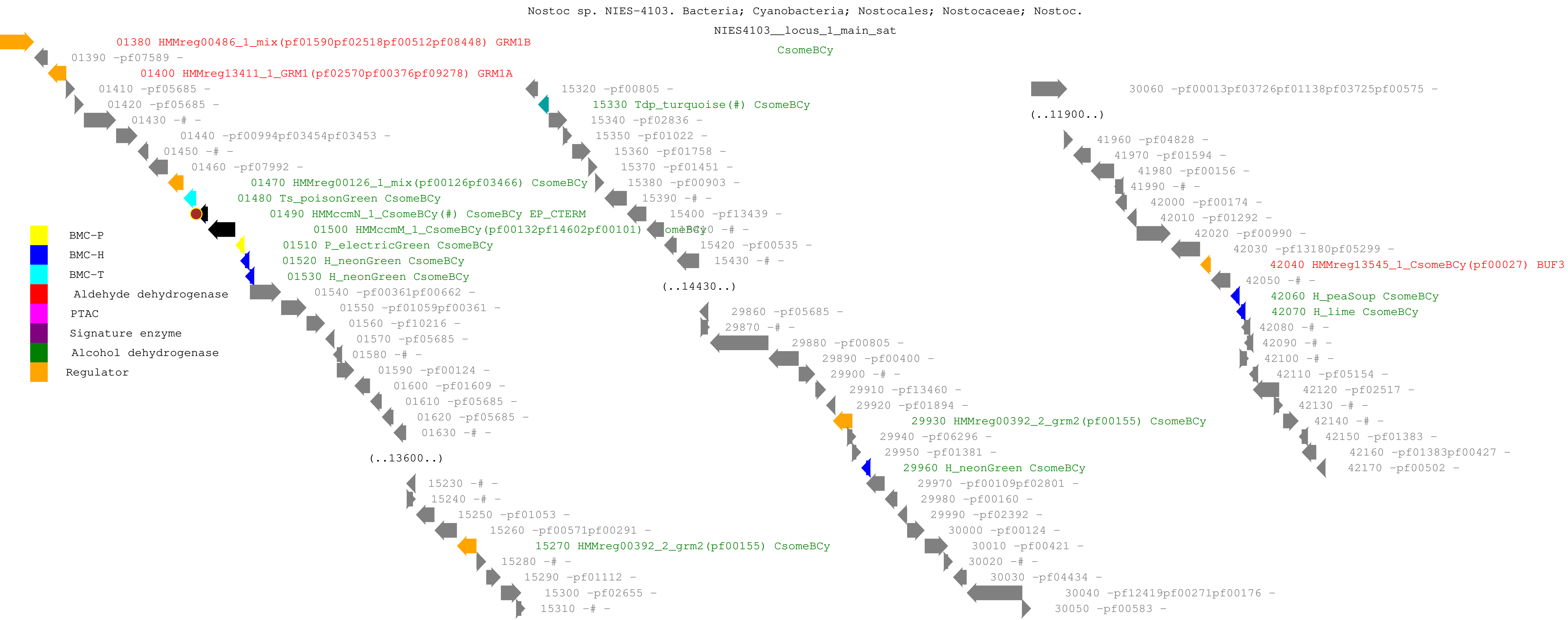

Nostoc sp. NIES-4103. Bacteria; Cyanobacteria; Nostocales; Nostocaceae; Nostoc.
NIES4103__locus_1_main_sat
CsomeBCy
01380 HMMreg00486_1_mix(pf01590pf02518pf00512pf08448) GRM1B
01390 -pf07589 -
01400 HMMreg13411_1_GRM1(pf02570pf00376pf09278) GRM1A
01410 -pf05685 -
01420 -pf05685 -
01430 -# -
01440 -pf00994pf03454pf03453 -
01450 -# -
01460 -pf07992 -
01470 HMMreg00126_1_mix(pf00126pf03466) CsomeBCy
01480 Ts_poisonGreen CsomeBCy
01490 HMMccmN_1_CsomeBCy(#) CsomeBCy EP_CTERM
01500 HMMccmM_1_CsomeBCy(pf00132pf14602pf00101) CsomeBCy
01510 P_electricGreen CsomeBCy
01520 H_neonGreen CsomeBCy
01530 H_neonGreen CsomeBCy
01540 -pf00361pf00662 -
01550 -pf01059pf00361 -
01560 -pf10216 -
01570 -pf05685 -
01580 -# -
01590 -pf00124 -
01600 -pf01609 -
01610 -pf05685 -
01620 -pf05685 -
01630 -# -
(..13600..)
15230 -# -
15240 -# -
15250 -pf01053 -
15260 -pf00571pf00291 -
15270 HMMreg00392_2_grm2(pf00155) CsomeBCy
15280 -# -
15290 -pf01112 -
15300 -pf02655 -
15310 -# -
15320 -pf00805 -
15330 Tdp_turquoise(#) CsomeBCy
15340 -pf02836 -
15350 -pf01022 -
15360 -pf01758 -
15370 -pf01451 -
15380 -pf00903 -
15390 -# -
15400 -pf13439 -
15410 -# -
15420 -pf00535 -
15430 -# -
(..14430..)
29860 -pf05685 -
29870 -# -
29880 -pf00805 -
29890 -pf00400 -
29900 -# -
29910 -pf13460 -
29920 -pf01894 -
29930 HMMreg00392_2_grm2(pf00155) CsomeBCy
29940 -pf06296 -
29950 -pf01381 -
29960 H_neonGreen CsomeBCy
29970 -pf00109pf02801 -
29980 -pf00160 -
29990 -pf02392 -
30000 -pf00124 -
30010 -pf00421 -
30020 -# -
30030 -pf04434 -
30040 -pf12419pf00271pf00176 -
30050 -pf00583 -
30060 -pf00013pf03726pf01138pf03725pf00575 -
(..11900..)
41960 -pf04828 -
41970 -pf01594 -
41980 -pf00156 -
41990 -# -
42000 -pf00174 -
42010 -pf01292 -
42020 -pf00990 -
42030 -pf13180pf05299 -
42040 HMMreg13545_1_CsomeBCy(pf00027) BUF3
42050 -# -
42060 H_peaSoup CsomeBCy
42070 H_lime CsomeBCy
42080 -# -
42090 -# -
42100 -# -
42110 -pf05154 -
42120 -pf02517 -
42130 -# -
42140 -# -
42150 -pf01383 -
42160 -pf01383pf00427 -
42170 -pf00502 -
BMC-P
BMC-H
BMC-T
Aldehyde dehydrogenase
PTAC
Signature enzyme
Alcohol dehydrogenase
Regulator

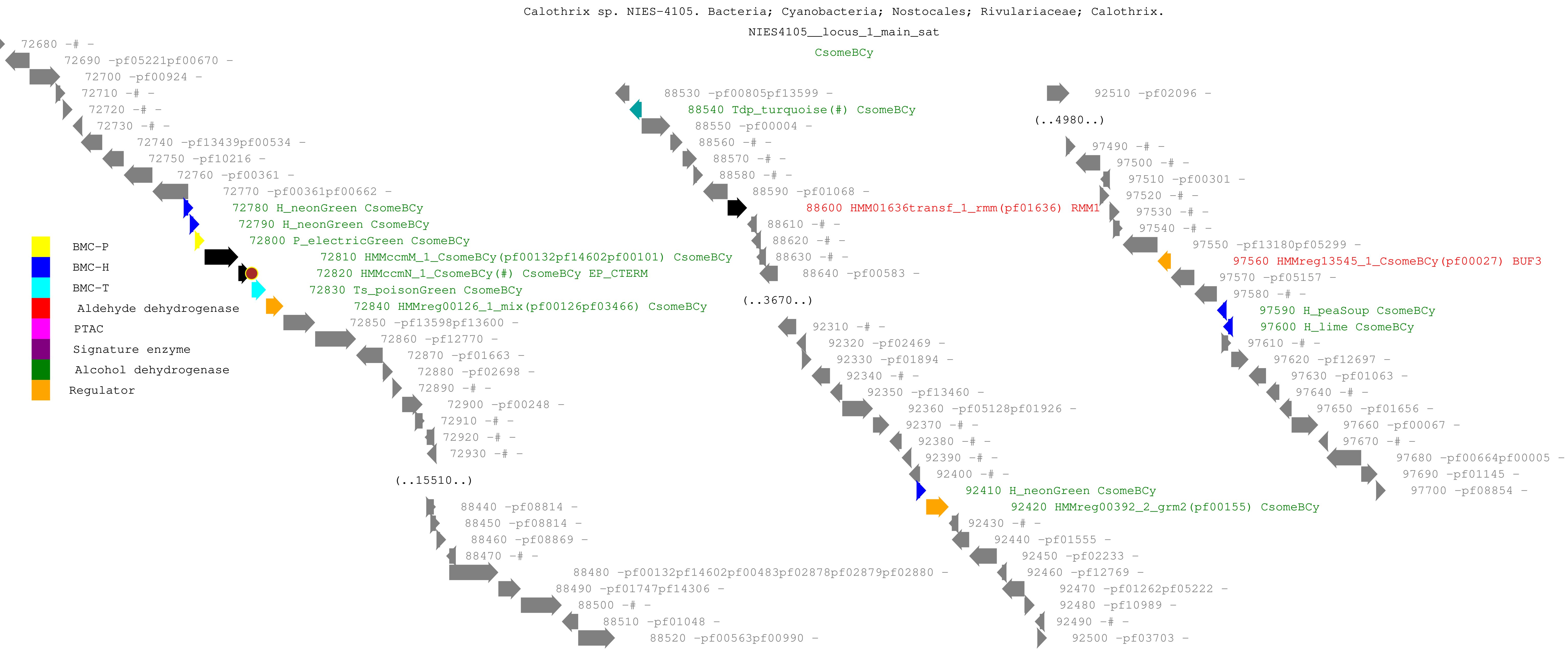

Calothrix sp. NIES-4105. Bacteria; Cyanobacteria; Nostocales; Rivulariaceae; Calothrix.
NIES4105__locus_1_main_sat
CsomeBCy
72680 -# -
72690 -pf05221pf00670 -
72700 -pf00924 -
72710 -# -
72720 -# -
72730 -# -
72740 -pf13439pf00534 -
72750 -pf10216 -
72760 -pf00361 -
72770 -pf00361pf00662 -
72780 H_neonGreen CsomeBCy
72790 H_neonGreen CsomeBCy
72800 P_electricGreen CsomeBCy
72810 HMMccmM_1_CsomeBCy(pf00132pf14602pf00101) CsomeBCy
72820 HMMccmN_1_CsomeBCy(#) CsomeBCy EP_CTERM
72830 Ts_poisonGreen CsomeBCy
72840 HMMreg00126_1_mix(pf00126pf03466) CsomeBCy
72850 -pf13598pf13600 -
72860 -pf12770 -
72870 -pf01663 -
72880 -pf02698 -
72890 -# -
72900 -pf00248 -
72910 -# -
72920 -# -
72930 -# -
(..15510..)
88440 -pf08814 -
88450 -pf08814 -
88460 -pf08869 -
88470 -# -
88480 -pf00132pf14602pf00483pf02878pf02879pf02880 -
88490 -pf01747pf14306 -
88500 -# -
88510 -pf01048 -
88520 -pf00563pf00990 -
88530 -pf00805pf13599 -
88540 Tdp_turquoise(#) CsomeBCy
88550 -pf00004 -
88560 -# -
88570 -# -
88580 -# -
88590 -pf01068 -
88600 HMM01636transf_1_rmm(pf01636) RMM1
88610 -# -
88620 -# -
88630 -# -
88640 -pf00583 -
(..3670..)
92310 -# -
92320 -pf02469 -
92330 -pf01894 -
92340 -# -
92350 -pf13460 -
92360 -pf05128pf01926 -
92370 -# -
92380 -# -
92390 -# -
92400 -# -
92410 H_neonGreen CsomeBCy
92420 HMMreg00392_2_grm2(pf00155) CsomeBCy
92430 -# -
92440 -pf01555 -
92450 -pf02233 -
92460 -pf12769 -
92470 -pf01262pf05222 -
92480 -pf10989 -
92490 -# -
92500 -pf03703 -
92510 -pf02096 -
(..4980..)
97490 -# -
97500 -# -
97510 -pf00301 -
97520 -# -
97530 -# -
97540 -# -
97550 -pf13180pf05299 -
97560 HMMreg13545_1_CsomeBCy(pf00027) BUF3
97570 -pf05157 -
97580 -# -
97590 H_peaSoup CsomeBCy
97600 H_lime CsomeBCy
97610 -# -
97620 -pf12697 -
97630 -pf01063 -
97640 -# -
97650 -pf01656 -
97660 -pf00067 -
97670 -# -
97680 -pf00664pf00005 -
97690 -pf01145 -
97700 -pf08854 -
BMC-P
BMC-H
BMC-T
Aldehyde dehydrogenase
PTAC
Signature enzyme
Alcohol dehydrogenase
Regulator

Fischerella sp. NIES-4106. Bacteria; Cyanobacteria; Nostocales; Hapalosiphonaceae; Fischerella.

NIES4106__locus_1_main_sat

CsomeBCy

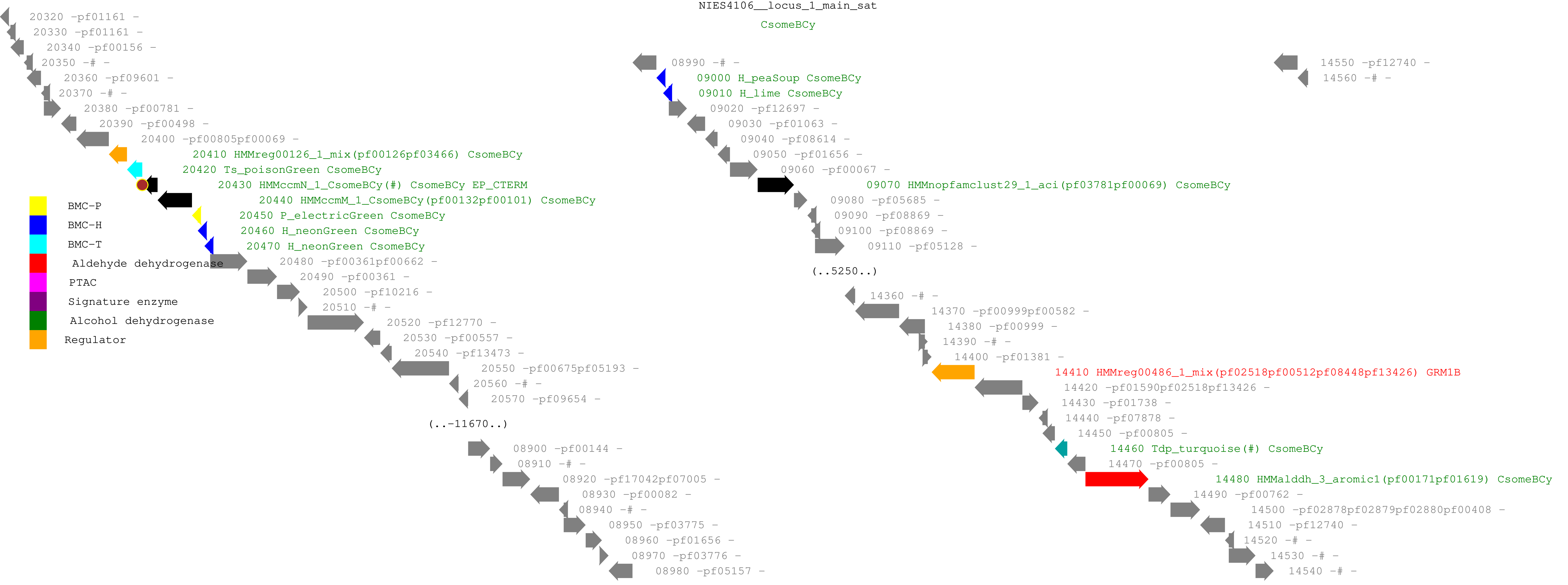

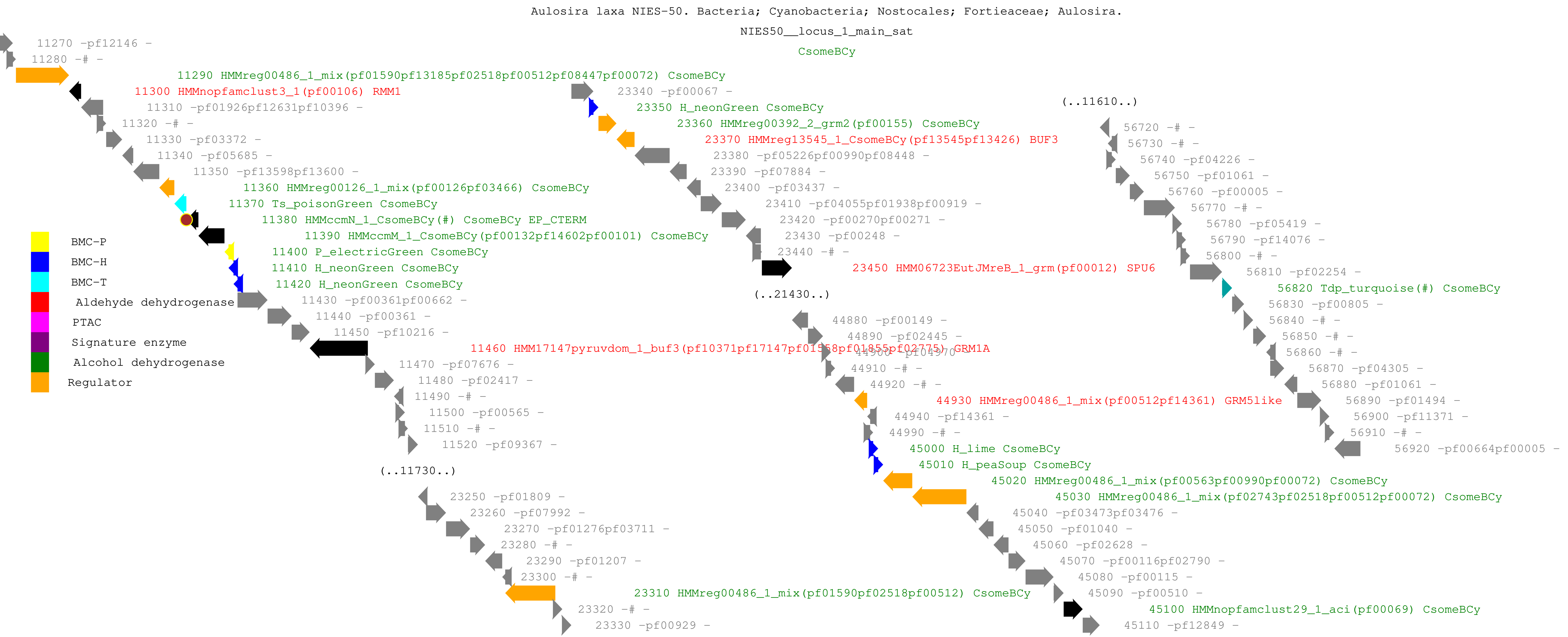

Aulosira laxa NIES-50. Bacteria; Cyanobacteria; Nostocales; Fortieaceae; Aulosira.
NIES50__locus_1_main_sat
CsomeBCy
11270 -pf12146 -
11280 -# -
11290 HMMreg00486_1_mix(pf01590pf13185pf02518pf00512pf08447pf00072) CsomeBCy
11300 HMMnopfamclust3_1(pf00106) RMM1
11310 -pf01926pf12631pf10396 -
11320 -# -
11330 -pf03372 -
11340 -pf05685 -
11350 -pf13598pf13600 -
11360 HMMreg00126_1_mix(pf00126pf03466) CsomeBCy
11370 Ts_poisonGreen CsomeBCy
11380 HMMccmN_1_CsomeBCy(#) CsomeBCy EP_CTERM
11390 HMMccmM_1_CsomeBCy(pf00132pf14602pf00101) CsomeBCy
11400 P_electricGreen CsomeBCy
11410 H_neonGreen CsomeBCy
11420 H_neonGreen CsomeBCy
11430 -pf00361pf00662 -
11440 -pf00361 -
11450 -pf10216 -
11460 HMM17147pyruvdom_1_buf3(pf10371pf17147pf01558pf01855pf02775) GRM1A
11470 -pf07676 -
11480 -pf02417 -
11490 -# -
11500 -pf00565 -
11510 -# -
11520 -pf09367 -
(..11730..)
23250 -pf01809 -
23260 -pf07992 -
23270 -pf01276pf03711 -
23280 -# -
23290 -pf01207 -
23300 -# -
23310 HMMreg00486_1_mix(pf01590pf02518pf00512) CsomeBCy
23320 -# -
23330 -pf00929 -
23340 -pf00067 -
23350 H_neonGreen CsomeBCy
23360 HMMreg00392_2_grm2(pf00155) CsomeBCy
23370 HMMreg13545_1_CsomeBCy(pf13545pf13426) BUF3
23380 -pf05226pf00990pf08448 -
23390 -pf07884 -
23400 -pf03437 -
23410 -pf04055pf01938pf00919 -
23420 -pf00270pf00271 -
23430 -pf00248 -
23440 -# -
23450 HMM06723EutJMreB_1_grm(pf00012) SPU6
(..21430..)
44880 -pf00149 -
44890 -pf02445 -
44900 -pf04970 -
44910 -# -
44920 -# -
44930 HMMreg00486_1_mix(pf00512pf14361) GRM5like
44940 -pf14361 -
44990 -# -
45000 H_lime CsomeBCy
45010 H_peaSoup CsomeBCy
45020 HMMreg00486_1_mix(pf00563pf00990pf00072) CsomeBCy
45030 HMMreg00486_1_mix(pf02743pf02518pf00512pf00072) CsomeBCy
45040 -pf03473pf03476 -
45050 -pf01040 -
45060 -pf02628 -
45070 -pf00116pf02790 -
45080 -pf00115 -
45090 -pf00510 -
45100 HMMnopfamclust29_1_aci(pf00069) CsomeBCy
45110 -pf12849 -
(..11610..)
56720 -# -
56730 -# -
56740 -pf04226 -
56750 -pf01061 -
56760 -pf00005 -
56770 -# -
56780 -pf05419 -
56790 -pf14076 -
56800 -# -
56810 -pf02254 -
56820 Tdp_turquoise(#) CsomeBCy
56830 -pf00805 -
56840 -# -
56850 -# -
56860 -# -
56870 -pf04305 -
56880 -pf01061 -
56890 -pf01494 -
56900 -pf11371 -
56910 -# -
56920 -pf00664pf00005 -
BMC-P
BMC-H
BMC-T
Aldehyde dehydrogenase
PTAC
Signature enzyme
Alcohol dehydrogenase
Regulator

Fischerella major NIES-592. Bacteria; Cyanobacteria; Nostocales; Hapalosiphonaceae; Fischerella.

NIES592__locus_1_main_sat

CsomeBCy

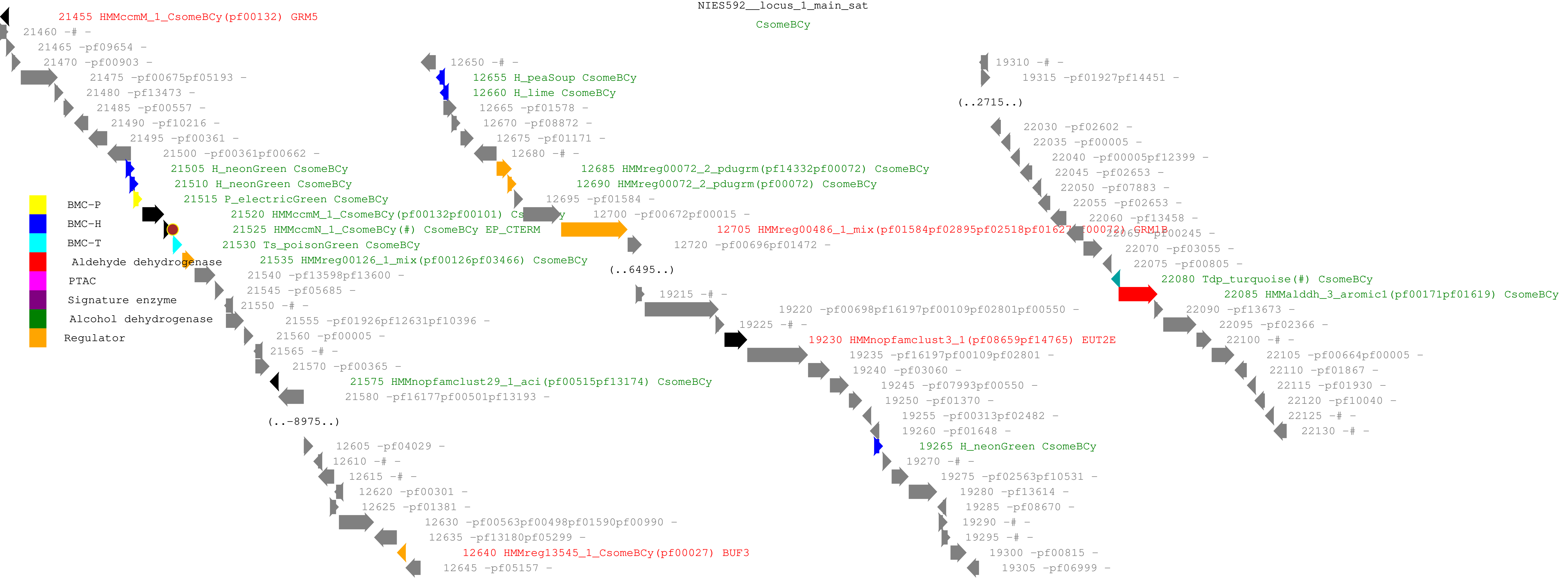

Hydrococcus rivularis NIES-593. Bacteria; Cyanobacteria; Pleurocapsales; Hydrococcaceae; Hydrococcus.

NIES593__locus_1_main_sat

CsomeBCy

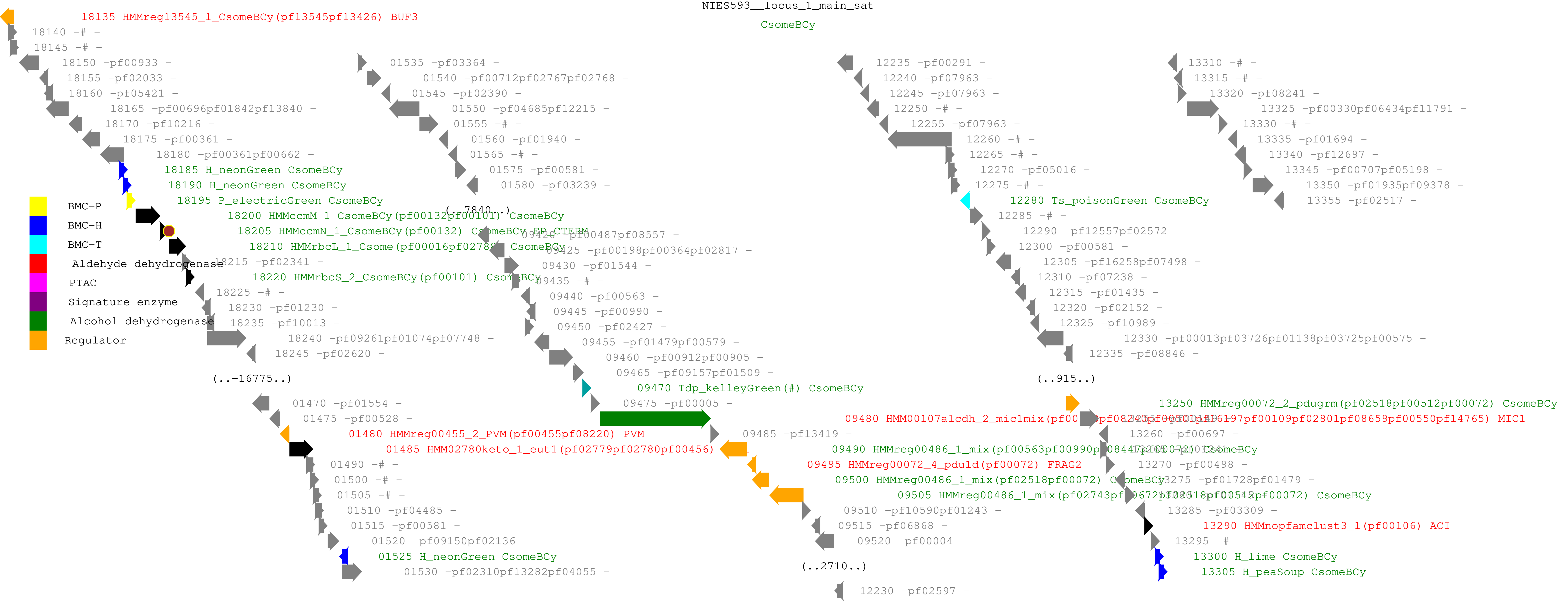

Sphaerospermopsis kisseleviana NIES-73. Bacteria; Cyanobacteria; Nostocales; Aphanizomenonaceae;

NIES73__locus_1_main_sat

CsomeBCy

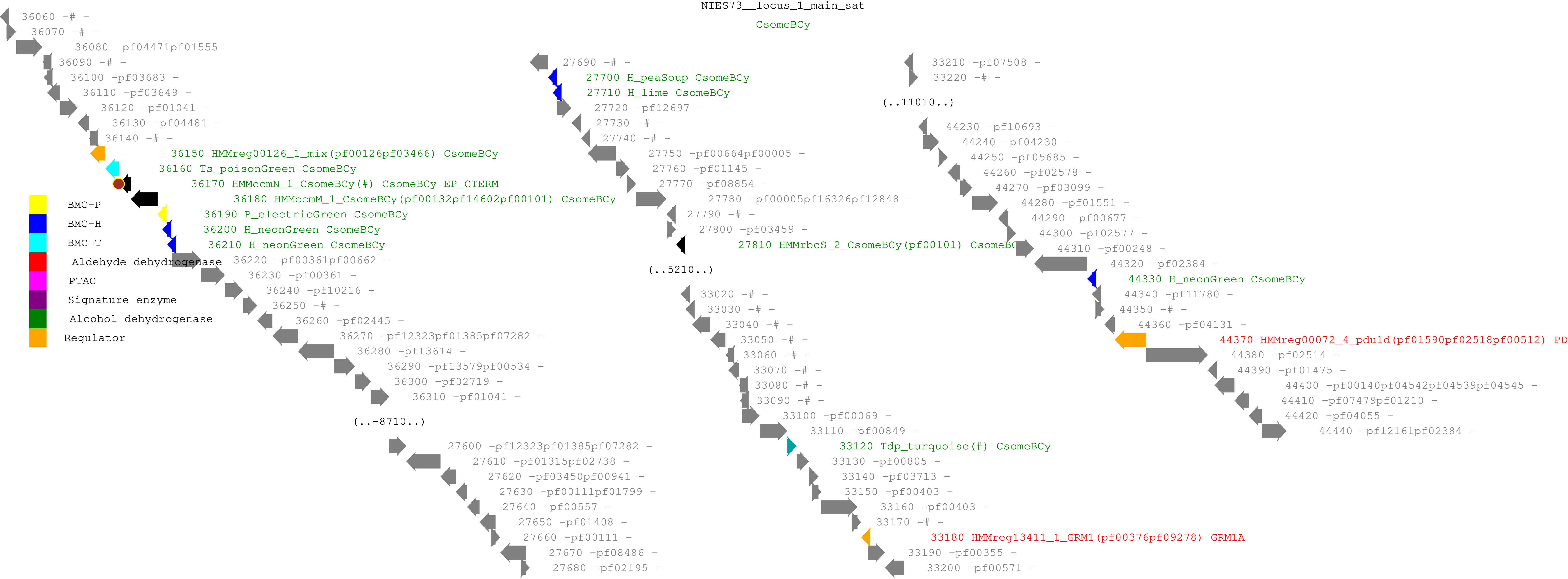

Dolichospermum compactum NIES-806. Bacteria; Cyanobacteria; Nostocales; Aphanizomenonaceae;

NIES806__locus_1_main_sat

CsomeBCy

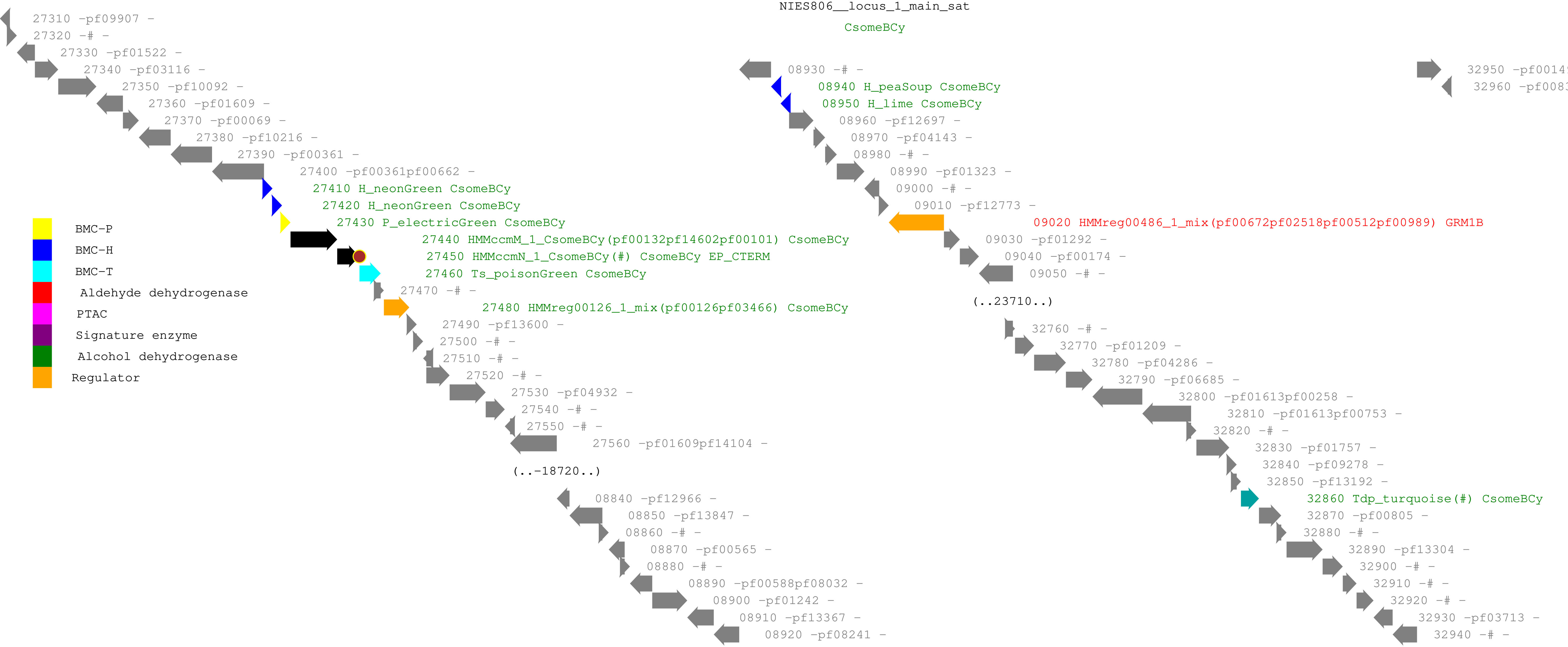

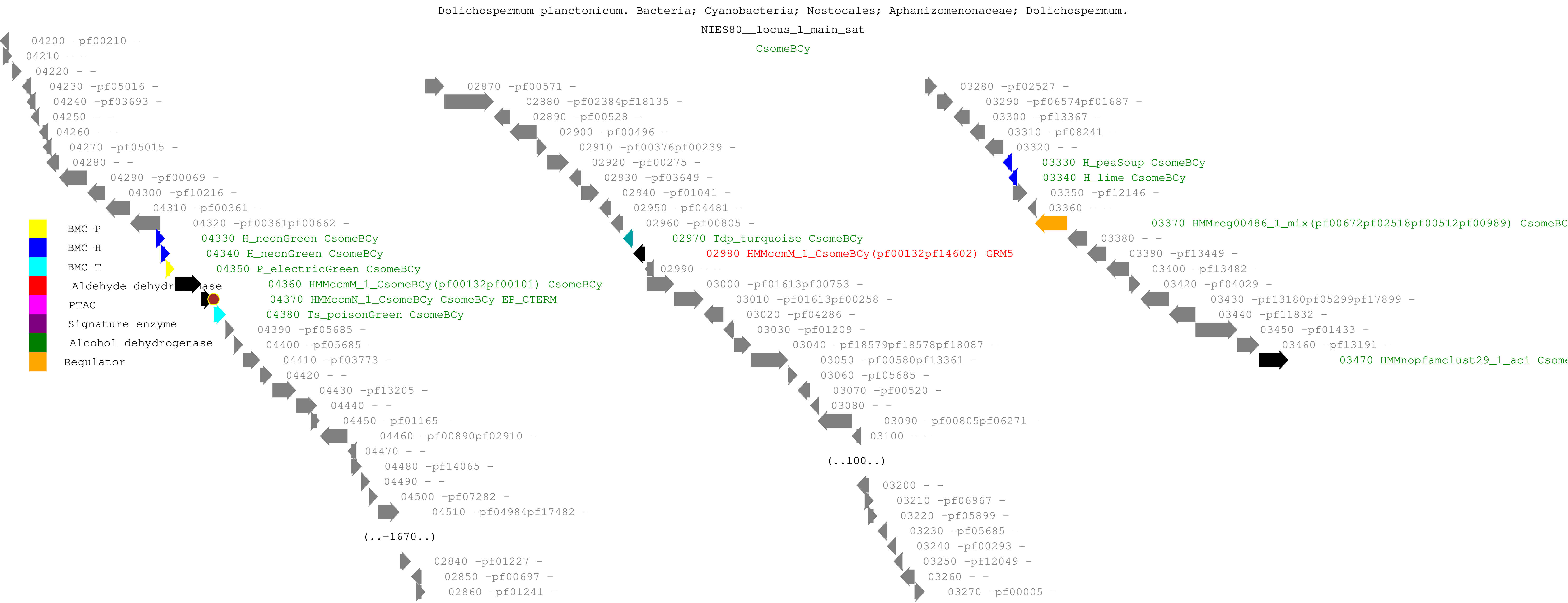
Dolichospermum planctonicum. Bacteria; Cyanobacteria; Nostocales; Aphanizomenonaceae; Dolichospermum.
NIES80__locus_1_main_sat
CsomeBCy
BMC-P
BMC-H
BMC-T
Aldehyde dehydrogenase
PTAC
Signature enzyme
Alcohol dehydrogenase
Regulator
04200 -pf00210 -
04210 - -
04220 - -
04230 -pf05016 -
04240 -pf03693 -
04250 - -
04260 - -
04270 -pf05015 -
04280 - -
04290 -pf00069 -
04300 -pf10216 -
04310 -pf00361 -
04320 -pf00361pf00662 -
04330 H_neonGreen CsomeBCy
04340 H_neonGreen CsomeBCy
04350 P_electricGreen CsomeBCy
04360 HMMccmM_1_CsomeBCy(pf00132pf00101) CsomeBCy
04370 HMMccmN_1_CsomeBCy CsomeBCy EP_CTERM
04380 Ts_poisonGreen CsomeBCy
04390 -pf05685 -
04400 -pf05685 -
04410 -pf03773 -
04420 - -
04430 -pf13205 -
04440 - -
04450 -pf01165 -
04460 -pf00890pf02910 -
04470 - -
04480 -pf14065 -
04490 - -
04500 -pf07282 -
04510 -pf04984pf17482 -
(..-1670..)
02840 -pf01227 -
02850 -pf00697 -
02860 -pf01241 -
02870 -pf00571 -
02880 -pf02384pf18135 -
02890 -pf00528 -
02900 -pf00496 -
02910 -pf00376pf00239 -
02920 -pf00275 -
02930 -pf03649 -
02940 -pf01041 -
02950 -pf04481 -
02960 -pf00805 -
02970 Tdp_turquoise CsomeBCy
02980 HMMccmM_1_CsomeBCy(pf00132pf14602) GRM5
02990 - -
03000 -pf01613pf00753 -
03010 -pf01613pf00258 -
03020 -pf04286 -
03030 -pf01209 -
03040 -pf18579pf18578pf18087 -
03050 -pf00580pf13361 -
03060 -pf05685 -
03070 -pf00520 -
03080 - -
03090 -pf00805pf06271 -
03100 - -
(..100..)
03200 - -
03210 -pf06967 -
03220 -pf05899 -
03230 -pf05685 -
03240 -pf00293 -
03250 -pf12049 -
03260 - -
03270 -pf00005 -
03280 -pf02527 -
03290 -pf06574pf01687 -
03300 -pf13367 -
03310 -pf08241 -
03320 - -
03330 H_peaSoup CsomeBCy
03340 H_lime CsomeBCy
03350 -pf12146 -
03360 - -
03370 HMMreg00486_1_mix(pf00672pf02518pf00512pf00989) CsomeBC
03380 - -
03390 -pf13449 -
03400 -pf13482 -
03420 -pf04029 -
03430 -pf13180pf05299pf17899 -
03440 -pf11832 -
03450 -pf01433 -
03460 -pf13191 -
03470 HMMnopfamclust29_1_aci Csome

Raphidiopsis curvata NIES-932. Bacteria; Cyanobacteria; Nostocales; Aphanizomenonaceae; Raphidiopsis.

NIES932__locus_1_main_sat

CsomeBCy

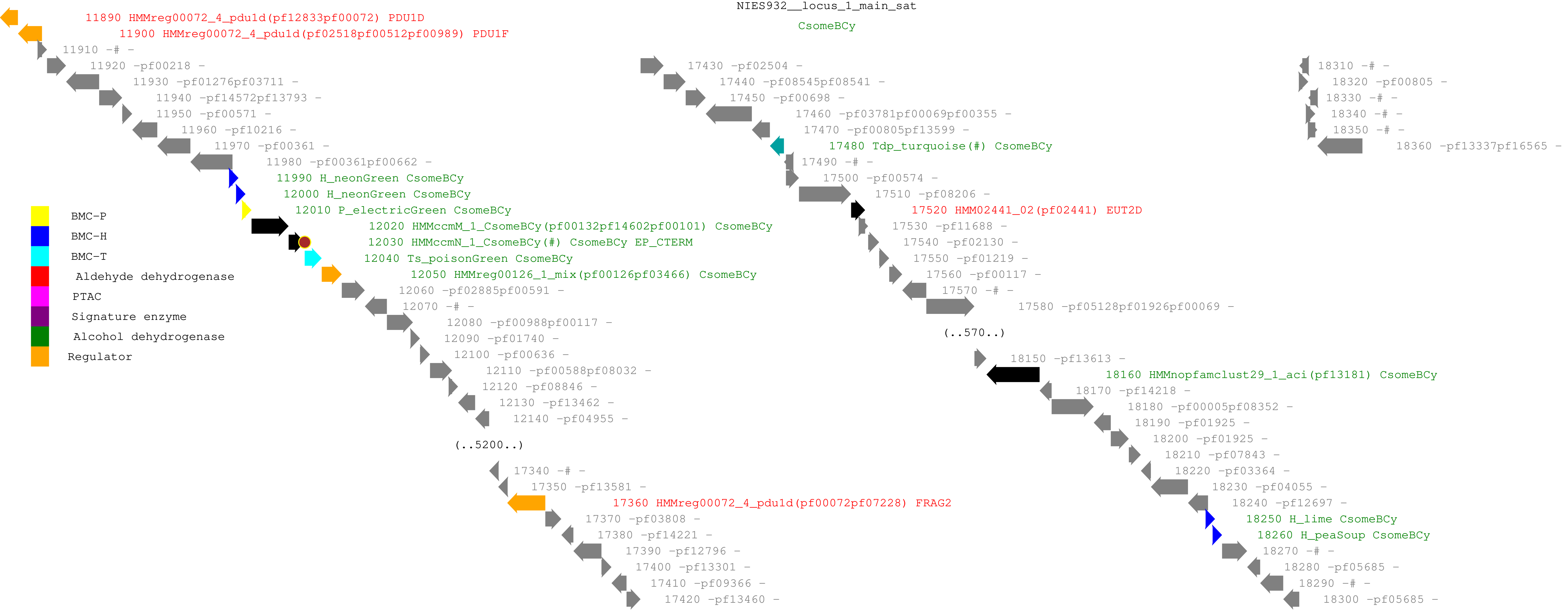

Synechococcus sp. NIES-970. Bacteria; Cyanobacteria; Synechococcales; Synechococcaceae;

NIES970__locus_1_main_sat

CsomeBCy

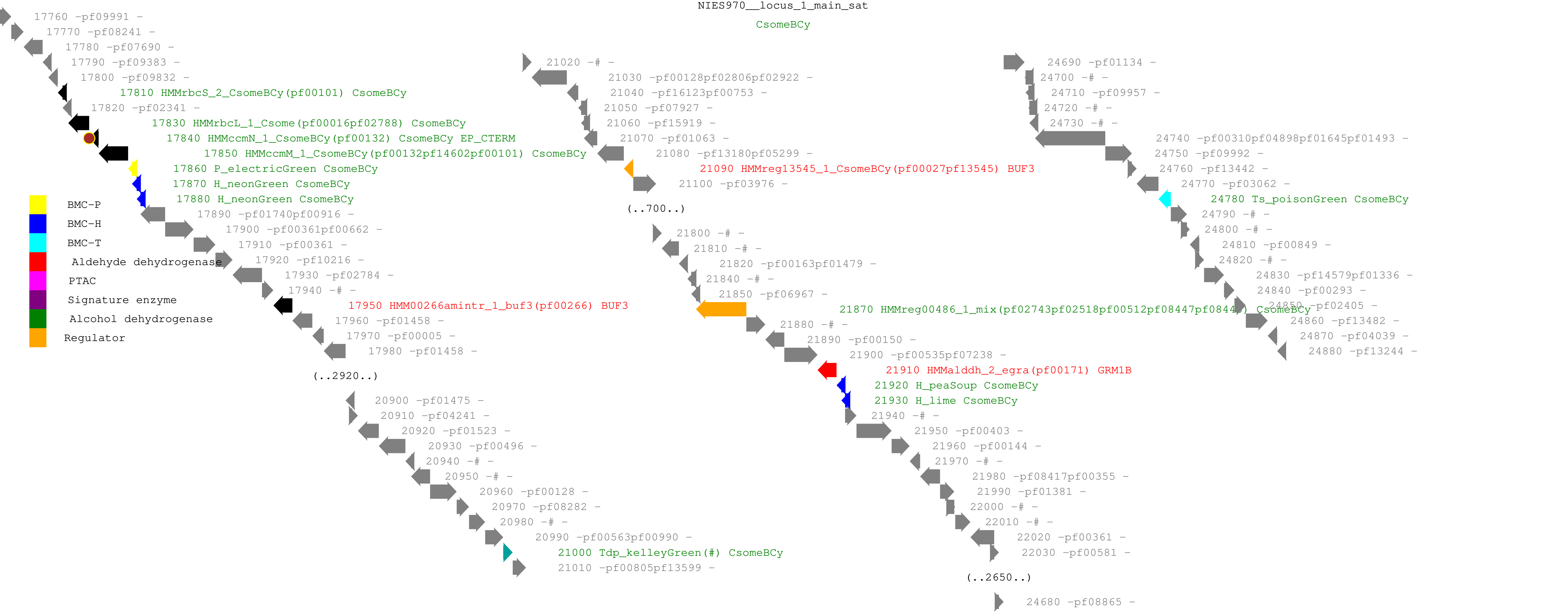

Thermosynechococcus sp. NK55a. Bacteria; Cyanobacteria; Synechococcales; Synechococcaceae;

NK55__locus_1_main_sat

CsomeBCy

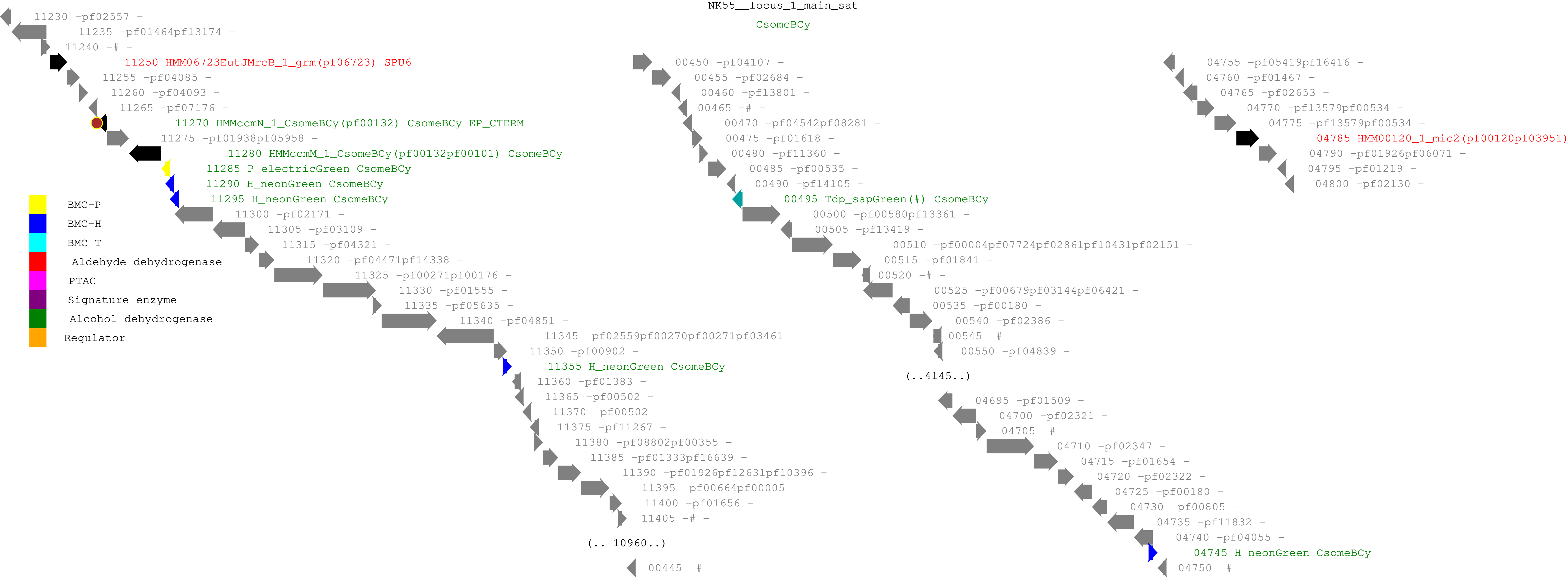

Nostoc sp. 'Lobaria pulmonaria (5183) cyanobiont'. Bacteria; Cyanobacteria; Nostocales; Nostocaceae; Nostoc.

NLP__locus_1_main_sat

CsomeBCy

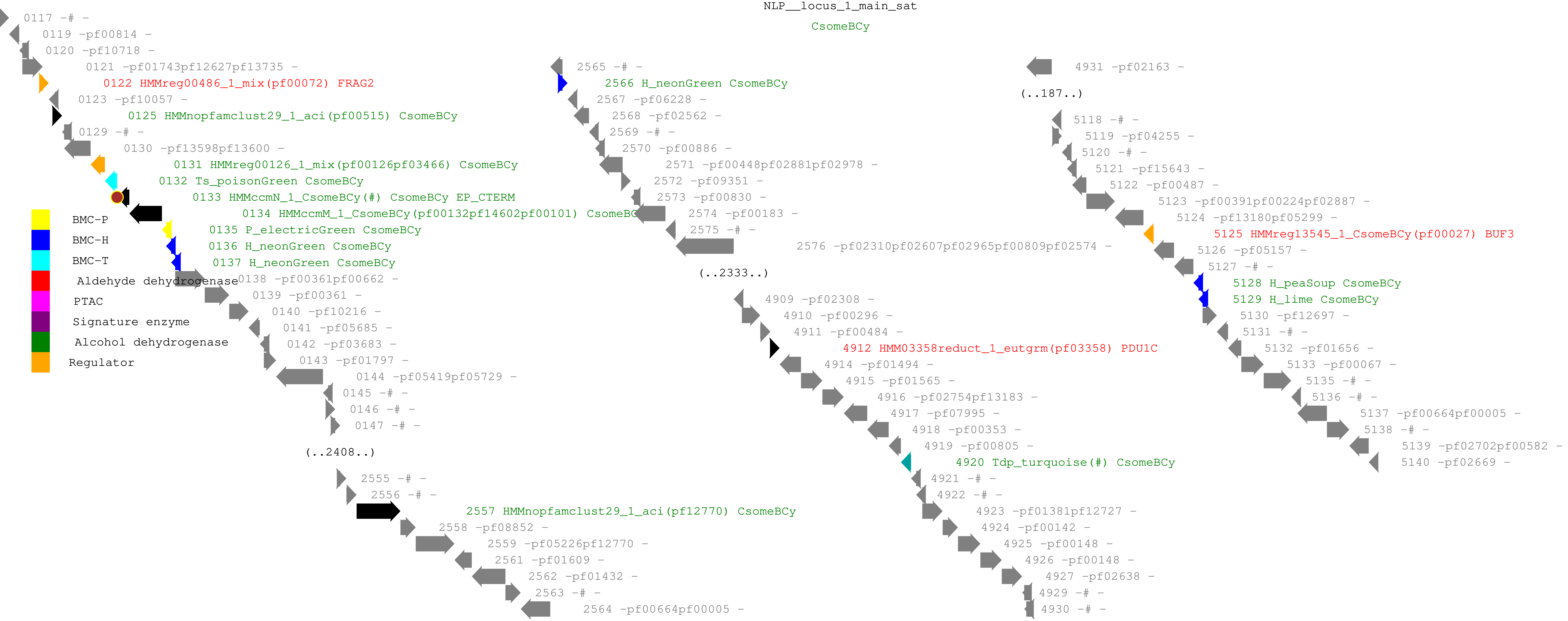

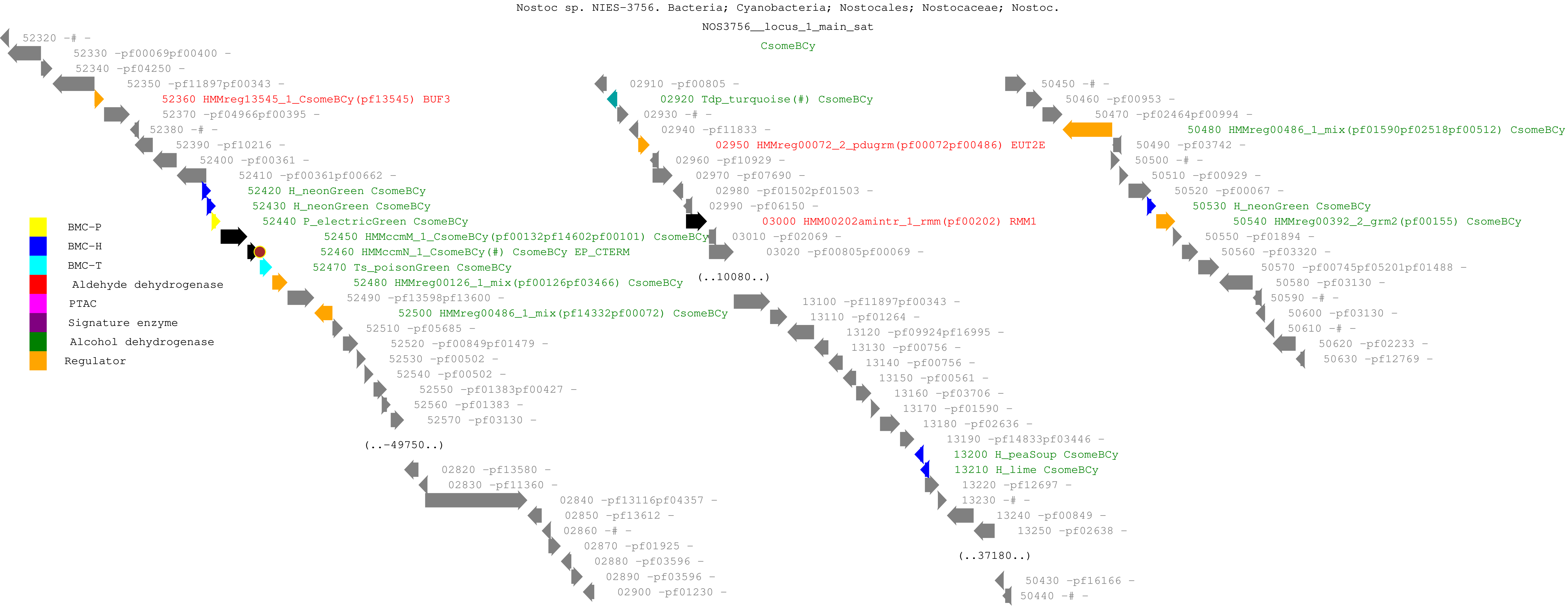

Nostoc sp. NIES-3756. Bacteria; Cyanobacteria; Nostocales; Nostocaceae; Nostoc.
NOS3756__locus_1_main_sat
CsomeBCy
52320 -# -
52330 -pf00069pf00400 -
52340 -pf04250 -
52350 -pf11897pf00343 -
52360 HMMreg13545_1_CsomeBCy(pf13545) BUF3
52370 -pf04966pf00395 -
52380 -# -
52390 -pf10216 -
52400 -pf00361 -
52410 -pf00361pf00662 -
52420 H_neonGreen CsomeBCy
52430 H_neonGreen CsomeBCy
52440 P_electricGreen CsomeBCy
52450 HMMccmM_1_CsomeBCy(pf00132pf14602pf00101) CsomeBCy
52460 HMMccmN_1_CsomeBCy(#) CsomeBCy EP_CTERM
52470 Ts_poisonGreen CsomeBCy
52480 HMMreg00126_1_mix(pf00126pf03466) CsomeBCy
52490 -pf13598pf13600 -
52500 HMMreg00486_1_mix(pf14332pf00072) CsomeBCy
52510 -pf05685 -
52520 -pf00849pf01479 -
52530 -pf00502 -
52540 -pf00502 -
52550 -pf01383pf00427 -
52560 -pf01383 -
52570 -pf03130 -
(..-49750..)
02820 -pf13580 -
02830 -pf11360 -
02840 -pf13116pf04357 -
02850 -pf13612 -
02860 -# -
02870 -pf01925 -
02880 -pf03596 -
02890 -pf03596 -
02900 -pf01230 -
02910 -pf00805 -
02920 Tdp_turquoise(#) CsomeBCy
02930 -# -
02940 -pf11833 -
02950 HMMreg00072_2_pdugrm(pf00072pf00486) EUT2E
02960 -pf10929 -
02970 -pf07690 -
02980 -pf01502pf01503 -
02990 -pf06150 -
03000 HMM00202amintr_1_rmm(pf00202) RMM1
03010 -pf02069 -
03020 -pf00805pf00069 -
(..10080..)
13100 -pf11897pf00343 -
13110 -pf01264 -
13120 -pf09924pf16995 -
13130 -pf00756 -
13140 -pf00756 -
13150 -pf00561 -
13160 -pf03706 -
13170 -pf01590 -
13180 -pf02636 -
13190 -pf14833pf03446 -
13200 H_peaSoup CsomeBCy
13210 H_lime CsomeBCy
13220 -pf12697 -
13230 -# -
13240 -pf00849 -
13250 -pf02638 -
(..37180..)
50430 -pf16166 -
50440 -# -
50450 -# -
50460 -pf00953 -
50470 -pf02464pf00994 -
50480 HMMreg00486_1_mix(pf01590pf02518pf00512) CsomeBCy
50490 -pf03742 -
50500 -# -
50510 -pf00929 -
50520 -pf00067 -
50530 H_neonGreen CsomeBCy
50540 HMMreg00392_2_grm2(pf00155) CsomeBCy
50550 -pf01894 -
50560 -pf03320 -
50570 -pf00745pf05201pf01488 -
50580 -pf03130 -
50590 -# -
50600 -pf03130 -
50610 -# -
50620 -pf02233 -
50630 -pf12769 -
BMC-P
BMC-H
BMC-T
Aldehyde dehydrogenase
PTAC
Signature enzyme
Alcohol dehydrogenase
Regulator

Nostoc sp. PCC 7107. Bacteria; Cyanobacteria; Nostocales; Nostocaceae; Nostoc.

Nos7107__locus_1_main_sat

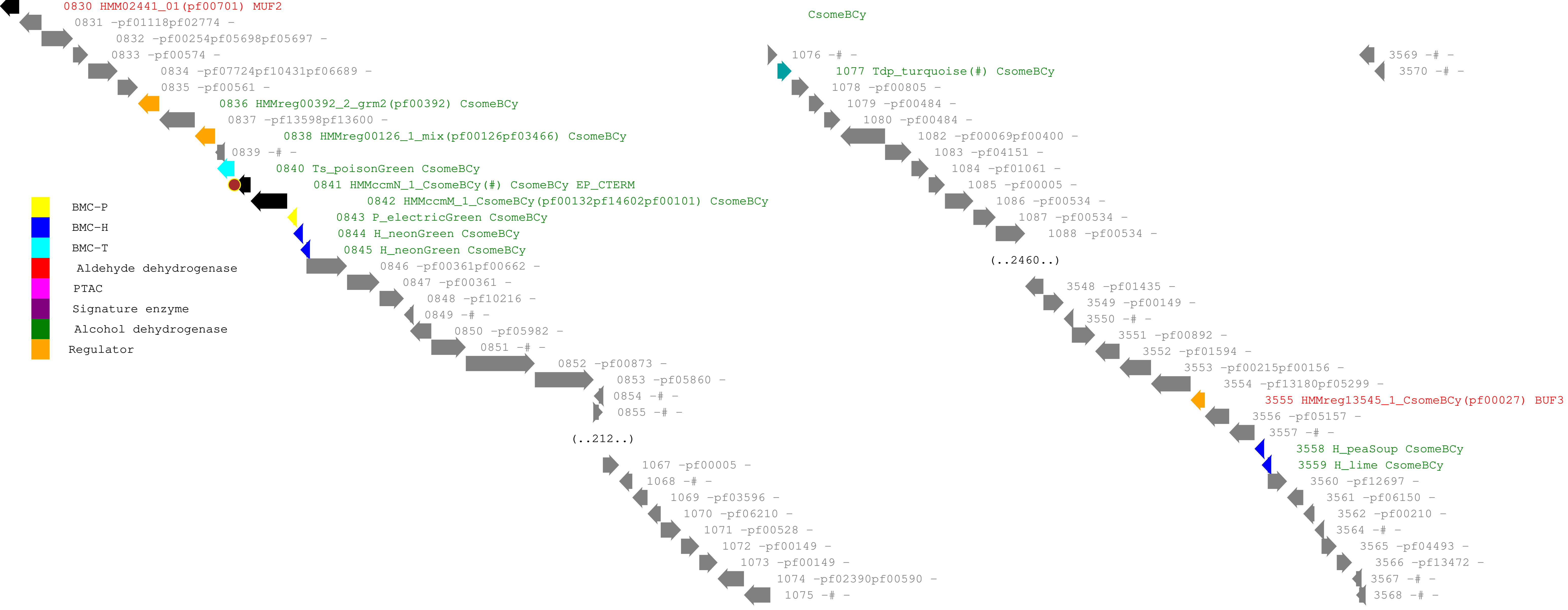

Nostoc sp. (strain ATCC 29411 / PCC 7524). Bacteria; Cyanobacteria; Nostocales; Nostocaceae; Nostoc.

Nos7524__locus_1_main_sat

CsomeBCy

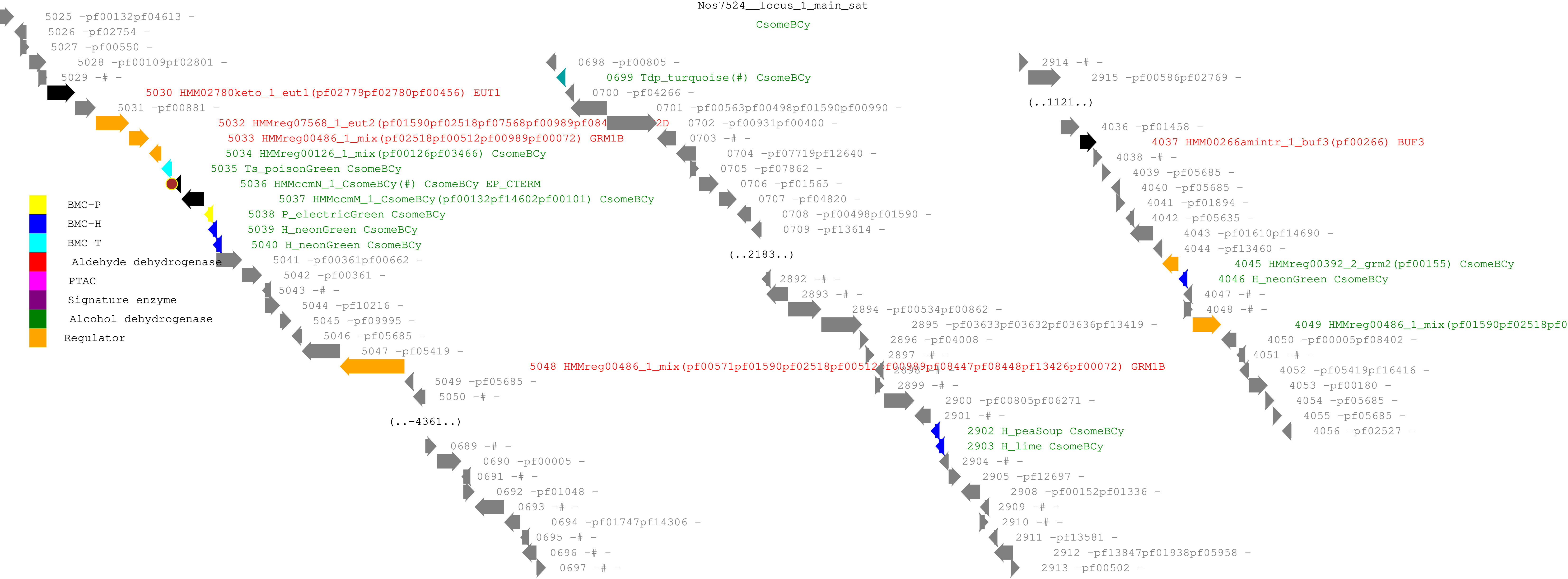

Nostoc sp. 'Peltigera membranacea cyanobiont N6'. Bacteria; Cyanobacteria; Nostocales; Nostocaceae; Nostoc.

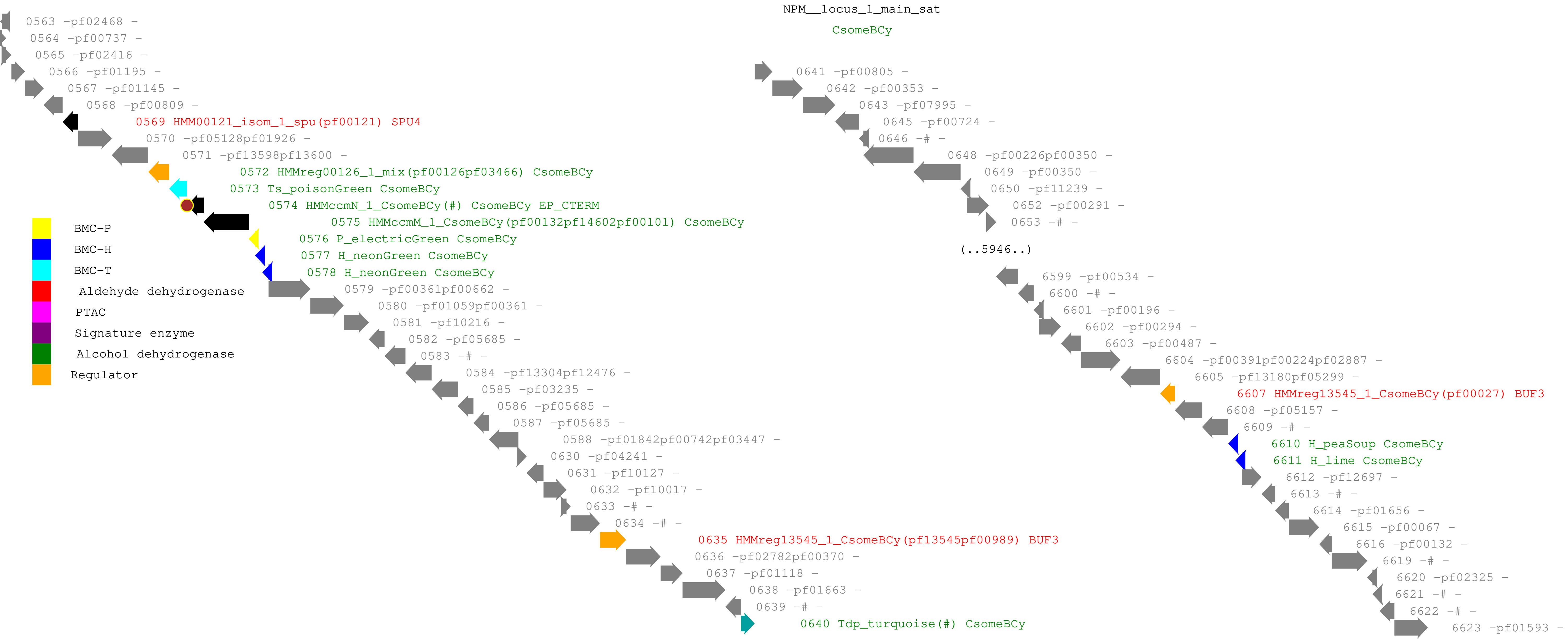

Nostoc punctiforme (strain ATCC 29133 / PCC 73102). Bacteria; Cyanobacteria; Nostocales; Nostocaceae; Nostoc.

Npun_F_locus_1_main_sat

CsomeBCy

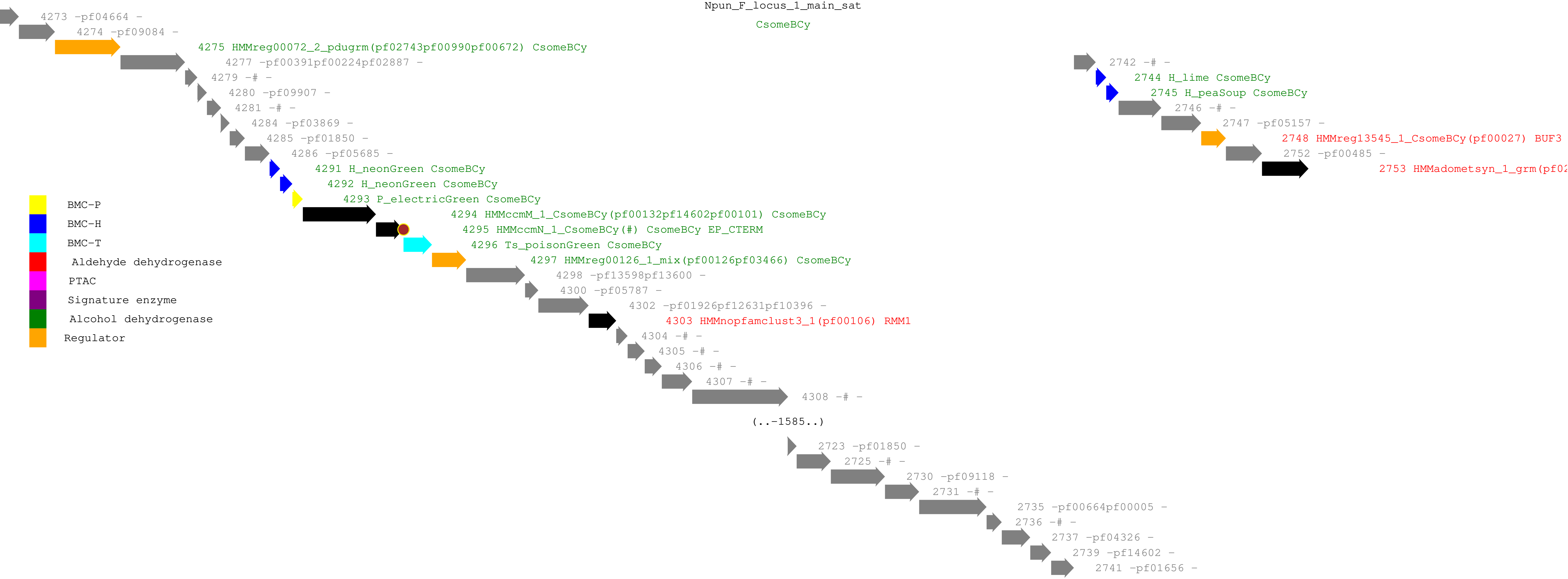

Nodularia spumigena CCY9414. Bacteria; Cyanobacteria; Nostocales; Aphanizomenonaceae; Nodularia.

NSP__locus_1_main_sat

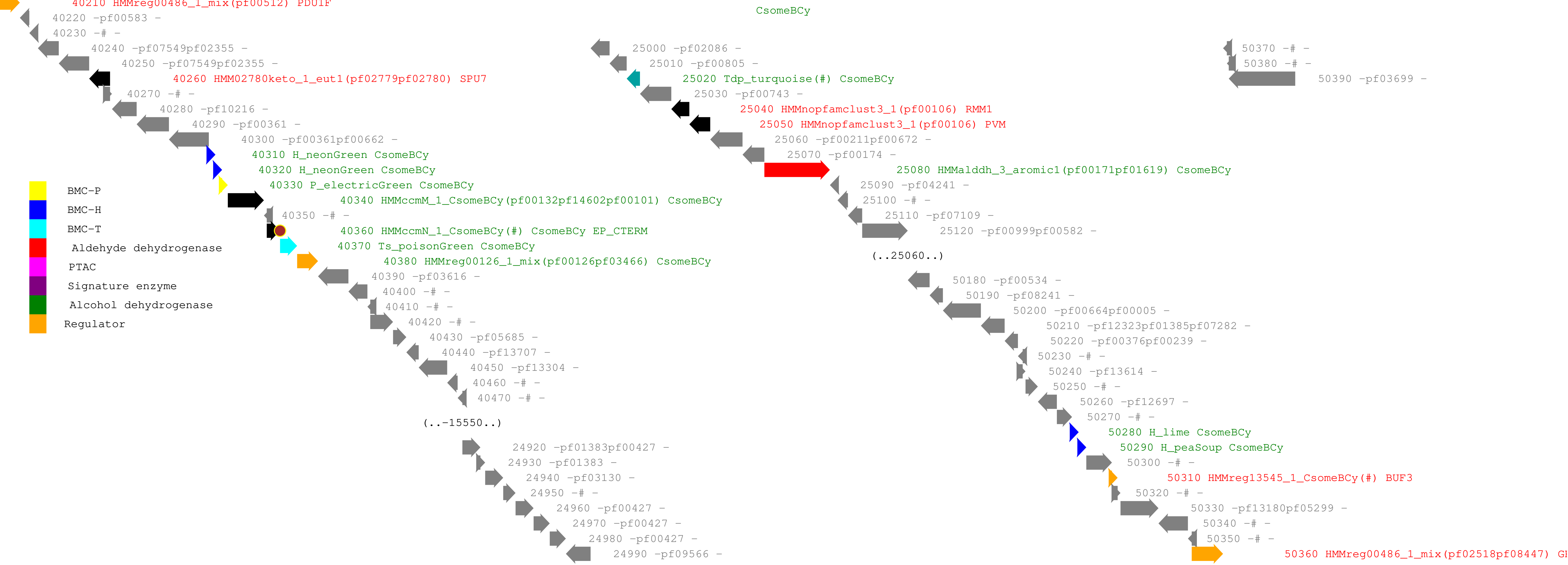

Microcystis aeruginosa TAIHU98. Bacteria; Cyanobacteria; Oscillatoriophycideae; Chroococcales;

O53__locus_fragmented

CsomeBCy

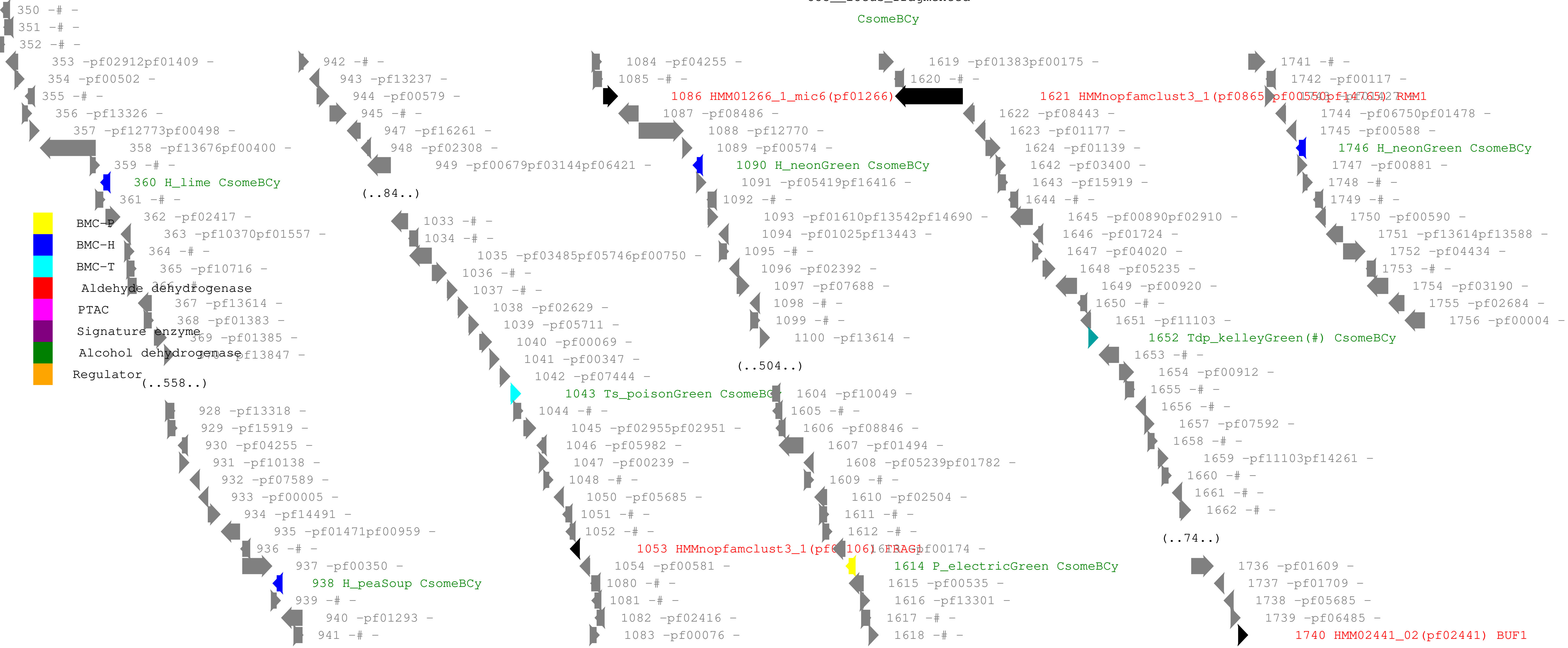

Leptolyngbya sp. O-77. Bacteria; Cyanobacteria; Synechococcales; Leptolyngbyaceae;

O77CONTIG1__locus_1_main_sat

CsomeBCy

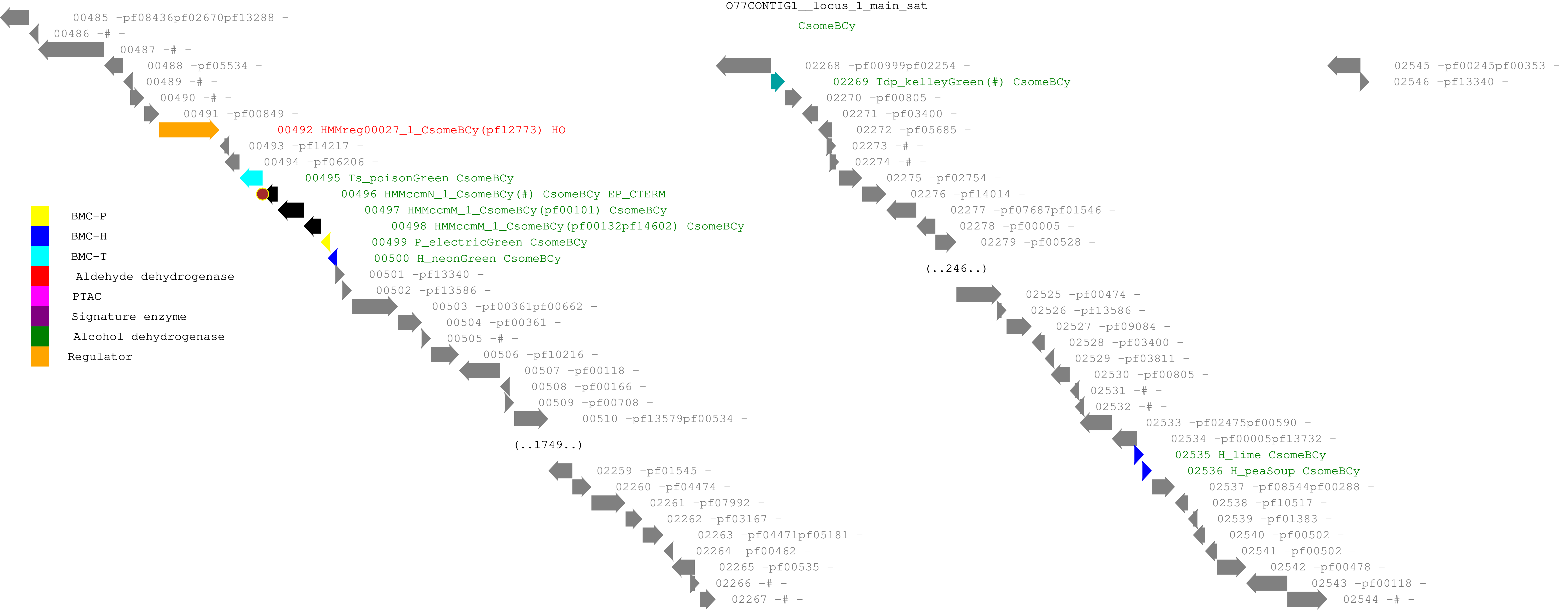

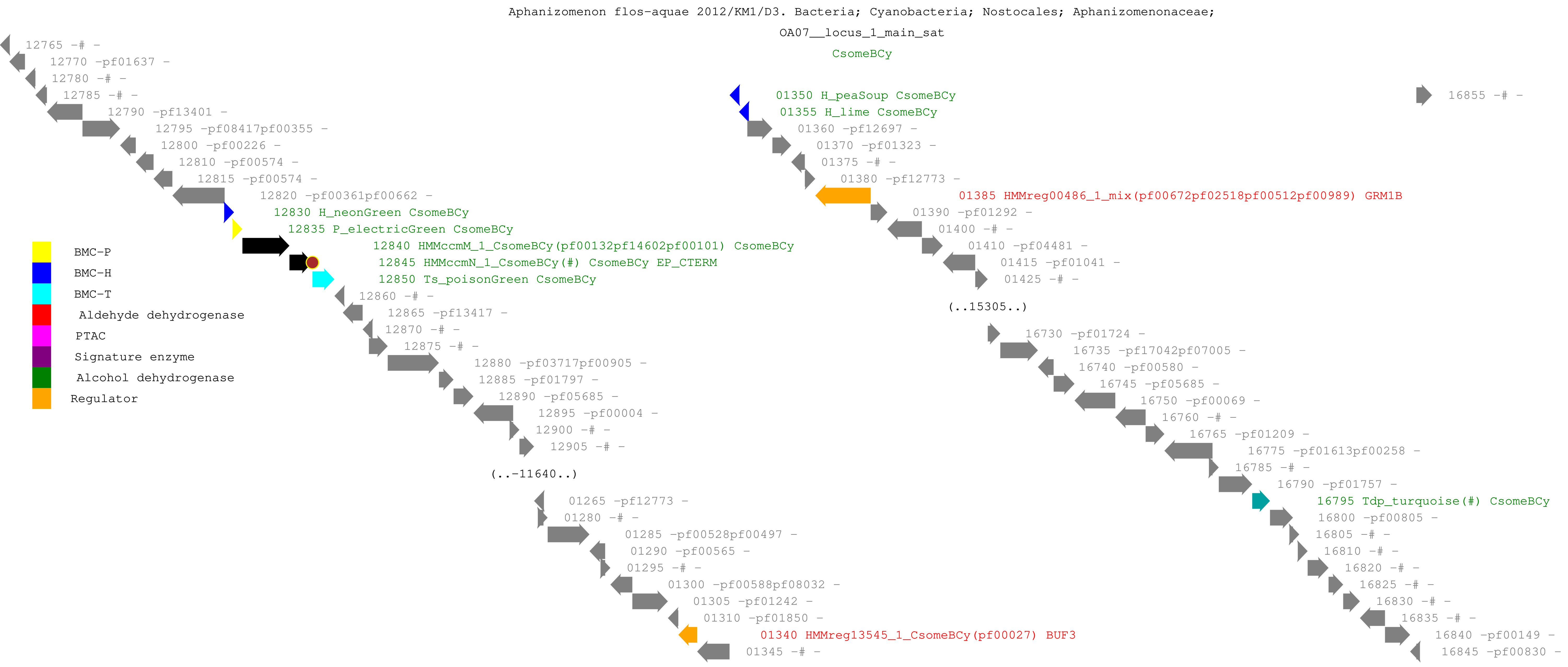
Aphanizomenon flos-aquae 2012/KM1/D3. Bacteria; Cyanobacteria; Nostocales; Aphanizomenonaceae;
OA07__locus_1_main_sat
CsomeBCy
12765 -# -
12770 -pf01637 -
12780 -# -
12785 -# -
12790 -pf13401 -
12795 -pf08417pf00355 -
12800 -pf00226 -
12810 -pf00574 -
12815 -pf00574 -
12820 -pf00361pf00662 -
12830 H_neonGreen CsomeBCy
12835 P_electricGreen CsomeBCy
12840 HMMccmM_1_CsomeBCy(pf00132pf14602pf00101) CsomeBCy
12845 HMMccmN_1_CsomeBCy(#) CsomeBCy EP_CTERM
12850 Ts_poisonGreen CsomeBCy
12860 -# -
12865 -pf13417 -
12870 -# -
12875 -# -
12880 -pf03717pf00905 -
12885 -pf01797 -
12890 -pf05685 -
12895 -pf00004 -
12900 -# -
12905 -# -
(..-11640..)
01265 -pf12773 -
01280 -# -
01285 -pf00528pf00497 -
01290 -pf00565 -
01295 -# -
01300 -pf00588pf08032 -
01305 -pf01242 -
01310 -pf01850 -
01340 HMMreg13545_1_CsomeBCy(pf00027) BUF3
01345 -# -
BMC-P
BMC-H
BMC-T
Aldehyde dehydrogenase
PTAC
Signature enzyme
Alcohol dehydrogenase
Regulator
01350 H_peaSoup CsomeBCy
01355 H_lime CsomeBCy
01360 -pf12697 -
01370 -pf01323 -
01375 -# -
01380 -pf12773 -
01385 HMMreg00486_1_mix(pf00672pf02518pf00512pf00989) GRM1B
01390 -pf01292 -
01400 -# -
01410 -pf04481 -
01415 -pf01041 -
01425 -# -
(..15305..)
16730 -pf01724 -
16735 -pf17042pf07005 -
16740 -pf00580 -
16745 -pf05685 -
16750 -pf00069 -
16760 -# -
16765 -pf01209 -
16775 -pf01613pf00258 -
16785 -# -
16790 -pf01757 -
16795 Tdp_turquoise(#) CsomeBCy
16800 -pf00805 -
16805 -# -
16810 -# -
16820 -# -
16825 -# -
16830 -# -
16835 -# -
16840 -pf00149 -
16845 -pf00830 -
16855 -# -

Microcystis aeruginosa NIES-88. Bacteria; Cyanobacteria; Oscillatoriophycideae; Chroococcales;

OA58__locus_1_main_sat

CsomeBCy

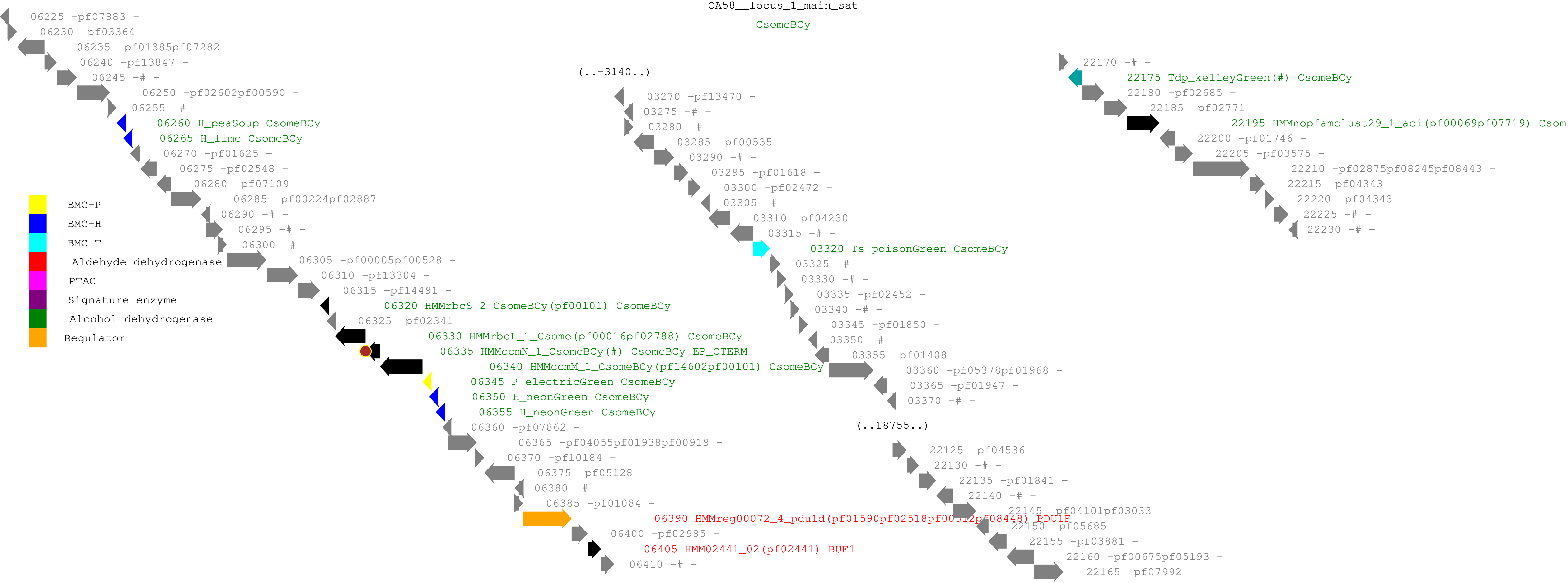

Oscillatoria nigro-viridis PCC 7112. Bacteria; Cyanobacteria; Oscillatoriophycideae; Oscillatoriales;

Osc7112__locus_1_main_sat

CsomeBCy

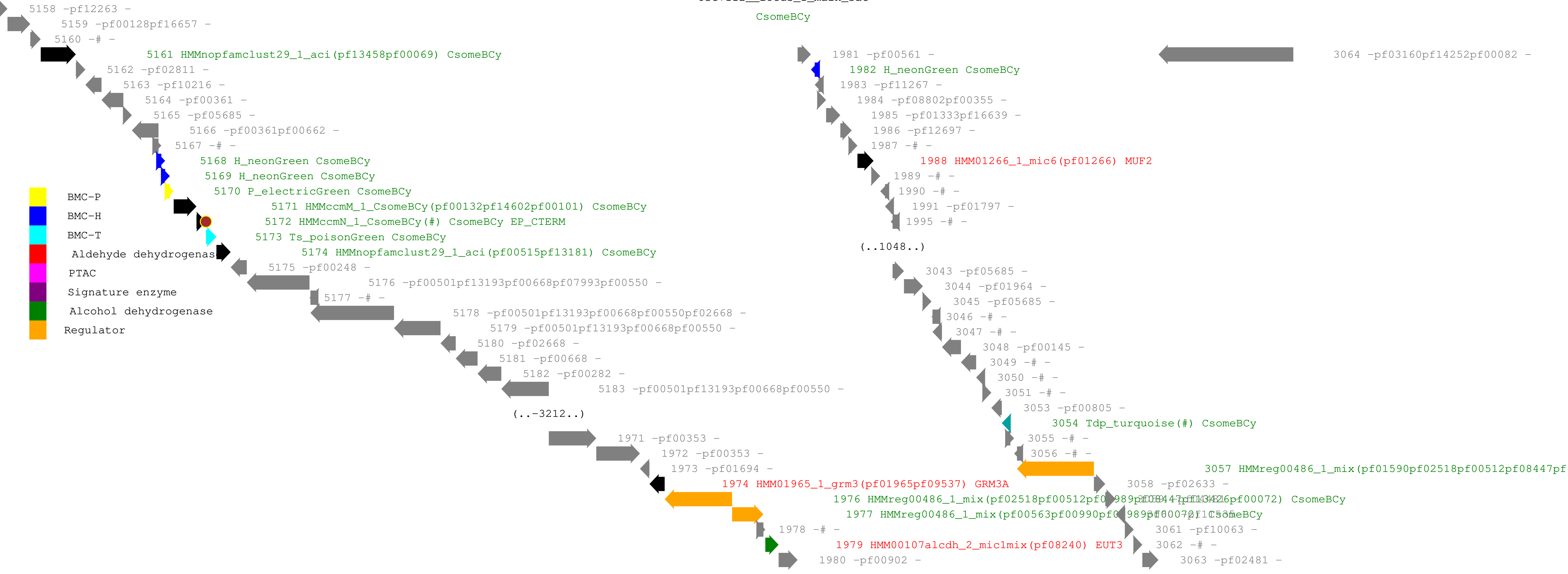

Oscillatoriales cyanobacterium JSC-12. Bacteria; Cyanobacteria; Oscillatoriophycideae; Oscillatoriales;

OsccyDRAFT__locus_1_main_sat

CsomeBCy

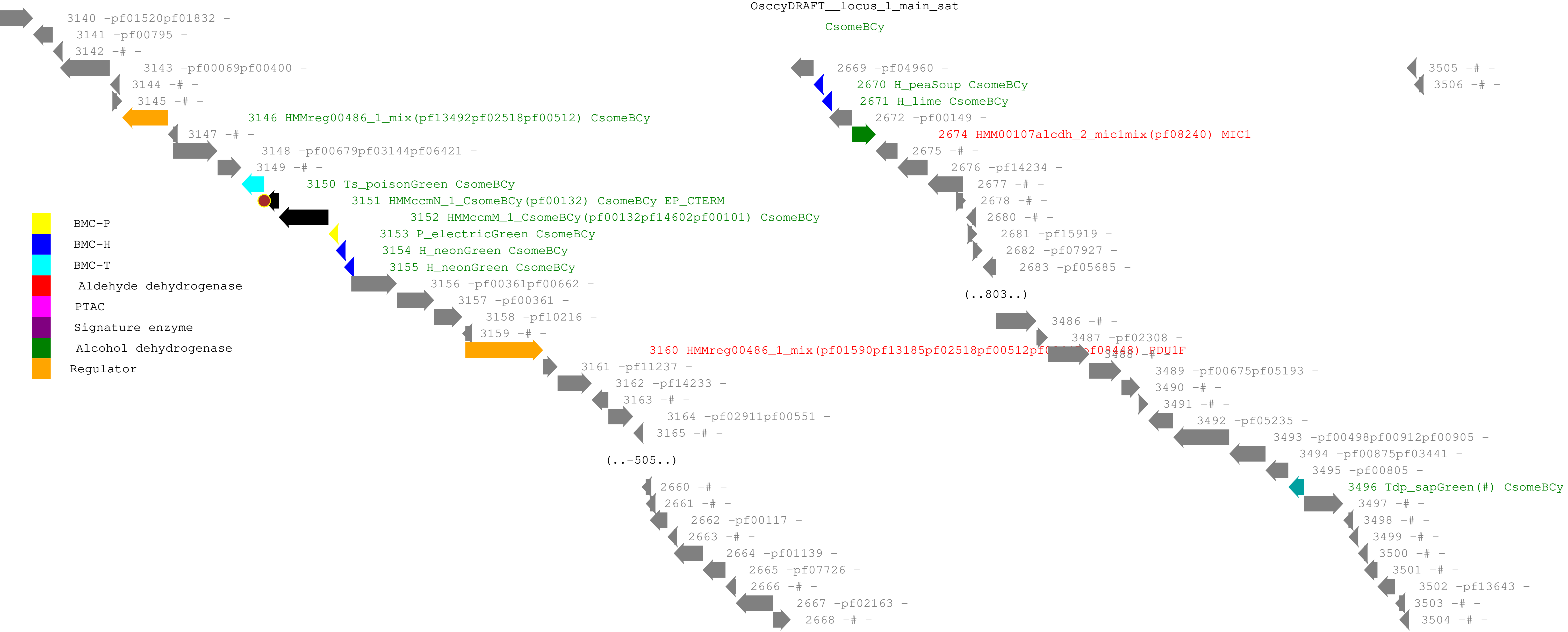

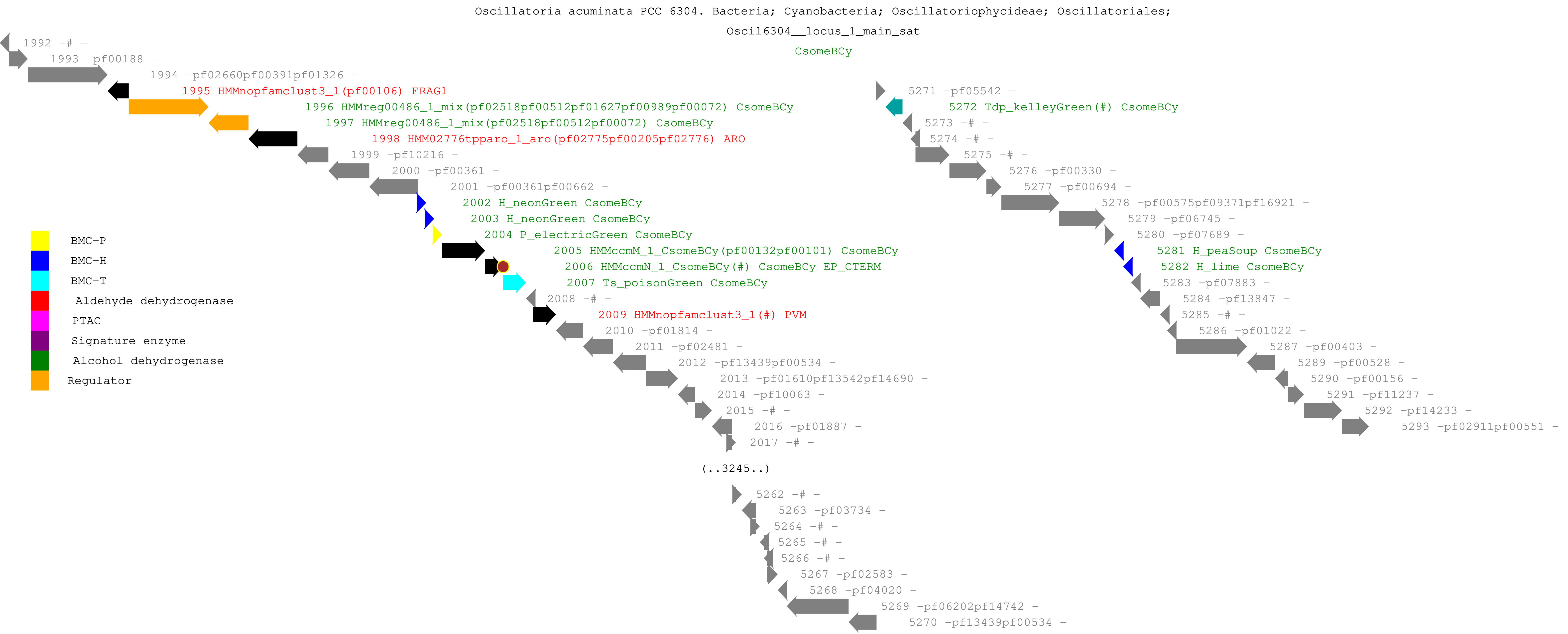

Oscillatoria acuminata PCC 6304. Bacteria; Cyanobacteria; Oscillatoriophycideae; Oscillatoriales;
Oscil6304__locus_1_main_sat
CsomeBCy
1992 -# -
1993 -pf00188 -
1994 -pf02660pf00391pf01326 -
1995 HMMnopfamclust3_1(pf00106) FRAG1
1996 HMMreg00486_1_mix(pf02518pf00512pf01627pf00989pf00072) CsomeBCy
1997 HMMreg00486_1_mix(pf02518pf00512pf00072) CsomeBCy
1998 HMM02776tpparo_1_aro(pf02775pf00205pf02776) ARO
1999 -pf10216 -
2000 -pf00361 -
2001 -pf00361pf00662 -
2002 H_neonGreen CsomeBCy
2003 H_neonGreen CsomeBCy
2004 P_electricGreen CsomeBCy
2005 HMMccmM_1_CsomeBCy(pf00132pf00101) CsomeBCy
2006 HMMccmN_1_CsomeBCy(#) CsomeBCy EP_CTERM
2007 Ts_poisonGreen CsomeBCy
2008 -# -
2009 HMMnopfamclust3_1(#) PVM
2010 -pf01814 -
2011 -pf02481 -
2012 -pf13439pf00534 -
2013 -pf01610pf13542pf14690 -
2014 -pf10063 -
2015 -# -
2016 -pf01887 -
2017 -# -
(..3245..)
5262 -# -
5263 -pf03734 -
5264 -# -
5265 -# -
5266 -# -
5267 -pf02583 -
5268 -pf04020 -
5269 -pf06202pf14742 -
5270 -pf13439pf00534 -
5271 -pf05542 -
5272 Tdp_kelleyGreen(#) CsomeBCy
5273 -# -
5274 -# -
5275 -# -
5276 -pf00330 -
5277 -pf00694 -
5278 -pf00575pf09371pf16921 -
5279 -pf06745 -
5280 -pf07689 -
5281 H_peaSoup CsomeBCy
5282 H_lime CsomeBCy
5283 -pf07883 -
5284 -pf13847 -
5285 -# -
5286 -pf01022 -
5287 -pf00403 -
5289 -pf00528 -
5290 -pf00156 -
5291 -pf11237 -
5292 -pf14233 -
5293 -pf02911pf00551 -
BMC-P
BMC-H
BMC-T
Aldehyde dehydrogenase
PTAC
Signature enzyme
Alcohol dehydrogenase
Regulator

Oscillatoria sp. PCC 6506. Bacteria; Cyanobacteria; Oscillatoriophycideae; Oscillatoriales;

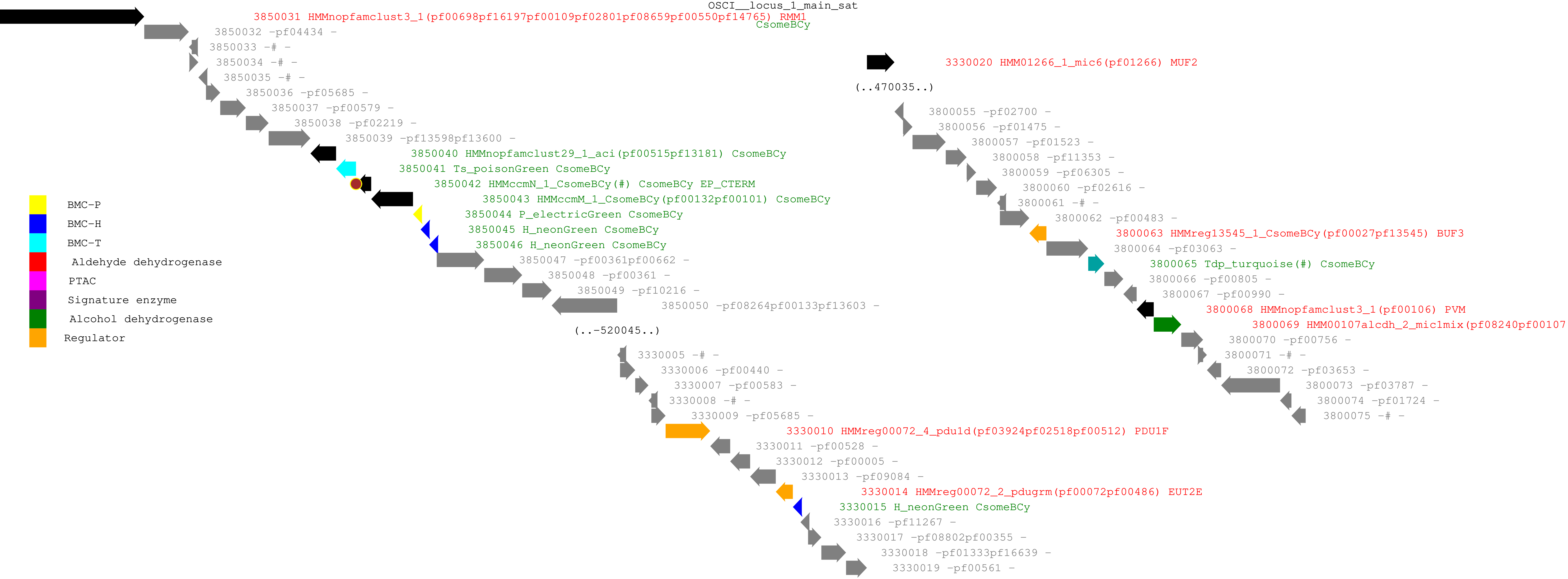

Planktothrix agardhii CCAP 1459/11A. Bacteria; Cyanobacteria; Oscillatoriophycideae; Oscillatoriales;

PA905__locus_1_main_sat

CsomeBCy

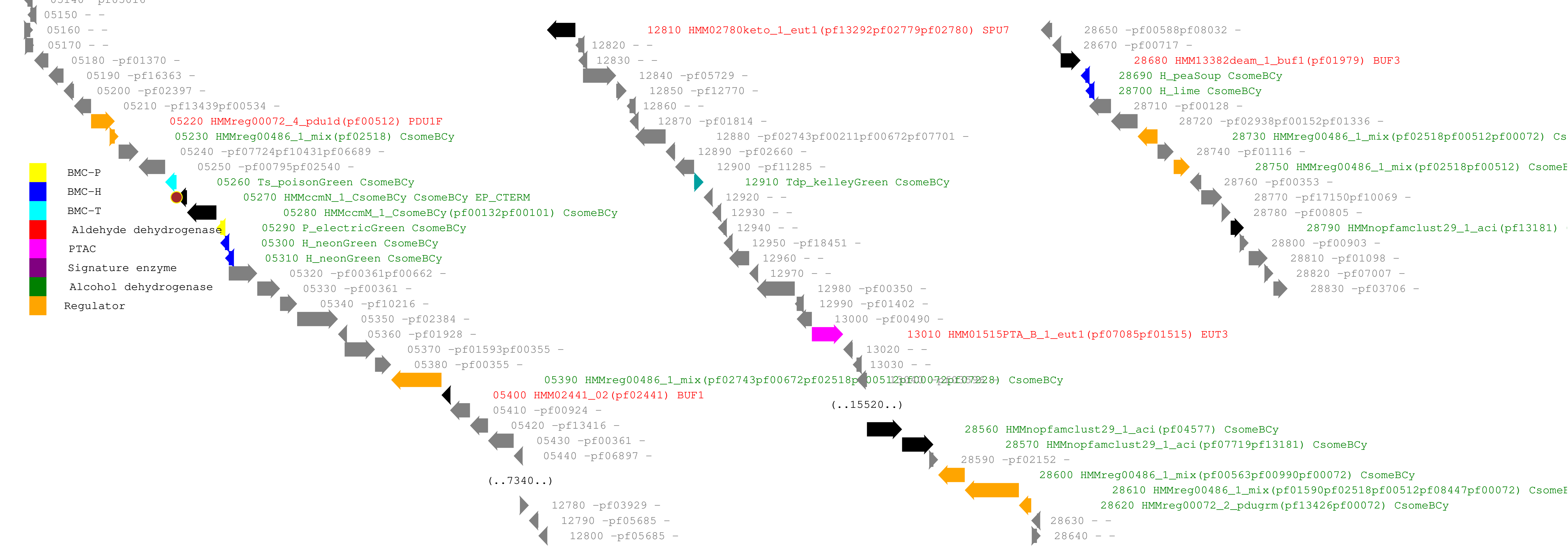

Chlorogloeopsis fritschii PCC 6912. Bacteria; Cyanobacteria; Nostocales; Chlorogloeopsidaceae; Chlorogloeopsis.

PCC6912__locus_1_main_sat

CsomeBCy

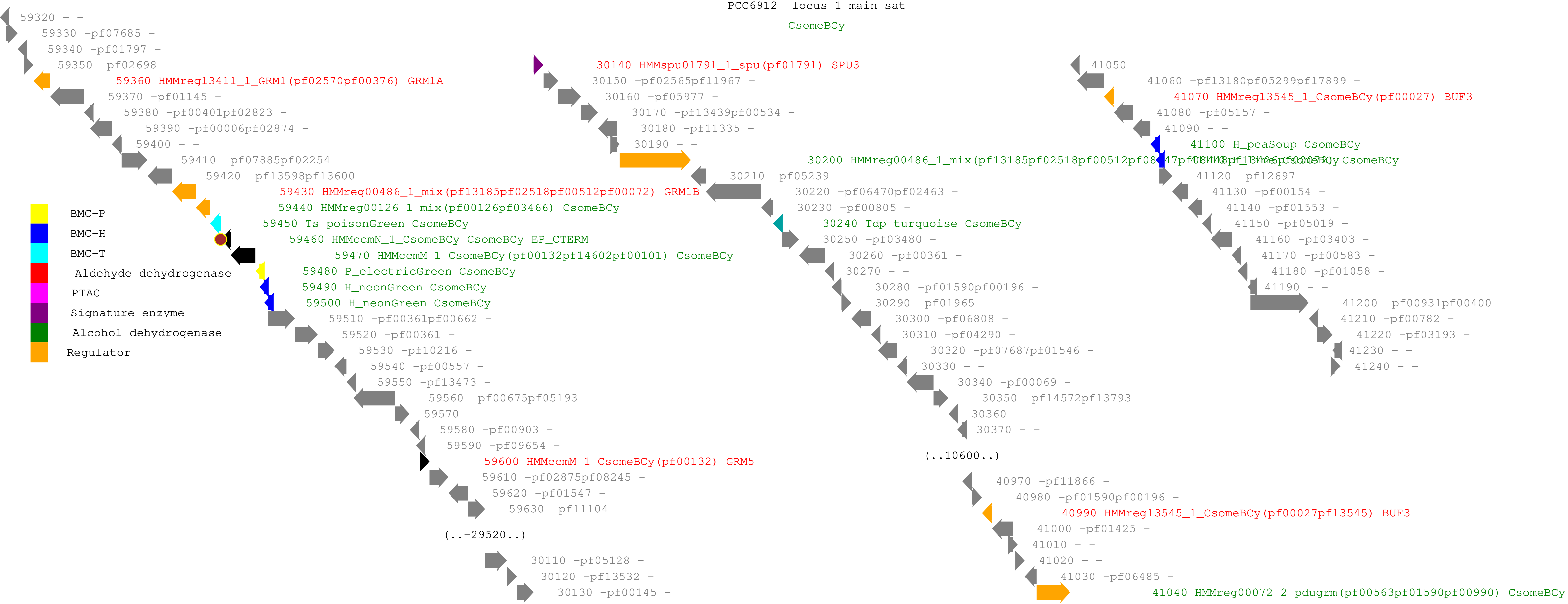

Halothece sp. (strain PCC 7418) (Synechococcus sp. (strain PCC 7418)). Bacteria; Cyanobacteria; Oscillatoriophycideae; Chroococcales;

PCC7418__locus_1_main_sat

CsomeBCy

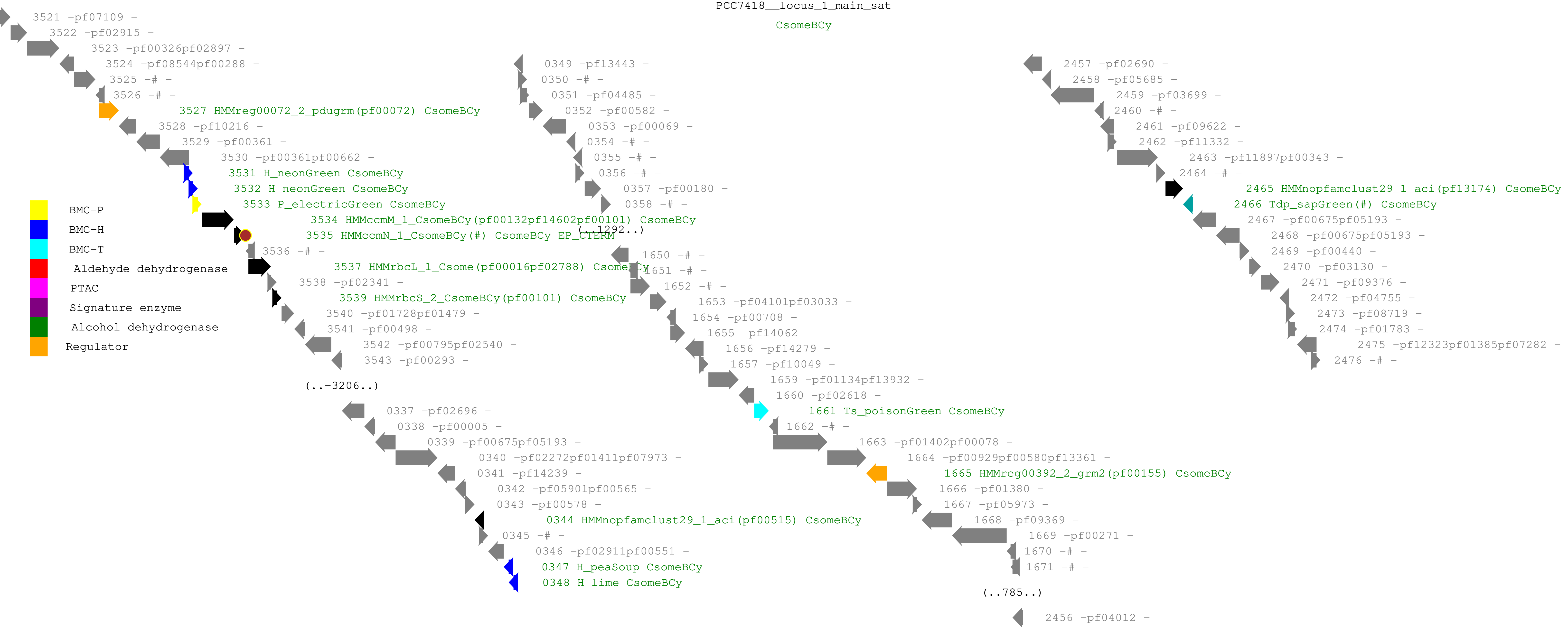

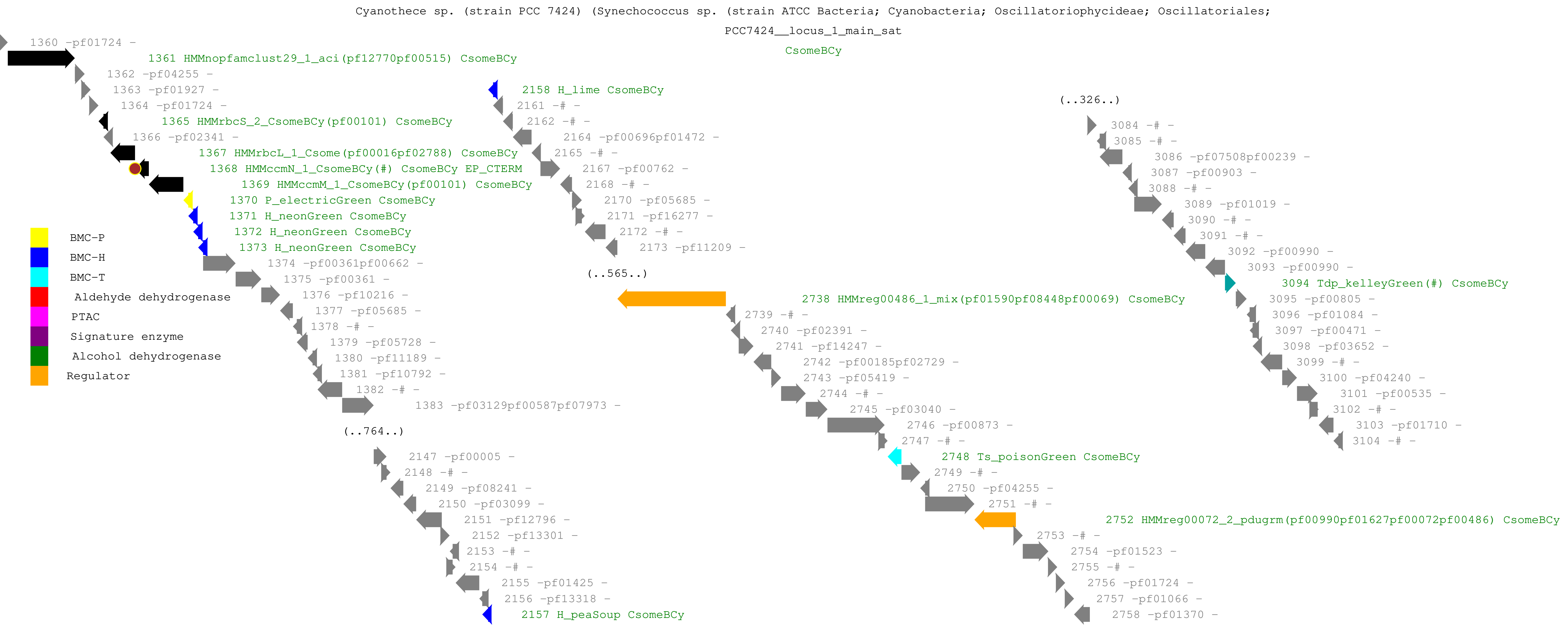

Cyanothece sp. (strain PCC 7424) (Synechococcus sp. (strain ATCC Bacteria; Cyanobacteria; Oscillatoriophycideae; Oscillatoriales;
PCC7424__locus_1_main_sat
CsomeBCy
1360 -pf01724 -
1361 HMMnopfamclust29_1_aci(pf12770pf00515) CsomeBCy
1362 -pf04255 -
1363 -pf01927 -
1364 -pf01724 -
1365 HMMrbcS_2_CsomeBCy(pf00101) CsomeBCy
1366 -pf02341 -
1367 HMMrbcL_1_Csome(pf00016pf02788) CsomeBCy
1368 HMMccmN_1_CsomeBCy(#) CsomeBCy EP_CTERM
1369 HMMccmM_1_CsomeBCy(pf00101) CsomeBCy
1370 P_electricGreen CsomeBCy
1371 H_neonGreen CsomeBCy
1372 H_neonGreen CsomeBCy
1373 H_neonGreen CsomeBCy
1374 -pf00361pf00662 -
1375 -pf00361 -
1376 -pf10216 -
1377 -pf05685 -
1378 -# -
1379 -pf05728 -
1380 -pf11189 -
1381 -pf10792 -
1382 -# -
1383 -pf03129pf00587pf07973 -
(..764..)
2147 -pf00005 -
2148 -# -
2149 -pf08241 -
2150 -pf03099 -
2151 -pf12796 -
2152 -pf13301 -
2153 -# -
2154 -# -
2155 -pf01425 -
2156 -pf13318 -
2157 H_peaSoup CsomeBCy
2158 H_lime CsomeBCy
2161 -# -
2162 -# -
2164 -pf00696pf01472 -
2165 -# -
2167 -pf00762 -
2168 -# -
2170 -pf05685 -
2171 -pf16277 -
2172 -# -
2173 -pf11209 -
(..565..)
2738 HMMreg00486_1_mix(pf01590pf08448pf00069) CsomeBCy
2739 -# -
2740 -pf02391 -
2741 -pf14247 -
2742 -pf00185pf02729 -
2743 -pf05419 -
2744 -# -
2745 -pf03040 -
2746 -pf00873 -
2747 -# -
2748 Ts_poisonGreen CsomeBCy
2749 -# -
2750 -pf04255 -
2751 -# -
2752 HMMreg00072_2_pdugrm(pf00990pf01627pf00072pf00486) CsomeBCy
2753 -# -
2754 -pf01523 -
2755 -# -
2756 -pf01724 -
2757 -pf01066 -
2758 -pf01370 -
(..326..)
3084 -# -
3085 -# -
3086 -pf07508pf00239 -
3087 -pf00903 -
3088 -# -
3089 -pf01019 -
3090 -# -
3091 -# -
3092 -pf00990 -
3093 -pf00990 -
3094 Tdp_kelleyGreen(#) CsomeBCy
3095 -pf00805 -
3096 -pf01084 -
3097 -pf00471 -
3098 -pf03652 -
3099 -# -
3100 -pf04240 -
3101 -pf00535 -
3102 -# -
3103 -pf01710 -
3104 -# -
BMC-P
BMC-H
BMC-T
Aldehyde dehydrogenase
PTAC
Signature enzyme
Alcohol dehydrogenase
Regulator

Cyanothece sp. (strain PCC 8801) (Synechococcus sp. (strain PCC 8801 / Bacteria; Cyanobacteria; Oscillatoriophycideae; Oscillatoriales;

PCC8801__locus_1_main_sat

CsomeBCy

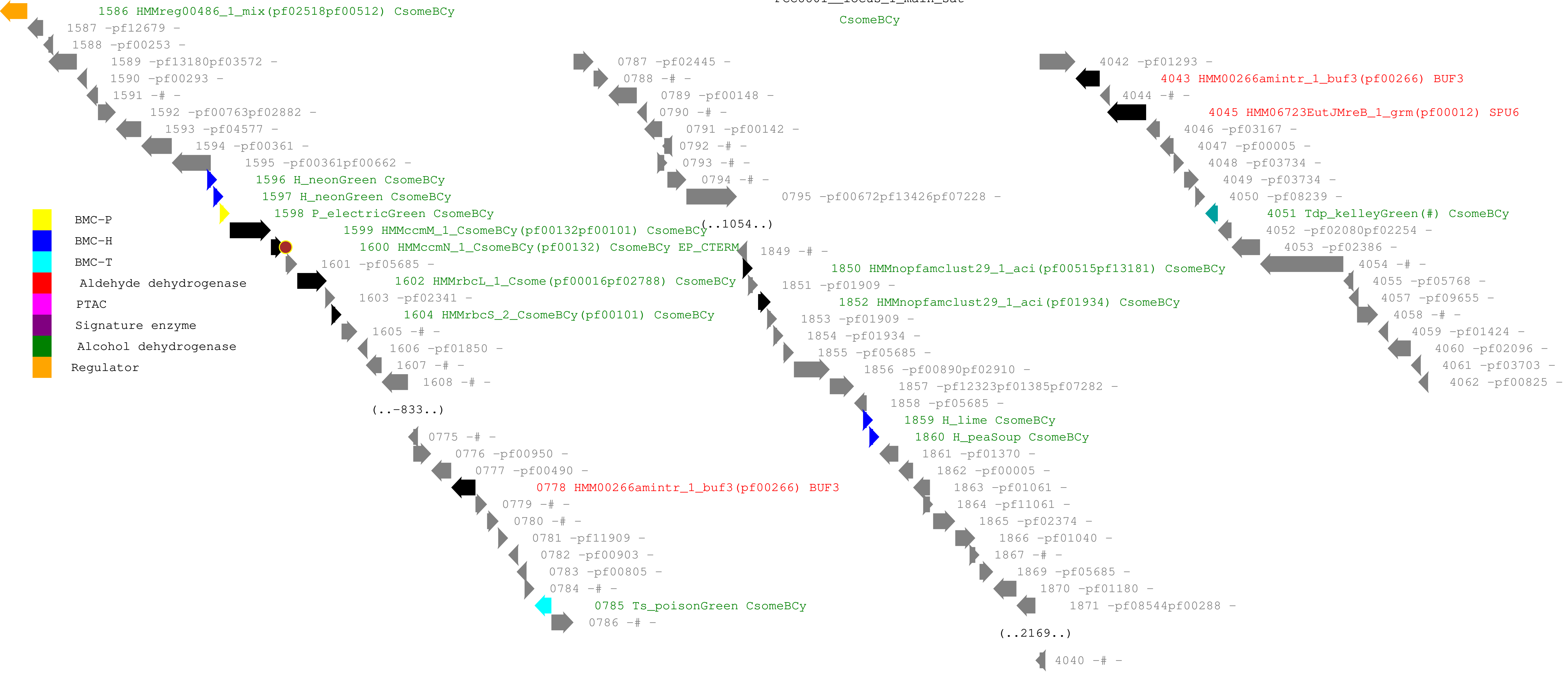

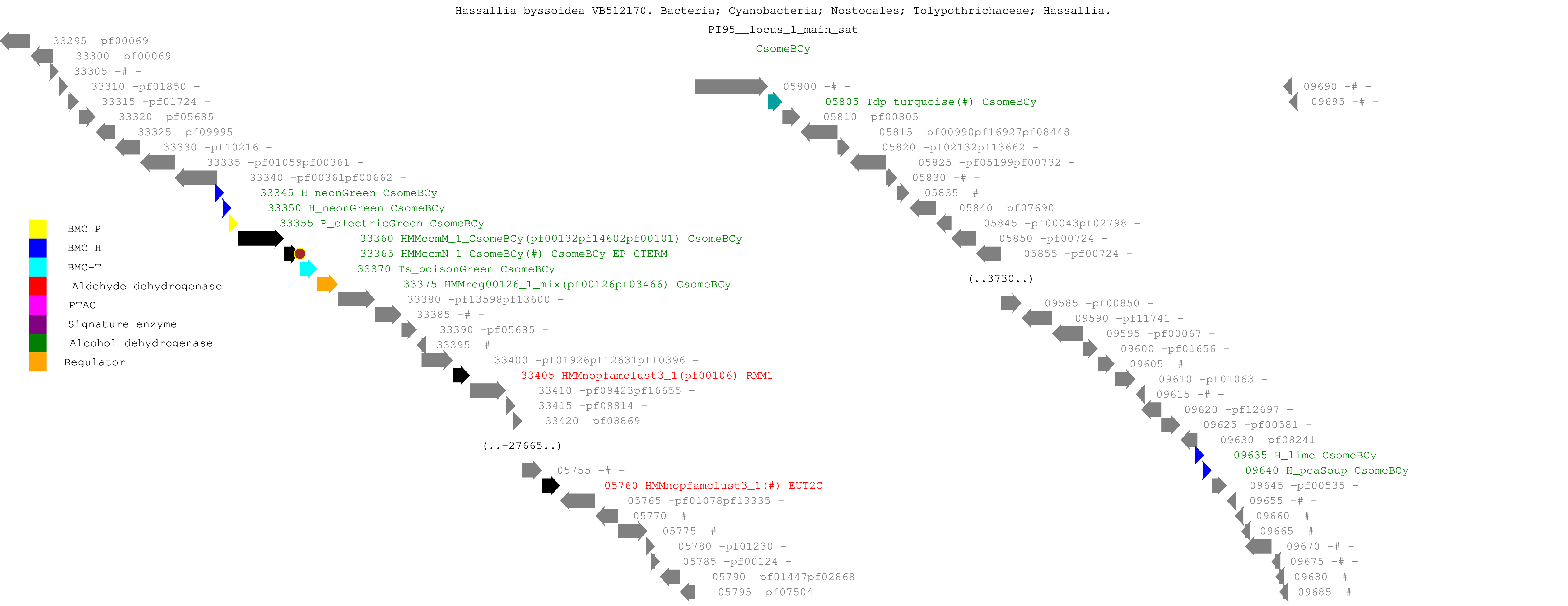

Hassallia byssoidea VB512170. Bacteria; Cyanobacteria; Nostocales; Tolypothrichaceae; Hassallia.
PI95__locus_1_main_sat
CsomeBCy
33295 -pf00069 -
33300 -pf00069 -
33305 -# -
33310 -pf01850 -
33315 -pf01724 -
33320 -pf05685 -
33325 -pf09995 -
33330 -pf10216 -
33335 -pf01059pf00361 -
33340 -pf00361pf00662 -
33345 H_neonGreen CsomeBCy
33350 H_neonGreen CsomeBCy
33355 P_electricGreen CsomeBCy
33360 HMMccmM_1_CsomeBCy(pf00132pf14602pf00101) CsomeBCy
33365 HMMccmN_1_CsomeBCy(#) CsomeBCy EP_CTERM
33370 Ts_poisonGreen CsomeBCy
33375 HMMreg00126_1_mix(pf00126pf03466) CsomeBCy
33380 -pf13598pf13600 -
33385 -# -
33390 -pf05685 -
33395 -# -
33400 -pf01926pf12631pf10396 -
33405 HMMnopfamclust3_1(pf00106) RMM1
33410 -pf09423pf16655 -
33415 -pf08814 -
33420 -pf08869 -
(..-27665..)
05755 -# -
05760 HMMnopfamclust3_1(#) EUT2C
05765 -pf01078pf13335 -
05770 -# -
05775 -# -
05780 -pf01230 -
05785 -pf00124 -
05790 -pf01447pf02868 -
05795 -pf07504 -
05800 -# -
05805 Tdp_turquoise(#) CsomeBCy
05810 -pf00805 -
05815 -pf00990pf16927pf08448 -
05820 -pf02132pf13662 -
05825 -pf05199pf00732 -
05830 -# -
05835 -# -
05840 -pf07690 -
05845 -pf00043pf02798 -
05850 -pf00724 -
05855 -pf00724 -
(..3730..)
09585 -pf00850 -
09590 -pf11741 -
09595 -pf00067 -
09600 -pf01656 -
09605 -# -
09610 -pf01063 -
09615 -# -
09620 -pf12697 -
09625 -pf00581 -
09630 -pf08241 -
09635 H_lime CsomeBCy
09640 H_peaSoup CsomeBCy
09645 -pf00535 -
09655 -# -
09660 -# -
09665 -# -
09670 -# -
09675 -# -
09680 -# -
09685 -# -
09690 -# -
09695 -# -
BMC-P
BMC-H
BMC-T
Aldehyde dehydrogenase
PTAC
Signature enzyme
Alcohol dehydrogenase
Regulator

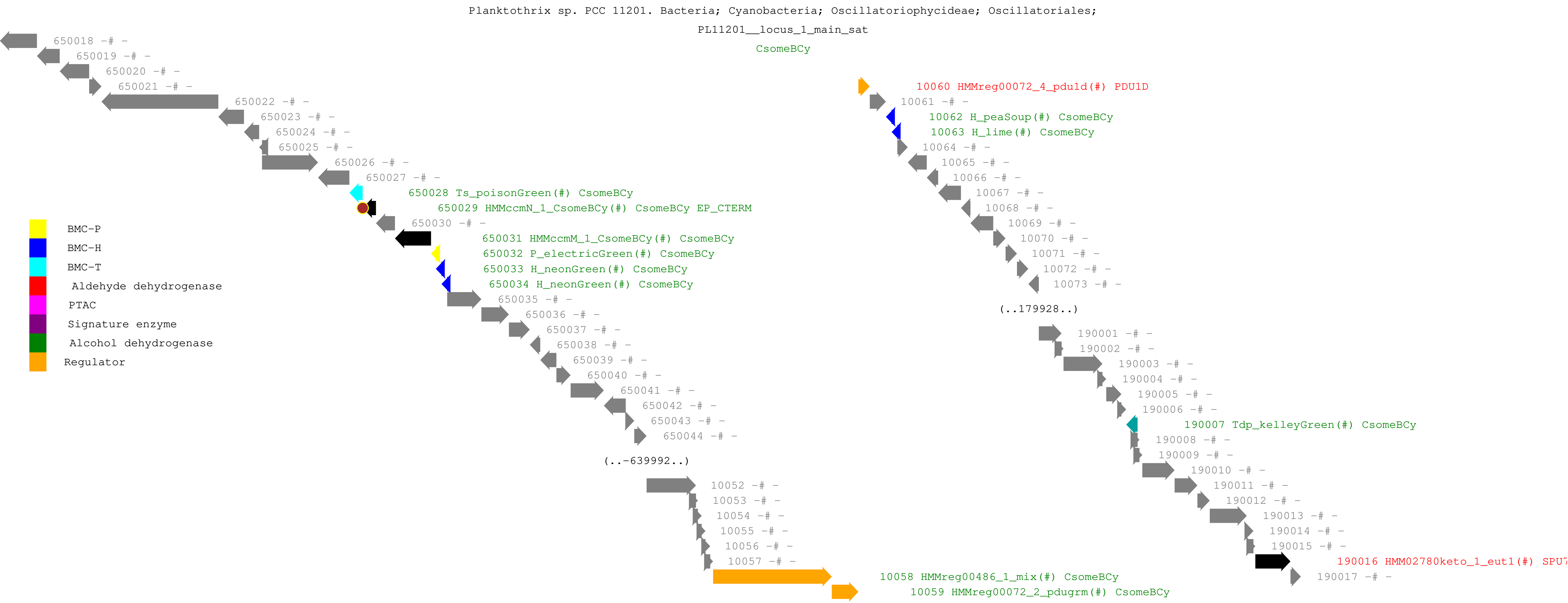

Planktothrix sp. PCC 11201. Bacteria; Cyanobacteria; Oscillatoriophycideae; Oscillatoriales;
PL11201__locus_1_main_sat
CsomeBCy
650018 -# -
650019 -# -
650020 -# -
650021 -# -
650022 -# -
650023 -# -
650024 -# -
650025 -# -
650026 -# -
650027 -# -
650028 Ts_poisonGreen(#) CsomeBCy
650029 HMMccmN_1_CsomeBCy(#) CsomeBCy EP_CTERM
650030 -# -
650031 HMMccmM_1_CsomeBCy(#) CsomeBCy
650032 P_electricGreen(#) CsomeBCy
650033 H_neonGreen(#) CsomeBCy
650034 H_neonGreen(#) CsomeBCy
650035 -# -
650036 -# -
650037 -# -
650038 -# -
650039 -# -
650040 -# -
650041 -# -
650042 -# -
650043 -# -
650044 -# -
(..-639992..)
10052 -# -
10053 -# -
10054 -# -
10055 -# -
10056 -# -
10057 -# -
10058 HMMreg00486_1_mix(#) CsomeBCy
10059 HMMreg00072_2_pdugrm(#) CsomeBCy
10060 HMMreg00072_4_pdu1d(#) PDU1D
10061 -# -
10062 H_peaSoup(#) CsomeBCy
10063 H_lime(#) CsomeBCy
10064 -# -
10065 -# -
10066 -# -
10067 -# -
10068 -# -
10069 -# -
10070 -# -
10071 -# -
10072 -# -
10073 -# -
(..179928..)
190001 -# -
190002 -# -
190003 -# -
190004 -# -
190005 -# -
190006 -# -
190007 Tdp_kelleyGreen(#) CsomeBCy
190008 -# -
190009 -# -
190010 -# -
190011 -# -
190012 -# -
190013 -# -
190014 -# -
190015 -# -
190016 HMM02780keto_1_eut1(#) SPU7
190017 -# -
BMC-P
BMC-H
BMC-T
Aldehyde dehydrogenase
PTAC
Signature enzyme
Alcohol dehydrogenase
Regulator

Planktothrix serta PCC 8927. Bacteria; Cyanobacteria; Oscillatoriophycideae; Oscillatoriales;

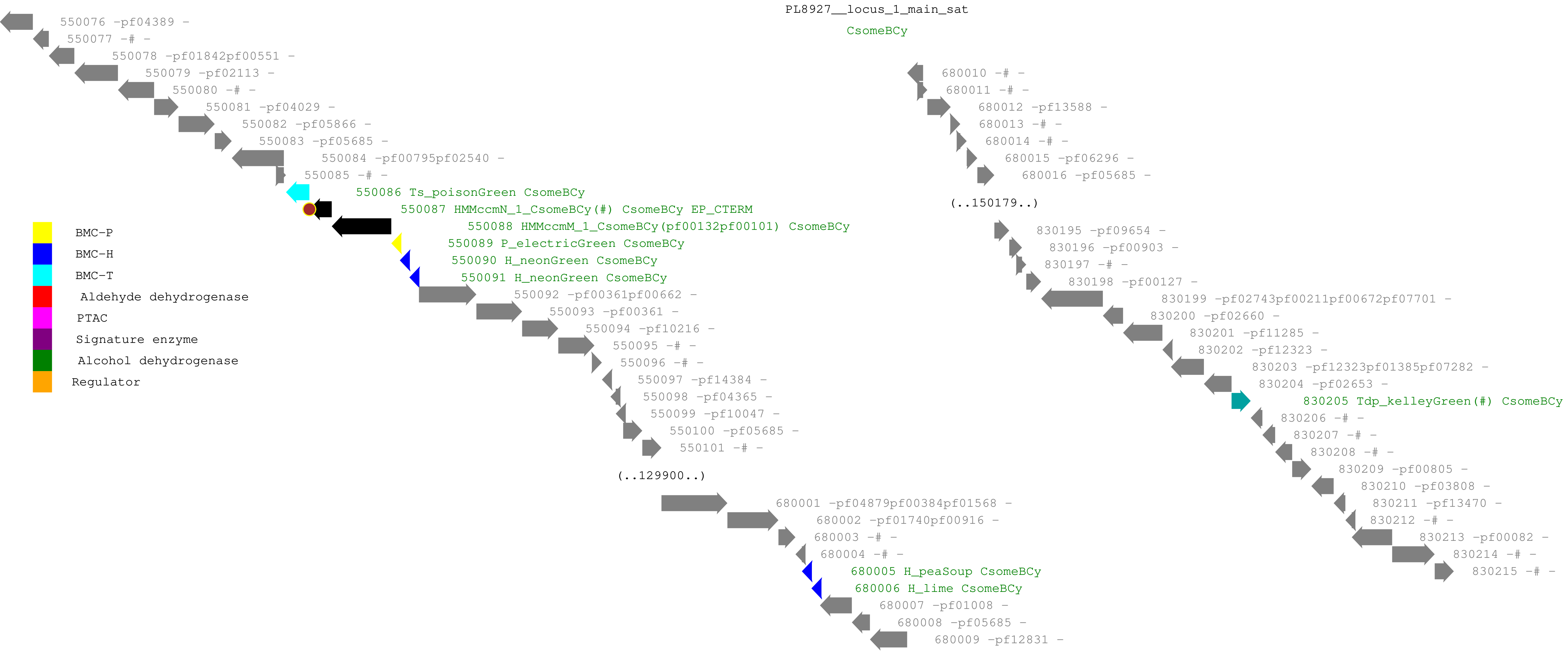

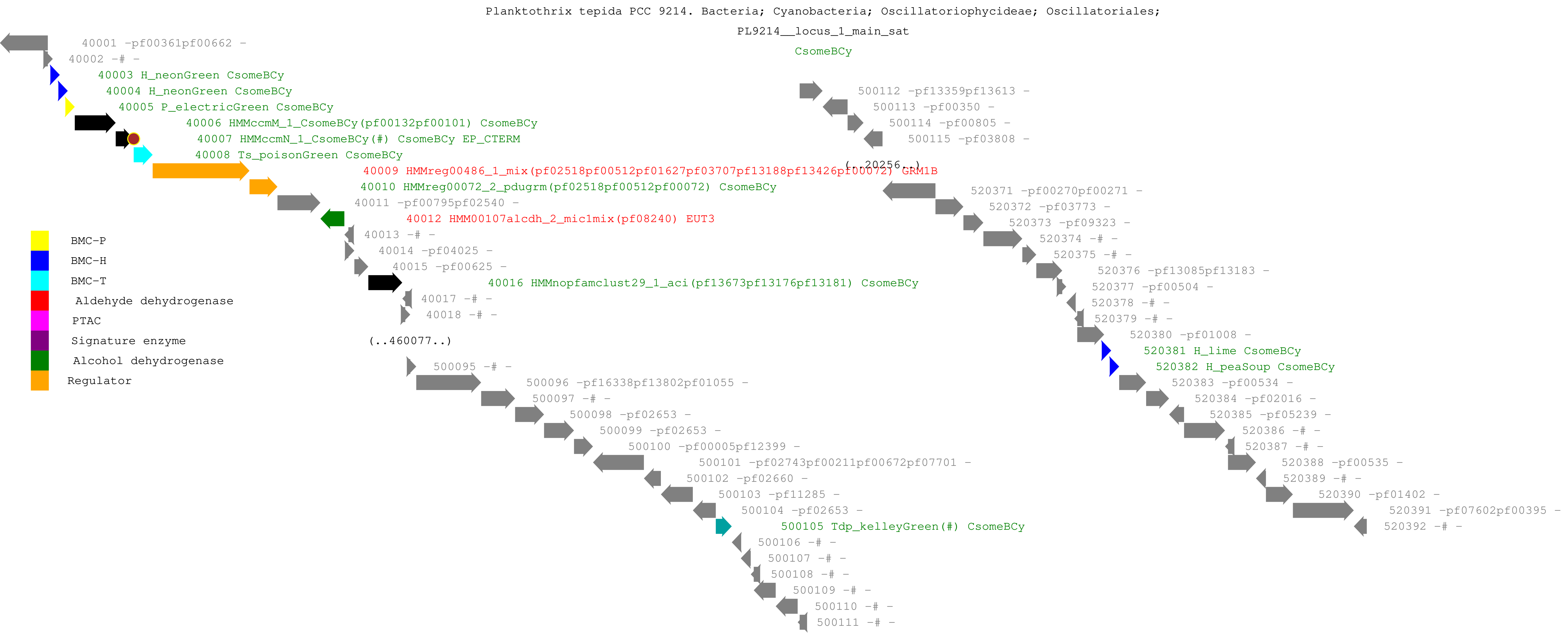

Planktothrix tepida PCC 9214. Bacteria; Cyanobacteria; Oscillatoriophycideae; Oscillatoriales;
PL9214__locus_1_main_sat
CsomeBCy
40001 -pf00361pf00662 -
40002 -# -
40003 H_neonGreen CsomeBCy
40004 H_neonGreen CsomeBCy
40005 P_electricGreen CsomeBCy
40006 HMMccmM_1_CsomeBCy(pf00132pf00101) CsomeBCy
40007 HMMccmN_1_CsomeBCy(#) CsomeBCy EP_CTERM
40008 Ts_poisonGreen CsomeBCy
40009 HMMreg00486_1_mix(pf02518pf00512pf01627pf03707pf13188pf13426pf00072) GRM1B
40010 HMMreg00072_2_pdugrm(pf02518pf00512pf00072) CsomeBCy
40011 -pf00795pf02540 -
40012 HMM00107alcdh_2_mic1mix(pf08240) EUT3
40013 -# -
40014 -pf04025 -
40015 -pf00625 -
40016 HMMnopfamclust29_1_aci(pf13673pf13176pf13181) CsomeBCy
40017 -# -
40018 -# -
(..460077..)
500095 -# -
500096 -pf16338pf13802pf01055 -
500097 -# -
500098 -pf02653 -
500099 -pf02653 -
500100 -pf00005pf12399 -
500101 -pf02743pf00211pf00672pf07701 -
500102 -pf02660 -
500103 -pf11285 -
500104 -pf02653 -
500105 Tdp_kelleyGreen(#) CsomeBCy
500106 -# -
500107 -# -
500108 -# -
500109 -# -
500110 -# -
500111 -# -
500112 -pf13359pf13613 -
500113 -pf00350 -
500114 -pf00805 -
500115 -pf03808 -
(..20256..)
520371 -pf00270pf00271 -
520372 -pf03773 -
520373 -pf09323 -
520374 -# -
520375 -# -
520376 -pf13085pf13183 -
520377 -pf00504 -
520378 -# -
520379 -# -
520380 -pf01008 -
520381 H_lime CsomeBCy
520382 H_peaSoup CsomeBCy
520383 -pf00534 -
520384 -pf02016 -
520385 -pf05239 -
520386 -# -
520387 -# -
520388 -pf00535 -
520389 -# -
520390 -pf01402 -
520391 -pf07602pf00395 -
520392 -# -
BMC-P
BMC-H
BMC-T
Aldehyde dehydrogenase
PTAC
Signature enzyme
Alcohol dehydrogenase
Regulator

Planktothrix paucivesiculata PCC 9631. Bacteria; Cyanobacteria; Oscillatoriophycideae; Oscillatoriales;

PL9631__locus_1_main_sat

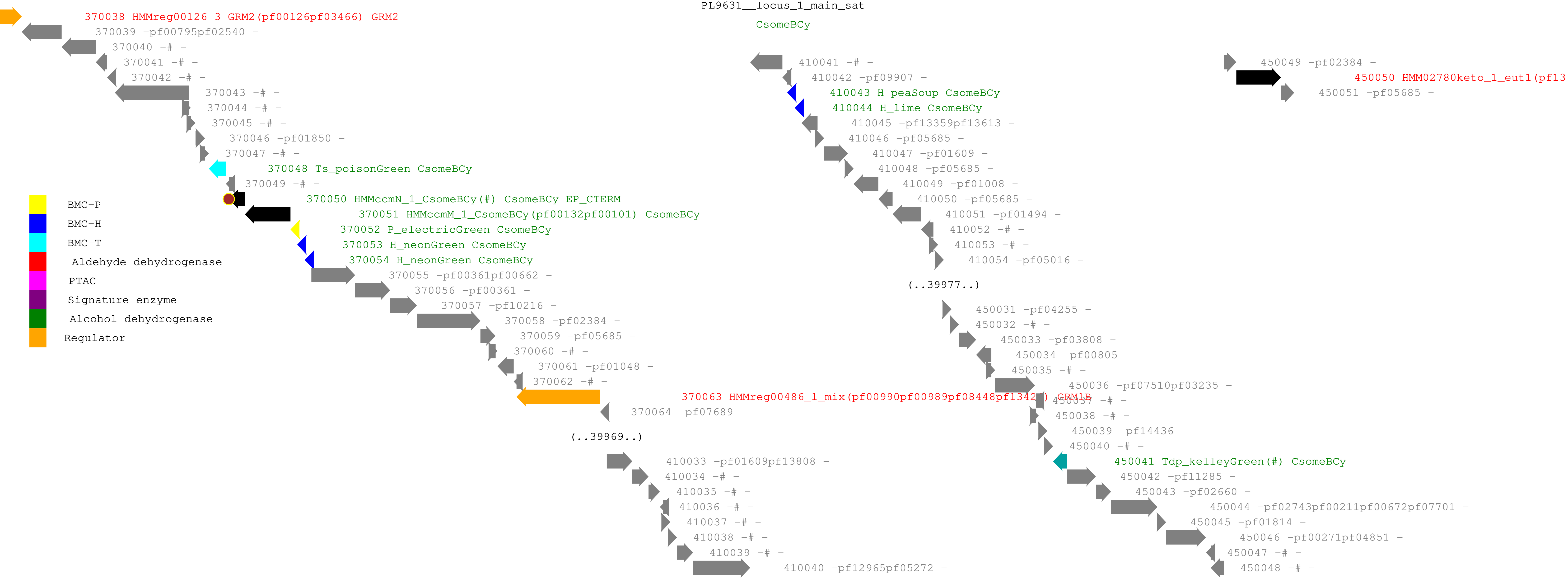

Planktothrix agardhii (Oscillatoria agardhii). Bacteria; Cyanobacteria; Oscillatoriophycideae; Oscillatoriales;

PLAM__locus_1_main_sat

CsomeBCy

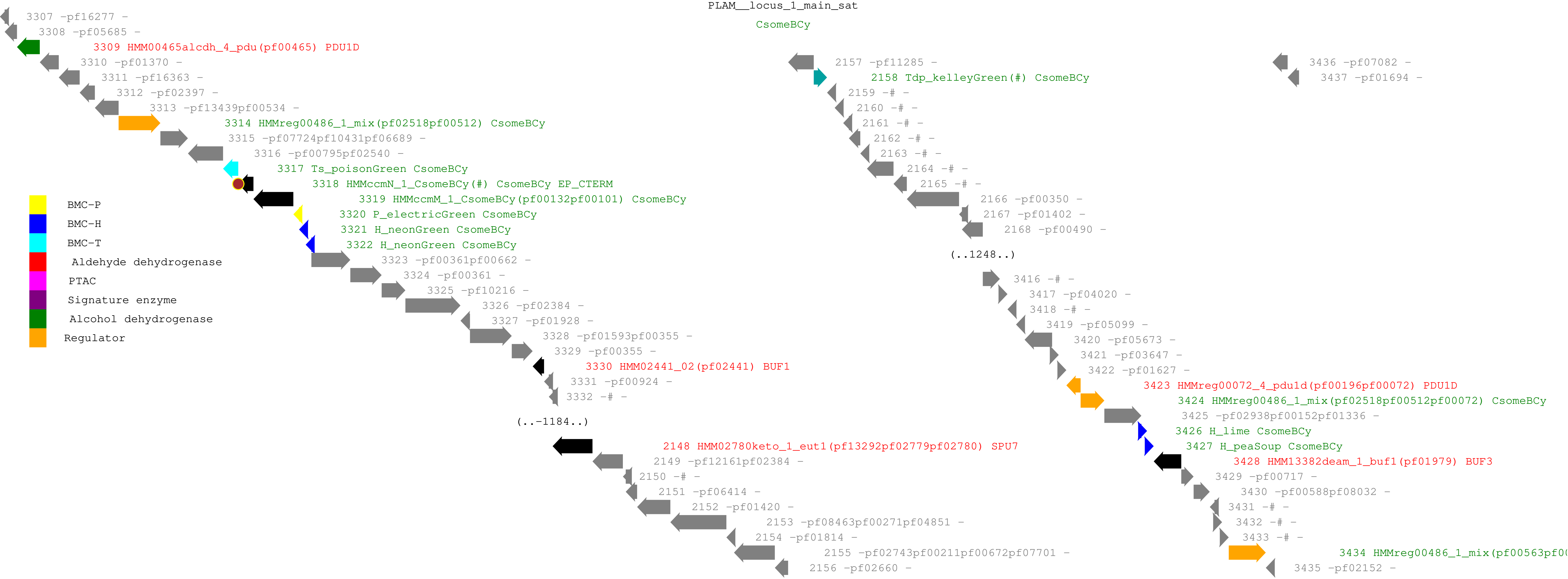

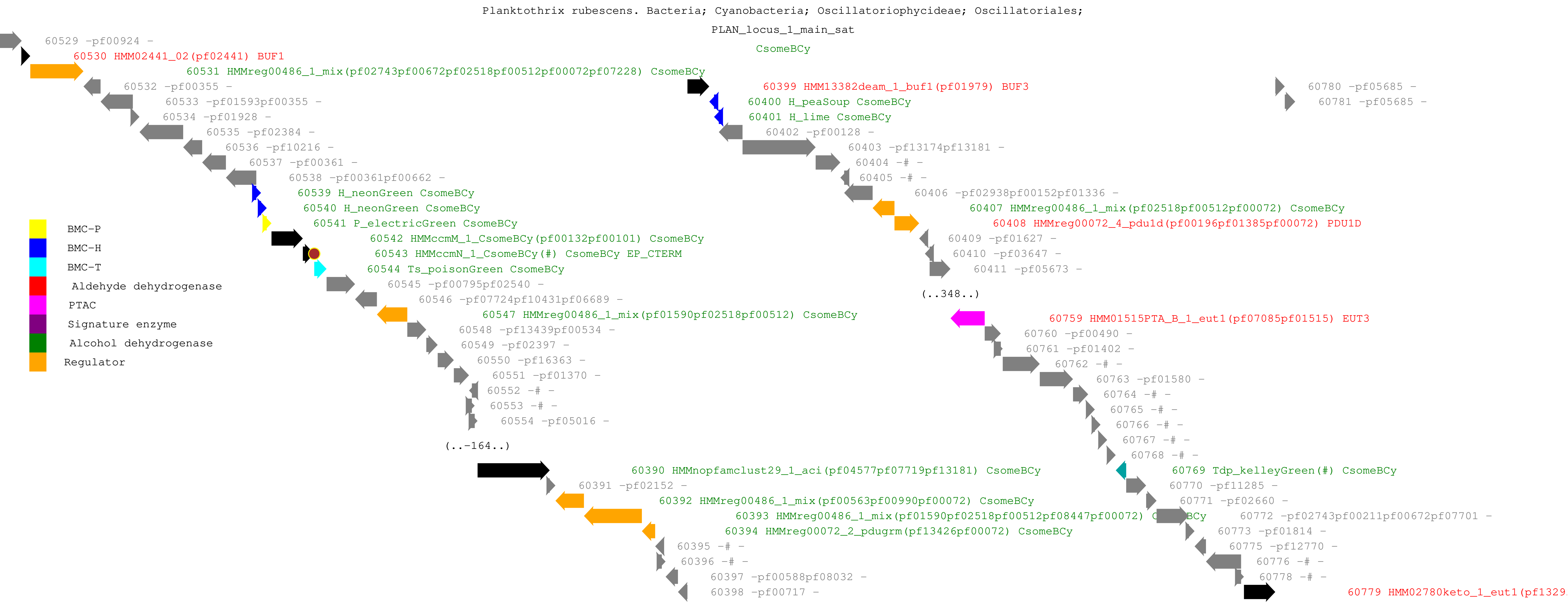
Planktothrix rubescens. Bacteria; Cyanobacteria; Oscillatoriophycideae; Oscillatoriales;
PLAN_locus_1_main_sat
CsomeBCy
60529 -pf00924 -
60530 HMM02441_02(pf02441) BUF1
60531 HMMreg00486_1_mix(pf02743pf00672pf02518pf00512pf00072pf07228) CsomeBCy
60532 -pf00355 -
60533 -pf01593pf00355 -
60534 -pf01928 -
60535 -pf02384 -
60536 -pf10216 -
60537 -pf00361 -
60538 -pf00361pf00662 -
60539 H_neonGreen CsomeBCy
60540 H_neonGreen CsomeBCy
60541 P_electricGreen CsomeBCy
60542 HMMccmM_1_CsomeBCy(pf00132pf00101) CsomeBCy
60543 HMMccmN_1_CsomeBCy(#) CsomeBCy EP_CTERM
60544 Ts_poisonGreen CsomeBCy
60545 -pf00795pf02540 -
60546 -pf07724pf10431pf06689 -
60547 HMMreg00486_1_mix(pf01590pf02518pf00512) CsomeBCy
60548 -pf13439pf00534 -
60549 -pf02397 -
60550 -pf16363 -
60551 -pf01370 -
60552 -# -
60553 -# -
60554 -pf05016 -
(..-164..)
60390 HMMnopfamclust29_1_aci(pf04577pf07719pf13181) CsomeBCy
60391 -pf02152 -
60392 HMMreg00486_1_mix(pf00563pf00990pf00072) CsomeBCy
60393 HMMreg00486_1_mix(pf01590pf02518pf00512pf08447pf00072) C BCy
60394 HMMreg00072_2_pdugrm(pf13426pf00072) CsomeBCy
60395 -# -
60396 -# -
60397 -pf00588pf08032 -
60398 -pf00717 -
BMC-P
BMC-H
BMC-T
Aldehyde dehydrogenase
PTAC
Signature enzyme
Alcohol dehydrogenase
Regulator
60399 HMM13382deam_1_buf1(pf01979) BUF3
60400 H_peaSoup CsomeBCy
60401 H_lime CsomeBCy
60402 -pf00128 -
60403 -pf13174pf13181 -
60404 -# -
60405 -# -
60406 -pf02938pf00152pf01336 -
60407 HMMreg00486_1_mix(pf02518pf00512pf00072) CsomeBCy
60408 HMMreg00072_4_pdu1d(pf00196pf01385pf00072) PDU1D
60409 -pf01627 -
60410 -pf03647 -
60411 -pf05673 -
(..348..)
60759 HMM01515PTA_B_1_eut1(pf07085pf01515) EUT3
60760 -pf00490 -
60761 -pf01402 -
60762 -# -
60763 -pf01580 -
60764 -# -
60765 -# -
60766 -# -
60767 -# -
60768 -# -
60769 Tdp_kelleyGreen(#) CsomeBCy
60770 -pf11285 -
60771 -pf02660 -
60772 -pf02743pf00211pf00672pf07701 -
60773 -pf01814 -
60775 -pf12770 -
60776 -# -
60778 -# -
60779 HMM02780keto_1_eut1(pf1329
60780 -pf05685 -
60781 -pf05685 -

Pleurocapsa sp. PCC 7327. Bacteria; Cyanobacteria; Pleurocapsales; Hyellaceae; Pleurocapsa.

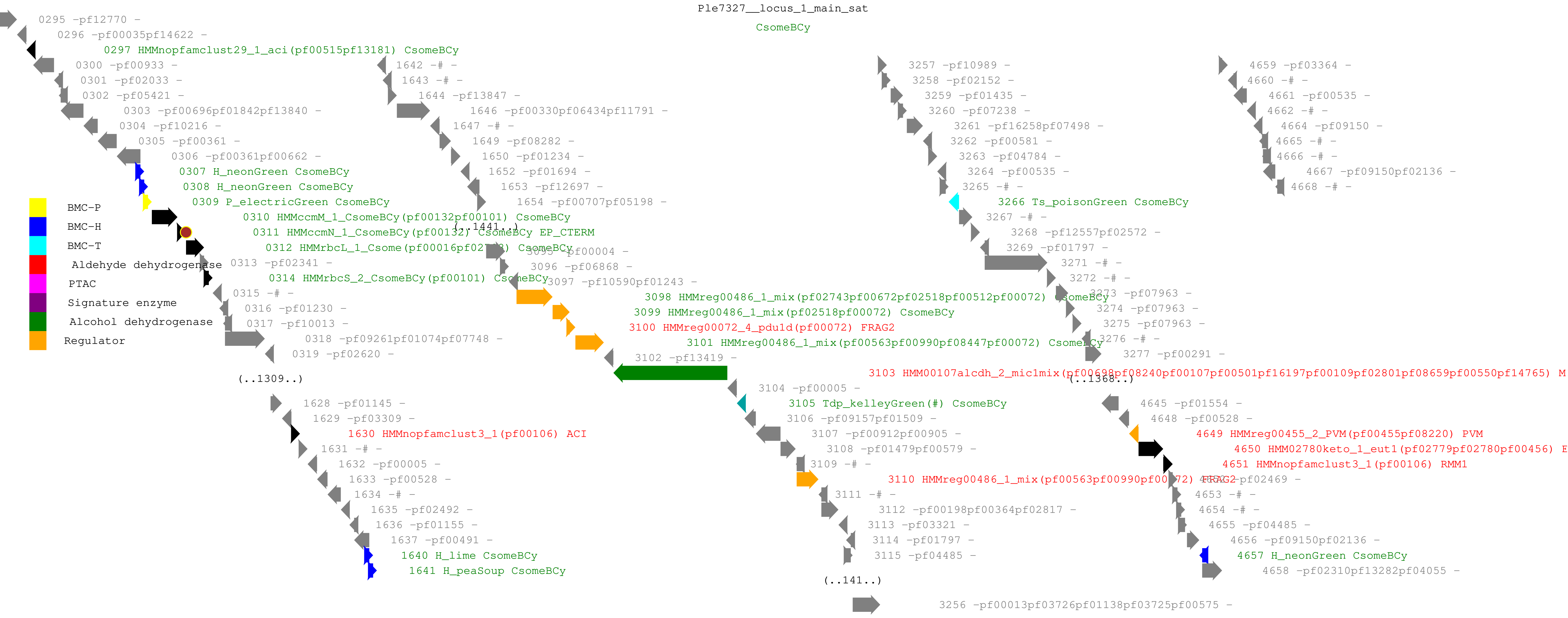

Prochlorothrix hollandica PCC 9006 = CALU 1027. Bacteria; Cyanobacteria; Synechococcales; Pseudanabaenaceae;

PROH__locus_1_main_sat

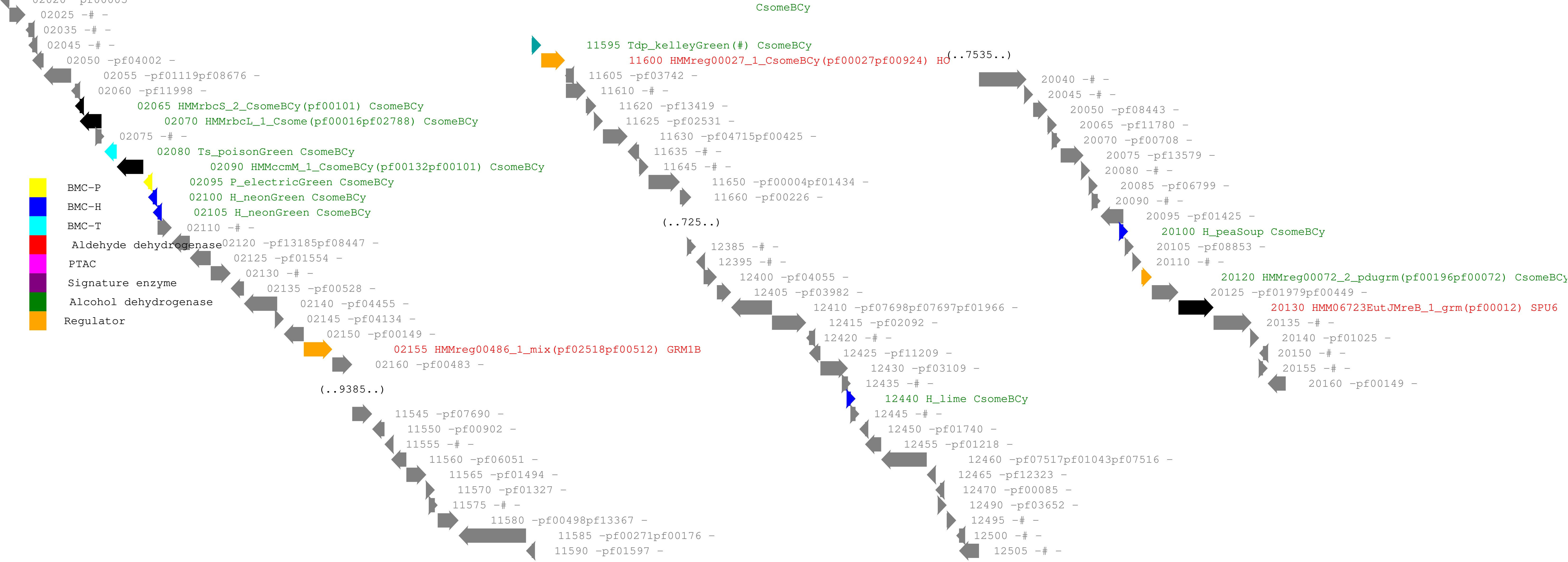

Pseudanabaena sp. PCC 7367. Bacteria; Cyanobacteria; Synechococcales; Pseudanabaenaceae;

Pse7367__locus_1_main_sat

CsomeBCy

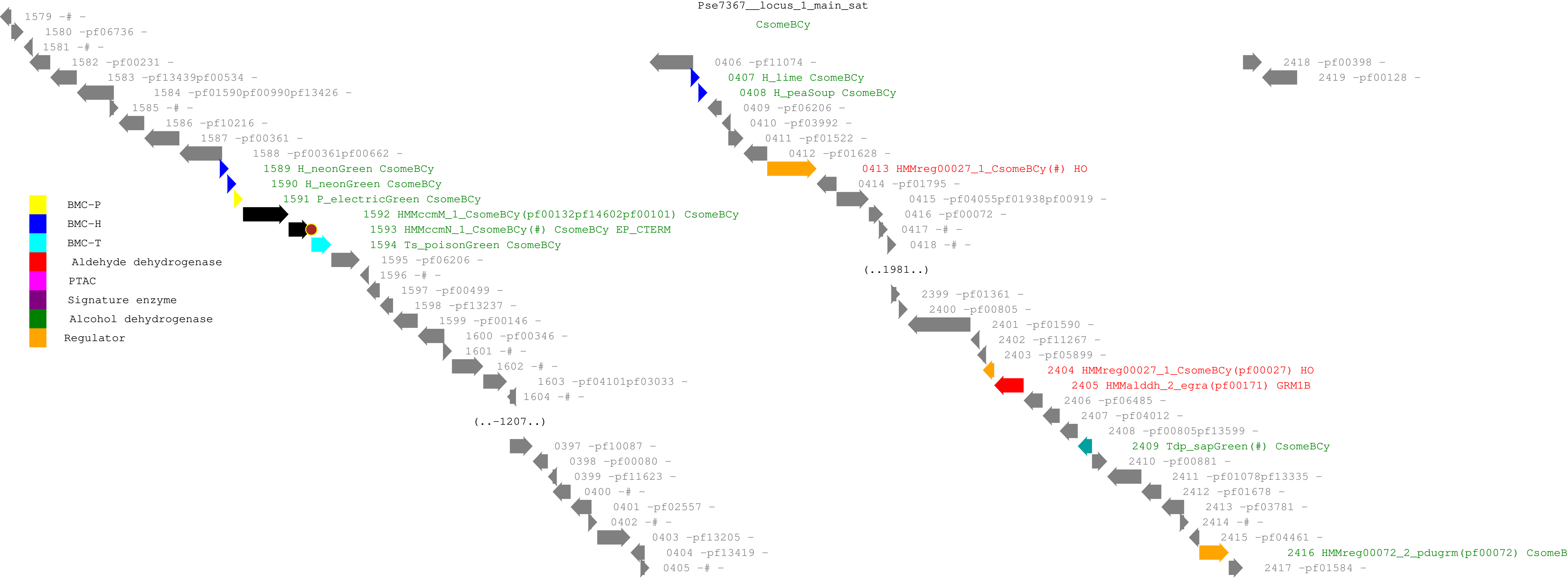

Pseudanabaena biceps PCC 7429. Bacteria; Cyanobacteria; Synechococcales; Pseudanabaenaceae;

Pse7429DRAFT__locus_1_main_sat

CsomeBCy

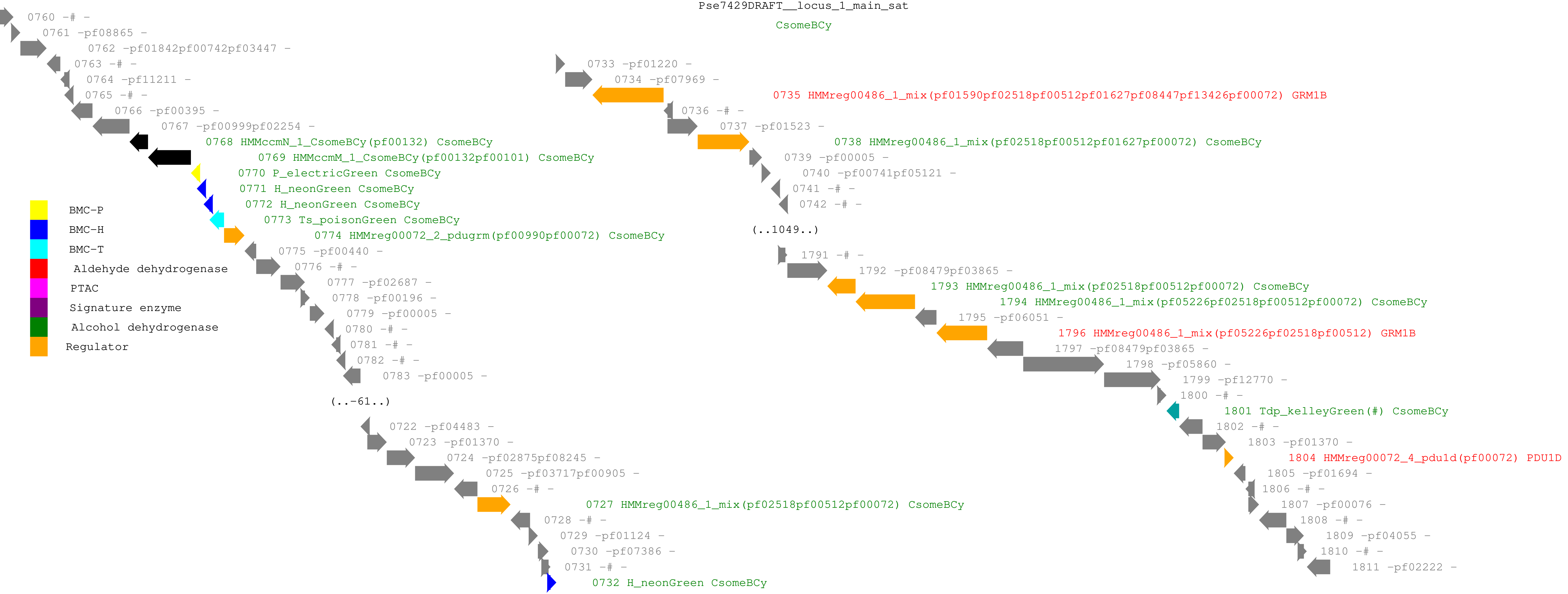

Scytonema millei VB511283. Bacteria; Cyanobacteria; Nostocales; Scytonemataceae; Scytonema.

QH73__locus_1_main_sat

CsomeBCy

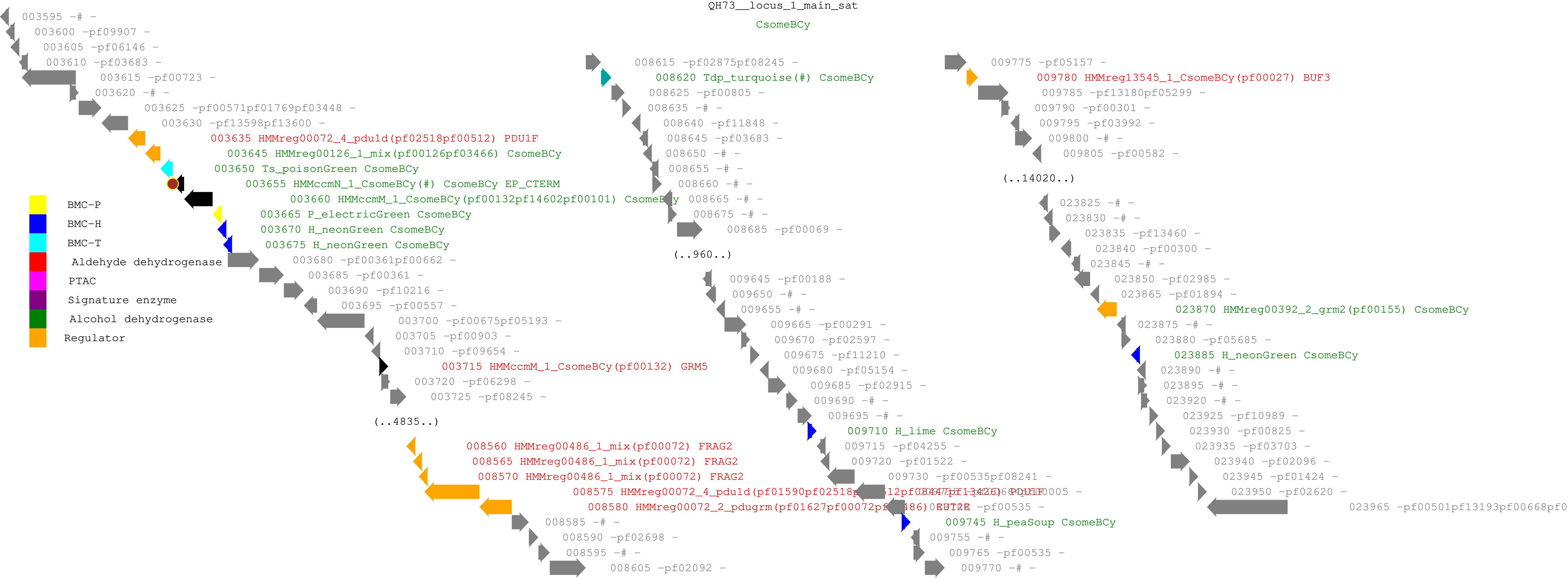

Aphanocapsa montana BDHKU210001. Bacteria; Cyanobacteria; Synechococcales; Merismopediaceae;

QP59__locus_1_main_sat

CsomeBCy

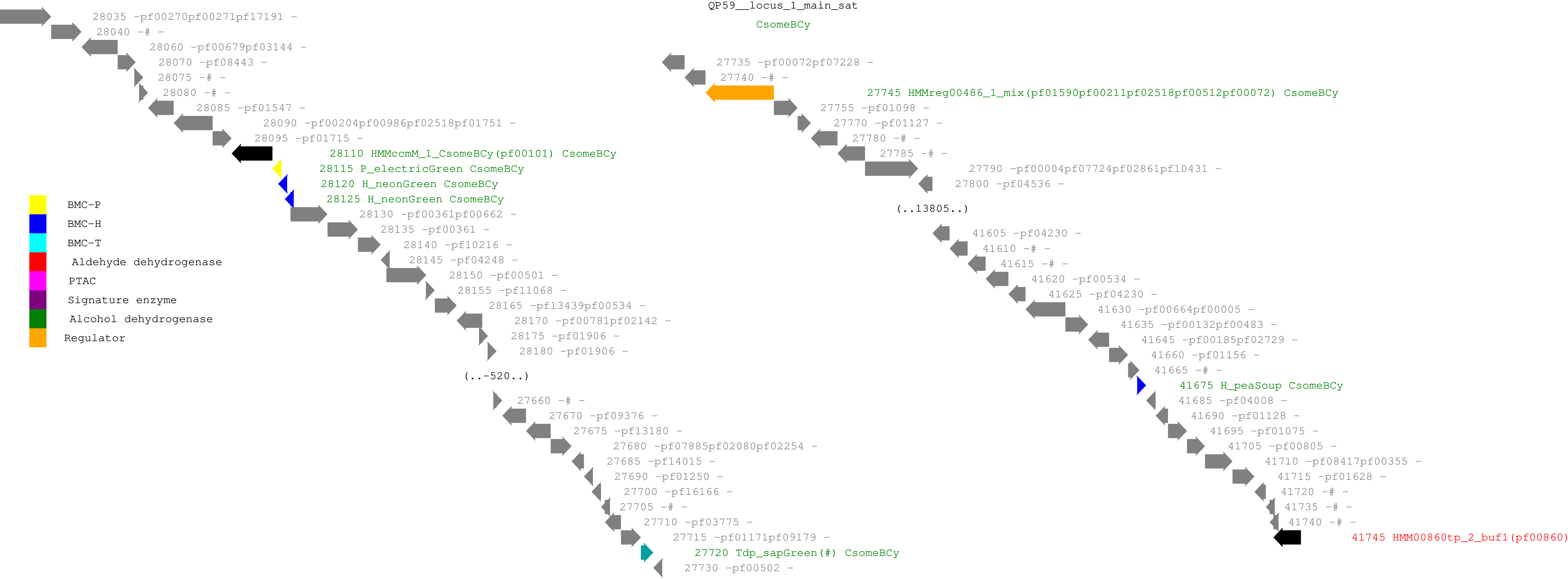

Lyngbya confervoides BDU141951. Bacteria; Cyanobacteria; Oscillatoriophycideae; Oscillatoriales;

QQ91__locus_1_main_sat

CsomeBCy

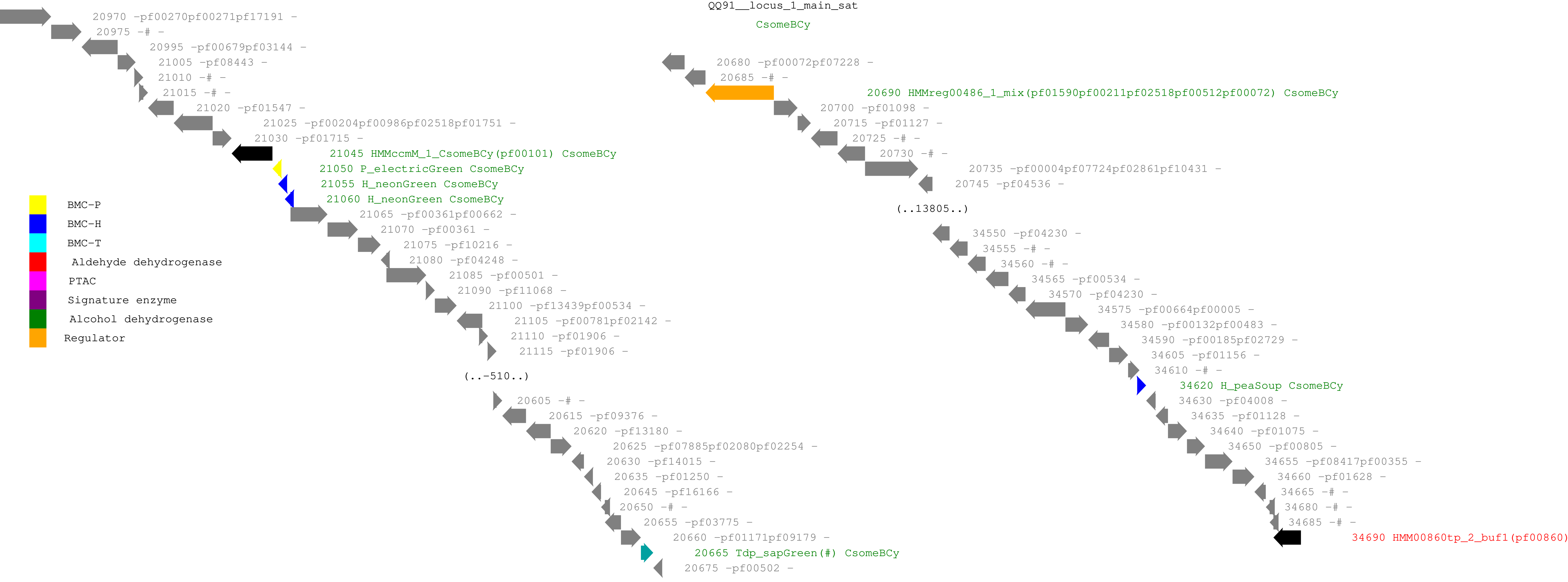

Richelia intracellularis HH01. Bacteria; Cyanobacteria; Nostocales; Nostocaceae; Richelia.

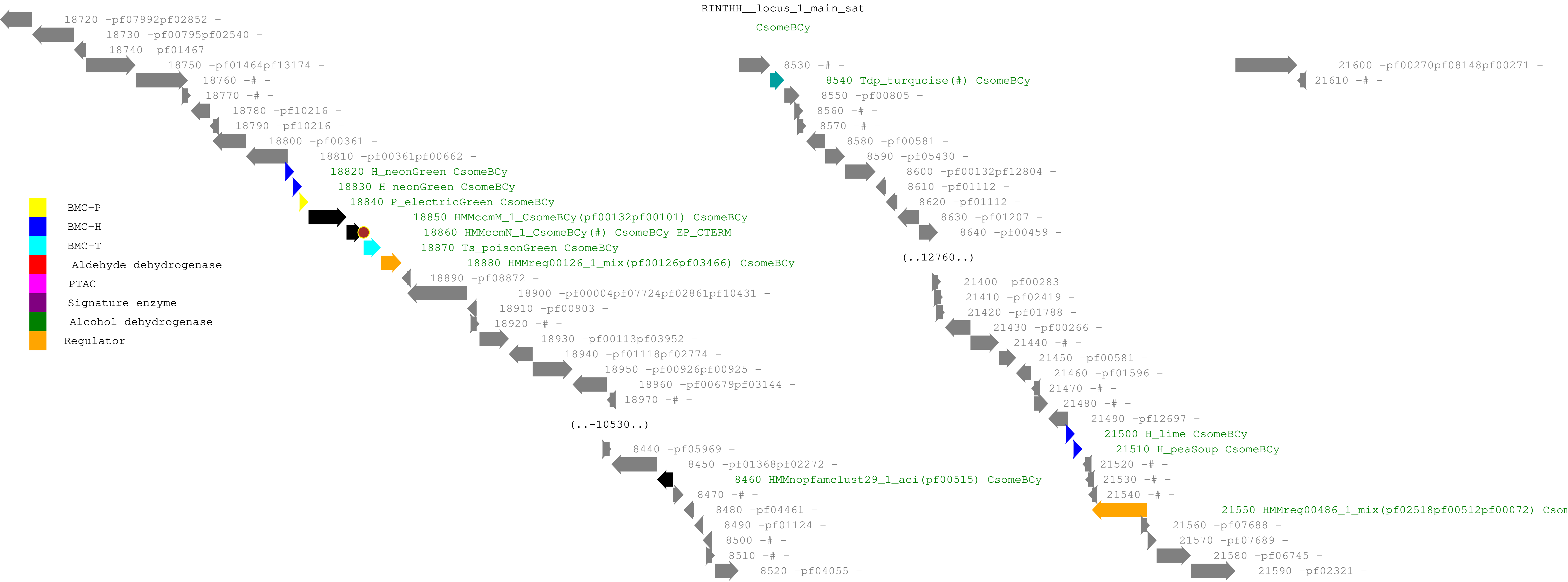

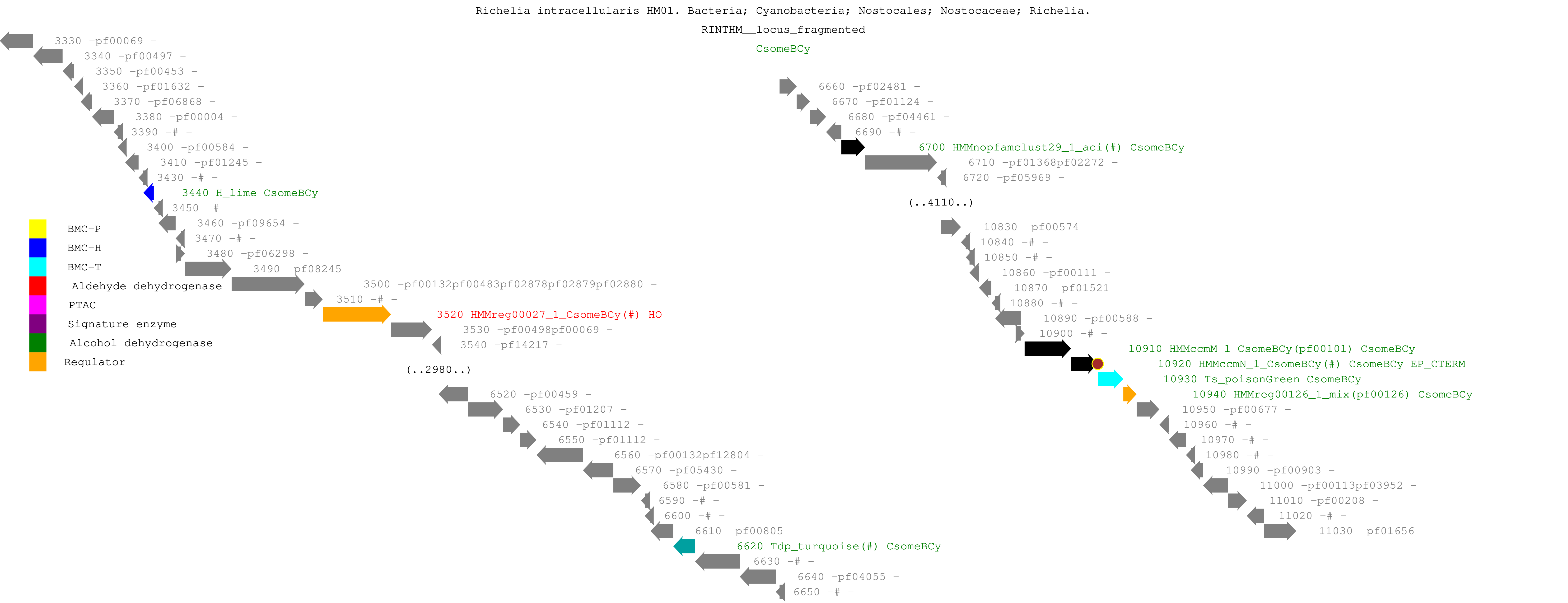

Richelia intracellularis HM01. Bacteria; Cyanobacteria; Nostocales; Nostocaceae; Richelia.
RINTHM__locus_fragmented
CsomeBCy
3330 -pf00069 -
3340 -pf00497 -
3350 -pf00453 -
3360 -pf01632 -
3370 -pf06868 -
3380 -pf00004 -
3390 -# -
3400 -pf00584 -
3410 -pf01245 -
3430 -# -
3440 H_lime CsomeBCy
3450 -# -
3460 -pf09654 -
3470 -# -
3480 -pf06298 -
3490 -pf08245 -
3500 -pf00132pf00483pf02878pf02879pf02880 -
3510 -# -
3520 HMMreg00027_1_CsomeBCy(#) HO
3530 -pf00498pf00069 -
3540 -pf14217 -
(..2980..)
6520 -pf00459 -
6530 -pf01207 -
6540 -pf01112 -
6550 -pf01112 -
6560 -pf00132pf12804 -
6570 -pf05430 -
6580 -pf00581 -
6590 -# -
6600 -# -
6610 -pf00805 -
6620 Tdp_turquoise(#) CsomeBCy
6630 -# -
6640 -pf04055 -
6650 -# -
6660 -pf02481 -
6670 -pf01124 -
6680 -pf04461 -
6690 -# -
6700 HMMnopfamclust29_1_aci(#) CsomeBCy
6710 -pf01368pf02272 -
6720 -pf05969 -
(..4110..)
10830 -pf00574 -
10840 -# -
10850 -# -
10860 -pf00111 -
10870 -pf01521 -
10880 -# -
10890 -pf00588 -
10900 -# -
10910 HMMccmM_1_CsomeBCy(pf00101) CsomeBCy
10920 HMMccmN_1_CsomeBCy(#) CsomeBCy EP_CTERM
10930 Ts_poisonGreen CsomeBCy
10940 HMMreg00126_1_mix(pf00126) CsomeBCy
10950 -pf00677 -
10960 -# -
10970 -# -
10980 -# -
10990 -pf00903 -
11000 -pf00113pf03952 -
11010 -pf00208 -
11020 -# -
11030 -pf01656 -
BMC-P
BMC-H
BMC-T
Aldehyde dehydrogenase
PTAC
Signature enzyme
Alcohol dehydrogenase
Regulator

Richelia intracellularis. Bacteria; Cyanobacteria; Nostocales; Nostocaceae; Richelia.

RintRC__locus_1_main_sat

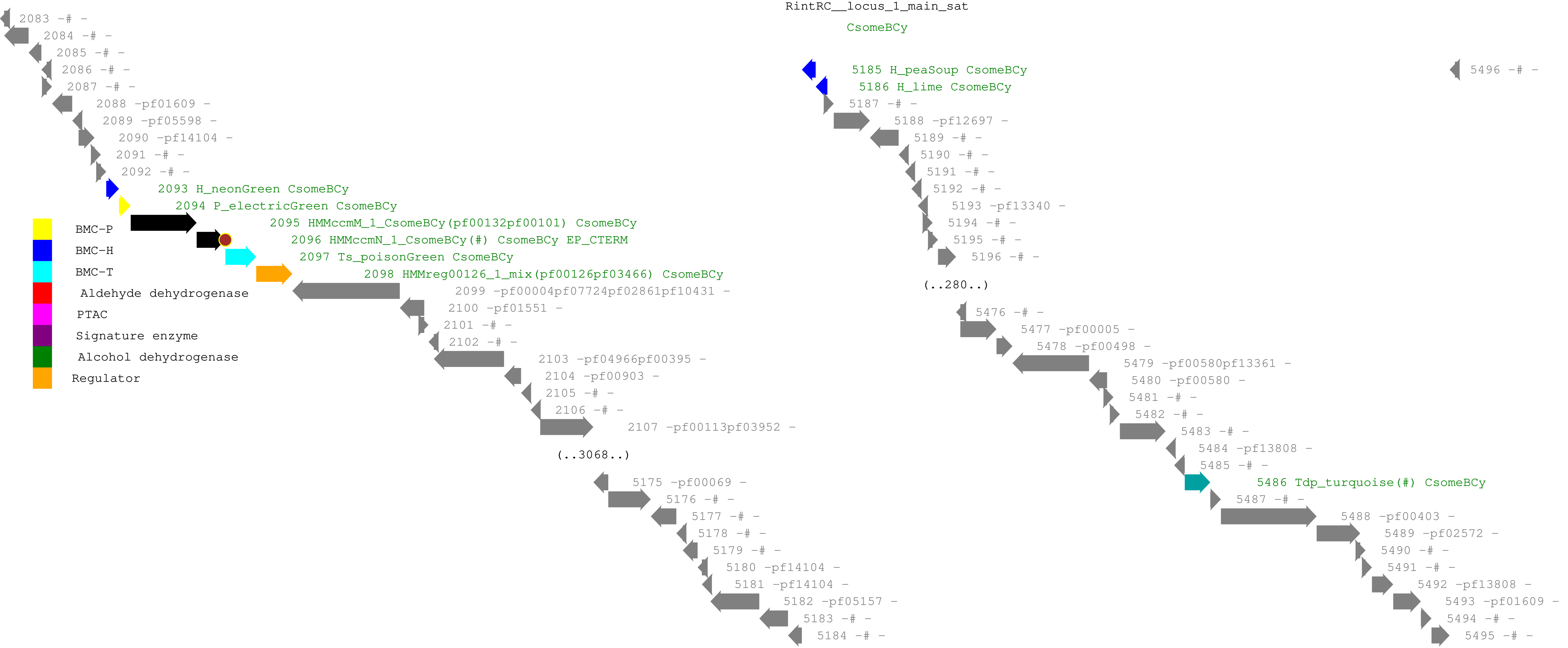

Rivularia sp. PCC 7116. Bacteria; Cyanobacteria; Nostocales; Rivulariaceae; Rivularia.

Riv7116__locus_1_main_sat

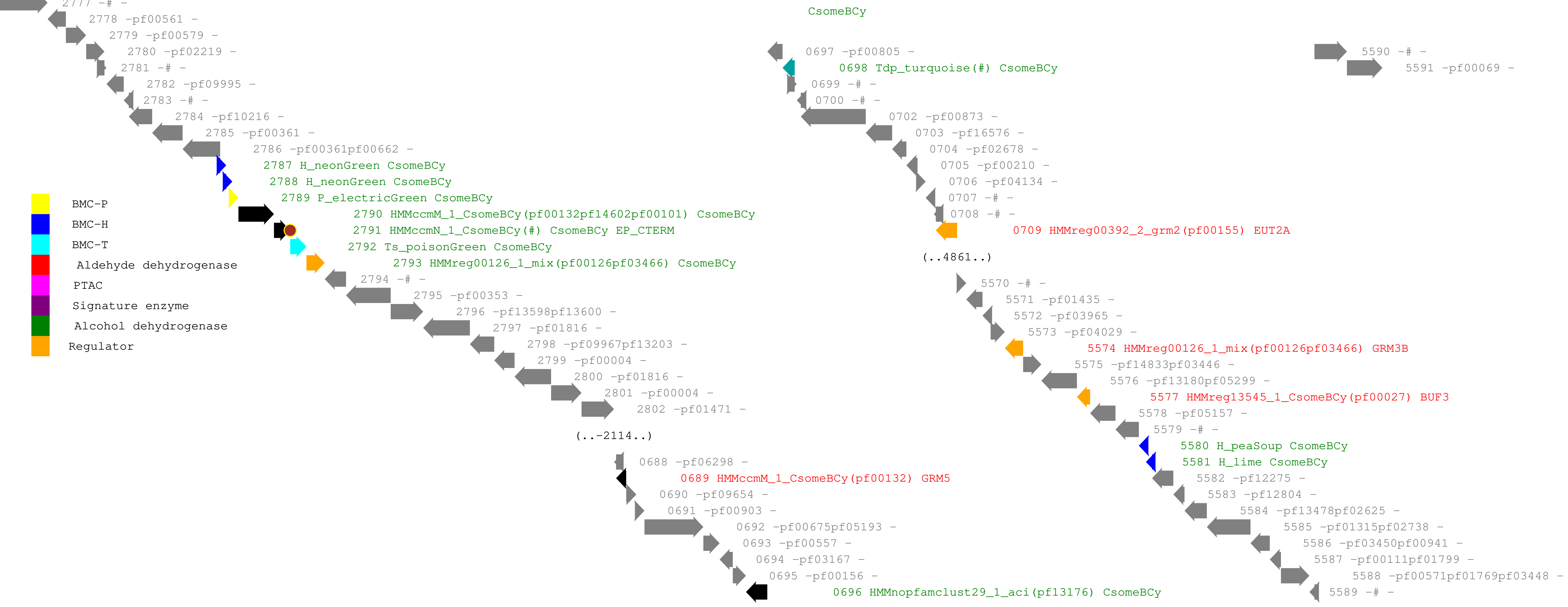

Scytonema sp. HK-05. Bacteria; Cyanobacteria; Nostocales; Scytonemataceae; Scytonema.

SAMD00079811__locus_1_main_sat

CsomeBCy

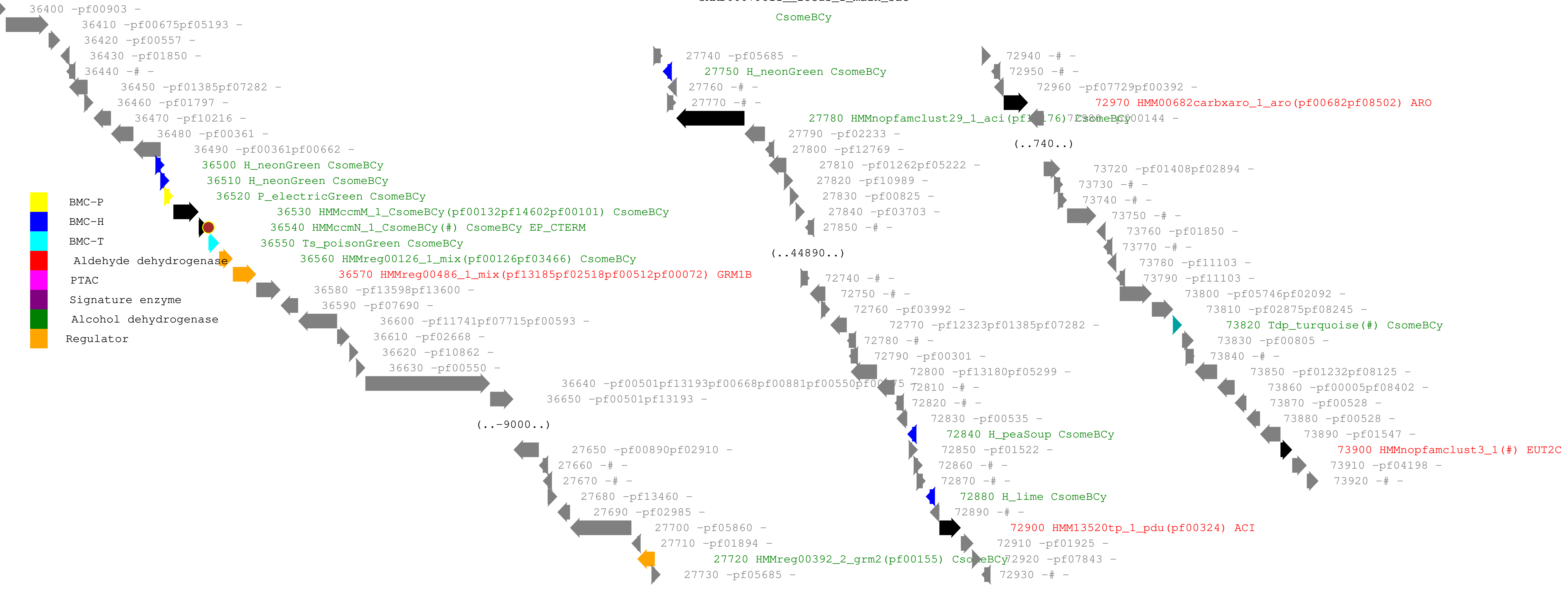

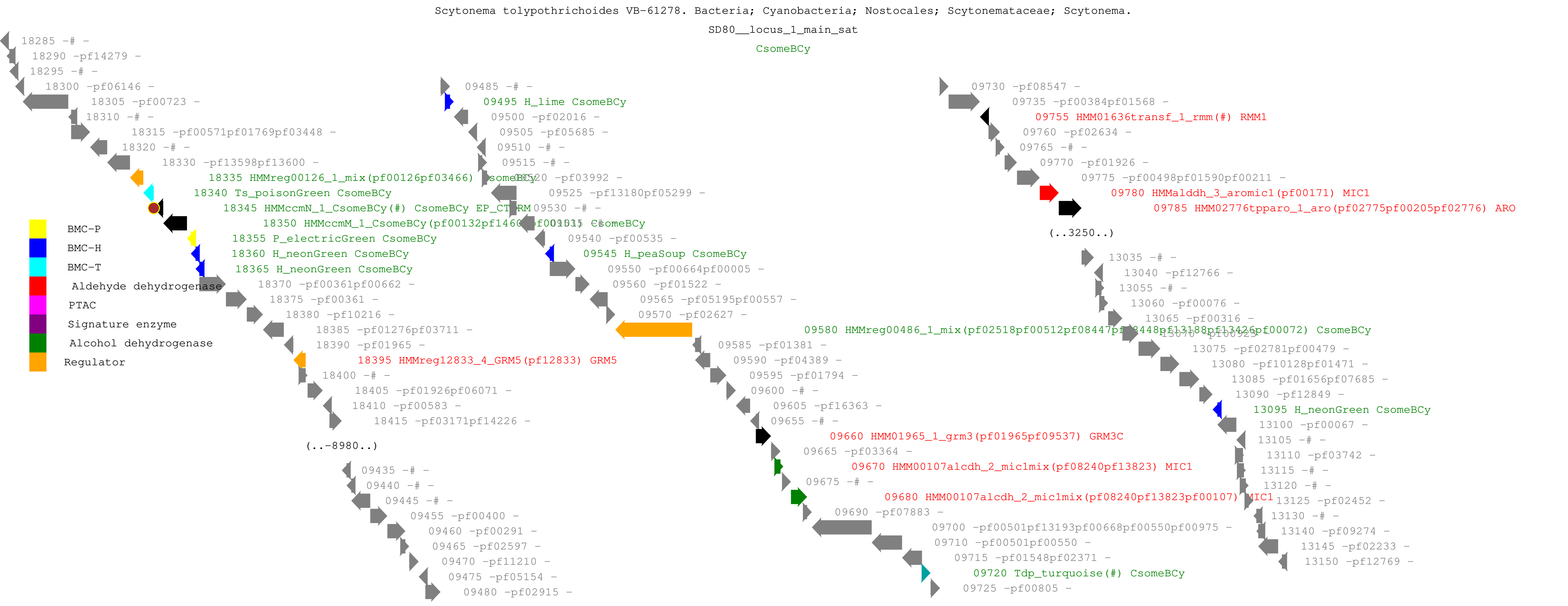
Scytonema tolypothrichoides VB-61278. Bacteria; Cyanobacteria; Nostocales; Scytonemataceae; Scytonema.
SD80__locus_1_main_sat
CsomeBCy
18285 -# -
18290 -pf14279 -
18295 -# -
18300 -pf06146 -
18305 -pf00723 -
18310 -# -
18315 -pf00571pf01769pf03448 -
18320 -# -
18330 -pf13598pf13600 -
18335 HMMreg00126_1_mix(pf00126pf03466) CsomeBCy
18340 Ts_poisonGreen CsomeBCy
18345 HMMccmN_1_CsomeBCy(#) CsomeBCy EP_CTERM
18350 HMMccmM_1_CsomeBCy(pf00132pf14602pf00101) CsomeBCy
18355 P_electricGreen CsomeBCy
18360 H_neonGreen CsomeBCy
18365 H_neonGreen CsomeBCy
18370 -pf00361pf00662 -
18375 -pf00361 -
18380 -pf10216 -
18385 -pf01276pf03711 -
18390 -pf01965 -
18395 HMMreg12833_4_GRM5(pf12833) GRM5
18400 -# -
18405 -pf01926pf06071 -
18410 -pf00583 -
18415 -pf03171pf14226 -
(..-8980..)
09435 -# -
09440 -# -
09445 -# -
09455 -pf00400 -
09460 -pf00291 -
09465 -pf02597 -
09470 -pf11210 -
09475 -pf05154 -
09480 -pf02915 -
09485 -# -
09495 H_lime CsomeBCy
09500 -pf02016 -
09505 -pf05685 -
09510 -# -
09515 -# -
09520 -pf03992 -
09525 -pf13180pf05299 -
09530 -# -
09535 CsomeBCy
09540 -pf00535 -
09545 H_peaSoup CsomeBCy
09550 -pf00664pf00005 -
09560 -pf01522 -
09565 -pf05195pf00557 -
09570 -pf02627 -
09580 HMMreg00486_1_mix(pf02518pf00512pf08447pf08448pf13188pf13426pf00072) CsomeBCy
09585 -pf01381 -
09590 -pf04389 -
09595 -pf01794 -
09600 -# -
09605 -pf16363 -
09655 -# -
09660 HMM01965_1_grm3(pf01965pf09537) GRM3C
09665 -pf03364 -
09670 HMM00107alcdh_2_mic1mix(pf08240pf13823) MIC1
09675 -# -
09680 HMM00107alcdh_2_mic1mix(pf08240pf13823pf00107) MIC1
09690 -pf07883 -
09700 -pf00501pf13193pf00668pf00550pf00975 -
09710 -pf00501pf00550 -
09715 -pf01548pf02371 -
09720 Tdp_turquoise(#) CsomeBCy
09725 -pf00805 -
09730 -pf08547 -
09735 -pf00384pf01568 -
09755 HMM01636transf_1_rmm(#) RMM1
09760 -pf02634 -
09765 -# -
09770 -pf01926 -
09775 -pf00498pf01590pf00211 -
09780 HMMaldd h_3_aromic1(pf00171) MIC1
09785 HMM02776tpparo_1_aro(pf02775pf00205pf02776) ARO
(..3250..)
13035 -# -
13040 -pf12766 -
13055 -# -
13060 -pf00076 -
13065 -pf00316 -
13070 -pf00923 -
13075 -pf02781pf00479 -
13080 -pf10128pf01471 -
13085 -pf01656pf07685 -
13090 -pf12849 -
13095 H_neonGreen CsomeBCy
13100 -pf00067 -
13105 -# -
13110 -pf03742 -
13115 -# -
13120 -# -
13125 -pf02452 -
13130 -# -
13140 -pf09274 -
13145 -pf02233 -
13150 -pf12769 -
BMC-P
BMC-H
BMC-T
Aldehyde dehydrogenase
PTAC
Signature enzyme
Alcohol dehydrogenase
Regulator

Tolypothrix campylonemoides VB511288. Bacteria; Cyanobacteria; Nostocales; Tolypothrichaceae; Tolypothrix.

SD81__locus_1_main_sat

CsomeBCy

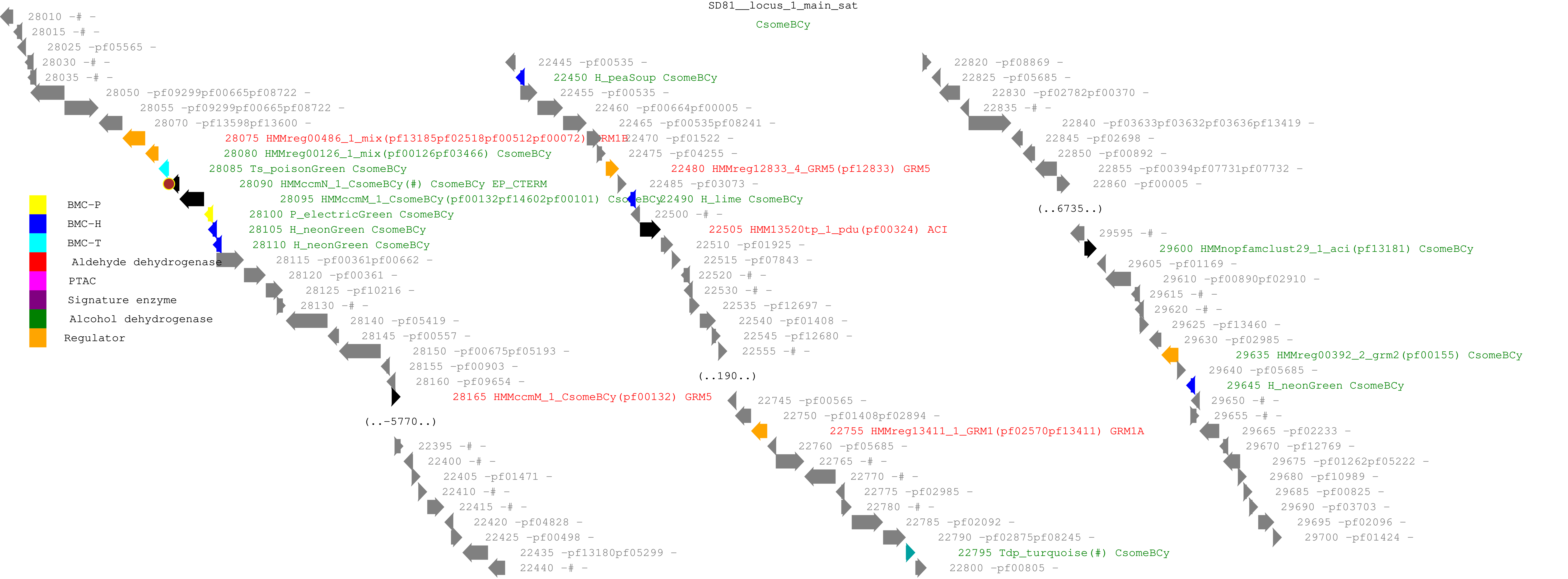

Synechocystis sp. (strain PCC 6803 / Kazusa). Bacteria; Cyanobacteria; Synechococcales; Merismopediaceae;

slr_locus_fragmented

CsomeBCy

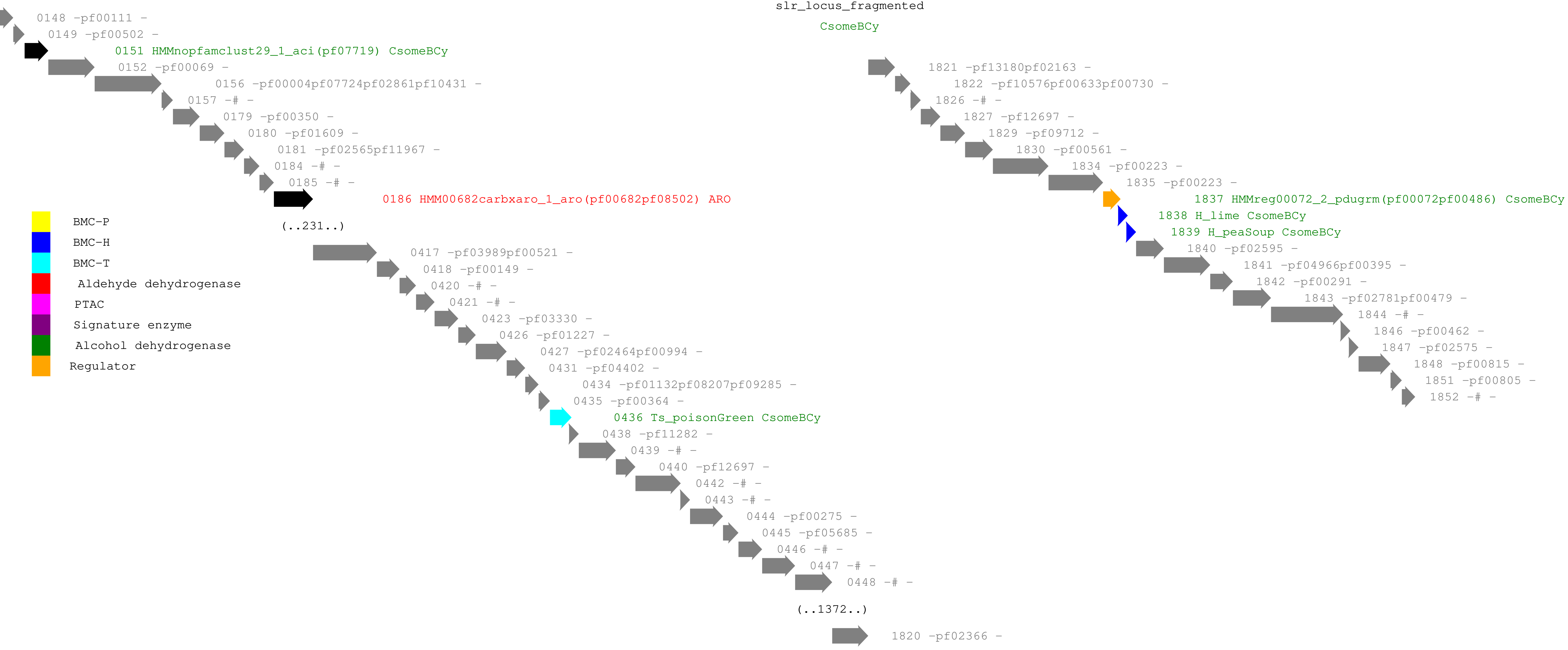

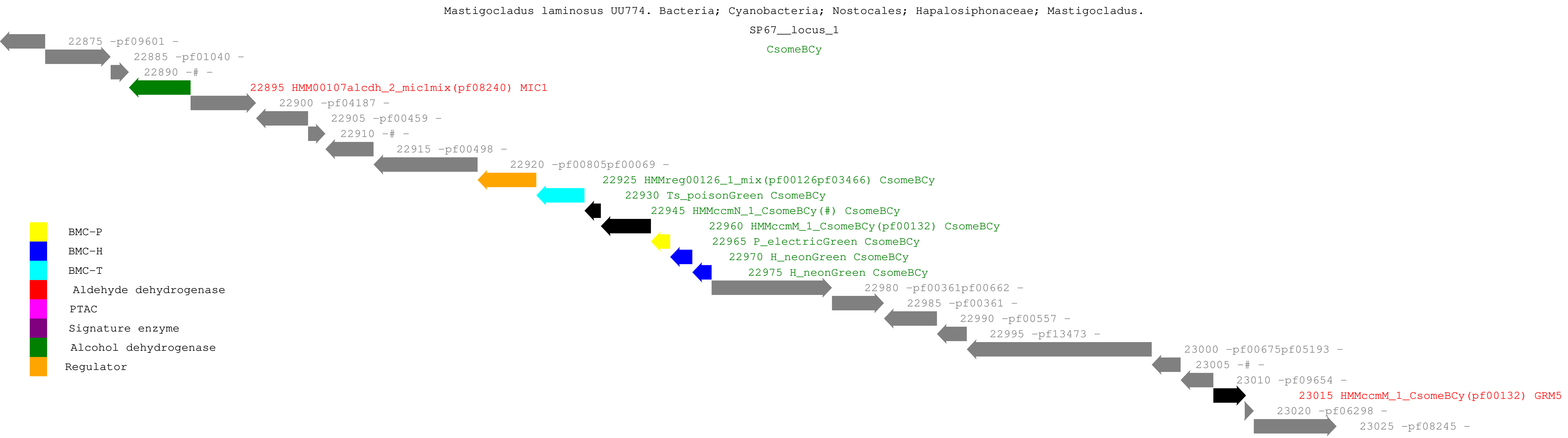

Mastigocladus laminosus UU774. Bacteria; Cyanobacteria; Nostocales; Hapalosiphonaceae; Mastigocladus.
SP67__locus_1
CsomeBCy
22875 -pf09601 -
22885 -pf01040 -
22890 -# -
22895 HMM00107alcdh_2_mic1mix(pf08240) MIC1
22900 -pf04187 -
22905 -pf00459 -
22910 -# -
22915 -pf00498 -
22920 -pf00805pf00069 -
22925 HMMreg00126_1_mix(pf00126pf03466) CsomeBCy
22930 Ts_poisonGreen CsomeBCy
22945 HMMccmN_1_CsomeBCy(#) CsomeBCy
22960 HMMccmM_1_CsomeBCy(pf00132) CsomeBCy
22965 P_electricGreen CsomeBCy
22970 H_neonGreen CsomeBCy
22975 H_neonGreen CsomeBCy
22980 -pf00361pf00662 -
22985 -pf00361 -
22990 -pf00557 -
22995 -pf13473 -
23000 -pf00675pf05193 -
23005 -# -
23010 -pf09654 -
23015 HMMccmM_1_CsomeBCy(pf00132) GRM5
23020 -pf06298 -
23025 -pf08245 -
BMC-P
BMC-H
BMC-T
Aldehyde dehydrogenase
PTAC
Signature enzyme
Alcohol dehydrogenase
Regulator

Arthrospira platensis C1. Bacteria; Cyanobacteria; Oscillatoriophycideae; Oscillatoriales;

SPLC1_S_locus_1_main_sat

CsomeBCy

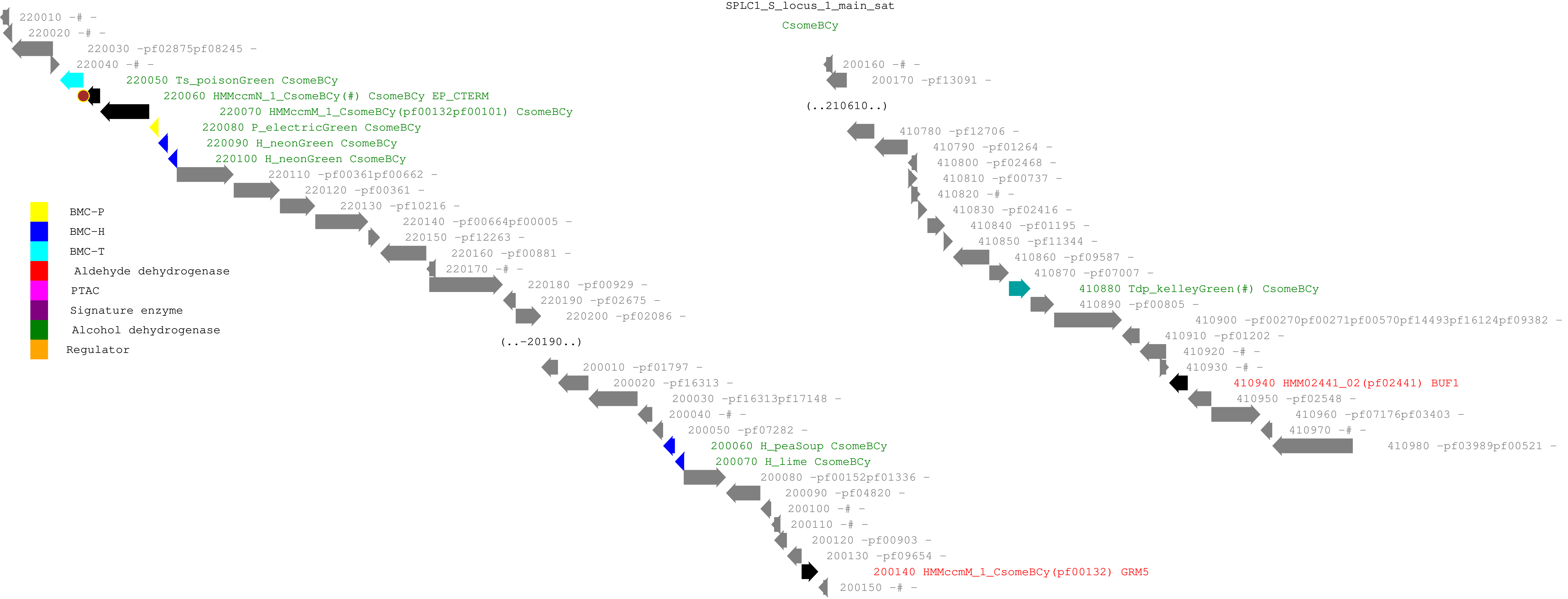

Sphaerospermopsis reniformis. Bacteria; Cyanobacteria; Nostocales; Aphanizomenonaceae; Sphaerospermopsis.

SR1949__locus_1_main_sat

CsomeBCy

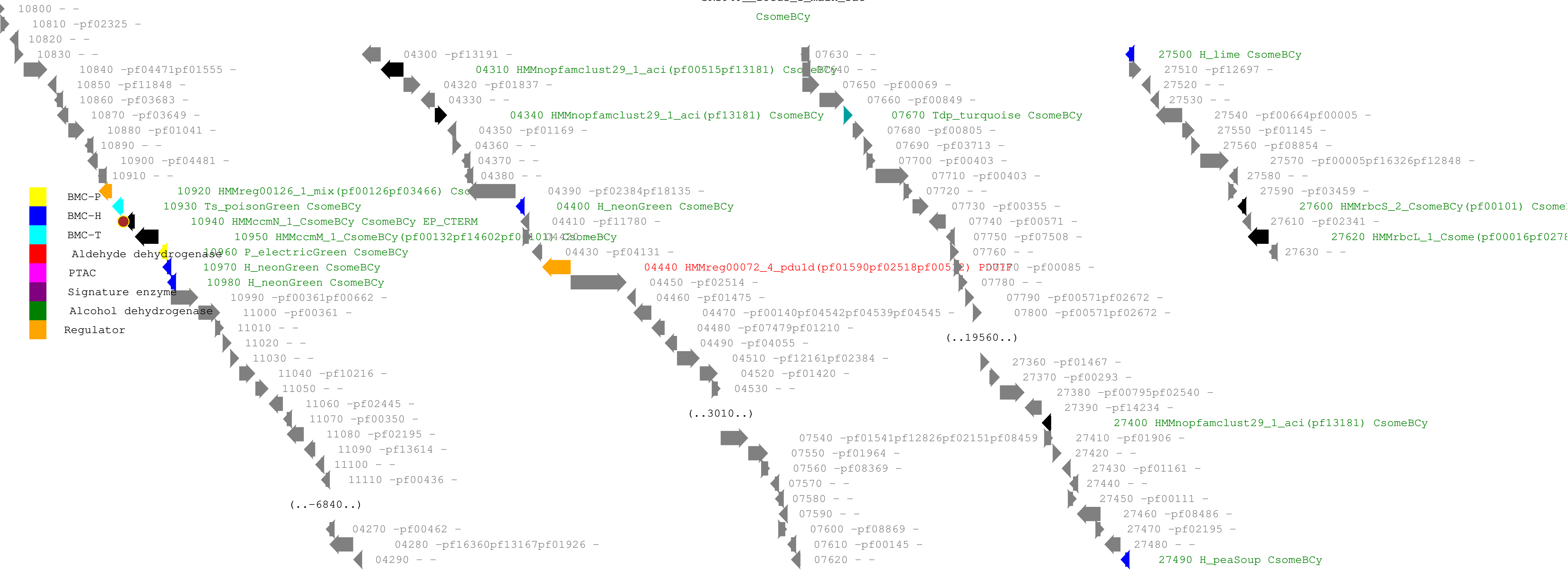

Stanieria sp. NIES-3757. Bacteria; Cyanobacteria; Pleurocapsales; Dermocarpellaceae; Stanieria.

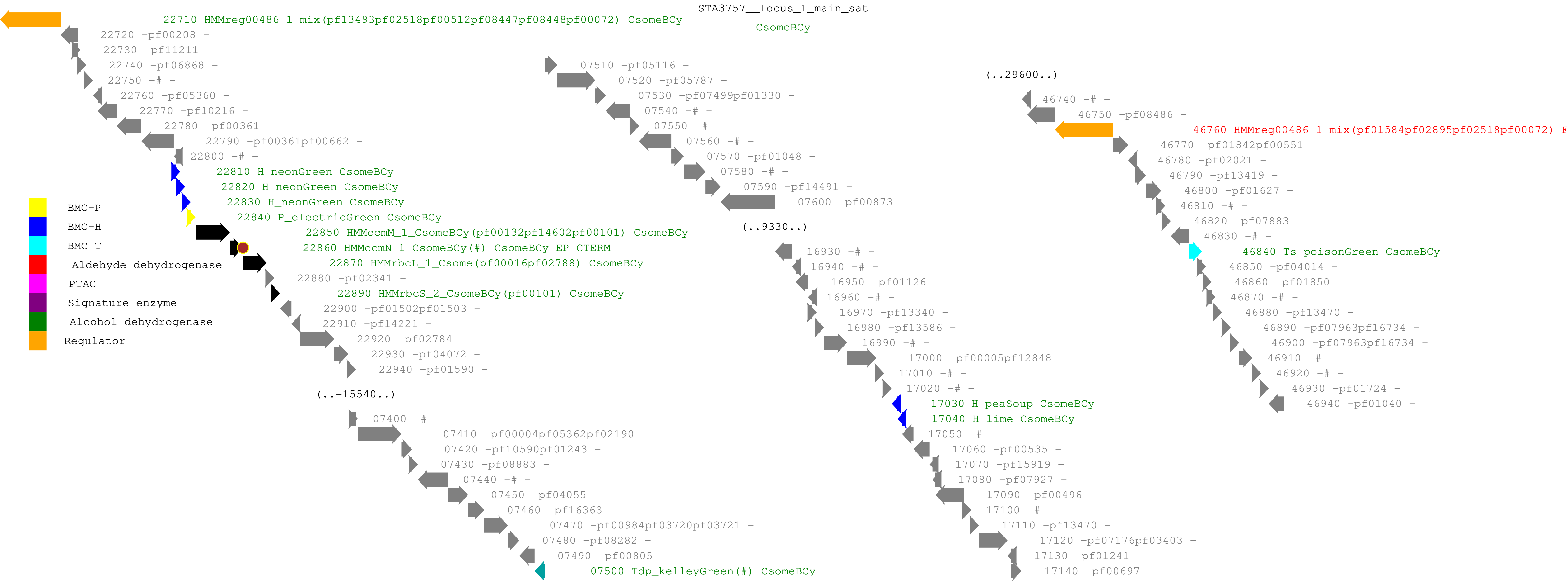

Stanieria cyanosphaera (strain ATCC 29371 / PCC 7437). Bacteria; Cyanobacteria; Pleurocapsales; Dermocarpellaceae; Stanieria.

Sta7437__locus_1_main_sat

CsomeBCy

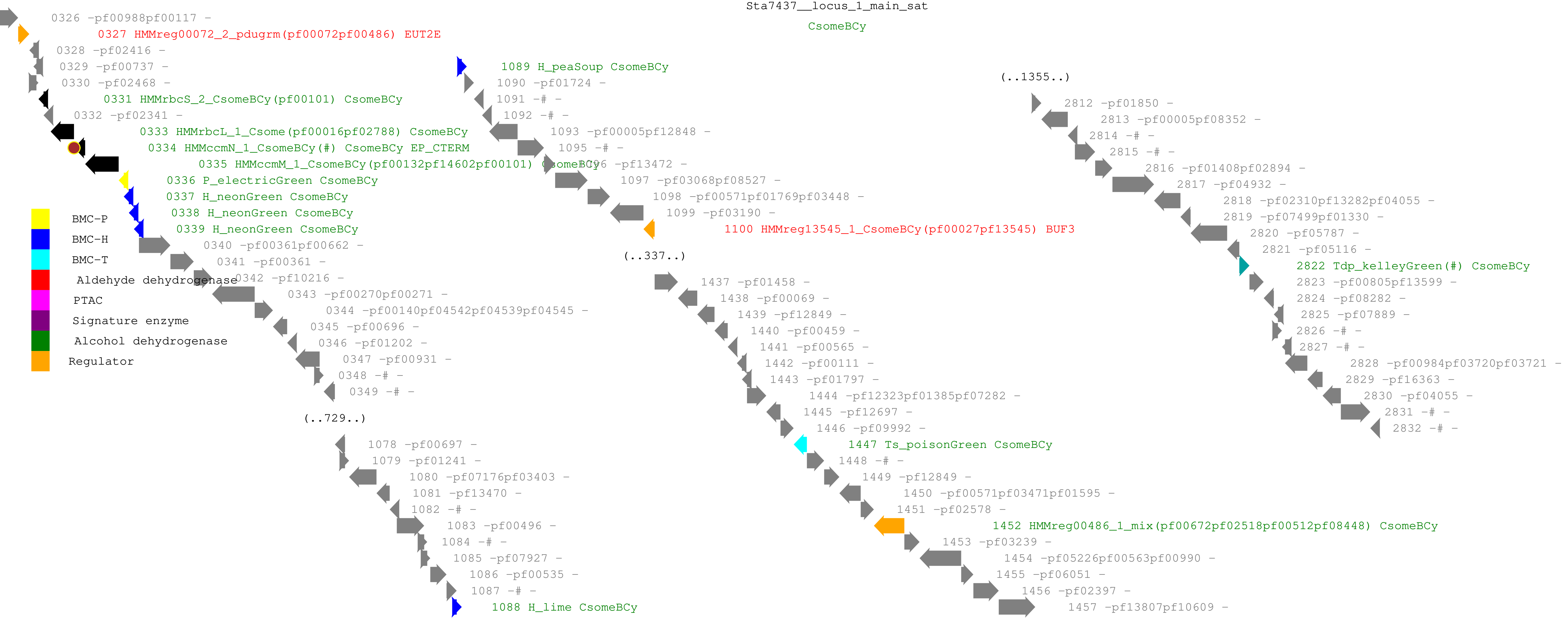

Synechococcus sp. 60AY4M2. Bacteria; Cyanobacteria; Synechococcales; Synechococcaceae;

SYN60AY4M2__locus_1_main_sat

CsomeBCy

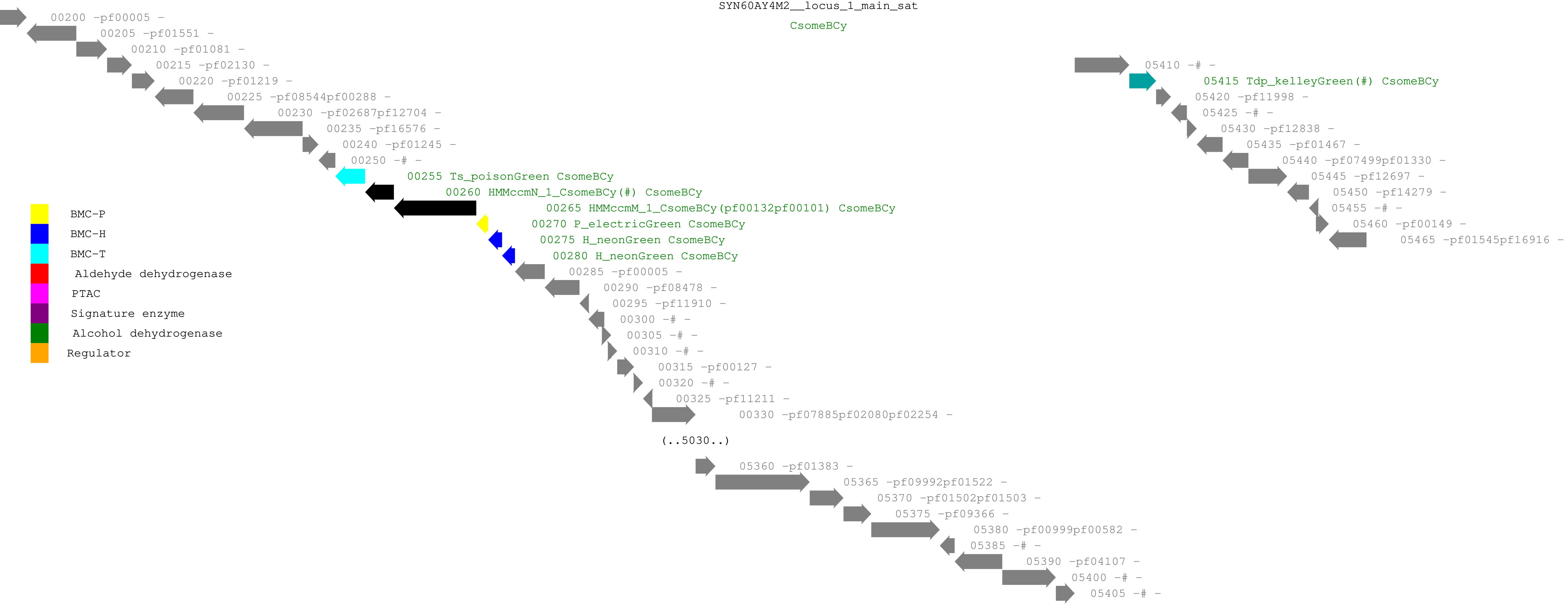

Synechococcus sp. (strain ATCC 27167 / PCC 6312). Bacteria; Cyanobacteria; Synechococcales; Synechococcaceae;

Syn6312__locus_1_main_sat

CsomeBCy

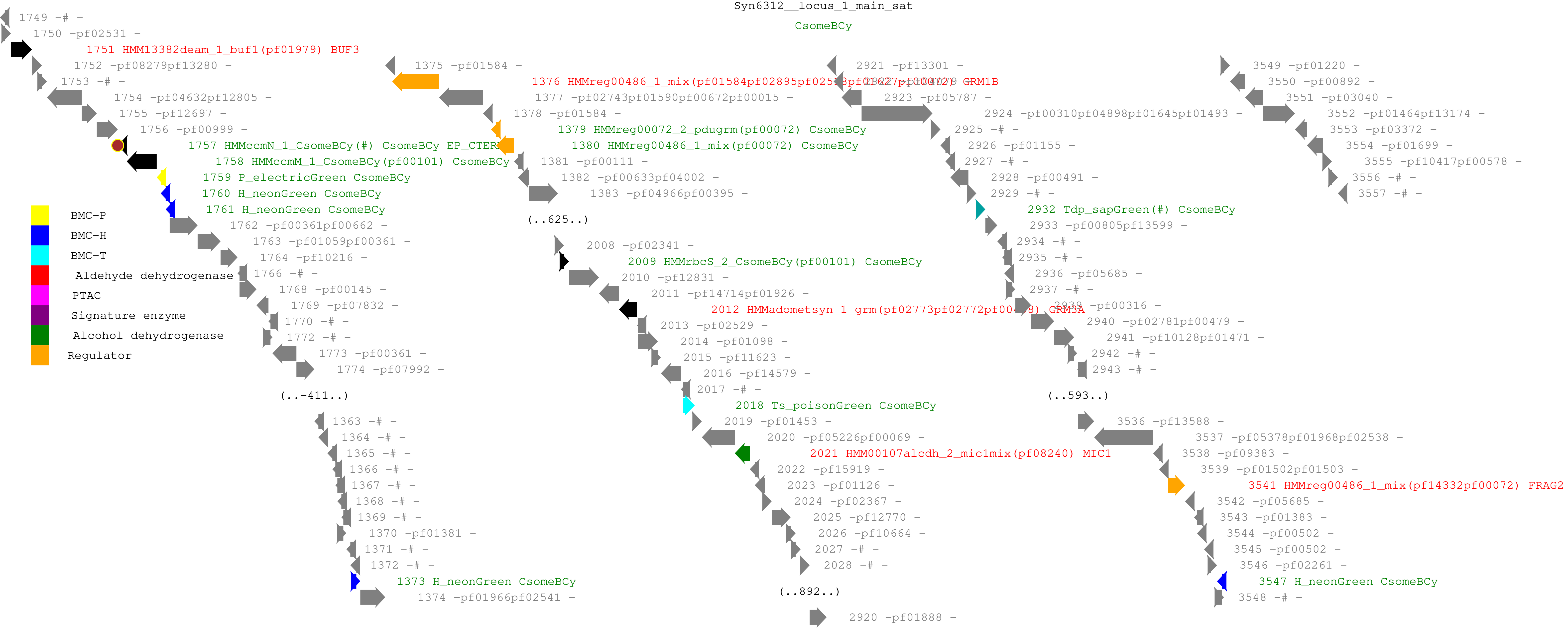

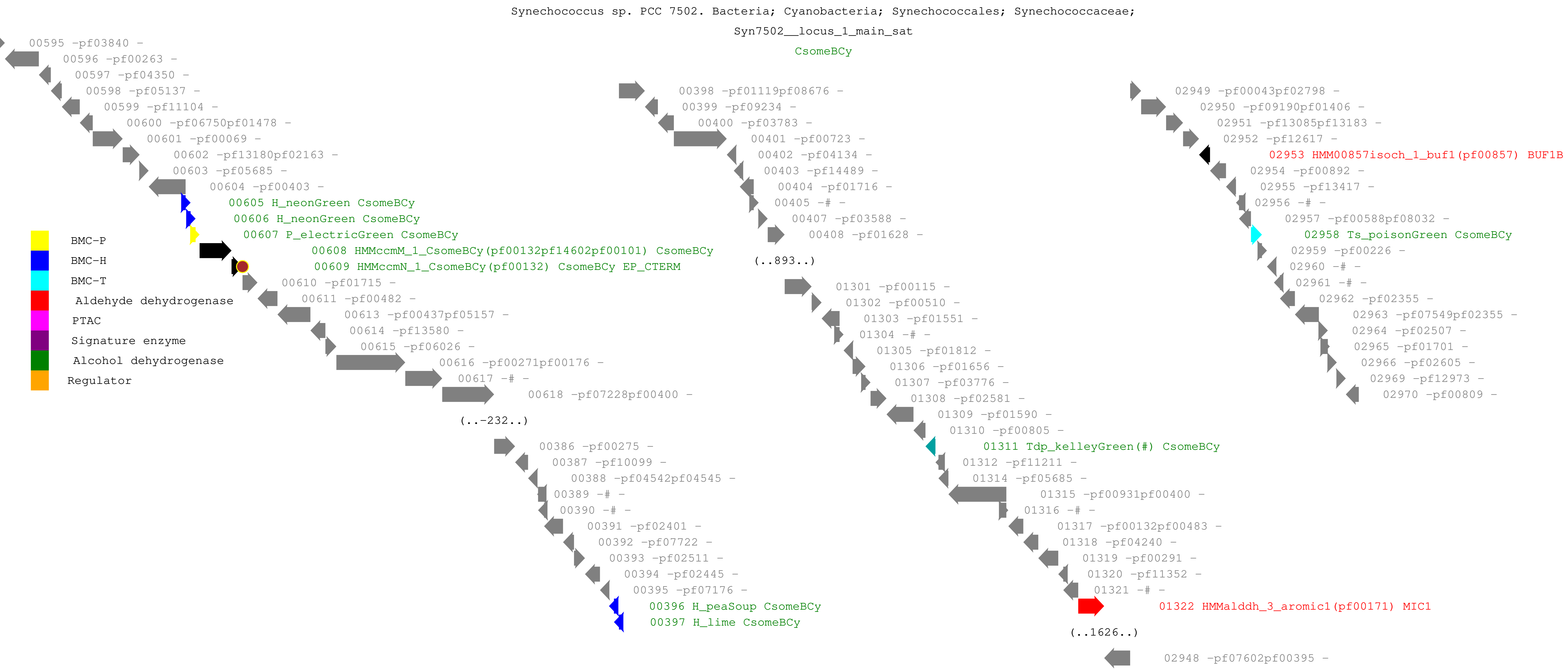

Synechococcus sp. PCC 7502. Bacteria; Cyanobacteria; Synechococcales; Synechococcaceae;
Syn7502__locus_1_main_sat
CsomeBCy
00595 -pf03840 -
00596 -pf00263 -
00597 -pf04350 -
00598 -pf05137 -
00599 -pf11104 -
00600 -pf06750pf01478 -
00601 -pf00069 -
00602 -pf13180pf02163 -
00603 -pf05685 -
00604 -pf00403 -
00605 H_neonGreen CsomeBCy
00606 H_neonGreen CsomeBCy
00607 P_electricGreen CsomeBCy
00608 HMMccmM_1_CsomeBCy(pf00132pf14602pf00101) CsomeBCy
00609 HMMccmN_1_CsomeBCy(pf00132) CsomeBCy EP_CTERM
00610 -pf01715 -
00611 -pf00482 -
00613 -pf00437pf05157 -
00614 -pf13580 -
00615 -pf06026 -
00616 -pf00271pf00176 -
00617 -# -
00618 -pf07228pf00400 -
(..-232..)
00386 -pf00275 -
00387 -pf10099 -
00388 -pf04542pf04545 -
00389 -# -
00390 -# -
00391 -pf02401 -
00392 -pf07722 -
00393 -pf02511 -
00394 -pf02445 -
00395 -pf07176 -
00396 H_peaSoup CsomeBCy
00397 H_lime CsomeBCy
00398 -pf01119pf08676 -
00399 -pf09234 -
00400 -pf03783 -
00401 -pf00723 -
00402 -pf04134 -
00403 -pf14489 -
00404 -pf01716 -
00405 -# -
00407 -pf03588 -
00408 -pf01628 -
(..893..)
01301 -pf00115 -
01302 -pf00510 -
01303 -pf01551 -
01304 -# -
01305 -pf01812 -
01306 -pf01656 -
01307 -pf03776 -
01308 -pf02581 -
01309 -pf01590 -
01310 -pf00805 -
01311 Tdp_kelleyGreen(#) CsomeBCy
01312 -pf11211 -
01314 -pf05685 -
01315 -pf00931pf00400 -
01316 -# -
01317 -pf00132pf00483 -
01318 -pf04240 -
01319 -pf00291 -
01320 -pf11352 -
01321 -# -
01322 HMMaldh_3_aromic1(pf00171) MIC1
(..1626..)
02948 -pf07602pf00395 -
02949 -pf00043pf02798 -
02950 -pf09190pf01406 -
02951 -pf13085pf13183 -
02952 -pf12617 -
02953 HMM00857isoch_1_buf1(pf00857) BUF1B
02954 -pf00892 -
02955 -pf13417 -
02956 -# -
02957 -pf00588pf08032 -
02958 Ts_poisonGreen CsomeBCy
02959 -pf00226 -
02960 -# -
02961 -# -
02962 -pf02355 -
02963 -pf07549pf02355 -
02964 -pf02507 -
02965 -pf01701 -
02966 -pf02605 -
02969 -pf12973 -
02970 -pf00809 -
BMC-P
BMC-H
BMC-T
Aldehyde dehydrogenase
PTAC
Signature enzyme
Alcohol dehydrogenase
Regulator

Synechococcus sp. (strain ATCC 27264 / PCC 7002 / PR-6) (Agmenellum Bacteria; Cyanobacteria; Synechococcales; Synechococcaceae;

SYNPCC7002_A_locus_1_main_sat

CsomeBCy

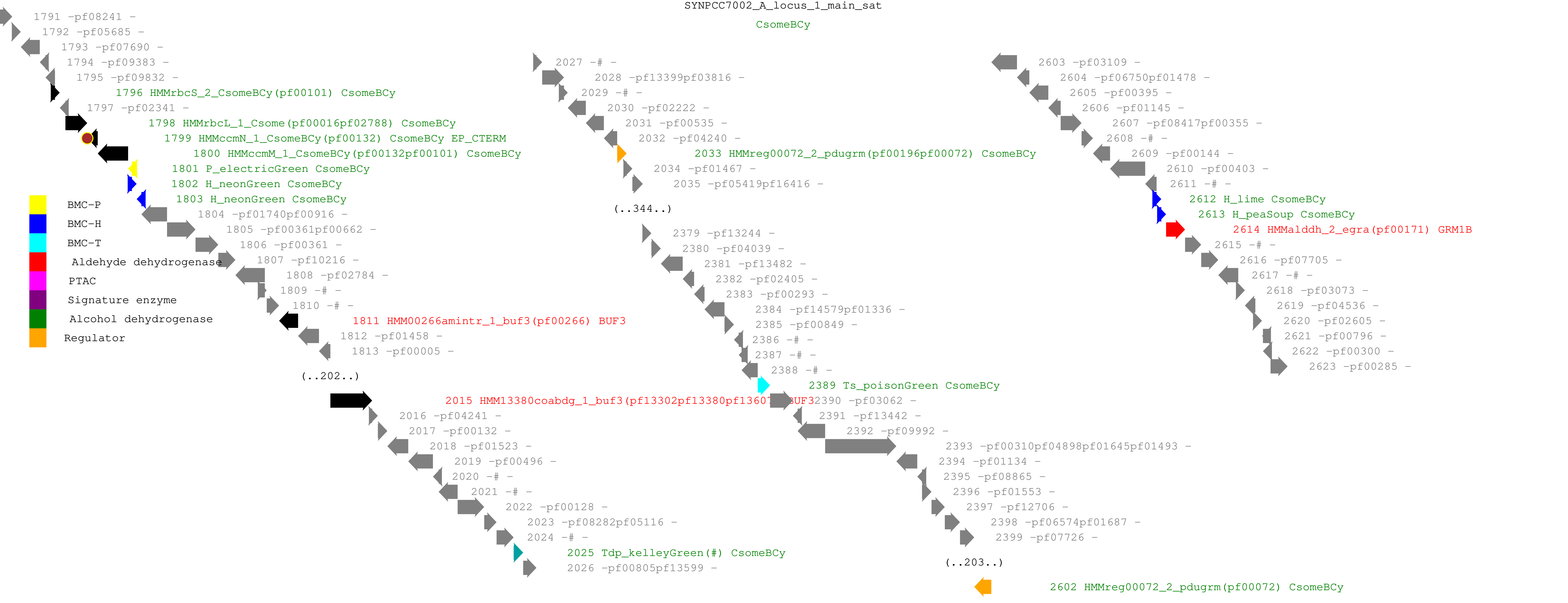

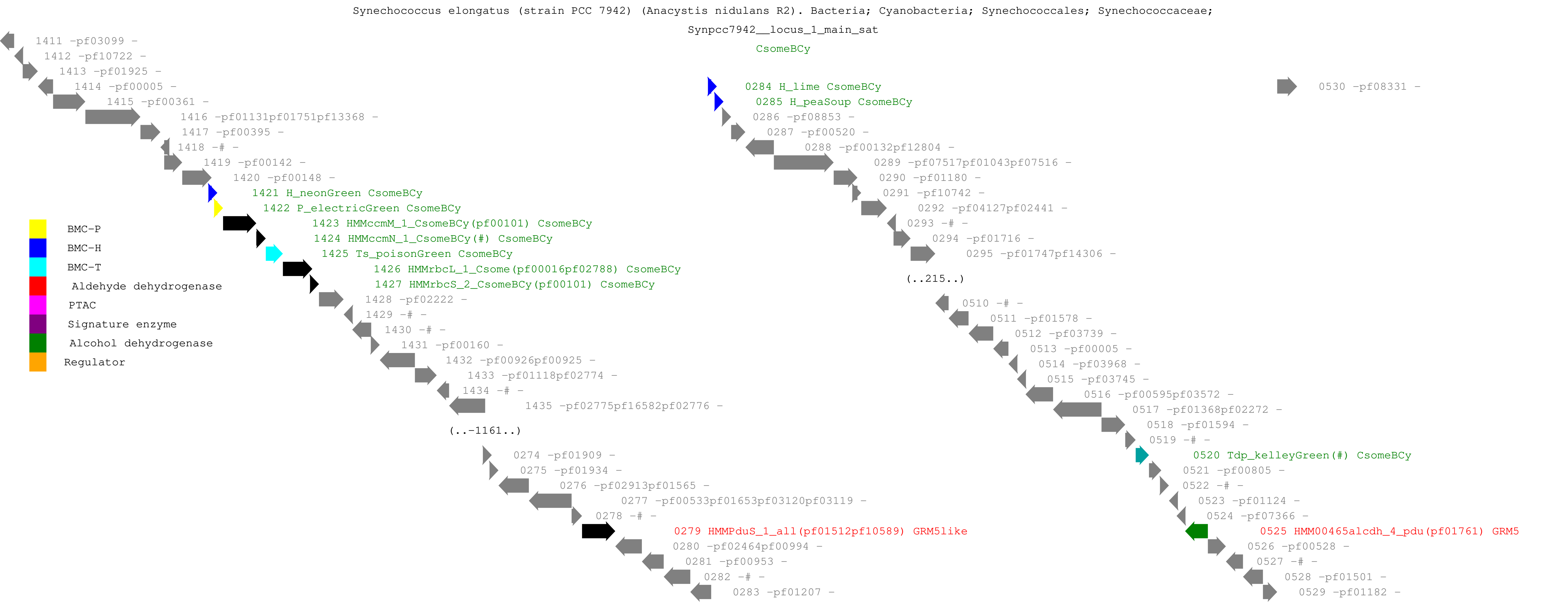

Synechococcus elongatus (strain PCC 7942) (Anacystis nidulans R2). Bacteria; Cyanobacteria; Synechococcales; Synechococcaceae;
Synpcc7942__locus_1_main_sat
CsomeBCy
1411 -pf03099 -
1412 -pf10722 -
1413 -pf01925 -
1414 -pf00005 -
1415 -pf00361 -
1416 -pf01131pf01751pf13368 -
1417 -pf00395 -
1418 -# -
1419 -pf00142 -
1420 -pf00148 -
1421 H_neonGreen CsomeBCy
1422 P_electricGreen CsomeBCy
1423 HMMccmM_1_CsomeBCy(pf00101) CsomeBCy
1424 HMMccmN_1_CsomeBCy(#) CsomeBCy
1425 Ts_poisonGreen CsomeBCy
1426 HMMrbcL_1_Csome(pf00016pf02788) CsomeBCy
1427 HMMrbcS_2_CsomeBCy(pf00101) CsomeBCy
1428 -pf02222 -
1429 -# -
1430 -# -
1431 -pf00160 -
1432 -pf00926pf00925 -
1433 -pf01118pf02774 -
1434 -# -
1435 -pf02775pf16582pf02776 -
(..-1161..)
0274 -pf01909 -
0275 -pf01934 -
0276 -pf02913pf01565 -
0277 -pf00533pf01653pf03120pf03119 -
0278 -# -
0279 HMMPduS_1_all(pf01512pf10589) GRM5like
0280 -pf02464pf00994 -
0281 -pf00953 -
0282 -# -
0283 -pf01207 -
0284 H_lime CsomeBCy
0285 H_peaSoup CsomeBCy
0286 -pf08853 -
0287 -pf00520 -
0288 -pf00132pf12804 -
0289 -pf07517pf01043pf07516 -
0290 -pf01180 -
0291 -pf10742 -
0292 -pf04127pf02441 -
0293 -# -
0294 -pf01716 -
0295 -pf01747pf14306 -
(..215..)
0510 -# -
0511 -pf01578 -
0512 -pf03739 -
0513 -pf00005 -
0514 -pf03968 -
0515 -pf03745 -
0516 -pf00595pf03572 -
0517 -pf01368pf02272 -
0518 -pf01594 -
0519 -# -
0520 Tdp_kelleyGreen(#) CsomeBCy
0521 -pf00805 -
0522 -# -
0523 -pf01124 -
0524 -pf07366 -
0525 HMM00465alcdh_4_pdu(pf01761) GRM5
0526 -pf00528 -
0527 -# -
0528 -pf01501 -
0529 -pf01182 -
0530 -pf08331 -
BMC-P
BMC-H
BMC-T
Aldehyde dehydrogenase
PTAC
Signature enzyme
Alcohol dehydrogenase
Regulator

Trichodesmium erythraeum (strain IMS101). Bacteria; Cyanobacteria; Oscillatoriophycideae; Oscillatoriales;

Tery__locus_1_main_sat

CsomeBCy

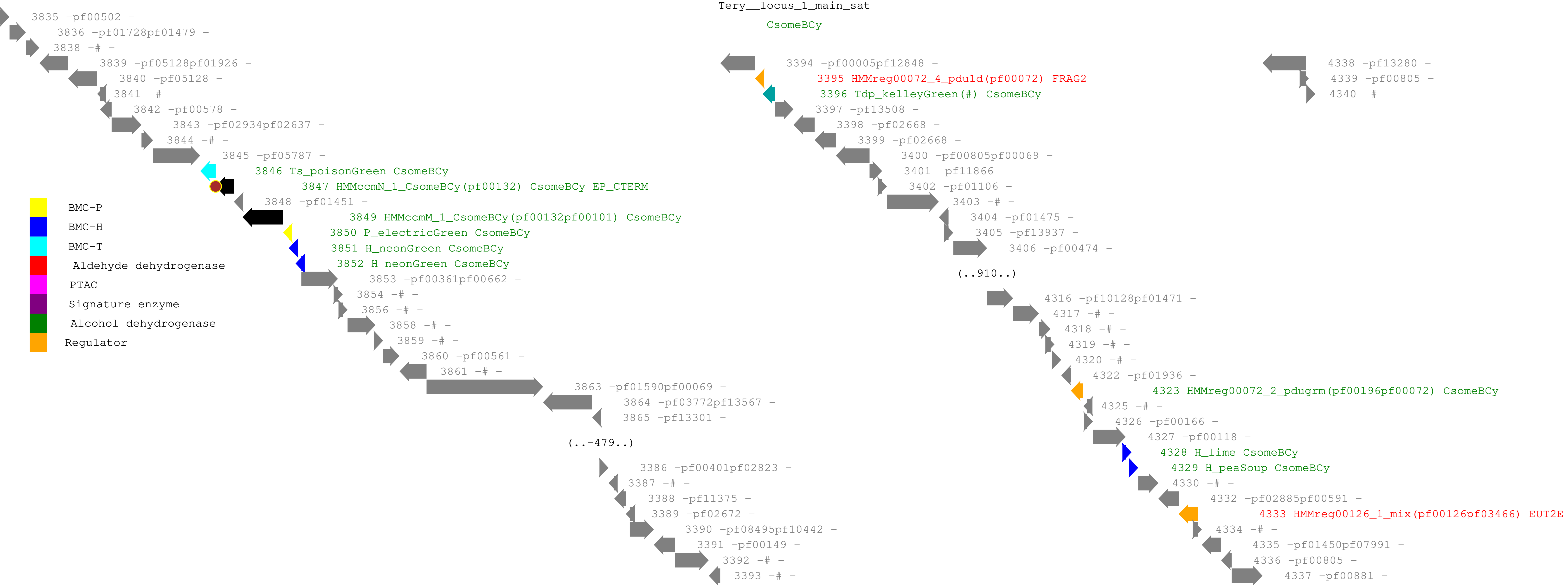

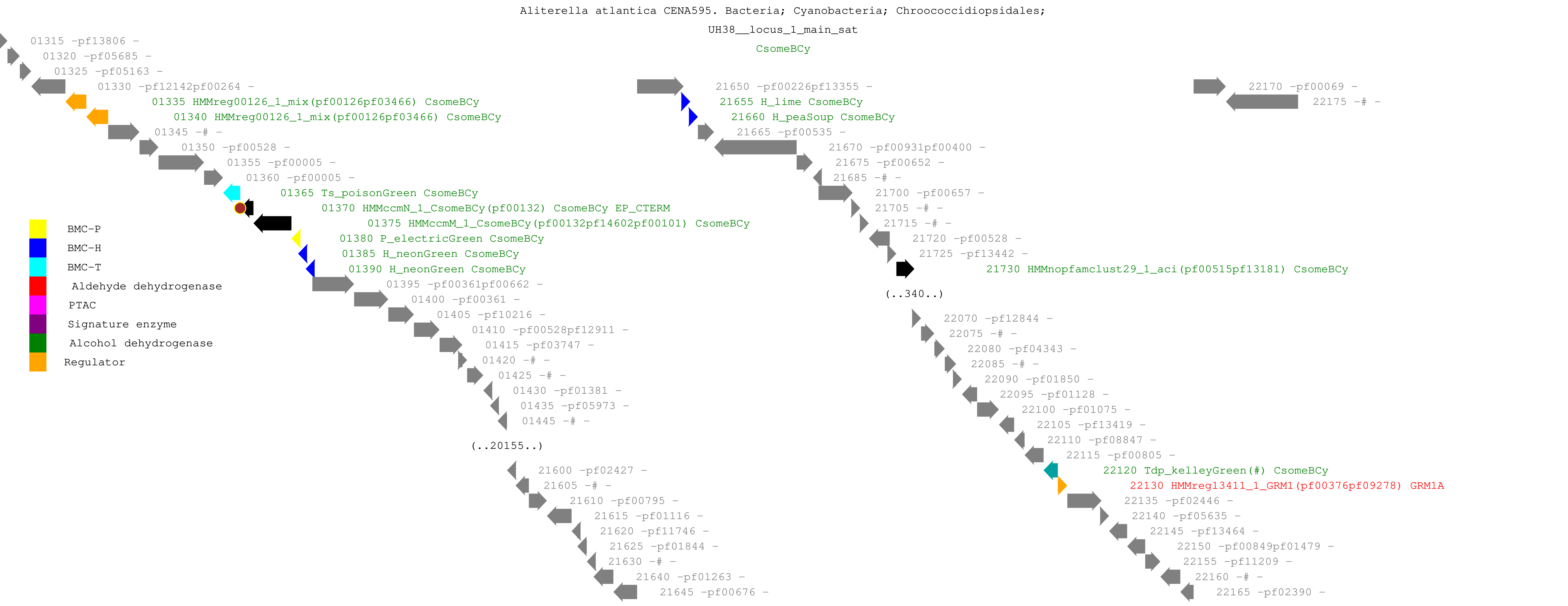
Aliterella atlantica CENA595. Bacteria; Cyanobacteria; Chroococcidiopsidales;
UH38__locus_1_main_sat
CsomeBCy
01315 -pf13806 -
01320 -pf05685 -
01325 -pf05163 -
01330 -pf12142pf00264 -
01335 HMMreg00126_1_mix(pf00126pf03466) CsomeBCy
01340 HMMreg00126_1_mix(pf00126pf03466) CsomeBCy
01345 -# -
01350 -pf00528 -
01355 -pf00005 -
01360 -pf00005 -
01365 Ts_poisonGreen CsomeBCy
01370 HMMccmN_1_CsomeBCy(pf00132) CsomeBCy EP_CTERM
01375 HMMccmM_1_CsomeBCy(pf00132pf14602pf00101) CsomeBCy
01380 P_electricGreen CsomeBCy
01385 H_neonGreen CsomeBCy
01390 H_neonGreen CsomeBCy
01395 -pf00361pf00662 -
01400 -pf00361 -
01405 -pf10216 -
01410 -pf00528pf12911 -
01415 -pf03747 -
01420 -# -
01425 -# -
01430 -pf01381 -
01435 -pf05973 -
01445 -# -
(..20155..)
21600 -pf02427 -
21605 -# -
21610 -pf00795 -
21615 -pf01116 -
21620 -pf11746 -
21625 -pf01844 -
21630 -# -
21640 -pf01263 -
21645 -pf00676 -
21650 -pf00226pf13355 -
21655 H_lime CsomeBCy
21660 H_peaSoup CsomeBCy
21665 -pf00535 -
21670 -pf00931pf00400 -
21675 -pf00652 -
21685 -# -
21700 -pf00657 -
21705 -# -
21715 -# -
21720 -pf00528 -
21725 -pf13442 -
21730 HMMnopfamclust29_1_aci(pf00515pf13181) CsomeBCy
(..340..)
22070 -pf12844 -
22075 -# -
22080 -pf04343 -
22085 -# -
22090 -pf01850 -
22095 -pf01128 -
22100 -pf01075 -
22105 -pf13419 -
22110 -pf08847 -
22115 -pf00805 -
22120 Tdp_kelleyGreen(#) CsomeBCy
22130 HMMreg13411_1_GRM1(pf00376pf09278) GRM1A
22135 -pf02446 -
22140 -pf05635 -
22145 -pf13464 -
22150 -pf00849pf01479 -
22155 -pf11209 -
22160 -# -
22165 -pf02390 -
22170 -pf00069 -
22175 -# -
BMC-P
BMC-H
BMC-T
Aldehyde dehydrogenase
PTAC
Signature enzyme
Alcohol dehydrogenase
Regulator

Nostoc linckia z16. Bacteria; Cyanobacteria; Nostocales; Nostocaceae; Nostoc.

VF13__locus_1_main_sat

CsomeBCy

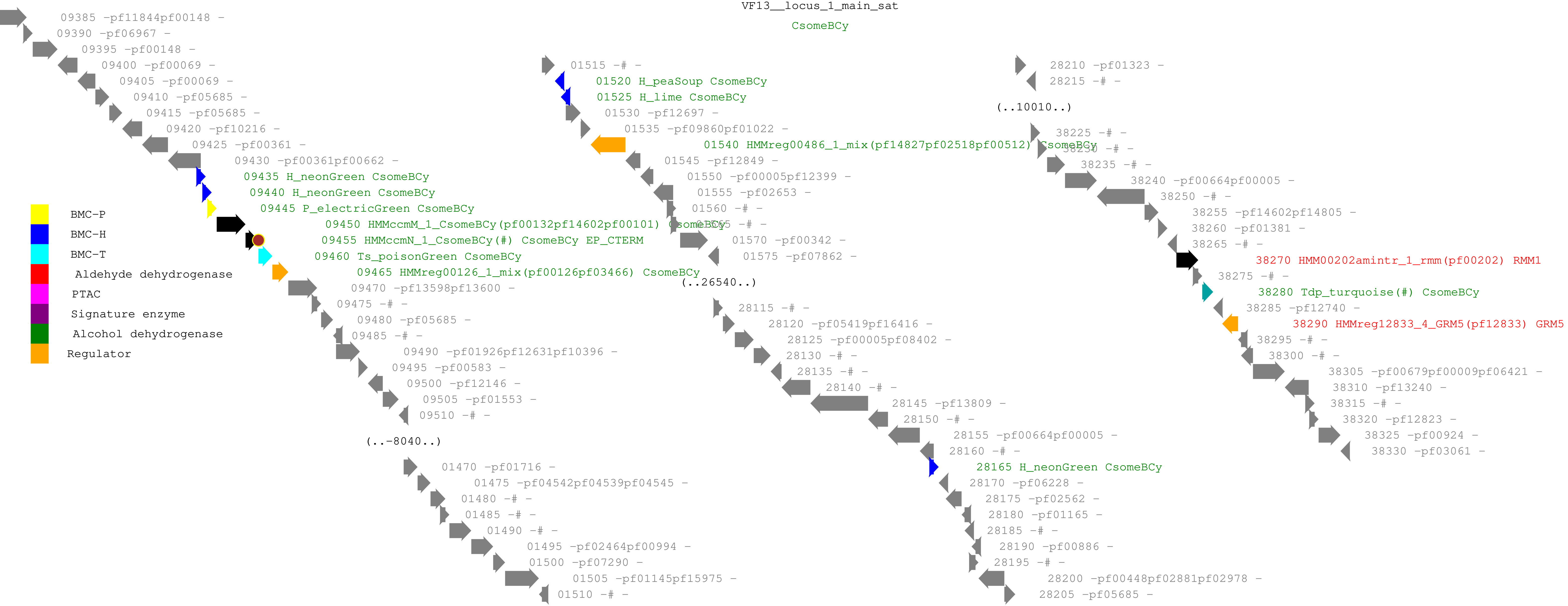

Microcystis panniformis FACHB-1757. Bacteria; Cyanobacteria; Oscillatoriophycideae; Chroococcales;

VL20__locus_1_main_sat

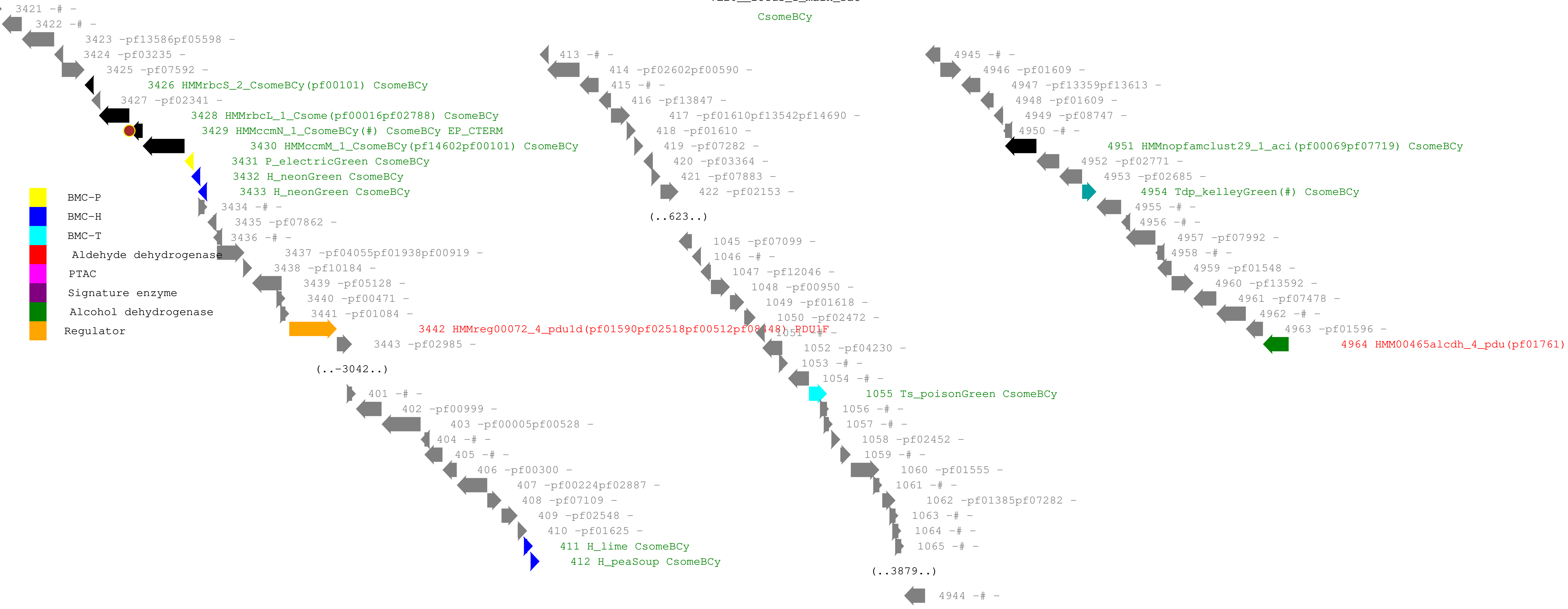

Scytonema hofmannii PCC 7110. Bacteria; Cyanobacteria; Nostocales; Scytonemataceae; Scytonema.

WA1__locus_1_main_sat

CsomeBCy

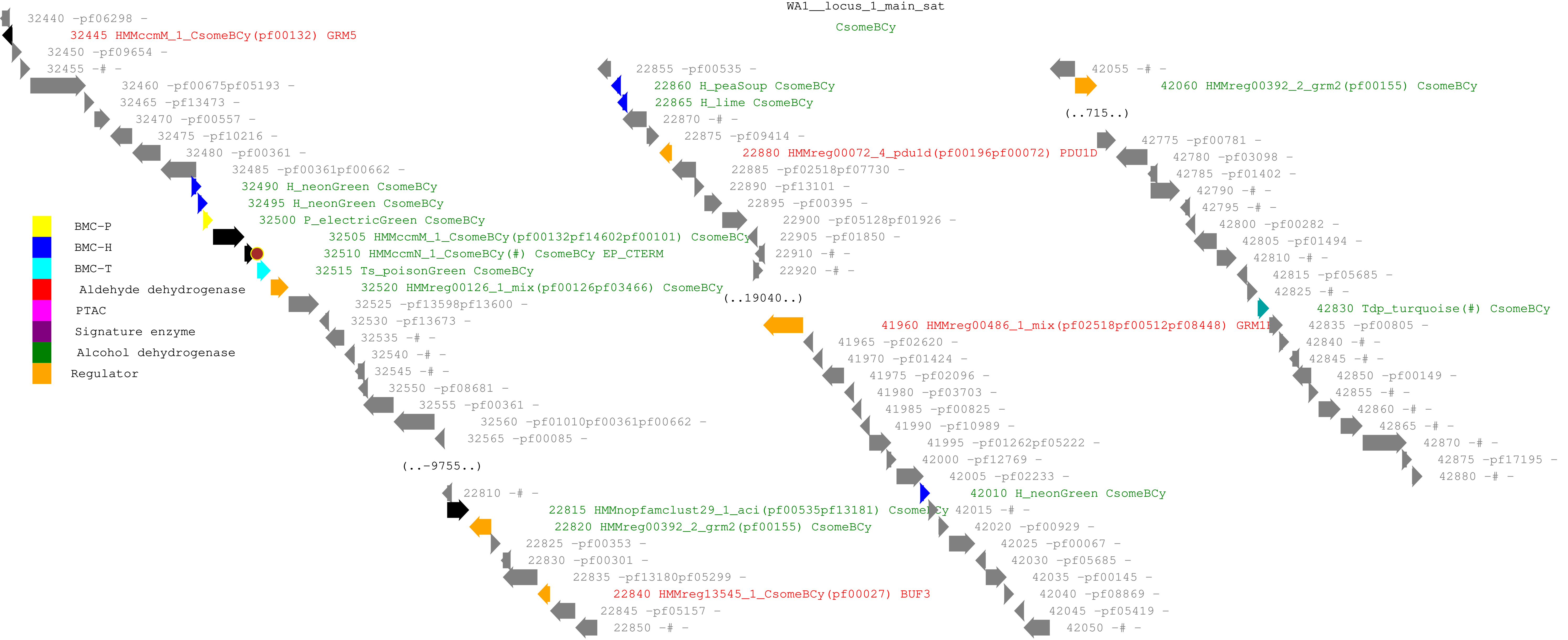

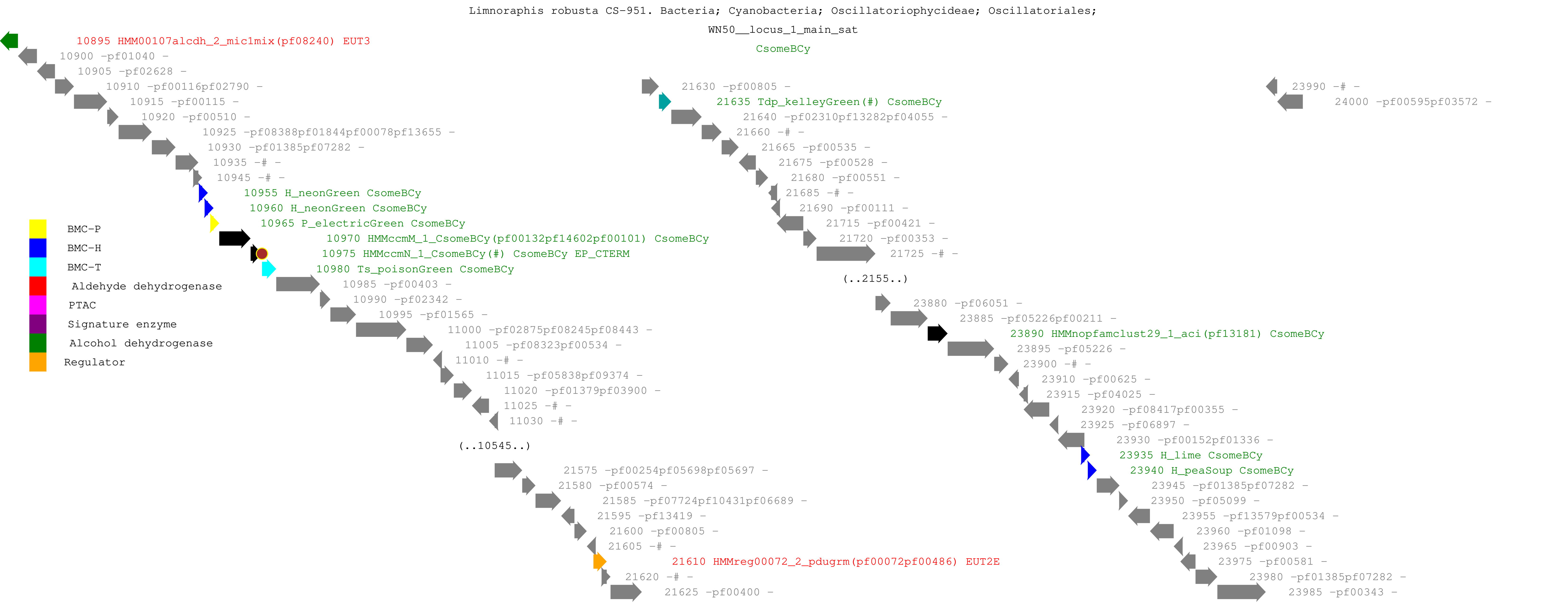
Limnoraphis robusta CS-951. Bacteria; Cyanobacteria; Oscillatoriophycideae; Oscillatoriales;
WN50__locus_1_main_sat
CsomeBCy
10895 HMM00107alcdh_2_mic1mix(pf08240) EUT3
10900 -pf01040 -
10905 -pf02628 -
10910 -pf00116pf02790 -
10915 -pf00115 -
10920 -pf00510 -
10925 -pf08388pf01844pf00078pf13655 -
10930 -pf01385pf07282 -
10935 -# -
10945 -# -
10955 H_neonGreen CsomeBCy
10960 H_neonGreen CsomeBCy
10965 P_electricGreen CsomeBCy
10970 HMMccmM_1_CsomeBCy(pf00132pf14602pf00101) CsomeBCy
10975 HMMccmN_1_CsomeBCy(#) CsomeBCy EP_CTERM
10980 Ts_poisonGreen CsomeBCy
10985 -pf00403 -
10990 -pf02342 -
10995 -pf01565 -
11000 -pf02875pf08245pf08443 -
11005 -pf08323pf00534 -
11010 -# -
11015 -pf05838pf09374 -
11020 -pf01379pf03900 -
11025 -# -
11030 -# -
(..10545..)
21575 -pf00254pf05698pf05697 -
21580 -pf00574 -
21585 -pf07724pf10431pf06689 -
21595 -pf13419 -
21600 -pf00805 -
21605 -# -
21610 HMMreg00072_2_pdugrm(pf00072pf00486) EUT2E
21620 -# -
21625 -pf00400 -
21630 -pf00805 -
21635 Tdp_kelleyGreen(#) CsomeBCy
21640 -pf02310pf13282pf04055 -
21660 -# -
21665 -pf00535 -
21675 -pf00528 -
21680 -pf00551 -
21685 -# -
21690 -pf00111 -
21715 -pf00421 -
21720 -pf00353 -
21725 -# -
(..2155..)
23880 -pf06051 -
23885 -pf05226pf00211 -
23890 HMMnopfamclust29_1_aci(pf13181) CsomeBCy
23895 -pf05226 -
23900 -# -
23910 -pf00625 -
23915 -pf04025 -
23920 -pf08417pf00355 -
23925 -pf06897 -
23930 -pf00152pf01336 -
23935 H_lime CsomeBCy
23940 H_peaSoup CsomeBCy
23945 -pf01385pf07282 -
23950 -pf05099 -
23955 -pf13579pf00534 -
23960 -pf01098 -
23965 -pf00903 -
23975 -pf00581 -
23980 -pf01385pf07282 -
23985 -pf00343 -
23990 -# -
24000 -pf00595pf03572 -
BMC-P
BMC-H
BMC-T
Aldehyde dehydrogenase
PTAC
Signature enzyme
Alcohol dehydrogenase
Regulator

Xenococcus sp. PCC 7305. Bacteria; Cyanobacteria; Pleurocapsales; Xenococcaceae; Xenococcus.

Xen7305DRAFT__locus_1_main_sat

CsomeBCy

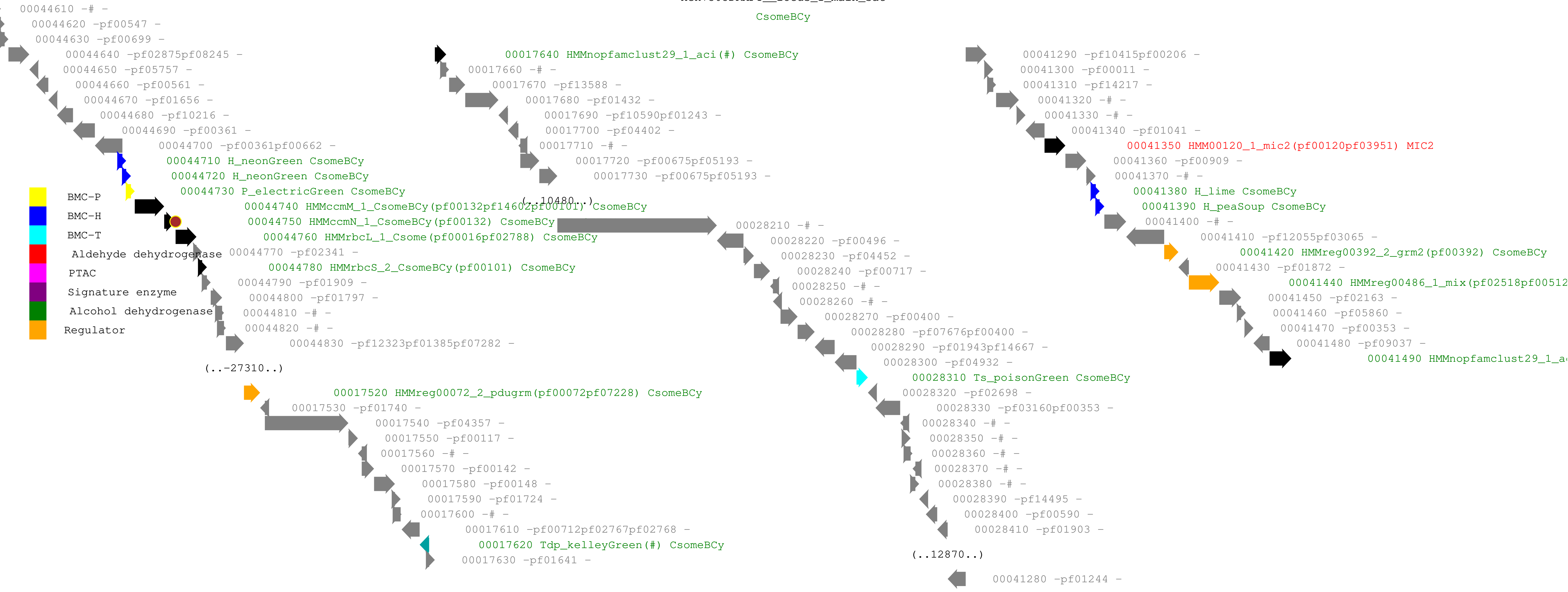

Halomicronema hongdechloris C2206. Bacteria; Cyanobacteria; Synechococcales; Leptolyngbyaceae;

XM38__locus_1_main_sat

CsomeBCy

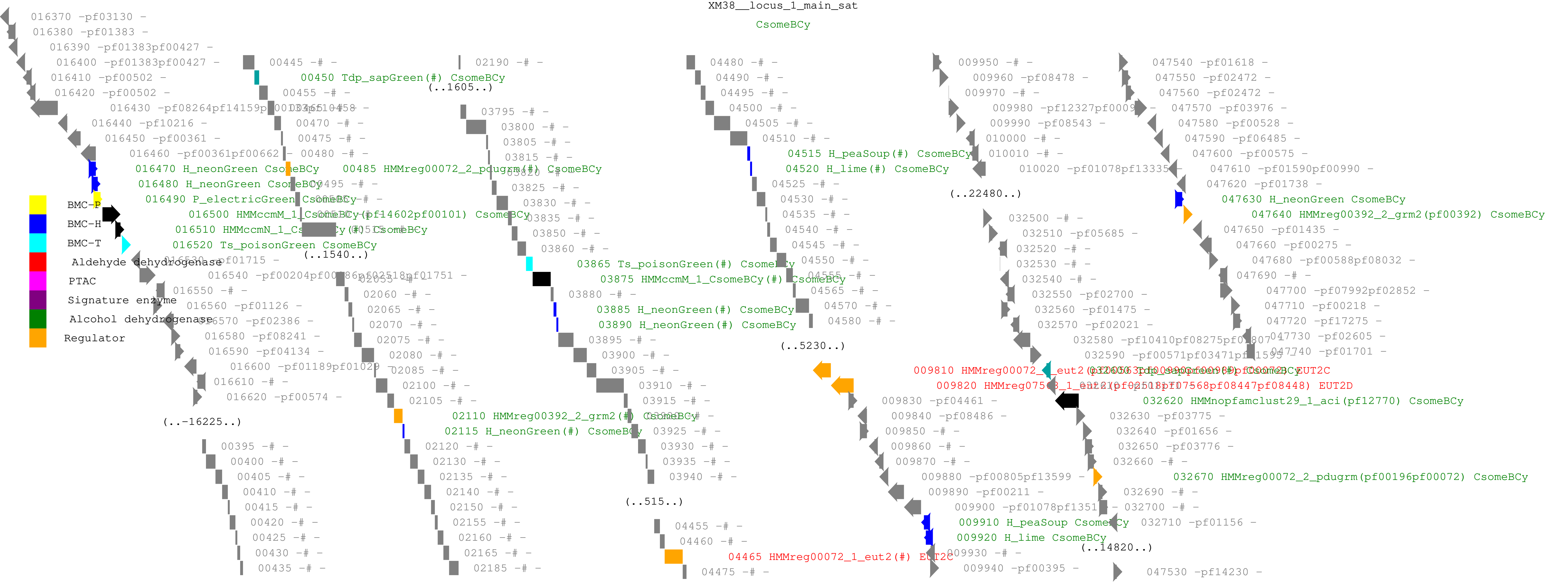

Clostridium kluyveri. Bacteria; Firmicutes; Clostridia; Clostridiales; Clostridiaceae;

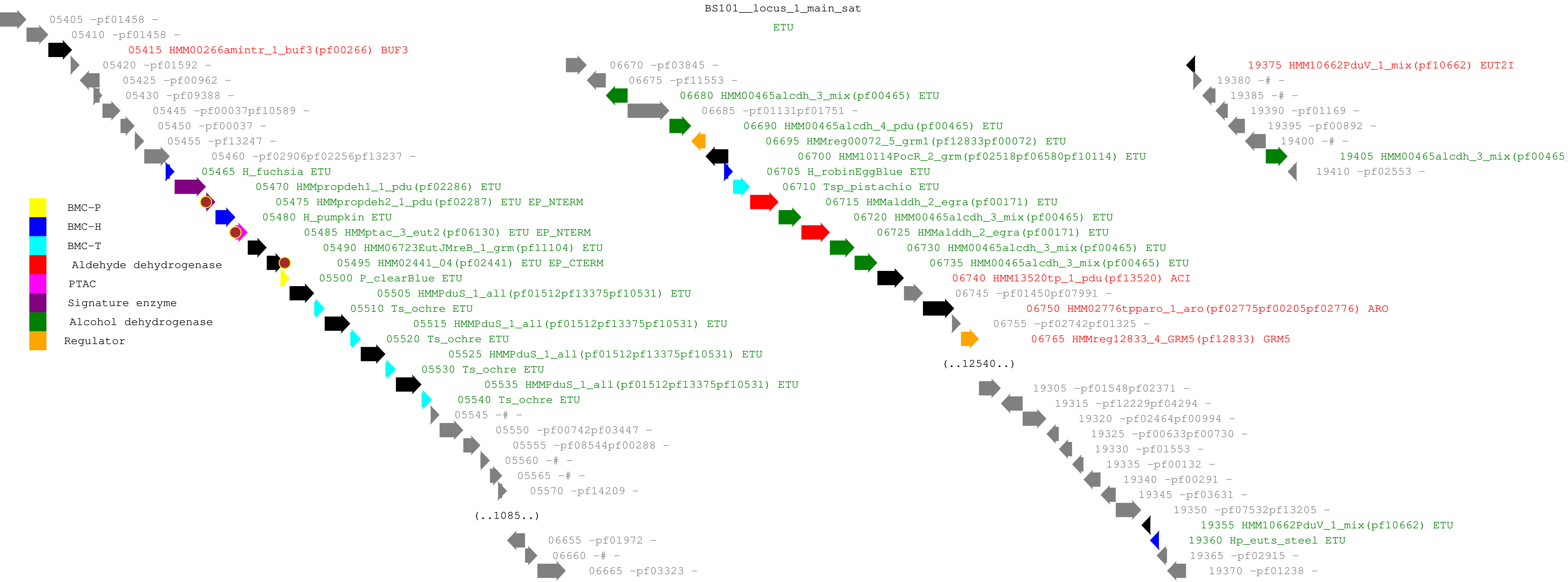

Clostridium kluyveri (strain ATCC 8527 / DSM 555 / NCIMB 10680). Bacteria; Firmicutes; Clostridia; Clostridiales; Clostridiaceae;

CKL__locus_1_main_sat

ETU

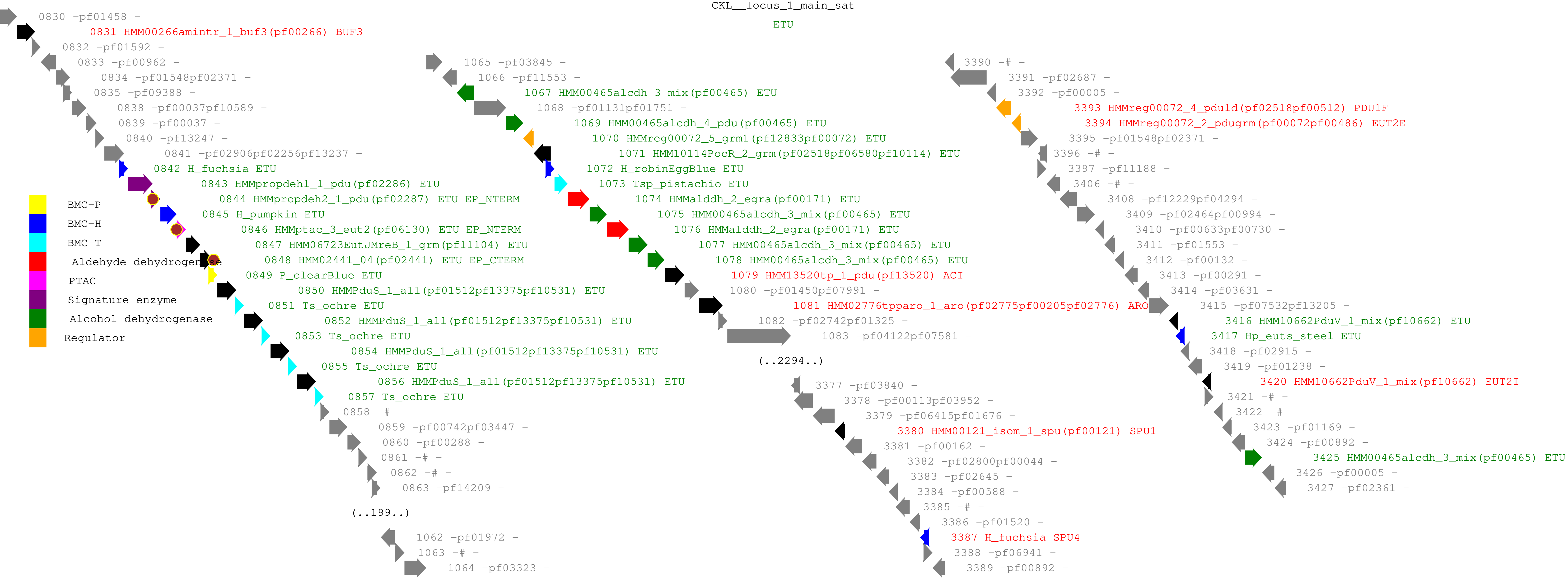

Clostridium kluyveri (strain NBRC 12016). Bacteria; Firmicutes; Clostridia; Clostridiales; Clostridiaceae;

CKR__locus_1_main_sat

ETU

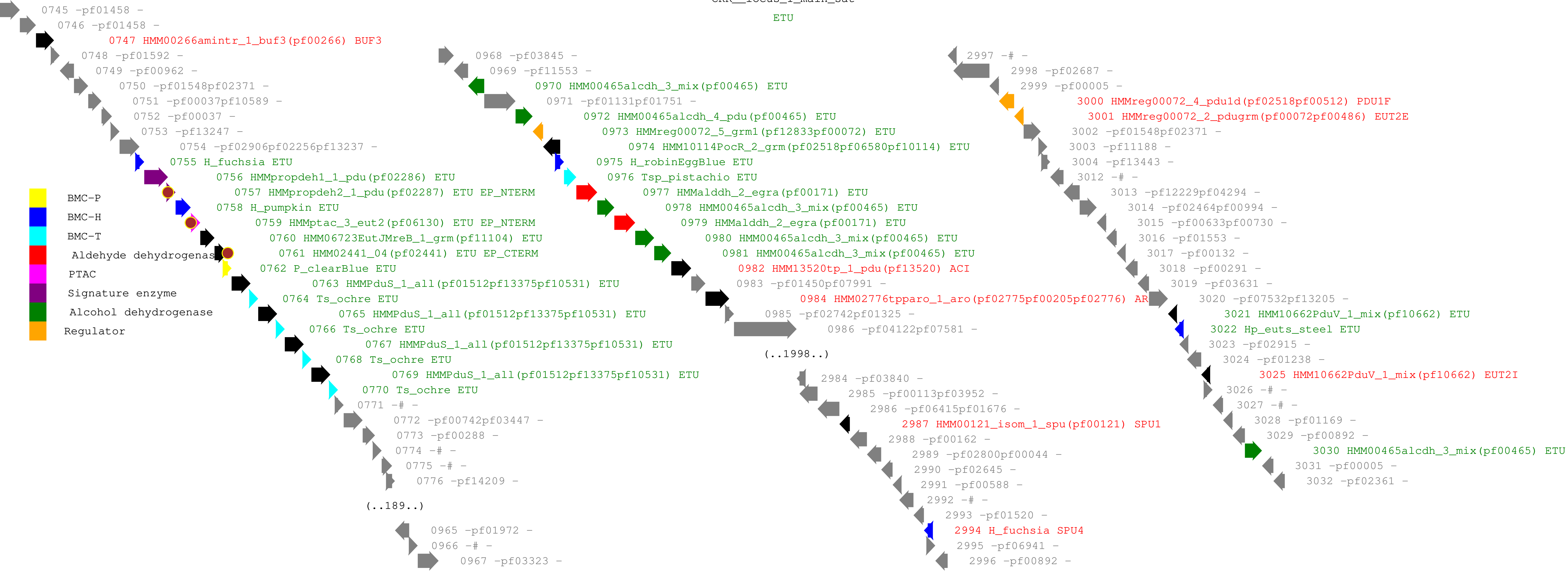

Shigella sp. PAMC 28760. Bacteria; Proteobacteria; Gammaproteobacteria; Enterobacterales;

A0259__locus_1

EUT1

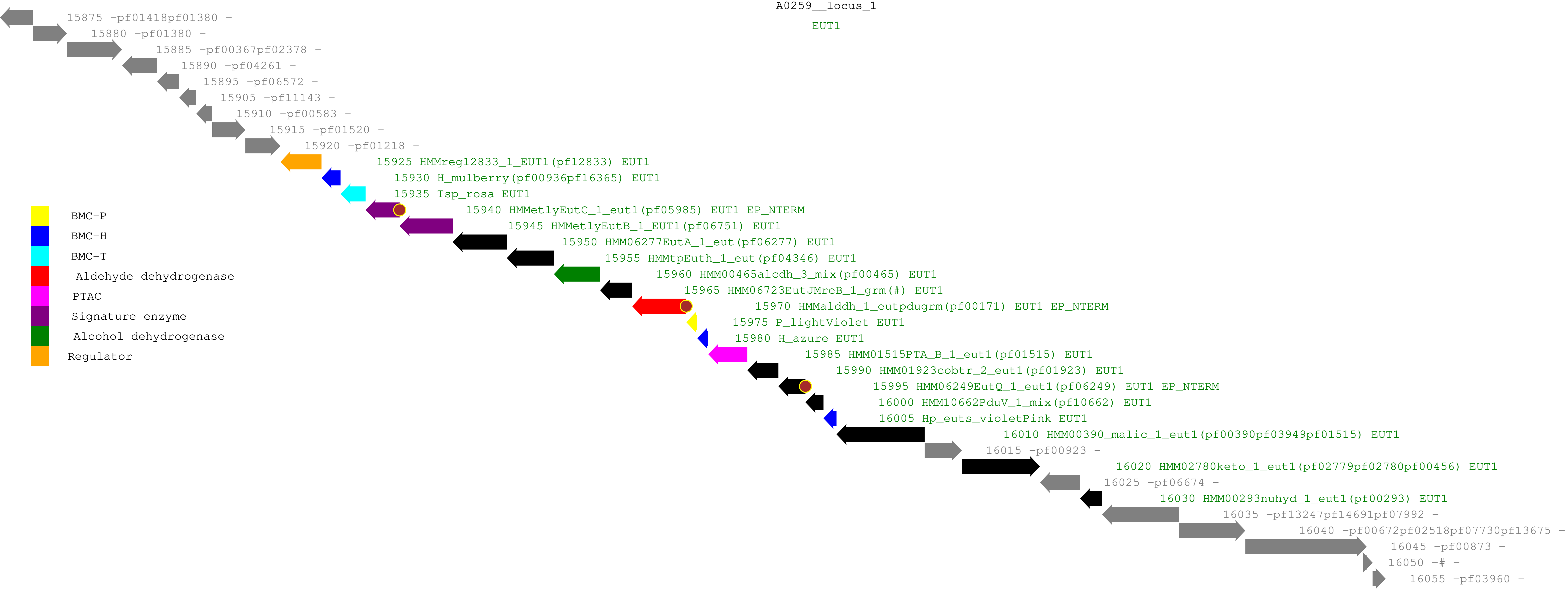

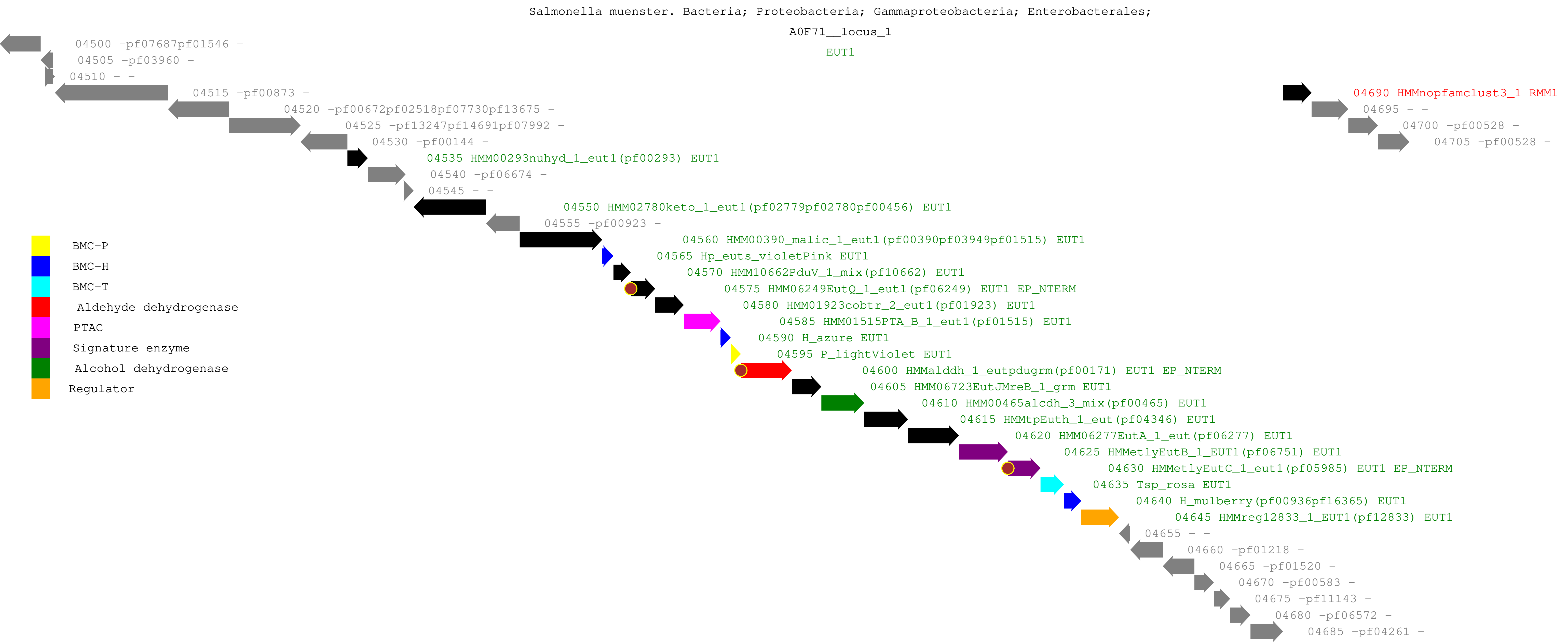

Salmonella muenster. Bacteria; Proteobacteria; Gammaproteobacteria; Enterobacterales;
A0F71__locus_1
EUT1
04500 -pf07687pf01546 -
04505 -pf03960 -
04510 - -
04515 -pf00873 -
04520 -pf00672pf02518pf07730pf13675 -
04525 -pf13247pf14691pf07992 -
04530 -pf00144 -
04535 HMM00293nuhyd_1_eut1(pf00293) EUT1
04540 -pf06674 -
04545 - -
04550 HMM02780keto_1_eut1(pf02779pf02780pf00456) EUT1
04555 -pf00923 -
04560 HMM00390_malic_1_eut1(pf00390pf03949pf01515) EUT1
04565 Hp_euts_violetPink EUT1
04570 HMM10662PduV_1_mix(pf10662) EUT1
04575 HMM06249EutQ_1_eut1(pf06249) EUT1 EP_NTERM
04580 HMM01923cobtr_2_eut1(pf01923) EUT1
04585 HMM01515PTA_B_1_eut1(pf01515) EUT1
04590 H_azure EUT1
04595 P_lightViolet EUT1
04600 HMMalddh_1_eutpdugrm(pf00171) EUT1 EP_NTERM
04605 HMM06723EutJMreB_1_grm EUT1
04610 HMM00465alcdh_3_mix(pf00465) EUT1
04615 HMMtpEuth_1_eut(pf04346) EUT1
04620 HMM06277EutA_1_eut(pf06277) EUT1
04625 HMMetlyEutB_1_EUT1(pf06751) EUT1
04630 HMMetlyEutC_1_eut1(pf05985) EUT1 EP_NTERM
04635 Tsp_rosa EUT1
04640 H_mulberry(pf00936pf16365) EUT1
04645 HMMreg12833_1_EUT1(pf12833) EUT1
04655 - -
04660 -pf01218 -
04665 -pf01520 -
04670 -pf00583 -
04675 -pf11143 -
04680 -pf06572 -
04685 -pf04261 -
04690 HMMnopfamclust3_1 RMM1
04695 - -
04700 -pf00528 -
04705 -pf00528 -
BMC-P
BMC-H
BMC-T
Aldehyde dehydrogenase
PTAC
Signature enzyme
Alcohol dehydrogenase
Regulator

Escherichia coli KTE182. Bacteria; Proteobacteria; Gammaproteobacteria; Enterobacterales;

A13A__locus_1

EUT1

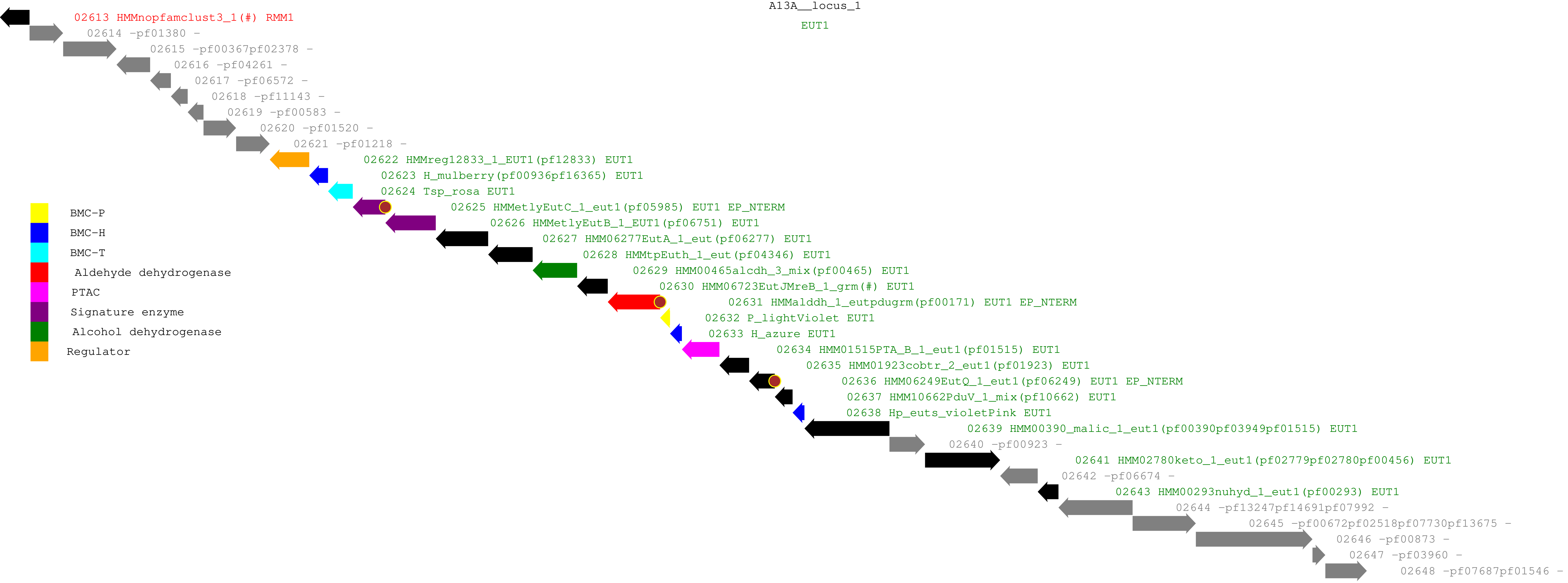

Escherichia coli KTE193. Bacteria; Proteobacteria; Gammaproteobacteria; Enterobacterales;

A13W__locus_1

EUT1

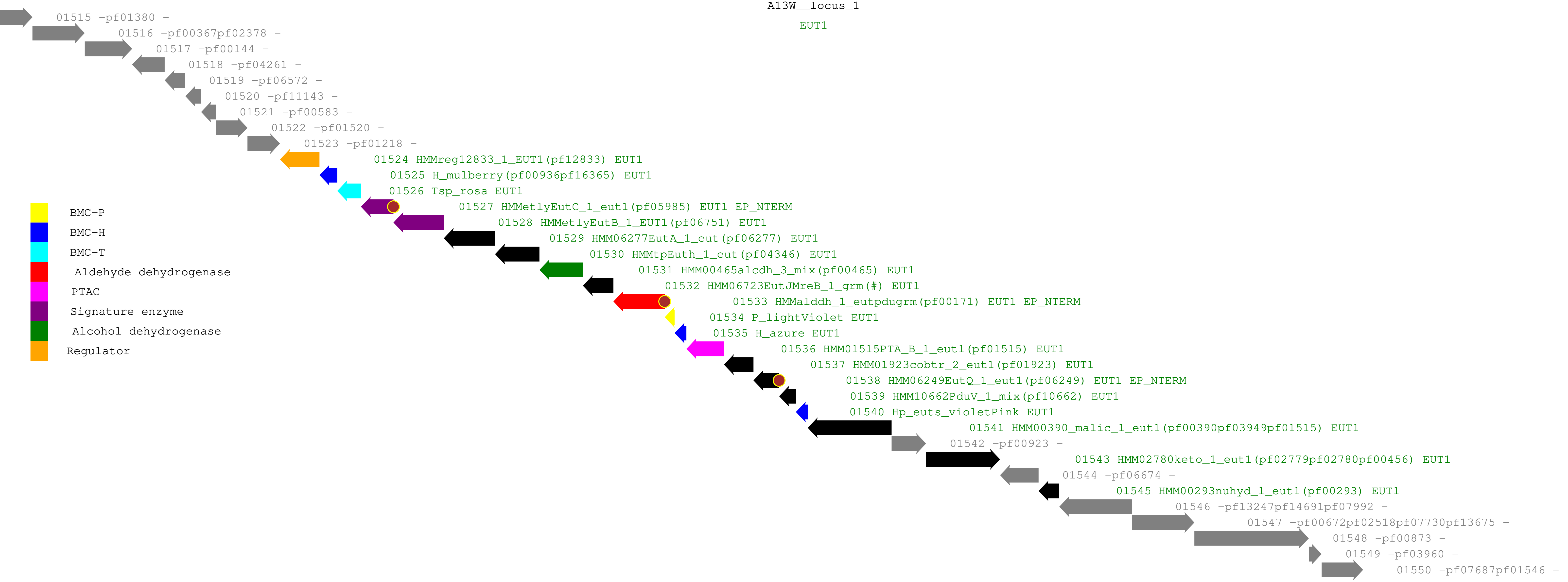

Escherichia sp. KTE52. Bacteria; Proteobacteria; Gammaproteobacteria; Enterobacterales;

A1SC__locus_2

EUT1

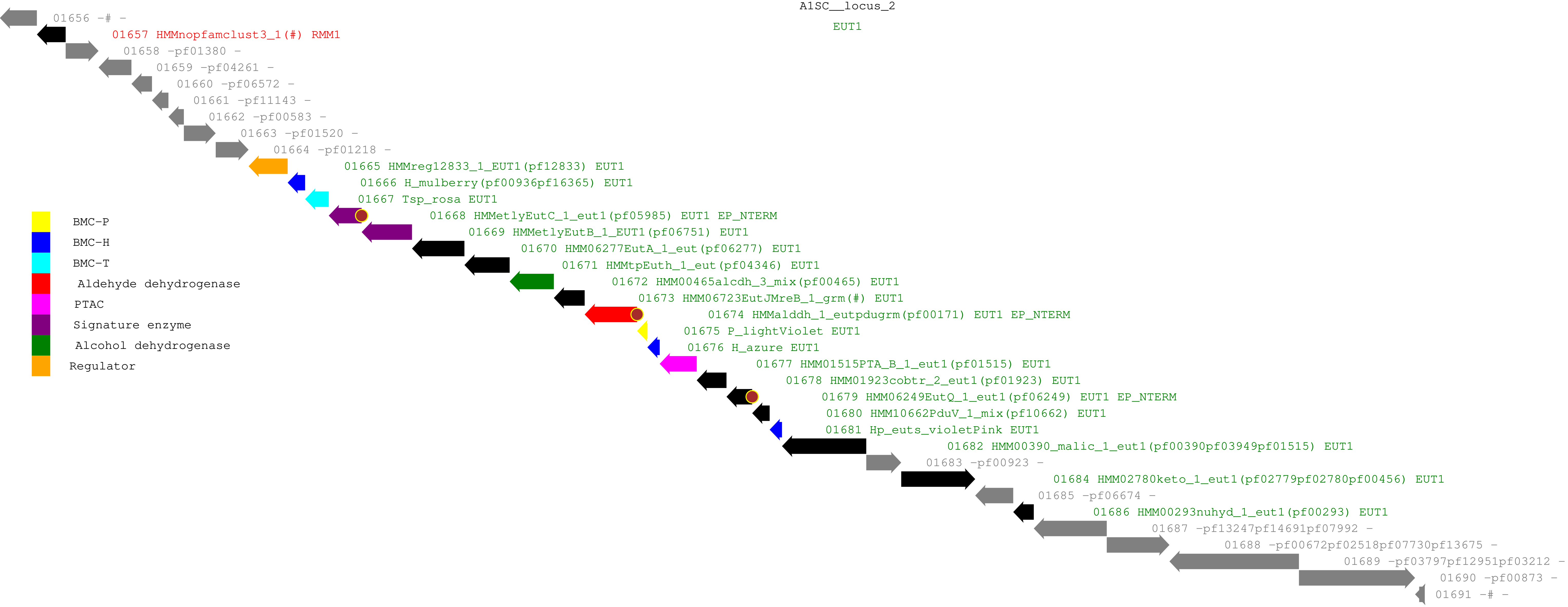

Escherichia coli KTE64. Bacteria; Proteobacteria; Gammaproteobacteria; Enterobacterales;

A1U1__locus_1

EUT1

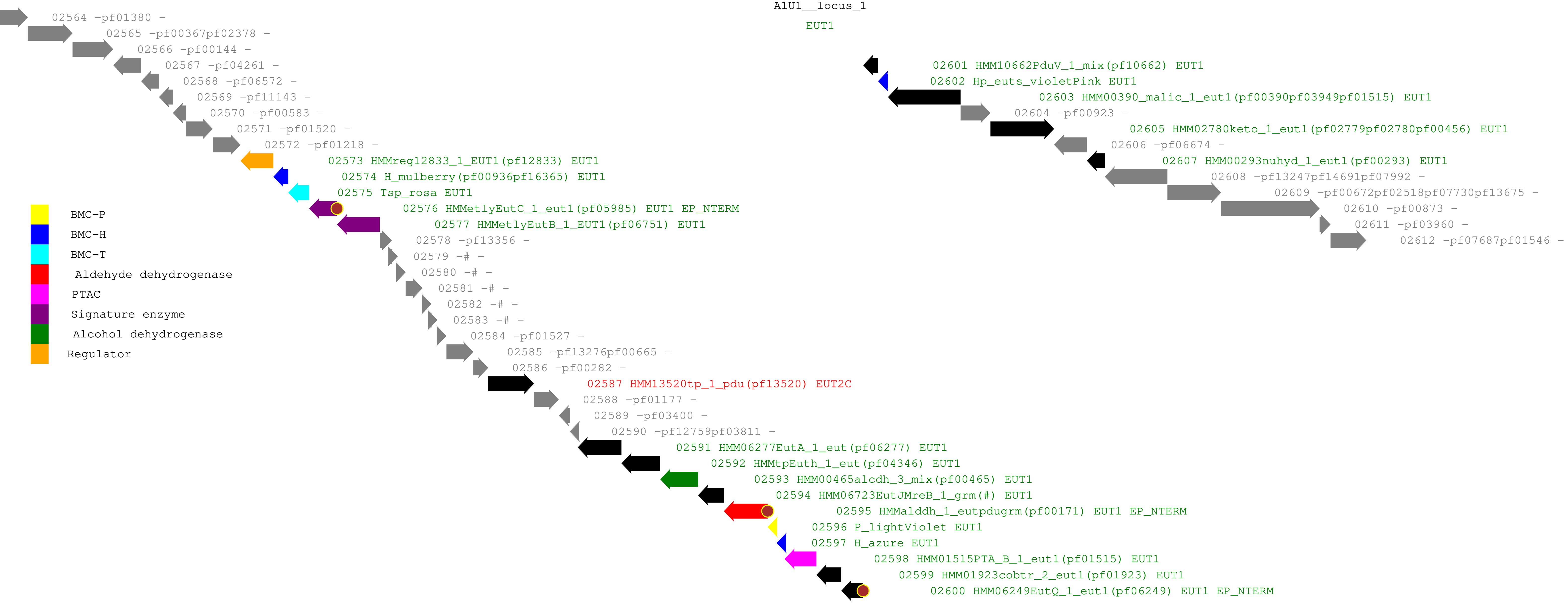

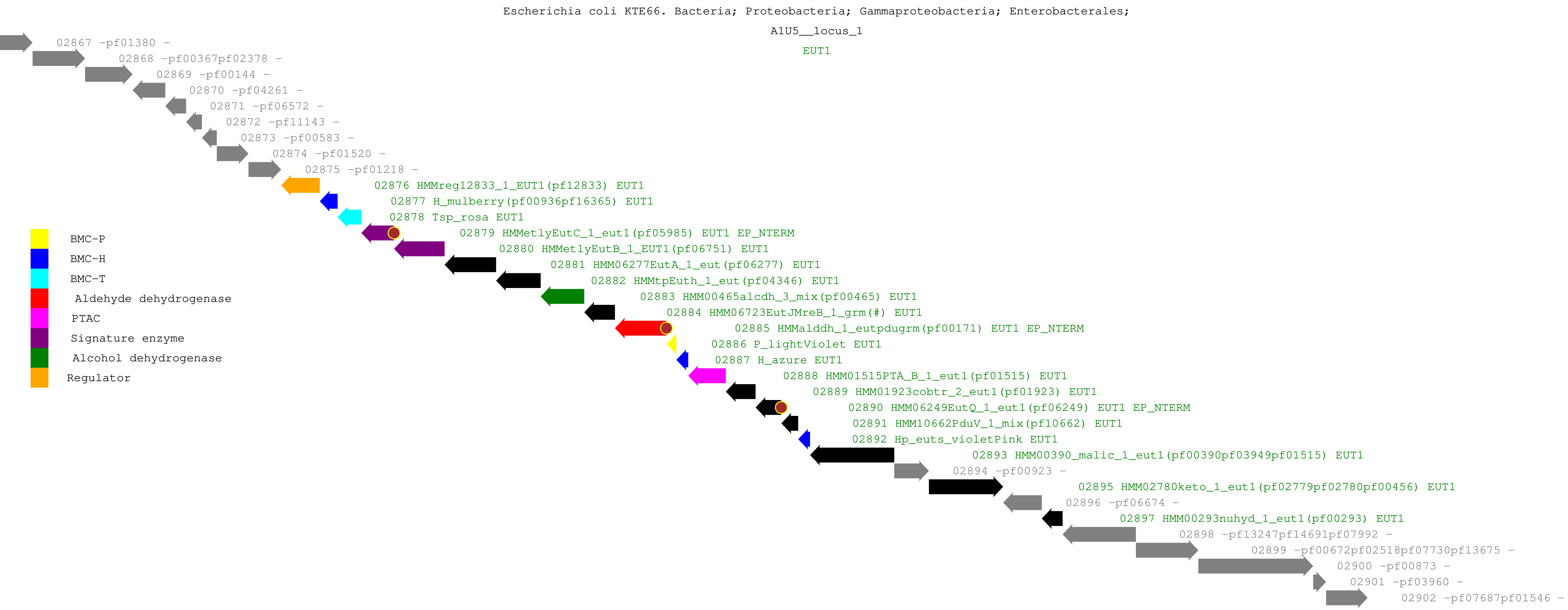
Escherichia coli KTE66. Bacteria; Proteobacteria; Gammaproteobacteria; Enterobacterales;
A1U5__locus_1
EUT1
02867 -pf01380 -
02868 -pf00367pf02378 -
02869 -pf00144 -
02870 -pf04261 -
02871 -pf06572 -
02872 -pf11143 -
02873 -pf00583 -
02874 -pf01520 -
02875 -pf01218 -
02876 HMMreg12833_1_EUT1(pf12833) EUT1
02877 H_mulberry(pf00936pf16365) EUT1
02878 Tsp_rosa EUT1
02879 HMMetlyEutC_1_eut1(pf05985) EUT1 EP_NTERM
02880 HMMetlyEutB_1_EUT1(pf06751) EUT1
02881 HMM06277EutA_1_eut(pf06277) EUT1
02882 HMMtpEuth_1_eut(pf04346) EUT1
02883 HMM00465alcdh_3_mix(pf00465) EUT1
02884 HMM06723EutJMreB_1_grm(#) EUT1
02885 HMMalddh_1_eutpdugrm(pf00171) EUT1 EP_NTERM
02886 P_lightViolet EUT1
02887 H_azure EUT1
02888 HMM01515PTA_B_1_eut1(pf01515) EUT1
02889 HMM01923cobtr_2_eut1(pf01923) EUT1
02890 HMM06249EutQ_1_eut1(pf06249) EUT1 EP_NTERM
02891 HMM10662PduV_1_mix(pf10662) EUT1
02892 Hp_euts_violetPink EUT1
02893 HMM00390_malic_1_eut1(pf00390pf03949pf01515) EUT1
02894 -pf00923 -
02895 HMM02780keto_1_eut1(pf02779pf02780pf00456) EUT1
02896 -pf06674 -
02897 HMM00293nuhyd_1_eut1(pf00293) EUT1
02898 -pf13247pf14691pf07992 -
02899 -pf00672pf02518pf07730pf13675 -
02900 -pf00873 -
02901 -pf03960 -
02902 -pf07687pf01546 -
BMC-P
BMC-H
BMC-T
Aldehyde dehydrogenase
PTAC
Signature enzyme
Alcohol dehydrogenase
Regulator

Escherichia coli KTE73. Bacteria; Proteobacteria; Gammaproteobacteria; Enterobacterales;

A1UI__locus_1

EUT1

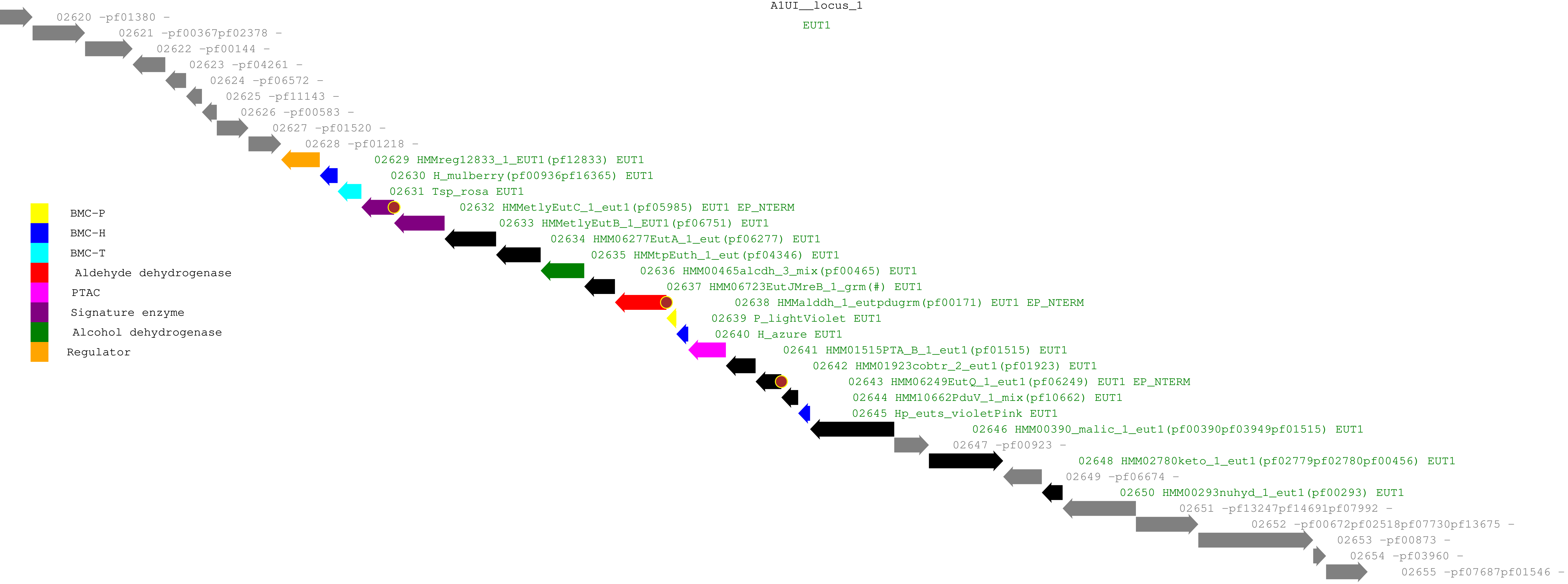

Escherichia coli KTE75. Bacteria; Proteobacteria; Gammaproteobacteria; Enterobacterales;

A1UM__locus_2

EUT1

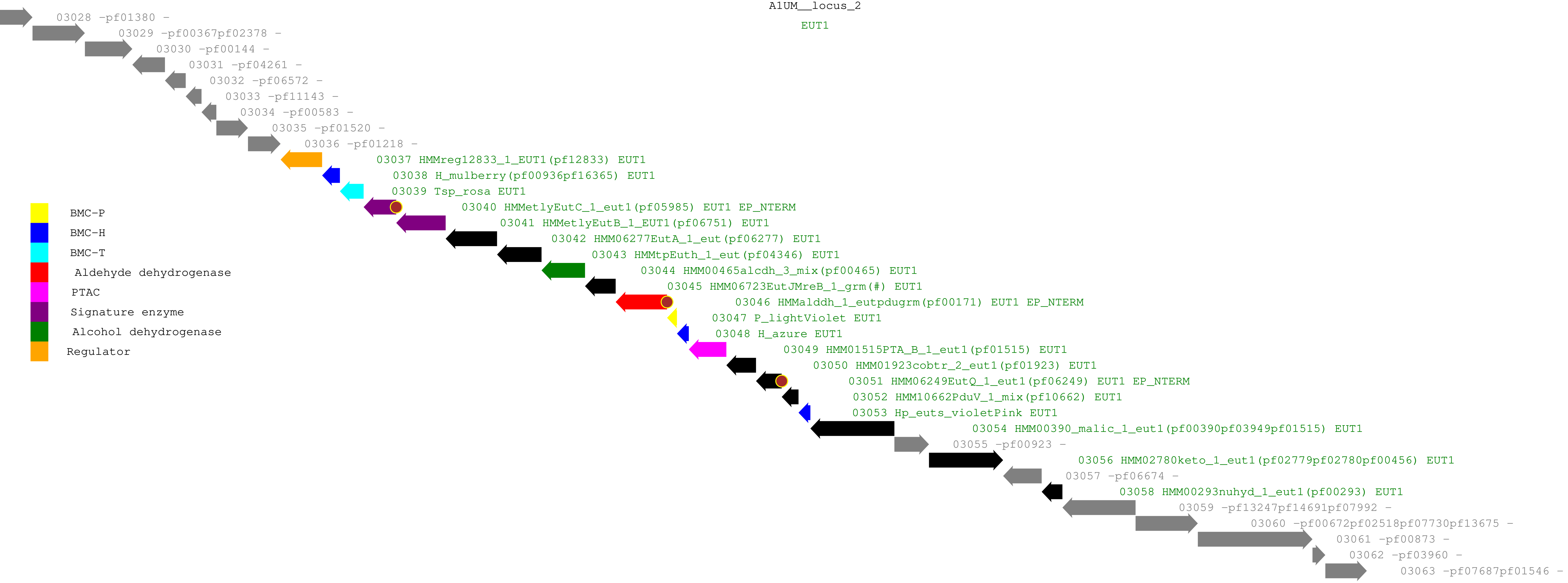

Escherichia coli KTE76. Bacteria; Proteobacteria; Gammaproteobacteria; Enterobacterales;

A1UO__locus_1

EUT1

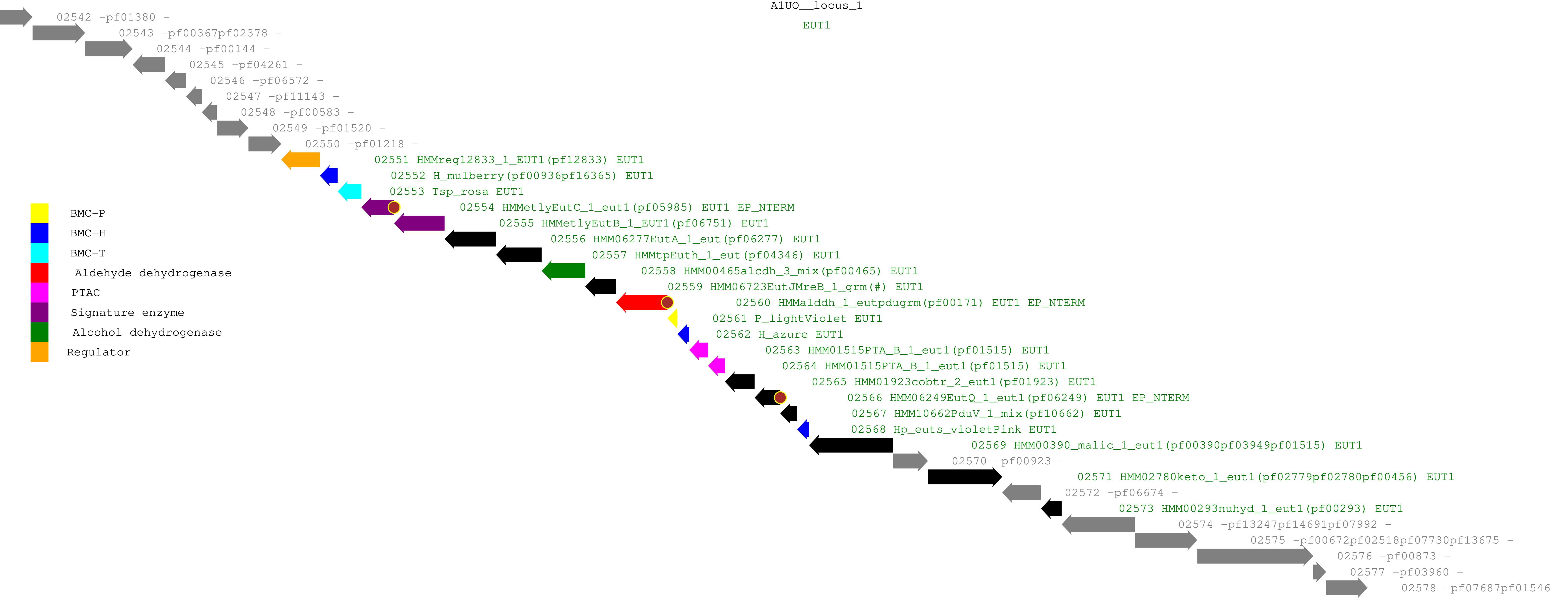

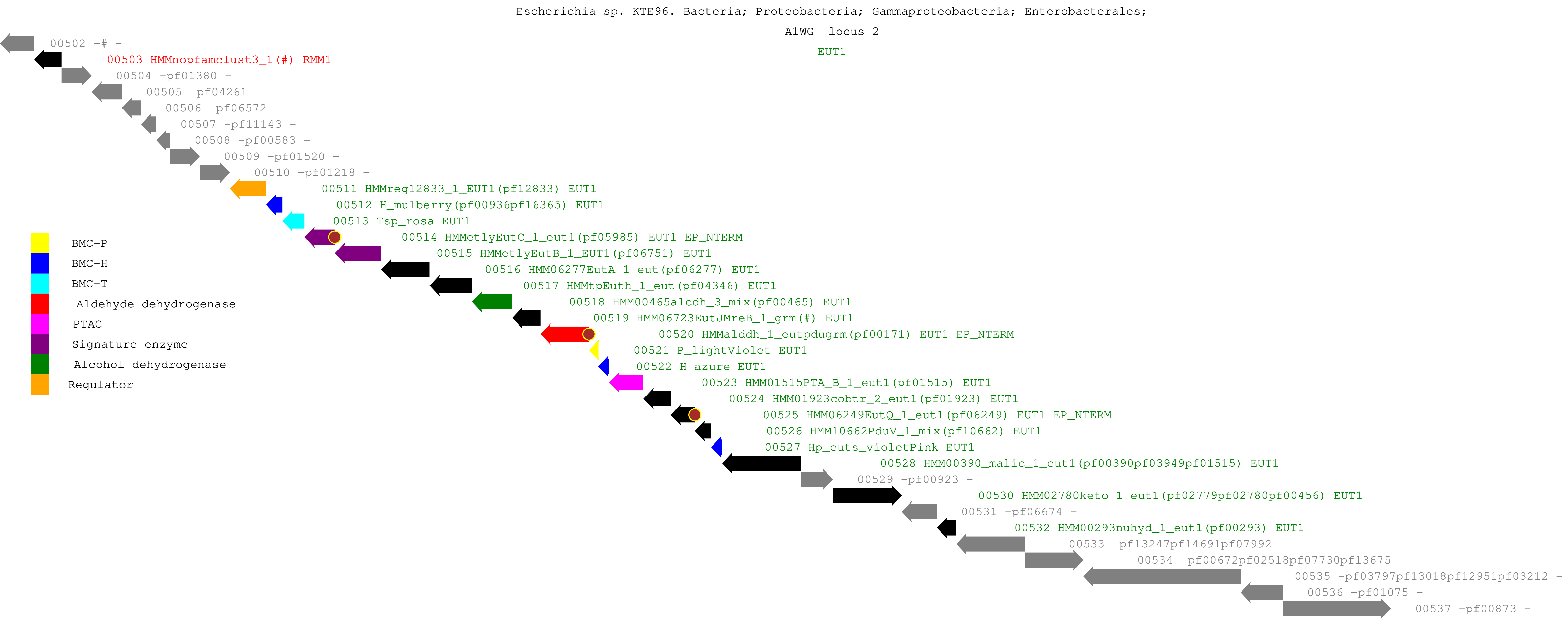
Escherichia sp. KTE96. Bacteria; Proteobacteria; Gammaproteobacteria; Enterobacterales;
A1WG__locus_2
EUT1
00502 -# -
00503 HMMnopfamclust3_1(#) RMM1
00504 -pf01380 -
00505 -pf04261 -
00506 -pf06572 -
00507 -pf11143 -
00508 -pf00583 -
00509 -pf01520 -
00510 -pf01218 -
00511 HMMreg12833_1_EUT1(pf12833) EUT1
00512 H_mulberry(pf00936pf16365) EUT1
00513 Tsp_rosa EUT1
00514 HMMetlyEutC_1_eut1(pf05985) EUT1 EP_NTERM
00515 HMMetlyEutB_1_EUT1(pf06751) EUT1
00516 HMM06277EutA_1_eut(pf06277) EUT1
00517 HMMtpEuth_1_eut(pf04346) EUT1
00518 HMM00465alcdh_3_mix(pf00465) EUT1
00519 HMM06723EutJMreB_1_grm(#) EUT1
00520 HMMalddh_1_eutpdugrm(pf00171) EUT1 EP_NTERM
00521 P_lightViolet EUT1
00522 H_azure EUT1
00523 HMM01515PTA_B_1_eut1(pf01515) EUT1
00524 HMM01923cobtr_2_eut1(pf01923) EUT1
00525 HMM06249EutQ_1_eut1(pf06249) EUT1 EP_NTERM
00526 HMM10662PduV_1_mix(pf10662) EUT1
00527 Hp_euts_violetPink EUT1
00528 HMM00390_malic_1_eut1(pf00390pf03949pf01515) EUT1
00529 -pf00923 -
00530 HMM02780keto_1_eut1(pf02779pf02780pf00456) EUT1
00531 -pf06674 -
00532 HMM00293nuhyd_1_eut1(pf00293) EUT1
00533 -pf13247pf14691pf07992 -
00534 -pf00672pf02518pf07730pf13675 -
00535 -pf03797pf13018pf12951pf03212 -
00536 -pf01075 -
00537 -pf00873 -
BMC-P
BMC-H
BMC-T
Aldehyde dehydrogenase
PTAC
Signature enzyme
Alcohol dehydrogenase
Regulator

Escherichia coli KTE100. Bacteria; Proteobacteria; Gammaproteobacteria; Enterobacterales;

A1WK__locus_1

EUT1

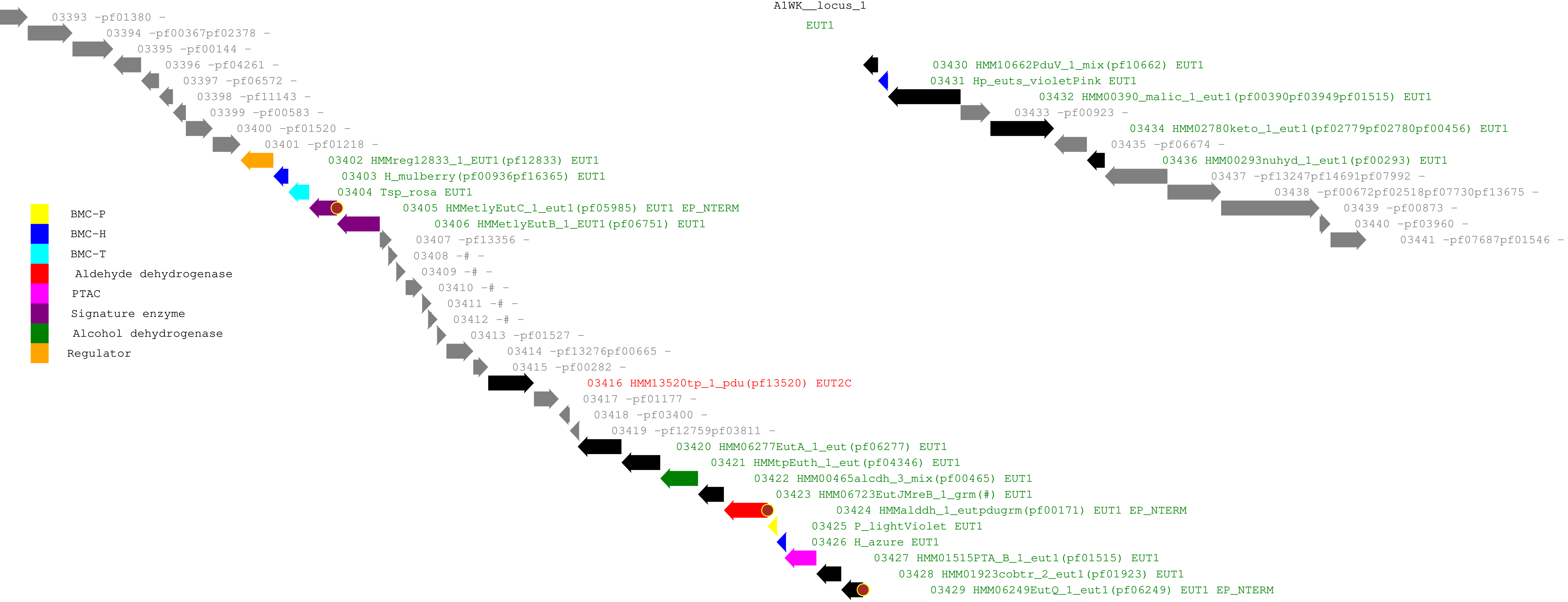

Escherichia coli KTE107. Bacteria; Proteobacteria; Gammaproteobacteria; Enterobacterales;

A1WS__locus_1

EUT1

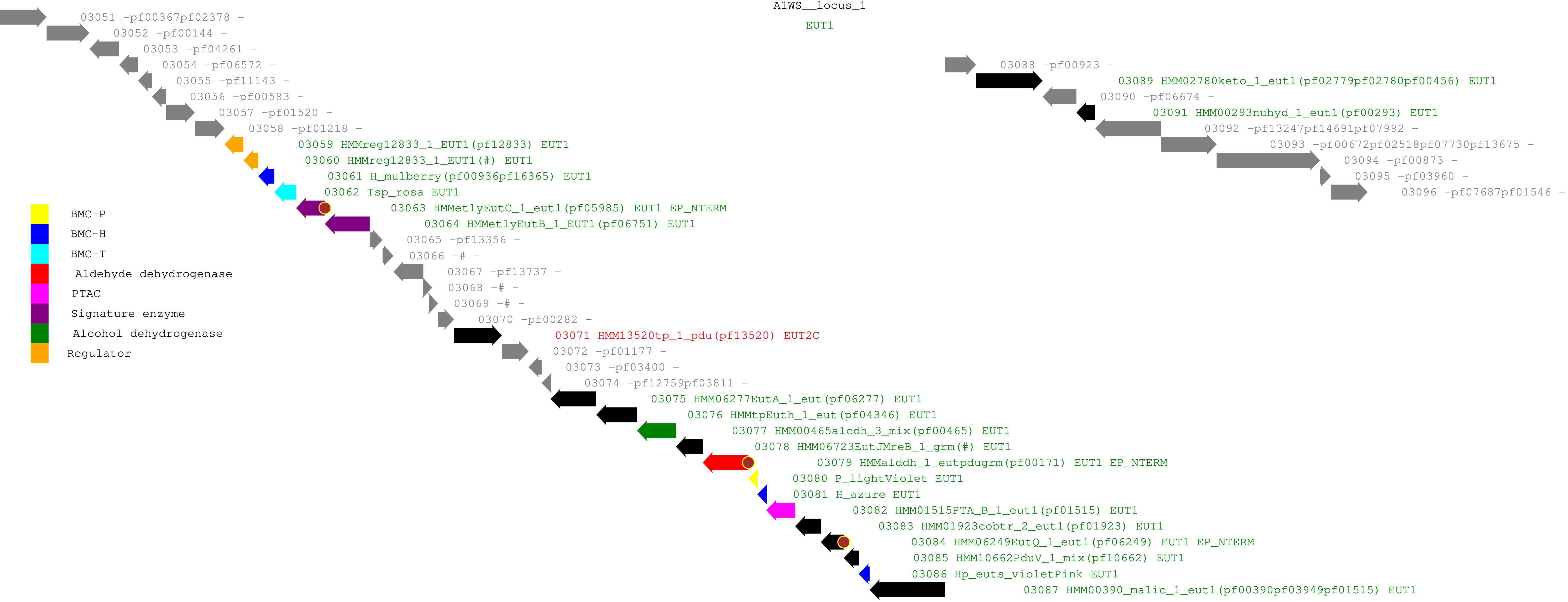

Escherichia coli KTE108. Bacteria; Proteobacteria; Gammaproteobacteria; Enterobacterales;

A1WU__locus_1

EUT1

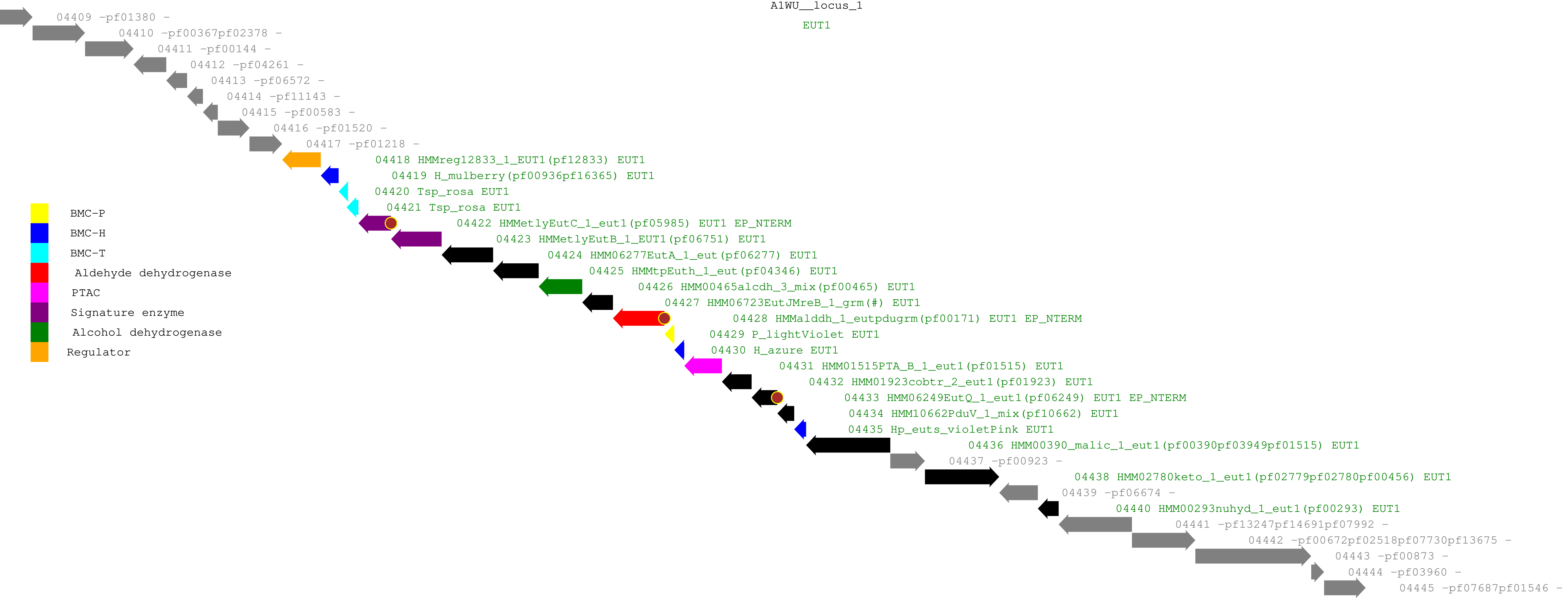

Klebsiella michiganensis E718. Bacteria; Proteobacteria; Gammaproteobacteria; Enterobacterales;

A225__locus_2

EUT1

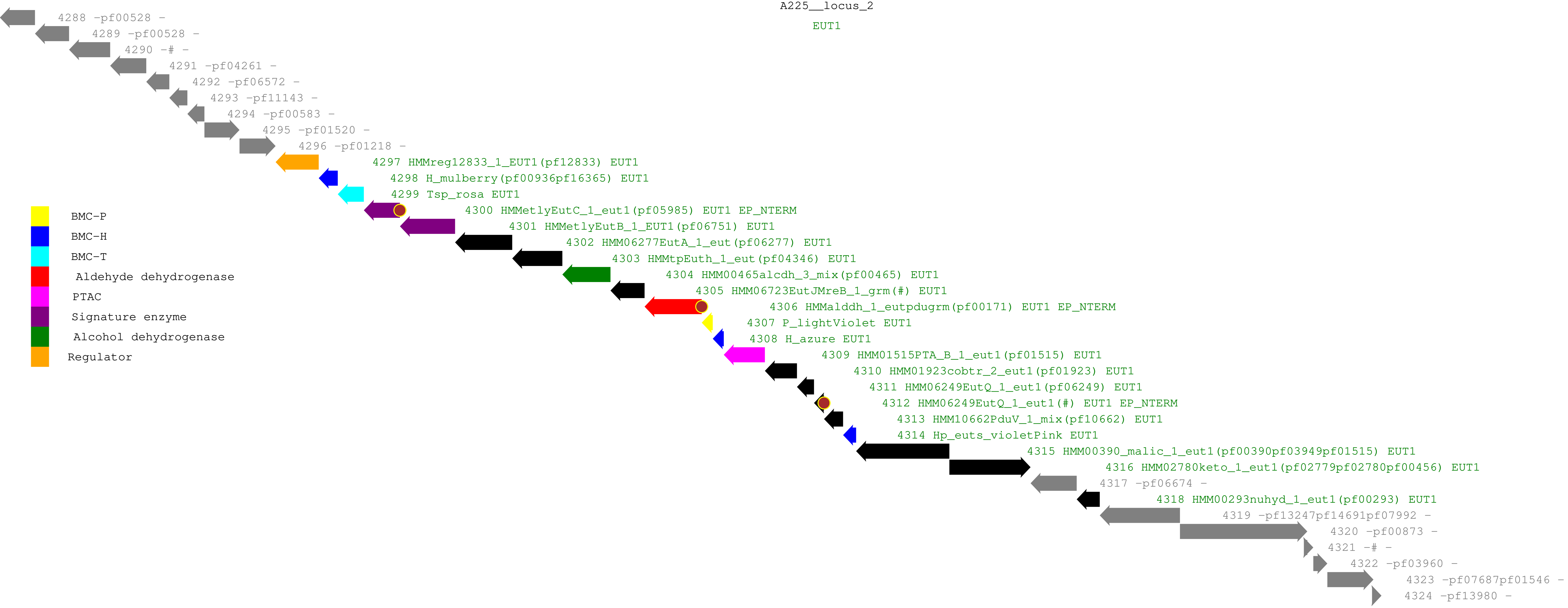

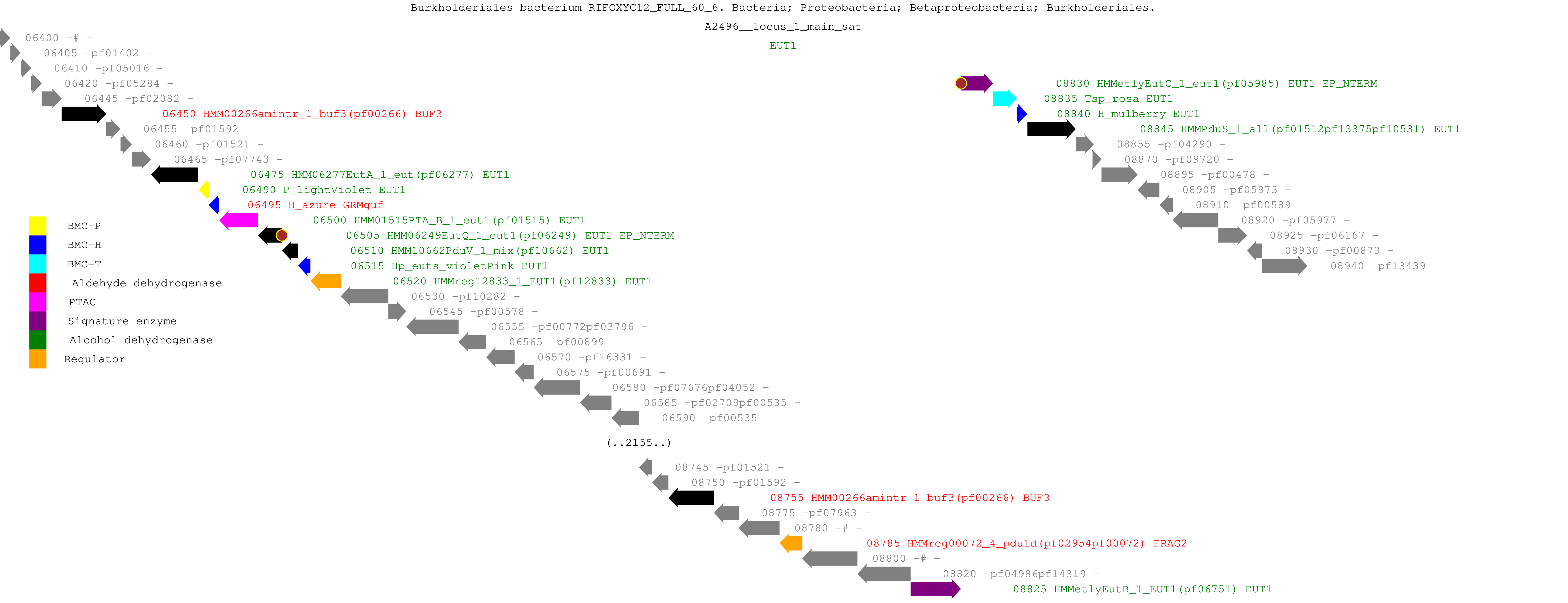
Burkholderiales bacterium RIFOXYC12_FULL_60_6. Bacteria; Proteobacteria; Betaproteobacteria; Burkholderiales.
A2496__locus_1_main_sat
EUT1
06400 -# -
06405 -pf01402 -
06410 -pf05016 -
06420 -pf05284 -
06445 -pf02082 -
06450 HMM00266amintr_1_buf3(pf00266) BUF3
06455 -pf01592 -
06460 -pf01521 -
06465 -pf07743 -
06475 HMM06277EutA_1_eut(pf06277) EUT1
06490 P_lightViolet EUT1
06495 H_azure GRMguf
06500 HMM01515PTA_B_1_eut1(pf01515) EUT1
06505 HMM06249EutQ_1_eut1(pf06249) EUT1 EP_NTERM
06510 HMM10662PduV_1_mix(pf10662) EUT1
06515 Hp_euts_violetPink EUT1
06520 HMMreg12833_1_EUT1(pf12833) EUT1
06530 -pf10282 -
06545 -pf00578 -
06555 -pf00772pf03796 -
06565 -pf00899 -
06570 -pf16331 -
06575 -pf00691 -
06580 -pf07676pf04052 -
06585 -pf02709pf00535 -
06590 -pf00535 -
(..2155..)
08745 -pf01521 -
08750 -pf01592 -
08755 HMM00266amintr_1_buf3(pf00266) BUF3
08775 -pf07963 -
08780 -# -
08785 HMMreg00072_4_pdu1d(pf02954pf00072) FRAG2
08800 -# -
08820 -pf04986pf14319 -
08825 HMMetlyEutB_1_EUT1(pf06751) EUT1
08830 HMMetlyEutC_1_eut1(pf05985) EUT1 EP_NTERM
08835 Tsp_rosa EUT1
08840 H_mulberry EUT1
08845 HMMPduS_1_all(pf01512pf13375pf10531) EUT1
08855 -pf04290 -
08870 -pf09720 -
08895 -pf00478 -
08905 -pf05973 -
08910 -pf00589 -
08920 -pf05977 -
08925 -pf06167 -
08930 -pf00873 -
08940 -pf13439 -
BMC-P
BMC-H
BMC-T
Aldehyde dehydrogenase
PTAC
Signature enzyme
Alcohol dehydrogenase
Regulator

Burkholderiales bacterium RIFOXYD12_FULL_59_19. Bacteria; Proteobacteria; Betaproteobacteria; Burkholderiales.

A2503__locus_1

EUT1

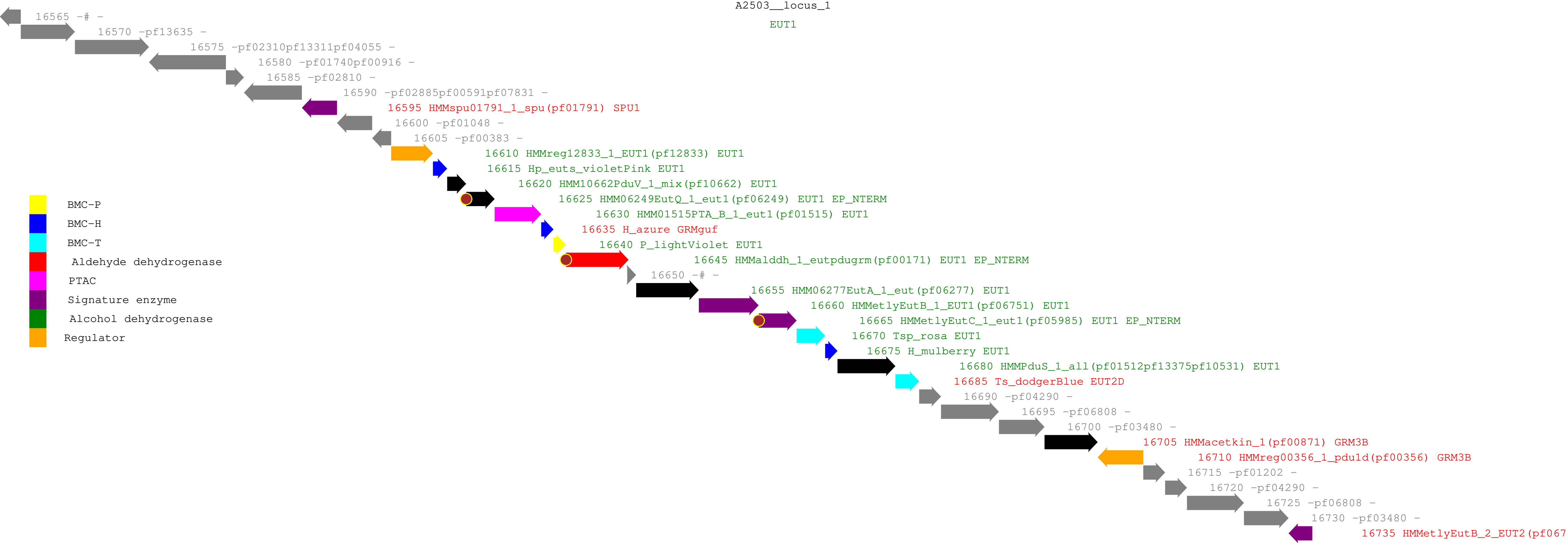

Escherichia coli KTE146. Bacteria; Proteobacteria; Gammaproteobacteria; Enterobacterales;

A311__locus_2

EUT1

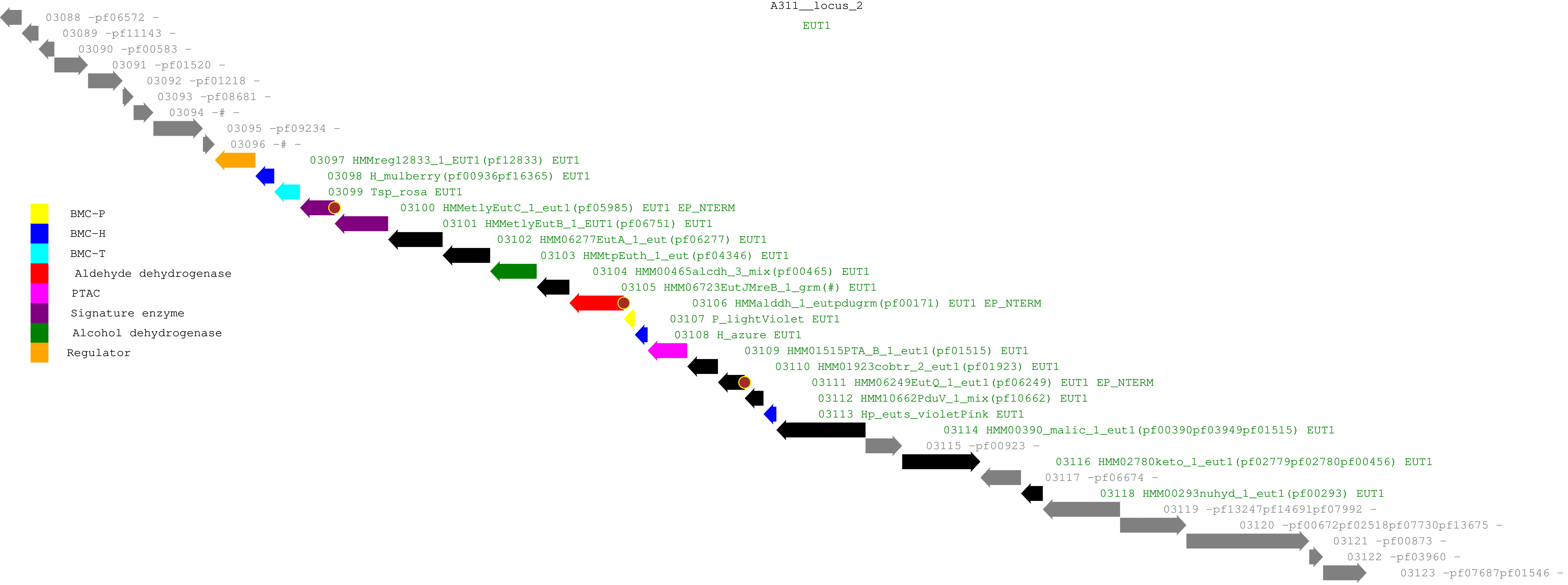

Escherichia sp. KTE159. Bacteria; Proteobacteria; Gammaproteobacteria; Enterobacterales;

A31E__locus_2

EUT1

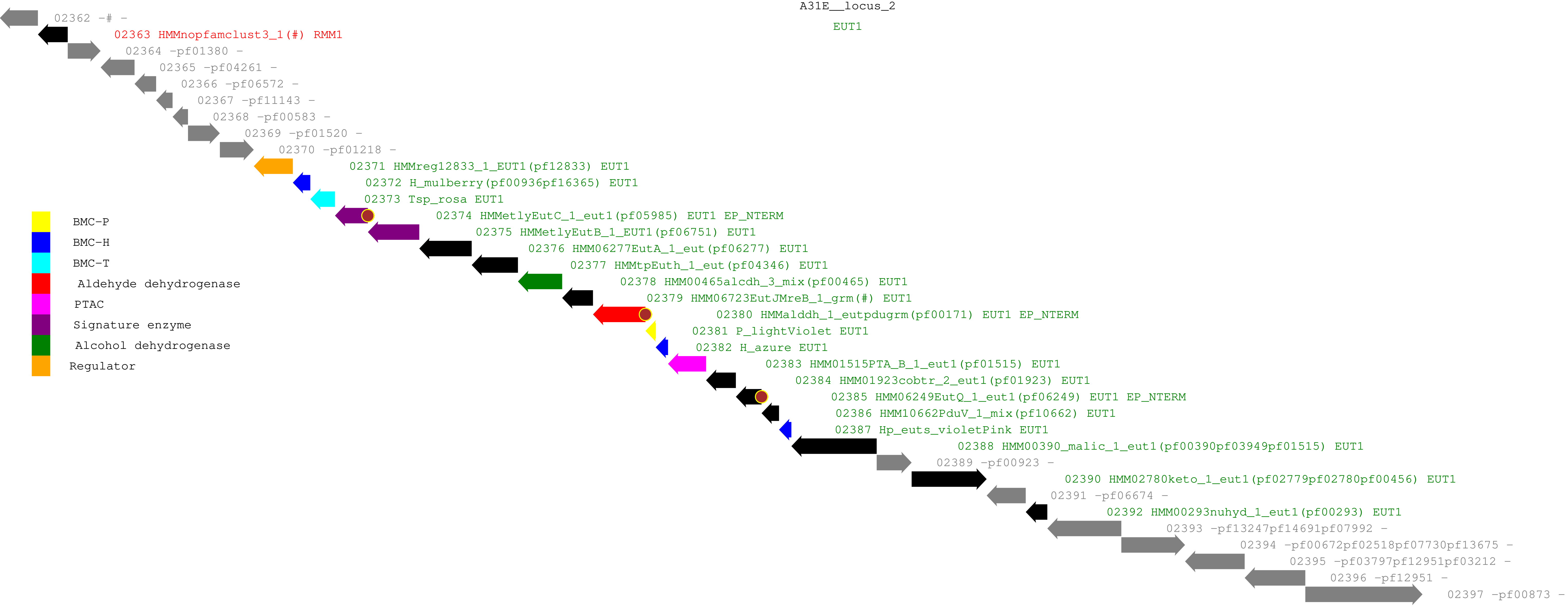

Salmonella paratyphi A. Bacteria; Proteobacteria; Gammaproteobacteria; Enterobacterales;

A3456__locus_1

EUT1

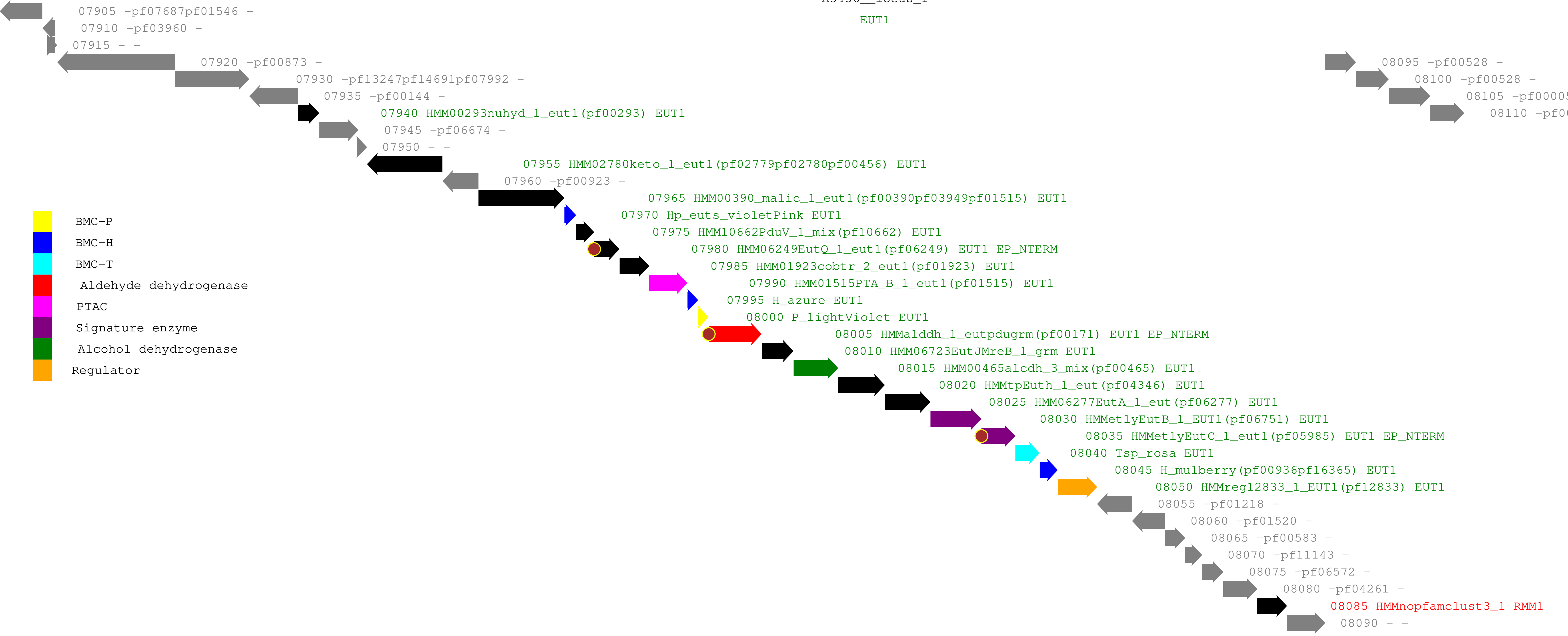

Burkholderiales bacterium RIFCSPLOWO2_12_FULL_61_40. Bacteria; Proteobacteria; Betaproteobacteria; Burkholderiales.

A3F78__locus_1_main_sat

EUT1

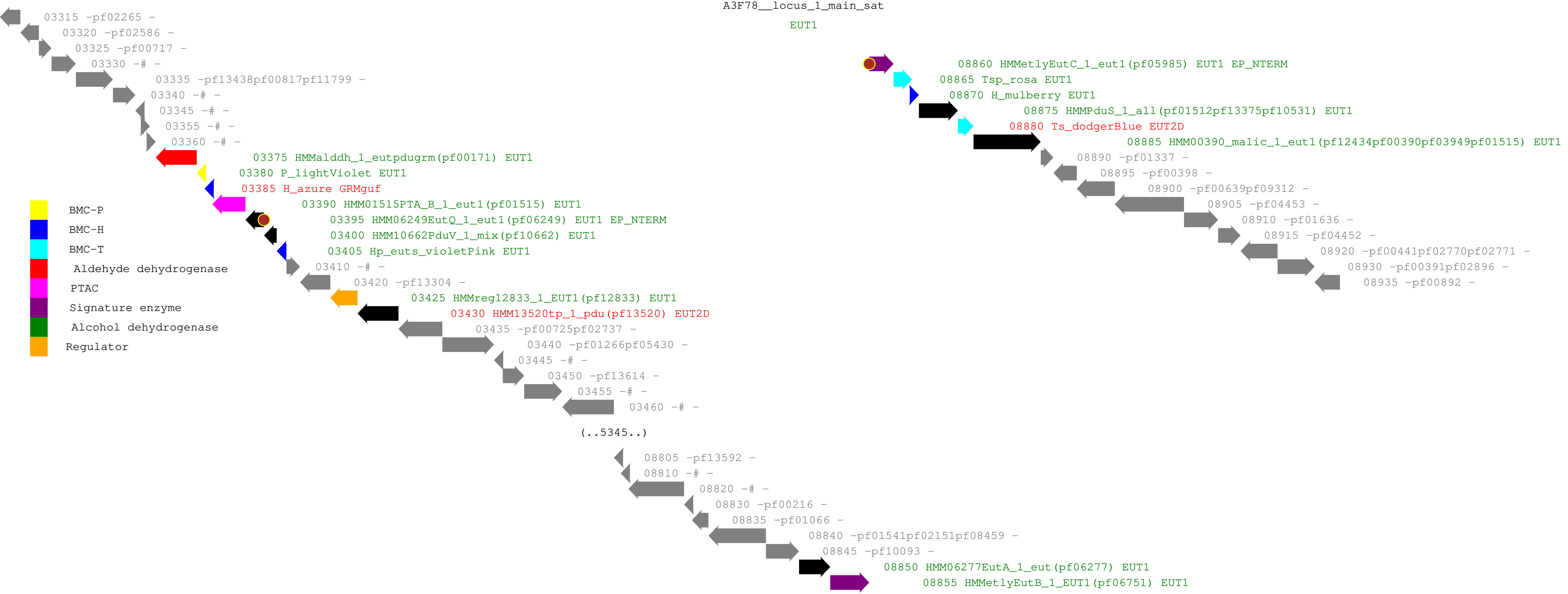

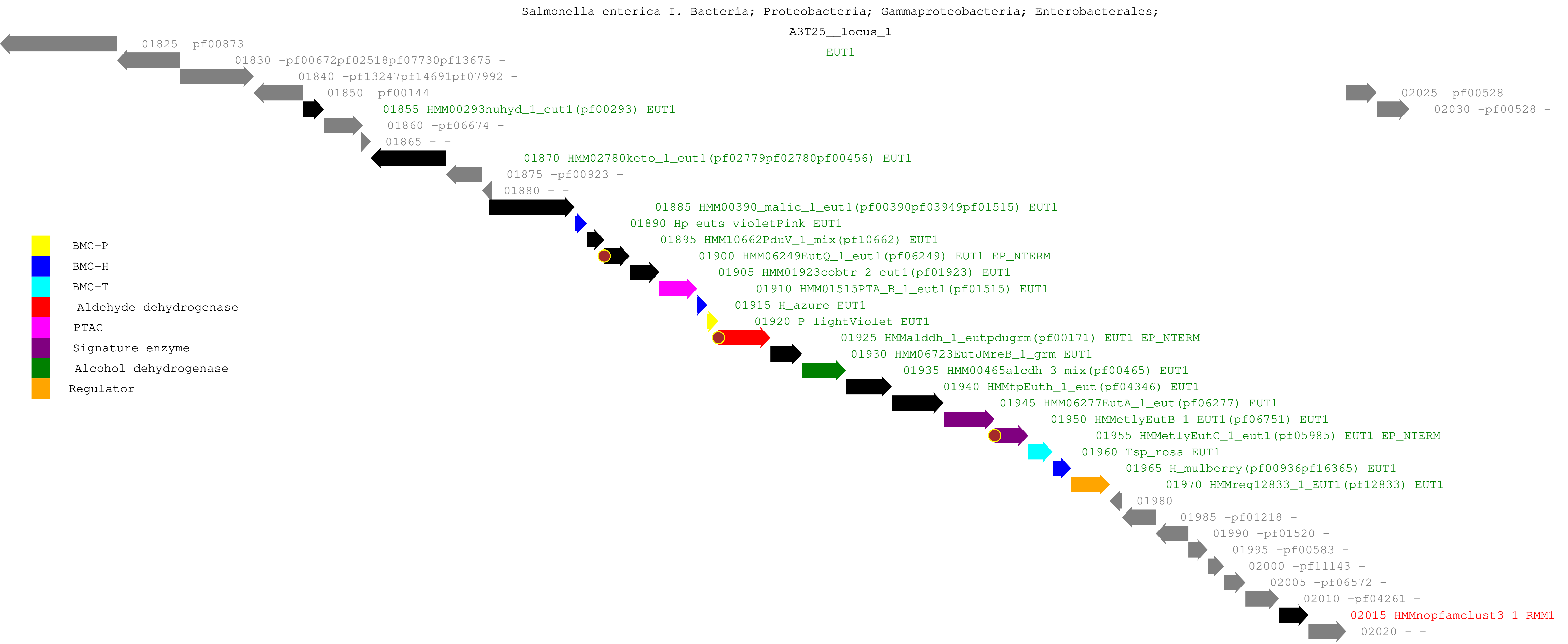
Salmonella enterica I. Bacteria; Proteobacteria; Gammaproteobacteria; Enterobacterales;
A3T25__locus_1
EUT1
01825 -pf00873 -
01830 -pf00672pf02518pf07730pf13675 -
01840 -pf13247pf14691pf07992 -
01850 -pf00144 -
01855 HMM00293nuhyd_1_eut1(pf00293) EUT1
01860 -pf06674 -
01865 - -
01870 HMM02780keto_1_eut1(pf02779pf02780pf00456) EUT1
01875 -pf00923 -
01880 - -
01885 HMM00390_malic_1_eut1(pf00390pf03949pf01515) EUT1
01890 Hp_euts_violetPink EUT1
01895 HMM10662PduV_1_mix(pf10662) EUT1
01900 HMM06249EutQ_1_eut1(pf06249) EUT1 EP_NTERM
01905 HMM01923cobtr_2_eut1(pf01923) EUT1
01910 HMM01515PTA_B_1_eut1(pf01515) EUT1
01915 H_azure EUT1
01920 P_lightViolet EUT1
01925 HMMalddh_1_eutpdugrm(pf00171) EUT1 EP_NTERM
01930 HMM06723EutJMreB_1_grm EUT1
01935 HMM00465alcdh_3_mix(pf00465) EUT1
01940 HMMtpEuth_1_eut(pf04346) EUT1
01945 HMM06277EutA_1_eut(pf06277) EUT1
01950 HMMetlyEutB_1_EUT1(pf06751) EUT1
01955 HMMetlyEutC_1_eut1(pf05985) EUT1 EP_NTERM
01960 Tsp_rosa EUT1
01965 H_mulberry(pf00936pf16365) EUT1
01970 HMMreg12833_1_EUT1(pf12833) EUT1
01980 - -
01985 -pf01218 -
01990 -pf01520 -
01995 -pf00583 -
02000 -pf11143 -
02005 -pf06572 -
02010 -pf04261 -
02015 HMMnopfamclust3_1 RMM1
02020 - -
02025 -pf00528 -
02030 -pf00528 -
BMC-P
BMC-H
BMC-T
Aldehyde dehydrogenase
PTAC
Signature enzyme
Alcohol dehydrogenase
Regulator

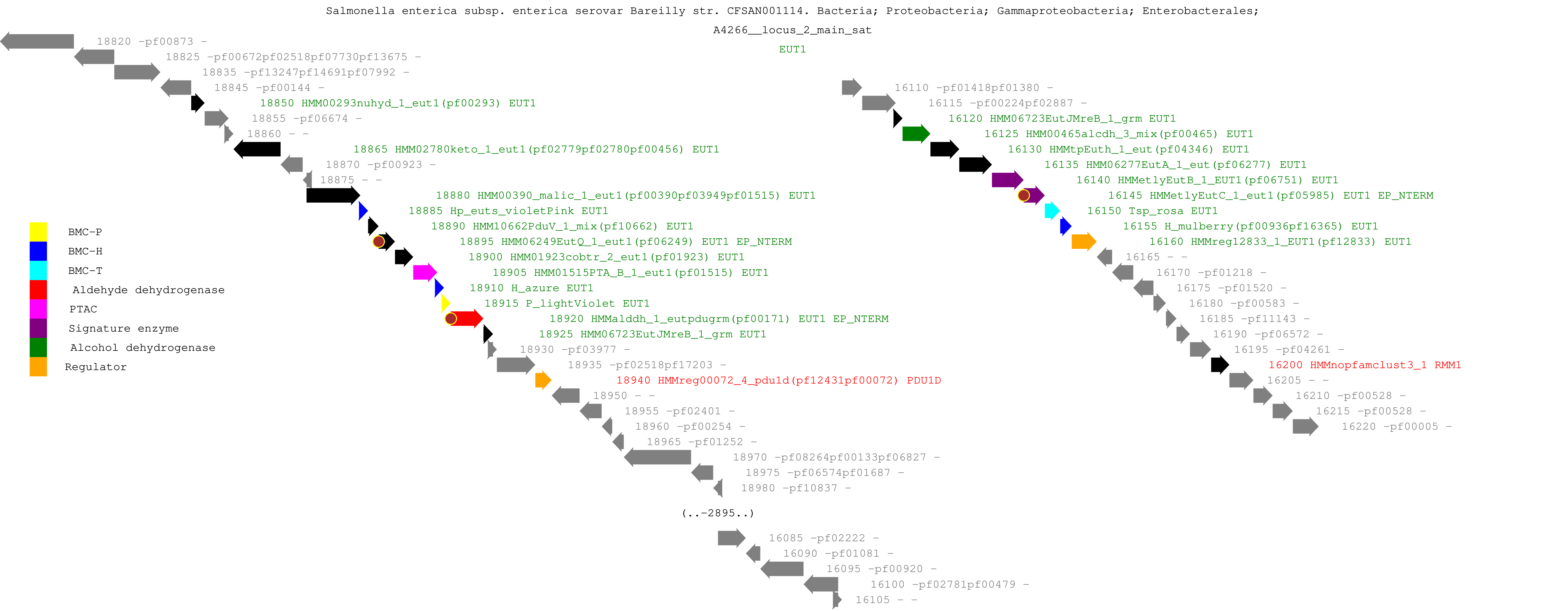
Salmonella enterica subsp. enterica serovar Bareilly str. CFSAN001114. Bacteria; Proteobacteria; Gammaproteobacteria; Enterobacterales;
A4266__locus_2_main_sat
EUT1
18820 -pf00873 -
18825 -pf00672pf02518pf07730pf13675 -
18835 -pf13247pf14691pf07992 -
18845 -pf00144 -
18850 HMM00293nuhyd_1_eut1(pf00293) EUT1
18855 -pf06674 -
18860 - -
18865 HMM02780keto_1_eut1(pf02779pf02780pf00456) EUT1
18870 -pf00923 -
18875 - -
18880 HMM00390_malic_1_eut1(pf00390pf03949pf01515) EUT1
18885 Hp_euts_violetPink EUT1
18890 HMM10662PduV_1_mix(pf10662) EUT1
18895 HMM06249EutQ_1_eut1(pf06249) EUT1 EP_NTERM
18900 HMM01923cobtr_2_eut1(pf01923) EUT1
18905 HMM01515PTA_B_1_eut1(pf01515) EUT1
18910 H_azure EUT1
18915 P_lightViolet EUT1
18920 HMMalddh_1_eutpdugrm(pf00171) EUT1 EP_NTERM
18925 HMM06723EutJMreB_1_grm EUT1
18930 -pf03977 -
18935 -pf02518pf17203 -
18940 HMMreg00072_4_pdu1d(pf12431pf00072) PDU1D
18950 - -
18955 -pf02401 -
18960 -pf00254 -
18965 -pf01252 -
18970 -pf08264pf00133pf06827 -
18975 -pf06574pf01687 -
18980 -pf10837 -
(..-2895..)
16085 -pf02222 -
16090 -pf01081 -
16095 -pf00920 -
16100 -pf02781pf00479 -
16105 - -
16110 -pf01418pf01380 -
16115 -pf00224pf02887 -
16120 HMM06723EutJMreB_1_grm EUT1
16125 HMM00465alcdh_3_mix(pf00465) EUT1
16130 HMMtpEuth_1_eut(pf04346) EUT1
16135 HMM06277EutA_1_eut(pf06277) EUT1
16140 HMMetlyEutB_1_EUT1(pf06751) EUT1
16145 HMMetlyEutC_1_eut1(pf05985) EUT1 EP_NTERM
16150 Tsp_rosa EUT1
16155 H_mulberry(pf00936pf16365) EUT1
16160 HMMreg12833_1_EUT1(pf12833) EUT1
16165 - -
16170 -pf01218 -
16175 -pf01520 -
16180 -pf00583 -
16185 -pf11143 -
16190 -pf06572 -
16195 -pf04261 -
16200 HMMnopfamclust3_1 RMM1
16205 - -
16210 -pf00528 -
16215 -pf00528 -
16220 -pf00005 -
BMC-P
BMC-H
BMC-T
Aldehyde dehydrogenase
PTAC
Signature enzyme
Alcohol dehydrogenase
Regulator

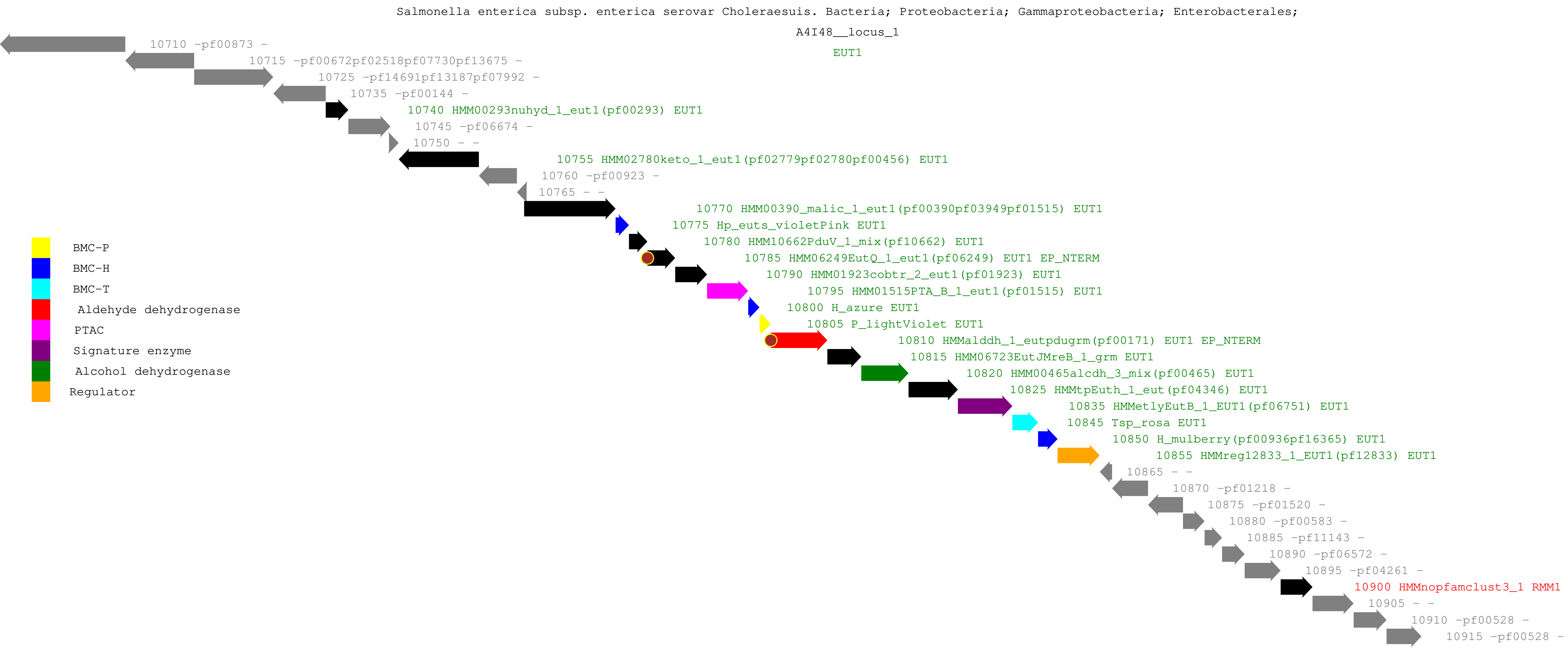

Salmonella enterica subsp. enterica serovar Choleraesuis. Bacteria; Proteobacteria; Gammaproteobacteria; Enterobacterales;
A4I48__locus_1
EUT1
10710 -pf00873 -
10715 -pf00672pf02518pf07730pf13675 -
10725 -pf14691pf13187pf07992 -
10735 -pf00144 -
10740 HMM00293nuhyd_1_eut1(pf00293) EUT1
10745 -pf06674 -
10750 - -
10755 HMM02780keto_1_eut1(pf02779pf02780pf00456) EUT1
10760 -pf00923 -
10765 - -
10770 HMM00390_malic_1_eut1(pf00390pf03949pf01515) EUT1
10775 Hp_euts_violetPink EUT1
10780 HMM10662PduV_1_mix(pf10662) EUT1
10785 HMM06249EutQ_1_eut1(pf06249) EUT1 EP_NTERM
10790 HMM01923cobtr_2_eut1(pf01923) EUT1
10795 HMM01515PTA_B_1_eut1(pf01515) EUT1
10800 H_azure EUT1
10805 P_lightViolet EUT1
10810 HMMalddh_1_eutpdugrm(pf00171) EUT1 EP_NTERM
10815 HMM06723EutJMreB_1_grm EUT1
10820 HMM00465alcdh_3_mix(pf00465) EUT1
10825 HMMtpEuth_1_eut(pf04346) EUT1
10835 HMMetlyEutB_1_EUT1(pf06751) EUT1
10845 Tsp_rosa EUT1
10850 H_mulberry(pf00936pf16365) EUT1
10855 HMMreg12833_1_EUT1(pf12833) EUT1
10865 - -
10870 -pf01218 -
10875 -pf01520 -
10880 -pf00583 -
10885 -pf11143 -
10890 -pf06572 -
10895 -pf04261 -
10900 HMMnopfamclust3_1 RMM1
10905 - -
10910 -pf00528 -
10915 -pf00528 -
BMC-P
BMC-H
BMC-T
Aldehyde dehydrogenase
PTAC
Signature enzyme
Alcohol dehydrogenase
Regulator

Salmonella enterica subsp. enterica serovar Panama. Bacteria; Proteobacteria; Gammaproteobacteria; Enterobacterales;

A4I59__locus_1

EUT1

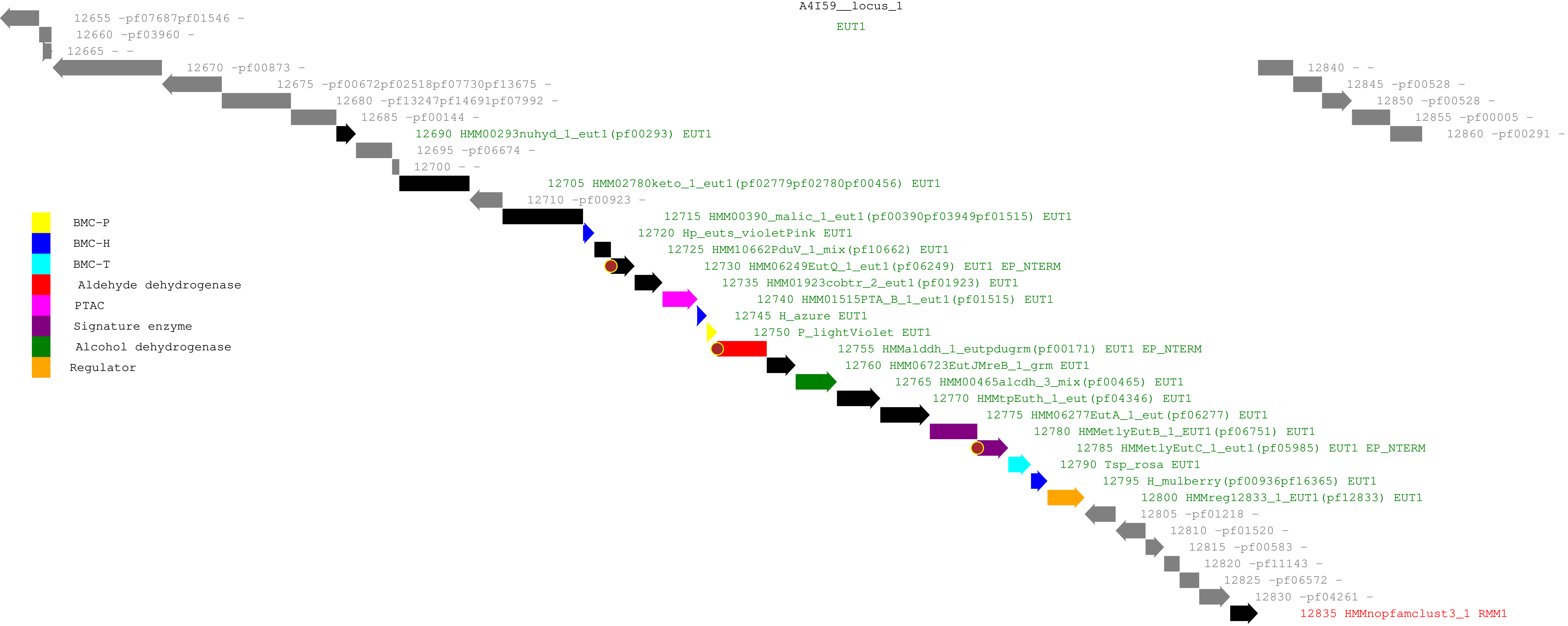

Salmonella enterica subsp. enterica serovar Richmond. Bacteria; Proteobacteria; Gammaproteobacteria; Enterobacterales;

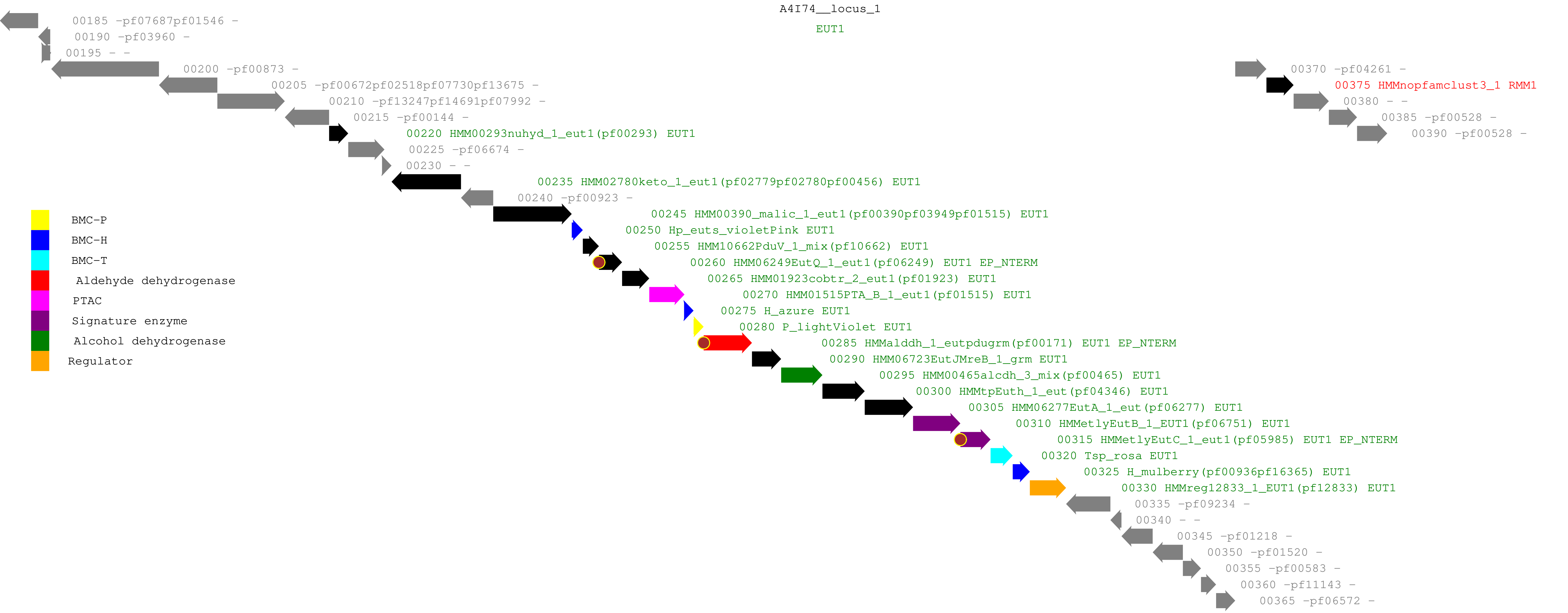

Salmonella enterica subsp. enterica serovar Kedougou. Bacteria; Proteobacteria; Gammaproteobacteria; Enterobacterales;

A4I75__locus_2

EUT1

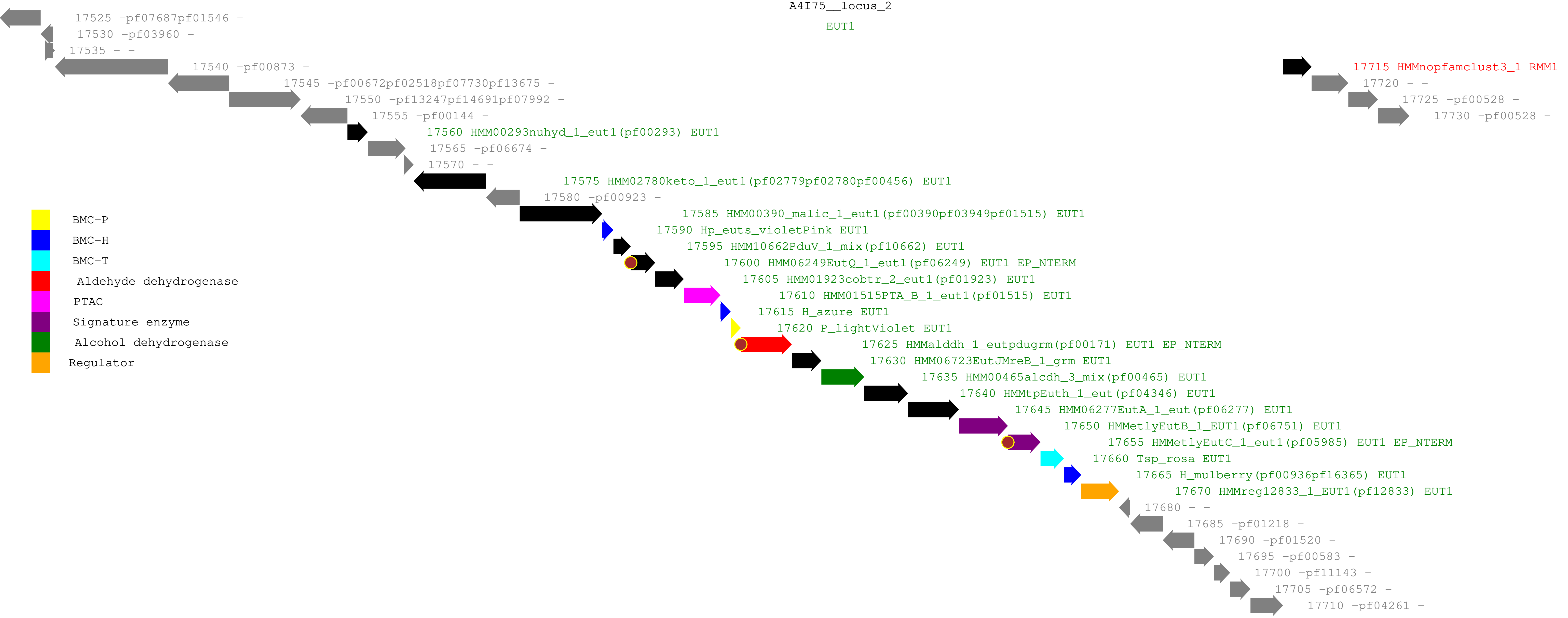

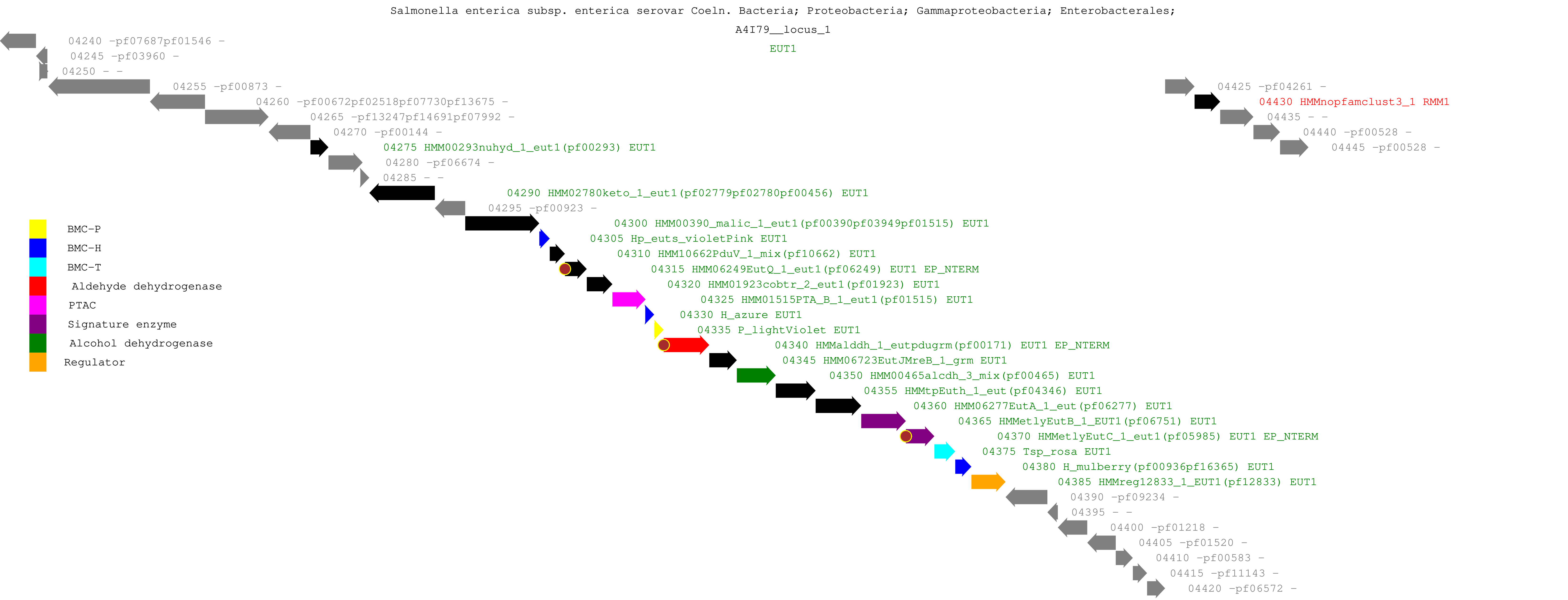
Salmonella enterica subsp. enterica serovar Coeln. Bacteria; Proteobacteria; Gammaproteobacteria; Enterobacterales;
A4I79__locus_1
EUT1
04240 -pf07687pf01546 -
04245 -pf03960 -
04250 - -
04255 -pf00873 -
04260 -pf00672pf02518pf07730pf13675 -
04265 -pf13247pf14691pf07992 -
04270 -pf00144 -
04275 HMM00293nuhyd_1_eut1(pf00293) EUT1
04280 -pf06674 -
04285 - -
04290 HMM02780keto_1_eut1(pf02779pf02780pf00456) EUT1
04295 -pf00923 -
04300 HMM00390_malic_1_eut1(pf00390pf03949pf01515) EUT1
04305 Hp_euts_violetPink EUT1
04310 HMM10662PduV_1_mix(pf10662) EUT1
04315 HMM06249EutQ_1_eut1(pf06249) EUT1 EP_NTERM
04320 HMM01923cobtr_2_eut1(pf01923) EUT1
04325 HMM01515PTA_B_1_eut1(pf01515) EUT1
04330 H_azure EUT1
04335 P_lightViolet EUT1
04340 HMMalddh_1_eutpdugrm(pf00171) EUT1 EP_NTERM
04345 HMM06723EutJMreB_1_grm EUT1
04350 HMM00465alcdh_3_mix(pf00465) EUT1
04355 HMMtpEuth_1_eut(pf04346) EUT1
04360 HMM06277EutA_1_eut(pf06277) EUT1
04365 HMMetlyEutB_1_EUT1(pf06751) EUT1
04370 HMMetlyEutC_1_eut1(pf05985) EUT1 EP_NTERM
04375 Tsp_rosa EUT1
04380 H_mulberry(pf00936pf16365) EUT1
04385 HMMreg12833_1_EUT1(pf12833) EUT1
04390 -pf09234 -
04395 - -
04400 -pf01218 -
04405 -pf01520 -
04410 -pf00583 -
04415 -pf11143 -
04420 -pf06572 -
04425 -pf04261 -
04430 HMMnopfamclust3_1 RMM1
04435 - -
04440 -pf00528 -
04445 -pf00528 -
BMC-P
BMC-H
BMC-T
Aldehyde dehydrogenase
PTAC
Signature enzyme
Alcohol dehydrogenase
Regulator

Salmonella oranienberg. Bacteria; Proteobacteria; Gammaproteobacteria; Enterobacterales;

A4I87__locus_1

EUT1

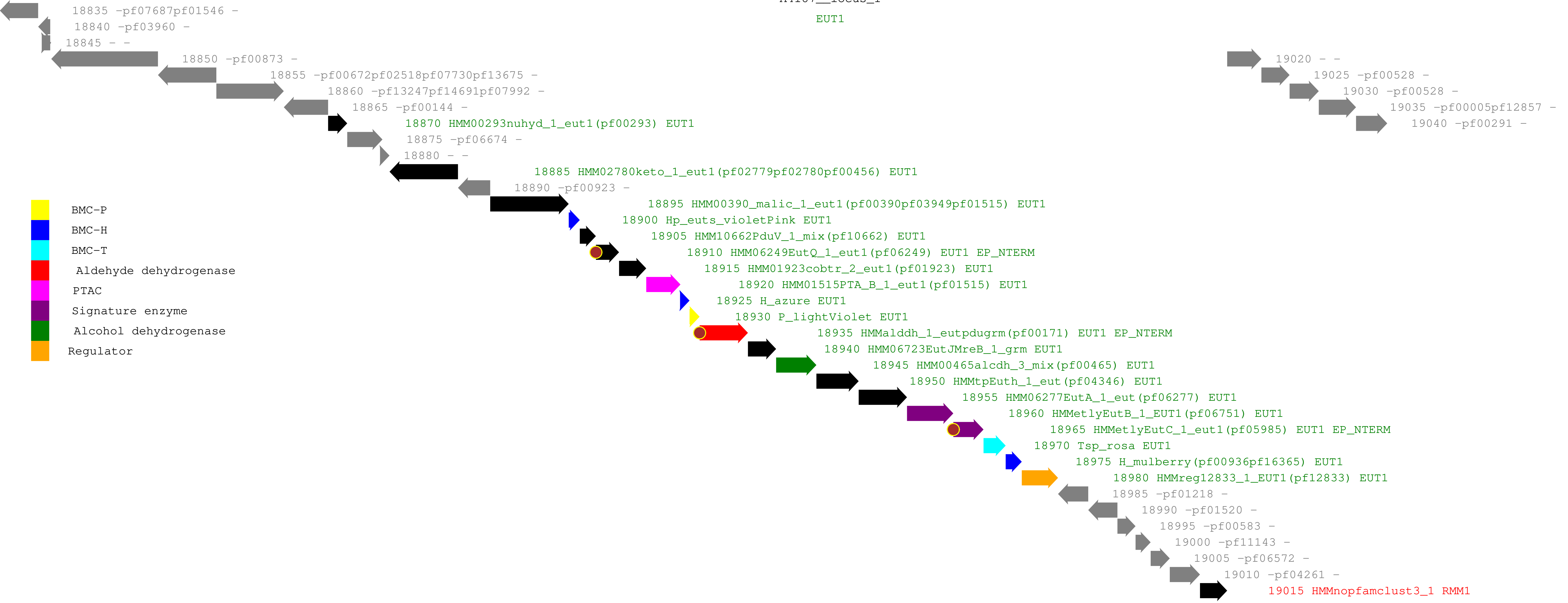

Salmonella enterica subsp. enterica serovar London. Bacteria; Proteobacteria; Gammaproteobacteria; Enterobacterales;

A4I94__locus_2

EUT1

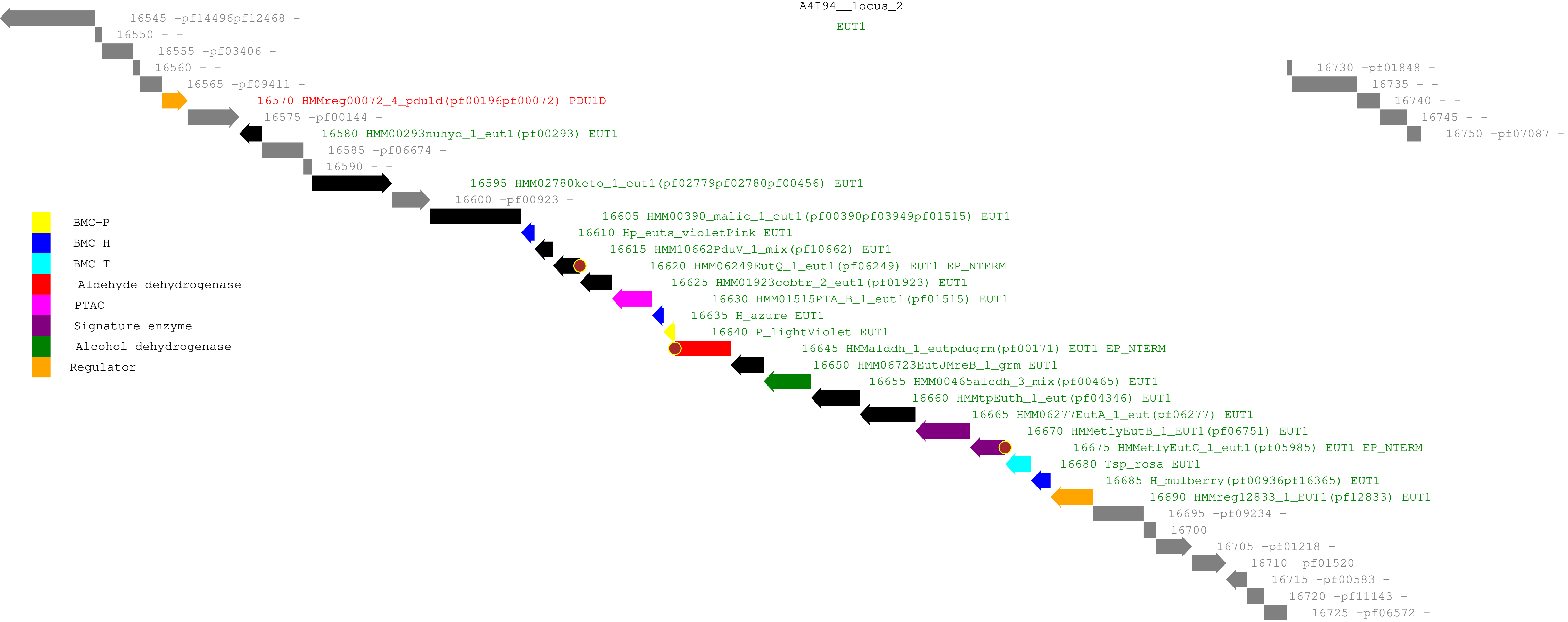

Salmonella enterica subsp. enterica serovar Sandiego. Bacteria; Proteobacteria; Gammaproteobacteria; Enterobacterales;

A4J53__locus_2

EUT1

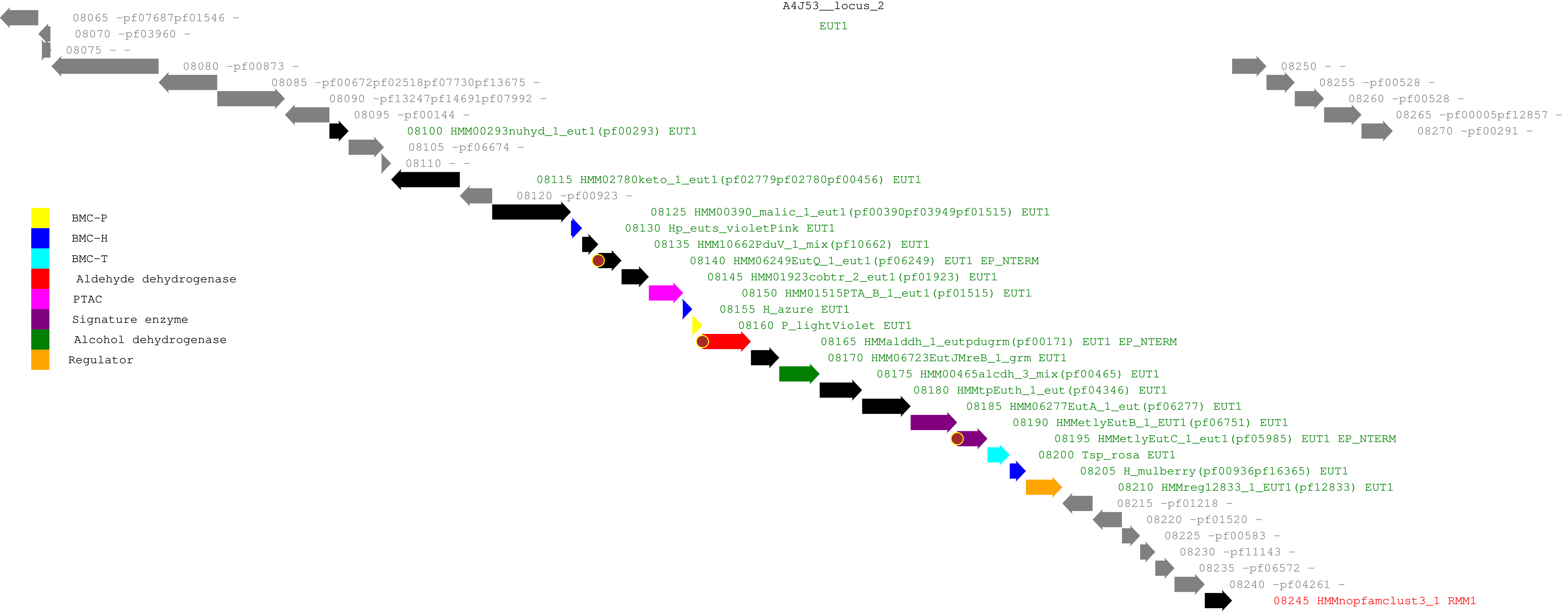

Salmonella enterica subsp. enterica serovar Joal. Bacteria; Proteobacteria; Gammaproteobacteria; Enterobacterales;

A4J61__locus_1

EUT1

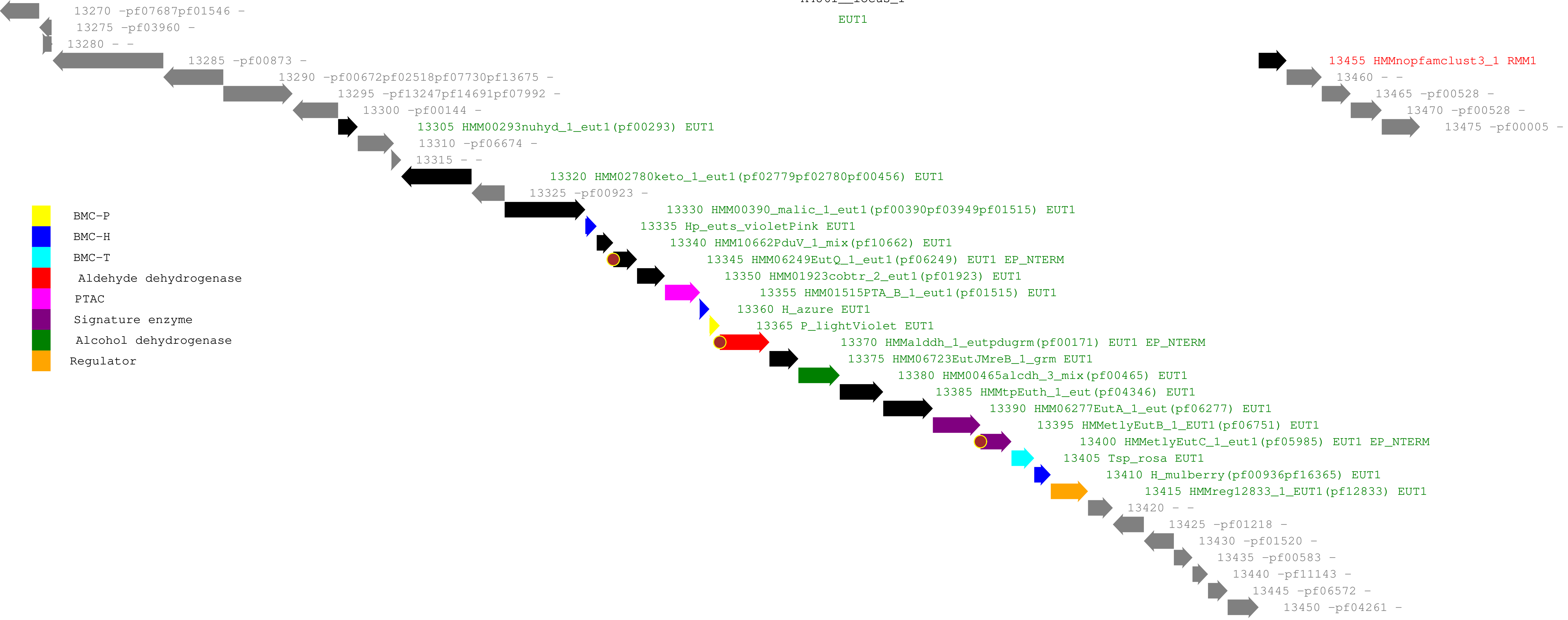

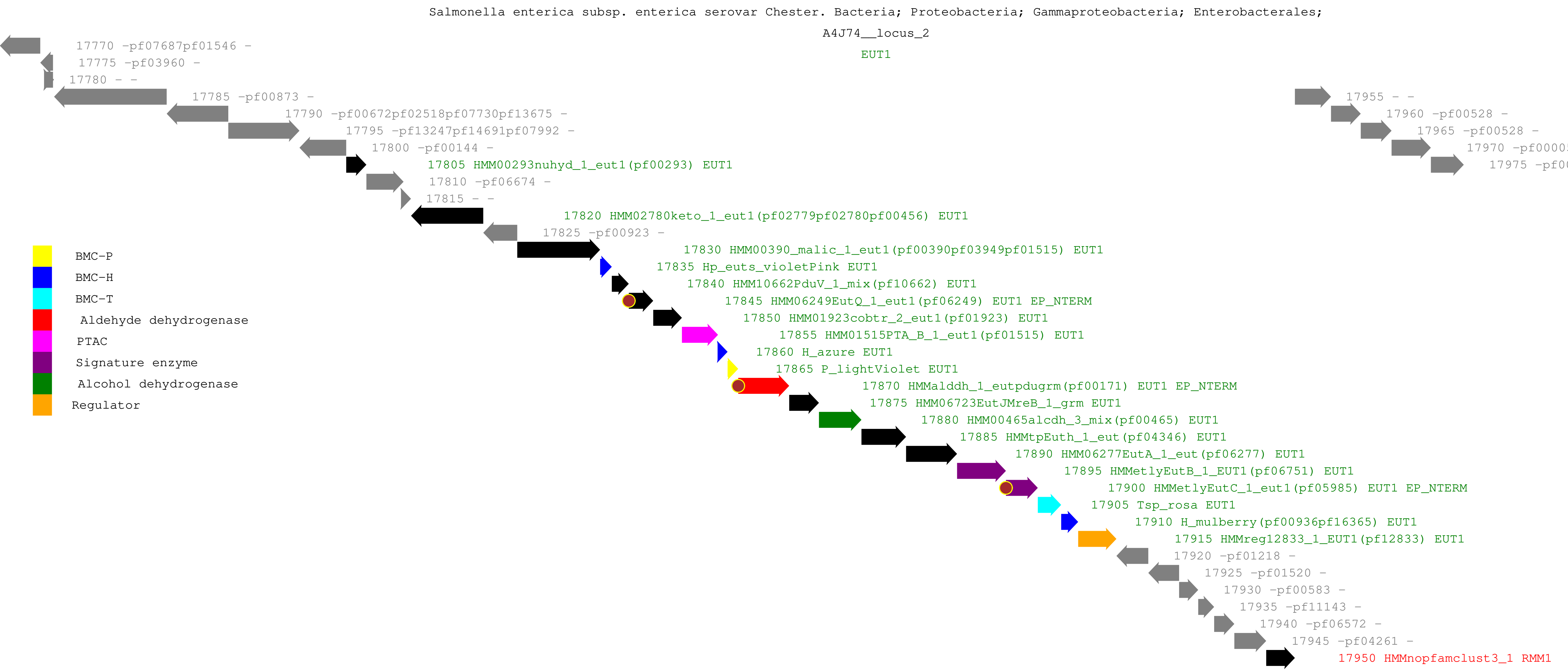
Salmonella enterica subsp. enterica serovar Chester. Bacteria; Proteobacteria; Gammaproteobacteria; Enterobacterales;
A4J74__locus_2
EUT1
17770 -pf07687pf01546 -
17775 -pf03960 -
17780 - -
17785 -pf00873 -
17790 -pf00672pf02518pf07730pf13675 -
17795 -pf13247pf14691pf07992 -
17800 -pf00144 -
17805 HMM00293nuhyd_1_eut1(pf00293) EUT1
17810 -pf06674 -
17815 - -
17820 HMM02780keto_1_eut1(pf02779pf02780pf00456) EUT1
17825 -pf00923 -
17830 HMM00390_malic_1_eut1(pf00390pf03949pf01515) EUT1
17835 Hp_euts_violetPink EUT1
17840 HMM10662PduV_1_mix(pf10662) EUT1
17845 HMM06249EutQ_1_eut1(pf06249) EUT1 EP_NTERM
17850 HMM01923cobtr_2_eut1(pf01923) EUT1
17855 HMM01515PTA_B_1_eut1(pf01515) EUT1
17860 H_azure EUT1
17865 P_lightViolet EUT1
17870 HMMalddh_1_eutpdugrm(pf00171) EUT1 EP_NTERM
17875 HMM06723EutJMreB_1_grm EUT1
17880 HMM00465alcdh_3_mix(pf00465) EUT1
17885 HMMtpEuth_1_eut(pf04346) EUT1
17890 HMM06277EutA_1_eut(pf06277) EUT1
17895 HMMetlyEutB_1_EUT1(pf06751) EUT1
17900 HMMetlyEutC_1_eut1(pf05985) EUT1 EP_NTERM
17905 Tsp_rosa EUT1
17910 H_mulberry(pf00936pf16365) EUT1
17915 HMMreg12833_1_EUT1(pf12833) EUT1
17920 -pf01218 -
17925 -pf01520 -
17930 -pf00583 -
17935 -pf11143 -
17940 -pf06572 -
17945 -pf04261 -
17950 HMMnopfamclust3_1 RMM1
17955 - -
17960 -pf00528 -
17965 -pf00528 -
17970 -pf00005pf12857 -
17975 -pf00291 -
BMC-P
BMC-H
BMC-T
Aldehyde dehydrogenase
PTAC
Signature enzyme
Alcohol dehydrogenase
Regulator

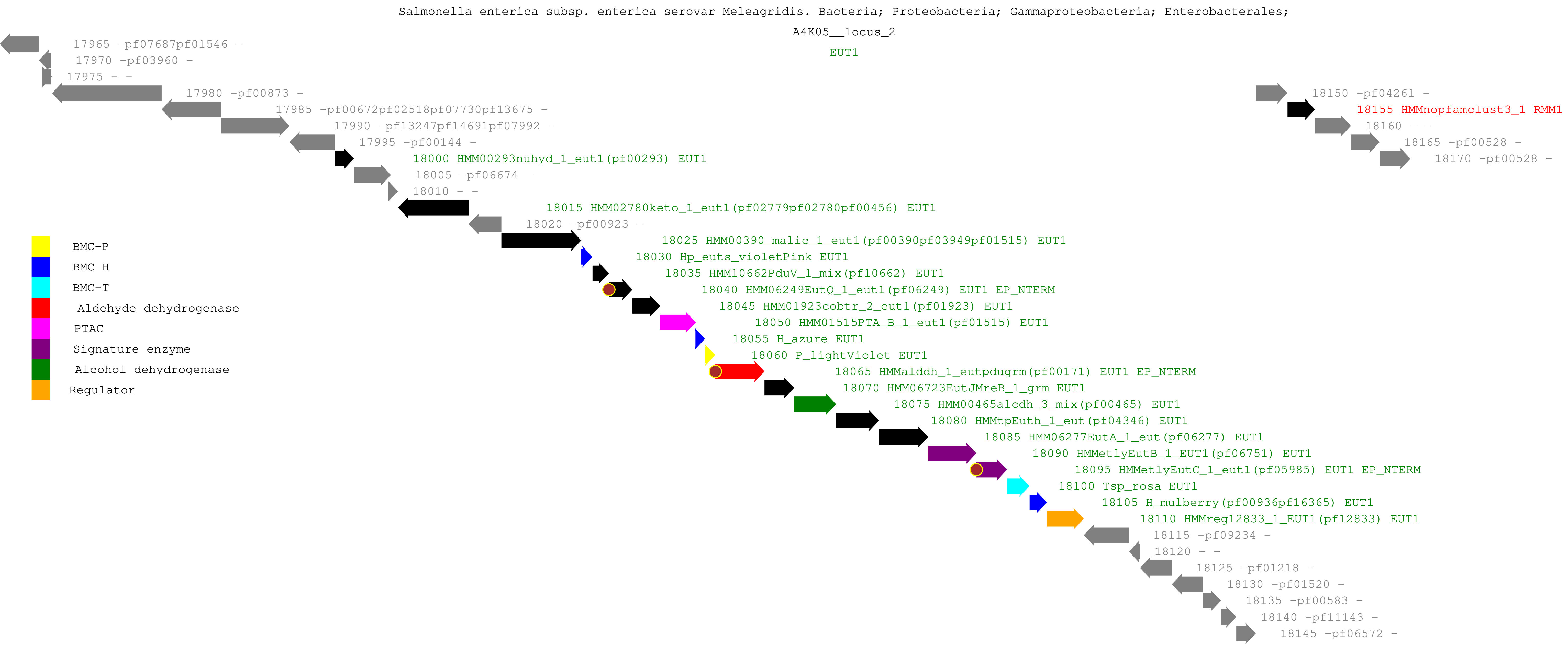
Salmonella enterica subsp. enterica serovar Meleagridis. Bacteria; Proteobacteria; Gammaproteobacteria; Enterobacterales;
A4K05__locus_2
EUT1
17965 -pf07687pf01546 -
17970 -pf03960 -
17975 - -
17980 -pf00873 -
17985 -pf00672pf02518pf07730pf13675 -
17990 -pf13247pf14691pf07992 -
17995 -pf00144 -
18000 HMM00293nuhyd_1_eut1(pf00293) EUT1
18005 -pf06674 -
18010 - -
18015 HMM02780keto_1_eut1(pf02779pf02780pf00456) EUT1
18020 -pf00923 -
18025 HMM00390_malic_1_eut1(pf00390pf03949pf01515) EUT1
18030 Hp_euts_violetPink EUT1
18035 HMM10662PduV_1_mix(pf10662) EUT1
18040 HMM06249EutQ_1_eut1(pf06249) EUT1 EP_NTERM
18045 HMM01923cobtr_2_eut1(pf01923) EUT1
18050 HMM01515PTA_B_1_eut1(pf01515) EUT1
18055 H_azure EUT1
18060 P_lightViolet EUT1
18065 HMMalddh_1_eutpdugrm(pf00171) EUT1 EP_NTERM
18070 HMM06723EutJMreB_1_grm EUT1
18075 HMM00465alcdh_3_mix(pf00465) EUT1
18080 HMMtpEuth_1_eut(pf04346) EUT1
18085 HMM06277EutA_1_eut(pf06277) EUT1
18090 HMMetlyEutB_1_EUT1(pf06751) EUT1
18095 HMMetlyEutC_1_eut1(pf05985) EUT1 EP_NTERM
18100 Tsp_rosa EUT1
18105 H_mulberry(pf00936pf16365) EUT1
18110 HMMreg12833_1_EUT1(pf12833) EUT1
18115 -pf09234 -
18120 - -
18125 -pf01218 -
18130 -pf01520 -
18135 -pf00583 -
18140 -pf11143 -
18145 -pf06572 -
18150 -pf04261 -
18155 HMMnopfamclust3_1 RMM1
18160 - -
18165 -pf00528 -
18170 -pf00528 -
BMC-P
BMC-H
BMC-T
Aldehyde dehydrogenase
PTAC
Signature enzyme
Alcohol dehydrogenase
Regulator

Salmonella enterica subsp. enterica serovar Bakau. Bacteria; Proteobacteria; Gammaproteobacteria; Enterobacterales;

A4K08__locus_2

EUT1

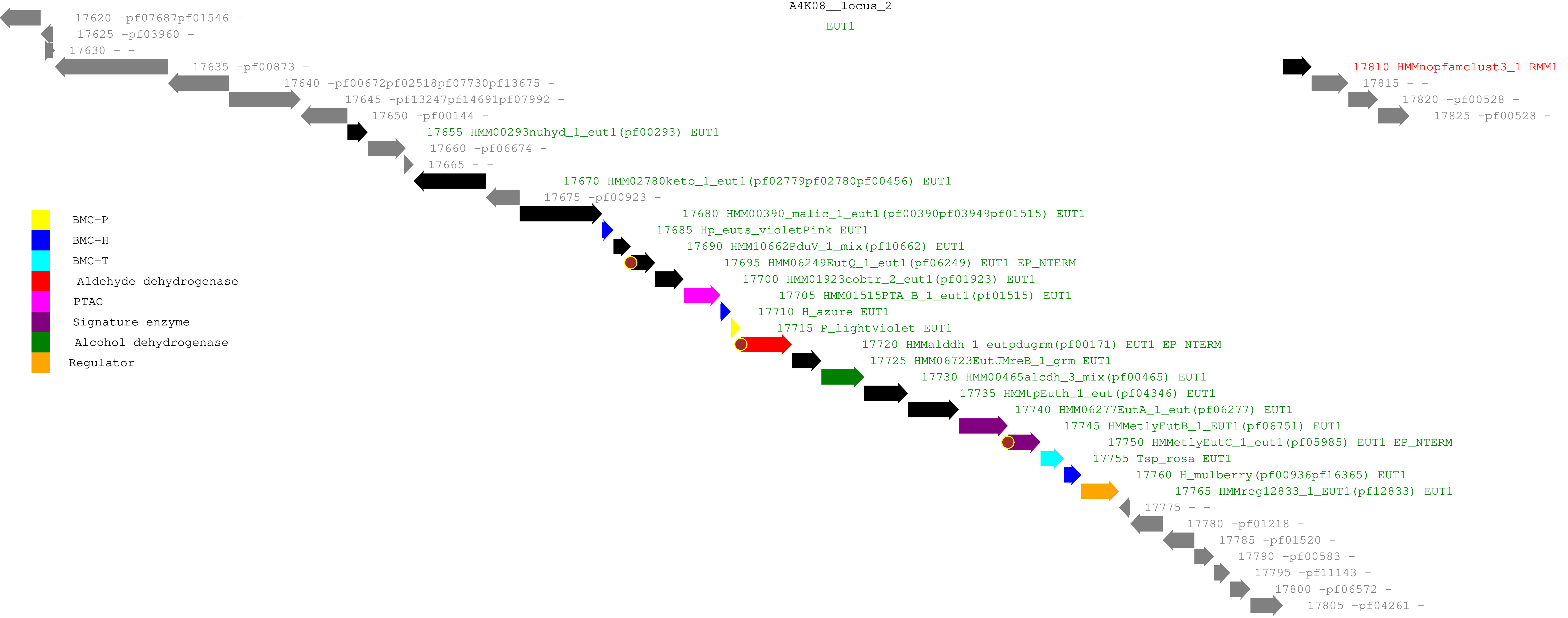

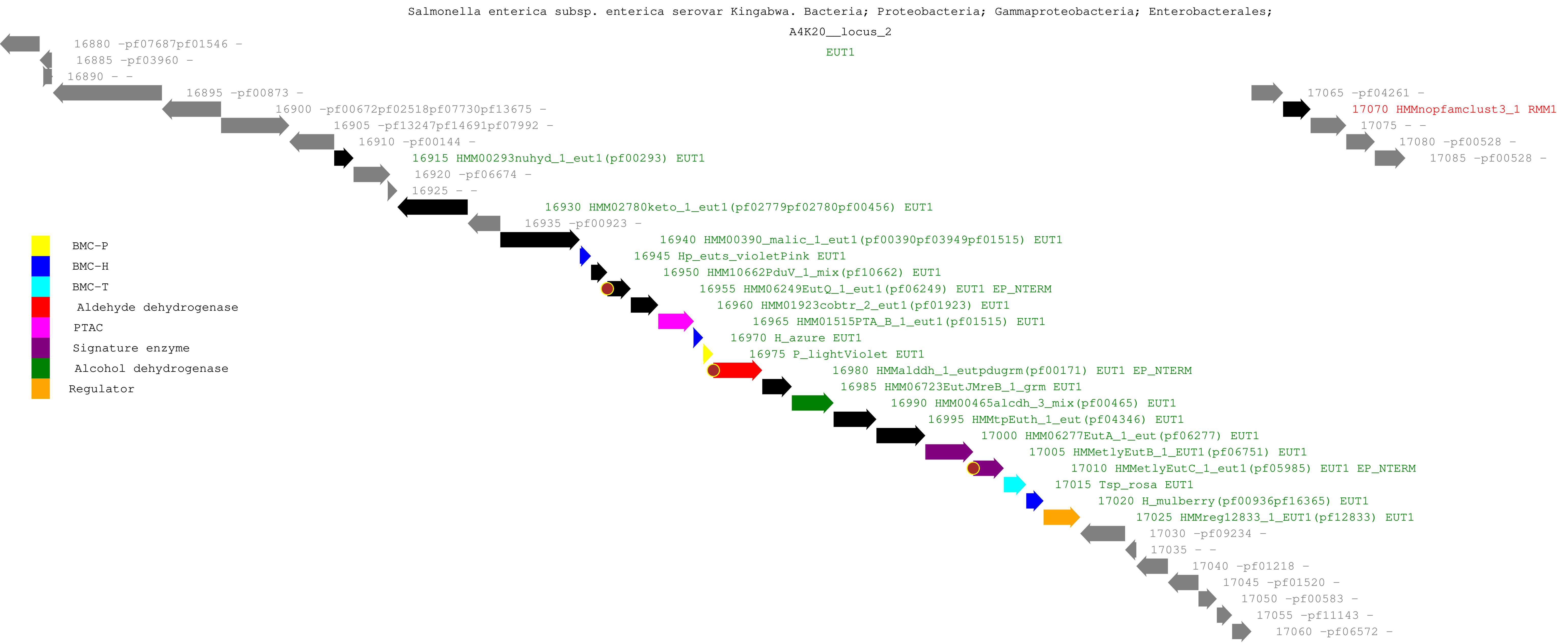
Salmonella enterica subsp. enterica serovar Kingabwa. Bacteria; Proteobacteria; Gammaproteobacteria; Enterobacterales;
A4K20__locus_2
EUT1
16880 -pf07687pf01546 -
16885 -pf03960 -
16890 - -
16895 -pf00873 -
16900 -pf00672pf02518pf07730pf13675 -
16905 -pf13247pf14691pf07992 -
16910 -pf00144 -
16915 HMM00293nuhyd_1_eut1(pf00293) EUT1
16920 -pf06674 -
16925 - -
16930 HMM02780keto_1_eut1(pf02779pf02780pf00456) EUT1
16935 -pf00923 -
16940 HMM00390_malic_1_eut1(pf00390pf03949pf01515) EUT1
16945 Hp_euts_violetPink EUT1
16950 HMM10662PduV_1_mix(pf10662) EUT1
16955 HMM06249EutQ_1_eut1(pf06249) EUT1 EP_NTERM
16960 HMM01923cobtr_2_eut1(pf01923) EUT1
16965 HMM01515PTA_B_1_eut1(pf01515) EUT1
16970 H_azure EUT1
16975 P_lightViolet EUT1
16980 HMMalddh_1_eutpdugrm(pf00171) EUT1 EP_NTERM
16985 HMM06723EutJMreB_1_grm EUT1
16990 HMM00465alcdh_3_mix(pf00465) EUT1
16995 HMMtpEuth_1_eut(pf04346) EUT1
17000 HMM06277EutA_1_eut(pf06277) EUT1
17005 HMMetlyEutB_1_EUT1(pf06751) EUT1
17010 HMMetlyEutC_1_eut1(pf05985) EUT1 EP_NTERM
17015 Tsp_rosa EUT1
17020 H_mulberry(pf00936pf16365) EUT1
17025 HMMreg12833_1_EUT1(pf12833) EUT1
17030 -pf09234 -
17035 - -
17040 -pf01218 -
17045 -pf01520 -
17050 -pf00583 -
17055 -pf11143 -
17060 -pf06572 -
17065 -pf04261 -
17070 HMMnopfamclust3_1 RMM1
17075 - -
17080 -pf00528 -
17085 -pf00528 -
BMC-P
BMC-H
BMC-T
Aldehyde dehydrogenase
PTAC
Signature enzyme
Alcohol dehydrogenase
Regulator

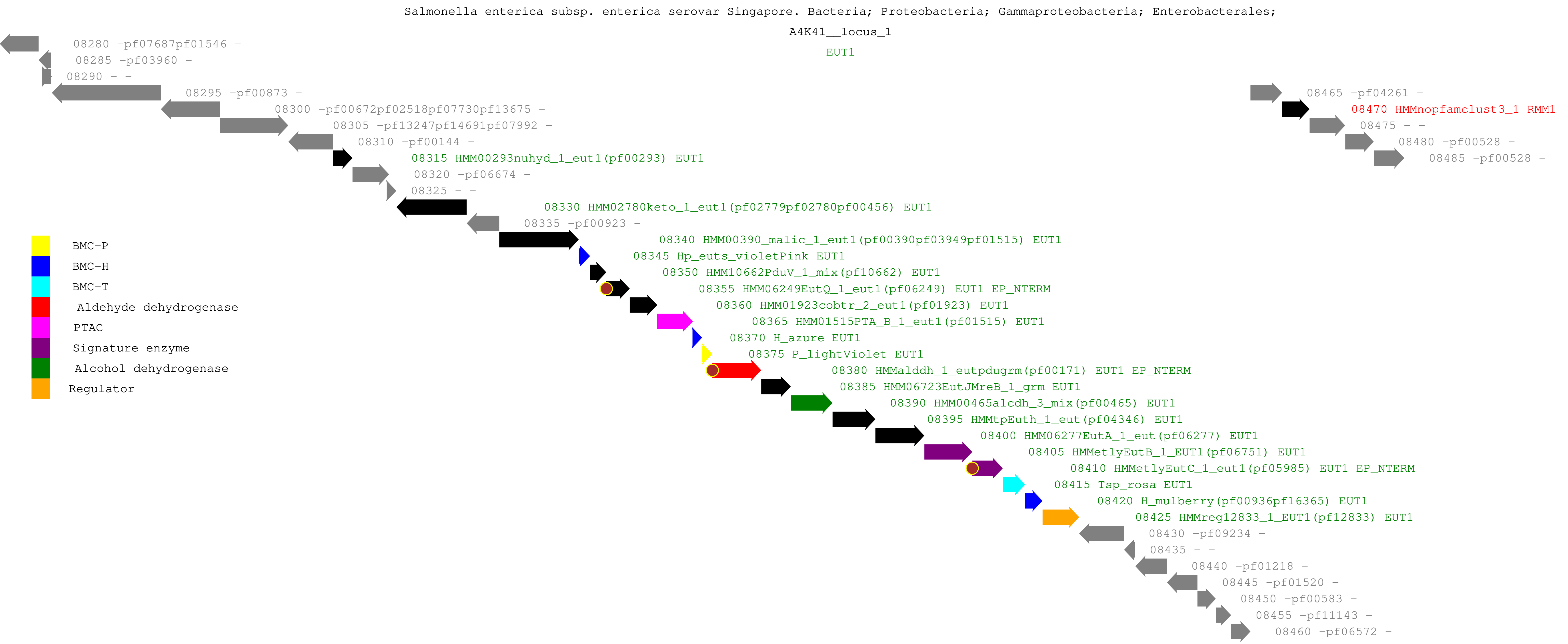
Salmonella enterica subsp. enterica serovar Singapore. Bacteria; Proteobacteria; Gammaproteobacteria; Enterobacterales;
A4K41__locus_1
EUT1
08280 -pf07687pf01546 -
08285 -pf03960 -
08290 - -
08295 -pf00873 -
08300 -pf00672pf02518pf07730pf13675 -
08305 -pf13247pf14691pf07992 -
08310 -pf00144 -
08315 HMM00293nuhyd_1_eut1(pf00293) EUT1
08320 -pf06674 -
08325 - -
08330 HMM02780keto_1_eut1(pf02779pf02780pf00456) EUT1
08335 -pf00923 -
08340 HMM00390_malic_1_eut1(pf00390pf03949pf01515) EUT1
08345 Hp_euts_violetPink EUT1
08350 HMM10662PduV_1_mix(pf10662) EUT1
08355 HMM06249EutQ_1_eut1(pf06249) EUT1 EP_NTERM
08360 HMM01923cobtr_2_eut1(pf01923) EUT1
08365 HMM01515PTA_B_1_eut1(pf01515) EUT1
08370 H_azure EUT1
08375 P_lightViolet EUT1
08380 HMMalddh_1_eutpdugrm(pf00171) EUT1 EP_NTERM
08385 HMM06723EutJMreB_1_grm EUT1
08390 HMM00465alcdh_3_mix(pf00465) EUT1
08395 HMMtpEuth_1_eut(pf04346) EUT1
08400 HMM06277EutA_1_eut(pf06277) EUT1
08405 HMMetlyEutB_1_EUT1(pf06751) EUT1
08410 HMMetlyEutC_1_eut1(pf05985) EUT1 EP_NTERM
08415 Tsp_rosa EUT1
08420 H_mulberry(pf00936pf16365) EUT1
08425 HMMreg12833_1_EUT1(pf12833) EUT1
08430 -pf09234 -
08435 - -
08440 -pf01218 -
08445 -pf01520 -
08450 -pf00583 -
08455 -pf11143 -
08460 -pf06572 -
08465 -pf04261 -
08470 HMMnopfamclust3_1 RMM1
08475 - -
08480 -pf00528 -
08485 -pf00528 -
BMC-P
BMC-H
BMC-T
Aldehyde dehydrogenase
PTAC
Signature enzyme
Alcohol dehydrogenase
Regulator

Salmonella enterica subsp. enterica serovar Mississippi. Bacteria; Proteobacteria; Gammaproteobacteria; Enterobacterales;

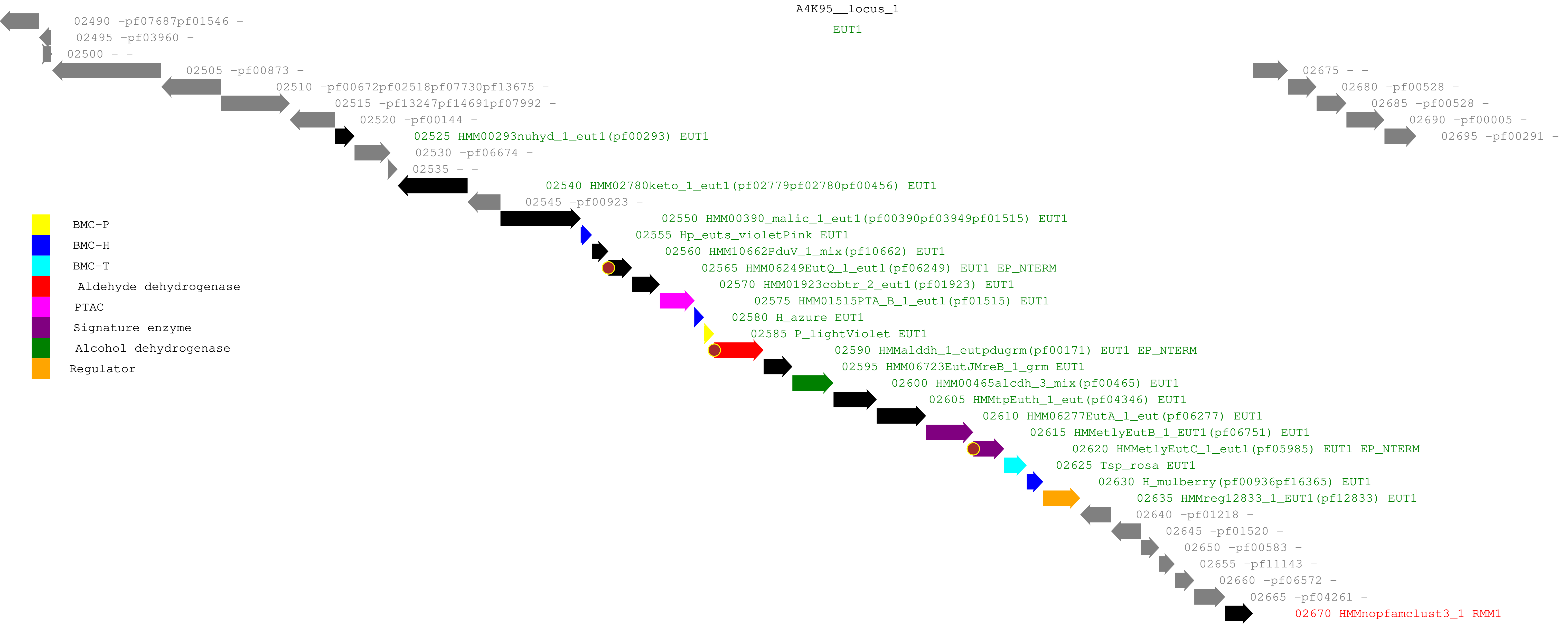

Salmonella dublin. Bacteria; Proteobacteria; Gammaproteobacteria; Enterobacterales;

A4L74__locus_2

EUT1

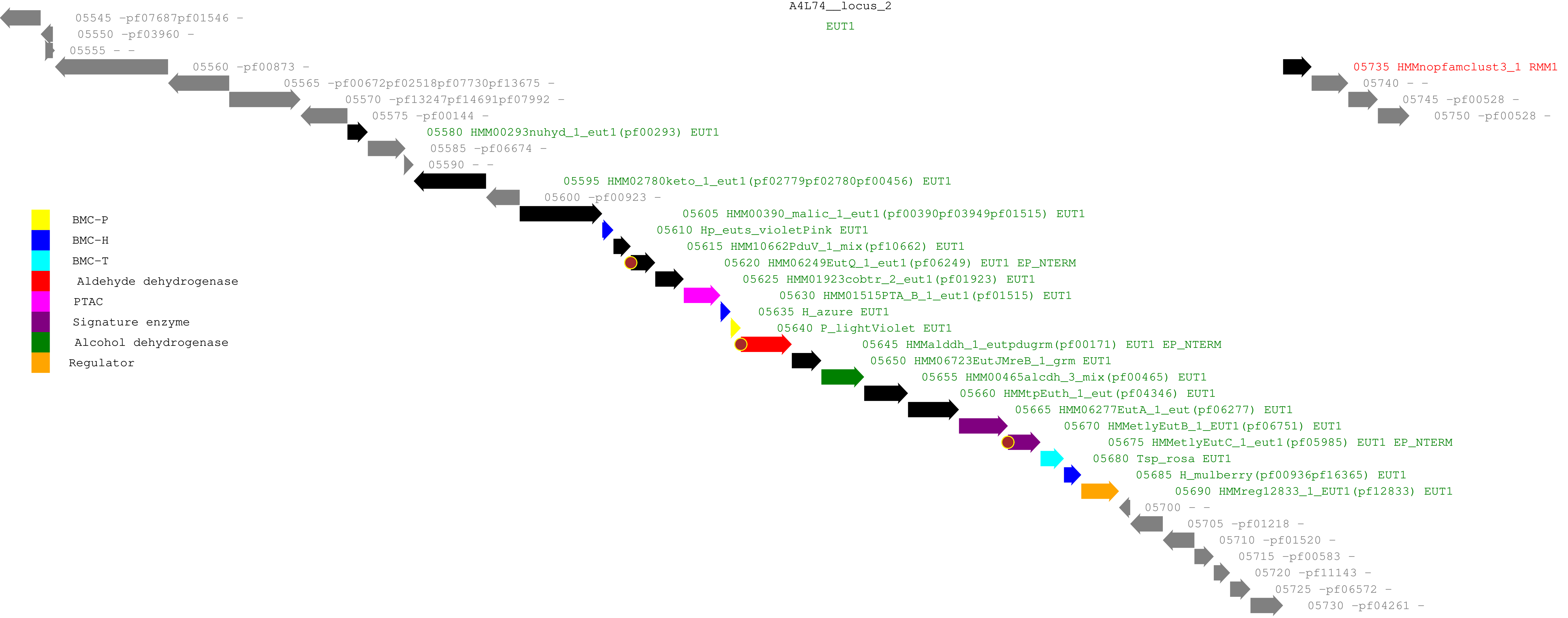

Salmonella virchow. Bacteria; Proteobacteria; Gammaproteobacteria; Enterobacterales;

A4L79__locus_1

EUT1

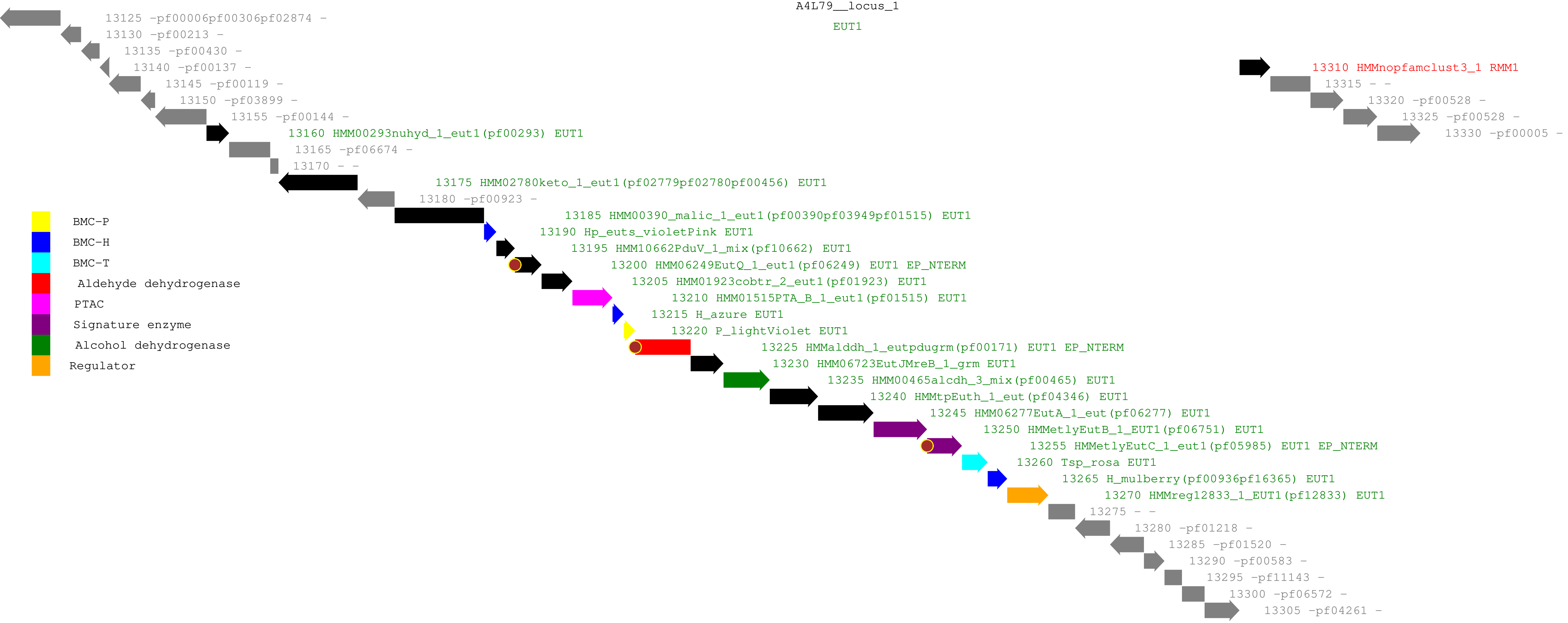

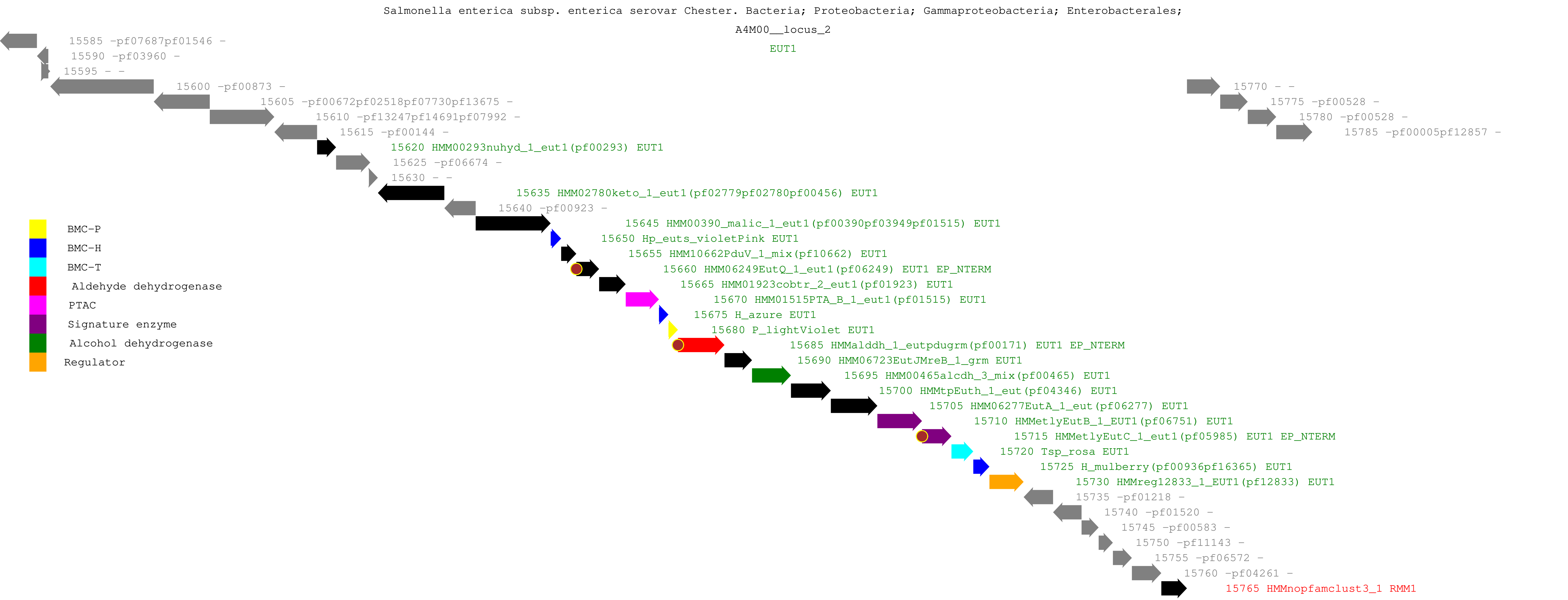

Salmonella enterica subsp. enterica serovar Chester. Bacteria; Proteobacteria; Gammaproteobacteria; Enterobacterales;
A4M00__locus_2
EUT1
15585 -pf07687pf01546 -
15590 -pf03960 -
15595 - -
15600 -pf00873 -
15605 -pf00672pf02518pf07730pf13675 -
15610 -pf13247pf14691pf07992 -
15615 -pf00144 -
15620 HMM00293nuhyd_1_eut1(pf00293) EUT1
15625 -pf06674 -
15630 - -
15635 HMM02780keto_1_eut1(pf02779pf02780pf00456) EUT1
15640 -pf00923 -
15645 HMM00390_malic_1_eut1(pf00390pf03949pf01515) EUT1
15650 Hp_euts_violetPink EUT1
15655 HMM10662PduV_1_mix(pf10662) EUT1
15660 HMM06249EutQ_1_eut1(pf06249) EUT1 EP_NTERM
15665 HMM01923cobtr_2_eut1(pf01923) EUT1
15670 HMM01515PTA_B_1_eut1(pf01515) EUT1
15675 H_azure EUT1
15680 P_lightViolet EUT1
15685 HMMalddh_1_eutpdugrm(pf00171) EUT1 EP_NTERM
15690 HMM06723EutJMreB_1_grm EUT1
15695 HMM00465alcdh_3_mix(pf00465) EUT1
15700 HMMtpEuth_1_eut(pf04346) EUT1
15705 HMM06277EutA_1_eut(pf06277) EUT1
15710 HMMetlyEutB_1_EUT1(pf06751) EUT1
15715 HMMetlyEutC_1_eut1(pf05985) EUT1 EP_NTERM
15720 Tsp_rosa EUT1
15725 H_mulberry(pf00936pf16365) EUT1
15730 HMMreg12833_1_EUT1(pf12833) EUT1
15735 -pf01218 -
15740 -pf01520 -
15745 -pf00583 -
15750 -pf11143 -
15755 -pf06572 -
15760 -pf04261 -
15765 HMMnopfamclust3_1 RMM1
15770 - -
15775 -pf00528 -
15780 -pf00528 -
15785 -pf00005pf12857 -
BMC-P
BMC-H
BMC-T
Aldehyde dehydrogenase
PTAC
Signature enzyme
Alcohol dehydrogenase
Regulator

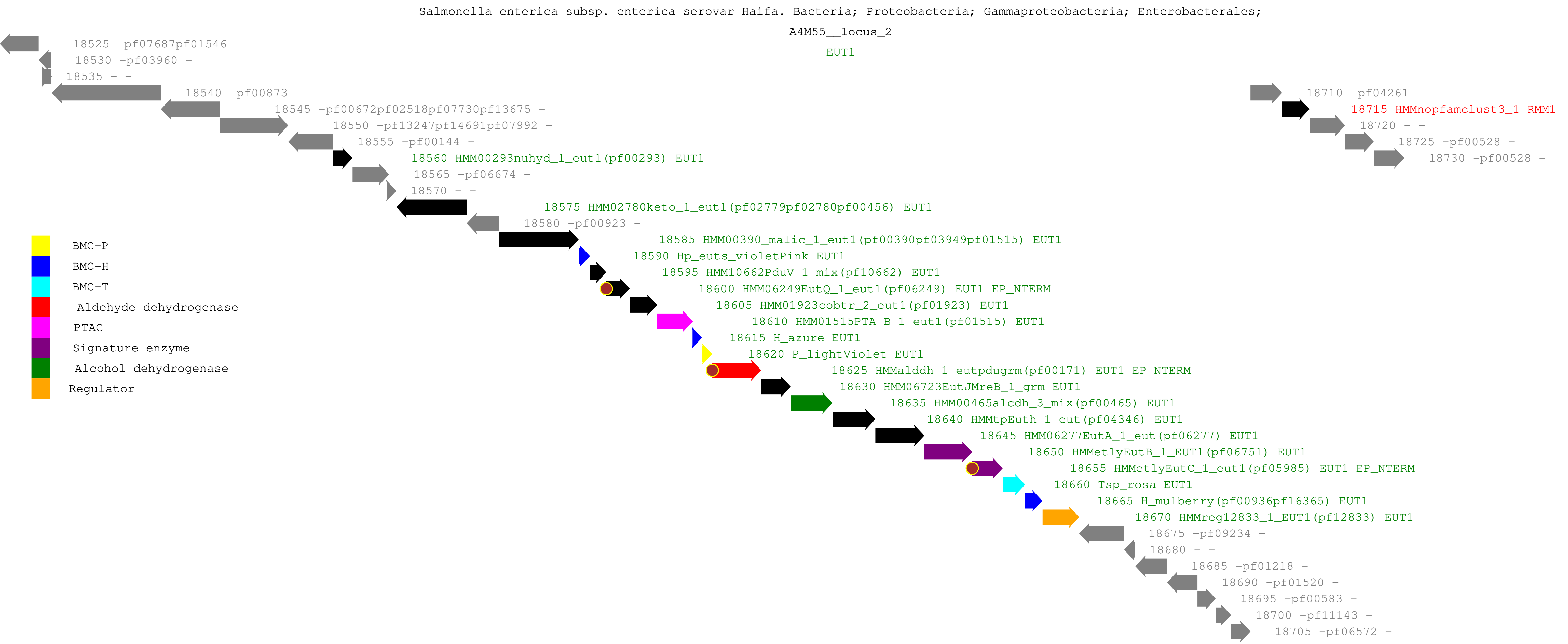
Salmonella enterica subsp. enterica serovar Haifa. Bacteria; Proteobacteria; Gammaproteobacteria; Enterobacterales;
A4M55__locus_2
EUT1
18525 -pf07687pf01546 -
18530 -pf03960 -
18535 - -
18540 -pf00873 -
18545 -pf00672pf02518pf07730pf13675 -
18550 -pf13247pf14691pf07992 -
18555 -pf00144 -
18560 HMM00293nuhyd_1_eut1(pf00293) EUT1
18565 -pf06674 -
18570 - -
18575 HMM02780keto_1_eut1(pf02779pf02780pf00456) EUT1
18580 -pf00923 -
18585 HMM00390_malic_1_eut1(pf00390pf03949pf01515) EUT1
18590 Hp_euts_violetPink EUT1
18595 HMM10662PduV_1_mix(pf10662) EUT1
18600 HMM06249EutQ_1_eut1(pf06249) EUT1 EP_NTERM
18605 HMM01923cobtr_2_eut1(pf01923) EUT1
18610 HMM01515PTA_B_1_eut1(pf01515) EUT1
18615 H_azure EUT1
18620 P_lightViolet EUT1
18625 HMMalddh_1_eutpdugrm(pf00171) EUT1 EP_NTERM
18630 HMM06723EutJMreB_1_grm EUT1
18635 HMM00465alcdh_3_mix(pf00465) EUT1
18640 HMMtpEuth_1_eut(pf04346) EUT1
18645 HMM06277EutA_1_eut(pf06277) EUT1
18650 HMMetlyEutB_1_EUT1(pf06751) EUT1
18655 HMMetlyEutC_1_eut1(pf05985) EUT1 EP_NTERM
18660 Tsp_rosa EUT1
18665 H_mulberry(pf00936pf16365) EUT1
18670 HMMreg12833_1_EUT1(pf12833) EUT1
18675 -pf09234 -
18680 - -
18685 -pf01218 -
18690 -pf01520 -
18695 -pf00583 -
18700 -pf11143 -
18705 -pf06572 -
18710 -pf04261 -
18715 HMMnopfamclust3_1 RMM1
18720 - -
18725 -pf00528 -
18730 -pf00528 -
BMC-P
BMC-H
BMC-T
Aldehyde dehydrogenase
PTAC
Signature enzyme
Alcohol dehydrogenase
Regulator

Salmonella enterica subsp. enterica serovar Braenderup. Bacteria; Proteobacteria; Gammaproteobacteria; Enterobacterales;

A4N09__locus_2

EUT1

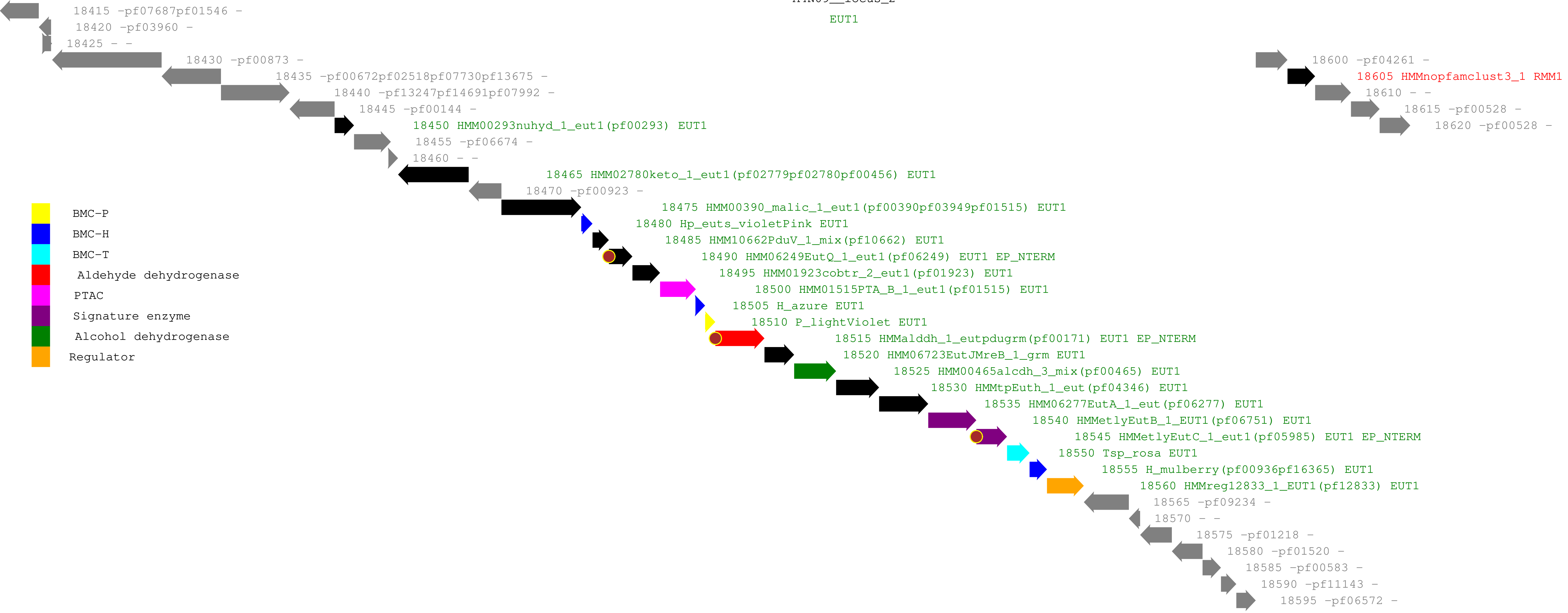

Salmonella enterica subsp. enterica serovar Isangi. Bacteria; Proteobacteria; Gammaproteobacteria; Enterobacterales;

A4N97__locus_1

EUT1

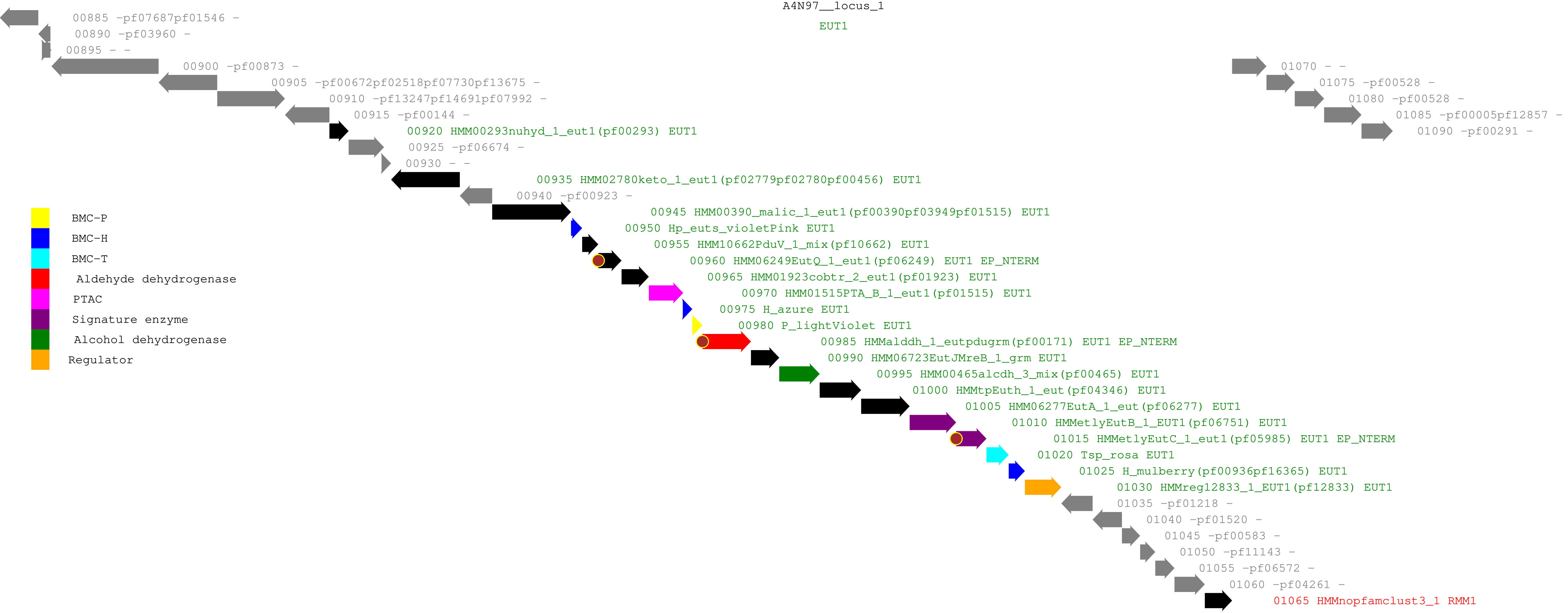

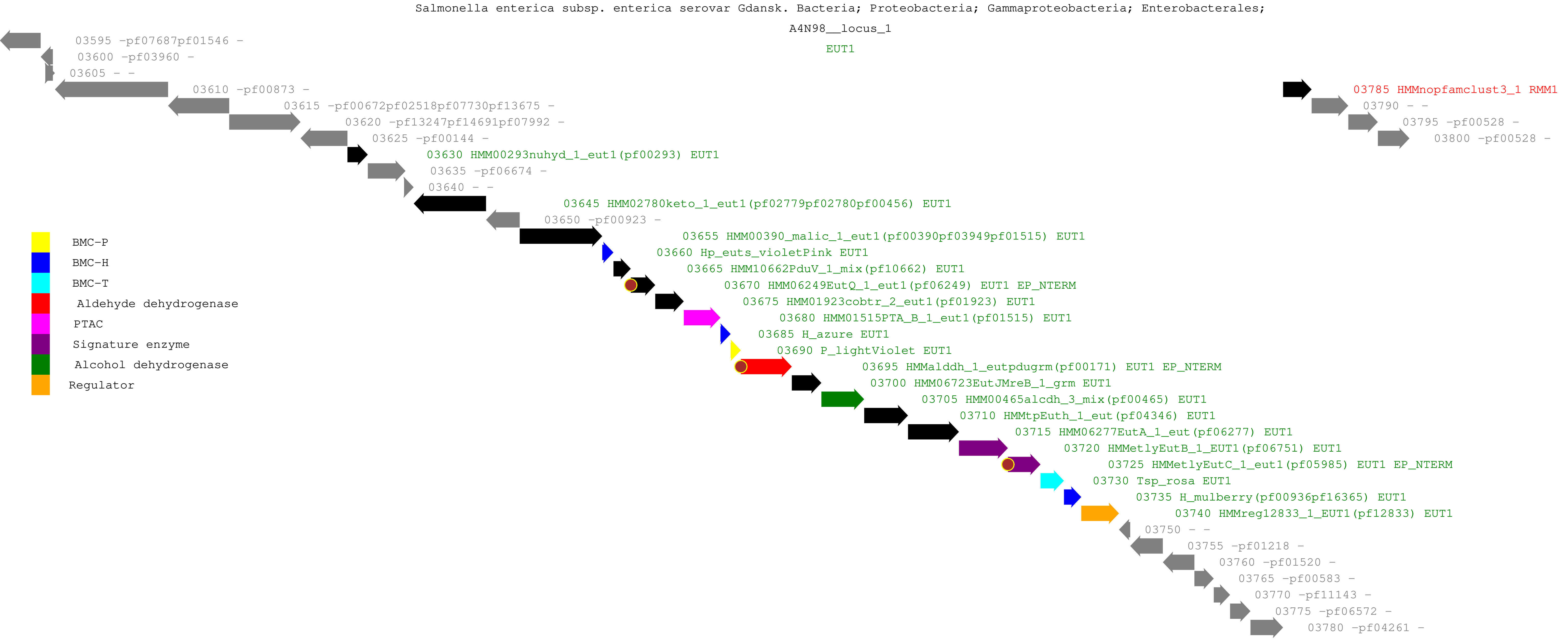

Salmonella enterica subsp. enterica serovar Gdansk. Bacteria; Proteobacteria; Gammaproteobacteria; Enterobacterales;
A4N98__locus_1
EUT1
03595 -pf07687pf01546 -
03600 -pf03960 -
03605 - -
03610 -pf00873 -
03615 -pf00672pf02518pf07730pf13675 -
03620 -pf13247pf14691pf07992 -
03625 -pf00144 -
03630 HMM00293nuhyd_1_eut1(pf00293) EUT1
03635 -pf06674 -
03640 - -
03645 HMM02780keto_1_eut1(pf02779pf02780pf00456) EUT1
03650 -pf00923 -
03655 HMM00390_malic_1_eut1(pf00390pf03949pf01515) EUT1
03660 Hp_euts_violetPink EUT1
03665 HMM10662PduV_1_mix(pf10662) EUT1
03670 HMM06249EutQ_1_eut1(pf06249) EUT1 EP_NTERM
03675 HMM01923cobtr_2_eut1(pf01923) EUT1
03680 HMM01515PTA_B_1_eut1(pf01515) EUT1
03685 H_azure EUT1
03690 P_lightViolet EUT1
03695 HMMalddh_1_eutpdugrm(pf00171) EUT1 EP_NTERM
03700 HMM06723EutJMreB_1_grm EUT1
03705 HMM00465alcdh_3_mix(pf00465) EUT1
03710 HMMtpEuth_1_eut(pf04346) EUT1
03715 HMM06277EutA_1_eut(pf06277) EUT1
03720 HMMetlyEutB_1_EUT1(pf06751) EUT1
03725 HMMetlyEutC_1_eut1(pf05985) EUT1 EP_NTERM
03730 Tsp_rosa EUT1
03735 H_mulberry(pf00936pf16365) EUT1
03740 HMMreg12833_1_EUT1(pf12833) EUT1
03750 - -
03755 -pf01218 -
03760 -pf01520 -
03765 -pf00583 -
03770 -pf11143 -
03775 -pf06572 -
03780 -pf04261 -
03785 HMMnopfamclust3_1 RMM1
03790 - -
03795 -pf00528 -
03800 -pf00528 -
BMC-P
BMC-H
BMC-T
Aldehyde dehydrogenase
PTAC
Signature enzyme
Alcohol dehydrogenase
Regulator

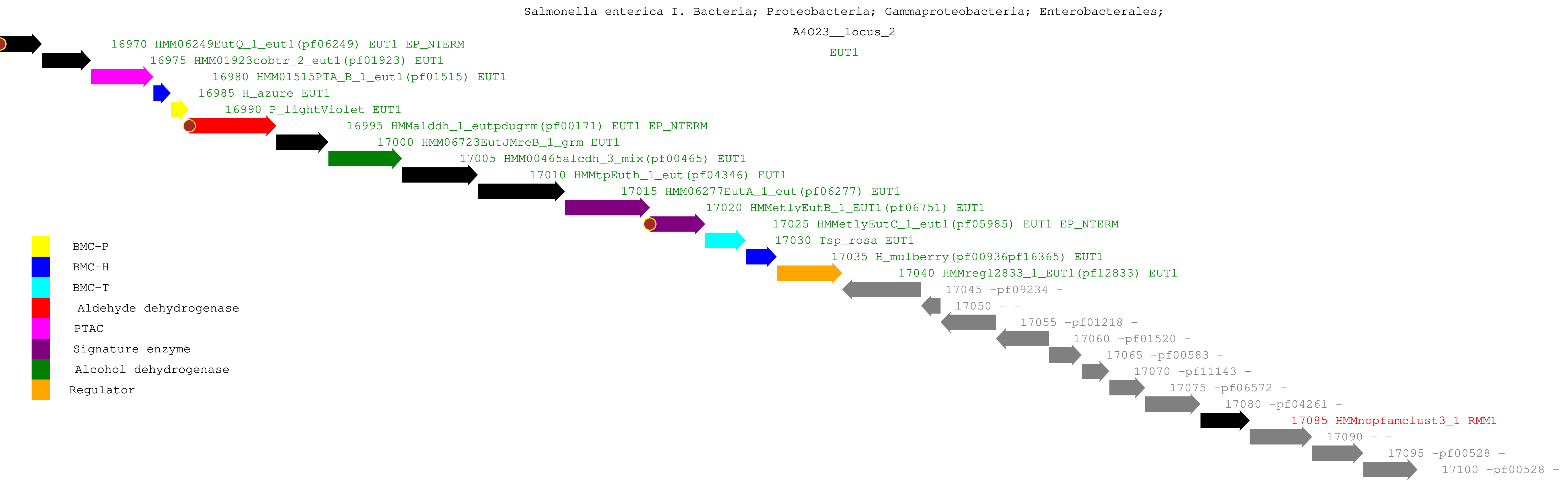
Salmonella enterica I. Bacteria; Proteobacteria; Gammaproteobacteria; Enterobacterales;
A4023__locus_2
EUT1
16970 HMM06249EutQ_1_eut1(pf06249) EUT1 EP_NTERM
16975 HMM01923cobtr_2_eut1(pf01923) EUT1
16980 HMM01515PTA_B_1_eut1(pf01515) EUT1
16985 H_azure EUT1
16990 P_lightViolet EUT1
16995 HMMalddh_1_eutpdugrm(pf00171) EUT1 EP_NTERM
17000 HMM06723EutJMreB_1_grm EUT1
17005 HMM00465alcdh_3_mix(pf00465) EUT1
17010 HMMtpEuth_1_eut(pf04346) EUT1
17015 HMM06277EutA_1_eut(pf06277) EUT1
17020 HMMetlyEutB_1_EUT1(pf06751) EUT1
17025 HMMetlyEutC_1_eut1(pf05985) EUT1 EP_NTERM
17030 Tsp_rosa EUT1
17035 H_mulberry(pf00936pf16365) EUT1
17040 HMMreg12833_1_EUT1(pf12833) EUT1
17045 -pf09234 -
17050 - -
17055 -pf01218 -
17060 -pf01520 -
17065 -pf00583 -
17070 -pf11143 -
17075 -pf06572 -
17080 -pf04261 -
17085 HMMnopfamclust3_1 RMM1
17090 - -
17095 -pf00528 -
17100 -pf00528 -
BMC-P
BMC-H
BMC-T
Aldehyde dehydrogenase
PTAC
Signature enzyme
Alcohol dehydrogenase
Regulator

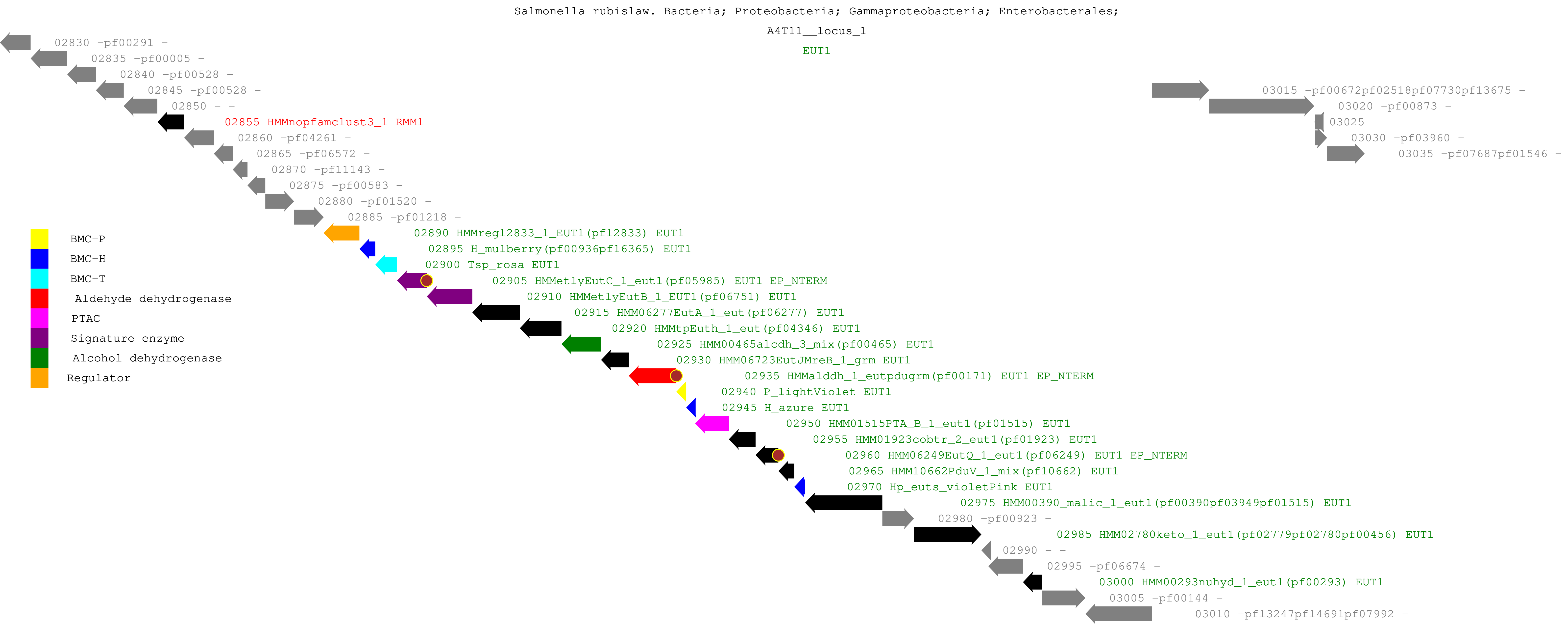

Salmonella rubislaw. Bacteria; Proteobacteria; Gammaproteobacteria; Enterobacterales;
A4T11__locus_1
EUT1
02830 -pf00291 -
02835 -pf00005 -
02840 -pf00528 -
02845 -pf00528 -
02850 - -
02855 HMMnopfamclust3_1 RMM1
02860 -pf04261 -
02865 -pf06572 -
02870 -pf11143 -
02875 -pf00583 -
02880 -pf01520 -
02885 -pf01218 -
02890 HMMreg12833_1_EUT1(pf12833) EUT1
02895 H_mulberry(pf00936pf16365) EUT1
02900 Tsp_rosa EUT1
02905 HMMetlyEutC_1_eut1(pf05985) EUT1 EP_NTERM
02910 HMMetlyEutB_1_EUT1(pf06751) EUT1
02915 HMM06277EutA_1_eut(pf06277) EUT1
02920 HMMtpEuth_1_eut(pf04346) EUT1
02925 HMM00465alcdh_3_mix(pf00465) EUT1
02930 HMM06723EutJMreB_1_grm EUT1
02935 HMMalddh_1_eutpdugrm(pf00171) EUT1 EP_NTERM
02940 P_lightViolet EUT1
02945 H_azure EUT1
02950 HMM01515PTA_B_1_eut1(pf01515) EUT1
02955 HMM01923cobtr_2_eut1(pf01923) EUT1
02960 HMM06249EutQ_1_eut1(pf06249) EUT1 EP_NTERM
02965 HMM10662PduV_1_mix(pf10662) EUT1
02970 Hp_euts_violetPink EUT1
02975 HMM00390_malic_1_eut1(pf00390pf03949pf01515) EUT1
02980 -pf00923 -
02985 HMM02780keto_1_eut1(pf02779pf02780pf00456) EUT1
02990 - -
02995 -pf06674 -
03000 HMM00293nuhyd_1_eut1(pf00293) EUT1
03005 -pf00144 -
03010 -pf13247pf14691pf07992 -
03015 -pf00672pf02518pf07730pf13675 -
03020 -pf00873 -
03025 - -
03030 -pf03960 -
03035 -pf07687pf01546 -
BMC-P
BMC-H
BMC-T
Aldehyde dehydrogenase
PTAC
Signature enzyme
Alcohol dehydrogenase
Regulator

Escherichia coli. Bacteria; Proteobacteria; Gammaproteobacteria; Enterobacterales;

A4T40__locus_1

EUT1

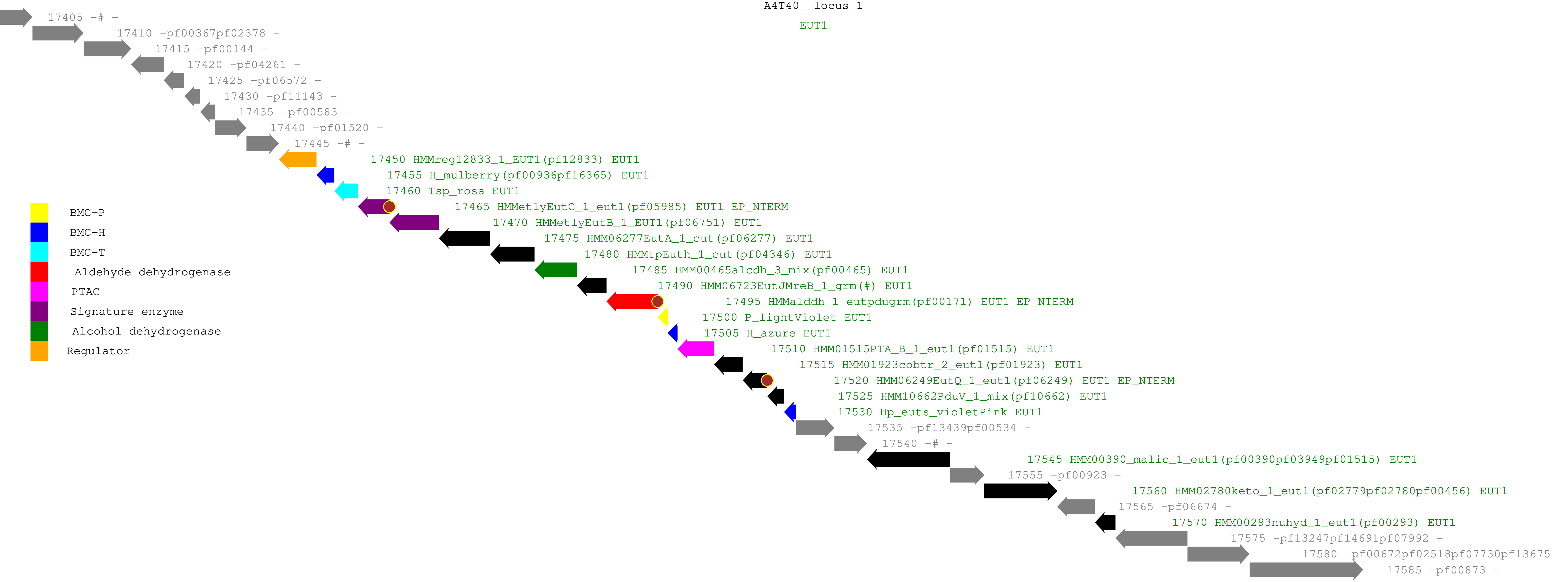

Escherichia coli. Bacteria; Proteobacteria; Gammaproteobacteria; Enterobacterales;

A4T50__locus_1

EUT1

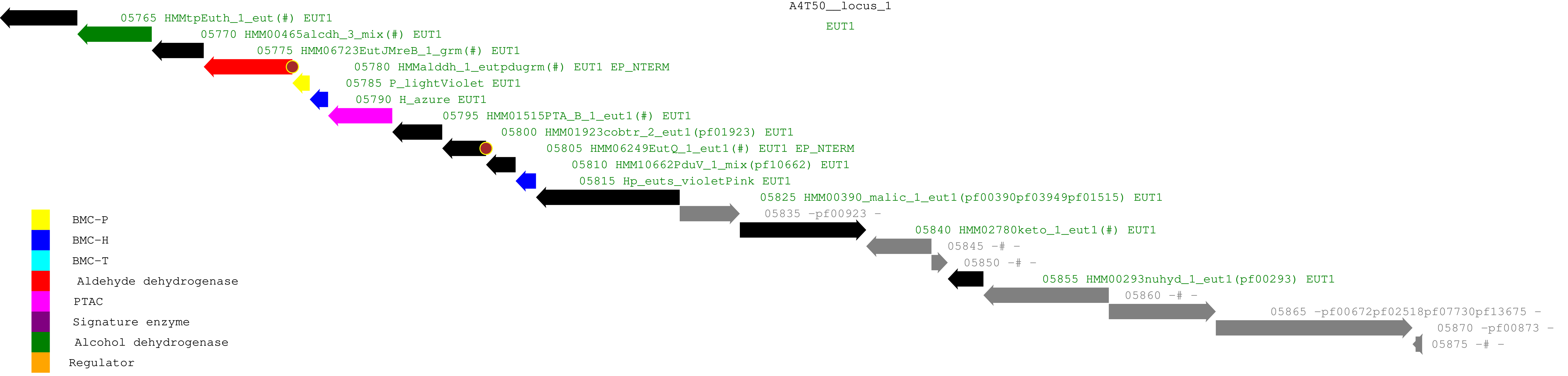

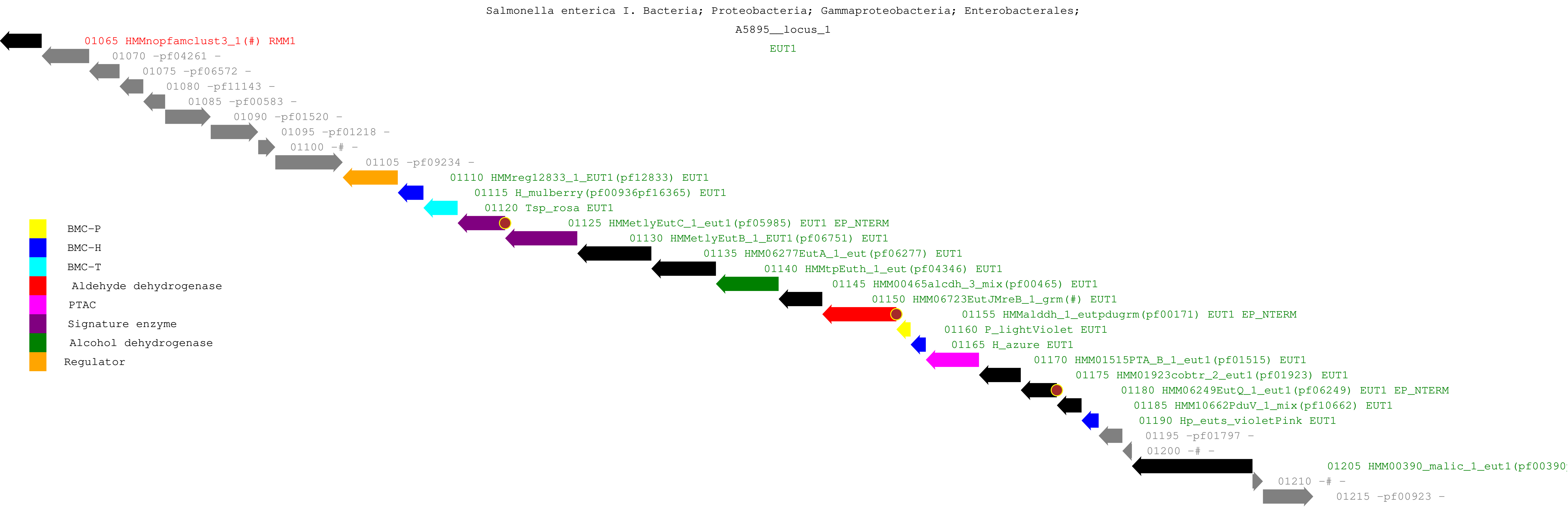
Salmonella enterica I. Bacteria; Proteobacteria; Gammaproteobacteria; Enterobacterales;
A5895__locus_1
EUT1
01065 HMMnopfamclust3_1(#) RMM1
01070 -pf04261 -
01075 -pf06572 -
01080 -pf11143 -
01085 -pf00583 -
01090 -pf01520 -
01095 -pf01218 -
01100 -# -
01105 -pf09234 -
01110 HMMreg12833_1_EUT1(pf12833) EUT1
01115 H_mulberry(pf00936pf16365) EUT1
01120 Tsp_rosa EUT1
01125 HMMetlyEutC_1_eut1(pf05985) EUT1 EP_NTERM
01130 HMMetlyEutB_1_EUT1(pf06751) EUT1
01135 HMM06277EutA_1_eut(pf06277) EUT1
01140 HMMtpEuth_1_eut(pf04346) EUT1
01145 HMM00465alcdh_3_mix(pf00465) EUT1
01150 HMM06723EutJMreB_1_grm(#) EUT1
01155 HMMalddh_1_eutpdugrm(pf00171) EUT1 EP_NTERM
01160 P_lightViolet EUT1
01165 H_azure EUT1
01170 HMM01515PTA_B_1_eut1(pf01515) EUT1
01175 HMM01923cobtr_2_eut1(pf01923) EUT1
01180 HMM06249EutQ_1_eut1(pf06249) EUT1 EP_NTERM
01185 HMM10662PduV_1_mix(pf10662) EUT1
01190 Hp_euts_violetPink EUT1
01195 -pf01797 -
01200 -# -
01205 HMM00390_malic_1_eut1(pf00390
01210 -# -
01215 -pf00923 -
BMC-P
BMC-H
BMC-T
Aldehyde dehydrogenase
PTAC
Signature enzyme
Alcohol dehydrogenase
Regulator

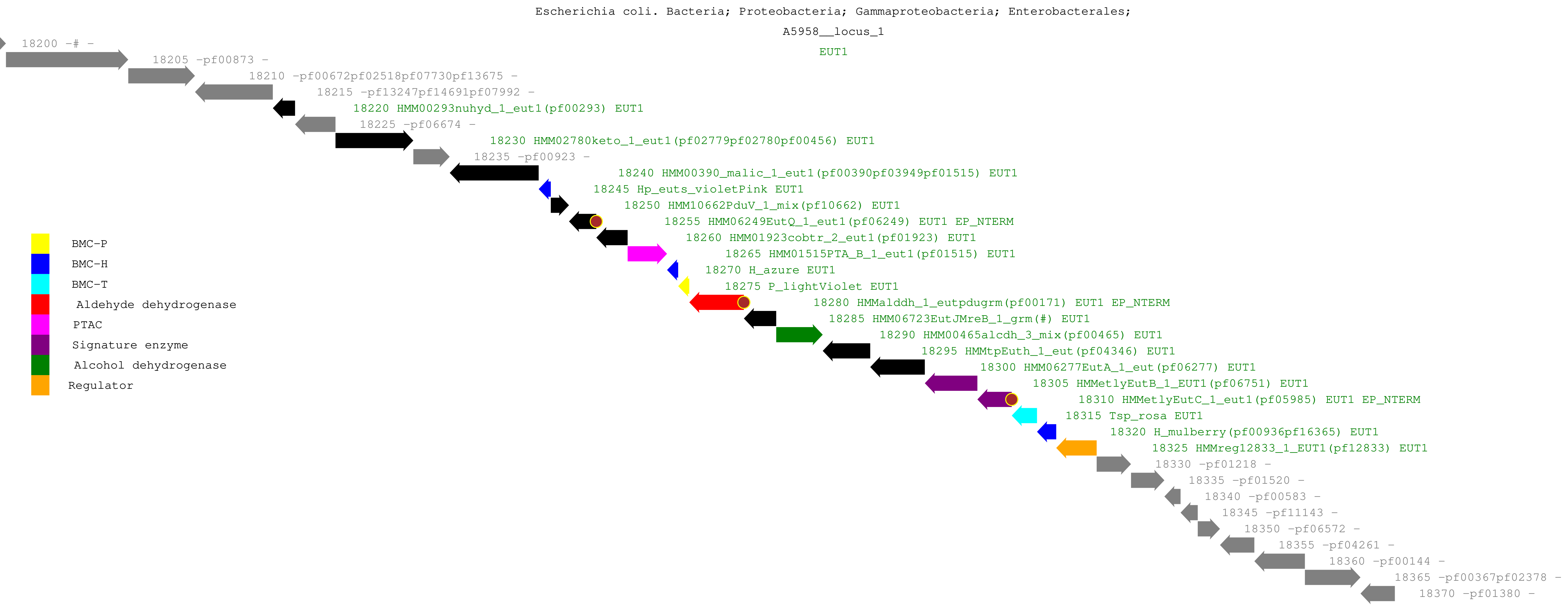

Escherichia coli. Bacteria; Proteobacteria; Gammaproteobacteria; Enterobacterales;
A5958__locus_1
EUT1
18200 -# -
18205 -pf00873 -
18210 -pf00672pf02518pf07730pf13675 -
18215 -pf13247pf14691pf07992 -
18220 HMM00293nuhyd_1_eut1(pf00293) EUT1
18225 -pf06674 -
18230 HMM02780keto_1_eut1(pf02779pf02780pf00456) EUT1
18235 -pf00923 -
18240 HMM00390_malic_1_eut1(pf00390pf03949pf01515) EUT1
18245 Hp_euts_violetPink EUT1
18250 HMM10662PduV_1_mix(pf10662) EUT1
18255 HMM06249EutQ_1_eut1(pf06249) EUT1 EP_NTERM
18260 HMM01923cobtr_2_eut1(pf01923) EUT1
18265 HMM01515PTA_B_1_eut1(pf01515) EUT1
18270 H_azure EUT1
18275 P_lightViolet EUT1
18280 HMMalddh_1_eutpdugrm(pf00171) EUT1 EP_NTERM
18285 HMM06723EutJMreB_1_grm(#) EUT1
18290 HMM00465alcdh_3_mix(pf00465) EUT1
18295 HMMtpEuth_1_eut(pf04346) EUT1
18300 HMM06277EutA_1_eut(pf06277) EUT1
18305 HMMetlyEutB_1_EUT1(pf06751) EUT1
18310 HMMetlyEutC_1_eut1(pf05985) EUT1 EP_NTERM
18315 Tsp_rosa EUT1
18320 H_mulberry(pf00936pf16365) EUT1
18325 HMMreg12833_1_EUT1(pf12833) EUT1
18330 -pf01218 -
18335 -pf01520 -
18340 -pf00583 -
18345 -pf11143 -
18350 -pf06572 -
18355 -pf04261 -
18360 -pf00144 -
18365 -pf00367pf02378 -
18370 -pf01380 -
BMC-P
BMC-H
BMC-T
Aldehyde dehydrogenase
PTAC
Signature enzyme
Alcohol dehydrogenase
Regulator

Escherichia coli NCCP15648. Bacteria; Proteobacteria; Gammaproteobacteria; Enterobacterales;

A610__locus_1

EUT1

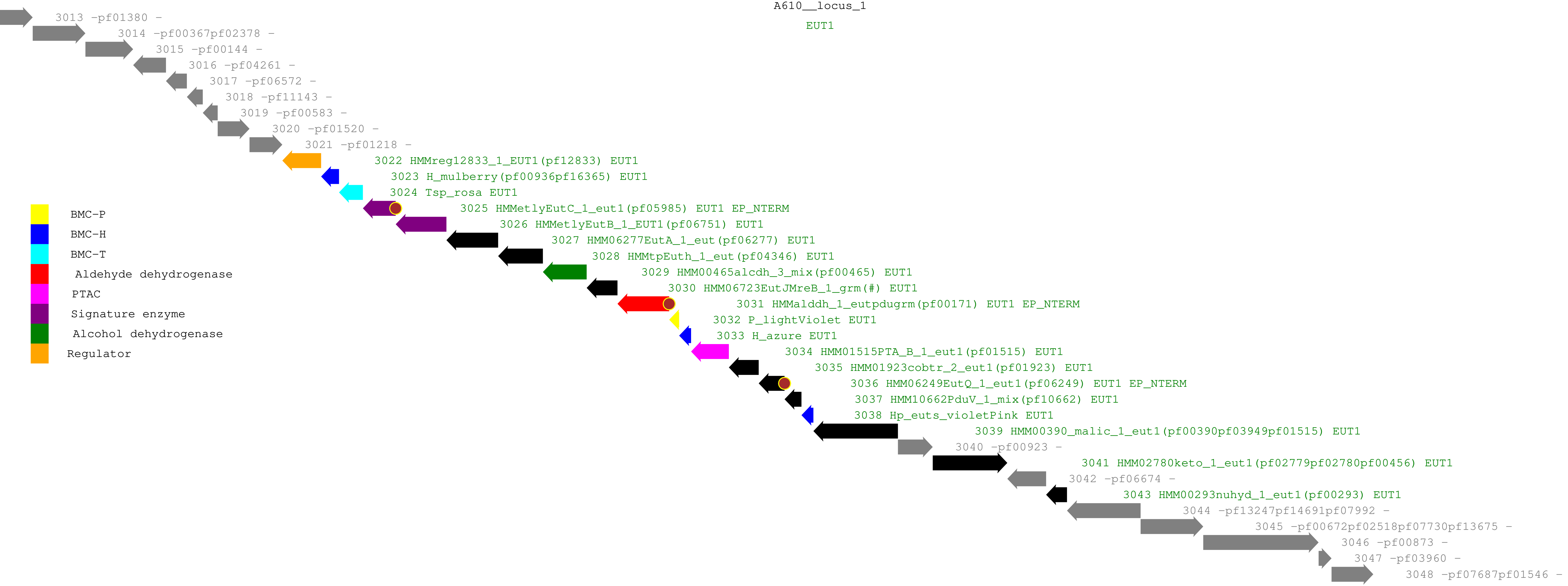

Salmonella enterica subsp. enterica serovar Cubana str. 76814. Bacteria; Proteobacteria; Gammaproteobacteria; Enterobacterales;

A628__locus_1

EUT1

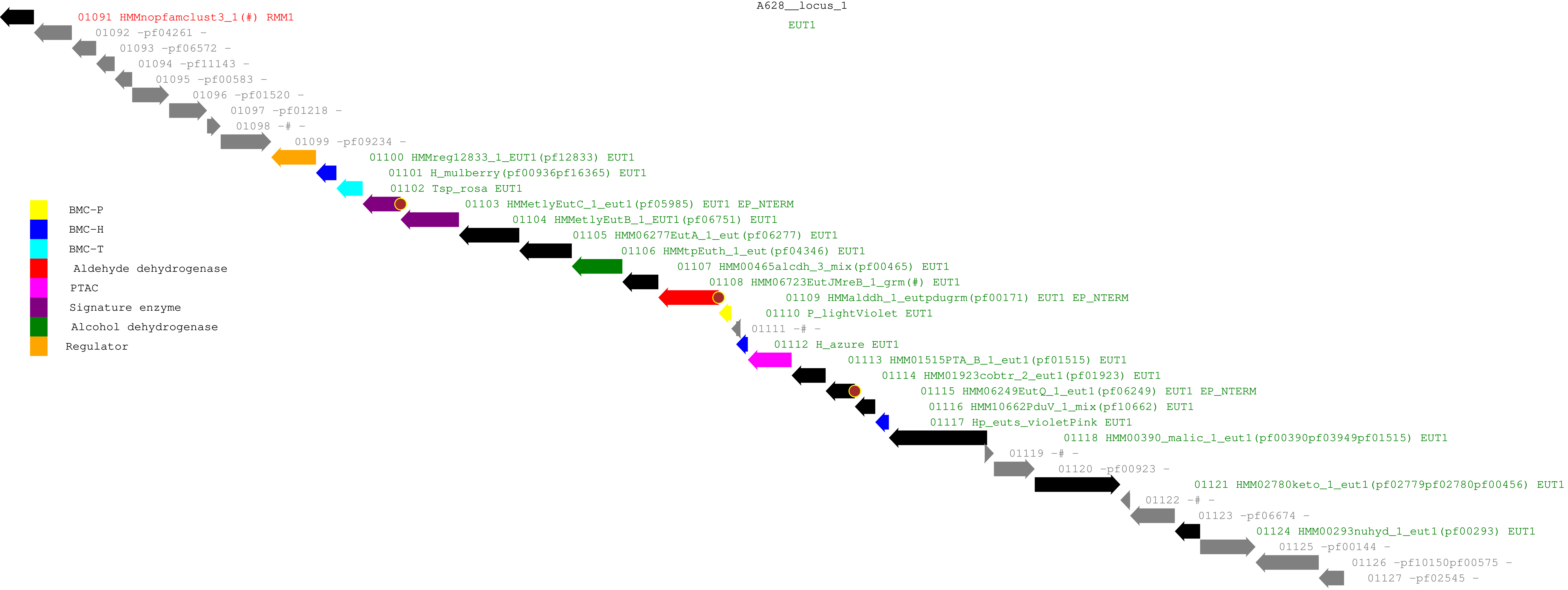

Salmonella enterica subsp. enterica serovar Dublin str. UC16. Bacteria; Proteobacteria; Gammaproteobacteria; Enterobacterales;

A670__locus_1

EUT1

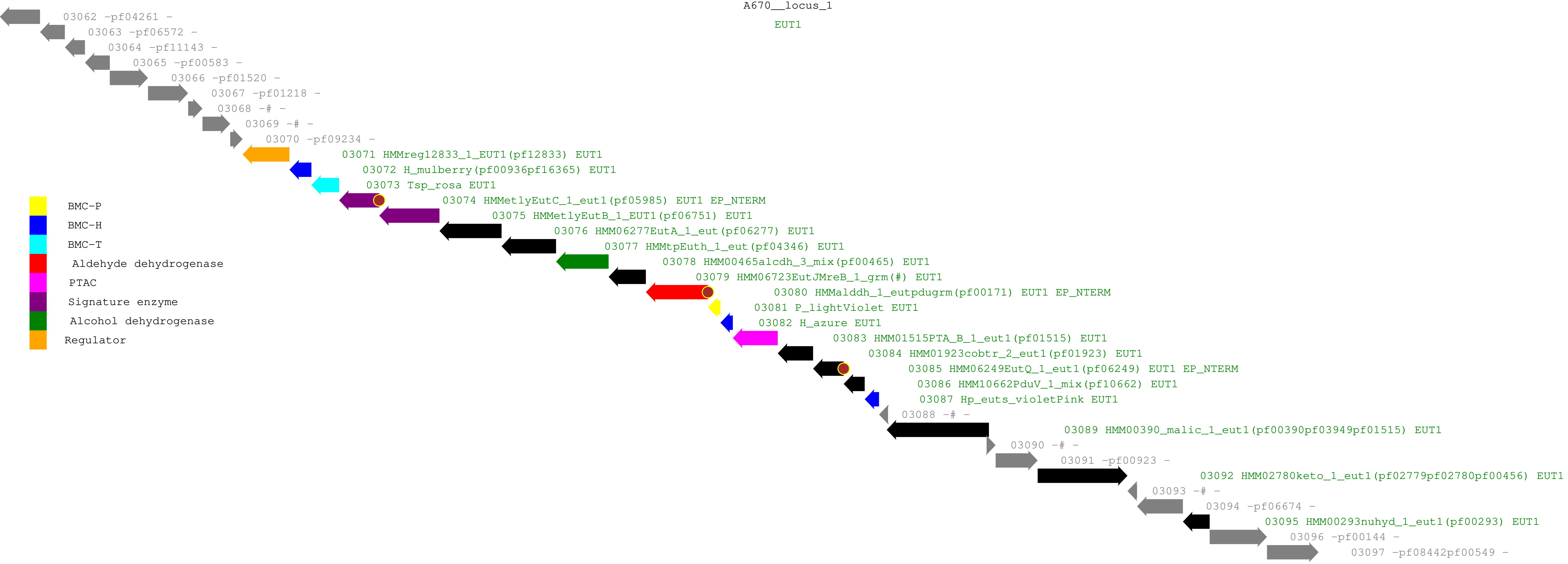

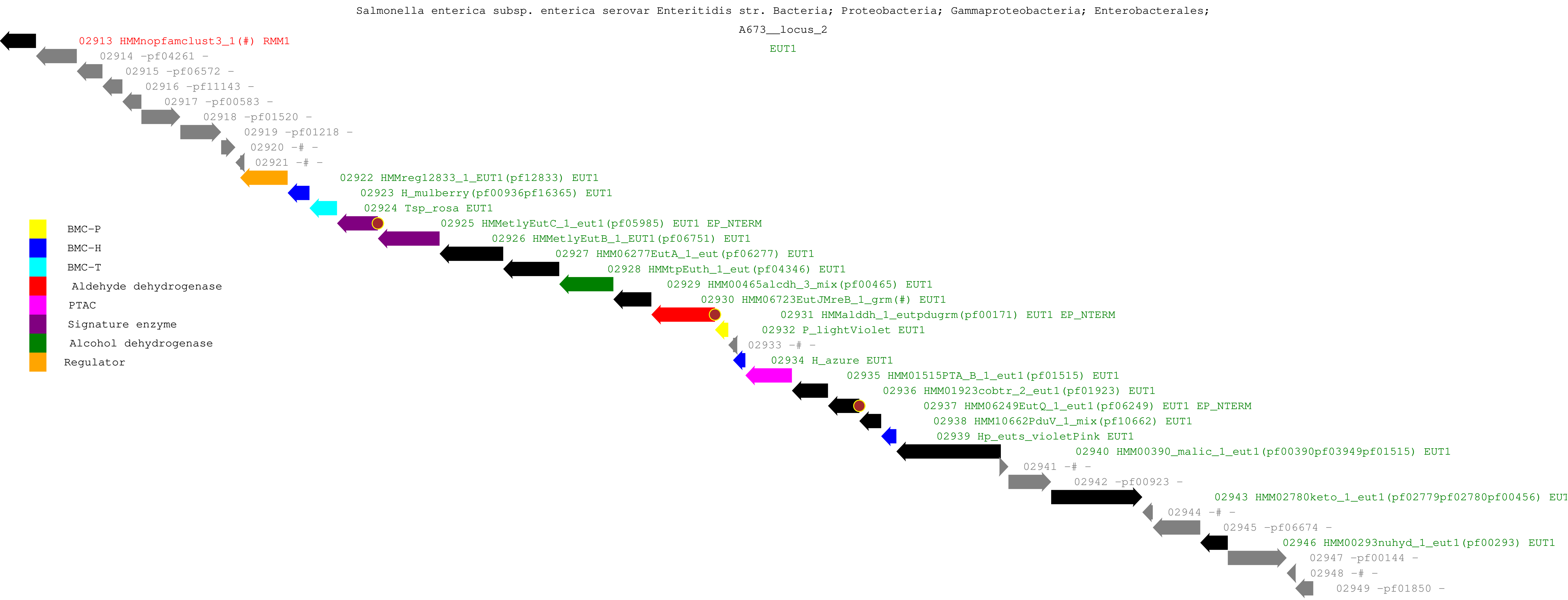
Salmonella enterica subsp. enterica serovar Enteritidis str. Bacteria; Proteobacteria; Gammaproteobacteria; Enterobacterales;
A673__locus_2
EUT1
02913 HMMnopfamclust3_1(#) RMM1
02914 -pf04261 -
02915 -pf06572 -
02916 -pf11143 -
02917 -pf00583 -
02918 -pf01520 -
02919 -pf01218 -
02920 -# -
02921 -# -
02922 HMMreg12833_1_EUT1(pf12833) EUT1
02923 H_mulberry(pf00936pf16365) EUT1
02924 Tsp_rosa EUT1
02925 HMMetlyEutC_1_eut1(pf05985) EUT1 EP_NTERM
02926 HMMetlyEutB_1_EUT1(pf06751) EUT1
02927 HMM06277EutA_1_eut(pf06277) EUT1
02928 HMMtpEuth_1_eut(pf04346) EUT1
02929 HMM00465alcdh_3_mix(pf00465) EUT1
02930 HMM06723EutJMreB_1_grm(#) EUT1
02931 HMMalddh_1_eutpdugrm(pf00171) EUT1 EP_NTERM
02932 P_lightViolet EUT1
02933 -# -
02934 H_azure EUT1
02935 HMM01515PTA_B_1_eut1(pf01515) EUT1
02936 HMM01923cobtr_2_eut1(pf01923) EUT1
02937 HMM06249EutQ_1_eut1(pf06249) EUT1 EP_NTERM
02938 HMM10662PduV_1_mix(pf10662) EUT1
02939 Hp_euts_violetPink EUT1
02940 HMM00390_malic_1_eut1(pf00390pf03949pf01515) EUT1
02941 -# -
02942 -pf00923 -
02943 HMM02780keto_1_eut1(pf02779pf02780pf00456) EUT
02944 -# -
02945 -pf06674 -
02946 HMM00293nuhyd_1_eut1(pf00293) EUT1
02947 -pf00144 -
02948 -# -
02949 -pf01850 -
BMC-P
BMC-H
BMC-T
Aldehyde dehydrogenase
PTAC
Signature enzyme
Alcohol dehydrogenase
Regulator

Klebsiella michiganensis. Bacteria; Proteobacteria; Gammaproteobacteria; Enterobacterales;

A6A30__locus_2

EUT1

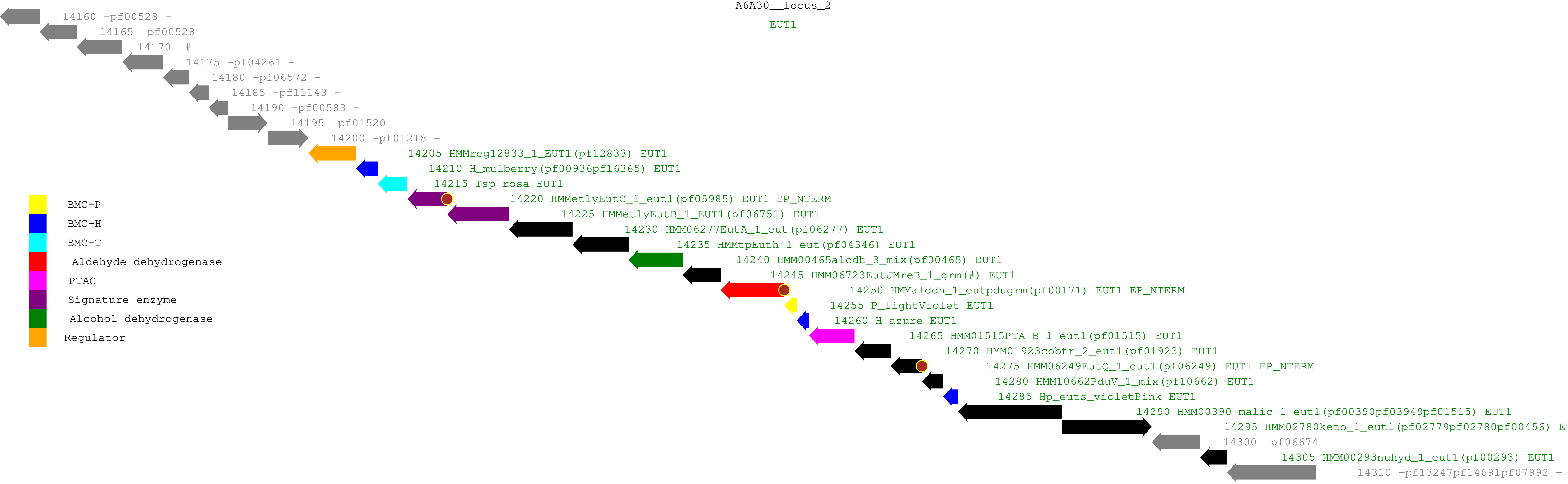

Salmonella enterica subsp. enterica serovar Rissen. Bacteria; Proteobacteria; Gammaproteobacteria; Enterobacterales;

A6S79__locus_1

EUT1

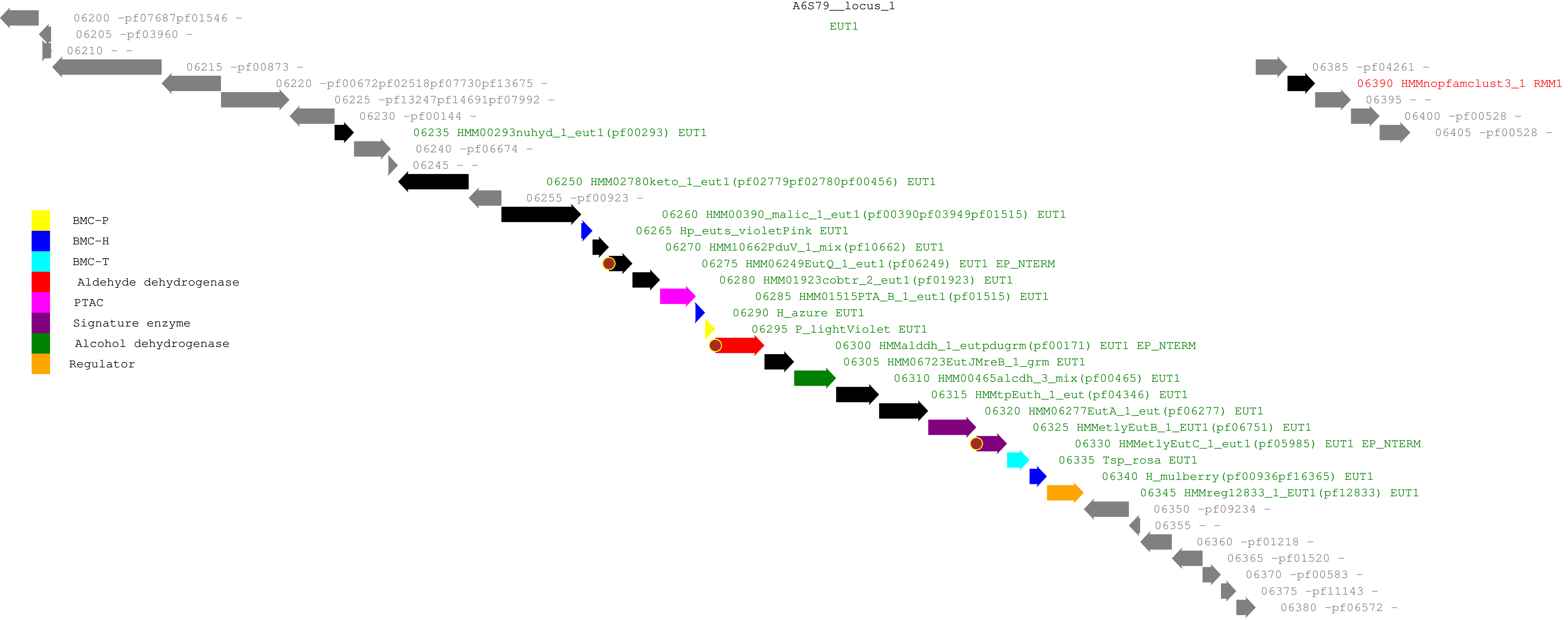

Salmonella rubislaw. Bacteria; Proteobacteria; Gammaproteobacteria; Enterobacterales;

A7C81__locus_1

EUT1

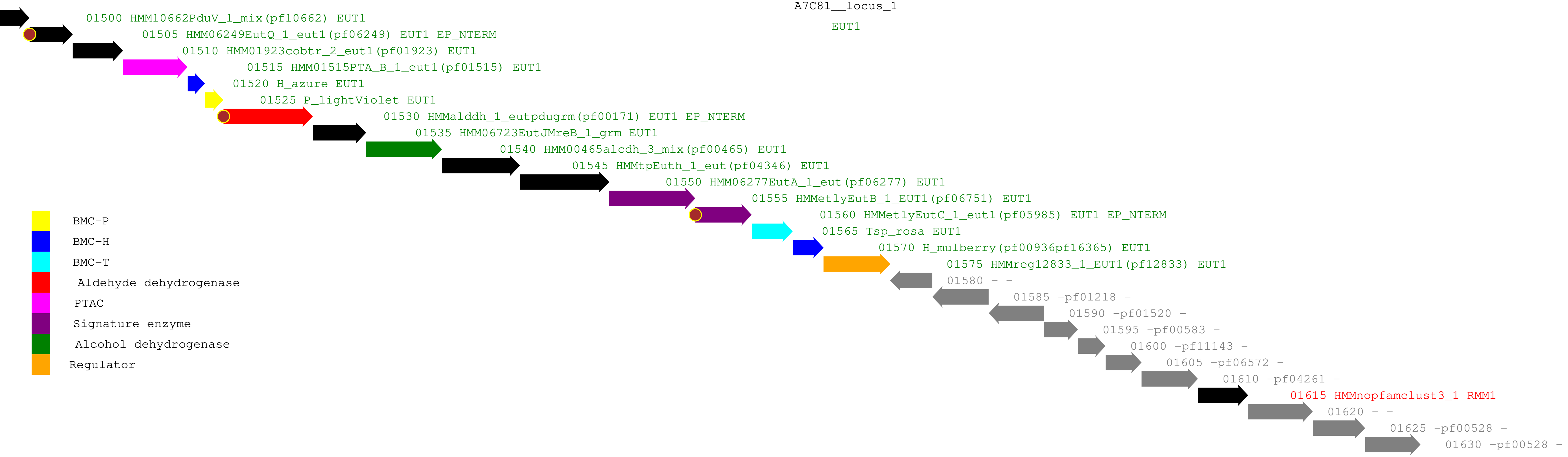

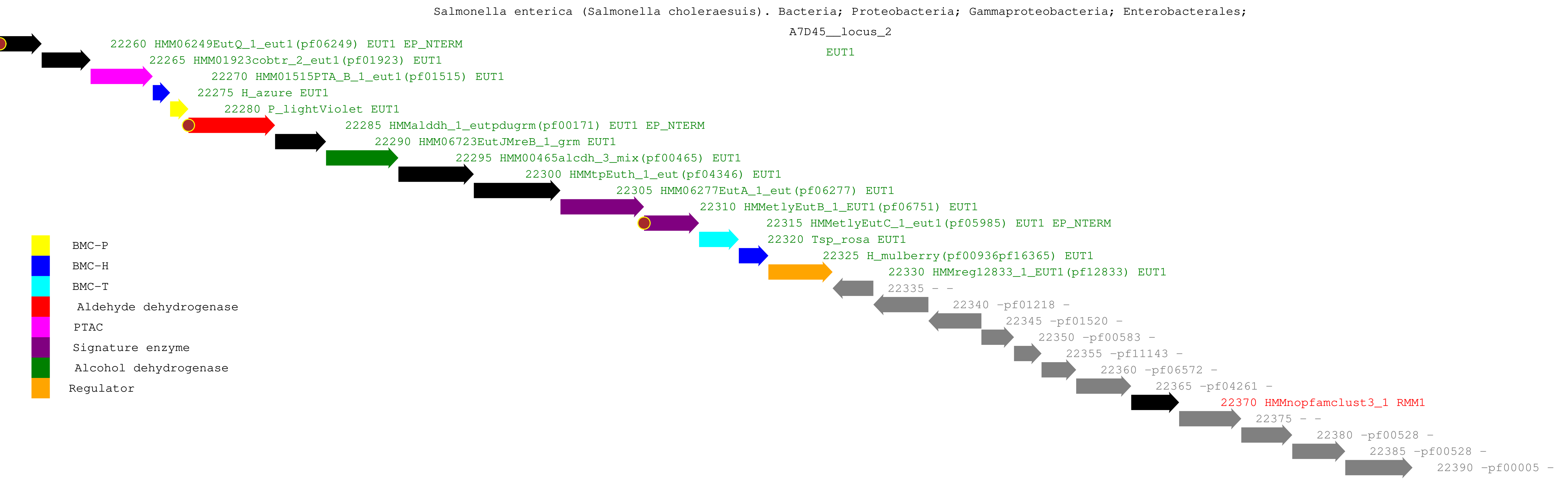
Salmonella enterica (Salmonella choleraesuis). Bacteria; Proteobacteria; Gammaproteobacteria; Enterobacterales;
A7D45__locus_2
EUT1
22260 HMM06249EutQ_1_eut1(pf06249) EUT1 EP_NTERM
22265 HMM01923cobtr_2_eut1(pf01923) EUT1
22270 HMM01515PTA_B_1_eut1(pf01515) EUT1
22275 H_azure EUT1
22280 P_lightViolet EUT1
22285 HMMalddh_1_eutpdugrm(pf00171) EUT1 EP_NTERM
22290 HMM06723EutJMreB_1_grm EUT1
22295 HMM00465alcdh_3_mix(pf00465) EUT1
22300 HMMtpEuth_1_eut(pf04346) EUT1
22305 HMM06277EutA_1_eut(pf06277) EUT1
22310 HMMetlyEutB_1_EUT1(pf06751) EUT1
22315 HMMetlyEutC_1_eut1(pf05985) EUT1 EP_NTERM
22320 Tsp_rosa EUT1
22325 H_mulberry(pf00936pf16365) EUT1
22330 HMMreg12833_1_EUT1(pf12833) EUT1
22335 - -
22340 -pf01218 -
22345 -pf01520 -
22350 -pf00583 -
22355 -pf11143 -
22360 -pf06572 -
22365 -pf04261 -
22370 HMMnopfamclust3_1 RMM1
22375 - -
22380 -pf00528 -
22385 -pf00528 -
22390 -pf00005 -
BMC-P
BMC-H
BMC-T
Aldehyde dehydrogenase
PTAC
Signature enzyme
Alcohol dehydrogenase
Regulator

Salmonella enterica (Salmonella choleraesuis). Bacteria; Proteobacteria; Gammaproteobacteria; Enterobacterales;

A7K84__locus_2

EUT1

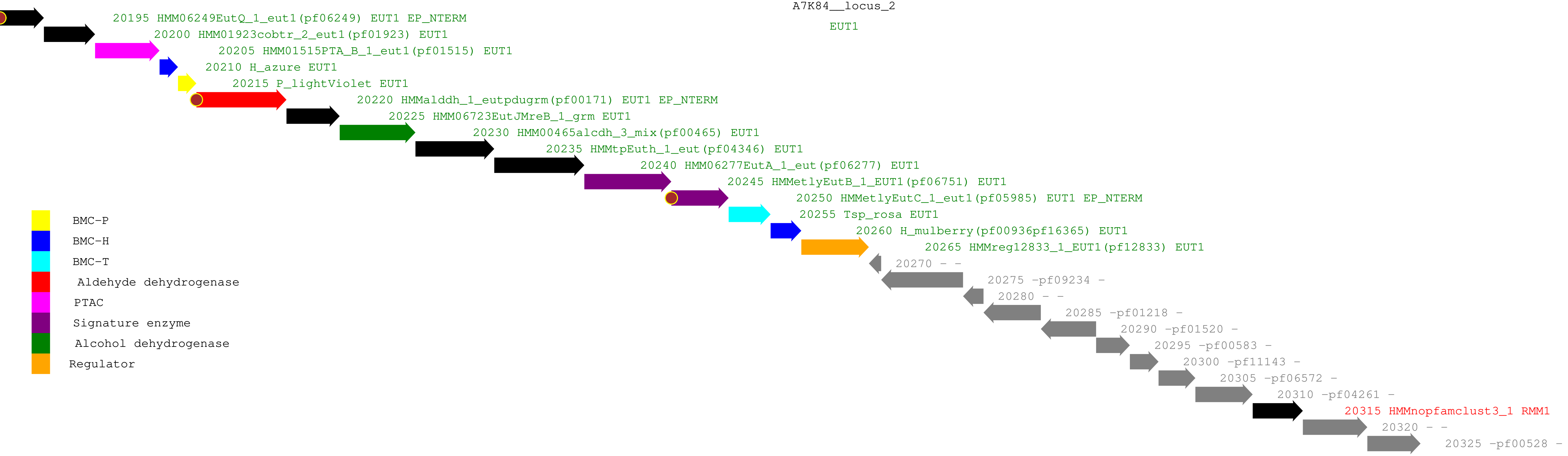

Salmonella choleraesuis. Bacteria; Proteobacteria; Gammaproteobacteria; Enterobacterales;

A7R90__locus_1

EUT1

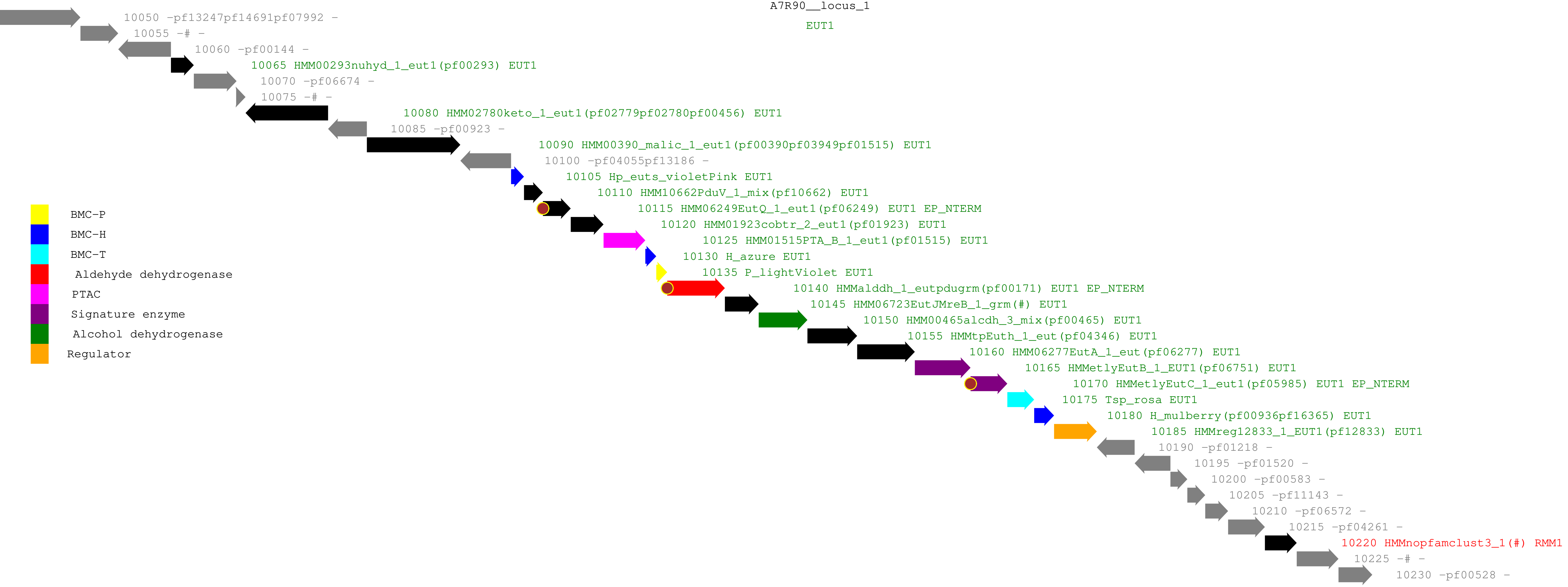

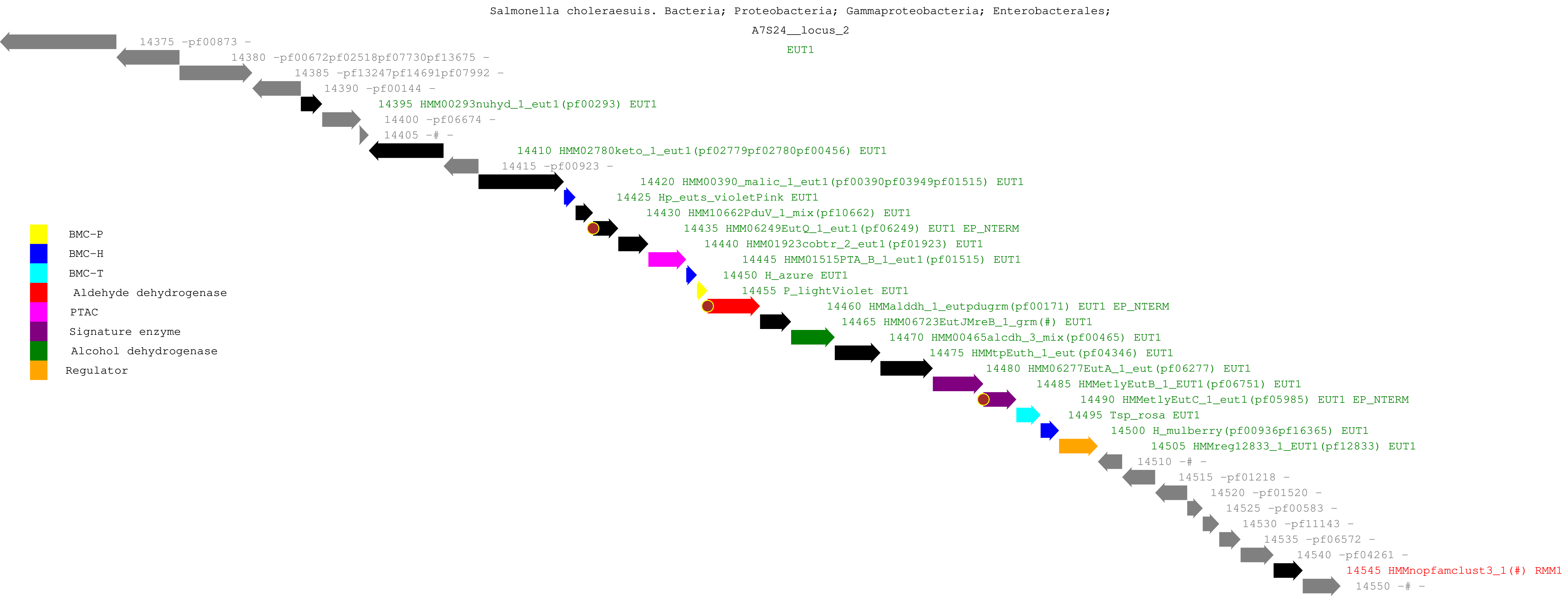

Salmonella choleraesuis. Bacteria; Proteobacteria; Gammaproteobacteria; Enterobacterales;
A7S24__locus_2
EUT1
14375 -pf00873 -
14380 -pf00672pf02518pf07730pf13675 -
14385 -pf13247pf14691pf07992 -
14390 -pf00144 -
14395 HMM00293nuhyd_1_eut1(pf00293) EUT1
14400 -pf06674 -
14405 -# -
14410 HMM02780keto_1_eut1(pf02779pf02780pf00456) EUT1
14415 -pf00923 -
14420 HMM00390_malic_1_eut1(pf00390pf03949pf01515) EUT1
14425 Hp_euts_violetPink EUT1
14430 HMM10662PduV_1_mix(pf10662) EUT1
14435 HMM06249EutQ_1_eut1(pf06249) EUT1 EP_NTERM
14440 HMM01923cobtr_2_eut1(pf01923) EUT1
14445 HMM01515PTA_B_1_eut1(pf01515) EUT1
14450 H_azure EUT1
14455 P_lightViolet EUT1
14460 HMMalddh_1_eutpdugrm(pf00171) EUT1 EP_NTERM
14465 HMM06723EutJMreB_1_grm(#) EUT1
14470 HMM00465alcdh_3_mix(pf00465) EUT1
14475 HMMtpEuth_1_eut(pf04346) EUT1
14480 HMM06277EutA_1_eut(pf06277) EUT1
14485 HMMetlyEutB_1_EUT1(pf06751) EUT1
14490 HMMetlyEutC_1_eut1(pf05985) EUT1 EP_NTERM
14495 Tsp_rosa EUT1
14500 H_mulberry(pf00936pf16365) EUT1
14505 HMMreg12833_1_EUT1(pf12833) EUT1
14510 -# -
14515 -pf01218 -
14520 -pf01520 -
14525 -pf00583 -
14530 -pf11143 -
14535 -pf06572 -
14540 -pf04261 -
14545 HMMnopfamclust3_1(#) RMM1
14550 -# -
BMC-P
BMC-H
BMC-T
Aldehyde dehydrogenase
PTAC
Signature enzyme
Alcohol dehydrogenase
Regulator

Salmonella enterica (Salmonella choleraesuis). Bacteria; Proteobacteria; Gammaproteobacteria; Enterobacterales;

A7S51__locus_1

EUT1

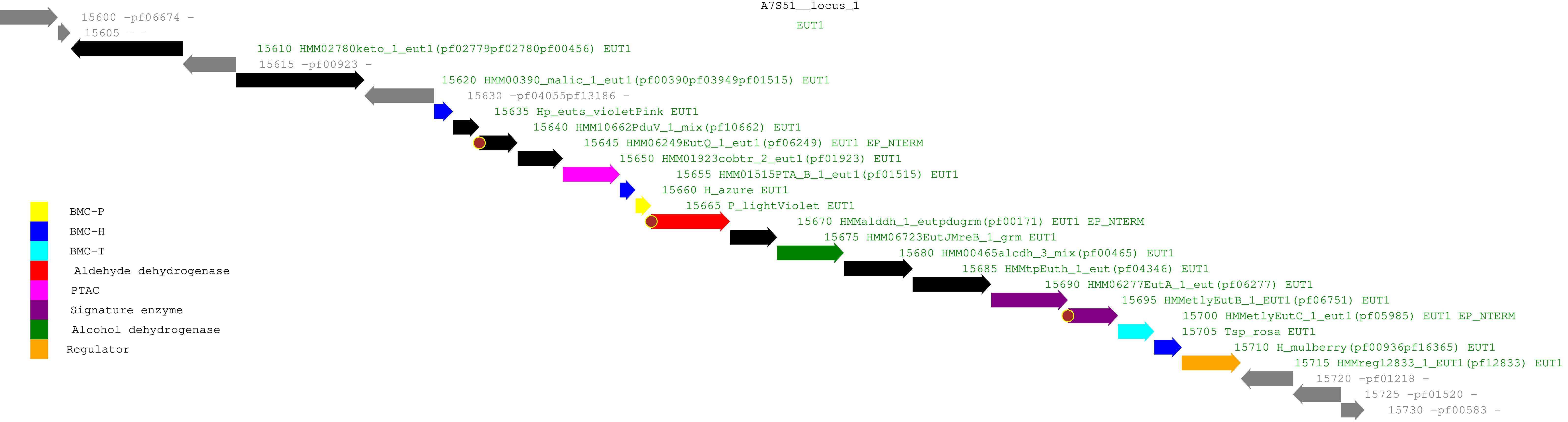

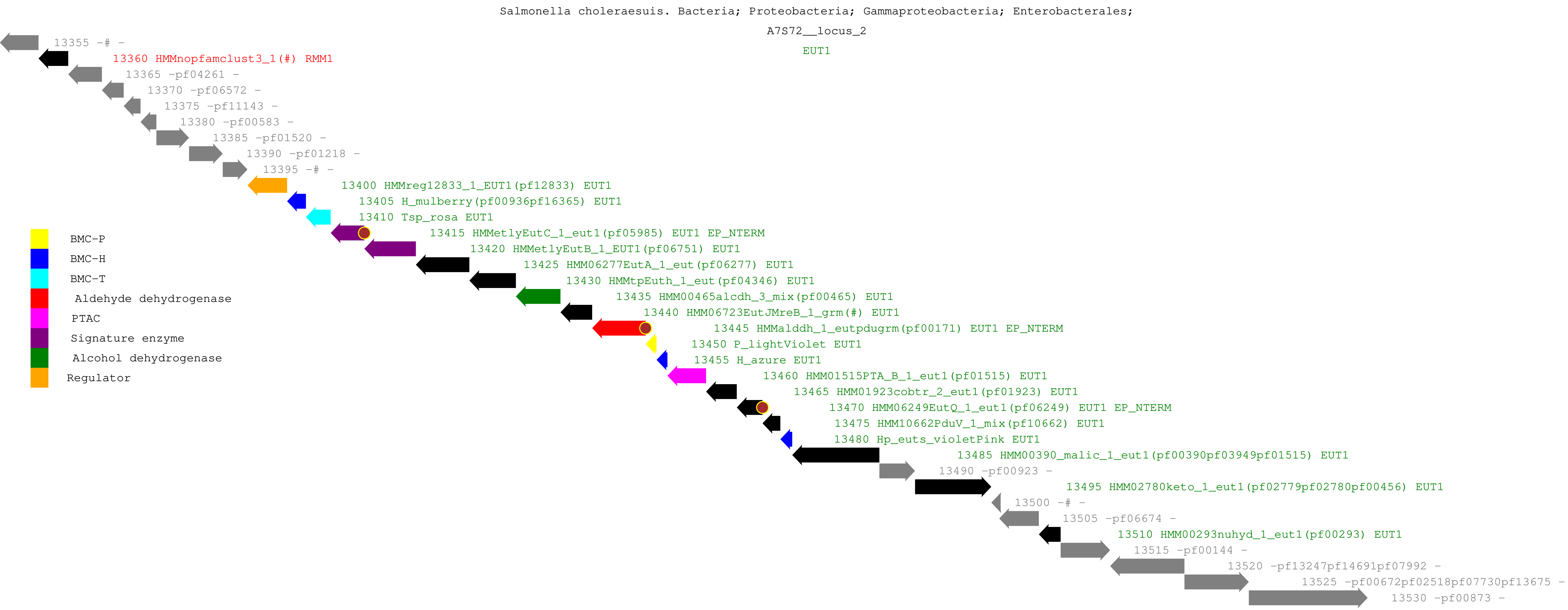
Salmonella choleraesuis. Bacteria; Proteobacteria; Gammaproteobacteria; Enterobacterales;
A7S72__locus_2
EUT1
13355 -# -
13360 HMMnopfamclust3_1(#) RMM1
13365 -pf04261 -
13370 -pf06572 -
13375 -pf11143 -
13380 -pf00583 -
13385 -pf01520 -
13390 -pf01218 -
13395 -# -
13400 HMMreg12833_1_EUT1(pf12833) EUT1
13405 H_mulberry(pf00936pf16365) EUT1
13410 Tsp_rosa EUT1
13415 HMMetlyEutC_1_eut1(pf05985) EUT1 EP_NTERM
13420 HMMetlyEutB_1_EUT1(pf06751) EUT1
13425 HMM06277EutA_1_eut(pf06277) EUT1
13430 HMMtpEuth_1_eut(pf04346) EUT1
13435 HMM00465alcdh_3_mix(pf00465) EUT1
13440 HMM06723EutJMreB_1_grm(#) EUT1
13445 HMMalddh_1_eutpdugrm(pf00171) EUT1 EP_NTERM
13450 P_lightViolet EUT1
13455 H_azure EUT1
13460 HMM01515PTA_B_1_eut1(pf01515) EUT1
13465 HMM01923cobtr_2_eut1(pf01923) EUT1
13470 HMM06249EutQ_1_eut1(pf06249) EUT1 EP_NTERM
13475 HMM10662PduV_1_mix(pf10662) EUT1
13480 Hp_euts_violetPink EUT1
13485 HMM00390_malic_1_eut1(pf00390pf03949pf01515) EUT1
13490 -pf00923 -
13495 HMM02780keto_1_eut1(pf02779pf02780pf00456) EUT1
13500 -# -
13505 -pf06674 -
13510 HMM00293nuhyd_1_eut1(pf00293) EUT1
13515 -pf00144 -
13520 -pf13247pf14691pf07992 -
13525 -pf00672pf02518pf07730pf13675 -
13530 -pf00873 -
BMC-P
BMC-H
BMC-T
Aldehyde dehydrogenase
PTAC
Signature enzyme
Alcohol dehydrogenase
Regulator

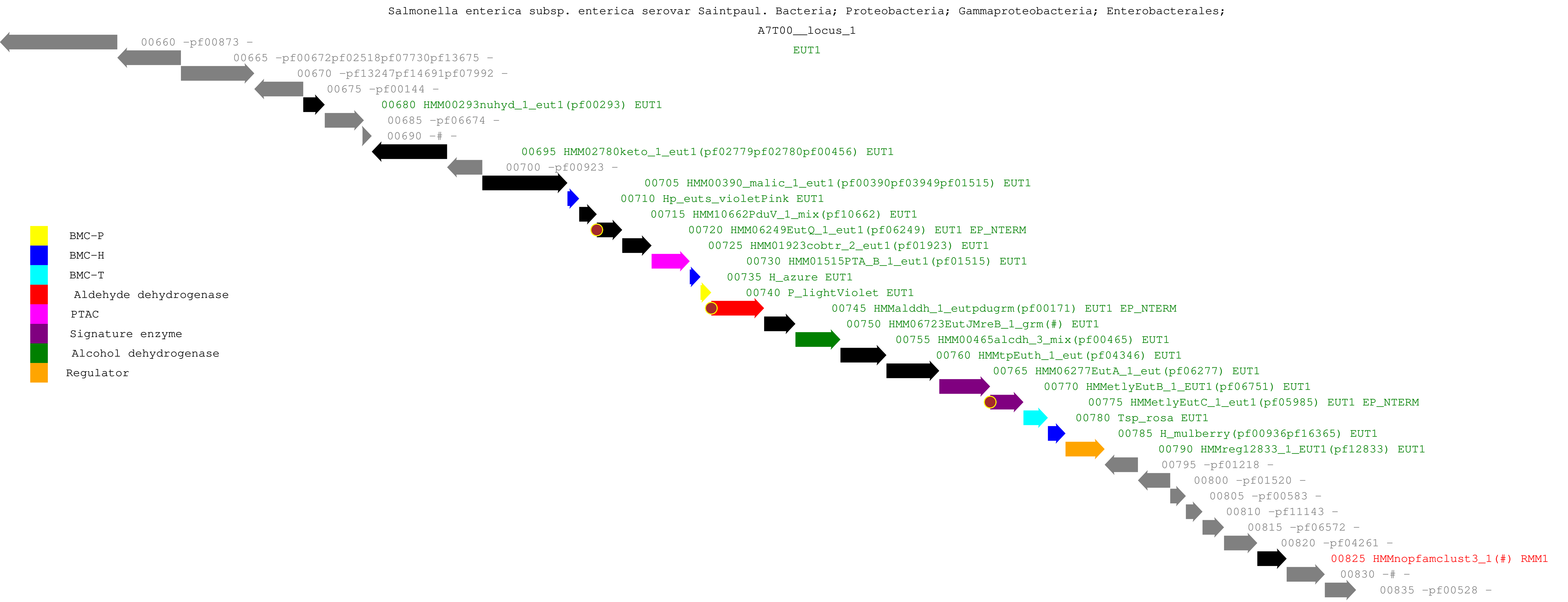
Salmonella enterica subsp. enterica serovar Saintpaul. Bacteria; Proteobacteria; Gammaproteobacteria; Enterobacterales;
A7T00__locus_1
EUT1
00660 -pf00873 -
00665 -pf00672pf02518pf07730pf13675 -
00670 -pf13247pf14691pf07992 -
00675 -pf00144 -
00680 HMM00293nuhyd_1_eut1(pf00293) EUT1
00685 -pf06674 -
00690 -# -
00695 HMM02780keto_1_eut1(pf02779pf02780pf00456) EUT1
00700 -pf00923 -
00705 HMM00390_malic_1_eut1(pf00390pf03949pf01515) EUT1
00710 Hp_euts_violetPink EUT1
00715 HMM10662PduV_1_mix(pf10662) EUT1
00720 HMM06249EutQ_1_eut1(pf06249) EUT1 EP_NTERM
00725 HMM01923cobtr_2_eut1(pf01923) EUT1
00730 HMM01515PTA_B_1_eut1(pf01515) EUT1
00735 H_azure EUT1
00740 P_lightViolet EUT1
00745 HMMalddh_1_eutpdugrm(pf00171) EUT1 EP_NTERM
00750 HMM06723EutJMreB_1_grm(#) EUT1
00755 HMM00465alcdh_3_mix(pf00465) EUT1
00760 HMMtpEuth_1_eut(pf04346) EUT1
00765 HMM06277EutA_1_eut(pf06277) EUT1
00770 HMMetlyEutB_1_EUT1(pf06751) EUT1
00775 HMMetlyEutC_1_eut1(pf05985) EUT1 EP_NTERM
00780 Tsp_rosa EUT1
00785 H_mulberry(pf00936pf16365) EUT1
00790 HMMreg12833_1_EUT1(pf12833) EUT1
00795 -pf01218 -
00800 -pf01520 -
00805 -pf00583 -
00810 -pf11143 -
00815 -pf06572 -
00820 -pf04261 -
00825 HMMnopfamclust3_1(#) RMM1
00830 -# -
00835 -pf00528 -
BMC-P
BMC-H
BMC-T
Aldehyde dehydrogenase
PTAC
Signature enzyme
Alcohol dehydrogenase
Regulator

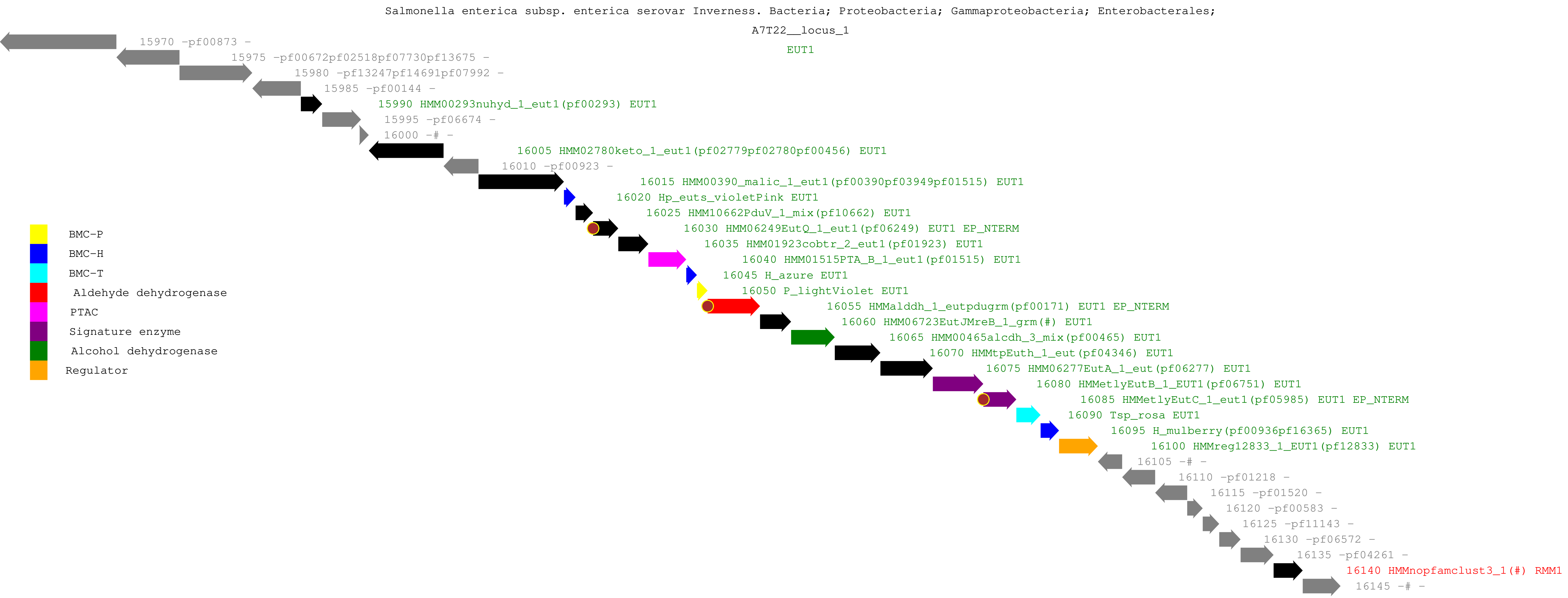
Salmonella enterica subsp. enterica serovar Inverness. Bacteria; Proteobacteria; Gammaproteobacteria; Enterobacterales;
A7T22__locus_1
EUT1
15970 -pf00873 -
15975 -pf00672pf02518pf07730pf13675 -
15980 -pf13247pf14691pf07992 -
15985 -pf00144 -
15990 HMM00293nuhyd_1_eut1(pf00293) EUT1
15995 -pf06674 -
16000 -# -
16005 HMM02780keto_1_eut1(pf02779pf02780pf00456) EUT1
16010 -pf00923 -
16015 HMM00390_malic_1_eut1(pf00390pf03949pf01515) EUT1
16020 Hp_euts_violetPink EUT1
16025 HMM10662PduV_1_mix(pf10662) EUT1
16030 HMM06249EutQ_1_eut1(pf06249) EUT1 EP_NTERM
16035 HMM01923cobtr_2_eut1(pf01923) EUT1
16040 HMM01515PTA_B_1_eut1(pf01515) EUT1
16045 H_azure EUT1
16050 P_lightViolet EUT1
16055 HMMalddh_1_eutpdugrm(pf00171) EUT1 EP_NTERM
16060 HMM06723EutJMreB_1_grm(#) EUT1
16065 HMM00465alcdh_3_mix(pf00465) EUT1
16070 HMMtpEuth_1_eut(pf04346) EUT1
16075 HMM06277EutA_1_eut(pf06277) EUT1
16080 HMMetlyEutB_1_EUT1(pf06751) EUT1
16085 HMMetlyEutC_1_eut1(pf05985) EUT1 EP_NTERM
16090 Tsp_rosa EUT1
16095 H_mulberry(pf00936pf16365) EUT1
16100 HMMreg12833_1_EUT1(pf12833) EUT1
16105 -# -
16110 -pf01218 -
16115 -pf01520 -
16120 -pf00583 -
16125 -pf11143 -
16130 -pf06572 -
16135 -pf04261 -
16140 HMMnopfamclust3_1(#) RMM1
16145 -# -
BMC-P
BMC-H
BMC-T
Aldehyde dehydrogenase
PTAC
Signature enzyme
Alcohol dehydrogenase
Regulator

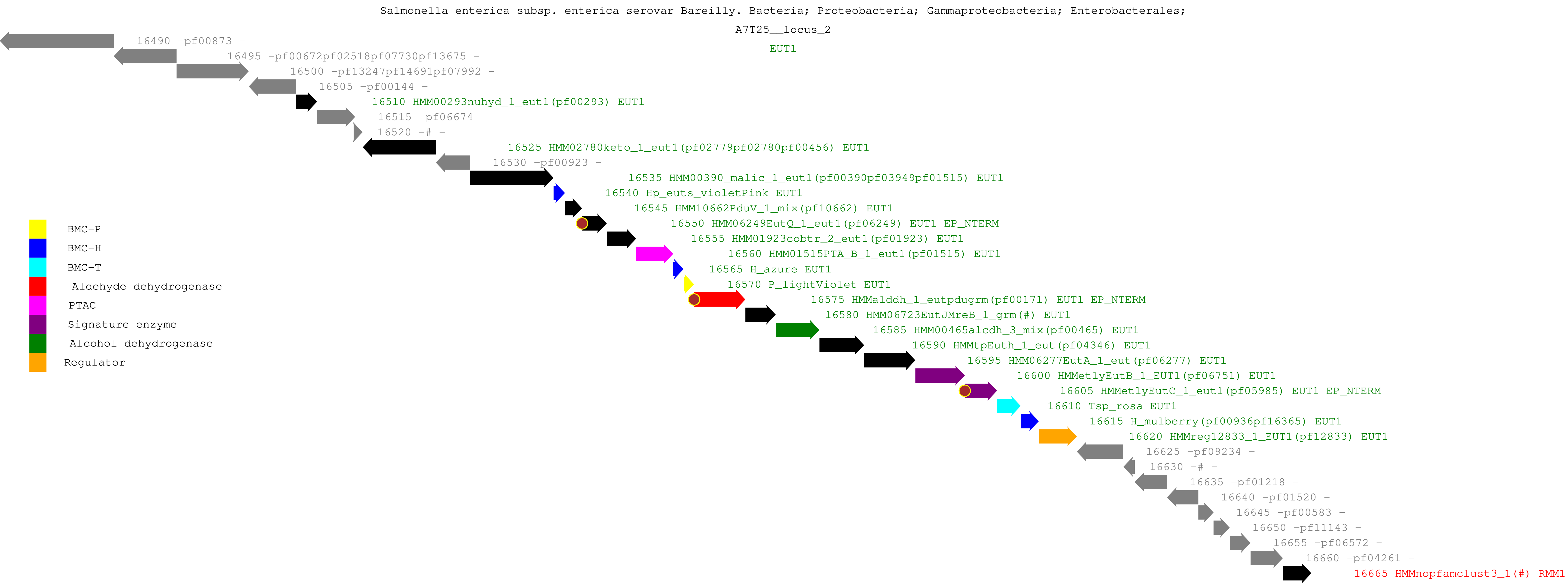
Salmonella enterica subsp. enterica serovar Bareilly. Bacteria; Proteobacteria; Gammaproteobacteria; Enterobacterales;
A7T25__locus_2
EUT1
16490 -pf00873 -
16495 -pf00672pf02518pf07730pf13675 -
16500 -pf13247pf14691pf07992 -
16505 -pf00144 -
16510 HMM00293nuhyd_1_eut1(pf00293) EUT1
16515 -pf06674 -
16520 -# -
16525 HMM02780keto_1_eut1(pf02779pf02780pf00456) EUT1
16530 -pf00923 -
16535 HMM00390_malic_1_eut1(pf00390pf03949pf01515) EUT1
16540 Hp_euts_violetPink EUT1
16545 HMM10662PduV_1_mix(pf10662) EUT1
16550 HMM06249EutQ_1_eut1(pf06249) EUT1 EP_NTERM
16555 HMM01923cobtr_2_eut1(pf01923) EUT1
16560 HMM01515PTA_B_1_eut1(pf01515) EUT1
16565 H_azure EUT1
16570 P_lightViolet EUT1
16575 HMMalddh_1_eutpdugrm(pf00171) EUT1 EP_NTERM
16580 HMM06723EutJMreB_1_grm(#) EUT1
16585 HMM00465alcdh_3_mix(pf00465) EUT1
16590 HMMtpEuth_1_eut(pf04346) EUT1
16595 HMM06277EutA_1_eut(pf06277) EUT1
16600 HMMetlyEutB_1_EUT1(pf06751) EUT1
16605 HMMetlyEutC_1_eut1(pf05985) EUT1 EP_NTERM
16610 Tsp_rosa EUT1
16615 H_mulberry(pf00936pf16365) EUT1
16620 HMMreg12833_1_EUT1(pf12833) EUT1
16625 -pf09234 -
16630 -# -
16635 -pf01218 -
16640 -pf01520 -
16645 -pf00583 -
16650 -pf11143 -
16655 -pf06572 -
16660 -pf04261 -
16665 HMMnopfamclust3_1(#) RMM1
BMC-P
BMC-H
BMC-T
Aldehyde dehydrogenase
PTAC
Signature enzyme
Alcohol dehydrogenase
Regulator

Salmonella anatum. Bacteria; Proteobacteria; Gammaproteobacteria; Enterobacterales;

A7T43__locus_2

EUT1

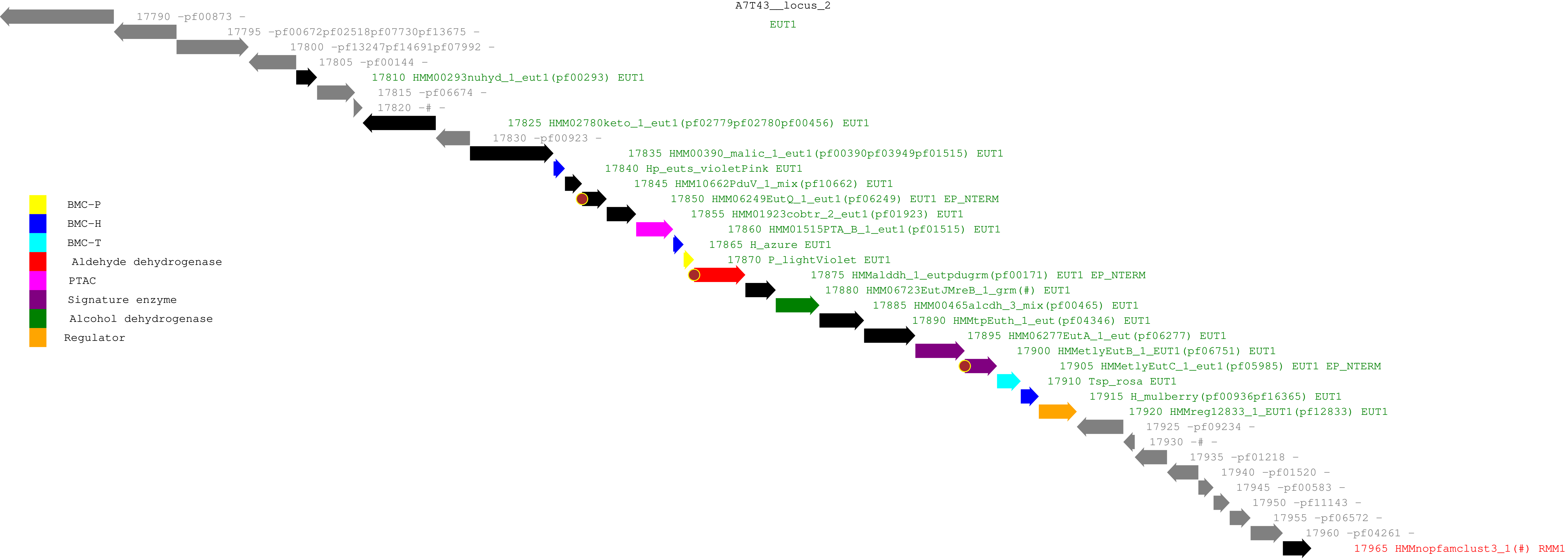

Salmonella enterica subsp. diarizonae serovar 16:z10:e,n,x,z15. Bacteria; Proteobacteria; Gammaproteobacteria; Enterobacterales;

A7T60__locus_1

EUT1

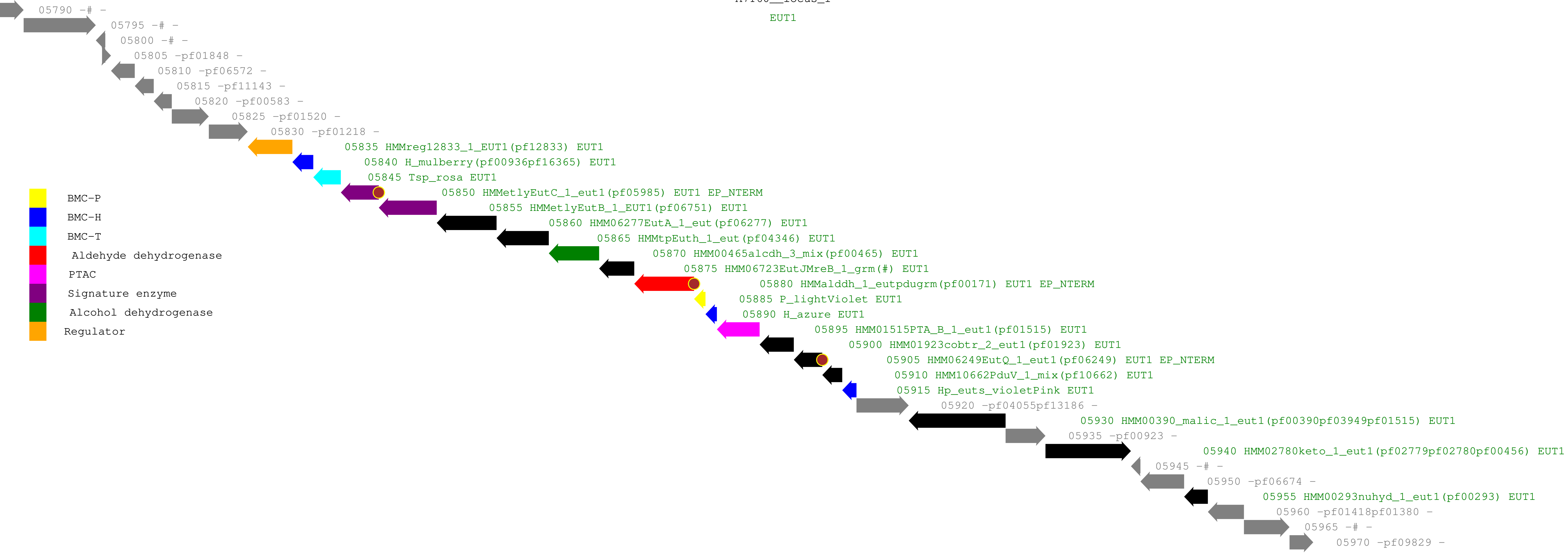

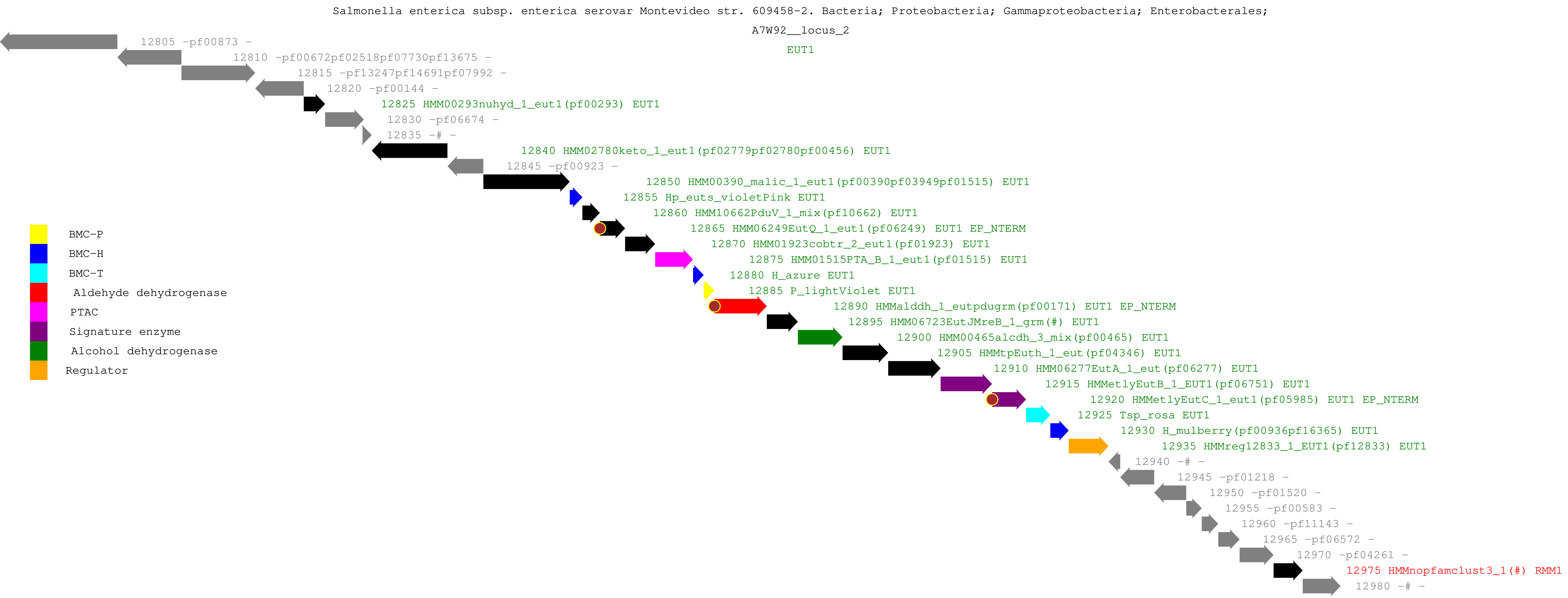
Salmonella enterica subsp. enterica serovar Montevideo str. 609458-2. Bacteria; Proteobacteria; Gammaproteobacteria; Enterobacterales;
A7W92__locus_2
EUT1
12805 -pf00873 -
12810 -pf00672pf02518pf07730pf13675 -
12815 -pf13247pf14691pf07992 -
12820 -pf00144 -
12825 HMM00293nuhyd_1_eut1(pf00293) EUT1
12830 -pf06674 -
12835 -# -
12840 HMM02780keto_1_eut1(pf02779pf02780pf00456) EUT1
12845 -pf00923 -
12850 HMM00390_malic_1_eut1(pf00390pf03949pf01515) EUT1
12855 Hp_euts_violetPink EUT1
12860 HMM10662PduV_1_mix(pf10662) EUT1
12865 HMM06249EutQ_1_eut1(pf06249) EUT1 EP_NTERM
12870 HMM01923cobtr_2_eut1(pf01923) EUT1
12875 HMM01515PTA_B_1_eut1(pf01515) EUT1
12880 H_azure EUT1
12885 P_lightViolet EUT1
12890 HMMalddh_1_eutpdugrm(pf00171) EUT1 EP_NTERM
12895 HMM06723EutJMreB_1_grm(#) EUT1
12900 HMM00465alcdh_3_mix(pf00465) EUT1
12905 HMMtpEuth_1_eut(pf04346) EUT1
12910 HMM06277EutA_1_eut(pf06277) EUT1
12915 HMMetlyEutB_1_EUT1(pf06751) EUT1
12920 HMMetlyEutC_1_eut1(pf05985) EUT1 EP_NTERM
12925 Tsp_rosa EUT1
12930 H_mulberry(pf00936pf16365) EUT1
12935 HMMreg12833_1_EUT1(pf12833) EUT1
12940 -# -
12945 -pf01218 -
12950 -pf01520 -
12955 -pf00583 -
12960 -pf11143 -
12965 -pf06572 -
12970 -pf04261 -
12975 HMMnopfamclust3_1(#) RMM1
12980 -# -
BMC-P
BMC-H
BMC-T
Aldehyde dehydrogenase
PTAC
Signature enzyme
Alcohol dehydrogenase
Regulator

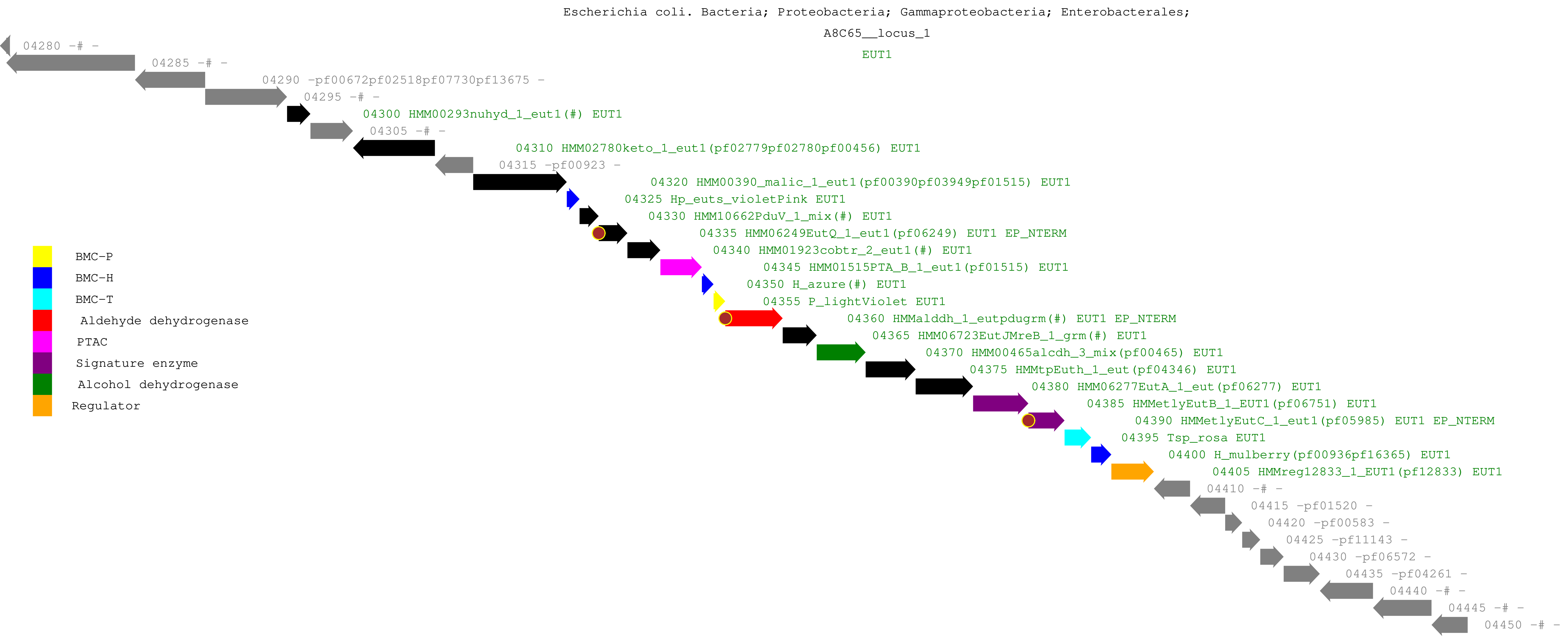
Escherichia coli. Bacteria; Proteobacteria; Gammaproteobacteria; Enterobacterales;
A8C65__locus_1
EUT1
04280 -# -
04285 -# -
04290 -pf00672pf02518pf07730pf13675 -
04295 -# -
04300 HMM00293nuhyd_1_eut1(#) EUT1
04305 -# -
04310 HMM02780keto_1_eut1(pf02779pf02780pf00456) EUT1
04315 -pf00923 -
04320 HMM00390_malic_1_eut1(pf00390pf03949pf01515) EUT1
04325 Hp_euts_violetPink EUT1
04330 HMM10662PduV_1_mix(#) EUT1
04335 HMM06249EutQ_1_eut1(pf06249) EUT1 EP_NTERM
04340 HMM01923cobtr_2_eut1(#) EUT1
04345 HMM01515PTA_B_1_eut1(pf01515) EUT1
04350 H_azure(#) EUT1
04355 P_lightViolet EUT1
04360 HMMalddh_1_eutpdugrm(#) EUT1 EP_NTERM
04365 HMM06723EutJMreB_1_grm(#) EUT1
04370 HMM00465alcdh_3_mix(pf00465) EUT1
04375 HMMtpEuth_1_eut(pf04346) EUT1
04380 HMM06277EutA_1_eut(pf06277) EUT1
04385 HMMetlyEutB_1_EUT1(pf06751) EUT1
04390 HMMetlyEutC_1_eut1(pf05985) EUT1 EP_NTERM
04395 Tsp_rosa EUT1
04400 H_mulberry(pf00936pf16365) EUT1
04405 HMMreg12833_1_EUT1(pf12833) EUT1
04410 -# -
04415 -pf01520 -
04420 -pf00583 -
04425 -pf11143 -
04430 -pf06572 -
04435 -pf04261 -
04440 -# -
04445 -# -
04450 -# -
BMC-P
BMC-H
BMC-T
Aldehyde dehydrogenase
PTAC
Signature enzyme
Alcohol dehydrogenase
Regulator

Escherichia coli. Bacteria; Proteobacteria; Gammaproteobacteria; Enterobacterales;

A8G17__locus_1

EUT1

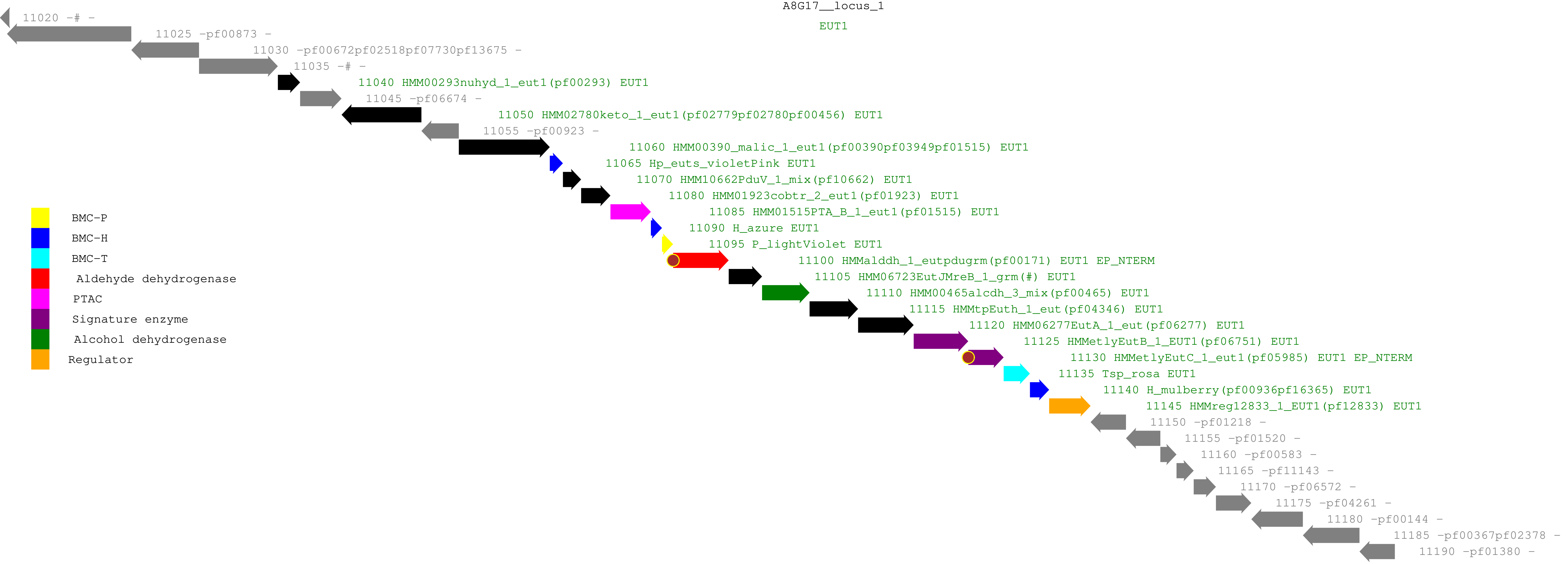

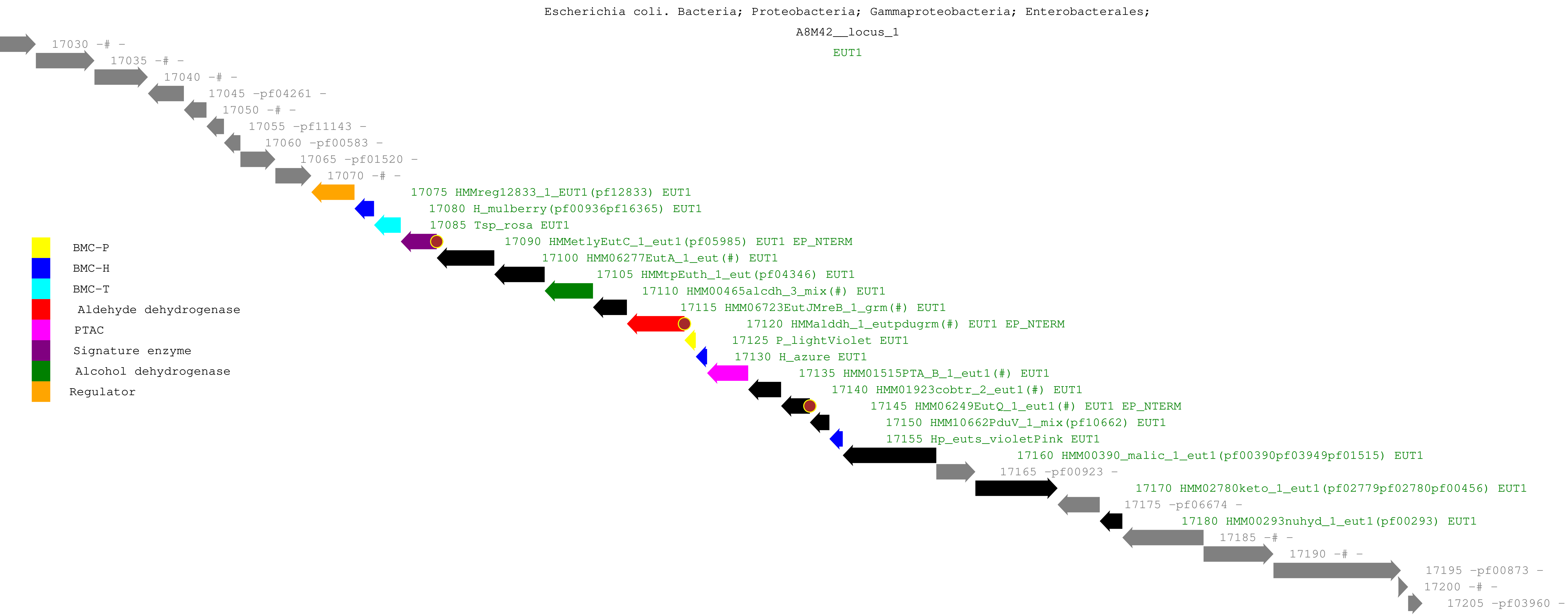

Escherichia coli. Bacteria; Proteobacteria; Gammaproteobacteria; Enterobacterales;
A8M42__locus_1
EUT1
17030 -# -
17035 -# -
17040 -# -
17045 -pf04261 -
17050 -# -
17055 -pf11143 -
17060 -pf00583 -
17065 -pf01520 -
17070 -# -
17075 HMMreg12833_1_EUT1(pf12833) EUT1
17080 H_mulberry(pf00936pf16365) EUT1
17085 Tsp_rosa EUT1
17090 HMMetlyEutC_1_eut1(pf05985) EUT1 EP_NTERM
17100 HMM06277EutA_1_eut(#) EUT1
17105 HMMtpEuth_1_eut(pf04346) EUT1
17110 HMM00465alcdh_3_mix(#) EUT1
17115 HMM06723EutJMreB_1_grm(#) EUT1
17120 HMMalddh_1_eutpdugrm(#) EUT1 EP_NTERM
17125 P_lightViolet EUT1
17130 H_azure EUT1
17135 HMM01515PTA_B_1_eut1(#) EUT1
17140 HMM01923cobtr_2_eut1(#) EUT1
17145 HMM06249EutQ_1_eut1(#) EUT1 EP_NTERM
17150 HMM10662PduV_1_mix(pf10662) EUT1
17155 Hp_euts_violetPink EUT1
17160 HMM00390_malic_1_eut1(pf00390pf03949pf01515) EUT1
17165 -pf00923 -
17170 HMM02780keto_1_eut1(pf02779pf02780pf00456) EUT1
17175 -pf06674 -
17180 HMM00293nuhyd_1_eut1(pf00293) EUT1
17185 -# -
17190 -# -
17195 -pf00873 -
17200 -# -
17205 -pf03960 -
BMC-P
BMC-H
BMC-T
Aldehyde dehydrogenase
PTAC
Signature enzyme
Alcohol dehydrogenase
Regulator

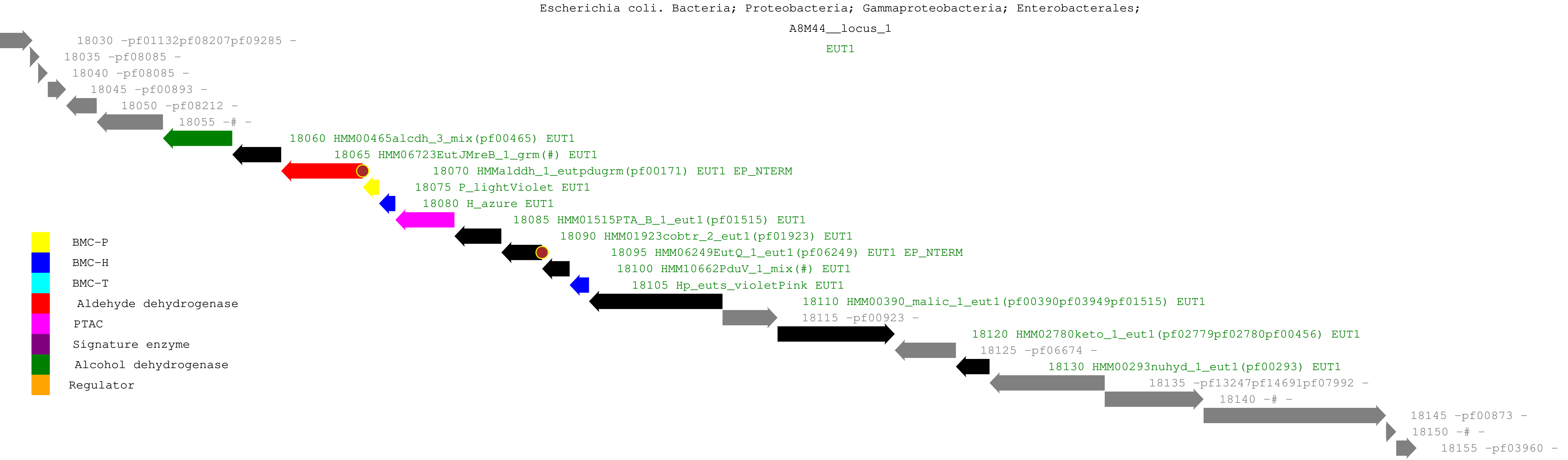

Escherichia coli. Bacteria; Proteobacteria; Gammaproteobacteria; Enterobacterales;
A8M44__locus_1
EUT1
18030 -pf01132pf08207pf09285 -
18035 -pf08085 -
18040 -pf08085 -
18045 -pf00893 -
18050 -pf08212 -
18055 -# -
18060 HMM00465alcdh_3_mix(pf00465) EUT1
18065 HMM06723EutJMreB_1_grm(#) EUT1
18070 HMMalddh_1_eutpdugrm(pf00171) EUT1 EP_NTERM
18075 P_lightViolet EUT1
18080 H_azure EUT1
18085 HMM01515PTA_B_1_eut1(pf01515) EUT1
18090 HMM01923cobtr_2_eut1(pf01923) EUT1
18095 HMM06249EutQ_1_eut1(pf06249) EUT1 EP_NTERM
18100 HMM10662PduV_1_mix(#) EUT1
18105 Hp_euts_violetPink EUT1
18110 HMM00390_malic_1_eut1(pf00390pf03949pf01515) EUT1
18115 -pf00923 -
18120 HMM02780keto_1_eut1(pf02779pf02780pf00456) EUT1
18125 -pf06674 -
18130 HMM00293nuhyd_1_eut1(pf00293) EUT1
18135 -pf13247pf14691pf07992 -
18140 -# -
18145 -pf00873 -
18150 -# -
18155 -pf03960 -
BMC-P
BMC-H
BMC-T
Aldehyde dehydrogenase
PTAC
Signature enzyme
Alcohol dehydrogenase
Regulator

Escherichia coli. Bacteria; Proteobacteria; Gammaproteobacteria; Enterobacterales;

A8M81__locus_1

EUT1

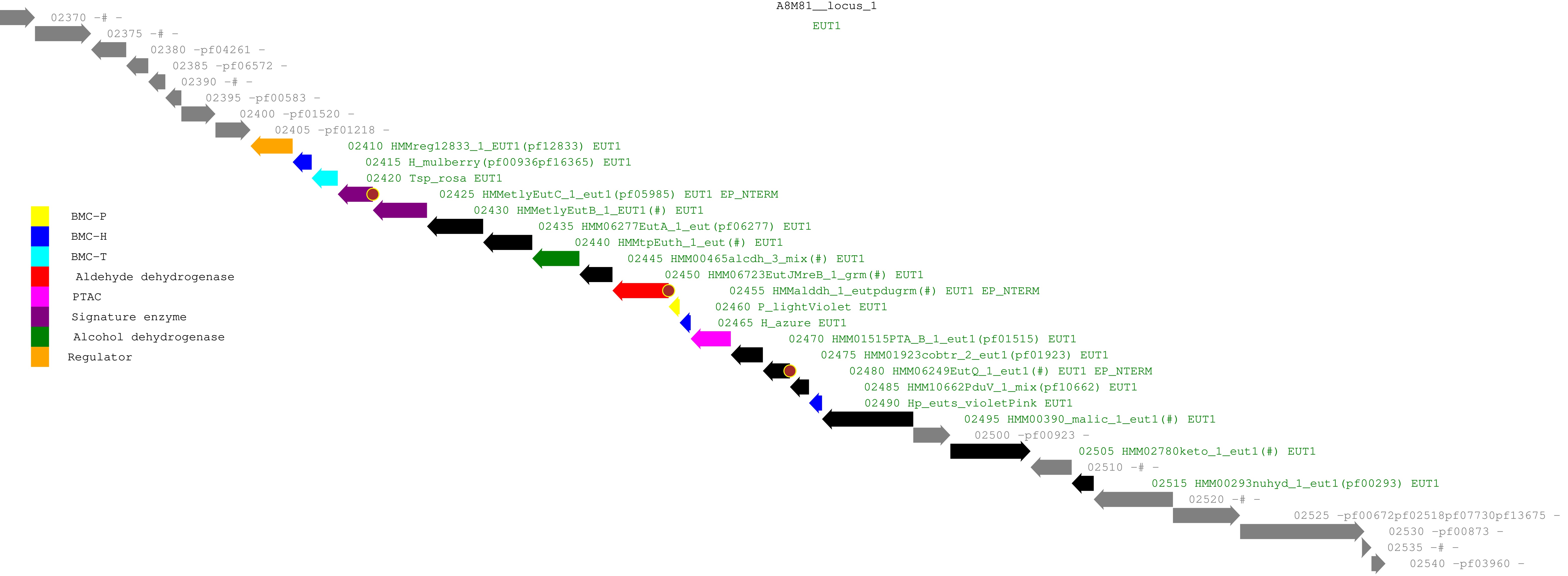

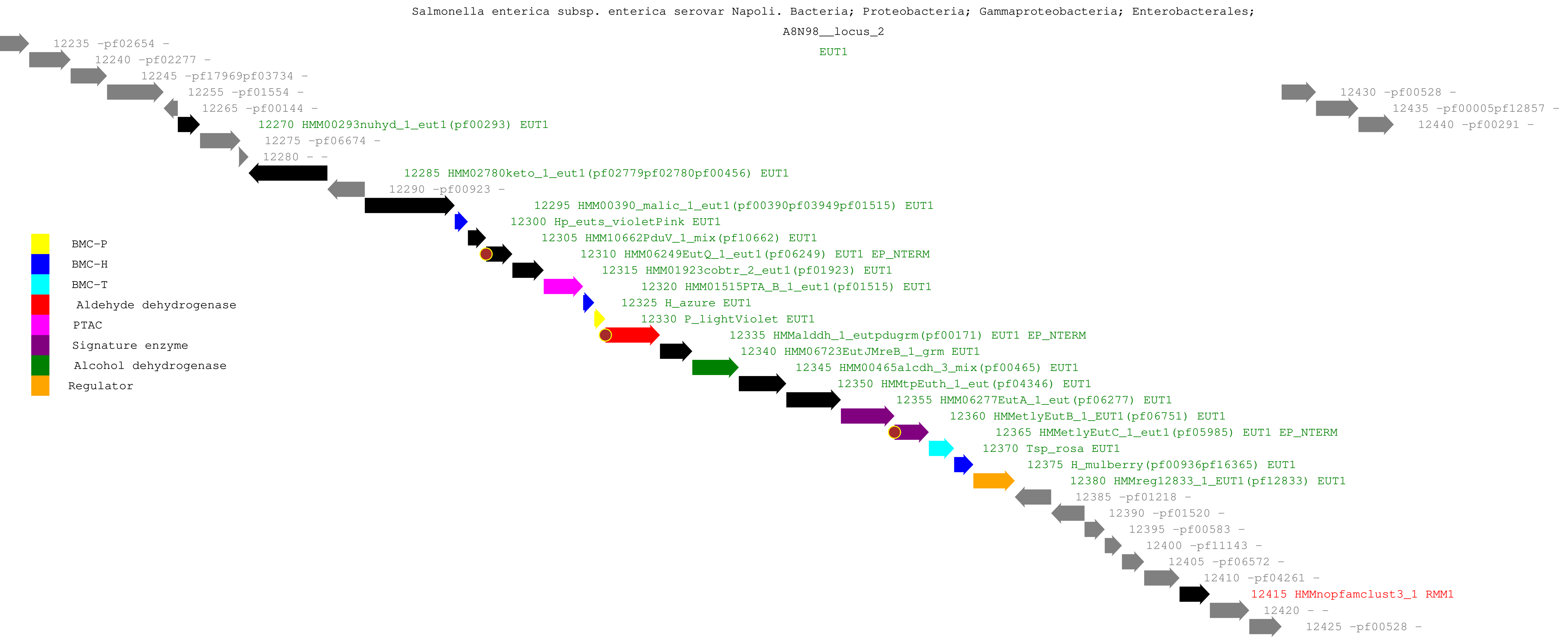
Salmonella enterica subsp. enterica serovar Napoli. Bacteria; Proteobacteria; Gammaproteobacteria; Enterobacterales;
A8N98__locus_2
EUT1
12235 -pf02654 -
12240 -pf02277 -
12245 -pf17969pf03734 -
12255 -pf01554 -
12265 -pf00144 -
12270 HMM00293nuhyd_1_eut1(pf00293) EUT1
12275 -pf06674 -
12280 - -
12285 HMM02780keto_1_eut1(pf02779pf02780pf00456) EUT1
12290 -pf00923 -
12295 HMM00390_malic_1_eut1(pf00390pf03949pf01515) EUT1
12300 Hp_euts_violetPink EUT1
12305 HMM10662PduV_1_mix(pf10662) EUT1
12310 HMM06249EutQ_1_eut1(pf06249) EUT1 EP_NTERM
12315 HMM01923cobtr_2_eut1(pf01923) EUT1
12320 HMM01515PTA_B_1_eut1(pf01515) EUT1
12325 H_azure EUT1
12330 P_lightViolet EUT1
12335 HMMalddh_1_eutpdugrm(pf00171) EUT1 EP_NTERM
12340 HMM06723EutJMreB_1_grm EUT1
12345 HMM00465alcdh_3_mix(pf00465) EUT1
12350 HMMtpEuth_1_eut(pf04346) EUT1
12355 HMM06277EutA_1_eut(pf06277) EUT1
12360 HMMetlyEutB_1_EUT1(pf06751) EUT1
12365 HMMetlyEutC_1_eut1(pf05985) EUT1 EP_NTERM
12370 Tsp_rosa EUT1
12375 H_mulberry(pf00936pf16365) EUT1
12380 HMMreg12833_1_EUT1(pf12833) EUT1
12385 -pf01218 -
12390 -pf01520 -
12395 -pf00583 -
12400 -pf11143 -
12405 -pf06572 -
12410 -pf04261 -
12415 HMMnopfamclust3_1 RMM1
12420 - -
12425 -pf00528 -
12430 -pf00528 -
12435 -pf00005pf12857 -
12440 -pf00291 -
BMC-P
BMC-H
BMC-T
Aldehyde dehydrogenase
PTAC
Signature enzyme
Alcohol dehydrogenase
Regulator

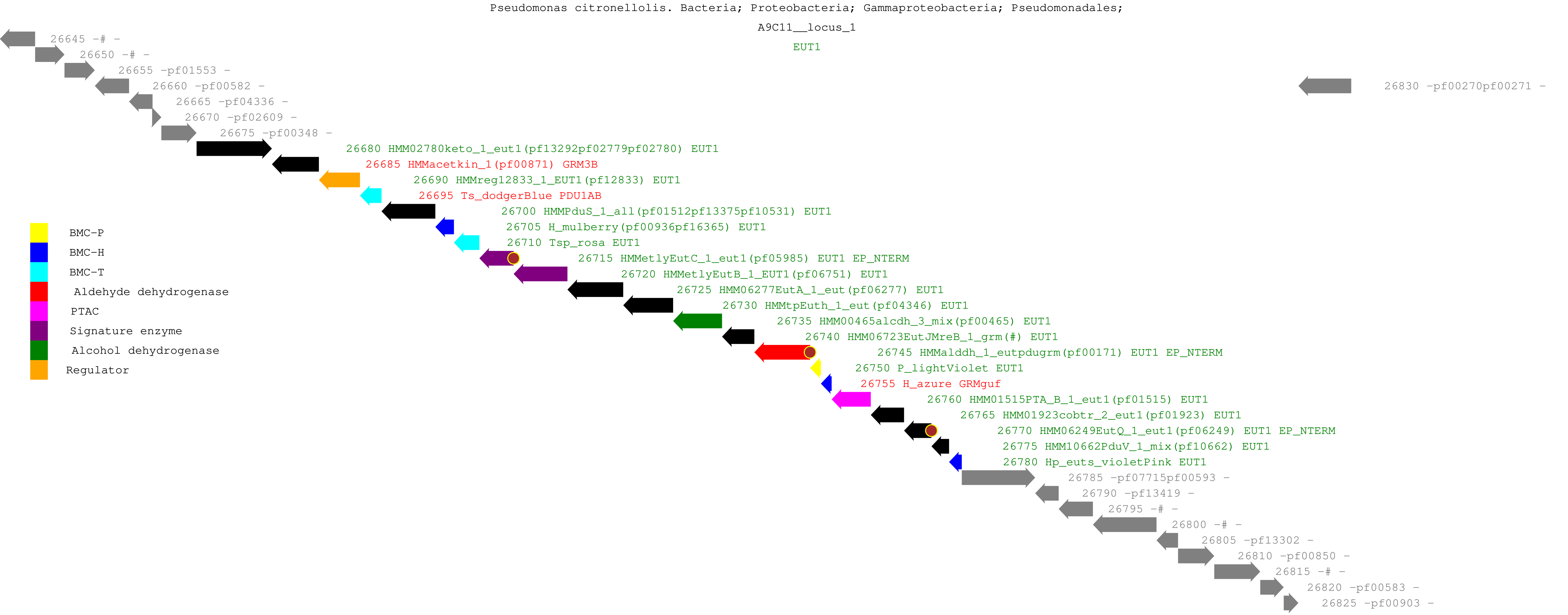

Pseudomonas citronellolis. Bacteria; Proteobacteria; Gammaproteobacteria; Pseudomonadales;
A9C11__locus_1
EUT1
26645 -# -
26650 -# -
26655 -pf01553 -
26660 -pf00582 -
26665 -pf04336 -
26670 -pf02609 -
26675 -pf00348 -
26680 HMM02780keto_1_eut1(pf13292pf02779pf02780) EUT1
26685 HMMacetkin_1(pf00871) GRM3B
26690 HMMreg12833_1_EUT1(pf12833) EUT1
26695 Ts_dodgerBlue PDU1AB
26700 HMMPduS_1_all(pf01512pf13375pf10531) EUT1
26705 H_mulberry(pf00936pf16365) EUT1
26710 Tsp_rosa EUT1
26715 HMMetlyEutC_1_eut1(pf05985) EUT1 EP_NTERM
26720 HMMetlyEutB_1_EUT1(pf06751) EUT1
26725 HMM06277EutA_1_eut(pf06277) EUT1
26730 HMMtpEuth_1_eut(pf04346) EUT1
26735 HMM00465alcdh_3_mix(pf00465) EUT1
26740 HMM06723EutJMreB_1_grm(#) EUT1
26745 HMMalddh_1_eutpdugrm(pf00171) EUT1 EP_NTERM
26750 P_lightViolet EUT1
26755 H_azure GRMguf
26760 HMM01515PTA_B_1_eut1(pf01515) EUT1
26765 HMM01923cobtr_2_eut1(pf01923) EUT1
26770 HMM06249EutQ_1_eut1(pf06249) EUT1 EP_NTERM
26775 HMM10662PduV_1_mix(pf10662) EUT1
26780 Hp_euts_violetPink EUT1
26785 -pf07715pf00593 -
26790 -pf13419 -
26795 -# -
26800 -# -
26805 -pf13302 -
26810 -pf00850 -
26815 -# -
26820 -pf00583 -
26825 -pf00903 -
26830 -pf00270pf00271 -
BMC-P
BMC-H
BMC-T
Aldehyde dehydrogenase
PTAC
Signature enzyme
Alcohol dehydrogenase
Regulator

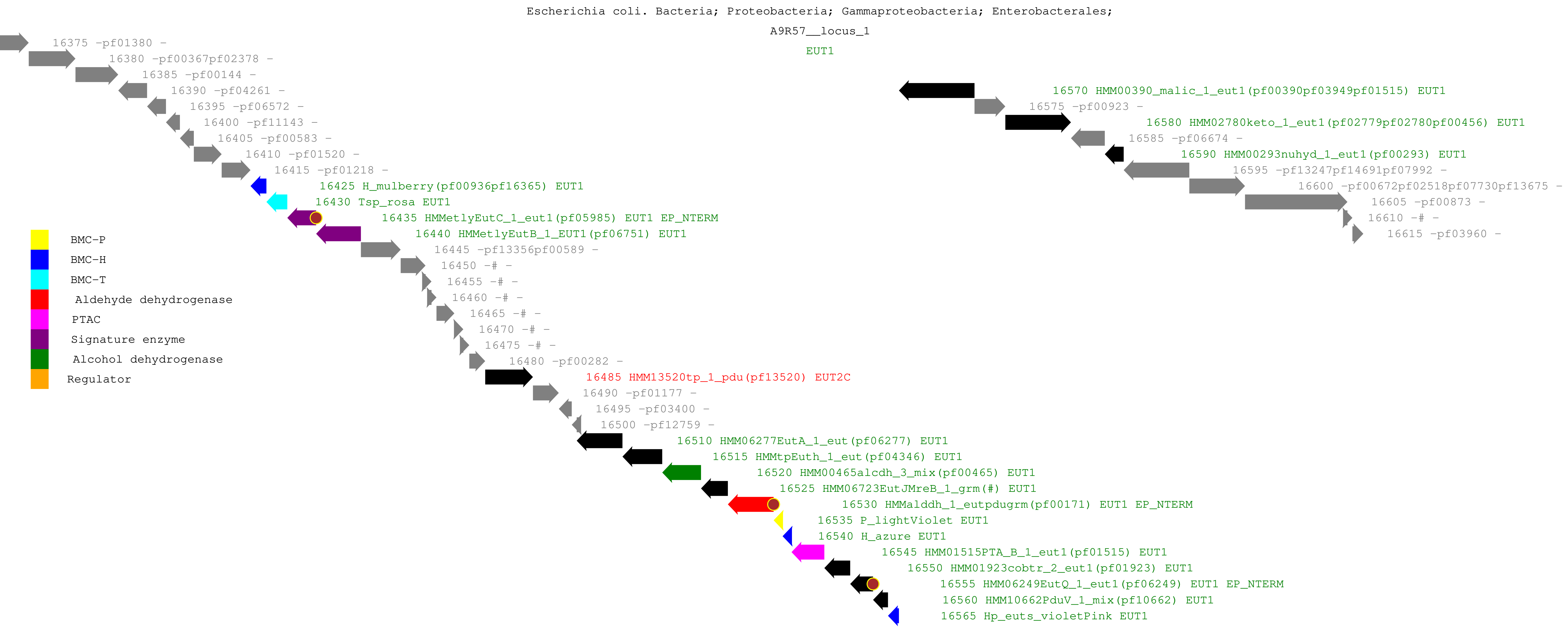
Escherichia coli. Bacteria; Proteobacteria; Gammaproteobacteria; Enterobacterales;
A9R57__locus_1
EUT1
16375 -pf01380 -
16380 -pf00367pf02378 -
16385 -pf00144 -
16390 -pf04261 -
16395 -pf06572 -
16400 -pf11143 -
16405 -pf00583 -
16410 -pf01520 -
16415 -pf01218 -
16425 H_mulberry(pf00936pf16365) EUT1
16430 Tsp_rosa EUT1
16435 HMMetlyEutC_1_eut1(pf05985) EUT1 EP_NTERM
16440 HMMetlyEutB_1_EUT1(pf06751) EUT1
16445 -pf13356pf00589 -
16450 -# -
16455 -# -
16460 -# -
16465 -# -
16470 -# -
16475 -# -
16480 -pf00282 -
16485 HMM13520tp_1_pdu(pf13520) EUT2C
16490 -pf01177 -
16495 -pf03400 -
16500 -pf12759 -
16510 HMM06277EutA_1_eut(pf06277) EUT1
16515 HMMtpEuth_1_eut(pf04346) EUT1
16520 HMM00465alcdh_3_mix(pf00465) EUT1
16525 HMM06723EutJMreB_1_grm(#) EUT1
16530 HMMalddh_1_eutpdugrm(pf00171) EUT1 EP_NTERM
16535 P_lightViolet EUT1
16540 H_azure EUT1
16545 HMM01515PTA_B_1_eut1(pf01515) EUT1
16550 HMM01923cobtr_2_eut1(pf01923) EUT1
16555 HMM06249EutQ_1_eut1(pf06249) EUT1 EP_NTERM
16560 HMM10662PduV_1_mix(pf10662) EUT1
16565 Hp_euts_violetPink EUT1
16570 HMM00390_malic_1_eut1(pf00390pf03949pf01515) EUT1
16575 -pf00923 -
16580 HMM02780keto_1_eut1(pf02779pf02780pf00456) EUT1
16585 -pf06674 -
16590 HMM00293nuhyd_1_eut1(pf00293) EUT1
16595 -pf13247pf14691pf07992 -
16600 -pf00672pf02518pf07730pf13675 -
16605 -pf00873 -
16610 -# -
16615 -pf03960 -
BMC-P
BMC-H
BMC-T
Aldehyde dehydrogenase
PTAC
Signature enzyme
Alcohol dehydrogenase
Regulator

Escherichia coli. Bacteria; Proteobacteria; Gammaproteobacteria; Enterobacterales;

A9X62__locus_1

EUT1

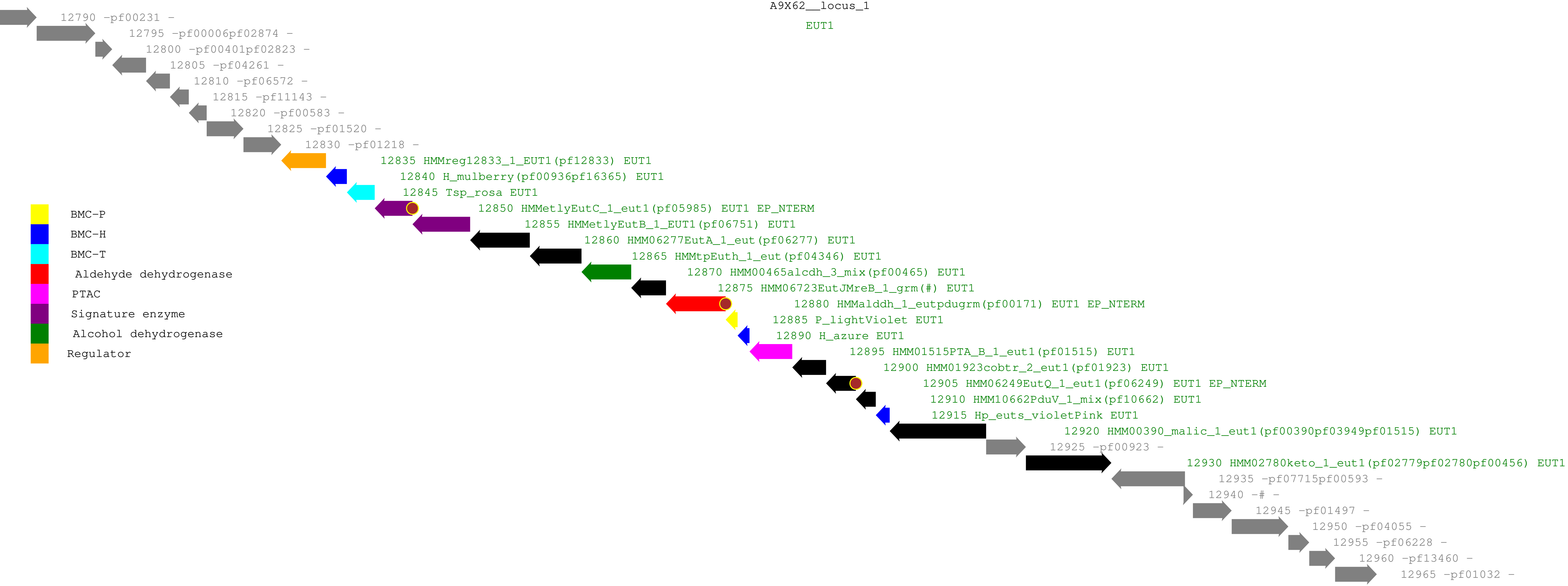

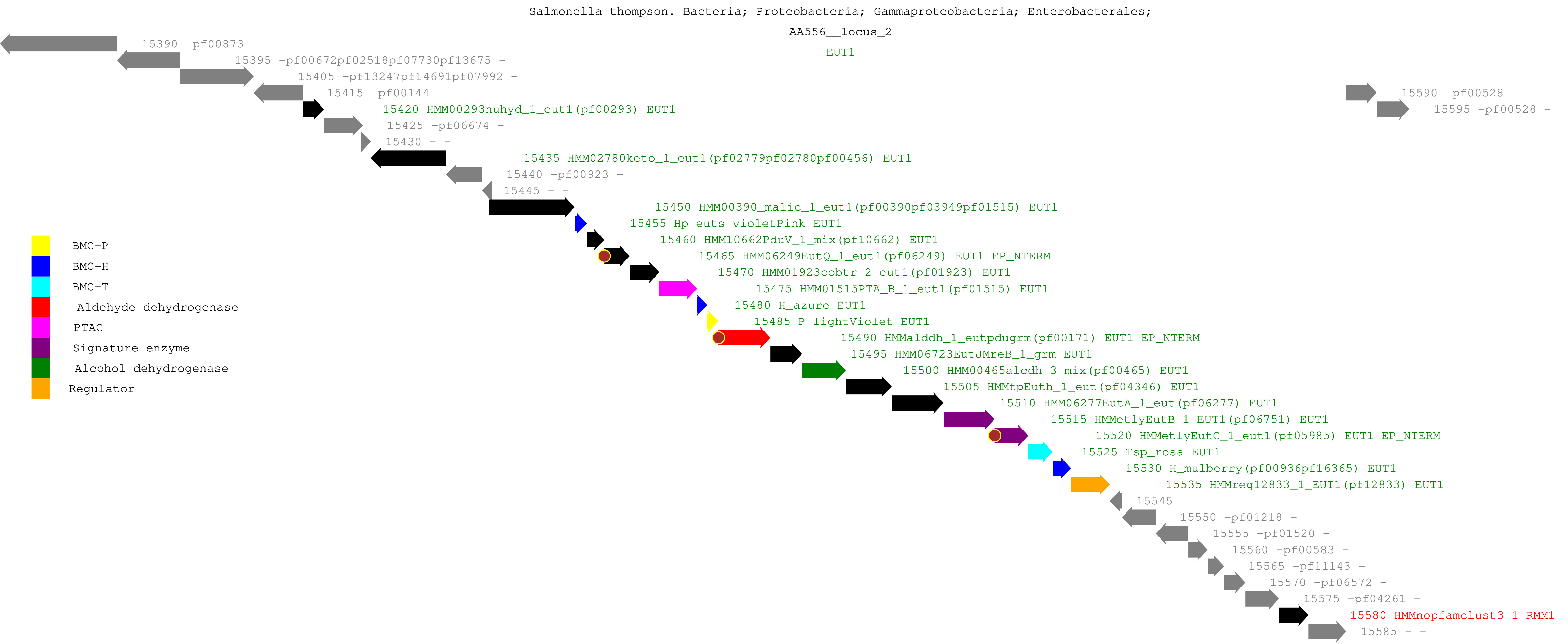
Salmonella thompson. Bacteria; Proteobacteria; Gammaproteobacteria; Enterobacterales;
AA556__locus_2
EUT1
15390 -pf00873 -
15395 -pf00672pf02518pf07730pf13675 -
15405 -pf13247pf14691pf07992 -
15415 -pf00144 -
15420 HMM00293nuhyd_1_eut1(pf00293) EUT1
15425 -pf06674 -
15430 - -
15435 HMM02780keto_1_eut1(pf02779pf02780pf00456) EUT1
15440 -pf00923 -
15445 - -
15450 HMM00390_malic_1_eut1(pf00390pf03949pf01515) EUT1
15455 Hp_euts_violetPink EUT1
15460 HMM10662PduV_1_mix(pf10662) EUT1
15465 HMM06249EutQ_1_eut1(pf06249) EUT1 EP_NTERM
15470 HMM01923cobtr_2_eut1(pf01923) EUT1
15475 HMM01515PTA_B_1_eut1(pf01515) EUT1
15480 H_azure EUT1
15485 P_lightViolet EUT1
15490 HMMalddh_1_eutpdugrm(pf00171) EUT1 EP_NTERM
15495 HMM06723EutJMreB_1_grm EUT1
15500 HMM00465alcdh_3_mix(pf00465) EUT1
15505 HMMtpEuth_1_eut(pf04346) EUT1
15510 HMM06277EutA_1_eut(pf06277) EUT1
15515 HMMetlyEutB_1_EUT1(pf06751) EUT1
15520 HMMetlyEutC_1_eut1(pf05985) EUT1 EP_NTERM
15525 Tsp_rosa EUT1
15530 H_mulberry(pf00936pf16365) EUT1
15535 HMMreg12833_1_EUT1(pf12833) EUT1
15545 - -
15550 -pf01218 -
15555 -pf01520 -
15560 -pf00583 -
15565 -pf11143 -
15570 -pf06572 -
15575 -pf04261 -
15580 HMMnopfamclust3_1 RMM1
15585 - -
15590 -pf00528 -
15595 -pf00528 -
BMC-P
BMC-H
BMC-T
Aldehyde dehydrogenase
PTAC
Signature enzyme
Alcohol dehydrogenase
Regulator

Salmonella enteritidis. Bacteria; Proteobacteria; Gammaproteobacteria; Enterobacterales;

AA689__locus_2

EUT1

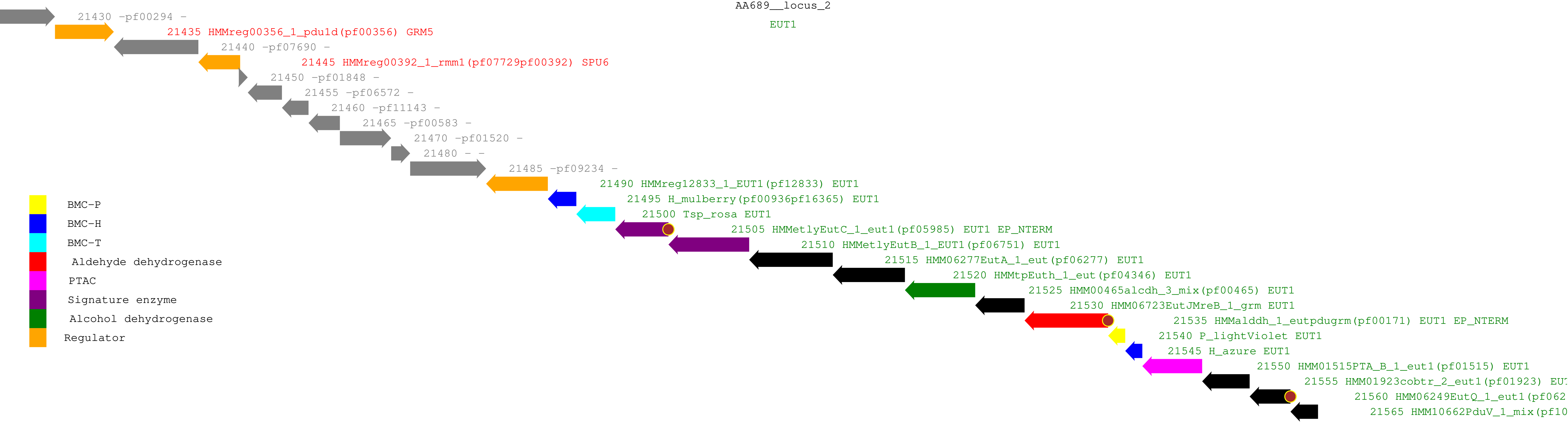

Escherichia coli 2-011-08_S1_C1. Bacteria; Proteobacteria; Gammaproteobacteria; Enterobacterales;

AA98__locus_1

EUT1

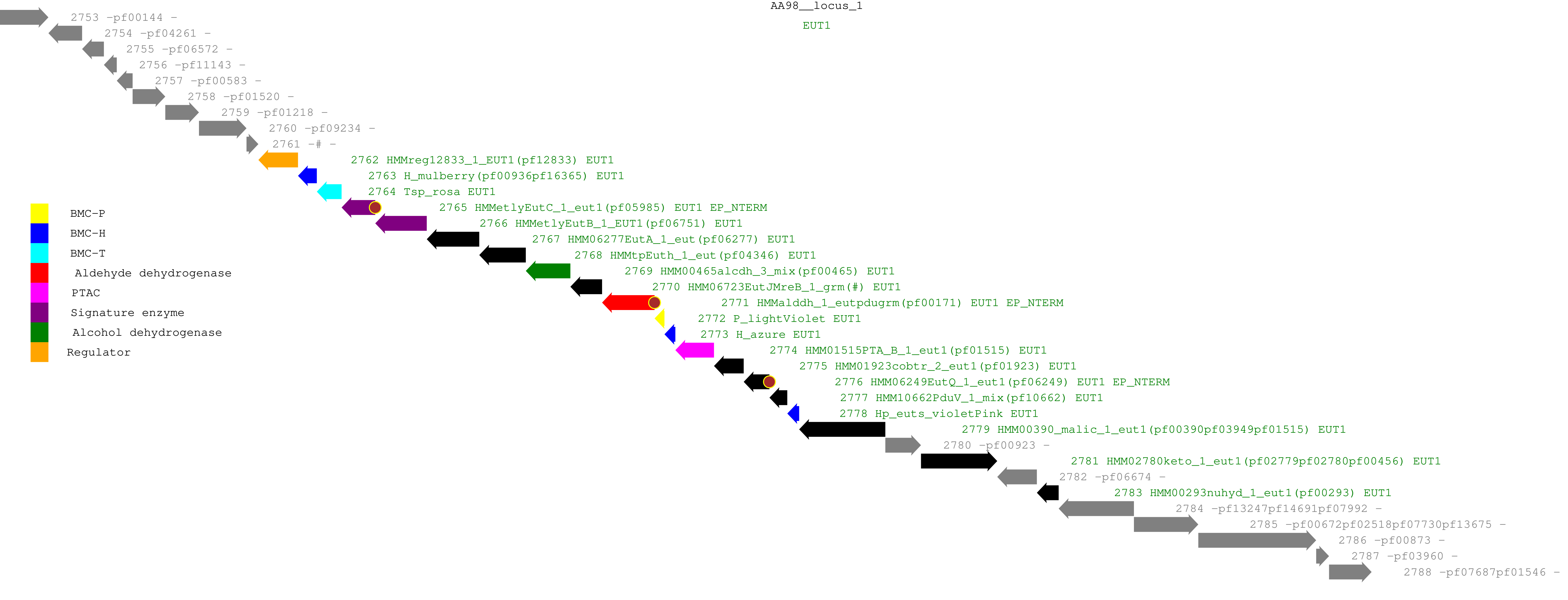

Salmonella newport. Bacteria; Proteobacteria; Gammaproteobacteria; Enterobacterales;

AAJ88__locus_2_main_sat

EUT1

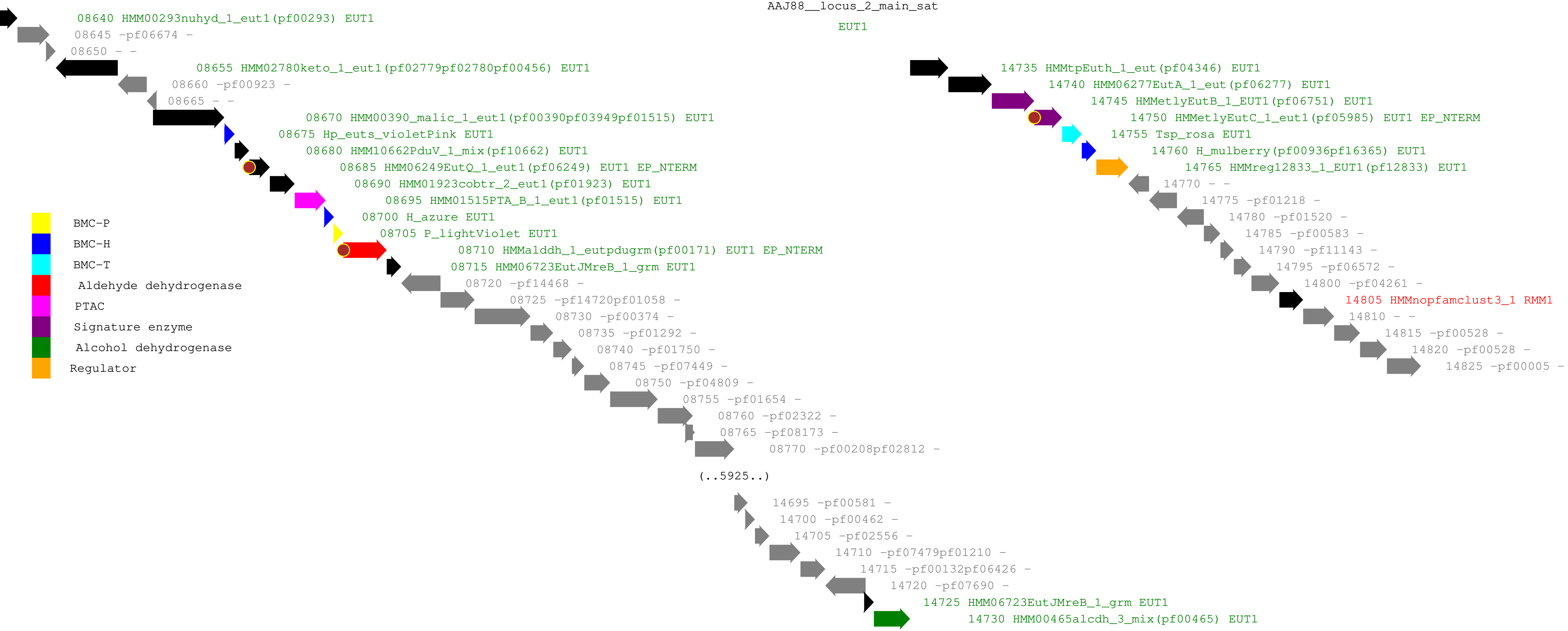

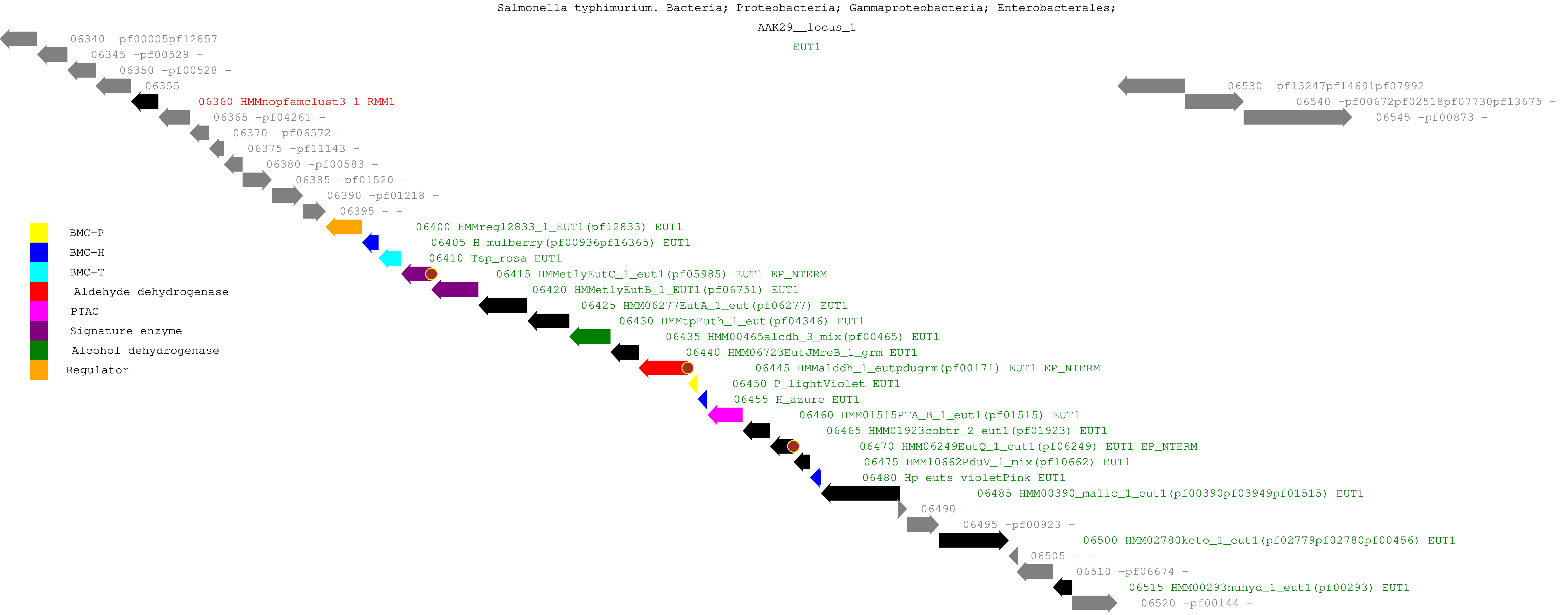

Salmonella typhimurium. Bacteria; Proteobacteria; Gammaproteobacteria; Enterobacterales;
AAK29__locus_1
EUT1
06340 -pf00005pf12857 -
06345 -pf00528 -
06350 -pf00528 -
06355 - -
06360 HMMnopfamclust3_1 RMM1
06365 -pf04261 -
06370 -pf06572 -
06375 -pf11143 -
06380 -pf00583 -
06385 -pf01520 -
06390 -pf01218 -
06395 - -
06400 HMMreg12833_1_EUT1(pf12833) EUT1
06405 H_mulberry(pf00936pf16365) EUT1
06410 Tsp_rosa EUT1
06415 HMMetlyEutC_1_eut1(pf05985) EUT1 EP_NTERM
06420 HMMetlyEutB_1_EUT1(pf06751) EUT1
06425 HMM06277EutA_1_eut(pf06277) EUT1
06430 HMMtpEuth_1_eut(pf04346) EUT1
06435 HMM00465alcdh_3_mix(pf00465) EUT1
06440 HMM06723EutJMreB_1_grm EUT1
06445 HMMalddh_1_eutpdugrm(pf00171) EUT1 EP_NTERM
06450 P_lightViolet EUT1
06455 H_azure EUT1
06460 HMM01515PTA_B_1_eut1(pf01515) EUT1
06465 HMM01923cobtr_2_eut1(pf01923) EUT1
06470 HMM06249EutQ_1_eut1(pf06249) EUT1 EP_NTERM
06475 HMM10662PduV_1_mix(pf10662) EUT1
06480 Hp_euts_violetPink EUT1
06485 HMM00390_malic_1_eut1(pf00390pf03949pf01515) EUT1
06490 - -
06495 -pf00923 -
06500 HMM02780keto_1_eut1(pf02779pf02780pf00456) EUT1
06505 - -
06510 -pf06674 -
06515 HMM00293nuhyd_1_eut1(pf00293) EUT1
06520 -pf00144 -
06530 -pf13247pf14691pf07992 -
06540 -pf00672pf02518pf07730pf13675 -
06545 -pf00873 -
BMC-P
BMC-H
BMC-T
Aldehyde dehydrogenase
PTAC
Signature enzyme
Alcohol dehydrogenase
Regulator

Salmonella enterica I. Bacteria; Proteobacteria; Gammaproteobacteria; Enterobacterales;

AAL03__locus_1

EUT1

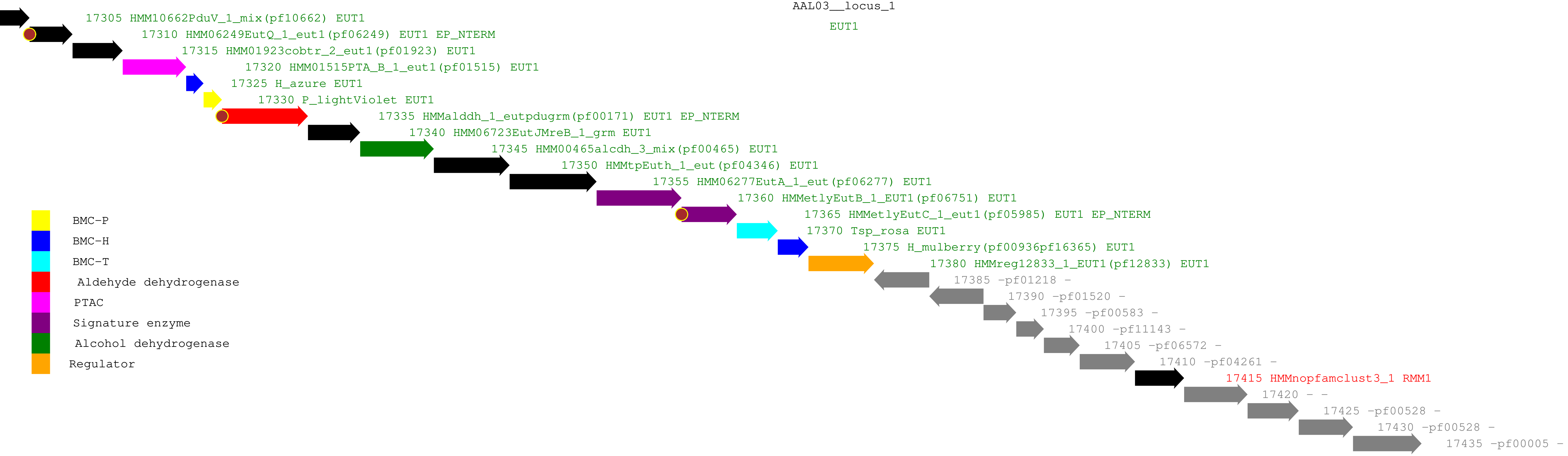

Salmonella enterica I. Bacteria; Proteobacteria; Gammaproteobacteria; Enterobacterales;

AAM89__locus_1

EUT1

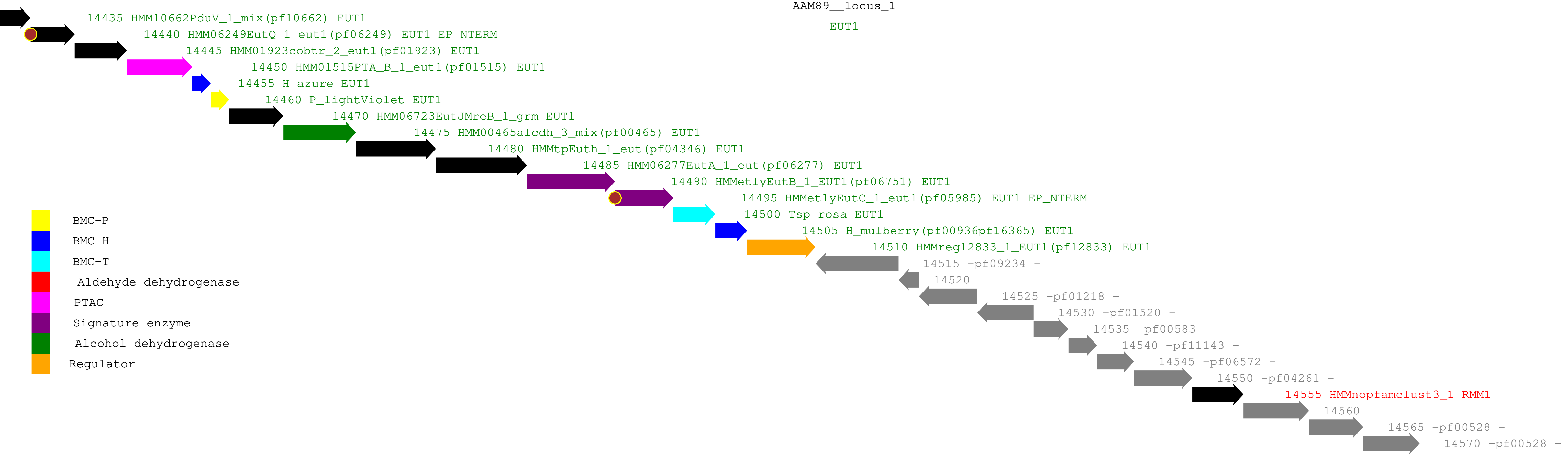

Salmonella montevideo. Bacteria; Proteobacteria; Gammaproteobacteria; Enterobacterales;

AAN00__locus_1

EUT1

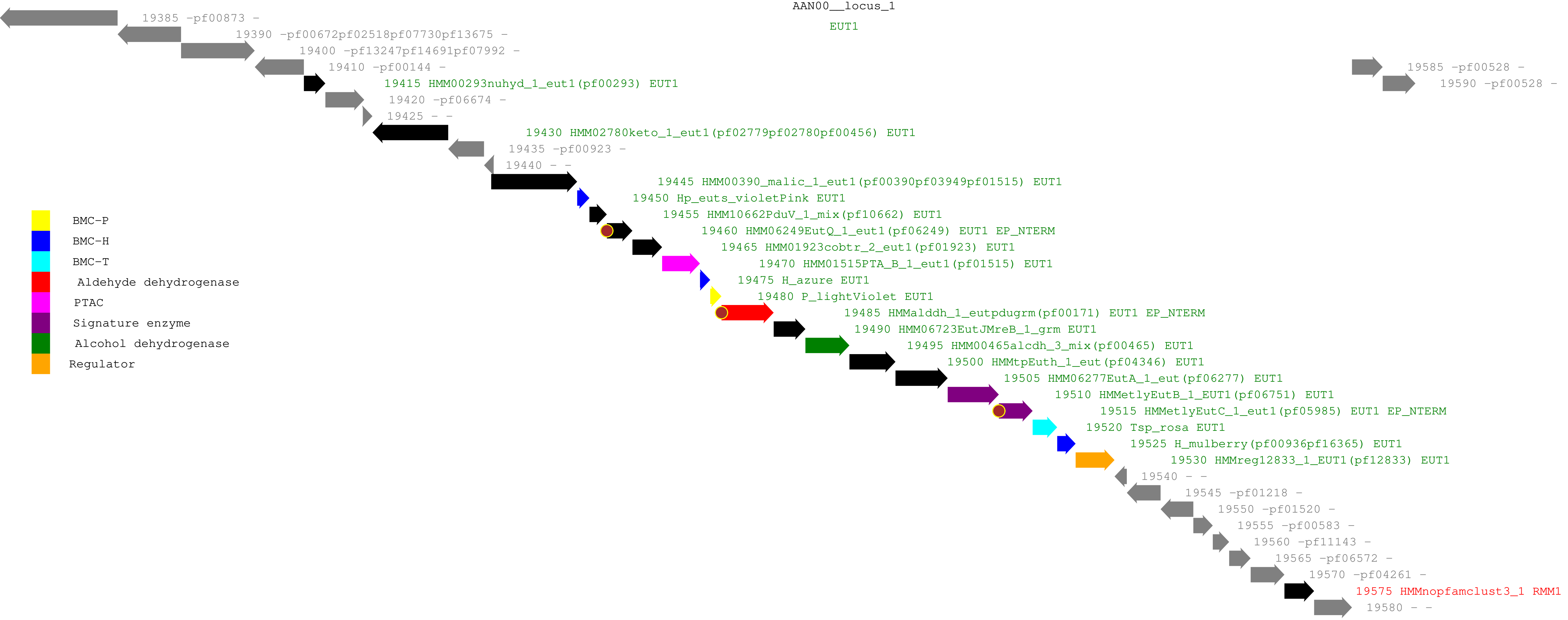

Salmonella derby. Bacteria; Proteobacteria; Gammaproteobacteria; Enterobacterales;

AAN63__locus_2

EUT1

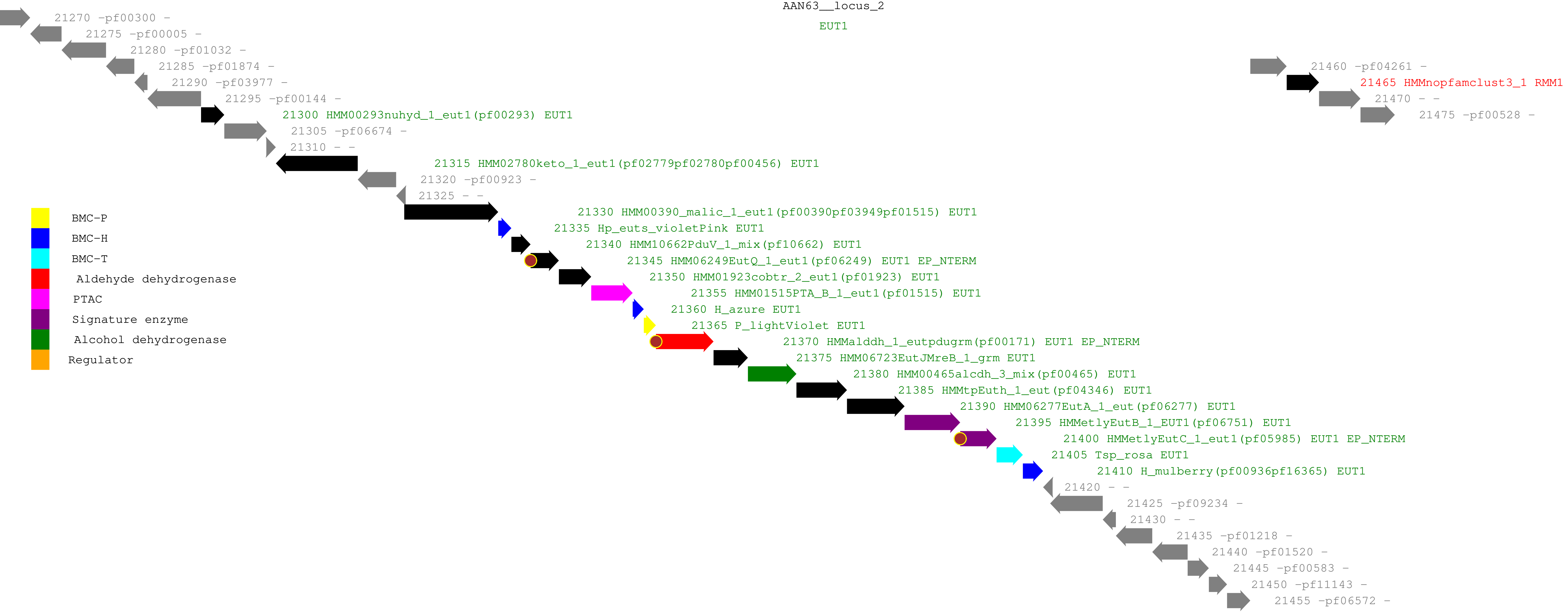

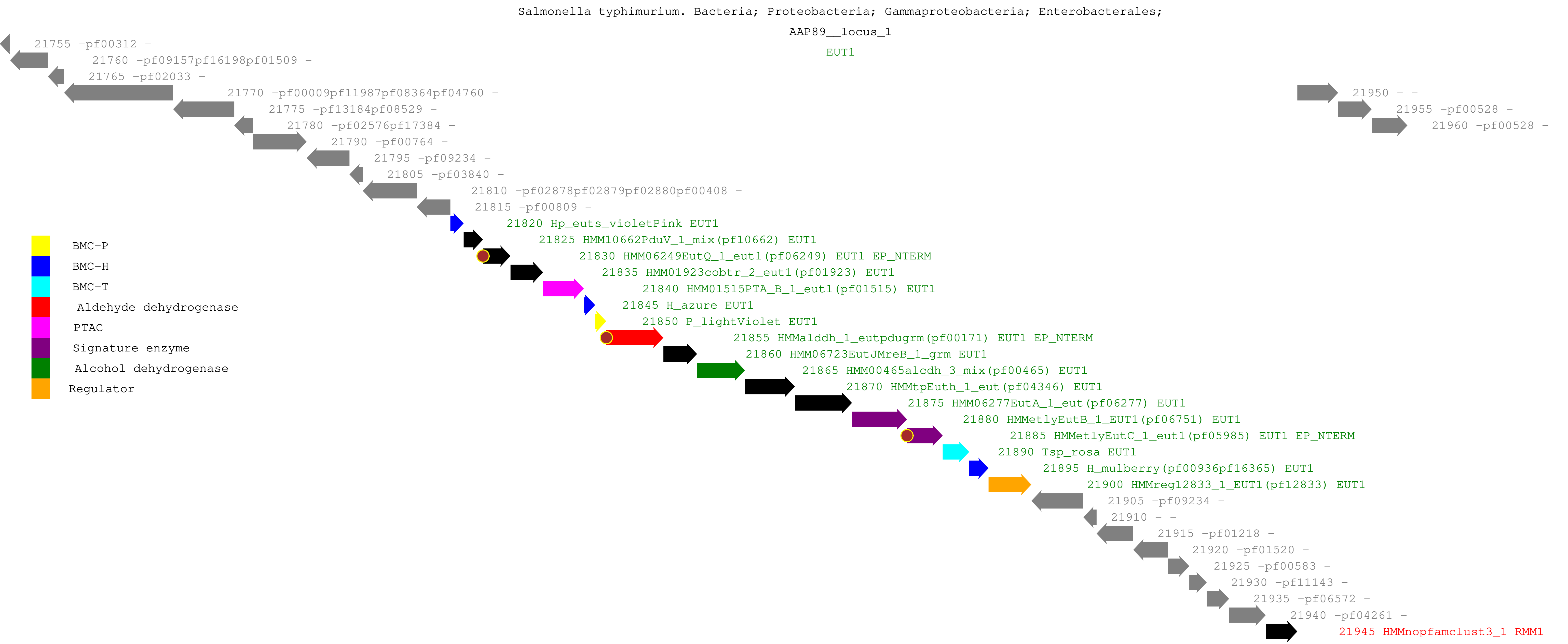
Salmonella typhimurium. Bacteria; Proteobacteria; Gammaproteobacteria; Enterobacterales;
AAP89__locus_1
EUT1
21755 -pf00312 -
21760 -pf09157pf16198pf01509 -
21765 -pf02033 -
21770 -pf00009pf11987pf08364pf04760 -
21775 -pf13184pf08529 -
21780 -pf02576pf17384 -
21790 -pf00764 -
21795 -pf09234 -
21805 -pf03840 -
21810 -pf02878pf02879pf02880pf00408 -
21815 -pf00809 -
21820 Hp_euts_violetPink EUT1
21825 HMM10662PduV_1_mix(pf10662) EUT1
21830 HMM06249EutQ_1_eut1(pf06249) EUT1 EP_NTERM
21835 HMM01923cobtr_2_eut1(pf01923) EUT1
21840 HMM01515PTA_B_1_eut1(pf01515) EUT1
21845 H_azure EUT1
21850 P_lightViolet EUT1
21855 HMMalddh_1_eutpdugrm(pf00171) EUT1 EP_NTERM
21860 HMM06723EutJMreB_1_grm EUT1
21865 HMM00465alcdh_3_mix(pf00465) EUT1
21870 HMMtpEuth_1_eut(pf04346) EUT1
21875 HMM06277EutA_1_eut(pf06277) EUT1
21880 HMMetlyEutB_1_EUT1(pf06751) EUT1
21885 HMMetlyEutC_1_eut1(pf05985) EUT1 EP_NTERM
21890 Tsp_rosa EUT1
21895 H_mulberry(pf00936pf16365) EUT1
21900 HMMreg12833_1_EUT1(pf12833) EUT1
21905 -pf09234 -
21910 - -
21915 -pf01218 -
21920 -pf01520 -
21925 -pf00583 -
21930 -pf11143 -
21935 -pf06572 -
21940 -pf04261 -
21945 HMMnopfamclust3_1 RMM1
21950 - -
21955 -pf00528 -
21960 -pf00528 -
BMC-P
BMC-H
BMC-T
Aldehyde dehydrogenase
PTAC
Signature enzyme
Alcohol dehydrogenase
Regulator

Escherichia coli. Bacteria; Proteobacteria; Gammaproteobacteria; Enterobacterales;

AAW06__locus_2

EUT1

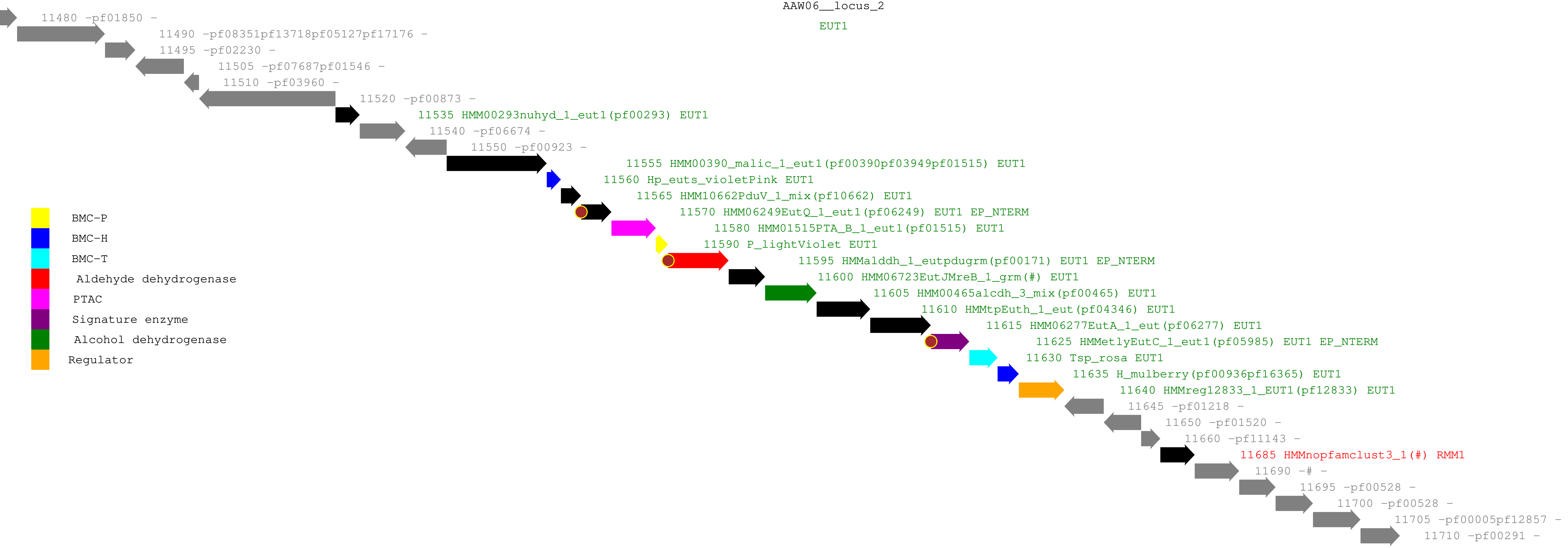

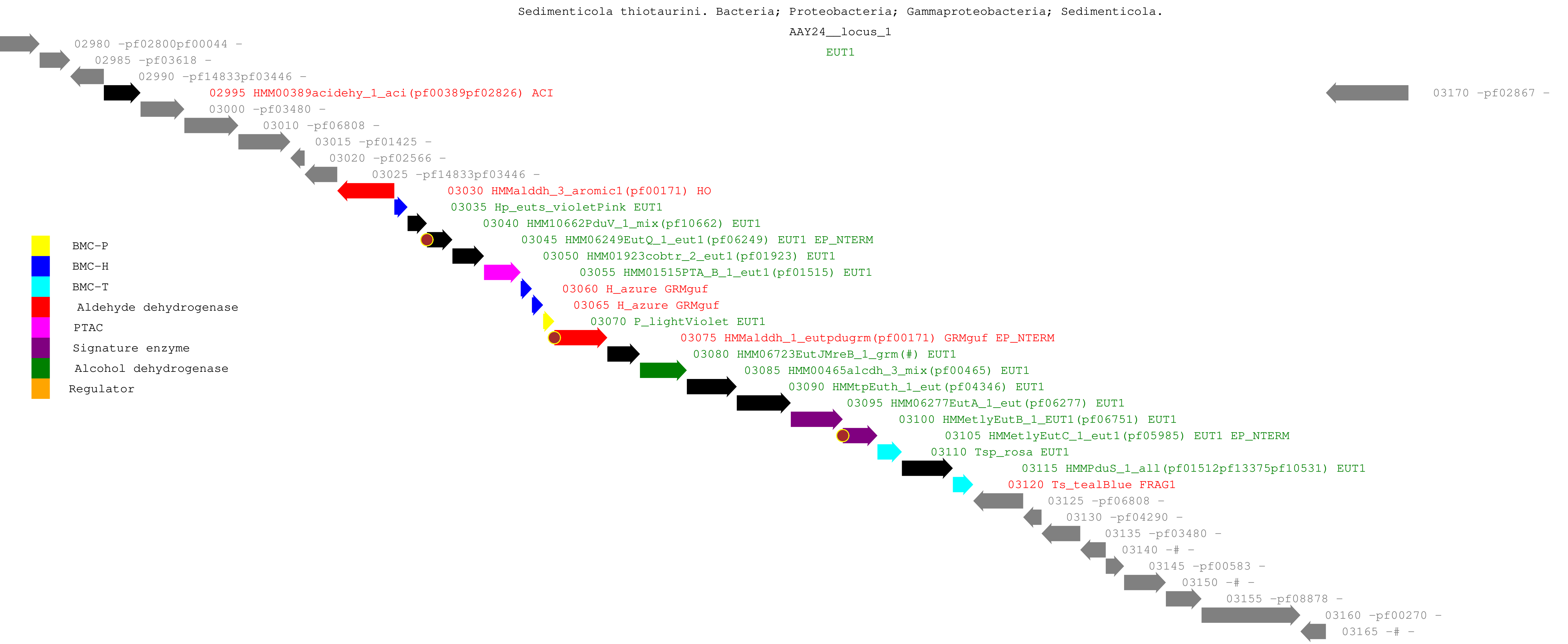

Sedimenticola thiotaurini. Bacteria; Proteobacteria; Gammaproteobacteria; Sedimenticola.
AAY24__locus_1
EUT1
02980 -pf02800pf00044 -
02985 -pf03618 -
02990 -pf14833pf03446 -
02995 HMM00389acidehy_1_aci(pf00389pf02826) ACI
03000 -pf03480 -
03010 -pf06808 -
03015 -pf01425 -
03020 -pf02566 -
03025 -pf14833pf03446 -
03030 HMMalddh_3_aromic1(pf00171) HO
03035 Hp_euts_violetPink EUT1
03040 HMM10662PduV_1_mix(pf10662) EUT1
03045 HMM06249EutQ_1_eut1(pf06249) EUT1 EP_NTERM
03050 HMM01923cobtr_2_eut1(pf01923) EUT1
03055 HMM01515PTA_B_1_eut1(pf01515) EUT1
03060 H_azure GRMguf
03065 H_azure GRMguf
03070 P_lightViolet EUT1
03075 HMMalddh_1_eutpdugrm(pf00171) GRMguf EP_NTERM
03080 HMM06723EutJMreB_1_grm(#) EUT1
03085 HMM00465alcdh_3_mix(pf00465) EUT1
03090 HMMtpEuth_1_eut(pf04346) EUT1
03095 HMM06277EutA_1_eut(pf06277) EUT1
03100 HMMetlyEutB_1_EUT1(pf06751) EUT1
03105 HMMetlyEutC_1_eut1(pf05985) EUT1 EP_NTERM
03110 Tsp_rosa EUT1
03115 HMMPduS_1_all(pf01512pf13375pf10531) EUT1
03120 Ts_tealBlue FRAG1
03125 -pf06808 -
03130 -pf04290 -
03135 -pf03480 -
03140 -# -
03145 -pf00583 -
03150 -# -
03155 -pf08878 -
03160 -pf00270 -
03165 -# -
03170 -pf02867 -
BMC-P
BMC-H
BMC-T
Aldehyde dehydrogenase
PTAC
Signature enzyme
Alcohol dehydrogenase
Regulator

Vibrio parahaemolyticus. Bacteria; Proteobacteria; Gammaproteobacteria; Vibrionales;

AAY51__locus_2

EUT1

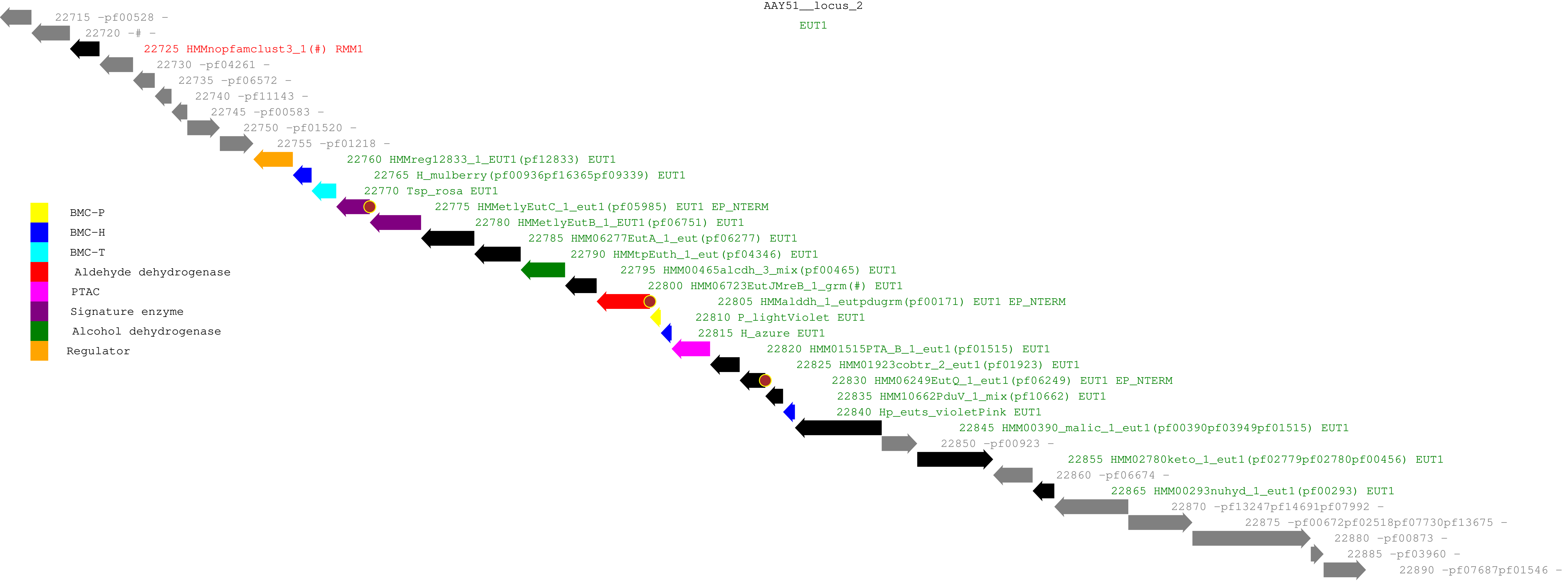

Escherichia coli 5-366-08_S1_C1. Bacteria; Proteobacteria; Gammaproteobacteria; Enterobacterales;

AB08__locus_1

EUT1

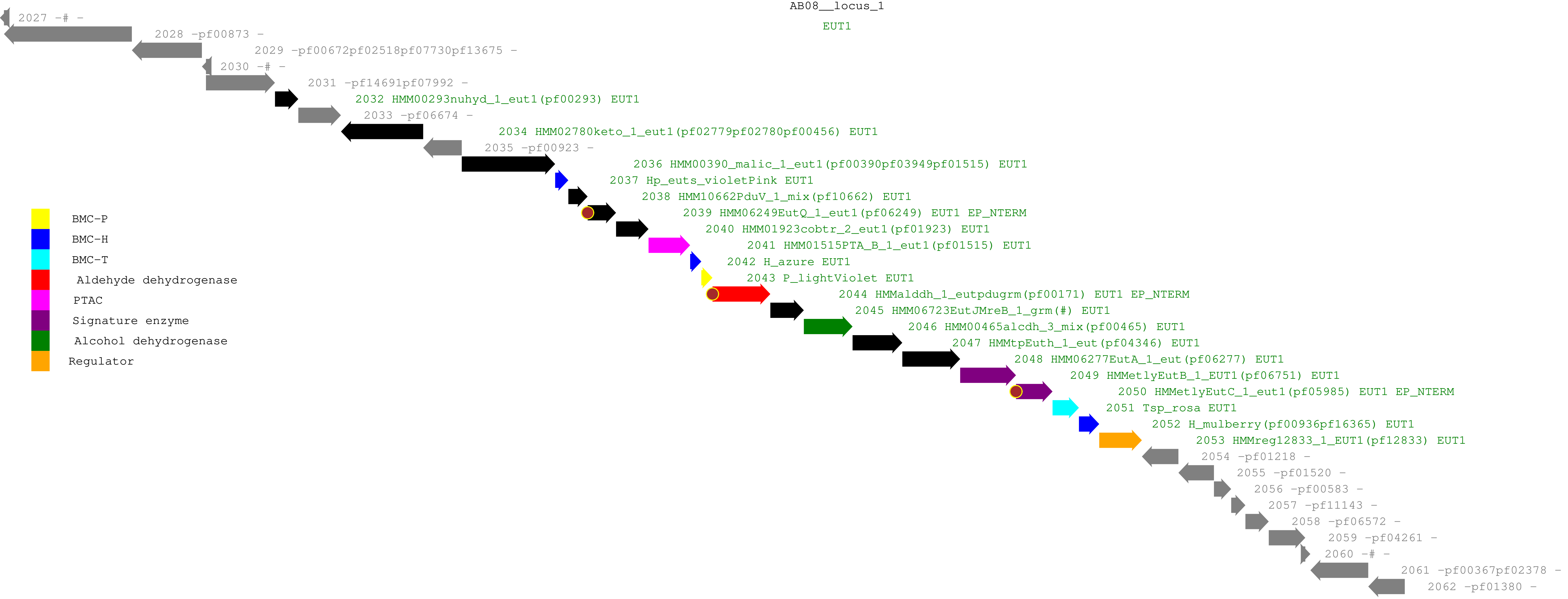

Klebsiella oxytoca. Bacteria; Proteobacteria; Gammaproteobacteria; Enterobacterales;

AB185__locus_2

EUT1

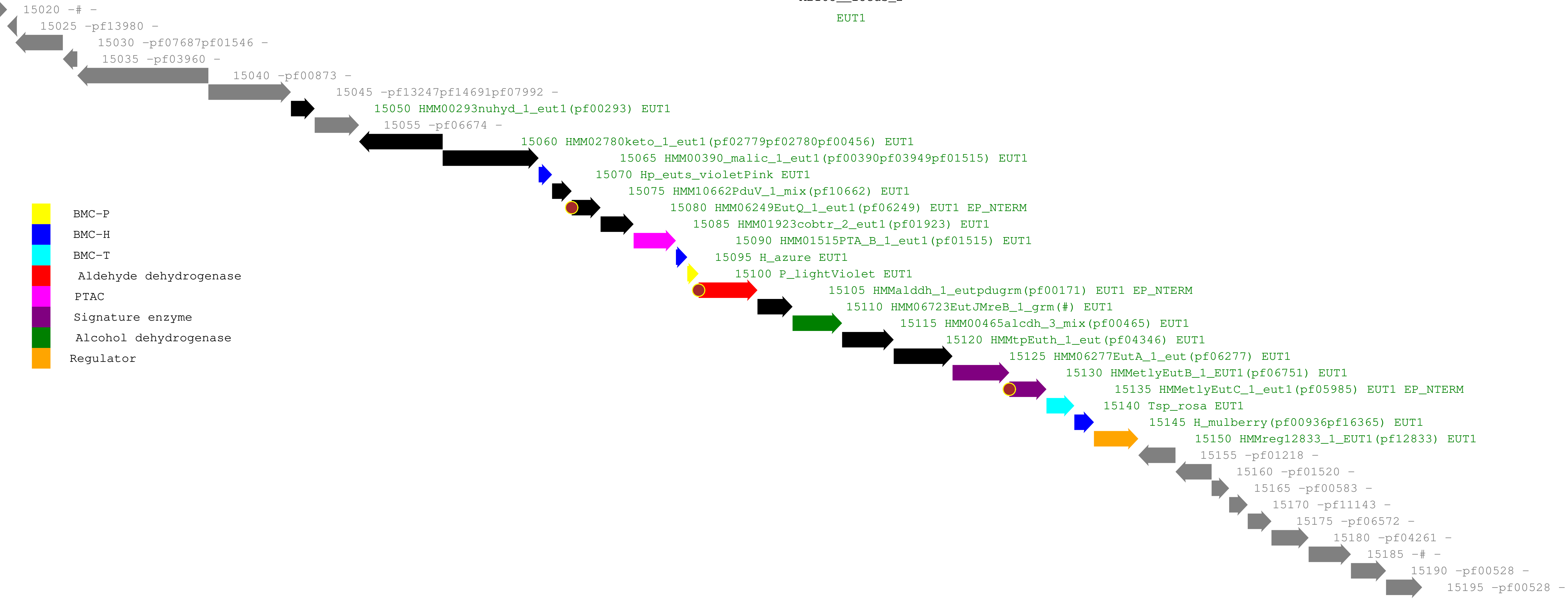

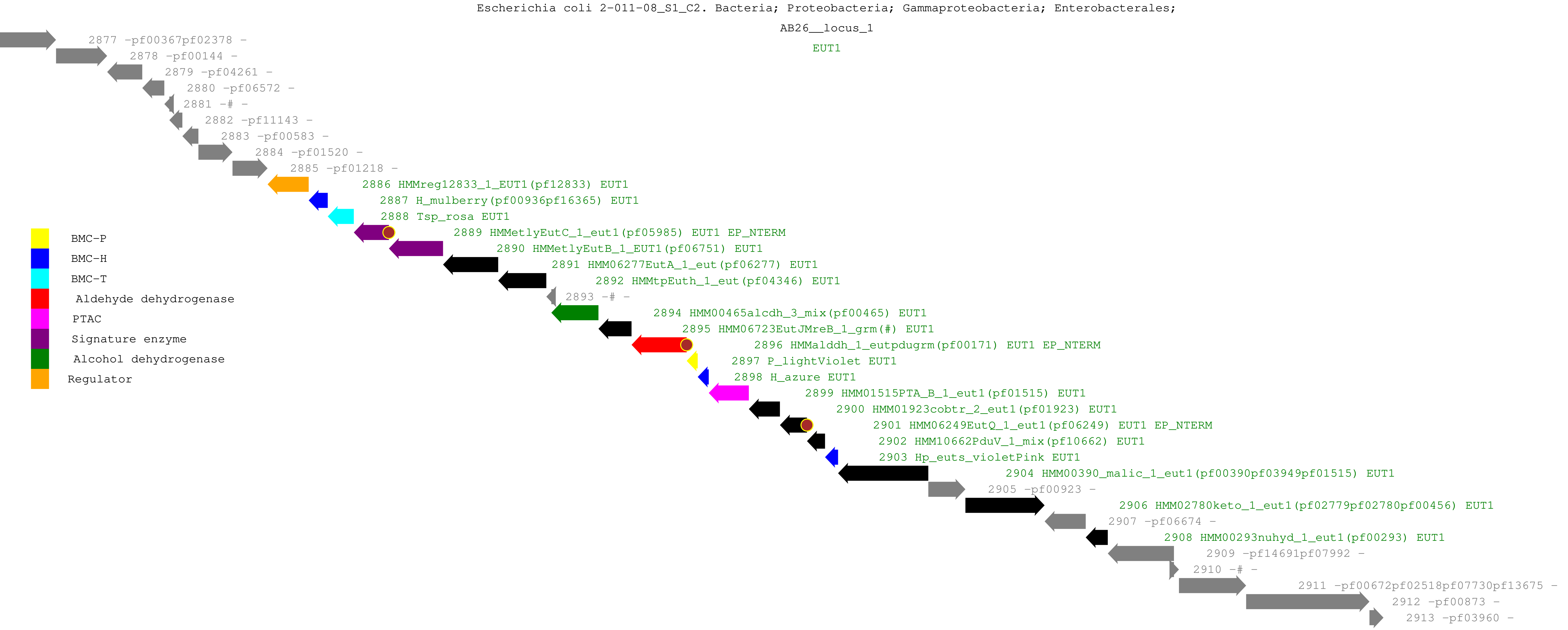

Escherichia coli 2-011-08_S1_C2. Bacteria; Proteobacteria; Gammaproteobacteria; Enterobacterales;
AB26__locus_1
EUT1
2877 -pf00367pf02378 -
2878 -pf00144 -
2879 -pf04261 -
2880 -pf06572 -
2881 -# -
2882 -pf11143 -
2883 -pf00583 -
2884 -pf01520 -
2885 -pf01218 -
2886 HMMreg12833_1_EUT1(pf12833) EUT1
2887 H_mulberry(pf00936pf16365) EUT1
2888 Tsp_rosa EUT1
2889 HMMetlyEutC_1_eut1(pf05985) EUT1 EP_NTERM
2890 HMMetlyEutB_1_EUT1(pf06751) EUT1
2891 HMM06277EutA_1_eut(pf06277) EUT1
2892 HMMtpEuth_1_eut(pf04346) EUT1
2893 -# -
2894 HMM00465alcdh_3_mix(pf00465) EUT1
2895 HMM06723EutJMreB_1_grm(#) EUT1
2896 HMMalddh_1_eutpdugrm(pf00171) EUT1 EP_NTERM
2897 P_lightViolet EUT1
2898 H_azure EUT1
2899 HMM01515PTA_B_1_eut1(pf01515) EUT1
2900 HMM01923cobtr_2_eut1(pf01923) EUT1
2901 HMM06249EutQ_1_eut1(pf06249) EUT1 EP_NTERM
2902 HMM10662PduV_1_mix(pf10662) EUT1
2903 Hp_euts_violetPink EUT1
2904 HMM00390_malic_1_eut1(pf00390pf03949pf01515) EUT1
2905 -pf00923 -
2906 HMM02780keto_1_eut1(pf02779pf02780pf00456) EUT1
2907 -pf06674 -
2908 HMM00293nuhyd_1_eut1(pf00293) EUT1
2909 -pf14691pf07992 -
2910 -# -
2911 -pf00672pf02518pf07730pf13675 -
2912 -pf00873 -
2913 -pf03960 -
BMC-P
BMC-H
BMC-T
Aldehyde dehydrogenase
PTAC
Signature enzyme
Alcohol dehydrogenase
Regulator

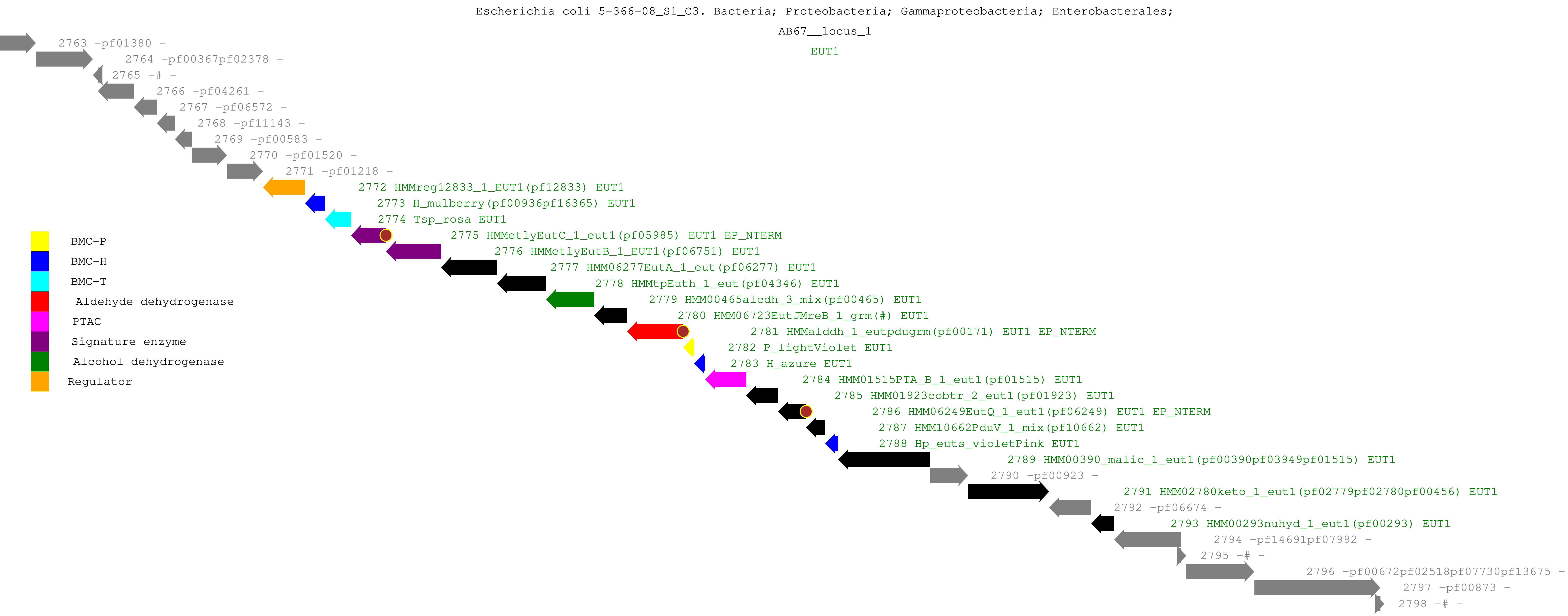

Escherichia coli 5-366-08_S1_C3. Bacteria; Proteobacteria; Gammaproteobacteria; Enterobacterales;
AB67__locus_1
EUT1
2763 -pf01380 -
2764 -pf00367pf02378 -
2765 -# -
2766 -pf04261 -
2767 -pf06572 -
2768 -pf11143 -
2769 -pf00583 -
2770 -pf01520 -
2771 -pf01218 -
2772 HMMreg12833_1_EUT1(pf12833) EUT1
2773 H_mulberry(pf00936pf16365) EUT1
2774 Tsp_rosa EUT1
2775 HMMetlyEutC_1_eut1(pf05985) EUT1 EP_NTERM
2776 HMMetlyEutB_1_EUT1(pf06751) EUT1
2777 HMM06277EutA_1_eut(pf06277) EUT1
2778 HMMtpEuth_1_eut(pf04346) EUT1
2779 HMM00465alcdh_3_mix(pf00465) EUT1
2780 HMM06723EutJMreB_1_grm(#) EUT1
2781 HMMalddh_1_eutpdugrm(pf00171) EUT1 EP_NTERM
2782 P_lightViolet EUT1
2783 H_azure EUT1
2784 HMM01515PTA_B_1_eut1(pf01515) EUT1
2785 HMM01923cobtr_2_eut1(pf01923) EUT1
2786 HMM06249EutQ_1_eut1(pf06249) EUT1 EP_NTERM
2787 HMM10662PduV_1_mix(pf10662) EUT1
2788 Hp_euts_violetPink EUT1
2789 HMM00390_malic_1_eut1(pf00390pf03949pf01515) EUT1
2790 -pf00923 -
2791 HMM02780keto_1_eut1(pf02779pf02780pf00456) EUT1
2792 -pf06674 -
2793 HMM00293nuhyd_1_eut1(pf00293) EUT1
2794 -pf14691pf07992 -
2795 -# -
2796 -pf00672pf02518pf07730pf13675 -
2797 -pf00873 -
2798 -# -
BMC-P
BMC-H
BMC-T
Aldehyde dehydrogenase
PTAC
Signature enzyme
Alcohol dehydrogenase
Regulator

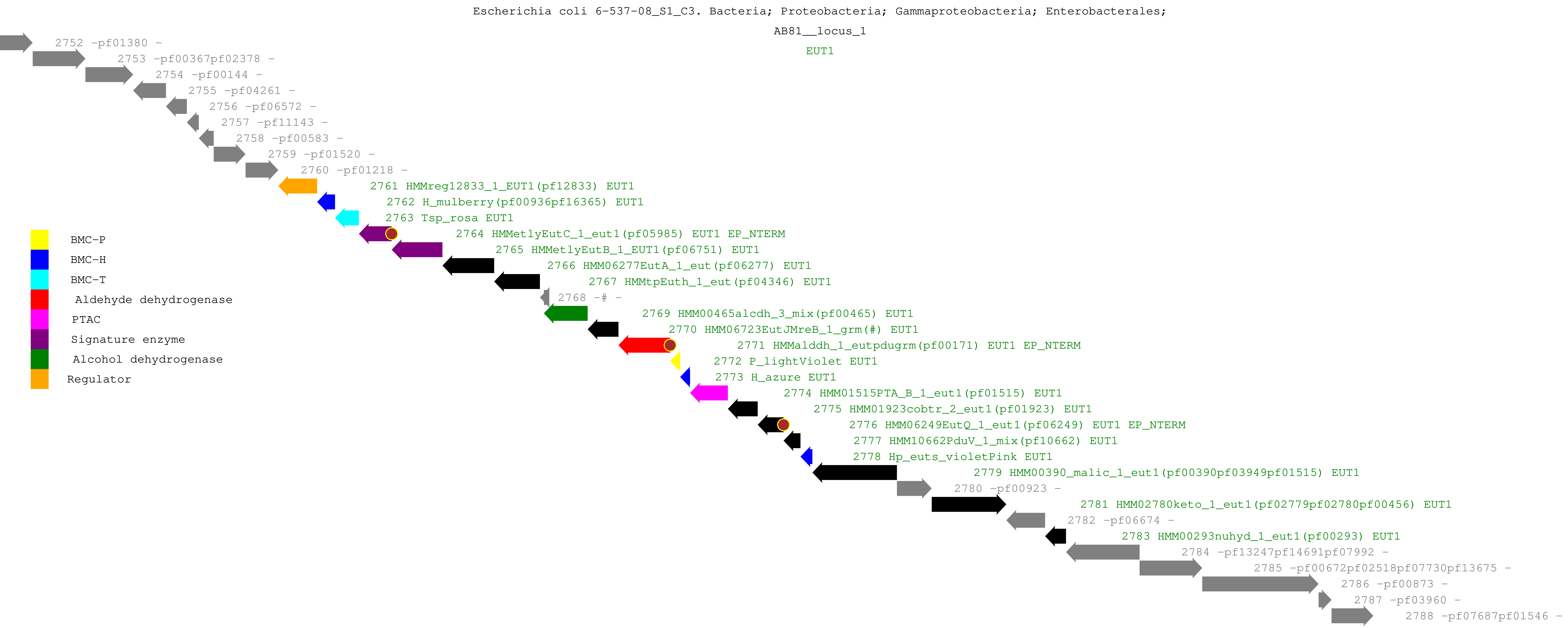
Escherichia coli 6-537-08_S1_C3. Bacteria; Proteobacteria; Gammaproteobacteria; Enterobacterales;
AB81__locus_1
EUT1
2752 -pf01380 -
2753 -pf00367pf02378 -
2754 -pf00144 -
2755 -pf04261 -
2756 -pf06572 -
2757 -pf11143 -
2758 -pf00583 -
2759 -pf01520 -
2760 -pf01218 -
2761 HMMreg12833_1_EUT1(pf12833) EUT1
2762 H_mulberry(pf00936pf16365) EUT1
2763 Tsp_rosa EUT1
2764 HMMetlyEutC_1_eut1(pf05985) EUT1 EP_NTERM
2765 HMMetlyEutB_1_EUT1(pf06751) EUT1
2766 HMM06277EutA_1_eut(pf06277) EUT1
2767 HMMtpEuth_1_eut(pf04346) EUT1
2768 -# -
2769 HMM00465alcdh_3_mix(pf00465) EUT1
2770 HMM06723EutJMreB_1_grm(#) EUT1
2771 HMMaldh_1_eutpdugrm(pf00171) EUT1 EP_NTERM
2772 P_lightViolet EUT1
2773 H_azure EUT1
2774 HMM01515PTA_B_1_eut1(pf01515) EUT1
2775 HMM01923cobtr_2_eut1(pf01923) EUT1
2776 HMM06249EutQ_1_eut1(pf06249) EUT1 EP_NTERM
2777 HMM10662PduV_1_mix(pf10662) EUT1
2778 Hp_euts_violetPink EUT1
2779 HMM00390_malic_1_eut1(pf00390pf03949pf01515) EUT1
2780 -pf00923 -
2781 HMM02780keto_1_eut1(pf02779pf02780pf00456) EUT1
2782 -pf06674 -
2783 HMM00293nuhyd_1_eut1(pf00293) EUT1
2784 -pf13247pf14691pf07992 -
2785 -pf00672pf02518pf07730pf13675 -
2786 -pf00873 -
2787 -pf03960 -
2788 -pf07687pf01546 -
BMC-P
BMC-H
BMC-T
Aldehyde dehydrogenase
PTAC
Signature enzyme
Alcohol dehydrogenase
Regulator

Escherichia coli 1-110-08_S3_C1. Bacteria; Proteobacteria; Gammaproteobacteria; Enterobacterales;

AB97__locus_1

EUT1

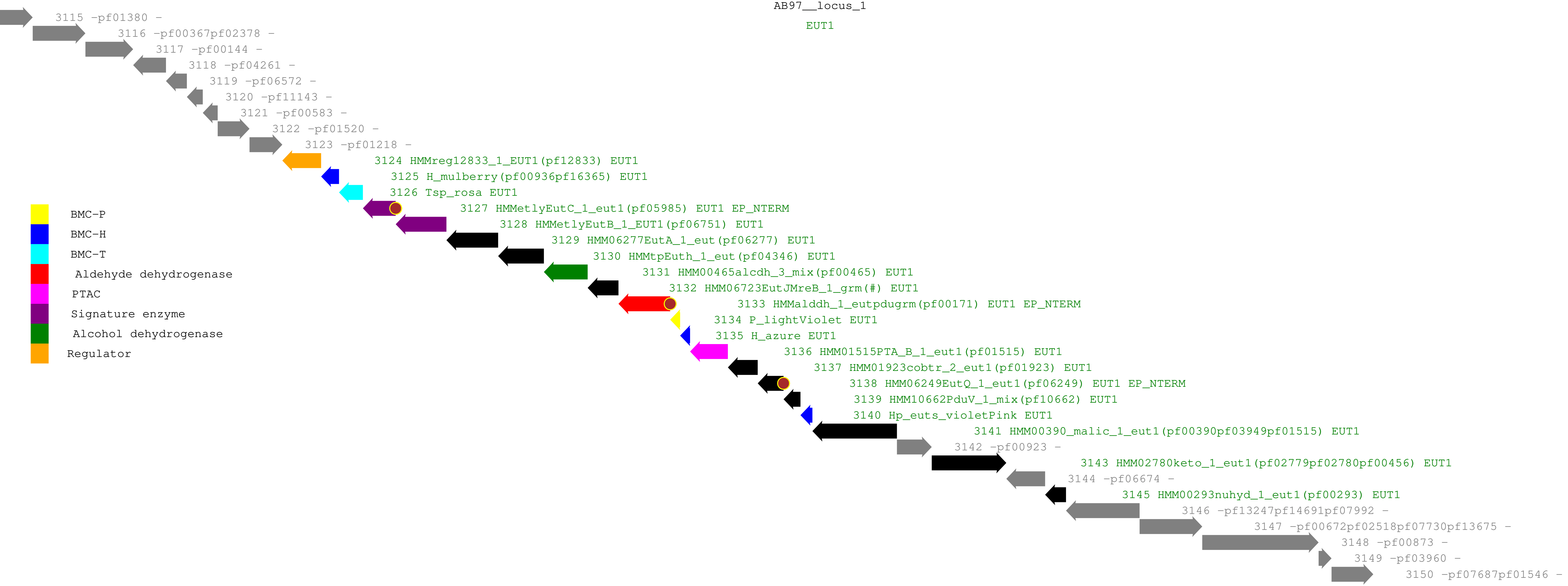

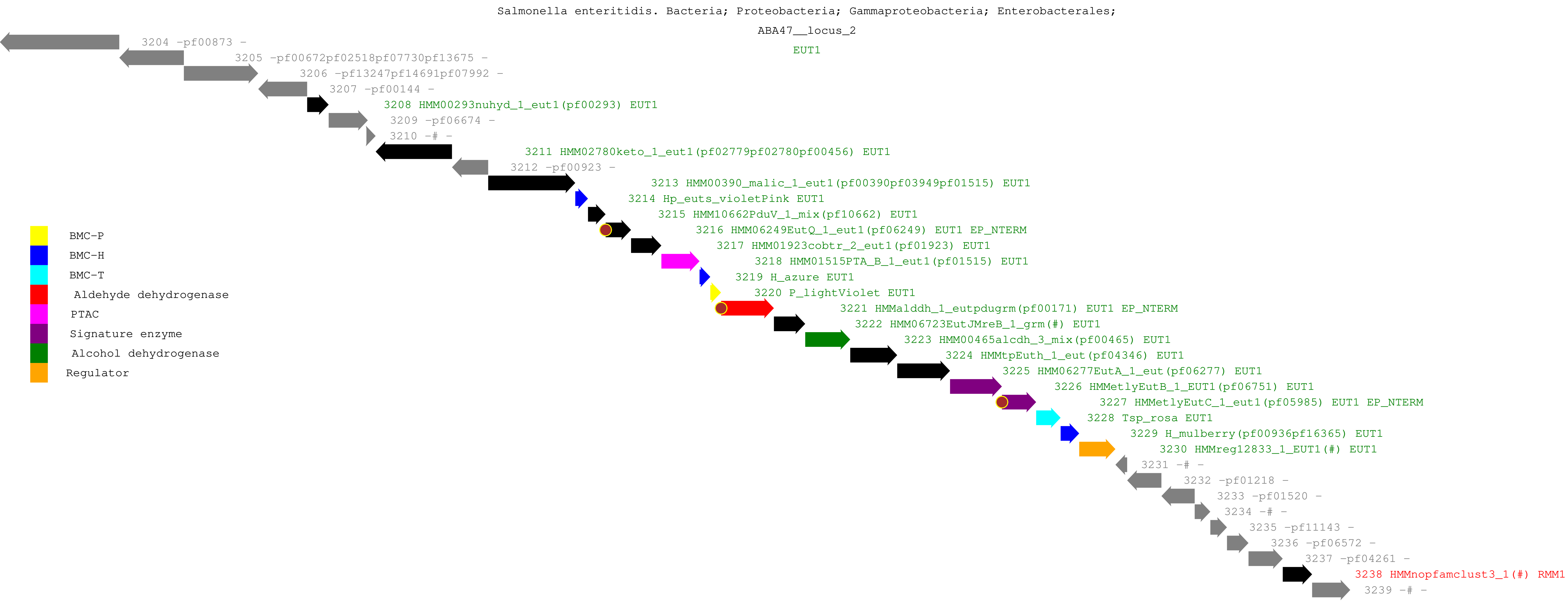
Salmonella enteritidis. Bacteria; Proteobacteria; Gammaproteobacteria; Enterobacterales;
ABA47__locus_2
EUT1
3204 -pf00873 -
3205 -pf00672pf02518pf07730pf13675 -
3206 -pf13247pf14691pf07992 -
3207 -pf00144 -
3208 HMM00293nuhyd_1_eut1(pf00293) EUT1
3209 -pf06674 -
3210 -# -
3211 HMM02780keto_1_eut1(pf02779pf02780pf00456) EUT1
3212 -pf00923 -
3213 HMM00390_malic_1_eut1(pf00390pf03949pf01515) EUT1
3214 Hp_euts_violetPink EUT1
3215 HMM10662PduV_1_mix(pf10662) EUT1
3216 HMM06249EutQ_1_eut1(pf06249) EUT1 EP_NTERM
3217 HMM01923cobtr_2_eut1(pf01923) EUT1
3218 HMM01515PTA_B_1_eut1(pf01515) EUT1
3219 H_azure EUT1
3220 P_lightViolet EUT1
3221 HMMalddh_1_eutpdugrm(pf00171) EUT1 EP_NTERM
3222 HMM06723EutJMreB_1_grm(#) EUT1
3223 HMM00465alcdh_3_mix(pf00465) EUT1
3224 HMMtpEuth_1_eut(pf04346) EUT1
3225 HMM06277EutA_1_eut(pf06277) EUT1
3226 HMMetlyEutB_1_EUT1(pf06751) EUT1
3227 HMMetlyEutC_1_eut1(pf05985) EUT1 EP_NTERM
3228 Tsp_rosa EUT1
3229 H_mulberry(pf00936pf16365) EUT1
3230 HMMreg12833_1_EUT1(#) EUT1
3231 -# -
3232 -pf01218 -
3233 -pf01520 -
3234 -# -
3235 -pf11143 -
3236 -pf06572 -
3237 -pf04261 -
3238 HMMnopfamclust3_1(#) RMM1
3239 -# -
BMC-P
BMC-H
BMC-T
Aldehyde dehydrogenase
PTAC
Signature enzyme
Alcohol dehydrogenase
Regulator

Shigella boydii. Bacteria; Proteobacteria; Gammaproteobacteria; Enterobacterales;

ABE81__locus_1

EUT1

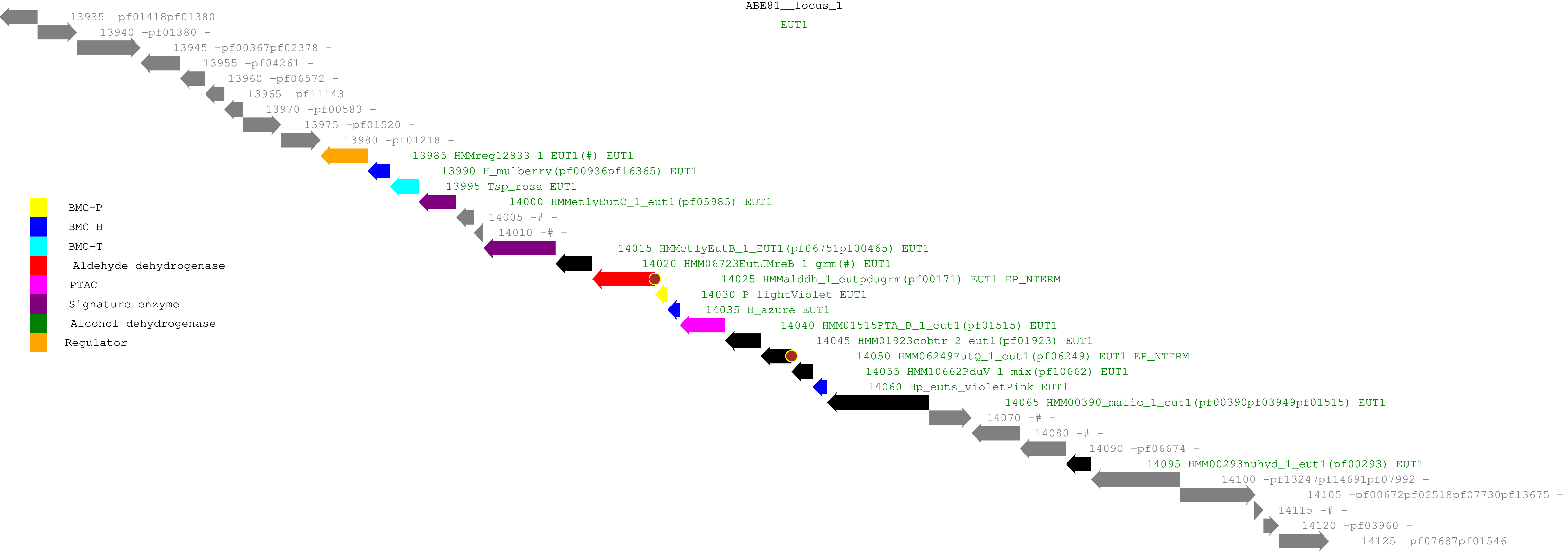

Salmonella enterica subsp. enterica serovar Mbandaka. Bacteria; Proteobacteria; Gammaproteobacteria; Enterobacterales;

ABJ60__locus_2_main_sat

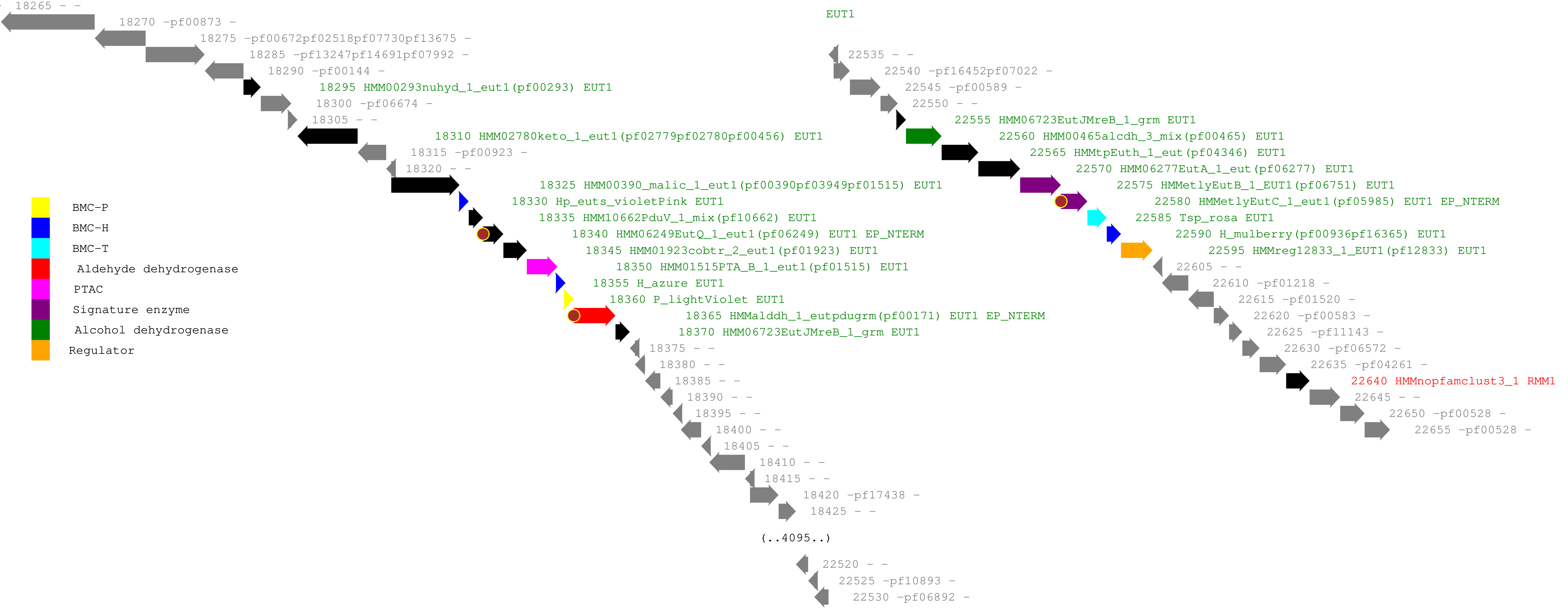

Salmonella senftenberg. Bacteria; Proteobacteria; Gammaproteobacteria; Enterobacterales;

ABM45__locus_2_main_sat

EUT1

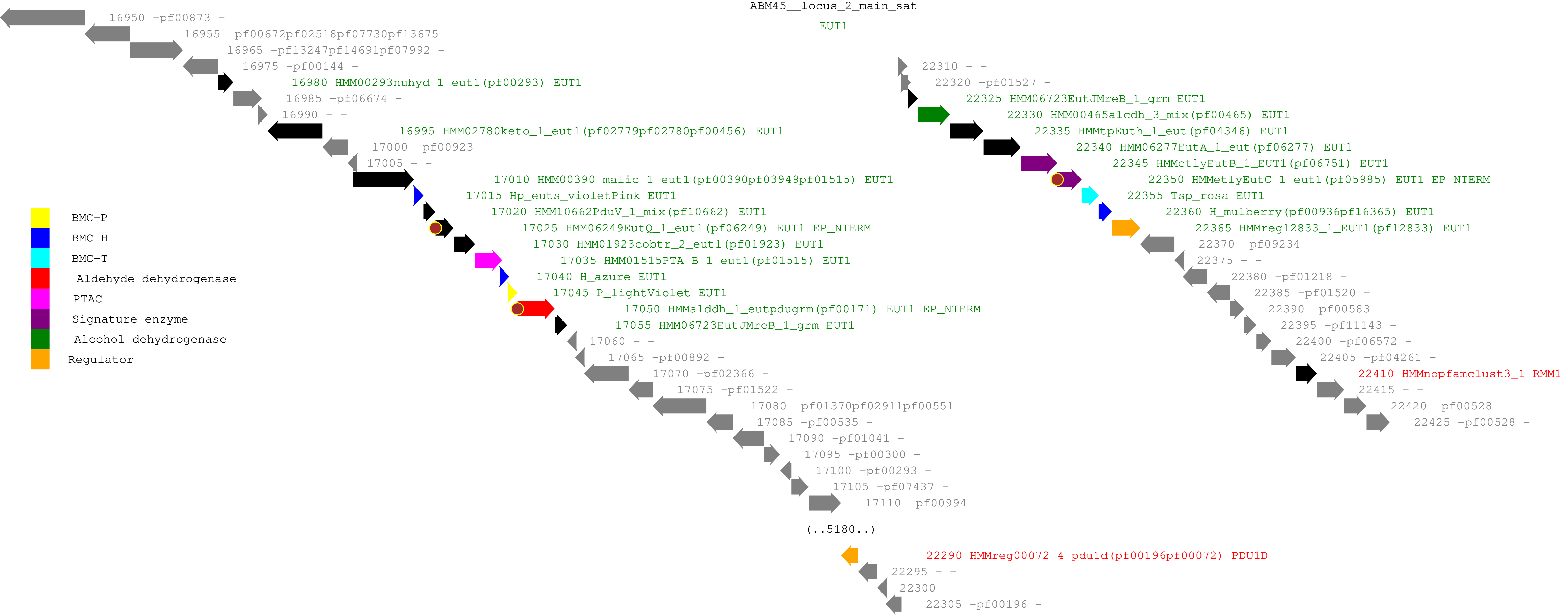

Salmonella typhi. Bacteria; Proteobacteria; Gammaproteobacteria; Enterobacterales;

ABP05__locus_1_main_sat

EUT1

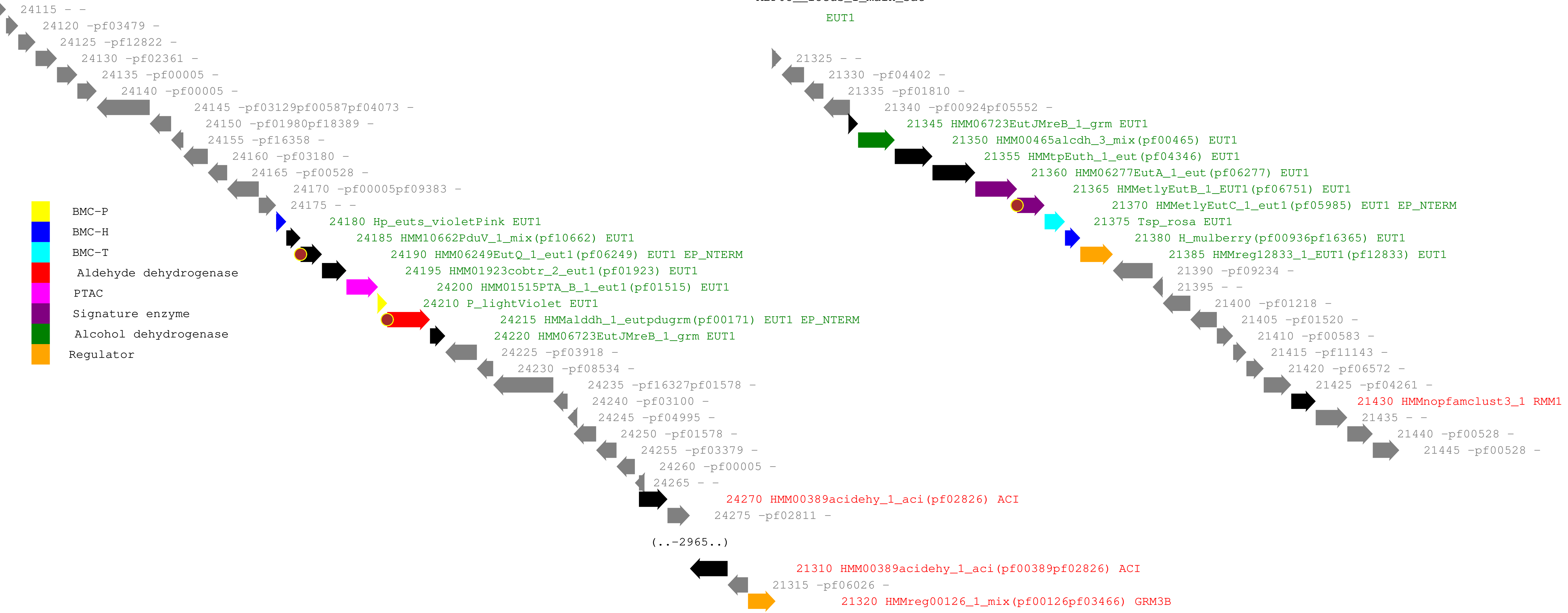

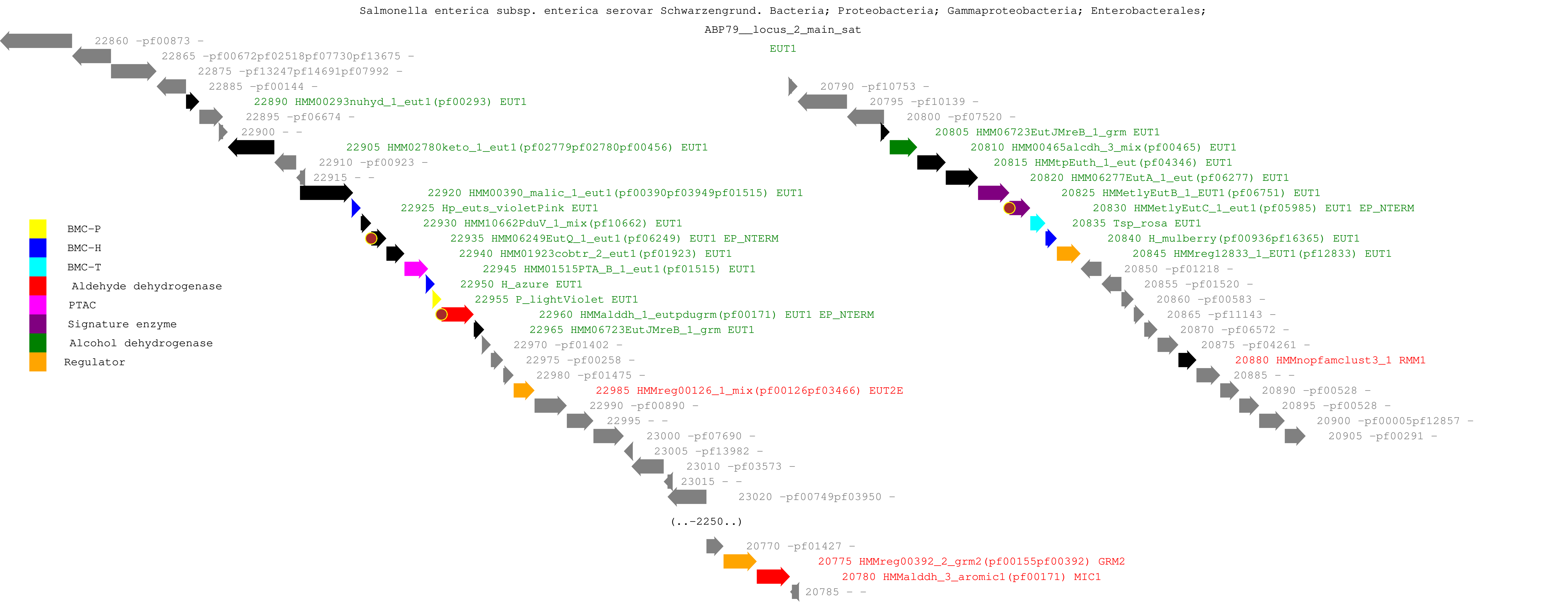
Salmonella enterica subsp. enterica serovar Schwarzengrund. Bacteria; Proteobacteria; Gammaproteobacteria; Enterobacterales;
ABP79__locus_2_main_sat
EUT1
22860 -pf00873 -
22865 -pf00672pf02518pf07730pf13675 -
22875 -pf13247pf14691pf07992 -
22885 -pf00144 -
22890 HMM00293nuhyd_1_eut1(pf00293) EUT1
22895 -pf06674 -
22900 - -
22905 HMM02780keto_1_eut1(pf02779pf02780pf00456) EUT1
22910 -pf00923 -
22915 - -
22920 HMM00390_malic_1_eut1(pf00390pf03949pf01515) EUT1
22925 Hp_euts_violetPink EUT1
22930 HMM10662PduV_1_mix(pf10662) EUT1
22935 HMM06249EutQ_1_eut1(pf06249) EUT1 EP_NTERM
22940 HMM01923cobtr_2_eut1(pf01923) EUT1
22945 HMM01515PTA_B_1_eut1(pf01515) EUT1
22950 H_azure EUT1
22955 P_lightViolet EUT1
22960 HMMalddh_1_eutpdugrm(pf00171) EUT1 EP_NTERM
22965 HMM06723EutJMreB_1_grm EUT1
22970 -pf01402 -
22975 -pf00258 -
22980 -pf01475 -
22985 HMMreg00126_1_mix(pf00126pf03466) EUT2E
22990 -pf00890 -
22995 - -
23000 -pf07690 -
23005 -pf13982 -
23010 -pf03573 -
23015 - -
23020 -pf00749pf03950 -
(..-2250..)
20770 -pf01427 -
20775 HMMreg00392_2_grm2(pf00155pf00392) GRM2
20780 HMMaldh_3_aromic1(pf00171) MIC1
20785 - -
20790 -pf10753 -
20795 -pf10139 -
20800 -pf07520 -
20805 HMM06723EutJMreB_1_grm EUT1
20810 HMM00465alcdh_3_mix(pf00465) EUT1
20815 HMMtpEuth_1_eut(pf04346) EUT1
20820 HMM06277EutA_1_eut(pf06277) EUT1
20825 HMMetlyEutB_1_EUT1(pf06751) EUT1
20830 HMMetlyEutC_1_eut1(pf05985) EUT1 EP_NTERM
20835 Tsp_rosa EUT1
20840 H_mulberry(pf00936pf16365) EUT1
20845 HMMreg12833_1_EUT1(pf12833) EUT1
20850 -pf01218 -
20855 -pf01520 -
20860 -pf00583 -
20865 -pf11143 -
20870 -pf06572 -
20875 -pf04261 -
20880 HMMnopfamclust3_1 RMM1
20885 - -
20890 -pf00528 -
20895 -pf00528 -
20900 -pf00005pf12857 -
20905 -pf00291 -
BMC-P
BMC-H
BMC-T
Aldehyde dehydrogenase
PTAC
Signature enzyme
Alcohol dehydrogenase
Regulator

Enterobacter sp. GN02600. Bacteria; Proteobacteria; Gammaproteobacteria; Enterobacterales;

ABR34__locus_2

EUT1

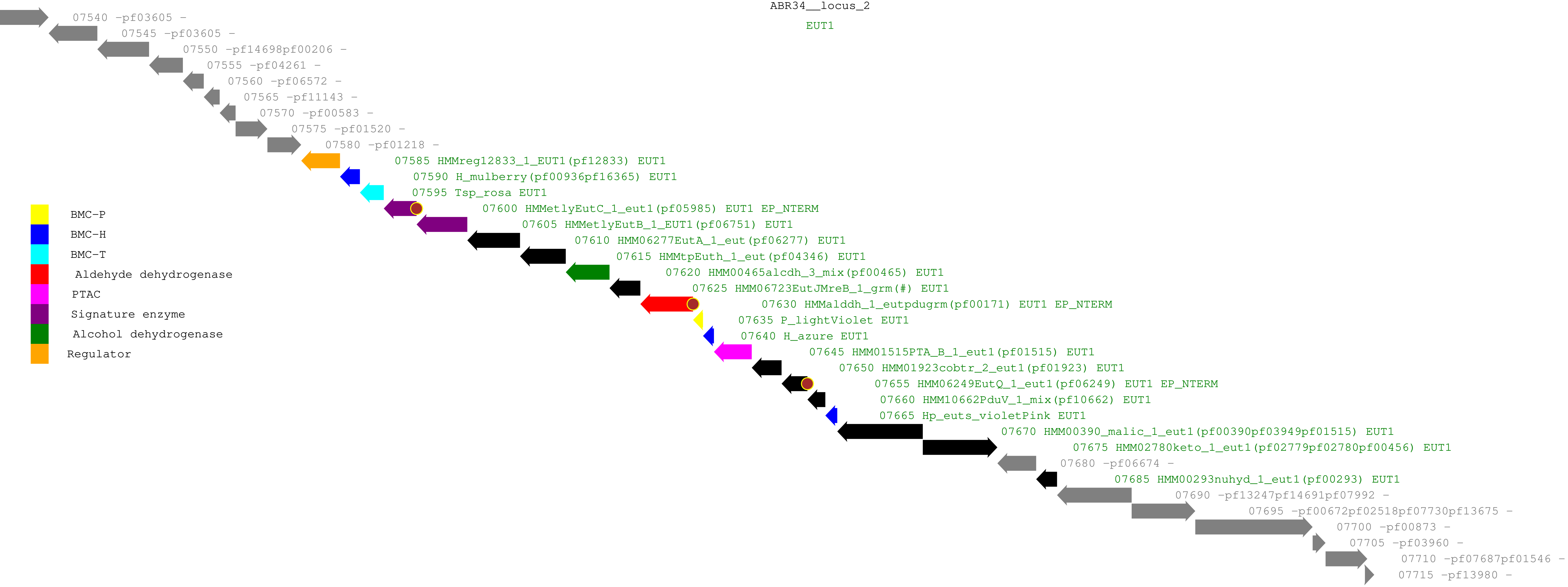

Photobacterium aquae. Bacteria; Proteobacteria; Gammaproteobacteria; Vibrionales;

ABT56__locus_1

EUT1

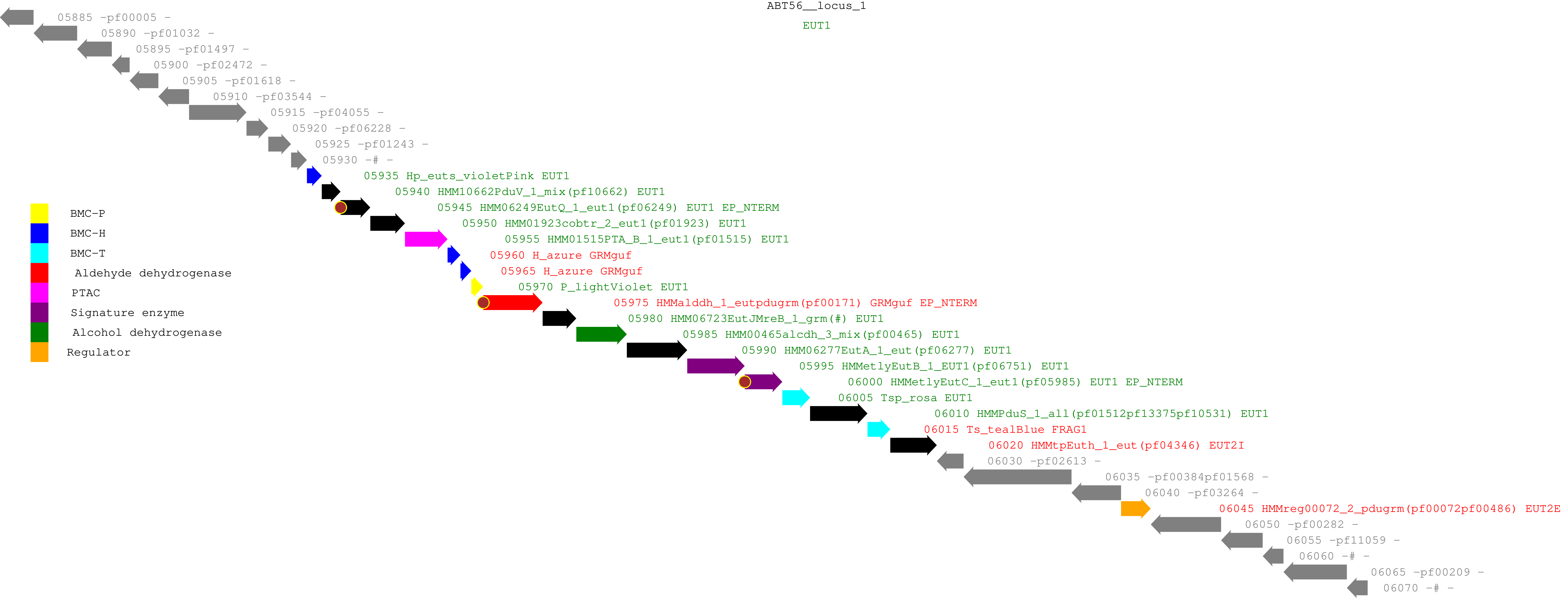

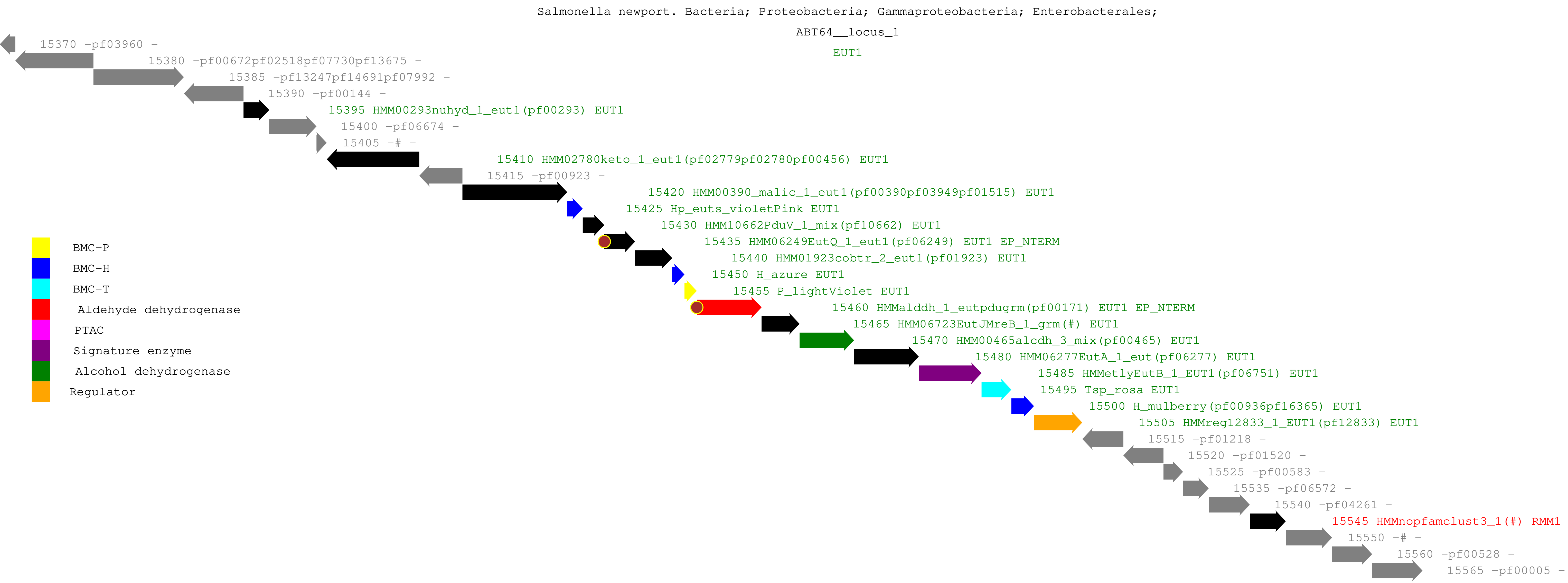

Salmonella newport. Bacteria; Proteobacteria; Gammaproteobacteria; Enterobacterales;
ABT64__locus_1
EUT1
15370 -pf03960 -
15380 -pf00672pf02518pf07730pf13675 -
15385 -pf13247pf14691pf07992 -
15390 -pf00144 -
15395 HMM00293nuhyd_1_eut1(pf00293) EUT1
15400 -pf06674 -
15405 -# -
15410 HMM02780keto_1_eut1(pf02779pf02780pf00456) EUT1
15415 -pf00923 -
15420 HMM00390_malic_1_eut1(pf00390pf03949pf01515) EUT1
15425 Hp_euts_violetPink EUT1
15430 HMM10662PduV_1_mix(pf10662) EUT1
15435 HMM06249EutQ_1_eut1(pf06249) EUT1 EP_NTERM
15440 HMM01923cobtr_2_eut1(pf01923) EUT1
15450 H_azure EUT1
15455 P_lightViolet EUT1
15460 HMMalddh_1_eutpdugrm(pf00171) EUT1 EP_NTERM
15465 HMM06723EutJMreB_1_grm(#) EUT1
15470 HMM00465alcdh_3_mix(pf00465) EUT1
15480 HMM06277EutA_1_eut(pf06277) EUT1
15485 HMMetlyEutB_1_EUT1(pf06751) EUT1
15495 Tsp_rosa EUT1
15500 H_mulberry(pf00936pf16365) EUT1
15505 HMMreg12833_1_EUT1(pf12833) EUT1
15515 -pf01218 -
15520 -pf01520 -
15525 -pf00583 -
15535 -pf06572 -
15540 -pf04261 -
15545 HMMnopfamclust3_1(#) RMM1
15550 -# -
15560 -pf00528 -
15565 -pf00005 -
BMC-P
BMC-H
BMC-T
Aldehyde dehydrogenase
PTAC
Signature enzyme
Alcohol dehydrogenase
Regulator

Pluralibacter gergoviae (Enterobacter gergoviae). Bacteria; Proteobacteria; Gammaproteobacteria; Enterobacterales;

ABW06__locus_1

EUT1

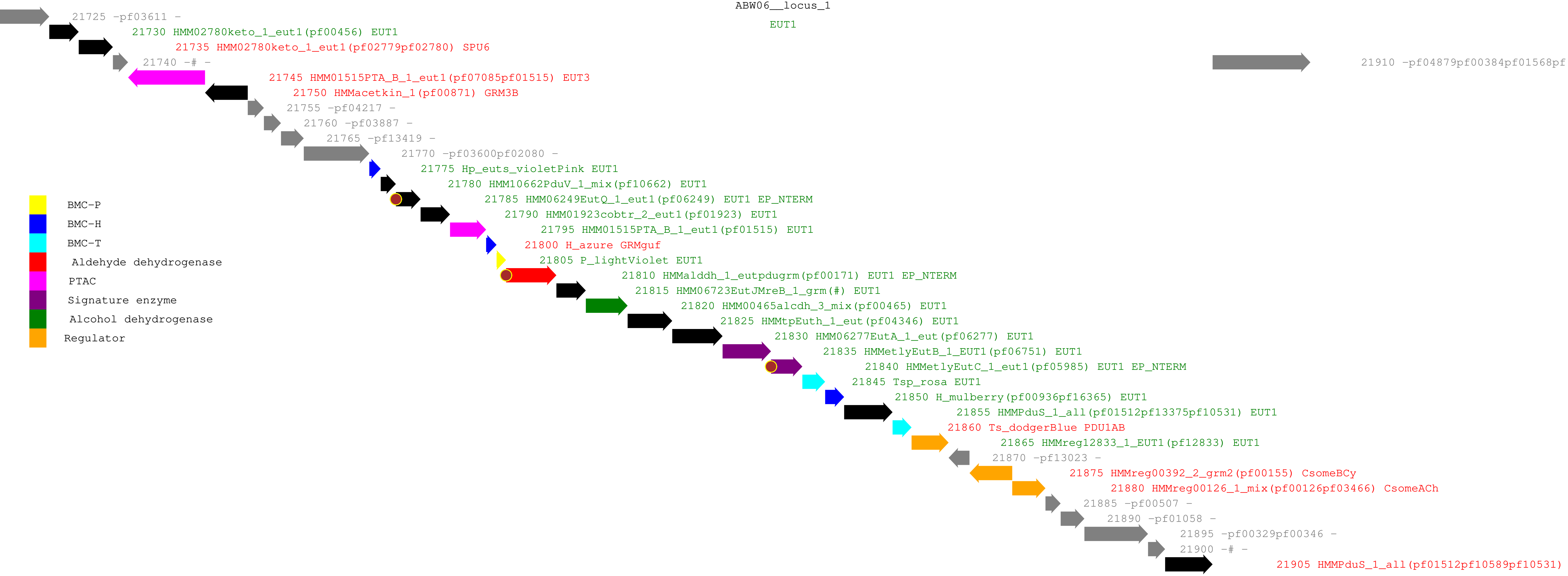

Escherichia coli. Bacteria; Proteobacteria; Gammaproteobacteria; Enterobacterales;

AC067__locus_1

EUT1

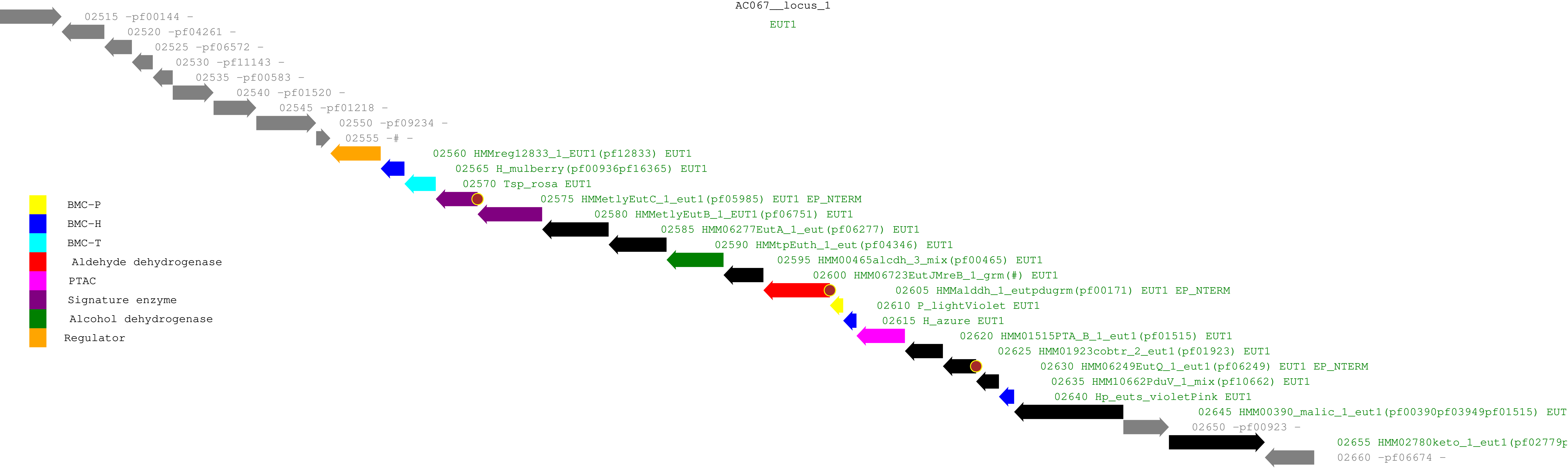

Escherichia coli 2-177-06_S3_C2. Bacteria; Proteobacteria; Gammaproteobacteria; Enterobacterales;

AC16__locus_1

EUT1

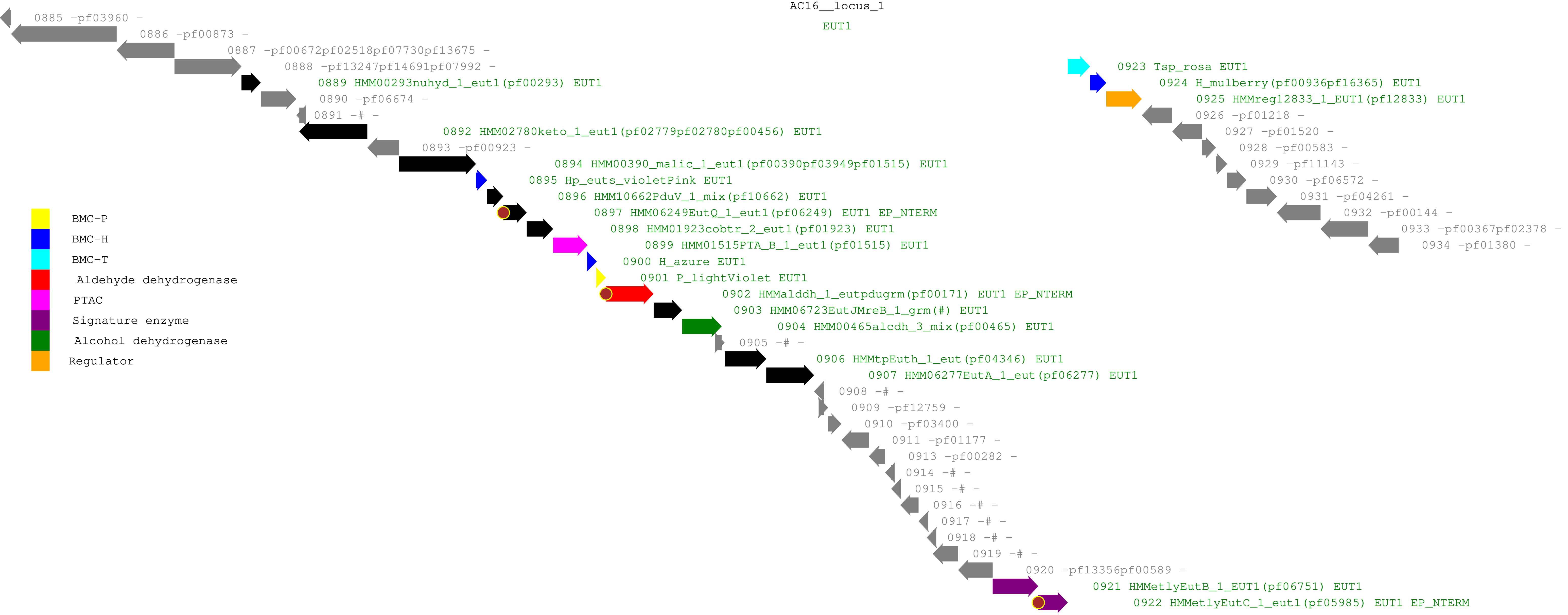

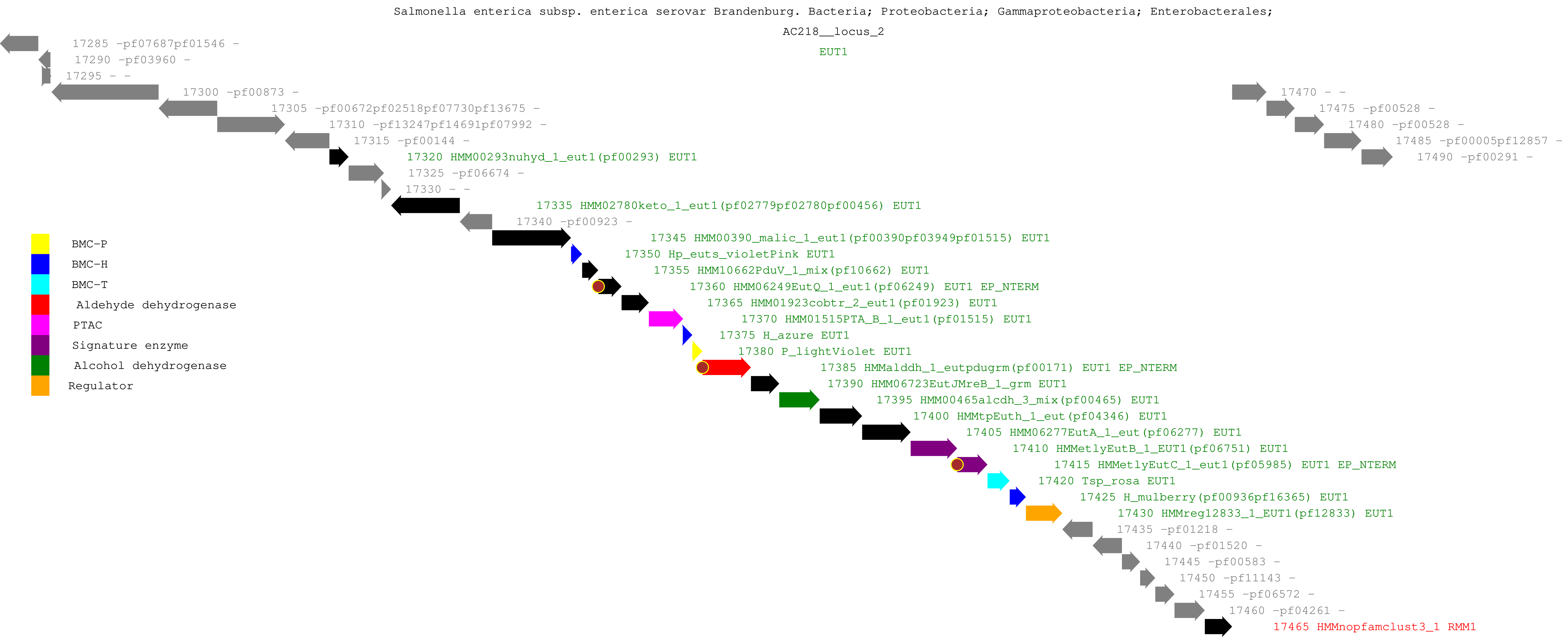
Salmonella enterica subsp. enterica serovar Brandenburg. Bacteria; Proteobacteria; Gammaproteobacteria; Enterobacterales;
AC218__locus_2
EUT1
17285 -pf07687pf01546 -
17290 -pf03960 -
17295 - -
17300 -pf00873 -
17305 -pf00672pf02518pf07730pf13675 -
17310 -pf13247pf14691pf07992 -
17315 -pf00144 -
17320 HMM00293nuhyd_1_eut1(pf00293) EUT1
17325 -pf06674 -
17330 - -
17335 HMM02780keto_1_eut1(pf02779pf02780pf00456) EUT1
17340 -pf00923 -
17345 HMM00390_malic_1_eut1(pf00390pf03949pf01515) EUT1
17350 Hp_euts_violetPink EUT1
17355 HMM10662PduV_1_mix(pf10662) EUT1
17360 HMM06249EutQ_1_eut1(pf06249) EUT1 EP_NTERM
17365 HMM01923cobtr_2_eut1(pf01923) EUT1
17370 HMM01515PTA_B_1_eut1(pf01515) EUT1
17375 H_azure EUT1
17380 P_lightViolet EUT1
17385 HMMalddh_1_eutpdugrm(pf00171) EUT1 EP_NTERM
17390 HMM06723EutJMreB_1_grm EUT1
17395 HMM00465alcdh_3_mix(pf00465) EUT1
17400 HMMtpEuth_1_eut(pf04346) EUT1
17405 HMM06277EutA_1_eut(pf06277) EUT1
17410 HMMetlyEutB_1_EUT1(pf06751) EUT1
17415 HMMetlyEutC_1_eut1(pf05985) EUT1 EP_NTERM
17420 Tsp_rosa EUT1
17425 H_mulberry(pf00936pf16365) EUT1
17430 HMMreg12833_1_EUT1(pf12833) EUT1
17435 -pf01218 -
17440 -pf01520 -
17445 -pf00583 -
17450 -pf11143 -
17455 -pf06572 -
17460 -pf04261 -
17465 HMMnopfamclust3_1 RMM1
17470 - -
17475 -pf00528 -
17480 -pf00528 -
17485 -pf00005pf12857 -
17490 -pf00291 -
BMC-P
BMC-H
BMC-T
Aldehyde dehydrogenase
PTAC
Signature enzyme
Alcohol dehydrogenase
Regulator

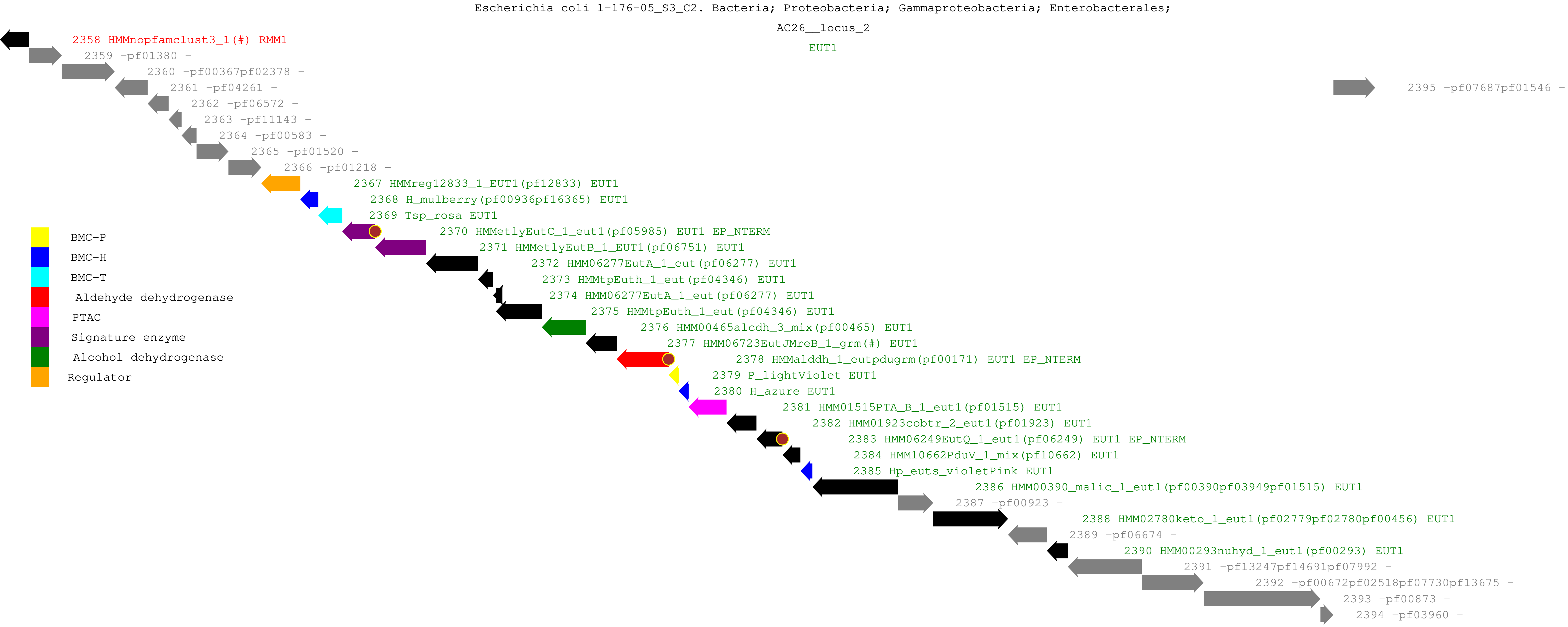
Escherichia coli 1-176-05_S3_C2. Bacteria; Proteobacteria; Gammaproteobacteria; Enterobacterales;
AC26__locus_2
EUT1
2358 HMMnopfamclust3_1(#) RMM1
2359 -pf01380 -
2360 -pf00367pf02378 -
2361 -pf04261 -
2362 -pf06572 -
2363 -pf11143 -
2364 -pf00583 -
2365 -pf01520 -
2366 -pf01218 -
2367 HMMreg12833_1_EUT1(pf12833) EUT1
2368 H_mulberry(pf00936pf16365) EUT1
2369 Tsp_rosa EUT1
2370 HMMetlyEutC_1_eut1(pf05985) EUT1 EP_NTERM
2371 HMMetlyEutB_1_EUT1(pf06751) EUT1
2372 HMM06277EutA_1_eut(pf06277) EUT1
2373 HMMtpEuth_1_eut(pf04346) EUT1
2374 HMM06277EutA_1_eut(pf06277) EUT1
2375 HMMtpEuth_1_eut(pf04346) EUT1
2376 HMM00465alcdh_3_mix(pf00465) EUT1
2377 HMM06723EutJMreB_1_grm(#) EUT1
2378 HMMalddh_1_eutpdugrm(pf00171) EUT1 EP_NTERM
2379 P_lightViolet EUT1
2380 H_azure EUT1
2381 HMM01515PTA_B_1_eut1(pf01515) EUT1
2382 HMM01923cobtr_2_eut1(pf01923) EUT1
2383 HMM06249EutQ_1_eut1(pf06249) EUT1 EP_NTERM
2384 HMM10662PduV_1_mix(pf10662) EUT1
2385 Hp_euts_violetPink EUT1
2386 HMM00390_malic_1_eut1(pf00390pf03949pf01515) EUT1
2387 -pf00923 -
2388 HMM02780keto_1_eut1(pf02779pf02780pf00456) EUT1
2389 -pf06674 -
2390 HMM00293nuhyd_1_eut1(pf00293) EUT1
2391 -pf13247pf14691pf07992 -
2392 -pf00672pf02518pf07730pf13675 -
2393 -pf00873 -
2394 -pf03960 -
2395 -pf07687pf01546 -
BMC-P
BMC-H
BMC-T
Aldehyde dehydrogenase
PTAC
Signature enzyme
Alcohol dehydrogenase
Regulator

Salmonella enterica subsp. salamae. Bacteria; Proteobacteria; Gammaproteobacteria; Enterobacterales;

AC322__locus_1

EUT1

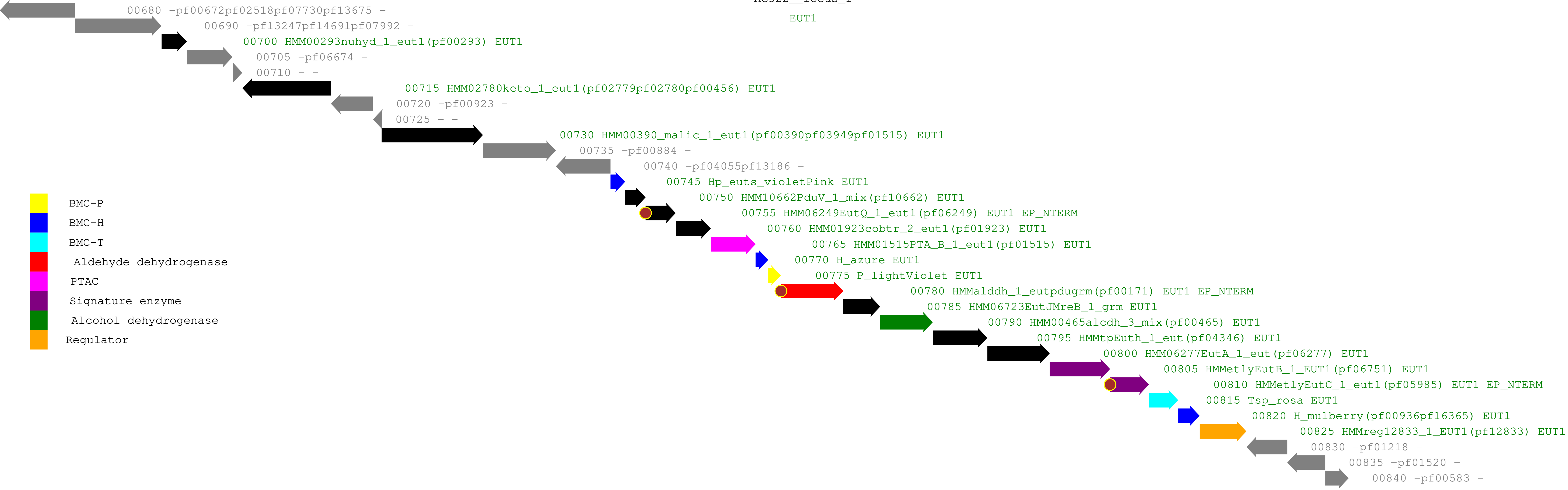

Escherichia coli 2-177-06_S3_C3. Bacteria; Proteobacteria; Gammaproteobacteria; Enterobacterales;

AC44__locus_1

EUT1

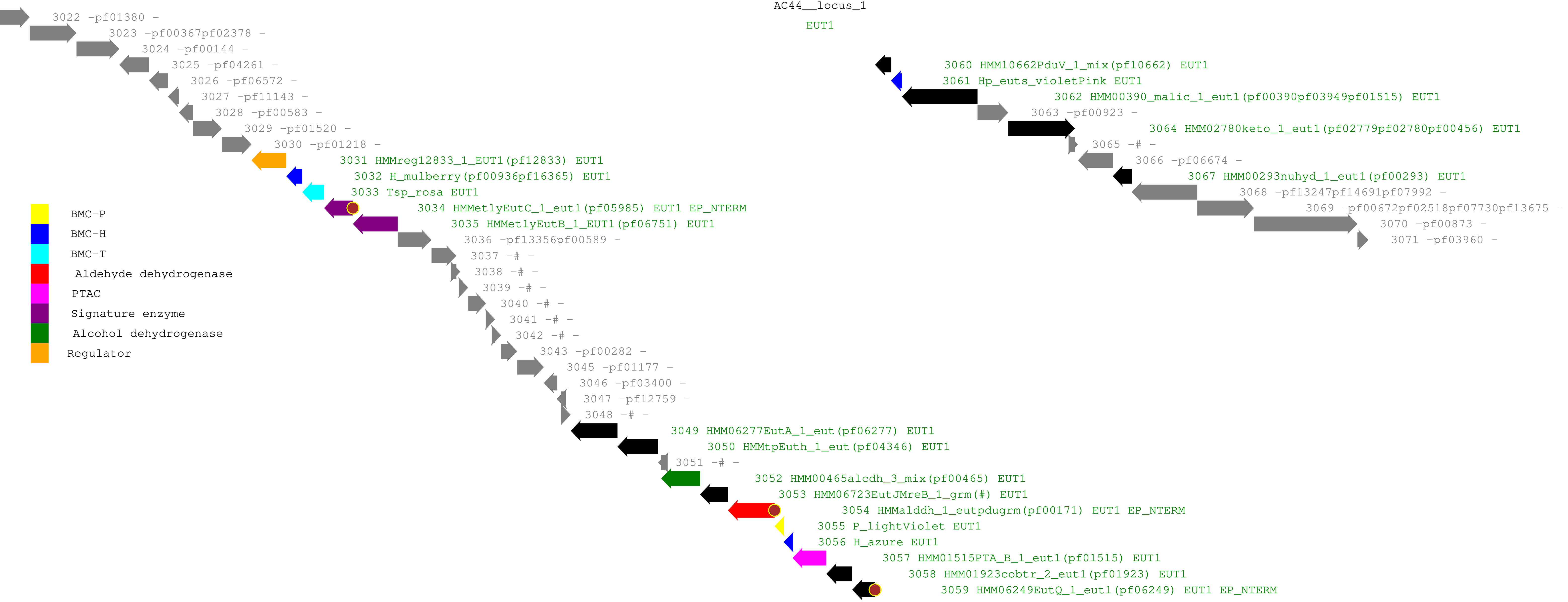

Escherichia coli 2-210-07_S3_C3. Bacteria; Proteobacteria; Gammaproteobacteria; Enterobacterales;

AC45__locus_1

EUT1

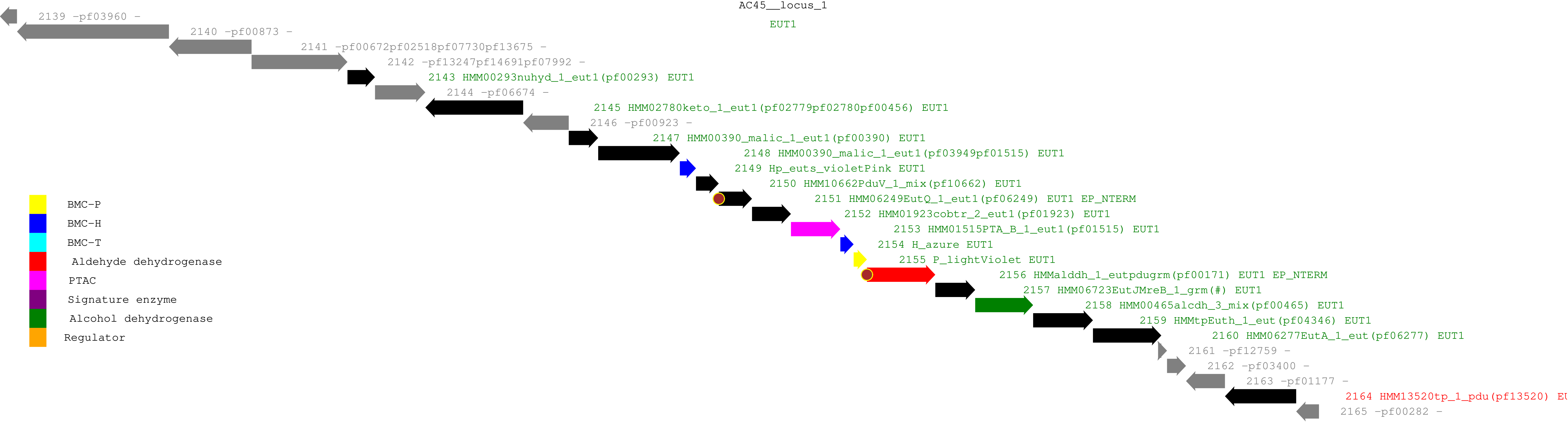

Escherichia coli 1-182-04_S3_C3. Bacteria; Proteobacteria; Gammaproteobacteria; Enterobacterales;

AC56__locus_1

EUT1

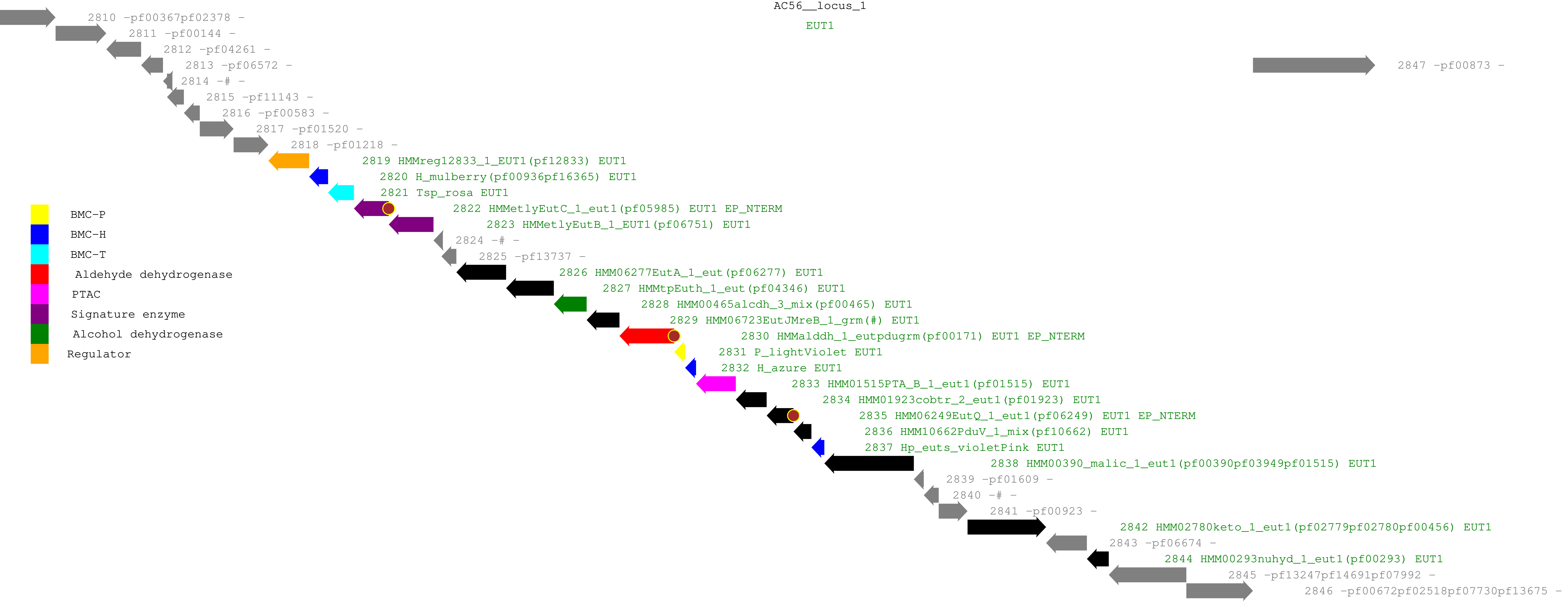

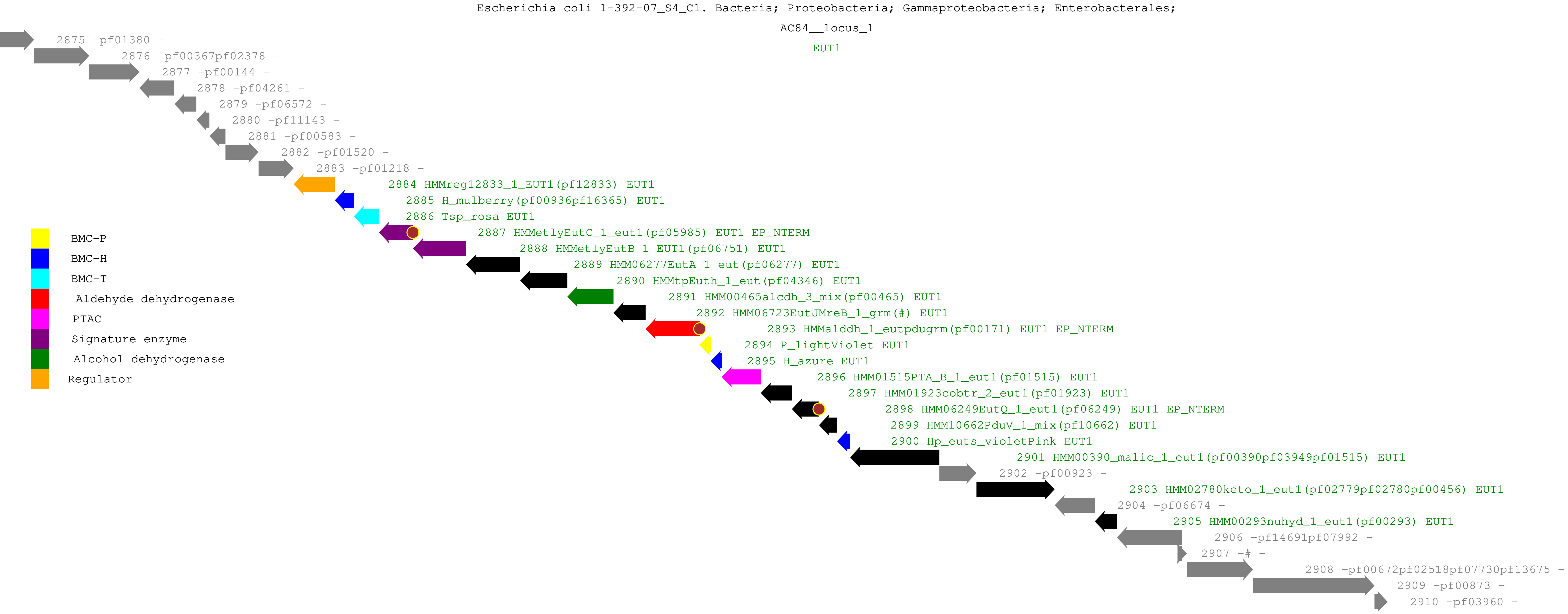

Escherichia coli 1-392-07_S4_C1. Bacteria; Proteobacteria; Gammaproteobacteria; Enterobacterales;
AC84__locus_1
EUT1
2875 -pf01380 -
2876 -pf00367pf02378 -
2877 -pf00144 -
2878 -pf04261 -
2879 -pf06572 -
2880 -pf11143 -
2881 -pf00583 -
2882 -pf01520 -
2883 -pf01218 -
2884 HMMreg12833_1_EUT1(pf12833) EUT1
2885 H_mulberry(pf00936pf16365) EUT1
2886 Tsp_rosa EUT1
2887 HMMetlyEutC_1_eut1(pf05985) EUT1 EP_NTERM
2888 HMMetlyEutB_1_EUT1(pf06751) EUT1
2889 HMM06277EutA_1_eut(pf06277) EUT1
2890 HMMtpEuth_1_eut(pf04346) EUT1
2891 HMM00465alcdh_3_mix(pf00465) EUT1
2892 HMM06723EutJMreB_1_grm(#) EUT1
2893 HMMaldh_1_eutpdugrm(pf00171) EUT1 EP_NTERM
2894 P_lightViolet EUT1
2895 H_azure EUT1
2896 HMM01515PTA_B_1_eut1(pf01515) EUT1
2897 HMM01923cobtr_2_eut1(pf01923) EUT1
2898 HMM06249EutQ_1_eut1(pf06249) EUT1 EP_NTERM
2899 HMM10662PduV_1_mix(pf10662) EUT1
2900 Hp_euts_violetPink EUT1
2901 HMM00390_malic_1_eut1(pf00390pf03949pf01515) EUT1
2902 -pf00923 -
2903 HMM02780keto_1_eut1(pf02779pf02780pf00456) EUT1
2904 -pf06674 -
2905 HMM00293nuhyd_1_eut1(pf00293) EUT1
2906 -pf14691pf07992 -
2907 -# -
2908 -pf00672pf02518pf07730pf13675 -
2909 -pf00873 -
2910 -pf03960 -
BMC-P
BMC-H
BMC-T
Aldehyde dehydrogenase
PTAC
Signature enzyme
Alcohol dehydrogenase
Regulator

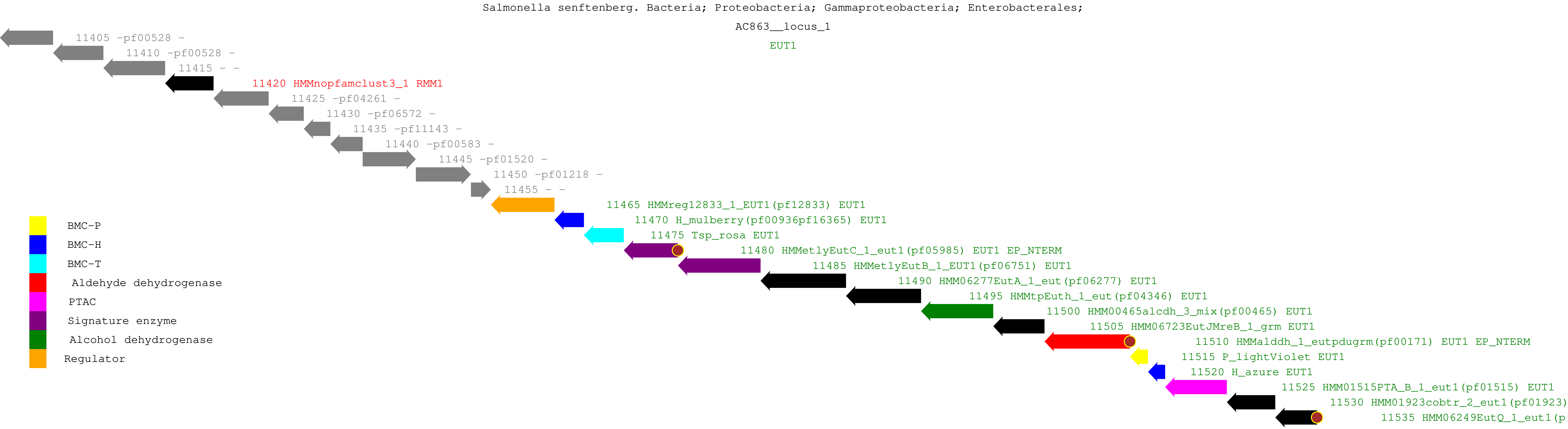
Salmonella senftenberg. Bacteria; Proteobacteria; Gammaproteobacteria; Enterobacterales;
AC863__locus_1
EUT1
11405 -pf00528 -
11410 -pf00528 -
11415 - -
11420 HMMnopfamclust3_1 RMM1
11425 -pf04261 -
11430 -pf06572 -
11435 -pf11143 -
11440 -pf00583 -
11445 -pf01520 -
11450 -pf01218 -
11455 - -
11465 HMMreg12833_1_EUT1(pf12833) EUT1
11470 H_mulberry(pf00936pf16365) EUT1
11475 Tsp_rosa EUT1
11480 HMMetlyEutC_1_eut1(pf05985) EUT1 EP_NTERM
11485 HMMetlyEutB_1_EUT1(pf06751) EUT1
11490 HMM06277EutA_1_eut(pf06277) EUT1
11495 HMMtpEuth_1_eut(pf04346) EUT1
11500 HMM00465alcdh_3_mix(pf00465) EUT1
11505 HMM06723EutJMreB_1_grm EUT1
11510 HMMaldh_1_eutpdugrm(pf00171) EUT1 EP_NTERM
11515 P_lightViolet EUT1
11520 H_azure EUT1
11525 HMM01515PTA_B_1_eut1(pf01515) EUT1
11530 HMM01923cobtr_2_eut1(pf01923)
11535 HMM06249EutQ_1_eut1(p
BMC-P
BMC-H
BMC-T
Aldehyde dehydrogenase
PTAC
Signature enzyme
Alcohol dehydrogenase
Regulator

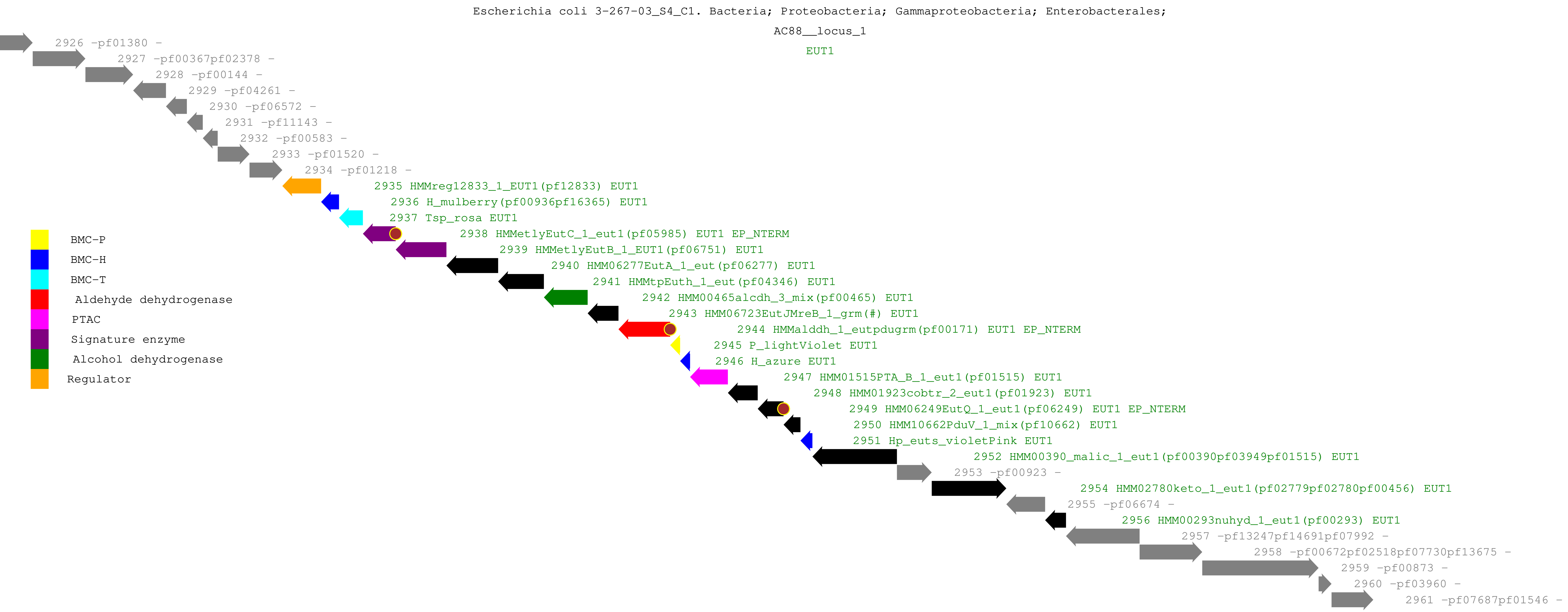

Escherichia coli 3-267-03_S4_C1. Bacteria; Proteobacteria; Gammaproteobacteria; Enterobacterales;
AC88__locus_1
EUT1
2926 -pf01380 -
2927 -pf00367pf02378 -
2928 -pf00144 -
2929 -pf04261 -
2930 -pf06572 -
2931 -pf11143 -
2932 -pf00583 -
2933 -pf01520 -
2934 -pf01218 -
2935 HMMreg12833_1_EUT1(pf12833) EUT1
2936 H_mulberry(pf00936pf16365) EUT1
2937 Tsp_rosa EUT1
2938 HMMetlyEutC_1_eut1(pf05985) EUT1 EP_NTERM
2939 HMMetlyEutB_1_EUT1(pf06751) EUT1
2940 HMM06277EutA_1_eut(pf06277) EUT1
2941 HMMtpEuth_1_eut(pf04346) EUT1
2942 HMM00465alcdh_3_mix(pf00465) EUT1
2943 HMM06723EutJMreB_1_grm(#) EUT1
2944 HMMaldh_1_eutpdugrm(pf00171) EUT1 EP_NTERM
2945 P_lightViolet EUT1
2946 H_azure EUT1
2947 HMM01515PTA_B_1_eut1(pf01515) EUT1
2948 HMM01923cobtr_2_eut1(pf01923) EUT1
2949 HMM06249EutQ_1_eut1(pf06249) EUT1 EP_NTERM
2950 HMM10662PduV_1_mix(pf10662) EUT1
2951 Hp_euts_violetPink EUT1
2952 HMM00390_malic_1_eut1(pf00390pf03949pf01515) EUT1
2953 -pf00923 -
2954 HMM02780keto_1_eut1(pf02779pf02780pf00456) EUT1
2955 -pf06674 -
2956 HMM00293nuhyd_1_eut1(pf00293) EUT1
2957 -pf13247pf14691pf07992 -
2958 -pf00672pf02518pf07730pf13675 -
2959 -pf00873 -
2960 -pf03960 -
2961 -pf07687pf01546 -
BMC-P
BMC-H
BMC-T
Aldehyde dehydrogenase
PTAC
Signature enzyme
Alcohol dehydrogenase
Regulator

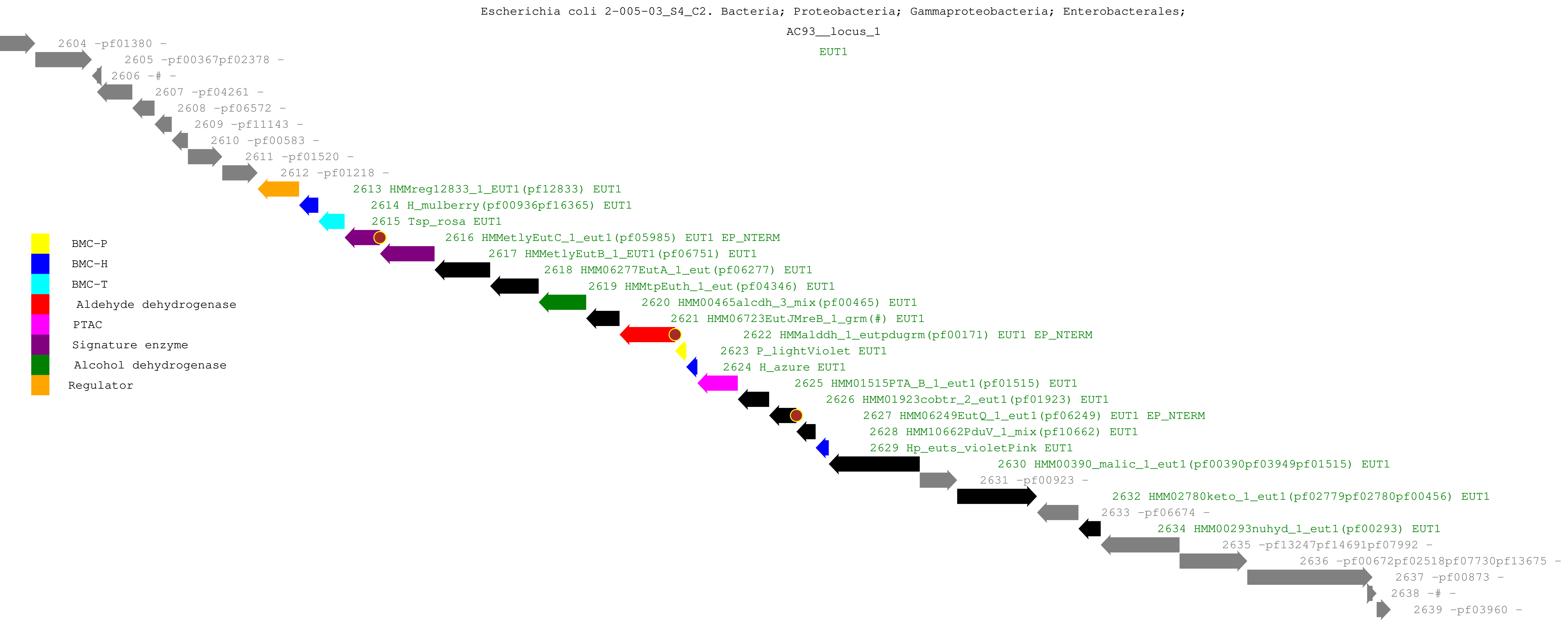
Escherichia coli 2-005-03_S4_C2. Bacteria; Proteobacteria; Gammaproteobacteria; Enterobacterales;
AC93__locus_1
EUT1
2604 -pf01380 -
2605 -pf00367pf02378 -
2606 -# -
2607 -pf04261 -
2608 -pf06572 -
2609 -pf11143 -
2610 -pf00583 -
2611 -pf01520 -
2612 -pf01218 -
2613 HMMreg12833_1_EUT1(pf12833) EUT1
2614 H_mulberry(pf00936pf16365) EUT1
2615 Tsp_rosa EUT1
2616 HMMetlyEutC_1_eut1(pf05985) EUT1 EP_NTERM
2617 HMMetlyEutB_1_EUT1(pf06751) EUT1
2618 HMM06277EutA_1_eut(pf06277) EUT1
2619 HMMtpEuth_1_eut(pf04346) EUT1
2620 HMM00465alcdh_3_mix(pf00465) EUT1
2621 HMM06723EutJMreB_1_grm(#) EUT1
2622 HMMalddh_1_eutpdugrm(pf00171) EUT1 EP_NTERM
2623 P_lightViolet EUT1
2624 H_azure EUT1
2625 HMM01515PTA_B_1_eut1(pf01515) EUT1
2626 HMM01923cobtr_2_eut1(pf01923) EUT1
2627 HMM06249EutQ_1_eut1(pf06249) EUT1 EP_NTERM
2628 HMM10662PduV_1_mix(pf10662) EUT1
2629 Hp_euts_violetPink EUT1
2630 HMM00390_malic_1_eut1(pf00390pf03949pf01515) EUT1
2631 -pf00923 -
2632 HMM02780keto_1_eut1(pf02779pf02780pf00456) EUT1
2633 -pf06674 -
2634 HMM00293nuhyd_1_eut1(pf00293) EUT1
2635 -pf13247pf14691pf07992 -
2636 -pf00672pf02518pf07730pf13675 -
2637 -pf00873 -
2638 -# -
2639 -pf03960 -
BMC-P
BMC-H
BMC-T
Aldehyde dehydrogenase
PTAC
Signature enzyme
Alcohol dehydrogenase
Regulator

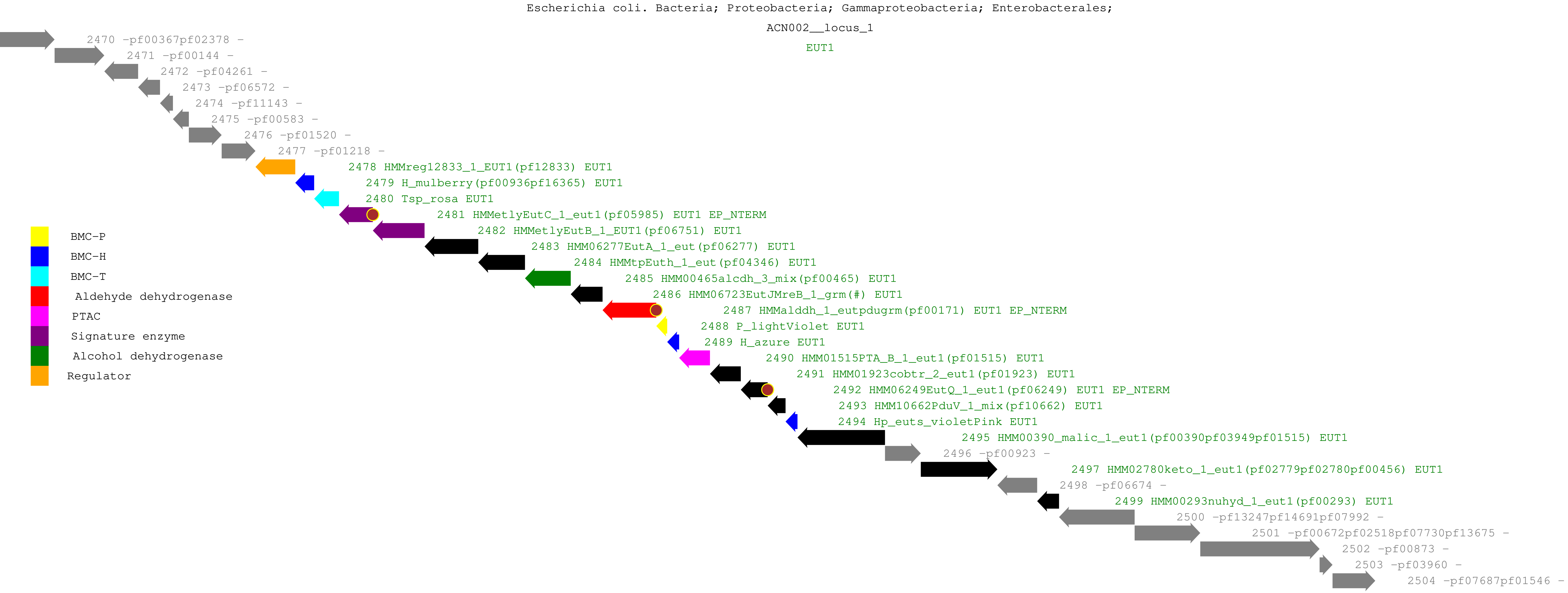

Escherichia coli. Bacteria; Proteobacteria; Gammaproteobacteria; Enterobacterales;
ACN002__locus_1
EUT1
2470 -pf00367pf02378 -
2471 -pf00144 -
2472 -pf04261 -
2473 -pf06572 -
2474 -pf11143 -
2475 -pf00583 -
2476 -pf01520 -
2477 -pf01218 -
2478 HMMreg12833_1_EUT1(pf12833) EUT1
2479 H_mulberry(pf00936pf16365) EUT1
2480 Tsp_rosa EUT1
2481 HMMetlyEutC_1_eut1(pf05985) EUT1 EP_NTERM
2482 HMMetlyEutB_1_EUT1(pf06751) EUT1
2483 HMM06277EutA_1_eut(pf06277) EUT1
2484 HMMtpEuth_1_eut(pf04346) EUT1
2485 HMM00465alcdh_3_mix(pf00465) EUT1
2486 HMM06723EutJMreB_1_grm(#) EUT1
2487 HMMalddh_1_eutpdugrm(pf00171) EUT1 EP_NTERM
2488 P_lightViolet EUT1
2489 H_azure EUT1
2490 HMM01515PTA_B_1_eut1(pf01515) EUT1
2491 HMM01923cobtr_2_eut1(pf01923) EUT1
2492 HMM06249EutQ_1_eut1(pf06249) EUT1 EP_NTERM
2493 HMM10662PduV_1_mix(pf10662) EUT1
2494 Hp_euts_violetPink EUT1
2495 HMM00390_malic_1_eut1(pf00390pf03949pf01515) EUT1
2496 -pf00923 -
2497 HMM02780keto_1_eut1(pf02779pf02780pf00456) EUT1
2498 -pf06674 -
2499 HMM00293nuhyd_1_eut1(pf00293) EUT1
2500 -pf13247pf14691pf07992 -
2501 -pf00672pf02518pf07730pf13675 -
2502 -pf00873 -
2503 -pf03960 -
2504 -pf07687pf01546 -
BMC-P
BMC-H
BMC-T
Aldehyde dehydrogenase
PTAC
Signature enzyme
Alcohol dehydrogenase
Regulator

Escherichia coli. Bacteria; Proteobacteria; Gammaproteobacteria; Enterobacterales;

ACU57__locus_1

EUT1

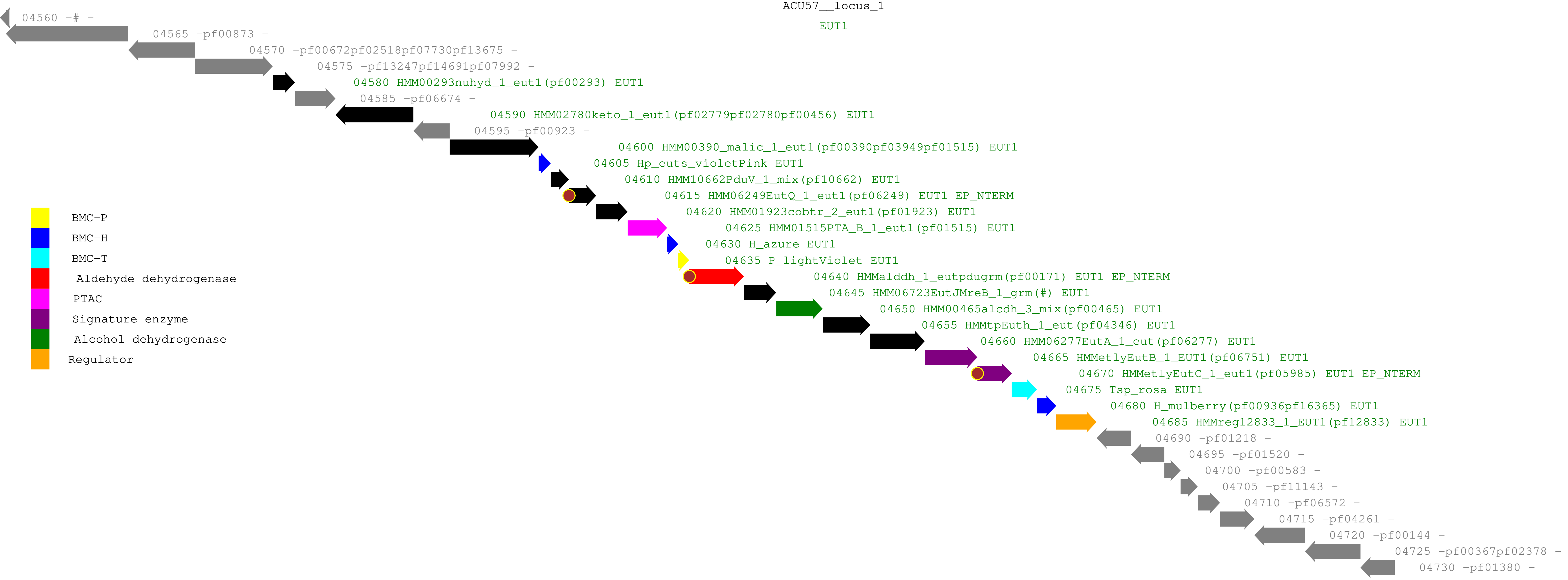

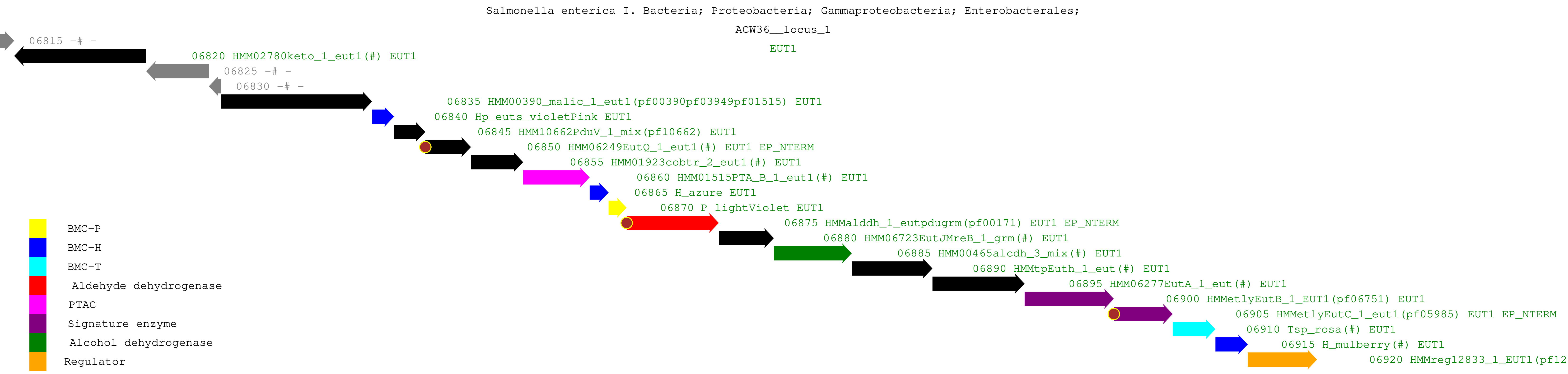
Salmonella enterica I. Bacteria; Proteobacteria; Gammaproteobacteria; Enterobacterales;
ACW36__locus_1
EUT1
06815 -# -
06820 HMM02780keto_1_eut1(#) EUT1
06825 -# -
06830 -# -
06835 HMM00390_malic_1_eut1(pf00390pf03949pf01515) EUT1
06840 Hp_euts_violetPink EUT1
06845 HMM10662PduV_1_mix(pf10662) EUT1
06850 HMM06249EutQ_1_eut1(#) EUT1 EP_NTERM
06855 HMM01923cobtr_2_eut1(#) EUT1
06860 HMM01515PTA_B_1_eut1(#) EUT1
06865 H_azure EUT1
06870 P_lightViolet EUT1
06875 HMMalddh_1_eutpdugrm(pf00171) EUT1 EP_NTERM
06880 HMM06723EutJMreB_1_grm(#) EUT1
06885 HMM00465alcdh_3_mix(#) EUT1
06890 HMMtpEuth_1_eut(#) EUT1
06895 HMM06277EutA_1_eut(#) EUT1
06900 HMMetlyEutB_1_EUT1(pf06751) EUT1
06905 HMMetlyEutC_1_eut1(pf05985) EUT1 EP_NTERM
06910 Tsp_rosa(#) EUT1
06915 H_mulberry(#) EUT1
06920 HMMreg12833_1_EUT1(pf12
BMC-P
BMC-H
BMC-T
Aldehyde dehydrogenase
PTAC
Signature enzyme
Alcohol dehydrogenase
Regulator

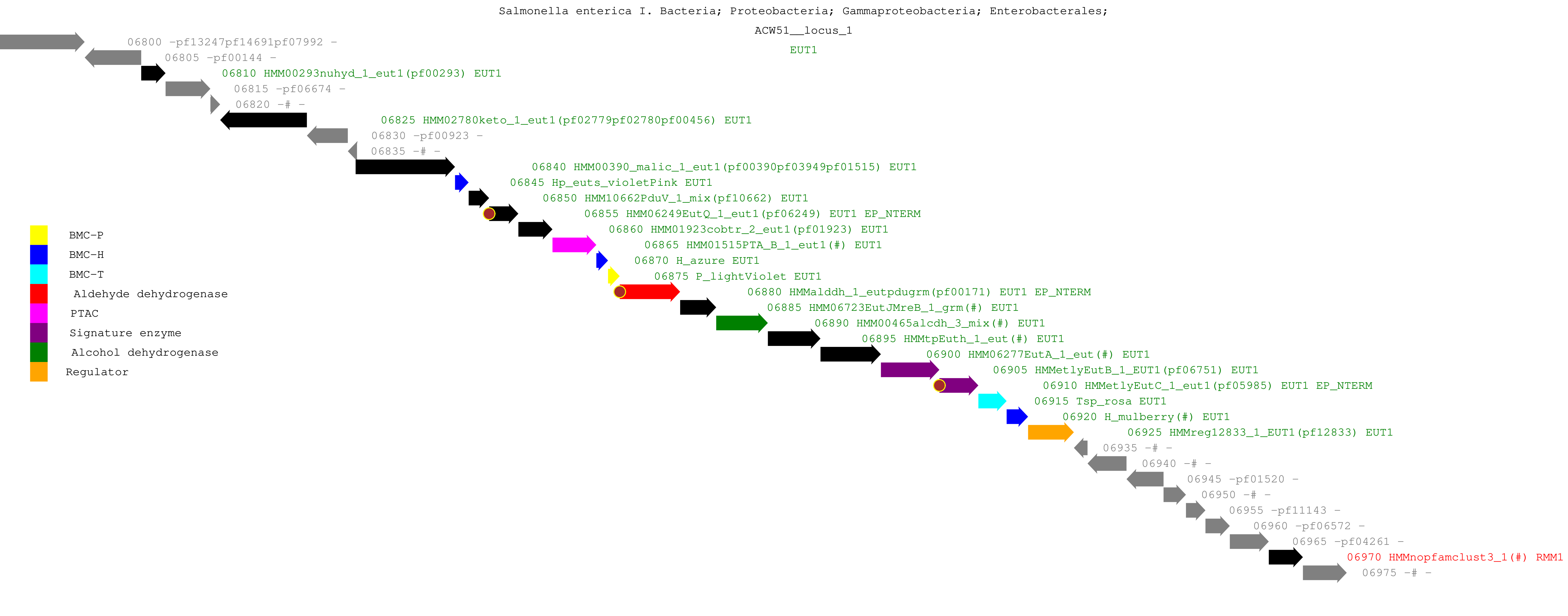
Salmonella enterica I. Bacteria; Proteobacteria; Gammaproteobacteria; Enterobacterales;
ACW51__locus_1
EUT1
06800 -pf13247pf14691pf07992 -
06805 -pf00144 -
06810 HMM00293nuhyd_1_eut1(pf00293) EUT1
06815 -pf06674 -
06820 -# -
06825 HMM02780keto_1_eut1(pf02779pf02780pf00456) EUT1
06830 -pf00923 -
06835 -# -
06840 HMM00390_malic_1_eut1(pf00390pf03949pf01515) EUT1
06845 Hp_euts_violetPink EUT1
06850 HMM10662PduV_1_mix(pf10662) EUT1
06855 HMM06249EutQ_1_eut1(pf06249) EUT1 EP_NTERM
06860 HMM01923cobtr_2_eut1(pf01923) EUT1
06865 HMM01515PTA_B_1_eut1(#) EUT1
06870 H_azure EUT1
06875 P_lightViolet EUT1
06880 HMMalddh_1_eutpdugrm(pf00171) EUT1 EP_NTERM
06885 HMM06723EutJMreB_1_grm(#) EUT1
06890 HMM00465alcdh_3_mix(#) EUT1
06895 HMMtpEuth_1_eut(#) EUT1
06900 HMM06277EutA_1_eut(#) EUT1
06905 HMMetlyEutB_1_EUT1(pf06751) EUT1
06910 HMMetlyEutC_1_eut1(pf05985) EUT1 EP_NTERM
06915 Tsp_rosa EUT1
06920 H_mulberry(#) EUT1
06925 HMMreg12833_1_EUT1(pf12833) EUT1
06935 -# -
06940 -# -
06945 -pf01520 -
06950 -# -
06955 -pf11143 -
06960 -pf06572 -
06965 -pf04261 -
06970 HMMnopfamclust3_1(#) RMM1
06975 -# -
BMC-P
BMC-H
BMC-T
Aldehyde dehydrogenase
PTAC
Signature enzyme
Alcohol dehydrogenase
Regulator

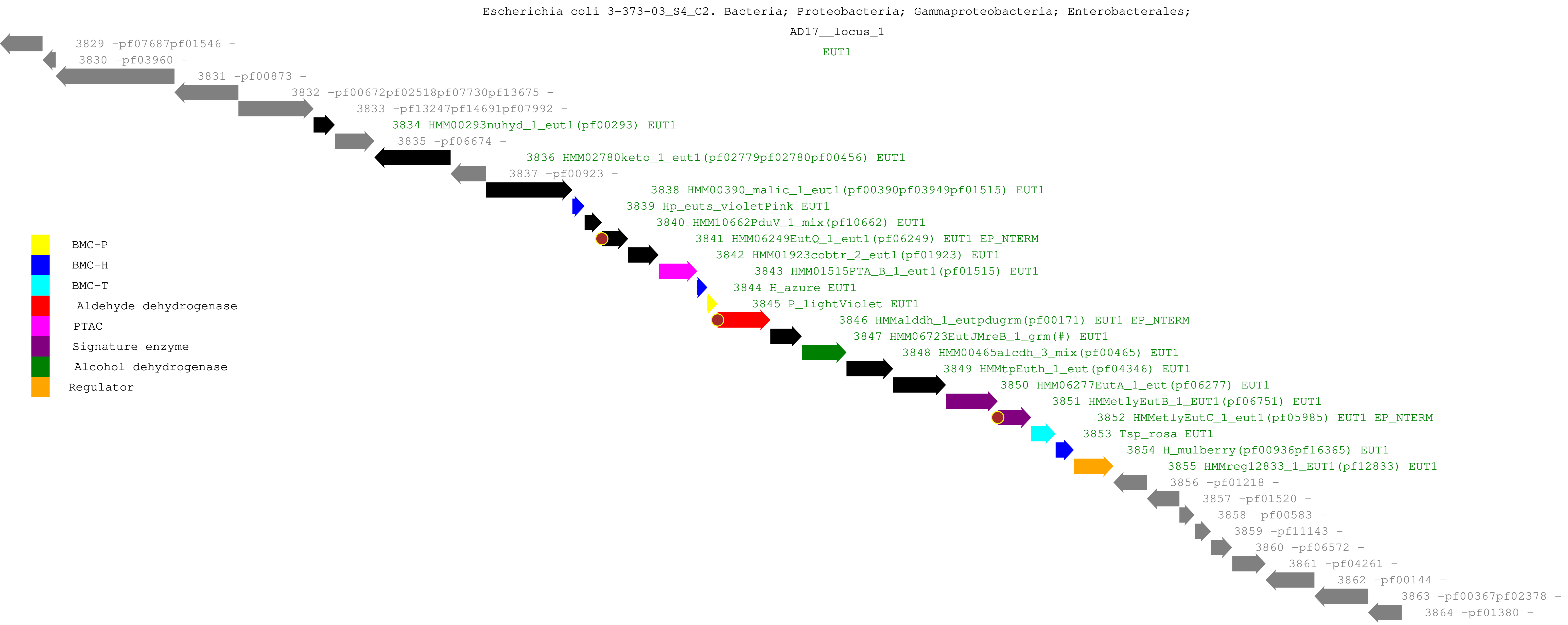
Escherichia coli 3-373-03_S4_C2. Bacteria; Proteobacteria; Gammaproteobacteria; Enterobacterales;
AD17__locus_1
EUT1
3829 -pf07687pf01546 -
3830 -pf03960 -
3831 -pf00873 -
3832 -pf00672pf02518pf07730pf13675 -
3833 -pf13247pf14691pf07992 -
3834 HMM00293nuhyd_1_eut1(pf00293) EUT1
3835 -pf06674 -
3836 HMM02780keto_1_eut1(pf02779pf02780pf00456) EUT1
3837 -pf00923 -
3838 HMM00390_malic_1_eut1(pf00390pf03949pf01515) EUT1
3839 Hp_euts_violetPink EUT1
3840 HMM10662PduV_1_mix(pf10662) EUT1
3841 HMM06249EutQ_1_eut1(pf06249) EUT1 EP_NTERM
3842 HMM01923cobtr_2_eut1(pf01923) EUT1
3843 HMM01515PTA_B_1_eut1(pf01515) EUT1
3844 H_azure EUT1
3845 P_lightViolet EUT1
3846 HMMalddh_1_eutpdugrm(pf00171) EUT1 EP_NTERM
3847 HMM06723EutJMreB_1_grm(#) EUT1
3848 HMM00465alcdh_3_mix(pf00465) EUT1
3849 HMMtpEuth_1_eut(pf04346) EUT1
3850 HMM06277EutA_1_eut(pf06277) EUT1
3851 HMMetlyEutB_1_EUT1(pf06751) EUT1
3852 HMMetlyEutC_1_eut1(pf05985) EUT1 EP_NTERM
3853 Tsp_rosa EUT1
3854 H_mulberry(pf00936pf16365) EUT1
3855 HMMreg12833_1_EUT1(pf12833) EUT1
3856 -pf01218 -
3857 -pf01520 -
3858 -pf00583 -
3859 -pf11143 -
3860 -pf06572 -
3861 -pf04261 -
3862 -pf00144 -
3863 -pf00367pf02378 -
3864 -pf01380 -
BMC-P
BMC-H
BMC-T
Aldehyde dehydrogenase
PTAC
Signature enzyme
Alcohol dehydrogenase
Regulator

Escherichia coli 2-005-03_S4_C3. Bacteria; Proteobacteria; Gammaproteobacteria; Enterobacterales;

AD23__locus_1

EUT1

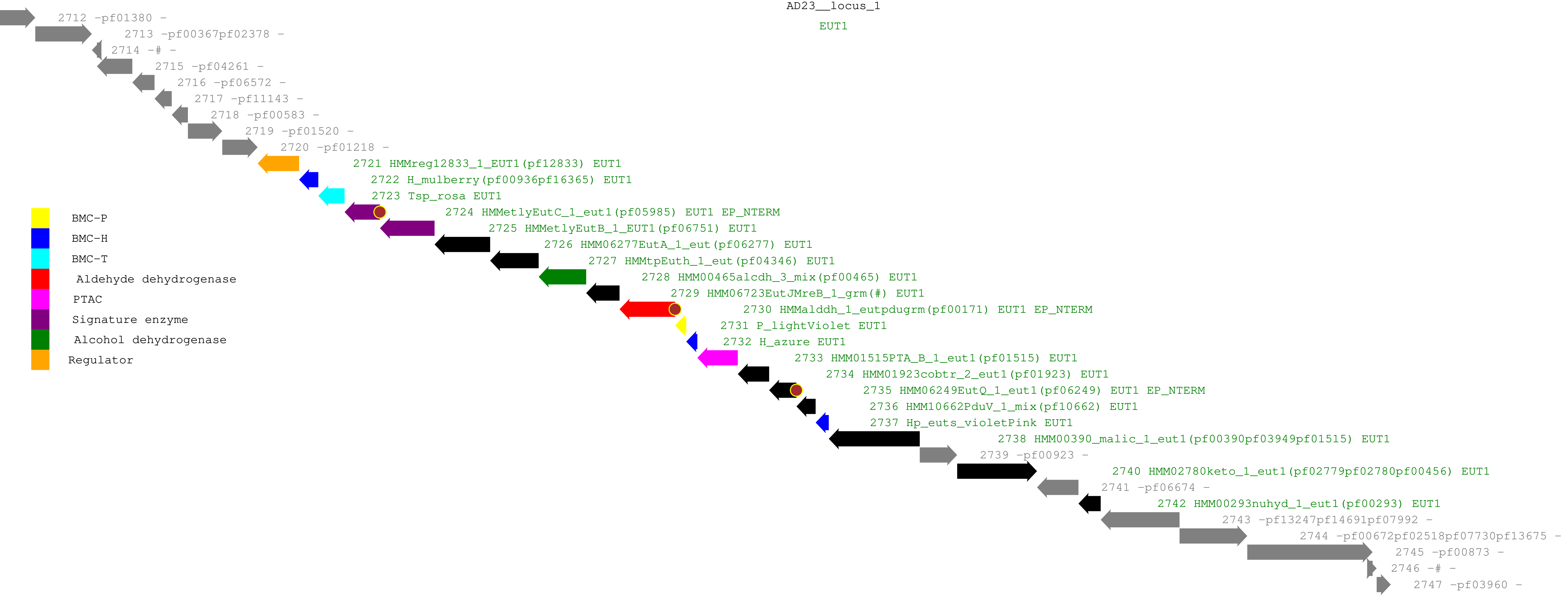

Escherichia coli 2-427-07_S4_C3. Bacteria; Proteobacteria; Gammaproteobacteria; Enterobacterales;

AD31__locus_1

EUT1

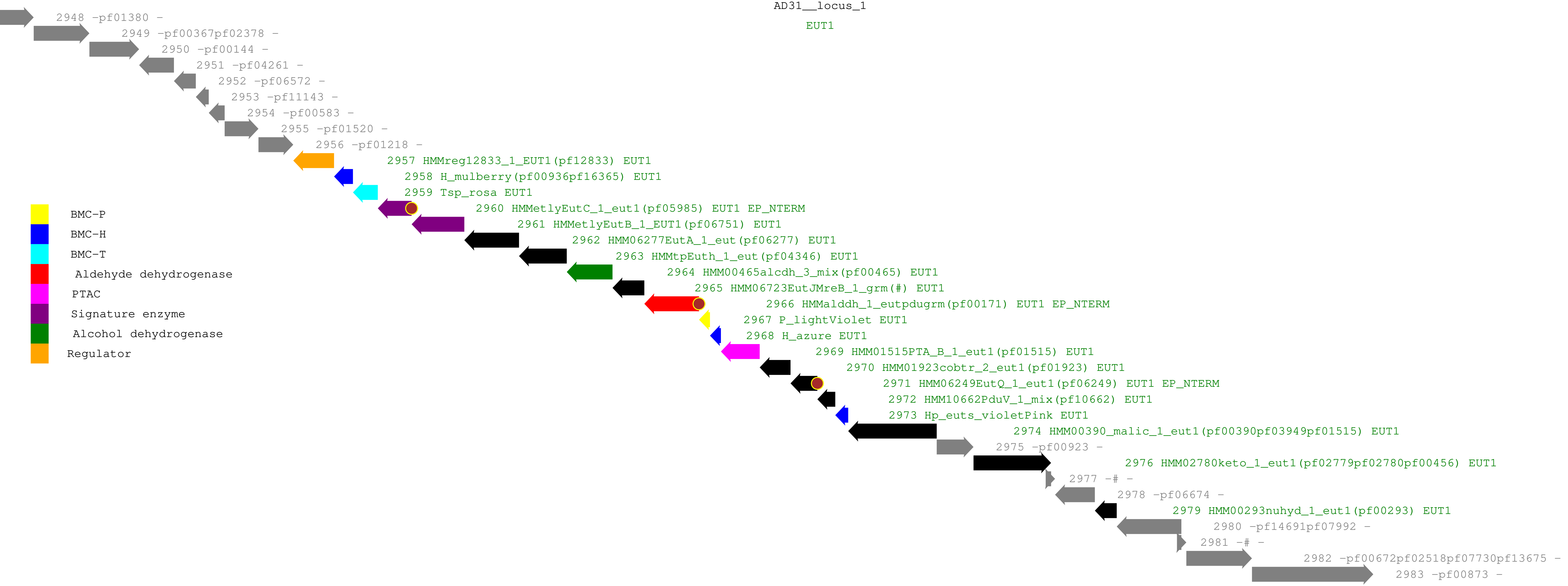

Escherichia coli 1-392-07_S4_C3. Bacteria; Proteobacteria; Gammaproteobacteria; Enterobacterales;

AD40__locus_1

EUT1

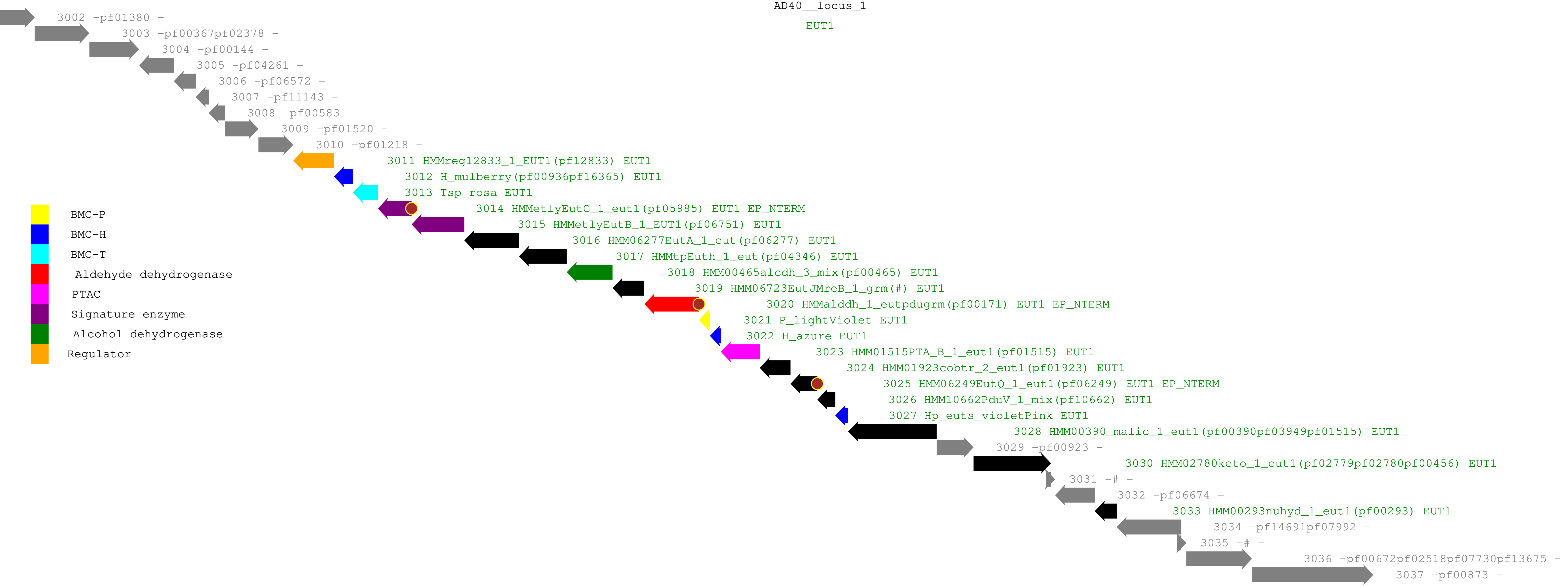

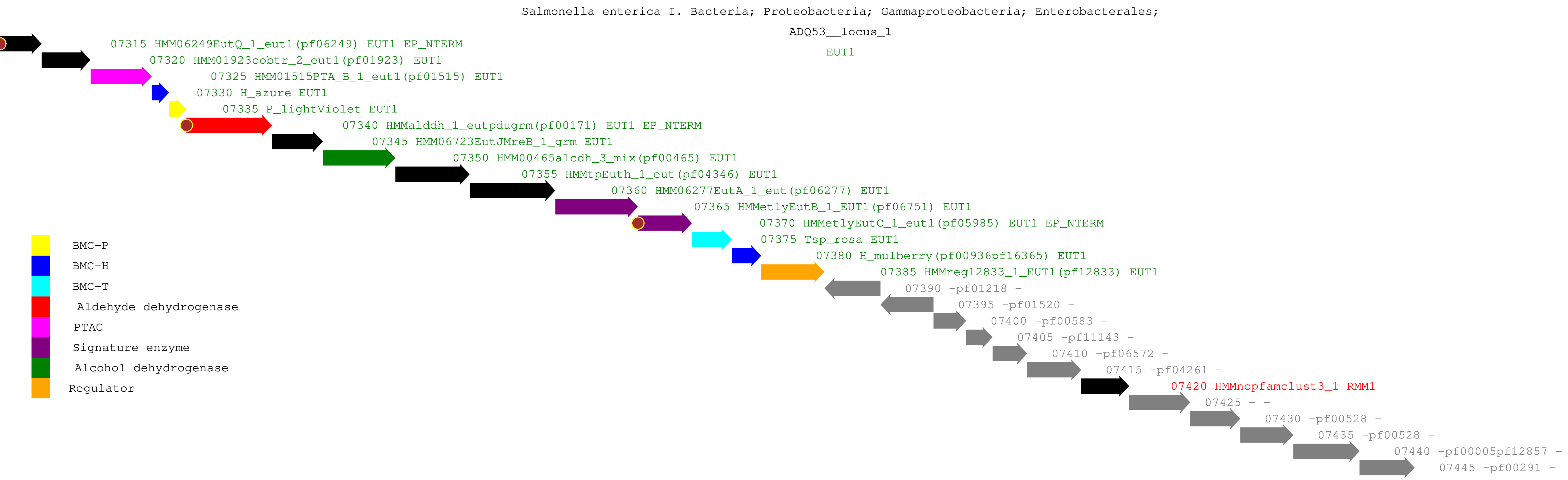
Salmonella enterica I. Bacteria; Proteobacteria; Gammaproteobacteria; Enterobacterales;
ADQ53__locus_1
EUT1
07315 HMM06249EutQ_1_eut1(pf06249) EUT1 EP_NTERM
07320 HMM01923cobtr_2_eut1(pf01923) EUT1
07325 HMM01515PTA_B_1_eut1(pf01515) EUT1
07330 H_azure EUT1
07335 P_lightViolet EUT1
07340 HMMalddh_1_eutpdugrm(pf00171) EUT1 EP_NTERM
07345 HMM06723EutJMreB_1_grm EUT1
07350 HMM00465alcdh_3_mix(pf00465) EUT1
07355 HMMtpEuth_1_eut(pf04346) EUT1
07360 HMM06277EutA_1_eut(pf06277) EUT1
07365 HMMetlyEutB_1_EUT1(pf06751) EUT1
07370 HMMetlyEutC_1_eut1(pf05985) EUT1 EP_NTERM
07375 Tsp_rosa EUT1
07380 H_mulberry(pf00936pf16365) EUT1
07385 HMMreg12833_1_EUT1(pf12833) EUT1
07390 -pf01218 -
07395 -pf01520 -
07400 -pf00583 -
07405 -pf11143 -
07410 -pf06572 -
07415 -pf04261 -
07420 HMMnopfamclust3_1 RMM1
07425 - -
07430 -pf00528 -
07435 -pf00528 -
07440 -pf00005pf12857 -
07445 -pf00291 -
BMC-P
BMC-H
BMC-T
Aldehyde dehydrogenase
PTAC
Signature enzyme
Alcohol dehydrogenase
Regulator

Salmonella enterica (Salmonella choleraesuis). Bacteria; Proteobacteria; Gammaproteobacteria; Enterobacterales;

ADQ66__locus_2

EUT1

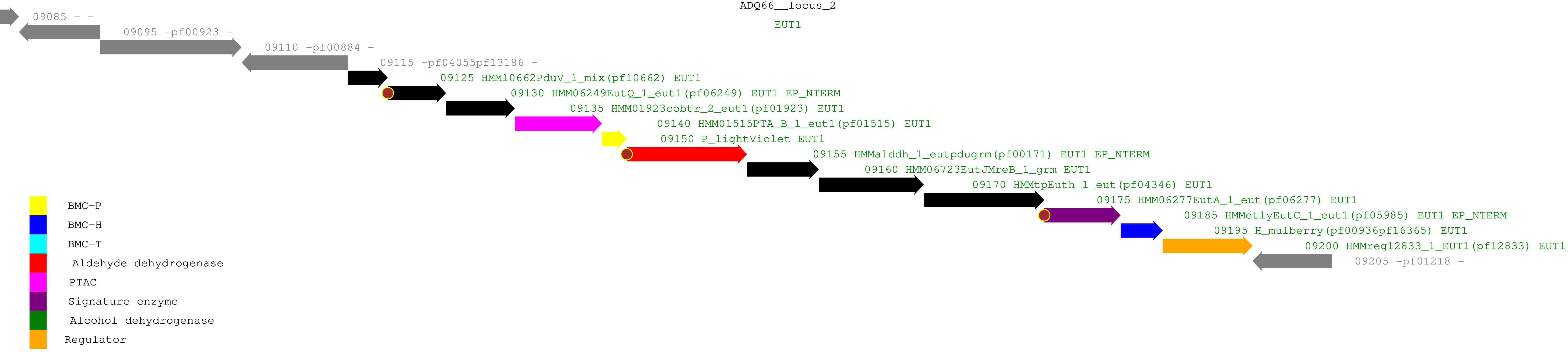

Escherichia coli UCI 65. Bacteria; Proteobacteria; Gammaproteobacteria; Enterobacterales;

AE24__locus_1

EUT1

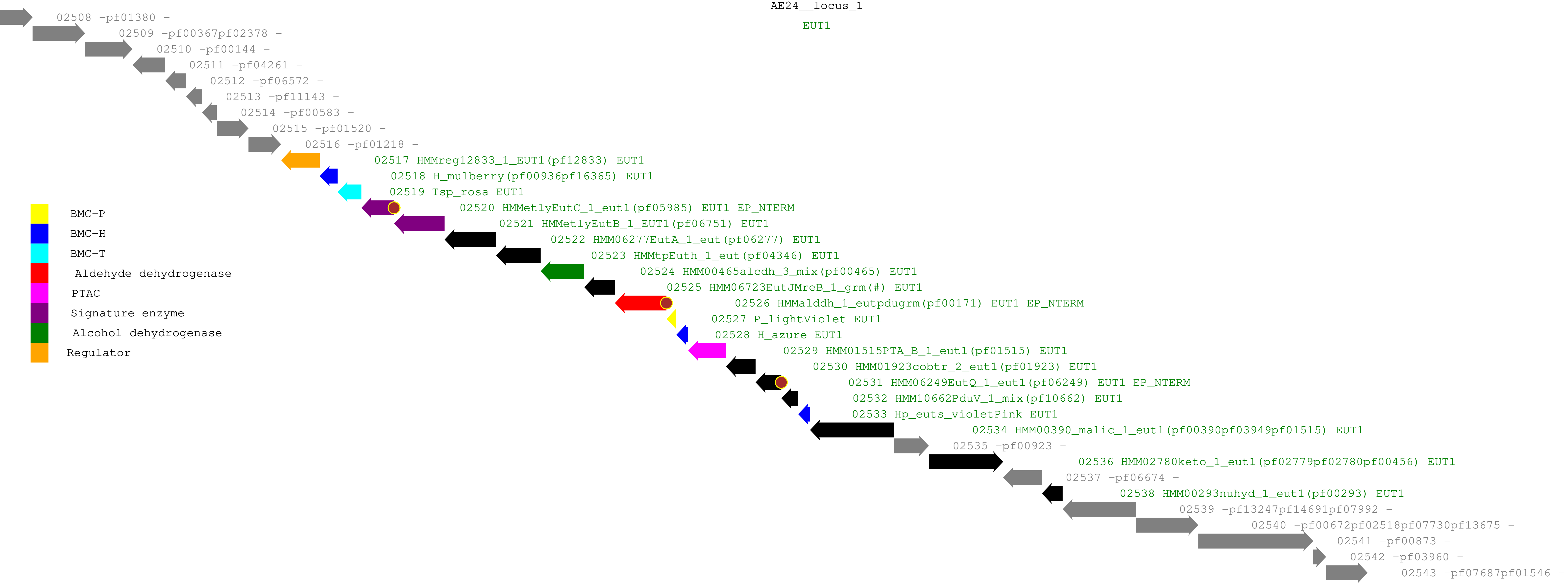

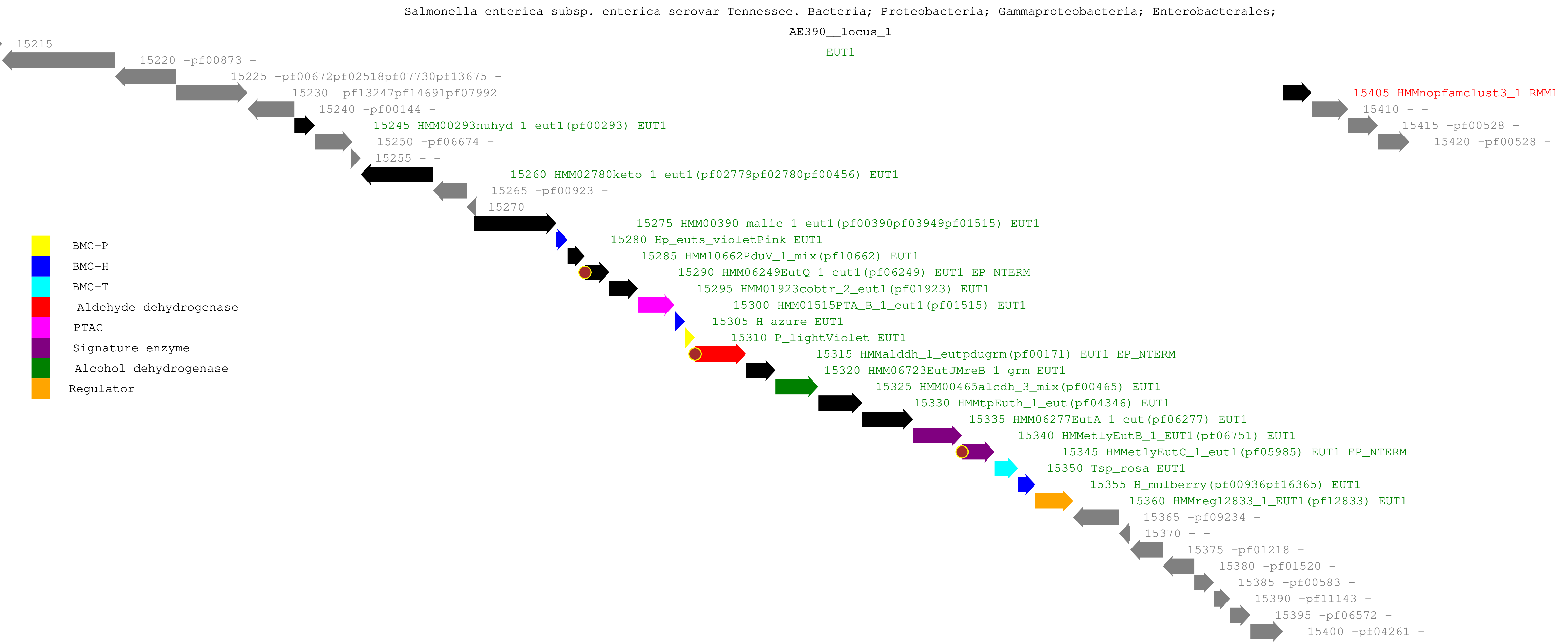
Salmonella enterica subsp. enterica serovar Tennessee. Bacteria; Proteobacteria; Gammaproteobacteria; Enterobacterales;
AE390__locus_1
EUT1
15215 - -
15220 -pf00873 -
15225 -pf00672pf02518pf07730pf13675 -
15230 -pf13247pf14691pf07992 -
15240 -pf00144 -
15245 HMM00293nuhyd_1_eut1(pf00293) EUT1
15250 -pf06674 -
15255 - -
15260 HMM02780keto_1_eut1(pf02779pf02780pf00456) EUT1
15265 -pf00923 -
15270 - -
15275 HMM00390_malic_1_eut1(pf00390pf03949pf01515) EUT1
15280 Hp_euts_violetPink EUT1
15285 HMM10662PduV_1_mix(pf10662) EUT1
15290 HMM06249EutQ_1_eut1(pf06249) EUT1 EP_NTERM
15295 HMM01923cobtr_2_eut1(pf01923) EUT1
15300 HMM01515PTA_B_1_eut1(pf01515) EUT1
15305 H_azure EUT1
15310 P_lightViolet EUT1
15315 HMMaldh_1_eutpdugrm(pf00171) EUT1 EP_NTERM
15320 HMM06723EutJMreB_1_grm EUT1
15325 HMM00465alcdh_3_mix(pf00465) EUT1
15330 HMMtpEuth_1_eut(pf04346) EUT1
15335 HMM06277EutA_1_eut(pf06277) EUT1
15340 HMMetlyEutB_1_EUT1(pf06751) EUT1
15345 HMMetlyEutC_1_eut1(pf05985) EUT1 EP_NTERM
15350 Tsp_rosa EUT1
15355 H_mulberry(pf00936pf16365) EUT1
15360 HMMreg12833_1_EUT1(pf12833) EUT1
15365 -pf09234 -
15370 - -
15375 -pf01218 -
15380 -pf01520 -
15385 -pf00583 -
15390 -pf11143 -
15395 -pf06572 -
15400 -pf04261 -
15405 HMMnopfamclust3_1 RMM1
15410 - -
15415 -pf00528 -
15420 -pf00528 -
BMC-P
BMC-H
BMC-T
Aldehyde dehydrogenase
PTAC
Signature enzyme
Alcohol dehydrogenase
Regulator

Salmonella enterica subsp. enterica serovar Heidelberg. Bacteria; Proteobacteria; Gammaproteobacteria; Enterobacterales;

AEV26__locus_1

EUT1

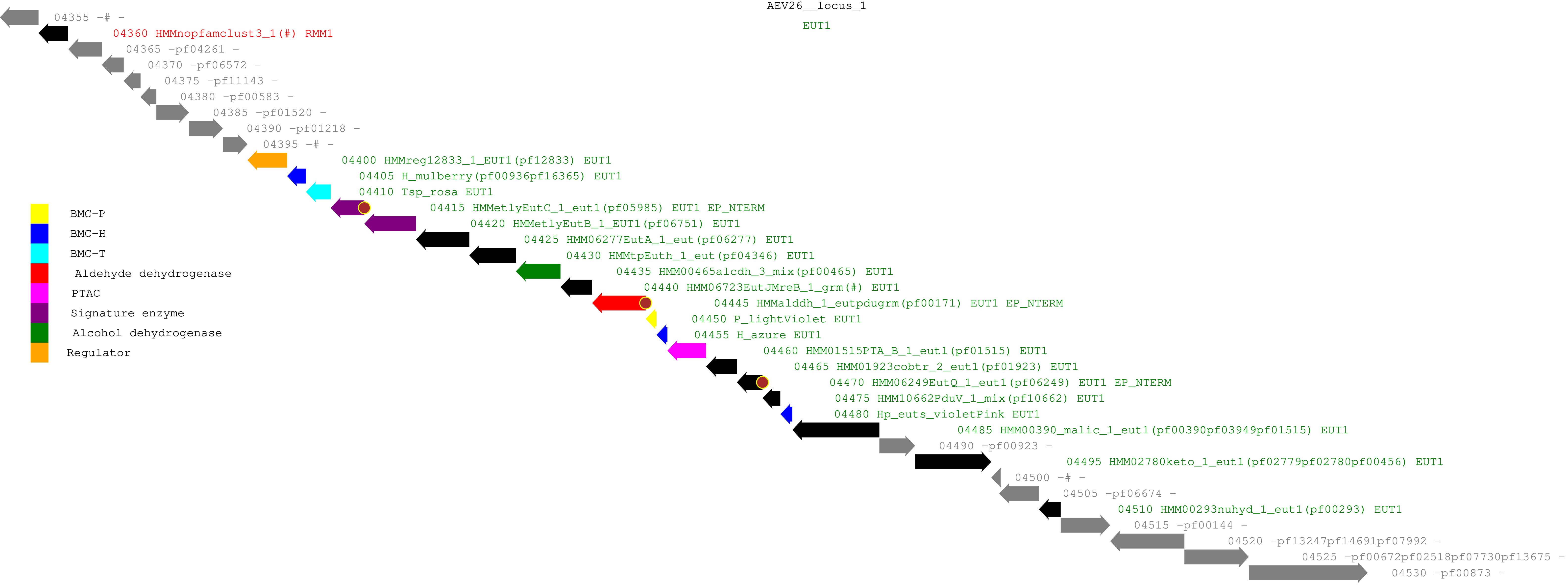

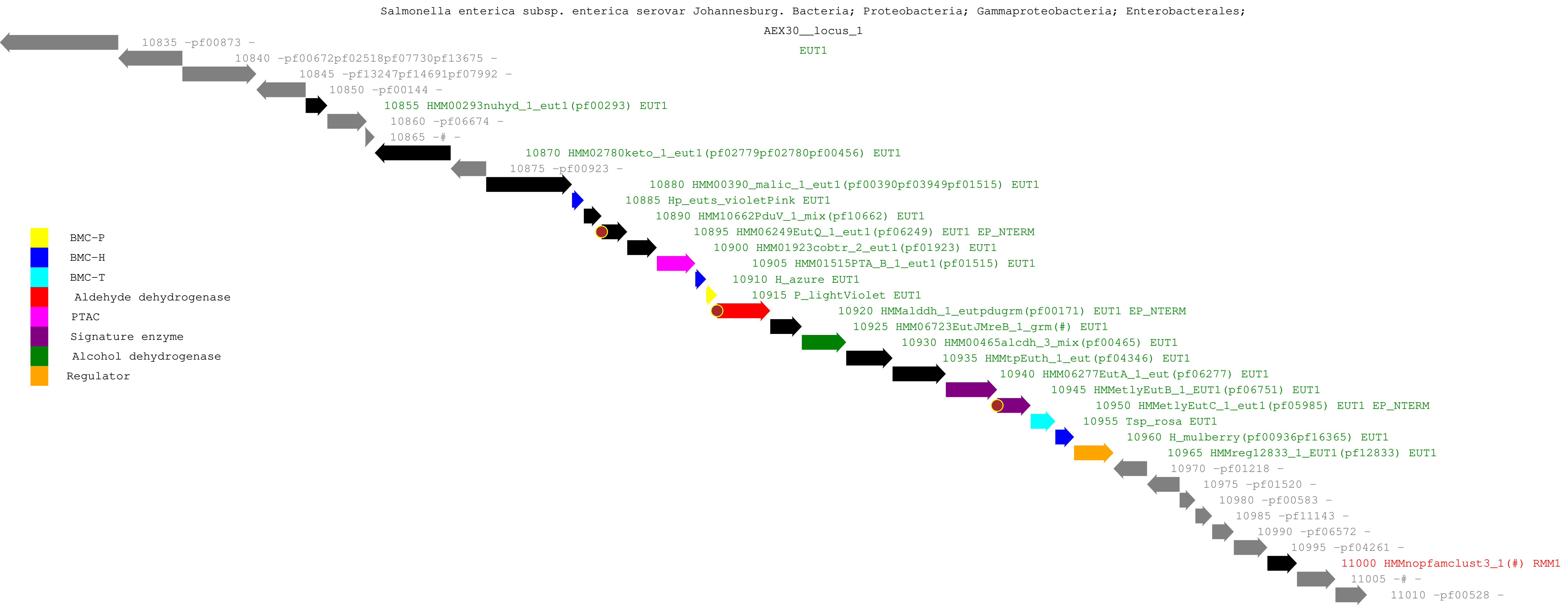
Salmonella enterica subsp. enterica serovar Johannesburg. Bacteria; Proteobacteria; Gammaproteobacteria; Enterobacterales;
AEX30__locus_1
EUT1
10835 -pf00873 -
10840 -pf00672pf02518pf07730pf13675 -
10845 -pf13247pf14691pf07992 -
10850 -pf00144 -
10855 HMM00293nuhyd_1_eut1(pf00293) EUT1
10860 -pf06674 -
10865 -# -
10870 HMM02780keto_1_eut1(pf02779pf02780pf00456) EUT1
10875 -pf00923 -
10880 HMM00390_malic_1_eut1(pf00390pf03949pf01515) EUT1
10885 Hp_euts_violetPink EUT1
10890 HMM10662PduV_1_mix(pf10662) EUT1
10895 HMM06249EutQ_1_eut1(pf06249) EUT1 EP_NTERM
10900 HMM01923cobtr_2_eut1(pf01923) EUT1
10905 HMM01515PTA_B_1_eut1(pf01515) EUT1
10910 H_azure EUT1
10915 P_lightViolet EUT1
10920 HMMalddh_1_eutpdugrm(pf00171) EUT1 EP_NTERM
10925 HMM06723EutJMreB_1_grm(#) EUT1
10930 HMM00465alcdh_3_mix(pf00465) EUT1
10935 HMMtpEuth_1_eut(pf04346) EUT1
10940 HMM06277EutA_1_eut(pf06277) EUT1
10945 HMMetlyEutB_1_EUT1(pf06751) EUT1
10950 HMMetlyEutC_1_eut1(pf05985) EUT1 EP_NTERM
10955 Tsp_rosa EUT1
10960 H_mulberry(pf00936pf16365) EUT1
10965 HMMreg12833_1_EUT1(pf12833) EUT1
10970 -pf01218 -
10975 -pf01520 -
10980 -pf00583 -
10985 -pf11143 -
10990 -pf06572 -
10995 -pf04261 -
11000 HMMnopfamclust3_1(#) RMM1
11005 -# -
11010 -pf00528 -
BMC-P
BMC-H
BMC-T
Aldehyde dehydrogenase
PTAC
Signature enzyme
Alcohol dehydrogenase
Regulator

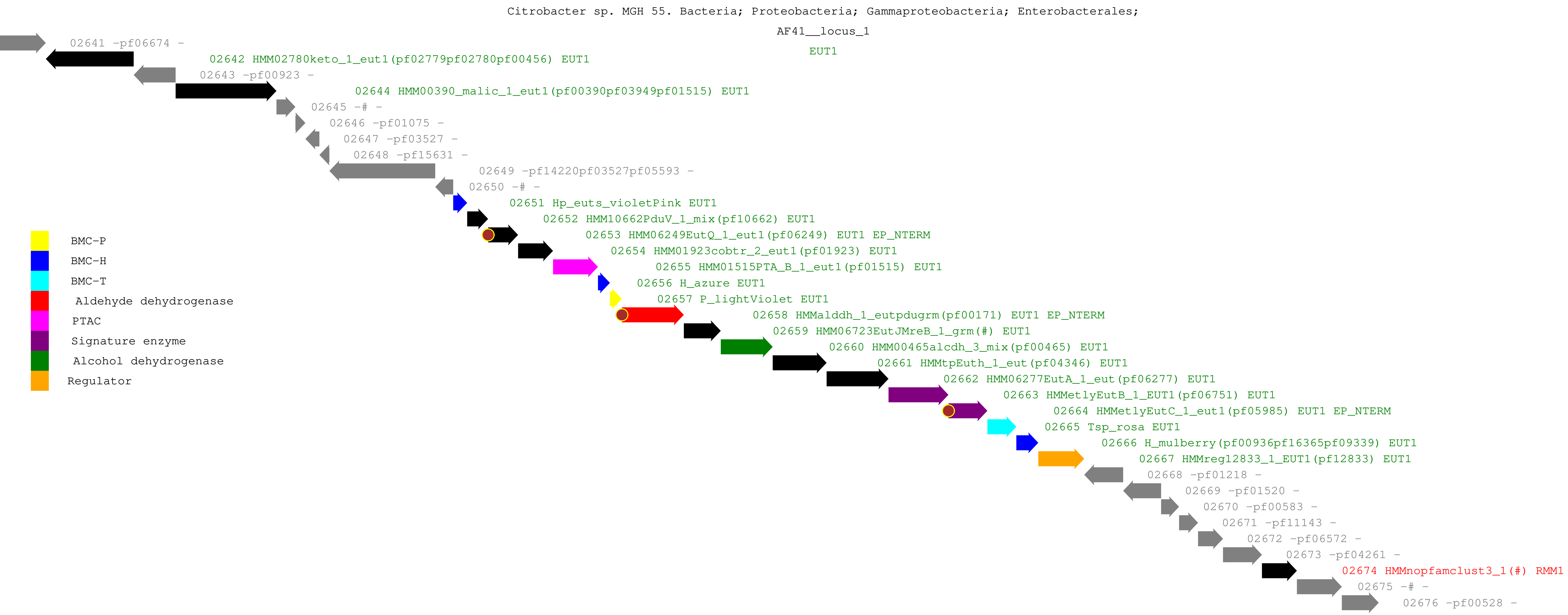
Citrobacter sp. MGH 55. Bacteria; Proteobacteria; Gammaproteobacteria; Enterobacterales;
AF41__locus_1
EUT1
02641 -pf06674 -
02642 HMM02780keto_1_eut1(pf02779pf02780pf00456) EUT1
02643 -pf00923 -
02644 HMM00390_malic_1_eut1(pf00390pf03949pf01515) EUT1
02645 -# -
02646 -pf01075 -
02647 -pf03527 -
02648 -pf15631 -
02649 -pf14220pf03527pf05593 -
02650 -# -
02651 Hp_euts_violetPink EUT1
02652 HMM10662PduV_1_mix(pf10662) EUT1
02653 HMM06249EutQ_1_eut1(pf06249) EUT1 EP_NTERM
02654 HMM01923cobtr_2_eut1(pf01923) EUT1
02655 HMM01515PTA_B_1_eut1(pf01515) EUT1
02656 H_azure EUT1
02657 P_lightViolet EUT1
02658 HMMalddh_1_eutpdugrm(pf00171) EUT1 EP_NTERM
02659 HMM06723EutJMreB_1_grm(#) EUT1
02660 HMM00465alcdh_3_mix(pf00465) EUT1
02661 HMMtpEuth_1_eut(pf04346) EUT1
02662 HMM06277EutA_1_eut(pf06277) EUT1
02663 HMMetlyEutB_1_EUT1(pf06751) EUT1
02664 HMMetlyEutC_1_eut1(pf05985) EUT1 EP_NTERM
02665 Tsp_rosa EUT1
02666 H_mulberry(pf00936pf16365pf09339) EUT1
02667 HMMreg12833_1_EUT1(pf12833) EUT1
02668 -pf01218 -
02669 -pf01520 -
02670 -pf00583 -
02671 -pf11143 -
02672 -pf06572 -
02673 -pf04261 -
02674 HMMnopfamclust3_1(#) RMM1
02675 -# -
02676 -pf00528 -
BMC-P
BMC-H
BMC-T
Aldehyde dehydrogenase
PTAC
Signature enzyme
Alcohol dehydrogenase
Regulator

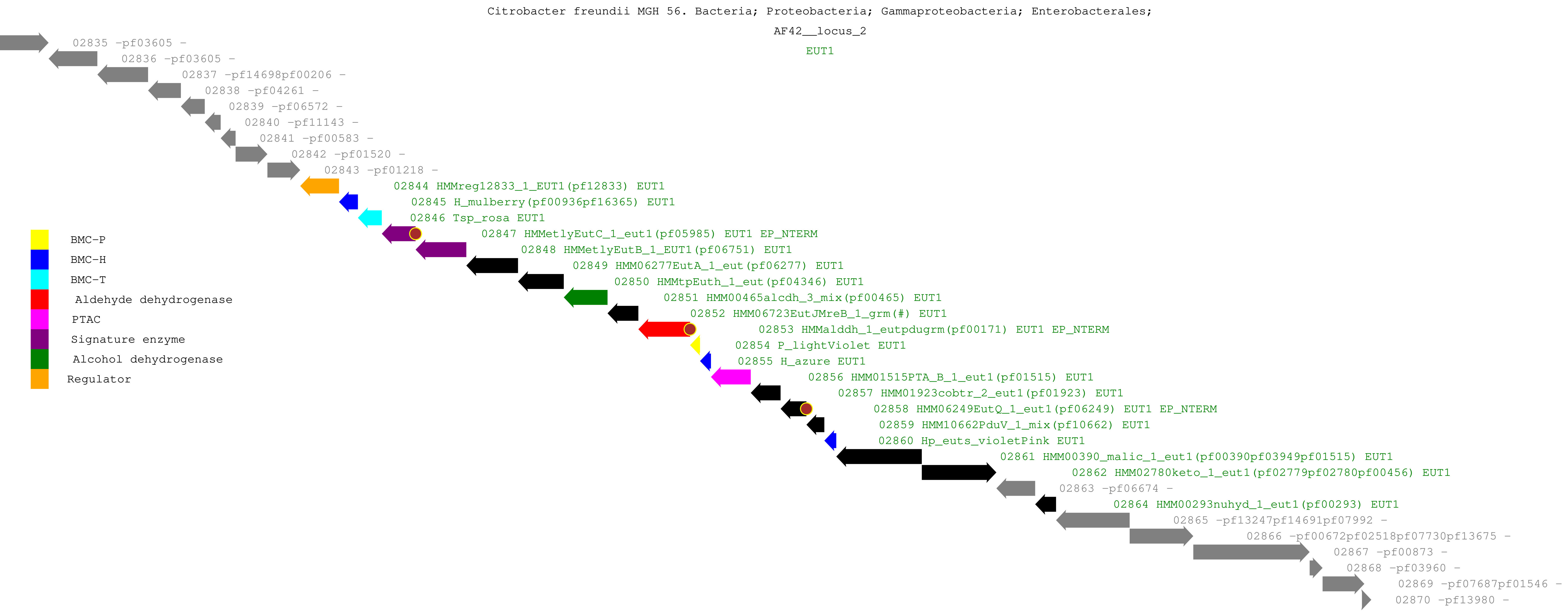
Citrobacter freundii MGH 56. Bacteria; Proteobacteria; Gammaproteobacteria; Enterobacterales;
AF42__locus_2
EUT1
02835 -pf03605 -
02836 -pf03605 -
02837 -pf14698pf00206 -
02838 -pf04261 -
02839 -pf06572 -
02840 -pf11143 -
02841 -pf00583 -
02842 -pf01520 -
02843 -pf01218 -
02844 HMMreg12833_1_EUT1(pf12833) EUT1
02845 H_mulberry(pf00936pf16365) EUT1
02846 Tsp_rosa EUT1
02847 HMMetlyEutC_1_eut1(pf05985) EUT1 EP_NTERM
02848 HMMetlyEutB_1_EUT1(pf06751) EUT1
02849 HMM06277EutA_1_eut(pf06277) EUT1
02850 HMMtpEuth_1_eut(pf04346) EUT1
02851 HMM00465alcdh_3_mix(pf00465) EUT1
02852 HMM06723EutJMreB_1_grm(#) EUT1
02853 HMMalddh_1_eutpdugrm(pf00171) EUT1 EP_NTERM
02854 P_lightViolet EUT1
02855 H_azure EUT1
02856 HMM01515PTA_B_1_eut1(pf01515) EUT1
02857 HMM01923cobtr_2_eut1(pf01923) EUT1
02858 HMM06249EutQ_1_eut1(pf06249) EUT1 EP_NTERM
02859 HMM10662PduV_1_mix(pf10662) EUT1
02860 Hp_euts_violetPink EUT1
02861 HMM00390_malic_1_eut1(pf00390pf03949pf01515) EUT1
02862 HMM02780keto_1_eut1(pf02779pf02780pf00456) EUT1
02863 -pf06674 -
02864 HMM00293nuhyd_1_eut1(pf00293) EUT1
02865 -pf13247pf14691pf07992 -
02866 -pf00672pf02518pf07730pf13675 -
02867 -pf00873 -
02868 -pf03960 -
02869 -pf07687pf01546 -
02870 -pf13980 -
BMC-P
BMC-H
BMC-T
Aldehyde dehydrogenase
PTAC
Signature enzyme
Alcohol dehydrogenase
Regulator

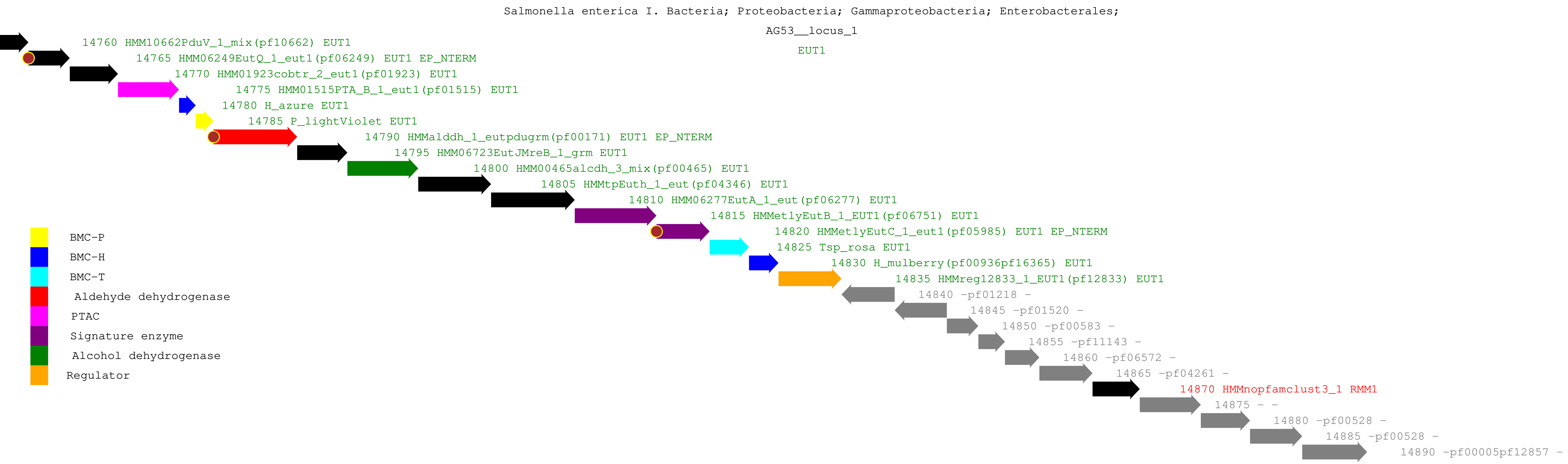
Salmonella enterica I. Bacteria; Proteobacteria; Gammaproteobacteria; Enterobacterales;
AG53__locus_1
EUT1
14760 HMM10662PduV_1_mix(pf10662) EUT1
14765 HMM06249EutQ_1_eut1(pf06249) EUT1 EP_NTERM
14770 HMM01923cobtr_2_eut1(pf01923) EUT1
14775 HMM01515PTA_B_1_eut1(pf01515) EUT1
14780 H_azure EUT1
14785 P_lightViolet EUT1
14790 HMMalddh_1_eutpdugrm(pf00171) EUT1 EP_NTERM
14795 HMM06723EutJMreB_1_grm EUT1
14800 HMM00465alcdh_3_mix(pf00465) EUT1
14805 HMMtpEuth_1_eut(pf04346) EUT1
14810 HMM06277EutA_1_eut(pf06277) EUT1
14815 HMMetlyEutB_1_EUT1(pf06751) EUT1
14820 HMMetlyEutC_1_eut1(pf05985) EUT1 EP_NTERM
14825 Tsp_rosa EUT1
14830 H_mulberry(pf00936pf16365) EUT1
14835 HMMreg12833_1_EUT1(pf12833) EUT1
14840 -pf01218 -
14845 -pf01520 -
14850 -pf00583 -
14855 -pf11143 -
14860 -pf06572 -
14865 -pf04261 -
14870 HMMnopfamclust3_1 RMM1
14875 - -
14880 -pf00528 -
14885 -pf00528 -
14890 -pf00005pf12857 -
BMC-P
BMC-H
BMC-T
Aldehyde dehydrogenase
PTAC
Signature enzyme
Alcohol dehydrogenase
Regulator

Klebsiella pneumoniae. Bacteria; Proteobacteria; Gammaproteobacteria; Enterobacterales;

AGG09__locus_2

EUT1

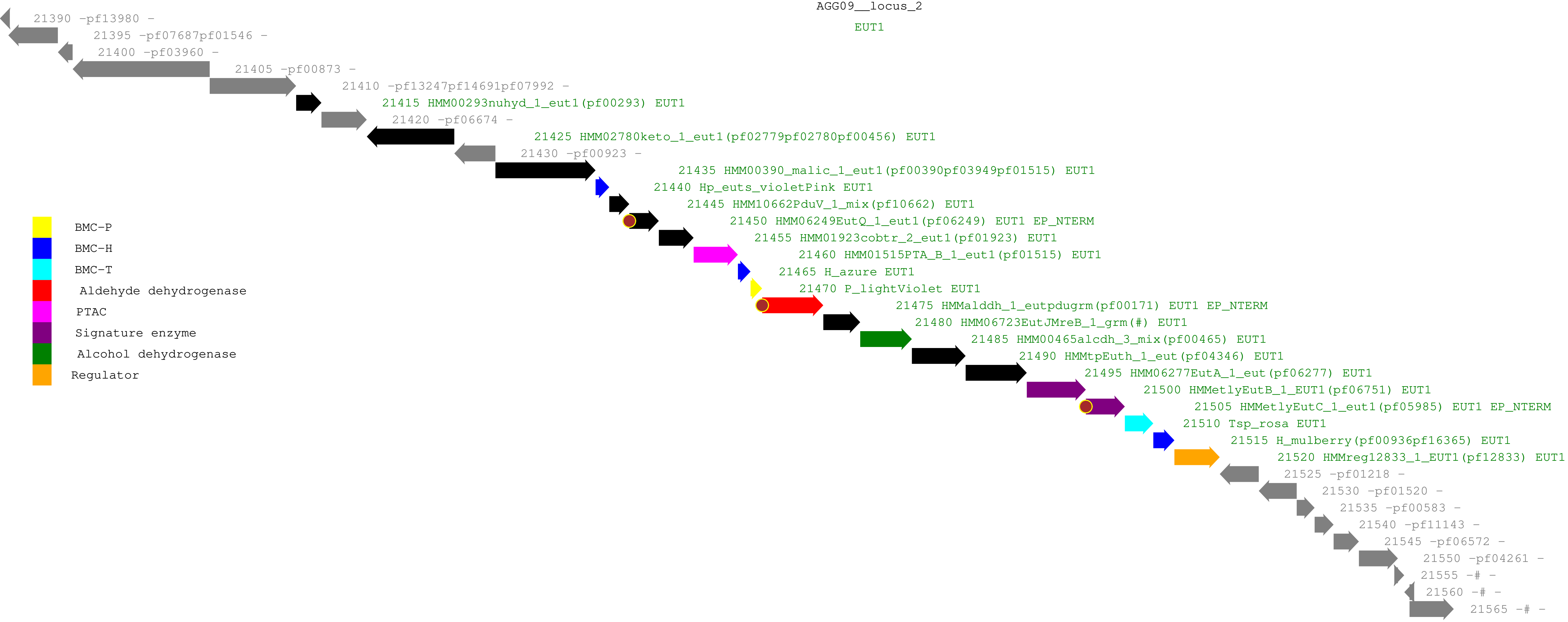

Klebsiella michiganensis. Bacteria; Proteobacteria; Gammaproteobacteria; Enterobacterales;

AGG97__locus_2

EUT1

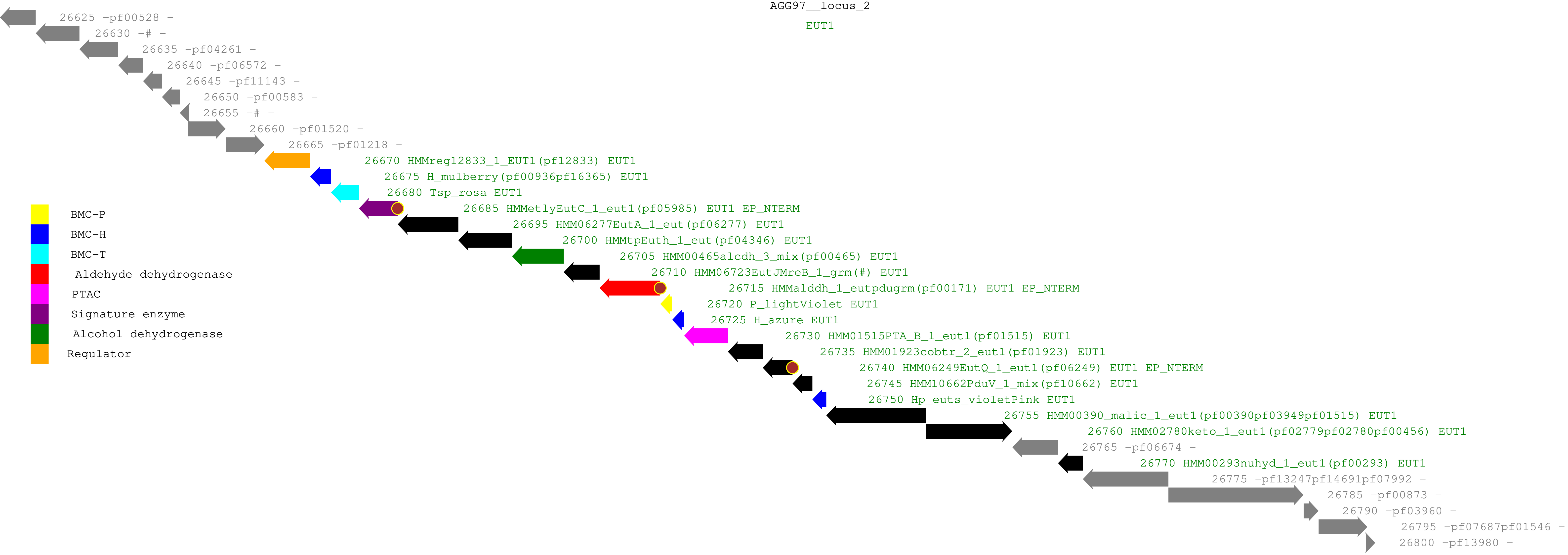

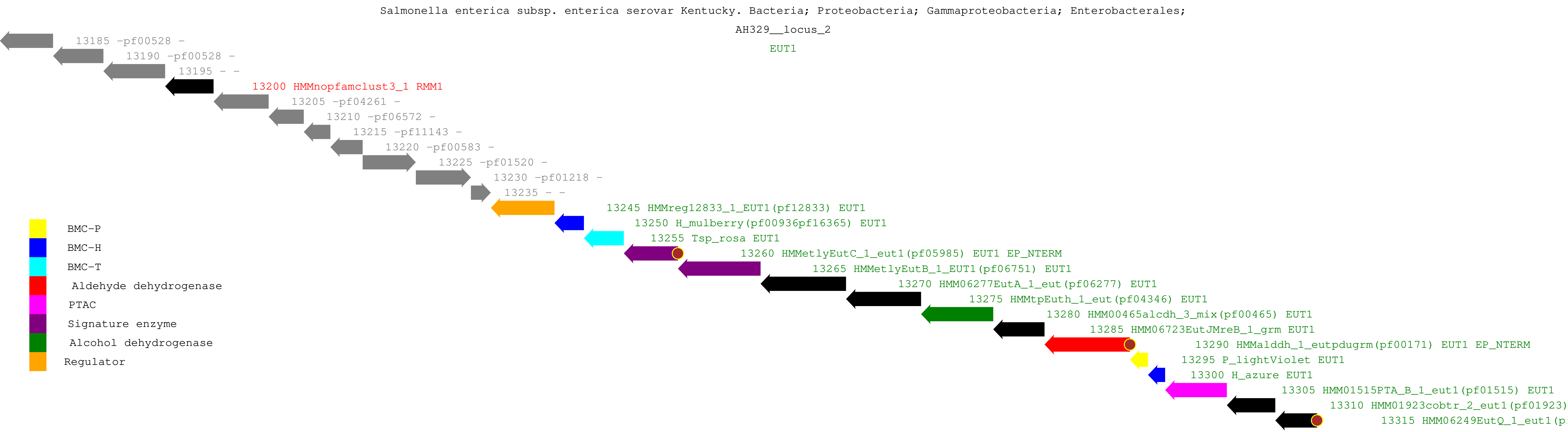
Salmonella enterica subsp. enterica serovar Kentucky. Bacteria; Proteobacteria; Gammaproteobacteria; Enterobacterales;
AH329__locus_2
EUT1
13185 -pf00528 -
13190 -pf00528 -
13195 - -
13200 HMMnopfamclust3_1 RMM1
13205 -pf04261 -
13210 -pf06572 -
13215 -pf11143 -
13220 -pf00583 -
13225 -pf01520 -
13230 -pf01218 -
13235 - -
13245 HMMreg12833_1_EUT1(pf12833) EUT1
13250 H_mulberry(pf00936pf16365) EUT1
13255 Tsp_rosa EUT1
13260 HMMetlyEutC_1_eut1(pf05985) EUT1 EP_NTERM
13265 HMMetlyEutB_1_EUT1(pf06751) EUT1
13270 HMM06277EutA_1_eut(pf06277) EUT1
13275 HMMtpEuth_1_eut(pf04346) EUT1
13280 HMM00465alcdh_3_mix(pf00465) EUT1
13285 HMM06723EutJMreB_1_grm EUT1
13290 HMMalddh_1_eutpdugrm(pf00171) EUT1 EP_NTERM
13295 P_lightViolet EUT1
13300 H_azure EUT1
13305 HMM01515PTA_B_1_eut1(pf01515) EUT1
13310 HMM01923cobtr_2_eut1(pf01923)
13315 HMM06249EutQ_1_eut1(p
BMC-P
BMC-H
BMC-T
Aldehyde dehydrogenase
PTAC
Signature enzyme
Alcohol dehydrogenase
Regulator

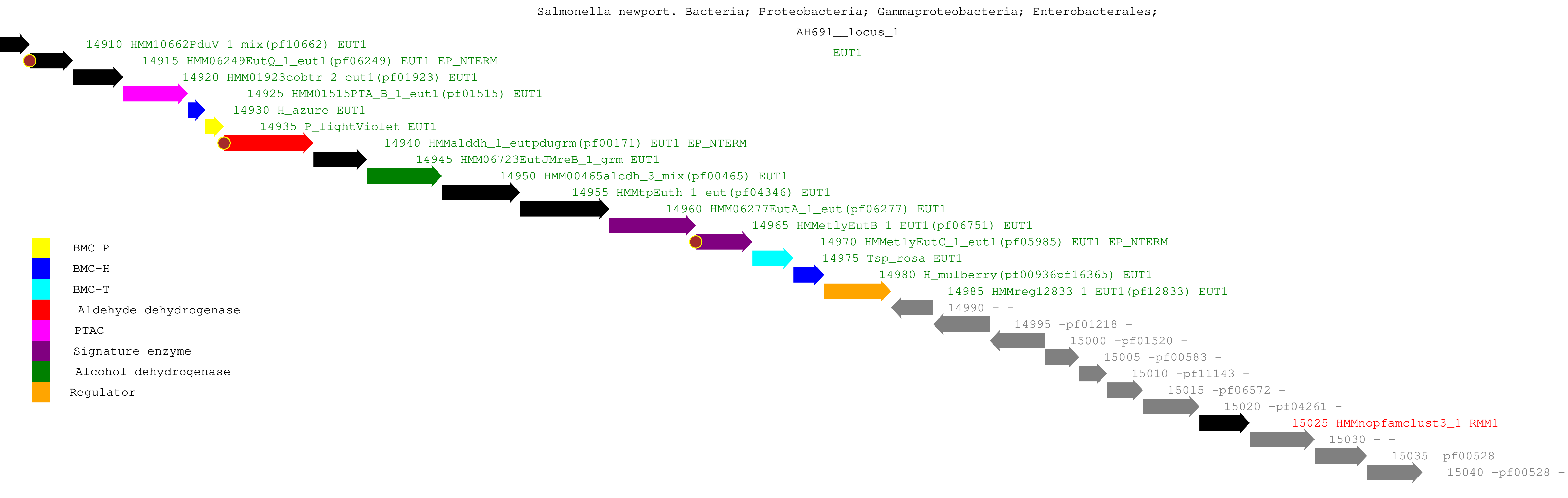
Salmonella newport. Bacteria; Proteobacteria; Gammaproteobacteria; Enterobacterales;
AH691__locus_1
EUT1
14910 HMM10662PduV_1_mix(pf10662) EUT1
14915 HMM06249EutQ_1_eut1(pf06249) EUT1 EP_NTERM
14920 HMM01923cobtr_2_eut1(pf01923) EUT1
14925 HMM01515PTA_B_1_eut1(pf01515) EUT1
14930 H_azure EUT1
14935 P_lightViolet EUT1
14940 HMMalddh_1_eutpdugrm(pf00171) EUT1 EP_NTERM
14945 HMM06723EutJMreB_1_grm EUT1
14950 HMM00465alcdh_3_mix(pf00465) EUT1
14955 HMMtpEuth_1_eut(pf04346) EUT1
14960 HMM06277EutA_1_eut(pf06277) EUT1
14965 HMMetlyEutB_1_EUT1(pf06751) EUT1
14970 HMMetlyEutC_1_eut1(pf05985) EUT1 EP_NTERM
14975 Tsp_rosa EUT1
14980 H_mulberry(pf00936pf16365) EUT1
14985 HMMreg12833_1_EUT1(pf12833) EUT1
14990 - -
14995 -pf01218 -
15000 -pf01520 -
15005 -pf00583 -
15010 -pf11143 -
15015 -pf06572 -
15020 -pf04261 -
15025 HMMnopfamclust3_1 RMM1
15030 - -
15035 -pf00528 -
15040 -pf00528 -
BMC-P
BMC-H
BMC-T
Aldehyde dehydrogenase
PTAC
Signature enzyme
Alcohol dehydrogenase
Regulator

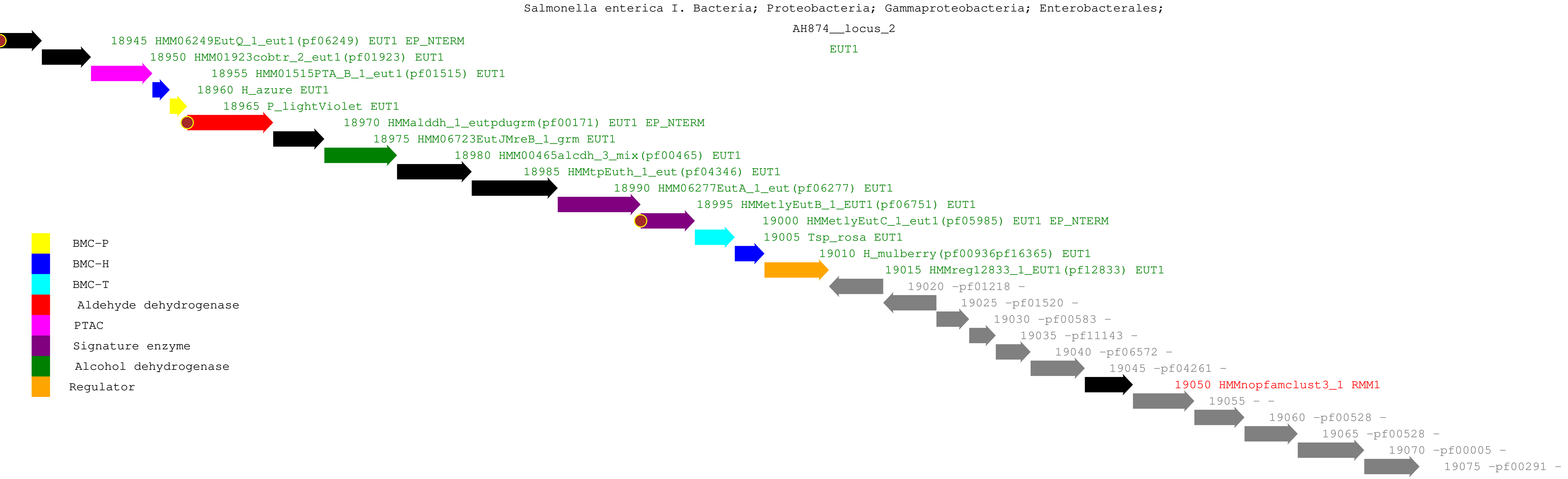
Salmonella enterica I. Bacteria; Proteobacteria; Gammaproteobacteria; Enterobacterales;
AH874__locus_2
EUT1
18945 HMM06249EutQ_1_eut1(pf06249) EUT1 EP_NTERM
18950 HMM01923cobtr_2_eut1(pf01923) EUT1
18955 HMM01515PTA_B_1_eut1(pf01515) EUT1
18960 H_azure EUT1
18965 P_lightViolet EUT1
18970 HMMalddh_1_eutpdugrm(pf00171) EUT1 EP_NTERM
18975 HMM06723EutJMreB_1_grm EUT1
18980 HMM00465alcdh_3_mix(pf00465) EUT1
18985 HMMtpEuth_1_eut(pf04346) EUT1
18990 HMM06277EutA_1_eut(pf06277) EUT1
18995 HMMetlyEutB_1_EUT1(pf06751) EUT1
19000 HMMetlyEutC_1_eut1(pf05985) EUT1 EP_NTERM
19005 Tsp_rosa EUT1
19010 H_mulberry(pf00936pf16365) EUT1
19015 HMMreg12833_1_EUT1(pf12833) EUT1
19020 -pf01218 -
19025 -pf01520 -
19030 -pf00583 -
19035 -pf11143 -
19040 -pf06572 -
19045 -pf04261 -
19050 HMMnopfamclust3_1 RMM1
19055 - -
19060 -pf00528 -
19065 -pf00528 -
19070 -pf00005 -
19075 -pf00291 -
BMC-P
BMC-H
BMC-T
Aldehyde dehydrogenase
PTAC
Signature enzyme
Alcohol dehydrogenase
Regulator

Salmonella enterica subsp. enterica serovar Havana. Bacteria; Proteobacteria; Gammaproteobacteria; Enterobacterales;

AH902__locus_1

EUT1

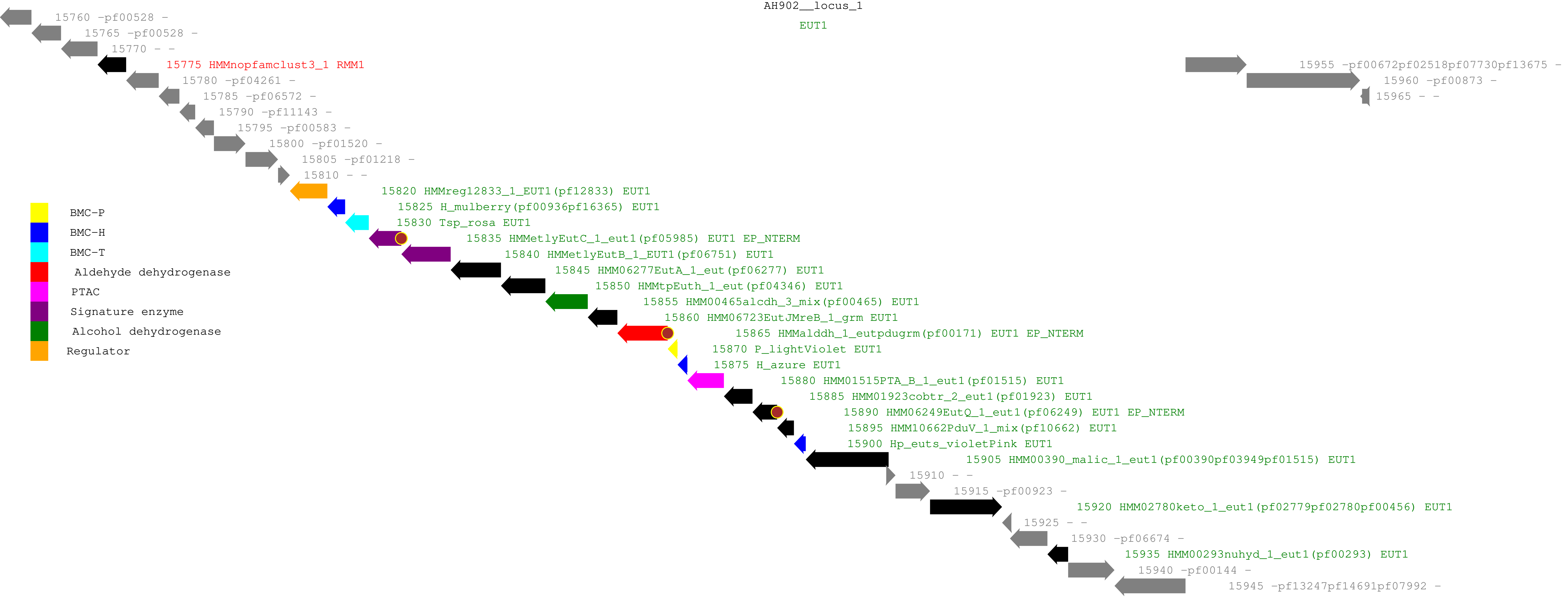

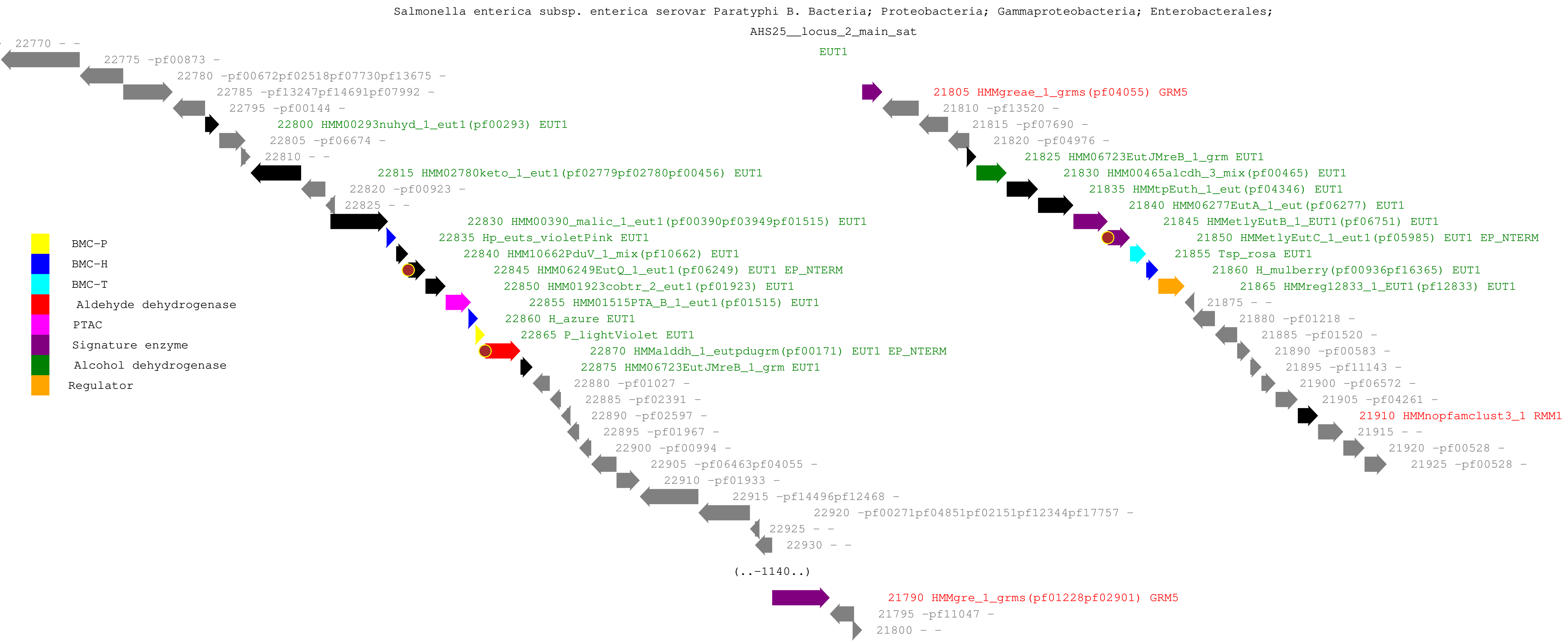
Salmonella enterica subsp. enterica serovar Paratyphi B. Bacteria; Proteobacteria; Gammaproteobacteria; Enterobacterales;
AHS25__locus_2_main_sat
EUT1
22770 - -
22775 -pf00873 -
22780 -pf00672pf02518pf07730pf13675 -
22785 -pf13247pf14691pf07992 -
22795 -pf00144 -
22800 HMM00293nuhyd_1_eut1(pf00293) EUT1
22805 -pf06674 -
22810 - -
22815 HMM02780keto_1_eut1(pf02779pf02780pf00456) EUT1
22820 -pf00923 -
22825 - -
22830 HMM00390_malic_1_eut1(pf00390pf03949pf01515) EUT1
22835 Hp_euts_violetPink EUT1
22840 HMM10662PduV_1_mix(pf10662) EUT1
22845 HMM06249EutQ_1_eut1(pf06249) EUT1 EP_NTERM
22850 HMM01923cobtr_2_eut1(pf01923) EUT1
22855 HMM01515PTA_B_1_eut1(pf01515) EUT1
22860 H_azure EUT1
22865 P_lightViolet EUT1
22870 HMMalddh_1_eutpdugrm(pf00171) EUT1 EP_NTERM
22875 HMM06723EutJMreB_1_grm EUT1
22880 -pf01027 -
22885 -pf02391 -
22890 -pf02597 -
22895 -pf01967 -
22900 -pf00994 -
22905 -pf06463pf04055 -
22910 -pf01933 -
22915 -pf14496pf12468 -
22920 -pf00271pf04851pf02151pf12344pf17757 -
22925 - -
22930 - -
(..-1140..)
21790 HMMgre_1_grms(pf01228pf02901) GRM5
21795 -pf11047 -
21800 - -
21805 HMMgreae_1_grms(pf04055) GRM5
21810 -pf13520 -
21815 -pf07690 -
21820 -pf04976 -
21825 HMM06723EutJMreB_1_grm EUT1
21830 HMM00465alcdh_3_mix(pf00465) EUT1
21835 HMMtpEuth_1_eut(pf04346) EUT1
21840 HMM06277EutA_1_eut(pf06277) EUT1
21845 HMMetlyEutB_1_EUT1(pf06751) EUT1
21850 HMMetlyEutC_1_eut1(pf05985) EUT1 EP_NTERM
21855 Tsp_rosa EUT1
21860 H_mulberry(pf00936pf16365) EUT1
21865 HMMreg12833_1_EUT1(pf12833) EUT1
21875 - -
21880 -pf01218 -
21885 -pf01520 -
21890 -pf00583 -
21895 -pf11143 -
21900 -pf06572 -
21905 -pf04261 -
21910 HMMnopfamclust3_1 RMM1
21915 - -
21920 -pf00528 -
21925 -pf00528 -
BMC-P
BMC-H
BMC-T
Aldehyde dehydrogenase
PTAC
Signature enzyme
Alcohol dehydrogenase
Regulator

Salmonella newport. Bacteria; Proteobacteria; Gammaproteobacteria; Enterobacterales;

AHY32__locus_1

EUT1

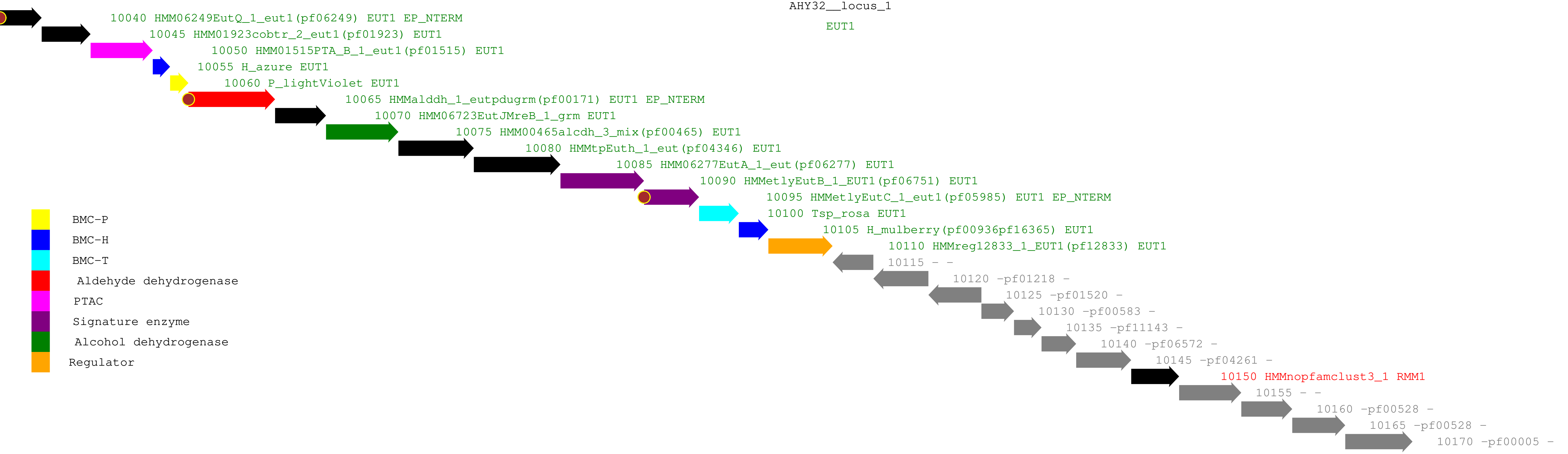

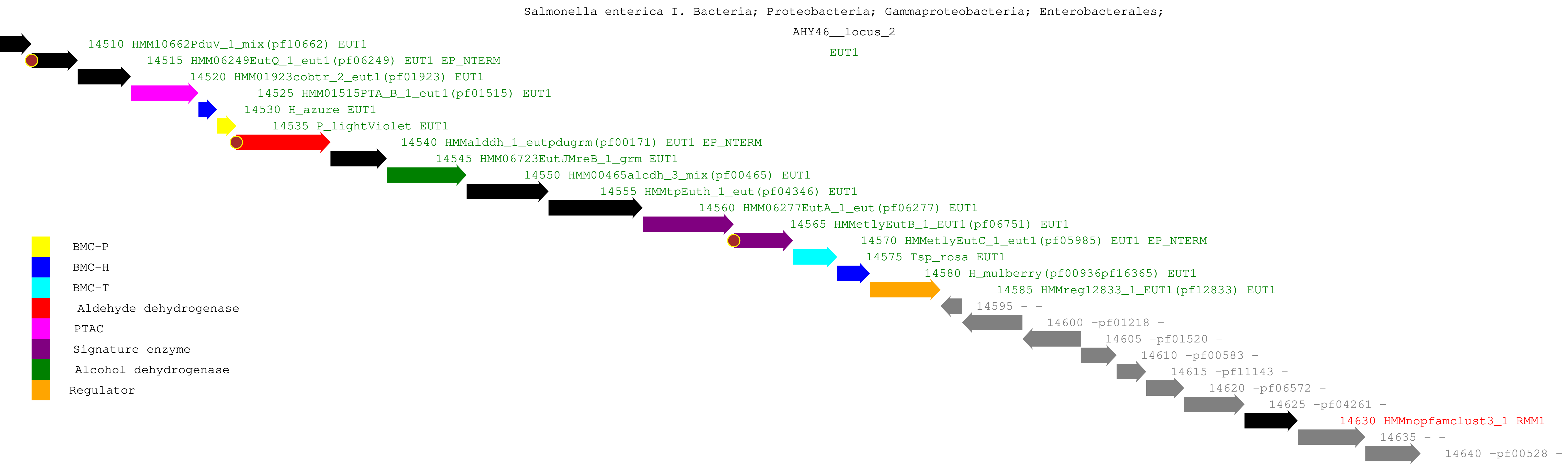
Salmonella enterica I. Bacteria; Proteobacteria; Gammaproteobacteria; Enterobacterales;
AHY46__locus_2
EUT1
14510 HMM10662PduV_1_mix(pf10662) EUT1
14515 HMM06249EutQ_1_eut1(pf06249) EUT1 EP_NTERM
14520 HMM01923cobtr_2_eut1(pf01923) EUT1
14525 HMM01515PTA_B_1_eut1(pf01515) EUT1
14530 H_azure EUT1
14535 P_lightViolet EUT1
14540 HMMaldh_1_eutpdugrm(pf00171) EUT1 EP_NTERM
14545 HMM06723EutJMreB_1_grm EUT1
14550 HMM00465alcdh_3_mix(pf00465) EUT1
14555 HMMtpEuth_1_eut(pf04346) EUT1
14560 HMM06277EutA_1_eut(pf06277) EUT1
14565 HMMetlyEutB_1_EUT1(pf06751) EUT1
14570 HMMetlyEutC_1_eut1(pf05985) EUT1 EP_NTERM
14575 Tsp_rosa EUT1
14580 H_mulberry(pf00936pf16365) EUT1
14585 HMMreg12833_1_EUT1(pf12833) EUT1
14595 - -
14600 -pf01218 -
14605 -pf01520 -
14610 -pf00583 -
14615 -pf11143 -
14620 -pf06572 -
14625 -pf04261 -
14630 HMMnopfamclust3_1 RMM1
14635 - -
14640 -pf00528 -
BMC-P
BMC-H
BMC-T
Aldehyde dehydrogenase
PTAC
Signature enzyme
Alcohol dehydrogenase
Regulator

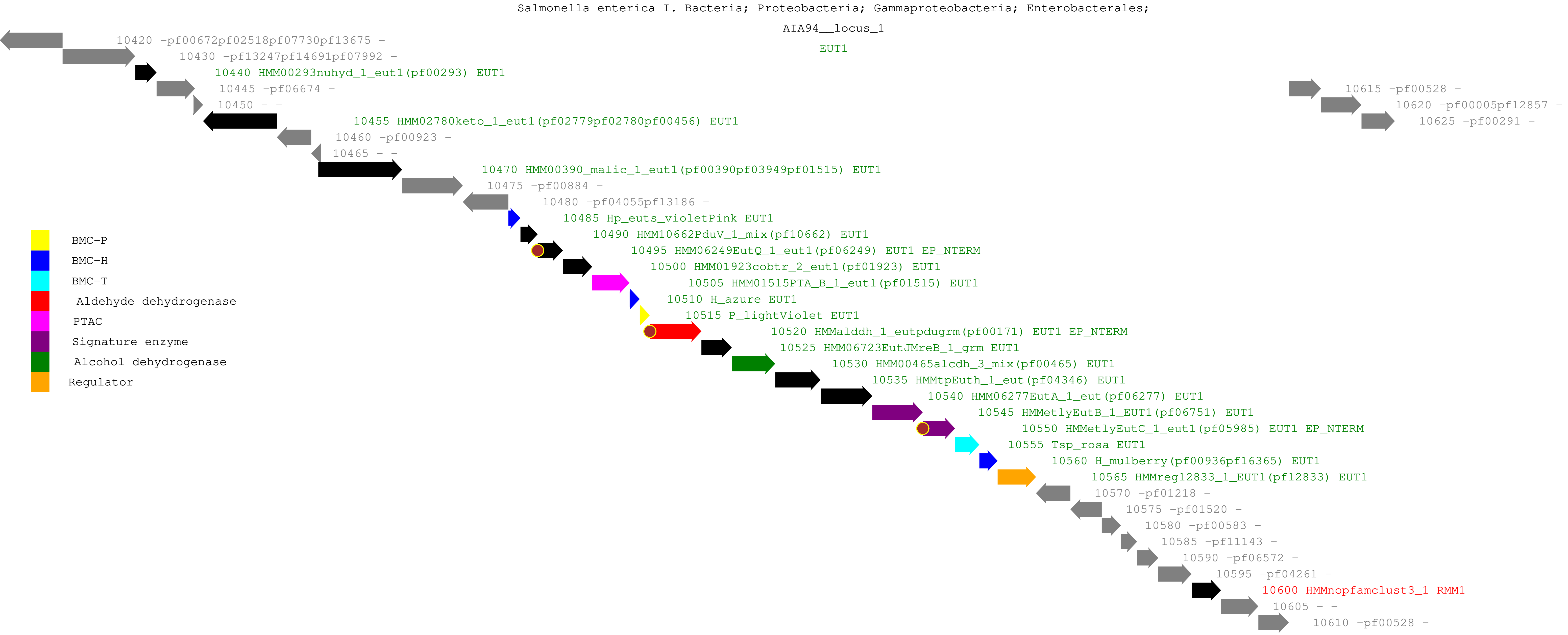

Salmonella enterica I. Bacteria; Proteobacteria; Gammaproteobacteria; Enterobacterales;
AIA94__locus_1
EUT1
10420 -pf00672pf02518pf07730pf13675 -
10430 -pf13247pf14691pf07992 -
10440 HMM00293nuhyd_1_eut1(pf00293) EUT1
10445 -pf06674 -
10450 - -
10455 HMM02780keto_1_eut1(pf02779pf02780pf00456) EUT1
10460 -pf00923 -
10465 - -
10470 HMM00390_malic_1_eut1(pf00390pf03949pf01515) EUT1
10475 -pf00884 -
10480 -pf04055pf13186 -
10485 Hp_euts_violetPink EUT1
10490 HMM10662PduV_1_mix(pf10662) EUT1
10495 HMM06249EutQ_1_eut1(pf06249) EUT1 EP_NTERM
10500 HMM01923cobtr_2_eut1(pf01923) EUT1
10505 HMM01515PTA_B_1_eut1(pf01515) EUT1
10510 H_azure EUT1
10515 P_lightViolet EUT1
10520 HMMalddh_1_eutpdugrm(pf00171) EUT1 EP_NTERM
10525 HMM06723EutJMreB_1_grm EUT1
10530 HMM00465alcdh_3_mix(pf00465) EUT1
10535 HMMtpEuth_1_eut(pf04346) EUT1
10540 HMM06277EutA_1_eut(pf06277) EUT1
10545 HMMetlyEutB_1_EUT1(pf06751) EUT1
10550 HMMetlyEutC_1_eut1(pf05985) EUT1 EP_NTERM
10555 Tsp_rosa EUT1
10560 H_mulberry(pf00936pf16365) EUT1
10565 HMMreg12833_1_EUT1(pf12833) EUT1
10570 -pf01218 -
10575 -pf01520 -
10580 -pf00583 -
10585 -pf11143 -
10590 -pf06572 -
10595 -pf04261 -
10600 HMMnopfamclust3_1 RMM1
10605 - -
10610 -pf00528 -
10615 -pf00528 -
10620 -pf00005pf12857 -
10625 -pf00291 -
BMC-P
BMC-H
BMC-T
Aldehyde dehydrogenase
PTAC
Signature enzyme
Alcohol dehydrogenase
Regulator

Salmonella thompson. Bacteria; Proteobacteria; Gammaproteobacteria; Enterobacterales;

AIG11__locus_1

EUT1

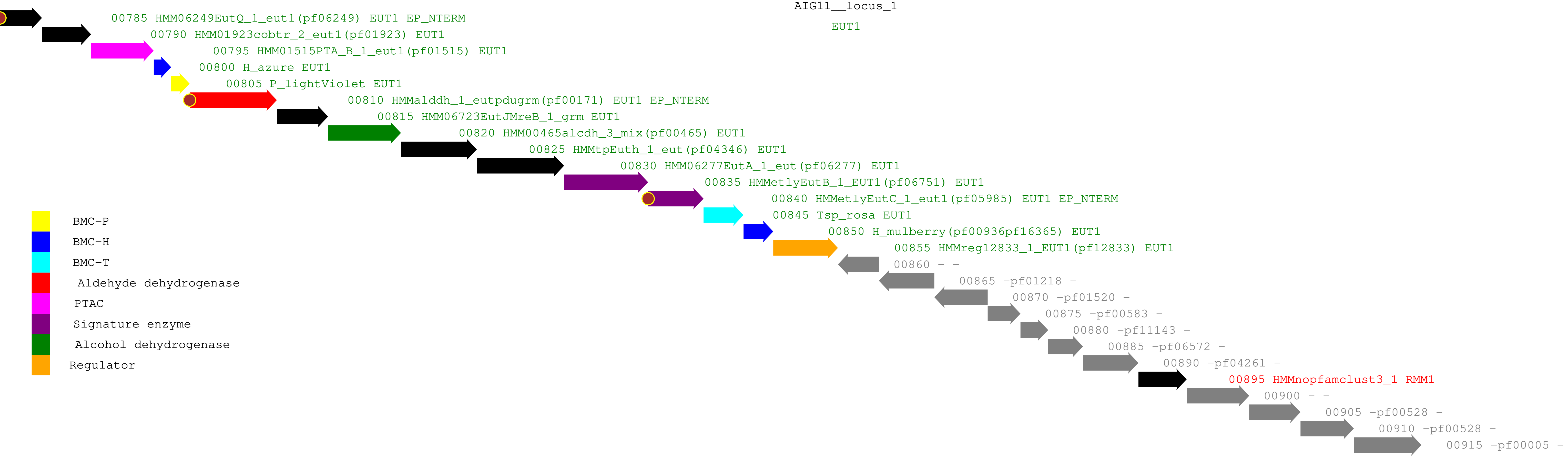

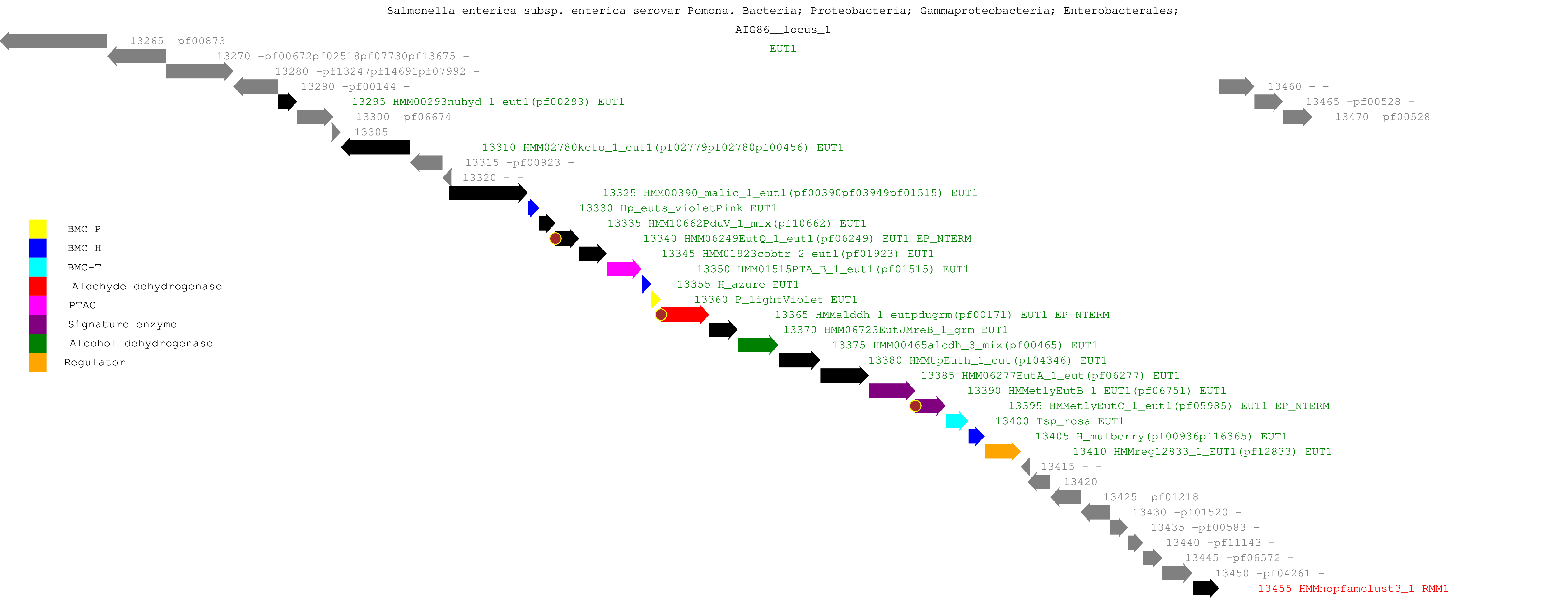
Salmonella enterica subsp. enterica serovar Pomona. Bacteria; Proteobacteria; Gammaproteobacteria; Enterobacterales;
AIG86__locus_1
EUT1
13265 -pf00873 -
13270 -pf00672pf02518pf07730pf13675 -
13280 -pf13247pf14691pf07992 -
13290 -pf00144 -
13295 HMM00293nuhyd_1_eut1(pf00293) EUT1
13300 -pf06674 -
13305 - -
13310 HMM02780keto_1_eut1(pf02779pf02780pf00456) EUT1
13315 -pf00923 -
13320 - -
13325 HMM00390_malic_1_eut1(pf00390pf03949pf01515) EUT1
13330 Hp_euts_violetPink EUT1
13335 HMM10662PduV_1_mix(pf10662) EUT1
13340 HMM06249EutQ_1_eut1(pf06249) EUT1 EP_NTERM
13345 HMM01923cobtr_2_eut1(pf01923) EUT1
13350 HMM01515PTA_B_1_eut1(pf01515) EUT1
13355 H_azure EUT1
13360 P_lightViolet EUT1
13365 HMMalddh_1_eutpdugrm(pf00171) EUT1 EP_NTERM
13370 HMM06723EutJMreB_1_grm EUT1
13375 HMM00465alcdh_3_mix(pf00465) EUT1
13380 HMMtpEuth_1_eut(pf04346) EUT1
13385 HMM06277EutA_1_eut(pf06277) EUT1
13390 HMMetlyEutB_1_EUT1(pf06751) EUT1
13395 HMMetlyEutC_1_eut1(pf05985) EUT1 EP_NTERM
13400 Tsp_rosa EUT1
13405 H_mulberry(pf00936pf16365) EUT1
13410 HMMreg12833_1_EUT1(pf12833) EUT1
13415 - -
13420 - -
13425 -pf01218 -
13430 -pf01520 -
13435 -pf00583 -
13440 -pf11143 -
13445 -pf06572 -
13450 -pf04261 -
13455 HMMnopfamclust3_1 RMM1
13460 - -
13465 -pf00528 -
13470 -pf00528 -
BMC-P
BMC-H
BMC-T
Aldehyde dehydrogenase
PTAC
Signature enzyme
Alcohol dehydrogenase
Regulator

Salmonella enterica subsp. enterica serovar Havana. Bacteria; Proteobacteria; Gammaproteobacteria; Enterobacterales;

AIH56__locus_2

EUT1

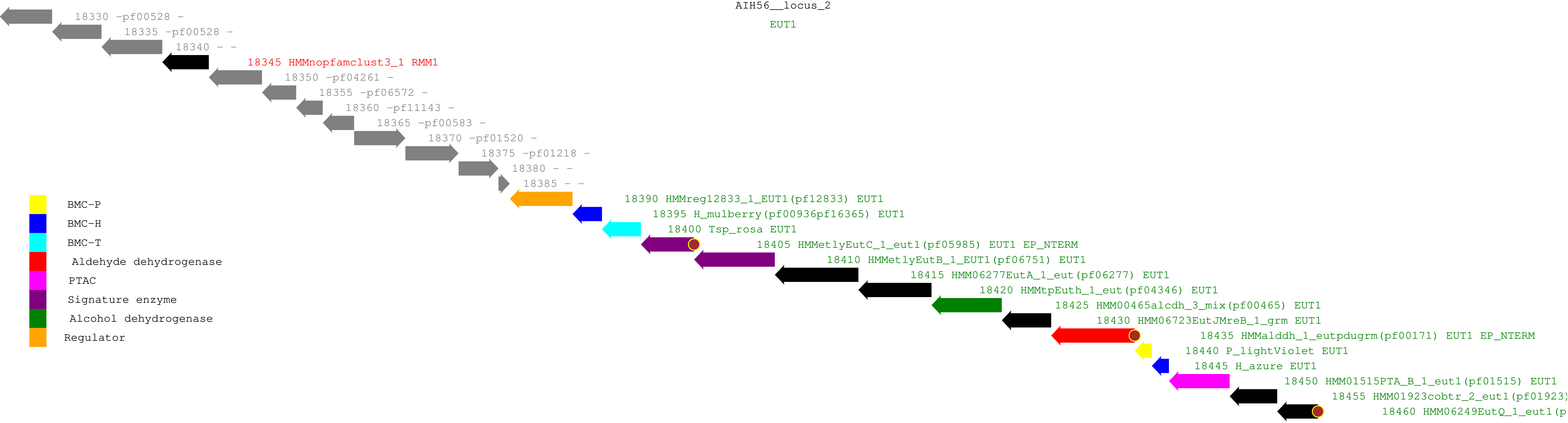

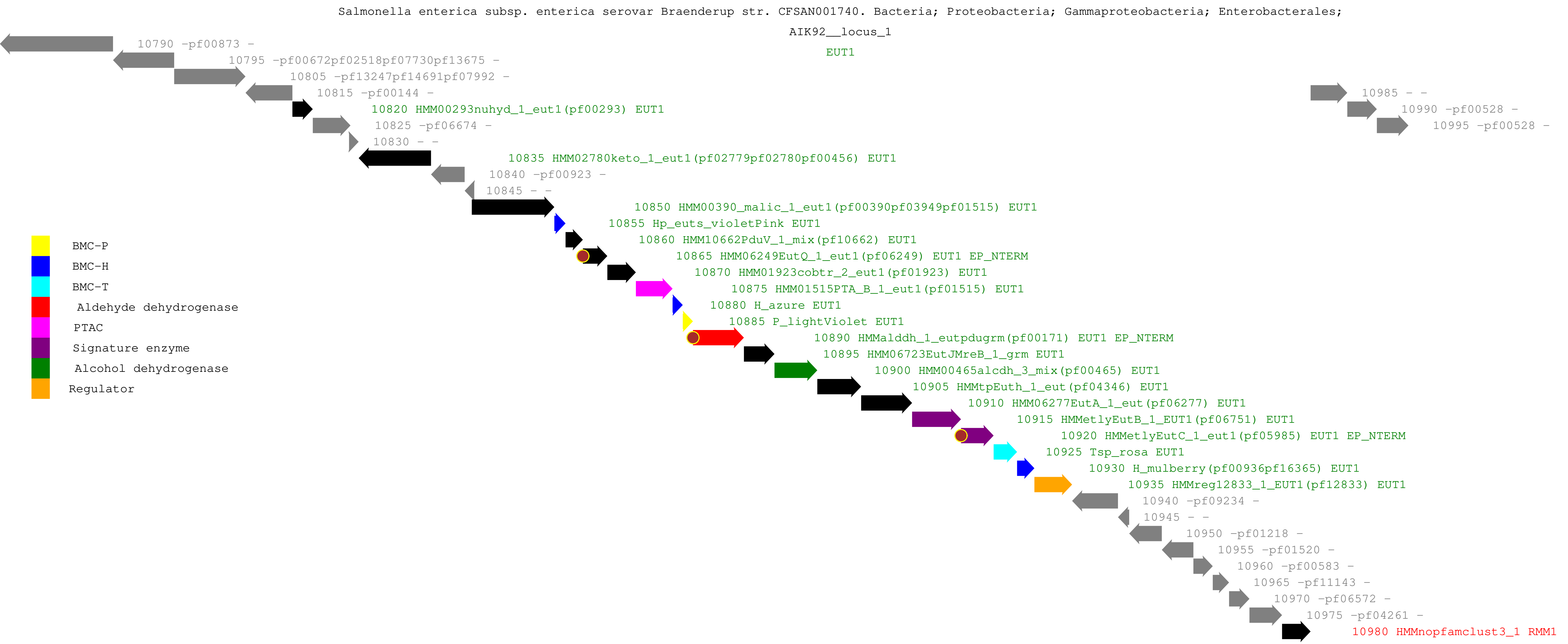

Salmonella enterica subsp. enterica serovar Braenderup str. CFSAN001740. Bacteria; Proteobacteria; Gammaproteobacteria; Enterobacterales;
AIK92__locus_1
EUT1
10790 -pf00873 -
10795 -pf00672pf02518pf07730pf13675 -
10805 -pf13247pf14691pf07992 -
10815 -pf00144 -
10820 HMM00293nuhyd_1_eut1(pf00293) EUT1
10825 -pf06674 -
10830 - -
10835 HMM02780keto_1_eut1(pf02779pf02780pf00456) EUT1
10840 -pf00923 -
10845 - -
10850 HMM00390_malic_1_eut1(pf00390pf03949pf01515) EUT1
10855 Hp_euts_violetPink EUT1
10860 HMM10662PduV_1_mix(pf10662) EUT1
10865 HMM06249EutQ_1_eut1(pf06249) EUT1 EP_NTERM
10870 HMM01923cobtr_2_eut1(pf01923) EUT1
10875 HMM01515PTA_B_1_eut1(pf01515) EUT1
10880 H_azure EUT1
10885 P_lightViolet EUT1
10890 HMMaldh_1_eutpdugrm(pf00171) EUT1 EP_NTERM
10895 HMM06723EutJMreB_1_grm EUT1
10900 HMM00465alcdh_3_mix(pf00465) EUT1
10905 HMMtpEuth_1_eut(pf04346) EUT1
10910 HMM06277EutA_1_eut(pf06277) EUT1
10915 HMMetlyEutB_1_EUT1(pf06751) EUT1
10920 HMMetlyEutC_1_eut1(pf05985) EUT1 EP_NTERM
10925 Tsp_rosa EUT1
10930 H_mulberry(pf00936pf16365) EUT1
10935 HMMreg12833_1_EUT1(pf12833) EUT1
10940 -pf09234 -
10945 - -
10950 -pf01218 -
10955 -pf01520 -
10960 -pf00583 -
10965 -pf11143 -
10970 -pf06572 -
10975 -pf04261 -
10980 HMMnopfamclust3_1 RMM1
10985 - -
10990 -pf00528 -
10995 -pf00528 -
BMC-P
BMC-H
BMC-T
Aldehyde dehydrogenase
PTAC
Signature enzyme
Alcohol dehydrogenase
Regulator

Salmonella enterica subsp. arizonae serovar 41:z4,z23:- str. 01-0089. Bacteria; Proteobacteria; Gammaproteobacteria; Enterobacterales;

AIT14__locus_1

EUT1

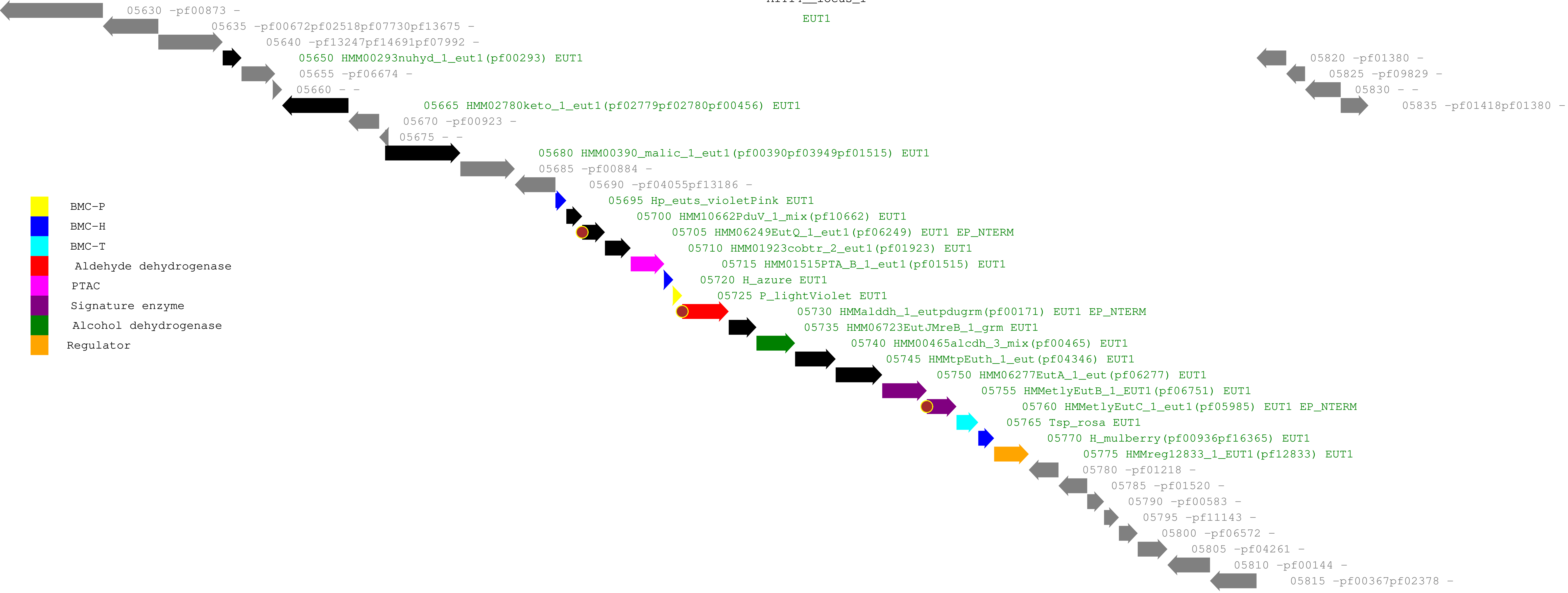

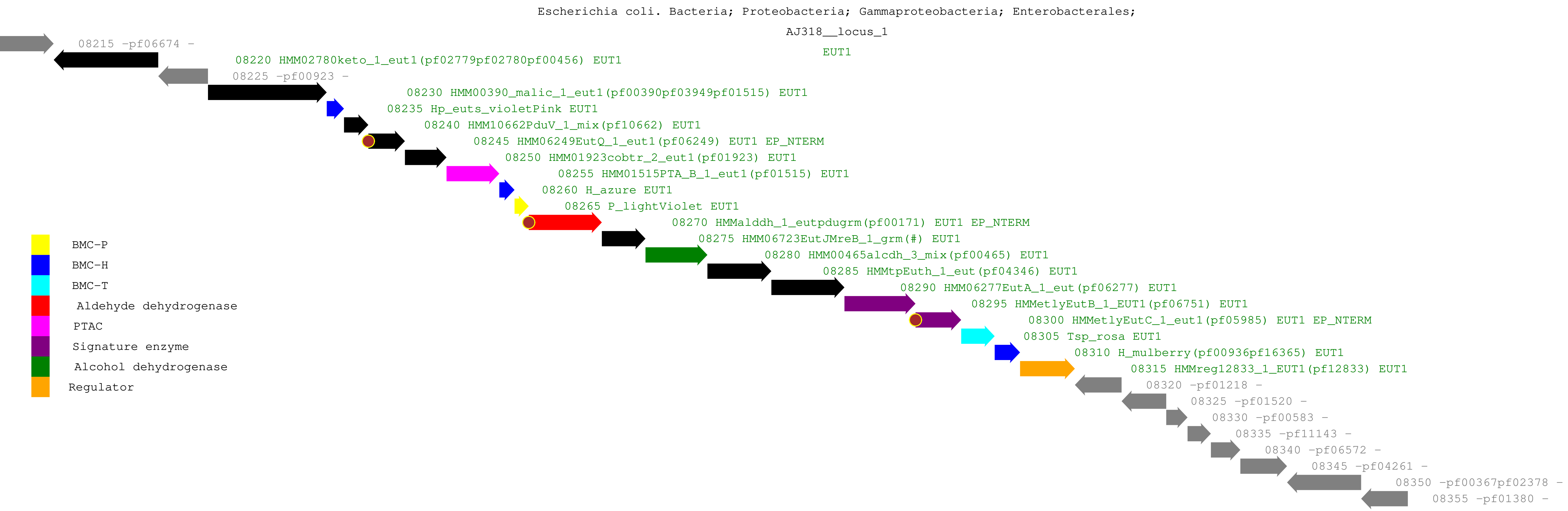
Escherichia coli. Bacteria; Proteobacteria; Gammaproteobacteria; Enterobacterales;
AJ318__locus_1
EUT1
08215 -pf06674 -
08220 HMM02780keto_1_eut1(pf02779pf02780pf00456) EUT1
08225 -pf00923 -
08230 HMM00390_malic_1_eut1(pf00390pf03949pf01515) EUT1
08235 Hp_euts_violetPink EUT1
08240 HMM10662PduV_1_mix(pf10662) EUT1
08245 HMM06249EutQ_1_eut1(pf06249) EUT1 EP_NTERM
08250 HMM01923cobtr_2_eut1(pf01923) EUT1
08255 HMM01515PTA_B_1_eut1(pf01515) EUT1
08260 H_azure EUT1
08265 P_lightViolet EUT1
08270 HMMalddh_1_eutpdugrm(pf00171) EUT1 EP_NTERM
08275 HMM06723EutJMreB_1_grm(#) EUT1
08280 HMM00465alcdh_3_mix(pf00465) EUT1
08285 HMMtpEuth_1_eut(pf04346) EUT1
08290 HMM06277EutA_1_eut(pf06277) EUT1
08295 HMMetlyEutB_1_EUT1(pf06751) EUT1
08300 HMMetlyEutC_1_eut1(pf05985) EUT1 EP_NTERM
08305 Tsp_rosa EUT1
08310 H_mulberry(pf00936pf16365) EUT1
08315 HMMreg12833_1_EUT1(pf12833) EUT1
08320 -pf01218 -
08325 -pf01520 -
08330 -pf00583 -
08335 -pf11143 -
08340 -pf06572 -
08345 -pf04261 -
08350 -pf00367pf02378 -
08355 -pf01380 -
BMC-P
BMC-H
BMC-T
Aldehyde dehydrogenase
PTAC
Signature enzyme
Alcohol dehydrogenase
Regulator

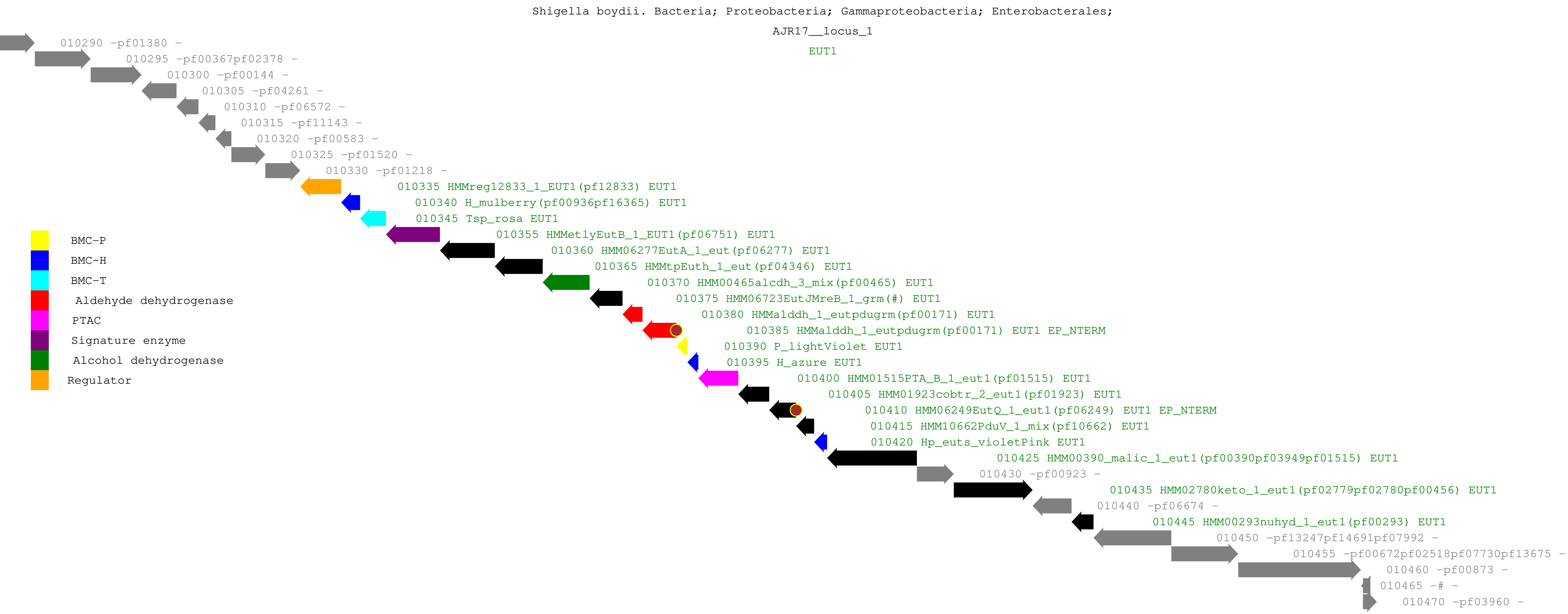
Shigella boydii. Bacteria; Proteobacteria; Gammaproteobacteria; Enterobacterales;
AJR17__locus_1
EUT1
010290 -pf01380 -
010295 -pf00367pf02378 -
010300 -pf00144 -
010305 -pf04261 -
010310 -pf06572 -
010315 -pf11143 -
010320 -pf00583 -
010325 -pf01520 -
010330 -pf01218 -
010335 HMMreg12833_1_EUT1(pf12833) EUT1
010340 H_mulberry(pf00936pf16365) EUT1
010345 Tsp_rosa EUT1
010355 HMMetlyEutB_1_EUT1(pf06751) EUT1
010360 HMM06277EutA_1_eut(pf06277) EUT1
010365 HMMtpEuth_1_eut(pf04346) EUT1
010370 HMM00465alcdh_3_mix(pf00465) EUT1
010375 HMM06723EutJMreB_1_grm(#) EUT1
010380 HMMalddh_1_eutpdugrm(pf00171) EUT1
010385 HMMalddh_1_eutpdugrm(pf00171) EUT1 EP_NTERM
010390 P_lightViolet EUT1
010395 H_azure EUT1
010400 HMM01515PTA_B_1_eut1(pf01515) EUT1
010405 HMM01923cobtr_2_eut1(pf01923) EUT1
010410 HMM06249EutQ_1_eut1(pf06249) EUT1 EP_NTERM
010415 HMM10662PduV_1_mix(pf10662) EUT1
010420 Hp_euts_violetPink EUT1
010425 HMM00390_malic_1_eut1(pf00390pf03949pf01515) EUT1
010430 -pf00923 -
010435 HMM02780keto_1_eut1(pf02779pf02780pf00456) EUT1
010440 -pf06674 -
010445 HMM00293nuhyd_1_eut1(pf00293) EUT1
010450 -pf13247pf14691pf07992 -
010455 -pf00672pf02518pf07730pf13675 -
010460 -pf00873 -
010465 -# -
010470 -pf03960 -
BMC-P
BMC-H
BMC-T
Aldehyde dehydrogenase
PTAC
Signature enzyme
Alcohol dehydrogenase
Regulator

Shigella boydii. Bacteria; Proteobacteria; Gammaproteobacteria; Enterobacterales;

AJR18__locus_1_main_sat

EUT1

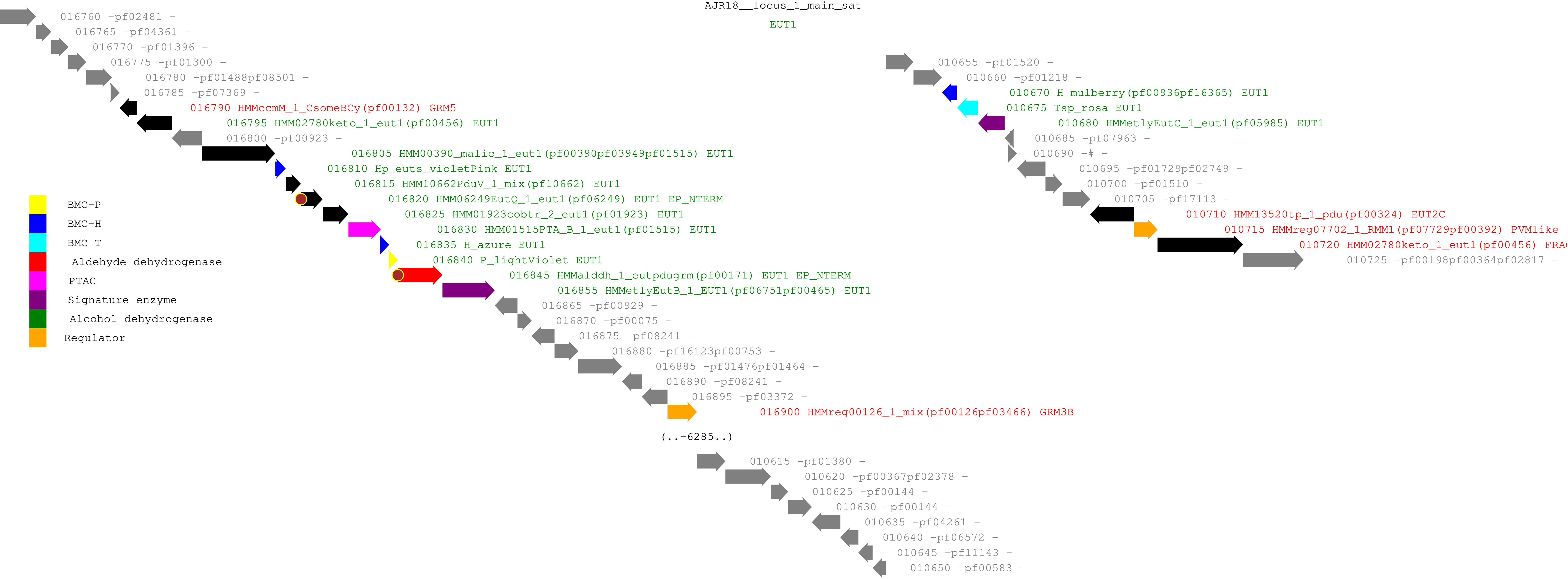

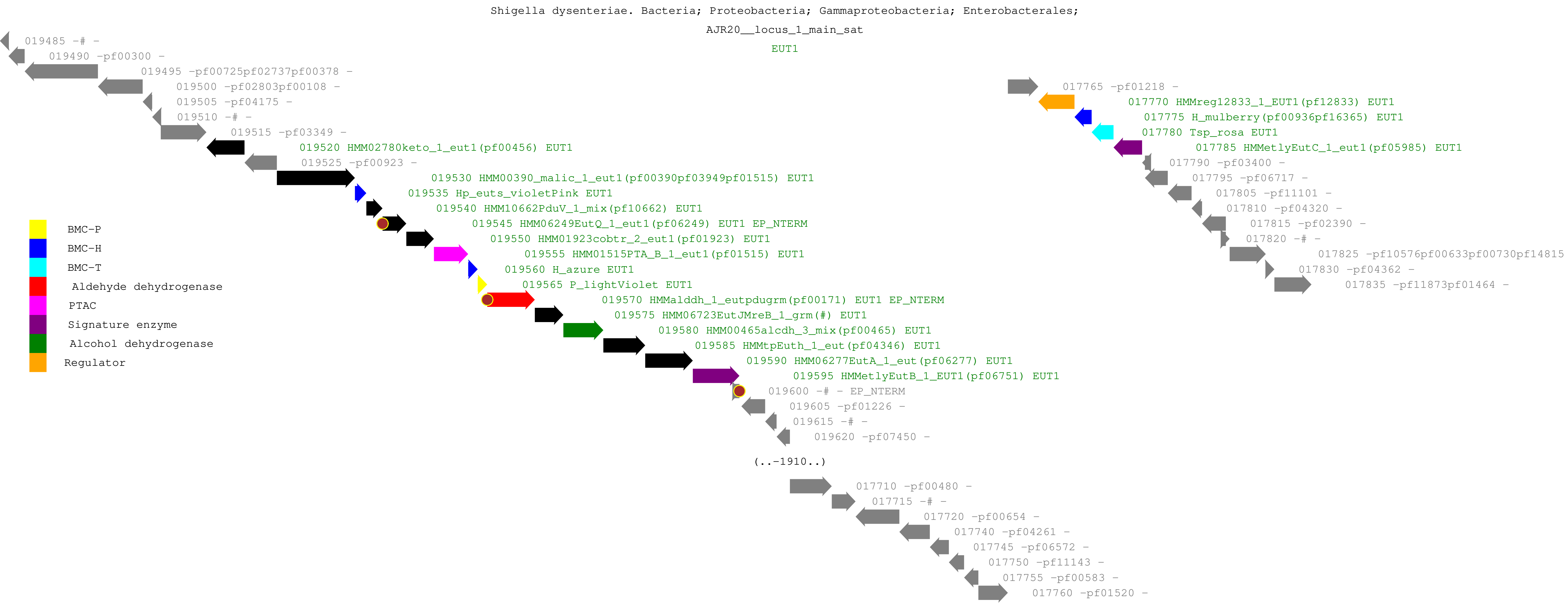
Shigella dysenteriae. Bacteria; Proteobacteria; Gammaproteobacteria; Enterobacterales;
AJR20__locus_1_main_sat
EUT1
019485 -# -
019490 -pf00300 -
019495 -pf00725pf02737pf00378 -
019500 -pf02803pf00108 -
019505 -pf04175 -
019510 -# -
019515 -pf03349 -
019520 HMM02780keto_1_eut1(pf00456) EUT1
019525 -pf00923 -
019530 HMM00390_malic_1_eut1(pf00390pf03949pf01515) EUT1
019535 Hp_euts_violetPink EUT1
019540 HMM10662PduV_1_mix(pf10662) EUT1
019545 HMM06249EutQ_1_eut1(pf06249) EUT1 EP_NTERM
019550 HMM01923cobtr_2_eut1(pf01923) EUT1
019555 HMM01515PTA_B_1_eut1(pf01515) EUT1
019560 H_azure EUT1
019565 P_lightViolet EUT1
019570 HMMaldh_1_eutpdugrm(pf00171) EUT1 EP_NTERM
019575 HMM06723EutJMreB_1_grm(#) EUT1
019580 HMM00465alcdh_3_mix(pf00465) EUT1
019585 HMMtpEuth_1_eut(pf04346) EUT1
019590 HMM06277EutA_1_eut(pf06277) EUT1
019595 HMMetlyEutB_1_EUT1(pf06751) EUT1
019600 -# - EP_NTERM
019605 -pf01226 -
019615 -# -
019620 -pf07450 -
(..-1910..)
017710 -pf00480 -
017715 -# -
017720 -pf00654 -
017740 -pf04261 -
017745 -pf06572 -
017750 -pf11143 -
017755 -pf00583 -
017760 -pf01520 -
017765 -pf01218 -
017770 HMMreg12833_1_EUT1(pf12833) EUT1
017775 H_mulberry(pf00936pf16365) EUT1
017780 Tsp_rosa EUT1
017785 HMMetlyEutC_1_eut1(pf05985) EUT1
017790 -pf03400 -
017795 -pf06717 -
017805 -pf11101 -
017810 -pf04320 -
017815 -pf02390 -
017820 -# -
017825 -pf10576pf00633pf00730pf14815
017830 -pf04362 -
017835 -pf11873pf01464 -
BMC-P
BMC-H
BMC-T
Aldehyde dehydrogenase
PTAC
Signature enzyme
Alcohol dehydrogenase
Regulator

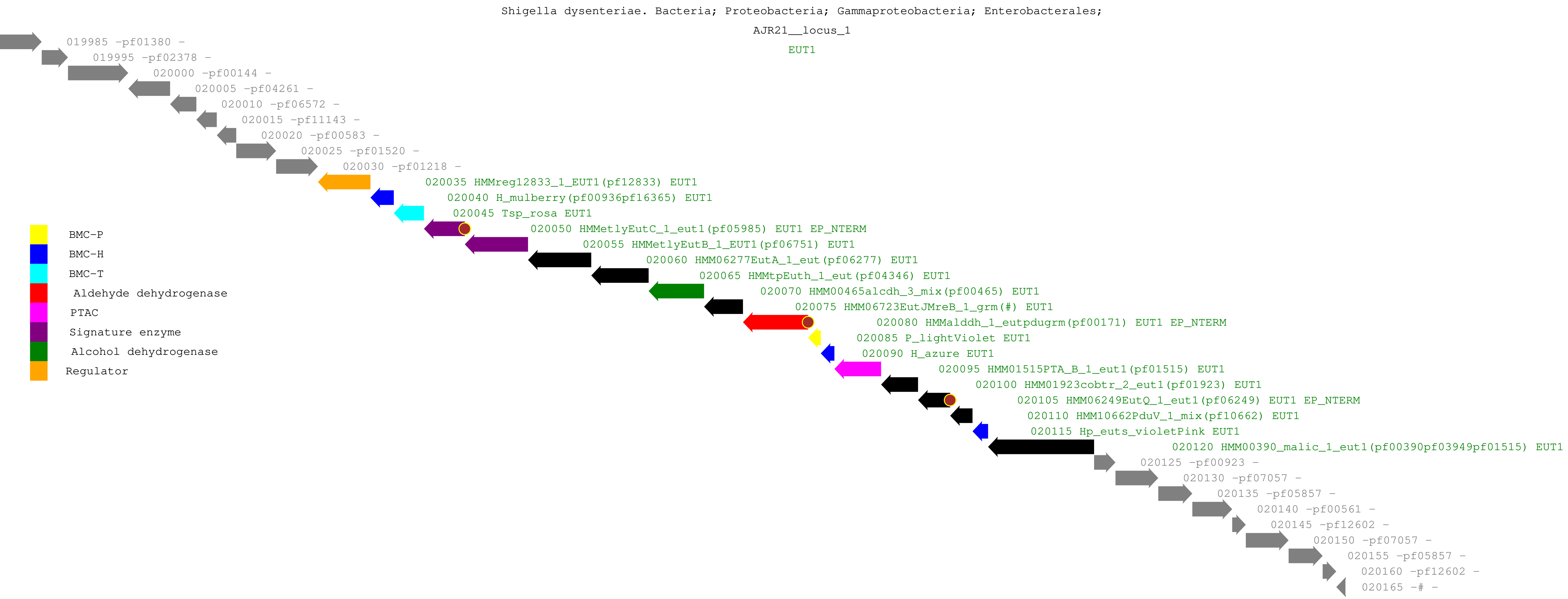
Shigella dysenteriae. Bacteria; Proteobacteria; Gammaproteobacteria; Enterobacterales;
AJR21__locus_1
EUT1
019985 -pf01380 -
019995 -pf02378 -
020000 -pf00144 -
020005 -pf04261 -
020010 -pf06572 -
020015 -pf11143 -
020020 -pf00583 -
020025 -pf01520 -
020030 -pf01218 -
020035 HMMreg12833_1_EUT1(pf12833) EUT1
020040 H_mulberry(pf00936pf16365) EUT1
020045 Tsp_rosa EUT1
020050 HMMetlyEutC_1_eut1(pf05985) EUT1 EP_NTERM
020055 HMMetlyEutB_1_EUT1(pf06751) EUT1
020060 HMM06277EutA_1_eut(pf06277) EUT1
020065 HMMtpEuth_1_eut(pf04346) EUT1
020070 HMM00465alcdh_3_mix(pf00465) EUT1
020075 HMM06723EutJMreB_1_grm(#) EUT1
020080 HMMalddh_1_eutpdugrm(pf00171) EUT1 EP_NTERM
020085 P_lightViolet EUT1
020090 H_azure EUT1
020095 HMM01515PTA_B_1_eut1(pf01515) EUT1
020100 HMM01923cobtr_2_eut1(pf01923) EUT1
020105 HMM06249EutQ_1_eut1(pf06249) EUT1 EP_NTERM
020110 HMM10662PduV_1_mix(pf10662) EUT1
020115 Hp_euts_violetPink EUT1
020120 HMM00390_malic_1_eut1(pf00390pf03949pf01515) EUT1
020125 -pf00923 -
020130 -pf07057 -
020135 -pf05857 -
020140 -pf00561 -
020145 -pf12602 -
020150 -pf07057 -
020155 -pf05857 -
020160 -pf12602 -
020165 -# -
BMC-P
BMC-H
BMC-T
Aldehyde dehydrogenase
PTAC
Signature enzyme
Alcohol dehydrogenase
Regulator

Shigella flexneri. Bacteria; Proteobacteria; Gammaproteobacteria; Enterobacterales;

AJR26__locus_1_main_sat

EUT1

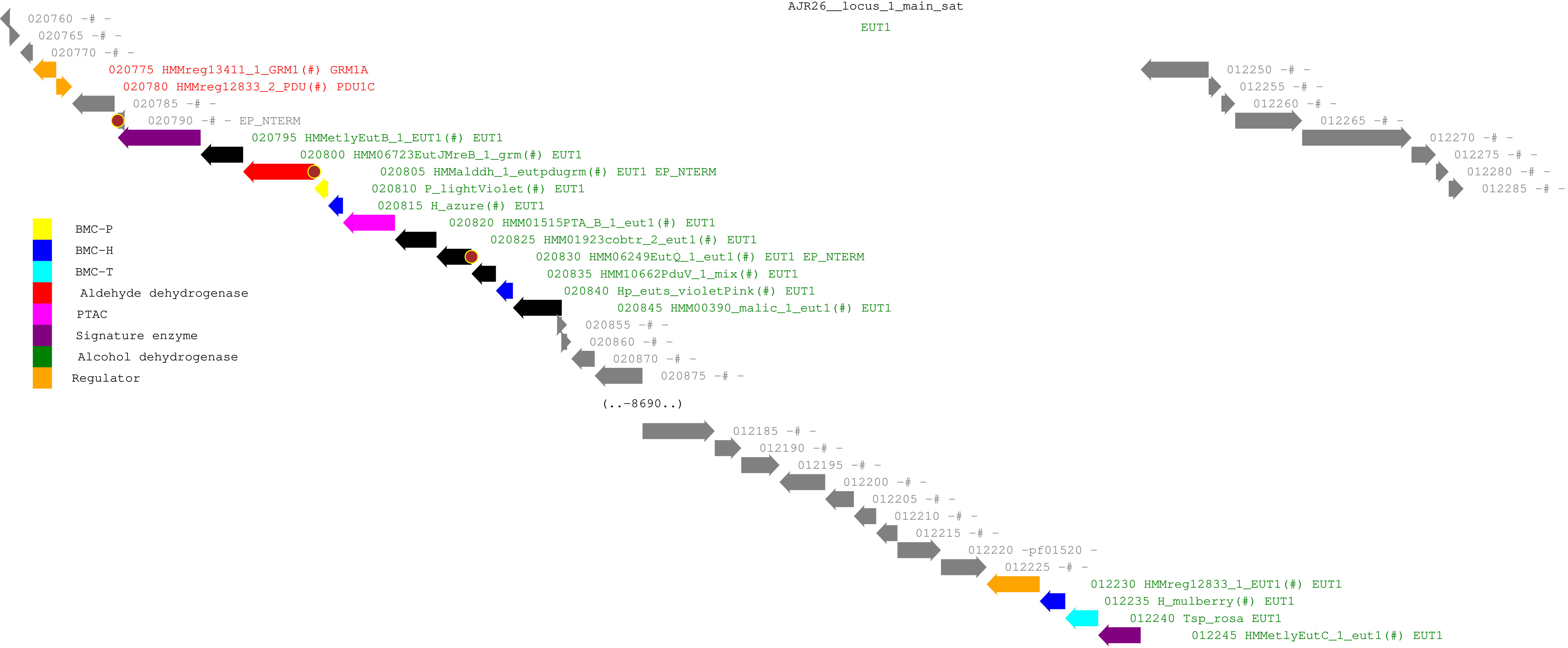

Shigella sonnei. Bacteria; Proteobacteria; Gammaproteobacteria; Enterobacterales;

AJR32__locus_2

EUT1

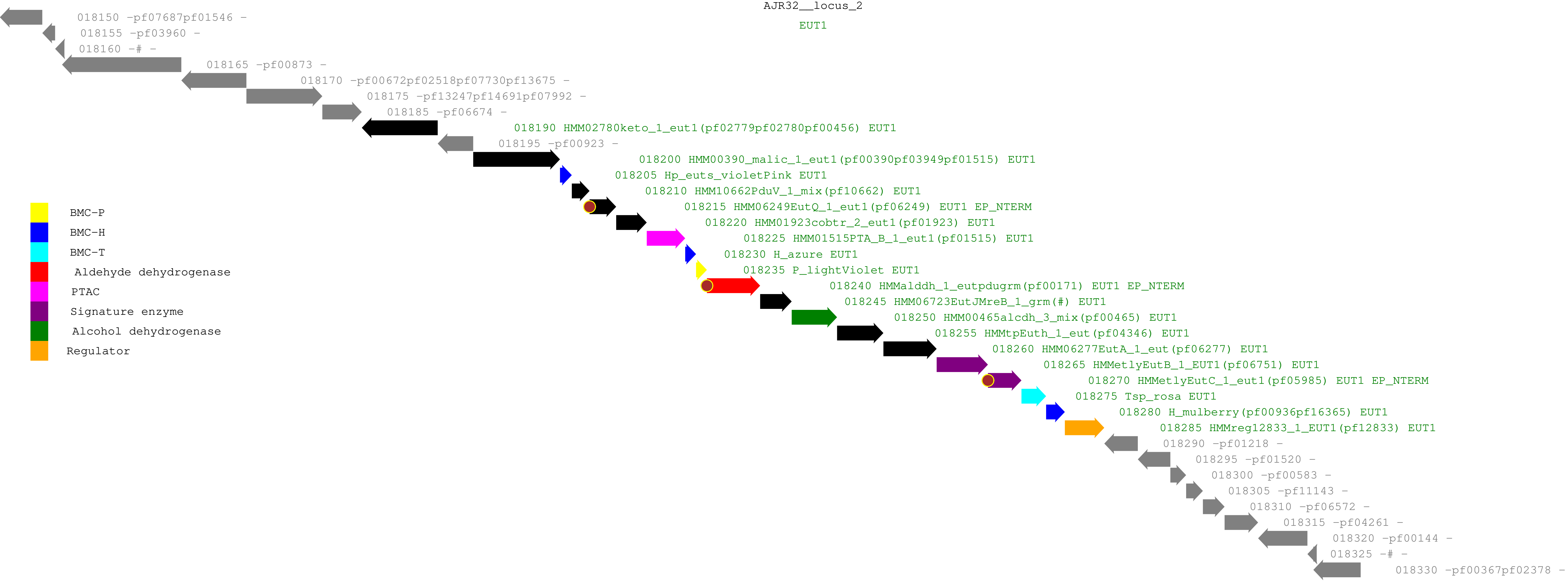

Escherichia coli. Bacteria; Proteobacteria; Gammaproteobacteria; Enterobacterales;

AKG99__locus_1

EUT1

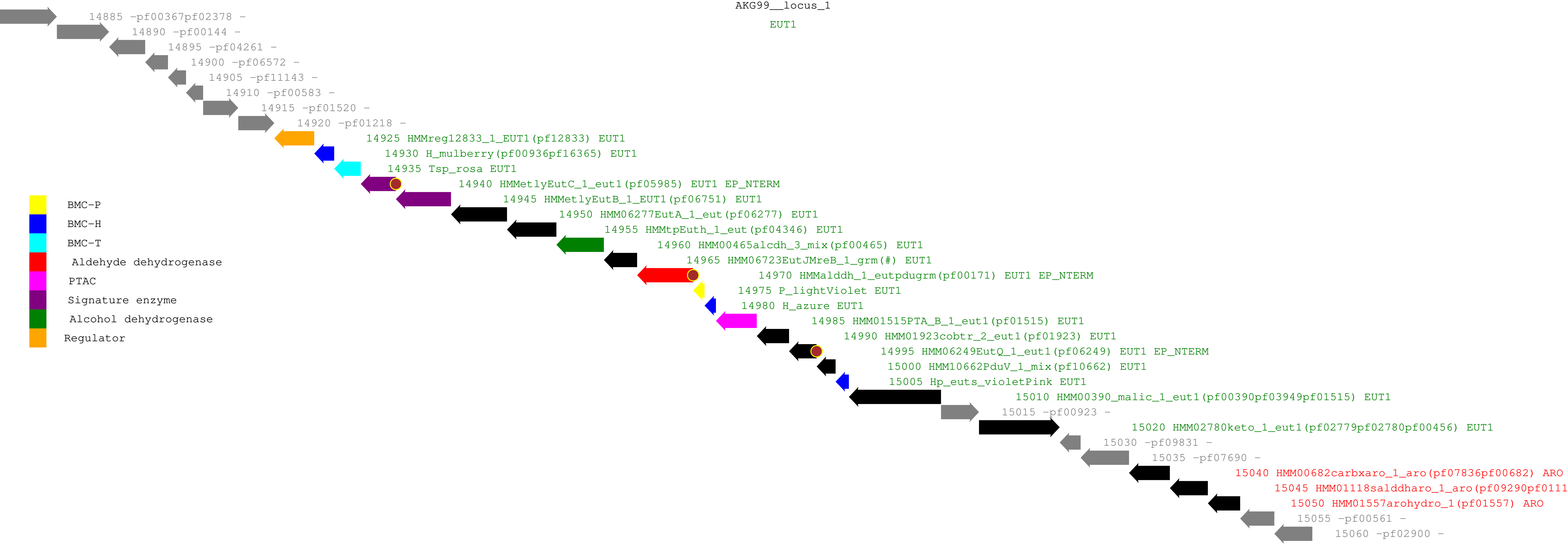

Escherichia coli. Bacteria; Proteobacteria; Gammaproteobacteria; Enterobacterales;

AKN40__locus_1

EUT1

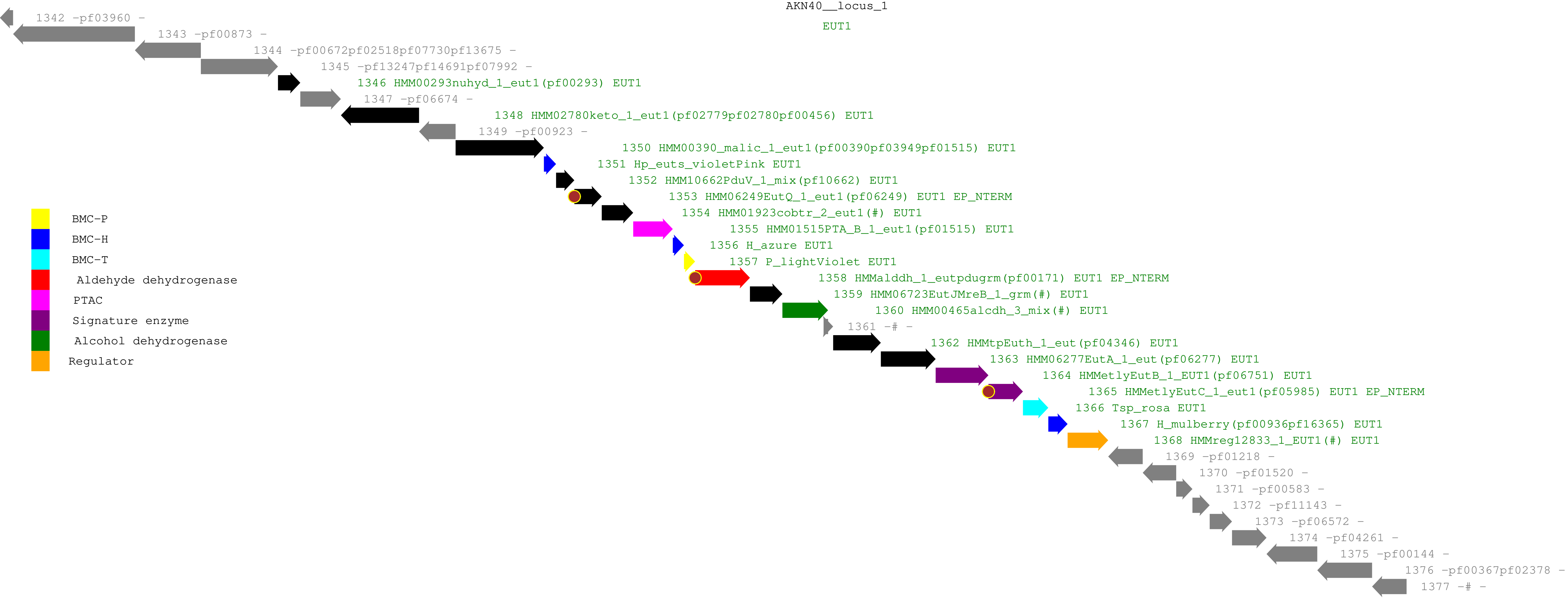

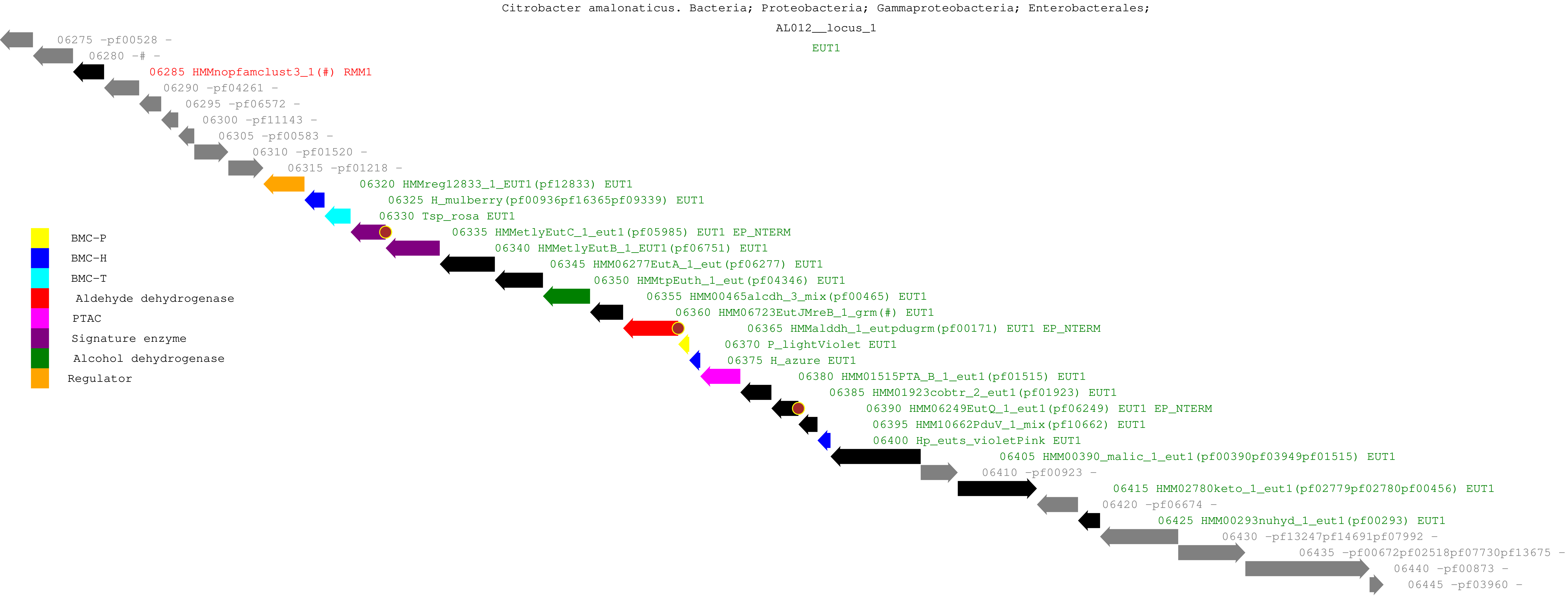
Citrobacter amalonaticus. Bacteria; Proteobacteria; Gammaproteobacteria; Enterobacterales;
AL012__locus_1
EUT1
06275 -pf00528 -
06280 -# -
06285 HMMnopfamclust3_1(#) RMM1
06290 -pf04261 -
06295 -pf06572 -
06300 -pf11143 -
06305 -pf00583 -
06310 -pf01520 -
06315 -pf01218 -
06320 HMMreg12833_1_EUT1(pf12833) EUT1
06325 H_mulberry(pf00936pf16365pf09339) EUT1
06330 Tsp_rosa EUT1
06335 HMMetlyEutC_1_eut1(pf05985) EUT1 EP_NTERM
06340 HMMetlyEutB_1_EUT1(pf06751) EUT1
06345 HMM06277EutA_1_eut(pf06277) EUT1
06350 HMMtpEuth_1_eut(pf04346) EUT1
06355 HMM00465alcdh_3_mix(pf00465) EUT1
06360 HMM06723EutJMreB_1_grm(#) EUT1
06365 HMMalddh_1_eutpdugrm(pf00171) EUT1 EP_NTERM
06370 P_lightViolet EUT1
06375 H_azure EUT1
06380 HMM01515PTA_B_1_eut1(pf01515) EUT1
06385 HMM01923cobtr_2_eut1(pf01923) EUT1
06390 HMM06249EutQ_1_eut1(pf06249) EUT1 EP_NTERM
06395 HMM10662PduV_1_mix(pf10662) EUT1
06400 Hp_euts_violetPink EUT1
06405 HMM00390_malic_1_eut1(pf00390pf03949pf01515) EUT1
06410 -pf00923 -
06415 HMM02780keto_1_eut1(pf02779pf02780pf00456) EUT1
06420 -pf06674 -
06425 HMM00293nuhyd_1_eut1(pf00293) EUT1
06430 -pf13247pf14691pf07992 -
06435 -pf00672pf02518pf07730pf13675 -
06440 -pf00873 -
06445 -pf03960 -
BMC-P
BMC-H
BMC-T
Aldehyde dehydrogenase
PTAC
Signature enzyme
Alcohol dehydrogenase
Regulator

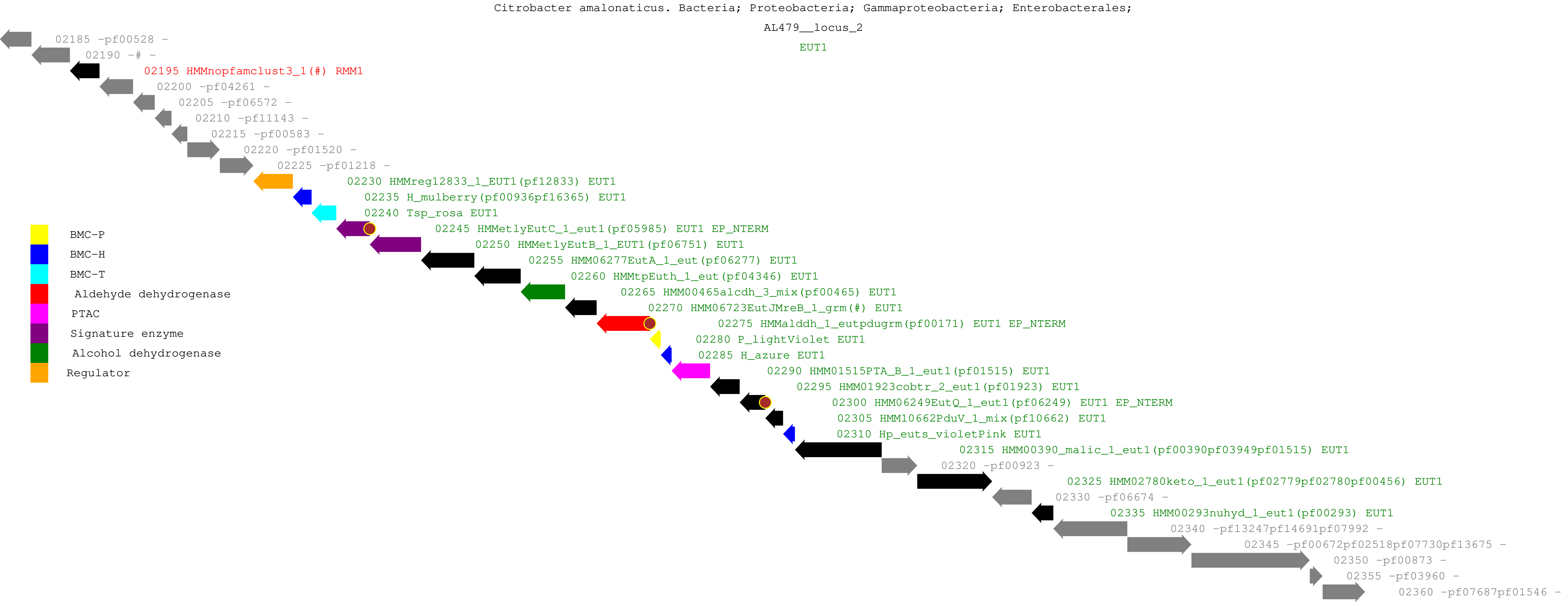

Citrobacter amalonaticus. Bacteria; Proteobacteria; Gammaproteobacteria; Enterobacterales;
AL479__locus_2
EUT1
02185 -pf00528 -
02190 -# -
02195 HMMnopfamclust3_1(#) RMM1
02200 -pf04261 -
02205 -pf06572 -
02210 -pf11143 -
02215 -pf00583 -
02220 -pf01520 -
02225 -pf01218 -
02230 HMMreg12833_1_EUT1(pf12833) EUT1
02235 H_mulberry(pf00936pf16365) EUT1
02240 Tsp_rosa EUT1
02245 HMMetlyEutC_1_eut1(pf05985) EUT1 EP_NTERM
02250 HMMetlyEutB_1_EUT1(pf06751) EUT1
02255 HMM06277EutA_1_eut(pf06277) EUT1
02260 HMMtpEuth_1_eut(pf04346) EUT1
02265 HMM00465alcdh_3_mix(pf00465) EUT1
02270 HMM06723EutJMreB_1_grm(#) EUT1
02275 HMMalddh_1_eutpdugrm(pf00171) EUT1 EP_NTERM
02280 P_lightViolet EUT1
02285 H_azure EUT1
02290 HMM01515PTA_B_1_eut1(pf01515) EUT1
02295 HMM01923cobtr_2_eut1(pf01923) EUT1
02300 HMM06249EutQ_1_eut1(pf06249) EUT1 EP_NTERM
02305 HMM10662PduV_1_mix(pf10662) EUT1
02310 Hp_euts_violetPink EUT1
02315 HMM00390_malic_1_eut1(pf00390pf03949pf01515) EUT1
02320 -pf00923 -
02325 HMM02780keto_1_eut1(pf02779pf02780pf00456) EUT1
02330 -pf06674 -
02335 HMM00293nuhyd_1_eut1(pf00293) EUT1
02340 -pf13247pf14691pf07992 -
02345 -pf00672pf02518pf07730pf13675 -
02350 -pf00873 -
02355 -pf03960 -
02360 -pf07687pf01546 -
BMC-P
BMC-H
BMC-T
Aldehyde dehydrogenase
PTAC
Signature enzyme
Alcohol dehydrogenase
Regulator

Escherichia fergusonii. Bacteria; Proteobacteria; Gammaproteobacteria; Enterobacterales;

AL530__locus_1

EUT1

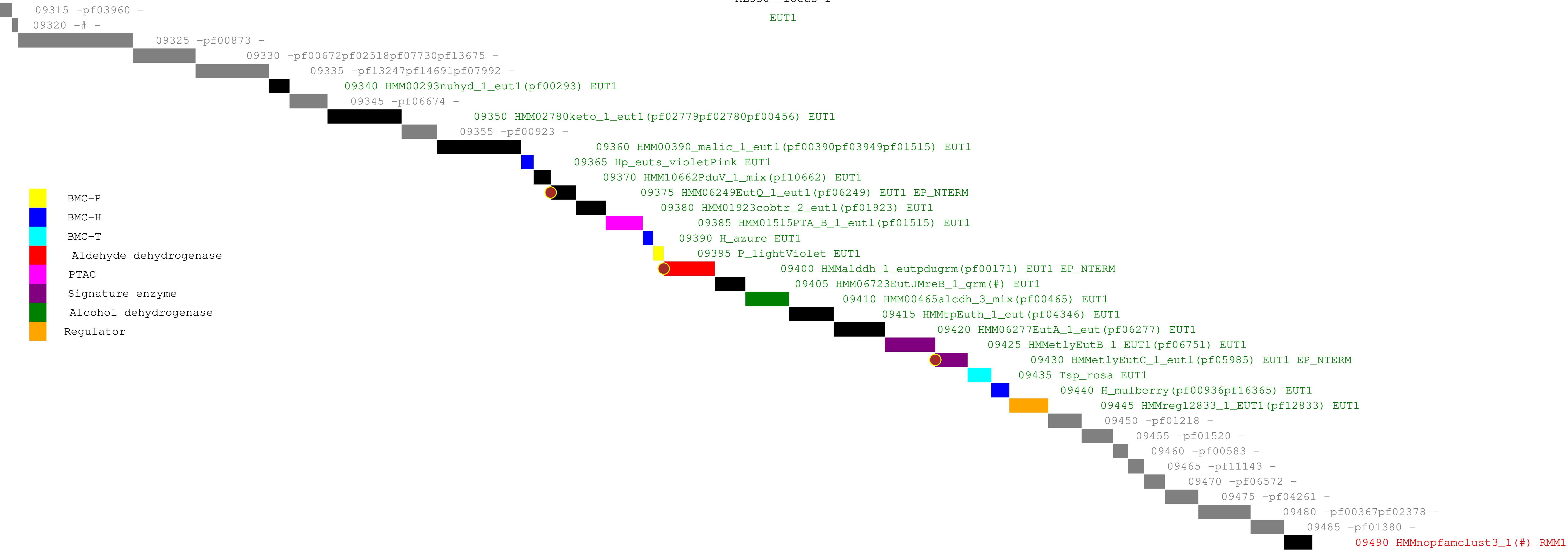

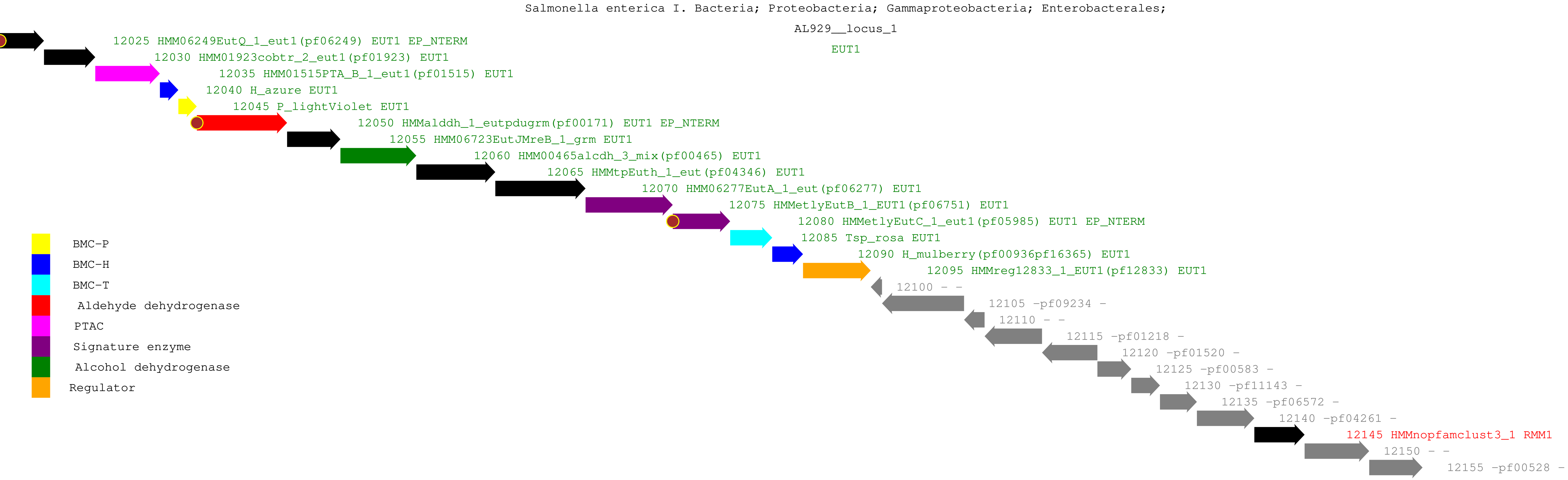

Salmonella enterica I. Bacteria; Proteobacteria; Gammaproteobacteria; Enterobacterales;
AL929__locus_1
EUT1
12025 HMM06249EutQ_1_eut1(pf06249) EUT1 EP_NTERM
12030 HMM01923cobtr_2_eut1(pf01923) EUT1
12035 HMM01515PTA_B_1_eut1(pf01515) EUT1
12040 H_azure EUT1
12045 P_lightViolet EUT1
12050 HMMalddh_1_eutpdugrm(pf00171) EUT1 EP_NTERM
12055 HMM06723EutJMreB_1_grm EUT1
12060 HMM00465alcdh_3_mix(pf00465) EUT1
12065 HMMtpEuth_1_eut(pf04346) EUT1
12070 HMM06277EutA_1_eut(pf06277) EUT1
12075 HMMetlyEutB_1_EUT1(pf06751) EUT1
12080 HMMetlyEutC_1_eut1(pf05985) EUT1 EP_NTERM
12085 Tsp_rosa EUT1
12090 H_mulberry(pf00936pf16365) EUT1
12095 HMMreg12833_1_EUT1(pf12833) EUT1
12100 - -
12105 -pf09234 -
12110 - -
12115 -pf01218 -
12120 -pf01520 -
12125 -pf00583 -
12130 -pf11143 -
12135 -pf06572 -
12140 -pf04261 -
12145 HMMnopfamclust3_1 RMM1
12150 - -
12155 -pf00528 -
BMC-P
BMC-H
BMC-T
Aldehyde dehydrogenase
PTAC
Signature enzyme
Alcohol dehydrogenase
Regulator

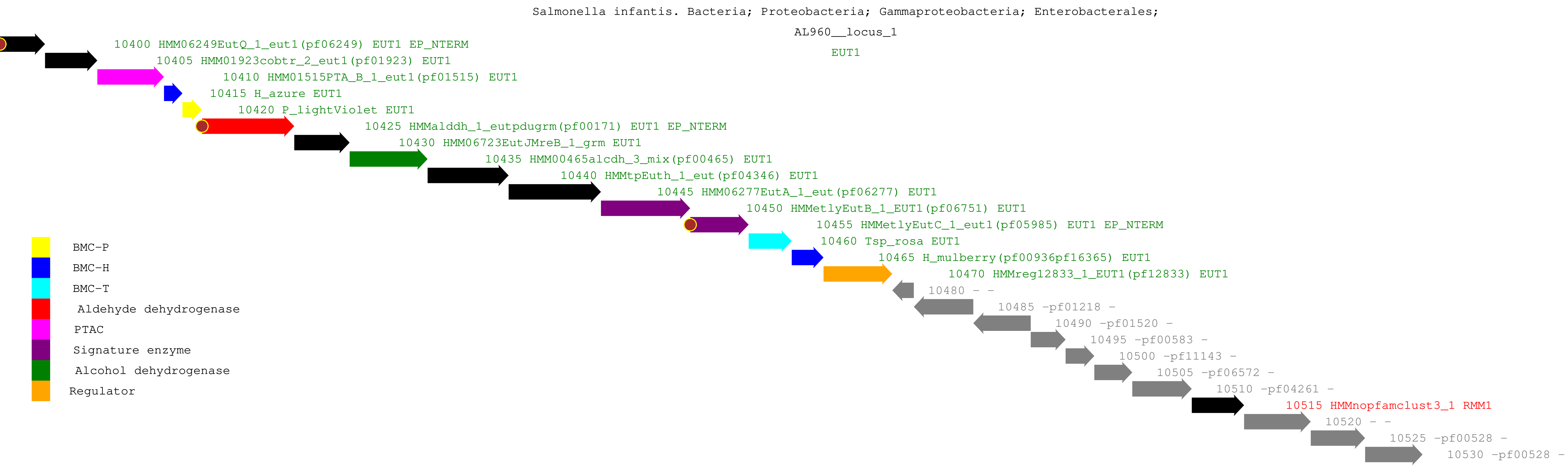

Salmonella infantis. Bacteria; Proteobacteria; Gammaproteobacteria; Enterobacterales;
AL960__locus_1
EUT1
10400 HMM06249EutQ_1_eut1(pf06249) EUT1 EP_NTERM
10405 HMM01923cobtr_2_eut1(pf01923) EUT1
10410 HMM01515PTA_B_1_eut1(pf01515) EUT1
10415 H_azure EUT1
10420 P_lightViolet EUT1
10425 HMMalddh_1_eutpdugrm(pf00171) EUT1 EP_NTERM
10430 HMM06723EutJMreB_1_grm EUT1
10435 HMM00465alcdh_3_mix(pf00465) EUT1
10440 HMMtpEuth_1_eut(pf04346) EUT1
10445 HMM06277EutA_1_eut(pf06277) EUT1
10450 HMMetlyEutB_1_EUT1(pf06751) EUT1
10455 HMMetlyEutC_1_eut1(pf05985) EUT1 EP_NTERM
10460 Tsp_rosa EUT1
10465 H_mulberry(pf00936pf16365) EUT1
10470 HMMreg12833_1_EUT1(pf12833) EUT1
10480 - -
10485 -pf01218 -
10490 -pf01520 -
10495 -pf00583 -
10500 -pf11143 -
10505 -pf06572 -
10510 -pf04261 -
10515 HMMnopfamclust3_1 RMM1
10520 - -
10525 -pf00528 -
10530 -pf00528 -
BMC-P
BMC-H
BMC-T
Aldehyde dehydrogenase
PTAC
Signature enzyme
Alcohol dehydrogenase
Regulator

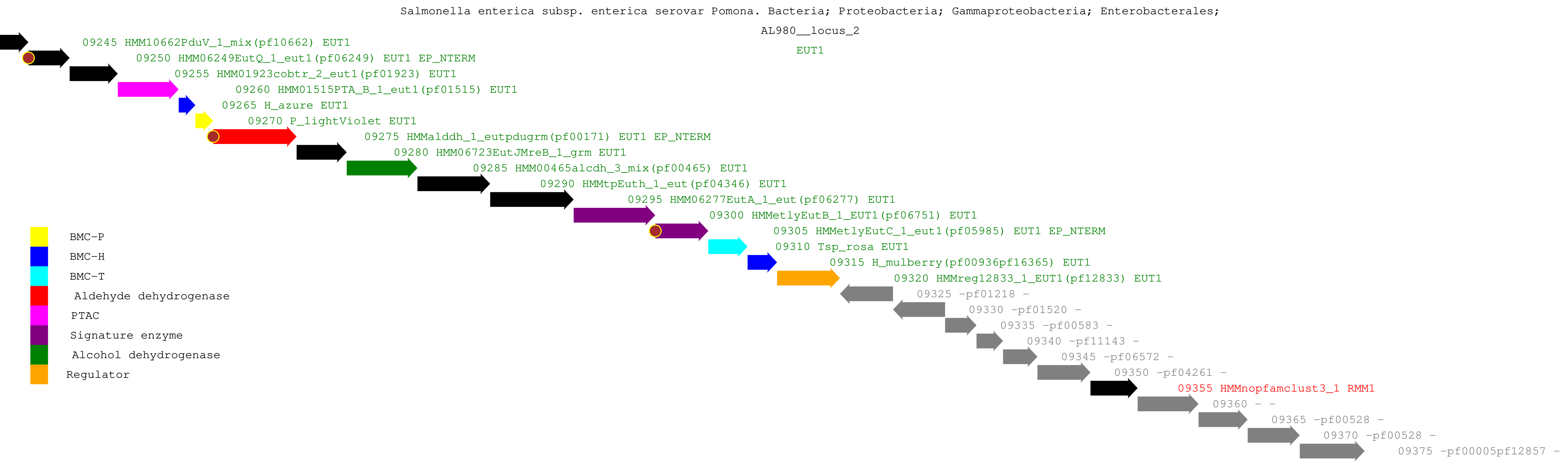
Salmonella enterica subsp. enterica serovar Pomona. Bacteria; Proteobacteria; Gammaproteobacteria; Enterobacterales;
AL980__locus_2
EUT1
09245 HMM10662PduV_1_mix(pf10662) EUT1
09250 HMM06249EutQ_1_eut1(pf06249) EUT1 EP_NTERM
09255 HMM01923cobtr_2_eut1(pf01923) EUT1
09260 HMM01515PTA_B_1_eut1(pf01515) EUT1
09265 H_azure EUT1
09270 P_lightViolet EUT1
09275 HMMalddh_1_eutpdugrm(pf00171) EUT1 EP_NTERM
09280 HMM06723EutJMreB_1_grm EUT1
09285 HMM00465alcdh_3_mix(pf00465) EUT1
09290 HMMtpEuth_1_eut(pf04346) EUT1
09295 HMM06277EutA_1_eut(pf06277) EUT1
09300 HMMetlyEutB_1_EUT1(pf06751) EUT1
09305 HMMetlyEutC_1_eut1(pf05985) EUT1 EP_NTERM
09310 Tsp_rosa EUT1
09315 H_mulberry(pf00936pf16365) EUT1
09320 HMMreg12833_1_EUT1(pf12833) EUT1
09325 -pf01218 -
09330 -pf01520 -
09335 -pf00583 -
09340 -pf11143 -
09345 -pf06572 -
09350 -pf04261 -
09355 HMMnopfamclust3_1 RMM1
09360 - -
09365 -pf00528 -
09370 -pf00528 -
09375 -pf00005pf12857 -
BMC-P
BMC-H
BMC-T
Aldehyde dehydrogenase
PTAC
Signature enzyme
Alcohol dehydrogenase
Regulator

Salmonella typhimurium. Bacteria; Proteobacteria; Gammaproteobacteria; Enterobacterales;

AM358__locus_1

EUT1

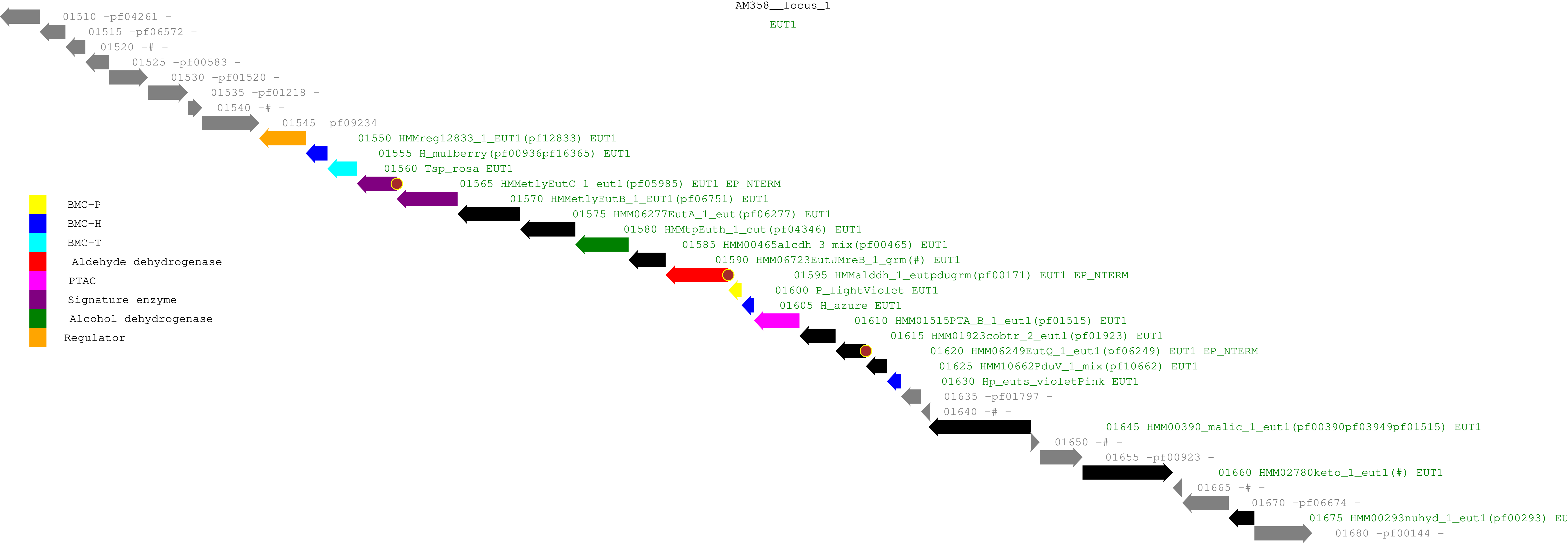

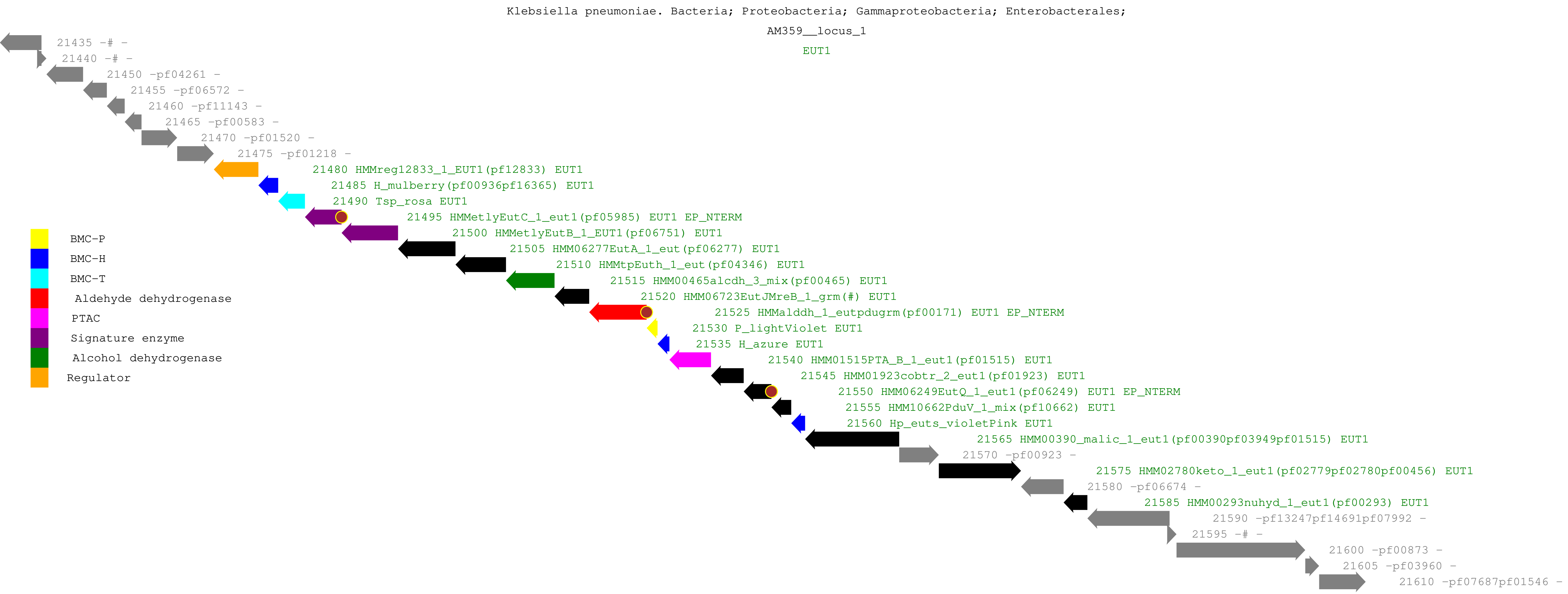
Klebsiella pneumoniae. Bacteria; Proteobacteria; Gammaproteobacteria; Enterobacterales;
AM359__locus_1
EUT1
21435 -# -
21440 -# -
21450 -pf04261 -
21455 -pf06572 -
21460 -pf11143 -
21465 -pf00583 -
21470 -pf01520 -
21475 -pf01218 -
21480 HMMreg12833_1_EUT1(pf12833) EUT1
21485 H_mulberry(pf00936pf16365) EUT1
21490 Tsp_rosa EUT1
21495 HMMetlyEutC_1_eut1(pf05985) EUT1 EP_NTERM
21500 HMMetlyEutB_1_EUT1(pf06751) EUT1
21505 HMM06277EutA_1_eut(pf06277) EUT1
21510 HMMtpEuth_1_eut(pf04346) EUT1
21515 HMM00465alcdh_3_mix(pf00465) EUT1
21520 HMM06723EutJMreB_1_grm(#) EUT1
21525 HMMalddh_1_eutpdugrm(pf00171) EUT1 EP_NTERM
21530 P_lightViolet EUT1
21535 H_azure EUT1
21540 HMM01515PTA_B_1_eut1(pf01515) EUT1
21545 HMM01923cobtr_2_eut1(pf01923) EUT1
21550 HMM06249EutQ_1_eut1(pf06249) EUT1 EP_NTERM
21555 HMM10662PduV_1_mix(pf10662) EUT1
21560 Hp_euts_violetPink EUT1
21565 HMM00390_malic_1_eut1(pf00390pf03949pf01515) EUT1
21570 -pf00923 -
21575 HMM02780keto_1_eut1(pf02779pf02780pf00456) EUT1
21580 -pf06674 -
21585 HMM00293nuhyd_1_eut1(pf00293) EUT1
21590 -pf13247pf14691pf07992 -
21595 -# -
21600 -pf00873 -
21605 -pf03960 -
21610 -pf07687pf01546 -
BMC-P
BMC-H
BMC-T
Aldehyde dehydrogenase
PTAC
Signature enzyme
Alcohol dehydrogenase
Regulator

Morganella morganii (Proteus morganii). Bacteria; Proteobacteria; Gammaproteobacteria; Enterobacterales;

AM380__locus_1

EUT1

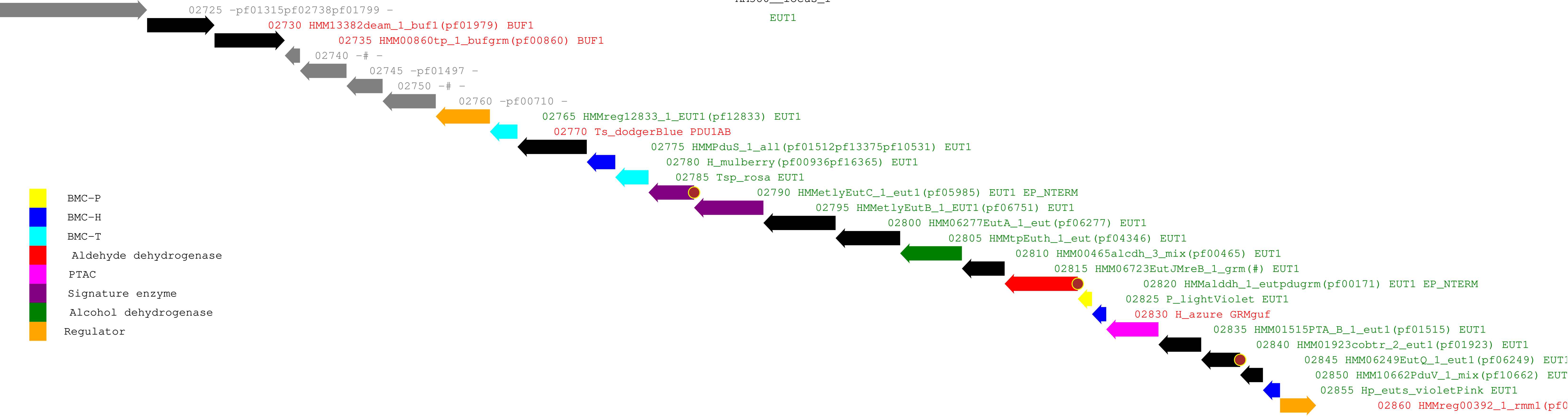

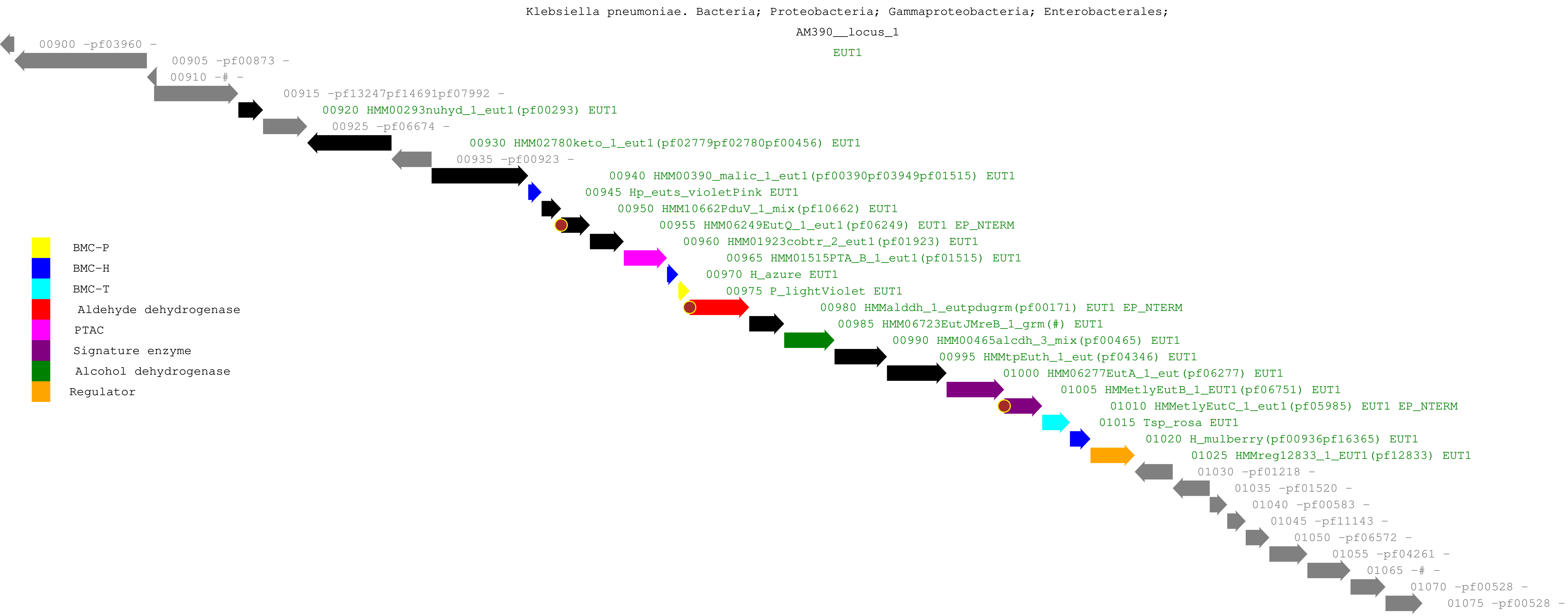
Klebsiella pneumoniae. Bacteria; Proteobacteria; Gammaproteobacteria; Enterobacterales;
AM390__locus_1
EUT1
00900 -pf03960 -
00905 -pf00873 -
00910 -# -
00915 -pf13247pf14691pf07992 -
00920 HMM00293nuhyd_1_eut1(pf00293) EUT1
00925 -pf06674 -
00930 HMM02780keto_1_eut1(pf02779pf02780pf00456) EUT1
00935 -pf00923 -
00940 HMM00390_malic_1_eut1(pf00390pf03949pf01515) EUT1
00945 Hp_euts_violetPink EUT1
00950 HMM10662PduV_1_mix(pf10662) EUT1
00955 HMM06249EutQ_1_eut1(pf06249) EUT1 EP_NTERM
00960 HMM01923cobtr_2_eut1(pf01923) EUT1
00965 HMM01515PTA_B_1_eut1(pf01515) EUT1
00970 H_azure EUT1
00975 P_lightViolet EUT1
00980 HMMalddh_1_eutpdugrm(pf00171) EUT1 EP_NTERM
00985 HMM06723EutJMreB_1_grm(#) EUT1
00990 HMM00465alcdh_3_mix(pf00465) EUT1
00995 HMMtpEuth_1_eut(pf04346) EUT1
01000 HMM06277EutA_1_eut(pf06277) EUT1
01005 HMMetlyEutB_1_EUT1(pf06751) EUT1
01010 HMMetlyEutC_1_eut1(pf05985) EUT1 EP_NTERM
01015 Tsp_rosa EUT1
01020 H_mulberry(pf00936pf16365) EUT1
01025 HMMreg12833_1_EUT1(pf12833) EUT1
01030 -pf01218 -
01035 -pf01520 -
01040 -pf00583 -
01045 -pf11143 -
01050 -pf06572 -
01055 -pf04261 -
01065 -# -
01070 -pf00528 -
01075 -pf00528 -
BMC-P
BMC-H
BMC-T
Aldehyde dehydrogenase
PTAC
Signature enzyme
Alcohol dehydrogenase
Regulator

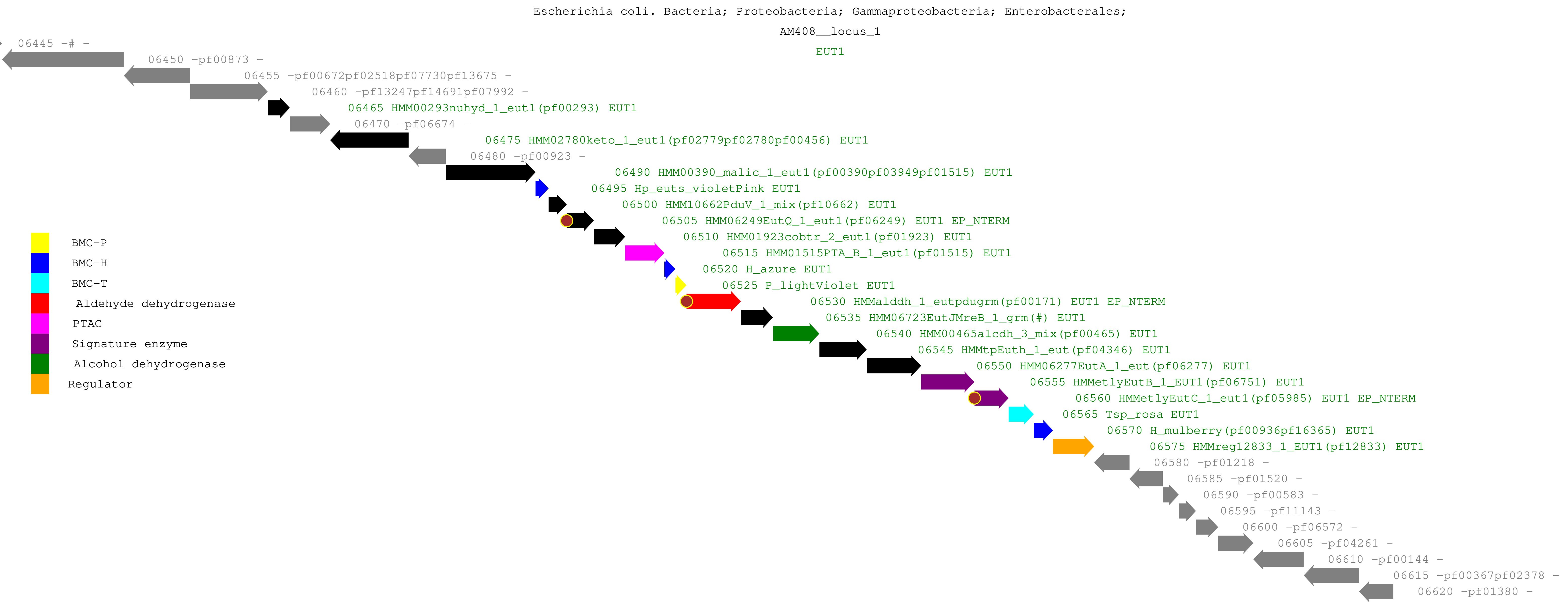
Escherichia coli. Bacteria; Proteobacteria; Gammaproteobacteria; Enterobacterales;
AM408__locus_1
EUT1
06445 -# -
06450 -pf00873 -
06455 -pf00672pf02518pf07730pf13675 -
06460 -pf13247pf14691pf07992 -
06465 HMM00293nuhyd_1_eut1(pf00293) EUT1
06470 -pf06674 -
06475 HMM02780keto_1_eut1(pf02779pf02780pf00456) EUT1
06480 -pf00923 -
06490 HMM00390_malic_1_eut1(pf00390pf03949pf01515) EUT1
06495 Hp_euts_violetPink EUT1
06500 HMM10662PduV_1_mix(pf10662) EUT1
06505 HMM06249EutQ_1_eut1(pf06249) EUT1 EP_NTERM
06510 HMM01923cobtr_2_eut1(pf01923) EUT1
06515 HMM01515PTA_B_1_eut1(pf01515) EUT1
06520 H_azure EUT1
06525 P_lightViolet EUT1
06530 HMMalddh_1_eutpdugrm(pf00171) EUT1 EP_NTERM
06535 HMM06723EutJMreB_1_grm(#) EUT1
06540 HMM00465alcdh_3_mix(pf00465) EUT1
06545 HMMtpEuth_1_eut(pf04346) EUT1
06550 HMM06277EutA_1_eut(pf06277) EUT1
06555 HMMetlyEutB_1_EUT1(pf06751) EUT1
06560 HMMetlyEutC_1_eut1(pf05985) EUT1 EP_NTERM
06565 Tsp_rosa EUT1
06570 H_mulberry(pf00936pf16365) EUT1
06575 HMMreg12833_1_EUT1(pf12833) EUT1
06580 -pf01218 -
06585 -pf01520 -
06590 -pf00583 -
06595 -pf11143 -
06600 -pf06572 -
06605 -pf04261 -
06610 -pf00144 -
06615 -pf00367pf02378 -
06620 -pf01380 -
BMC-P
BMC-H
BMC-T
Aldehyde dehydrogenase
PTAC
Signature enzyme
Alcohol dehydrogenase
Regulator

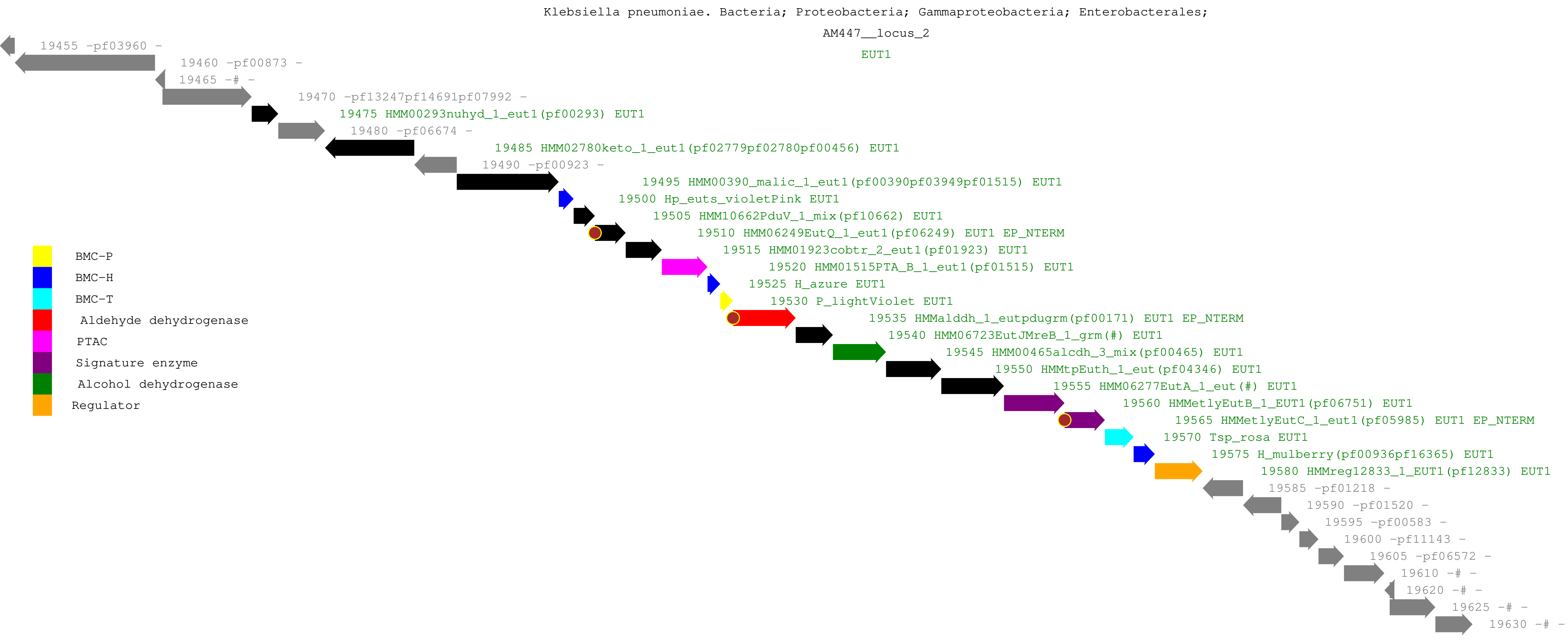
Klebsiella pneumoniae. Bacteria; Proteobacteria; Gammaproteobacteria; Enterobacterales;
AM447__locus_2
EUT1
19455 -pf03960 -
19460 -pf00873 -
19465 -# -
19470 -pf13247pf14691pf07992 -
19475 HMM00293nuhyd_1_eut1(pf00293) EUT1
19480 -pf06674 -
19485 HMM02780keto_1_eut1(pf02779pf02780pf00456) EUT1
19490 -pf00923 -
19495 HMM00390_malic_1_eut1(pf00390pf03949pf01515) EUT1
19500 Hp_euts_violetPink EUT1
19505 HMM10662PduV_1_mix(pf10662) EUT1
19510 HMM06249EutQ_1_eut1(pf06249) EUT1 EP_NTERM
19515 HMM01923cobtr_2_eut1(pf01923) EUT1
19520 HMM01515PTA_B_1_eut1(pf01515) EUT1
19525 H_azure EUT1
19530 P_lightViolet EUT1
19535 HMMalddh_1_eutpdugrm(pf00171) EUT1 EP_NTERM
19540 HMM06723EutJMreB_1_grm(#) EUT1
19545 HMM00465alcdh_3_mix(pf00465) EUT1
19550 HMMtpEuth_1_eut(pf04346) EUT1
19555 HMM06277EutA_1_eut(#) EUT1
19560 HMMetlyEutB_1_EUT1(pf06751) EUT1
19565 HMMetlyEutC_1_eut1(pf05985) EUT1 EP_NTERM
19570 Tsp_rosa EUT1
19575 H_mulberry(pf00936pf16365) EUT1
19580 HMMreg12833_1_EUT1(pf12833) EUT1
19585 -pf01218 -
19590 -pf01520 -
19595 -pf00583 -
19600 -pf11143 -
19605 -pf06572 -
19610 -# -
19620 -# -
19625 -# -
19630 -# -
BMC-P
BMC-H
BMC-T
Aldehyde dehydrogenase
PTAC
Signature enzyme
Alcohol dehydrogenase
Regulator

Salmonella enterica subsp. enterica serovar Brandenburg. Bacteria; Proteobacteria; Gammaproteobacteria; Enterobacterales;

AMA33__locus_1

EUT1

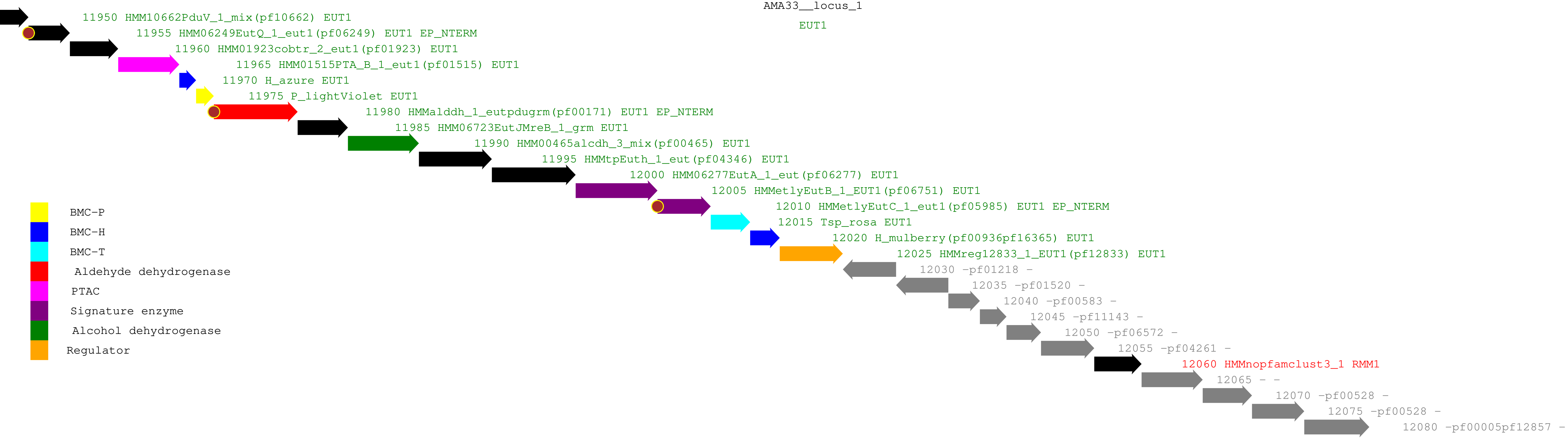

Escherichia coli. Bacteria; Proteobacteria; Gammaproteobacteria; Enterobacterales;

AML35__locus_1

EUT1

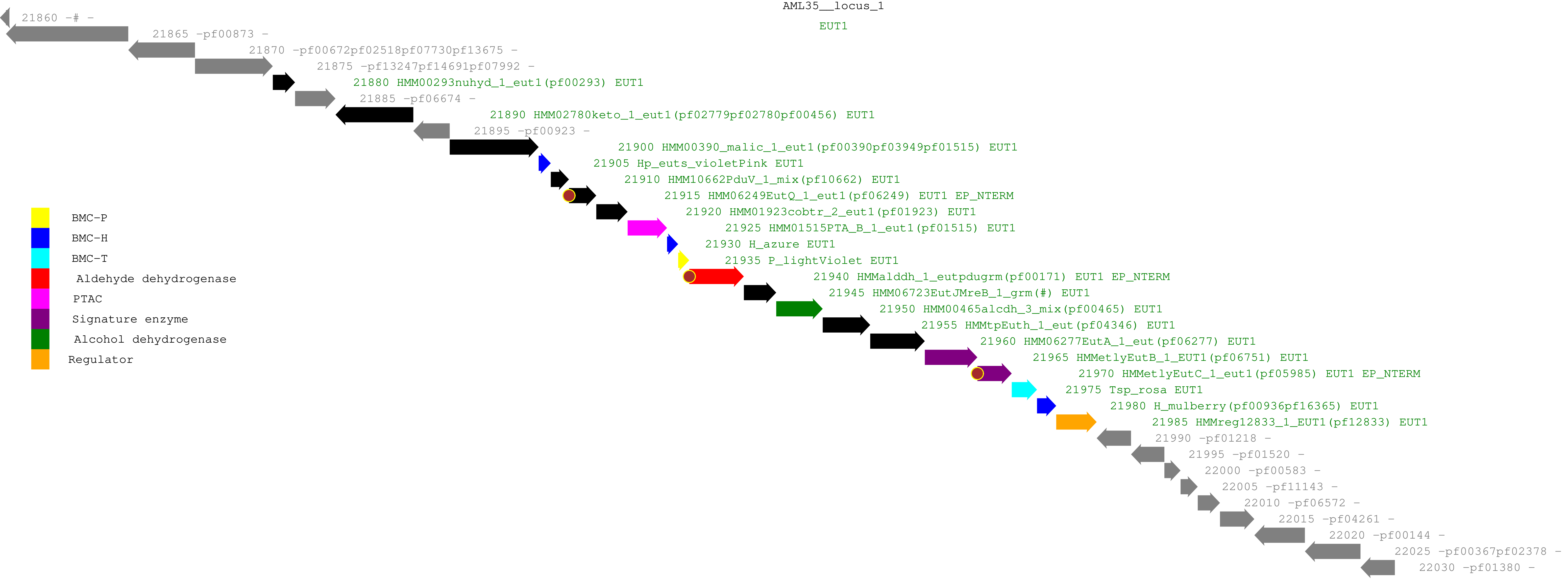

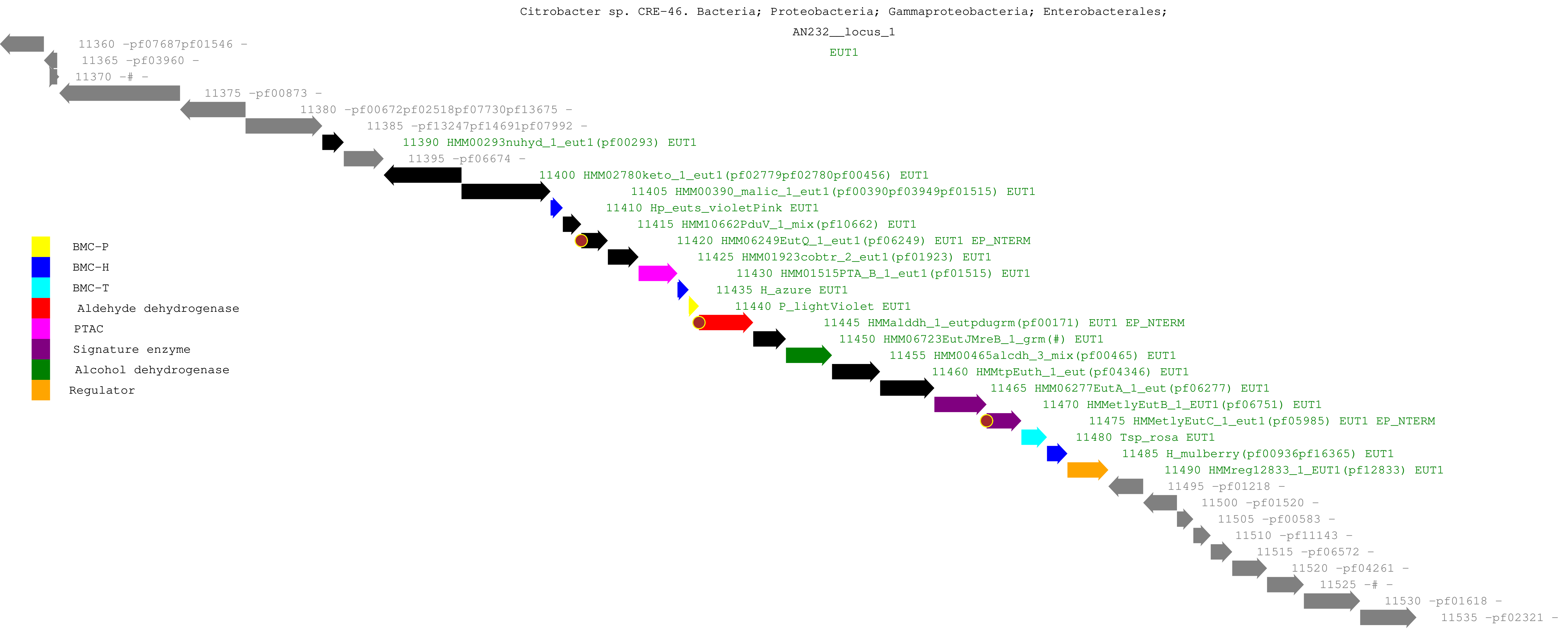
Citrobacter sp. CRE-46. Bacteria; Proteobacteria; Gammaproteobacteria; Enterobacterales;
AN232__locus_1
EUT1
11360 -pf07687pf01546 -
11365 -pf03960 -
11370 -# -
11375 -pf00873 -
11380 -pf00672pf02518pf07730pf13675 -
11385 -pf13247pf14691pf07992 -
11390 HMM00293nuhyd_1_eut1(pf00293) EUT1
11395 -pf06674 -
11400 HMM02780keto_1_eut1(pf02779pf02780pf00456) EUT1
11405 HMM00390_malic_1_eut1(pf00390pf03949pf01515) EUT1
11410 Hp_euts_violetPink EUT1
11415 HMM10662PduV_1_mix(pf10662) EUT1
11420 HMM06249EutQ_1_eut1(pf06249) EUT1 EP_NTERM
11425 HMM01923cobtr_2_eut1(pf01923) EUT1
11430 HMM01515PTA_B_1_eut1(pf01515) EUT1
11435 H_azure EUT1
11440 P_lightViolet EUT1
11445 HMMalddh_1_eutpdugrm(pf00171) EUT1 EP_NTERM
11450 HMM06723EutJMreB_1_grm(#) EUT1
11455 HMM00465alcdh_3_mix(pf00465) EUT1
11460 HMMtpEuth_1_eut(pf04346) EUT1
11465 HMM06277EutA_1_eut(pf06277) EUT1
11470 HMMetlyEutB_1_EUT1(pf06751) EUT1
11475 HMMetlyEutC_1_eut1(pf05985) EUT1 EP_NTERM
11480 Tsp_rosa EUT1
11485 H_mulberry(pf00936pf16365) EUT1
11490 HMMreg12833_1_EUT1(pf12833) EUT1
11495 -pf01218 -
11500 -pf01520 -
11505 -pf00583 -
11510 -pf11143 -
11515 -pf06572 -
11520 -pf04261 -
11525 -# -
11530 -pf01618 -
11535 -pf02321 -
BMC-P
BMC-H
BMC-T
Aldehyde dehydrogenase
PTAC
Signature enzyme
Alcohol dehydrogenase
Regulator

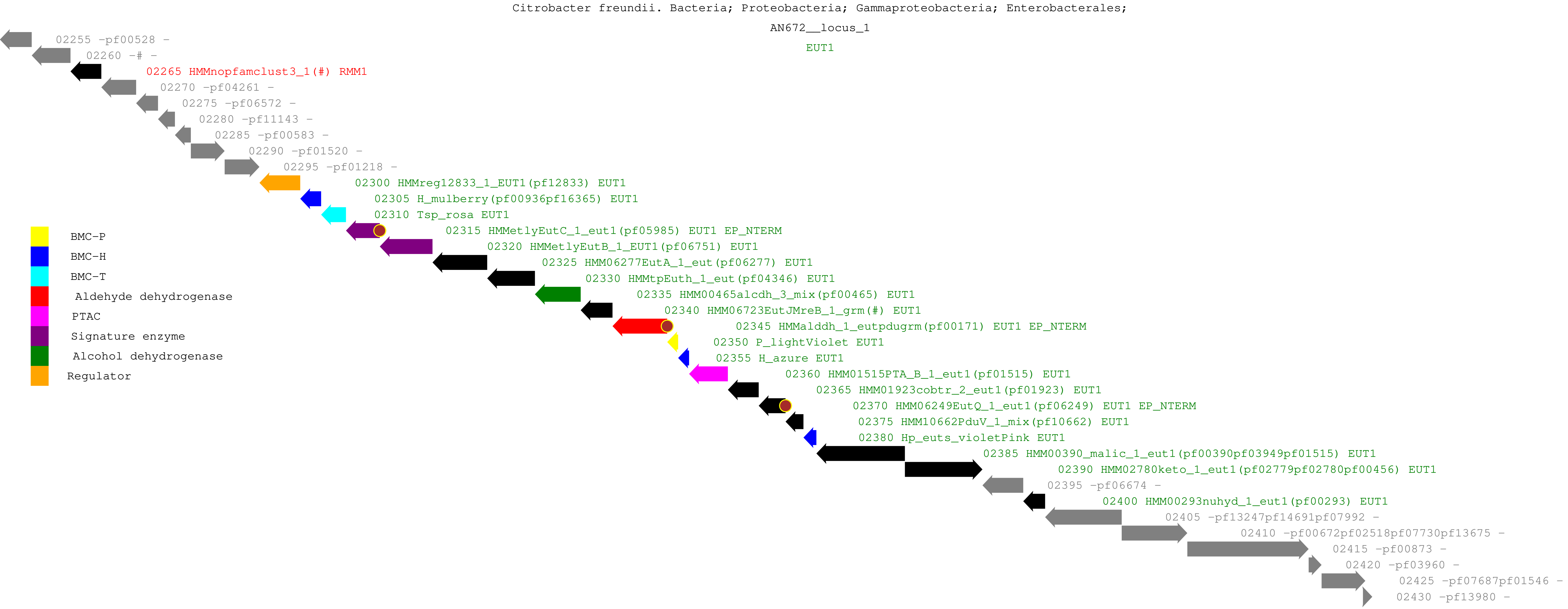

Citrobacter freundii. Bacteria; Proteobacteria; Gammaproteobacteria; Enterobacterales;
AN672__locus_1
EUT1
02255 -pf00528 -
02260 -# -
02265 HMMnopfamclust3_1(#) RMM1
02270 -pf04261 -
02275 -pf06572 -
02280 -pf11143 -
02285 -pf00583 -
02290 -pf01520 -
02295 -pf01218 -
02300 HMMreg12833_1_EUT1(pf12833) EUT1
02305 H_mulberry(pf00936pf16365) EUT1
02310 Tsp_rosa EUT1
02315 HMMetlyEutC_1_eut1(pf05985) EUT1 EP_NTERM
02320 HMMetlyEutB_1_EUT1(pf06751) EUT1
02325 HMM06277EutA_1_eut(pf06277) EUT1
02330 HMMtpEuth_1_eut(pf04346) EUT1
02335 HMM00465alcdh_3_mix(pf00465) EUT1
02340 HMM06723EutJMreB_1_grm(#) EUT1
02345 HMMalddh_1_eutpdugrm(pf00171) EUT1 EP_NTERM
02350 P_lightViolet EUT1
02355 H_azure EUT1
02360 HMM01515PTA_B_1_eut1(pf01515) EUT1
02365 HMM01923cobtr_2_eut1(pf01923) EUT1
02370 HMM06249EutQ_1_eut1(pf06249) EUT1 EP_NTERM
02375 HMM10662PduV_1_mix(pf10662) EUT1
02380 Hp_euts_violetPink EUT1
02385 HMM00390_malic_1_eut1(pf00390pf03949pf01515) EUT1
02390 HMM02780keto_1_eut1(pf02779pf02780pf00456) EUT1
02395 -pf06674 -
02400 HMM00293nuhyd_1_eut1(pf00293) EUT1
02405 -pf13247pf14691pf07992 -
02410 -pf00672pf02518pf07730pf13675 -
02415 -pf00873 -
02420 -pf03960 -
02425 -pf07687pf01546 -
02430 -pf13980 -
BMC-P
BMC-H
BMC-T
Aldehyde dehydrogenase
PTAC
Signature enzyme
Alcohol dehydrogenase
Regulator

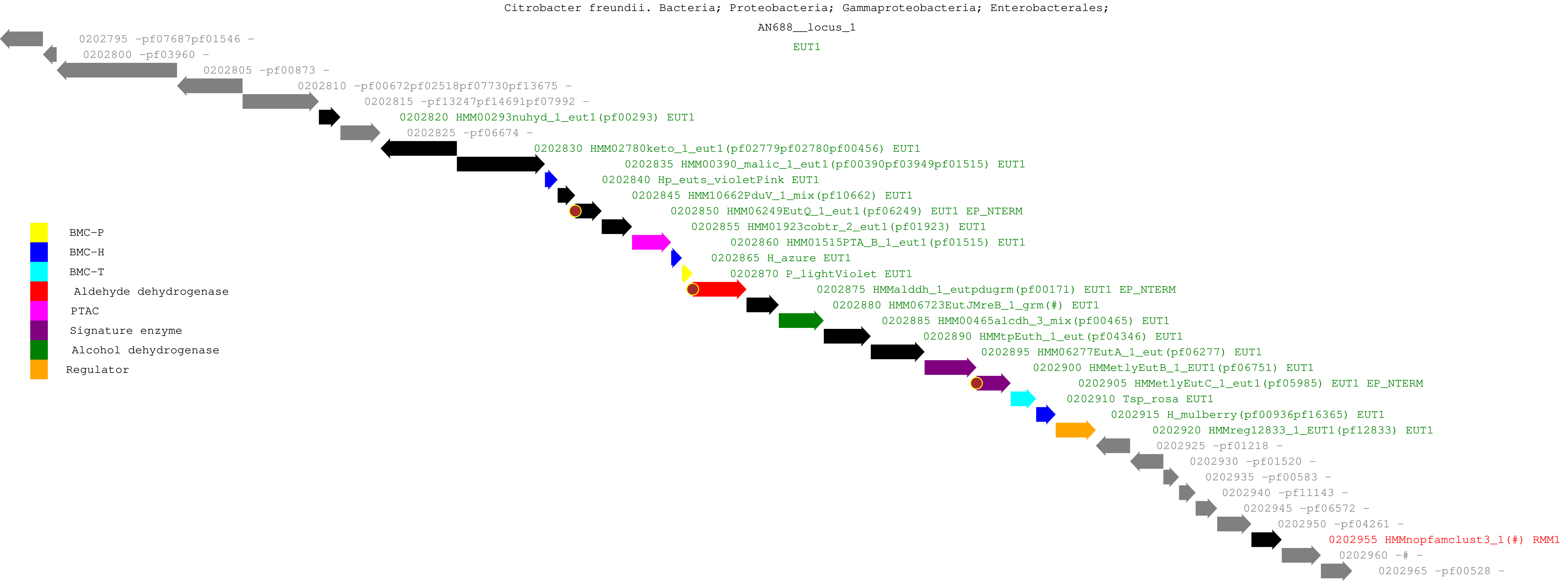

Citrobacter freundii. Bacteria; Proteobacteria; Gammaproteobacteria; Enterobacterales;
AN688__locus_1
EUT1
0202795 -pf07687pf01546 -
0202800 -pf03960 -
0202805 -pf00873 -
0202810 -pf00672pf02518pf07730pf13675 -
0202815 -pf13247pf14691pf07992 -
0202820 HMM00293nuhyd_1_eut1(pf00293) EUT1
0202825 -pf06674 -
0202830 HMM02780keto_1_eut1(pf02779pf02780pf00456) EUT1
0202835 HMM00390_malic_1_eut1(pf00390pf03949pf01515) EUT1
0202840 Hp_euts_violetPink EUT1
0202845 HMM10662PduV_1_mix(pf10662) EUT1
0202850 HMM06249EutQ_1_eut1(pf06249) EUT1 EP_NTERM
0202855 HMM01923cobtr_2_eut1(pf01923) EUT1
0202860 HMM01515PTA_B_1_eut1(pf01515) EUT1
0202865 H_azure EUT1
0202870 P_lightViolet EUT1
0202875 HMMalddh_1_eutpdugrm(pf00171) EUT1 EP_NTERM
0202880 HMM06723EutJMreB_1_grm(#) EUT1
0202885 HMM00465alcdh_3_mix(pf00465) EUT1
0202890 HMMtpEuth_1_eut(pf04346) EUT1
0202895 HMM06277EutA_1_eut(pf06277) EUT1
0202900 HMMetlyEutB_1_EUT1(pf06751) EUT1
0202905 HMMetlyEutC_1_eut1(pf05985) EUT1 EP_NTERM
0202910 Tsp_rosa EUT1
0202915 H_mulberry(pf00936pf16365) EUT1
0202920 HMMreg12833_1_EUT1(pf12833) EUT1
0202925 -pf01218 -
0202930 -pf01520 -
0202935 -pf00583 -
0202940 -pf11143 -
0202945 -pf06572 -
0202950 -pf04261 -
0202955 HMMnopfamclust3_1(#) RMM1
0202960 -# -
0202965 -pf00528 -
BMC-P
BMC-H
BMC-T
Aldehyde dehydrogenase
PTAC
Signature enzyme
Alcohol dehydrogenase
Regulator

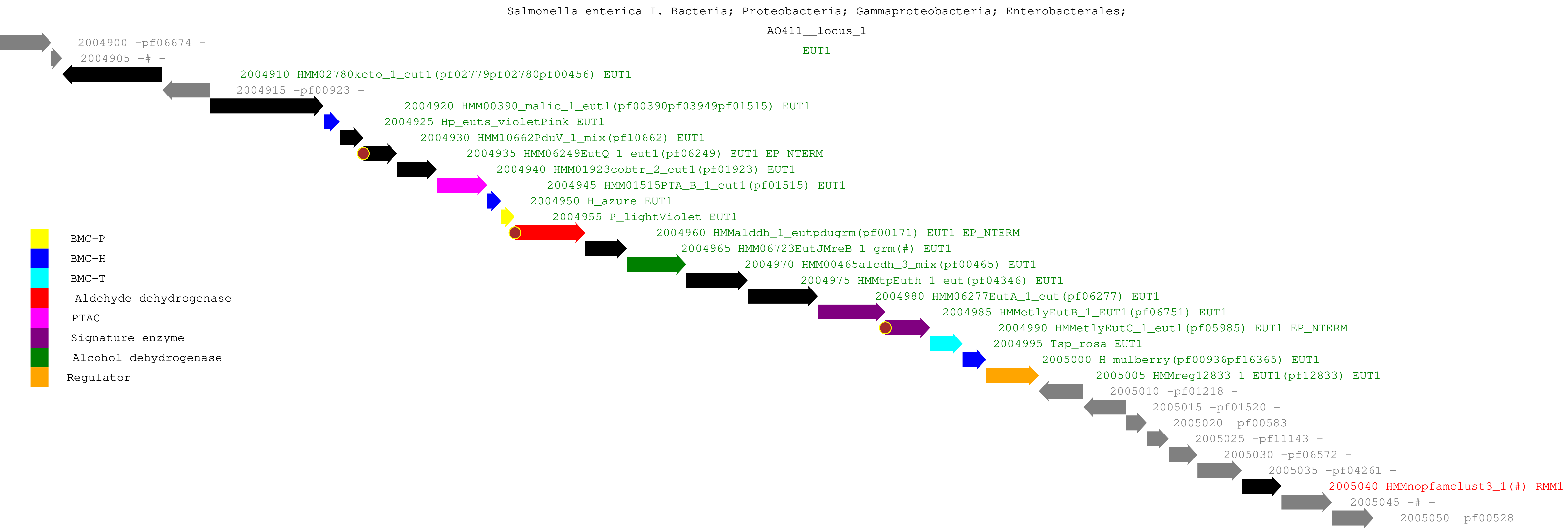

Salmonella enterica I. Bacteria; Proteobacteria; Gammaproteobacteria; Enterobacterales;
AO411__locus_1
EUT1
2004900 -pf06674 -
2004905 -# -
2004910 HMM02780keto_1_eut1(pf02779pf02780pf00456) EUT1
2004915 -pf00923 -
2004920 HMM00390_malic_1_eut1(pf00390pf03949pf01515) EUT1
2004925 Hp_euts_violetPink EUT1
2004930 HMM10662PduV_1_mix(pf10662) EUT1
2004935 HMM06249EutQ_1_eut1(pf06249) EUT1 EP_NTERM
2004940 HMM01923cobtr_2_eut1(pf01923) EUT1
2004945 HMM01515PTA_B_1_eut1(pf01515) EUT1
2004950 H_azure EUT1
2004955 P_lightViolet EUT1
2004960 HMMaldh_1_eutpdugrm(pf00171) EUT1 EP_NTERM
2004965 HMM06723EutJMreB_1_grm(#) EUT1
2004970 HMM00465alcdh_3_mix(pf00465) EUT1
2004975 HMMtpEuth_1_eut(pf04346) EUT1
2004980 HMM06277EutA_1_eut(pf06277) EUT1
2004985 HMMetlyEutB_1_EUT1(pf06751) EUT1
2004990 HMMetlyEutC_1_eut1(pf05985) EUT1 EP_NTERM
2004995 Tsp_rosa EUT1
2005000 H_mulberry(pf00936pf16365) EUT1
2005005 HMMreg12833_1_EUT1(pf12833) EUT1
2005010 -pf01218 -
2005015 -pf01520 -
2005020 -pf00583 -
2005025 -pf11143 -
2005030 -pf06572 -
2005035 -pf04261 -
2005040 HMMnopfamclust3_1(#) RMM1
2005045 -# -
2005050 -pf00528 -
BMC-P
BMC-H
BMC-T
Aldehyde dehydrogenase
PTAC
Signature enzyme
Alcohol dehydrogenase
Regulator

Pseudomonas sp. (strain ADP). Bacteria; Proteobacteria.

AOX63__locus_1

EUT1

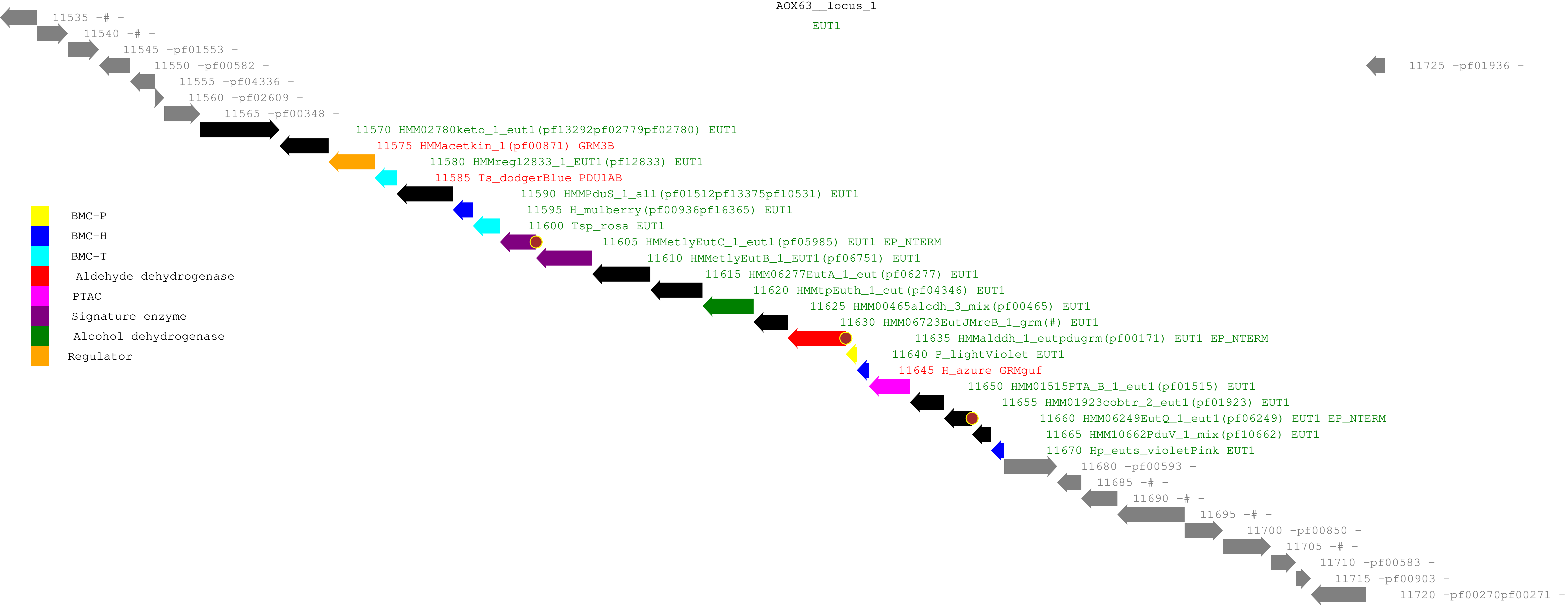

Escherichia coli APEC O18. Bacteria; Proteobacteria; Gammaproteobacteria; Enterobacterales;

APECO18__locus_1

EUT1

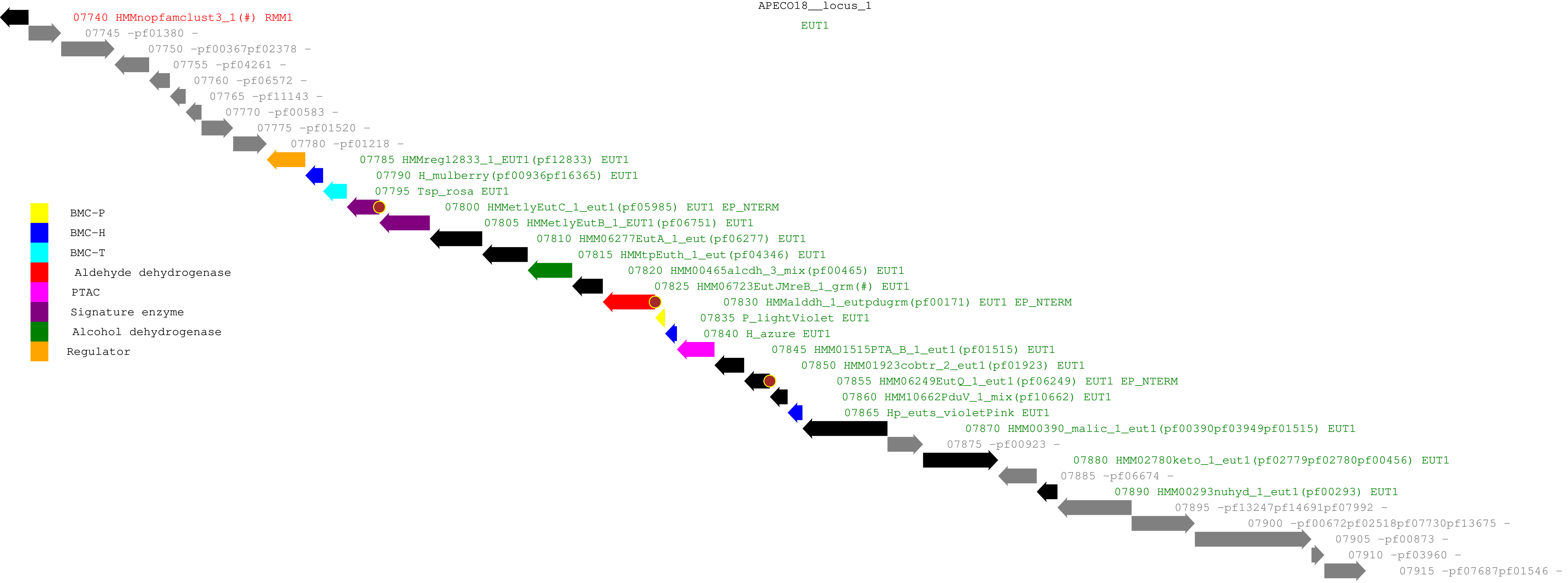

Escherichia coli O1:K1 / APEC. Bacteria; Proteobacteria; Gammaproteobacteria; Enterobacterales;

APECO1__locus_2

EUT1

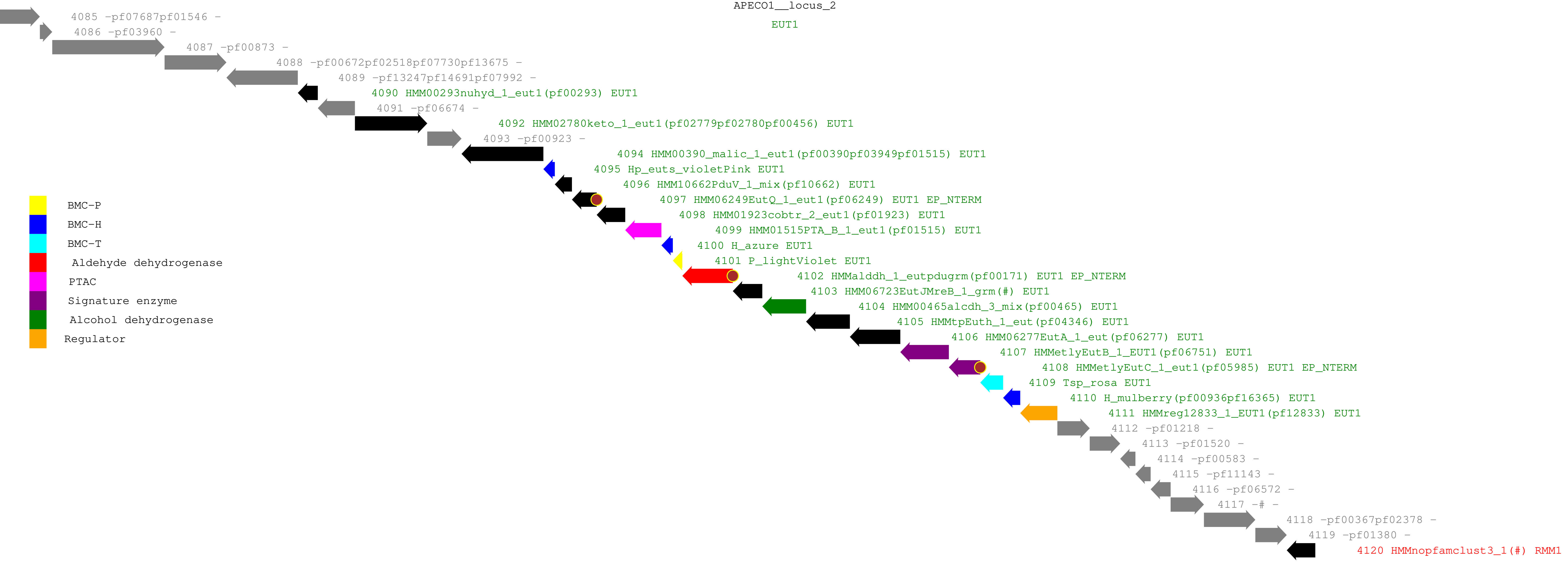

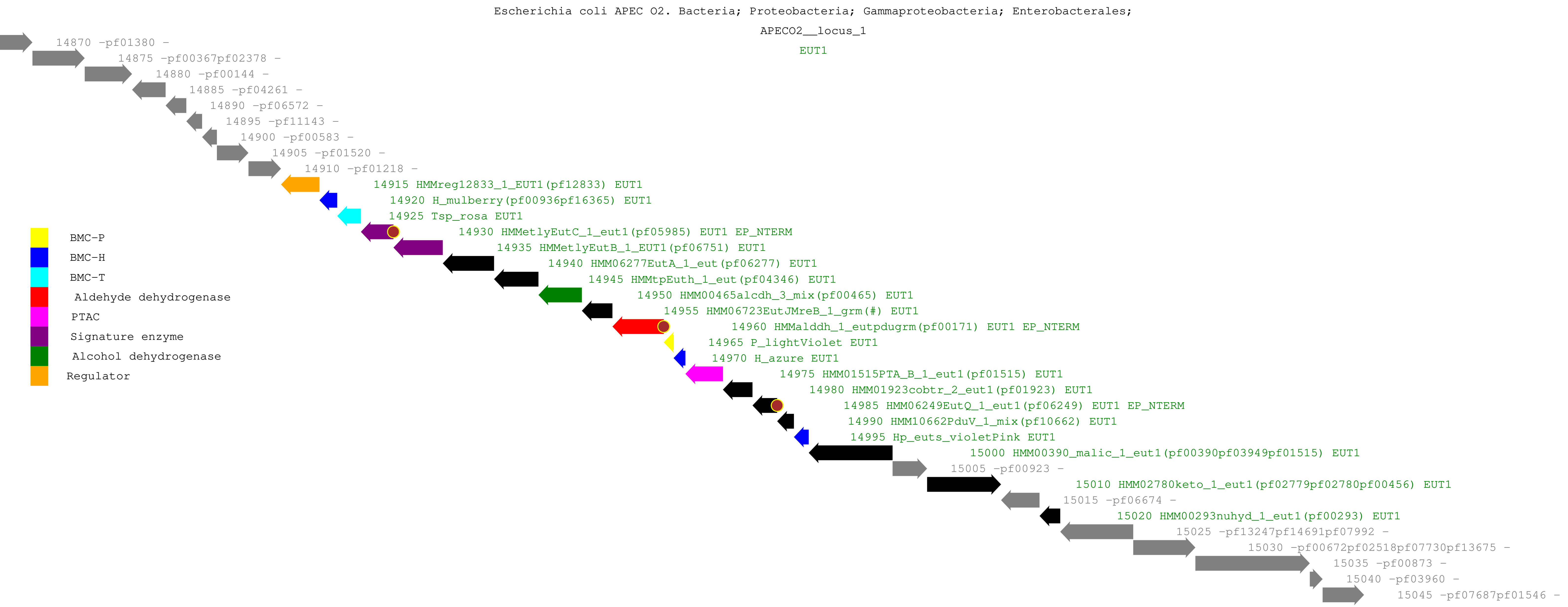
Escherichia coli APEC O2. Bacteria; Proteobacteria; Gammaproteobacteria; Enterobacterales;
APECO2__locus_1
EUT1
14870 -pf01380 -
14875 -pf00367pf02378 -
14880 -pf00144 -
14885 -pf04261 -
14890 -pf06572 -
14895 -pf11143 -
14900 -pf00583 -
14905 -pf01520 -
14910 -pf01218 -
14915 HMMreg12833_1_EUT1(pf12833) EUT1
14920 H_mulberry(pf00936pf16365) EUT1
14925 Tsp_rosa EUT1
14930 HMMetlyEutC_1_eut1(pf05985) EUT1 EP_NTERM
14935 HMMetlyEutB_1_EUT1(pf06751) EUT1
14940 HMM06277EutA_1_eut(pf06277) EUT1
14945 HMMtpEuth_1_eut(pf04346) EUT1
14950 HMM00465alcdh_3_mix(pf00465) EUT1
14955 HMM06723EutJMreB_1_grm(#) EUT1
14960 HMMalddh_1_eutpdugrm(pf00171) EUT1 EP_NTERM
14965 P_lightViolet EUT1
14970 H_azure EUT1
14975 HMM01515PTA_B_1_eut1(pf01515) EUT1
14980 HMM01923cobtr_2_eut1(pf01923) EUT1
14985 HMM06249EutQ_1_eut1(pf06249) EUT1 EP_NTERM
14990 HMM10662PduV_1_mix(pf10662) EUT1
14995 Hp_euts_violetPink EUT1
15000 HMM00390_malic_1_eut1(pf00390pf03949pf01515) EUT1
15005 -pf00923 -
15010 HMM02780keto_1_eut1(pf02779pf02780pf00456) EUT1
15015 -pf06674 -
15020 HMM00293nuhyd_1_eut1(pf00293) EUT1
15025 -pf13247pf14691pf07992 -
15030 -pf00672pf02518pf07730pf13675 -
15035 -pf00873 -
15040 -pf03960 -
15045 -pf07687pf01546 -
BMC-P
BMC-H
BMC-T
Aldehyde dehydrogenase
PTAC
Signature enzyme
Alcohol dehydrogenase
Regulator

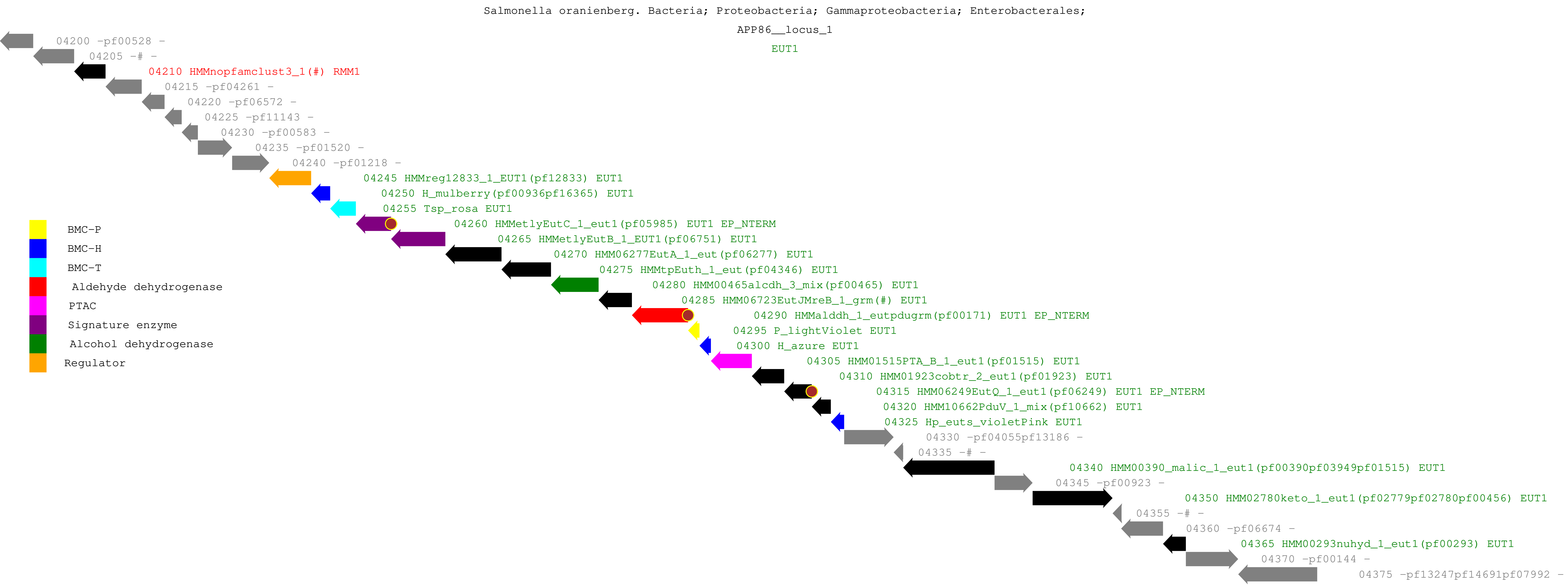
Salmonella oranienberg. Bacteria; Proteobacteria; Gammaproteobacteria; Enterobacterales;
APP86__locus_1
EUT1
04200 -pf00528 -
04205 -# -
04210 HMMnopfamclust3_1(#) RMM1
04215 -pf04261 -
04220 -pf06572 -
04225 -pf11143 -
04230 -pf00583 -
04235 -pf01520 -
04240 -pf01218 -
04245 HMMreg12833_1_EUT1(pf12833) EUT1
04250 H_mulberry(pf00936pf16365) EUT1
04255 Tsp_rosa EUT1
04260 HMMetlyEutC_1_eut1(pf05985) EUT1 EP_NTERM
04265 HMMetlyEutB_1_EUT1(pf06751) EUT1
04270 HMM06277EutA_1_eut(pf06277) EUT1
04275 HMMtpEuth_1_eut(pf04346) EUT1
04280 HMM00465alcdh_3_mix(pf00465) EUT1
04285 HMM06723EutJMreB_1_grm(#) EUT1
04290 HMMalddh_1_eutpdugrm(pf00171) EUT1 EP_NTERM
04295 P_lightViolet EUT1
04300 H_azure EUT1
04305 HMM01515PTA_B_1_eut1(pf01515) EUT1
04310 HMM01923cobtr_2_eut1(pf01923) EUT1
04315 HMM06249EutQ_1_eut1(pf06249) EUT1 EP_NTERM
04320 HMM10662PduV_1_mix(pf10662) EUT1
04325 Hp_euts_violetPink EUT1
04330 -pf04055pf13186 -
04335 -# -
04340 HMM00390_malic_1_eut1(pf00390pf03949pf01515) EUT1
04345 -pf00923 -
04350 HMM02780keto_1_eut1(pf02779pf02780pf00456) EUT1
04355 -# -
04360 -pf06674 -
04365 HMM00293nuhyd_1_eut1(pf00293) EUT1
04370 -pf00144 -
04375 -pf13247pf14691pf07992 -
BMC-P
BMC-H
BMC-T
Aldehyde dehydrogenase
PTAC
Signature enzyme
Alcohol dehydrogenase
Regulator

Escherichia coli. Bacteria; Proteobacteria; Gammaproteobacteria; Enterobacterales;

APT94__locus_1

EUT1

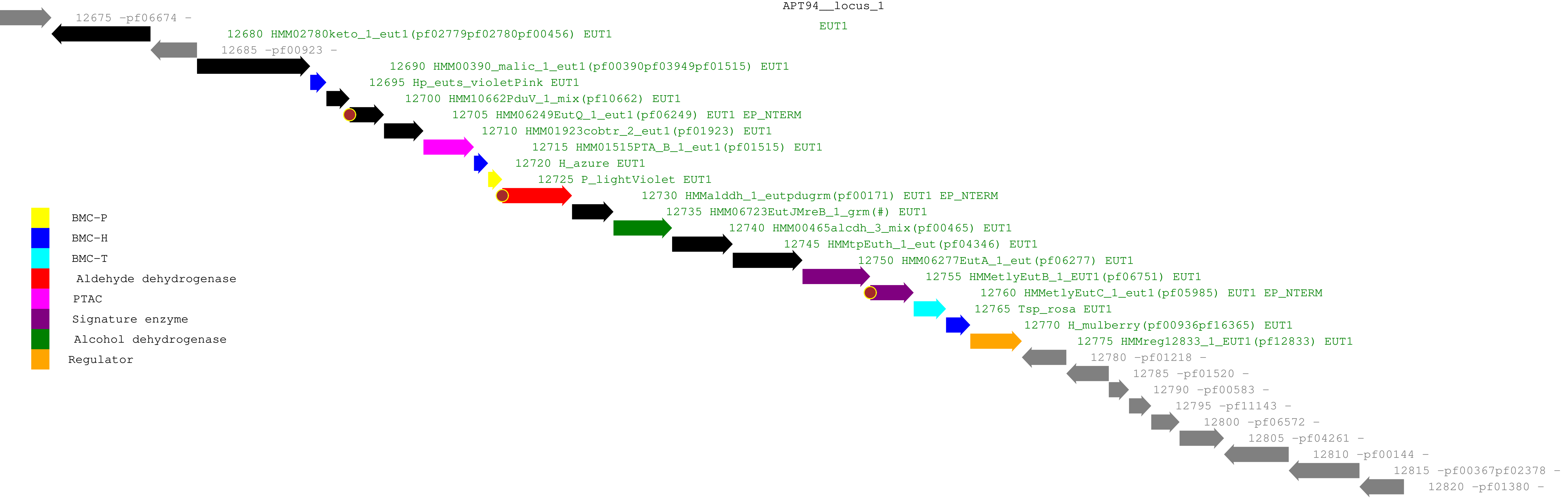

Citrobacter sp. 50677481. Bacteria; Proteobacteria; Gammaproteobacteria; Enterobacterales;

APU02__locus_1

EUT1

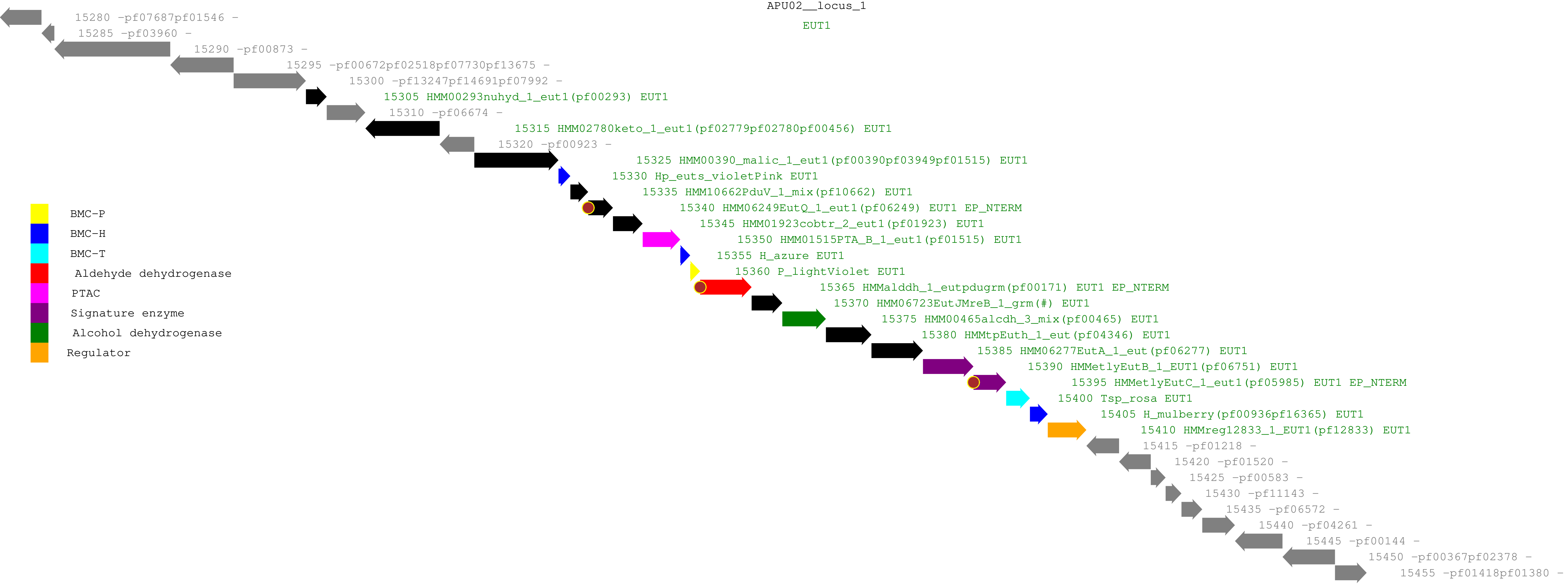

Escherichia coli. Bacteria; Proteobacteria; Gammaproteobacteria; Enterobacterales;

APU18__locus_1

EUT1

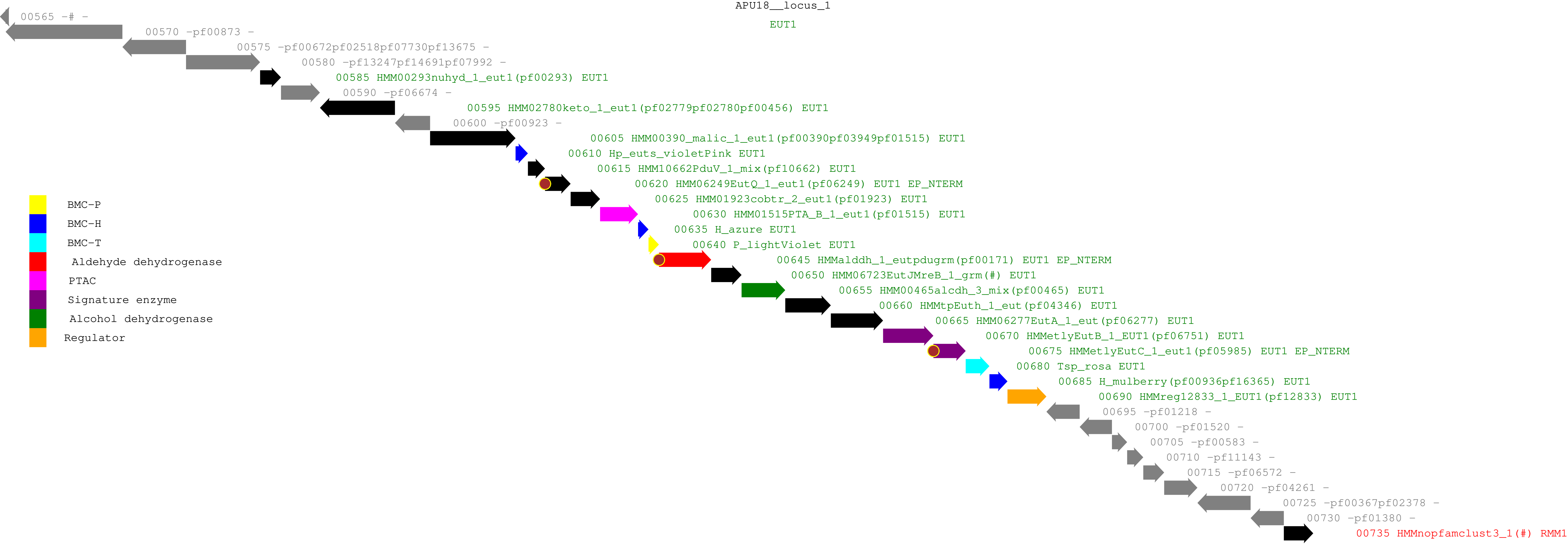

Salmonella enterica subsp. enterica serovar 4,12:d:-. Bacteria; Proteobacteria; Gammaproteobacteria; Enterobacterales;

AR931__locus_1

EUT1

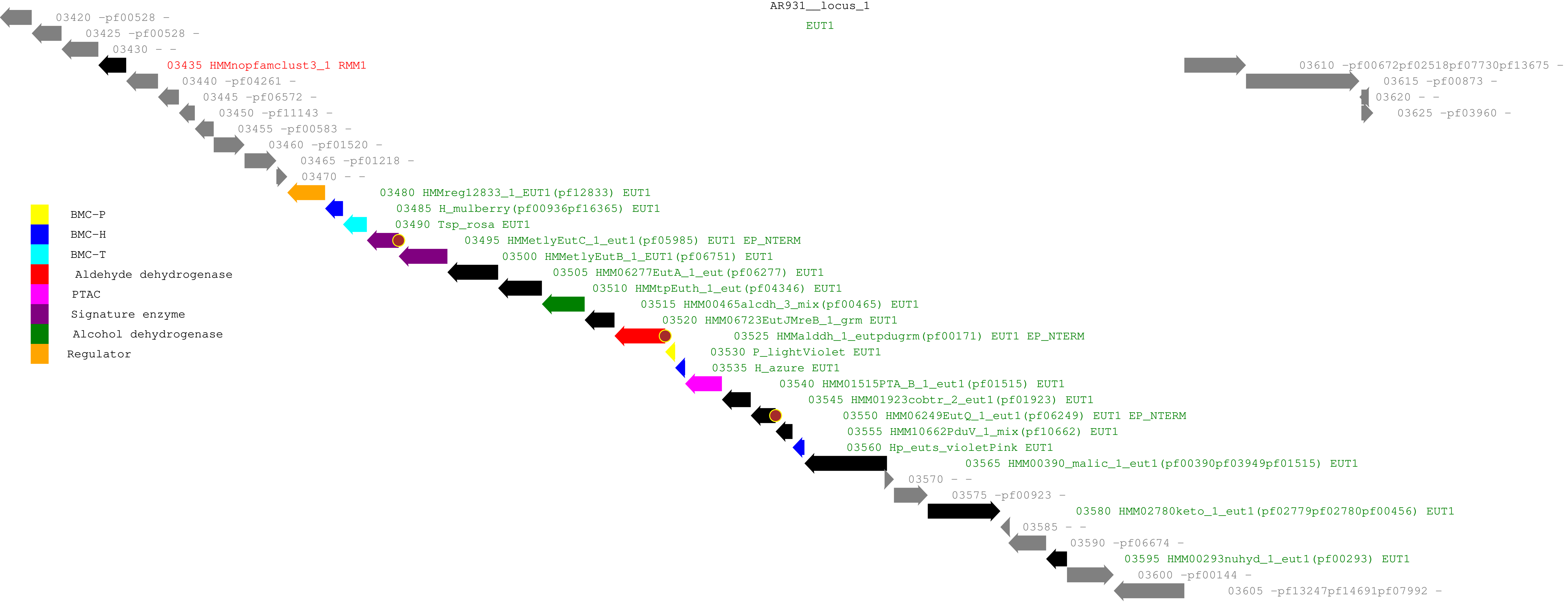

Escherichia coli. Bacteria; Proteobacteria; Gammaproteobacteria; Enterobacterales;

ARC77__locus_1

EUT1

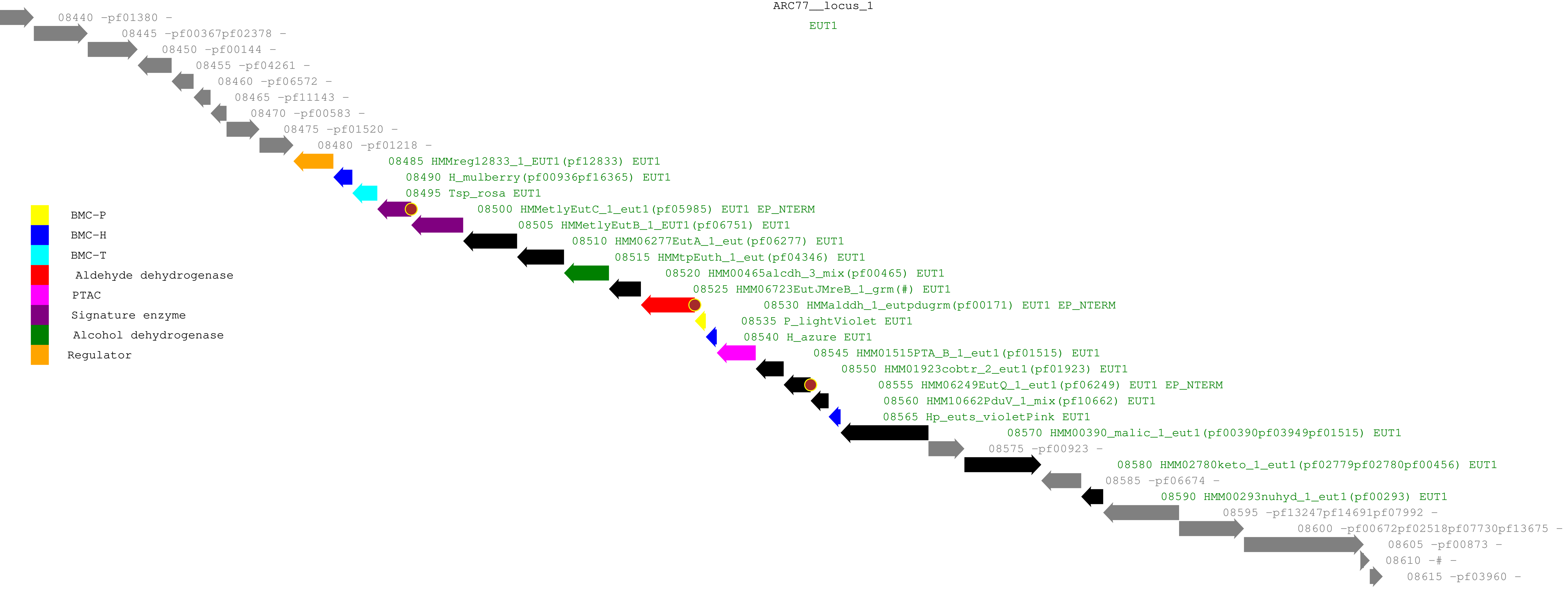

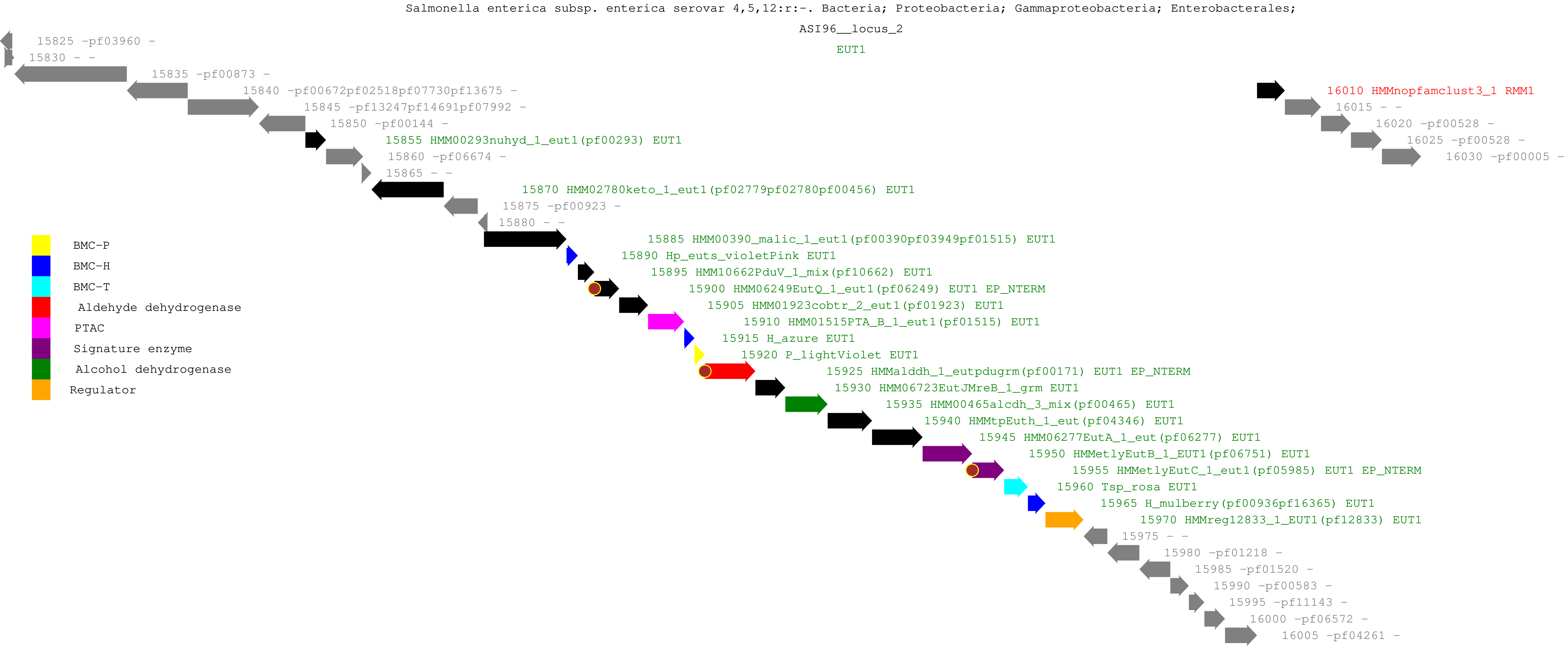
Salmonella enterica subsp. enterica serovar 4,5,12:r:-. Bacteria; Proteobacteria; Gammaproteobacteria; Enterobacterales;
ASI96__locus_2
EUT1
15825 -pf03960 -
15830 - -
15835 -pf00873 -
15840 -pf00672pf02518pf07730pf13675 -
15845 -pf13247pf14691pf07992 -
15850 -pf00144 -
15855 HMM00293nuhyd_1_eut1(pf00293) EUT1
15860 -pf06674 -
15865 - -
15870 HMM02780keto_1_eut1(pf02779pf02780pf00456) EUT1
15875 -pf00923 -
15880 - -
15885 HMM00390_malic_1_eut1(pf00390pf03949pf01515) EUT1
15890 Hp_euts_violetPink EUT1
15895 HMM10662PduV_1_mix(pf10662) EUT1
15900 HMM06249EutQ_1_eut1(pf06249) EUT1 EP_NTERM
15905 HMM01923cobtr_2_eut1(pf01923) EUT1
15910 HMM01515PTA_B_1_eut1(pf01515) EUT1
15915 H_azure EUT1
15920 P_lightViolet EUT1
15925 HMMalddh_1_eutpdugrm(pf00171) EUT1 EP_NTERM
15930 HMM06723EutJMreB_1_grm EUT1
15935 HMM00465alcdh_3_mix(pf00465) EUT1
15940 HMMtpEuth_1_eut(pf04346) EUT1
15945 HMM06277EutA_1_eut(pf06277) EUT1
15950 HMMetlyEutB_1_EUT1(pf06751) EUT1
15955 HMMetlyEutC_1_eut1(pf05985) EUT1 EP_NTERM
15960 Tsp_rosa EUT1
15965 H_mulberry(pf00936pf16365) EUT1
15970 HMMreg12833_1_EUT1(pf12833) EUT1
15975 - -
15980 -pf01218 -
15985 -pf01520 -
15990 -pf00583 -
15995 -pf11143 -
16000 -pf06572 -
16005 -pf04261 -
16010 HMMnopfamclust3_1 RMM1
16015 - -
16020 -pf00528 -
16025 -pf00528 -
16030 -pf00005 -
BMC-P
BMC-H
BMC-T
Aldehyde dehydrogenase
PTAC
Signature enzyme
Alcohol dehydrogenase
Regulator

Hafnia alvei FB1. Bacteria; Proteobacteria; Gammaproteobacteria; Enterobacterales;

AT03__locus_1

EUT1

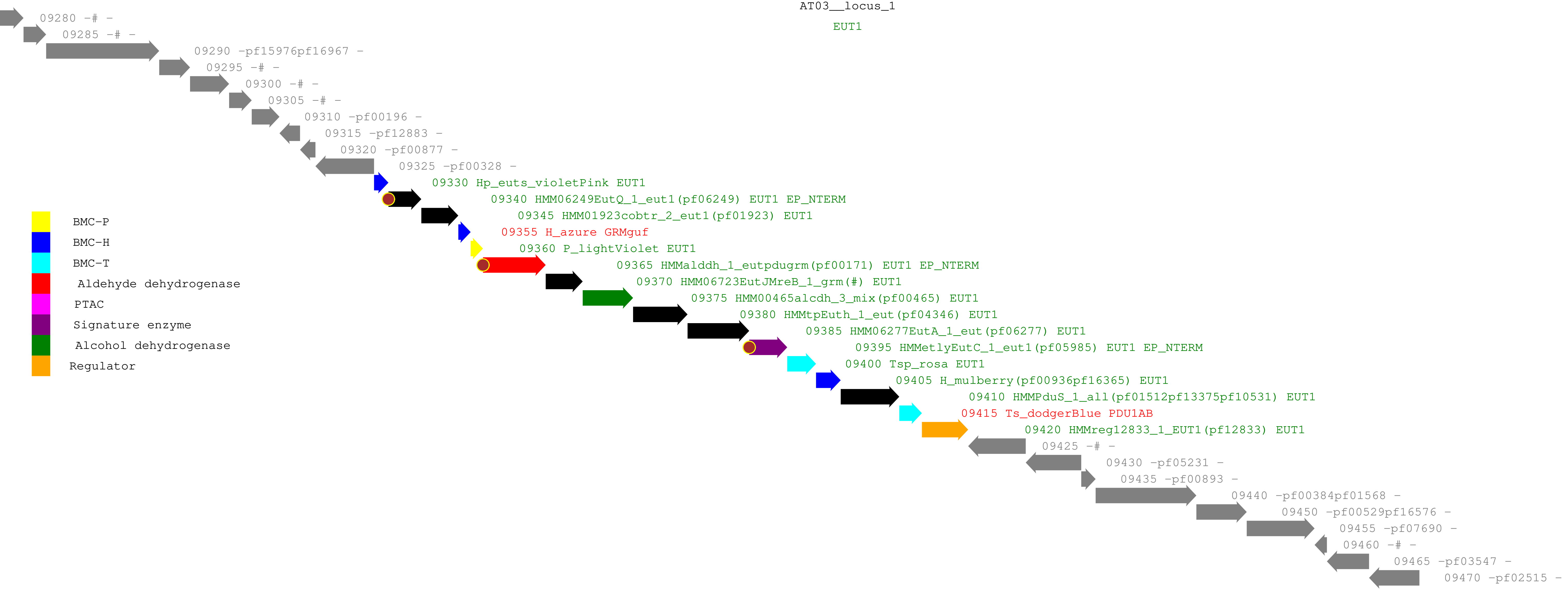

Marinobacter sp. LV10R520-4. Bacteria; Proteobacteria; Gammaproteobacteria; Alteromonadales;

ATG98__locus_1

EUT1

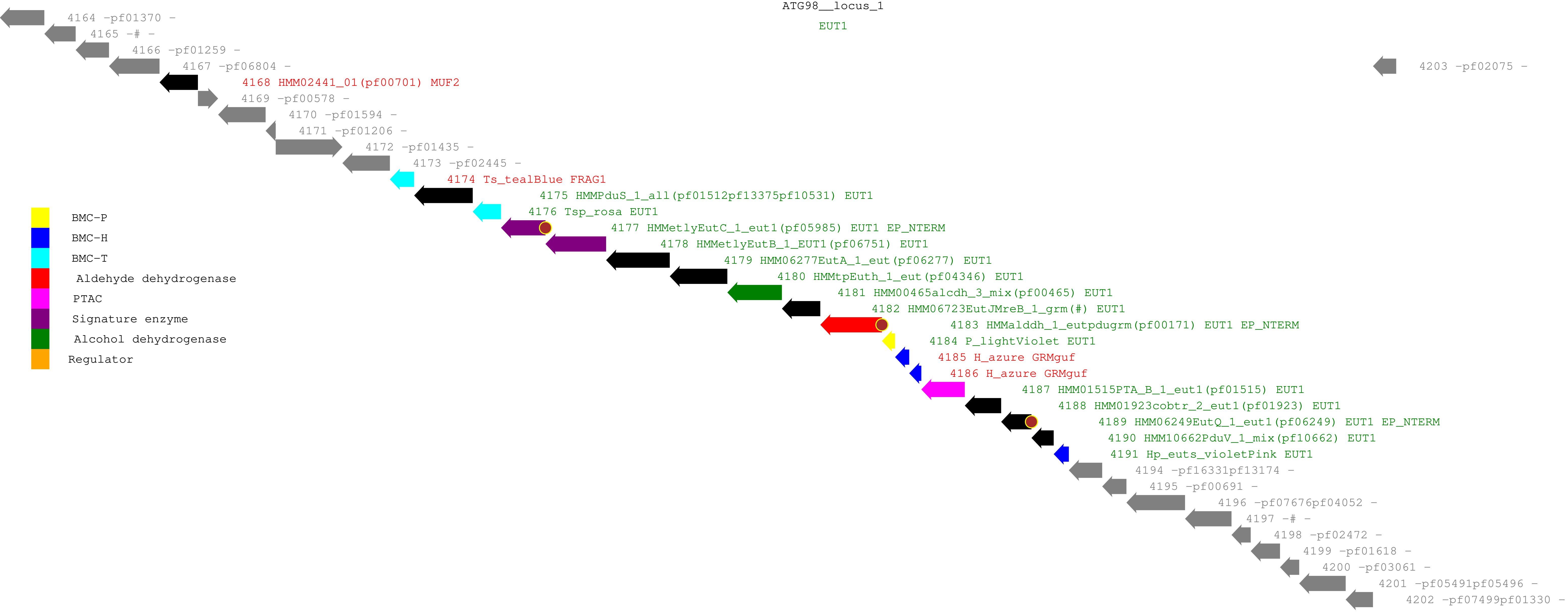

Salmonella enterica (Salmonella choleraesuis). Bacteria; Proteobacteria; Gammaproteobacteria; Enterobacterales;

ATP91__locus_1

EUT1

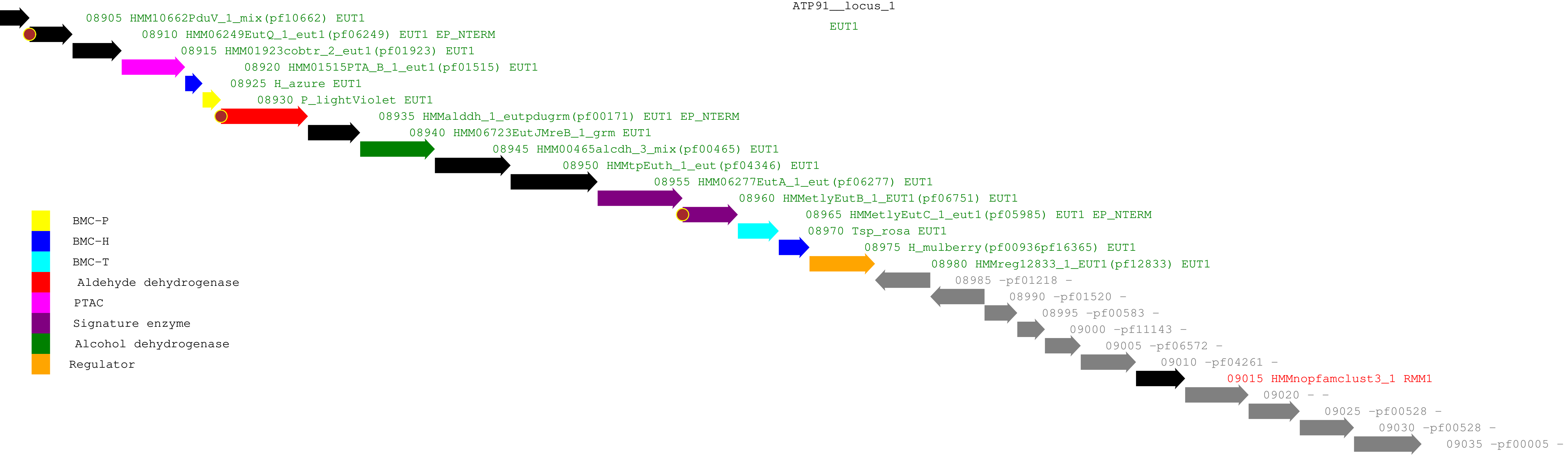

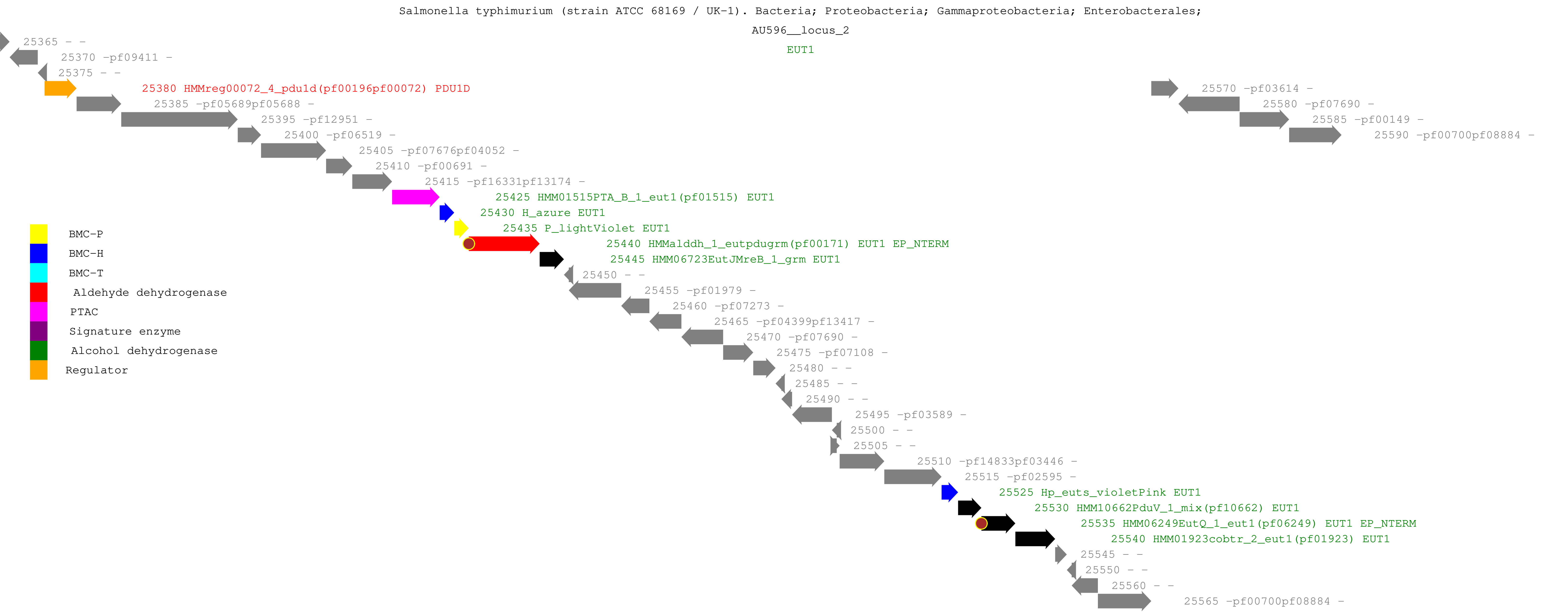
Salmonella typhimurium (strain ATCC 68169 / UK-1). Bacteria; Proteobacteria; Gammaproteobacteria; Enterobacterales;
AU596__locus_2
EUT1
25365 - -
25370 -pf09411 -
25375 - -
25380 HMMreg00072_4_pdu1d(pf00196pf00072) PDU1D
25385 -pf05689pf05688 -
25395 -pf12951 -
25400 -pf06519 -
25405 -pf07676pf04052 -
25410 -pf00691 -
25415 -pf16331pf13174 -
25425 HMM01515PTA_B_1_eut1(pf01515) EUT1
25430 H_azure EUT1
25435 P_lightViolet EUT1
25440 HMMaldh_1_eutpdugrm(pf00171) EUT1 EP_NTERM
25445 HMM06723EutJMreB_1_grm EUT1
25450 - -
25455 -pf01979 -
25460 -pf07273 -
25465 -pf04399pf13417 -
25470 -pf07690 -
25475 -pf07108 -
25480 - -
25485 - -
25490 - -
25495 -pf03589 -
25500 - -
25505 - -
25510 -pf14833pf03446 -
25515 -pf02595 -
25525 Hp_euts_violetPink EUT1
25530 HMM10662PduV_1_mix(pf10662) EUT1
25535 HMM06249EutQ_1_eut1(pf06249) EUT1 EP_NTERM
25540 HMM01923cobtr_2_eut1(pf01923) EUT1
25545 - -
25550 - -
25560 - -
25565 -pf00700pf08884 -
25570 -pf03614 -
25580 -pf07690 -
25585 -pf00149 -
25590 -pf00700pf08884 -
BMC-P
BMC-H
BMC-T
Aldehyde dehydrogenase
PTAC
Signature enzyme
Alcohol dehydrogenase
Regulator

Salmonella enterica subsp. enterica serovar Sekondi. Bacteria; Proteobacteria; Gammaproteobacteria; Enterobacterales;

AU626__locus_3_main_sat

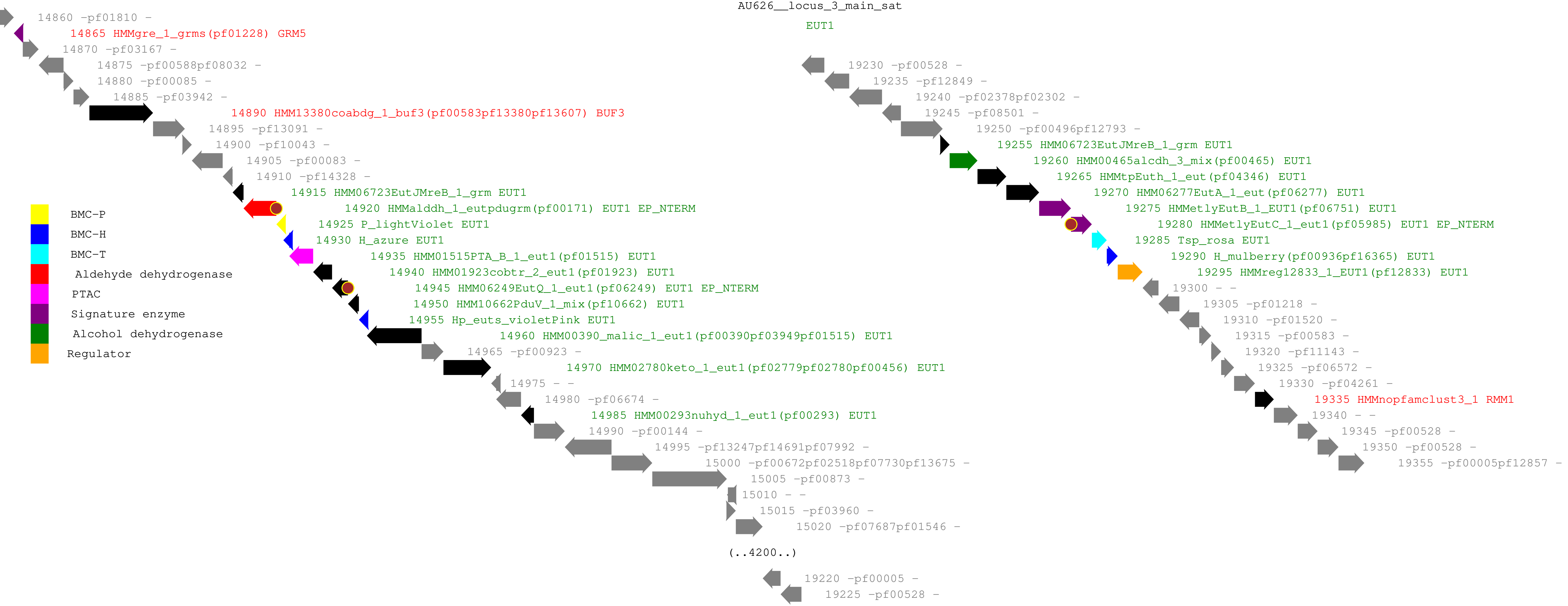

Salmonella enterica subsp. enterica serovar Ridge. Bacteria; Proteobacteria; Gammaproteobacteria; Enterobacterales;

AU648__locus_2

EUT1

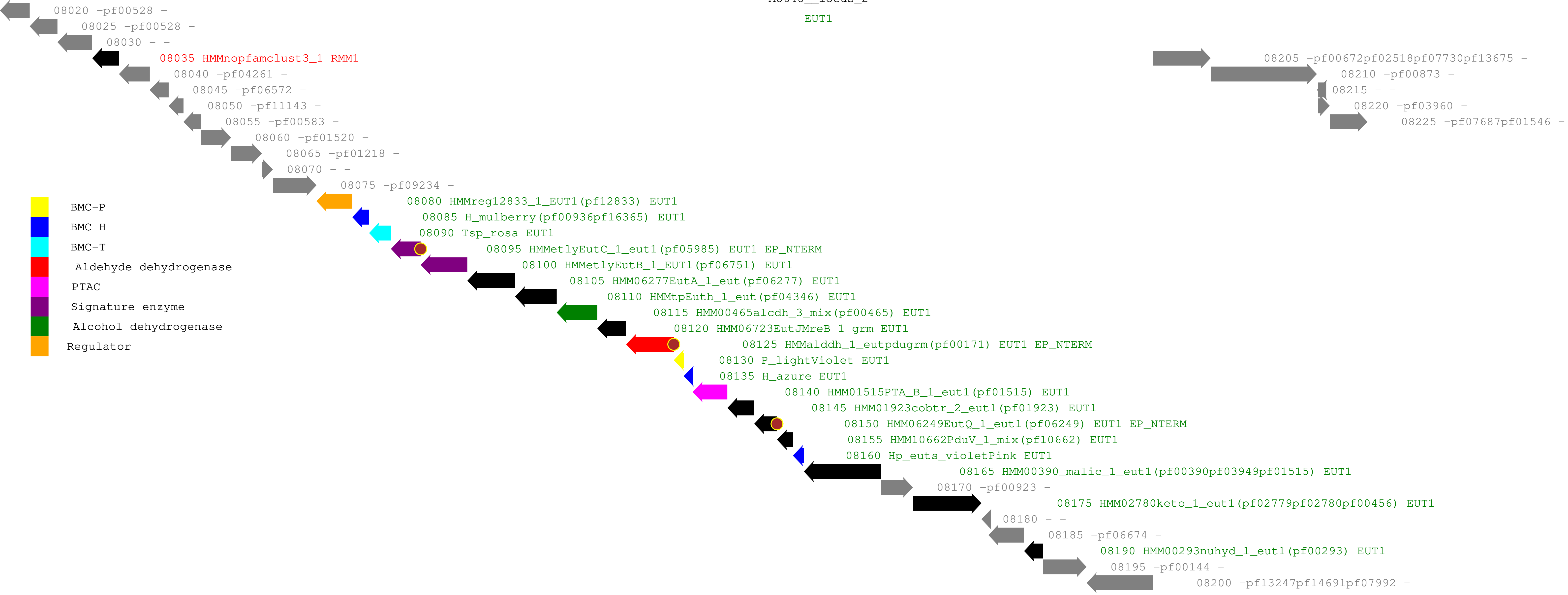

Salmonella abony. Bacteria; Proteobacteria; Gammaproteobacteria; Enterobacterales;

AU743__locus_2_main_sat

EUT1

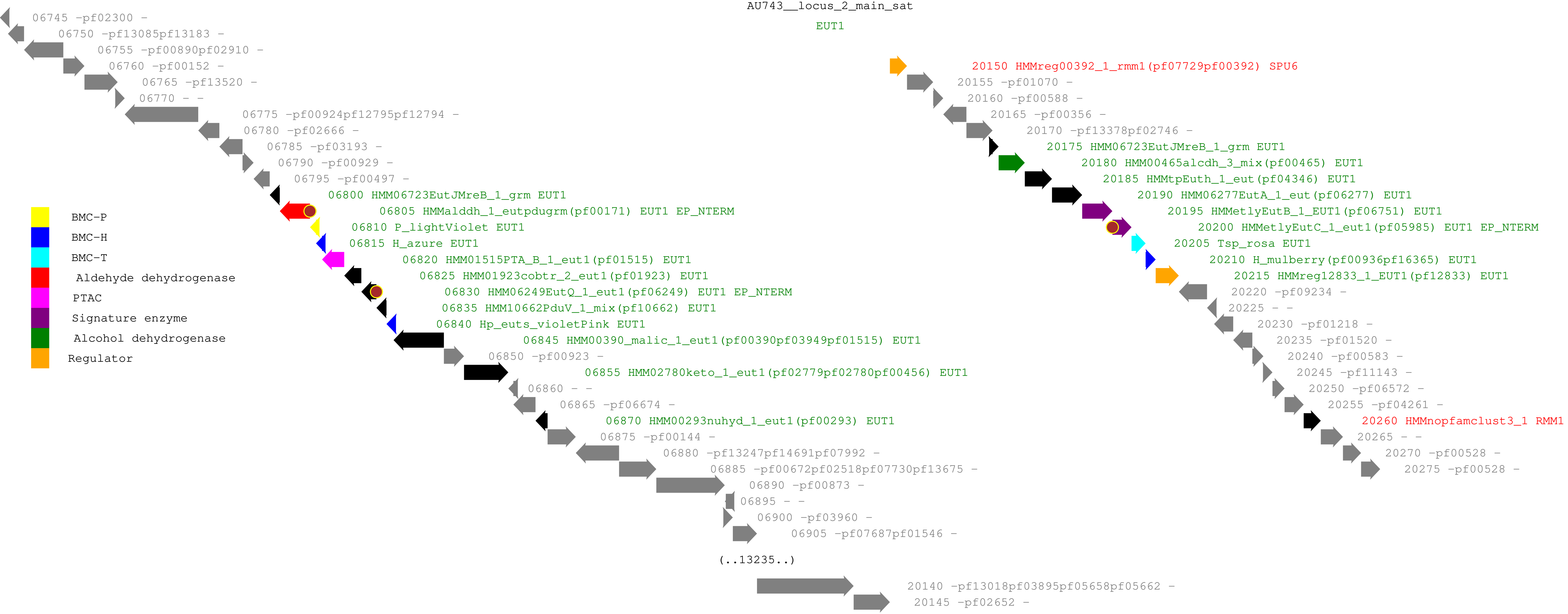

Salmonella enterica subsp. enterica serovar Curacao. Bacteria; Proteobacteria; Gammaproteobacteria; Enterobacterales;

AU769__locus_2_main_sat

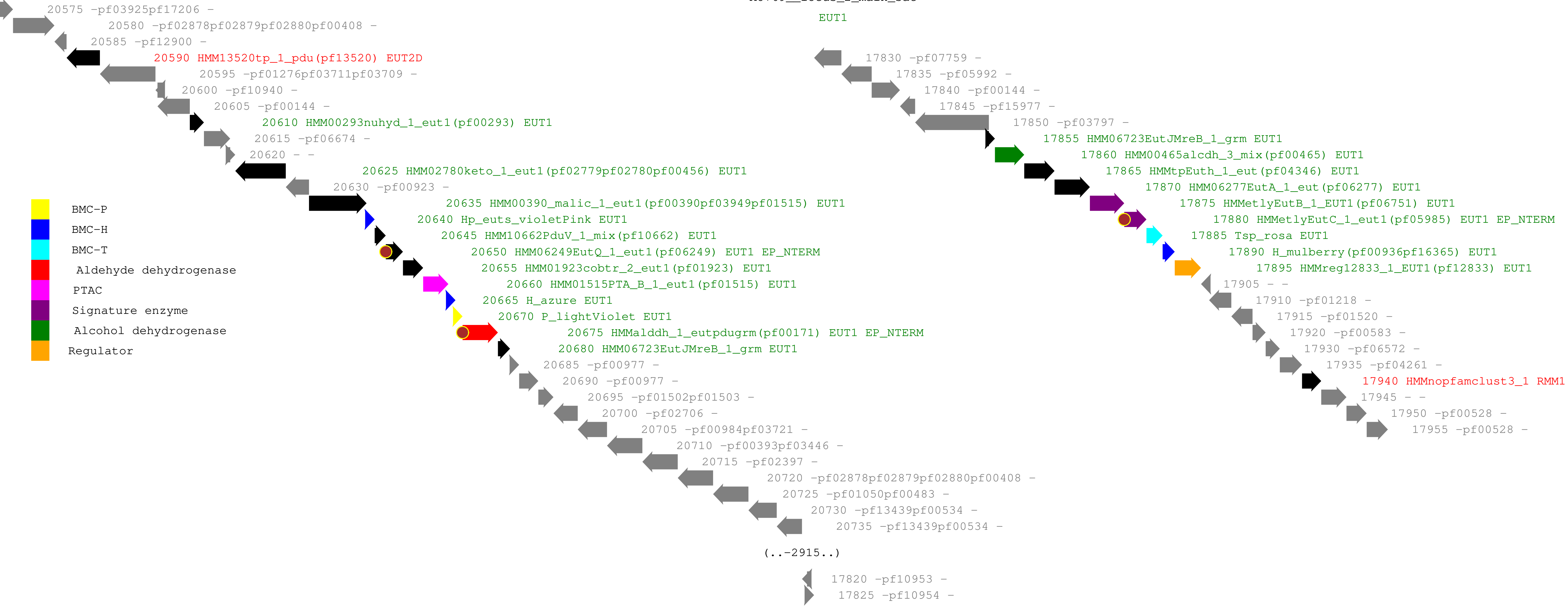

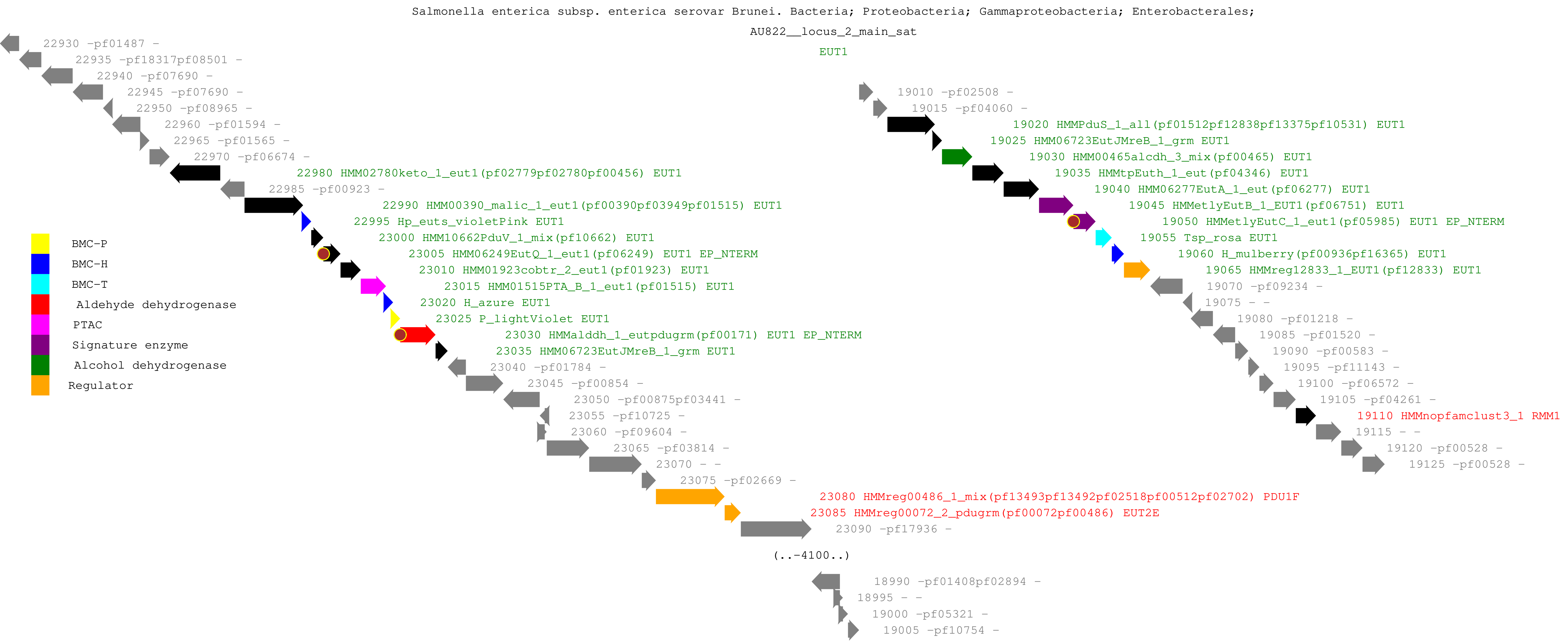
Salmonella enterica subsp. enterica serovar Brunei. Bacteria; Proteobacteria; Gammaproteobacteria; Enterobacterales;
AU822__locus_2_main_sat
EUT1
22930 -pf01487 -
22935 -pf18317pf08501 -
22940 -pf07690 -
22945 -pf07690 -
22950 -pf08965 -
22960 -pf01594 -
22965 -pf01565 -
22970 -pf06674 -
22980 HMM02780keto_1_eut1(pf02779pf02780pf00456) EUT1
22985 -pf00923 -
22990 HMM00390_malic_1_eut1(pf00390pf03949pf01515) EUT1
22995 Hp_euts_violetPink EUT1
23000 HMM10662PduV_1_mix(pf10662) EUT1
23005 HMM06249EutQ_1_eut1(pf06249) EUT1 EP_NTERM
23010 HMM01923cobtr_2_eut1(pf01923) EUT1
23015 HMM01515PTA_B_1_eut1(pf01515) EUT1
23020 H_azure EUT1
23025 P_lightViolet EUT1
23030 HMMalddh_1_eutpdugrm(pf00171) EUT1 EP_NTERM
23035 HMM06723EutJMreB_1_grm EUT1
23040 -pf01784 -
23045 -pf00854 -
23050 -pf00875pf03441 -
23055 -pf10725 -
23060 -pf09604 -
23065 -pf03814 -
23070 - -
23075 -pf02669 -
23080 HMMreg00486_1_mix(pf13493pf13492pf02518pf00512pf02702) PDU1F
23085 HMMreg00072_2_pdugrm(pf00072pf00486) EUT2E
23090 -pf17936 -
(..-4100..)
18990 -pf01408pf02894 -
18995 - -
19000 -pf05321 -
19005 -pf10754 -
19010 -pf02508 -
19015 -pf04060 -
19020 HMMPduS_1_all(pf01512pf12838pf13375pf10531) EUT1
19025 HMM06723EutJMreB_1_grm EUT1
19030 HMM00465alcdh_3_mix(pf00465) EUT1
19035 HMMtpEuth_1_eut(pf04346) EUT1
19040 HMM06277EutA_1_eut(pf06277) EUT1
19045 HMMetlyEutB_1_EUT1(pf06751) EUT1
19050 HMMetlyEutC_1_eut1(pf05985) EUT1 EP_NTERM
19055 Tsp_rosa EUT1
19060 H_mulberry(pf00936pf16365) EUT1
19065 HMMreg12833_1_EUT1(pf12833) EUT1
19070 -pf09234 -
19075 - -
19080 -pf01218 -
19085 -pf01520 -
19090 -pf00583 -
19095 -pf11143 -
19100 -pf06572 -
19105 -pf04261 -
19110 HMMnopfamclust3_1 RMM1
19115 - -
19120 -pf00528 -
19125 -pf00528 -
BMC-P
BMC-H
BMC-T
Aldehyde dehydrogenase
PTAC
Signature enzyme
Alcohol dehydrogenase
Regulator

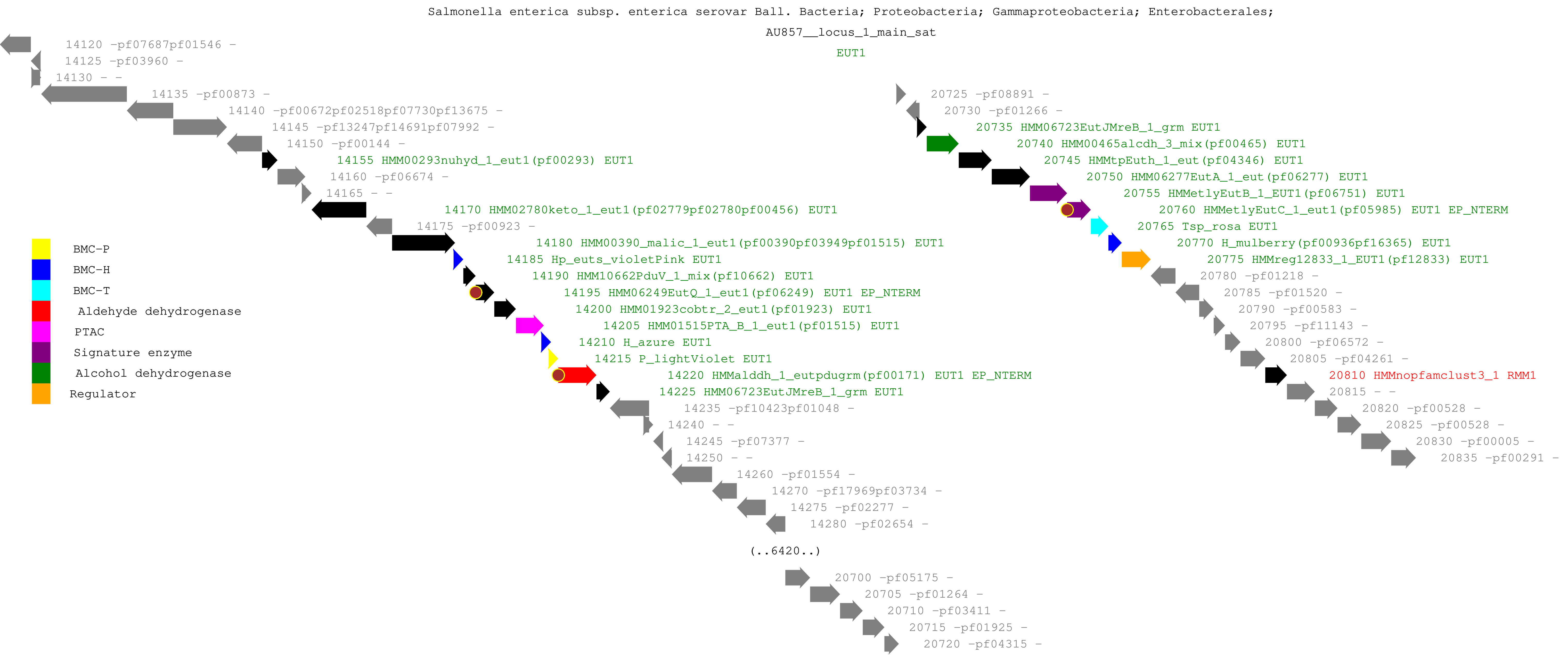
Salmonella enterica subsp. enterica serovar Ball. Bacteria; Proteobacteria; Gammaproteobacteria; Enterobacterales;
AU857__locus_1_main_sat
EUT1
14120 -pf07687pf01546 -
14125 -pf03960 -
14130 - -
14135 -pf00873 -
14140 -pf00672pf02518pf07730pf13675 -
14145 -pf13247pf14691pf07992 -
14150 -pf00144 -
14155 HMM00293nuhyd_1_eut1(pf00293) EUT1
14160 -pf06674 -
14165 - -
14170 HMM02780keto_1_eut1(pf02779pf02780pf00456) EUT1
14175 -pf00923 -
14180 HMM00390_malic_1_eut1(pf00390pf03949pf01515) EUT1
14185 Hp_euts_violetPink EUT1
14190 HMM10662PduV_1_mix(pf10662) EUT1
14195 HMM06249EutQ_1_eut1(pf06249) EUT1 EP_NTERM
14200 HMM01923cobtr_2_eut1(pf01923) EUT1
14205 HMM01515PTA_B_1_eut1(pf01515) EUT1
14210 H_azure EUT1
14215 P_lightViolet EUT1
14220 HMMalddh_1_eutpdugrm(pf00171) EUT1 EP_NTERM
14225 HMM06723EutJMreB_1_grm EUT1
14235 -pf10423pf01048 -
14240 - -
14245 -pf07377 -
14250 - -
14260 -pf01554 -
14270 -pf17969pf03734 -
14275 -pf02277 -
14280 -pf02654 -
(..6420..)
20700 -pf05175 -
20705 -pf01264 -
20710 -pf03411 -
20715 -pf01925 -
20720 -pf04315 -
20725 -pf08891 -
20730 -pf01266 -
20735 HMM06723EutJMreB_1_grm EUT1
20740 HMM00465alcdh_3_mix(pf00465) EUT1
20745 HMMtpEuth_1_eut(pf04346) EUT1
20750 HMM06277EutA_1_eut(pf06277) EUT1
20755 HMMetlyEutB_1_EUT1(pf06751) EUT1
20760 HMMetlyEutC_1_eut1(pf05985) EUT1 EP_NTERM
20765 Tsp_rosa EUT1
20770 H_mulberry(pf00936pf16365) EUT1
20775 HMMreg12833_1_EUT1(pf12833) EUT1
20780 -pf01218 -
20785 -pf01520 -
20790 -pf00583 -
20795 -pf11143 -
20800 -pf06572 -
20805 -pf04261 -
20810 HMMnopfamclust3_1 RMM1
20815 - -
20820 -pf00528 -
20825 -pf00528 -
20830 -pf00005 -
20835 -pf00291 -
BMC-P
BMC-H
BMC-T
Aldehyde dehydrogenase
PTAC
Signature enzyme
Alcohol dehydrogenase
Regulator

Salmonella enterica subsp. enterica serovar Ibadan. Bacteria; Proteobacteria; Gammaproteobacteria; Enterobacterales;

AU867__locus_2

EUT1

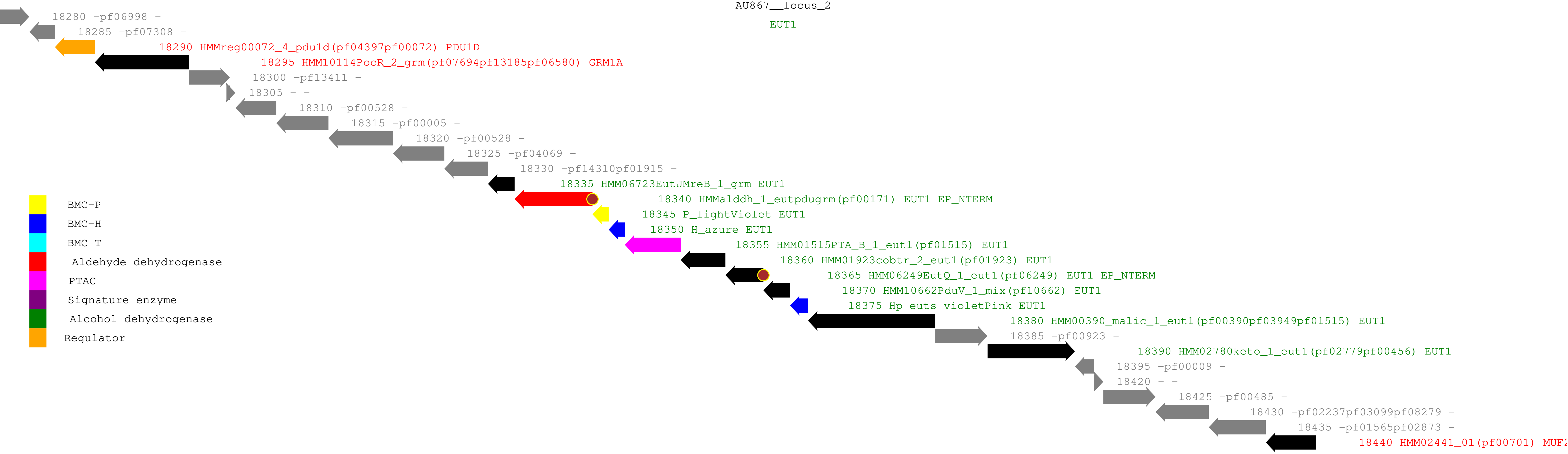

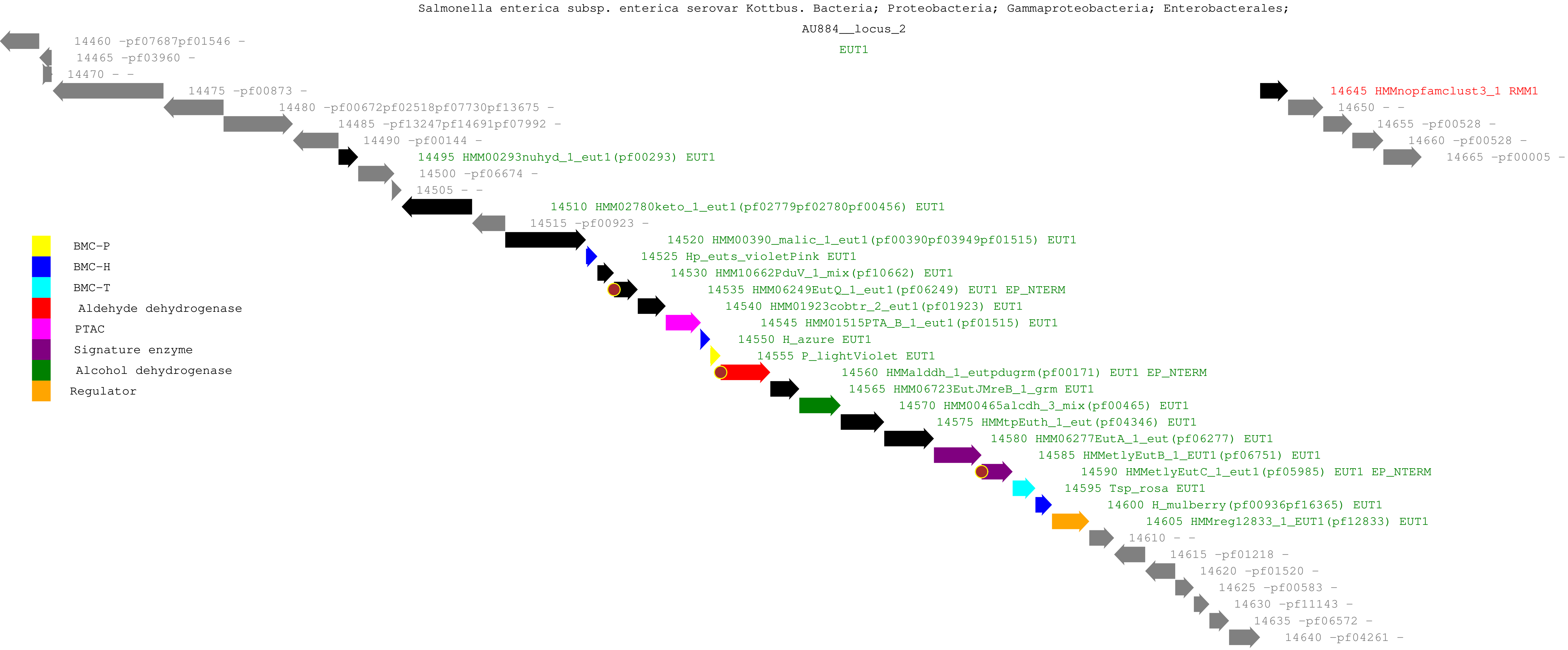

Salmonella enterica subsp. enterica serovar Kottbus. Bacteria; Proteobacteria; Gammaproteobacteria; Enterobacterales;
AU884__locus_2
EUT1
14460 -pf07687pf01546 -
14465 -pf03960 -
14470 - -
14475 -pf00873 -
14480 -pf00672pf02518pf07730pf13675 -
14485 -pf13247pf14691pf07992 -
14490 -pf00144 -
14495 HMM00293nuhyd_1_eut1(pf00293) EUT1
14500 -pf06674 -
14505 - -
14510 HMM02780keto_1_eut1(pf02779pf02780pf00456) EUT1
14515 -pf00923 -
14520 HMM00390_malic_1_eut1(pf00390pf03949pf01515) EUT1
14525 Hp_euts_violetPink EUT1
14530 HMM10662PduV_1_mix(pf10662) EUT1
14535 HMM06249EutQ_1_eut1(pf06249) EUT1 EP_NTERM
14540 HMM01923cobtr_2_eut1(pf01923) EUT1
14545 HMM01515PTA_B_1_eut1(pf01515) EUT1
14550 H_azure EUT1
14555 P_lightViolet EUT1
14560 HMMalddh_1_eutpdugrm(pf00171) EUT1 EP_NTERM
14565 HMM06723EutJMreB_1_grm EUT1
14570 HMM00465alcdh_3_mix(pf00465) EUT1
14575 HMMtpEuth_1_eut(pf04346) EUT1
14580 HMM06277EutA_1_eut(pf06277) EUT1
14585 HMMetlyEutB_1_EUT1(pf06751) EUT1
14590 HMMetlyEutC_1_eut1(pf05985) EUT1 EP_NTERM
14595 Tsp_rosa EUT1
14600 H_mulberry(pf00936pf16365) EUT1
14605 HMMreg12833_1_EUT1(pf12833) EUT1
14610 - -
14615 -pf01218 -
14620 -pf01520 -
14625 -pf00583 -
14630 -pf11143 -
14635 -pf06572 -
14640 -pf04261 -
14645 HMMnopfamclust3_1 RMM1
14650 - -
14655 -pf00528 -
14660 -pf00528 -
14665 -pf00005 -
BMC-P
BMC-H
BMC-T
Aldehyde dehydrogenase
PTAC
Signature enzyme
Alcohol dehydrogenase
Regulator

Salmonella enterica subsp. enterica serovar Ago. Bacteria; Proteobacteria; Gammaproteobacteria; Enterobacterales;

AU897__locus_2_main_sat

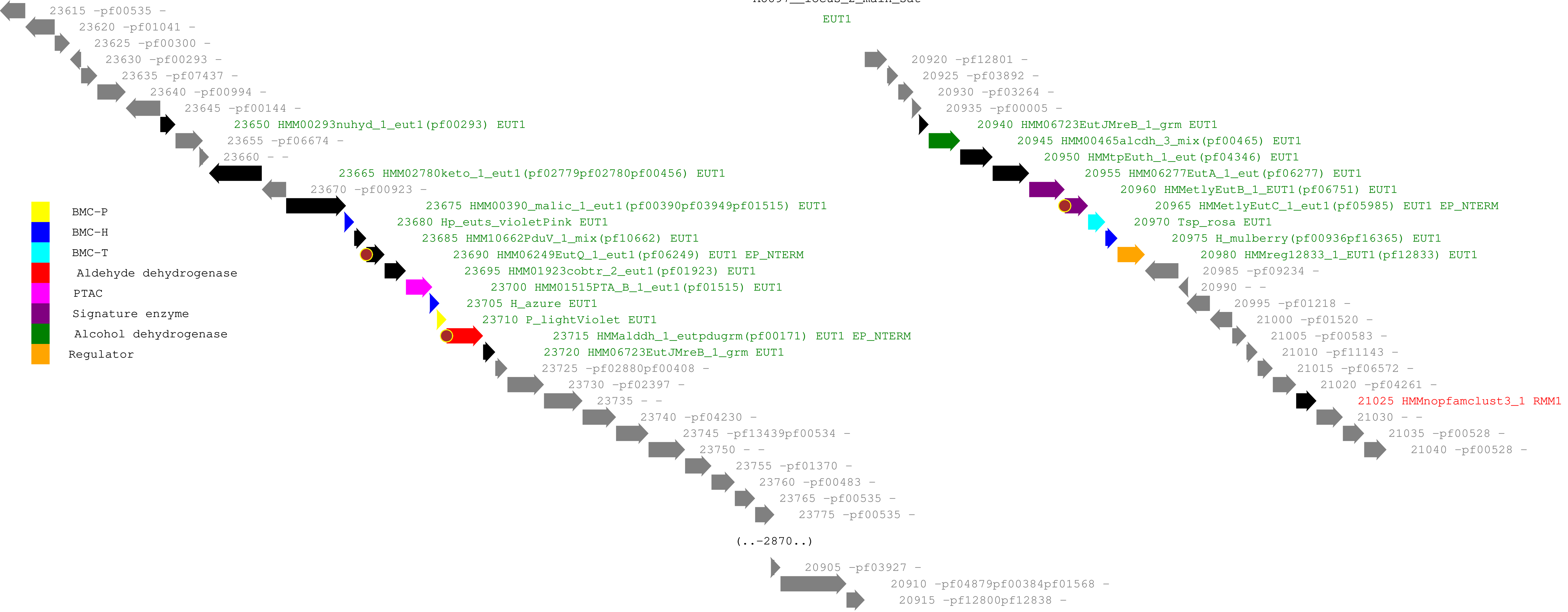

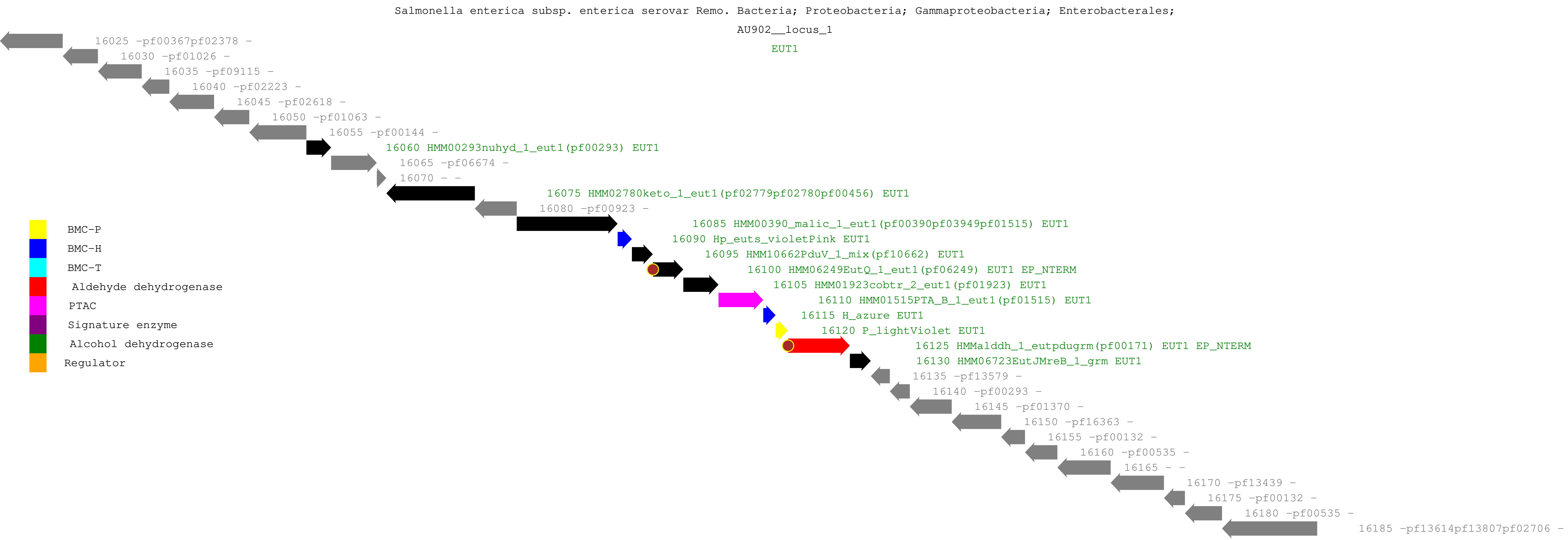
Salmonella enterica subsp. enterica serovar Remo. Bacteria; Proteobacteria; Gammaproteobacteria; Enterobacterales;
AU902__locus_1
EUT1
16025 -pf00367pf02378 -
16030 -pf01026 -
16035 -pf09115 -
16040 -pf02223 -
16045 -pf02618 -
16050 -pf01063 -
16055 -pf00144 -
16060 HMM00293nuhyd_1_eut1(pf00293) EUT1
16065 -pf06674 -
16070 - -
16075 HMM02780keto_1_eut1(pf02779pf02780pf00456) EUT1
16080 -pf00923 -
16085 HMM00390_malic_1_eut1(pf00390pf03949pf01515) EUT1
16090 Hp_euts_violetPink EUT1
16095 HMM10662PduV_1_mix(pf10662) EUT1
16100 HMM06249EutQ_1_eut1(pf06249) EUT1 EP_NTERM
16105 HMM01923cobtr_2_eut1(pf01923) EUT1
16110 HMM01515PTA_B_1_eut1(pf01515) EUT1
16115 H_azure EUT1
16120 P_lightViolet EUT1
16125 HMMalddh_1_eutpdugrm(pf00171) EUT1 EP_NTERM
16130 HMM06723EutJMreB_1_grm EUT1
16135 -pf13579 -
16140 -pf00293 -
16145 -pf01370 -
16150 -pf16363 -
16155 -pf00132 -
16160 -pf00535 -
16165 - -
16170 -pf13439 -
16175 -pf00132 -
16180 -pf00535 -
16185 -pf13614pf13807pf02706 -
BMC-P
BMC-H
BMC-T
Aldehyde dehydrogenase
PTAC
Signature enzyme
Alcohol dehydrogenase
Regulator

Salmonella enterica subsp. enterica serovar Eastbourne. Bacteria; Proteobacteria; Gammaproteobacteria; Enterobacterales;

AU911__locus_2

EUT1

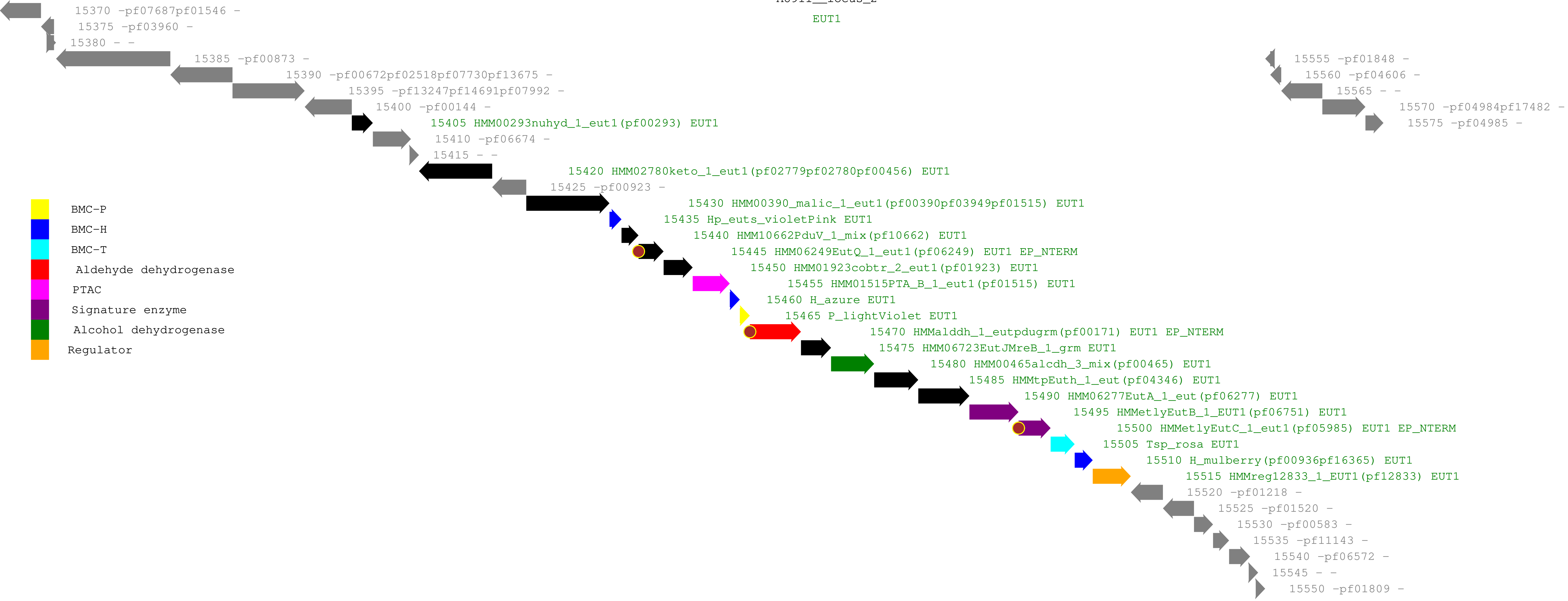

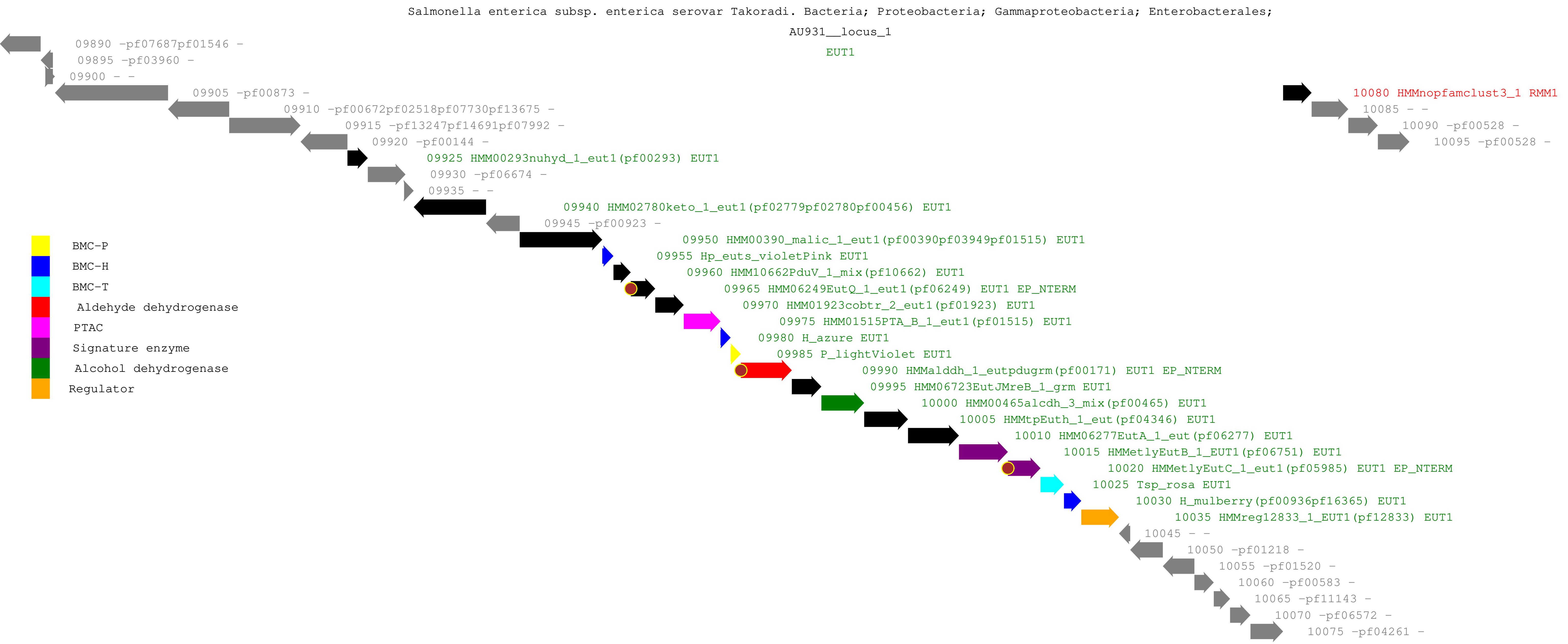
Salmonella enterica subsp. enterica serovar Takoradi. Bacteria; Proteobacteria; Gammaproteobacteria; Enterobacterales;
AU931__locus_1
EUT1
09890 -pf07687pf01546 -
09895 -pf03960 -
09900 - -
09905 -pf00873 -
09910 -pf00672pf02518pf07730pf13675 -
09915 -pf13247pf14691pf07992 -
09920 -pf00144 -
09925 HMM00293nuhyd_1_eut1(pf00293) EUT1
09930 -pf06674 -
09935 - -
09940 HMM02780keto_1_eut1(pf02779pf02780pf00456) EUT1
09945 -pf00923 -
09950 HMM00390_malic_1_eut1(pf00390pf03949pf01515) EUT1
09955 Hp_euts_violetPink EUT1
09960 HMM10662PduV_1_mix(pf10662) EUT1
09965 HMM06249EutQ_1_eut1(pf06249) EUT1 EP_NTERM
09970 HMM01923cobtr_2_eut1(pf01923) EUT1
09975 HMM01515PTA_B_1_eut1(pf01515) EUT1
09980 H_azure EUT1
09985 P_lightViolet EUT1
09990 HMMalddh_1_eutpdugrm(pf00171) EUT1 EP_NTERM
09995 HMM06723EutJMreB_1_grm EUT1
10000 HMM00465alcdh_3_mix(pf00465) EUT1
10005 HMMtpEuth_1_eut(pf04346) EUT1
10010 HMM06277EutA_1_eut(pf06277) EUT1
10015 HMMetlyEutB_1_EUT1(pf06751) EUT1
10020 HMMetlyEutC_1_eut1(pf05985) EUT1 EP_NTERM
10025 Tsp_rosa EUT1
10030 H_mulberry(pf00936pf16365) EUT1
10035 HMMreg12833_1_EUT1(pf12833) EUT1
10045 - -
10050 -pf01218 -
10055 -pf01520 -
10060 -pf00583 -
10065 -pf11143 -
10070 -pf06572 -
10075 -pf04261 -
10080 HMMnopfamclust3_1 RMM1
10085 - -
10090 -pf00528 -
10095 -pf00528 -
BMC-P
BMC-H
BMC-T
Aldehyde dehydrogenase
PTAC
Signature enzyme
Alcohol dehydrogenase
Regulator

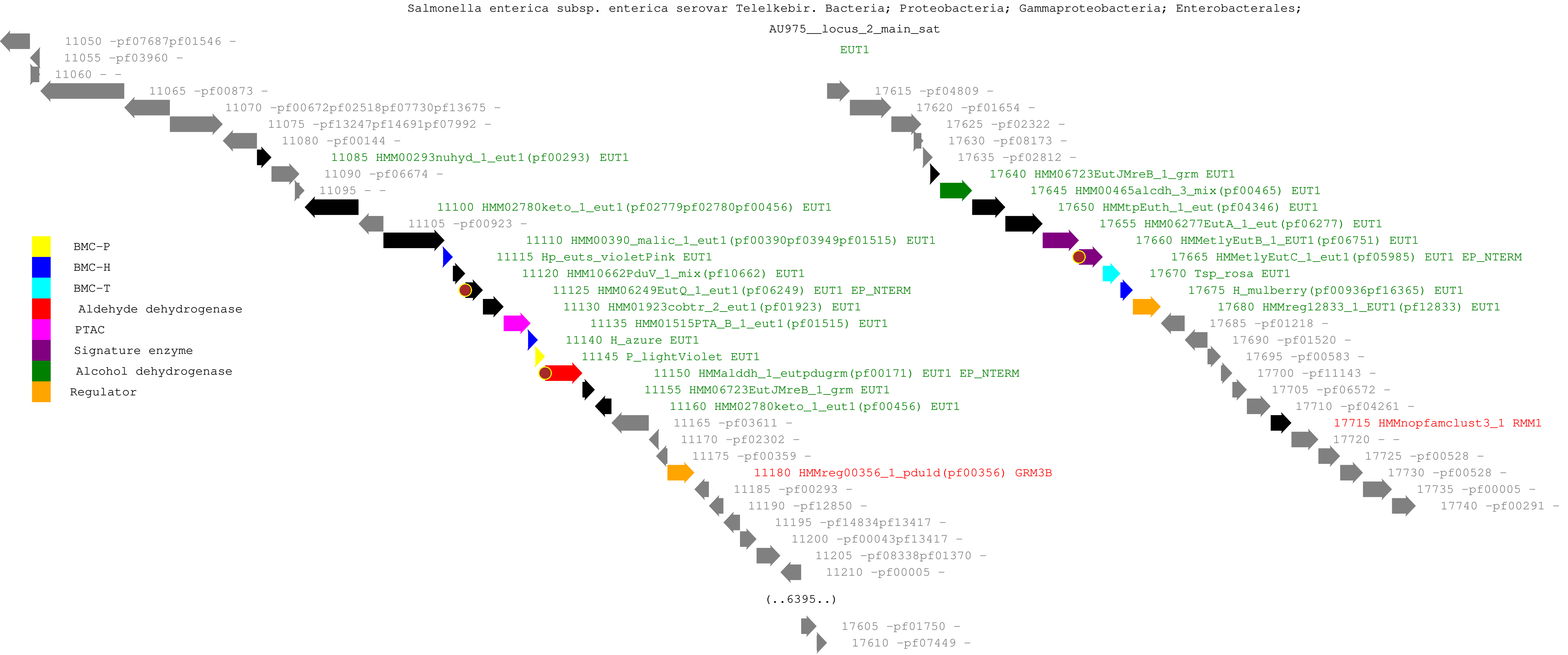
Salmonella enterica subsp. enterica serovar Telelkebir. Bacteria; Proteobacteria; Gammaproteobacteria; Enterobacterales;
AU975__locus_2_main_sat
EUT1
11050 -pf07687pf01546 -
11055 -pf03960 -
11060 - -
11065 -pf00873 -
11070 -pf00672pf02518pf07730pf13675 -
11075 -pf13247pf14691pf07992 -
11080 -pf00144 -
11085 HMM00293nuhyd_1_eut1(pf00293) EUT1
11090 -pf06674 -
11095 - -
11100 HMM02780keto_1_eut1(pf02779pf02780pf00456) EUT1
11105 -pf00923 -
11110 HMM00390_malic_1_eut1(pf00390pf03949pf01515) EUT1
11115 Hp_euts_violetPink EUT1
11120 HMM10662PduV_1_mix(pf10662) EUT1
11125 HMM06249EutQ_1_eut1(pf06249) EUT1 EP_NTERM
11130 HMM01923cobtr_2_eut1(pf01923) EUT1
11135 HMM01515PTA_B_1_eut1(pf01515) EUT1
11140 H_azure EUT1
11145 P_lightViolet EUT1
11150 HMMaldh_1_eutpdugrm(pf00171) EUT1 EP_NTERM
11155 HMM06723EutJMreB_1_grm EUT1
11160 HMM02780keto_1_eut1(pf00456) EUT1
11165 -pf03611 -
11170 -pf02302 -
11175 -pf00359 -
11180 HMMreg00356_1_pdu1d(pf00356) GRM3B
11185 -pf00293 -
11190 -pf12850 -
11195 -pf14834pf13417 -
11200 -pf00043pf13417 -
11205 -pf08338pf01370 -
11210 -pf00005 -
(..6395..)
17605 -pf01750 -
17610 -pf07449 -
17615 -pf04809 -
17620 -pf01654 -
17625 -pf02322 -
17630 -pf08173 -
17635 -pf02812 -
17640 HMM06723EutJMreB_1_grm EUT1
17645 HMM00465alcdh_3_mix(pf00465) EUT1
17650 HMMtpEuth_1_eut(pf04346) EUT1
17655 HMM06277EutA_1_eut(pf06277) EUT1
17660 HMMetlyEutB_1_EUT1(pf06751) EUT1
17665 HMMetlyEutC_1_eut1(pf05985) EUT1 EP_NTERM
17670 Tsp_rosa EUT1
17675 H_mulberry(pf00936pf16365) EUT1
17680 HMMreg12833_1_EUT1(pf12833) EUT1
17685 -pf01218 -
17690 -pf01520 -
17695 -pf00583 -
17700 -pf11143 -
17705 -pf06572 -
17710 -pf04261 -
17715 HMMnopfamclust3_1 RMM1
17720 - -
17725 -pf00528 -
17730 -pf00528 -
17735 -pf00005 -
17740 -pf00291 -
BMC-P
BMC-H
BMC-T
Aldehyde dehydrogenase
PTAC
Signature enzyme
Alcohol dehydrogenase
Regulator

Salmonella paratyphi B (strain ATCC BAA-1250 / SPB7). Bacteria; Proteobacteria; Gammaproteobacteria; Enterobacterales;

AU980__locus_2

EUT1

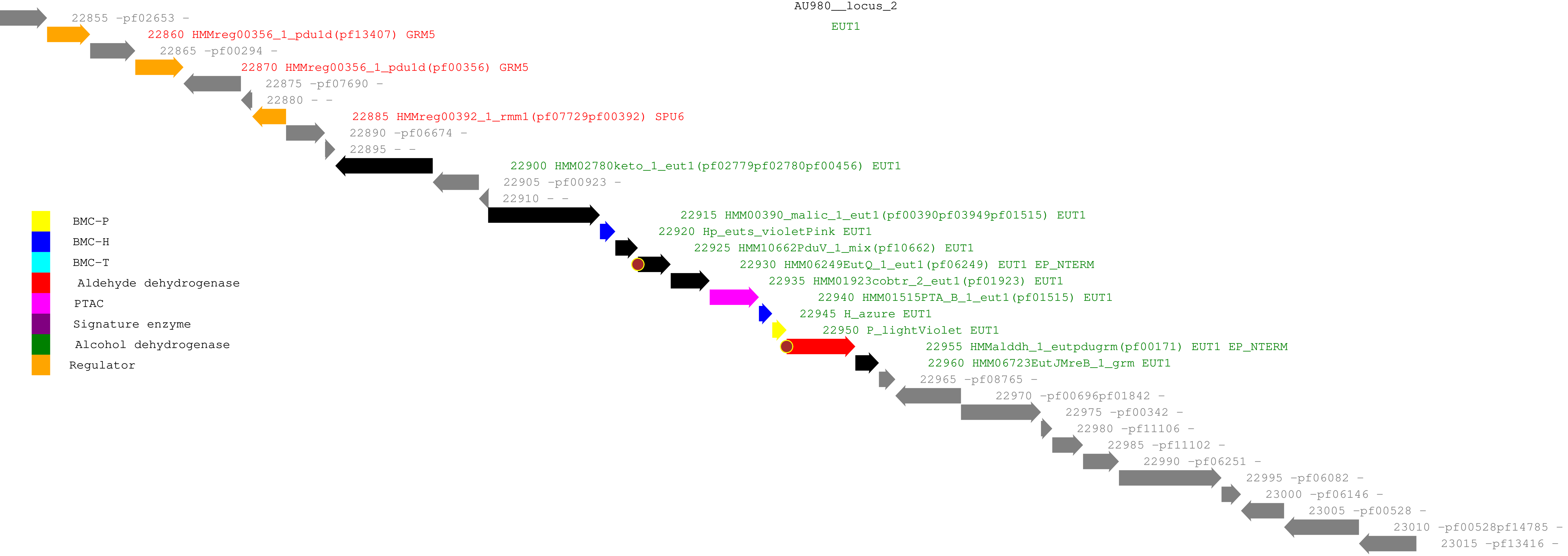

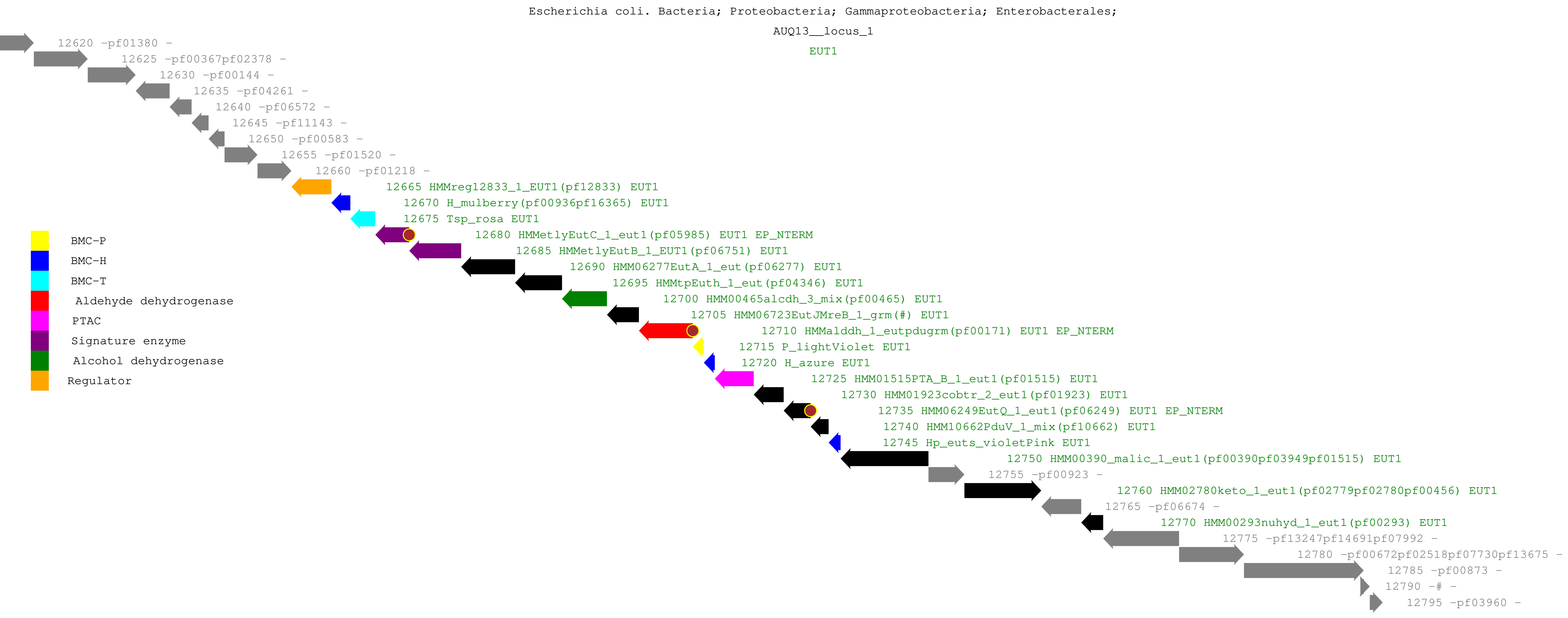

Escherichia coli. Bacteria; Proteobacteria; Gammaproteobacteria; Enterobacterales;
AUQ13__locus_1
EUT1
12620 -pf01380 -
12625 -pf00367pf02378 -
12630 -pf00144 -
12635 -pf04261 -
12640 -pf06572 -
12645 -pf11143 -
12650 -pf00583 -
12655 -pf01520 -
12660 -pf01218 -
12665 HMMreg12833_1_EUT1(pf12833) EUT1
12670 H_mulberry(pf00936pf16365) EUT1
12675 Tsp_rosa EUT1
12680 HMMetlyEutC_1_eut1(pf05985) EUT1 EP_NTERM
12685 HMMetlyEutB_1_EUT1(pf06751) EUT1
12690 HMM06277EutA_1_eut(pf06277) EUT1
12695 HMMtpEuth_1_eut(pf04346) EUT1
12700 HMM00465alcdh_3_mix(pf00465) EUT1
12705 HMM06723EutJMreB_1_grm(#) EUT1
12710 HMMalddh_1_eutpdugrm(pf00171) EUT1 EP_NTERM
12715 P_lightViolet EUT1
12720 H_azure EUT1
12725 HMM01515PTA_B_1_eut1(pf01515) EUT1
12730 HMM01923cobtr_2_eut1(pf01923) EUT1
12735 HMM06249EutQ_1_eut1(pf06249) EUT1 EP_NTERM
12740 HMM10662PduV_1_mix(pf10662) EUT1
12745 Hp_euts_violetPink EUT1
12750 HMM00390_malic_1_eut1(pf00390pf03949pf01515) EUT1
12755 -pf00923 -
12760 HMM02780keto_1_eut1(pf02779pf02780pf00456) EUT1
12765 -pf06674 -
12770 HMM00293nuhyd_1_eut1(pf00293) EUT1
12775 -pf13247pf14691pf07992 -
12780 -pf00672pf02518pf07730pf13675 -
12785 -pf00873 -
12790 -# -
12795 -pf03960 -
BMC-P
BMC-H
BMC-T
Aldehyde dehydrogenase
PTAC
Signature enzyme
Alcohol dehydrogenase
Regulator

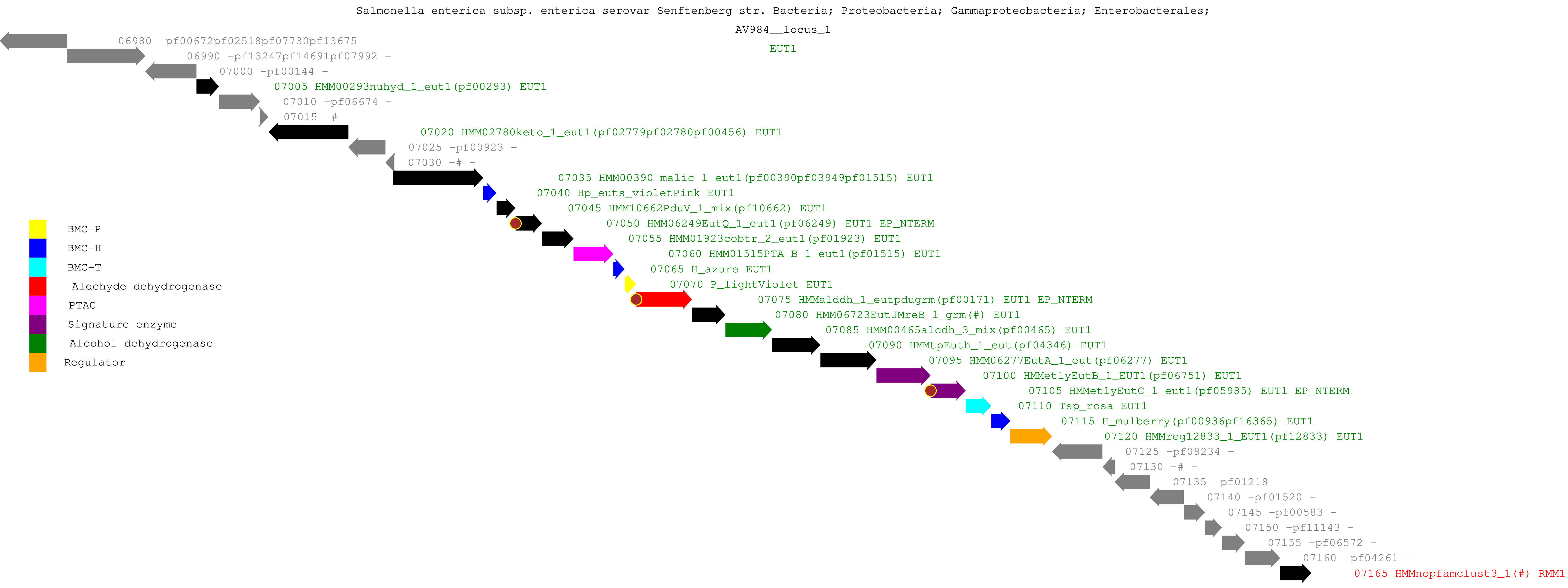
Salmonella enterica subsp. enterica serovar Senftenberg str. Bacteria; Proteobacteria; Gammaproteobacteria; Enterobacterales;
AV984__locus_1
EUT1
06980 -pf00672pf02518pf07730pf13675 -
06990 -pf13247pf14691pf07992 -
07000 -pf00144 -
07005 HMM00293nuhyd_1_eut1(pf00293) EUT1
07010 -pf06674 -
07015 -# -
07020 HMM02780keto_1_eut1(pf02779pf02780pf00456) EUT1
07025 -pf00923 -
07030 -# -
07035 HMM00390_malic_1_eut1(pf00390pf03949pf01515) EUT1
07040 Hp_euts_violetPink EUT1
07045 HMM10662PduV_1_mix(pf10662) EUT1
07050 HMM06249EutQ_1_eut1(pf06249) EUT1 EP_NTERM
07055 HMM01923cobtr_2_eut1(pf01923) EUT1
07060 HMM01515PTA_B_1_eut1(pf01515) EUT1
07065 H_azure EUT1
07070 P_lightViolet EUT1
07075 HMMalddh_1_eutpdugrm(pf00171) EUT1 EP_NTERM
07080 HMM06723EutJMreB_1_grm(#) EUT1
07085 HMM00465alcdh_3_mix(pf00465) EUT1
07090 HMMtpEuth_1_eut(pf04346) EUT1
07095 HMM06277EutA_1_eut(pf06277) EUT1
07100 HMMetlyEutB_1_EUT1(pf06751) EUT1
07105 HMMetlyEutC_1_eut1(pf05985) EUT1 EP_NTERM
07110 Tsp_rosa EUT1
07115 H_mulberry(pf00936pf16365) EUT1
07120 HMMreg12833_1_EUT1(pf12833) EUT1
07125 -pf09234 -
07130 -# -
07135 -pf01218 -
07140 -pf01520 -
07145 -pf00583 -
07150 -pf11143 -
07155 -pf06572 -
07160 -pf04261 -
07165 HMMnopfamclust3_1(#) RMM1
BMC-P
BMC-H
BMC-T
Aldehyde dehydrogenase
PTAC
Signature enzyme
Alcohol dehydrogenase
Regulator

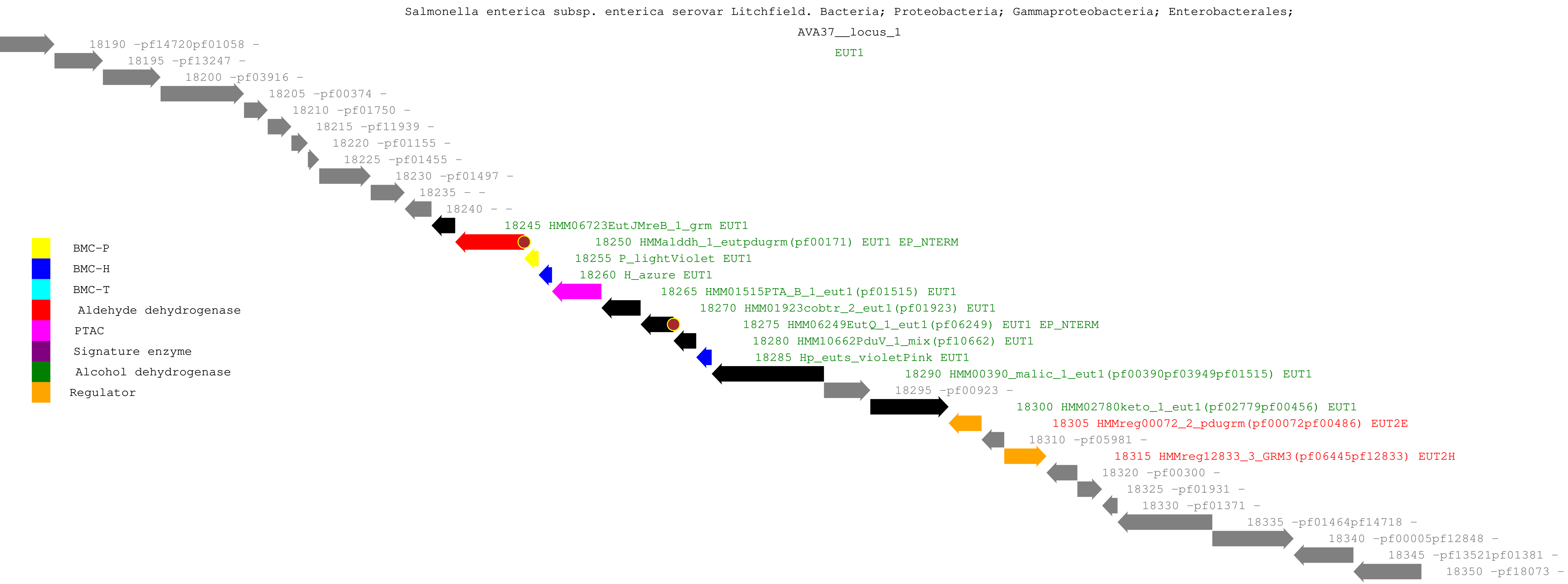
Salmonella enterica subsp. enterica serovar Litchfield. Bacteria; Proteobacteria; Gammaproteobacteria; Enterobacterales;
AVA37__locus_1
EUT1
18190 -pf14720pf01058 -
18195 -pf13247 -
18200 -pf03916 -
18205 -pf00374 -
18210 -pf01750 -
18215 -pf11939 -
18220 -pf01155 -
18225 -pf01455 -
18230 -pf01497 -
18235 - -
18240 - -
18245 HMM06723EutJMreB_1_grm EUT1
18250 HMMalddh_1_eutpdugrm(pf00171) EUT1 EP_NTERM
18255 P_lightViolet EUT1
18260 H_azure EUT1
18265 HMM01515PTA_B_1_eut1(pf01515) EUT1
18270 HMM01923cobtr_2_eut1(pf01923) EUT1
18275 HMM06249EutQ_1_eut1(pf06249) EUT1 EP_NTERM
18280 HMM10662PduV_1_mix(pf10662) EUT1
18285 Hp_euts_violetPink EUT1
18290 HMM00390_malic_1_eut1(pf00390pf03949pf01515) EUT1
18295 -pf00923 -
18300 HMM02780keto_1_eut1(pf02779pf00456) EUT1
18305 HMMreg00072_2_pdugrm(pf00072pf00486) EUT2E
18310 -pf05981 -
18315 HMMreg12833_3_GRM3(pf06445pf12833) EUT2H
18320 -pf00300 -
18325 -pf01931 -
18330 -pf01371 -
18335 -pf01464pf14718 -
18340 -pf00005pf12848 -
18345 -pf13521pf01381 -
18350 -pf18073 -
BMC-P
BMC-H
BMC-T
Aldehyde dehydrogenase
PTAC
Signature enzyme
Alcohol dehydrogenase
Regulator

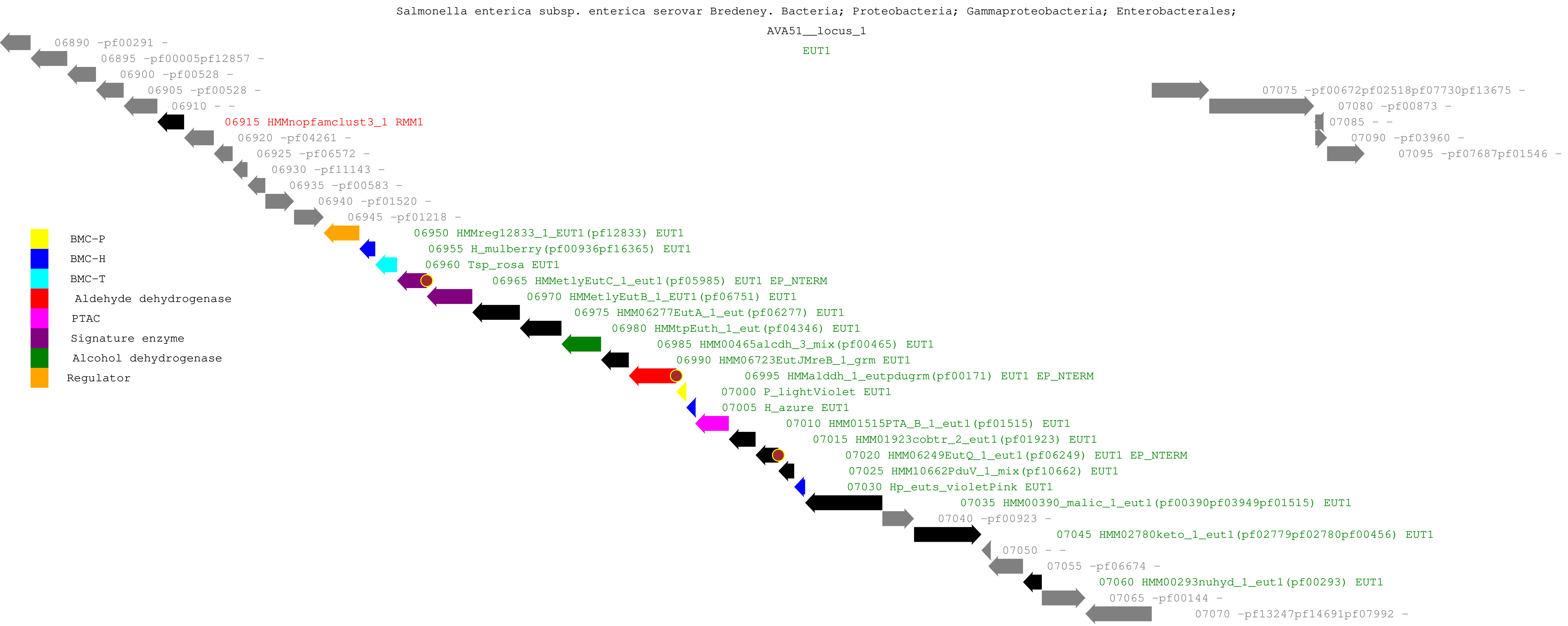

Salmonella enterica subsp. enterica serovar Bredeney. Bacteria; Proteobacteria; Gammaproteobacteria; Enterobacterales;
AVA51__locus_1
EUT1
06890 -pf00291 -
06895 -pf00005pf12857 -
06900 -pf00528 -
06905 -pf00528 -
06910 - -
06915 HMMnopfamclust3_1 RMM1
06920 -pf04261 -
06925 -pf06572 -
06930 -pf11143 -
06935 -pf00583 -
06940 -pf01520 -
06945 -pf01218 -
06950 HMMreg12833_1_EUT1(pf12833) EUT1
06955 H_mulberry(pf00936pf16365) EUT1
06960 Tsp_rosa EUT1
06965 HMMetlyEutC_1_eut1(pf05985) EUT1 EP_NTERM
06970 HMMetlyEutB_1_EUT1(pf06751) EUT1
06975 HMM06277EutA_1_eut(pf06277) EUT1
06980 HMMtpEuth_1_eut(pf04346) EUT1
06985 HMM00465alcdh_3_mix(pf00465) EUT1
06990 HMM06723EutJMreB_1_grm EUT1
06995 HMMalddh_1_eutpdugrm(pf00171) EUT1 EP_NTERM
07000 P_lightViolet EUT1
07005 H_azure EUT1
07010 HMM01515PTA_B_1_eut1(pf01515) EUT1
07015 HMM01923cobtr_2_eut1(pf01923) EUT1
07020 HMM06249EutQ_1_eut1(pf06249) EUT1 EP_NTERM
07025 HMM10662PduV_1_mix(pf10662) EUT1
07030 Hp_euts_violetPink EUT1
07035 HMM00390_malic_1_eut1(pf00390pf03949pf01515) EUT1
07040 -pf00923 -
07045 HMM02780keto_1_eut1(pf02779pf02780pf00456) EUT1
07050 - -
07055 -pf06674 -
07060 HMM00293nuhyd_1_eut1(pf00293) EUT1
07065 -pf00144 -
07070 -pf13247pf14691pf07992 -
07075 -pf00672pf02518pf07730pf13675 -
07080 -pf00873 -
07085 - -
07090 -pf03960 -
07095 -pf07687pf01546 -
BMC-P
BMC-H
BMC-T
Aldehyde dehydrogenase
PTAC
Signature enzyme
Alcohol dehydrogenase
Regulator

Salmonella enterica subsp. enterica serovar Altona. Bacteria; Proteobacteria; Gammaproteobacteria; Enterobacterales;

AVA57__locus_3_main_sat

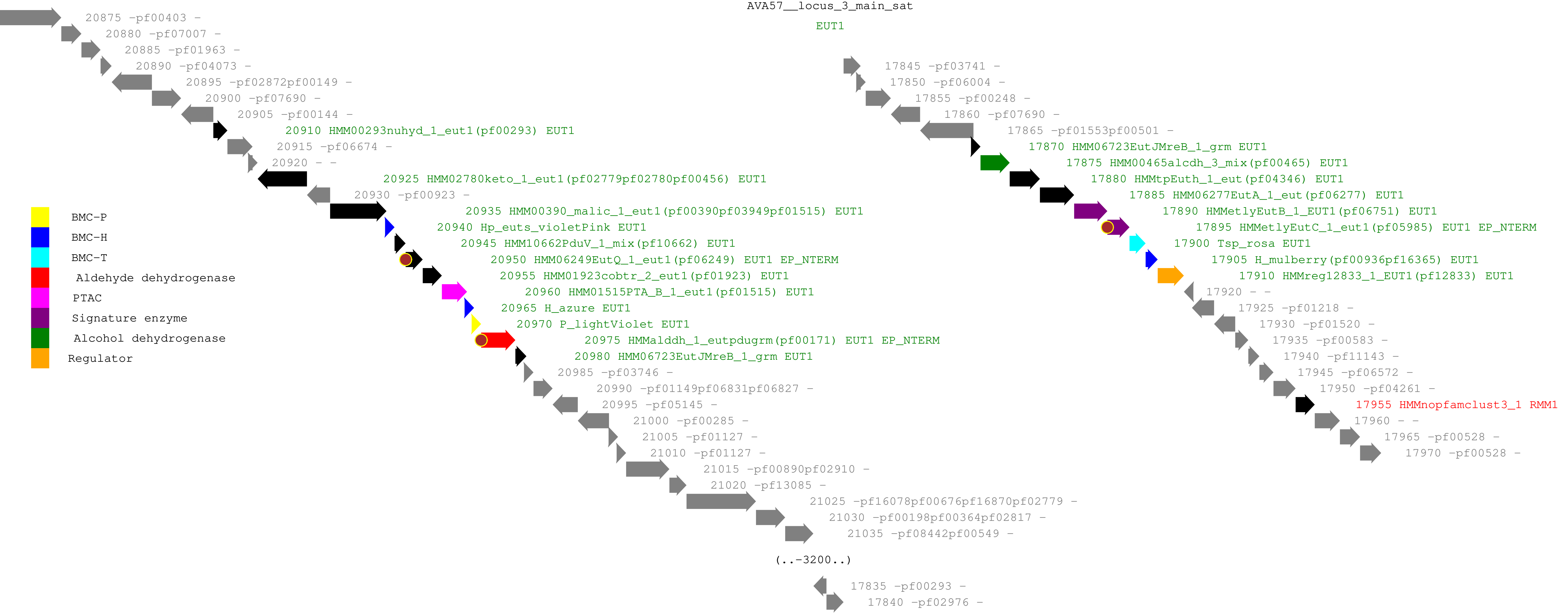

Salmonella enterica subsp. enterica serovar Nima. Bacteria; Proteobacteria; Gammaproteobacteria; Enterobacterales;

AVA78__locus_2_main_sat

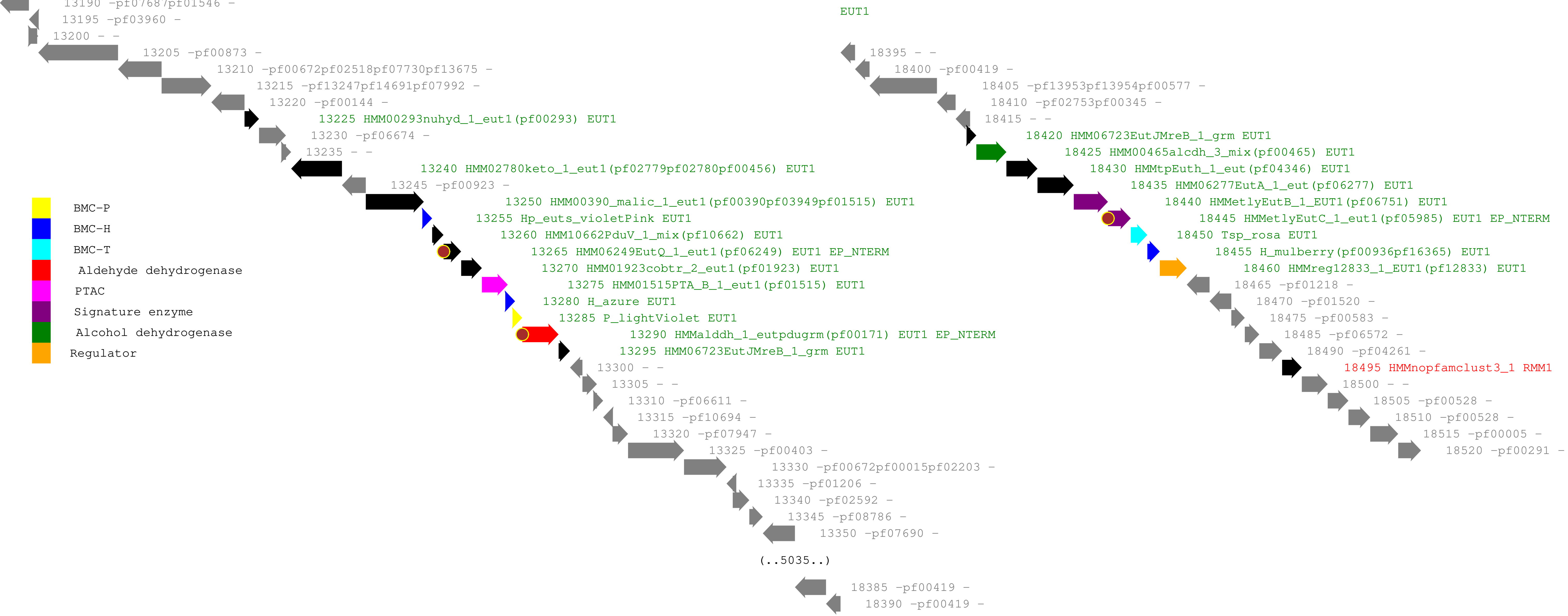

Salmonella enterica subsp. enterica serovar Livingstone. Bacteria; Proteobacteria; Gammaproteobacteria; Enterobacterales;

AVA86__locus_2_main_sat

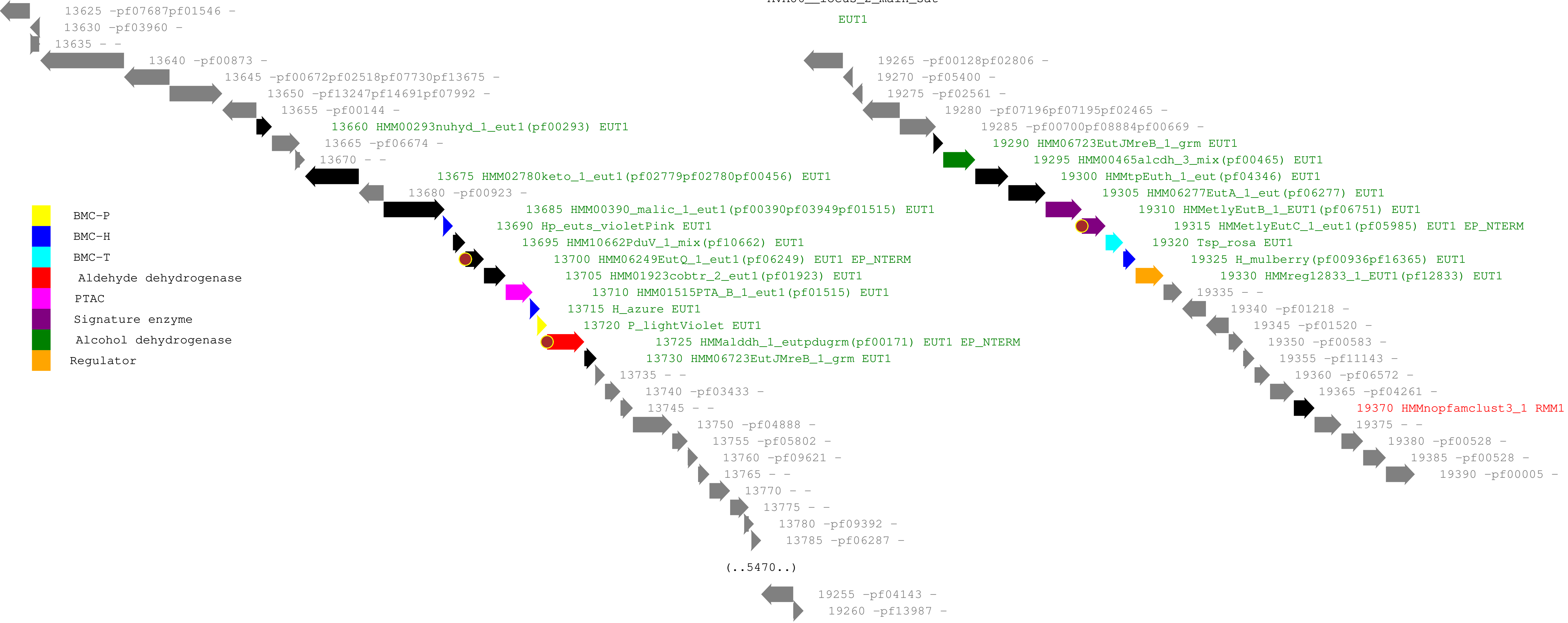

Salmonella enterica subsp. enterica serovar Hvittingfoss. Bacteria; Proteobacteria; Gammaproteobacteria; Enterobacterales;

AVB75__locus_2_main_sat

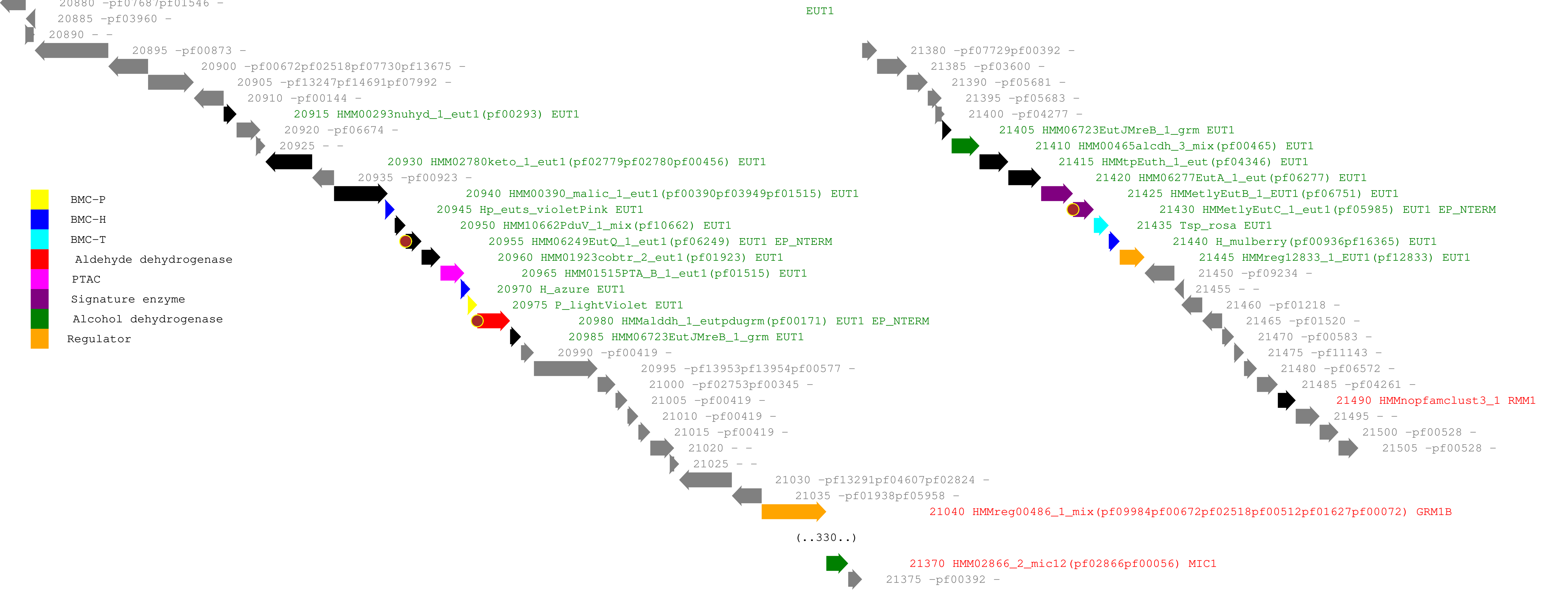

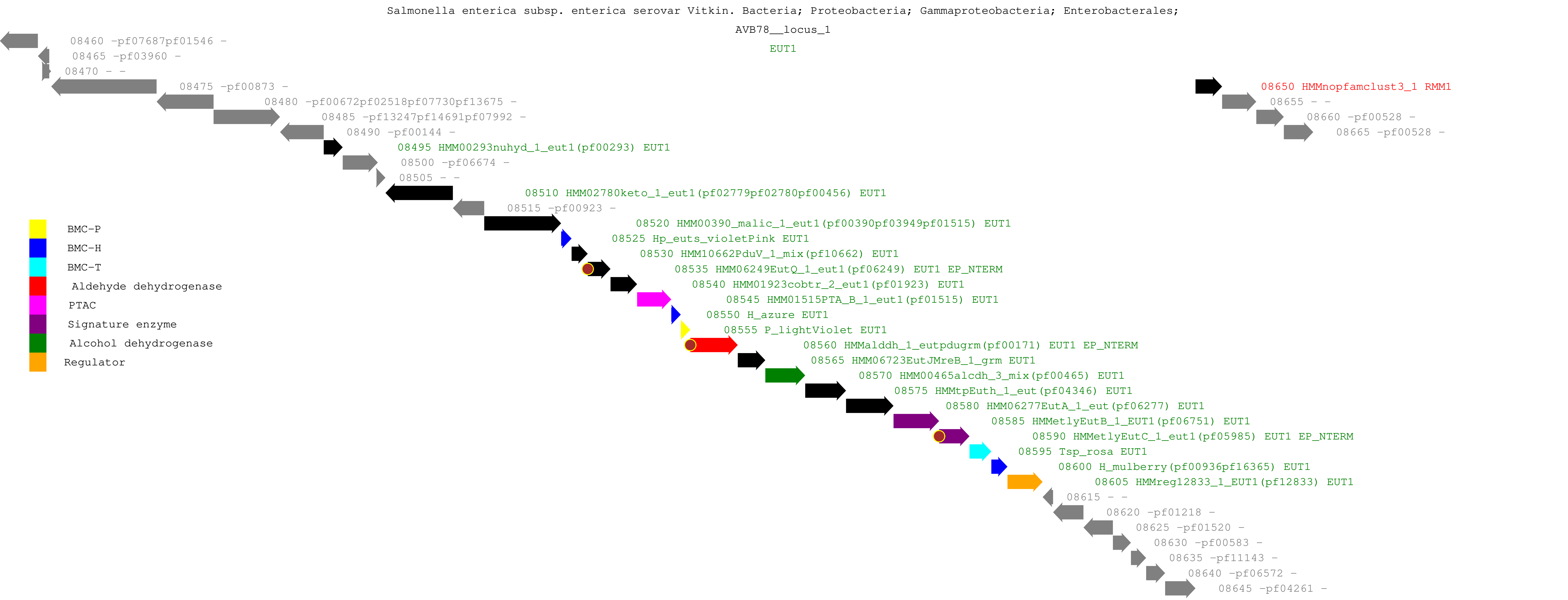

Salmonella enterica subsp. enterica serovar Vitkin. Bacteria; Proteobacteria; Gammaproteobacteria; Enterobacterales;
AVB78__locus_1
EUT1
08460 -pf07687pf01546 -
08465 -pf03960 -
08470 - -
08475 -pf00873 -
08480 -pf00672pf02518pf07730pf13675 -
08485 -pf13247pf14691pf07992 -
08490 -pf00144 -
08495 HMM00293nuhyd_1_eut1(pf00293) EUT1
08500 -pf06674 -
08505 - -
08510 HMM02780keto_1_eut1(pf02779pf02780pf00456) EUT1
08515 -pf00923 -
08520 HMM00390_malic_1_eut1(pf00390pf03949pf01515) EUT1
08525 Hp_euts_violetPink EUT1
08530 HMM10662PduV_1_mix(pf10662) EUT1
08535 HMM06249EutQ_1_eut1(pf06249) EUT1 EP_NTERM
08540 HMM01923cobtr_2_eut1(pf01923) EUT1
08545 HMM01515PTA_B_1_eut1(pf01515) EUT1
08550 H_azure EUT1
08555 P_lightViolet EUT1
08560 HMMalddh_1_eutpdugrm(pf00171) EUT1 EP_NTERM
08565 HMM06723EutJMreB_1_grm EUT1
08570 HMM00465alcdh_3_mix(pf00465) EUT1
08575 HMMtpEuth_1_eut(pf04346) EUT1
08580 HMM06277EutA_1_eut(pf06277) EUT1
08585 HMMetlyEutB_1_EUT1(pf06751) EUT1
08590 HMMetlyEutC_1_eut1(pf05985) EUT1 EP_NTERM
08595 Tsp_rosa EUT1
08600 H_mulberry(pf00936pf16365) EUT1
08605 HMMreg12833_1_EUT1(pf12833) EUT1
08615 - -
08620 -pf01218 -
08625 -pf01520 -
08630 -pf00583 -
08635 -pf11143 -
08640 -pf06572 -
08645 -pf04261 -
08650 HMMnopfamclust3_1 RMM1
08655 - -
08660 -pf00528 -
08665 -pf00528 -
BMC-P
BMC-H
BMC-T
Aldehyde dehydrogenase
PTAC
Signature enzyme
Alcohol dehydrogenase
Regulator

Salmonella enterica subsp. enterica serovar Ajiobo. Bacteria; Proteobacteria; Gammaproteobacteria; Enterobacterales;

AVB95__locus_2

EUT1

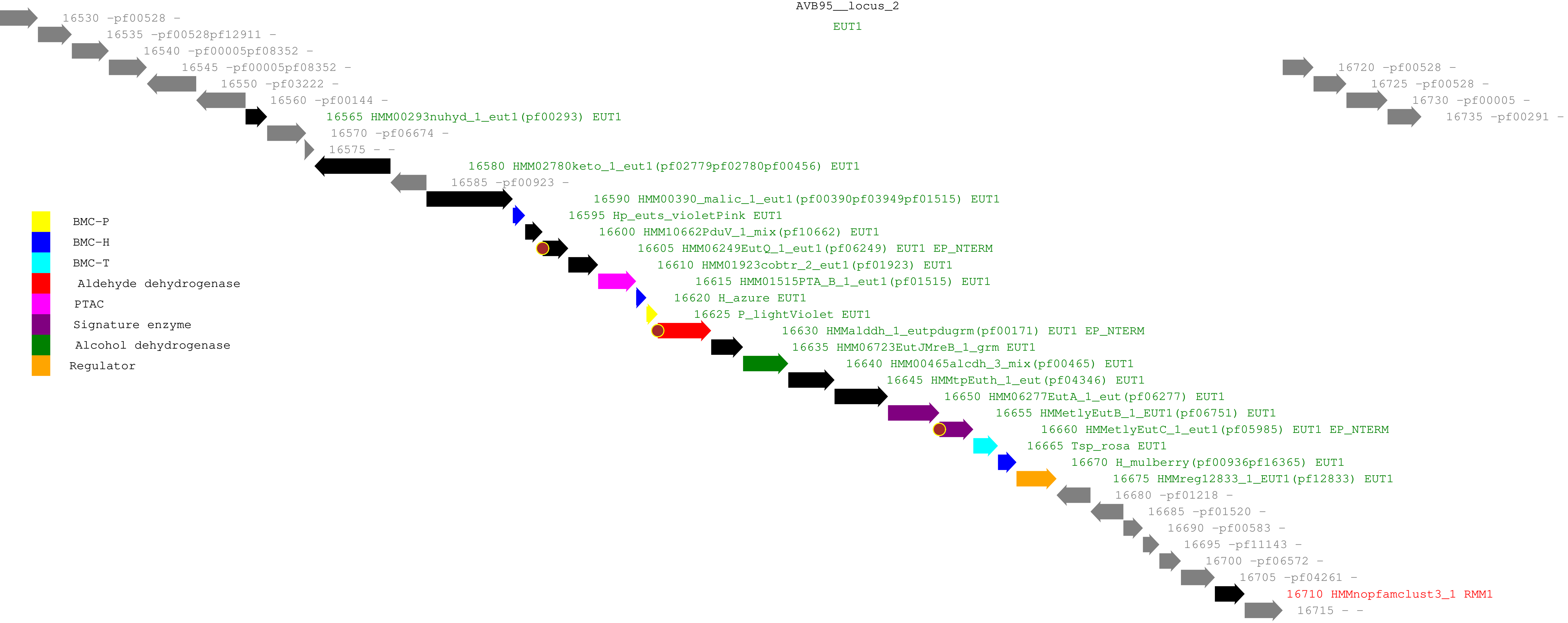

Salmonella enterica subsp. enterica serovar Kambole. Bacteria; Proteobacteria; Gammaproteobacteria; Enterobacterales;

AVC51__locus_1_main_sat

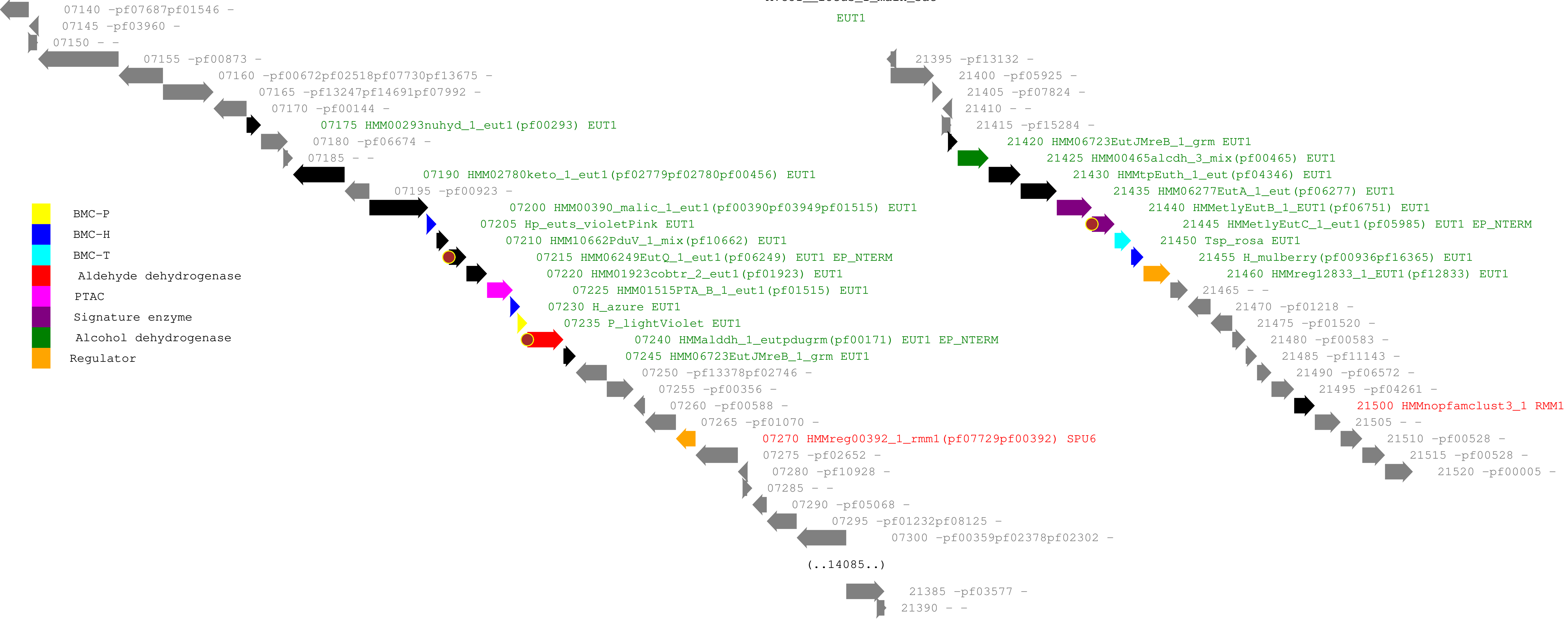

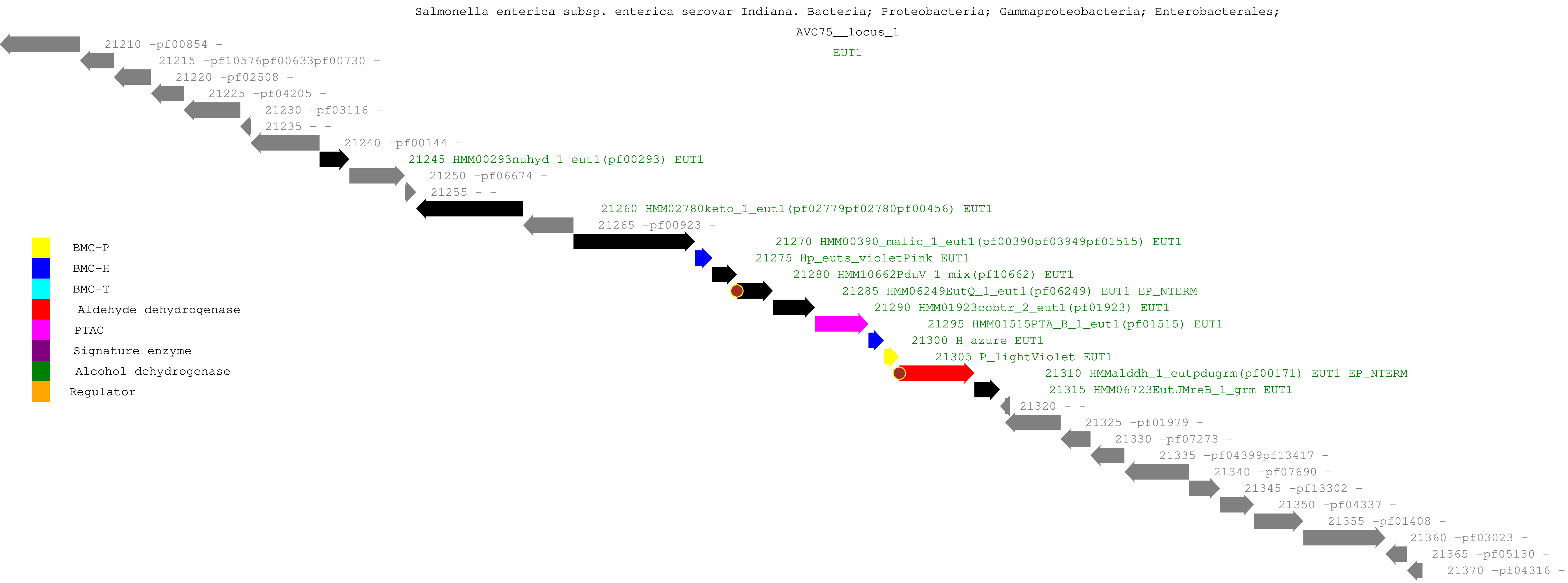
Salmonella enterica subsp. enterica serovar Indiana. Bacteria; Proteobacteria; Gammaproteobacteria; Enterobacterales;
AVC75__locus_1
EUT1
21210 -pf00854 -
21215 -pf10576pf00633pf00730 -
21220 -pf02508 -
21225 -pf04205 -
21230 -pf03116 -
21235 - -
21240 -pf00144 -
21245 HMM00293nuhyd_1_eut1(pf00293) EUT1
21250 -pf06674 -
21255 - -
21260 HMM02780keto_1_eut1(pf02779pf02780pf00456) EUT1
21265 -pf00923 -
21270 HMM00390_malic_1_eut1(pf00390pf03949pf01515) EUT1
21275 Hp_euts_violetPink EUT1
21280 HMM10662PduV_1_mix(pf10662) EUT1
21285 HMM06249EutQ_1_eut1(pf06249) EUT1 EP_NTERM
21290 HMM01923cobtr_2_eut1(pf01923) EUT1
21295 HMM01515PTA_B_1_eut1(pf01515) EUT1
21300 H_azure EUT1
21305 P_lightViolet EUT1
21310 HMMalddh_1_eutpdugrm(pf00171) EUT1 EP_NTERM
21315 HMM06723EutJMreB_1_grm EUT1
21320 - -
21325 -pf01979 -
21330 -pf07273 -
21335 -pf04399pf13417 -
21340 -pf07690 -
21345 -pf13302 -
21350 -pf04337 -
21355 -pf01408 -
21360 -pf03023 -
21365 -pf05130 -
21370 -pf04316 -
BMC-P
BMC-H
BMC-T
Aldehyde dehydrogenase
PTAC
Signature enzyme
Alcohol dehydrogenase
Regulator

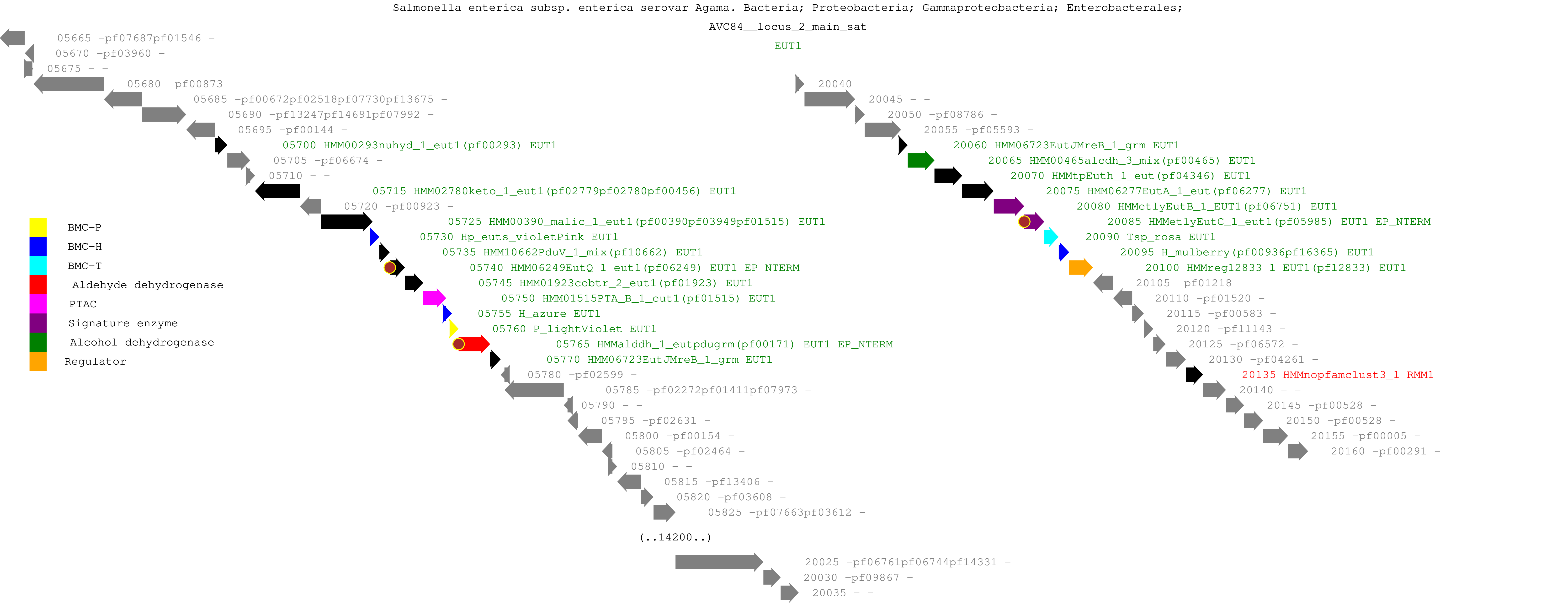
Salmonella enterica subsp. enterica serovar Agama. Bacteria; Proteobacteria; Gammaproteobacteria; Enterobacterales;
AVC84__locus_2_main_sat
EUT1
05665 -pf07687pf01546 -
05670 -pf03960 -
05675 - -
05680 -pf00873 -
05685 -pf00672pf02518pf07730pf13675 -
05690 -pf13247pf14691pf07992 -
05695 -pf00144 -
05700 HMM00293nuhyd_1_eut1(pf00293) EUT1
05705 -pf06674 -
05710 - -
05715 HMM02780keto_1_eut1(pf02779pf02780pf00456) EUT1
05720 -pf00923 -
05725 HMM00390_malic_1_eut1(pf00390pf03949pf01515) EUT1
05730 Hp_euts_violetPink EUT1
05735 HMM10662PduV_1_mix(pf10662) EUT1
05740 HMM06249EutQ_1_eut1(pf06249) EUT1 EP_NTERM
05745 HMM01923cobtr_2_eut1(pf01923) EUT1
05750 HMM01515PTA_B_1_eut1(pf01515) EUT1
05755 H_azure EUT1
05760 P_lightViolet EUT1
05765 HMMalddh_1_eutpdugrm(pf00171) EUT1 EP_NTERM
05770 HMM06723EutJMreB_1_grm EUT1
05780 -pf02599 -
05785 -pf02272pf01411pf07973 -
05790 - -
05795 -pf02631 -
05800 -pf00154 -
05805 -pf02464 -
05810 - -
05815 -pf13406 -
05820 -pf03608 -
05825 -pf07663pf03612 -
(..14200..)
20025 -pf06761pf06744pf14331 -
20030 -pf09867 -
20035 - -
20040 - -
20045 - -
20050 -pf08786 -
20055 -pf05593 -
20060 HMM06723EutJMreB_1_grm EUT1
20065 HMM00465alcdh_3_mix(pf00465) EUT1
20070 HMMtpEuth_1_eut(pf04346) EUT1
20075 HMM06277EutA_1_eut(pf06277) EUT1
20080 HMMetlyEutB_1_EUT1(pf06751) EUT1
20085 HMMetlyEutC_1_eut1(pf05985) EUT1 EP_NTERM
20090 Tsp_rosa EUT1
20095 H_mulberry(pf00936pf16365) EUT1
20100 HMMreg12833_1_EUT1(pf12833) EUT1
20105 -pf01218 -
20110 -pf01520 -
20115 -pf00583 -
20120 -pf11143 -
20125 -pf06572 -
20130 -pf04261 -
20135 HMMnopfamclust3_1 RMM1
20140 - -
20145 -pf00528 -
20150 -pf00528 -
20155 -pf00005 -
20160 -pf00291 -
BMC-P
BMC-H
BMC-T
Aldehyde dehydrogenase
PTAC
Signature enzyme
Alcohol dehydrogenase
Regulator

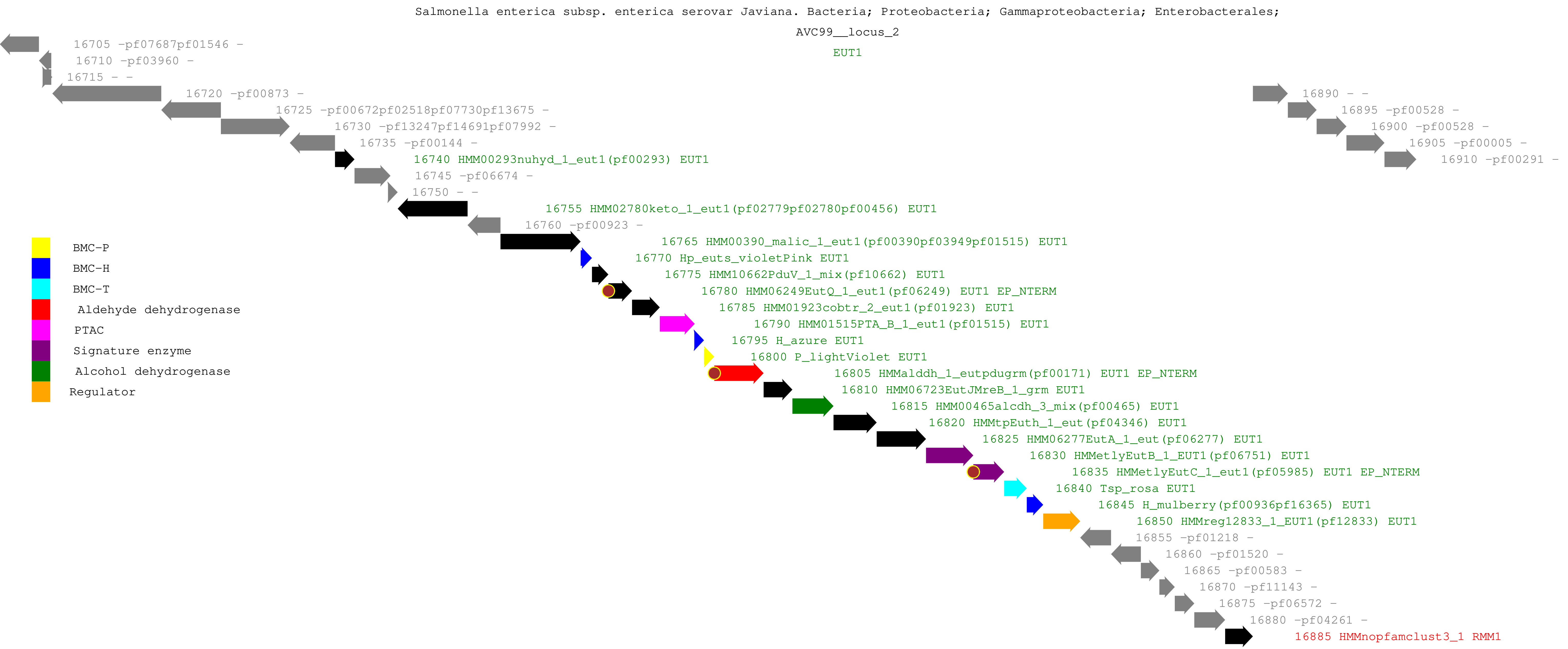
Salmonella enterica subsp. enterica serovar Javiana. Bacteria; Proteobacteria; Gammaproteobacteria; Enterobacterales;
AVC99__locus_2
EUT1
16705 -pf07687pf01546 -
16710 -pf03960 -
16715 - -
16720 -pf00873 -
16725 -pf00672pf02518pf07730pf13675 -
16730 -pf13247pf14691pf07992 -
16735 -pf00144 -
16740 HMM00293nuhyd_1_eut1(pf00293) EUT1
16745 -pf06674 -
16750 - -
16755 HMM02780keto_1_eut1(pf02779pf02780pf00456) EUT1
16760 -pf00923 -
16765 HMM00390_malic_1_eut1(pf00390pf03949pf01515) EUT1
16770 Hp_euts_violetPink EUT1
16775 HMM10662PduV_1_mix(pf10662) EUT1
16780 HMM06249EutQ_1_eut1(pf06249) EUT1 EP_NTERM
16785 HMM01923cobtr_2_eut1(pf01923) EUT1
16790 HMM01515PTA_B_1_eut1(pf01515) EUT1
16795 H_azure EUT1
16800 P_lightViolet EUT1
16805 HMMalddh_1_eutpdugrm(pf00171) EUT1 EP_NTERM
16810 HMM06723EutJMreB_1_grm EUT1
16815 HMM00465alcdh_3_mix(pf00465) EUT1
16820 HMMtpEuth_1_eut(pf04346) EUT1
16825 HMM06277EutA_1_eut(pf06277) EUT1
16830 HMMetlyEutB_1_EUT1(pf06751) EUT1
16835 HMMetlyEutC_1_eut1(pf05985) EUT1 EP_NTERM
16840 Tsp_rosa EUT1
16845 H_mulberry(pf00936pf16365) EUT1
16850 HMMreg12833_1_EUT1(pf12833) EUT1
16855 -pf01218 -
16860 -pf01520 -
16865 -pf00583 -
16870 -pf11143 -
16875 -pf06572 -
16880 -pf04261 -
16885 HMMnopfamclust3_1 RMM1
16890 - -
16895 -pf00528 -
16900 -pf00528 -
16905 -pf00005 -
16910 -pf00291 -
BMC-P
BMC-H
BMC-T
Aldehyde dehydrogenase
PTAC
Signature enzyme
Alcohol dehydrogenase
Regulator

Salmonella enterica subsp. enterica serovar Alachua. Bacteria; Proteobacteria; Gammaproteobacteria; Enterobacterales;

AVD57__locus_2

EUT1

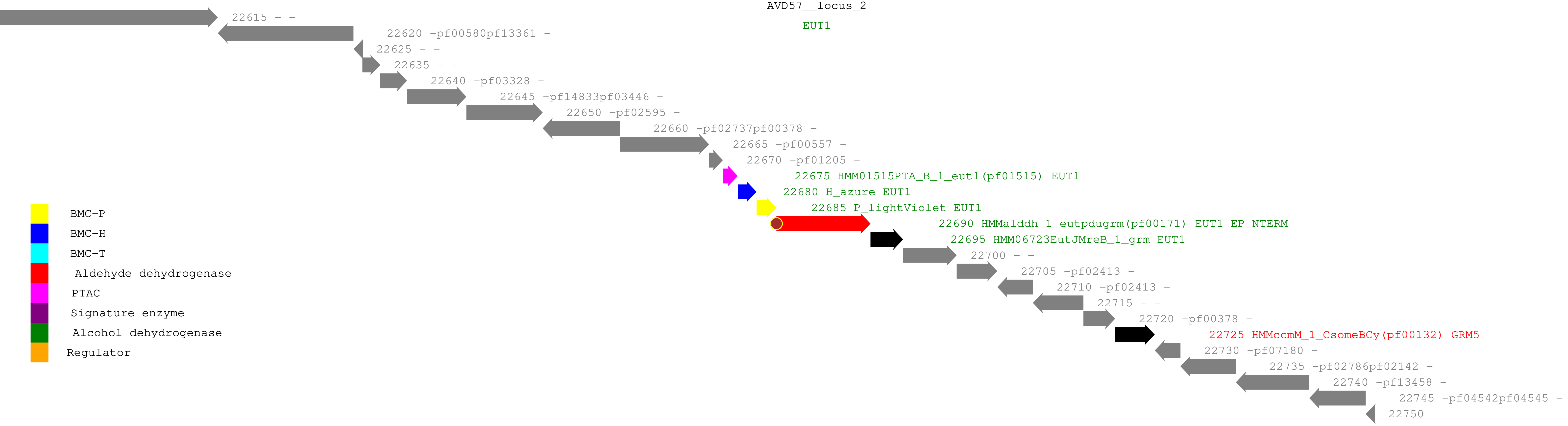

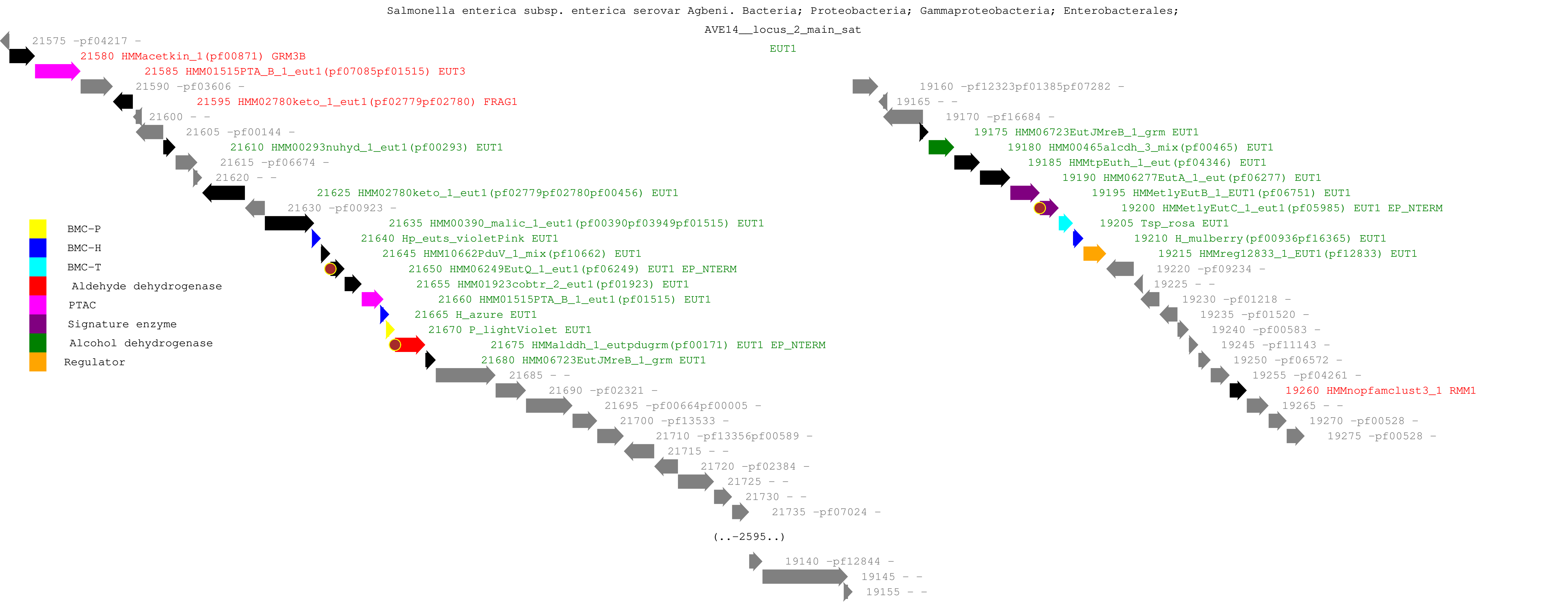
Salmonella enterica subsp. enterica serovar Agbeni. Bacteria; Proteobacteria; Gammaproteobacteria; Enterobacterales;
AVE14__locus_2_main_sat
EUT1
21575 -pf04217 -
21580 HMMacetkin_1(pf00871) GRM3B
21585 HMM01515PTA_B_1_eut1(pf07085pf01515) EUT3
21590 -pf03606 -
21595 HMM02780keto_1_eut1(pf02779pf02780) FRAG1
21600 - -
21605 -pf00144 -
21610 HMM00293nuhyd_1_eut1(pf00293) EUT1
21615 -pf06674 -
21620 - -
21625 HMM02780keto_1_eut1(pf02779pf02780pf00456) EUT1
21630 -pf00923 -
21635 HMM00390_malic_1_eut1(pf00390pf03949pf01515) EUT1
21640 Hp_euts_violetPink EUT1
21645 HMM10662PduV_1_mix(pf10662) EUT1
21650 HMM06249EutQ_1_eut1(pf06249) EUT1 EP_NTERM
21655 HMM01923cobtr_2_eut1(pf01923) EUT1
21660 HMM01515PTA_B_1_eut1(pf01515) EUT1
21665 H_azure EUT1
21670 P_lightViolet EUT1
21675 HMMalddh_1_eutpdugrm(pf00171) EUT1 EP_NTERM
21680 HMM06723EutJMreB_1_grm EUT1
21685 - -
21690 -pf02321 -
21695 -pf00664pf00005 -
21700 -pf13533 -
21710 -pf13356pf00589 -
21715 - -
21720 -pf02384 -
21725 - -
21730 - -
21735 -pf07024 -
(..-2595..)
19140 -pf12844 -
19145 - -
19155 - -
19160 -pf12323pf01385pf07282 -
19165 - -
19170 -pf16684 -
19175 HMM06723EutJMreB_1_grm EUT1
19180 HMM00465alcdh_3_mix(pf00465) EUT1
19185 HMMtpEuth_1_eut(pf04346) EUT1
19190 HMM06277EutA_1_eut(pf06277) EUT1
19195 HMMetlyEutB_1_EUT1(pf06751) EUT1
19200 HMMetlyEutC_1_eut1(pf05985) EUT1 EP_NTERM
19205 Tsp_rosa EUT1
19210 H_mulberry(pf00936pf16365) EUT1
19215 HMMreg12833_1_EUT1(pf12833) EUT1
19220 -pf09234 -
19225 - -
19230 -pf01218 -
19235 -pf01520 -
19240 -pf00583 -
19245 -pf11143 -
19250 -pf06572 -
19255 -pf04261 -
19260 HMMnopfamclust3_1 RMM1
19265 - -
19270 -pf00528 -
19275 -pf00528 -
BMC-P
BMC-H
BMC-T
Aldehyde dehydrogenase
PTAC
Signature enzyme
Alcohol dehydrogenase
Regulator

Salmonella enterica subsp. enterica serovar Havana. Bacteria; Proteobacteria; Gammaproteobacteria; Enterobacterales;

AVF28__locus_2

EUT1

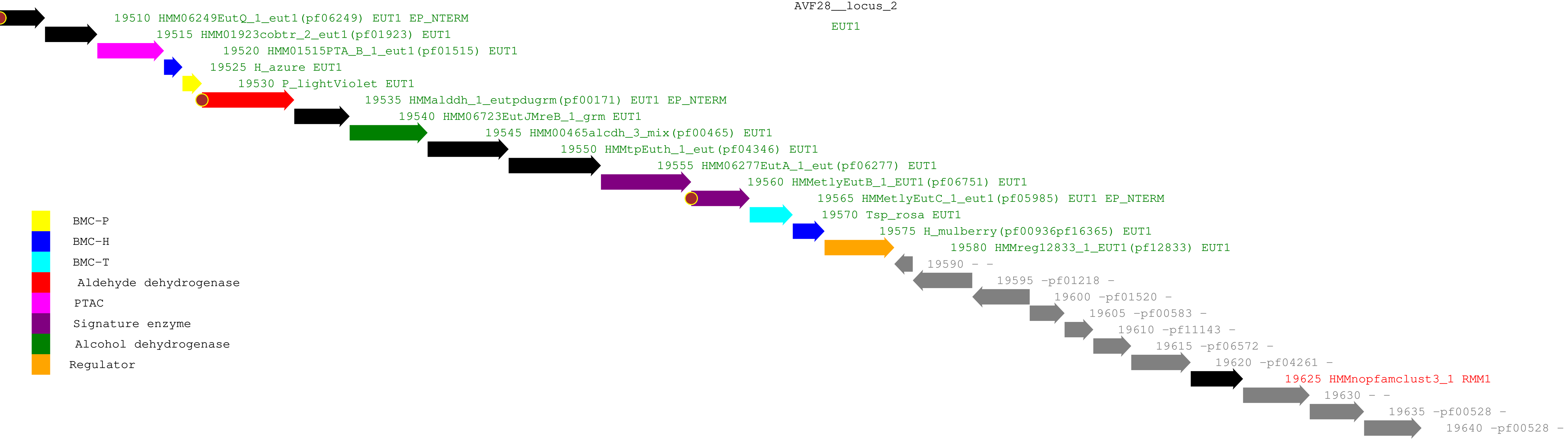

Salmonella enterica subsp. enterica serovar Mikawasima. Bacteria; Proteobacteria; Gammaproteobacteria; Enterobacterales;

AVF35__locus_2_main_sat

EUT1

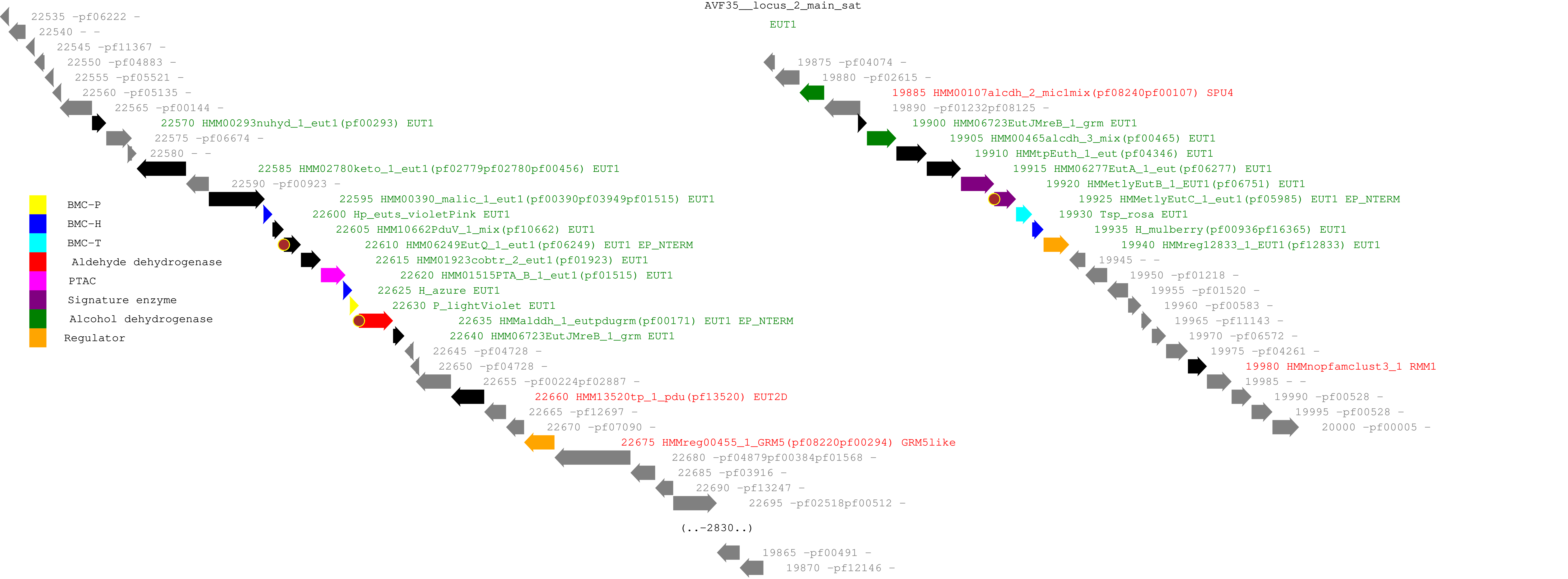

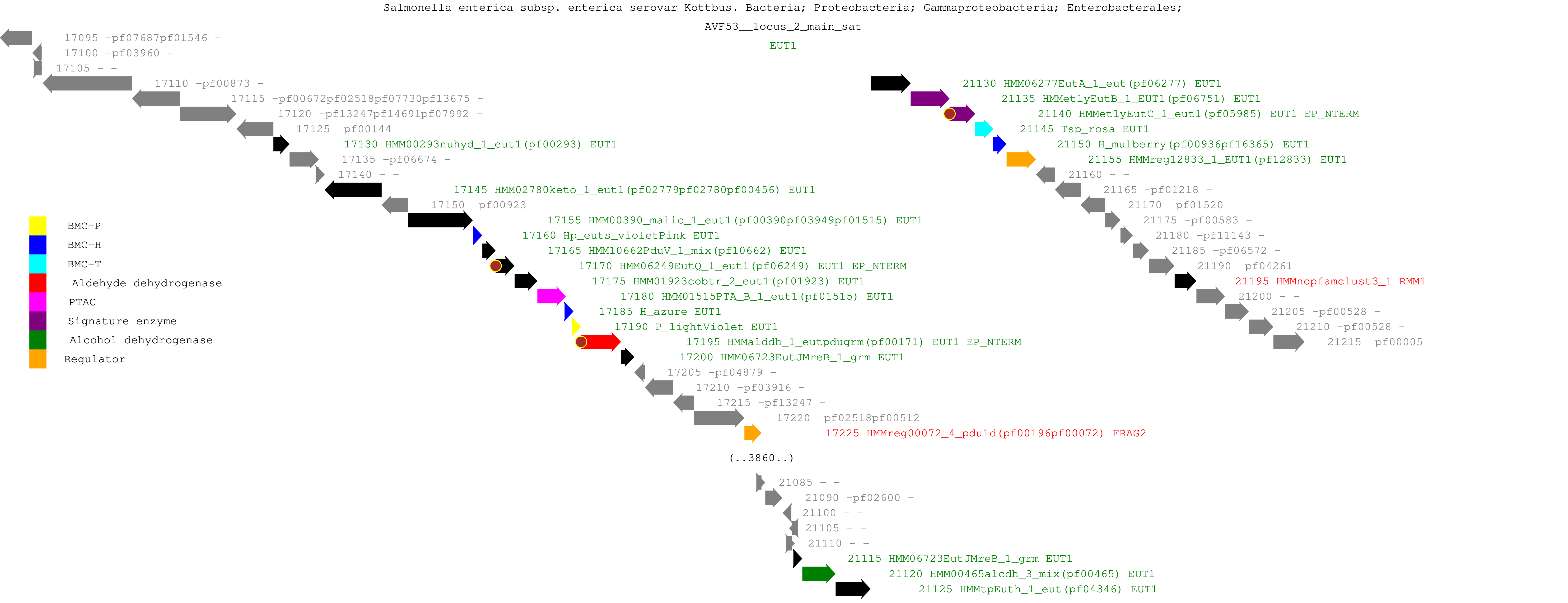
Salmonella enterica subsp. enterica serovar Kottbus. Bacteria; Proteobacteria; Gammaproteobacteria; Enterobacterales;
AVF53__locus_2_main_sat
EUT1
17095 -pf07687pf01546 -
17100 -pf03960 -
17105 - -
17110 -pf00873 -
17115 -pf00672pf02518pf07730pf13675 -
17120 -pf13247pf14691pf07992 -
17125 -pf00144 -
17130 HMM00293nuhyd_1_eut1(pf00293) EUT1
17135 -pf06674 -
17140 - -
17145 HMM02780keto_1_eut1(pf02779pf02780pf00456) EUT1
17150 -pf00923 -
17155 HMM00390_malic_1_eut1(pf00390pf03949pf01515) EUT1
17160 Hp_euts_violetPink EUT1
17165 HMM10662PduV_1_mix(pf10662) EUT1
17170 HMM06249EutQ_1_eut1(pf06249) EUT1 EP_NTERM
17175 HMM01923cobtr_2_eut1(pf01923) EUT1
17180 HMM01515PTA_B_1_eut1(pf01515) EUT1
17185 H_azure EUT1
17190 P_lightViolet EUT1
17195 HMMalddh_1_eutpdugrm(pf00171) EUT1 EP_NTERM
17200 HMM06723EutJMreB_1_grm EUT1
17205 -pf04879 -
17210 -pf03916 -
17215 -pf13247 -
17220 -pf02518pf00512 -
17225 HMMreg00072_4_pdu1d(pf00196pf00072) FRAG2
(..3860..)
21085 - -
21090 -pf02600 -
21100 - -
21105 - -
21110 - -
21115 HMM06723EutJMreB_1_grm EUT1
21120 HMM00465alcdh_3_mix(pf00465) EUT1
21125 HMMtpEuth_1_eut(pf04346) EUT1
21130 HMM06277EutA_1_eut(pf06277) EUT1
21135 HMMetlyEutB_1_EUT1(pf06751) EUT1
21140 HMMetlyEutC_1_eut1(pf05985) EUT1 EP_NTERM
21145 Tsp_rosa EUT1
21150 H_mulberry(pf00936pf16365) EUT1
21155 HMMreg12833_1_EUT1(pf12833) EUT1
21160 - -
21165 -pf01218 -
21170 -pf01520 -
21175 -pf00583 -
21180 -pf11143 -
21185 -pf06572 -
21190 -pf04261 -
21195 HMMnopfamclust3_1 RMM1
21200 - -
21205 -pf00528 -
21210 -pf00528 -
21215 -pf00005 -
BMC-P
BMC-H
BMC-T
Aldehyde dehydrogenase
PTAC
Signature enzyme
Alcohol dehydrogenase
Regulator

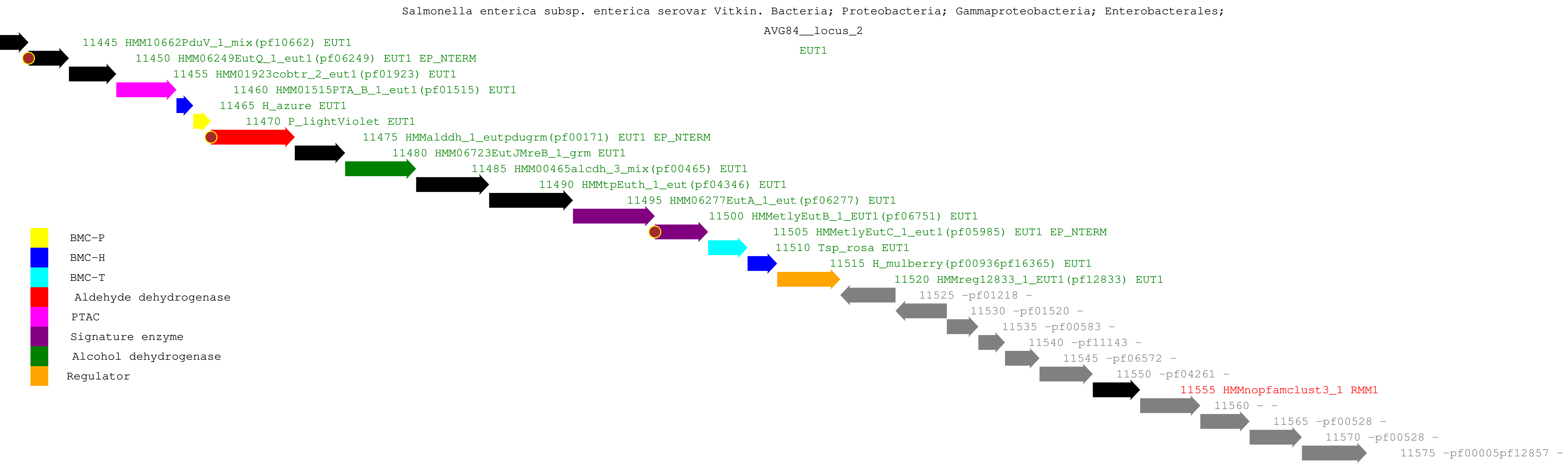
Salmonella enterica subsp. enterica serovar Vitkin. Bacteria; Proteobacteria; Gammaproteobacteria; Enterobacterales;
AVG84__locus_2
EUT1
11445 HMM10662PduV_1_mix(pf10662) EUT1
11450 HMM06249EutQ_1_eut1(pf06249) EUT1 EP_NTERM
11455 HMM01923cobtr_2_eut1(pf01923) EUT1
11460 HMM01515PTA_B_1_eut1(pf01515) EUT1
11465 H_azure EUT1
11470 P_lightViolet EUT1
11475 HMMalddh_1_eutpdugrm(pf00171) EUT1 EP_NTERM
11480 HMM06723EutJMreB_1_grm EUT1
11485 HMM00465alcdh_3_mix(pf00465) EUT1
11490 HMMtpEuth_1_eut(pf04346) EUT1
11495 HMM06277EutA_1_eut(pf06277) EUT1
11500 HMMetlyEutB_1_EUT1(pf06751) EUT1
11505 HMMetlyEutC_1_eut1(pf05985) EUT1 EP_NTERM
11510 Tsp_rosa EUT1
11515 H_mulberry(pf00936pf16365) EUT1
11520 HMMreg12833_1_EUT1(pf12833) EUT1
11525 -pf01218 -
11530 -pf01520 -
11535 -pf00583 -
11540 -pf11143 -
11545 -pf06572 -
11550 -pf04261 -
11555 HMMnopfamclust3_1 RMM1
11560 - -
11565 -pf00528 -
11570 -pf00528 -
11575 -pf00005pf12857 -
BMC-P
BMC-H
BMC-T
Aldehyde dehydrogenase
PTAC
Signature enzyme
Alcohol dehydrogenase
Regulator

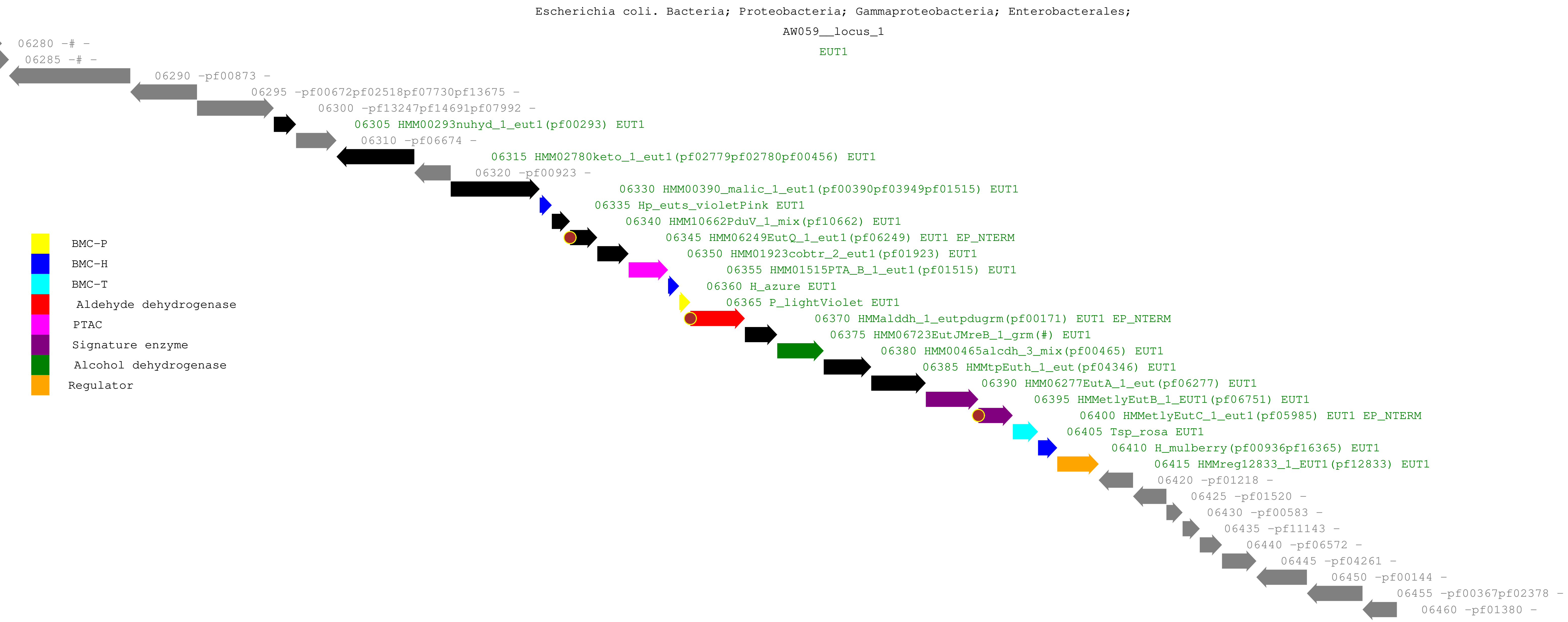
Escherichia coli. Bacteria; Proteobacteria; Gammaproteobacteria; Enterobacterales;
AW059__locus_1
EUT1
06280 -# -
06285 -# -
06290 -pf00873 -
06295 -pf00672pf02518pf07730pf13675 -
06300 -pf13247pf14691pf07992 -
06305 HMM00293nuhyd_1_eut1(pf00293) EUT1
06310 -pf06674 -
06315 HMM02780keto_1_eut1(pf02779pf02780pf00456) EUT1
06320 -pf00923 -
06330 HMM00390_malic_1_eut1(pf00390pf03949pf01515) EUT1
06335 Hp_euts_violetPink EUT1
06340 HMM10662PduV_1_mix(pf10662) EUT1
06345 HMM06249EutQ_1_eut1(pf06249) EUT1 EP_NTERM
06350 HMM01923cobtr_2_eut1(pf01923) EUT1
06355 HMM01515PTA_B_1_eut1(pf01515) EUT1
06360 H_azure EUT1
06365 P_lightViolet EUT1
06370 HMMalddh_1_eutpdugrm(pf00171) EUT1 EP_NTERM
06375 HMM06723EutJMreB_1_grm(#) EUT1
06380 HMM00465alcdh_3_mix(pf00465) EUT1
06385 HMMtpEuth_1_eut(pf04346) EUT1
06390 HMM06277EutA_1_eut(pf06277) EUT1
06395 HMMetlyEutB_1_EUT1(pf06751) EUT1
06400 HMMetlyEutC_1_eut1(pf05985) EUT1 EP_NTERM
06405 Tsp_rosa EUT1
06410 H_mulberry(pf00936pf16365) EUT1
06415 HMMreg12833_1_EUT1(pf12833) EUT1
06420 -pf01218 -
06425 -pf01520 -
06430 -pf00583 -
06435 -pf11143 -
06440 -pf06572 -
06445 -pf04261 -
06450 -pf00144 -
06455 -pf00367pf02378 -
06460 -pf01380 -
BMC-P
BMC-H
BMC-T
Aldehyde dehydrogenase
PTAC
Signature enzyme
Alcohol dehydrogenase
Regulator

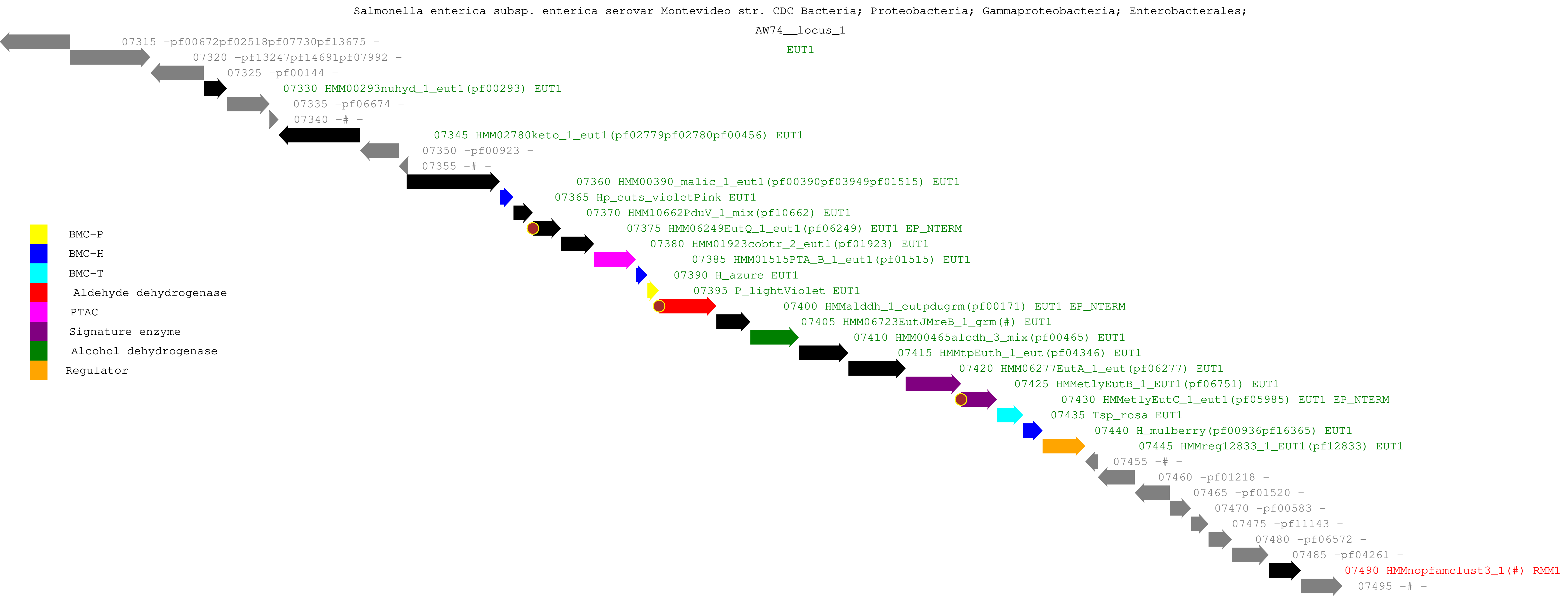

Salmonella enterica subsp. enterica serovar Montevideo str. CDC Bacteria; Proteobacteria; Gammaproteobacteria; Enterobacterales;
AW74__locus_1
EUT1
07315 -pf00672pf02518pf07730pf13675 -
07320 -pf13247pf14691pf07992 -
07325 -pf00144 -
07330 HMM00293nuhyd_1_eut1(pf00293) EUT1
07335 -pf06674 -
07340 -# -
07345 HMM02780keto_1_eut1(pf02779pf02780pf00456) EUT1
07350 -pf00923 -
07355 -# -
07360 HMM00390_malic_1_eut1(pf00390pf03949pf01515) EUT1
07365 Hp_euts_violetPink EUT1
07370 HMM10662PduV_1_mix(pf10662) EUT1
07375 HMM06249EutQ_1_eut1(pf06249) EUT1 EP_NTERM
07380 HMM01923cobtr_2_eut1(pf01923) EUT1
07385 HMM01515PTA_B_1_eut1(pf01515) EUT1
07390 H_azure EUT1
07395 P_lightViolet EUT1
07400 HMMalddh_1_eutpdugrm(pf00171) EUT1 EP_NTERM
07405 HMM06723EutJMreB_1_grm(#) EUT1
07410 HMM00465alcdh_3_mix(pf00465) EUT1
07415 HMMtpEuth_1_eut(pf04346) EUT1
07420 HMM06277EutA_1_eut(pf06277) EUT1
07425 HMMetlyEutB_1_EUT1(pf06751) EUT1
07430 HMMetlyEutC_1_eut1(pf05985) EUT1 EP_NTERM
07435 Tsp_rosa EUT1
07440 H_mulberry(pf00936pf16365) EUT1
07445 HMMreg12833_1_EUT1(pf12833) EUT1
07455 -# -
07460 -pf01218 -
07465 -pf01520 -
07470 -pf00583 -
07475 -pf11143 -
07480 -pf06572 -
07485 -pf04261 -
07490 HMMnopfamclust3_1(#) RMM1
07495 -# -
BMC-P
BMC-H
BMC-T
Aldehyde dehydrogenase
PTAC
Signature enzyme
Alcohol dehydrogenase
Regulator

Escherichia coli. Bacteria; Proteobacteria; Gammaproteobacteria; Enterobacterales;

AWF22__locus_1

EUT1

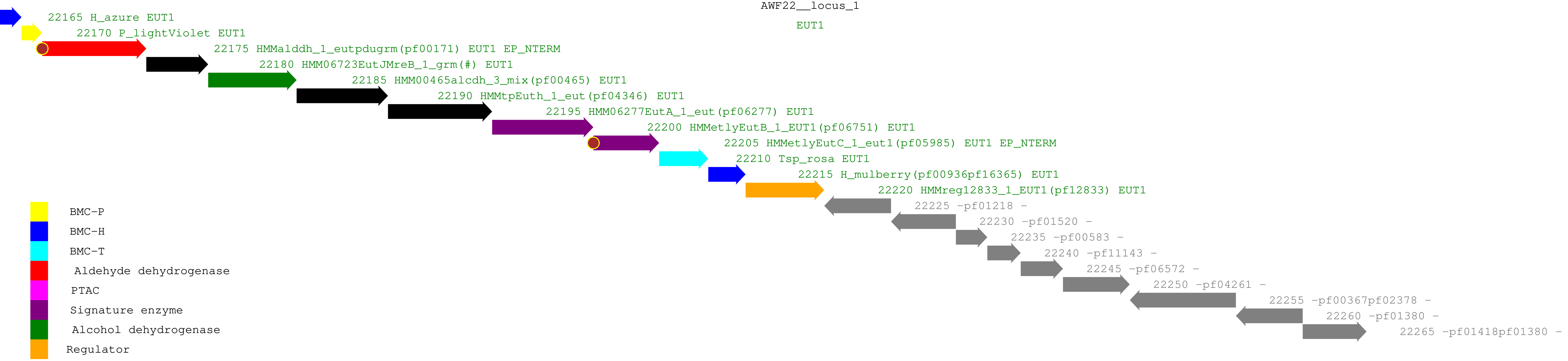

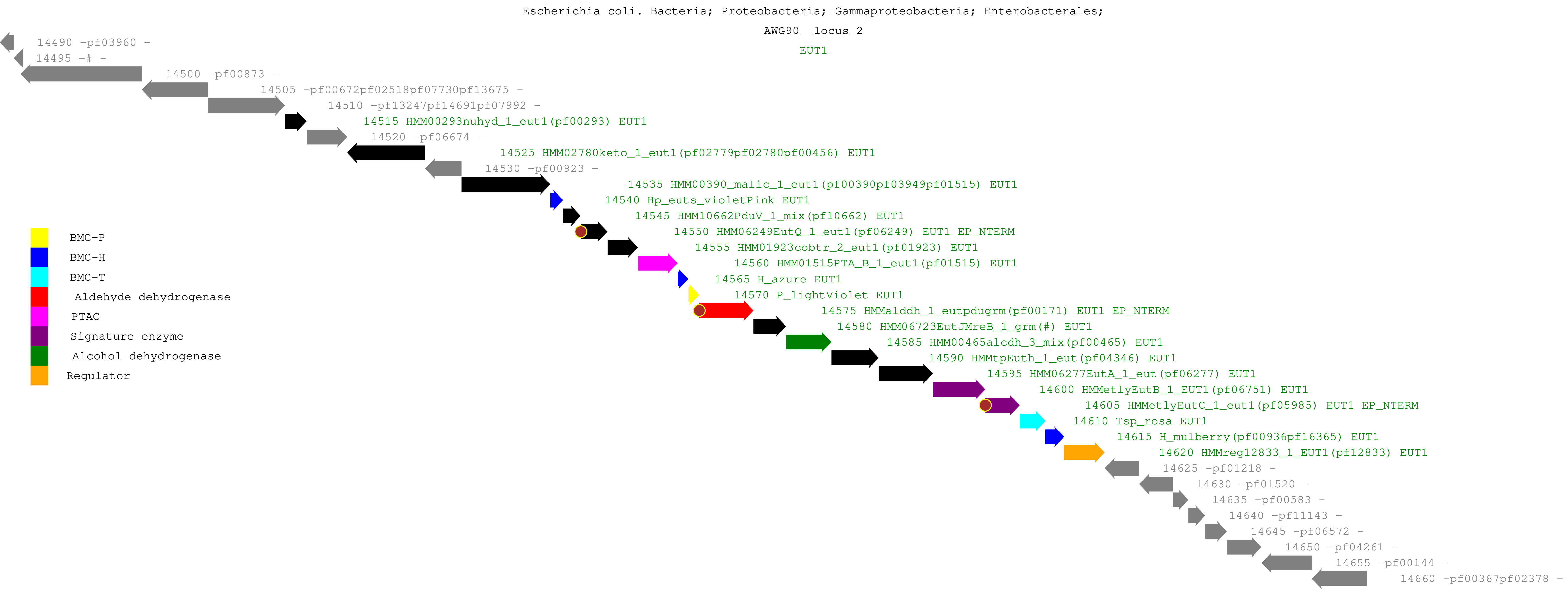

Escherichia coli. Bacteria; Proteobacteria; Gammaproteobacteria; Enterobacterales;
AWG90__locus_2
EUT1
14490 -pf03960 -
14495 -# -
14500 -pf00873 -
14505 -pf00672pf02518pf07730pf13675 -
14510 -pf13247pf14691pf07992 -
14515 HMM00293nuhyd_1_eut1(pf00293) EUT1
14520 -pf06674 -
14525 HMM02780keto_1_eut1(pf02779pf02780pf00456) EUT1
14530 -pf00923 -
14535 HMM00390_malic_1_eut1(pf00390pf03949pf01515) EUT1
14540 Hp_euts_violetPink EUT1
14545 HMM10662PduV_1_mix(pf10662) EUT1
14550 HMM06249EutQ_1_eut1(pf06249) EUT1 EP_NTERM
14555 HMM01923cobtr_2_eut1(pf01923) EUT1
14560 HMM01515PTA_B_1_eut1(pf01515) EUT1
14565 H_azure EUT1
14570 P_lightViolet EUT1
14575 HMMalddh_1_eutpdugrm(pf00171) EUT1 EP_NTERM
14580 HMM06723EutJMreB_1_grm(#) EUT1
14585 HMM00465alcdh_3_mix(pf00465) EUT1
14590 HMMtpEuth_1_eut(pf04346) EUT1
14595 HMM06277EutA_1_eut(pf06277) EUT1
14600 HMMetlyEutB_1_EUT1(pf06751) EUT1
14605 HMMetlyEutC_1_eut1(pf05985) EUT1 EP_NTERM
14610 Tsp_rosa EUT1
14615 H_mulberry(pf00936pf16365) EUT1
14620 HMMreg12833_1_EUT1(pf12833) EUT1
14625 -pf01218 -
14630 -pf01520 -
14635 -pf00583 -
14640 -pf11143 -
14645 -pf06572 -
14650 -pf04261 -
14655 -pf00144 -
14660 -pf00367pf02378 -
BMC-P
BMC-H
BMC-T
Aldehyde dehydrogenase
PTAC
Signature enzyme
Alcohol dehydrogenase
Regulator

Gammaproteobacteria bacterium. Bacteria; Proteobacteria; Gammaproteobacteria.

AwWohl__locus_1_main_sat

EUT1

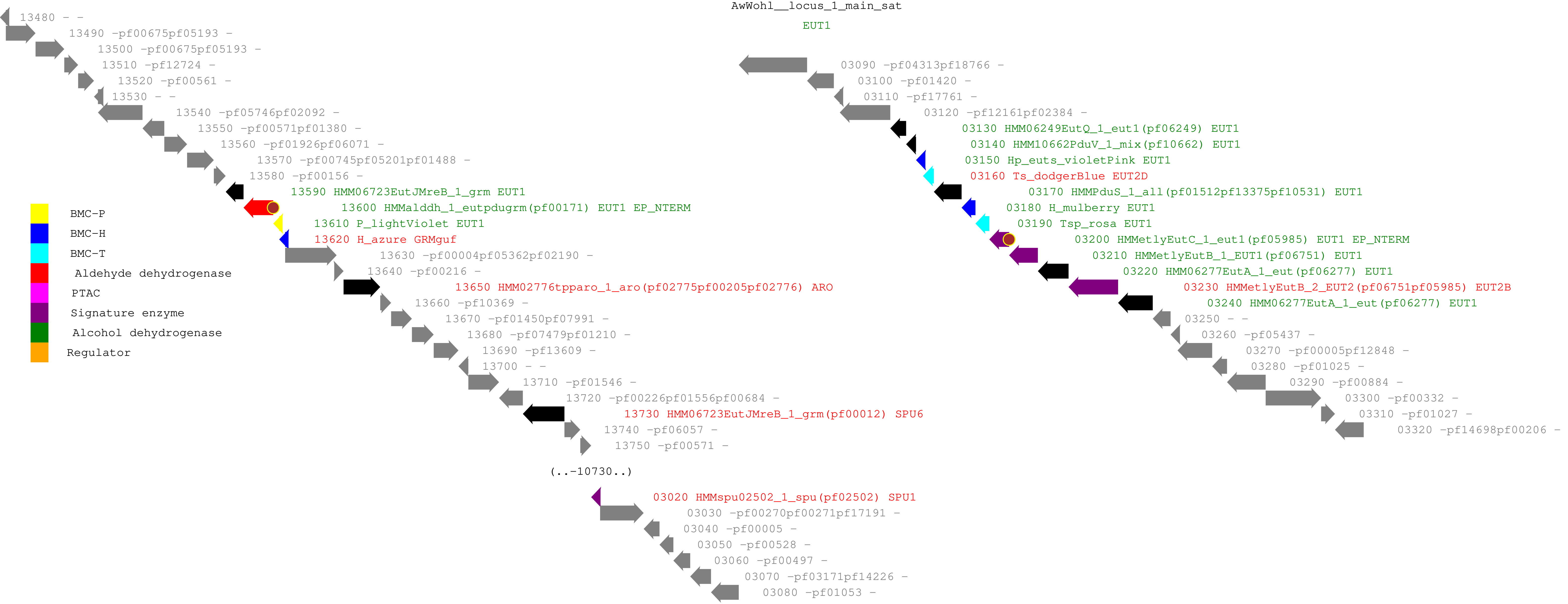

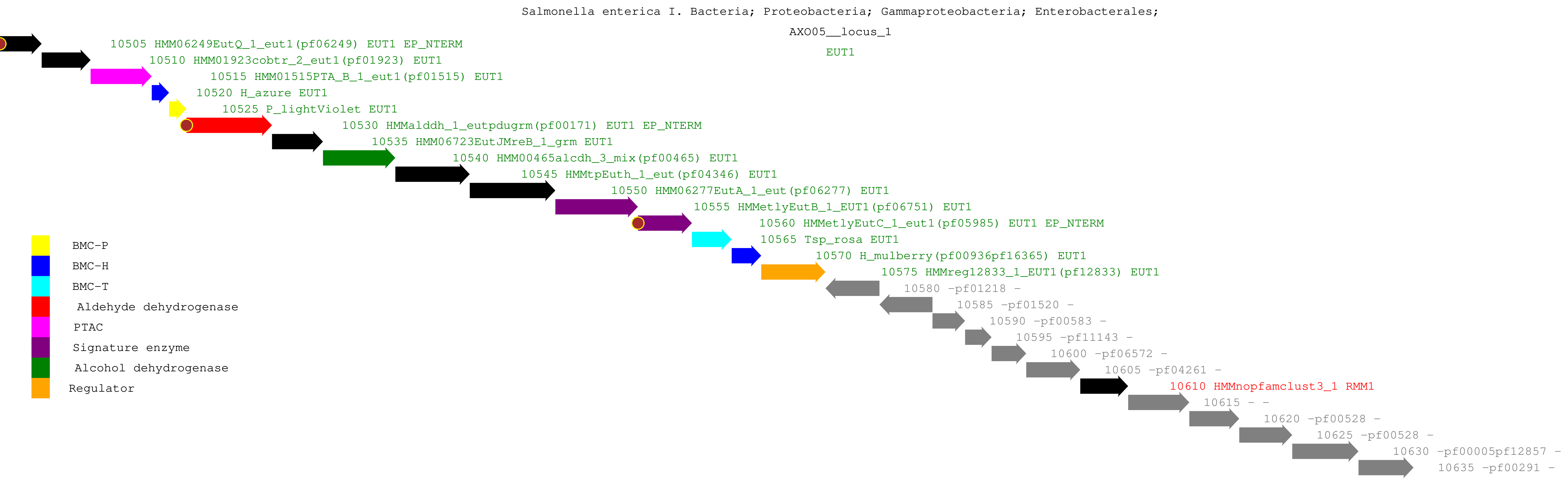

Salmonella enterica I. Bacteria; Proteobacteria; Gammaproteobacteria; Enterobacterales;
AXO05__locus_1
EUT1
10505 HMM06249EutQ_1_eut1(pf06249) EUT1 EP_NTERM
10510 HMM01923cobtr_2_eut1(pf01923) EUT1
10515 HMM01515PTA_B_1_eut1(pf01515) EUT1
10520 H_azure EUT1
10525 P_lightViolet EUT1
10530 HMMalddh_1_eutpdugrm(pf00171) EUT1 EP_NTERM
10535 HMM06723EutJMreB_1_grm EUT1
10540 HMM00465alcdh_3_mix(pf00465) EUT1
10545 HMMtpEuth_1_eut(pf04346) EUT1
10550 HMM06277EutA_1_eut(pf06277) EUT1
10555 HMMetlyEutB_1_EUT1(pf06751) EUT1
10560 HMMetlyEutC_1_eut1(pf05985) EUT1 EP_NTERM
10565 Tsp_rosa EUT1
10570 H_mulberry(pf00936pf16365) EUT1
10575 HMMreg12833_1_EUT1(pf12833) EUT1
10580 -pf01218 -
10585 -pf01520 -
10590 -pf00583 -
10595 -pf11143 -
10600 -pf06572 -
10605 -pf04261 -
10610 HMMnopfamclust3_1 RMM1
10615 - -
10620 -pf00528 -
10625 -pf00528 -
10630 -pf00005pf12857 -
10635 -pf00291 -
BMC-P
BMC-H
BMC-T
Aldehyde dehydrogenase
PTAC
Signature enzyme
Alcohol dehydrogenase
Regulator

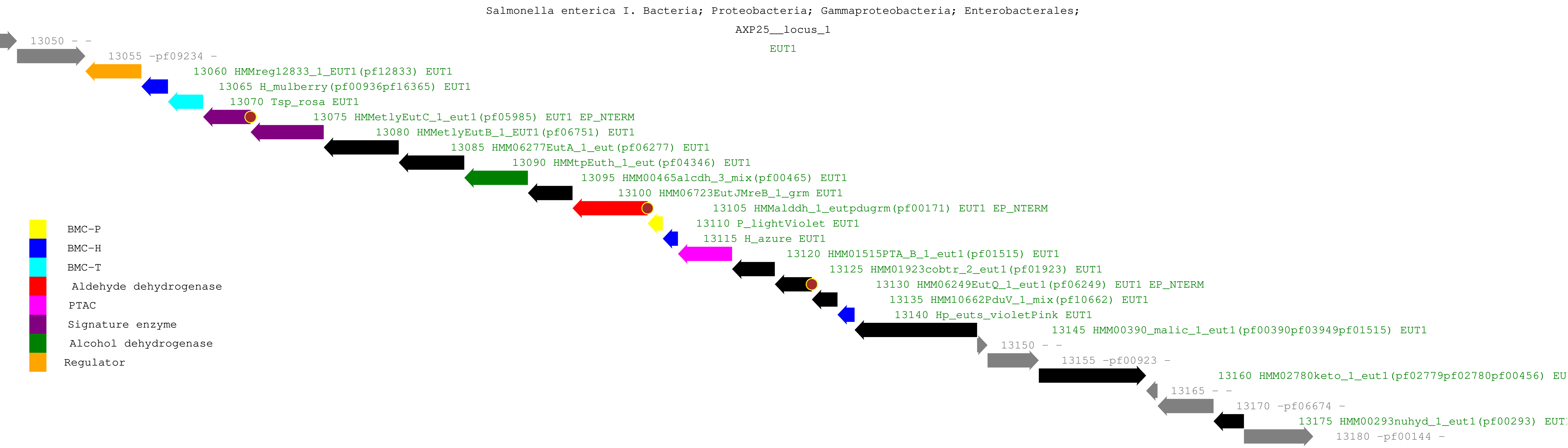
Salmonella enterica I. Bacteria; Proteobacteria; Gammaproteobacteria; Enterobacterales;
AXP25__locus_1
EUT1
13050 - -
13055 -pf09234 -
13060 HMMreg12833_1_EUT1(pf12833) EUT1
13065 H_mulberry(pf00936pf16365) EUT1
13070 Tsp_rosa EUT1
13075 HMMetlyEutC_1_eut1(pf05985) EUT1 EP_NTERM
13080 HMMetlyEutB_1_EUT1(pf06751) EUT1
13085 HMM06277EutA_1_eut(pf06277) EUT1
13090 HMMtpEuth_1_eut(pf04346) EUT1
13095 HMM00465alcdh_3_mix(pf00465) EUT1
13100 HMM06723EutJMreB_1_grm EUT1
13105 HMMalddh_1_eutpdugrm(pf00171) EUT1 EP_NTERM
13110 P_lightViolet EUT1
13115 H_azure EUT1
13120 HMM01515PTA_B_1_eut1(pf01515) EUT1
13125 HMM01923cobtr_2_eut1(pf01923) EUT1
13130 HMM06249EutQ_1_eut1(pf06249) EUT1 EP_NTERM
13135 HMM10662PduV_1_mix(pf10662) EUT1
13140 Hp_euts_violetPink EUT1
13145 HMM00390_malic_1_eut1(pf00390pf03949pf01515) EUT1
13150 - -
13155 -pf00923 -
13160 HMM02780keto_1_eut1(pf02779pf02780pf00456) EU
13165 - -
13170 -pf06674 -
13175 HMM00293nuhyd_1_eut1(pf00293) EUT
13180 -pf00144 -
BMC-P
BMC-H
BMC-T
Aldehyde dehydrogenase
PTAC
Signature enzyme
Alcohol dehydrogenase
Regulator

Salmonella montevideo. Bacteria; Proteobacteria; Gammaproteobacteria; Enterobacterales;

AXU21__locus_1

EUT1

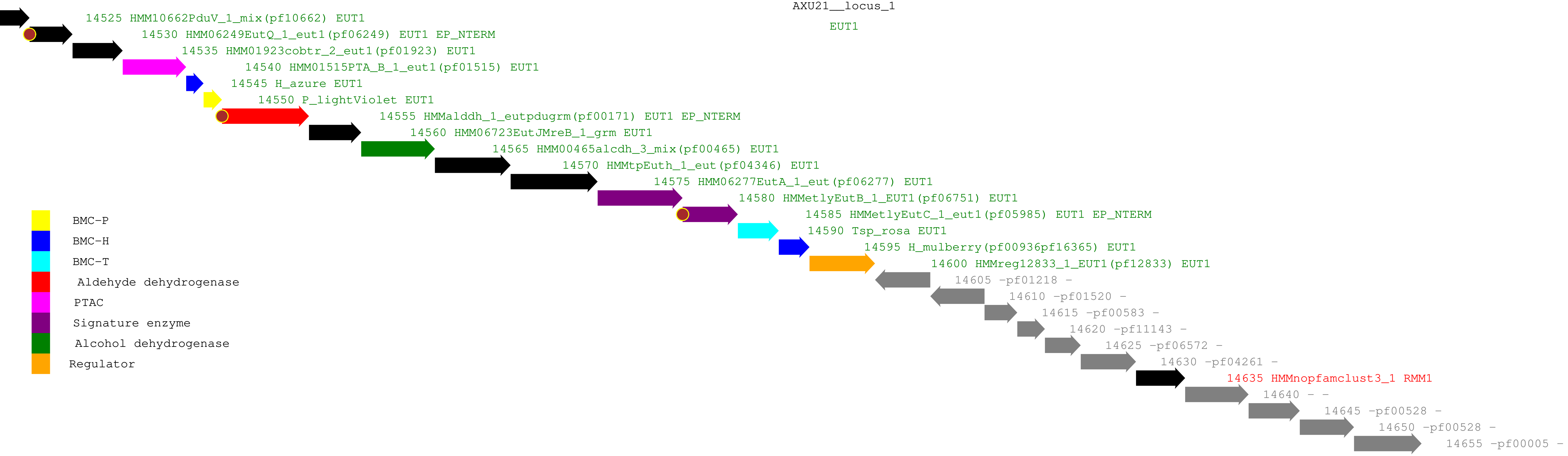

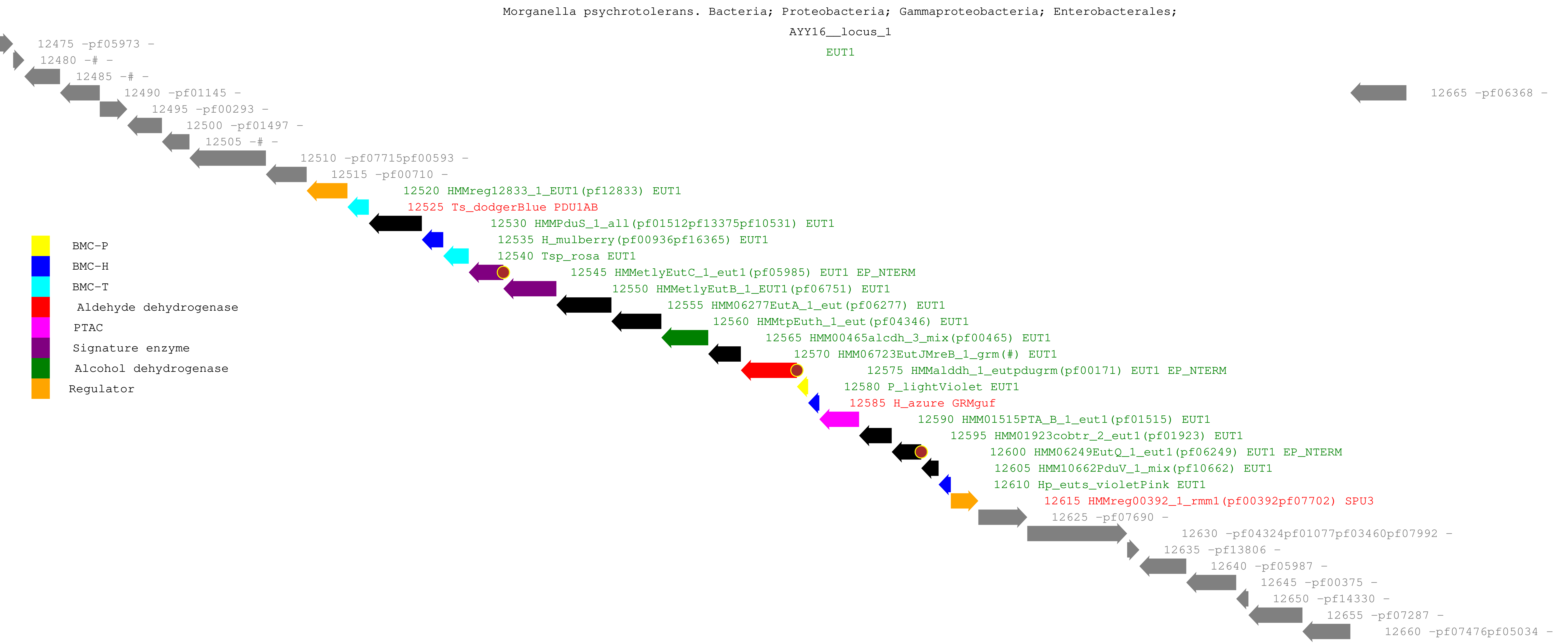

Morganella psychrotolerans. Bacteria; Proteobacteria; Gammaproteobacteria; Enterobacterales;
AYY16__locus_1
EUT1
12475 -pf05973 -
12480 -# -
12485 -# -
12490 -pf01145 -
12495 -pf00293 -
12500 -pf01497 -
12505 -# -
12510 -pf07715pf00593 -
12515 -pf00710 -
12520 HMMreg12833_1_EUT1(pf12833) EUT1
12525 Ts_dodgerBlue PDU1AB
12530 HMMPduS_1_all(pf01512pf13375pf10531) EUT1
12535 H_mulberry(pf00936pf16365) EUT1
12540 Tsp_rosa EUT1
12545 HMMetlyEutC_1_eut1(pf05985) EUT1 EP_NTERM
12550 HMMetlyEutB_1_EUT1(pf06751) EUT1
12555 HMM06277EutA_1_eut(pf06277) EUT1
12560 HMMtpEuth_1_eut(pf04346) EUT1
12565 HMM00465alcdh_3_mix(pf00465) EUT1
12570 HMM06723EutJMreB_1_grm(#) EUT1
12575 HMMalddh_1_eutpdugrm(pf00171) EUT1 EP_NTERM
12580 P_lightViolet EUT1
12585 H_azure GRMguf
12590 HMM01515PTA_B_1_eut1(pf01515) EUT1
12595 HMM01923cobtr_2_eut1(pf01923) EUT1
12600 HMM06249EutQ_1_eut1(pf06249) EUT1 EP_NTERM
12605 HMM10662PduV_1_mix(pf10662) EUT1
12610 Hp_euts_violetPink EUT1
12615 HMMreg00392_1_rmm1(pf00392pf07702) SPU3
12625 -pf07690 -
12630 -pf04324pf01077pf03460pf07992 -
12635 -pf13806 -
12640 -pf05987 -
12645 -pf00375 -
12650 -pf14330 -
12655 -pf07287 -
12660 -pf07476pf05034 -
12665 -pf06368 -
BMC-P
BMC-H
BMC-T
Aldehyde dehydrogenase
PTAC
Signature enzyme
Alcohol dehydrogenase
Regulator

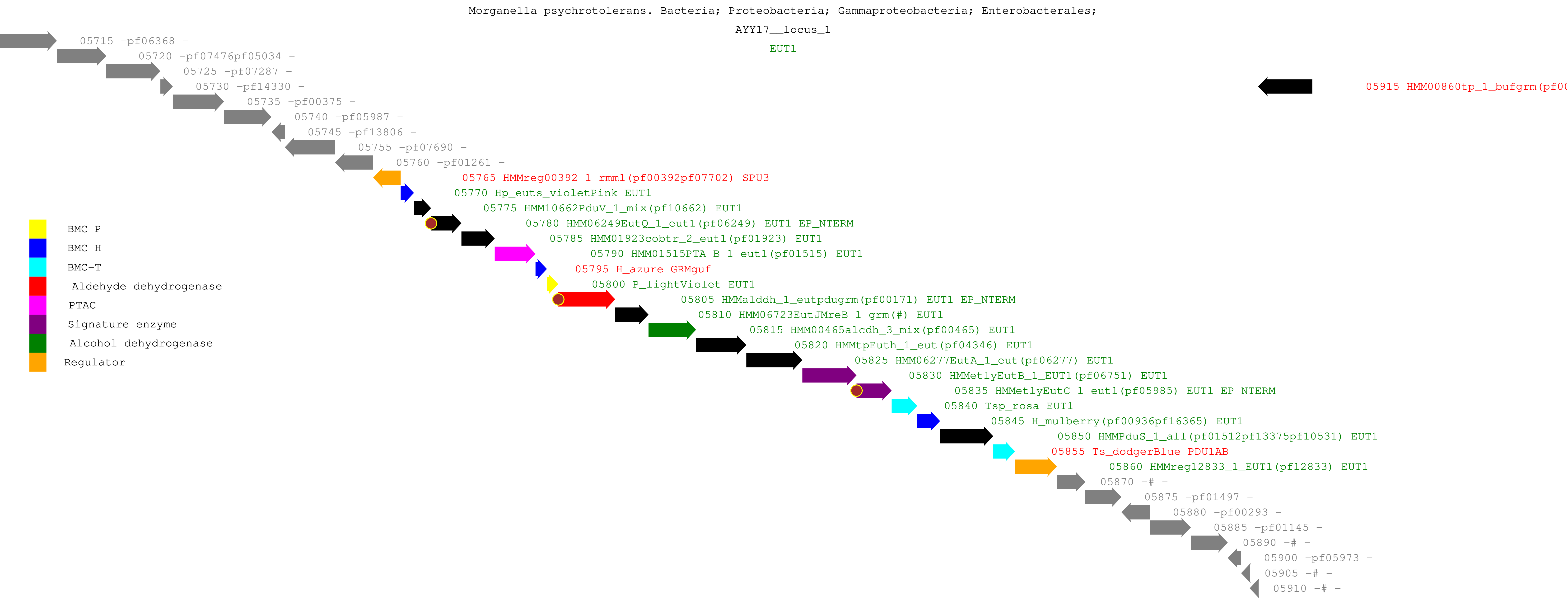
Morganella psychrotolerans. Bacteria; Proteobacteria; Gammaproteobacteria; Enterobacterales;
AYY17__locus_1
EUT1
05715 -pf06368 -
05720 -pf07476pf05034 -
05725 -pf07287 -
05730 -pf14330 -
05735 -pf00375 -
05740 -pf05987 -
05745 -pf13806 -
05755 -pf07690 -
05760 -pf01261 -
05765 HMMreg00392_1_rmm1(pf00392pf07702) SPU3
05770 Hp_euts_violetPink EUT1
05775 HMM10662PduV_1_mix(pf10662) EUT1
05780 HMM06249EutQ_1_eut1(pf06249) EUT1 EP_NTERM
05785 HMM01923cobtr_2_eut1(pf01923) EUT1
05790 HMM01515PTA_B_1_eut1(pf01515) EUT1
05795 H_azure GRMguf
05800 P_lightViolet EUT1
05805 HMMalddh_1_eutpdugrm(pf00171) EUT1 EP_NTERM
05810 HMM06723EutJMreB_1_grm(#) EUT1
05815 HMM00465alcdh_3_mix(pf00465) EUT1
05820 HMMtpEuth_1_eut(pf04346) EUT1
05825 HMM06277EutA_1_eut(pf06277) EUT1
05830 HMMetlyEutB_1_EUT1(pf06751) EUT1
05835 HMMetlyEutC_1_eut1(pf05985) EUT1 EP_NTERM
05840 Tsp_rosa EUT1
05845 H_mulberry(pf00936pf16365) EUT1
05850 HMMPduS_1_all(pf01512pf13375pf10531) EUT1
05855 Ts_dodgerBlue PDU1AB
05860 HMMreg12833_1_EUT1(pf12833) EUT1
05870 -# -
05875 -pf01497 -
05880 -pf00293 -
05885 -pf01145 -
05890 -# -
05900 -pf05973 -
05905 -# -
05910 -# -
05915 HMM00860tp_1_bufgrm(pf0
BMC-P
BMC-H
BMC-T
Aldehyde dehydrogenase
PTAC
Signature enzyme
Alcohol dehydrogenase
Regulator

Morganella psychrotolerans. Bacteria; Proteobacteria; Gammaproteobacteria; Enterobacterales;

AYY18__locus_1

EUT1

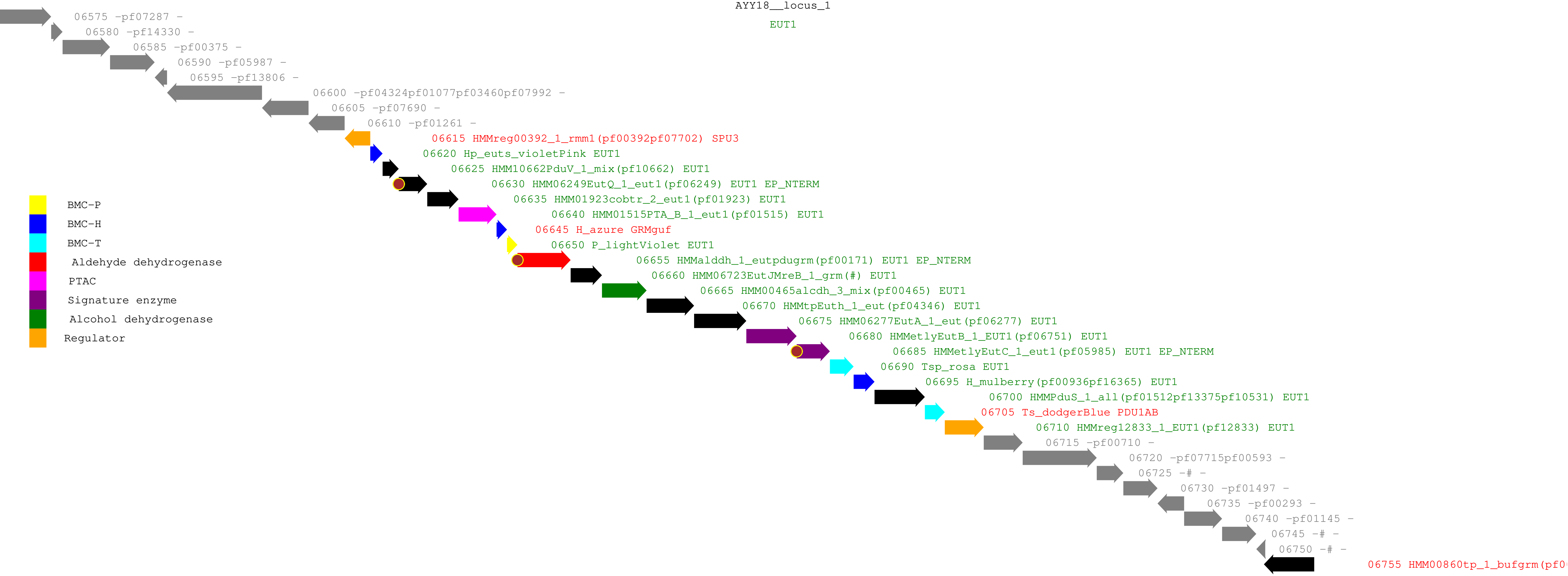

Citrobacter freundii. Bacteria; Proteobacteria; Gammaproteobacteria; Enterobacterales;

AZ002__locus_1

EUT1

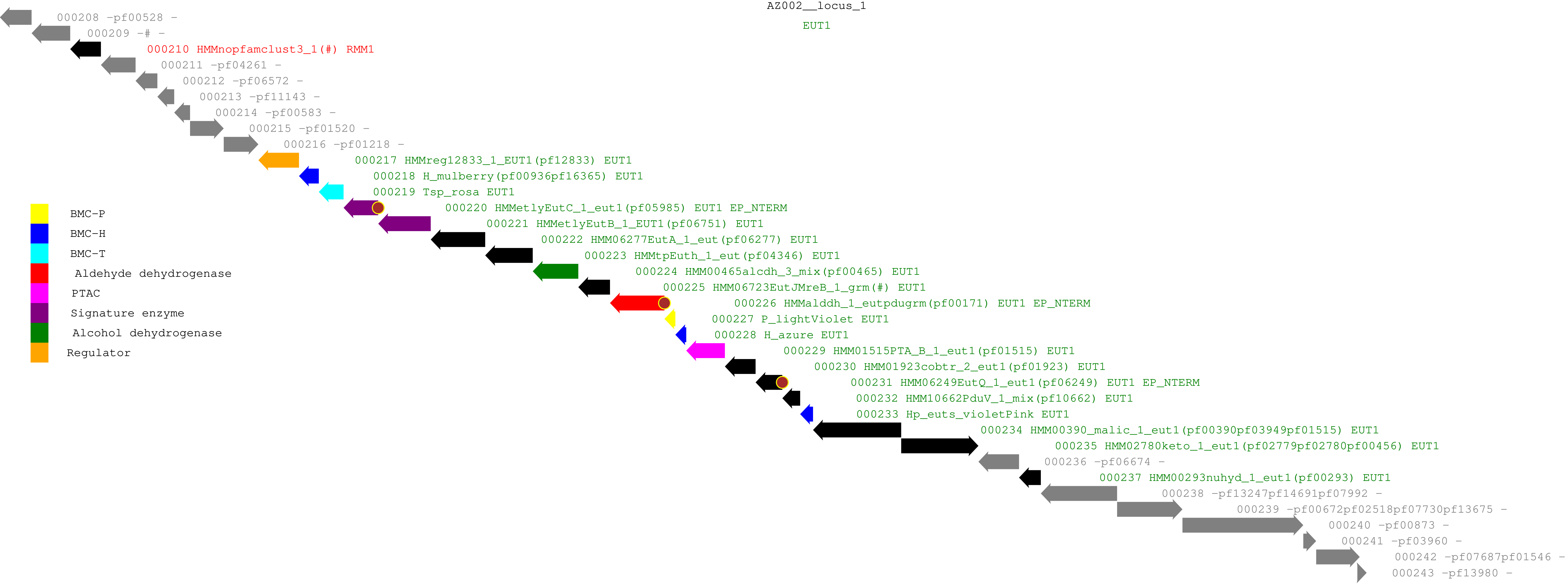

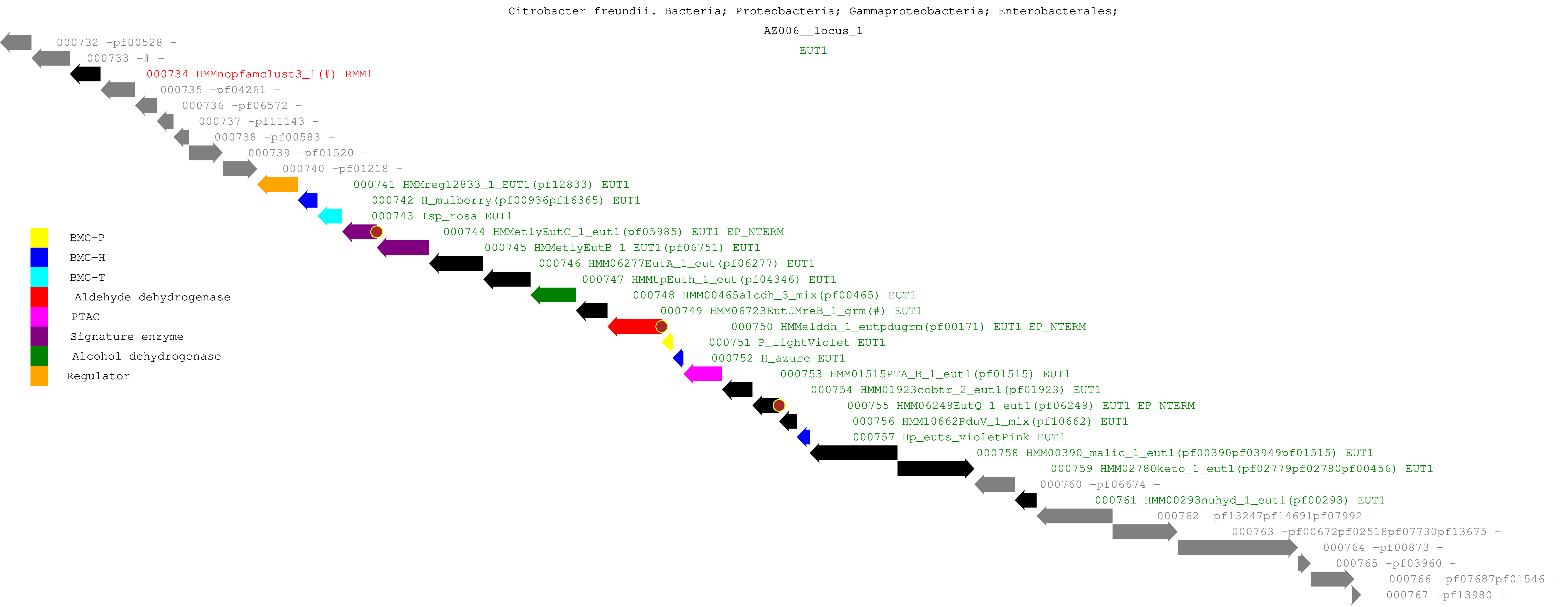
Citrobacter freundii. Bacteria; Proteobacteria; Gammaproteobacteria; Enterobacterales;
AZ006__locus_1
EUT1
000732 -pf00528 -
000733 -# -
000734 HMMnopfamclust3_1(#) RMM1
000735 -pf04261 -
000736 -pf06572 -
000737 -pf11143 -
000738 -pf00583 -
000739 -pf01520 -
000740 -pf01218 -
000741 HMMreg12833_1_EUT1(pf12833) EUT1
000742 H_mulberry(pf00936pf16365) EUT1
000743 Tsp_rosa EUT1
000744 HMMetlyEutC_1_eut1(pf05985) EUT1 EP_NTERM
000745 HMMetlyEutB_1_EUT1(pf06751) EUT1
000746 HMM06277EutA_1_eut(pf06277) EUT1
000747 HMMtpEuth_1_eut(pf04346) EUT1
000748 HMM00465alcdh_3_mix(pf00465) EUT1
000749 HMM06723EutJMreB_1_grm(#) EUT1
000750 HMMalddh_1_eutpdugrm(pf00171) EUT1 EP_NTERM
000751 P_lightViolet EUT1
000752 H_azure EUT1
000753 HMM01515PTA_B_1_eut1(pf01515) EUT1
000754 HMM01923cobtr_2_eut1(pf01923) EUT1
000755 HMM06249EutQ_1_eut1(pf06249) EUT1 EP_NTERM
000756 HMM10662PduV_1_mix(pf10662) EUT1
000757 Hp_euts_violetPink EUT1
000758 HMM00390_malic_1_eut1(pf00390pf03949pf01515) EUT1
000759 HMM02780keto_1_eut1(pf02779pf02780pf00456) EUT1
000760 -pf06674 -
000761 HMM00293nuhyd_1_eut1(pf00293) EUT1
000762 -pf13247pf14691pf07992 -
000763 -pf00672pf02518pf07730pf13675 -
000764 -pf00873 -
000765 -pf03960 -
000766 -pf07687pf01546 -
000767 -pf13980 -
BMC-P
BMC-H
BMC-T
Aldehyde dehydrogenase
PTAC
Signature enzyme
Alcohol dehydrogenase
Regulator

Citrobacter freundii. Bacteria; Proteobacteria; Gammaproteobacteria; Enterobacterales;

AZ013__locus_1

EUT1

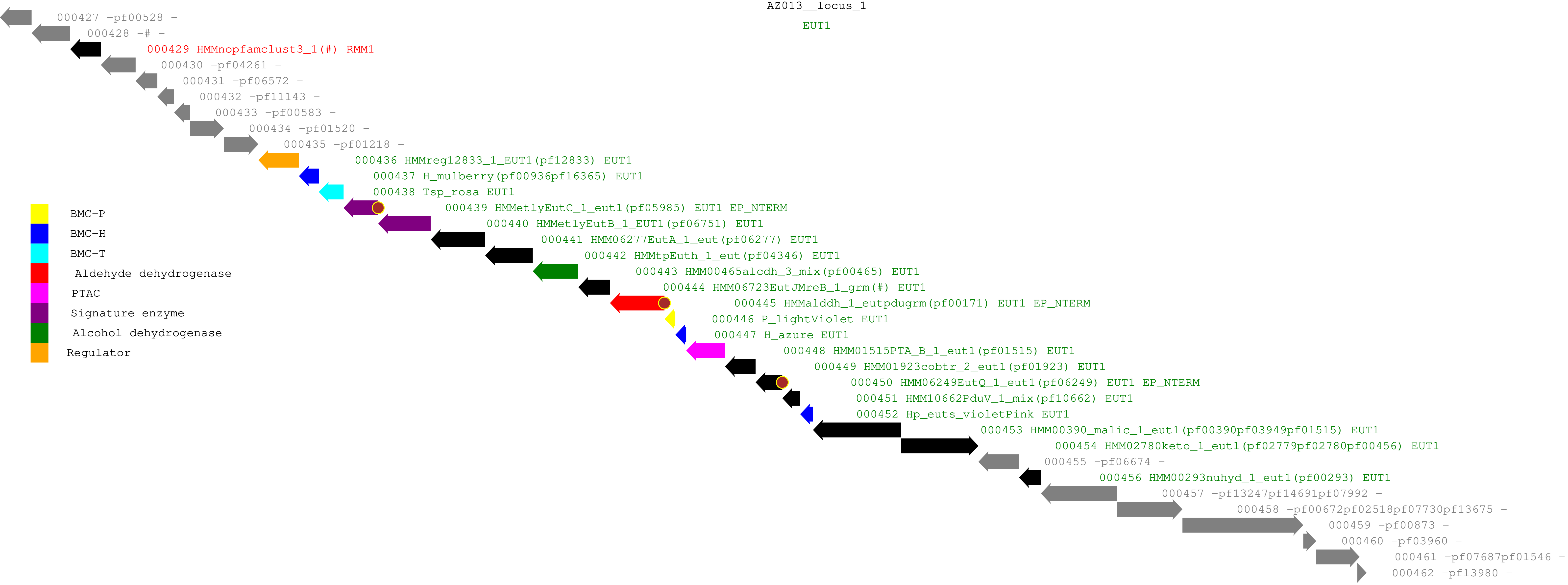

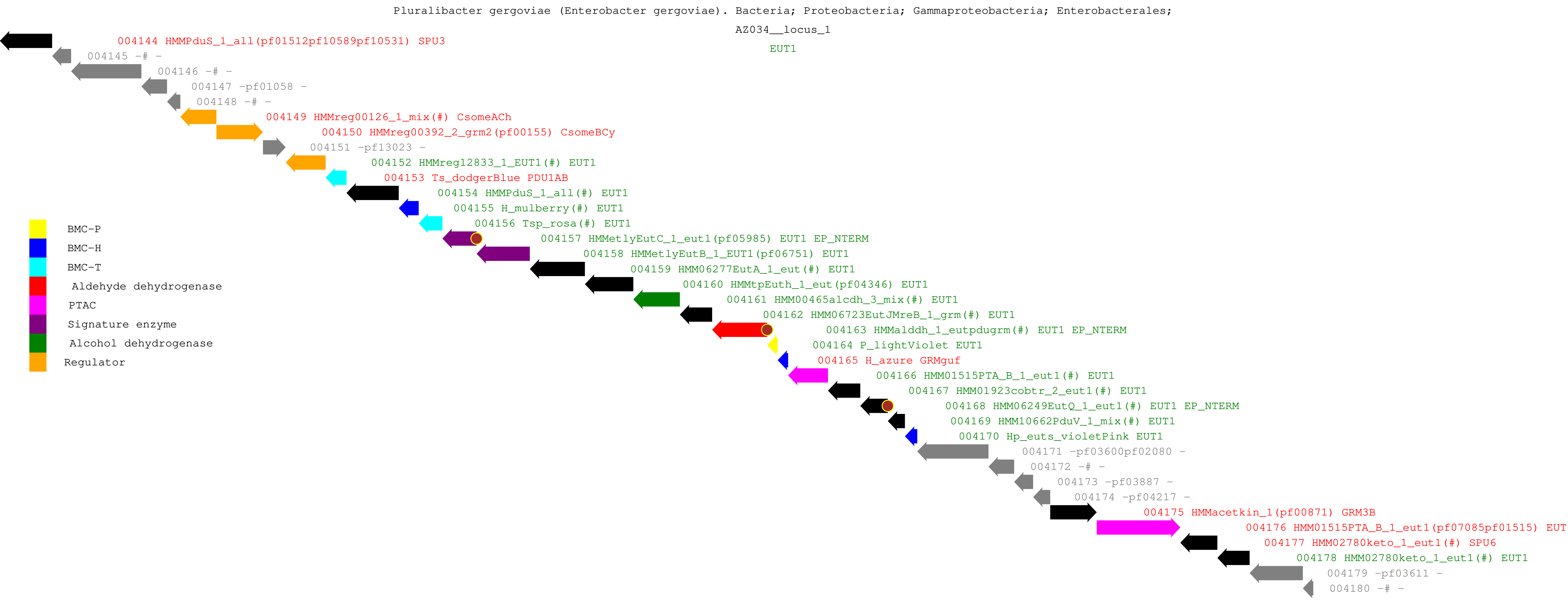
Pluralibacter gergoviae (Enterobacter gergoviae). Bacteria; Proteobacteria; Gammaproteobacteria; Enterobacterales;
AZ034__locus_1
EUT1
004144 HMMPduS_1_all(pf01512pf10589pf10531) SPU3
004145 -# -
004146 -# -
004147 -pf01058 -
004148 -# -
004149 HMMreg00126_1_mix(#) CsomeACh
004150 HMMreg00392_2_grm2(pf00155) CsomeBCy
004151 -pf13023 -
004152 HMMreg12833_1_EUT1(#) EUT1
004153 Ts_dodgerBlue PDU1AB
004154 HMMPduS_1_all(#) EUT1
004155 H_mulberry(#) EUT1
004156 Tsp_rosa(#) EUT1
004157 HMMetlyEutC_1_eut1(pf05985) EUT1 EP_NTERM
004158 HMMetlyEutB_1_EUT1(pf06751) EUT1
004159 HMM06277EutA_1_eut(#) EUT1
004160 HMMtpEuth_1_eut(pf04346) EUT1
004161 HMM00465alcdh_3_mix(#) EUT1
004162 HMM06723EutJMreB_1_grm(#) EUT1
004163 HMMalddh_1_eutpdugrm(#) EUT1 EP_NTERM
004164 P_lightViolet EUT1
004165 H_azure GRMguf
004166 HMM01515PTA_B_1_eut1(#) EUT1
004167 HMM01923cobtr_2_eut1(#) EUT1
004168 HMM06249EutQ_1_eut1(#) EUT1 EP_NTERM
004169 HMM10662PduV_1_mix(#) EUT1
004170 Hp_euts_violetPink EUT1
004171 -pf03600pf02080 -
004172 -# -
004173 -pf03887 -
004174 -pf04217 -
004175 HMMacetkin_1(pf00871) GRM3B
004176 HMM01515PTA_B_1_eut1(pf07085pf01515) EUT
004177 HMM02780keto_1_eut1(#) SPU6
004178 HMM02780keto_1_eut1(#) EUT1
004179 -pf03611 -
004180 -# -
BMC-P
BMC-H
BMC-T
Aldehyde dehydrogenase
PTAC
Signature enzyme
Alcohol dehydrogenase
Regulator

Escherichia coli NU14. Bacteria; Proteobacteria; Gammaproteobacteria; Enterobacterales;

B0908__locus_1

EUT1

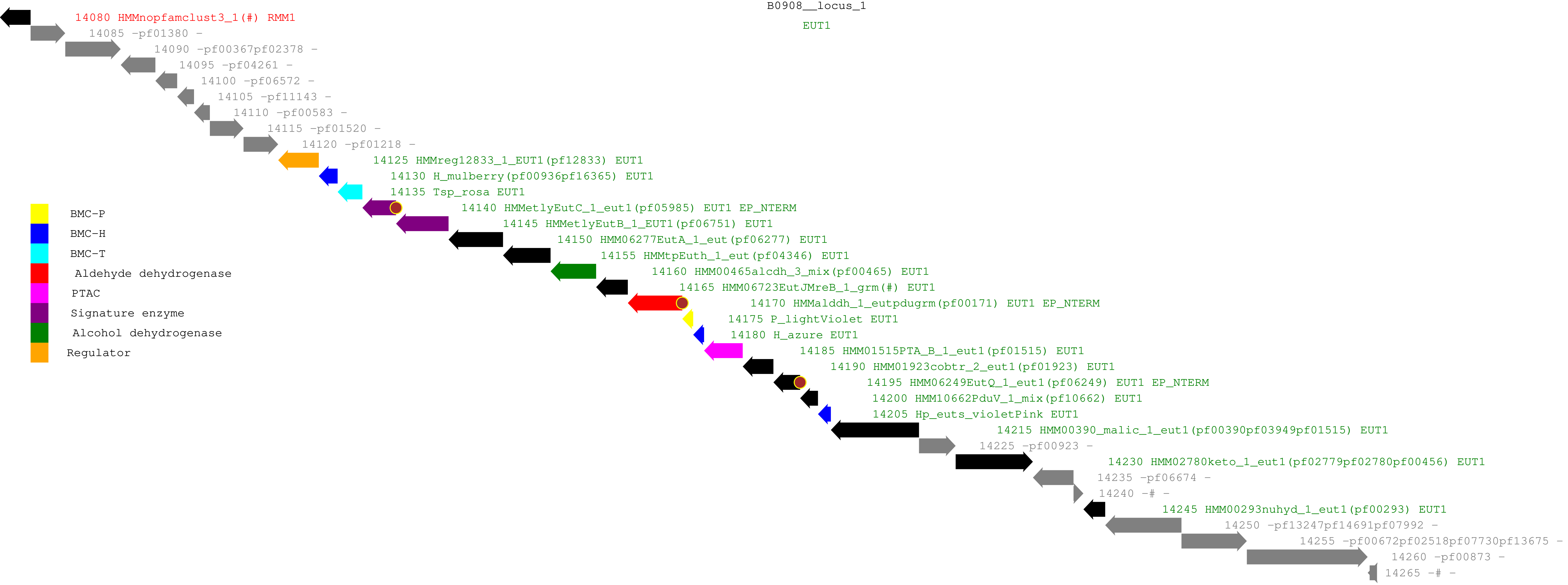

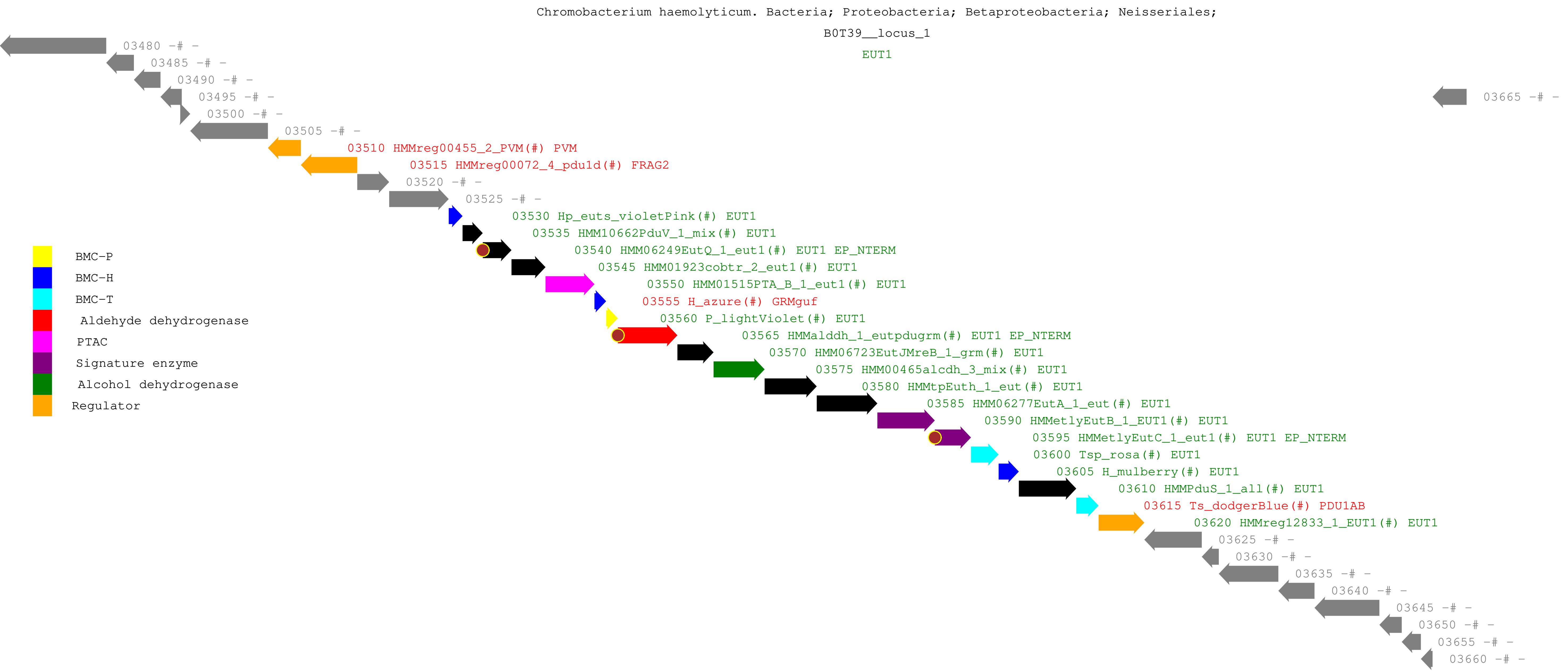
Chromobacterium haemolyticum. Bacteria; Proteobacteria; Betaproteobacteria; Neisseriales;
B0T39__locus_1
EUT1
03480 -# -
03485 -# -
03490 -# -
03495 -# -
03500 -# -
03505 -# -
03510 HMMreg00455_2_PVM(#) PVM
03515 HMMreg00072_4_pdu1d(#) FRAG2
03520 -# -
03525 -# -
03530 Hp_euts_violetPink(#) EUT1
03535 HMM10662PduV_1_mix(#) EUT1
03540 HMM06249EutQ_1_eut1(#) EUT1 EP_NTERM
03545 HMM01923cobtr_2_eut1(#) EUT1
03550 HMM01515PTA_B_1_eut1(#) EUT1
03555 H_azure(#) GRMguf
03560 P_lightViolet(#) EUT1
03565 HMMalddh_1_eutpdugrm(#) EUT1 EP_NTERM
03570 HMM06723EutJMreB_1_grm(#) EUT1
03575 HMM00465alcdh_3_mix(#) EUT1
03580 HMMtpEuth_1_eut(#) EUT1
03585 HMM06277EutA_1_eut(#) EUT1
03590 HMMetlyEutB_1_EUT1(#) EUT1
03595 HMMetlyEutC_1_eut1(#) EUT1 EP_NTERM
03600 Tsp_rosa(#) EUT1
03605 H_mulberry(#) EUT1
03610 HMMPduS_1_all(#) EUT1
03615 Ts_dodgerBlue(#) PDU1AB
03620 HMMreg12833_1_EUT1(#) EUT1
03625 -# -
03630 -# -
03635 -# -
03640 -# -
03645 -# -
03650 -# -
03655 -# -
03660 -# -
03665 -# -
BMC-P
BMC-H
BMC-T
Aldehyde dehydrogenase
PTAC
Signature enzyme
Alcohol dehydrogenase
Regulator

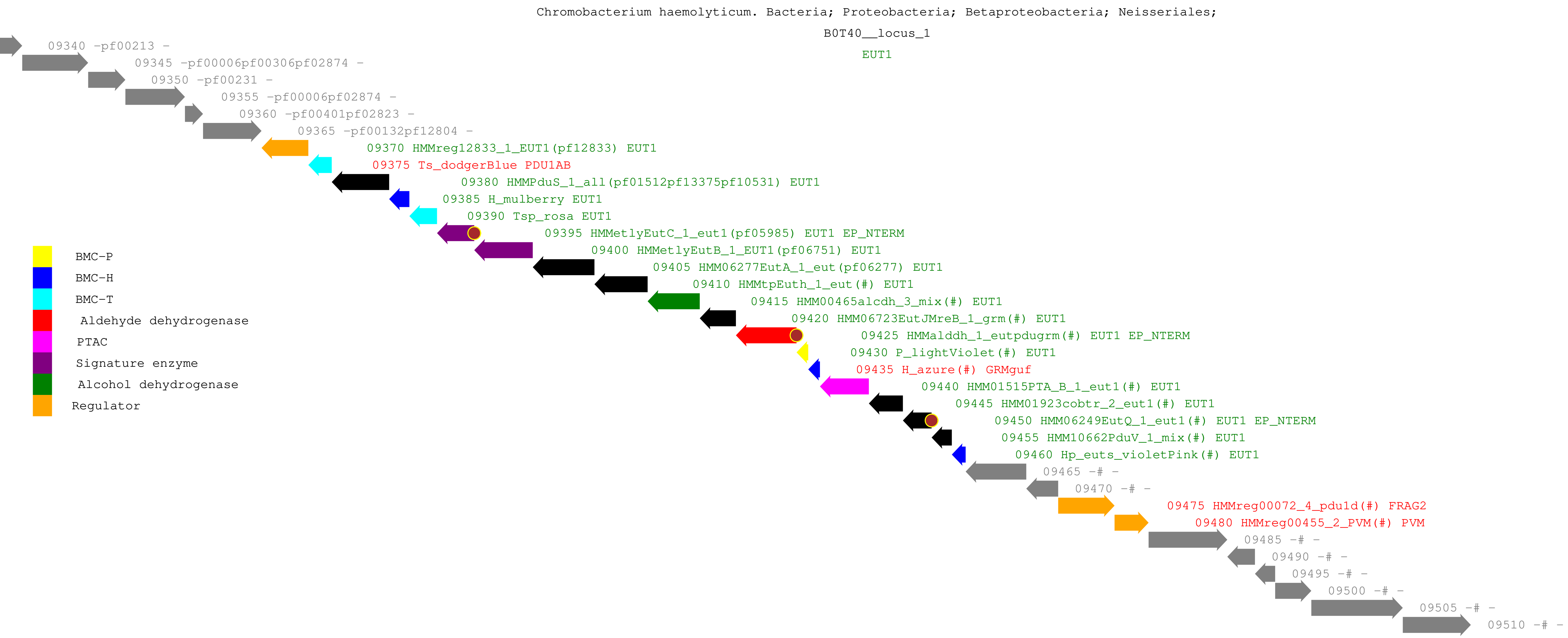
Chromobacterium haemolyticum. Bacteria; Proteobacteria; Betaproteobacteria; Neisseriales;
B0T40__locus_1
EUT1
09340 -pf00213 -
09345 -pf00006pf00306pf02874 -
09350 -pf00231 -
09355 -pf00006pf02874 -
09360 -pf00401pf02823 -
09365 -pf00132pf12804 -
09370 HMMreg12833_1_EUT1(pf12833) EUT1
09375 Ts_dodgerBlue PDU1AB
09380 HMMPduS_1_all(pf01512pf13375pf10531) EUT1
09385 H_mulberry EUT1
09390 Tsp_rosa EUT1
09395 HMMetlyEutC_1_eut1(pf05985) EUT1 EP_NTERM
09400 HMMetlyEutB_1_EUT1(pf06751) EUT1
09405 HMM06277EutA_1_eut(pf06277) EUT1
09410 HMMtpEuth_1_eut(#) EUT1
09415 HMM00465alcdh_3_mix(#) EUT1
09420 HMM06723EutJMreB_1_grm(#) EUT1
09425 HMMalddh_1_eutpdugrm(#) EUT1 EP_NTERM
09430 P_lightViolet(#) EUT1
09435 H_azure(#) GRMguf
09440 HMM01515PTA_B_1_eut1(#) EUT1
09445 HMM01923cobtr_2_eut1(#) EUT1
09450 HMM06249EutQ_1_eut1(#) EUT1 EP_NTERM
09455 HMM10662PduV_1_mix(#) EUT1
09460 Hp_euts_violetPink(#) EUT1
09465 -# -
09470 -# -
09475 HMMreg00072_4_pdu1d(#) FRAG2
09480 HMMreg00455_2_PVM(#) PVM
09485 -# -
09490 -# -
09495 -# -
09500 -# -
09505 -# -
09510 -# -
BMC-P
BMC-H
BMC-T
Aldehyde dehydrogenase
PTAC
Signature enzyme
Alcohol dehydrogenase
Regulator

Salmonella enterica subsp. enterica serovar Irumu. Bacteria; Proteobacteria; Gammaproteobacteria; Enterobacterales;

B0T94__locus_2_main_sat

EUT1

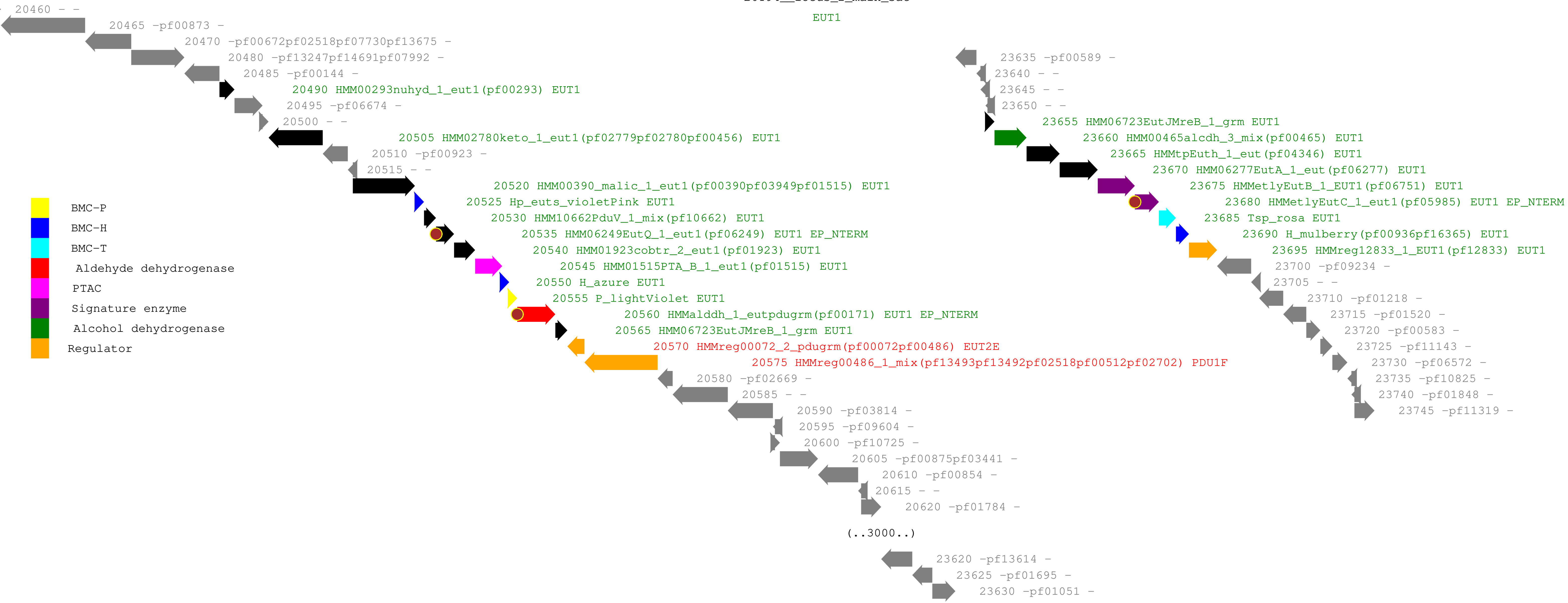

Escherichia coli. Bacteria; Proteobacteria; Gammaproteobacteria; Enterobacterales;

B1K96__locus_2

EUT1

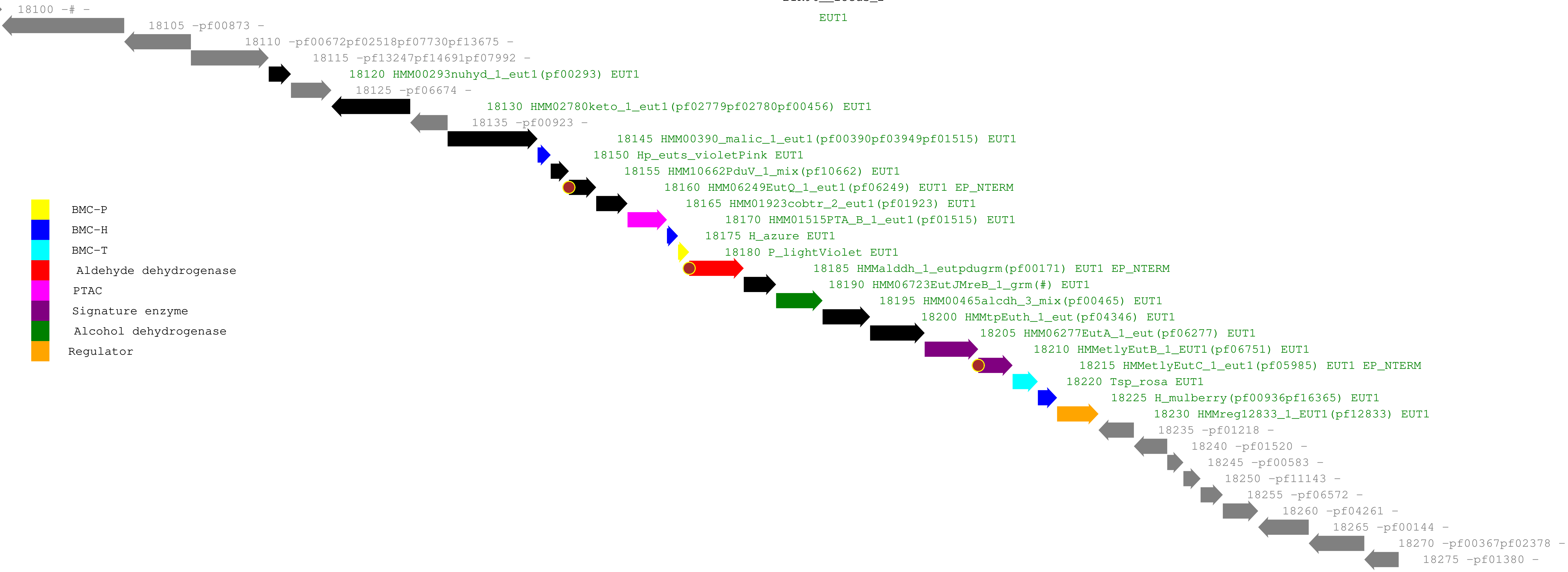

Escherichia coli M8. Bacteria; Proteobacteria; Gammaproteobacteria; Enterobacterales;

B2H83__locus_2

EUT1

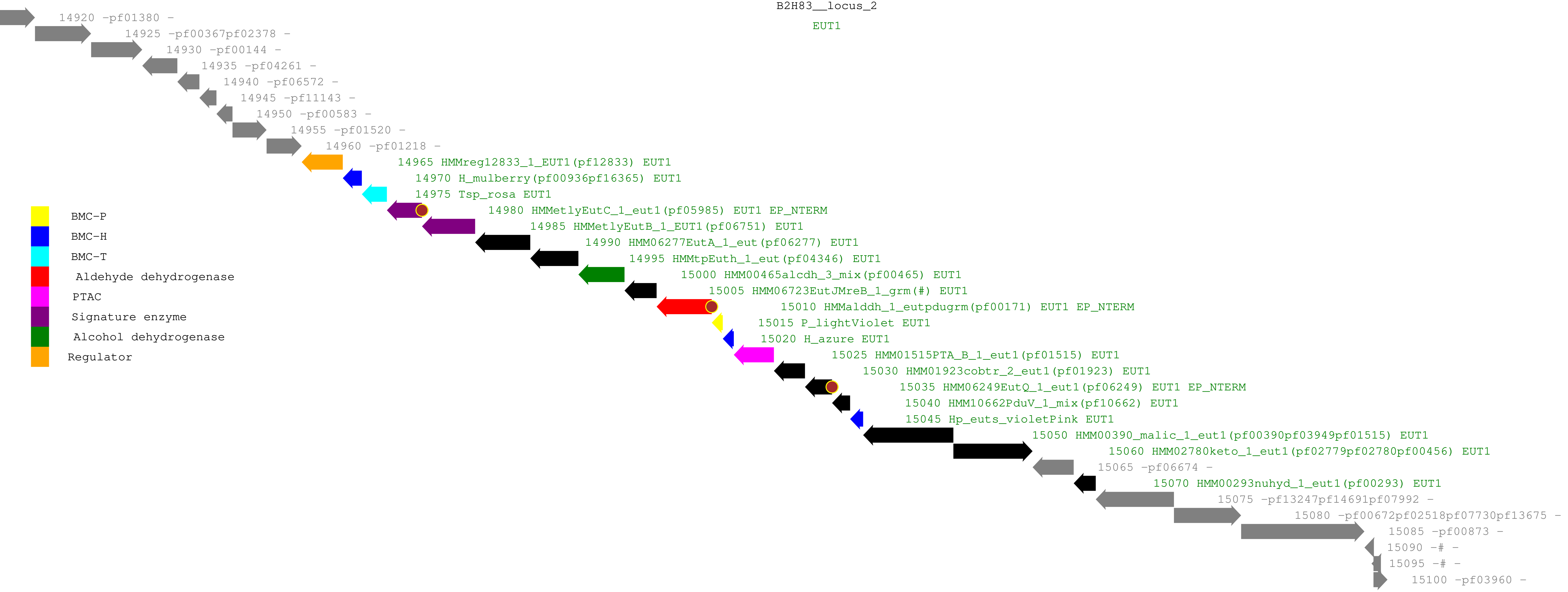

Klebsiella pneumoniae. Bacteria; Proteobacteria; Gammaproteobacteria; Enterobacterales;

B4U22__locus_1

EUT1

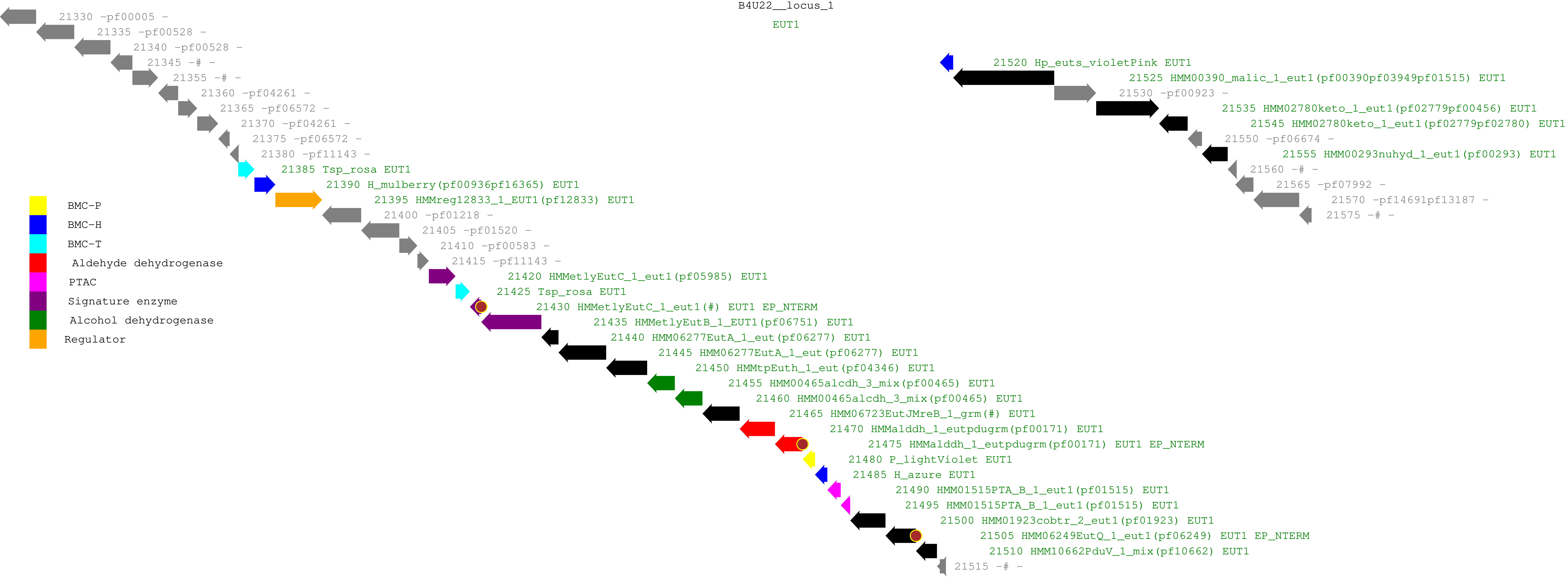

Klebsiella pneumoniae. Bacteria; Proteobacteria; Gammaproteobacteria; Enterobacterales;

B4U25__locus_1

EUT1

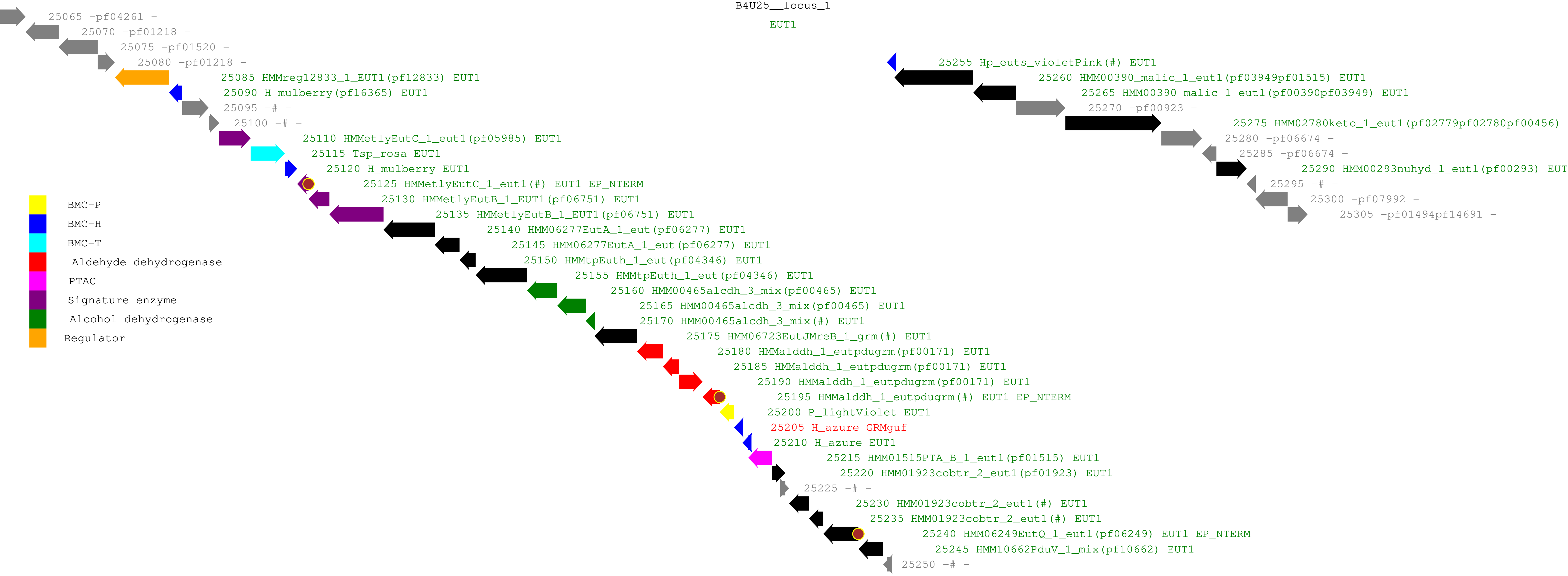

Klebsiella pneumoniae. Bacteria; Proteobacteria; Gammaproteobacteria; Enterobacterales;

B4U27__locus_1

EUT1

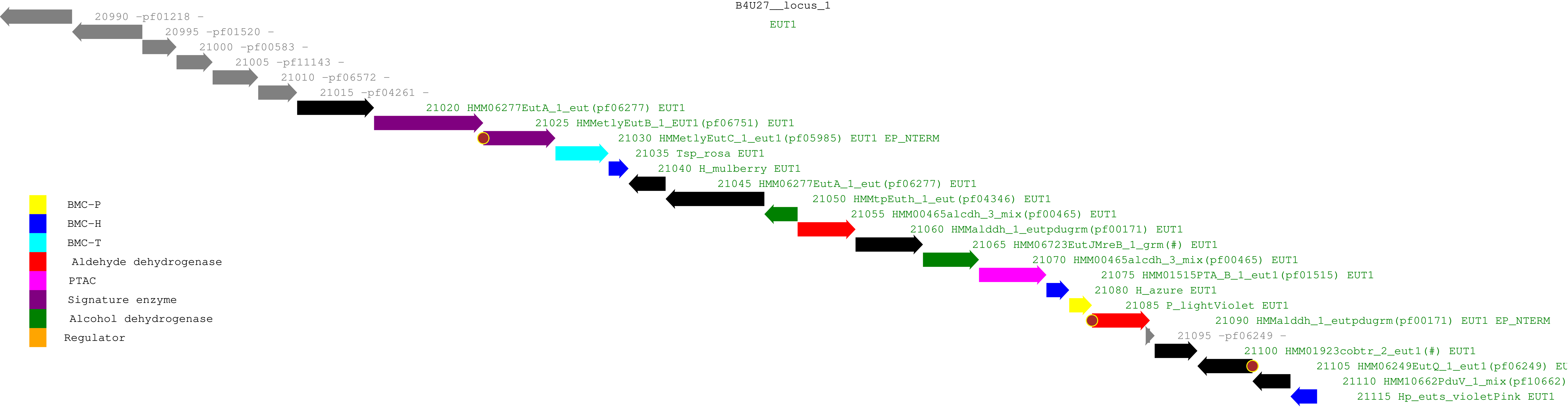

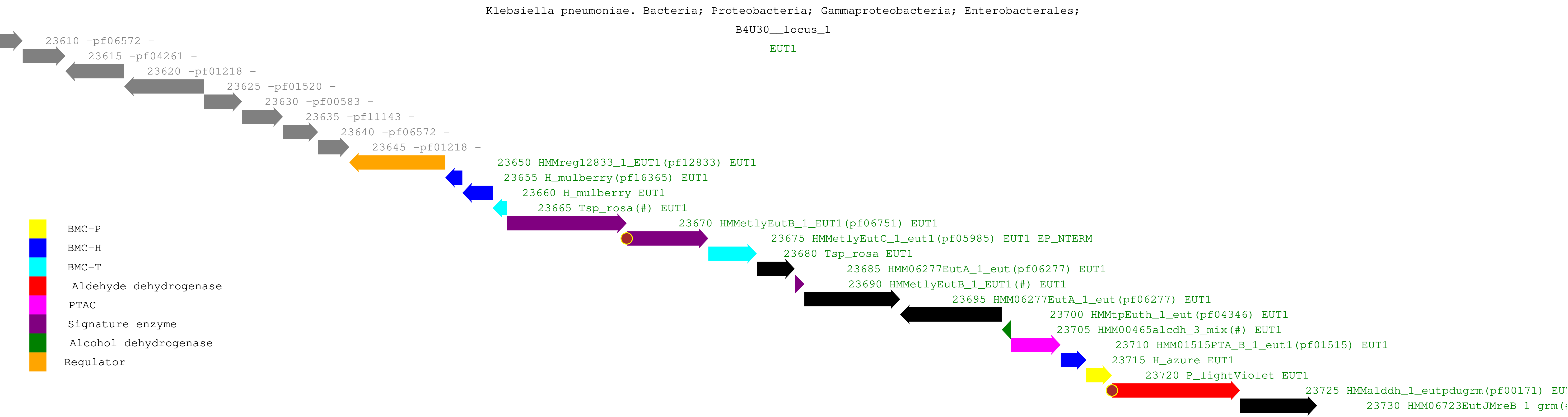
Klebsiella pneumoniae. Bacteria; Proteobacteria; Gammaproteobacteria; Enterobacterales;
B4U30__locus_1
EUT1
23610 -pf06572 -
23615 -pf04261 -
23620 -pf01218 -
23625 -pf01520 -
23630 -pf00583 -
23635 -pf11143 -
23640 -pf06572 -
23645 -pf01218 -
23650 HMMreg12833_1_EUT1(pf12833) EUT1
23655 H_mulberry(pf16365) EUT1
23660 H_mulberry EUT1
23665 Tsp_rosa(#) EUT1
23670 HMMetlyEutB_1_EUT1(pf06751) EUT1
23675 HMMetlyEutC_1_eut1(pf05985) EUT1 EP_NTERM
23680 Tsp_rosa EUT1
23685 HMM06277EutA_1_eut(pf06277) EUT1
23690 HMMetlyEutB_1_EUT1(#) EUT1
23695 HMM06277EutA_1_eut(pf06277) EUT1
23700 HMMtpEuth_1_eut(pf04346) EUT1
23705 HMM00465alcdh_3_mix(#) EUT1
23710 HMM01515PTA_B_1_eut1(pf01515) EUT1
23715 H_azure EUT1
23720 P_lightViolet EUT1
23725 HMMalddh_1_eutpdugrm(pf00171) EU
23730 HMM06723EutJMreB_1_grm(
BMC-P
BMC-H
BMC-T
Aldehyde dehydrogenase
PTAC
Signature enzyme
Alcohol dehydrogenase
Regulator

Klebsiella pneumoniae. Bacteria; Proteobacteria; Gammaproteobacteria; Enterobacterales;

B4U61__locus_1

EUT1

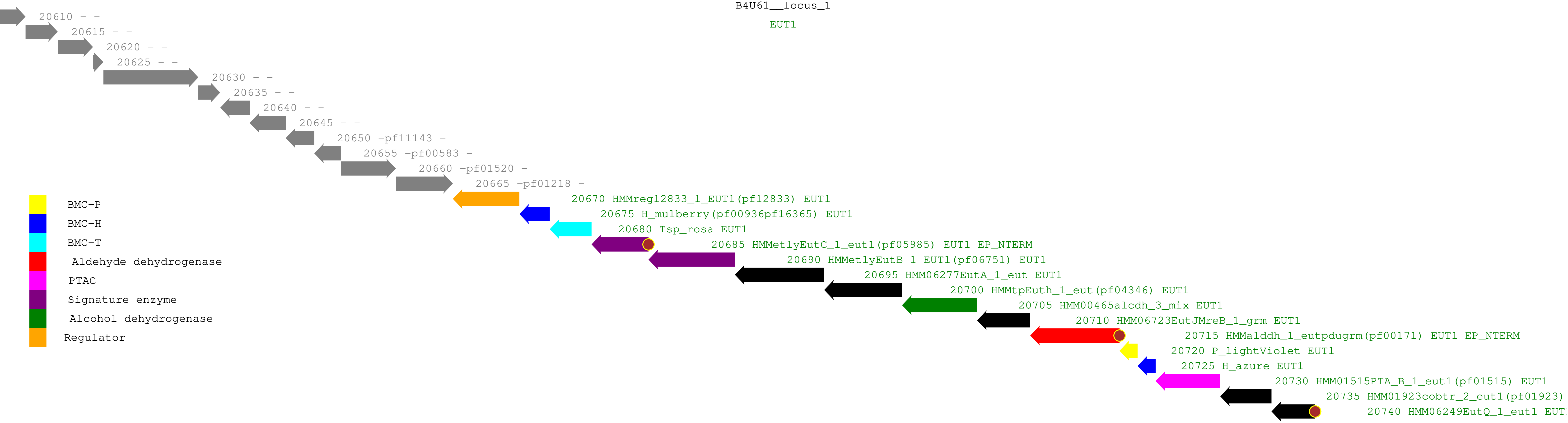

Salmonella enteritidis. Bacteria; Proteobacteria; Gammaproteobacteria; Enterobacterales;

B5567__locus_2

EUT1

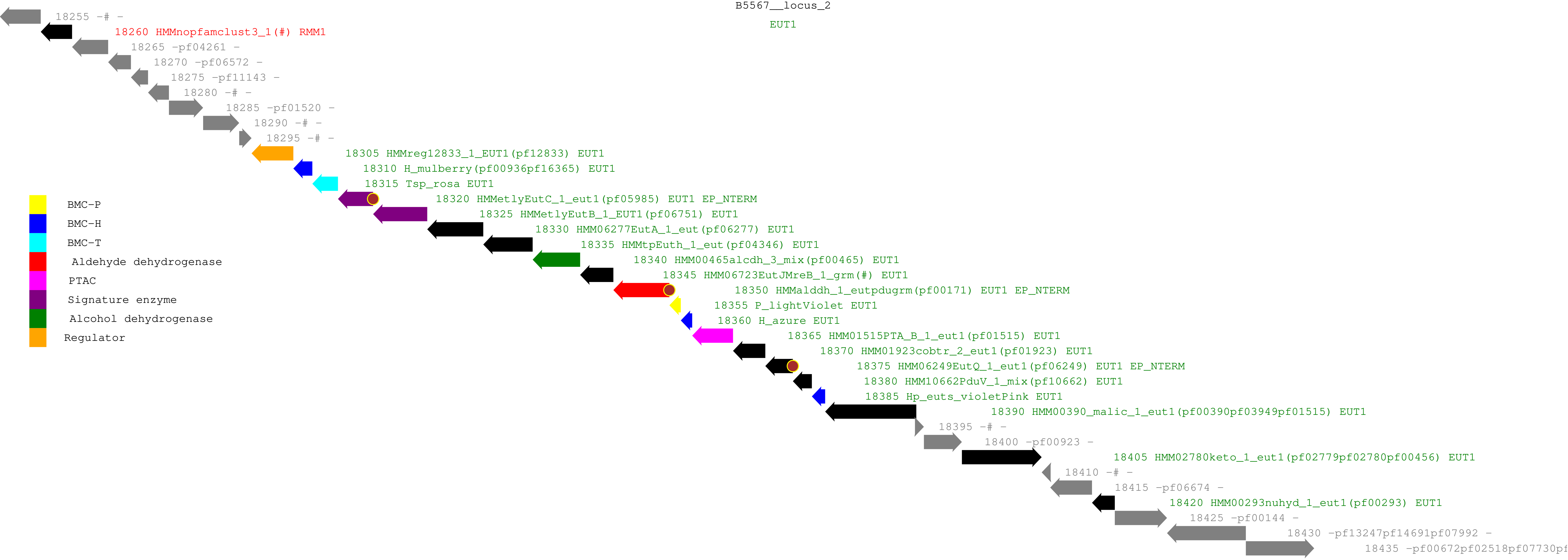

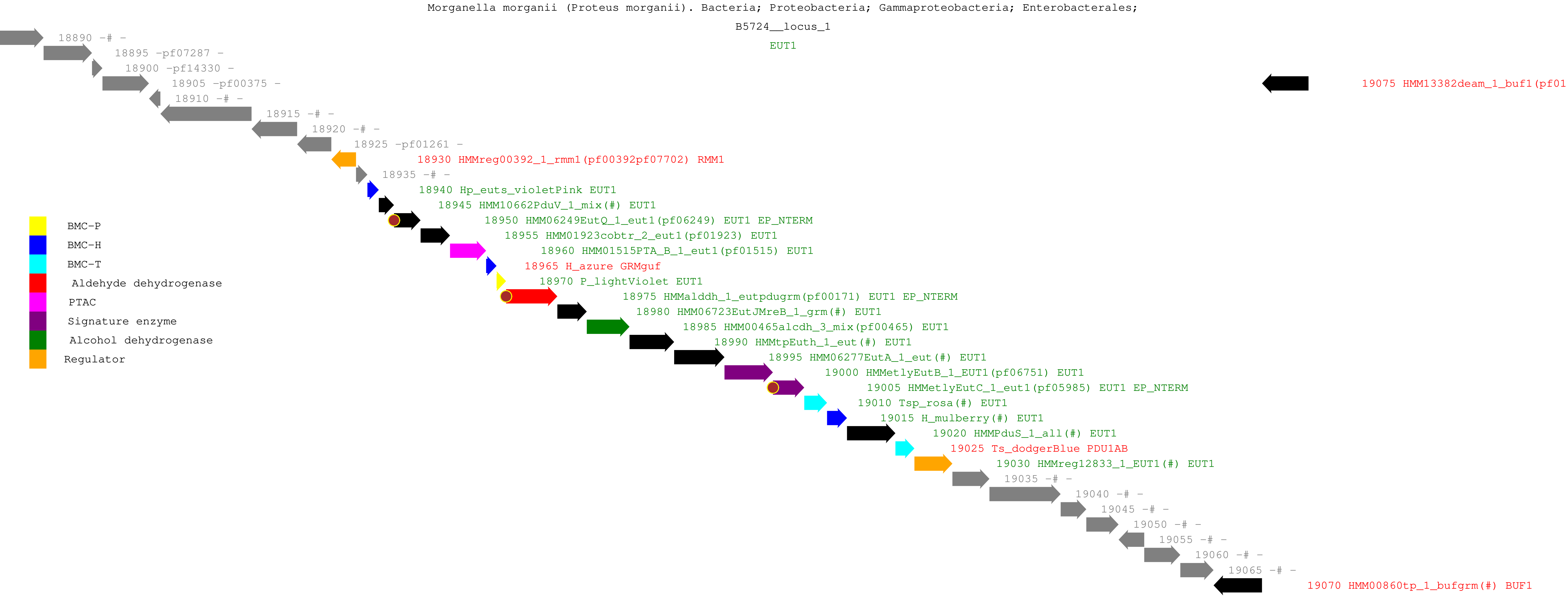

Morganella morganii (Proteus morganii). Bacteria; Proteobacteria; Gammaproteobacteria; Enterobacterales;
B5724__locus_1
EUT1
18890 -# -
18895 -pf07287 -
18900 -pf14330 -
18905 -pf00375 -
18910 -# -
18915 -# -
18920 -# -
18925 -pf01261 -
18930 HMMreg00392_1_rmm1(pf00392pf07702) RMM1
18935 -# -
18940 Hp_euts_violetPink EUT1
18945 HMM10662PduV_1_mix(#) EUT1
18950 HMM06249EutQ_1_eut1(pf06249) EUT1 EP_NTERM
18955 HMM01923cobtr_2_eut1(pf01923) EUT1
18960 HMM01515PTA_B_1_eut1(pf01515) EUT1
18965 H_azure GRMguf
18970 P_lightViolet EUT1
18975 HMMalddh_1_eutpdugrm(pf00171) EUT1 EP_NTERM
18980 HMM06723EutJMreB_1_grm(#) EUT1
18985 HMM00465alcdh_3_mix(pf00465) EUT1
18990 HMMtpEuth_1_eut(#) EUT1
18995 HMM06277EutA_1_eut(#) EUT1
19000 HMMetlyEutB_1_EUT1(pf06751) EUT1
19005 HMMetlyEutC_1_eut1(pf05985) EUT1 EP_NTERM
19010 Tsp_rosa(#) EUT1
19015 H_mulberry(#) EUT1
19020 HMMPduS_1_all(#) EUT1
19025 Ts_dodgerBlue PDU1AB
19030 HMMreg12833_1_EUT1(#) EUT1
19035 -# -
19040 -# -
19045 -# -
19050 -# -
19055 -# -
19060 -# -
19065 -# -
19070 HMM00860tp_1_bufgrm(#) BUF1
19075 HMM13382deam_1_buf1(pf01
BMC-P
BMC-H
BMC-T
Aldehyde dehydrogenase
PTAC
Signature enzyme
Alcohol dehydrogenase
Regulator

Klebsiella pneumoniae. Bacteria; Proteobacteria; Gammaproteobacteria; Enterobacterales;

B5G58__locus_2

EUT1

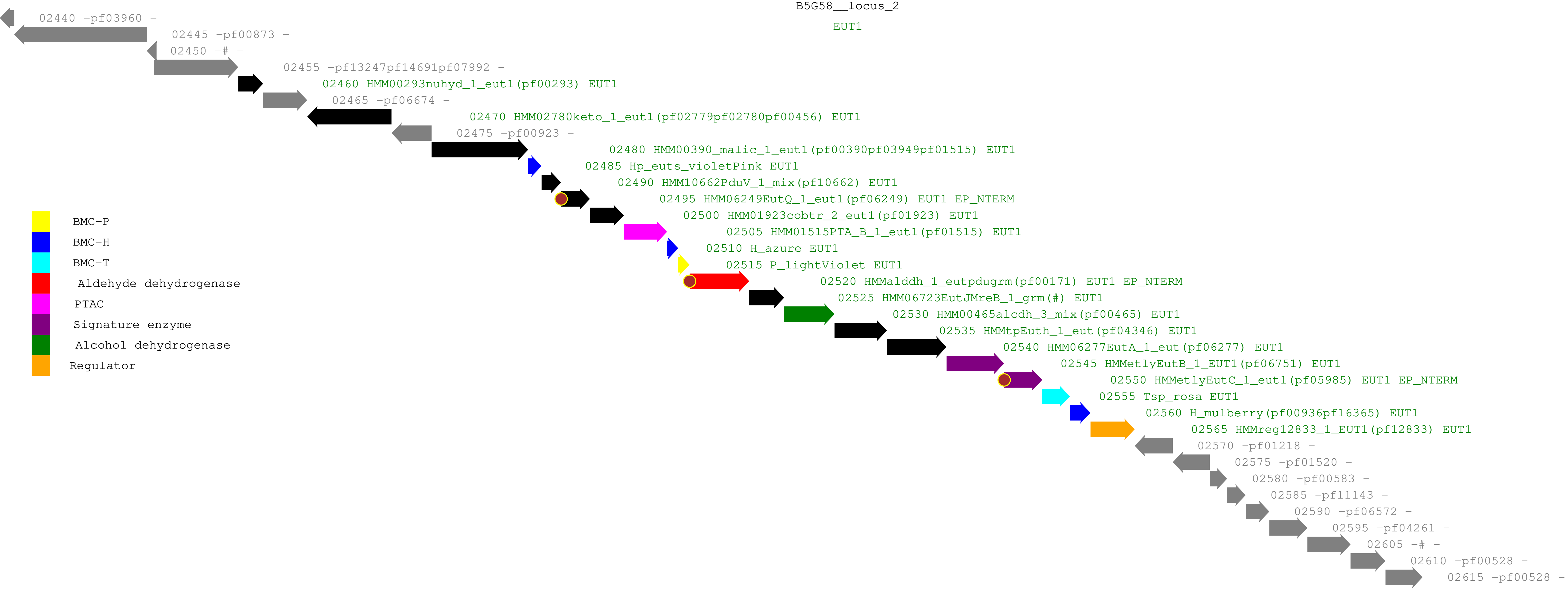

Klebsiella pneumoniae subsp. pneumoniae. Bacteria; Proteobacteria; Gammaproteobacteria; Enterobacterales;

B5K12__locus_1

EUT1

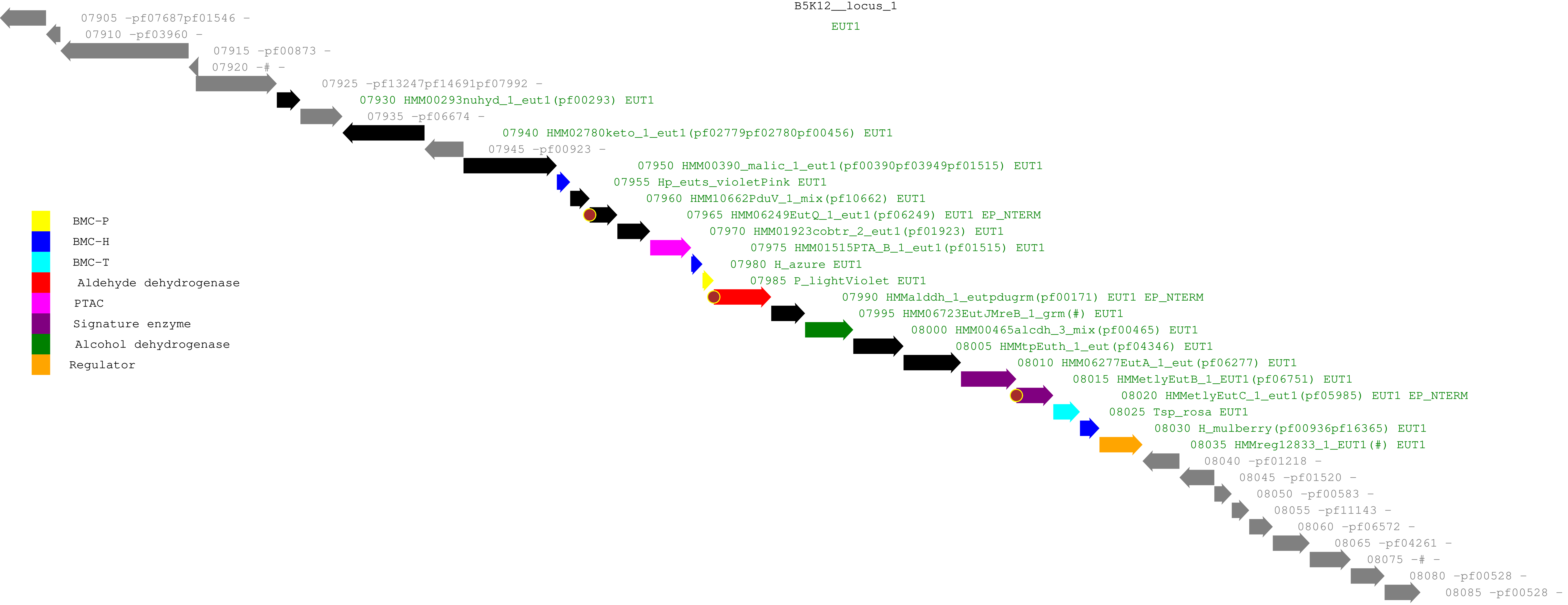

Pluralibacter gergoviae (Enterobacter gergoviae). Bacteria; Proteobacteria; Gammaproteobacteria; Enterobacterales;

B5M10__locus_1_main_sat

EUT1

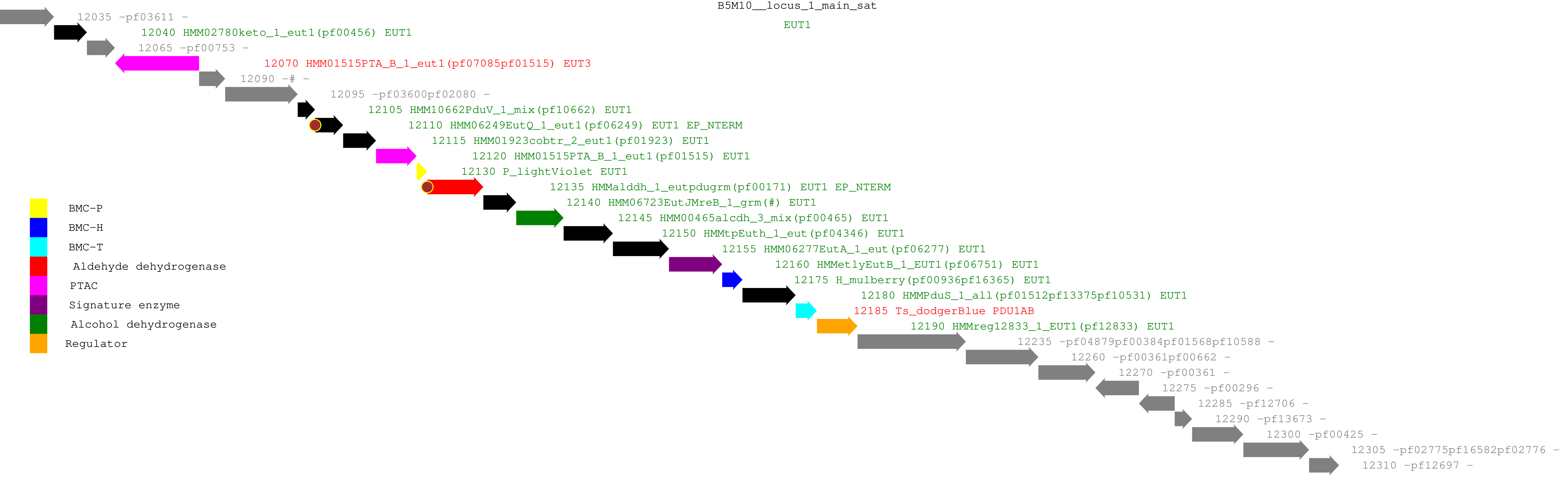

Morganella morganii (Proteus morganii). Bacteria; Proteobacteria; Gammaproteobacteria; Enterobacterales;

B5S45__locus_1

EUT1

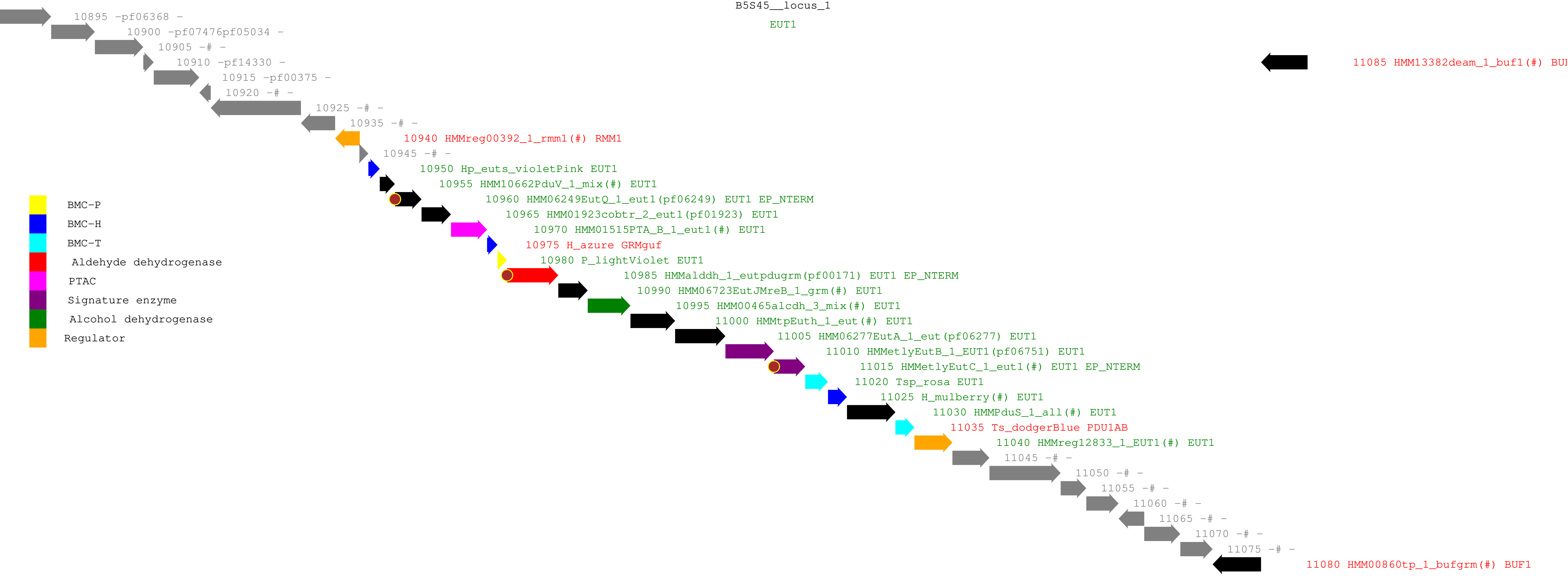

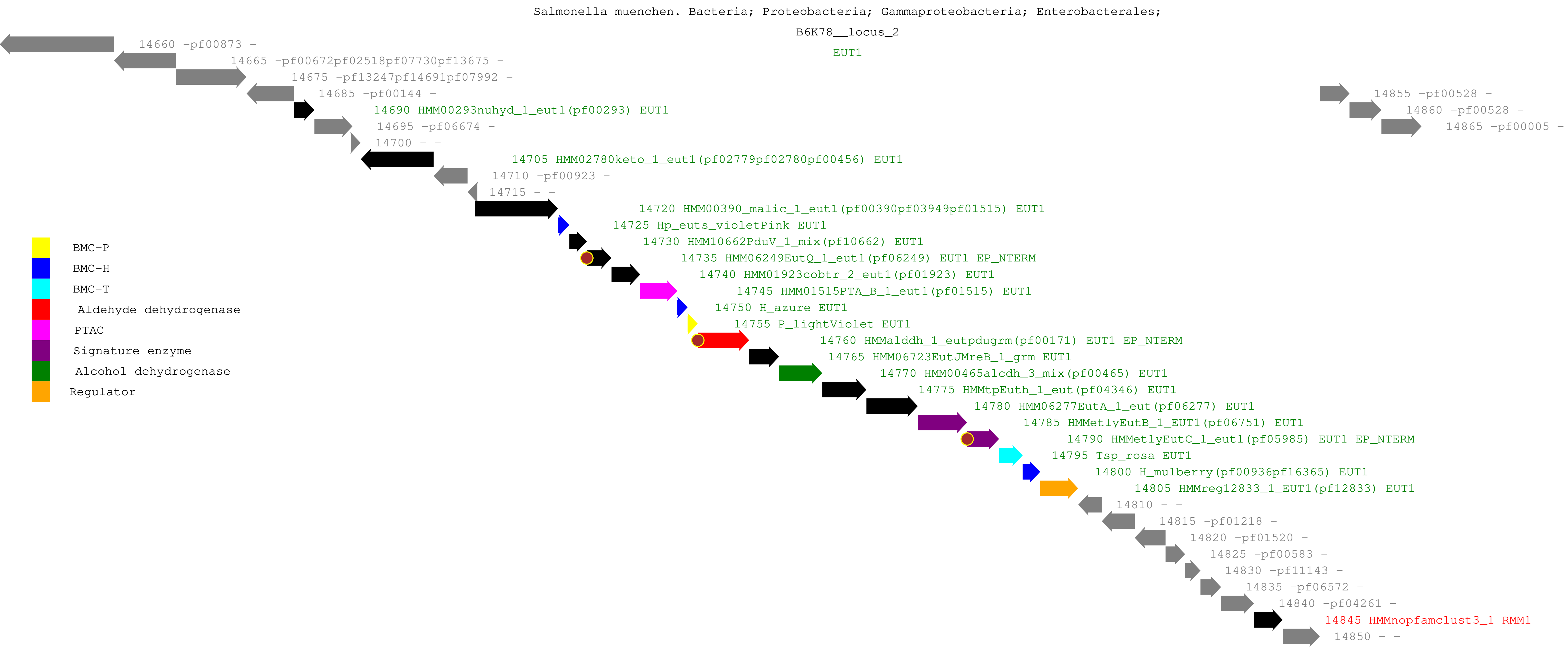
Salmonella muenchen. Bacteria; Proteobacteria; Gammaproteobacteria; Enterobacterales;
B6K78__locus_2
EUT1
14660 -pf00873 -
14665 -pf00672pf02518pf07730pf13675 -
14675 -pf13247pf14691pf07992 -
14685 -pf00144 -
14690 HMM00293nuhyd_1_eut1(pf00293) EUT1
14695 -pf06674 -
14700 - -
14705 HMM02780keto_1_eut1(pf02779pf02780pf00456) EUT1
14710 -pf00923 -
14715 - -
14720 HMM00390_malic_1_eut1(pf00390pf03949pf01515) EUT1
14725 Hp_euts_violetPink EUT1
14730 HMM10662PduV_1_mix(pf10662) EUT1
14735 HMM06249EutQ_1_eut1(pf06249) EUT1 EP_NTERM
14740 HMM01923cobtr_2_eut1(pf01923) EUT1
14745 HMM01515PTA_B_1_eut1(pf01515) EUT1
14750 H_azure EUT1
14755 P_lightViolet EUT1
14760 HMMalddh_1_eutpdugrm(pf00171) EUT1 EP_NTERM
14765 HMM06723EutJMreB_1_grm EUT1
14770 HMM00465alcdh_3_mix(pf00465) EUT1
14775 HMMtpEuth_1_eut(pf04346) EUT1
14780 HMM06277EutA_1_eut(pf06277) EUT1
14785 HMMetlyEutB_1_EUT1(pf06751) EUT1
14790 HMMetlyEutC_1_eut1(pf05985) EUT1 EP_NTERM
14795 Tsp_rosa EUT1
14800 H_mulberry(pf00936pf16365) EUT1
14805 HMMreg12833_1_EUT1(pf12833) EUT1
14810 - -
14815 -pf01218 -
14820 -pf01520 -
14825 -pf00583 -
14830 -pf11143 -
14835 -pf06572 -
14840 -pf04261 -
14845 HMMnopfamclust3_1 RMM1
14850 - -
14855 -pf00528 -
14860 -pf00528 -
14865 -pf00005 -
BMC-P
BMC-H
BMC-T
Aldehyde dehydrogenase
PTAC
Signature enzyme
Alcohol dehydrogenase
Regulator

Escherichia coli. Bacteria; Proteobacteria; Gammaproteobacteria; Enterobacterales;

B7C53__locus_1

EUT1

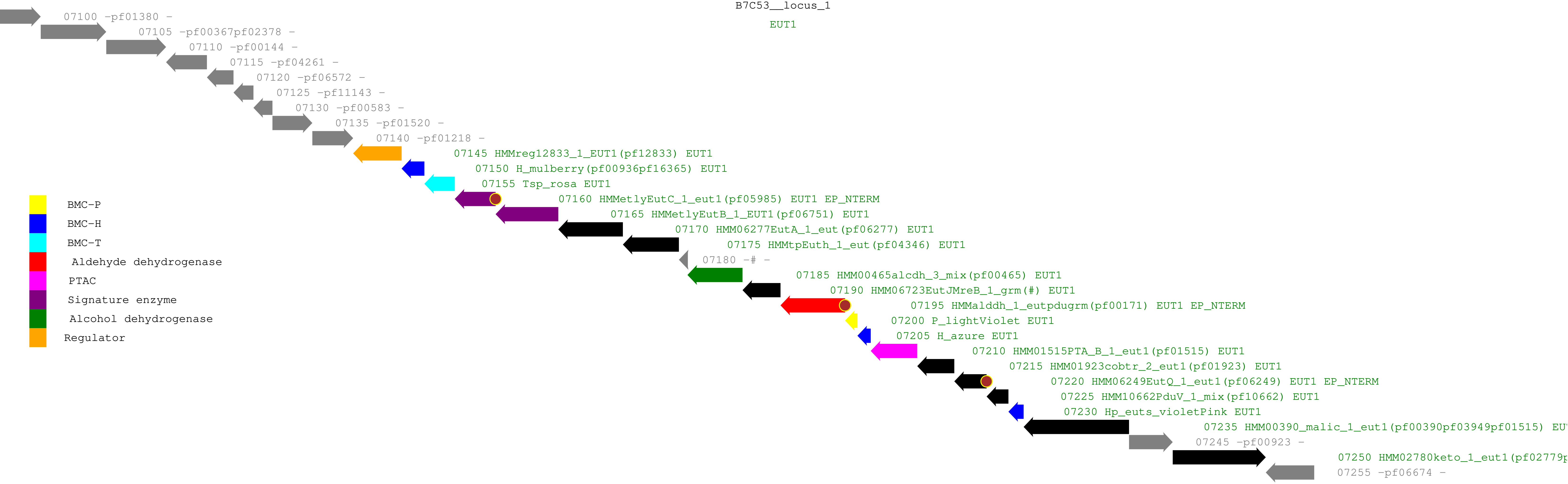

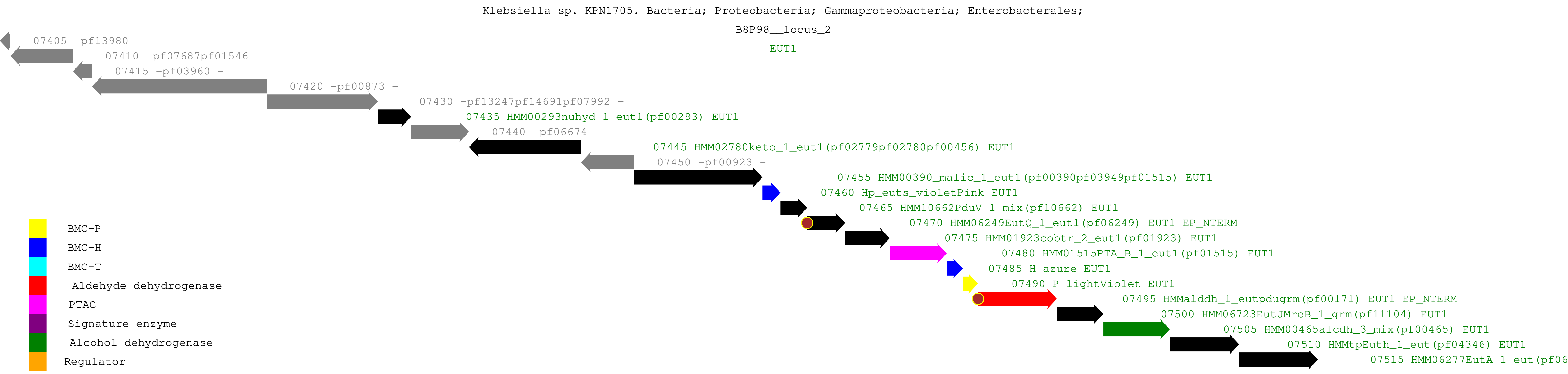

Klebsiella sp. KPN1705. Bacteria; Proteobacteria; Gammaproteobacteria; Enterobacterales;
B8P98__locus_2
EUT1
07405 -pf13980 -
07410 -pf07687pf01546 -
07415 -pf03960 -
07420 -pf00873 -
07430 -pf13247pf14691pf07992 -
07435 HMM00293nuhyd_1_eut1(pf00293) EUT1
07440 -pf06674 -
07445 HMM02780keto_1_eut1(pf02779pf02780pf00456) EUT1
07450 -pf00923 -
07455 HMM00390_malic_1_eut1(pf00390pf03949pf01515) EUT1
07460 Hp_euts_violetPink EUT1
07465 HMM10662PduV_1_mix(pf10662) EUT1
07470 HMM06249EutQ_1_eut1(pf06249) EUT1 EP_NTERM
07475 HMM01923cobtr_2_eut1(pf01923) EUT1
07480 HMM01515PTA_B_1_eut1(pf01515) EUT1
07485 H_azure EUT1
07490 P_lightViolet EUT1
07495 HMMalddh_1_eutpdugrm(pf00171) EUT1 EP_NTERM
07500 HMM06723EutJMreB_1_grm(pf11104) EUT1
07505 HMM00465alcdh_3_mix(pf00465) EUT1
07510 HMMtpEuth_1_eut(pf04346) EUT1
07515 HMM06277EutA_1_eut(pf06
BMC-P
BMC-H
BMC-T
Aldehyde dehydrogenase
PTAC
Signature enzyme
Alcohol dehydrogenase
Regulator

Citrobacter werkmanii. Bacteria; Proteobacteria; Gammaproteobacteria; Enterobacterales;

B9081__locus_2

EUT1

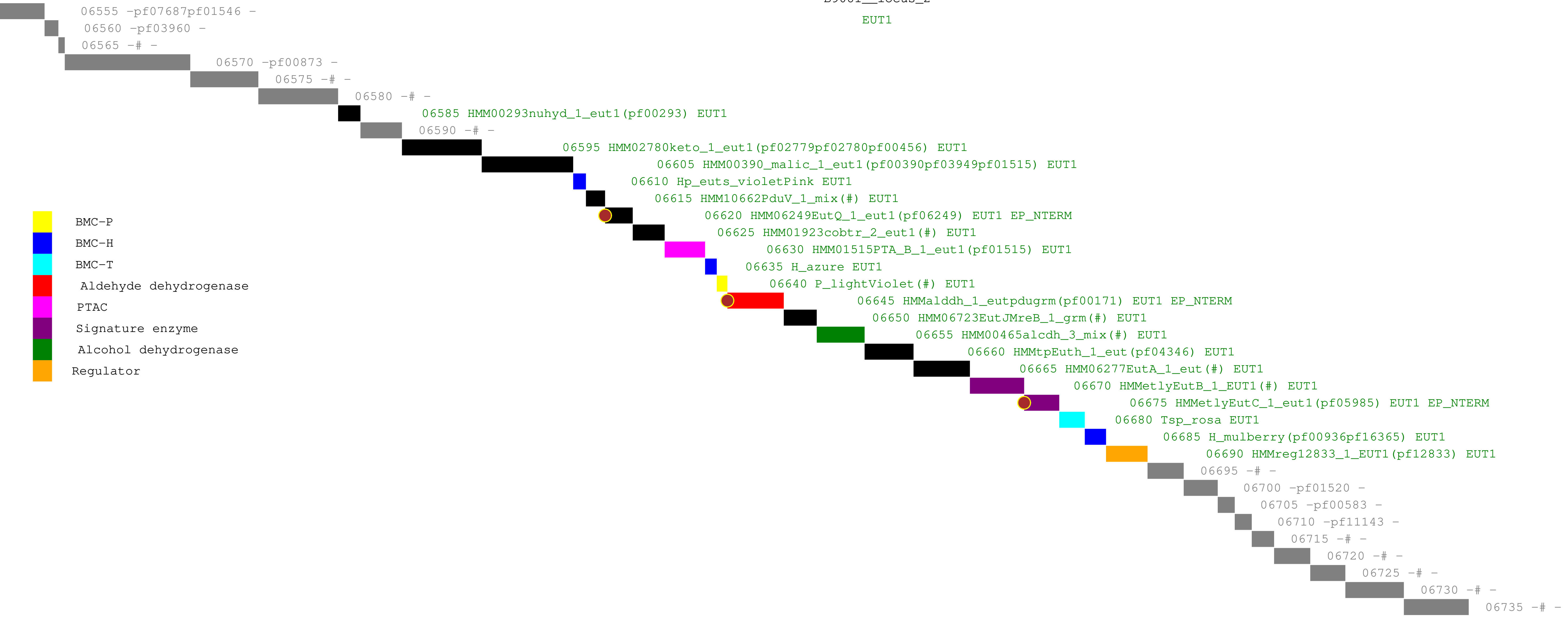

Citrobacter sp. L55. Bacteria; Proteobacteria; Gammaproteobacteria; Enterobacterales;

B9P82__locus_2

EUT1

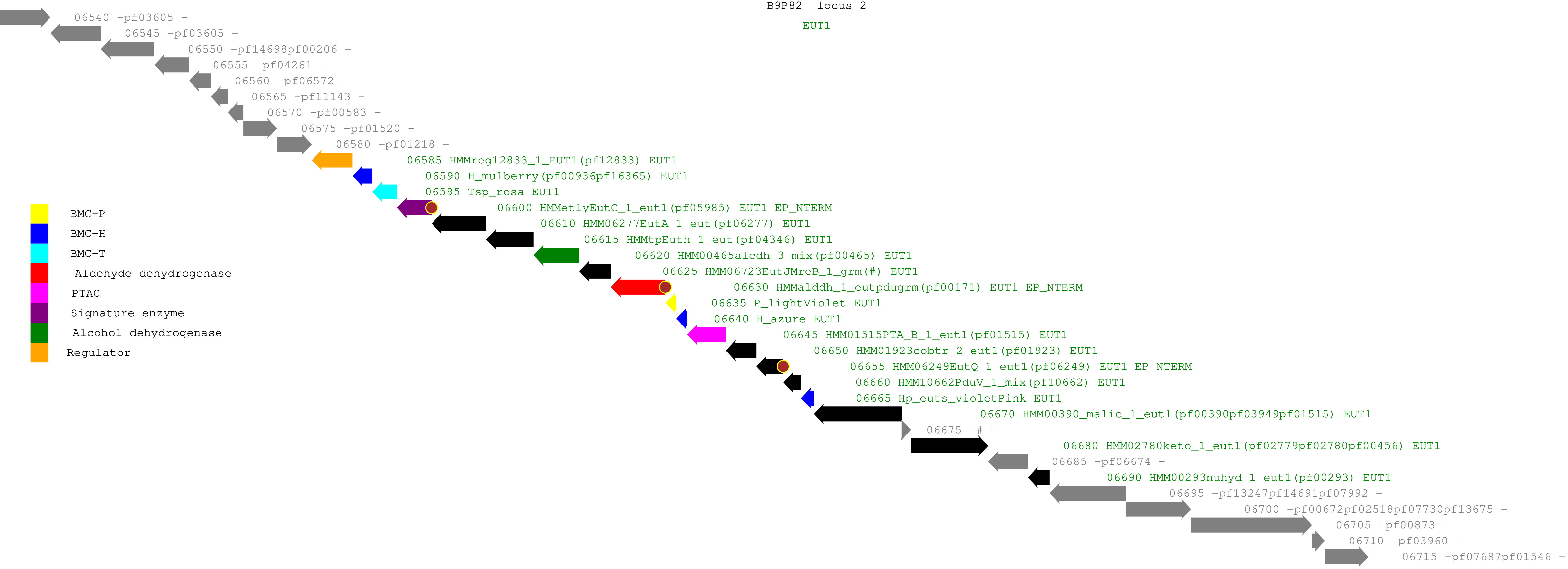

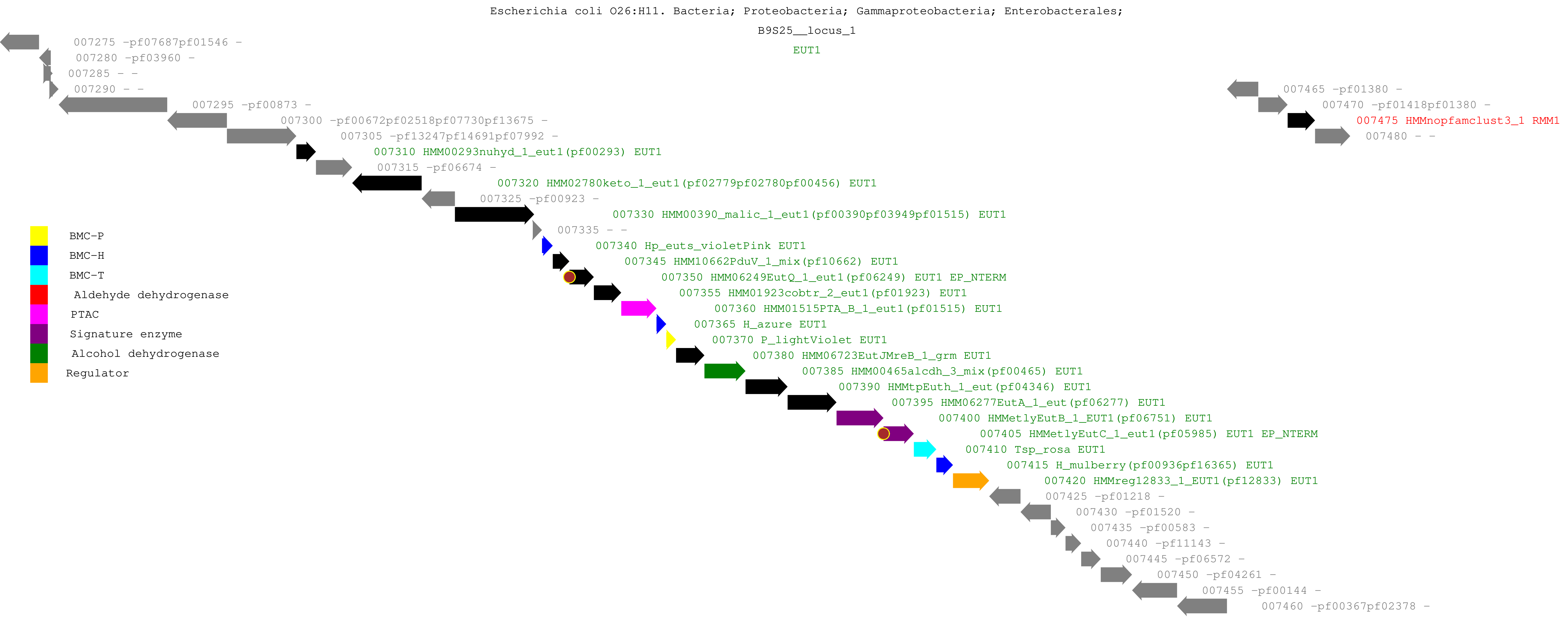
Escherichia coli O26:H11. Bacteria; Proteobacteria; Gammaproteobacteria; Enterobacterales;
B9S25__locus_1
EUT1
007275 -pf07687pf01546 -
007280 -pf03960 -
007285 - -
007290 - -
007295 -pf00873 -
007300 -pf00672pf02518pf07730pf13675 -
007305 -pf13247pf14691pf07992 -
007310 HMM00293nuhyd_1_eut1(pf00293) EUT1
007315 -pf06674 -
007320 HMM02780keto_1_eut1(pf02779pf02780pf00456) EUT1
007325 -pf00923 -
007330 HMM00390_malic_1_eut1(pf00390pf03949pf01515) EUT1
007335 - -
007340 Hp_euts_violetPink EUT1
007345 HMM10662PduV_1_mix(pf10662) EUT1
007350 HMM06249EutQ_1_eut1(pf06249) EUT1 EP_NTERM
007355 HMM01923cobtr_2_eut1(pf01923) EUT1
007360 HMM01515PTA_B_1_eut1(pf01515) EUT1
007365 H_azure EUT1
007370 P_lightViolet EUT1
007380 HMM06723EutJMreB_1_grm EUT1
007385 HMM00465alcdh_3_mix(pf00465) EUT1
007390 HMMtpEuth_1_eut(pf04346) EUT1
007395 HMM06277EutA_1_eut(pf06277) EUT1
007400 HMMetlyEutB_1_EUT1(pf06751) EUT1
007405 HMMetlyEutC_1_eut1(pf05985) EUT1 EP_NTERM
007410 Tsp_rosa EUT1
007415 H_mulberry(pf00936pf16365) EUT1
007420 HMMreg12833_1_EUT1(pf12833) EUT1
007425 -pf01218 -
007430 -pf01520 -
007435 -pf00583 -
007440 -pf11143 -
007445 -pf06572 -
007450 -pf04261 -
007455 -pf00144 -
007460 -pf00367pf02378 -
007465 -pf01380 -
007470 -pf01418pf01380 -
007475 HMMnopfamclust3_1 RMM1
007480 - -
BMC-P
BMC-H
BMC-T
Aldehyde dehydrogenase
PTAC
Signature enzyme
Alcohol dehydrogenase
Regulator

Escherichia coli O121:H19. Bacteria; Proteobacteria; Gammaproteobacteria; Enterobacterales;

B9X35__locus_1

EUT1

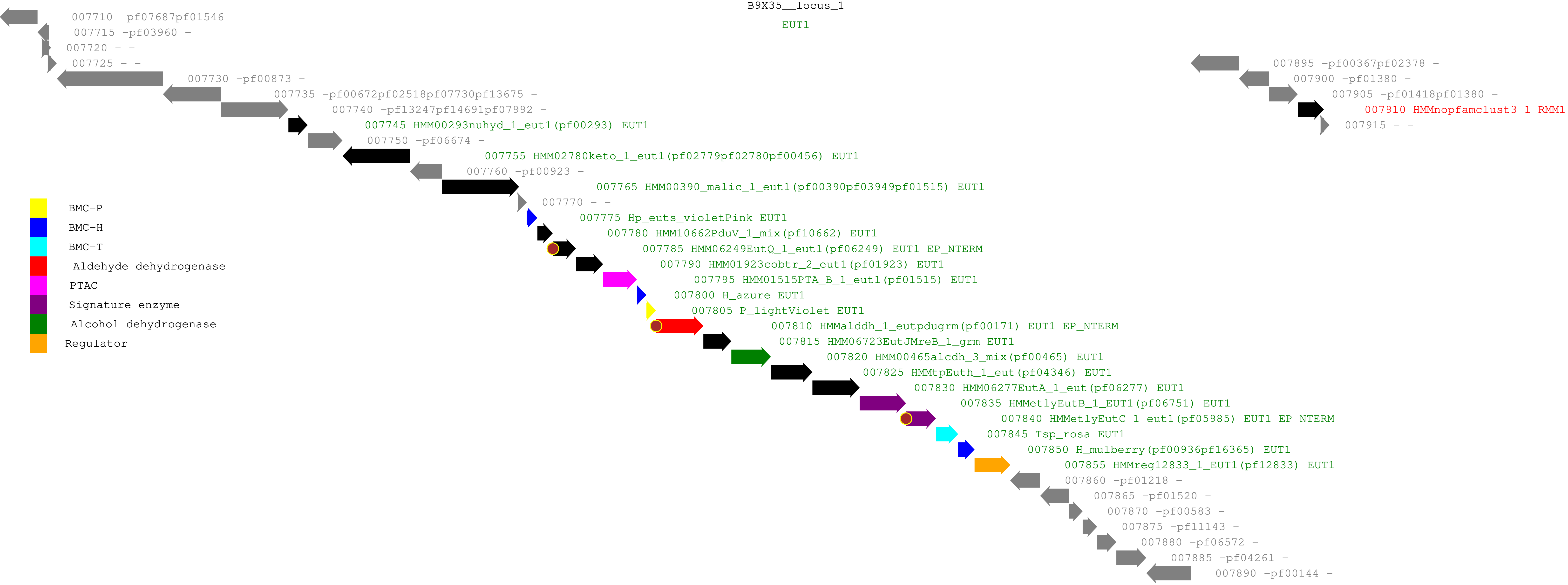

Morganella morganii subsp. morganii. Bacteria; Proteobacteria; Gammaproteobacteria; Enterobacterales;

B9Z91__locus_1

EUT1

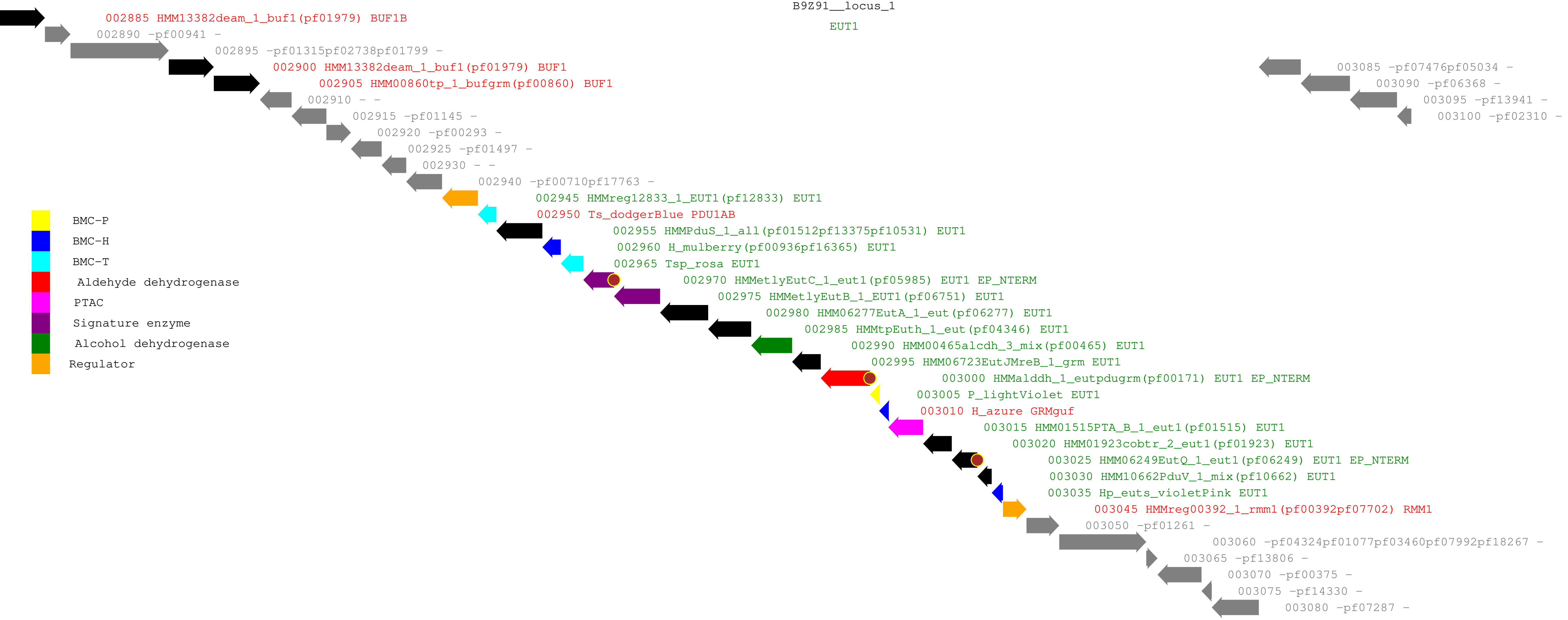

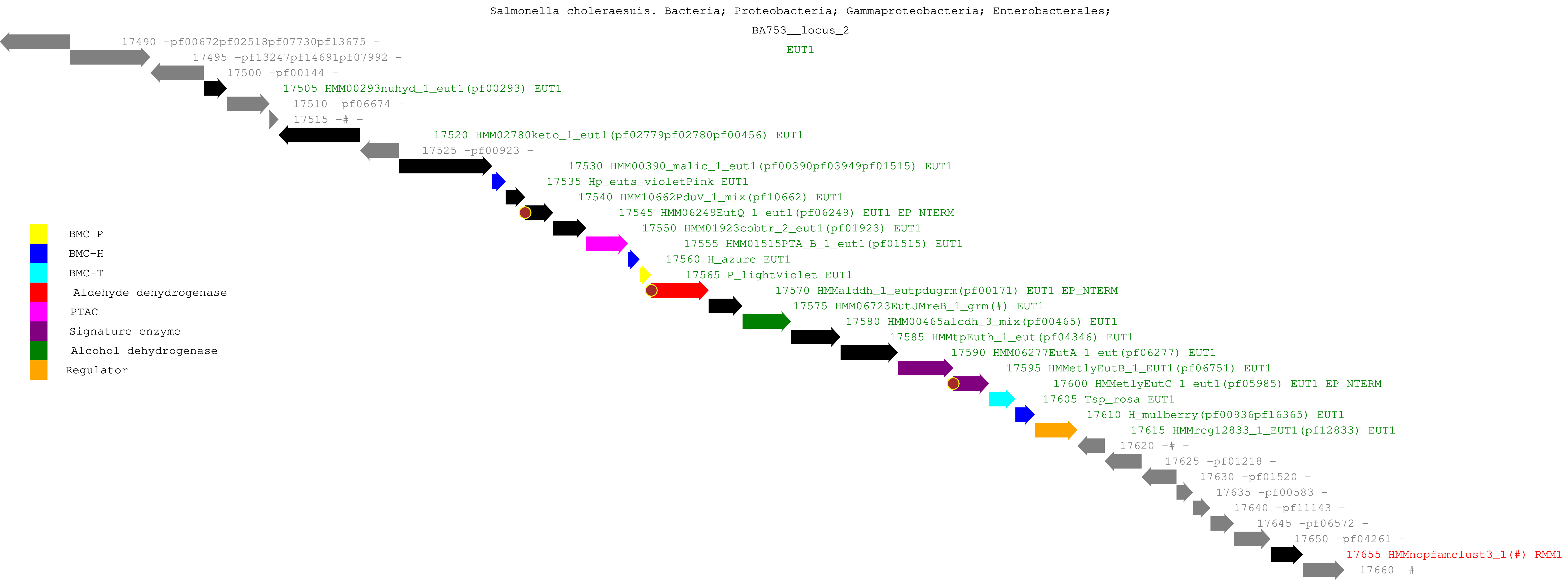
Salmonella choleraesuis. Bacteria; Proteobacteria; Gammaproteobacteria; Enterobacterales;
BA753__locus_2
EUT1
17490 -pf00672pf02518pf07730pf13675 -
17495 -pf13247pf14691pf07992 -
17500 -pf00144 -
17505 HMM00293nuhyd_1_eut1(pf00293) EUT1
17510 -pf06674 -
17515 -# -
17520 HMM02780keto_1_eut1(pf02779pf02780pf00456) EUT1
17525 -pf00923 -
17530 HMM00390_malic_1_eut1(pf00390pf03949pf01515) EUT1
17535 Hp_euts_violetPink EUT1
17540 HMM10662PduV_1_mix(pf10662) EUT1
17545 HMM06249EutQ_1_eut1(pf06249) EUT1 EP_NTERM
17550 HMM01923cobtr_2_eut1(pf01923) EUT1
17555 HMM01515PTA_B_1_eut1(pf01515) EUT1
17560 H_azure EUT1
17565 P_lightViolet EUT1
17570 HMMalddh_1_eutpdugrm(pf00171) EUT1 EP_NTERM
17575 HMM06723EutJMreB_1_grm(#) EUT1
17580 HMM00465alcdh_3_mix(pf00465) EUT1
17585 HMMtpEuth_1_eut(pf04346) EUT1
17590 HMM06277EutA_1_eut(pf06277) EUT1
17595 HMMetlyEutB_1_EUT1(pf06751) EUT1
17600 HMMetlyEutC_1_eut1(pf05985) EUT1 EP_NTERM
17605 Tsp_rosa EUT1
17610 H_mulberry(pf00936pf16365) EUT1
17615 HMMreg12833_1_EUT1(pf12833) EUT1
17620 -# -
17625 -pf01218 -
17630 -pf01520 -
17635 -pf00583 -
17640 -pf11143 -
17645 -pf06572 -
17650 -pf04261 -
17655 HMMnopfamclust3_1(#) RMM1
17660 -# -
BMC-P
BMC-H
BMC-T
Aldehyde dehydrogenase
PTAC
Signature enzyme
Alcohol dehydrogenase
Regulator

Pseudomonas sp. EGD-AKN5. Bacteria; Proteobacteria; Gammaproteobacteria; Pseudomonadales;

BAE52__locus_1

EUT1

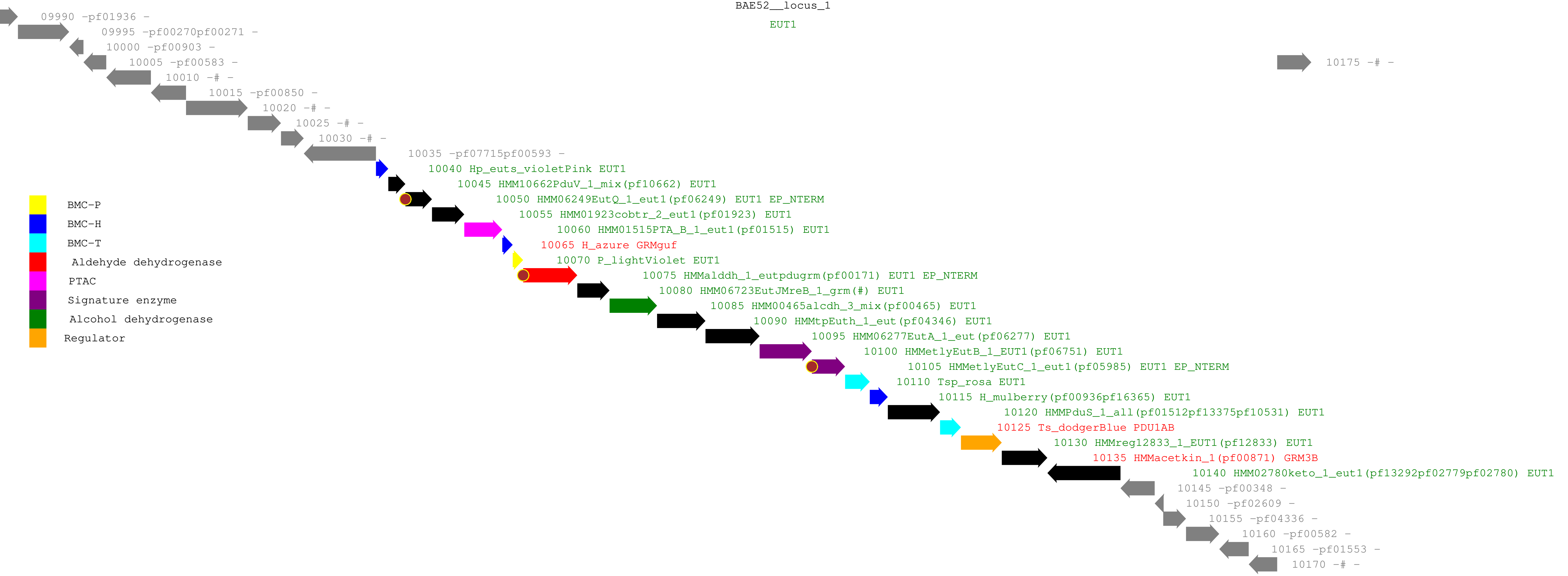

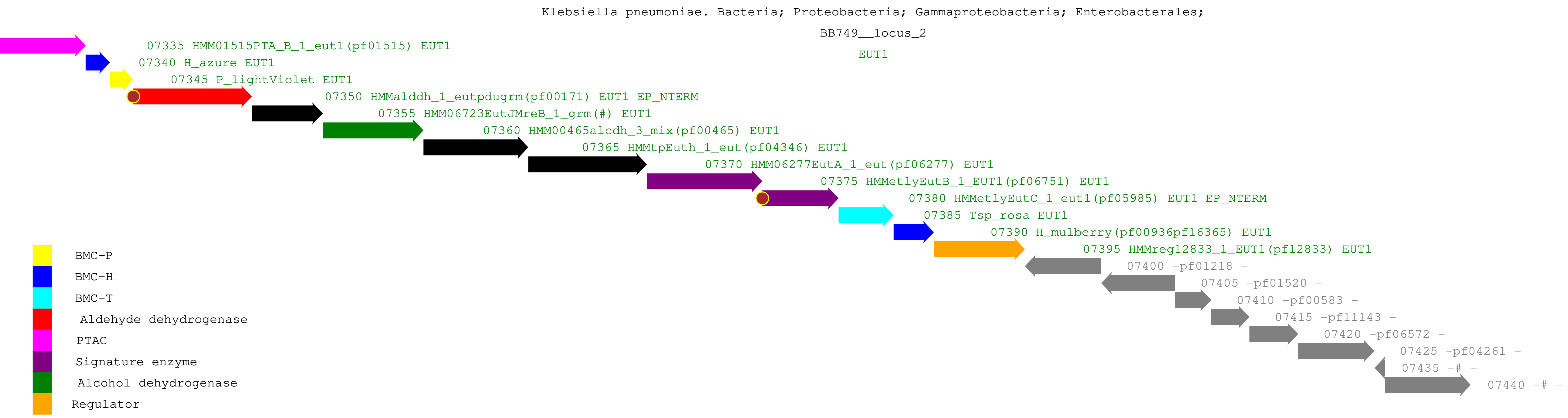
Klebsiella pneumoniae. Bacteria; Proteobacteria; Gammaproteobacteria; Enterobacterales;
BB749__locus_2
EUT1
07335 HMM01515PTA_B_1_eut1(pf01515) EUT1
07340 H_azure EUT1
07345 P_lightViolet EUT1
07350 HMMalddh_1_eutpdugrm(pf00171) EUT1 EP_NTERM
07355 HMM06723EutJMreB_1_grm(#) EUT1
07360 HMM00465alcdh_3_mix(pf00465) EUT1
07365 HMMtpEuth_1_eut(pf04346) EUT1
07370 HMM06277EutA_1_eut(pf06277) EUT1
07375 HMMetlyEutB_1_EUT1(pf06751) EUT1
07380 HMMetlyEutC_1_eut1(pf05985) EUT1 EP_NTERM
07385 Tsp_rosa EUT1
07390 H_mulberry(pf00936pf16365) EUT1
07395 HMMreg12833_1_EUT1(pf12833) EUT1
07400 -pf01218 -
07405 -pf01520 -
07410 -pf00583 -
07415 -pf11143 -
07420 -pf06572 -
07425 -pf04261 -
07435 -# -
07440 -# -
BMC-P
BMC-H
BMC-T
Aldehyde dehydrogenase
PTAC
Signature enzyme
Alcohol dehydrogenase
Regulator

Pluralibacter gergoviae (Enterobacter gergoviae). Bacteria; Proteobacteria; Gammaproteobacteria; Enterobacterales;

BB778__locus_1

EUT1

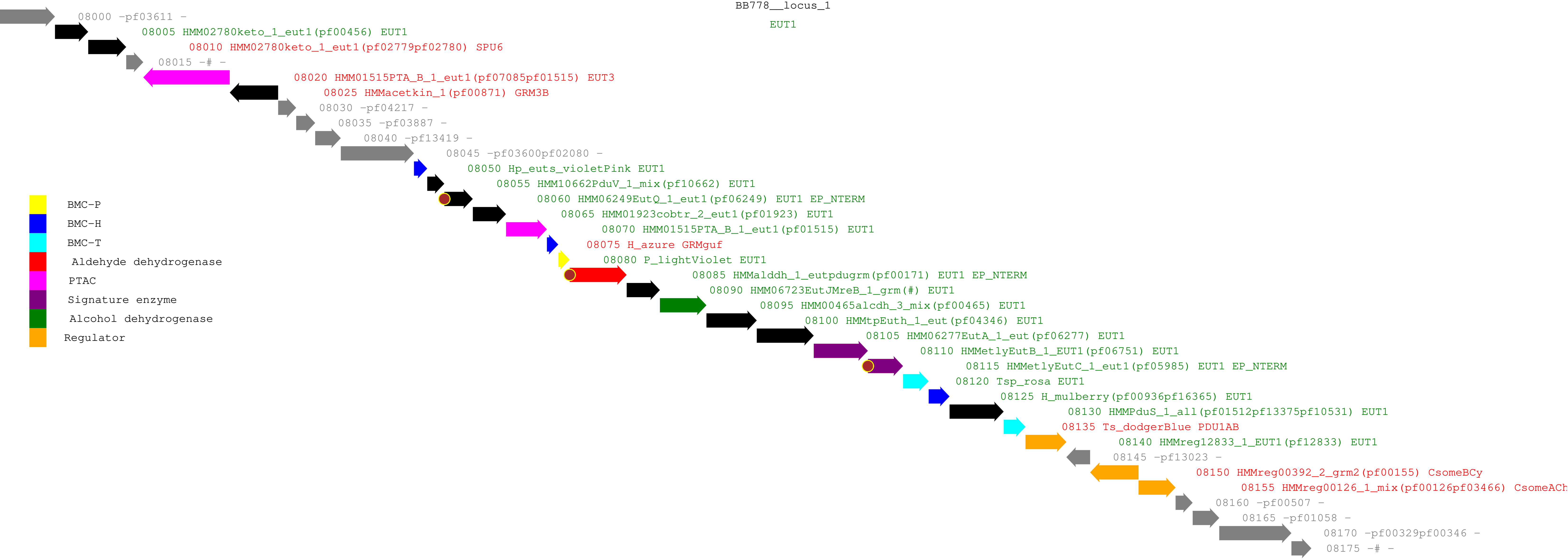

Salmonella enterica subsp. enterica serovar Javiana str. CFSAN000904. Bacteria; Proteobacteria; Gammaproteobacteria; Enterobacterales;

BBD49__locus_2

EUT1

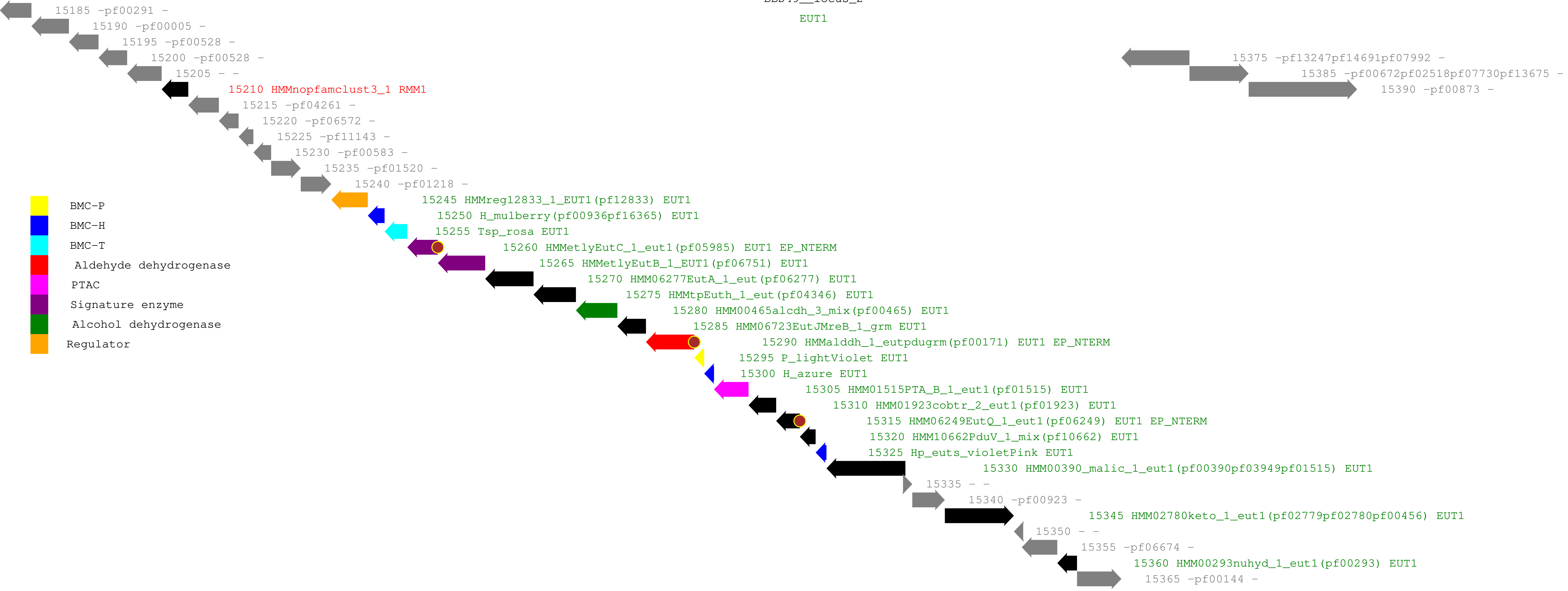

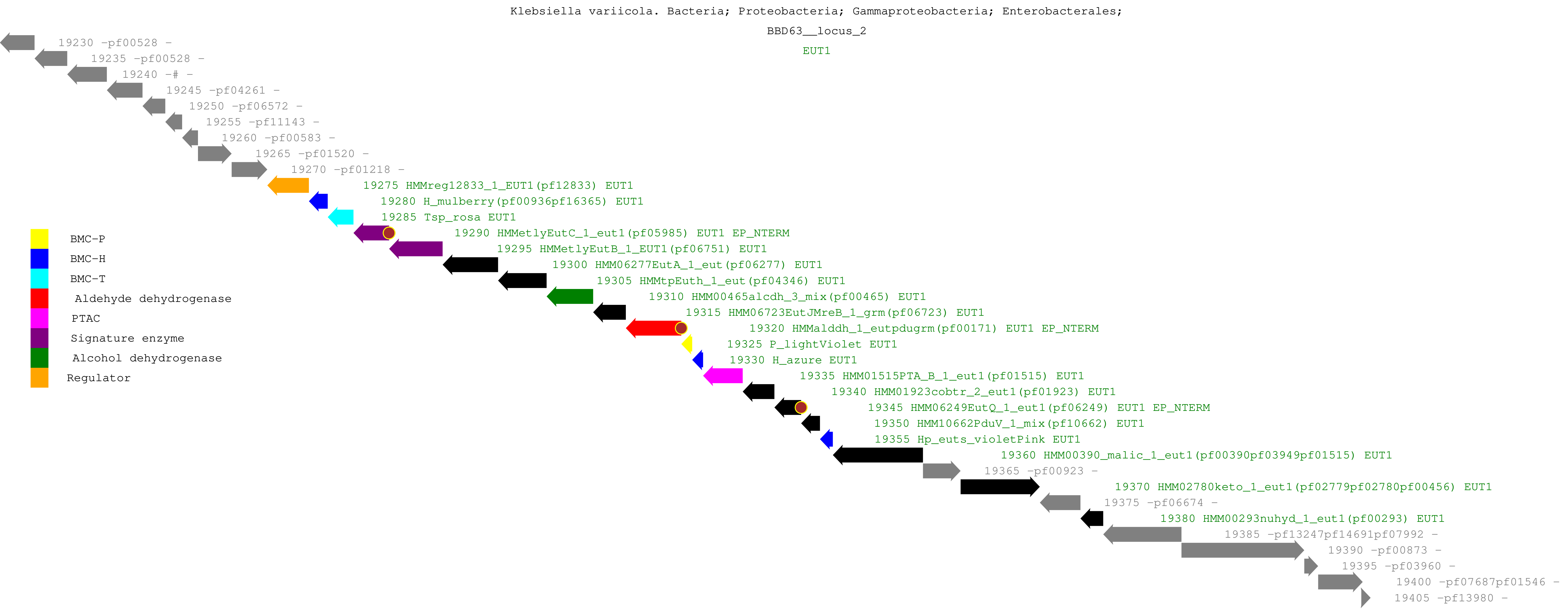
Klebsiella variicola. Bacteria; Proteobacteria; Gammaproteobacteria; Enterobacterales;
BBD63__locus_2
EUT1
19230 -pf00528 -
19235 -pf00528 -
19240 -# -
19245 -pf04261 -
19250 -pf06572 -
19255 -pf11143 -
19260 -pf00583 -
19265 -pf01520 -
19270 -pf01218 -
19275 HMMreg12833_1_EUT1(pf12833) EUT1
19280 H_mulberry(pf00936pf16365) EUT1
19285 Tsp_rosa EUT1
19290 HMMetlyEutC_1_eut1(pf05985) EUT1 EP_NTERM
19295 HMMetlyEutB_1_EUT1(pf06751) EUT1
19300 HMM06277EutA_1_eut(pf06277) EUT1
19305 HMMtpEuth_1_eut(pf04346) EUT1
19310 HMM00465alcdh_3_mix(pf00465) EUT1
19315 HMM06723EutJMreB_1_grm(pf06723) EUT1
19320 HMMaldh_1_eutpdugrm(pf00171) EUT1 EP_NTERM
19325 P_lightViolet EUT1
19330 H_azure EUT1
19335 HMM01515PTA_B_1_eut1(pf01515) EUT1
19340 HMM01923cobtr_2_eut1(pf01923) EUT1
19345 HMM06249EutQ_1_eut1(pf06249) EUT1 EP_NTERM
19350 HMM10662PduV_1_mix(pf10662) EUT1
19355 Hp_euts_violetPink EUT1
19360 HMM00390_malic_1_eut1(pf00390pf03949pf01515) EUT1
19365 -pf00923 -
19370 HMM02780keto_1_eut1(pf02779pf02780pf00456) EUT1
19375 -pf06674 -
19380 HMM00293nuhyd_1_eut1(pf00293) EUT1
19385 -pf13247pf14691pf07992 -
19390 -pf00873 -
19395 -pf03960 -
19400 -pf07687pf01546 -
19405 -pf13980 -
BMC-P
BMC-H
BMC-T
Aldehyde dehydrogenase
PTAC
Signature enzyme
Alcohol dehydrogenase
Regulator

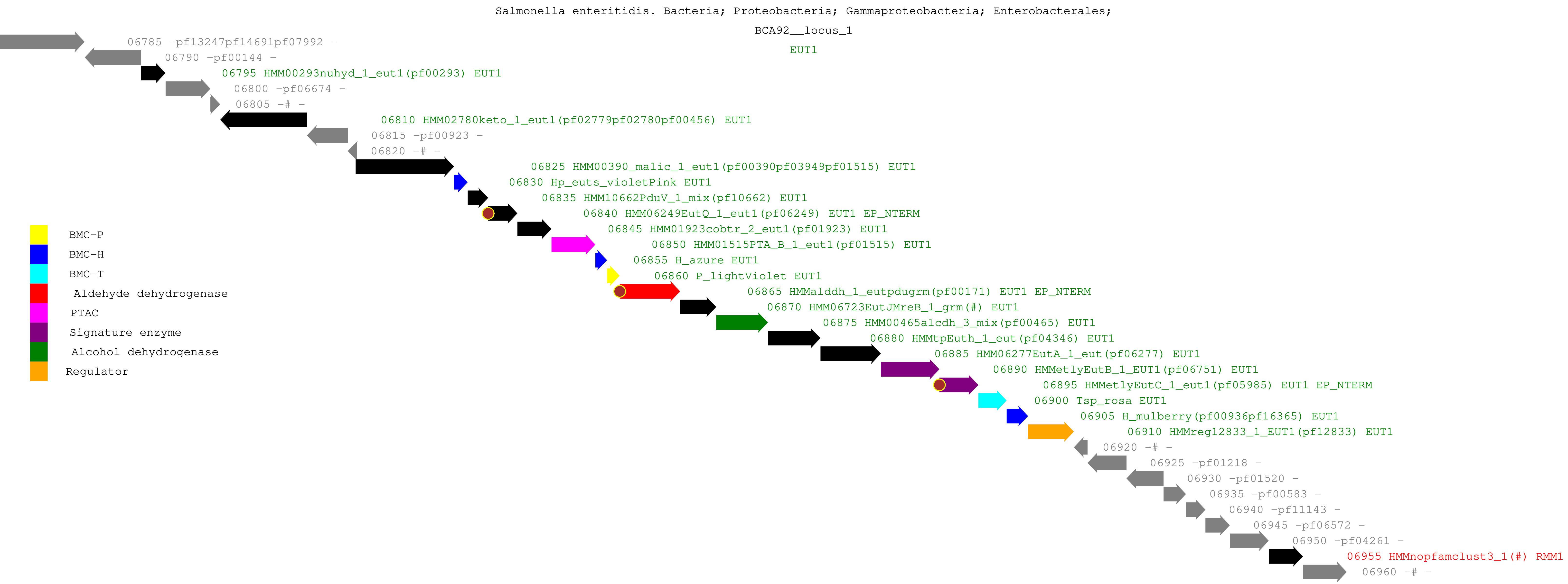
Salmonella enteritidis. Bacteria; Proteobacteria; Gammaproteobacteria; Enterobacterales;
BCA92__locus_1
EUT1
06785 -pf13247pf14691pf07992 -
06790 -pf00144 -
06795 HMM00293nuhyd_1_eut1(pf00293) EUT1
06800 -pf06674 -
06805 -# -
06810 HMM02780keto_1_eut1(pf02779pf02780pf00456) EUT1
06815 -pf00923 -
06820 -# -
06825 HMM00390_malic_1_eut1(pf00390pf03949pf01515) EUT1
06830 Hp_euts_violetPink EUT1
06835 HMM10662PduV_1_mix(pf10662) EUT1
06840 HMM06249EutQ_1_eut1(pf06249) EUT1 EP_NTERM
06845 HMM01923cobtr_2_eut1(pf01923) EUT1
06850 HMM01515PTA_B_1_eut1(pf01515) EUT1
06855 H_azure EUT1
06860 P_lightViolet EUT1
06865 HMMalddh_1_eutpdugrm(pf00171) EUT1 EP_NTERM
06870 HMM06723EutJMreB_1_grm(#) EUT1
06875 HMM00465alcdh_3_mix(pf00465) EUT1
06880 HMMtpEuth_1_eut(pf04346) EUT1
06885 HMM06277EutA_1_eut(pf06277) EUT1
06890 HMMetlyEutB_1_EUT1(pf06751) EUT1
06895 HMMetlyEutC_1_eut1(pf05985) EUT1 EP_NTERM
06900 Tsp_rosa EUT1
06905 H_mulberry(pf00936pf16365) EUT1
06910 HMMreg12833_1_EUT1(pf12833) EUT1
06920 -# -
06925 -pf01218 -
06930 -pf01520 -
06935 -pf00583 -
06940 -pf11143 -
06945 -pf06572 -
06950 -pf04261 -
06955 HMMnopfamclust3_1(#) RMM1
06960 -# -
BMC-P
BMC-H
BMC-T
Aldehyde dehydrogenase
PTAC
Signature enzyme
Alcohol dehydrogenase
Regulator

Klebsiella pneumoniae. Bacteria; Proteobacteria; Gammaproteobacteria; Enterobacterales;

BCB67__locus_2

EUT1

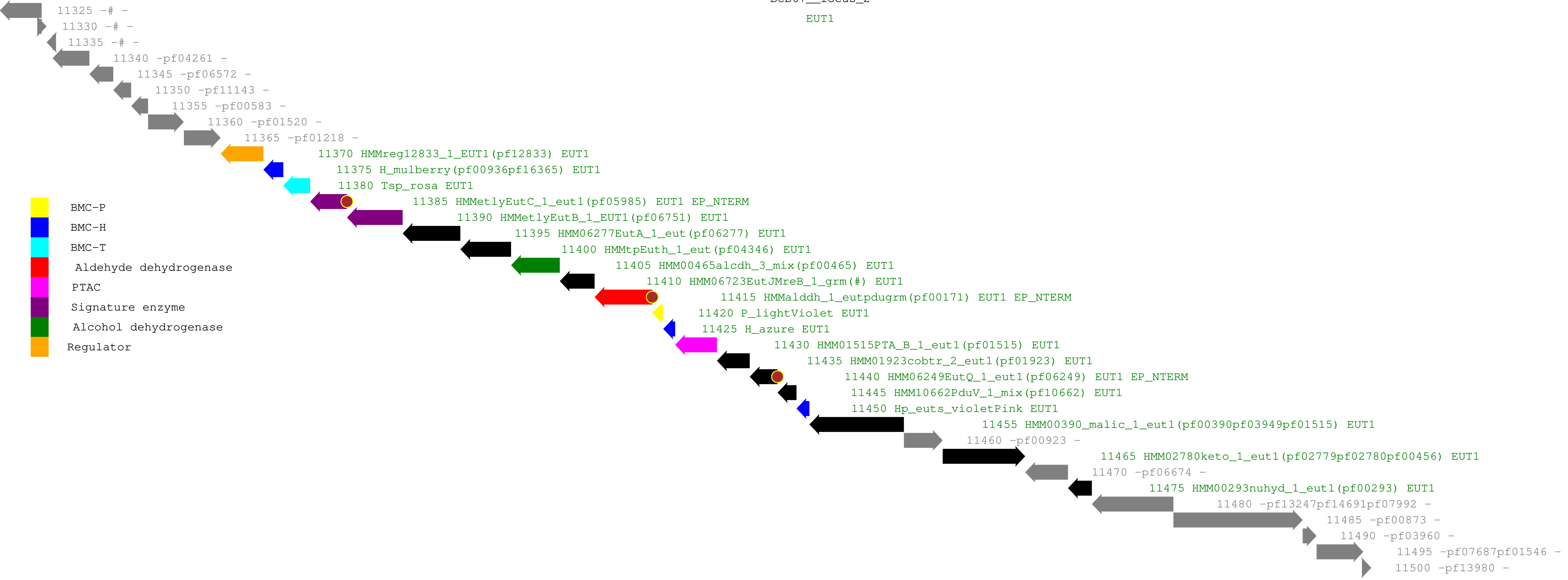

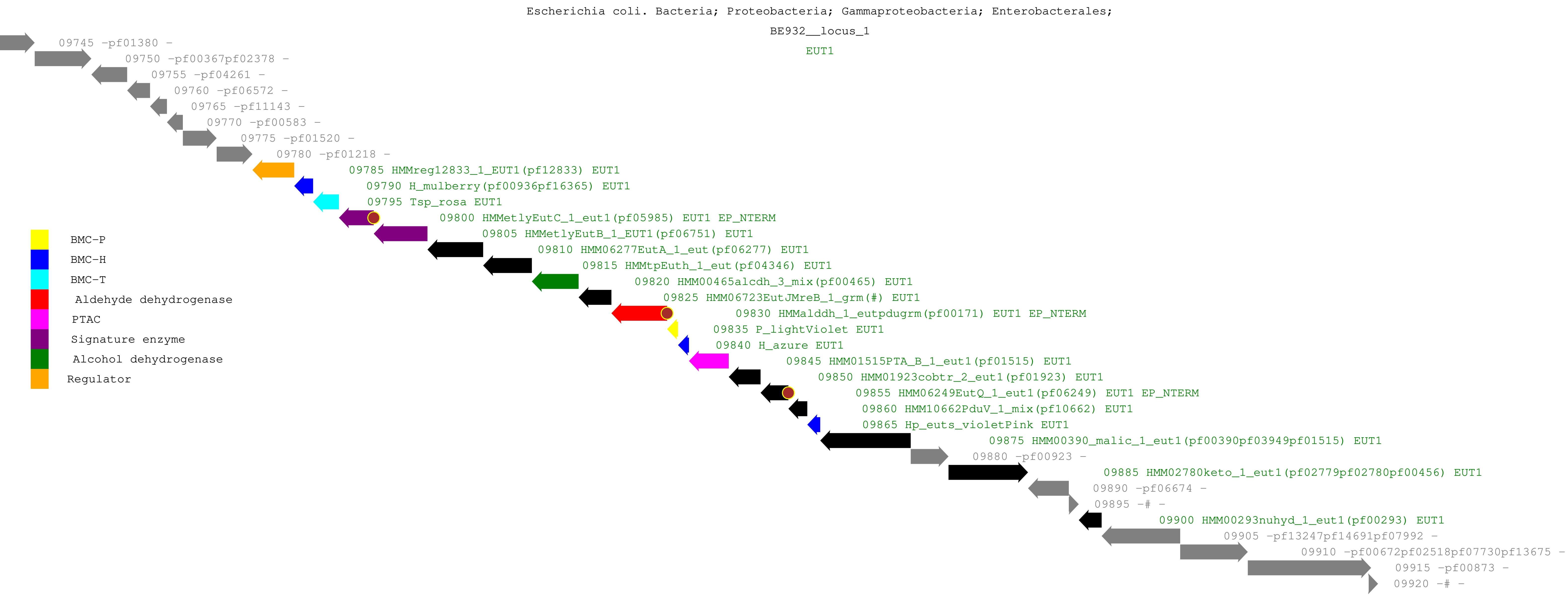

Escherichia coli. Bacteria; Proteobacteria; Gammaproteobacteria; Enterobacterales;
BE932__locus_1
EUT1
09745 -pf01380 -
09750 -pf00367pf02378 -
09755 -pf04261 -
09760 -pf06572 -
09765 -pf11143 -
09770 -pf00583 -
09775 -pf01520 -
09780 -pf01218 -
09785 HMMreg12833_1_EUT1(pf12833) EUT1
09790 H_mulberry(pf00936pf16365) EUT1
09795 Tsp_rosa EUT1
09800 HMMetlyEutC_1_eut1(pf05985) EUT1 EP_NTERM
09805 HMMetlyEutB_1_EUT1(pf06751) EUT1
09810 HMM06277EutA_1_eut(pf06277) EUT1
09815 HMMtpEuth_1_eut(pf04346) EUT1
09820 HMM00465alcdh_3_mix(pf00465) EUT1
09825 HMM06723EutJMreB_1_grm(#) EUT1
09830 HMMaldddh_1_eutpdugrm(pf00171) EUT1 EP_NTERM
09835 P_lightViolet EUT1
09840 H_azure EUT1
09845 HMM01515PTA_B_1_eut1(pf01515) EUT1
09850 HMM01923cobtr_2_eut1(pf01923) EUT1
09855 HMM06249EutQ_1_eut1(pf06249) EUT1 EP_NTERM
09860 HMM10662PduV_1_mix(pf10662) EUT1
09865 Hp_euts_violetPink EUT1
09875 HMM00390_malic_1_eut1(pf00390pf03949pf01515) EUT1
09880 -pf00923 -
09885 HMM02780keto_1_eut1(pf02779pf02780pf00456) EUT1
09890 -pf06674 -
09895 -# -
09900 HMM00293nuhyd_1_eut1(pf00293) EUT1
09905 -pf13247pf14691pf07992 -
09910 -pf00672pf02518pf07730pf13675 -
09915 -pf00873 -
09920 -# -
BMC-P
BMC-H
BMC-T
Aldehyde dehydrogenase
PTAC
Signature enzyme
Alcohol dehydrogenase
Regulator

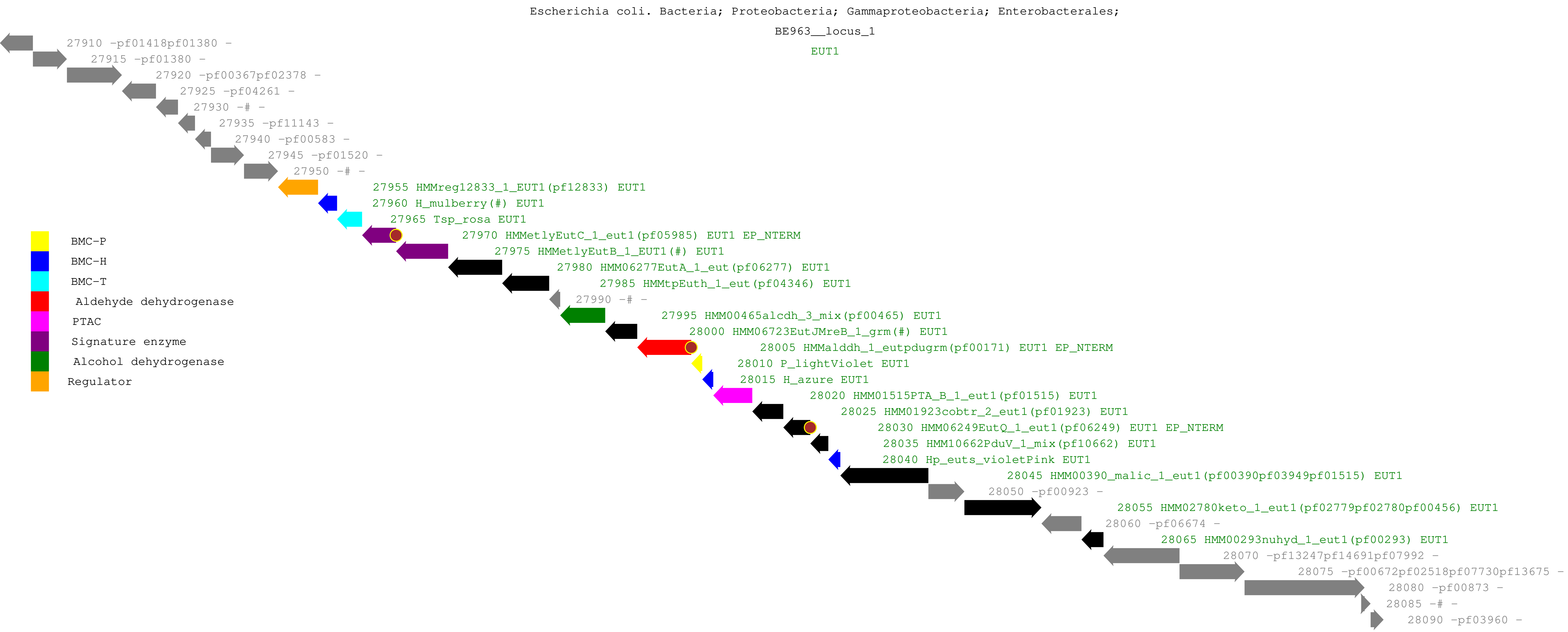

Escherichia coli. Bacteria; Proteobacteria; Gammaproteobacteria; Enterobacterales;
BE963__locus_1
EUT1
27910 -pf01418pf01380 -
27915 -pf01380 -
27920 -pf00367pf02378 -
27925 -pf04261 -
27930 -# -
27935 -pf11143 -
27940 -pf00583 -
27945 -pf01520 -
27950 -# -
27955 HMMreg12833_1_EUT1(pf12833) EUT1
27960 H_mulberry(#) EUT1
27965 Tsp_rosa EUT1
27970 HMMetlyEutC_1_eut1(pf05985) EUT1 EP_NTERM
27975 HMMetlyEutB_1_EUT1(#) EUT1
27980 HMM06277EutA_1_eut(pf06277) EUT1
27985 HMMtpEuth_1_eut(pf04346) EUT1
27990 -# -
27995 HMM00465alcdh_3_mix(pf00465) EUT1
28000 HMM06723EutJMreB_1_grm(#) EUT1
28005 HMMalddh_1_eutpdugrm(pf00171) EUT1 EP_NTERM
28010 P_lightViolet EUT1
28015 H_azure EUT1
28020 HMM01515PTA_B_1_eut1(pf01515) EUT1
28025 HMM01923cobtr_2_eut1(pf01923) EUT1
28030 HMM06249EutQ_1_eut1(pf06249) EUT1 EP_NTERM
28035 HMM10662PduV_1_mix(pf10662) EUT1
28040 Hp_euts_violetPink EUT1
28045 HMM00390_malic_1_eut1(pf00390pf03949pf01515) EUT1
28050 -pf00923 -
28055 HMM02780keto_1_eut1(pf02779pf02780pf00456) EUT1
28060 -pf06674 -
28065 HMM00293nuhyd_1_eut1(pf00293) EUT1
28070 -pf13247pf14691pf07992 -
28075 -pf00672pf02518pf07730pf13675 -
28080 -pf00873 -
28085 -# -
28090 -pf03960 -
BMC-P
BMC-H
BMC-T
Aldehyde dehydrogenase
PTAC
Signature enzyme
Alcohol dehydrogenase
Regulator

Citrobacter freundii. Bacteria; Proteobacteria; Gammaproteobacteria; Enterobacterales;

BED30__locus_1

EUT1

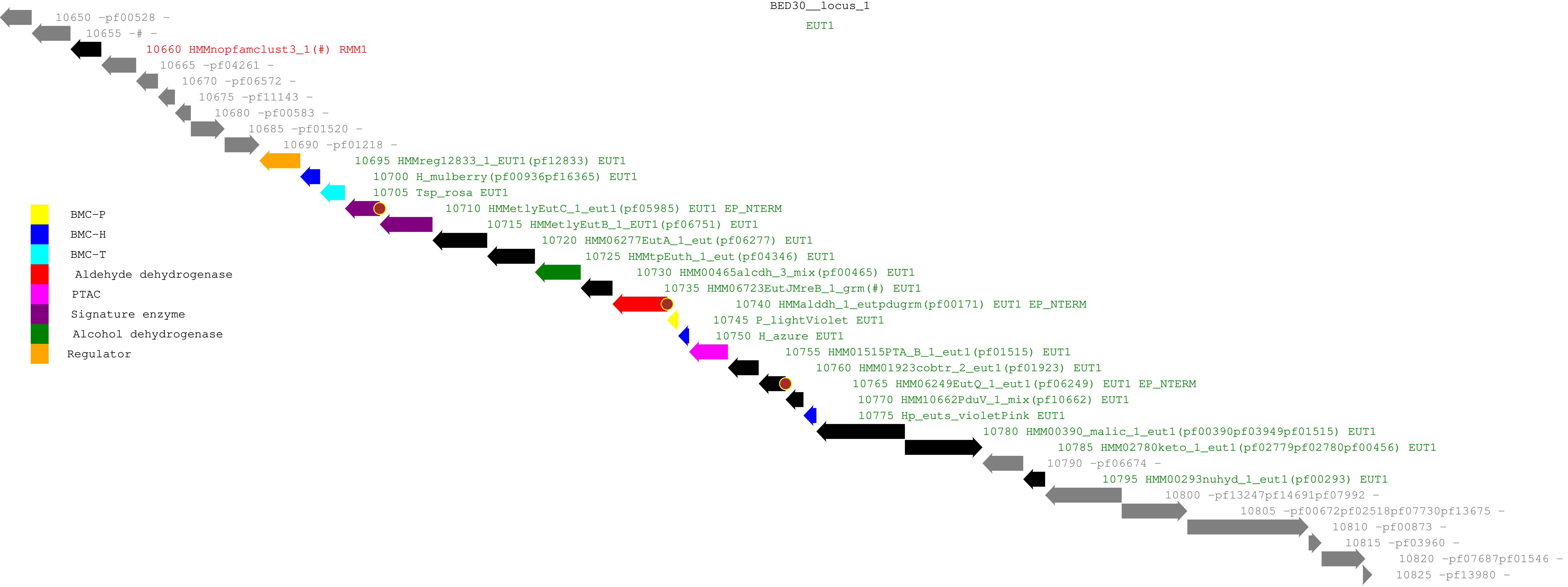

Citrobacter freundii. Bacteria; Proteobacteria; Gammaproteobacteria; Enterobacterales;

BED45__locus_2

EUT1

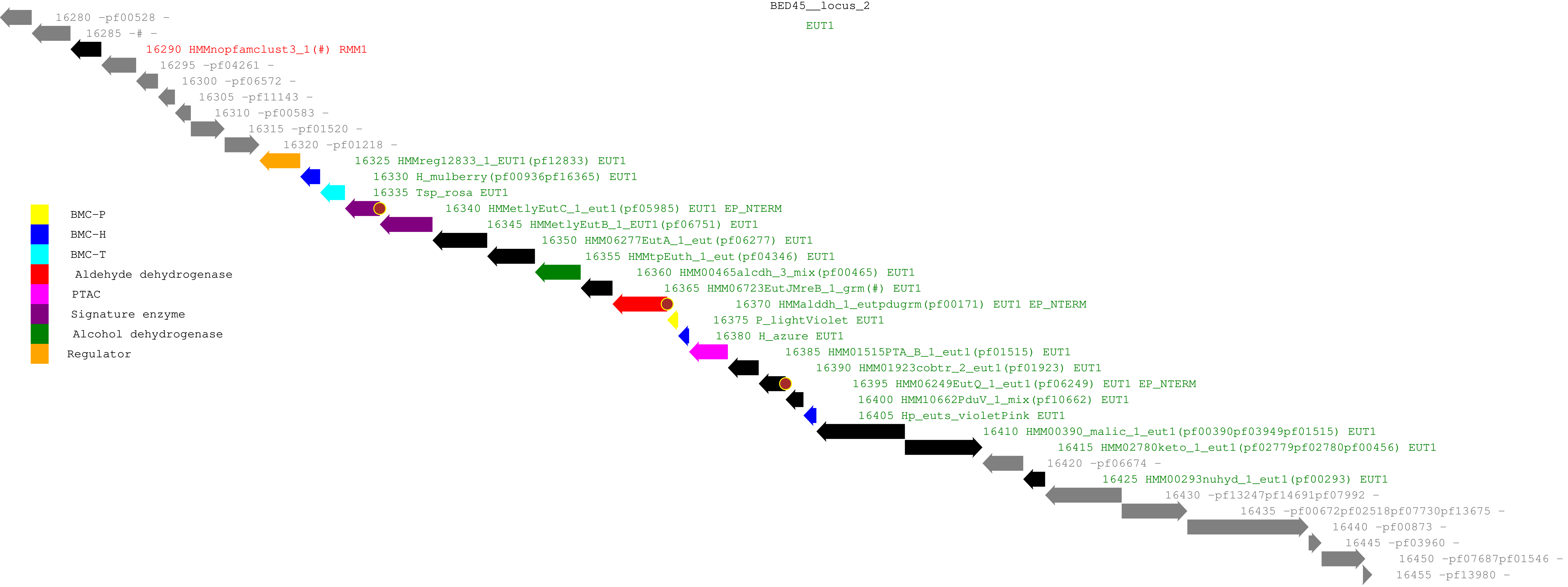

Salmonella enterica subsp. enterica serovar Grumpensis. Bacteria; Proteobacteria; Gammaproteobacteria; Enterobacterales;

BEL51__locus_1

EUT1

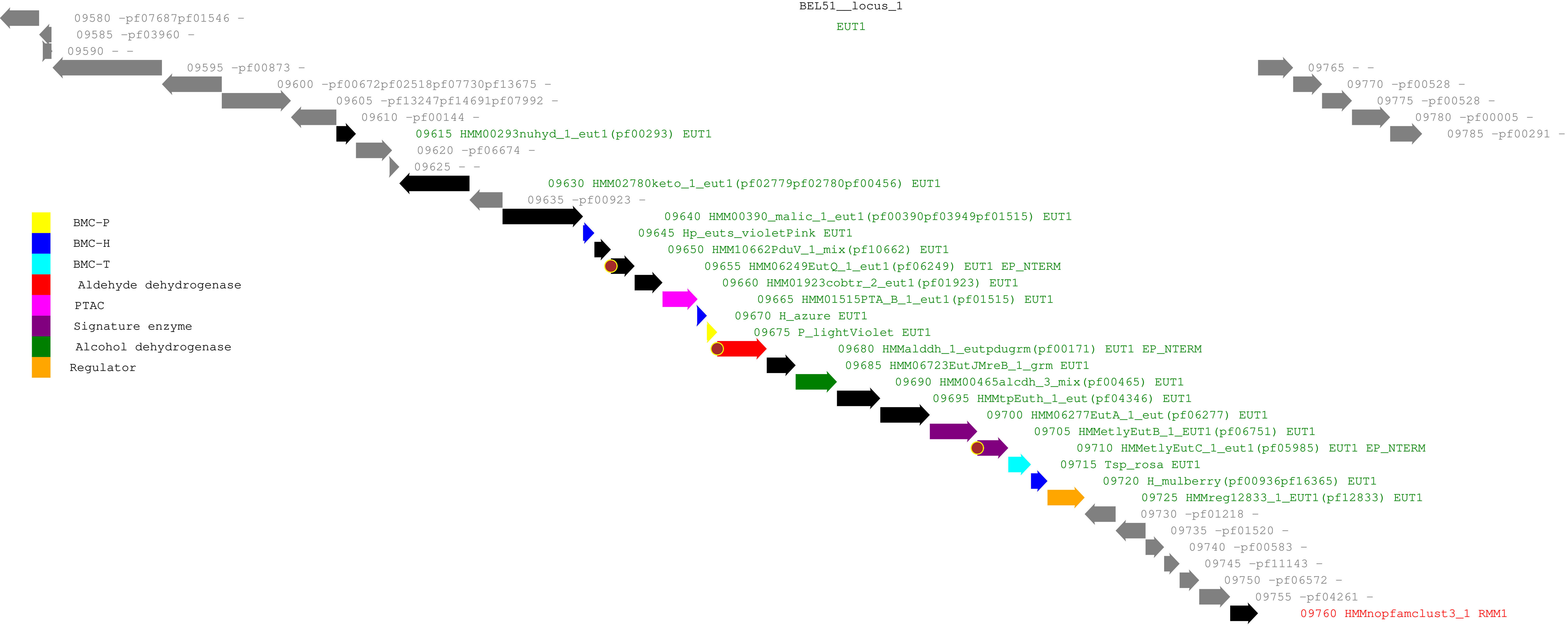

Escherichia sp. MOD1-EC5451. Bacteria; Proteobacteria; Gammaproteobacteria; Enterobacterales;

BEM40__locus_1

EUT1

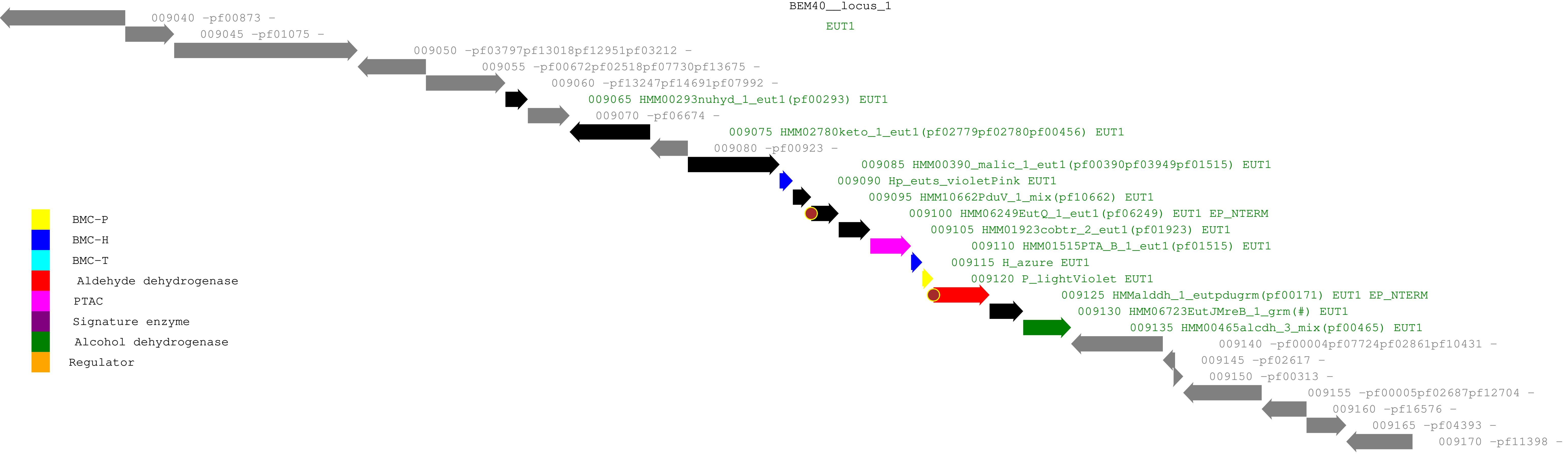

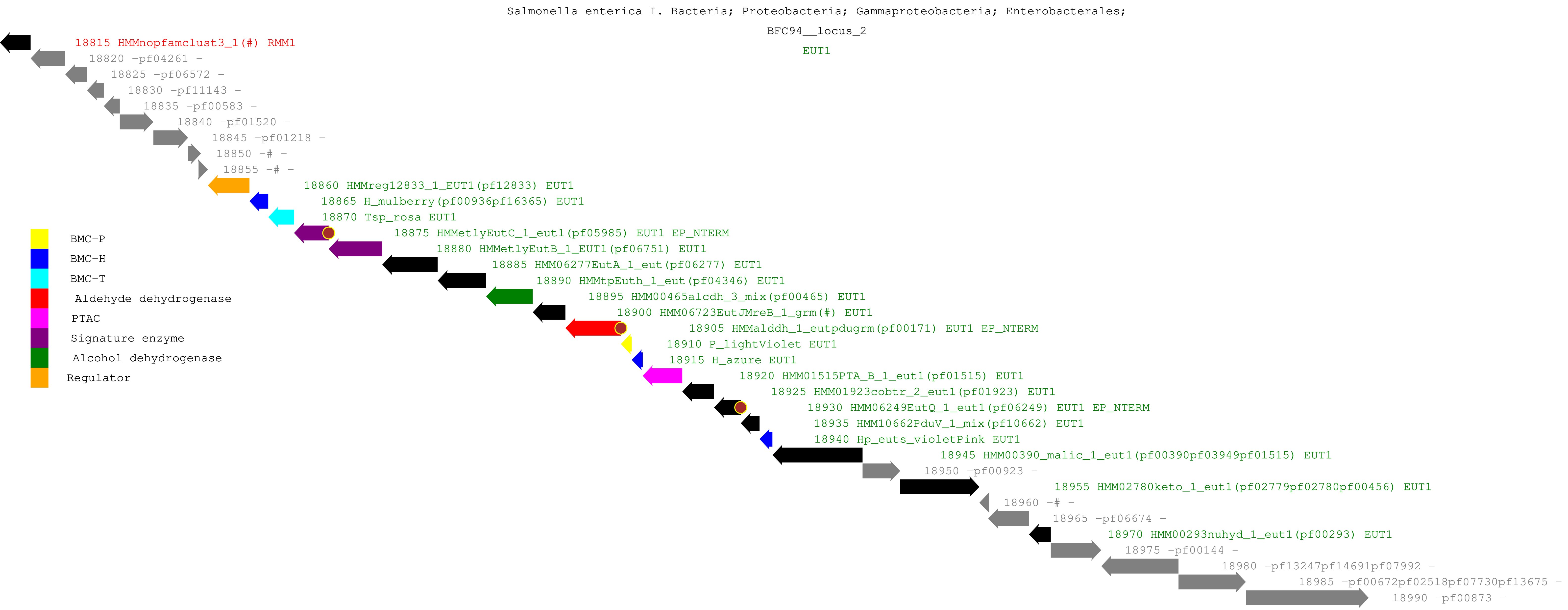

Salmonella enterica I. Bacteria; Proteobacteria; Gammaproteobacteria; Enterobacterales;
BFC94__locus_2
EUT1
18815 HMMnopfamclust3_1(#) RMM1
18820 -pf04261 -
18825 -pf06572 -
18830 -pf11143 -
18835 -pf00583 -
18840 -pf01520 -
18845 -pf01218 -
18850 -# -
18855 -# -
18860 HMMreg12833_1_EUT1(pf12833) EUT1
18865 H_mulberry(pf00936pf16365) EUT1
18870 Tsp_rosa EUT1
18875 HMMetlyEutC_1_eut1(pf05985) EUT1 EP_NTERM
18880 HMMetlyEutB_1_EUT1(pf06751) EUT1
18885 HMM06277EutA_1_eut(pf06277) EUT1
18890 HMMtpEuth_1_eut(pf04346) EUT1
18895 HMM00465alcdh_3_mix(pf00465) EUT1
18900 HMM06723EutJMreB_1_grm(#) EUT1
18905 HMMalddh_1_eutpdugrm(pf00171) EUT1 EP_NTERM
18910 P_lightViolet EUT1
18915 H_azure EUT1
18920 HMM01515PTA_B_1_eut1(pf01515) EUT1
18925 HMM01923cobtr_2_eut1(pf01923) EUT1
18930 HMM06249EutQ_1_eut1(pf06249) EUT1 EP_NTERM
18935 HMM10662PduV_1_mix(pf10662) EUT1
18940 Hp_euts_violetPink EUT1
18945 HMM00390_malic_1_eut1(pf00390pf03949pf01515) EUT1
18950 -pf00923 -
18955 HMM02780keto_1_eut1(pf02779pf02780pf00456) EUT1
18960 -# -
18965 -pf06674 -
18970 HMM00293nuhyd_1_eut1(pf00293) EUT1
18975 -pf00144 -
18980 -pf13247pf14691pf07992 -
18985 -pf00672pf02518pf07730pf13675 -
18990 -pf00873 -
BMC-P
BMC-H
BMC-T
Aldehyde dehydrogenase
PTAC
Signature enzyme
Alcohol dehydrogenase
Regulator

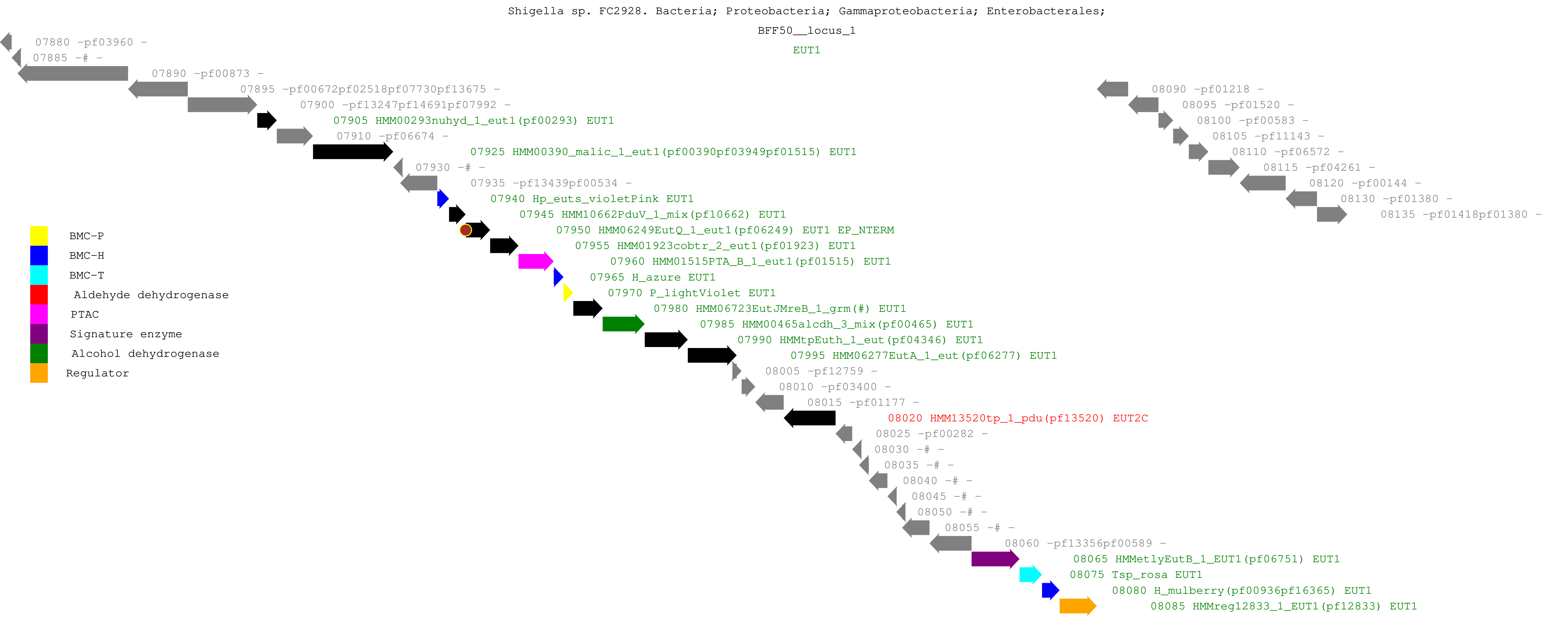
Shigella sp. FC2928. Bacteria; Proteobacteria; Gammaproteobacteria; Enterobacterales;
BFF50__locus_1
EUT1
07880 -pf03960 -
07885 -# -
07890 -pf00873 -
07895 -pf00672pf02518pf07730pf13675 -
07900 -pf13247pf14691pf07992 -
07905 HMM00293nuhyd_1_eut1(pf00293) EUT1
07910 -pf06674 -
07925 HMM00390_malic_1_eut1(pf00390pf03949pf01515) EUT1
07930 -# -
07935 -pf13439pf00534 -
07940 Hp_euts_violetPink EUT1
07945 HMM10662PduV_1_mix(pf10662) EUT1
07950 HMM06249EutQ_1_eut1(pf06249) EUT1 EP_NTERM
07955 HMM01923cobtr_2_eut1(pf01923) EUT1
07960 HMM01515PTA_B_1_eut1(pf01515) EUT1
07965 H_azure EUT1
07970 P_lightViolet EUT1
07980 HMM06723EutJMreB_1_grm(#) EUT1
07985 HMM00465alcdh_3_mix(pf00465) EUT1
07990 HMMtpEuth_1_eut(pf04346) EUT1
07995 HMM06277EutA_1_eut(pf06277) EUT1
08005 -pf12759 -
08010 -pf03400 -
08015 -pf01177 -
08020 HMM13520tp_1_pdu(pf13520) EUT2C
08025 -pf00282 -
08030 -# -
08035 -# -
08040 -# -
08045 -# -
08050 -# -
08055 -# -
08060 -pf13356pf00589 -
08065 HMMetlyEutB_1_EUT1(pf06751) EUT1
08075 Tsp_rosa EUT1
08080 H_mulberry(pf00936pf16365) EUT1
08085 HMMreg12833_1_EUT1(pf12833) EUT1
08090 -pf01218 -
08095 -pf01520 -
08100 -pf00583 -
08105 -pf11143 -
08110 -pf06572 -
08115 -pf04261 -
08120 -pf00144 -
08130 -pf01380 -
08135 -pf01418pf01380 -
BMC-P
BMC-H
BMC-T
Aldehyde dehydrogenase
PTAC
Signature enzyme
Alcohol dehydrogenase
Regulator

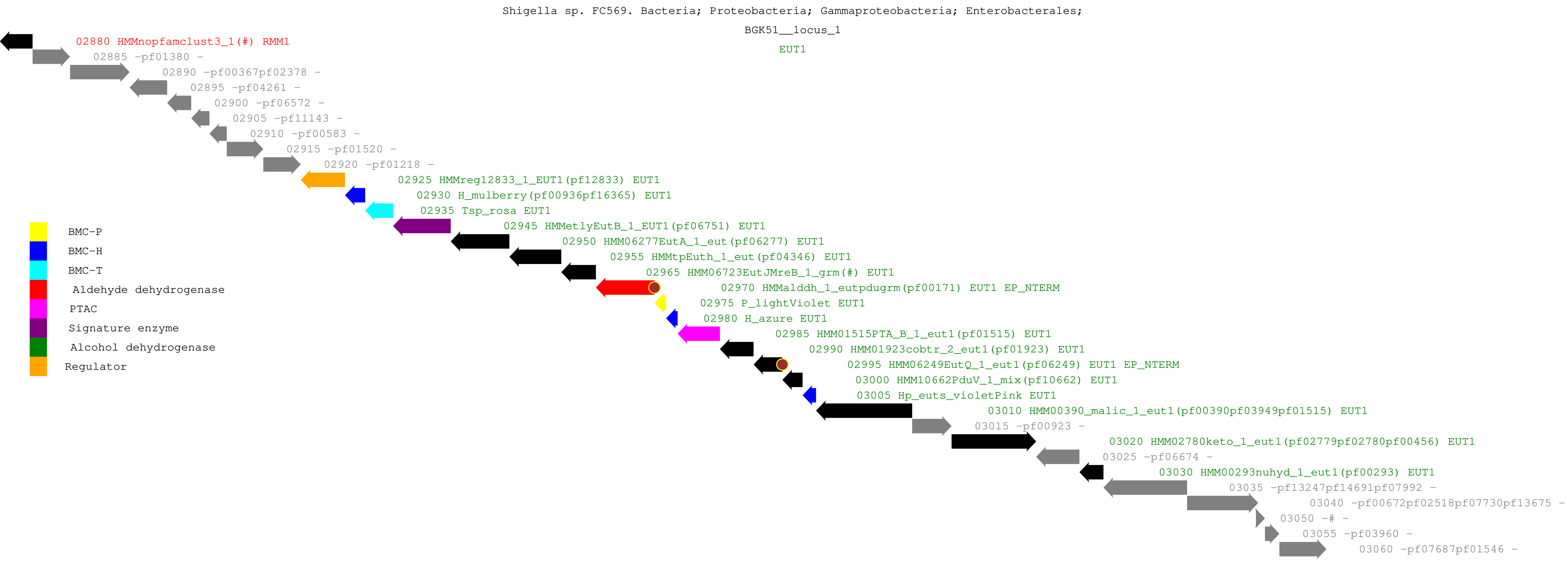
Shigella sp. FC569. Bacteria; Proteobacteria; Gammaproteobacteria; Enterobacterales;
BGK51__locus_1
EUT1
02880 HMMnopfamclust3_1(#) RMM1
02885 -pf01380 -
02890 -pf00367pf02378 -
02895 -pf04261 -
02900 -pf06572 -
02905 -pf11143 -
02910 -pf00583 -
02915 -pf01520 -
02920 -pf01218 -
02925 HMMreg12833_1_EUT1(pf12833) EUT1
02930 H_mulberry(pf00936pf16365) EUT1
02935 Tsp_rosa EUT1
02945 HMMetlyEutB_1_EUT1(pf06751) EUT1
02950 HMM06277EutA_1_eut(pf06277) EUT1
02955 HMMtpEuth_1_eut(pf04346) EUT1
02965 HMM06723EutJMreB_1_grm(#) EUT1
02970 HMMalddh_1_eutpdugrm(pf00171) EUT1 EP_NTERM
02975 P_lightViolet EUT1
02980 H_azure EUT1
02985 HMM01515PTA_B_1_eut1(pf01515) EUT1
02990 HMM01923cobtr_2_eut1(pf01923) EUT1
02995 HMM06249EutQ_1_eut1(pf06249) EUT1 EP_NTERM
03000 HMM10662PduV_1_mix(pf10662) EUT1
03005 Hp_euts_violetPink EUT1
03010 HMM00390_malic_1_eut1(pf00390pf03949pf01515) EUT1
03015 -pf00923 -
03020 HMM02780keto_1_eut1(pf02779pf02780pf00456) EUT1
03025 -pf06674 -
03030 HMM00293nuhyd_1_eut1(pf00293) EUT1
03035 -pf13247pf14691pf07992 -
03040 -pf00672pf02518pf07730pf13675 -
03050 -# -
03055 -pf03960 -
03060 -pf07687pf01546 -
BMC-P
BMC-H
BMC-T
Aldehyde dehydrogenase
PTAC
Signature enzyme
Alcohol dehydrogenase
Regulator

Salmonella enterica (Salmonella choleraesuis). Bacteria; Proteobacteria; Gammaproteobacteria; Enterobacterales;

BH006__locus_1

EUT1

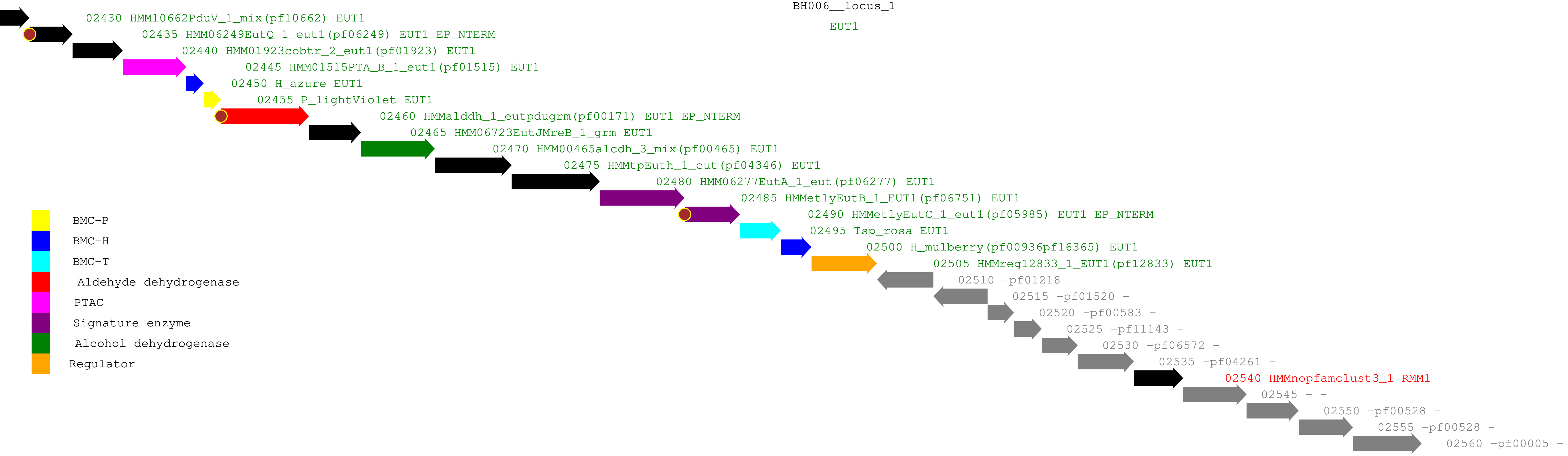

Escherichia coli. Bacteria; Proteobacteria; Gammaproteobacteria; Enterobacterales;

BIQ87__locus_1

EUT1

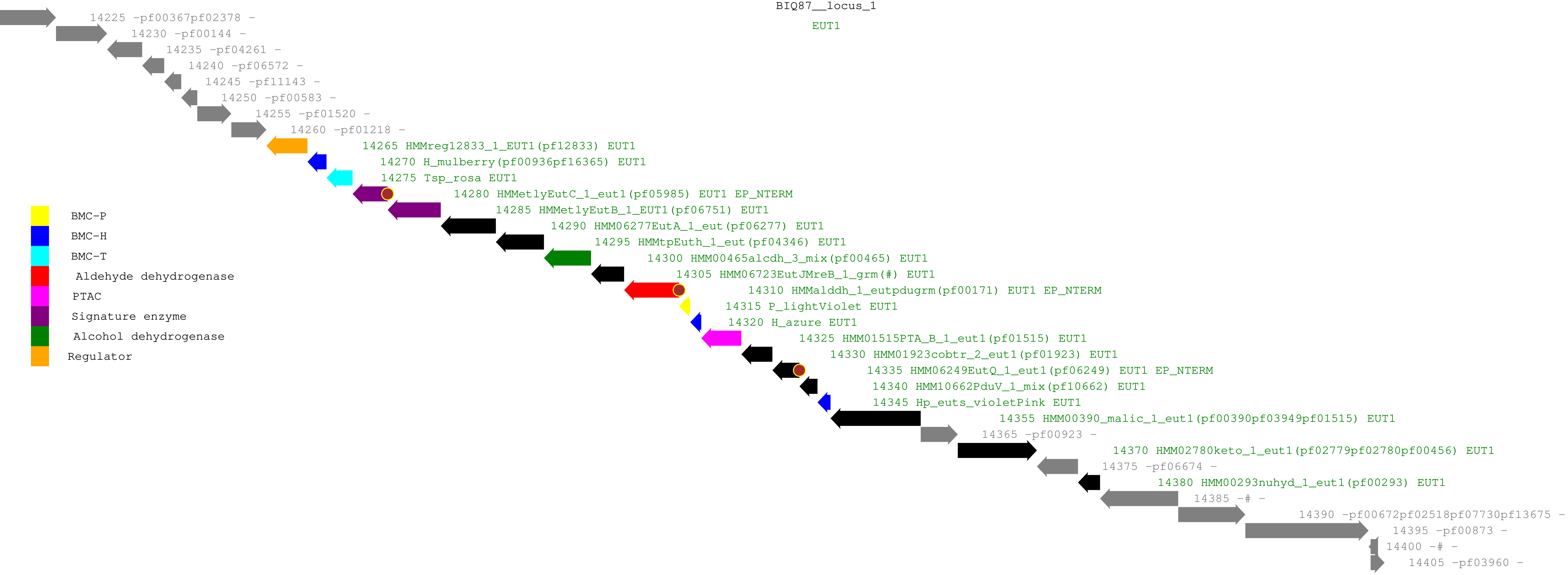

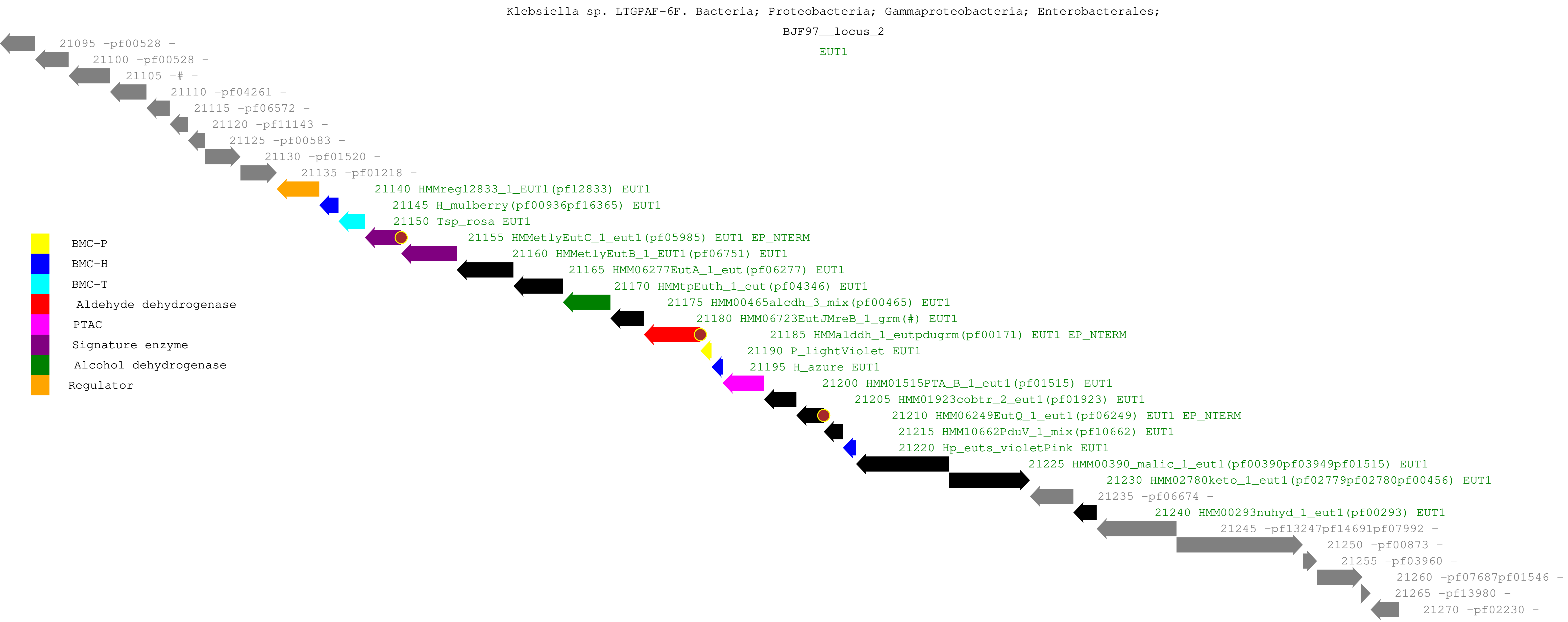
Klebsiella sp. LTGPAF-6F. Bacteria; Proteobacteria; Gammaproteobacteria; Enterobacterales;
BJF97__locus_2
EUT1
21095 -pf00528 -
21100 -pf00528 -
21105 -# -
21110 -pf04261 -
21115 -pf06572 -
21120 -pf11143 -
21125 -pf00583 -
21130 -pf01520 -
21135 -pf01218 -
21140 HMMreg12833_1_EUT1(pf12833) EUT1
21145 H_mulberry(pf00936pf16365) EUT1
21150 Tsp_rosa EUT1
21155 HMMetlyEutC_1_eut1(pf05985) EUT1 EP_NTERM
21160 HMMetlyEutB_1_EUT1(pf06751) EUT1
21165 HMM06277EutA_1_eut(pf06277) EUT1
21170 HMMtpEuth_1_eut(pf04346) EUT1
21175 HMM00465alcdh_3_mix(pf00465) EUT1
21180 HMM06723EutJMreB_1_grm(#) EUT1
21185 HMMalddh_1_eutpdugrm(pf00171) EUT1 EP_NTERM
21190 P_lightViolet EUT1
21195 H_azure EUT1
21200 HMM01515PTA_B_1_eut1(pf01515) EUT1
21205 HMM01923cobtr_2_eut1(pf01923) EUT1
21210 HMM06249EutQ_1_eut1(pf06249) EUT1 EP_NTERM
21215 HMM10662PduV_1_mix(pf10662) EUT1
21220 Hp_euts_violetPink EUT1
21225 HMM00390_malic_1_eut1(pf00390pf03949pf01515) EUT1
21230 HMM02780keto_1_eut1(pf02779pf02780pf00456) EUT1
21235 -pf06674 -
21240 HMM00293nuhyd_1_eut1(pf00293) EUT1
21245 -pf13247pf14691pf07992 -
21250 -pf00873 -
21255 -pf03960 -
21260 -pf07687pf01546 -
21265 -pf13980 -
21270 -pf02230 -
BMC-P
BMC-H
BMC-T
Aldehyde dehydrogenase
PTAC
Signature enzyme
Alcohol dehydrogenase
Regulator

Escherichia coli. Bacteria; Proteobacteria; Gammaproteobacteria; Enterobacterales;

BK373__locus_1

EUT1

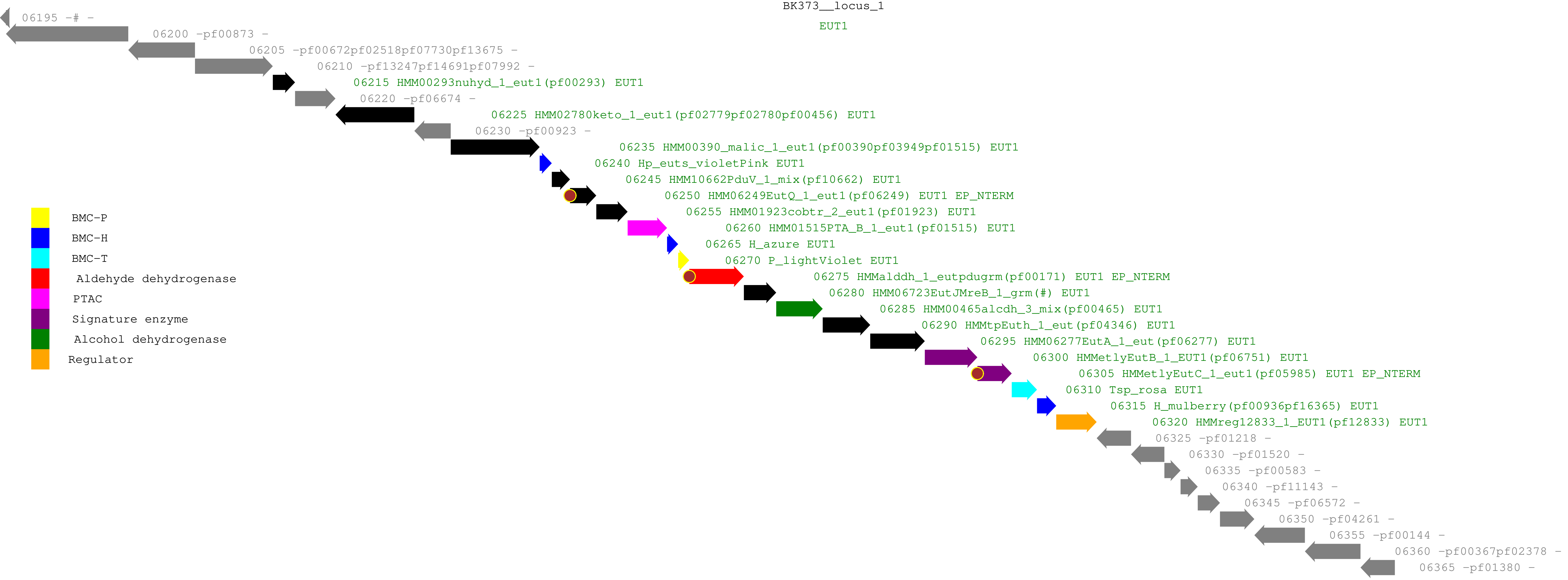

Escherichia coli. Bacteria; Proteobacteria; Gammaproteobacteria; Enterobacterales;

BK375__locus_1

EUT1

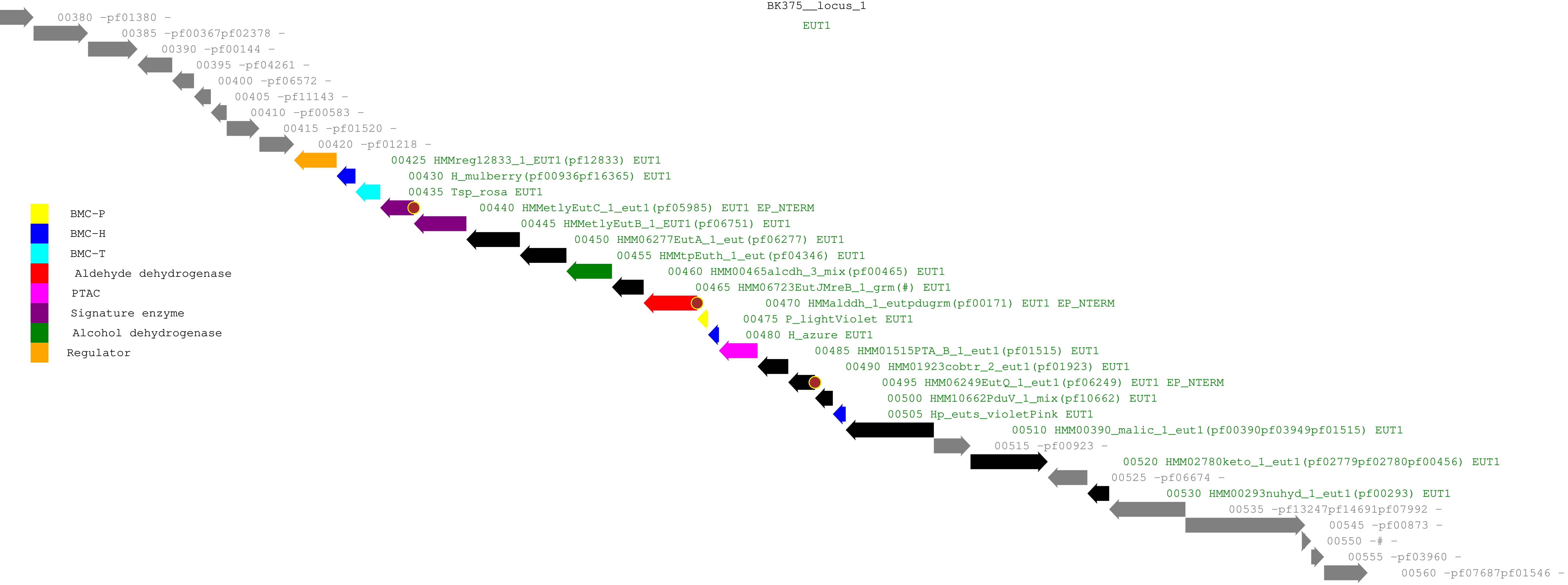

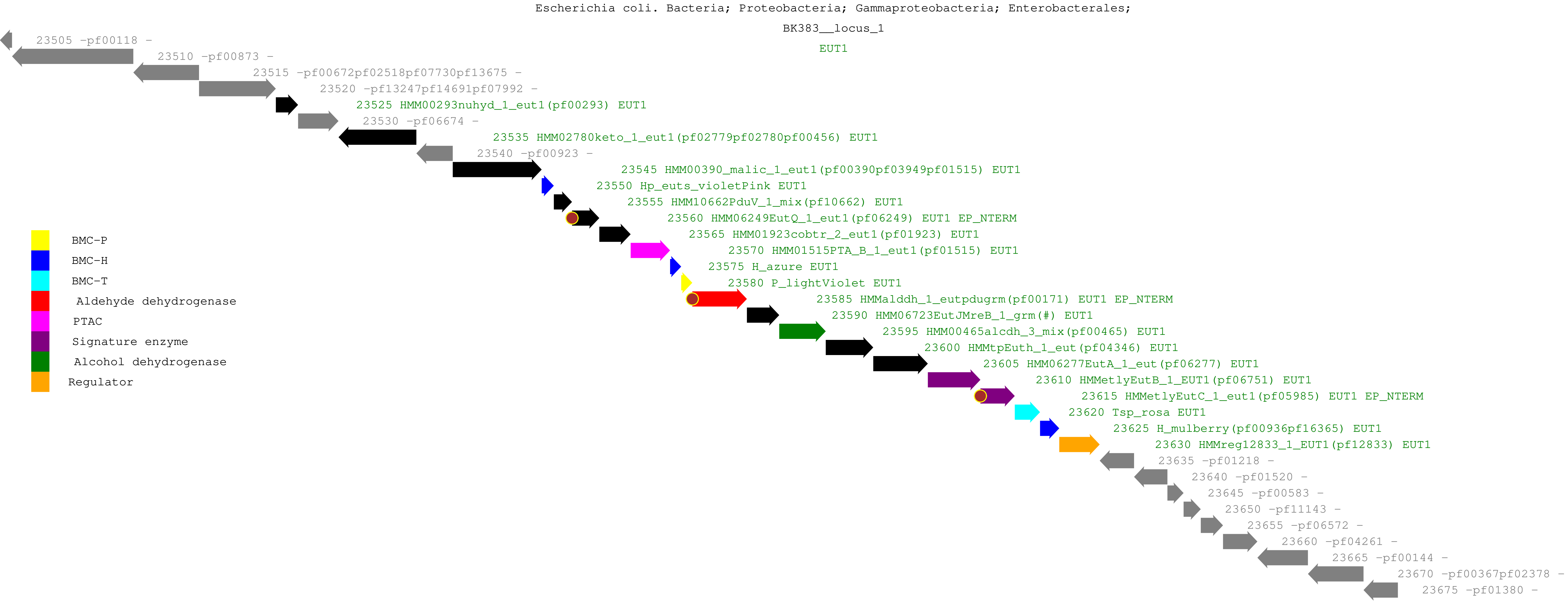
Escherichia coli. Bacteria; Proteobacteria; Gammaproteobacteria; Enterobacterales;
BK383__locus_1
EUT1
23505 -pf00118 -
23510 -pf00873 -
23515 -pf00672pf02518pf07730pf13675 -
23520 -pf13247pf14691pf07992 -
23525 HMM00293nuhyd_1_eut1(pf00293) EUT1
23530 -pf06674 -
23535 HMM02780keto_1_eut1(pf02779pf02780pf00456) EUT1
23540 -pf00923 -
23545 HMM00390_malic_1_eut1(pf00390pf03949pf01515) EUT1
23550 Hp_euts_violetPink EUT1
23555 HMM10662PduV_1_mix(pf10662) EUT1
23560 HMM06249EutQ_1_eut1(pf06249) EUT1 EP_NTERM
23565 HMM01923cobtr_2_eut1(pf01923) EUT1
23570 HMM01515PTA_B_1_eut1(pf01515) EUT1
23575 H_azure EUT1
23580 P_lightViolet EUT1
23585 HMMalddh_1_eutpdugrm(pf00171) EUT1 EP_NTERM
23590 HMM06723EutJMreB_1_grm(#) EUT1
23595 HMM00465alcdh_3_mix(pf00465) EUT1
23600 HMMtpEuth_1_eut(pf04346) EUT1
23605 HMM06277EutA_1_eut(pf06277) EUT1
23610 HMMetlyEutB_1_EUT1(pf06751) EUT1
23615 HMMetlyEutC_1_eut1(pf05985) EUT1 EP_NTERM
23620 Tsp_rosa EUT1
23625 H_mulberry(pf00936pf16365) EUT1
23630 HMMreg12833_1_EUT1(pf12833) EUT1
23635 -pf01218 -
23640 -pf01520 -
23645 -pf00583 -
23650 -pf11143 -
23655 -pf06572 -
23660 -pf04261 -
23665 -pf00144 -
23670 -pf00367pf02378 -
23675 -pf01380 -
BMC-P
BMC-H
BMC-T
Aldehyde dehydrogenase
PTAC
Signature enzyme
Alcohol dehydrogenase
Regulator

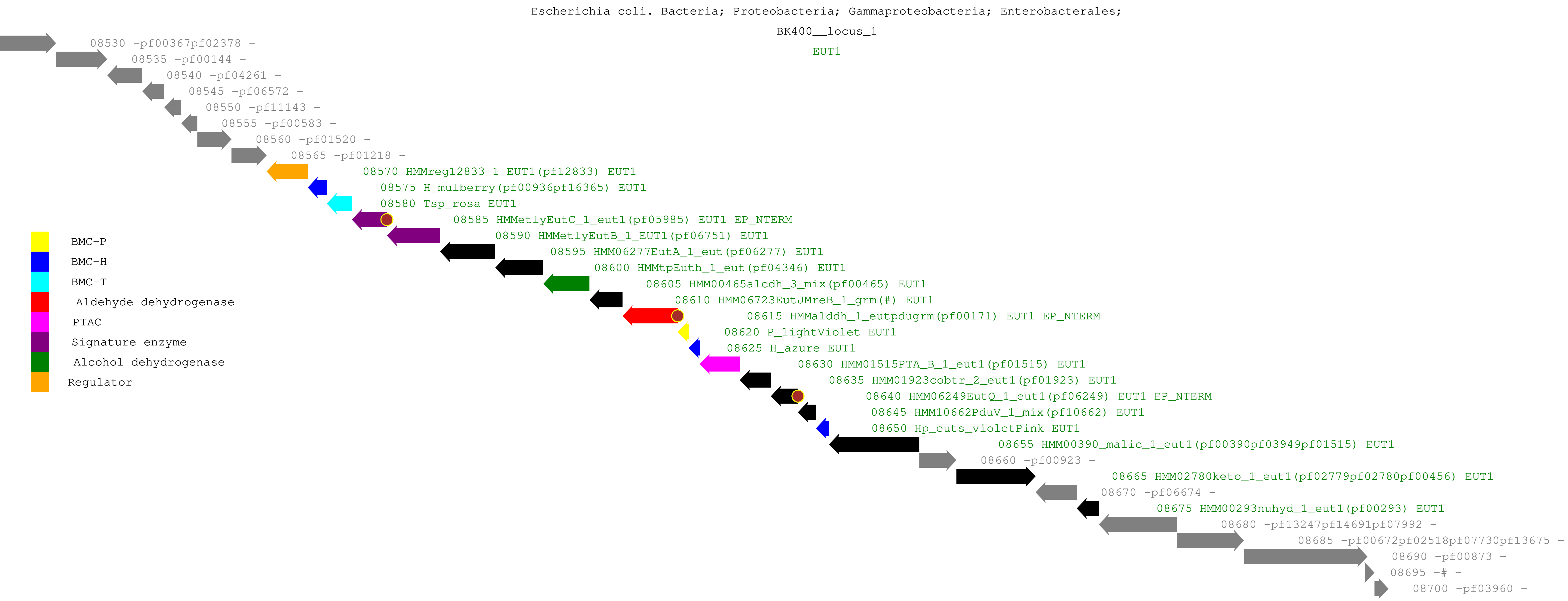
Escherichia coli. Bacteria; Proteobacteria; Gammaproteobacteria; Enterobacterales;
BK400__locus_1
EUT1
08530 -pf00367pf02378 -
08535 -pf00144 -
08540 -pf04261 -
08545 -pf06572 -
08550 -pf11143 -
08555 -pf00583 -
08560 -pf01520 -
08565 -pf01218 -
08570 HMMreg12833_1_EUT1(pf12833) EUT1
08575 H_mulberry(pf00936pf16365) EUT1
08580 Tsp_rosa EUT1
08585 HMMetlyEutC_1_eut1(pf05985) EUT1 EP_NTERM
08590 HMMetlyEutB_1_EUT1(pf06751) EUT1
08595 HMM06277EutA_1_eut(pf06277) EUT1
08600 HMMtpEuth_1_eut(pf04346) EUT1
08605 HMM00465alcdh_3_mix(pf00465) EUT1
08610 HMM06723EutJMreB_1_grm(#) EUT1
08615 HMMalddh_1_eutpdugrm(pf00171) EUT1 EP_NTERM
08620 P_lightViolet EUT1
08625 H_azure EUT1
08630 HMM01515PTA_B_1_eut1(pf01515) EUT1
08635 HMM01923cobtr_2_eut1(pf01923) EUT1
08640 HMM06249EutQ_1_eut1(pf06249) EUT1 EP_NTERM
08645 HMM10662PduV_1_mix(pf10662) EUT1
08650 Hp_euts_violetPink EUT1
08655 HMM00390_malic_1_eut1(pf00390pf03949pf01515) EUT1
08660 -pf00923 -
08665 HMM02780keto_1_eut1(pf02779pf02780pf00456) EUT1
08670 -pf06674 -
08675 HMM00293nuhyd_1_eut1(pf00293) EUT1
08680 -pf13247pf14691pf07992 -
08685 -pf00672pf02518pf07730pf13675 -
08690 -pf00873 -
08695 -# -
08700 -pf03960 -
BMC-P
BMC-H
BMC-T
Aldehyde dehydrogenase
PTAC
Signature enzyme
Alcohol dehydrogenase
Regulator

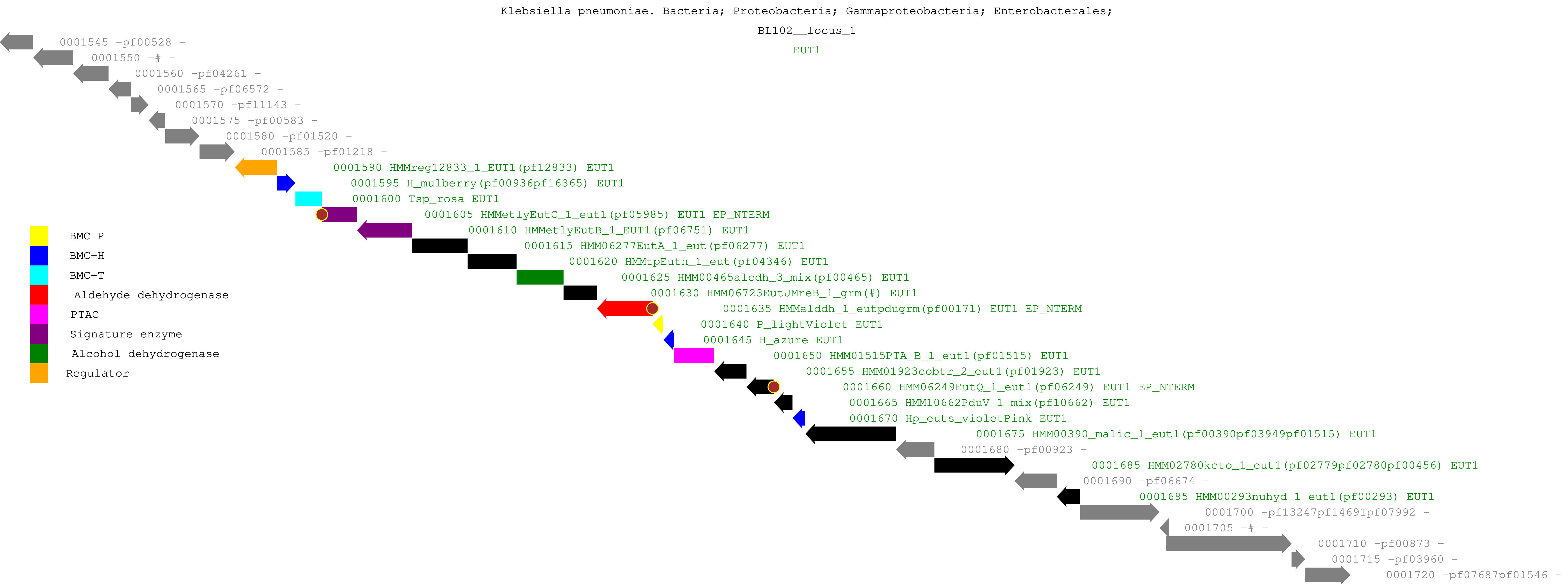
Klebsiella pneumoniae. Bacteria; Proteobacteria; Gammaproteobacteria; Enterobacterales;
BL102__locus_1
EUT1
0001545 -pf00528 -
0001550 -# -
0001560 -pf04261 -
0001565 -pf06572 -
0001570 -pf11143 -
0001575 -pf00583 -
0001580 -pf01520 -
0001585 -pf01218 -
0001590 HMMreg12833_1_EUT1(pf12833) EUT1
0001595 H_mulberry(pf00936pf16365) EUT1
0001600 Tsp_rosa EUT1
0001605 HMMetlyEutC_1_eut1(pf05985) EUT1 EP_NTERM
0001610 HMMetlyEutB_1_EUT1(pf06751) EUT1
0001615 HMM06277EutA_1_eut(pf06277) EUT1
0001620 HMMtpEuth_1_eut(pf04346) EUT1
0001625 HMM00465alcdh_3_mix(pf00465) EUT1
0001630 HMM06723EutJMreB_1_grm(#) EUT1
0001635 HMMalddh_1_eutpdugrm(pf00171) EUT1 EP_NTERM
0001640 P_lightViolet EUT1
0001645 H_azure EUT1
0001650 HMM01515PTA_B_1_eut1(pf01515) EUT1
0001655 HMM01923cobtr_2_eut1(pf01923) EUT1
0001660 HMM06249EutQ_1_eut1(pf06249) EUT1 EP_NTERM
0001665 HMM10662PduV_1_mix(pf10662) EUT1
0001670 Hp_euts_violetPink EUT1
0001675 HMM00390_malic_1_eut1(pf00390pf03949pf01515) EUT1
0001680 -pf00923 -
0001685 HMM02780keto_1_eut1(pf02779pf02780pf00456) EUT1
0001690 -pf06674 -
0001695 HMM00293nuhyd_1_eut1(pf00293) EUT1
0001700 -pf13247pf14691pf07992 -
0001705 -# -
0001710 -pf00873 -
0001715 -pf03960 -
0001720 -pf07687pf01546 -
BMC-P
BMC-H
BMC-T
Aldehyde dehydrogenase
PTAC
Signature enzyme
Alcohol dehydrogenase
Regulator

Klebsiella sp. PO552. Bacteria; Proteobacteria; Gammaproteobacteria; Enterobacterales;

BMD99__locus_2

EUT1

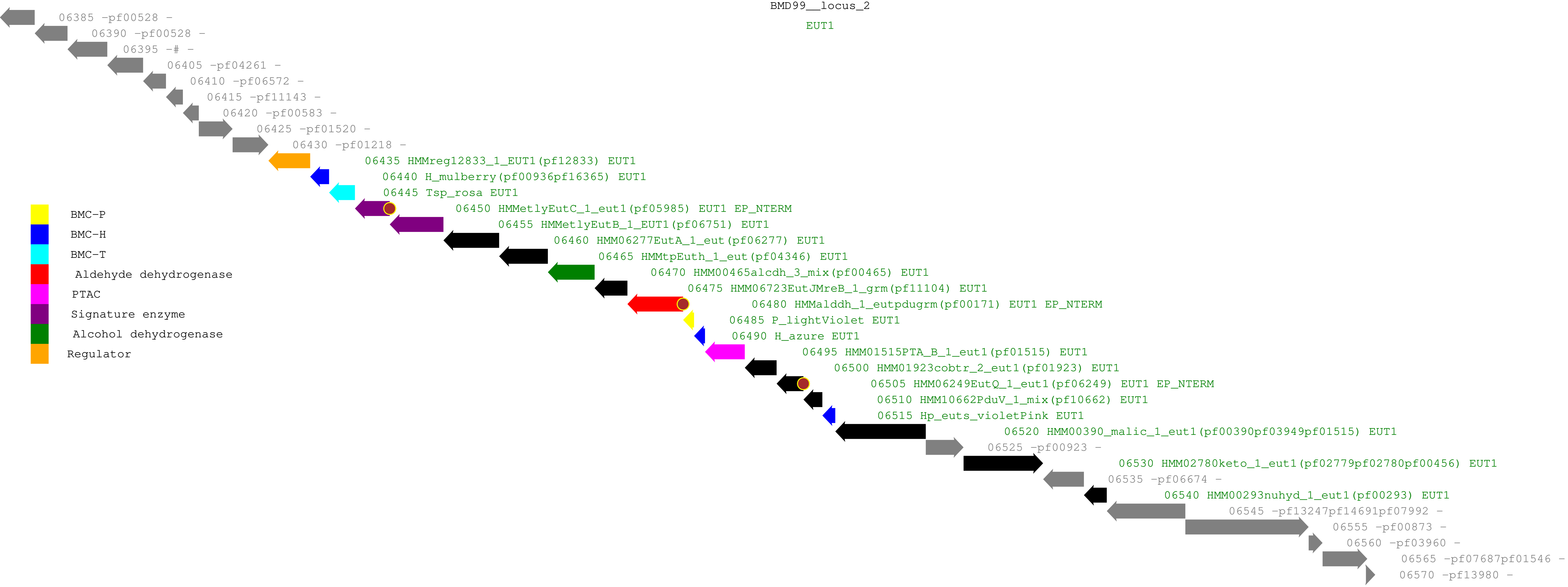

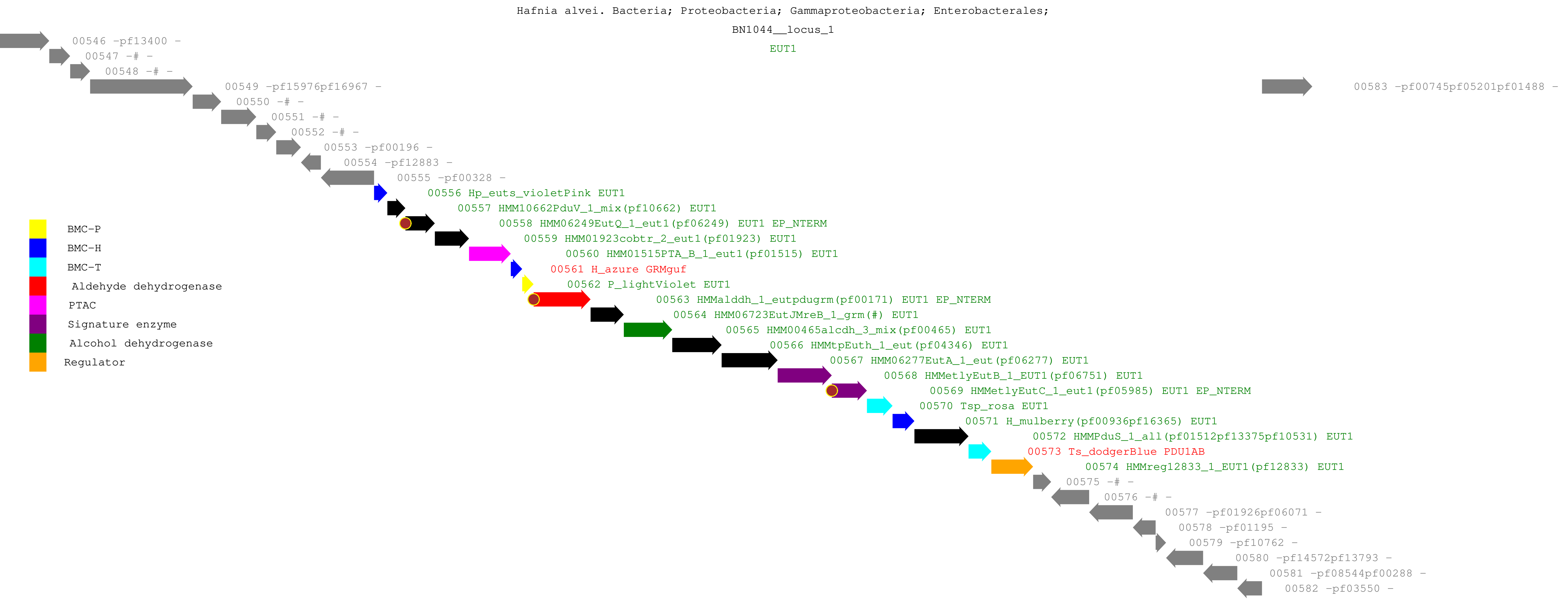
Hafnia alvei. Bacteria; Proteobacteria; Gammaproteobacteria; Enterobacterales;
BN1044__locus_1
EUT1
00546 -pf13400 -
00547 -# -
00548 -# -
00549 -pf15976pf16967 -
00550 -# -
00551 -# -
00552 -# -
00553 -pf00196 -
00554 -pf12883 -
00555 -pf00328 -
00556 Hp_euts_violetPink EUT1
00557 HMM10662PduV_1_mix(pf10662) EUT1
00558 HMM06249EutQ_1_eut1(pf06249) EUT1 EP_NTERM
00559 HMM01923cobtr_2_eut1(pf01923) EUT1
00560 HMM01515PTA_B_1_eut1(pf01515) EUT1
00561 H_azure GRMguf
00562 P_lightViolet EUT1
00563 HMMalddh_1_eutpdugrm(pf00171) EUT1 EP_NTERM
00564 HMM06723EutJMreB_1_grm(#) EUT1
00565 HMM00465alcdh_3_mix(pf00465) EUT1
00566 HMMtpEuth_1_eut(pf04346) EUT1
00567 HMM06277EutA_1_eut(pf06277) EUT1
00568 HMMetlyEutB_1_EUT1(pf06751) EUT1
00569 HMMetlyEutC_1_eut1(pf05985) EUT1 EP_NTERM
00570 Tsp_rosa EUT1
00571 H_mulberry(pf00936pf16365) EUT1
00572 HMMPduS_1_all(pf01512pf13375pf10531) EUT1
00573 Ts_dodgerBlue PDU1AB
00574 HMMreg12833_1_EUT1(pf12833) EUT1
00575 -# -
00576 -# -
00577 -pf01926pf06071 -
00578 -pf01195 -
00579 -pf10762 -
00580 -pf14572pf13793 -
00581 -pf08544pf00288 -
00582 -pf03550 -
00583 -pf00745pf05201pf01488 -
BMC-P
BMC-H
BMC-T
Aldehyde dehydrogenase
PTAC
Signature enzyme
Alcohol dehydrogenase
Regulator

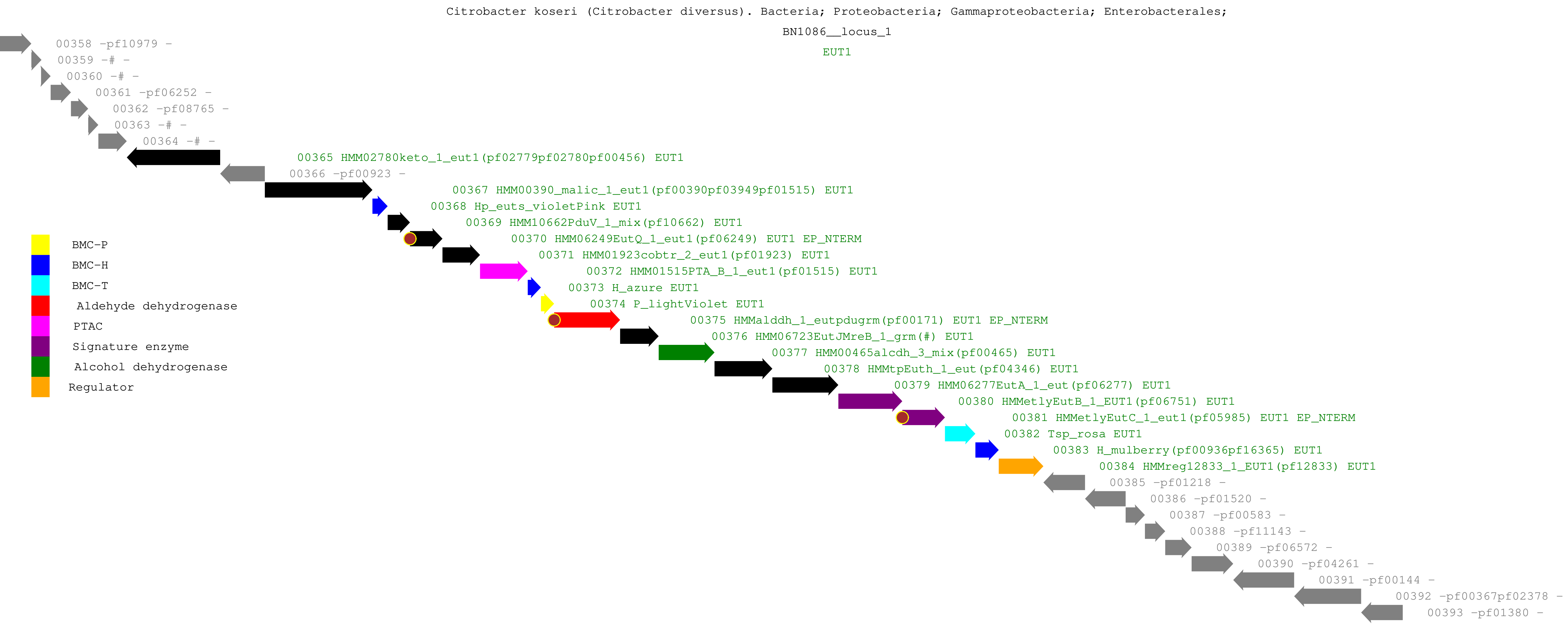
Citrobacter koseri (Citrobacter diversus). Bacteria; Proteobacteria; Gammaproteobacteria; Enterobacterales;
BN1086__locus_1
EUT1
00358 -pf10979 -
00359 -# -
00360 -# -
00361 -pf06252 -
00362 -pf08765 -
00363 -# -
00364 -# -
00365 HMM02780keto_1_eut1(pf02779pf02780pf00456) EUT1
00366 -pf00923 -
00367 HMM00390_malic_1_eut1(pf00390pf03949pf01515) EUT1
00368 Hp_euts_violetPink EUT1
00369 HMM10662PduV_1_mix(pf10662) EUT1
00370 HMM06249EutQ_1_eut1(pf06249) EUT1 EP_NTERM
00371 HMM01923cobtr_2_eut1(pf01923) EUT1
00372 HMM01515PTA_B_1_eut1(pf01515) EUT1
00373 H_azure EUT1
00374 P_lightViolet EUT1
00375 HMMalddh_1_eutpdugrm(pf00171) EUT1 EP_NTERM
00376 HMM06723EutJMreB_1_grm(#) EUT1
00377 HMM00465alcdh_3_mix(pf00465) EUT1
00378 HMMtpEuth_1_eut(pf04346) EUT1
00379 HMM06277EutA_1_eut(pf06277) EUT1
00380 HMMetlyEutB_1_EUT1(pf06751) EUT1
00381 HMMetlyEutC_1_eut1(pf05985) EUT1 EP_NTERM
00382 Tsp_rosa EUT1
00383 H_mulberry(pf00936pf16365) EUT1
00384 HMMreg12833_1_EUT1(pf12833) EUT1
00385 -pf01218 -
00386 -pf01520 -
00387 -pf00583 -
00388 -pf11143 -
00389 -pf06572 -
00390 -pf04261 -
00391 -pf00144 -
00392 -pf00367pf02378 -
00393 -pf01380 -
BMC-P
BMC-H
BMC-T
Aldehyde dehydrogenase
PTAC
Signature enzyme
Alcohol dehydrogenase
Regulator

Escherichia coli chi7122. Bacteria; Proteobacteria; Gammaproteobacteria; Enterobacterales;

BN16__locus_1

EUT1

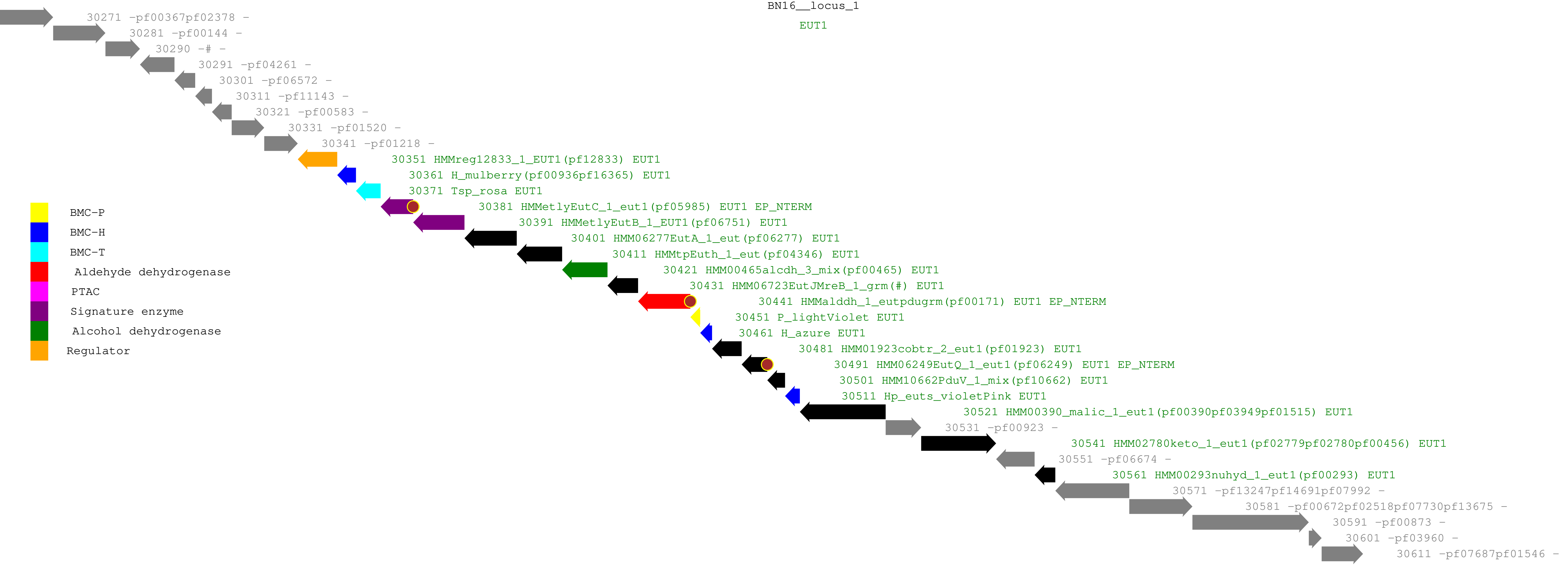

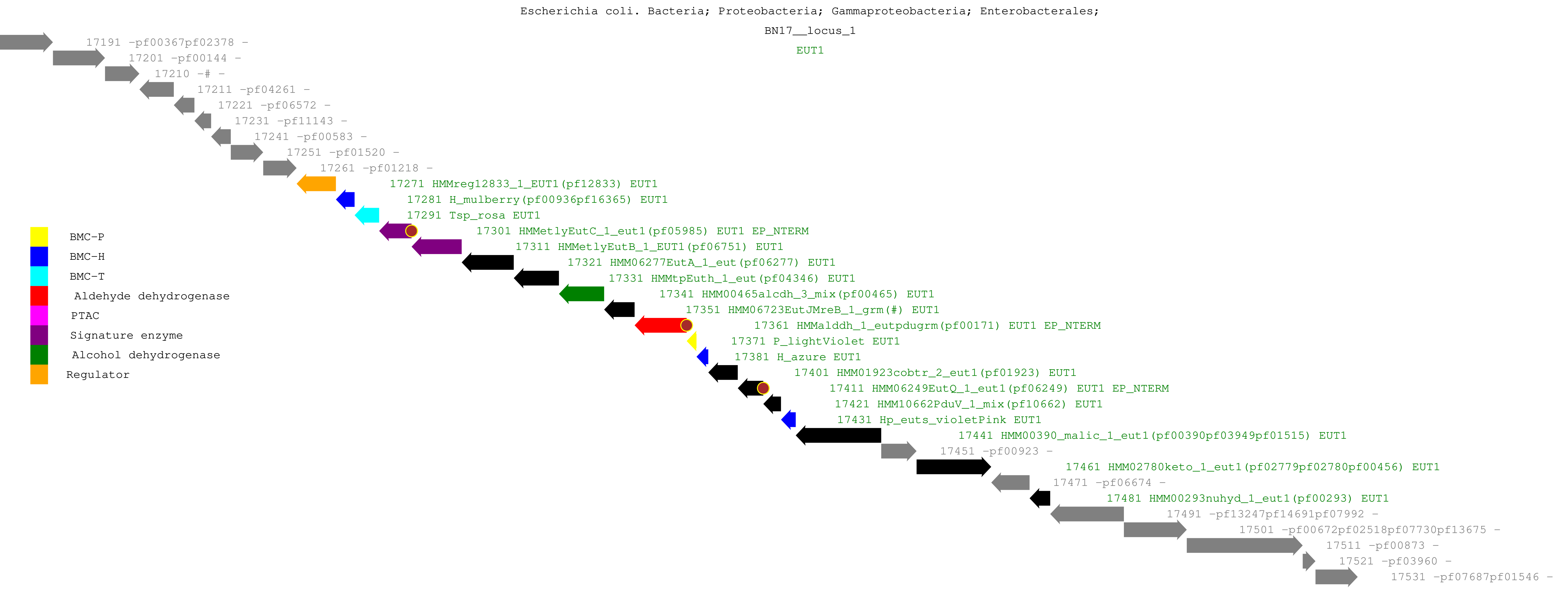

Escherichia coli. Bacteria; Proteobacteria; Gammaproteobacteria; Enterobacterales;
BN17__locus_1
EUT1
17191 -pf00367pf02378 -
17201 -pf00144 -
17210 -# -
17211 -pf04261 -
17221 -pf06572 -
17231 -pf11143 -
17241 -pf00583 -
17251 -pf01520 -
17261 -pf01218 -
17271 HMMreg12833_1_EUT1(pf12833) EUT1
17281 H_mulberry(pf00936pf16365) EUT1
17291 Tsp_rosa EUT1
17301 HMMetlyEutC_1_eut1(pf05985) EUT1 EP_NTERM
17311 HMMetlyEutB_1_EUT1(pf06751) EUT1
17321 HMM06277EutA_1_eut(pf06277) EUT1
17331 HMMtpEuth_1_eut(pf04346) EUT1
17341 HMM00465alcdh_3_mix(pf00465) EUT1
17351 HMM06723EutJMreB_1_grm(#) EUT1
17361 HMMaldh_1_eutpdugrm(pf00171) EUT1 EP_NTERM
17371 P_lightViolet EUT1
17381 H_azure EUT1
17401 HMM01923cobtr_2_eut1(pf01923) EUT1
17411 HMM06249EutQ_1_eut1(pf06249) EUT1 EP_NTERM
17421 HMM10662PduV_1_mix(pf10662) EUT1
17431 Hp_euts_violetPink EUT1
17441 HMM00390_malic_1_eut1(pf00390pf03949pf01515) EUT1
17451 -pf00923 -
17461 HMM02780keto_1_eut1(pf02779pf02780pf00456) EUT1
17471 -pf06674 -
17481 HMM00293nuhyd_1_eut1(pf00293) EUT1
17491 -pf13247pf14691pf07992 -
17501 -pf00672pf02518pf07730pf13675 -
17511 -pf00873 -
17521 -pf03960 -
17531 -pf07687pf01546 -
BMC-P
BMC-H
BMC-T
Aldehyde dehydrogenase
PTAC
Signature enzyme
Alcohol dehydrogenase
Regulator

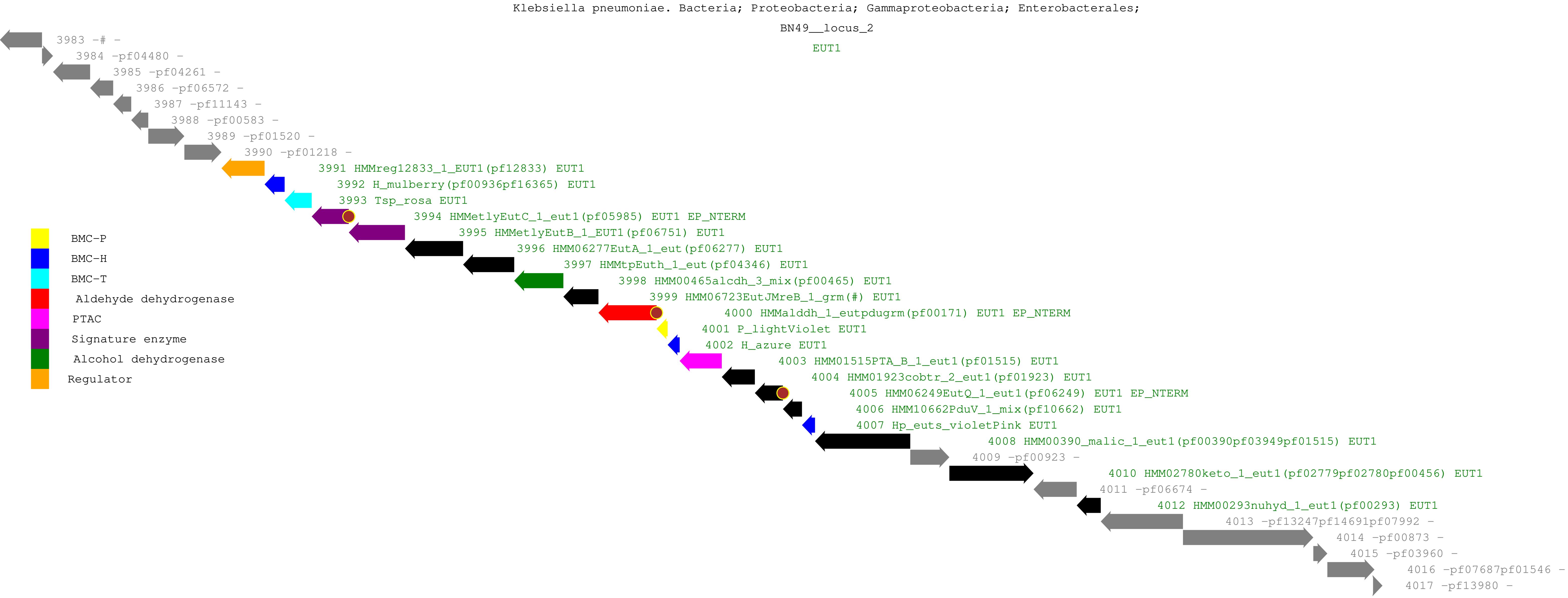
Klebsiella pneumoniae. Bacteria; Proteobacteria; Gammaproteobacteria; Enterobacterales;
BN49__locus_2
EUT1
3983 -# -
3984 -pf04480 -
3985 -pf04261 -
3986 -pf06572 -
3987 -pf11143 -
3988 -pf00583 -
3989 -pf01520 -
3990 -pf01218 -
3991 HMMreg12833_1_EUT1(pf12833) EUT1
3992 H_mulberry(pf00936pf16365) EUT1
3993 Tsp_rosa EUT1
3994 HMMetlyEutC_1_eut1(pf05985) EUT1 EP_NTERM
3995 HMMetlyEutB_1_EUT1(pf06751) EUT1
3996 HMM06277EutA_1_eut(pf06277) EUT1
3997 HMMtpEuth_1_eut(pf04346) EUT1
3998 HMM00465alcdh_3_mix(pf00465) EUT1
3999 HMM06723EutJMreB_1_grm(#) EUT1
4000 HMMalddh_1_eutpdugrm(pf00171) EUT1 EP_NTERM
4001 P_lightViolet EUT1
4002 H_azure EUT1
4003 HMM01515PTA_B_1_eut1(pf01515) EUT1
4004 HMM01923cobtr_2_eut1(pf01923) EUT1
4005 HMM06249EutQ_1_eut1(pf06249) EUT1 EP_NTERM
4006 HMM10662PduV_1_mix(pf10662) EUT1
4007 Hp_euts_violetPink EUT1
4008 HMM00390_malic_1_eut1(pf00390pf03949pf01515) EUT1
4009 -pf00923 -
4010 HMM02780keto_1_eut1(pf02779pf02780pf00456) EUT1
4011 -pf06674 -
4012 HMM00293nuhyd_1_eut1(pf00293) EUT1
4013 -pf13247pf14691pf07992 -
4014 -pf00873 -
4015 -pf03960 -
4016 -pf07687pf01546 -
4017 -pf13980 -
BMC-P
BMC-H
BMC-T
Aldehyde dehydrogenase
PTAC
Signature enzyme
Alcohol dehydrogenase
Regulator

Escherichia coli O127:H6. Bacteria; Proteobacteria; Gammaproteobacteria; Enterobacterales;

BQ9544__locus_2

EUT1

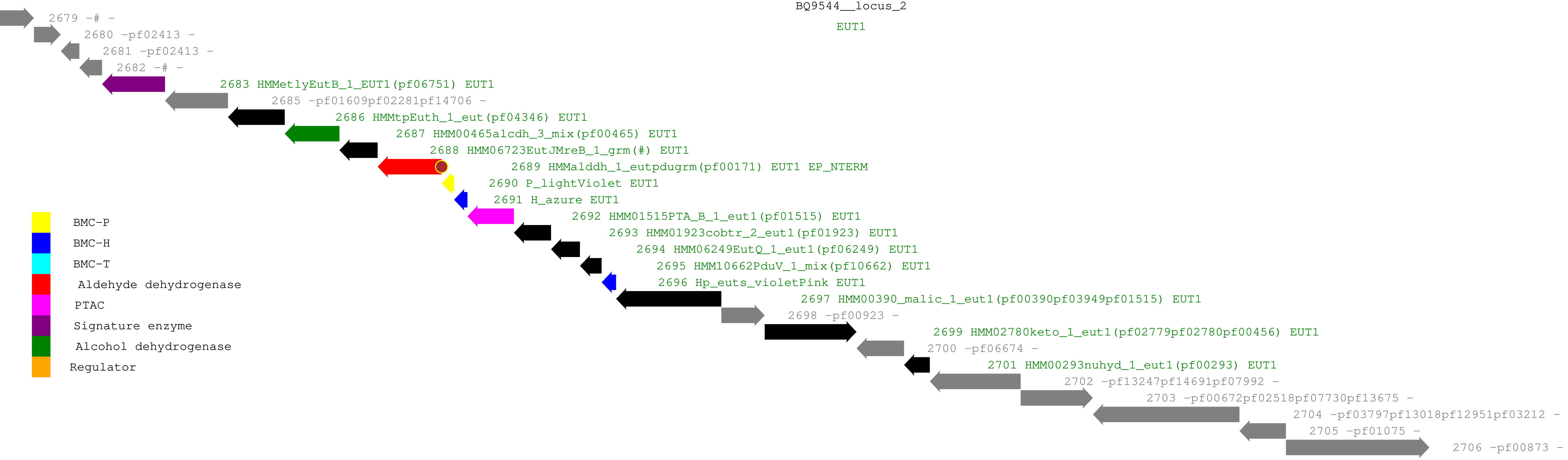

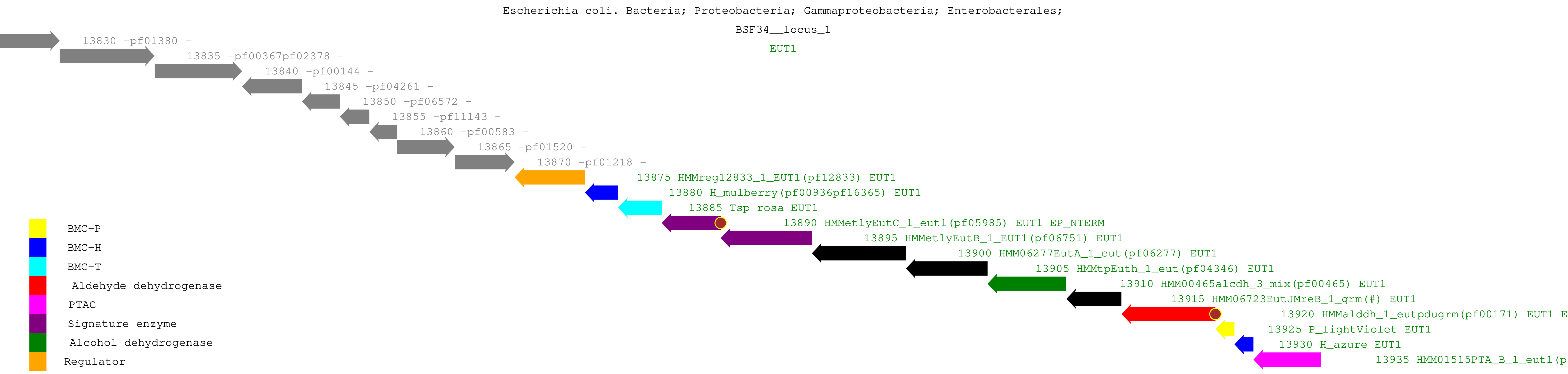
Escherichia coli. Bacteria; Proteobacteria; Gammaproteobacteria; Enterobacterales;
BSF34__locus_1
EUT1
13830 -pf01380 -
13835 -pf00367pf02378 -
13840 -pf00144 -
13845 -pf04261 -
13850 -pf06572 -
13855 -pf11143 -
13860 -pf00583 -
13865 -pf01520 -
13870 -pf01218 -
13875 HMMreg12833_1_EUT1(pf12833) EUT1
13880 H_mulberry(pf00936pf16365) EUT1
13885 Tsp_rosa EUT1
13890 HMMetlyEutC_1_eut1(pf05985) EUT1 EP_NTERM
13895 HMMetlyEutB_1_EUT1(pf06751) EUT1
13900 HMM06277EutA_1_eut(pf06277) EUT1
13905 HMMtpEuth_1_eut(pf04346) EUT1
13910 HMM00465alcdh_3_mix(pf00465) EUT1
13915 HMM06723EutJMreB_1_grm(#) EUT1
13920 HMMaldh_1_eutpdugrm(pf00171) EUT1 E
13925 P_lightViolet EUT1
13930 H_azure EUT1
13935 HMM01515PTA_B_1_eut1(p
BMC-P
BMC-H
BMC-T
Aldehyde dehydrogenase
PTAC
Signature enzyme
Alcohol dehydrogenase
Regulator

Buttiauxella sp. A111. Bacteria; Proteobacteria; Gammaproteobacteria; Enterobacterales;

BSPA111__locus_1

EUT1

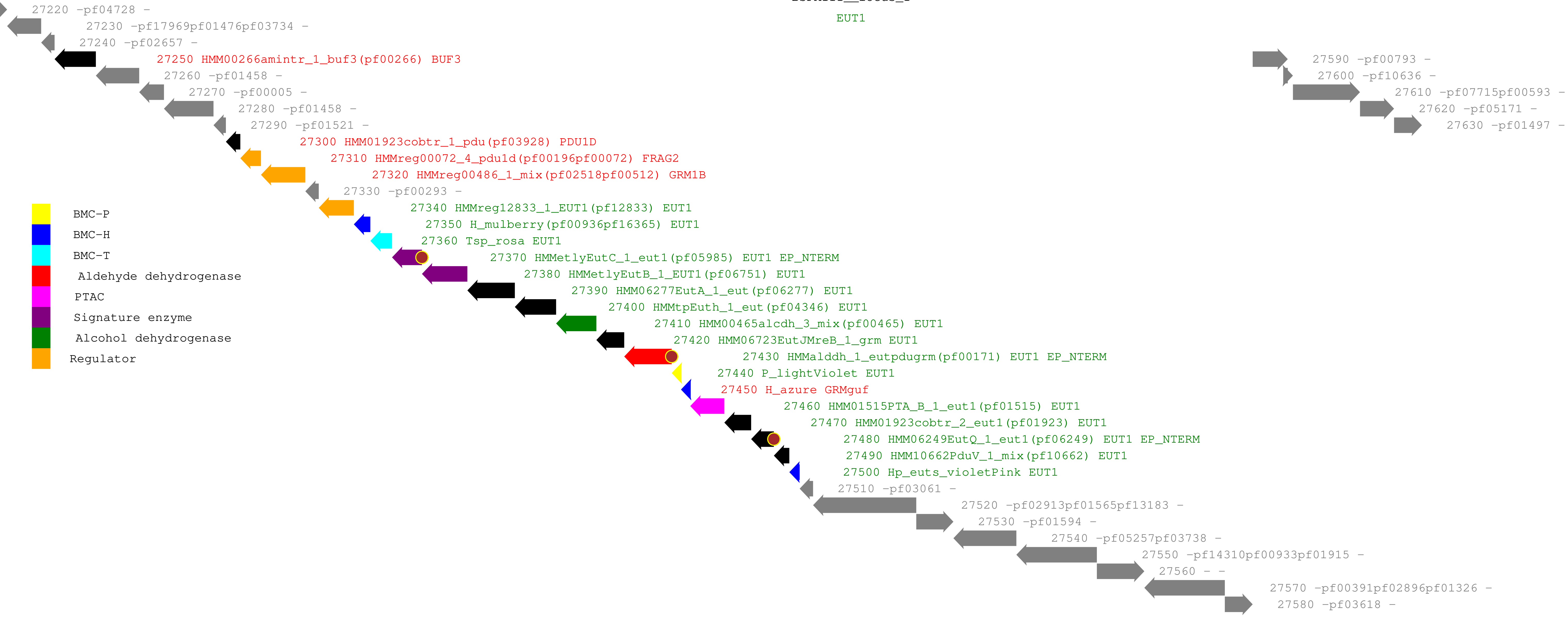

Salmonella enteritidis. Bacteria; Proteobacteria; Gammaproteobacteria; Enterobacterales;

BTN74__locus_2

EUT1

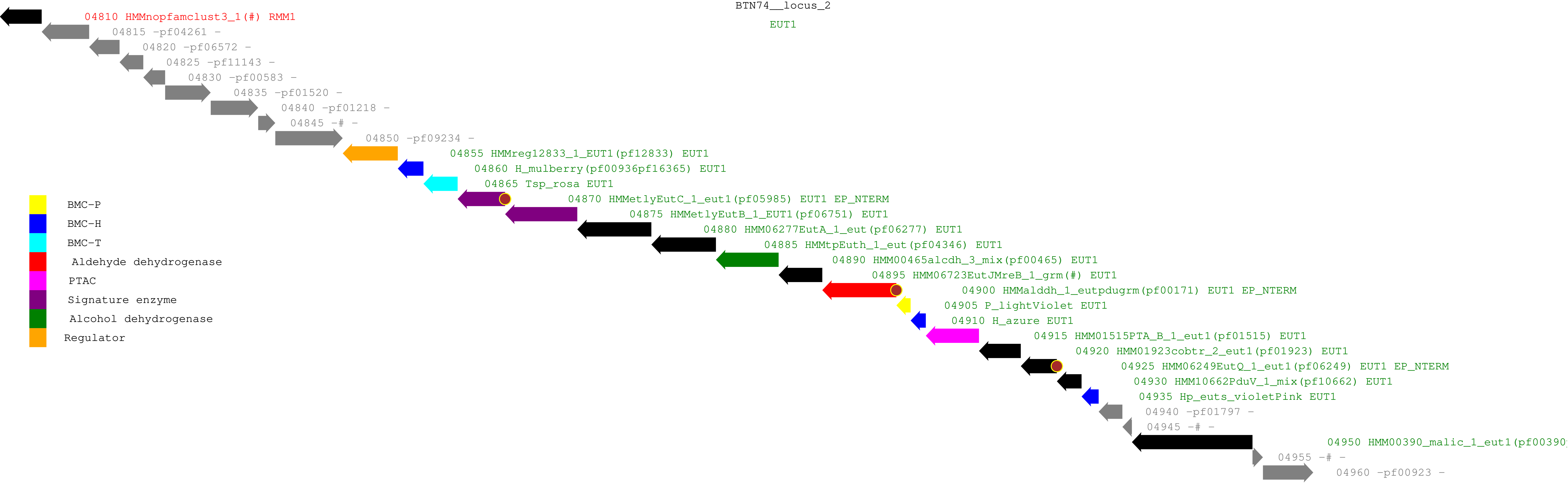

Escherichia coli O146:H21 str. 2010C-3325. Bacteria; Proteobacteria; Gammaproteobacteria; Enterobacterales;

BU55__locus_1

EUT1

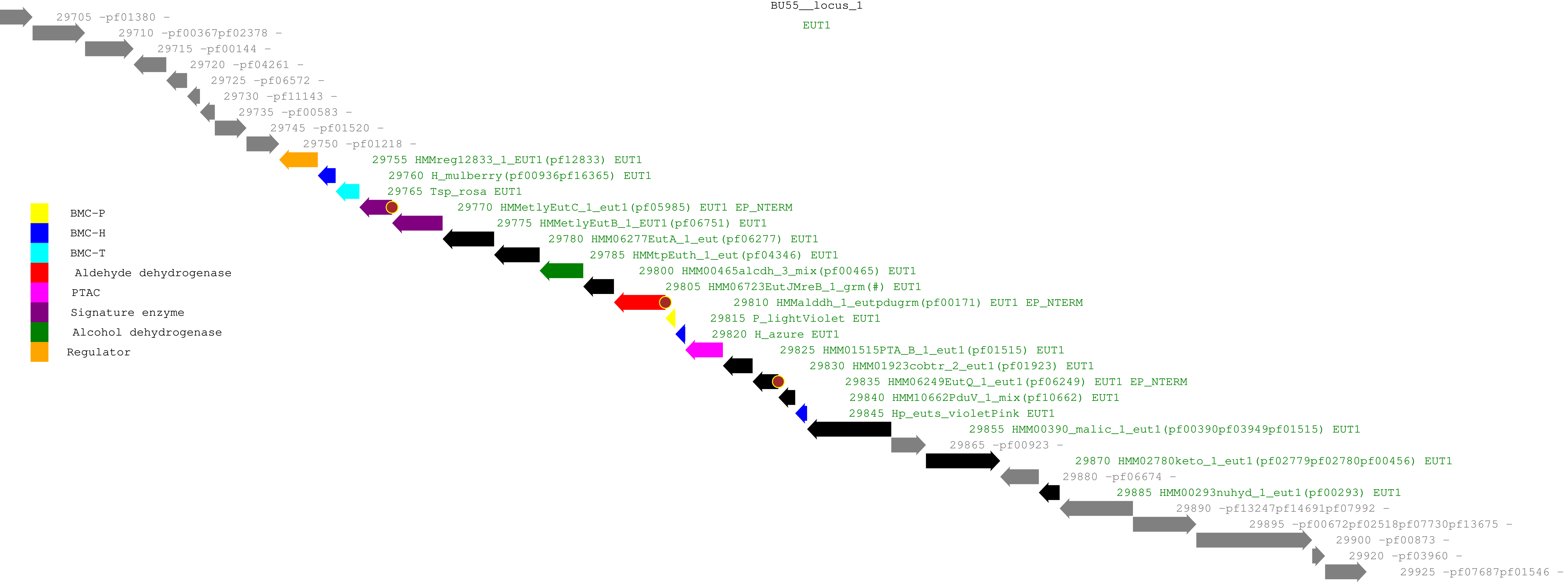

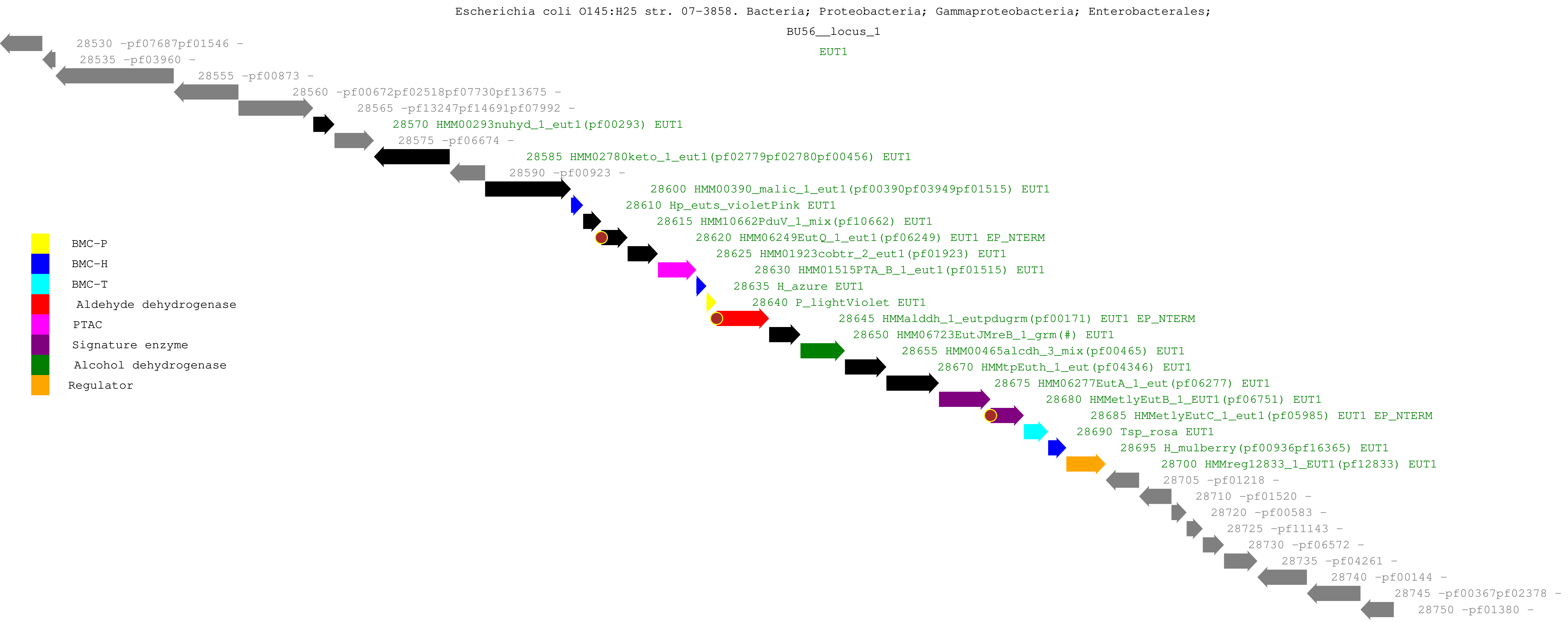
Escherichia coli O145:H25 str. 07-3858. Bacteria; Proteobacteria; Gammaproteobacteria; Enterobacterales;
BU56__locus_1
EUT1
28530 -pf07687pf01546 -
28535 -pf03960 -
28555 -pf00873 -
28560 -pf00672pf02518pf07730pf13675 -
28565 -pf13247pf14691pf07992 -
28570 HMM00293nuhyd_1_eut1(pf00293) EUT1
28575 -pf06674 -
28585 HMM02780keto_1_eut1(pf02779pf02780pf00456) EUT1
28590 -pf00923 -
28600 HMM00390_malic_1_eut1(pf00390pf03949pf01515) EUT1
28610 Hp_euts_violetPink EUT1
28615 HMM10662PduV_1_mix(pf10662) EUT1
28620 HMM06249EutQ_1_eut1(pf06249) EUT1 EP_NTERM
28625 HMM01923cobtr_2_eut1(pf01923) EUT1
28630 HMM01515PTA_B_1_eut1(pf01515) EUT1
28635 H_azure EUT1
28640 P_lightViolet EUT1
28645 HMMalddh_1_eutpdugrm(pf00171) EUT1 EP_NTERM
28650 HMM06723EutJMreB_1_grm(#) EUT1
28655 HMM00465alcdh_3_mix(pf00465) EUT1
28670 HMMtpEuth_1_eut(pf04346) EUT1
28675 HMM06277EutA_1_eut(pf06277) EUT1
28680 HMMetlyEutB_1_EUT1(pf06751) EUT1
28685 HMMetlyEutC_1_eut1(pf05985) EUT1 EP_NTERM
28690 Tsp_rosa EUT1
28695 H_mulberry(pf00936pf16365) EUT1
28700 HMMreg12833_1_EUT1(pf12833) EUT1
28705 -pf01218 -
28710 -pf01520 -
28720 -pf00583 -
28725 -pf11143 -
28730 -pf06572 -
28735 -pf04261 -
28740 -pf00144 -
28745 -pf00367pf02378 -
28750 -pf01380 -
BMC-P
BMC-H
BMC-T
Aldehyde dehydrogenase
PTAC
Signature enzyme
Alcohol dehydrogenase
Regulator

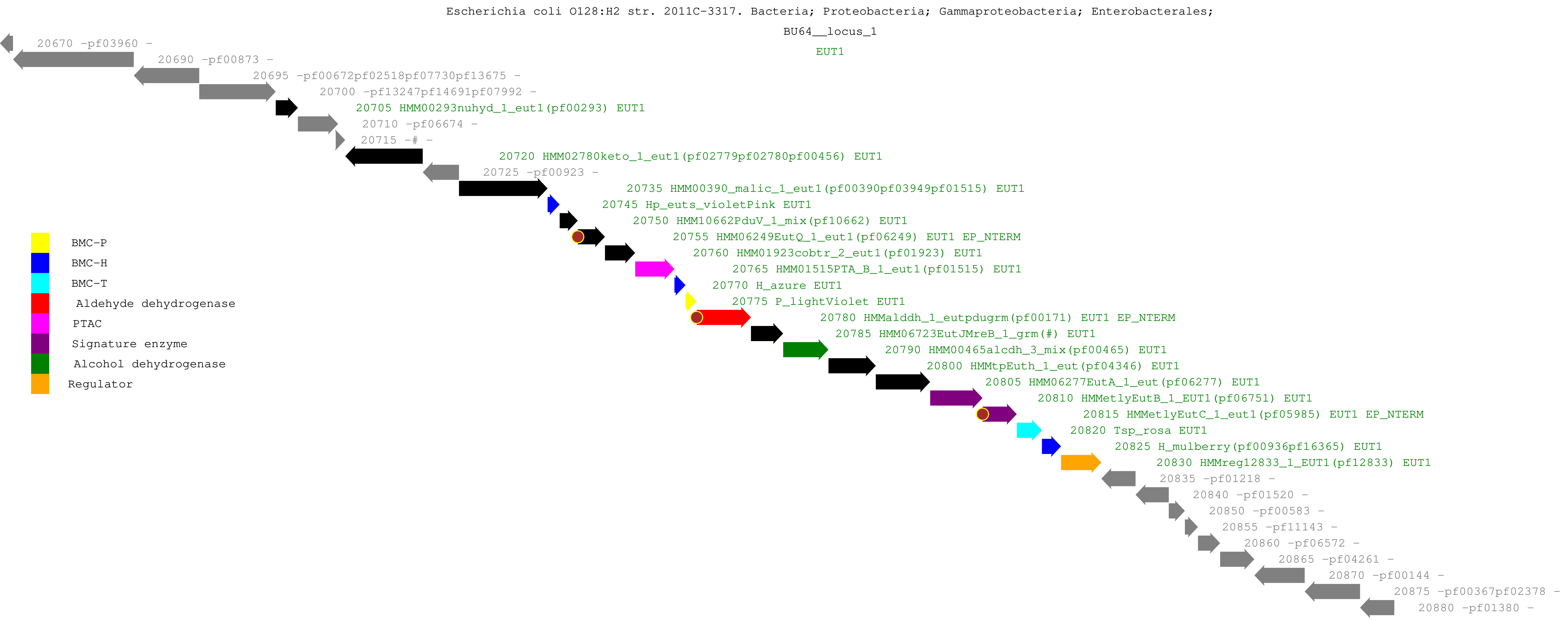
Escherichia coli O128:H2 str. 2011C-3317. Bacteria; Proteobacteria; Gammaproteobacteria; Enterobacterales;
BU64__locus_1
EUT1
20670 -pf03960 -
20690 -pf00873 -
20695 -pf00672pf02518pf07730pf13675 -
20700 -pf13247pf14691pf07992 -
20705 HMM00293nuhyd_1_eut1(pf00293) EUT1
20710 -pf06674 -
20715 -# -
20720 HMM02780keto_1_eut1(pf02779pf02780pf00456) EUT1
20725 -pf00923 -
20735 HMM00390_malic_1_eut1(pf00390pf03949pf01515) EUT1
20745 Hp_euts_violetPink EUT1
20750 HMM10662PduV_1_mix(pf10662) EUT1
20755 HMM06249EutQ_1_eut1(pf06249) EUT1 EP_NTERM
20760 HMM01923cobtr_2_eut1(pf01923) EUT1
20765 HMM01515PTA_B_1_eut1(pf01515) EUT1
20770 H_azure EUT1
20775 P_lightViolet EUT1
20780 HMMalddh_1_eutpdugrm(pf00171) EUT1 EP_NTERM
20785 HMM06723EutJMreB_1_grm(#) EUT1
20790 HMM00465alcdh_3_mix(pf00465) EUT1
20800 HMMtpEuth_1_eut(pf04346) EUT1
20805 HMM06277EutA_1_eut(pf06277) EUT1
20810 HMMetlyEutB_1_EUT1(pf06751) EUT1
20815 HMMetlyEutC_1_eut1(pf05985) EUT1 EP_NTERM
20820 Tsp_rosa EUT1
20825 H_mulberry(pf00936pf16365) EUT1
20830 HMMreg12833_1_EUT1(pf12833) EUT1
20835 -pf01218 -
20840 -pf01520 -
20850 -pf00583 -
20855 -pf11143 -
20860 -pf06572 -
20865 -pf04261 -
20870 -pf00144 -
20875 -pf00367pf02378 -
20880 -pf01380 -
BMC-P
BMC-H
BMC-T
Aldehyde dehydrogenase
PTAC
Signature enzyme
Alcohol dehydrogenase
Regulator

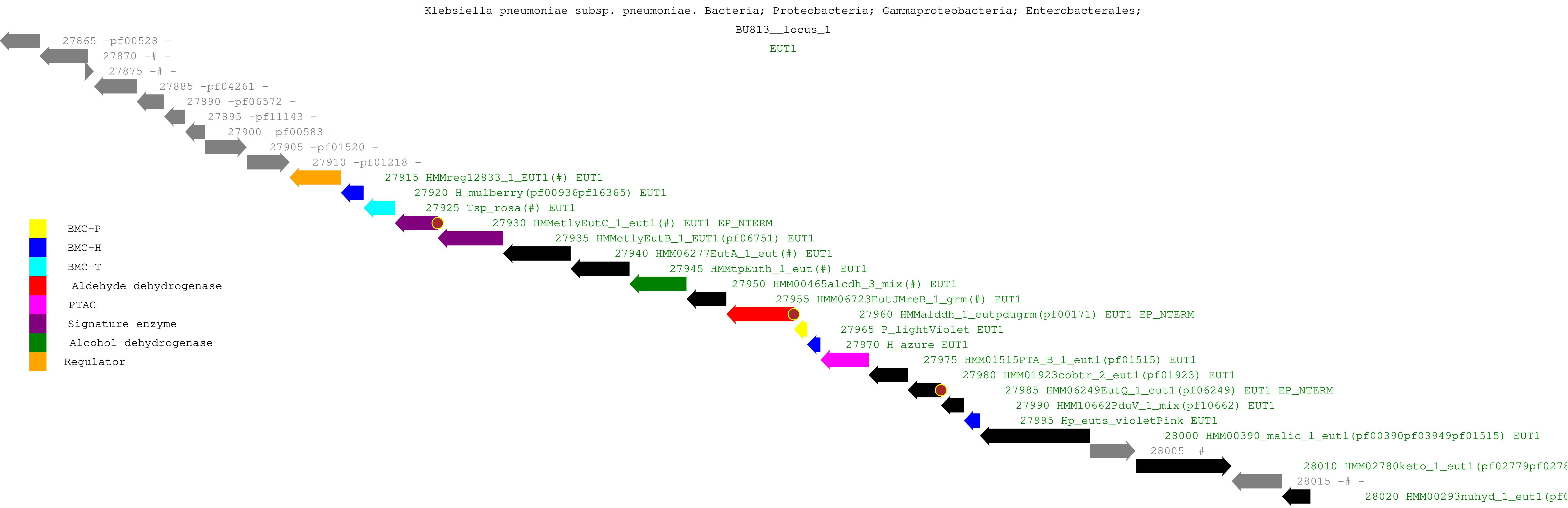

Klebsiella pneumoniae subsp. pneumoniae. Bacteria; Proteobacteria; Gammaproteobacteria; Enterobacterales;
BU813__locus_1
EUT1
27865 -pf00528 -
27870 -# -
27875 -# -
27885 -pf04261 -
27890 -pf06572 -
27895 -pf11143 -
27900 -pf00583 -
27905 -pf01520 -
27910 -pf01218 -
27915 HMMreg12833_1_EUT1(#) EUT1
27920 H_mulberry(pf00936pf16365) EUT1
27925 Tsp_rosa(#) EUT1
27930 HMMetlyEutC_1_eut1(#) EUT1 EP_NTERM
27935 HMMetlyEutB_1_EUT1(pf06751) EUT1
27940 HMM06277EutA_1_eut(#) EUT1
27945 HMMtpEuth_1_eut(#) EUT1
27950 HMM00465alcdh_3_mix(#) EUT1
27955 HMM06723EutJMreB_1_grm(#) EUT1
27960 HMMalddh_1_eutpdugrm(pf00171) EUT1 EP_NTERM
27965 P_lightViolet EUT1
27970 H_azure EUT1
27975 HMM01515PTA_B_1_eut1(pf01515) EUT1
27980 HMM01923cobtr_2_eut1(pf01923) EUT1
27985 HMM06249EutQ_1_eut1(pf06249) EUT1 EP_NTERM
27990 HMM10662PduV_1_mix(pf10662) EUT1
27995 Hp_euts_violetPink EUT1
28000 HMM00390_malic_1_eut1(pf00390pf03949pf01515) EUT1
28005 -# -
28010 HMM02780keto_1_eut1(pf02779pf0278
28015 -# -
28020 HMM00293nuhyd_1_eut1(pf0
BMC-P
BMC-H
BMC-T
Aldehyde dehydrogenase
PTAC
Signature enzyme
Alcohol dehydrogenase
Regulator

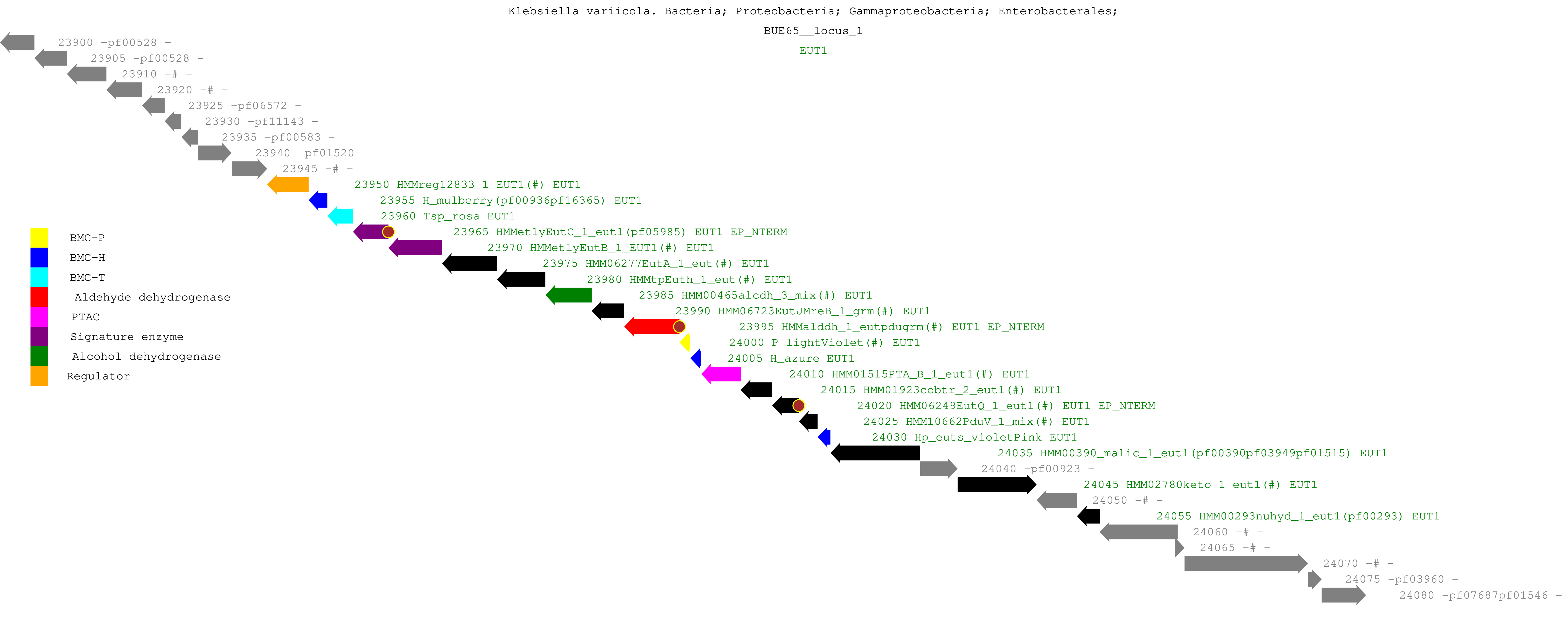
Klebsiella variicola. Bacteria; Proteobacteria; Gammaproteobacteria; Enterobacterales;
BUE65__locus_1
EUT1
23900 -pf00528 -
23905 -pf00528 -
23910 -# -
23920 -# -
23925 -pf06572 -
23930 -pf11143 -
23935 -pf00583 -
23940 -pf01520 -
23945 -# -
23950 HMMreg12833_1_EUT1(#) EUT1
23955 H_mulberry(pf00936pf16365) EUT1
23960 Tsp_rosa EUT1
23965 HMMetlyEutC_1_eut1(pf05985) EUT1 EP_NTERM
23970 HMMetlyEutB_1_EUT1(#) EUT1
23975 HMM06277EutA_1_eut(#) EUT1
23980 HMMtpEuth_1_eut(#) EUT1
23985 HMM00465alcdh_3_mix(#) EUT1
23990 HMM06723EutJMreB_1_grm(#) EUT1
23995 HMMalddh_1_eutpdugrm(#) EUT1 EP_NTERM
24000 P_lightViolet(#) EUT1
24005 H_azure EUT1
24010 HMM01515PTA_B_1_eut1(#) EUT1
24015 HMM01923cobtr_2_eut1(#) EUT1
24020 HMM06249EutQ_1_eut1(#) EUT1 EP_NTERM
24025 HMM10662PduV_1_mix(#) EUT1
24030 Hp_euts_violetPink EUT1
24035 HMM00390_malic_1_eut1(pf00390pf03949pf01515) EUT1
24040 -pf00923 -
24045 HMM02780keto_1_eut1(#) EUT1
24050 -# -
24055 HMM00293nuhyd_1_eut1(pf00293) EUT1
24060 -# -
24065 -# -
24070 -# -
24075 -pf03960 -
24080 -pf07687pf01546 -
BMC-P
BMC-H
BMC-T
Aldehyde dehydrogenase
PTAC
Signature enzyme
Alcohol dehydrogenase
Regulator

Escherichia coli. Bacteria; Proteobacteria; Gammaproteobacteria; Enterobacterales;

BUE81__locus_1

EUT1

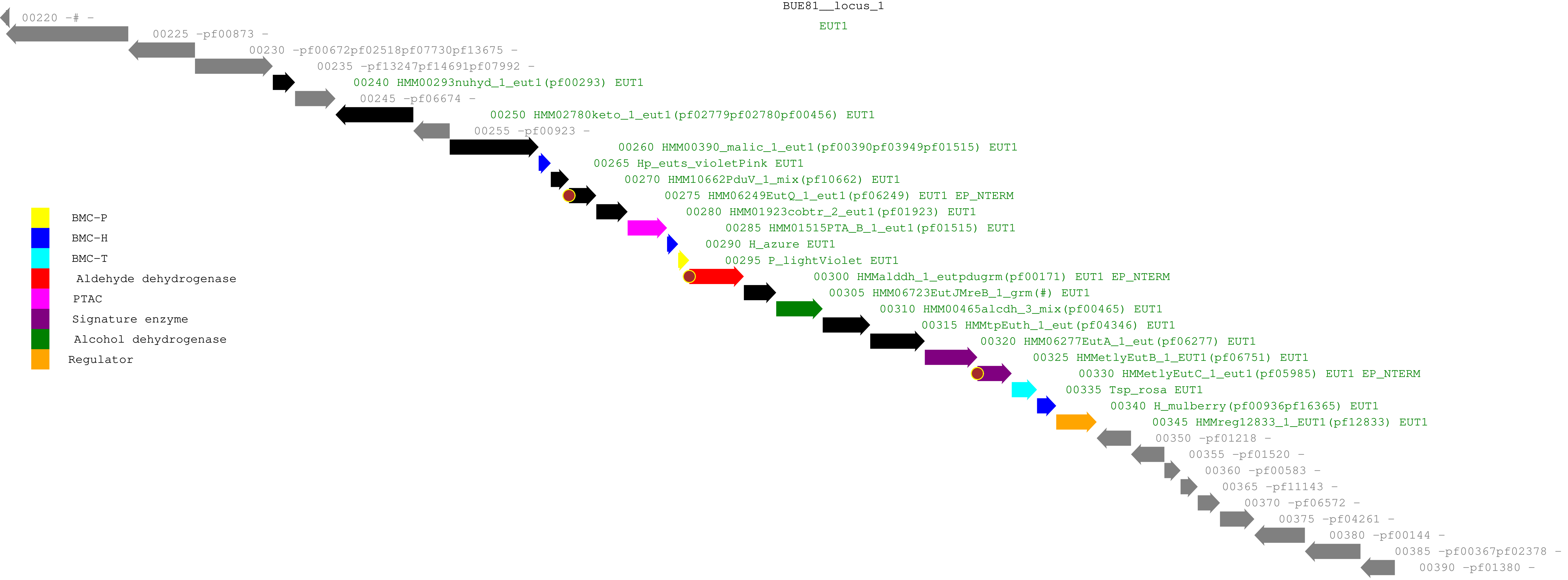

Salmonella enterica subsp. diarizonae serovar 60:r:e,n,x,z15. Bacteria; Proteobacteria; Gammaproteobacteria; Enterobacterales;

BUJ13__locus_1

EUT1

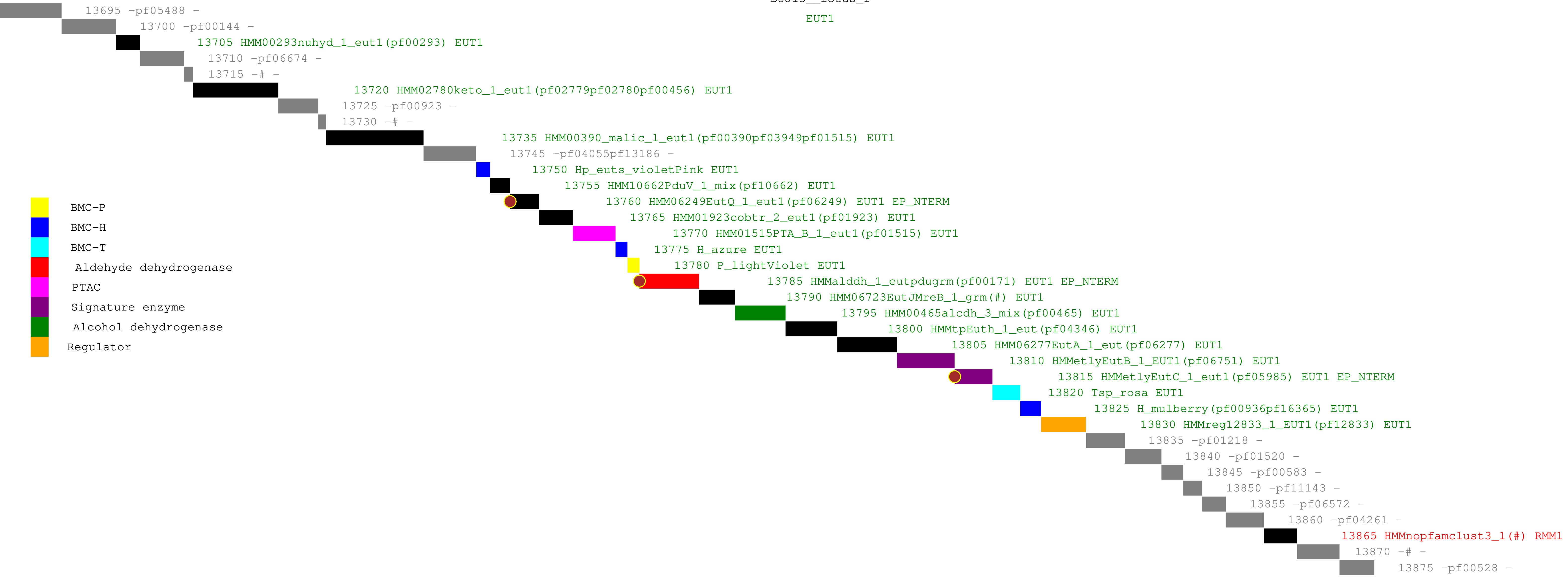

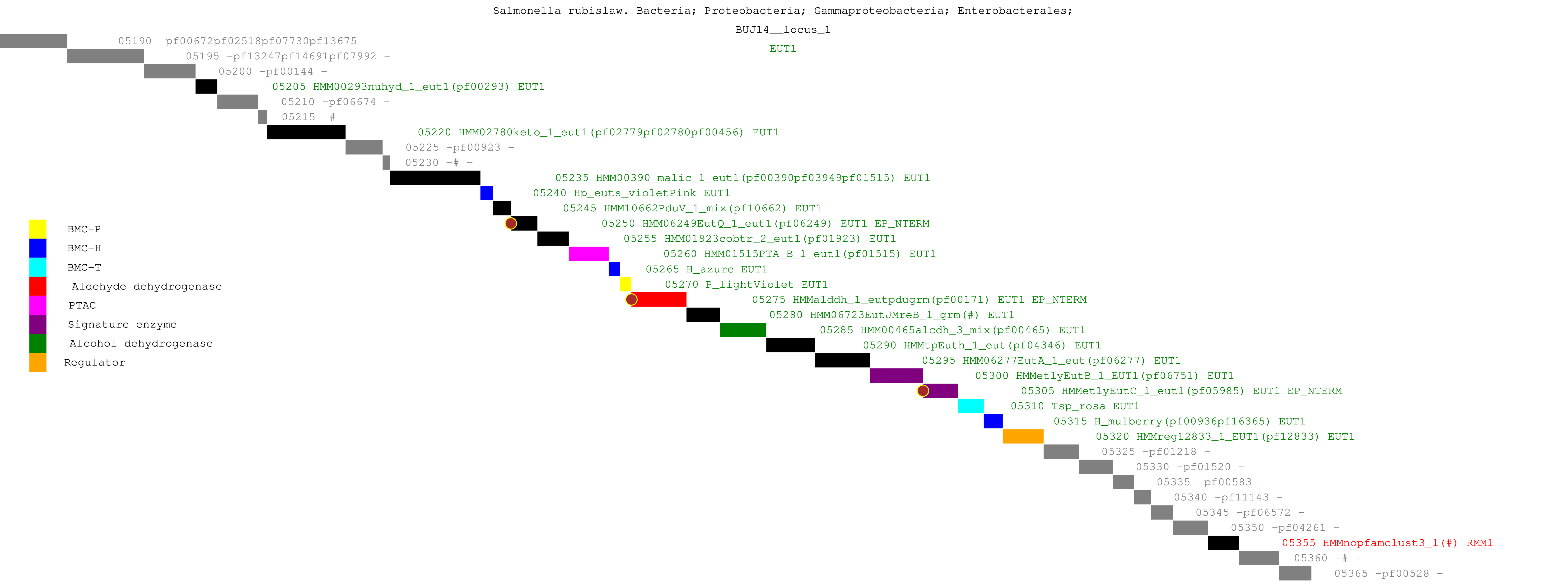
Salmonella rubislaw. Bacteria; Proteobacteria; Gammaproteobacteria; Enterobacterales;
BUJ14__locus_1
EUT1
05190 -pf00672pf02518pf07730pf13675 -
05195 -pf13247pf14691pf07992 -
05200 -pf00144 -
05205 HMM00293nuhyd_1_eut1(pf00293) EUT1
05210 -pf06674 -
05215 -# -
05220 HMM02780keto_1_eut1(pf02779pf02780pf00456) EUT1
05225 -pf00923 -
05230 -# -
05235 HMM00390_malic_1_eut1(pf00390pf03949pf01515) EUT1
05240 Hp_euts_violetPink EUT1
05245 HMM10662PduV_1_mix(pf10662) EUT1
05250 HMM06249EutQ_1_eut1(pf06249) EUT1 EP_NTERM
05255 HMM01923cobtr_2_eut1(pf01923) EUT1
05260 HMM01515PTA_B_1_eut1(pf01515) EUT1
05265 H_azure EUT1
05270 P_lightViolet EUT1
05275 HMMalddh_1_eutpdugrm(pf00171) EUT1 EP_NTERM
05280 HMM06723EutJMreB_1_grm(#) EUT1
05285 HMM00465alcdh_3_mix(pf00465) EUT1
05290 HMMtpEuth_1_eut(pf04346) EUT1
05295 HMM06277EutA_1_eut(pf06277) EUT1
05300 HMMetlyEutB_1_EUT1(pf06751) EUT1
05305 HMMetlyEutC_1_eut1(pf05985) EUT1 EP_NTERM
05310 Tsp_rosa EUT1
05315 H_mulberry(pf00936pf16365) EUT1
05320 HMMreg12833_1_EUT1(pf12833) EUT1
05325 -pf01218 -
05330 -pf01520 -
05335 -pf00583 -
05340 -pf11143 -
05345 -pf06572 -
05350 -pf04261 -
05355 HMMnopfamclust3_1(#) RMM1
05360 -# -
05365 -pf00528 -
BMC-P
BMC-H
BMC-T
Aldehyde dehydrogenase
PTAC
Signature enzyme
Alcohol dehydrogenase
Regulator

Salmonella newport. Bacteria; Proteobacteria; Gammaproteobacteria; Enterobacterales;

BUJ15__locus_1

EUT1

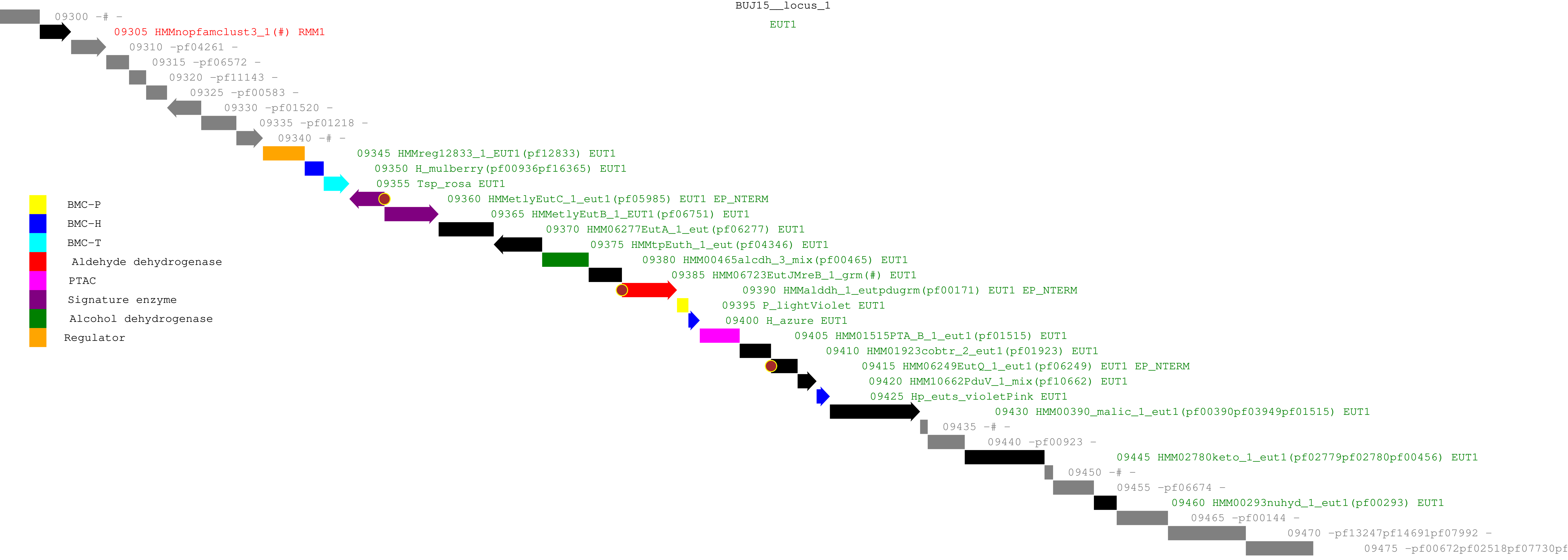

Salmonella choleraesuis. Bacteria; Proteobacteria; Gammaproteobacteria; Enterobacterales;

BUJ19__locus_1

EUT1

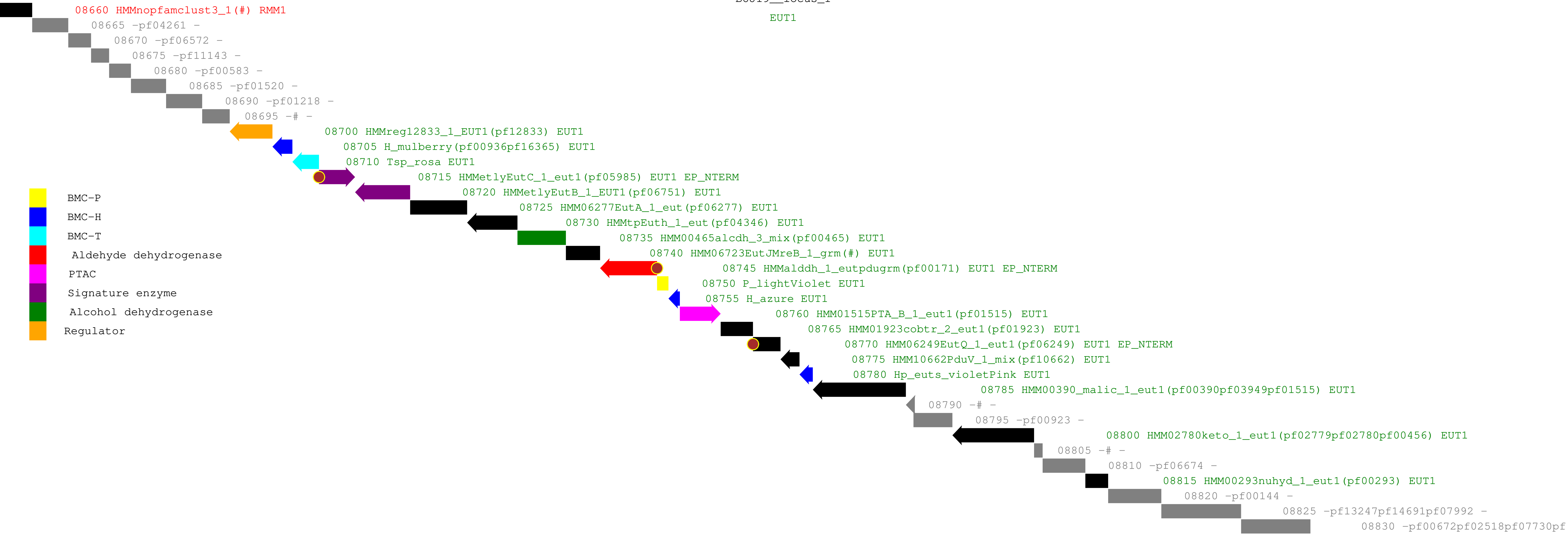

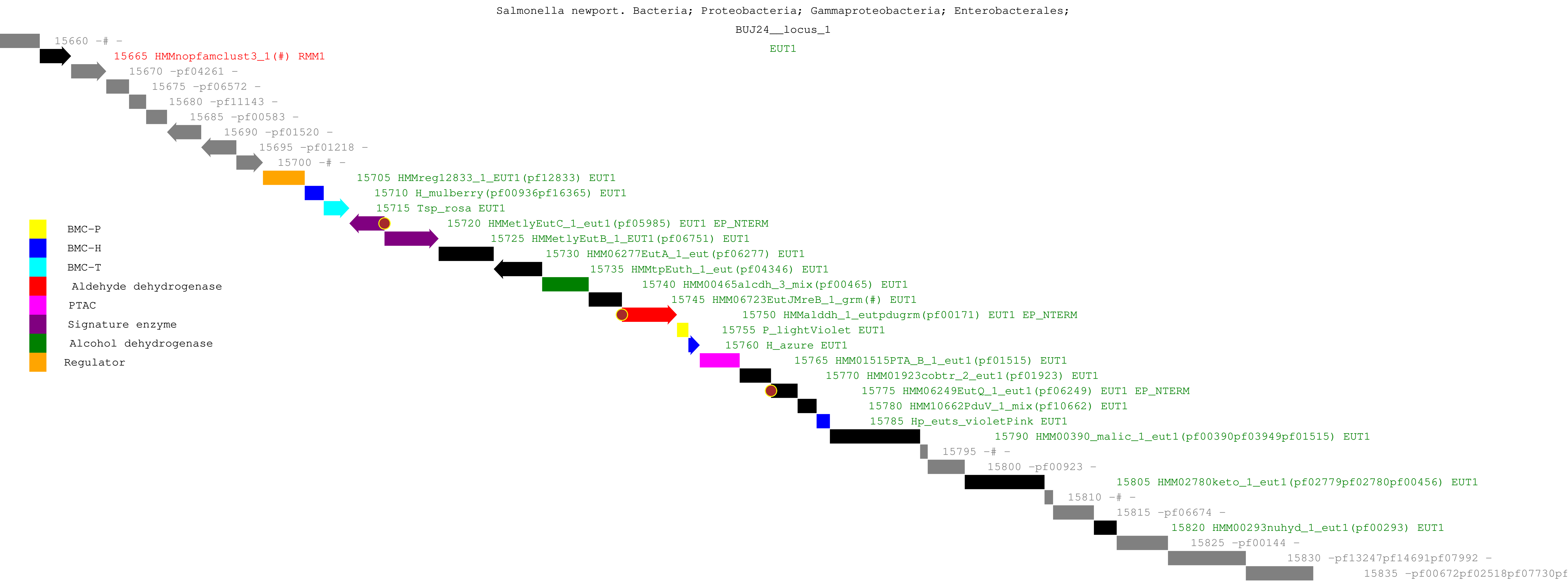
Salmonella newport. Bacteria; Proteobacteria; Gammaproteobacteria; Enterobacterales;
BUJ24__locus_1
EUT1
15660 -# -
15665 HMMnopfamclust3_1(#) RMM1
15670 -pf04261 -
15675 -pf06572 -
15680 -pf11143 -
15685 -pf00583 -
15690 -pf01520 -
15695 -pf01218 -
15700 -# -
15705 HMMreg12833_1_EUT1(pf12833) EUT1
15710 H_mulberry(pf00936pf16365) EUT1
15715 Tsp_rosa EUT1
15720 HMMetlyEutC_1_eut1(pf05985) EUT1 EP_NTERM
15725 HMMetlyEutB_1_EUT1(pf06751) EUT1
15730 HMM06277EutA_1_eut(pf06277) EUT1
15735 HMMtpEuth_1_eut(pf04346) EUT1
15740 HMM00465alcdh_3_mix(pf00465) EUT1
15745 HMM06723EutJMreB_1_grm(#) EUT1
15750 HMMalddh_1_eutpdugrm(pf00171) EUT1 EP_NTERM
15755 P_lightViolet EUT1
15760 H_azure EUT1
15765 HMM01515PTA_B_1_eut1(pf01515) EUT1
15770 HMM01923cobtr_2_eut1(pf01923) EUT1
15775 HMM06249EutQ_1_eut1(pf06249) EUT1 EP_NTERM
15780 HMM10662PduV_1_mix(pf10662) EUT1
15785 Hp_euts_violetPink EUT1
15790 HMM00390_malic_1_eut1(pf00390pf03949pf01515) EUT1
15795 -# -
15800 -pf00923 -
15805 HMM02780keto_1_eut1(pf02779pf02780pf00456) EUT1
15810 -# -
15815 -pf06674 -
15820 HMM00293nuhyd_1_eut1(pf00293) EUT1
15825 -pf00144 -
15830 -pf13247pf14691pf07992 -
15835 -pf00672pf02518pf07730pf
BMC-P
BMC-H
BMC-T
Aldehyde dehydrogenase
PTAC
Signature enzyme
Alcohol dehydrogenase
Regulator

Salmonella enterica subsp. enterica serovar Bardo. Bacteria; Proteobacteria; Gammaproteobacteria; Enterobacterales;

BUJ26__locus_1

EUT1

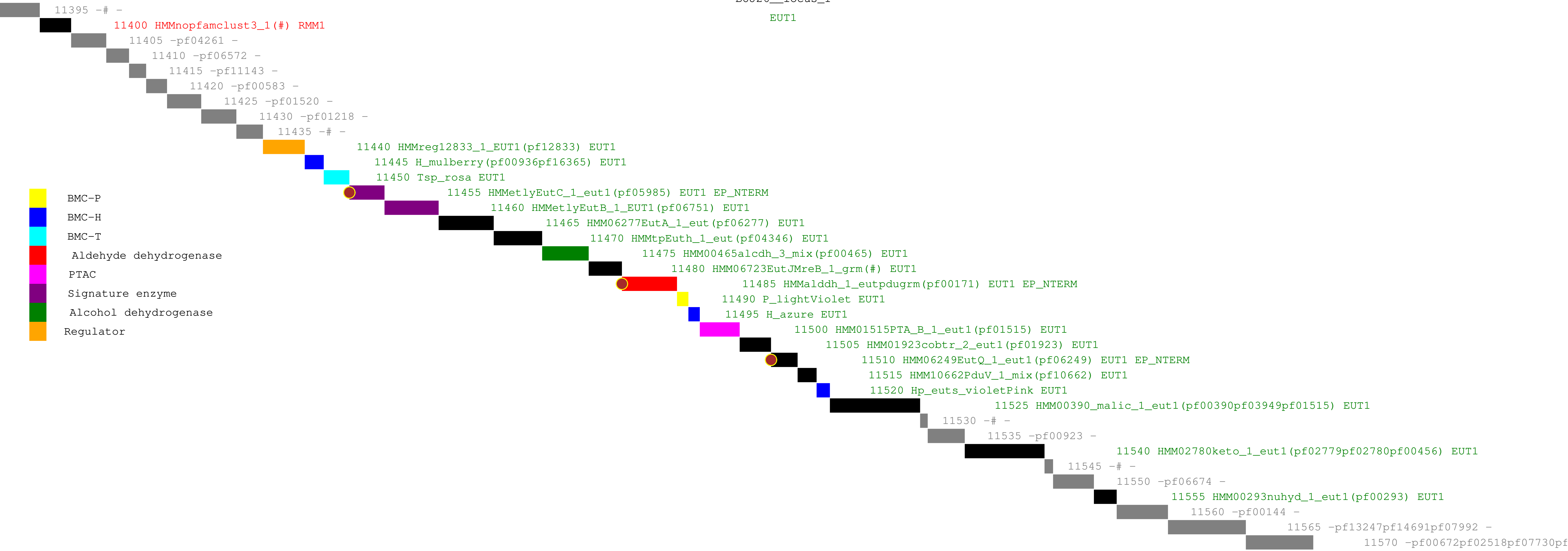

Salmonella enterica subsp. enterica serovar Hartford. Bacteria; Proteobacteria; Gammaproteobacteria; Enterobacterales;

BUJ27__locus_1

EUT1

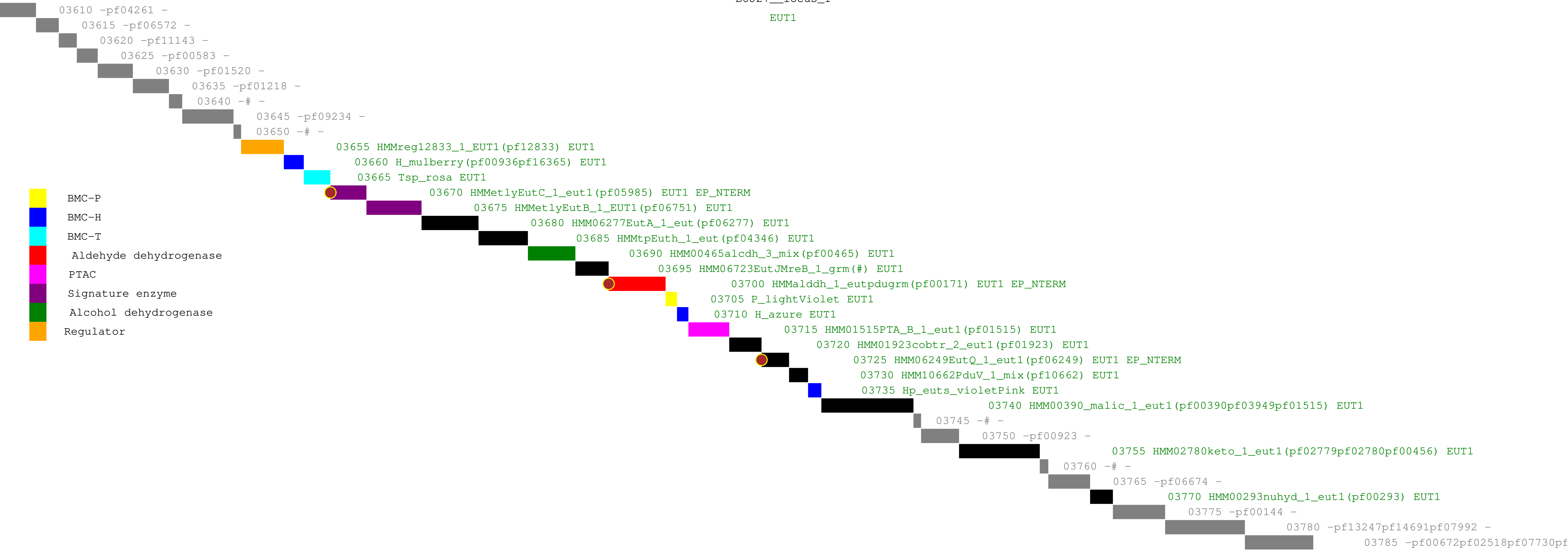

Salmonella muenchen. Bacteria; Proteobacteria; Gammaproteobacteria; Enterobacterales;

BUJ29__locus_2

EUT1

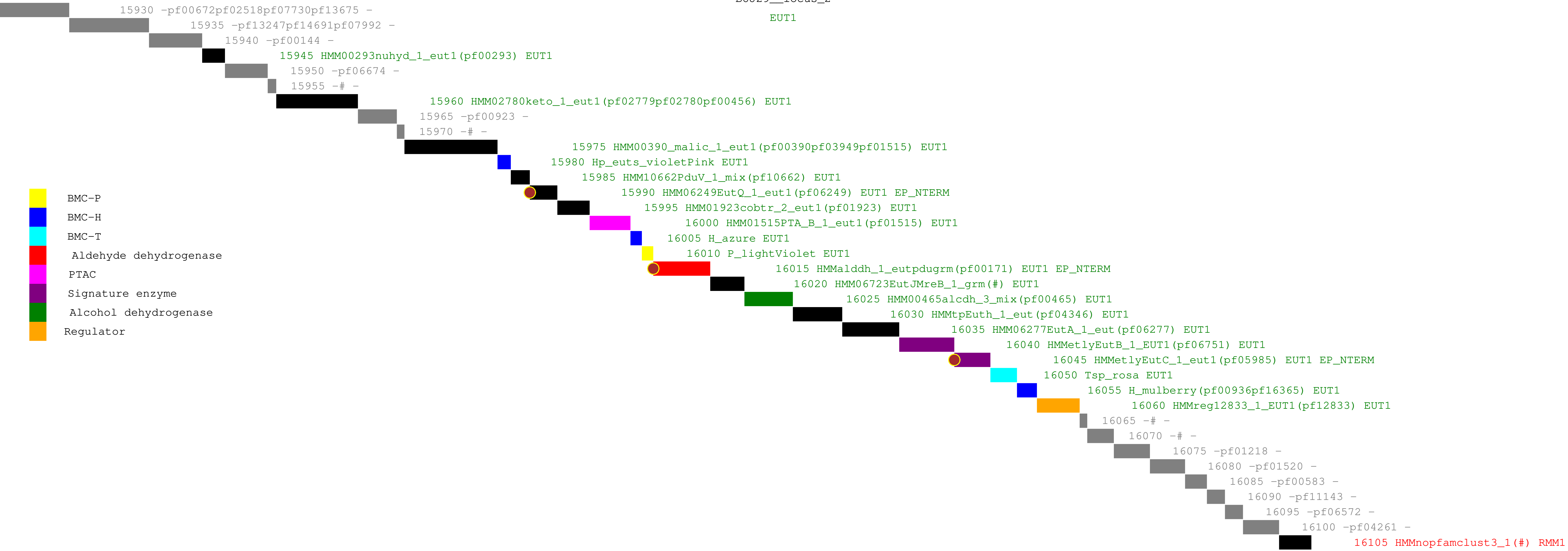

Escherichia coli. Bacteria; Proteobacteria; Gammaproteobacteria; Enterobacterales;

BvCmsHHP056__locus_1

EUT1

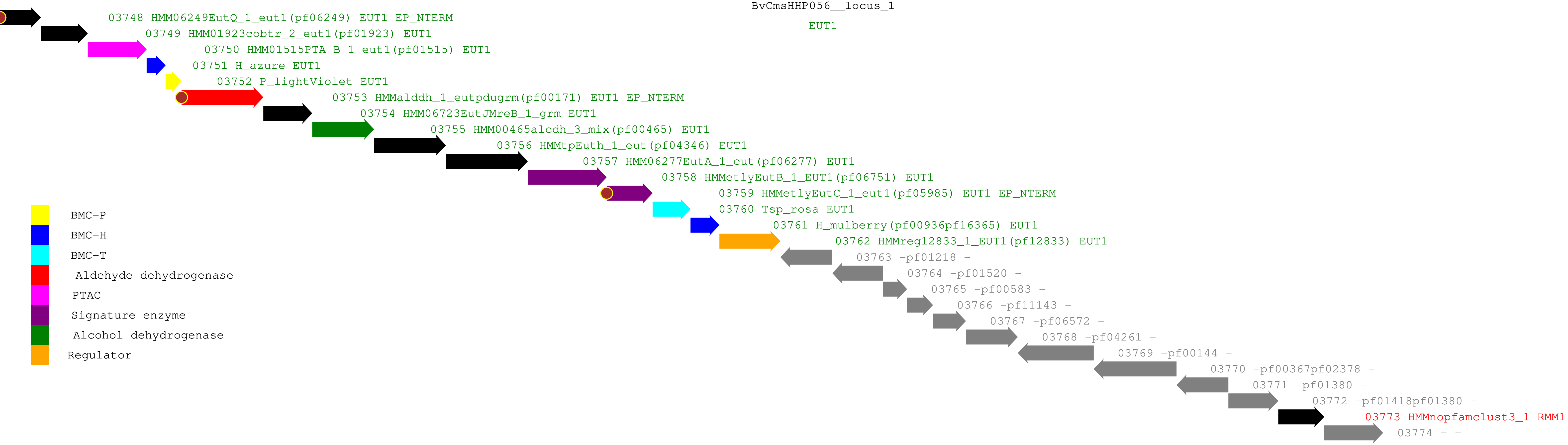

Escherichia coli. Bacteria; Proteobacteria; Gammaproteobacteria; Enterobacterales;

BvCmsKSP026__locus_1

EUT1

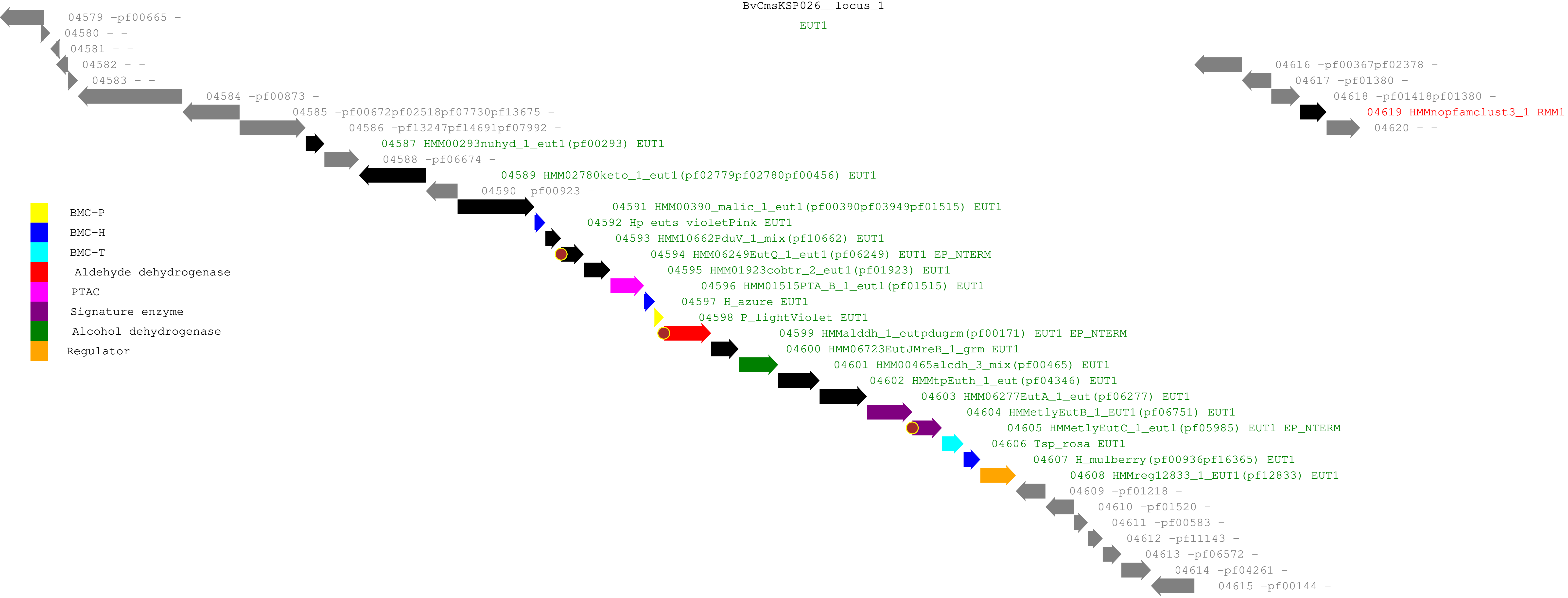

Escherichia coli. Bacteria; Proteobacteria; Gammaproteobacteria; Enterobacterales;

BvCmsKSP058__locus_1

EUT1

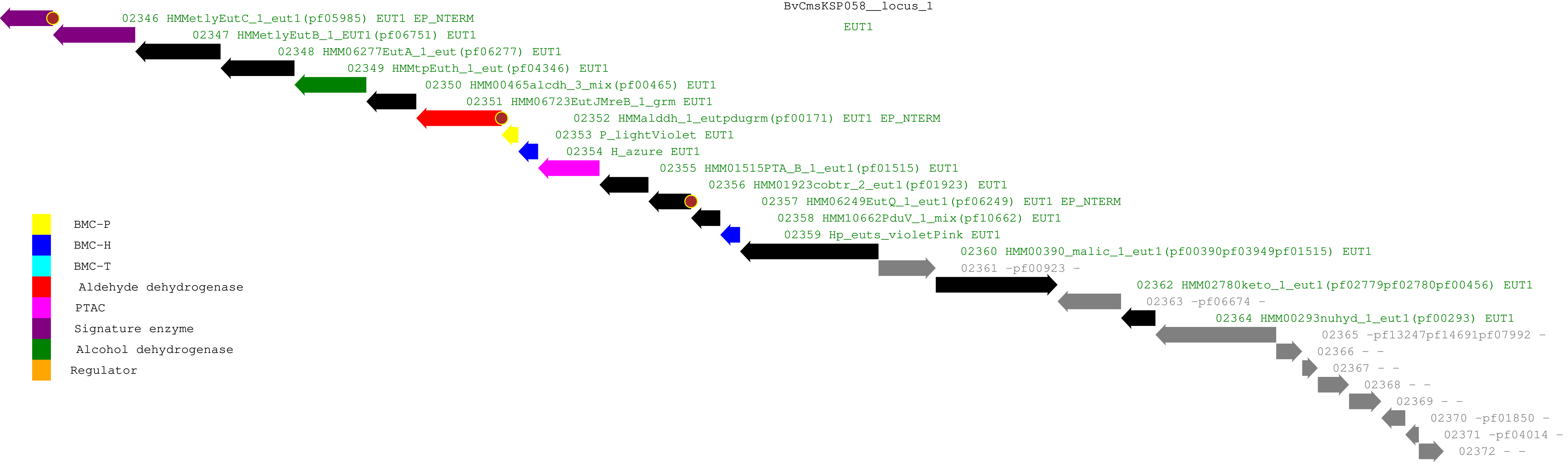

Escherichia coli. Bacteria; Proteobacteria; Gammaproteobacteria; Enterobacterales;

BVL39__locus_1

EUT1

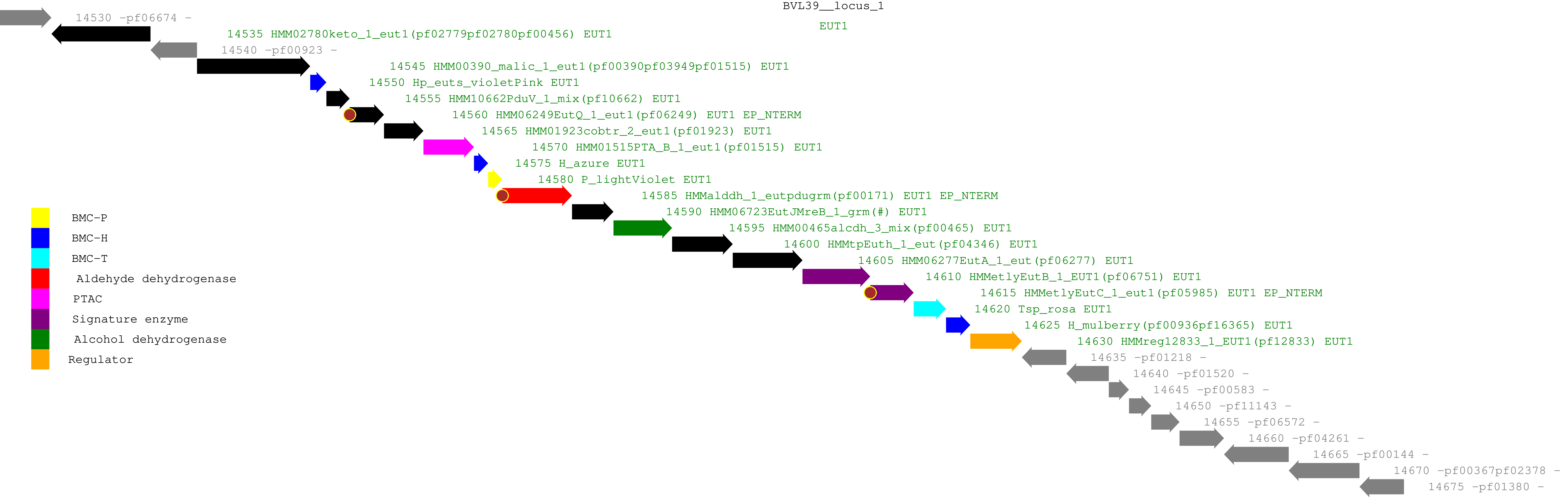

Escherichia coli O174:H8 str. 04-3038. Bacteria; Proteobacteria; Gammaproteobacteria; Enterobacterales;

BW70__locus_1

EUT1

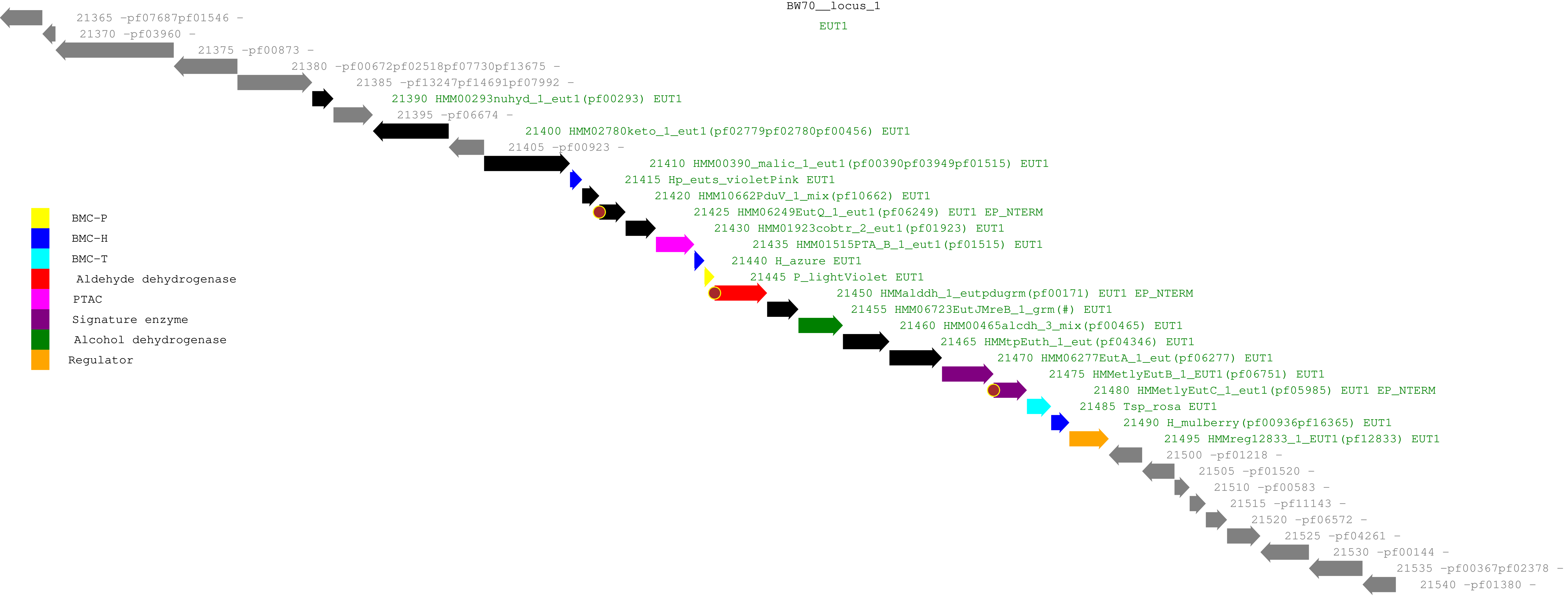

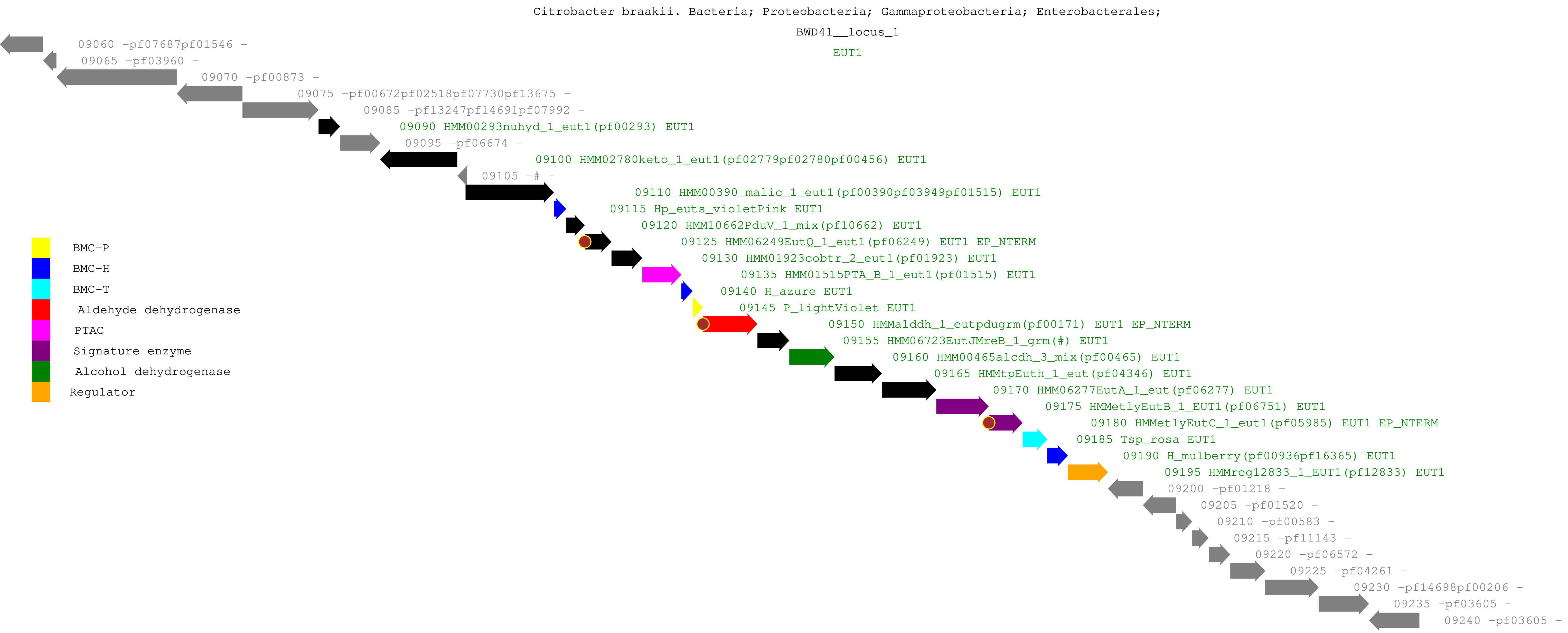
Citrobacter braakii. Bacteria; Proteobacteria; Gammaproteobacteria; Enterobacterales;
BWD41__locus_1
EUT1
09060 -pf07687pf01546 -
09065 -pf03960 -
09070 -pf00873 -
09075 -pf00672pf02518pf07730pf13675 -
09085 -pf13247pf14691pf07992 -
09090 HMM00293nuhyd_1_eut1(pf00293) EUT1
09095 -pf06674 -
09100 HMM02780keto_1_eut1(pf02779pf02780pf00456) EUT1
09105 -# -
09110 HMM00390_malic_1_eut1(pf00390pf03949pf01515) EUT1
09115 Hp_euts_violetPink EUT1
09120 HMM10662PduV_1_mix(pf10662) EUT1
09125 HMM06249EutQ_1_eut1(pf06249) EUT1 EP_NTERM
09130 HMM01923cobtr_2_eut1(pf01923) EUT1
09135 HMM01515PTA_B_1_eut1(pf01515) EUT1
09140 H_azure EUT1
09145 P_lightViolet EUT1
09150 HMMalddh_1_eutpdugrm(pf00171) EUT1 EP_NTERM
09155 HMM06723EutJMreB_1_grm(#) EUT1
09160 HMM00465alcdh_3_mix(pf00465) EUT1
09165 HMMtpEuth_1_eut(pf04346) EUT1
09170 HMM06277EutA_1_eut(pf06277) EUT1
09175 HMMetlyEutB_1_EUT1(pf06751) EUT1
09180 HMMetlyEutC_1_eut1(pf05985) EUT1 EP_NTERM
09185 Tsp_rosa EUT1
09190 H_mulberry(pf00936pf16365) EUT1
09195 HMMreg12833_1_EUT1(pf12833) EUT1
09200 -pf01218 -
09205 -pf01520 -
09210 -pf00583 -
09215 -pf11143 -
09220 -pf06572 -
09225 -pf04261 -
09230 -pf14698pf00206 -
09235 -pf03605 -
09240 -pf03605 -
BMC-P
BMC-H
BMC-T
Aldehyde dehydrogenase
PTAC
Signature enzyme
Alcohol dehydrogenase
Regulator

Salmonella enterica subsp. enterica serovar Adelaide. Bacteria; Proteobacteria; Gammaproteobacteria; Enterobacterales;

BWP08__locus_2

EUT1

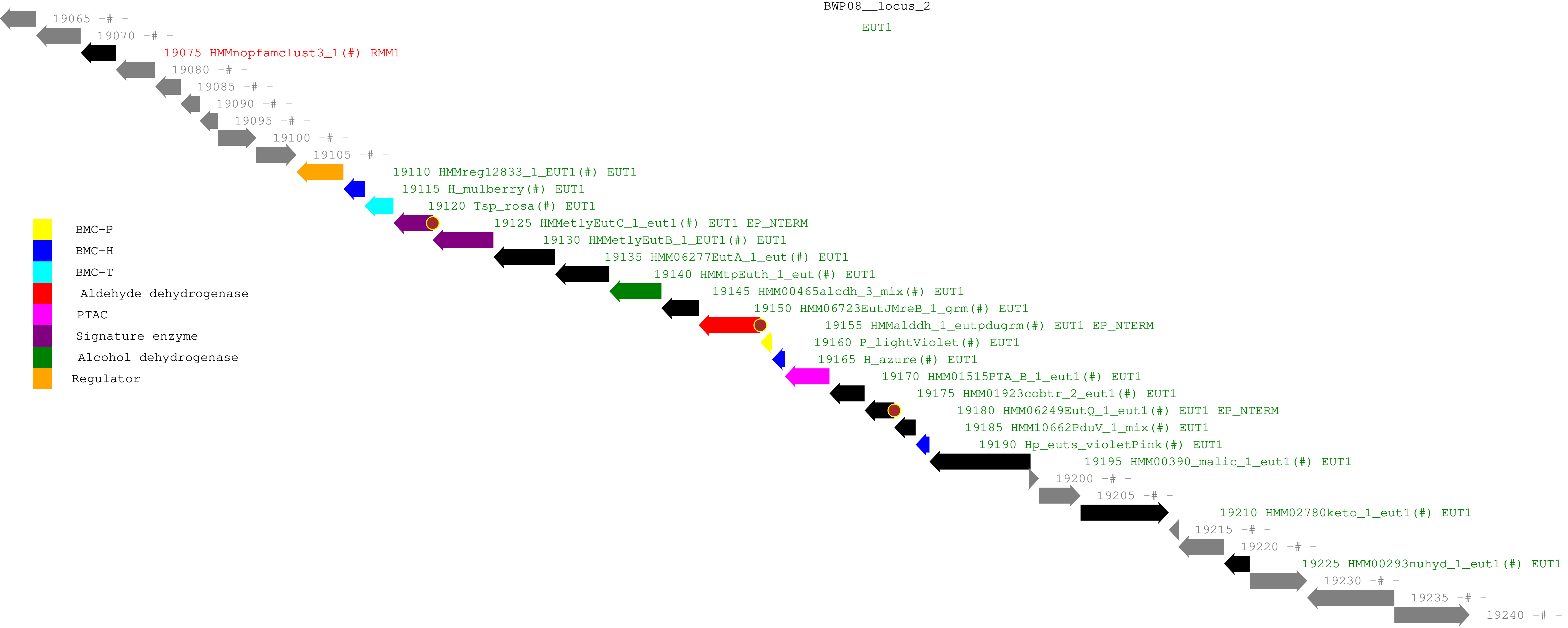

Salmonella enterica subsp. enterica serovar Worthington. Bacteria; Proteobacteria; Gammaproteobacteria; Enterobacterales;

BWP16__locus_1

EUT1

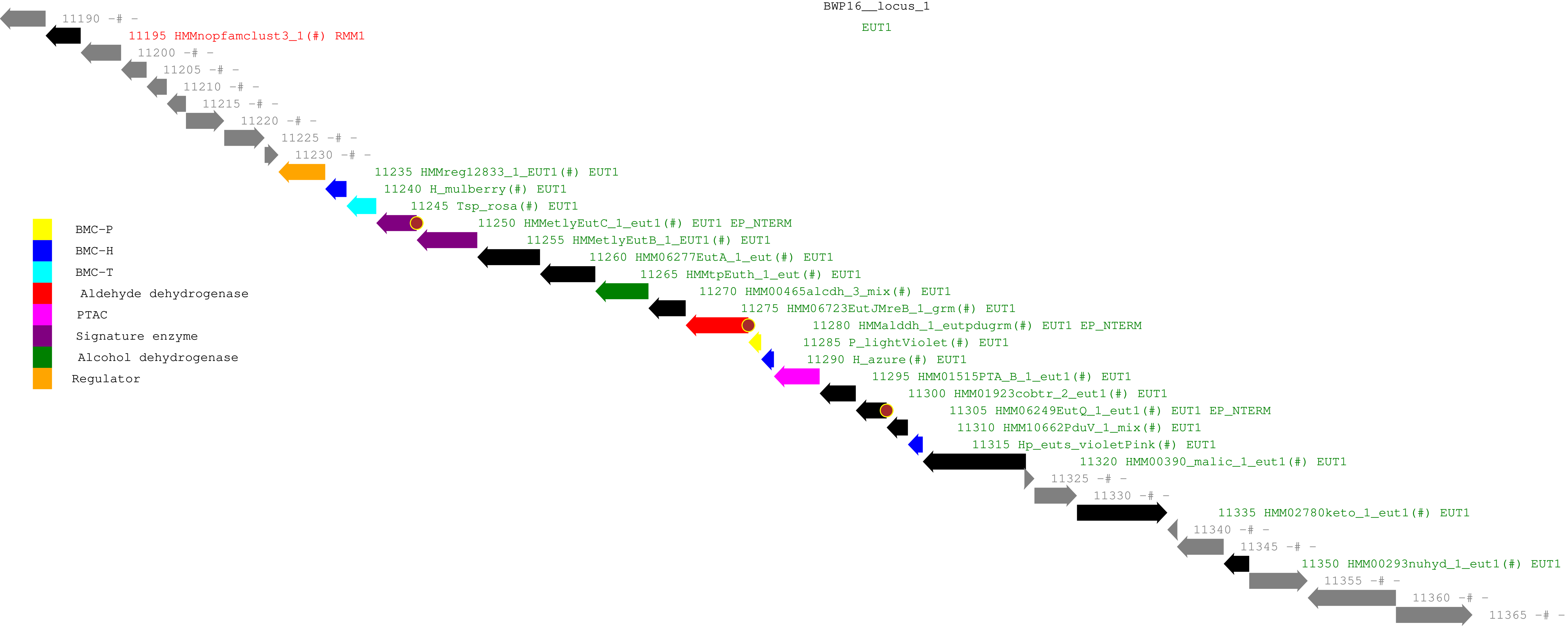

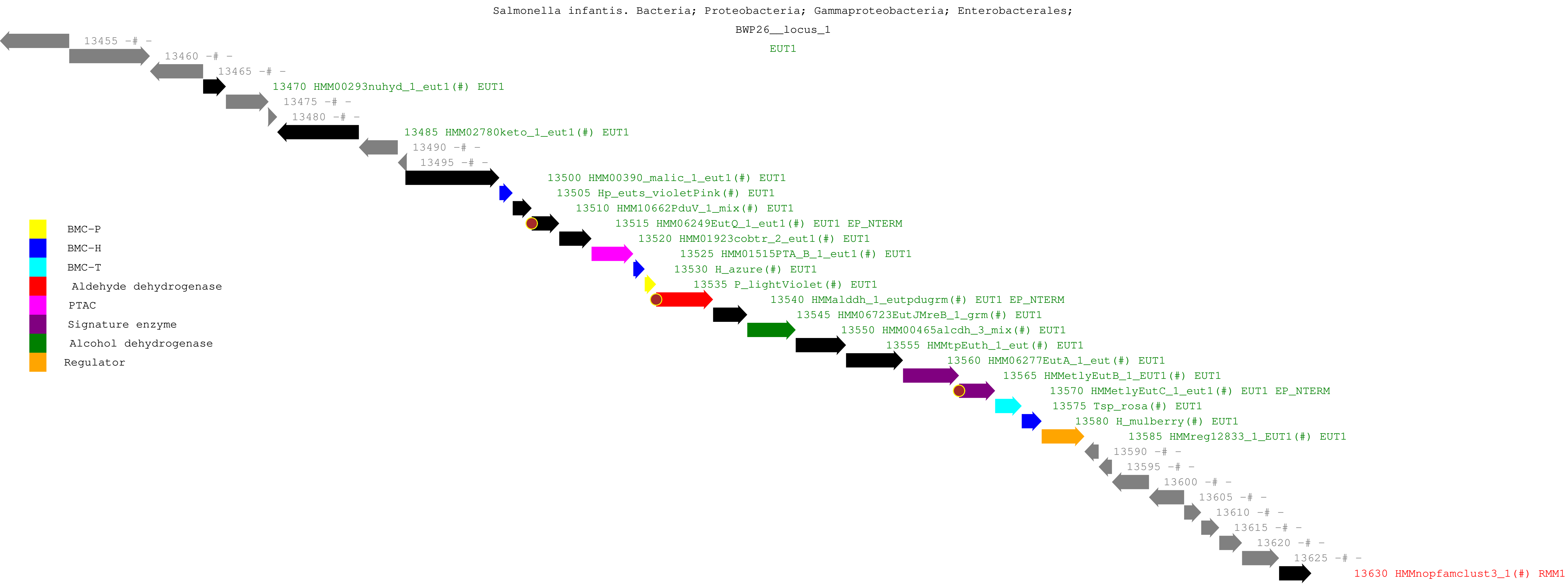
Salmonella infantis. Bacteria; Proteobacteria; Gammaproteobacteria; Enterobacterales;
BWP26__locus_1
EUT1
13455 -# -
13460 -# -
13465 -# -
13470 HMM00293nuhyd_1_eut1(#) EUT1
13475 -# -
13480 -# -
13485 HMM02780keto_1_eut1(#) EUT1
13490 -# -
13495 -# -
13500 HMM00390_malic_1_eut1(#) EUT1
13505 Hp_euts_violetPink(#) EUT1
13510 HMM10662PduV_1_mix(#) EUT1
13515 HMM06249EutQ_1_eut1(#) EUT1 EP_NTERM
13520 HMM01923cobtr_2_eut1(#) EUT1
13525 HMM01515PTA_B_1_eut1(#) EUT1
13530 H_azure(#) EUT1
13535 P_lightViolet(#) EUT1
13540 HMMaldh_1_eutpdugrm(#) EUT1 EP_NTERM
13545 HMM06723EutJMreB_1_grm(#) EUT1
13550 HMM00465alcdh_3_mix(#) EUT1
13555 HMMtpEuth_1_eut(#) EUT1
13560 HMM06277EutA_1_eut(#) EUT1
13565 HMMetlyEutB_1_EUT1(#) EUT1
13570 HMMetlyEutC_1_eut1(#) EUT1 EP_NTERM
13575 Tsp_rosa(#) EUT1
13580 H_mulberry(#) EUT1
13585 HMMreg12833_1_EUT1(#) EUT1
13590 -# -
13595 -# -
13600 -# -
13605 -# -
13610 -# -
13615 -# -
13620 -# -
13625 -# -
13630 HMMnopfamclust3_1(#) RMM1
BMC-P
BMC-H
BMC-T
Aldehyde dehydrogenase
PTAC
Signature enzyme
Alcohol dehydrogenase
Regulator

Escherichia coli O69:H11 str. 08-4661. Bacteria; Proteobacteria; Gammaproteobacteria; Enterobacterales;

BX06__locus_1

EUT1

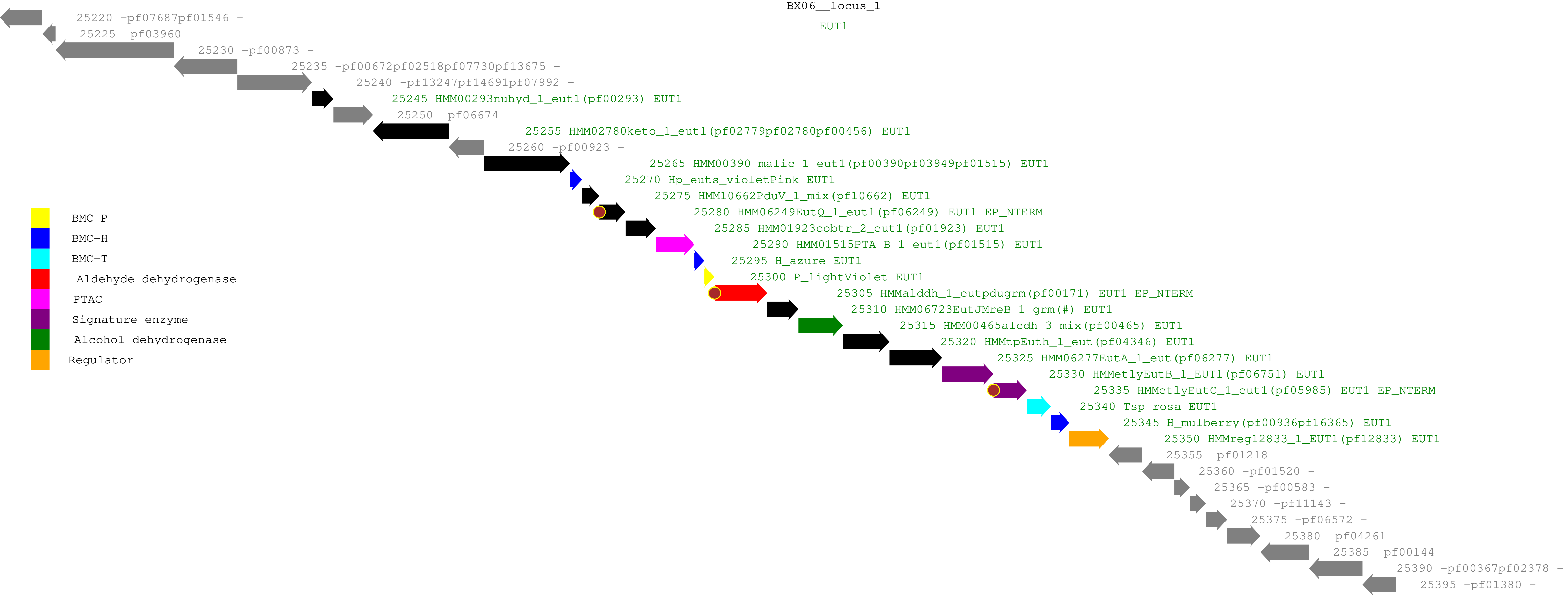

Escherichia coli O118:H16 str. 2009C-4446. Bacteria; Proteobacteria; Gammaproteobacteria; Enterobacterales;

BX23__locus_1

EUT1

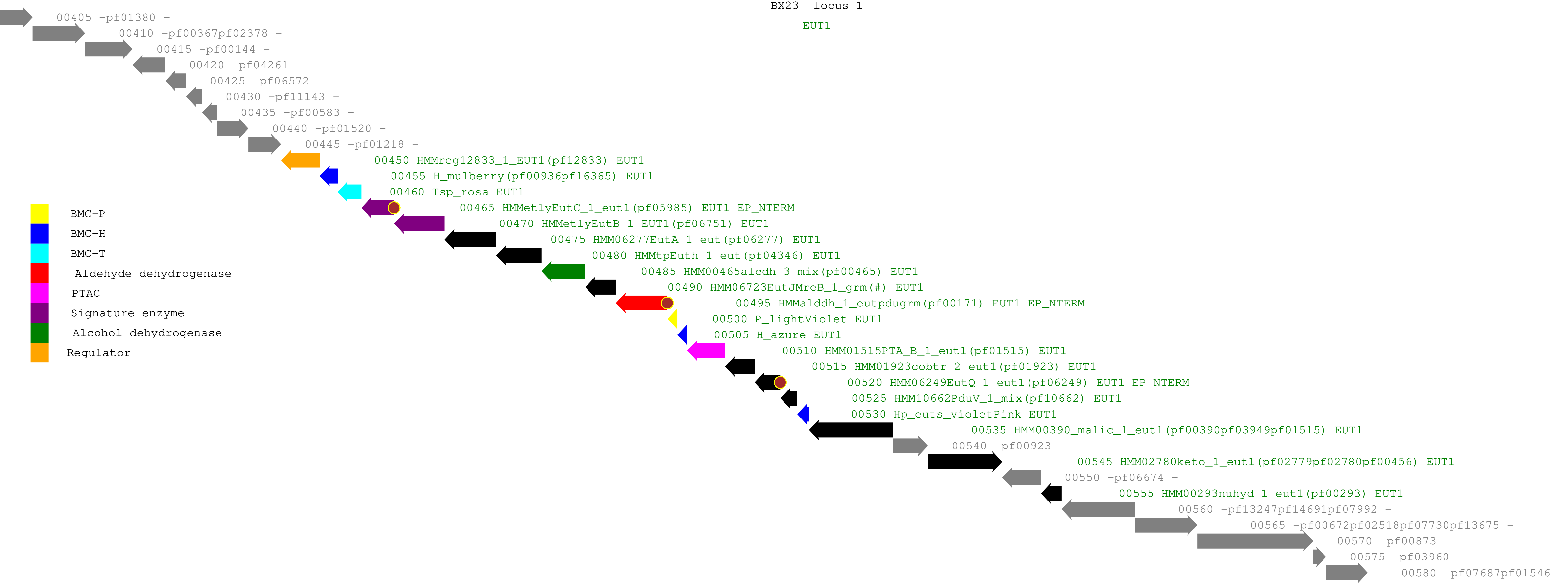

Escherichia coli O145:NM str. 2010C-3526. Bacteria; Proteobacteria; Gammaproteobacteria; Enterobacterales;

BX51__locus_1

EUT1

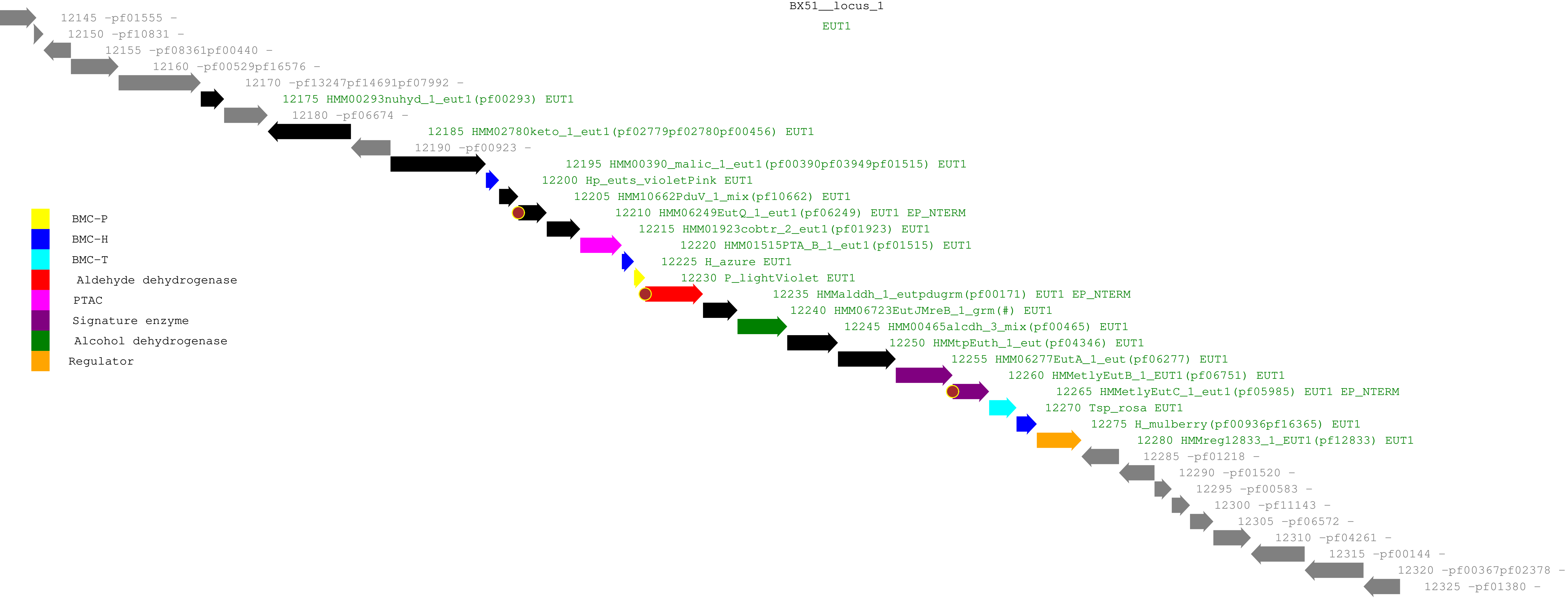

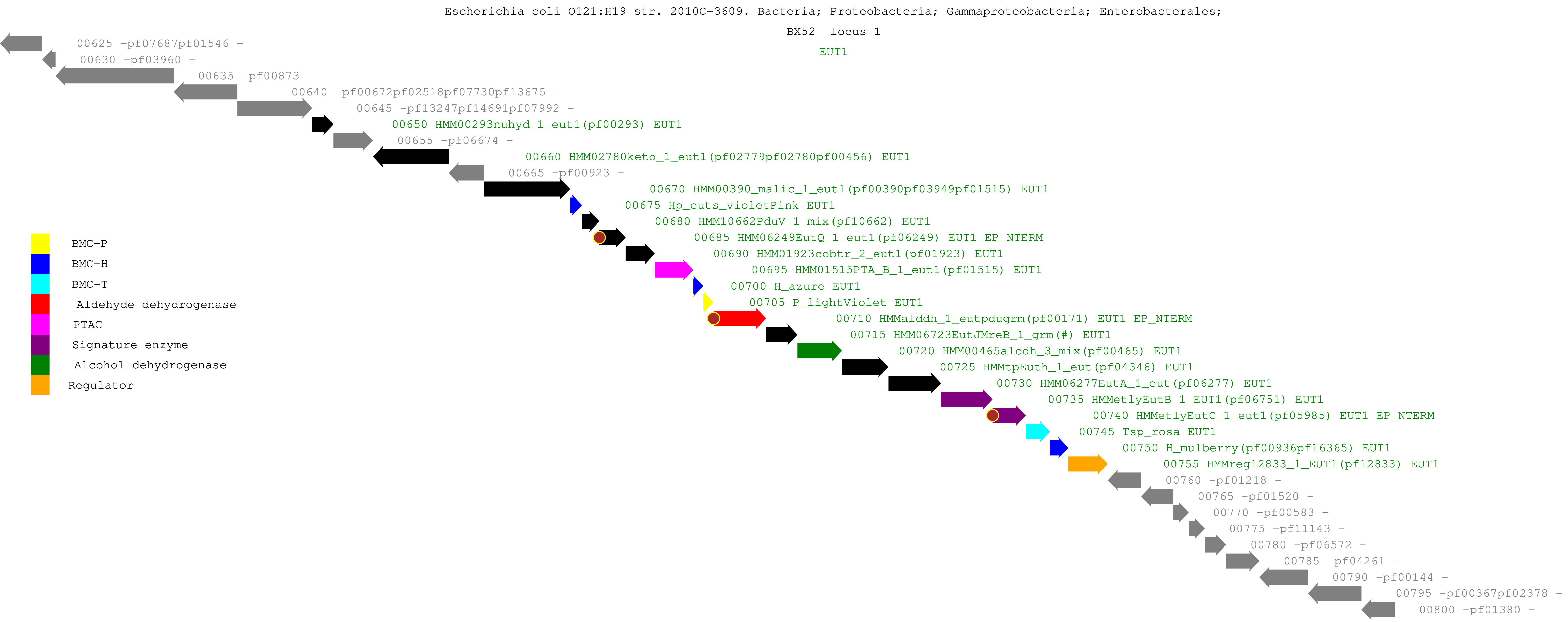

Escherichia coli O121:H19 str. 2010C-3609. Bacteria; Proteobacteria; Gammaproteobacteria; Enterobacterales;
BX52__locus_1
EUT1
00625 -pf07687pf01546 -
00630 -pf03960 -
00635 -pf00873 -
00640 -pf00672pf02518pf07730pf13675 -
00645 -pf13247pf14691pf07992 -
00650 HMM00293nuhyd_1_eut1(pf00293) EUT1
00655 -pf06674 -
00660 HMM02780keto_1_eut1(pf02779pf02780pf00456) EUT1
00665 -pf00923 -
00670 HMM00390_malic_1_eut1(pf00390pf03949pf01515) EUT1
00675 Hp_euts_violetPink EUT1
00680 HMM10662PduV_1_mix(pf10662) EUT1
00685 HMM06249EutQ_1_eut1(pf06249) EUT1 EP_NTERM
00690 HMM01923cobtr_2_eut1(pf01923) EUT1
00695 HMM01515PTA_B_1_eut1(pf01515) EUT1
00700 H_azure EUT1
00705 P_lightViolet EUT1
00710 HMMalddh_1_eutpdugrm(pf00171) EUT1 EP_NTERM
00715 HMM06723EutJMreB_1_grm(#) EUT1
00720 HMM00465alcdh_3_mix(pf00465) EUT1
00725 HMMtpEuth_1_eut(pf04346) EUT1
00730 HMM06277EutA_1_eut(pf06277) EUT1
00735 HMMetlyEutB_1_EUT1(pf06751) EUT1
00740 HMMetlyEutC_1_eut1(pf05985) EUT1 EP_NTERM
00745 Tsp_rosa EUT1
00750 H_mulberry(pf00936pf16365) EUT1
00755 HMMreg12833_1_EUT1(pf12833) EUT1
00760 -pf01218 -
00765 -pf01520 -
00770 -pf00583 -
00775 -pf11143 -
00780 -pf06572 -
00785 -pf04261 -
00790 -pf00144 -
00795 -pf00367pf02378 -
00800 -pf01380 -
BMC-P
BMC-H
BMC-T
Aldehyde dehydrogenase
PTAC
Signature enzyme
Alcohol dehydrogenase
Regulator

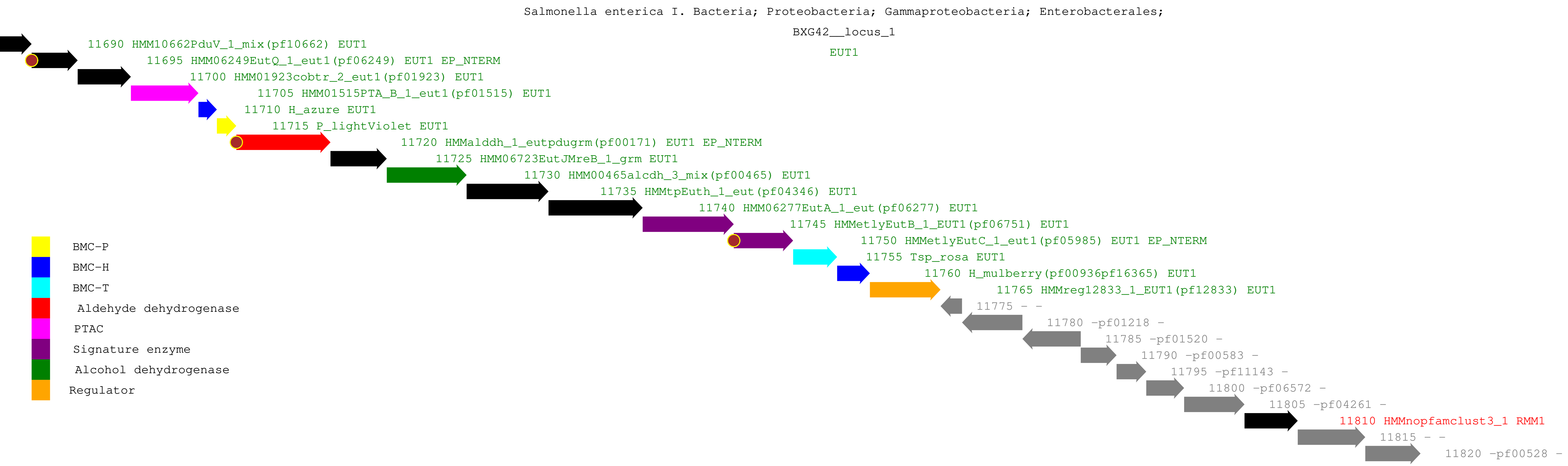
Salmonella enterica I. Bacteria; Proteobacteria; Gammaproteobacteria; Enterobacterales;
BXG42__locus_1
EUT1
11690 HMM10662PduV_1_mix(pf10662) EUT1
11695 HMM06249EutQ_1_eut1(pf06249) EUT1 EP_NTERM
11700 HMM01923cobtr_2_eut1(pf01923) EUT1
11705 HMM01515PTA_B_1_eut1(pf01515) EUT1
11710 H_azure EUT1
11715 P_lightViolet EUT1
11720 HMMaldh_1_eutpdugrm(pf00171) EUT1 EP_NTERM
11725 HMM06723EutJMreB_1_grm EUT1
11730 HMM00465alcdh_3_mix(pf00465) EUT1
11735 HMMtpEuth_1_eut(pf04346) EUT1
11740 HMM06277EutA_1_eut(pf06277) EUT1
11745 HMMetlyEutB_1_EUT1(pf06751) EUT1
11750 HMMetlyEutC_1_eut1(pf05985) EUT1 EP_NTERM
11755 Tsp_rosa EUT1
11760 H_mulberry(pf00936pf16365) EUT1
11765 HMMreg12833_1_EUT1(pf12833) EUT1
11775 - -
11780 -pf01218 -
11785 -pf01520 -
11790 -pf00583 -
11795 -pf11143 -
11800 -pf06572 -
11805 -pf04261 -
11810 HMMnopfamclust3_1 RMM1
11815 - -
11820 -pf00528 -
BMC-P
BMC-H
BMC-T
Aldehyde dehydrogenase
PTAC
Signature enzyme
Alcohol dehydrogenase
Regulator

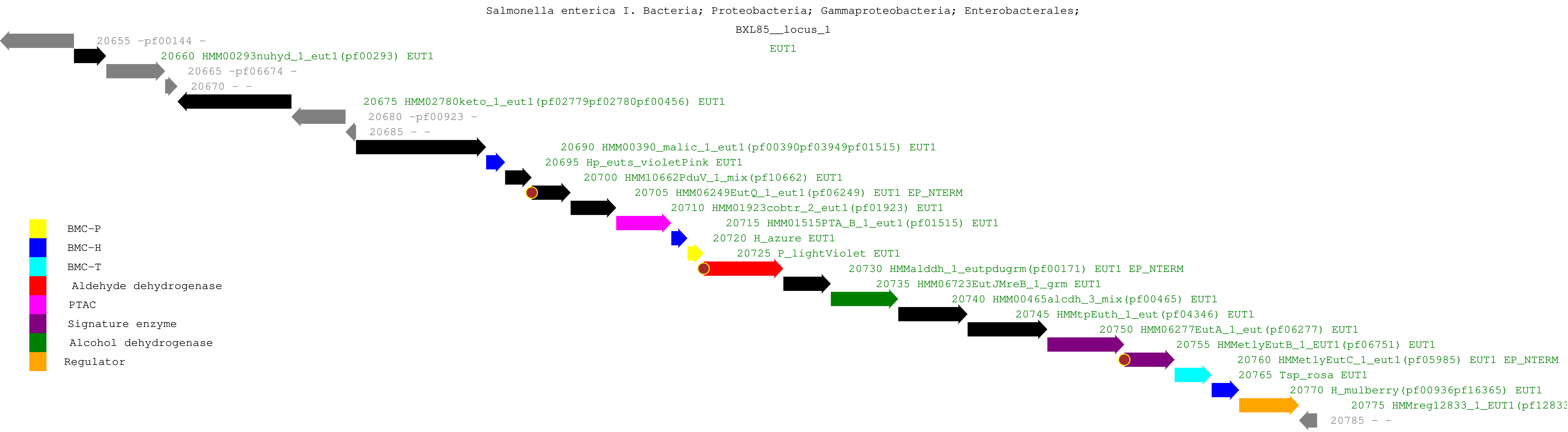
Salmonella enterica I. Bacteria; Proteobacteria; Gammaproteobacteria; Enterobacterales;
BXL85__locus_1
EUT1
20655 -pf00144 -
20660 HMM00293nuhyd_1_eut1(pf00293) EUT1
20665 -pf06674 -
20670 - -
20675 HMM02780keto_1_eut1(pf02779pf02780pf00456) EUT1
20680 -pf00923 -
20685 - -
20690 HMM00390_malic_1_eut1(pf00390pf03949pf01515) EUT1
20695 Hp_euts_violetPink EUT1
20700 HMM10662PduV_1_mix(pf10662) EUT1
20705 HMM06249EutQ_1_eut1(pf06249) EUT1 EP_NTERM
20710 HMM01923cobtr_2_eut1(pf01923) EUT1
20715 HMM01515PTA_B_1_eut1(pf01515) EUT1
20720 H_azure EUT1
20725 P_lightViolet EUT1
20730 HMMalddh_1_eutpdugrm(pf00171) EUT1 EP_NTERM
20735 HMM06723EutJMreB_1_grm EUT1
20740 HMM00465alcdh_3_mix(pf00465) EUT1
20745 HMMtpEuth_1_eut(pf04346) EUT1
20750 HMM06277EutA_1_eut(pf06277) EUT1
20755 HMMetlyEutB_1_EUT1(pf06751) EUT1
20760 HMMetlyEutC_1_eut1(pf05985) EUT1 EP_NTERM
20765 Tsp_rosa EUT1
20770 H_mulberry(pf00936pf16365) EUT1
20775 HMMreg12833_1_EUT1(pf1283
20785 - -
BMC-P
BMC-H
BMC-T
Aldehyde dehydrogenase
PTAC
Signature enzyme
Alcohol dehydrogenase
Regulator

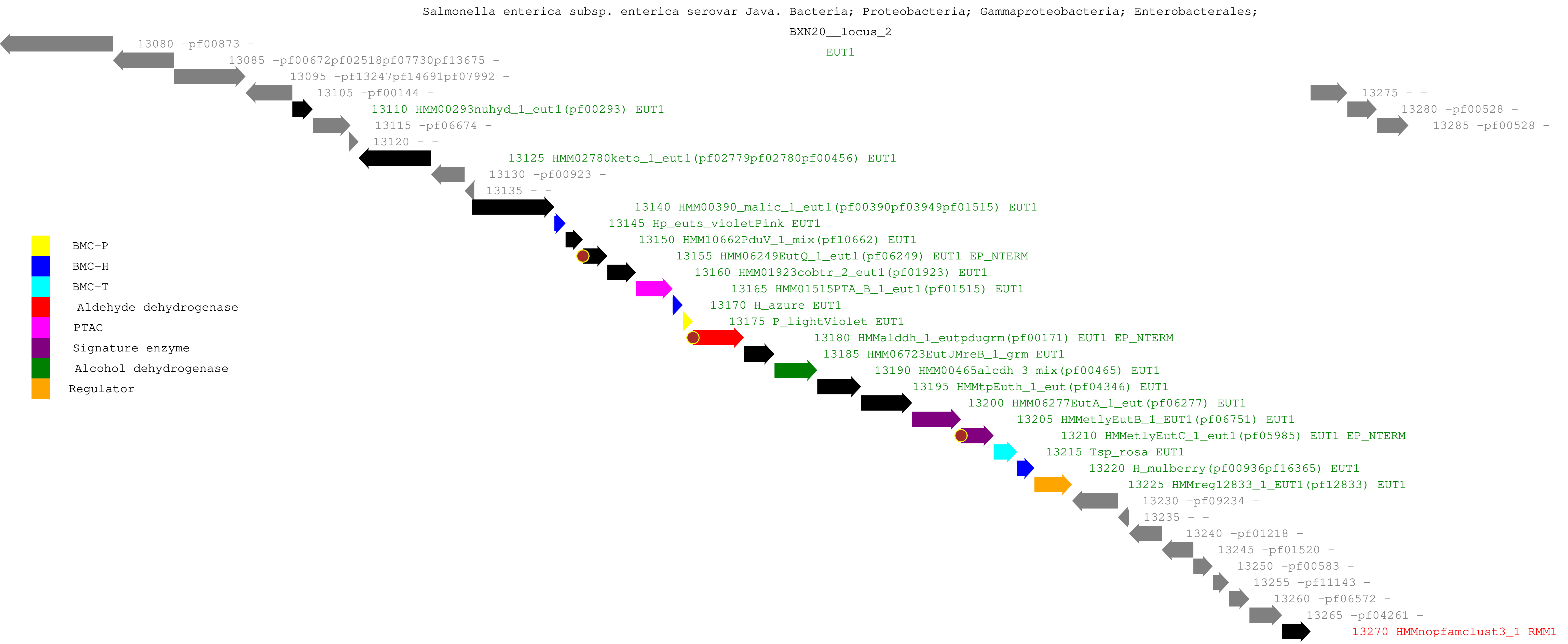
Salmonella enterica subsp. enterica serovar Java. Bacteria; Proteobacteria; Gammaproteobacteria; Enterobacterales;
BXN20__locus_2
EUT1
13080 -pf00873 -
13085 -pf00672pf02518pf07730pf13675 -
13095 -pf13247pf14691pf07992 -
13105 -pf00144 -
13110 HMM00293nuhyd_1_eut1(pf00293) EUT1
13115 -pf06674 -
13120 - -
13125 HMM02780keto_1_eut1(pf02779pf02780pf00456) EUT1
13130 -pf00923 -
13135 - -
13140 HMM00390_malic_1_eut1(pf00390pf03949pf01515) EUT1
13145 Hp_euts_violetPink EUT1
13150 HMM10662PduV_1_mix(pf10662) EUT1
13155 HMM06249EutQ_1_eut1(pf06249) EUT1 EP_NTERM
13160 HMM01923cobtr_2_eut1(pf01923) EUT1
13165 HMM01515PTA_B_1_eut1(pf01515) EUT1
13170 H_azure EUT1
13175 P_lightViolet EUT1
13180 HMMaldh_1_eutpdugrm(pf00171) EUT1 EP_NTERM
13185 HMM06723EutJMreB_1_grm EUT1
13190 HMM00465alcdh_3_mix(pf00465) EUT1
13195 HMMtpEuth_1_eut(pf04346) EUT1
13200 HMM06277EutA_1_eut(pf06277) EUT1
13205 HMMetlyEutB_1_EUT1(pf06751) EUT1
13210 HMMetlyEutC_1_eut1(pf05985) EUT1 EP_NTERM
13215 Tsp_rosa EUT1
13220 H_mulberry(pf00936pf16365) EUT1
13225 HMMreg12833_1_EUT1(pf12833) EUT1
13230 -pf09234 -
13235 - -
13240 -pf01218 -
13245 -pf01520 -
13250 -pf00583 -
13255 -pf11143 -
13260 -pf06572 -
13265 -pf04261 -
13270 HMMnopfamclust3_1 RMM1
13275 - -
13280 -pf00528 -
13285 -pf00528 -
BMC-P
BMC-H
BMC-T
Aldehyde dehydrogenase
PTAC
Signature enzyme
Alcohol dehydrogenase
Regulator

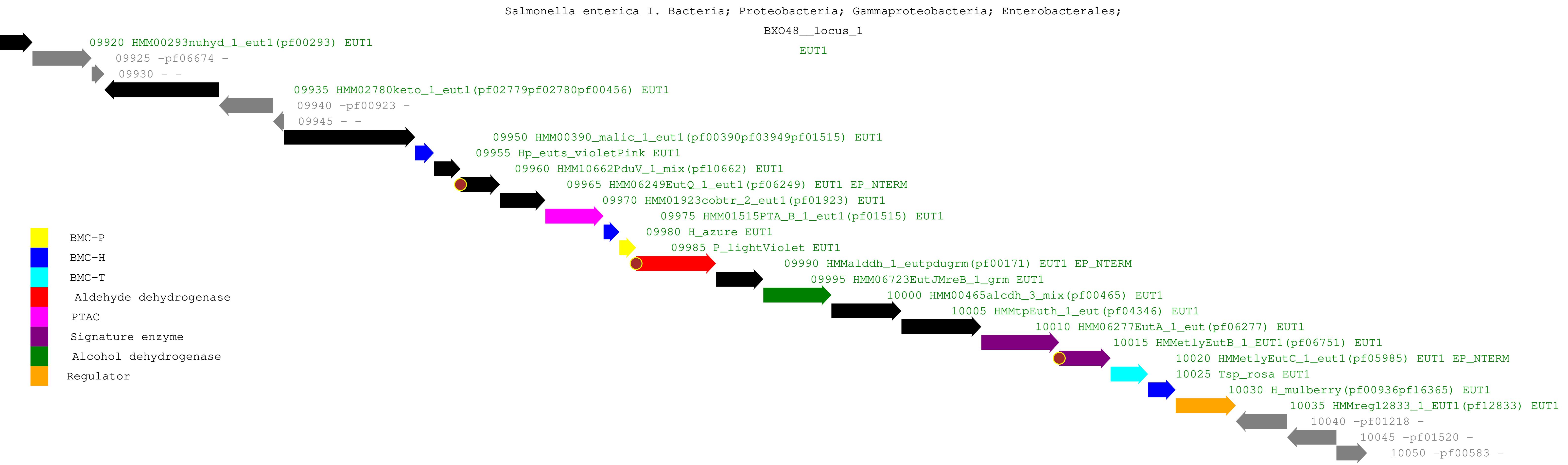
Salmonella enterica I. Bacteria; Proteobacteria; Gammaproteobacteria; Enterobacterales;
BX048__locus_1
EUT1
09920 HMM00293nuhyd_1_eut1(pf00293) EUT1
09925 -pf06674 -
09930 - -
09935 HMM02780keto_1_eut1(pf02779pf02780pf00456) EUT1
09940 -pf00923 -
09945 - -
09950 HMM00390_malic_1_eut1(pf00390pf03949pf01515) EUT1
09955 Hp_euts_violetPink EUT1
09960 HMM10662PduV_1_mix(pf10662) EUT1
09965 HMM06249EutQ_1_eut1(pf06249) EUT1 EP_NTERM
09970 HMM01923cobtr_2_eut1(pf01923) EUT1
09975 HMM01515PTA_B_1_eut1(pf01515) EUT1
09980 H_azure EUT1
09985 P_lightViolet EUT1
09990 HMMalddh_1_eutpdugrm(pf00171) EUT1 EP_NTERM
09995 HMM06723EutJMreB_1_grm EUT1
10000 HMM00465alcdh_3_mix(pf00465) EUT1
10005 HMMtpEuth_1_eut(pf04346) EUT1
10010 HMM06277EutA_1_eut(pf06277) EUT1
10015 HMMetlyEutB_1_EUT1(pf06751) EUT1
10020 HMMetlyEutC_1_eut1(pf05985) EUT1 EP_NTERM
10025 Tsp_rosa EUT1
10030 H_mulberry(pf00936pf16365) EUT1
10035 HMMreg12833_1_EUT1(pf12833) EUT1
10040 -pf01218 -
10045 -pf01520 -
10050 -pf00583 -
BMC-P
BMC-H
BMC-T
Aldehyde dehydrogenase
PTAC
Signature enzyme
Alcohol dehydrogenase
Regulator

Escherichia coli O111:NM str. K6722. Bacteria; Proteobacteria; Gammaproteobacteria; Enterobacterales;

BY96__locus_1

EUT1

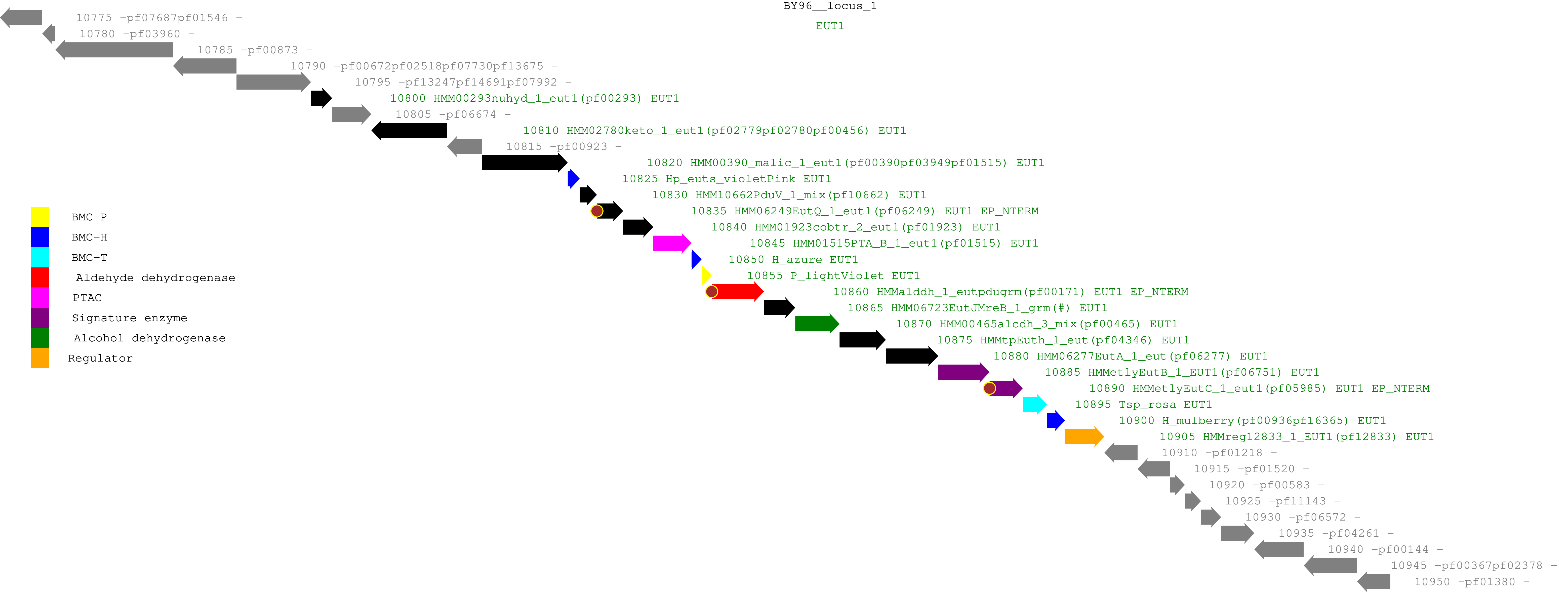

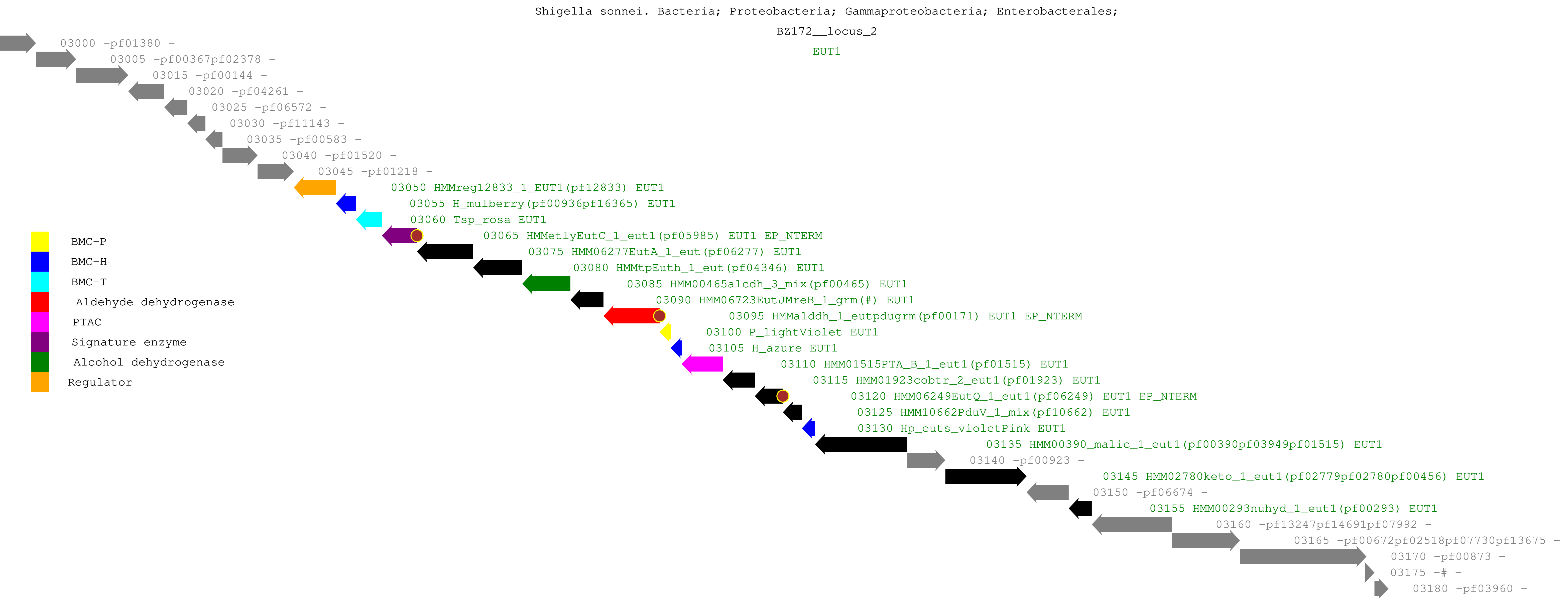

Shigella sonnei. Bacteria; Proteobacteria; Gammaproteobacteria; Enterobacterales;
BZ172__locus_2
EUT1
03000 -pf01380 -
03005 -pf00367pf02378 -
03015 -pf00144 -
03020 -pf04261 -
03025 -pf06572 -
03030 -pf11143 -
03035 -pf00583 -
03040 -pf01520 -
03045 -pf01218 -
03050 HMMreg12833_1_EUT1(pf12833) EUT1
03055 H_mulberry(pf00936pf16365) EUT1
03060 Tsp_rosa EUT1
03065 HMMetlyEutC_1_eut1(pf05985) EUT1 EP_NTERM
03075 HMM06277EutA_1_eut(pf06277) EUT1
03080 HMMtpEuth_1_eut(pf04346) EUT1
03085 HMM00465alcdh_3_mix(pf00465) EUT1
03090 HMM06723EutJMreB_1_grm(#) EUT1
03095 HMMalddh_1_eutpdugrm(pf00171) EUT1 EP_NTERM
03100 P_lightViolet EUT1
03105 H_azure EUT1
03110 HMM01515PTA_B_1_eut1(pf01515) EUT1
03115 HMM01923cobtr_2_eut1(pf01923) EUT1
03120 HMM06249EutQ_1_eut1(pf06249) EUT1 EP_NTERM
03125 HMM10662PduV_1_mix(pf10662) EUT1
03130 Hp_euts_violetPink EUT1
03135 HMM00390_malic_1_eut1(pf00390pf03949pf01515) EUT1
03140 -pf00923 -
03145 HMM02780keto_1_eut1(pf02779pf02780pf00456) EUT1
03150 -pf06674 -
03155 HMM00293nuhyd_1_eut1(pf00293) EUT1
03160 -pf13247pf14691pf07992 -
03165 -pf00672pf02518pf07730pf13675 -
03170 -pf00873 -
03175 -# -
03180 -pf03960 -
BMC-P
BMC-H
BMC-T
Aldehyde dehydrogenase
PTAC
Signature enzyme
Alcohol dehydrogenase
Regulator

Citrobacter sp. A316. Bacteria; Proteobacteria; Gammaproteobacteria; Enterobacterales;

BZK41__locus_1

EUT1

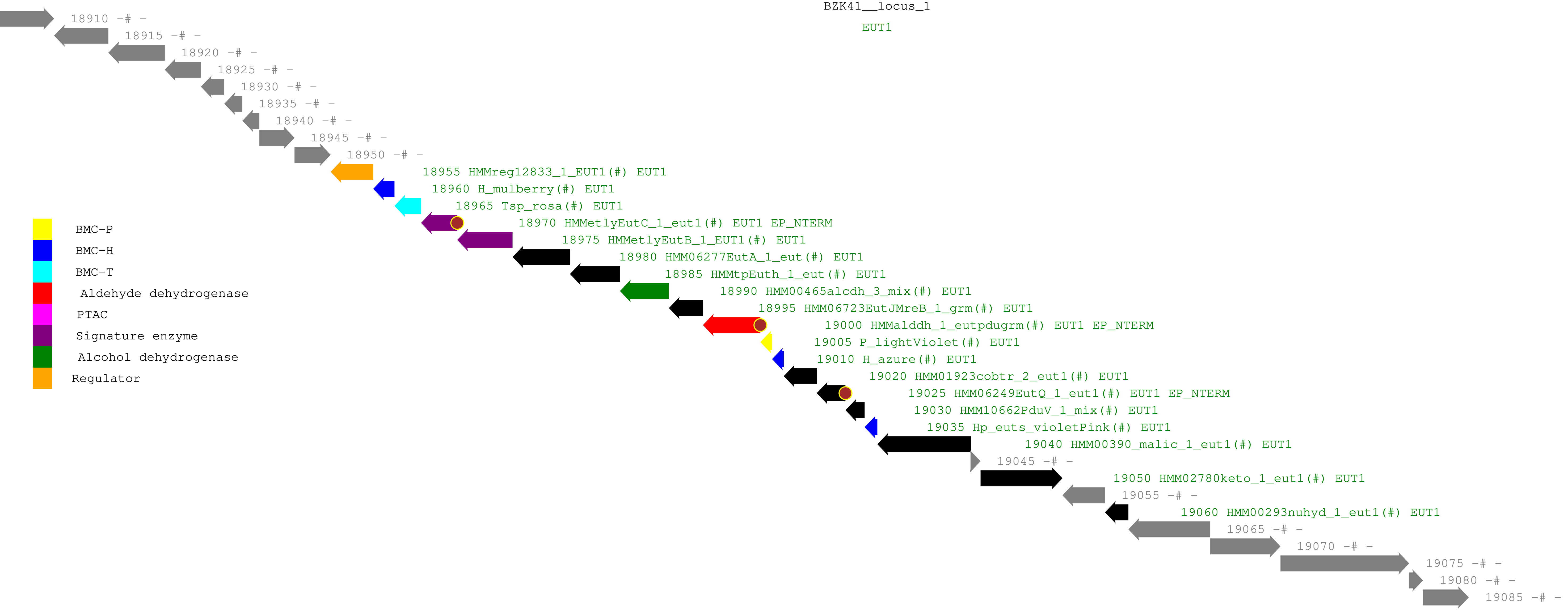

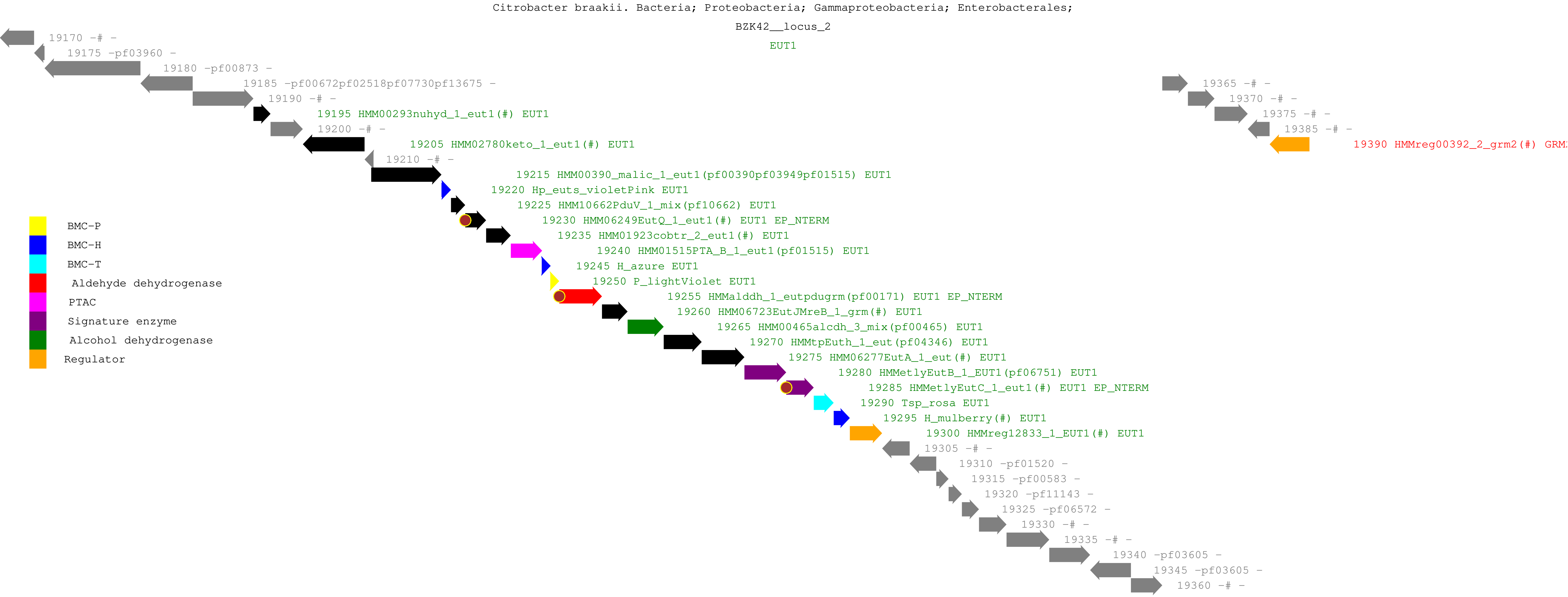

Citrobacter braakii. Bacteria; Proteobacteria; Gammaproteobacteria; Enterobacterales;
BZK42__locus_2
EUT1
19170 -# -
19175 -pf03960 -
19180 -pf00873 -
19185 -pf00672pf02518pf07730pf13675 -
19190 -# -
19195 HMM00293nuhyd_1_eut1(#) EUT1
19200 -# -
19205 HMM02780keto_1_eut1(#) EUT1
19210 -# -
19215 HMM00390_malic_1_eut1(pf00390pf03949pf01515) EUT1
19220 Hp_euts_violetPink EUT1
19225 HMM10662PduV_1_mix(pf10662) EUT1
19230 HMM06249EutQ_1_eut1(#) EUT1 EP_NTERM
19235 HMM01923cobtr_2_eut1(#) EUT1
19240 HMM01515PTA_B_1_eut1(pf01515) EUT1
19245 H_azure EUT1
19250 P_lightViolet EUT1
19255 HMMaldh_1_eutpdugrm(pf00171) EUT1 EP_NTERM
19260 HMM06723EutJMreB_1_grm(#) EUT1
19265 HMM00465alcdh_3_mix(pf00465) EUT1
19270 HMMtpEuth_1_eut(pf04346) EUT1
19275 HMM06277EutA_1_eut(#) EUT1
19280 HMMetlyEutB_1_EUT1(pf06751) EUT1
19285 HMMetlyEutC_1_eut1(#) EUT1 EP_NTERM
19290 Tsp_rosa EUT1
19295 H_mulberry(#) EUT1
19300 HMMreg12833_1_EUT1(#) EUT1
19305 -# -
19310 -pf01520 -
19315 -pf00583 -
19320 -pf11143 -
19325 -pf06572 -
19330 -# -
19335 -# -
19340 -pf03605 -
19345 -pf03605 -
19360 -# -
19365 -# -
19370 -# -
19375 -# -
19385 -# -
19390 HMMreg00392_2_grm2(#) GRM
BMC-P
BMC-H
BMC-T
Aldehyde dehydrogenase
PTAC
Signature enzyme
Alcohol dehydrogenase
Regulator

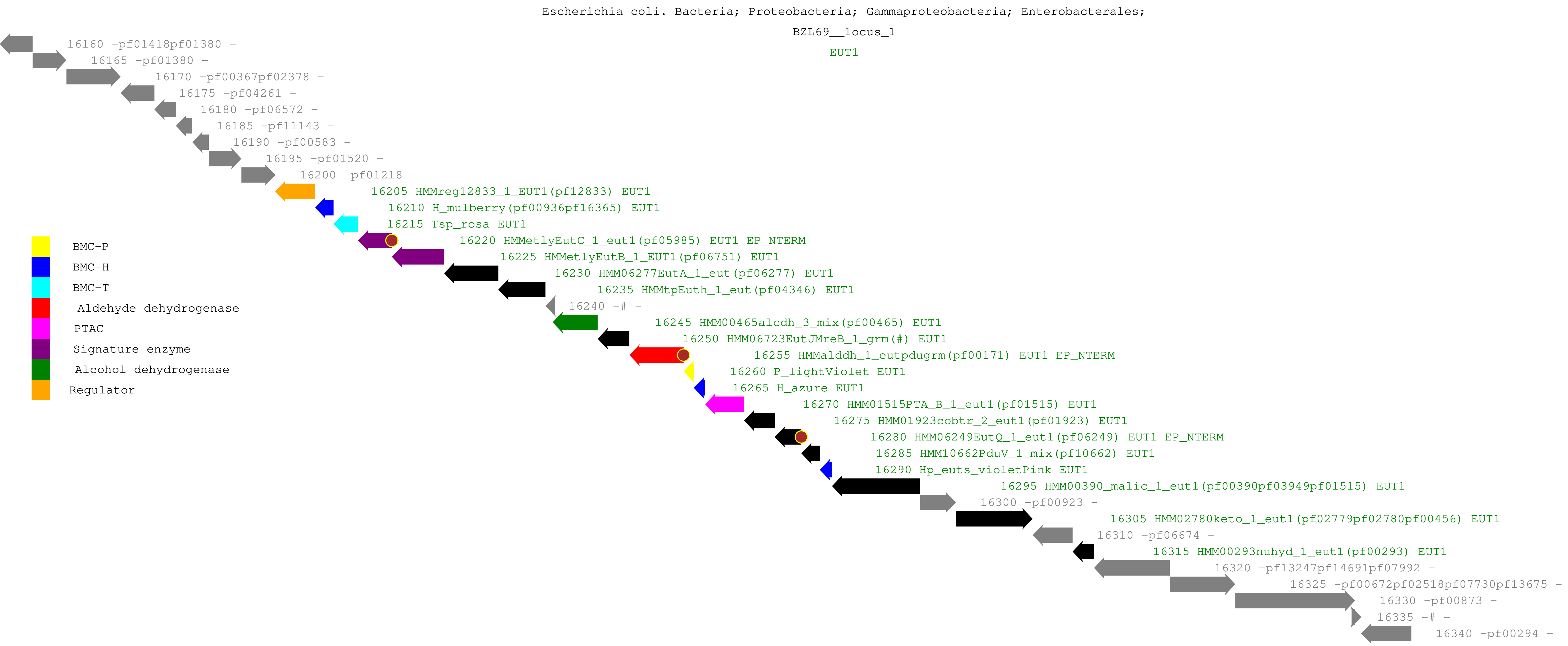
Escherichia coli. Bacteria; Proteobacteria; Gammaproteobacteria; Enterobacterales;
BZL69__locus_1
EUT1
16160 -pf01418pf01380 -
16165 -pf01380 -
16170 -pf00367pf02378 -
16175 -pf04261 -
16180 -pf06572 -
16185 -pf11143 -
16190 -pf00583 -
16195 -pf01520 -
16200 -pf01218 -
16205 HMMreg12833_1_EUT1(pf12833) EUT1
16210 H_mulberry(pf00936pf16365) EUT1
16215 Tsp_rosa EUT1
16220 HMMetlyEutC_1_eut1(pf05985) EUT1 EP_NTERM
16225 HMMetlyEutB_1_EUT1(pf06751) EUT1
16230 HMM06277EutA_1_eut(pf06277) EUT1
16235 HMMtpEuth_1_eut(pf04346) EUT1
16240 -# -
16245 HMM00465alcdh_3_mix(pf00465) EUT1
16250 HMM06723EutJMreB_1_grm(#) EUT1
16255 HMMalddh_1_eutpdugrm(pf00171) EUT1 EP_NTERM
16260 P_lightViolet EUT1
16265 H_azure EUT1
16270 HMM01515PTA_B_1_eut1(pf01515) EUT1
16275 HMM01923cobtr_2_eut1(pf01923) EUT1
16280 HMM06249EutQ_1_eut1(pf06249) EUT1 EP_NTERM
16285 HMM10662PduV_1_mix(pf10662) EUT1
16290 Hp_euts_violetPink EUT1
16295 HMM00390_malic_1_eut1(pf00390pf03949pf01515) EUT1
16300 -pf00923 -
16305 HMM02780keto_1_eut1(pf02779pf02780pf00456) EUT1
16310 -pf06674 -
16315 HMM00293nuhyd_1_eut1(pf00293) EUT1
16320 -pf13247pf14691pf07992 -
16325 -pf00672pf02518pf07730pf13675 -
16330 -pf00873 -
16335 -# -
16340 -pf00294 -
BMC-P
BMC-H
BMC-T
Aldehyde dehydrogenase
PTAC
Signature enzyme
Alcohol dehydrogenase
Regulator

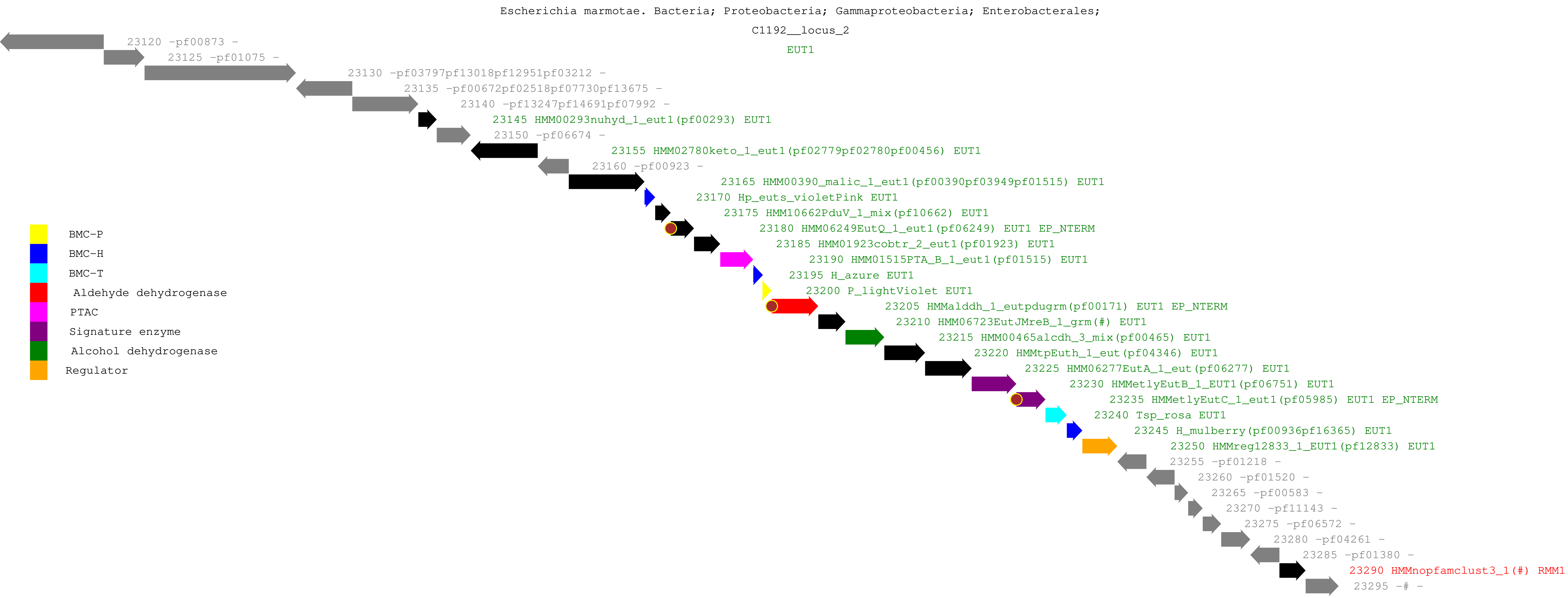

Escherichia marmotae. Bacteria; Proteobacteria; Gammaproteobacteria; Enterobacterales;
C1192__locus_2
EUT1
23120 -pf00873 -
23125 -pf01075 -
23130 -pf03797pf13018pf12951pf03212 -
23135 -pf00672pf02518pf07730pf13675 -
23140 -pf13247pf14691pf07992 -
23145 HMM00293nuhyd_1_eut1(pf00293) EUT1
23150 -pf06674 -
23155 HMM02780keto_1_eut1(pf02779pf02780pf00456) EUT1
23160 -pf00923 -
23165 HMM00390_malic_1_eut1(pf00390pf03949pf01515) EUT1
23170 Hp_euts_violetPink EUT1
23175 HMM10662PduV_1_mix(pf10662) EUT1
23180 HMM06249EutQ_1_eut1(pf06249) EUT1 EP_NTERM
23185 HMM01923cobtr_2_eut1(pf01923) EUT1
23190 HMM01515PTA_B_1_eut1(pf01515) EUT1
23195 H_azure EUT1
23200 P_lightViolet EUT1
23205 HMMalddh_1_eutpdugrm(pf00171) EUT1 EP_NTERM
23210 HMM06723EutJMreB_1_grm(#) EUT1
23215 HMM00465alcdh_3_mix(pf00465) EUT1
23220 HMMtpEuth_1_eut(pf04346) EUT1
23225 HMM06277EutA_1_eut(pf06277) EUT1
23230 HMMetlyEutB_1_EUT1(pf06751) EUT1
23235 HMMetlyEutC_1_eut1(pf05985) EUT1 EP_NTERM
23240 Tsp_rosa EUT1
23245 H_mulberry(pf00936pf16365) EUT1
23250 HMMreg12833_1_EUT1(pf12833) EUT1
23255 -pf01218 -
23260 -pf01520 -
23265 -pf00583 -
23270 -pf11143 -
23275 -pf06572 -
23280 -pf04261 -
23285 -pf01380 -
23290 HMMnopfamclust3_1(#) RMM1
23295 -# -
BMC-P
BMC-H
BMC-T
Aldehyde dehydrogenase
PTAC
Signature enzyme
Alcohol dehydrogenase
Regulator

Klebsiella pneumoniae. Bacteria; Proteobacteria; Gammaproteobacteria; Enterobacterales;

C1459__locus_1

EUT1

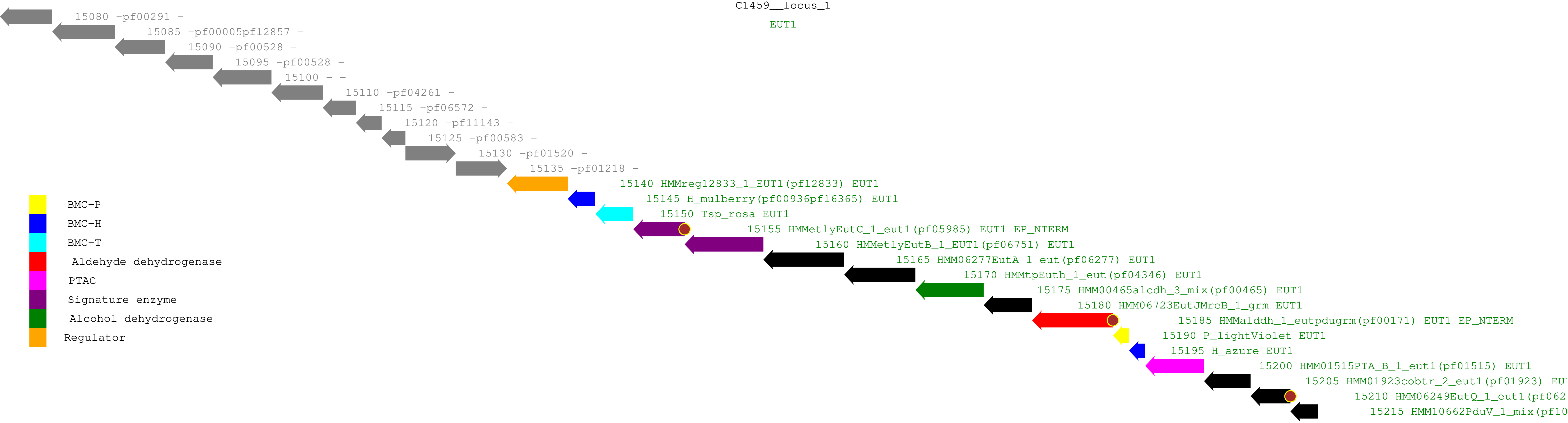

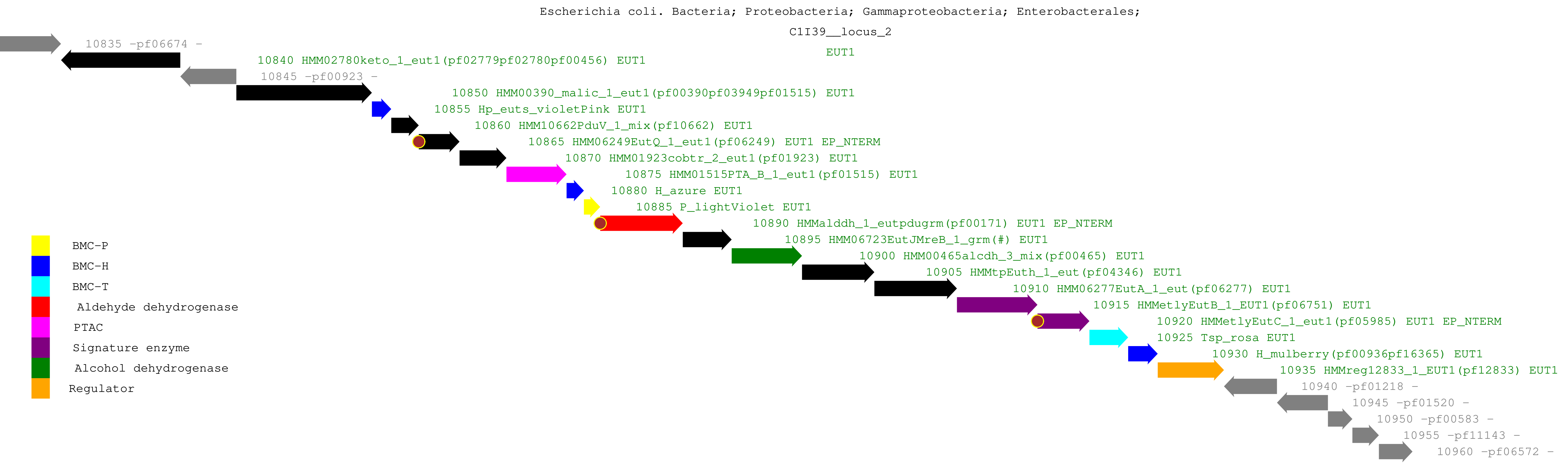
Escherichia coli. Bacteria; Proteobacteria; Gammaproteobacteria; Enterobacterales;
C1I39__locus_2
EUT1
10835 -pf06674 -
10840 HMM02780keto_1_eut1(pf02779pf02780pf00456) EUT1
10845 -pf00923 -
10850 HMM00390_malic_1_eut1(pf00390pf03949pf01515) EUT1
10855 Hp_euts_violetPink EUT1
10860 HMM10662PduV_1_mix(pf10662) EUT1
10865 HMM06249EutQ_1_eut1(pf06249) EUT1 EP_NTERM
10870 HMM01923cobtr_2_eut1(pf01923) EUT1
10875 HMM01515PTA_B_1_eut1(pf01515) EUT1
10880 H_azure EUT1
10885 P_lightViolet EUT1
10890 HMMalddh_1_eutpdugrm(pf00171) EUT1 EP_NTERM
10895 HMM06723EutJMreB_1_grm(#) EUT1
10900 HMM00465alcdh_3_mix(pf00465) EUT1
10905 HMMtpEuth_1_eut(pf04346) EUT1
10910 HMM06277EutA_1_eut(pf06277) EUT1
10915 HMMetlyEutB_1_EUT1(pf06751) EUT1
10920 HMMetlyEutC_1_eut1(pf05985) EUT1 EP_NTERM
10925 Tsp_rosa EUT1
10930 H_mulberry(pf00936pf16365) EUT1
10935 HMMreg12833_1_EUT1(pf12833) EUT1
10940 -pf01218 -
10945 -pf01520 -
10950 -pf00583 -
10955 -pf11143 -
10960 -pf06572 -
BMC-P
BMC-H
BMC-T
Aldehyde dehydrogenase
PTAC
Signature enzyme
Alcohol dehydrogenase
Regulator

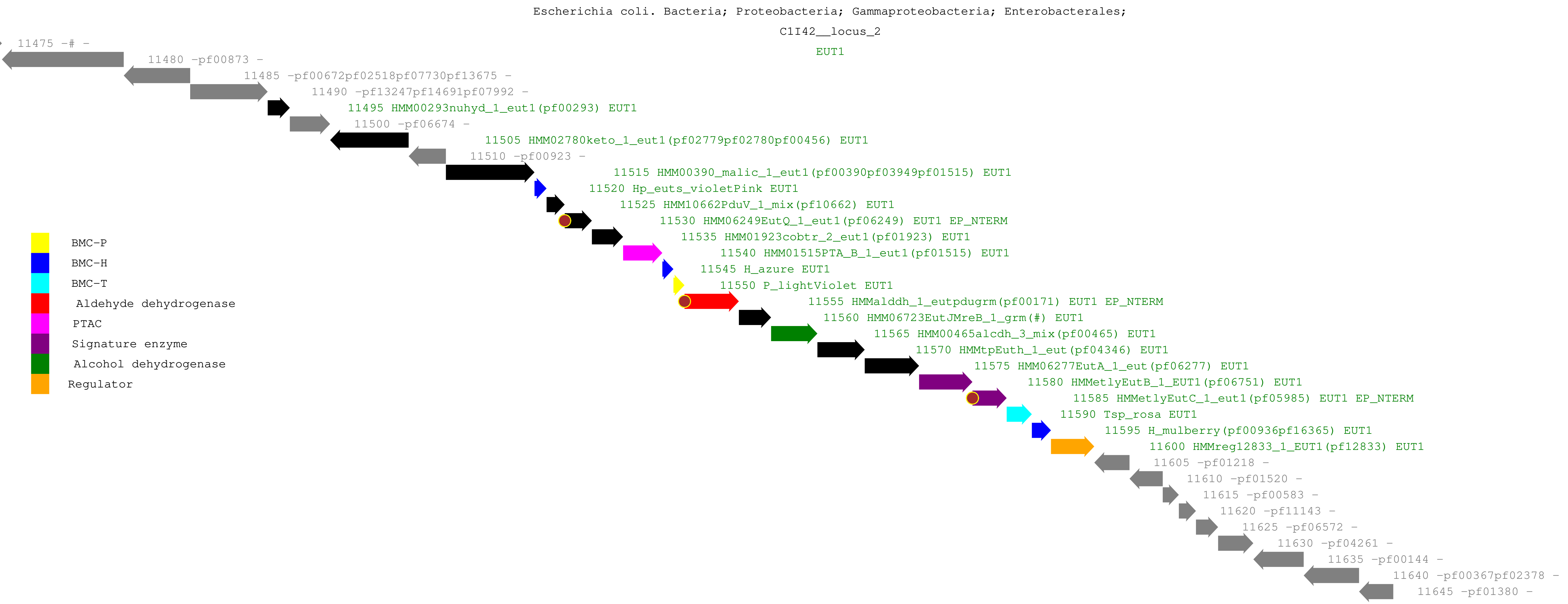

Escherichia coli. Bacteria; Proteobacteria; Gammaproteobacteria; Enterobacterales;
C1I42__locus_2
EUT1
11475 -# -
11480 -pf00873 -
11485 -pf00672pf02518pf07730pf13675 -
11490 -pf13247pf14691pf07992 -
11495 HMM00293nuhyd_1_eut1(pf00293) EUT1
11500 -pf06674 -
11505 HMM02780keto_1_eut1(pf02779pf02780pf00456) EUT1
11510 -pf00923 -
11515 HMM00390_malic_1_eut1(pf00390pf03949pf01515) EUT1
11520 Hp_euts_violetPink EUT1
11525 HMM10662PduV_1_mix(pf10662) EUT1
11530 HMM06249EutQ_1_eut1(pf06249) EUT1 EP_NTERM
11535 HMM01923cobtr_2_eut1(pf01923) EUT1
11540 HMM01515PTA_B_1_eut1(pf01515) EUT1
11545 H_azure EUT1
11550 P_lightViolet EUT1
11555 HMMaldh_1_eutpdugrm(pf00171) EUT1 EP_NTERM
11560 HMM06723EutJMreB_1_grm(#) EUT1
11565 HMM00465alcdh_3_mix(pf00465) EUT1
11570 HMMtpEuth_1_eut(pf04346) EUT1
11575 HMM06277EutA_1_eut(pf06277) EUT1
11580 HMMetlyEutB_1_EUT1(pf06751) EUT1
11585 HMMetlyEutC_1_eut1(pf05985) EUT1 EP_NTERM
11590 Tsp_rosa EUT1
11595 H_mulberry(pf00936pf16365) EUT1
11600 HMMreg12833_1_EUT1(pf12833) EUT1
11605 -pf01218 -
11610 -pf01520 -
11615 -pf00583 -
11620 -pf11143 -
11625 -pf06572 -
11630 -pf04261 -
11635 -pf00144 -
11640 -pf00367pf02378 -
11645 -pf01380 -
BMC-P
BMC-H
BMC-T
Aldehyde dehydrogenase
PTAC
Signature enzyme
Alcohol dehydrogenase
Regulator

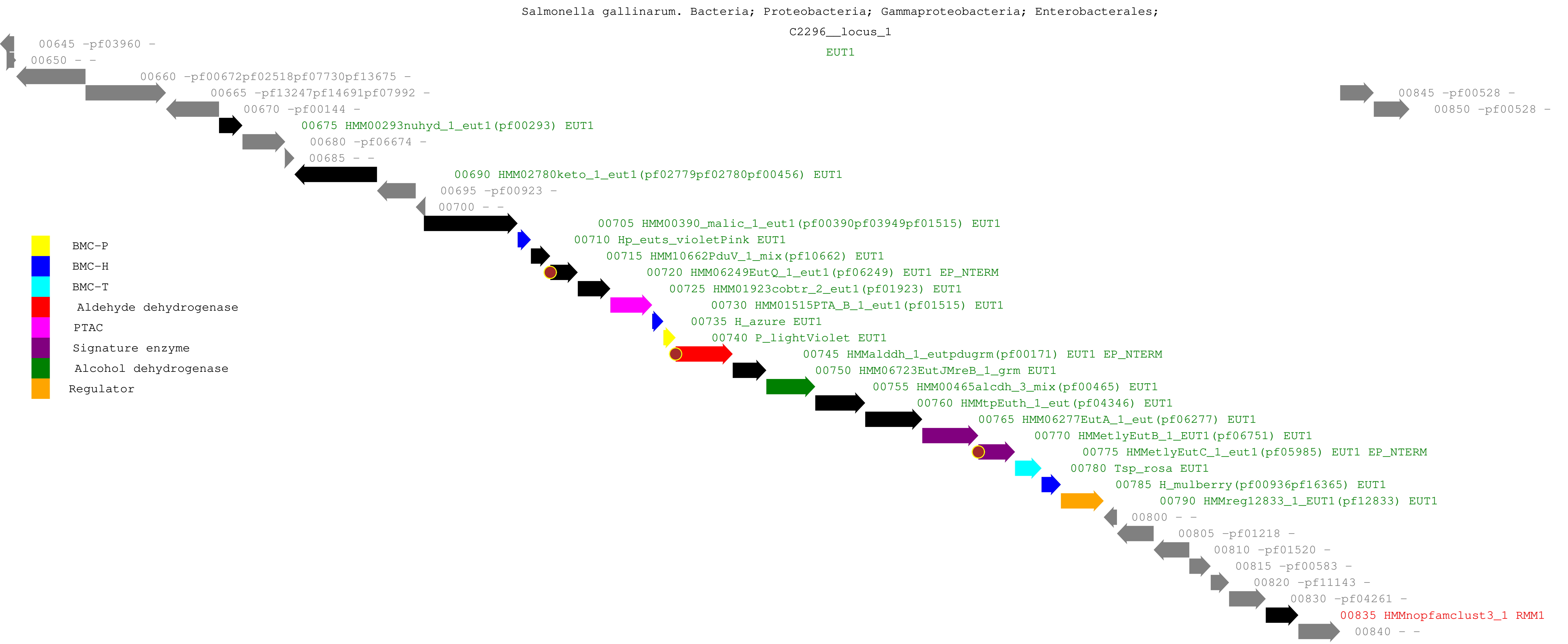
Salmonella gallinarum. Bacteria; Proteobacteria; Gammaproteobacteria; Enterobacterales;
C2296__locus_1
EUT1
00645 -pf03960 -
00650 - -
00660 -pf00672pf02518pf07730pf13675 -
00665 -pf13247pf14691pf07992 -
00670 -pf00144 -
00675 HMM00293nuhyd_1_eut1(pf00293) EUT1
00680 -pf06674 -
00685 - -
00690 HMM02780keto_1_eut1(pf02779pf02780pf00456) EUT1
00695 -pf00923 -
00700 - -
00705 HMM00390_malic_1_eut1(pf00390pf03949pf01515) EUT1
00710 Hp_euts_violetPink EUT1
00715 HMM10662PduV_1_mix(pf10662) EUT1
00720 HMM06249EutQ_1_eut1(pf06249) EUT1 EP_NTERM
00725 HMM01923cobtr_2_eut1(pf01923) EUT1
00730 HMM01515PTA_B_1_eut1(pf01515) EUT1
00735 H_azure EUT1
00740 P_lightViolet EUT1
00745 HMMalddh_1_eutpdugrm(pf00171) EUT1 EP_NTERM
00750 HMM06723EutJMreB_1_grm EUT1
00755 HMM00465alcdh_3_mix(pf00465) EUT1
00760 HMMtpEuth_1_eut(pf04346) EUT1
00765 HMM06277EutA_1_eut(pf06277) EUT1
00770 HMMetlyEutB_1_EUT1(pf06751) EUT1
00775 HMMetlyEutC_1_eut1(pf05985) EUT1 EP_NTERM
00780 Tsp_rosa EUT1
00785 H_mulberry(pf00936pf16365) EUT1
00790 HMMreg12833_1_EUT1(pf12833) EUT1
00800 - -
00805 -pf01218 -
00810 -pf01520 -
00815 -pf00583 -
00820 -pf11143 -
00830 -pf04261 -
00835 HMMnopfamclust3_1 RMM1
00840 - -
00845 -pf00528 -
00850 -pf00528 -
BMC-P
BMC-H
BMC-T
Aldehyde dehydrogenase
PTAC
Signature enzyme
Alcohol dehydrogenase
Regulator

Citrobacter freundii complex sp. CFNIH3. Bacteria; Proteobacteria; Gammaproteobacteria; Enterobacterales;

C2U38__locus_2

EUT1

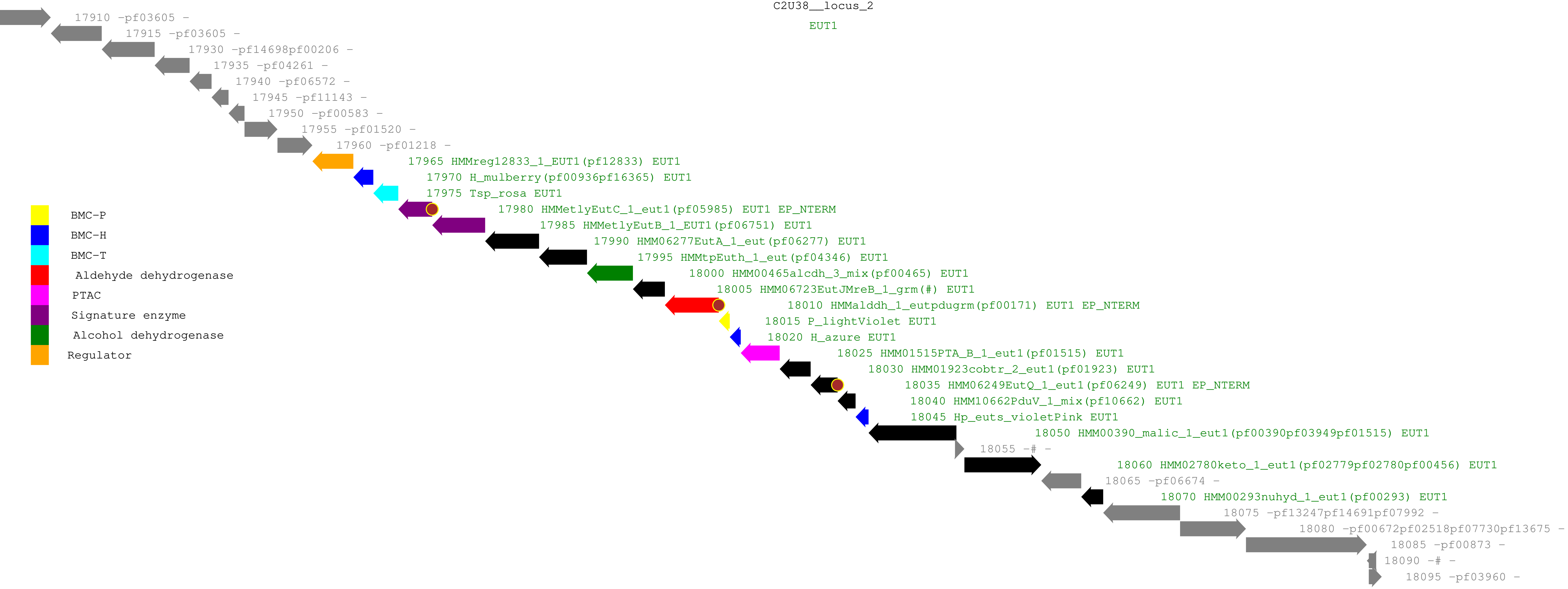

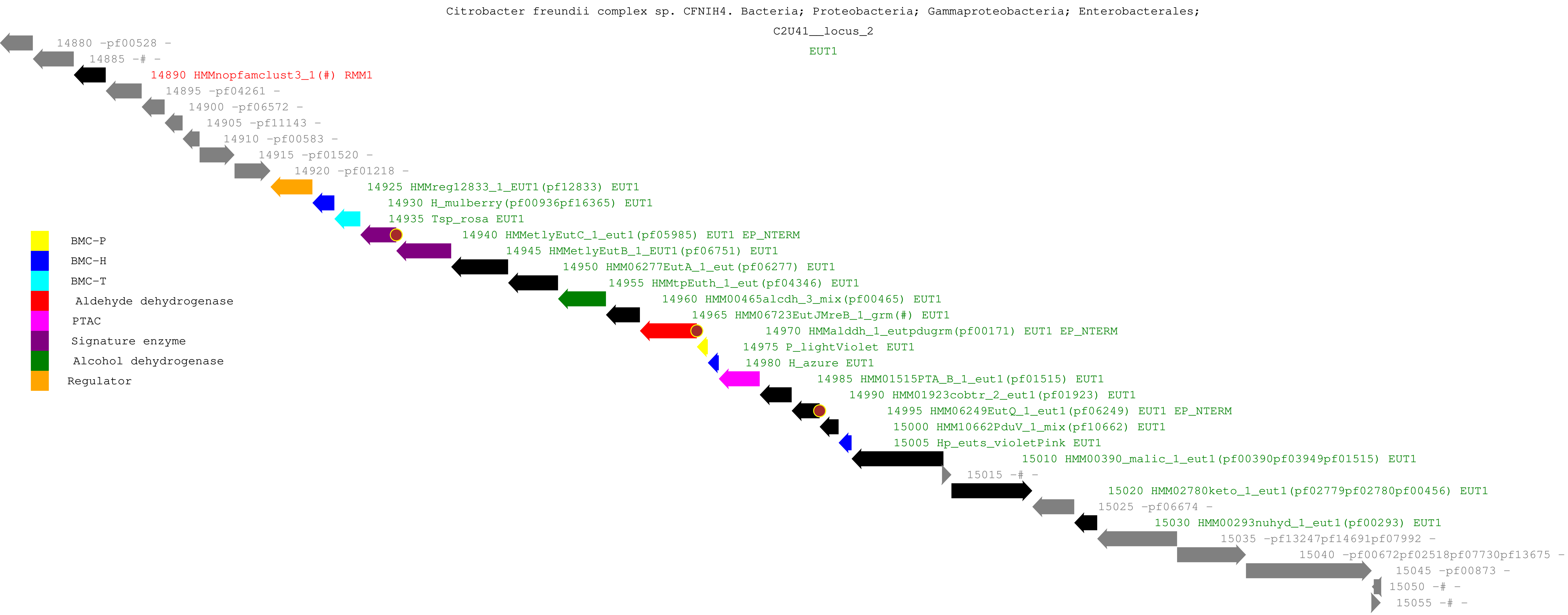
Citrobacter freundii complex sp. CFNIH4. Bacteria; Proteobacteria; Gammaproteobacteria; Enterobacterales;
C2U41__locus_2
EUT1
14880 -pf00528 -
14885 -# -
14890 HMMnopfamclust3_1(#) RMM1
14895 -pf04261 -
14900 -pf06572 -
14905 -pf11143 -
14910 -pf00583 -
14915 -pf01520 -
14920 -pf01218 -
14925 HMMreg12833_1_EUT1(pf12833) EUT1
14930 H_mulberry(pf00936pf16365) EUT1
14935 Tsp_rosa EUT1
14940 HMMetlyEutC_1_eut1(pf05985) EUT1 EP_NTERM
14945 HMMetlyEutB_1_EUT1(pf06751) EUT1
14950 HMM06277EutA_1_eut(pf06277) EUT1
14955 HMMtpEuth_1_eut(pf04346) EUT1
14960 HMM00465alcdh_3_mix(pf00465) EUT1
14965 HMM06723EutJMreB_1_grm(#) EUT1
14970 HMMalddh_1_eutpdugrm(pf00171) EUT1 EP_NTERM
14975 P_lightViolet EUT1
14980 H_azure EUT1
14985 HMM01515PTA_B_1_eut1(pf01515) EUT1
14990 HMM01923cobtr_2_eut1(pf01923) EUT1
14995 HMM06249EutQ_1_eut1(pf06249) EUT1 EP_NTERM
15000 HMM10662PduV_1_mix(pf10662) EUT1
15005 Hp_euts_violetPink EUT1
15010 HMM00390_malic_1_eut1(pf00390pf03949pf01515) EUT1
15015 -# -
15020 HMM02780keto_1_eut1(pf02779pf02780pf00456) EUT1
15025 -pf06674 -
15030 HMM00293nuhyd_1_eut1(pf00293) EUT1
15035 -pf13247pf14691pf07992 -
15040 -pf00672pf02518pf07730pf13675 -
15045 -pf00873 -
15050 -# -
15055 -# -
BMC-P
BMC-H
BMC-T
Aldehyde dehydrogenase
PTAC
Signature enzyme
Alcohol dehydrogenase
Regulator

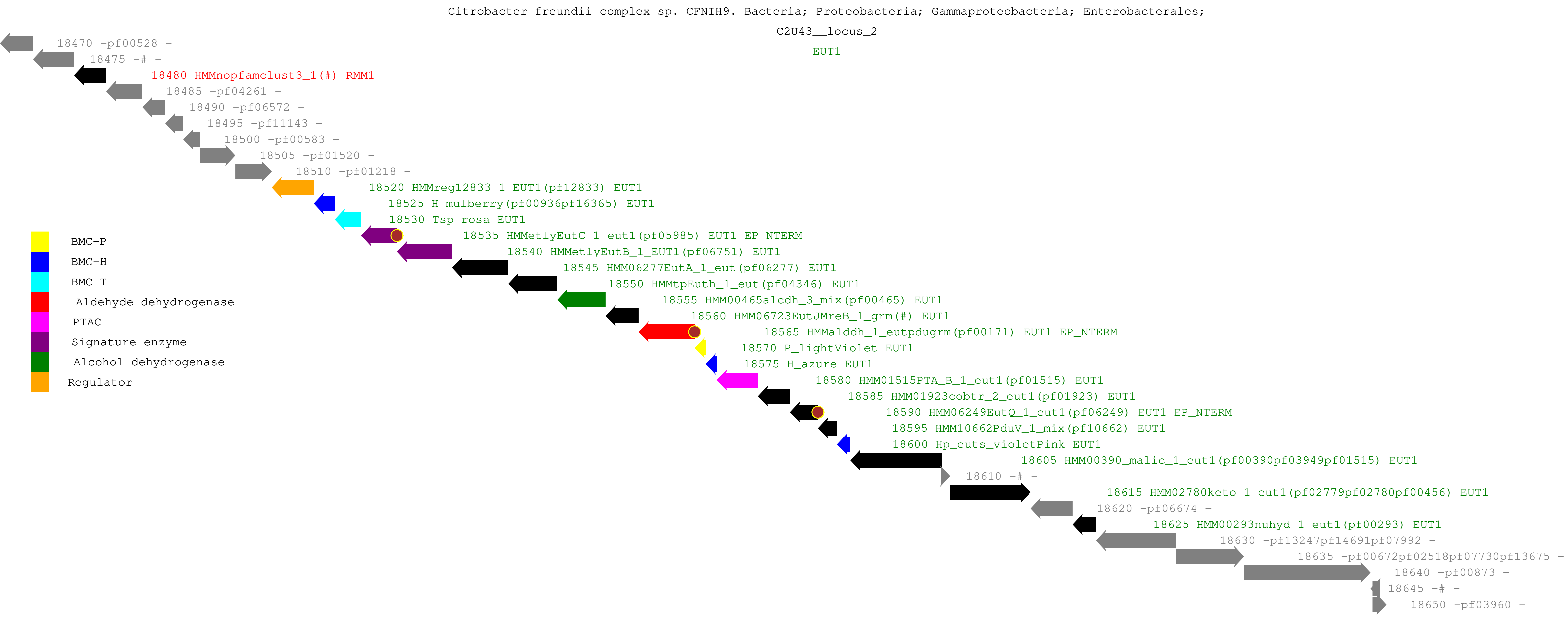
Citrobacter freundii complex sp. CFNIH9. Bacteria; Proteobacteria; Gammaproteobacteria; Enterobacterales;
C2U43__locus_2
EUT1
18470 -pf00528 -
18475 -# -
18480 HMMnopfamclust3_1(#) RMM1
18485 -pf04261 -
18490 -pf06572 -
18495 -pf11143 -
18500 -pf00583 -
18505 -pf01520 -
18510 -pf01218 -
18520 HMMreg12833_1_EUT1(pf12833) EUT1
18525 H_mulberry(pf00936pf16365) EUT1
18530 Tsp_rosa EUT1
18535 HMMetlyEutC_1_eut1(pf05985) EUT1 EP_NTERM
18540 HMMetlyEutB_1_EUT1(pf06751) EUT1
18545 HMM06277EutA_1_eut(pf06277) EUT1
18550 HMMtpEuth_1_eut(pf04346) EUT1
18555 HMM00465alcdh_3_mix(pf00465) EUT1
18560 HMM06723EutJMreB_1_grm(#) EUT1
18565 HMMalddh_1_eutpdugrm(pf00171) EUT1 EP_NTERM
18570 P_lightViolet EUT1
18575 H_azure EUT1
18580 HMM01515PTA_B_1_eut1(pf01515) EUT1
18585 HMM01923cobtr_2_eut1(pf01923) EUT1
18590 HMM06249EutQ_1_eut1(pf06249) EUT1 EP_NTERM
18595 HMM10662PduV_1_mix(pf10662) EUT1
18600 Hp_euts_violetPink EUT1
18605 HMM00390_malic_1_eut1(pf00390pf03949pf01515) EUT1
18610 -# -
18615 HMM02780keto_1_eut1(pf02779pf02780pf00456) EUT1
18620 -pf06674 -
18625 HMM00293nuhyd_1_eut1(pf00293) EUT1
18630 -pf13247pf14691pf07992 -
18635 -pf00672pf02518pf07730pf13675 -
18640 -pf00873 -
18645 -# -
18650 -pf03960 -
BMC-P
BMC-H
BMC-T
Aldehyde dehydrogenase
PTAC
Signature enzyme
Alcohol dehydrogenase
Regulator

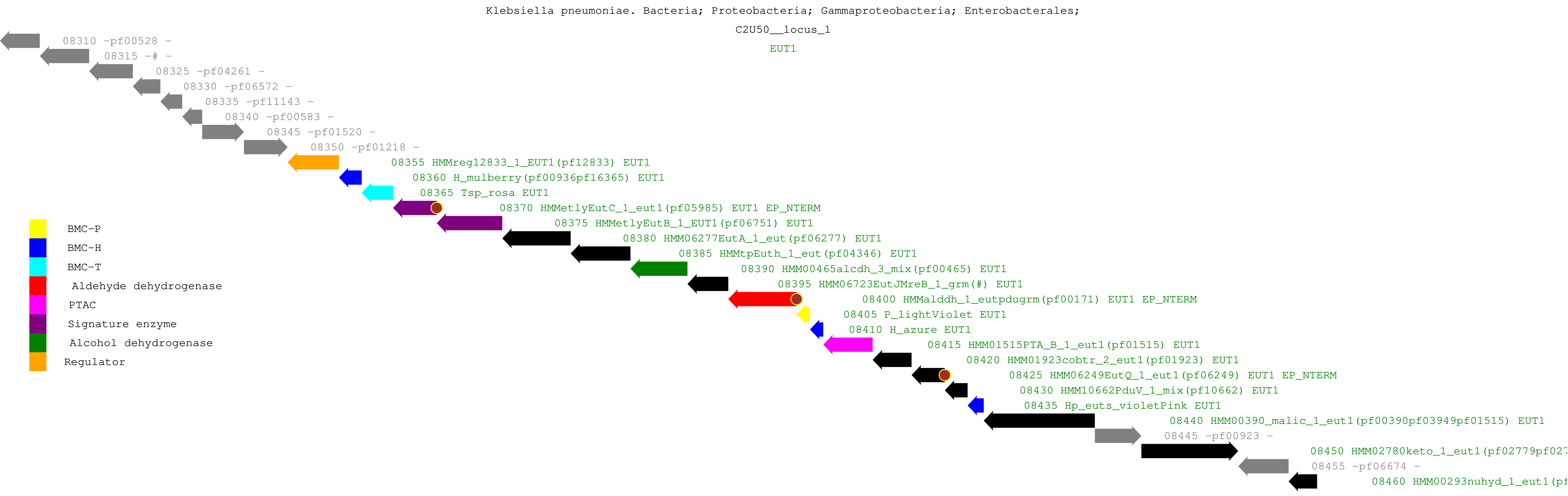
Klebsiella pneumoniae. Bacteria; Proteobacteria; Gammaproteobacteria; Enterobacterales;
C2U50__locus_1
EUT1
08310 -pf00528 -
08315 -# -
08325 -pf04261 -
08330 -pf06572 -
08335 -pf11143 -
08340 -pf00583 -
08345 -pf01520 -
08350 -pf01218 -
08355 HMMreg12833_1_EUT1(pf12833) EUT1
08360 H_mulberry(pf00936pf16365) EUT1
08365 Tsp_rosa EUT1
08370 HMMetlyEutC_1_eut1(pf05985) EUT1 EP_NTERM
08375 HMMetlyEutB_1_EUT1(pf06751) EUT1
08380 HMM06277EutA_1_eut(pf06277) EUT1
08385 HMMtpEuth_1_eut(pf04346) EUT1
08390 HMM00465alcdh_3_mix(pf00465) EUT1
08395 HMM06723EutJMreB_1_grm(#) EUT1
08400 HMMalddh_1_eutpdugrm(pf00171) EUT1 EP_NTERM
08405 P_lightViolet EUT1
08410 H_azure EUT1
08415 HMM01515PTA_B_1_eut1(pf01515) EUT1
08420 HMM01923cobtr_2_eut1(pf01923) EUT1
08425 HMM06249EutQ_1_eut1(pf06249) EUT1 EP_NTERM
08430 HMM10662PduV_1_mix(pf10662) EUT1
08435 Hp_euts_violetPink EUT1
08440 HMM00390_malic_1_eut1(pf00390pf03949pf01515) EUT1
08445 -pf00923 -
08450 HMM02780keto_1_eut1(pf02779pf027
08455 -pf06674 -
08460 HMM00293nuhyd_1_eut1(pf
BMC-P
BMC-H
BMC-T
Aldehyde dehydrogenase
PTAC
Signature enzyme
Alcohol dehydrogenase
Regulator

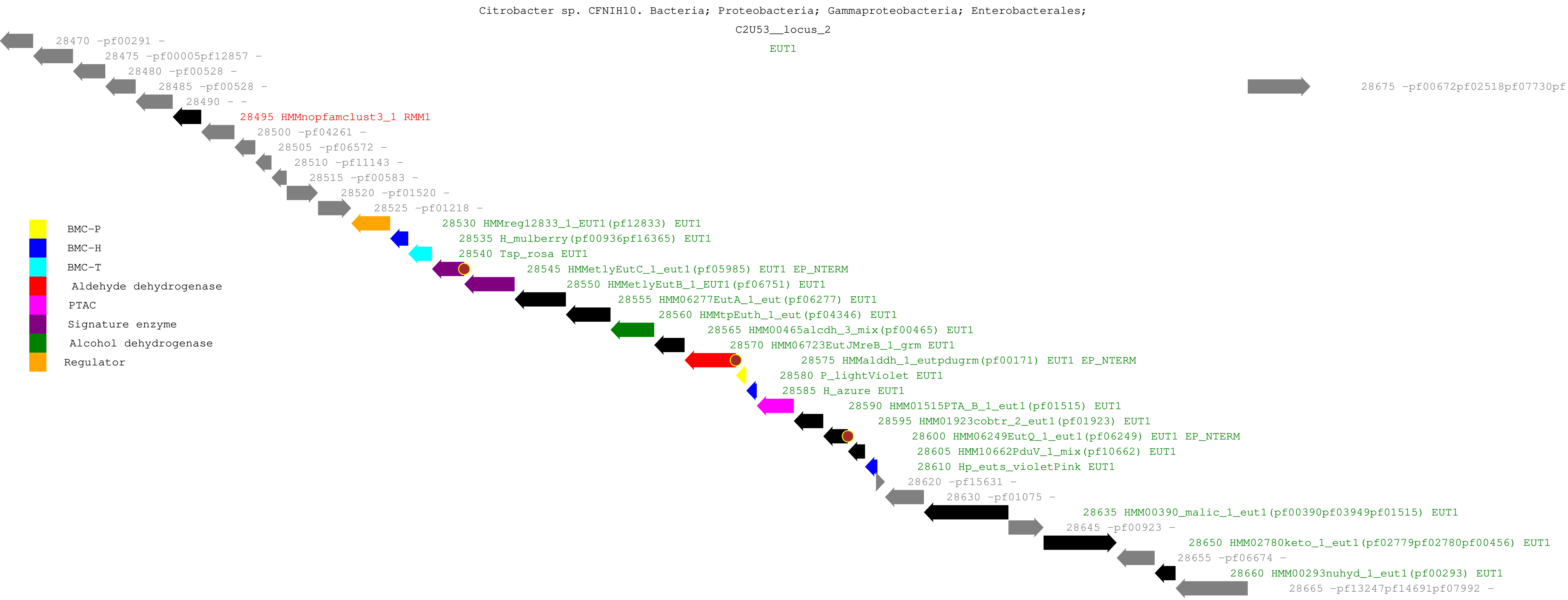
Citrobacter sp. CFNIH10. Bacteria; Proteobacteria; Gammaproteobacteria; Enterobacterales;
C2U53__locus_2
EUT1
28470 -pf00291 -
28475 -pf00005pf12857 -
28480 -pf00528 -
28485 -pf00528 -
28490 - -
28495 HMMnopfamclust3_1 RMM1
28500 -pf04261 -
28505 -pf06572 -
28510 -pf11143 -
28515 -pf00583 -
28520 -pf01520 -
28525 -pf01218 -
28530 HMMreg12833_1_EUT1(pf12833) EUT1
28535 H_mulberry(pf00936pf16365) EUT1
28540 Tsp_rosa EUT1
28545 HMMetlyEutC_1_eut1(pf05985) EUT1 EP_NTERM
28550 HMMetlyEutB_1_EUT1(pf06751) EUT1
28555 HMM06277EutA_1_eut(pf06277) EUT1
28560 HMMtpEuth_1_eut(pf04346) EUT1
28565 HMM00465alcdh_3_mix(pf00465) EUT1
28570 HMM06723EutJMreB_1_grm EUT1
28575 HMMaldh_1_eutpdugrm(pf00171) EUT1 EP_NTERM
28580 P_lightViolet EUT1
28585 H_azure EUT1
28590 HMM01515PTA_B_1_eut1(pf01515) EUT1
28595 HMM01923cobtr_2_eut1(pf01923) EUT1
28600 HMM06249EutQ_1_eut1(pf06249) EUT1 EP_NTERM
28605 HMM10662PduV_1_mix(pf10662) EUT1
28610 Hp_euts_violetPink EUT1
28620 -pf15631 -
28630 -pf01075 -
28635 HMM00390_malic_1_eut1(pf00390pf03949pf01515) EUT1
28645 -pf00923 -
28650 HMM02780keto_1_eut1(pf02779pf02780pf00456) EUT1
28655 -pf06674 -
28660 HMM00293nuhyd_1_eut1(pf00293) EUT1
28665 -pf13247pf14691pf07992 -
28675 -pf00672pf02518pf07730pf
BMC-P
BMC-H
BMC-T
Aldehyde dehydrogenase
PTAC
Signature enzyme
Alcohol dehydrogenase
Regulator

Citrobacter amalonaticus. Bacteria; Proteobacteria; Gammaproteobacteria; Enterobacterales;

C3430__locus_2

EUT1

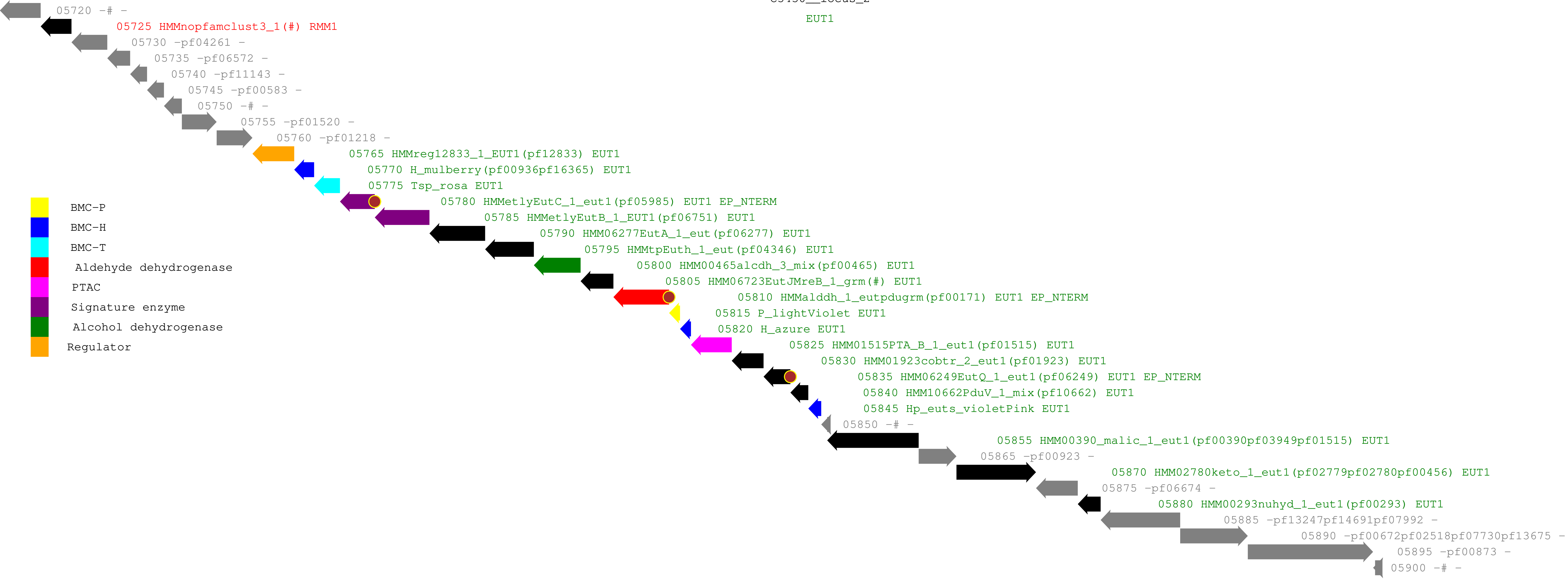

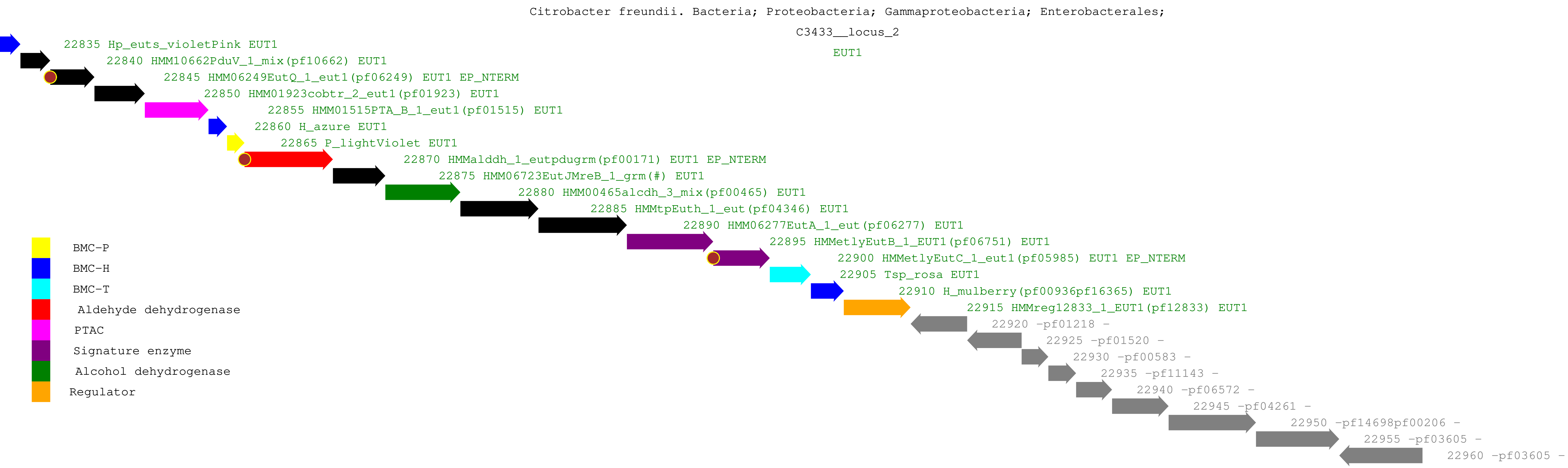
Citrobacter freundii. Bacteria; Proteobacteria; Gammaproteobacteria; Enterobacterales;
C3433__locus_2
EUT1
22835 Hp_euts_violetPink EUT1
22840 HMM10662PduV_1_mix(pf10662) EUT1
22845 HMM06249EutQ_1_eut1(pf06249) EUT1 EP_NTERM
22850 HMM01923cobtr_2_eut1(pf01923) EUT1
22855 HMM01515PTA_B_1_eut1(pf01515) EUT1
22860 H_azure EUT1
22865 P_lightViolet EUT1
22870 HMMaldh_1_eutpdugrm(pf00171) EUT1 EP_NTERM
22875 HMM06723EutJMreB_1_grm(#) EUT1
22880 HMM00465alcdh_3_mix(pf00465) EUT1
22885 HMMtpEuth_1_eut(pf04346) EUT1
22890 HMM06277EutA_1_eut(pf06277) EUT1
22895 HMMetlyEutB_1_EUT1(pf06751) EUT1
22900 HMMetlyEutC_1_eut1(pf05985) EUT1 EP_NTERM
22905 Tsp_rosa EUT1
22910 H_mulberry(pf00936pf16365) EUT1
22915 HMMreg12833_1_EUT1(pf12833) EUT1
22920 -pf01218 -
22925 -pf01520 -
22930 -pf00583 -
22935 -pf11143 -
22940 -pf06572 -
22945 -pf04261 -
22950 -pf14698pf00206 -
22955 -pf03605 -
22960 -pf03605 -
BMC-P
BMC-H
BMC-T
Aldehyde dehydrogenase
PTAC
Signature enzyme
Alcohol dehydrogenase
Regulator

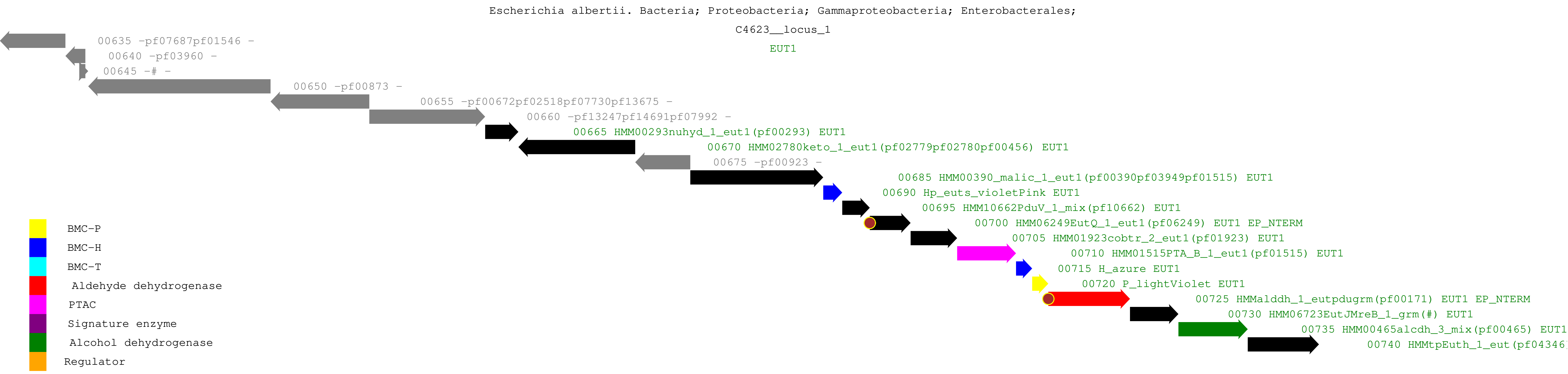
Escherichia albertii. Bacteria; Proteobacteria; Gammaproteobacteria; Enterobacterales;
C4623__locus_1
EUT1
00635 -pf07687pf01546 -
00640 -pf03960 -
00645 -# -
00650 -pf00873 -
00655 -pf00672pf02518pf07730pf13675 -
00660 -pf13247pf14691pf07992 -
00665 HMM00293nuhyd_1_eut1(pf00293) EUT1
00670 HMM02780keto_1_eut1(pf02779pf02780pf00456) EUT1
00675 -pf00923 -
00685 HMM00390_malic_1_eut1(pf00390pf03949pf01515) EUT1
00690 Hp_euts_violetPink EUT1
00695 HMM10662PduV_1_mix(pf10662) EUT1
00700 HMM06249EutQ_1_eut1(pf06249) EUT1 EP_NTERM
00705 HMM01923cobtr_2_eut1(pf01923) EUT1
00710 HMM01515PTA_B_1_eut1(pf01515) EUT1
00715 H_azure EUT1
00720 P_lightViolet EUT1
00725 HMMaldh_1_eutpdugrm(pf00171) EUT1 EP_NTERM
00730 HMM06723EutJMreB_1_grm(#) EUT1
00735 HMM00465alcdh_3_mix(pf00465) EUT1
00740 HMMtpEuth_1_eut(pf04346
BMC-P
BMC-H
BMC-T
Aldehyde dehydrogenase
PTAC
Signature enzyme
Alcohol dehydrogenase
Regulator

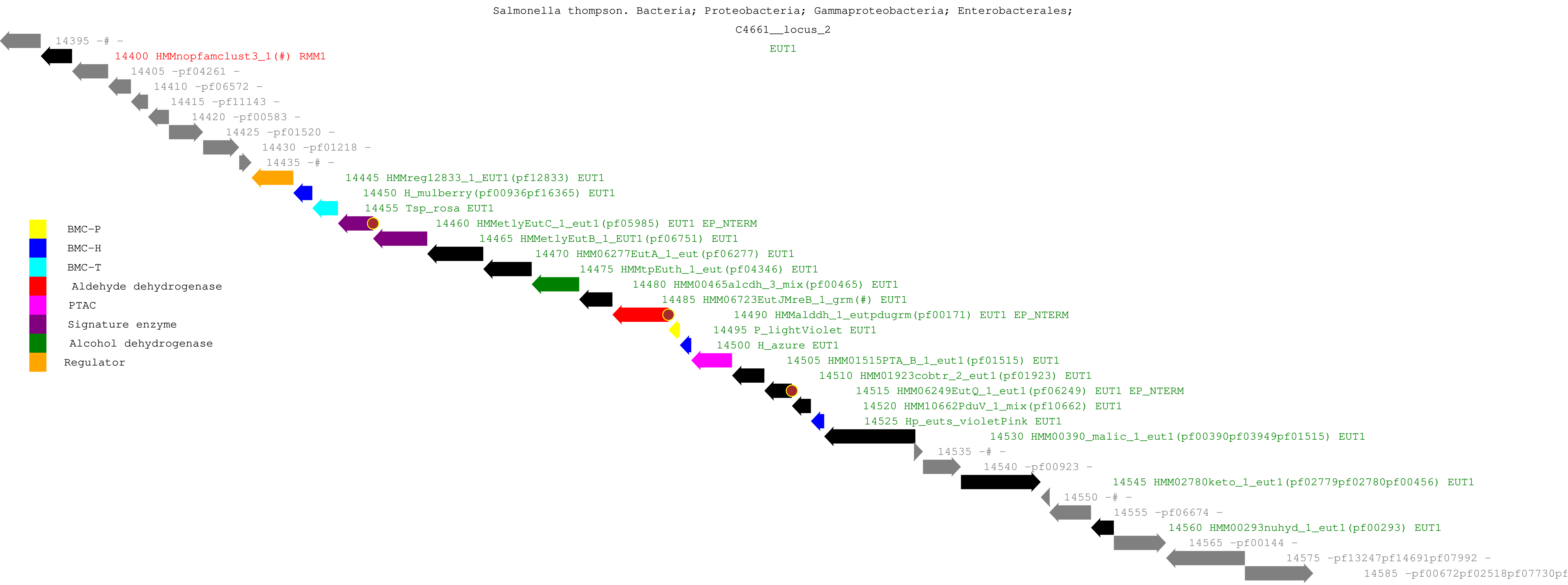

Salmonella thompson. Bacteria; Proteobacteria; Gammaproteobacteria; Enterobacterales;
C4661__locus_2
EUT1
14395 -# -
14400 HMMnopfamclust3_1(#) RMM1
14405 -pf04261 -
14410 -pf06572 -
14415 -pf11143 -
14420 -pf00583 -
14425 -pf01520 -
14430 -pf01218 -
14435 -# -
14445 HMMreg12833_1_EUT1(pf12833) EUT1
14450 H_mulberry(pf00936pf16365) EUT1
14455 Tsp_rosa EUT1
14460 HMMetlyEutC_1_eut1(pf05985) EUT1 EP_NTERM
14465 HMMetlyEutB_1_EUT1(pf06751) EUT1
14470 HMM06277EutA_1_eut(pf06277) EUT1
14475 HMMtpEuth_1_eut(pf04346) EUT1
14480 HMM00465alcdh_3_mix(pf00465) EUT1
14485 HMM06723EutJMreB_1_grm(#) EUT1
14490 HMMalddh_1_eutpdugrm(pf00171) EUT1 EP_NTERM
14495 P_lightViolet EUT1
14500 H_azure EUT1
14505 HMM01515PTA_B_1_eut1(pf01515) EUT1
14510 HMM01923cobtr_2_eut1(pf01923) EUT1
14515 HMM06249EutQ_1_eut1(pf06249) EUT1 EP_NTERM
14520 HMM10662PduV_1_mix(pf10662) EUT1
14525 Hp_euts_violetPink EUT1
14530 HMM00390_malic_1_eut1(pf00390pf03949pf01515) EUT1
14535 -# -
14540 -pf00923 -
14545 HMM02780keto_1_eut1(pf02779pf02780pf00456) EUT1
14550 -# -
14555 -pf06674 -
14560 HMM00293nuhyd_1_eut1(pf00293) EUT1
14565 -pf00144 -
14575 -pf13247pf14691pf07992 -
14585 -pf00672pf02518pf07730pf
BMC-P
BMC-H
BMC-T
Aldehyde dehydrogenase
PTAC
Signature enzyme
Alcohol dehydrogenase
Regulator

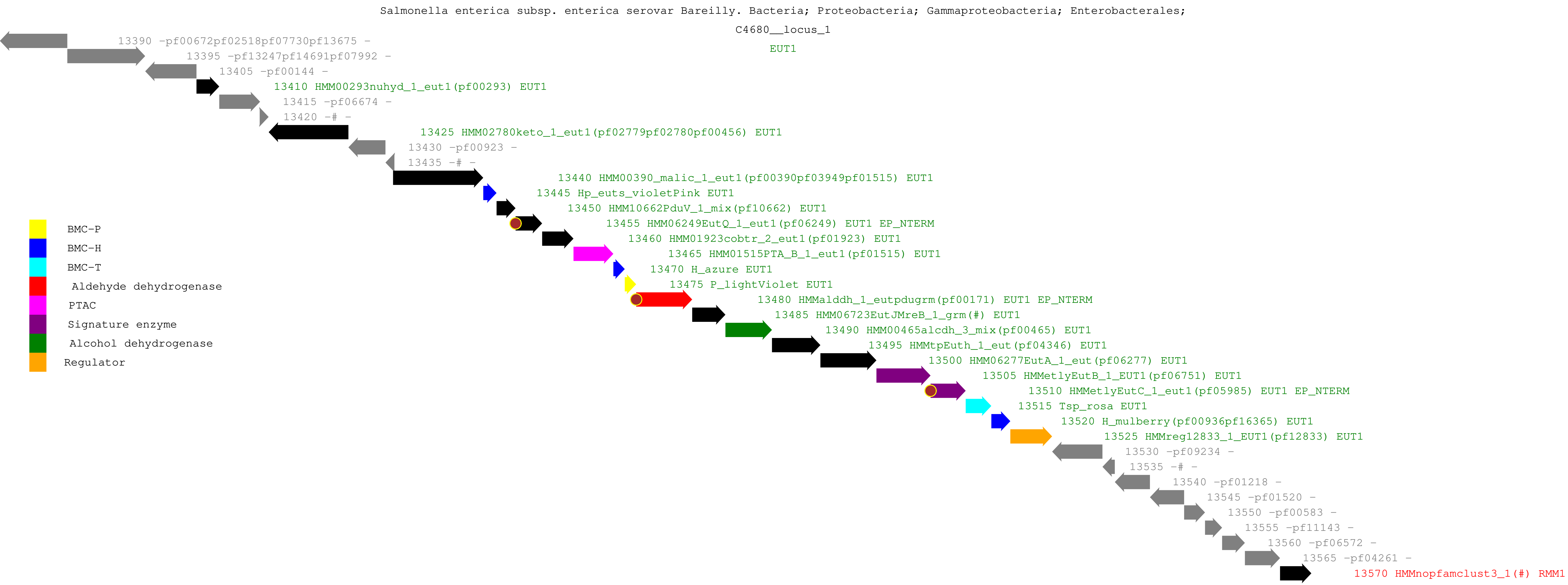
Salmonella enterica subsp. enterica serovar Bareilly. Bacteria; Proteobacteria; Gammaproteobacteria; Enterobacterales;
C4680__locus_1
EUT1
13390 -pf00672pf02518pf07730pf13675 -
13395 -pf13247pf14691pf07992 -
13405 -pf00144 -
13410 HMM00293nuhyd_1_eut1(pf00293) EUT1
13415 -pf06674 -
13420 -# -
13425 HMM02780keto_1_eut1(pf02779pf02780pf00456) EUT1
13430 -pf00923 -
13435 -# -
13440 HMM00390_malic_1_eut1(pf00390pf03949pf01515) EUT1
13445 Hp_euts_violetPink EUT1
13450 HMM10662PduV_1_mix(pf10662) EUT1
13455 HMM06249EutQ_1_eut1(pf06249) EUT1 EP_NTERM
13460 HMM01923cobtr_2_eut1(pf01923) EUT1
13465 HMM01515PTA_B_1_eut1(pf01515) EUT1
13470 H_azure EUT1
13475 P_lightViolet EUT1
13480 HMMalddh_1_eutpdugrm(pf00171) EUT1 EP_NTERM
13485 HMM06723EutJMreB_1_grm(#) EUT1
13490 HMM00465alcdh_3_mix(pf00465) EUT1
13495 HMMtpEuth_1_eut(pf04346) EUT1
13500 HMM06277EutA_1_eut(pf06277) EUT1
13505 HMMetlyEutB_1_EUT1(pf06751) EUT1
13510 HMMetlyEutC_1_eut1(pf05985) EUT1 EP_NTERM
13515 Tsp_rosa EUT1
13520 H_mulberry(pf00936pf16365) EUT1
13525 HMMreg12833_1_EUT1(pf12833) EUT1
13530 -pf09234 -
13535 -# -
13540 -pf01218 -
13545 -pf01520 -
13550 -pf00583 -
13555 -pf11143 -
13560 -pf06572 -
13565 -pf04261 -
13570 HMMnopfamclust3_1(#) RMM1
BMC-P
BMC-H
BMC-T
Aldehyde dehydrogenase
PTAC
Signature enzyme
Alcohol dehydrogenase
Regulator

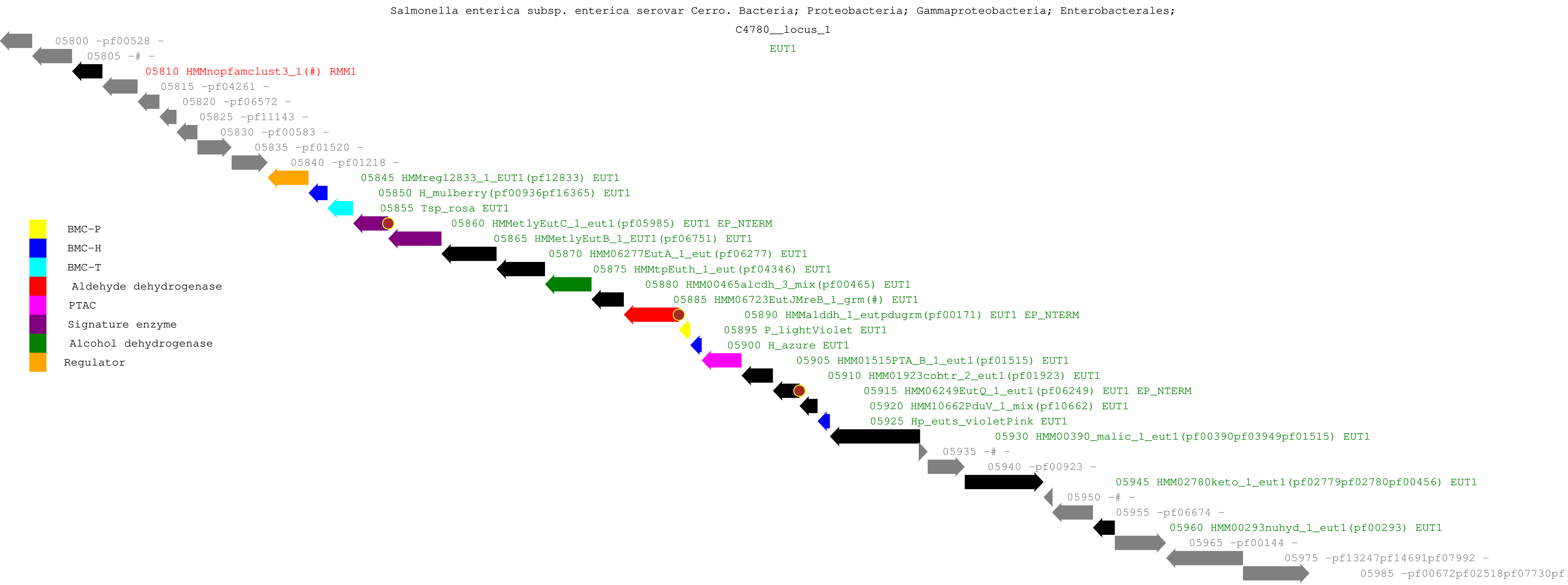
Salmonella enterica subsp. enterica serovar Cerro. Bacteria; Proteobacteria; Gammaproteobacteria; Enterobacterales;
C4780__locus_1
EUT1
05800 -pf00528 -
05805 -# -
05810 HMMnopfamclust3_1(#) RMM1
05815 -pf04261 -
05820 -pf06572 -
05825 -pf11143 -
05830 -pf00583 -
05835 -pf01520 -
05840 -pf01218 -
05845 HMMreg12833_1_EUT1(pf12833) EUT1
05850 H_mulberry(pf00936pf16365) EUT1
05855 Tsp_rosa EUT1
05860 HMMetlyEutC_1_eut1(pf05985) EUT1 EP_NTERM
05865 HMMetlyEutB_1_EUT1(pf06751) EUT1
05870 HMM06277EutA_1_eut(pf06277) EUT1
05875 HMMtpEuth_1_eut(pf04346) EUT1
05880 HMM00465alcdh_3_mix(pf00465) EUT1
05885 HMM06723EutJMreB_1_grm(#) EUT1
05890 HMMalddh_1_eutpdugrm(pf00171) EUT1 EP_NTERM
05895 P_lightViolet EUT1
05900 H_azure EUT1
05905 HMM01515PTA_B_1_eut1(pf01515) EUT1
05910 HMM01923cobtr_2_eut1(pf01923) EUT1
05915 HMM06249EutQ_1_eut1(pf06249) EUT1 EP_NTERM
05920 HMM10662PduV_1_mix(pf10662) EUT1
05925 Hp_euts_violetPink EUT1
05930 HMM00390_malic_1_eut1(pf00390pf03949pf01515) EUT1
05935 -# -
05940 -pf00923 -
05945 HMM02780keto_1_eut1(pf02779pf02780pf00456) EUT1
05950 -# -
05955 -pf06674 -
05960 HMM00293nuhyd_1_eut1(pf00293) EUT1
05965 -pf00144 -
05975 -pf13247pf14691pf07992 -
05985 -pf00672pf02518pf07730pf
BMC-P
BMC-H
BMC-T
Aldehyde dehydrogenase
PTAC
Signature enzyme
Alcohol dehydrogenase
Regulator

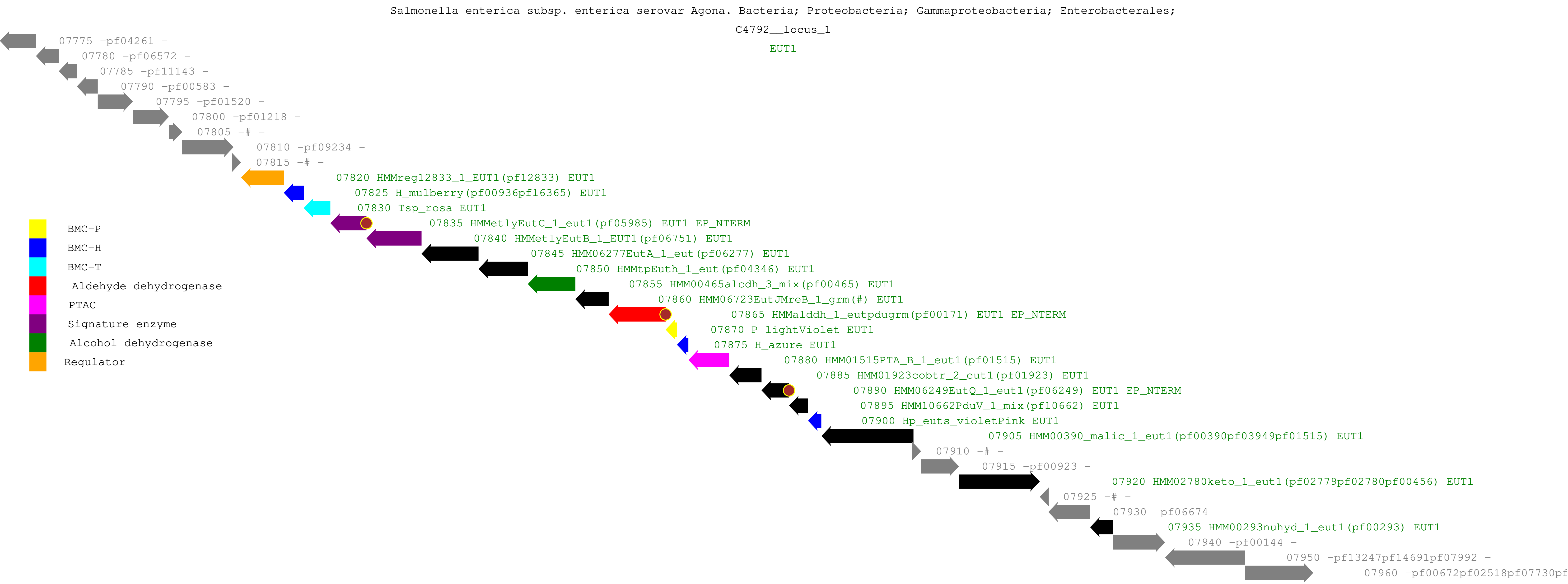
Salmonella enterica subsp. enterica serovar Agona. Bacteria; Proteobacteria; Gammaproteobacteria; Enterobacterales;
C4792__locus_1
EUT1
07775 -pf04261 -
07780 -pf06572 -
07785 -pf11143 -
07790 -pf00583 -
07795 -pf01520 -
07800 -pf01218 -
07805 -# -
07810 -pf09234 -
07815 -# -
07820 HMMreg12833_1_EUT1(pf12833) EUT1
07825 H_mulberry(pf00936pf16365) EUT1
07830 Tsp_rosa EUT1
07835 HMMetlyEutC_1_eut1(pf05985) EUT1 EP_NTERM
07840 HMMetlyEutB_1_EUT1(pf06751) EUT1
07845 HMM06277EutA_1_eut(pf06277) EUT1
07850 HMMtpEuth_1_eut(pf04346) EUT1
07855 HMM00465alcdh_3_mix(pf00465) EUT1
07860 HMM06723EutJMreB_1_grm(#) EUT1
07865 HMMalddh_1_eutpdugrm(pf00171) EUT1 EP_NTERM
07870 P_lightViolet EUT1
07875 H_azure EUT1
07880 HMM01515PTA_B_1_eut1(pf01515) EUT1
07885 HMM01923cobtr_2_eut1(pf01923) EUT1
07890 HMM06249EutQ_1_eut1(pf06249) EUT1 EP_NTERM
07895 HMM10662PduV_1_mix(pf10662) EUT1
07900 Hp_euts_violetPink EUT1
07905 HMM00390_malic_1_eut1(pf00390pf03949pf01515) EUT1
07910 -# -
07915 -pf00923 -
07920 HMM02780keto_1_eut1(pf02779pf02780pf00456) EUT1
07925 -# -
07930 -pf06674 -
07935 HMM00293nuhyd_1_eut1(pf00293) EUT1
07940 -pf00144 -
07950 -pf13247pf14691pf07992 -
07960 -pf00672pf02518pf07730pf
BMC-P
BMC-H
BMC-T
Aldehyde dehydrogenase
PTAC
Signature enzyme
Alcohol dehydrogenase
Regulator

Salmonella choleraesuis. Bacteria; Proteobacteria; Gammaproteobacteria; Enterobacterales;

C4801__locus_1

EUT1

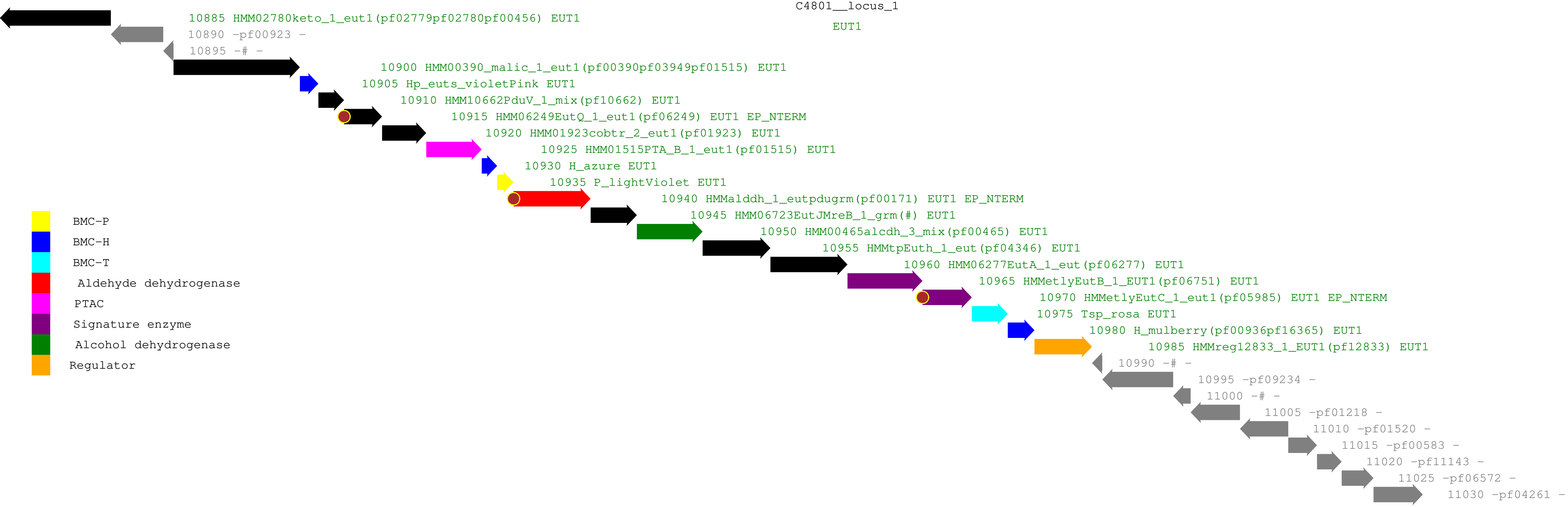

Salmonella enterica subsp. arizonae serovar 51:g,z51:-. Bacteria; Proteobacteria; Gammaproteobacteria; Enterobacterales;

C4803__locus_2

EUT1

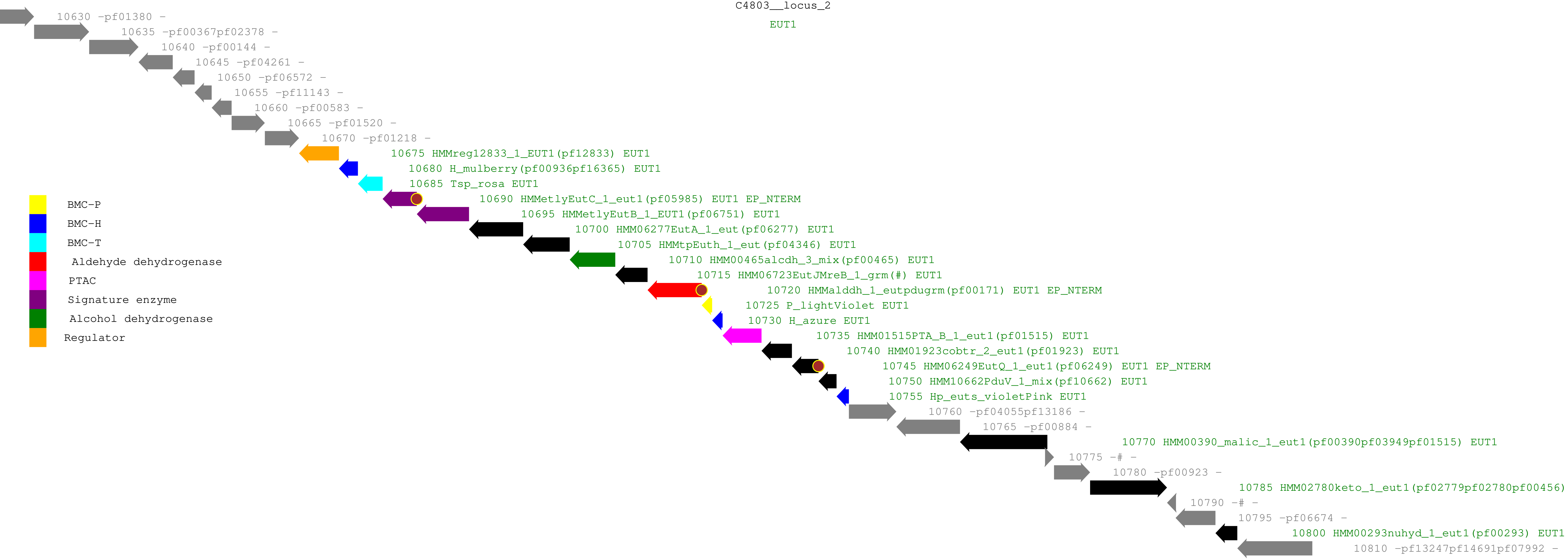

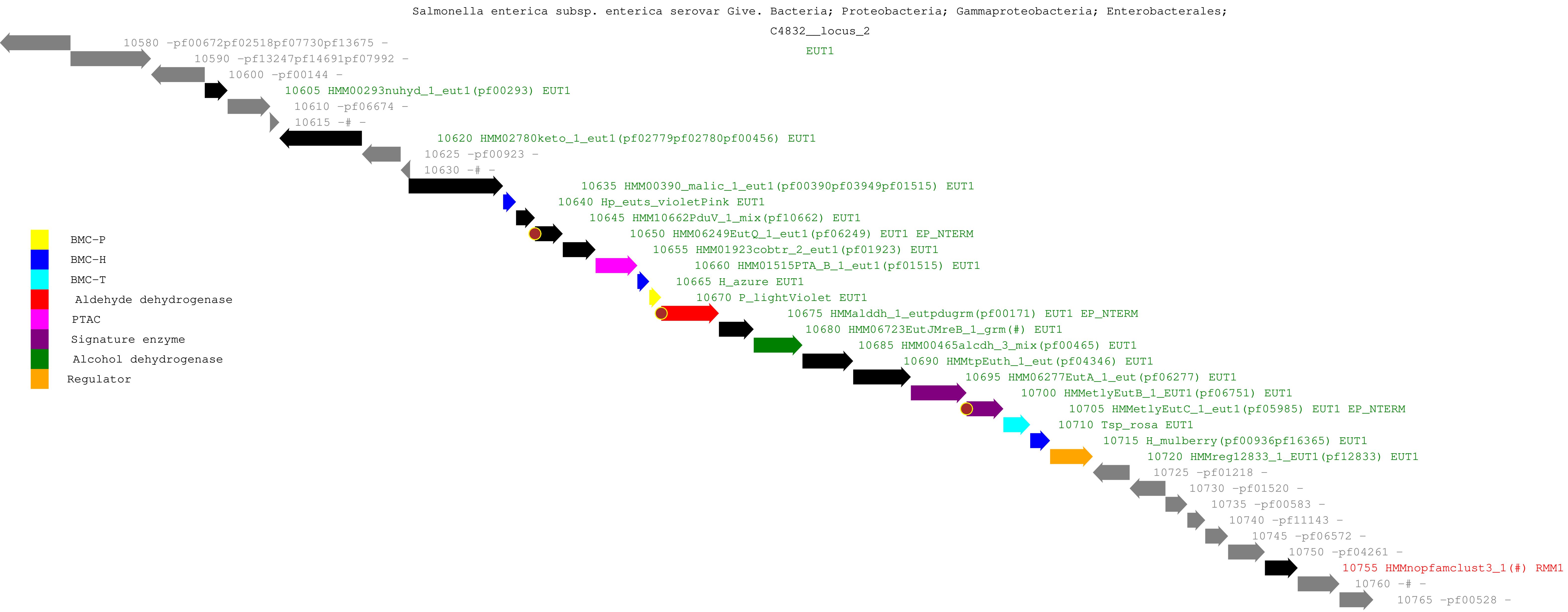

Salmonella enterica subsp. enterica serovar Give. Bacteria; Proteobacteria; Gammaproteobacteria; Enterobacterales;
C4832__locus_2
EUT1
10580 -pf00672pf02518pf07730pf13675 -
10590 -pf13247pf14691pf07992 -
10600 -pf00144 -
10605 HMM00293nuhyd_1_eut1(pf00293) EUT1
10610 -pf06674 -
10615 -# -
10620 HMM02780keto_1_eut1(pf02779pf02780pf00456) EUT1
10625 -pf00923 -
10630 -# -
10635 HMM00390_malic_1_eut1(pf00390pf03949pf01515) EUT1
10640 Hp_euts_violetPink EUT1
10645 HMM10662PduV_1_mix(pf10662) EUT1
10650 HMM06249EutQ_1_eut1(pf06249) EUT1 EP_NTERM
10655 HMM01923cobtr_2_eut1(pf01923) EUT1
10660 HMM01515PTA_B_1_eut1(pf01515) EUT1
10665 H_azure EUT1
10670 P_lightViolet EUT1
10675 HMMalddh_1_eutpdugrm(pf00171) EUT1 EP_NTERM
10680 HMM06723EutJMreB_1_grm(#) EUT1
10685 HMM00465alcdh_3_mix(pf00465) EUT1
10690 HMMtpEuth_1_eut(pf04346) EUT1
10695 HMM06277EutA_1_eut(pf06277) EUT1
10700 HMMetlyEutB_1_EUT1(pf06751) EUT1
10705 HMMetlyEutC_1_eut1(pf05985) EUT1 EP_NTERM
10710 Tsp_rosa EUT1
10715 H_mulberry(pf00936pf16365) EUT1
10720 HMMreg12833_1_EUT1(pf12833) EUT1
10725 -pf01218 -
10730 -pf01520 -
10735 -pf00583 -
10740 -pf11143 -
10745 -pf06572 -
10750 -pf04261 -
10755 HMMnopfamclust3_1(#) RMM1
10760 -# -
10765 -pf00528 -
BMC-P
BMC-H
BMC-T
Aldehyde dehydrogenase
PTAC
Signature enzyme
Alcohol dehydrogenase
Regulator

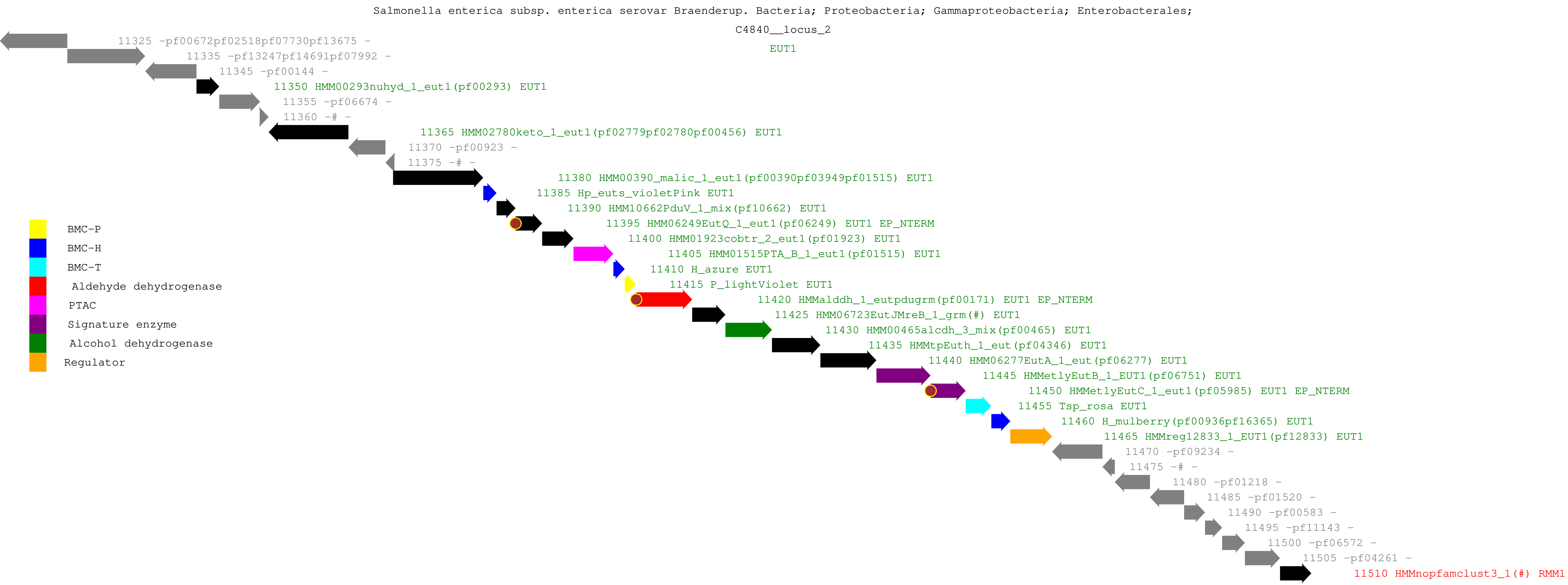
Salmonella enterica subsp. enterica serovar Braenderup. Bacteria; Proteobacteria; Gammaproteobacteria; Enterobacterales;
C4840__locus_2
EUT1
11325 -pf00672pf02518pf07730pf13675 -
11335 -pf13247pf14691pf07992 -
11345 -pf00144 -
11350 HMM00293nuhyd_1_eut1(pf00293) EUT1
11355 -pf06674 -
11360 -# -
11365 HMM02780keto_1_eut1(pf02779pf02780pf00456) EUT1
11370 -pf00923 -
11375 -# -
11380 HMM00390_malic_1_eut1(pf00390pf03949pf01515) EUT1
11385 Hp_euts_violetPink EUT1
11390 HMM10662PduV_1_mix(pf10662) EUT1
11395 HMM06249EutQ_1_eut1(pf06249) EUT1 EP_NTERM
11400 HMM01923cobtr_2_eut1(pf01923) EUT1
11405 HMM01515PTA_B_1_eut1(pf01515) EUT1
11410 H_azure EUT1
11415 P_lightViolet EUT1
11420 HMMalddh_1_eutpdugrm(pf00171) EUT1 EP_NTERM
11425 HMM06723EutJMreB_1_grm(#) EUT1
11430 HMM00465alcdh_3_mix(pf00465) EUT1
11435 HMMtpEuth_1_eut(pf04346) EUT1
11440 HMM06277EutA_1_eut(pf06277) EUT1
11445 HMMetlyEutB_1_EUT1(pf06751) EUT1
11450 HMMetlyEutC_1_eut1(pf05985) EUT1 EP_NTERM
11455 Tsp_rosa EUT1
11460 H_mulberry(pf00936pf16365) EUT1
11465 HMMreg12833_1_EUT1(pf12833) EUT1
11470 -pf09234 -
11475 -# -
11480 -pf01218 -
11485 -pf01520 -
11490 -pf00583 -
11495 -pf11143 -
11500 -pf06572 -
11505 -pf04261 -
11510 HMMnopfamclust3_1(#) RMM1
BMC-P
BMC-H
BMC-T
Aldehyde dehydrogenase
PTAC
Signature enzyme
Alcohol dehydrogenase
Regulator

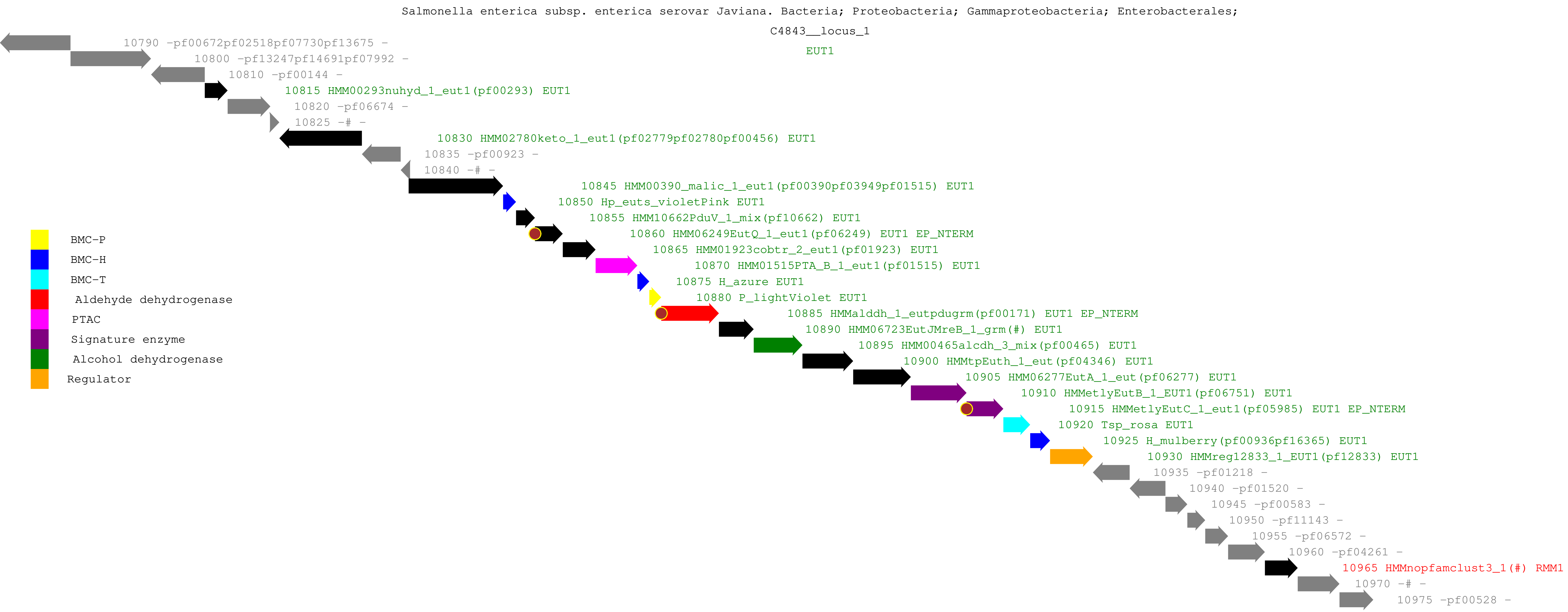

Salmonella enterica subsp. enterica serovar Javiana. Bacteria; Proteobacteria; Gammaproteobacteria; Enterobacterales;
C4843__locus_1
EUT1
10790 -pf00672pf02518pf07730pf13675 -
10800 -pf13247pf14691pf07992 -
10810 -pf00144 -
10815 HMM00293nuhyd_1_eut1(pf00293) EUT1
10820 -pf06674 -
10825 -# -
10830 HMM02780keto_1_eut1(pf02779pf02780pf00456) EUT1
10835 -pf00923 -
10840 -# -
10845 HMM00390_malic_1_eut1(pf00390pf03949pf01515) EUT1
10850 Hp_euts_violetPink EUT1
10855 HMM10662PduV_1_mix(pf10662) EUT1
10860 HMM06249EutQ_1_eut1(pf06249) EUT1 EP_NTERM
10865 HMM01923cobtr_2_eut1(pf01923) EUT1
10870 HMM01515PTA_B_1_eut1(pf01515) EUT1
10875 H_azure EUT1
10880 P_lightViolet EUT1
10885 HMMalddh_1_eutpdugrm(pf00171) EUT1 EP_NTERM
10890 HMM06723EutJMreB_1_grm(#) EUT1
10895 HMM00465alcdh_3_mix(pf00465) EUT1
10900 HMMtpEuth_1_eut(pf04346) EUT1
10905 HMM06277EutA_1_eut(pf06277) EUT1
10910 HMMetlyEutB_1_EUT1(pf06751) EUT1
10915 HMMetlyEutC_1_eut1(pf05985) EUT1 EP_NTERM
10920 Tsp_rosa EUT1
10925 H_mulberry(pf00936pf16365) EUT1
10930 HMMreg12833_1_EUT1(pf12833) EUT1
10935 -pf01218 -
10940 -pf01520 -
10945 -pf00583 -
10950 -pf11143 -
10955 -pf06572 -
10960 -pf04261 -
10965 HMMnopfamclust3_1(#) RMM1
10970 -# -
10975 -pf00528 -
BMC-P
BMC-H
BMC-T
Aldehyde dehydrogenase
PTAC
Signature enzyme
Alcohol dehydrogenase
Regulator

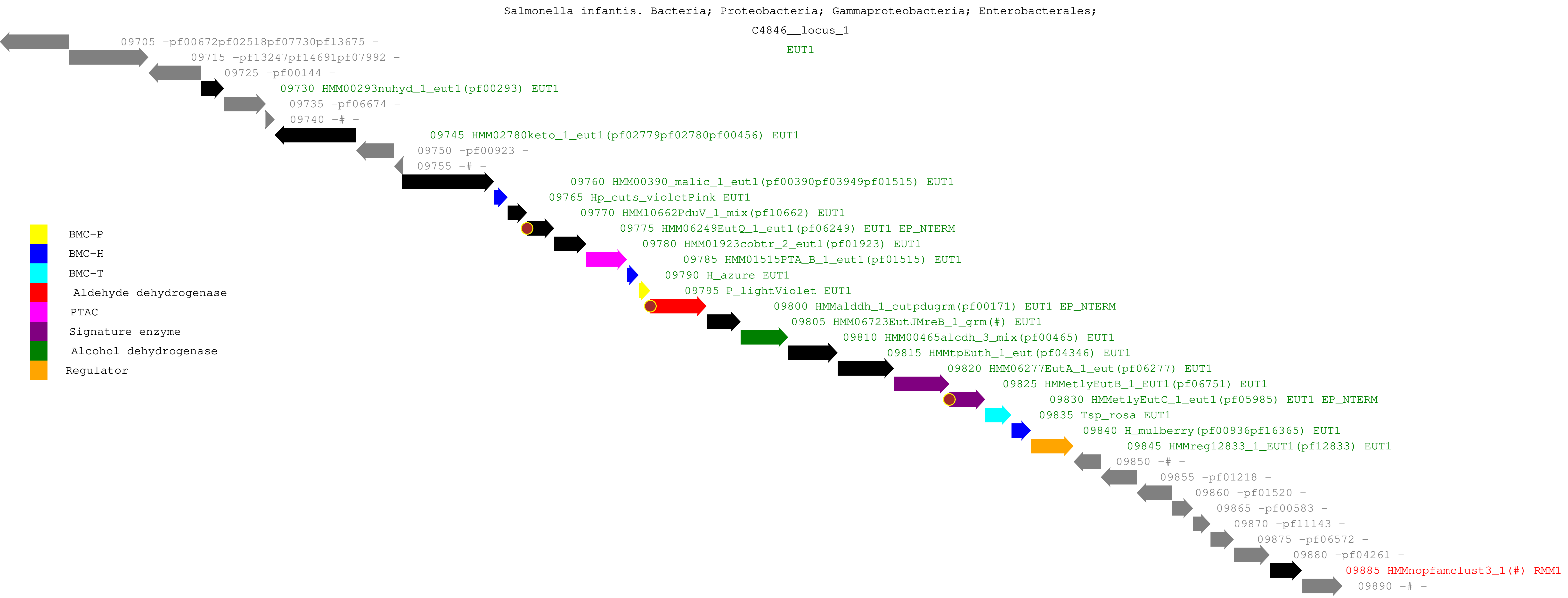
Salmonella infantis. Bacteria; Proteobacteria; Gammaproteobacteria; Enterobacterales;
C4846__locus_1
EUT1
09705 -pf00672pf02518pf07730pf13675 -
09715 -pf13247pf14691pf07992 -
09725 -pf00144 -
09730 HMM00293nuhyd_1_eut1(pf00293) EUT1
09735 -pf06674 -
09740 -# -
09745 HMM02780keto_1_eut1(pf02779pf02780pf00456) EUT1
09750 -pf00923 -
09755 -# -
09760 HMM00390_malic_1_eut1(pf00390pf03949pf01515) EUT1
09765 Hp_euts_violetPink EUT1
09770 HMM10662PduV_1_mix(pf10662) EUT1
09775 HMM06249EutQ_1_eut1(pf06249) EUT1 EP_NTERM
09780 HMM01923cobtr_2_eut1(pf01923) EUT1
09785 HMM01515PTA_B_1_eut1(pf01515) EUT1
09790 H_azure EUT1
09795 P_lightViolet EUT1
09800 HMMaldh_1_eutpdugrm(pf00171) EUT1 EP_NTERM
09805 HMM06723EutJMreB_1_grm(#) EUT1
09810 HMM00465alcdh_3_mix(pf00465) EUT1
09815 HMMtpEuth_1_eut(pf04346) EUT1
09820 HMM06277EutA_1_eut(pf06277) EUT1
09825 HMMetlyEutB_1_EUT1(pf06751) EUT1
09830 HMMetlyEutC_1_eut1(pf05985) EUT1 EP_NTERM
09835 Tsp_rosa EUT1
09840 H_mulberry(pf00936pf16365) EUT1
09845 HMMreg12833_1_EUT1(pf12833) EUT1
09850 -# -
09855 -pf01218 -
09860 -pf01520 -
09865 -pf00583 -
09870 -pf11143 -
09875 -pf06572 -
09880 -pf04261 -
09885 HMMnopfamclust3_1(#) RMM1
09890 -# -
BMC-P
BMC-H
BMC-T
Aldehyde dehydrogenase
PTAC
Signature enzyme
Alcohol dehydrogenase
Regulator

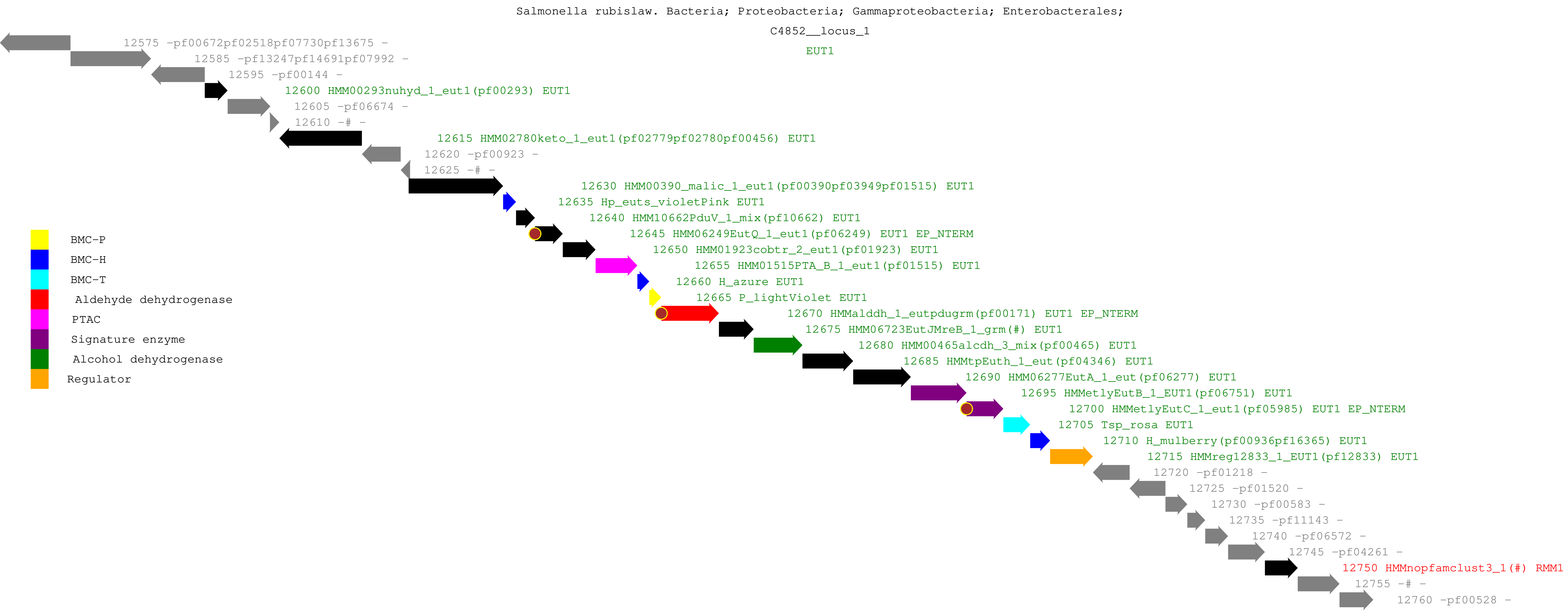

Salmonella rubislaw. Bacteria; Proteobacteria; Gammaproteobacteria; Enterobacterales;
C4852__locus_1
EUT1
12575 -pf00672pf02518pf07730pf13675 -
12585 -pf13247pf14691pf07992 -
12595 -pf00144 -
12600 HMM00293nuhyd_1_eut1(pf00293) EUT1
12605 -pf06674 -
12610 -# -
12615 HMM02780keto_1_eut1(pf02779pf02780pf00456) EUT1
12620 -pf00923 -
12625 -# -
12630 HMM00390_malic_1_eut1(pf00390pf03949pf01515) EUT1
12635 Hp_euts_violetPink EUT1
12640 HMM10662PduV_1_mix(pf10662) EUT1
12645 HMM06249EutQ_1_eut1(pf06249) EUT1 EP_NTERM
12650 HMM01923cobtr_2_eut1(pf01923) EUT1
12655 HMM01515PTA_B_1_eut1(pf01515) EUT1
12660 H_azure EUT1
12665 P_lightViolet EUT1
12670 HMMalddh_1_eutpdugrm(pf00171) EUT1 EP_NTERM
12675 HMM06723EutJMreB_1_grm(#) EUT1
12680 HMM00465alcdh_3_mix(pf00465) EUT1
12685 HMMtpEuth_1_eut(pf04346) EUT1
12690 HMM06277EutA_1_eut(pf06277) EUT1
12695 HMMetlyEutB_1_EUT1(pf06751) EUT1
12700 HMMetlyEutC_1_eut1(pf05985) EUT1 EP_NTERM
12705 Tsp_rosa EUT1
12710 H_mulberry(pf00936pf16365) EUT1
12715 HMMreg12833_1_EUT1(pf12833) EUT1
12720 -pf01218 -
12725 -pf01520 -
12730 -pf00583 -
12735 -pf11143 -
12740 -pf06572 -
12745 -pf04261 -
12750 HMMnopfamclust3_1(#) RMM1
12755 -# -
12760 -pf00528 -
BMC-P
BMC-H
BMC-T
Aldehyde dehydrogenase
PTAC
Signature enzyme
Alcohol dehydrogenase
Regulator

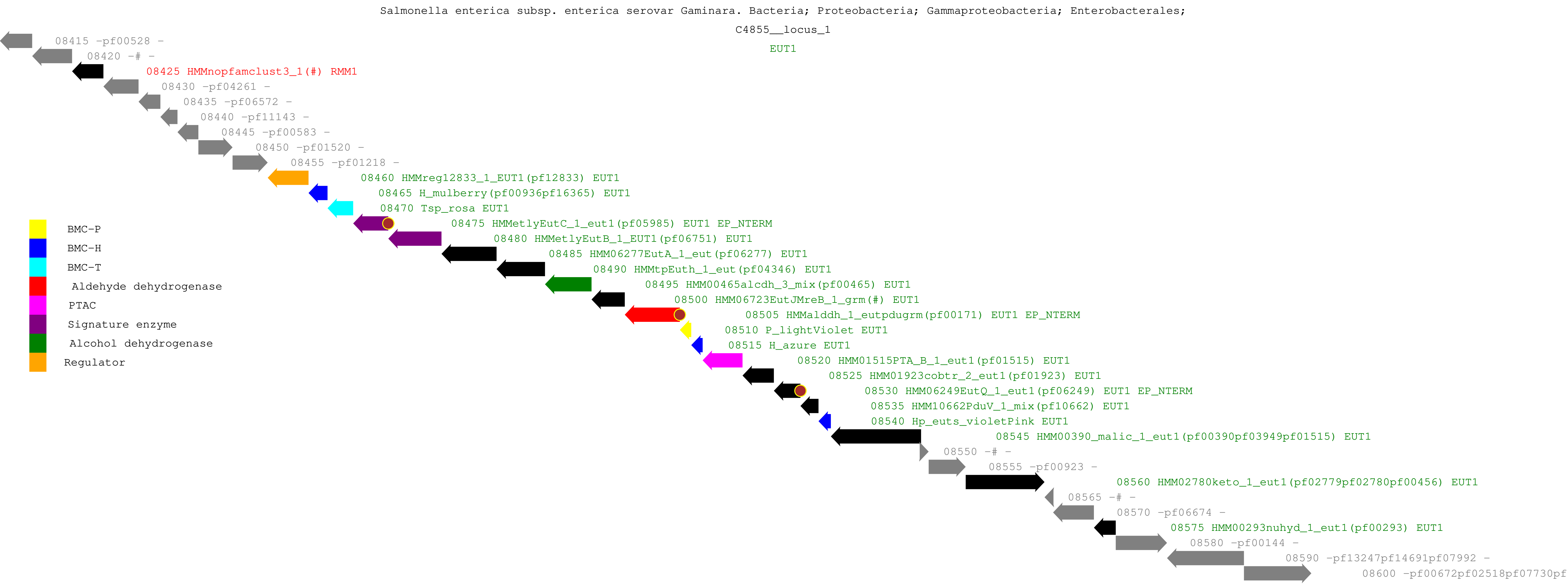

Salmonella enterica subsp. enterica serovar Gaminara. Bacteria; Proteobacteria; Gammaproteobacteria; Enterobacterales;
C4855__locus_1
EUT1
08415 -pf00528 -
08420 -# -
08425 HMMnopfamclust3_1(#) RMM1
08430 -pf04261 -
08435 -pf06572 -
08440 -pf11143 -
08445 -pf00583 -
08450 -pf01520 -
08455 -pf01218 -
08460 HMMreg12833_1_EUT1(pf12833) EUT1
08465 H_mulberry(pf00936pf16365) EUT1
08470 Tsp_rosa EUT1
08475 HMMetlyEutC_1_eut1(pf05985) EUT1 EP_NTERM
08480 HMMetlyEutB_1_EUT1(pf06751) EUT1
08485 HMM06277EutA_1_eut(pf06277) EUT1
08490 HMMtpEuth_1_eut(pf04346) EUT1
08495 HMM00465alcdh_3_mix(pf00465) EUT1
08500 HMM06723EutJMreB_1_grm(#) EUT1
08505 HMMalddh_1_eutpdugrm(pf00171) EUT1 EP_NTERM
08510 P_lightViolet EUT1
08515 H_azure EUT1
08520 HMM01515PTA_B_1_eut1(pf01515) EUT1
08525 HMM01923cobtr_2_eut1(pf01923) EUT1
08530 HMM06249EutQ_1_eut1(pf06249) EUT1 EP_NTERM
08535 HMM10662PduV_1_mix(pf10662) EUT1
08540 Hp_euts_violetPink EUT1
08545 HMM00390_malic_1_eut1(pf00390pf03949pf01515) EUT1
08550 -# -
08555 -pf00923 -
08560 HMM02780keto_1_eut1(pf02779pf02780pf00456) EUT1
08565 -# -
08570 -pf06674 -
08575 HMM00293nuhyd_1_eut1(pf00293) EUT1
08580 -pf00144 -
08590 -pf13247pf14691pf07992 -
08600 -pf00672pf02518pf07730pf
BMC-P
BMC-H
BMC-T
Aldehyde dehydrogenase
PTAC
Signature enzyme
Alcohol dehydrogenase
Regulator

Salmonella enterica subsp. enterica serovar 4,12:i:-. Bacteria; Proteobacteria; Gammaproteobacteria; Enterobacterales;

C4866__locus_1

EUT1

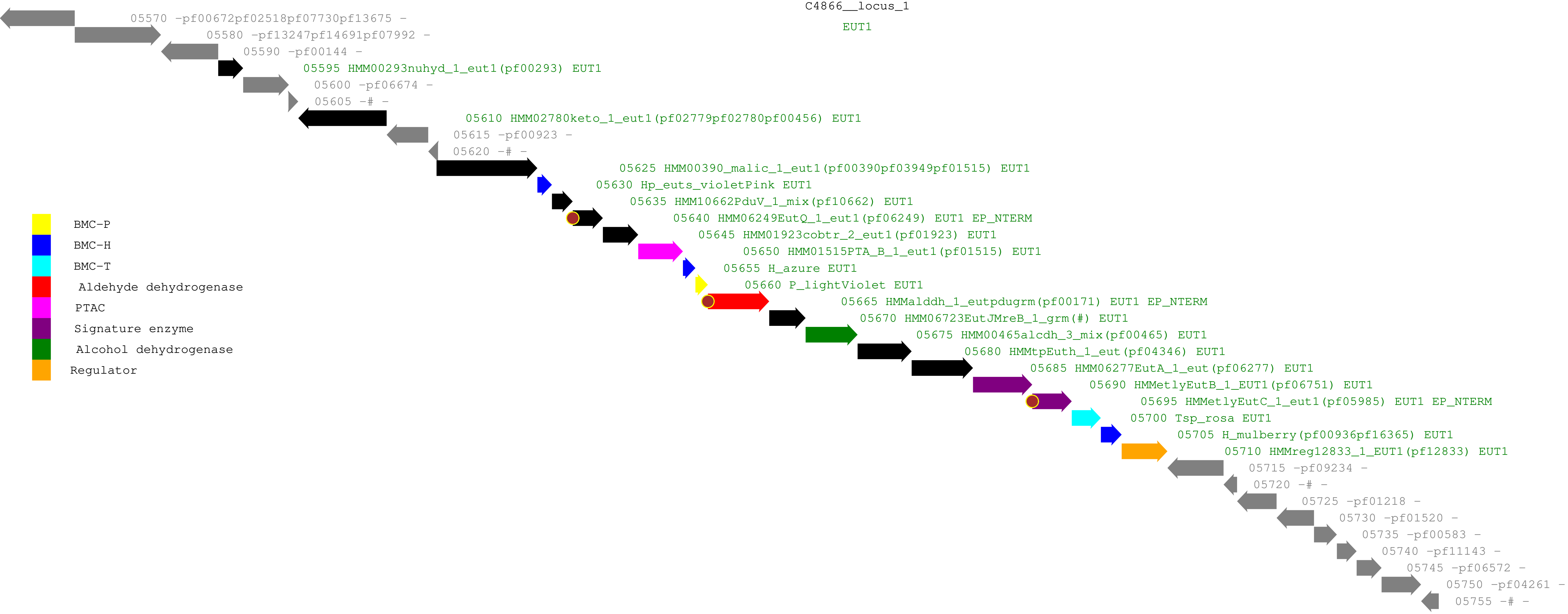

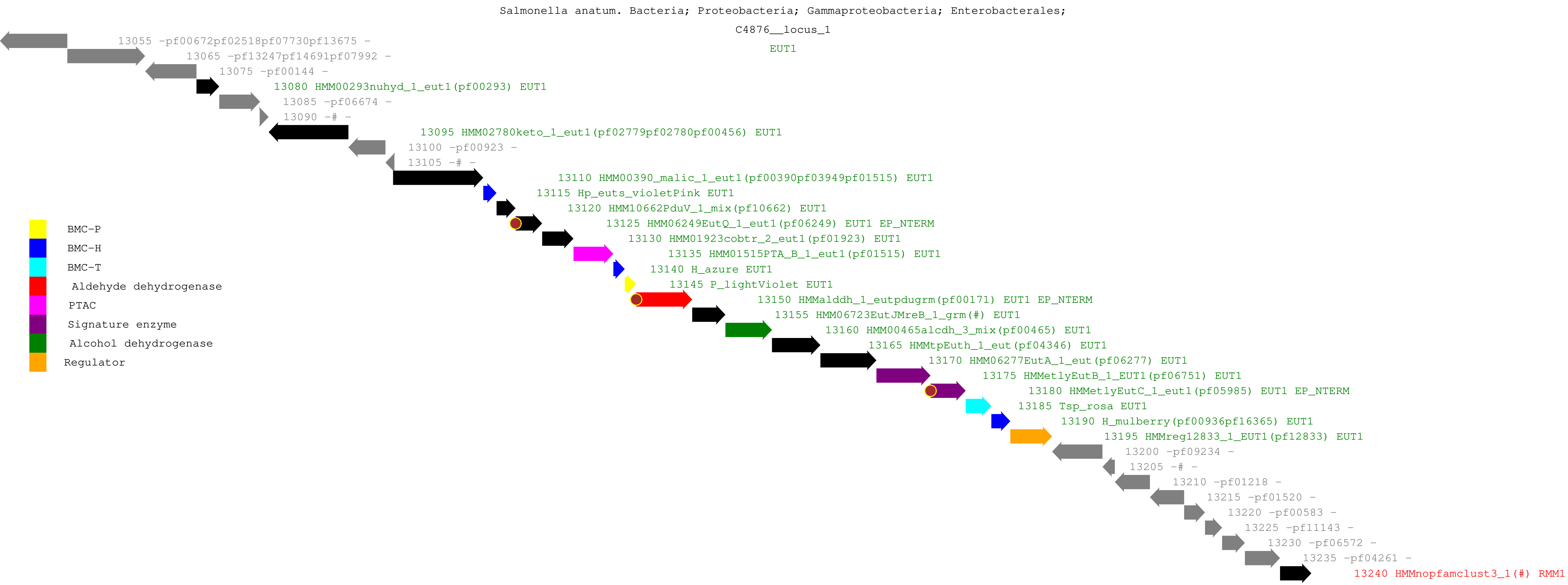
Salmonella anatum. Bacteria; Proteobacteria; Gammaproteobacteria; Enterobacterales;
C4876__locus_1
EUT1
13055 -pf00672pf02518pf07730pf13675 -
13065 -pf13247pf14691pf07992 -
13075 -pf00144 -
13080 HMM00293nuhyd_1_eut1(pf00293) EUT1
13085 -pf06674 -
13090 -# -
13095 HMM02780keto_1_eut1(pf02779pf02780pf00456) EUT1
13100 -pf00923 -
13105 -# -
13110 HMM00390_malic_1_eut1(pf00390pf03949pf01515) EUT1
13115 Hp_euts_violetPink EUT1
13120 HMM10662PduV_1_mix(pf10662) EUT1
13125 HMM06249EutQ_1_eut1(pf06249) EUT1 EP_NTERM
13130 HMM01923cobtr_2_eut1(pf01923) EUT1
13135 HMM01515PTA_B_1_eut1(pf01515) EUT1
13140 H_azure EUT1
13145 P_lightViolet EUT1
13150 HMMalddh_1_eutpdugrm(pf00171) EUT1 EP_NTERM
13155 HMM06723EutJMreB_1_grm(#) EUT1
13160 HMM00465alcdh_3_mix(pf00465) EUT1
13165 HMMtpEuth_1_eut(pf04346) EUT1
13170 HMM06277EutA_1_eut(pf06277) EUT1
13175 HMMetlyEutB_1_EUT1(pf06751) EUT1
13180 HMMetlyEutC_1_eut1(pf05985) EUT1 EP_NTERM
13185 Tsp_rosa EUT1
13190 H_mulberry(pf00936pf16365) EUT1
13195 HMMreg12833_1_EUT1(pf12833) EUT1
13200 -pf09234 -
13205 -# -
13210 -pf01218 -
13215 -pf01520 -
13220 -pf00583 -
13225 -pf11143 -
13230 -pf06572 -
13235 -pf04261 -
13240 HMMnopfamclust3_1(#) RMM1
BMC-P
BMC-H
BMC-T
Aldehyde dehydrogenase
PTAC
Signature enzyme
Alcohol dehydrogenase
Regulator

Salmonella enterica subsp. enterica serovar Kentucky. Bacteria; Proteobacteria; Gammaproteobacteria; Enterobacterales;

C4878__locus_2

EUT1

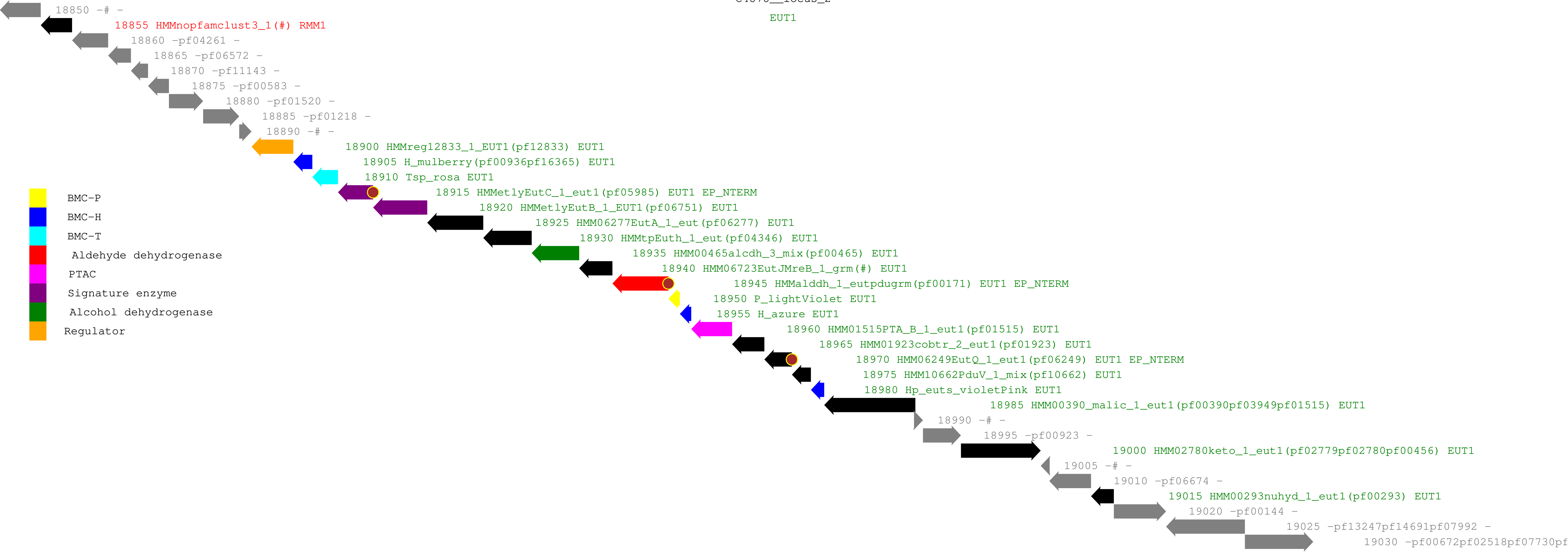

Escherichia marmotae. Bacteria; Proteobacteria; Gammaproteobacteria; Enterobacterales;

C4A13__locus_1_main_sat

EUT1

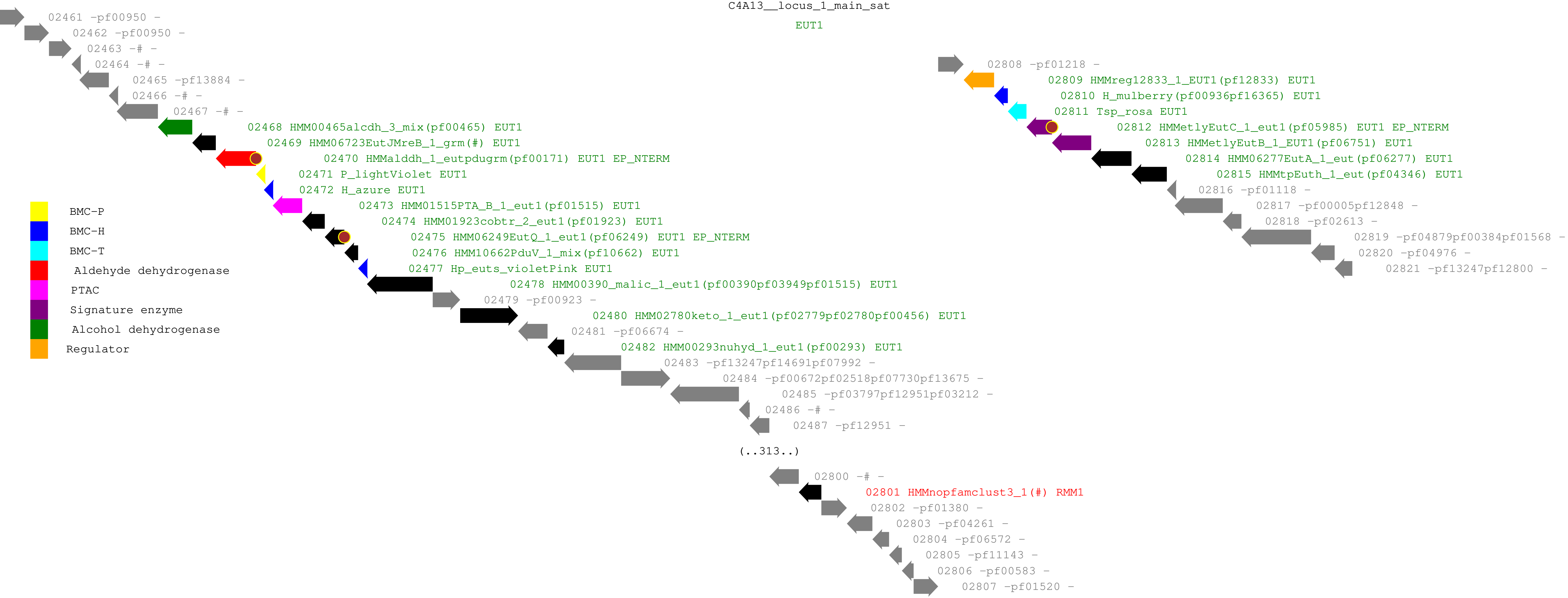

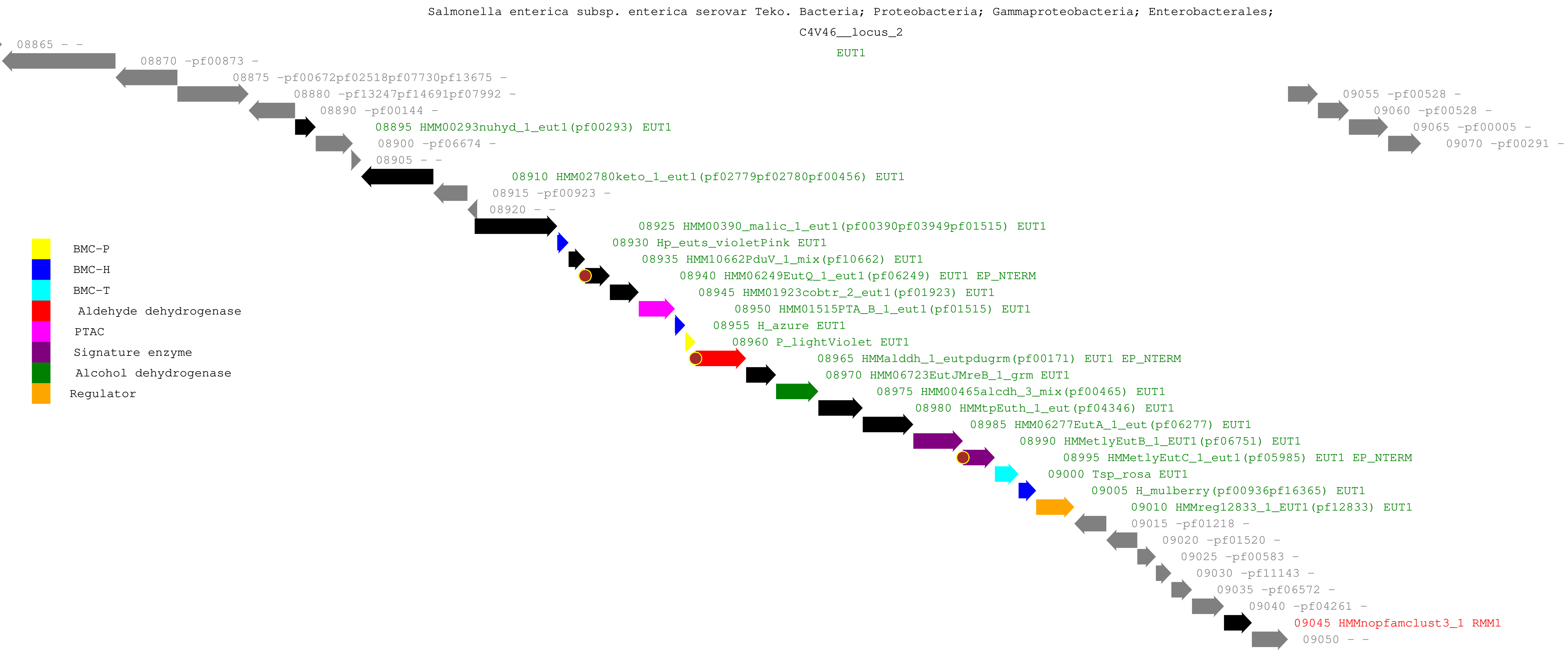
Salmonella enterica subsp. enterica serovar Teko. Bacteria; Proteobacteria; Gammaproteobacteria; Enterobacterales;
C4V46__locus_2
EUT1
08865 - -
08870 -pf00873 -
08875 -pf00672pf02518pf07730pf13675 -
08880 -pf13247pf14691pf07992 -
08890 -pf00144 -
08895 HMM00293nuhyd_1_eut1(pf00293) EUT1
08900 -pf06674 -
08905 - -
08910 HMM02780keto_1_eut1(pf02779pf02780pf00456) EUT1
08915 -pf00923 -
08920 - -
08925 HMM00390_malic_1_eut1(pf00390pf03949pf01515) EUT1
08930 Hp_euts_violetPink EUT1
08935 HMM10662PduV_1_mix(pf10662) EUT1
08940 HMM06249EutQ_1_eut1(pf06249) EUT1 EP_NTERM
08945 HMM01923cobtr_2_eut1(pf01923) EUT1
08950 HMM01515PTA_B_1_eut1(pf01515) EUT1
08955 H_azure EUT1
08960 P_lightViolet EUT1
08965 HMMalddh_1_eutpdugrm(pf00171) EUT1 EP_NTERM
08970 HMM06723EutJMreB_1_grm EUT1
08975 HMM00465alcdh_3_mix(pf00465) EUT1
08980 HMMtpEuth_1_eut(pf04346) EUT1
08985 HMM06277EutA_1_eut(pf06277) EUT1
08990 HMMetlyEutB_1_EUT1(pf06751) EUT1
08995 HMMetlyEutC_1_eut1(pf05985) EUT1 EP_NTERM
09000 Tsp_rosa EUT1
09005 H_mulberry(pf00936pf16365) EUT1
09010 HMMreg12833_1_EUT1(pf12833) EUT1
09015 -pf01218 -
09020 -pf01520 -
09025 -pf00583 -
09030 -pf11143 -
09035 -pf06572 -
09040 -pf04261 -
09045 HMMnopfamclust3_1 RMM1
09050 - -
09055 -pf00528 -
09060 -pf00528 -
09065 -pf00005 -
09070 -pf00291 -
BMC-P
BMC-H
BMC-T
Aldehyde dehydrogenase
PTAC
Signature enzyme
Alcohol dehydrogenase
Regulator

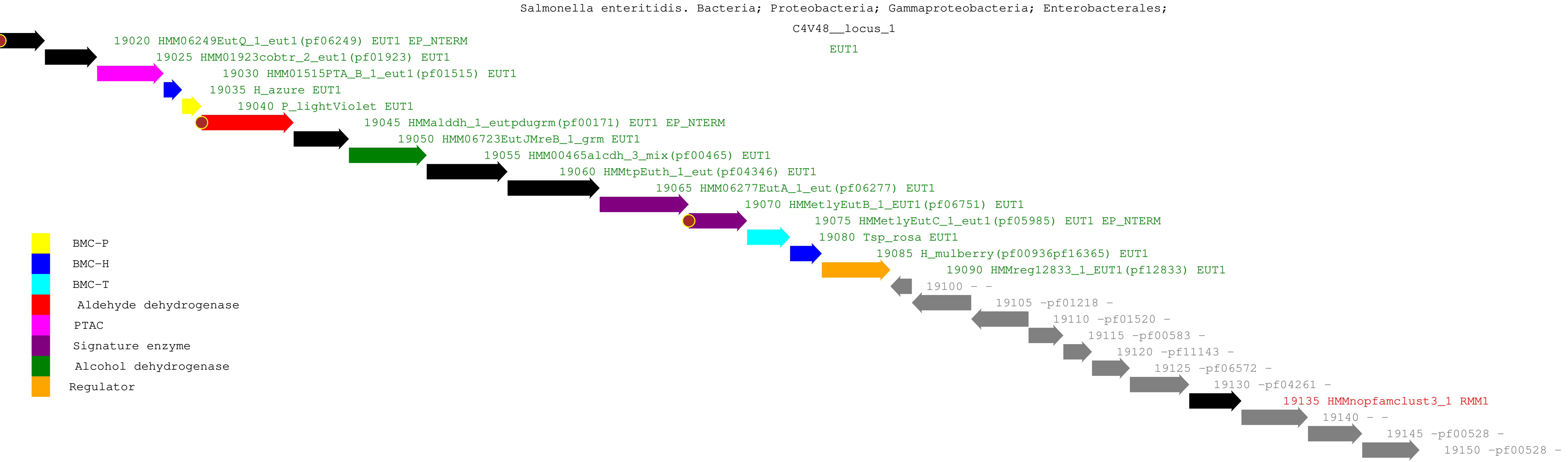

Salmonella enteritidis. Bacteria; Proteobacteria; Gammaproteobacteria; Enterobacterales;
C4V48__locus_1
EUT1
19020 HMM06249EutQ_1_eut1(pf06249) EUT1 EP_NTERM
19025 HMM01923cobtr_2_eut1(pf01923) EUT1
19030 HMM01515PTA_B_1_eut1(pf01515) EUT1
19035 H_azure EUT1
19040 P_lightViolet EUT1
19045 HMMalddh_1_eutpdugrm(pf00171) EUT1 EP_NTERM
19050 HMM06723EutJMreB_1_grm EUT1
19055 HMM00465alcdh_3_mix(pf00465) EUT1
19060 HMMtpEuth_1_eut(pf04346) EUT1
19065 HMM06277EutA_1_eut(pf06277) EUT1
19070 HMMetlyEutB_1_EUT1(pf06751) EUT1
19075 HMMetlyEutC_1_eut1(pf05985) EUT1 EP_NTERM
19080 Tsp_rosa EUT1
19085 H_mulberry(pf00936pf16365) EUT1
19090 HMMreg12833_1_EUT1(pf12833) EUT1
19100 - -
19105 -pf01218 -
19110 -pf01520 -
19115 -pf00583 -
19120 -pf11143 -
19125 -pf06572 -
19130 -pf04261 -
19135 HMMnopfamclust3_1 RMM1
19140 - -
19145 -pf00528 -
19150 -pf00528 -
BMC-P
BMC-H
BMC-T
Aldehyde dehydrogenase
PTAC
Signature enzyme
Alcohol dehydrogenase
Regulator

Klebsiella pneumoniae subsp. pneumoniae. Bacteria; Proteobacteria; Gammaproteobacteria; Enterobacterales;

C4Y59__locus_2

EUT1

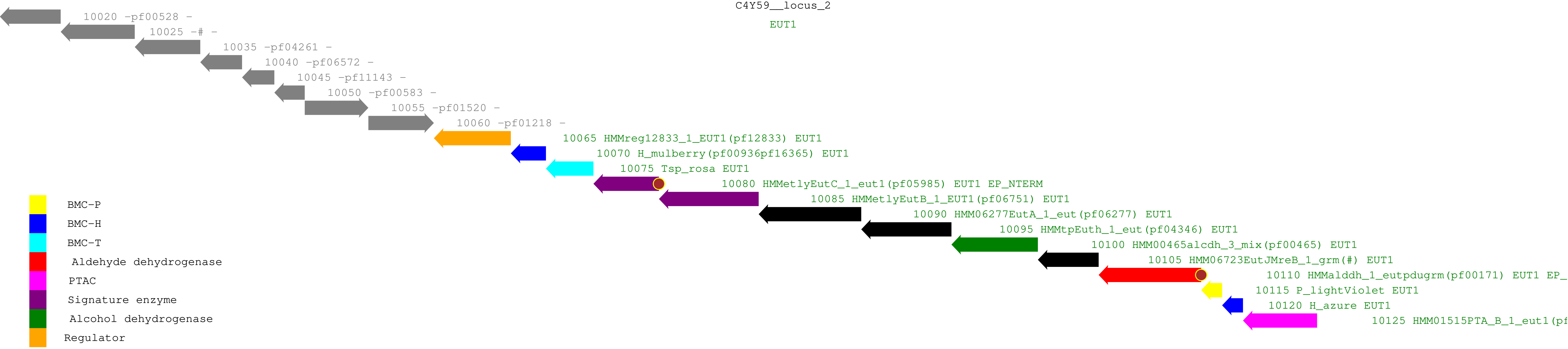

Klebsiella pneumoniae subsp. pneumoniae. Bacteria; Proteobacteria; Gammaproteobacteria; Enterobacterales;

C4Y86__locus_1

EUT1

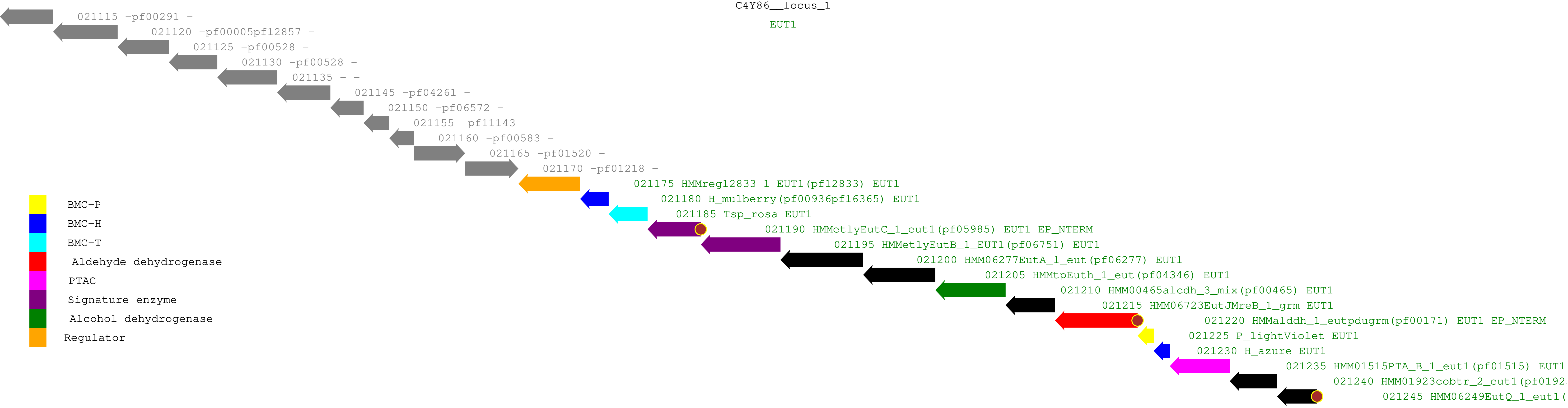

Klebsiella pneumoniae subsp. pneumoniae. Bacteria; Proteobacteria; Gammaproteobacteria; Enterobacterales;

C4Z38__locus_2

EUT1

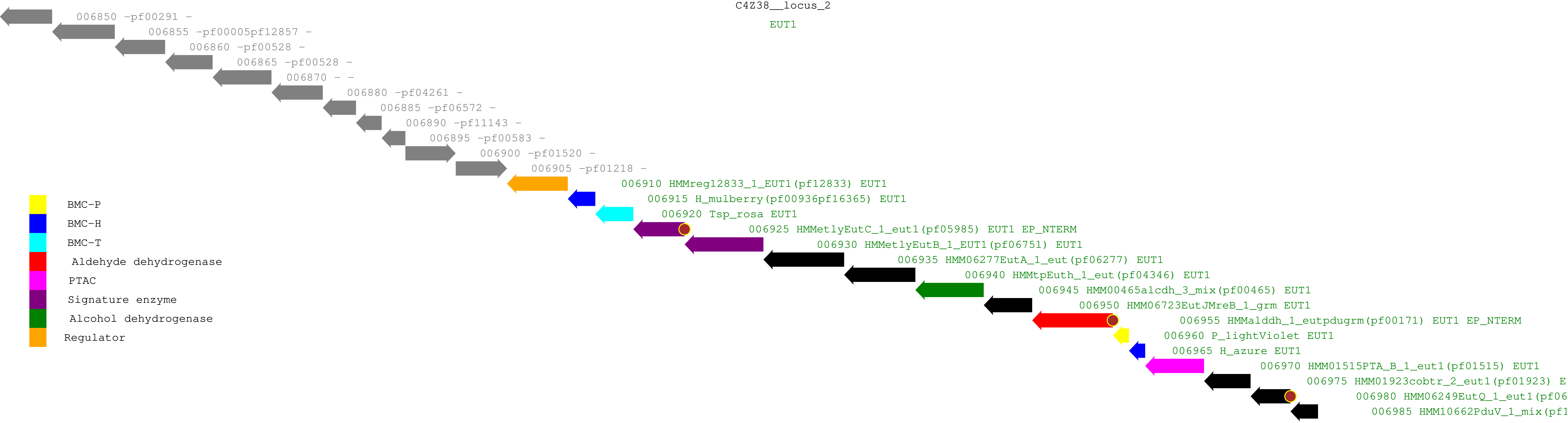

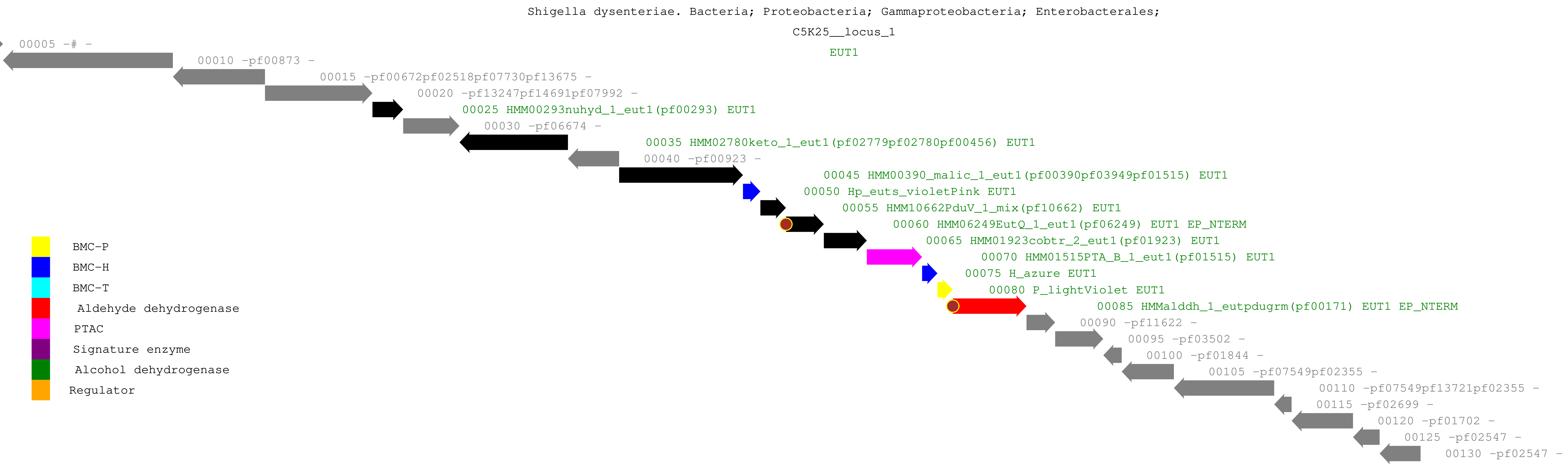
Shigella dysenteriae. Bacteria; Proteobacteria; Gammaproteobacteria; Enterobacterales;
C5K25__locus_1
EUT1
00005 -# -
00010 -pf00873 -
00015 -pf00672pf02518pf07730pf13675 -
00020 -pf13247pf14691pf07992 -
00025 HMM00293nuhyd_1_eut1(pf00293) EUT1
00030 -pf06674 -
00035 HMM02780keto_1_eut1(pf02779pf02780pf00456) EUT1
00040 -pf00923 -
00045 HMM00390_malic_1_eut1(pf00390pf03949pf01515) EUT1
00050 Hp_euts_violetPink EUT1
00055 HMM10662PduV_1_mix(pf10662) EUT1
00060 HMM06249EutQ_1_eut1(pf06249) EUT1 EP_NTERM
00065 HMM01923cobtr_2_eut1(pf01923) EUT1
00070 HMM01515PTA_B_1_eut1(pf01515) EUT1
00075 H_azure EUT1
00080 P_lightViolet EUT1
00085 HMMalddh_1_eutpdugrm(pf00171) EUT1 EP_NTERM
00090 -pf11622 -
00095 -pf03502 -
00100 -pf01844 -
00105 -pf07549pf02355 -
00110 -pf07549pf13721pf02355 -
00115 -pf02699 -
00120 -pf01702 -
00125 -pf02547 -
00130 -pf02547 -
BMC-P
BMC-H
BMC-T
Aldehyde dehydrogenase
PTAC
Signature enzyme
Alcohol dehydrogenase
Regulator

Salmonella enterica subsp. enterica serovar Concord. Bacteria; Proteobacteria; Gammaproteobacteria; Enterobacterales;

C6651__locus_1

EUT1

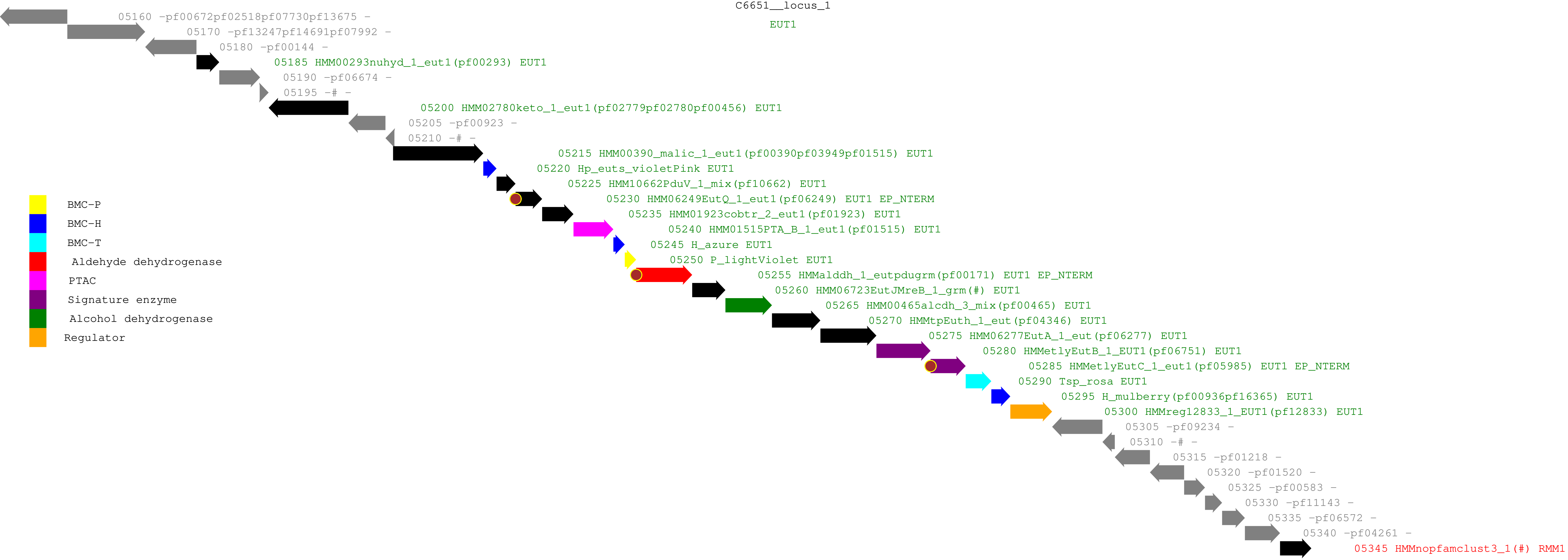

Thauera linaloolentis 47Lol = DSM 12138. Bacteria; Proteobacteria; Betaproteobacteria; Rhodocyclales;

C666__locus_1_main_sat

EUT1

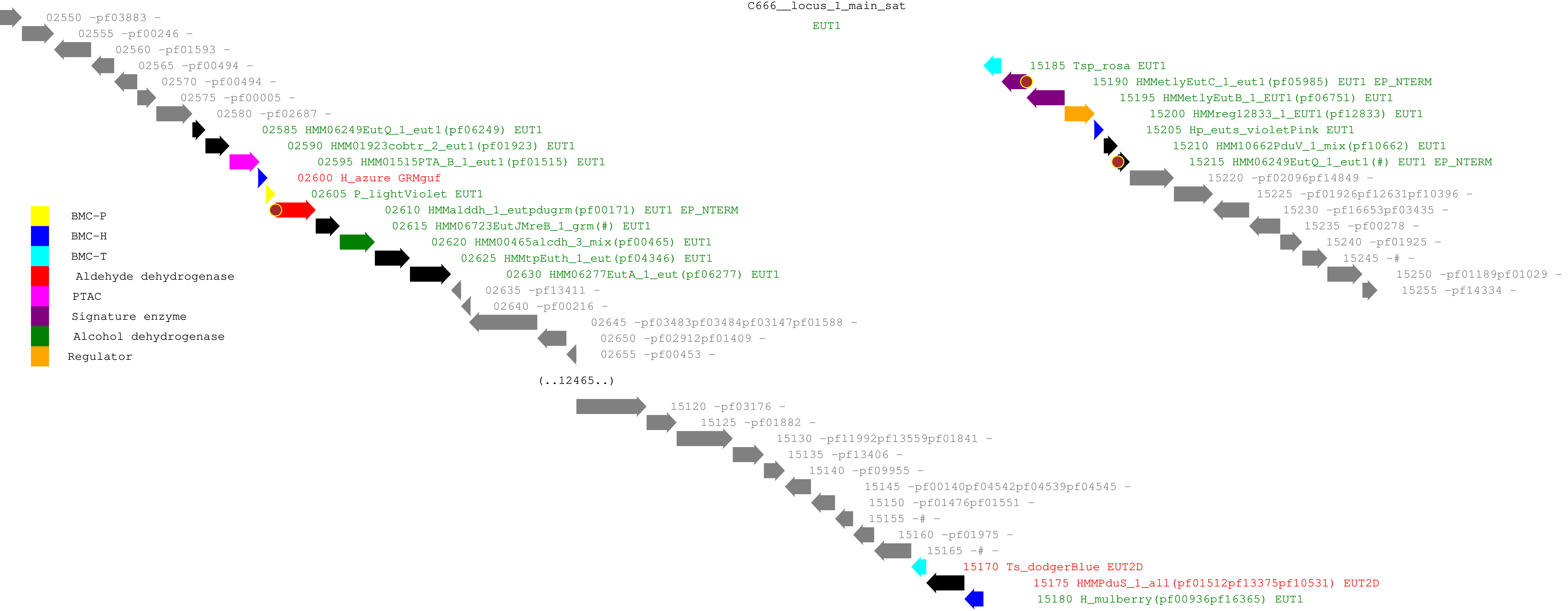

Escherichia coli. Bacteria; Proteobacteria; Gammaproteobacteria; Enterobacterales;

C6985__locus_2

EUT1

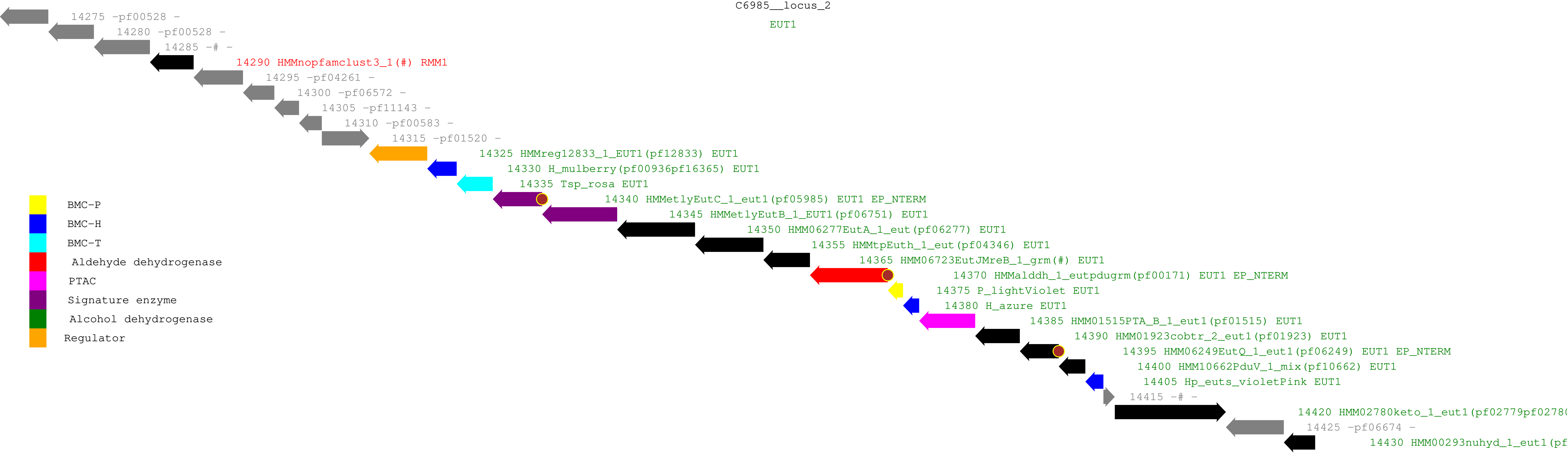

Escherichia coli. Bacteria; Proteobacteria; Gammaproteobacteria; Enterobacterales;

C6B13__locus_1

EUT1

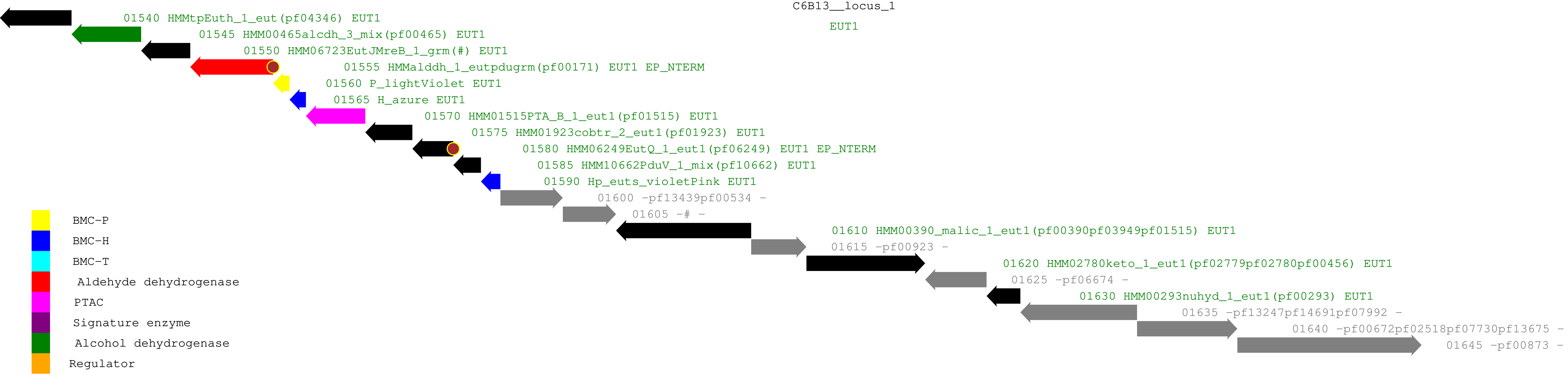

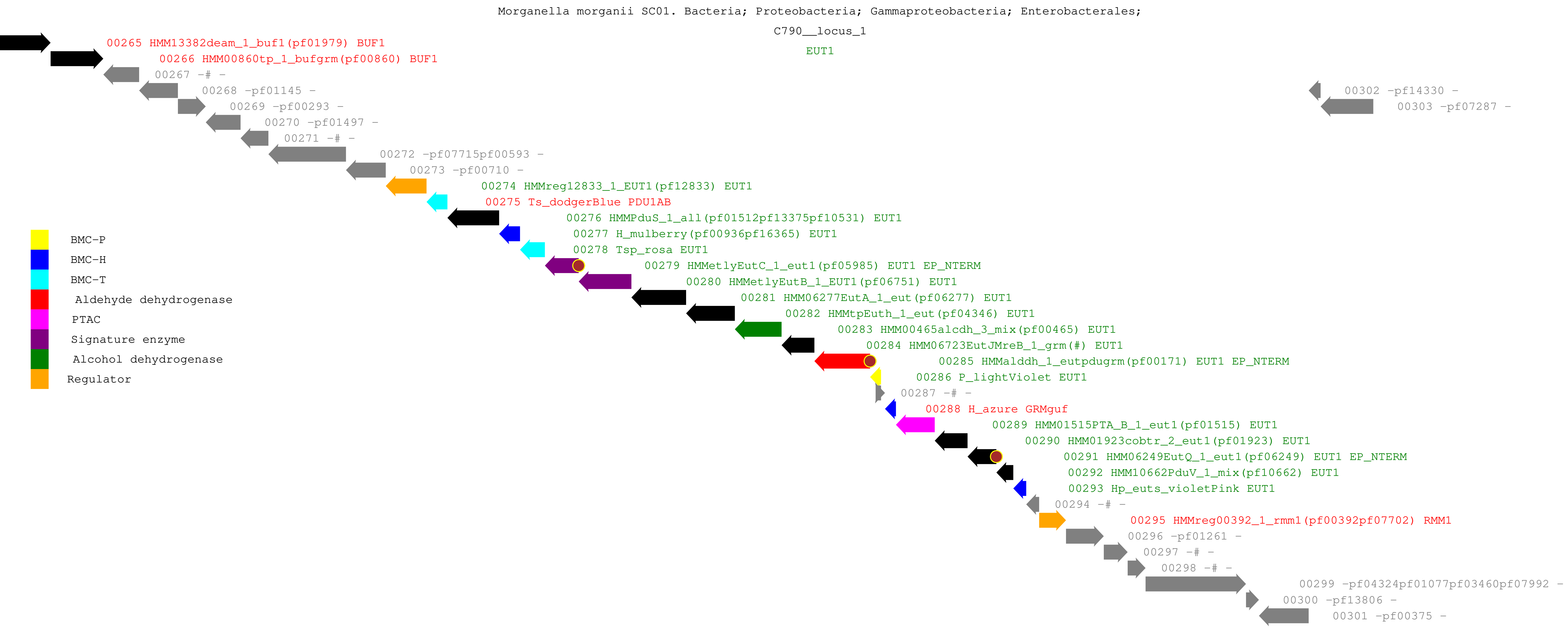

Morganella morganii SC01. Bacteria; Proteobacteria; Gammaproteobacteria; Enterobacterales;
C790__locus_1
EUT1
00265 HMM13382deam_1_buf1(pf01979) BUF1
00266 HMM00860tp_1_bufgrm(pf00860) BUF1
00267 -# -
00268 -pf01145 -
00269 -pf00293 -
00270 -pf01497 -
00271 -# -
00272 -pf07715pf00593 -
00273 -pf00710 -
00274 HMMreg12833_1_EUT1(pf12833) EUT1
00275 Ts_dodgerBlue PDU1AB
00276 HMMPduS_1_all(pf01512pf13375pf10531) EUT1
00277 H_mulberry(pf00936pf16365) EUT1
00278 Tsp_rosa EUT1
00279 HMMetlyEutC_1_eut1(pf05985) EUT1 EP_NTERM
00280 HMMetlyEutB_1_EUT1(pf06751) EUT1
00281 HMM06277EutA_1_eut(pf06277) EUT1
00282 HMMtpEuth_1_eut(pf04346) EUT1
00283 HMM00465alcdh_3_mix(pf00465) EUT1
00284 HMM06723EutJMreB_1_grm(#) EUT1
00285 HMMalddh_1_eutpdugrm(pf00171) EUT1 EP_NTERM
00286 P_lightViolet EUT1
00287 -# -
00288 H_azure GRMguf
00289 HMM01515PTA_B_1_eut1(pf01515) EUT1
00290 HMM01923cobtr_2_eut1(pf01923) EUT1
00291 HMM06249EutQ_1_eut1(pf06249) EUT1 EP_NTERM
00292 HMM10662PduV_1_mix(pf10662) EUT1
00293 Hp_euts_violetPink EUT1
00294 -# -
00295 HMMreg00392_1_rmm1(pf00392pf07702) RMM1
00296 -pf01261 -
00297 -# -
00298 -# -
00299 -pf04324pf01077pf03460pf07992 -
00300 -pf13806 -
00301 -pf00375 -
00302 -pf14330 -
00303 -pf07287 -
BMC-P
BMC-H
BMC-T
Aldehyde dehydrogenase
PTAC
Signature enzyme
Alcohol dehydrogenase
Regulator

Escherichia sp. 4726-5. Bacteria; Proteobacteria; Gammaproteobacteria; Enterobacterales;

C7B04__locus_2

EUT1

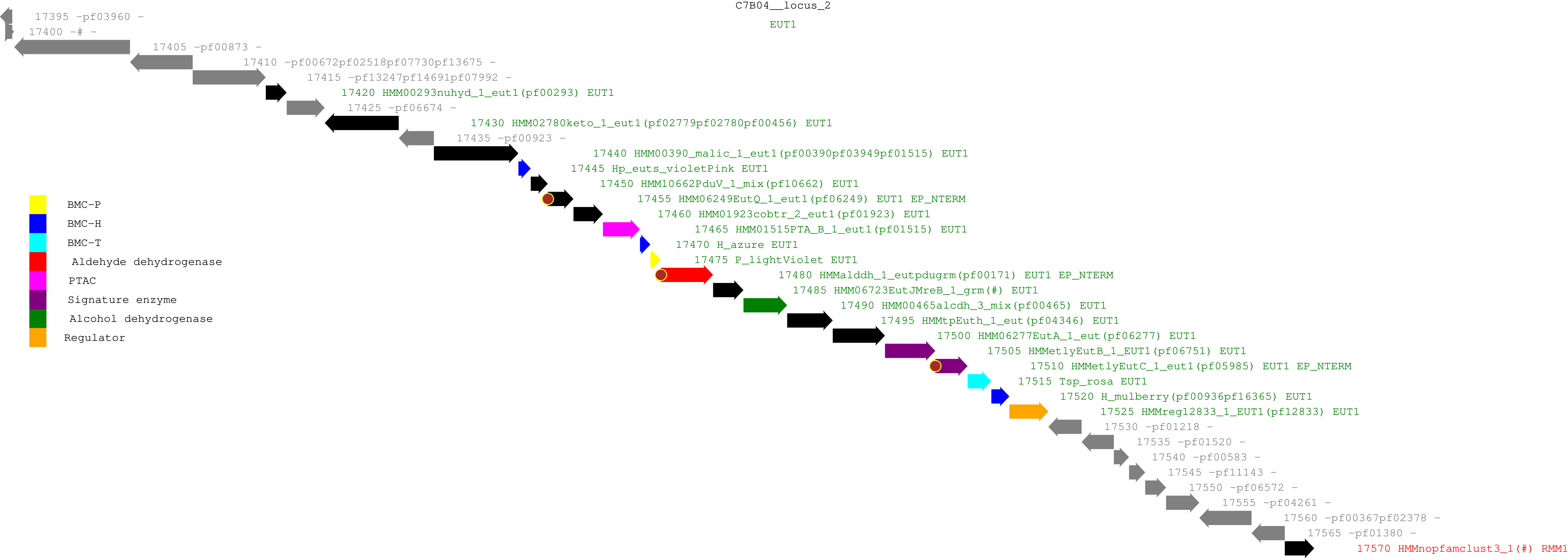

Escherichia albertii. Bacteria; Proteobacteria; Gammaproteobacteria; Enterobacterales;

C7B09__locus_2

EUT1

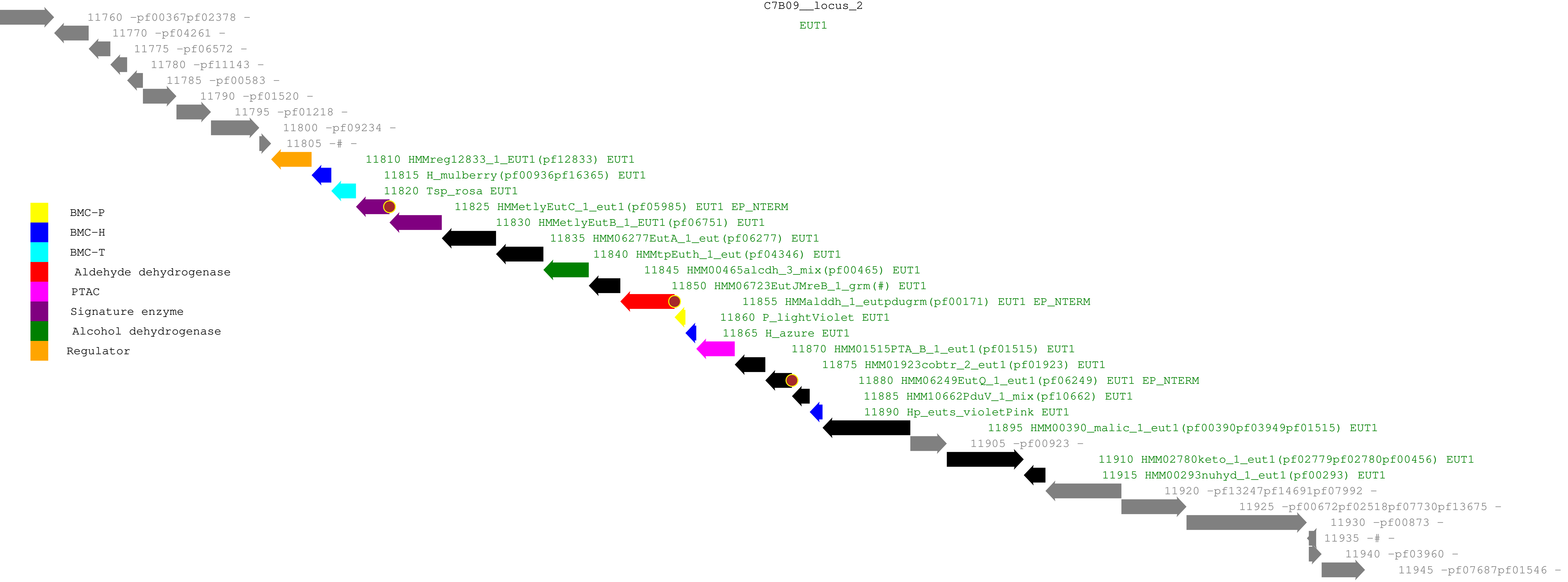

Salmonella enterica subsp. enterica serovar 4,5,12:b:-. Bacteria; Proteobacteria; Gammaproteobacteria; Enterobacterales;

C7S56__locus_2

EUT1

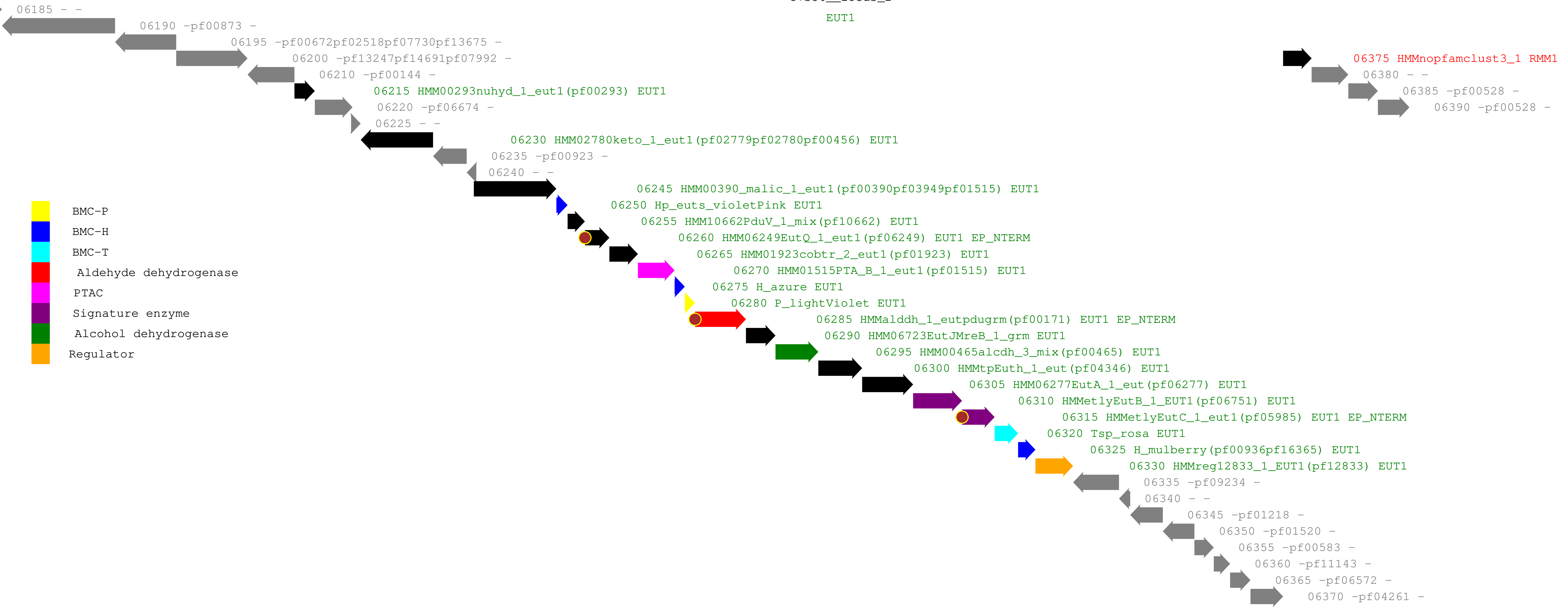

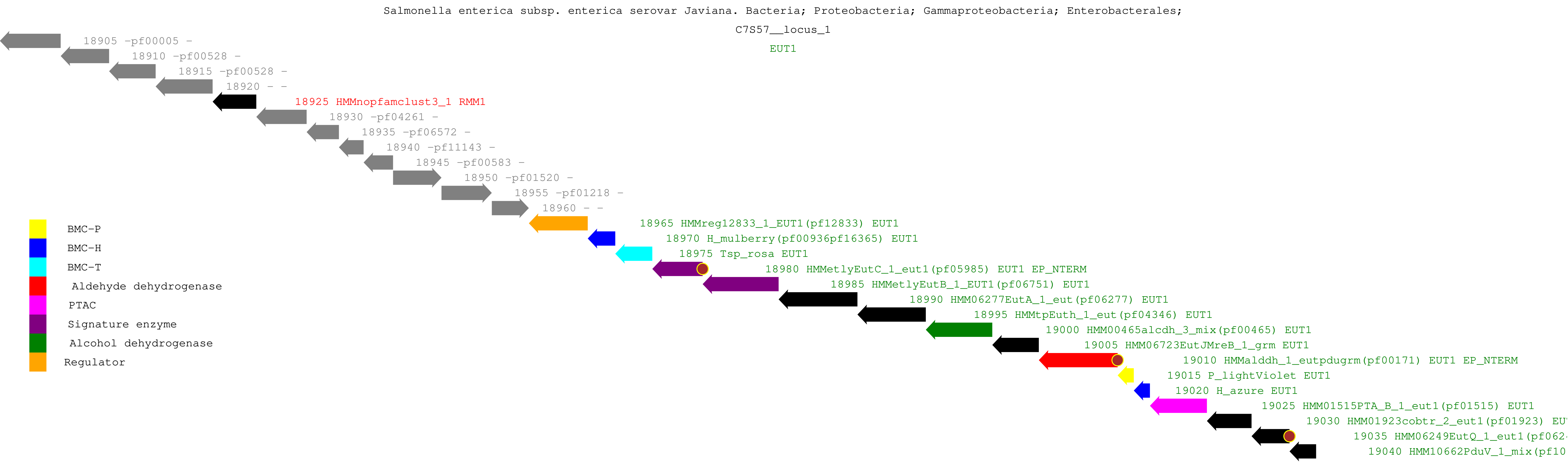
Salmonella enterica subsp. enterica serovar Javiana. Bacteria; Proteobacteria; Gammaproteobacteria; Enterobacterales;
C7S57__locus_1
EUT1
18905 -pf00005 -
18910 -pf00528 -
18915 -pf00528 -
18920 - -
18925 HMMnopfamclust3_1 RMM1
18930 -pf04261 -
18935 -pf06572 -
18940 -pf11143 -
18945 -pf00583 -
18950 -pf01520 -
18955 -pf01218 -
18960 - -
18965 HMMreg12833_1_EUT1(pf12833) EUT1
18970 H_mulberry(pf00936pf16365) EUT1
18975 Tsp_rosa EUT1
18980 HMMetlyEutC_1_eut1(pf05985) EUT1 EP_NTERM
18985 HMMetlyEutB_1_EUT1(pf06751) EUT1
18990 HMM06277EutA_1_eut(pf06277) EUT1
18995 HMMtpEuth_1_eut(pf04346) EUT1
19000 HMM00465alcdh_3_mix(pf00465) EUT1
19005 HMM06723EutJMreB_1_grm EUT1
19010 HMMalddh_1_eutpdugrm(pf00171) EUT1 EP_NTERM
19015 P_lightViolet EUT1
19020 H_azure EUT1
19025 HMM01515PTA_B_1_eut1(pf01515) EUT1
19030 HMM01923cobtr_2_eut1(pf01923) EU
19035 HMM06249EutQ_1_eut1(pf062
19040 HMM10662PduV_1_mix(pf10
BMC-P
BMC-H
BMC-T
Aldehyde dehydrogenase
PTAC
Signature enzyme
Alcohol dehydrogenase
Regulator

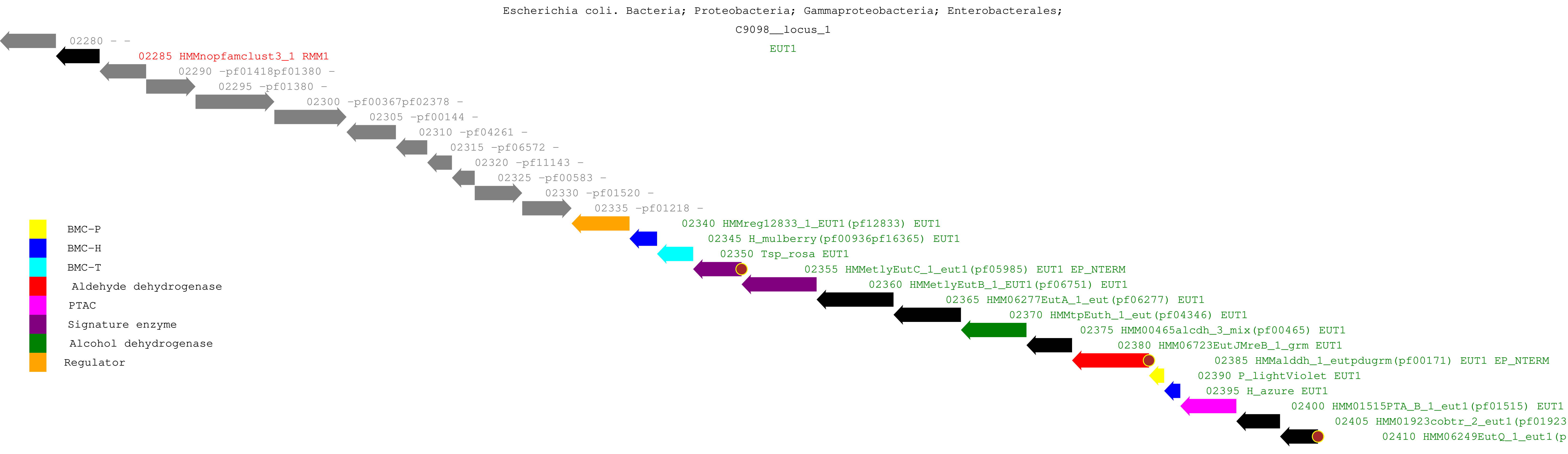
Escherichia coli. Bacteria; Proteobacteria; Gammaproteobacteria; Enterobacterales;
C9098__locus_1
EUT1
02280 - -
02285 HMMnopfamclust3_1 RMM1
02290 -pf01418pf01380 -
02295 -pf01380 -
02300 -pf00367pf02378 -
02305 -pf00144 -
02310 -pf04261 -
02315 -pf06572 -
02320 -pf11143 -
02325 -pf00583 -
02330 -pf01520 -
02335 -pf01218 -
02340 HMMreg12833_1_EUT1(pf12833) EUT1
02345 H_mulberry(pf00936pf16365) EUT1
02350 Tsp_rosa EUT1
02355 HMMetlyEutC_1_eut1(pf05985) EUT1 EP_NTERM
02360 HMMetlyEutB_1_EUT1(pf06751) EUT1
02365 HMM06277EutA_1_eut(pf06277) EUT1
02370 HMMtpEuth_1_eut(pf04346) EUT1
02375 HMM00465alcdh_3_mix(pf00465) EUT1
02380 HMM06723EutJMreB_1_grm EUT1
02385 HMMalddh_1_eutpdugrm(pf00171) EUT1 EP_NTERM
02390 P_lightViolet EUT1
02395 H_azure EUT1
02400 HMM01515PTA_B_1_eut1(pf01515) EUT1
02405 HMM01923cobtr_2_eut1(pf01923
02410 HMM06249EutQ_1_eut1(p
BMC-P
BMC-H
BMC-T
Aldehyde dehydrogenase
PTAC
Signature enzyme
Alcohol dehydrogenase
Regulator

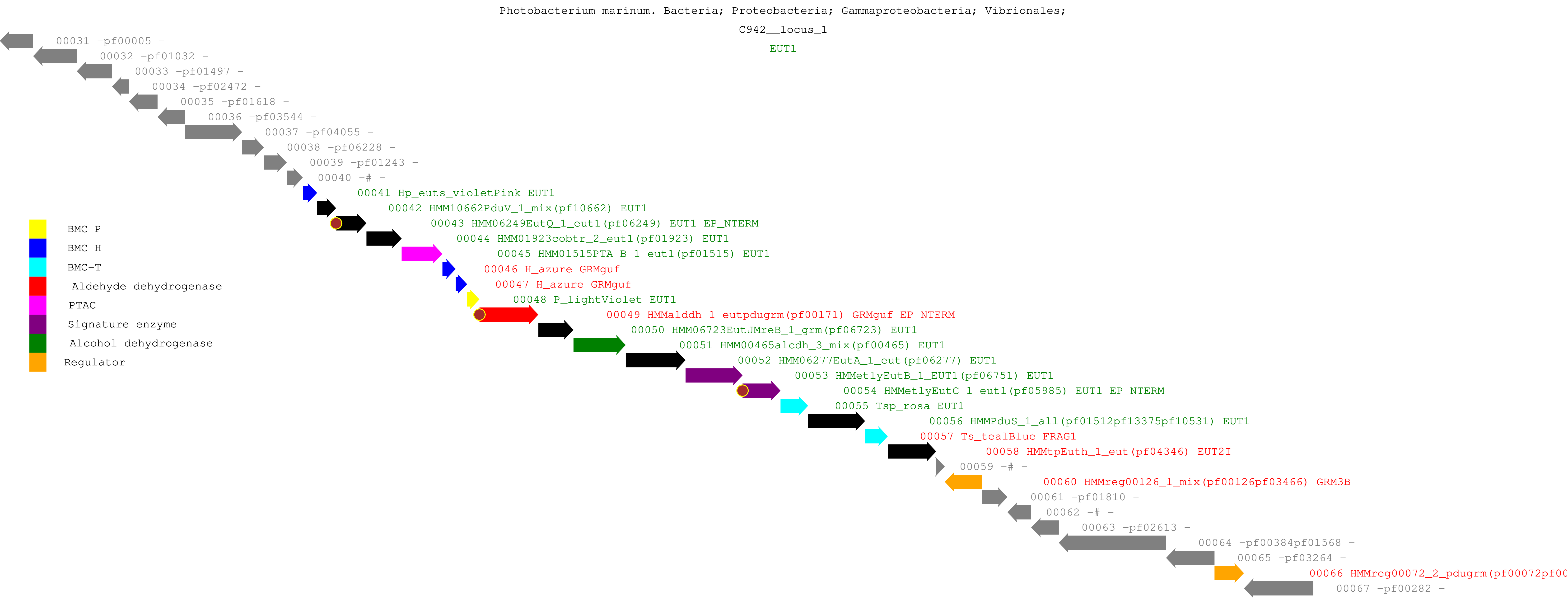

Photobacterium marinum. Bacteria; Proteobacteria; Gammaproteobacteria; Vibrionales;
C942__locus_1
EUT1
00031 -pf00005 -
00032 -pf01032 -
00033 -pf01497 -
00034 -pf02472 -
00035 -pf01618 -
00036 -pf03544 -
00037 -pf04055 -
00038 -pf06228 -
00039 -pf01243 -
00040 -# -
00041 Hp_euts_violetPink EUT1
00042 HMM10662PduV_1_mix(pf10662) EUT1
00043 HMM06249EutQ_1_eut1(pf06249) EUT1 EP_NTERM
00044 HMM01923cobtr_2_eut1(pf01923) EUT1
00045 HMM01515PTA_B_1_eut1(pf01515) EUT1
00046 H_azure GRMguf
00047 H_azure GRMguf
00048 P_lightViolet EUT1
00049 HMMaldh_1_eutpdugrm(pf00171) GRMguf EP_NTERM
00050 HMM06723EutJMreB_1_grm(pf06723) EUT1
00051 HMM00465alcdh_3_mix(pf00465) EUT1
00052 HMM06277EutA_1_eut(pf06277) EUT1
00053 HMMetlyEutB_1_EUT1(pf06751) EUT1
00054 HMMetlyEutC_1_eut1(pf05985) EUT1 EP_NTERM
00055 Tsp_rosa EUT1
00056 HMMPduS_1_all(pf01512pf13375pf10531) EUT1
00057 Ts_tealBlue FRAG1
00058 HMMtpEuth_1_eut(pf04346) EUT2I
00059 -# -
00060 HMMreg00126_1_mix(pf00126pf03466) GRM3B
00061 -pf01810 -
00062 -# -
00063 -pf02613 -
00064 -pf00384pf01568 -
00065 -pf03264 -
00066 HMMreg00072_2_pdugrm(pf00072pf00
00067 -pf00282 -
BMC-P
BMC-H
BMC-T
Aldehyde dehydrogenase
PTAC
Signature enzyme
Alcohol dehydrogenase
Regulator

Photobacterium lipolyticum. Bacteria; Proteobacteria; Gammaproteobacteria; Vibrionales;

C9I89__locus_1

EUT1

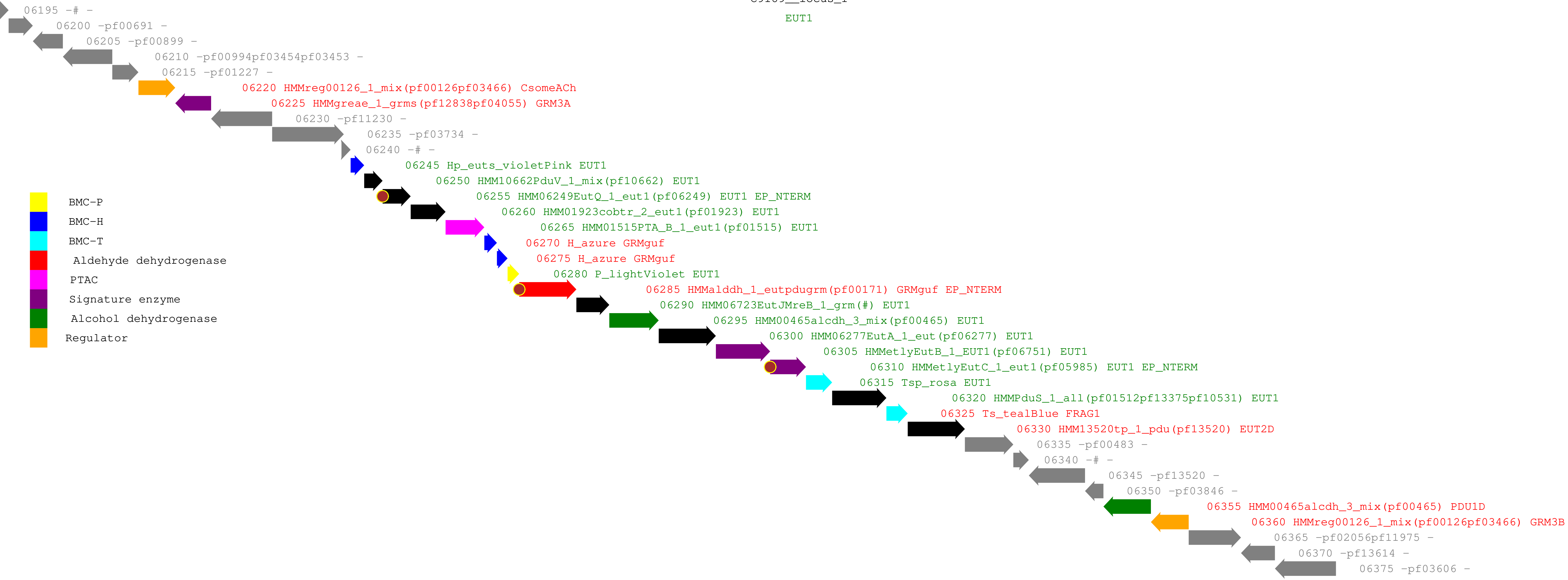

Photobacterium lutimaris. Bacteria; Proteobacteria; Gammaproteobacteria; Vibrionales;

C9I99__locus_1

EUT1

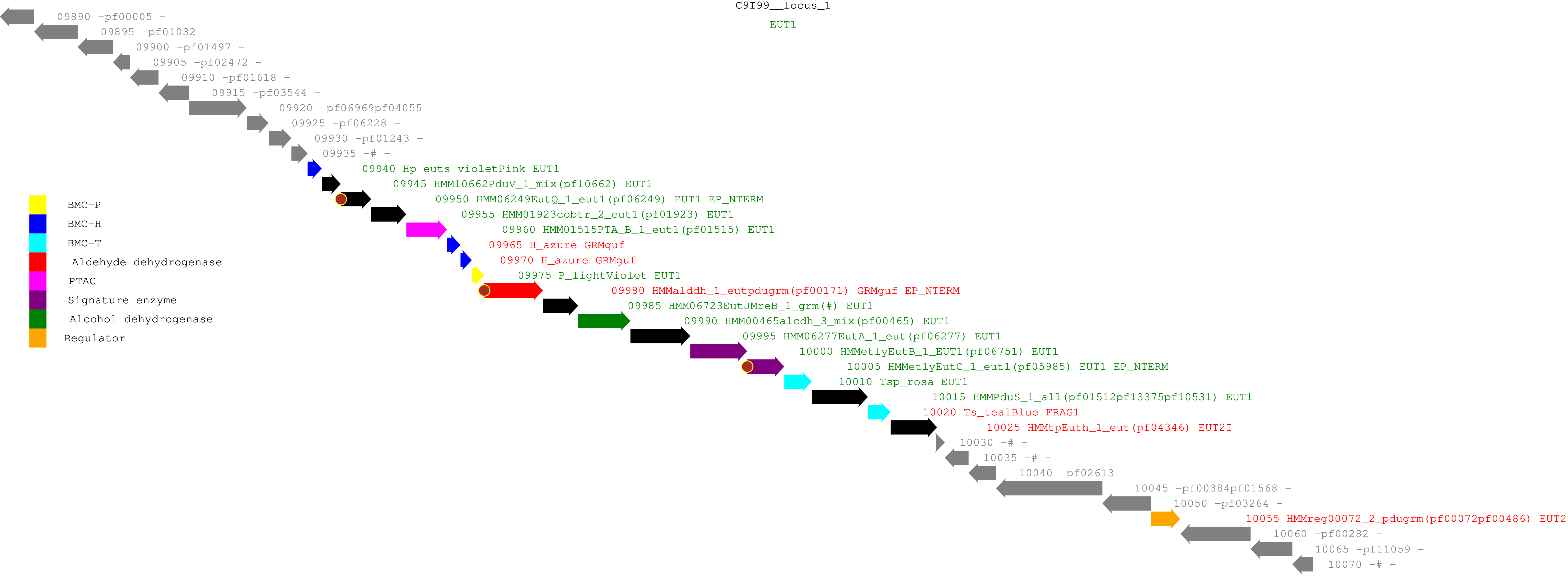

Photobacterium indicum. Bacteria; Proteobacteria; Gammaproteobacteria; Vibrionales;

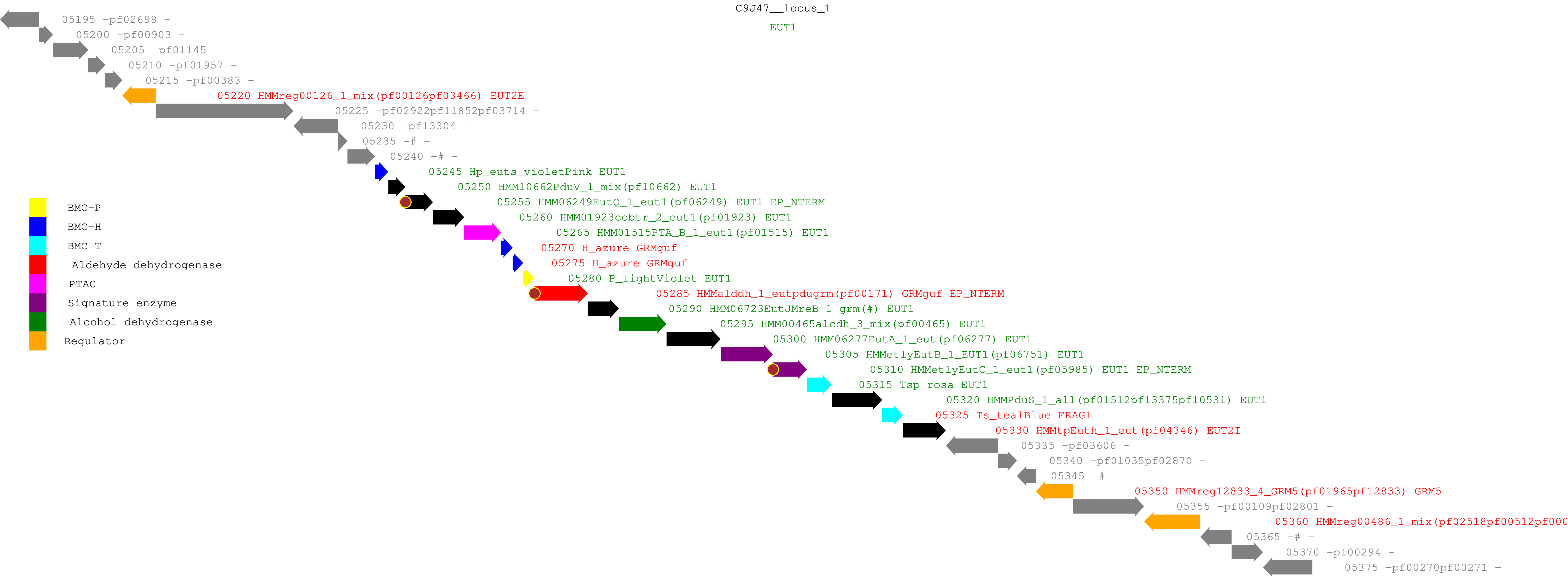

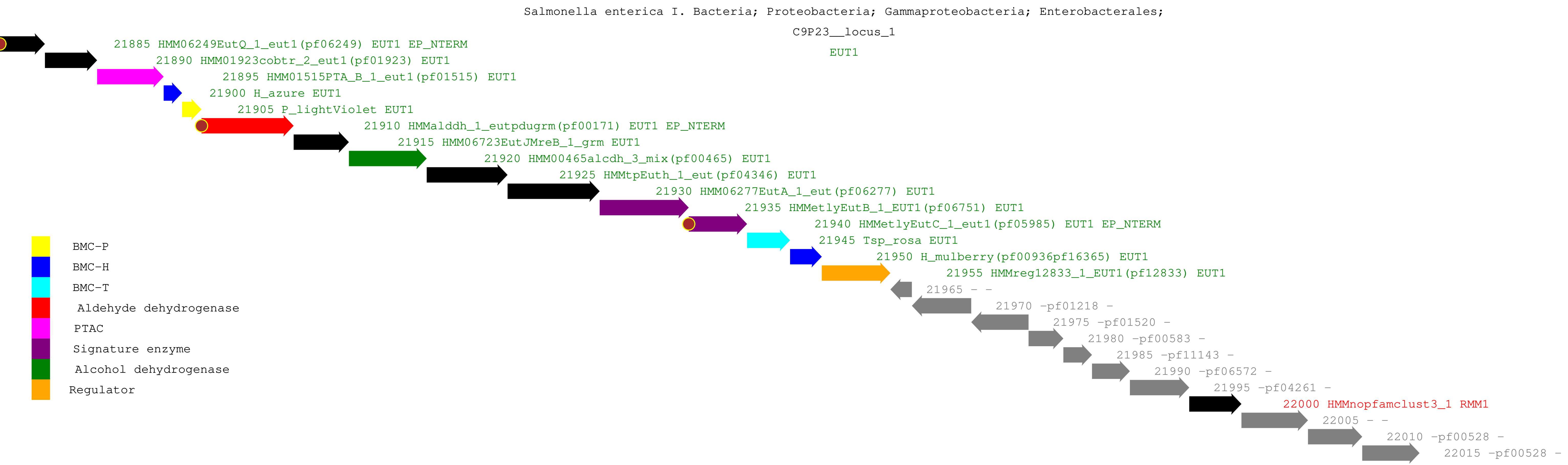
Salmonella enterica I. Bacteria; Proteobacteria; Gammaproteobacteria; Enterobacterales;
C9P23__locus_1
EUT1
21885 HMM06249EutQ_1_eut1(pf06249) EUT1 EP_NTERM
21890 HMM01923cobtr_2_eut1(pf01923) EUT1
21895 HMM01515PTA_B_1_eut1(pf01515) EUT1
21900 H_azure EUT1
21905 P_lightViolet EUT1
21910 HMMalddh_1_eutpdugrm(pf00171) EUT1 EP_NTERM
21915 HMM06723EutJMreB_1_grm EUT1
21920 HMM00465alcdh_3_mix(pf00465) EUT1
21925 HMMtpEuth_1_eut(pf04346) EUT1
21930 HMM06277EutA_1_eut(pf06277) EUT1
21935 HMMetlyEutB_1_EUT1(pf06751) EUT1
21940 HMMetlyEutC_1_eut1(pf05985) EUT1 EP_NTERM
21945 Tsp_rosa EUT1
21950 H_mulberry(pf00936pf16365) EUT1
21955 HMMreg12833_1_EUT1(pf12833) EUT1
21965 - -
21970 -pf01218 -
21975 -pf01520 -
21980 -pf00583 -
21985 -pf11143 -
21990 -pf06572 -
21995 -pf04261 -
22000 HMMnopfamclust3_1 RMM1
22005 - -
22010 -pf00528 -
22015 -pf00528 -
BMC-P
BMC-H
BMC-T
Aldehyde dehydrogenase
PTAC
Signature enzyme
Alcohol dehydrogenase
Regulator

Salmonella enteritidis. Bacteria; Proteobacteria; Gammaproteobacteria; Enterobacterales;

CA285__locus_2

EUT1

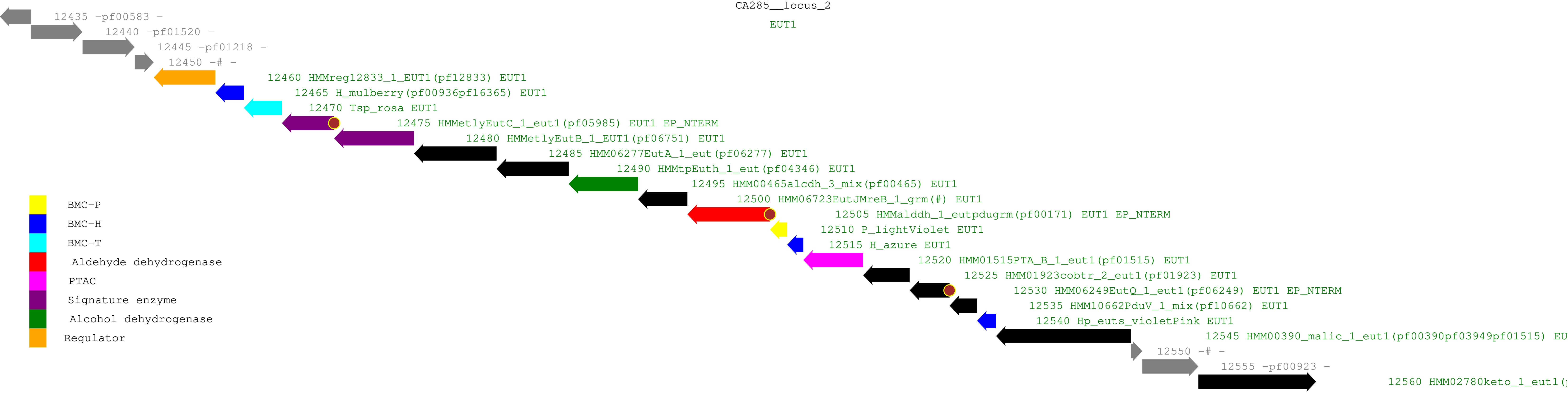

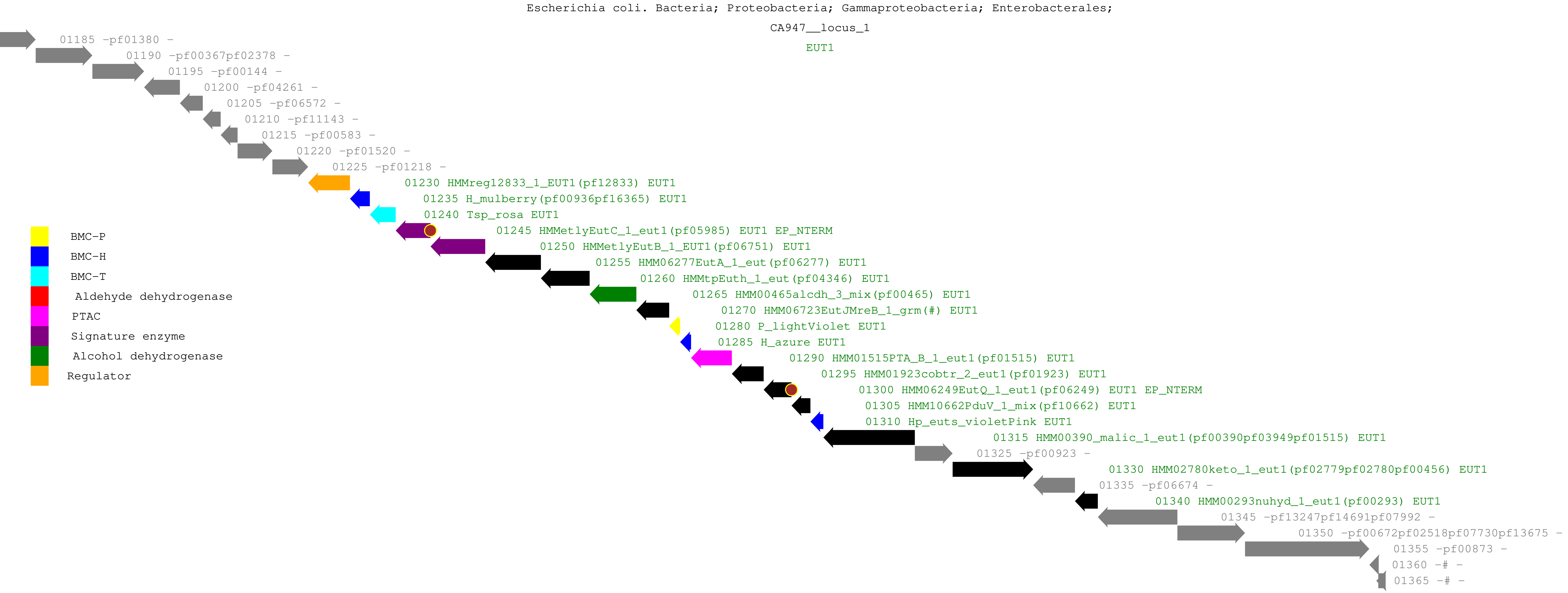
Escherichia coli. Bacteria; Proteobacteria; Gammaproteobacteria; Enterobacterales;
CA947__locus_1
EUT1
01185 -pf01380 -
01190 -pf00367pf02378 -
01195 -pf00144 -
01200 -pf04261 -
01205 -pf06572 -
01210 -pf11143 -
01215 -pf00583 -
01220 -pf01520 -
01225 -pf01218 -
01230 HMMreg12833_1_EUT1(pf12833) EUT1
01235 H_mulberry(pf00936pf16365) EUT1
01240 Tsp_rosa EUT1
01245 HMMetlyEutC_1_eut1(pf05985) EUT1 EP_NTERM
01250 HMMetlyEutB_1_EUT1(pf06751) EUT1
01255 HMM06277EutA_1_eut(pf06277) EUT1
01260 HMMtpEuth_1_eut(pf04346) EUT1
01265 HMM00465alcdh_3_mix(pf00465) EUT1
01270 HMM06723EutJMreB_1_grm(#) EUT1
01280 P_lightViolet EUT1
01285 H_azure EUT1
01290 HMM01515PTA_B_1_eut1(pf01515) EUT1
01295 HMM01923cobtr_2_eut1(pf01923) EUT1
01300 HMM06249EutQ_1_eut1(pf06249) EUT1 EP_NTERM
01305 HMM10662PduV_1_mix(pf10662) EUT1
01310 Hp_euts_violetPink EUT1
01315 HMM00390_malic_1_eut1(pf00390pf03949pf01515) EUT1
01325 -pf00923 -
01330 HMM02780keto_1_eut1(pf02779pf02780pf00456) EUT1
01335 -pf06674 -
01340 HMM00293nuhyd_1_eut1(pf00293) EUT1
01345 -pf13247pf14691pf07992 -
01350 -pf00672pf02518pf07730pf13675 -
01355 -pf00873 -
01360 -# -
01365 -# -
BMC-P
BMC-H
BMC-T
Aldehyde dehydrogenase
PTAC
Signature enzyme
Alcohol dehydrogenase
Regulator

Salmonella enterica subsp. enterica serovar Paratyphi B str. SARA61. Bacteria; Proteobacteria; Gammaproteobacteria; Enterobacterales;

CAH10__locus_1

EUT1

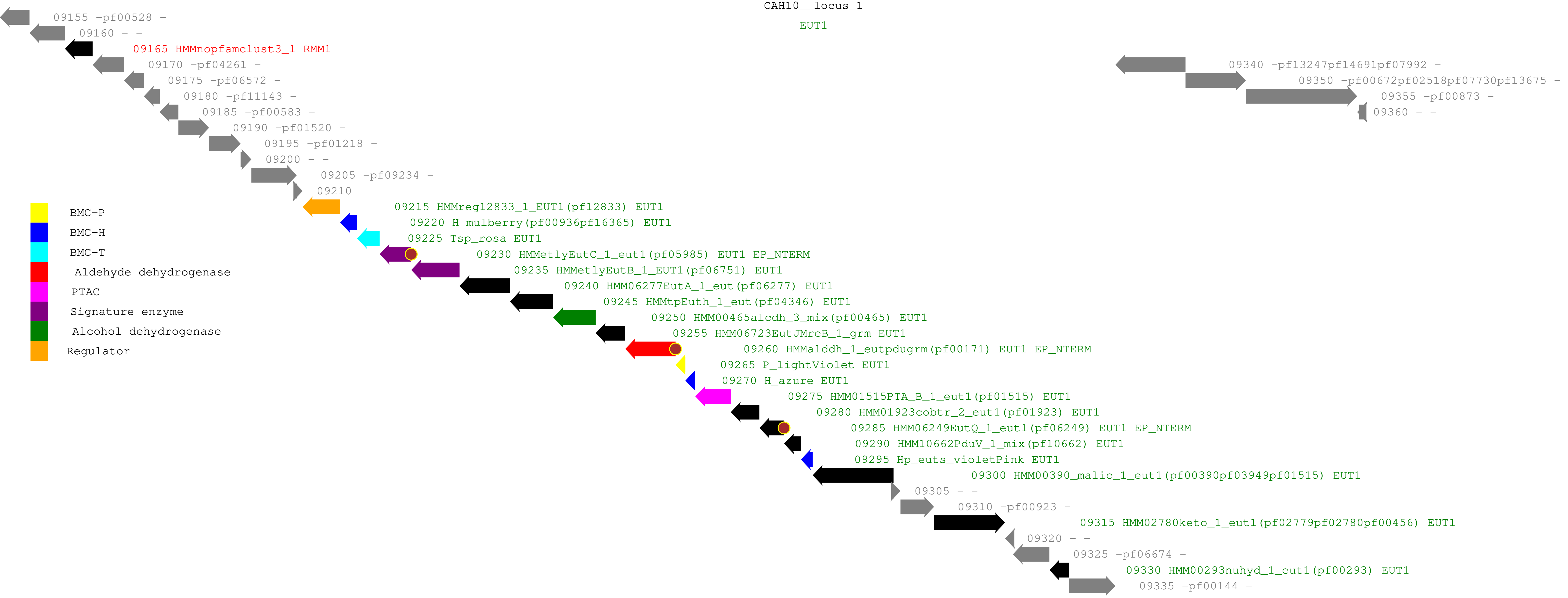

Salmonella enterica subsp. enterica serovar Typhimurium str. CFSAN000648. Bacteria; Proteobacteria; Gammaproteobacteria; Enterobacterales;

CAI81__locus_2

EUT1

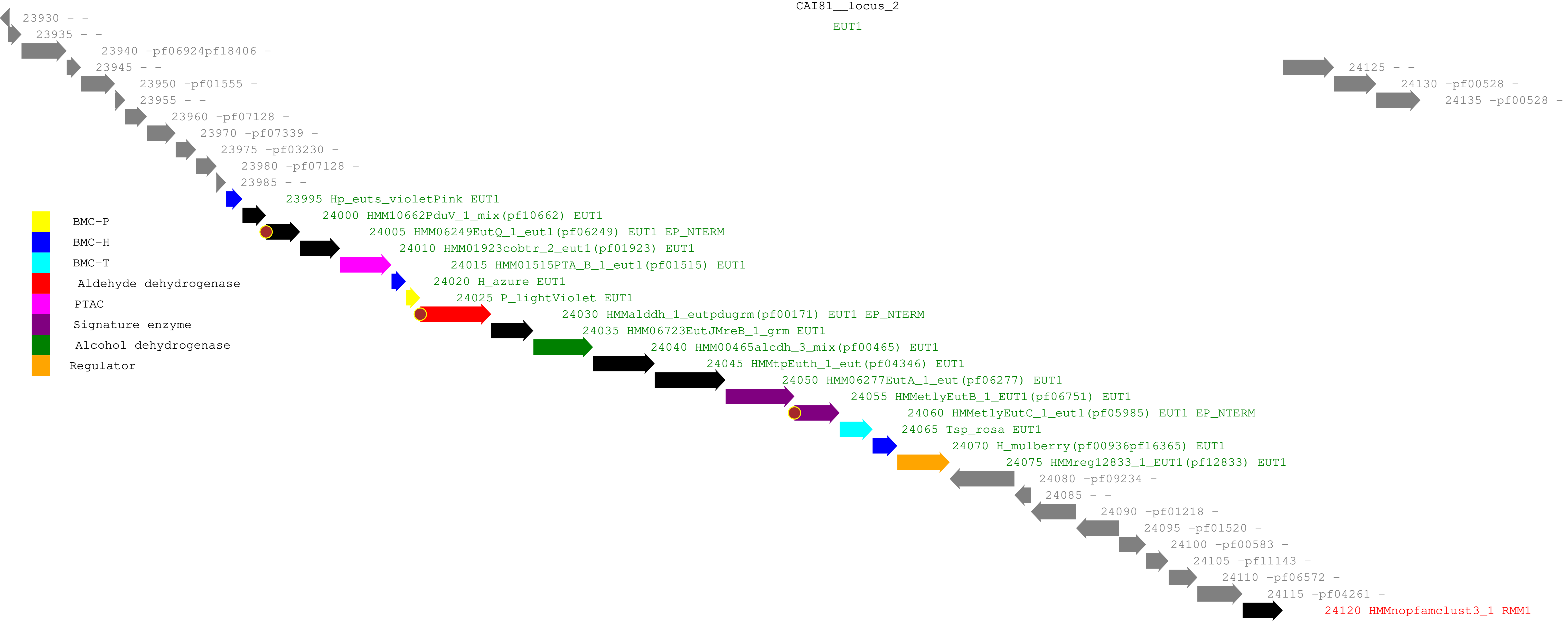

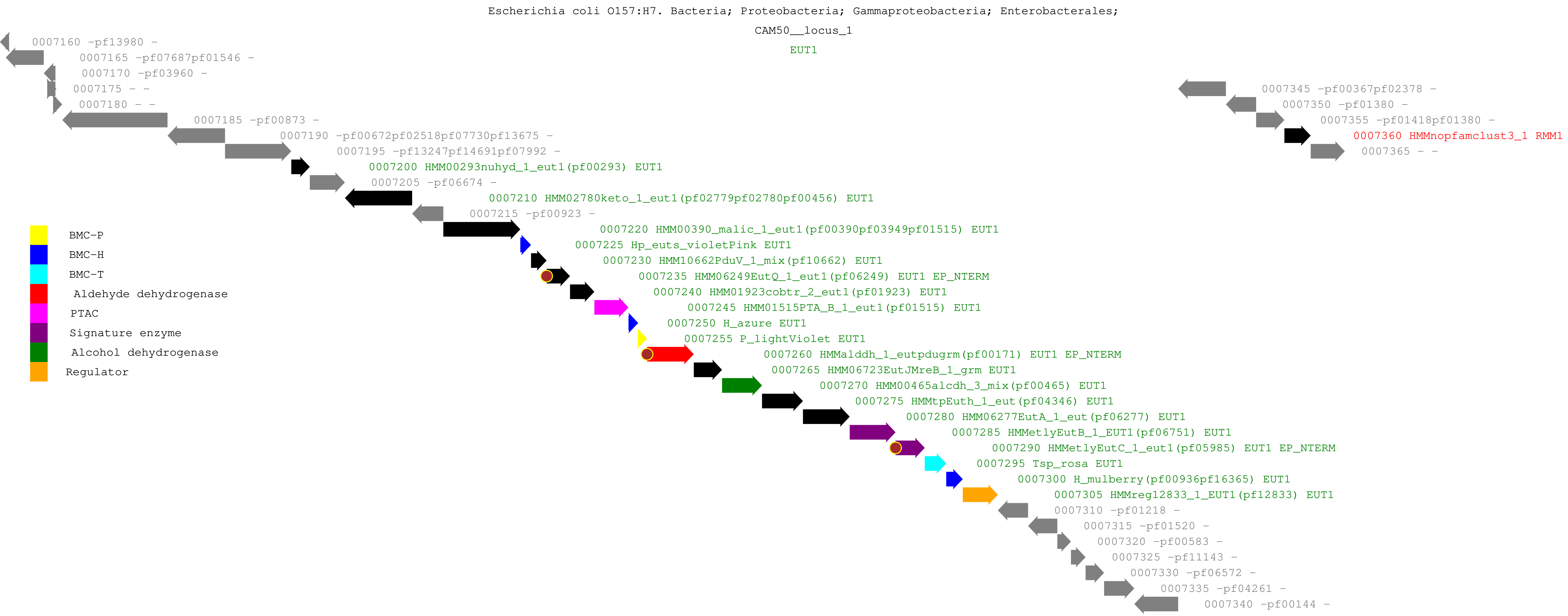
Escherichia coli O157:H7. Bacteria; Proteobacteria; Gammaproteobacteria; Enterobacterales;
CAM50__locus_1
EUT1
0007160 -pf13980 -
0007165 -pf07687pf01546 -
0007170 -pf03960 -
0007175 - -
0007180 - -
0007185 -pf00873 -
0007190 -pf00672pf02518pf07730pf13675 -
0007195 -pf13247pf14691pf07992 -
0007200 HMM00293nuhyd_1_eut1(pf00293) EUT1
0007205 -pf06674 -
0007210 HMM02780keto_1_eut1(pf02779pf02780pf00456) EUT1
0007215 -pf00923 -
0007220 HMM00390_malic_1_eut1(pf00390pf03949pf01515) EUT1
0007225 Hp_euts_violetPink EUT1
0007230 HMM10662PduV_1_mix(pf10662) EUT1
0007235 HMM06249EutQ_1_eut1(pf06249) EUT1 EP_NTERM
0007240 HMM01923cobtr_2_eut1(pf01923) EUT1
0007245 HMM01515PTA_B_1_eut1(pf01515) EUT1
0007250 H_azure EUT1
0007255 P_lightViolet EUT1
0007260 HMMalddh_1_eutpdugrm(pf00171) EUT1 EP_NTERM
0007265 HMM06723EutJMreB_1_grm EUT1
0007270 HMM00465alcdh_3_mix(pf00465) EUT1
0007275 HMMtpEuth_1_eut(pf04346) EUT1
0007280 HMM06277EutA_1_eut(pf06277) EUT1
0007285 HMMetlyEutB_1_EUT1(pf06751) EUT1
0007290 HMMetlyEutC_1_eut1(pf05985) EUT1 EP_NTERM
0007295 Tsp_rosa EUT1
0007300 H_mulberry(pf00936pf16365) EUT1
0007305 HMMreg12833_1_EUT1(pf12833) EUT1
0007310 -pf01218 -
0007315 -pf01520 -
0007320 -pf00583 -
0007325 -pf11143 -
0007330 -pf06572 -
0007335 -pf04261 -
0007340 -pf00144 -
0007345 -pf00367pf02378 -
0007350 -pf01380 -
0007355 -pf01418pf01380 -
0007360 HMMnopfamclust3_1 RMM1
0007365 - -
BMC-P
BMC-H
BMC-T
Aldehyde dehydrogenase
PTAC
Signature enzyme
Alcohol dehydrogenase
Regulator

Salmonella choleraesuis. Bacteria; Proteobacteria; Gammaproteobacteria; Enterobacterales;

CBI64__locus_1

EUT1

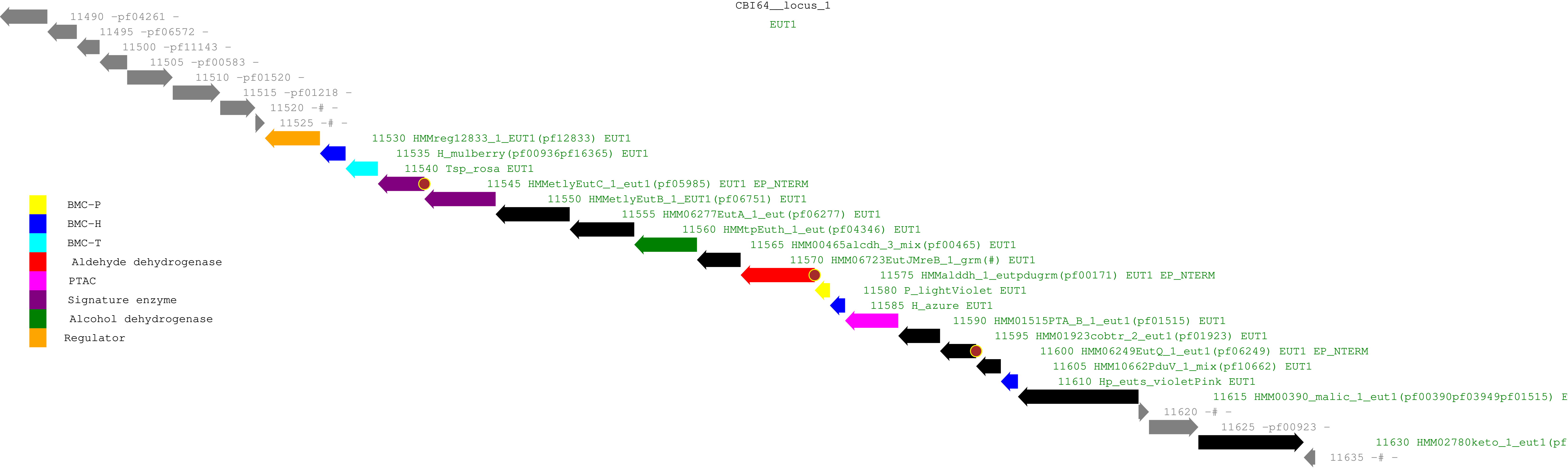

Shigella flexneri. Bacteria; Proteobacteria; Gammaproteobacteria; Enterobacterales;

CBL21__locus_1

EUT1

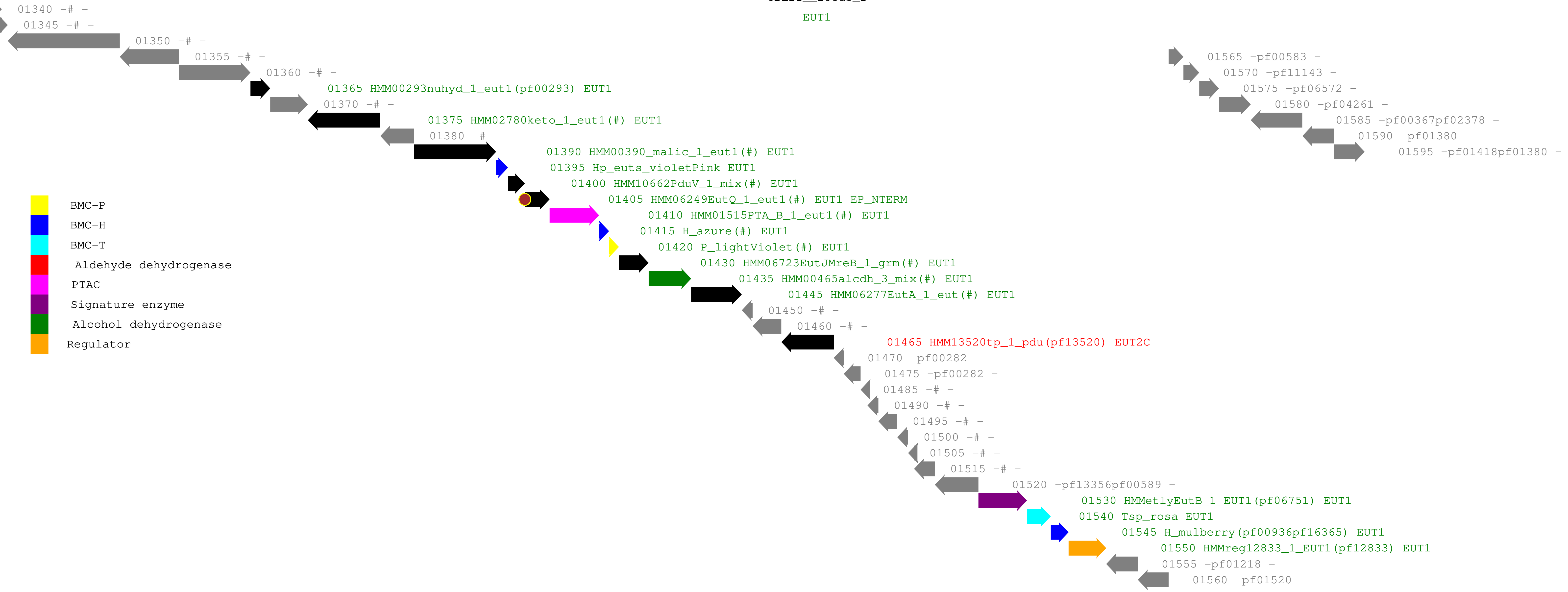

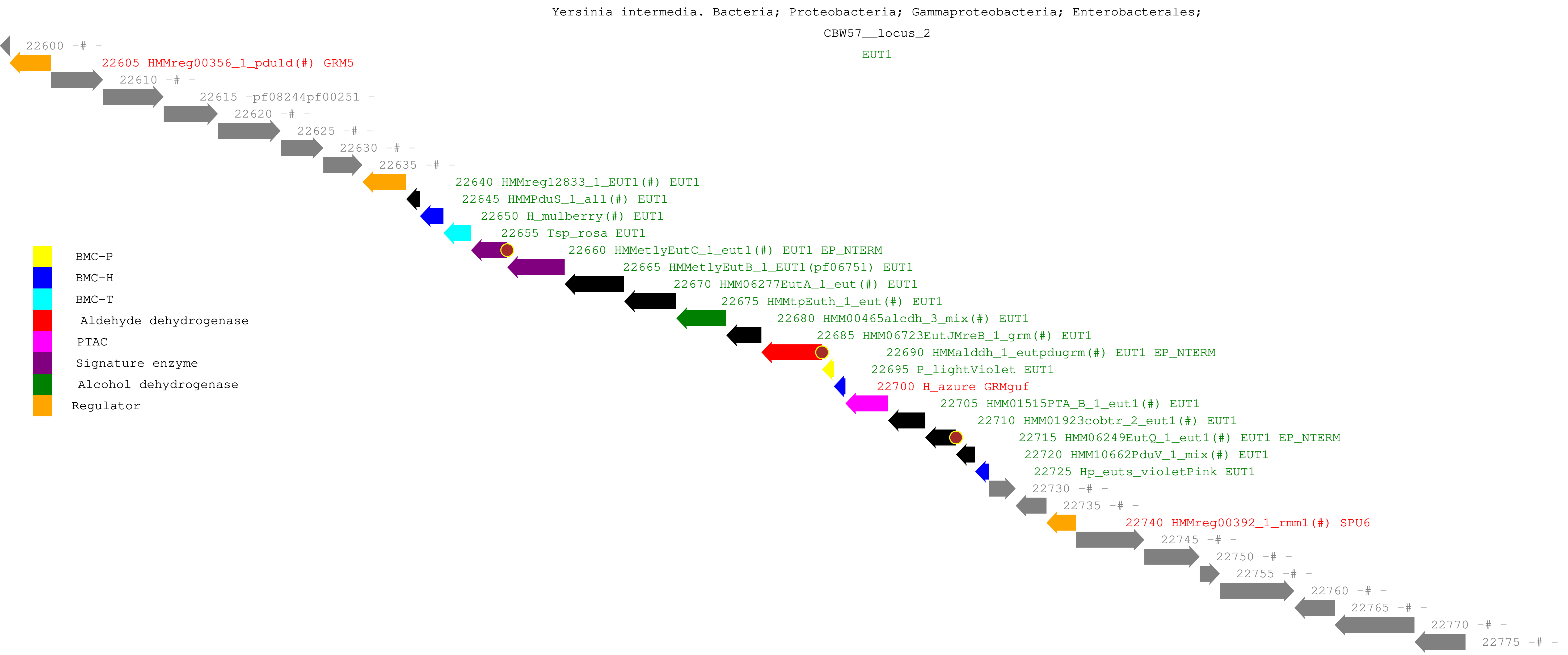
Yersinia intermedia. Bacteria; Proteobacteria; Gammaproteobacteria; Enterobacterales;
CBW57__locus_2
EUT1
22600 -# -
22605 HMMreg00356_1_pdu1d(#) GRM5
22610 -# -
22615 -pf08244pf00251 -
22620 -# -
22625 -# -
22630 -# -
22635 -# -
22640 HMMreg12833_1_EUT1(#) EUT1
22645 HMMPduS_1_all(#) EUT1
22650 H_mulberry(#) EUT1
22655 Tsp_rosa EUT1
22660 HMMetlyEutC_1_eut1(#) EUT1 EP_NTERM
22665 HMMetlyEutB_1_EUT1(pf06751) EUT1
22670 HMM06277EutA_1_eut(#) EUT1
22675 HMMtpEuth_1_eut(#) EUT1
22680 HMM00465alcdh_3_mix(#) EUT1
22685 HMM06723EutJMreB_1_grm(#) EUT1
22690 HMMalddh_1_eutpdugrm(#) EUT1 EP_NTERM
22695 P_lightViolet EUT1
22700 H_azure GRMguf
22705 HMM01515PTA_B_1_eut1(#) EUT1
22710 HMM01923cobtr_2_eut1(#) EUT1
22715 HMM06249EutQ_1_eut1(#) EUT1 EP_NTERM
22720 HMM10662PduV_1_mix(#) EUT1
22725 Hp_euts_violetPink EUT1
22730 -# -
22735 -# -
22740 HMMreg00392_1_rmm1(#) SPU6
22745 -# -
22750 -# -
22755 -# -
22760 -# -
22765 -# -
22770 -# -
22775 -# -
BMC-P
BMC-H
BMC-T
Aldehyde dehydrogenase
PTAC
Signature enzyme
Alcohol dehydrogenase
Regulator

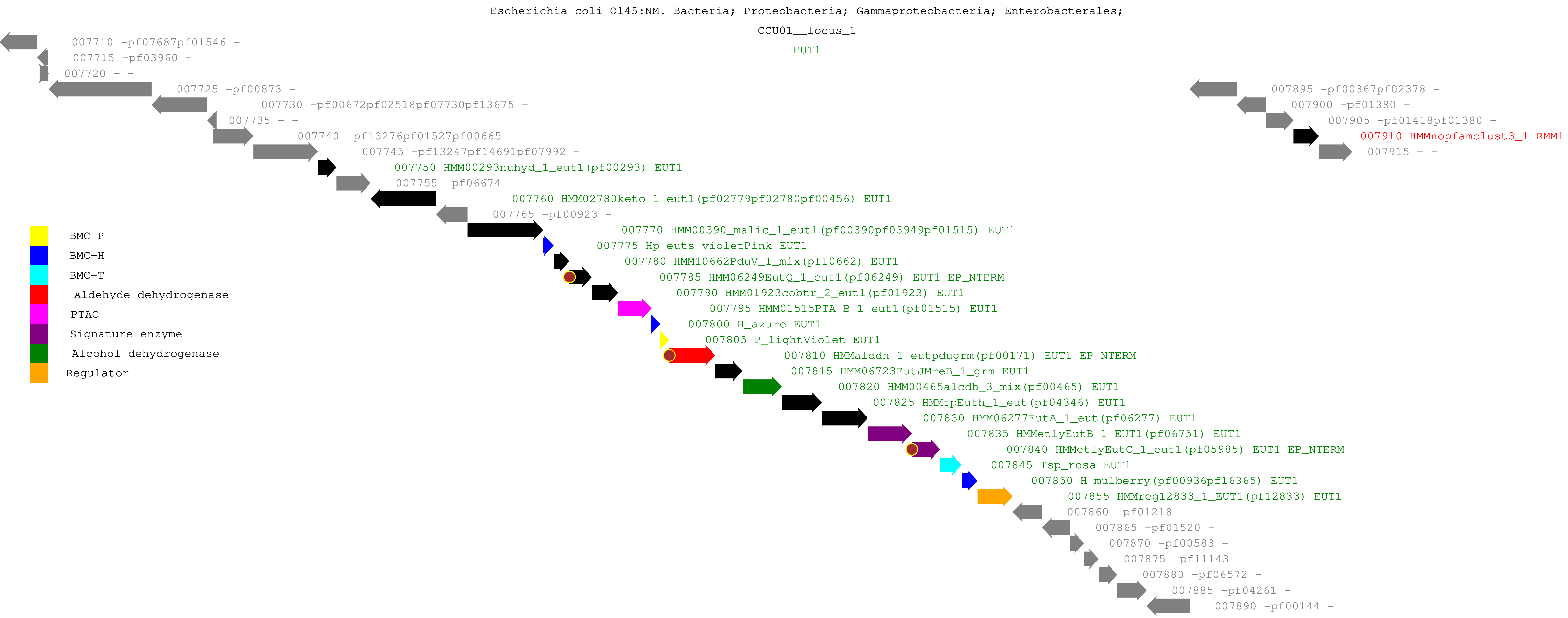
Escherichia coli O145:NM. Bacteria; Proteobacteria; Gammaproteobacteria; Enterobacterales;
CCU01__locus_1
EUT1
007710 -pf07687pf01546 -
007715 -pf03960 -
007720 - -
007725 -pf00873 -
007730 -pf00672pf02518pf07730pf13675 -
007735 - -
007740 -pf13276pf01527pf00665 -
007745 -pf13247pf14691pf07992 -
007750 HMM00293nuhyd_1_eut1(pf00293) EUT1
007755 -pf06674 -
007760 HMM02780keto_1_eut1(pf02779pf02780pf00456) EUT1
007765 -pf00923 -
007770 HMM00390_malic_1_eut1(pf00390pf03949pf01515) EUT1
007775 Hp_euts_violetPink EUT1
007780 HMM10662PduV_1_mix(pf10662) EUT1
007785 HMM06249EutQ_1_eut1(pf06249) EUT1 EP_NTERM
007790 HMM01923cobtr_2_eut1(pf01923) EUT1
007795 HMM01515PTA_B_1_eut1(pf01515) EUT1
007800 H_azure EUT1
007805 P_lightViolet EUT1
007810 HMMalddh_1_eutpdugrm(pf00171) EUT1 EP_NTERM
007815 HMM06723EutJMreB_1_grm EUT1
007820 HMM00465alcdh_3_mix(pf00465) EUT1
007825 HMMtpEuth_1_eut(pf04346) EUT1
007830 HMM06277EutA_1_eut(pf06277) EUT1
007835 HMMetlyEutB_1_EUT1(pf06751) EUT1
007840 HMMetlyEutC_1_eut1(pf05985) EUT1 EP_NTERM
007845 Tsp_rosa EUT1
007850 H_mulberry(pf00936pf16365) EUT1
007855 HMMreg12833_1_EUT1(pf12833) EUT1
007860 -pf01218 -
007865 -pf01520 -
007870 -pf00583 -
007875 -pf11143 -
007880 -pf06572 -
007885 -pf04261 -
007890 -pf00144 -
007895 -pf00367pf02378 -
007900 -pf01380 -
007905 -pf01418pf01380 -
007910 HMMnopfamclust3_1 RMM1
007915 - -
BMC-P
BMC-H
BMC-T
Aldehyde dehydrogenase
PTAC
Signature enzyme
Alcohol dehydrogenase
Regulator

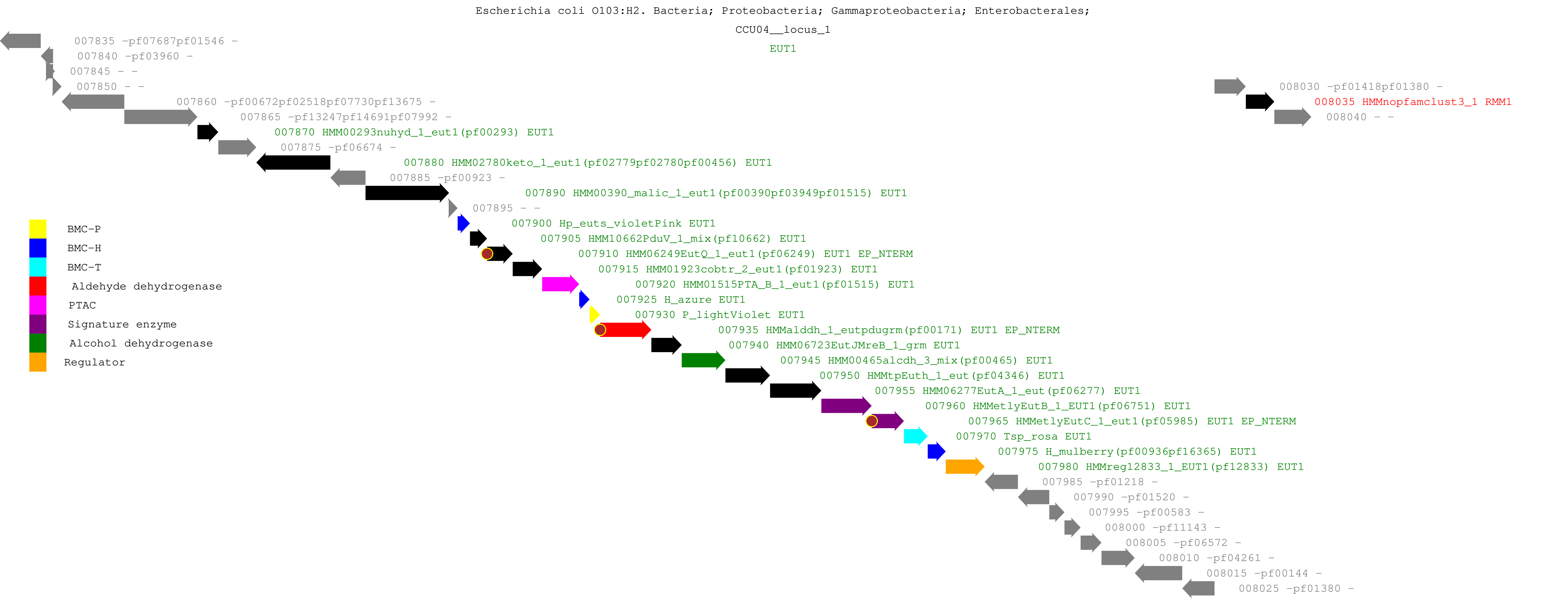
Escherichia coli O103:H2. Bacteria; Proteobacteria; Gammaproteobacteria; Enterobacterales;
CCU04__locus_1
EUT1
007835 -pf07687pf01546 -
007840 -pf03960 -
007845 - -
007850 - -
007860 -pf00672pf02518pf07730pf13675 -
007865 -pf13247pf14691pf07992 -
007870 HMM00293nuhyd_1_eut1(pf00293) EUT1
007875 -pf06674 -
007880 HMM02780keto_1_eut1(pf02779pf02780pf00456) EUT1
007885 -pf00923 -
007890 HMM00390_malic_1_eut1(pf00390pf03949pf01515) EUT1
007895 - -
007900 Hp_euts_violetPink EUT1
007905 HMM10662PduV_1_mix(pf10662) EUT1
007910 HMM06249EutQ_1_eut1(pf06249) EUT1 EP_NTERM
007915 HMM01923cobtr_2_eut1(pf01923) EUT1
007920 HMM01515PTA_B_1_eut1(pf01515) EUT1
007925 H_azure EUT1
007930 P_lightViolet EUT1
007935 HMMaldh_1_eutpdugrm(pf00171) EUT1 EP_NTERM
007940 HMM06723EutJMreB_1_grm EUT1
007945 HMM00465alcdh_3_mix(pf00465) EUT1
007950 HMMtpEuth_1_eut(pf04346) EUT1
007955 HMM06277EutA_1_eut(pf06277) EUT1
007960 HMMetlyEutB_1_EUT1(pf06751) EUT1
007965 HMMetlyEutC_1_eut1(pf05985) EUT1 EP_NTERM
007970 Tsp_rosa EUT1
007975 H_mulberry(pf00936pf16365) EUT1
007980 HMMreg12833_1_EUT1(pf12833) EUT1
007985 -pf01218 -
007990 -pf01520 -
007995 -pf00583 -
008000 -pf11143 -
008005 -pf06572 -
008010 -pf04261 -
008015 -pf00144 -
008025 -pf01380 -
008030 -pf01418pf01380 -
008035 HMMnopfamclust3_1 RMM1
008040 - -
BMC-P
BMC-H
BMC-T
Aldehyde dehydrogenase
PTAC
Signature enzyme
Alcohol dehydrogenase
Regulator

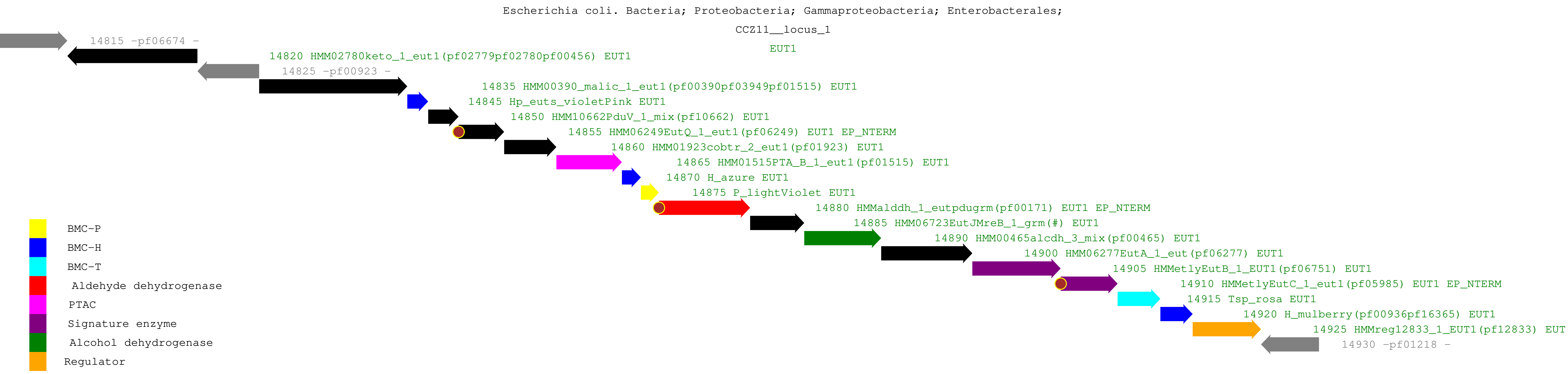

Escherichia coli. Bacteria; Proteobacteria; Gammaproteobacteria; Enterobacterales;
CCZ11__locus_1
EUT1
14815 -pf06674 -
14820 HMM02780keto_1_eut1(pf02779pf02780pf00456) EUT1
14825 -pf00923 -
14835 HMM00390_malic_1_eut1(pf00390pf03949pf01515) EUT1
14845 Hp_euts_violetPink EUT1
14850 HMM10662PduV_1_mix(pf10662) EUT1
14855 HMM06249EutQ_1_eut1(pf06249) EUT1 EP_NTERM
14860 HMM01923cobtr_2_eut1(pf01923) EUT1
14865 HMM01515PTA_B_1_eut1(pf01515) EUT1
14870 H_azure EUT1
14875 P_lightViolet EUT1
14880 HMMalddh_1_eutpdugrm(pf00171) EUT1 EP_NTERM
14885 HMM06723EutJMreB_1_grm(#) EUT1
14890 HMM00465alcdh_3_mix(pf00465) EUT1
14900 HMM06277EutA_1_eut(pf06277) EUT1
14905 HMMetlyEutB_1_EUT1(pf06751) EUT1
14910 HMMetlyEutC_1_eut1(pf05985) EUT1 EP_NTERM
14915 Tsp_rosa EUT1
14920 H_mulberry(pf00936pf16365) EUT1
14925 HMMreg12833_1_EUT1(pf12833) EUT
14930 -pf01218 -
BMC-P
BMC-H
BMC-T
Aldehyde dehydrogenase
PTAC
Signature enzyme
Alcohol dehydrogenase
Regulator

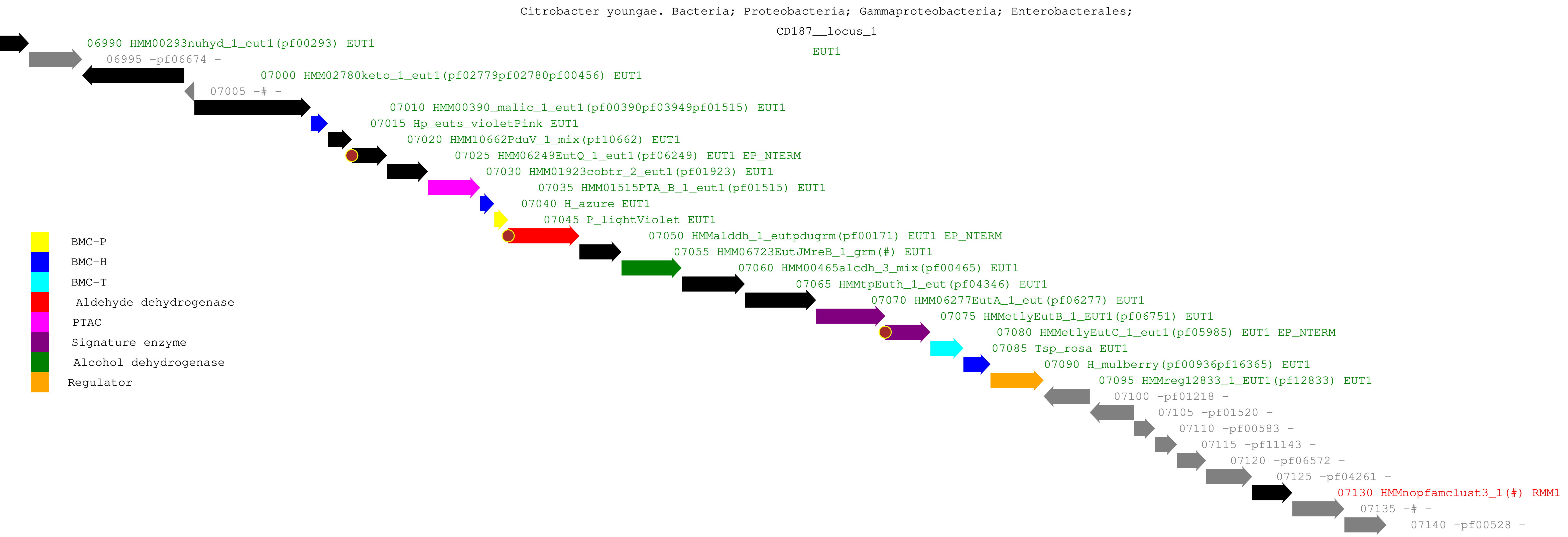
Citrobacter youngae. Bacteria; Proteobacteria; Gammaproteobacteria; Enterobacterales;
CD187__locus_1
EUT1
06990 HMM00293nuhyd_1_eut1(pf00293) EUT1
06995 -pf06674 -
07000 HMM02780keto_1_eut1(pf02779pf02780pf00456) EUT1
07005 -# -
07010 HMM00390_malic_1_eut1(pf00390pf03949pf01515) EUT1
07015 Hp_euts_violetPink EUT1
07020 HMM10662PduV_1_mix(pf10662) EUT1
07025 HMM06249EutQ_1_eut1(pf06249) EUT1 EP_NTERM
07030 HMM01923cobtr_2_eut1(pf01923) EUT1
07035 HMM01515PTA_B_1_eut1(pf01515) EUT1
07040 H_azure EUT1
07045 P_lightViolet EUT1
07050 HMMalddh_1_eutpdugrm(pf00171) EUT1 EP_NTERM
07055 HMM06723EutJMreB_1_grm(#) EUT1
07060 HMM00465alcdh_3_mix(pf00465) EUT1
07065 HMMtpEuth_1_eut(pf04346) EUT1
07070 HMM06277EutA_1_eut(pf06277) EUT1
07075 HMMetlyEutB_1_EUT1(pf06751) EUT1
07080 HMMetlyEutC_1_eut1(pf05985) EUT1 EP_NTERM
07085 Tsp_rosa EUT1
07090 H_mulberry(pf00936pf16365) EUT1
07095 HMMreg12833_1_EUT1(pf12833) EUT1
07100 -pf01218 -
07105 -pf01520 -
07110 -pf00583 -
07115 -pf11143 -
07120 -pf06572 -
07125 -pf04261 -
07130 HMMnopfamclust3_1(#) RMM1
07135 -# -
07140 -pf00528 -
BMC-P
BMC-H
BMC-T
Aldehyde dehydrogenase
PTAC
Signature enzyme
Alcohol dehydrogenase
Regulator

Salmonella senftenberg. Bacteria; Proteobacteria; Gammaproteobacteria; Enterobacterales;

CD793__locus_2

EUT1

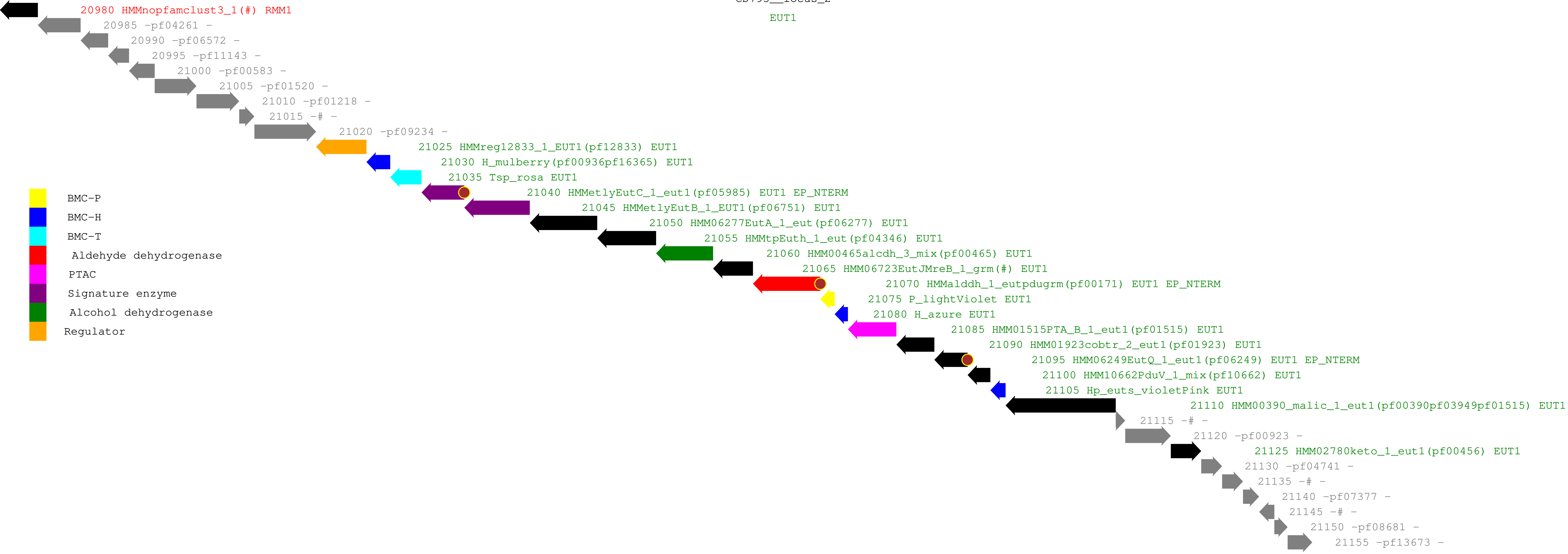

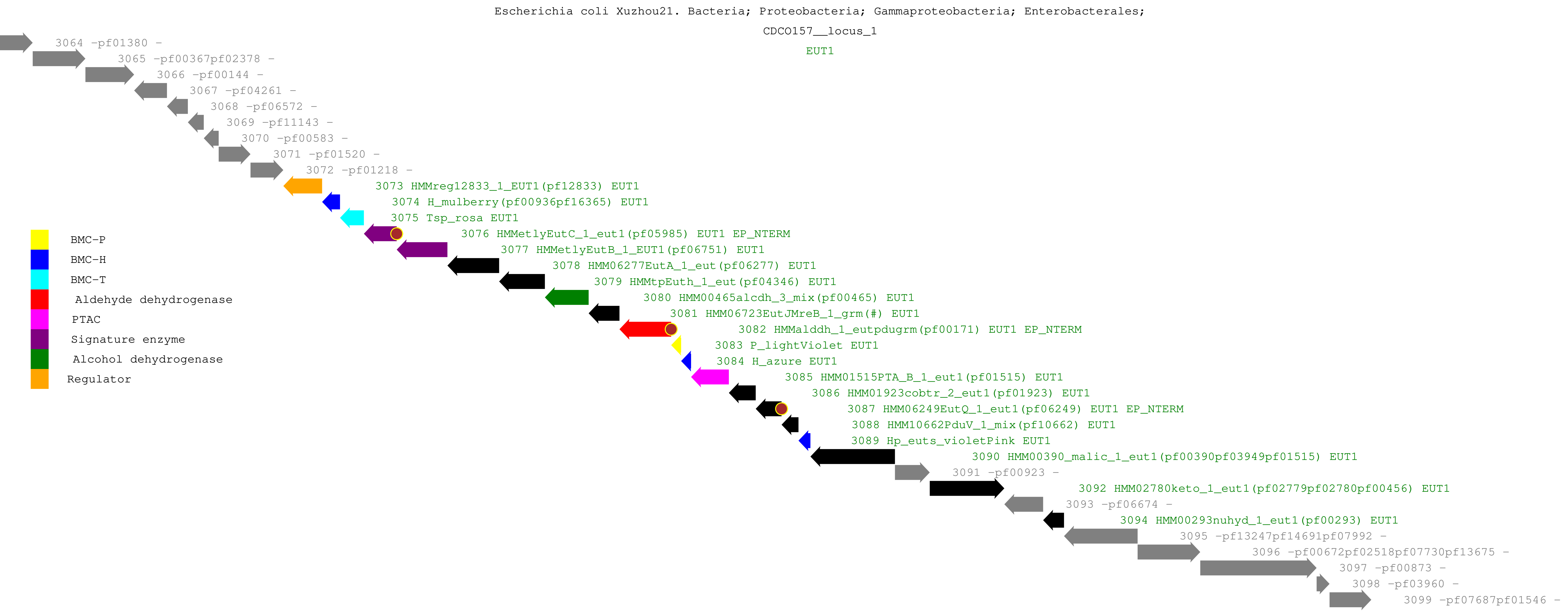
Escherichia coli Xuzhou21. Bacteria; Proteobacteria; Gammaproteobacteria; Enterobacterales;
CDCO157__locus_1
EUT1
3064 -pf01380 -
3065 -pf00367pf02378 -
3066 -pf00144 -
3067 -pf04261 -
3068 -pf06572 -
3069 -pf11143 -
3070 -pf00583 -
3071 -pf01520 -
3072 -pf01218 -
3073 HMMreg12833_1_EUT1(pf12833) EUT1
3074 H_mulberry(pf00936pf16365) EUT1
3075 Tsp_rosa EUT1
3076 HMMetlyEutC_1_eut1(pf05985) EUT1 EP_NTERM
3077 HMMetlyEutB_1_EUT1(pf06751) EUT1
3078 HMM06277EutA_1_eut(pf06277) EUT1
3079 HMMtpEuth_1_eut(pf04346) EUT1
3080 HMM00465alcdh_3_mix(pf00465) EUT1
3081 HMM06723EutJMreB_1_grm(#) EUT1
3082 HMMalddh_1_eutpdugrm(pf00171) EUT1 EP_NTERM
3083 P_lightViolet EUT1
3084 H_azure EUT1
3085 HMM01515PTA_B_1_eut1(pf01515) EUT1
3086 HMM01923cobtr_2_eut1(pf01923) EUT1
3087 HMM06249EutQ_1_eut1(pf06249) EUT1 EP_NTERM
3088 HMM10662PduV_1_mix(pf10662) EUT1
3089 Hp_euts_violetPink EUT1
3090 HMM00390_malic_1_eut1(pf00390pf03949pf01515) EUT1
3091 -pf00923 -
3092 HMM02780keto_1_eut1(pf02779pf02780pf00456) EUT1
3093 -pf06674 -
3094 HMM00293nuhyd_1_eut1(pf00293) EUT1
3095 -pf13247pf14691pf07992 -
3096 -pf00672pf02518pf07730pf13675 -
3097 -pf00873 -
3098 -pf03960 -
3099 -pf07687pf01546 -
BMC-P
BMC-H
BMC-T
Aldehyde dehydrogenase
PTAC
Signature enzyme
Alcohol dehydrogenase
Regulator

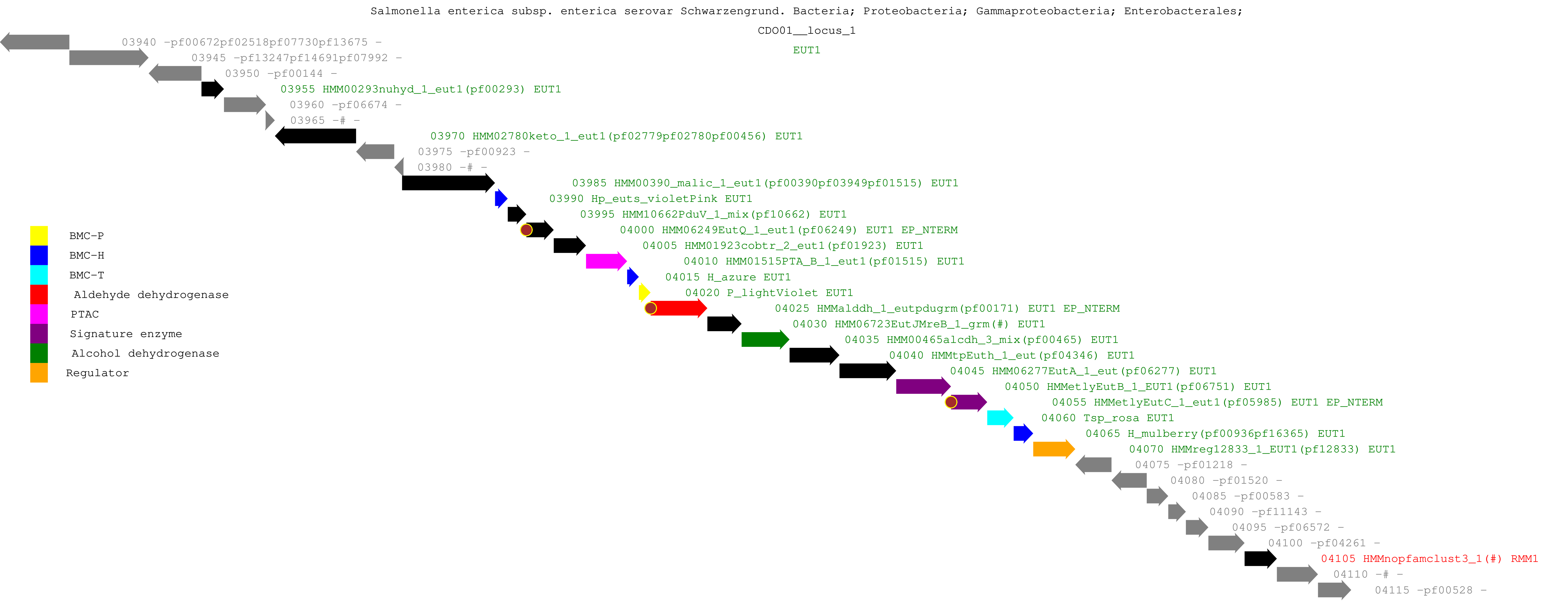
Salmonella enterica subsp. enterica serovar Schwarzengrund. Bacteria; Proteobacteria; Gammaproteobacteria; Enterobacterales;
CDO01__locus_1
EUT1
03940 -pf00672pf02518pf07730pf13675 -
03945 -pf13247pf14691pf07992 -
03950 -pf00144 -
03955 HMM00293nuhyd_1_eut1(pf00293) EUT1
03960 -pf06674 -
03965 -# -
03970 HMM02780keto_1_eut1(pf02779pf02780pf00456) EUT1
03975 -pf00923 -
03980 -# -
03985 HMM00390_malic_1_eut1(pf00390pf03949pf01515) EUT1
03990 Hp_euts_violetPink EUT1
03995 HMM10662PduV_1_mix(pf10662) EUT1
04000 HMM06249EutQ_1_eut1(pf06249) EUT1 EP_NTERM
04005 HMM01923cobtr_2_eut1(pf01923) EUT1
04010 HMM01515PTA_B_1_eut1(pf01515) EUT1
04015 H_azure EUT1
04020 P_lightViolet EUT1
04025 HMMalddh_1_eutpdugrm(pf00171) EUT1 EP_NTERM
04030 HMM06723EutJMreB_1_grm(#) EUT1
04035 HMM00465alcdh_3_mix(pf00465) EUT1
04040 HMMtpEuth_1_eut(pf04346) EUT1
04045 HMM06277EutA_1_eut(pf06277) EUT1
04050 HMMetlyEutB_1_EUT1(pf06751) EUT1
04055 HMMetlyEutC_1_eut1(pf05985) EUT1 EP_NTERM
04060 Tsp_rosa EUT1
04065 H_mulberry(pf00936pf16365) EUT1
04070 HMMreg12833_1_EUT1(pf12833) EUT1
04075 -pf01218 -
04080 -pf01520 -
04085 -pf00583 -
04090 -pf11143 -
04095 -pf06572 -
04100 -pf04261 -
04105 HMMnopfamclust3_1(#) RMM1
04110 -# -
04115 -pf00528 -
BMC-P
BMC-H
BMC-T
Aldehyde dehydrogenase
PTAC
Signature enzyme
Alcohol dehydrogenase
Regulator

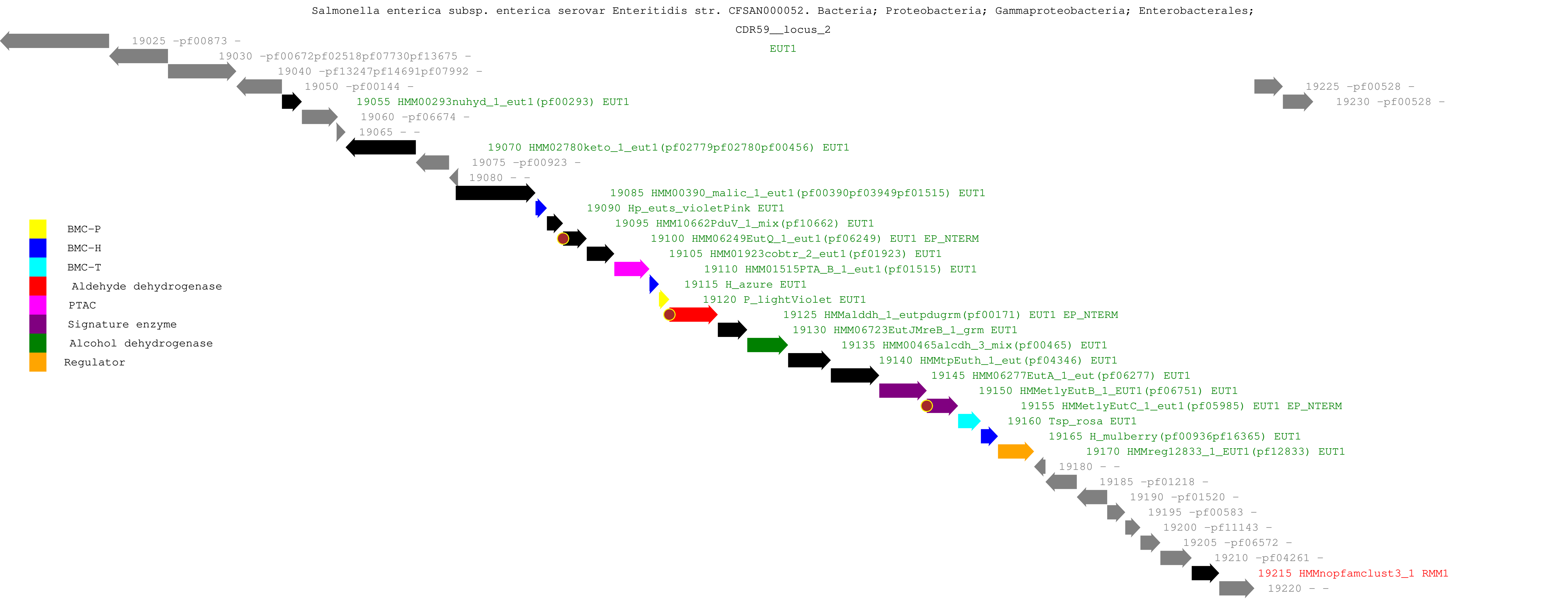
Salmonella enterica subsp. enterica serovar Enteritidis str. CFSAN000052. Bacteria; Proteobacteria; Gammaproteobacteria; Enterobacterales;
CDR59__locus_2
EUT1
19025 -pf00873 -
19030 -pf00672pf02518pf07730pf13675 -
19040 -pf13247pf14691pf07992 -
19050 -pf00144 -
19055 HMM00293nuhyd_1_eut1(pf00293) EUT1
19060 -pf06674 -
19065 - -
19070 HMM02780keto_1_eut1(pf02779pf02780pf00456) EUT1
19075 -pf00923 -
19080 - -
19085 HMM00390_malic_1_eut1(pf00390pf03949pf01515) EUT1
19090 Hp_euts_violetPink EUT1
19095 HMM10662PduV_1_mix(pf10662) EUT1
19100 HMM06249EutQ_1_eut1(pf06249) EUT1 EP_NTERM
19105 HMM01923cobtr_2_eut1(pf01923) EUT1
19110 HMM01515PTA_B_1_eut1(pf01515) EUT1
19115 H_azure EUT1
19120 P_lightViolet EUT1
19125 HMMaldh_1_eutpdugrm(pf00171) EUT1 EP_NTERM
19130 HMM06723EutJMreB_1_grm EUT1
19135 HMM00465alcdh_3_mix(pf00465) EUT1
19140 HMMtpEuth_1_eut(pf04346) EUT1
19145 HMM06277EutA_1_eut(pf06277) EUT1
19150 HMMetlyEutB_1_EUT1(pf06751) EUT1
19155 HMMetlyEutC_1_eut1(pf05985) EUT1 EP_NTERM
19160 Tsp_rosa EUT1
19165 H_mulberry(pf00936pf16365) EUT1
19170 HMMreg12833_1_EUT1(pf12833) EUT1
19180 - -
19185 -pf01218 -
19190 -pf01520 -
19195 -pf00583 -
19200 -pf11143 -
19205 -pf06572 -
19210 -pf04261 -
19215 HMMnopfamclust3_1 RMM1
19220 - -
19225 -pf00528 -
19230 -pf00528 -
BMC-P
BMC-H
BMC-T
Aldehyde dehydrogenase
PTAC
Signature enzyme
Alcohol dehydrogenase
Regulator

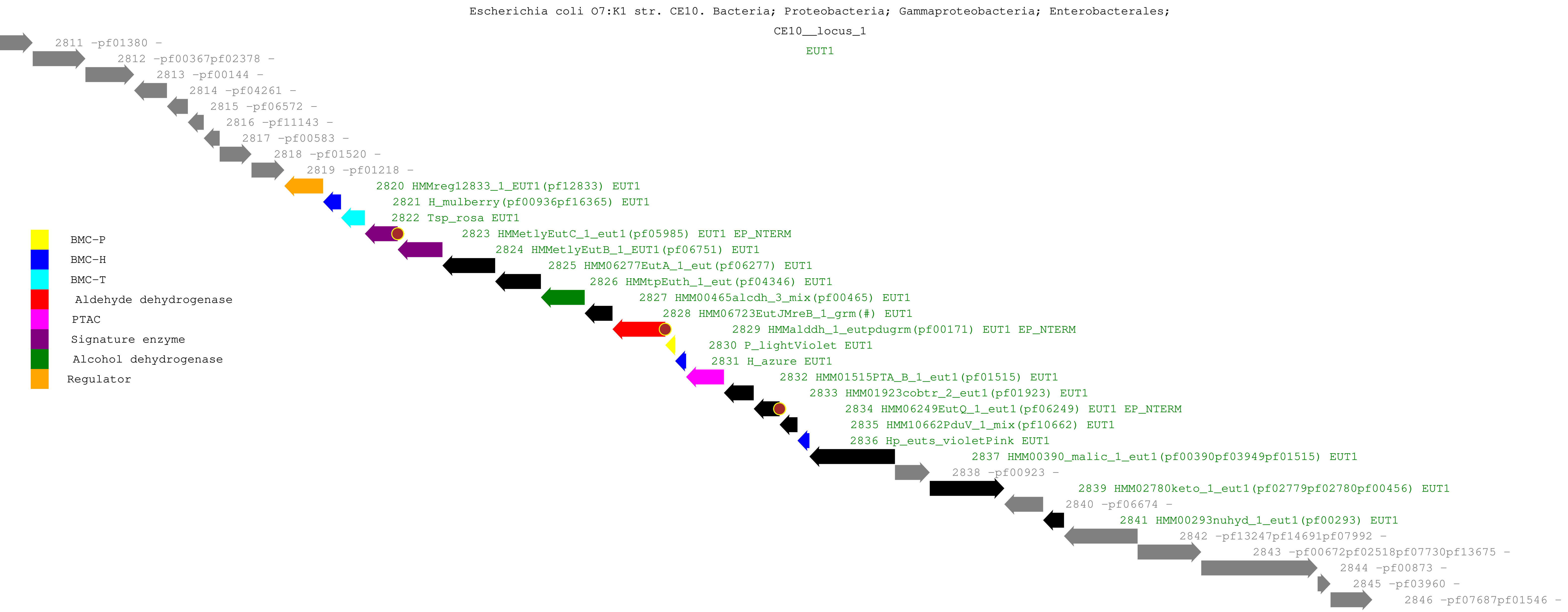

Escherichia coli O7:K1 str. CE10. Bacteria; Proteobacteria; Gammaproteobacteria; Enterobacterales;
CE10__locus_1
EUT1
2811 -pf01380 -
2812 -pf00367pf02378 -
2813 -pf00144 -
2814 -pf04261 -
2815 -pf06572 -
2816 -pf11143 -
2817 -pf00583 -
2818 -pf01520 -
2819 -pf01218 -
2820 HMMreg12833_1_EUT1(pf12833) EUT1
2821 H_mulberry(pf00936pf16365) EUT1
2822 Tsp_rosa EUT1
2823 HMMetlyEutC_1_eut1(pf05985) EUT1 EP_NTERM
2824 HMMetlyEutB_1_EUT1(pf06751) EUT1
2825 HMM06277EutA_1_eut(pf06277) EUT1
2826 HMMtpEuth_1_eut(pf04346) EUT1
2827 HMM00465alcdh_3_mix(pf00465) EUT1
2828 HMM06723EutJMreB_1_grm(#) EUT1
2829 HMMaldh_1_eutpdugrm(pf00171) EUT1 EP_NTERM
2830 P_lightViolet EUT1
2831 H_azure EUT1
2832 HMM01515PTA_B_1_eut1(pf01515) EUT1
2833 HMM01923cobtr_2_eut1(pf01923) EUT1
2834 HMM06249EutQ_1_eut1(pf06249) EUT1 EP_NTERM
2835 HMM10662PduV_1_mix(pf10662) EUT1
2836 Hp_euts_violetPink EUT1
2837 HMM00390_malic_1_eut1(pf00390pf03949pf01515) EUT1
2838 -pf00923 -
2839 HMM02780keto_1_eut1(pf02779pf02780pf00456) EUT1
2840 -pf06674 -
2841 HMM00293nuhyd_1_eut1(pf00293) EUT1
2842 -pf13247pf14691pf07992 -
2843 -pf00672pf02518pf07730pf13675 -
2844 -pf00873 -
2845 -pf03960 -
2846 -pf07687pf01546 -
BMC-P
BMC-H
BMC-T
Aldehyde dehydrogenase
PTAC
Signature enzyme
Alcohol dehydrogenase
Regulator

Klebsiella pneumoniae subsp. pneumoniae. Bacteria; Proteobacteria; Gammaproteobacteria; Enterobacterales;

CEB43__locus_1

EUT1

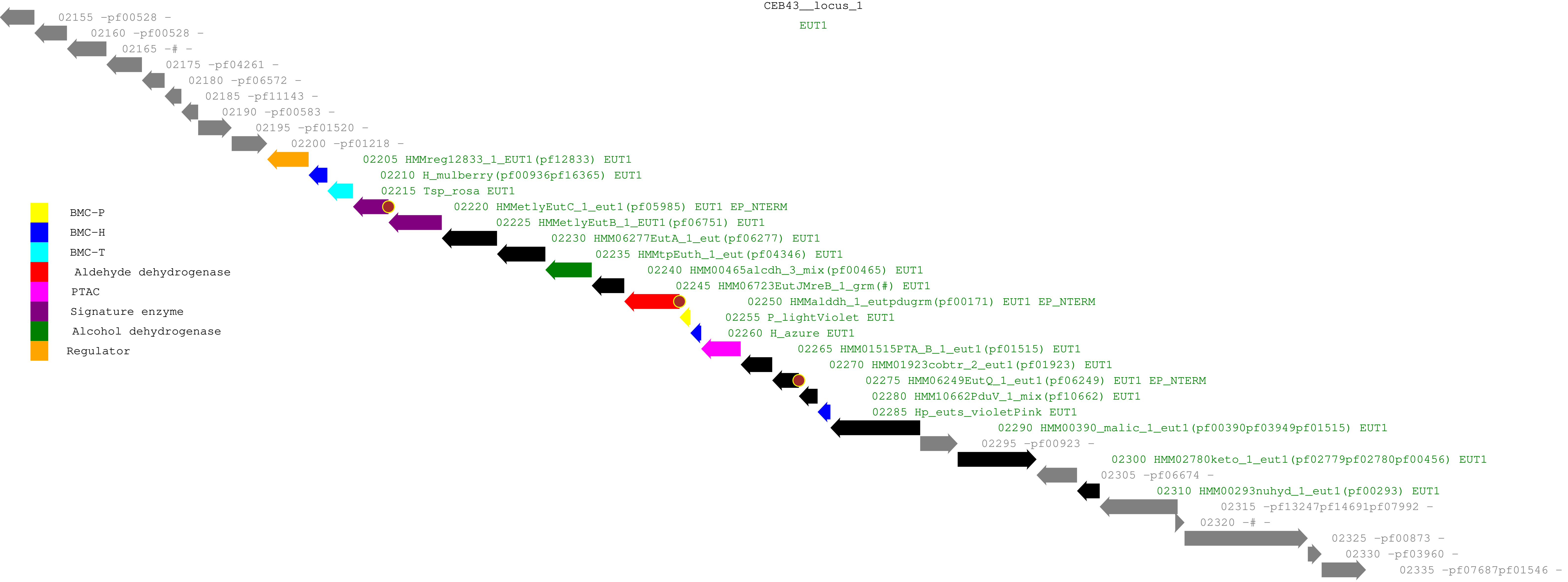

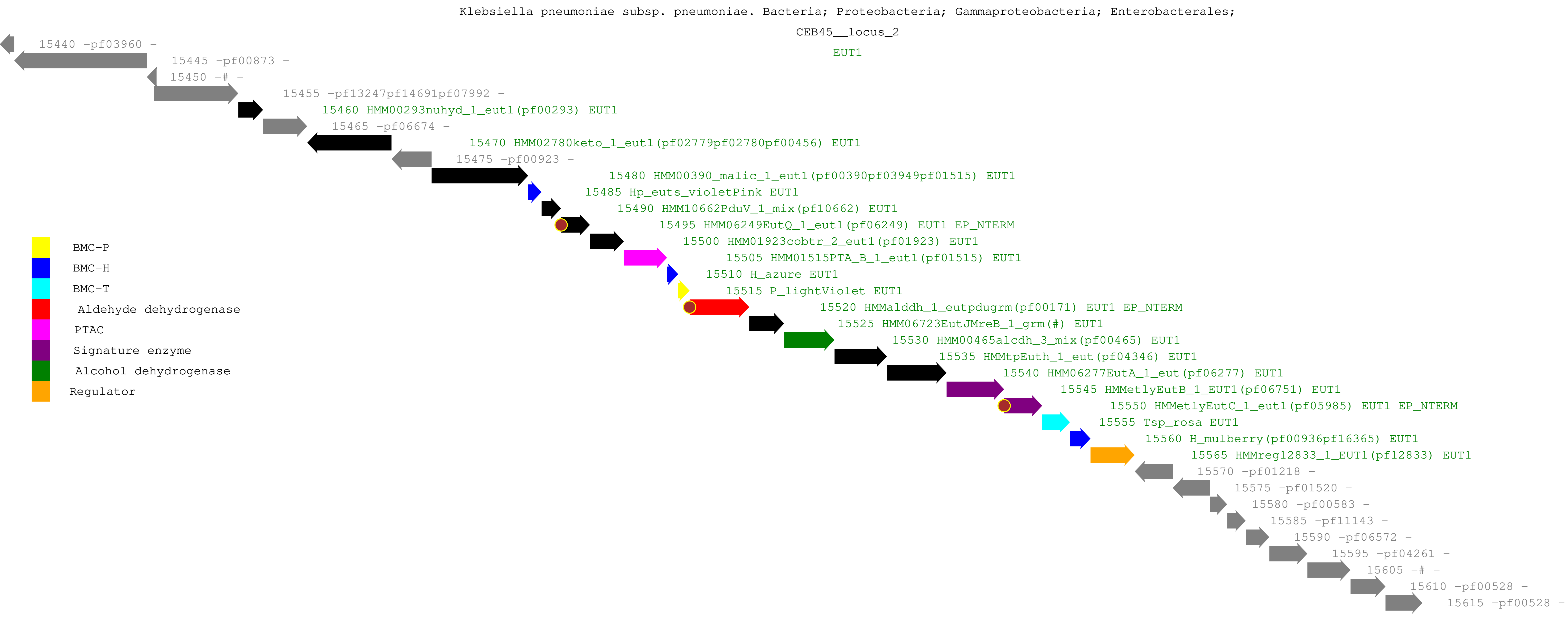
Klebsiella pneumoniae subsp. pneumoniae. Bacteria; Proteobacteria; Gammaproteobacteria; Enterobacterales;
CEB45__locus_2
EUT1
15440 -pf03960 -
15445 -pf00873 -
15450 -# -
15455 -pf13247pf14691pf07992 -
15460 HMM00293nuhyd_1_eut1(pf00293) EUT1
15465 -pf06674 -
15470 HMM02780keto_1_eut1(pf02779pf02780pf00456) EUT1
15475 -pf00923 -
15480 HMM00390_malic_1_eut1(pf00390pf03949pf01515) EUT1
15485 Hp_euts_violetPink EUT1
15490 HMM10662PduV_1_mix(pf10662) EUT1
15495 HMM06249EutQ_1_eut1(pf06249) EUT1 EP_NTERM
15500 HMM01923cobtr_2_eut1(pf01923) EUT1
15505 HMM01515PTA_B_1_eut1(pf01515) EUT1
15510 H_azure EUT1
15515 P_lightViolet EUT1
15520 HMMalddh_1_eutpdugrm(pf00171) EUT1 EP_NTERM
15525 HMM06723EutJMreB_1_grm(#) EUT1
15530 HMM00465alcdh_3_mix(pf00465) EUT1
15535 HMMtpEuth_1_eut(pf04346) EUT1
15540 HMM06277EutA_1_eut(pf06277) EUT1
15545 HMMetlyEutB_1_EUT1(pf06751) EUT1
15550 HMMetlyEutC_1_eut1(pf05985) EUT1 EP_NTERM
15555 Tsp_rosa EUT1
15560 H_mulberry(pf00936pf16365) EUT1
15565 HMMreg12833_1_EUT1(pf12833) EUT1
15570 -pf01218 -
15575 -pf01520 -
15580 -pf00583 -
15585 -pf11143 -
15590 -pf06572 -
15595 -pf04261 -
15605 -# -
15610 -pf00528 -
15615 -pf00528 -
BMC-P
BMC-H
BMC-T
Aldehyde dehydrogenase
PTAC
Signature enzyme
Alcohol dehydrogenase
Regulator

Klebsiella pneumoniae subsp. pneumoniae. Bacteria; Proteobacteria; Gammaproteobacteria; Enterobacterales;

CEB46__locus_2

EUT1

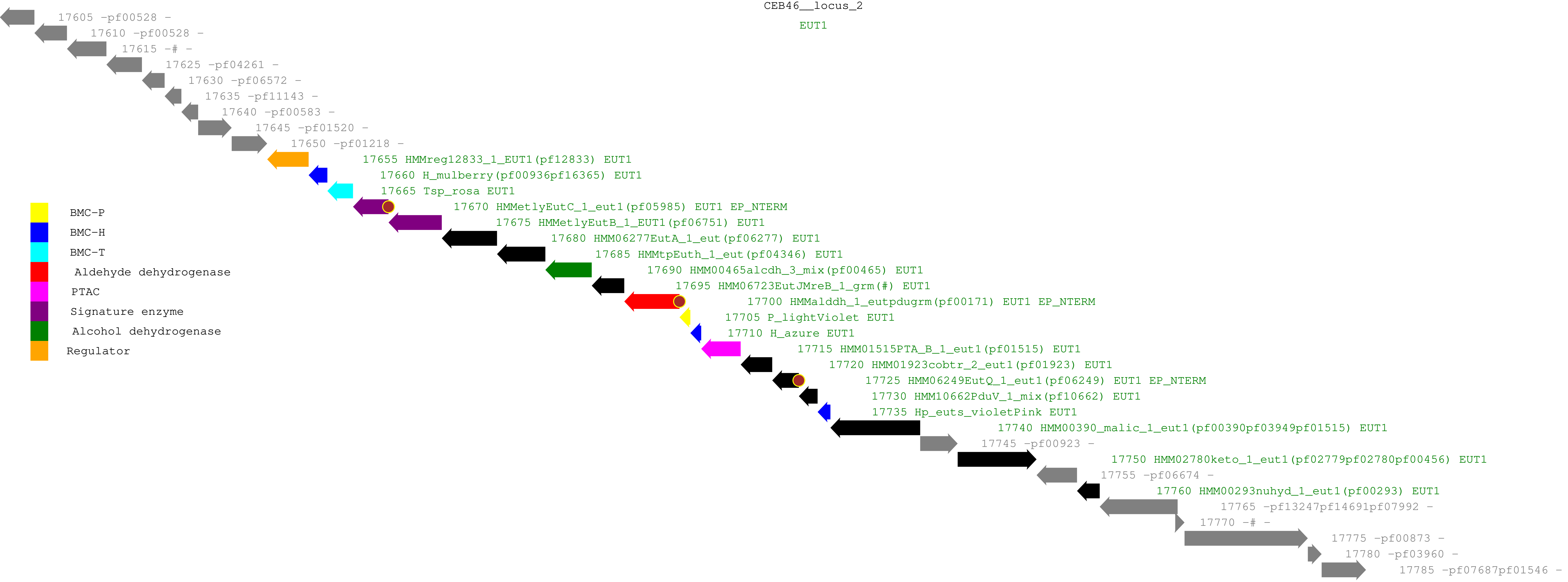

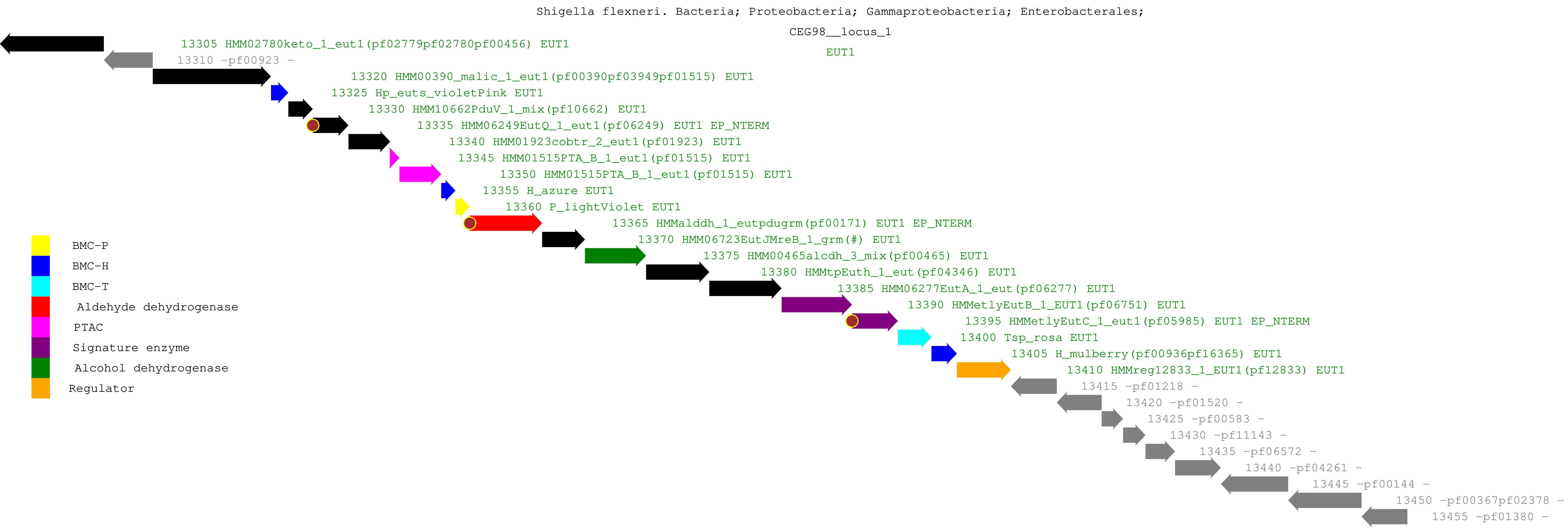
Shigella flexneri. Bacteria; Proteobacteria; Gammaproteobacteria; Enterobacterales;
CEG98__locus_1
EUT1
13305 HMM02780keto_1_eut1(pf02779pf02780pf00456) EUT1
13310 -pf00923 -
13320 HMM00390_malic_1_eut1(pf00390pf03949pf01515) EUT1
13325 Hp_euts_violetPink EUT1
13330 HMM10662PduV_1_mix(pf10662) EUT1
13335 HMM06249EutQ_1_eut1(pf06249) EUT1 EP_NTERM
13340 HMM01923cobtr_2_eut1(pf01923) EUT1
13345 HMM01515PTA_B_1_eut1(pf01515) EUT1
13350 HMM01515PTA_B_1_eut1(pf01515) EUT1
13355 H_azure EUT1
13360 P_lightViolet EUT1
13365 HMMalddh_1_eutpdugrm(pf00171) EUT1 EP_NTERM
13370 HMM06723EutJMreB_1_grm(#) EUT1
13375 HMM00465alcdh_3_mix(pf00465) EUT1
13380 HMMtpEuth_1_eut(pf04346) EUT1
13385 HMM06277EutA_1_eut(pf06277) EUT1
13390 HMMetlyEutB_1_EUT1(pf06751) EUT1
13395 HMMetlyEutC_1_eut1(pf05985) EUT1 EP_NTERM
13400 Tsp_rosa EUT1
13405 H_mulberry(pf00936pf16365) EUT1
13410 HMMreg12833_1_EUT1(pf12833) EUT1
13415 -pf01218 -
13420 -pf01520 -
13425 -pf00583 -
13430 -pf11143 -
13435 -pf06572 -
13440 -pf04261 -
13445 -pf00144 -
13450 -pf00367pf02378 -
13455 -pf01380 -
BMC-P
BMC-H
BMC-T
Aldehyde dehydrogenase
PTAC
Signature enzyme
Alcohol dehydrogenase
Regulator

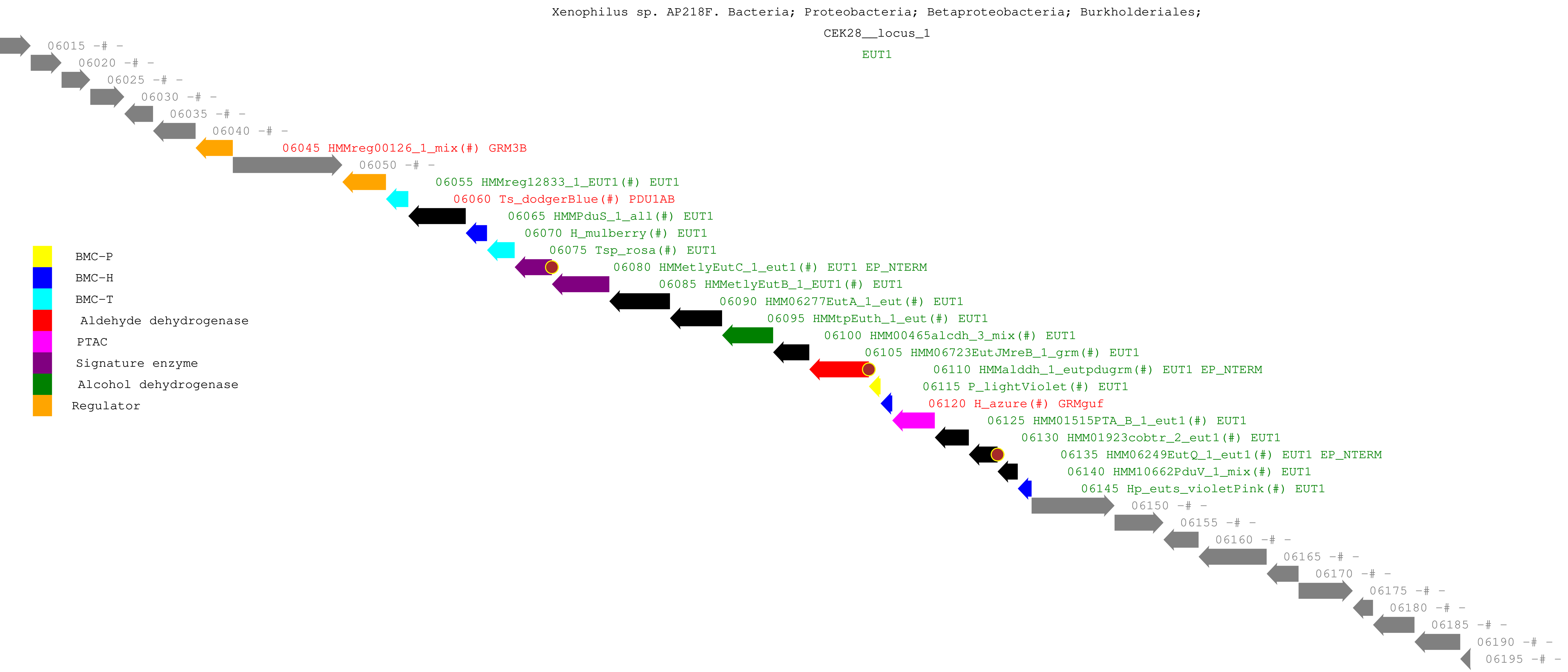

Xenophilus sp. AP218F. Bacteria; Proteobacteria; Betaproteobacteria; Burkholderiales;
CEK28__locus_1
EUT1
06015 -# -
06020 -# -
06025 -# -
06030 -# -
06035 -# -
06040 -# -
06045 HMMreg00126_1_mix(#) GRM3B
06050 -# -
06055 HMMreg12833_1_EUT1(#) EUT1
06060 Ts_dodgerBlue(#) PDU1AB
06065 HMMPduS_1_all(#) EUT1
06070 H_mulberry(#) EUT1
06075 Tsp_rosa(#) EUT1
06080 HMMetlyEutC_1_eut1(#) EUT1 EP_NTERM
06085 HMMetlyEutB_1_EUT1(#) EUT1
06090 HMM06277EutA_1_eut(#) EUT1
06095 HMMtpEuth_1_eut(#) EUT1
06100 HMM00465alcdh_3_mix(#) EUT1
06105 HMM06723EutJMreB_1_grm(#) EUT1
06110 HMMaldh_1_eutpdugrm(#) EUT1 EP_NTERM
06115 P_lightViolet(#) EUT1
06120 H_azure(#) GRMguf
06125 HMM01515PTA_B_1_eut1(#) EUT1
06130 HMM01923cobtr_2_eut1(#) EUT1
06135 HMM06249EutQ_1_eut1(#) EUT1 EP_NTERM
06140 HMM10662PduV_1_mix(#) EUT1
06145 Hp_euts_violetPink(#) EUT1
06150 -# -
06155 -# -
06160 -# -
06165 -# -
06170 -# -
06175 -# -
06180 -# -
06185 -# -
06190 -# -
06195 -# -
BMC-P
BMC-H
BMC-T
Aldehyde dehydrogenase
PTAC
Signature enzyme
Alcohol dehydrogenase
Regulator

Escherichia coli O157. Bacteria; Proteobacteria; Gammaproteobacteria; Enterobacterales;

CEP72__locus_1

EUT1

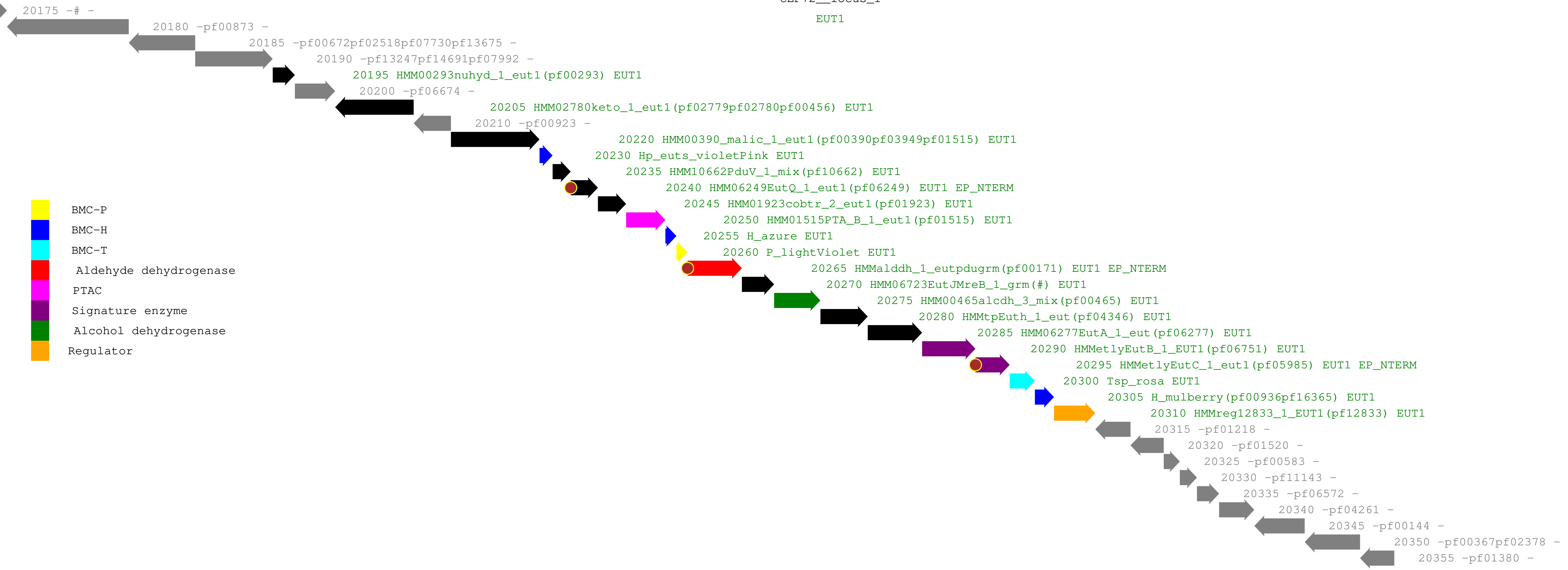

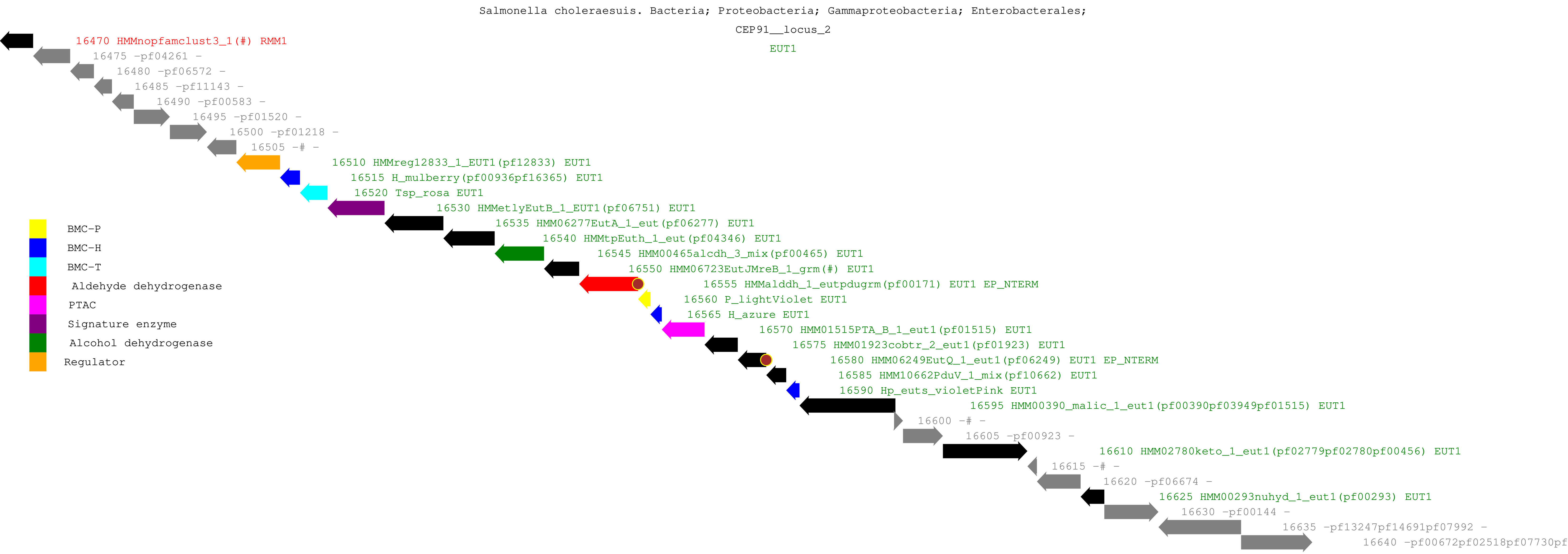
Salmonella choleraesuis. Bacteria; Proteobacteria; Gammaproteobacteria; Enterobacterales;
CEP91__locus_2
EUT1
16470 HMMnopfamclust3_1(#) RMM1
16475 -pf04261 -
16480 -pf06572 -
16485 -pf11143 -
16490 -pf00583 -
16495 -pf01520 -
16500 -pf01218 -
16505 -# -
16510 HMMreg12833_1_EUT1(pf12833) EUT1
16515 H_mulberry(pf00936pf16365) EUT1
16520 Tsp_rosa EUT1
16530 HMMetlyEutB_1_EUT1(pf06751) EUT1
16535 HMM06277EutA_1_eut(pf06277) EUT1
16540 HMMtpEuth_1_eut(pf04346) EUT1
16545 HMM00465alcdh_3_mix(pf00465) EUT1
16550 HMM06723EutJMreB_1_grm(#) EUT1
16555 HMMalddh_1_eutpdugrm(pf00171) EUT1 EP_NTERM
16560 P_lightViolet EUT1
16565 H_azure EUT1
16570 HMM01515PTA_B_1_eut1(pf01515) EUT1
16575 HMM01923cobtr_2_eut1(pf01923) EUT1
16580 HMM06249EutQ_1_eut1(pf06249) EUT1 EP_NTERM
16585 HMM10662PduV_1_mix(pf10662) EUT1
16590 Hp_euts_violetPink EUT1
16595 HMM00390_malic_1_eut1(pf00390pf03949pf01515) EUT1
16600 -# -
16605 -pf00923 -
16610 HMM02780keto_1_eut1(pf02779pf02780pf00456) EUT1
16615 -# -
16620 -pf06674 -
16625 HMM00293nuhyd_1_eut1(pf00293) EUT1
16630 -pf00144 -
16635 -pf13247pf14691pf07992 -
16640 -pf00672pf02518pf07730pf
BMC-P
BMC-H
BMC-T
Aldehyde dehydrogenase
PTAC
Signature enzyme
Alcohol dehydrogenase
Regulator

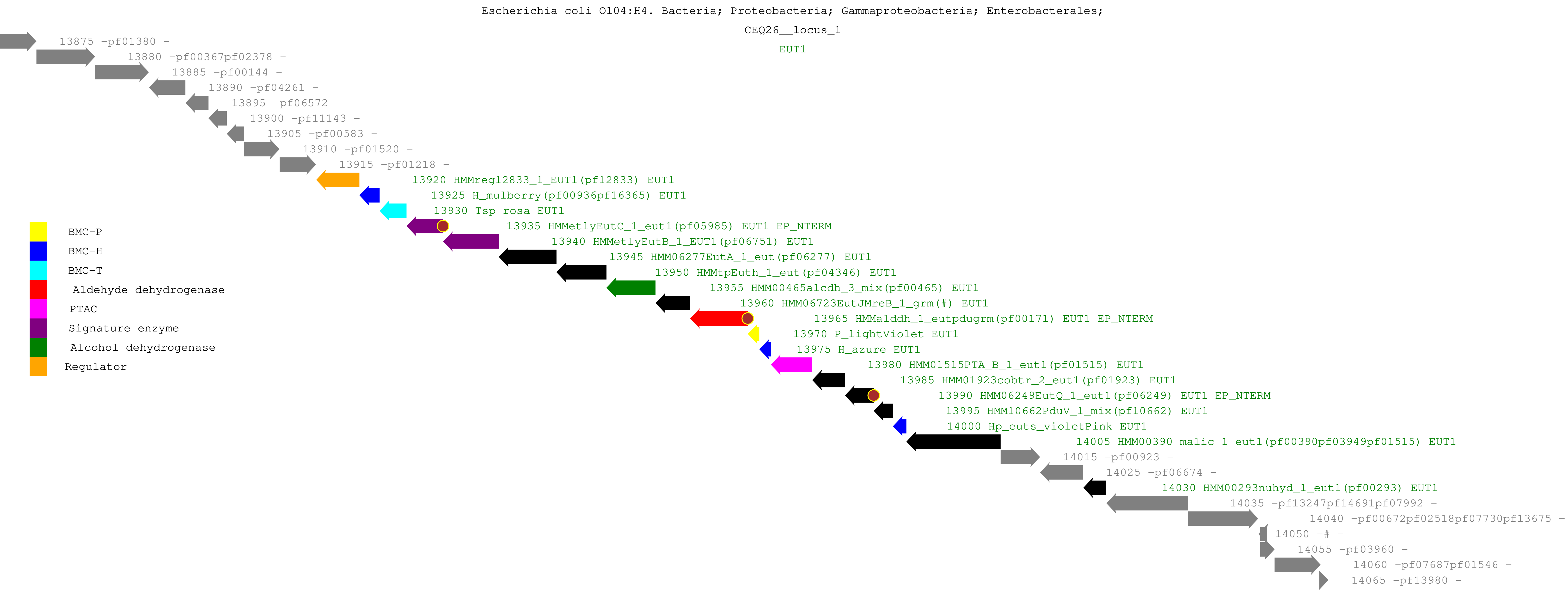
Escherichia coli O104:H4. Bacteria; Proteobacteria; Gammaproteobacteria; Enterobacterales;
CEQ26__locus_1
EUT1
13875 -pf01380 -
13880 -pf00367pf02378 -
13885 -pf00144 -
13890 -pf04261 -
13895 -pf06572 -
13900 -pf11143 -
13905 -pf00583 -
13910 -pf01520 -
13915 -pf01218 -
13920 HMMreg12833_1_EUT1(pf12833) EUT1
13925 H_mulberry(pf00936pf16365) EUT1
13930 Tsp_rosa EUT1
13935 HMMetlyEutC_1_eut1(pf05985) EUT1 EP_NTERM
13940 HMMetlyEutB_1_EUT1(pf06751) EUT1
13945 HMM06277EutA_1_eut(pf06277) EUT1
13950 HMMtpEuth_1_eut(pf04346) EUT1
13955 HMM00465alcdh_3_mix(pf00465) EUT1
13960 HMM06723EutJMreB_1_grm(#) EUT1
13965 HMMaldddh_1_eutpdugrm(pf00171) EUT1 EP_NTERM
13970 P_lightViolet EUT1
13975 H_azure EUT1
13980 HMM01515PTA_B_1_eut1(pf01515) EUT1
13985 HMM01923cobtr_2_eut1(pf01923) EUT1
13990 HMM06249EutQ_1_eut1(pf06249) EUT1 EP_NTERM
13995 HMM10662PduV_1_mix(pf10662) EUT1
14000 Hp_euts_violetPink EUT1
14005 HMM00390_malic_1_eut1(pf00390pf03949pf01515) EUT1
14015 -pf00923 -
14025 -pf06674 -
14030 HMM00293nuhyd_1_eut1(pf00293) EUT1
14035 -pf13247pf14691pf07992 -
14040 -pf00672pf02518pf07730pf13675 -
14050 -# -
14055 -pf03960 -
14060 -pf07687pf01546 -
14065 -pf13980 -
BMC-P
BMC-H
BMC-T
Aldehyde dehydrogenase
PTAC
Signature enzyme
Alcohol dehydrogenase
Regulator

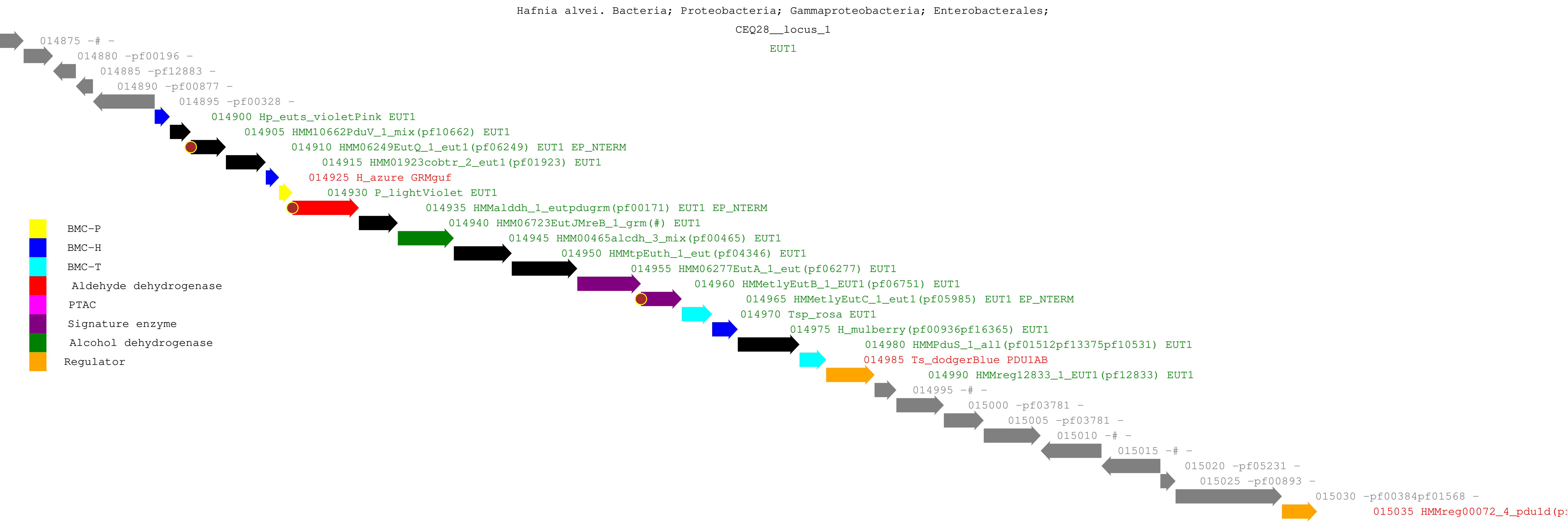
Hafnia alvei. Bacteria; Proteobacteria; Gammaproteobacteria; Enterobacterales;
CEQ28__locus_1
EUT1
014875 -# -
014880 -pf00196 -
014885 -pf12883 -
014890 -pf00877 -
014895 -pf00328 -
014900 Hp_euts_violetPink EUT1
014905 HMM10662PduV_1_mix(pf10662) EUT1
014910 HMM06249EutQ_1_eut1(pf06249) EUT1 EP_NTERM
014915 HMM01923cobtr_2_eut1(pf01923) EUT1
014925 H_azure GRMguf
014930 P_lightViolet EUT1
014935 HMMalddh_1_eutpdugrm(pf00171) EUT1 EP_NTERM
014940 HMM06723EutJMreB_1_grm(#) EUT1
014945 HMM00465alcdh_3_mix(pf00465) EUT1
014950 HMMtpEuth_1_eut(pf04346) EUT1
014955 HMM06277EutA_1_eut(pf06277) EUT1
014960 HMMetlyEutB_1_EUT1(pf06751) EUT1
014965 HMMetlyEutC_1_eut1(pf05985) EUT1 EP_NTERM
014970 Tsp_rosa EUT1
014975 H_mulberry(pf00936pf16365) EUT1
014980 HMMPduS_1_all(pf01512pf13375pf10531) EUT1
014985 Ts_dodgerBlue PDU1AB
014990 HMMreg12833_1_EUT1(pf12833) EUT1
014995 -# -
015000 -pf03781 -
015005 -pf03781 -
015010 -# -
015015 -# -
015020 -pf05231 -
015025 -pf00893 -
015030 -pf00384pf01568 -
015035 HMMreg00072_4_pdu1d(p
BMC-P
BMC-H
BMC-T
Aldehyde dehydrogenase
PTAC
Signature enzyme
Alcohol dehydrogenase
Regulator

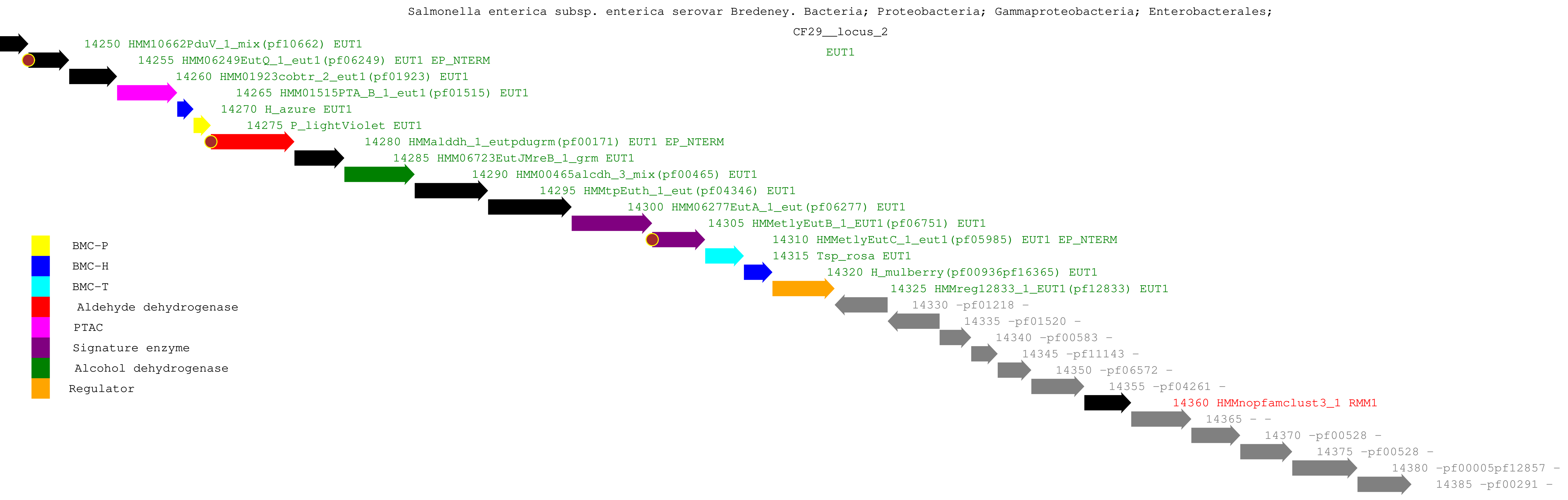
Salmonella enterica subsp. enterica serovar Bredeney. Bacteria; Proteobacteria; Gammaproteobacteria; Enterobacterales;
CF29__locus_2
EUT1
14250 HMM10662PduV_1_mix(pf10662) EUT1
14255 HMM06249EutQ_1_eut1(pf06249) EUT1 EP_NTERM
14260 HMM01923cobtr_2_eut1(pf01923) EUT1
14265 HMM01515PTA_B_1_eut1(pf01515) EUT1
14270 H_azure EUT1
14275 P_lightViolet EUT1
14280 HMMalddh_1_eutpdugrm(pf00171) EUT1 EP_NTERM
14285 HMM06723EutJMreB_1_grm EUT1
14290 HMM00465alcdh_3_mix(pf00465) EUT1
14295 HMMtpEuth_1_eut(pf04346) EUT1
14300 HMM06277EutA_1_eut(pf06277) EUT1
14305 HMMetlyEutB_1_EUT1(pf06751) EUT1
14310 HMMetlyEutC_1_eut1(pf05985) EUT1 EP_NTERM
14315 Tsp_rosa EUT1
14320 H_mulberry(pf00936pf16365) EUT1
14325 HMMreg12833_1_EUT1(pf12833) EUT1
14330 -pf01218 -
14335 -pf01520 -
14340 -pf00583 -
14345 -pf11143 -
14350 -pf06572 -
14355 -pf04261 -
14360 HMMnopfamclust3_1 RMM1
14365 - -
14370 -pf00528 -
14375 -pf00528 -
14380 -pf00005pf12857 -
14385 -pf00291 -
BMC-P
BMC-H
BMC-T
Aldehyde dehydrogenase
PTAC
Signature enzyme
Alcohol dehydrogenase
Regulator

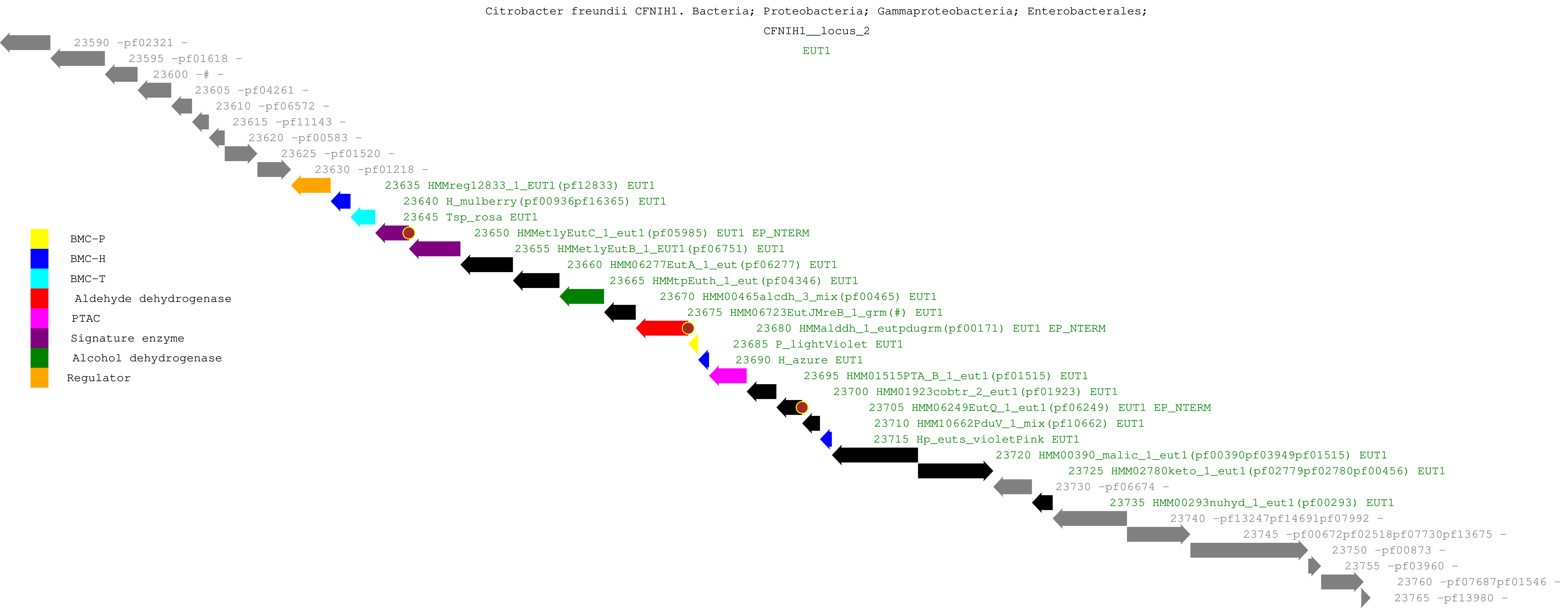
Citrobacter freundii CFNIH1. Bacteria; Proteobacteria; Gammaproteobacteria; Enterobacterales;
CFNIH1__locus_2
EUT1
23590 -pf02321 -
23595 -pf01618 -
23600 -# -
23605 -pf04261 -
23610 -pf06572 -
23615 -pf11143 -
23620 -pf00583 -
23625 -pf01520 -
23630 -pf01218 -
23635 HMMreg12833_1_EUT1(pf12833) EUT1
23640 H_mulberry(pf00936pf16365) EUT1
23645 Tsp_rosa EUT1
23650 HMMetlyEutC_1_eut1(pf05985) EUT1 EP_NTERM
23655 HMMetlyEutB_1_EUT1(pf06751) EUT1
23660 HMM06277EutA_1_eut(pf06277) EUT1
23665 HMMtpEuth_1_eut(pf04346) EUT1
23670 HMM00465alcdh_3_mix(pf00465) EUT1
23675 HMM06723EutJMreB_1_grm(#) EUT1
23680 HMMalddh_1_eutpdugrm(pf00171) EUT1 EP_NTERM
23685 P_lightViolet EUT1
23690 H_azure EUT1
23695 HMM01515PTA_B_1_eut1(pf01515) EUT1
23700 HMM01923cobtr_2_eut1(pf01923) EUT1
23705 HMM06249EutQ_1_eut1(pf06249) EUT1 EP_NTERM
23710 HMM10662PduV_1_mix(pf10662) EUT1
23715 Hp_euts_violetPink EUT1
23720 HMM00390_malic_1_eut1(pf00390pf03949pf01515) EUT1
23725 HMM02780keto_1_eut1(pf02779pf02780pf00456) EUT1
23730 -pf06674 -
23735 HMM00293nuhyd_1_eut1(pf00293) EUT1
23740 -pf13247pf14691pf07992 -
23745 -pf00672pf02518pf07730pf13675 -
23750 -pf00873 -
23755 -pf03960 -
23760 -pf07687pf01546 -
23765 -pf13980 -
BMC-P
BMC-H
BMC-T
Aldehyde dehydrogenase
PTAC
Signature enzyme
Alcohol dehydrogenase
Regulator

Salmonella enterica subsp. enterica serovar Choleraesuis str. CFSAN000514. Bacteria; Proteobacteria; Gammaproteobacteria; Enterobacterales;

CFSAN000514__locus_1

EUT1

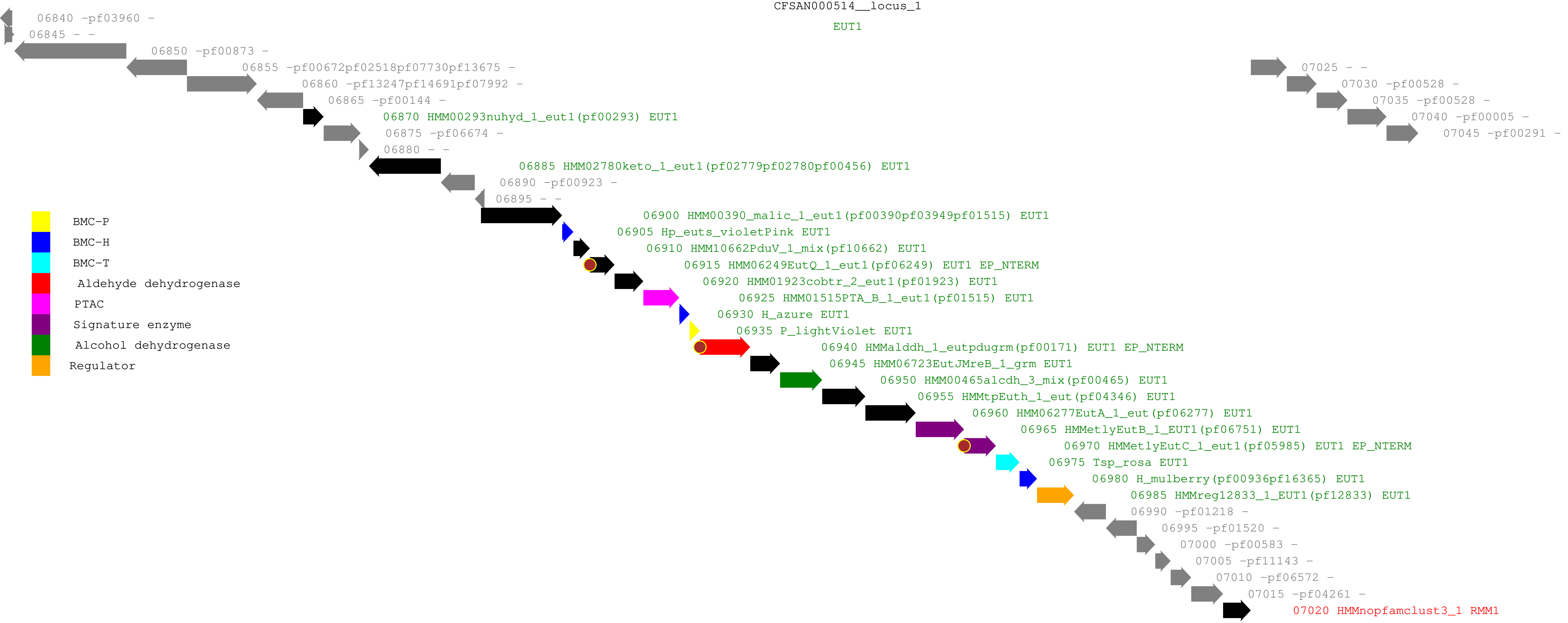

Salmonella enterica subsp. houtenae str. CFSAN000557. Bacteria; Proteobacteria; Gammaproteobacteria; Enterobacterales;

CFSAN000557__locus_1

EUT1

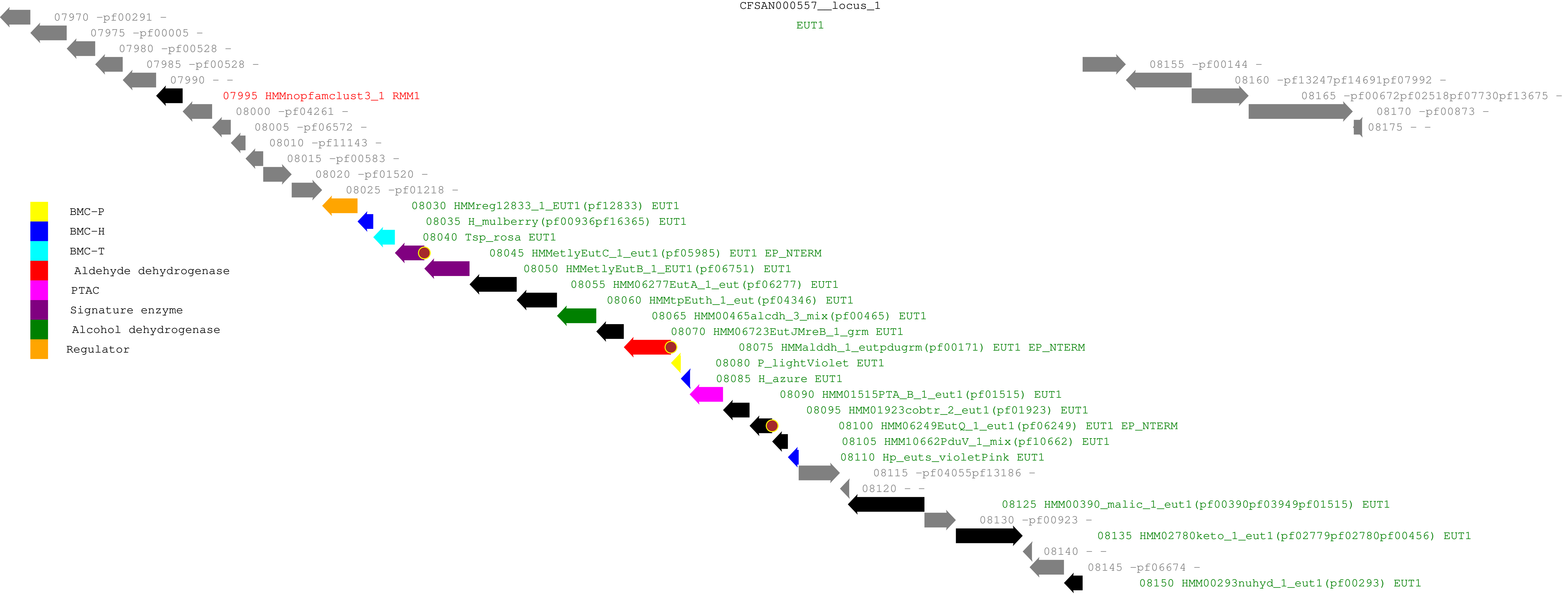

Salmonella enterica subsp. salamae str. CFSAN000559. Bacteria; Proteobacteria; Gammaproteobacteria; Enterobacterales;

CFSAN000559__locus_1

EUT1

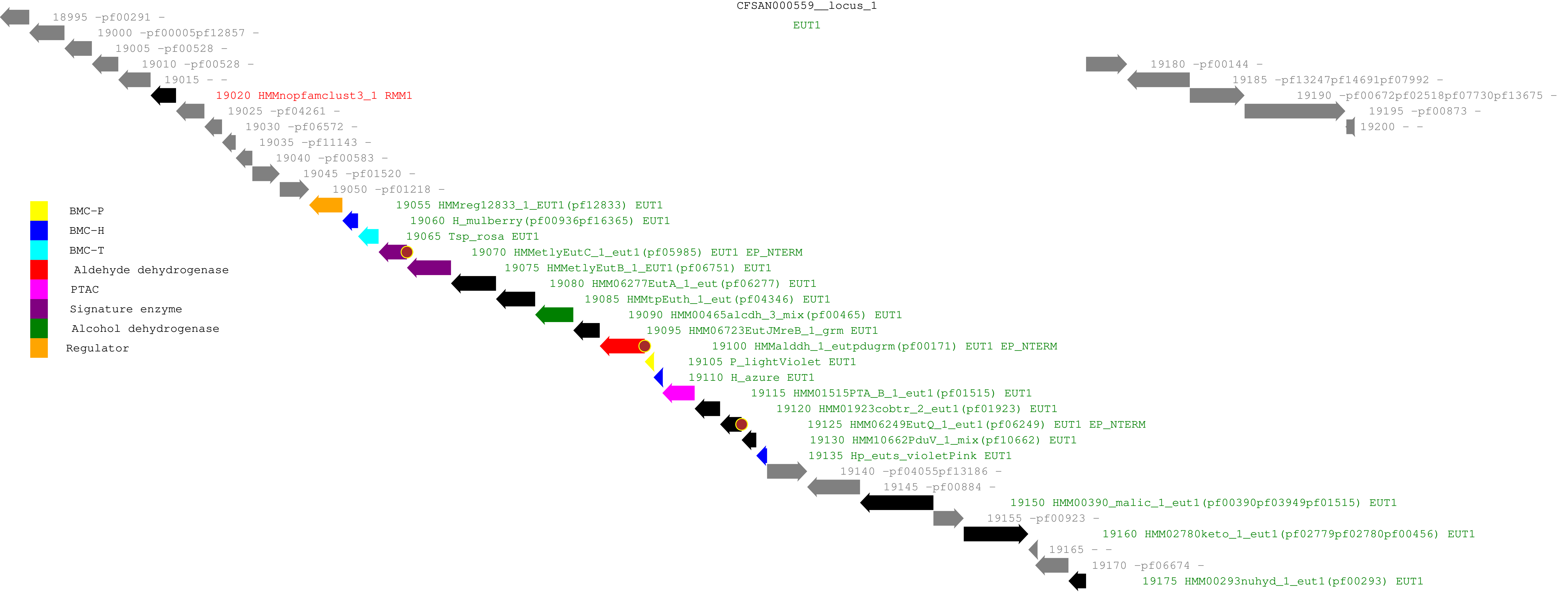

Salmonella enterica subsp. arizonae str. CFSAN000560. Bacteria; Proteobacteria; Gammaproteobacteria; Enterobacterales;

CFSAN000560__locus_1

EUT1

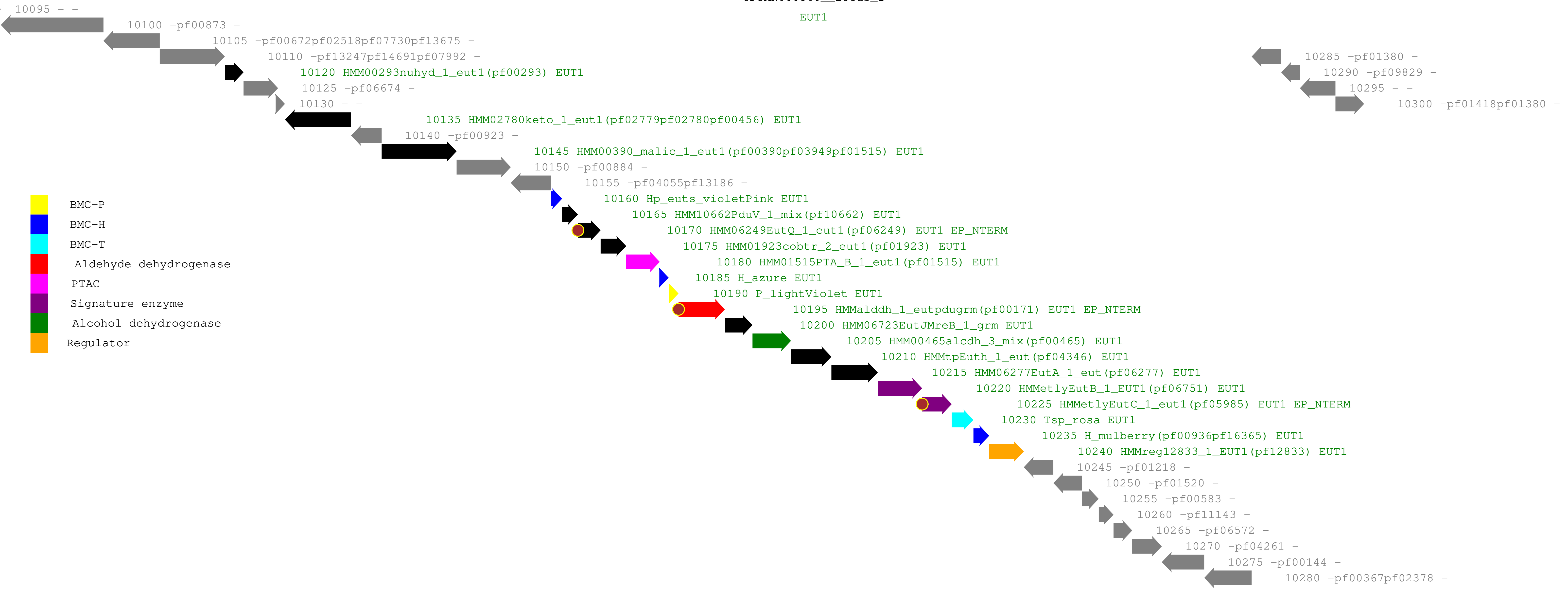

Salmonella enterica subsp. enterica serovar Gallinarum str. CFSAN000571. Bacteria; Proteobacteria; Gammaproteobacteria; Enterobacterales;

CFSAN000571__locus_2

EUT1

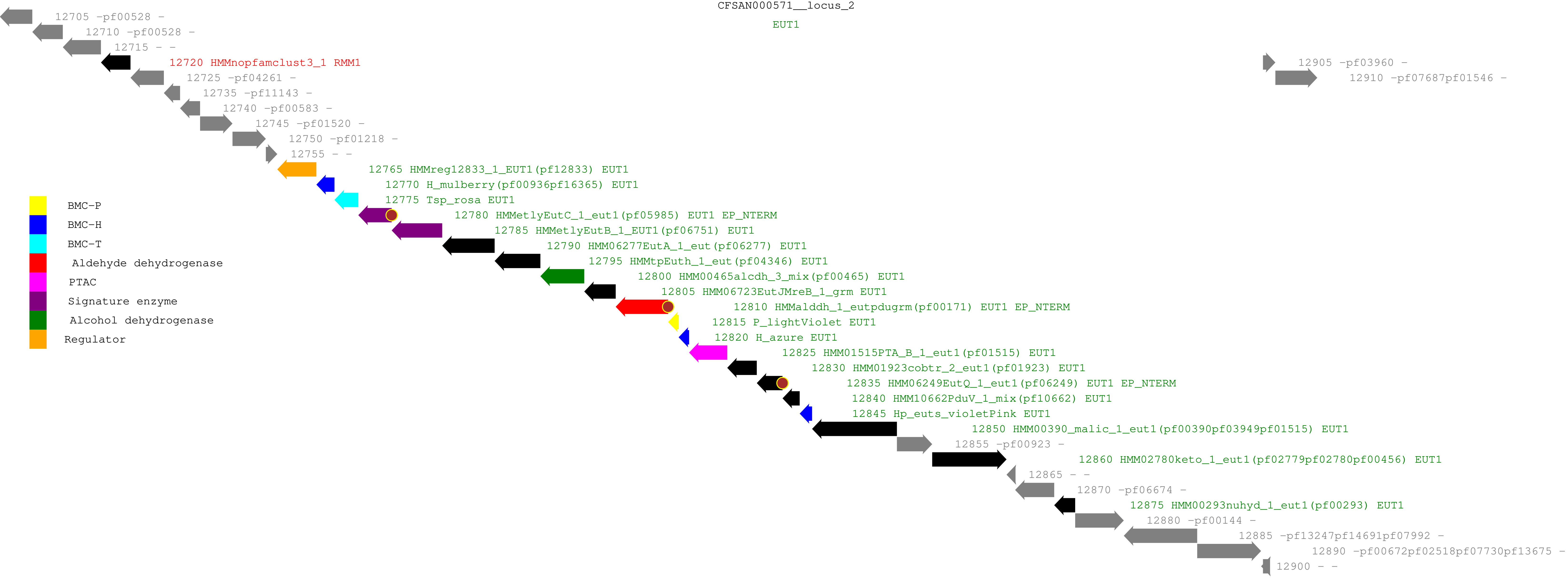

Salmonella enterica subsp. enterica serovar Muenchen str. CFSAN000593. Bacteria; Proteobacteria; Gammaproteobacteria; Enterobacterales;

CFSAN000593__locus_2

EUT1

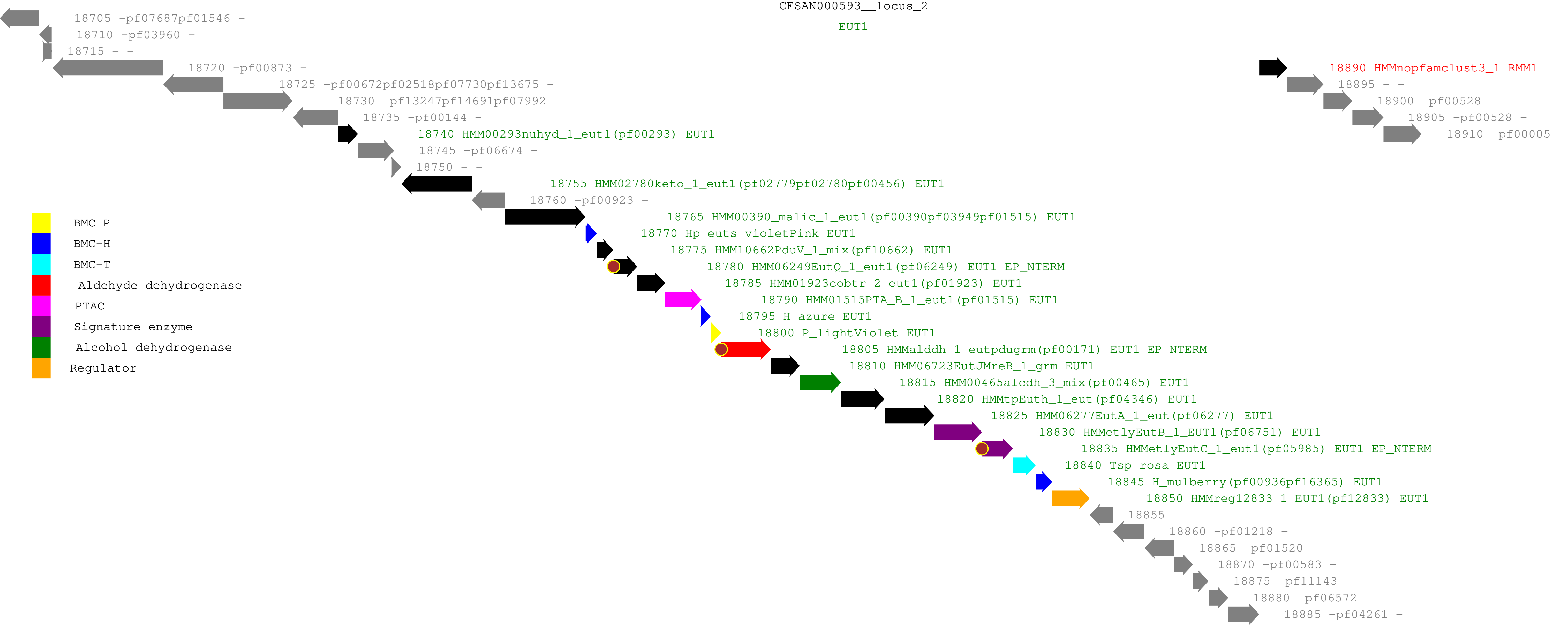

Salmonella enterica subsp. enterica serovar Muenchen str. CFSAN000596. Bacteria; Proteobacteria; Gammaproteobacteria; Enterobacterales;

CFSAN000596__locus_1

EUT1

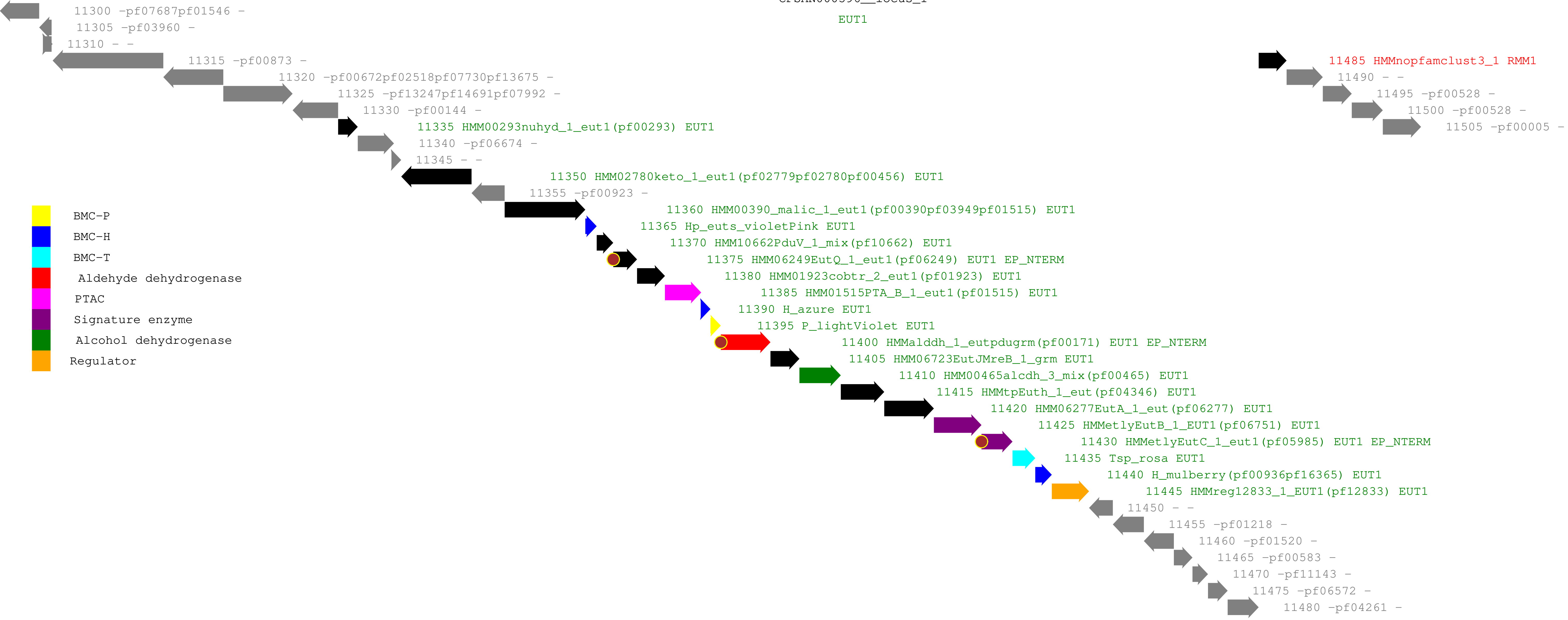

Salmonella enterica subsp. enterica serovar Panama str. CFSAN000600. Bacteria; Proteobacteria; Gammaproteobacteria; Enterobacterales;

CFSAN000600__locus_2

EUT1

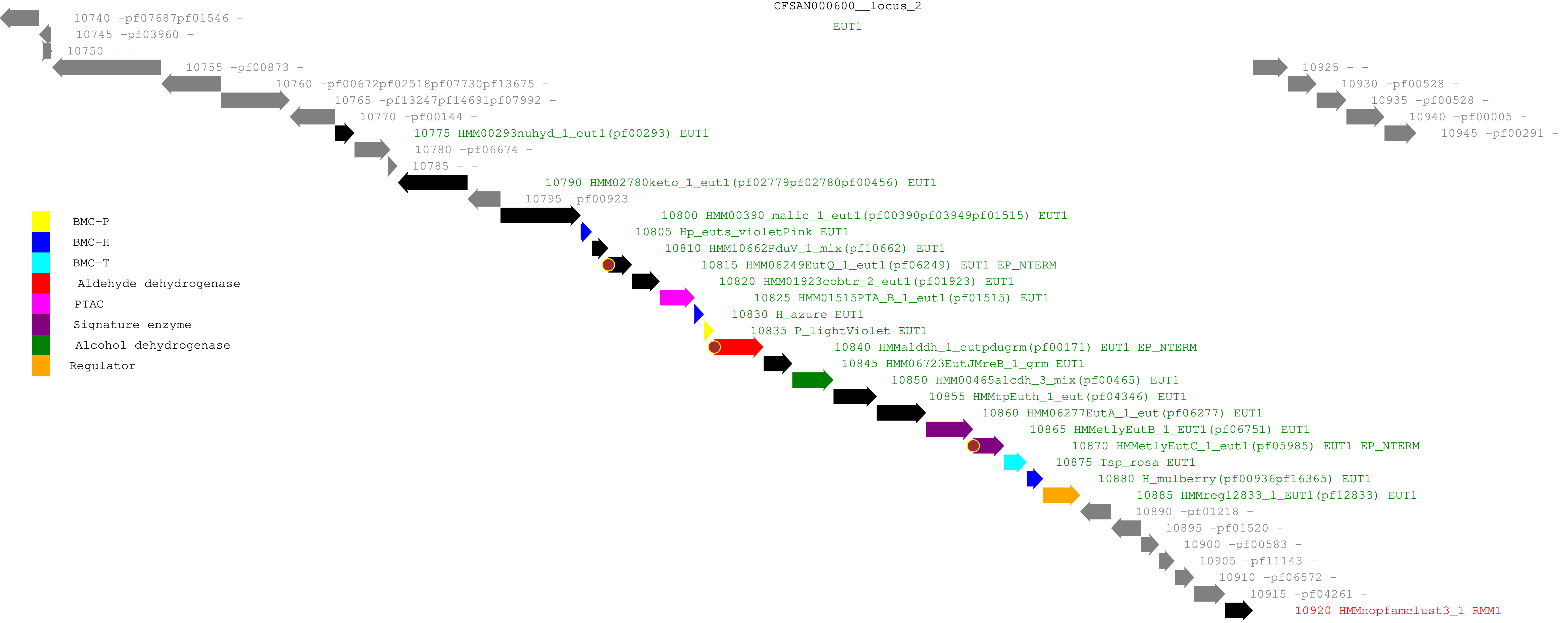

Salmonella enterica subsp. enterica serovar Saintpaul str. CFSAN000618. Bacteria; Proteobacteria; Gammaproteobacteria; Enterobacterales;

CFSAN000618__locus_1

EUT1

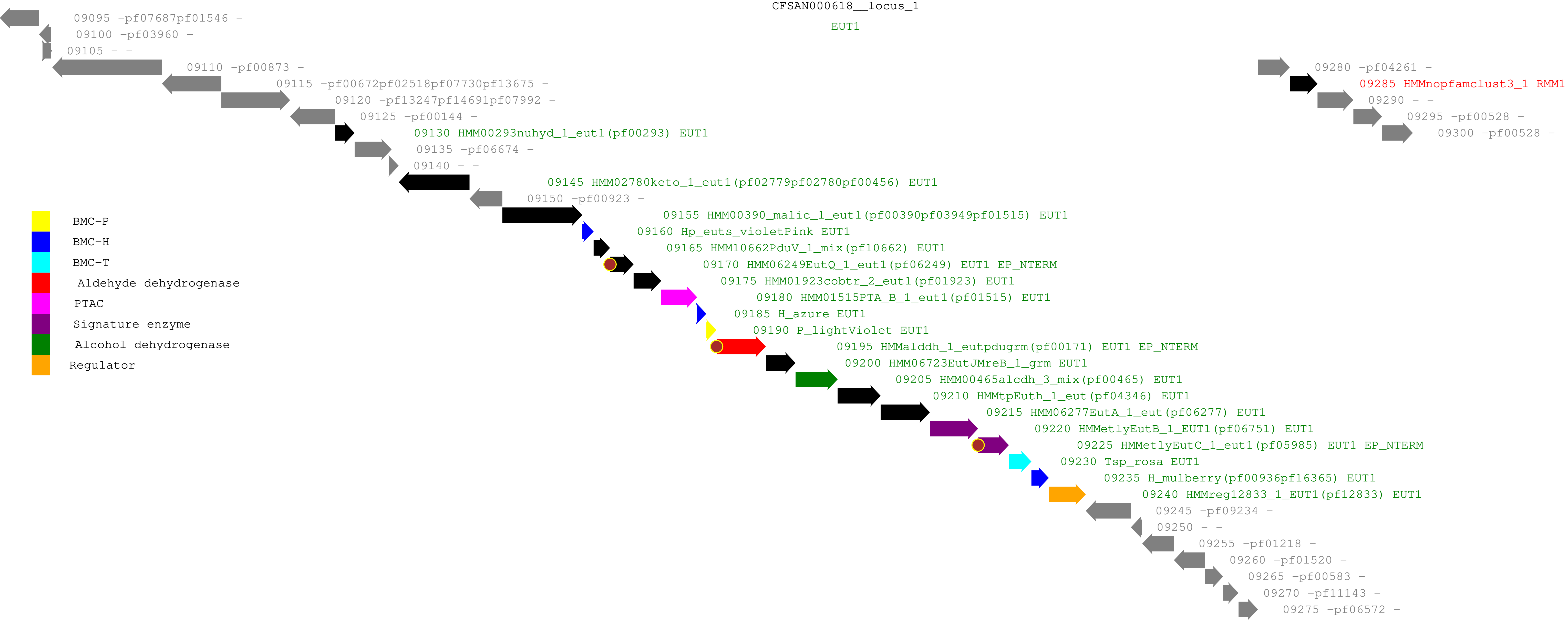

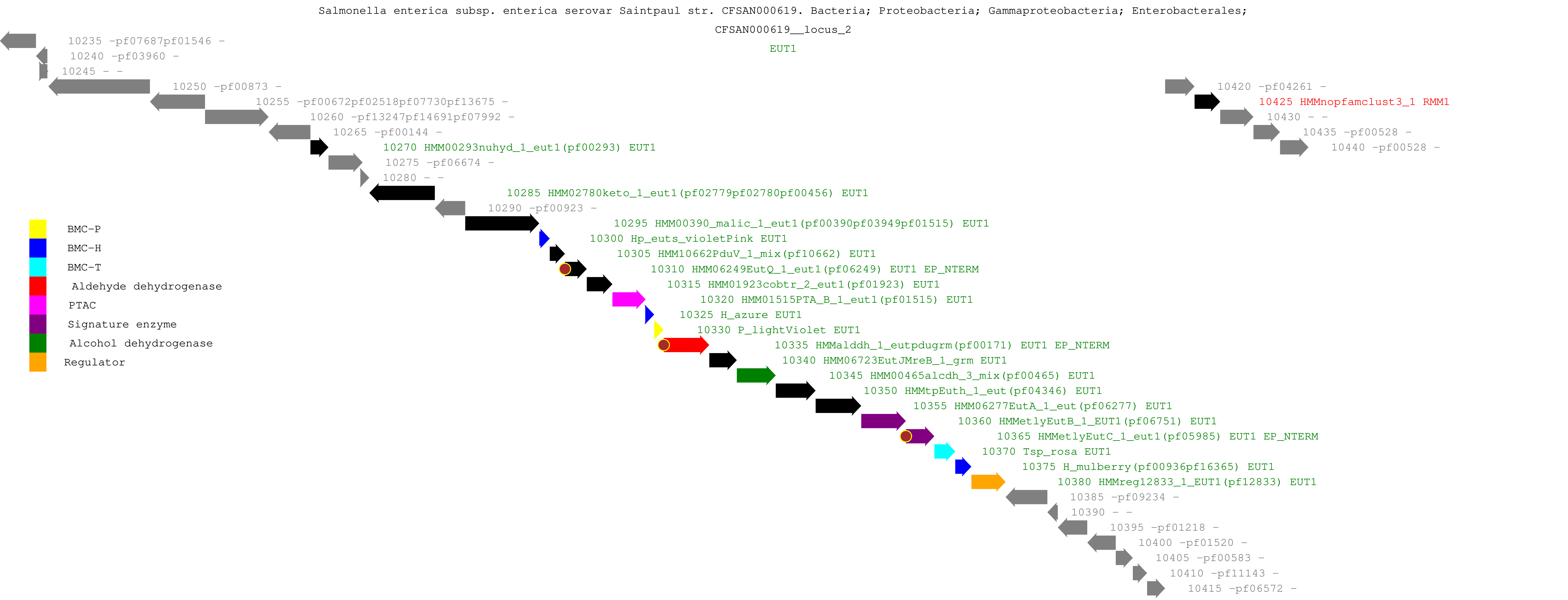
Salmonella enterica subsp. enterica serovar Saintpaul str. CFSAN000619. Bacteria; Proteobacteria; Gammaproteobacteria; Enterobacterales;
CFSAN000619__locus_2
EUT1
10235 -pf07687pf01546 -
10240 -pf03960 -
10245 - -
10250 -pf00873 -
10255 -pf00672pf02518pf07730pf13675 -
10260 -pf13247pf14691pf07992 -
10265 -pf00144 -
10270 HMM00293nuhyd_1_eut1(pf00293) EUT1
10275 -pf06674 -
10280 - -
10285 HMM02780keto_1_eut1(pf02779pf02780pf00456) EUT1
10290 -pf00923 -
10295 HMM00390_malic_1_eut1(pf00390pf03949pf01515) EUT1
10300 Hp_euts_violetPink EUT1
10305 HMM10662PduV_1_mix(pf10662) EUT1
10310 HMM06249EutQ_1_eut1(pf06249) EUT1 EP_NTERM
10315 HMM01923cobtr_2_eut1(pf01923) EUT1
10320 HMM01515PTA_B_1_eut1(pf01515) EUT1
10325 H_azure EUT1
10330 P_lightViolet EUT1
10335 HMMalddh_1_eutpdugrm(pf00171) EUT1 EP_NTERM
10340 HMM06723EutJMreB_1_grm EUT1
10345 HMM00465alcdh_3_mix(pf00465) EUT1
10350 HMMtpEuth_1_eut(pf04346) EUT1
10355 HMM06277EutA_1_eut(pf06277) EUT1
10360 HMMetlyEutB_1_EUT1(pf06751) EUT1
10365 HMMetlyEutC_1_eut1(pf05985) EUT1 EP_NTERM
10370 Tsp_rosa EUT1
10375 H_mulberry(pf00936pf16365) EUT1
10380 HMMreg12833_1_EUT1(pf12833) EUT1
10385 -pf09234 -
10390 - -
10395 -pf01218 -
10400 -pf01520 -
10405 -pf00583 -
10410 -pf11143 -
10415 -pf06572 -
10420 -pf04261 -
10425 HMMnopfamclust3_1 RMM1
10430 - -
10435 -pf00528 -
10440 -pf00528 -
BMC-P
BMC-H
BMC-T
Aldehyde dehydrogenase
PTAC
Signature enzyme
Alcohol dehydrogenase
Regulator

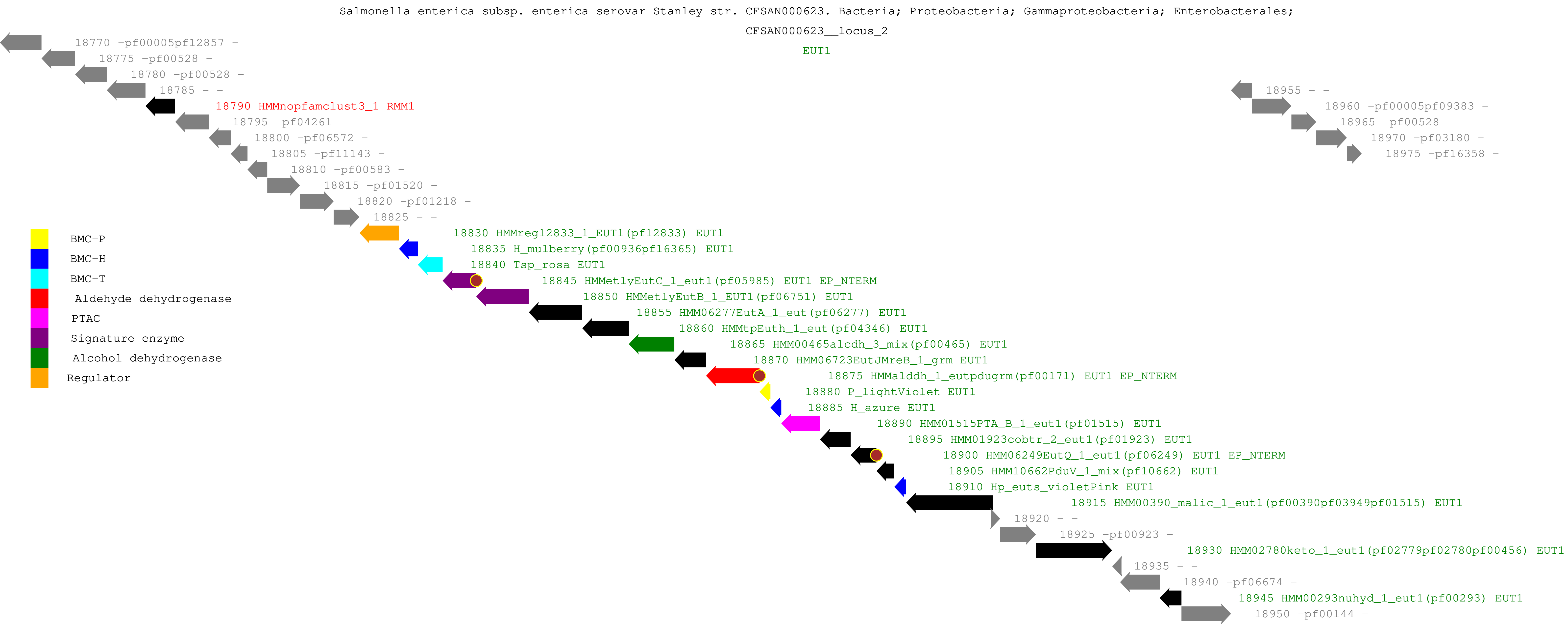
Salmonella enterica subsp. enterica serovar Stanley str. CFSAN000623. Bacteria; Proteobacteria; Gammaproteobacteria; Enterobacterales;
CFSAN000623__locus_2
EUT1
18770 -pf00005pf12857 -
18775 -pf00528 -
18780 -pf00528 -
18785 - -
18790 HMMnopfamclust3_1 RMM1
18795 -pf04261 -
18800 -pf06572 -
18805 -pf11143 -
18810 -pf00583 -
18815 -pf01520 -
18820 -pf01218 -
18825 - -
18830 HMMreg12833_1_EUT1(pf12833) EUT1
18835 H_mulberry(pf00936pf16365) EUT1
18840 Tsp_rosa EUT1
18845 HMMetlyEutC_1_eut1(pf05985) EUT1 EP_NTERM
18850 HMMetlyEutB_1_EUT1(pf06751) EUT1
18855 HMM06277EutA_1_eut(pf06277) EUT1
18860 HMMtpEuth_1_eut(pf04346) EUT1
18865 HMM00465alcdh_3_mix(pf00465) EUT1
18870 HMM06723EutJMreB_1_grm EUT1
18875 HMMalddh_1_eutpdugrm(pf00171) EUT1 EP_NTERM
18880 P_lightViolet EUT1
18885 H_azure EUT1
18890 HMM01515PTA_B_1_eut1(pf01515) EUT1
18895 HMM01923cobtr_2_eut1(pf01923) EUT1
18900 HMM06249EutQ_1_eut1(pf06249) EUT1 EP_NTERM
18905 HMM10662PduV_1_mix(pf10662) EUT1
18910 Hp_euts_violetPink EUT1
18915 HMM00390_malic_1_eut1(pf00390pf03949pf01515) EUT1
18920 - -
18925 -pf00923 -
18930 HMM02780keto_1_eut1(pf02779pf02780pf00456) EUT1
18935 - -
18940 -pf06674 -
18945 HMM00293nuhyd_1_eut1(pf00293) EUT1
18950 -pf00144 -
18955 - -
18960 -pf00005pf09383 -
18965 -pf00528 -
18970 -pf03180 -
18975 -pf16358 -
BMC-P
BMC-H
BMC-T
Aldehyde dehydrogenase
PTAC
Signature enzyme
Alcohol dehydrogenase
Regulator

Salmonella enterica subsp. enterica serovar Thompson str. CFSAN000625. Bacteria; Proteobacteria; Gammaproteobacteria; Enterobacterales;

CFSAN000625__locus_2

EUT1

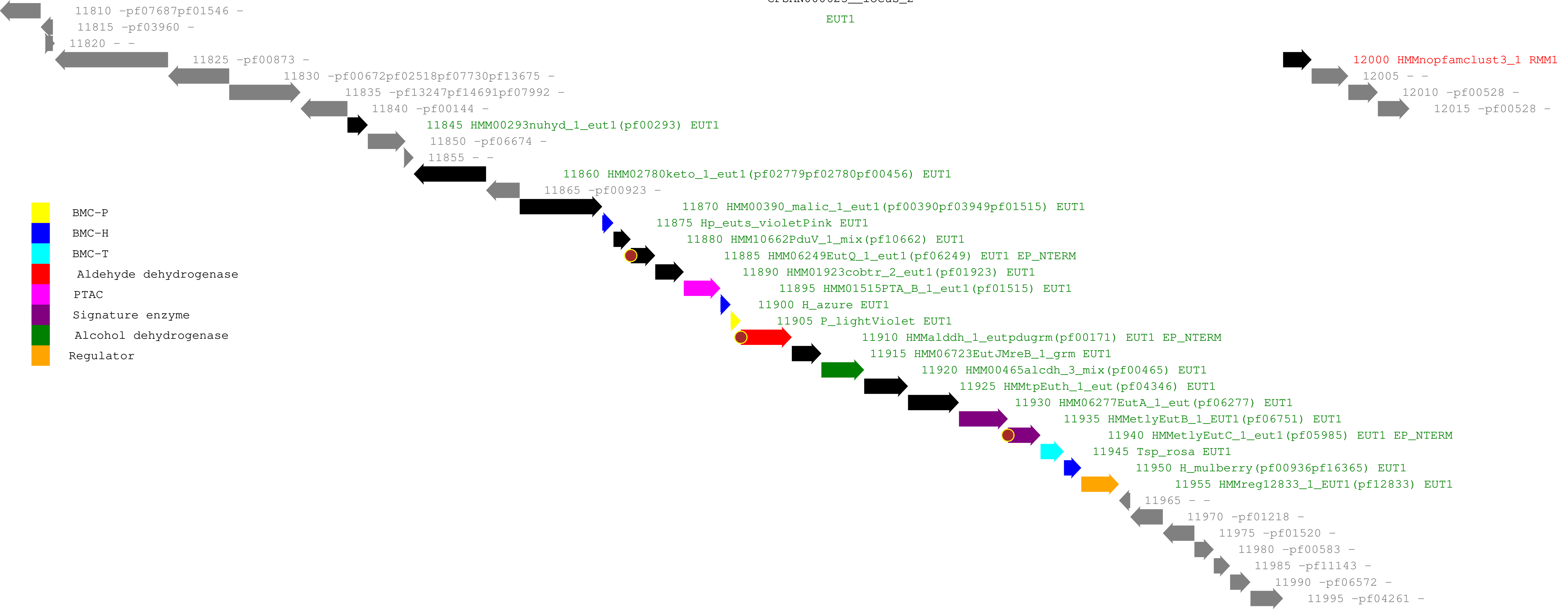

Salmonella enterica subsp. enterica serovar Cerro str. CFSAN001590. Bacteria; Proteobacteria; Gammaproteobacteria; Enterobacterales;

CFSAN001590__locus_2

EUT1

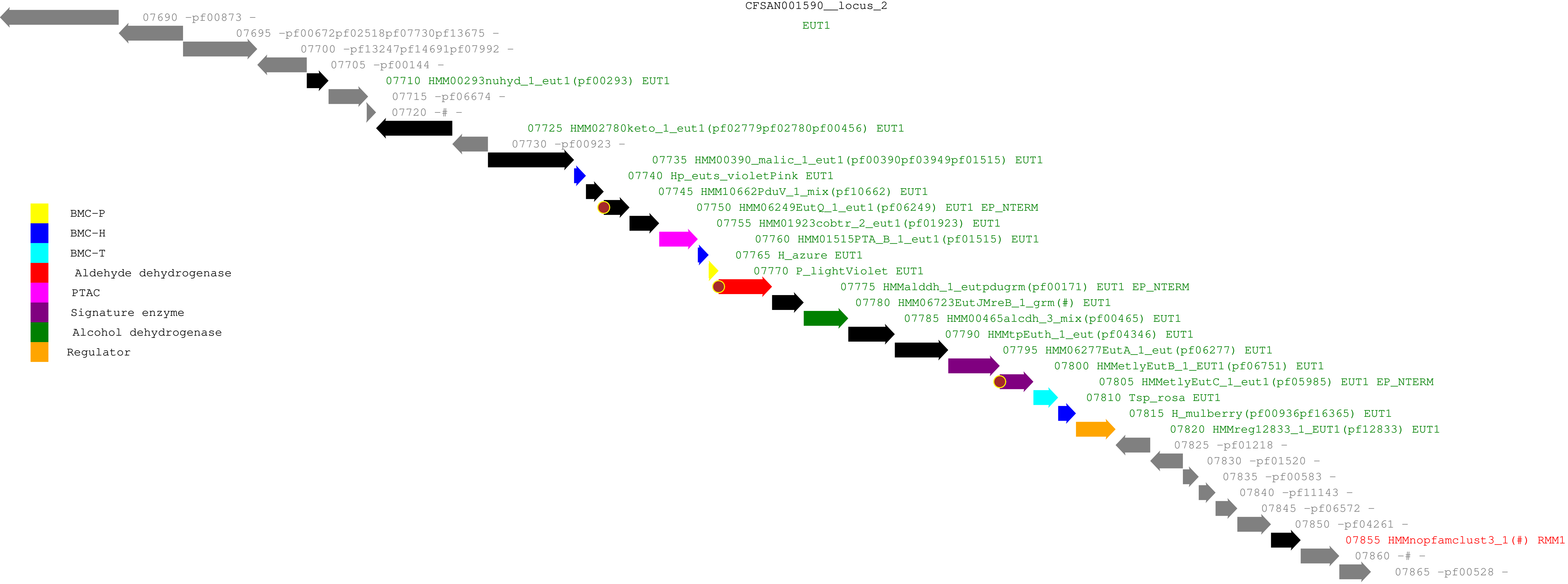

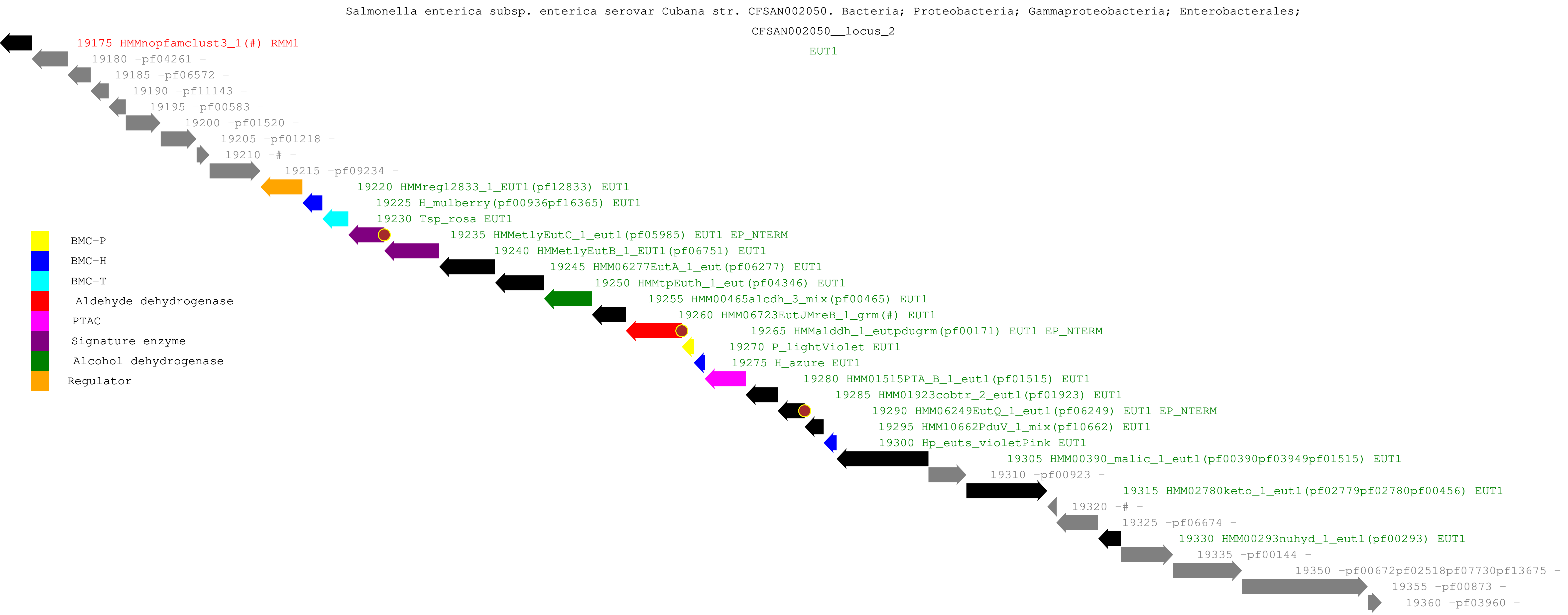
Salmonella enterica subsp. enterica serovar Cubana str. CFSAN002050. Bacteria; Proteobacteria; Gammaproteobacteria; Enterobacterales;
CFSAN002050__locus_2
EUT1
19175 HMMnopfamclust3_1(#) RMM1
19180 -pf04261 -
19185 -pf06572 -
19190 -pf11143 -
19195 -pf00583 -
19200 -pf01520 -
19205 -pf01218 -
19210 -# -
19215 -pf09234 -
19220 HMMreg12833_1_EUT1(pf12833) EUT1
19225 H_mulberry(pf00936pf16365) EUT1
19230 Tsp_rosa EUT1
19235 HMMetlyEutC_1_eut1(pf05985) EUT1 EP_NTERM
19240 HMMetlyEutB_1_EUT1(pf06751) EUT1
19245 HMM06277EutA_1_eut(pf06277) EUT1
19250 HMMtpEuth_1_eut(pf04346) EUT1
19255 HMM00465alcdh_3_mix(pf00465) EUT1
19260 HMM06723EutJMreB_1_grm(#) EUT1
19265 HMMalddh_1_eutpdugrm(pf00171) EUT1 EP_NTERM
19270 P_lightViolet EUT1
19275 H_azure EUT1
19280 HMM01515PTA_B_1_eut1(pf01515) EUT1
19285 HMM01923cobtr_2_eut1(pf01923) EUT1
19290 HMM06249EutQ_1_eut1(pf06249) EUT1 EP_NTERM
19295 HMM10662PduV_1_mix(pf10662) EUT1
19300 Hp_euts_violetPink EUT1
19305 HMM00390_malic_1_eut1(pf00390pf03949pf01515) EUT1
19310 -pf00923 -
19315 HMM02780keto_1_eut1(pf02779pf02780pf00456) EUT1
19320 -# -
19325 -pf06674 -
19330 HMM00293nuhyd_1_eut1(pf00293) EUT1
19335 -pf00144 -
19350 -pf00672pf02518pf07730pf13675 -
19355 -pf00873 -
19360 -pf03960 -
BMC-P
BMC-H
BMC-T
Aldehyde dehydrogenase
PTAC
Signature enzyme
Alcohol dehydrogenase
Regulator

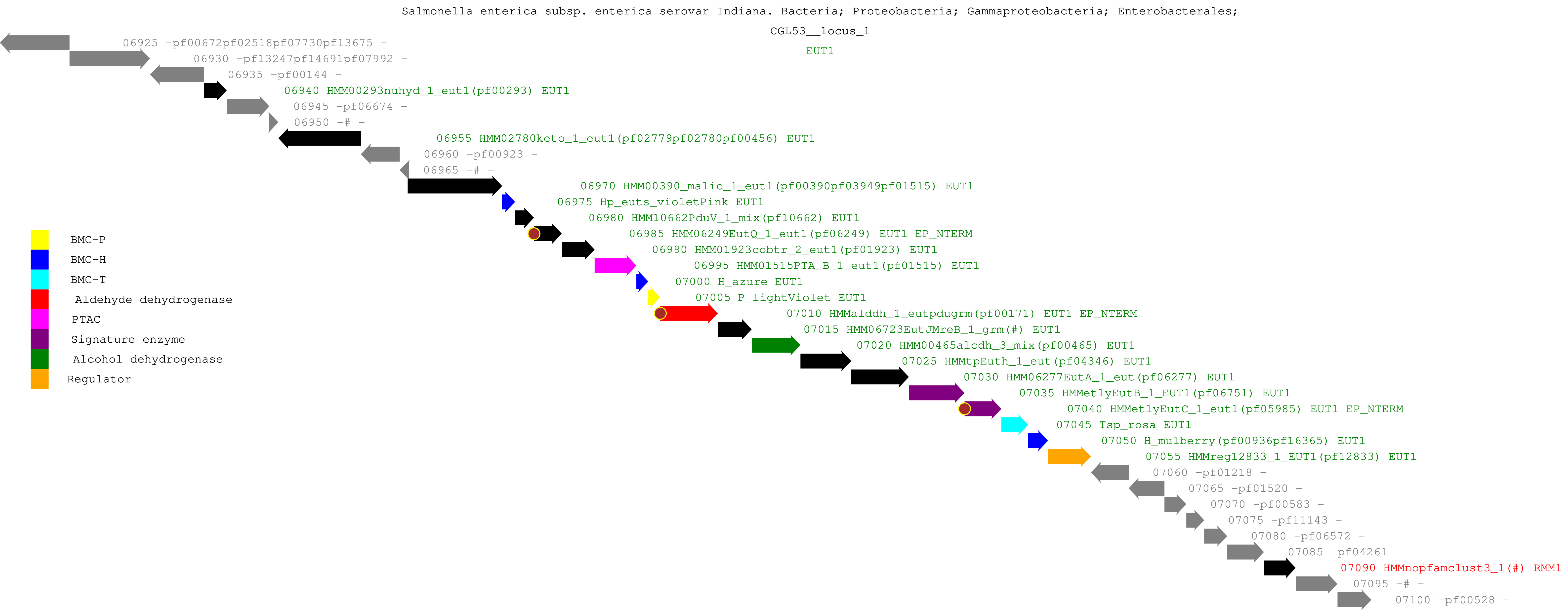

Salmonella enterica subsp. enterica serovar Indiana. Bacteria; Proteobacteria; Gammaproteobacteria; Enterobacterales;
CGL53__locus_1
EUT1
06925 -pf00672pf02518pf07730pf13675 -
06930 -pf13247pf14691pf07992 -
06935 -pf00144 -
06940 HMM00293nuhyd_1_eut1(pf00293) EUT1
06945 -pf06674 -
06950 -# -
06955 HMM02780keto_1_eut1(pf02779pf02780pf00456) EUT1
06960 -pf00923 -
06965 -# -
06970 HMM00390_malic_1_eut1(pf00390pf03949pf01515) EUT1
06975 Hp_euts_violetPink EUT1
06980 HMM10662PduV_1_mix(pf10662) EUT1
06985 HMM06249EutQ_1_eut1(pf06249) EUT1 EP_NTERM
06990 HMM01923cobtr_2_eut1(pf01923) EUT1
06995 HMM01515PTA_B_1_eut1(pf01515) EUT1
07000 H_azure EUT1
07005 P_lightViolet EUT1
07010 HMMalddh_1_eutpdugrm(pf00171) EUT1 EP_NTERM
07015 HMM06723EutJMreB_1_grm(#) EUT1
07020 HMM00465alcdh_3_mix(pf00465) EUT1
07025 HMMtpEuth_1_eut(pf04346) EUT1
07030 HMM06277EutA_1_eut(pf06277) EUT1
07035 HMMetlyEutB_1_EUT1(pf06751) EUT1
07040 HMMetlyEutC_1_eut1(pf05985) EUT1 EP_NTERM
07045 Tsp_rosa EUT1
07050 H_mulberry(pf00936pf16365) EUT1
07055 HMMreg12833_1_EUT1(pf12833) EUT1
07060 -pf01218 -
07065 -pf01520 -
07070 -pf00583 -
07075 -pf11143 -
07080 -pf06572 -
07085 -pf04261 -
07090 HMMnopfamclust3_1(#) RMM1
07095 -# -
07100 -pf00528 -
BMC-P
BMC-H
BMC-T
Aldehyde dehydrogenase
PTAC
Signature enzyme
Alcohol dehydrogenase
Regulator

Salmonella enterica subsp. enterica serovar Heidelberg. Bacteria; Proteobacteria; Gammaproteobacteria; Enterobacterales;

CH228__locus_1

EUT1

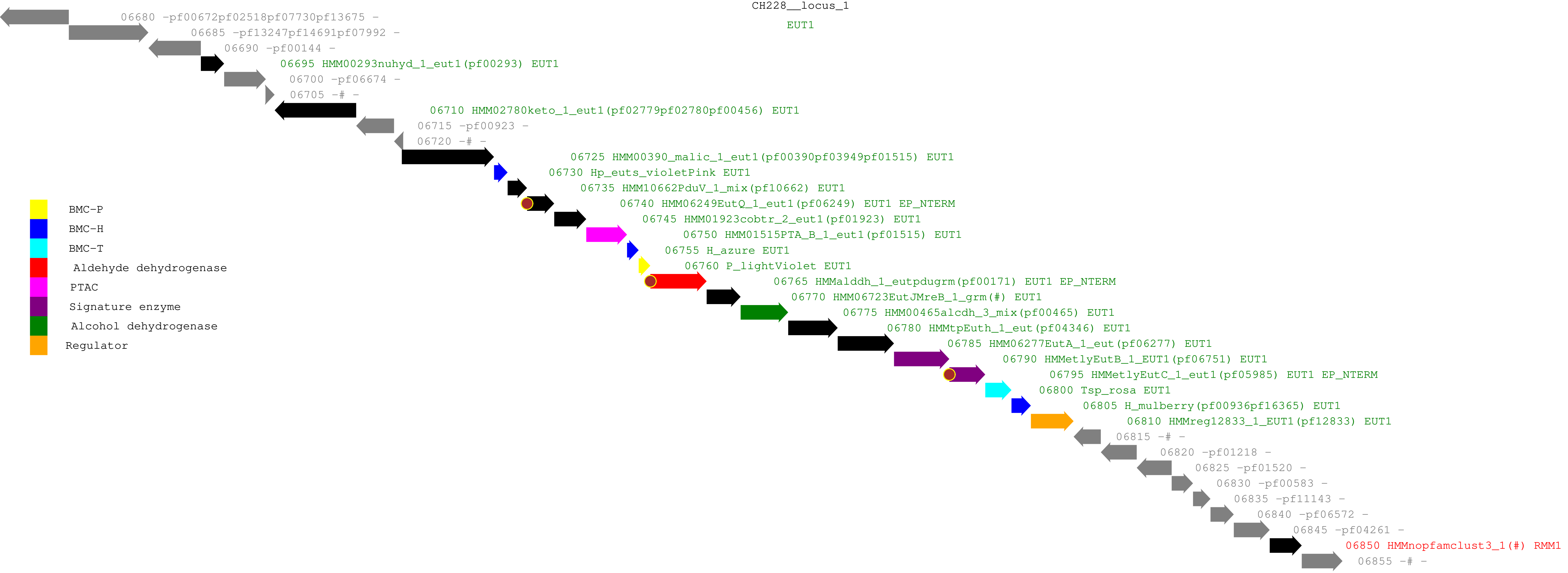

Salmonella choleraesuis. Bacteria; Proteobacteria; Gammaproteobacteria; Enterobacterales;

CHC34__locus_1

EUT1

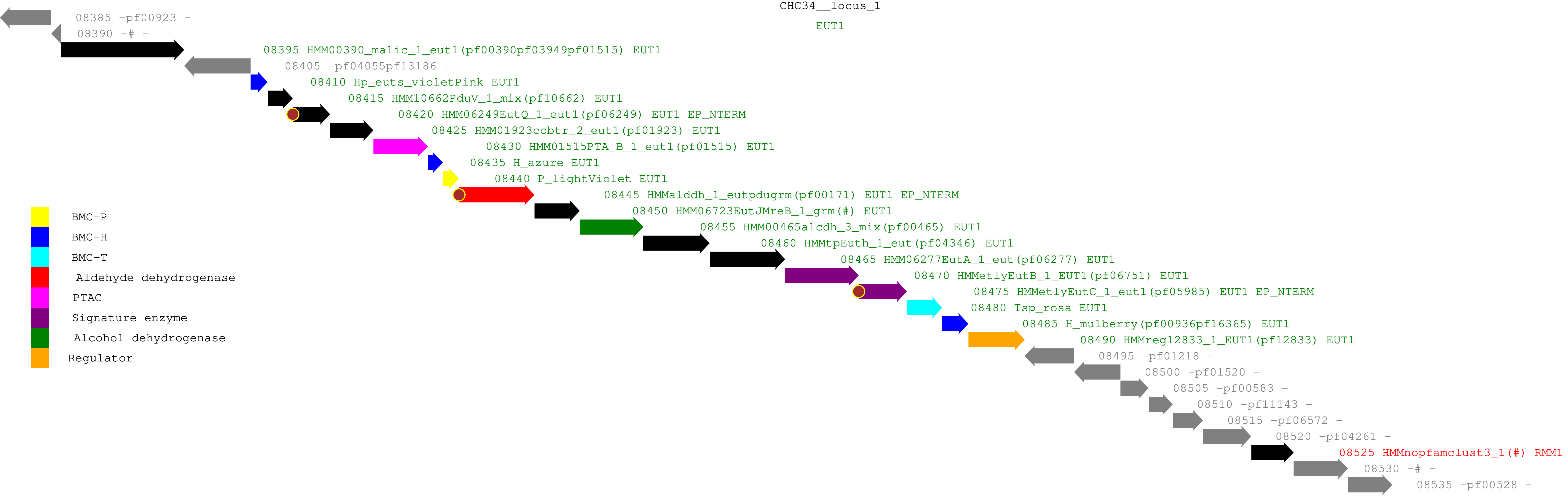

Salmonella choleraesuis. Bacteria; Proteobacteria; Gammaproteobacteria; Enterobacterales;

CHD23__locus_1

EUT1

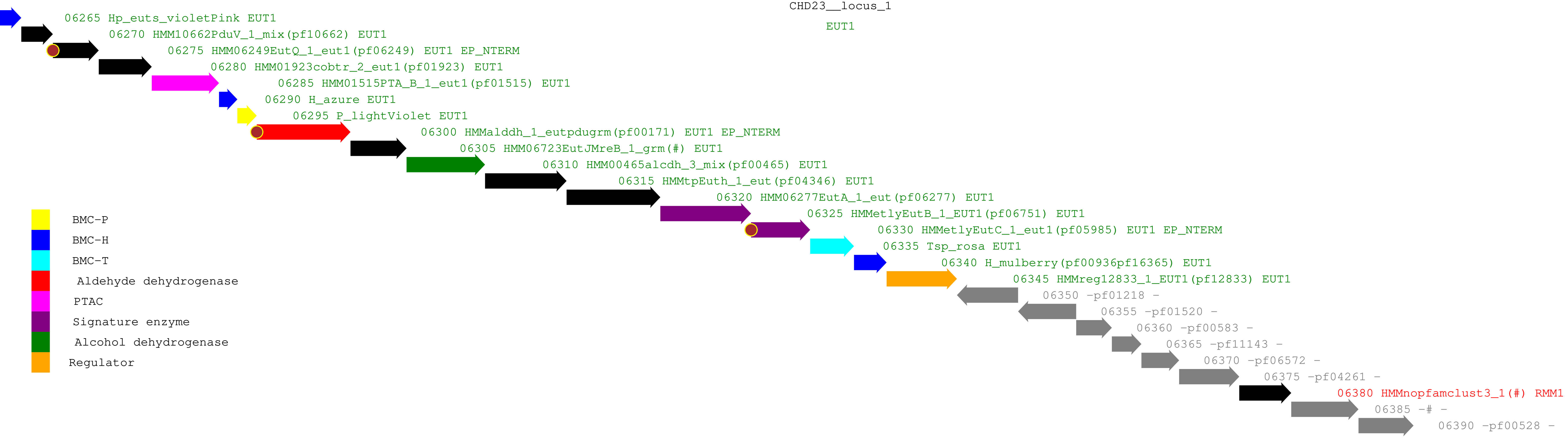

Klebsiella pneumoniae subsp. pneumoniae. Bacteria; Proteobacteria; Gammaproteobacteria; Enterobacterales;

CI103__locus_2

EUT1

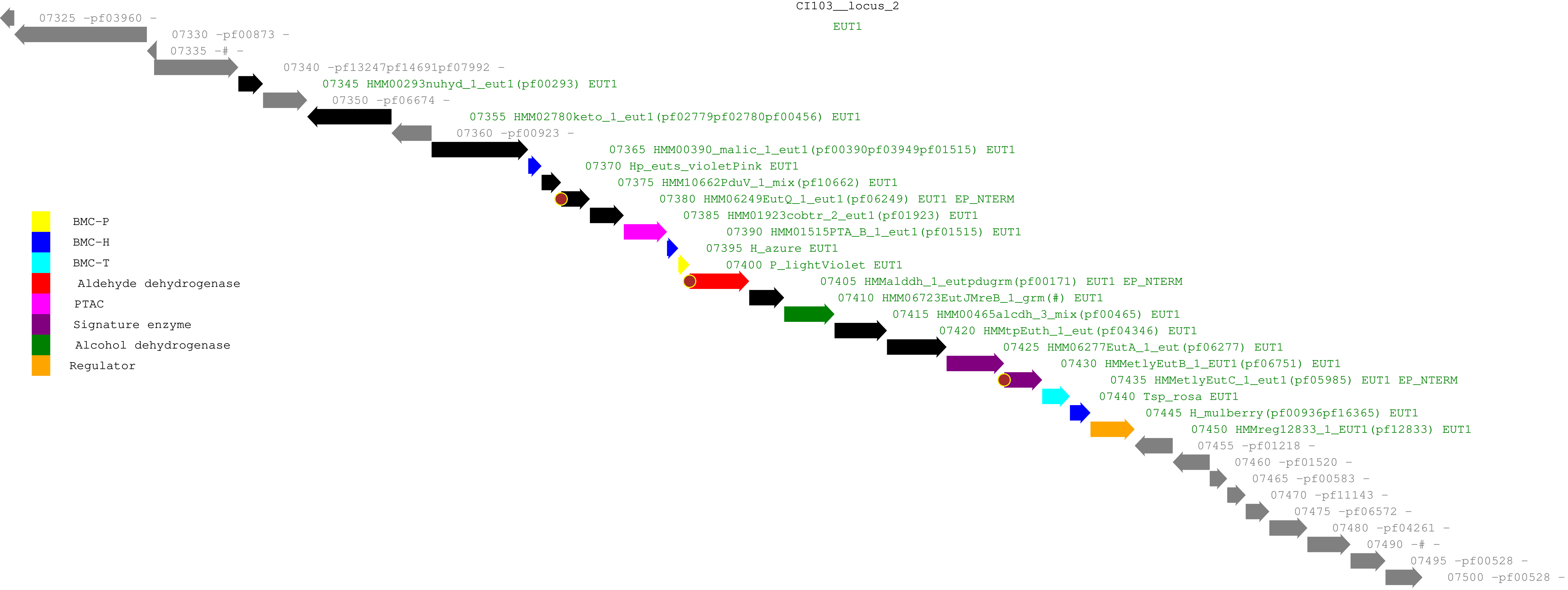

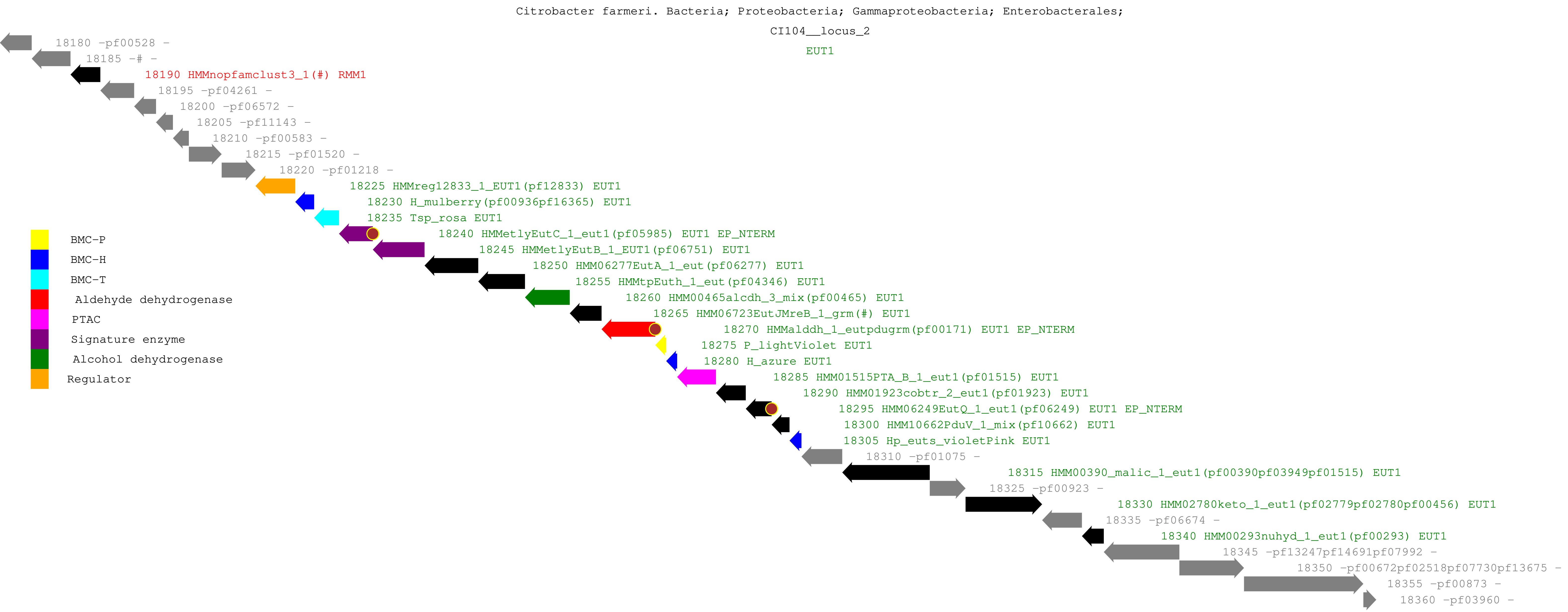
Citrobacter farmeri. Bacteria; Proteobacteria; Gammaproteobacteria; Enterobacterales;
CI104__locus_2
EUT1
18180 -pf00528 -
18185 -# -
18190 HMMnopfamclust3_1(#) RMM1
18195 -pf04261 -
18200 -pf06572 -
18205 -pf11143 -
18210 -pf00583 -
18215 -pf01520 -
18220 -pf01218 -
18225 HMMreg12833_1_EUT1(pf12833) EUT1
18230 H_mulberry(pf00936pf16365) EUT1
18235 Tsp_rosa EUT1
18240 HMMetlyEutC_1_eut1(pf05985) EUT1 EP_NTERM
18245 HMMetlyEutB_1_EUT1(pf06751) EUT1
18250 HMM06277EutA_1_eut(pf06277) EUT1
18255 HMMtpEuth_1_eut(pf04346) EUT1
18260 HMM00465alcdh_3_mix(pf00465) EUT1
18265 HMM06723EutJMreB_1_grm(#) EUT1
18270 HMMalddh_1_eutpdugrm(pf00171) EUT1 EP_NTERM
18275 P_lightViolet EUT1
18280 H_azure EUT1
18285 HMM01515PTA_B_1_eut1(pf01515) EUT1
18290 HMM01923cobtr_2_eut1(pf01923) EUT1
18295 HMM06249EutQ_1_eut1(pf06249) EUT1 EP_NTERM
18300 HMM10662PduV_1_mix(pf10662) EUT1
18305 Hp_euts_violetPink EUT1
18310 -pf01075 -
18315 HMM00390_malic_1_eut1(pf00390pf03949pf01515) EUT1
18325 -pf00923 -
18330 HMM02780keto_1_eut1(pf02779pf02780pf00456) EUT1
18335 -pf06674 -
18340 HMM00293nuhyd_1_eut1(pf00293) EUT1
18345 -pf13247pf14691pf07992 -
18350 -pf00672pf02518pf07730pf13675 -
18355 -pf00873 -
18360 -pf03960 -
BMC-P
BMC-H
BMC-T
Aldehyde dehydrogenase
PTAC
Signature enzyme
Alcohol dehydrogenase
Regulator

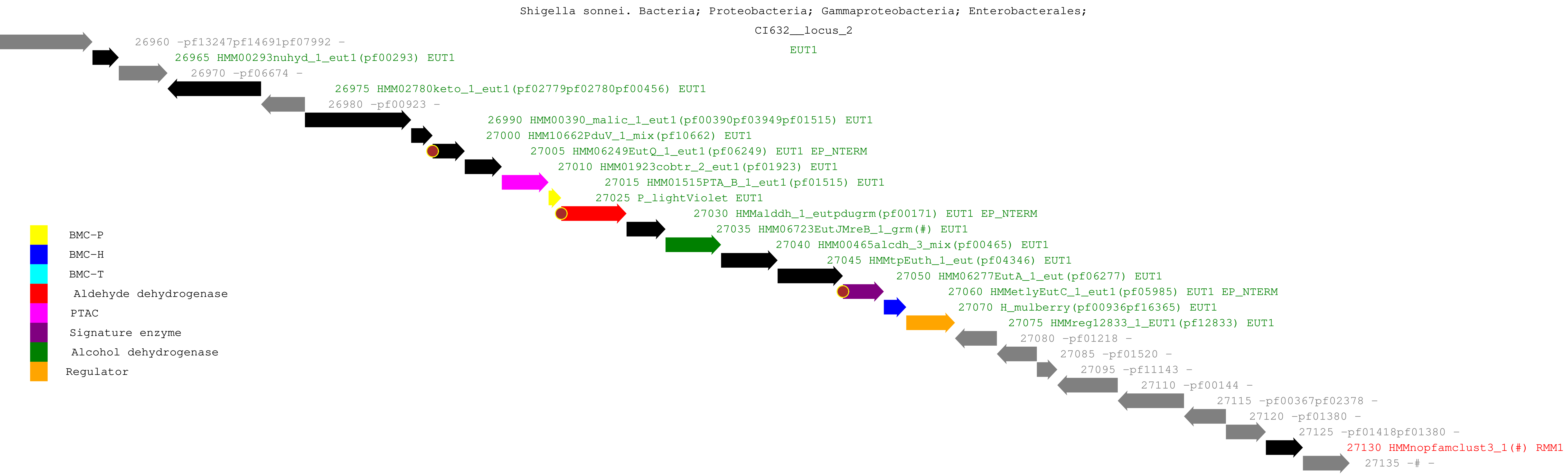
Shigella sonnei. Bacteria; Proteobacteria; Gammaproteobacteria; Enterobacterales;
CI632__locus_2
EUT1
26960 -pf13247pf14691pf07992 -
26965 HMM00293nuhyd_1_eut1(pf00293) EUT1
26970 -pf06674 -
26975 HMM02780keto_1_eut1(pf02779pf02780pf00456) EUT1
26980 -pf00923 -
26990 HMM00390_malic_1_eut1(pf00390pf03949pf01515) EUT1
27000 HMM10662PduV_1_mix(pf10662) EUT1
27005 HMM06249EutQ_1_eut1(pf06249) EUT1 EP_NTERM
27010 HMM01923cobtr_2_eut1(pf01923) EUT1
27015 HMM01515PTA_B_1_eut1(pf01515) EUT1
27025 P_lightViolet EUT1
27030 HMMaldh_1_eutpdugrm(pf00171) EUT1 EP_NTERM
27035 HMM06723EutJMreB_1_grm(#) EUT1
27040 HMM00465alcdh_3_mix(pf00465) EUT1
27045 HMMtpEuth_1_eut(pf04346) EUT1
27050 HMM06277EutA_1_eut(pf06277) EUT1
27060 HMMetlyEutC_1_eut1(pf05985) EUT1 EP_NTERM
27070 H_mulberry(pf00936pf16365) EUT1
27075 HMMreg12833_1_EUT1(pf12833) EUT1
27080 -pf01218 -
27085 -pf01520 -
27095 -pf11143 -
27110 -pf00144 -
27115 -pf00367pf02378 -
27120 -pf01380 -
27125 -pf01418pf01380 -
27130 HMMnopfamclust3_1(#) RMM1
27135 -# -
BMC-P
BMC-H
BMC-T
Aldehyde dehydrogenase
PTAC
Signature enzyme
Alcohol dehydrogenase
Regulator

Shigella sonnei. Bacteria; Proteobacteria; Gammaproteobacteria; Enterobacterales;

CI641__locus_2

EUT1

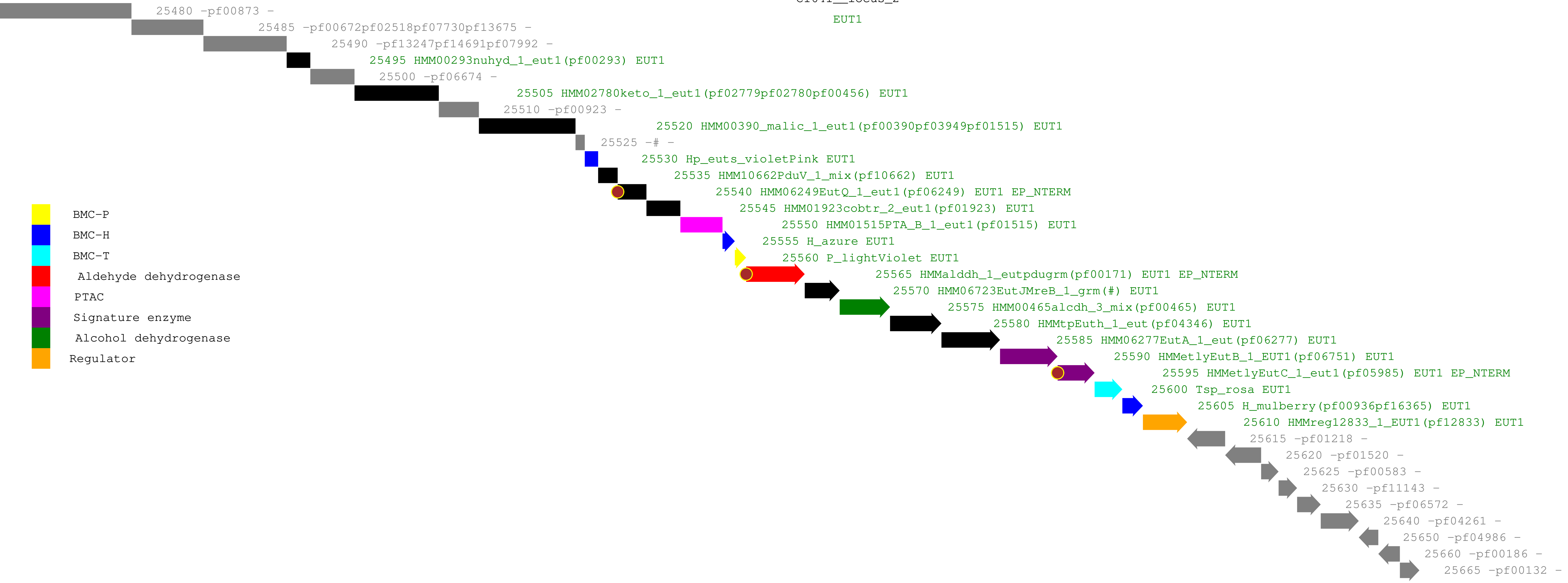

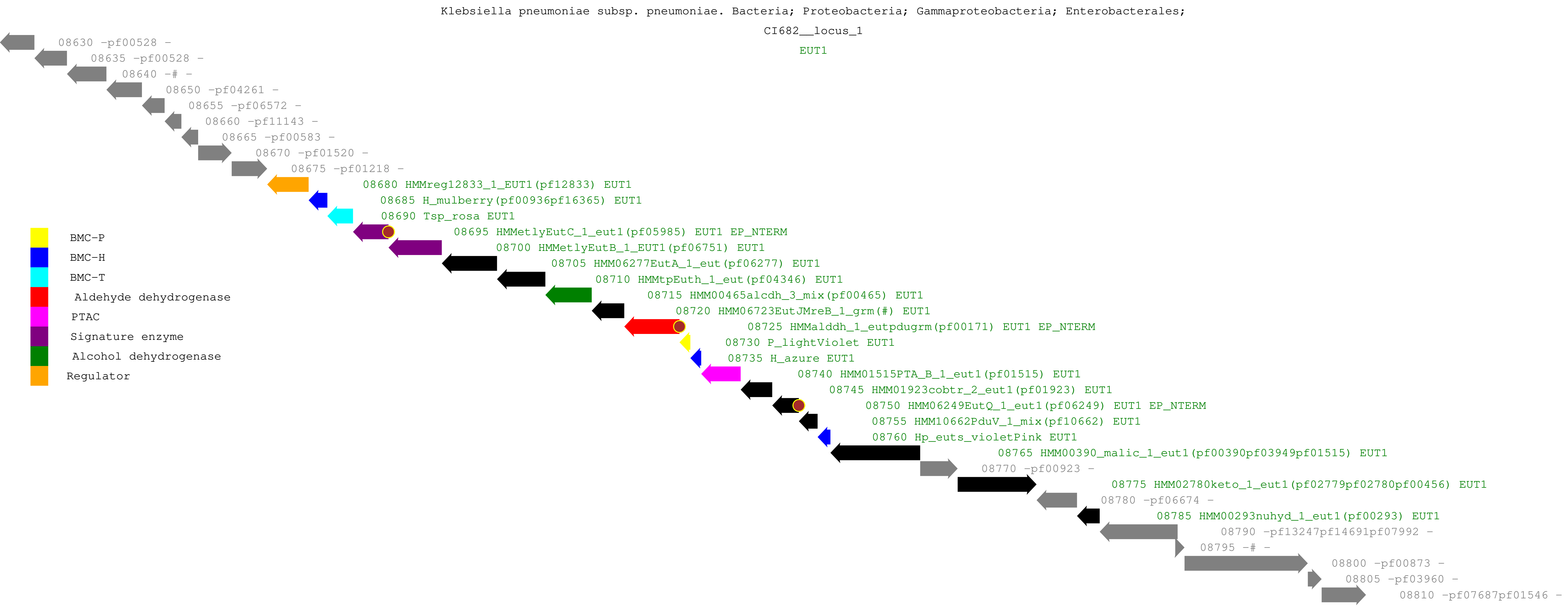
Klebsiella pneumoniae subsp. pneumoniae. Bacteria; Proteobacteria; Gammaproteobacteria; Enterobacterales;
CI682__locus_1
EUT1
08630 -pf00528 -
08635 -pf00528 -
08640 -# -
08650 -pf04261 -
08655 -pf06572 -
08660 -pf11143 -
08665 -pf00583 -
08670 -pf01520 -
08675 -pf01218 -
08680 HMMreg12833_1_EUT1(pf12833) EUT1
08685 H_mulberry(pf00936pf16365) EUT1
08690 Tsp_rosa EUT1
08695 HMMetlyEutC_1_eut1(pf05985) EUT1 EP_NTERM
08700 HMMetlyEutB_1_EUT1(pf06751) EUT1
08705 HMM06277EutA_1_eut(pf06277) EUT1
08710 HMMtpEuth_1_eut(pf04346) EUT1
08715 HMM00465alcdh_3_mix(pf00465) EUT1
08720 HMM06723EutJMreB_1_grm(#) EUT1
08725 HMMalddh_1_eutpdugrm(pf00171) EUT1 EP_NTERM
08730 P_lightViolet EUT1
08735 H_azure EUT1
08740 HMM01515PTA_B_1_eut1(pf01515) EUT1
08745 HMM01923cobtr_2_eut1(pf01923) EUT1
08750 HMM06249EutQ_1_eut1(pf06249) EUT1 EP_NTERM
08755 HMM10662PduV_1_mix(pf10662) EUT1
08760 Hp_euts_violetPink EUT1
08765 HMM00390_malic_1_eut1(pf00390pf03949pf01515) EUT1
08770 -pf00923 -
08775 HMM02780keto_1_eut1(pf02779pf02780pf00456) EUT1
08780 -pf06674 -
08785 HMM00293nuhyd_1_eut1(pf00293) EUT1
08790 -pf13247pf14691pf07992 -
08795 -# -
08800 -pf00873 -
08805 -pf03960 -
08810 -pf07687pf01546 -
BMC-P
BMC-H
BMC-T
Aldehyde dehydrogenase
PTAC
Signature enzyme
Alcohol dehydrogenase
Regulator

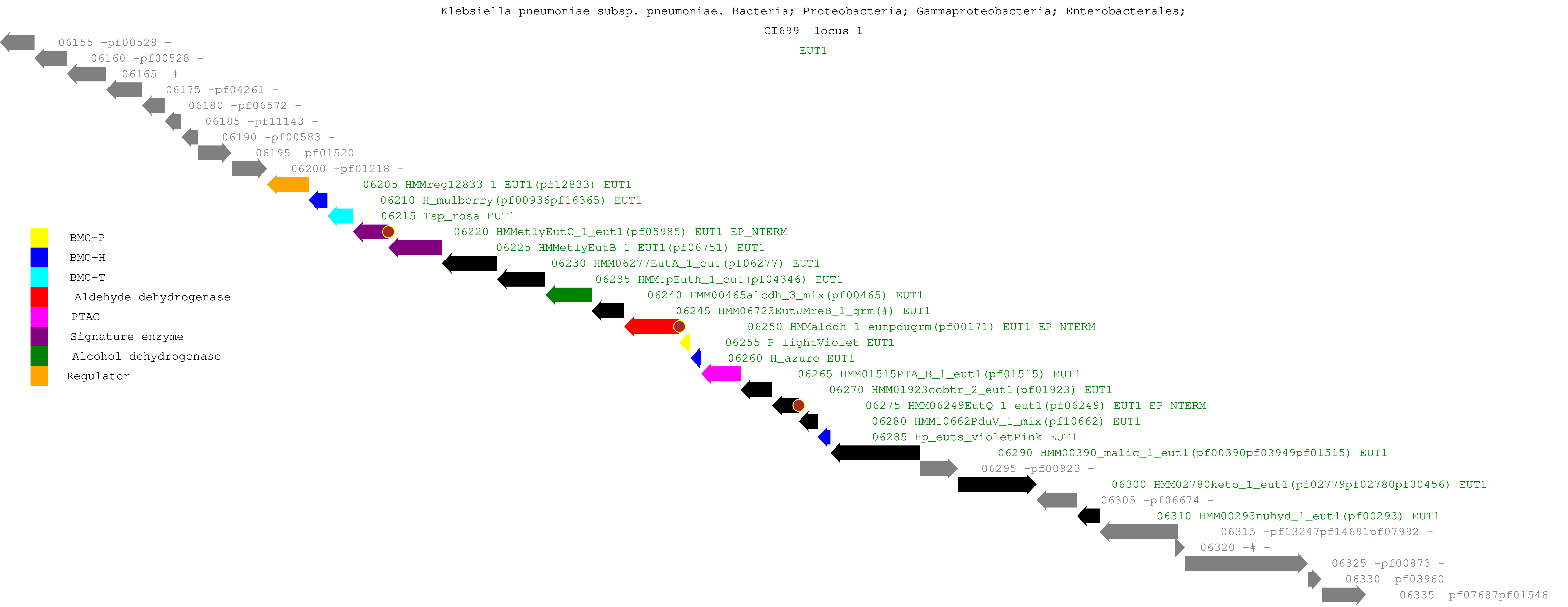
Klebsiella pneumoniae subsp. pneumoniae. Bacteria; Proteobacteria; Gammaproteobacteria; Enterobacterales;
CI699__locus_1
EUT1
06155 -pf00528 -
06160 -pf00528 -
06165 -# -
06175 -pf04261 -
06180 -pf06572 -
06185 -pf11143 -
06190 -pf00583 -
06195 -pf01520 -
06200 -pf01218 -
06205 HMMreg12833_1_EUT1(pf12833) EUT1
06210 H_mulberry(pf00936pf16365) EUT1
06215 Tsp_rosa EUT1
06220 HMMetlyEutC_1_eut1(pf05985) EUT1 EP_NTERM
06225 HMMetlyEutB_1_EUT1(pf06751) EUT1
06230 HMM06277EutA_1_eut(pf06277) EUT1
06235 HMMtpEuth_1_eut(pf04346) EUT1
06240 HMM00465alcdh_3_mix(pf00465) EUT1
06245 HMM06723EutJMreB_1_grm(#) EUT1
06250 HMMalddh_1_eutpdugrm(pf00171) EUT1 EP_NTERM
06255 P_lightViolet EUT1
06260 H_azure EUT1
06265 HMM01515PTA_B_1_eut1(pf01515) EUT1
06270 HMM01923cobtr_2_eut1(pf01923) EUT1
06275 HMM06249EutQ_1_eut1(pf06249) EUT1 EP_NTERM
06280 HMM10662PduV_1_mix(pf10662) EUT1
06285 Hp_euts_violetPink EUT1
06290 HMM00390_malic_1_eut1(pf00390pf03949pf01515) EUT1
06295 -pf00923 -
06300 HMM02780keto_1_eut1(pf02779pf02780pf00456) EUT1
06305 -pf06674 -
06310 HMM00293nuhyd_1_eut1(pf00293) EUT1
06315 -pf13247pf14691pf07992 -
06320 -# -
06325 -pf00873 -
06330 -pf03960 -
06335 -pf07687pf01546 -
BMC-P
BMC-H
BMC-T
Aldehyde dehydrogenase
PTAC
Signature enzyme
Alcohol dehydrogenase
Regulator

Shigella boydii. Bacteria; Proteobacteria; Gammaproteobacteria; Enterobacterales;

CI719__locus_1

EUT1

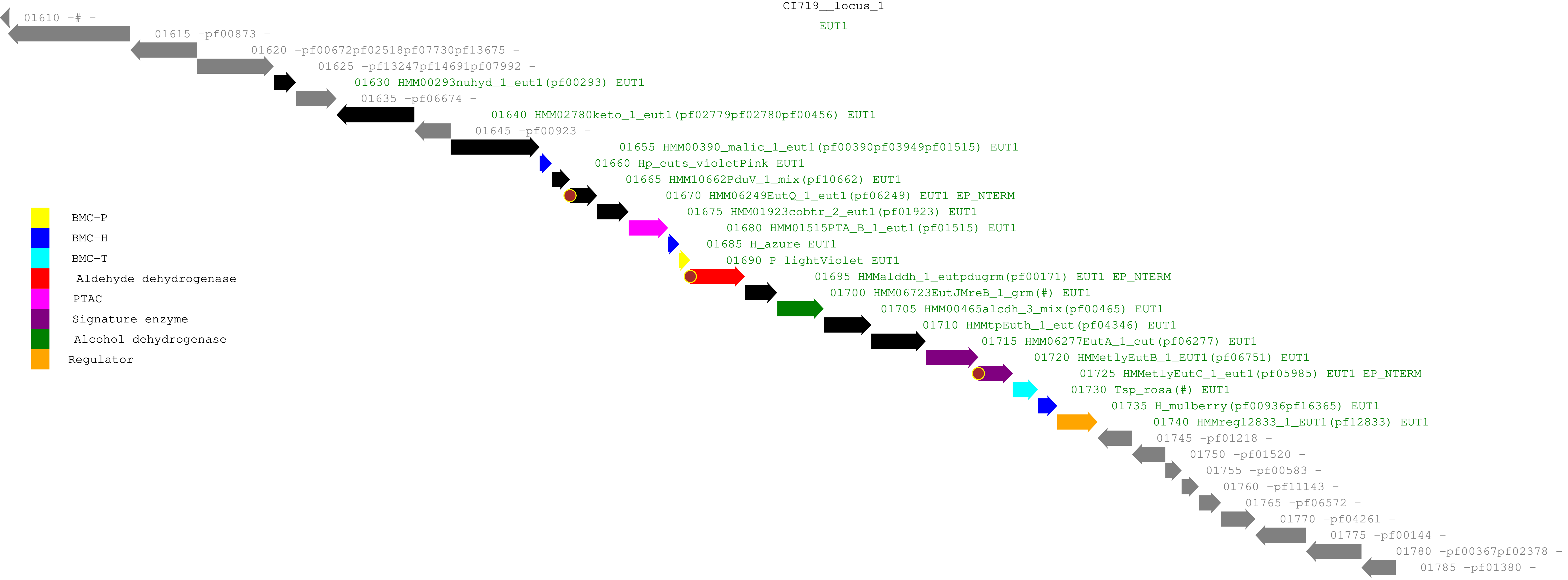

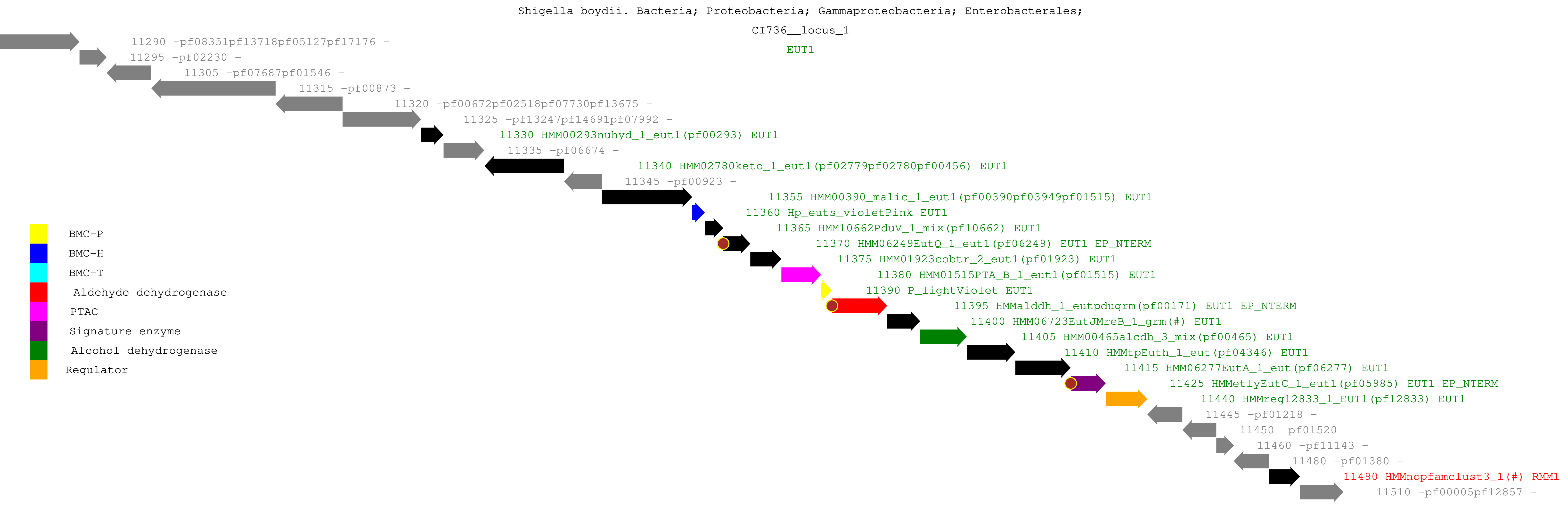
Shigella boydii. Bacteria; Proteobacteria; Gammaproteobacteria; Enterobacterales;
CI736__locus_1
EUT1
11290 -pf08351pf13718pf05127pf17176 -
11295 -pf02230 -
11305 -pf07687pf01546 -
11315 -pf00873 -
11320 -pf00672pf02518pf07730pf13675 -
11325 -pf13247pf14691pf07992 -
11330 HMM00293nuhyd_1_eut1(pf00293) EUT1
11335 -pf06674 -
11340 HMM02780keto_1_eut1(pf02779pf02780pf00456) EUT1
11345 -pf00923 -
11355 HMM00390_malic_1_eut1(pf00390pf03949pf01515) EUT1
11360 Hp_euts_violetPink EUT1
11365 HMM10662PduV_1_mix(pf10662) EUT1
11370 HMM06249EutQ_1_eut1(pf06249) EUT1 EP_NTERM
11375 HMM01923cobtr_2_eut1(pf01923) EUT1
11380 HMM01515PTA_B_1_eut1(pf01515) EUT1
11390 P_lightViolet EUT1
11395 HMMalddh_1_eutpdugrm(pf00171) EUT1 EP_NTERM
11400 HMM06723EutJMreB_1_grm(#) EUT1
11405 HMM00465alcdh_3_mix(pf00465) EUT1
11410 HMMtpEuth_1_eut(pf04346) EUT1
11415 HMM06277EutA_1_eut(pf06277) EUT1
11425 HMMetlyEutC_1_eut1(pf05985) EUT1 EP_NTERM
11440 HMMreg12833_1_EUT1(pf12833) EUT1
11445 -pf01218 -
11450 -pf01520 -
11460 -pf11143 -
11480 -pf01380 -
11490 HMMnopfamclust3_1(#) RMM1
11510 -pf00005pf12857 -
BMC-P
BMC-H
BMC-T
Aldehyde dehydrogenase
PTAC
Signature enzyme
Alcohol dehydrogenase
Regulator

Klebsiella pneumoniae subsp. pneumoniae. Bacteria; Proteobacteria; Gammaproteobacteria; Enterobacterales;

CI751__locus_2

EUT1

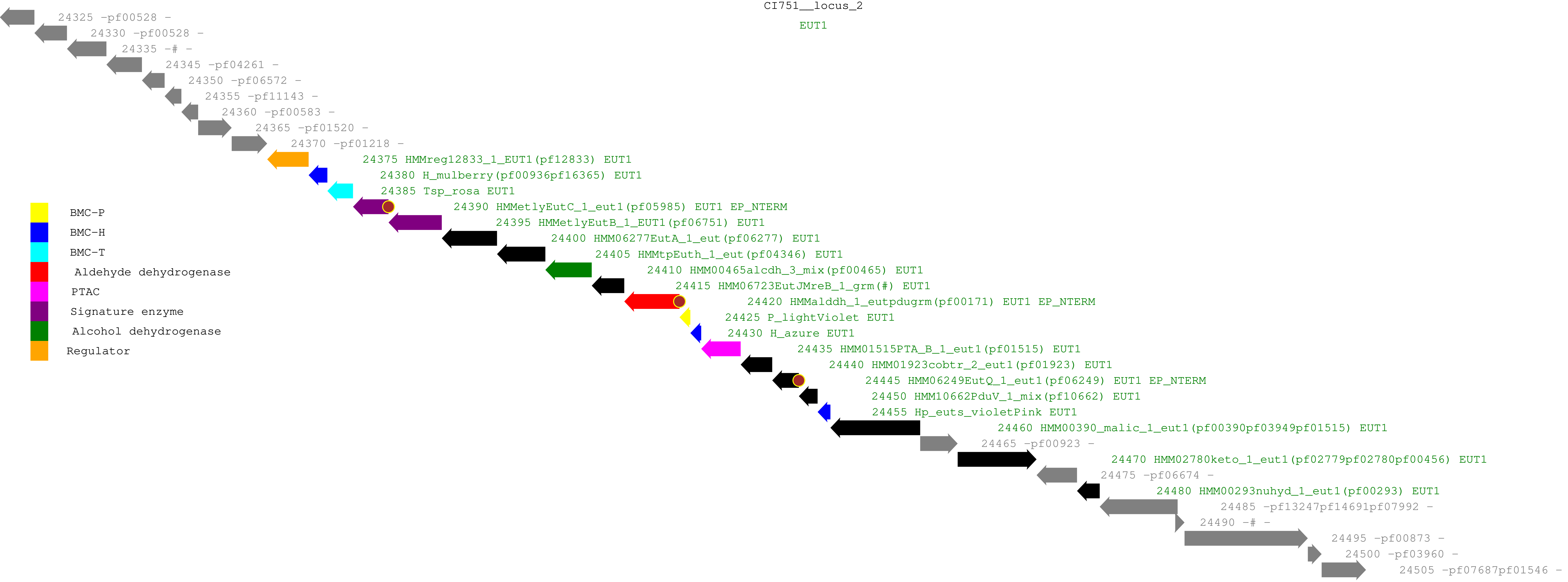

Klebsiella pneumoniae subsp. pneumoniae. Bacteria; Proteobacteria; Gammaproteobacteria; Enterobacterales;

CI752__locus_1

EUT1

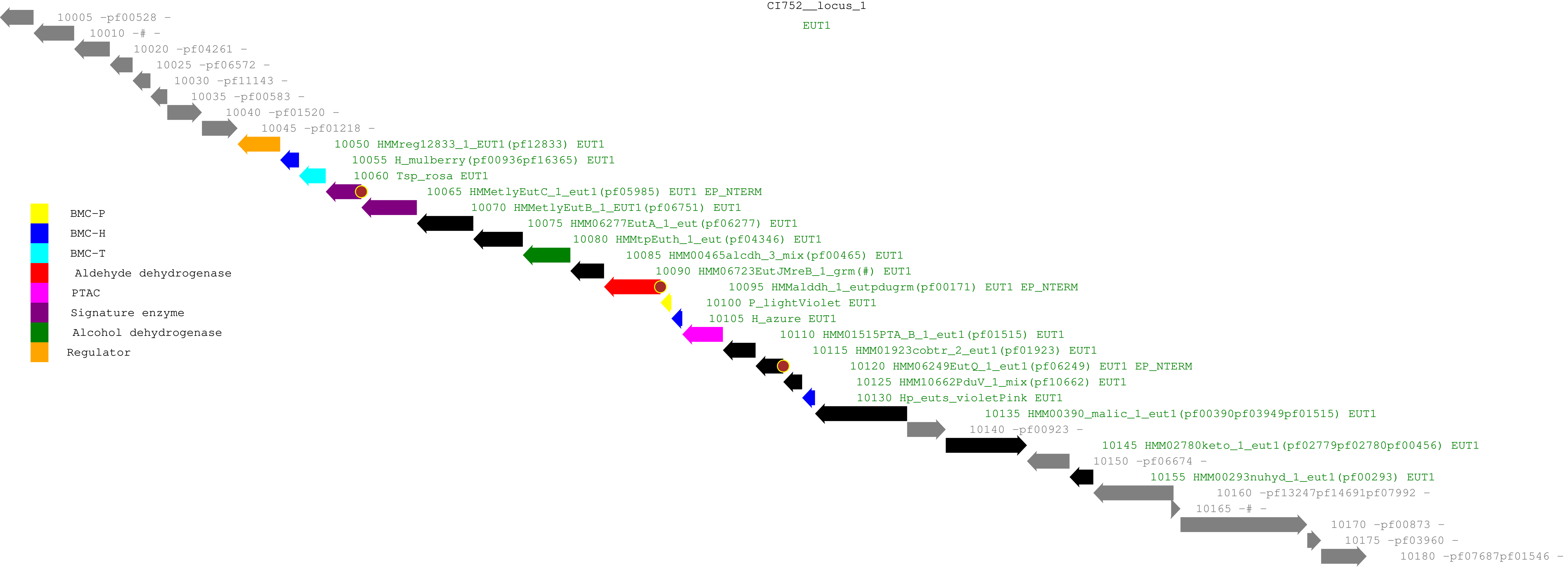

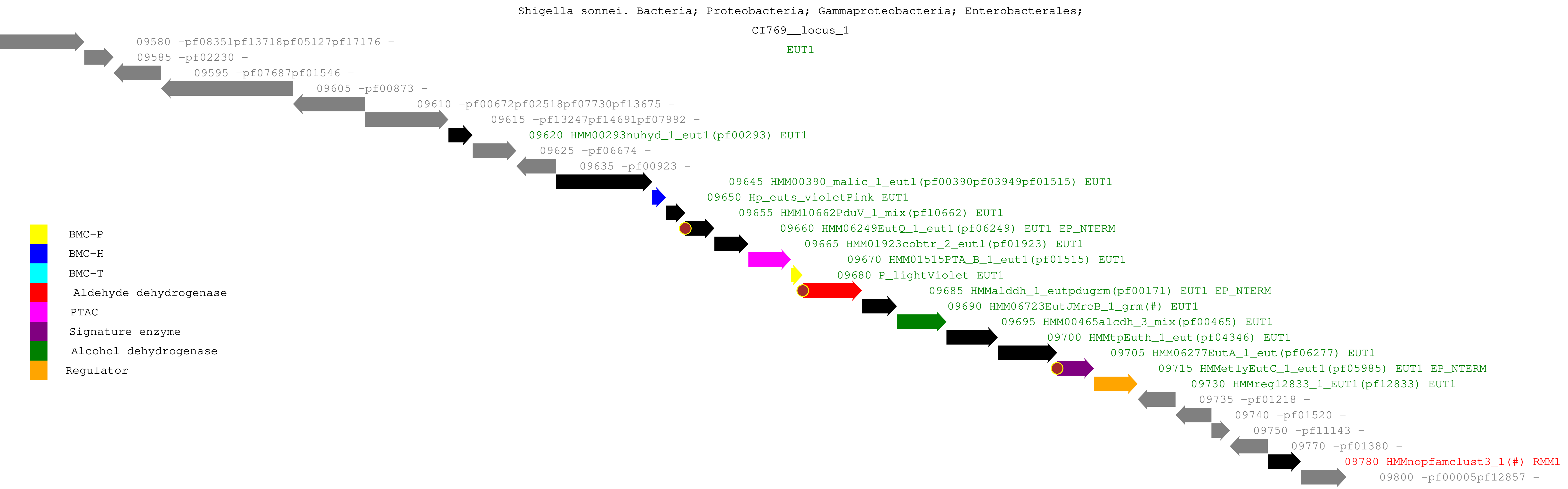
Shigella sonnei. Bacteria; Proteobacteria; Gammaproteobacteria; Enterobacterales;
CI769__locus_1
EUT1
09580 -pf08351pf13718pf05127pf17176 -
09585 -pf02230 -
09595 -pf07687pf01546 -
09605 -pf00873 -
09610 -pf00672pf02518pf07730pf13675 -
09615 -pf13247pf14691pf07992 -
09620 HMM00293nuhyd_1_eut1(pf00293) EUT1
09625 -pf06674 -
09635 -pf00923 -
09645 HMM00390_malic_1_eut1(pf00390pf03949pf01515) EUT1
09650 Hp_euts_violetPink EUT1
09655 HMM10662PduV_1_mix(pf10662) EUT1
09660 HMM06249EutQ_1_eut1(pf06249) EUT1 EP_NTERM
09665 HMM01923cobtr_2_eut1(pf01923) EUT1
09670 HMM01515PTA_B_1_eut1(pf01515) EUT1
09680 P_lightViolet EUT1
09685 HMMalddh_1_eutpdugrm(pf00171) EUT1 EP_NTERM
09690 HMM06723EutJMreB_1_grm(#) EUT1
09695 HMM00465alcdh_3_mix(pf00465) EUT1
09700 HMMtpEuth_1_eut(pf04346) EUT1
09705 HMM06277EutA_1_eut(pf06277) EUT1
09715 HMMetlyEutC_1_eut1(pf05985) EUT1 EP_NTERM
09730 HMMreg12833_1_EUT1(pf12833) EUT1
09735 -pf01218 -
09740 -pf01520 -
09750 -pf11143 -
09770 -pf01380 -
09780 HMMnopfamclust3_1(#) RMM1
09800 -pf00005pf12857 -
BMC-P
BMC-H
BMC-T
Aldehyde dehydrogenase
PTAC
Signature enzyme
Alcohol dehydrogenase
Regulator

Citrobacter farmeri GTC 1319. Bacteria; Proteobacteria; Gammaproteobacteria; Enterobacterales;

CIFAM_01__locus_1

EUT1

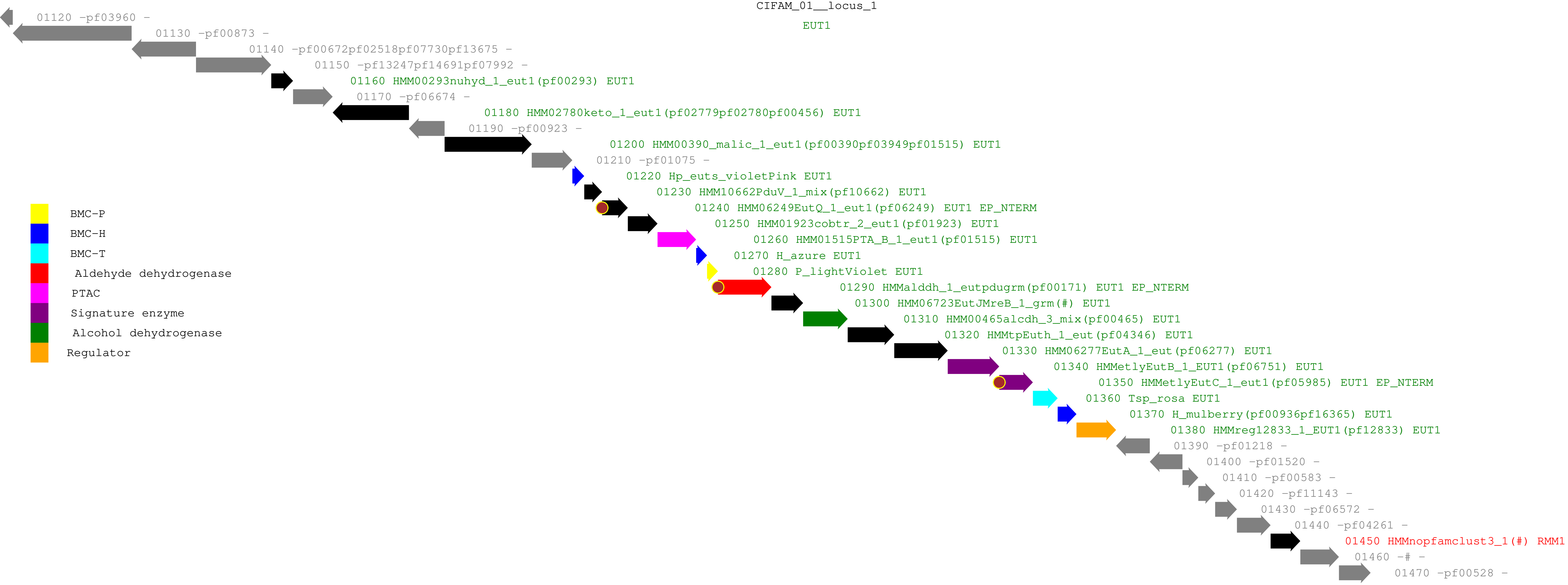

Citrobacter youngae ATCC 29220. Bacteria; Proteobacteria; Gammaproteobacteria; Enterobacterales;

CIT292__locus_1

EUT1

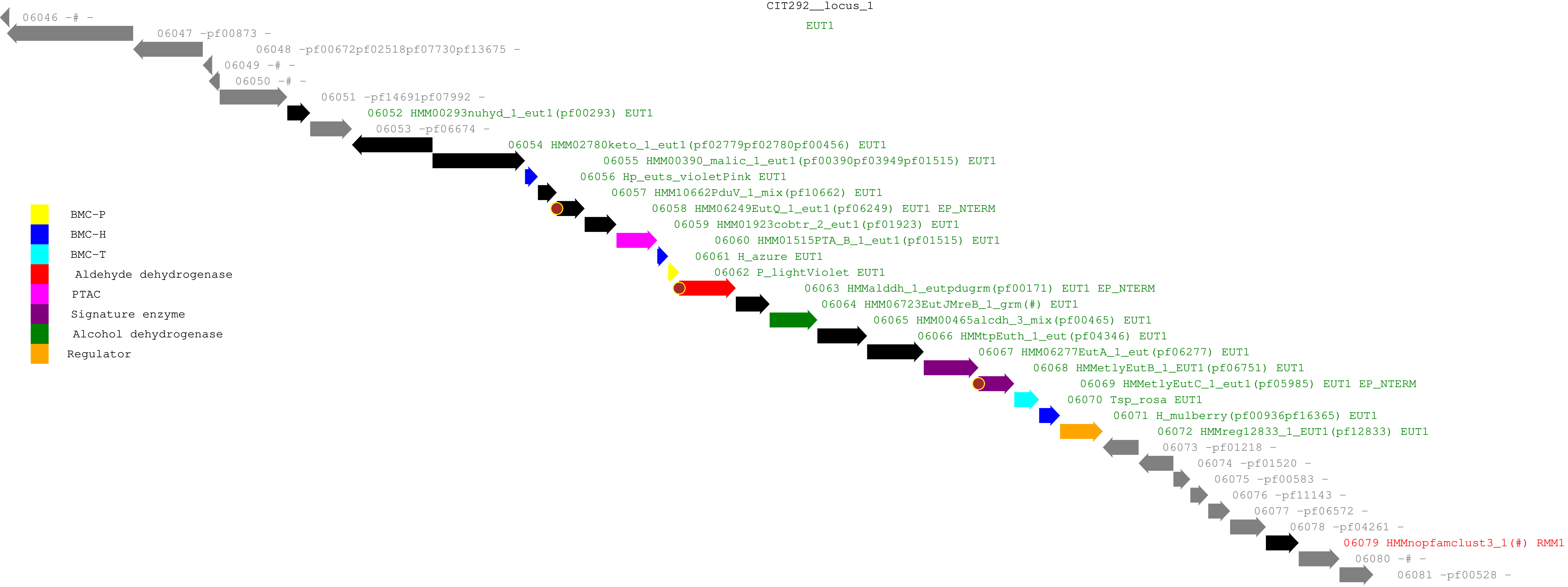

Citrobacter sp. 86. Bacteria; Proteobacteria; Gammaproteobacteria; Enterobacterales;

CITRO86__locus_1

EUT1

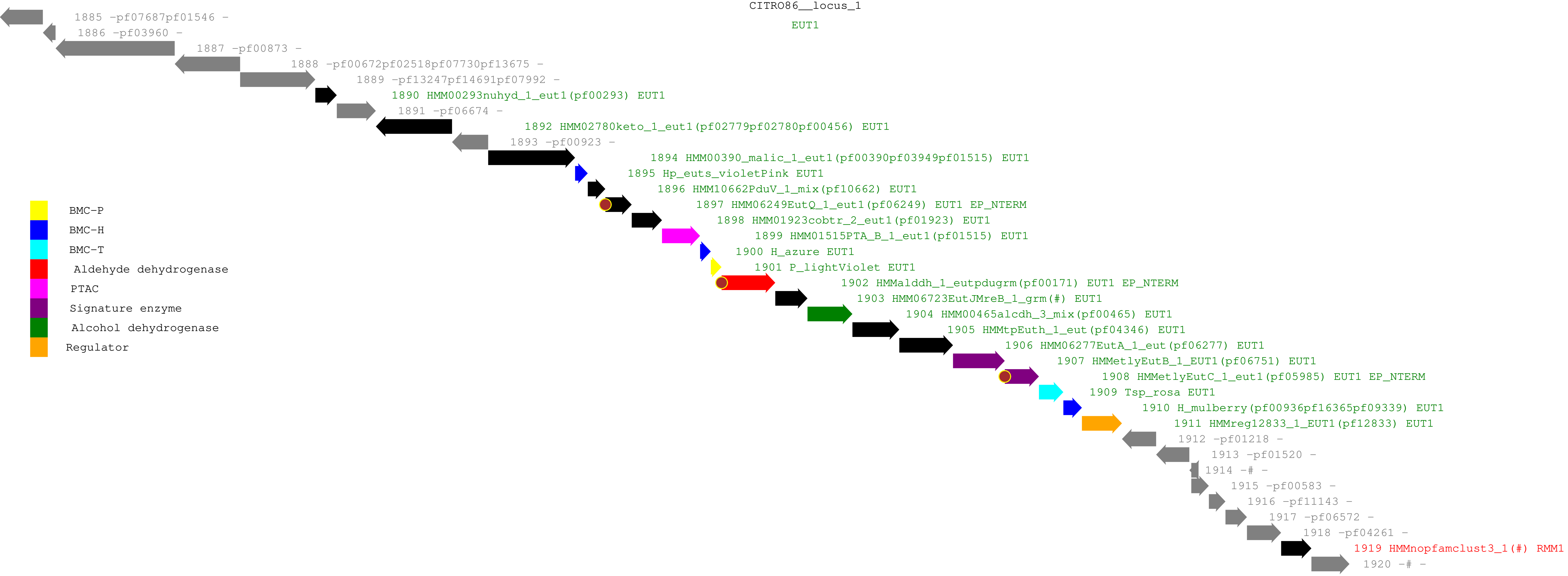

Citrobacter sp. 92. Bacteria; Proteobacteria; Gammaproteobacteria; Enterobacterales;

CITRO92__locus_2

EUT1

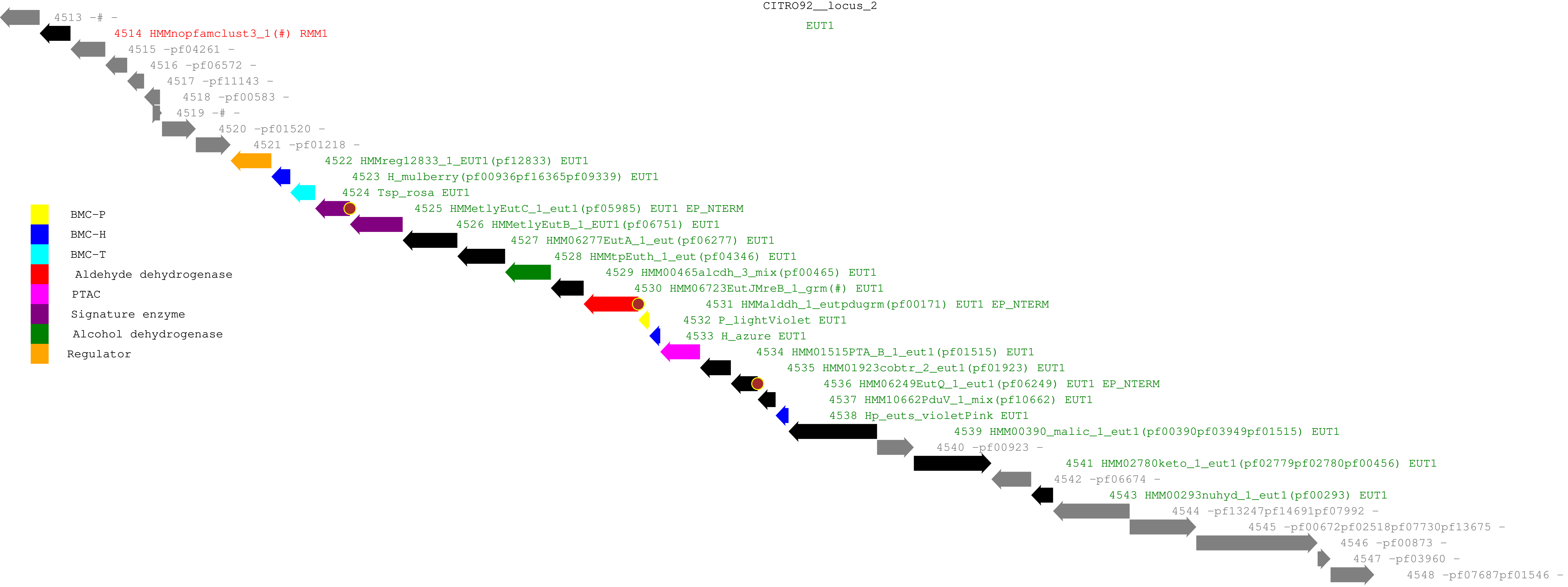

Citrobacter werkmanii NBRC 105721. Bacteria; Proteobacteria; Gammaproteobacteria; Enterobacterales;

CIWKM_02__locus_1

EUT1

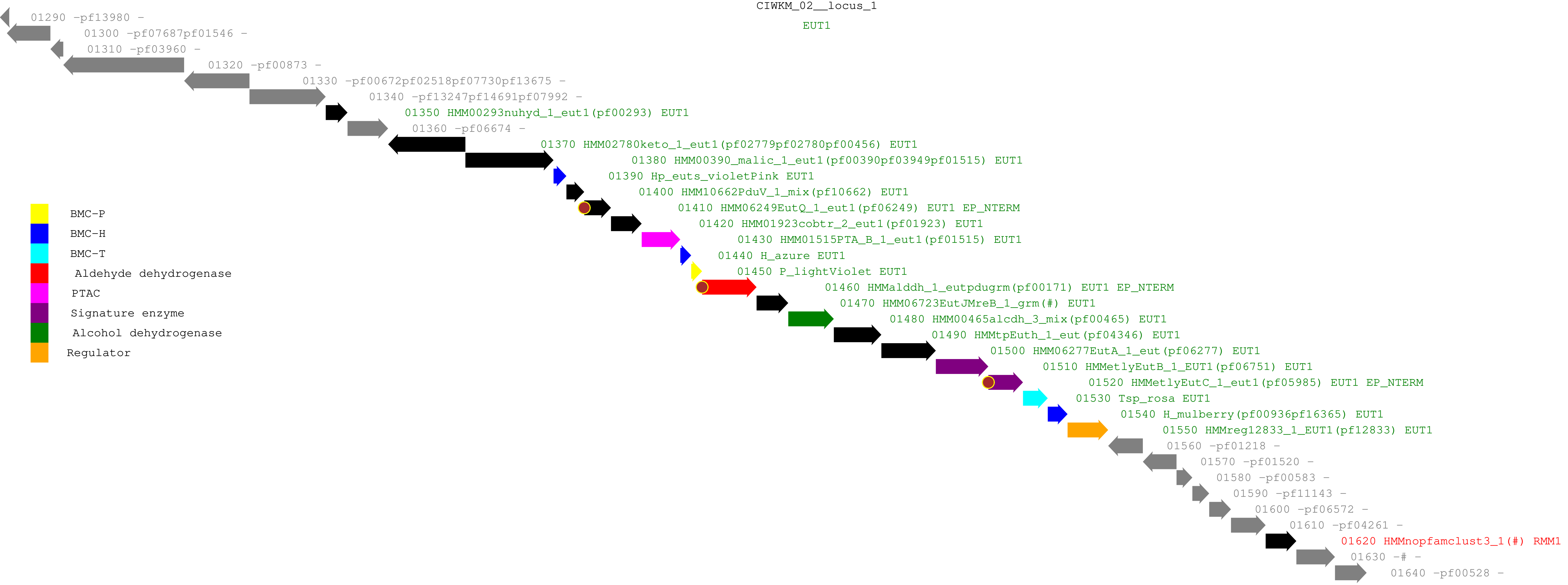

Salmonella enterica subsp. enterica serovar Schwarzengrund. Bacteria; Proteobacteria; Gammaproteobacteria; Enterobacterales;

CJS39__locus_2

EUT1

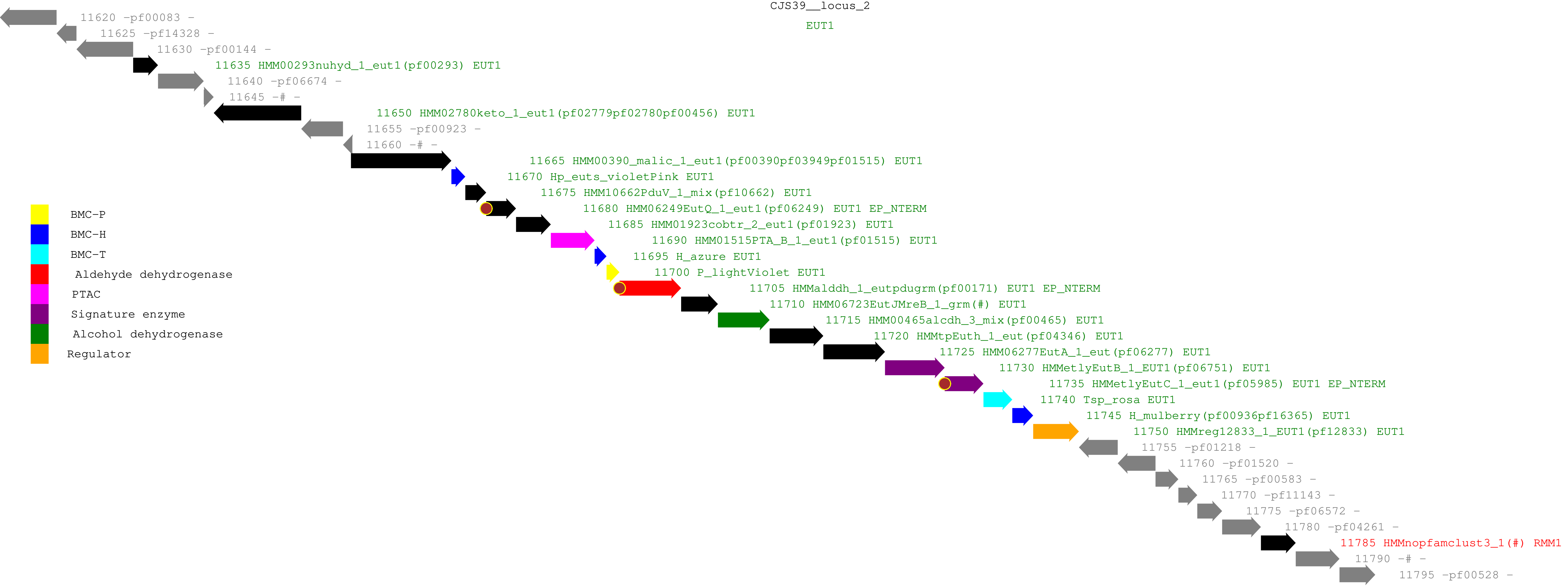

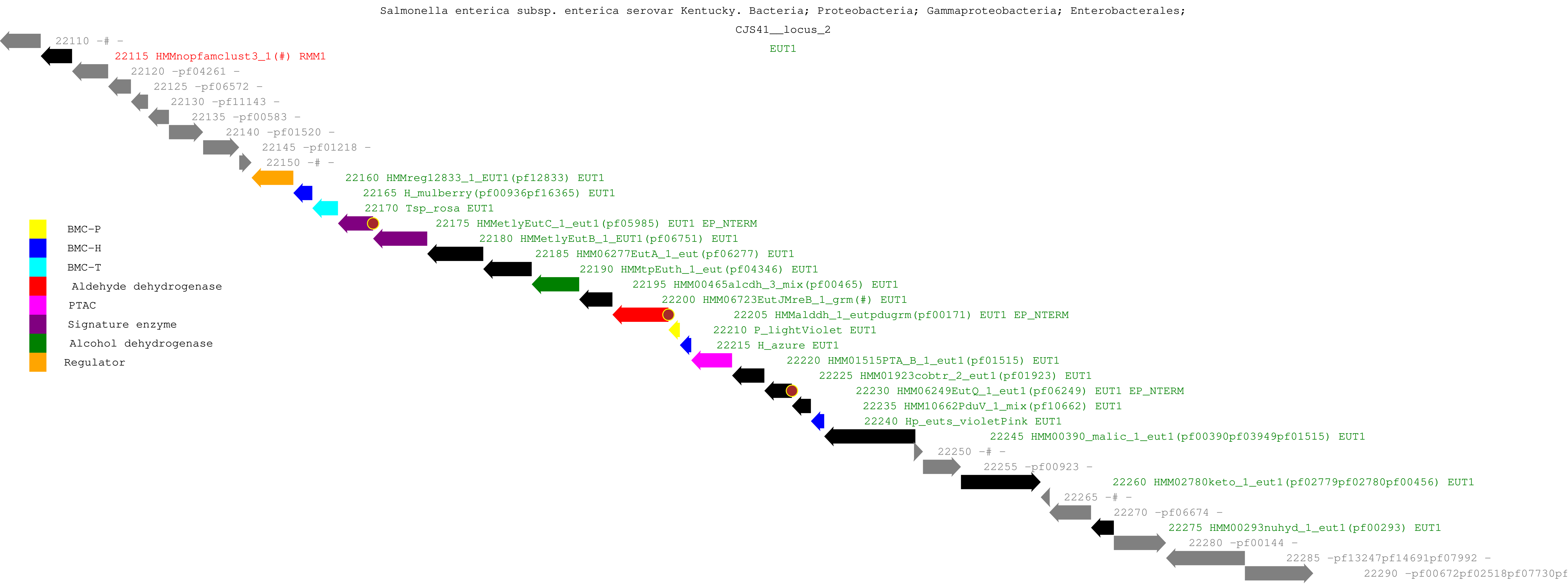
Salmonella enterica subsp. enterica serovar Kentucky. Bacteria; Proteobacteria; Gammaproteobacteria; Enterobacterales;
CJS41__locus_2
EUT1
22110 -# -
22115 HMMnopfamclust3_1(#) RMM1
22120 -pf04261 -
22125 -pf06572 -
22130 -pf11143 -
22135 -pf00583 -
22140 -pf01520 -
22145 -pf01218 -
22150 -# -
22160 HMMreg12833_1_EUT1(pf12833) EUT1
22165 H_mulberry(pf00936pf16365) EUT1
22170 Tsp_rosa EUT1
22175 HMMetlyEutC_1_eut1(pf05985) EUT1 EP_NTERM
22180 HMMetlyEutB_1_EUT1(pf06751) EUT1
22185 HMM06277EutA_1_eut(pf06277) EUT1
22190 HMMtpEuth_1_eut(pf04346) EUT1
22195 HMM00465alcdh_3_mix(pf00465) EUT1
22200 HMM06723EutJMreB_1_grm(#) EUT1
22205 HMMalddh_1_eutpdugrm(pf00171) EUT1 EP_NTERM
22210 P_lightViolet EUT1
22215 H_azure EUT1
22220 HMM01515PTA_B_1_eut1(pf01515) EUT1
22225 HMM01923cobtr_2_eut1(pf01923) EUT1
22230 HMM06249EutQ_1_eut1(pf06249) EUT1 EP_NTERM
22235 HMM10662PduV_1_mix(pf10662) EUT1
22240 Hp_euts_violetPink EUT1
22245 HMM00390_malic_1_eut1(pf00390pf03949pf01515) EUT1
22250 -# -
22255 -pf00923 -
22260 HMM02780keto_1_eut1(pf02779pf02780pf00456) EUT1
22265 -# -
22270 -pf06674 -
22275 HMM00293nuhyd_1_eut1(pf00293) EUT1
22280 -pf00144 -
22285 -pf13247pf14691pf07992 -
22290 -pf00672pf02518pf07730pf
BMC-P
BMC-H
BMC-T
Aldehyde dehydrogenase
PTAC
Signature enzyme
Alcohol dehydrogenase
Regulator

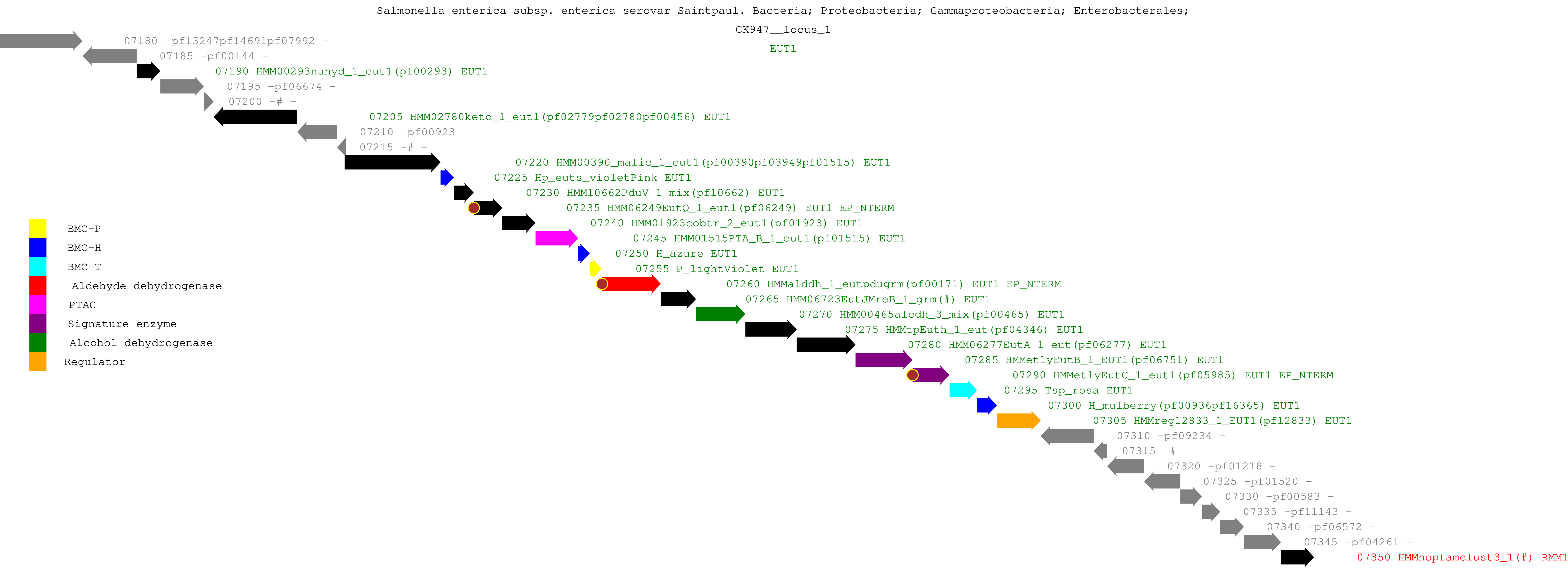
Salmonella enterica subsp. enterica serovar Saintpaul. Bacteria; Proteobacteria; Gammaproteobacteria; Enterobacterales;
CK947__locus_1
EUT1
07180 -pf13247pf14691pf07992 -
07185 -pf00144 -
07190 HMM00293nuhyd_1_eut1(pf00293) EUT1
07195 -pf06674 -
07200 -# -
07205 HMM02780keto_1_eut1(pf02779pf02780pf00456) EUT1
07210 -pf00923 -
07215 -# -
07220 HMM00390_malic_1_eut1(pf00390pf03949pf01515) EUT1
07225 Hp_euts_violetPink EUT1
07230 HMM10662PduV_1_mix(pf10662) EUT1
07235 HMM06249EutQ_1_eut1(pf06249) EUT1 EP_NTERM
07240 HMM01923cobtr_2_eut1(pf01923) EUT1
07245 HMM01515PTA_B_1_eut1(pf01515) EUT1
07250 H_azure EUT1
07255 P_lightViolet EUT1
07260 HMMalddh_1_eutpdugrm(pf00171) EUT1 EP_NTERM
07265 HMM06723EutJMreB_1_grm(#) EUT1
07270 HMM00465alcdh_3_mix(pf00465) EUT1
07275 HMMtpEuth_1_eut(pf04346) EUT1
07280 HMM06277EutA_1_eut(pf06277) EUT1
07285 HMMetlyEutB_1_EUT1(pf06751) EUT1
07290 HMMetlyEutC_1_eut1(pf05985) EUT1 EP_NTERM
07295 Tsp_rosa EUT1
07300 H_mulberry(pf00936pf16365) EUT1
07305 HMMreg12833_1_EUT1(pf12833) EUT1
07310 -pf09234 -
07315 -# -
07320 -pf01218 -
07325 -pf01520 -
07330 -pf00583 -
07335 -pf11143 -
07340 -pf06572 -
07345 -pf04261 -
07350 HMMnopfamclust3_1(#) RMM1
BMC-P
BMC-H
BMC-T
Aldehyde dehydrogenase
PTAC
Signature enzyme
Alcohol dehydrogenase
Regulator

Morganella morganii (Proteus morganii). Bacteria; Proteobacteria; Gammaproteobacteria; Enterobacterales;

CKG00__locus_1

EUT1

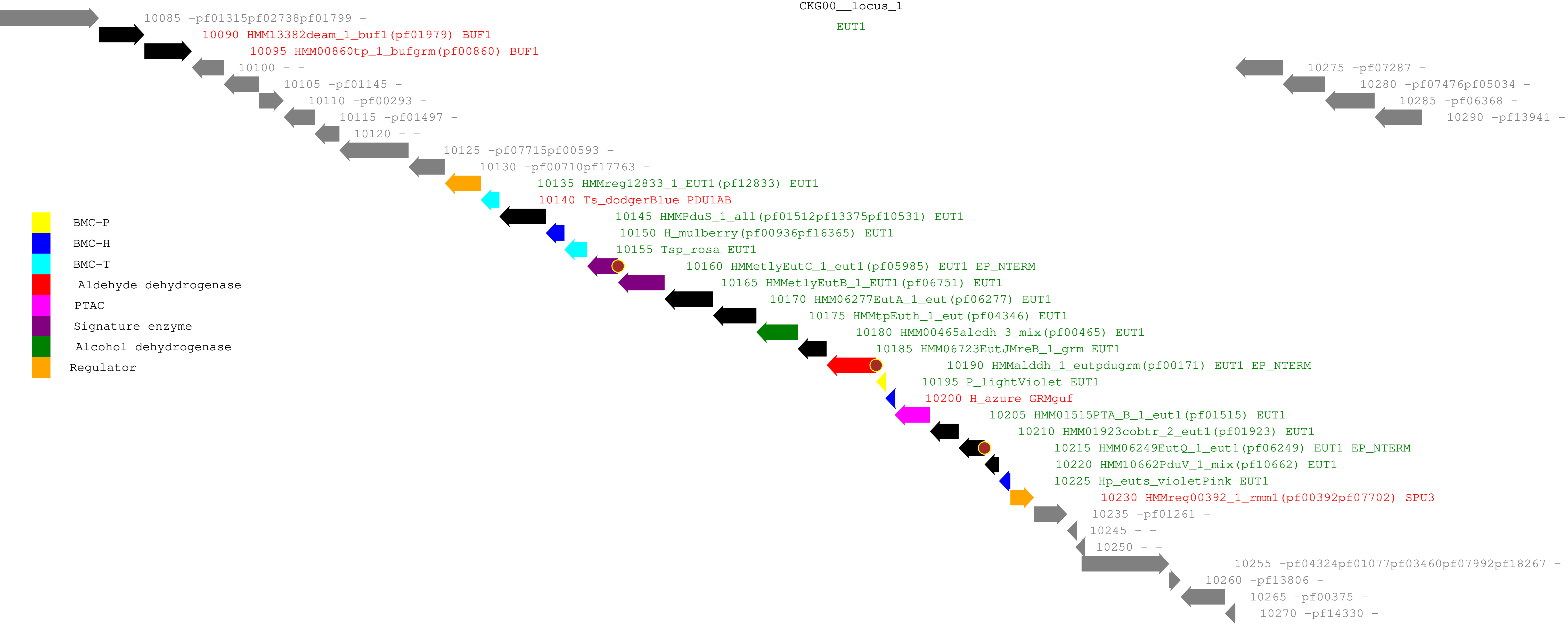

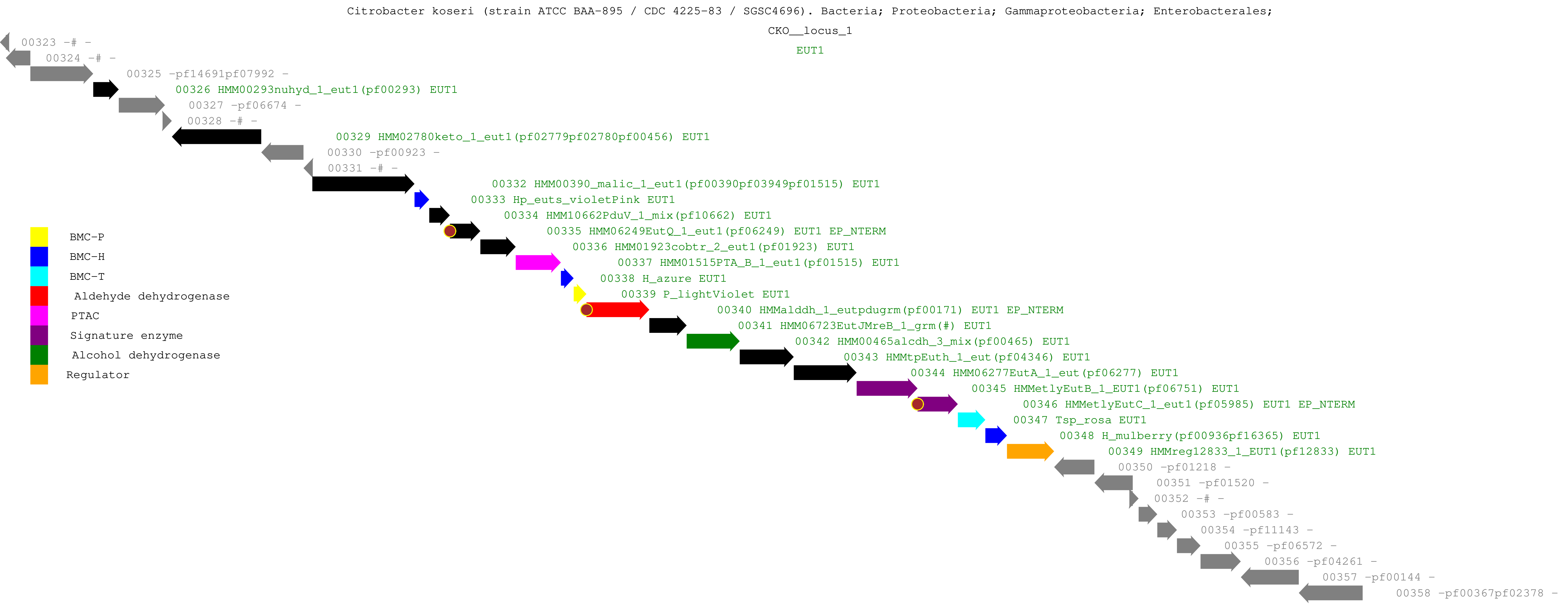
Citrobacter koseri (strain ATCC BAA-895 / CDC 4225-83 / SGSC4696). Bacteria; Proteobacteria; Gammaproteobacteria; Enterobacterales;
CKO__locus_1
EUT1
00323 -# -
00324 -# -
00325 -pf14691pf07992 -
00326 HMM00293nuhyd_1_eut1(pf00293) EUT1
00327 -pf06674 -
00328 -# -
00329 HMM02780keto_1_eut1(pf02779pf02780pf00456) EUT1
00330 -pf00923 -
00331 -# -
00332 HMM00390_malic_1_eut1(pf00390pf03949pf01515) EUT1
00333 Hp_euts_violetPink EUT1
00334 HMM10662PduV_1_mix(pf10662) EUT1
00335 HMM06249EutQ_1_eut1(pf06249) EUT1 EP_NTERM
00336 HMM01923cobtr_2_eut1(pf01923) EUT1
00337 HMM01515PTA_B_1_eut1(pf01515) EUT1
00338 H_azure EUT1
00339 P_lightViolet EUT1
00340 HMMalddh_1_eutpdugrm(pf00171) EUT1 EP_NTERM
00341 HMM06723EutJMreB_1_grm(#) EUT1
00342 HMM00465alcdh_3_mix(pf00465) EUT1
00343 HMMtpEuth_1_eut(pf04346) EUT1
00344 HMM06277EutA_1_eut(pf06277) EUT1
00345 HMMetlyEutB_1_EUT1(pf06751) EUT1
00346 HMMetlyEutC_1_eut1(pf05985) EUT1 EP_NTERM
00347 Tsp_rosa EUT1
00348 H_mulberry(pf00936pf16365) EUT1
00349 HMMreg12833_1_EUT1(pf12833) EUT1
00350 -pf01218 -
00351 -pf01520 -
00352 -# -
00353 -pf00583 -
00354 -pf11143 -
00355 -pf06572 -
00356 -pf04261 -
00357 -pf00144 -
00358 -pf00367pf02378 -
BMC-P
BMC-H
BMC-T
Aldehyde dehydrogenase
PTAC
Signature enzyme
Alcohol dehydrogenase
Regulator

Escherichia coli O6:H1 (strain CFT073 / ATCC 700928 / UPEC). Bacteria; Proteobacteria; Gammaproteobacteria; Enterobacterales;

c_locus_1

EUT1

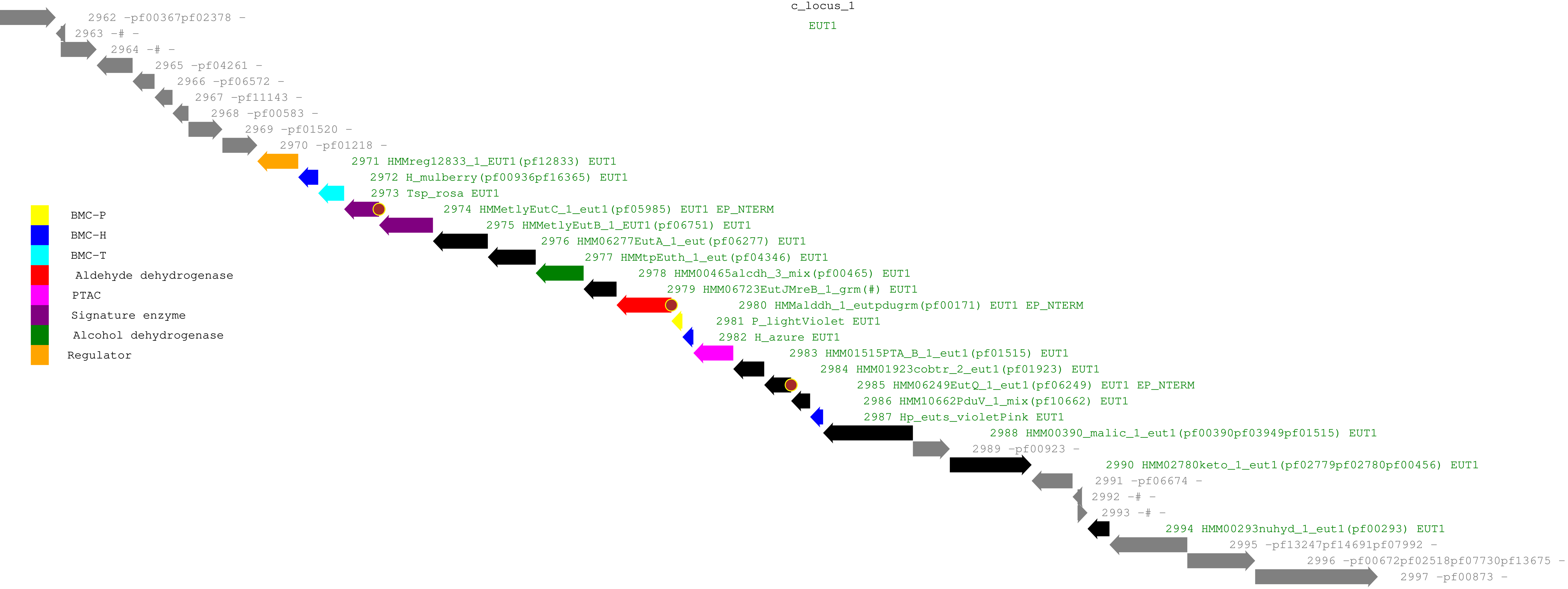

Salmonella montevideo. Bacteria; Proteobacteria; Gammaproteobacteria; Enterobacterales;

CM22__locus_1

EUT1

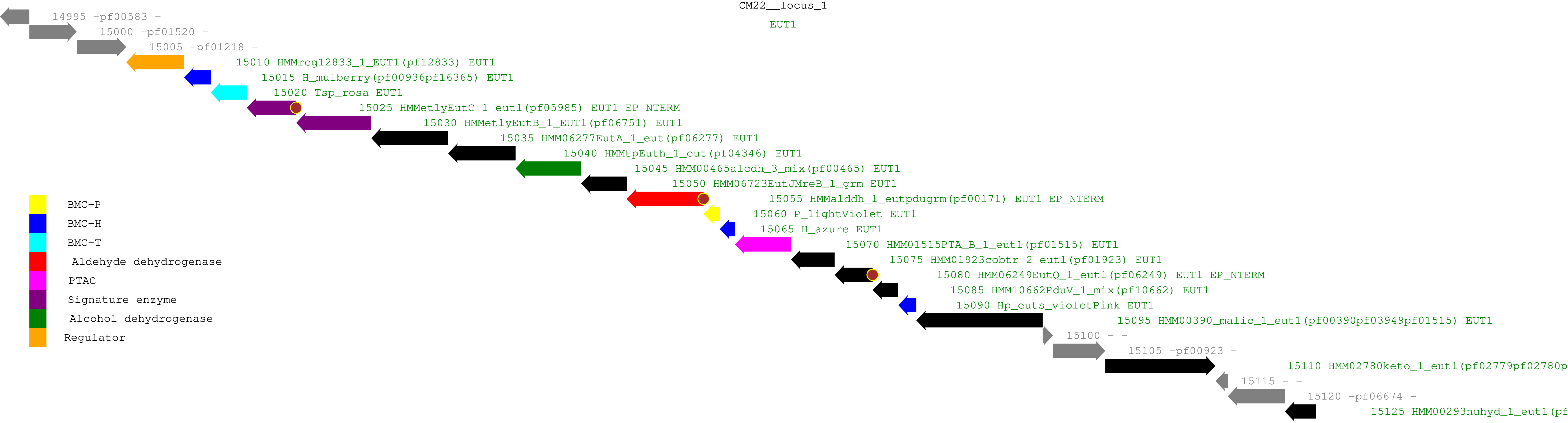

Salmonella diarizonae. Bacteria; Proteobacteria; Gammaproteobacteria; Enterobacterales;

CNQ75__locus_1

EUT1

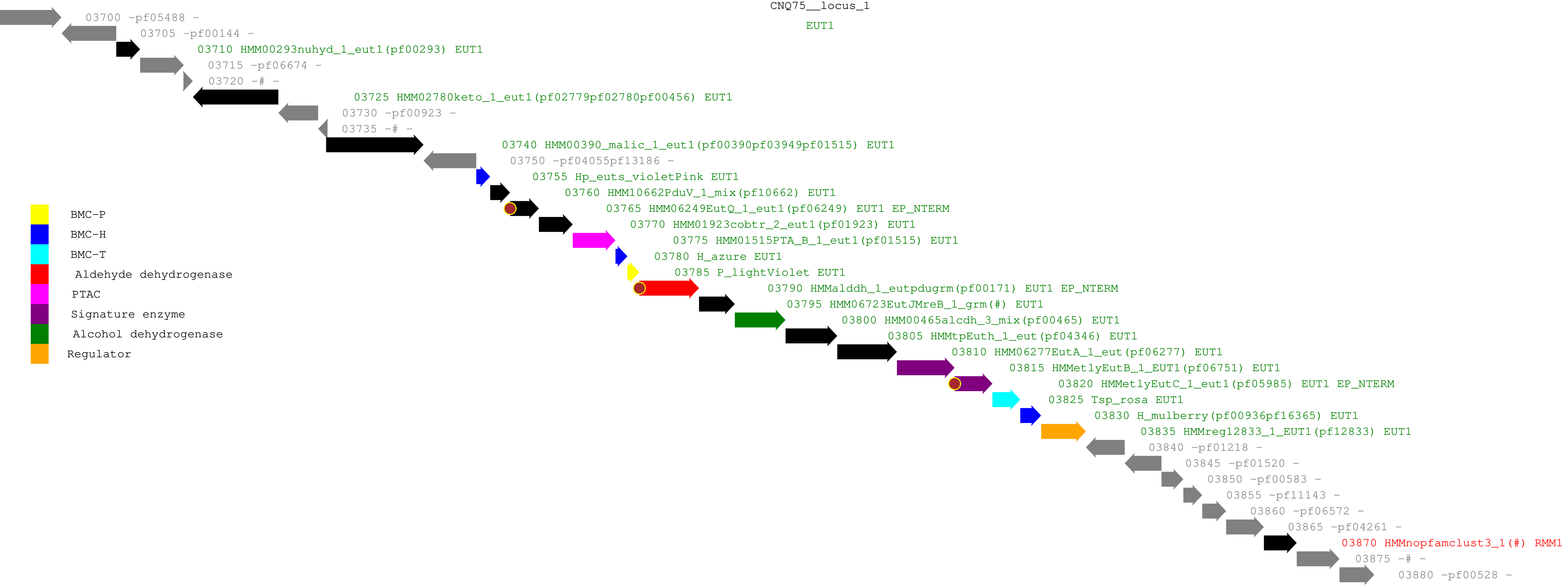

Citrobacter werkmanii. Bacteria; Proteobacteria; Gammaproteobacteria; Enterobacterales;

CO701__locus_1

EUT1

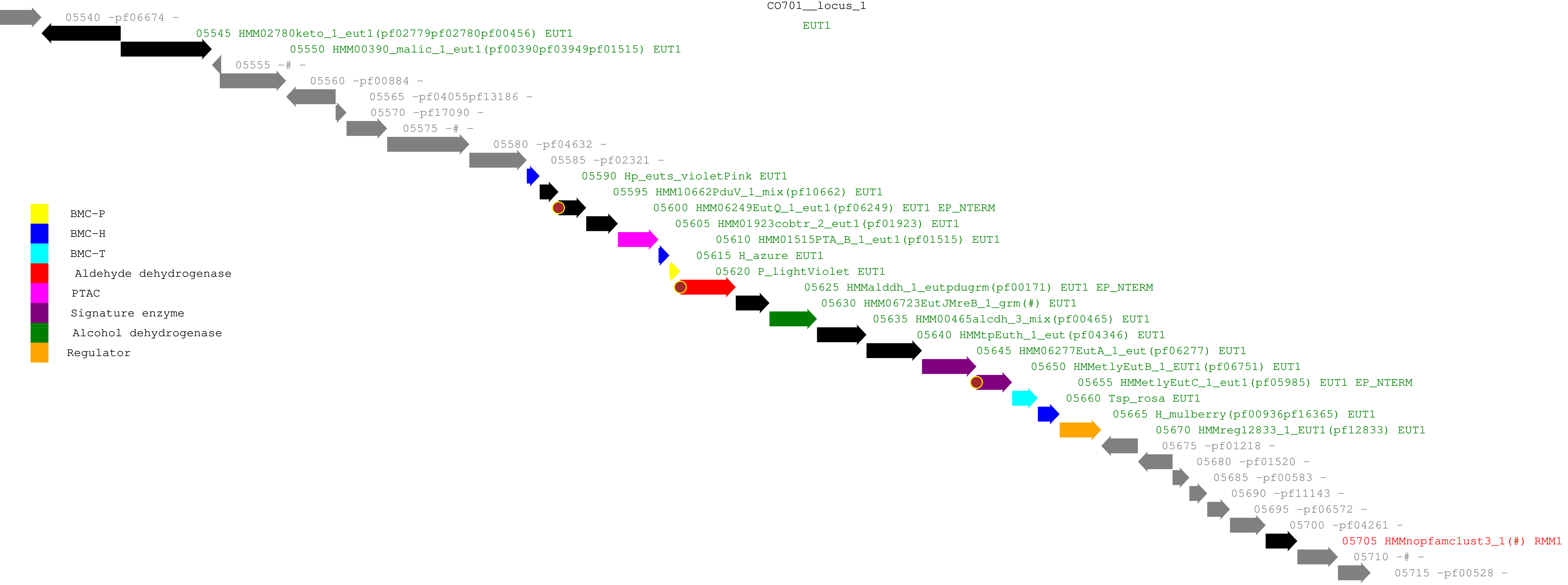

Oceanospirillales bacterium. Bacteria; Proteobacteria; Gammaproteobacteria; Oceanospirillales;

COB25__locus_1_main_sat

EUT1

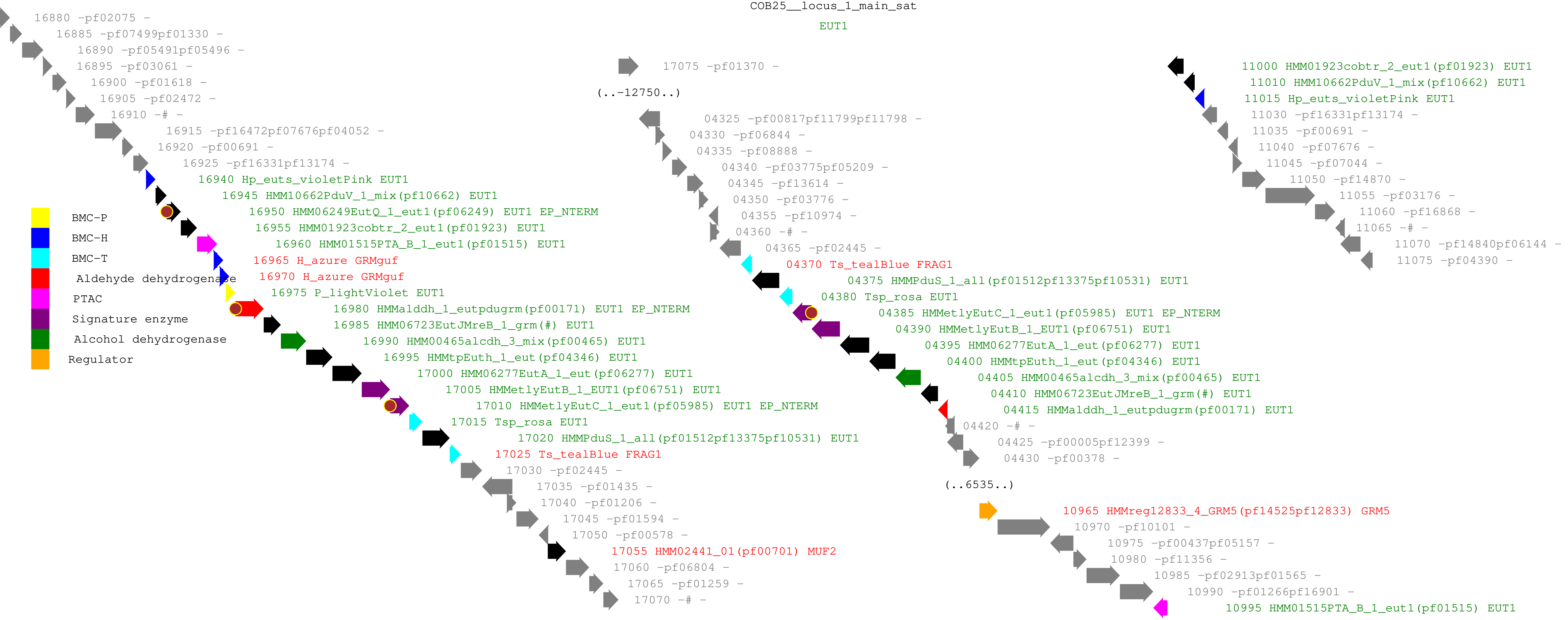

Marinobacter sp. ANT_B65. Bacteria; Proteobacteria; Gammaproteobacteria; Alteromonadales;

CPA50__locus_1

EUT1

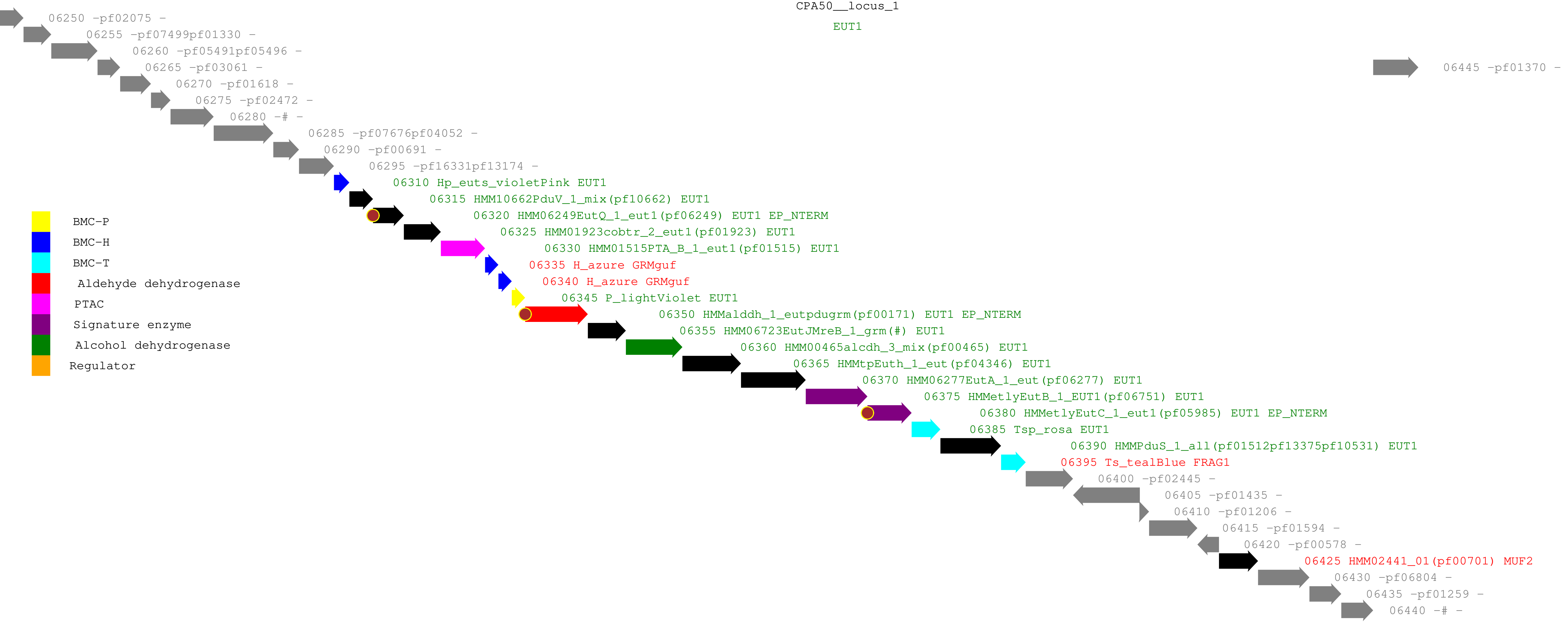

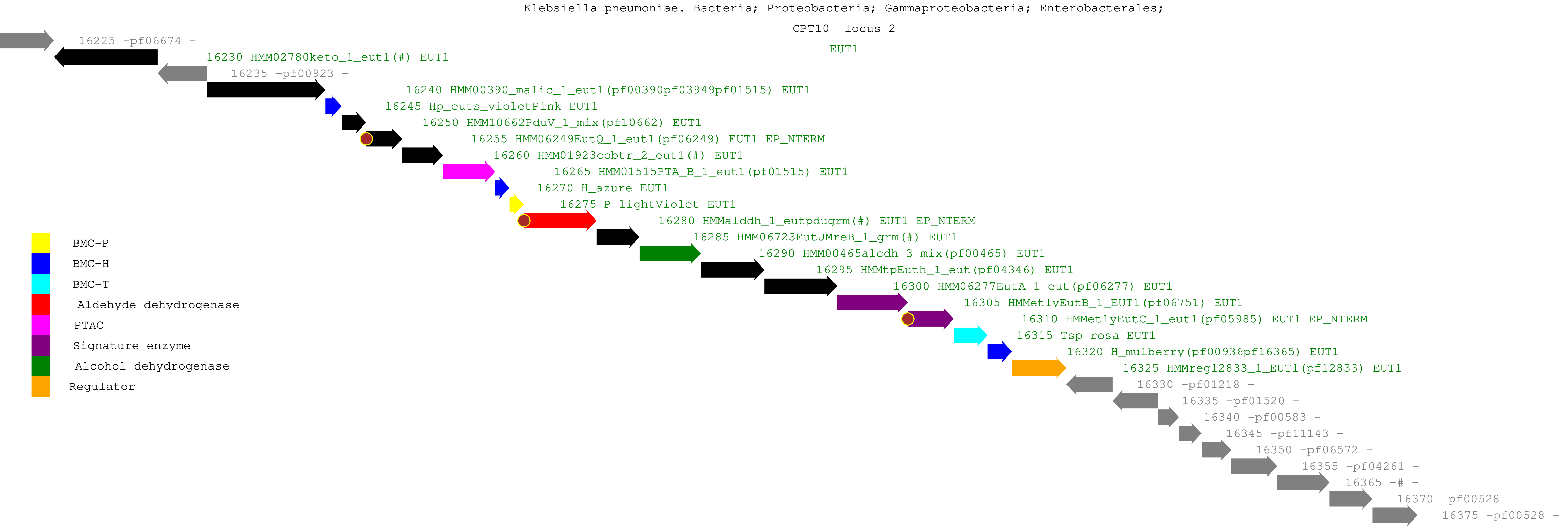

Klebsiella pneumoniae. Bacteria; Proteobacteria; Gammaproteobacteria; Enterobacterales;
CPT10__locus_2
EUT1
16225 -pf06674 -
16230 HMM02780keto_1_eut1(#) EUT1
16235 -pf00923 -
16240 HMM00390_malic_1_eut1(pf00390pf03949pf01515) EUT1
16245 Hp_euts_violetPink EUT1
16250 HMM10662PduV_1_mix(pf10662) EUT1
16255 HMM06249EutQ_1_eut1(pf06249) EUT1 EP_NTERM
16260 HMM01923cobtr_2_eut1(#) EUT1
16265 HMM01515PTA_B_1_eut1(pf01515) EUT1
16270 H_azure EUT1
16275 P_lightViolet EUT1
16280 HMMalddh_1_eutpdugrm(#) EUT1 EP_NTERM
16285 HMM06723EutJMreB_1_grm(#) EUT1
16290 HMM00465alcdh_3_mix(pf00465) EUT1
16295 HMMtpEuth_1_eut(pf04346) EUT1
16300 HMM06277EutA_1_eut(pf06277) EUT1
16305 HMMetlyEutB_1_EUT1(pf06751) EUT1
16310 HMMetlyEutC_1_eut1(pf05985) EUT1 EP_NTERM
16315 Tsp_rosa EUT1
16320 H_mulberry(pf00936pf16365) EUT1
16325 HMMreg12833_1_EUT1(pf12833) EUT1
16330 -pf01218 -
16335 -pf01520 -
16340 -pf00583 -
16345 -pf11143 -
16350 -pf06572 -
16355 -pf04261 -
16365 -# -
16370 -pf00528 -
16375 -pf00528 -
BMC-P
BMC-H
BMC-T
Aldehyde dehydrogenase
PTAC
Signature enzyme
Alcohol dehydrogenase
Regulator

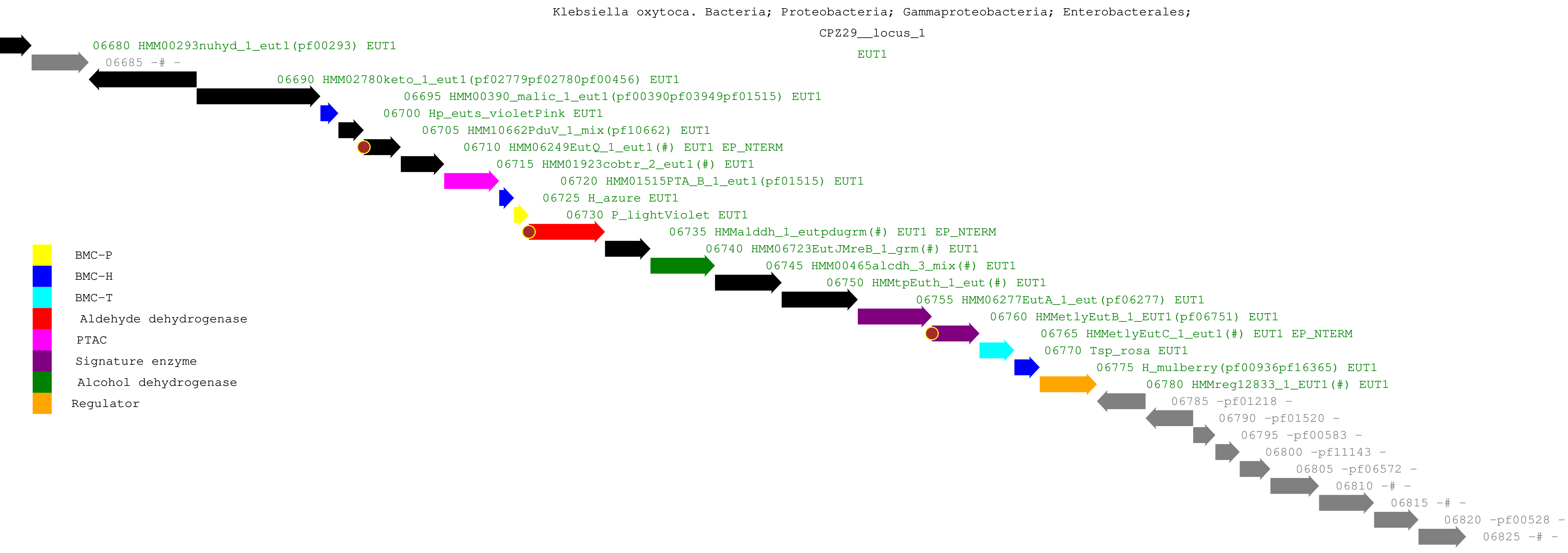
Klebsiella oxytoca. Bacteria; Proteobacteria; Gammaproteobacteria; Enterobacterales;
CPZ29__locus_1
EUT1
06680 HMM00293nuhyd_1_eut1(pf00293) EUT1
06685 -# -
06690 HMM02780keto_1_eut1(pf02779pf02780pf00456) EUT1
06695 HMM00390_malic_1_eut1(pf00390pf03949pf01515) EUT1
06700 Hp_euts_violetPink EUT1
06705 HMM10662PduV_1_mix(pf10662) EUT1
06710 HMM06249EutQ_1_eut1(#) EUT1 EP_NTERM
06715 HMM01923cobtr_2_eut1(#) EUT1
06720 HMM01515PTA_B_1_eut1(pf01515) EUT1
06725 H_azure EUT1
06730 P_lightViolet EUT1
06735 HMMalddh_1_eutpdugrm(#) EUT1 EP_NTERM
06740 HMM06723EutJMreB_1_grm(#) EUT1
06745 HMM00465alcdh_3_mix(#) EUT1
06750 HMMtpEuth_1_eut(#) EUT1
06755 HMM06277EutA_1_eut(pf06277) EUT1
06760 HMMetlyEutB_1_EUT1(pf06751) EUT1
06765 HMMetlyEutC_1_eut1(#) EUT1 EP_NTERM
06770 Tsp_rosa EUT1
06775 H_mulberry(pf00936pf16365) EUT1
06780 HMMreg12833_1_EUT1(#) EUT1
06785 -pf01218 -
06790 -pf01520 -
06795 -pf00583 -
06800 -pf11143 -
06805 -pf06572 -
06810 -# -
06815 -# -
06820 -pf00528 -
06825 -# -
BMC-P
BMC-H
BMC-T
Aldehyde dehydrogenase
PTAC
Signature enzyme
Alcohol dehydrogenase
Regulator

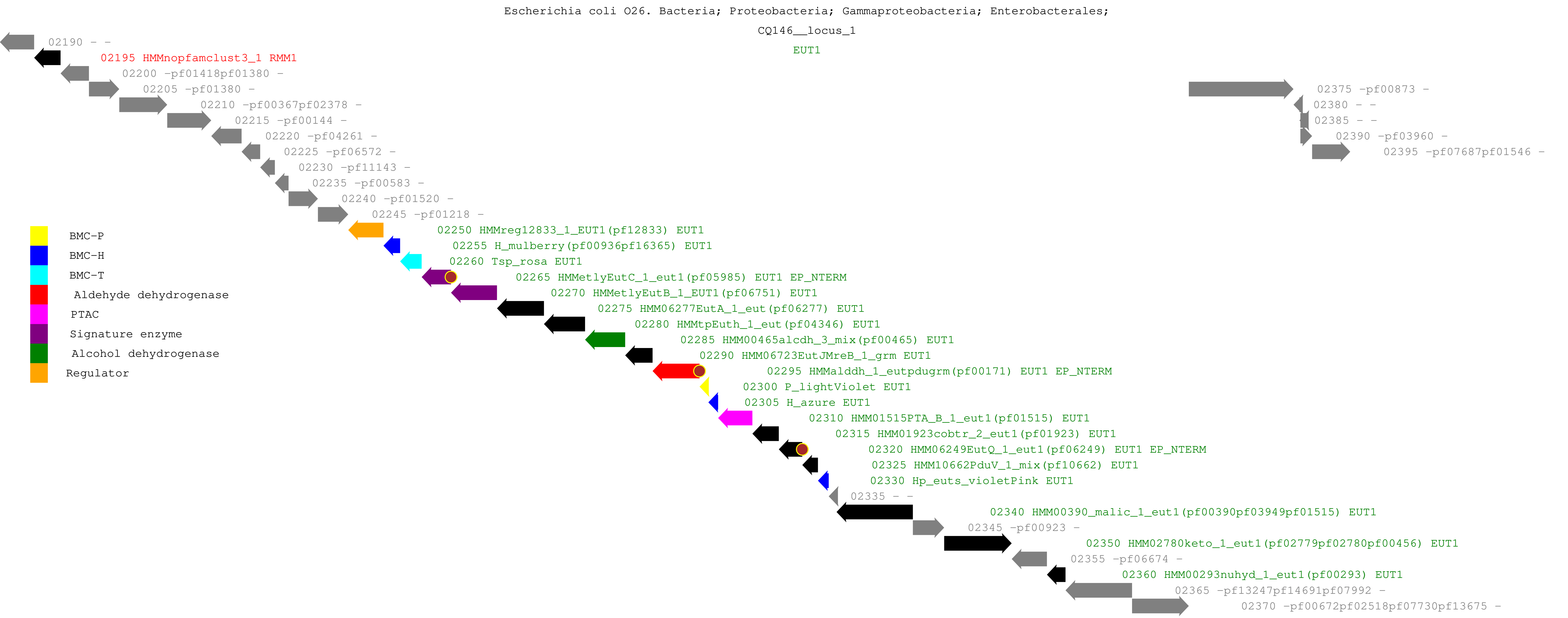
Escherichia coli O26. Bacteria; Proteobacteria; Gammaproteobacteria; Enterobacterales;
CQ146__locus_1
EUT1
02190 - -
02195 HMMnopfamclust3_1 RMM1
02200 -pf01418pf01380 -
02205 -pf01380 -
02210 -pf00367pf02378 -
02215 -pf00144 -
02220 -pf04261 -
02225 -pf06572 -
02230 -pf11143 -
02235 -pf00583 -
02240 -pf01520 -
02245 -pf01218 -
02250 HMMreg12833_1_EUT1(pf12833) EUT1
02255 H_mulberry(pf00936pf16365) EUT1
02260 Tsp_rosa EUT1
02265 HMMetlyEutC_1_eut1(pf05985) EUT1 EP_NTERM
02270 HMMetlyEutB_1_EUT1(pf06751) EUT1
02275 HMM06277EutA_1_eut(pf06277) EUT1
02280 HMMtpEuth_1_eut(pf04346) EUT1
02285 HMM00465alcdh_3_mix(pf00465) EUT1
02290 HMM06723EutJMreB_1_grm EUT1
02295 HMMalddh_1_eutpdugrm(pf00171) EUT1 EP_NTERM
02300 P_lightViolet EUT1
02305 H_azure EUT1
02310 HMM01515PTA_B_1_eut1(pf01515) EUT1
02315 HMM01923cobtr_2_eut1(pf01923) EUT1
02320 HMM06249EutQ_1_eut1(pf06249) EUT1 EP_NTERM
02325 HMM10662PduV_1_mix(pf10662) EUT1
02330 Hp_euts_violetPink EUT1
02335 - -
02340 HMM00390_malic_1_eut1(pf00390pf03949pf01515) EUT1
02345 -pf00923 -
02350 HMM02780keto_1_eut1(pf02779pf02780pf00456) EUT1
02355 -pf06674 -
02360 HMM00293nuhyd_1_eut1(pf00293) EUT1
02365 -pf13247pf14691pf07992 -
02370 -pf00672pf02518pf07730pf13675 -
02375 -pf00873 -
02380 - -
02385 - -
02390 -pf03960 -
02395 -pf07687pf01546 -
BMC-P
BMC-H
BMC-T
Aldehyde dehydrogenase
PTAC
Signature enzyme
Alcohol dehydrogenase
Regulator

Escherichia coli. Bacteria; Proteobacteria; Gammaproteobacteria; Enterobacterales;

CQB02__locus_2

EUT1

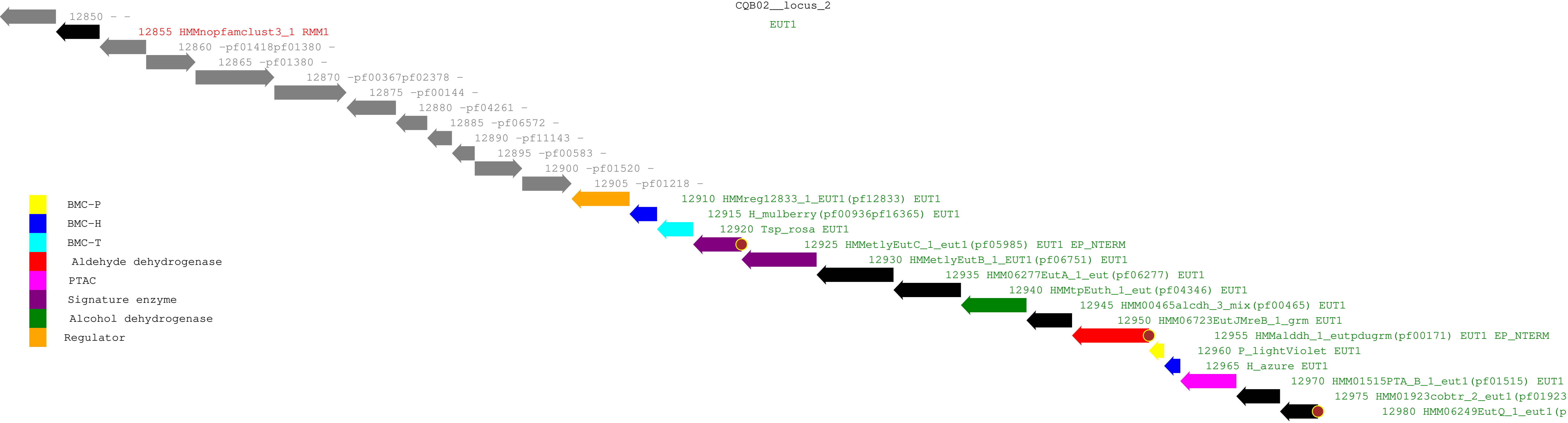

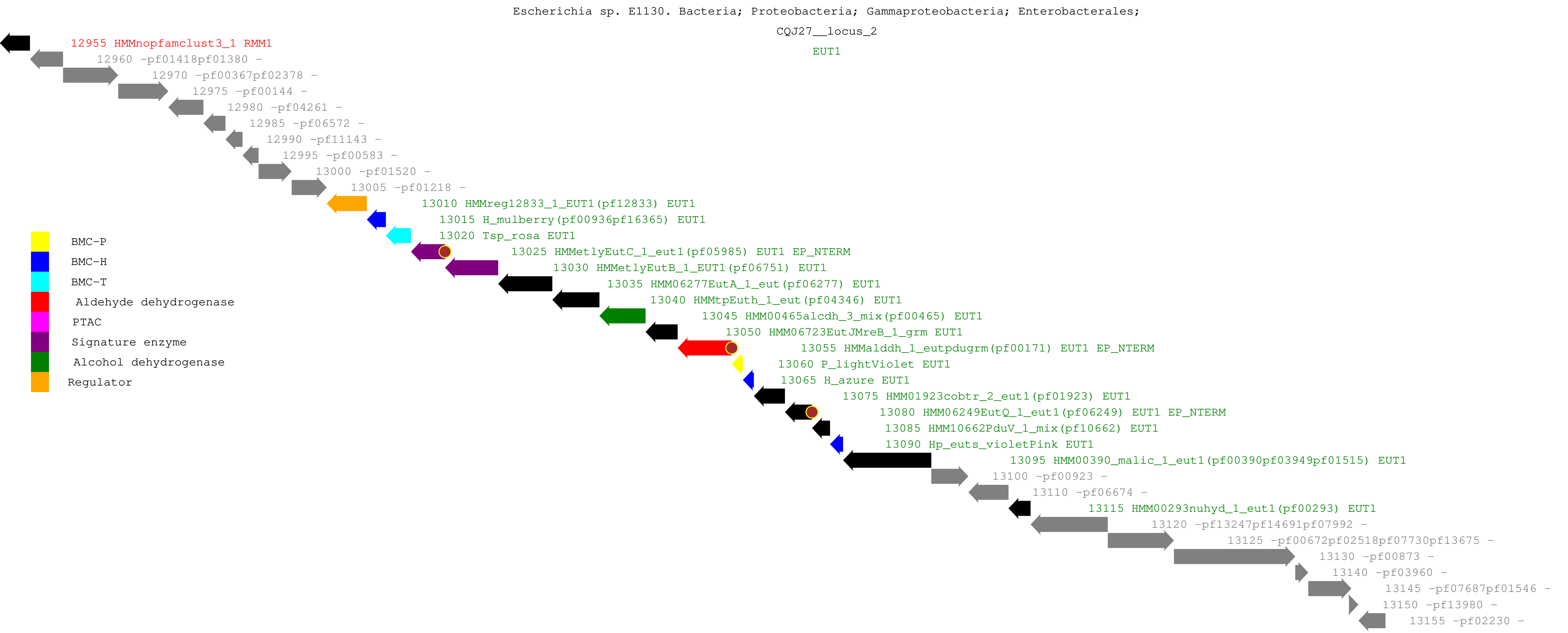

Escherichia sp. E1130. Bacteria; Proteobacteria; Gammaproteobacteria; Enterobacterales;
CQJ27__locus_2
EUT1
12955 HMMnopfamclust3_1 RMM1
12960 -pf01418pf01380 -
12970 -pf00367pf02378 -
12975 -pf00144 -
12980 -pf04261 -
12985 -pf06572 -
12990 -pf11143 -
12995 -pf00583 -
13000 -pf01520 -
13005 -pf01218 -
13010 HMMreg12833_1_EUT1(pf12833) EUT1
13015 H_mulberry(pf00936pf16365) EUT1
13020 Tsp_rosa EUT1
13025 HMMetlyEutC_1_eut1(pf05985) EUT1 EP_NTERM
13030 HMMetlyEutB_1_EUT1(pf06751) EUT1
13035 HMM06277EutA_1_eut(pf06277) EUT1
13040 HMMtpEuth_1_eut(pf04346) EUT1
13045 HMM00465alcdh_3_mix(pf00465) EUT1
13050 HMM06723EutJMreB_1_grm EUT1
13055 HMMalddh_1_eutpdugrm(pf00171) EUT1 EP_NTERM
13060 P_lightViolet EUT1
13065 H_azure EUT1
13075 HMM01923cobtr_2_eut1(pf01923) EUT1
13080 HMM06249EutQ_1_eut1(pf06249) EUT1 EP_NTERM
13085 HMM10662PduV_1_mix(pf10662) EUT1
13090 Hp_euts_violetPink EUT1
13095 HMM00390_malic_1_eut1(pf00390pf03949pf01515) EUT1
13100 -pf00923 -
13110 -pf06674 -
13115 HMM00293nuhyd_1_eut1(pf00293) EUT1
13120 -pf13247pf14691pf07992 -
13125 -pf00672pf02518pf07730pf13675 -
13130 -pf00873 -
13140 -pf03960 -
13145 -pf07687pf01546 -
13150 -pf13980 -
13155 -pf02230 -
BMC-P
BMC-H
BMC-T
Aldehyde dehydrogenase
PTAC
Signature enzyme
Alcohol dehydrogenase
Regulator

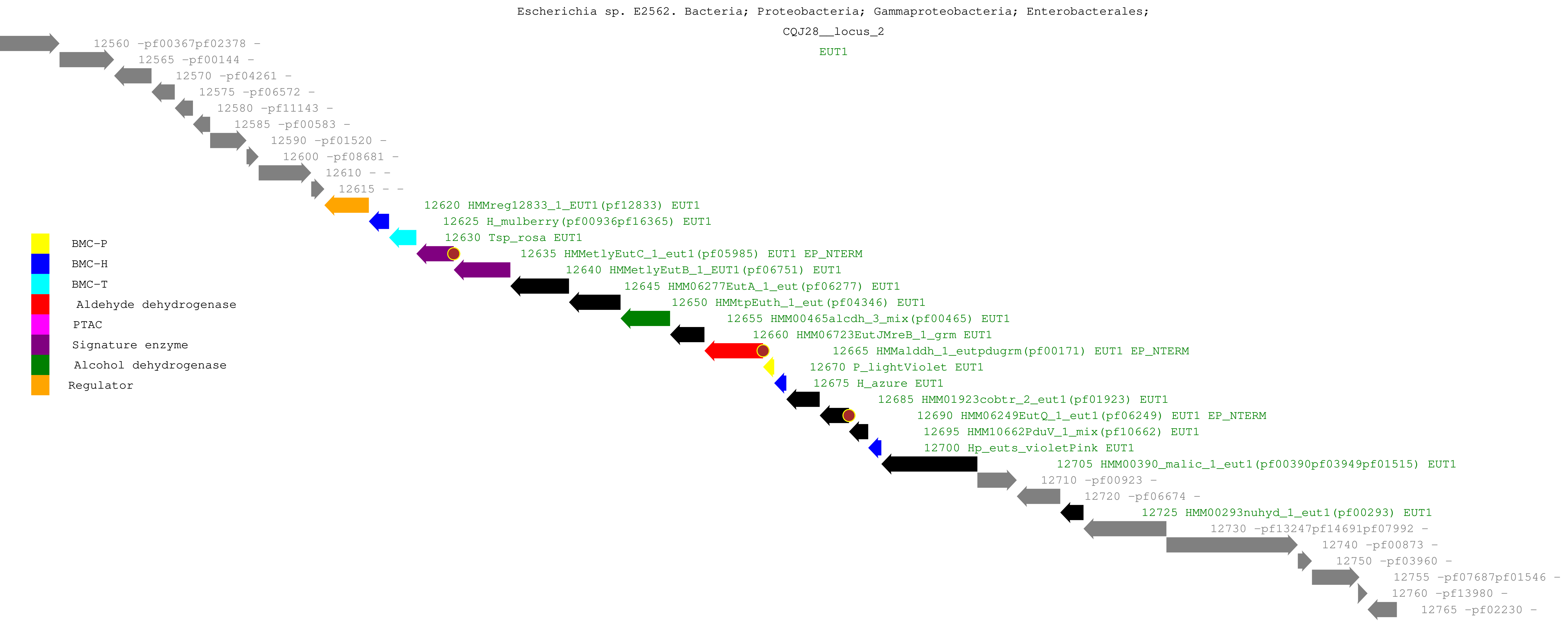
Escherichia sp. E2562. Bacteria; Proteobacteria; Gammaproteobacteria; Enterobacterales;
CQJ28__locus_2
EUT1
12560 -pf00367pf02378 -
12565 -pf00144 -
12570 -pf04261 -
12575 -pf06572 -
12580 -pf11143 -
12585 -pf00583 -
12590 -pf01520 -
12600 -pf08681 -
12610 - -
12615 - -
12620 HMMreg12833_1_EUT1(pf12833) EUT1
12625 H_mulberry(pf00936pf16365) EUT1
12630 Tsp_rosa EUT1
12635 HMMetlyEutC_1_eut1(pf05985) EUT1 EP_NTERM
12640 HMMetlyEutB_1_EUT1(pf06751) EUT1
12645 HMM06277EutA_1_eut(pf06277) EUT1
12650 HMMtpEuth_1_eut(pf04346) EUT1
12655 HMM00465alcdh_3_mix(pf00465) EUT1
12660 HMM06723EutJMreB_1_grm EUT1
12665 HMMalddh_1_eutpdugrm(pf00171) EUT1 EP_NTERM
12670 P_lightViolet EUT1
12675 H_azure EUT1
12685 HMM01923cobtr_2_eut1(pf01923) EUT1
12690 HMM06249EutQ_1_eut1(pf06249) EUT1 EP_NTERM
12695 HMM10662PduV_1_mix(pf10662) EUT1
12700 Hp_euts_violetPink EUT1
12705 HMM00390_malic_1_eut1(pf00390pf03949pf01515) EUT1
12710 -pf00923 -
12720 -pf06674 -
12725 HMM00293nuhyd_1_eut1(pf00293) EUT1
12730 -pf13247pf14691pf07992 -
12740 -pf00873 -
12750 -pf03960 -
12755 -pf07687pf01546 -
12760 -pf13980 -
12765 -pf02230 -
BMC-P
BMC-H
BMC-T
Aldehyde dehydrogenase
PTAC
Signature enzyme
Alcohol dehydrogenase
Regulator

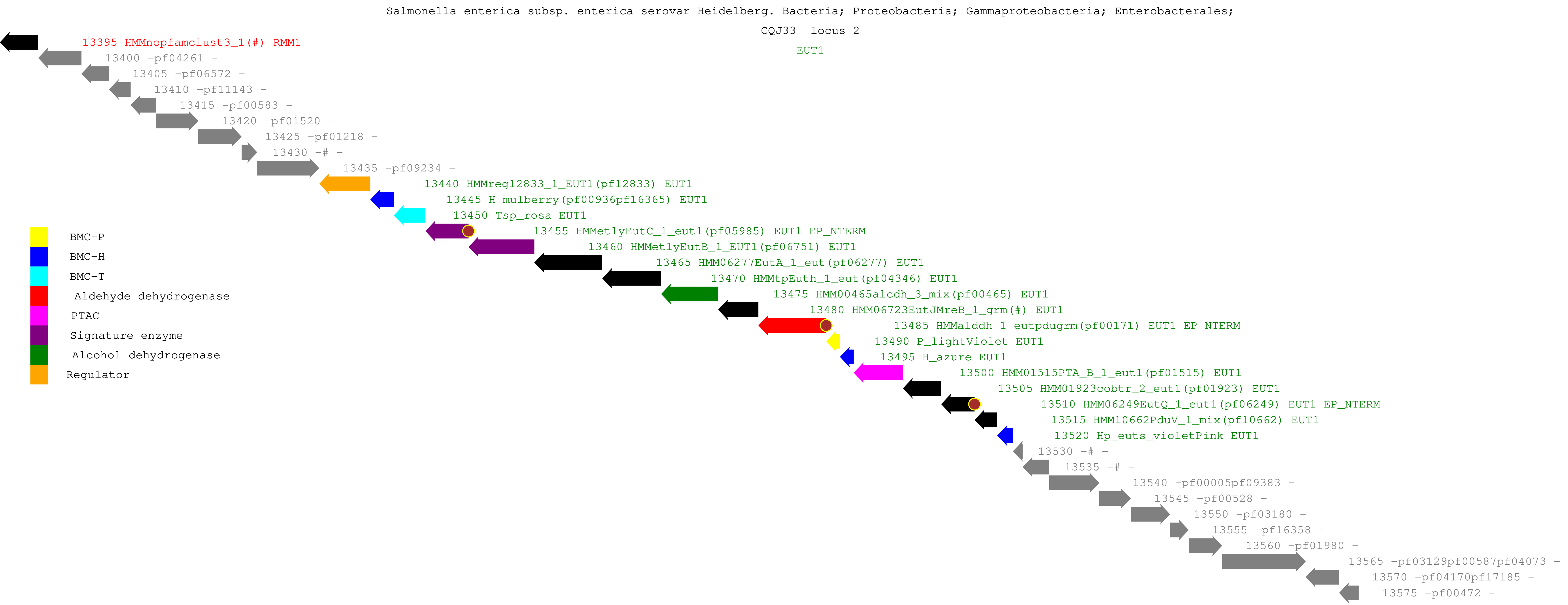

Salmonella enterica subsp. enterica serovar Heidelberg. Bacteria; Proteobacteria; Gammaproteobacteria; Enterobacterales;
CQJ33__locus_2
EUT1
13395 HMMnopfamclust3_1(#) RMM1
13400 -pf04261 -
13405 -pf06572 -
13410 -pf11143 -
13415 -pf00583 -
13420 -pf01520 -
13425 -pf01218 -
13430 -# -
13435 -pf09234 -
13440 HMMreg12833_1_EUT1(pf12833) EUT1
13445 H_mulberry(pf00936pf16365) EUT1
13450 Tsp_rosa EUT1
13455 HMMetlyEutC_1_eut1(pf05985) EUT1 EP_NTERM
13460 HMMetlyEutB_1_EUT1(pf06751) EUT1
13465 HMM06277EutA_1_eut(pf06277) EUT1
13470 HMMtpEuth_1_eut(pf04346) EUT1
13475 HMM00465alcdh_3_mix(pf00465) EUT1
13480 HMM06723EutJMreB_1_grm(#) EUT1
13485 HMMalddh_1_eutpdugrm(pf00171) EUT1 EP_NTERM
13490 P_lightViolet EUT1
13495 H_azure EUT1
13500 HMM01515PTA_B_1_eut1(pf01515) EUT1
13505 HMM01923cobtr_2_eut1(pf01923) EUT1
13510 HMM06249EutQ_1_eut1(pf06249) EUT1 EP_NTERM
13515 HMM10662PduV_1_mix(pf10662) EUT1
13520 Hp_euts_violetPink EUT1
13530 -# -
13535 -# -
13540 -pf00005pf09383 -
13545 -pf00528 -
13550 -pf03180 -
13555 -pf16358 -
13560 -pf01980 -
13565 -pf03129pf00587pf04073 -
13570 -pf04170pf17185 -
13575 -pf00472 -
BMC-P
BMC-H
BMC-T
Aldehyde dehydrogenase
PTAC
Signature enzyme
Alcohol dehydrogenase
Regulator

Salmonella enterica subsp. enterica serovar Ouakam. Bacteria; Proteobacteria; Gammaproteobacteria; Enterobacterales;

CQU62__locus_1

EUT1

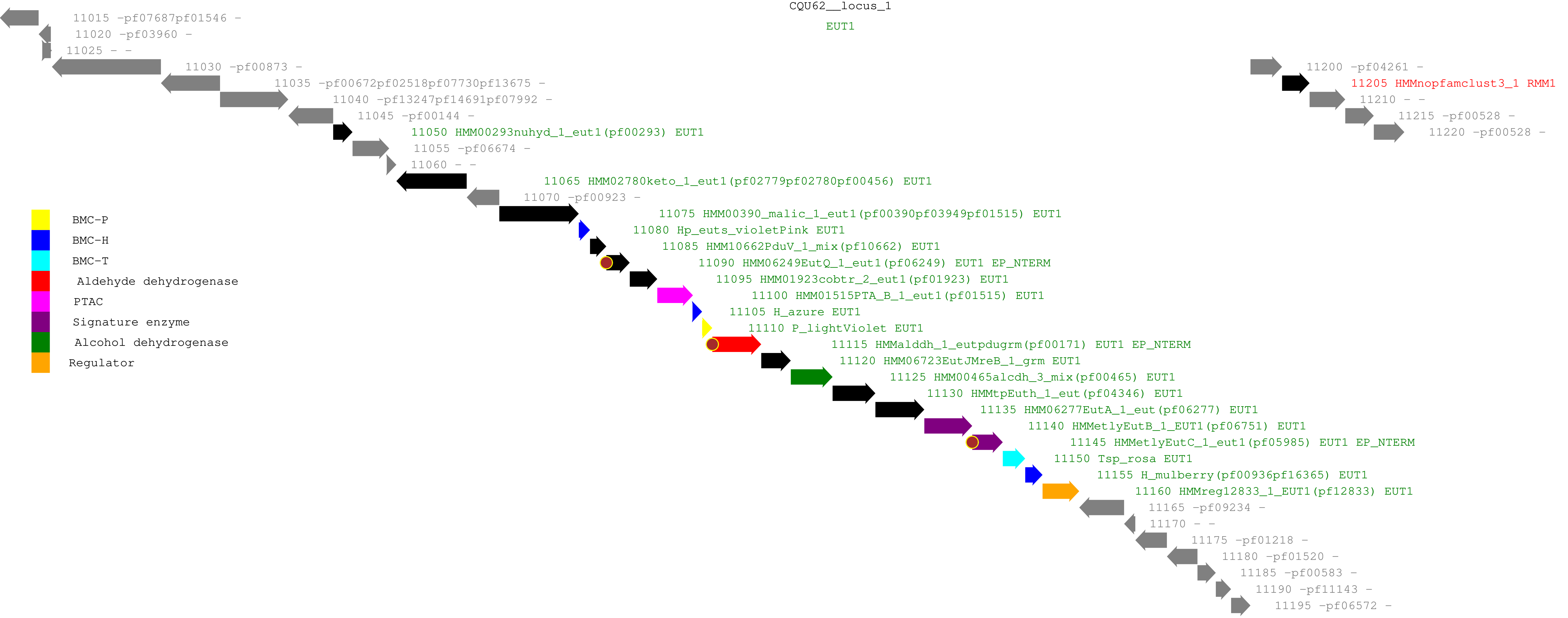

Salmonella enterica (Salmonella choleraesuis). Bacteria; Proteobacteria; Gammaproteobacteria; Enterobacterales;

CRE05__locus_1

EUT1

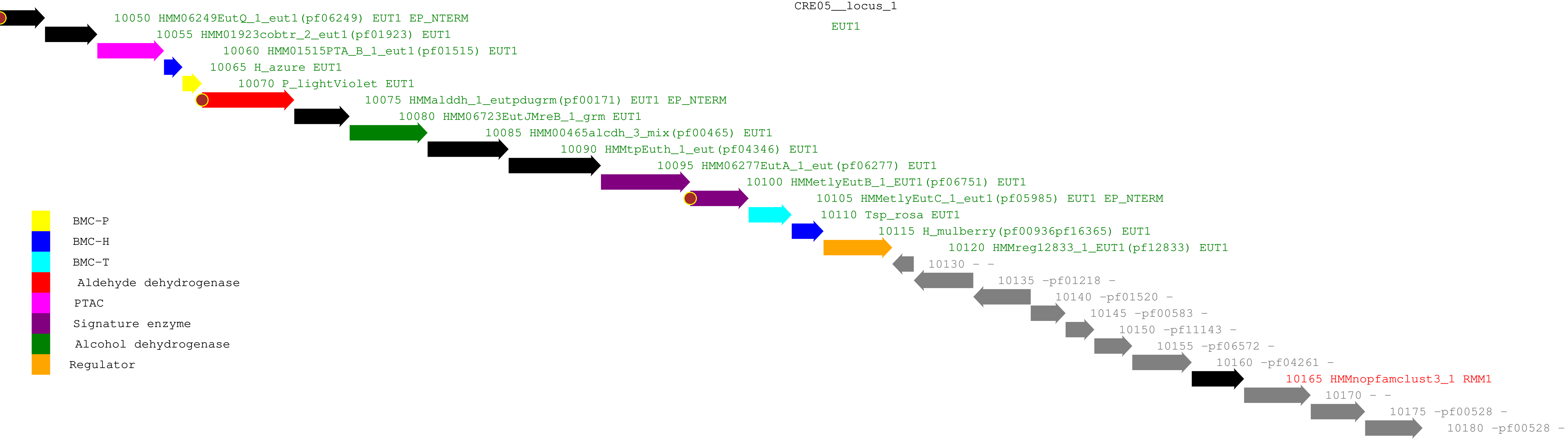

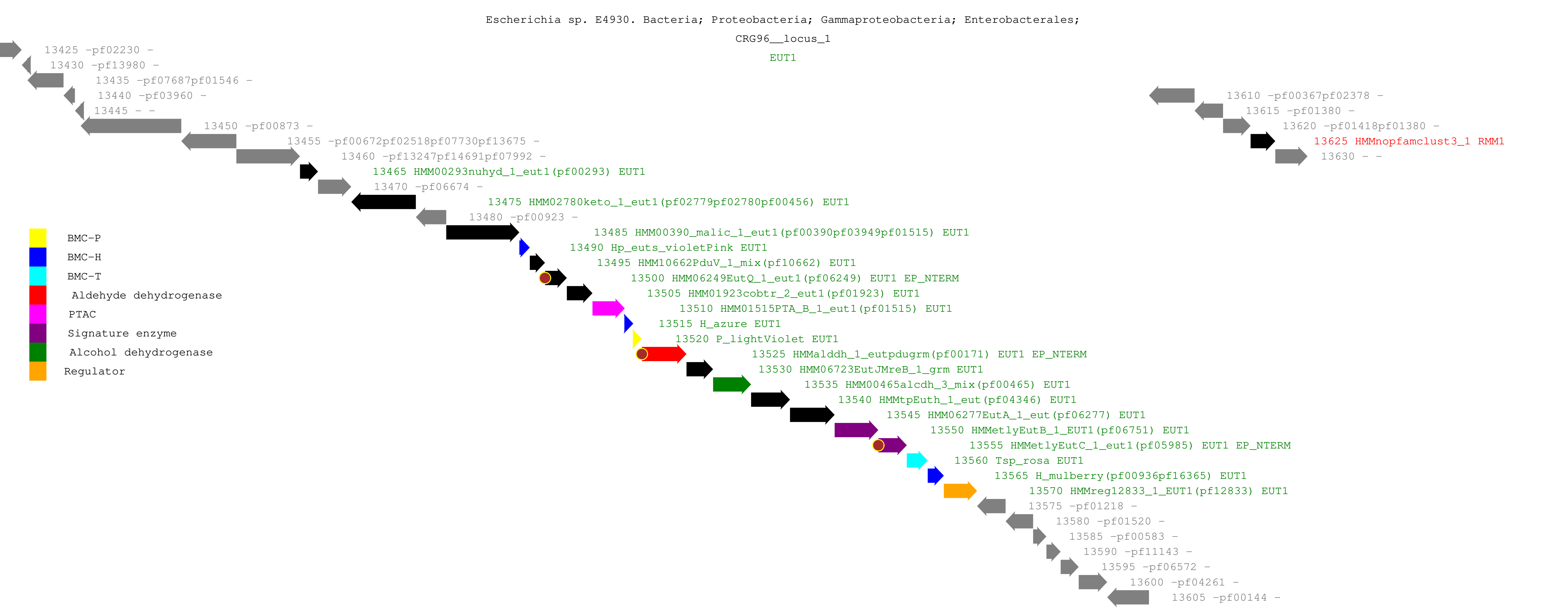
Escherichia sp. E4930. Bacteria; Proteobacteria; Gammaproteobacteria; Enterobacterales;
CRG96__locus_1
EUT1
13425 -pf02230 -
13430 -pf13980 -
13435 -pf07687pf01546 -
13440 -pf03960 -
13445 - -
13450 -pf00873 -
13455 -pf00672pf02518pf07730pf13675 -
13460 -pf13247pf14691pf07992 -
13465 HMM00293nuhyd_1_eut1(pf00293) EUT1
13470 -pf06674 -
13475 HMM02780keto_1_eut1(pf02779pf02780pf00456) EUT1
13480 -pf00923 -
13485 HMM00390_malic_1_eut1(pf00390pf03949pf01515) EUT1
13490 Hp_euts_violetPink EUT1
13495 HMM10662PduV_1_mix(pf10662) EUT1
13500 HMM06249EutQ_1_eut1(pf06249) EUT1 EP_NTERM
13505 HMM01923cobtr_2_eut1(pf01923) EUT1
13510 HMM01515PTA_B_1_eut1(pf01515) EUT1
13515 H_azure EUT1
13520 P_lightViolet EUT1
13525 HMMalddh_1_eutpdugrm(pf00171) EUT1 EP_NTERM
13530 HMM06723EutJMreB_1_grm EUT1
13535 HMM00465alcdh_3_mix(pf00465) EUT1
13540 HMMtpEuth_1_eut(pf04346) EUT1
13545 HMM06277EutA_1_eut(pf06277) EUT1
13550 HMMetlyEutB_1_EUT1(pf06751) EUT1
13555 HMMetlyEutC_1_eut1(pf05985) EUT1 EP_NTERM
13560 Tsp_rosa EUT1
13565 H_mulberry(pf00936pf16365) EUT1
13570 HMMreg12833_1_EUT1(pf12833) EUT1
13575 -pf01218 -
13580 -pf01520 -
13585 -pf00583 -
13590 -pf11143 -
13595 -pf06572 -
13600 -pf04261 -
13605 -pf00144 -
13610 -pf00367pf02378 -
13615 -pf01380 -
13620 -pf01418pf01380 -
13625 HMMnopfamclust3_1 RMM1
13630 - -
BMC-P
BMC-H
BMC-T
Aldehyde dehydrogenase
PTAC
Signature enzyme
Alcohol dehydrogenase
Regulator

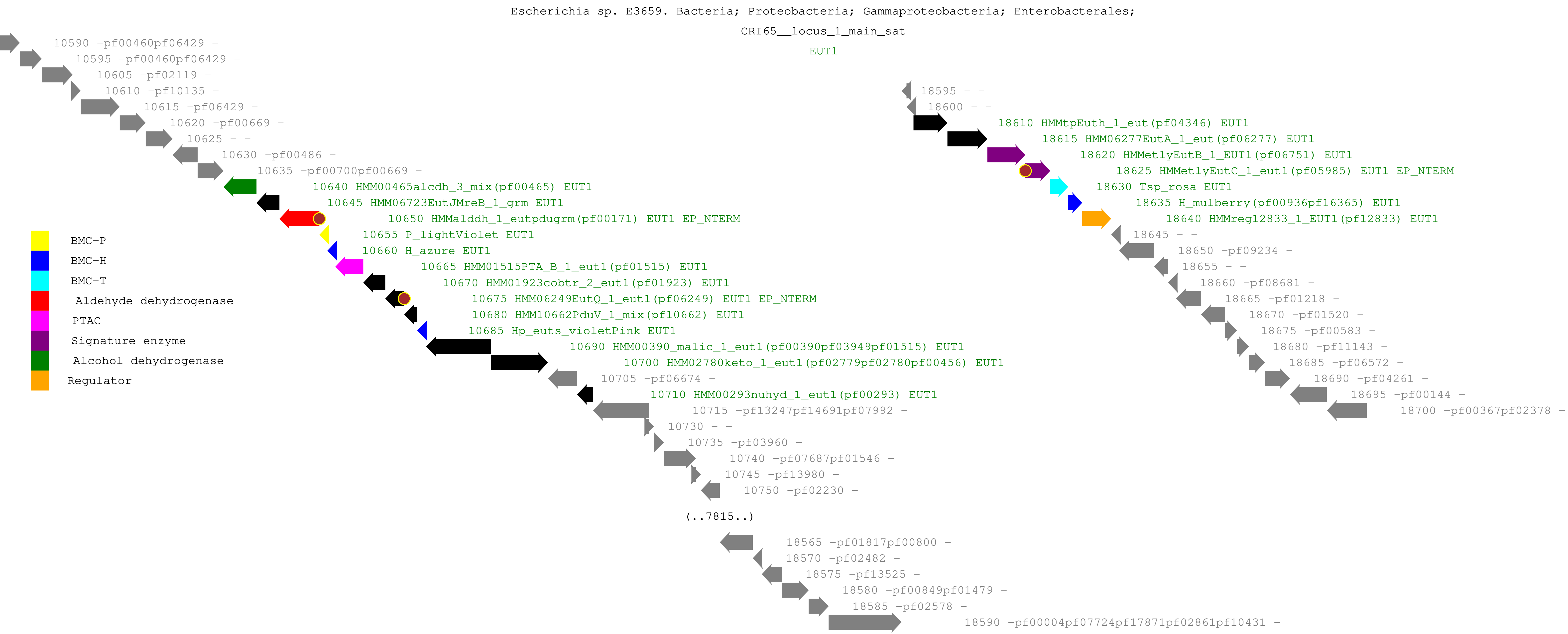
Escherichia sp. E3659. Bacteria; Proteobacteria; Gammaproteobacteria; Enterobacterales;
CRI65__locus_1_main_sat
EUT1
10590 -pf00460pf06429 -
10595 -pf00460pf06429 -
10605 -pf02119 -
10610 -pf10135 -
10615 -pf06429 -
10620 -pf00669 -
10625 - -
10630 -pf00486 -
10635 -pf00700pf00669 -
10640 HMM00465alcdh_3_mix(pf00465) EUT1
10645 HMM06723EutJMreB_1_grm EUT1
10650 HMMalddh_1_eutpdugrm(pf00171) EUT1 EP_NTERM
10655 P_lightViolet EUT1
10660 H_azure EUT1
10665 HMM01515PTA_B_1_eut1(pf01515) EUT1
10670 HMM01923cobtr_2_eut1(pf01923) EUT1
10675 HMM06249EutQ_1_eut1(pf06249) EUT1 EP_NTERM
10680 HMM10662PduV_1_mix(pf10662) EUT1
10685 Hp_euts_violetPink EUT1
10690 HMM00390_malic_1_eut1(pf00390pf03949pf01515) EUT1
10700 HMM02780keto_1_eut1(pf02779pf02780pf00456) EUT1
10705 -pf06674 -
10710 HMM00293nuhyd_1_eut1(pf00293) EUT1
10715 -pf13247pf14691pf07992 -
10730 - -
10735 -pf03960 -
10740 -pf07687pf01546 -
10745 -pf13980 -
10750 -pf02230 -
(..7815..)
18565 -pf01817pf00800 -
18570 -pf02482 -
18575 -pf13525 -
18580 -pf00849pf01479 -
18585 -pf02578 -
18590 -pf00004pf07724pf17871pf02861pf10431 -
18595 - -
18600 - -
18610 HMMtpEuth_1_eut(pf04346) EUT1
18615 HMM06277EutA_1_eut(pf06277) EUT1
18620 HMMetlyEutB_1_EUT1(pf06751) EUT1
18625 HMMetlyEutC_1_eut1(pf05985) EUT1 EP_NTERM
18630 Tsp_rosa EUT1
18635 H_mulberry(pf00936pf16365) EUT1
18640 HMMreg12833_1_EUT1(pf12833) EUT1
18645 - -
18650 -pf09234 -
18655 - -
18660 -pf08681 -
18665 -pf01218 -
18670 -pf01520 -
18675 -pf00583 -
18680 -pf11143 -
18685 -pf06572 -
18690 -pf04261 -
18695 -pf00144 -
18700 -pf00367pf02378 -
BMC-P
BMC-H
BMC-T
Aldehyde dehydrogenase
PTAC
Signature enzyme
Alcohol dehydrogenase
Regulator

Escherichia sp. E4702. Bacteria; Proteobacteria; Gammaproteobacteria; Enterobacterales;

CRI67__locus_1

EUT1

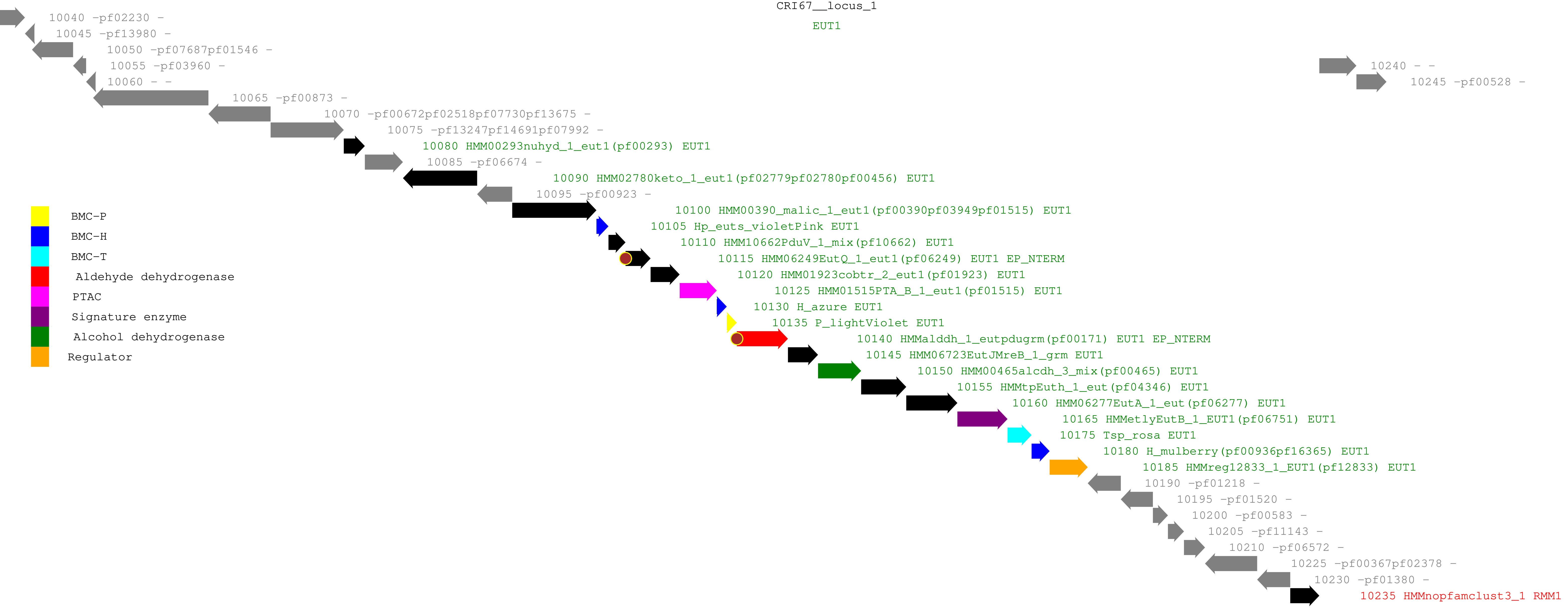

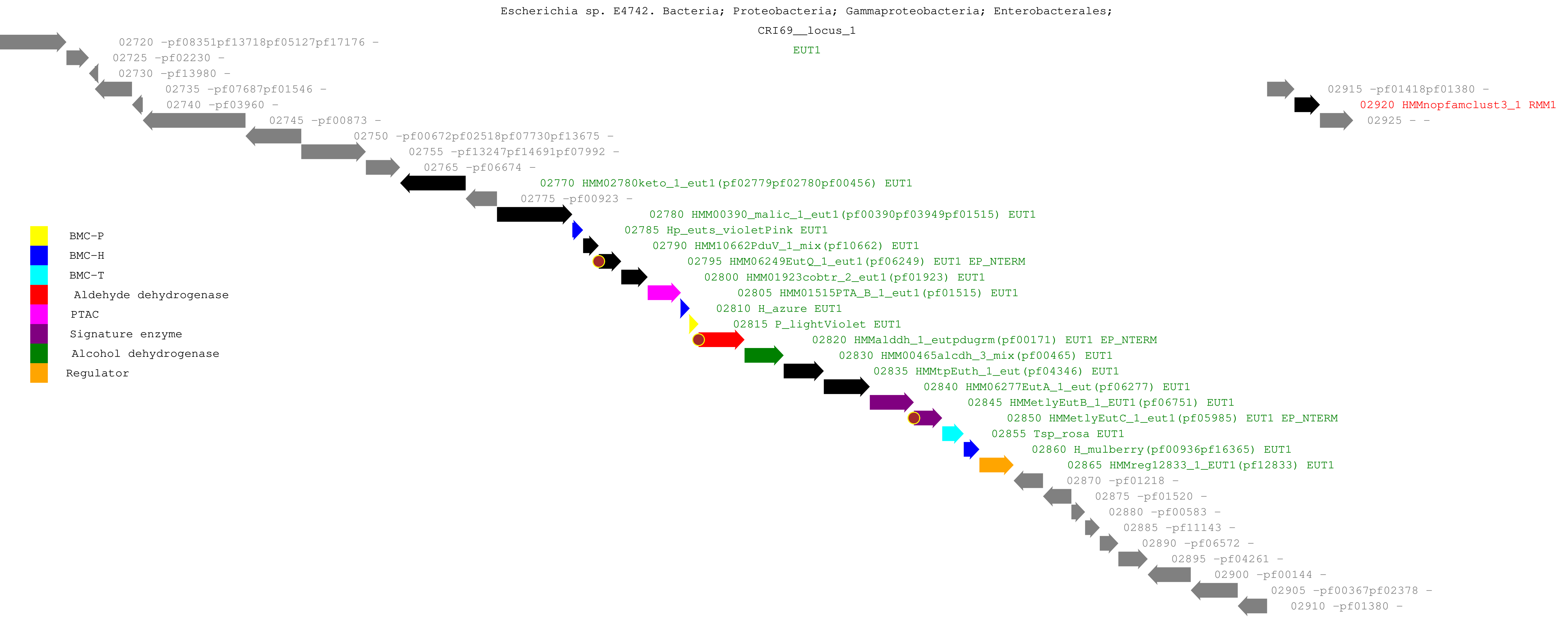

Escherichia sp. E4742. Bacteria; Proteobacteria; Gammaproteobacteria; Enterobacterales;
CRI69__locus_1
EUT1
02720 -pf08351pf13718pf05127pf17176 -
02725 -pf02230 -
02730 -pf13980 -
02735 -pf07687pf01546 -
02740 -pf03960 -
02745 -pf00873 -
02750 -pf00672pf02518pf07730pf13675 -
02755 -pf13247pf14691pf07992 -
02765 -pf06674 -
02770 HMM02780keto_1_eut1(pf02779pf02780pf00456) EUT1
02775 -pf00923 -
02780 HMM00390_malic_1_eut1(pf00390pf03949pf01515) EUT1
02785 Hp_euts_violetPink EUT1
02790 HMM10662PduV_1_mix(pf10662) EUT1
02795 HMM06249EutQ_1_eut1(pf06249) EUT1 EP_NTERM
02800 HMM01923cobtr_2_eut1(pf01923) EUT1
02805 HMM01515PTA_B_1_eut1(pf01515) EUT1
02810 H_azure EUT1
02815 P_lightViolet EUT1
02820 HMMaldddh_1_eutpdugrm(pf00171) EUT1 EP_NTERM
02830 HMM00465alcdh_3_mix(pf00465) EUT1
02835 HMMtpEuth_1_eut(pf04346) EUT1
02840 HMM06277EutA_1_eut(pf06277) EUT1
02845 HMMetlyEutB_1_EUT1(pf06751) EUT1
02850 HMMetlyEutC_1_eut1(pf05985) EUT1 EP_NTERM
02855 Tsp_rosa EUT1
02860 H_mulberry(pf00936pf16365) EUT1
02865 HMMreg12833_1_EUT1(pf12833) EUT1
02870 -pf01218 -
02875 -pf01520 -
02880 -pf00583 -
02885 -pf11143 -
02890 -pf06572 -
02895 -pf04261 -
02900 -pf00144 -
02905 -pf00367pf02378 -
02910 -pf01380 -
02915 -pf01418pf01380 -
02920 HMMnopfamclust3_1 RMM1
02925 - -
BMC-P
BMC-H
BMC-T
Aldehyde dehydrogenase
PTAC
Signature enzyme
Alcohol dehydrogenase
Regulator

Escherichia sp. E5028. Bacteria; Proteobacteria; Gammaproteobacteria; Enterobacterales;

CRT22__locus_2

EUT1

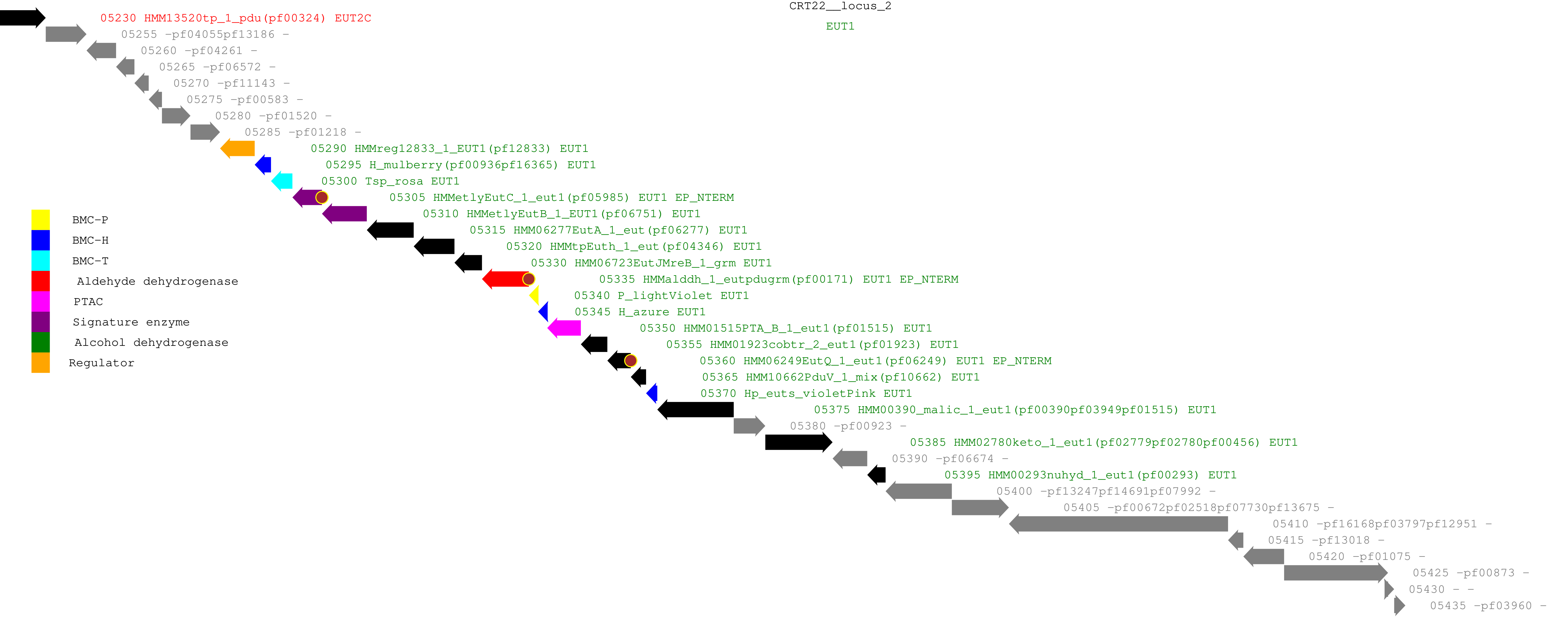

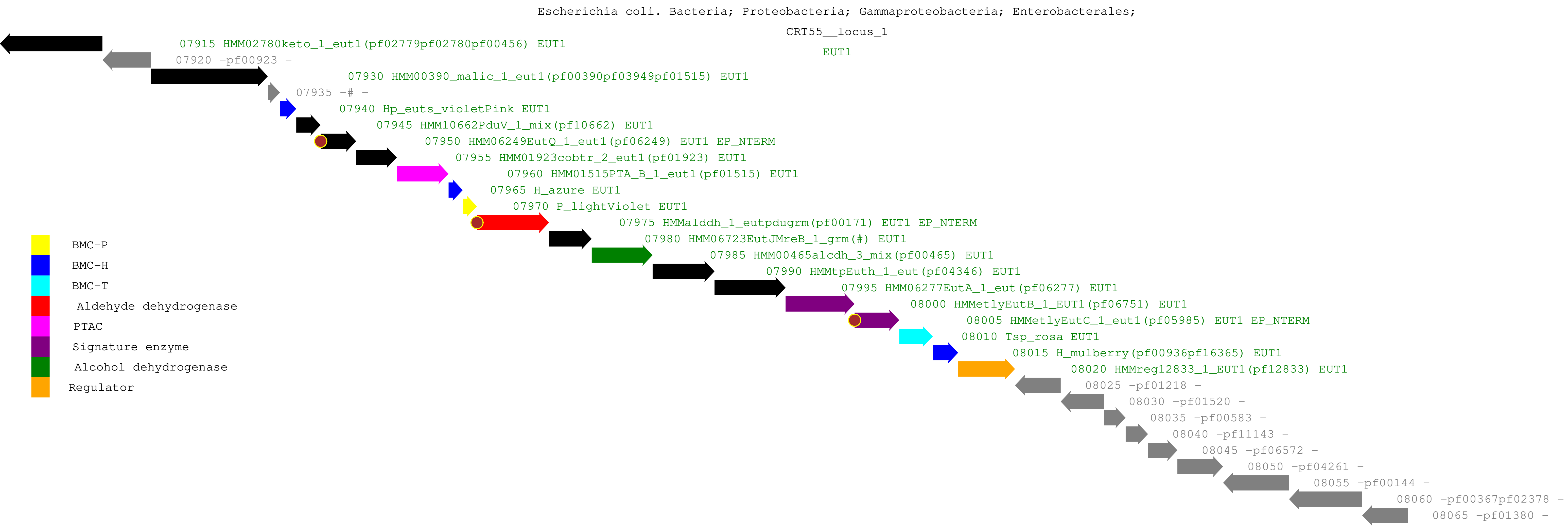
Escherichia coli. Bacteria; Proteobacteria; Gammaproteobacteria; Enterobacterales;
CRT55__locus_1
EUT1
07915 HMM02780keto_1_eut1(pf02779pf02780pf00456) EUT1
07920 -pf00923 -
07930 HMM00390_malic_1_eut1(pf00390pf03949pf01515) EUT1
07935 -# -
07940 Hp_euts_violetPink EUT1
07945 HMM10662PduV_1_mix(pf10662) EUT1
07950 HMM06249EutQ_1_eut1(pf06249) EUT1 EP_NTERM
07955 HMM01923cobtr_2_eut1(pf01923) EUT1
07960 HMM01515PTA_B_1_eut1(pf01515) EUT1
07965 H_azure EUT1
07970 P_lightViolet EUT1
07975 HMMalddh_1_eutpdugrm(pf00171) EUT1 EP_NTERM
07980 HMM06723EutJMreB_1_grm(#) EUT1
07985 HMM00465alcdh_3_mix(pf00465) EUT1
07990 HMMtpEuth_1_eut(pf04346) EUT1
07995 HMM06277EutA_1_eut(pf06277) EUT1
08000 HMMetlyEutB_1_EUT1(pf06751) EUT1
08005 HMMetlyEutC_1_eut1(pf05985) EUT1 EP_NTERM
08010 Tsp_rosa EUT1
08015 H_mulberry(pf00936pf16365) EUT1
08020 HMMreg12833_1_EUT1(pf12833) EUT1
08025 -pf01218 -
08030 -pf01520 -
08035 -pf00583 -
08040 -pf11143 -
08045 -pf06572 -
08050 -pf04261 -
08055 -pf00144 -
08060 -pf00367pf02378 -
08065 -pf01380 -
BMC-P
BMC-H
BMC-T
Aldehyde dehydrogenase
PTAC
Signature enzyme
Alcohol dehydrogenase
Regulator

Escherichia sp. E4385. Bacteria; Proteobacteria; Gammaproteobacteria; Enterobacterales;

CRU79__locus_2

EUT1

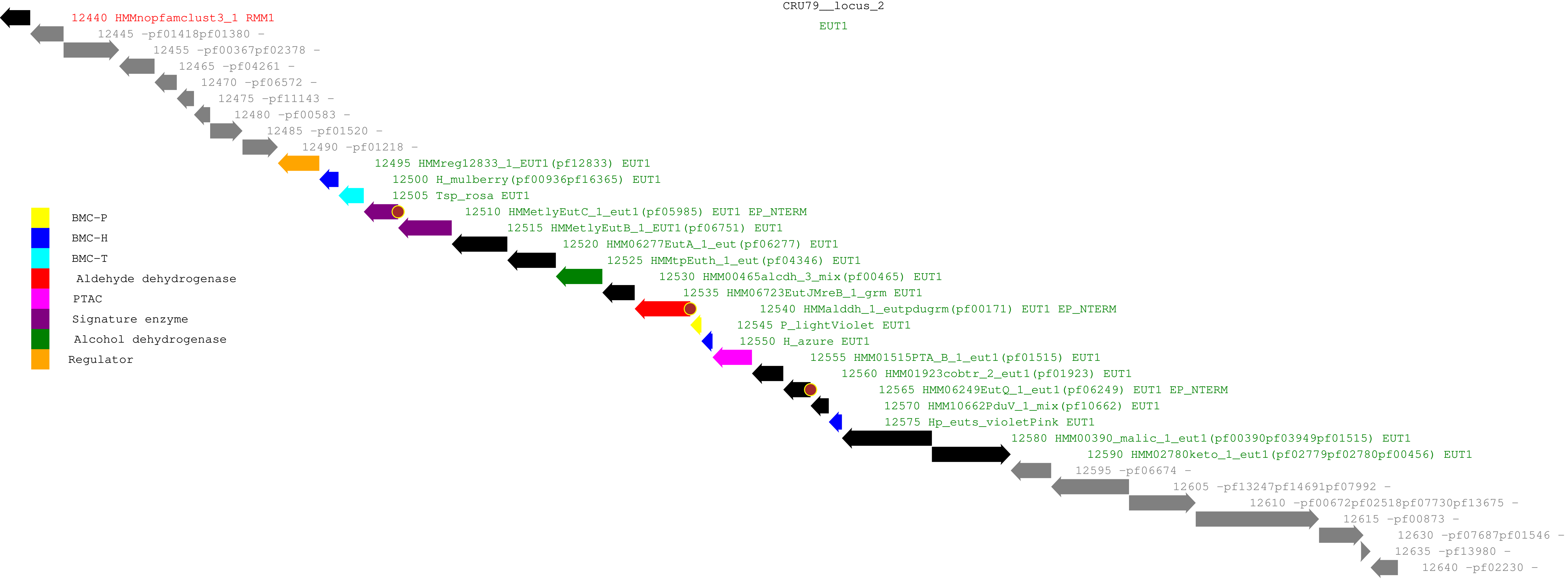

Morganella morganii (Proteus morganii). Bacteria; Proteobacteria; Gammaproteobacteria; Enterobacterales;

CRX48__locus_1

EUT1

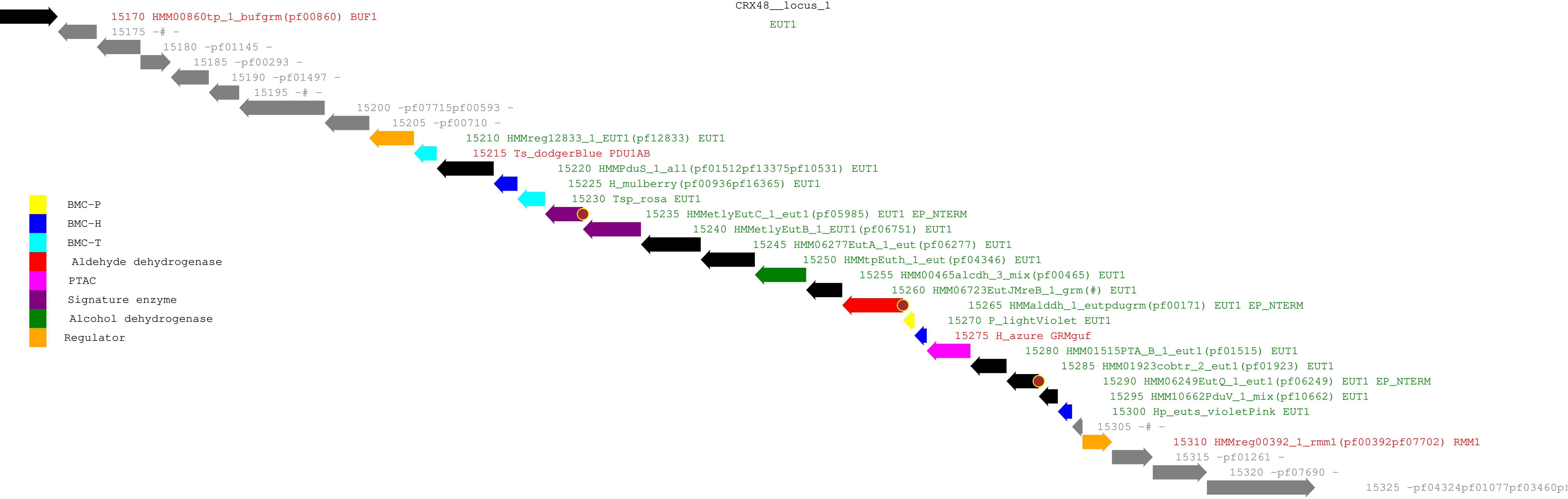

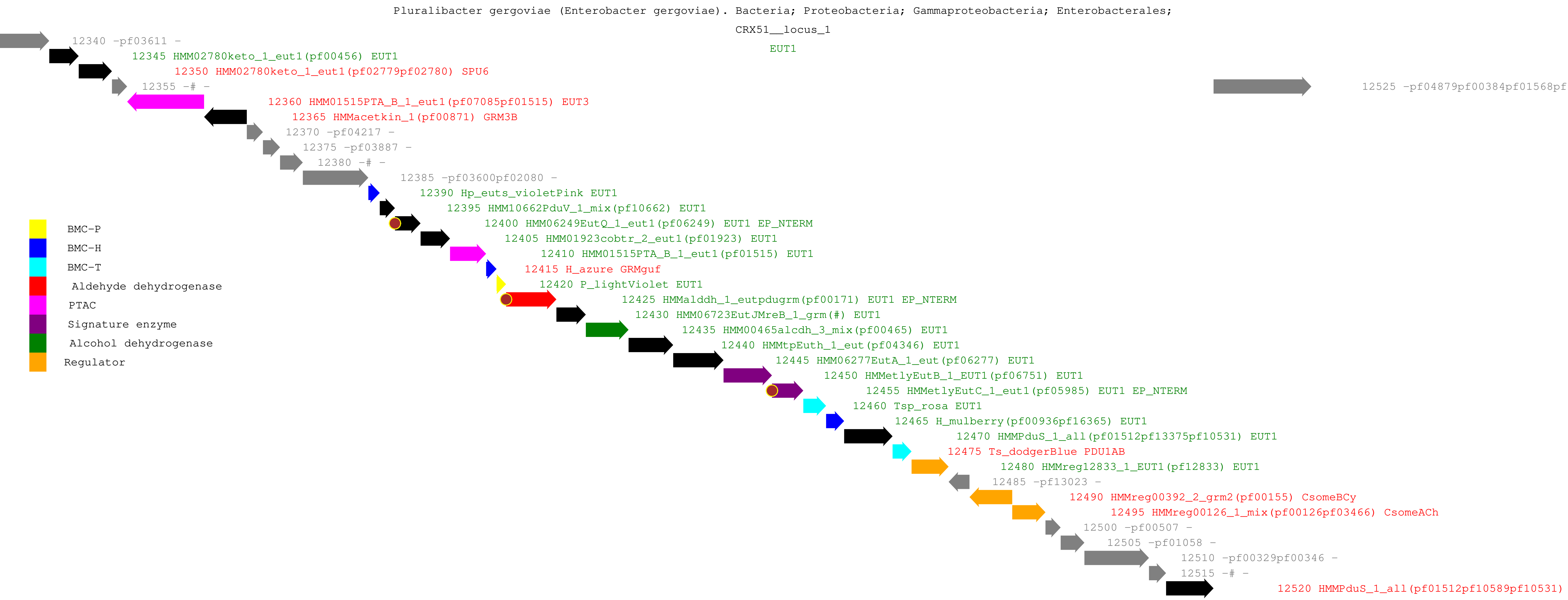

Pluralibacter gergoviae (Enterobacter gergoviae). Bacteria; Proteobacteria; Gammaproteobacteria; Enterobacterales;
CRX51__locus_1
EUT1
12340 -pf03611 -
12345 HMM02780keto_1_eut1(pf00456) EUT1
12350 HMM02780keto_1_eut1(pf02779pf02780) SPU6
12355 -# -
12525 -pf04879pf00384pf01568pf
12360 HMM01515PTA_B_1_eut1(pf07085pf01515) EUT3
12365 HMMacetkin_1(pf00871) GRM3B
12370 -pf04217 -
12375 -pf03887 -
12380 -# -
12385 -pf03600pf02080 -
12390 Hp_euts_violetPink EUT1
12395 HMM10662PduV_1_mix(pf10662) EUT1
12400 HMM06249EutQ_1_eut1(pf06249) EUT1 EP_NTERM
12405 HMM01923cobtr_2_eut1(pf01923) EUT1
12410 HMM01515PTA_B_1_eut1(pf01515) EUT1
12415 H_azure GRMguf
12420 P_lightViolet EUT1
12425 HMMalddh_1_eutpdugrm(pf00171) EUT1 EP_NTERM
12430 HMM06723EutJMreB_1_grm(#) EUT1
12435 HMM00465alcdh_3_mix(pf00465) EUT1
12440 HMMtpEuth_1_eut(pf04346) EUT1
12445 HMM06277EutA_1_eut(pf06277) EUT1
12450 HMMetlyEutB_1_EUT1(pf06751) EUT1
12455 HMMetlyEutC_1_eut1(pf05985) EUT1 EP_NTERM
12460 Tsp_rosa EUT1
12465 H_mulberry(pf00936pf16365) EUT1
12470 HMMPduS_1_all(pf01512pf13375pf10531) EUT1
12475 Ts_dodgerBlue PDU1AB
12480 HMMreg12833_1_EUT1(pf12833) EUT1
12485 -pf13023 -
12490 HMMreg00392_2_grm2(pf00155) CsomeBCy
12495 HMMreg00126_1_mix(pf00126pf03466) CsomeACh
12500 -pf00507 -
12505 -pf01058 -
12510 -pf00329pf00346 -
12515 -# -
12520 HMMPduS_1_all(pf01512pf10589pf10531)
BMC-P
BMC-H
BMC-T
Aldehyde dehydrogenase
PTAC
Signature enzyme
Alcohol dehydrogenase
Regulator

Escherichia coli O103. Bacteria; Proteobacteria; Gammaproteobacteria; Enterobacterales;

CRY61__locus_1

EUT1

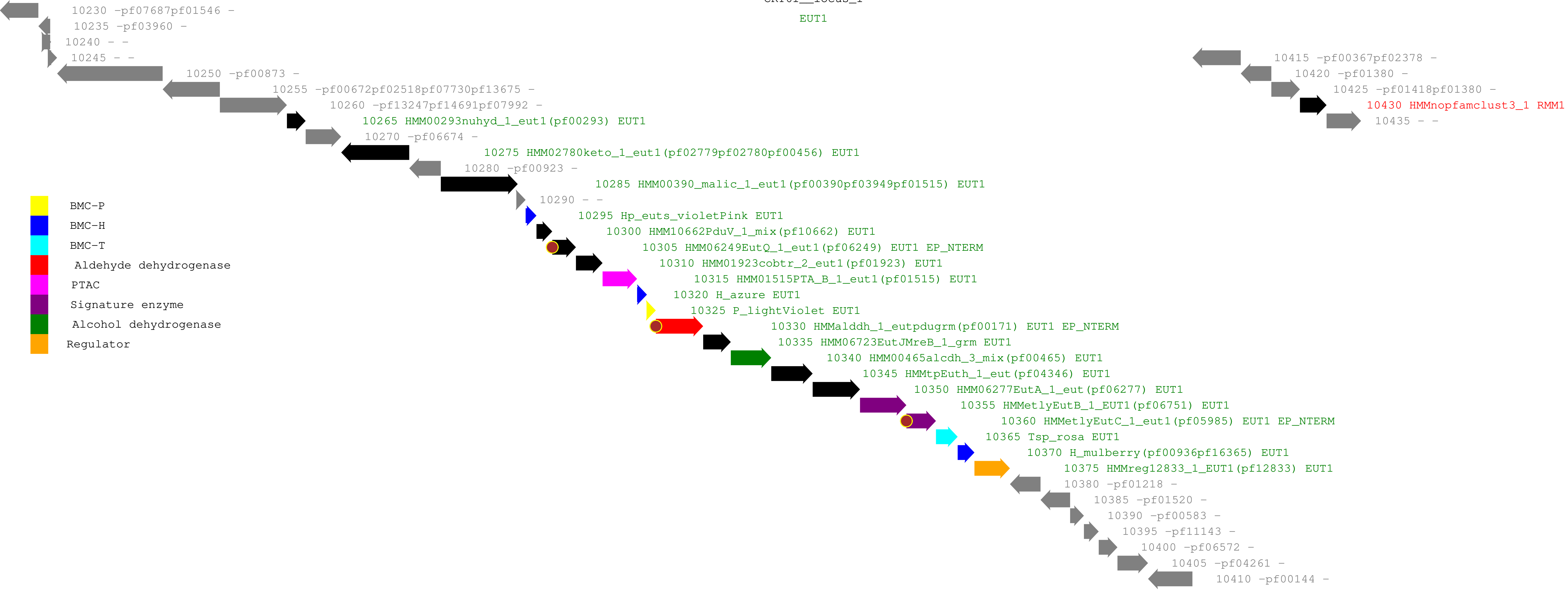

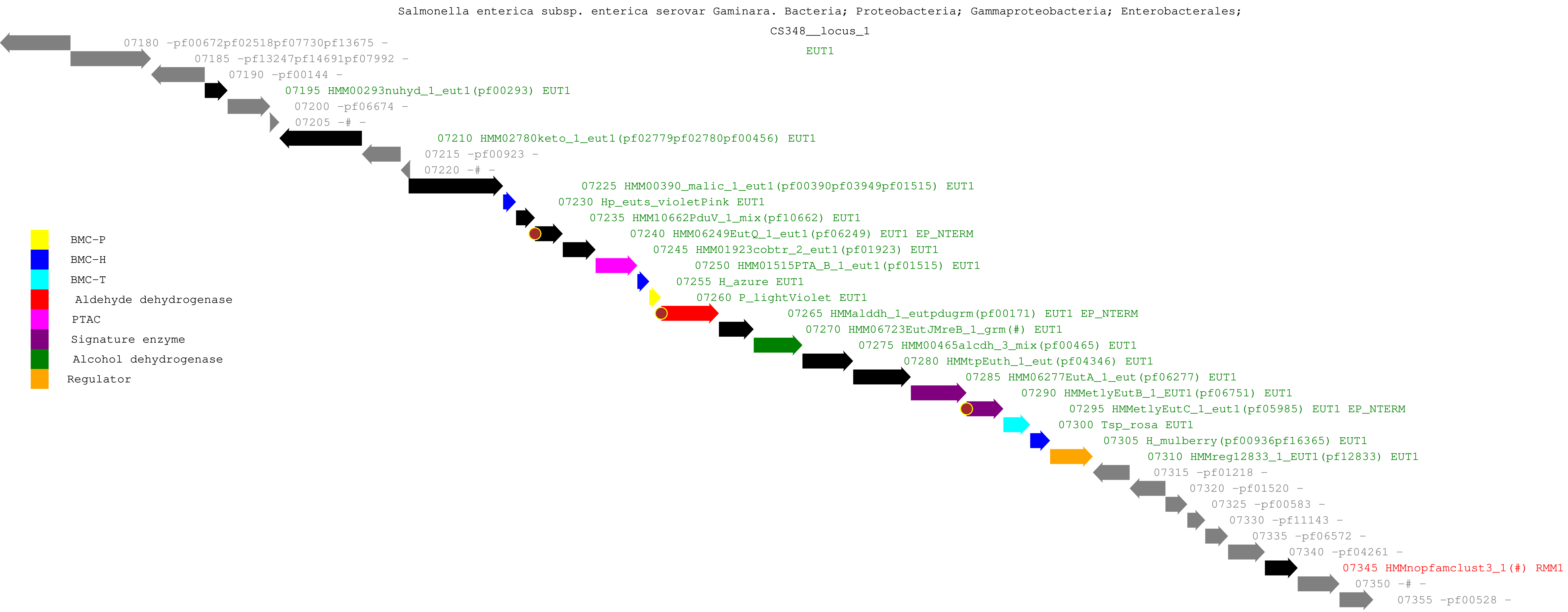
Salmonella enterica subsp. enterica serovar Gaminara. Bacteria; Proteobacteria; Gammaproteobacteria; Enterobacterales;
CS348__locus_1
EUT1
07180 -pf00672pf02518pf07730pf13675 -
07185 -pf13247pf14691pf07992 -
07190 -pf00144 -
07195 HMM00293nuhyd_1_eut1(pf00293) EUT1
07200 -pf06674 -
07205 -# -
07210 HMM02780keto_1_eut1(pf02779pf02780pf00456) EUT1
07215 -pf00923 -
07220 -# -
07225 HMM00390_malic_1_eut1(pf00390pf03949pf01515) EUT1
07230 Hp_euts_violetPink EUT1
07235 HMM10662PduV_1_mix(pf10662) EUT1
07240 HMM06249EutQ_1_eut1(pf06249) EUT1 EP_NTERM
07245 HMM01923cobtr_2_eut1(pf01923) EUT1
07250 HMM01515PTA_B_1_eut1(pf01515) EUT1
07255 H_azure EUT1
07260 P_lightViolet EUT1
07265 HMMalddh_1_eutpdugrm(pf00171) EUT1 EP_NTERM
07270 HMM06723EutJMreB_1_grm(#) EUT1
07275 HMM00465alcdh_3_mix(pf00465) EUT1
07280 HMMtpEuth_1_eut(pf04346) EUT1
07285 HMM06277EutA_1_eut(pf06277) EUT1
07290 HMMetlyEutB_1_EUT1(pf06751) EUT1
07295 HMMetlyEutC_1_eut1(pf05985) EUT1 EP_NTERM
07300 Tsp_rosa EUT1
07305 H_mulberry(pf00936pf16365) EUT1
07310 HMMreg12833_1_EUT1(pf12833) EUT1
07315 -pf01218 -
07320 -pf01520 -
07325 -pf00583 -
07330 -pf11143 -
07335 -pf06572 -
07340 -pf04261 -
07345 HMMnopfamclust3_1(#) RMM1
07350 -# -
07355 -pf00528 -
BMC-P
BMC-H
BMC-T
Aldehyde dehydrogenase
PTAC
Signature enzyme
Alcohol dehydrogenase
Regulator

Salmonella enterica subsp. enterica serovar Tennessee. Bacteria; Proteobacteria; Gammaproteobacteria; Enterobacterales;

CS351__locus_1

EUT1

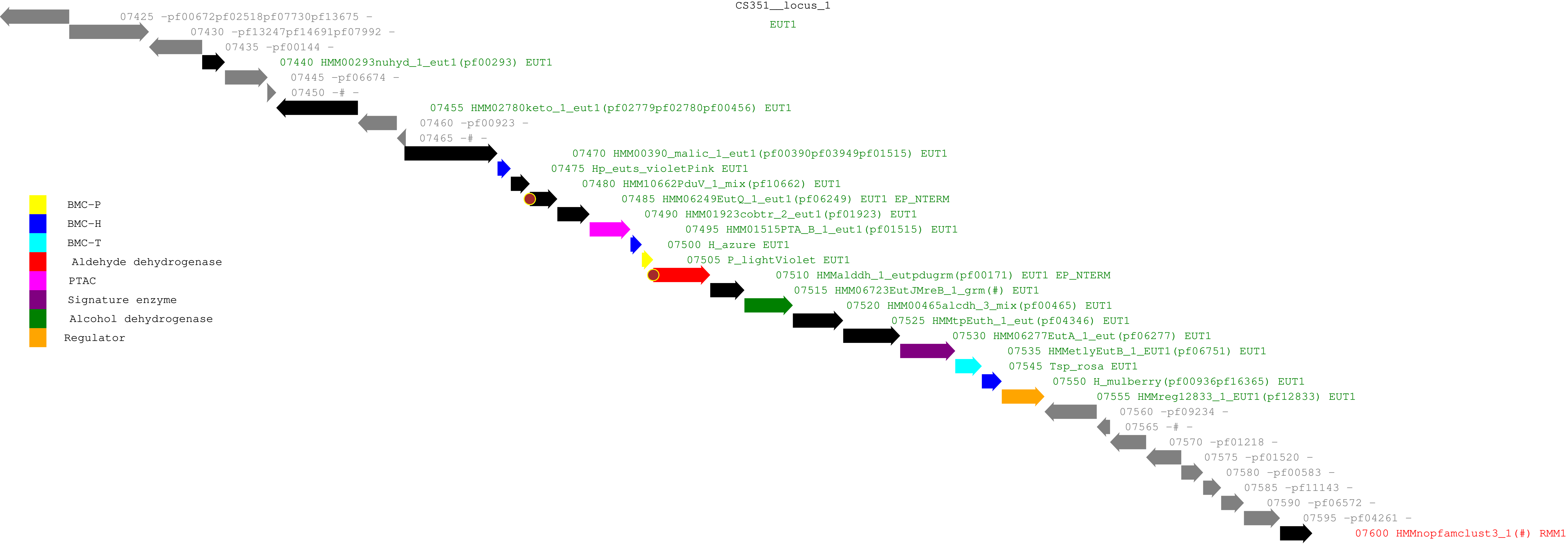

Proteobacteria bacterium. Bacteria; Proteobacteria.

CSB48__locus_1

EUT1

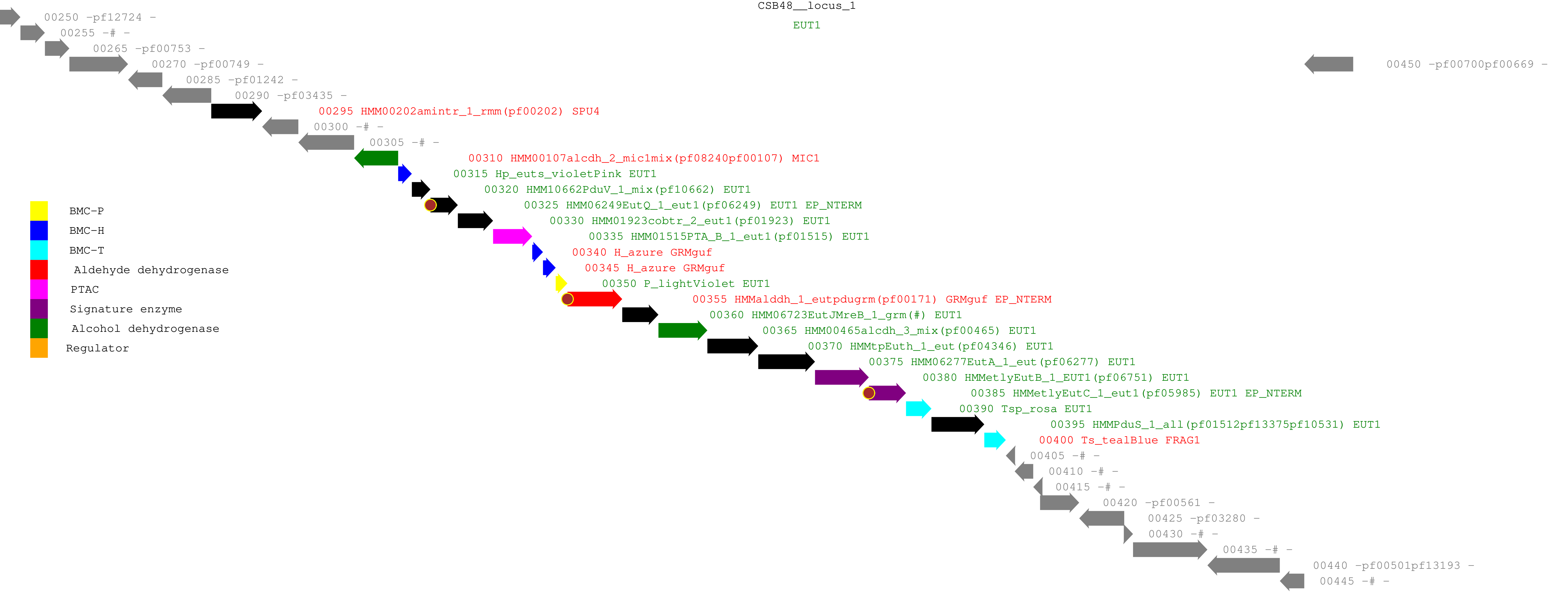

Klebsiella pneumoniae. Bacteria; Proteobacteria; Gammaproteobacteria; Enterobacterales;

CSC88__locus_1_main_sat

EUT1

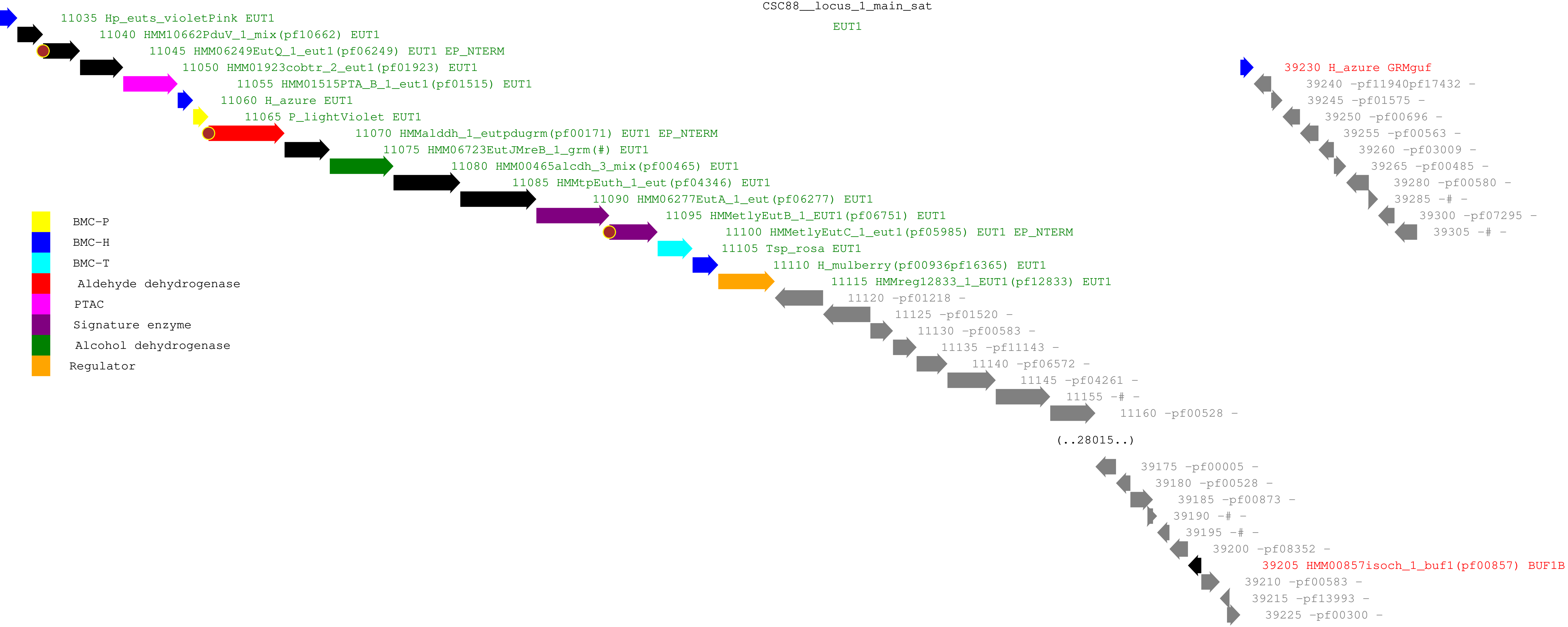

Salmonella enterica subsp. arizonae. Bacteria; Proteobacteria; Gammaproteobacteria; Enterobacterales;

CSM93__locus_1

EUT1

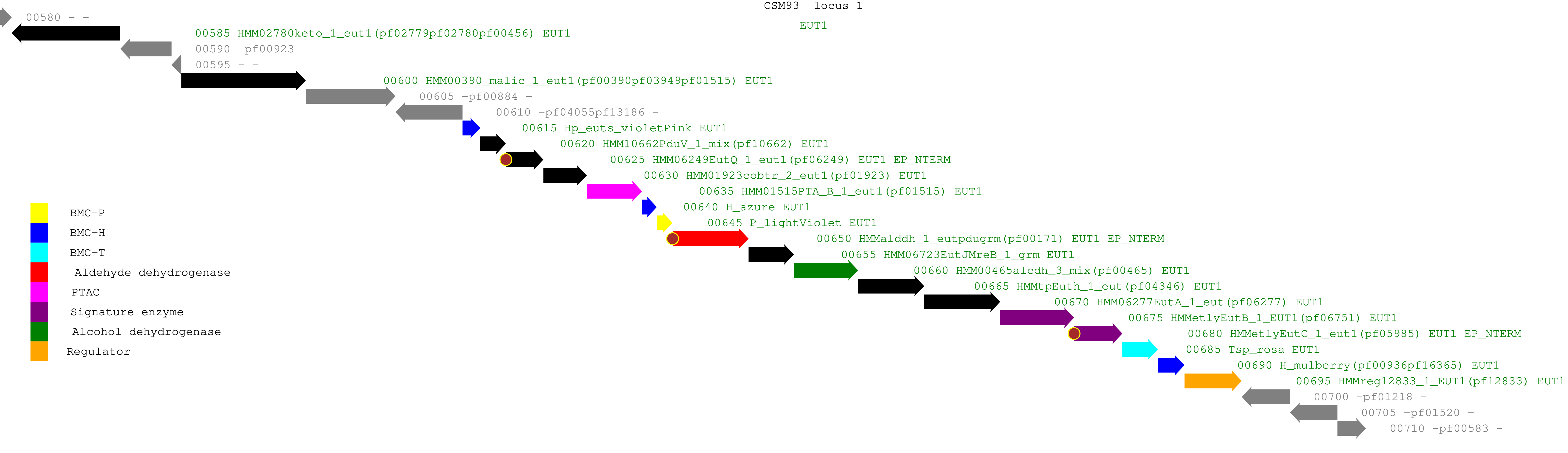

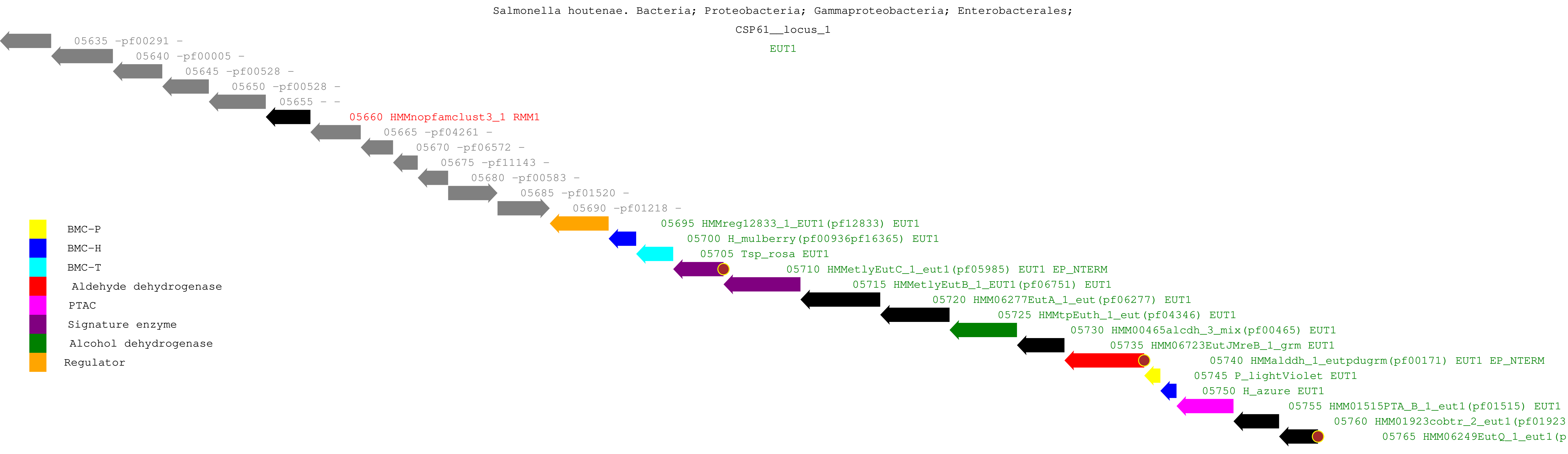
Salmonella houtenae. Bacteria; Proteobacteria; Gammaproteobacteria; Enterobacterales;
CSP61__locus_1
EUT1
05635 -pf00291 -
05640 -pf00005 -
05645 -pf00528 -
05650 -pf00528 -
05655 - -
05660 HMMnopfamclust3_1 RMM1
05665 -pf04261 -
05670 -pf06572 -
05675 -pf11143 -
05680 -pf00583 -
05685 -pf01520 -
05690 -pf01218 -
05695 HMMreg12833_1_EUT1(pf12833) EUT1
05700 H_mulberry(pf00936pf16365) EUT1
05705 Tsp_rosa EUT1
05710 HMMetlyEutC_1_eut1(pf05985) EUT1 EP_NTERM
05715 HMMetlyEutB_1_EUT1(pf06751) EUT1
05720 HMM06277EutA_1_eut(pf06277) EUT1
05725 HMMtpEuth_1_eut(pf04346) EUT1
05730 HMM00465alcdh_3_mix(pf00465) EUT1
05735 HMM06723EutJMreB_1_grm EUT1
05740 HMMalddh_1_eutpdugrm(pf00171) EUT1 EP_NTERM
05745 P_lightViolet EUT1
05750 H_azure EUT1
05755 HMM01515PTA_B_1_eut1(pf01515) EUT1
05760 HMM01923cobtr_2_eut1(pf01923
05765 HMM06249EutQ_1_eut1(p
BMC-P
BMC-H
BMC-T
Aldehyde dehydrogenase
PTAC
Signature enzyme
Alcohol dehydrogenase
Regulator

Iodobacter sp. BJB302. Bacteria; Proteobacteria; Betaproteobacteria; Neisseriales;

CSQ88__locus_1

EUT1

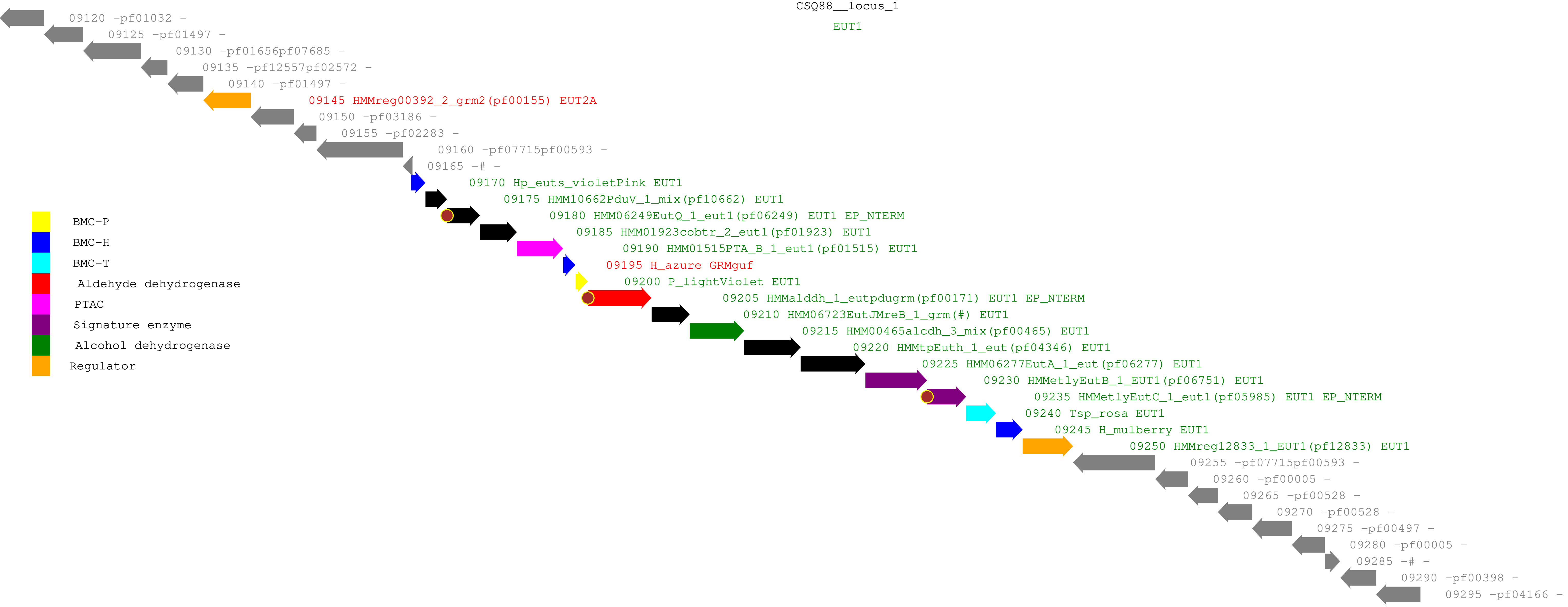

Salmonella enterica subsp. enterica serovar Weltevreden. Bacteria; Proteobacteria; Gammaproteobacteria; Enterobacterales;

CSR62__locus_1

EUT1

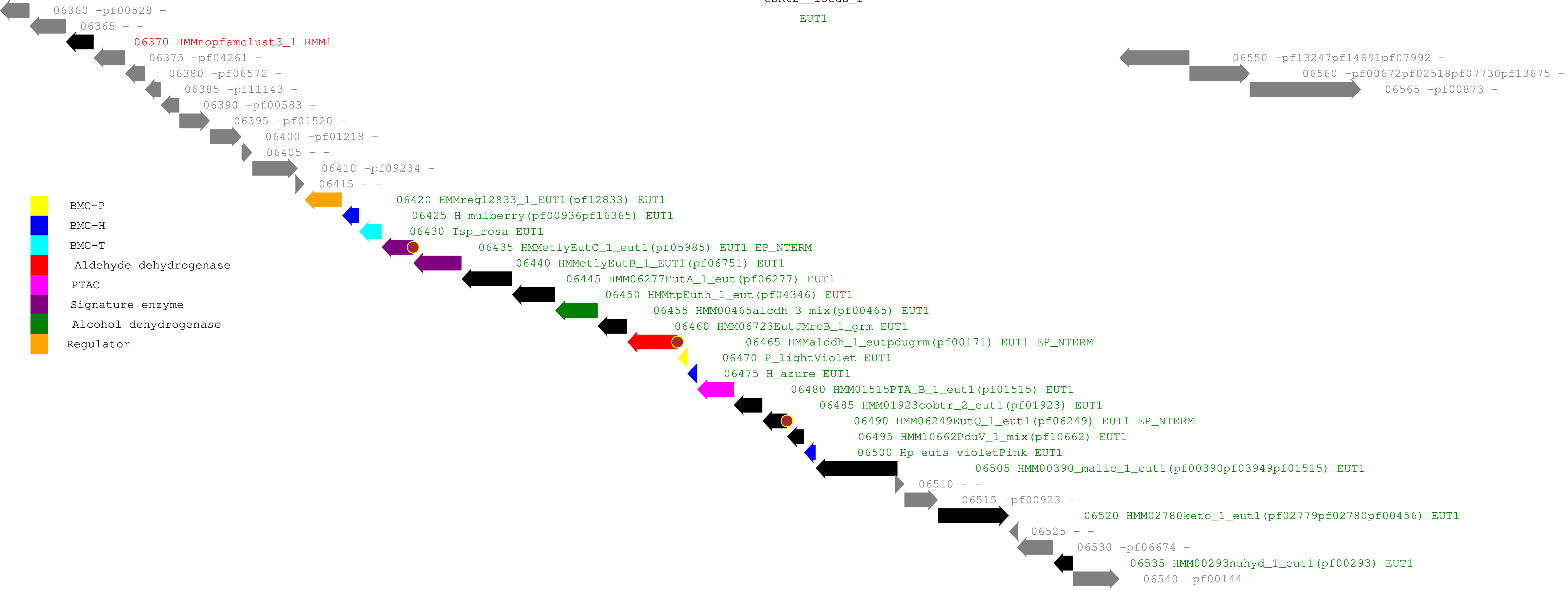

Salmonella diarizonae. Bacteria; Proteobacteria; Gammaproteobacteria; Enterobacterales;

CSU32__locus_1

EUT1

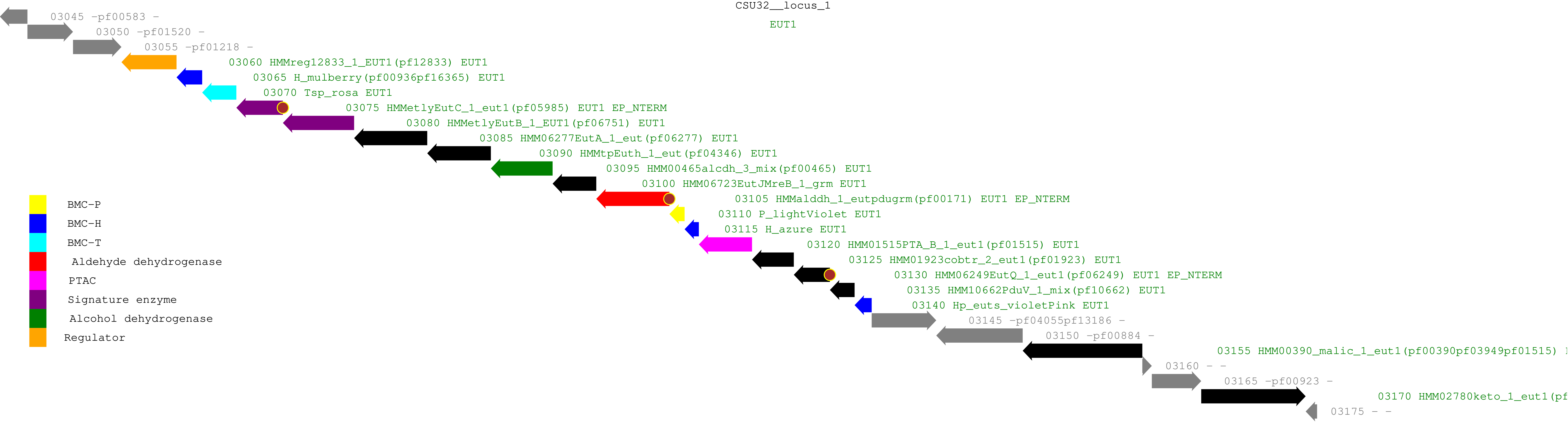

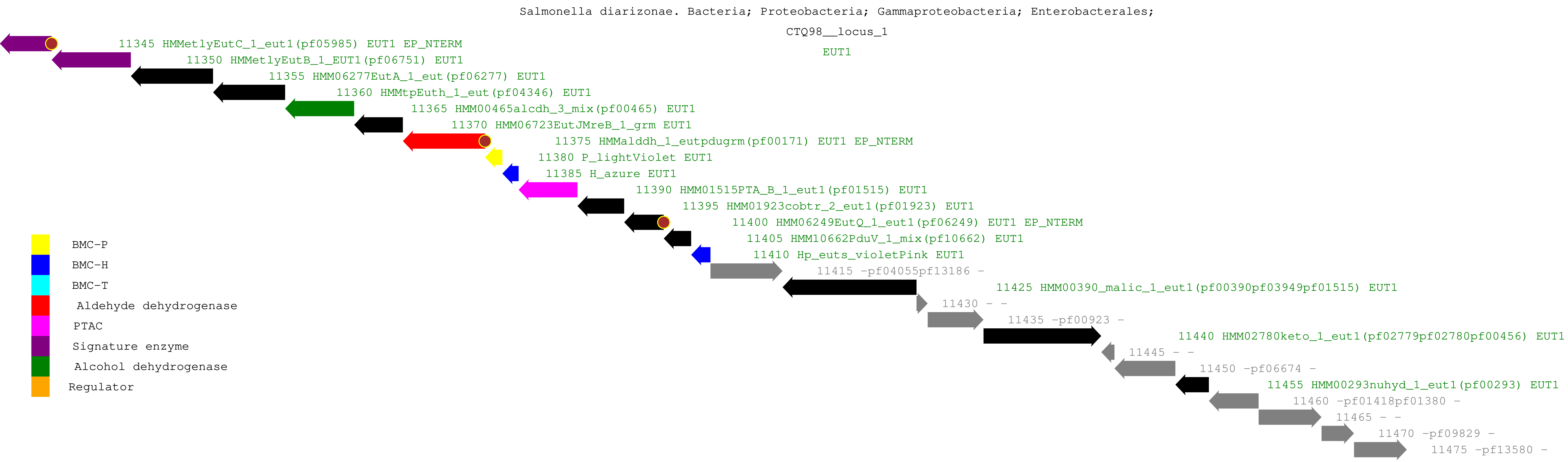

Salmonella diarizonae. Bacteria; Proteobacteria; Gammaproteobacteria; Enterobacterales;
CTQ98__locus_1
EUT1
11345 HMMetlyEutC_1_eut1(pf05985) EUT1 EP_NTERM
11350 HMMetlyEutB_1_EUT1(pf06751) EUT1
11355 HMM06277EutA_1_eut(pf06277) EUT1
11360 HMMtpEuth_1_eut(pf04346) EUT1
11365 HMM00465alcdh_3_mix(pf00465) EUT1
11370 HMM06723EutJMreB_1_grm EUT1
11375 HMMaldh_1_eutpdugrm(pf00171) EUT1 EP_NTERM
11380 P_lightViolet EUT1
11385 H_azure EUT1
11390 HMM01515PTA_B_1_eut1(pf01515) EUT1
11395 HMM01923cobtr_2_eut1(pf01923) EUT1
11400 HMM06249EutQ_1_eut1(pf06249) EUT1 EP_NTERM
11405 HMM10662PduV_1_mix(pf10662) EUT1
11410 Hp_euts_violetPink EUT1
11415 -pf04055pf13186 -
11425 HMM00390_malic_1_eut1(pf00390pf03949pf01515) EUT1
11430 - -
11435 -pf00923 -
11440 HMM02780keto_1_eut1(pf02779pf02780pf00456) EUT1
11445 - -
11450 -pf06674 -
11455 HMM00293nuhyd_1_eut1(pf00293) EUT1
11460 -pf01418pf01380 -
11465 - -
11470 -pf09829 -
11475 -pf13580 -
BMC-P
BMC-H
BMC-T
Aldehyde dehydrogenase
PTAC
Signature enzyme
Alcohol dehydrogenase
Regulator

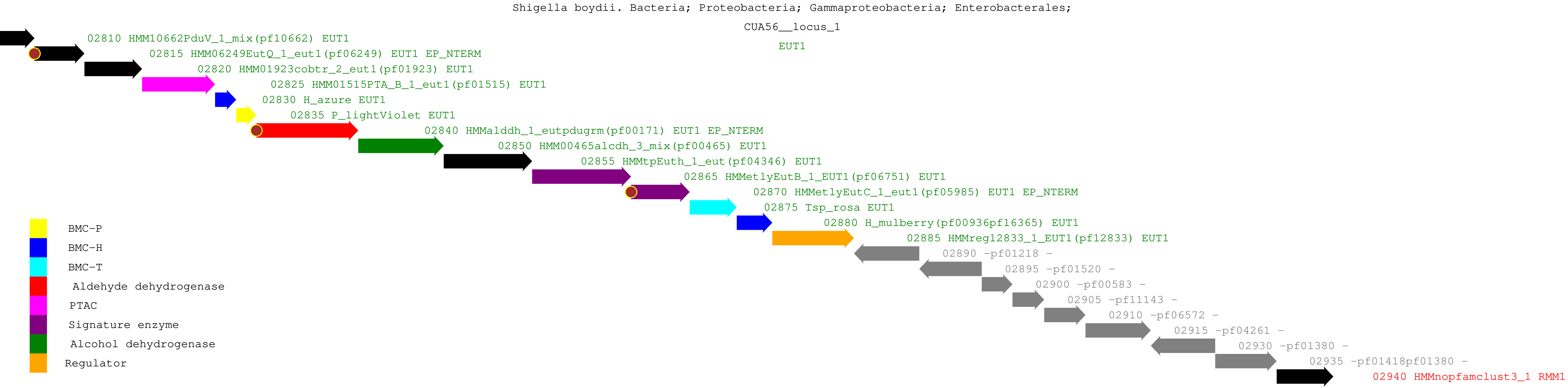

Shigella boydii. Bacteria; Proteobacteria; Gammaproteobacteria; Enterobacterales;
CUA56__locus_1
EUT1
02810 HMM10662PduV_1_mix(pf10662) EUT1
02815 HMM06249EutQ_1_eut1(pf06249) EUT1 EP_NTERM
02820 HMM01923cobtr_2_eut1(pf01923) EUT1
02825 HMM01515PTA_B_1_eut1(pf01515) EUT1
02830 H_azure EUT1
02835 P_lightViolet EUT1
02840 HMMalddh_1_eutpdugrm(pf00171) EUT1 EP_NTERM
02850 HMM00465alcdh_3_mix(pf00465) EUT1
02855 HMMtpEuth_1_eut(pf04346) EUT1
02865 HMMetlyEutB_1_EUT1(pf06751) EUT1
02870 HMMetlyEutC_1_eut1(pf05985) EUT1 EP_NTERM
02875 Tsp_rosa EUT1
02880 H_mulberry(pf00936pf16365) EUT1
02885 HMMreg12833_1_EUT1(pf12833) EUT1
02890 -pf01218 -
02895 -pf01520 -
02900 -pf00583 -
02905 -pf11143 -
02910 -pf06572 -
02915 -pf04261 -
02930 -pf01380 -
02935 -pf01418pf01380 -
02940 HMMnopfamclust3_1 RMM1
BMC-P
BMC-H
BMC-T
Aldehyde dehydrogenase
PTAC
Signature enzyme
Alcohol dehydrogenase
Regulator

Salmonella muenster. Bacteria; Proteobacteria; Gammaproteobacteria; Enterobacterales;

CUR71__locus_2

EUT1

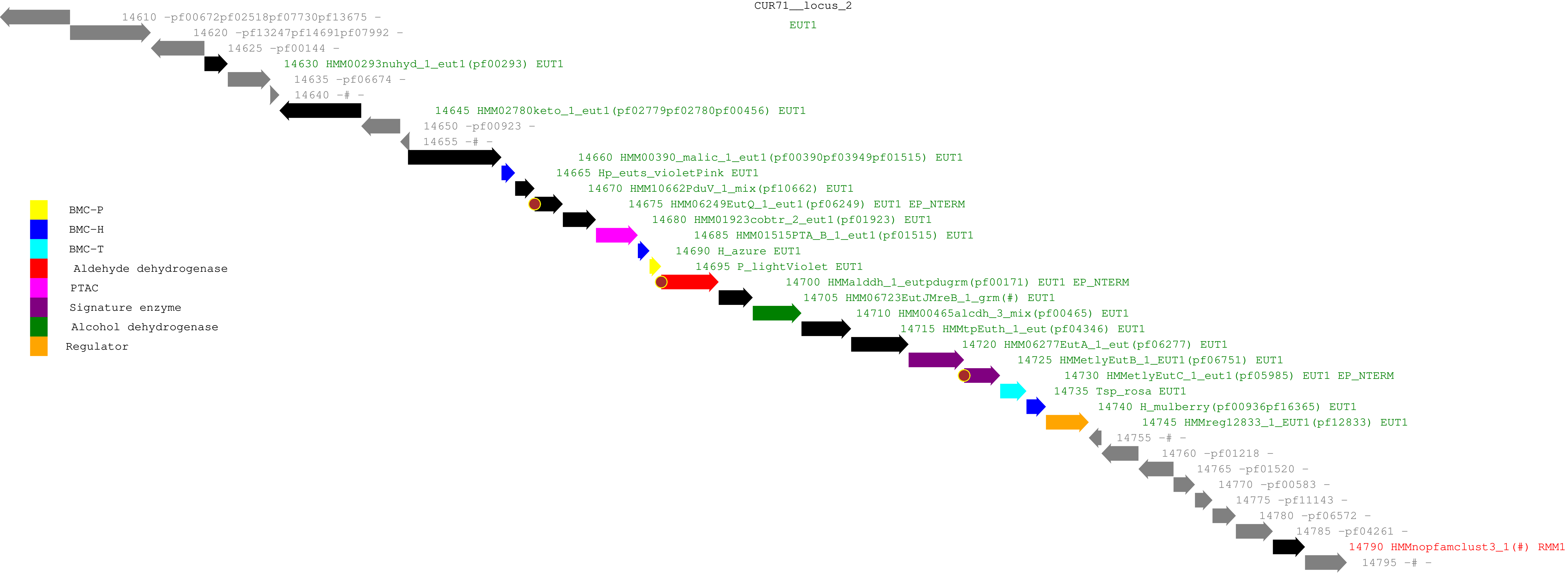

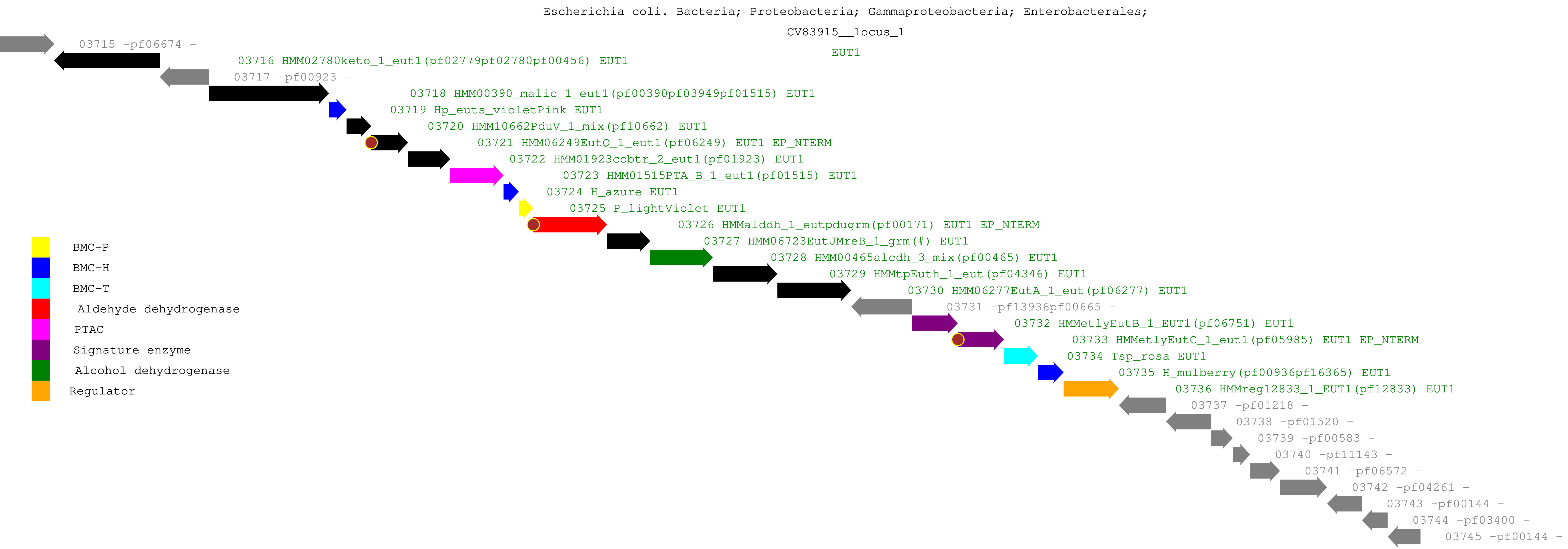
Escherichia coli. Bacteria; Proteobacteria; Gammaproteobacteria; Enterobacterales;
CV83915__locus_1
EUT1
03715 -pf06674 -
03716 HMM02780keto_1_eut1(pf02779pf02780pf00456) EUT1
03717 -pf00923 -
03718 HMM00390_malic_1_eut1(pf00390pf03949pf01515) EUT1
03719 Hp_euts_violetPink EUT1
03720 HMM10662PduV_1_mix(pf10662) EUT1
03721 HMM06249EutQ_1_eut1(pf06249) EUT1 EP_NTERM
03722 HMM01923cobtr_2_eut1(pf01923) EUT1
03723 HMM01515PTA_B_1_eut1(pf01515) EUT1
03724 H_azure EUT1
03725 P_lightViolet EUT1
03726 HMMalddh_1_eutpdugrm(pf00171) EUT1 EP_NTERM
03727 HMM06723EutJMreB_1_grm(#) EUT1
03728 HMM00465alcdh_3_mix(pf00465) EUT1
03729 HMMtpEuth_1_eut(pf04346) EUT1
03730 HMM06277EutA_1_eut(pf06277) EUT1
03731 -pf13936pf00665 -
03732 HMMetlyEutB_1_EUT1(pf06751) EUT1
03733 HMMetlyEutC_1_eut1(pf05985) EUT1 EP_NTERM
03734 Tsp_rosa EUT1
03735 H_mulberry(pf00936pf16365) EUT1
03736 HMMreg12833_1_EUT1(pf12833) EUT1
03737 -pf01218 -
03738 -pf01520 -
03739 -pf00583 -
03740 -pf11143 -
03741 -pf06572 -
03742 -pf04261 -
03743 -pf00144 -
03744 -pf03400 -
03745 -pf00144 -
BMC-P
BMC-H
BMC-T
Aldehyde dehydrogenase
PTAC
Signature enzyme
Alcohol dehydrogenase
Regulator

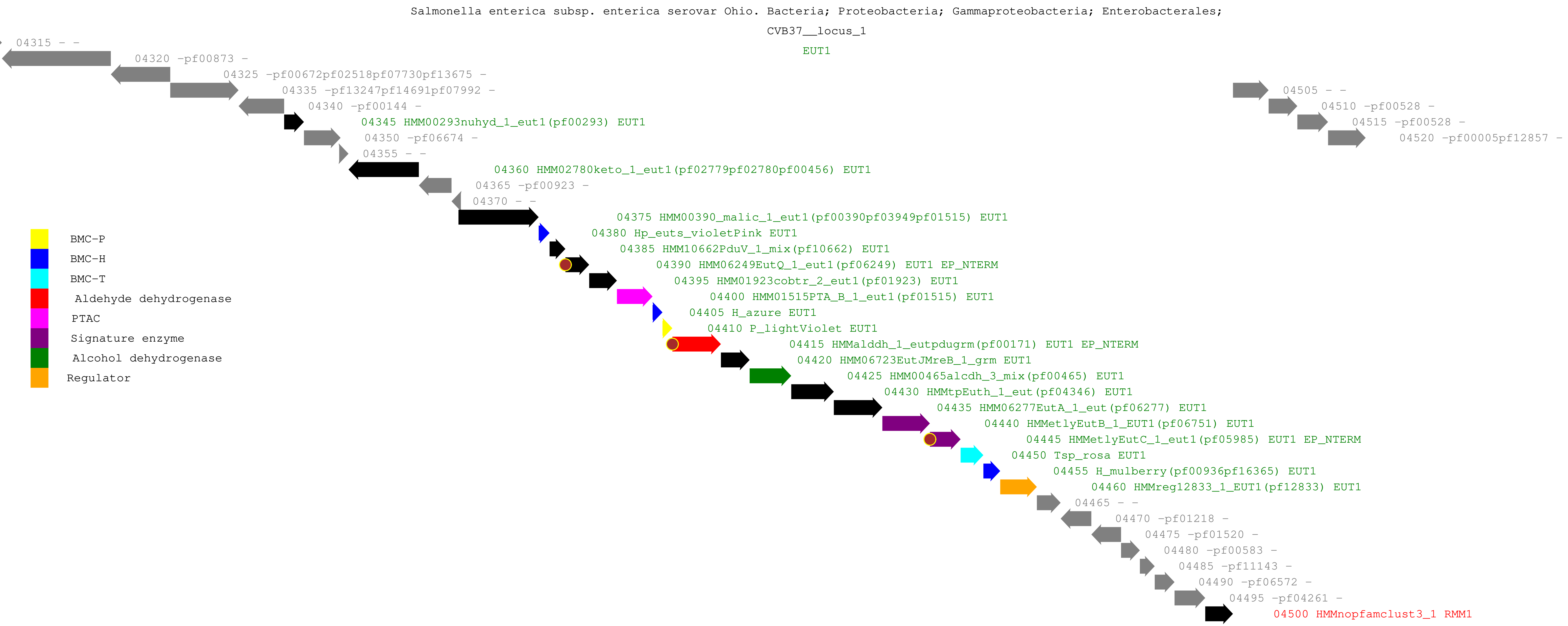
Salmonella enterica subsp. enterica serovar Ohio. Bacteria; Proteobacteria; Gammaproteobacteria; Enterobacterales;
CVB37__locus_1
EUT1
04315 - -
04320 -pf00873 -
04325 -pf00672pf02518pf07730pf13675 -
04335 -pf13247pf14691pf07992 -
04340 -pf00144 -
04345 HMM00293nuhyd_1_eut1(pf00293) EUT1
04350 -pf06674 -
04355 - -
04360 HMM02780keto_1_eut1(pf02779pf02780pf00456) EUT1
04365 -pf00923 -
04370 - -
04375 HMM00390_malic_1_eut1(pf00390pf03949pf01515) EUT1
04380 Hp_euts_violetPink EUT1
04385 HMM10662PduV_1_mix(pf10662) EUT1
04390 HMM06249EutQ_1_eut1(pf06249) EUT1 EP_NTERM
04395 HMM01923cobtr_2_eut1(pf01923) EUT1
04400 HMM01515PTA_B_1_eut1(pf01515) EUT1
04405 H_azure EUT1
04410 P_lightViolet EUT1
04415 HMMalddh_1_eutpdugrm(pf00171) EUT1 EP_NTERM
04420 HMM06723EutJMreB_1_grm EUT1
04425 HMM00465alcdh_3_mix(pf00465) EUT1
04430 HMMtpEuth_1_eut(pf04346) EUT1
04435 HMM06277EutA_1_eut(pf06277) EUT1
04440 HMMetlyEutB_1_EUT1(pf06751) EUT1
04445 HMMetlyEutC_1_eut1(pf05985) EUT1 EP_NTERM
04450 Tsp_rosa EUT1
04455 H_mulberry(pf00936pf16365) EUT1
04460 HMMreg12833_1_EUT1(pf12833) EUT1
04465 - -
04470 -pf01218 -
04475 -pf01520 -
04480 -pf00583 -
04485 -pf11143 -
04490 -pf06572 -
04495 -pf04261 -
04500 HMMnopfamclust3_1 RMM1
04505 - -
04510 -pf00528 -
04515 -pf00528 -
04520 -pf00005pf12857 -
BMC-P
BMC-H
BMC-T
Aldehyde dehydrogenase
PTAC
Signature enzyme
Alcohol dehydrogenase
Regulator

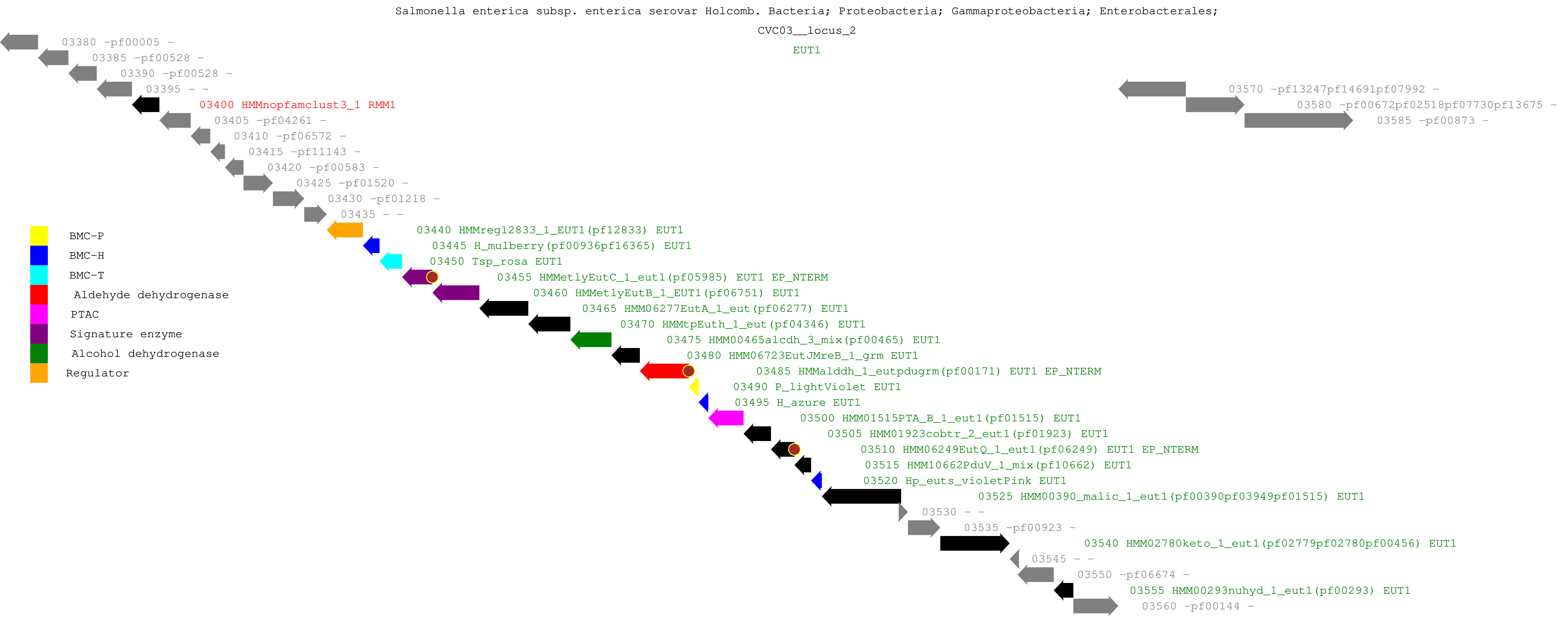
Salmonella enterica subsp. enterica serovar Holcomb. Bacteria; Proteobacteria; Gammaproteobacteria; Enterobacterales;
CVC03__locus_2
EUT1
03380 -pf00005 -
03385 -pf00528 -
03390 -pf00528 -
03395 - -
03400 HMMnopfamclust3_1 RMM1
03405 -pf04261 -
03410 -pf06572 -
03415 -pf11143 -
03420 -pf00583 -
03425 -pf01520 -
03430 -pf01218 -
03435 - -
03440 HMMreg12833_1_EUT1(pf12833) EUT1
03445 H_mulberry(pf00936pf16365) EUT1
03450 Tsp_rosa EUT1
03455 HMMetlyEutC_1_eut1(pf05985) EUT1 EP_NTERM
03460 HMMetlyEutB_1_EUT1(pf06751) EUT1
03465 HMM06277EutA_1_eut(pf06277) EUT1
03470 HMMtpEuth_1_eut(pf04346) EUT1
03475 HMM00465alcdh_3_mix(pf00465) EUT1
03480 HMM06723EutJMreB_1_grm EUT1
03485 HMMalddh_1_eutpdugrm(pf00171) EUT1 EP_NTERM
03490 P_lightViolet EUT1
03495 H_azure EUT1
03500 HMM01515PTA_B_1_eut1(pf01515) EUT1
03505 HMM01923cobtr_2_eut1(pf01923) EUT1
03510 HMM06249EutQ_1_eut1(pf06249) EUT1 EP_NTERM
03515 HMM10662PduV_1_mix(pf10662) EUT1
03520 Hp_euts_violetPink EUT1
03525 HMM00390_malic_1_eut1(pf00390pf03949pf01515) EUT1
03530 - -
03535 -pf00923 -
03540 HMM02780keto_1_eut1(pf02779pf02780pf00456) EUT1
03545 - -
03550 -pf06674 -
03555 HMM00293nuhyd_1_eut1(pf00293) EUT1
03560 -pf00144 -
03570 -pf13247pf14691pf07992 -
03580 -pf00672pf02518pf07730pf13675 -
03585 -pf00873 -
BMC-P
BMC-H
BMC-T
Aldehyde dehydrogenase
PTAC
Signature enzyme
Alcohol dehydrogenase
Regulator

Pseudomonas citronellolis. Bacteria; Proteobacteria; Gammaproteobacteria; Pseudomonadales;

CW310__locus_1

EUT1

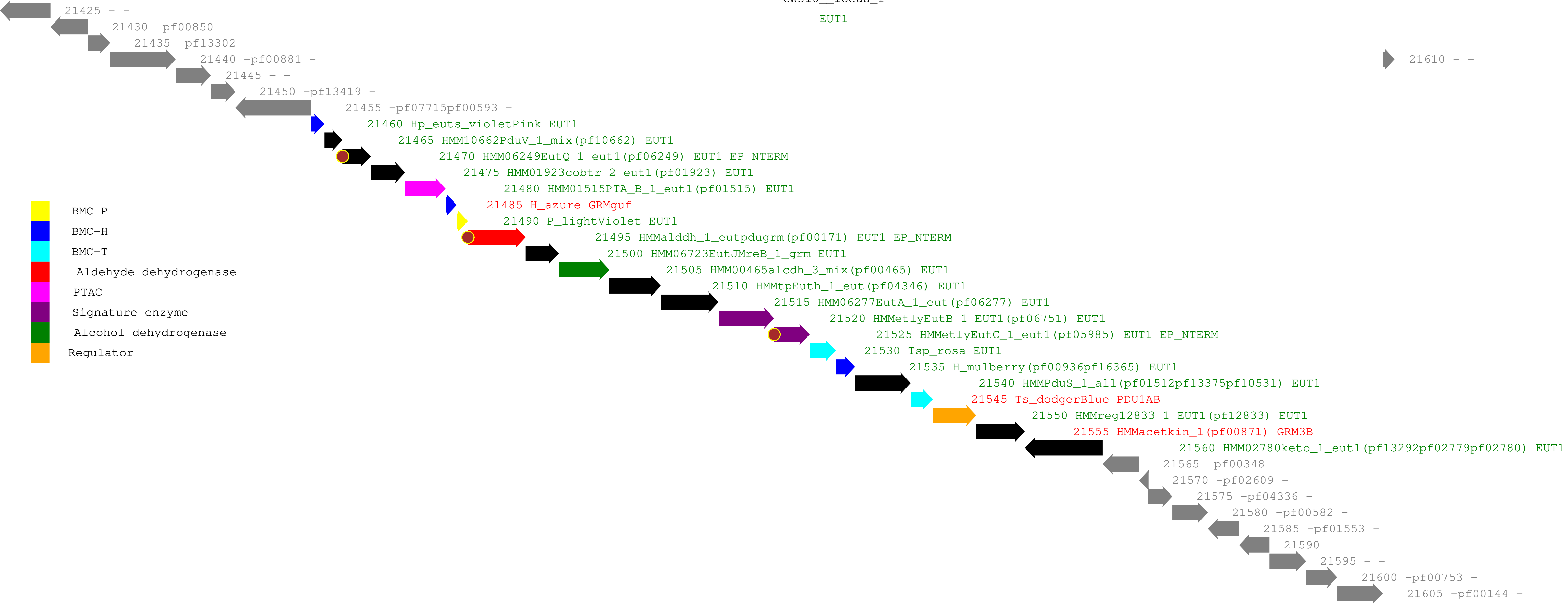

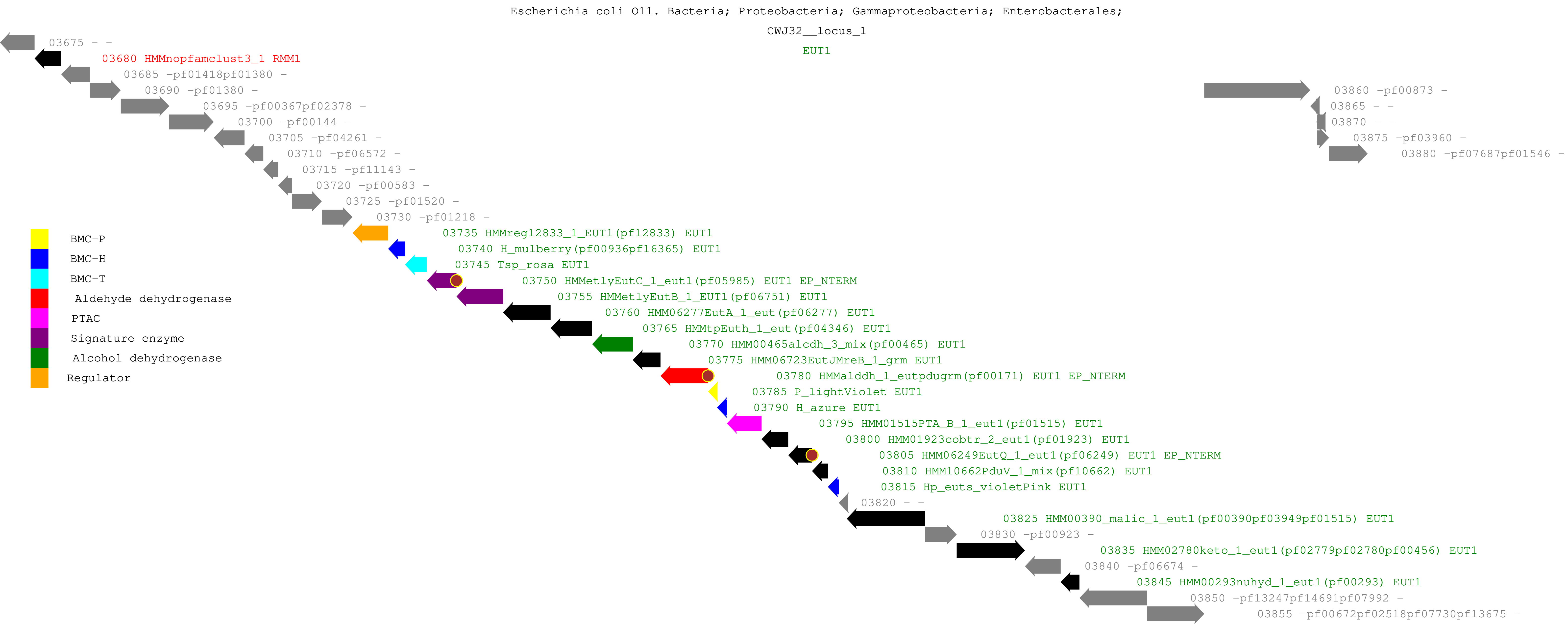
Escherichia coli O11. Bacteria; Proteobacteria; Gammaproteobacteria; Enterobacterales;
CWJ32__locus_1
EUT1
03675 - -
03680 HMMnopfamclust3_1 RMM1
03685 -pf01418pf01380 -
03690 -pf01380 -
03695 -pf00367pf02378 -
03700 -pf00144 -
03705 -pf04261 -
03710 -pf06572 -
03715 -pf11143 -
03720 -pf00583 -
03725 -pf01520 -
03730 -pf01218 -
03735 HMMreg12833_1_EUT1(pf12833) EUT1
03740 H_mulberry(pf00936pf16365) EUT1
03745 Tsp_rosa EUT1
03750 HMMetlyEutC_1_eut1(pf05985) EUT1 EP_NTERM
03755 HMMetlyEutB_1_EUT1(pf06751) EUT1
03760 HMM06277EutA_1_eut(pf06277) EUT1
03765 HMMtpEuth_1_eut(pf04346) EUT1
03770 HMM00465alcdh_3_mix(pf00465) EUT1
03775 HMM06723EutJMreB_1_grm EUT1
03780 HMMalddh_1_eutpdugrm(pf00171) EUT1 EP_NTERM
03785 P_lightViolet EUT1
03790 H_azure EUT1
03795 HMM01515PTA_B_1_eut1(pf01515) EUT1
03800 HMM01923cobtr_2_eut1(pf01923) EUT1
03805 HMM06249EutQ_1_eut1(pf06249) EUT1 EP_NTERM
03810 HMM10662PduV_1_mix(pf10662) EUT1
03815 Hp_euts_violetPink EUT1
03820 - -
03825 HMM00390_malic_1_eut1(pf00390pf03949pf01515) EUT1
03830 -pf00923 -
03835 HMM02780keto_1_eut1(pf02779pf02780pf00456) EUT1
03840 -pf06674 -
03845 HMM00293nuhyd_1_eut1(pf00293) EUT1
03850 -pf13247pf14691pf07992 -
03855 -pf00672pf02518pf07730pf13675 -
03860 -pf00873 -
03865 - -
03870 - -
03875 -pf03960 -
03880 -pf07687pf01546 -
BMC-P
BMC-H
BMC-T
Aldehyde dehydrogenase
PTAC
Signature enzyme
Alcohol dehydrogenase
Regulator

Morganella morganii (Proteus morganii). Bacteria; Proteobacteria; Gammaproteobacteria; Enterobacterales;

CXB74__locus_1

EUT1

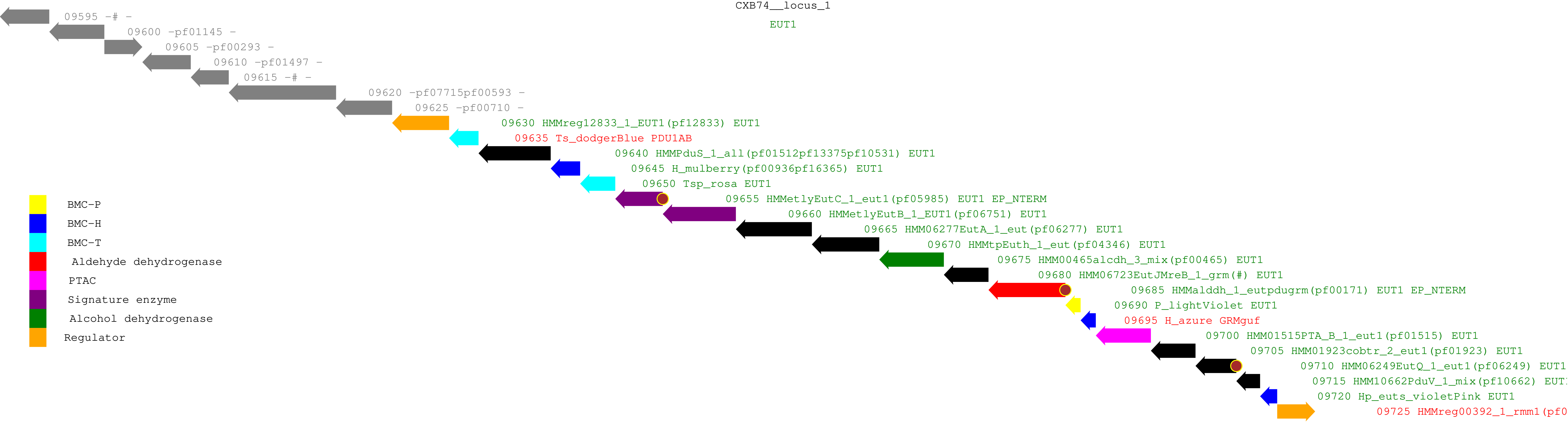

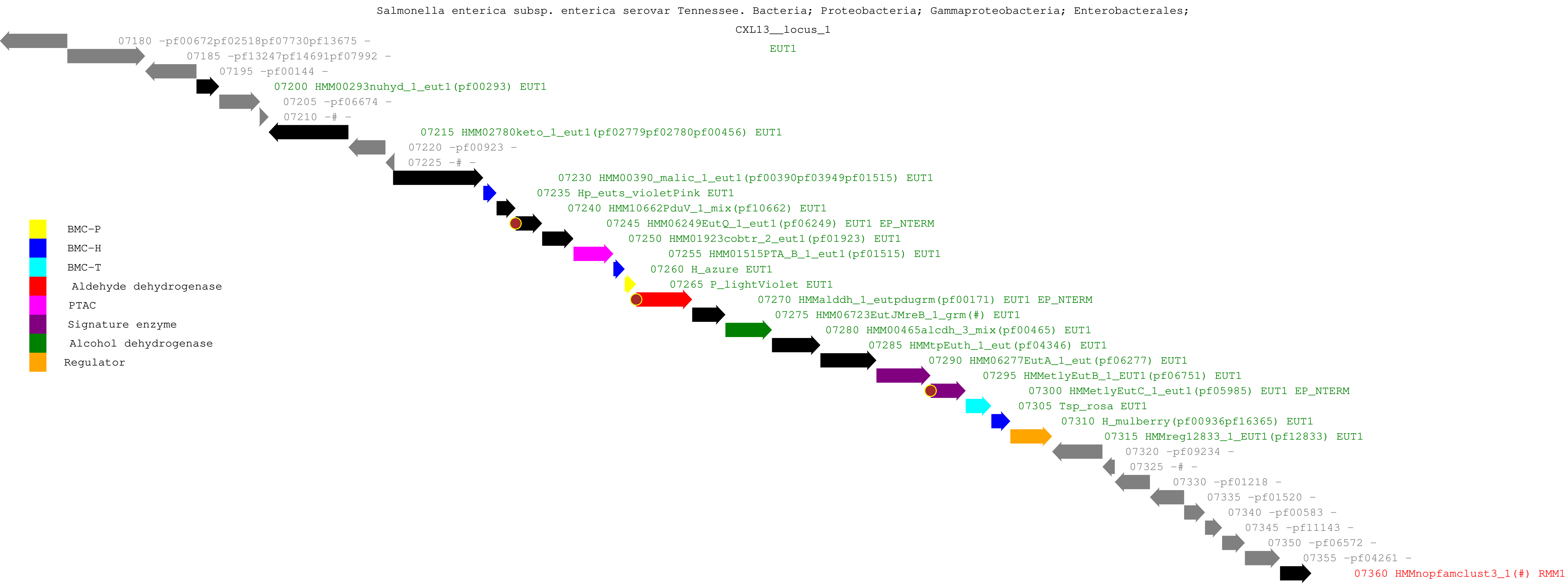

Salmonella enterica subsp. enterica serovar Tennessee. Bacteria; Proteobacteria; Gammaproteobacteria; Enterobacterales;
CXL13__locus_1
EUT1
07180 -pf00672pf02518pf07730pf13675 -
07185 -pf13247pf14691pf07992 -
07195 -pf00144 -
07200 HMM00293nuhyd_1_eut1(pf00293) EUT1
07205 -pf06674 -
07210 -# -
07215 HMM02780keto_1_eut1(pf02779pf02780pf00456) EUT1
07220 -pf00923 -
07225 -# -
07230 HMM00390_malic_1_eut1(pf00390pf03949pf01515) EUT1
07235 Hp_euts_violetPink EUT1
07240 HMM10662PduV_1_mix(pf10662) EUT1
07245 HMM06249EutQ_1_eut1(pf06249) EUT1 EP_NTERM
07250 HMM01923cobtr_2_eut1(pf01923) EUT1
07255 HMM01515PTA_B_1_eut1(pf01515) EUT1
07260 H_azure EUT1
07265 P_lightViolet EUT1
07270 HMMalddh_1_eutpdugrm(pf00171) EUT1 EP_NTERM
07275 HMM06723EutJMreB_1_grm(#) EUT1
07280 HMM00465alcdh_3_mix(pf00465) EUT1
07285 HMMtpEuth_1_eut(pf04346) EUT1
07290 HMM06277EutA_1_eut(pf06277) EUT1
07295 HMMetlyEutB_1_EUT1(pf06751) EUT1
07300 HMMetlyEutC_1_eut1(pf05985) EUT1 EP_NTERM
07305 Tsp_rosa EUT1
07310 H_mulberry(pf00936pf16365) EUT1
07315 HMMreg12833_1_EUT1(pf12833) EUT1
07320 -pf09234 -
07325 -# -
07330 -pf01218 -
07335 -pf01520 -
07340 -pf00583 -
07345 -pf11143 -
07350 -pf06572 -
07355 -pf04261 -
07360 HMMnopfamclust3_1(#) RMM1
BMC-P
BMC-H
BMC-T
Aldehyde dehydrogenase
PTAC
Signature enzyme
Alcohol dehydrogenase
Regulator

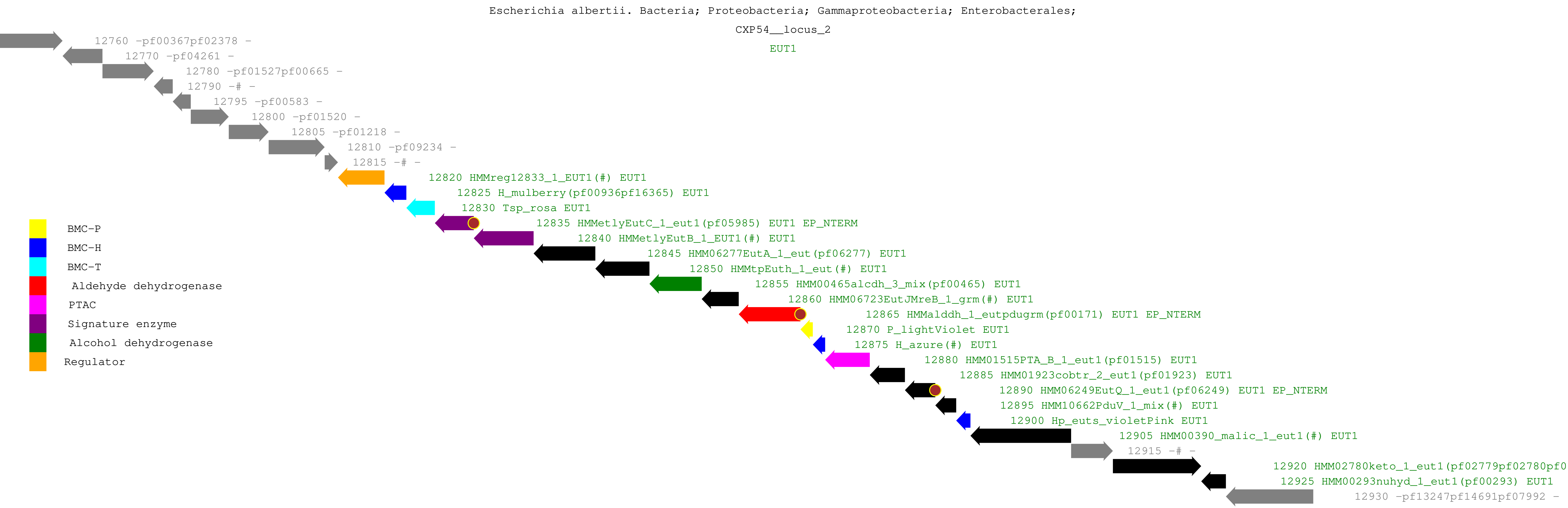

Escherichia albertii. Bacteria; Proteobacteria; Gammaproteobacteria; Enterobacterales;
CXP54__locus_2
EUT1
12760 -pf00367pf02378 -
12770 -pf04261 -
12780 -pf01527pf00665 -
12790 -# -
12795 -pf00583 -
12800 -pf01520 -
12805 -pf01218 -
12810 -pf09234 -
12815 -# -
12820 HMMreg12833_1_EUT1(#) EUT1
12825 H_mulberry(pf00936pf16365) EUT1
12830 Tsp_rosa EUT1
12835 HMMetlyEutC_1_eut1(pf05985) EUT1 EP_NTERM
12840 HMMetlyEutB_1_EUT1(#) EUT1
12845 HMM06277EutA_1_eut(pf06277) EUT1
12850 HMMtpEuth_1_eut(#) EUT1
12855 HMM00465alcdh_3_mix(pf00465) EUT1
12860 HMM06723EutJMreB_1_grm(#) EUT1
12865 HMMalddh_1_eutpdugrm(pf00171) EUT1 EP_NTERM
12870 P_lightViolet EUT1
12875 H_azure(#) EUT1
12880 HMM01515PTA_B_1_eut1(pf01515) EUT1
12885 HMM01923cobtr_2_eut1(pf01923) EUT1
12890 HMM06249EutQ_1_eut1(pf06249) EUT1 EP_NTERM
12895 HMM10662PduV_1_mix(#) EUT1
12900 Hp_euts_violetPink EUT1
12905 HMM00390_malic_1_eut1(#) EUT1
12915 -# -
12920 HMM02780keto_1_eut1(pf02779pf02780pf0
12925 HMM00293nuhyd_1_eut1(pf00293) EUT1
12930 -pf13247pf14691pf07992 -
BMC-P
BMC-H
BMC-T
Aldehyde dehydrogenase
PTAC
Signature enzyme
Alcohol dehydrogenase
Regulator

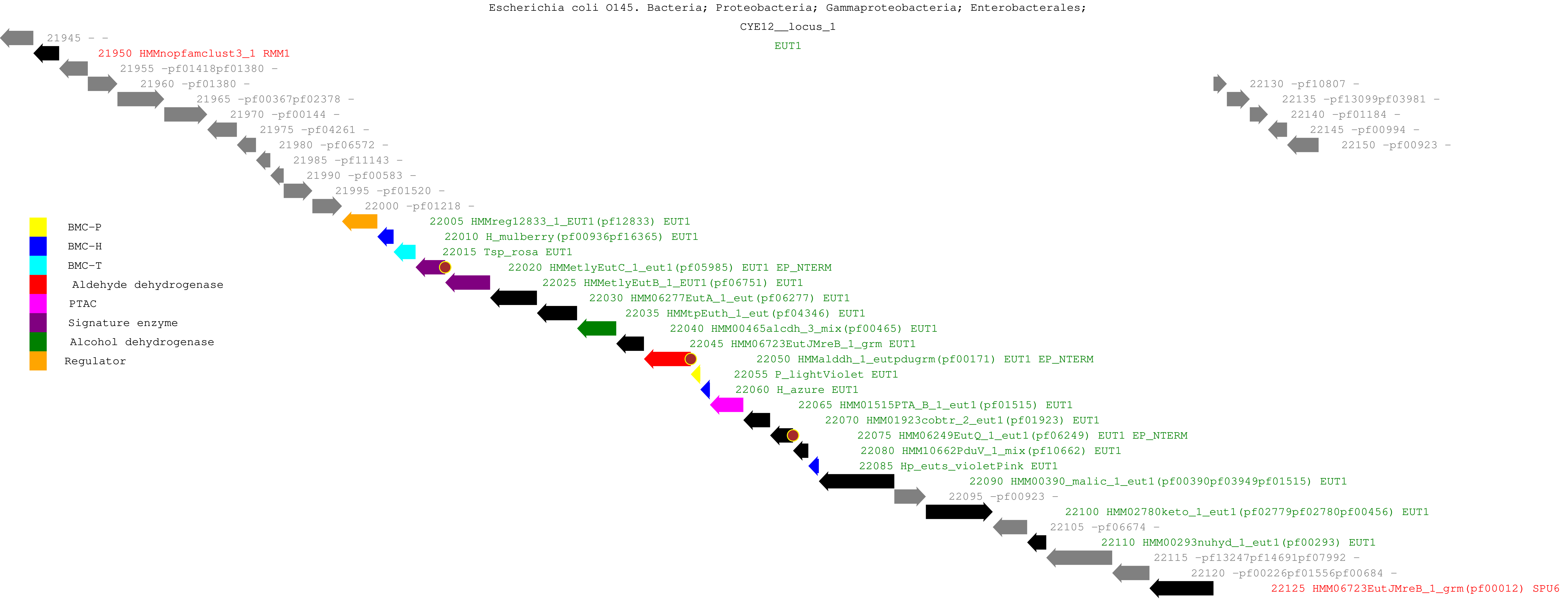
Escherichia coli O145. Bacteria; Proteobacteria; Gammaproteobacteria; Enterobacterales;
CYE12__locus_1
EUT1
21945 - -
21950 HMMnopfamclust3_1 RMM1
21955 -pf01418pf01380 -
21960 -pf01380 -
21965 -pf00367pf02378 -
21970 -pf00144 -
21975 -pf04261 -
21980 -pf06572 -
21985 -pf11143 -
21990 -pf00583 -
21995 -pf01520 -
22000 -pf01218 -
22005 HMMreg12833_1_EUT1(pf12833) EUT1
22010 H_mulberry(pf00936pf16365) EUT1
22015 Tsp_rosa EUT1
22020 HMMetlyEutC_1_eut1(pf05985) EUT1 EP_NTERM
22025 HMMetlyEutB_1_EUT1(pf06751) EUT1
22030 HMM06277EutA_1_eut(pf06277) EUT1
22035 HMMtpEuth_1_eut(pf04346) EUT1
22040 HMM00465alcdh_3_mix(pf00465) EUT1
22045 HMM06723EutJMreB_1_grm EUT1
22050 HMMalddh_1_eutpdugrm(pf00171) EUT1 EP_NTERM
22055 P_lightViolet EUT1
22060 H_azure EUT1
22065 HMM01515PTA_B_1_eut1(pf01515) EUT1
22070 HMM01923cobtr_2_eut1(pf01923) EUT1
22075 HMM06249EutQ_1_eut1(pf06249) EUT1 EP_NTERM
22080 HMM10662PduV_1_mix(pf10662) EUT1
22085 Hp_euts_violetPink EUT1
22090 HMM00390_malic_1_eut1(pf00390pf03949pf01515) EUT1
22095 -pf00923 -
22100 HMM02780keto_1_eut1(pf02779pf02780pf00456) EUT1
22105 -pf06674 -
22110 HMM00293nuhyd_1_eut1(pf00293) EUT1
22115 -pf13247pf14691pf07992 -
22120 -pf00226pf01556pf00684 -
22125 HMM06723EutJMreB_1_grm(pf00012) SPU6
22130 -pf10807 -
22135 -pf13099pf03981 -
22140 -pf01184 -
22145 -pf00994 -
22150 -pf00923 -
BMC-P
BMC-H
BMC-T
Aldehyde dehydrogenase
PTAC
Signature enzyme
Alcohol dehydrogenase
Regulator

Salmonella enterica subsp. houtenae serovar 16:z4,z32:-- str. RKS3027. Bacteria; Proteobacteria; Gammaproteobacteria; Enterobacterales;

D088__locus_1

EUT1

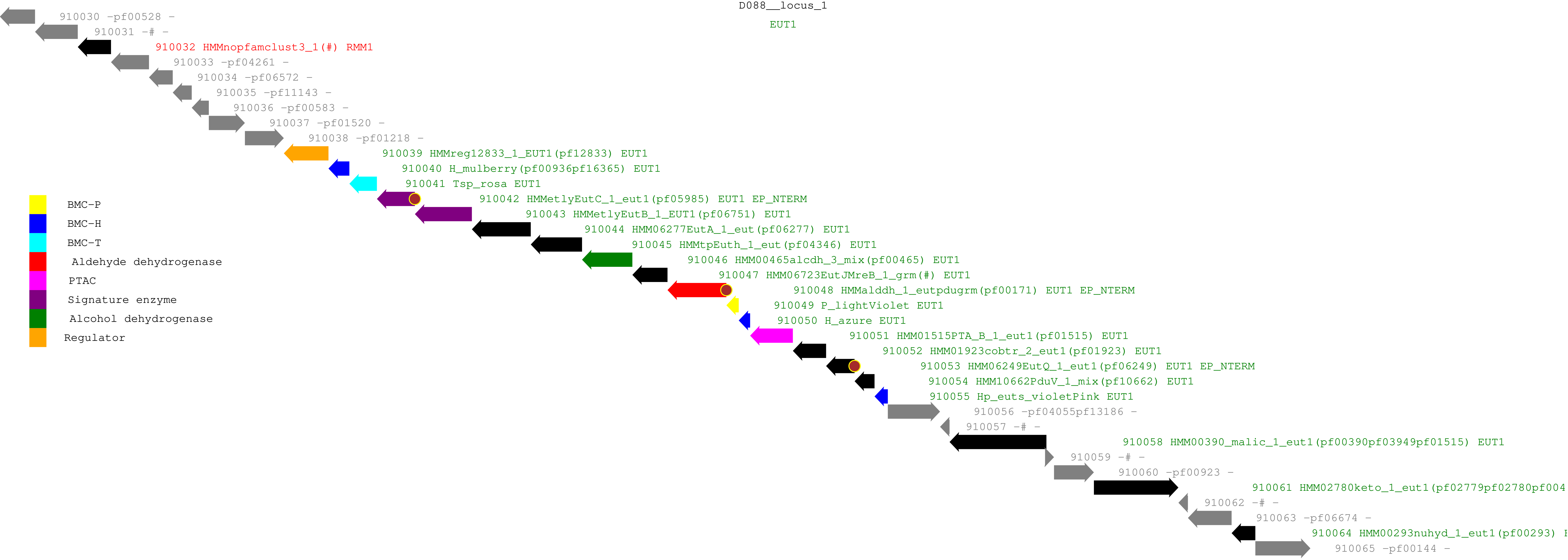

Salmonella enterica subsp. enterica serovar Saintpaul. Bacteria; Proteobacteria; Gammaproteobacteria; Enterobacterales;

D1118__locus_1

EUT1

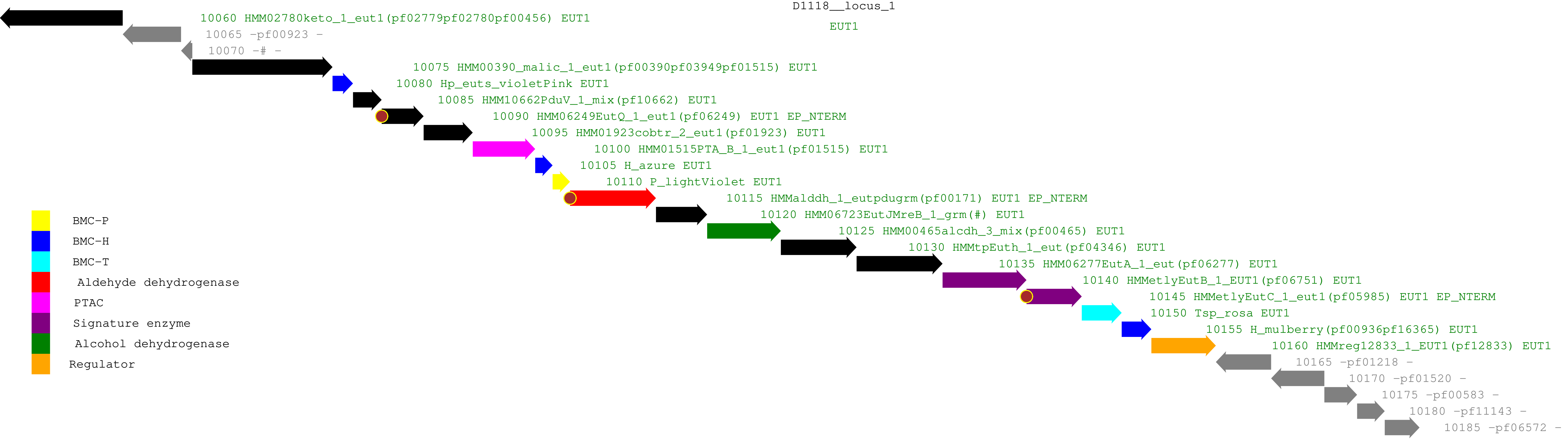

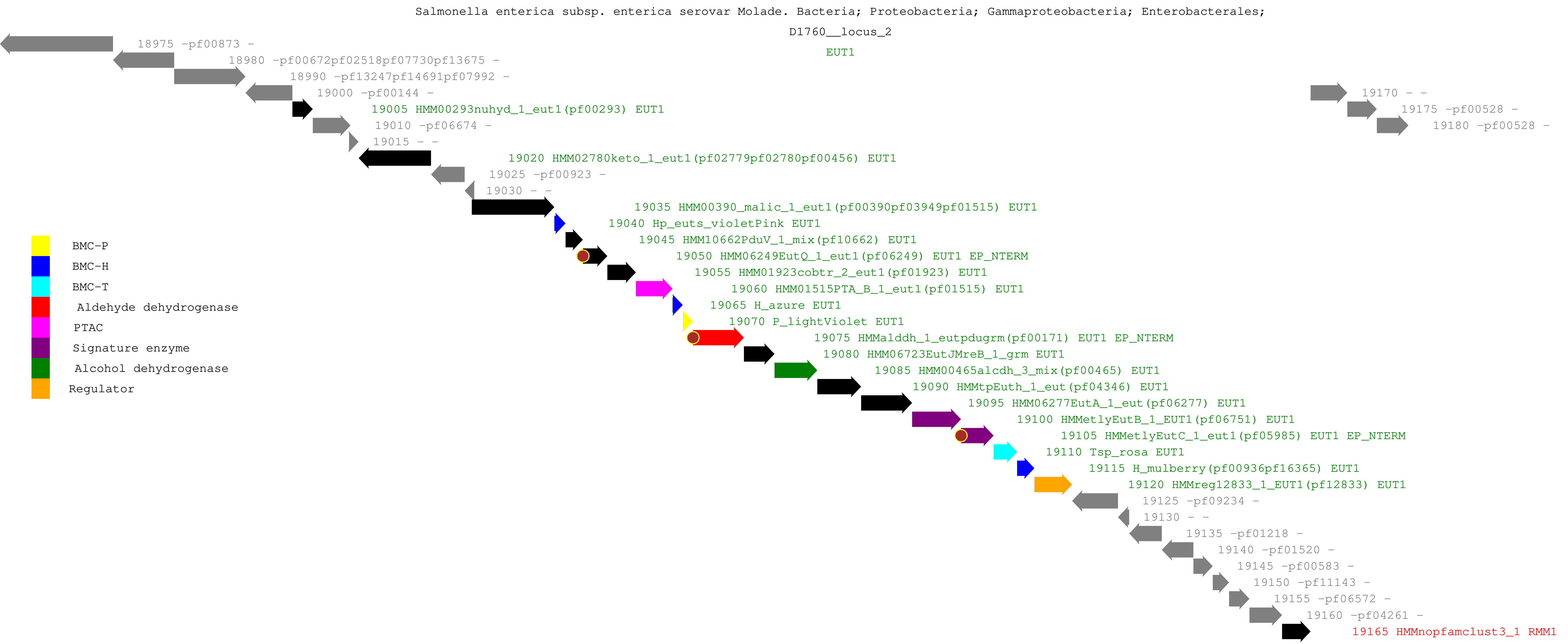
Salmonella enterica subsp. enterica serovar Molade. Bacteria; Proteobacteria; Gammaproteobacteria; Enterobacterales;
D1760__locus_2
EUT1
18975 -pf00873 -
18980 -pf00672pf02518pf07730pf13675 -
18990 -pf13247pf14691pf07992 -
19000 -pf00144 -
19005 HMM00293nuhyd_1_eut1(pf00293) EUT1
19010 -pf06674 -
19015 - -
19020 HMM02780keto_1_eut1(pf02779pf02780pf00456) EUT1
19025 -pf00923 -
19030 - -
19035 HMM00390_malic_1_eut1(pf00390pf03949pf01515) EUT1
19040 Hp_euts_violetPink EUT1
19045 HMM10662PduV_1_mix(pf10662) EUT1
19050 HMM06249EutQ_1_eut1(pf06249) EUT1 EP_NTERM
19055 HMM01923cobtr_2_eut1(pf01923) EUT1
19060 HMM01515PTA_B_1_eut1(pf01515) EUT1
19065 H_azure EUT1
19070 P_lightViolet EUT1
19075 HMMalddh_1_eutpdugrm(pf00171) EUT1 EP_NTERM
19080 HMM06723EutJMreB_1_grm EUT1
19085 HMM00465alcdh_3_mix(pf00465) EUT1
19090 HMMtpEuth_1_eut(pf04346) EUT1
19095 HMM06277EutA_1_eut(pf06277) EUT1
19100 HMMetlyEutB_1_EUT1(pf06751) EUT1
19105 HMMetlyEutC_1_eut1(pf05985) EUT1 EP_NTERM
19110 Tsp_rosa EUT1
19115 H_mulberry(pf00936pf16365) EUT1
19120 HMMreg12833_1_EUT1(pf12833) EUT1
19125 -pf09234 -
19130 - -
19135 -pf01218 -
19140 -pf01520 -
19145 -pf00583 -
19150 -pf11143 -
19155 -pf06572 -
19160 -pf04261 -
19165 HMMnopfamclust3_1 RMM1
19170 - -
19175 -pf00528 -
19180 -pf00528 -
BMC-P
BMC-H
BMC-T
Aldehyde dehydrogenase
PTAC
Signature enzyme
Alcohol dehydrogenase
Regulator

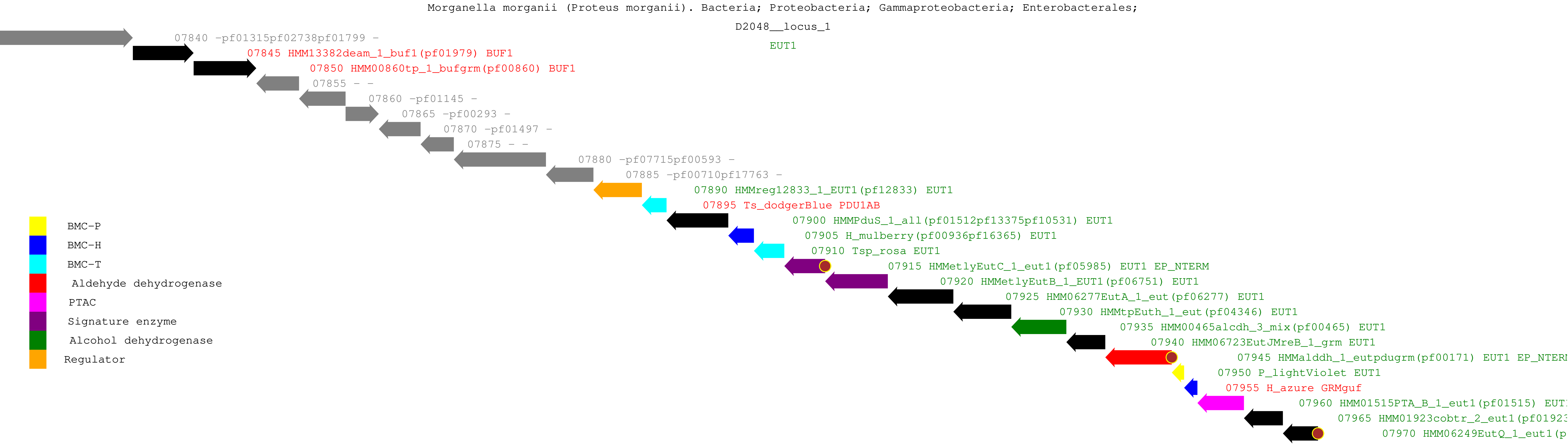
Morganella morganii (Proteus morganii). Bacteria; Proteobacteria; Gammaproteobacteria; Enterobacterales;
D2048__locus_1
EUT1
07840 -pf01315pf02738pf01799 -
07845 HMM13382deam_1_buf1(pf01979) BUF1
07850 HMM00860tp_1_bufgrm(pf00860) BUF1
07855 - -
07860 -pf01145 -
07865 -pf00293 -
07870 -pf01497 -
07875 - -
07880 -pf07715pf00593 -
07885 -pf00710pf17763 -
07890 HMMreg12833_1_EUT1(pf12833) EUT1
07895 Ts_dodgerBlue PDU1AB
07900 HMMPduS_1_all(pf01512pf13375pf10531) EUT1
07905 H_mulberry(pf00936pf16365) EUT1
07910 Tsp_rosa EUT1
07915 HMMetlyEutC_1_eut1(pf05985) EUT1 EP_NTERM
07920 HMMetlyEutB_1_EUT1(pf06751) EUT1
07925 HMM06277EutA_1_eut(pf06277) EUT1
07930 HMMtpEuth_1_eut(pf04346) EUT1
07935 HMM00465alcdh_3_mix(pf00465) EUT1
07940 HMM06723EutJMreB_1_grm EUT1
07945 HMMaldh_1_eutpdugrm(pf00171) EUT1 EP_NTERM
07950 P_lightViolet EUT1
07955 H_azure GRMguf
07960 HMM01515PTA_B_1_eut1(pf01515) EUT
07965 HMM01923cobtr_2_eut1(pf0192
07970 HMM06249EutQ_1_eut1(p
BMC-P
BMC-H
BMC-T
Aldehyde dehydrogenase
PTAC
Signature enzyme
Alcohol dehydrogenase
Regulator

Morganella morganii (Proteus morganii). Bacteria; Proteobacteria; Gammaproteobacteria; Enterobacterales;

D2048__locus_3

EUT1

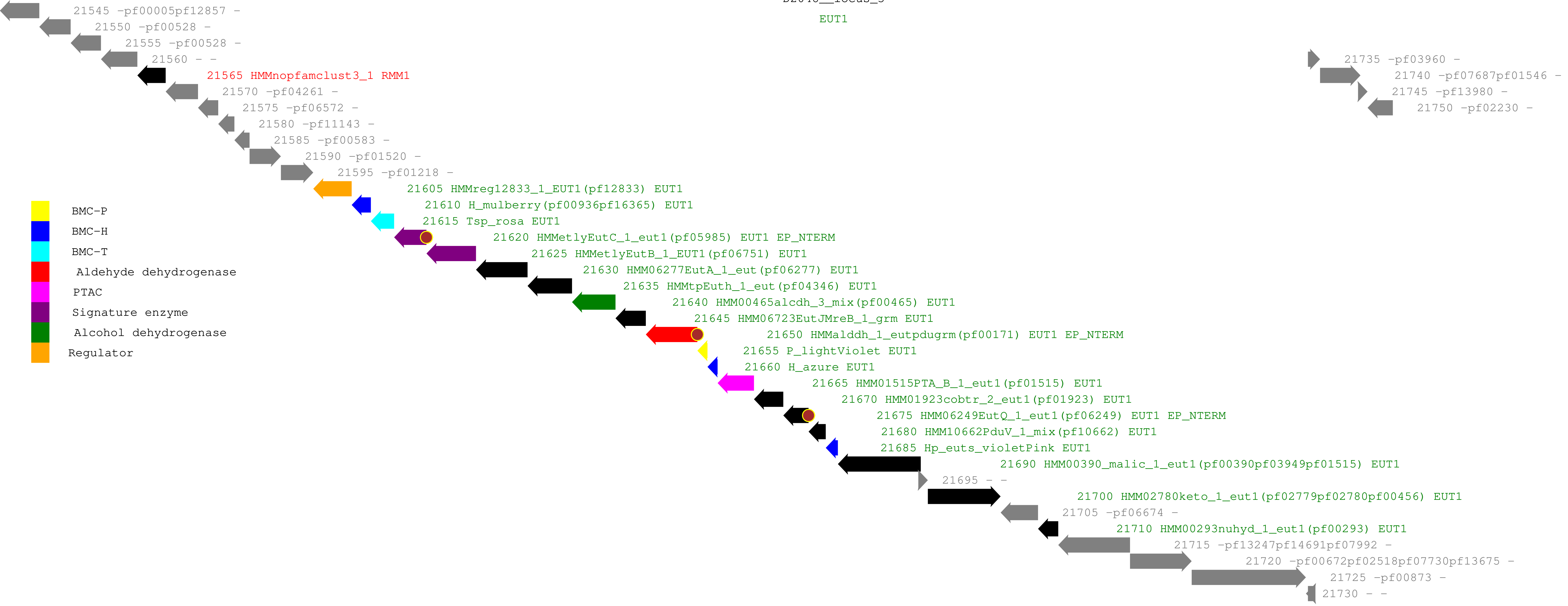

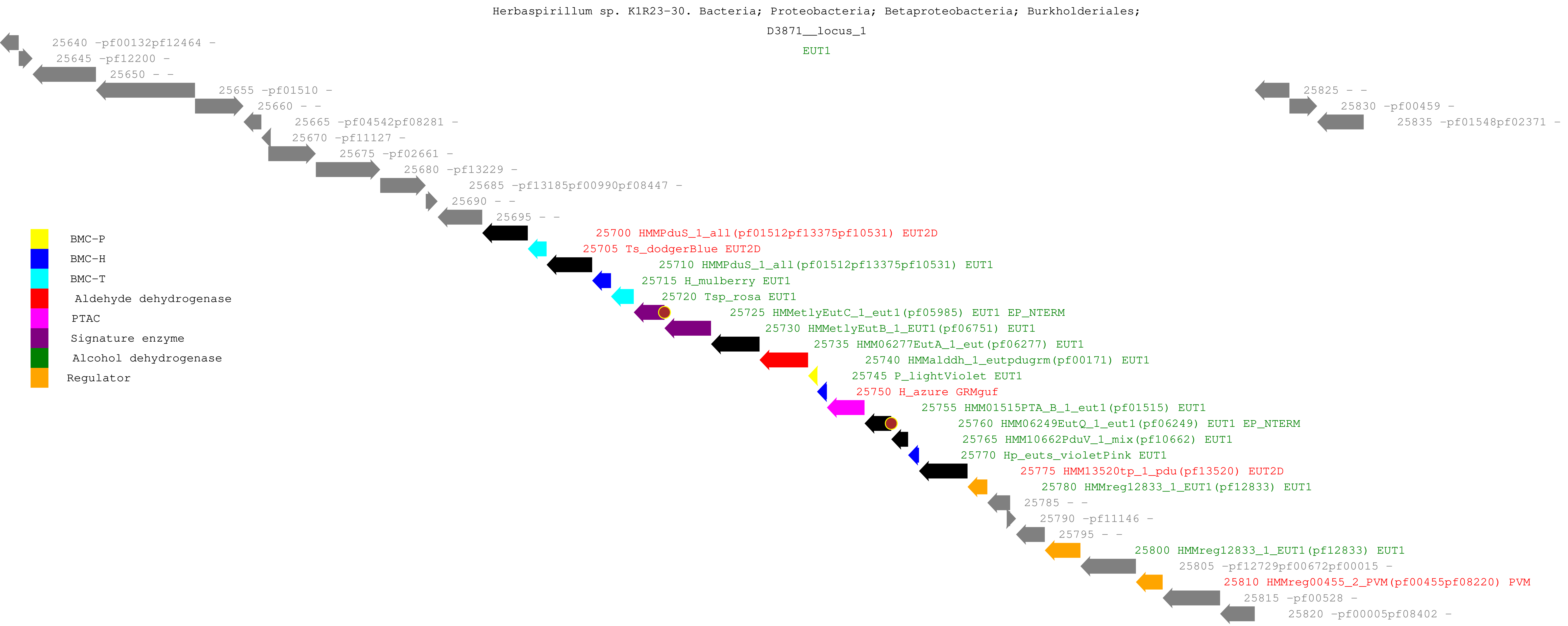
Herbaspirillum sp. K1R23-30. Bacteria; Proteobacteria; Betaproteobacteria; Burkholderiales;
D3871__locus_1
EUT1
25640 -pf00132pf12464 -
25645 -pf12200 -
25650 - -
25655 -pf01510 -
25660 - -
25665 -pf04542pf08281 -
25670 -pf11127 -
25675 -pf02661 -
25680 -pf13229 -
25685 -pf13185pf00990pf08447 -
25690 - -
25695 - -
25700 HMMPduS_1_all(pf01512pf13375pf10531) EUT2D
25705 Ts_dodgerBlue EUT2D
25710 HMMPduS_1_all(pf01512pf13375pf10531) EUT1
25715 H_mulberry EUT1
25720 Tsp_rosa EUT1
25725 HMMetlyEutC_1_eut1(pf05985) EUT1 EP_NTERM
25730 HMMetlyEutB_1_EUT1(pf06751) EUT1
25735 HMM06277EutA_1_eut(pf06277) EUT1
25740 HMMalddh_1_eutpdugrm(pf00171) EUT1
25745 P_lightViolet EUT1
25750 H_azure GRMguf
25755 HMM01515PTA_B_1_eut1(pf01515) EUT1
25760 HMM06249EutQ_1_eut1(pf06249) EUT1 EP_NTERM
25765 HMM10662PduV_1_mix(pf10662) EUT1
25770 Hp_euts_violetPink EUT1
25775 HMM13520tp_1_pdu(pf13520) EUT2D
25780 HMMreg12833_1_EUT1(pf12833) EUT1
25785 - -
25790 -pf11146 -
25795 - -
25800 HMMreg12833_1_EUT1(pf12833) EUT1
25805 -pf12729pf00672pf00015 -
25810 HMMreg00455_2_PVM(pf00455pf08220) PVM
25815 -pf00528 -
25820 -pf00005pf08402 -
25825 - -
25830 -pf00459 -
25835 -pf01548pf02371 -
BMC-P
BMC-H
BMC-T
Aldehyde dehydrogenase
PTAC
Signature enzyme
Alcohol dehydrogenase
Regulator

Escherichia coli. Bacteria; Proteobacteria; Gammaproteobacteria; Enterobacterales;

D3O91__locus_1

EUT1

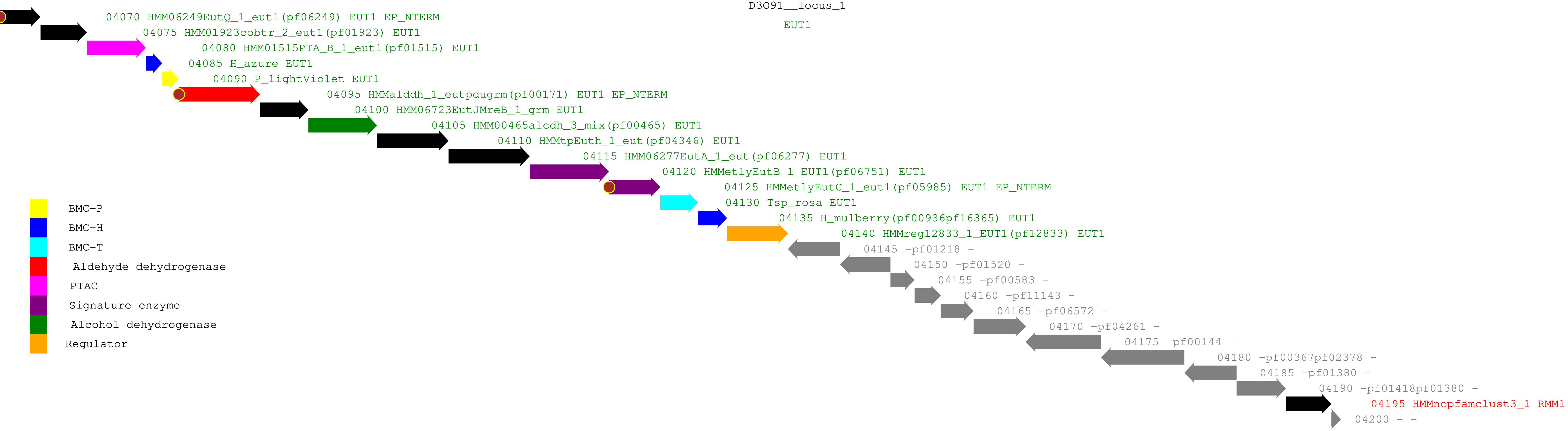

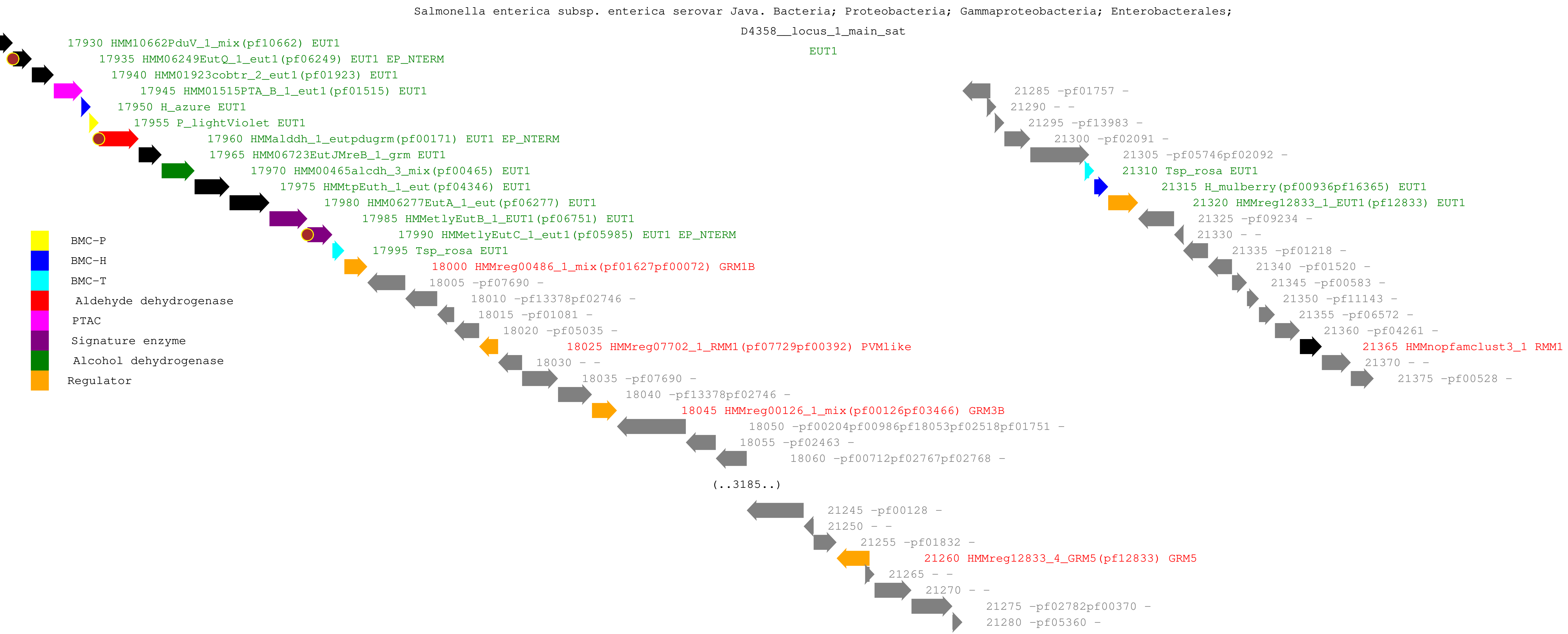
Salmonella enterica subsp. enterica serovar Java. Bacteria; Proteobacteria; Gammaproteobacteria; Enterobacterales;
D4358__locus_1_main_sat
EUT1
17930 HMM10662PduV_1_mix(pf10662) EUT1
17935 HMM06249EutQ_1_eut1(pf06249) EUT1 EP_NTERM
17940 HMM01923cobtr_2_eut1(pf01923) EUT1
17945 HMM01515PTA_B_1_eut1(pf01515) EUT1
17950 H_azure EUT1
17955 P_lightViolet EUT1
17960 HMMaldh_1_eutpdugrm(pf00171) EUT1 EP_NTERM
17965 HMM06723EutJMreB_1_grm EUT1
17970 HMM00465alcdh_3_mix(pf00465) EUT1
17975 HMMtpEuth_1_eut(pf04346) EUT1
17980 HMM06277EutA_1_eut(pf06277) EUT1
17985 HMMetlyEutB_1_EUT1(pf06751) EUT1
17990 HMMetlyEutC_1_eut1(pf05985) EUT1 EP_NTERM
17995 Tsp_rosa EUT1
18000 HMMreg00486_1_mix(pf01627pf00072) GRM1B
18005 -pf07690 -
18010 -pf13378pf02746 -
18015 -pf01081 -
18020 -pf05035 -
18025 HMMreg07702_1_RMM1(pf07729pf00392) PVMlike
18030 - -
18035 -pf07690 -
18040 -pf13378pf02746 -
18045 HMMreg00126_1_mix(pf00126pf03466) GRM3B
18050 -pf00204pf00986pf18053pf02518pf01751 -
18055 -pf02463 -
18060 -pf00712pf02767pf02768 -
(..3185..)
21245 -pf00128 -
21250 - -
21255 -pf01832 -
21260 HMMreg12833_4_GRM5(pf12833) GRM5
21265 - -
21270 - -
21275 -pf02782pf00370 -
21280 -pf05360 -
21285 -pf01757 -
21290 - -
21295 -pf13983 -
21300 -pf02091 -
21305 -pf05746pf02092 -
21310 Tsp_rosa EUT1
21315 H_mulberry(pf00936pf16365) EUT1
21320 HMMreg12833_1_EUT1(pf12833) EUT1
21325 -pf09234 -
21330 - -
21335 -pf01218 -
21340 -pf01520 -
21345 -pf00583 -
21350 -pf11143 -
21355 -pf06572 -
21360 -pf04261 -
21365 HMMnopfamclust3_1 RMM1
21370 - -
21375 -pf00528 -
BMC-P
BMC-H
BMC-T
Aldehyde dehydrogenase
PTAC
Signature enzyme
Alcohol dehydrogenase
Regulator

Salmonella typhimurium. Bacteria; Proteobacteria; Gammaproteobacteria; Enterobacterales;

D4380__locus_1

EUT1

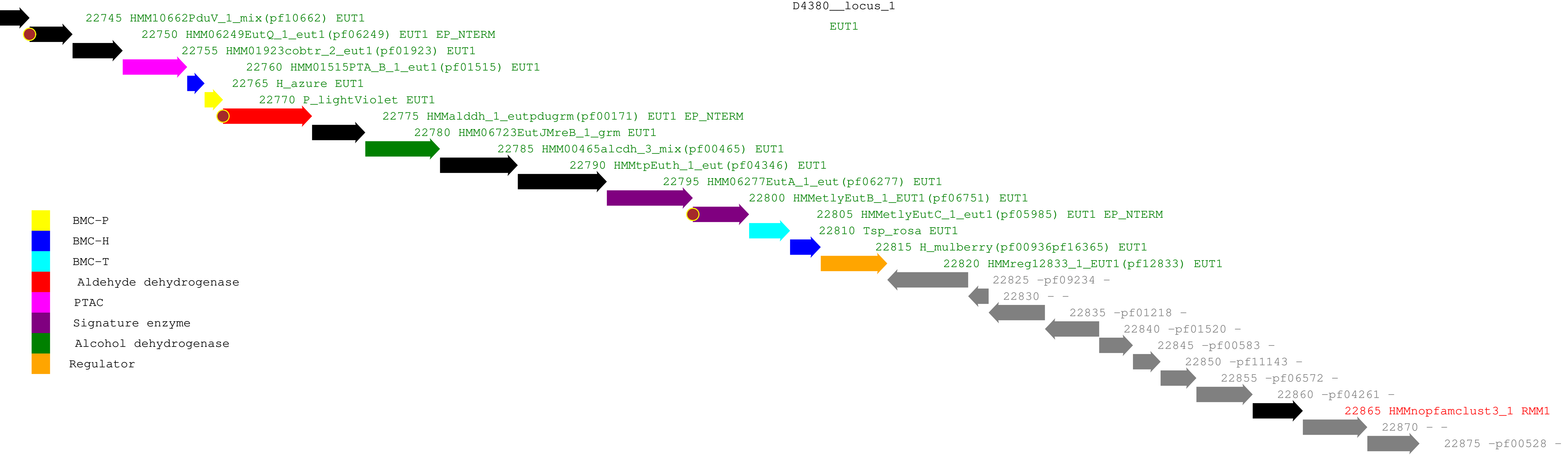

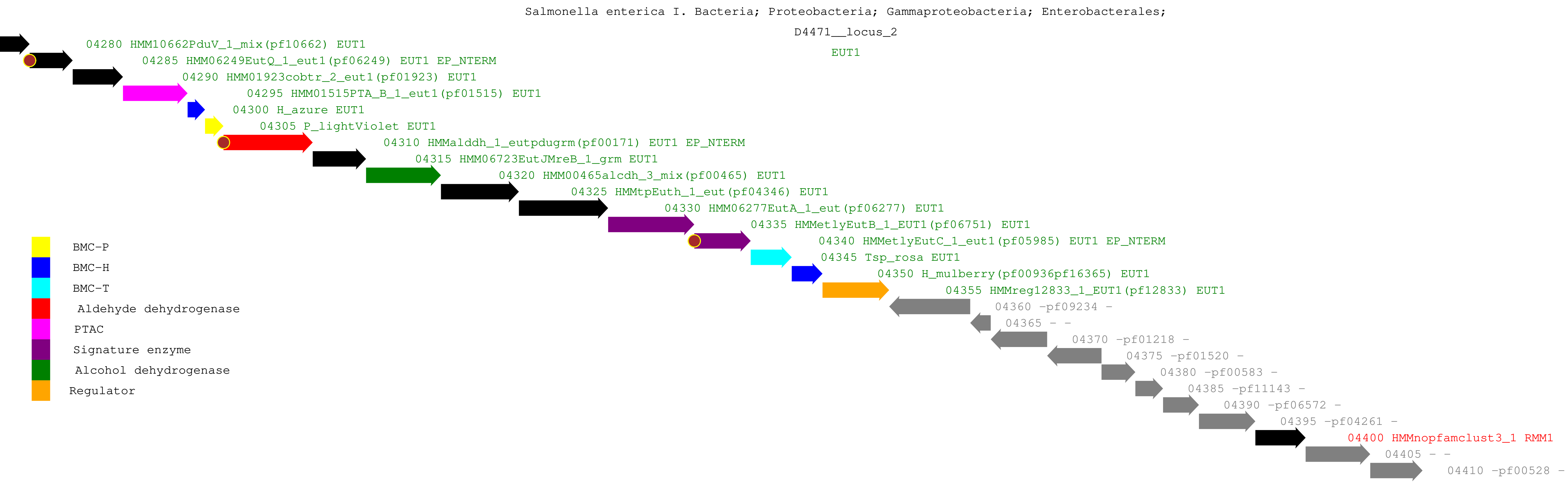
Salmonella enterica I. Bacteria; Proteobacteria; Gammaproteobacteria; Enterobacterales;
D4471__locus_2
EUT1
04280 HMM10662PduV_1_mix(pf10662) EUT1
04285 HMM06249EutQ_1_eut1(pf06249) EUT1 EP_NTERM
04290 HMM01923cobtr_2_eut1(pf01923) EUT1
04295 HMM01515PTA_B_1_eut1(pf01515) EUT1
04300 H_azure EUT1
04305 P_lightViolet EUT1
04310 HMMalddh_1_eutpdugrm(pf00171) EUT1 EP_NTERM
04315 HMM06723EutJMreB_1_grm EUT1
04320 HMM00465alcdh_3_mix(pf00465) EUT1
04325 HMMtpEuth_1_eut(pf04346) EUT1
04330 HMM06277EutA_1_eut(pf06277) EUT1
04335 HMMetlyEutB_1_EUT1(pf06751) EUT1
04340 HMMetlyEutC_1_eut1(pf05985) EUT1 EP_NTERM
04345 Tsp_rosa EUT1
04350 H_mulberry(pf00936pf16365) EUT1
04355 HMMreg12833_1_EUT1(pf12833) EUT1
04360 -pf09234 -
04365 - -
04370 -pf01218 -
04375 -pf01520 -
04380 -pf00583 -
04385 -pf11143 -
04390 -pf06572 -
04395 -pf04261 -
04400 HMMnopfamclust3_1 RMM1
04405 - -
04410 -pf00528 -
BMC-P
BMC-H
BMC-T
Aldehyde dehydrogenase
PTAC
Signature enzyme
Alcohol dehydrogenase
Regulator

Salmonella enterica subsp. enterica serovar Kentucky. Bacteria; Proteobacteria; Gammaproteobacteria; Enterobacterales;

D4523__locus_2

EUT1

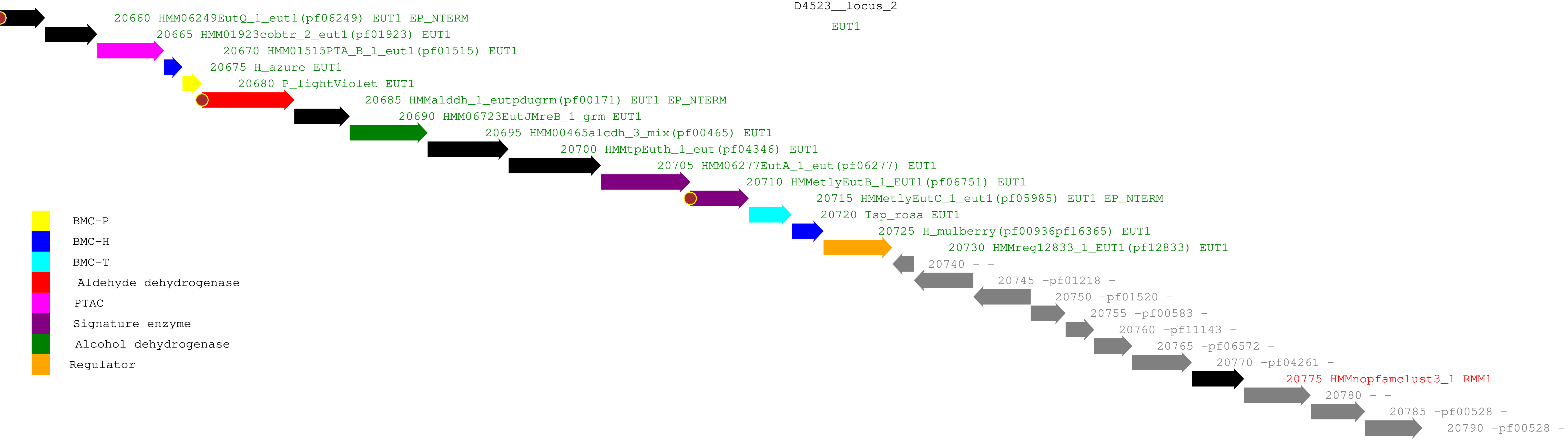

Salmonella enterica I. Bacteria; Proteobacteria; Gammaproteobacteria; Enterobacterales;

D4E70__locus_1

EUT1

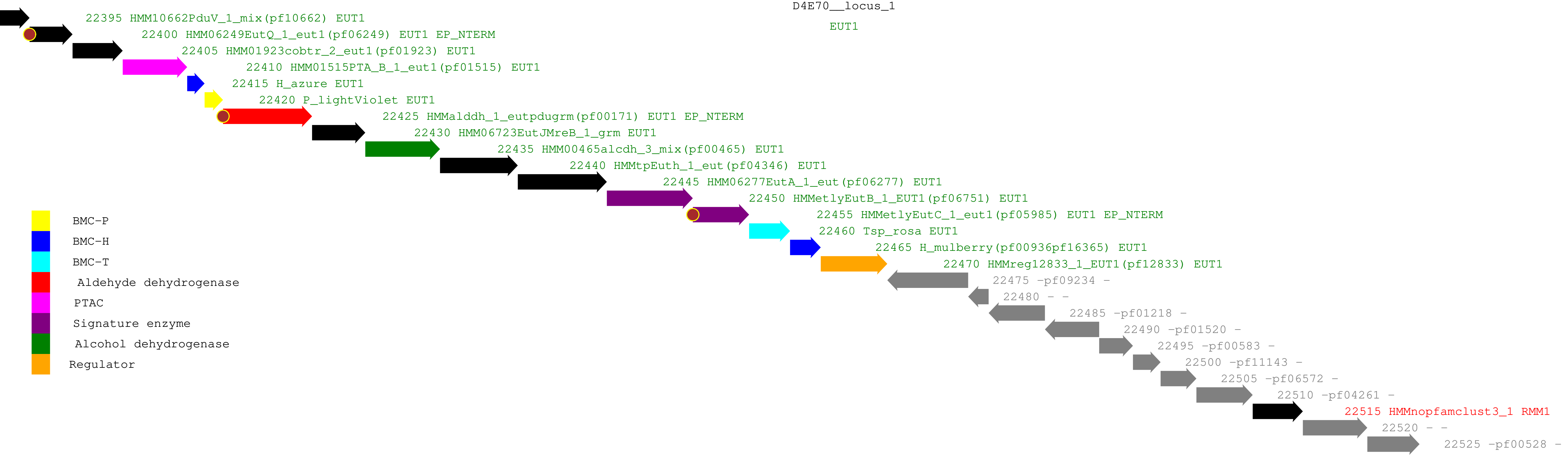

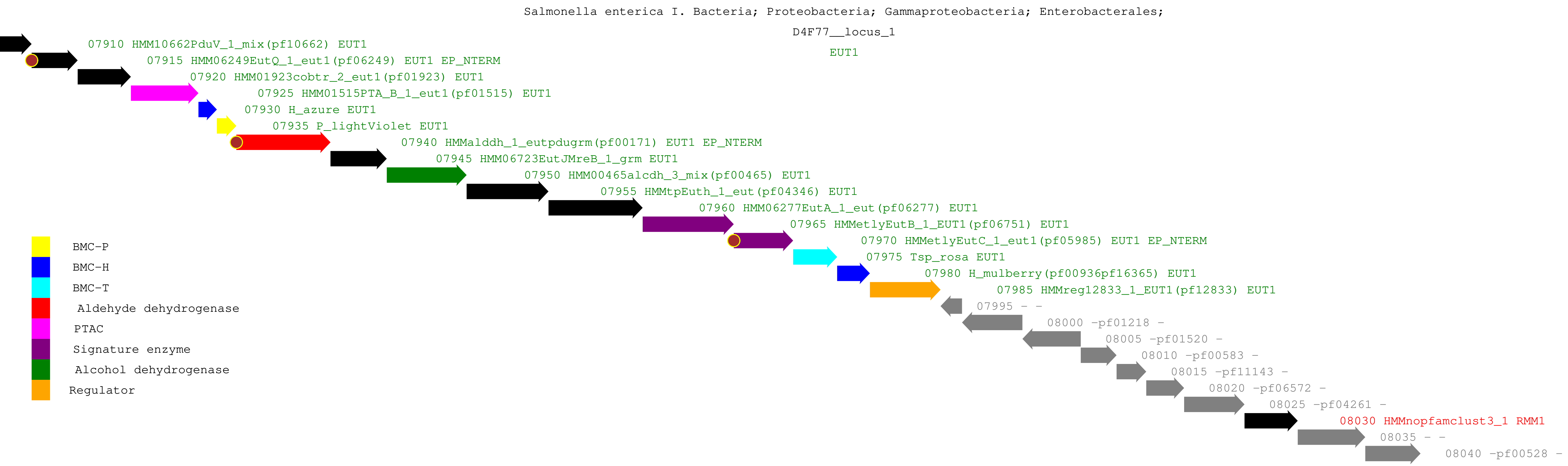

Salmonella enterica I. Bacteria; Proteobacteria; Gammaproteobacteria; Enterobacterales;
D4F77__locus_1
EUT1
07910 HMM10662PduV_1_mix(pf10662) EUT1
07915 HMM06249EutQ_1_eut1(pf06249) EUT1 EP_NTERM
07920 HMM01923cobtr_2_eut1(pf01923) EUT1
07925 HMM01515PTA_B_1_eut1(pf01515) EUT1
07930 H_azure EUT1
07935 P_lightViolet EUT1
07940 HMMaldddh_1_eutpdugrm(pf00171) EUT1 EP_NTERM
07945 HMM06723EutJMreB_1_grm EUT1
07950 HMM00465alcdh_3_mix(pf00465) EUT1
07955 HMMtpEuth_1_eut(pf04346) EUT1
07960 HMM06277EutA_1_eut(pf06277) EUT1
07965 HMMetlyEutB_1_EUT1(pf06751) EUT1
07970 HMMetlyEutC_1_eut1(pf05985) EUT1 EP_NTERM
07975 Tsp_rosa EUT1
07980 H_mulberry(pf00936pf16365) EUT1
07985 HMMreg12833_1_EUT1(pf12833) EUT1
07995 - -
08000 -pf01218 -
08005 -pf01520 -
08010 -pf00583 -
08015 -pf11143 -
08020 -pf06572 -
08025 -pf04261 -
08030 HMMnopfamclust3_1 RMM1
08035 - -
08040 -pf00528 -
BMC-P
BMC-H
BMC-T
Aldehyde dehydrogenase
PTAC
Signature enzyme
Alcohol dehydrogenase
Regulator

Salmonella enterica I. Bacteria; Proteobacteria; Gammaproteobacteria; Enterobacterales;

D4G09__locus_1

EUT1

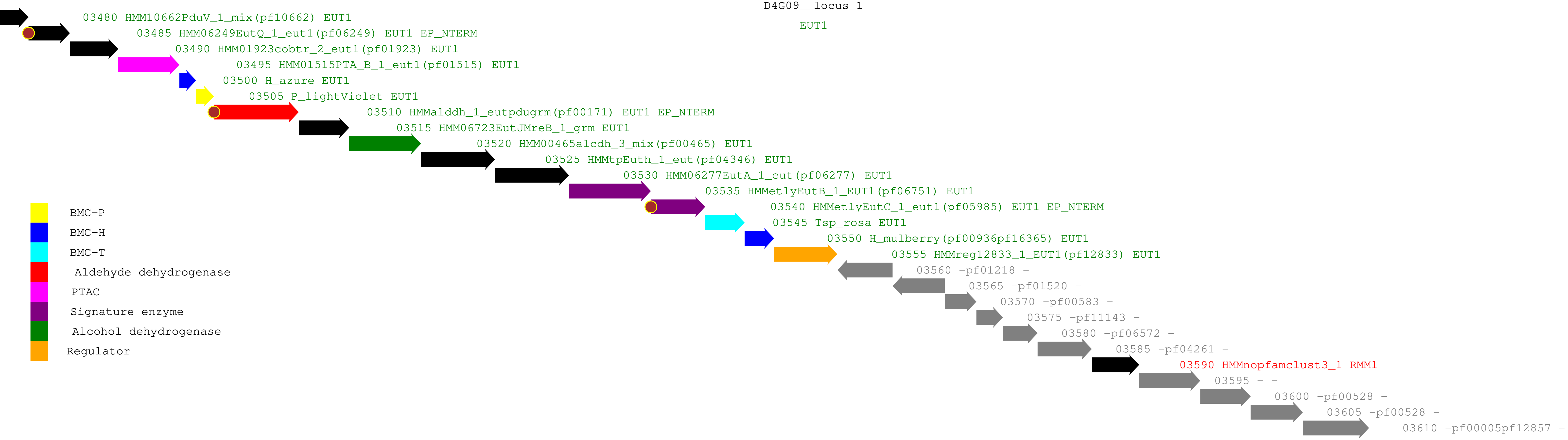

Salmonella enterica I. Bacteria; Proteobacteria; Gammaproteobacteria; Enterobacterales;

D4Y27__locus_1

EUT1

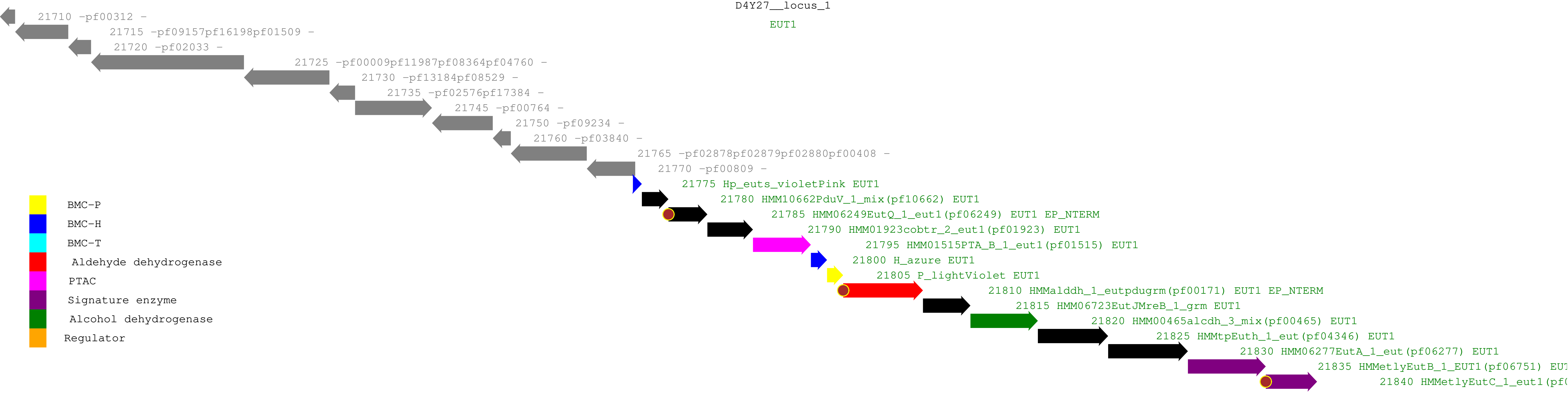

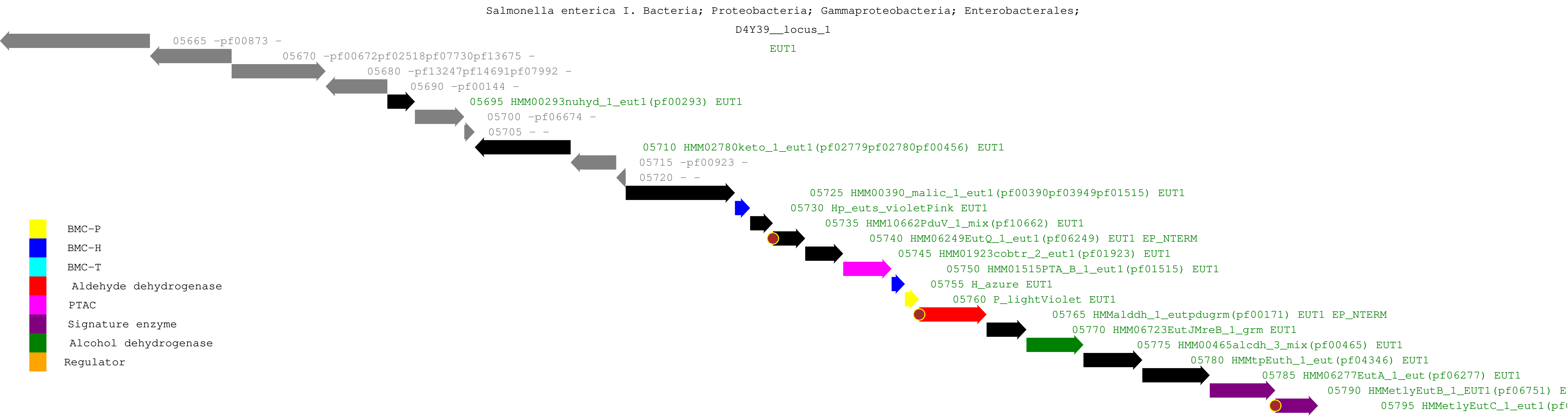
Salmonella enterica I. Bacteria; Proteobacteria; Gammaproteobacteria; Enterobacterales;
D4Y39__locus_1
EUT1
05665 -pf00873 -
05670 -pf00672pf02518pf07730pf13675 -
05680 -pf13247pf14691pf07992 -
05690 -pf00144 -
05695 HMM00293nuhyd_1_eut1(pf00293) EUT1
05700 -pf06674 -
05705 - -
05710 HMM02780keto_1_eut1(pf02779pf02780pf00456) EUT1
05715 -pf00923 -
05720 - -
05725 HMM00390_malic_1_eut1(pf00390pf03949pf01515) EUT1
05730 Hp_euts_violetPink EUT1
05735 HMM10662PduV_1_mix(pf10662) EUT1
05740 HMM06249EutQ_1_eut1(pf06249) EUT1 EP_NTERM
05745 HMM01923cobtr_2_eut1(pf01923) EUT1
05750 HMM01515PTA_B_1_eut1(pf01515) EUT1
05755 H_azure EUT1
05760 P_lightViolet EUT1
05765 HMMalddh_1_eutpdugrm(pf00171) EUT1 EP_NTERM
05770 HMM06723EutJMreB_1_grm EUT1
05775 HMM00465alcdh_3_mix(pf00465) EUT1
05780 HMMtpEuth_1_eut(pf04346) EUT1
05785 HMM06277EutA_1_eut(pf06277) EUT1
05790 HMMetlyEutB_1_EUT1(pf06751)
05795 HMMetlyEutC_1_eut1(pf
BMC-P
BMC-H
BMC-T
Aldehyde dehydrogenase
PTAC
Signature enzyme
Alcohol dehydrogenase
Regulator

Salmonella enterica I. Bacteria; Proteobacteria; Gammaproteobacteria; Enterobacterales;

D4Z58__locus_2

EUT1

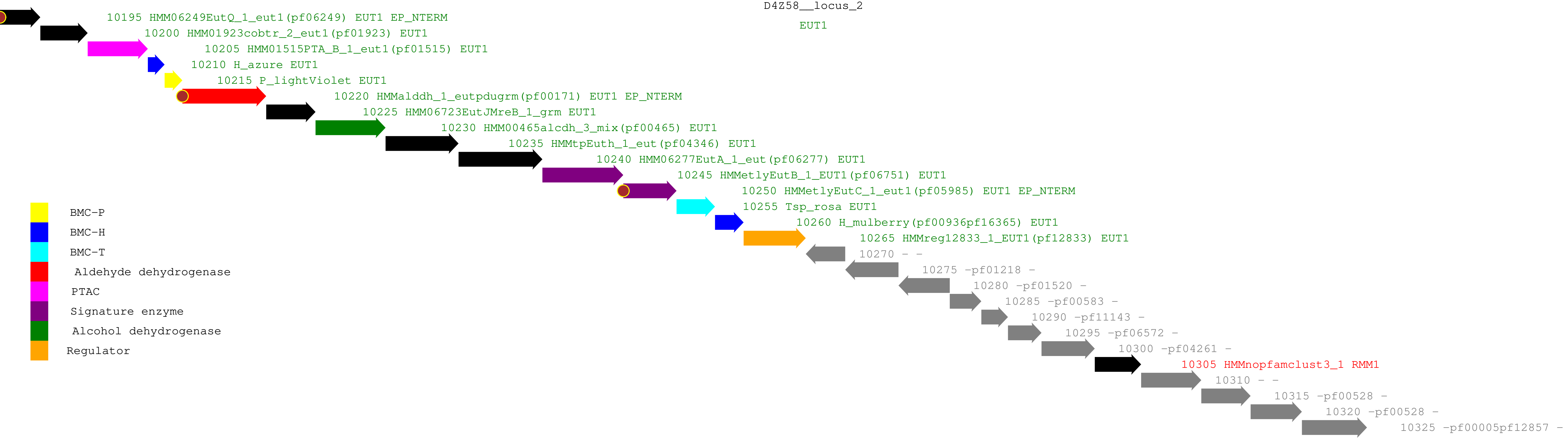

Salmonella enterica I. Bacteria; Proteobacteria; Gammaproteobacteria; Enterobacterales;

D5810__locus_1

EUT1

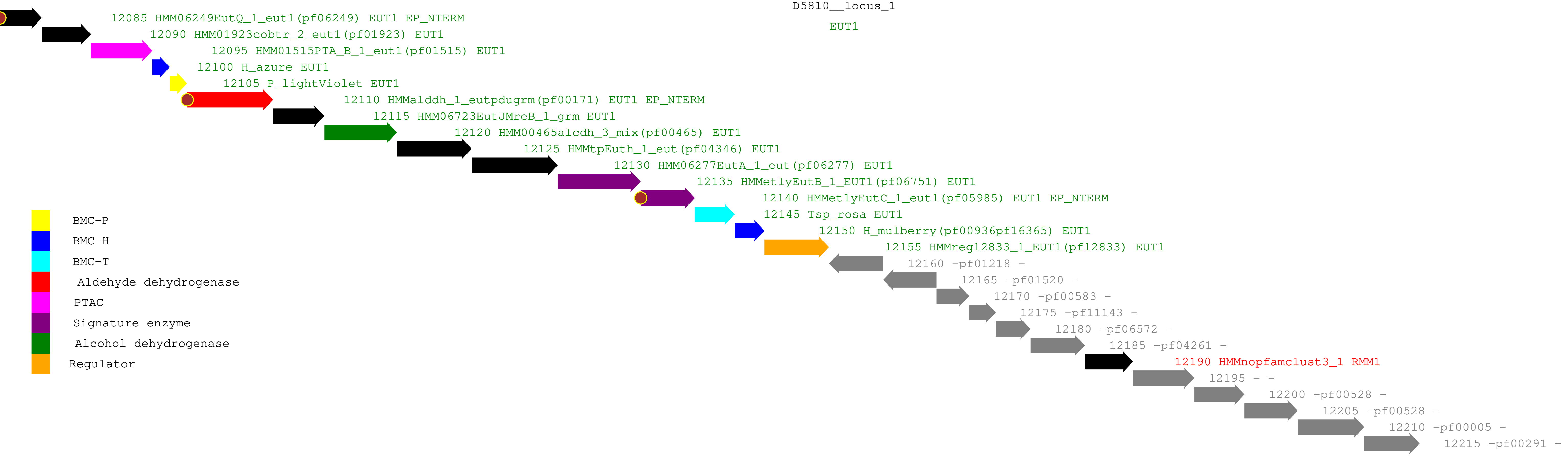

Salmonella enterica I. Bacteria; Proteobacteria; Gammaproteobacteria; Enterobacterales;

D5848__locus_1

EUT1

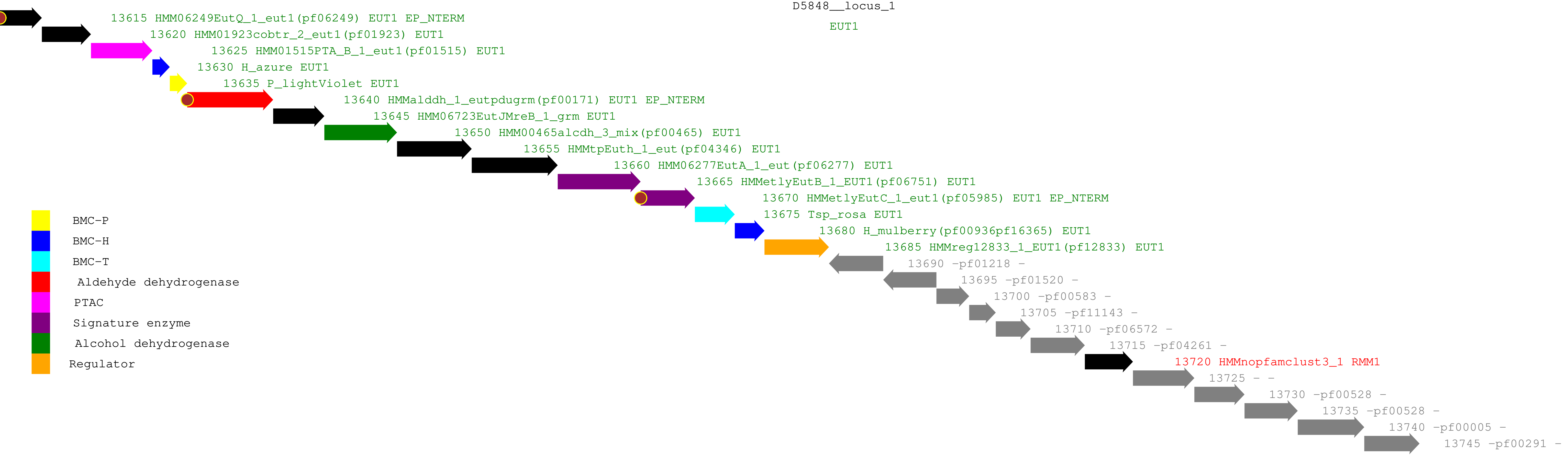

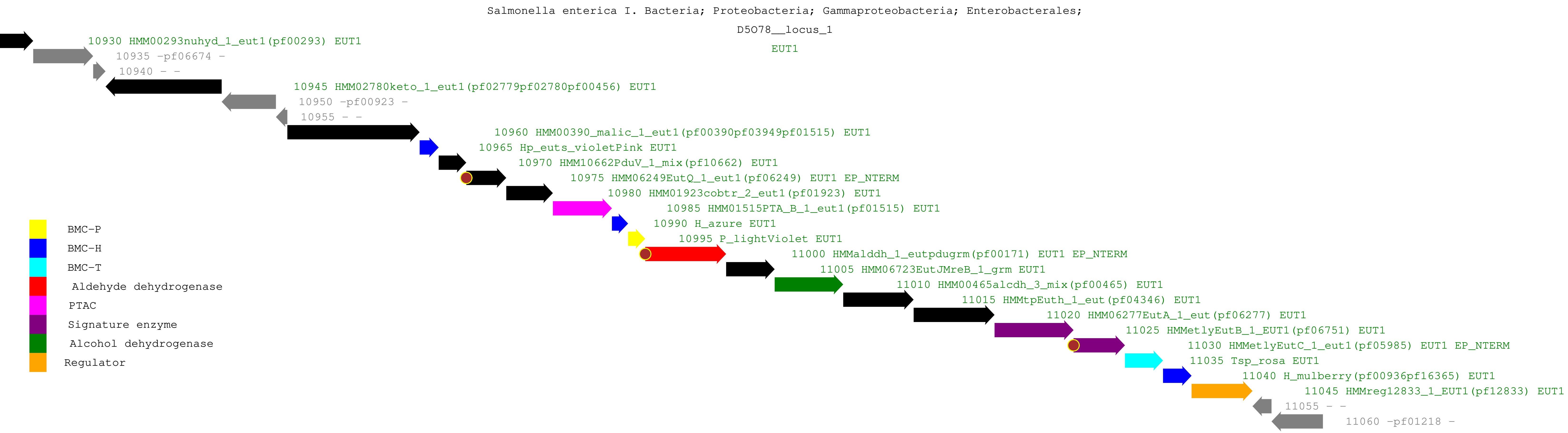
Salmonella enterica I. Bacteria; Proteobacteria; Gammaproteobacteria; Enterobacterales;
D5078__locus_1
EUT1
10930 HMM00293nuhyd_1_eut1(pf00293) EUT1
10935 -pf06674 -
10940 - -
10945 HMM02780keto_1_eut1(pf02779pf02780pf00456) EUT1
10950 -pf00923 -
10955 - -
10960 HMM00390_malic_1_eut1(pf00390pf03949pf01515) EUT1
10965 Hp_euts_violetPink EUT1
10970 HMM10662PduV_1_mix(pf10662) EUT1
10975 HMM06249EutQ_1_eut1(pf06249) EUT1 EP_NTERM
10980 HMM01923cobtr_2_eut1(pf01923) EUT1
10985 HMM01515PTA_B_1_eut1(pf01515) EUT1
10990 H_azure EUT1
10995 P_lightViolet EUT1
11000 HMMalddh_1_eutpdugrm(pf00171) EUT1 EP_NTERM
11005 HMM06723EutJMreB_1_grm EUT1
11010 HMM00465alcdh_3_mix(pf00465) EUT1
11015 HMMtpEuth_1_eut(pf04346) EUT1
11020 HMM06277EutA_1_eut(pf06277) EUT1
11025 HMMetlyEutB_1_EUT1(pf06751) EUT1
11030 HMMetlyEutC_1_eut1(pf05985) EUT1 EP_NTERM
11035 Tsp_rosa EUT1
11040 H_mulberry(pf00936pf16365) EUT1
11045 HMMreg12833_1_EUT1(pf12833) EUT1
11055 - -
11060 -pf01218 -
BMC-P
BMC-H
BMC-T
Aldehyde dehydrogenase
PTAC
Signature enzyme
Alcohol dehydrogenase
Regulator

Salmonella enterica I. Bacteria; Proteobacteria; Gammaproteobacteria; Enterobacterales;

D5Y01__locus_1

EUT1

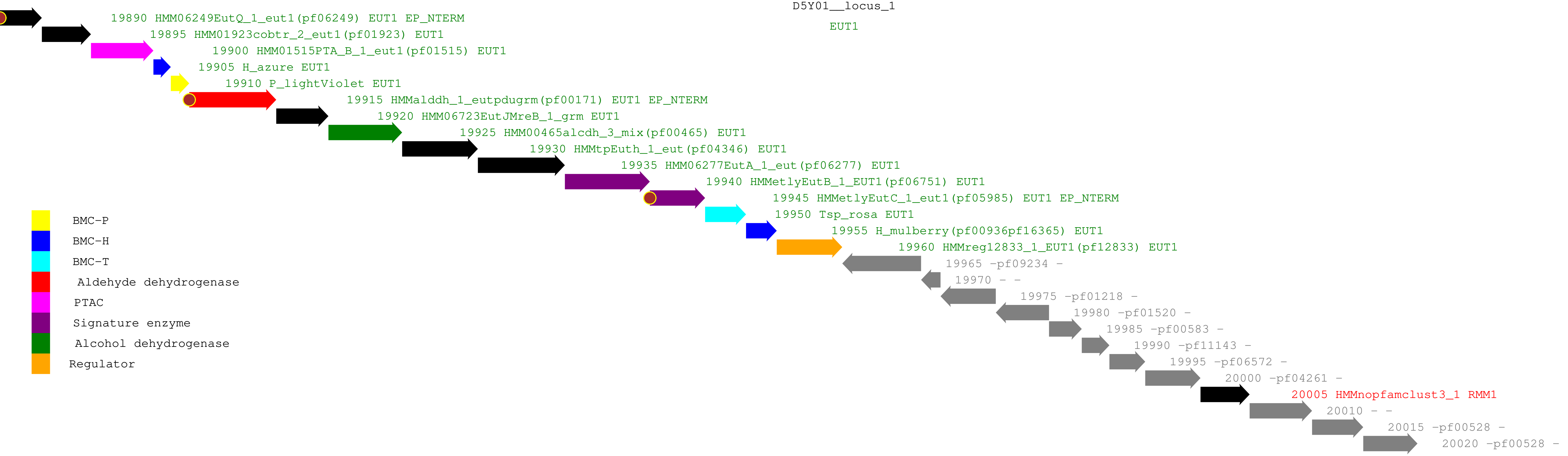

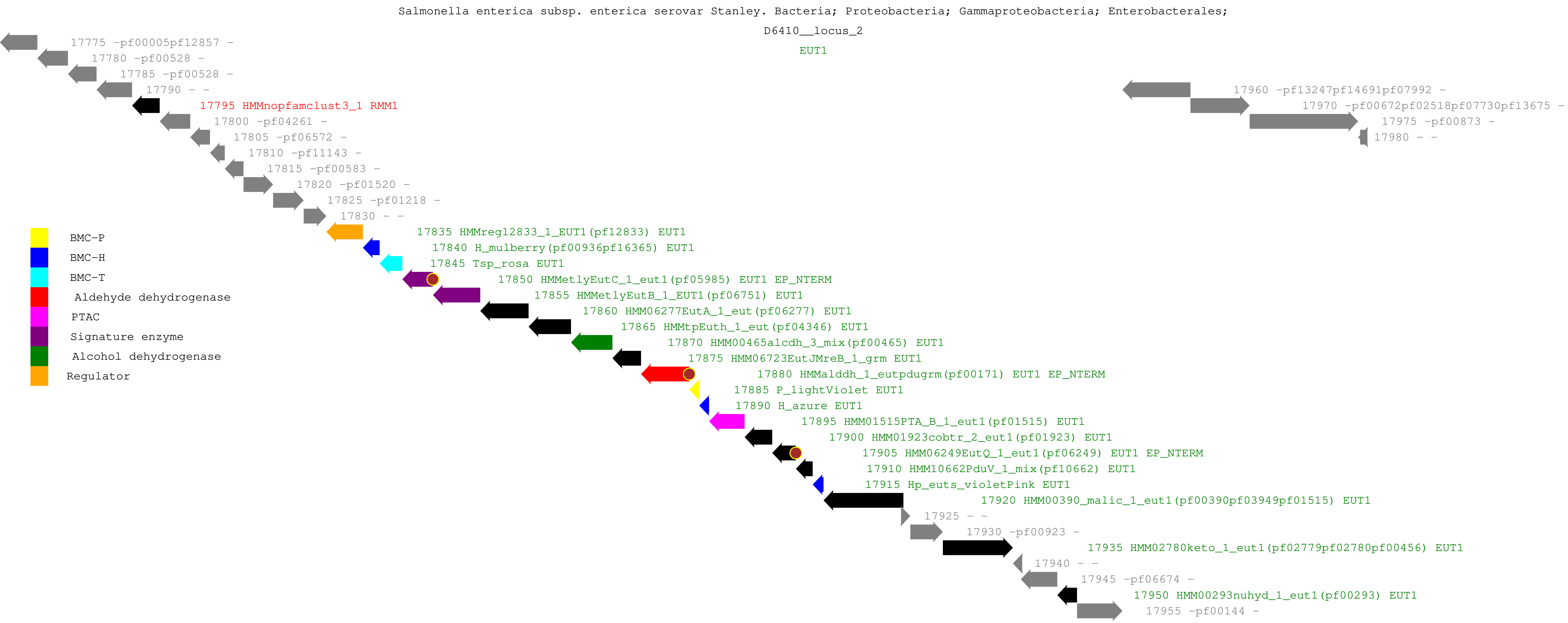
Salmonella enterica subsp. enterica serovar Stanley. Bacteria; Proteobacteria; Gammaproteobacteria; Enterobacterales;
D6410__locus_2
EUT1
17775 -pf00005pf12857 -
17780 -pf00528 -
17785 -pf00528 -
17790 - -
17795 HMMnopfamclust3_1 RMM1
17800 -pf04261 -
17805 -pf06572 -
17810 -pf11143 -
17815 -pf00583 -
17820 -pf01520 -
17825 -pf01218 -
17830 - -
17835 HMMreg12833_1_EUT1(pf12833) EUT1
17840 H_mulberry(pf00936pf16365) EUT1
17845 Tsp_rosa EUT1
17850 HMMetlyEutC_1_eut1(pf05985) EUT1 EP_NTERM
17855 HMMetlyEutB_1_EUT1(pf06751) EUT1
17860 HMM06277EutA_1_eut(pf06277) EUT1
17865 HMMtpEuth_1_eut(pf04346) EUT1
17870 HMM00465alcdh_3_mix(pf00465) EUT1
17875 HMM06723EutJMreB_1_grm EUT1
17880 HMMalddh_1_eutpdugrm(pf00171) EUT1 EP_NTERM
17885 P_lightViolet EUT1
17890 H_azure EUT1
17895 HMM01515PTA_B_1_eut1(pf01515) EUT1
17900 HMM01923cobtr_2_eut1(pf01923) EUT1
17905 HMM06249EutQ_1_eut1(pf06249) EUT1 EP_NTERM
17910 HMM10662PduV_1_mix(pf10662) EUT1
17915 Hp_euts_violetPink EUT1
17920 HMM00390_malic_1_eut1(pf00390pf03949pf01515) EUT1
17925 - -
17930 -pf00923 -
17935 HMM02780keto_1_eut1(pf02779pf02780pf00456) EUT1
17940 - -
17945 -pf06674 -
17950 HMM00293nuhyd_1_eut1(pf00293) EUT1
17955 -pf00144 -
17960 -pf13247pf14691pf07992 -
17970 -pf00672pf02518pf07730pf13675 -
17975 -pf00873 -
17980 - -
BMC-P
BMC-H
BMC-T
Aldehyde dehydrogenase
PTAC
Signature enzyme
Alcohol dehydrogenase
Regulator

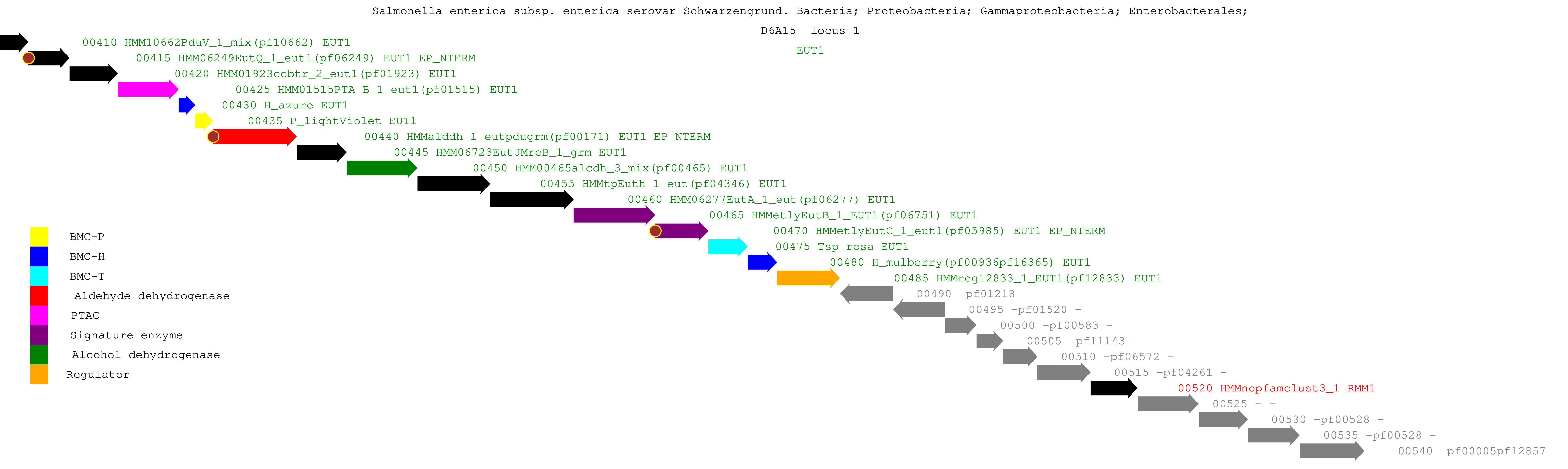
Salmonella enterica subsp. enterica serovar Schwarzengrund. Bacteria; Proteobacteria; Gammaproteobacteria; Enterobacterales;
D6A15__locus_1
EUT1
00410 HMM10662PduV_1_mix(pf10662) EUT1
00415 HMM06249EutQ_1_eut1(pf06249) EUT1 EP_NTERM
00420 HMM01923cobtr_2_eut1(pf01923) EUT1
00425 HMM01515PTA_B_1_eut1(pf01515) EUT1
00430 H_azure EUT1
00435 P_lightViolet EUT1
00440 HMMaldh_1_eutpdugrm(pf00171) EUT1 EP_NTERM
00445 HMM06723EutJMreB_1_grm EUT1
00450 HMM00465alcdh_3_mix(pf00465) EUT1
00455 HMMtpEuth_1_eut(pf04346) EUT1
00460 HMM06277EutA_1_eut(pf06277) EUT1
00465 HMMetlyEutB_1_EUT1(pf06751) EUT1
00470 HMMetlyEutC_1_eut1(pf05985) EUT1 EP_NTERM
00475 Tsp_rosa EUT1
00480 H_mulberry(pf00936pf16365) EUT1
00485 HMMreg12833_1_EUT1(pf12833) EUT1
00490 -pf01218 -
00495 -pf01520 -
00500 -pf00583 -
00505 -pf11143 -
00510 -pf06572 -
00515 -pf04261 -
00520 HMMnopfamclust3_1 RMM1
00525 - -
00530 -pf00528 -
00535 -pf00528 -
00540 -pf00005pf12857 -
BMC-P
BMC-H
BMC-T
Aldehyde dehydrogenase
PTAC
Signature enzyme
Alcohol dehydrogenase
Regulator

Salmonella enterica I. Bacteria; Proteobacteria; Gammaproteobacteria; Enterobacterales;

D6J43__locus_2

EUT1

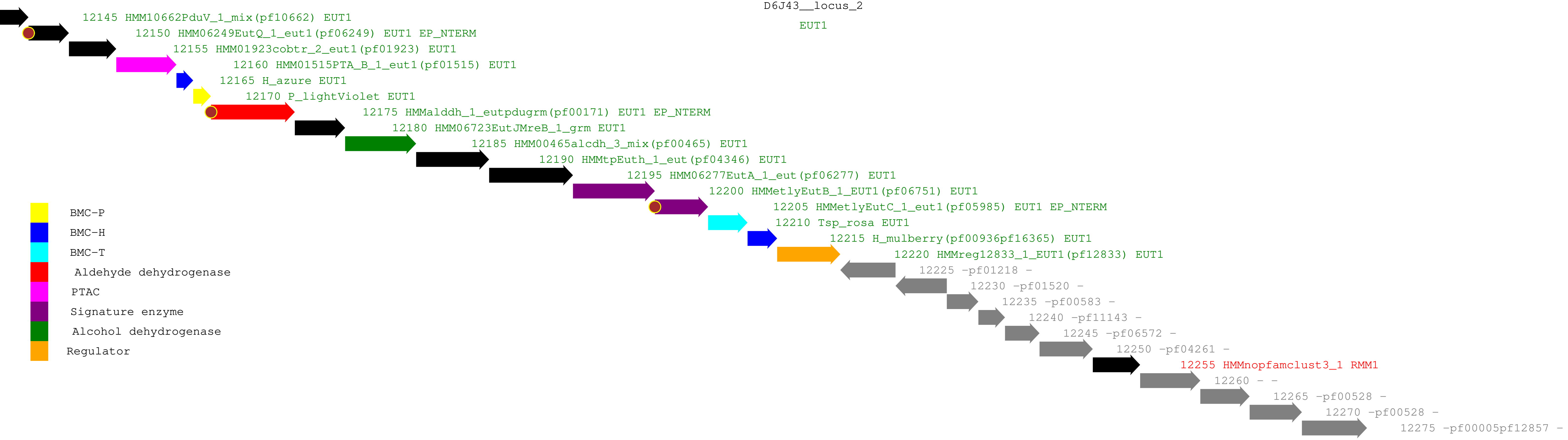

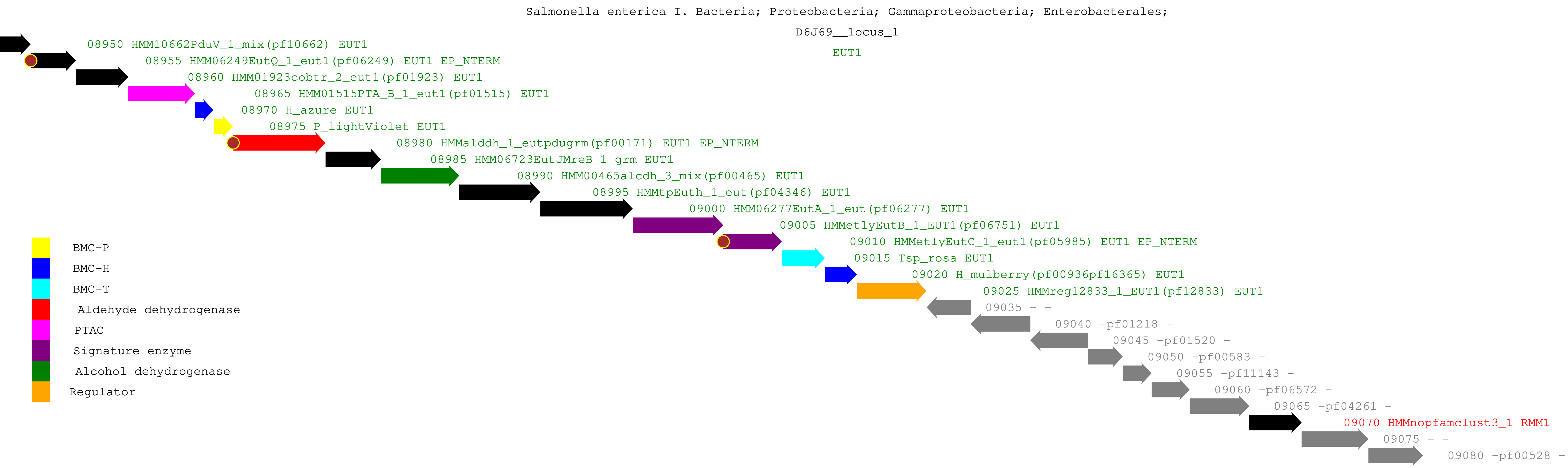
Salmonella enterica I. Bacteria; Proteobacteria; Gammaproteobacteria; Enterobacterales;
D6J69__locus_1
EUT1
08950 HMM10662PduV_1_mix(pf10662) EUT1
08955 HMM06249EutQ_1_eut1(pf06249) EUT1 EP_NTERM
08960 HMM01923cobtr_2_eut1(pf01923) EUT1
08965 HMM01515PTA_B_1_eut1(pf01515) EUT1
08970 H_azure EUT1
08975 P_lightViolet EUT1
08980 HMMalddh_1_eutpdugrm(pf00171) EUT1 EP_NTERM
08985 HMM06723EutJMreB_1_grm EUT1
08990 HMM00465alcdh_3_mix(pf00465) EUT1
08995 HMMtpEuth_1_eut(pf04346) EUT1
09000 HMM06277EutA_1_eut(pf06277) EUT1
09005 HMMetlyEutB_1_EUT1(pf06751) EUT1
09010 HMMetlyEutC_1_eut1(pf05985) EUT1 EP_NTERM
09015 Tsp_rosa EUT1
09020 H_mulberry(pf00936pf16365) EUT1
09025 HMMreg12833_1_EUT1(pf12833) EUT1
09035 - -
09040 -pf01218 -
09045 -pf01520 -
09050 -pf00583 -
09055 -pf11143 -
09060 -pf06572 -
09065 -pf04261 -
09070 HMMnopfamclust3_1 RMM1
09075 - -
09080 -pf00528 -
BMC-P
BMC-H
BMC-T
Aldehyde dehydrogenase
PTAC
Signature enzyme
Alcohol dehydrogenase
Regulator

Salmonella enterica subsp. enterica serovar Java. Bacteria; Proteobacteria; Gammaproteobacteria; Enterobacterales;

D6K54__locus_1

EUT1

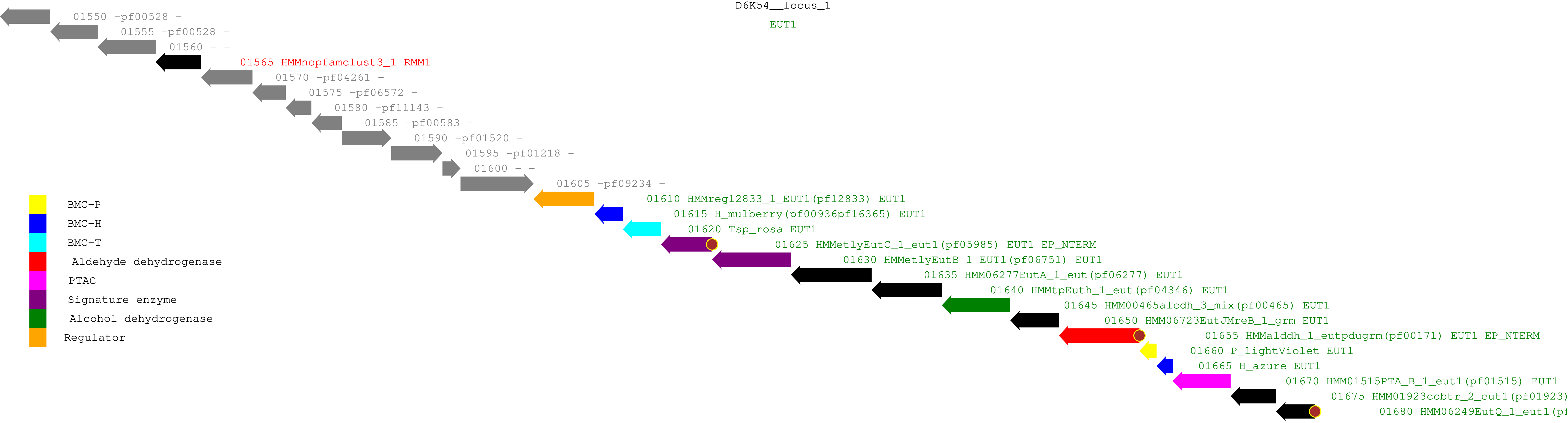

Salmonella enterica I. Bacteria; Proteobacteria; Gammaproteobacteria; Enterobacterales;

D6K86__locus_1

EUT1

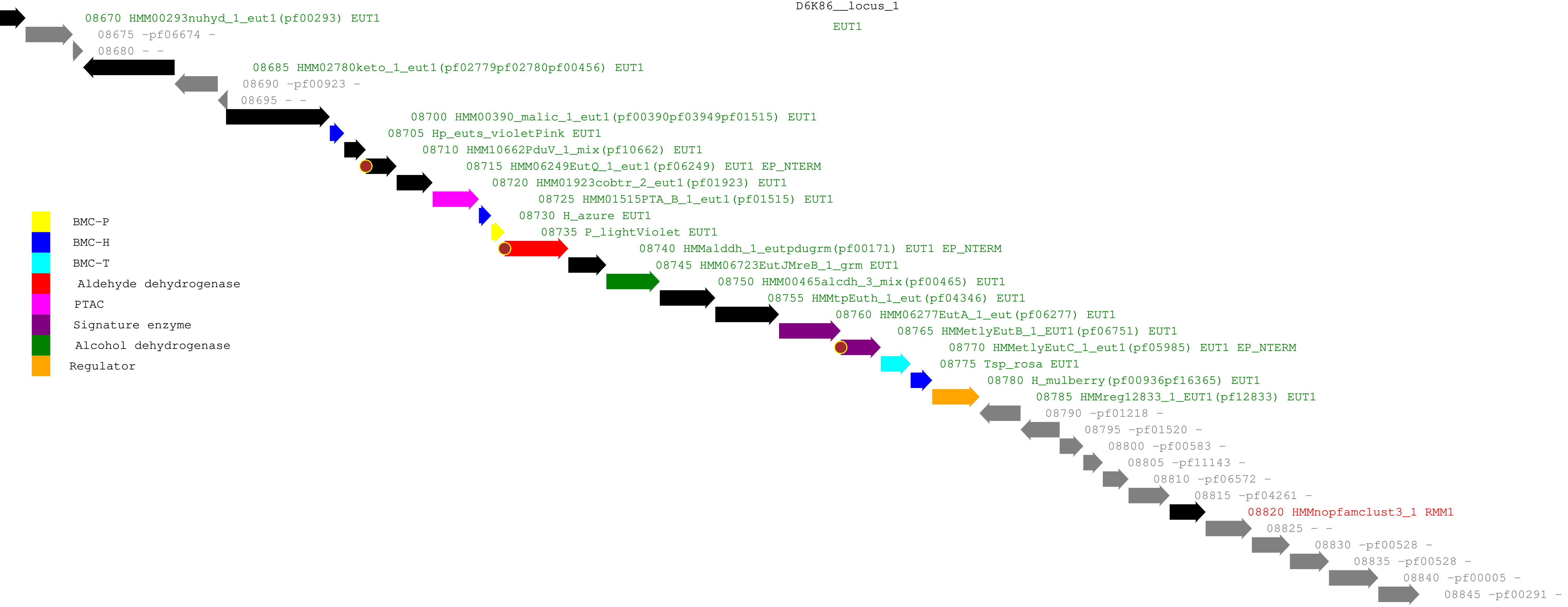

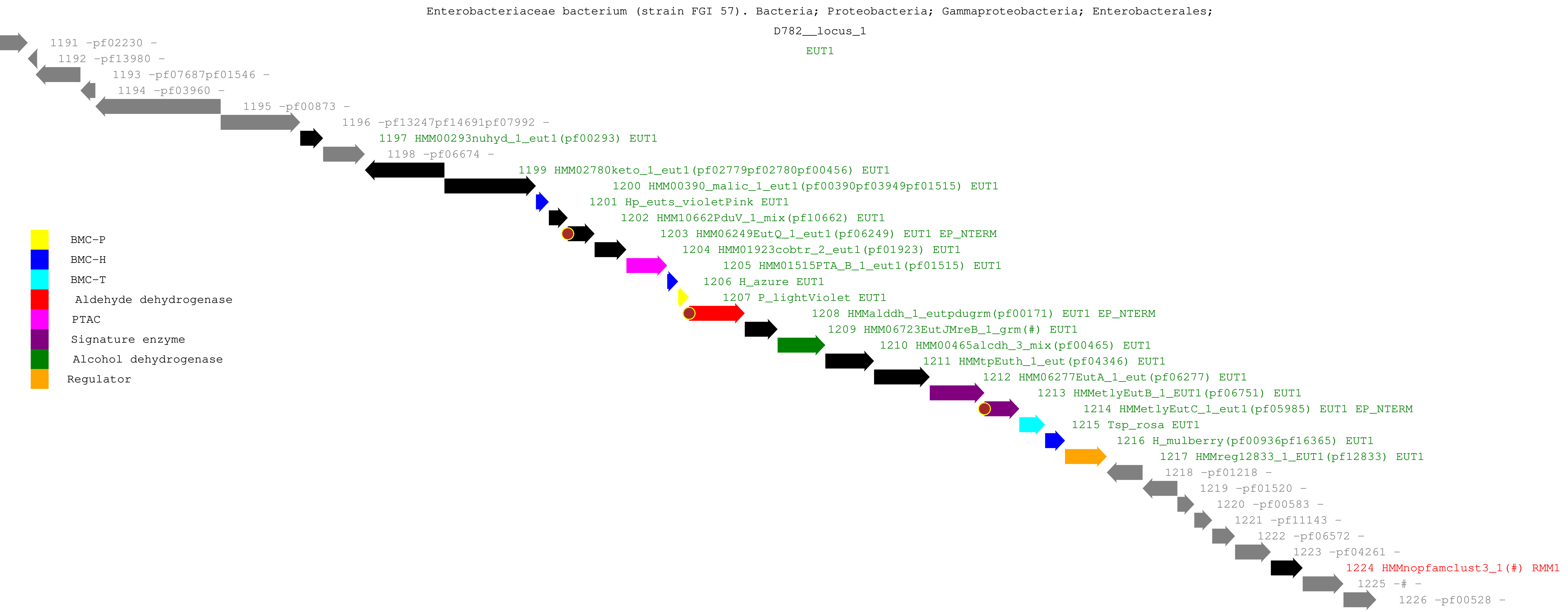
Enterobacteriaceae bacterium (strain FGI 57). Bacteria; Proteobacteria; Gammaproteobacteria; Enterobacterales;
D782__locus_1
EUT1
1191 -pf02230 -
1192 -pf13980 -
1193 -pf07687pf01546 -
1194 -pf03960 -
1195 -pf00873 -
1196 -pf13247pf14691pf07992 -
1197 HMM00293nuhyd_1_eut1(pf00293) EUT1
1198 -pf06674 -
1199 HMM02780keto_1_eut1(pf02779pf02780pf00456) EUT1
1200 HMM00390_malic_1_eut1(pf00390pf03949pf01515) EUT1
1201 Hp_euts_violetPink EUT1
1202 HMM10662PduV_1_mix(pf10662) EUT1
1203 HMM06249EutQ_1_eut1(pf06249) EUT1 EP_NTERM
1204 HMM01923cobtr_2_eut1(pf01923) EUT1
1205 HMM01515PTA_B_1_eut1(pf01515) EUT1
1206 H_azure EUT1
1207 P_lightViolet EUT1
1208 HMMaldh_1_eutpdugrm(pf00171) EUT1 EP_NTERM
1209 HMM06723EutJMreB_1_grm(#) EUT1
1210 HMM00465alcdh_3_mix(pf00465) EUT1
1211 HMMtpEuth_1_eut(pf04346) EUT1
1212 HMM06277EutA_1_eut(pf06277) EUT1
1213 HMMetlyEutB_1_EUT1(pf06751) EUT1
1214 HMMetlyEutC_1_eut1(pf05985) EUT1 EP_NTERM
1215 Tsp_rosa EUT1
1216 H_mulberry(pf00936pf16365) EUT1
1217 HMMreg12833_1_EUT1(pf12833) EUT1
1218 -pf01218 -
1219 -pf01520 -
1220 -pf00583 -
1221 -pf11143 -
1222 -pf06572 -
1223 -pf04261 -
1224 HMMnopfamclust3_1(#) RMM1
1225 -# -
1226 -pf00528 -
BMC-P
BMC-H
BMC-T
Aldehyde dehydrogenase
PTAC
Signature enzyme
Alcohol dehydrogenase
Regulator

Citrobacter sp. MH181794. Bacteria; Proteobacteria; Gammaproteobacteria; Enterobacterales;

D7I40__locus_2

EUT1

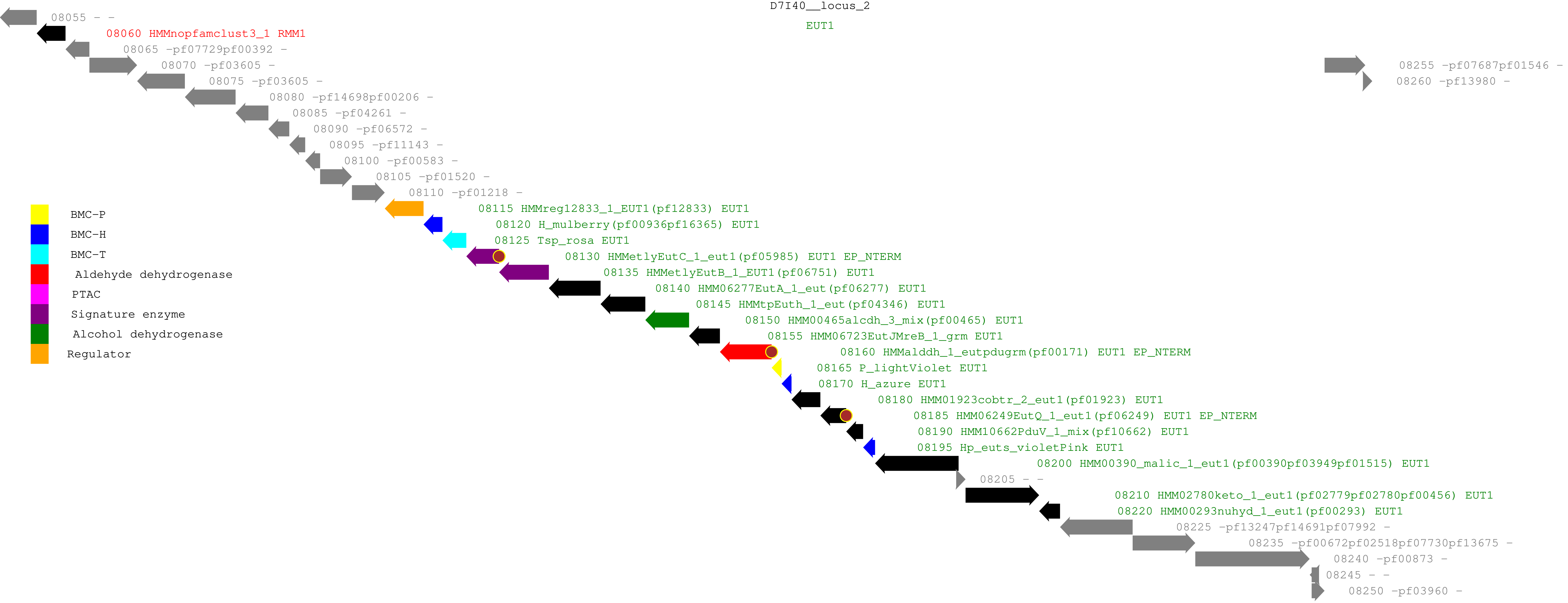

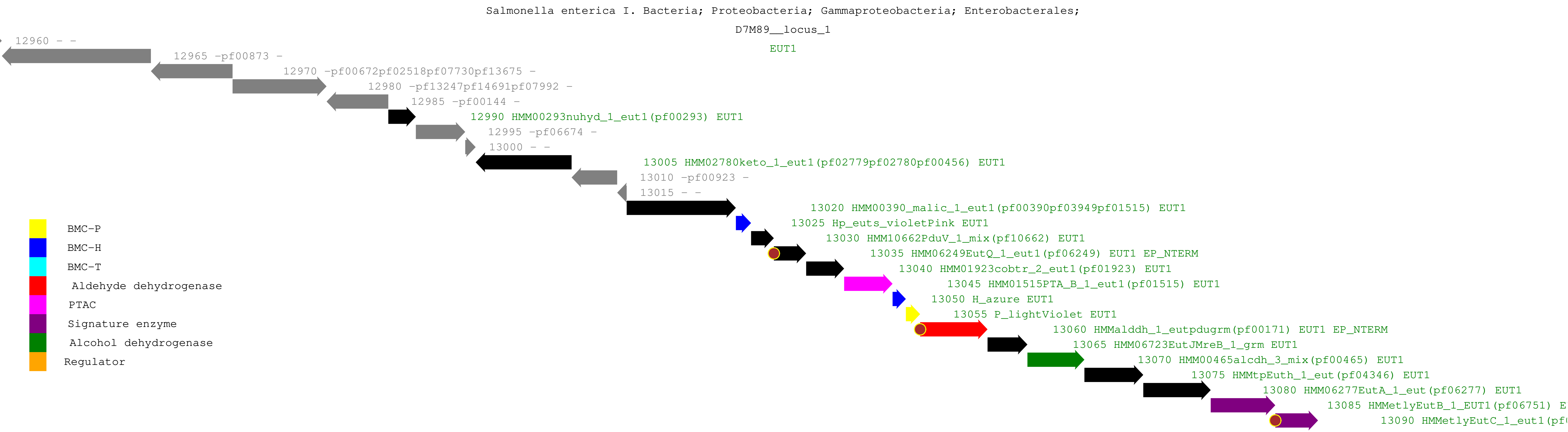
Salmonella enterica I. Bacteria; Proteobacteria; Gammaproteobacteria; Enterobacterales;
D7M89__locus_1
EUT1
12960 - -
12965 -pf00873 -
12970 -pf00672pf02518pf07730pf13675 -
12980 -pf13247pf14691pf07992 -
12985 -pf00144 -
12990 HMM00293nuhyd_1_eut1(pf00293) EUT1
12995 -pf06674 -
13000 - -
13005 HMM02780keto_1_eut1(pf02779pf02780pf00456) EUT1
13010 -pf00923 -
13015 - -
13020 HMM00390_malic_1_eut1(pf00390pf03949pf01515) EUT1
13025 Hp_euts_violetPink EUT1
13030 HMM10662PduV_1_mix(pf10662) EUT1
13035 HMM06249EutQ_1_eut1(pf06249) EUT1 EP_NTERM
13040 HMM01923cobtr_2_eut1(pf01923) EUT1
13045 HMM01515PTA_B_1_eut1(pf01515) EUT1
13050 H_azure EUT1
13055 P_lightViolet EUT1
13060 HMMalddh_1_eutpdugrm(pf00171) EUT1 EP_NTERM
13065 HMM06723EutJMreB_1_grm EUT1
13070 HMM00465alcdh_3_mix(pf00465) EUT1
13075 HMMtpEuth_1_eut(pf04346) EUT1
13080 HMM06277EutA_1_eut(pf06277) EUT1
13085 HMMetlyEutB_1_EUT1(pf06751) E
13090 HMMetlyEutC_1_eut1(pf
BMC-P
BMC-H
BMC-T
Aldehyde dehydrogenase
PTAC
Signature enzyme
Alcohol dehydrogenase
Regulator

Salmonella hadar. Bacteria; Proteobacteria; Gammaproteobacteria; Enterobacterales;

D7N39__locus_1

EUT1

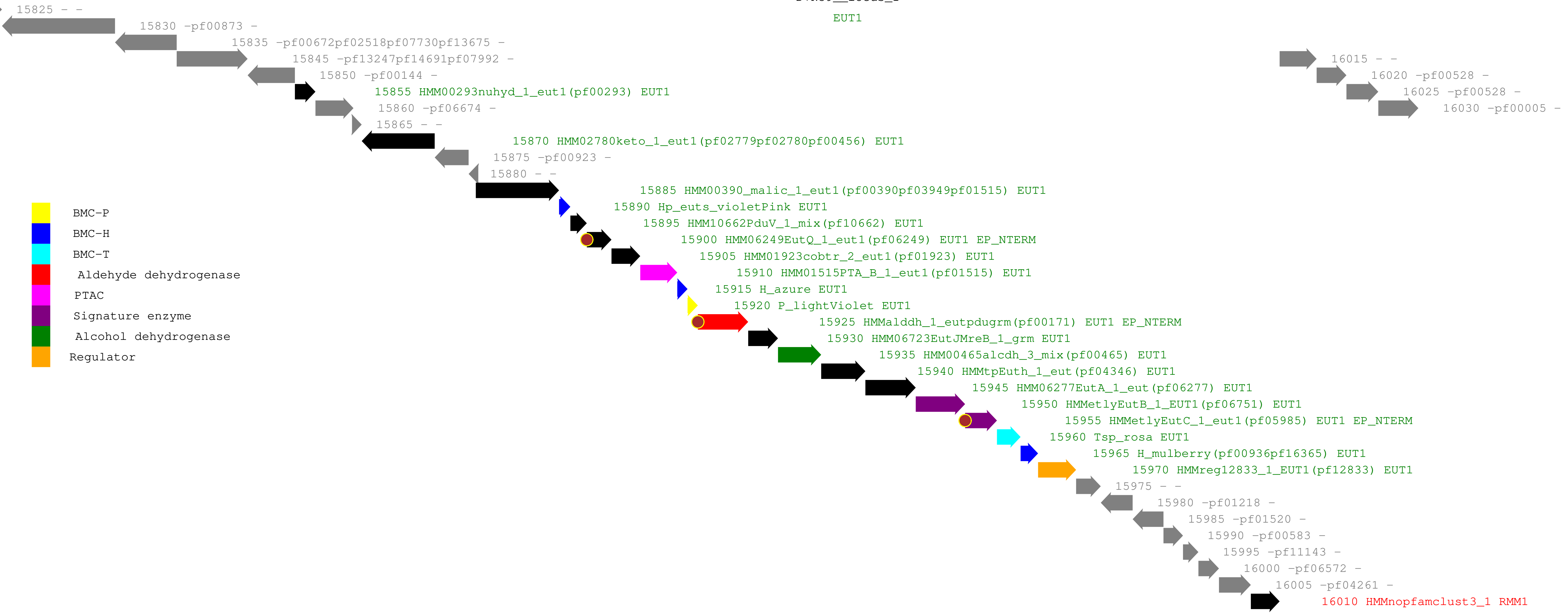

Salmonella enterica I. Bacteria; Proteobacteria; Gammaproteobacteria; Enterobacterales;

D7N51__locus_2

EUT1

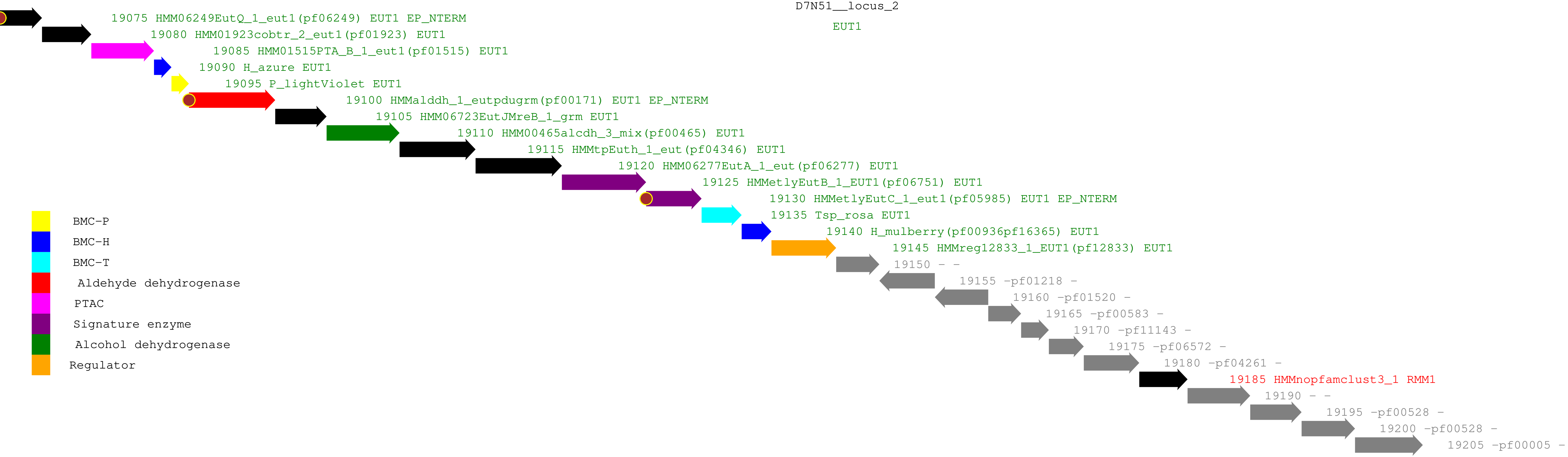

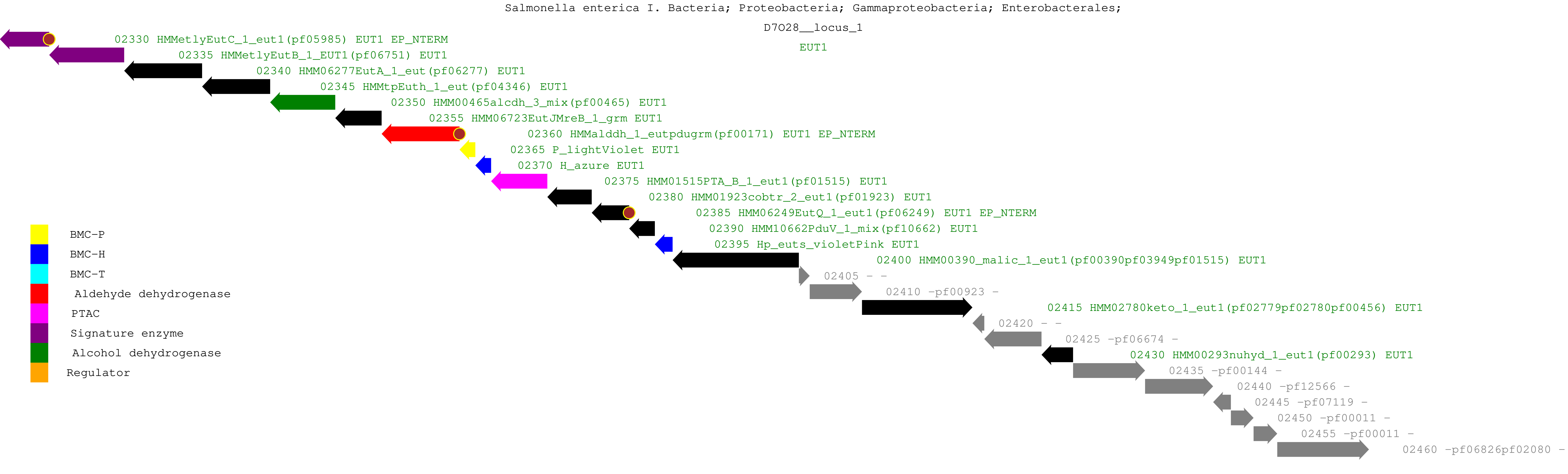
Salmonella enterica I. Bacteria; Proteobacteria; Gammaproteobacteria; Enterobacterales;
D7028__locus_1
EUT1
02330 HMMetlyEutC_1_eut1(pf05985) EUT1 EP_NTERM
02335 HMMetlyEutB_1_EUT1(pf06751) EUT1
02340 HMM06277EutA_1_eut(pf06277) EUT1
02345 HMMtpEuth_1_eut(pf04346) EUT1
02350 HMM00465alcdh_3_mix(pf00465) EUT1
02355 HMM06723EutJMreB_1_grm EUT1
02360 HMMalddh_1_eutpdugrm(pf00171) EUT1 EP_NTERM
02365 P_lightViolet EUT1
02370 H_azure EUT1
02375 HMM01515PTA_B_1_eut1(pf01515) EUT1
02380 HMM01923cobtr_2_eut1(pf01923) EUT1
02385 HMM06249EutQ_1_eut1(pf06249) EUT1 EP_NTERM
02390 HMM10662PduV_1_mix(pf10662) EUT1
02395 Hp_euts_violetPink EUT1
02400 HMM00390_malic_1_eut1(pf00390pf03949pf01515) EUT1
02405 - -
02410 -pf00923 -
02415 HMM02780keto_1_eut1(pf02779pf02780pf00456) EUT1
02420 - -
02425 -pf06674 -
02430 HMM00293nuhyd_1_eut1(pf00293) EUT1
02435 -pf00144 -
02440 -pf12566 -
02445 -pf07119 -
02450 -pf00011 -
02455 -pf00011 -
02460 -pf06826pf02080 -
BMC-P
BMC-H
BMC-T
Aldehyde dehydrogenase
PTAC
Signature enzyme
Alcohol dehydrogenase
Regulator

Salmonella enterica subsp. enterica serovar Saintpaul. Bacteria; Proteobacteria; Gammaproteobacteria; Enterobacterales;

D7O55__locus_1

EUT1

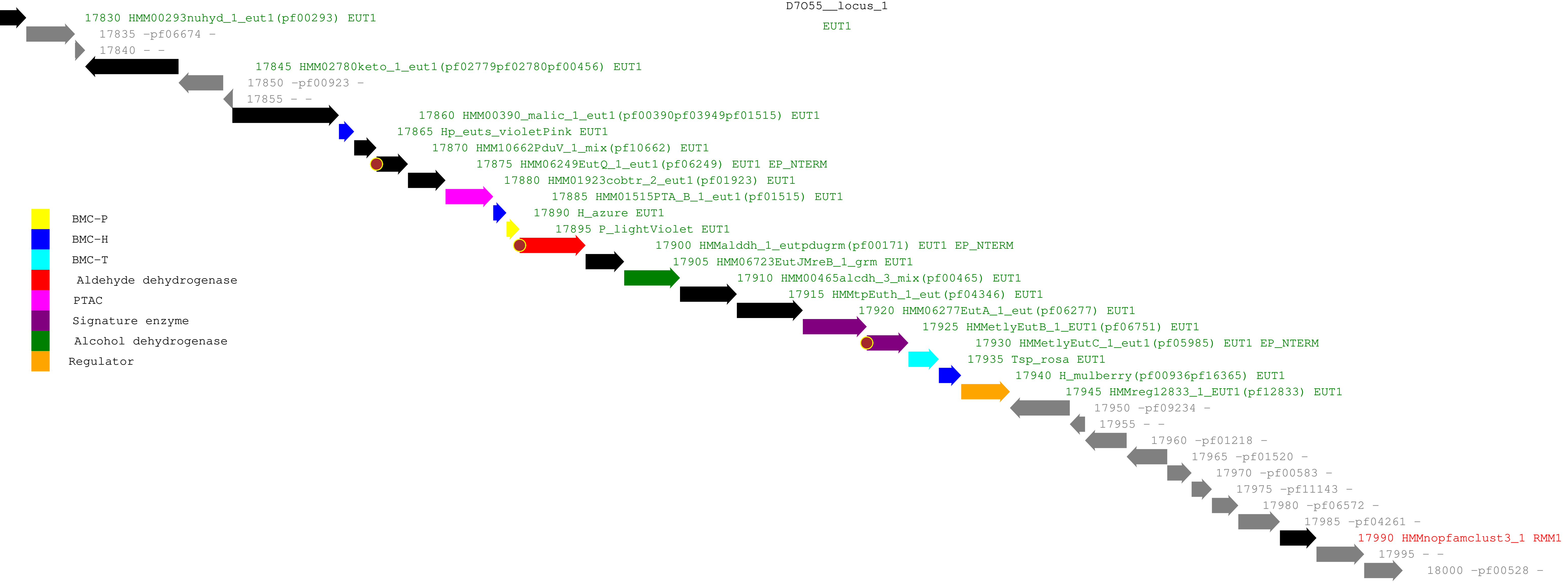

Buttiauxella sp. 3AFRM03. Bacteria; Proteobacteria; Gammaproteobacteria; Enterobacterales;

D8682__locus_2

EUT1

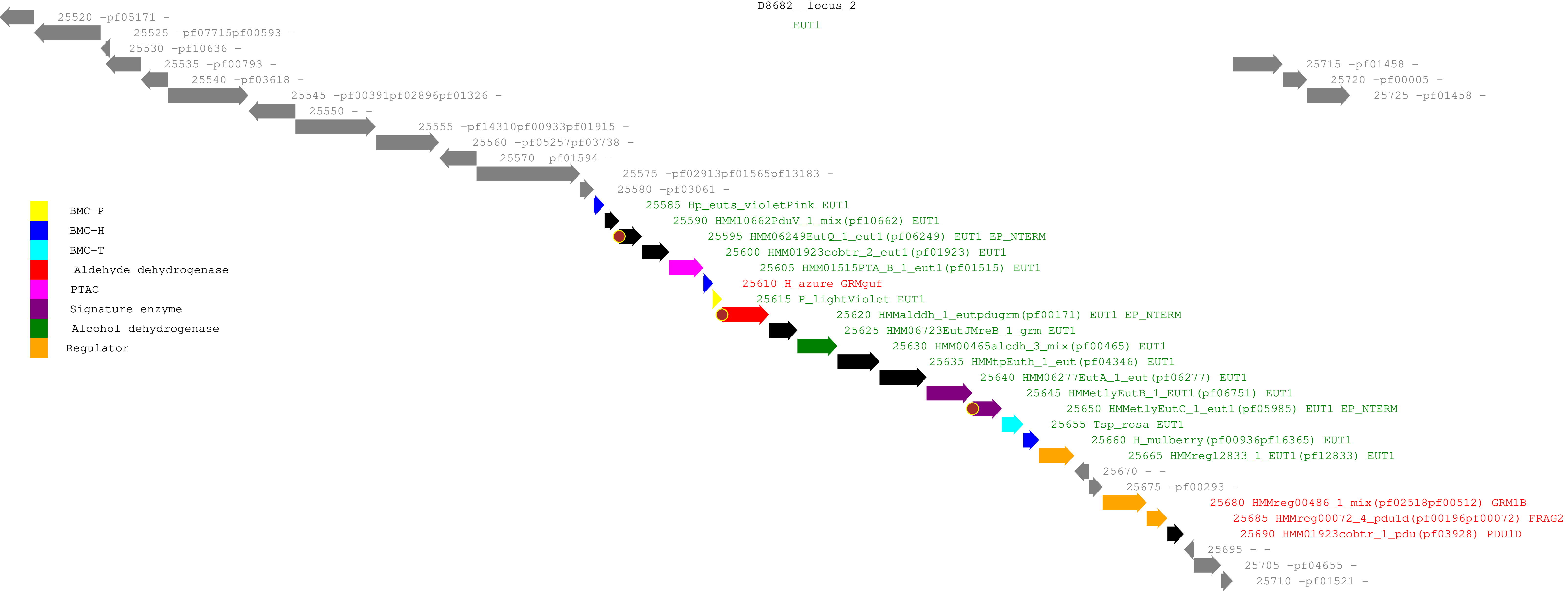

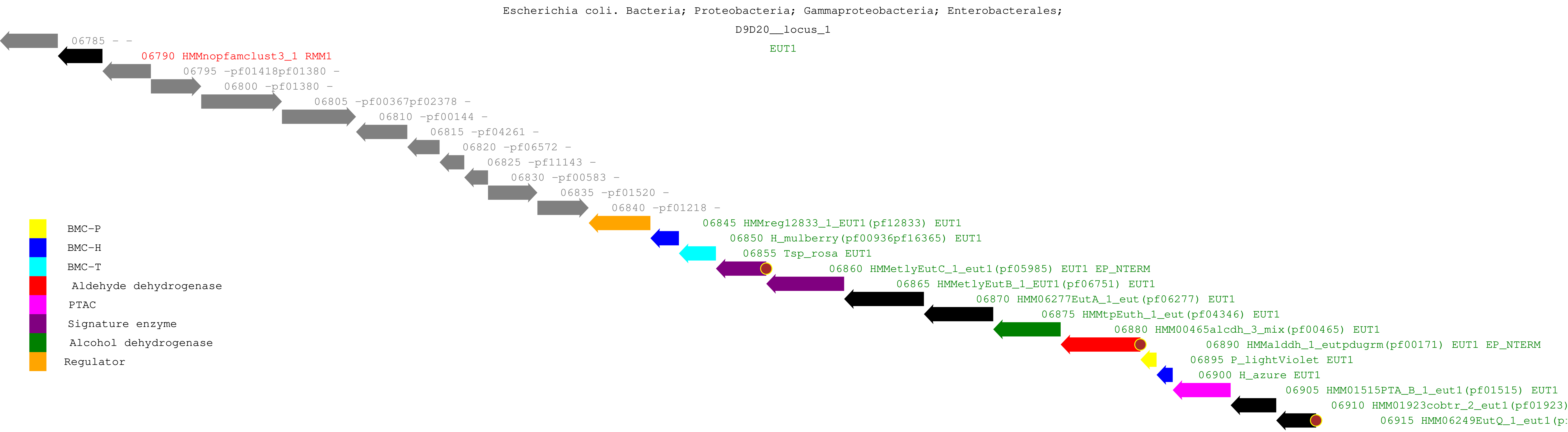
Escherichia coli. Bacteria; Proteobacteria; Gammaproteobacteria; Enterobacterales;
D9D20__locus_1
EUT1
06785 - -
06790 HMMnopfamclust3_1 RMM1
06795 -pf01418pf01380 -
06800 -pf01380 -
06805 -pf00367pf02378 -
06810 -pf00144 -
06815 -pf04261 -
06820 -pf06572 -
06825 -pf11143 -
06830 -pf00583 -
06835 -pf01520 -
06840 -pf01218 -
06845 HMMreg12833_1_EUT1(pf12833) EUT1
06850 H_mulberry(pf00936pf16365) EUT1
06855 Tsp_rosa EUT1
06860 HMMetlyEutC_1_eut1(pf05985) EUT1 EP_NTERM
06865 HMMetlyEutB_1_EUT1(pf06751) EUT1
06870 HMM06277EutA_1_eut(pf06277) EUT1
06875 HMMtpEuth_1_eut(pf04346) EUT1
06880 HMM00465alcdh_3_mix(pf00465) EUT1
06890 HMMaldh_1_eutpdugrm(pf00171) EUT1 EP_NTERM
06895 P_lightViolet EUT1
06900 H_azure EUT1
06905 HMM01515PTA_B_1_eut1(pf01515) EUT1
06910 HMM01923cobtr_2_eut1(pf01923)
06915 HMM06249EutQ_1_eut1(p
BMC-P
BMC-H
BMC-T
Aldehyde dehydrogenase
PTAC
Signature enzyme
Alcohol dehydrogenase
Regulator

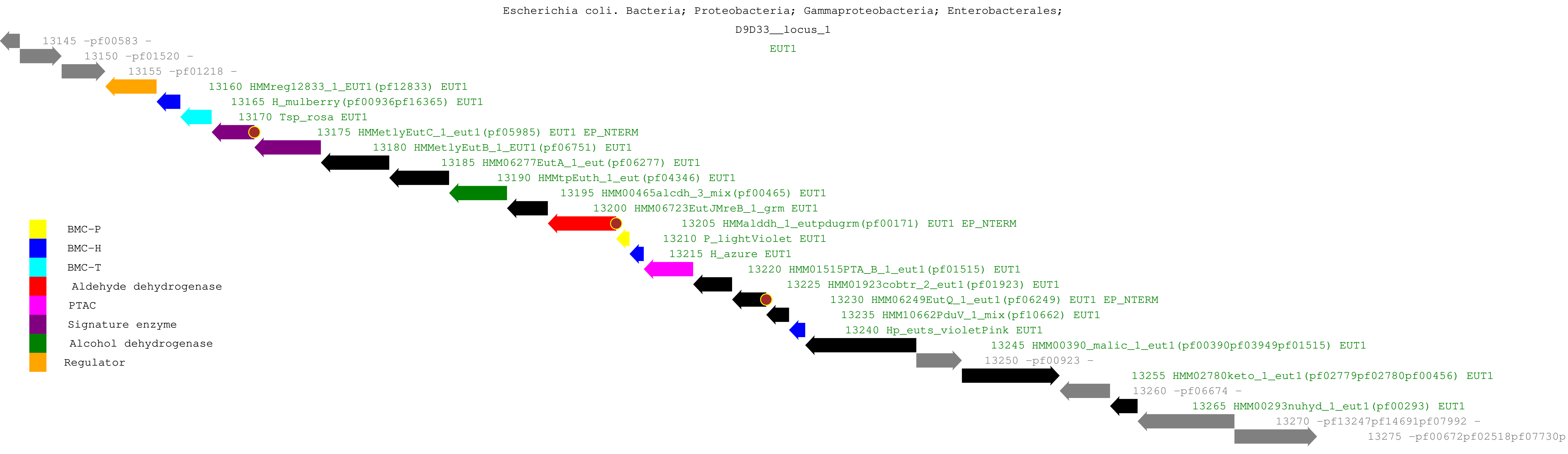
Escherichia coli. Bacteria; Proteobacteria; Gammaproteobacteria; Enterobacterales;
D9D33__locus_1
EUT1
13145 -pf00583 -
13150 -pf01520 -
13155 -pf01218 -
13160 HMMreg12833_1_EUT1(pf12833) EUT1
13165 H_mulberry(pf00936pf16365) EUT1
13170 Tsp_rosa EUT1
13175 HMMetlyEutC_1_eut1(pf05985) EUT1 EP_NTERM
13180 HMMetlyEutB_1_EUT1(pf06751) EUT1
13185 HMM06277EutA_1_eut(pf06277) EUT1
13190 HMMtpEuth_1_eut(pf04346) EUT1
13195 HMM00465alcdh_3_mix(pf00465) EUT1
13200 HMM06723EutJMreB_1_grm EUT1
13205 HMMalddh_1_eutpdugrm(pf00171) EUT1 EP_NTERM
13210 P_lightViolet EUT1
13215 H_azure EUT1
13220 HMM01515PTA_B_1_eut1(pf01515) EUT1
13225 HMM01923cobtr_2_eut1(pf01923) EUT1
13230 HMM06249EutQ_1_eut1(pf06249) EUT1 EP_NTERM
13235 HMM10662PduV_1_mix(pf10662) EUT1
13240 Hp_euts_violetPink EUT1
13245 HMM00390_malic_1_eut1(pf00390pf03949pf01515) EUT1
13250 -pf00923 -
13255 HMM02780keto_1_eut1(pf02779pf02780pf00456) EUT1
13260 -pf06674 -
13265 HMM00293nuhyd_1_eut1(pf00293) EUT1
13270 -pf13247pf14691pf07992 -
13275 -pf00672pf02518pf07730p
BMC-P
BMC-H
BMC-T
Aldehyde dehydrogenase
PTAC
Signature enzyme
Alcohol dehydrogenase
Regulator

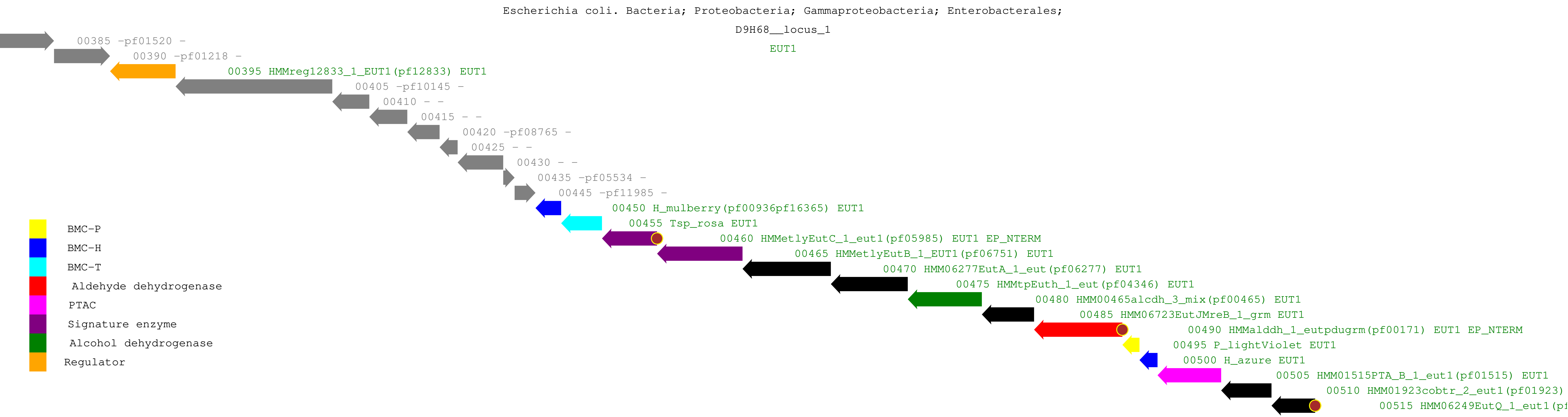

Escherichia coli. Bacteria; Proteobacteria; Gammaproteobacteria; Enterobacterales;
D9H68__locus_1
EUT1
00385 -pf01520 -
00390 -pf01218 -
00395 HMMreg12833_1_EUT1(pf12833) EUT1
00405 -pf10145 -
00410 - -
00415 - -
00420 -pf08765 -
00425 - -
00430 - -
00435 -pf05534 -
00445 -pf11985 -
00450 H_mulberry(pf00936pf16365) EUT1
00455 Tsp_rosa EUT1
00460 HMMetlyEutC_1_eut1(pf05985) EUT1 EP_NTERM
00465 HMMetlyEutB_1_EUT1(pf06751) EUT1
00470 HMM06277EutA_1_eut(pf06277) EUT1
00475 HMMtpEuth_1_eut(pf04346) EUT1
00480 HMM00465alcdh_3_mix(pf00465) EUT1
00485 HMM06723EutJMreB_1_grm EUT1
00490 HMMalddh_1_eutpdugrm(pf00171) EUT1 EP_NTERM
00495 P_lightViolet EUT1
00500 H_azure EUT1
00505 HMM01515PTA_B_1_eut1(pf01515) EUT1
00510 HMM01923cobtr_2_eut1(pf01923)
00515 HMM06249EutQ_1_eut1(p
BMC-P
BMC-H
BMC-T
Aldehyde dehydrogenase
PTAC
Signature enzyme
Alcohol dehydrogenase
Regulator

Escherichia coli. Bacteria; Proteobacteria; Gammaproteobacteria; Enterobacterales;

D9I18__locus_1

EUT1

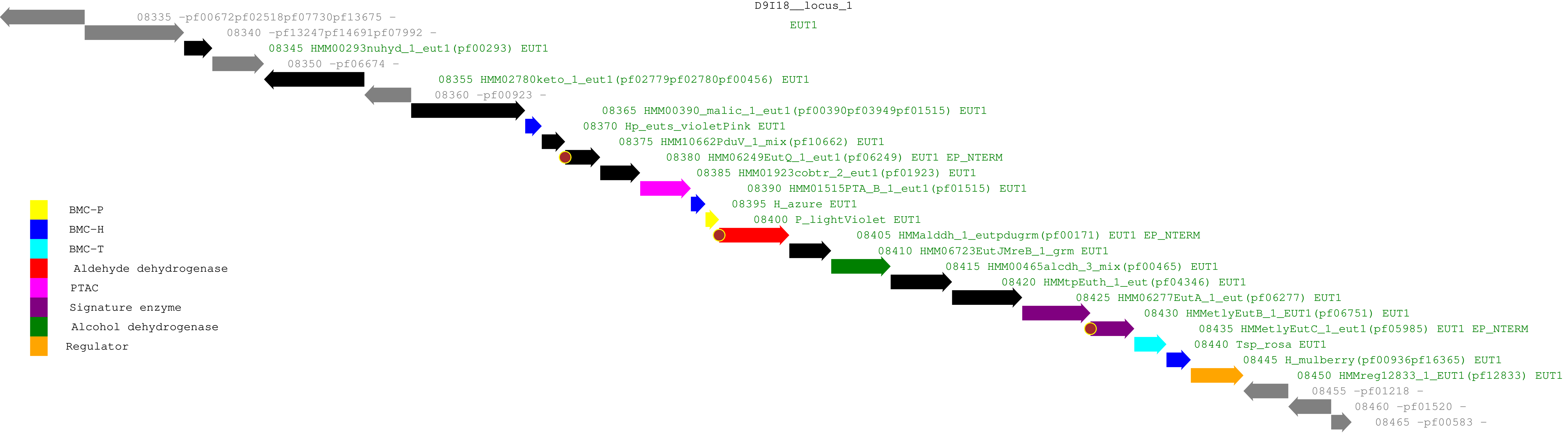

Escherichia coli. Bacteria; Proteobacteria; Gammaproteobacteria; Enterobacterales;

D9I20__locus_1

EUT1

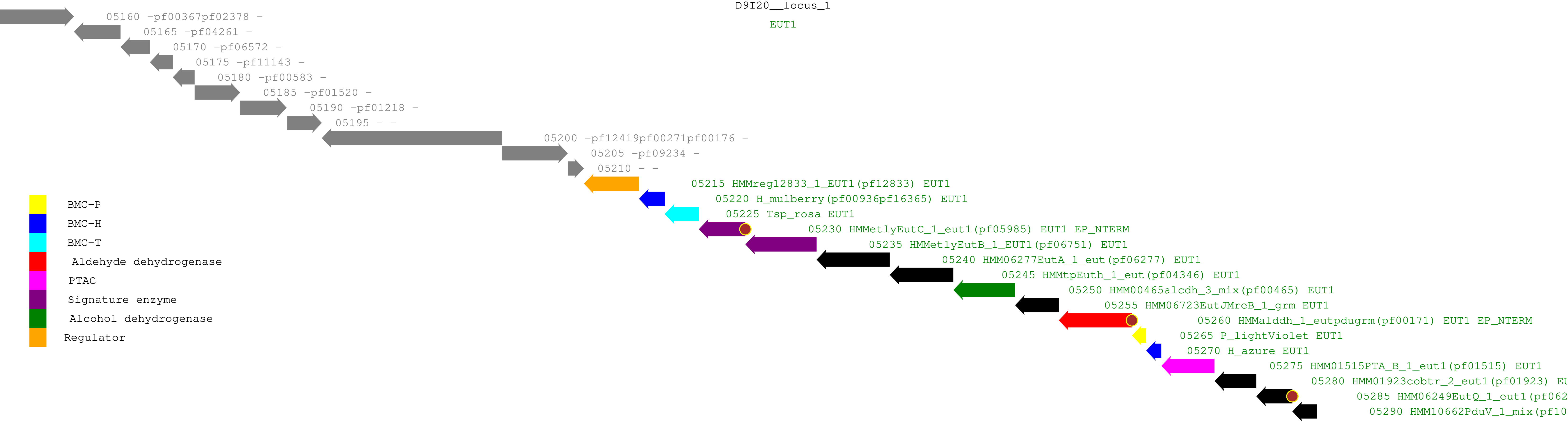

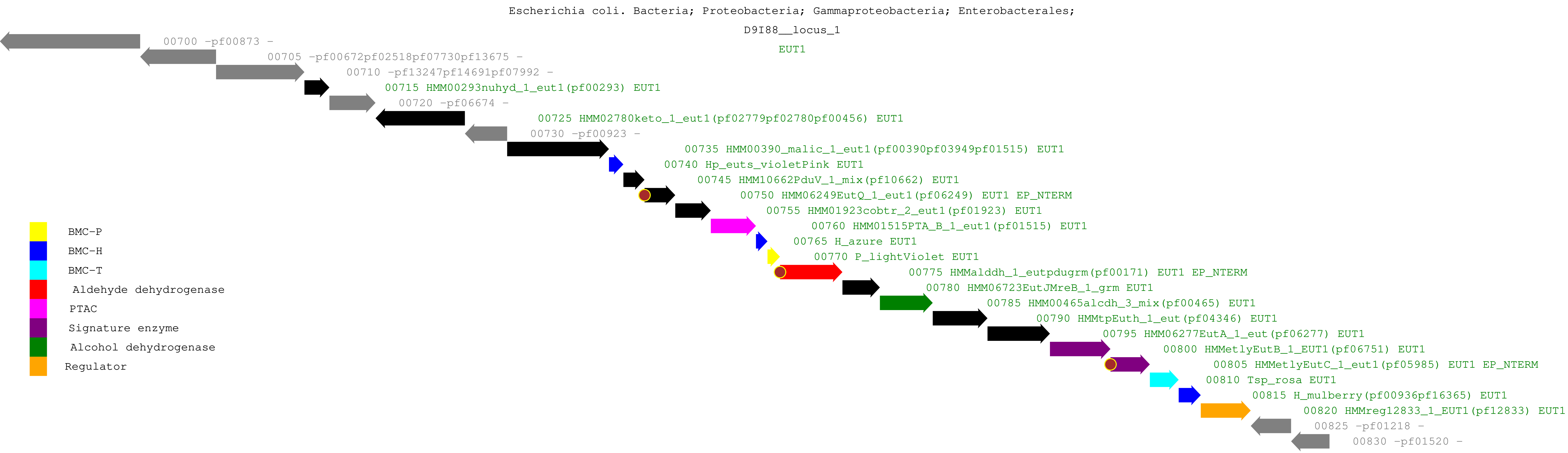
Escherichia coli. Bacteria; Proteobacteria; Gammaproteobacteria; Enterobacterales;
D9I88__locus_1
EUT1
00700 -pf00873 -
00705 -pf00672pf02518pf07730pf13675 -
00710 -pf13247pf14691pf07992 -
00715 HMM00293nuhyd_1_eut1(pf00293) EUT1
00720 -pf06674 -
00725 HMM02780keto_1_eut1(pf02779pf02780pf00456) EUT1
00730 -pf00923 -
00735 HMM00390_malic_1_eut1(pf00390pf03949pf01515) EUT1
00740 Hp_euts_violetPink EUT1
00745 HMM10662PduV_1_mix(pf10662) EUT1
00750 HMM06249EutQ_1_eut1(pf06249) EUT1 EP_NTERM
00755 HMM01923cobtr_2_eut1(pf01923) EUT1
00760 HMM01515PTA_B_1_eut1(pf01515) EUT1
00765 H_azure EUT1
00770 P_lightViolet EUT1
00775 HMMaldh_1_eutpdugrm(pf00171) EUT1 EP_NTERM
00780 HMM06723EutJMreB_1_grm EUT1
00785 HMM00465alcdh_3_mix(pf00465) EUT1
00790 HMMtpEuth_1_eut(pf04346) EUT1
00795 HMM06277EutA_1_eut(pf06277) EUT1
00800 HMMetlyEutB_1_EUT1(pf06751) EUT1
00805 HMMetlyEutC_1_eut1(pf05985) EUT1 EP_NTERM
00810 Tsp_rosa EUT1
00815 H_mulberry(pf00936pf16365) EUT1
00820 HMMreg12833_1_EUT1(pf12833) EUT1
00825 -pf01218 -
00830 -pf01520 -
BMC-P
BMC-H
BMC-T
Aldehyde dehydrogenase
PTAC
Signature enzyme
Alcohol dehydrogenase
Regulator

Escherichia coli. Bacteria; Proteobacteria; Gammaproteobacteria; Enterobacterales;

D9J44__locus_1

EUT1

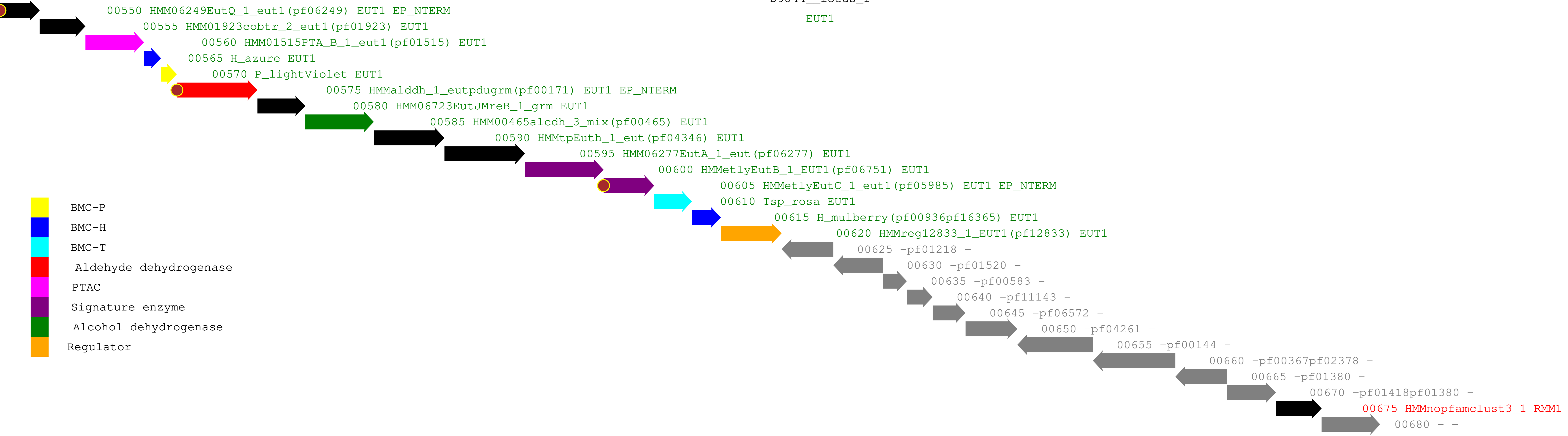

Salmonella enterica (Salmonella choleraesuis). Bacteria; Proteobacteria; Gammaproteobacteria; Enterobacterales;

D9031__locus_2

EUT1

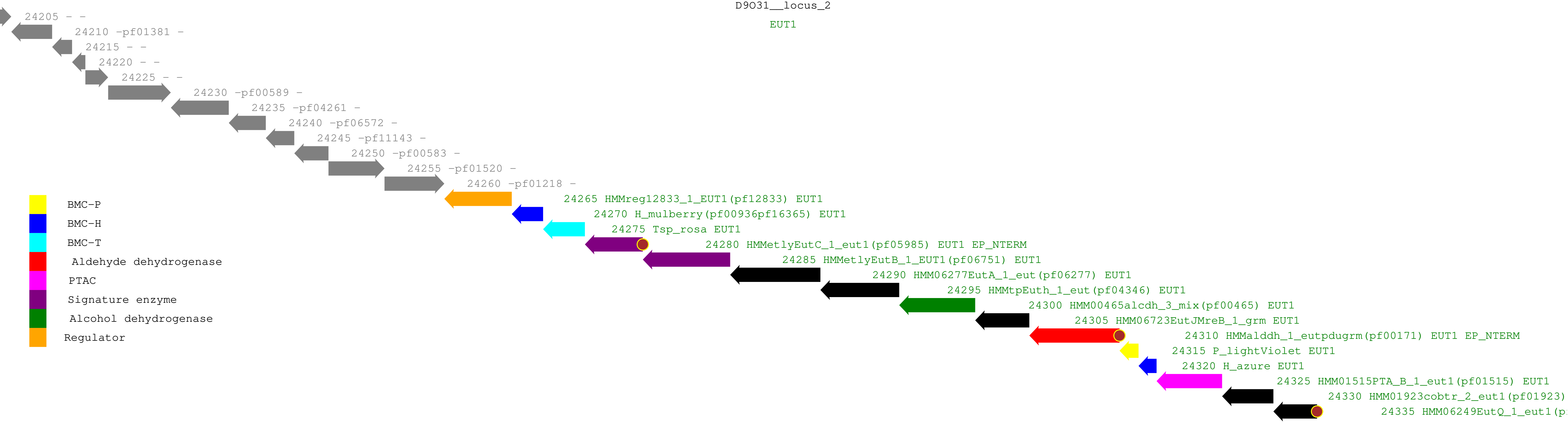

Salmonella enterica subsp. enterica serovar Bareilly. Bacteria; Proteobacteria; Gammaproteobacteria; Enterobacterales;

D9Q14__locus_2

EUT1

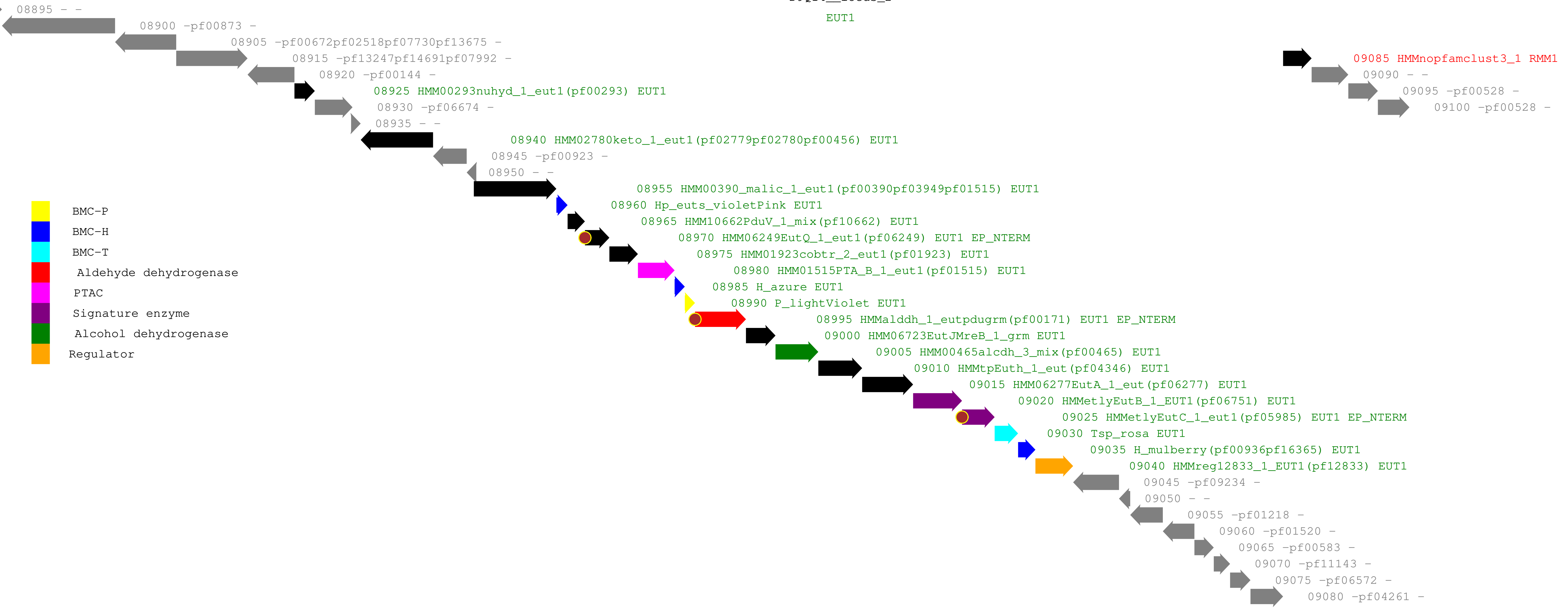

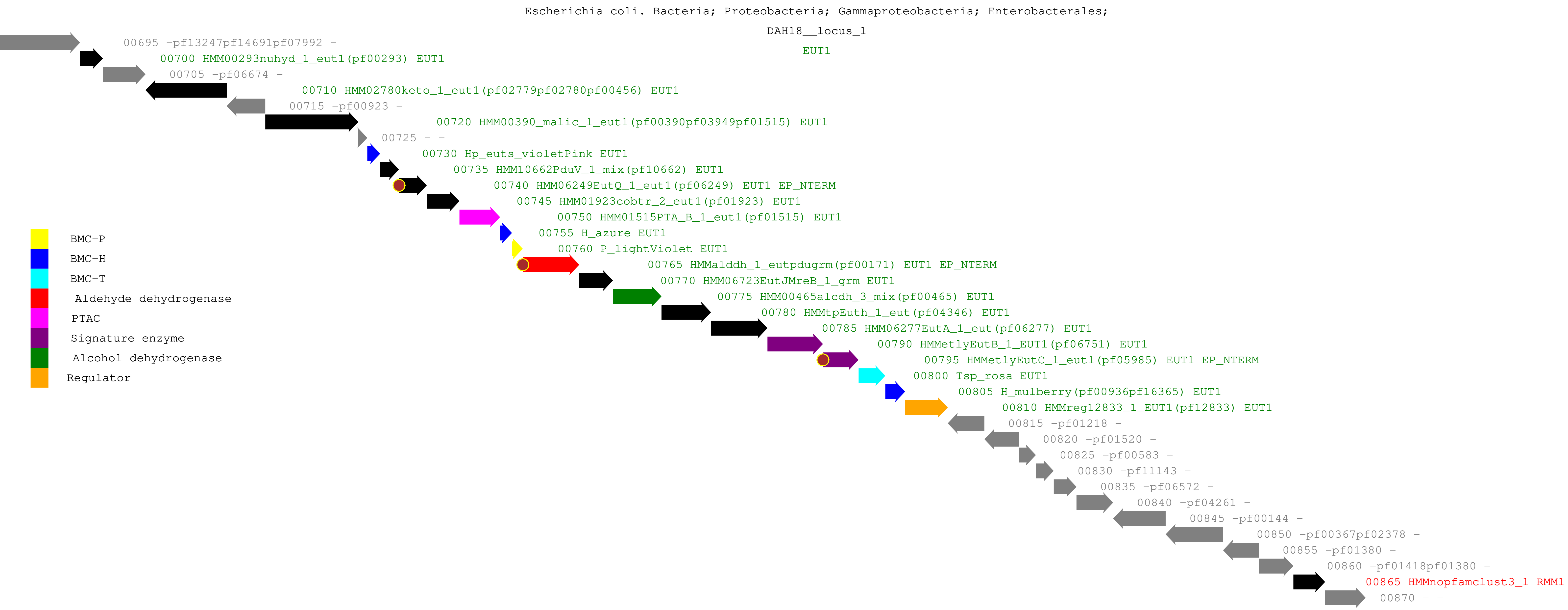
Escherichia coli. Bacteria; Proteobacteria; Gammaproteobacteria; Enterobacterales;
DAH18__locus_1
EUT1
00695 -pf13247pf14691pf07992 -
00700 HMM00293nuhyd_1_eut1(pf00293) EUT1
00705 -pf06674 -
00710 HMM02780keto_1_eut1(pf02779pf02780pf00456) EUT1
00715 -pf00923 -
00720 HMM00390_malic_1_eut1(pf00390pf03949pf01515) EUT1
00725 - -
00730 Hp_euts_violetPink EUT1
00735 HMM10662PduV_1_mix(pf10662) EUT1
00740 HMM06249EutQ_1_eut1(pf06249) EUT1 EP_NTERM
00745 HMM01923cobtr_2_eut1(pf01923) EUT1
00750 HMM01515PTA_B_1_eut1(pf01515) EUT1
00755 H_azure EUT1
00760 P_lightViolet EUT1
00765 HMMalddh_1_eutpdugrm(pf00171) EUT1 EP_NTERM
00770 HMM06723EutJMreB_1_grm EUT1
00775 HMM00465alcdh_3_mix(pf00465) EUT1
00780 HMMtpEuth_1_eut(pf04346) EUT1
00785 HMM06277EutA_1_eut(pf06277) EUT1
00790 HMMetlyEutB_1_EUT1(pf06751) EUT1
00795 HMMetlyEutC_1_eut1(pf05985) EUT1 EP_NTERM
00800 Tsp_rosa EUT1
00805 H_mulberry(pf00936pf16365) EUT1
00810 HMMreg12833_1_EUT1(pf12833) EUT1
00815 -pf01218 -
00820 -pf01520 -
00825 -pf00583 -
00830 -pf11143 -
00835 -pf06572 -
00840 -pf04261 -
00845 -pf00144 -
00850 -pf00367pf02378 -
00855 -pf01380 -
00860 -pf01418pf01380 -
00865 HMMnopfamclust3_1 RMM1
00870 - -
BMC-P
BMC-H
BMC-T
Aldehyde dehydrogenase
PTAC
Signature enzyme
Alcohol dehydrogenase
Regulator

Salmonella enterica I. Bacteria; Proteobacteria; Gammaproteobacteria; Enterobacterales;

DAX84__locus_1

EUT1

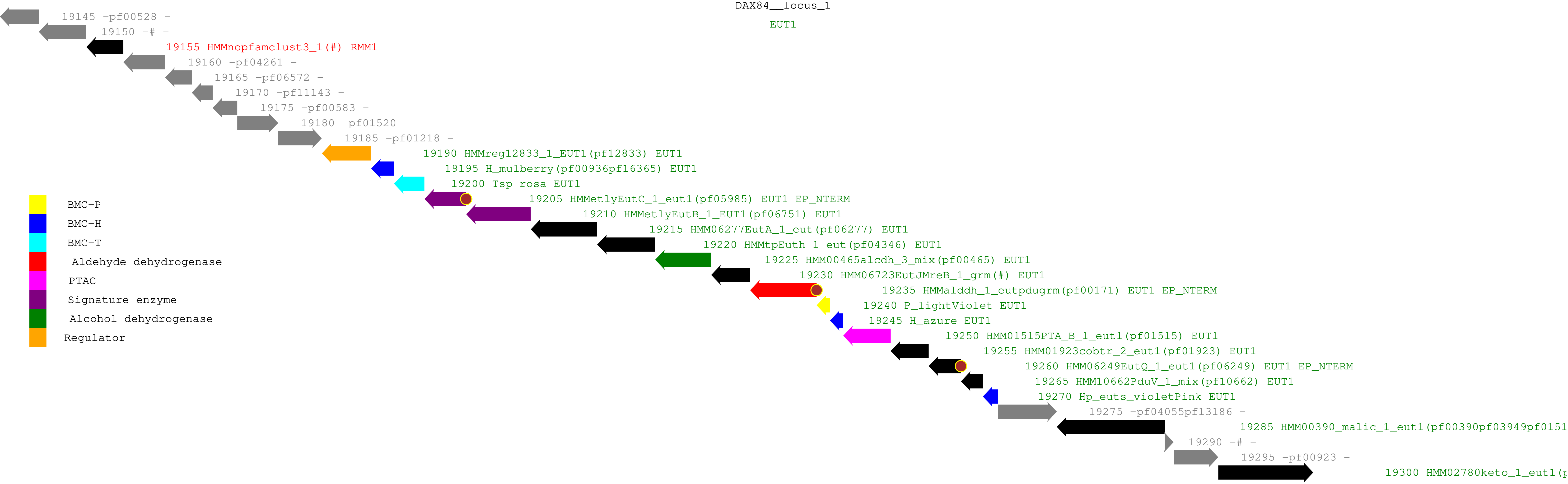

Salmonella enterica I. Bacteria; Proteobacteria; Gammaproteobacteria; Enterobacterales;

DAX91__locus_1

EUT1

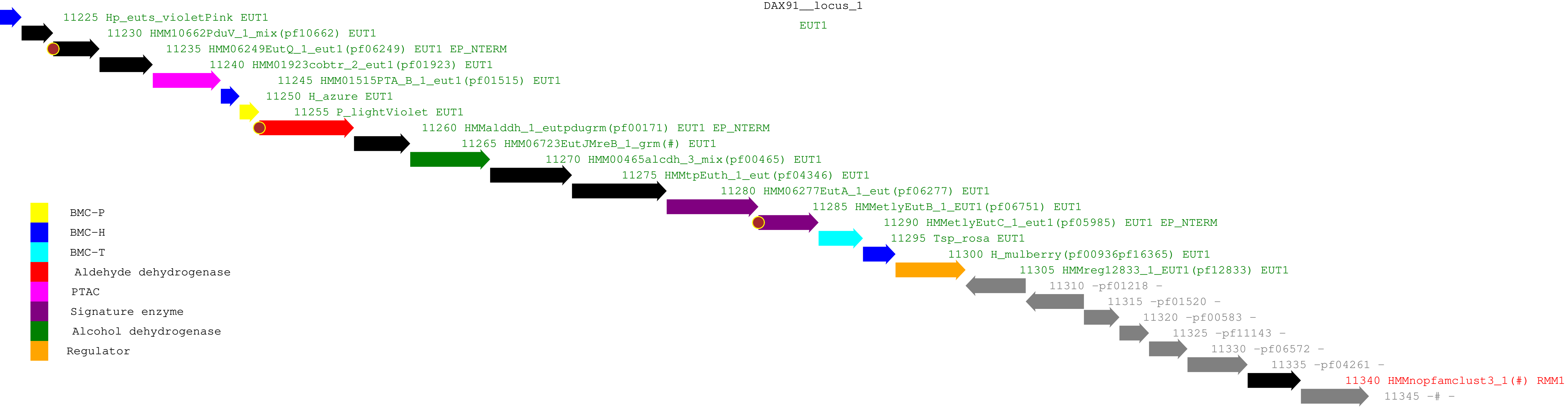

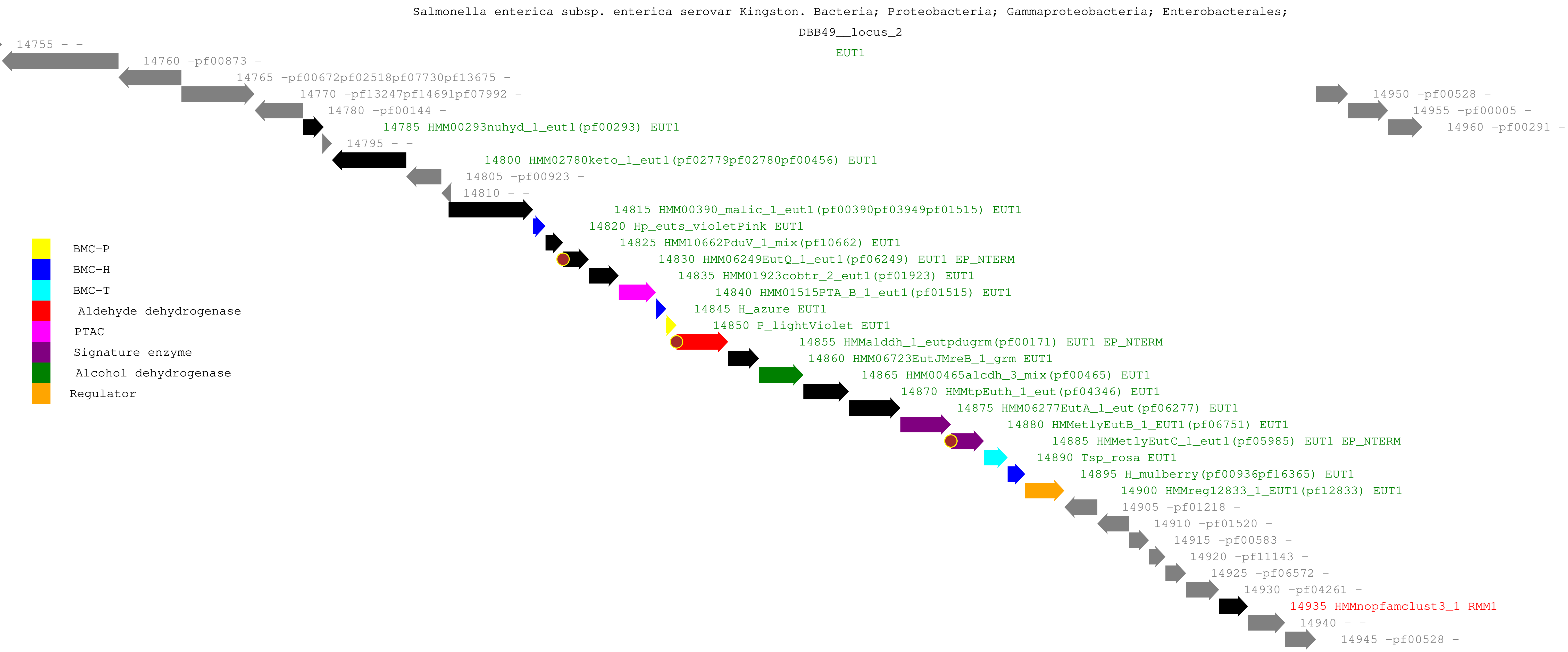
Salmonella enterica subsp. enterica serovar Kingston. Bacteria; Proteobacteria; Gammaproteobacteria; Enterobacterales;
DBB49__locus_2
EUT1
14755 - -
14760 -pf00873 -
14765 -pf00672pf02518pf07730pf13675 -
14770 -pf13247pf14691pf07992 -
14780 -pf00144 -
14785 HMM00293nuhyd_1_eut1(pf00293) EUT1
14795 - -
14800 HMM02780keto_1_eut1(pf02779pf02780pf00456) EUT1
14805 -pf00923 -
14810 - -
14815 HMM00390_malic_1_eut1(pf00390pf03949pf01515) EUT1
14820 Hp_euts_violetPink EUT1
14825 HMM10662PduV_1_mix(pf10662) EUT1
14830 HMM06249EutQ_1_eut1(pf06249) EUT1 EP_NTERM
14835 HMM01923cobtr_2_eut1(pf01923) EUT1
14840 HMM01515PTA_B_1_eut1(pf01515) EUT1
14845 H_azure EUT1
14850 P_lightViolet EUT1
14855 HMMalddh_1_eutpdugrm(pf00171) EUT1 EP_NTERM
14860 HMM06723EutJMreB_1_grm EUT1
14865 HMM00465alcdh_3_mix(pf00465) EUT1
14870 HMMtpEuth_1_eut(pf04346) EUT1
14875 HMM06277EutA_1_eut(pf06277) EUT1
14880 HMMetlyEutB_1_EUT1(pf06751) EUT1
14885 HMMetlyEutC_1_eut1(pf05985) EUT1 EP_NTERM
14890 Tsp_rosa EUT1
14895 H_mulberry(pf00936pf16365) EUT1
14900 HMMreg12833_1_EUT1(pf12833) EUT1
14905 -pf01218 -
14910 -pf01520 -
14915 -pf00583 -
14920 -pf11143 -
14925 -pf06572 -
14930 -pf04261 -
14935 HMMnopfamclust3_1 RMM1
14940 - -
14945 -pf00528 -
14950 -pf00528 -
14955 -pf00005 -
14960 -pf00291 -
BMC-P
BMC-H
BMC-T
Aldehyde dehydrogenase
PTAC
Signature enzyme
Alcohol dehydrogenase
Regulator

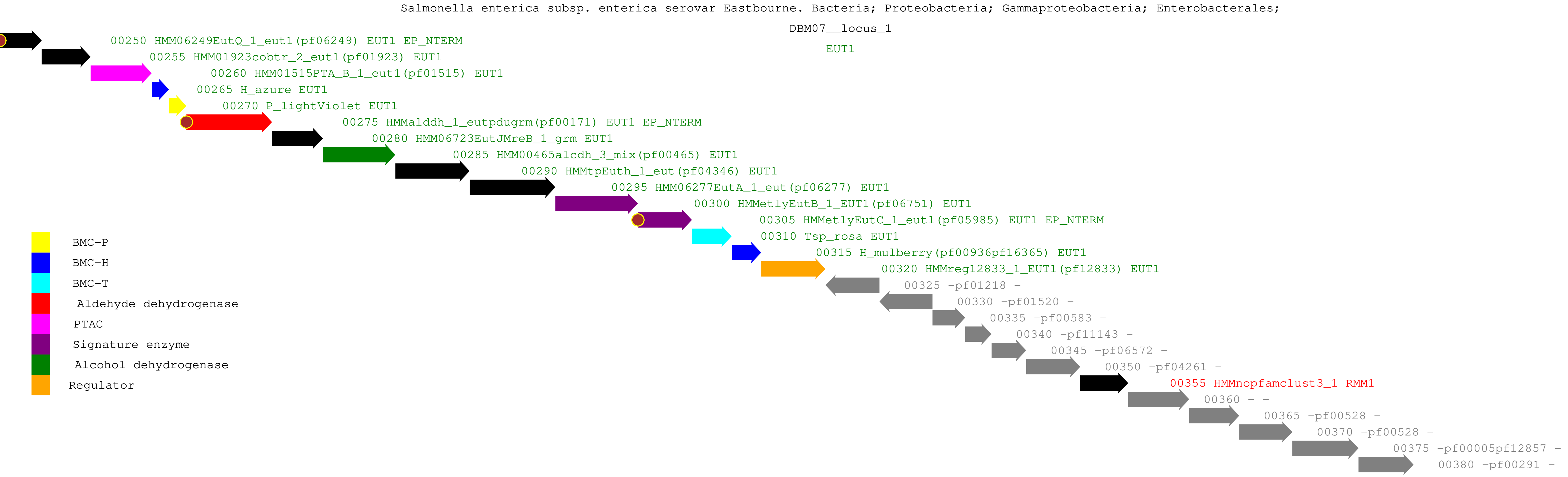
Salmonella enterica subsp. enterica serovar Eastbourne. Bacteria; Proteobacteria; Gammaproteobacteria; Enterobacterales;
DBM07__locus_1
EUT1
00250 HMM06249EutQ_1_eut1(pf06249) EUT1 EP_NTERM
00255 HMM01923cobtr_2_eut1(pf01923) EUT1
00260 HMM01515PTA_B_1_eut1(pf01515) EUT1
00265 H_azure EUT1
00270 P_lightViolet EUT1
00275 HMMalddh_1_eutpdugrm(pf00171) EUT1 EP_NTERM
00280 HMM06723EutJMreB_1_grm EUT1
00285 HMM00465alcdh_3_mix(pf00465) EUT1
00290 HMMtpEuth_1_eut(pf04346) EUT1
00295 HMM06277EutA_1_eut(pf06277) EUT1
00300 HMMetlyEutB_1_EUT1(pf06751) EUT1
00305 HMMetlyEutC_1_eut1(pf05985) EUT1 EP_NTERM
00310 Tsp_rosa EUT1
00315 H_mulberry(pf00936pf16365) EUT1
00320 HMMreg12833_1_EUT1(pf12833) EUT1
00325 -pf01218 -
00330 -pf01520 -
00335 -pf00583 -
00340 -pf11143 -
00345 -pf06572 -
00350 -pf04261 -
00355 HMMnopfamclust3_1 RMM1
00360 - -
00365 -pf00528 -
00370 -pf00528 -
00375 -pf00005pf12857 -
00380 -pf00291 -
BMC-P
BMC-H
BMC-T
Aldehyde dehydrogenase
PTAC
Signature enzyme
Alcohol dehydrogenase
Regulator

Enterobacter sp. EC-NT1. Bacteria; Proteobacteria; Gammaproteobacteria; Enterobacterales;

DBP23__locus_1

EUT1

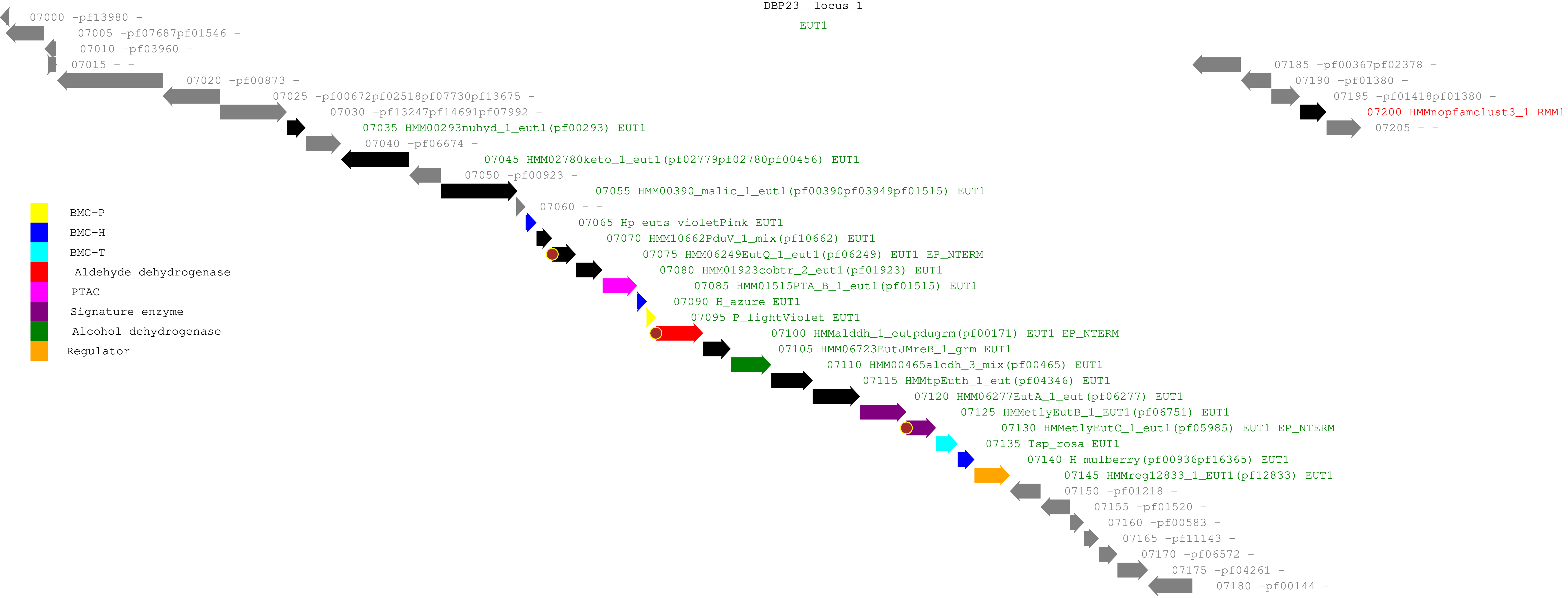

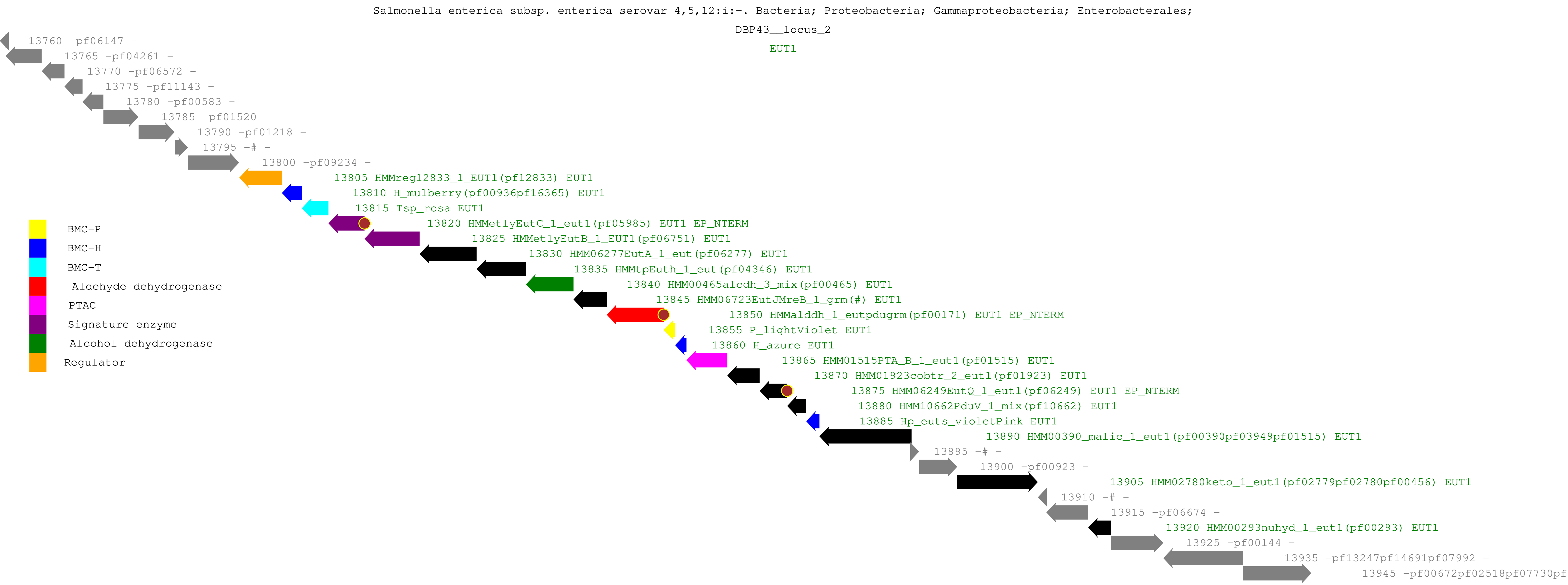
Salmonella enterica subsp. enterica serovar 4,5,12:i:-. Bacteria; Proteobacteria; Gammaproteobacteria; Enterobacterales;
DBP43__locus_2
EUT1
13760 -pf06147 -
13765 -pf04261 -
13770 -pf06572 -
13775 -pf11143 -
13780 -pf00583 -
13785 -pf01520 -
13790 -pf01218 -
13795 -# -
13800 -pf09234 -
13805 HMMreg12833_1_EUT1(pf12833) EUT1
13810 H_mulberry(pf00936pf16365) EUT1
13815 Tsp_rosa EUT1
13820 HMMetlyEutC_1_eut1(pf05985) EUT1 EP_NTERM
13825 HMMetlyEutB_1_EUT1(pf06751) EUT1
13830 HMM06277EutA_1_eut(pf06277) EUT1
13835 HMMtpEuth_1_eut(pf04346) EUT1
13840 HMM00465alcdh_3_mix(pf00465) EUT1
13845 HMM06723EutJMreB_1_grm(#) EUT1
13850 HMMalddh_1_eutpdugrm(pf00171) EUT1 EP_NTERM
13855 P_lightViolet EUT1
13860 H_azure EUT1
13865 HMM01515PTA_B_1_eut1(pf01515) EUT1
13870 HMM01923cobtr_2_eut1(pf01923) EUT1
13875 HMM06249EutQ_1_eut1(pf06249) EUT1 EP_NTERM
13880 HMM10662PduV_1_mix(pf10662) EUT1
13885 Hp_euts_violetPink EUT1
13890 HMM00390_malic_1_eut1(pf00390pf03949pf01515) EUT1
13895 -# -
13900 -pf00923 -
13905 HMM02780keto_1_eut1(pf02779pf02780pf00456) EUT1
13910 -# -
13915 -pf06674 -
13920 HMM00293nuhyd_1_eut1(pf00293) EUT1
13925 -pf00144 -
13935 -pf13247pf14691pf07992 -
13945 -pf00672pf02518pf07730pf
BMC-P
BMC-H
BMC-T
Aldehyde dehydrogenase
PTAC
Signature enzyme
Alcohol dehydrogenase
Regulator

Salmonella enterica subsp. enterica serovar Albany. Bacteria; Proteobacteria; Gammaproteobacteria; Enterobacterales;

DBZ74__locus_2

EUT1

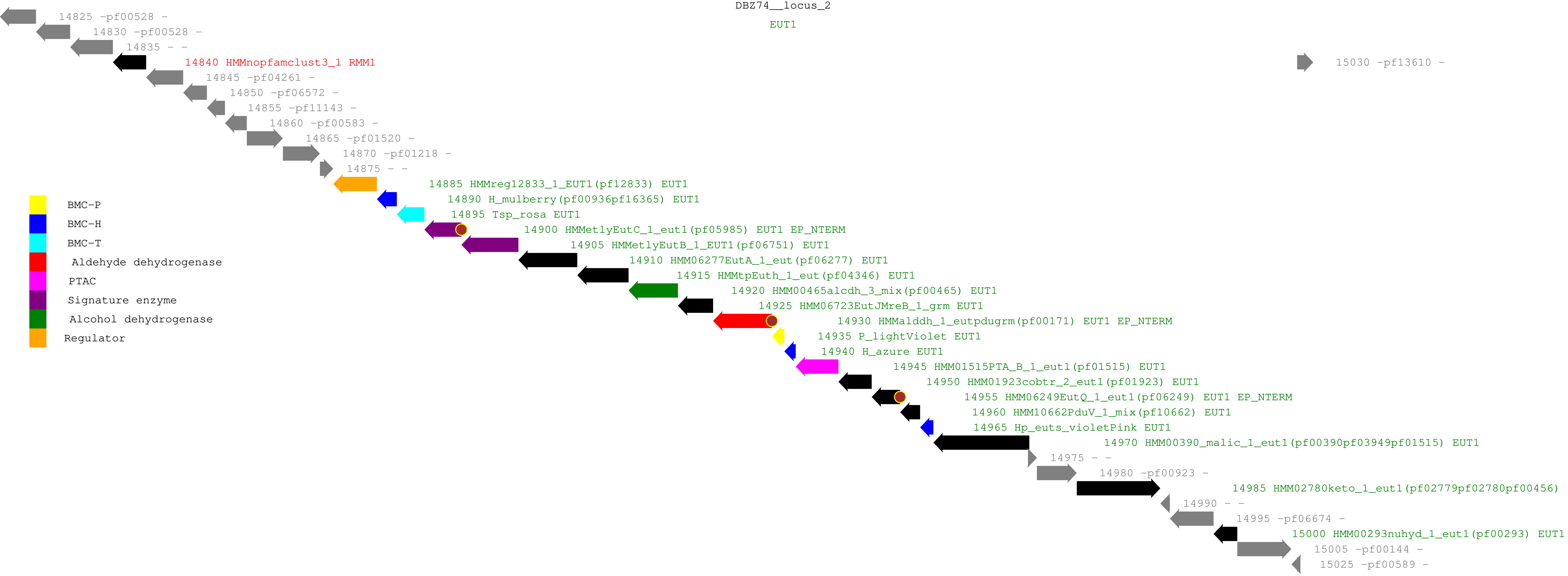

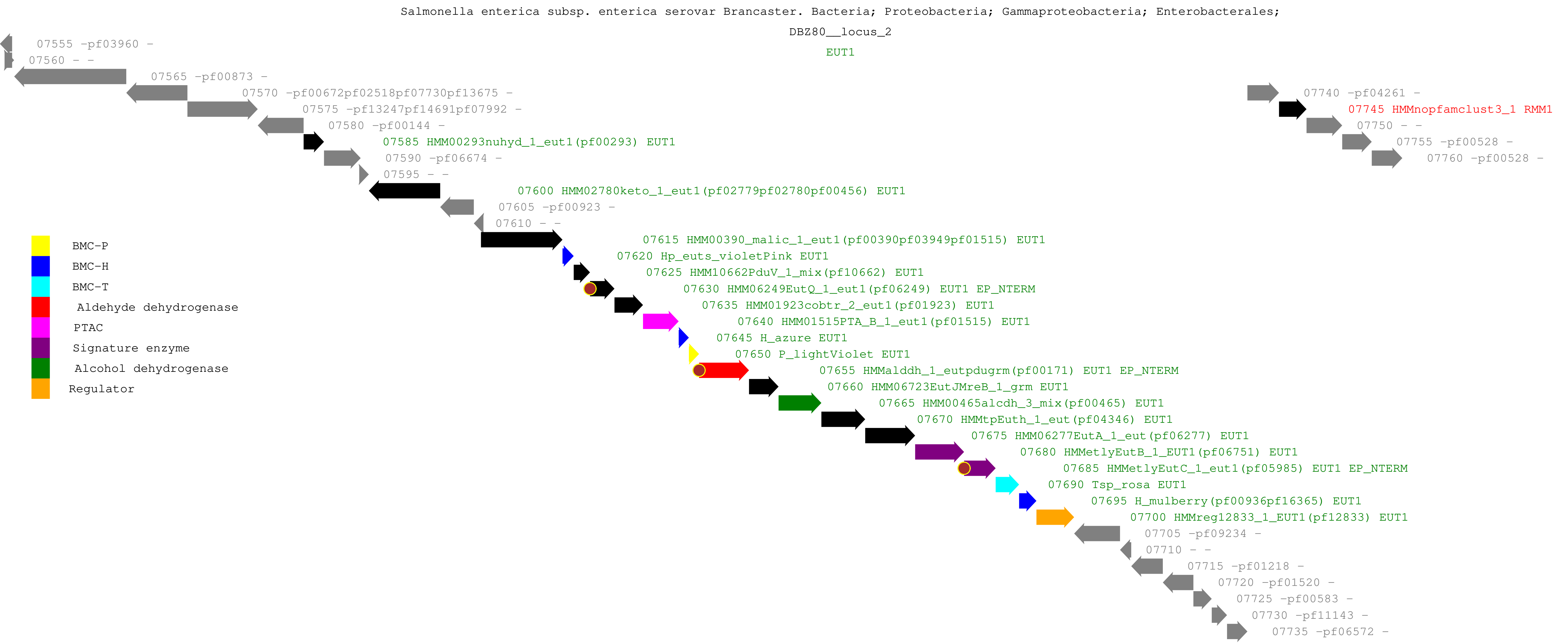
Salmonella enterica subsp. enterica serovar Brancaster. Bacteria; Proteobacteria; Gammaproteobacteria; Enterobacterales;
DBZ80__locus_2
EUT1
07555 -pf03960 -
07560 - -
07565 -pf00873 -
07570 -pf00672pf02518pf07730pf13675 -
07575 -pf13247pf14691pf07992 -
07580 -pf00144 -
07585 HMM00293nuhyd_1_eut1(pf00293) EUT1
07590 -pf06674 -
07595 - -
07600 HMM02780keto_1_eut1(pf02779pf02780pf00456) EUT1
07605 -pf00923 -
07610 - -
07615 HMM00390_malic_1_eut1(pf00390pf03949pf01515) EUT1
07620 Hp_euts_violetPink EUT1
07625 HMM10662PduV_1_mix(pf10662) EUT1
07630 HMM06249EutQ_1_eut1(pf06249) EUT1 EP_NTERM
07635 HMM01923cobtr_2_eut1(pf01923) EUT1
07640 HMM01515PTA_B_1_eut1(pf01515) EUT1
07645 H_azure EUT1
07650 P_lightViolet EUT1
07655 HMMalddh_1_eutpdugrm(pf00171) EUT1 EP_NTERM
07660 HMM06723EutJMreB_1_grm EUT1
07665 HMM00465alcdh_3_mix(pf00465) EUT1
07670 HMMtpEuth_1_eut(pf04346) EUT1
07675 HMM06277EutA_1_eut(pf06277) EUT1
07680 HMMetlyEutB_1_EUT1(pf06751) EUT1
07685 HMMetlyEutC_1_eut1(pf05985) EUT1 EP_NTERM
07690 Tsp_rosa EUT1
07695 H_mulberry(pf00936pf16365) EUT1
07700 HMMreg12833_1_EUT1(pf12833) EUT1
07705 -pf09234 -
07710 - -
07715 -pf01218 -
07720 -pf01520 -
07725 -pf00583 -
07730 -pf11143 -
07735 -pf06572 -
07740 -pf04261 -
07745 HMMnopfamclust3_1 RMM1
07750 - -
07755 -pf00528 -
07760 -pf00528 -
BMC-P
BMC-H
BMC-T
Aldehyde dehydrogenase
PTAC
Signature enzyme
Alcohol dehydrogenase
Regulator

Ignatzschineria ureiclastica. Bacteria; Proteobacteria; Gammaproteobacteria; Ignatzschineria.

DC083__locus_1

EUT1

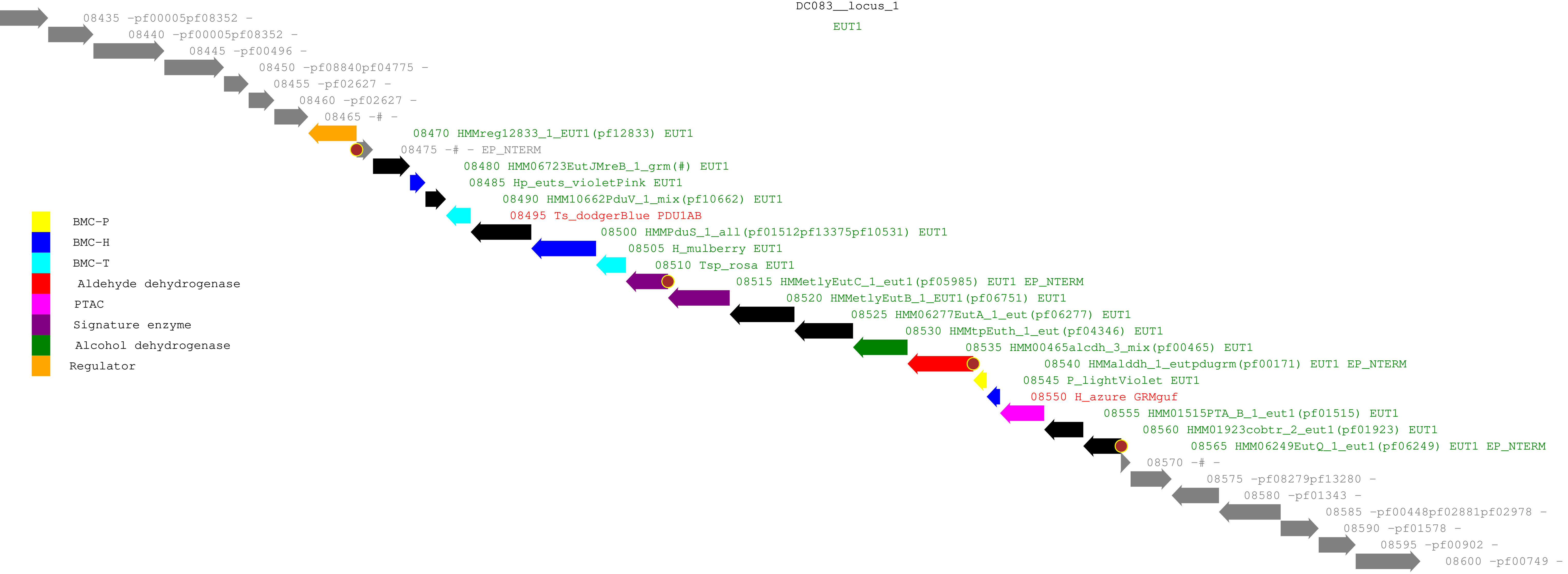

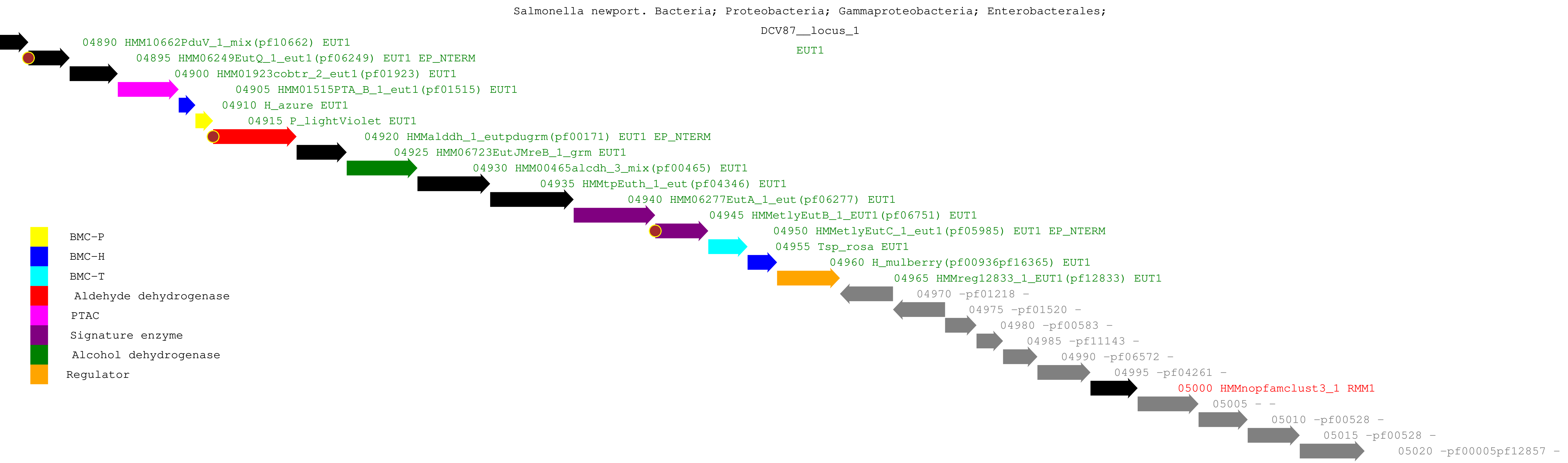

Salmonella newport. Bacteria; Proteobacteria; Gammaproteobacteria; Enterobacterales;
DCV87__locus_1
EUT1
04890 HMM10662PduV_1_mix(pf10662) EUT1
04895 HMM06249EutQ_1_eut1(pf06249) EUT1 EP_NTERM
04900 HMM01923cobtr_2_eut1(pf01923) EUT1
04905 HMM01515PTA_B_1_eut1(pf01515) EUT1
04910 H_azure EUT1
04915 P_lightViolet EUT1
04920 HMMalddh_1_eutpdugrm(pf00171) EUT1 EP_NTERM
04925 HMM06723EutJMreB_1_grm EUT1
04930 HMM00465alcdh_3_mix(pf00465) EUT1
04935 HMMtpEuth_1_eut(pf04346) EUT1
04940 HMM06277EutA_1_eut(pf06277) EUT1
04945 HMMetlyEutB_1_EUT1(pf06751) EUT1
04950 HMMetlyEutC_1_eut1(pf05985) EUT1 EP_NTERM
04955 Tsp_rosa EUT1
04960 H_mulberry(pf00936pf16365) EUT1
04965 HMMreg12833_1_EUT1(pf12833) EUT1
04970 -pf01218 -
04975 -pf01520 -
04980 -pf00583 -
04985 -pf11143 -
04990 -pf06572 -
04995 -pf04261 -
05000 HMMnopfamclust3_1 RMM1
05005 - -
05010 -pf00528 -
05015 -pf00528 -
05020 -pf00005pf12857 -
BMC-P
BMC-H
BMC-T
Aldehyde dehydrogenase
PTAC
Signature enzyme
Alcohol dehydrogenase
Regulator

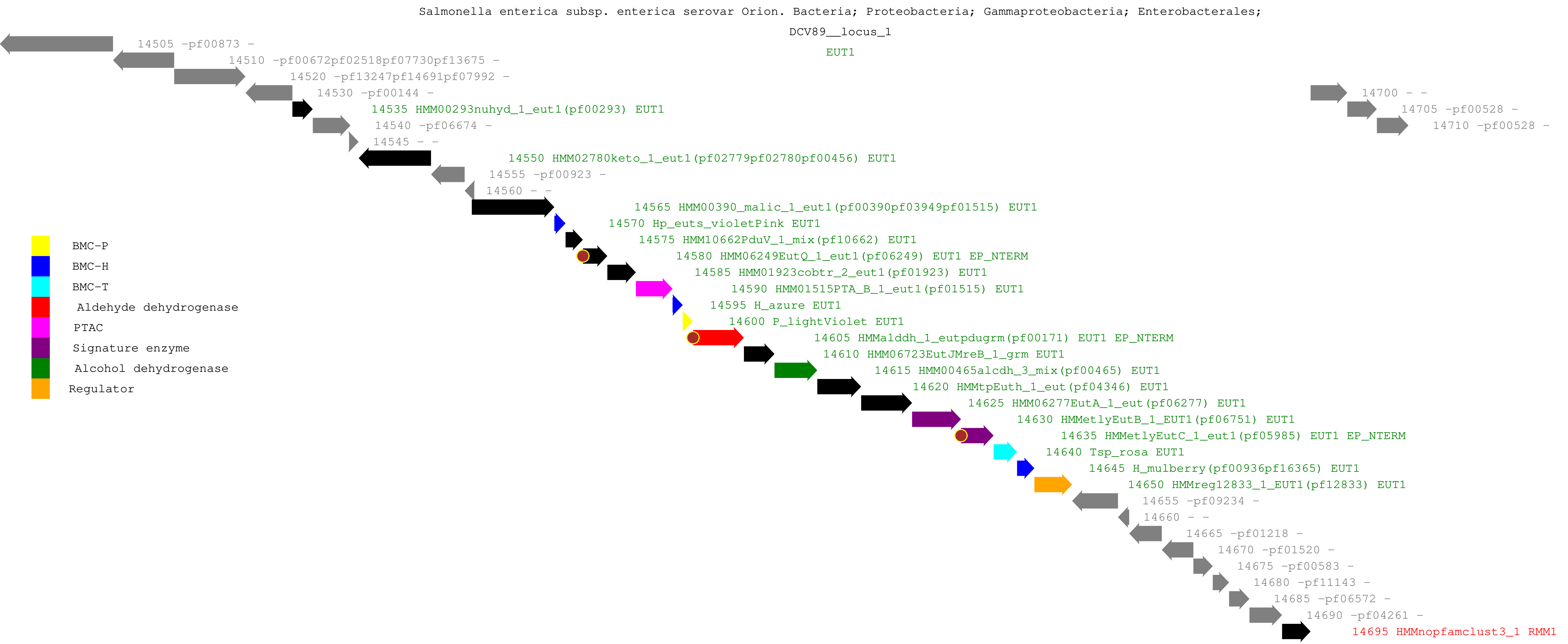

Salmonella enterica subsp. enterica serovar Orion. Bacteria; Proteobacteria; Gammaproteobacteria; Enterobacterales;
DCV89__locus_1
EUT1
14505 -pf00873 -
14510 -pf00672pf02518pf07730pf13675 -
14520 -pf13247pf14691pf07992 -
14530 -pf00144 -
14535 HMM00293nuhyd_1_eut1(pf00293) EUT1
14540 -pf06674 -
14545 - -
14550 HMM02780keto_1_eut1(pf02779pf02780pf00456) EUT1
14555 -pf00923 -
14560 - -
14565 HMM00390_malic_1_eut1(pf00390pf03949pf01515) EUT1
14570 Hp_euts_violetPink EUT1
14575 HMM10662PduV_1_mix(pf10662) EUT1
14580 HMM06249EutQ_1_eut1(pf06249) EUT1 EP_NTERM
14585 HMM01923cobtr_2_eut1(pf01923) EUT1
14590 HMM01515PTA_B_1_eut1(pf01515) EUT1
14595 H_azure EUT1
14600 P_lightViolet EUT1
14605 HMMaldh_1_eutpdugrm(pf00171) EUT1 EP_NTERM
14610 HMM06723EutJMreB_1_grm EUT1
14615 HMM00465alcdh_3_mix(pf00465) EUT1
14620 HMMtpEuth_1_eut(pf04346) EUT1
14625 HMM06277EutA_1_eut(pf06277) EUT1
14630 HMMetlyEutB_1_EUT1(pf06751) EUT1
14635 HMMetlyEutC_1_eut1(pf05985) EUT1 EP_NTERM
14640 Tsp_rosa EUT1
14645 H_mulberry(pf00936pf16365) EUT1
14650 HMMreg12833_1_EUT1(pf12833) EUT1
14655 -pf09234 -
14660 - -
14665 -pf01218 -
14670 -pf01520 -
14675 -pf00583 -
14680 -pf11143 -
14685 -pf06572 -
14690 -pf04261 -
14695 HMMnopfamclust3_1 RMM1
14700 - -
14705 -pf00528 -
14710 -pf00528 -
BMC-P
BMC-H
BMC-T
Aldehyde dehydrogenase
PTAC
Signature enzyme
Alcohol dehydrogenase
Regulator

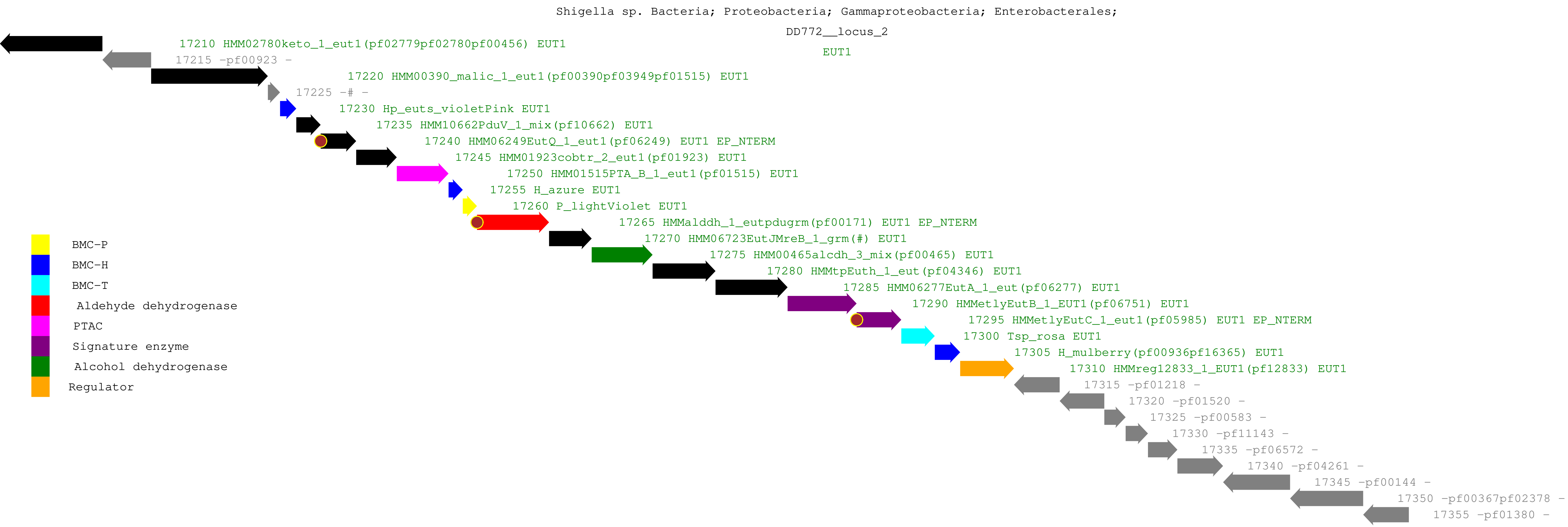
Shigella sp. Bacteria; Proteobacteria; Gammaproteobacteria; Enterobacterales;
DD772__locus_2
EUT1
17210 HMM02780keto_1_eut1(pf02779pf02780pf00456) EUT1
17215 -pf00923 -
17220 HMM00390_malic_1_eut1(pf00390pf03949pf01515) EUT1
17225 -# -
17230 Hp_euts_violetPink EUT1
17235 HMM10662PduV_1_mix(pf10662) EUT1
17240 HMM06249EutQ_1_eut1(pf06249) EUT1 EP_NTERM
17245 HMM01923cobtr_2_eut1(pf01923) EUT1
17250 HMM01515PTA_B_1_eut1(pf01515) EUT1
17255 H_azure EUT1
17260 P_lightViolet EUT1
17265 HMMalddh_1_eutpdugrm(pf00171) EUT1 EP_NTERM
17270 HMM06723EutJMreB_1_grm(#) EUT1
17275 HMM00465alcdh_3_mix(pf00465) EUT1
17280 HMMtpEuth_1_eut(pf04346) EUT1
17285 HMM06277EutA_1_eut(pf06277) EUT1
17290 HMMetlyEutB_1_EUT1(pf06751) EUT1
17295 HMMetlyEutC_1_eut1(pf05985) EUT1 EP_NTERM
17300 Tsp_rosa EUT1
17305 H_mulberry(pf00936pf16365) EUT1
17310 HMMreg12833_1_EUT1(pf12833) EUT1
17315 -pf01218 -
17320 -pf01520 -
17325 -pf00583 -
17330 -pf11143 -
17335 -pf06572 -
17340 -pf04261 -
17345 -pf00144 -
17350 -pf00367pf02378 -
17355 -pf01380 -
BMC-P
BMC-H
BMC-T
Aldehyde dehydrogenase
PTAC
Signature enzyme
Alcohol dehydrogenase
Regulator

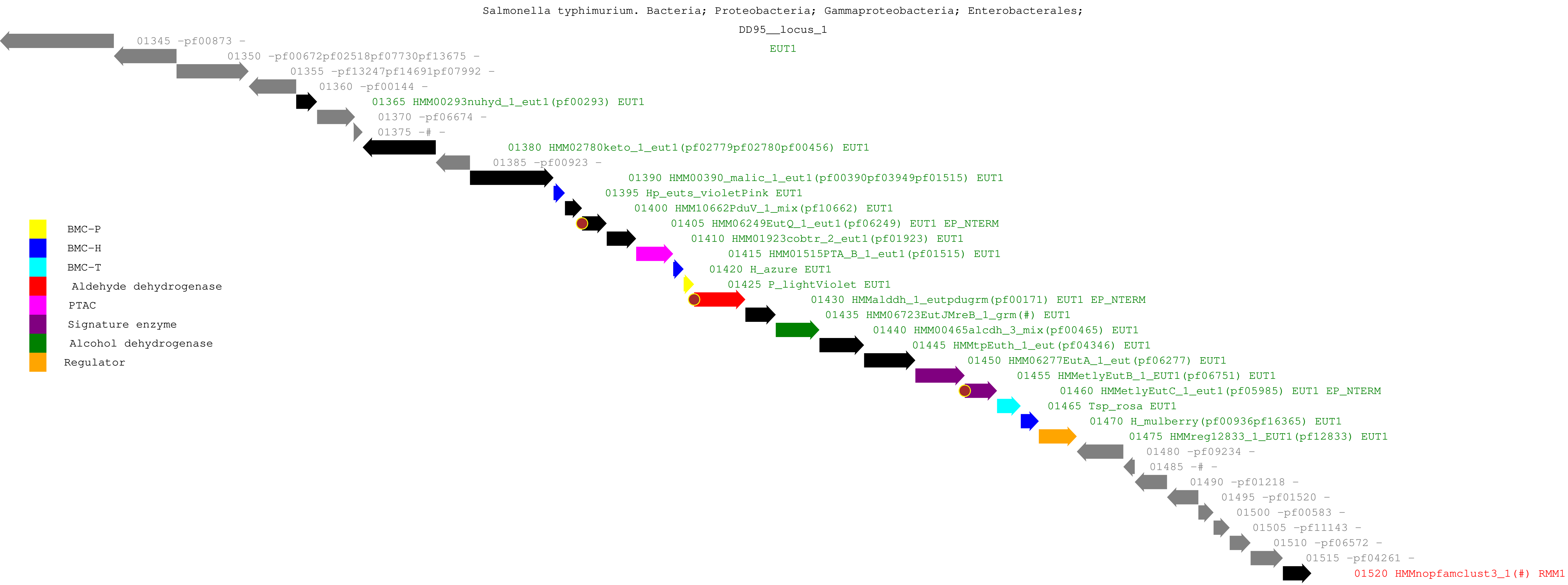
Salmonella typhimurium. Bacteria; Proteobacteria; Gammaproteobacteria; Enterobacterales;
DD95__locus_1
EUT1
01345 -pf00873 -
01350 -pf00672pf02518pf07730pf13675 -
01355 -pf13247pf14691pf07992 -
01360 -pf00144 -
01365 HMM00293nuhyd_1_eut1(pf00293) EUT1
01370 -pf06674 -
01375 -# -
01380 HMM02780keto_1_eut1(pf02779pf02780pf00456) EUT1
01385 -pf00923 -
01390 HMM00390_malic_1_eut1(pf00390pf03949pf01515) EUT1
01395 Hp_euts_violetPink EUT1
01400 HMM10662PduV_1_mix(pf10662) EUT1
01405 HMM06249EutQ_1_eut1(pf06249) EUT1 EP_NTERM
01410 HMM01923cobtr_2_eut1(pf01923) EUT1
01415 HMM01515PTA_B_1_eut1(pf01515) EUT1
01420 H_azure EUT1
01425 P_lightViolet EUT1
01430 HMMalddh_1_eutpdugrm(pf00171) EUT1 EP_NTERM
01435 HMM06723EutJMreB_1_grm(#) EUT1
01440 HMM00465alcdh_3_mix(pf00465) EUT1
01445 HMMtpEuth_1_eut(pf04346) EUT1
01450 HMM06277EutA_1_eut(pf06277) EUT1
01455 HMMetlyEutB_1_EUT1(pf06751) EUT1
01460 HMMetlyEutC_1_eut1(pf05985) EUT1 EP_NTERM
01465 Tsp_rosa EUT1
01470 H_mulberry(pf00936pf16365) EUT1
01475 HMMreg12833_1_EUT1(pf12833) EUT1
01480 -pf09234 -
01485 -# -
01490 -pf01218 -
01495 -pf01520 -
01500 -pf00583 -
01505 -pf11143 -
01510 -pf06572 -
01515 -pf04261 -
01520 HMMnopfamclust3_1(#) RMM1
BMC-P
BMC-H
BMC-T
Aldehyde dehydrogenase
PTAC
Signature enzyme
Alcohol dehydrogenase
Regulator

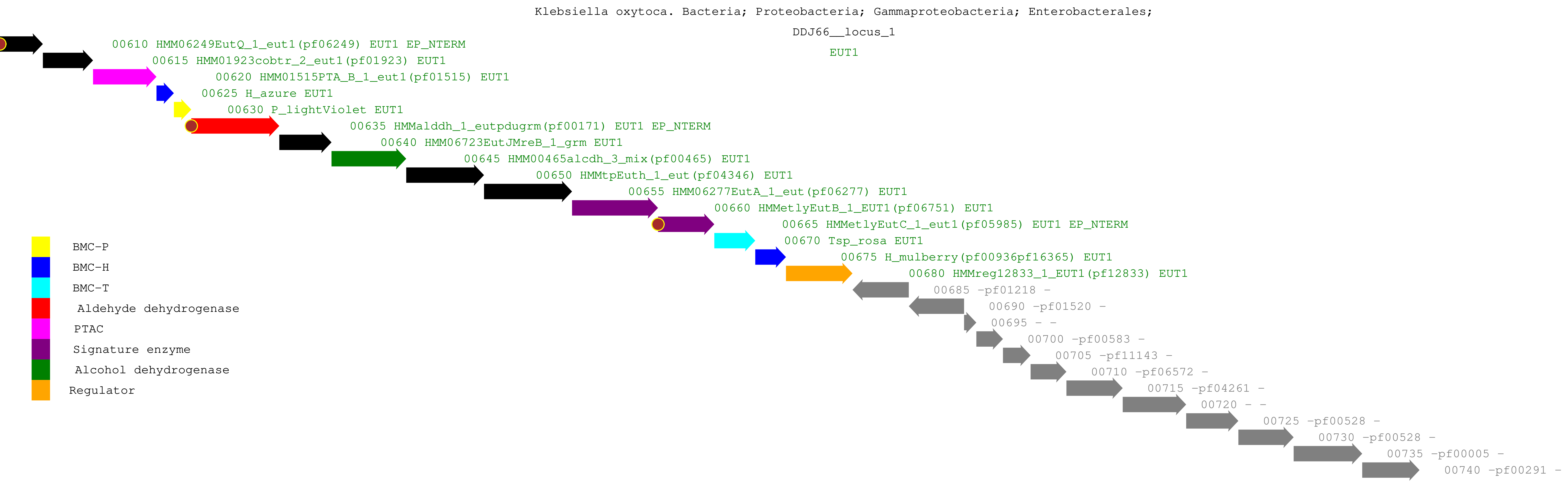

Klebsiella oxytoca. Bacteria; Proteobacteria; Gammaproteobacteria; Enterobacterales;
DDJ66__locus_1
EUT1
00610 HMM06249EutQ_1_eut1(pf06249) EUT1 EP_NTERM
00615 HMM01923cobtr_2_eut1(pf01923) EUT1
00620 HMM01515PTA_B_1_eut1(pf01515) EUT1
00625 H_azure EUT1
00630 P_lightViolet EUT1
00635 HMMalddh_1_eutpdugrm(pf00171) EUT1 EP_NTERM
00640 HMM06723EutJMreB_1_grm EUT1
00645 HMM00465alcdh_3_mix(pf00465) EUT1
00650 HMMtpEuth_1_eut(pf04346) EUT1
00655 HMM06277EutA_1_eut(pf06277) EUT1
00660 HMMetlyEutB_1_EUT1(pf06751) EUT1
00665 HMMetlyEutC_1_eut1(pf05985) EUT1 EP_NTERM
00670 Tsp_rosa EUT1
00675 H_mulberry(pf00936pf16365) EUT1
00680 HMMreg12833_1_EUT1(pf12833) EUT1
00685 -pf01218 -
00690 -pf01520 -
00695 - -
00700 -pf00583 -
00705 -pf11143 -
00710 -pf06572 -
00715 -pf04261 -
00720 - -
00725 -pf00528 -
00730 -pf00528 -
00735 -pf00005 -
00740 -pf00291 -
BMC-P
BMC-H
BMC-T
Aldehyde dehydrogenase
PTAC
Signature enzyme
Alcohol dehydrogenase
Regulator

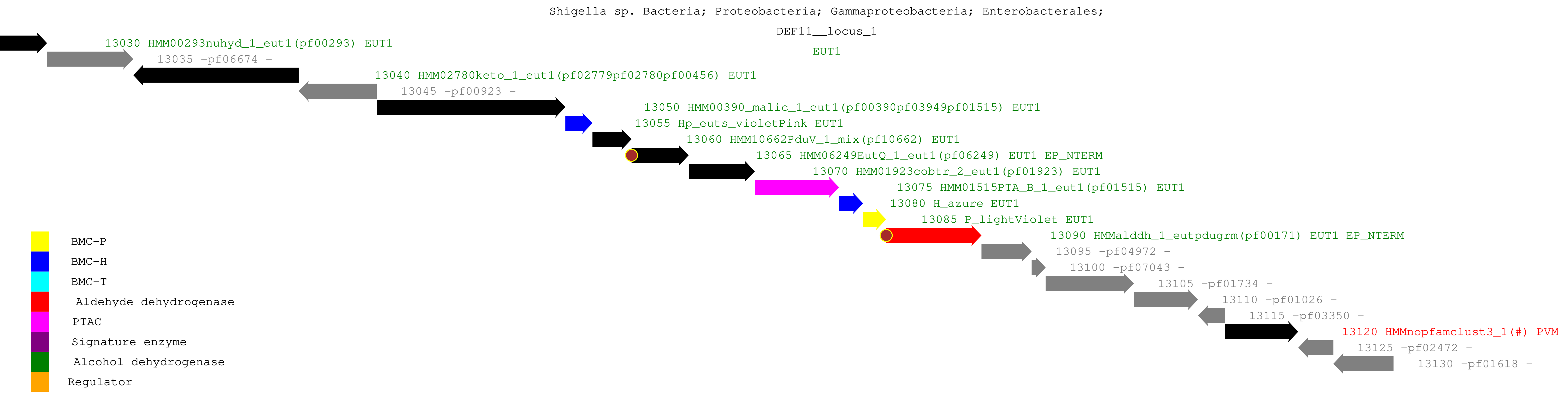

Shigella sp. Bacteria; Proteobacteria; Gammaproteobacteria; Enterobacterales;
DEF11__locus_1
EUT1
13030 HMM00293nuhyd_1_eut1(pf00293) EUT1
13035 -pf06674 -
13040 HMM02780keto_1_eut1(pf02779pf02780pf00456) EUT1
13045 -pf00923 -
13050 HMM00390_malic_1_eut1(pf00390pf03949pf01515) EUT1
13055 Hp_euts_violetPink EUT1
13060 HMM10662PduV_1_mix(pf10662) EUT1
13065 HMM06249EutQ_1_eut1(pf06249) EUT1 EP_NTERM
13070 HMM01923cobtr_2_eut1(pf01923) EUT1
13075 HMM01515PTA_B_1_eut1(pf01515) EUT1
13080 H_azure EUT1
13085 P_lightViolet EUT1
13090 HMMalddh_1_eutpdugrm(pf00171) EUT1 EP_NTERM
13095 -pf04972 -
13100 -pf07043 -
13105 -pf01734 -
13110 -pf01026 -
13115 -pf03350 -
13120 HMMnopfamclust3_1(#) PVM
13125 -pf02472 -
13130 -pf01618 -
BMC-P
BMC-H
BMC-T
Aldehyde dehydrogenase
PTAC
Signature enzyme
Alcohol dehydrogenase
Regulator

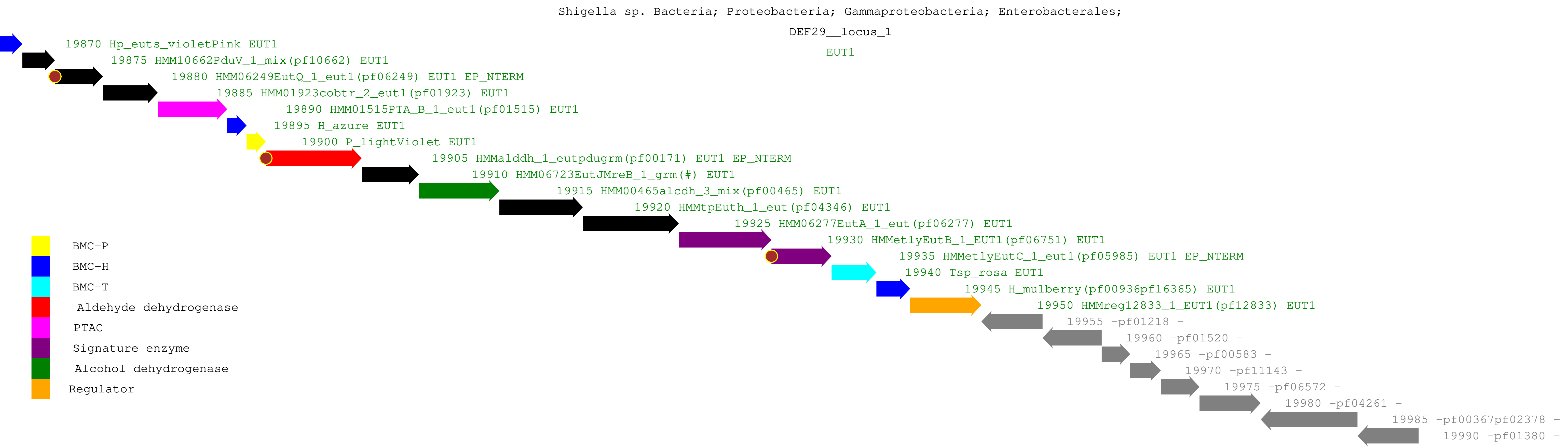
Shigella sp. Bacteria; Proteobacteria; Gammaproteobacteria; Enterobacterales;
DEF29__locus_1
EUT1
19870 Hp_euts_violetPink EUT1
19875 HMM10662PduV_1_mix(pf10662) EUT1
19880 HMM06249EutQ_1_eut1(pf06249) EUT1 EP_NTERM
19885 HMM01923cobtr_2_eut1(pf01923) EUT1
19890 HMM01515PTA_B_1_eut1(pf01515) EUT1
19895 H_azure EUT1
19900 P_lightViolet EUT1
19905 HMMalddh_1_eutpdugrm(pf00171) EUT1 EP_NTERM
19910 HMM06723EutJMreB_1_grm(#) EUT1
19915 HMM00465alcdh_3_mix(pf00465) EUT1
19920 HMMtpEuth_1_eut(pf04346) EUT1
19925 HMM06277EutA_1_eut(pf06277) EUT1
19930 HMMetlyEutB_1_EUT1(pf06751) EUT1
19935 HMMetlyEutC_1_eut1(pf05985) EUT1 EP_NTERM
19940 Tsp_rosa EUT1
19945 H_mulberry(pf00936pf16365) EUT1
19950 HMMreg12833_1_EUT1(pf12833) EUT1
19955 -pf01218 -
19960 -pf01520 -
19965 -pf00583 -
19970 -pf11143 -
19975 -pf06572 -
19980 -pf04261 -
19985 -pf00367pf02378 -
19990 -pf01380 -
BMC-P
BMC-H
BMC-T
Aldehyde dehydrogenase
PTAC
Signature enzyme
Alcohol dehydrogenase
Regulator

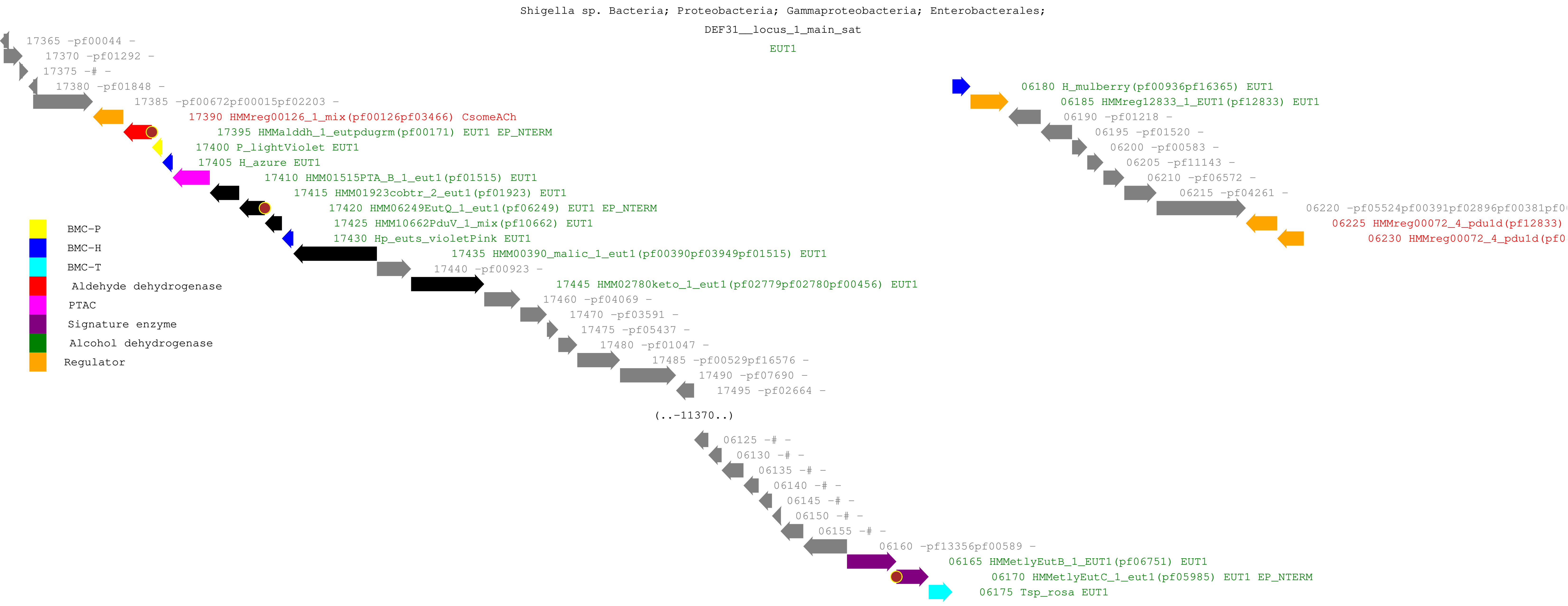
Shigella sp. Bacteria; Proteobacteria; Gammaproteobacteria; Enterobacterales;
DEF31__locus_1_main_sat
EUT1
17365 -pf00044 -
17370 -pf01292 -
17375 -# -
17380 -pf01848 -
17385 -pf00672pf00015pf02203 -
17390 HMMreg00126_1_mix(pf00126pf03466) CsomeACh
17395 HMMalddh_1_eutpdugrm(pf00171) EUT1 EP_NTERM
17400 P_lightViolet EUT1
17405 H_azure EUT1
17410 HMM01515PTA_B_1_eut1(pf01515) EUT1
17415 HMM01923cobtr_2_eut1(pf01923) EUT1
17420 HMM06249EutQ_1_eut1(pf06249) EUT1 EP_NTERM
17425 HMM10662PduV_1_mix(pf10662) EUT1
17430 Hp_euts_violetPink EUT1
17435 HMM00390_malic_1_eut1(pf00390pf03949pf01515) EUT1
17440 -pf00923 -
17445 HMM02780keto_1_eut1(pf02779pf02780pf00456) EUT1
17460 -pf04069 -
17470 -pf03591 -
17475 -pf05437 -
17480 -pf01047 -
17485 -pf00529pf16576 -
17490 -pf07690 -
17495 -pf02664 -
(..-11370..)
06125 -# -
06130 -# -
06135 -# -
06140 -# -
06145 -# -
06150 -# -
06155 -# -
06160 -pf13356pf00589 -
06165 HMMetlyEutB_1_EUT1(pf06751) EUT1
06170 HMMetlyEutC_1_eut1(pf05985) EUT1 EP_NTERM
06175 Tsp_rosa EUT1
06180 H_mulberry(pf00936pf16365) EUT1
06185 HMMreg12833_1_EUT1(pf12833) EUT1
06190 -pf01218 -
06195 -pf01520 -
06200 -pf00583 -
06205 -pf11143 -
06210 -pf06572 -
06215 -pf04261 -
06220 -pf05524pf00391pf02896pf00381pf0
06225 HMMreg00072_4_pdu1d(pf12833)
06230 HMMreg00072_4_pdu1d(pf0
BMC-P
BMC-H
BMC-T
Aldehyde dehydrogenase
PTAC
Signature enzyme
Alcohol dehydrogenase
Regulator

Klebsiella oxytoca. Bacteria; Proteobacteria; Gammaproteobacteria; Enterobacterales;

DET57__locus_3

EUT1

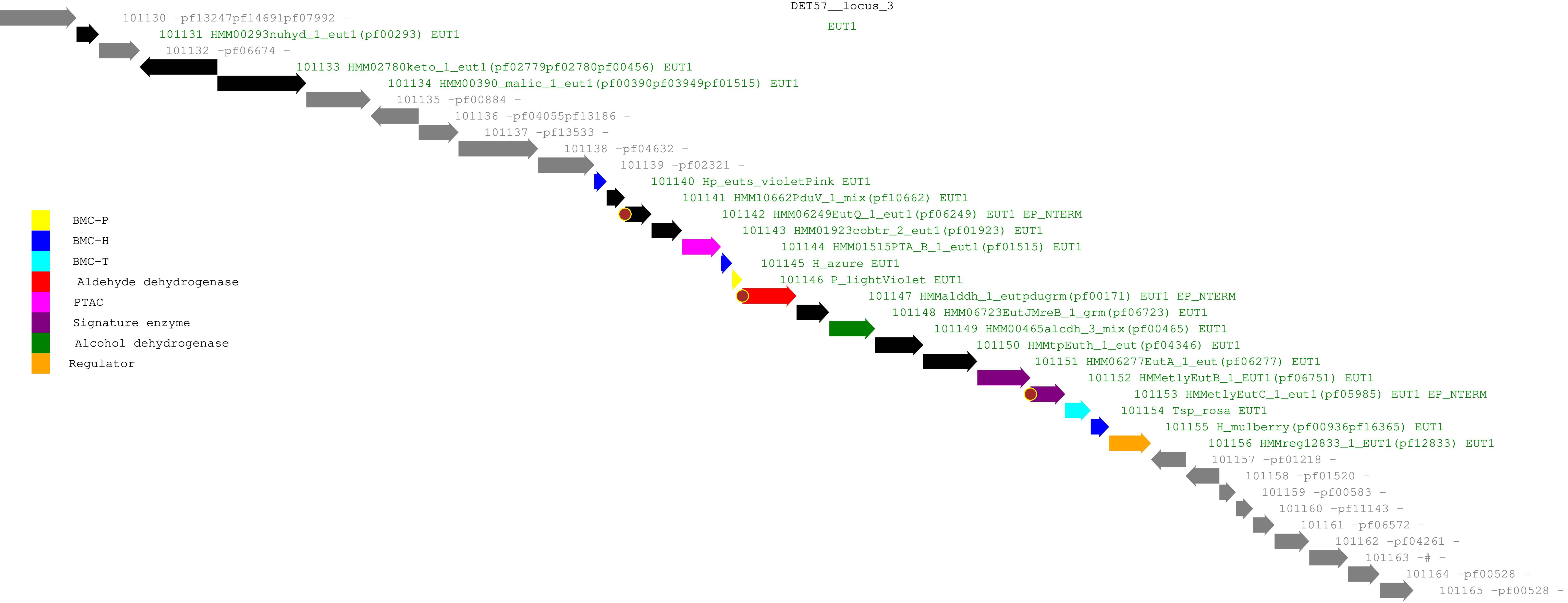

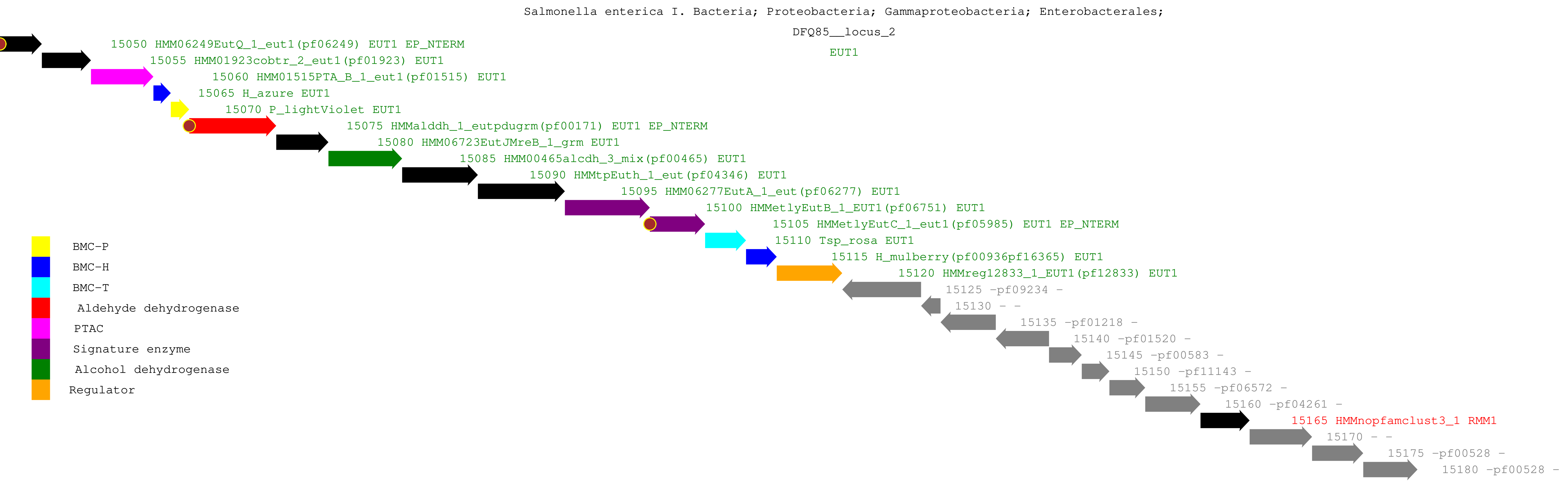
Salmonella enterica I. Bacteria; Proteobacteria; Gammaproteobacteria; Enterobacterales;
DFQ85__locus_2
EUT1
15050 HMM06249EutQ_1_eut1(pf06249) EUT1 EP_NTERM
15055 HMM01923cobtr_2_eut1(pf01923) EUT1
15060 HMM01515PTA_B_1_eut1(pf01515) EUT1
15065 H_azure EUT1
15070 P_lightViolet EUT1
15075 HMMalddh_1_eutpdugrm(pf00171) EUT1 EP_NTERM
15080 HMM06723EutJMreB_1_grm EUT1
15085 HMM00465alcdh_3_mix(pf00465) EUT1
15090 HMMtpEuth_1_eut(pf04346) EUT1
15095 HMM06277EutA_1_eut(pf06277) EUT1
15100 HMMetlyEutB_1_EUT1(pf06751) EUT1
15105 HMMetlyEutC_1_eut1(pf05985) EUT1 EP_NTERM
15110 Tsp_rosa EUT1
15115 H_mulberry(pf00936pf16365) EUT1
15120 HMMreg12833_1_EUT1(pf12833) EUT1
15125 -pf09234 -
15130 - -
15135 -pf01218 -
15140 -pf01520 -
15145 -pf00583 -
15150 -pf11143 -
15155 -pf06572 -
15160 -pf04261 -
15165 HMMnopfamclust3_1 RMM1
15170 - -
15175 -pf00528 -
15180 -pf00528 -
BMC-P
BMC-H
BMC-T
Aldehyde dehydrogenase
PTAC
Signature enzyme
Alcohol dehydrogenase
Regulator

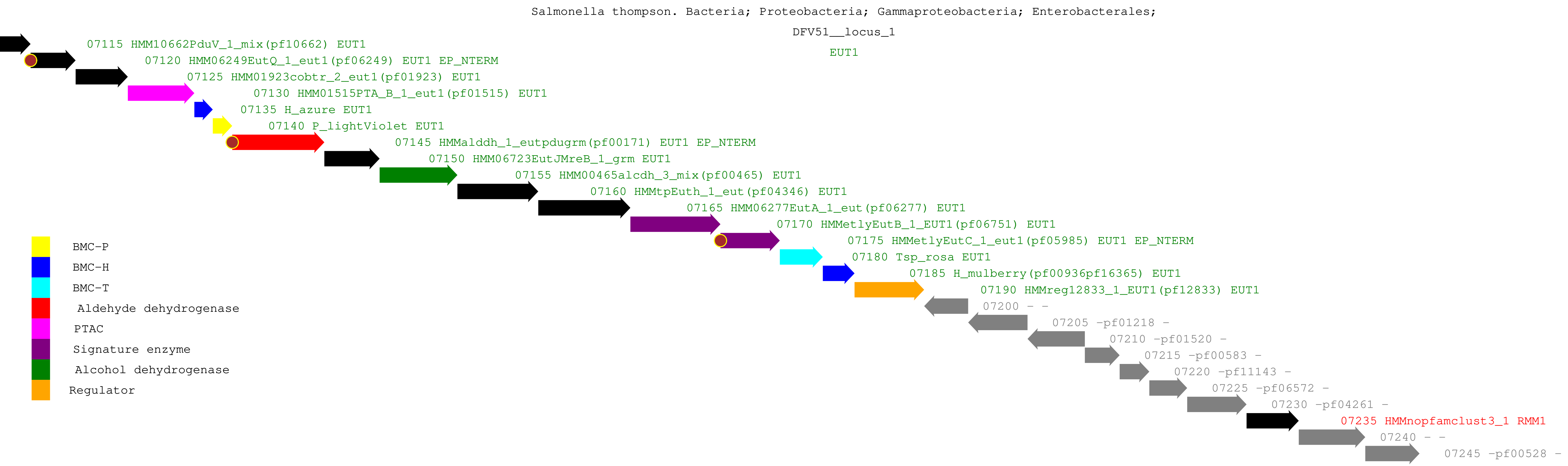
Salmonella thompson. Bacteria; Proteobacteria; Gammaproteobacteria; Enterobacterales;
DFV51__locus_1
EUT1
07115 HMM10662PduV_1_mix(pf10662) EUT1
07120 HMM06249EutQ_1_eut1(pf06249) EUT1 EP_NTERM
07125 HMM01923cobtr_2_eut1(pf01923) EUT1
07130 HMM01515PTA_B_1_eut1(pf01515) EUT1
07135 H_azure EUT1
07140 P_lightViolet EUT1
07145 HMMalddh_1_eutpdugrm(pf00171) EUT1 EP_NTERM
07150 HMM06723EutJMreB_1_grm EUT1
07155 HMM00465alcdh_3_mix(pf00465) EUT1
07160 HMMtpEuth_1_eut(pf04346) EUT1
07165 HMM06277EutA_1_eut(pf06277) EUT1
07170 HMMetlyEutB_1_EUT1(pf06751) EUT1
07175 HMMetlyEutC_1_eut1(pf05985) EUT1 EP_NTERM
07180 Tsp_rosa EUT1
07185 H_mulberry(pf00936pf16365) EUT1
07190 HMMreg12833_1_EUT1(pf12833) EUT1
07200 - -
07205 -pf01218 -
07210 -pf01520 -
07215 -pf00583 -
07220 -pf11143 -
07225 -pf06572 -
07230 -pf04261 -
07235 HMMnopfamclust3_1 RMM1
07240 - -
07245 -pf00528 -
BMC-P
BMC-H
BMC-T
Aldehyde dehydrogenase
PTAC
Signature enzyme
Alcohol dehydrogenase
Regulator

Morganella sp. (in: Bacteria). Bacteria; Proteobacteria; Gammaproteobacteria; Enterobacterales;

DIT05__locus_1

EUT1

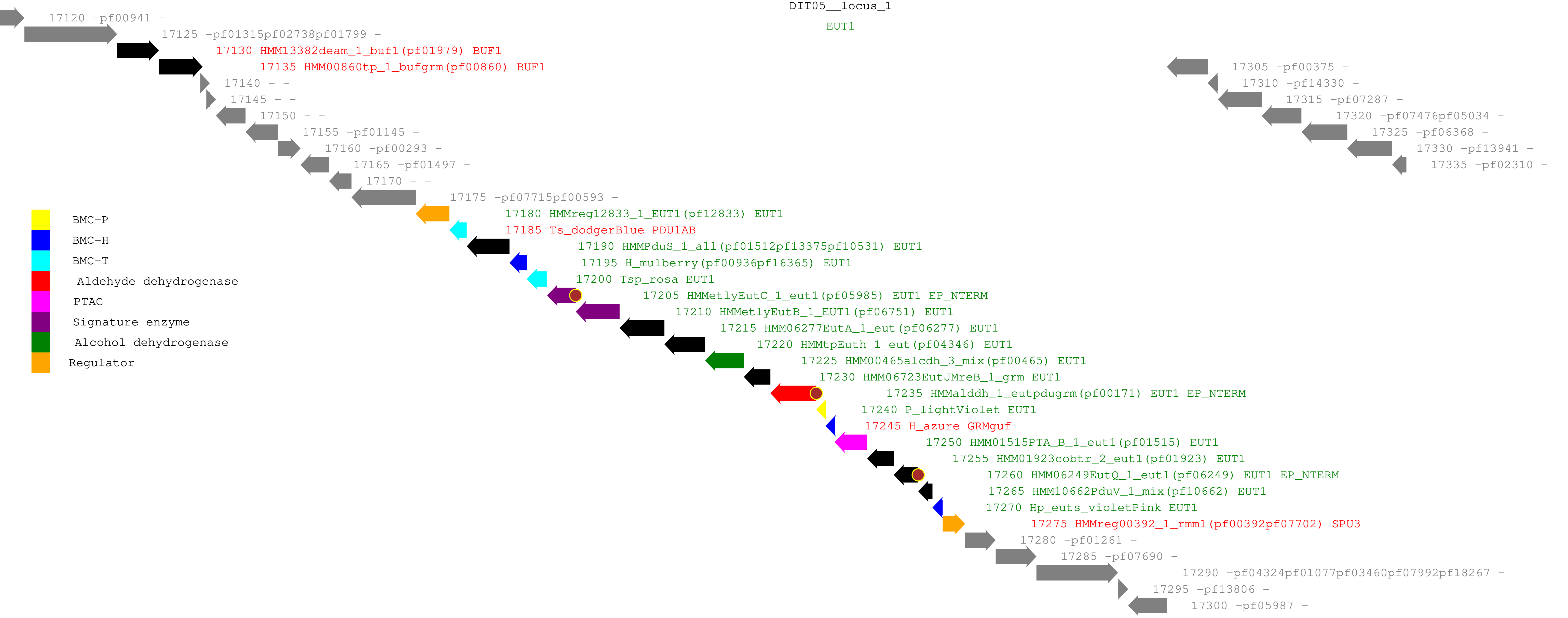

Salmonella enterica I. Bacteria; Proteobacteria; Gammaproteobacteria; Enterobacterales;

DJ273__locus_2

EUT1

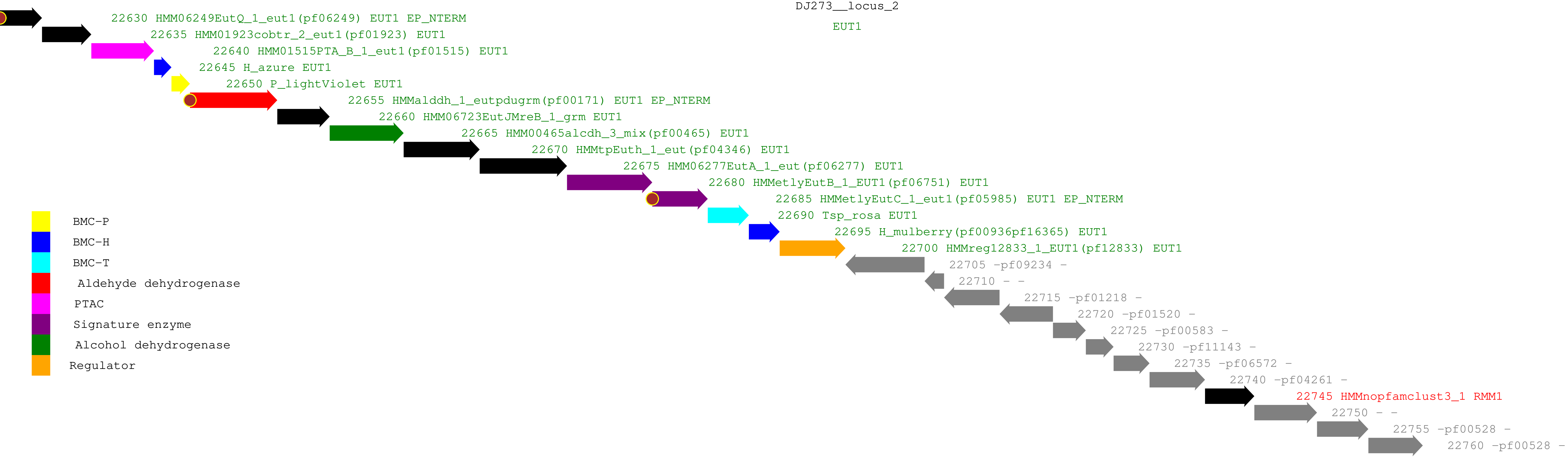

Escherichia coli. Bacteria; Proteobacteria; Gammaproteobacteria; Enterobacterales;

DJ503__locus_1

EUT1

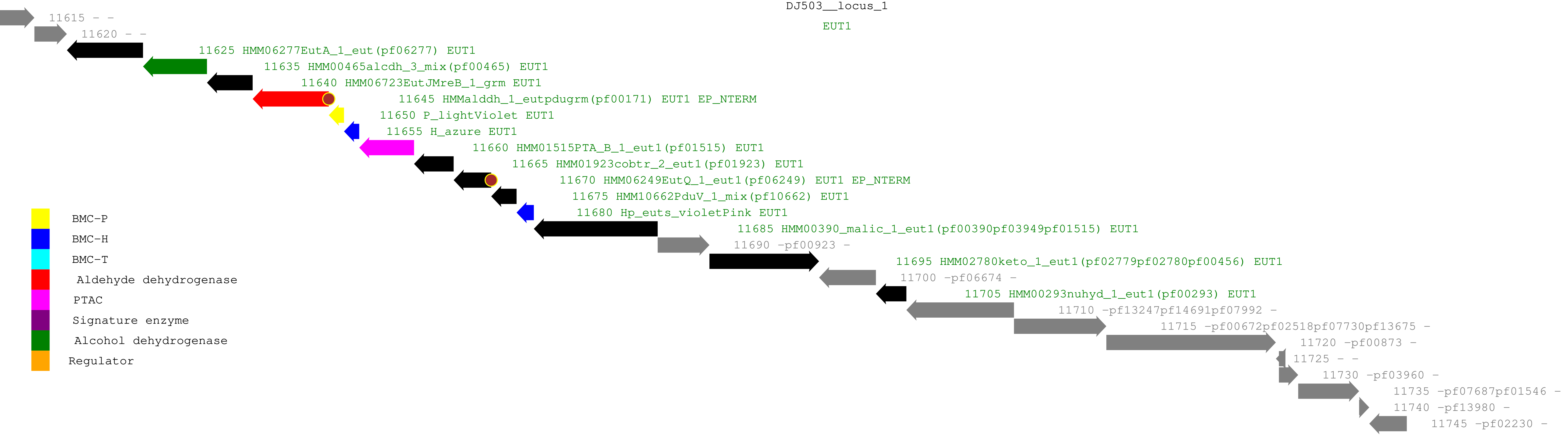

Klebsiella grimontii. Bacteria; Proteobacteria; Gammaproteobacteria; Enterobacterales;

DJ542__locus_2

EUT1

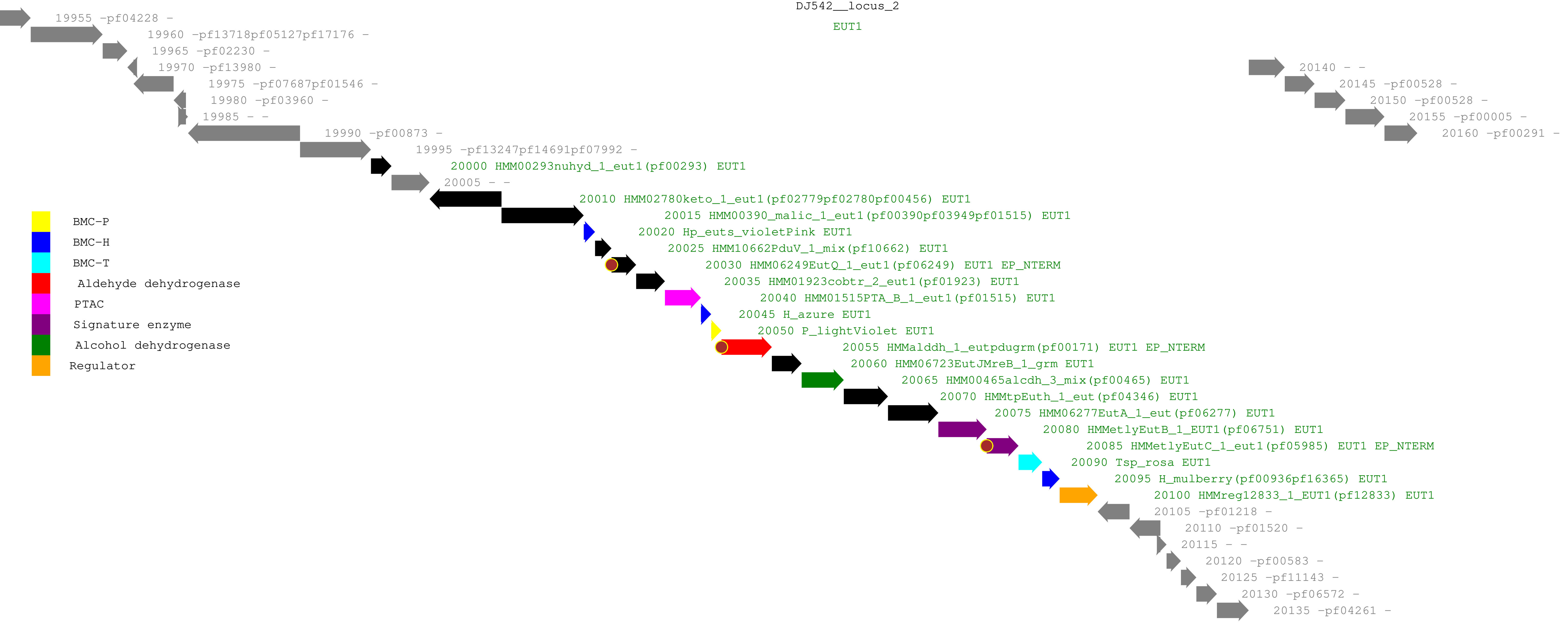

Salmonella enteritidis. Bacteria; Proteobacteria; Gammaproteobacteria; Enterobacterales;

DK092__locus_2

EUT1

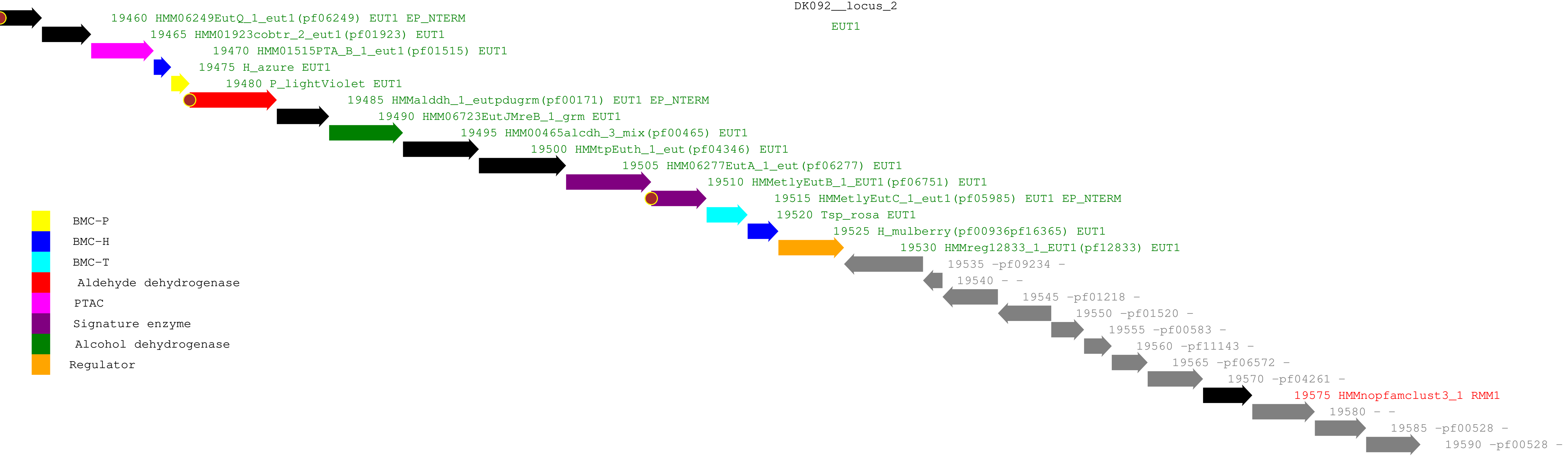

Pseudomonas sp. RW407. Bacteria; Proteobacteria; Gammaproteobacteria; Pseudomonadales;

DK254__locus_1_main_sat

EUT1

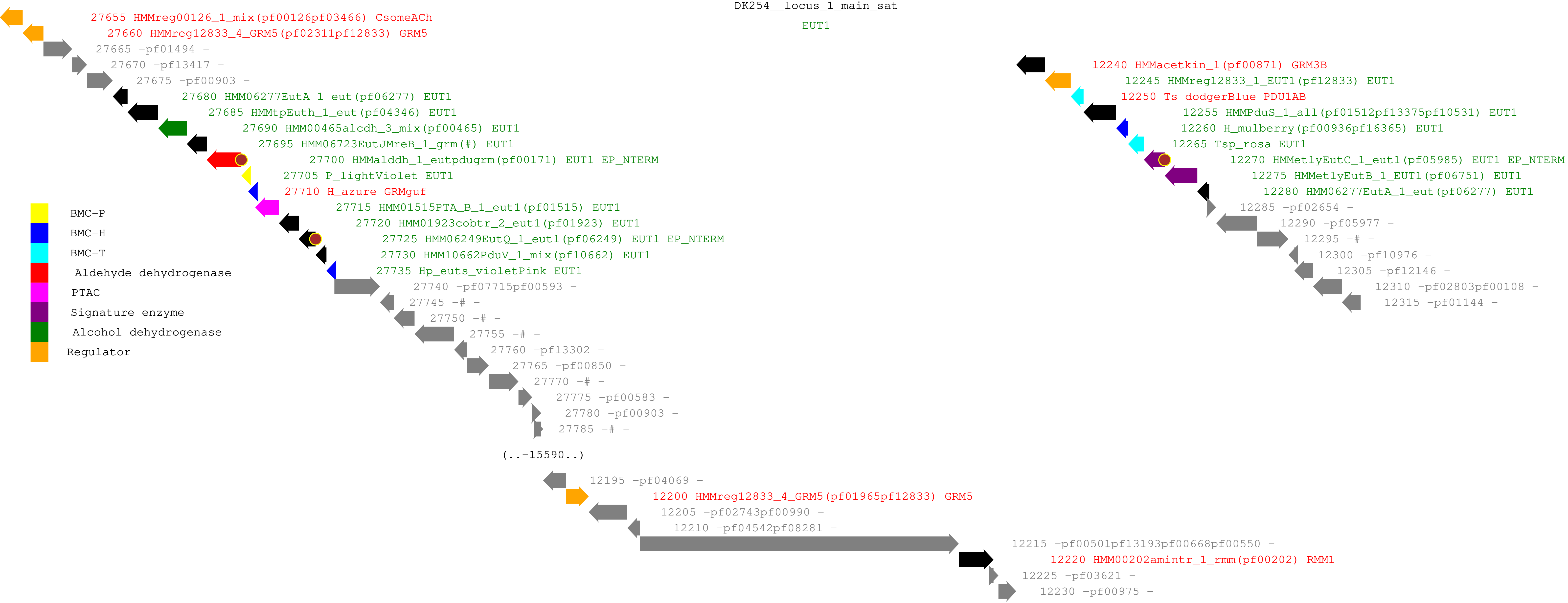

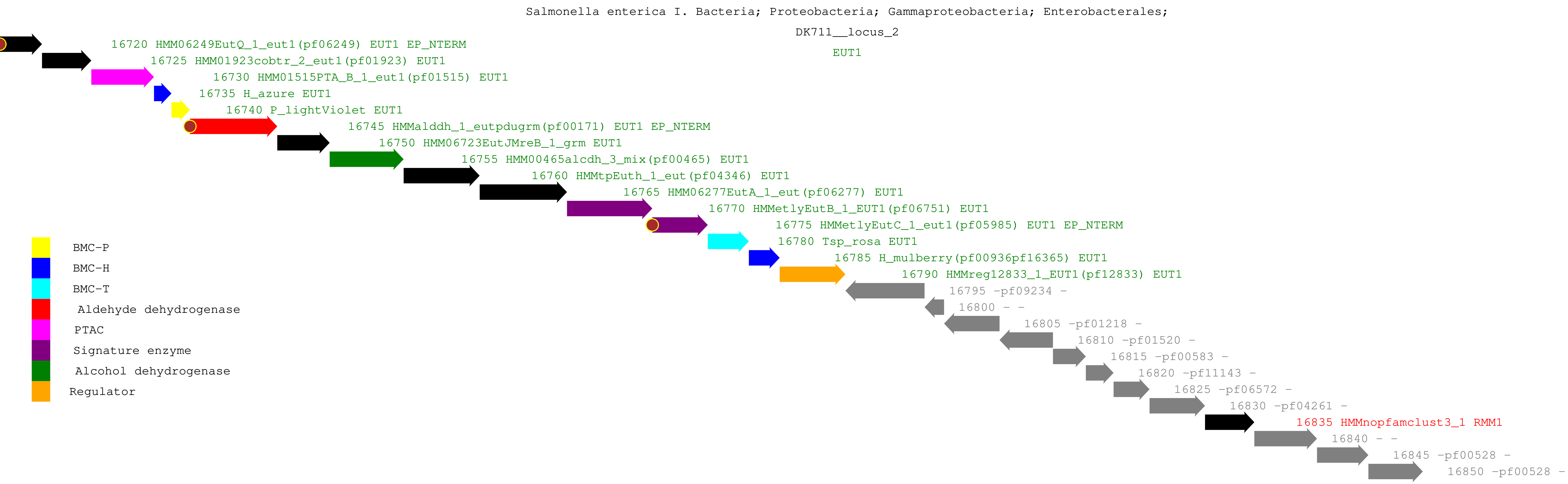
Salmonella enterica I. Bacteria; Proteobacteria; Gammaproteobacteria; Enterobacterales;
DK711__locus_2
EUT1
16720 HMM06249EutQ_1_eut1(pf06249) EUT1 EP_NTERM
16725 HMM01923cobtr_2_eut1(pf01923) EUT1
16730 HMM01515PTA_B_1_eut1(pf01515) EUT1
16735 H_azure EUT1
16740 P_lightViolet EUT1
16745 HMMalddh_1_eutpdugrm(pf00171) EUT1 EP_NTERM
16750 HMM06723EutJMreB_1_grm EUT1
16755 HMM00465alcdh_3_mix(pf00465) EUT1
16760 HMMtpEuth_1_eut(pf04346) EUT1
16765 HMM06277EutA_1_eut(pf06277) EUT1
16770 HMMetlyEutB_1_EUT1(pf06751) EUT1
16775 HMMetlyEutC_1_eut1(pf05985) EUT1 EP_NTERM
16780 Tsp_rosa EUT1
16785 H_mulberry(pf00936pf16365) EUT1
16790 HMMreg12833_1_EUT1(pf12833) EUT1
16795 -pf09234 -
16800 - -
16805 -pf01218 -
16810 -pf01520 -
16815 -pf00583 -
16820 -pf11143 -
16825 -pf06572 -
16830 -pf04261 -
16835 HMMnopfamclust3_1 RMM1
16840 - -
16845 -pf00528 -
16850 -pf00528 -
BMC-P
BMC-H
BMC-T
Aldehyde dehydrogenase
PTAC
Signature enzyme
Alcohol dehydrogenase
Regulator

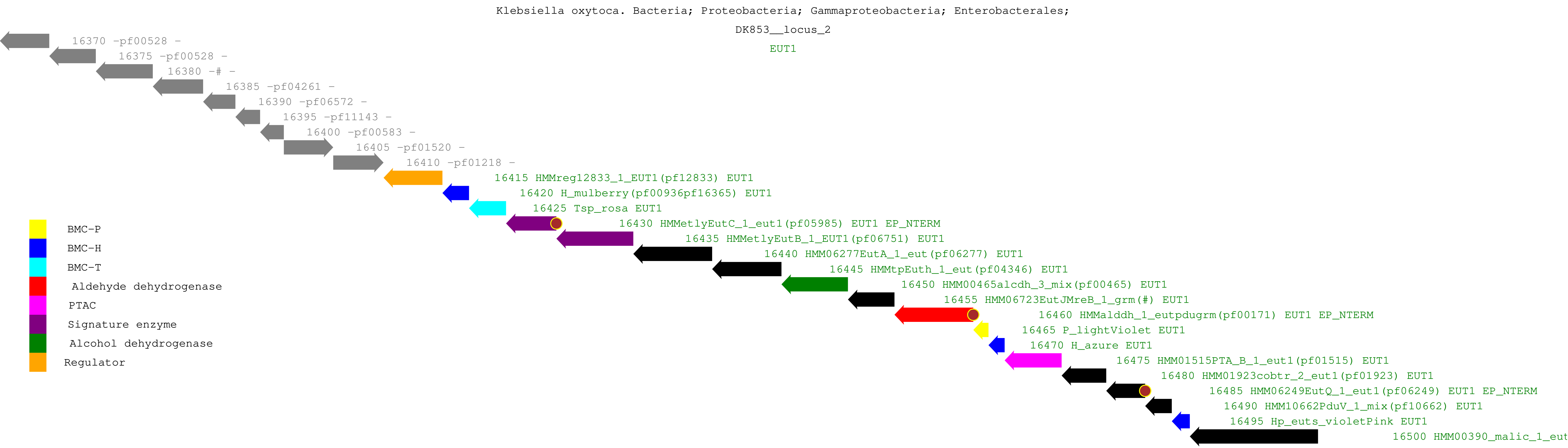

Klebsiella oxytoca. Bacteria; Proteobacteria; Gammaproteobacteria; Enterobacterales;
DK853__locus_2
EUT1
16370 -pf00528 -
16375 -pf00528 -
16380 -# -
16385 -pf04261 -
16390 -pf06572 -
16395 -pf11143 -
16400 -pf00583 -
16405 -pf01520 -
16410 -pf01218 -
16415 HMMreg12833_1_EUT1(pf12833) EUT1
16420 H_mulberry(pf00936pf16365) EUT1
16425 Tsp_rosa EUT1
16430 HMMetlyEutC_1_eut1(pf05985) EUT1 EP_NTERM
16435 HMMetlyEutB_1_EUT1(pf06751) EUT1
16440 HMM06277EutA_1_eut(pf06277) EUT1
16445 HMMtpEuth_1_eut(pf04346) EUT1
16450 HMM00465alcdh_3_mix(pf00465) EUT1
16455 HMM06723EutJMreB_1_grm(#) EUT1
16460 HMMalddh_1_eutpdugrm(pf00171) EUT1 EP_NTERM
16465 P_lightViolet EUT1
16470 H_azure EUT1
16475 HMM01515PTA_B_1_eut1(pf01515) EUT1
16480 HMM01923cobtr_2_eut1(pf01923) EUT1
16485 HMM06249EutQ_1_eut1(pf06249) EUT1 EP_NTERM
16490 HMM10662PduV_1_mix(pf10662) EUT1
16495 Hp_euts_violetPink EUT1
16500 HMM00390_malic_1_eut
BMC-P
BMC-H
BMC-T
Aldehyde dehydrogenase
PTAC
Signature enzyme
Alcohol dehydrogenase
Regulator

Salmonella enterica subsp. enterica serovar Mishmarhaemek. Bacteria; Proteobacteria; Gammaproteobacteria; Enterobacterales;

DKA99__locus_1

EUT1

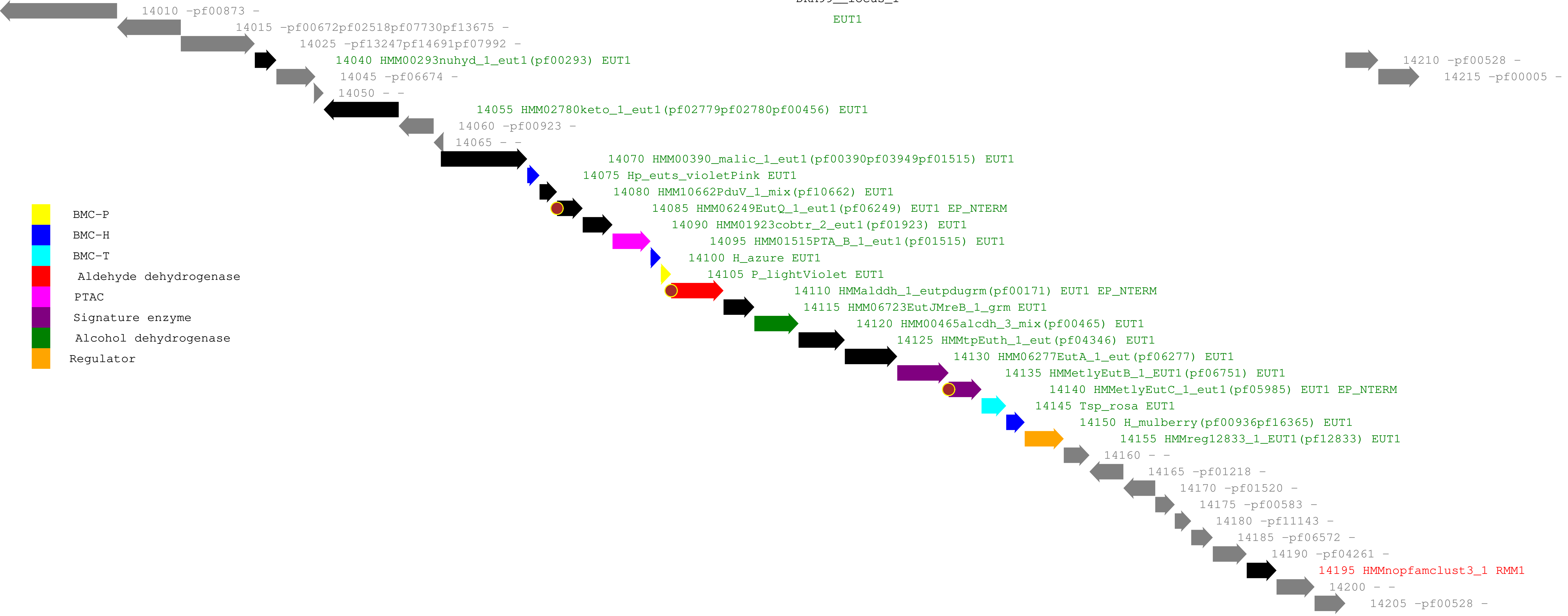

Salmonella enterica subsp. enterica serovar Lexington. Bacteria; Proteobacteria; Gammaproteobacteria; Enterobacterales;

DKG82__locus_2

EUT1

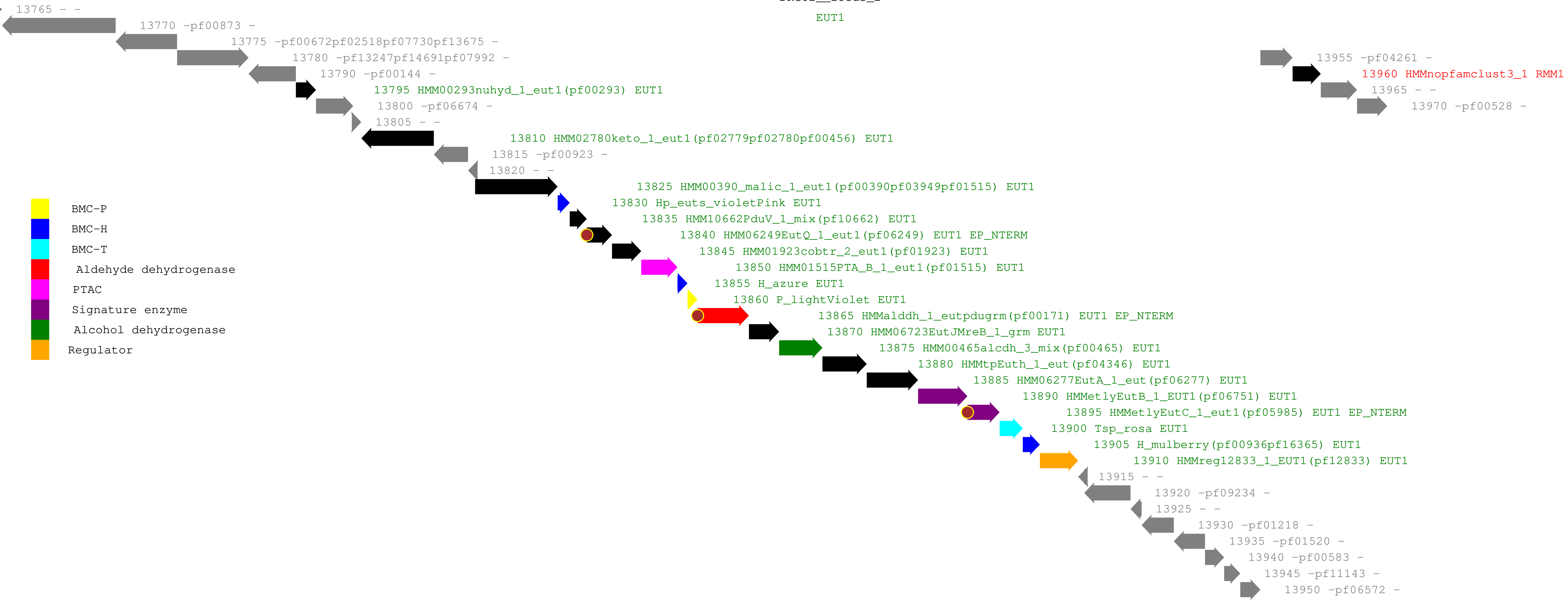

Salmonella typhimurium. Bacteria; Proteobacteria; Gammaproteobacteria; Enterobacterales;

DKJ11__locus_1

EUT1

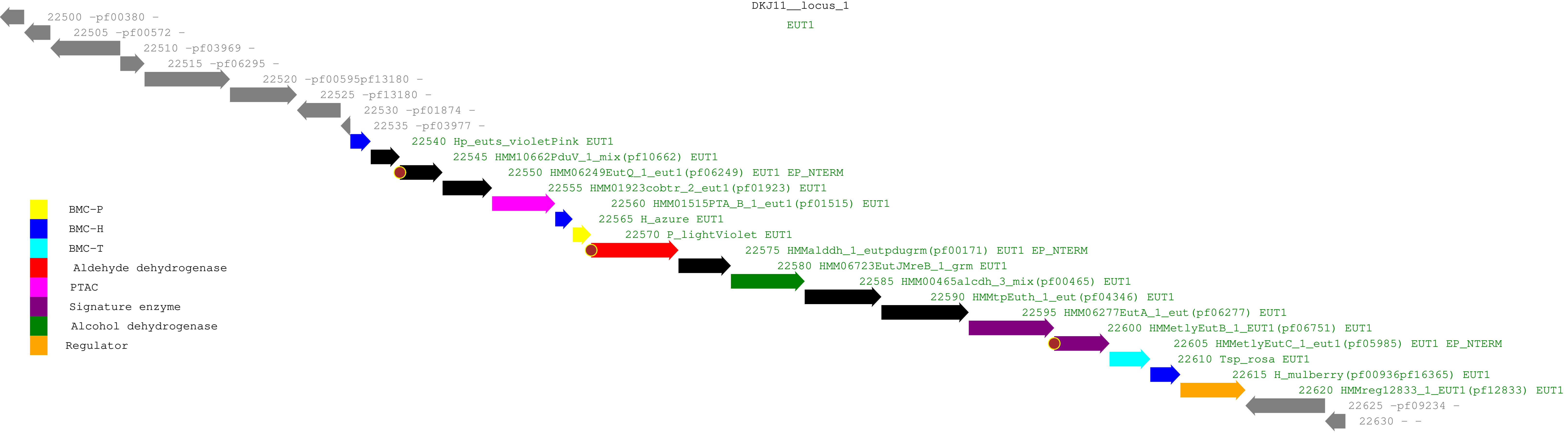

Gammaproteobacteria bacterium ESL0073. Bacteria; Proteobacteria; Gammaproteobacteria.

DKL61__locus_1

EUT1

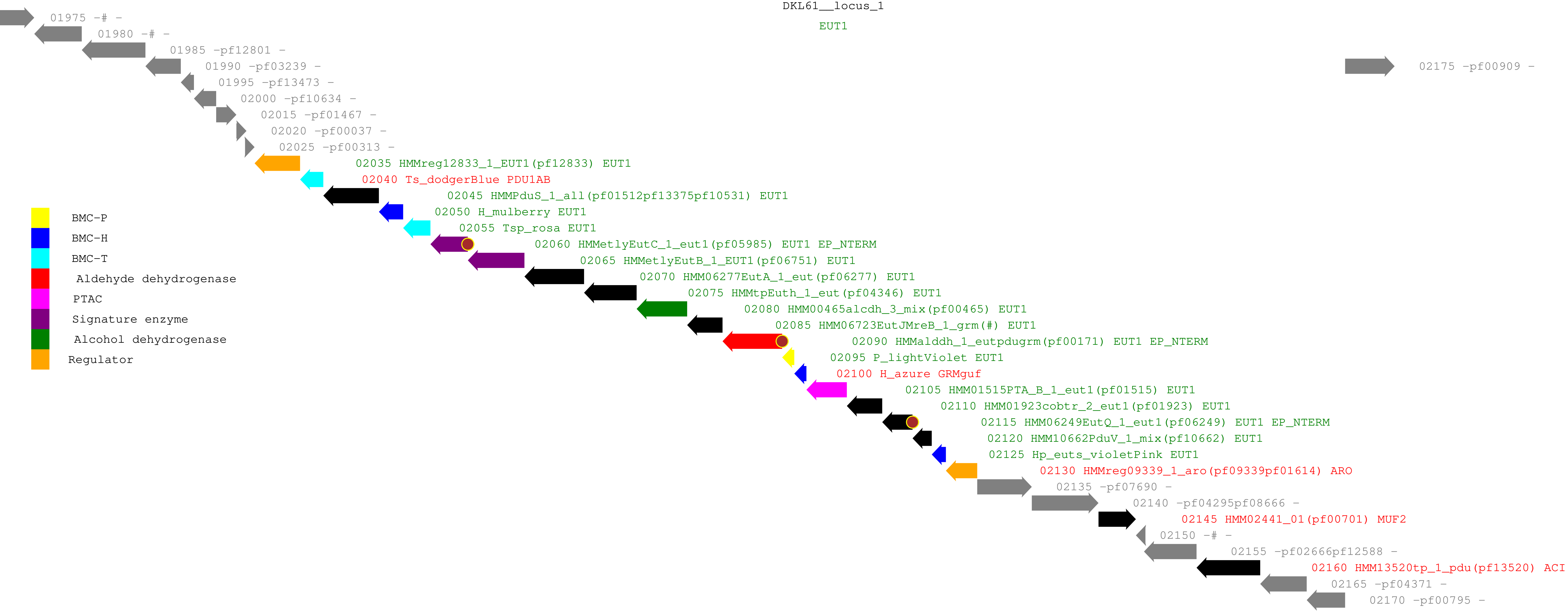

Salmonella diarizonae. Bacteria; Proteobacteria; Gammaproteobacteria; Enterobacterales;

DKP45__locus_1

EUT1

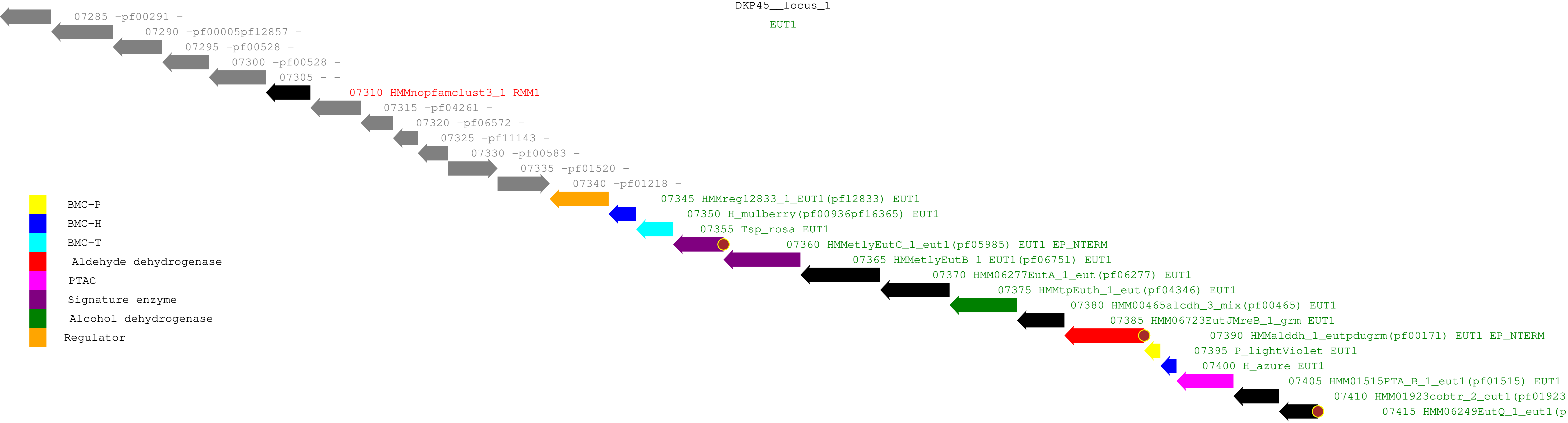

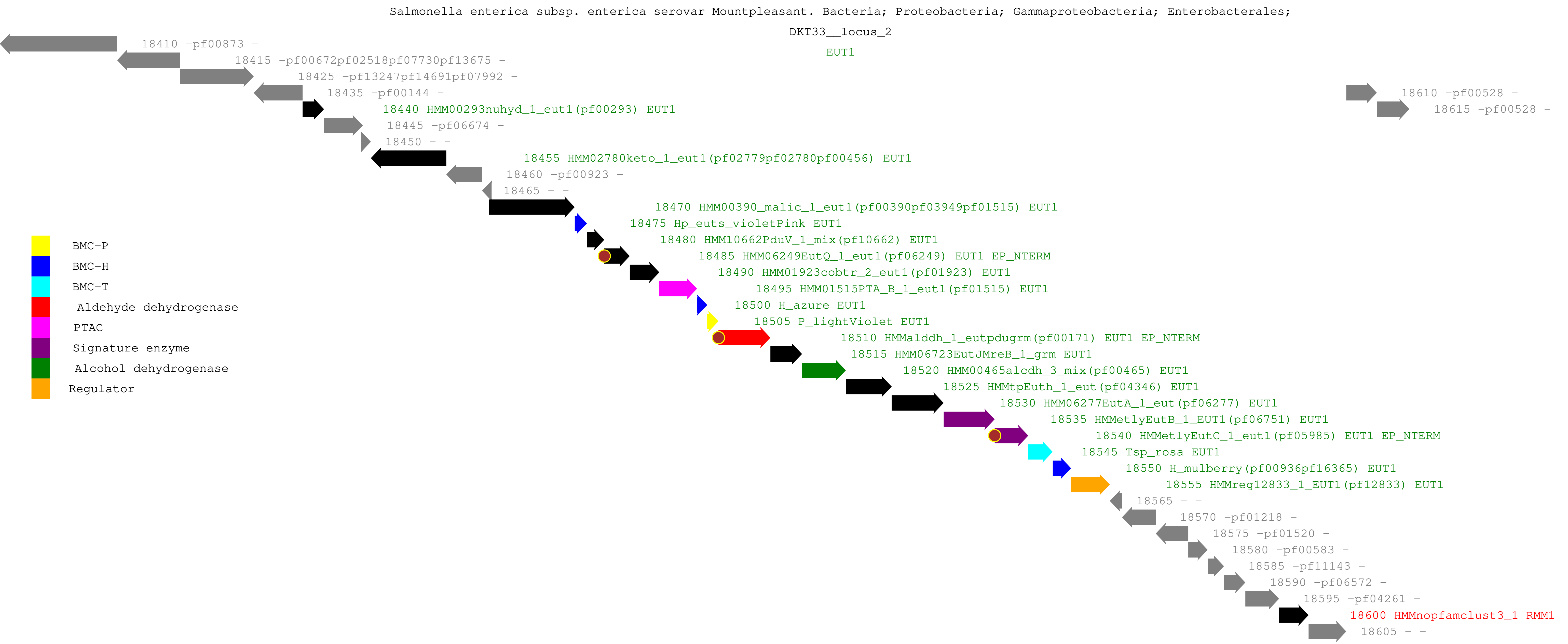
Salmonella enterica subsp. enterica serovar Mountpleasant. Bacteria; Proteobacteria; Gammaproteobacteria; Enterobacterales;
DKT33__locus_2
EUT1
18410 -pf00873 -
18415 -pf00672pf02518pf07730pf13675 -
18425 -pf13247pf14691pf07992 -
18435 -pf00144 -
18440 HMM00293nuhyd_1_eut1(pf00293) EUT1
18445 -pf06674 -
18450 - -
18455 HMM02780keto_1_eut1(pf02779pf02780pf00456) EUT1
18460 -pf00923 -
18465 - -
18470 HMM00390_malic_1_eut1(pf00390pf03949pf01515) EUT1
18475 Hp_euts_violetPink EUT1
18480 HMM10662PduV_1_mix(pf10662) EUT1
18485 HMM06249EutQ_1_eut1(pf06249) EUT1 EP_NTERM
18490 HMM01923cobtr_2_eut1(pf01923) EUT1
18495 HMM01515PTA_B_1_eut1(pf01515) EUT1
18500 H_azure EUT1
18505 P_lightViolet EUT1
18510 HMMalddh_1_eutpdugrm(pf00171) EUT1 EP_NTERM
18515 HMM06723EutJMreB_1_grm EUT1
18520 HMM00465alcdh_3_mix(pf00465) EUT1
18525 HMMtpEuth_1_eut(pf04346) EUT1
18530 HMM06277EutA_1_eut(pf06277) EUT1
18535 HMMetlyEutB_1_EUT1(pf06751) EUT1
18540 HMMetlyEutC_1_eut1(pf05985) EUT1 EP_NTERM
18545 Tsp_rosa EUT1
18550 H_mulberry(pf00936pf16365) EUT1
18555 HMMreg12833_1_EUT1(pf12833) EUT1
18565 - -
18570 -pf01218 -
18575 -pf01520 -
18580 -pf00583 -
18585 -pf11143 -
18590 -pf06572 -
18595 -pf04261 -
18600 HMMnopfamclust3_1 RMM1
18605 - -
18610 -pf00528 -
18615 -pf00528 -
BMC-P
BMC-H
BMC-T
Aldehyde dehydrogenase
PTAC
Signature enzyme
Alcohol dehydrogenase
Regulator

Salmonella enterica I. Bacteria; Proteobacteria; Gammaproteobacteria; Enterobacterales;

DL097__locus_2

EUT1

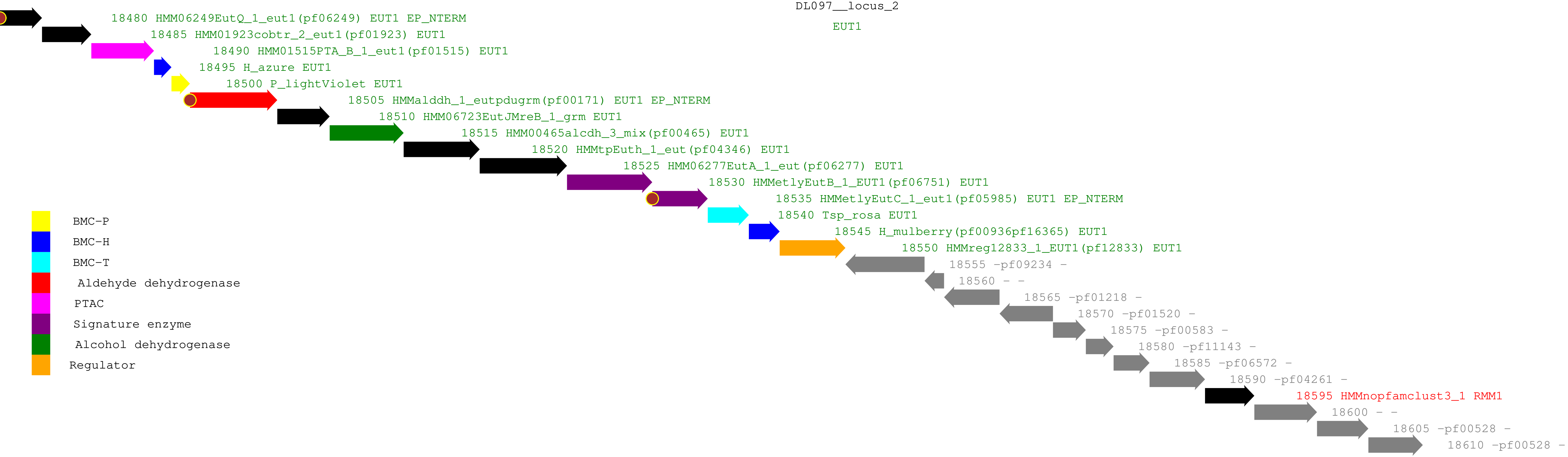

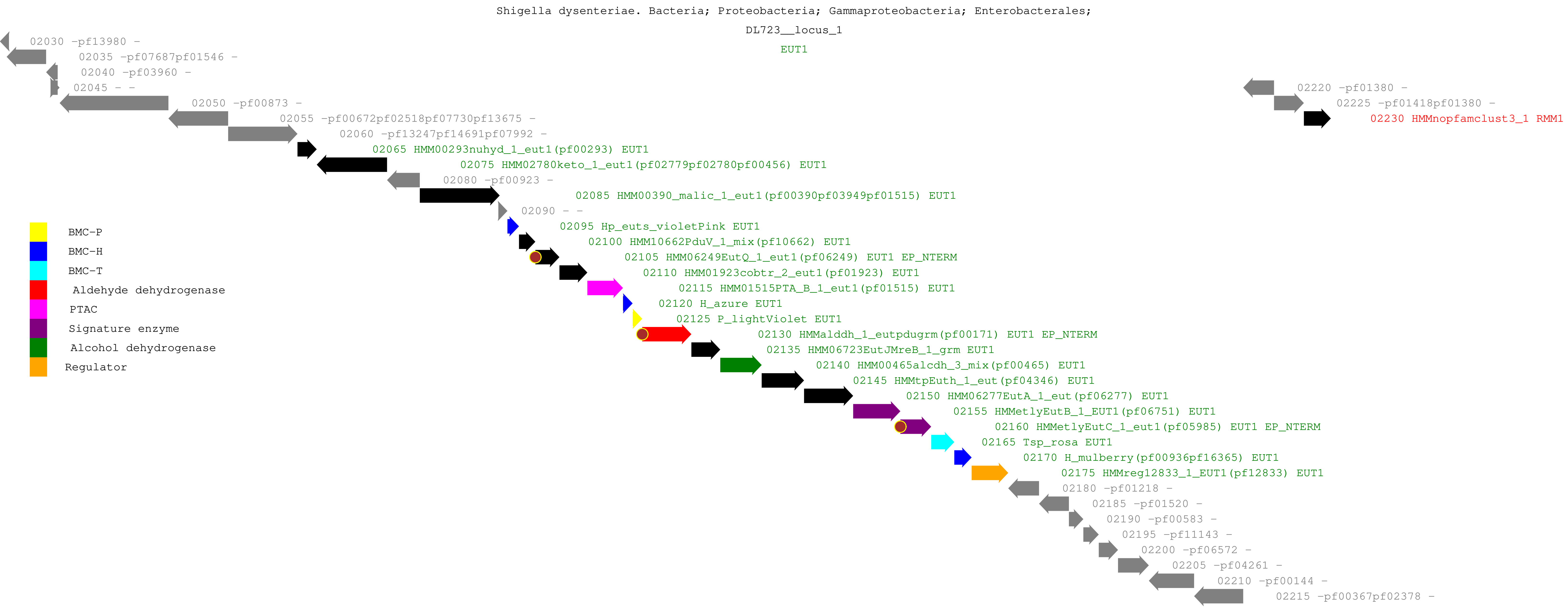
Shigella dysenteriae. Bacteria; Proteobacteria; Gammaproteobacteria; Enterobacterales;
DL723__locus_1
EUT1
02030 -pf13980 -
02035 -pf07687pf01546 -
02040 -pf03960 -
02045 - -
02050 -pf00873 -
02055 -pf00672pf02518pf07730pf13675 -
02060 -pf13247pf14691pf07992 -
02065 HMM00293nuhyd_1_eut1(pf00293) EUT1
02075 HMM02780keto_1_eut1(pf02779pf02780pf00456) EUT1
02080 -pf00923 -
02085 HMM00390_malic_1_eut1(pf00390pf03949pf01515) EUT1
02090 - -
02095 Hp_euts_violetPink EUT1
02100 HMM10662PduV_1_mix(pf10662) EUT1
02105 HMM06249EutQ_1_eut1(pf06249) EUT1 EP_NTERM
02110 HMM01923cobtr_2_eut1(pf01923) EUT1
02115 HMM01515PTA_B_1_eut1(pf01515) EUT1
02120 H_azure EUT1
02125 P_lightViolet EUT1
02130 HMMalddh_1_eutpdugrm(pf00171) EUT1 EP_NTERM
02135 HMM06723EutJMreB_1_grm EUT1
02140 HMM00465alcdh_3_mix(pf00465) EUT1
02145 HMMtpEuth_1_eut(pf04346) EUT1
02150 HMM06277EutA_1_eut(pf06277) EUT1
02155 HMMetlyEutB_1_EUT1(pf06751) EUT1
02160 HMMetlyEutC_1_eut1(pf05985) EUT1 EP_NTERM
02165 Tsp_rosa EUT1
02170 H_mulberry(pf00936pf16365) EUT1
02175 HMMreg12833_1_EUT1(pf12833) EUT1
02180 -pf01218 -
02185 -pf01520 -
02190 -pf00583 -
02195 -pf11143 -
02200 -pf06572 -
02205 -pf04261 -
02210 -pf00144 -
02215 -pf00367pf02378 -
02220 -pf01380 -
02225 -pf01418pf01380 -
02230 HMMnopfamclust3_1 RMM1
BMC-P
BMC-H
BMC-T
Aldehyde dehydrogenase
PTAC
Signature enzyme
Alcohol dehydrogenase
Regulator

Salmonella typhimurium. Bacteria; Proteobacteria; Gammaproteobacteria; Enterobacterales;

DLM31__locus_1

EUT1

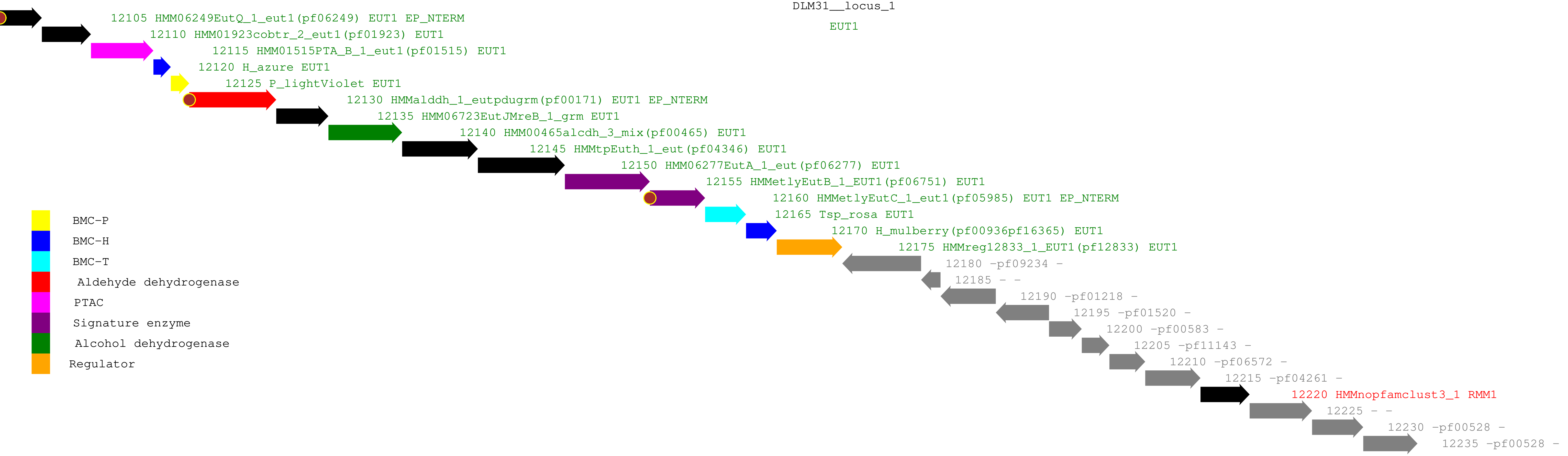

Klebsiella pneumoniae. Bacteria; Proteobacteria; Gammaproteobacteria; Enterobacterales;

DM059__locus_1

EUT1

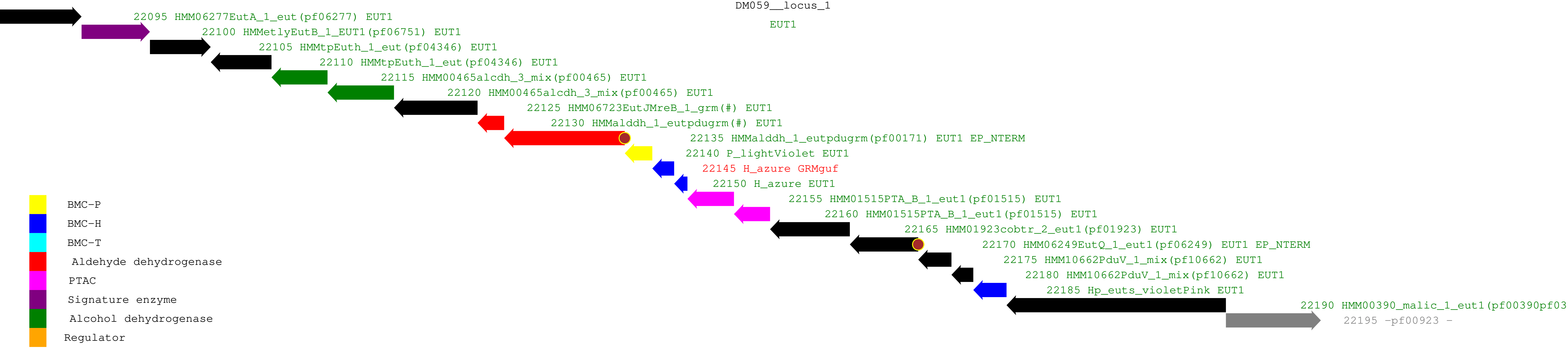

Klebsiella pneumoniae. Bacteria; Proteobacteria; Gammaproteobacteria; Enterobacterales;

DM060__locus_1

EUT1

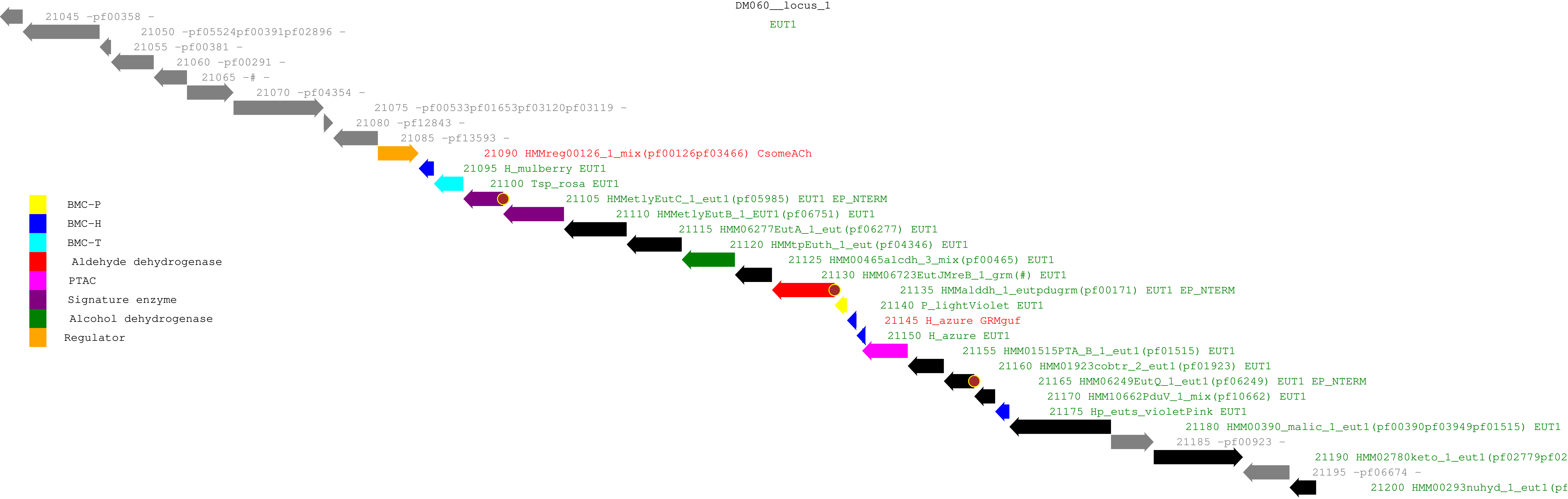

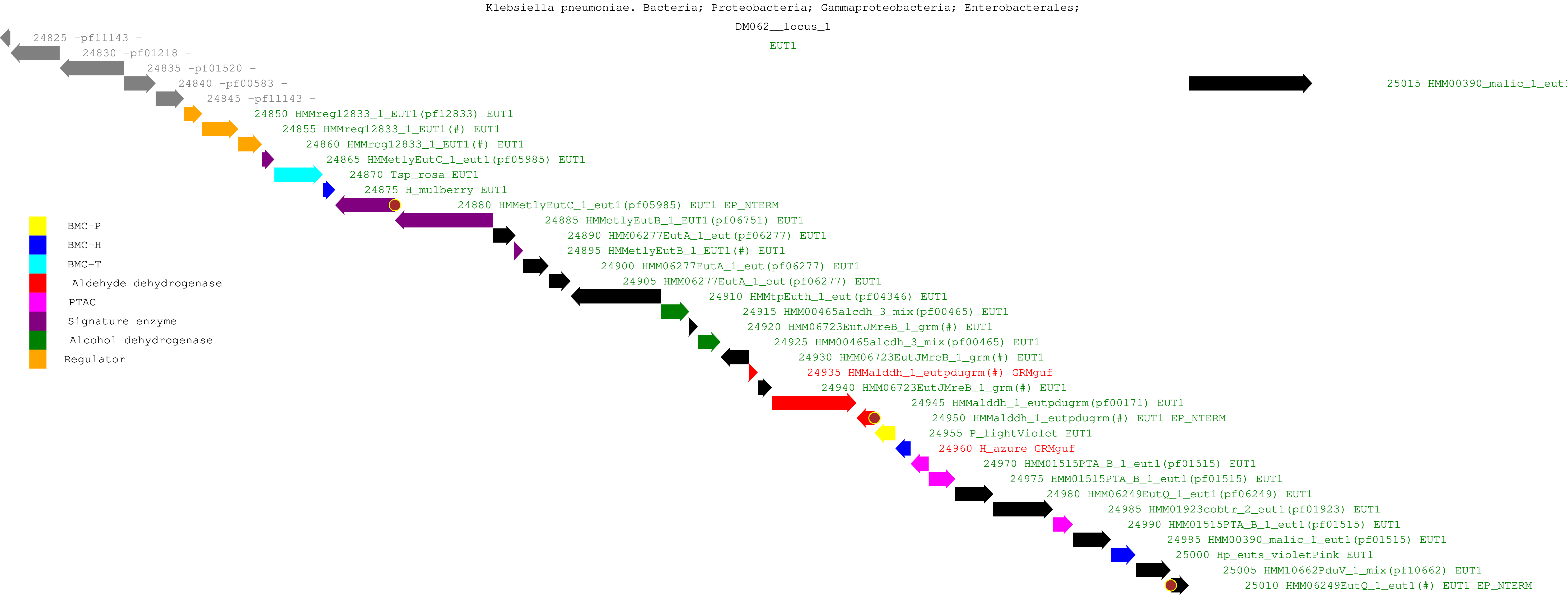

Klebsiella pneumoniae. Bacteria; Proteobacteria; Gammaproteobacteria; Enterobacterales;
DM062__locus_1
EUT1
24825 -pf11143 -
24830 -pf01218 -
24835 -pf01520 -
24840 -pf00583 -
24845 -pf11143 -
24850 HMMreg12833_1_EUT1(pf12833) EUT1
24855 HMMreg12833_1_EUT1(#) EUT1
24860 HMMreg12833_1_EUT1(#) EUT1
24865 HMMetlyEutC_1_eut1(pf05985) EUT1
24870 Tsp_rosa EUT1
24875 H_mulberry EUT1
24880 HMMetlyEutC_1_eut1(pf05985) EUT1 EP_NTERM
24885 HMMetlyEutB_1_EUT1(pf06751) EUT1
24890 HMM06277EutA_1_eut(pf06277) EUT1
24895 HMMetlyEutB_1_EUT1(#) EUT1
24900 HMM06277EutA_1_eut(pf06277) EUT1
24905 HMM06277EutA_1_eut(pf06277) EUT1
24910 HMMtpEuth_1_eut(pf04346) EUT1
24915 HMM00465alcdh_3_mix(pf00465) EUT1
24920 HMM06723EutJMreB_1_grm(#) EUT1
24925 HMM00465alcdh_3_mix(pf00465) EUT1
24930 HMM06723EutJMreB_1_grm(#) EUT1
24935 HMMalddh_1_eutpdugrm(#) GRMguf
24940 HMM06723EutJMreB_1_grm(#) EUT1
24945 HMMalddh_1_eutpdugrm(pf00171) EUT1
24950 HMMalddh_1_eutpdugrm(#) EUT1 EP_NTERM
24955 P_lightViolet EUT1
24960 H_azure GRMguf
24970 HMM01515PTA_B_1_eut1(pf01515) EUT1
24975 HMM01515PTA_B_1_eut1(pf01515) EUT1
24980 HMM06249EutQ_1_eut1(pf06249) EUT1
24985 HMM01923cobtr_2_eut1(pf01923) EUT1
24990 HMM01515PTA_B_1_eut1(pf01515) EUT1
24995 HMM00390_malic_1_eut1(pf01515) EUT1
25000 Hp_euts_violetPink EUT1
25005 HMM10662PduV_1_mix(pf10662) EUT1
25010 HMM06249EutQ_1_eut1(#) EUT1 EP_NTERM
25015 HMM00390_malic_1_eut1
BMC-P
BMC-H
BMC-T
Aldehyde dehydrogenase
PTAC
Signature enzyme
Alcohol dehydrogenase
Regulator

Escherichia coli. Bacteria; Proteobacteria; Gammaproteobacteria; Enterobacterales;

DM102__locus_1

EUT1

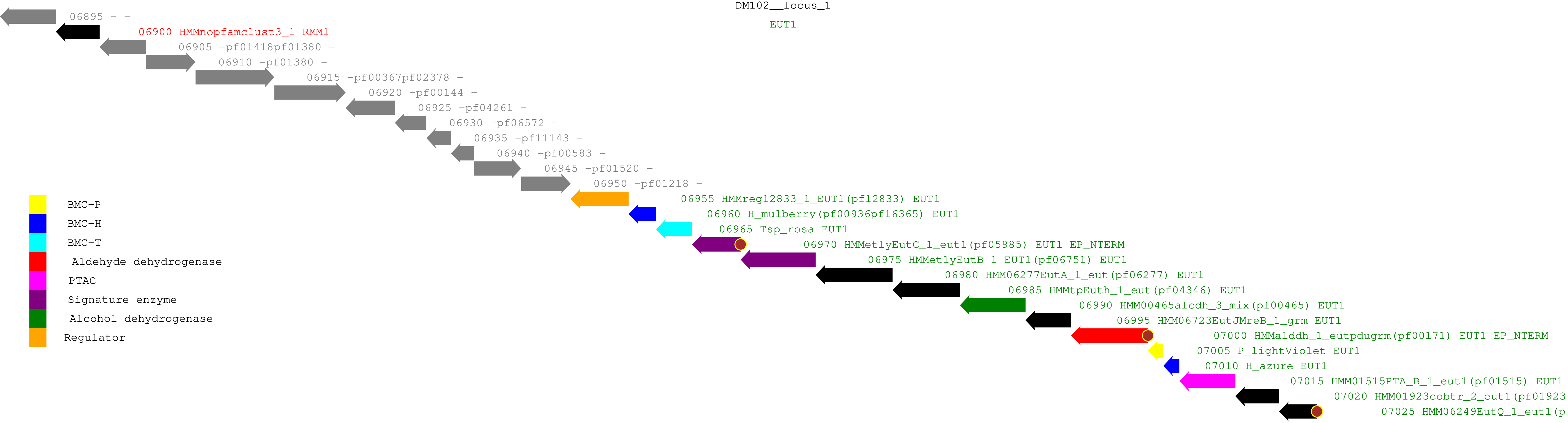

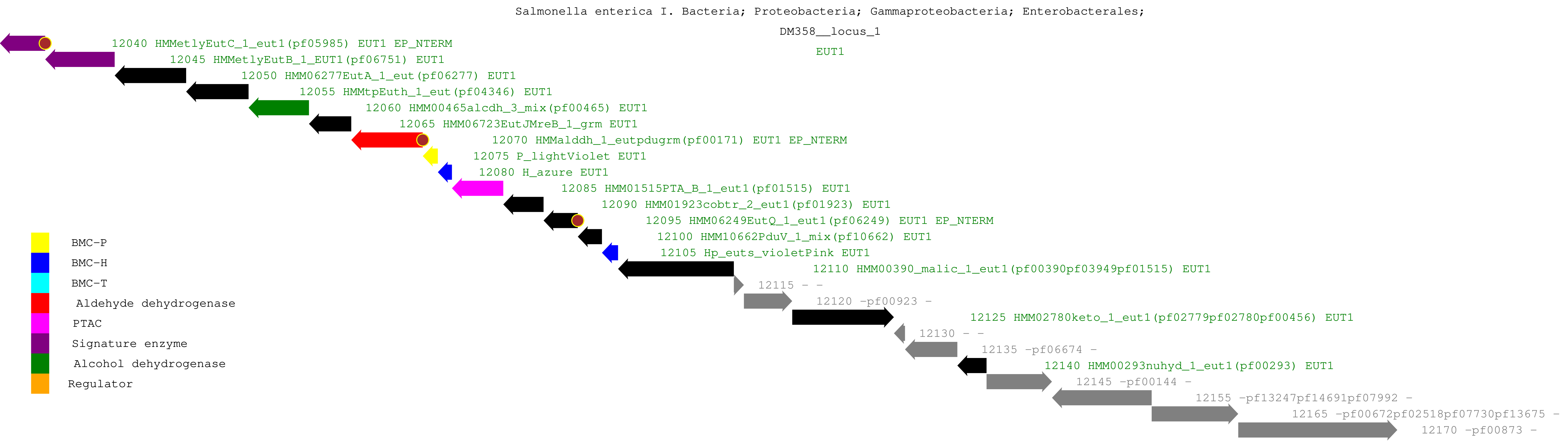
Salmonella enterica I. Bacteria; Proteobacteria; Gammaproteobacteria; Enterobacterales;
DM358__locus_1
EUT1
12040 HMMetlyEutC_1_eut1(pf05985) EUT1 EP_NTERM
12045 HMMetlyEutB_1_EUT1(pf06751) EUT1
12050 HMM06277EutA_1_eut(pf06277) EUT1
12055 HMMtpEuth_1_eut(pf04346) EUT1
12060 HMM00465alcdh_3_mix(pf00465) EUT1
12065 HMM06723EutJMreB_1_grm EUT1
12070 HMMalddh_1_eutpdugrm(pf00171) EUT1 EP_NTERM
12075 P_lightViolet EUT1
12080 H_azure EUT1
12085 HMM01515PTA_B_1_eut1(pf01515) EUT1
12090 HMM01923cobtr_2_eut1(pf01923) EUT1
12095 HMM06249EutQ_1_eut1(pf06249) EUT1 EP_NTERM
12100 HMM10662PduV_1_mix(pf10662) EUT1
12105 Hp_euts_violetPink EUT1
12110 HMM00390_malic_1_eut1(pf00390pf03949pf01515) EUT1
12115 - -
12120 -pf00923 -
12125 HMM02780keto_1_eut1(pf02779pf02780pf00456) EUT1
12130 - -
12135 -pf06674 -
12140 HMM00293nuhyd_1_eut1(pf00293) EUT1
12145 -pf00144 -
12155 -pf13247pf14691pf07992 -
12165 -pf00672pf02518pf07730pf13675 -
12170 -pf00873 -
BMC-P
BMC-H
BMC-T
Aldehyde dehydrogenase
PTAC
Signature enzyme
Alcohol dehydrogenase
Regulator

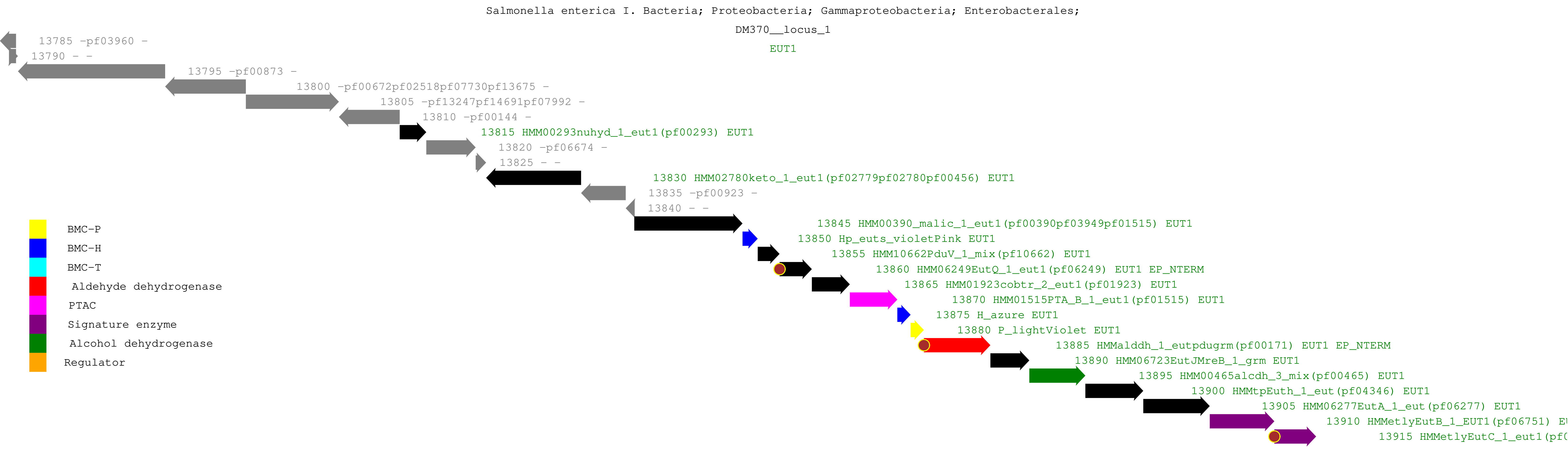
Salmonella enterica I. Bacteria; Proteobacteria; Gammaproteobacteria; Enterobacterales;
DM370__locus_1
EUT1
13785 -pf03960 -
13790 - -
13795 -pf00873 -
13800 -pf00672pf02518pf07730pf13675 -
13805 -pf13247pf14691pf07992 -
13810 -pf00144 -
13815 HMM00293nuhyd_1_eut1(pf00293) EUT1
13820 -pf06674 -
13825 - -
13830 HMM02780keto_1_eut1(pf02779pf02780pf00456) EUT1
13835 -pf00923 -
13840 - -
13845 HMM00390_malic_1_eut1(pf00390pf03949pf01515) EUT1
13850 Hp_euts_violetPink EUT1
13855 HMM10662PduV_1_mix(pf10662) EUT1
13860 HMM06249EutQ_1_eut1(pf06249) EUT1 EP_NTERM
13865 HMM01923cobtr_2_eut1(pf01923) EUT1
13870 HMM01515PTA_B_1_eut1(pf01515) EUT1
13875 H_azure EUT1
13880 P_lightViolet EUT1
13885 HMMalddh_1_eutpdugrm(pf00171) EUT1 EP_NTERM
13890 HMM06723EutJMreB_1_grm EUT1
13895 HMM00465alcdh_3_mix(pf00465) EUT1
13900 HMMtpEuth_1_eut(pf04346) EUT1
13905 HMM06277EutA_1_eut(pf06277) EUT1
13910 HMMetlyEutB_1_EUT1(pf06751)
13915 HMMetlyEutC_1_eut1
BMC-P
BMC-H
BMC-T
Aldehyde dehydrogenase
PTAC
Signature enzyme
Alcohol dehydrogenase
Regulator

Salmonella enterica subsp. enterica serovar Telhashomer. Bacteria; Proteobacteria; Gammaproteobacteria; Enterobacterales;

DM732__locus_2

EUT1

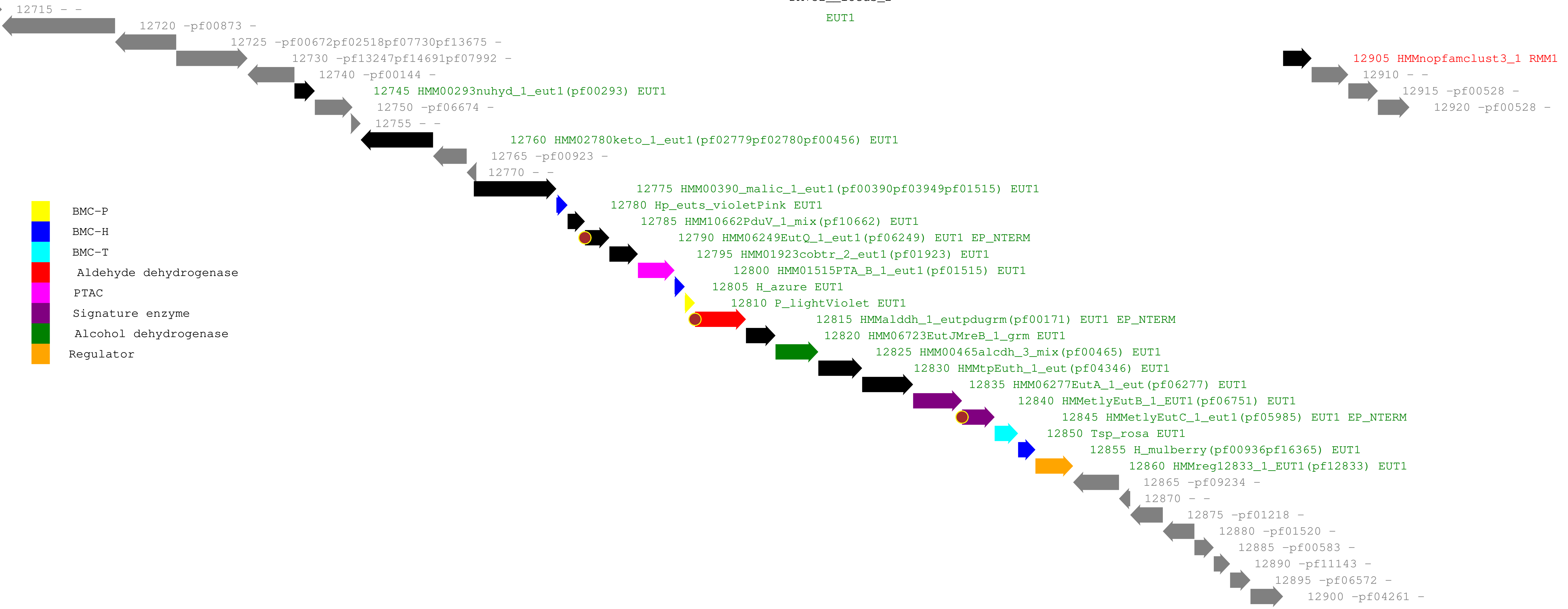

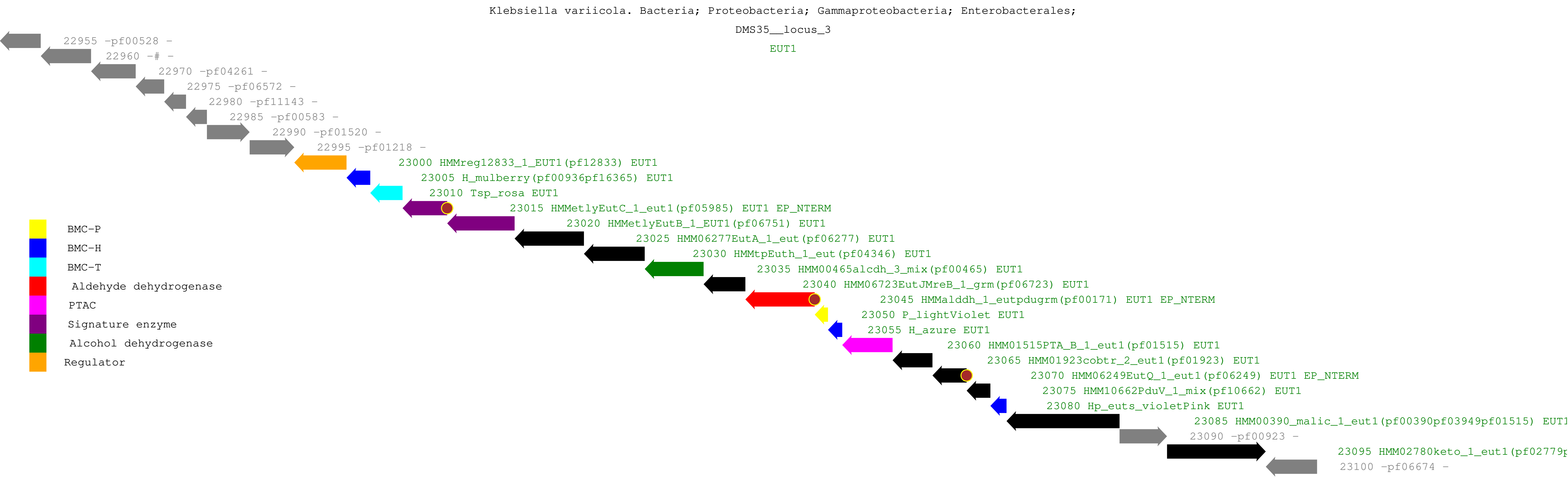
Klebsiella variicola. Bacteria; Proteobacteria; Gammaproteobacteria; Enterobacterales;
DMS35__locus_3
EUT1
22955 -pf00528 -
22960 -# -
22970 -pf04261 -
22975 -pf06572 -
22980 -pf11143 -
22985 -pf00583 -
22990 -pf01520 -
22995 -pf01218 -
23000 HMMreg12833_1_EUT1(pf12833) EUT1
23005 H_mulberry(pf00936pf16365) EUT1
23010 Tsp_rosa EUT1
23015 HMMetlyEutC_1_eut1(pf05985) EUT1 EP_NTERM
23020 HMMetlyEutB_1_EUT1(pf06751) EUT1
23025 HMM06277EutA_1_eut(pf06277) EUT1
23030 HMMtpEuth_1_eut(pf04346) EUT1
23035 HMM00465alcdh_3_mix(pf00465) EUT1
23040 HMM06723EutJMreB_1_grm(pf06723) EUT1
23045 HMMalddh_1_eutpdugrm(pf00171) EUT1 EP_NTERM
23050 P_lightViolet EUT1
23055 H_azure EUT1
23060 HMM01515PTA_B_1_eut1(pf01515) EUT1
23065 HMM01923cobtr_2_eut1(pf01923) EUT1
23070 HMM06249EutQ_1_eut1(pf06249) EUT1 EP_NTERM
23075 HMM10662PduV_1_mix(pf10662) EUT1
23080 Hp_euts_violetPink EUT1
23085 HMM00390_malic_1_eut1(pf00390pf03949pf01515) EUT
23090 -pf00923 -
23095 HMM02780keto_1_eut1(pf02779p
23100 -pf06674 -
BMC-P
BMC-H
BMC-T
Aldehyde dehydrogenase
PTAC
Signature enzyme
Alcohol dehydrogenase
Regulator

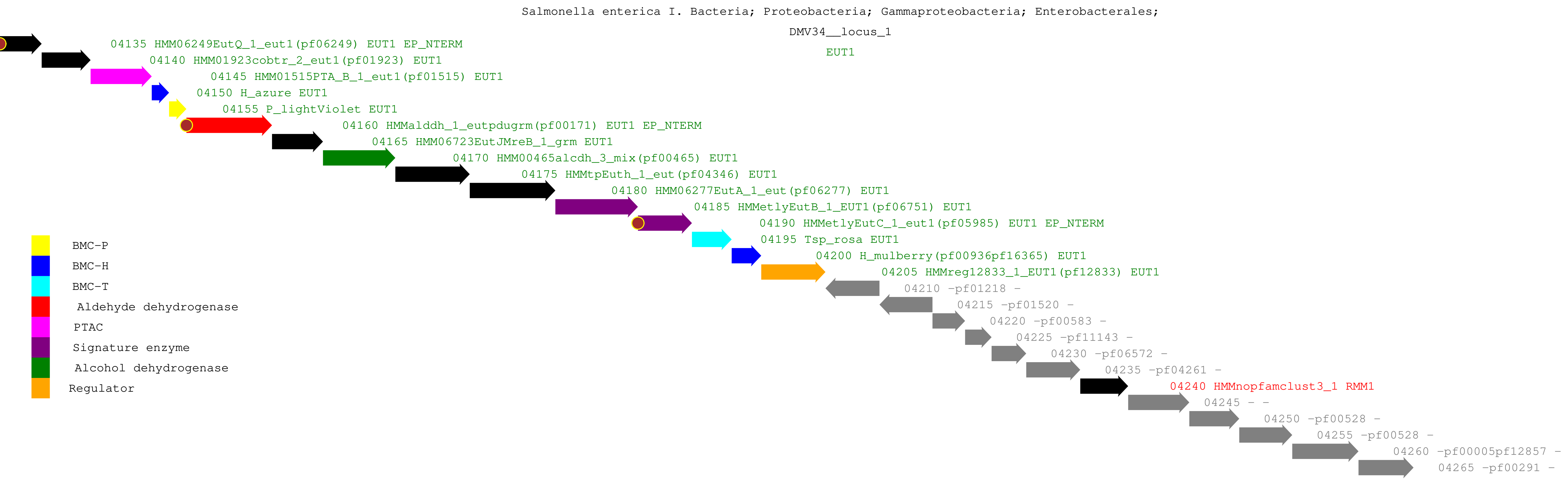
Salmonella enterica I. Bacteria; Proteobacteria; Gammaproteobacteria; Enterobacterales;
DMV34__locus_1
EUT1
04135 HMM06249EutQ_1_eut1(pf06249) EUT1 EP_NTERM
04140 HMM01923cobtr_2_eut1(pf01923) EUT1
04145 HMM01515PTA_B_1_eut1(pf01515) EUT1
04150 H_azure EUT1
04155 P_lightViolet EUT1
04160 HMMalddh_1_eutpdugrm(pf00171) EUT1 EP_NTERM
04165 HMM06723EutJMreB_1_grm EUT1
04170 HMM00465alcdh_3_mix(pf00465) EUT1
04175 HMMtpEuth_1_eut(pf04346) EUT1
04180 HMM06277EutA_1_eut(pf06277) EUT1
04185 HMMetlyEutB_1_EUT1(pf06751) EUT1
04190 HMMetlyEutC_1_eut1(pf05985) EUT1 EP_NTERM
04195 Tsp_rosa EUT1
04200 H_mulberry(pf00936pf16365) EUT1
04205 HMMreg12833_1_EUT1(pf12833) EUT1
04210 -pf01218 -
04215 -pf01520 -
04220 -pf00583 -
04225 -pf11143 -
04230 -pf06572 -
04235 -pf04261 -
04240 HMMnopfamclust3_1 RMM1
04245 - -
04250 -pf00528 -
04255 -pf00528 -
04260 -pf00005pf12857 -
04265 -pf00291 -
BMC-P
BMC-H
BMC-T
Aldehyde dehydrogenase
PTAC
Signature enzyme
Alcohol dehydrogenase
Regulator

Salmonella enterica I. Bacteria; Proteobacteria; Gammaproteobacteria; Enterobacterales;

DN322__locus_1

EUT1

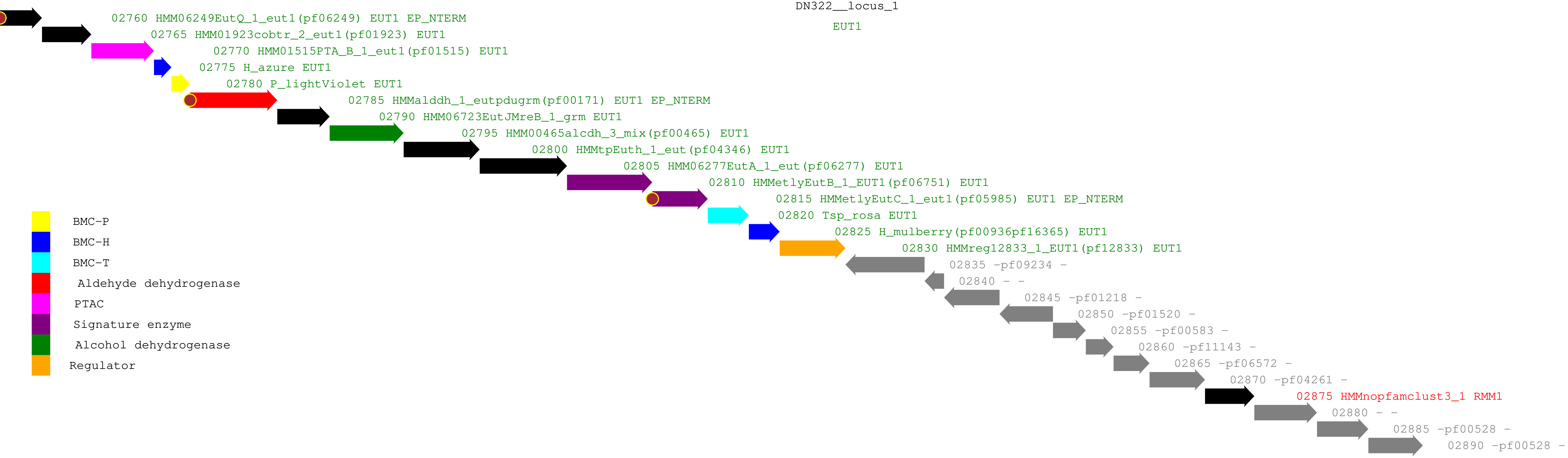

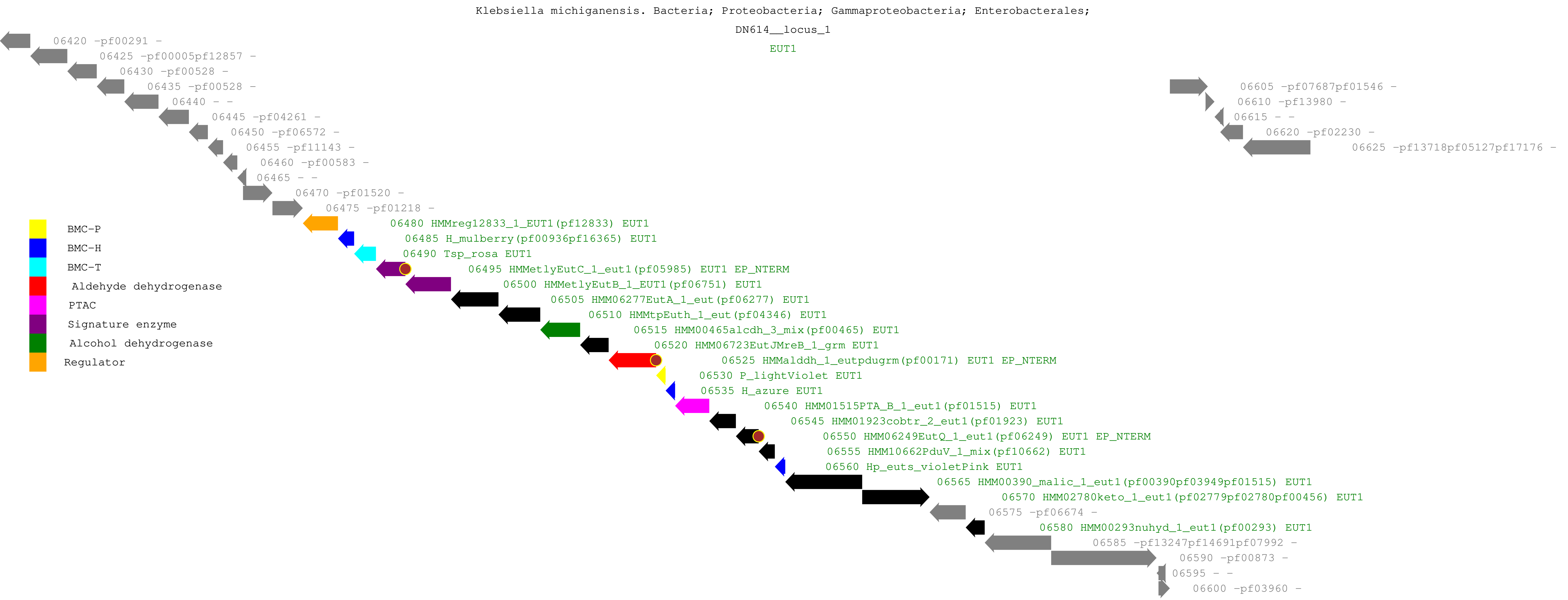

Klebsiella michiganensis. Bacteria; Proteobacteria; Gammaproteobacteria; Enterobacterales;
DN614__locus_1
EUT1
06420 -pf00291 -
06425 -pf00005pf12857 -
06430 -pf00528 -
06435 -pf00528 -
06440 - -
06445 -pf04261 -
06450 -pf06572 -
06455 -pf11143 -
06460 -pf00583 -
06465 - -
06470 -pf01520 -
06475 -pf01218 -
06480 HMMreg12833_1_EUT1(pf12833) EUT1
06485 H_mulberry(pf00936pf16365) EUT1
06490 Tsp_rosa EUT1
06495 HMMetlyEutC_1_eut1(pf05985) EUT1 EP_NTERM
06500 HMMetlyEutB_1_EUT1(pf06751) EUT1
06505 HMM06277EutA_1_eut(pf06277) EUT1
06510 HMMtpEuth_1_eut(pf04346) EUT1
06515 HMM00465alcdh_3_mix(pf00465) EUT1
06520 HMM06723EutJMreB_1_grm EUT1
06525 HMMalddh_1_eutpdugrm(pf00171) EUT1 EP_NTERM
06530 P_lightViolet EUT1
06535 H_azure EUT1
06540 HMM01515PTA_B_1_eut1(pf01515) EUT1
06545 HMM01923cobtr_2_eut1(pf01923) EUT1
06550 HMM06249EutQ_1_eut1(pf06249) EUT1 EP_NTERM
06555 HMM10662PduV_1_mix(pf10662) EUT1
06560 Hp_euts_violetPink EUT1
06565 HMM00390_malic_1_eut1(pf00390pf03949pf01515) EUT1
06570 HMM02780keto_1_eut1(pf02779pf02780pf00456) EUT1
06575 -pf06674 -
06580 HMM00293nuhyd_1_eut1(pf00293) EUT1
06585 -pf13247pf14691pf07992 -
06590 -pf00873 -
06595 - -
06600 -pf03960 -
06605 -pf07687pf01546 -
06610 -pf13980 -
06615 - -
06620 -pf02230 -
06625 -pf13718pf05127pf17176 -
BMC-P
BMC-H
BMC-T
Aldehyde dehydrogenase
PTAC
Signature enzyme
Alcohol dehydrogenase
Regulator

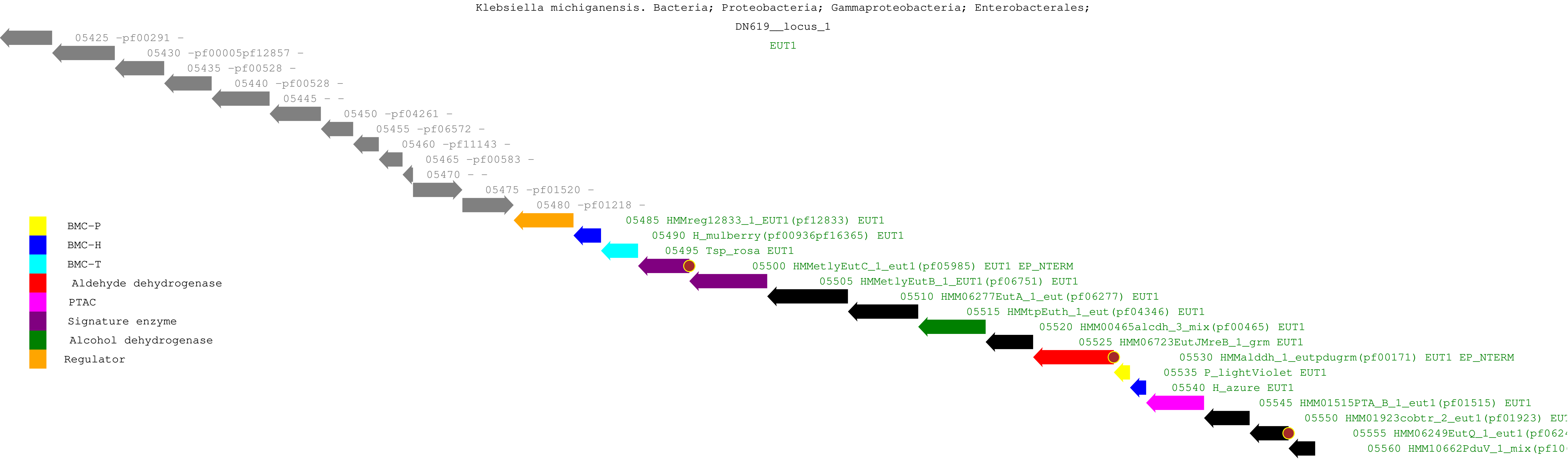
Klebsiella michiganensis. Bacteria; Proteobacteria; Gammaproteobacteria; Enterobacterales;
DN619__locus_1
EUT1
05425 -pf00291 -
05430 -pf00005pf12857 -
05435 -pf00528 -
05440 -pf00528 -
05445 - -
05450 -pf04261 -
05455 -pf06572 -
05460 -pf11143 -
05465 -pf00583 -
05470 - -
05475 -pf01520 -
05480 -pf01218 -
05485 HMMreg12833_1_EUT1(pf12833) EUT1
05490 H_mulberry(pf00936pf16365) EUT1
05495 Tsp_rosa EUT1
05500 HMMetlyEutC_1_eut1(pf05985) EUT1 EP_NTERM
05505 HMMetlyEutB_1_EUT1(pf06751) EUT1
05510 HMM06277EutA_1_eut(pf06277) EUT1
05515 HMMtpEuth_1_eut(pf04346) EUT1
05520 HMM00465alcdh_3_mix(pf00465) EUT1
05525 HMM06723EutJMreB_1_grm EUT1
05530 HMMalddh_1_eutpdugrm(pf00171) EUT1 EP_NTERM
05535 P_lightViolet EUT1
05540 H_azure EUT1
05545 HMM01515PTA_B_1_eut1(pf01515) EUT1
05550 HMM01923cobtr_2_eut1(pf01923) EU
05555 HMM06249EutQ_1_eut1(pf062
05560 HMM10662PduV_1_mix(pf10
BMC-P
BMC-H
BMC-T
Aldehyde dehydrogenase
PTAC
Signature enzyme
Alcohol dehydrogenase
Regulator

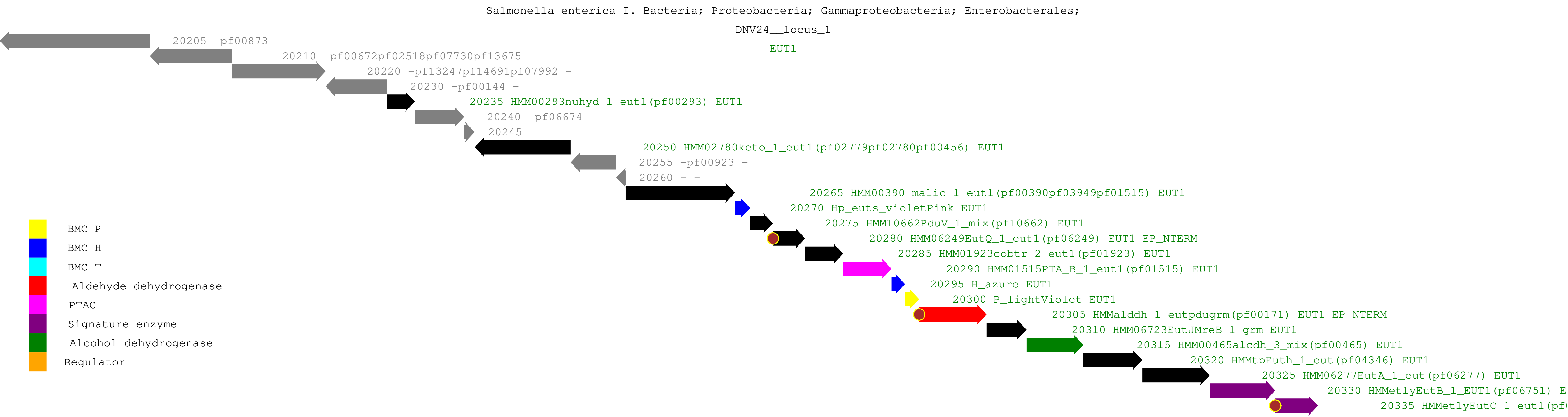
Salmonella enterica I. Bacteria; Proteobacteria; Gammaproteobacteria; Enterobacterales;
DNV24__locus_1
EUT1
20205 -pf00873 -
20210 -pf00672pf02518pf07730pf13675 -
20220 -pf13247pf14691pf07992 -
20230 -pf00144 -
20235 HMM00293nuhyd_1_eut1(pf00293) EUT1
20240 -pf06674 -
20245 - -
20250 HMM02780keto_1_eut1(pf02779pf02780pf00456) EUT1
20255 -pf00923 -
20260 - -
20265 HMM00390_malic_1_eut1(pf00390pf03949pf01515) EUT1
20270 Hp_euts_violetPink EUT1
20275 HMM10662PduV_1_mix(pf10662) EUT1
20280 HMM06249EutQ_1_eut1(pf06249) EUT1 EP_NTERM
20285 HMM01923cobtr_2_eut1(pf01923) EUT1
20290 HMM01515PTA_B_1_eut1(pf01515) EUT1
20295 H_azure EUT1
20300 P_lightViolet EUT1
20305 HMMalddh_1_eutpdugrm(pf00171) EUT1 EP_NTERM
20310 HMM06723EutJMreB_1_grm EUT1
20315 HMM00465alcdh_3_mix(pf00465) EUT1
20320 HMMtpEuth_1_eut(pf04346) EUT1
20325 HMM06277EutA_1_eut(pf06277) EUT1
20330 HMMetlyEutB_1_EUT1(pf06751) E
20335 HMMetlyEutC_1_eut1(pf
BMC-P
BMC-H
BMC-T
Aldehyde dehydrogenase
PTAC
Signature enzyme
Alcohol dehydrogenase
Regulator

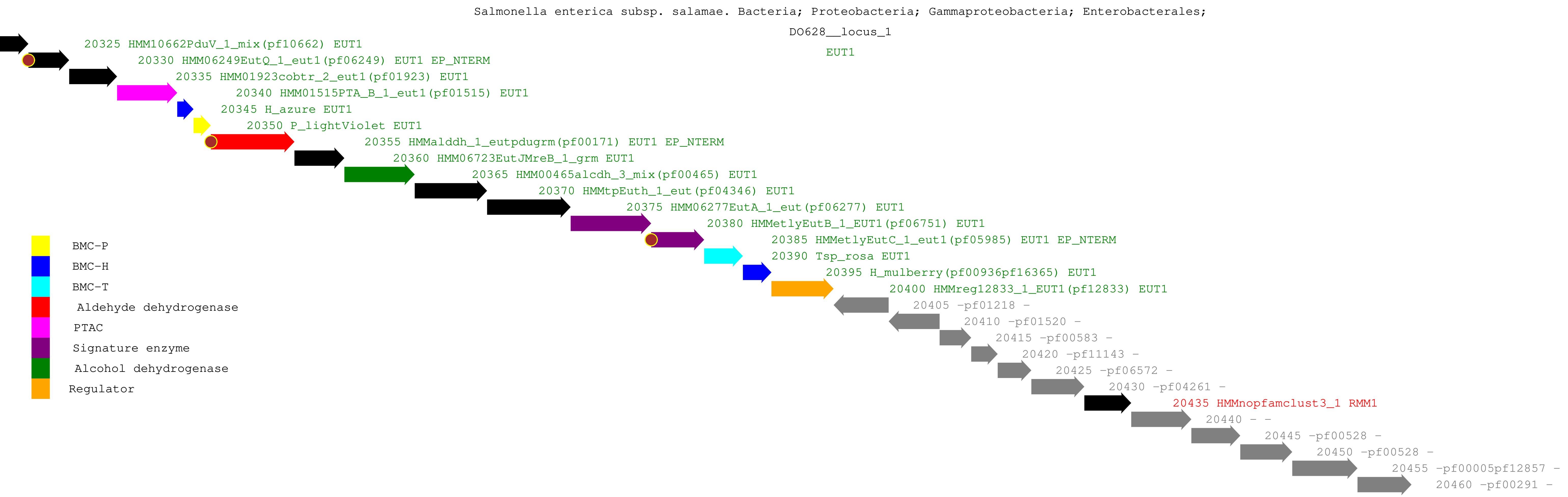
Salmonella enterica subsp. salamae. Bacteria; Proteobacteria; Gammaproteobacteria; Enterobacterales;
DO628__locus_1
EUT1
20325 HMM10662PduV_1_mix(pf10662) EUT1
20330 HMM06249EutQ_1_eut1(pf06249) EUT1 EP_NTERM
20335 HMM01923cobtr_2_eut1(pf01923) EUT1
20340 HMM01515PTA_B_1_eut1(pf01515) EUT1
20345 H_azure EUT1
20350 P_lightViolet EUT1
20355 HMMalddh_1_eutpdugrm(pf00171) EUT1 EP_NTERM
20360 HMM06723EutJMreB_1_grm EUT1
20365 HMM00465alcdh_3_mix(pf00465) EUT1
20370 HMMtpEuth_1_eut(pf04346) EUT1
20375 HMM06277EutA_1_eut(pf06277) EUT1
20380 HMMetlyEutB_1_EUT1(pf06751) EUT1
20385 HMMetlyEutC_1_eut1(pf05985) EUT1 EP_NTERM
20390 Tsp_rosa EUT1
20395 H_mulberry(pf00936pf16365) EUT1
20400 HMMreg12833_1_EUT1(pf12833) EUT1
20405 -pf01218 -
20410 -pf01520 -
20415 -pf00583 -
20420 -pf11143 -
20425 -pf06572 -
20430 -pf04261 -
20435 HMMnopfamclust3_1 RMM1
20440 - -
20445 -pf00528 -
20450 -pf00528 -
20455 -pf00005pf12857 -
20460 -pf00291 -
BMC-P
BMC-H
BMC-T
Aldehyde dehydrogenase
PTAC
Signature enzyme
Alcohol dehydrogenase
Regulator

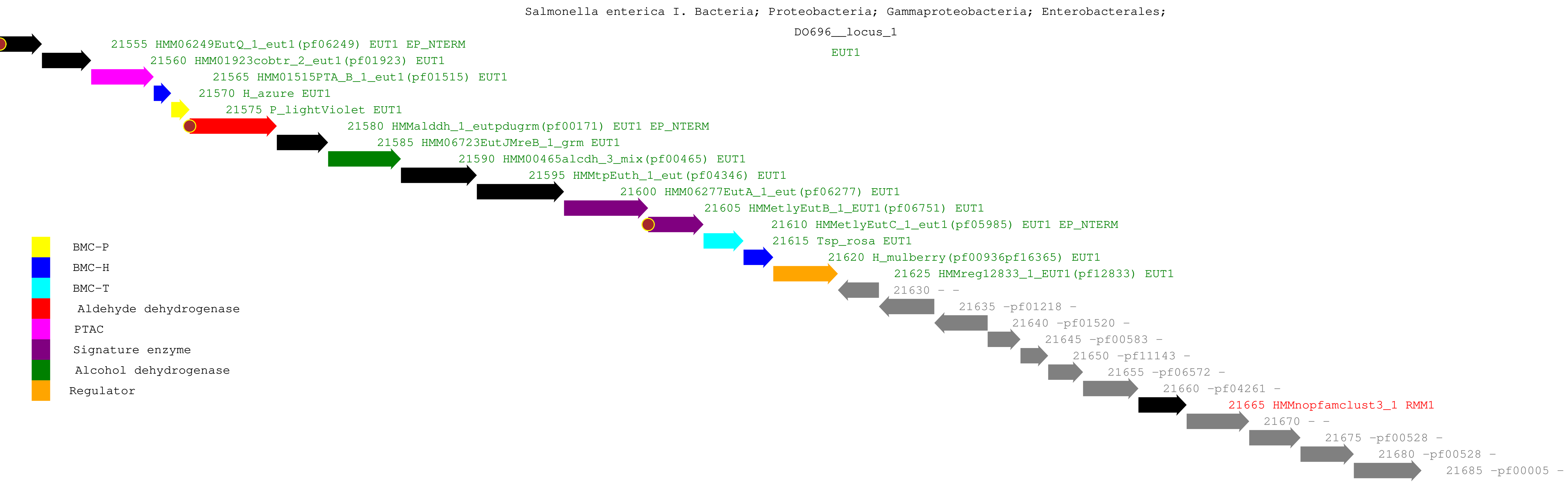
Salmonella enterica I. Bacteria; Proteobacteria; Gammaproteobacteria; Enterobacterales;
DO696__locus_1
EUT1
21555 HMM06249EutQ_1_eut1(pf06249) EUT1 EP_NTERM
21560 HMM01923cobtr_2_eut1(pf01923) EUT1
21565 HMM01515PTA_B_1_eut1(pf01515) EUT1
21570 H_azure EUT1
21575 P_lightViolet EUT1
21580 HMMalddh_1_eutpdugrm(pf00171) EUT1 EP_NTERM
21585 HMM06723EutJMreB_1_grm EUT1
21590 HMM00465alcdh_3_mix(pf00465) EUT1
21595 HMMtpEuth_1_eut(pf04346) EUT1
21600 HMM06277EutA_1_eut(pf06277) EUT1
21605 HMMetlyEutB_1_EUT1(pf06751) EUT1
21610 HMMetlyEutC_1_eut1(pf05985) EUT1 EP_NTERM
21615 Tsp_rosa EUT1
21620 H_mulberry(pf00936pf16365) EUT1
21625 HMMreg12833_1_EUT1(pf12833) EUT1
21630 - -
21635 -pf01218 -
21640 -pf01520 -
21645 -pf00583 -
21650 -pf11143 -
21655 -pf06572 -
21660 -pf04261 -
21665 HMMnopfamclust3_1 RMM1
21670 - -
21675 -pf00528 -
21680 -pf00528 -
21685 -pf00005 -
BMC-P
BMC-H
BMC-T
Aldehyde dehydrogenase
PTAC
Signature enzyme
Alcohol dehydrogenase
Regulator

Salmonella enteritidis. Bacteria; Proteobacteria; Gammaproteobacteria; Enterobacterales;

DO961__locus_1

EUT1

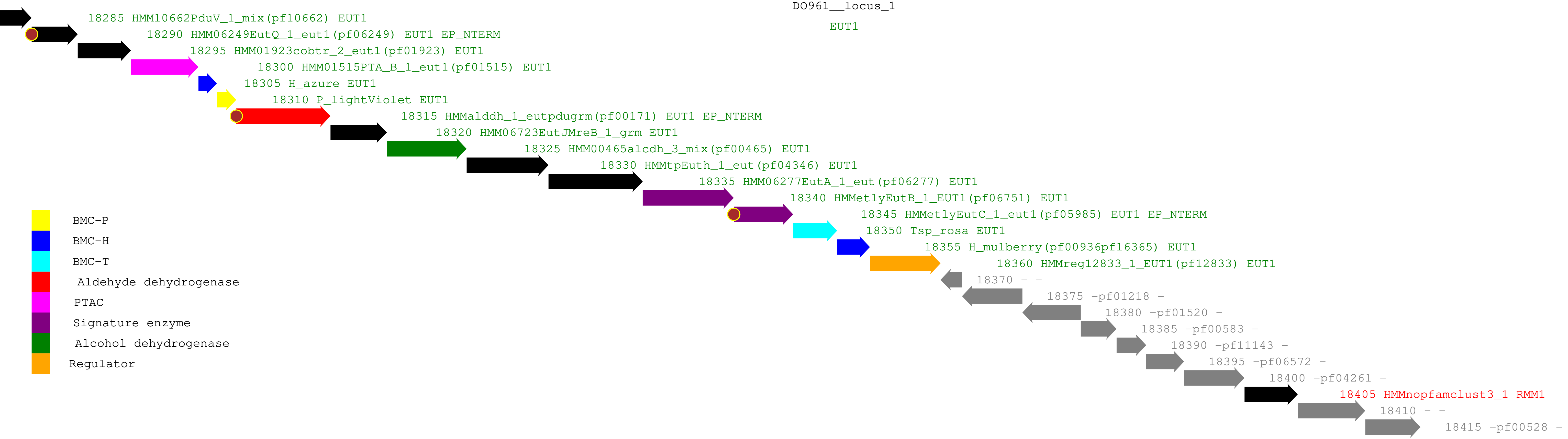

Salmonella enterica I. Bacteria; Proteobacteria; Gammaproteobacteria; Enterobacterales;

DOE57__locus_1

EUT1

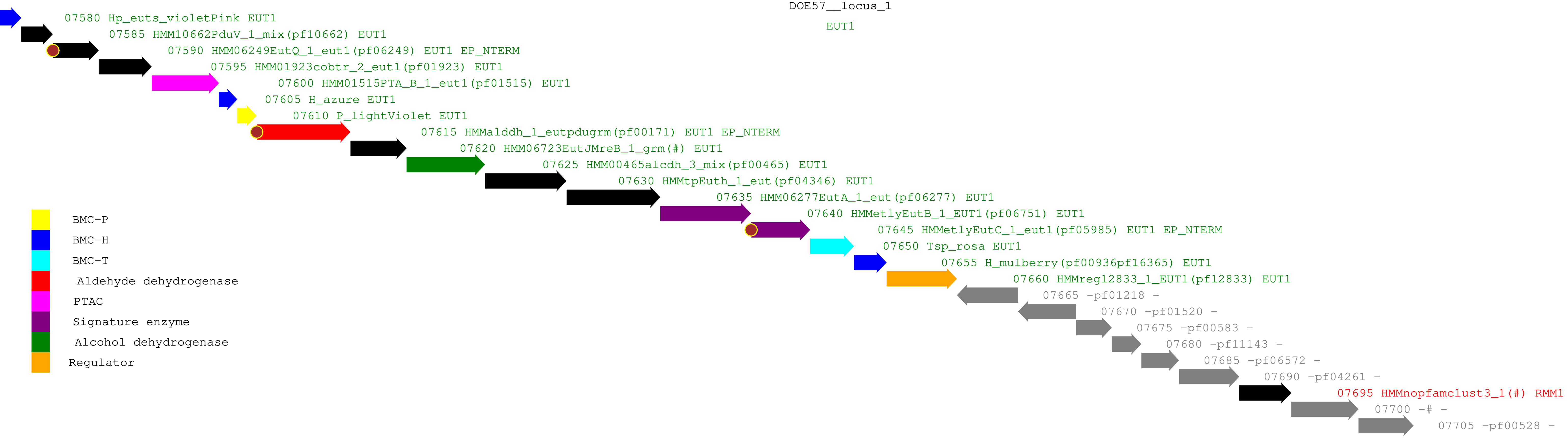

Salmonella enterica I. Bacteria; Proteobacteria; Gammaproteobacteria; Enterobacterales;

DOE63__locus_2

EUT1

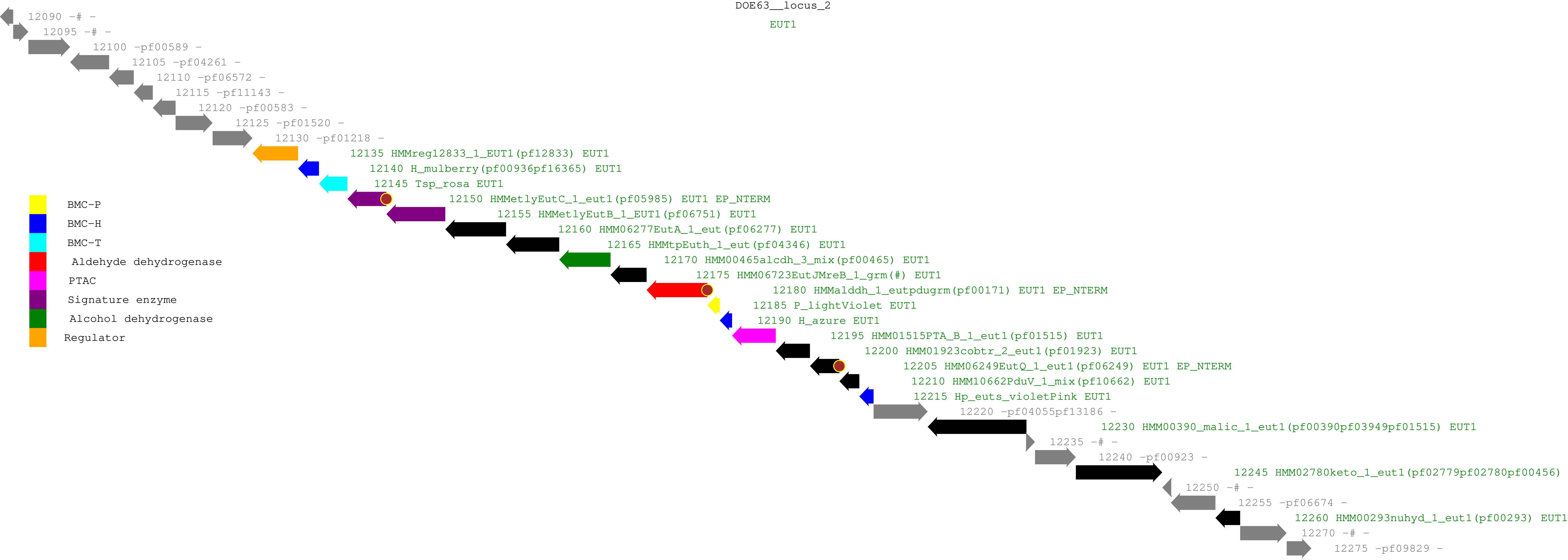

Salmonella enterica I. Bacteria; Proteobacteria; Gammaproteobacteria; Enterobacterales;

DOG94__locus_1

EUT1

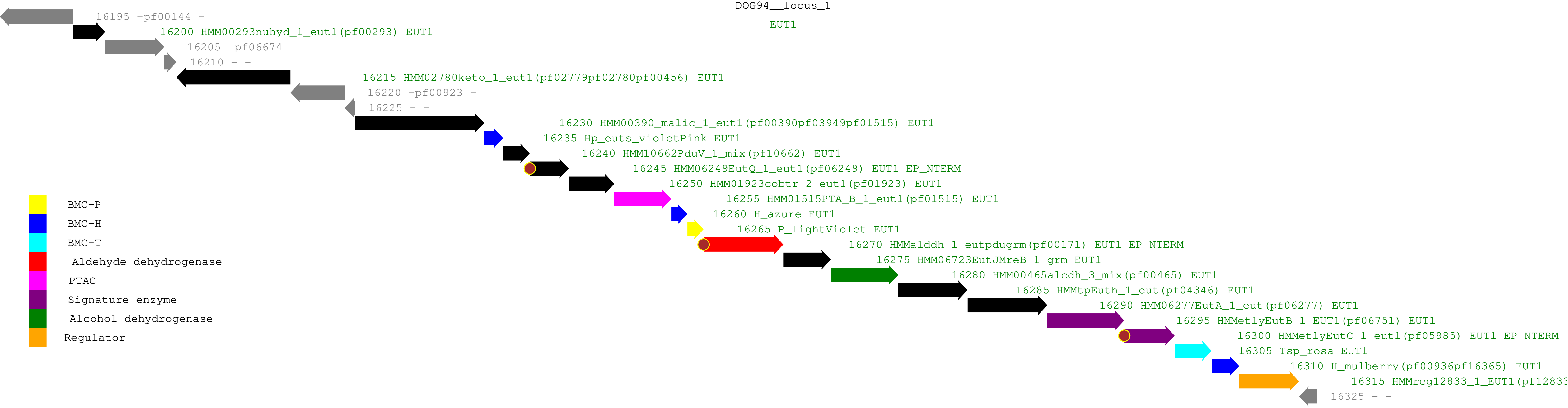

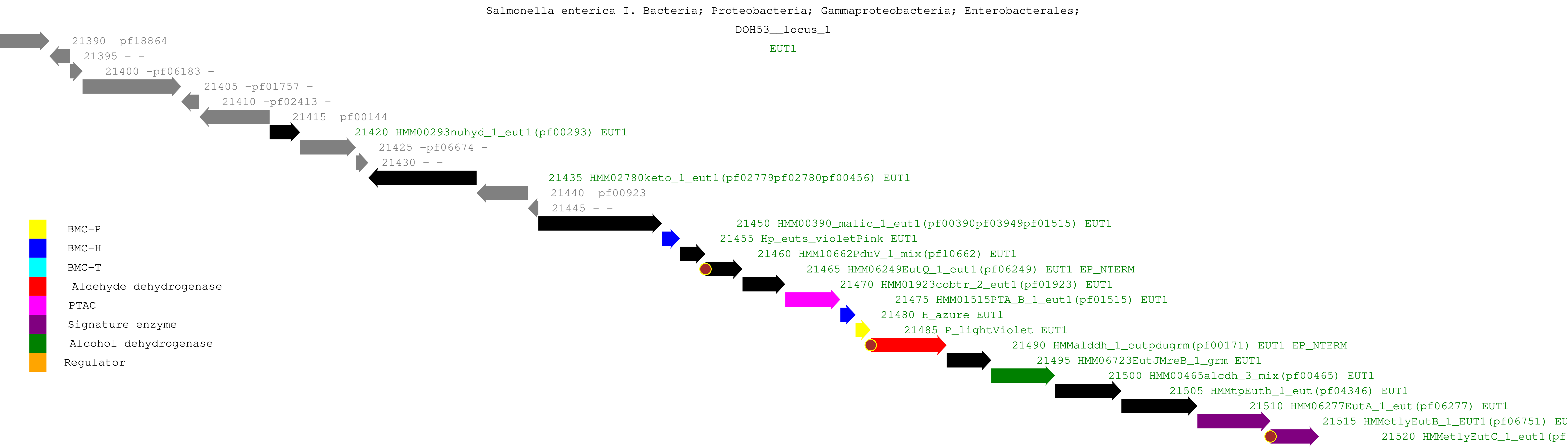

Salmonella enterica I. Bacteria; Proteobacteria; Gammaproteobacteria; Enterobacterales;
DOH53__locus_1
EUT1
21390 -pf18864 -
21395 - -
21400 -pf06183 -
21405 -pf01757 -
21410 -pf02413 -
21415 -pf00144 -
21420 HMM00293nuhyd_1_eut1(pf00293) EUT1
21425 -pf06674 -
21430 - -
21435 HMM02780keto_1_eut1(pf02779pf02780pf00456) EUT1
21440 -pf00923 -
21445 - -
21450 HMM00390_malic_1_eut1(pf00390pf03949pf01515) EUT1
21455 Hp_euts_violetPink EUT1
21460 HMM10662PduV_1_mix(pf10662) EUT1
21465 HMM06249EutQ_1_eut1(pf06249) EUT1 EP_NTERM
21470 HMM01923cobtr_2_eut1(pf01923) EUT1
21475 HMM01515PTA_B_1_eut1(pf01515) EUT1
21480 H_azure EUT1
21485 P_lightViolet EUT1
21490 HMMalddh_1_eutpdugrm(pf00171) EUT1 EP_NTERM
21495 HMM06723EutJMreB_1_grm EUT1
21500 HMM00465alcdh_3_mix(pf00465) EUT1
21505 HMMtpEuth_1_eut(pf04346) EUT1
21510 HMM06277EutA_1_eut(pf06277) EUT1
21515 HMMetlyEutB_1_EUT1(pf06751) EU
21520 HMMetlyEutC_1_eut1(pf
BMC-P
BMC-H
BMC-T
Aldehyde dehydrogenase
PTAC
Signature enzyme
Alcohol dehydrogenase
Regulator

Salmonella newport. Bacteria; Proteobacteria; Gammaproteobacteria; Enterobacterales;

DOI40__locus_2

EUT1

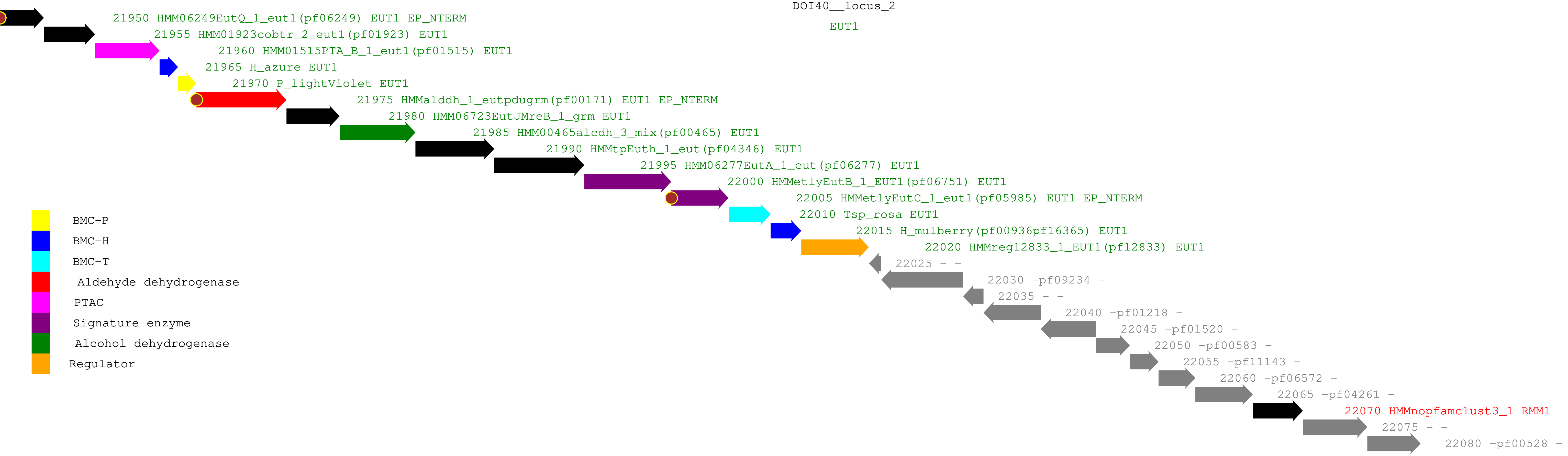

Shigella flexneri. Bacteria; Proteobacteria; Gammaproteobacteria; Enterobacterales;

DOU91__locus_1_main_sat

EUT1

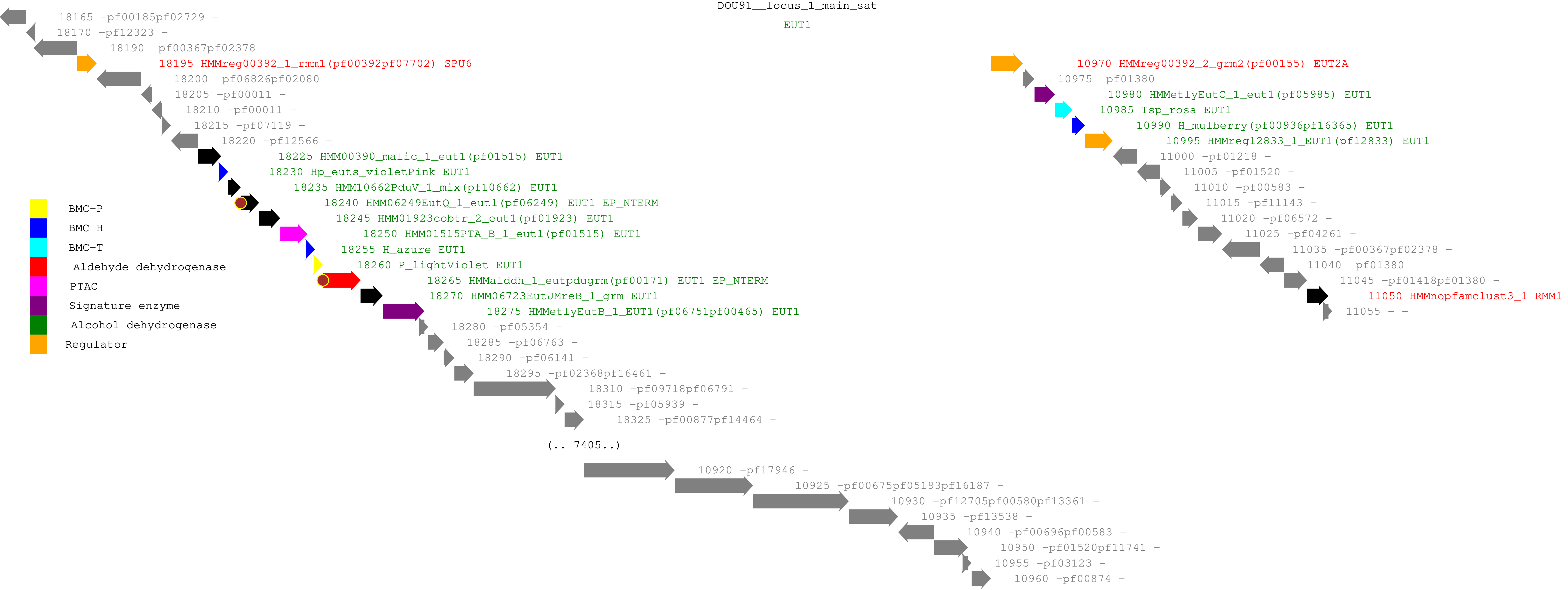

Salmonella enterica subsp. enterica serovar Give. Bacteria; Proteobacteria; Gammaproteobacteria; Enterobacterales;

DOW88__locus_2

EUT1

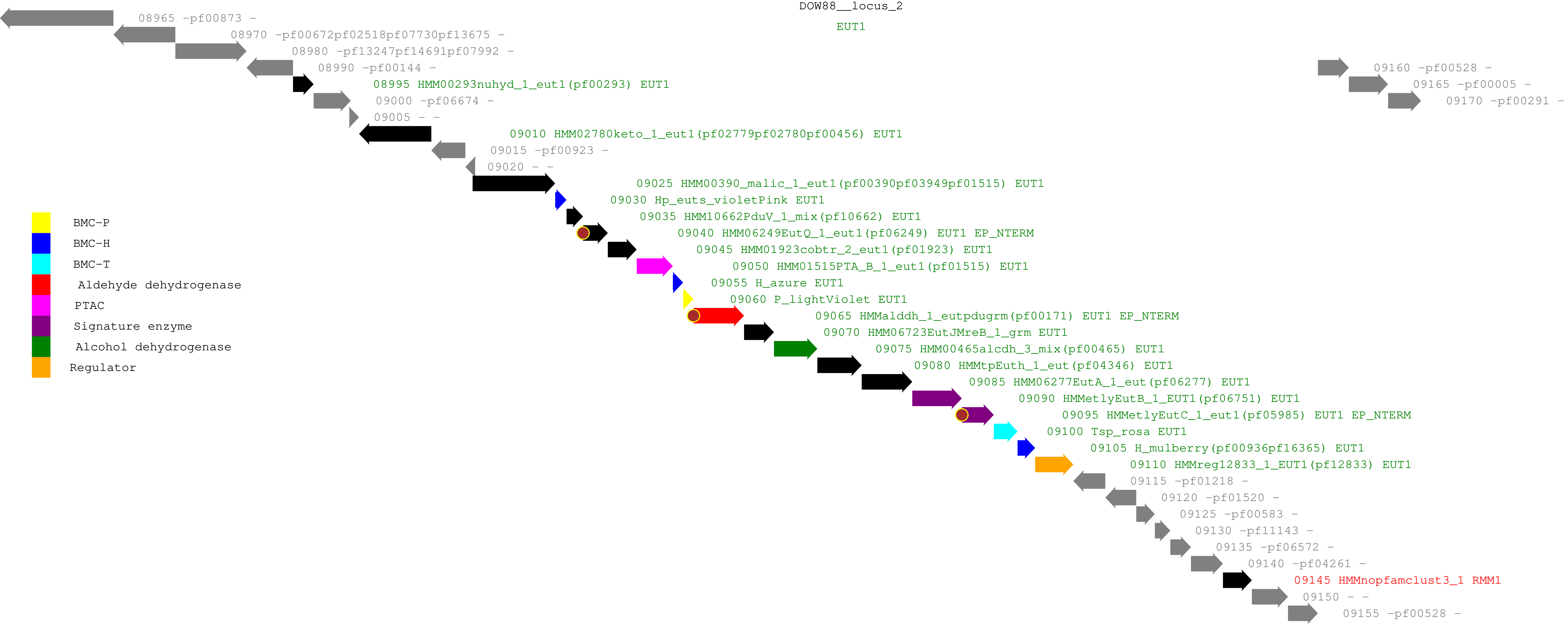

Escherichia coli. Bacteria; Proteobacteria; Gammaproteobacteria; Enterobacterales;

DP277__locus_1

EUT1

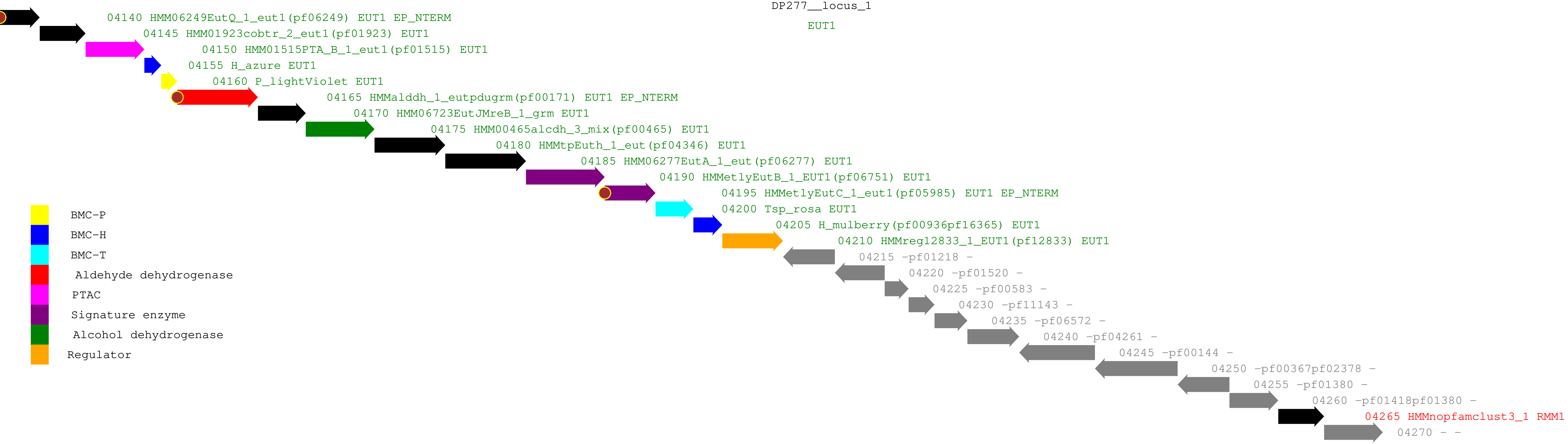

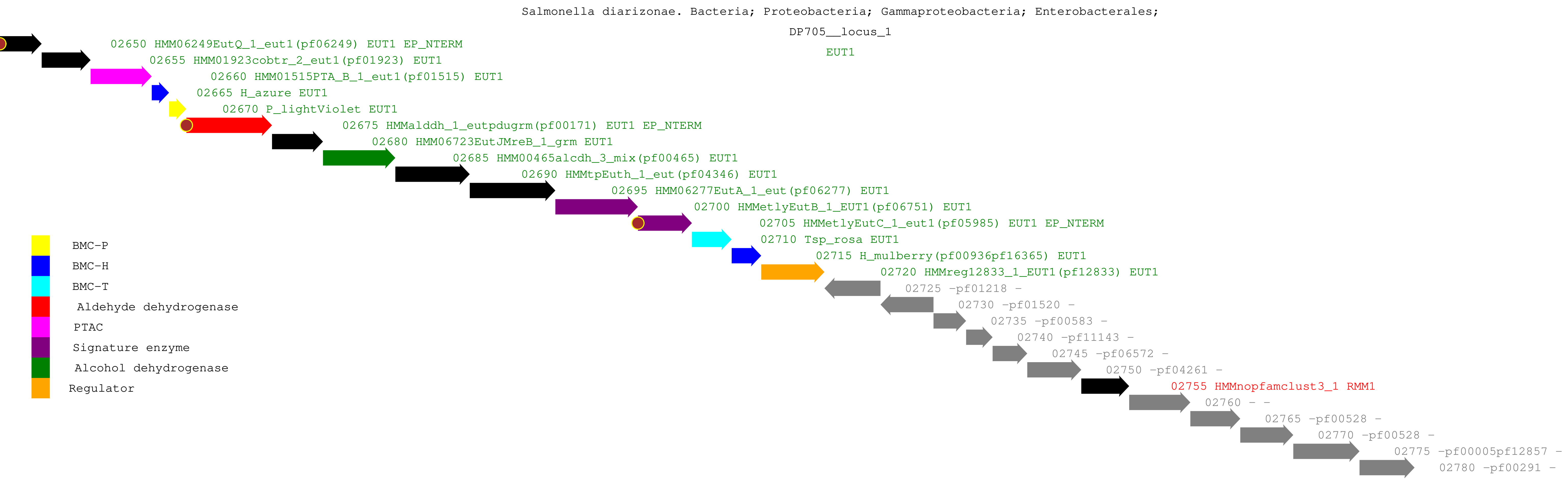

Salmonella diarizonae. Bacteria; Proteobacteria; Gammaproteobacteria; Enterobacterales;
DP705__locus_1
EUT1
02650 HMM06249EutQ_1_eut1(pf06249) EUT1 EP_NTERM
02655 HMM01923cobtr_2_eut1(pf01923) EUT1
02660 HMM01515PTA_B_1_eut1(pf01515) EUT1
02665 H_azure EUT1
02670 P_lightViolet EUT1
02675 HMMaldh_1_eutpdugrm(pf00171) EUT1 EP_NTERM
02680 HMM06723EutJMreB_1_grm EUT1
02685 HMM00465alcdh_3_mix(pf00465) EUT1
02690 HMMtpEuth_1_eut(pf04346) EUT1
02695 HMM06277EutA_1_eut(pf06277) EUT1
02700 HMMetlyEutB_1_EUT1(pf06751) EUT1
02705 HMMetlyEutC_1_eut1(pf05985) EUT1 EP_NTERM
02710 Tsp_rosa EUT1
02715 H_mulberry(pf00936pf16365) EUT1
02720 HMMreg12833_1_EUT1(pf12833) EUT1
02725 -pf01218 -
02730 -pf01520 -
02735 -pf00583 -
02740 -pf11143 -
02745 -pf06572 -
02750 -pf04261 -
02755 HMMnopfamclust3_1 RMM1
02760 - -
02765 -pf00528 -
02770 -pf00528 -
02775 -pf00005pf12857 -
02780 -pf00291 -
BMC-P
BMC-H
BMC-T
Aldehyde dehydrogenase
PTAC
Signature enzyme
Alcohol dehydrogenase
Regulator

Salmonella enterica I. Bacteria; Proteobacteria; Gammaproteobacteria; Enterobacterales;

DP727__locus_1

EUT1

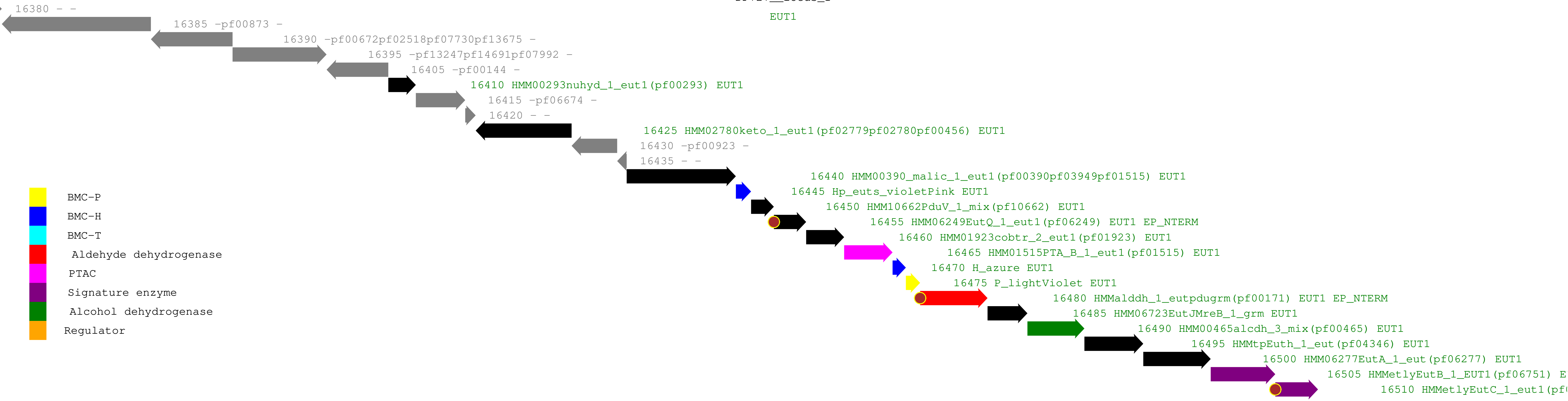

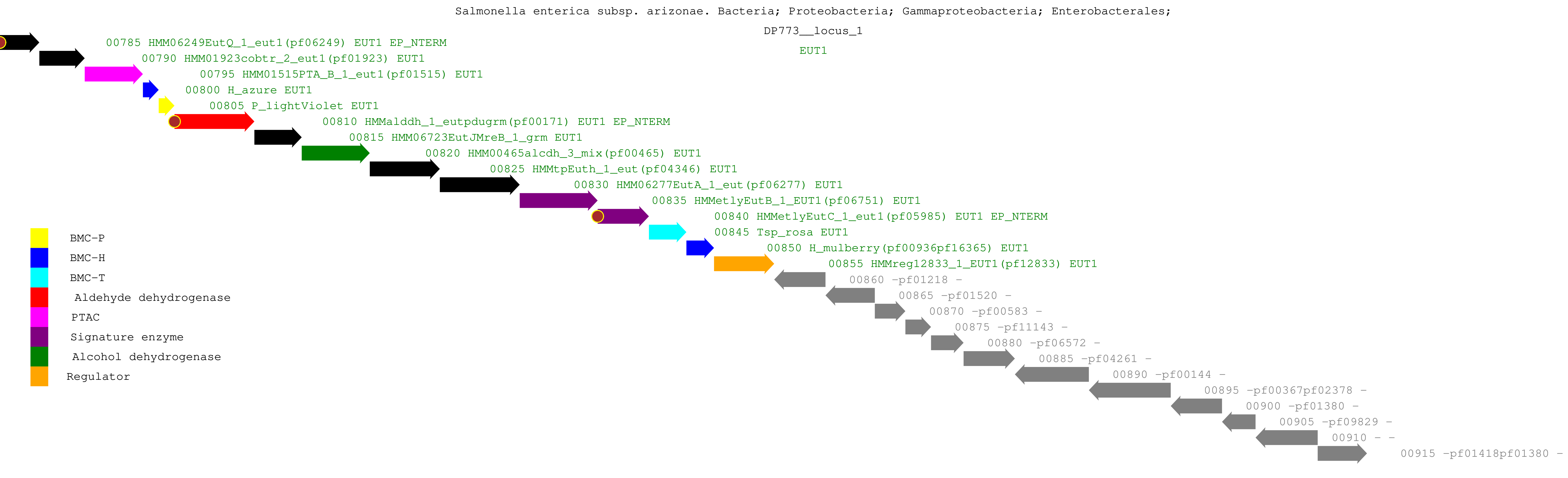
Salmonella enterica subsp. arizonae. Bacteria; Proteobacteria; Gammaproteobacteria; Enterobacterales;
DP773__locus_1
EUT1
00785 HMM06249EutQ_1_eut1(pf06249) EUT1 EP_NTERM
00790 HMM01923cobtr_2_eut1(pf01923) EUT1
00795 HMM01515PTA_B_1_eut1(pf01515) EUT1
00800 H_azure EUT1
00805 P_lightViolet EUT1
00810 HMMalddh_1_eutpdugrm(pf00171) EUT1 EP_NTERM
00815 HMM06723EutJMreB_1_grm EUT1
00820 HMM00465alcdh_3_mix(pf00465) EUT1
00825 HMMtpEuth_1_eut(pf04346) EUT1
00830 HMM06277EutA_1_eut(pf06277) EUT1
00835 HMMetlyEutB_1_EUT1(pf06751) EUT1
00840 HMMetlyEutC_1_eut1(pf05985) EUT1 EP_NTERM
00845 Tsp_rosa EUT1
00850 H_mulberry(pf00936pf16365) EUT1
00855 HMMreg12833_1_EUT1(pf12833) EUT1
00860 -pf01218 -
00865 -pf01520 -
00870 -pf00583 -
00875 -pf11143 -
00880 -pf06572 -
00885 -pf04261 -
00890 -pf00144 -
00895 -pf00367pf02378 -
00900 -pf01380 -
00905 -pf09829 -
00910 - -
00915 -pf01418pf01380 -
BMC-P
BMC-H
BMC-T
Aldehyde dehydrogenase
PTAC
Signature enzyme
Alcohol dehydrogenase
Regulator

Salmonella enterica I. Bacteria; Proteobacteria; Gammaproteobacteria; Enterobacterales;

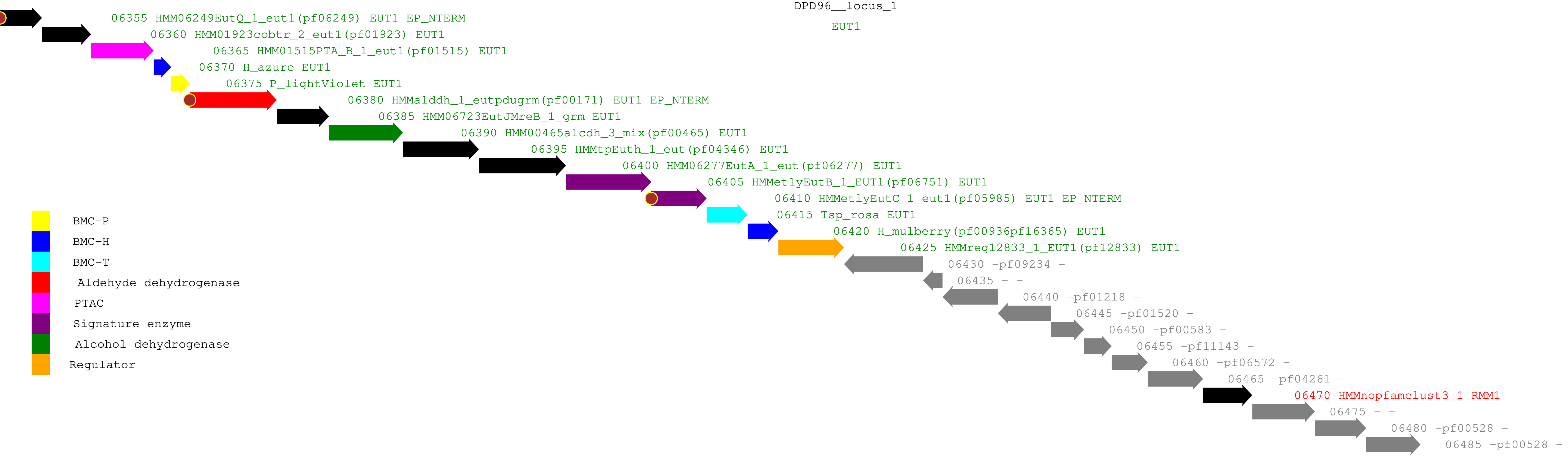

Salmonella diarizonae. Bacteria; Proteobacteria; Gammaproteobacteria; Enterobacterales;

DPF13__locus_1

EUT1

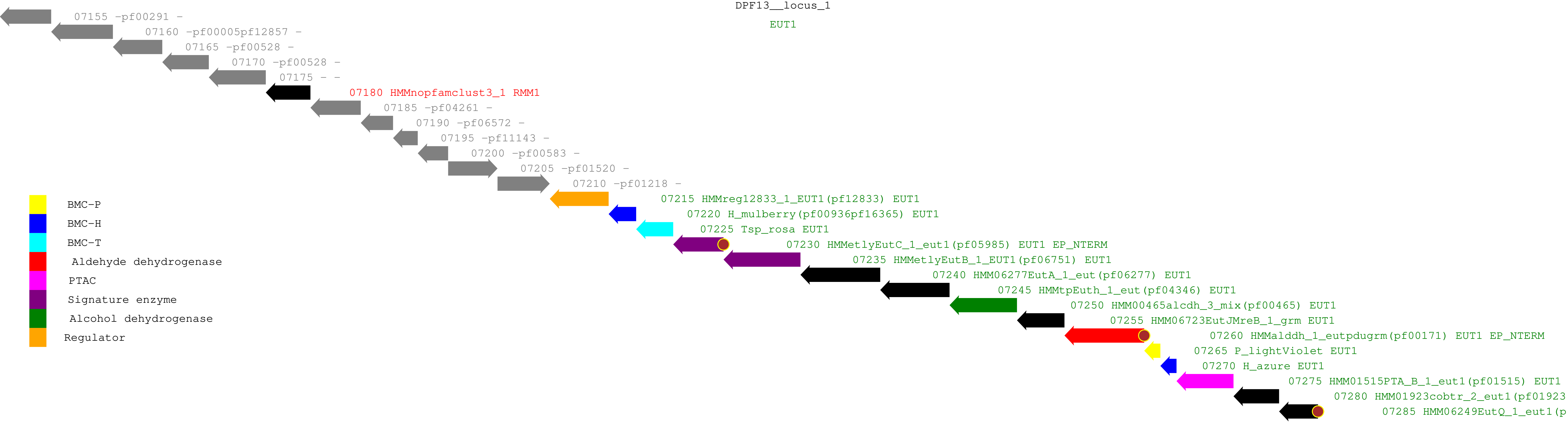

Salmonella enterica I. Bacteria; Proteobacteria; Gammaproteobacteria; Enterobacterales;

DPK49__locus_1

EUT1

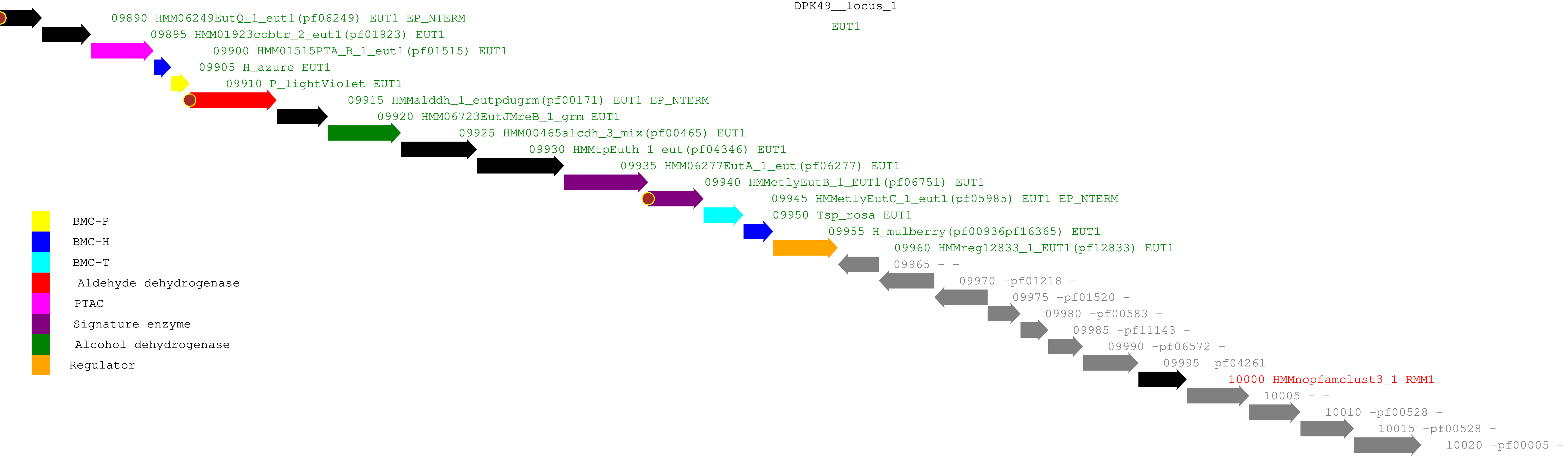

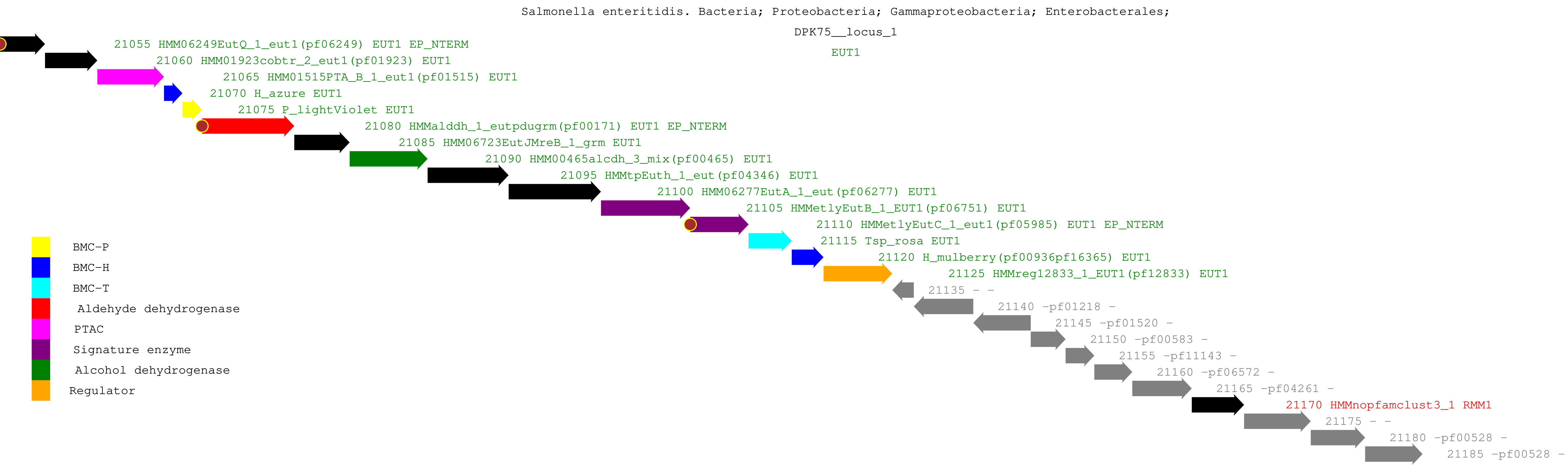
Salmonella enteritidis. Bacteria; Proteobacteria; Gammaproteobacteria; Enterobacterales;
DPK75__locus_1
EUT1
21055 HMM06249EutQ_1_eut1(pf06249) EUT1 EP_NTERM
21060 HMM01923cobtr_2_eut1(pf01923) EUT1
21065 HMM01515PTA_B_1_eut1(pf01515) EUT1
21070 H_azure EUT1
21075 P_lightViolet EUT1
21080 HMMalddh_1_eutpdugrm(pf00171) EUT1 EP_NTERM
21085 HMM06723EutJMreB_1_grm EUT1
21090 HMM00465alcdh_3_mix(pf00465) EUT1
21095 HMMtpEuth_1_eut(pf04346) EUT1
21100 HMM06277EutA_1_eut(pf06277) EUT1
21105 HMMetlyEutB_1_EUT1(pf06751) EUT1
21110 HMMetlyEutC_1_eut1(pf05985) EUT1 EP_NTERM
21115 Tsp_rosa EUT1
21120 H_mulberry(pf00936pf16365) EUT1
21125 HMMreg12833_1_EUT1(pf12833) EUT1
21135 - -
21140 -pf01218 -
21145 -pf01520 -
21150 -pf00583 -
21155 -pf11143 -
21160 -pf06572 -
21165 -pf04261 -
21170 HMMnopfamclust3_1 RMM1
21175 - -
21180 -pf00528 -
21185 -pf00528 -
BMC-P
BMC-H
BMC-T
Aldehyde dehydrogenase
PTAC
Signature enzyme
Alcohol dehydrogenase
Regulator

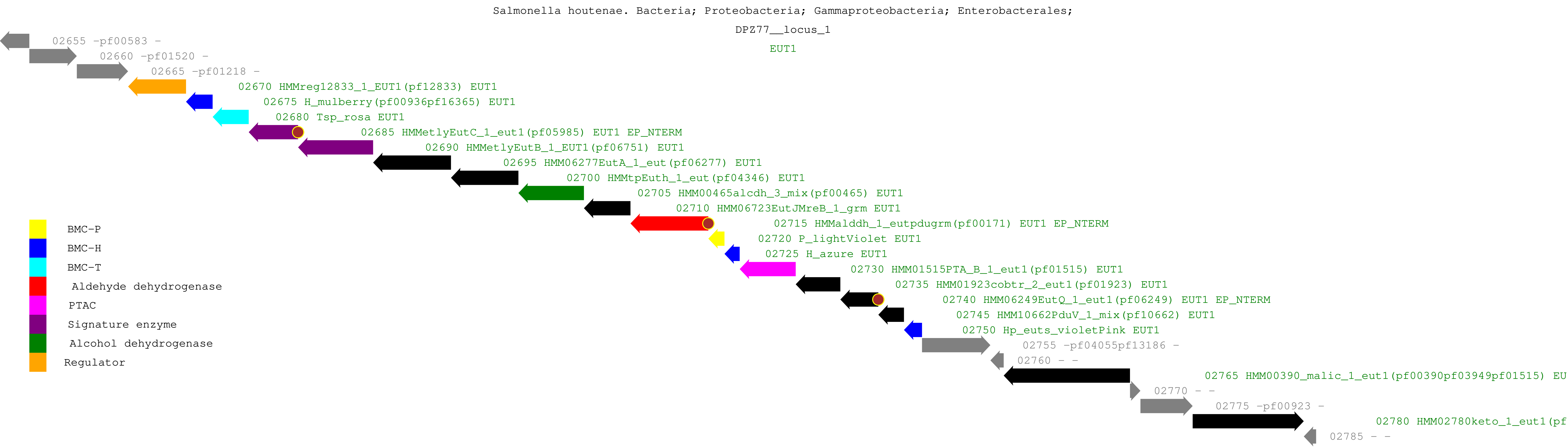
Salmonella houtenae. Bacteria; Proteobacteria; Gammaproteobacteria; Enterobacterales;
DPZ77__locus_1
EUT1
02655 -pf00583 -
02660 -pf01520 -
02665 -pf01218 -
02670 HMMreg12833_1_EUT1(pf12833) EUT1
02675 H_mulberry(pf00936pf16365) EUT1
02680 Tsp_rosa EUT1
02685 HMMetlyEutC_1_eut1(pf05985) EUT1 EP_NTERM
02690 HMMetlyEutB_1_EUT1(pf06751) EUT1
02695 HMM06277EutA_1_eut(pf06277) EUT1
02700 HMMtpEuth_1_eut(pf04346) EUT1
02705 HMM00465alcdh_3_mix(pf00465) EUT1
02710 HMM06723EutJMreB_1_grm EUT1
02715 HMMaldh_1_eutpdugrm(pf00171) EUT1 EP_NTERM
02720 P_lightViolet EUT1
02725 H_azure EUT1
02730 HMM01515PTA_B_1_eut1(pf01515) EUT1
02735 HMM01923cobtr_2_eut1(pf01923) EUT1
02740 HMM06249EutQ_1_eut1(pf06249) EUT1 EP_NTERM
02745 HMM10662PduV_1_mix(pf10662) EUT1
02750 Hp_euts_violetPink EUT1
02755 -pf04055pf13186 -
02760 - -
02765 HMM00390_malic_1_eut1(pf00390pf03949pf01515) EU
02770 - -
02775 -pf00923 -
02780 HMM02780keto_1_eut1(pf
02785 - -
BMC-P
BMC-H
BMC-T
Aldehyde dehydrogenase
PTAC
Signature enzyme
Alcohol dehydrogenase
Regulator

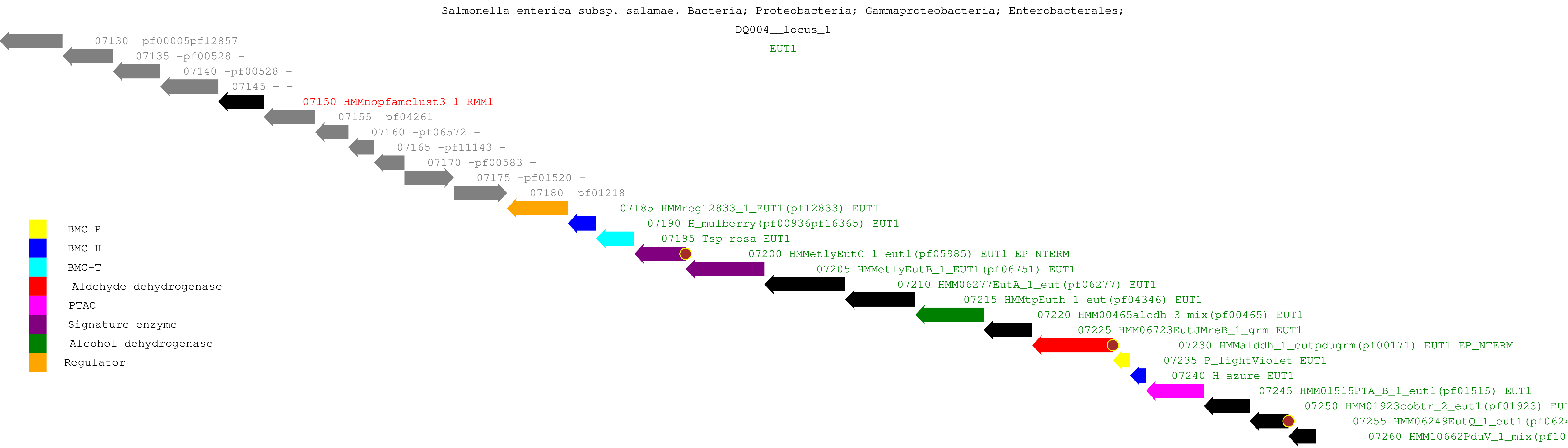
Salmonella enterica subsp. salamae. Bacteria; Proteobacteria; Gammaproteobacteria; Enterobacterales;
DQ004__locus_1
EUT1
07130 -pf00005pf12857 -
07135 -pf00528 -
07140 -pf00528 -
07145 - -
07150 HMMnopfamclust3_1 RMM1
07155 -pf04261 -
07160 -pf06572 -
07165 -pf11143 -
07170 -pf00583 -
07175 -pf01520 -
07180 -pf01218 -
07185 HMMreg12833_1_EUT1(pf12833) EUT1
07190 H_mulberry(pf00936pf16365) EUT1
07195 Tsp_rosa EUT1
07200 HMMetlyEutC_1_eut1(pf05985) EUT1 EP_NTERM
07205 HMMetlyEutB_1_EUT1(pf06751) EUT1
07210 HMM06277EutA_1_eut(pf06277) EUT1
07215 HMMtpEuth_1_eut(pf04346) EUT1
07220 HMM00465alcdh_3_mix(pf00465) EUT1
07225 HMM06723EutJMreB_1_grm EUT1
07230 HMMalddh_1_eutpdugrm(pf00171) EUT1 EP_NTERM
07235 P_lightViolet EUT1
07240 H_azure EUT1
07245 HMM01515PTA_B_1_eut1(pf01515) EUT1
07250 HMM01923cobtr_2_eut1(pf01923) EU
07255 HMM06249EutQ_1_eut1(pf062
07260 HMM10662PduV_1_mix(pf10
BMC-P
BMC-H
BMC-T
Aldehyde dehydrogenase
PTAC
Signature enzyme
Alcohol dehydrogenase
Regulator

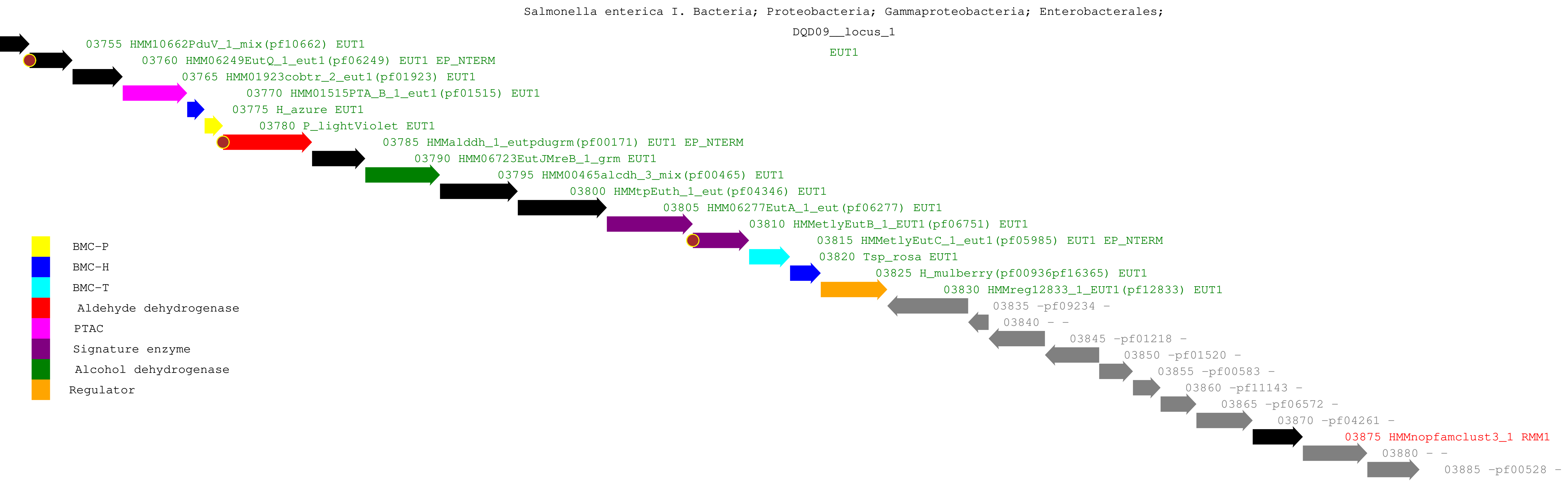
Salmonella enterica I. Bacteria; Proteobacteria; Gammaproteobacteria; Enterobacterales;
DQD09__locus_1
EUT1
03755 HMM10662PduV_1_mix(pf10662) EUT1
03760 HMM06249EutQ_1_eut1(pf06249) EUT1 EP_NTERM
03765 HMM01923cobtr_2_eut1(pf01923) EUT1
03770 HMM01515PTA_B_1_eut1(pf01515) EUT1
03775 H_azure EUT1
03780 P_lightViolet EUT1
03785 HMMalddh_1_eutpdugrm(pf00171) EUT1 EP_NTERM
03790 HMM06723EutJMreB_1_grm EUT1
03795 HMM00465alcdh_3_mix(pf00465) EUT1
03800 HMMtpEuth_1_eut(pf04346) EUT1
03805 HMM06277EutA_1_eut(pf06277) EUT1
03810 HMMetlyEutB_1_EUT1(pf06751) EUT1
03815 HMMetlyEutC_1_eut1(pf05985) EUT1 EP_NTERM
03820 Tsp_rosa EUT1
03825 H_mulberry(pf00936pf16365) EUT1
03830 HMMreg12833_1_EUT1(pf12833) EUT1
03835 -pf09234 -
03840 - -
03845 -pf01218 -
03850 -pf01520 -
03855 -pf00583 -
03860 -pf11143 -
03865 -pf06572 -
03870 -pf04261 -
03875 HMMnopfamclust3_1 RMM1
03880 - -
03885 -pf00528 -
BMC-P
BMC-H
BMC-T
Aldehyde dehydrogenase
PTAC
Signature enzyme
Alcohol dehydrogenase
Regulator

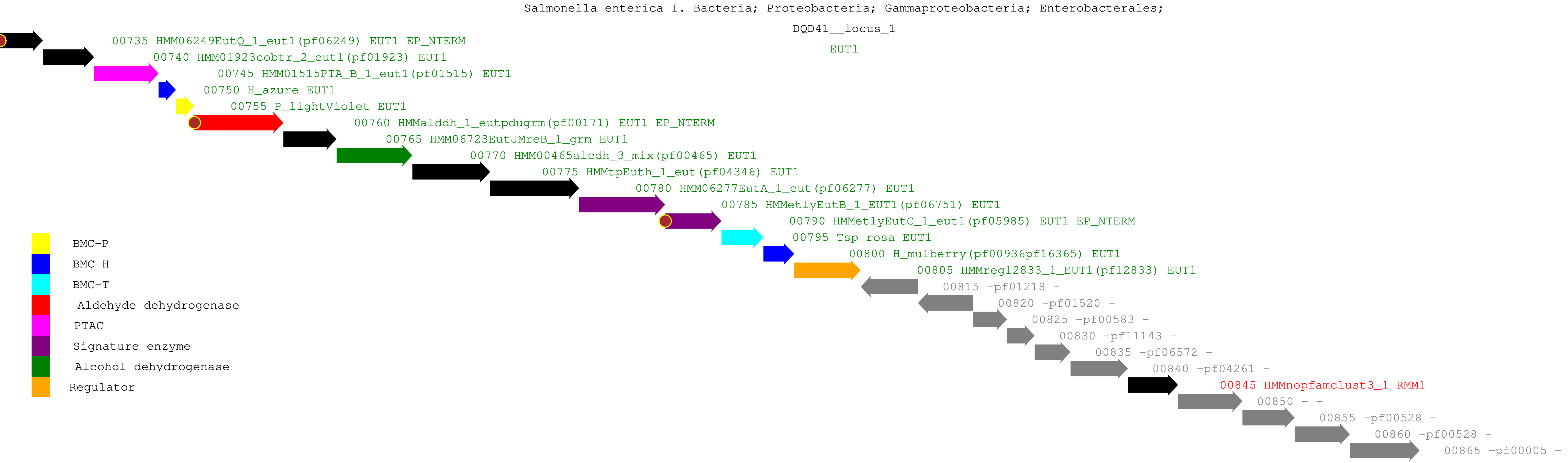
Salmonella enterica I. Bacteria; Proteobacteria; Gammaproteobacteria; Enterobacterales;
DQD41__locus_1
EUT1
00735 HMM06249EutQ_1_eut1(pf06249) EUT1 EP_NTERM
00740 HMM01923cobtr_2_eut1(pf01923) EUT1
00745 HMM01515PTA_B_1_eut1(pf01515) EUT1
00750 H_azure EUT1
00755 P_lightViolet EUT1
00760 HMMaldh_1_eutpdugrm(pf00171) EUT1 EP_NTERM
00765 HMM06723EutJMreB_1_grm EUT1
00770 HMM00465alcdh_3_mix(pf00465) EUT1
00775 HMMtpEuth_1_eut(pf04346) EUT1
00780 HMM06277EutA_1_eut(pf06277) EUT1
00785 HMMetlyEutB_1_EUT1(pf06751) EUT1
00790 HMMetlyEutC_1_eut1(pf05985) EUT1 EP_NTERM
00795 Tsp_rosa EUT1
00800 H_mulberry(pf00936pf16365) EUT1
00805 HMMreg12833_1_EUT1(pf12833) EUT1
00815 -pf01218 -
00820 -pf01520 -
00825 -pf00583 -
00830 -pf11143 -
00835 -pf06572 -
00840 -pf04261 -
00845 HMMnopfamclust3_1 RMM1
00850 - -
00855 -pf00528 -
00860 -pf00528 -
00865 -pf00005 -
BMC-P
BMC-H
BMC-T
Aldehyde dehydrogenase
PTAC
Signature enzyme
Alcohol dehydrogenase
Regulator

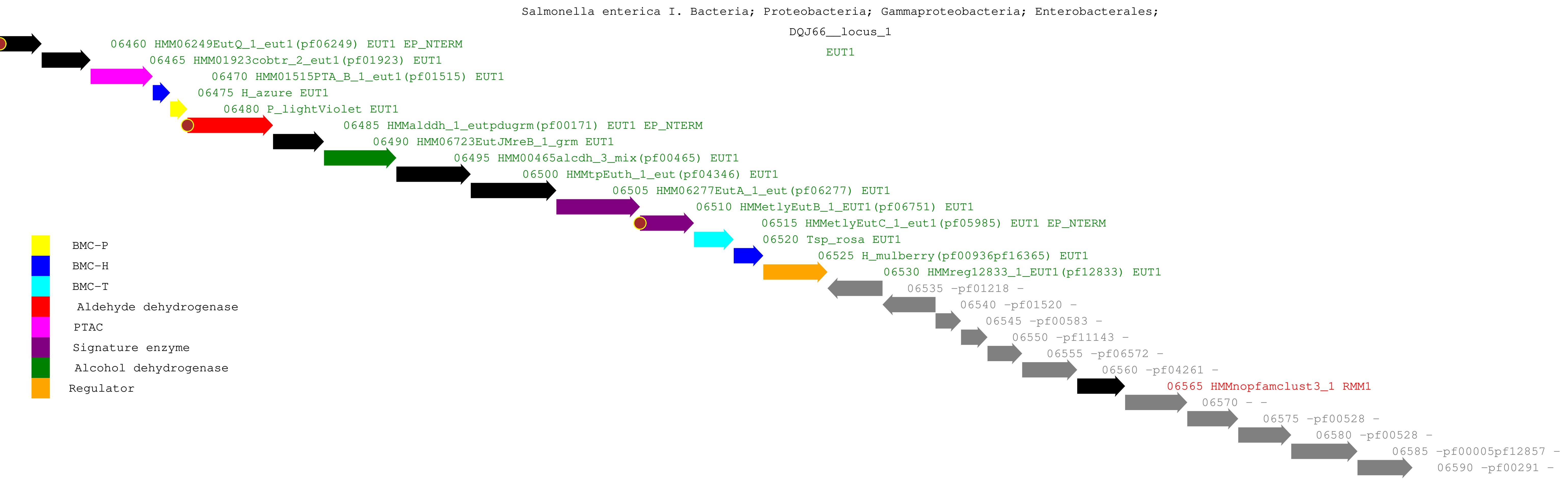
Salmonella enterica I. Bacteria; Proteobacteria; Gammaproteobacteria; Enterobacterales;
DQJ66__locus_1
EUT1
06460 HMM06249EutQ_1_eut1(pf06249) EUT1 EP_NTERM
06465 HMM01923cobtr_2_eut1(pf01923) EUT1
06470 HMM01515PTA_B_1_eut1(pf01515) EUT1
06475 H_azure EUT1
06480 P_lightViolet EUT1
06485 HMMaldh_1_eutpdugrm(pf00171) EUT1 EP_NTERM
06490 HMM06723EutJMreB_1_grm EUT1
06495 HMM00465alcdh_3_mix(pf00465) EUT1
06500 HMMtpEuth_1_eut(pf04346) EUT1
06505 HMM06277EutA_1_eut(pf06277) EUT1
06510 HMMetlyEutB_1_EUT1(pf06751) EUT1
06515 HMMetlyEutC_1_eut1(pf05985) EUT1 EP_NTERM
06520 Tsp_rosa EUT1
06525 H_mulberry(pf00936pf16365) EUT1
06530 HMMreg12833_1_EUT1(pf12833) EUT1
06535 -pf01218 -
06540 -pf01520 -
06545 -pf00583 -
06550 -pf11143 -
06555 -pf06572 -
06560 -pf04261 -
06565 HMMnopfamclust3_1 RMM1
06570 - -
06575 -pf00528 -
06580 -pf00528 -
06585 -pf00005pf12857 -
06590 -pf00291 -
BMC-P
BMC-H
BMC-T
Aldehyde dehydrogenase
PTAC
Signature enzyme
Alcohol dehydrogenase
Regulator

Salmonella enterica I. Bacteria; Proteobacteria; Gammaproteobacteria; Enterobacterales;

DQJ71__locus_1

EUT1

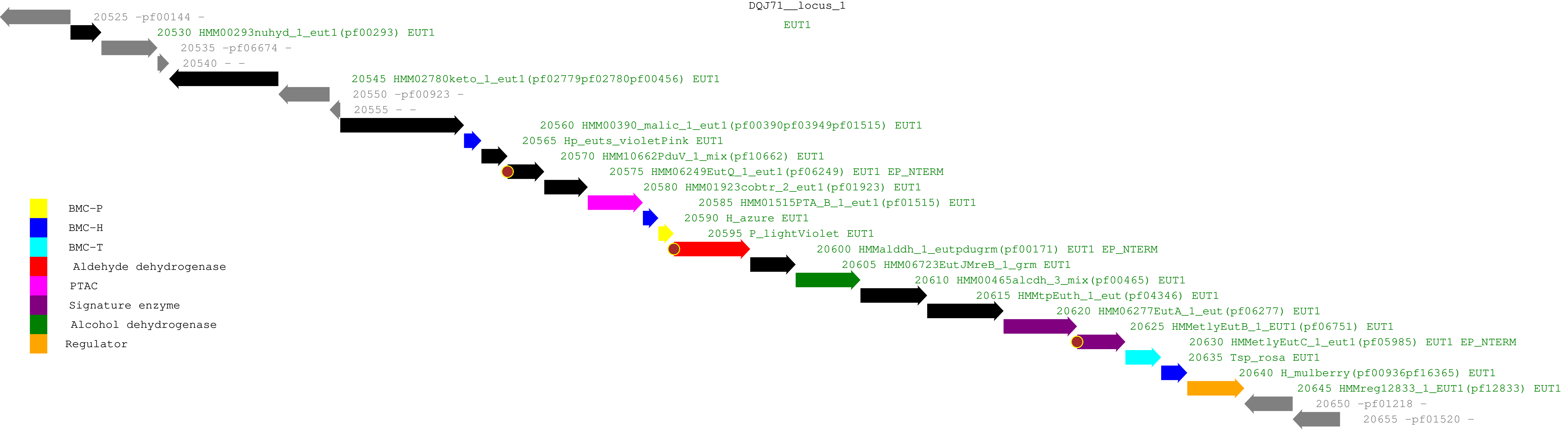

Salmonella enterica subsp. enterica serovar Havana. Bacteria; Proteobacteria; Gammaproteobacteria; Enterobacterales;

DQK02__locus_1

EUT1

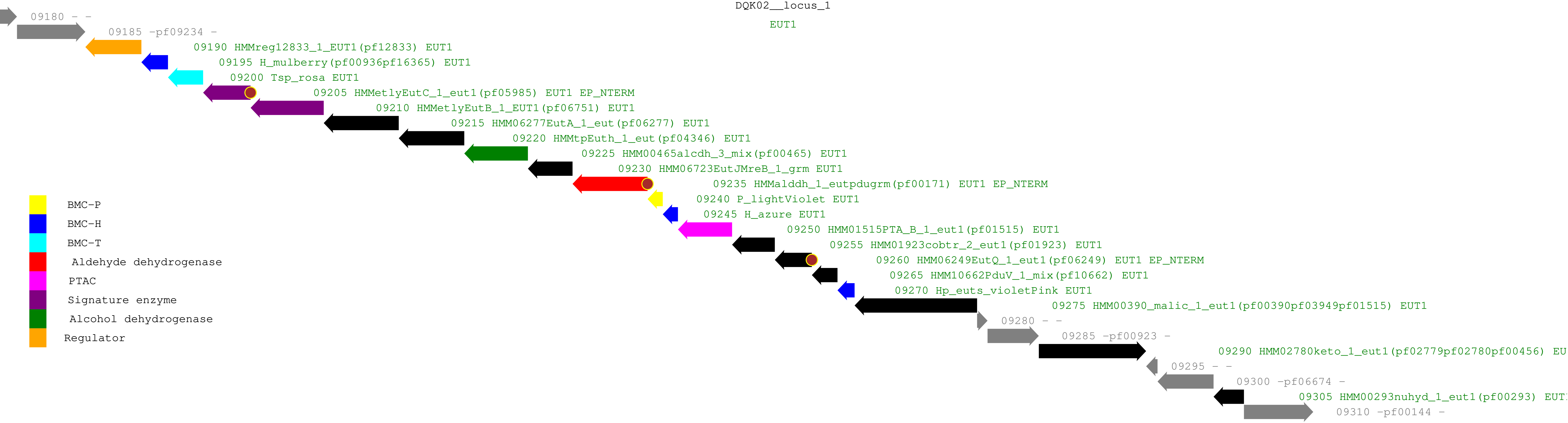

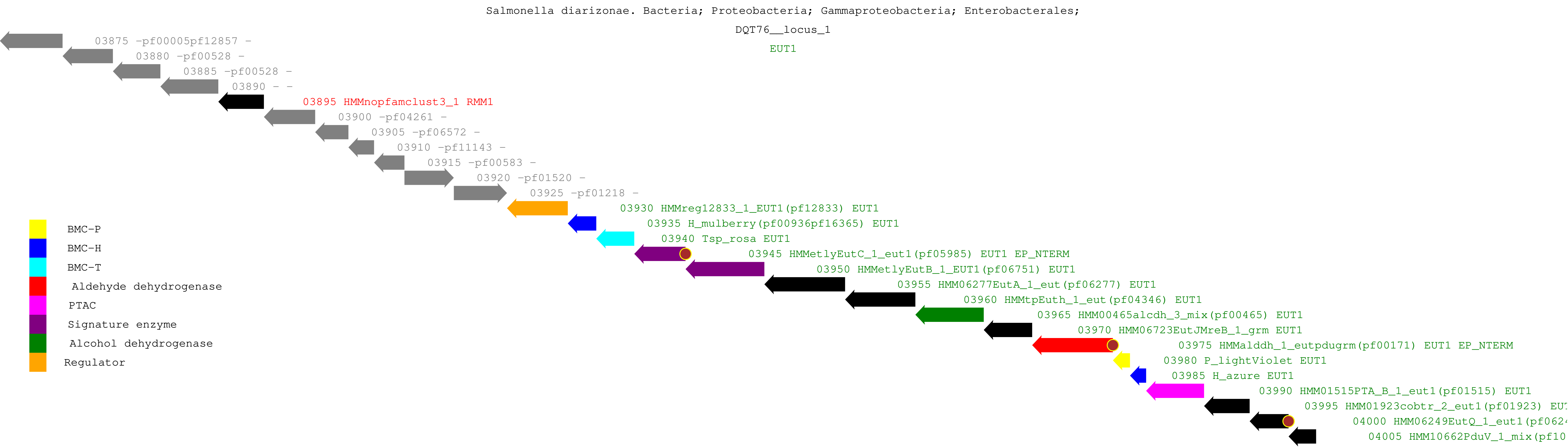
Salmonella diarizonae. Bacteria; Proteobacteria; Gammaproteobacteria; Enterobacterales;
DQT76__locus_1
EUT1
03875 -pf00005pf12857 -
03880 -pf00528 -
03885 -pf00528 -
03890 - -
03895 HMMnopfamclust3_1 RMM1
03900 -pf04261 -
03905 -pf06572 -
03910 -pf11143 -
03915 -pf00583 -
03920 -pf01520 -
03925 -pf01218 -
03930 HMMreg12833_1_EUT1(pf12833) EUT1
03935 H_mulberry(pf00936pf16365) EUT1
03940 Tsp_rosa EUT1
03945 HMMetlyEutC_1_eut1(pf05985) EUT1 EP_NTERM
03950 HMMetlyEutB_1_EUT1(pf06751) EUT1
03955 HMM06277EutA_1_eut(pf06277) EUT1
03960 HMMtpEuth_1_eut(pf04346) EUT1
03965 HMM00465alcdh_3_mix(pf00465) EUT1
03970 HMM06723EutJMreB_1_grm EUT1
03975 HMMalddh_1_eutpdugrm(pf00171) EUT1 EP_NTERM
03980 P_lightViolet EUT1
03985 H_azure EUT1
03990 HMM01515PTA_B_1_eut1(pf01515) EUT1
03995 HMM01923cobtr_2_eut1(pf01923) EU
04000 HMM06249EutQ_1_eut1(pf062
04005 HMM10662PduV_1_mix(pf10
BMC-P
BMC-H
BMC-T
Aldehyde dehydrogenase
PTAC
Signature enzyme
Alcohol dehydrogenase
Regulator

Escherichia coli O111:NM. Bacteria; Proteobacteria; Gammaproteobacteria; Enterobacterales;

DQW49__locus_1

EUT1

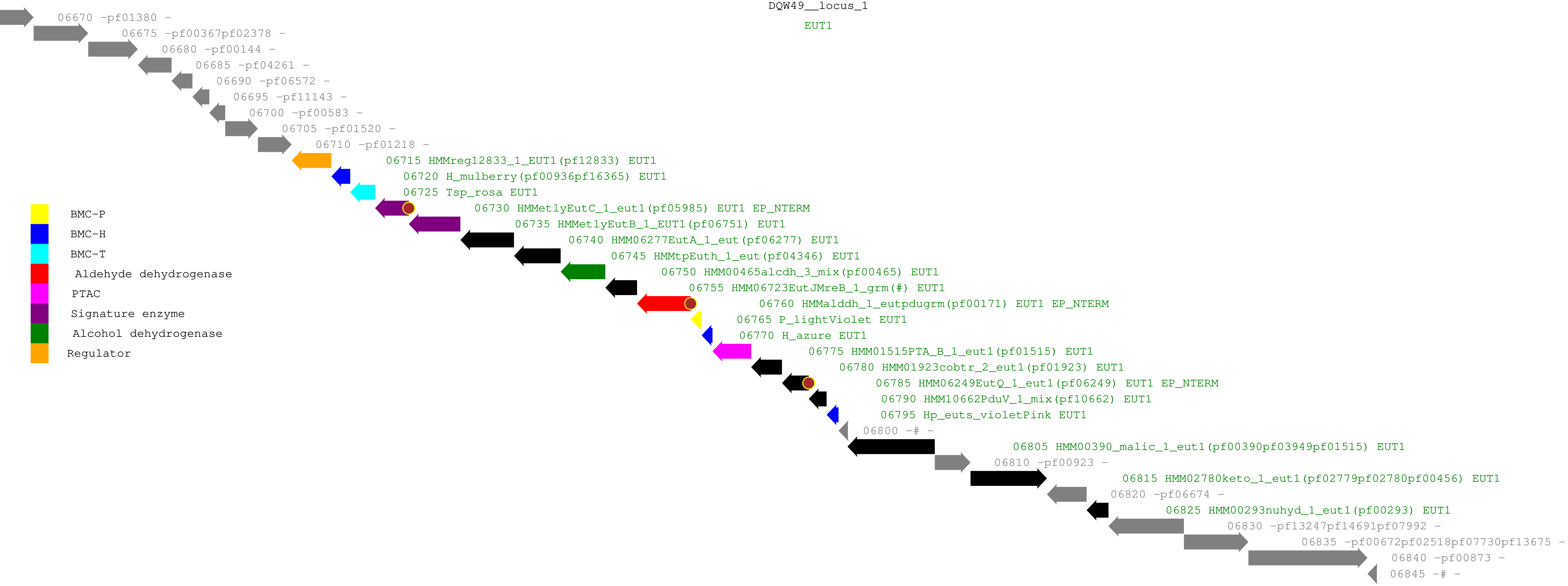

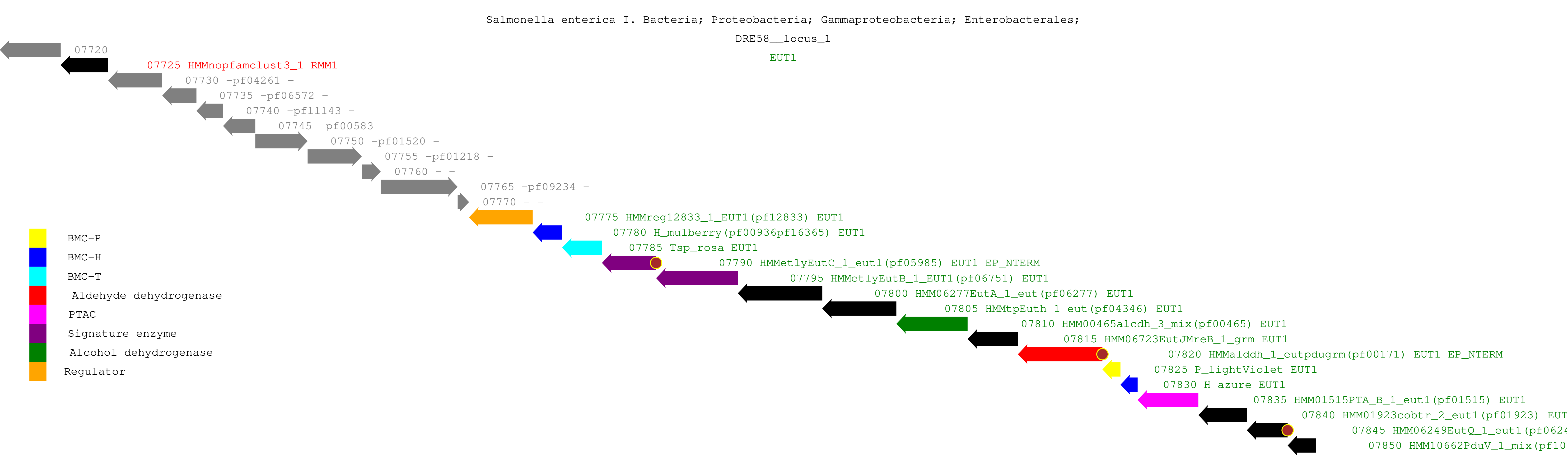

Salmonella enterica I. Bacteria; Proteobacteria; Gammaproteobacteria; Enterobacterales;
DRE58__locus_1
EUT1
07720 - -
07725 HMMnopfamclust3_1 RMM1
07730 -pf04261 -
07735 -pf06572 -
07740 -pf11143 -
07745 -pf00583 -
07750 -pf01520 -
07755 -pf01218 -
07760 - -
07765 -pf09234 -
07770 - -
07775 HMMreg12833_1_EUT1(pf12833) EUT1
07780 H_mulberry(pf00936pf16365) EUT1
07785 Tsp_rosa EUT1
07790 HMMetlyEutC_1_eut1(pf05985) EUT1 EP_NTERM
07795 HMMetlyEutB_1_EUT1(pf06751) EUT1
07800 HMM06277EutA_1_eut(pf06277) EUT1
07805 HMMtpEuth_1_eut(pf04346) EUT1
07810 HMM00465alcdh_3_mix(pf00465) EUT1
07815 HMM06723EutJMreB_1_grm EUT1
07820 HMMalddh_1_eutpdugrm(pf00171) EUT1 EP_NTERM
07825 P_lightViolet EUT1
07830 H_azure EUT1
07835 HMM01515PTA_B_1_eut1(pf01515) EUT1
07840 HMM01923cobtr_2_eut1(pf01923) EUT
07845 HMM06249EutQ_1_eut1(pf0624
07850 HMM10662PduV_1_mix(pf10
BMC-P
BMC-H
BMC-T
Aldehyde dehydrogenase
PTAC
Signature enzyme
Alcohol dehydrogenase
Regulator

Salmonella enterica I. Bacteria; Proteobacteria; Gammaproteobacteria; Enterobacterales;

DRL00__locus_1

EUT1

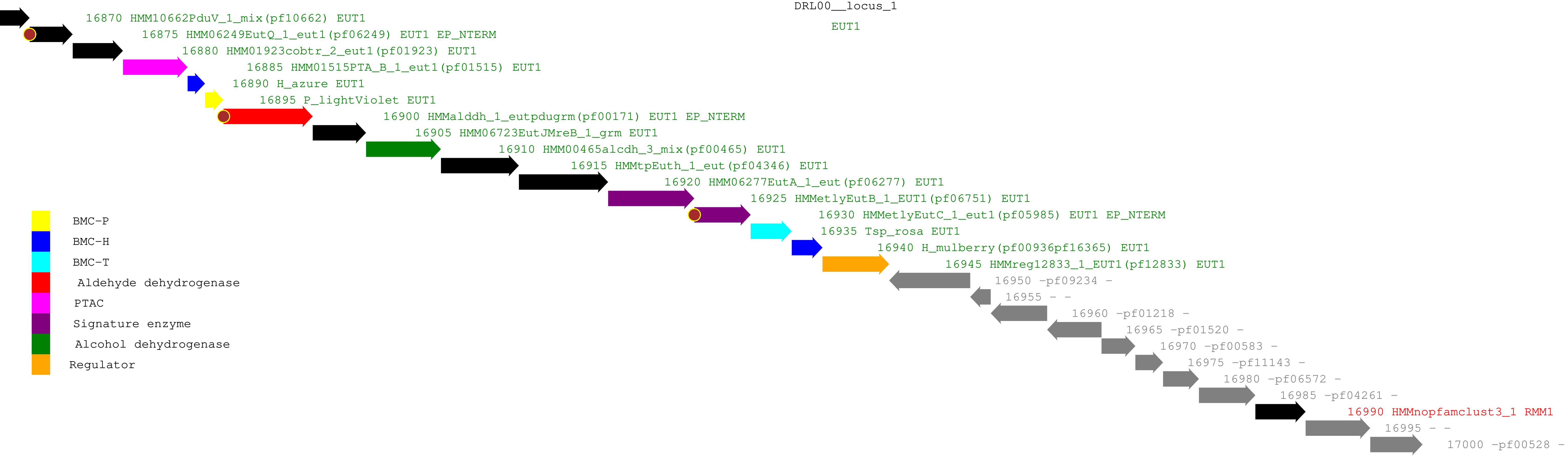

Shigella dysenteriae. Bacteria; Proteobacteria; Gammaproteobacteria; Enterobacterales;

DRW31__locus_1_main_sat

EUT1

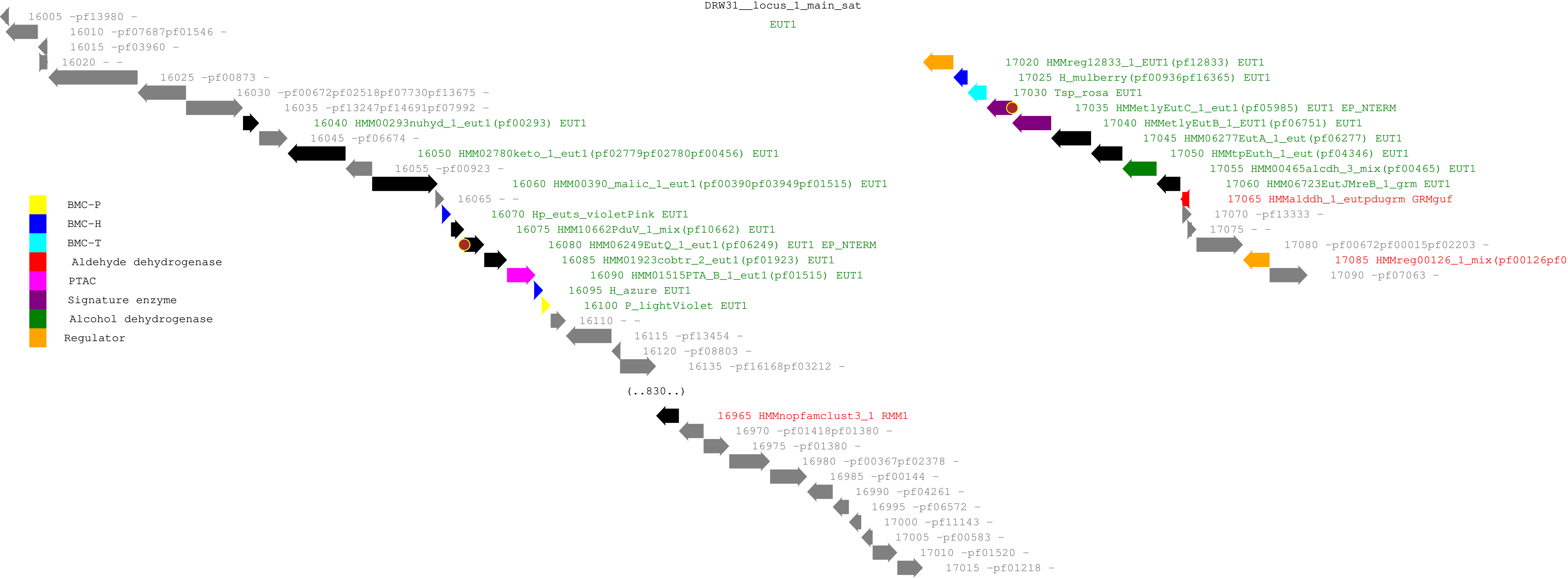

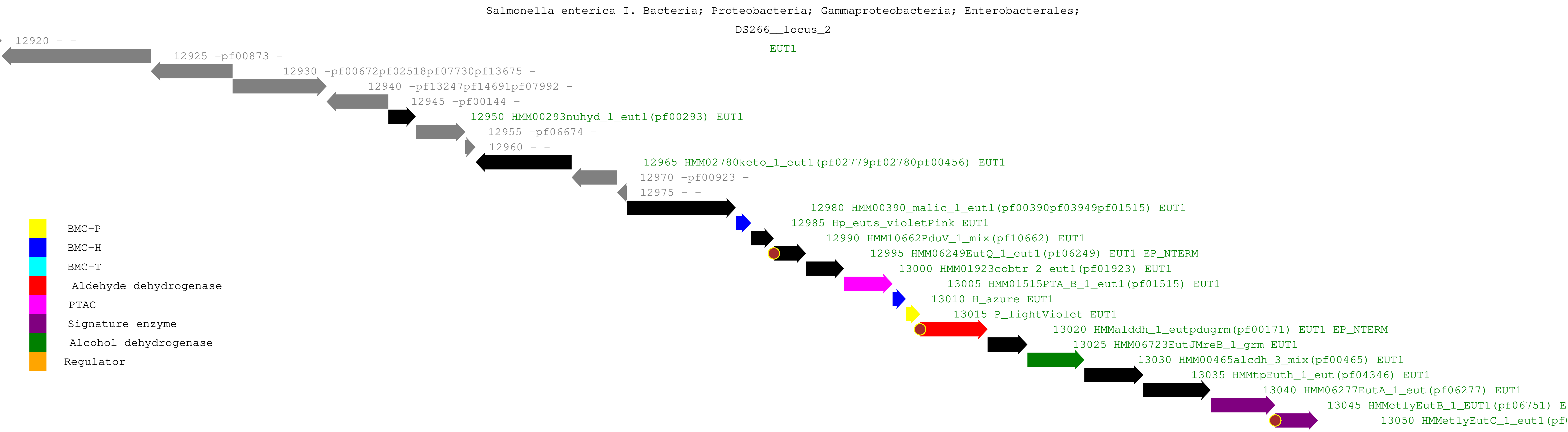
Salmonella enterica I. Bacteria; Proteobacteria; Gammaproteobacteria; Enterobacterales;
DS266__locus_2
EUT1
12920 - -
12925 -pf00873 -
12930 -pf00672pf02518pf07730pf13675 -
12940 -pf13247pf14691pf07992 -
12945 -pf00144 -
12950 HMM00293nuhyd_1_eut1(pf00293) EUT1
12955 -pf06674 -
12960 - -
12965 HMM02780keto_1_eut1(pf02779pf02780pf00456) EUT1
12970 -pf00923 -
12975 - -
12980 HMM00390_malic_1_eut1(pf00390pf03949pf01515) EUT1
12985 Hp_euts_violetPink EUT1
12990 HMM10662PduV_1_mix(pf10662) EUT1
12995 HMM06249EutQ_1_eut1(pf06249) EUT1 EP_NTERM
13000 HMM01923cobtr_2_eut1(pf01923) EUT1
13005 HMM01515PTA_B_1_eut1(pf01515) EUT1
13010 H_azure EUT1
13015 P_lightViolet EUT1
13020 HMMalddh_1_eutpdugrm(pf00171) EUT1 EP_NTERM
13025 HMM06723EutJMreB_1_grm EUT1
13030 HMM00465alcdh_3_mix(pf00465) EUT1
13035 HMMtpEuth_1_eut(pf04346) EUT1
13040 HMM06277EutA_1_eut(pf06277) EUT1
13045 HMMetlyEutB_1_EUT1(pf06751) E
13050 HMMetlyEutC_1_eut1(pf
BMC-P
BMC-H
BMC-T
Aldehyde dehydrogenase
PTAC
Signature enzyme
Alcohol dehydrogenase
Regulator

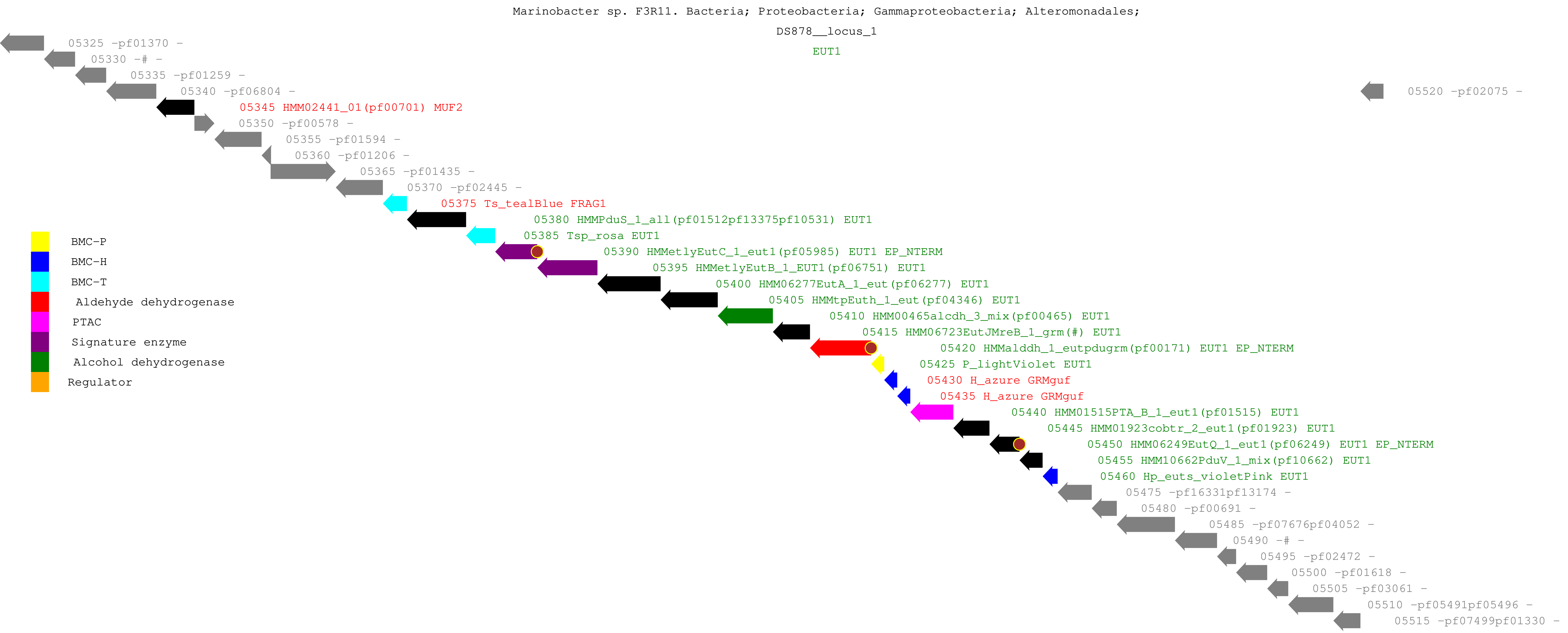

Marinobacter sp. F3R11. Bacteria; Proteobacteria; Gammaproteobacteria; Alteromonadales;
DS878__locus_1
EUT1
05325 -pf01370 -
05330 -# -
05335 -pf01259 -
05340 -pf06804 -
05345 HMM02441_01(pf00701) MUF2
05350 -pf00578 -
05355 -pf01594 -
05360 -pf01206 -
05365 -pf01435 -
05370 -pf02445 -
05375 Ts_tealBlue FRAG1
05380 HMMPduS_1_all(pf01512pf13375pf10531) EUT1
05385 Tsp_rosa EUT1
05390 HMMetlyEutC_1_eut1(pf05985) EUT1 EP_NTERM
05395 HMMetlyEutB_1_EUT1(pf06751) EUT1
05400 HMM06277EutA_1_eut(pf06277) EUT1
05405 HMMtpEuth_1_eut(pf04346) EUT1
05410 HMM00465alcdh_3_mix(pf00465) EUT1
05415 HMM06723EutJMreB_1_grm(#) EUT1
05420 HMMalddh_1_eutpdugrm(pf00171) EUT1 EP_NTERM
05425 P_lightViolet EUT1
05430 H_azure GRMguf
05435 H_azure GRMguf
05440 HMM01515PTA_B_1_eut1(pf01515) EUT1
05445 HMM01923cobtr_2_eut1(pf01923) EUT1
05450 HMM06249EutQ_1_eut1(pf06249) EUT1 EP_NTERM
05455 HMM10662PduV_1_mix(pf10662) EUT1
05460 Hp_euts_violetPink EUT1
05475 -pf16331pf13174 -
05480 -pf00691 -
05485 -pf07676pf04052 -
05490 -# -
05495 -pf02472 -
05500 -pf01618 -
05505 -pf03061 -
05510 -pf05491pf05496 -
05515 -pf07499pf01330 -
05520 -pf02075 -
BMC-P
BMC-H
BMC-T
Aldehyde dehydrogenase
PTAC
Signature enzyme
Alcohol dehydrogenase
Regulator

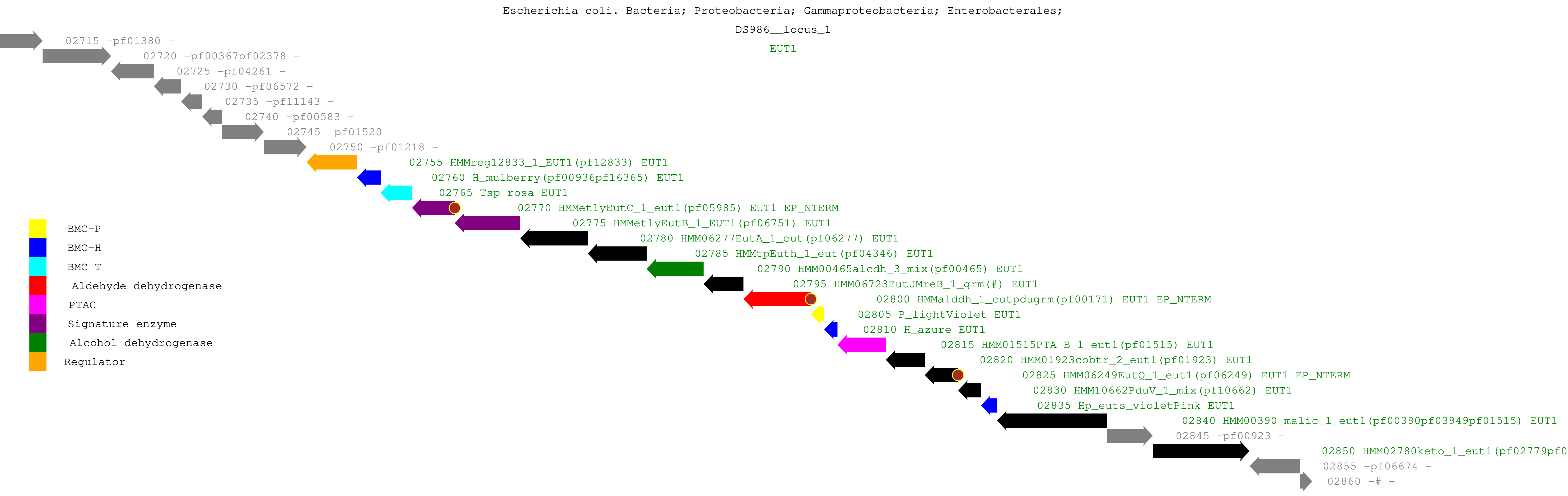

Escherichia coli. Bacteria; Proteobacteria; Gammaproteobacteria; Enterobacterales;
DS986__locus_1
EUT1
02715 -pf01380 -
02720 -pf00367pf02378 -
02725 -pf04261 -
02730 -pf06572 -
02735 -pf11143 -
02740 -pf00583 -
02745 -pf01520 -
02750 -pf01218 -
02755 HMMreg12833_1_EUT1(pf12833) EUT1
02760 H_mulberry(pf00936pf16365) EUT1
02765 Tsp_rosa EUT1
02770 HMMetlyEutC_1_eut1(pf05985) EUT1 EP_NTERM
02775 HMMetlyEutB_1_EUT1(pf06751) EUT1
02780 HMM06277EutA_1_eut(pf06277) EUT1
02785 HMMtpEuth_1_eut(pf04346) EUT1
02790 HMM00465alcdh_3_mix(pf00465) EUT1
02795 HMM06723EutJMreB_1_grm(#) EUT1
02800 HMMalddh_1_eutpdugrm(pf00171) EUT1 EP_NTERM
02805 P_lightViolet EUT1
02810 H_azure EUT1
02815 HMM01515PTA_B_1_eut1(pf01515) EUT1
02820 HMM01923cobtr_2_eut1(pf01923) EUT1
02825 HMM06249EutQ_1_eut1(pf06249) EUT1 EP_NTERM
02830 HMM10662PduV_1_mix(pf10662) EUT1
02835 Hp_euts_violetPink EUT1
02840 HMM00390_malic_1_eut1(pf00390pf03949pf01515) EUT1
02845 -pf00923 -
02850 HMM02780keto_1_eut1(pf02779pf0
02855 -pf06674 -
02860 -# -
BMC-P
BMC-H
BMC-T
Aldehyde dehydrogenase
PTAC
Signature enzyme
Alcohol dehydrogenase
Regulator

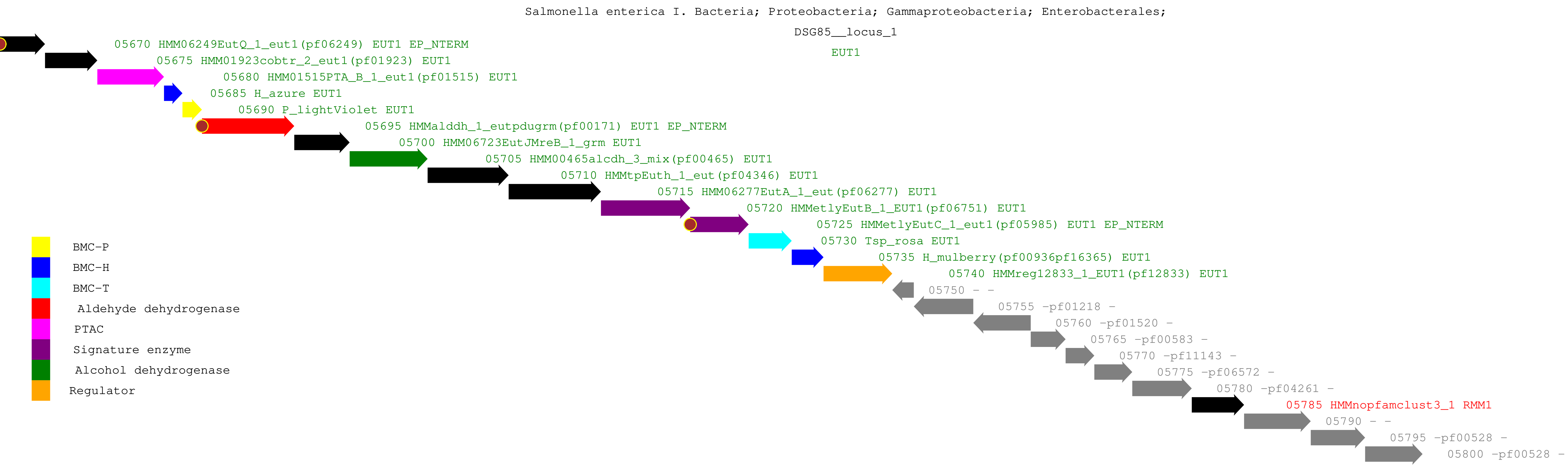

Salmonella enterica I. Bacteria; Proteobacteria; Gammaproteobacteria; Enterobacterales;
DSG85__locus_1
EUT1
05670 HMM06249EutQ_1_eut1(pf06249) EUT1 EP_NTERM
05675 HMM01923cobtr_2_eut1(pf01923) EUT1
05680 HMM01515PTA_B_1_eut1(pf01515) EUT1
05685 H_azure EUT1
05690 P_lightViolet EUT1
05695 HMMalddh_1_eutpdugrm(pf00171) EUT1 EP_NTERM
05700 HMM06723EutJMreB_1_grm EUT1
05705 HMM00465alcdh_3_mix(pf00465) EUT1
05710 HMMtpEuth_1_eut(pf04346) EUT1
05715 HMM06277EutA_1_eut(pf06277) EUT1
05720 HMMetlyEutB_1_EUT1(pf06751) EUT1
05725 HMMetlyEutC_1_eut1(pf05985) EUT1 EP_NTERM
05730 Tsp_rosa EUT1
05735 H_mulberry(pf00936pf16365) EUT1
05740 HMMreg12833_1_EUT1(pf12833) EUT1
05750 - -
05755 -pf01218 -
05760 -pf01520 -
05765 -pf00583 -
05770 -pf11143 -
05775 -pf06572 -
05780 -pf04261 -
05785 HMMnopfamclust3_1 RMM1
05790 - -
05795 -pf00528 -
05800 -pf00528 -
BMC-P
BMC-H
BMC-T
Aldehyde dehydrogenase
PTAC
Signature enzyme
Alcohol dehydrogenase
Regulator

Salmonella enterica I. Bacteria; Proteobacteria; Gammaproteobacteria; Enterobacterales;

DT150__locus_1

EUT1

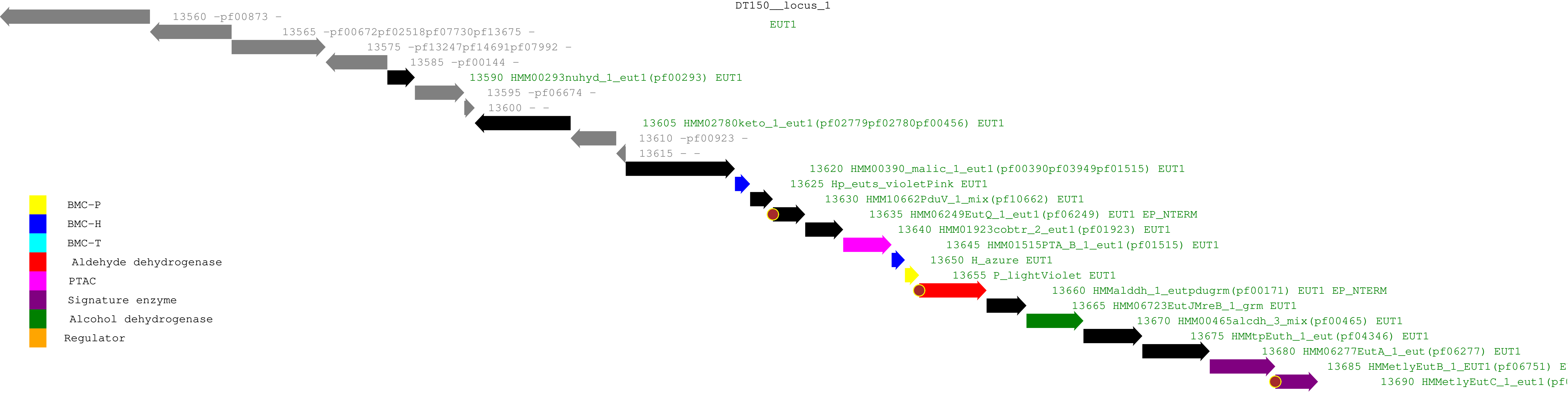

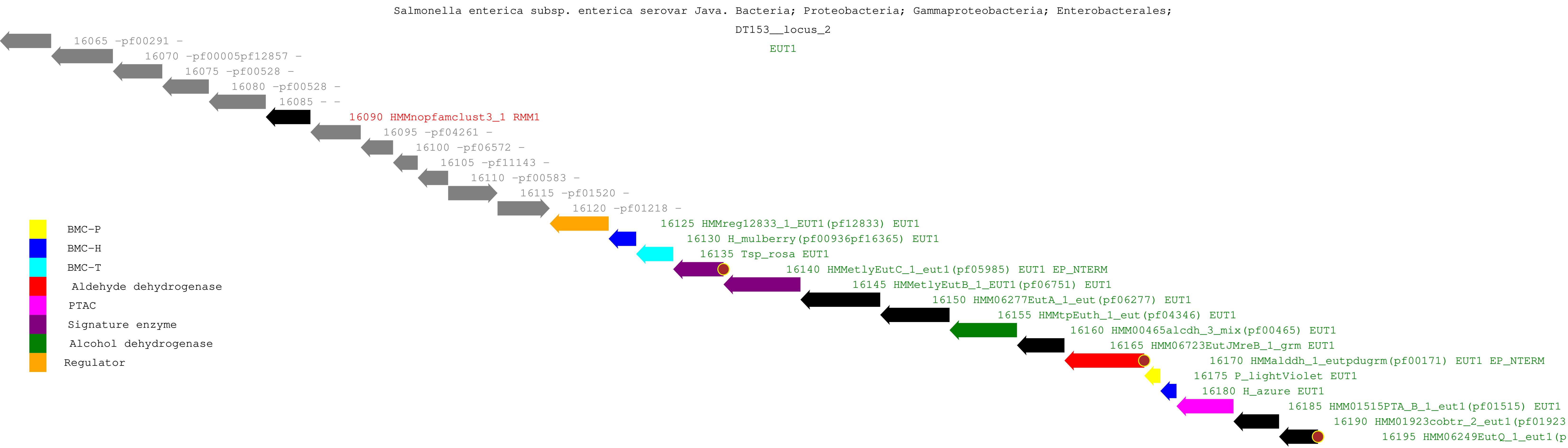

Salmonella enterica subsp. enterica serovar Java. Bacteria; Proteobacteria; Gammaproteobacteria; Enterobacterales;
DT153__locus_2
EUT1
16065 -pf00291 -
16070 -pf00005pf12857 -
16075 -pf00528 -
16080 -pf00528 -
16085 - -
16090 HMMnopfamclust3_1 RMM1
16095 -pf04261 -
16100 -pf06572 -
16105 -pf11143 -
16110 -pf00583 -
16115 -pf01520 -
16120 -pf01218 -
16125 HMMreg12833_1_EUT1(pf12833) EUT1
16130 H_mulberry(pf00936pf16365) EUT1
16135 Tsp_rosa EUT1
16140 HMMetlyEutC_1_eut1(pf05985) EUT1 EP_NTERM
16145 HMMetlyEutB_1_EUT1(pf06751) EUT1
16150 HMM06277EutA_1_eut(pf06277) EUT1
16155 HMMtpEuth_1_eut(pf04346) EUT1
16160 HMM00465alcdh_3_mix(pf00465) EUT1
16165 HMM06723EutJMreB_1_grm EUT1
16170 HMMaldh_1_eutpdugrm(pf00171) EUT1 EP_NTERM
16175 P_lightViolet EUT1
16180 H_azure EUT1
16185 HMM01515PTA_B_1_eut1(pf01515) EUT1
16190 HMM01923cobtr_2_eut1(pf01923
16195 HMM06249EutQ_1_eut1(p
BMC-P
BMC-H
BMC-T
Aldehyde dehydrogenase
PTAC
Signature enzyme
Alcohol dehydrogenase
Regulator

Klebsiella sp. P1CD1. Bacteria; Proteobacteria; Gammaproteobacteria; Enterobacterales;

DTA24__locus_2

EUT1

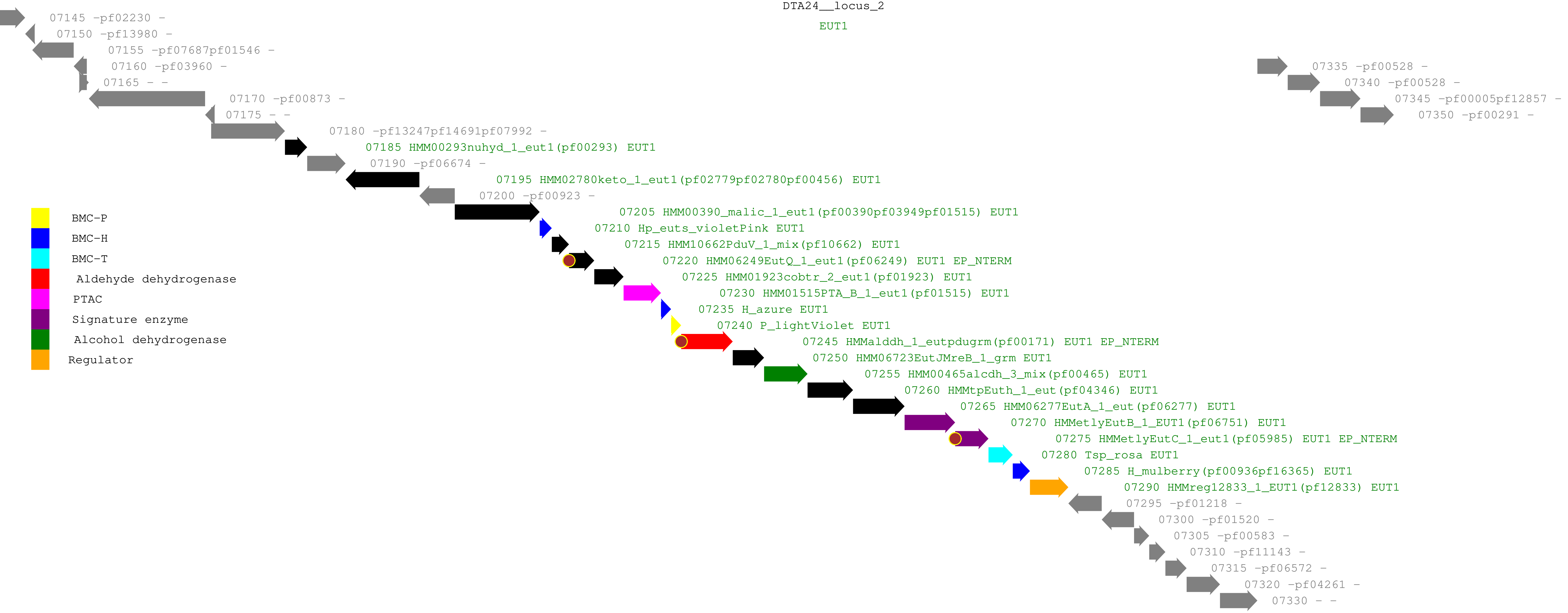

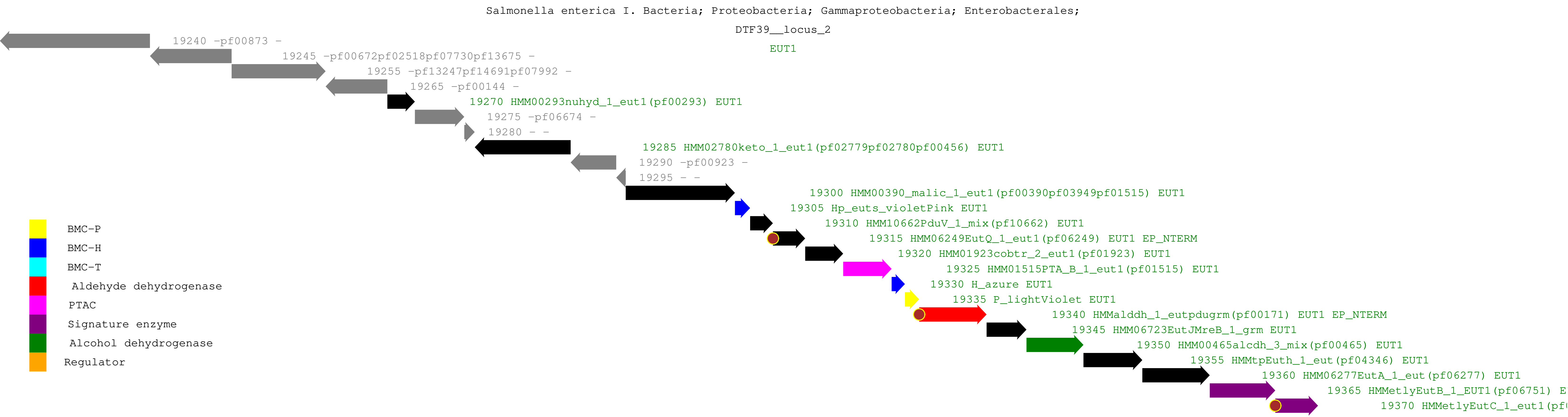

Salmonella enterica I. Bacteria; Proteobacteria; Gammaproteobacteria; Enterobacterales;
DTF39__locus_2
EUT1
19240 -pf00873 -
19245 -pf00672pf02518pf07730pf13675 -
19255 -pf13247pf14691pf07992 -
19265 -pf00144 -
19270 HMM00293nuhyd_1_eut1(pf00293) EUT1
19275 -pf06674 -
19280 - -
19285 HMM02780keto_1_eut1(pf02779pf02780pf00456) EUT1
19290 -pf00923 -
19295 - -
19300 HMM00390_malic_1_eut1(pf00390pf03949pf01515) EUT1
19305 Hp_euts_violetPink EUT1
19310 HMM10662PduV_1_mix(pf10662) EUT1
19315 HMM06249EutQ_1_eut1(pf06249) EUT1 EP_NTERM
19320 HMM01923cobtr_2_eut1(pf01923) EUT1
19325 HMM01515PTA_B_1_eut1(pf01515) EUT1
19330 H_azure EUT1
19335 P_lightViolet EUT1
19340 HMMalddh_1_eutpdugrm(pf00171) EUT1 EP_NTERM
19345 HMM06723EutJMreB_1_grm EUT1
19350 HMM00465alcdh_3_mix(pf00465) EUT1
19355 HMMtpEuth_1_eut(pf04346) EUT1
19360 HMM06277EutA_1_eut(pf06277) EUT1
19365 HMMetlyEutB_1_EUT1(pf06751) E
19370 HMMetlyEutC_1_eut1(pf
BMC-P
BMC-H
BMC-T
Aldehyde dehydrogenase
PTAC
Signature enzyme
Alcohol dehydrogenase
Regulator

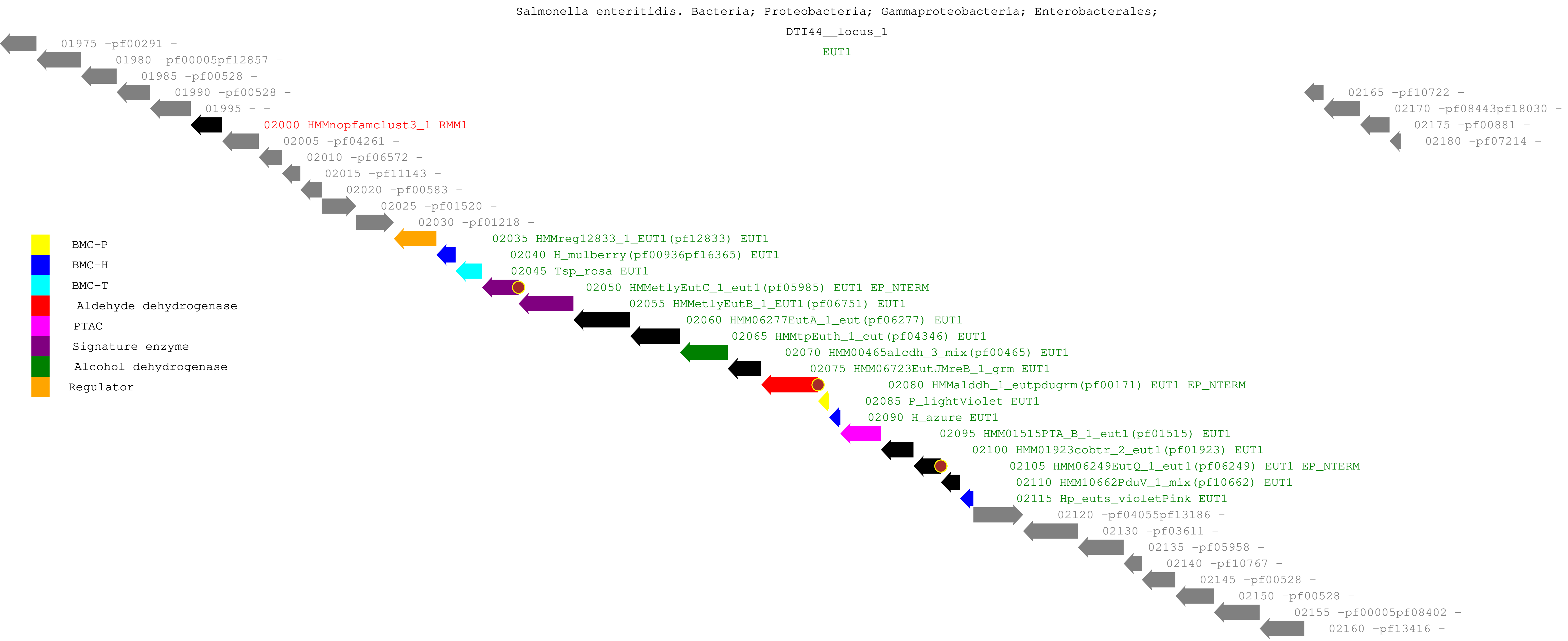

Salmonella enteritidis. Bacteria; Proteobacteria; Gammaproteobacteria; Enterobacterales;
DTI44__locus_1
EUT1
01975 -pf00291 -
01980 -pf00005pf12857 -
01985 -pf00528 -
01990 -pf00528 -
01995 - -
02000 HMMnopfamclust3_1 RMM1
02005 -pf04261 -
02010 -pf06572 -
02015 -pf11143 -
02020 -pf00583 -
02025 -pf01520 -
02030 -pf01218 -
02035 HMMreg12833_1_EUT1(pf12833) EUT1
02040 H_mulberry(pf00936pf16365) EUT1
02045 Tsp_rosa EUT1
02050 HMMetlyEutC_1_eut1(pf05985) EUT1 EP_NTERM
02055 HMMetlyEutB_1_EUT1(pf06751) EUT1
02060 HMM06277EutA_1_eut(pf06277) EUT1
02065 HMMtpEuth_1_eut(pf04346) EUT1
02070 HMM00465alcdh_3_mix(pf00465) EUT1
02075 HMM06723EutJMreB_1_grm EUT1
02080 HMMalddh_1_eutpdugrm(pf00171) EUT1 EP_NTERM
02085 P_lightViolet EUT1
02090 H_azure EUT1
02095 HMM01515PTA_B_1_eut1(pf01515) EUT1
02100 HMM01923cobtr_2_eut1(pf01923) EUT1
02105 HMM06249EutQ_1_eut1(pf06249) EUT1 EP_NTERM
02110 HMM10662PduV_1_mix(pf10662) EUT1
02115 Hp_euts_violetPink EUT1
02120 -pf04055pf13186 -
02130 -pf03611 -
02135 -pf05958 -
02140 -pf10767 -
02145 -pf00528 -
02150 -pf00528 -
02155 -pf00005pf08402 -
02160 -pf13416 -
02165 -pf10722 -
02170 -pf08443pf18030 -
02175 -pf00881 -
02180 -pf07214 -
BMC-P
BMC-H
BMC-T
Aldehyde dehydrogenase
PTAC
Signature enzyme
Alcohol dehydrogenase
Regulator

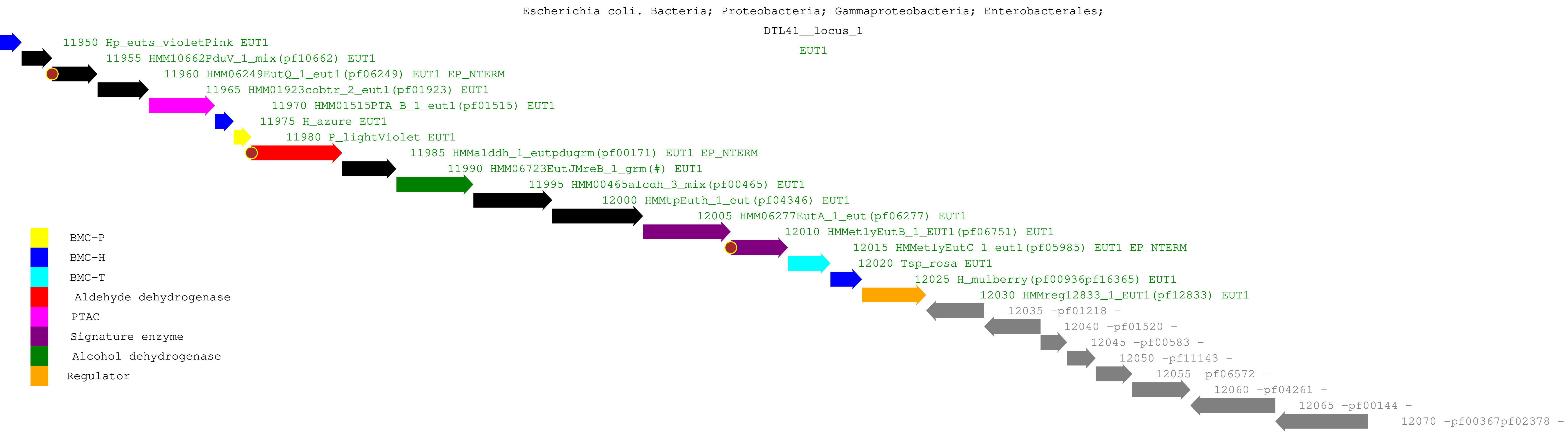

Escherichia coli. Bacteria; Proteobacteria; Gammaproteobacteria; Enterobacterales;
DTL41__locus_1
EUT1
11950 Hp_euts_violetPink EUT1
11955 HMM10662PduV_1_mix(pf10662) EUT1
11960 HMM06249EutQ_1_eut1(pf06249) EUT1 EP_NTERM
11965 HMM01923cobtr_2_eut1(pf01923) EUT1
11970 HMM01515PTA_B_1_eut1(pf01515) EUT1
11975 H_azure EUT1
11980 P_lightViolet EUT1
11985 HMMaldh_1_eutpdugrm(pf00171) EUT1 EP_NTERM
11990 HMM06723EutJMreB_1_grm(#) EUT1
11995 HMM00465alcdh_3_mix(pf00465) EUT1
12000 HMMtpEuth_1_eut(pf04346) EUT1
12005 HMM06277EutA_1_eut(pf06277) EUT1
12010 HMMetlyEutB_1_EUT1(pf06751) EUT1
12015 HMMetlyEutC_1_eut1(pf05985) EUT1 EP_NTERM
12020 Tsp_rosa EUT1
12025 H_mulberry(pf00936pf16365) EUT1
12030 HMMreg12833_1_EUT1(pf12833) EUT1
12035 -pf01218 -
12040 -pf01520 -
12045 -pf00583 -
12050 -pf11143 -
12055 -pf06572 -
12060 -pf04261 -
12065 -pf00144 -
12070 -pf00367pf02378 -
BMC-P
BMC-H
BMC-T
Aldehyde dehydrogenase
PTAC
Signature enzyme
Alcohol dehydrogenase
Regulator

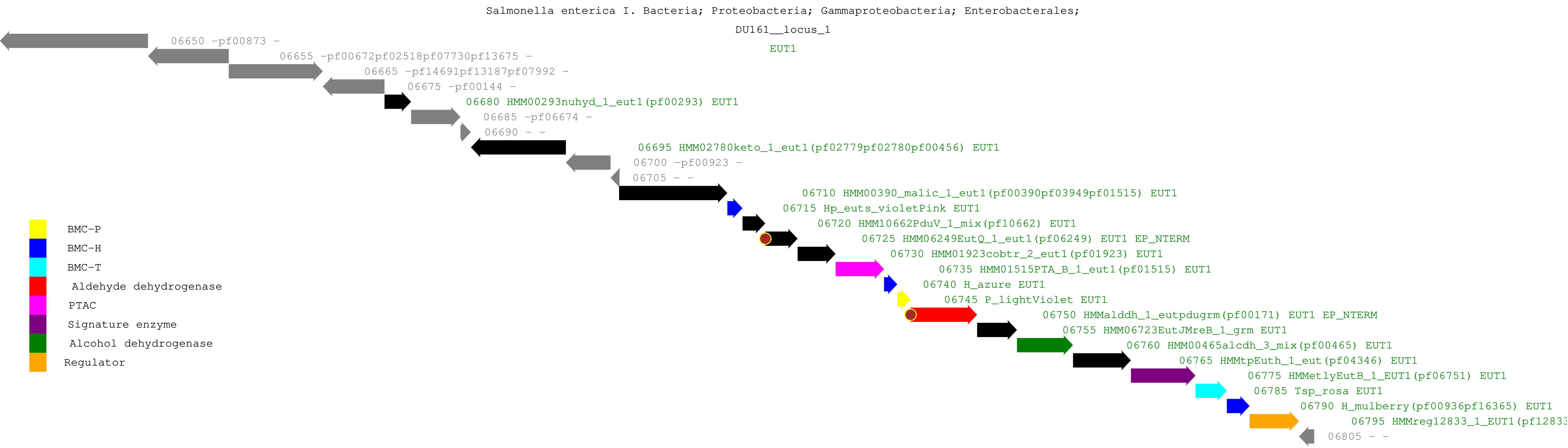

Salmonella enterica I. Bacteria; Proteobacteria; Gammaproteobacteria; Enterobacterales;
DU161__locus_1
EUT1
06650 -pf00873 -
06655 -pf00672pf02518pf07730pf13675 -
06665 -pf14691pf13187pf07992 -
06675 -pf00144 -
06680 HMM00293nuhyd_1_eut1(pf00293) EUT1
06685 -pf06674 -
06690 - -
06695 HMM02780keto_1_eut1(pf02779pf02780pf00456) EUT1
06700 -pf00923 -
06705 - -
06710 HMM00390_malic_1_eut1(pf00390pf03949pf01515) EUT1
06715 Hp_euts_violetPink EUT1
06720 HMM10662PduV_1_mix(pf10662) EUT1
06725 HMM06249EutQ_1_eut1(pf06249) EUT1 EP_NTERM
06730 HMM01923cobtr_2_eut1(pf01923) EUT1
06735 HMM01515PTA_B_1_eut1(pf01515) EUT1
06740 H_azure EUT1
06745 P_lightViolet EUT1
06750 HMMalddh_1_eutpdugrm(pf00171) EUT1 EP_NTERM
06755 HMM06723EutJMreB_1_grm EUT1
06760 HMM00465alcdh_3_mix(pf00465) EUT1
06765 HMMtpEuth_1_eut(pf04346) EUT1
06775 HMMetlyEutB_1_EUT1(pf06751) EUT1
06785 Tsp_rosa EUT1
06790 H_mulberry(pf00936pf16365) EUT1
06795 HMMreg12833_1_EUT1(pf1283
06805 - -
BMC-P
BMC-H
BMC-T
Aldehyde dehydrogenase
PTAC
Signature enzyme
Alcohol dehydrogenase
Regulator

Salmonella enterica I. Bacteria; Proteobacteria; Gammaproteobacteria; Enterobacterales;

DU777__locus_1

EUT1

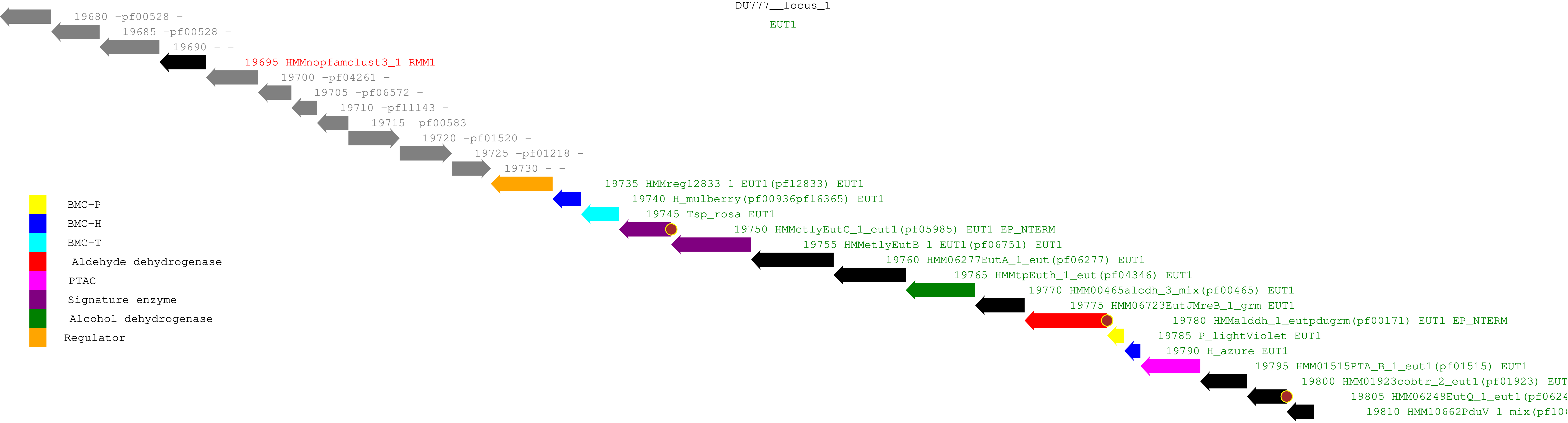

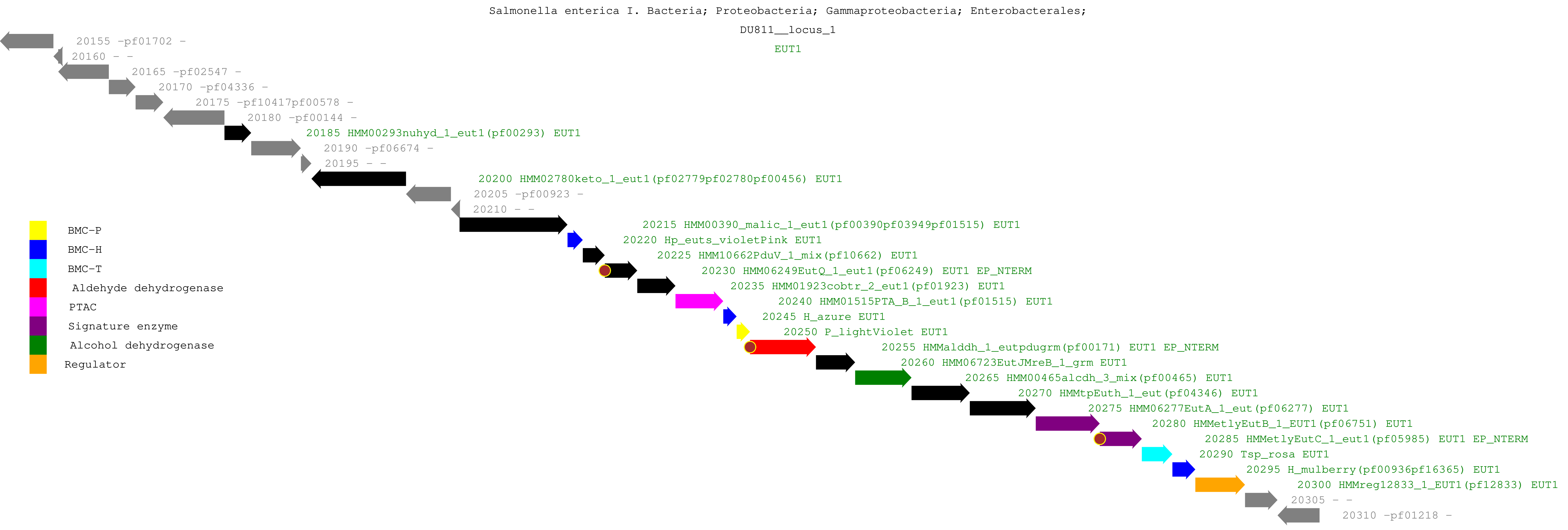

Salmonella enterica I. Bacteria; Proteobacteria; Gammaproteobacteria; Enterobacterales;
DU811__locus_1
EUT1
20155 -pf01702 -
20160 - -
20165 -pf02547 -
20170 -pf04336 -
20175 -pf10417pf00578 -
20180 -pf00144 -
20185 HMM00293nuhyd_1_eut1(pf00293) EUT1
20190 -pf06674 -
20195 - -
20200 HMM02780keto_1_eut1(pf02779pf02780pf00456) EUT1
20205 -pf00923 -
20210 - -
20215 HMM00390_malic_1_eut1(pf00390pf03949pf01515) EUT1
20220 Hp_euts_violetPink EUT1
20225 HMM10662PduV_1_mix(pf10662) EUT1
20230 HMM06249EutQ_1_eut1(pf06249) EUT1 EP_NTERM
20235 HMM01923cobtr_2_eut1(pf01923) EUT1
20240 HMM01515PTA_B_1_eut1(pf01515) EUT1
20245 H_azure EUT1
20250 P_lightViolet EUT1
20255 HMMalddh_1_eutpdugrm(pf00171) EUT1 EP_NTERM
20260 HMM06723EutJMreB_1_grm EUT1
20265 HMM00465alcdh_3_mix(pf00465) EUT1
20270 HMMtpEuth_1_eut(pf04346) EUT1
20275 HMM06277EutA_1_eut(pf06277) EUT1
20280 HMMetlyEutB_1_EUT1(pf06751) EUT1
20285 HMMetlyEutC_1_eut1(pf05985) EUT1 EP_NTERM
20290 Tsp_rosa EUT1
20295 H_mulberry(pf00936pf16365) EUT1
20300 HMMreg12833_1_EUT1(pf12833) EUT1
20305 - -
20310 -pf01218 -
BMC-P
BMC-H
BMC-T
Aldehyde dehydrogenase
PTAC
Signature enzyme
Alcohol dehydrogenase
Regulator

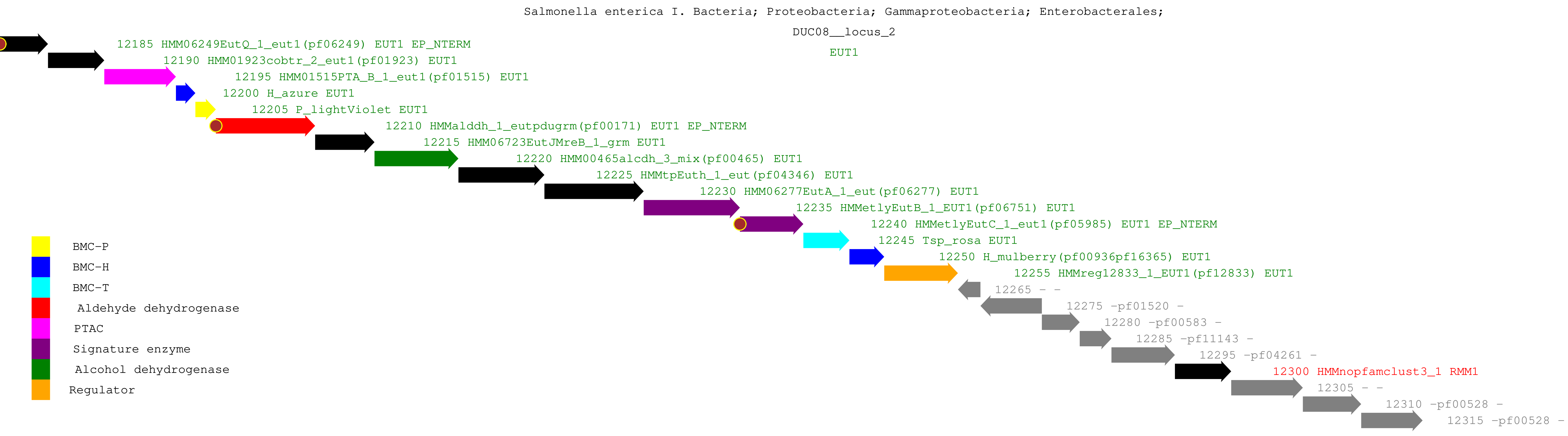
Salmonella enterica I. Bacteria; Proteobacteria; Gammaproteobacteria; Enterobacterales;
DUC08__locus_2
EUT1
12185 HMM06249EutQ_1_eut1(pf06249) EUT1 EP_NTERM
12190 HMM01923cobtr_2_eut1(pf01923) EUT1
12195 HMM01515PTA_B_1_eut1(pf01515) EUT1
12200 H_azure EUT1
12205 P_lightViolet EUT1
12210 HMMalddh_1_eutpdugrm(pf00171) EUT1 EP_NTERM
12215 HMM06723EutJMreB_1_grm EUT1
12220 HMM00465alcdh_3_mix(pf00465) EUT1
12225 HMMtpEuth_1_eut(pf04346) EUT1
12230 HMM06277EutA_1_eut(pf06277) EUT1
12235 HMMetlyEutB_1_EUT1(pf06751) EUT1
12240 HMMetlyEutC_1_eut1(pf05985) EUT1 EP_NTERM
12245 Tsp_rosa EUT1
12250 H_mulberry(pf00936pf16365) EUT1
12255 HMMreg12833_1_EUT1(pf12833) EUT1
12265 - -
12275 -pf01520 -
12280 -pf00583 -
12285 -pf11143 -
12295 -pf04261 -
12300 HMMnopfamclust3_1 RMM1
12305 - -
12310 -pf00528 -
12315 -pf00528 -
BMC-P
BMC-H
BMC-T
Aldehyde dehydrogenase
PTAC
Signature enzyme
Alcohol dehydrogenase
Regulator

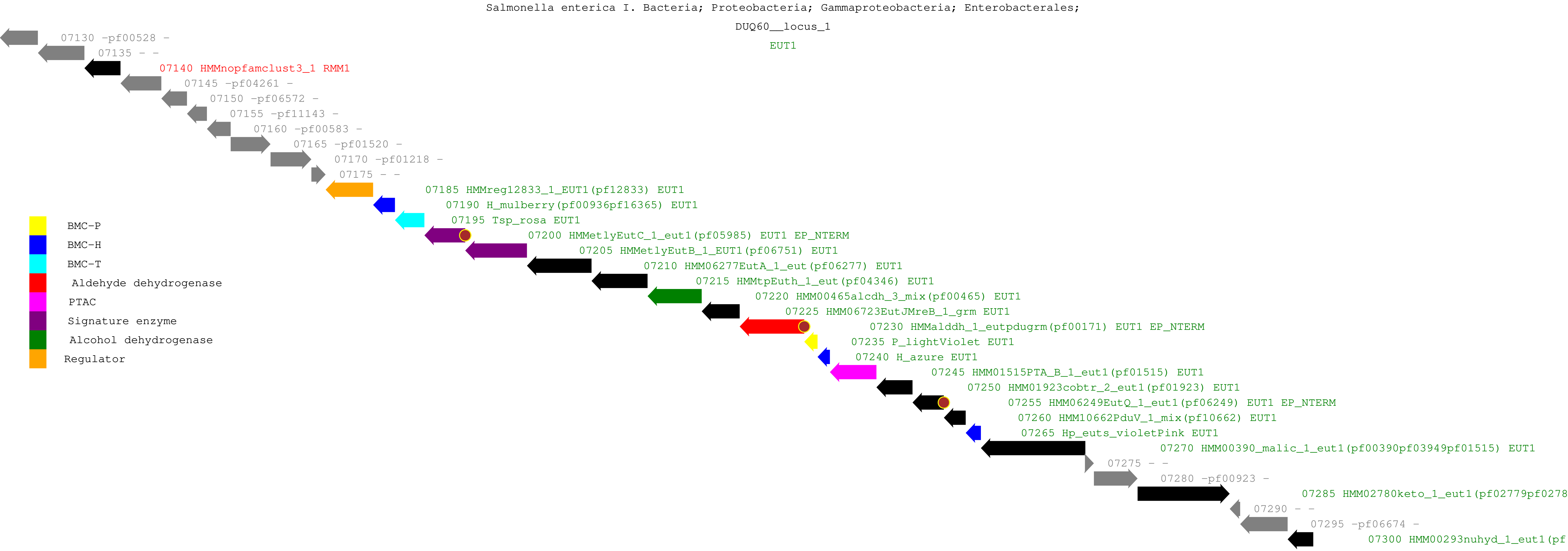

Salmonella enterica I. Bacteria; Proteobacteria; Gammaproteobacteria; Enterobacterales;
DUQ60__locus_1
EUT1
07130 -pf00528 -
07135 - -
07140 HMMnopfamclust3_1 RMM1
07145 -pf04261 -
07150 -pf06572 -
07155 -pf11143 -
07160 -pf00583 -
07165 -pf01520 -
07170 -pf01218 -
07175 - -
07185 HMMreg12833_1_EUT1(pf12833) EUT1
07190 H_mulberry(pf00936pf16365) EUT1
07195 Tsp_rosa EUT1
07200 HMMetlyEutC_1_eut1(pf05985) EUT1 EP_NTERM
07205 HMMetlyEutB_1_EUT1(pf06751) EUT1
07210 HMM06277EutA_1_eut(pf06277) EUT1
07215 HMMtpEuth_1_eut(pf04346) EUT1
07220 HMM00465alcdh_3_mix(pf00465) EUT1
07225 HMM06723EutJMreB_1_grm EUT1
07230 HMMalddh_1_eutpdugrm(pf00171) EUT1 EP_NTERM
07235 P_lightViolet EUT1
07240 H_azure EUT1
07245 HMM01515PTA_B_1_eut1(pf01515) EUT1
07250 HMM01923cobtr_2_eut1(pf01923) EUT1
07255 HMM06249EutQ_1_eut1(pf06249) EUT1 EP_NTERM
07260 HMM10662PduV_1_mix(pf10662) EUT1
07265 Hp_euts_violetPink EUT1
07270 HMM00390_malic_1_eut1(pf00390pf03949pf01515) EUT1
07275 - -
07280 -pf00923 -
07285 HMM02780keto_1_eut1(pf02779pf0278
07290 - -
07295 -pf06674 -
07300 HMM00293nuhyd_1_eut1(pf
BMC-P
BMC-H
BMC-T
Aldehyde dehydrogenase
PTAC
Signature enzyme
Alcohol dehydrogenase
Regulator

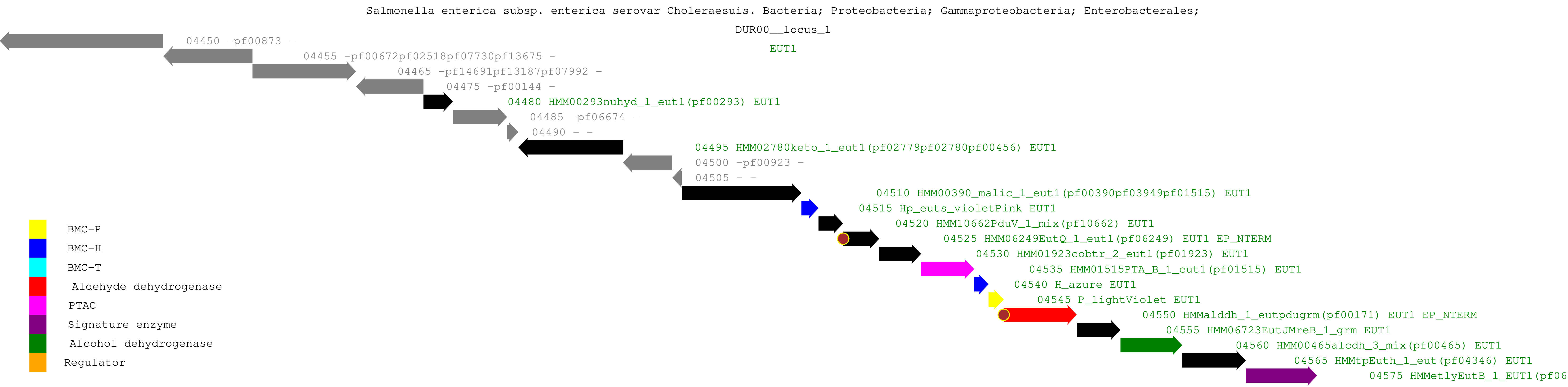
Salmonella enterica subsp. enterica serovar Choleraesuis. Bacteria; Proteobacteria; Gammaproteobacteria; Enterobacterales;
DUR00__locus_1
EUT1
04450 -pf00873 -
04455 -pf00672pf02518pf07730pf13675 -
04465 -pf14691pf13187pf07992 -
04475 -pf00144 -
04480 HMM00293nuhyd_1_eut1(pf00293) EUT1
04485 -pf06674 -
04490 - -
04495 HMM02780keto_1_eut1(pf02779pf02780pf00456) EUT1
04500 -pf00923 -
04505 - -
04510 HMM00390_malic_1_eut1(pf00390pf03949pf01515) EUT1
04515 Hp_euts_violetPink EUT1
04520 HMM10662PduV_1_mix(pf10662) EUT1
04525 HMM06249EutQ_1_eut1(pf06249) EUT1 EP_NTERM
04530 HMM01923cobtr_2_eut1(pf01923) EUT1
04535 HMM01515PTA_B_1_eut1(pf01515) EUT1
04540 H_azure EUT1
04545 P_lightViolet EUT1
04550 HMMaldh_1_eutpdugrm(pf00171) EUT1 EP_NTERM
04555 HMM06723EutJMreB_1_grm EUT1
04560 HMM00465alcdh_3_mix(pf00465) EUT1
04565 HMMtpEuth_1_eut(pf04346) EUT1
04575 HMMetlyEutB_1_EUT1(pf06
BMC-P
BMC-H
BMC-T
Aldehyde dehydrogenase
PTAC
Signature enzyme
Alcohol dehydrogenase
Regulator

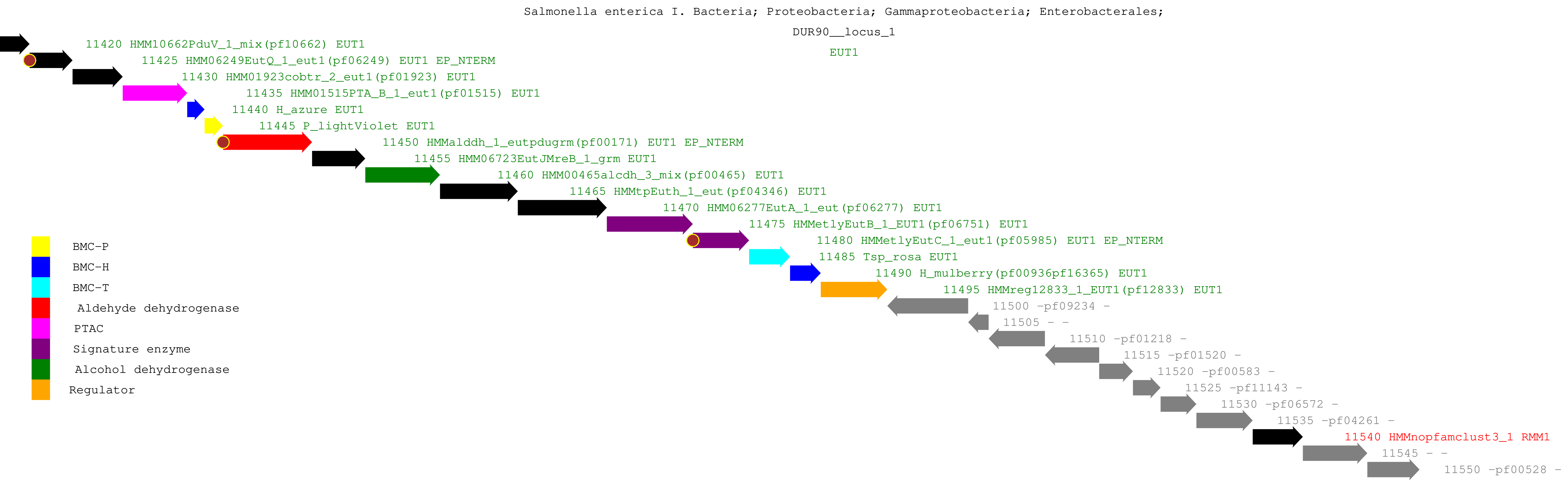
Salmonella enterica I. Bacteria; Proteobacteria; Gammaproteobacteria; Enterobacterales;
DUR90__locus_1
EUT1
11420 HMM10662PduV_1_mix(pf10662) EUT1
11425 HMM06249EutQ_1_eut1(pf06249) EUT1 EP_NTERM
11430 HMM01923cobtr_2_eut1(pf01923) EUT1
11435 HMM01515PTA_B_1_eut1(pf01515) EUT1
11440 H_azure EUT1
11445 P_lightViolet EUT1
11450 HMMalddh_1_eutpdugrm(pf00171) EUT1 EP_NTERM
11455 HMM06723EutJMreB_1_grm EUT1
11460 HMM00465alcdh_3_mix(pf00465) EUT1
11465 HMMtpEuth_1_eut(pf04346) EUT1
11470 HMM06277EutA_1_eut(pf06277) EUT1
11475 HMMetlyEutB_1_EUT1(pf06751) EUT1
11480 HMMetlyEutC_1_eut1(pf05985) EUT1 EP_NTERM
11485 Tsp_rosa EUT1
11490 H_mulberry(pf00936pf16365) EUT1
11495 HMMreg12833_1_EUT1(pf12833) EUT1
11500 -pf09234 -
11505 - -
11510 -pf01218 -
11515 -pf01520 -
11520 -pf00583 -
11525 -pf11143 -
11530 -pf06572 -
11535 -pf04261 -
11540 HMMnopfamclust3_1 RMM1
11545 - -
11550 -pf00528 -
BMC-P
BMC-H
BMC-T
Aldehyde dehydrogenase
PTAC
Signature enzyme
Alcohol dehydrogenase
Regulator

Klebsiella oxytoca. Bacteria; Proteobacteria; Gammaproteobacteria; Enterobacterales;

DVB85__locus_2

EUT1

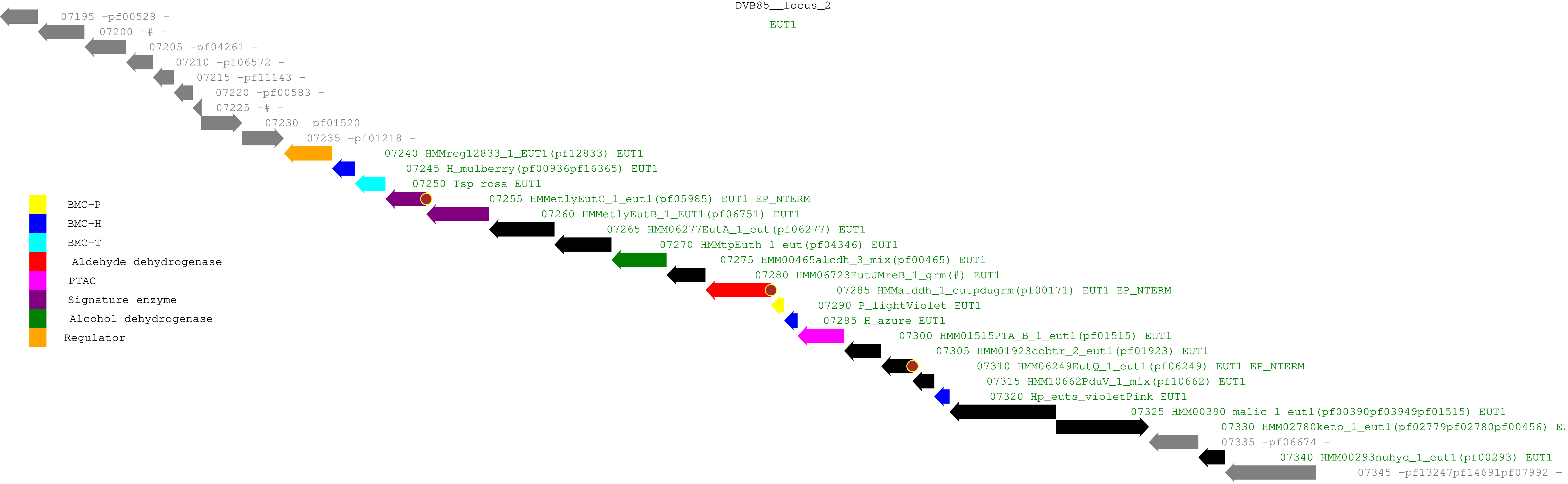

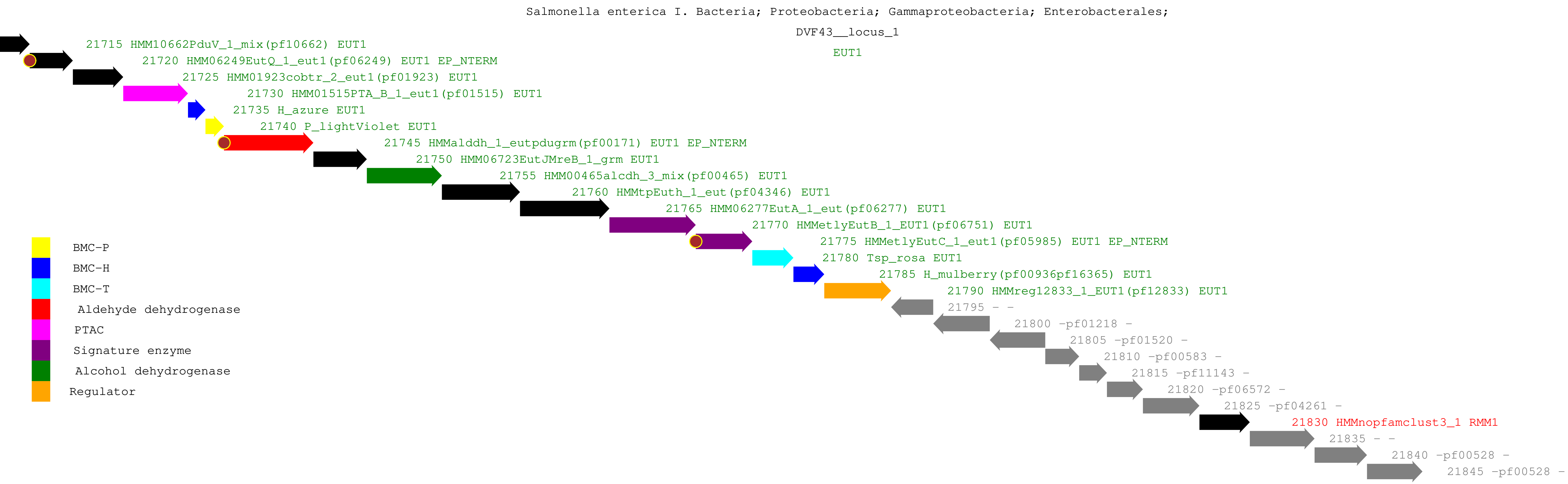
Salmonella enterica I. Bacteria; Proteobacteria; Gammaproteobacteria; Enterobacterales;
DVF43__locus_1
EUT1
21715 HMM10662PduV_1_mix(pf10662) EUT1
21720 HMM06249EutQ_1_eut1(pf06249) EUT1 EP_NTERM
21725 HMM01923cobtr_2_eut1(pf01923) EUT1
21730 HMM01515PTA_B_1_eut1(pf01515) EUT1
21735 H_azure EUT1
21740 P_lightViolet EUT1
21745 HMMalddh_1_eutpdugrm(pf00171) EUT1 EP_NTERM
21750 HMM06723EutJMreB_1_grm EUT1
21755 HMM00465alcdh_3_mix(pf00465) EUT1
21760 HMMtpEuth_1_eut(pf04346) EUT1
21765 HMM06277EutA_1_eut(pf06277) EUT1
21770 HMMetlyEutB_1_EUT1(pf06751) EUT1
21775 HMMetlyEutC_1_eut1(pf05985) EUT1 EP_NTERM
21780 Tsp_rosa EUT1
21785 H_mulberry(pf00936pf16365) EUT1
21790 HMMreg12833_1_EUT1(pf12833) EUT1
21795 - -
21800 -pf01218 -
21805 -pf01520 -
21810 -pf00583 -
21815 -pf11143 -
21820 -pf06572 -
21825 -pf04261 -
21830 HMMnopfamclust3_1 RMM1
21835 - -
21840 -pf00528 -
21845 -pf00528 -
BMC-P
BMC-H
BMC-T
Aldehyde dehydrogenase
PTAC
Signature enzyme
Alcohol dehydrogenase
Regulator

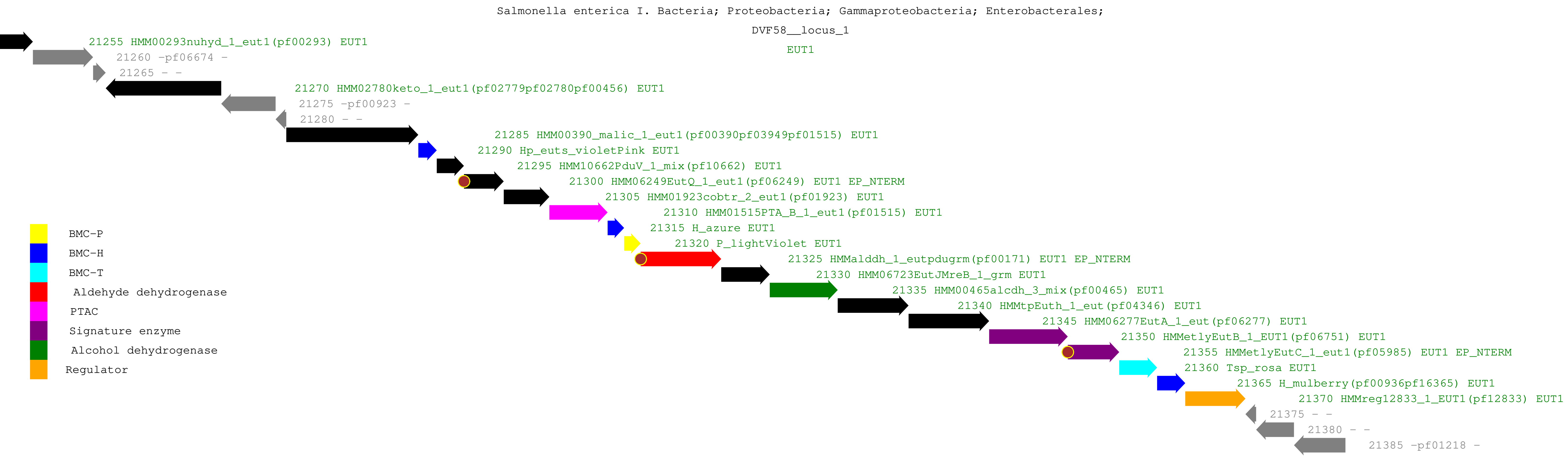

Salmonella enterica I. Bacteria; Proteobacteria; Gammaproteobacteria; Enterobacterales;
DVF58__locus_1
EUT1
21255 HMM00293nuhyd_1_eut1(pf00293) EUT1
21260 -pf06674 -
21265 - -
21270 HMM02780keto_1_eut1(pf02779pf02780pf00456) EUT1
21275 -pf00923 -
21280 - -
21285 HMM00390_malic_1_eut1(pf00390pf03949pf01515) EUT1
21290 Hp_euts_violetPink EUT1
21295 HMM10662PduV_1_mix(pf10662) EUT1
21300 HMM06249EutQ_1_eut1(pf06249) EUT1 EP_NTERM
21305 HMM01923cobtr_2_eut1(pf01923) EUT1
21310 HMM01515PTA_B_1_eut1(pf01515) EUT1
21315 H_azure EUT1
21320 P_lightViolet EUT1
21325 HMMalddh_1_eutpdugrm(pf00171) EUT1 EP_NTERM
21330 HMM06723EutJMreB_1_grm EUT1
21335 HMM00465alcdh_3_mix(pf00465) EUT1
21340 HMMtpEuth_1_eut(pf04346) EUT1
21345 HMM06277EutA_1_eut(pf06277) EUT1
21350 HMMetlyEutB_1_EUT1(pf06751) EUT1
21355 HMMetlyEutC_1_eut1(pf05985) EUT1 EP_NTERM
21360 Tsp_rosa EUT1
21365 H_mulberry(pf00936pf16365) EUT1
21370 HMMreg12833_1_EUT1(pf12833) EUT1
21375 - -
21380 - -
21385 -pf01218 -
BMC-P
BMC-H
BMC-T
Aldehyde dehydrogenase
PTAC
Signature enzyme
Alcohol dehydrogenase
Regulator

Citrobacter gillenii. Bacteria; Proteobacteria; Gammaproteobacteria; Enterobacterales;

DZA29__locus_2

EUT1

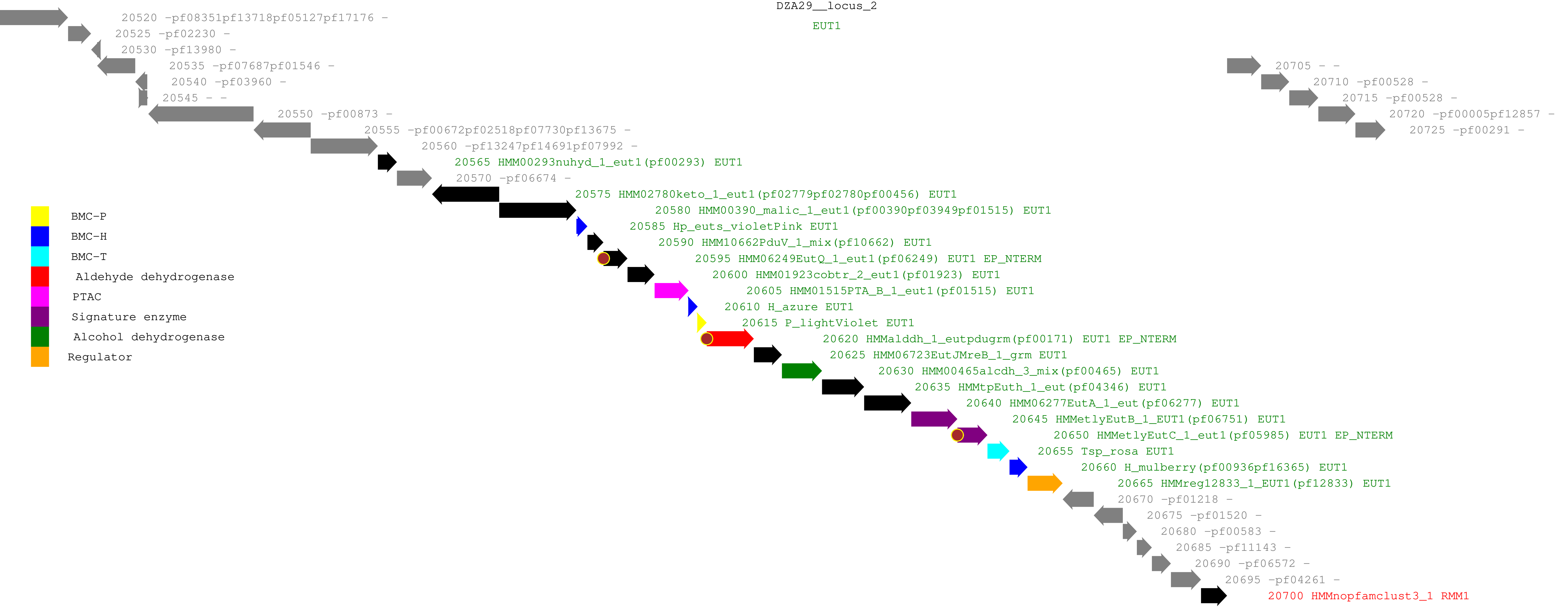

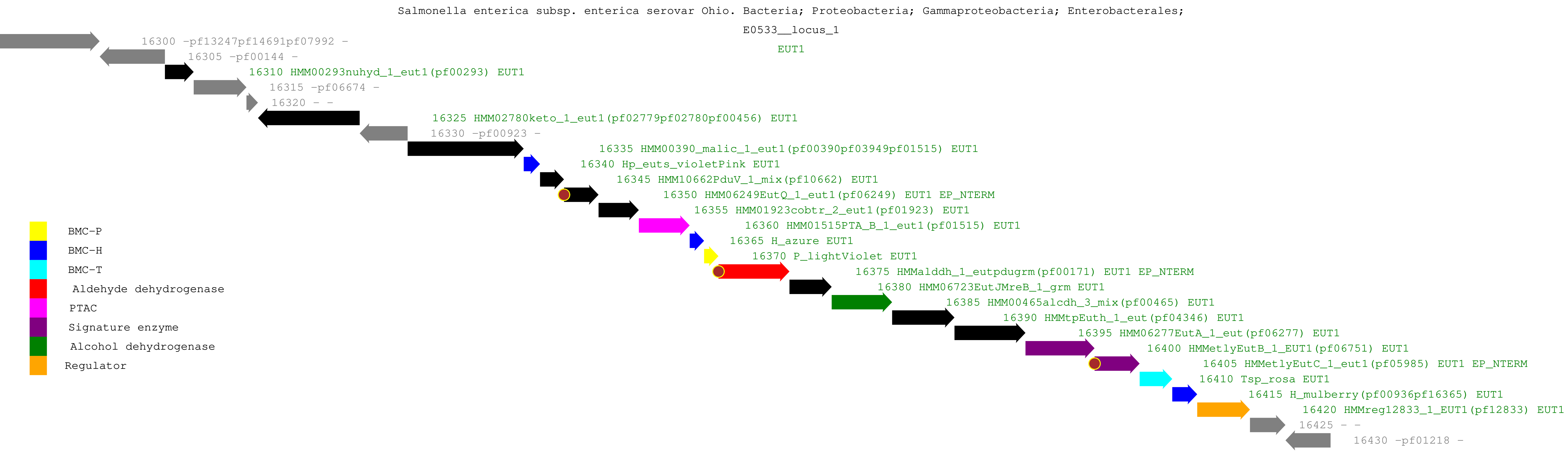
Salmonella enterica subsp. enterica serovar Ohio. Bacteria; Proteobacteria; Gammaproteobacteria; Enterobacterales;
E0533__locus_1
EUT1
16300 -pf13247pf14691pf07992 -
16305 -pf00144 -
16310 HMM00293nuhyd_1_eut1(pf00293) EUT1
16315 -pf06674 -
16320 - -
16325 HMM02780keto_1_eut1(pf02779pf02780pf00456) EUT1
16330 -pf00923 -
16335 HMM00390_malic_1_eut1(pf00390pf03949pf01515) EUT1
16340 Hp_euts_violetPink EUT1
16345 HMM10662PduV_1_mix(pf10662) EUT1
16350 HMM06249EutQ_1_eut1(pf06249) EUT1 EP_NTERM
16355 HMM01923cobtr_2_eut1(pf01923) EUT1
16360 HMM01515PTA_B_1_eut1(pf01515) EUT1
16365 H_azure EUT1
16370 P_lightViolet EUT1
16375 HMMalddh_1_eutpdugrm(pf00171) EUT1 EP_NTERM
16380 HMM06723EutJMreB_1_grm EUT1
16385 HMM00465alcdh_3_mix(pf00465) EUT1
16390 HMMtpEuth_1_eut(pf04346) EUT1
16395 HMM06277EutA_1_eut(pf06277) EUT1
16400 HMMetlyEutB_1_EUT1(pf06751) EUT1
16405 HMMetlyEutC_1_eut1(pf05985) EUT1 EP_NTERM
16410 Tsp_rosa EUT1
16415 H_mulberry(pf00936pf16365) EUT1
16420 HMMreg12833_1_EUT1(pf12833) EUT1
16425 - -
16430 -pf01218 -
BMC-P
BMC-H
BMC-T
Aldehyde dehydrogenase
PTAC
Signature enzyme
Alcohol dehydrogenase
Regulator

Salmonella enterica subsp. enterica serovar Tilene. Bacteria; Proteobacteria; Gammaproteobacteria; Enterobacterales;

E0539__locus_1

EUT1

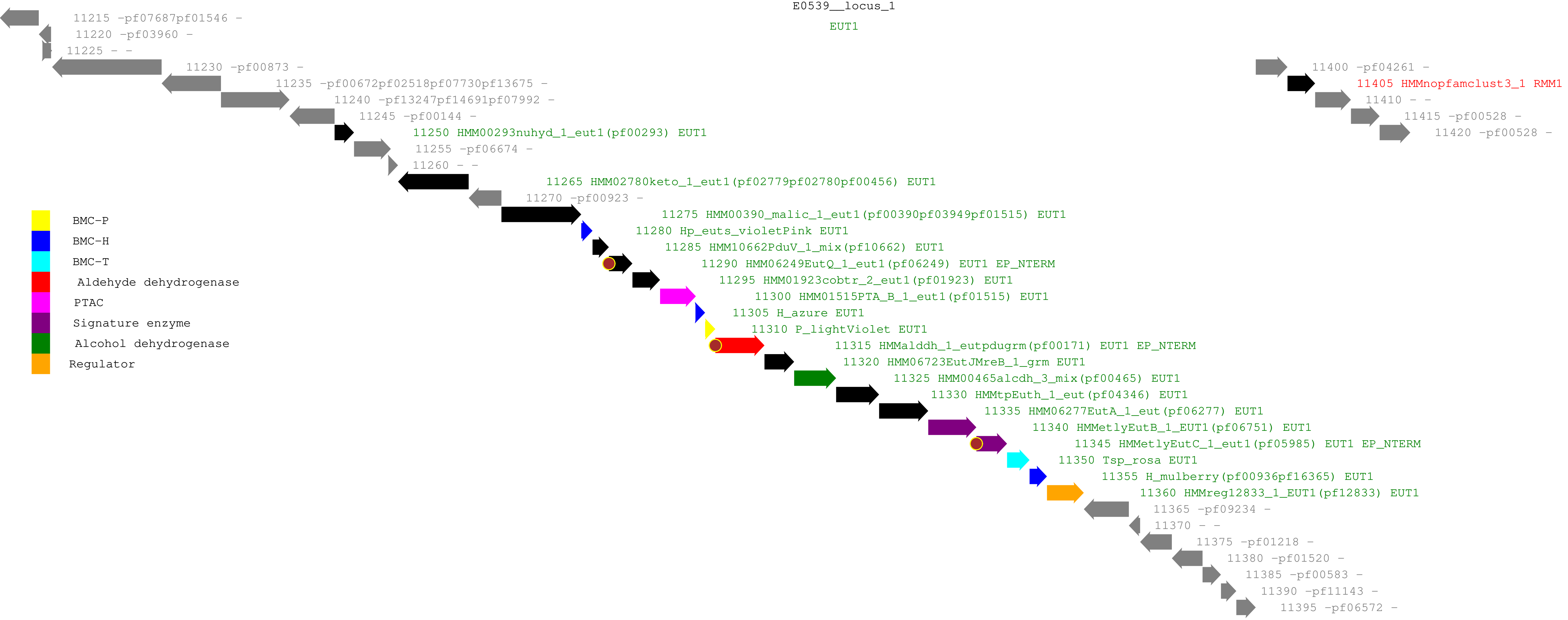

Salmonella enterica subsp. enterica serovar London. Bacteria; Proteobacteria; Gammaproteobacteria; Enterobacterales;

E0541__locus_2

EUT1

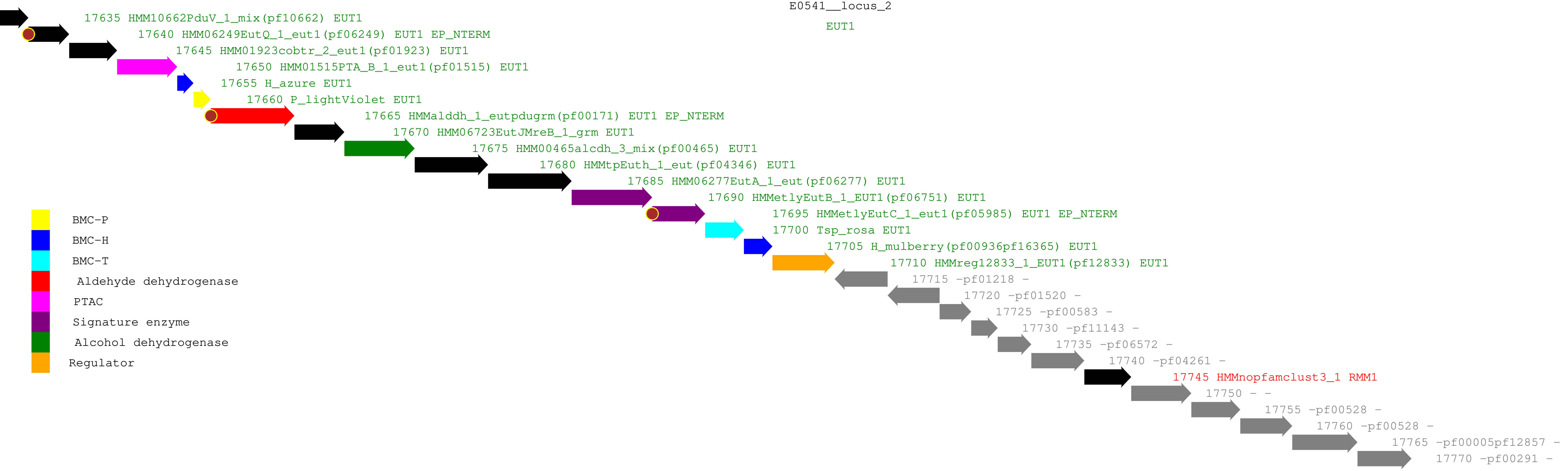

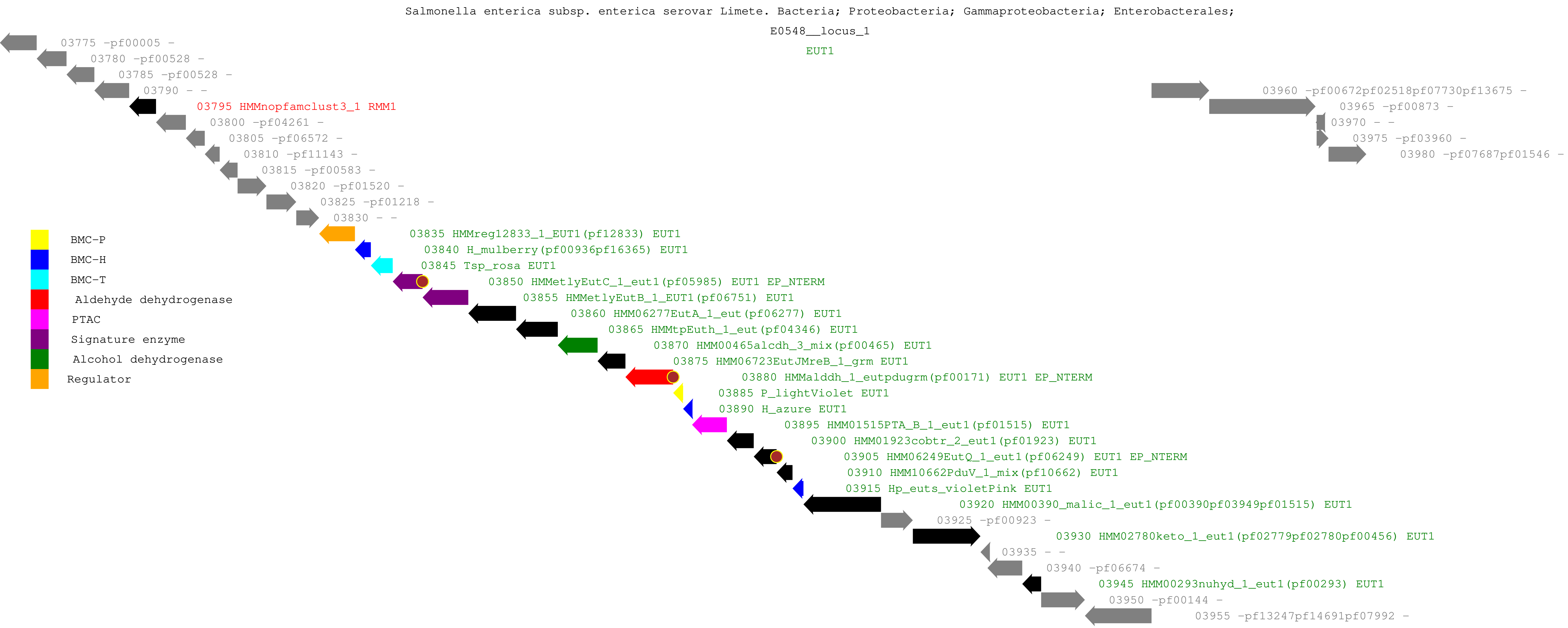
Salmonella enterica subsp. enterica serovar Limete. Bacteria; Proteobacteria; Gammaproteobacteria; Enterobacterales;
E0548__locus_1
EUT1
03775 -pf00005 -
03780 -pf00528 -
03785 -pf00528 -
03790 - -
03795 HMMnopfamclust3_1 RMM1
03800 -pf04261 -
03805 -pf06572 -
03810 -pf11143 -
03815 -pf00583 -
03820 -pf01520 -
03825 -pf01218 -
03830 - -
03835 HMMreg12833_1_EUT1(pf12833) EUT1
03840 H_mulberry(pf00936pf16365) EUT1
03845 Tsp_rosa EUT1
03850 HMMetlyEutC_1_eut1(pf05985) EUT1 EP_NTERM
03855 HMMetlyEutB_1_EUT1(pf06751) EUT1
03860 HMM06277EutA_1_eut(pf06277) EUT1
03865 HMMtpEuth_1_eut(pf04346) EUT1
03870 HMM00465alcdh_3_mix(pf00465) EUT1
03875 HMM06723EutJMreB_1_grm EUT1
03880 HMMalddh_1_eutpdugrm(pf00171) EUT1 EP_NTERM
03885 P_lightViolet EUT1
03890 H_azure EUT1
03895 HMM01515PTA_B_1_eut1(pf01515) EUT1
03900 HMM01923cobtr_2_eut1(pf01923) EUT1
03905 HMM06249EutQ_1_eut1(pf06249) EUT1 EP_NTERM
03910 HMM10662PduV_1_mix(pf10662) EUT1
03915 Hp_euts_violetPink EUT1
03920 HMM00390_malic_1_eut1(pf00390pf03949pf01515) EUT1
03925 -pf00923 -
03930 HMM02780keto_1_eut1(pf02779pf02780pf00456) EUT1
03935 - -
03940 -pf06674 -
03945 HMM00293nuhyd_1_eut1(pf00293) EUT1
03950 -pf00144 -
03955 -pf13247pf14691pf07992 -
03960 -pf00672pf02518pf07730pf13675 -
03965 -pf00873 -
03970 - -
03975 -pf03960 -
03980 -pf07687pf01546 -
BMC-P
BMC-H
BMC-T
Aldehyde dehydrogenase
PTAC
Signature enzyme
Alcohol dehydrogenase
Regulator

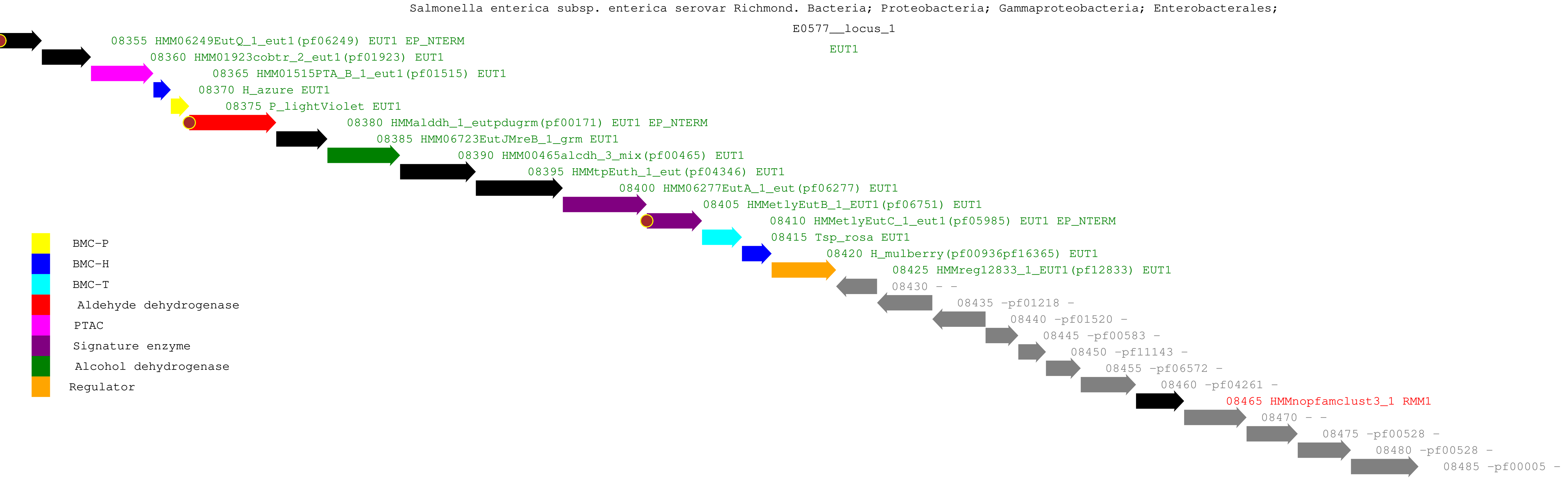
Salmonella enterica subsp. enterica serovar Richmond. Bacteria; Proteobacteria; Gammaproteobacteria; Enterobacterales;
E0577__locus_1
EUT1
08355 HMM06249EutQ_1_eut1(pf06249) EUT1 EP_NTERM
08360 HMM01923cobtr_2_eut1(pf01923) EUT1
08365 HMM01515PTA_B_1_eut1(pf01515) EUT1
08370 H_azure EUT1
08375 P_lightViolet EUT1
08380 HMMalddh_1_eutpdugrm(pf00171) EUT1 EP_NTERM
08385 HMM06723EutJMreB_1_grm EUT1
08390 HMM00465alcdh_3_mix(pf00465) EUT1
08395 HMMtpEuth_1_eut(pf04346) EUT1
08400 HMM06277EutA_1_eut(pf06277) EUT1
08405 HMMetlyEutB_1_EUT1(pf06751) EUT1
08410 HMMetlyEutC_1_eut1(pf05985) EUT1 EP_NTERM
08415 Tsp_rosa EUT1
08420 H_mulberry(pf00936pf16365) EUT1
08425 HMMreg12833_1_EUT1(pf12833) EUT1
08430 - -
08435 -pf01218 -
08440 -pf01520 -
08445 -pf00583 -
08450 -pf11143 -
08455 -pf06572 -
08460 -pf04261 -
08465 HMMnopfamclust3_1 RMM1
08470 - -
08475 -pf00528 -
08480 -pf00528 -
08485 -pf00005 -
BMC-P
BMC-H
BMC-T
Aldehyde dehydrogenase
PTAC
Signature enzyme
Alcohol dehydrogenase
Regulator

Salmonella enterica subsp. enterica serovar Carmel. Bacteria; Proteobacteria; Gammaproteobacteria; Enterobacterales;

E0601__locus_2

EUT1

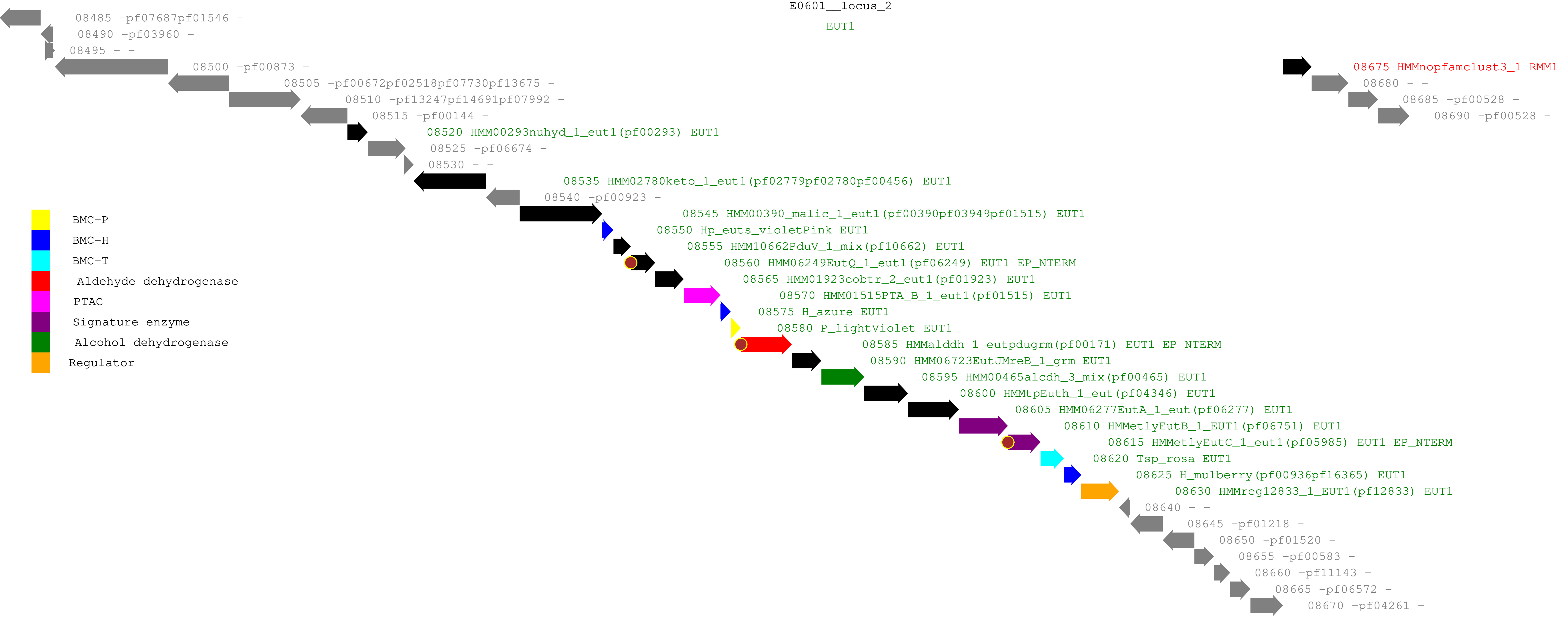

Salmonella abony. Bacteria; Proteobacteria; Gammaproteobacteria; Enterobacterales;

E0657__locus_2

EUT1

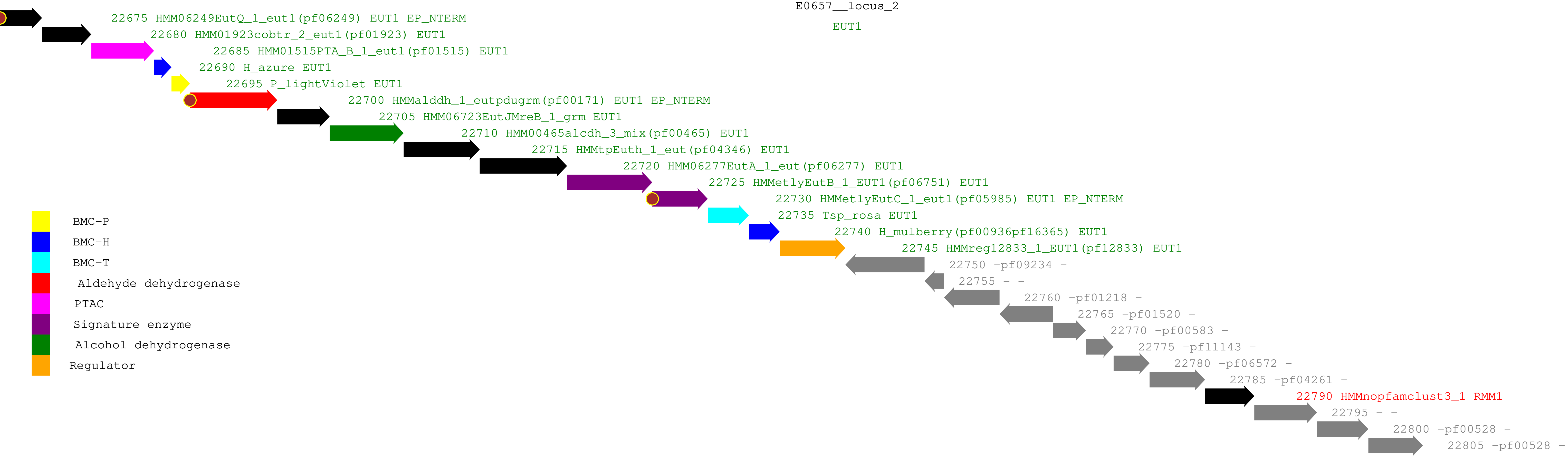

Salmonella enterica subsp. enterica serovar Oslo. Bacteria; Proteobacteria; Gammaproteobacteria; Enterobacterales;

E0660__locus_1

EUT1

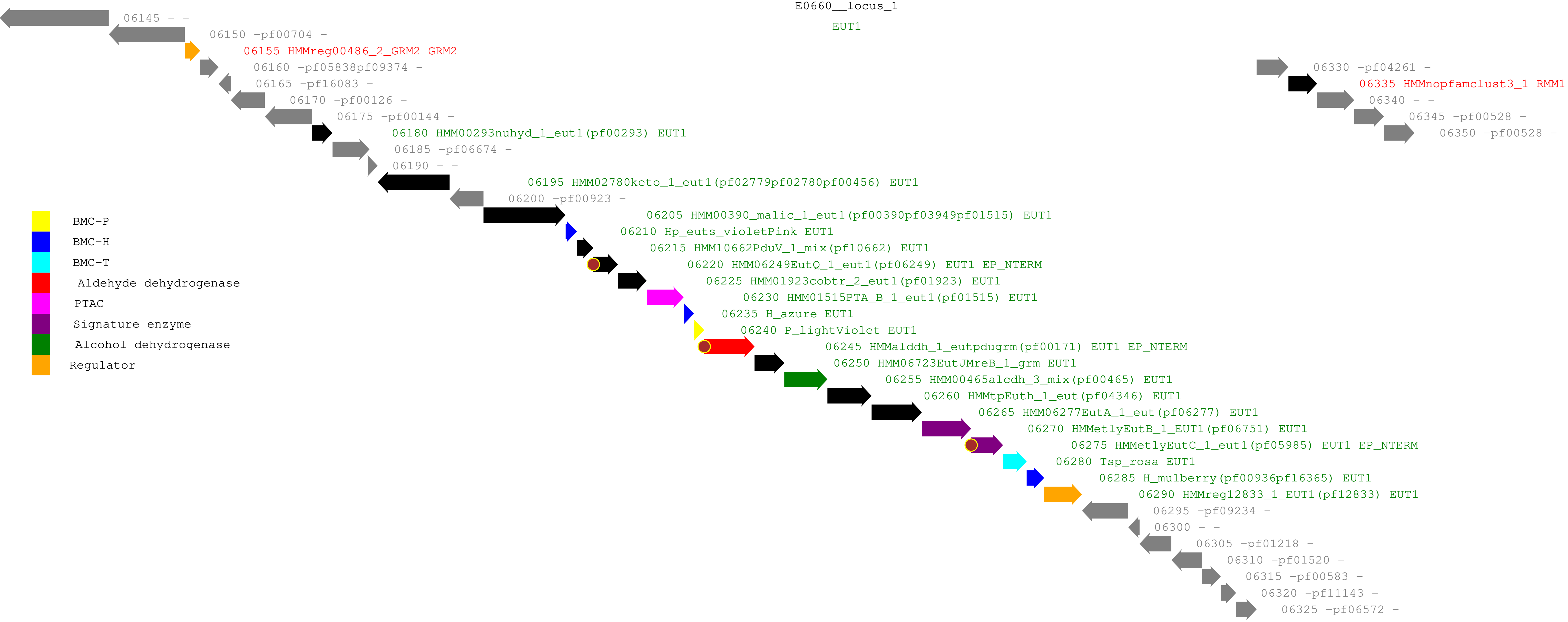

Salmonella berta. Bacteria; Proteobacteria; Gammaproteobacteria; Enterobacterales;

E0917__locus_2

EUT1

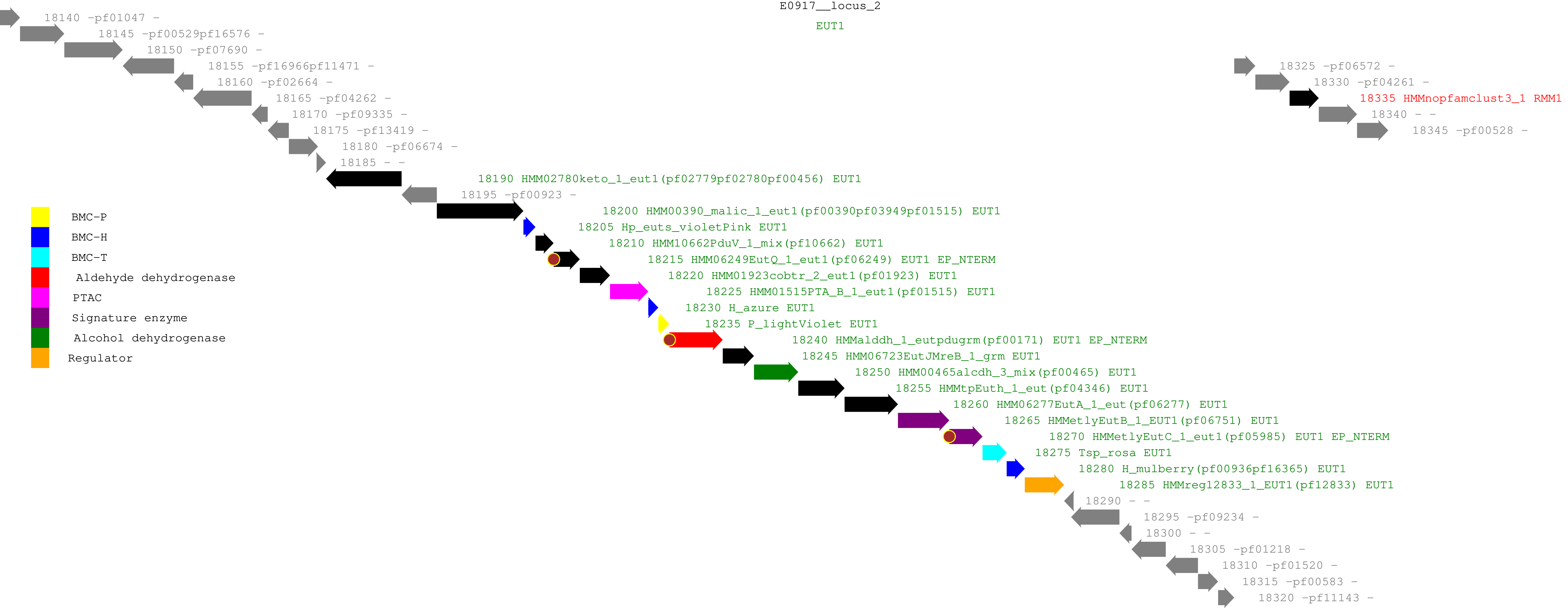

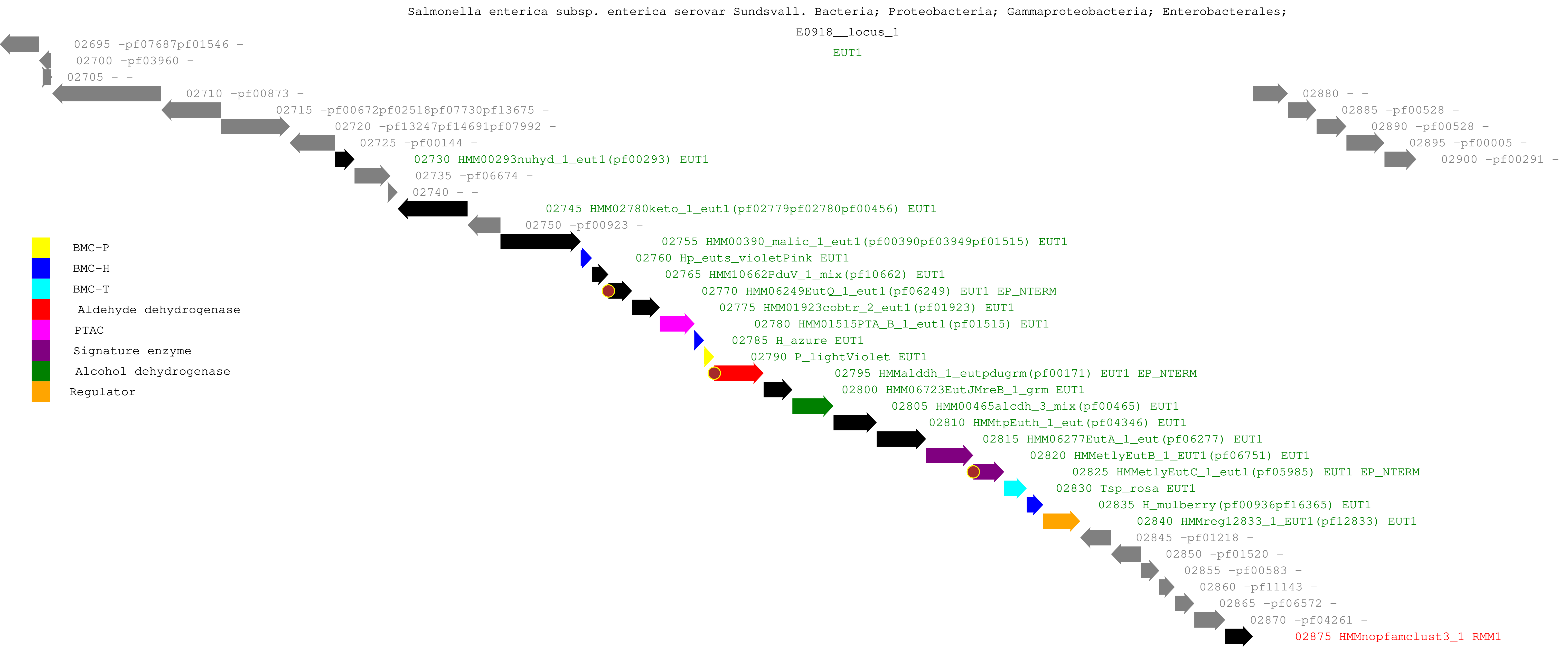
Salmonella enterica subsp. enterica serovar Sundsvall. Bacteria; Proteobacteria; Gammaproteobacteria; Enterobacterales;
E0918__locus_1
EUT1
02695 -pf07687pf01546 -
02700 -pf03960 -
02705 - -
02710 -pf00873 -
02715 -pf00672pf02518pf07730pf13675 -
02720 -pf13247pf14691pf07992 -
02725 -pf00144 -
02730 HMM00293nuhyd_1_eut1(pf00293) EUT1
02735 -pf06674 -
02740 - -
02745 HMM02780keto_1_eut1(pf02779pf02780pf00456) EUT1
02750 -pf00923 -
02755 HMM00390_malic_1_eut1(pf00390pf03949pf01515) EUT1
02760 Hp_euts_violetPink EUT1
02765 HMM10662PduV_1_mix(pf10662) EUT1
02770 HMM06249EutQ_1_eut1(pf06249) EUT1 EP_NTERM
02775 HMM01923cobtr_2_eut1(pf01923) EUT1
02780 HMM01515PTA_B_1_eut1(pf01515) EUT1
02785 H_azure EUT1
02790 P_lightViolet EUT1
02795 HMMalddh_1_eutpdugrm(pf00171) EUT1 EP_NTERM
02800 HMM06723EutJMreB_1_grm EUT1
02805 HMM00465alcdh_3_mix(pf00465) EUT1
02810 HMMtpEuth_1_eut(pf04346) EUT1
02815 HMM06277EutA_1_eut(pf06277) EUT1
02820 HMMetlyEutB_1_EUT1(pf06751) EUT1
02825 HMMetlyEutC_1_eut1(pf05985) EUT1 EP_NTERM
02830 Tsp_rosa EUT1
02835 H_mulberry(pf00936pf16365) EUT1
02840 HMMreg12833_1_EUT1(pf12833) EUT1
02845 -pf01218 -
02850 -pf01520 -
02855 -pf00583 -
02860 -pf11143 -
02865 -pf06572 -
02870 -pf04261 -
02875 HMMnopfamclust3_1 RMM1
02880 - -
02885 -pf00528 -
02890 -pf00528 -
02895 -pf00005 -
02900 -pf00291 -
BMC-P
BMC-H
BMC-T
Aldehyde dehydrogenase
PTAC
Signature enzyme
Alcohol dehydrogenase
Regulator

Salmonella enterica subsp. enterica serovar Szentes. Bacteria; Proteobacteria; Gammaproteobacteria; Enterobacterales;

E0931__locus_2

EUT1

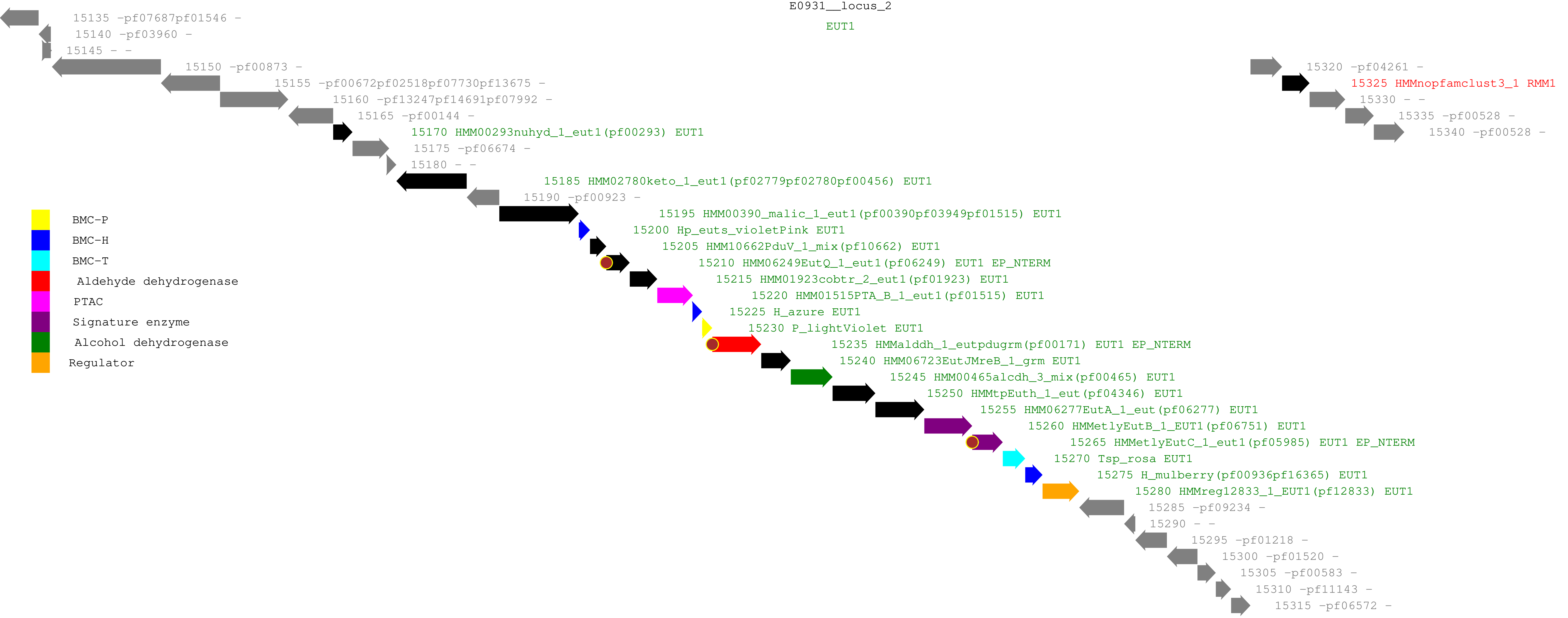

Salmonella enterica subsp. enterica serovar Mapo. Bacteria; Proteobacteria; Gammaproteobacteria; Enterobacterales;

E0940__locus_2

EUT1

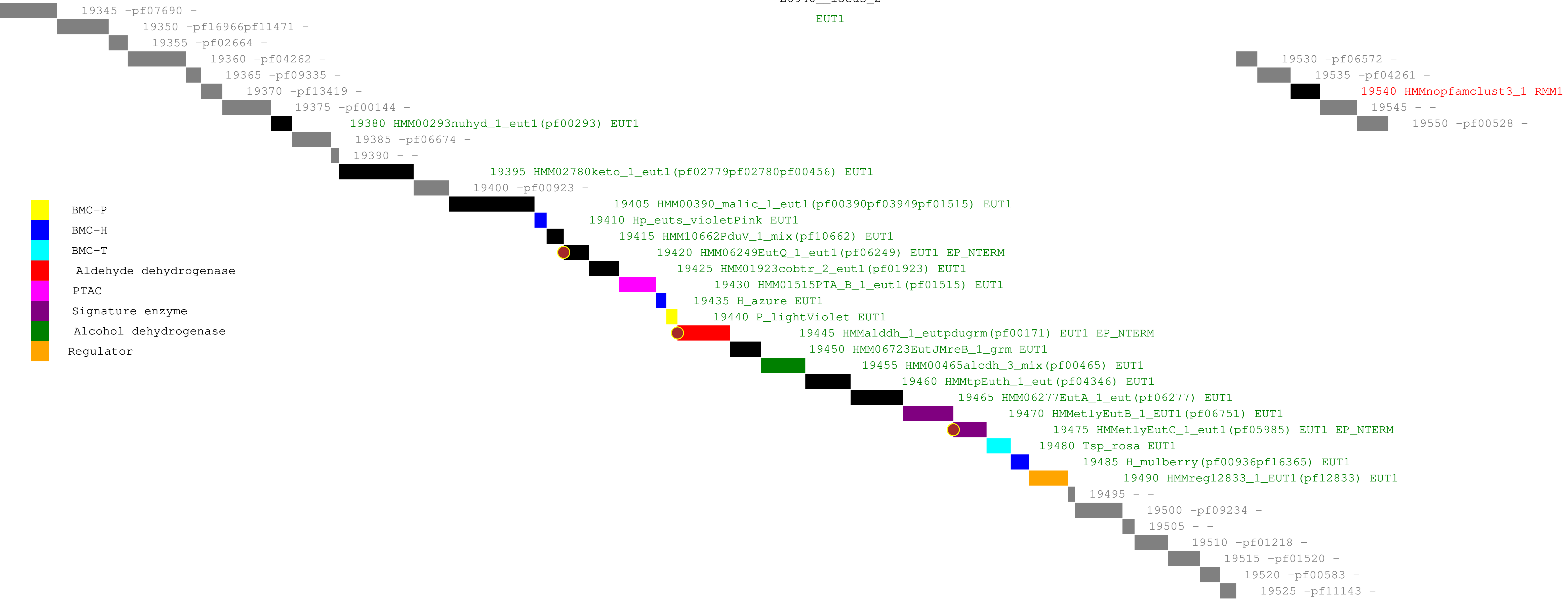

Salmonella enterica subsp. enterica serovar Manhattan. Bacteria; Proteobacteria; Gammaproteobacteria; Enterobacterales;

E0960__locus_2

EUT1

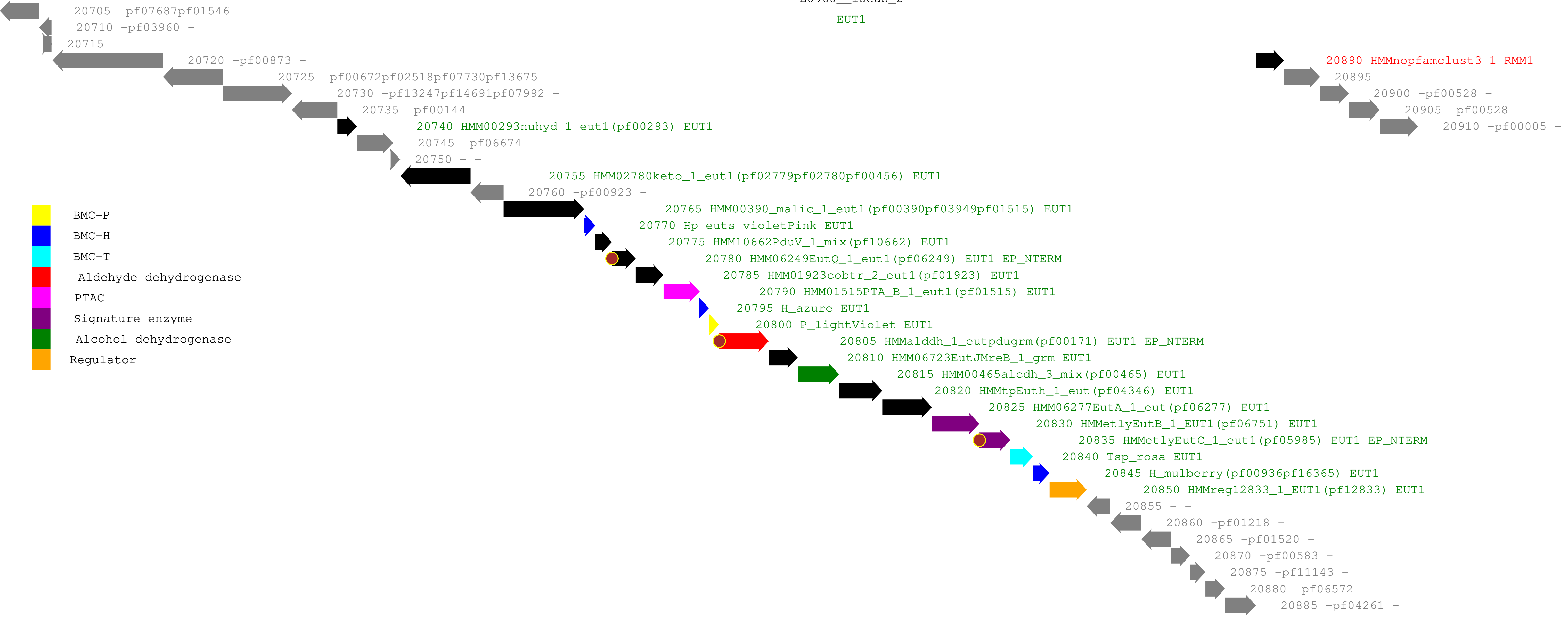

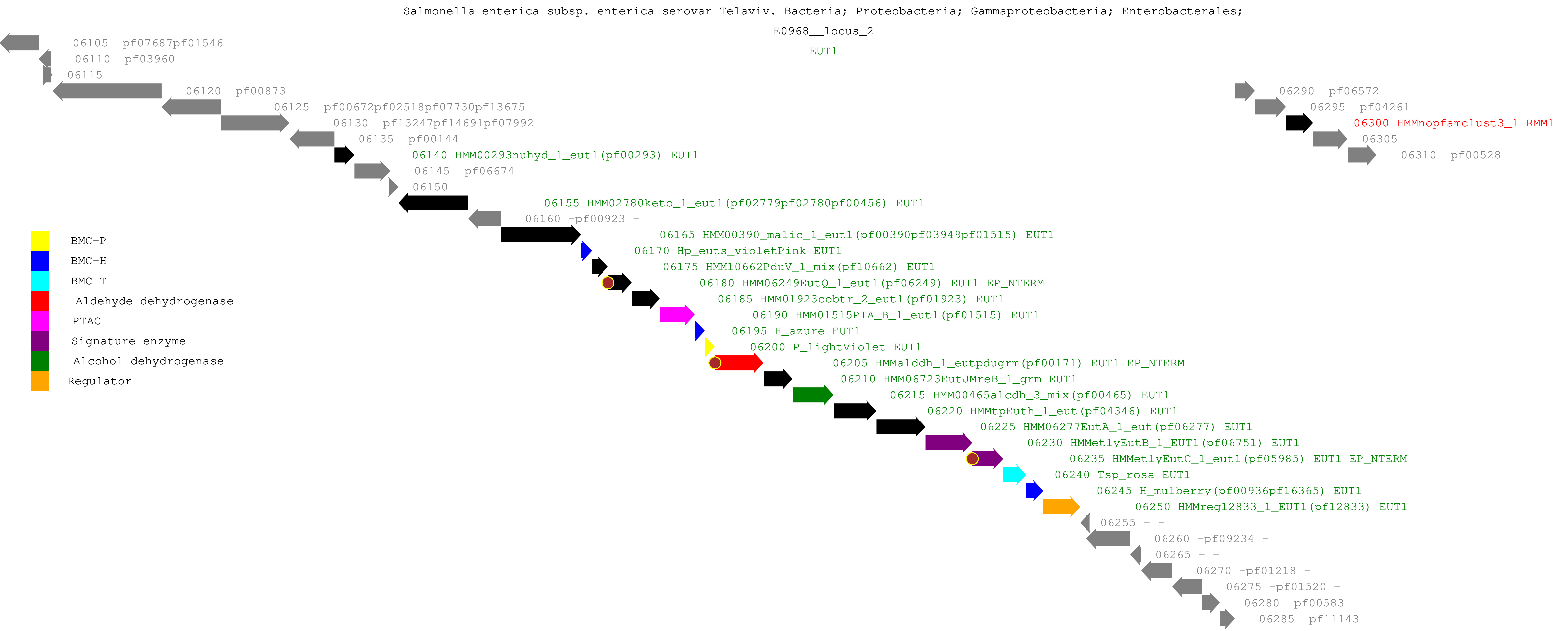
Salmonella enterica subsp. enterica serovar Telaviv. Bacteria; Proteobacteria; Gammaproteobacteria; Enterobacterales;
E0968__locus_2
EUT1
06105 -pf07687pf01546 -
06110 -pf03960 -
06115 - -
06120 -pf00873 -
06125 -pf00672pf02518pf07730pf13675 -
06130 -pf13247pf14691pf07992 -
06135 -pf00144 -
06140 HMM00293nuhyd_1_eut1(pf00293) EUT1
06145 -pf06674 -
06150 - -
06155 HMM02780keto_1_eut1(pf02779pf02780pf00456) EUT1
06160 -pf00923 -
06165 HMM00390_malic_1_eut1(pf00390pf03949pf01515) EUT1
06170 Hp_euts_violetPink EUT1
06175 HMM10662PduV_1_mix(pf10662) EUT1
06180 HMM06249EutQ_1_eut1(pf06249) EUT1 EP_NTERM
06185 HMM01923cobtr_2_eut1(pf01923) EUT1
06190 HMM01515PTA_B_1_eut1(pf01515) EUT1
06195 H_azure EUT1
06200 P_lightViolet EUT1
06205 HMMalddh_1_eutpdugrm(pf00171) EUT1 EP_NTERM
06210 HMM06723EutJMreB_1_grm EUT1
06215 HMM00465alcdh_3_mix(pf00465) EUT1
06220 HMMtpEuth_1_eut(pf04346) EUT1
06225 HMM06277EutA_1_eut(pf06277) EUT1
06230 HMMetlyEutB_1_EUT1(pf06751) EUT1
06235 HMMetlyEutC_1_eut1(pf05985) EUT1 EP_NTERM
06240 Tsp_rosa EUT1
06245 H_mulberry(pf00936pf16365) EUT1
06250 HMMreg12833_1_EUT1(pf12833) EUT1
06255 - -
06260 -pf09234 -
06265 - -
06270 -pf01218 -
06275 -pf01520 -
06280 -pf00583 -
06285 -pf11143 -
06290 -pf06572 -
06295 -pf04261 -
06300 HMMnopfamclust3_1 RMM1
06305 - -
06310 -pf00528 -
BMC-P
BMC-H
BMC-T
Aldehyde dehydrogenase
PTAC
Signature enzyme
Alcohol dehydrogenase
Regulator

Salmonella enterica subsp. enterica serovar Waycross. Bacteria; Proteobacteria; Gammaproteobacteria; Enterobacterales;

E0980__locus_2

EUT1

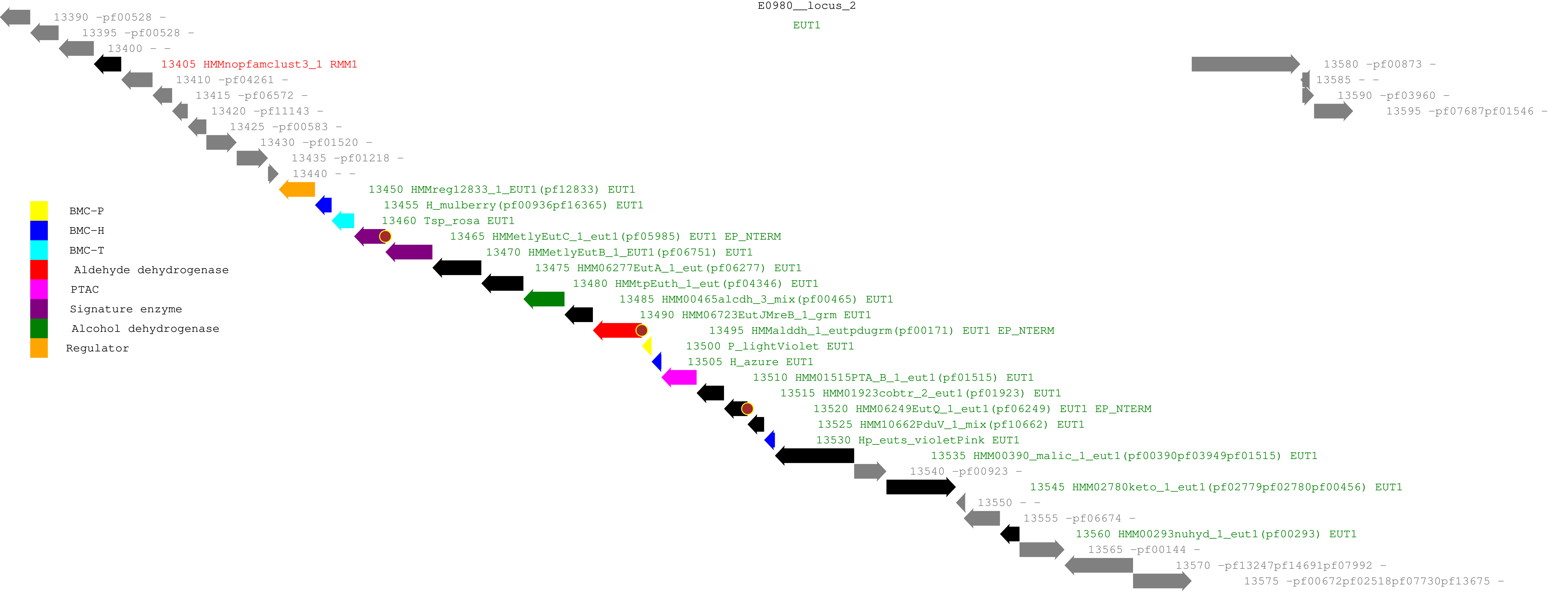

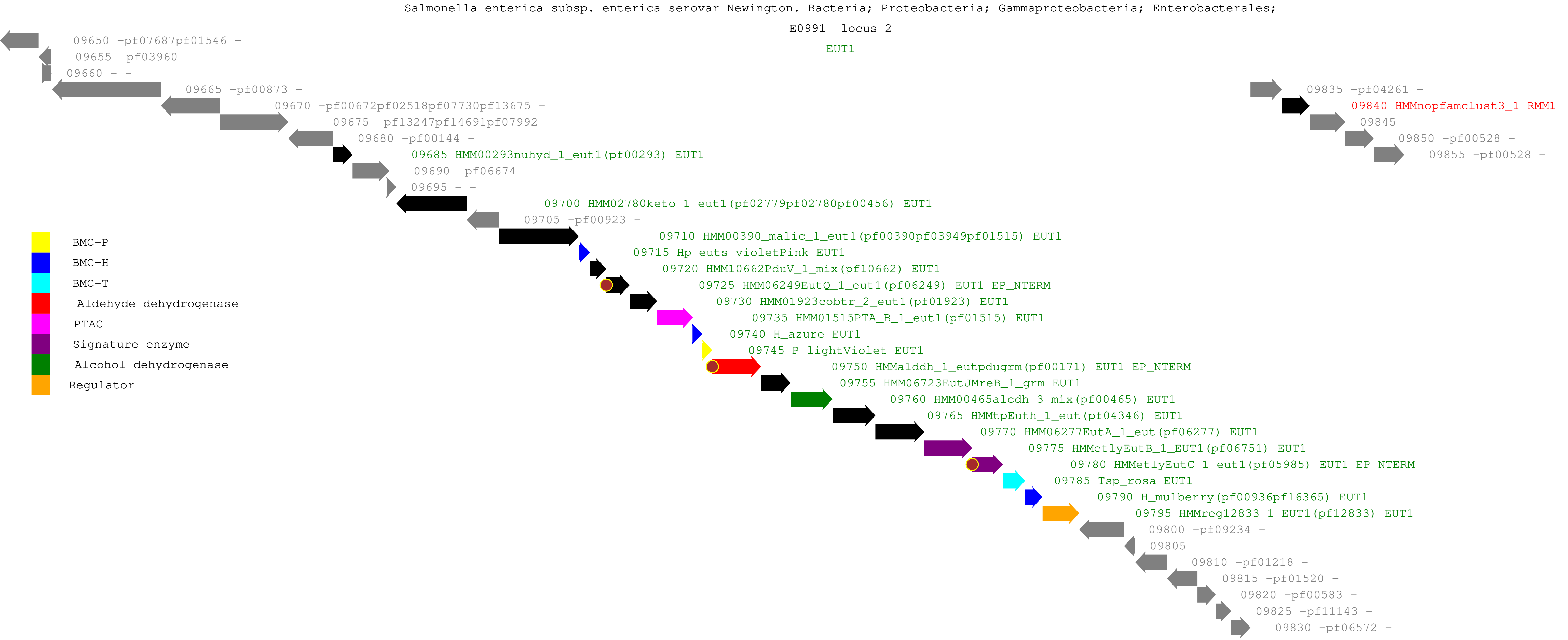

Salmonella enterica subsp. enterica serovar Newington. Bacteria; Proteobacteria; Gammaproteobacteria; Enterobacterales;
E0991__locus_2
EUT1
09650 -pf07687pf01546 -
09655 -pf03960 -
09660 - -
09665 -pf00873 -
09670 -pf00672pf02518pf07730pf13675 -
09675 -pf13247pf14691pf07992 -
09680 -pf00144 -
09685 HMM00293nuhyd_1_eut1(pf00293) EUT1
09690 -pf06674 -
09695 - -
09700 HMM02780keto_1_eut1(pf02779pf02780pf00456) EUT1
09705 -pf00923 -
09710 HMM00390_malic_1_eut1(pf00390pf03949pf01515) EUT1
09715 Hp_euts_violetPink EUT1
09720 HMM10662PduV_1_mix(pf10662) EUT1
09725 HMM06249EutQ_1_eut1(pf06249) EUT1 EP_NTERM
09730 HMM01923cobtr_2_eut1(pf01923) EUT1
09735 HMM01515PTA_B_1_eut1(pf01515) EUT1
09740 H_azure EUT1
09745 P_lightViolet EUT1
09750 HMMalddh_1_eutpdugrm(pf00171) EUT1 EP_NTERM
09755 HMM06723EutJMreB_1_grm EUT1
09760 HMM00465alcdh_3_mix(pf00465) EUT1
09765 HMMtpEuth_1_eut(pf04346) EUT1
09770 HMM06277EutA_1_eut(pf06277) EUT1
09775 HMMetlyEutB_1_EUT1(pf06751) EUT1
09780 HMMetlyEutC_1_eut1(pf05985) EUT1 EP_NTERM
09785 Tsp_rosa EUT1
09790 H_mulberry(pf00936pf16365) EUT1
09795 HMMreg12833_1_EUT1(pf12833) EUT1
09800 -pf09234 -
09805 - -
09810 -pf01218 -
09815 -pf01520 -
09820 -pf00583 -
09825 -pf11143 -
09830 -pf06572 -
09835 -pf04261 -
09840 HMMnopfamclust3_1 RMM1
09845 - -
09850 -pf00528 -
09855 -pf00528 -
BMC-P
BMC-H
BMC-T
Aldehyde dehydrogenase
PTAC
Signature enzyme
Alcohol dehydrogenase
Regulator

Salmonella enterica subsp. enterica serovar Urbana. Bacteria; Proteobacteria; Gammaproteobacteria; Enterobacterales;

E0999__locus_1

EUT1

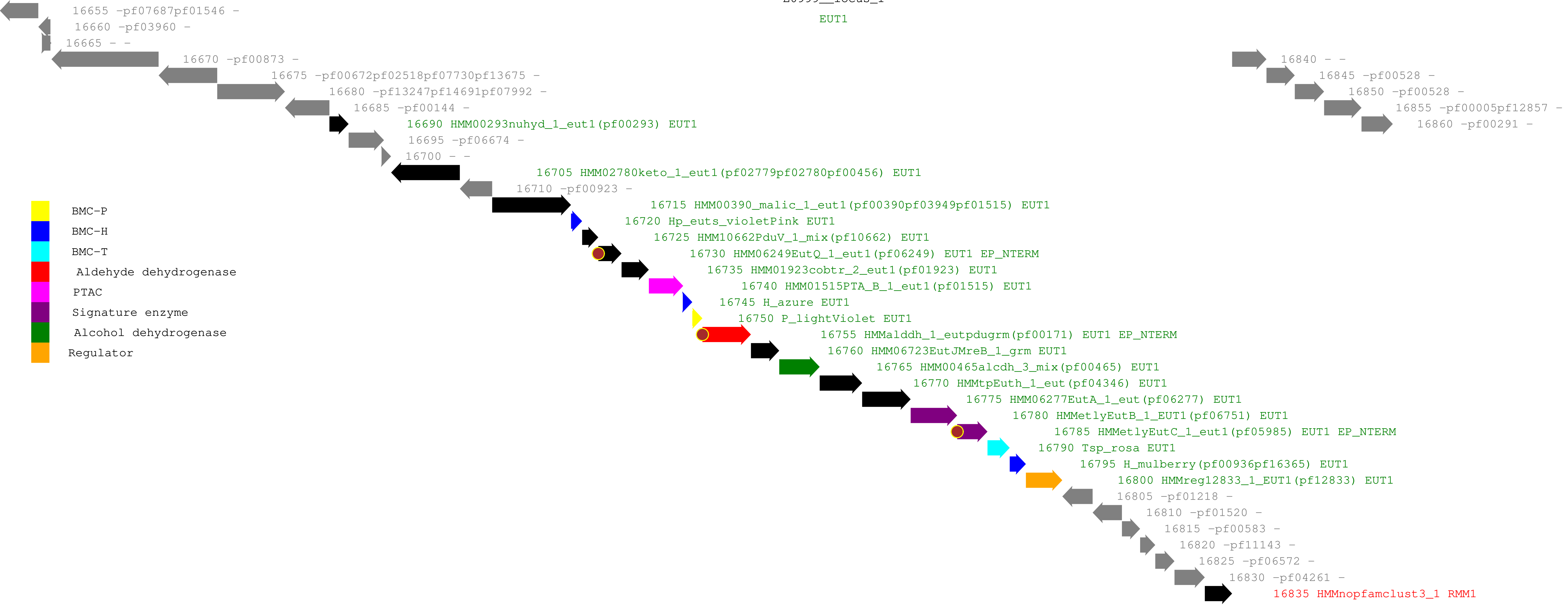

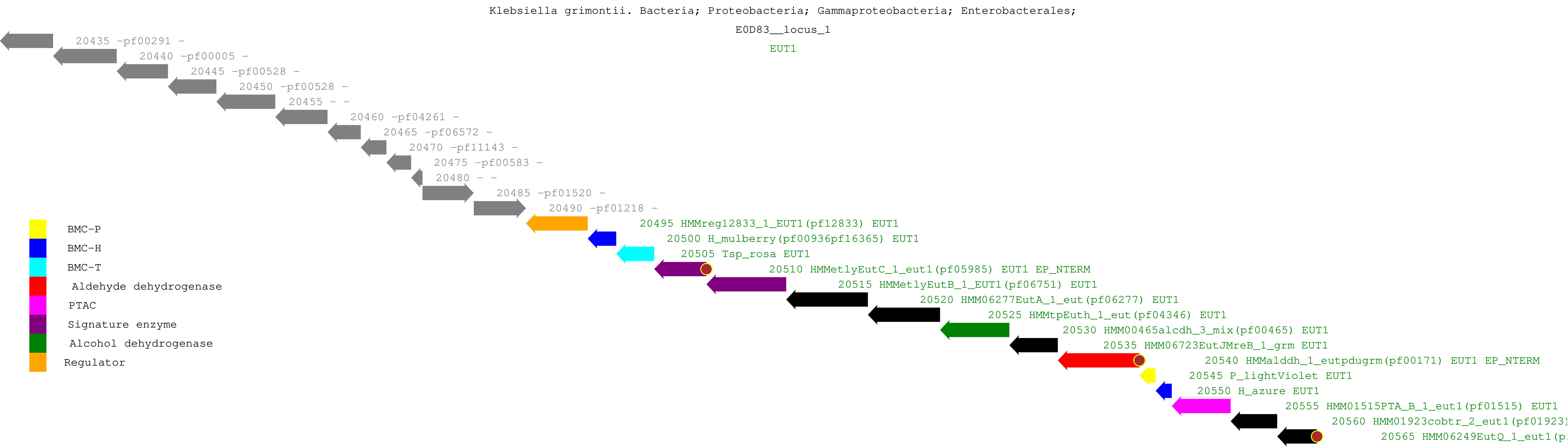
Klebsiella grimontii. Bacteria; Proteobacteria; Gammaproteobacteria; Enterobacterales;
E0D83__locus_1
EUT1
20435 -pf00291 -
20440 -pf00005 -
20445 -pf00528 -
20450 -pf00528 -
20455 - -
20460 -pf04261 -
20465 -pf06572 -
20470 -pf11143 -
20475 -pf00583 -
20480 - -
20485 -pf01520 -
20490 -pf01218 -
20495 HMMreg12833_1_EUT1(pf12833) EUT1
20500 H_mulberry(pf00936pf16365) EUT1
20505 Tsp_rosa EUT1
20510 HMMetlyEutC_1_eut1(pf05985) EUT1 EP_NTERM
20515 HMMetlyEutB_1_EUT1(pf06751) EUT1
20520 HMM06277EutA_1_eut(pf06277) EUT1
20525 HMMtpEuth_1_eut(pf04346) EUT1
20530 HMM00465alcdh_3_mix(pf00465) EUT1
20535 HMM06723EutJMreB_1_grm EUT1
20540 HMMalddh_1_eutpdugrm(pf00171) EUT1 EP_NTERM
20545 P_lightViolet EUT1
20550 H_azure EUT1
20555 HMM01515PTA_B_1_eut1(pf01515) EUT1
20560 HMM01923cobtr_2_eut1(pf01923)
20565 HMM06249EutQ_1_eut1(p
BMC-P
BMC-H
BMC-T
Aldehyde dehydrogenase
PTAC
Signature enzyme
Alcohol dehydrogenase
Regulator

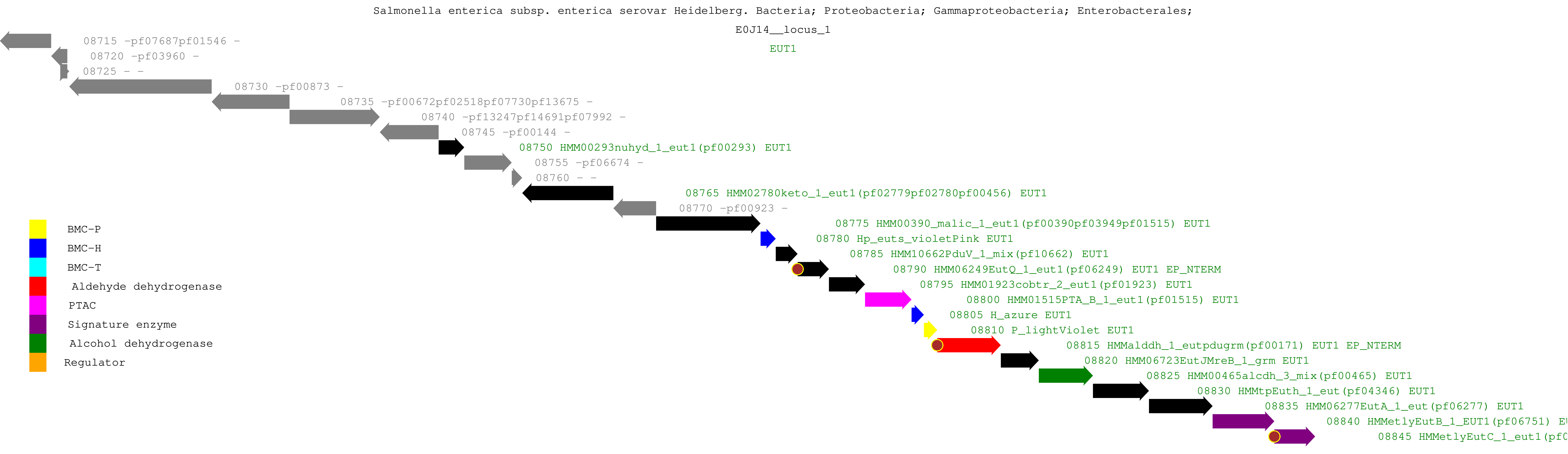
Salmonella enterica subsp. enterica serovar Heidelberg. Bacteria; Proteobacteria; Gammaproteobacteria; Enterobacterales;
E0J14__locus_1
EUT1
08715 -pf07687pf01546 -
08720 -pf03960 -
08725 - -
08730 -pf00873 -
08735 -pf00672pf02518pf07730pf13675 -
08740 -pf13247pf14691pf07992 -
08745 -pf00144 -
08750 HMM00293nuhyd_1_eut1(pf00293) EUT1
08755 -pf06674 -
08760 - -
08765 HMM02780keto_1_eut1(pf02779pf02780pf00456) EUT1
08770 -pf00923 -
08775 HMM00390_malic_1_eut1(pf00390pf03949pf01515) EUT1
08780 Hp_euts_violetPink EUT1
08785 HMM10662PduV_1_mix(pf10662) EUT1
08790 HMM06249EutQ_1_eut1(pf06249) EUT1 EP_NTERM
08795 HMM01923cobtr_2_eut1(pf01923) EUT1
08800 HMM01515PTA_B_1_eut1(pf01515) EUT1
08805 H_azure EUT1
08810 P_lightViolet EUT1
08815 HMMalddh_1_eutpdugrm(pf00171) EUT1 EP_NTERM
08820 HMM06723EutJMreB_1_grm EUT1
08825 HMM00465alcdh_3_mix(pf00465) EUT1
08830 HMMtpEuth_1_eut(pf04346) EUT1
08835 HMM06277EutA_1_eut(pf06277) EUT1
08840 HMMetlyEutB_1_EUT1(pf06751)
08845 HMMetlyEutC_1_eut1
BMC-P
BMC-H
BMC-T
Aldehyde dehydrogenase
PTAC
Signature enzyme
Alcohol dehydrogenase
Regulator

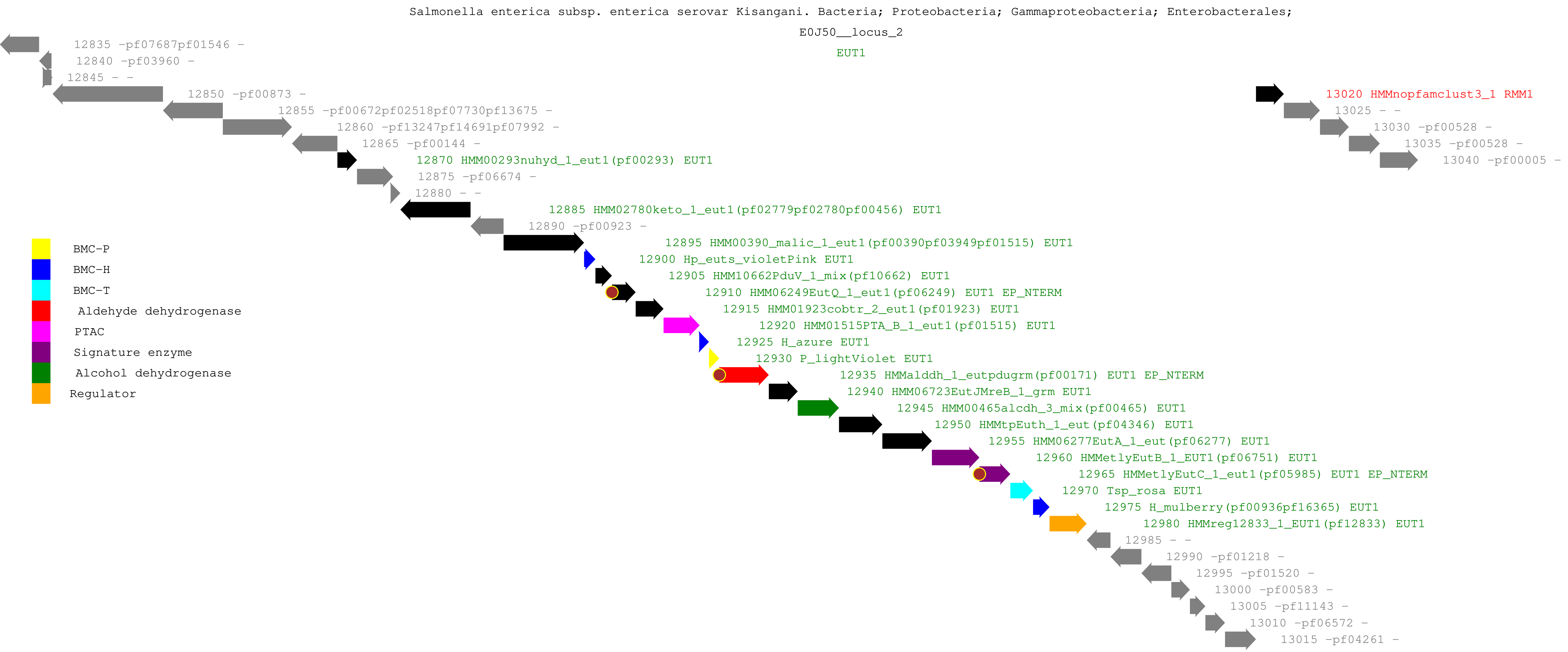
Salmonella enterica subsp. enterica serovar Kisangani. Bacteria; Proteobacteria; Gammaproteobacteria; Enterobacterales;
E0J50__locus_2
EUT1
12835 -pf07687pf01546 -
12840 -pf03960 -
12845 - -
12850 -pf00873 -
12855 -pf00672pf02518pf07730pf13675 -
12860 -pf13247pf14691pf07992 -
12865 -pf00144 -
12870 HMM00293nuhyd_1_eut1(pf00293) EUT1
12875 -pf06674 -
12880 - -
12885 HMM02780keto_1_eut1(pf02779pf02780pf00456) EUT1
12890 -pf00923 -
12895 HMM00390_malic_1_eut1(pf00390pf03949pf01515) EUT1
12900 Hp_euts_violetPink EUT1
12905 HMM10662PduV_1_mix(pf10662) EUT1
12910 HMM06249EutQ_1_eut1(pf06249) EUT1 EP_NTERM
12915 HMM01923cobtr_2_eut1(pf01923) EUT1
12920 HMM01515PTA_B_1_eut1(pf01515) EUT1
12925 H_azure EUT1
12930 P_lightViolet EUT1
12935 HMMalddh_1_eutpdugrm(pf00171) EUT1 EP_NTERM
12940 HMM06723EutJMreB_1_grm EUT1
12945 HMM00465alcdh_3_mix(pf00465) EUT1
12950 HMMtpEuth_1_eut(pf04346) EUT1
12955 HMM06277EutA_1_eut(pf06277) EUT1
12960 HMMetlyEutB_1_EUT1(pf06751) EUT1
12965 HMMetlyEutC_1_eut1(pf05985) EUT1 EP_NTERM
12970 Tsp_rosa EUT1
12975 H_mulberry(pf00936pf16365) EUT1
12980 HMMreg12833_1_EUT1(pf12833) EUT1
12985 - -
12990 -pf01218 -
12995 -pf01520 -
13000 -pf00583 -
13005 -pf11143 -
13010 -pf06572 -
13015 -pf04261 -
13020 HMMnopfamclust3_1 RMM1
13025 - -
13030 -pf00528 -
13035 -pf00528 -
13040 -pf00005 -
BMC-P
BMC-H
BMC-T
Aldehyde dehydrogenase
PTAC
Signature enzyme
Alcohol dehydrogenase
Regulator

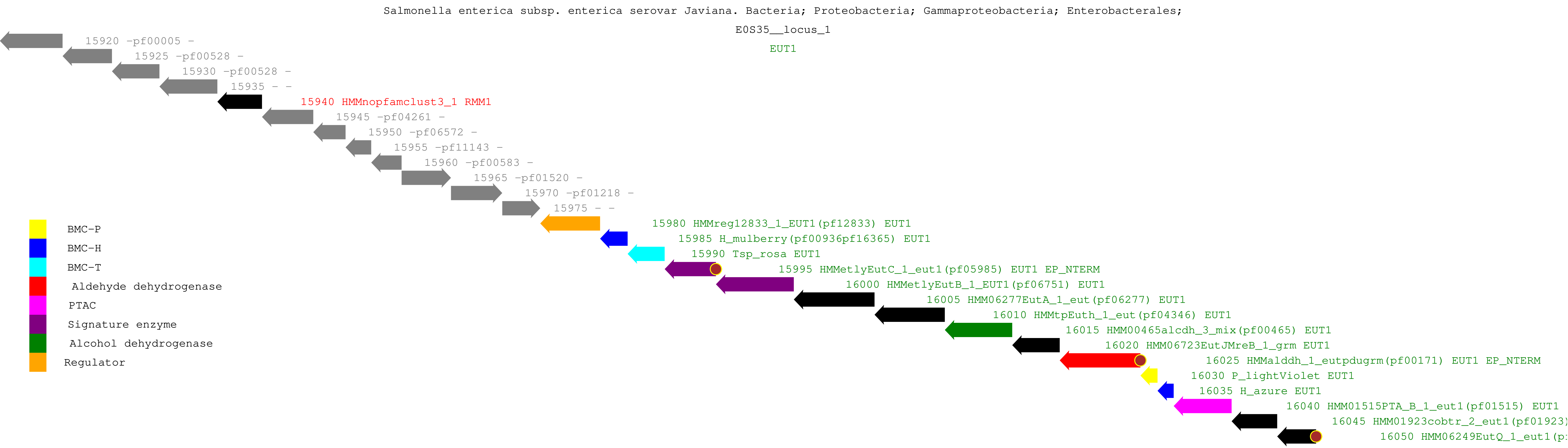
Salmonella enterica subsp. enterica serovar Javiana. Bacteria; Proteobacteria; Gammaproteobacteria; Enterobacterales;
E0S35__locus_1
EUT1
15920 -pf00005 -
15925 -pf00528 -
15930 -pf00528 -
15935 - -
15940 HMMnopfamclust3_1 RMM1
15945 -pf04261 -
15950 -pf06572 -
15955 -pf11143 -
15960 -pf00583 -
15965 -pf01520 -
15970 -pf01218 -
15975 - -
15980 HMMreg12833_1_EUT1(pf12833) EUT1
15985 H_mulberry(pf00936pf16365) EUT1
15990 Tsp_rosa EUT1
15995 HMMetlyEutC_1_eut1(pf05985) EUT1 EP_NTERM
16000 HMMetlyEutB_1_EUT1(pf06751) EUT1
16005 HMM06277EutA_1_eut(pf06277) EUT1
16010 HMMtpEuth_1_eut(pf04346) EUT1
16015 HMM00465alcdh_3_mix(pf00465) EUT1
16020 HMM06723EutJMreB_1_grm EUT1
16025 HMMaldh_1_eutpdugrm(pf00171) EUT1 EP_NTERM
16030 P_lightViolet EUT1
16035 H_azure EUT1
16040 HMM01515PTA_B_1_eut1(pf01515) EUT1
16045 HMM01923cobtr_2_eut1(pf01923)
16050 HMM06249EutQ_1_eut1(p
BMC-P
BMC-H
BMC-T
Aldehyde dehydrogenase
PTAC
Signature enzyme
Alcohol dehydrogenase
Regulator

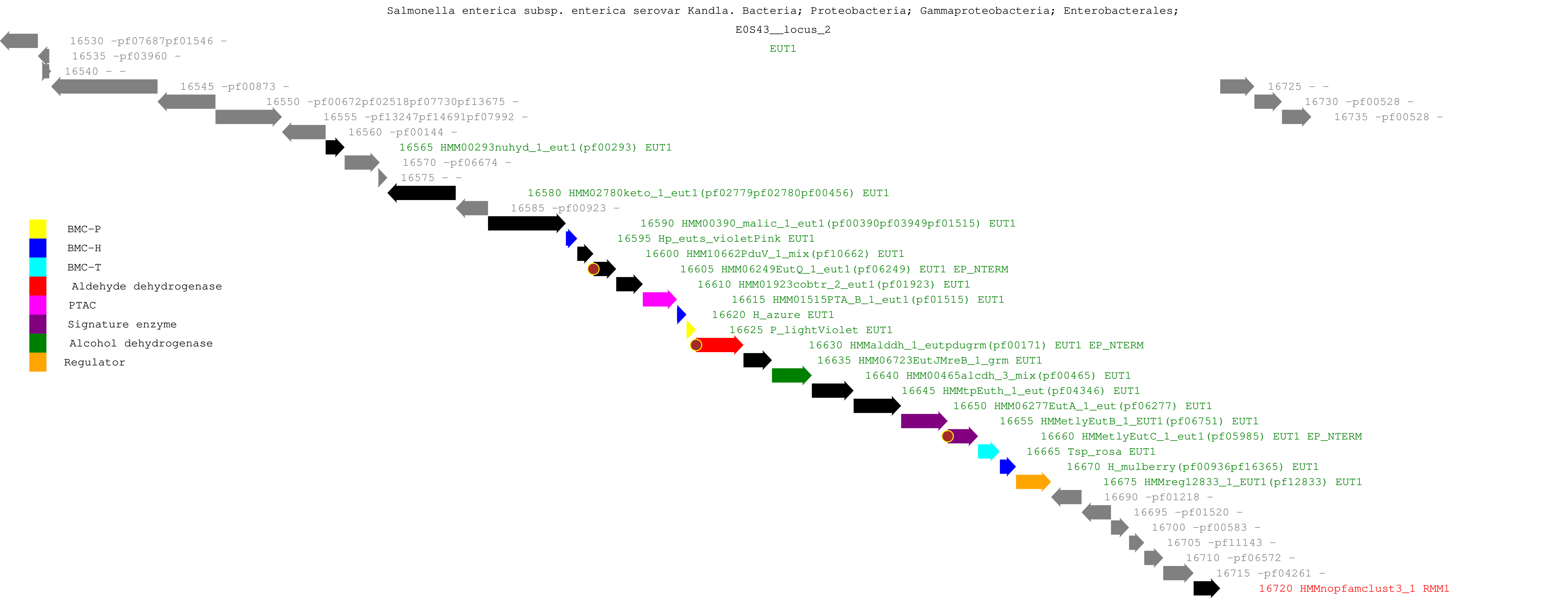

Salmonella enterica subsp. enterica serovar Kandla. Bacteria; Proteobacteria; Gammaproteobacteria; Enterobacterales;
E0S43__locus_2
EUT1
16530 -pf07687pf01546 -
16535 -pf03960 -
16540 - -
16545 -pf00873 -
16550 -pf00672pf02518pf07730pf13675 -
16555 -pf13247pf14691pf07992 -
16560 -pf00144 -
16565 HMM00293nuhyd_1_eut1(pf00293) EUT1
16570 -pf06674 -
16575 - -
16580 HMM02780keto_1_eut1(pf02779pf02780pf00456) EUT1
16585 -pf00923 -
16590 HMM00390_malic_1_eut1(pf00390pf03949pf01515) EUT1
16595 Hp_euts_violetPink EUT1
16600 HMM10662PduV_1_mix(pf10662) EUT1
16605 HMM06249EutQ_1_eut1(pf06249) EUT1 EP_NTERM
16610 HMM01923cobtr_2_eut1(pf01923) EUT1
16615 HMM01515PTA_B_1_eut1(pf01515) EUT1
16620 H_azure EUT1
16625 P_lightViolet EUT1
16630 HMMalddh_1_eutpdugrm(pf00171) EUT1 EP_NTERM
16635 HMM06723EutJMreB_1_grm EUT1
16640 HMM00465alcdh_3_mix(pf00465) EUT1
16645 HMMtpEuth_1_eut(pf04346) EUT1
16650 HMM06277EutA_1_eut(pf06277) EUT1
16655 HMMetlyEutB_1_EUT1(pf06751) EUT1
16660 HMMetlyEutC_1_eut1(pf05985) EUT1 EP_NTERM
16665 Tsp_rosa EUT1
16670 H_mulberry(pf00936pf16365) EUT1
16675 HMMreg12833_1_EUT1(pf12833) EUT1
16690 -pf01218 -
16695 -pf01520 -
16700 -pf00583 -
16705 -pf11143 -
16710 -pf06572 -
16715 -pf04261 -
16720 HMMnopfamclust3_1 RMM1
16725 - -
16730 -pf00528 -
16735 -pf00528 -
BMC-P
BMC-H
BMC-T
Aldehyde dehydrogenase
PTAC
Signature enzyme
Alcohol dehydrogenase
Regulator

Salmonella enterica subsp. enterica serovar Matopeni. Bacteria; Proteobacteria; Gammaproteobacteria; Enterobacterales;

E0S53__locus_1

EUT1

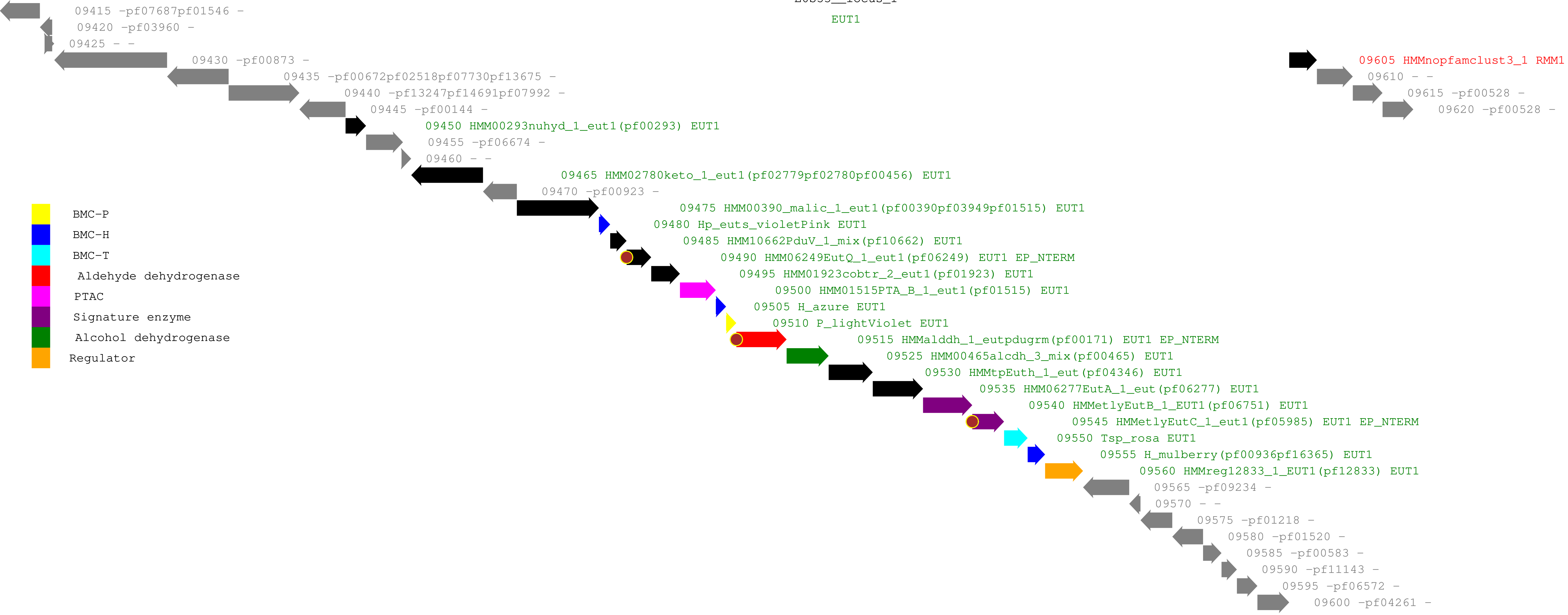

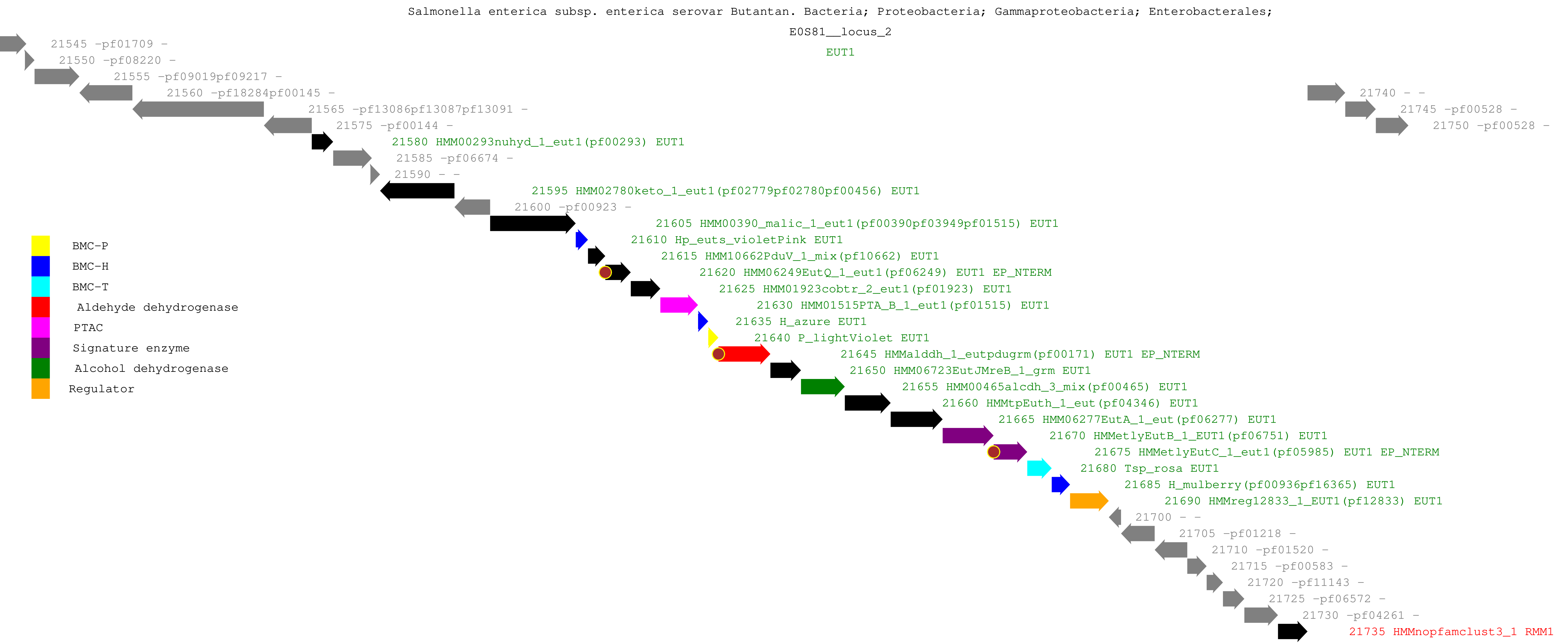
Salmonella enterica subsp. enterica serovar Butantan. Bacteria; Proteobacteria; Gammaproteobacteria; Enterobacterales;
E0S81__locus_2
EUT1
21545 -pf01709 -
21550 -pf08220 -
21555 -pf09019pf09217 -
21560 -pf18284pf00145 -
21565 -pf13086pf13087pf13091 -
21575 -pf00144 -
21580 HMM00293nuhyd_1_eut1(pf00293) EUT1
21585 -pf06674 -
21590 - -
21595 HMM02780keto_1_eut1(pf02779pf02780pf00456) EUT1
21600 -pf00923 -
21605 HMM00390_malic_1_eut1(pf00390pf03949pf01515) EUT1
21610 Hp_euts_violetPink EUT1
21615 HMM10662PduV_1_mix(pf10662) EUT1
21620 HMM06249EutQ_1_eut1(pf06249) EUT1 EP_NTERM
21625 HMM01923cobtr_2_eut1(pf01923) EUT1
21630 HMM01515PTA_B_1_eut1(pf01515) EUT1
21635 H_azure EUT1
21640 P_lightViolet EUT1
21645 HMMaldh_1_eutpdugrm(pf00171) EUT1 EP_NTERM
21650 HMM06723EutJMreB_1_grm EUT1
21655 HMM00465alcdh_3_mix(pf00465) EUT1
21660 HMMtpEuth_1_eut(pf04346) EUT1
21665 HMM06277EutA_1_eut(pf06277) EUT1
21670 HMMetlyEutB_1_EUT1(pf06751) EUT1
21675 HMMetlyEutC_1_eut1(pf05985) EUT1 EP_NTERM
21680 Tsp_rosa EUT1
21685 H_mulberry(pf00936pf16365) EUT1
21690 HMMreg12833_1_EUT1(pf12833) EUT1
21700 - -
21705 -pf01218 -
21710 -pf01520 -
21715 -pf00583 -
21720 -pf11143 -
21725 -pf06572 -
21730 -pf04261 -
21735 HMMnopfamclust3_1 RMM1
21740 - -
21745 -pf00528 -
21750 -pf00528 -
BMC-P
BMC-H
BMC-T
Aldehyde dehydrogenase
PTAC
Signature enzyme
Alcohol dehydrogenase
Regulator

Salmonella enterica subsp. enterica serovar Galiema. Bacteria; Proteobacteria; Gammaproteobacteria; Enterobacterales;

E0U28__locus_2

EUT1

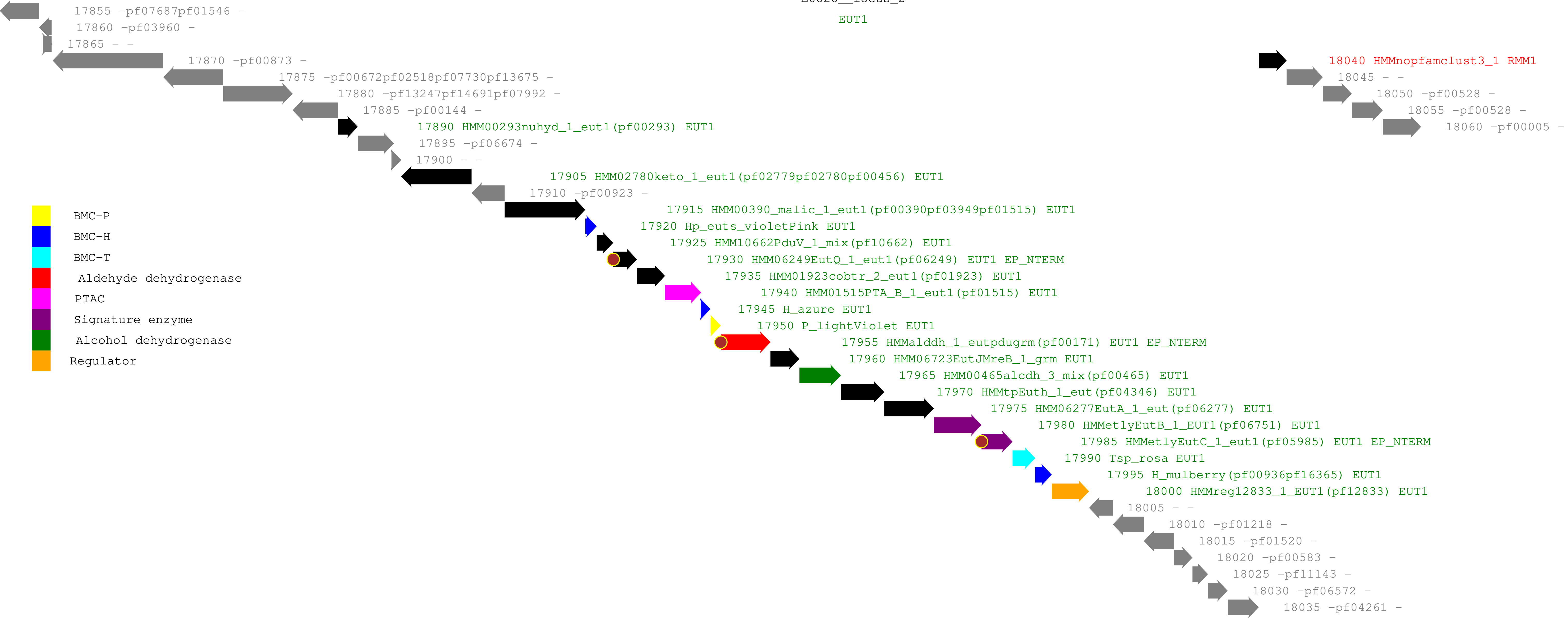

Salmonella enterica subsp. enterica serovar Cotham. Bacteria; Proteobacteria; Gammaproteobacteria; Enterobacterales;

E0U40__locus_2

EUT1

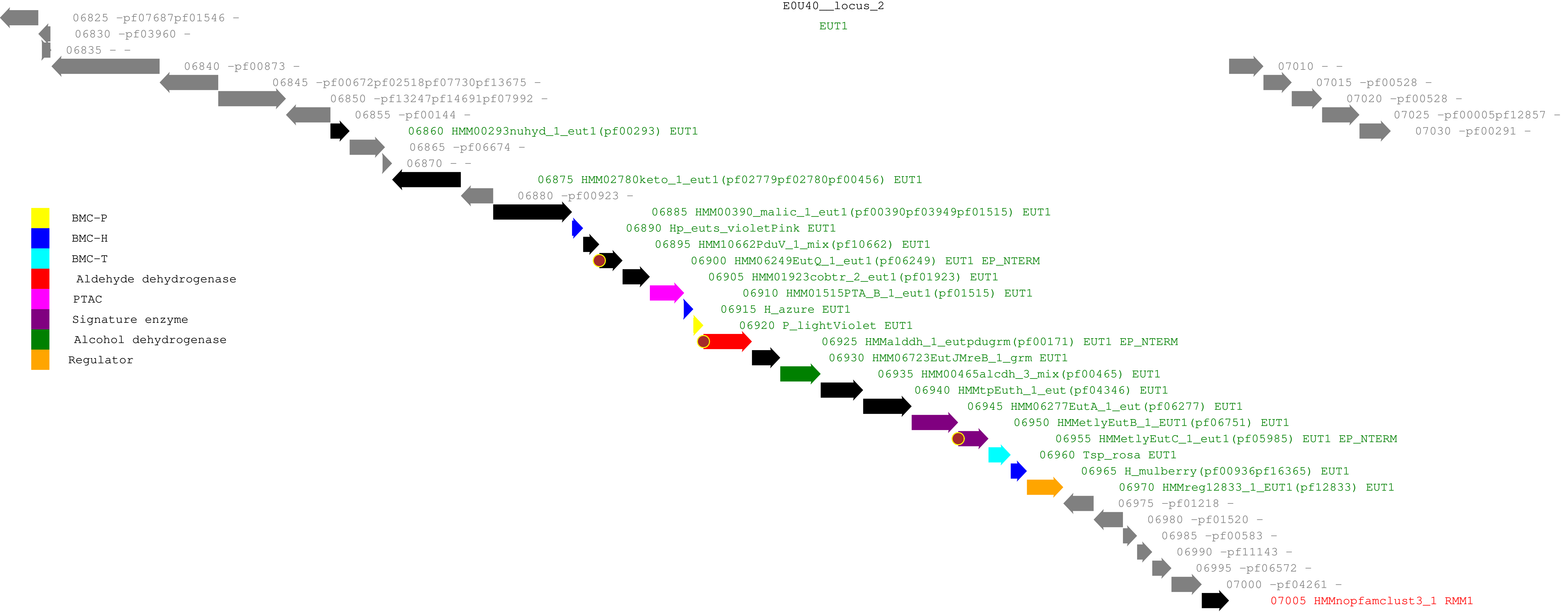

Salmonella enterica subsp. enterica serovar Durban. Bacteria; Proteobacteria; Gammaproteobacteria; Enterobacterales;

E0U47__locus_2

EUT1

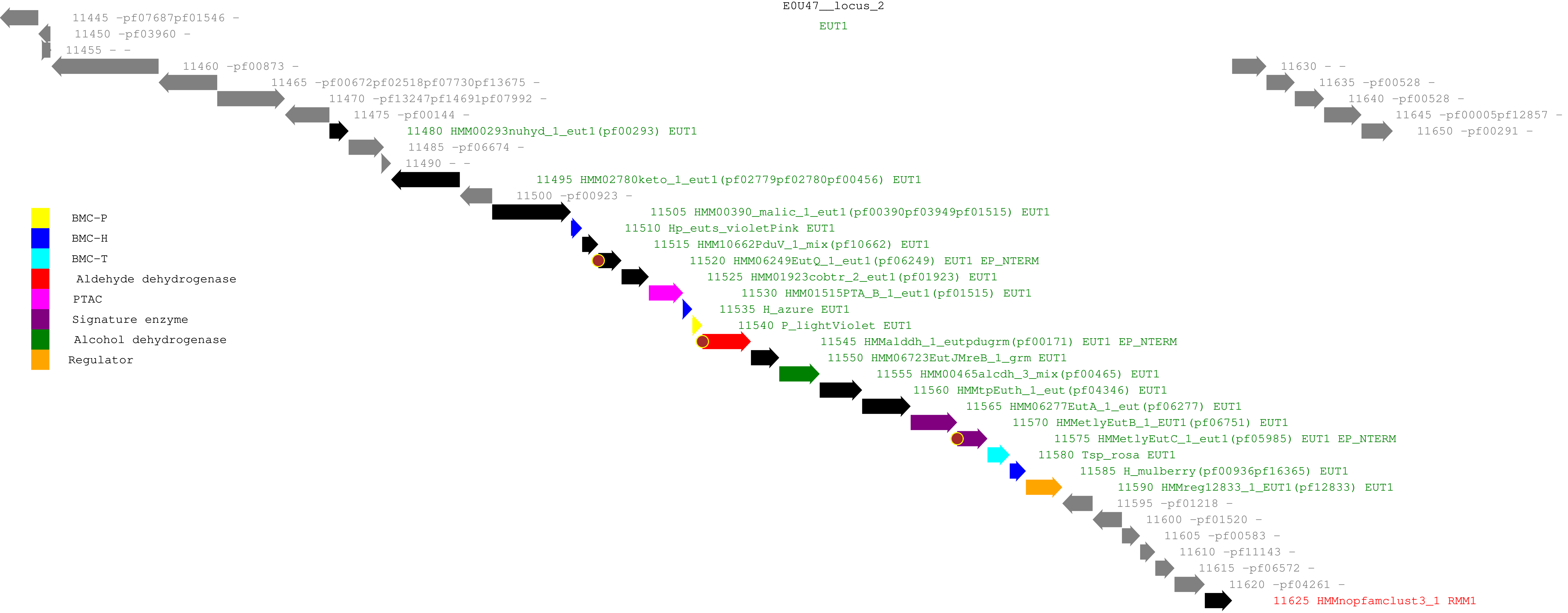

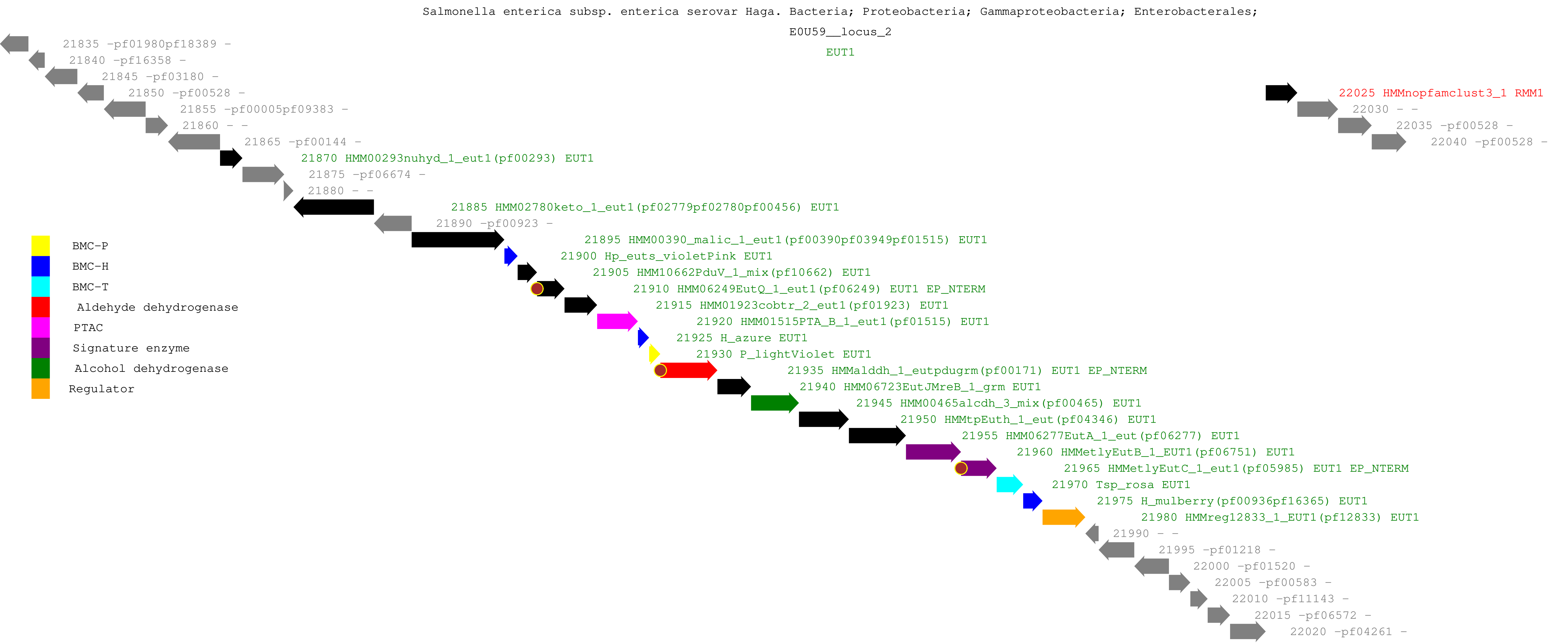
Salmonella enterica subsp. enterica serovar Haga. Bacteria; Proteobacteria; Gammaproteobacteria; Enterobacterales;
E0U59__locus_2
EUT1
21835 -pf01980pf18389 -
21840 -pf16358 -
21845 -pf03180 -
21850 -pf00528 -
21855 -pf00005pf09383 -
21860 - -
21865 -pf00144 -
21870 HMM00293nuhyd_1_eut1(pf00293) EUT1
21875 -pf06674 -
21880 - -
21885 HMM02780keto_1_eut1(pf02779pf02780pf00456) EUT1
21890 -pf00923 -
21895 HMM00390_malic_1_eut1(pf00390pf03949pf01515) EUT1
21900 Hp_euts_violetPink EUT1
21905 HMM10662PduV_1_mix(pf10662) EUT1
21910 HMM06249EutQ_1_eut1(pf06249) EUT1 EP_NTERM
21915 HMM01923cobtr_2_eut1(pf01923) EUT1
21920 HMM01515PTA_B_1_eut1(pf01515) EUT1
21925 H_azure EUT1
21930 P_lightViolet EUT1
21935 HMMaldh_1_eutpdugrm(pf00171) EUT1 EP_NTERM
21940 HMM06723EutJMreB_1_grm EUT1
21945 HMM00465alcdh_3_mix(pf00465) EUT1
21950 HMMtpEuth_1_eut(pf04346) EUT1
21955 HMM06277EutA_1_eut(pf06277) EUT1
21960 HMMetlyEutB_1_EUT1(pf06751) EUT1
21965 HMMetlyEutC_1_eut1(pf05985) EUT1 EP_NTERM
21970 Tsp_rosa EUT1
21975 H_mulberry(pf00936pf16365) EUT1
21980 HMMreg12833_1_EUT1(pf12833) EUT1
21990 - -
21995 -pf01218 -
22000 -pf01520 -
22005 -pf00583 -
22010 -pf11143 -
22015 -pf06572 -
22020 -pf04261 -
22025 HMMnopfamclust3_1 RMM1
22030 - -
22035 -pf00528 -
22040 -pf00528 -
BMC-P
BMC-H
BMC-T
Aldehyde dehydrogenase
PTAC
Signature enzyme
Alcohol dehydrogenase
Regulator

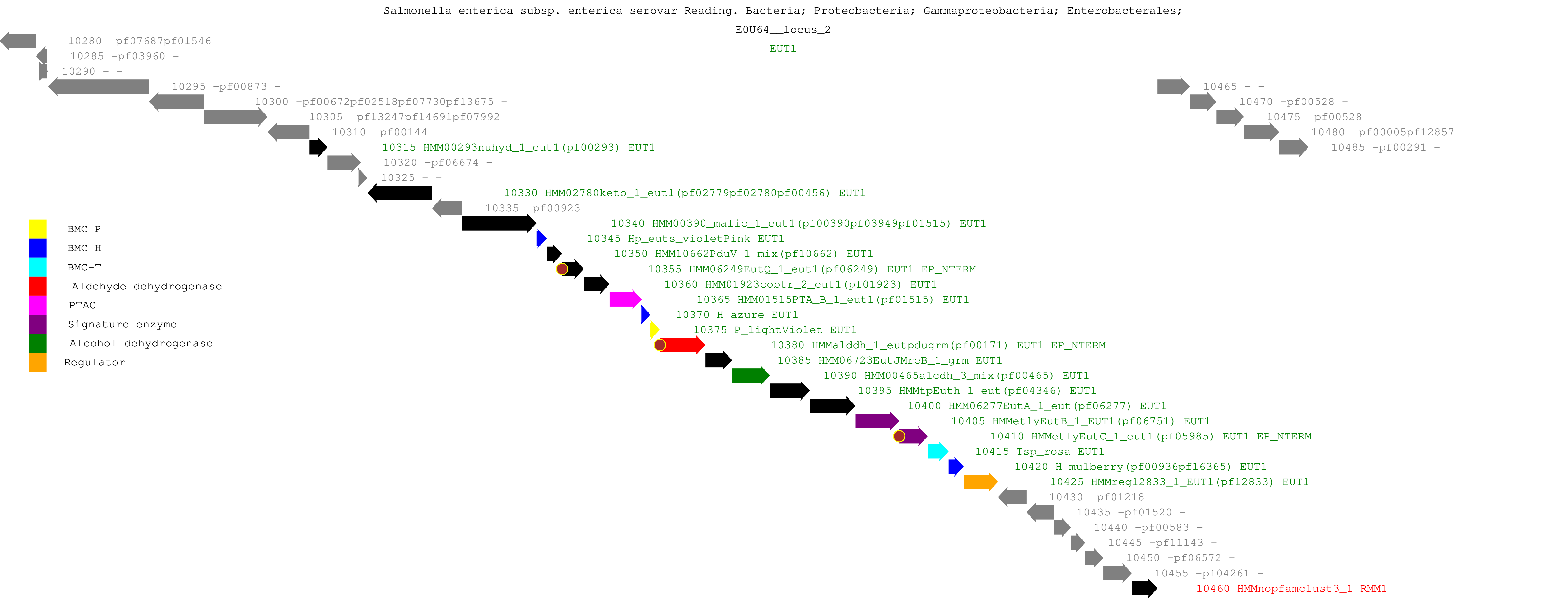
Salmonella enterica subsp. enterica serovar Reading. Bacteria; Proteobacteria; Gammaproteobacteria; Enterobacterales;
E0U64__locus_2
EUT1
10280 -pf07687pf01546 -
10285 -pf03960 -
10290 - -
10295 -pf00873 -
10300 -pf00672pf02518pf07730pf13675 -
10305 -pf13247pf14691pf07992 -
10310 -pf00144 -
10315 HMM00293nuhyd_1_eut1(pf00293) EUT1
10320 -pf06674 -
10325 - -
10330 HMM02780keto_1_eut1(pf02779pf02780pf00456) EUT1
10335 -pf00923 -
10340 HMM00390_malic_1_eut1(pf00390pf03949pf01515) EUT1
10345 Hp_euts_violetPink EUT1
10350 HMM10662PduV_1_mix(pf10662) EUT1
10355 HMM06249EutQ_1_eut1(pf06249) EUT1 EP_NTERM
10360 HMM01923cobtr_2_eut1(pf01923) EUT1
10365 HMM01515PTA_B_1_eut1(pf01515) EUT1
10370 H_azure EUT1
10375 P_lightViolet EUT1
10380 HMMalddh_1_eutpdugrm(pf00171) EUT1 EP_NTERM
10385 HMM06723EutJMreB_1_grm EUT1
10390 HMM00465alcdh_3_mix(pf00465) EUT1
10395 HMMtpEuth_1_eut(pf04346) EUT1
10400 HMM06277EutA_1_eut(pf06277) EUT1
10405 HMMetlyEutB_1_EUT1(pf06751) EUT1
10410 HMMetlyEutC_1_eut1(pf05985) EUT1 EP_NTERM
10415 Tsp_rosa EUT1
10420 H_mulberry(pf00936pf16365) EUT1
10425 HMMreg12833_1_EUT1(pf12833) EUT1
10430 -pf01218 -
10435 -pf01520 -
10440 -pf00583 -
10445 -pf11143 -
10450 -pf06572 -
10455 -pf04261 -
10460 HMMnopfamclust3_1 RMM1
10465 - -
10470 -pf00528 -
10475 -pf00528 -
10480 -pf00005pf12857 -
10485 -pf00291 -
BMC-P
BMC-H
BMC-T
Aldehyde dehydrogenase
PTAC
Signature enzyme
Alcohol dehydrogenase
Regulator

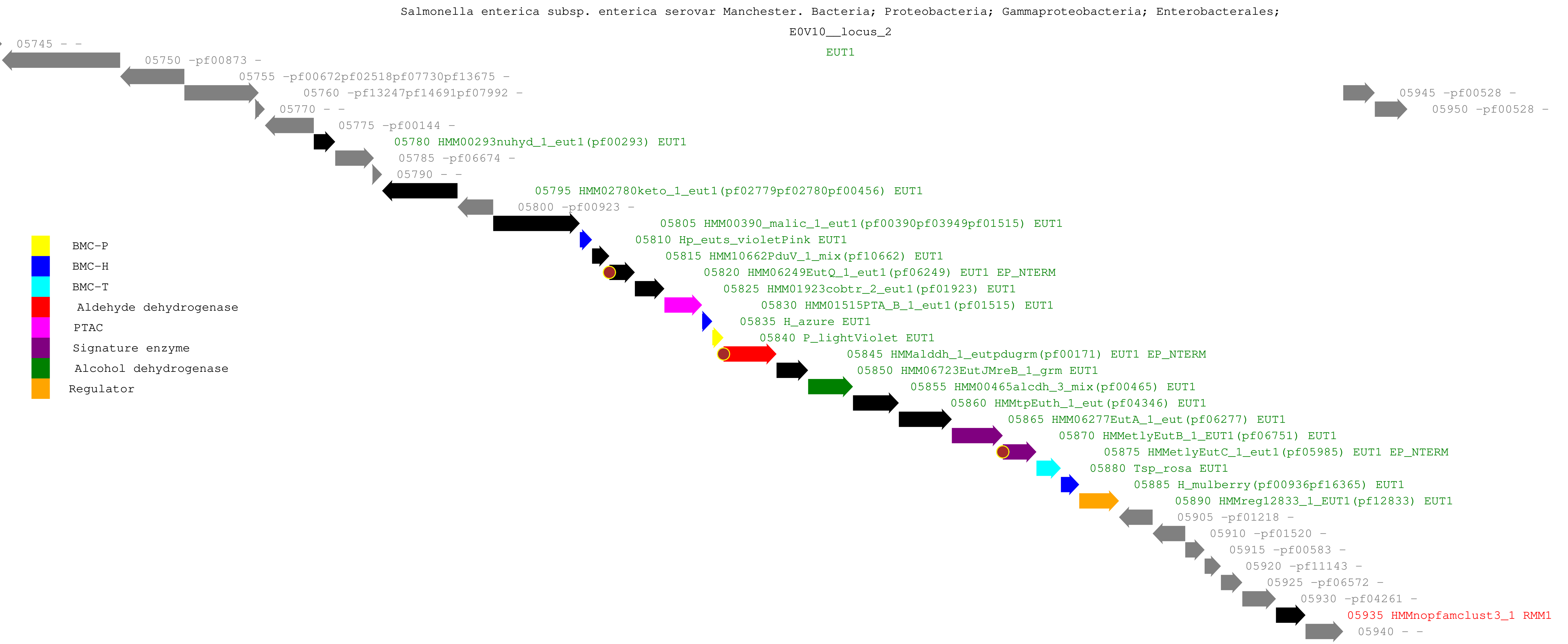

Salmonella enterica subsp. enterica serovar Manchester. Bacteria; Proteobacteria; Gammaproteobacteria; Enterobacterales;
E0V10__locus_2
EUT1
05745 - -
05750 -pf00873 -
05755 -pf00672pf02518pf07730pf13675 -
05760 -pf13247pf14691pf07992 -
05770 - -
05775 -pf00144 -
05780 HMM00293nuhyd_1_eut1(pf00293) EUT1
05785 -pf06674 -
05790 - -
05795 HMM02780keto_1_eut1(pf02779pf02780pf00456) EUT1
05800 -pf00923 -
05805 HMM00390_malic_1_eut1(pf00390pf03949pf01515) EUT1
05810 Hp_euts_violetPink EUT1
05815 HMM10662PduV_1_mix(pf10662) EUT1
05820 HMM06249EutQ_1_eut1(pf06249) EUT1 EP_NTERM
05825 HMM01923cobtr_2_eut1(pf01923) EUT1
05830 HMM01515PTA_B_1_eut1(pf01515) EUT1
05835 H_azure EUT1
05840 P_lightViolet EUT1
05845 HMMalddh_1_eutpdugrm(pf00171) EUT1 EP_NTERM
05850 HMM06723EutJMreB_1_grm EUT1
05855 HMM00465alcdh_3_mix(pf00465) EUT1
05860 HMMtpEuth_1_eut(pf04346) EUT1
05865 HMM06277EutA_1_eut(pf06277) EUT1
05870 HMMetlyEutB_1_EUT1(pf06751) EUT1
05875 HMMetlyEutC_1_eut1(pf05985) EUT1 EP_NTERM
05880 Tsp_rosa EUT1
05885 H_mulberry(pf00936pf16365) EUT1
05890 HMMreg12833_1_EUT1(pf12833) EUT1
05905 -pf01218 -
05910 -pf01520 -
05915 -pf00583 -
05920 -pf11143 -
05925 -pf06572 -
05930 -pf04261 -
05935 HMMnopfamclust3_1 RMM1
05940 - -
05945 -pf00528 -
05950 -pf00528 -
BMC-P
BMC-H
BMC-T
Aldehyde dehydrogenase
PTAC
Signature enzyme
Alcohol dehydrogenase
Regulator

Salmonella enterica subsp. enterica serovar Wangata. Bacteria; Proteobacteria; Gammaproteobacteria; Enterobacterales;

E0V16__locus_1

EUT1

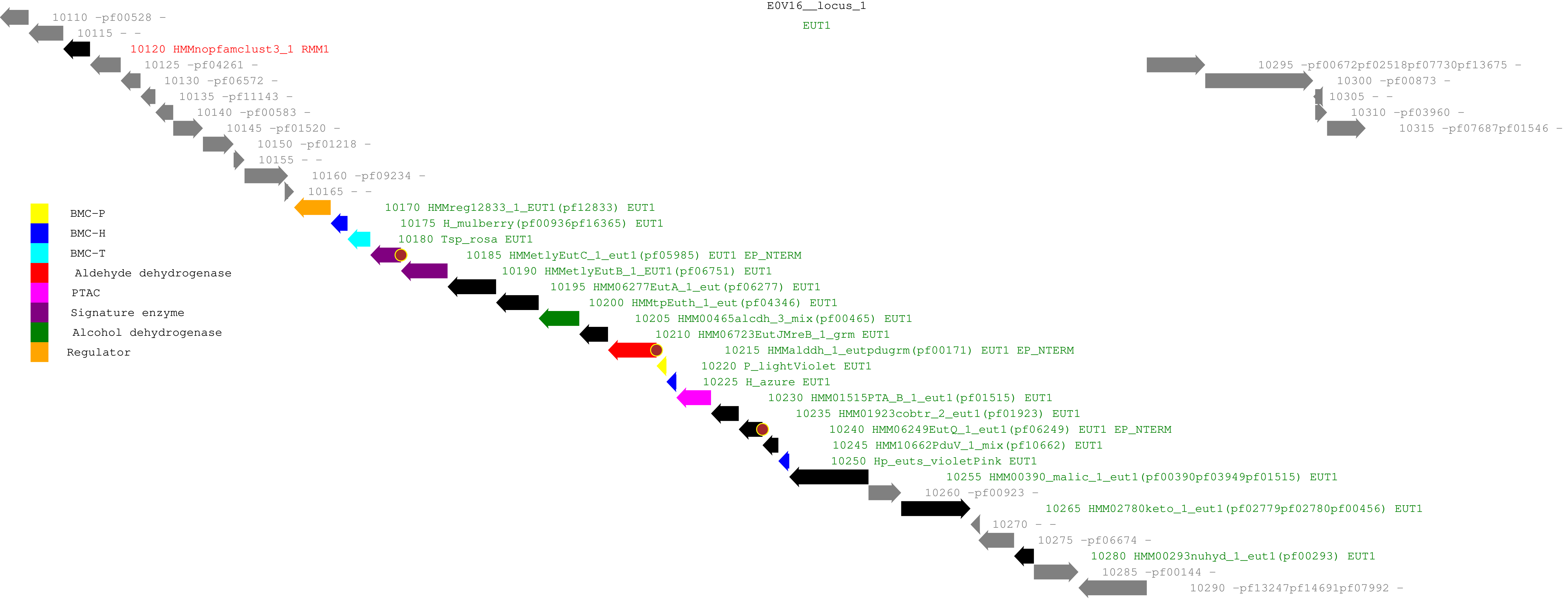

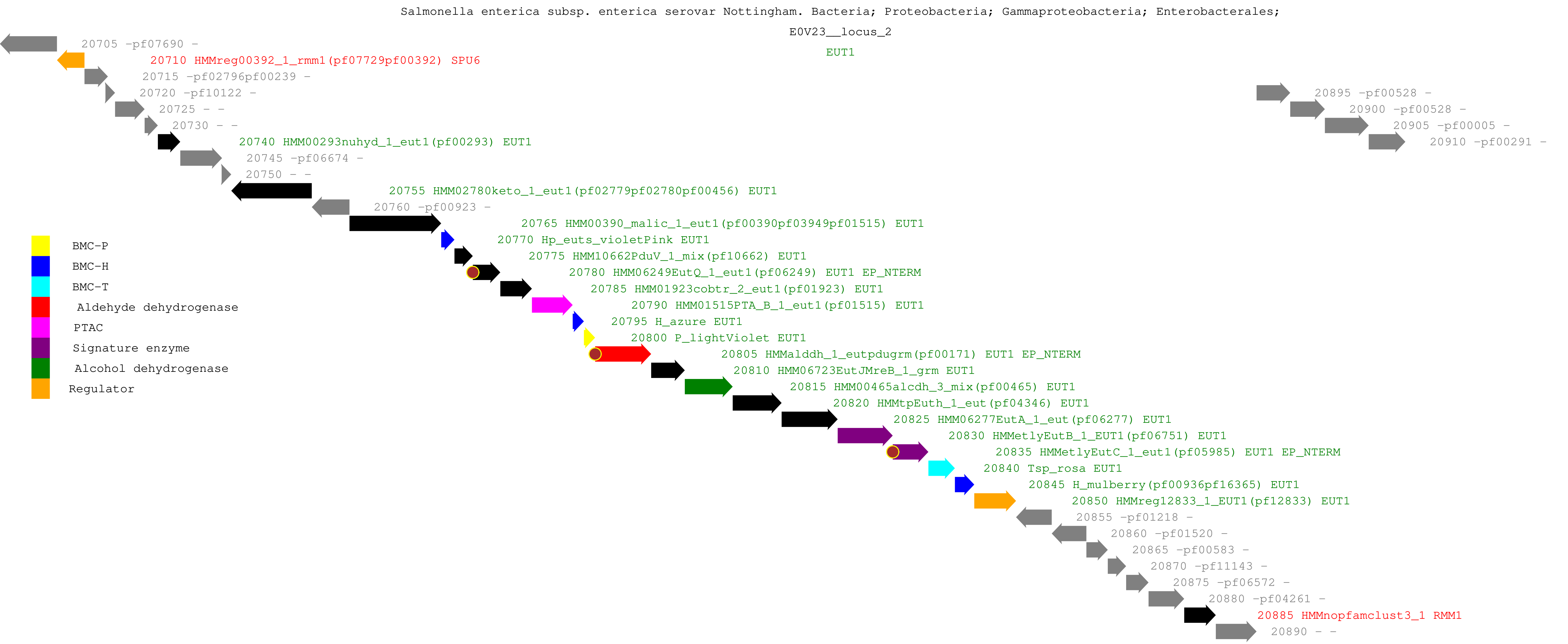
Salmonella enterica subsp. enterica serovar Nottingham. Bacteria; Proteobacteria; Gammaproteobacteria; Enterobacterales;
E0V23__locus_2
EUT1
20705 -pf07690 -
20710 HMMreg00392_1_rmm1(pf07729pf00392) SPU6
20715 -pf02796pf00239 -
20720 -pf10122 -
20725 - -
20730 - -
20740 HMM00293nuhyd_1_eut1(pf00293) EUT1
20745 -pf06674 -
20750 - -
20755 HMM02780keto_1_eut1(pf02779pf02780pf00456) EUT1
20760 -pf00923 -
20765 HMM00390_malic_1_eut1(pf00390pf03949pf01515) EUT1
20770 Hp_euts_violetPink EUT1
20775 HMM10662PduV_1_mix(pf10662) EUT1
20780 HMM06249EutQ_1_eut1(pf06249) EUT1 EP_NTERM
20785 HMM01923cobtr_2_eut1(pf01923) EUT1
20790 HMM01515PTA_B_1_eut1(pf01515) EUT1
20795 H_azure EUT1
20800 P_lightViolet EUT1
20805 HMMalddh_1_eutpdugrm(pf00171) EUT1 EP_NTERM
20810 HMM06723EutJMreB_1_grm EUT1
20815 HMM00465alcdh_3_mix(pf00465) EUT1
20820 HMMtpEuth_1_eut(pf04346) EUT1
20825 HMM06277EutA_1_eut(pf06277) EUT1
20830 HMMetlyEutB_1_EUT1(pf06751) EUT1
20835 HMMetlyEutC_1_eut1(pf05985) EUT1 EP_NTERM
20840 Tsp_rosa EUT1
20845 H_mulberry(pf00936pf16365) EUT1
20850 HMMreg12833_1_EUT1(pf12833) EUT1
20855 -pf01218 -
20860 -pf01520 -
20865 -pf00583 -
20870 -pf11143 -
20875 -pf06572 -
20880 -pf04261 -
20885 HMMnopfamclust3_1 RMM1
20890 - -
20895 -pf00528 -
20900 -pf00528 -
20905 -pf00005 -
20910 -pf00291 -
BMC-P
BMC-H
BMC-T
Aldehyde dehydrogenase
PTAC
Signature enzyme
Alcohol dehydrogenase
Regulator

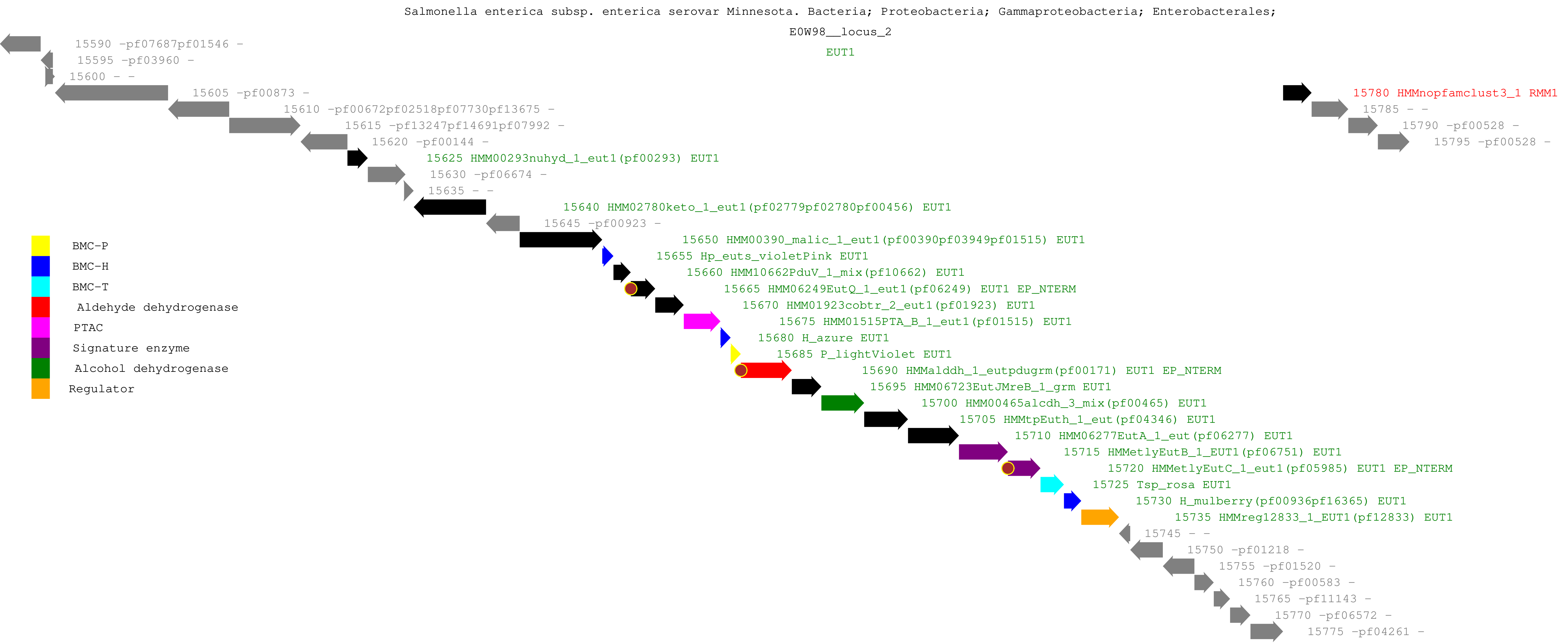
Salmonella enterica subsp. enterica serovar Minnesota. Bacteria; Proteobacteria; Gammaproteobacteria; Enterobacterales;
E0W98__locus_2
EUT1
15590 -pf07687pf01546 -
15595 -pf03960 -
15600 - -
15605 -pf00873 -
15610 -pf00672pf02518pf07730pf13675 -
15615 -pf13247pf14691pf07992 -
15620 -pf00144 -
15625 HMM00293nuhyd_1_eut1(pf00293) EUT1
15630 -pf06674 -
15635 - -
15640 HMM02780keto_1_eut1(pf02779pf02780pf00456) EUT1
15645 -pf00923 -
15650 HMM00390_malic_1_eut1(pf00390pf03949pf01515) EUT1
15655 Hp_euts_violetPink EUT1
15660 HMM10662PduV_1_mix(pf10662) EUT1
15665 HMM06249EutQ_1_eut1(pf06249) EUT1 EP_NTERM
15670 HMM01923cobtr_2_eut1(pf01923) EUT1
15675 HMM01515PTA_B_1_eut1(pf01515) EUT1
15680 H_azure EUT1
15685 P_lightViolet EUT1
15690 HMMalddh_1_eutpdugrm(pf00171) EUT1 EP_NTERM
15695 HMM06723EutJMreB_1_grm EUT1
15700 HMM00465alcdh_3_mix(pf00465) EUT1
15705 HMMtpEuth_1_eut(pf04346) EUT1
15710 HMM06277EutA_1_eut(pf06277) EUT1
15715 HMMetlyEutB_1_EUT1(pf06751) EUT1
15720 HMMetlyEutC_1_eut1(pf05985) EUT1 EP_NTERM
15725 Tsp_rosa EUT1
15730 H_mulberry(pf00936pf16365) EUT1
15735 HMMreg12833_1_EUT1(pf12833) EUT1
15745 - -
15750 -pf01218 -
15755 -pf01520 -
15760 -pf00583 -
15765 -pf11143 -
15770 -pf06572 -
15775 -pf04261 -
15780 HMMnopfamclust3_1 RMM1
15785 - -
15790 -pf00528 -
15795 -pf00528 -
BMC-P
BMC-H
BMC-T
Aldehyde dehydrogenase
PTAC
Signature enzyme
Alcohol dehydrogenase
Regulator

Salmonella enterica subsp. enterica serovar Tudu. Bacteria; Proteobacteria; Gammaproteobacteria; Enterobacterales;

E0X18__locus_2

EUT1

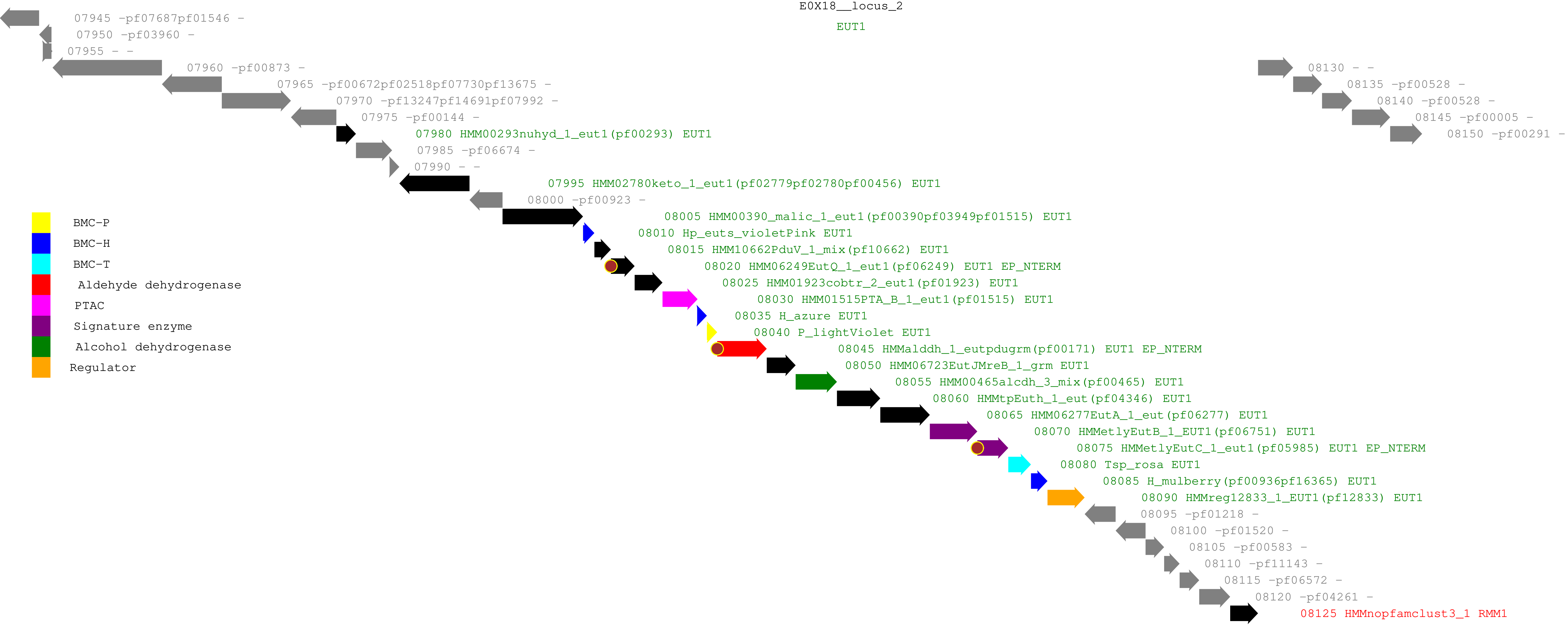

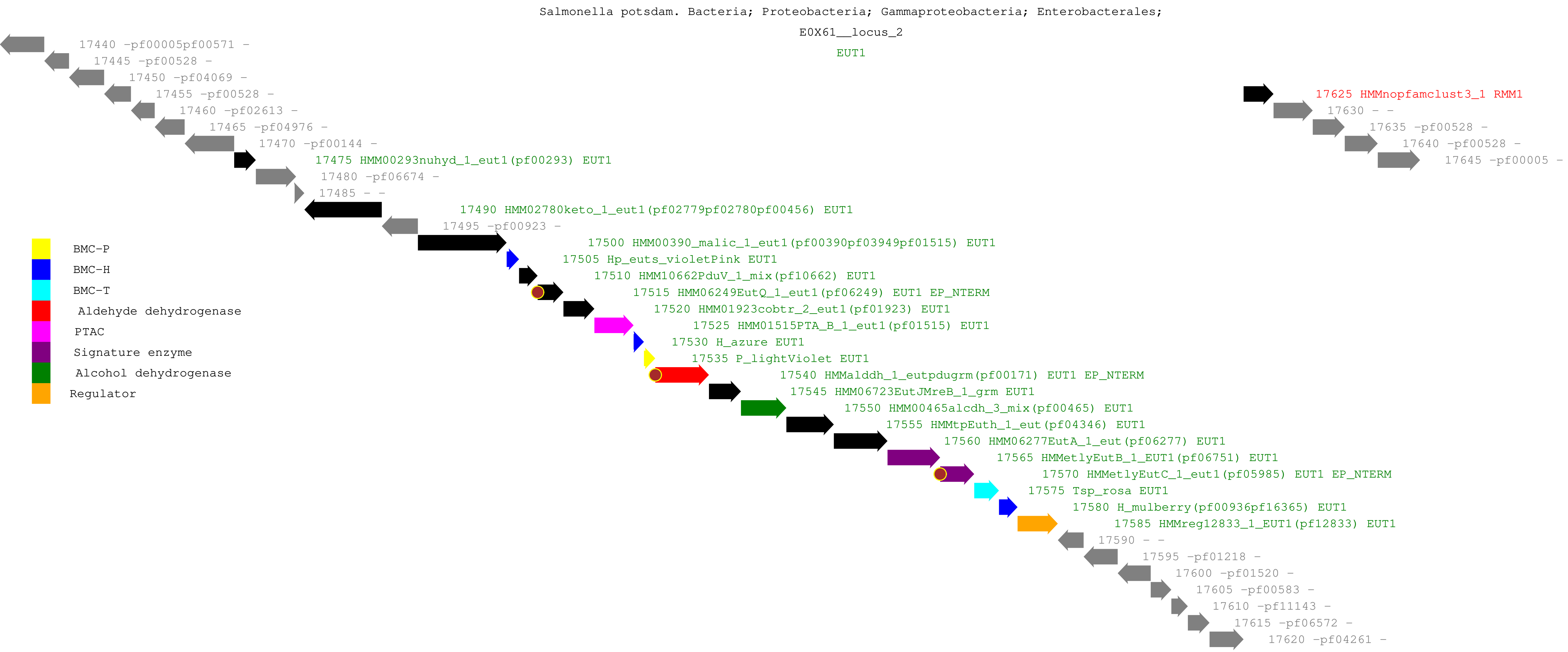

Salmonella potsdam. Bacteria; Proteobacteria; Gammaproteobacteria; Enterobacterales;
E0X61__locus_2
EUT1
17440 -pf00005pf00571 -
17445 -pf00528 -
17450 -pf04069 -
17455 -pf00528 -
17460 -pf02613 -
17465 -pf04976 -
17470 -pf00144 -
17475 HMM00293nuhyd_1_eut1(pf00293) EUT1
17480 -pf06674 -
17485 - -
17490 HMM02780keto_1_eut1(pf02779pf02780pf00456) EUT1
17495 -pf00923 -
17500 HMM00390_malic_1_eut1(pf00390pf03949pf01515) EUT1
17505 Hp_euts_violetPink EUT1
17510 HMM10662PduV_1_mix(pf10662) EUT1
17515 HMM06249EutQ_1_eut1(pf06249) EUT1 EP_NTERM
17520 HMM01923cobtr_2_eut1(pf01923) EUT1
17525 HMM01515PTA_B_1_eut1(pf01515) EUT1
17530 H_azure EUT1
17535 P_lightViolet EUT1
17540 HMMalddh_1_eutpdugrm(pf00171) EUT1 EP_NTERM
17545 HMM06723EutJMreB_1_grm EUT1
17550 HMM00465alcdh_3_mix(pf00465) EUT1
17555 HMMtpEuth_1_eut(pf04346) EUT1
17560 HMM06277EutA_1_eut(pf06277) EUT1
17565 HMMetlyEutB_1_EUT1(pf06751) EUT1
17570 HMMetlyEutC_1_eut1(pf05985) EUT1 EP_NTERM
17575 Tsp_rosa EUT1
17580 H_mulberry(pf00936pf16365) EUT1
17585 HMMreg12833_1_EUT1(pf12833) EUT1
17590 - -
17595 -pf01218 -
17600 -pf01520 -
17605 -pf00583 -
17610 -pf11143 -
17615 -pf06572 -
17620 -pf04261 -
17625 HMMnopfamclust3_1 RMM1
17630 - -
17635 -pf00528 -
17640 -pf00528 -
17645 -pf00005 -
BMC-P
BMC-H
BMC-T
Aldehyde dehydrogenase
PTAC
Signature enzyme
Alcohol dehydrogenase
Regulator

Salmonella enterica subsp. enterica serovar Miami. Bacteria; Proteobacteria; Gammaproteobacteria; Enterobacterales;

E0Y77__locus_2

EUT1

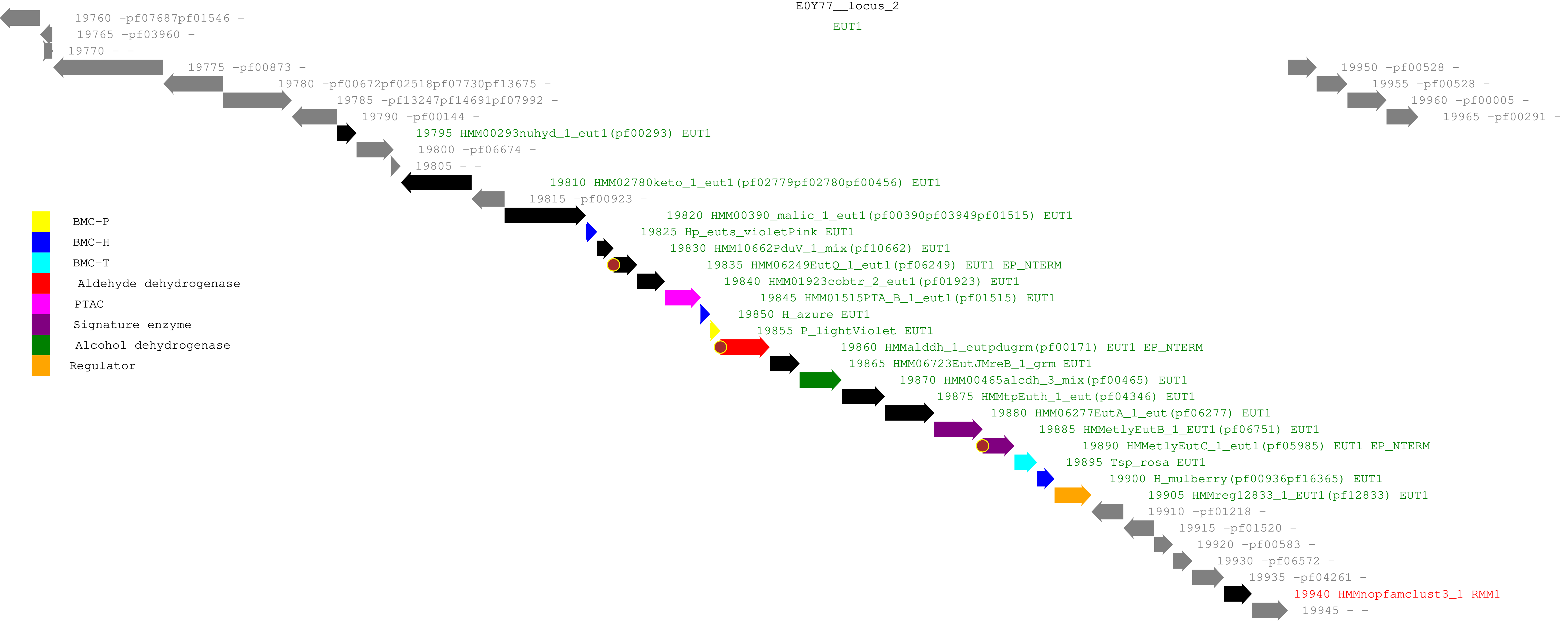

Salmonella enterica subsp. enterica serovar Wentworth. Bacteria; Proteobacteria; Gammaproteobacteria; Enterobacterales;

E1361__locus_1

EUT1

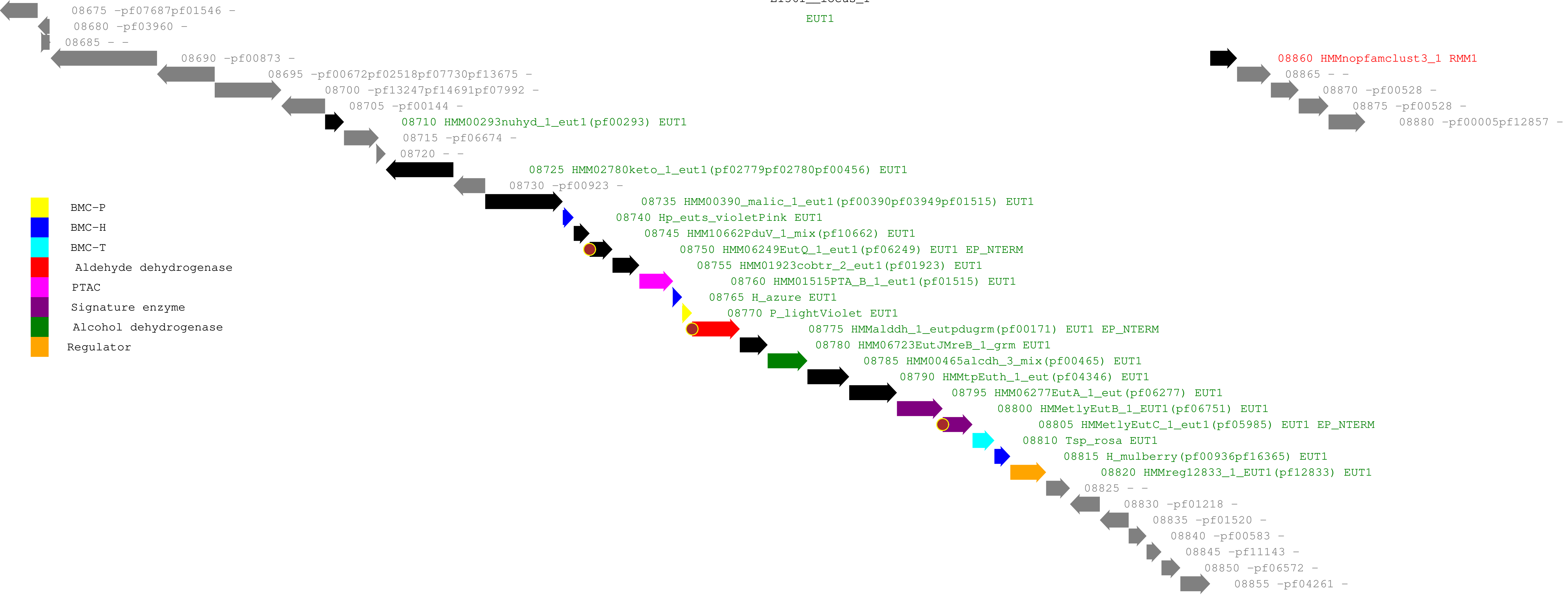

Salmonella enterica subsp. enterica serovar Falkensee. Bacteria; Proteobacteria; Gammaproteobacteria; Enterobacterales;

E1364__locus_1

EUT1

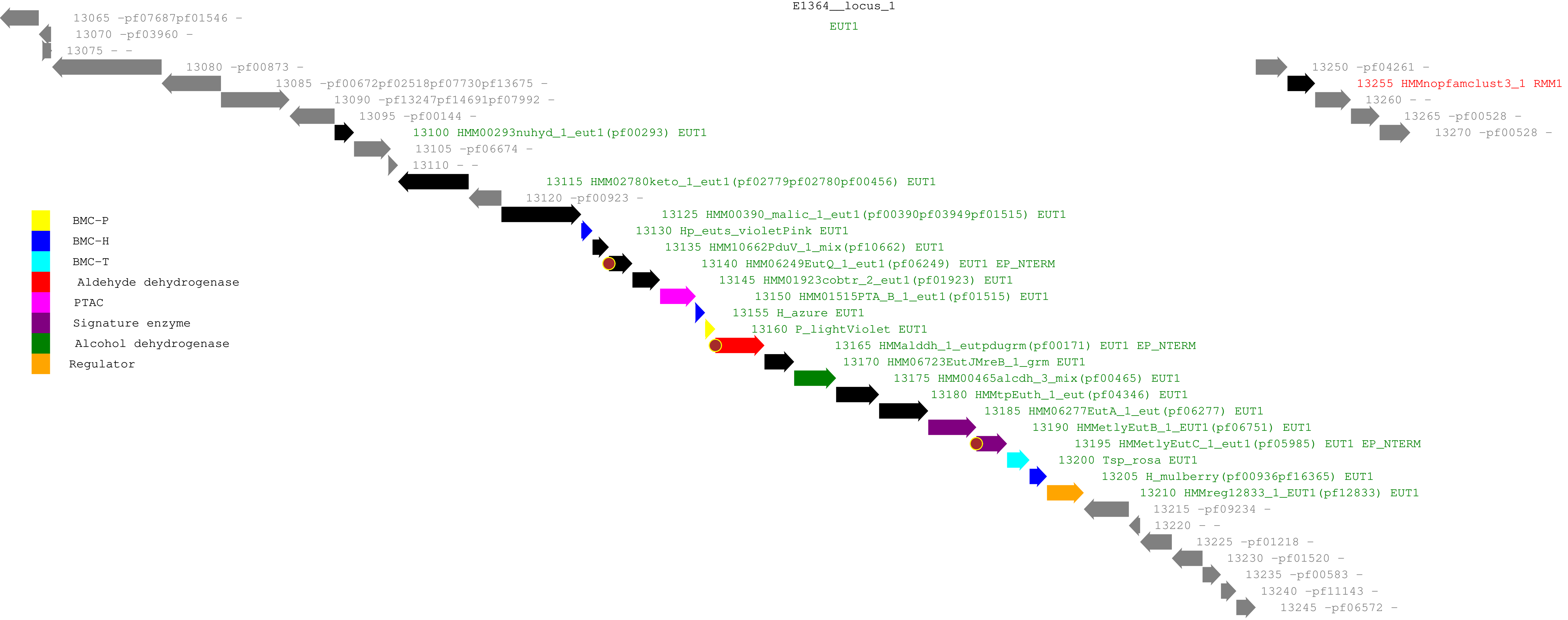

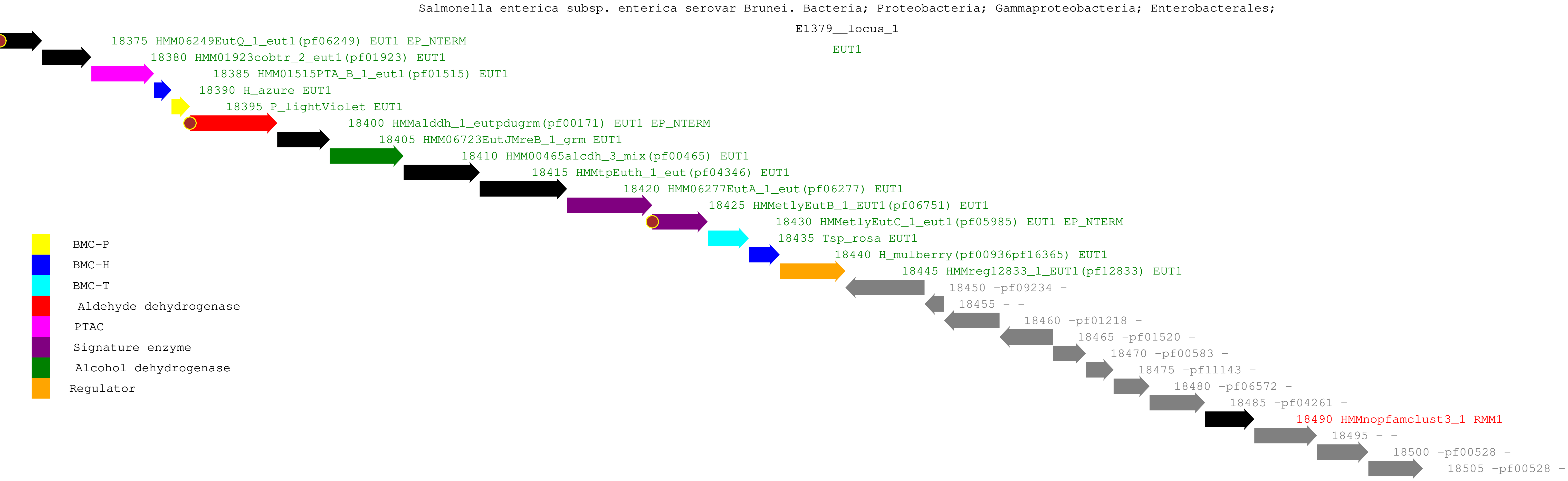
Salmonella enterica subsp. enterica serovar Brunei. Bacteria; Proteobacteria; Gammaproteobacteria; Enterobacterales;
E1379__locus_1
EUT1
18375 HMM06249EutQ_1_eut1(pf06249) EUT1 EP_NTERM
18380 HMM01923cobtr_2_eut1(pf01923) EUT1
18385 HMM01515PTA_B_1_eut1(pf01515) EUT1
18390 H_azure EUT1
18395 P_lightViolet EUT1
18400 HMMalddh_1_eutpdugrm(pf00171) EUT1 EP_NTERM
18405 HMM06723EutJMreB_1_grm EUT1
18410 HMM00465alcdh_3_mix(pf00465) EUT1
18415 HMMtpEuth_1_eut(pf04346) EUT1
18420 HMM06277EutA_1_eut(pf06277) EUT1
18425 HMMetlyEutB_1_EUT1(pf06751) EUT1
18430 HMMetlyEutC_1_eut1(pf05985) EUT1 EP_NTERM
18435 Tsp_rosa EUT1
18440 H_mulberry(pf00936pf16365) EUT1
18445 HMMreg12833_1_EUT1(pf12833) EUT1
18450 -pf09234 -
18455 - -
18460 -pf01218 -
18465 -pf01520 -
18470 -pf00583 -
18475 -pf11143 -
18480 -pf06572 -
18485 -pf04261 -
18490 HMMnopfamclust3_1 RMM1
18495 - -
18500 -pf00528 -
18505 -pf00528 -
BMC-P
BMC-H
BMC-T
Aldehyde dehydrogenase
PTAC
Signature enzyme
Alcohol dehydrogenase
Regulator

Salmonella enterica subsp. enterica serovar Anecho. Bacteria; Proteobacteria; Gammaproteobacteria; Enterobacterales;

E1393__locus_2

EUT1

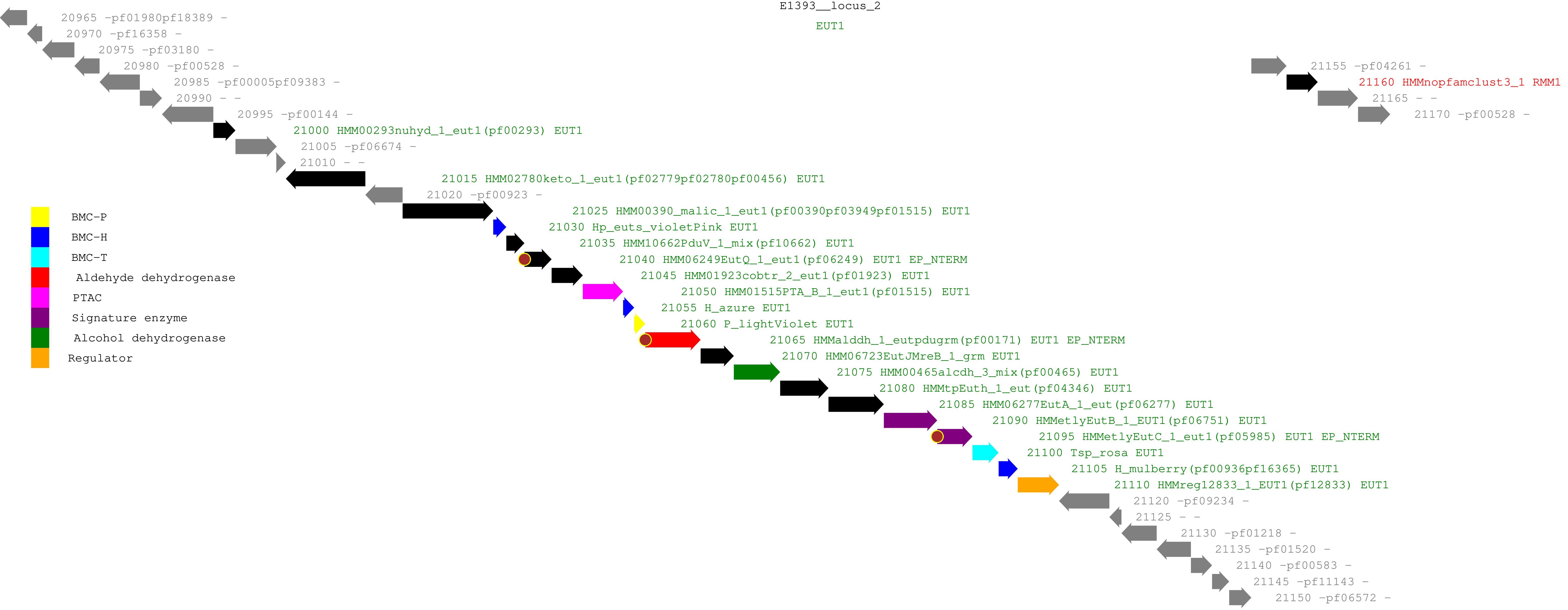

Salmonella enterica subsp. enterica serovar Lille. Bacteria; Proteobacteria; Gammaproteobacteria; Enterobacterales;

E1399__locus_1

EUT1

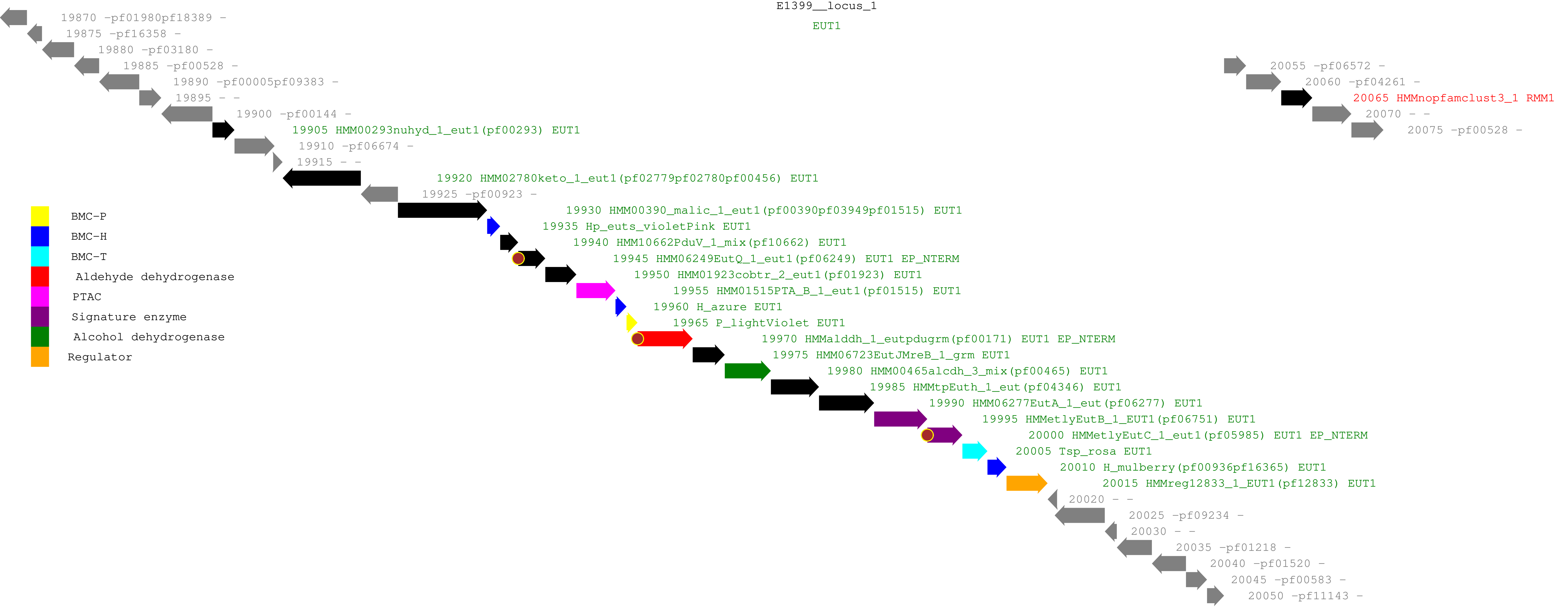

Salmonella enterica subsp. enterica serovar Augustenborg. Bacteria; Proteobacteria; Gammaproteobacteria; Enterobacterales;

E1429__locus_2

EUT1

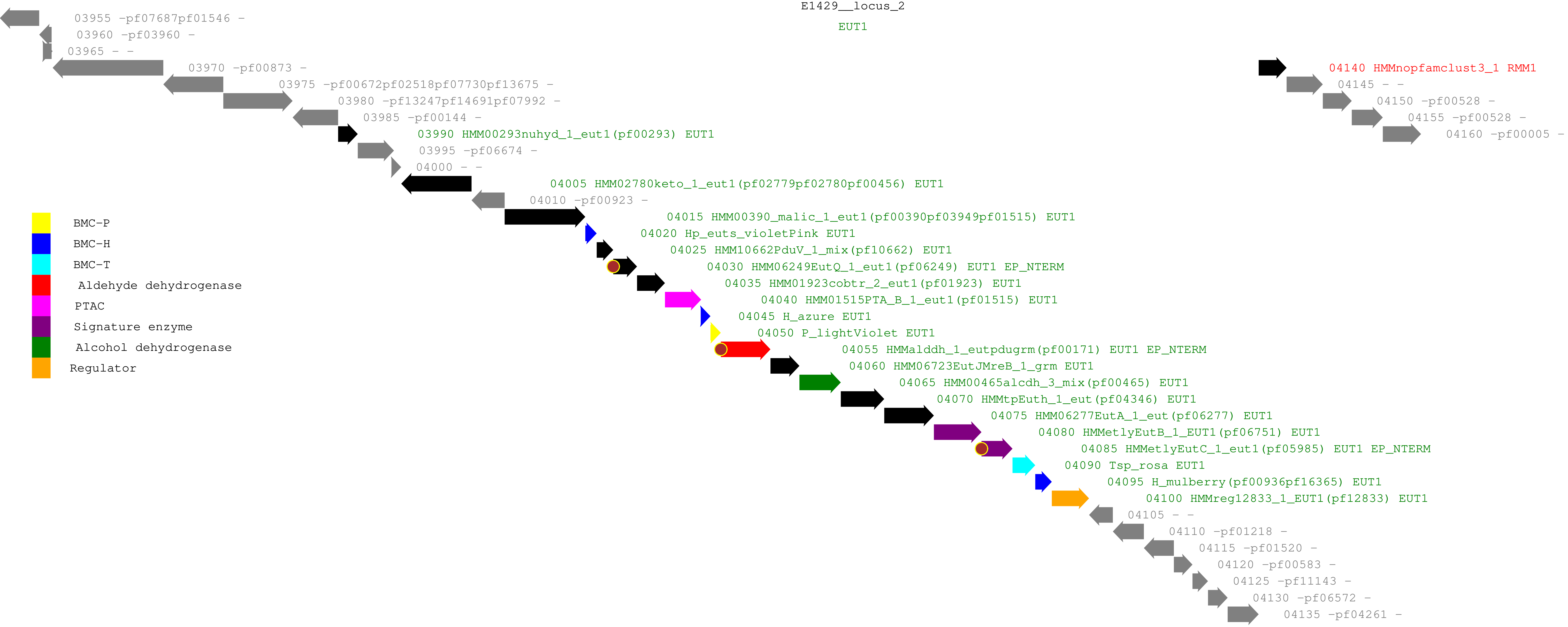

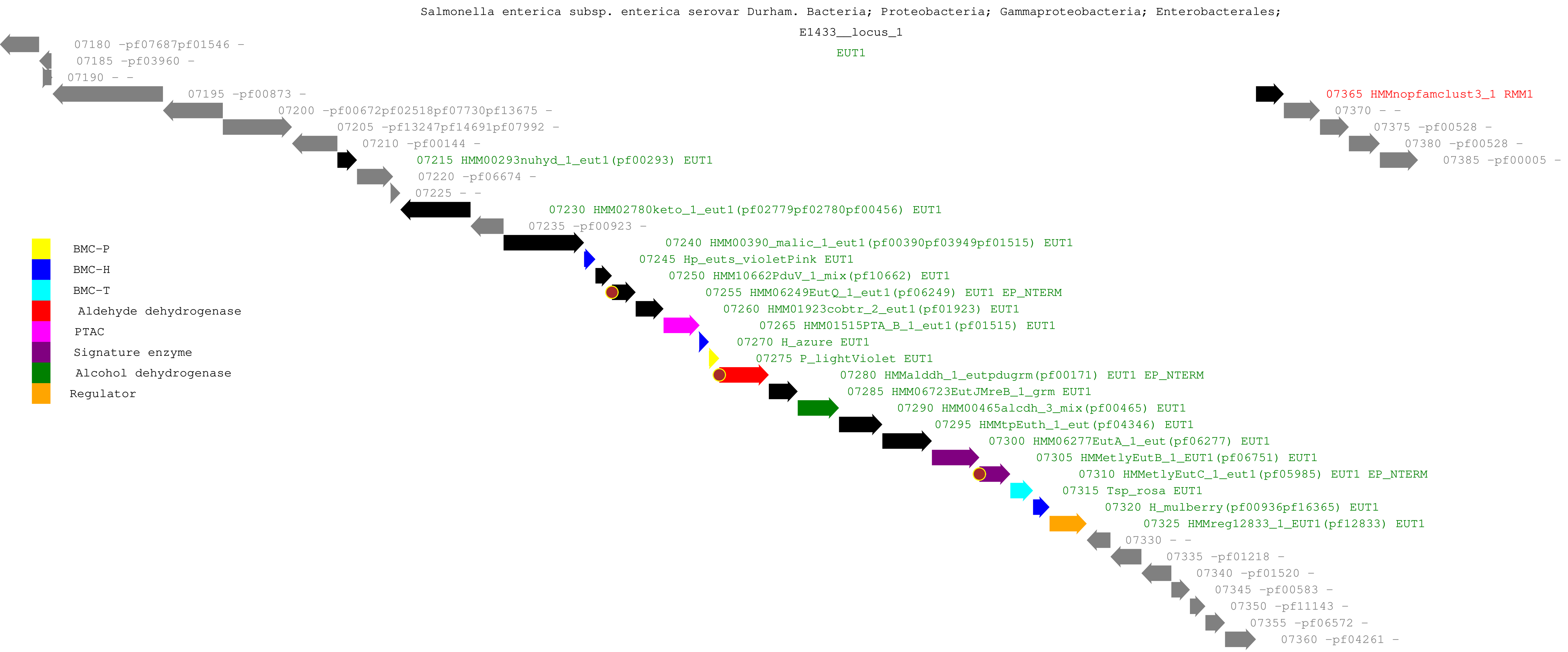

Salmonella enterica subsp. enterica serovar Durham. Bacteria; Proteobacteria; Gammaproteobacteria; Enterobacterales;
E1433__locus_1
EUT1
07180 -pf07687pf01546 -
07185 -pf03960 -
07190 - -
07195 -pf00873 -
07200 -pf00672pf02518pf07730pf13675 -
07205 -pf13247pf14691pf07992 -
07210 -pf00144 -
07215 HMM00293nuhyd_1_eut1(pf00293) EUT1
07220 -pf06674 -
07225 - -
07230 HMM02780keto_1_eut1(pf02779pf02780pf00456) EUT1
07235 -pf00923 -
07240 HMM00390_malic_1_eut1(pf00390pf03949pf01515) EUT1
07245 Hp_euts_violetPink EUT1
07250 HMM10662PduV_1_mix(pf10662) EUT1
07255 HMM06249EutQ_1_eut1(pf06249) EUT1 EP_NTERM
07260 HMM01923cobtr_2_eut1(pf01923) EUT1
07265 HMM01515PTA_B_1_eut1(pf01515) EUT1
07270 H_azure EUT1
07275 P_lightViolet EUT1
07280 HMMalddh_1_eutpdugrm(pf00171) EUT1 EP_NTERM
07285 HMM06723EutJMreB_1_grm EUT1
07290 HMM00465alcdh_3_mix(pf00465) EUT1
07295 HMMtpEuth_1_eut(pf04346) EUT1
07300 HMM06277EutA_1_eut(pf06277) EUT1
07305 HMMetlyEutB_1_EUT1(pf06751) EUT1
07310 HMMetlyEutC_1_eut1(pf05985) EUT1 EP_NTERM
07315 Tsp_rosa EUT1
07320 H_mulberry(pf00936pf16365) EUT1
07325 HMMreg12833_1_EUT1(pf12833) EUT1
07330 - -
07335 -pf01218 -
07340 -pf01520 -
07345 -pf00583 -
07350 -pf11143 -
07355 -pf06572 -
07360 -pf04261 -
07365 HMMnopfamclust3_1 RMM1
07370 - -
07375 -pf00528 -
07380 -pf00528 -
07385 -pf00005 -
BMC-P
BMC-H
BMC-T
Aldehyde dehydrogenase
PTAC
Signature enzyme
Alcohol dehydrogenase
Regulator

Salmonella enterica subsp. enterica serovar Kimuenza. Bacteria; Proteobacteria; Gammaproteobacteria; Enterobacterales;

E1444__locus_1

EUT1

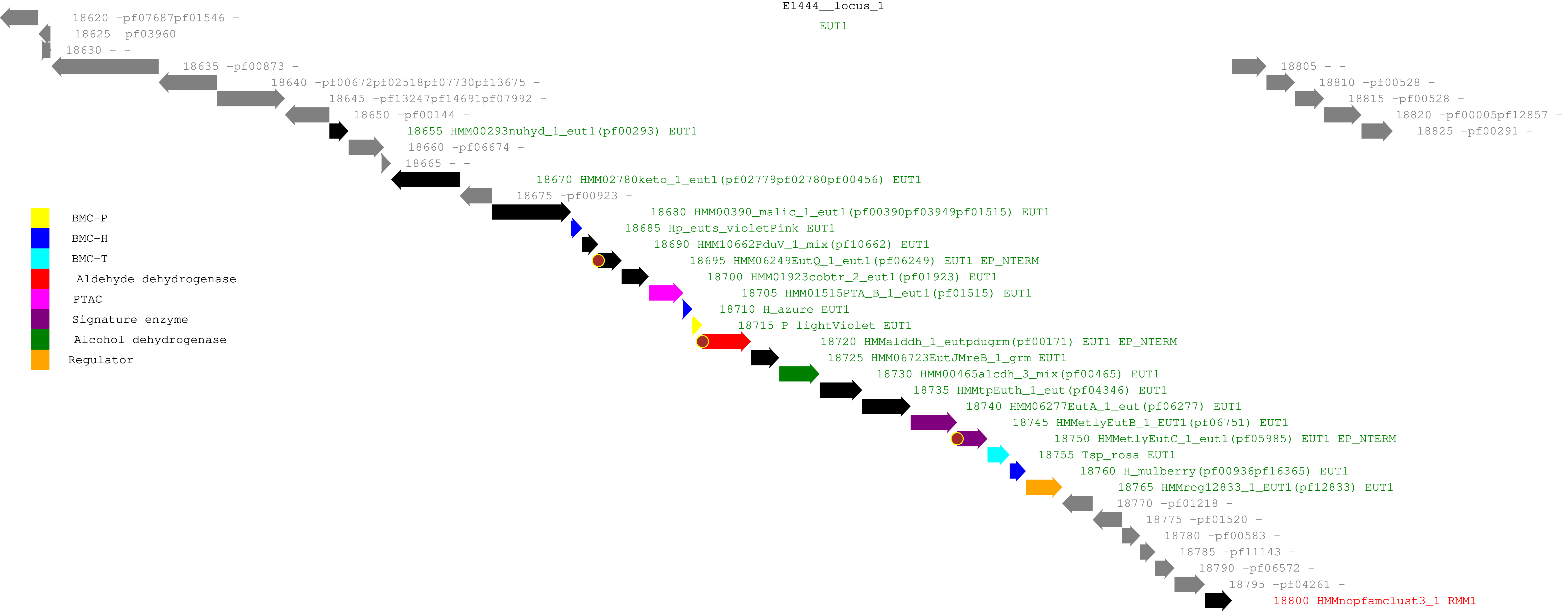

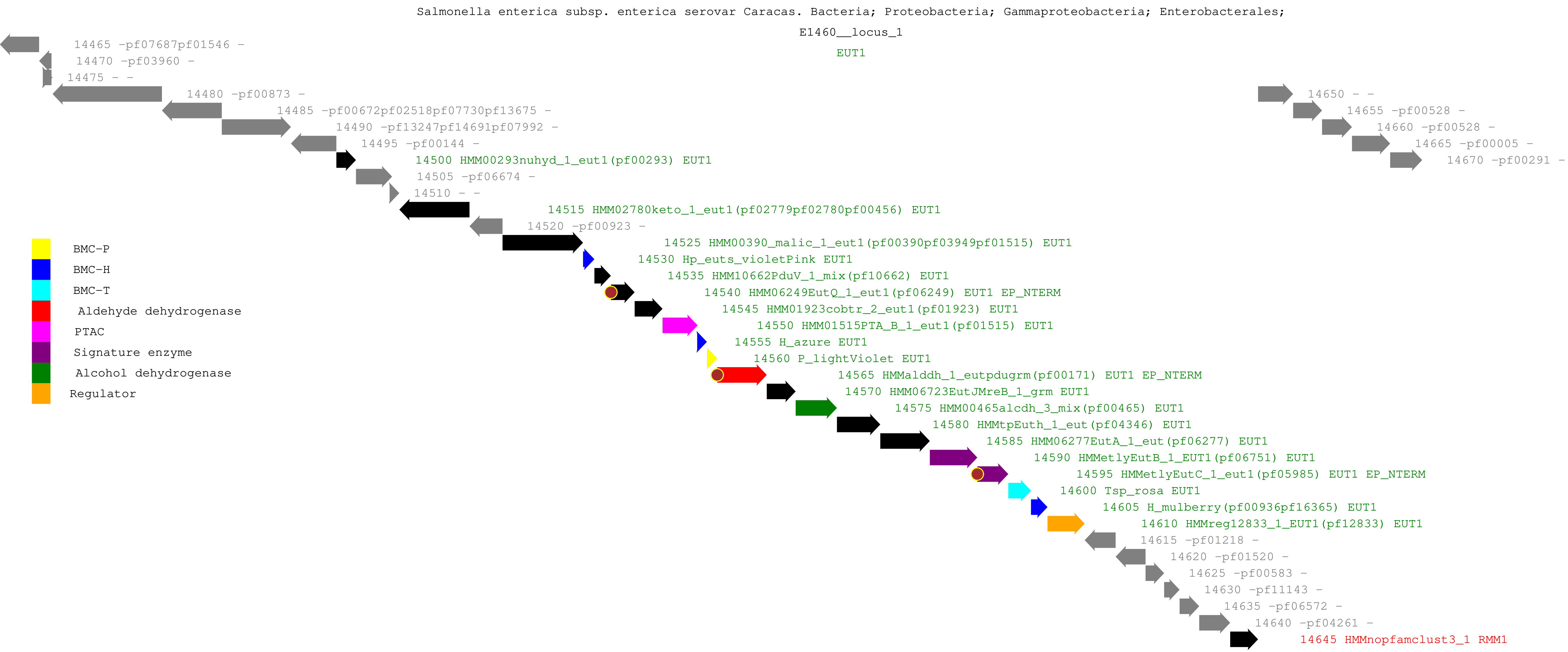
Salmonella enterica subsp. enterica serovar Caracas. Bacteria; Proteobacteria; Gammaproteobacteria; Enterobacterales;
E1460__locus_1
EUT1
14465 -pf07687pf01546 -
14470 -pf03960 -
14475 - -
14480 -pf00873 -
14485 -pf00672pf02518pf07730pf13675 -
14490 -pf13247pf14691pf07992 -
14495 -pf00144 -
14500 HMM00293nuhyd_1_eut1(pf00293) EUT1
14505 -pf06674 -
14510 - -
14515 HMM02780keto_1_eut1(pf02779pf02780pf00456) EUT1
14520 -pf00923 -
14525 HMM00390_malic_1_eut1(pf00390pf03949pf01515) EUT1
14530 Hp_euts_violetPink EUT1
14535 HMM10662PduV_1_mix(pf10662) EUT1
14540 HMM06249EutQ_1_eut1(pf06249) EUT1 EP_NTERM
14545 HMM01923cobtr_2_eut1(pf01923) EUT1
14550 HMM01515PTA_B_1_eut1(pf01515) EUT1
14555 H_azure EUT1
14560 P_lightViolet EUT1
14565 HMMalddh_1_eutpdugrm(pf00171) EUT1 EP_NTERM
14570 HMM06723EutJMreB_1_grm EUT1
14575 HMM00465alcdh_3_mix(pf00465) EUT1
14580 HMMtpEuth_1_eut(pf04346) EUT1
14585 HMM06277EutA_1_eut(pf06277) EUT1
14590 HMMetlyEutB_1_EUT1(pf06751) EUT1
14595 HMMetlyEutC_1_eut1(pf05985) EUT1 EP_NTERM
14600 Tsp_rosa EUT1
14605 H_mulberry(pf00936pf16365) EUT1
14610 HMMreg12833_1_EUT1(pf12833) EUT1
14615 -pf01218 -
14620 -pf01520 -
14625 -pf00583 -
14630 -pf11143 -
14635 -pf06572 -
14640 -pf04261 -
14645 HMMnopfamclust3_1 RMM1
14650 - -
14655 -pf00528 -
14660 -pf00528 -
14665 -pf00005 -
14670 -pf00291 -
BMC-P
BMC-H
BMC-T
Aldehyde dehydrogenase
PTAC
Signature enzyme
Alcohol dehydrogenase
Regulator

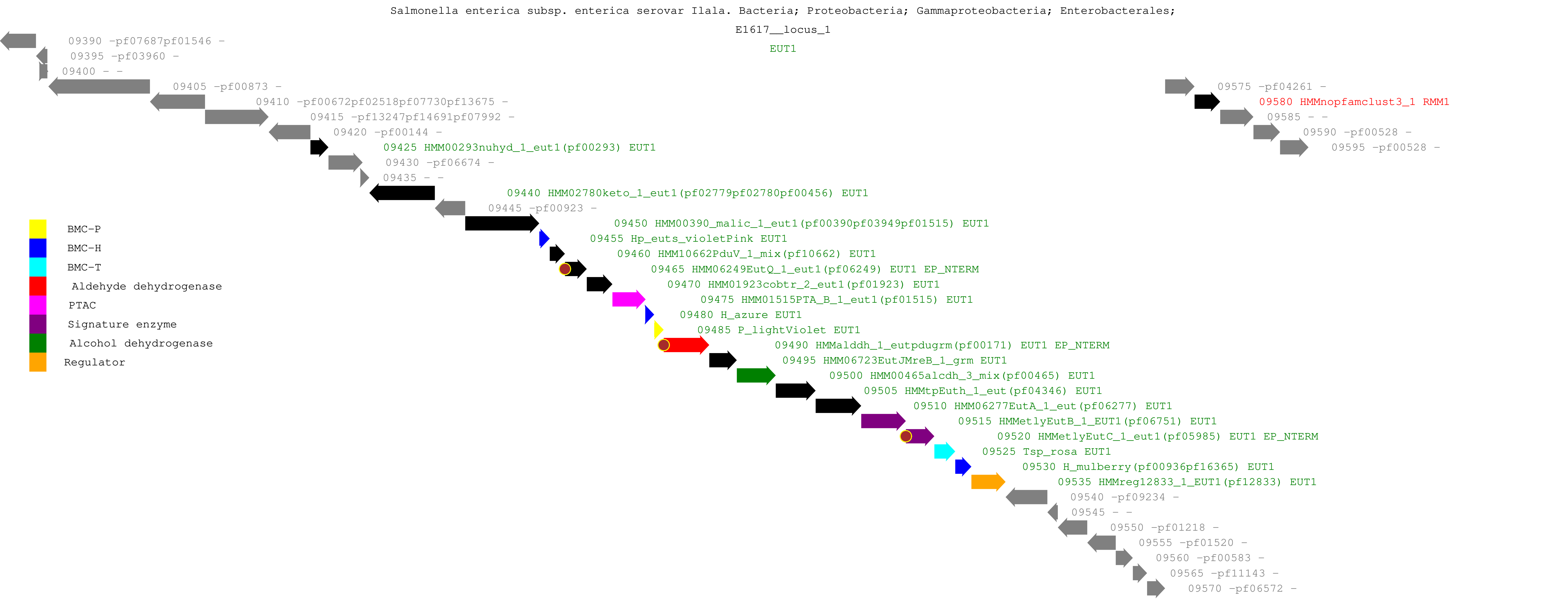
Salmonella enterica subsp. enterica serovar Ilala. Bacteria; Proteobacteria; Gammaproteobacteria; Enterobacterales;
E1617__locus_1
EUT1
09390 -pf07687pf01546 -
09395 -pf03960 -
09400 - -
09405 -pf00873 -
09410 -pf00672pf02518pf07730pf13675 -
09415 -pf13247pf14691pf07992 -
09420 -pf00144 -
09425 HMM00293nuhyd_1_eut1(pf00293) EUT1
09430 -pf06674 -
09435 - -
09440 HMM02780keto_1_eut1(pf02779pf02780pf00456) EUT1
09445 -pf00923 -
09450 HMM00390_malic_1_eut1(pf00390pf03949pf01515) EUT1
09455 Hp_euts_violetPink EUT1
09460 HMM10662PduV_1_mix(pf10662) EUT1
09465 HMM06249EutQ_1_eut1(pf06249) EUT1 EP_NTERM
09470 HMM01923cobtr_2_eut1(pf01923) EUT1
09475 HMM01515PTA_B_1_eut1(pf01515) EUT1
09480 H_azure EUT1
09485 P_lightViolet EUT1
09490 HMMalddh_1_eutpdugrm(pf00171) EUT1 EP_NTERM
09495 HMM06723EutJMreB_1_grm EUT1
09500 HMM00465alcdh_3_mix(pf00465) EUT1
09505 HMMtpEuth_1_eut(pf04346) EUT1
09510 HMM06277EutA_1_eut(pf06277) EUT1
09515 HMMetlyEutB_1_EUT1(pf06751) EUT1
09520 HMMetlyEutC_1_eut1(pf05985) EUT1 EP_NTERM
09525 Tsp_rosa EUT1
09530 H_mulberry(pf00936pf16365) EUT1
09535 HMMreg12833_1_EUT1(pf12833) EUT1
09540 -pf09234 -
09545 - -
09550 -pf01218 -
09555 -pf01520 -
09560 -pf00583 -
09565 -pf11143 -
09570 -pf06572 -
09575 -pf04261 -
09580 HMMnopfamclust3_1 RMM1
09585 - -
09590 -pf00528 -
09595 -pf00528 -
BMC-P
BMC-H
BMC-T
Aldehyde dehydrogenase
PTAC
Signature enzyme
Alcohol dehydrogenase
Regulator

Salmonella enterica subsp. enterica serovar Baguida. Bacteria; Proteobacteria; Gammaproteobacteria; Enterobacterales;

E1630__locus_1

EUT1

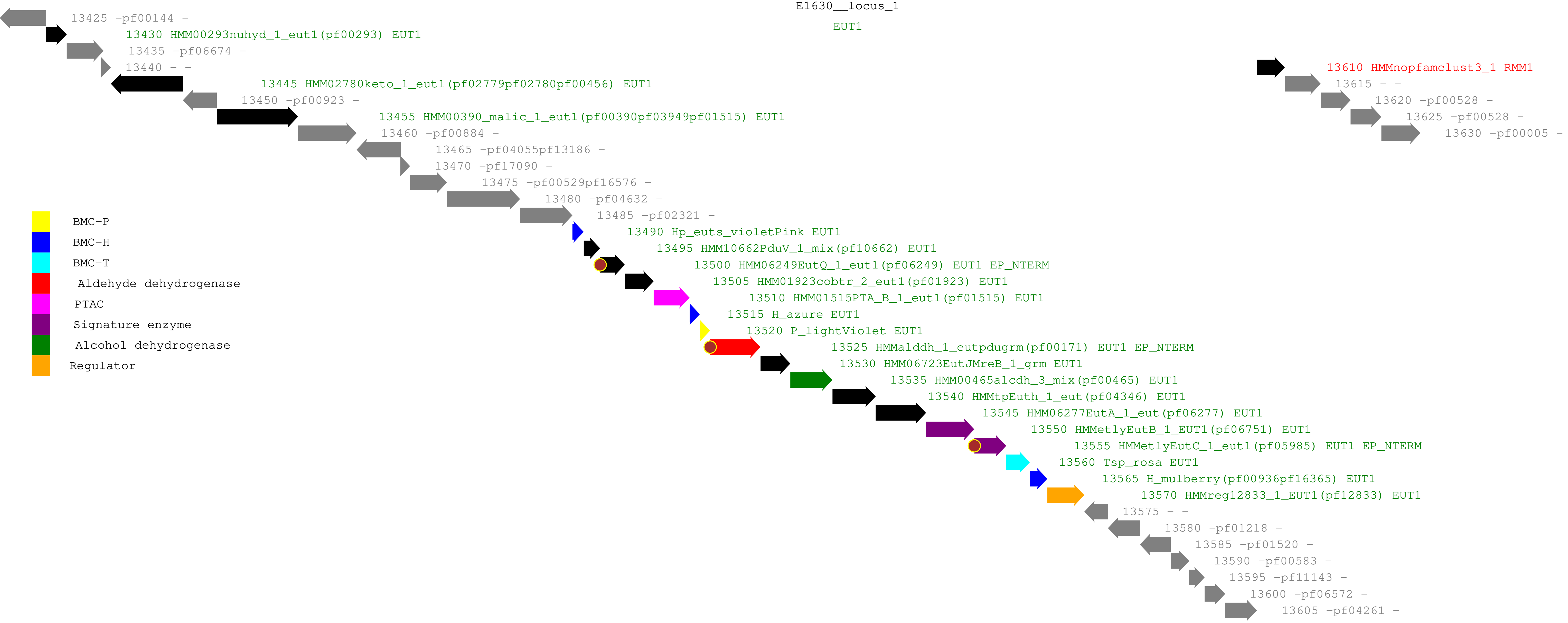

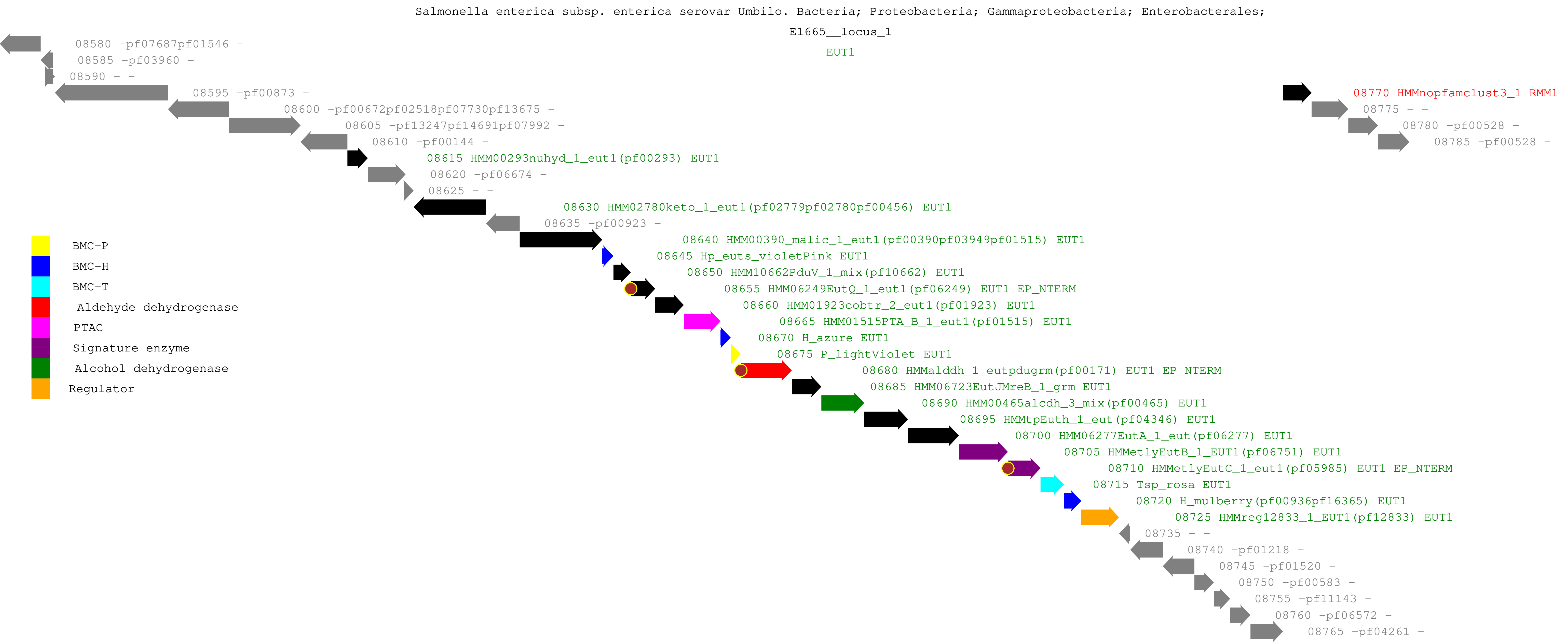

Salmonella enterica subsp. enterica serovar Umbilo. Bacteria; Proteobacteria; Gammaproteobacteria; Enterobacterales;
E1665__locus_1
EUT1
08580 -pf07687pf01546 -
08585 -pf03960 -
08590 - -
08595 -pf00873 -
08600 -pf00672pf02518pf07730pf13675 -
08605 -pf13247pf14691pf07992 -
08610 -pf00144 -
08615 HMM00293nuhyd_1_eut1(pf00293) EUT1
08620 -pf06674 -
08625 - -
08630 HMM02780keto_1_eut1(pf02779pf02780pf00456) EUT1
08635 -pf00923 -
08640 HMM00390_malic_1_eut1(pf00390pf03949pf01515) EUT1
08645 Hp_euts_violetPink EUT1
08650 HMM10662PduV_1_mix(pf10662) EUT1
08655 HMM06249EutQ_1_eut1(pf06249) EUT1 EP_NTERM
08660 HMM01923cobtr_2_eut1(pf01923) EUT1
08665 HMM01515PTA_B_1_eut1(pf01515) EUT1
08670 H_azure EUT1
08675 P_lightViolet EUT1
08680 HMMalddh_1_eutpdugrm(pf00171) EUT1 EP_NTERM
08685 HMM06723EutJMreB_1_grm EUT1
08690 HMM00465alcdh_3_mix(pf00465) EUT1
08695 HMMtpEuth_1_eut(pf04346) EUT1
08700 HMM06277EutA_1_eut(pf06277) EUT1
08705 HMMetlyEutB_1_EUT1(pf06751) EUT1
08710 HMMetlyEutC_1_eut1(pf05985) EUT1 EP_NTERM
08715 Tsp_rosa EUT1
08720 H_mulberry(pf00936pf16365) EUT1
08725 HMMreg12833_1_EUT1(pf12833) EUT1
08735 - -
08740 -pf01218 -
08745 -pf01520 -
08750 -pf00583 -
08755 -pf11143 -
08760 -pf06572 -
08765 -pf04261 -
08770 HMMnopfamclust3_1 RMM1
08775 - -
08780 -pf00528 -
08785 -pf00528 -
BMC-P
BMC-H
BMC-T
Aldehyde dehydrogenase
PTAC
Signature enzyme
Alcohol dehydrogenase
Regulator

Salmonella blockley. Bacteria; Proteobacteria; Gammaproteobacteria; Enterobacterales;

E1672__locus_2

EUT1

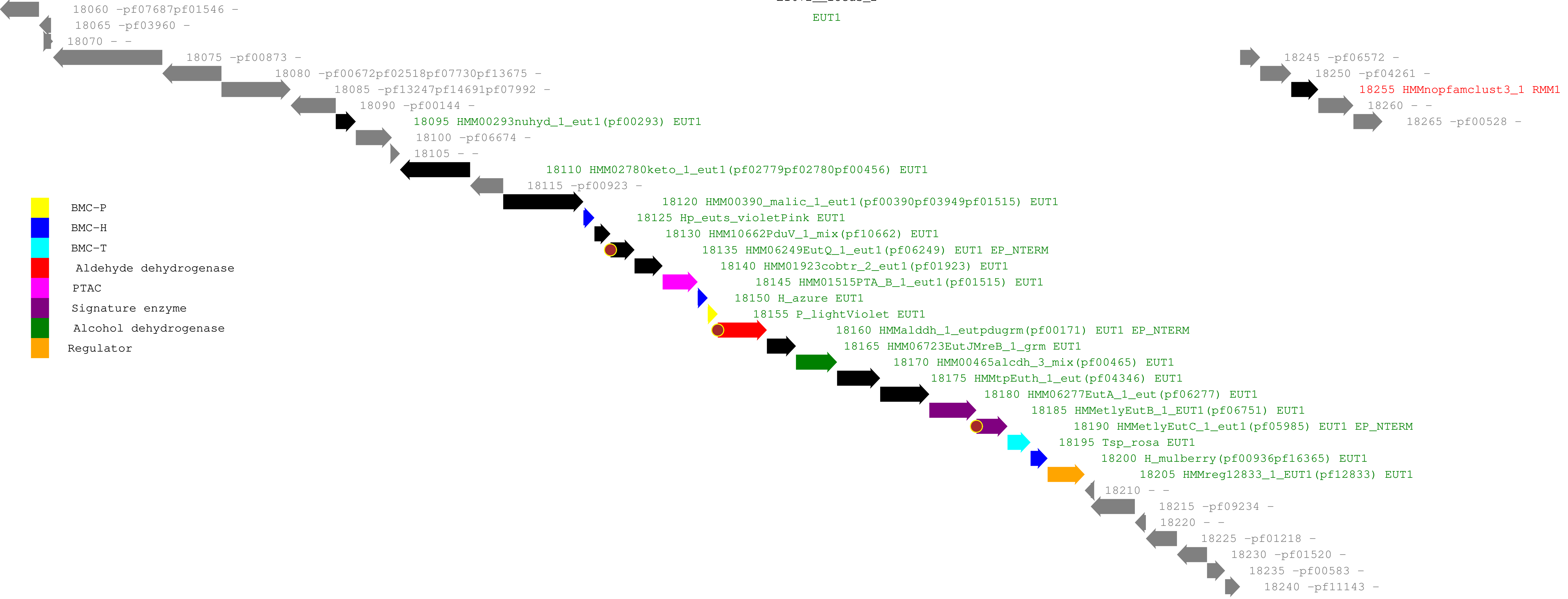

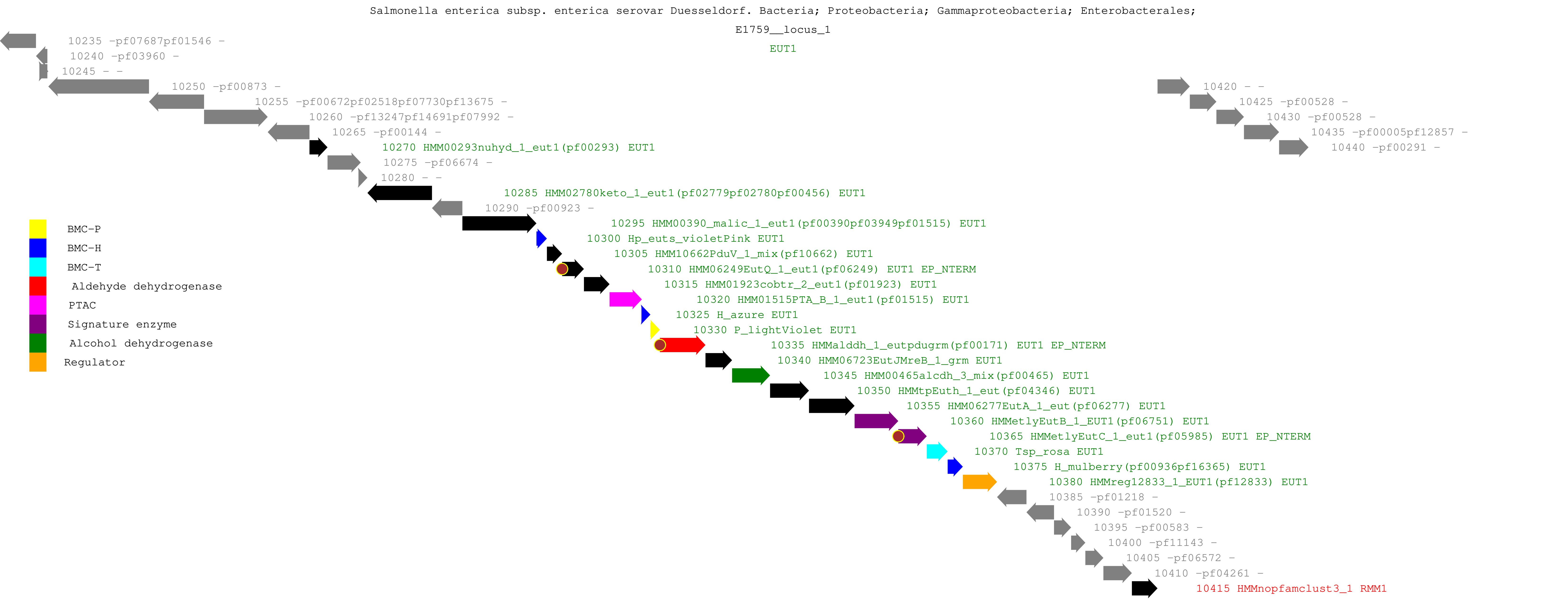
Salmonella enterica subsp. enterica serovar Duesseldorf. Bacteria; Proteobacteria; Gammaproteobacteria; Enterobacterales;
E1759__locus_1
EUT1
10235 -pf07687pf01546 -
10240 -pf03960 -
10245 - -
10250 -pf00873 -
10255 -pf00672pf02518pf07730pf13675 -
10260 -pf13247pf14691pf07992 -
10265 -pf00144 -
10270 HMM00293nuhyd_1_eut1(pf00293) EUT1
10275 -pf06674 -
10280 - -
10285 HMM02780keto_1_eut1(pf02779pf02780pf00456) EUT1
10290 -pf00923 -
10295 HMM00390_malic_1_eut1(pf00390pf03949pf01515) EUT1
10300 Hp_euts_violetPink EUT1
10305 HMM10662PduV_1_mix(pf10662) EUT1
10310 HMM06249EutQ_1_eut1(pf06249) EUT1 EP_NTERM
10315 HMM01923cobtr_2_eut1(pf01923) EUT1
10320 HMM01515PTA_B_1_eut1(pf01515) EUT1
10325 H_azure EUT1
10330 P_lightViolet EUT1
10335 HMMalddh_1_eutpdugrm(pf00171) EUT1 EP_NTERM
10340 HMM06723EutJMreB_1_grm EUT1
10345 HMM00465alcdh_3_mix(pf00465) EUT1
10350 HMMtpEuth_1_eut(pf04346) EUT1
10355 HMM06277EutA_1_eut(pf06277) EUT1
10360 HMMetlyEutB_1_EUT1(pf06751) EUT1
10365 HMMetlyEutC_1_eut1(pf05985) EUT1 EP_NTERM
10370 Tsp_rosa EUT1
10375 H_mulberry(pf00936pf16365) EUT1
10380 HMMreg12833_1_EUT1(pf12833) EUT1
10385 -pf01218 -
10390 -pf01520 -
10395 -pf00583 -
10400 -pf11143 -
10405 -pf06572 -
10410 -pf04261 -
10415 HMMnopfamclust3_1 RMM1
10420 - -
10425 -pf00528 -
10430 -pf00528 -
10435 -pf00005pf12857 -
10440 -pf00291 -
BMC-P
BMC-H
BMC-T
Aldehyde dehydrogenase
PTAC
Signature enzyme
Alcohol dehydrogenase
Regulator

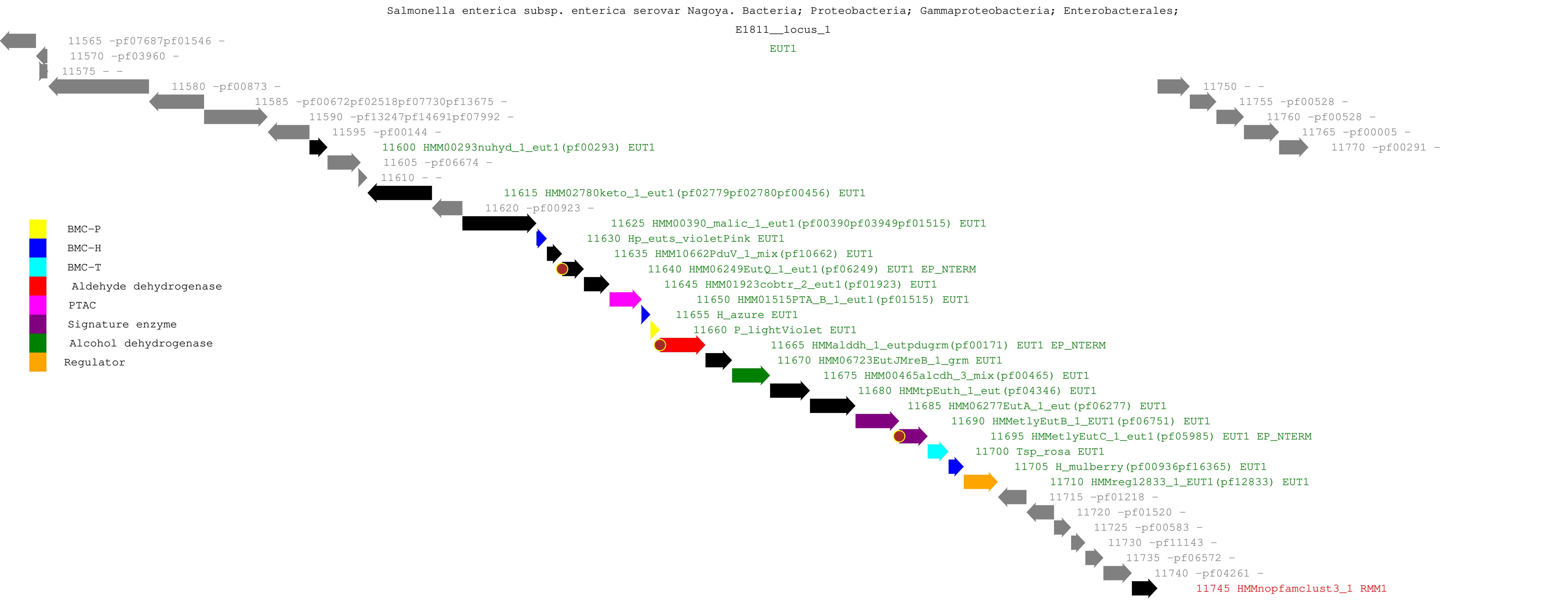
Salmonella enterica subsp. enterica serovar Nagoya. Bacteria; Proteobacteria; Gammaproteobacteria; Enterobacterales;
E1811__locus_1
EUT1
11565 -pf07687pf01546 -
11570 -pf03960 -
11575 - -
11580 -pf00873 -
11585 -pf00672pf02518pf07730pf13675 -
11590 -pf13247pf14691pf07992 -
11595 -pf00144 -
11600 HMM00293nuhyd_1_eut1(pf00293) EUT1
11605 -pf06674 -
11610 - -
11615 HMM02780keto_1_eut1(pf02779pf02780pf00456) EUT1
11620 -pf00923 -
11625 HMM00390_malic_1_eut1(pf00390pf03949pf01515) EUT1
11630 Hp_euts_violetPink EUT1
11635 HMM10662PduV_1_mix(pf10662) EUT1
11640 HMM06249EutQ_1_eut1(pf06249) EUT1 EP_NTERM
11645 HMM01923cobtr_2_eut1(pf01923) EUT1
11650 HMM01515PTA_B_1_eut1(pf01515) EUT1
11655 H_azure EUT1
11660 P_lightViolet EUT1
11665 HMMalddh_1_eutpdugrm(pf00171) EUT1 EP_NTERM
11670 HMM06723EutJMreB_1_grm EUT1
11675 HMM00465alcdh_3_mix(pf00465) EUT1
11680 HMMtpEuth_1_eut(pf04346) EUT1
11685 HMM06277EutA_1_eut(pf06277) EUT1
11690 HMMetlyEutB_1_EUT1(pf06751) EUT1
11695 HMMetlyEutC_1_eut1(pf05985) EUT1 EP_NTERM
11700 Tsp_rosa EUT1
11705 H_mulberry(pf00936pf16365) EUT1
11710 HMMreg12833_1_EUT1(pf12833) EUT1
11715 -pf01218 -
11720 -pf01520 -
11725 -pf00583 -
11730 -pf11143 -
11735 -pf06572 -
11740 -pf04261 -
11745 HMMnopfamclust3_1 RMM1
11750 - -
11755 -pf00528 -
11760 -pf00528 -
11765 -pf00005 -
11770 -pf00291 -
BMC-P
BMC-H
BMC-T
Aldehyde dehydrogenase
PTAC
Signature enzyme
Alcohol dehydrogenase
Regulator

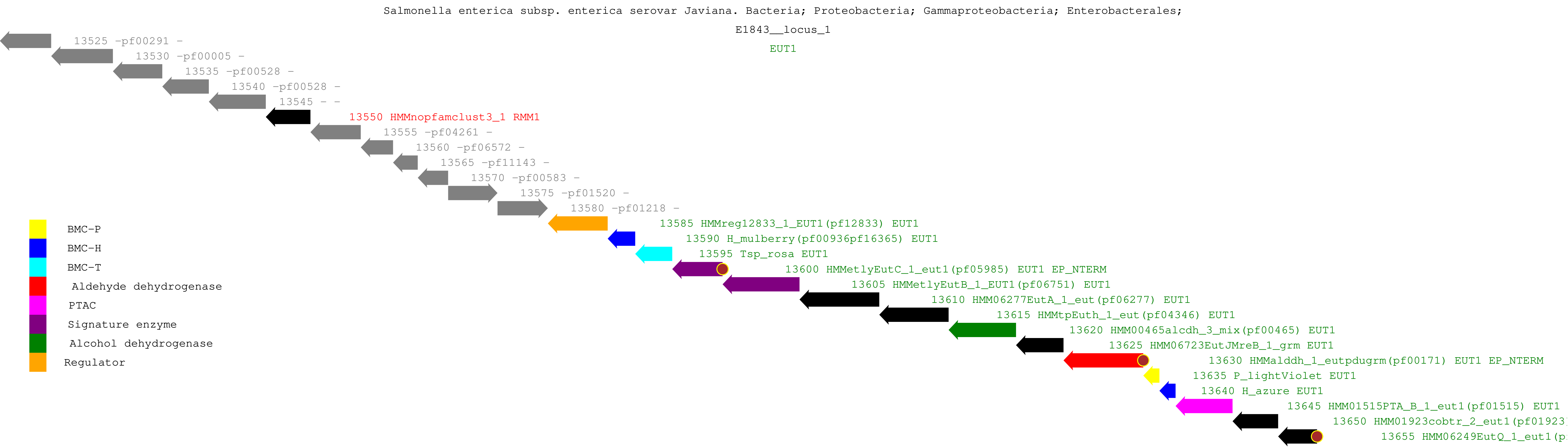

Salmonella enterica subsp. enterica serovar Javiana. Bacteria; Proteobacteria; Gammaproteobacteria; Enterobacterales;
E1843__locus_1
EUT1
13525 -pf00291 -
13530 -pf00005 -
13535 -pf00528 -
13540 -pf00528 -
13545 - -
13550 HMMnopfamclust3_1 RMM1
13555 -pf04261 -
13560 -pf06572 -
13565 -pf11143 -
13570 -pf00583 -
13575 -pf01520 -
13580 -pf01218 -
13585 HMMreg12833_1_EUT1(pf12833) EUT1
13590 H_mulberry(pf00936pf16365) EUT1
13595 Tsp_rosa EUT1
13600 HMMetlyEutC_1_eut1(pf05985) EUT1 EP_NTERM
13605 HMMetlyEutB_1_EUT1(pf06751) EUT1
13610 HMM06277EutA_1_eut(pf06277) EUT1
13615 HMMtpEuth_1_eut(pf04346) EUT1
13620 HMM00465alcdh_3_mix(pf00465) EUT1
13625 HMM06723EutJMreB_1_grm EUT1
13630 HMMalddh_1_eutpdugrm(pf00171) EUT1 EP_NTERM
13635 P_lightViolet EUT1
13640 H_azure EUT1
13645 HMM01515PTA_B_1_eut1(pf01515) EUT1
13650 HMM01923cobtr_2_eut1(pf01923
13655 HMM06249EutQ_1_eut1(p
BMC-P
BMC-H
BMC-T
Aldehyde dehydrogenase
PTAC
Signature enzyme
Alcohol dehydrogenase
Regulator

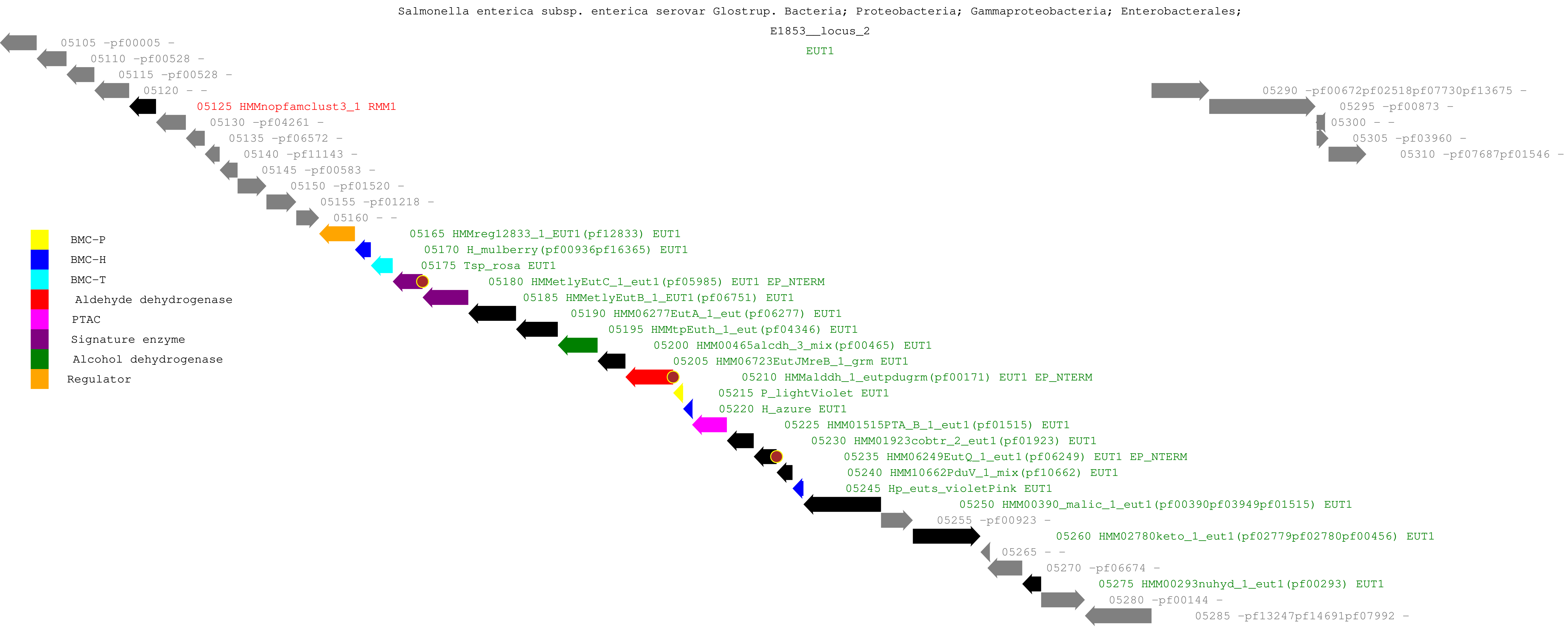
Salmonella enterica subsp. enterica serovar Glostrup. Bacteria; Proteobacteria; Gammaproteobacteria; Enterobacterales;
E1853__locus_2
EUT1
05105 -pf00005 -
05110 -pf00528 -
05115 -pf00528 -
05120 - -
05125 HMMnopfamclust3_1 RMM1
05130 -pf04261 -
05135 -pf06572 -
05140 -pf11143 -
05145 -pf00583 -
05150 -pf01520 -
05155 -pf01218 -
05160 - -
05165 HMMreg12833_1_EUT1(pf12833) EUT1
05170 H_mulberry(pf00936pf16365) EUT1
05175 Tsp_rosa EUT1
05180 HMMetlyEutC_1_eut1(pf05985) EUT1 EP_NTERM
05185 HMMetlyEutB_1_EUT1(pf06751) EUT1
05190 HMM06277EutA_1_eut(pf06277) EUT1
05195 HMMtpEuth_1_eut(pf04346) EUT1
05200 HMM00465alcdh_3_mix(pf00465) EUT1
05205 HMM06723EutJMreB_1_grm EUT1
05210 HMMalddh_1_eutpdugrm(pf00171) EUT1 EP_NTERM
05215 P_lightViolet EUT1
05220 H_azure EUT1
05225 HMM01515PTA_B_1_eut1(pf01515) EUT1
05230 HMM01923cobtr_2_eut1(pf01923) EUT1
05235 HMM06249EutQ_1_eut1(pf06249) EUT1 EP_NTERM
05240 HMM10662PduV_1_mix(pf10662) EUT1
05245 Hp_euts_violetPink EUT1
05250 HMM00390_malic_1_eut1(pf00390pf03949pf01515) EUT1
05255 -pf00923 -
05260 HMM02780keto_1_eut1(pf02779pf02780pf00456) EUT1
05265 - -
05270 -pf06674 -
05275 HMM00293nuhyd_1_eut1(pf00293) EUT1
05280 -pf00144 -
05285 -pf13247pf14691pf07992 -
05290 -pf00672pf02518pf07730pf13675 -
05295 -pf00873 -
05300 - -
05305 -pf03960 -
05310 -pf07687pf01546 -
BMC-P
BMC-H
BMC-T
Aldehyde dehydrogenase
PTAC
Signature enzyme
Alcohol dehydrogenase
Regulator

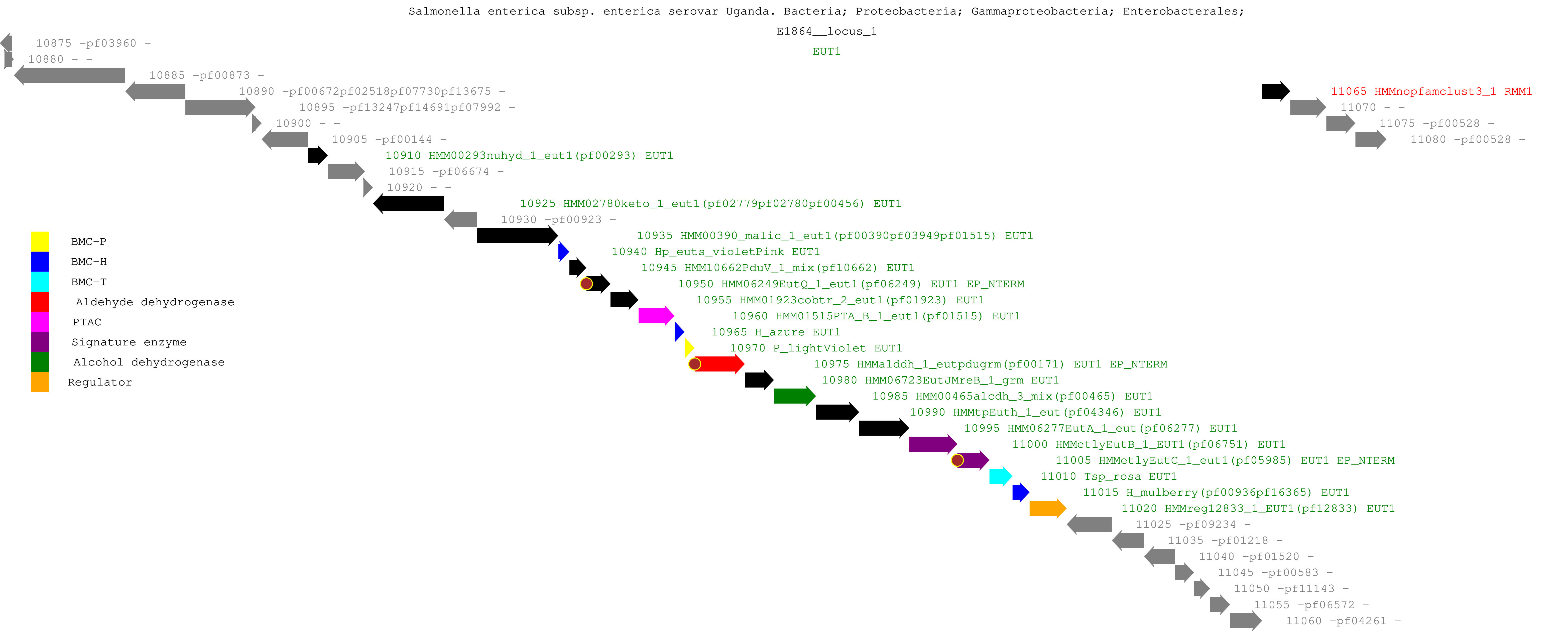
Salmonella enterica subsp. enterica serovar Uganda. Bacteria; Proteobacteria; Gammaproteobacteria; Enterobacterales;
E1864__locus_1
EUT1
10875 -pf03960 -
10880 - -
10885 -pf00873 -
10890 -pf00672pf02518pf07730pf13675 -
10895 -pf13247pf14691pf07992 -
10900 - -
10905 -pf00144 -
10910 HMM00293nuhyd_1_eut1(pf00293) EUT1
10915 -pf06674 -
10920 - -
10925 HMM02780keto_1_eut1(pf02779pf02780pf00456) EUT1
10930 -pf00923 -
10935 HMM00390_malic_1_eut1(pf00390pf03949pf01515) EUT1
10940 Hp_euts_violetPink EUT1
10945 HMM10662PduV_1_mix(pf10662) EUT1
10950 HMM06249EutQ_1_eut1(pf06249) EUT1 EP_NTERM
10955 HMM01923cobtr_2_eut1(pf01923) EUT1
10960 HMM01515PTA_B_1_eut1(pf01515) EUT1
10965 H_azure EUT1
10970 P_lightViolet EUT1
10975 HMMalddh_1_eutpdugrm(pf00171) EUT1 EP_NTERM
10980 HMM06723EutJMreB_1_grm EUT1
10985 HMM00465alcdh_3_mix(pf00465) EUT1
10990 HMMtpEuth_1_eut(pf04346) EUT1
10995 HMM06277EutA_1_eut(pf06277) EUT1
11000 HMMetlyEutB_1_EUT1(pf06751) EUT1
11005 HMMetlyEutC_1_eut1(pf05985) EUT1 EP_NTERM
11010 Tsp_rosa EUT1
11015 H_mulberry(pf00936pf16365) EUT1
11020 HMMreg12833_1_EUT1(pf12833) EUT1
11025 -pf09234 -
11035 -pf01218 -
11040 -pf01520 -
11045 -pf00583 -
11050 -pf11143 -
11055 -pf06572 -
11060 -pf04261 -
11065 HMMnopfamclust3_1 RMM1
11070 - -
11075 -pf00528 -
11080 -pf00528 -
BMC-P
BMC-H
BMC-T
Aldehyde dehydrogenase
PTAC
Signature enzyme
Alcohol dehydrogenase
Regulator

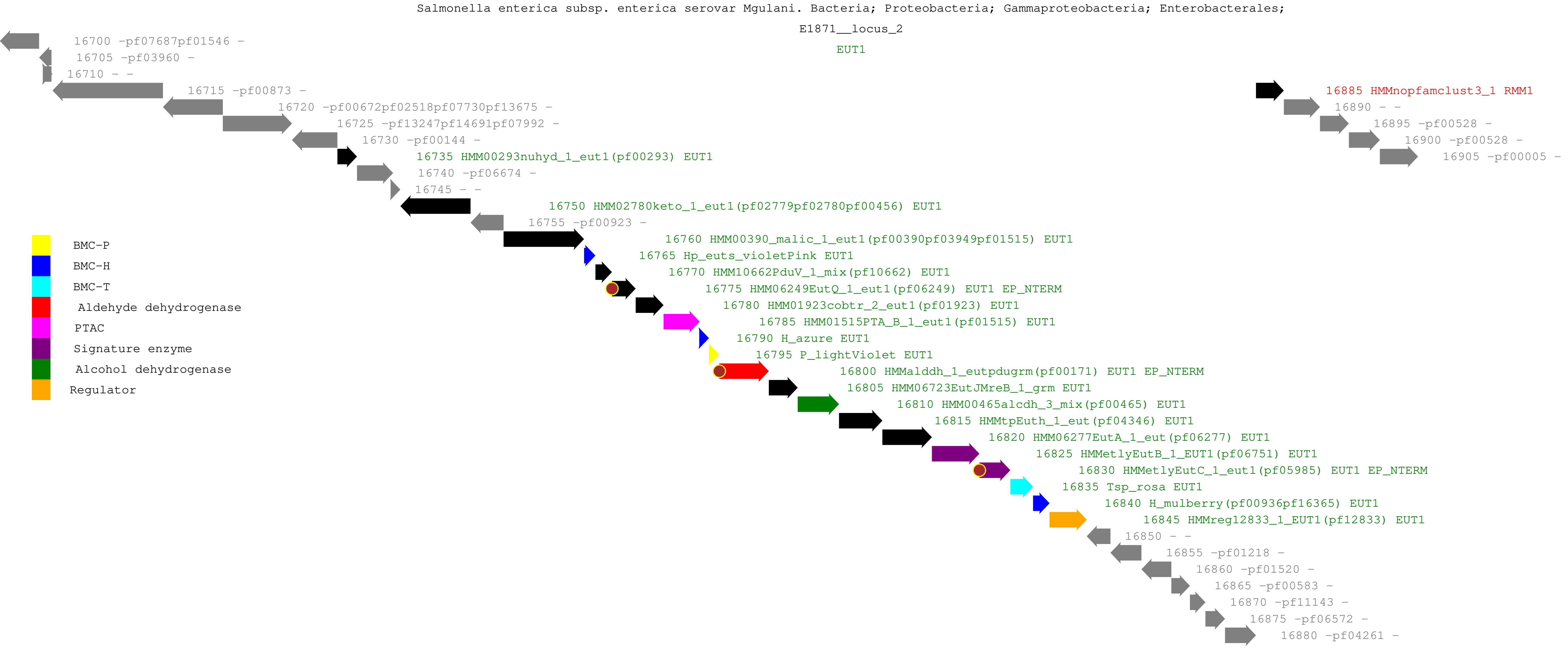
Salmonella enterica subsp. enterica serovar Mgulani. Bacteria; Proteobacteria; Gammaproteobacteria; Enterobacterales;
E1871__locus_2
EUT1
16700 -pf07687pf01546 -
16705 -pf03960 -
16710 - -
16715 -pf00873 -
16720 -pf00672pf02518pf07730pf13675 -
16725 -pf13247pf14691pf07992 -
16730 -pf00144 -
16735 HMM00293nuhyd_1_eut1(pf00293) EUT1
16740 -pf06674 -
16745 - -
16750 HMM02780keto_1_eut1(pf02779pf02780pf00456) EUT1
16755 -pf00923 -
16760 HMM00390_malic_1_eut1(pf00390pf03949pf01515) EUT1
16765 Hp_euts_violetPink EUT1
16770 HMM10662PduV_1_mix(pf10662) EUT1
16775 HMM06249EutQ_1_eut1(pf06249) EUT1 EP_NTERM
16780 HMM01923cobtr_2_eut1(pf01923) EUT1
16785 HMM01515PTA_B_1_eut1(pf01515) EUT1
16790 H_azure EUT1
16795 P_lightViolet EUT1
16800 HMMalddh_1_eutpdugrm(pf00171) EUT1 EP_NTERM
16805 HMM06723EutJMreB_1_grm EUT1
16810 HMM00465alcdh_3_mix(pf00465) EUT1
16815 HMMtpEuth_1_eut(pf04346) EUT1
16820 HMM06277EutA_1_eut(pf06277) EUT1
16825 HMMetlyEutB_1_EUT1(pf06751) EUT1
16830 HMMetlyEutC_1_eut1(pf05985) EUT1 EP_NTERM
16835 Tsp_rosa EUT1
16840 H_mulberry(pf00936pf16365) EUT1
16845 HMMreg12833_1_EUT1(pf12833) EUT1
16850 - -
16855 -pf01218 -
16860 -pf01520 -
16865 -pf00583 -
16870 -pf11143 -
16875 -pf06572 -
16880 -pf04261 -
16885 HMMnopfamclust3_1 RMM1
16890 - -
16895 -pf00528 -
16900 -pf00528 -
16905 -pf00005 -
BMC-P
BMC-H
BMC-T
Aldehyde dehydrogenase
PTAC
Signature enzyme
Alcohol dehydrogenase
Regulator

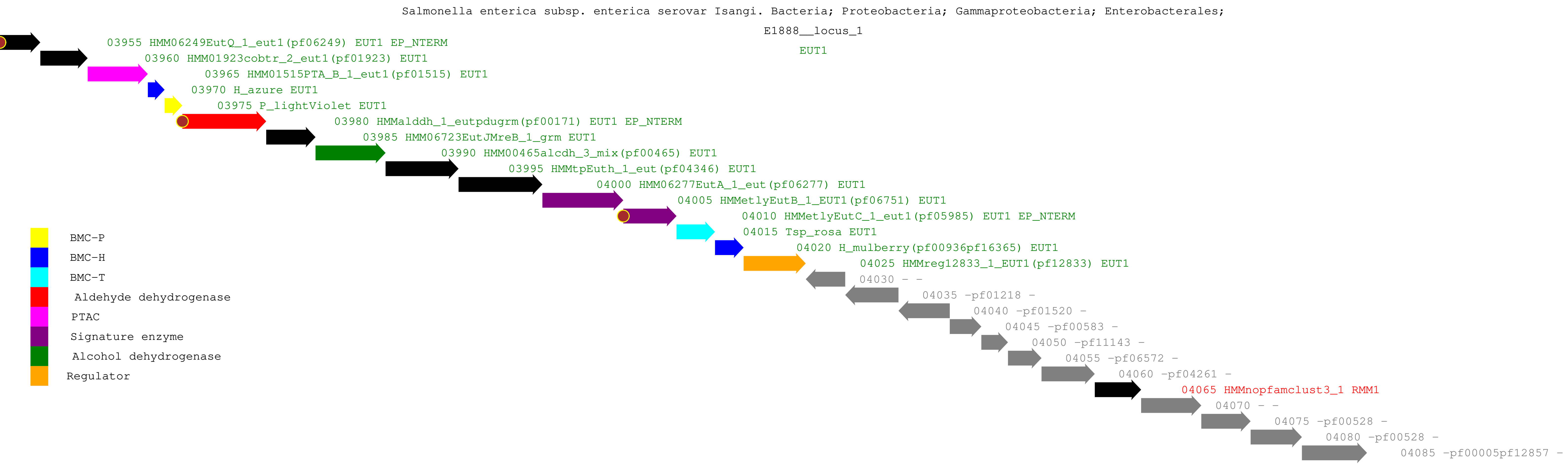
Salmonella enterica subsp. enterica serovar Isangi. Bacteria; Proteobacteria; Gammaproteobacteria; Enterobacterales;
E1888__locus_1
EUT1
03955 HMM06249EutQ_1_eut1(pf06249) EUT1 EP_NTERM
03960 HMM01923cobtr_2_eut1(pf01923) EUT1
03965 HMM01515PTA_B_1_eut1(pf01515) EUT1
03970 H_azure EUT1
03975 P_lightViolet EUT1
03980 HMMalddh_1_eutpdugrm(pf00171) EUT1 EP_NTERM
03985 HMM06723EutJMreB_1_grm EUT1
03990 HMM00465alcdh_3_mix(pf00465) EUT1
03995 HMMtpEuth_1_eut(pf04346) EUT1
04000 HMM06277EutA_1_eut(pf06277) EUT1
04005 HMMetlyEutB_1_EUT1(pf06751) EUT1
04010 HMMetlyEutC_1_eut1(pf05985) EUT1 EP_NTERM
04015 Tsp_rosa EUT1
04020 H_mulberry(pf00936pf16365) EUT1
04025 HMMreg12833_1_EUT1(pf12833) EUT1
04030 - -
04035 -pf01218 -
04040 -pf01520 -
04045 -pf00583 -
04050 -pf11143 -
04055 -pf06572 -
04060 -pf04261 -
04065 HMMnopfamclust3_1 RMM1
04070 - -
04075 -pf00528 -
04080 -pf00528 -
04085 -pf00005pf12857 -
BMC-P
BMC-H
BMC-T
Aldehyde dehydrogenase
PTAC
Signature enzyme
Alcohol dehydrogenase
Regulator

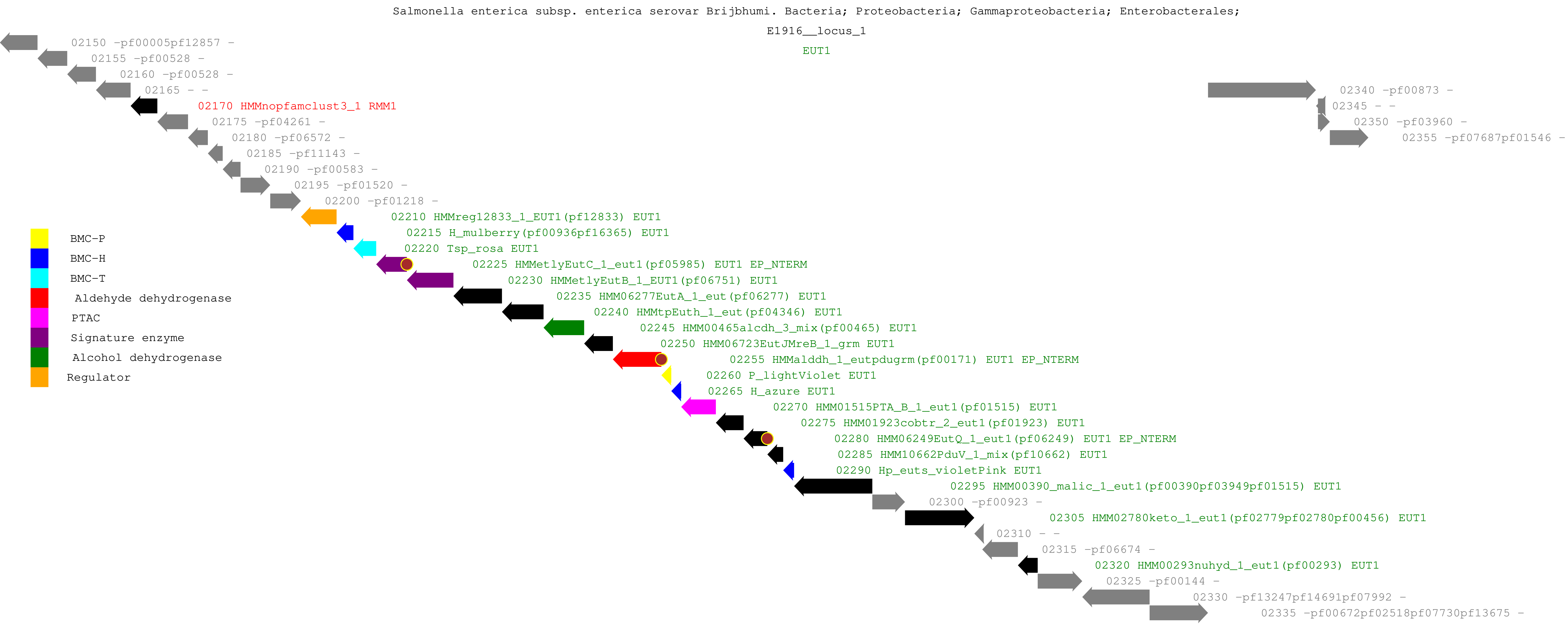
Salmonella enterica subsp. enterica serovar Brijbhumi. Bacteria; Proteobacteria; Gammaproteobacteria; Enterobacterales;
E1916__locus_1
EUT1
02150 -pf00005pf12857 -
02155 -pf00528 -
02160 -pf00528 -
02165 - -
02170 HMMnopfamclust3_1 RMM1
02175 -pf04261 -
02180 -pf06572 -
02185 -pf11143 -
02190 -pf00583 -
02195 -pf01520 -
02200 -pf01218 -
02210 HMMreg12833_1_EUT1(pf12833) EUT1
02215 H_mulberry(pf00936pf16365) EUT1
02220 Tsp_rosa EUT1
02225 HMMetlyEutC_1_eut1(pf05985) EUT1 EP_NTERM
02230 HMMetlyEutB_1_EUT1(pf06751) EUT1
02235 HMM06277EutA_1_eut(pf06277) EUT1
02240 HMMtpEuth_1_eut(pf04346) EUT1
02245 HMM00465alcdh_3_mix(pf00465) EUT1
02250 HMM06723EutJMreB_1_grm EUT1
02255 HMMalddh_1_eutpdugrm(pf00171) EUT1 EP_NTERM
02260 P_lightViolet EUT1
02265 H_azure EUT1
02270 HMM01515PTA_B_1_eut1(pf01515) EUT1
02275 HMM01923cobtr_2_eut1(pf01923) EUT1
02280 HMM06249EutQ_1_eut1(pf06249) EUT1 EP_NTERM
02285 HMM10662PduV_1_mix(pf10662) EUT1
02290 Hp_euts_violetPink EUT1
02295 HMM00390_malic_1_eut1(pf00390pf03949pf01515) EUT1
02300 -pf00923 -
02305 HMM02780keto_1_eut1(pf02779pf02780pf00456) EUT1
02310 - -
02315 -pf06674 -
02320 HMM00293nuhyd_1_eut1(pf00293) EUT1
02325 -pf00144 -
02330 -pf13247pf14691pf07992 -
02335 -pf00672pf02518pf07730pf13675 -
02340 -pf00873 -
02345 - -
02350 -pf03960 -
02355 -pf07687pf01546 -
BMC-P
BMC-H
BMC-T
Aldehyde dehydrogenase
PTAC
Signature enzyme
Alcohol dehydrogenase
Regulator

Salmonella enterica subsp. enterica serovar Ahuza. Bacteria; Proteobacteria; Gammaproteobacteria; Enterobacterales;

E1944__locus_2

EUT1

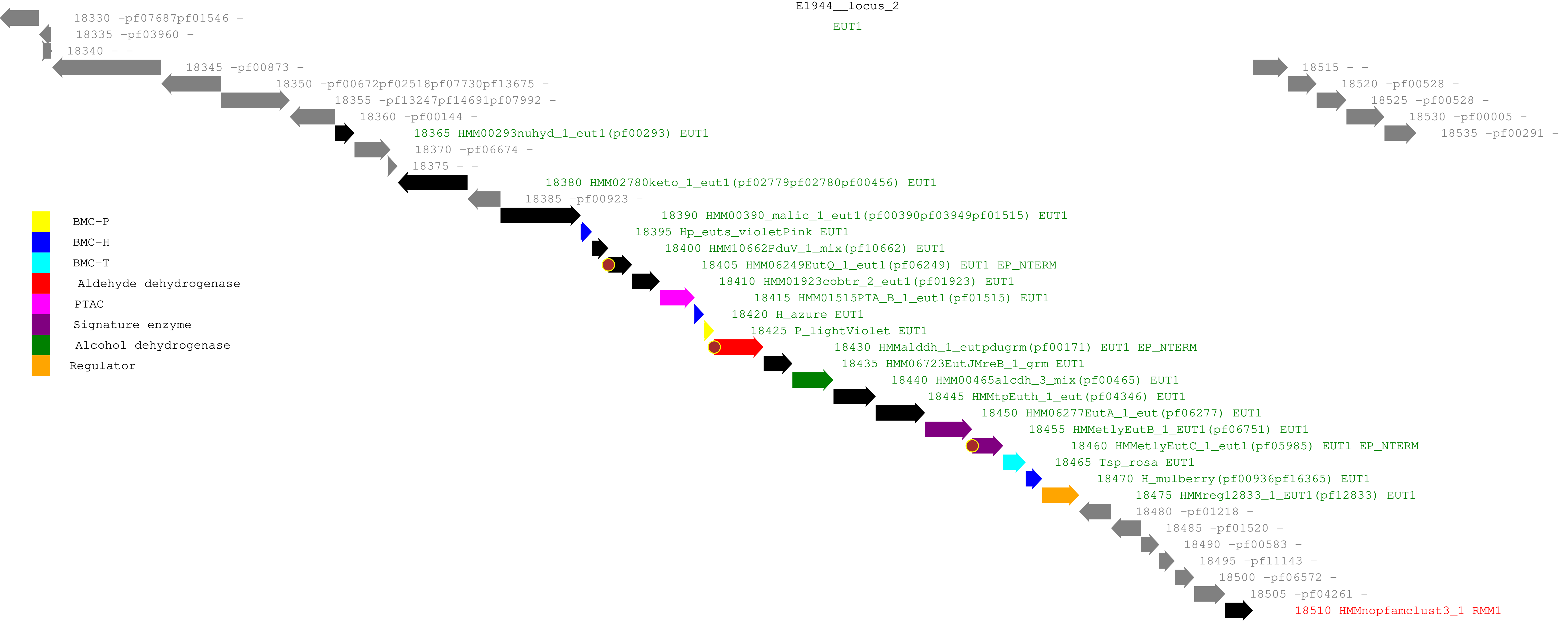

Salmonella enterica subsp. enterica serovar Cubana. Bacteria; Proteobacteria; Gammaproteobacteria; Enterobacterales;

E1A24__locus_2

EUT1

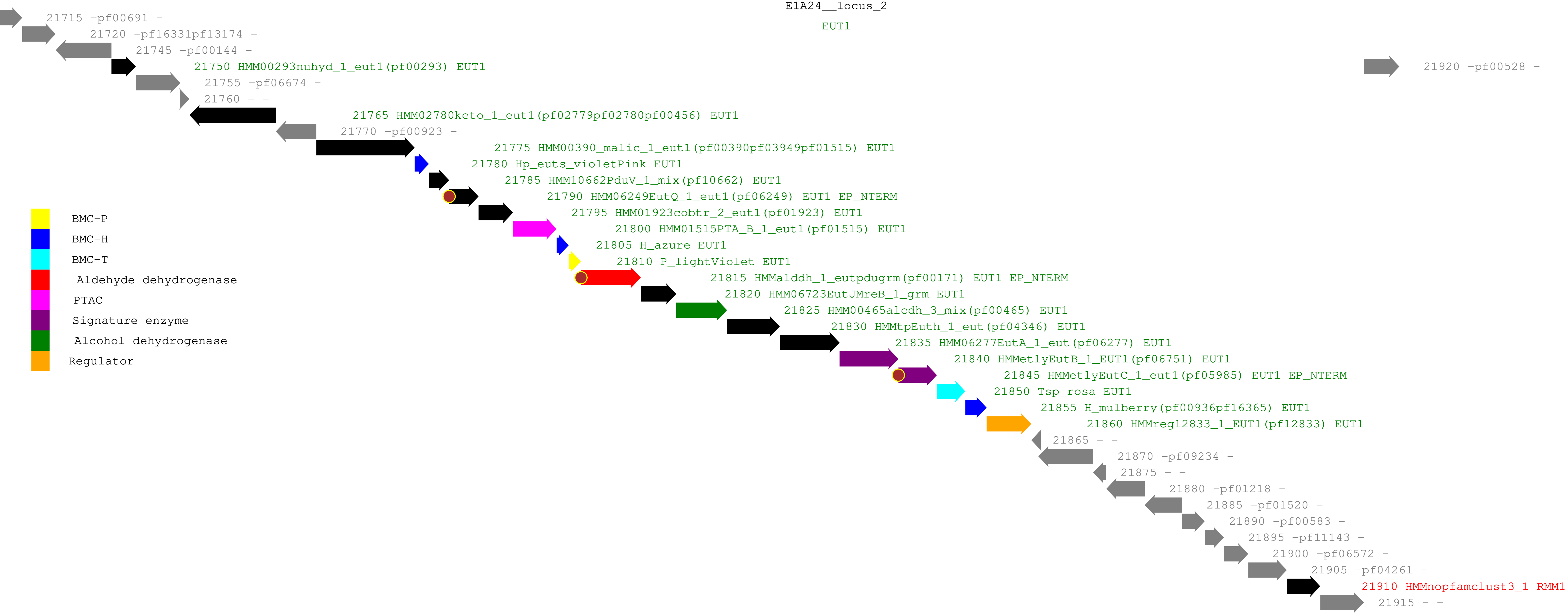

Citrobacter sp. LY-1. Bacteria; Proteobacteria; Gammaproteobacteria; Enterobacterales;

E1B03__locus_2

EUT1

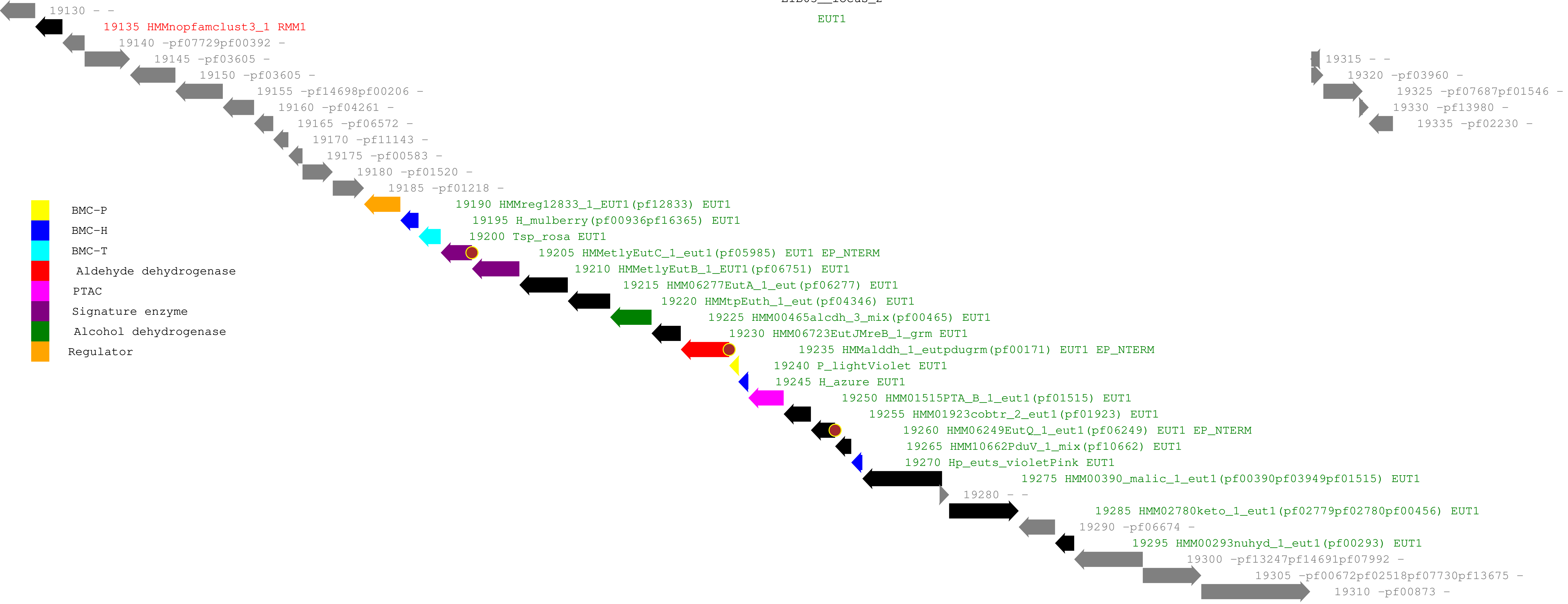

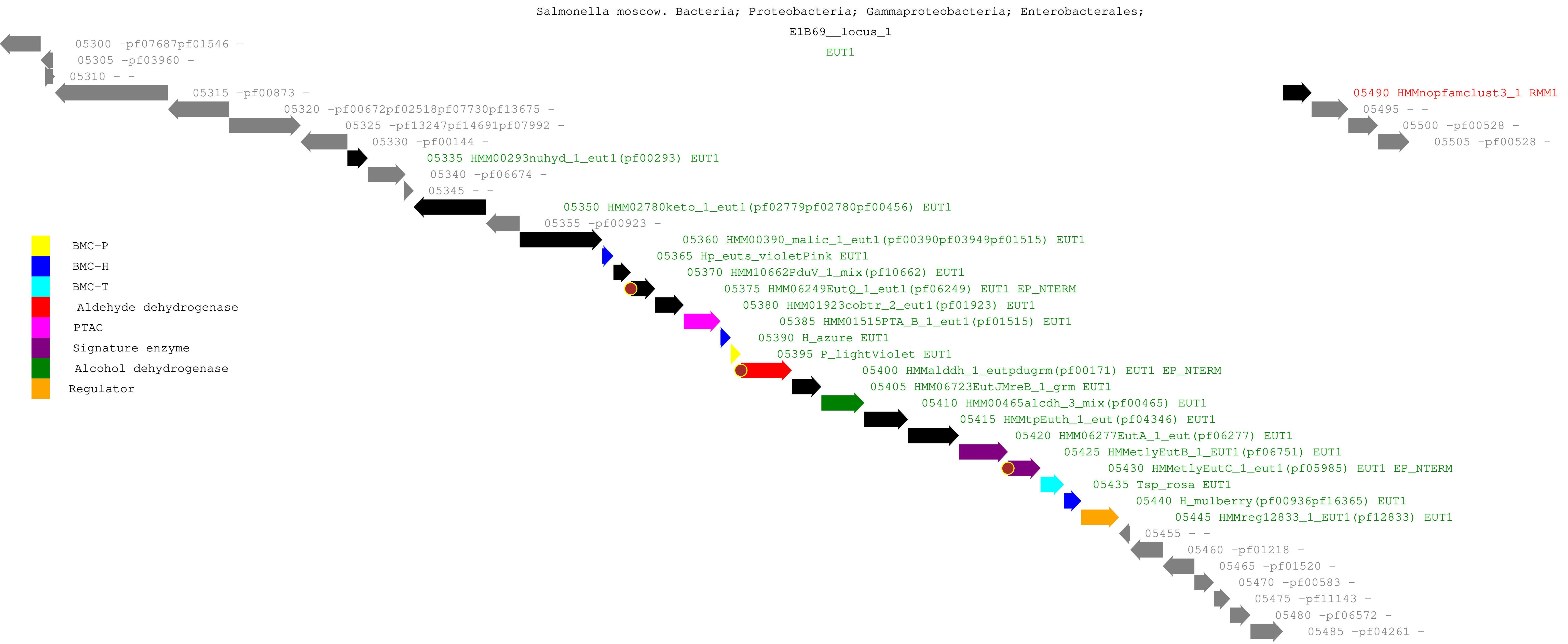
Salmonella moscow. Bacteria; Proteobacteria; Gammaproteobacteria; Enterobacterales;
E1B69__locus_1
EUT1
05300 -pf07687pf01546 -
05305 -pf03960 -
05310 - -
05315 -pf00873 -
05320 -pf00672pf02518pf07730pf13675 -
05325 -pf13247pf14691pf07992 -
05330 -pf00144 -
05335 HMM00293nuhyd_1_eut1(pf00293) EUT1
05340 -pf06674 -
05345 - -
05350 HMM02780keto_1_eut1(pf02779pf02780pf00456) EUT1
05355 -pf00923 -
05360 HMM00390_malic_1_eut1(pf00390pf03949pf01515) EUT1
05365 Hp_euts_violetPink EUT1
05370 HMM10662PduV_1_mix(pf10662) EUT1
05375 HMM06249EutQ_1_eut1(pf06249) EUT1 EP_NTERM
05380 HMM01923cobtr_2_eut1(pf01923) EUT1
05385 HMM01515PTA_B_1_eut1(pf01515) EUT1
05390 H_azure EUT1
05395 P_lightViolet EUT1
05400 HMMaldh_1_eutpdugrm(pf00171) EUT1 EP_NTERM
05405 HMM06723EutJMreB_1_grm EUT1
05410 HMM00465alcdh_3_mix(pf00465) EUT1
05415 HMMtpEuth_1_eut(pf04346) EUT1
05420 HMM06277EutA_1_eut(pf06277) EUT1
05425 HMMetlyEutB_1_EUT1(pf06751) EUT1
05430 HMMetlyEutC_1_eut1(pf05985) EUT1 EP_NTERM
05435 Tsp_rosa EUT1
05440 H_mulberry(pf00936pf16365) EUT1
05445 HMMreg12833_1_EUT1(pf12833) EUT1
05455 - -
05460 -pf01218 -
05465 -pf01520 -
05470 -pf00583 -
05475 -pf11143 -
05480 -pf06572 -
05485 -pf04261 -
05490 HMMnopfamclust3_1 RMM1
05495 - -
05500 -pf00528 -
05505 -pf00528 -
BMC-P
BMC-H
BMC-T
Aldehyde dehydrogenase
PTAC
Signature enzyme
Alcohol dehydrogenase
Regulator

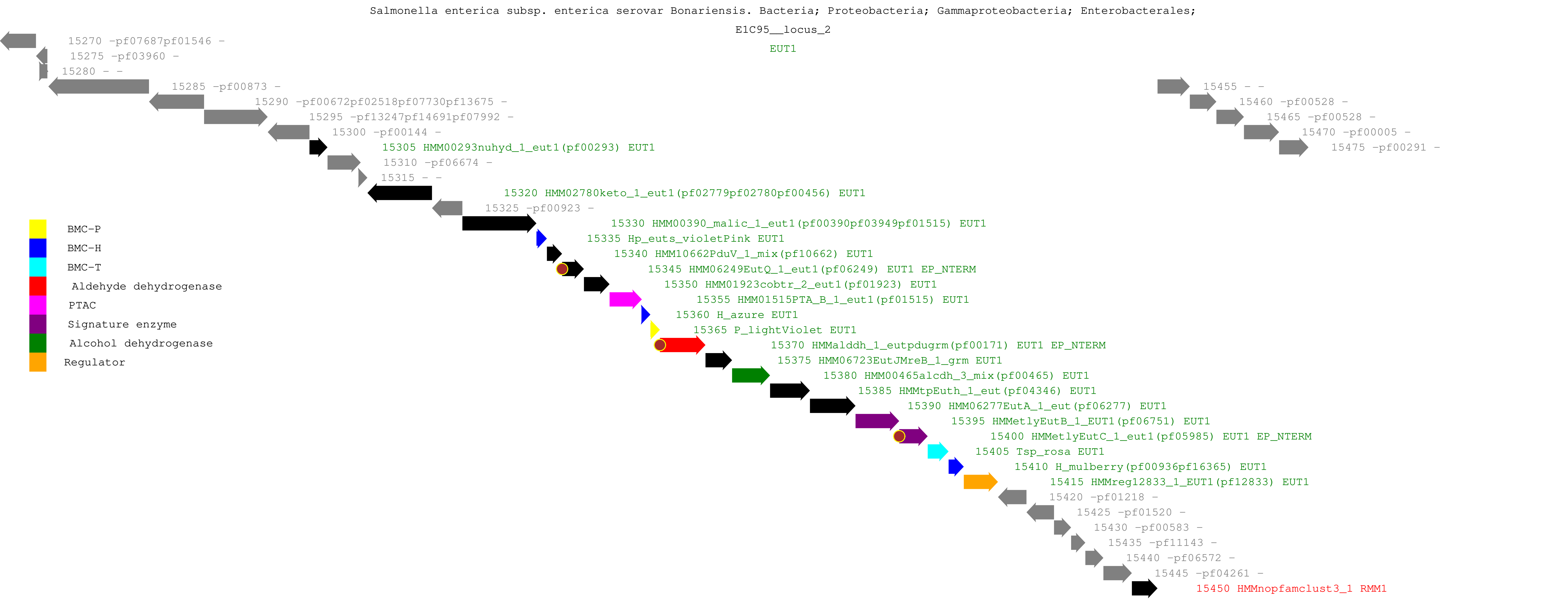

Salmonella enterica subsp. enterica serovar Bonariensis. Bacteria; Proteobacteria; Gammaproteobacteria; Enterobacterales;
E1C95__locus_2
EUT1
15270 -pf07687pf01546 -
15275 -pf03960 -
15280 - -
15285 -pf00873 -
15290 -pf00672pf02518pf07730pf13675 -
15295 -pf13247pf14691pf07992 -
15300 -pf00144 -
15305 HMM00293nuhyd_1_eut1(pf00293) EUT1
15310 -pf06674 -
15315 - -
15320 HMM02780keto_1_eut1(pf02779pf02780pf00456) EUT1
15325 -pf00923 -
15330 HMM00390_malic_1_eut1(pf00390pf03949pf01515) EUT1
15335 Hp_euts_violetPink EUT1
15340 HMM10662PduV_1_mix(pf10662) EUT1
15345 HMM06249EutQ_1_eut1(pf06249) EUT1 EP_NTERM
15350 HMM01923cobtr_2_eut1(pf01923) EUT1
15355 HMM01515PTA_B_1_eut1(pf01515) EUT1
15360 H_azure EUT1
15365 P_lightViolet EUT1
15370 HMMalddh_1_eutpdugrm(pf00171) EUT1 EP_NTERM
15375 HMM06723EutJMreB_1_grm EUT1
15380 HMM00465alcdh_3_mix(pf00465) EUT1
15385 HMMtpEuth_1_eut(pf04346) EUT1
15390 HMM06277EutA_1_eut(pf06277) EUT1
15395 HMMetlyEutB_1_EUT1(pf06751) EUT1
15400 HMMetlyEutC_1_eut1(pf05985) EUT1 EP_NTERM
15405 Tsp_rosa EUT1
15410 H_mulberry(pf00936pf16365) EUT1
15415 HMMreg12833_1_EUT1(pf12833) EUT1
15420 -pf01218 -
15425 -pf01520 -
15430 -pf00583 -
15435 -pf11143 -
15440 -pf06572 -
15445 -pf04261 -
15450 HMMnopfamclust3_1 RMM1
15455 - -
15460 -pf00528 -
15465 -pf00528 -
15470 -pf00005 -
15475 -pf00291 -
BMC-P
BMC-H
BMC-T
Aldehyde dehydrogenase
PTAC
Signature enzyme
Alcohol dehydrogenase
Regulator

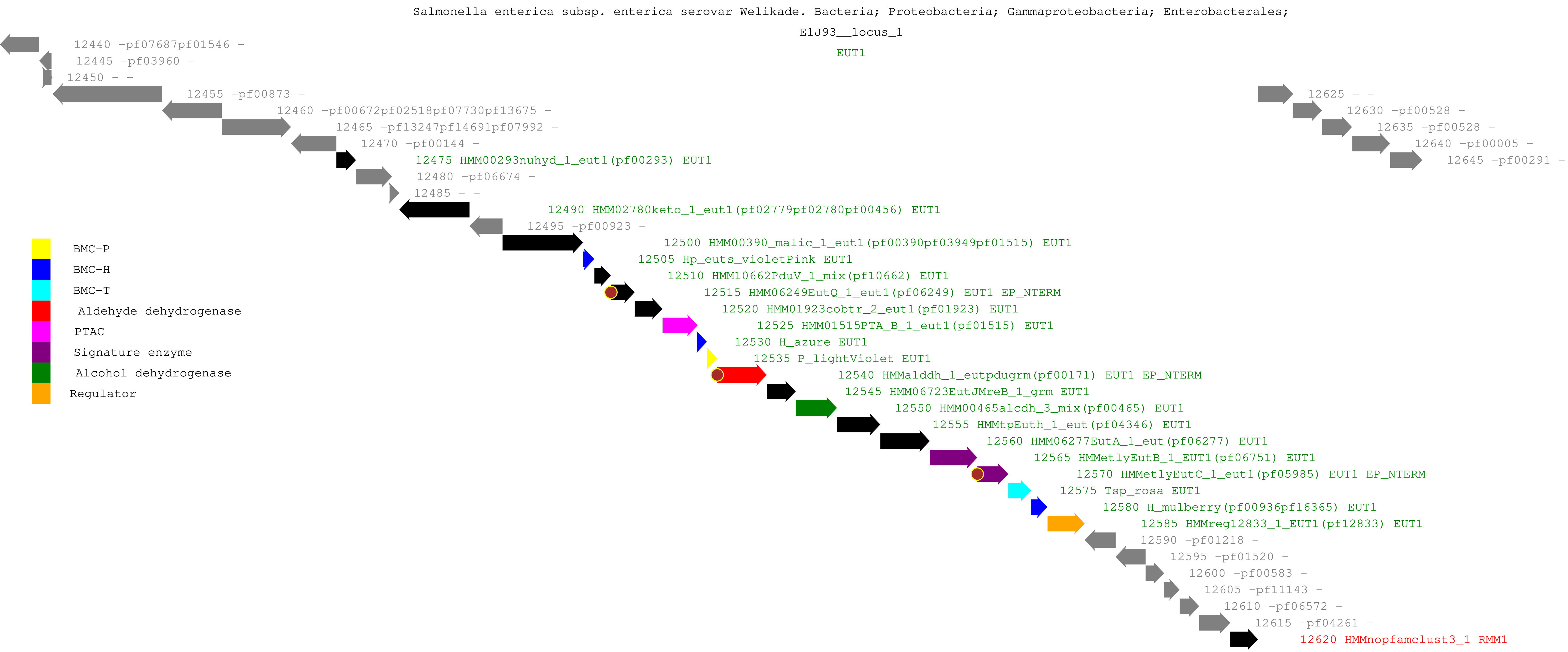
Salmonella enterica subsp. enterica serovar Welikade. Bacteria; Proteobacteria; Gammaproteobacteria; Enterobacterales;
E1J93__locus_1
EUT1
12440 -pf07687pf01546 -
12445 -pf03960 -
12450 - -
12455 -pf00873 -
12460 -pf00672pf02518pf07730pf13675 -
12465 -pf13247pf14691pf07992 -
12470 -pf00144 -
12475 HMM00293nuhyd_1_eut1(pf00293) EUT1
12480 -pf06674 -
12485 - -
12490 HMM02780keto_1_eut1(pf02779pf02780pf00456) EUT1
12495 -pf00923 -
12500 HMM00390_malic_1_eut1(pf00390pf03949pf01515) EUT1
12505 Hp_euts_violetPink EUT1
12510 HMM10662PduV_1_mix(pf10662) EUT1
12515 HMM06249EutQ_1_eut1(pf06249) EUT1 EP_NTERM
12520 HMM01923cobtr_2_eut1(pf01923) EUT1
12525 HMM01515PTA_B_1_eut1(pf01515) EUT1
12530 H_azure EUT1
12535 P_lightViolet EUT1
12540 HMMalddh_1_eutpdugrm(pf00171) EUT1 EP_NTERM
12545 HMM06723EutJMreB_1_grm EUT1
12550 HMM00465alcdh_3_mix(pf00465) EUT1
12555 HMMtpEuth_1_eut(pf04346) EUT1
12560 HMM06277EutA_1_eut(pf06277) EUT1
12565 HMMetlyEutB_1_EUT1(pf06751) EUT1
12570 HMMetlyEutC_1_eut1(pf05985) EUT1 EP_NTERM
12575 Tsp_rosa EUT1
12580 H_mulberry(pf00936pf16365) EUT1
12585 HMMreg12833_1_EUT1(pf12833) EUT1
12590 -pf01218 -
12595 -pf01520 -
12600 -pf00583 -
12605 -pf11143 -
12610 -pf06572 -
12615 -pf04261 -
12620 HMMnopfamclust3_1 RMM1
12625 - -
12630 -pf00528 -
12635 -pf00528 -
12640 -pf00005 -
12645 -pf00291 -
BMC-P
BMC-H
BMC-T
Aldehyde dehydrogenase
PTAC
Signature enzyme
Alcohol dehydrogenase
Regulator

Salmonella enterica subsp. enterica serovar Amager. Bacteria; Proteobacteria; Gammaproteobacteria; Enterobacterales;

E1K22__locus_2

EUT1

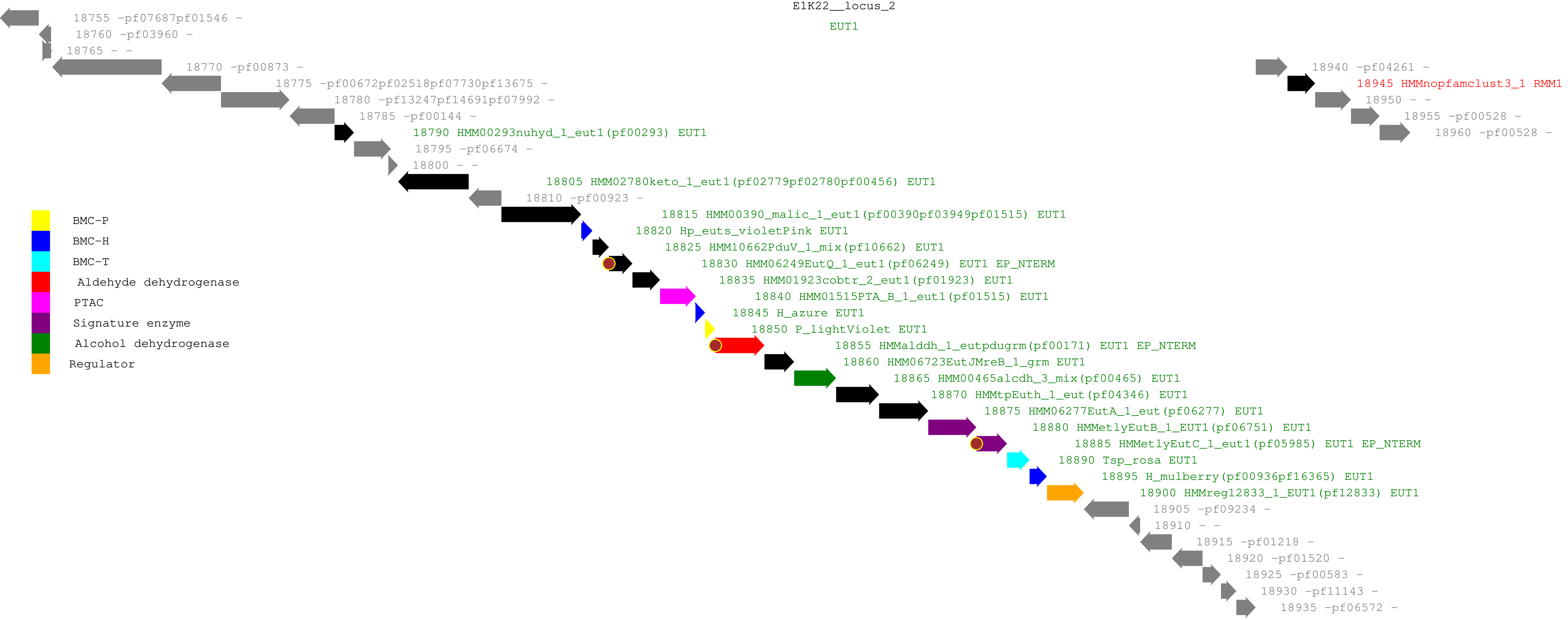

Salmonella enterica subsp. enterica serovar Ruiru. Bacteria; Proteobacteria; Gammaproteobacteria; Enterobacterales;

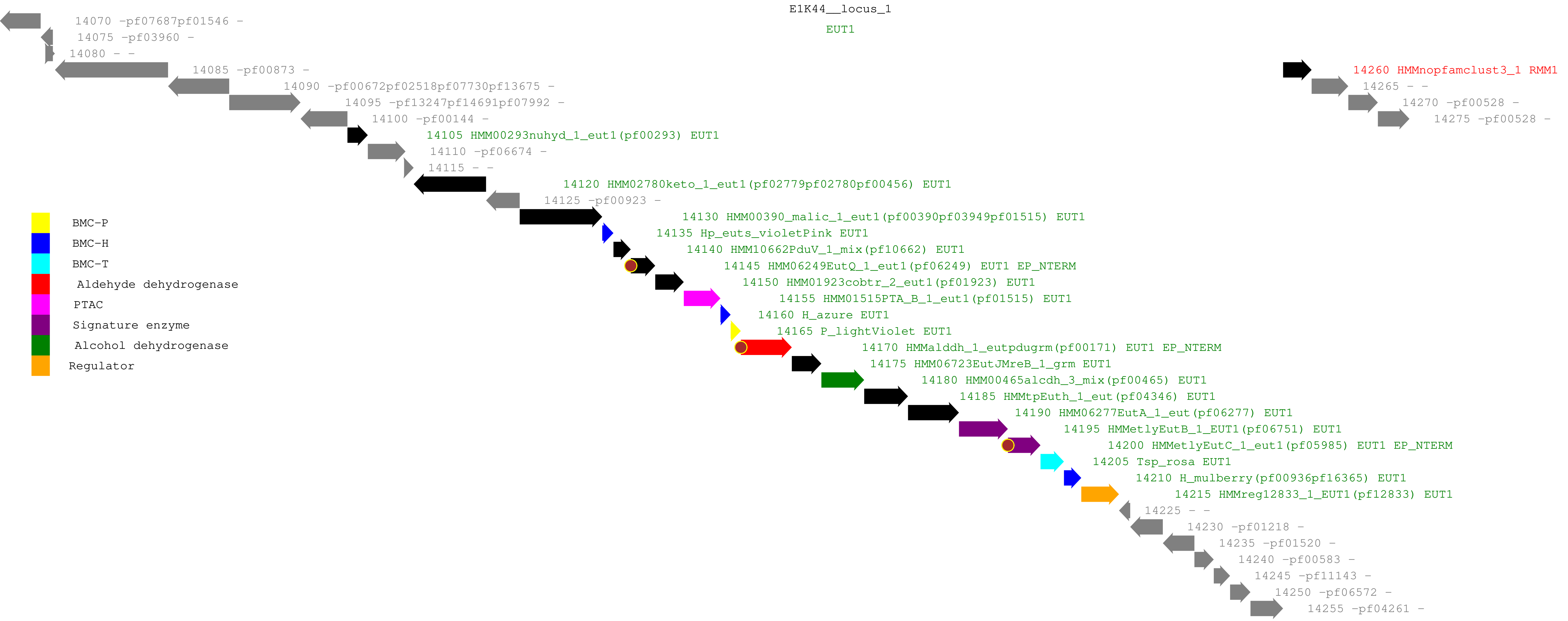

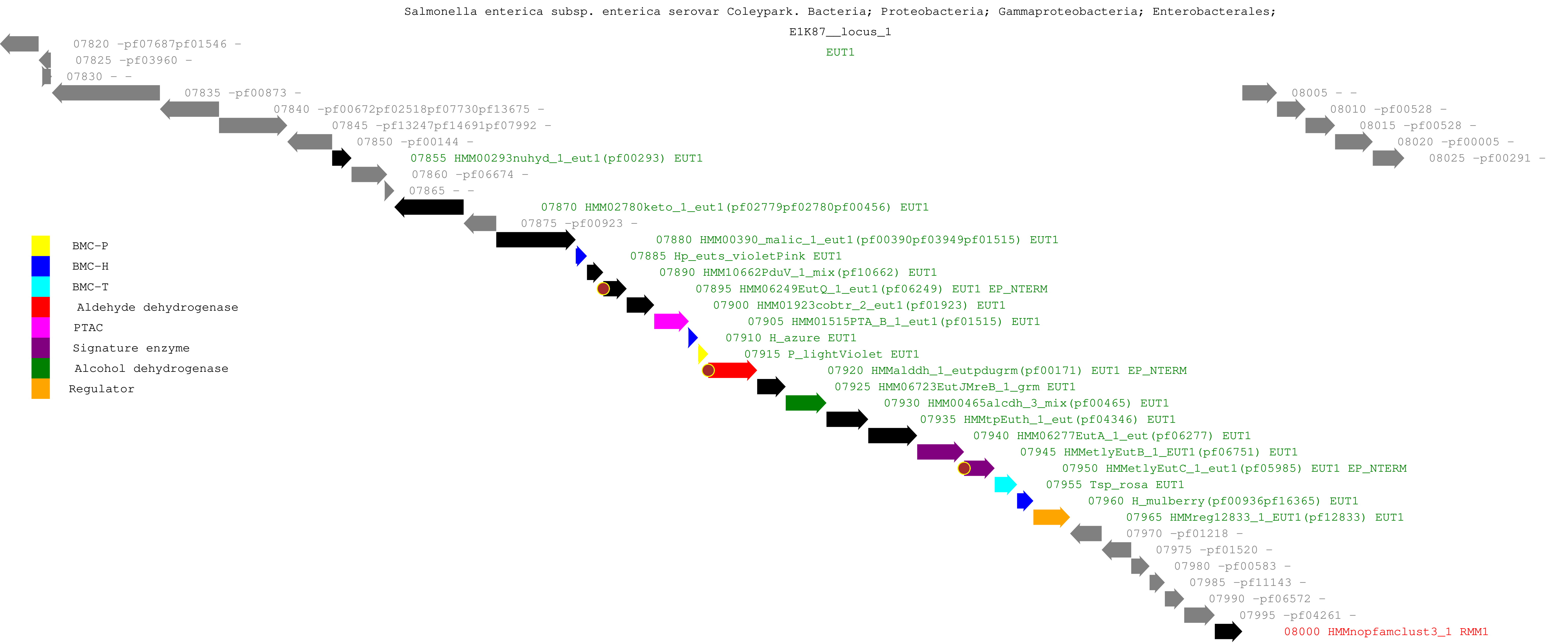
Salmonella enterica subsp. enterica serovar Coleypark. Bacteria; Proteobacteria; Gammaproteobacteria; Enterobacterales;
E1K87__locus_1
EUT1
07820 -pf07687pf01546 -
07825 -pf03960 -
07830 - -
07835 -pf00873 -
07840 -pf00672pf02518pf07730pf13675 -
07845 -pf13247pf14691pf07992 -
07850 -pf00144 -
07855 HMM00293nuhyd_1_eut1(pf00293) EUT1
07860 -pf06674 -
07865 - -
07870 HMM02780keto_1_eut1(pf02779pf02780pf00456) EUT1
07875 -pf00923 -
07880 HMM00390_malic_1_eut1(pf00390pf03949pf01515) EUT1
07885 Hp_euts_violetPink EUT1
07890 HMM10662PduV_1_mix(pf10662) EUT1
07895 HMM06249EutQ_1_eut1(pf06249) EUT1 EP_NTERM
07900 HMM01923cobtr_2_eut1(pf01923) EUT1
07905 HMM01515PTA_B_1_eut1(pf01515) EUT1
07910 H_azure EUT1
07915 P_lightViolet EUT1
07920 HMMalddh_1_eutpdugrm(pf00171) EUT1 EP_NTERM
07925 HMM06723EutJMreB_1_grm EUT1
07930 HMM00465alcdh_3_mix(pf00465) EUT1
07935 HMMtpEuth_1_eut(pf04346) EUT1
07940 HMM06277EutA_1_eut(pf06277) EUT1
07945 HMMetlyEutB_1_EUT1(pf06751) EUT1
07950 HMMetlyEutC_1_eut1(pf05985) EUT1 EP_NTERM
07955 Tsp_rosa EUT1
07960 H_mulberry(pf00936pf16365) EUT1
07965 HMMreg12833_1_EUT1(pf12833) EUT1
07970 -pf01218 -
07975 -pf01520 -
07980 -pf00583 -
07985 -pf11143 -
07990 -pf06572 -
07995 -pf04261 -
08000 HMMnopfamclust3_1 RMM1
08005 - -
08010 -pf00528 -
08015 -pf00528 -
08020 -pf00005 -
08025 -pf00291 -
BMC-P
BMC-H
BMC-T
Aldehyde dehydrogenase
PTAC
Signature enzyme
Alcohol dehydrogenase
Regulator

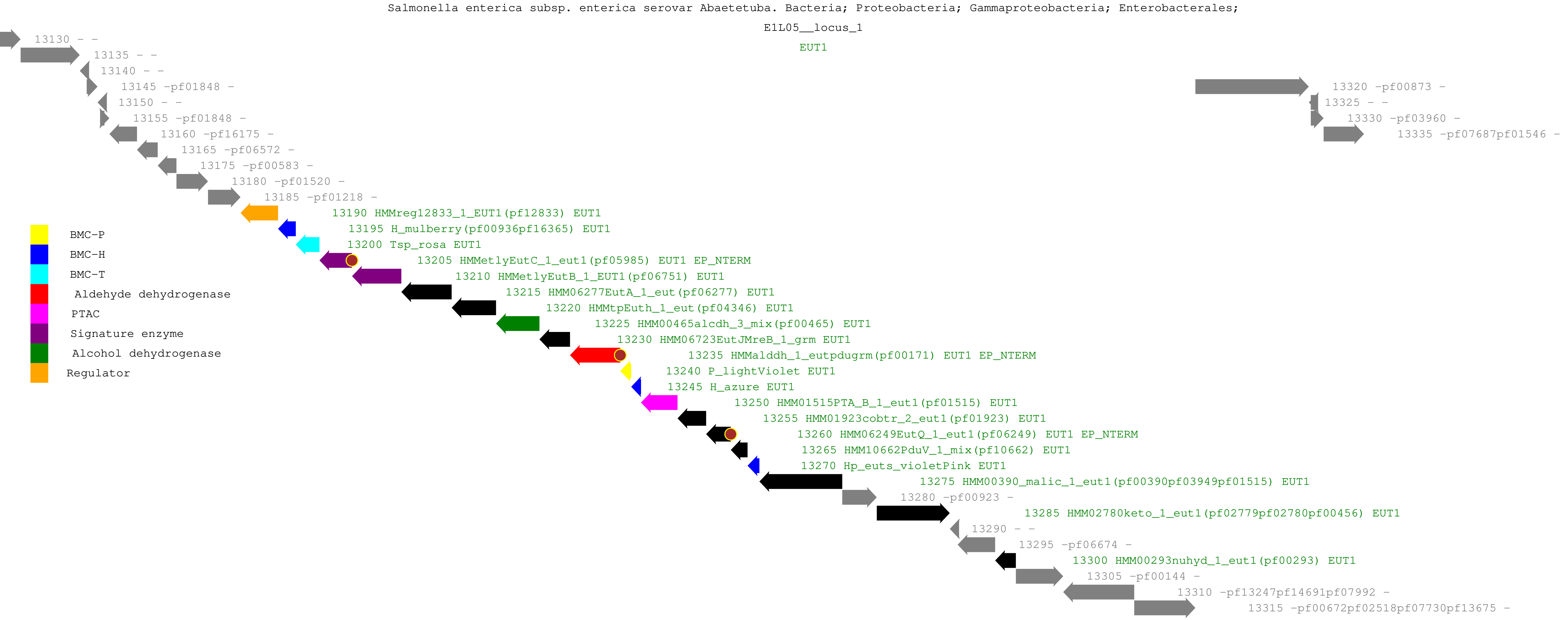

Salmonella enterica subsp. enterica serovar Abaetetuba. Bacteria; Proteobacteria; Gammaproteobacteria; Enterobacterales;
E1L05__locus_1
EUT1
13130 - -
13135 - -
13140 - -
13145 -pf01848 -
13150 - -
13155 -pf01848 -
13160 -pf16175 -
13165 -pf06572 -
13175 -pf00583 -
13180 -pf01520 -
13185 -pf01218 -
13190 HMMreg12833_1_EUT1(pf12833) EUT1
13195 H_mulberry(pf00936pf16365) EUT1
13200 Tsp_rosa EUT1
13205 HMMetlyEutC_1_eut1(pf05985) EUT1 EP_NTERM
13210 HMMetlyEutB_1_EUT1(pf06751) EUT1
13215 HMM06277EutA_1_eut(pf06277) EUT1
13220 HMMtpEuth_1_eut(pf04346) EUT1
13225 HMM00465alcdh_3_mix(pf00465) EUT1
13230 HMM06723EutJMreB_1_grm EUT1
13235 HMMalddh_1_eutpdugrm(pf00171) EUT1 EP_NTERM
13240 P_lightViolet EUT1
13245 H_azure EUT1
13250 HMM01515PTA_B_1_eut1(pf01515) EUT1
13255 HMM01923cobtr_2_eut1(pf01923) EUT1
13260 HMM06249EutQ_1_eut1(pf06249) EUT1 EP_NTERM
13265 HMM10662PduV_1_mix(pf10662) EUT1
13270 Hp_euts_violetPink EUT1
13275 HMM00390_malic_1_eut1(pf00390pf03949pf01515) EUT1
13280 -pf00923 -
13285 HMM02780keto_1_eut1(pf02779pf02780pf00456) EUT1
13290 - -
13295 -pf06674 -
13300 HMM00293nuhyd_1_eut1(pf00293) EUT1
13305 -pf00144 -
13310 -pf13247pf14691pf07992 -
13315 -pf00672pf02518pf07730pf13675 -
13320 -pf00873 -
13325 - -
13330 -pf03960 -
13335 -pf07687pf01546 -
BMC-P
BMC-H
BMC-T
Aldehyde dehydrogenase
PTAC
Signature enzyme
Alcohol dehydrogenase
Regulator

Salmonella enterica subsp. enterica serovar Everleigh. Bacteria; Proteobacteria; Gammaproteobacteria; Enterobacterales;

E1L13__locus_2

EUT1

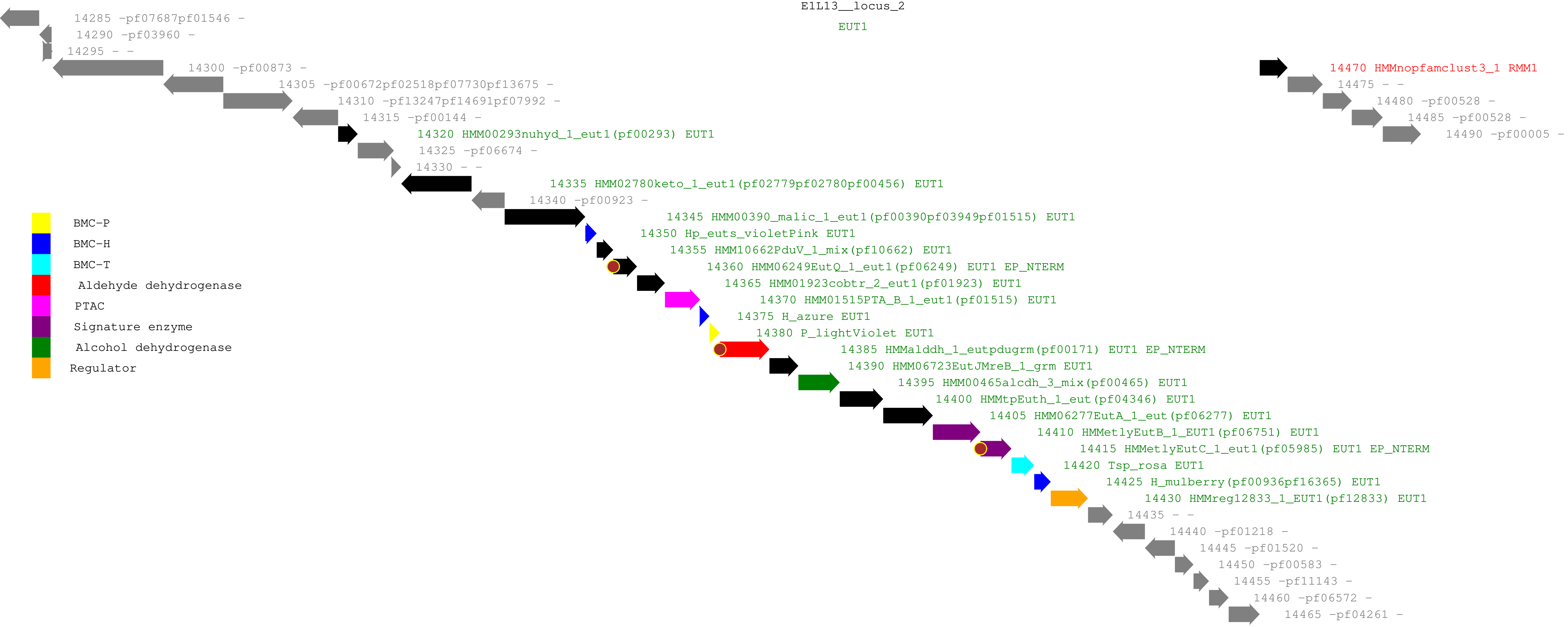

Salmonella enterica subsp. enterica serovar Wien. Bacteria; Proteobacteria; Gammaproteobacteria; Enterobacterales;

E2332__locus_2

EUT1

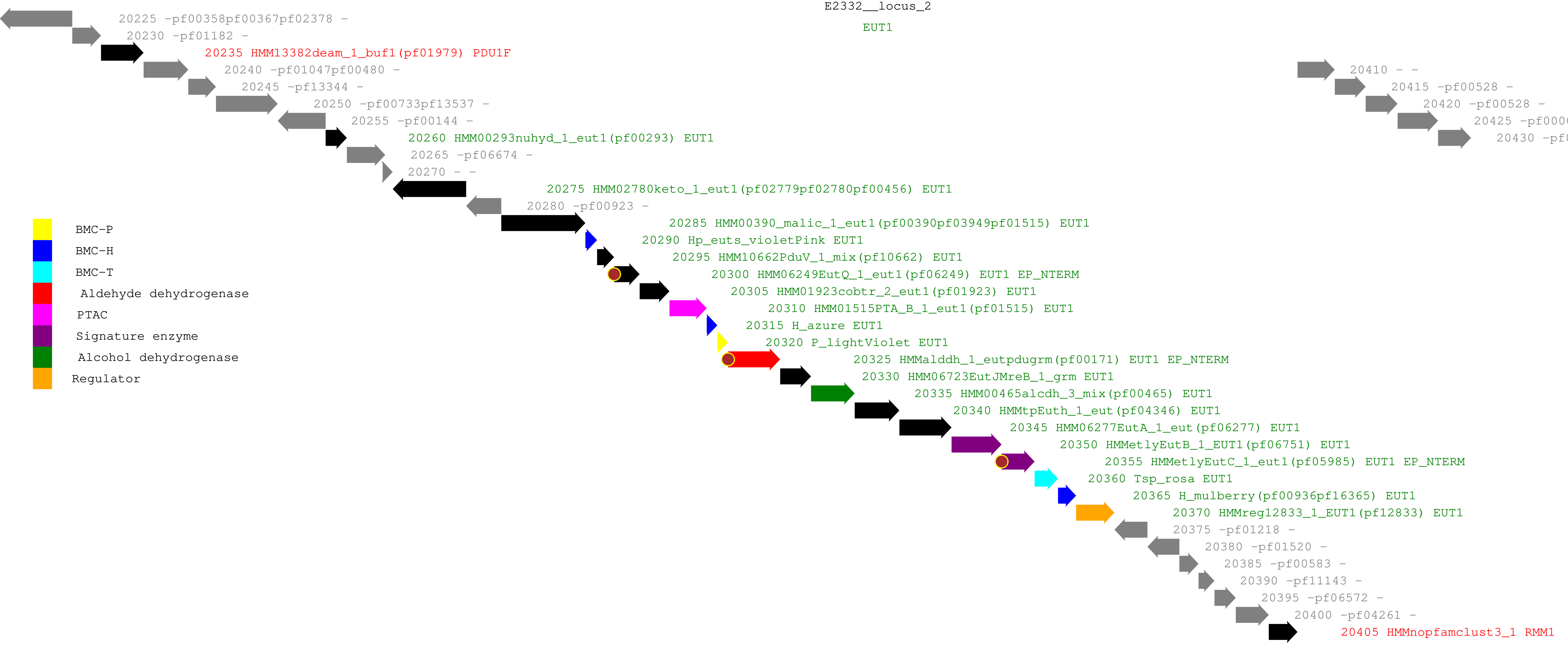

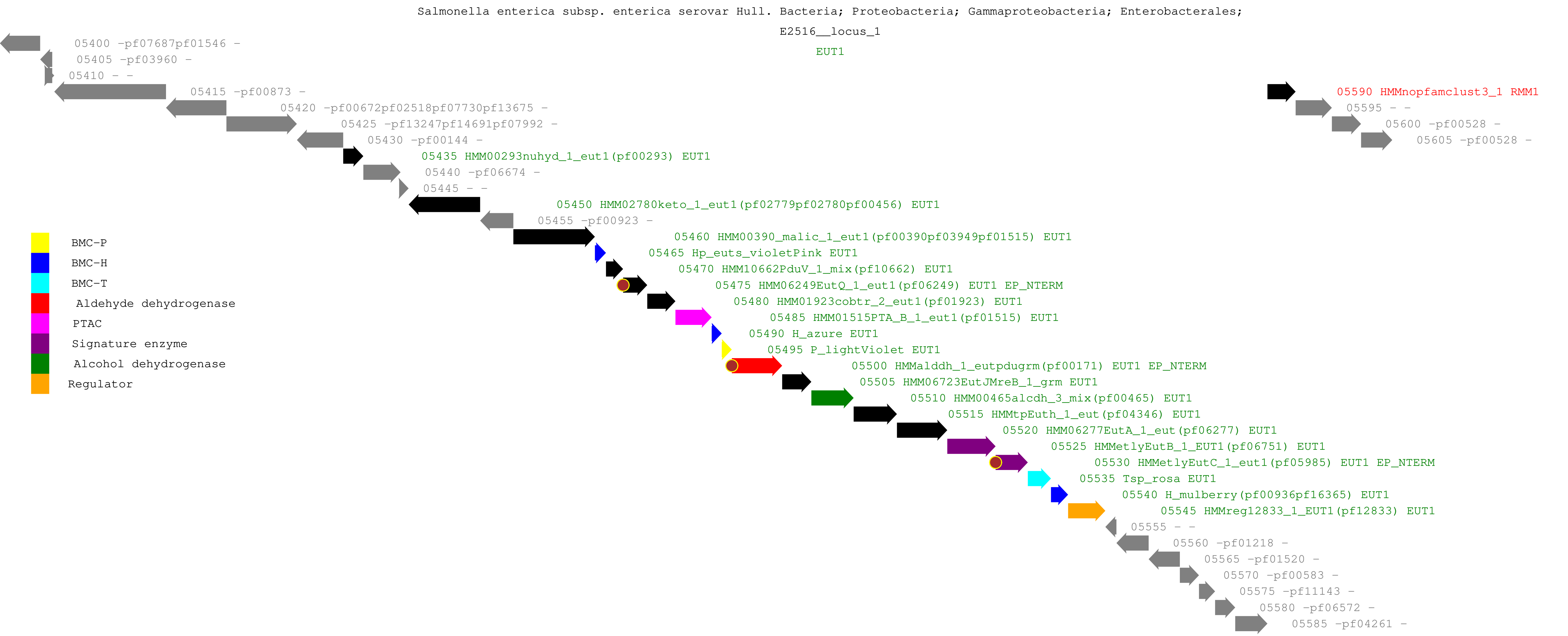
Salmonella enterica subsp. enterica serovar Hull. Bacteria; Proteobacteria; Gammaproteobacteria; Enterobacterales;
E2516__locus_1
EUT1
05400 -pf07687pf01546 -
05405 -pf03960 -
05410 - -
05415 -pf00873 -
05420 -pf00672pf02518pf07730pf13675 -
05425 -pf13247pf14691pf07992 -
05430 -pf00144 -
05435 HMM00293nuhyd_1_eut1(pf00293) EUT1
05440 -pf06674 -
05445 - -
05450 HMM02780keto_1_eut1(pf02779pf02780pf00456) EUT1
05455 -pf00923 -
05460 HMM00390_malic_1_eut1(pf00390pf03949pf01515) EUT1
05465 Hp_euts_violetPink EUT1
05470 HMM10662PduV_1_mix(pf10662) EUT1
05475 HMM06249EutQ_1_eut1(pf06249) EUT1 EP_NTERM
05480 HMM01923cobtr_2_eut1(pf01923) EUT1
05485 HMM01515PTA_B_1_eut1(pf01515) EUT1
05490 H_azure EUT1
05495 P_lightViolet EUT1
05500 HMMaldh_1_eutpdugrm(pf00171) EUT1 EP_NTERM
05505 HMM06723EutJMreB_1_grm EUT1
05510 HMM00465alcdh_3_mix(pf00465) EUT1
05515 HMMtpEuth_1_eut(pf04346) EUT1
05520 HMM06277EutA_1_eut(pf06277) EUT1
05525 HMMetlyEutB_1_EUT1(pf06751) EUT1
05530 HMMetlyEutC_1_eut1(pf05985) EUT1 EP_NTERM
05535 Tsp_rosa EUT1
05540 H_mulberry(pf00936pf16365) EUT1
05545 HMMreg12833_1_EUT1(pf12833) EUT1
05555 - -
05560 -pf01218 -
05565 -pf01520 -
05570 -pf00583 -
05575 -pf11143 -
05580 -pf06572 -
05585 -pf04261 -
05590 HMMnopfamclust3_1 RMM1
05595 - -
05600 -pf00528 -
05605 -pf00528 -
BMC-P
BMC-H
BMC-T
Aldehyde dehydrogenase
PTAC
Signature enzyme
Alcohol dehydrogenase
Regulator

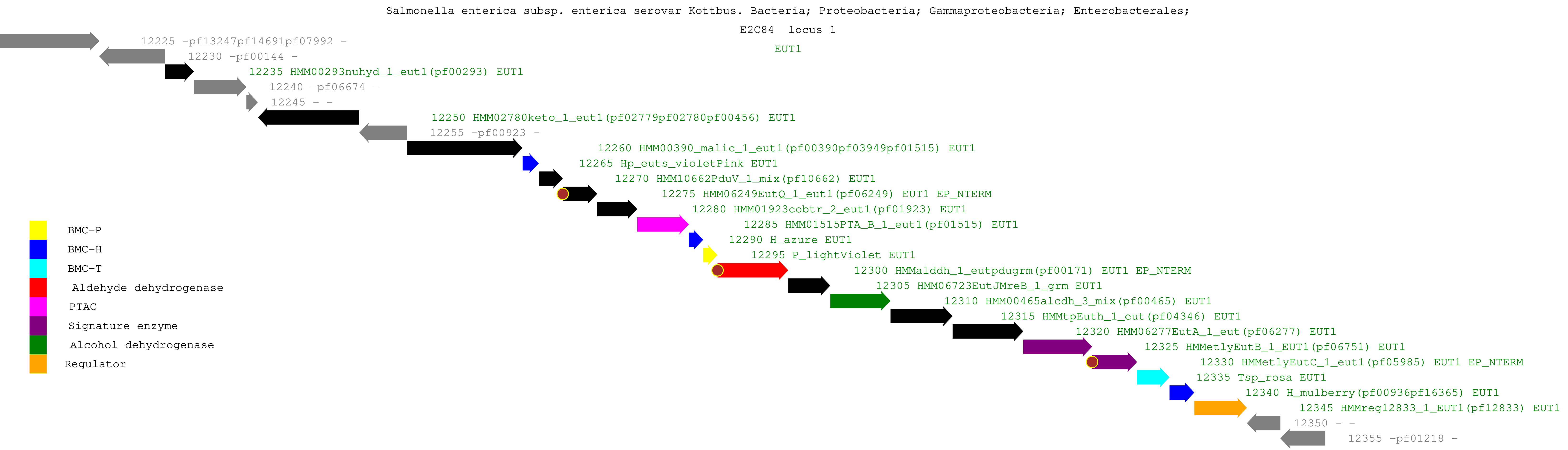

Salmonella enterica subsp. enterica serovar Kottbus. Bacteria; Proteobacteria; Gammaproteobacteria; Enterobacterales;
E2C84__locus_1
EUT1
12225 -pf13247pf14691pf07992 -
12230 -pf00144 -
12235 HMM00293nuhyd_1_eut1(pf00293) EUT1
12240 -pf06674 -
12245 - -
12250 HMM02780keto_1_eut1(pf02779pf02780pf00456) EUT1
12255 -pf00923 -
12260 HMM00390_malic_1_eut1(pf00390pf03949pf01515) EUT1
12265 Hp_euts_violetPink EUT1
12270 HMM10662PduV_1_mix(pf10662) EUT1
12275 HMM06249EutQ_1_eut1(pf06249) EUT1 EP_NTERM
12280 HMM01923cobtr_2_eut1(pf01923) EUT1
12285 HMM01515PTA_B_1_eut1(pf01515) EUT1
12290 H_azure EUT1
12295 P_lightViolet EUT1
12300 HMMaldh_1_eutpdugrm(pf00171) EUT1 EP_NTERM
12305 HMM06723EutJMreB_1_grm EUT1
12310 HMM00465alcdh_3_mix(pf00465) EUT1
12315 HMMtpEuth_1_eut(pf04346) EUT1
12320 HMM06277EutA_1_eut(pf06277) EUT1
12325 HMMetlyEutB_1_EUT1(pf06751) EUT1
12330 HMMetlyEutC_1_eut1(pf05985) EUT1 EP_NTERM
12335 Tsp_rosa EUT1
12340 H_mulberry(pf00936pf16365) EUT1
12345 HMMreg12833_1_EUT1(pf12833) EUT1
12350 - -
12355 -pf01218 -
BMC-P
BMC-H
BMC-T
Aldehyde dehydrogenase
PTAC
Signature enzyme
Alcohol dehydrogenase
Regulator

Salmonella enterica subsp. enterica serovar Miami. Bacteria; Proteobacteria; Gammaproteobacteria; Enterobacterales;

E2D53__locus_1

EUT1

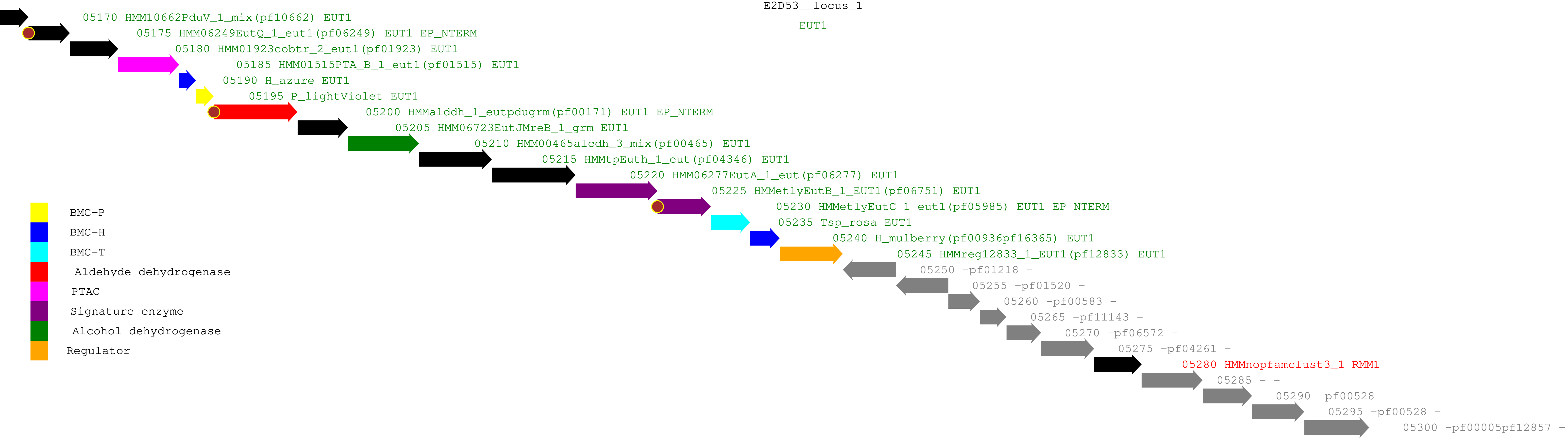

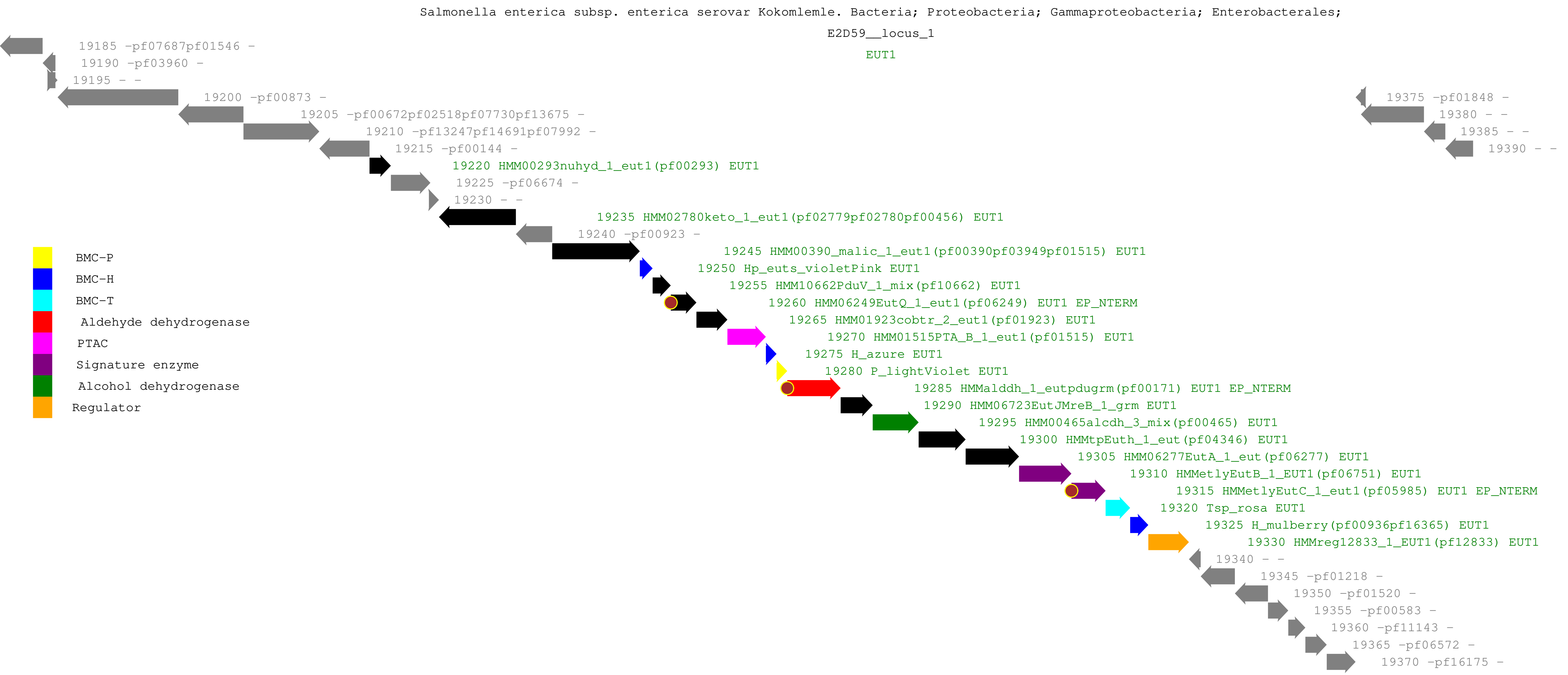
Salmonella enterica subsp. enterica serovar Kokomlemle. Bacteria; Proteobacteria; Gammaproteobacteria; Enterobacterales;
E2D59__locus_1
EUT1
19185 -pf07687pf01546 -
19190 -pf03960 -
19195 - -
19200 -pf00873 -
19205 -pf00672pf02518pf07730pf13675 -
19210 -pf13247pf14691pf07992 -
19215 -pf00144 -
19220 HMM00293nuhyd_1_eut1(pf00293) EUT1
19225 -pf06674 -
19230 - -
19235 HMM02780keto_1_eut1(pf02779pf02780pf00456) EUT1
19240 -pf00923 -
19245 HMM00390_malic_1_eut1(pf00390pf03949pf01515) EUT1
19250 Hp_euts_violetPink EUT1
19255 HMM10662PduV_1_mix(pf10662) EUT1
19260 HMM06249EutQ_1_eut1(pf06249) EUT1 EP_NTERM
19265 HMM01923cobtr_2_eut1(pf01923) EUT1
19270 HMM01515PTA_B_1_eut1(pf01515) EUT1
19275 H_azure EUT1
19280 P_lightViolet EUT1
19285 HMMaldh_1_eutpdugrm(pf00171) EUT1 EP_NTERM
19290 HMM06723EutJMreB_1_grm EUT1
19295 HMM00465alcdh_3_mix(pf00465) EUT1
19300 HMMtpEuth_1_eut(pf04346) EUT1
19305 HMM06277EutA_1_eut(pf06277) EUT1
19310 HMMetlyEutB_1_EUT1(pf06751) EUT1
19315 HMMetlyEutC_1_eut1(pf05985) EUT1 EP_NTERM
19320 Tsp_rosa EUT1
19325 H_mulberry(pf00936pf16365) EUT1
19330 HMMreg12833_1_EUT1(pf12833) EUT1
19340 - -
19345 -pf01218 -
19350 -pf01520 -
19355 -pf00583 -
19360 -pf11143 -
19365 -pf06572 -
19370 -pf16175 -
19375 -pf01848 -
19380 - -
19385 - -
19390 - -
BMC-P
BMC-H
BMC-T
Aldehyde dehydrogenase
PTAC
Signature enzyme
Alcohol dehydrogenase
Regulator

Salmonella enterica subsp. enterica serovar Weltevreden. Bacteria; Proteobacteria; Gammaproteobacteria; Enterobacterales;

E2E91__locus_1

EUT1

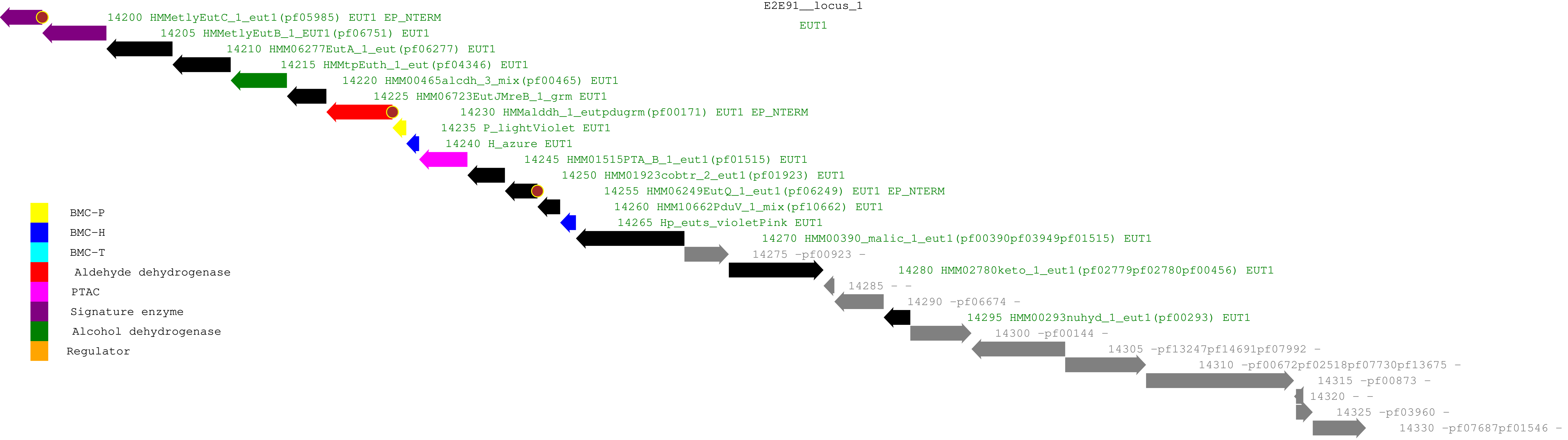

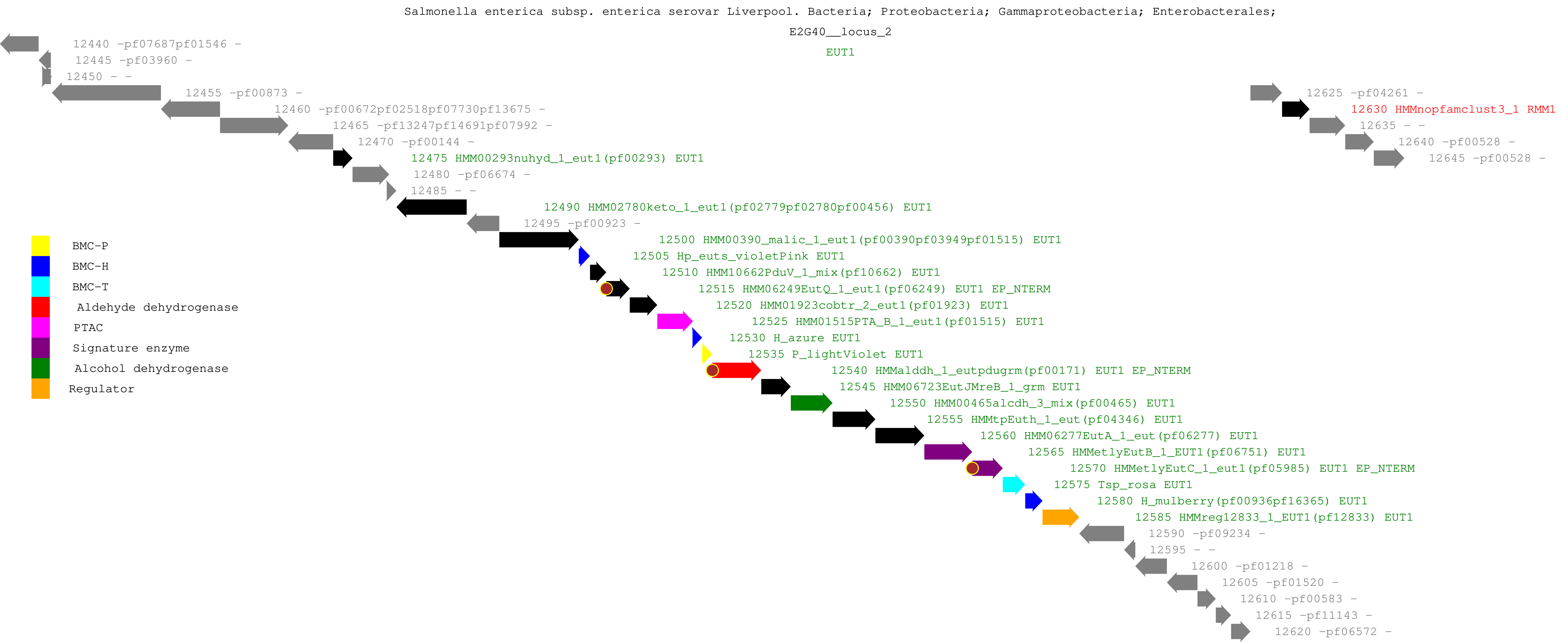
Salmonella enterica subsp. enterica serovar Liverpool. Bacteria; Proteobacteria; Gammaproteobacteria; Enterobacterales;
E2G40__locus_2
EUT1
12440 -pf07687pf01546 -
12445 -pf03960 -
12450 - -
12455 -pf00873 -
12460 -pf00672pf02518pf07730pf13675 -
12465 -pf13247pf14691pf07992 -
12470 -pf00144 -
12475 HMM00293nuhyd_1_eut1(pf00293) EUT1
12480 -pf06674 -
12485 - -
12490 HMM02780keto_1_eut1(pf02779pf02780pf00456) EUT1
12495 -pf00923 -
12500 HMM00390_malic_1_eut1(pf00390pf03949pf01515) EUT1
12505 Hp_euts_violetPink EUT1
12510 HMM10662PduV_1_mix(pf10662) EUT1
12515 HMM06249EutQ_1_eut1(pf06249) EUT1 EP_NTERM
12520 HMM01923cobtr_2_eut1(pf01923) EUT1
12525 HMM01515PTA_B_1_eut1(pf01515) EUT1
12530 H_azure EUT1
12535 P_lightViolet EUT1
12540 HMMalddh_1_eutpdugrm(pf00171) EUT1 EP_NTERM
12545 HMM06723EutJMreB_1_grm EUT1
12550 HMM00465alcdh_3_mix(pf00465) EUT1
12555 HMMtpEuth_1_eut(pf04346) EUT1
12560 HMM06277EutA_1_eut(pf06277) EUT1
12565 HMMetlyEutB_1_EUT1(pf06751) EUT1
12570 HMMetlyEutC_1_eut1(pf05985) EUT1 EP_NTERM
12575 Tsp_rosa EUT1
12580 H_mulberry(pf00936pf16365) EUT1
12585 HMMreg12833_1_EUT1(pf12833) EUT1
12590 -pf09234 -
12595 - -
12600 -pf01218 -
12605 -pf01520 -
12610 -pf00583 -
12615 -pf11143 -
12620 -pf06572 -
12625 -pf04261 -
12630 HMMnopfamclust3_1 RMM1
12635 - -
12640 -pf00528 -
12645 -pf00528 -
BMC-P
BMC-H
BMC-T
Aldehyde dehydrogenase
PTAC
Signature enzyme
Alcohol dehydrogenase
Regulator

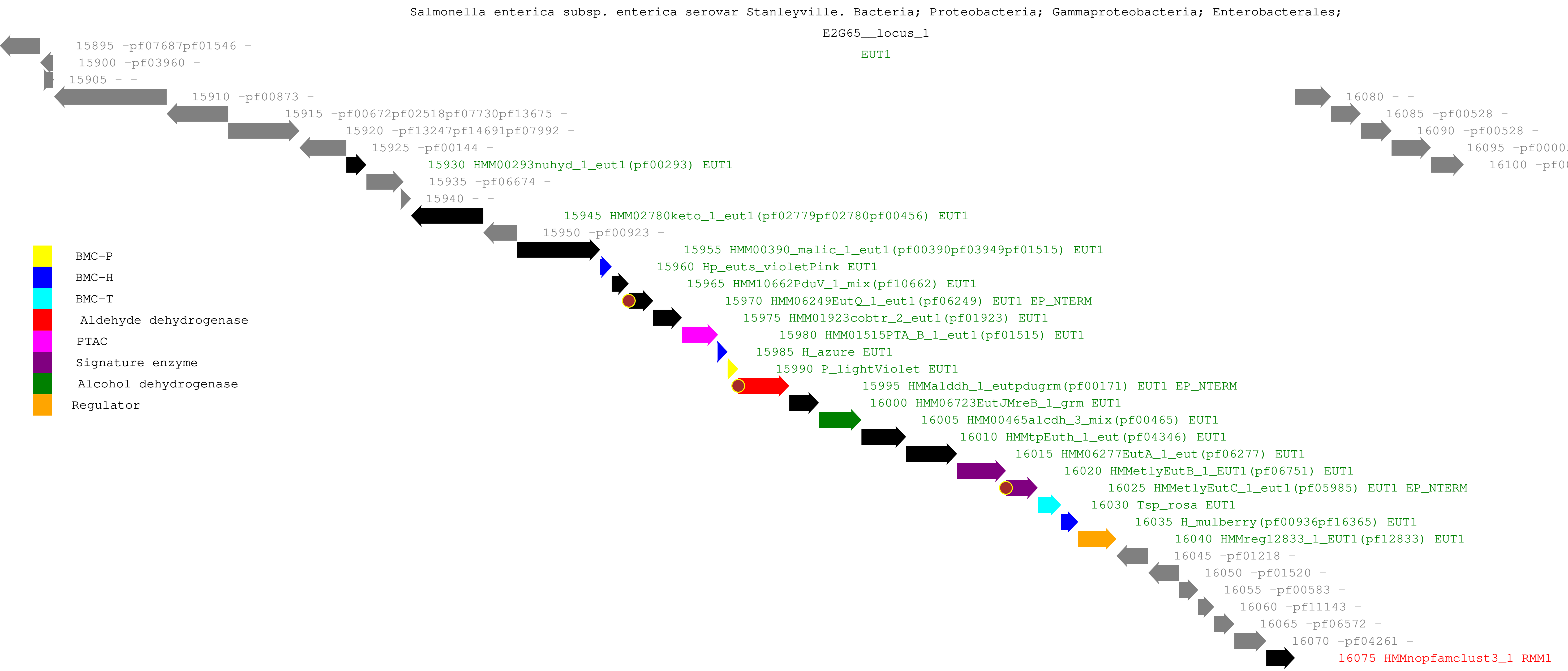

Salmonella enterica subsp. enterica serovar Stanleyville. Bacteria; Proteobacteria; Gammaproteobacteria; Enterobacterales;
E2G65__locus_1
EUT1
15895 -pf07687pf01546 -
15900 -pf03960 -
15905 - -
15910 -pf00873 -
15915 -pf00672pf02518pf07730pf13675 -
15920 -pf13247pf14691pf07992 -
15925 -pf00144 -
15930 HMM00293nuhyd_1_eut1(pf00293) EUT1
15935 -pf06674 -
15940 - -
15945 HMM02780keto_1_eut1(pf02779pf02780pf00456) EUT1
15950 -pf00923 -
15955 HMM00390_malic_1_eut1(pf00390pf03949pf01515) EUT1
15960 Hp_euts_violetPink EUT1
15965 HMM10662PduV_1_mix(pf10662) EUT1
15970 HMM06249EutQ_1_eut1(pf06249) EUT1 EP_NTERM
15975 HMM01923cobtr_2_eut1(pf01923) EUT1
15980 HMM01515PTA_B_1_eut1(pf01515) EUT1
15985 H_azure EUT1
15990 P_lightViolet EUT1
15995 HMMalddh_1_eutpdugrm(pf00171) EUT1 EP_NTERM
16000 HMM06723EutJMreB_1_grm EUT1
16005 HMM00465alcdh_3_mix(pf00465) EUT1
16010 HMMtpEuth_1_eut(pf04346) EUT1
16015 HMM06277EutA_1_eut(pf06277) EUT1
16020 HMMetlyEutB_1_EUT1(pf06751) EUT1
16025 HMMetlyEutC_1_eut1(pf05985) EUT1 EP_NTERM
16030 Tsp_rosa EUT1
16035 H_mulberry(pf00936pf16365) EUT1
16040 HMMreg12833_1_EUT1(pf12833) EUT1
16045 -pf01218 -
16050 -pf01520 -
16055 -pf00583 -
16060 -pf11143 -
16065 -pf06572 -
16070 -pf04261 -
16075 HMMnopfamclust3_1 RMM1
16080 - -
16085 -pf00528 -
16090 -pf00528 -
16095 -pf00005pf12857 -
16100 -pf00291 -
BMC-P
BMC-H
BMC-T
Aldehyde dehydrogenase
PTAC
Signature enzyme
Alcohol dehydrogenase
Regulator

Salmonella enterica subsp. enterica serovar Wien. Bacteria; Proteobacteria; Gammaproteobacteria; Enterobacterales;

E2G78__locus_1

EUT1

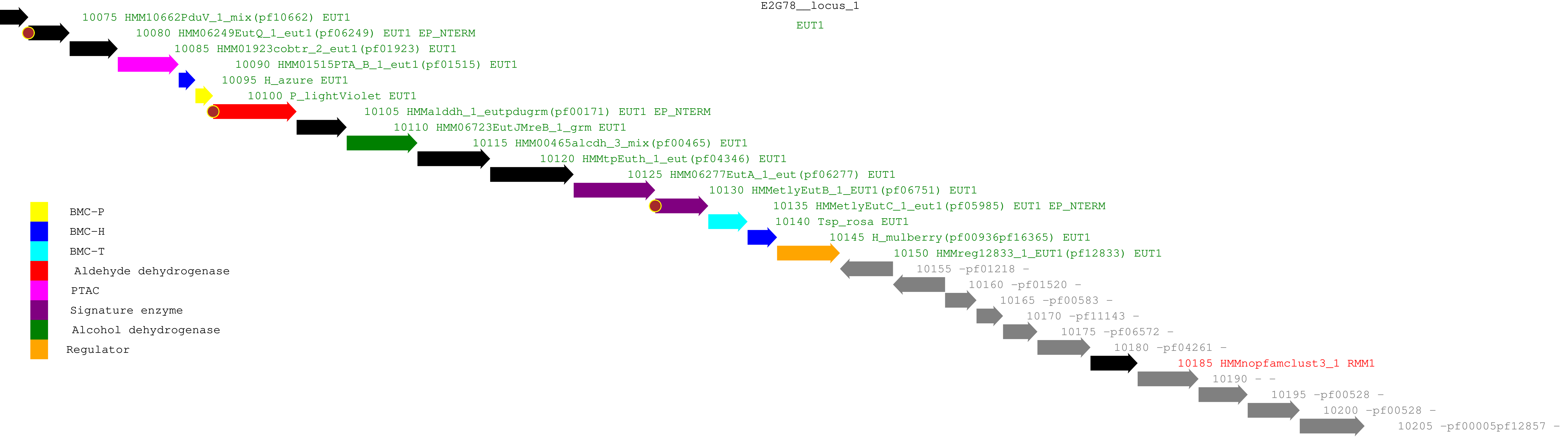

Salmonella enterica subsp. enterica serovar Ramatgan. Bacteria; Proteobacteria; Gammaproteobacteria; Enterobacterales;

E2G82__locus_1

EUT1

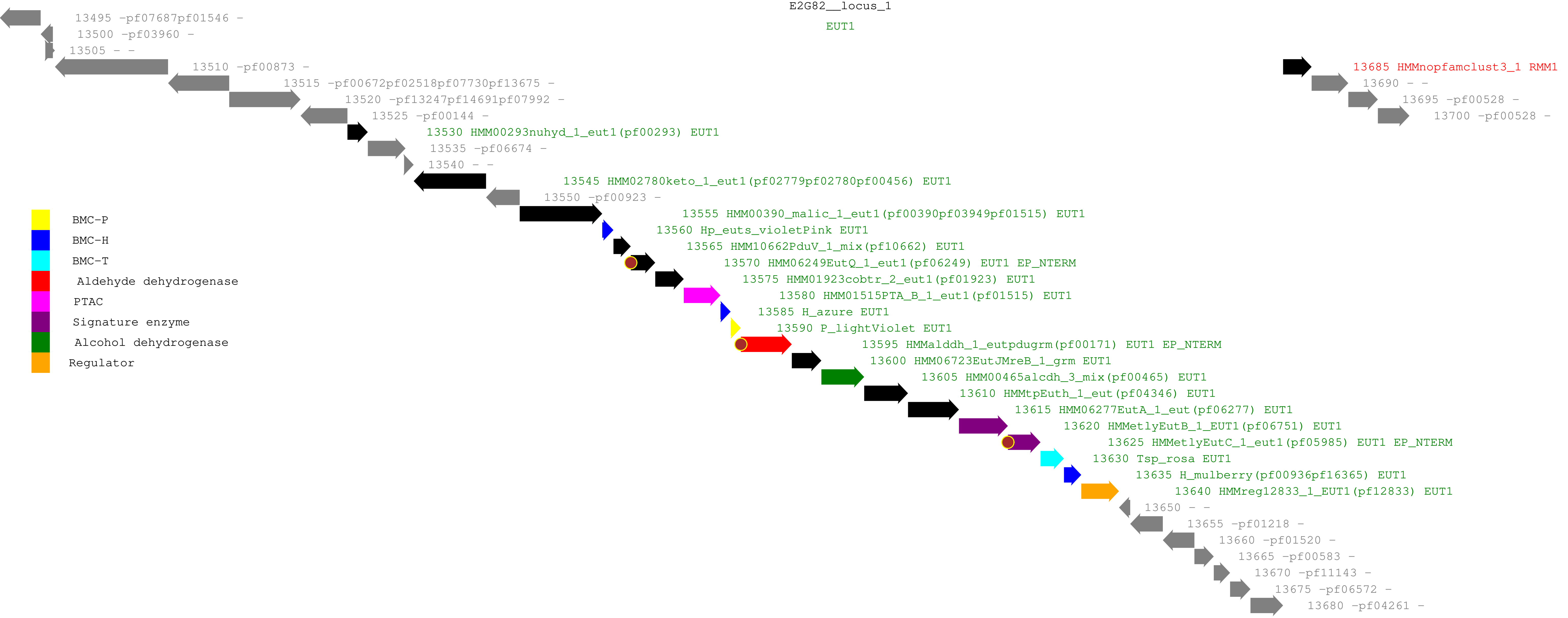

Salmonella enterica subsp. enterica serovar Volkmarsdorf. Bacteria; Proteobacteria; Gammaproteobacteria; Enterobacterales;

E2H20__locus_2

EUT1

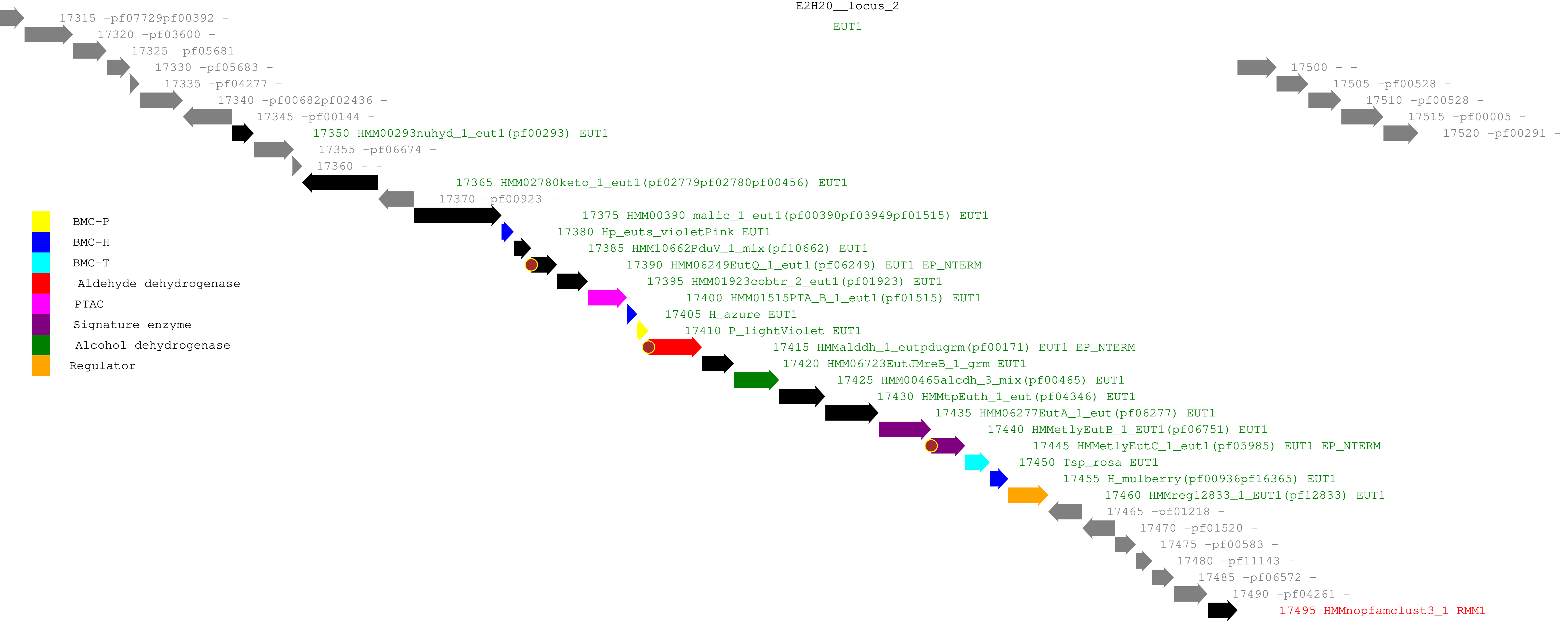

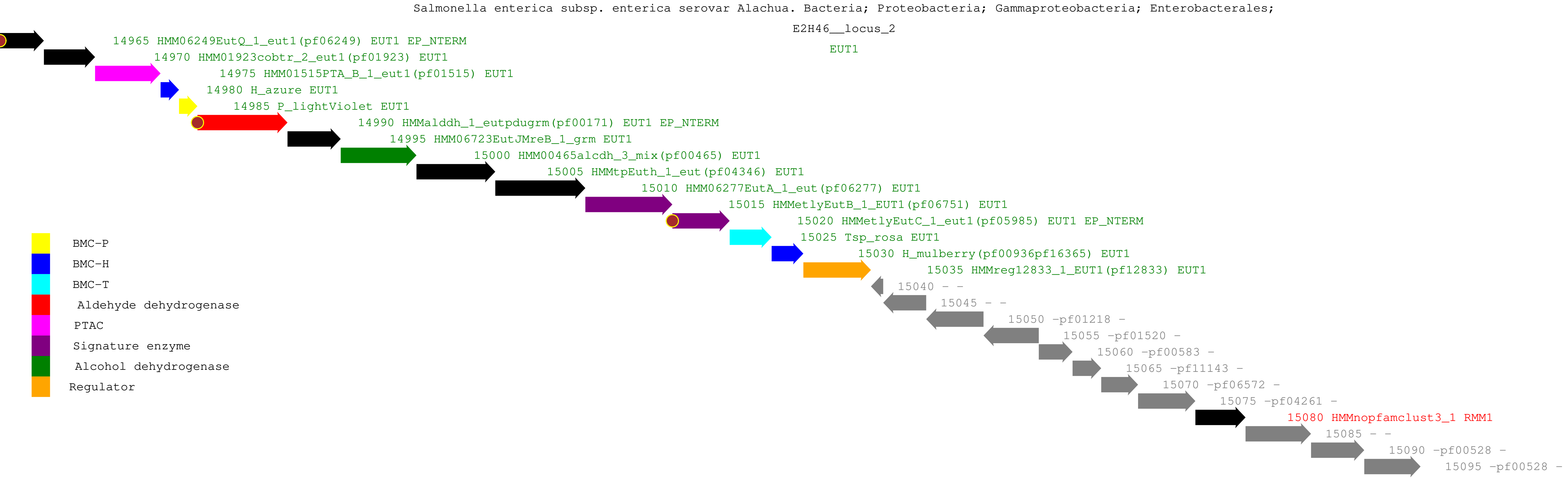

Salmonella enterica subsp. enterica serovar Alachua. Bacteria; Proteobacteria; Gammaproteobacteria; Enterobacterales;
E2H46__locus_2
EUT1
14965 HMM06249EutQ_1_eut1(pf06249) EUT1 EP_NTERM
14970 HMM01923cobtr_2_eut1(pf01923) EUT1
14975 HMM01515PTA_B_1_eut1(pf01515) EUT1
14980 H_azure EUT1
14985 P_lightViolet EUT1
14990 HMMalddh_1_eutpdugrm(pf00171) EUT1 EP_NTERM
14995 HMM06723EutJMreB_1_grm EUT1
15000 HMM00465alcdh_3_mix(pf00465) EUT1
15005 HMMtpEuth_1_eut(pf04346) EUT1
15010 HMM06277EutA_1_eut(pf06277) EUT1
15015 HMMetlyEutB_1_EUT1(pf06751) EUT1
15020 HMMetlyEutC_1_eut1(pf05985) EUT1 EP_NTERM
15025 Tsp_rosa EUT1
15030 H_mulberry(pf00936pf16365) EUT1
15035 HMMreg12833_1_EUT1(pf12833) EUT1
15040 - -
15045 - -
15050 -pf01218 -
15055 -pf01520 -
15060 -pf00583 -
15065 -pf11143 -
15070 -pf06572 -
15075 -pf04261 -
15080 HMMnopfamclust3_1 RMM1
15085 - -
15090 -pf00528 -
15095 -pf00528 -
BMC-P
BMC-H
BMC-T
Aldehyde dehydrogenase
PTAC
Signature enzyme
Alcohol dehydrogenase
Regulator

Salmonella enterica subsp. enterica serovar Krefeld. Bacteria; Proteobacteria; Gammaproteobacteria; Enterobacterales;

E2L63__locus_2

EUT1

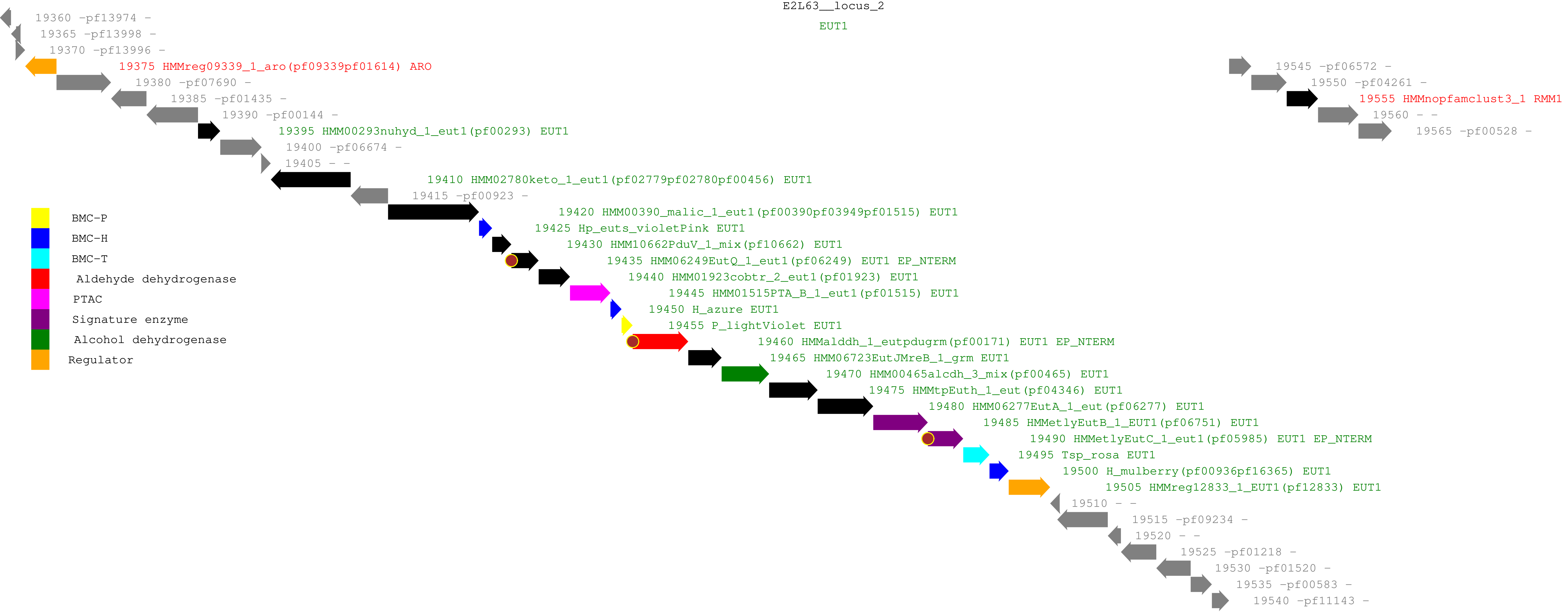

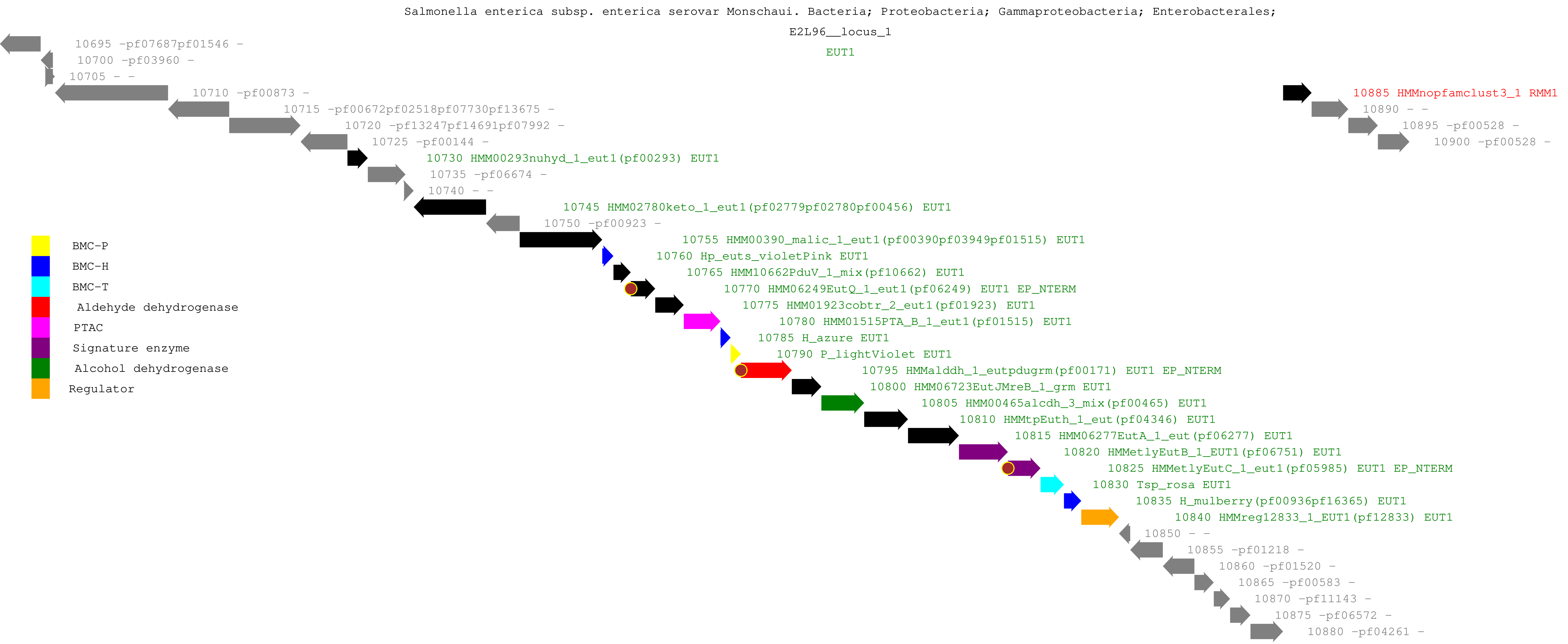
Salmonella enterica subsp. enterica serovar Monschaui. Bacteria; Proteobacteria; Gammaproteobacteria; Enterobacterales;
E2L96__locus_1
EUT1
10695 -pf07687pf01546 -
10700 -pf03960 -
10705 - -
10710 -pf00873 -
10715 -pf00672pf02518pf07730pf13675 -
10720 -pf13247pf14691pf07992 -
10725 -pf00144 -
10730 HMM00293nuhyd_1_eut1(pf00293) EUT1
10735 -pf06674 -
10740 - -
10745 HMM02780keto_1_eut1(pf02779pf02780pf00456) EUT1
10750 -pf00923 -
10755 HMM00390_malic_1_eut1(pf00390pf03949pf01515) EUT1
10760 Hp_euts_violetPink EUT1
10765 HMM10662PduV_1_mix(pf10662) EUT1
10770 HMM06249EutQ_1_eut1(pf06249) EUT1 EP_NTERM
10775 HMM01923cobtr_2_eut1(pf01923) EUT1
10780 HMM01515PTA_B_1_eut1(pf01515) EUT1
10785 H_azure EUT1
10790 P_lightViolet EUT1
10795 HMMalddh_1_eutpdugrm(pf00171) EUT1 EP_NTERM
10800 HMM06723EutJMreB_1_grm EUT1
10805 HMM00465alcdh_3_mix(pf00465) EUT1
10810 HMMtpEuth_1_eut(pf04346) EUT1
10815 HMM06277EutA_1_eut(pf06277) EUT1
10820 HMMetlyEutB_1_EUT1(pf06751) EUT1
10825 HMMetlyEutC_1_eut1(pf05985) EUT1 EP_NTERM
10830 Tsp_rosa EUT1
10835 H_mulberry(pf00936pf16365) EUT1
10840 HMMreg12833_1_EUT1(pf12833) EUT1
10850 - -
10855 -pf01218 -
10860 -pf01520 -
10865 -pf00583 -
10870 -pf11143 -
10875 -pf06572 -
10880 -pf04261 -
10885 HMMnopfamclust3_1 RMM1
10890 - -
10895 -pf00528 -
10900 -pf00528 -
BMC-P
BMC-H
BMC-T
Aldehyde dehydrogenase
PTAC
Signature enzyme
Alcohol dehydrogenase
Regulator

Salmonella enterica subsp. enterica serovar Arechavaleta. Bacteria; Proteobacteria; Gammaproteobacteria; Enterobacterales;

E2M22__locus_1_main_sat

EUT1

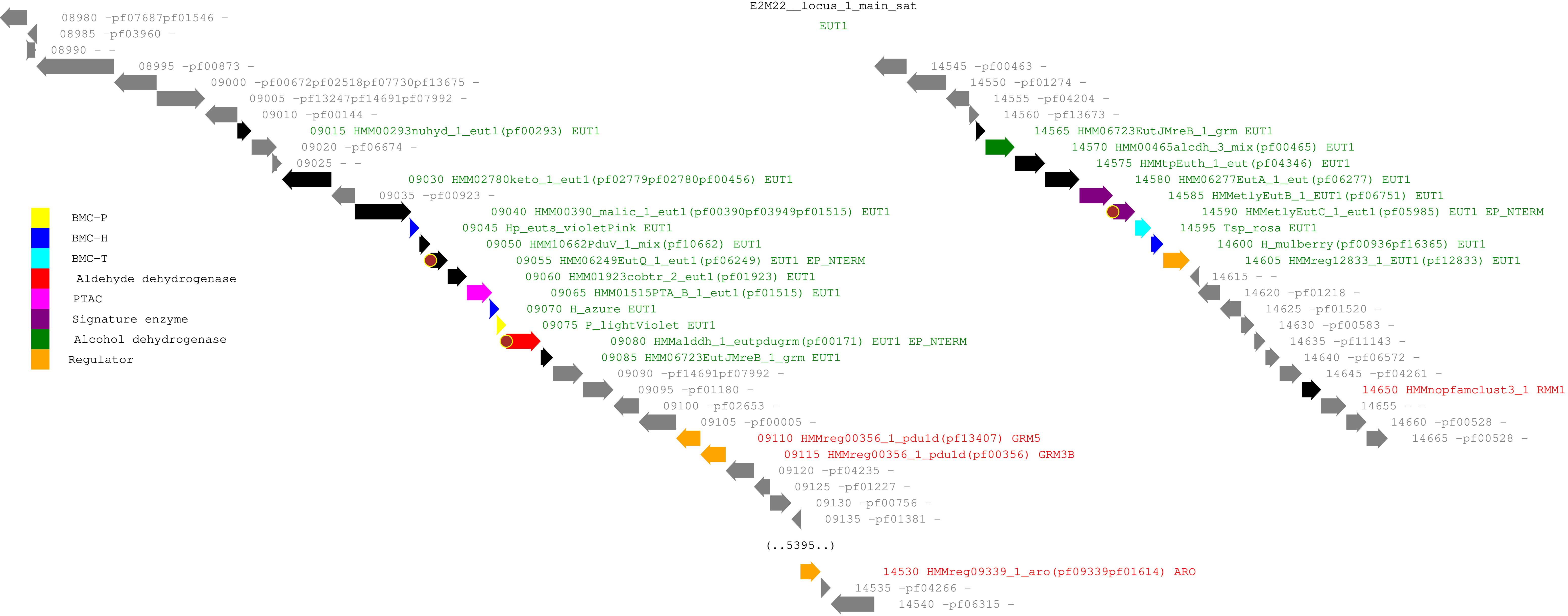

Salmonella enterica subsp. enterica serovar Nima. Bacteria; Proteobacteria; Gammaproteobacteria; Enterobacterales;

E2M36__locus_2

EUT1

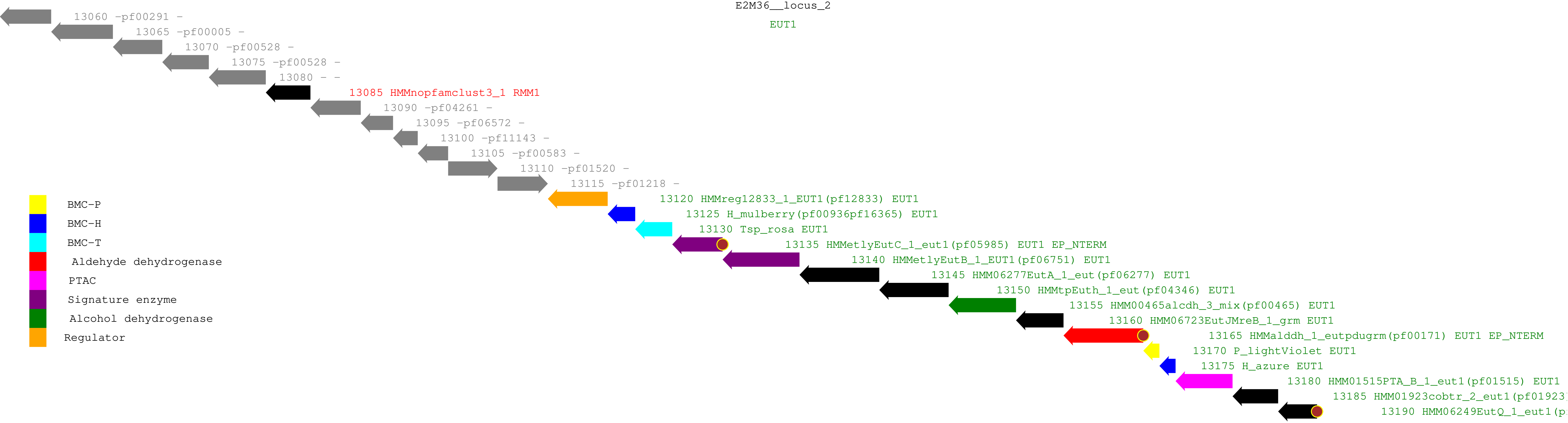

Salmonella enterica subsp. enterica serovar Othmarschen. Bacteria; Proteobacteria; Gammaproteobacteria; Enterobacterales;

E2M46__locus_2

EUT1

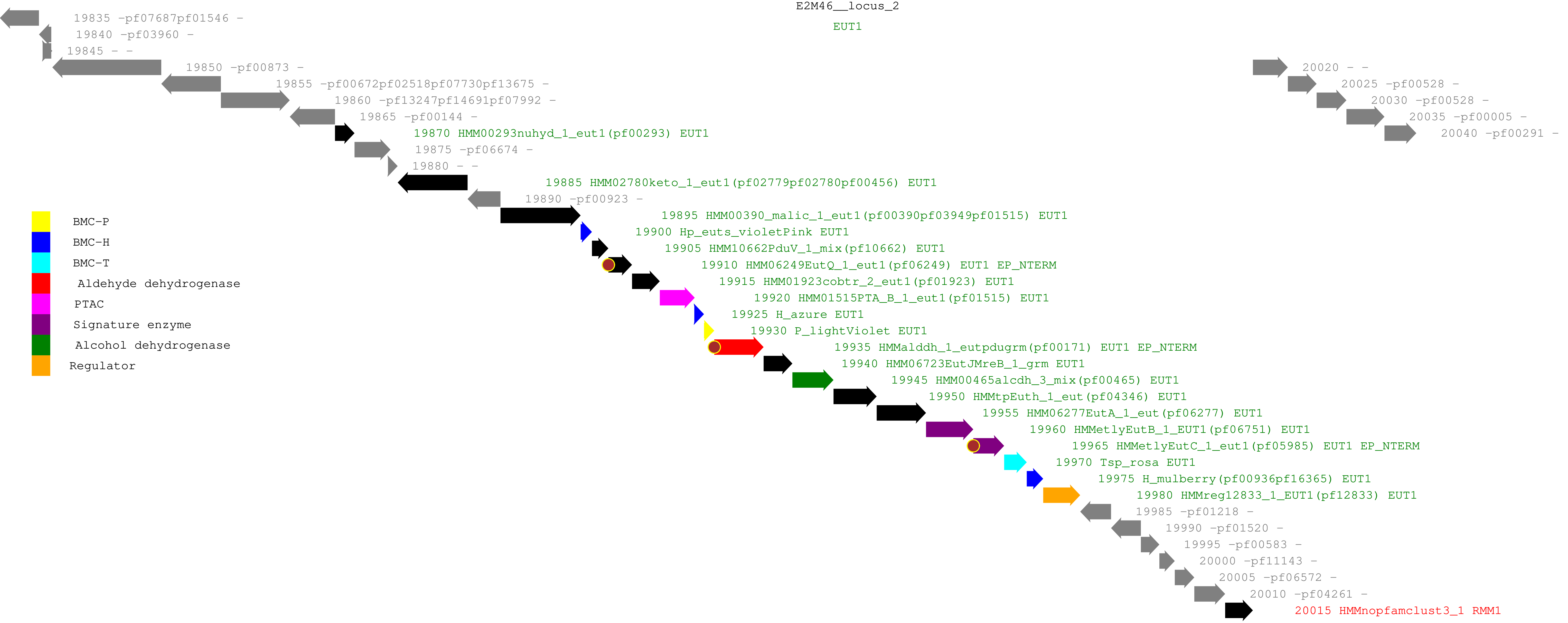

Citrobacter rodentium. Bacteria; Proteobacteria; Gammaproteobacteria; Enterobacterales;

E2R62__locus_2

EUT1

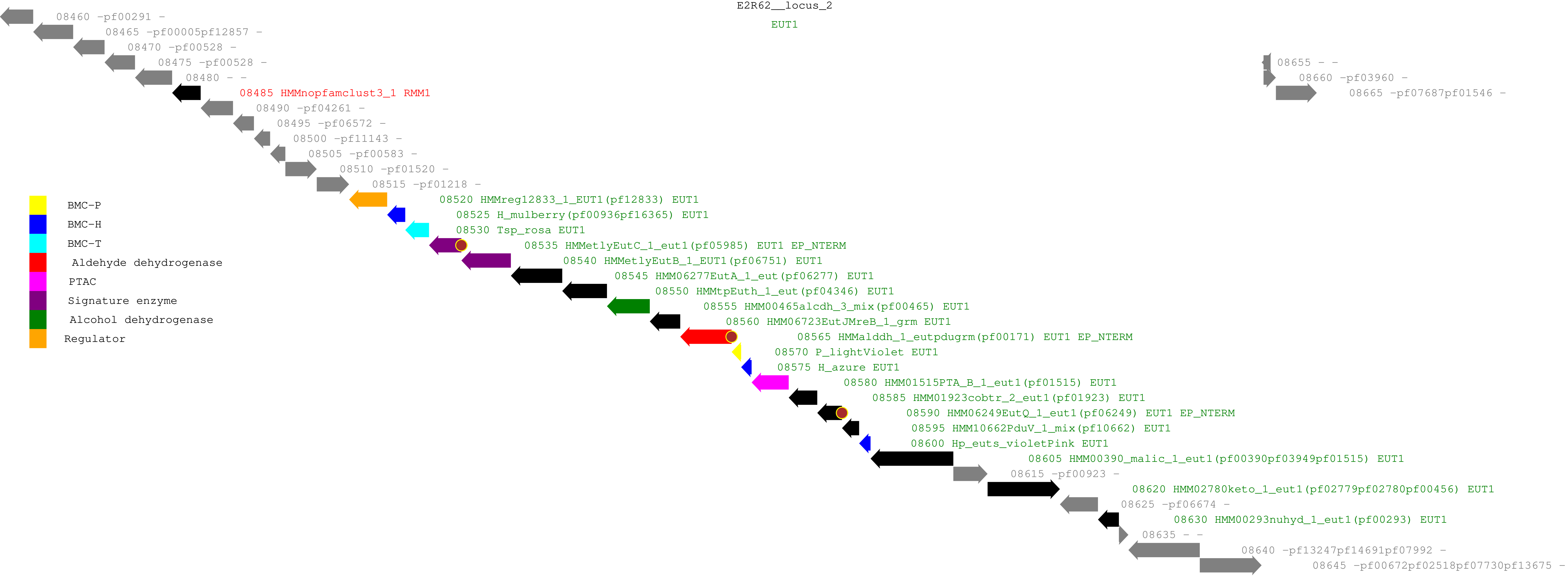

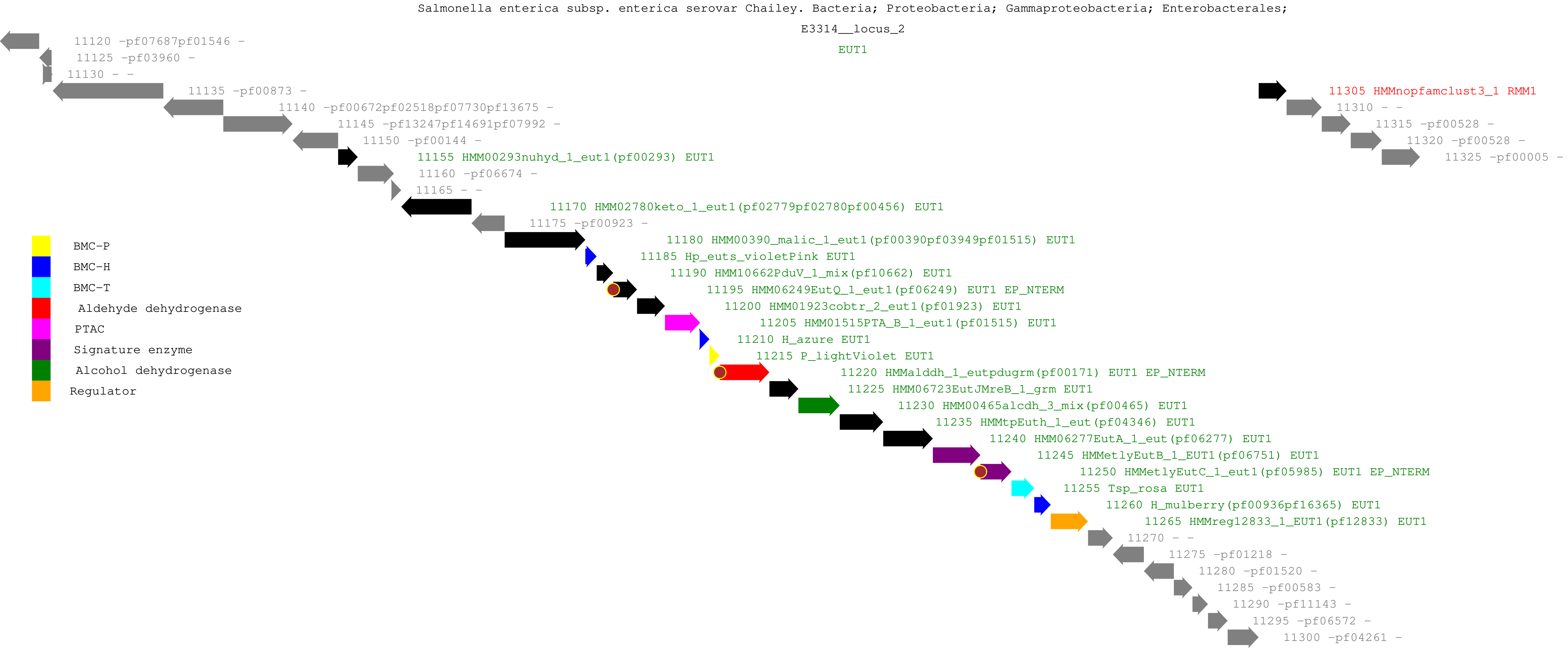

Salmonella enterica subsp. enterica serovar Chailey. Bacteria; Proteobacteria; Gammaproteobacteria; Enterobacterales;
E3314__locus_2
EUT1
11120 -pf07687pf01546 -
11125 -pf03960 -
11130 - -
11135 -pf00873 -
11140 -pf00672pf02518pf07730pf13675 -
11145 -pf13247pf14691pf07992 -
11150 -pf00144 -
11155 HMM00293nuhyd_1_eut1(pf00293) EUT1
11160 -pf06674 -
11165 - -
11170 HMM02780keto_1_eut1(pf02779pf02780pf00456) EUT1
11175 -pf00923 -
11180 HMM00390_malic_1_eut1(pf00390pf03949pf01515) EUT1
11185 Hp_euts_violetPink EUT1
11190 HMM10662PduV_1_mix(pf10662) EUT1
11195 HMM06249EutQ_1_eut1(pf06249) EUT1 EP_NTERM
11200 HMM01923cobtr_2_eut1(pf01923) EUT1
11205 HMM01515PTA_B_1_eut1(pf01515) EUT1
11210 H_azure EUT1
11215 P_lightViolet EUT1
11220 HMMalddh_1_eutpdugrm(pf00171) EUT1 EP_NTERM
11225 HMM06723EutJMreB_1_grm EUT1
11230 HMM00465alcdh_3_mix(pf00465) EUT1
11235 HMMtpEuth_1_eut(pf04346) EUT1
11240 HMM06277EutA_1_eut(pf06277) EUT1
11245 HMMetlyEutB_1_EUT1(pf06751) EUT1
11250 HMMetlyEutC_1_eut1(pf05985) EUT1 EP_NTERM
11255 Tsp_rosa EUT1
11260 H_mulberry(pf00936pf16365) EUT1
11265 HMMreg12833_1_EUT1(pf12833) EUT1
11270 - -
11275 -pf01218 -
11280 -pf01520 -
11285 -pf00583 -
11290 -pf11143 -
11295 -pf06572 -
11300 -pf04261 -
11305 HMMnopfamclust3_1 RMM1
11310 - -
11315 -pf00528 -
11320 -pf00528 -
11325 -pf00005 -
BMC-P
BMC-H
BMC-T
Aldehyde dehydrogenase
PTAC
Signature enzyme
Alcohol dehydrogenase
Regulator

Salmonella enterica subsp. enterica serovar Stourbridge. Bacteria; Proteobacteria; Gammaproteobacteria; Enterobacterales;

E3348__locus_2

EUT1

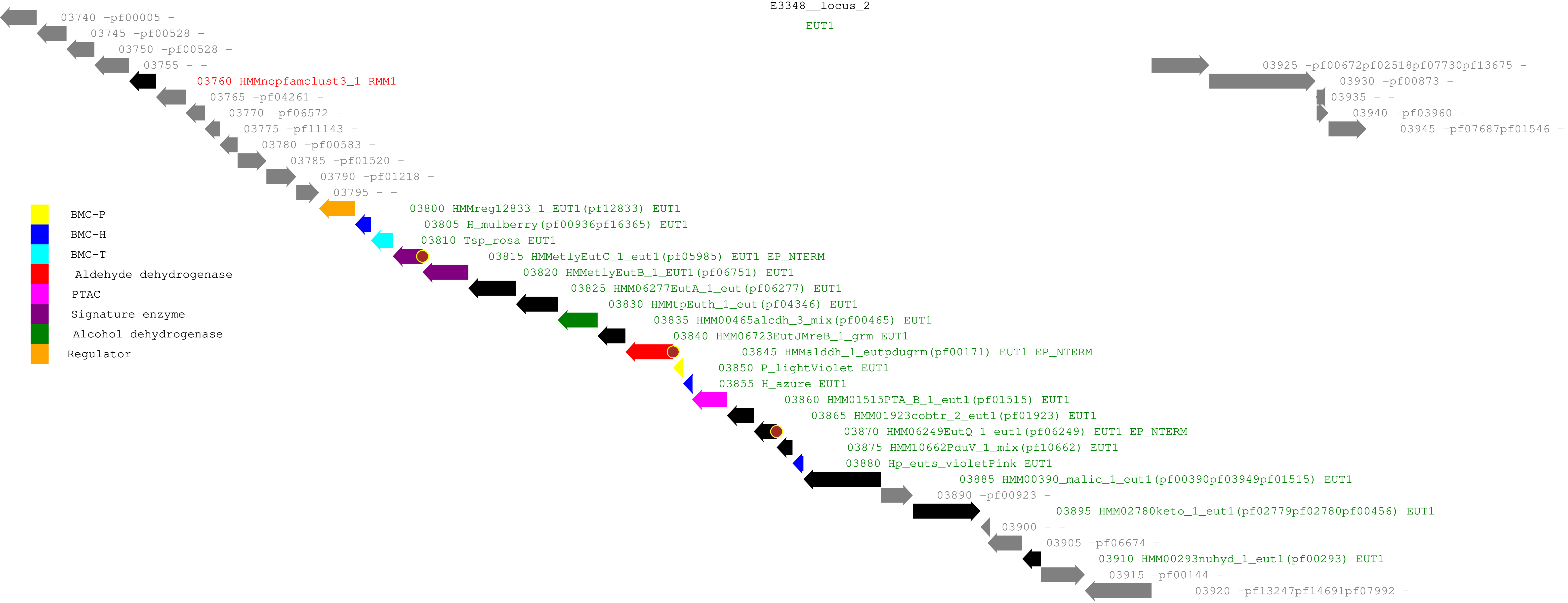

Lelliottia nimipressuralis. Bacteria; Proteobacteria; Gammaproteobacteria; Enterobacterales;

E3U32__locus_1

EUT1

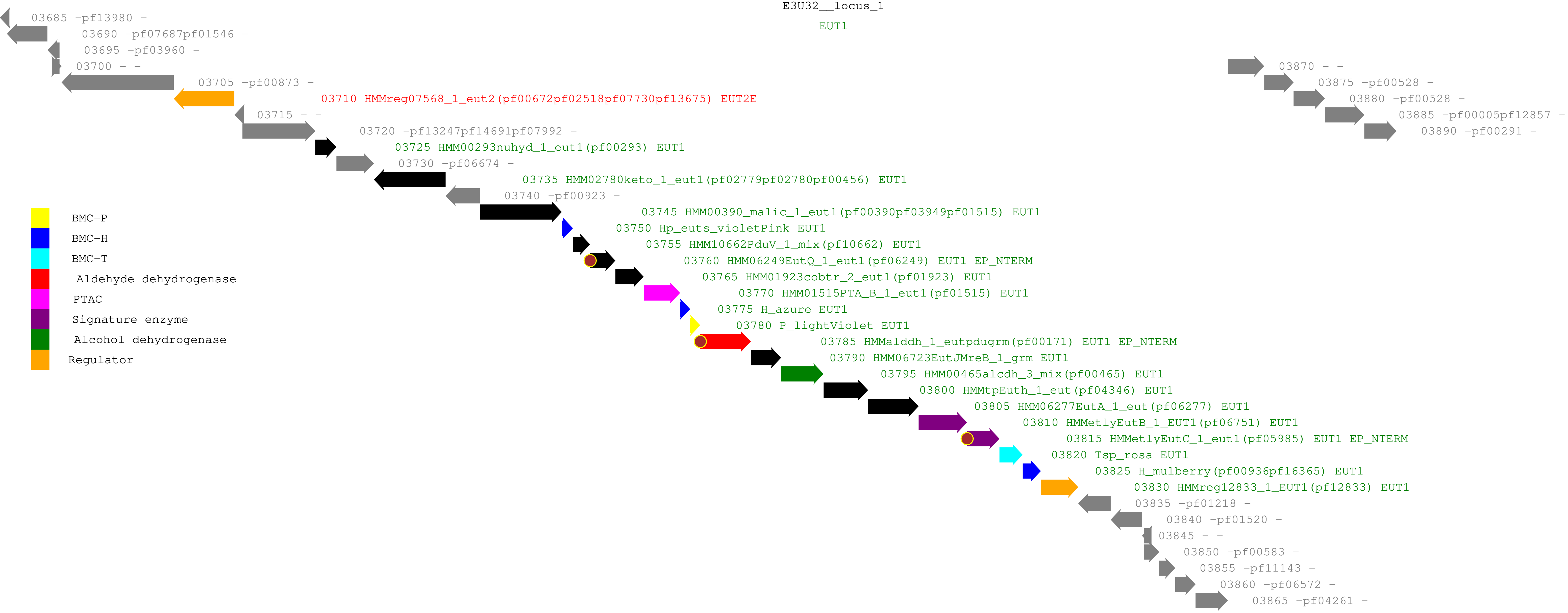

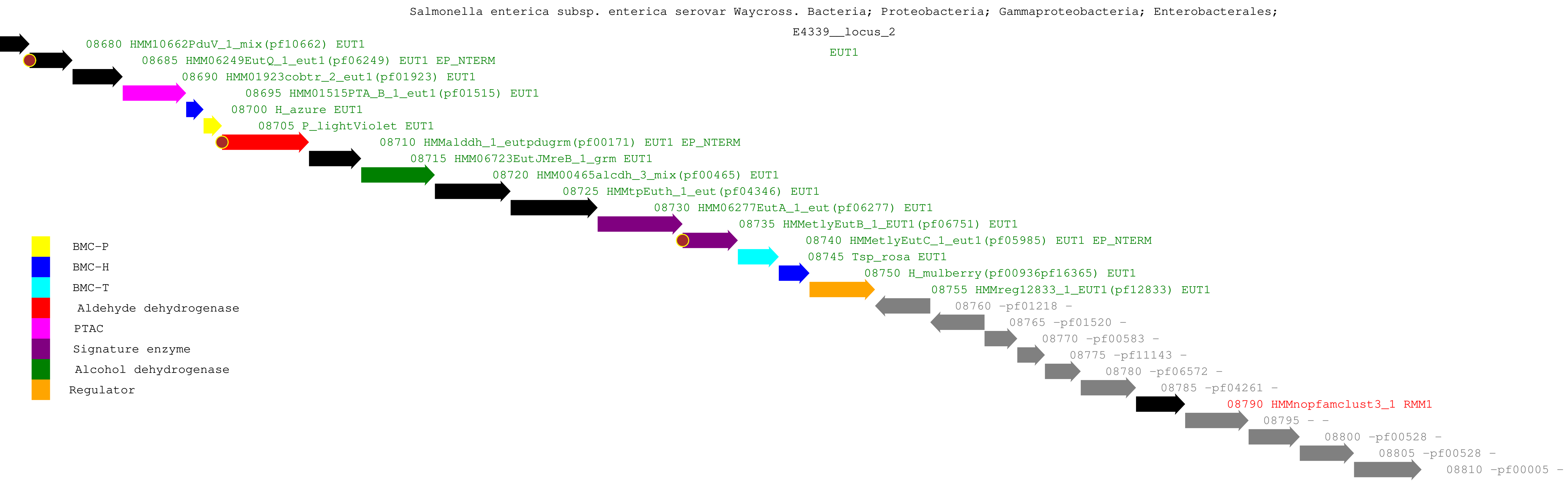
Salmonella enterica subsp. enterica serovar Waycross. Bacteria; Proteobacteria; Gammaproteobacteria; Enterobacterales;
E4339__locus_2
EUT1
08680 HMM10662PduV_1_mix(pf10662) EUT1
08685 HMM06249EutQ_1_eut1(pf06249) EUT1 EP_NTERM
08690 HMM01923cobtr_2_eut1(pf01923) EUT1
08695 HMM01515PTA_B_1_eut1(pf01515) EUT1
08700 H_azure EUT1
08705 P_lightViolet EUT1
08710 HMMaldh_1_eutpdugrm(pf00171) EUT1 EP_NTERM
08715 HMM06723EutJMreB_1_grm EUT1
08720 HMM00465alcdh_3_mix(pf00465) EUT1
08725 HMMtpEuth_1_eut(pf04346) EUT1
08730 HMM06277EutA_1_eut(pf06277) EUT1
08735 HMMetlyEutB_1_EUT1(pf06751) EUT1
08740 HMMetlyEutC_1_eut1(pf05985) EUT1 EP_NTERM
08745 Tsp_rosa EUT1
08750 H_mulberry(pf00936pf16365) EUT1
08755 HMMreg12833_1_EUT1(pf12833) EUT1
08760 -pf01218 -
08765 -pf01520 -
08770 -pf00583 -
08775 -pf11143 -
08780 -pf06572 -
08785 -pf04261 -
08790 HMMnopfamclust3_1 RMM1
08795 - -
08800 -pf00528 -
08805 -pf00528 -
08810 -pf00005 -
BMC-P
BMC-H
BMC-T
Aldehyde dehydrogenase
PTAC
Signature enzyme
Alcohol dehydrogenase
Regulator

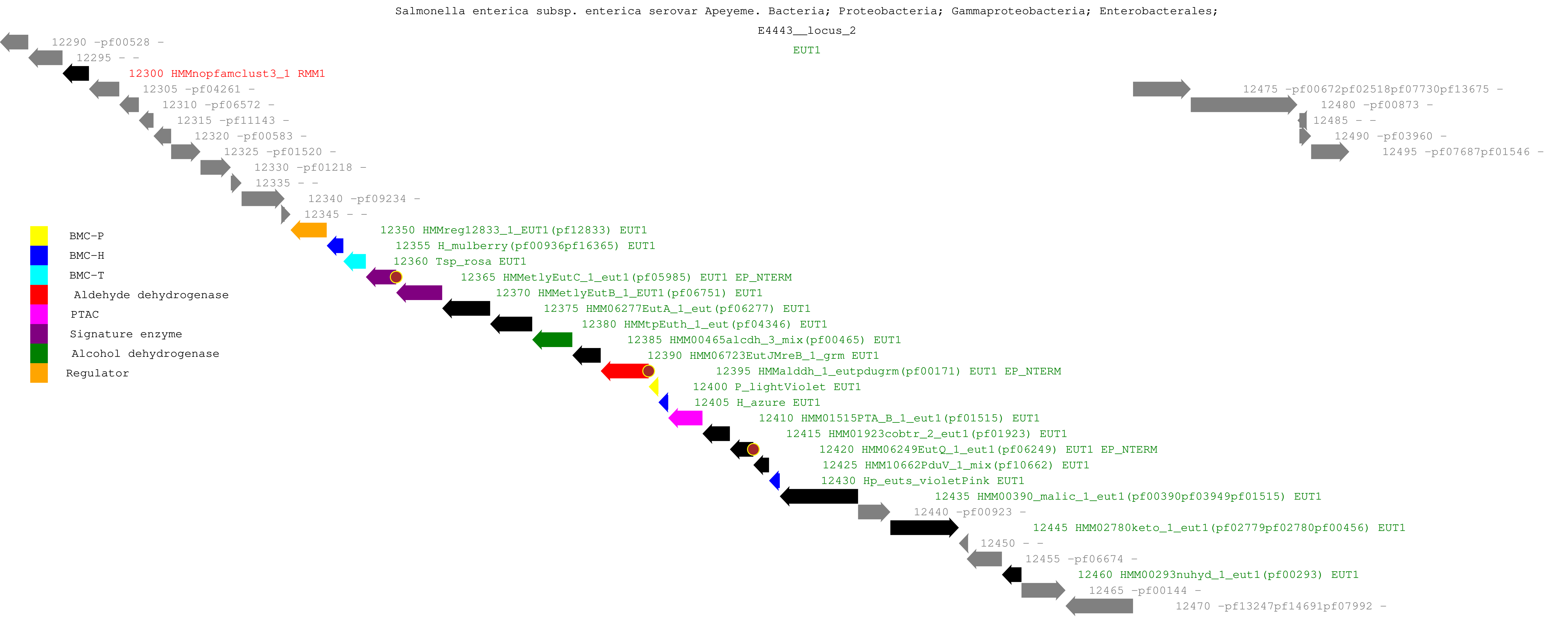

Salmonella enterica subsp. enterica serovar Apeyeme. Bacteria; Proteobacteria; Gammaproteobacteria; Enterobacterales;
E4443__locus_2
EUT1
12290 -pf00528 -
12295 - -
12300 HMMnopfamclust3_1 RMM1
12305 -pf04261 -
12310 -pf06572 -
12315 -pf11143 -
12320 -pf00583 -
12325 -pf01520 -
12330 -pf01218 -
12335 - -
12340 -pf09234 -
12345 - -
12350 HMMreg12833_1_EUT1(pf12833) EUT1
12355 H_mulberry(pf00936pf16365) EUT1
12360 Tsp_rosa EUT1
12365 HMMetlyEutC_1_eut1(pf05985) EUT1 EP_NTERM
12370 HMMetlyEutB_1_EUT1(pf06751) EUT1
12375 HMM06277EutA_1_eut(pf06277) EUT1
12380 HMMtpEuth_1_eut(pf04346) EUT1
12385 HMM00465alcdh_3_mix(pf00465) EUT1
12390 HMM06723EutJMreB_1_grm EUT1
12395 HMMalddh_1_eutpdugrm(pf00171) EUT1 EP_NTERM
12400 P_lightViolet EUT1
12405 H_azure EUT1
12410 HMM01515PTA_B_1_eut1(pf01515) EUT1
12415 HMM01923cobtr_2_eut1(pf01923) EUT1
12420 HMM06249EutQ_1_eut1(pf06249) EUT1 EP_NTERM
12425 HMM10662PduV_1_mix(pf10662) EUT1
12430 Hp_euts_violetPink EUT1
12435 HMM00390_malic_1_eut1(pf00390pf03949pf01515) EUT1
12440 -pf00923 -
12445 HMM02780keto_1_eut1(pf02779pf02780pf00456) EUT1
12450 - -
12455 -pf06674 -
12460 HMM00293nuhyd_1_eut1(pf00293) EUT1
12465 -pf00144 -
12470 -pf13247pf14691pf07992 -
12475 -pf00672pf02518pf07730pf13675 -
12480 -pf00873 -
12485 - -
12490 -pf03960 -
12495 -pf07687pf01546 -
BMC-P
BMC-H
BMC-T
Aldehyde dehydrogenase
PTAC
Signature enzyme
Alcohol dehydrogenase
Regulator

Salmonella enterica subsp. enterica serovar Ank. Bacteria; Proteobacteria; Gammaproteobacteria; Enterobacterales;

E4A63__locus_2

EUT1

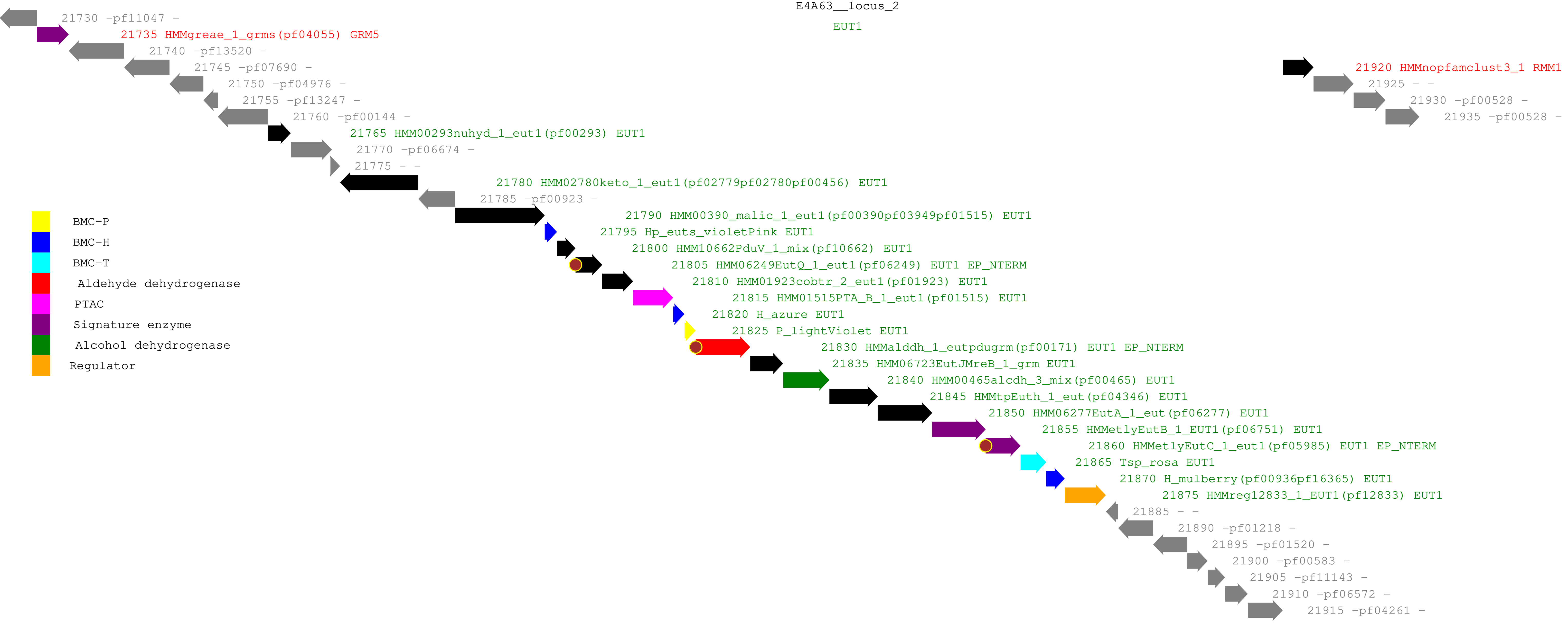

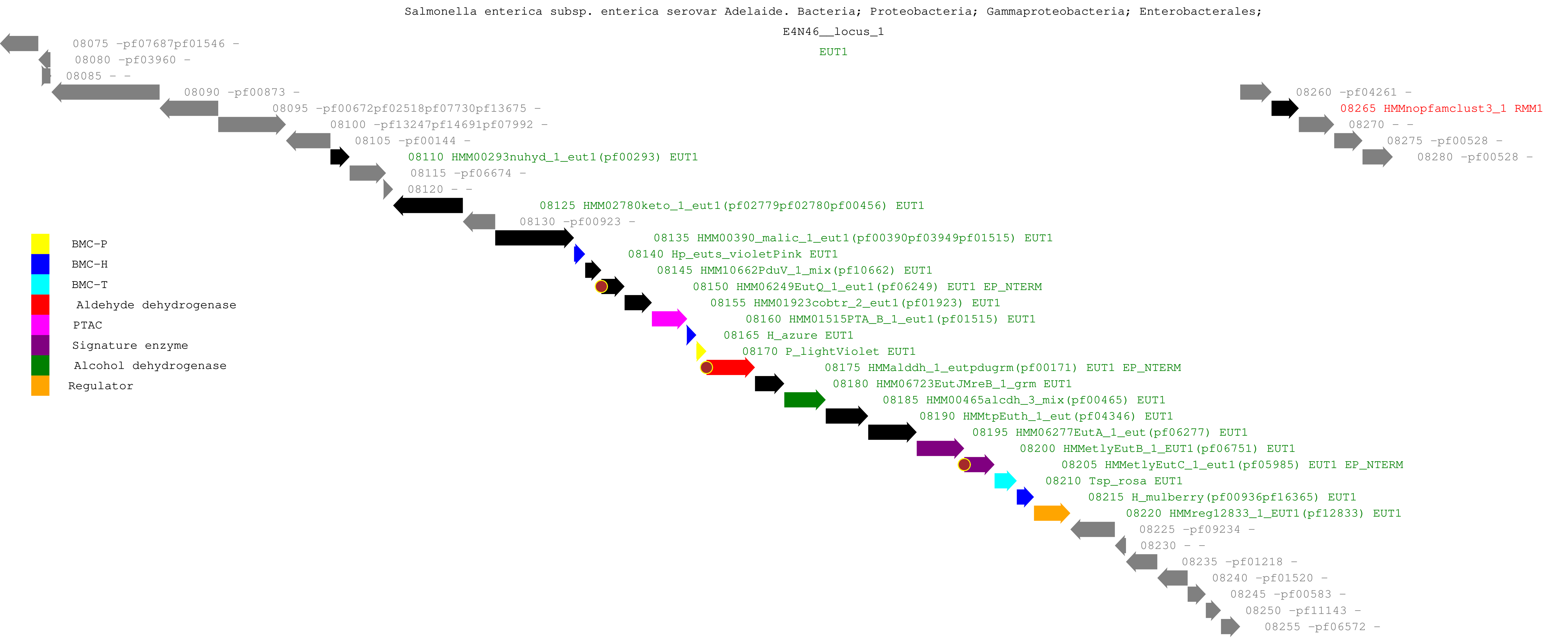
Salmonella enterica subsp. enterica serovar Adelaide. Bacteria; Proteobacteria; Gammaproteobacteria; Enterobacterales;
E4N46__locus_1
EUT1
08075 -pf07687pf01546 -
08080 -pf03960 -
08085 - -
08090 -pf00873 -
08095 -pf00672pf02518pf07730pf13675 -
08100 -pf13247pf14691pf07992 -
08105 -pf00144 -
08110 HMM00293nuhyd_1_eut1(pf00293) EUT1
08115 -pf06674 -
08120 - -
08125 HMM02780keto_1_eut1(pf02779pf02780pf00456) EUT1
08130 -pf00923 -
08135 HMM00390_malic_1_eut1(pf00390pf03949pf01515) EUT1
08140 Hp_euts_violetPink EUT1
08145 HMM10662PduV_1_mix(pf10662) EUT1
08150 HMM06249EutQ_1_eut1(pf06249) EUT1 EP_NTERM
08155 HMM01923cobtr_2_eut1(pf01923) EUT1
08160 HMM01515PTA_B_1_eut1(pf01515) EUT1
08165 H_azure EUT1
08170 P_lightViolet EUT1
08175 HMMalddh_1_eutpdugrm(pf00171) EUT1 EP_NTERM
08180 HMM06723EutJMreB_1_grm EUT1
08185 HMM00465alcdh_3_mix(pf00465) EUT1
08190 HMMtpEuth_1_eut(pf04346) EUT1
08195 HMM06277EutA_1_eut(pf06277) EUT1
08200 HMMetlyEutB_1_EUT1(pf06751) EUT1
08205 HMMetlyEutC_1_eut1(pf05985) EUT1 EP_NTERM
08210 Tsp_rosa EUT1
08215 H_mulberry(pf00936pf16365) EUT1
08220 HMMreg12833_1_EUT1(pf12833) EUT1
08225 -pf09234 -
08230 - -
08235 -pf01218 -
08240 -pf01520 -
08245 -pf00583 -
08250 -pf11143 -
08255 -pf06572 -
08260 -pf04261 -
08265 HMMnopfamclust3_1 RMM1
08270 - -
08275 -pf00528 -
08280 -pf00528 -
BMC-P
BMC-H
BMC-T
Aldehyde dehydrogenase
PTAC
Signature enzyme
Alcohol dehydrogenase
Regulator

Citrobacter sp. SNU WT2. Bacteria; Proteobacteria; Gammaproteobacteria; Enterobacterales;

E4Z61__locus_2

EUT1

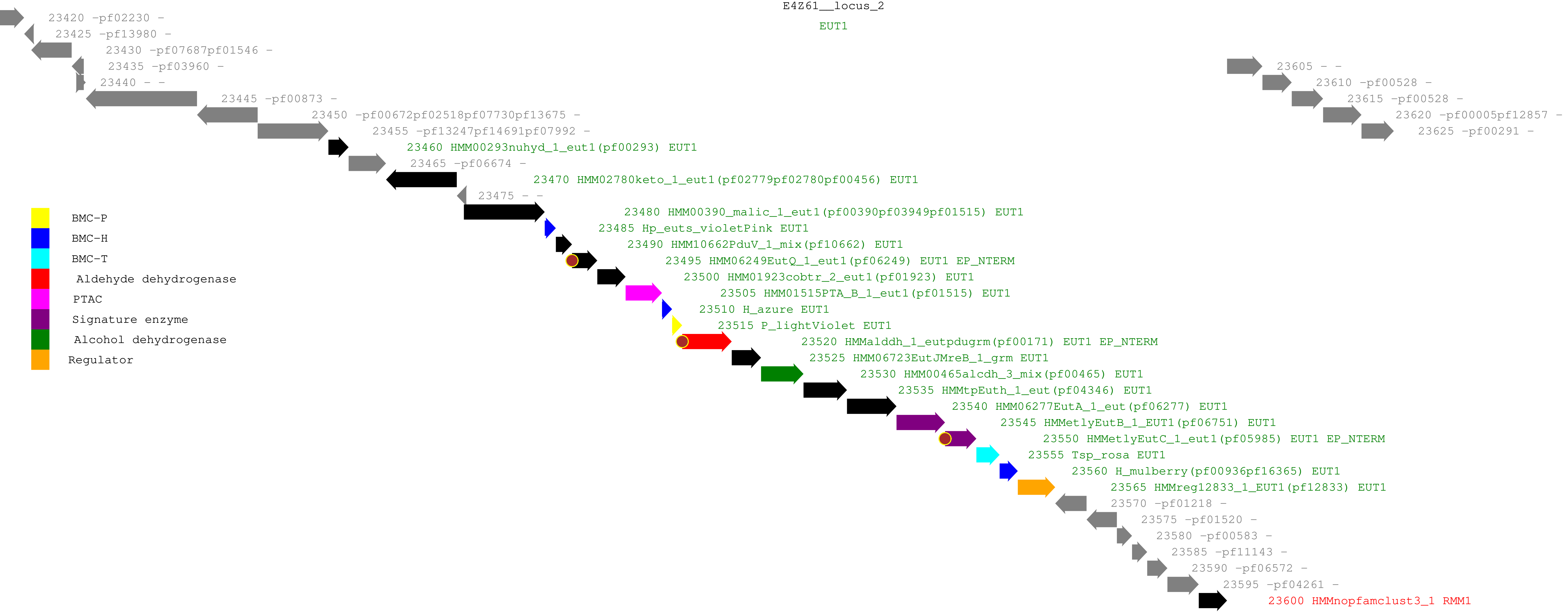

Salmonella enterica subsp. enterica serovar Florida. Bacteria; Proteobacteria; Gammaproteobacteria; Enterobacterales;

E5156__locus_2

EUT1

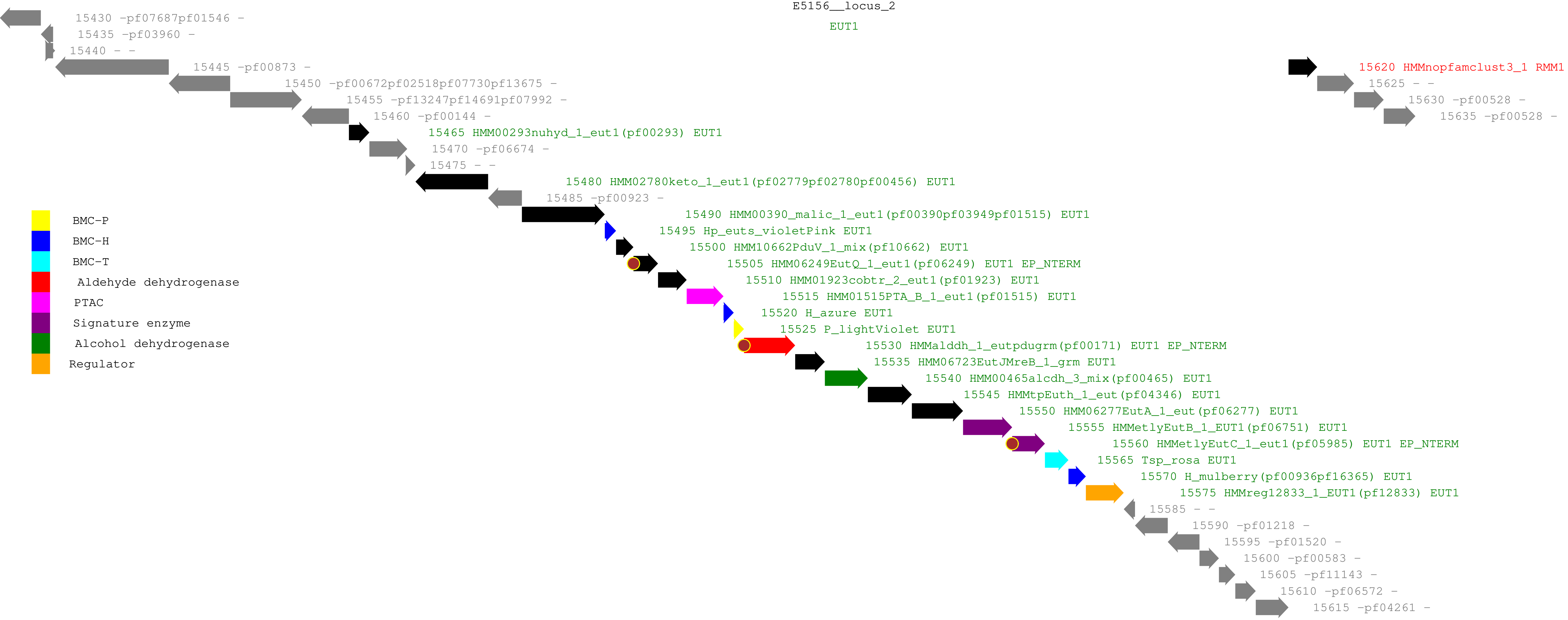

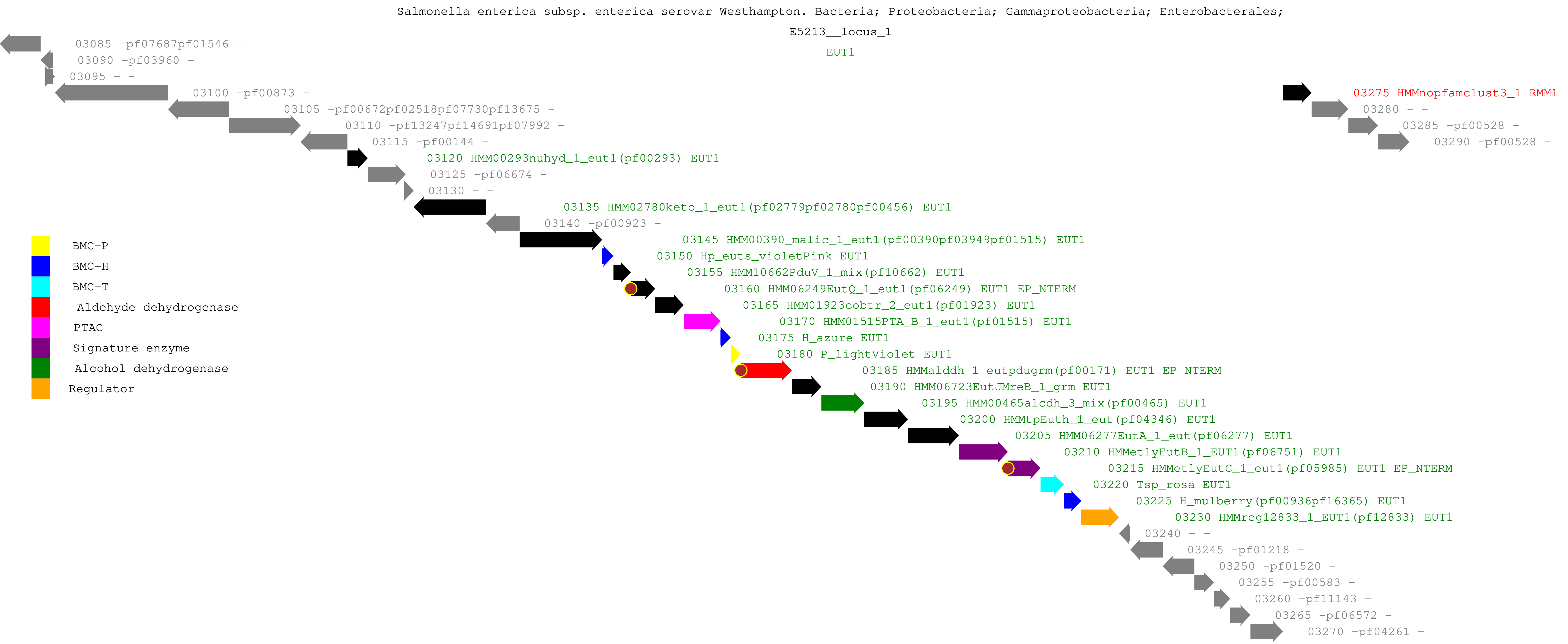

Salmonella enterica subsp. enterica serovar Westhampton. Bacteria; Proteobacteria; Gammaproteobacteria; Enterobacterales;
E5213__locus_1
EUT1
03085 -pf07687pf01546 -
03090 -pf03960 -
03095 - -
03100 -pf00873 -
03105 -pf00672pf02518pf07730pf13675 -
03110 -pf13247pf14691pf07992 -
03115 -pf00144 -
03120 HMM00293nuhyd_1_eut1(pf00293) EUT1
03125 -pf06674 -
03130 - -
03135 HMM02780keto_1_eut1(pf02779pf02780pf00456) EUT1
03140 -pf00923 -
03145 HMM00390_malic_1_eut1(pf00390pf03949pf01515) EUT1
03150 Hp_euts_violetPink EUT1
03155 HMM10662PduV_1_mix(pf10662) EUT1
03160 HMM06249EutQ_1_eut1(pf06249) EUT1 EP_NTERM
03165 HMM01923cobtr_2_eut1(pf01923) EUT1
03170 HMM01515PTA_B_1_eut1(pf01515) EUT1
03175 H_azure EUT1
03180 P_lightViolet EUT1
03185 HMMalddh_1_eutpdugrm(pf00171) EUT1 EP_NTERM
03190 HMM06723EutJMreB_1_grm EUT1
03195 HMM00465alcdh_3_mix(pf00465) EUT1
03200 HMMtpEuth_1_eut(pf04346) EUT1
03205 HMM06277EutA_1_eut(pf06277) EUT1
03210 HMMetlyEutB_1_EUT1(pf06751) EUT1
03215 HMMetlyEutC_1_eut1(pf05985) EUT1 EP_NTERM
03220 Tsp_rosa EUT1
03225 H_mulberry(pf00936pf16365) EUT1
03230 HMMreg12833_1_EUT1(pf12833) EUT1
03240 - -
03245 -pf01218 -
03250 -pf01520 -
03255 -pf00583 -
03260 -pf11143 -
03265 -pf06572 -
03270 -pf04261 -
03275 HMMnopfamclust3_1 RMM1
03280 - -
03285 -pf00528 -
03290 -pf00528 -
BMC-P
BMC-H
BMC-T
Aldehyde dehydrogenase
PTAC
Signature enzyme
Alcohol dehydrogenase
Regulator

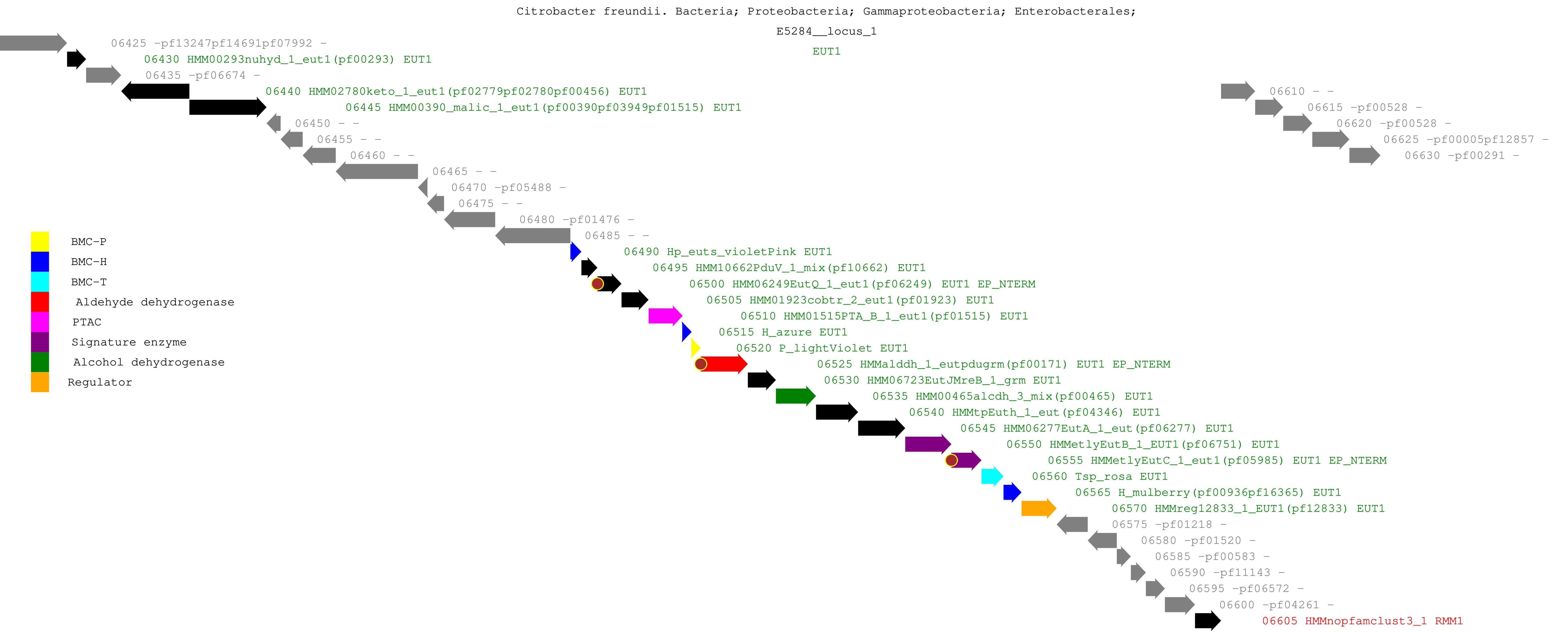
Citrobacter freundii. Bacteria; Proteobacteria; Gammaproteobacteria; Enterobacterales;
E5284__locus_1
EUT1
06425 -pf13247pf14691pf07992 -
06430 HMM00293nuhyd_1_eut1(pf00293) EUT1
06435 -pf06674 -
06440 HMM02780keto_1_eut1(pf02779pf02780pf00456) EUT1
06445 HMM00390_malic_1_eut1(pf00390pf03949pf01515) EUT1
06450 - -
06455 - -
06460 - -
06465 - -
06470 -pf05488 -
06475 - -
06480 -pf01476 -
06485 - -
06490 Hp_euts_violetPink EUT1
06495 HMM10662PduV_1_mix(pf10662) EUT1
06500 HMM06249EutQ_1_eut1(pf06249) EUT1 EP_NTERM
06505 HMM01923cobtr_2_eut1(pf01923) EUT1
06510 HMM01515PTA_B_1_eut1(pf01515) EUT1
06515 H_azure EUT1
06520 P_lightViolet EUT1
06525 HMMaldh_1_eutpdugrm(pf00171) EUT1 EP_NTERM
06530 HMM06723EutJMreB_1_grm EUT1
06535 HMM00465alcdh_3_mix(pf00465) EUT1
06540 HMMtpEuth_1_eut(pf04346) EUT1
06545 HMM06277EutA_1_eut(pf06277) EUT1
06550 HMMetlyEutB_1_EUT1(pf06751) EUT1
06555 HMMetlyEutC_1_eut1(pf05985) EUT1 EP_NTERM
06560 Tsp_rosa EUT1
06565 H_mulberry(pf00936pf16365) EUT1
06570 HMMreg12833_1_EUT1(pf12833) EUT1
06575 -pf01218 -
06580 -pf01520 -
06585 -pf00583 -
06590 -pf11143 -
06595 -pf06572 -
06600 -pf04261 -
06605 HMMnopfamclust3_1 RMM1
06610 - -
06615 -pf00528 -
06620 -pf00528 -
06625 -pf00005pf12857 -
06630 -pf00291 -
BMC-P
BMC-H
BMC-T
Aldehyde dehydrogenase
PTAC
Signature enzyme
Alcohol dehydrogenase
Regulator

Escherichia coli O157:H7. Bacteria; Proteobacteria; Gammaproteobacteria; Enterobacterales;

E5F07__locus_1

EUT1

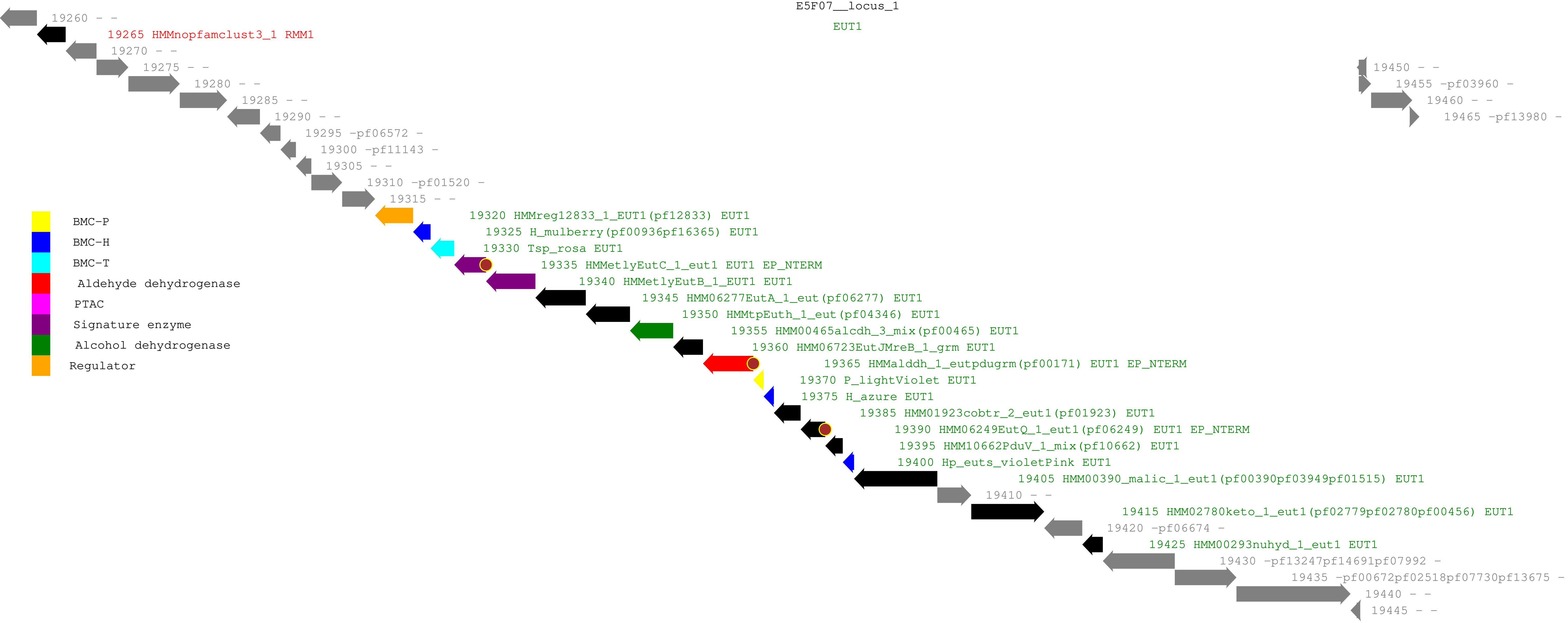

Salmonella enterica subsp. enterica serovar Lagos. Bacteria; Proteobacteria; Gammaproteobacteria; Enterobacterales;

E5J91__locus_2

EUT1

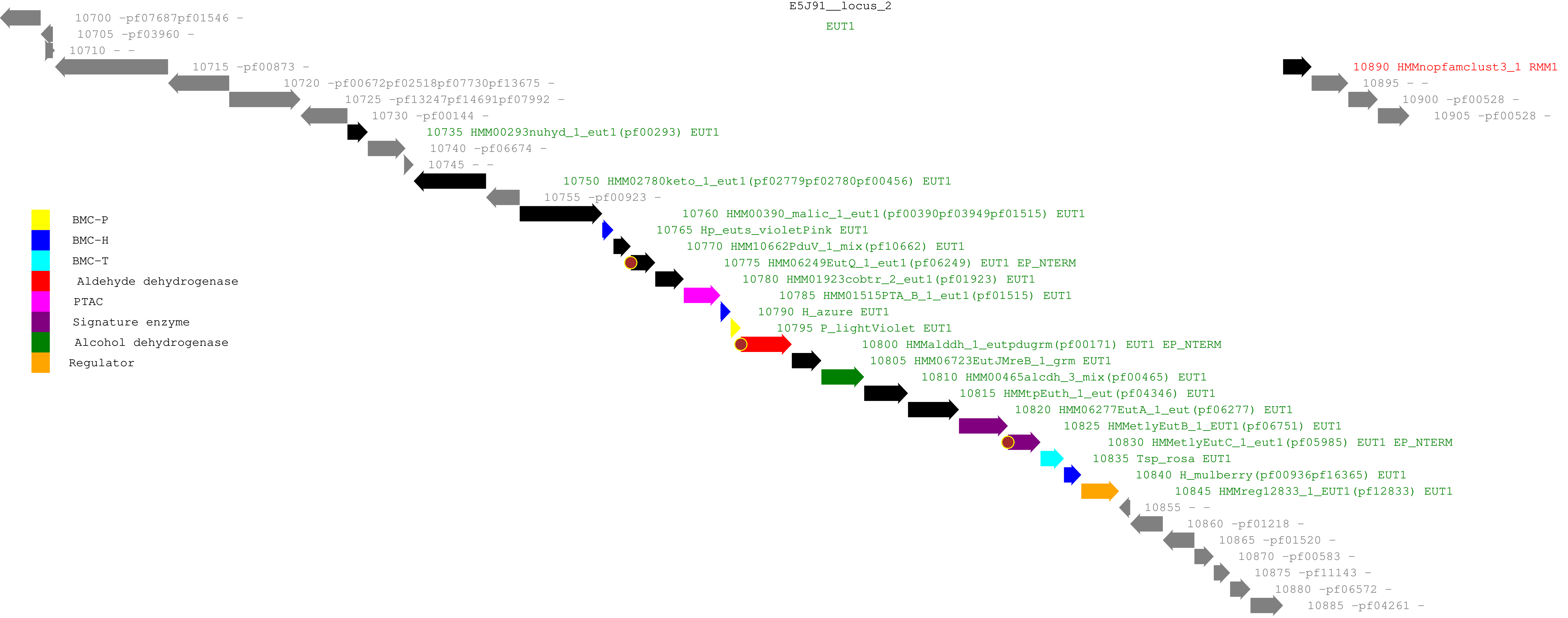

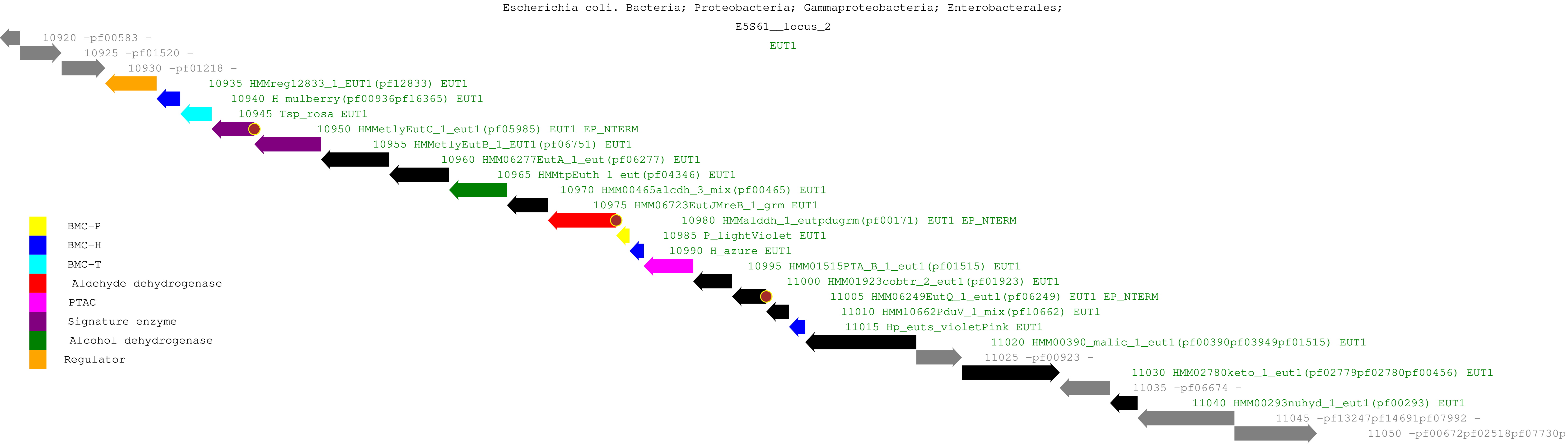
Escherichia coli. Bacteria; Proteobacteria; Gammaproteobacteria; Enterobacterales;
E5S61__locus_2
EUT1
10920 -pf00583 -
10925 -pf01520 -
10930 -pf01218 -
10935 HMMreg12833_1_EUT1(pf12833) EUT1
10940 H_mulberry(pf00936pf16365) EUT1
10945 Tsp_rosa EUT1
10950 HMMetlyEutC_1_eut1(pf05985) EUT1 EP_NTERM
10955 HMMetlyEutB_1_EUT1(pf06751) EUT1
10960 HMM06277EutA_1_eut(pf06277) EUT1
10965 HMMtpEuth_1_eut(pf04346) EUT1
10970 HMM00465alcdh_3_mix(pf00465) EUT1
10975 HMM06723EutJMreB_1_grm EUT1
10980 HMMalddh_1_eutpdugrm(pf00171) EUT1 EP_NTERM
10985 P_lightViolet EUT1
10990 H_azure EUT1
10995 HMM01515PTA_B_1_eut1(pf01515) EUT1
11000 HMM01923cobtr_2_eut1(pf01923) EUT1
11005 HMM06249EutQ_1_eut1(pf06249) EUT1 EP_NTERM
11010 HMM10662PduV_1_mix(pf10662) EUT1
11015 Hp_euts_violetPink EUT1
11020 HMM00390_malic_1_eut1(pf00390pf03949pf01515) EUT1
11025 -pf00923 -
11030 HMM02780keto_1_eut1(pf02779pf02780pf00456) EUT1
11035 -pf06674 -
11040 HMM00293nuhyd_1_eut1(pf00293) EUT1
11045 -pf13247pf14691pf07992 -
11050 -pf00672pf02518pf07730p
BMC-P
BMC-H
BMC-T
Aldehyde dehydrogenase
PTAC
Signature enzyme
Alcohol dehydrogenase
Regulator

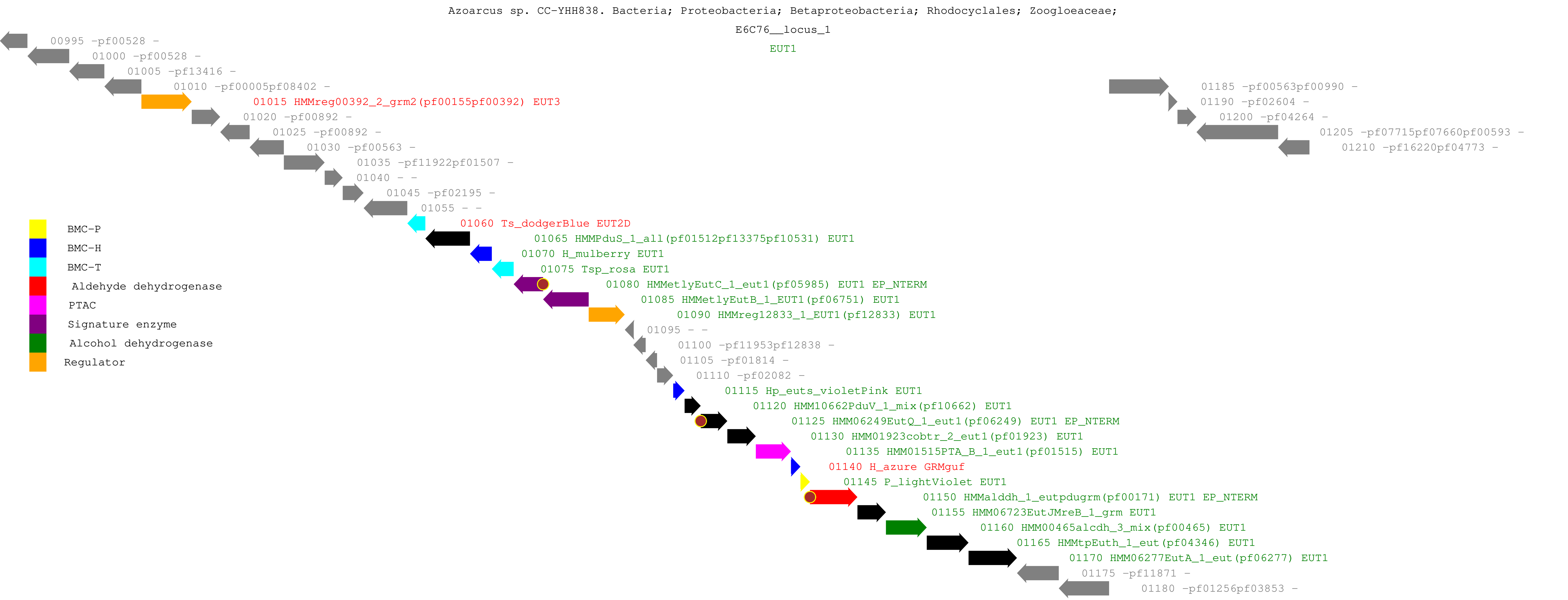
Azoarcus sp. CC-YHH838. Bacteria; Proteobacteria; Betaproteobacteria; Rhodocyclales; Zoogloeaceae;
E6C76__locus_1
EUT1
00995 -pf00528 -
01000 -pf00528 -
01005 -pf13416 -
01010 -pf00005pf08402 -
01015 HMMreg00392_2_grm2(pf00155pf00392) EUT3
01020 -pf00892 -
01025 -pf00892 -
01030 -pf00563 -
01035 -pf11922pf01507 -
01040 - -
01045 -pf02195 -
01055 - -
01060 Ts_dodgerBlue EUT2D
01065 HMMPduS_1_all(pf01512pf13375pf10531) EUT1
01070 H_mulberry EUT1
01075 Tsp_rosa EUT1
01080 HMMetlyEutC_1_eut1(pf05985) EUT1 EP_NTERM
01085 HMMetlyEutB_1_EUT1(pf06751) EUT1
01090 HMMreg12833_1_EUT1(pf12833) EUT1
01095 - -
01100 -pf11953pf12838 -
01105 -pf01814 -
01110 -pf02082 -
01115 Hp_euts_violetPink EUT1
01120 HMM10662PduV_1_mix(pf10662) EUT1
01125 HMM06249EutQ_1_eut1(pf06249) EUT1 EP_NTERM
01130 HMM01923cobtr_2_eut1(pf01923) EUT1
01135 HMM01515PTA_B_1_eut1(pf01515) EUT1
01140 H_azure GRMguf
01145 P_lightViolet EUT1
01150 HMMalddh_1_eutpdugrm(pf00171) EUT1 EP_NTERM
01155 HMM06723EutJMreB_1_grm EUT1
01160 HMM00465alcdh_3_mix(pf00465) EUT1
01165 HMMtpEuth_1_eut(pf04346) EUT1
01170 HMM06277EutA_1_eut(pf06277) EUT1
01175 -pf11871 -
01180 -pf01256pf03853 -
01185 -pf00563pf00990 -
01190 -pf02604 -
01200 -pf04264 -
01205 -pf07715pf07660pf00593 -
01210 -pf16220pf04773 -
BMC-P
BMC-H
BMC-T
Aldehyde dehydrogenase
PTAC
Signature enzyme
Alcohol dehydrogenase
Regulator

Azoarcus sp. CC-YHH848. Bacteria; Proteobacteria; Betaproteobacteria; Rhodocyclales; Zoogloeaceae;

E6O51__locus_1

EUT1

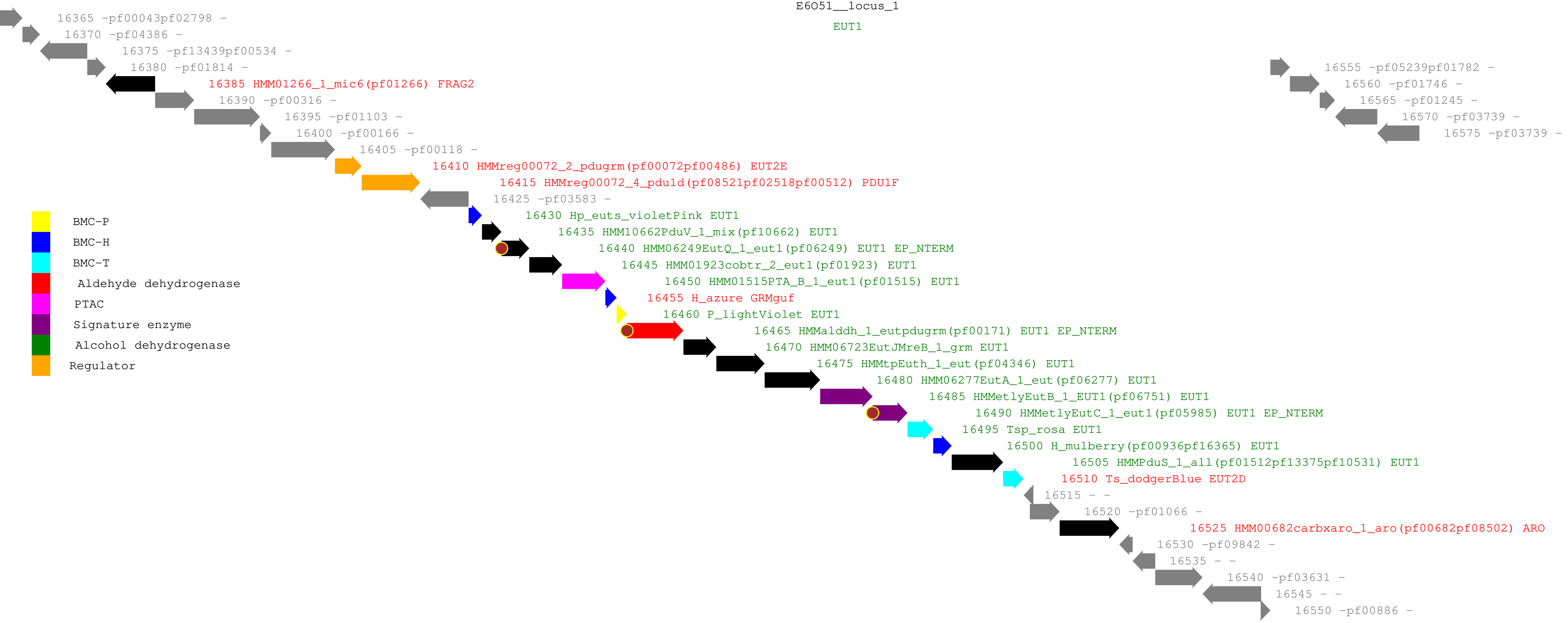

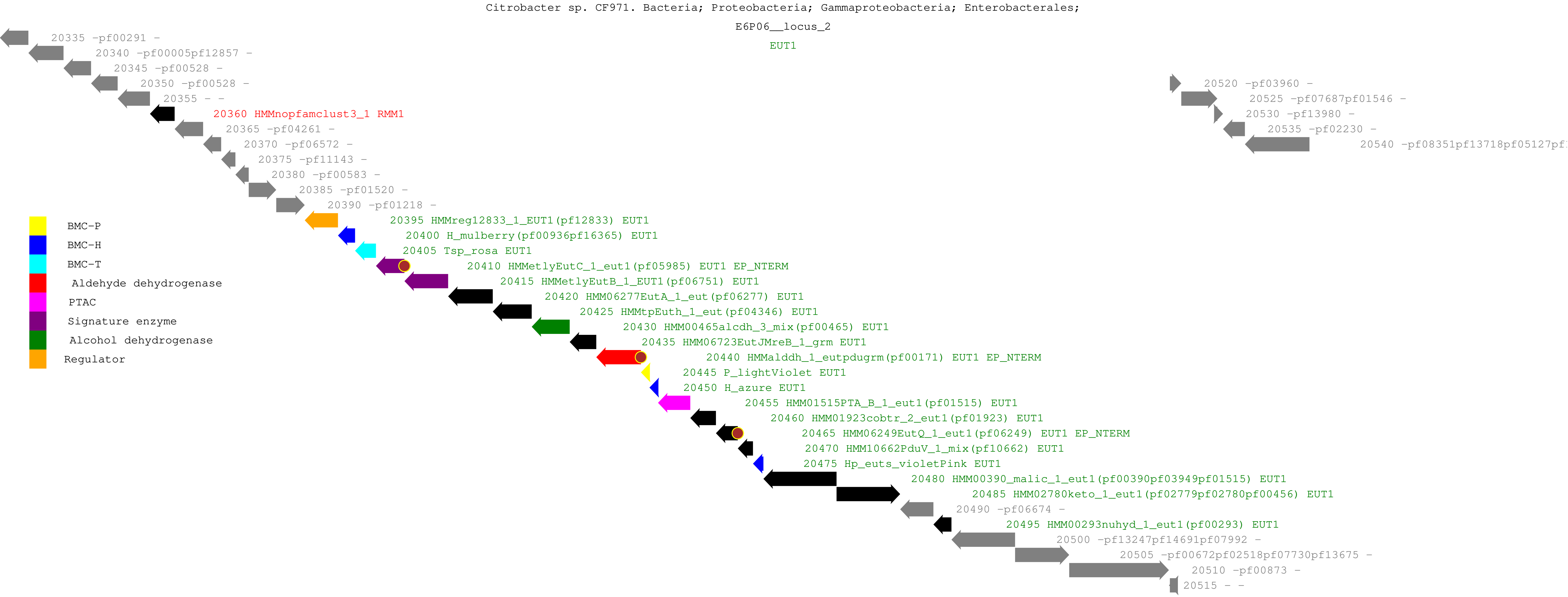
Citrobacter sp. CF971. Bacteria; Proteobacteria; Gammaproteobacteria; Enterobacterales;
E6P06__locus_2
EUT1
20335 -pf00291 -
20340 -pf00005pf12857 -
20345 -pf00528 -
20350 -pf00528 -
20355 - -
20360 HMMnopfamclust3_1 RMM1
20365 -pf04261 -
20370 -pf06572 -
20375 -pf11143 -
20380 -pf00583 -
20385 -pf01520 -
20390 -pf01218 -
20395 HMMreg12833_1_EUT1(pf12833) EUT1
20400 H_mulberry(pf00936pf16365) EUT1
20405 Tsp_rosa EUT1
20410 HMMetlyEutC_1_eut1(pf05985) EUT1 EP_NTERM
20415 HMMetlyEutB_1_EUT1(pf06751) EUT1
20420 HMM06277EutA_1_eut(pf06277) EUT1
20425 HMMtpEuth_1_eut(pf04346) EUT1
20430 HMM00465alcdh_3_mix(pf00465) EUT1
20435 HMM06723EutJMreB_1_grm EUT1
20440 HMMalddh_1_eutpdugrm(pf00171) EUT1 EP_NTERM
20445 P_lightViolet EUT1
20450 H_azure EUT1
20455 HMM01515PTA_B_1_eut1(pf01515) EUT1
20460 HMM01923cobtr_2_eut1(pf01923) EUT1
20465 HMM06249EutQ_1_eut1(pf06249) EUT1 EP_NTERM
20470 HMM10662PduV_1_mix(pf10662) EUT1
20475 Hp_euts_violetPink EUT1
20480 HMM00390_malic_1_eut1(pf00390pf03949pf01515) EUT1
20485 HMM02780keto_1_eut1(pf02779pf02780pf00456) EUT1
20490 -pf06674 -
20495 HMM00293nuhyd_1_eut1(pf00293) EUT1
20500 -pf13247pf14691pf07992 -
20505 -pf00672pf02518pf07730pf13675 -
20510 -pf00873 -
20515 - -
20520 -pf03960 -
20525 -pf07687pf01546 -
20530 -pf13980 -
20535 -pf02230 -
20540 -pf08351pf13718pf05127pf
BMC-P
BMC-H
BMC-T
Aldehyde dehydrogenase
PTAC
Signature enzyme
Alcohol dehydrogenase
Regulator

Salmonella enterica subsp. enterica serovar Berkeley. Bacteria; Proteobacteria; Gammaproteobacteria; Enterobacterales;

E6W86__locus_1

EUT1

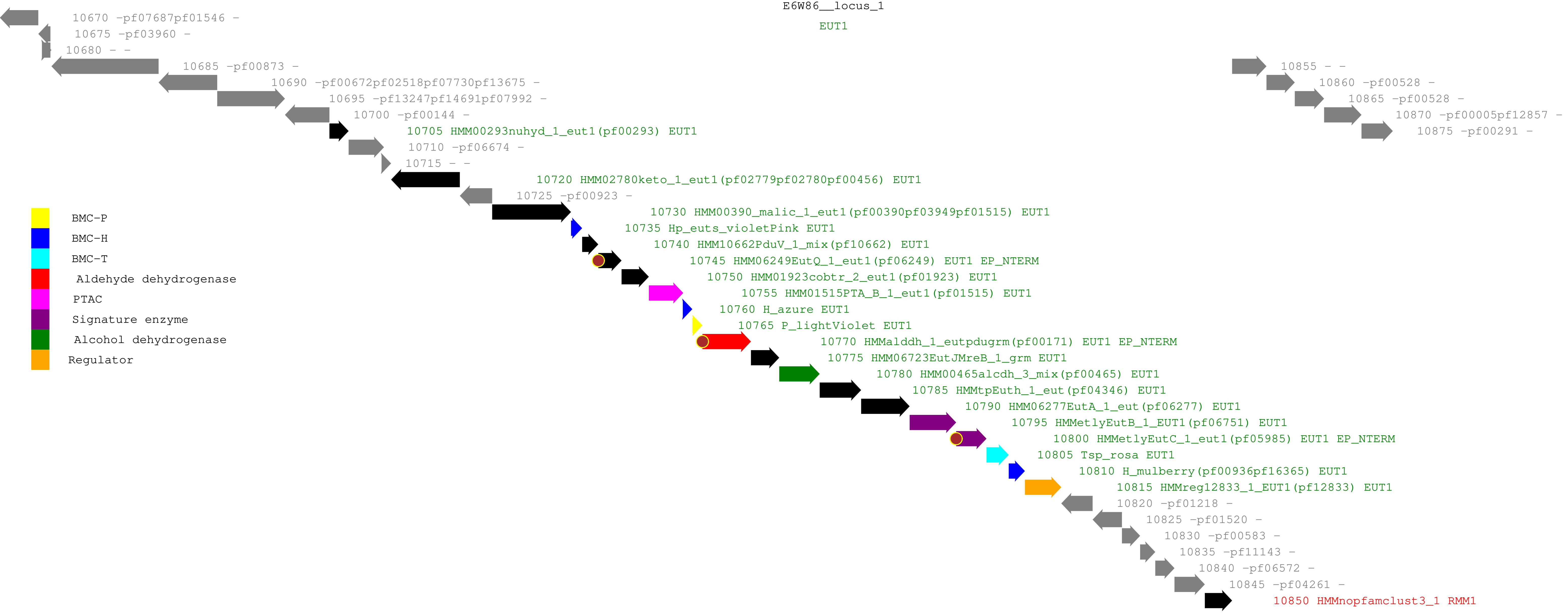

Salmonella enterica subsp. enterica serovar Uppsala. Bacteria; Proteobacteria; Gammaproteobacteria; Enterobacterales;

E6W93__locus_2

EUT1

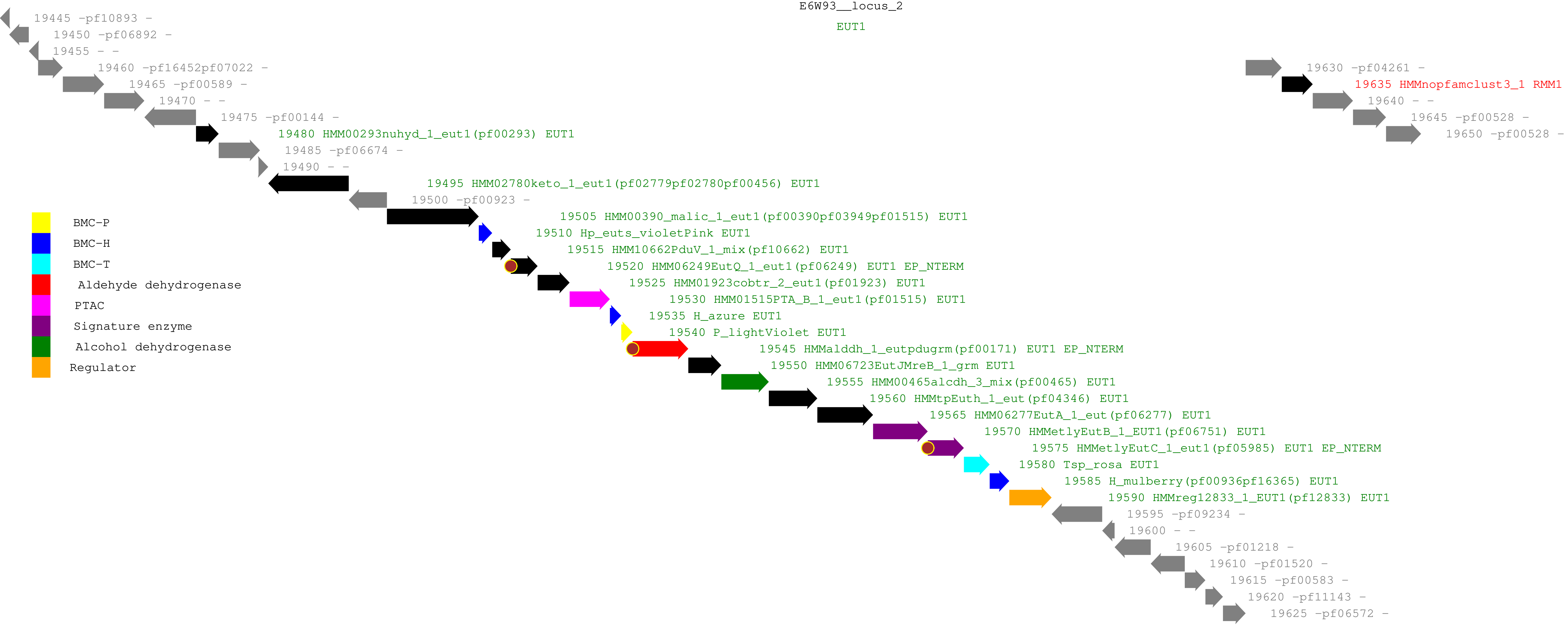

Salmonella enterica subsp. enterica serovar Poano. Bacteria; Proteobacteria; Gammaproteobacteria; Enterobacterales;

E6W94__locus_1

EUT1

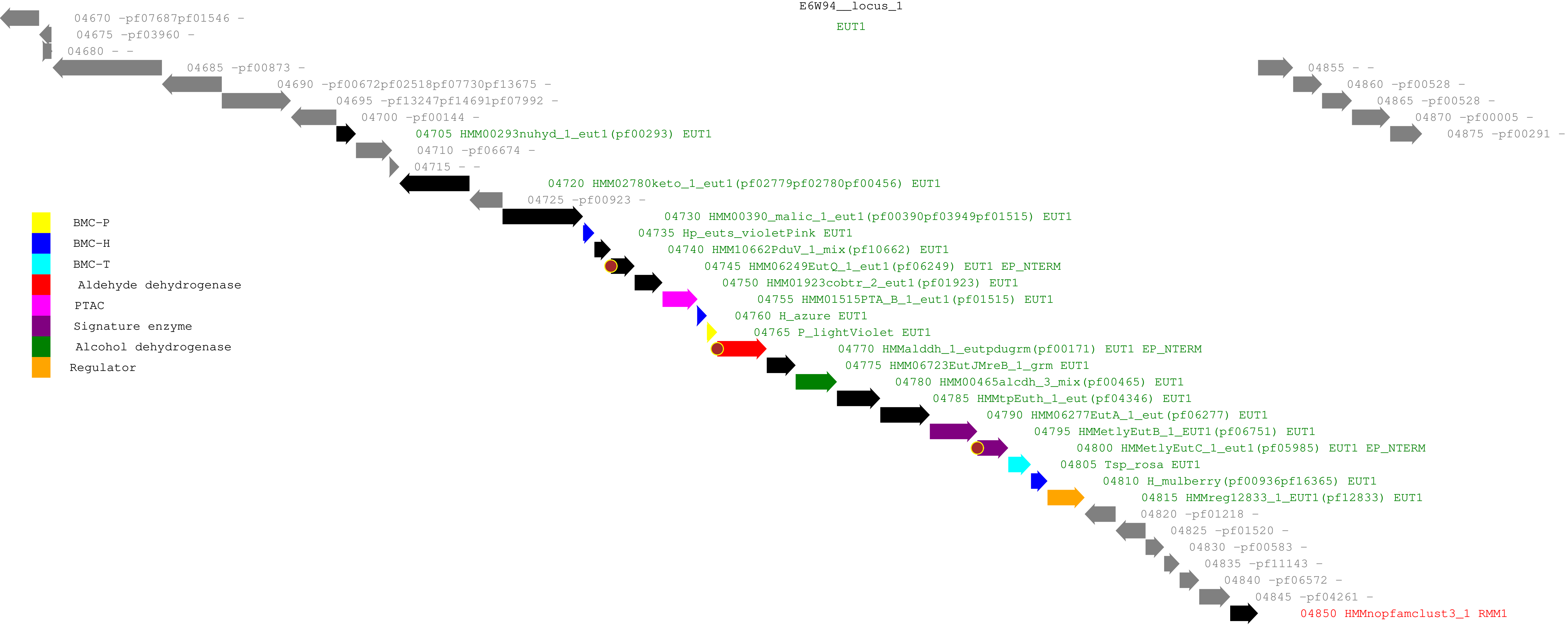

Salmonella enterica subsp. enterica serovar Tamale. Bacteria; Proteobacteria; Gammaproteobacteria; Enterobacterales;

E8C79__locus_1

EUT1

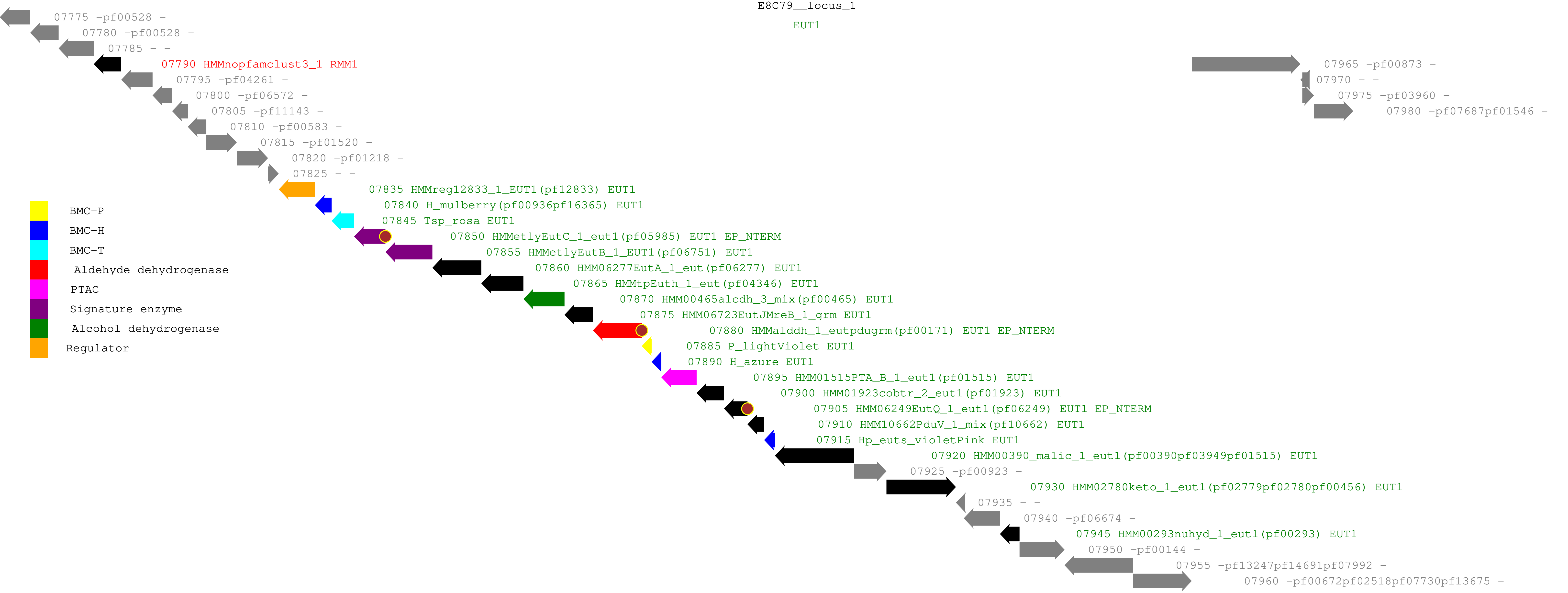

Salmonella enterica subsp. enterica serovar Amsterdam. Bacteria; Proteobacteria; Gammaproteobacteria; Enterobacterales;

E8Q13__locus_2

EUT1

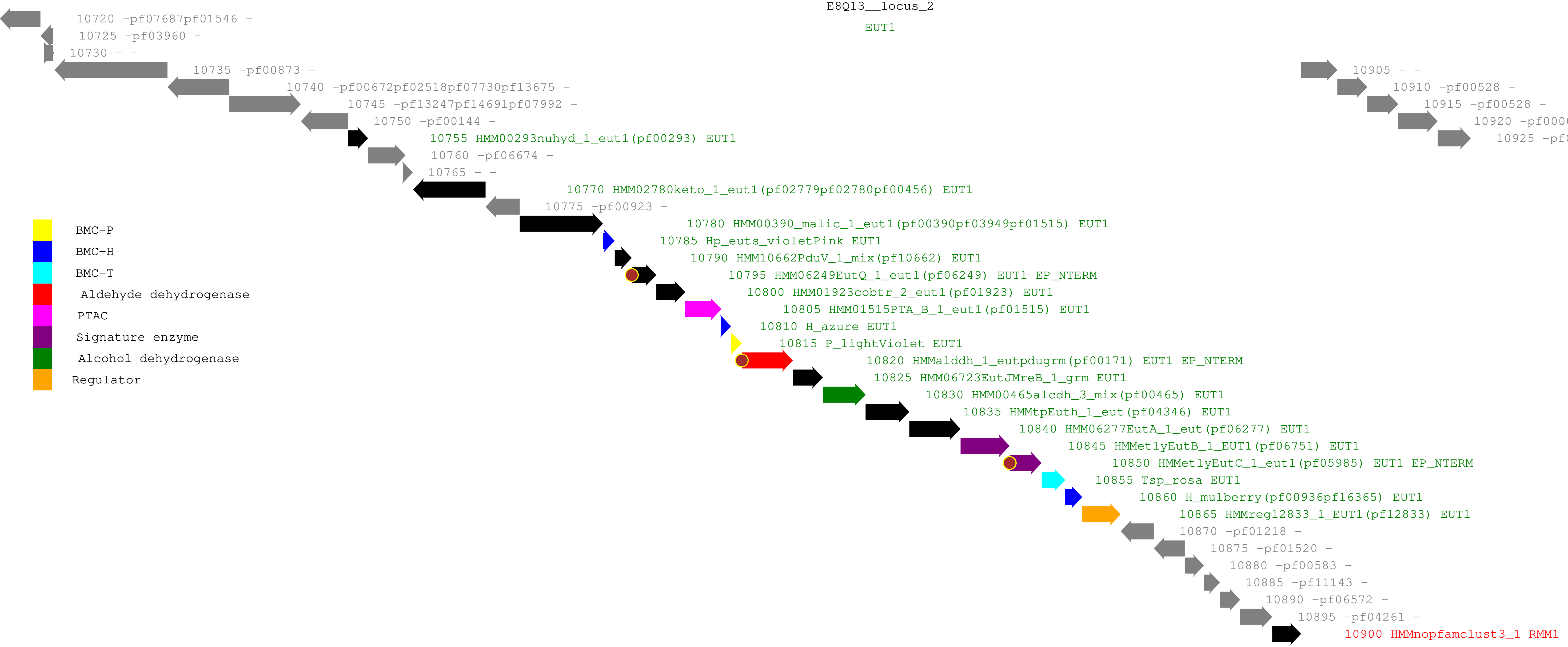

Salmonella enterica subsp. enterica serovar Goldcoast. Bacteria; Proteobacteria; Gammaproteobacteria; Enterobacterales;

E9126__locus_2

EUT1

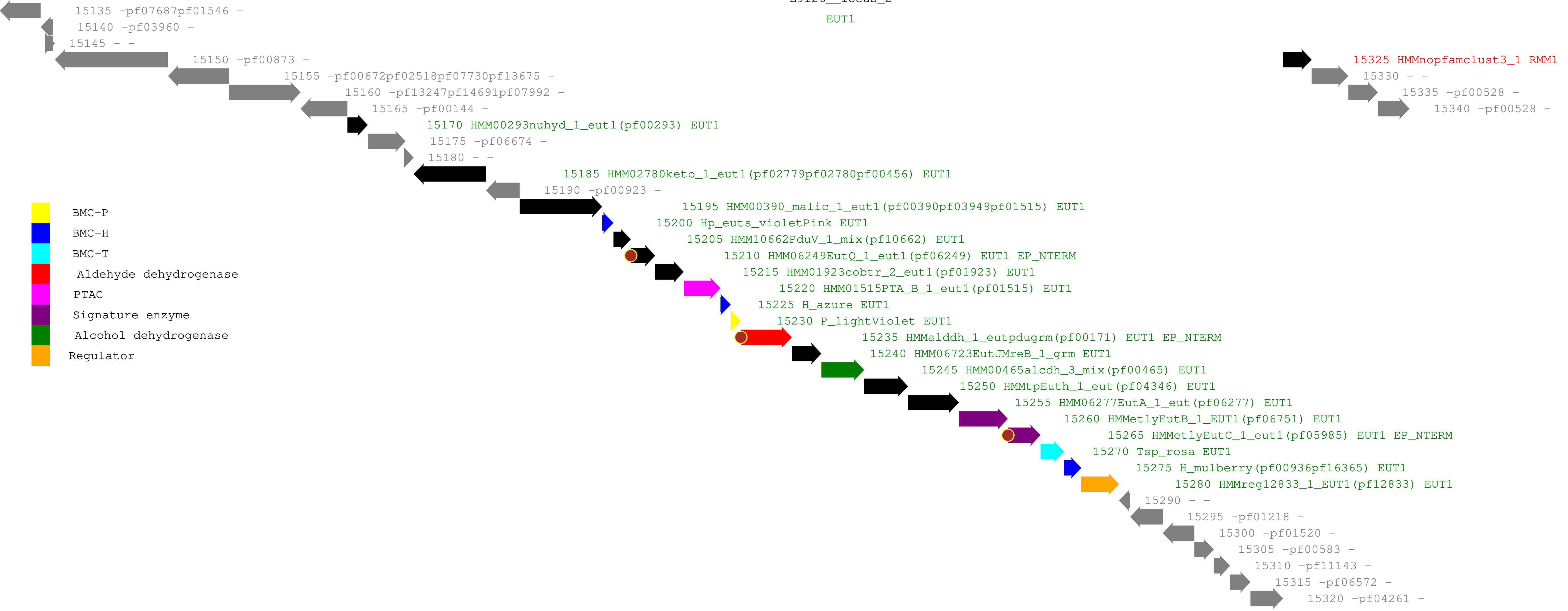

Klebsiella pneumoniae subsp. pneumoniae. Bacteria; Proteobacteria; Gammaproteobacteria; Enterobacterales;

E9161__locus_2

EUT1

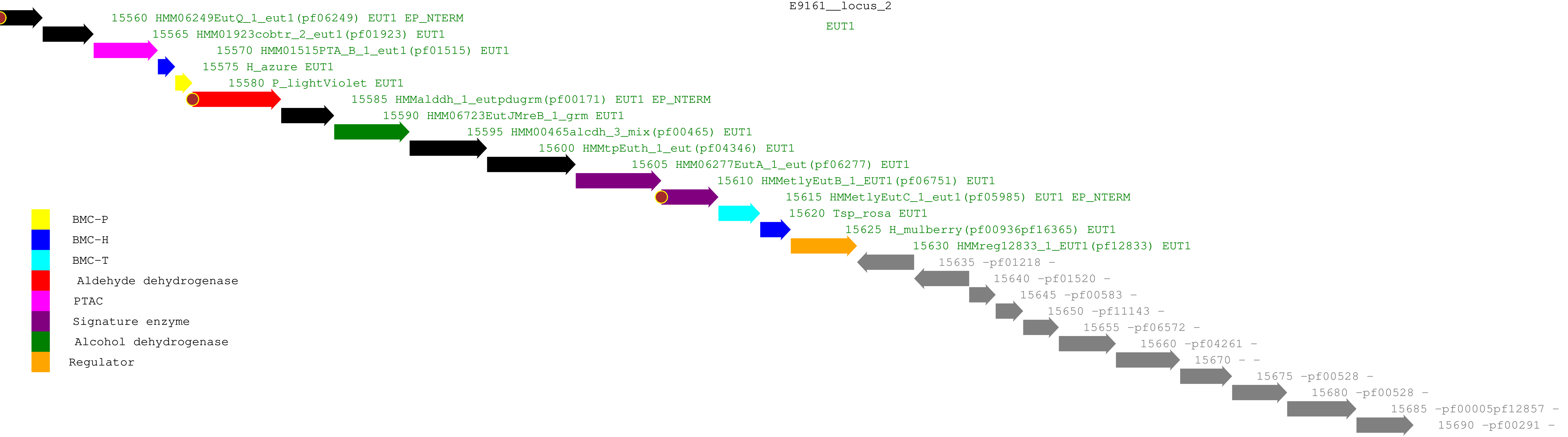

Escherichia albertii. Bacteria; Proteobacteria; Gammaproteobacteria; Enterobacterales;

EAB3F__locus_2

EUT1

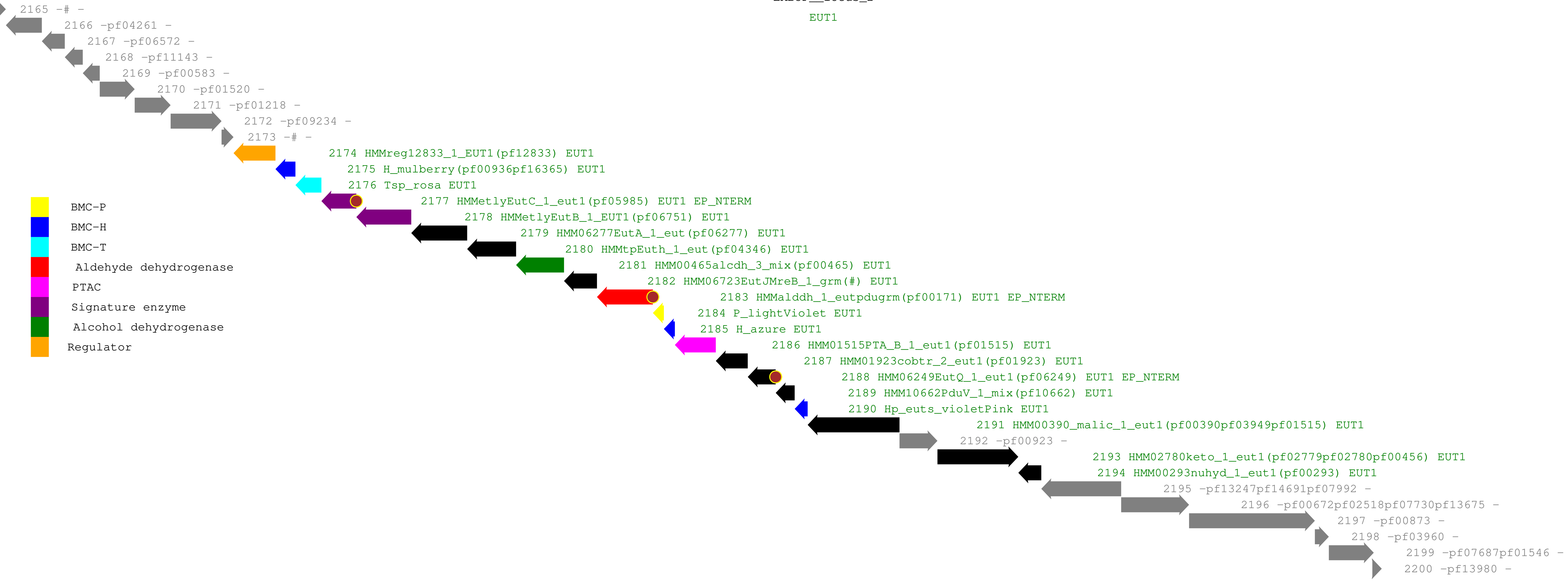

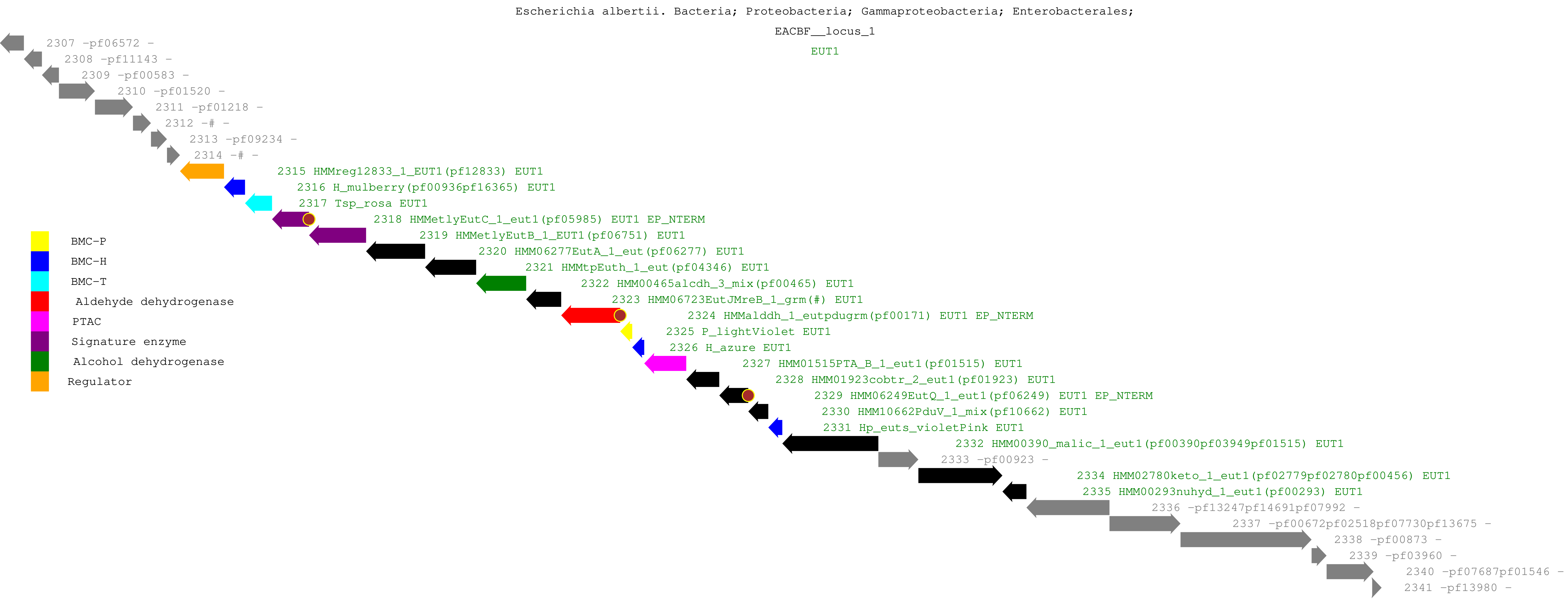

Escherichia albertii. Bacteria; Proteobacteria; Gammaproteobacteria; Enterobacterales;
EACBF__locus_1
EUT1
2307 -pf06572 -
2308 -pf11143 -
2309 -pf00583 -
2310 -pf01520 -
2311 -pf01218 -
2312 -# -
2313 -pf09234 -
2314 -# -
2315 HMMreg12833_1_EUT1(pf12833) EUT1
2316 H_mulberry(pf00936pf16365) EUT1
2317 Tsp_rosa EUT1
2318 HMMetlyEutC_1_eut1(pf05985) EUT1 EP_NTERM
2319 HMMetlyEutB_1_EUT1(pf06751) EUT1
2320 HMM06277EutA_1_eut(pf06277) EUT1
2321 HMMtpEuth_1_eut(pf04346) EUT1
2322 HMM00465alcdh_3_mix(pf00465) EUT1
2323 HMM06723EutJMreB_1_grm(#) EUT1
2324 HMMalddh_1_eutpdugrm(pf00171) EUT1 EP_NTERM
2325 P_lightViolet EUT1
2326 H_azure EUT1
2327 HMM01515PTA_B_1_eut1(pf01515) EUT1
2328 HMM01923cobtr_2_eut1(pf01923) EUT1
2329 HMM06249EutQ_1_eut1(pf06249) EUT1 EP_NTERM
2330 HMM10662PduV_1_mix(pf10662) EUT1
2331 Hp_euts_violetPink EUT1
2332 HMM00390_malic_1_eut1(pf00390pf03949pf01515) EUT1
2333 -pf00923 -
2334 HMM02780keto_1_eut1(pf02779pf02780pf00456) EUT1
2335 HMM00293nuhyd_1_eut1(pf00293) EUT1
2336 -pf13247pf14691pf07992 -
2337 -pf00672pf02518pf07730pf13675 -
2338 -pf00873 -
2339 -pf03960 -
2340 -pf07687pf01546 -
2341 -pf13980 -
BMC-P
BMC-H
BMC-T
Aldehyde dehydrogenase
PTAC
Signature enzyme
Alcohol dehydrogenase
Regulator

Escherichia coli E1114. Bacteria; Proteobacteria; Gammaproteobacteria; Enterobacterales;

EADG__locus_1

EUT1

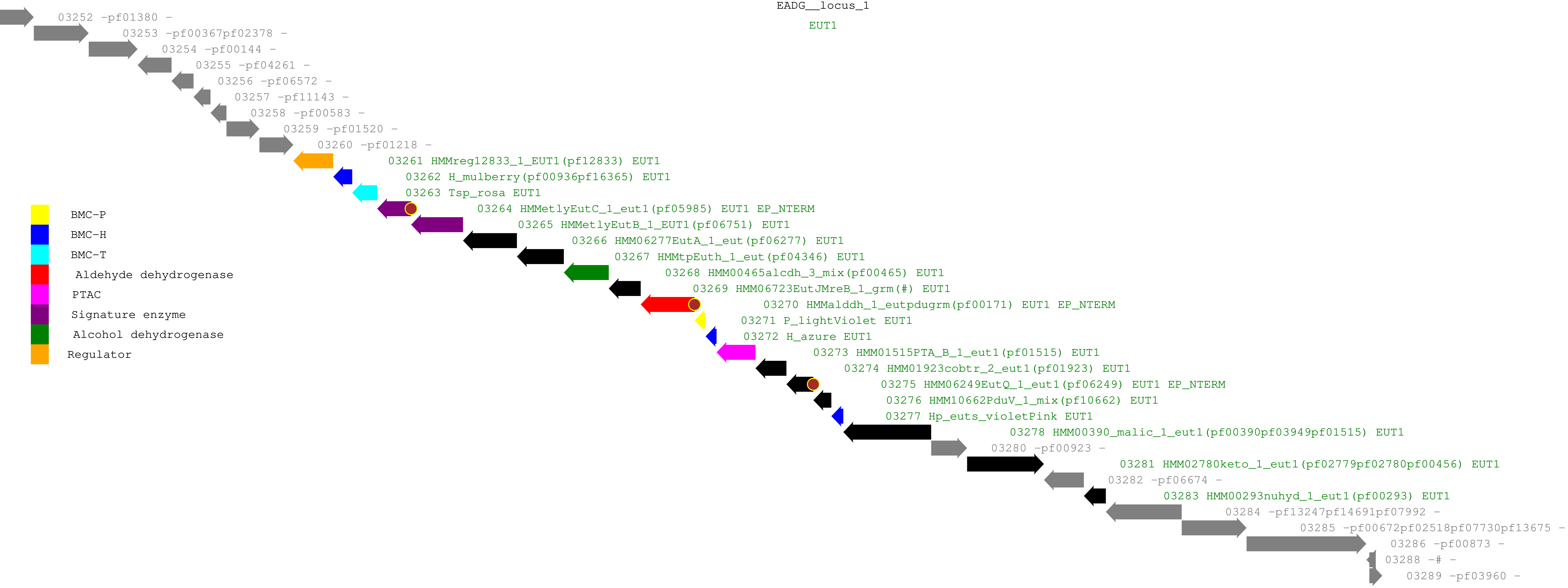

Escherichia coli. Bacteria; Proteobacteria; Gammaproteobacteria; Enterobacterales;

EAI42__locus_1

EUT1

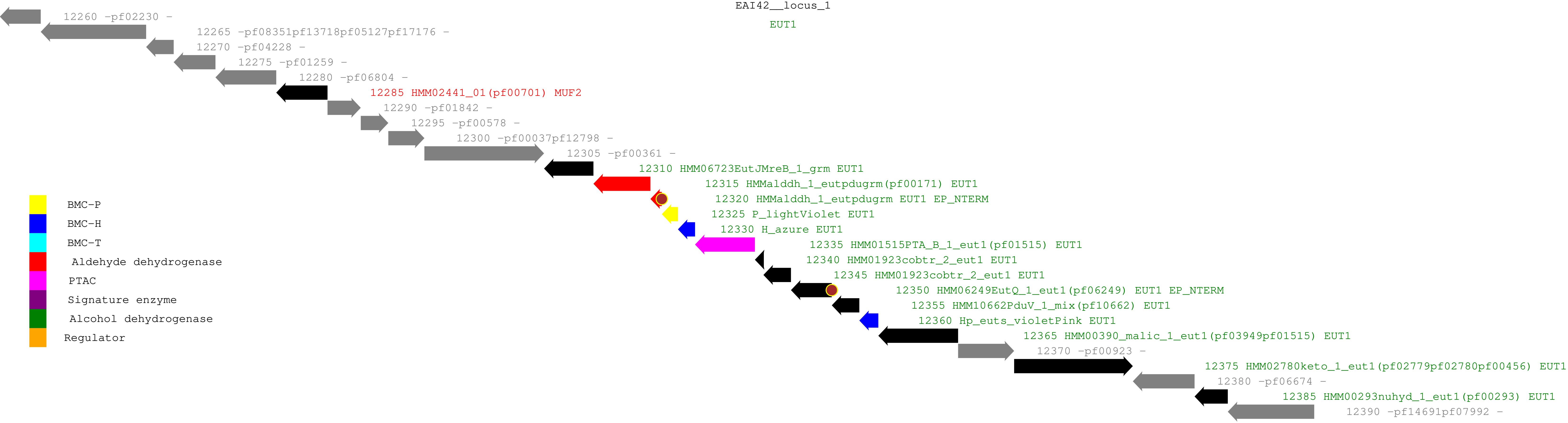

Escherichia albertii KF1. Bacteria; Proteobacteria; Gammaproteobacteria; Enterobacterales;

EAKF1_ch_locus_1

EUT1

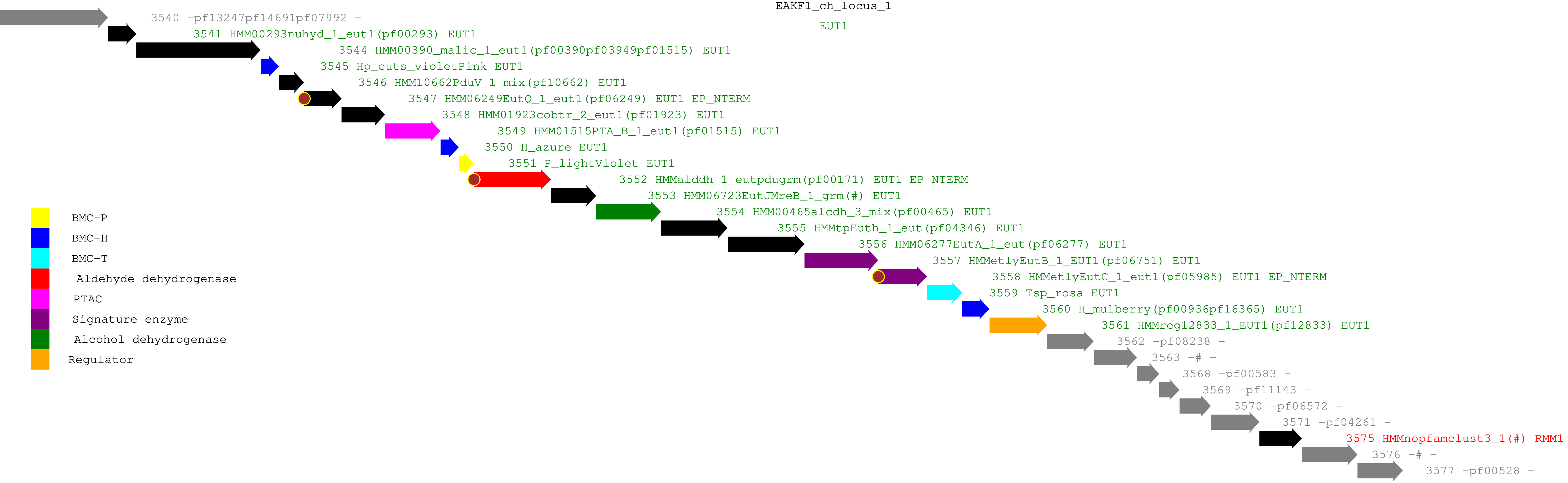

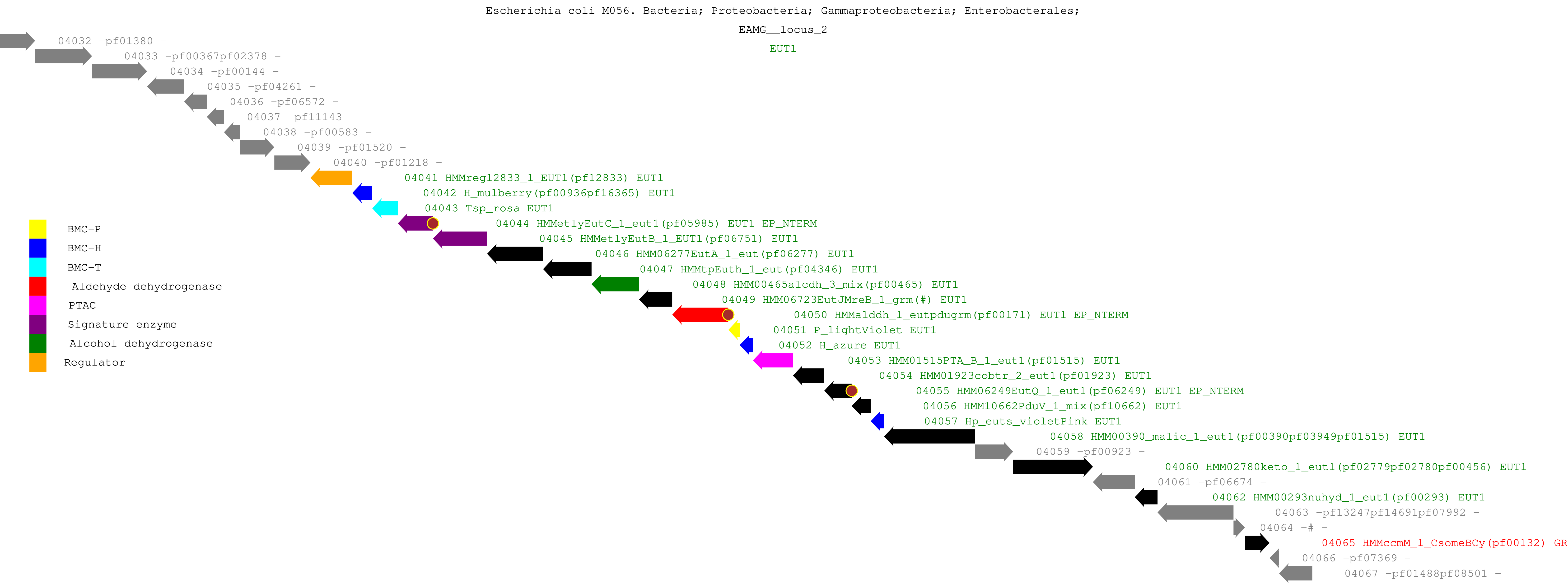

Escherichia coli M056. Bacteria; Proteobacteria; Gammaproteobacteria; Enterobacterales;
EAMG__locus_2
EUT1
04032 -pf01380 -
04033 -pf00367pf02378 -
04034 -pf00144 -
04035 -pf04261 -
04036 -pf06572 -
04037 -pf11143 -
04038 -pf00583 -
04039 -pf01520 -
04040 -pf01218 -
04041 HMMreg12833_1_EUT1(pf12833) EUT1
04042 H_mulberry(pf00936pf16365) EUT1
04043 Tsp_rosa EUT1
04044 HMMetlyEutC_1_eut1(pf05985) EUT1 EP_NTERM
04045 HMMetlyEutB_1_EUT1(pf06751) EUT1
04046 HMM06277EutA_1_eut(pf06277) EUT1
04047 HMMtpEuth_1_eut(pf04346) EUT1
04048 HMM00465alcdh_3_mix(pf00465) EUT1
04049 HMM06723EutJMreB_1_grm(#) EUT1
04050 HMMalddh_1_eutpdugrm(pf00171) EUT1 EP_NTERM
04051 P_lightViolet EUT1
04052 H_azure EUT1
04053 HMM01515PTA_B_1_eut1(pf01515) EUT1
04054 HMM01923cobtr_2_eut1(pf01923) EUT1
04055 HMM06249EutQ_1_eut1(pf06249) EUT1 EP_NTERM
04056 HMM10662PduV_1_mix(pf10662) EUT1
04057 Hp_euts_violetPink EUT1
04058 HMM00390_malic_1_eut1(pf00390pf03949pf01515) EUT1
04059 -pf00923 -
04060 HMM02780keto_1_eut1(pf02779pf02780pf00456) EUT1
04061 -pf06674 -
04062 HMM00293nuhyd_1_eut1(pf00293) EUT1
04063 -pf13247pf14691pf07992 -
04064 -# -
04065 HMMccmM_1_CsomeBCy(pf00132) GR
04066 -pf07369 -
04067 -pf01488pf08501 -
BMC-P
BMC-H
BMC-T
Aldehyde dehydrogenase
PTAC
Signature enzyme
Alcohol dehydrogenase
Regulator

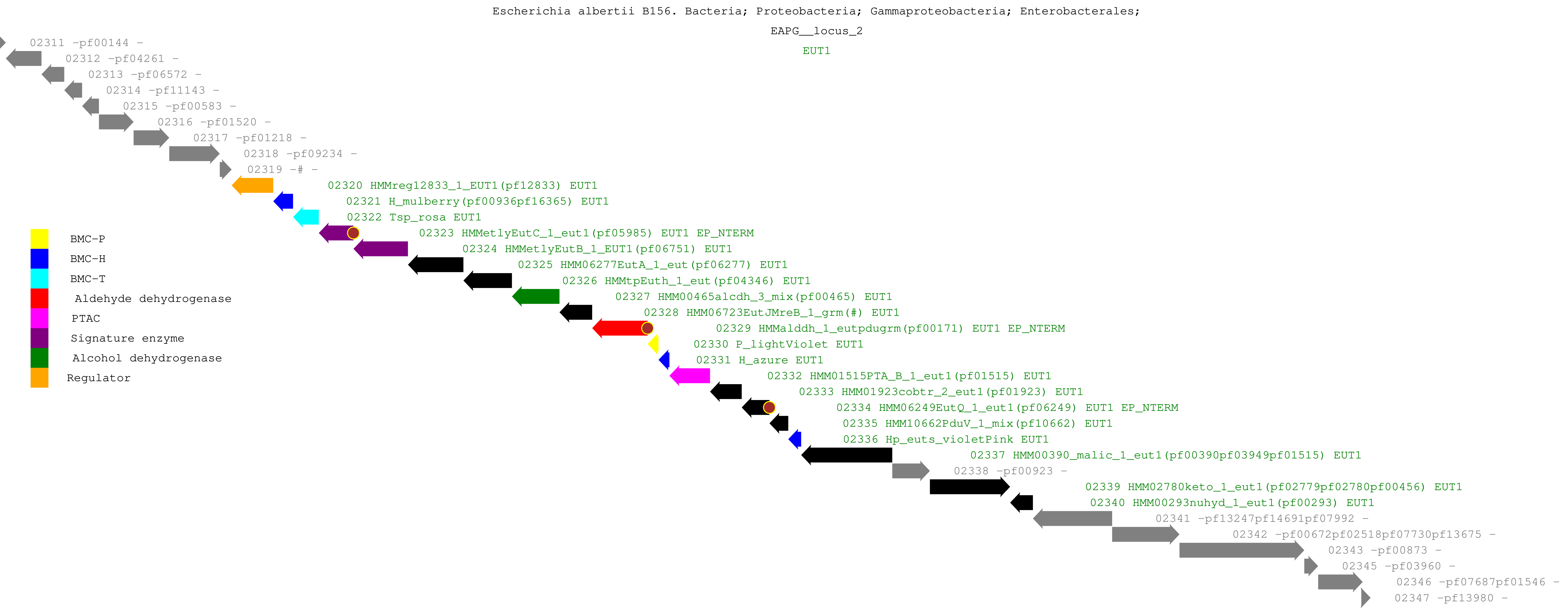

Escherichia albertii B156. Bacteria; Proteobacteria; Gammaproteobacteria; Enterobacterales;
EAPG__locus_2
EUT1
02311 -pf00144 -
02312 -pf04261 -
02313 -pf06572 -
02314 -pf11143 -
02315 -pf00583 -
02316 -pf01520 -
02317 -pf01218 -
02318 -pf09234 -
02319 -# -
02320 HMMreg12833_1_EUT1(pf12833) EUT1
02321 H_mulberry(pf00936pf16365) EUT1
02322 Tsp_rosa EUT1
02323 HMMetlyEutC_1_eut1(pf05985) EUT1 EP_NTERM
02324 HMMetlyEutB_1_EUT1(pf06751) EUT1
02325 HMM06277EutA_1_eut(pf06277) EUT1
02326 HMMtpEuth_1_eut(pf04346) EUT1
02327 HMM00465alcdh_3_mix(pf00465) EUT1
02328 HMM06723EutJMreB_1_grm(#) EUT1
02329 HMMalddh_1_eutpdugrm(pf00171) EUT1 EP_NTERM
02330 P_lightViolet EUT1
02331 H_azure EUT1
02332 HMM01515PTA_B_1_eut1(pf01515) EUT1
02333 HMM01923cobtr_2_eut1(pf01923) EUT1
02334 HMM06249EutQ_1_eut1(pf06249) EUT1 EP_NTERM
02335 HMM10662PduV_1_mix(pf10662) EUT1
02336 Hp_euts_violetPink EUT1
02337 HMM00390_malic_1_eut1(pf00390pf03949pf01515) EUT1
02338 -pf00923 -
02339 HMM02780keto_1_eut1(pf02779pf02780pf00456) EUT1
02340 HMM00293nuhyd_1_eut1(pf00293) EUT1
02341 -pf13247pf14691pf07992 -
02342 -pf00672pf02518pf07730pf13675 -
02343 -pf00873 -
02345 -pf03960 -
02346 -pf07687pf01546 -
02347 -pf13980 -
BMC-P
BMC-H
BMC-T
Aldehyde dehydrogenase
PTAC
Signature enzyme
Alcohol dehydrogenase
Regulator

Escherichia coli H461. Bacteria; Proteobacteria; Gammaproteobacteria; Enterobacterales;

EARG__locus_1

EUT1

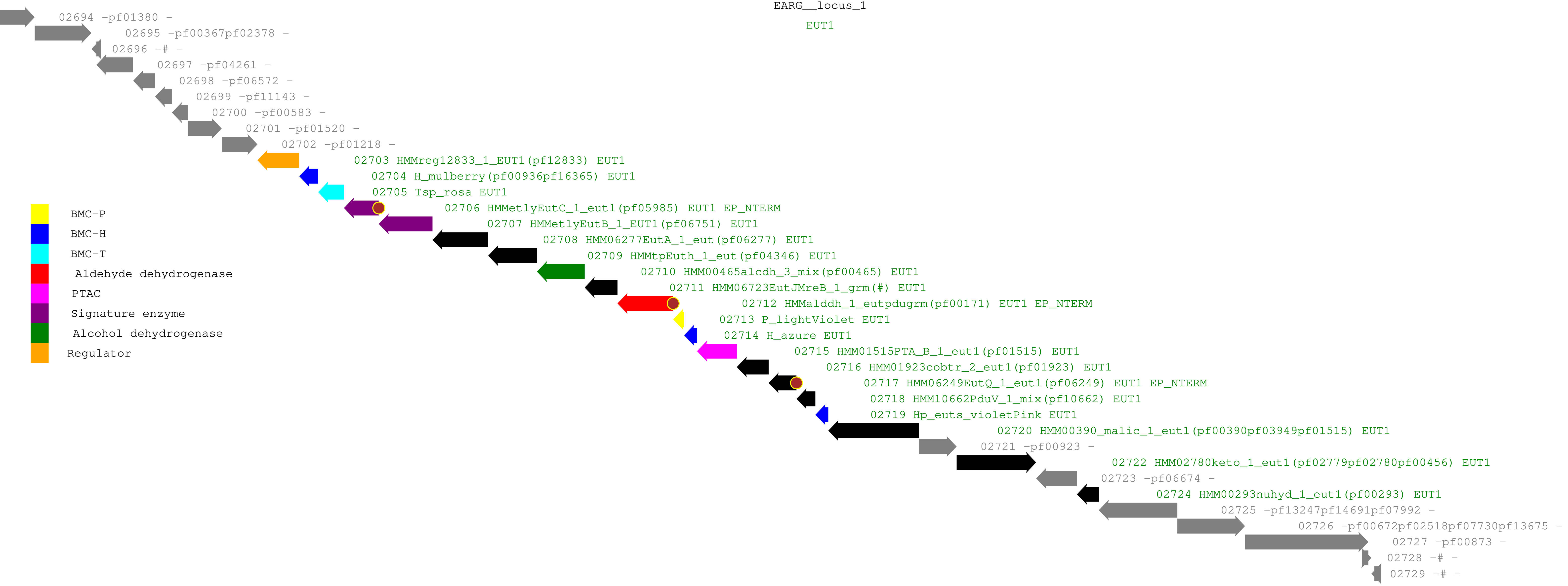

Escherichia coli H605. Bacteria; Proteobacteria; Gammaproteobacteria; Enterobacterales;

EATG__locus_1

EUT1

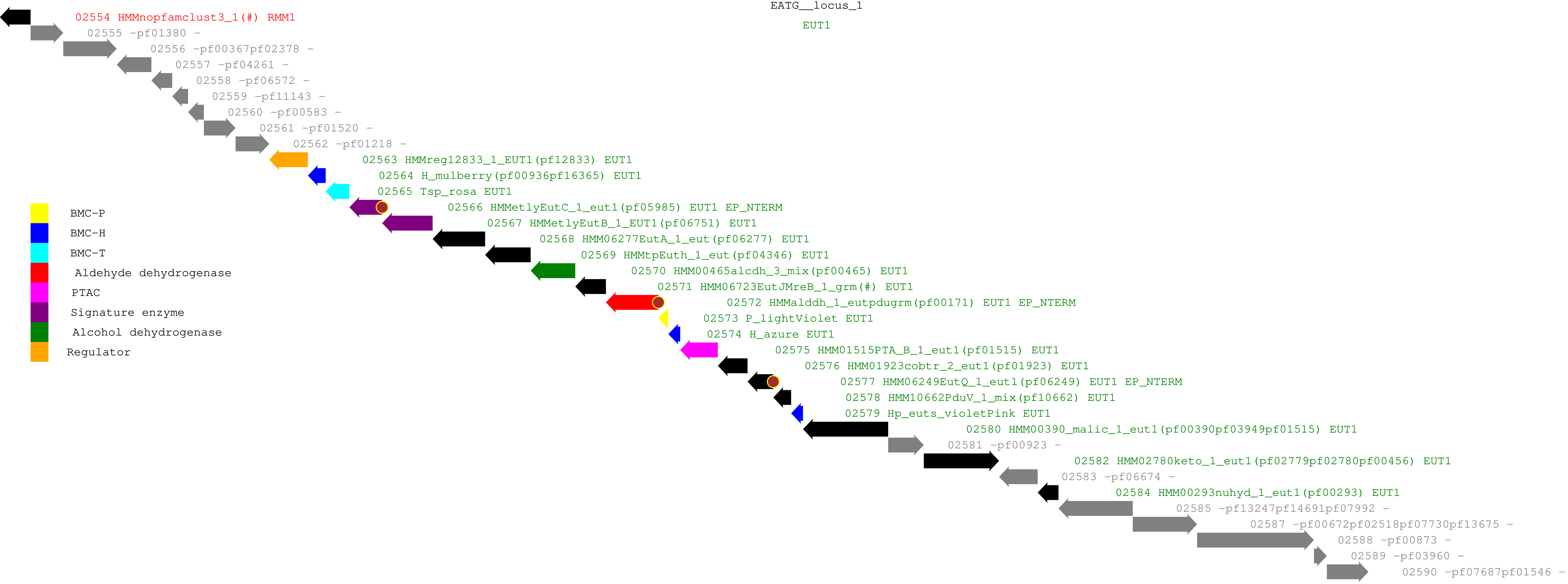

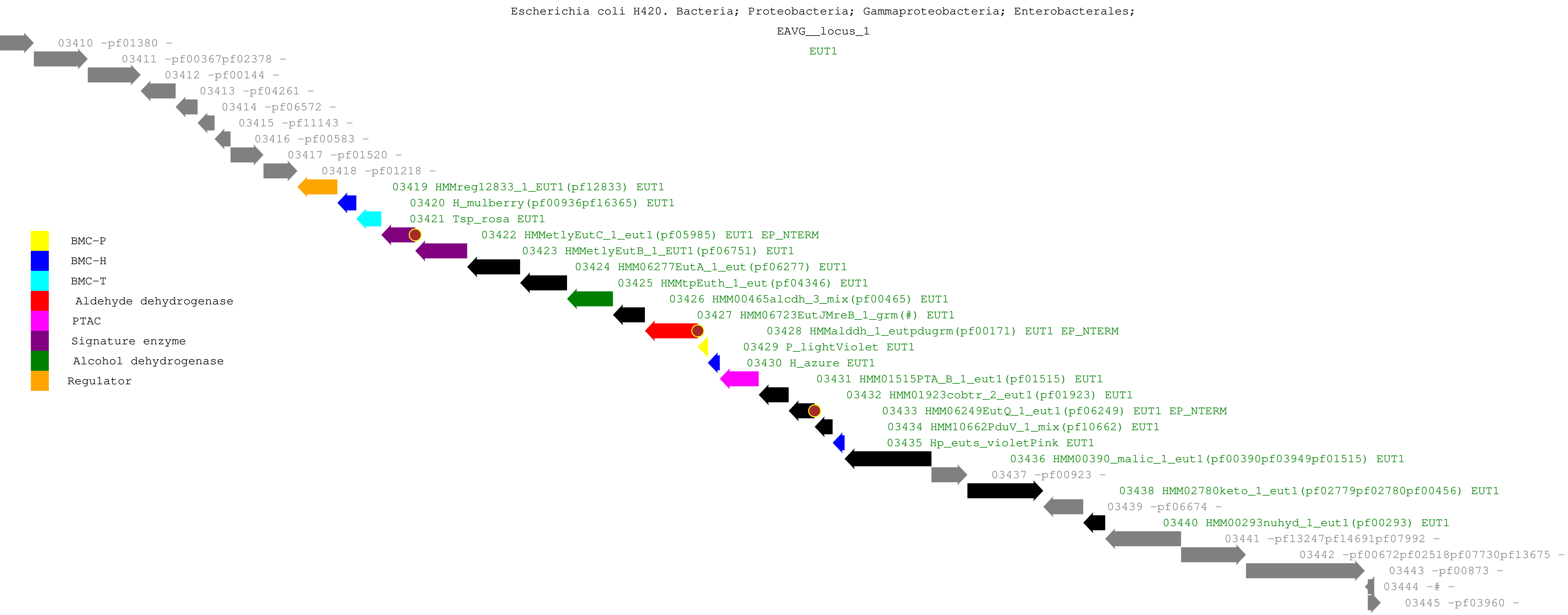
Escherichia coli H420. Bacteria; Proteobacteria; Gammaproteobacteria; Enterobacterales;
EAVG__locus_1
EUT1
03410 -pf01380 -
03411 -pf00367pf02378 -
03412 -pf00144 -
03413 -pf04261 -
03414 -pf06572 -
03415 -pf11143 -
03416 -pf00583 -
03417 -pf01520 -
03418 -pf01218 -
03419 HMMreg12833_1_EUT1(pf12833) EUT1
03420 H_mulberry(pf00936pf16365) EUT1
03421 Tsp_rosa EUT1
03422 HMMetlyEutC_1_eut1(pf05985) EUT1 EP_NTERM
03423 HMMetlyEutB_1_EUT1(pf06751) EUT1
03424 HMM06277EutA_1_eut(pf06277) EUT1
03425 HMMtpEuth_1_eut(pf04346) EUT1
03426 HMM00465alcdh_3_mix(pf00465) EUT1
03427 HMM06723EutJMreB_1_grm(#) EUT1
03428 HMMaldh_1_eutpdugrm(pf00171) EUT1 EP_NTERM
03429 P_lightViolet EUT1
03430 H_azure EUT1
03431 HMM01515PTA_B_1_eut1(pf01515) EUT1
03432 HMM01923cobtr_2_eut1(pf01923) EUT1
03433 HMM06249EutQ_1_eut1(pf06249) EUT1 EP_NTERM
03434 HMM10662PduV_1_mix(pf10662) EUT1
03435 Hp_euts_violetPink EUT1
03436 HMM00390_malic_1_eut1(pf00390pf03949pf01515) EUT1
03437 -pf00923 -
03438 HMM02780keto_1_eut1(pf02779pf02780pf00456) EUT1
03439 -pf06674 -
03440 HMM00293nuhyd_1_eut1(pf00293) EUT1
03441 -pf13247pf14691pf07992 -
03442 -pf00672pf02518pf07730pf13675 -
03443 -pf00873 -
03444 -# -
03445 -pf03960 -
BMC-P
BMC-H
BMC-T
Aldehyde dehydrogenase
PTAC
Signature enzyme
Alcohol dehydrogenase
Regulator

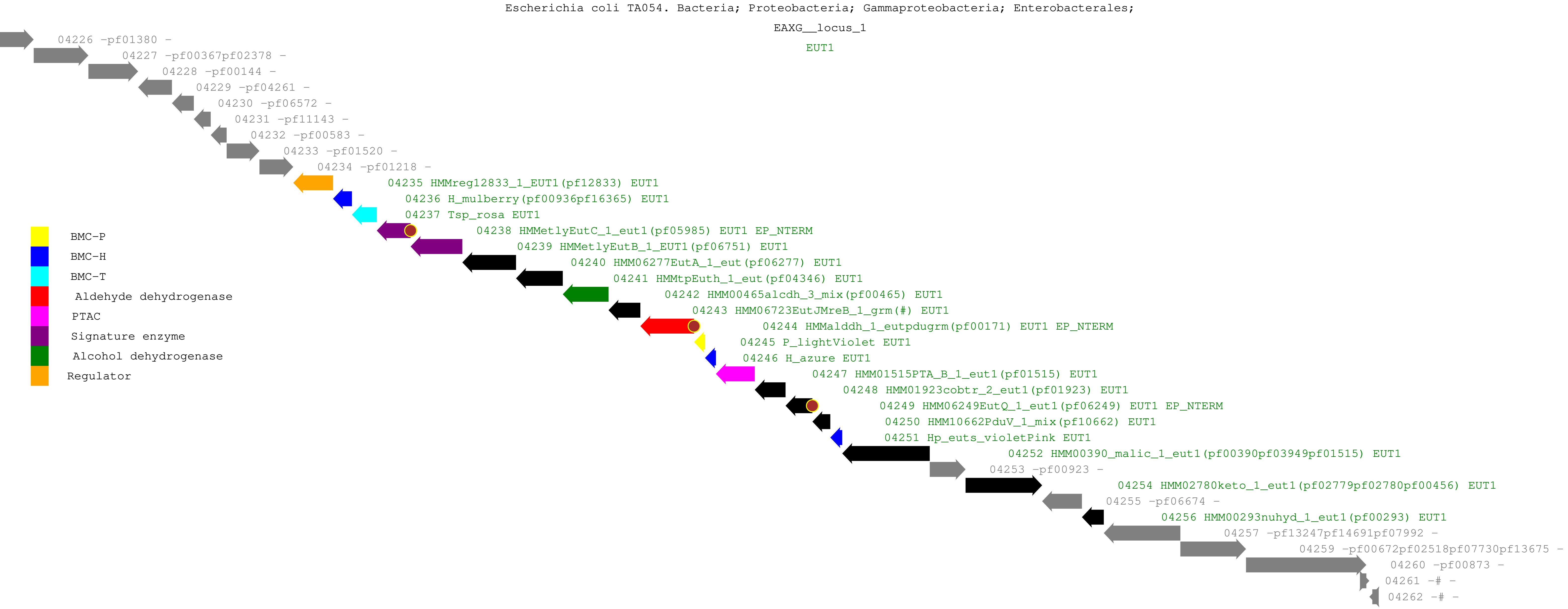

Escherichia coli TA054. Bacteria; Proteobacteria; Gammaproteobacteria; Enterobacterales;
EAXG__locus_1
EUT1
04226 -pf01380 -
04227 -pf00367pf02378 -
04228 -pf00144 -
04229 -pf04261 -
04230 -pf06572 -
04231 -pf11143 -
04232 -pf00583 -
04233 -pf01520 -
04234 -pf01218 -
04235 HMMreg12833_1_EUT1(pf12833) EUT1
04236 H_mulberry(pf00936pf16365) EUT1
04237 Tsp_rosa EUT1
04238 HMMetlyEutC_1_eut1(pf05985) EUT1 EP_NTERM
04239 HMMetlyEutB_1_EUT1(pf06751) EUT1
04240 HMM06277EutA_1_eut(pf06277) EUT1
04241 HMMtpEuth_1_eut(pf04346) EUT1
04242 HMM00465alcdh_3_mix(pf00465) EUT1
04243 HMM06723EutJMreB_1_grm(#) EUT1
04244 HMMalddh_1_eutpdugrm(pf00171) EUT1 EP_NTERM
04245 P_lightViolet EUT1
04246 H_azure EUT1
04247 HMM01515PTA_B_1_eut1(pf01515) EUT1
04248 HMM01923cobtr_2_eut1(pf01923) EUT1
04249 HMM06249EutQ_1_eut1(pf06249) EUT1 EP_NTERM
04250 HMM10662PduV_1_mix(pf10662) EUT1
04251 Hp_euts_violetPink EUT1
04252 HMM00390_malic_1_eut1(pf00390pf03949pf01515) EUT1
04253 -pf00923 -
04254 HMM02780keto_1_eut1(pf02779pf02780pf00456) EUT1
04255 -pf06674 -
04256 HMM00293nuhyd_1_eut1(pf00293) EUT1
04257 -pf13247pf14691pf07992 -
04259 -pf00672pf02518pf07730pf13675 -
04260 -pf00873 -
04261 -# -
04262 -# -
BMC-P
BMC-H
BMC-T
Aldehyde dehydrogenase
PTAC
Signature enzyme
Alcohol dehydrogenase
Regulator

Escherichia coli TA249. Bacteria; Proteobacteria; Gammaproteobacteria; Enterobacterales;

EAZG__locus_2

EUT1

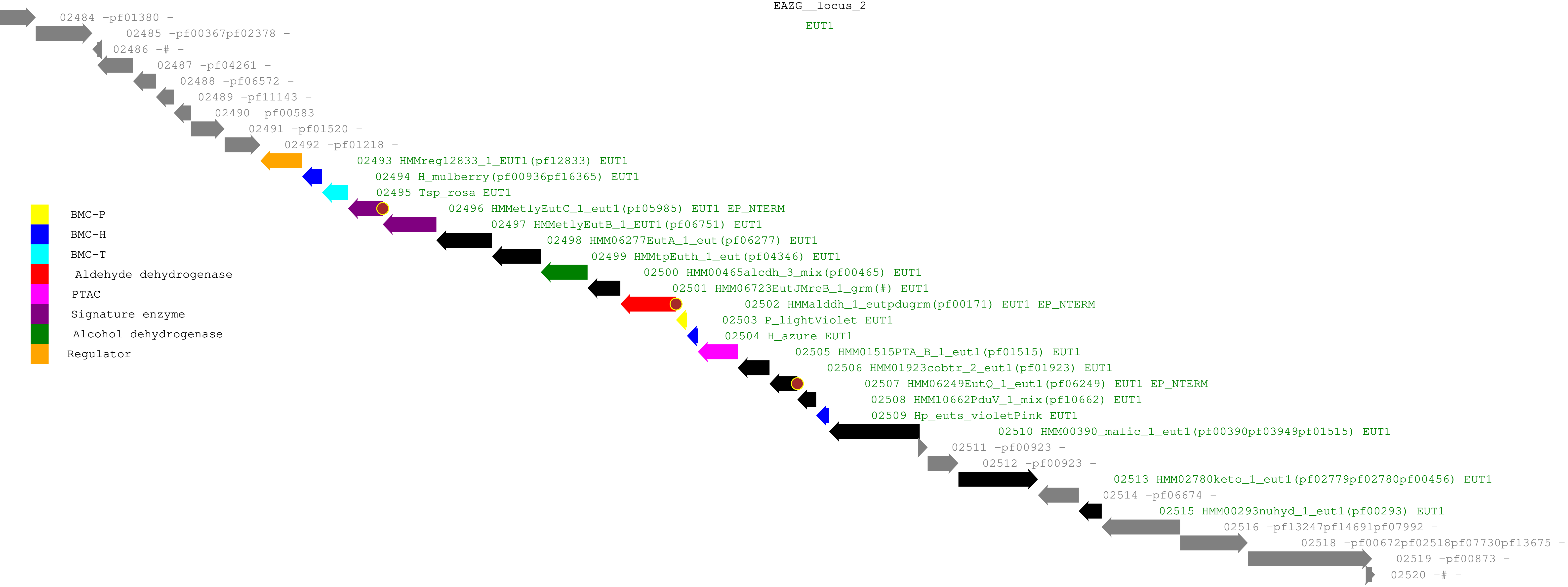

Escherichia coli E1118. Bacteria; Proteobacteria; Gammaproteobacteria; Enterobacterales;

EBCG__locus_2

EUT1

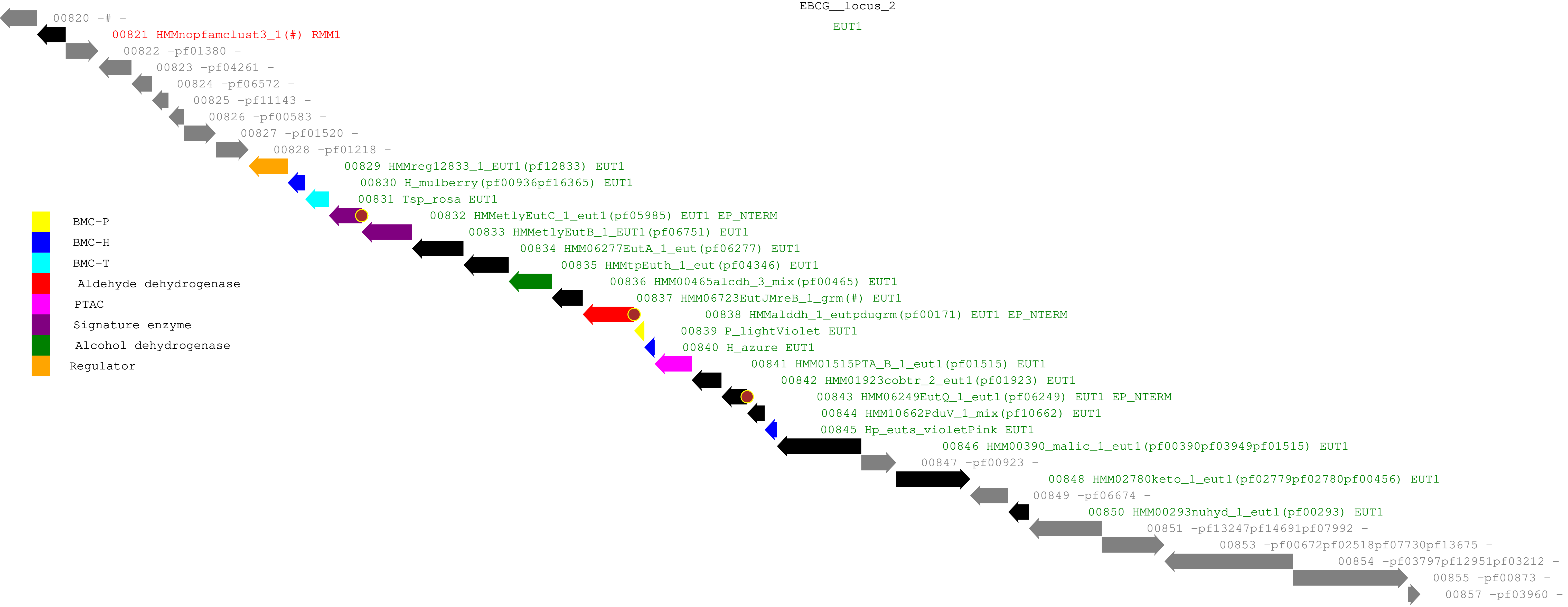

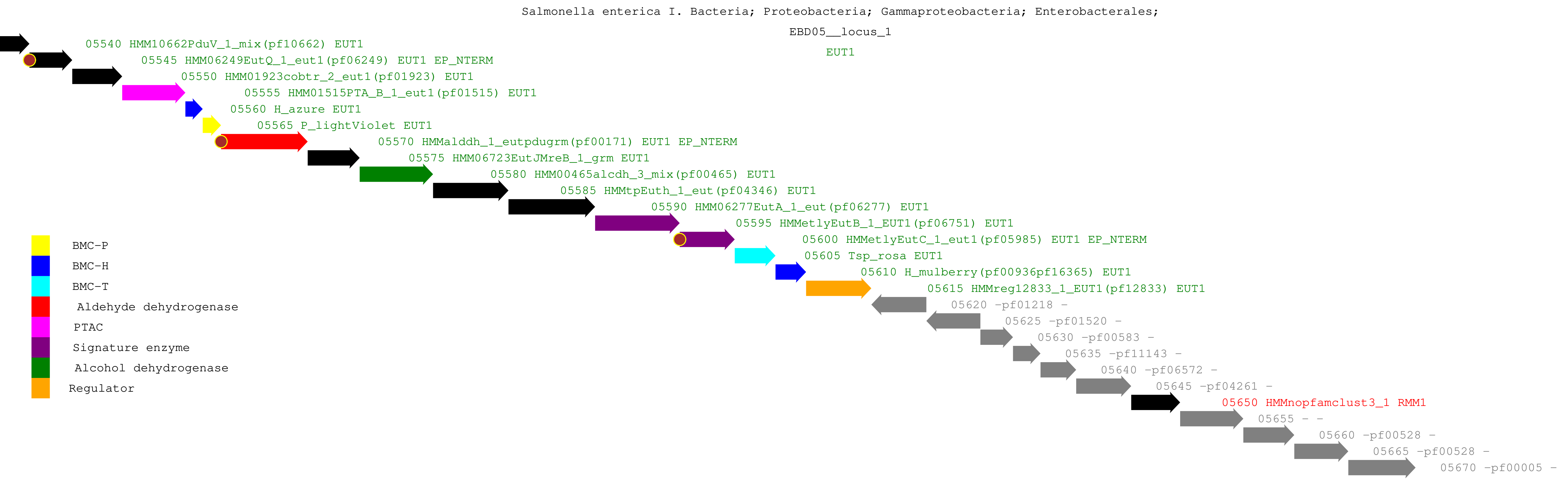

Salmonella enterica I. Bacteria; Proteobacteria; Gammaproteobacteria; Enterobacterales;
EBD05__locus_1
EUT1
05540 HMM10662PduV_1_mix(pf10662) EUT1
05545 HMM06249EutQ_1_eut1(pf06249) EUT1 EP_NTERM
05550 HMM01923cobtr_2_eut1(pf01923) EUT1
05555 HMM01515PTA_B_1_eut1(pf01515) EUT1
05560 H_azure EUT1
05565 P_lightViolet EUT1
05570 HMMalddh_1_eutpdugrm(pf00171) EUT1 EP_NTERM
05575 HMM06723EutJMreB_1_grm EUT1
05580 HMM00465alcdh_3_mix(pf00465) EUT1
05585 HMMtpEuth_1_eut(pf04346) EUT1
05590 HMM06277EutA_1_eut(pf06277) EUT1
05595 HMMetlyEutB_1_EUT1(pf06751) EUT1
05600 HMMetlyEutC_1_eut1(pf05985) EUT1 EP_NTERM
05605 Tsp_rosa EUT1
05610 H_mulberry(pf00936pf16365) EUT1
05615 HMMreg12833_1_EUT1(pf12833) EUT1
05620 -pf01218 -
05625 -pf01520 -
05630 -pf00583 -
05635 -pf11143 -
05640 -pf06572 -
05645 -pf04261 -
05650 HMMnopfamclust3_1 RMM1
05655 - -
05660 -pf00528 -
05665 -pf00528 -
05670 -pf00005 -
BMC-P
BMC-H
BMC-T
Aldehyde dehydrogenase
PTAC
Signature enzyme
Alcohol dehydrogenase
Regulator

Escherichia coli O44:H18 (strain 042 / EAEC). Bacteria; Proteobacteria; Gammaproteobacteria; Enterobacterales;

EC042__locus_1

EUT1

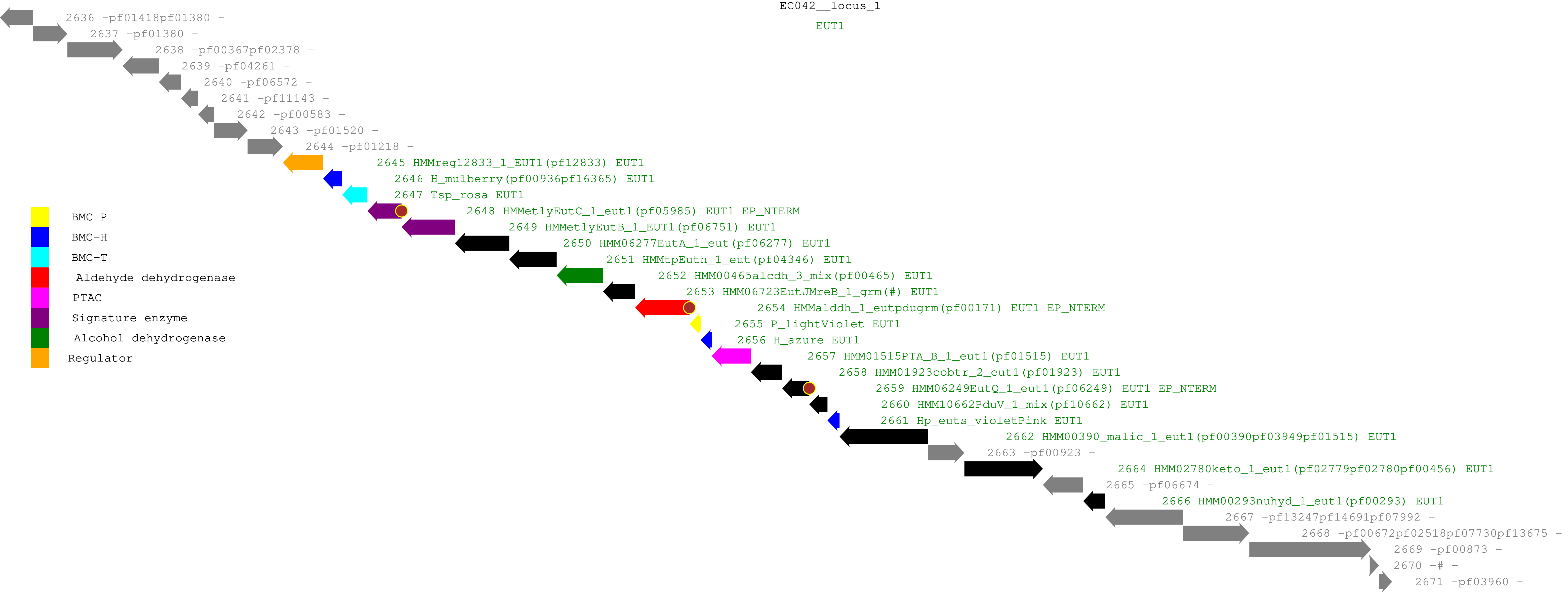

Escherichia coli O145:H28. Bacteria; Proteobacteria; Gammaproteobacteria; Enterobacterales;

EC10942__locus_1

EUT1

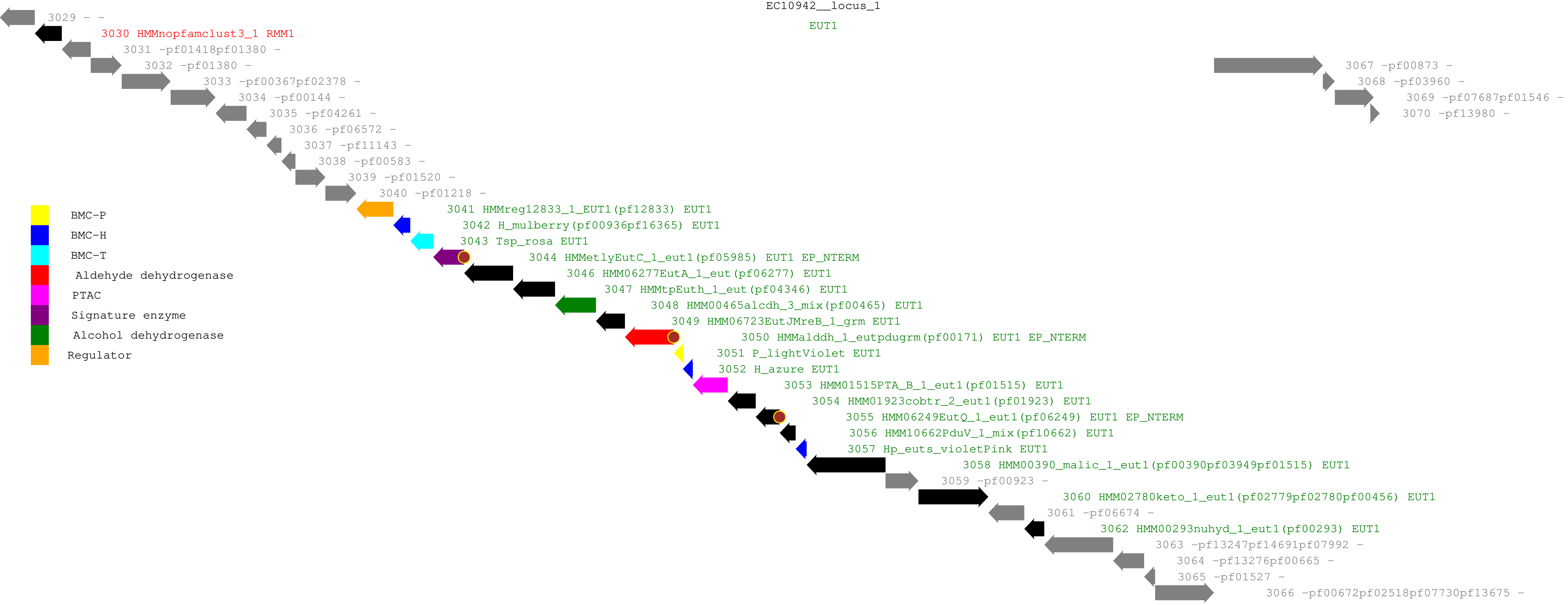

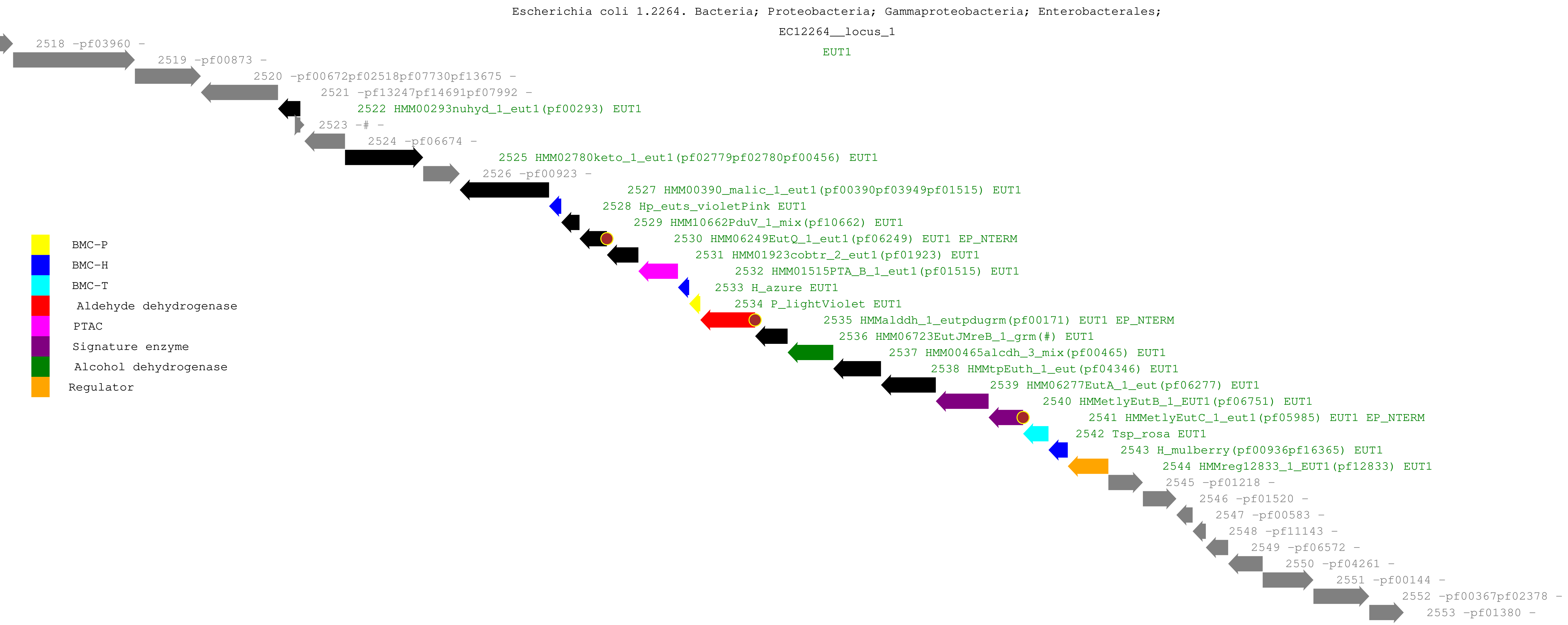
Escherichia coli 1.2264. Bacteria; Proteobacteria; Gammaproteobacteria; Enterobacterales;
EC12264__locus_1
EUT1
2518 -pf03960 -
2519 -pf00873 -
2520 -pf00672pf02518pf07730pf13675 -
2521 -pf13247pf14691pf07992 -
2522 HMM00293nuhyd_1_eut1(pf00293) EUT1
2523 -# -
2524 -pf06674 -
2525 HMM02780keto_1_eut1(pf02779pf02780pf00456) EUT1
2526 -pf00923 -
2527 HMM00390_malic_1_eut1(pf00390pf03949pf01515) EUT1
2528 Hp_euts_violetPink EUT1
2529 HMM10662PduV_1_mix(pf10662) EUT1
2530 HMM06249EutQ_1_eut1(pf06249) EUT1 EP_NTERM
2531 HMM01923cobtr_2_eut1(pf01923) EUT1
2532 HMM01515PTA_B_1_eut1(pf01515) EUT1
2533 H_azure EUT1
2534 P_lightViolet EUT1
2535 HMMalddh_1_eutpdugrm(pf00171) EUT1 EP_NTERM
2536 HMM06723EutJMreB_1_grm(#) EUT1
2537 HMM00465alcdh_3_mix(pf00465) EUT1
2538 HMMtpEuth_1_eut(pf04346) EUT1
2539 HMM06277EutA_1_eut(pf06277) EUT1
2540 HMMetlyEutB_1_EUT1(pf06751) EUT1
2541 HMMetlyEutC_1_eut1(pf05985) EUT1 EP_NTERM
2542 Tsp_rosa EUT1
2543 H_mulberry(pf00936pf16365) EUT1
2544 HMMreg12833_1_EUT1(pf12833) EUT1
2545 -pf01218 -
2546 -pf01520 -
2547 -pf00583 -
2548 -pf11143 -
2549 -pf06572 -
2550 -pf04261 -
2551 -pf00144 -
2552 -pf00367pf02378 -
2553 -pf01380 -
BMC-P
BMC-H
BMC-T
Aldehyde dehydrogenase
PTAC
Signature enzyme
Alcohol dehydrogenase
Regulator

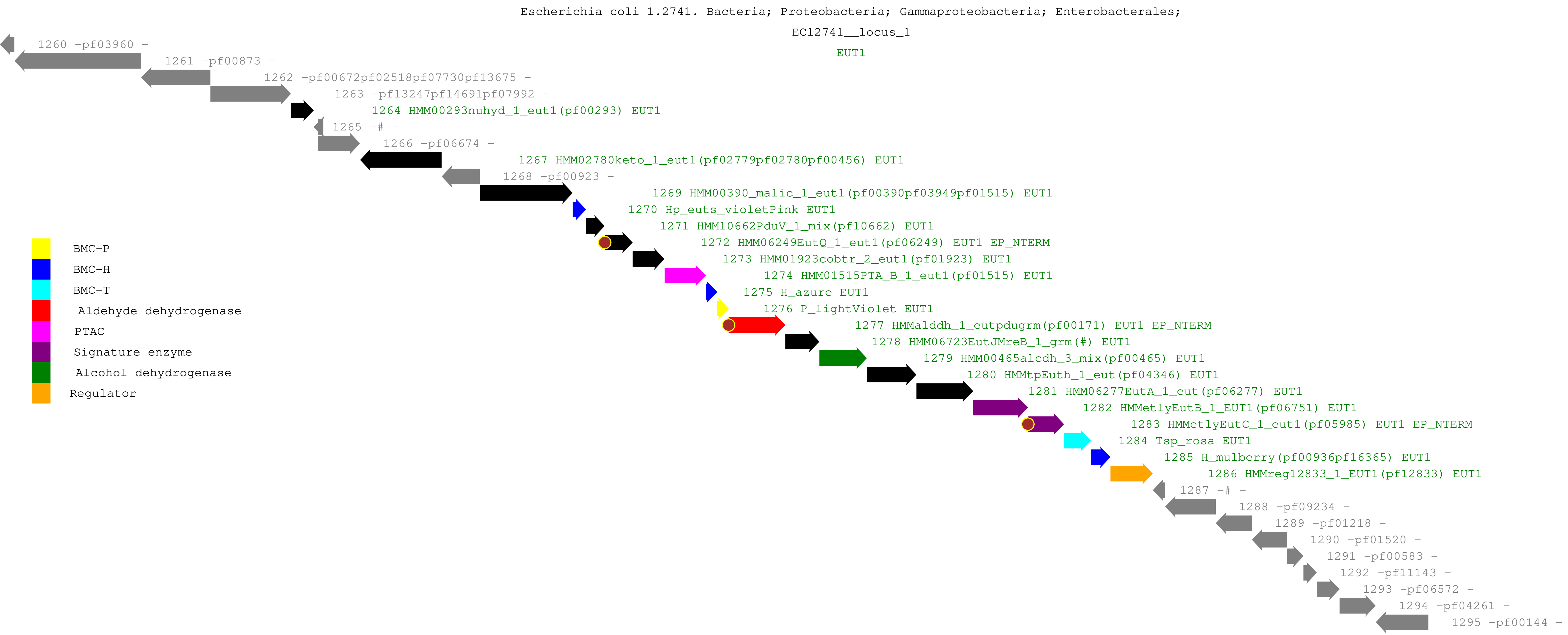
Escherichia coli 1.2741. Bacteria; Proteobacteria; Gammaproteobacteria; Enterobacterales;
EC12741__locus_1
EUT1
1260 -pf03960 -
1261 -pf00873 -
1262 -pf00672pf02518pf07730pf13675 -
1263 -pf13247pf14691pf07992 -
1264 HMM00293nuhyd_1_eut1(pf00293) EUT1
1265 -# -
1266 -pf06674 -
1267 HMM02780keto_1_eut1(pf02779pf02780pf00456) EUT1
1268 -pf00923 -
1269 HMM00390_malic_1_eut1(pf00390pf03949pf01515) EUT1
1270 Hp_euts_violetPink EUT1
1271 HMM10662PduV_1_mix(pf10662) EUT1
1272 HMM06249EutQ_1_eut1(pf06249) EUT1 EP_NTERM
1273 HMM01923cobtr_2_eut1(pf01923) EUT1
1274 HMM01515PTA_B_1_eut1(pf01515) EUT1
1275 H_azure EUT1
1276 P_lightViolet EUT1
1277 HMMaldddh_1_eutpdugrm(pf00171) EUT1 EP_NTERM
1278 HMM06723EutJMreB_1_grm(#) EUT1
1279 HMM00465alcdh_3_mix(pf00465) EUT1
1280 HMMtpEuth_1_eut(pf04346) EUT1
1281 HMM06277EutA_1_eut(pf06277) EUT1
1282 HMMetlyEutB_1_EUT1(pf06751) EUT1
1283 HMMetlyEutC_1_eut1(pf05985) EUT1 EP_NTERM
1284 Tsp_rosa EUT1
1285 H_mulberry(pf00936pf16365) EUT1
1286 HMMreg12833_1_EUT1(pf12833) EUT1
1287 -# -
1288 -pf09234 -
1289 -pf01218 -
1290 -pf01520 -
1291 -pf00583 -
1292 -pf11143 -
1293 -pf06572 -
1294 -pf04261 -
1295 -pf00144 -
BMC-P
BMC-H
BMC-T
Aldehyde dehydrogenase
PTAC
Signature enzyme
Alcohol dehydrogenase
Regulator

Escherichia coli 1303. Bacteria; Proteobacteria; Gammaproteobacteria; Enterobacterales;

EC1303_c_locus_1

EUT1

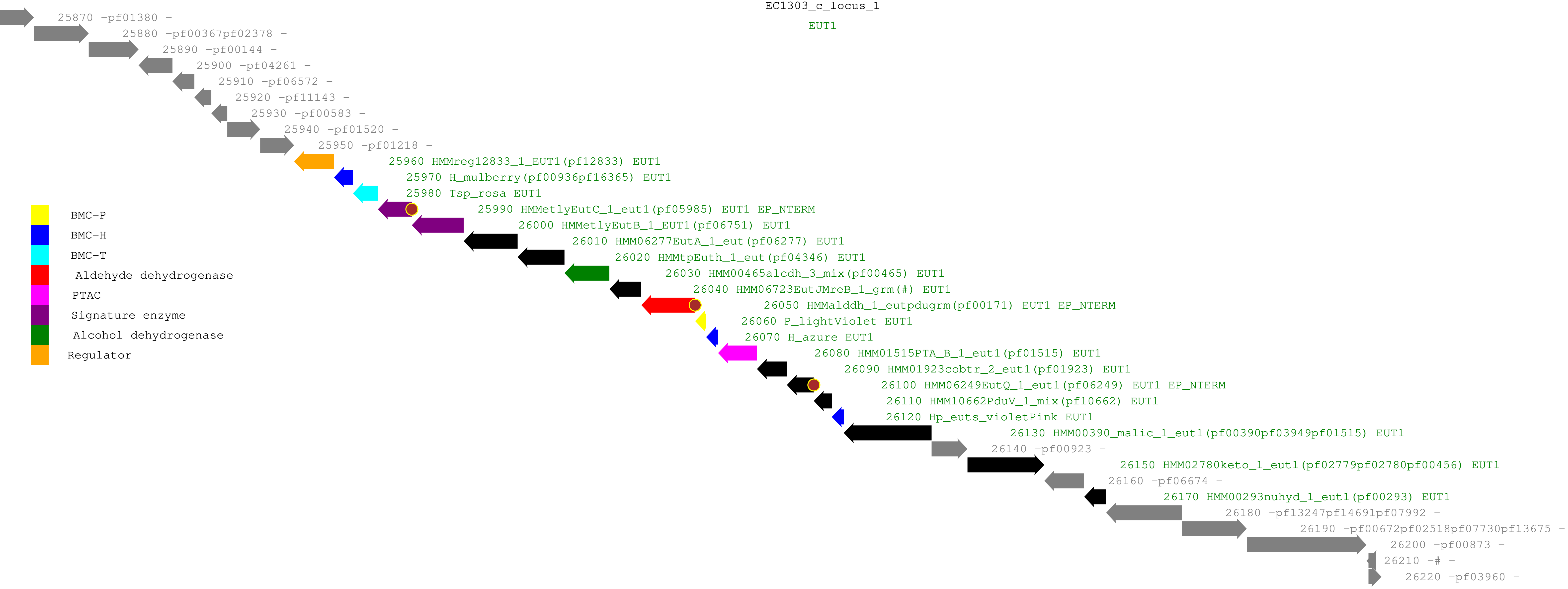

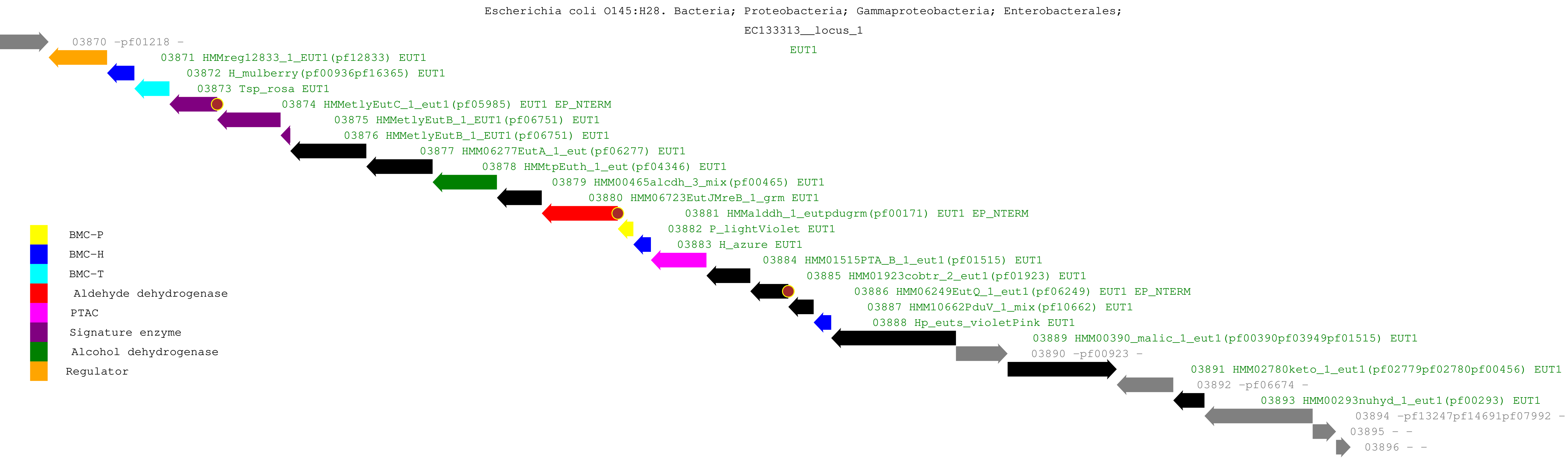
Escherichia coli O145:H28. Bacteria; Proteobacteria; Gammaproteobacteria; Enterobacterales;
EC133313__locus_1
EUT1
03870 -pf01218 -
03871 HMMreg12833_1_EUT1(pf12833) EUT1
03872 H_mulberry(pf00936pf16365) EUT1
03873 Tsp_rosa EUT1
03874 HMMetlyEutC_1_eut1(pf05985) EUT1 EP_NTERM
03875 HMMetlyEutB_1_EUT1(pf06751) EUT1
03876 HMMetlyEutB_1_EUT1(pf06751) EUT1
03877 HMM06277EutA_1_eut(pf06277) EUT1
03878 HMMtpEuth_1_eut(pf04346) EUT1
03879 HMM00465alcdh_3_mix(pf00465) EUT1
03880 HMM06723EutJMreB_1_grm EUT1
03881 HMMaldh_1_eutpdugrm(pf00171) EUT1 EP_NTERM
03882 P_lightViolet EUT1
03883 H_azure EUT1
03884 HMM01515PTA_B_1_eut1(pf01515) EUT1
03885 HMM01923cobtr_2_eut1(pf01923) EUT1
03886 HMM06249EutQ_1_eut1(pf06249) EUT1 EP_NTERM
03887 HMM10662PduV_1_mix(pf10662) EUT1
03888 Hp_euts_violetPink EUT1
03889 HMM00390_malic_1_eut1(pf00390pf03949pf01515) EUT1
03890 -pf00923 -
03891 HMM02780keto_1_eut1(pf02779pf02780pf00456) EUT1
03892 -pf06674 -
03893 HMM00293nuhyd_1_eut1(pf00293) EUT1
03894 -pf13247pf14691pf07992 -
03895 - -
03896 - -
BMC-P
BMC-H
BMC-T
Aldehyde dehydrogenase
PTAC
Signature enzyme
Alcohol dehydrogenase
Regulator

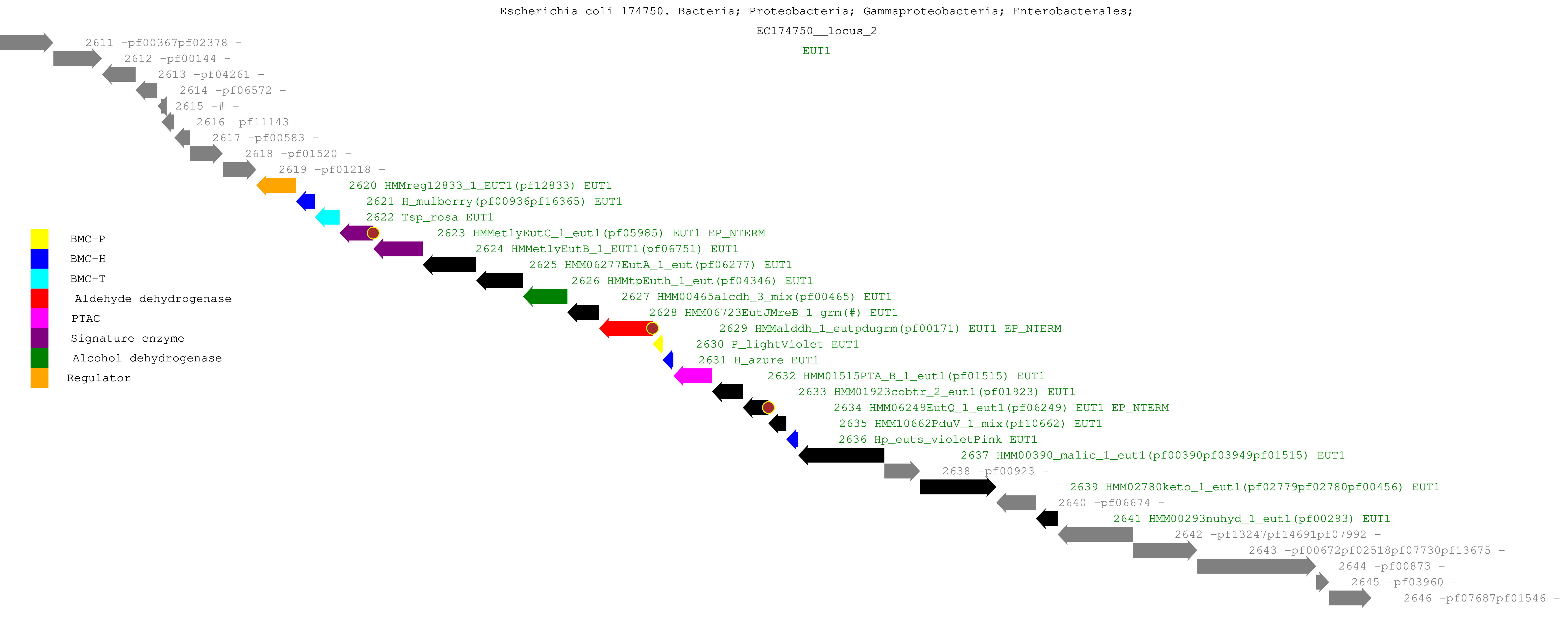
Escherichia coli 174750. Bacteria; Proteobacteria; Gammaproteobacteria; Enterobacterales;
EC174750__locus_2
EUT1
2611 -pf00367pf02378 -
2612 -pf00144 -
2613 -pf04261 -
2614 -pf06572 -
2615 -# -
2616 -pf11143 -
2617 -pf00583 -
2618 -pf01520 -
2619 -pf01218 -
2620 HMMreg12833_1_EUT1(pf12833) EUT1
2621 H_mulberry(pf00936pf16365) EUT1
2622 Tsp_rosa EUT1
2623 HMMetlyEutC_1_eut1(pf05985) EUT1 EP_NTERM
2624 HMMetlyEutB_1_EUT1(pf06751) EUT1
2625 HMM06277EutA_1_eut(pf06277) EUT1
2626 HMMtpEuth_1_eut(pf04346) EUT1
2627 HMM00465alcdh_3_mix(pf00465) EUT1
2628 HMM06723EutJMreB_1_grm(#) EUT1
2629 HMMalddh_1_eutpdugrm(pf00171) EUT1 EP_NTERM
2630 P_lightViolet EUT1
2631 H_azure EUT1
2632 HMM01515PTA_B_1_eut1(pf01515) EUT1
2633 HMM01923cobtr_2_eut1(pf01923) EUT1
2634 HMM06249EutQ_1_eut1(pf06249) EUT1 EP_NTERM
2635 HMM10662PduV_1_mix(pf10662) EUT1
2636 Hp_euts_violetPink EUT1
2637 HMM00390_malic_1_eut1(pf00390pf03949pf01515) EUT1
2638 -pf00923 -
2639 HMM02780keto_1_eut1(pf02779pf02780pf00456) EUT1
2640 -pf06674 -
2641 HMM00293nuhyd_1_eut1(pf00293) EUT1
2642 -pf13247pf14691pf07992 -
2643 -pf00672pf02518pf07730pf13675 -
2644 -pf00873 -
2645 -pf03960 -
2646 -pf07687pf01546 -
BMC-P
BMC-H
BMC-T
Aldehyde dehydrogenase
PTAC
Signature enzyme
Alcohol dehydrogenase
Regulator

Escherichia coli 2362-75. Bacteria; Proteobacteria; Gammaproteobacteria; Enterobacterales;

EC236275__locus_2

EUT1

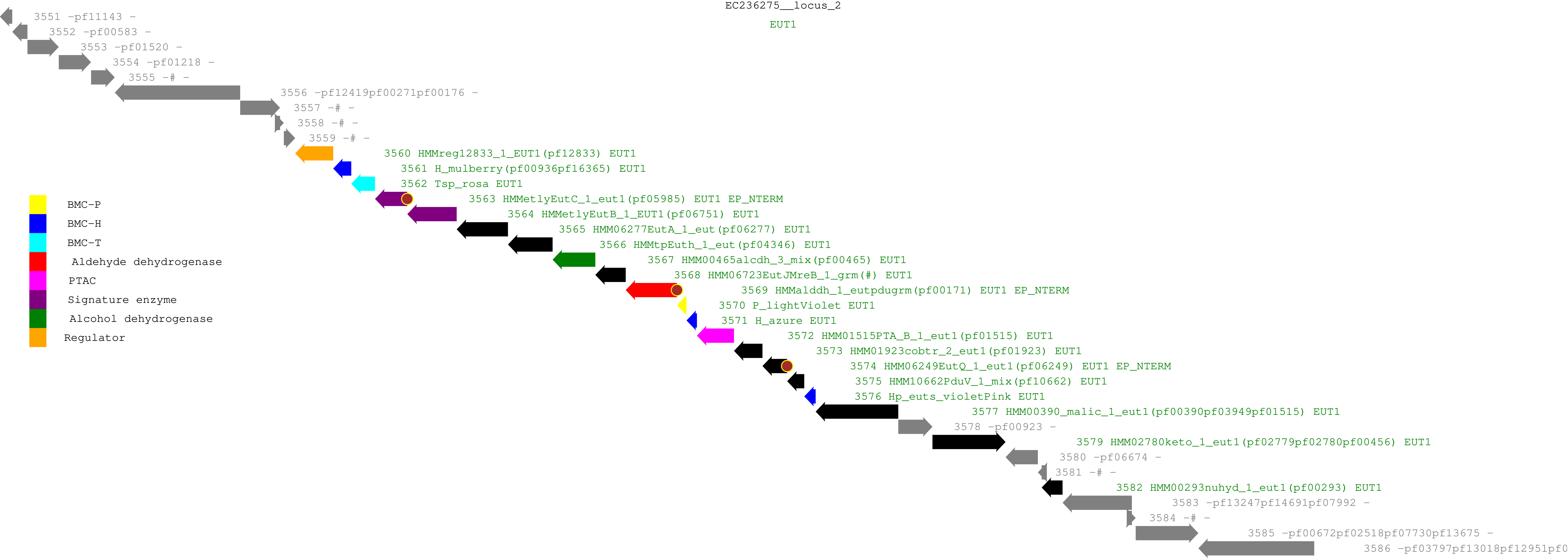

Escherichia coli 2.3916. Bacteria; Proteobacteria; Gammaproteobacteria; Enterobacterales;

EC23916__locus_1

EUT1

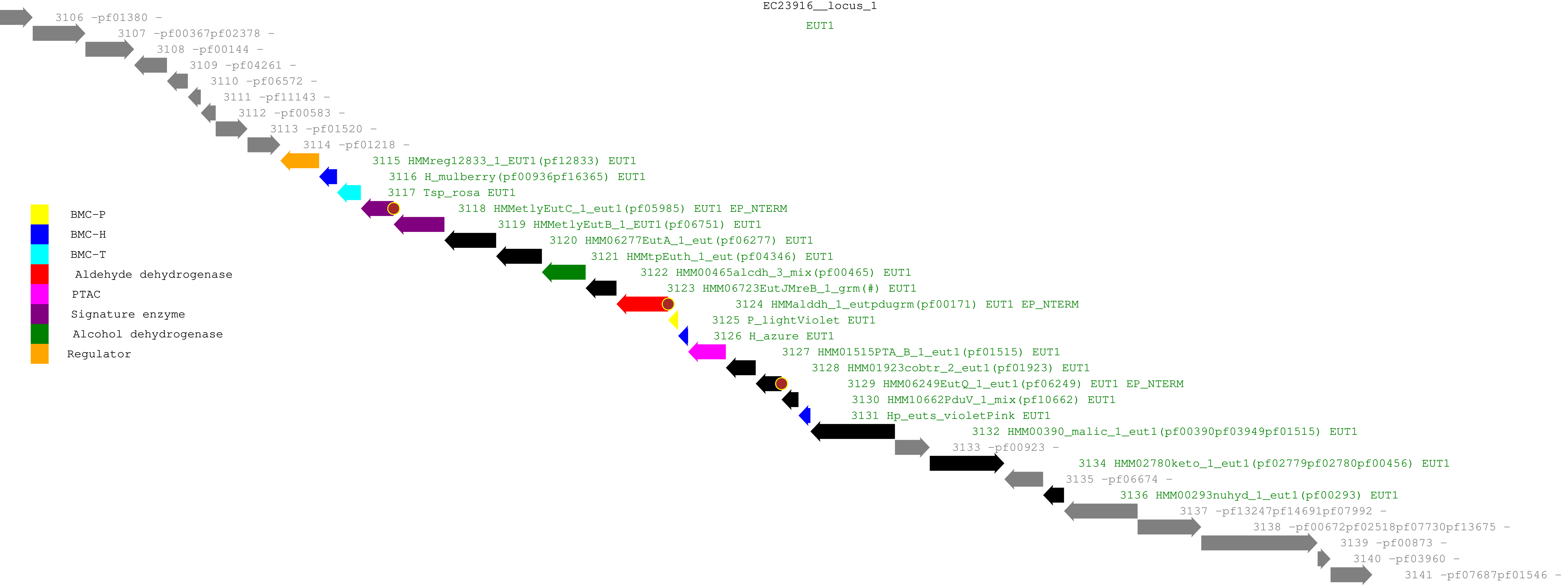

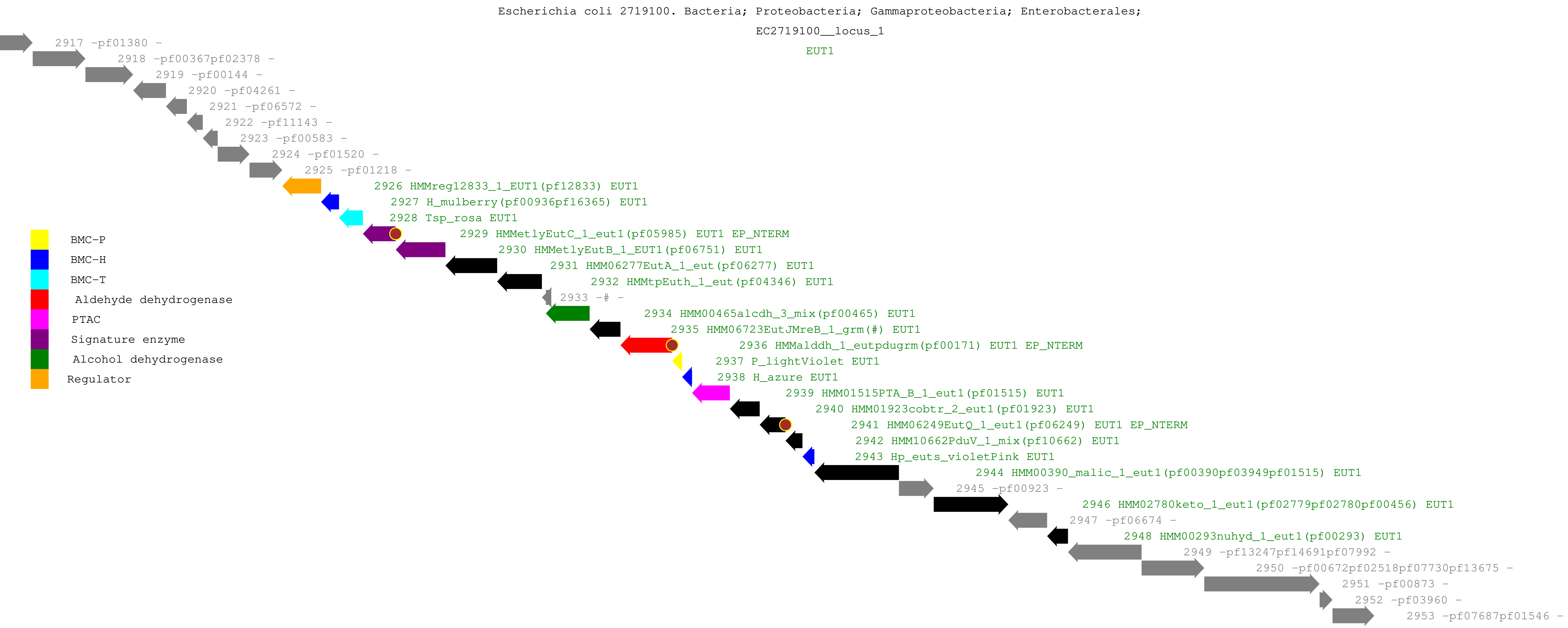
Escherichia coli 2719100. Bacteria; Proteobacteria; Gammaproteobacteria; Enterobacterales;
EC2719100__locus_1
EUT1
2917 -pf01380 -
2918 -pf00367pf02378 -
2919 -pf00144 -
2920 -pf04261 -
2921 -pf06572 -
2922 -pf11143 -
2923 -pf00583 -
2924 -pf01520 -
2925 -pf01218 -
2926 HMMreg12833_1_EUT1(pf12833) EUT1
2927 H_mulberry(pf00936pf16365) EUT1
2928 Tsp_rosa EUT1
2929 HMMetlyEutC_1_eut1(pf05985) EUT1 EP_NTERM
2930 HMMetlyEutB_1_EUT1(pf06751) EUT1
2931 HMM06277EutA_1_eut(pf06277) EUT1
2932 HMMtpEuth_1_eut(pf04346) EUT1
2933 -# -
2934 HMM00465alcdh_3_mix(pf00465) EUT1
2935 HMM06723EutJMreB_1_grm(#) EUT1
2936 HMMaldh_1_eutpdugrm(pf00171) EUT1 EP_NTERM
2937 P_lightViolet EUT1
2938 H_azure EUT1
2939 HMM01515PTA_B_1_eut1(pf01515) EUT1
2940 HMM01923cobtr_2_eut1(pf01923) EUT1
2941 HMM06249EutQ_1_eut1(pf06249) EUT1 EP_NTERM
2942 HMM10662PduV_1_mix(pf10662) EUT1
2943 Hp_euts_violetPink EUT1
2944 HMM00390_malic_1_eut1(pf00390pf03949pf01515) EUT1
2945 -pf00923 -
2946 HMM02780keto_1_eut1(pf02779pf02780pf00456) EUT1
2947 -pf06674 -
2948 HMM00293nuhyd_1_eut1(pf00293) EUT1
2949 -pf13247pf14691pf07992 -
2950 -pf00672pf02518pf07730pf13675 -
2951 -pf00873 -
2952 -pf03960 -
2953 -pf07687pf01546 -
BMC-P
BMC-H
BMC-T
Aldehyde dehydrogenase
PTAC
Signature enzyme
Alcohol dehydrogenase
Regulator

Escherichia coli 3.4880. Bacteria; Proteobacteria; Gammaproteobacteria; Enterobacterales;

EC34880__locus_1

EUT1

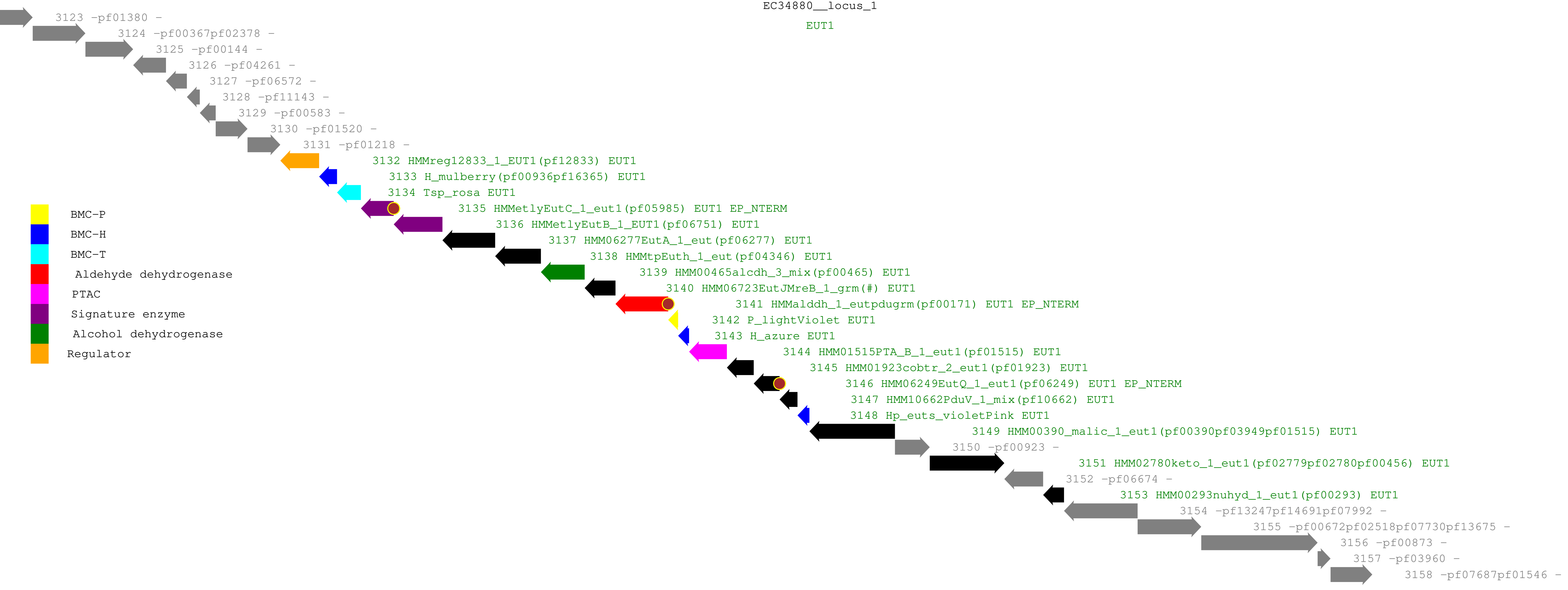

Escherichia coli 4.0522. Bacteria; Proteobacteria; Gammaproteobacteria; Enterobacterales;

EC40522__locus_1

EUT1

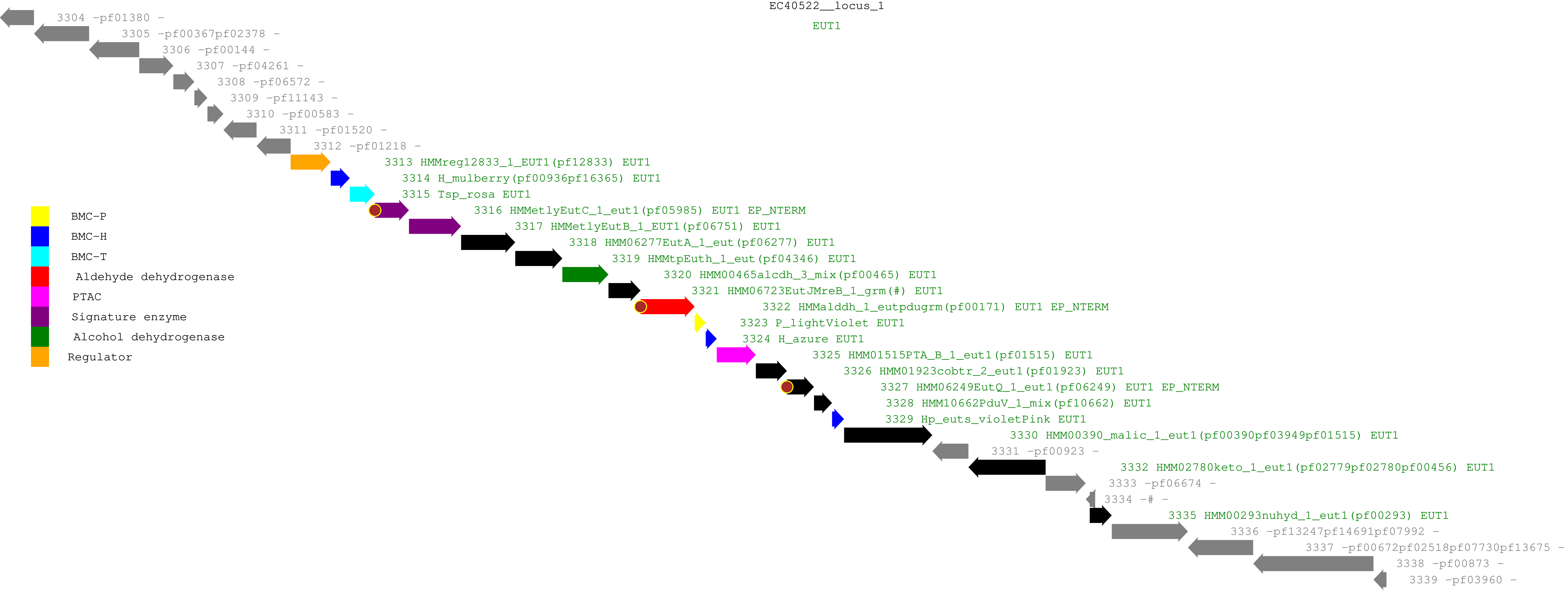

Escherichia coli 4.0967. Bacteria; Proteobacteria; Gammaproteobacteria; Enterobacterales;

EC40967__locus_1

EUT1

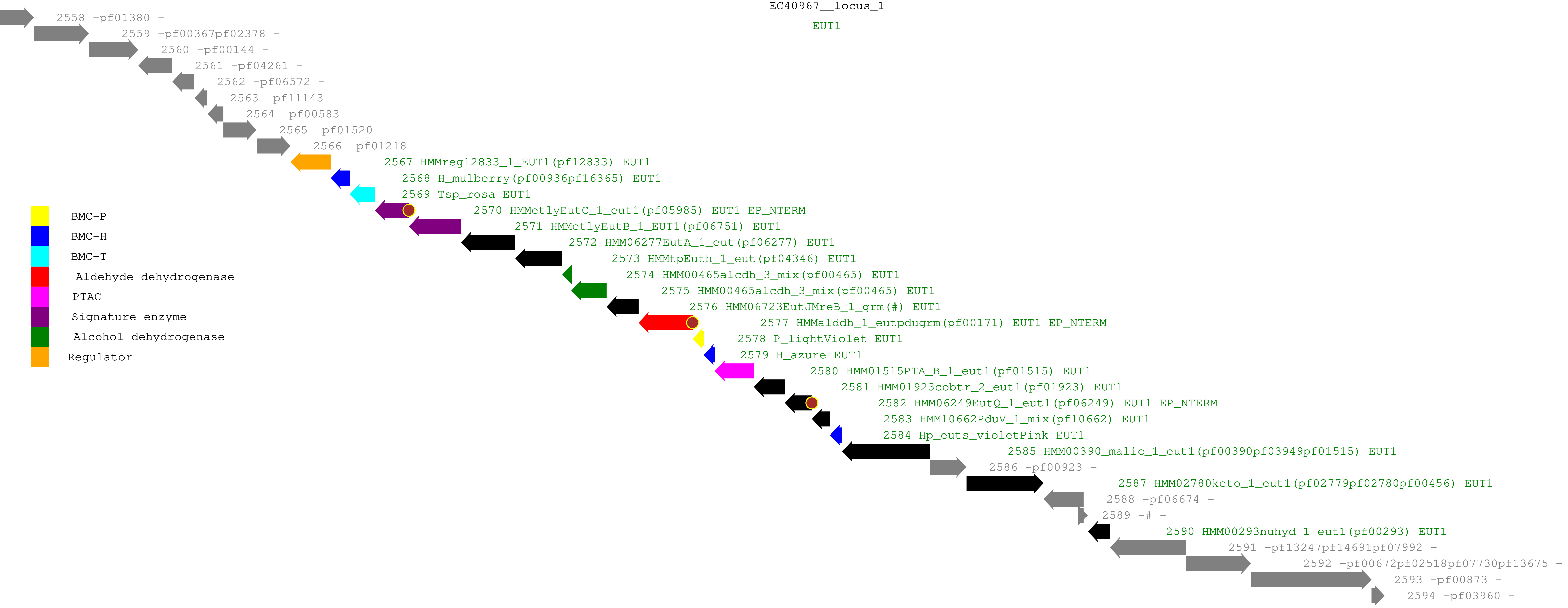

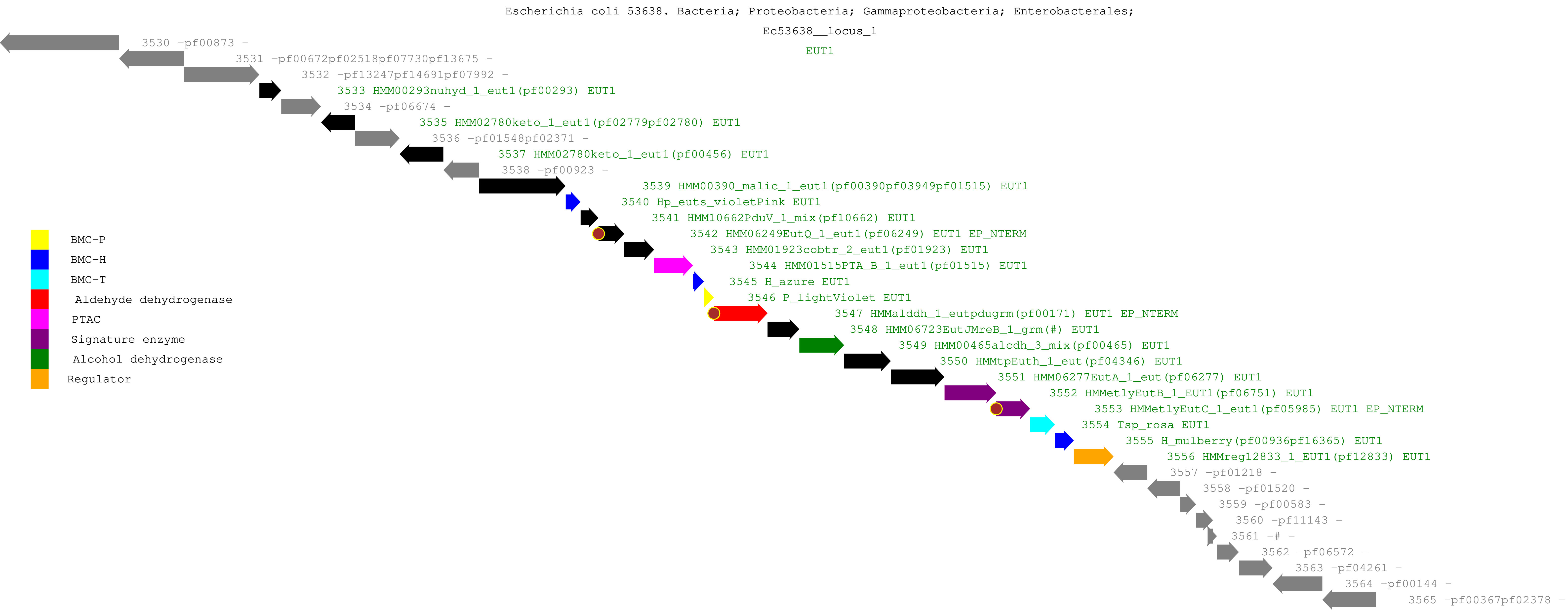

Escherichia coli 53638. Bacteria; Proteobacteria; Gammaproteobacteria; Enterobacterales;
Ec53638__locus_1
EUT1
3530 -pf00873 -
3531 -pf00672pf02518pf07730pf13675 -
3532 -pf13247pf14691pf07992 -
3533 HMM00293nuhyd_1_eut1(pf00293) EUT1
3534 -pf06674 -
3535 HMM02780keto_1_eut1(pf02779pf02780) EUT1
3536 -pf01548pf02371 -
3537 HMM02780keto_1_eut1(pf00456) EUT1
3538 -pf00923 -
3539 HMM00390_malic_1_eut1(pf00390pf03949pf01515) EUT1
3540 Hp_euts_violetPink EUT1
3541 HMM10662PduV_1_mix(pf10662) EUT1
3542 HMM06249EutQ_1_eut1(pf06249) EUT1 EP_NTERM
3543 HMM01923cobtr_2_eut1(pf01923) EUT1
3544 HMM01515PTA_B_1_eut1(pf01515) EUT1
3545 H_azure EUT1
3546 P_lightViolet EUT1
3547 HMMalddh_1_eutpdugrm(pf00171) EUT1 EP_NTERM
3548 HMM06723EutJMreB_1_grm(#) EUT1
3549 HMM00465alcdh_3_mix(pf00465) EUT1
3550 HMMtpEuth_1_eut(pf04346) EUT1
3551 HMM06277EutA_1_eut(pf06277) EUT1
3552 HMMetlyEutB_1_EUT1(pf06751) EUT1
3553 HMMetlyEutC_1_eut1(pf05985) EUT1 EP_NTERM
3554 Tsp_rosa EUT1
3555 H_mulberry(pf00936pf16365) EUT1
3556 HMMreg12833_1_EUT1(pf12833) EUT1
3557 -pf01218 -
3558 -pf01520 -
3559 -pf00583 -
3560 -pf11143 -
3561 -# -
3562 -pf06572 -
3563 -pf04261 -
3564 -pf00144 -
3565 -pf00367pf02378 -
BMC-P
BMC-H
BMC-T
Aldehyde dehydrogenase
PTAC
Signature enzyme
Alcohol dehydrogenase
Regulator

Escherichia coli 541-15. Bacteria; Proteobacteria; Gammaproteobacteria; Enterobacterales;

EC54115__locus_1

EUT1

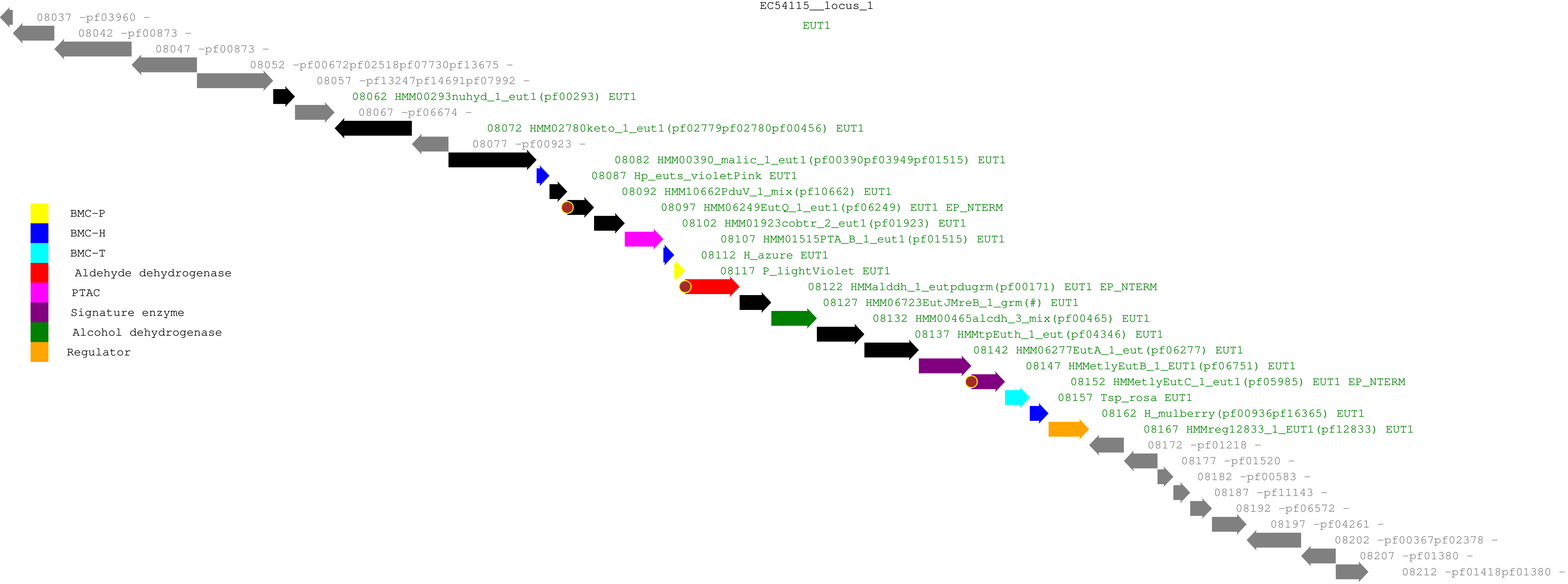

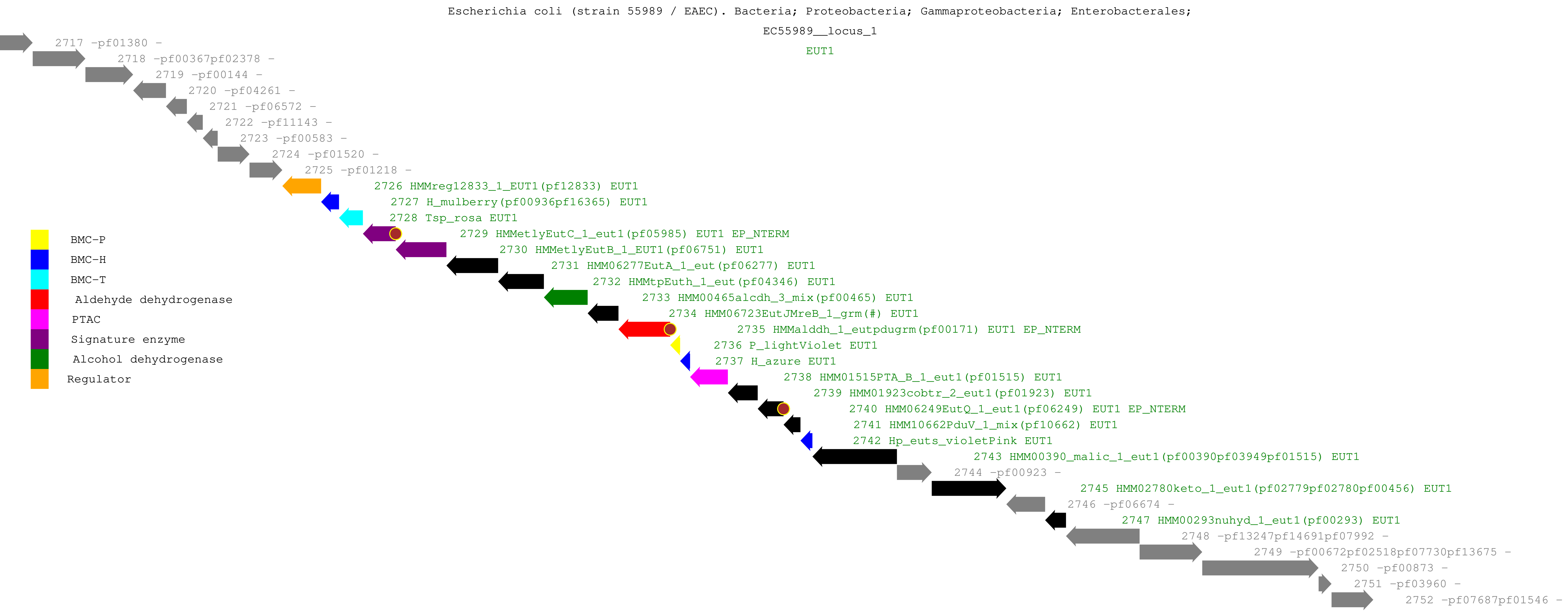
Escherichia coli (strain 55989 / EAEC). Bacteria; Proteobacteria; Gammaproteobacteria; Enterobacterales;
EC55989__locus_1
EUT1
2717 -pf01380 -
2718 -pf00367pf02378 -
2719 -pf00144 -
2720 -pf04261 -
2721 -pf06572 -
2722 -pf11143 -
2723 -pf00583 -
2724 -pf01520 -
2725 -pf01218 -
2726 HMMreg12833_1_EUT1(pf12833) EUT1
2727 H_mulberry(pf00936pf16365) EUT1
2728 Tsp_rosa EUT1
2729 HMMetlyEutC_1_eut1(pf05985) EUT1 EP_NTERM
2730 HMMetlyEutB_1_EUT1(pf06751) EUT1
2731 HMM06277EutA_1_eut(pf06277) EUT1
2732 HMMtpEuth_1_eut(pf04346) EUT1
2733 HMM00465alcdh_3_mix(pf00465) EUT1
2734 HMM06723EutJMreB_1_grm(#) EUT1
2735 HMMaldh_1_eutpdugrm(pf00171) EUT1 EP_NTERM
2736 P_lightViolet EUT1
2737 H_azure EUT1
2738 HMM01515PTA_B_1_eut1(pf01515) EUT1
2739 HMM01923cobtr_2_eut1(pf01923) EUT1
2740 HMM06249EutQ_1_eut1(pf06249) EUT1 EP_NTERM
2741 HMM10662PduV_1_mix(pf10662) EUT1
2742 Hp_euts_violetPink EUT1
2743 HMM00390_malic_1_eut1(pf00390pf03949pf01515) EUT1
2744 -pf00923 -
2745 HMM02780keto_1_eut1(pf02779pf02780pf00456) EUT1
2746 -pf06674 -
2747 HMM00293nuhyd_1_eut1(pf00293) EUT1
2748 -pf13247pf14691pf07992 -
2749 -pf00672pf02518pf07730pf13675 -
2750 -pf00873 -
2751 -pf03960 -
2752 -pf07687pf01546 -
BMC-P
BMC-H
BMC-T
Aldehyde dehydrogenase
PTAC
Signature enzyme
Alcohol dehydrogenase
Regulator

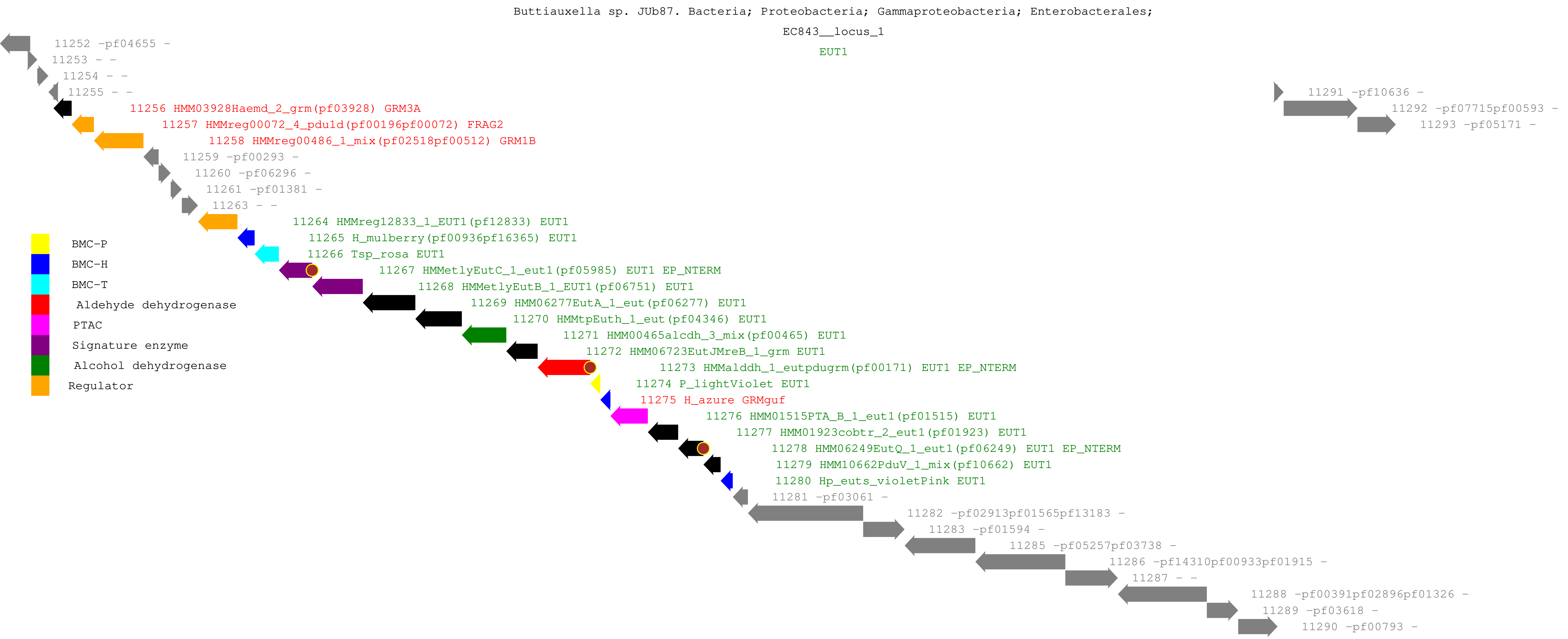
Buttiauxella sp. JUb87. Bacteria; Proteobacteria; Gammaproteobacteria; Enterobacterales;
EC843__locus_1
EUT1
11252 -pf04655 -
11253 - -
11254 - -
11255 - -
11256 HMM03928Haemd_2_grm(pf03928) GRM3A
11257 HMMreg00072_4_pdu1d(pf00196pf00072) FRAG2
11258 HMMreg00486_1_mix(pf02518pf00512) GRM1B
11259 -pf00293 -
11260 -pf06296 -
11261 -pf01381 -
11263 - -
11264 HMMreg12833_1_EUT1(pf12833) EUT1
11265 H_mulberry(pf00936pf16365) EUT1
11266 Tsp_rosa EUT1
11267 HMMetlyEutC_1_eut1(pf05985) EUT1 EP_NTERM
11268 HMMetlyEutB_1_EUT1(pf06751) EUT1
11269 HMM06277EutA_1_eut(pf06277) EUT1
11270 HMMtpEuth_1_eut(pf04346) EUT1
11271 HMM00465alcdh_3_mix(pf00465) EUT1
11272 HMM06723EutJMreB_1_grm EUT1
11273 HMMaldh_1_eutpdugrm(pf00171) EUT1 EP_NTERM
11274 P_lightViolet EUT1
11275 H_azure GRMguf
11276 HMM01515PTA_B_1_eut1(pf01515) EUT1
11277 HMM01923cobtr_2_eut1(pf01923) EUT1
11278 HMM06249EutQ_1_eut1(pf06249) EUT1 EP_NTERM
11279 HMM10662PduV_1_mix(pf10662) EUT1
11280 Hp_euts_violetPink EUT1
11281 -pf03061 -
11282 -pf02913pf01565pf13183 -
11283 -pf01594 -
11285 -pf05257pf03738 -
11286 -pf14310pf00933pf01915 -
11287 - -
11288 -pf00391pf02896pf01326 -
11289 -pf03618 -
11290 -pf00793 -
11291 -pf10636 -
11292 -pf07715pf00593 -
11293 -pf05171 -
BMC-P
BMC-H
BMC-T
Aldehyde dehydrogenase
PTAC
Signature enzyme
Alcohol dehydrogenase
Regulator

Escherichia coli 9.0111. Bacteria; Proteobacteria; Gammaproteobacteria; Enterobacterales;

EC90111__locus_1

EUT1

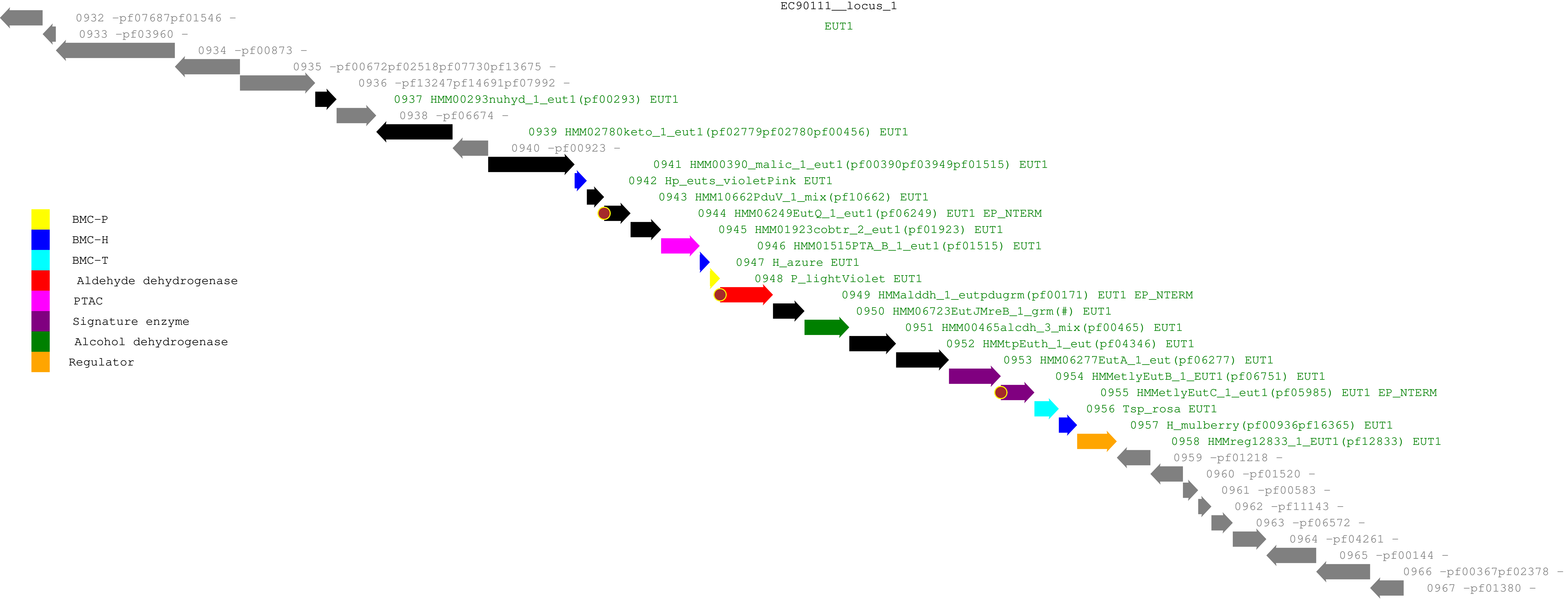

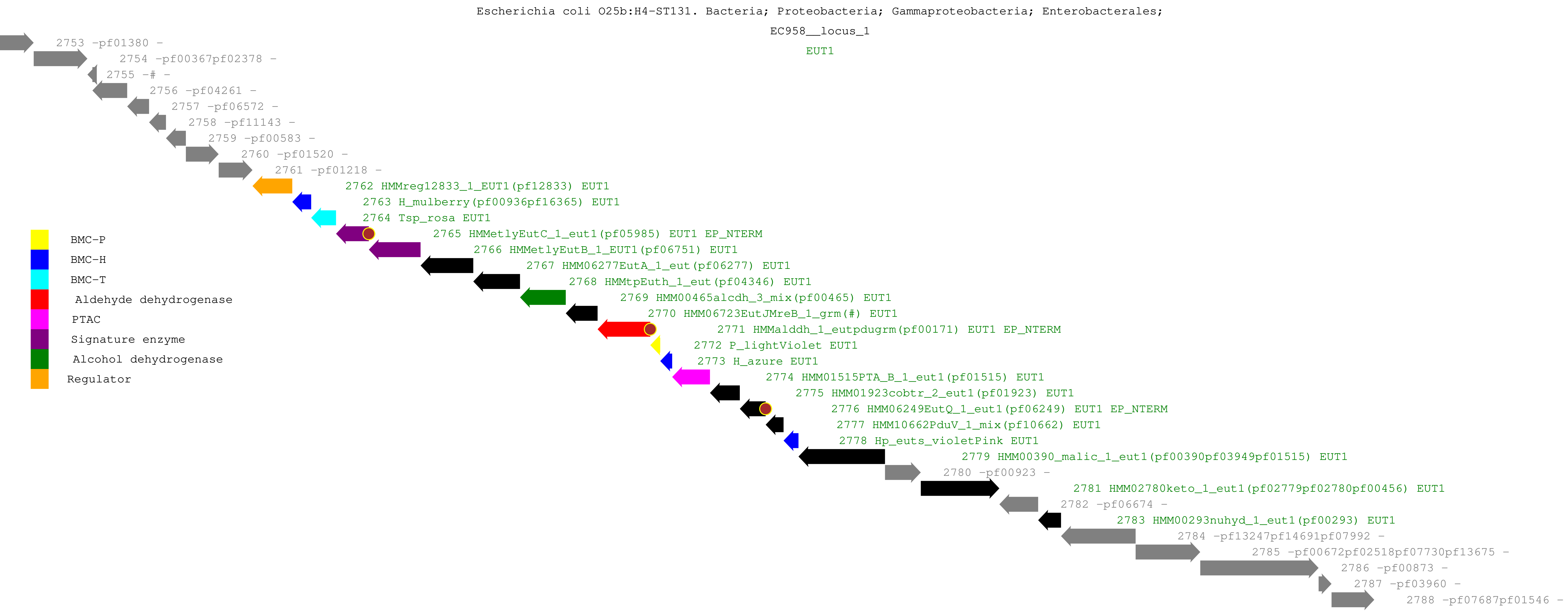
Escherichia coli O25b:H4-ST131. Bacteria; Proteobacteria; Gammaproteobacteria; Enterobacterales;
EC958__locus_1
EUT1
2753 -pf01380 -
2754 -pf00367pf02378 -
2755 -# -
2756 -pf04261 -
2757 -pf06572 -
2758 -pf11143 -
2759 -pf00583 -
2760 -pf01520 -
2761 -pf01218 -
2762 HMMreg12833_1_EUT1(pf12833) EUT1
2763 H_mulberry(pf00936pf16365) EUT1
2764 Tsp_rosa EUT1
2765 HMMetlyEutC_1_eut1(pf05985) EUT1 EP_NTERM
2766 HMMetlyEutB_1_EUT1(pf06751) EUT1
2767 HMM06277EutA_1_eut(pf06277) EUT1
2768 HMMtpEuth_1_eut(pf04346) EUT1
2769 HMM00465alcdh_3_mix(pf00465) EUT1
2770 HMM06723EutJMreB_1_grm(#) EUT1
2771 HMMalddh_1_eutpdugrm(pf00171) EUT1 EP_NTERM
2772 P_lightViolet EUT1
2773 H_azure EUT1
2774 HMM01515PTA_B_1_eut1(pf01515) EUT1
2775 HMM01923cobtr_2_eut1(pf01923) EUT1
2776 HMM06249EutQ_1_eut1(pf06249) EUT1 EP_NTERM
2777 HMM10662PduV_1_mix(pf10662) EUT1
2778 Hp_euts_violetPink EUT1
2779 HMM00390_malic_1_eut1(pf00390pf03949pf01515) EUT1
2780 -pf00923 -
2781 HMM02780keto_1_eut1(pf02779pf02780pf00456) EUT1
2782 -pf06674 -
2783 HMM00293nuhyd_1_eut1(pf00293) EUT1
2784 -pf13247pf14691pf07992 -
2785 -pf00672pf02518pf07730pf13675 -
2786 -pf00873 -
2787 -pf03960 -
2788 -pf07687pf01546 -
BMC-P
BMC-H
BMC-T
Aldehyde dehydrogenase
PTAC
Signature enzyme
Alcohol dehydrogenase
Regulator

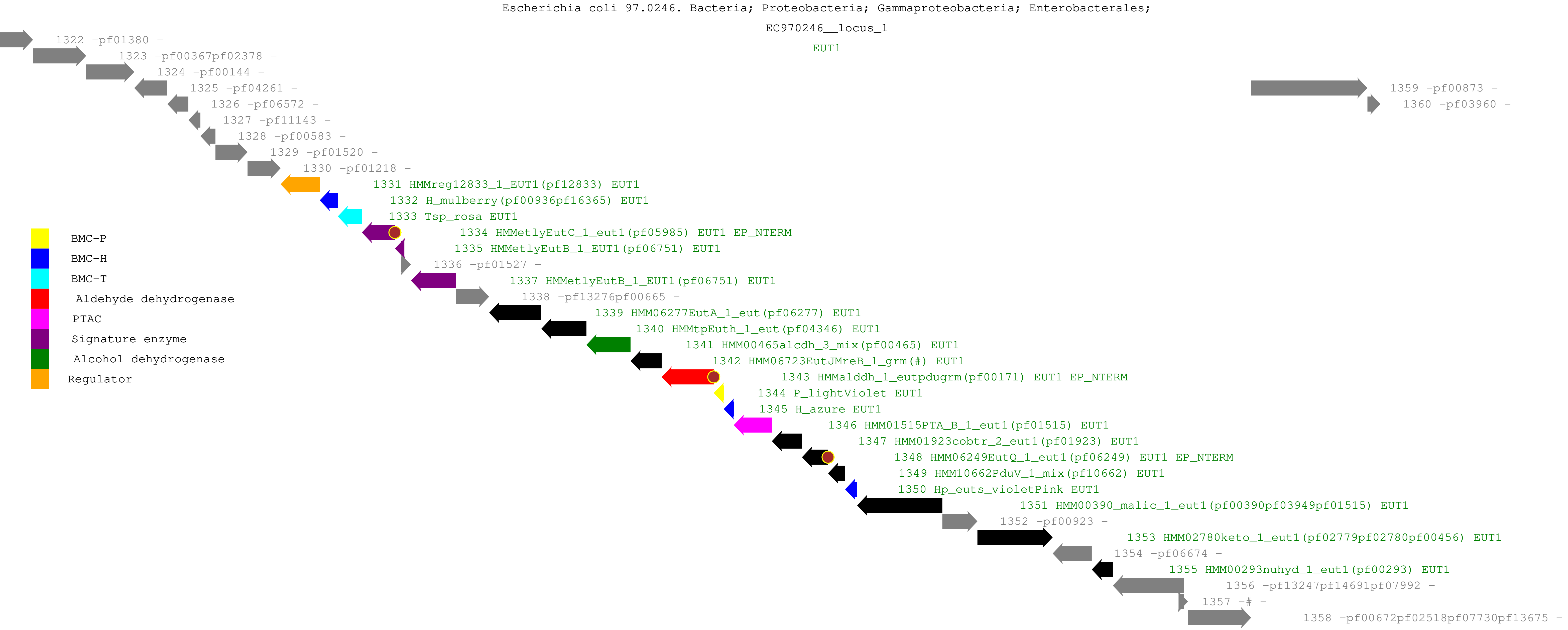

Escherichia coli 97.0246. Bacteria; Proteobacteria; Gammaproteobacteria; Enterobacterales;
EC970246__locus_1
EUT1
1322 -pf01380 -
1323 -pf00367pf02378 -
1324 -pf00144 -
1325 -pf04261 -
1326 -pf06572 -
1327 -pf11143 -
1328 -pf00583 -
1329 -pf01520 -
1330 -pf01218 -
1331 HMMreg12833_1_EUT1(pf12833) EUT1
1332 H_mulberry(pf00936pf16365) EUT1
1333 Tsp_rosa EUT1
1334 HMMetlyEutC_1_eut1(pf05985) EUT1 EP_NTERM
1335 HMMetlyEutB_1_EUT1(pf06751) EUT1
1336 -pf01527 -
1337 HMMetlyEutB_1_EUT1(pf06751) EUT1
1338 -pf13276pf00665 -
1339 HMM06277EutA_1_eut(pf06277) EUT1
1340 HMMtpEuth_1_eut(pf04346) EUT1
1341 HMM00465alcdh_3_mix(pf00465) EUT1
1342 HMM06723EutJMreB_1_grm(#) EUT1
1343 HMMalddh_1_eutpdugrm(pf00171) EUT1 EP_NTERM
1344 P_lightViolet EUT1
1345 H_azure EUT1
1346 HMM01515PTA_B_1_eut1(pf01515) EUT1
1347 HMM01923cobtr_2_eut1(pf01923) EUT1
1348 HMM06249EutQ_1_eut1(pf06249) EUT1 EP_NTERM
1349 HMM10662PduV_1_mix(pf10662) EUT1
1350 Hp_euts_violetPink EUT1
1351 HMM00390_malic_1_eut1(pf00390pf03949pf01515) EUT1
1352 -pf00923 -
1353 HMM02780keto_1_eut1(pf02779pf02780pf00456) EUT1
1354 -pf06674 -
1355 HMM00293nuhyd_1_eut1(pf00293) EUT1
1356 -pf13247pf14691pf07992 -
1357 -# -
1358 -pf00672pf02518pf07730pf13675 -
1359 -pf00873 -
1360 -pf03960 -
BMC-P
BMC-H
BMC-T
Aldehyde dehydrogenase
PTAC
Signature enzyme
Alcohol dehydrogenase
Regulator

Escherichia coli 99.0741. Bacteria; Proteobacteria; Gammaproteobacteria; Enterobacterales;

EC970259__locus_1

EUT1

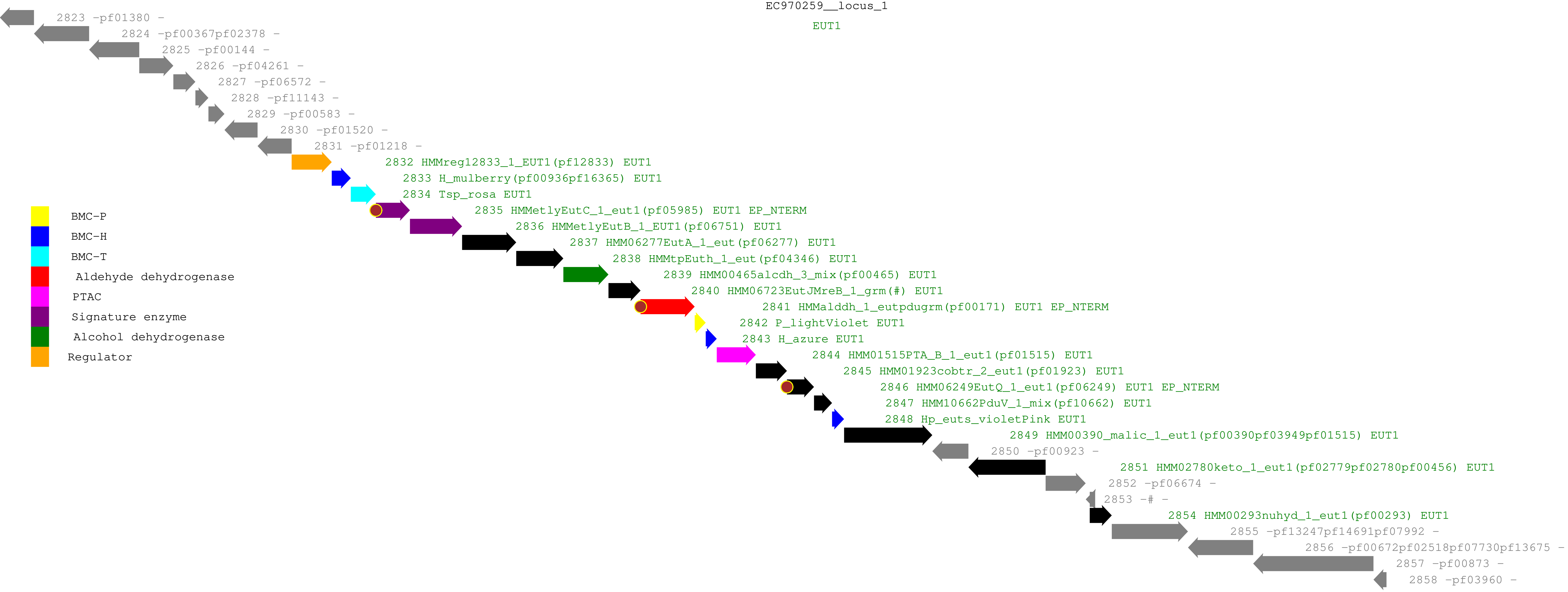

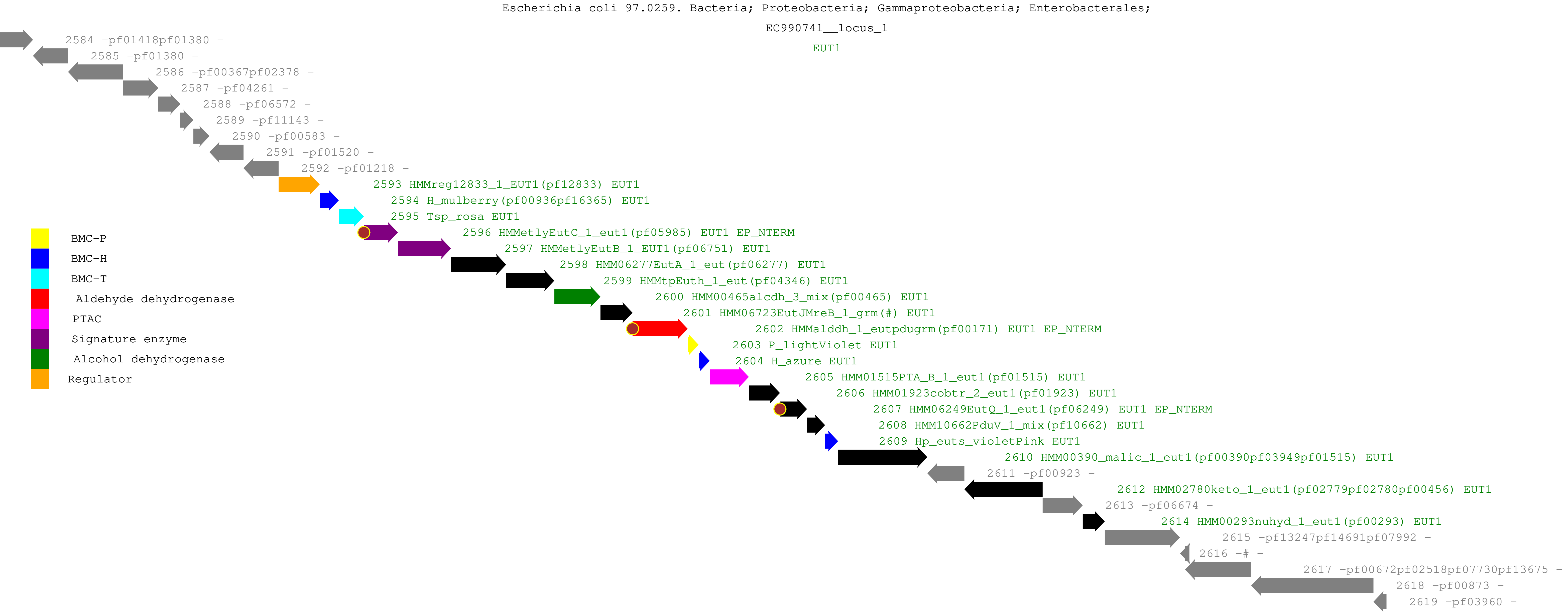

Escherichia coli 97.0259. Bacteria; Proteobacteria; Gammaproteobacteria; Enterobacterales;
EC990741__locus_1
EUT1
2584 -pf01418pf01380 -
2585 -pf01380 -
2586 -pf00367pf02378 -
2587 -pf04261 -
2588 -pf06572 -
2589 -pf11143 -
2590 -pf00583 -
2591 -pf01520 -
2592 -pf01218 -
2593 HMMreg12833_1_EUT1(pf12833) EUT1
2594 H_mulberry(pf00936pf16365) EUT1
2595 Tsp_rosa EUT1
2596 HMMetlyEutC_1_eut1(pf05985) EUT1 EP_NTERM
2597 HMMetlyEutB_1_EUT1(pf06751) EUT1
2598 HMM06277EutA_1_eut(pf06277) EUT1
2599 HMMtpEuth_1_eut(pf04346) EUT1
2600 HMM00465alcdh_3_mix(pf00465) EUT1
2601 HMM06723EutJMreB_1_grm(#) EUT1
2602 HMMaldh_1_eutpdugrm(pf00171) EUT1 EP_NTERM
2603 P_lightViolet EUT1
2604 H_azure EUT1
2605 HMM01515PTA_B_1_eut1(pf01515) EUT1
2606 HMM01923cobtr_2_eut1(pf01923) EUT1
2607 HMM06249EutQ_1_eut1(pf06249) EUT1 EP_NTERM
2608 HMM10662PduV_1_mix(pf10662) EUT1
2609 Hp_euts_violetPink EUT1
2610 HMM00390_malic_1_eut1(pf00390pf03949pf01515) EUT1
2611 -pf00923 -
2612 HMM02780keto_1_eut1(pf02779pf02780pf00456) EUT1
2613 -pf06674 -
2614 HMM00293nuhyd_1_eut1(pf00293) EUT1
2615 -pf13247pf14691pf07992 -
2616 -# -
2617 -pf00672pf02518pf07730pf13675 -
2618 -pf00873 -
2619 -pf03960 -
BMC-P
BMC-H
BMC-T
Aldehyde dehydrogenase
PTAC
Signature enzyme
Alcohol dehydrogenase
Regulator

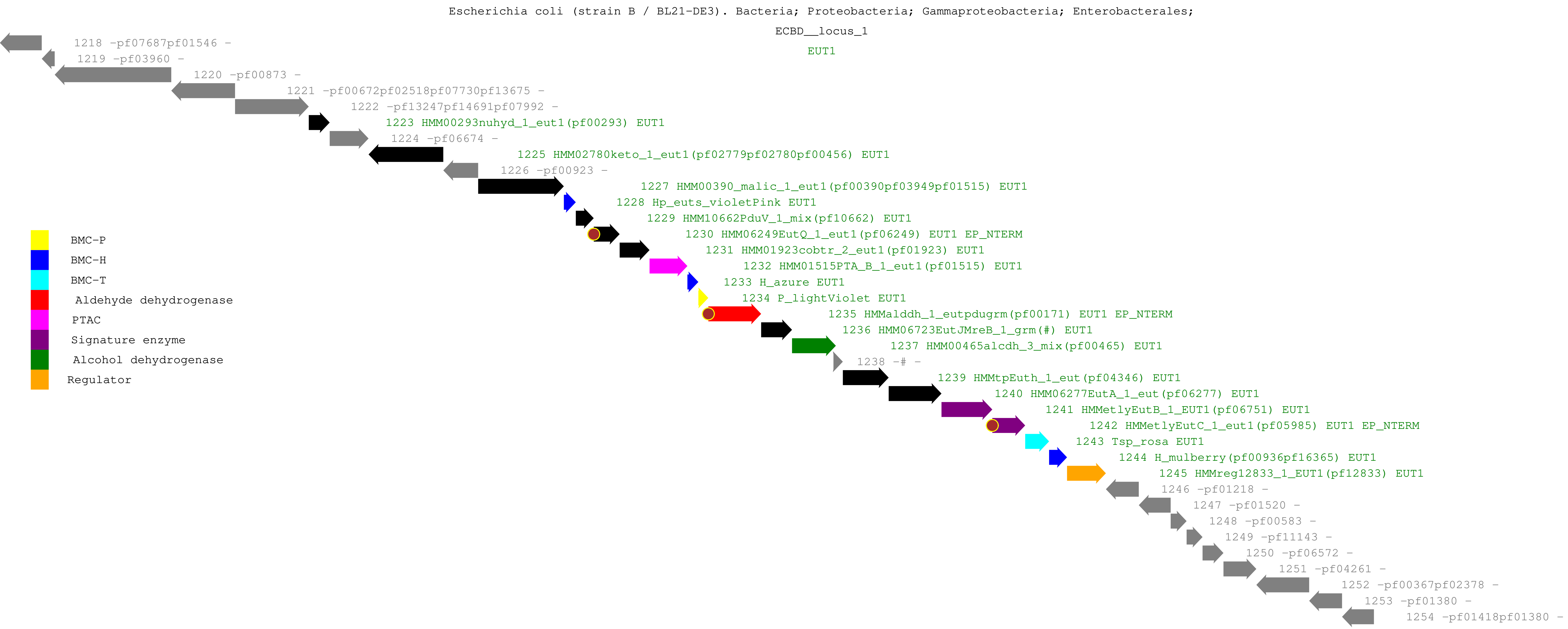

Escherichia coli (strain B / BL21-DE3). Bacteria; Proteobacteria; Gammaproteobacteria; Enterobacterales;
ECBD__locus_1
EUT1
1218 -pf07687pf01546 -
1219 -pf03960 -
1220 -pf00873 -
1221 -pf00672pf02518pf07730pf13675 -
1222 -pf13247pf14691pf07992 -
1223 HMM00293nuhyd_1_eut1(pf00293) EUT1
1224 -pf06674 -
1225 HMM02780keto_1_eut1(pf02779pf02780pf00456) EUT1
1226 -pf00923 -
1227 HMM00390_malic_1_eut1(pf00390pf03949pf01515) EUT1
1228 Hp_euts_violetPink EUT1
1229 HMM10662PduV_1_mix(pf10662) EUT1
1230 HMM06249EutQ_1_eut1(pf06249) EUT1 EP_NTERM
1231 HMM01923cobtr_2_eut1(pf01923) EUT1
1232 HMM01515PTA_B_1_eut1(pf01515) EUT1
1233 H_azure EUT1
1234 P_lightViolet EUT1
1235 HMMaldh_1_eutpdugrm(pf00171) EUT1 EP_NTERM
1236 HMM06723EutJMreB_1_grm(#) EUT1
1237 HMM00465alcdh_3_mix(pf00465) EUT1
1238 -# -
1239 HMMtpEuth_1_eut(pf04346) EUT1
1240 HMM06277EutA_1_eut(pf06277) EUT1
1241 HMMetlyEutB_1_EUT1(pf06751) EUT1
1242 HMMetlyEutC_1_eut1(pf05985) EUT1 EP_NTERM
1243 Tsp_rosa EUT1
1244 H_mulberry(pf00936pf16365) EUT1
1245 HMMreg12833_1_EUT1(pf12833) EUT1
1246 -pf01218 -
1247 -pf01520 -
1248 -pf00583 -
1249 -pf11143 -
1250 -pf06572 -
1251 -pf04261 -
1252 -pf00367pf02378 -
1253 -pf01380 -
1254 -pf01418pf01380 -
BMC-P
BMC-H
BMC-T
Aldehyde dehydrogenase
PTAC
Signature enzyme
Alcohol dehydrogenase
Regulator

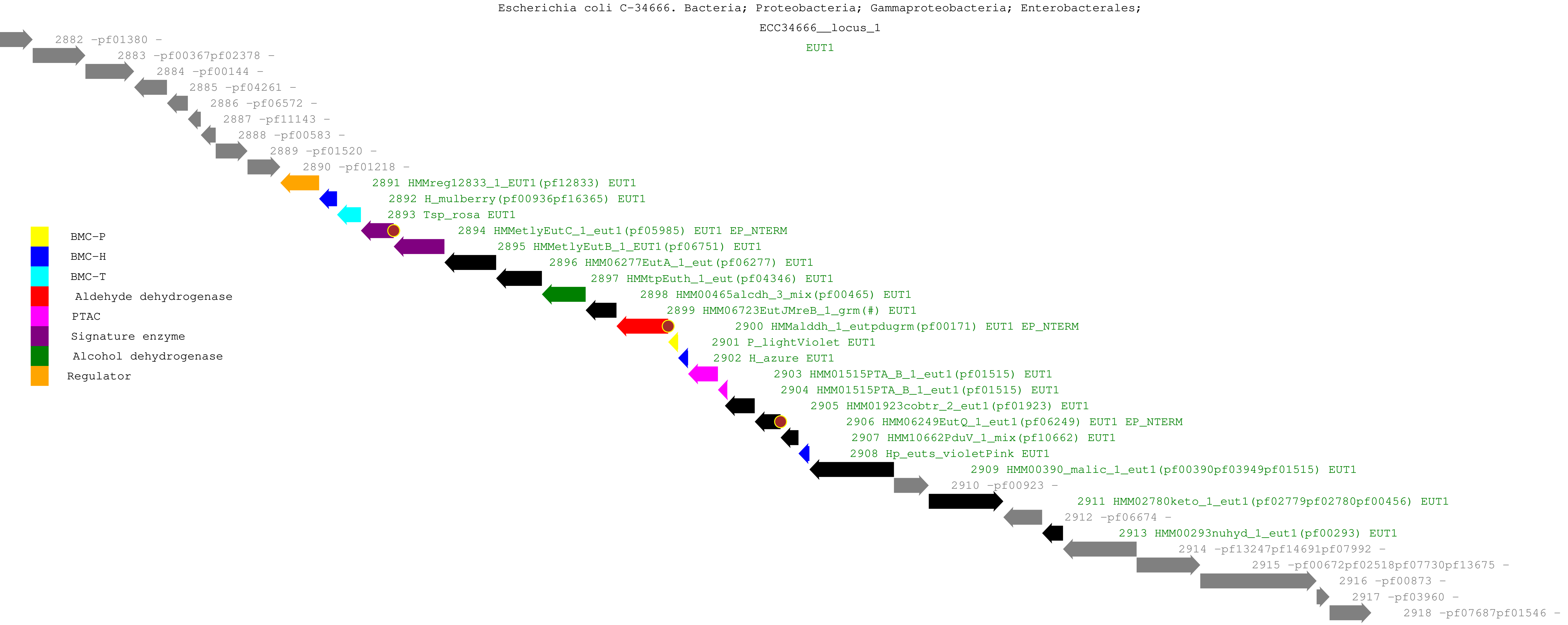
Escherichia coli C-34666. Bacteria; Proteobacteria; Gammaproteobacteria; Enterobacterales;
ECC34666__locus_1
EUT1
2882 -pf01380 -
2883 -pf00367pf02378 -
2884 -pf00144 -
2885 -pf04261 -
2886 -pf06572 -
2887 -pf11143 -
2888 -pf00583 -
2889 -pf01520 -
2890 -pf01218 -
2891 HMMreg12833_1_EUT1(pf12833) EUT1
2892 H_mulberry(pf00936pf16365) EUT1
2893 Tsp_rosa EUT1
2894 HMMetlyEutC_1_eut1(pf05985) EUT1 EP_NTERM
2895 HMMetlyEutB_1_EUT1(pf06751) EUT1
2896 HMM06277EutA_1_eut(pf06277) EUT1
2897 HMMtpEuth_1_eut(pf04346) EUT1
2898 HMM00465alcdh_3_mix(pf00465) EUT1
2899 HMM06723EutJMreB_1_grm(#) EUT1
2900 HMMalddh_1_eutpdugrm(pf00171) EUT1 EP_NTERM
2901 P_lightViolet EUT1
2902 H_azure EUT1
2903 HMM01515PTA_B_1_eut1(pf01515) EUT1
2904 HMM01515PTA_B_1_eut1(pf01515) EUT1
2905 HMM01923cobtr_2_eut1(pf01923) EUT1
2906 HMM06249EutQ_1_eut1(pf06249) EUT1 EP_NTERM
2907 HMM10662PduV_1_mix(pf10662) EUT1
2908 Hp_euts_violetPink EUT1
2909 HMM00390_malic_1_eut1(pf00390pf03949pf01515) EUT1
2910 -pf00923 -
2911 HMM02780keto_1_eut1(pf02779pf02780pf00456) EUT1
2912 -pf06674 -
2913 HMM00293nuhyd_1_eut1(pf00293) EUT1
2914 -pf13247pf14691pf07992 -
2915 -pf00672pf02518pf07730pf13675 -
2916 -pf00873 -
2917 -pf03960 -
2918 -pf07687pf01546 -
BMC-P
BMC-H
BMC-T
Aldehyde dehydrogenase
PTAC
Signature enzyme
Alcohol dehydrogenase
Regulator

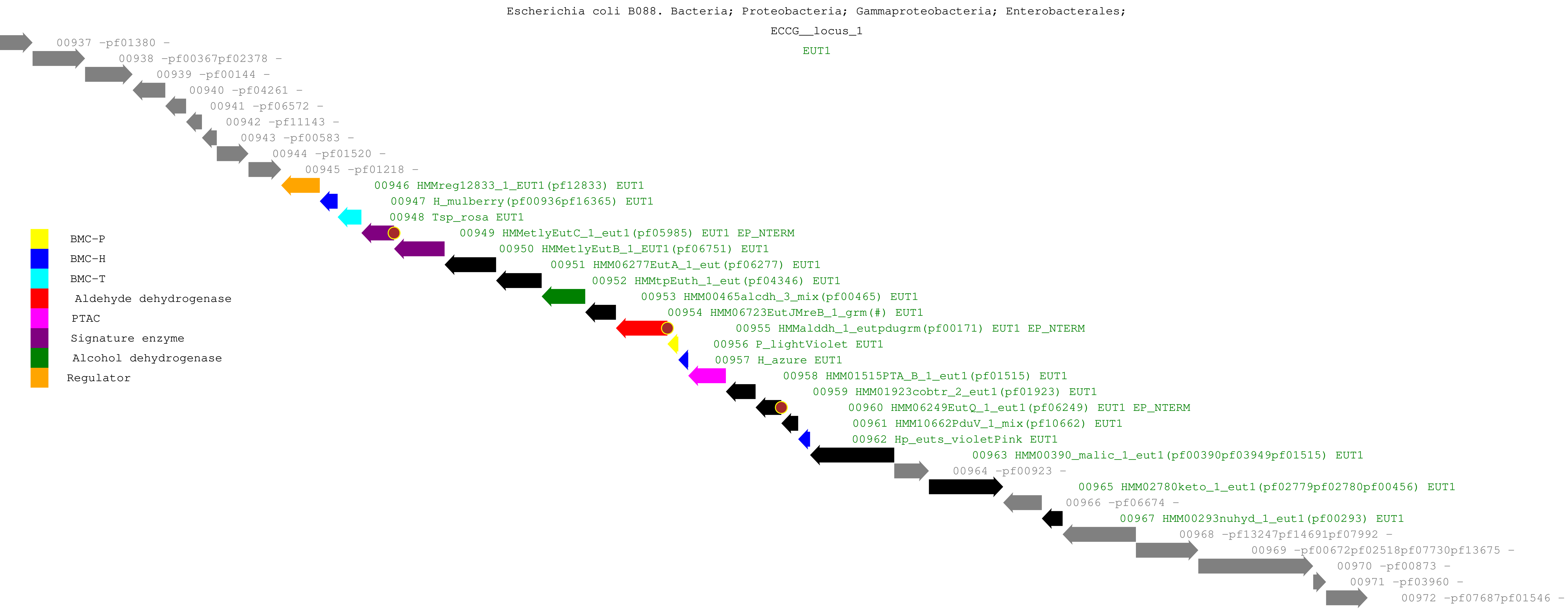

Escherichia coli B088. Bacteria; Proteobacteria; Gammaproteobacteria; Enterobacterales;
ECCG__locus_1
EUT1
00937 -pf01380 -
00938 -pf00367pf02378 -
00939 -pf00144 -
00940 -pf04261 -
00941 -pf06572 -
00942 -pf11143 -
00943 -pf00583 -
00944 -pf01520 -
00945 -pf01218 -
00946 HMMreg12833_1_EUT1(pf12833) EUT1
00947 H_mulberry(pf00936pf16365) EUT1
00948 Tsp_rosa EUT1
00949 HMMetlyEutC_1_eut1(pf05985) EUT1 EP_NTERM
00950 HMMetlyEutB_1_EUT1(pf06751) EUT1
00951 HMM06277EutA_1_eut(pf06277) EUT1
00952 HMMtpEuth_1_eut(pf04346) EUT1
00953 HMM00465alcdh_3_mix(pf00465) EUT1
00954 HMM06723EutJMreB_1_grm(#) EUT1
00955 HMMalddh_1_eutpdugrm(pf00171) EUT1 EP_NTERM
00956 P_lightViolet EUT1
00957 H_azure EUT1
00958 HMM01515PTA_B_1_eut1(pf01515) EUT1
00959 HMM01923cobtr_2_eut1(pf01923) EUT1
00960 HMM06249EutQ_1_eut1(pf06249) EUT1 EP_NTERM
00961 HMM10662PduV_1_mix(pf10662) EUT1
00962 Hp_euts_violetPink EUT1
00963 HMM00390_malic_1_eut1(pf00390pf03949pf01515) EUT1
00964 -pf00923 -
00965 HMM02780keto_1_eut1(pf02779pf02780pf00456) EUT1
00966 -pf06674 -
00967 HMM00293nuhyd_1_eut1(pf00293) EUT1
00968 -pf13247pf14691pf07992 -
00969 -pf00672pf02518pf07730pf13675 -
00970 -pf00873 -
00971 -pf03960 -
00972 -pf07687pf01546 -
BMC-P
BMC-H
BMC-T
Aldehyde dehydrogenase
PTAC
Signature enzyme
Alcohol dehydrogenase
Regulator

Escherichia coli DEC1B. Bacteria; Proteobacteria; Gammaproteobacteria; Enterobacterales;

ECDEC1B__locus_2

EUT1

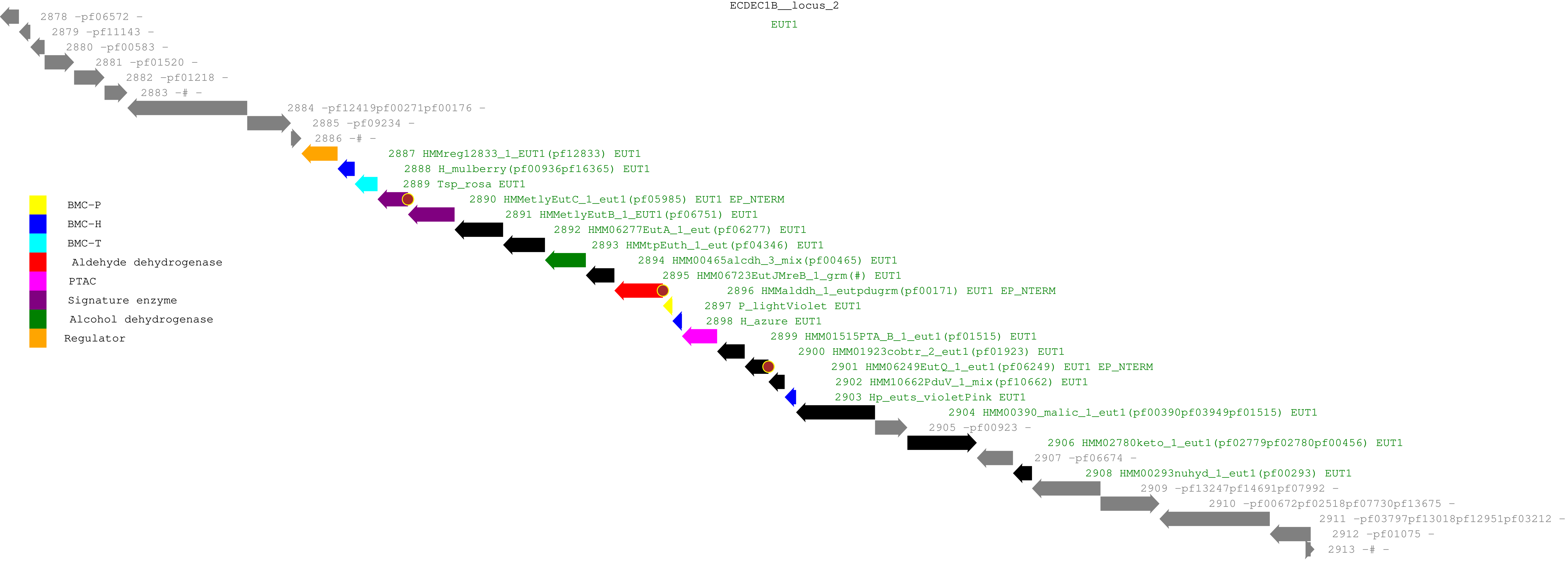

Escherichia coli DEC1C. Bacteria; Proteobacteria; Gammaproteobacteria; Enterobacterales;

ECDEC1C__locus_2

EUT1

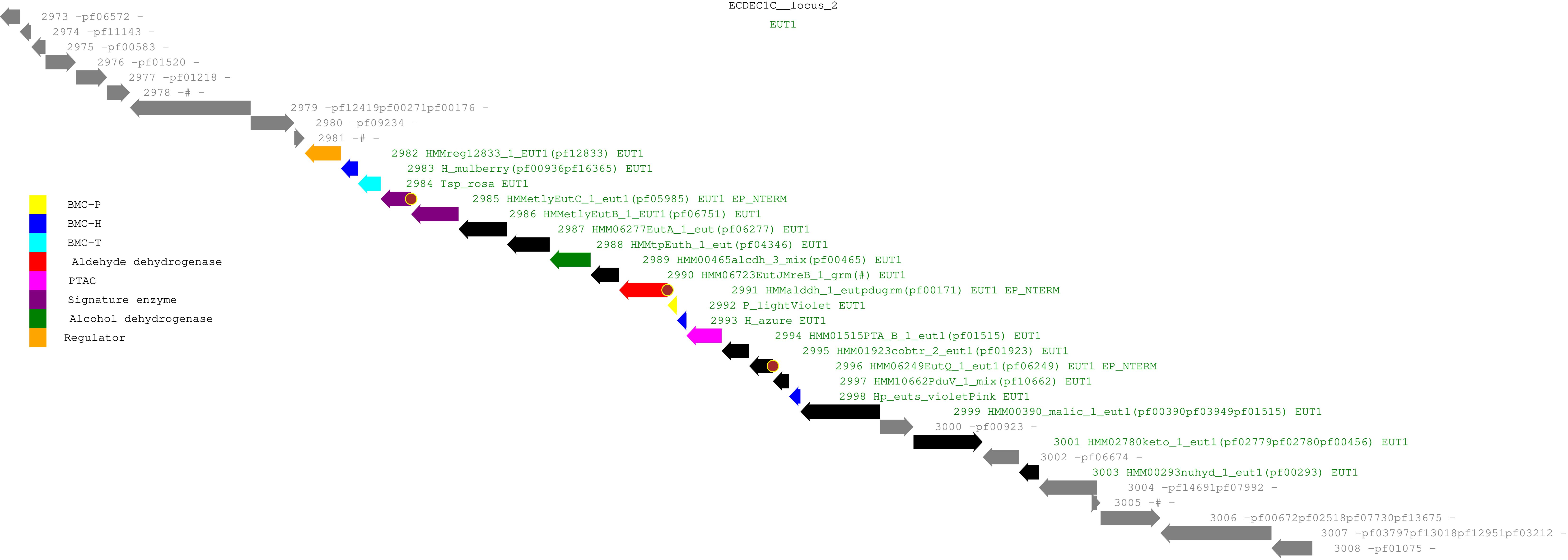

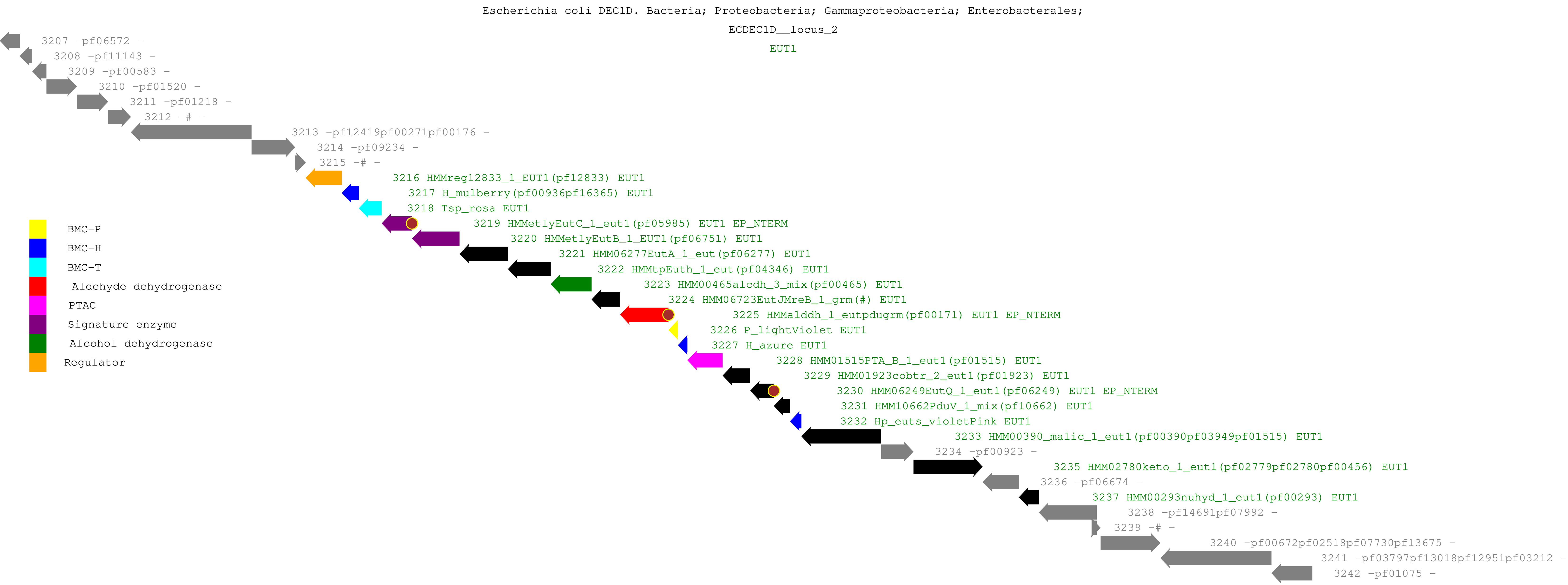
Escherichia coli DEC1D. Bacteria; Proteobacteria; Gammaproteobacteria; Enterobacterales;
ECDEC1D__locus_2
EUT1
3207 -pf06572 -
3208 -pf11143 -
3209 -pf00583 -
3210 -pf01520 -
3211 -pf01218 -
3212 -# -
3213 -pf12419pf00271pf00176 -
3214 -pf09234 -
3215 -# -
3216 HMMreg12833_1_EUT1(pf12833) EUT1
3217 H_mulberry(pf00936pf16365) EUT1
3218 Tsp_rosa EUT1
3219 HMMetlyEutC_1_eut1(pf05985) EUT1 EP_NTERM
3220 HMMetlyEutB_1_EUT1(pf06751) EUT1
3221 HMM06277EutA_1_eut(pf06277) EUT1
3222 HMMtpEuth_1_eut(pf04346) EUT1
3223 HMM00465alcdh_3_mix(pf00465) EUT1
3224 HMM06723EutJMreB_1_grm(#) EUT1
3225 HMMalddh_1_eutpdugrm(pf00171) EUT1 EP_NTERM
3226 P_lightViolet EUT1
3227 H_azure EUT1
3228 HMM01515PTA_B_1_eut1(pf01515) EUT1
3229 HMM01923cobtr_2_eut1(pf01923) EUT1
3230 HMM06249EutQ_1_eut1(pf06249) EUT1 EP_NTERM
3231 HMM10662PduV_1_mix(pf10662) EUT1
3232 Hp_euts_violetPink EUT1
3233 HMM00390_malic_1_eut1(pf00390pf03949pf01515) EUT1
3234 -pf00923 -
3235 HMM02780keto_1_eut1(pf02779pf02780pf00456) EUT1
3236 -pf06674 -
3237 HMM00293nuhyd_1_eut1(pf00293) EUT1
3238 -pf14691pf07992 -
3239 -# -
3240 -pf00672pf02518pf07730pf13675 -
3241 -pf03797pf13018pf12951pf03212 -
3242 -pf01075 -
BMC-P
BMC-H
BMC-T
Aldehyde dehydrogenase
PTAC
Signature enzyme
Alcohol dehydrogenase
Regulator

Escherichia coli DEC2C. Bacteria; Proteobacteria; Gammaproteobacteria; Enterobacterales;

ECDEC2C__locus_2

EUT1

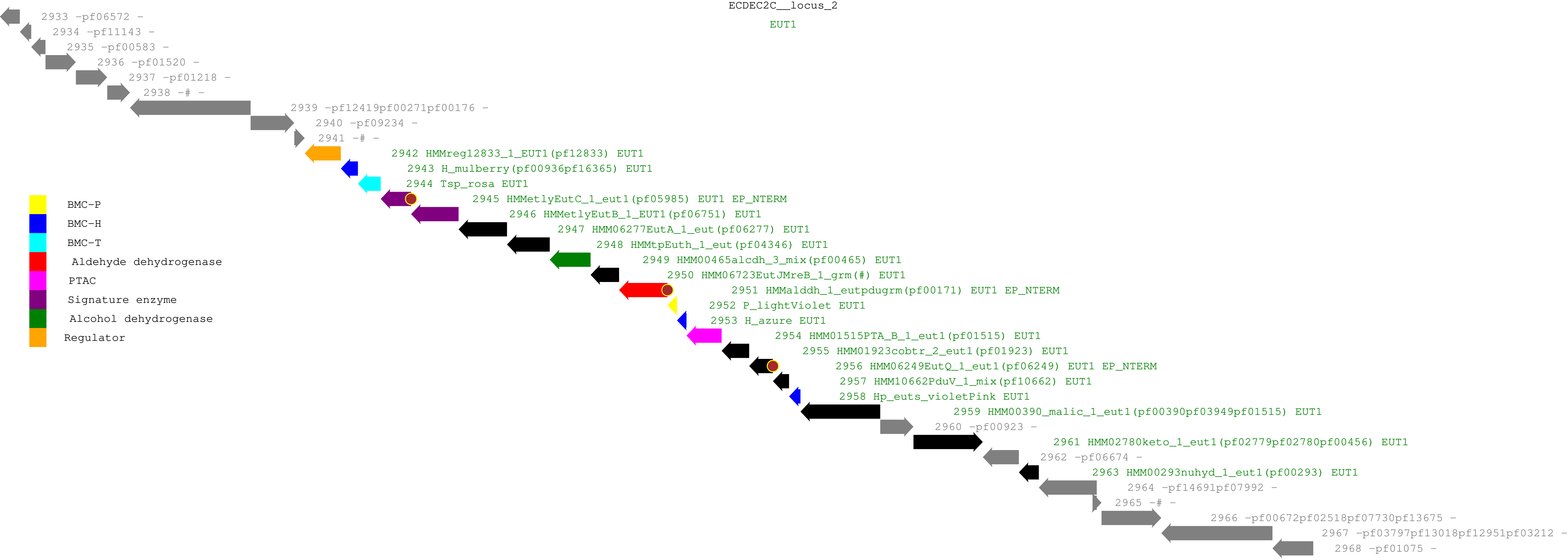

Escherichia coli DEC2E. Bacteria; Proteobacteria; Gammaproteobacteria; Enterobacterales;

ECDEC2E__locus_2

EUT1

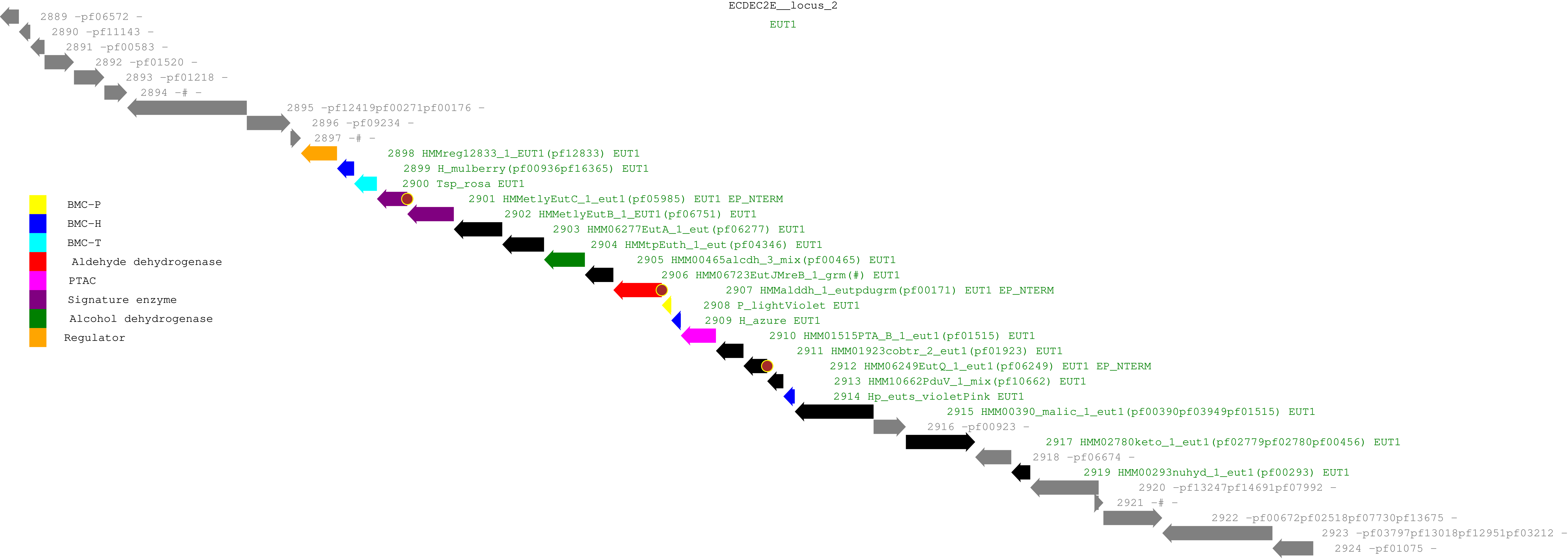

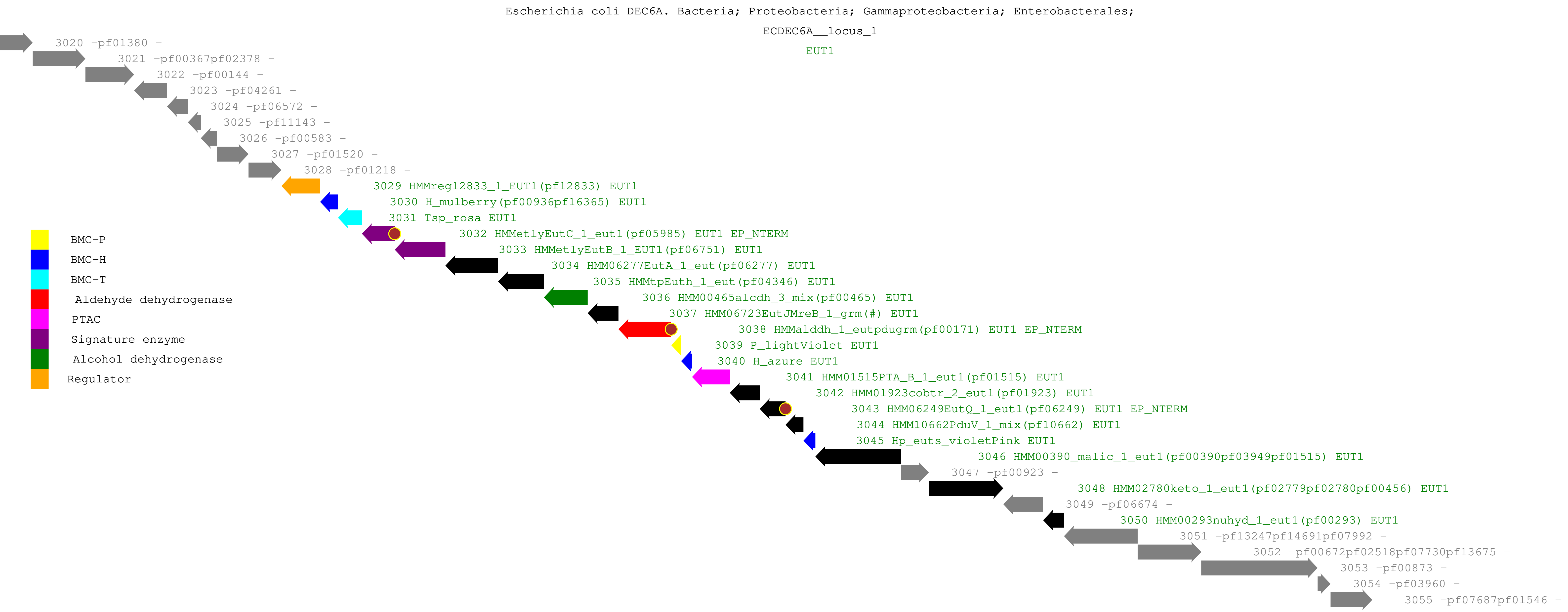
Escherichia coli DEC6A. Bacteria; Proteobacteria; Gammaproteobacteria; Enterobacterales;
ECDEC6A__locus_1
EUT1
3020 -pf01380 -
3021 -pf00367pf02378 -
3022 -pf00144 -
3023 -pf04261 -
3024 -pf06572 -
3025 -pf11143 -
3026 -pf00583 -
3027 -pf01520 -
3028 -pf01218 -
3029 HMMreg12833_1_EUT1(pf12833) EUT1
3030 H_mulberry(pf00936pf16365) EUT1
3031 Tsp_rosa EUT1
3032 HMMetlyEutC_1_eut1(pf05985) EUT1 EP_NTERM
3033 HMMetlyEutB_1_EUT1(pf06751) EUT1
3034 HMM06277EutA_1_eut(pf06277) EUT1
3035 HMMtpEuth_1_eut(pf04346) EUT1
3036 HMM00465alcdh_3_mix(pf00465) EUT1
3037 HMM06723EutJMreB_1_grm(#) EUT1
3038 HMMalddh_1_eutpdugrm(pf00171) EUT1 EP_NTERM
3039 P_lightViolet EUT1
3040 H_azure EUT1
3041 HMM01515PTA_B_1_eut1(pf01515) EUT1
3042 HMM01923cobtr_2_eut1(pf01923) EUT1
3043 HMM06249EutQ_1_eut1(pf06249) EUT1 EP_NTERM
3044 HMM10662PduV_1_mix(pf10662) EUT1
3045 Hp_euts_violetPink EUT1
3046 HMM00390_malic_1_eut1(pf00390pf03949pf01515) EUT1
3047 -pf00923 -
3048 HMM02780keto_1_eut1(pf02779pf02780pf00456) EUT1
3049 -pf06674 -
3050 HMM00293nuhyd_1_eut1(pf00293) EUT1
3051 -pf13247pf14691pf07992 -
3052 -pf00672pf02518pf07730pf13675 -
3053 -pf00873 -
3054 -pf03960 -
3055 -pf07687pf01546 -
BMC-P
BMC-H
BMC-T
Aldehyde dehydrogenase
PTAC
Signature enzyme
Alcohol dehydrogenase
Regulator

Escherichia coli B185. Bacteria; Proteobacteria; Gammaproteobacteria; Enterobacterales;

ECDG__locus_1

EUT1

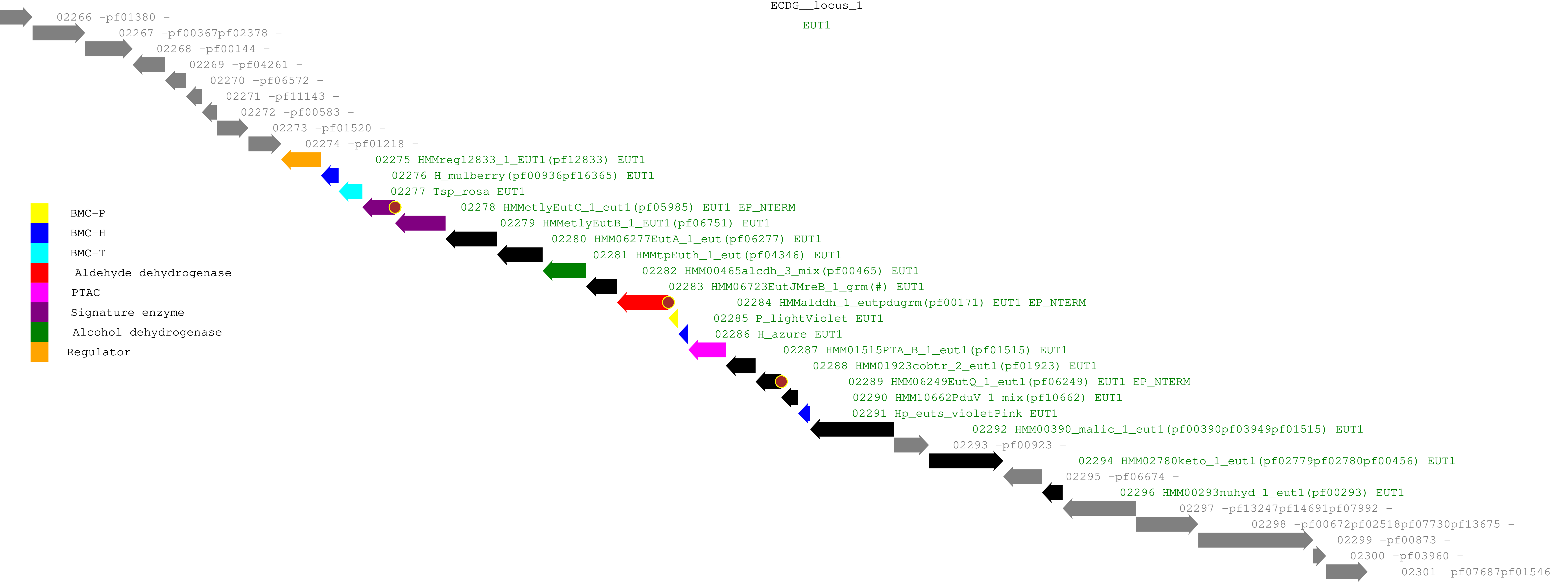

Escherichia coli O139:H28 (strain E24377A / ETEC). Bacteria; Proteobacteria; Gammaproteobacteria; Enterobacterales;

EcE24377A__locus_2

EUT1

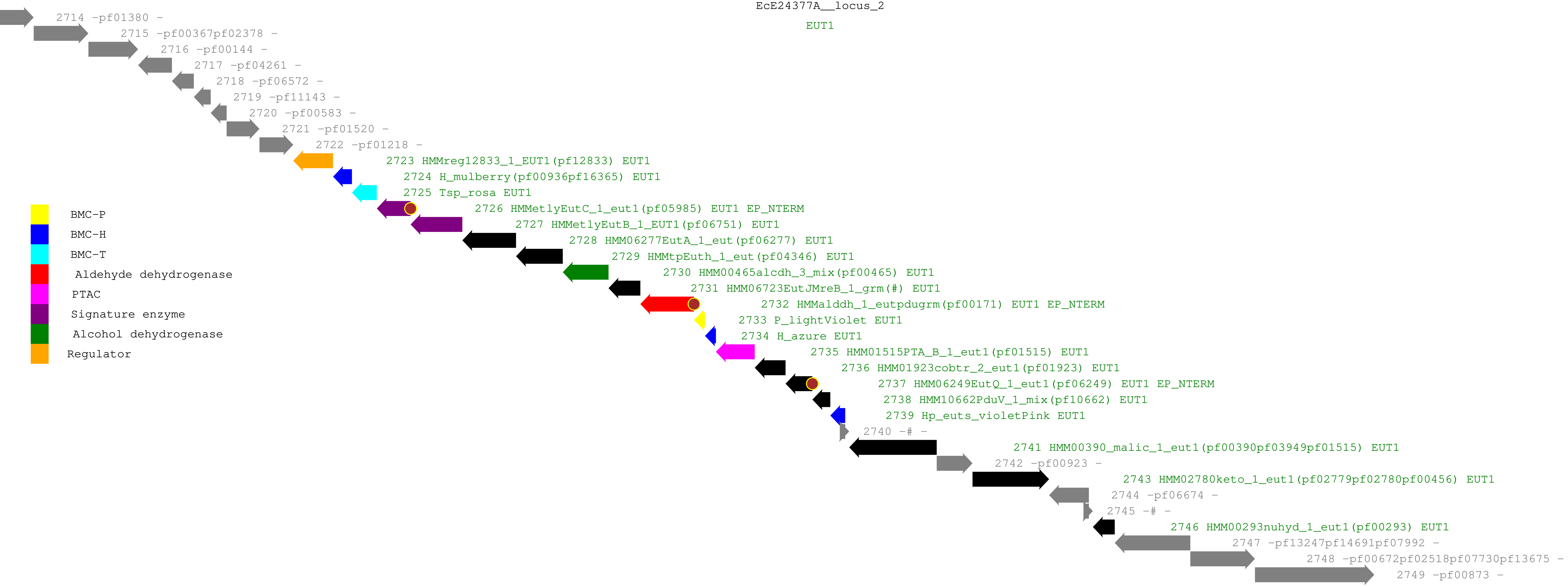

Escherichia coli O81 (strain ED1a). Bacteria; Proteobacteria; Gammaproteobacteria; Enterobacterales;

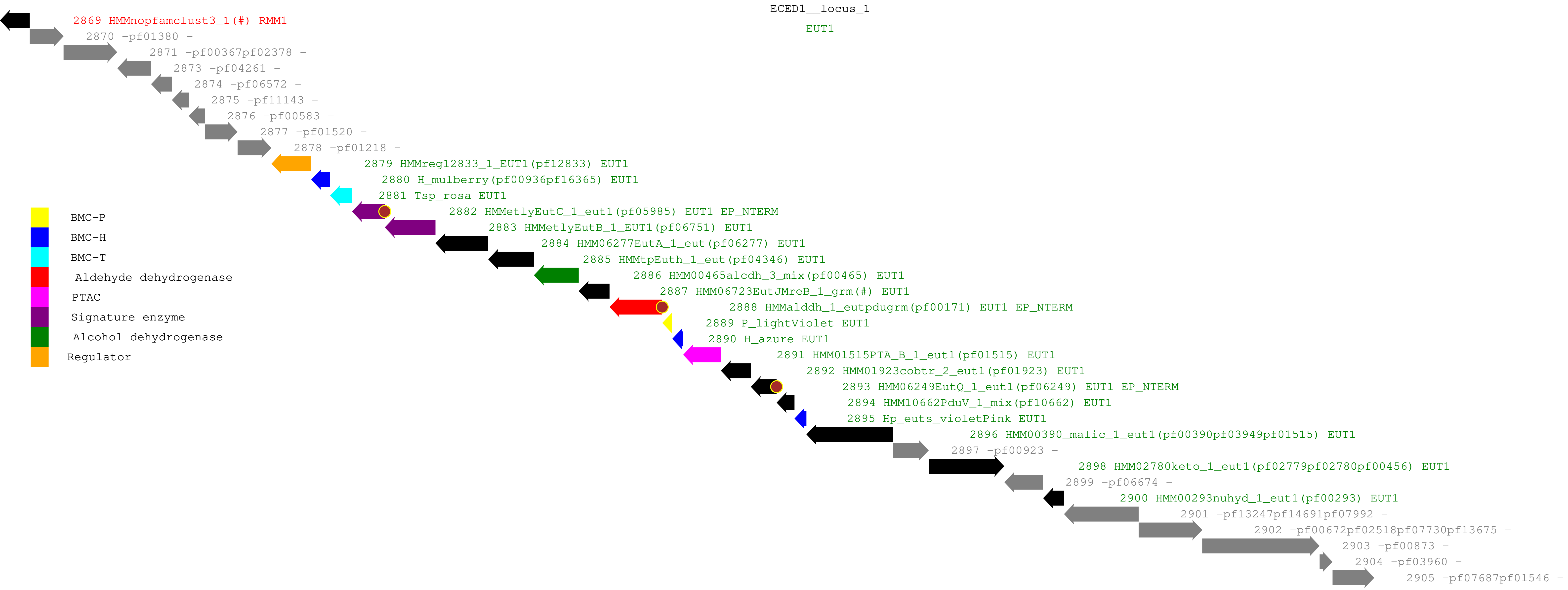

Escherichia coli B354. Bacteria; Proteobacteria; Gammaproteobacteria; Enterobacterales;

ECEG__locus_2

EUT1

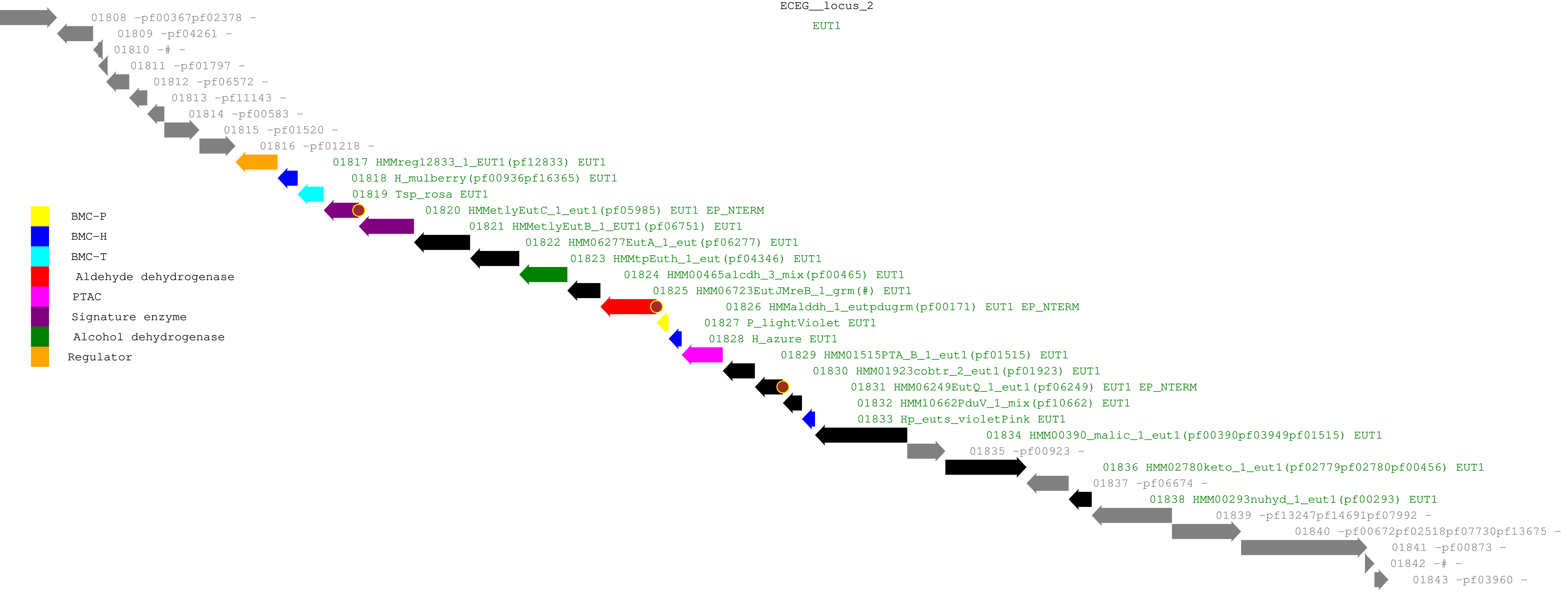

Escherichia coli O157:H7 (strain EC869). Bacteria; Proteobacteria; Gammaproteobacteria; Enterobacterales;

ECH7EC869__locus_1

EUT1

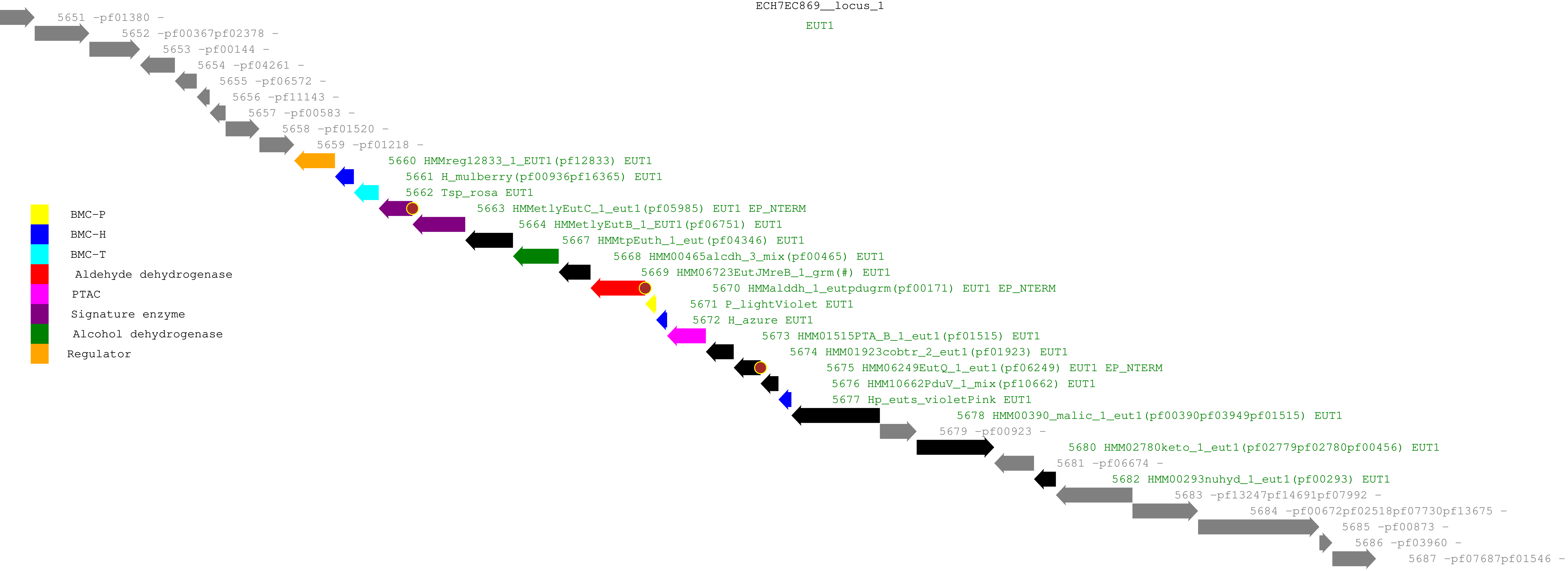

Escherichia coli H736. Bacteria; Proteobacteria; Gammaproteobacteria; Enterobacterales;

ECHG__locus_1

EUT1

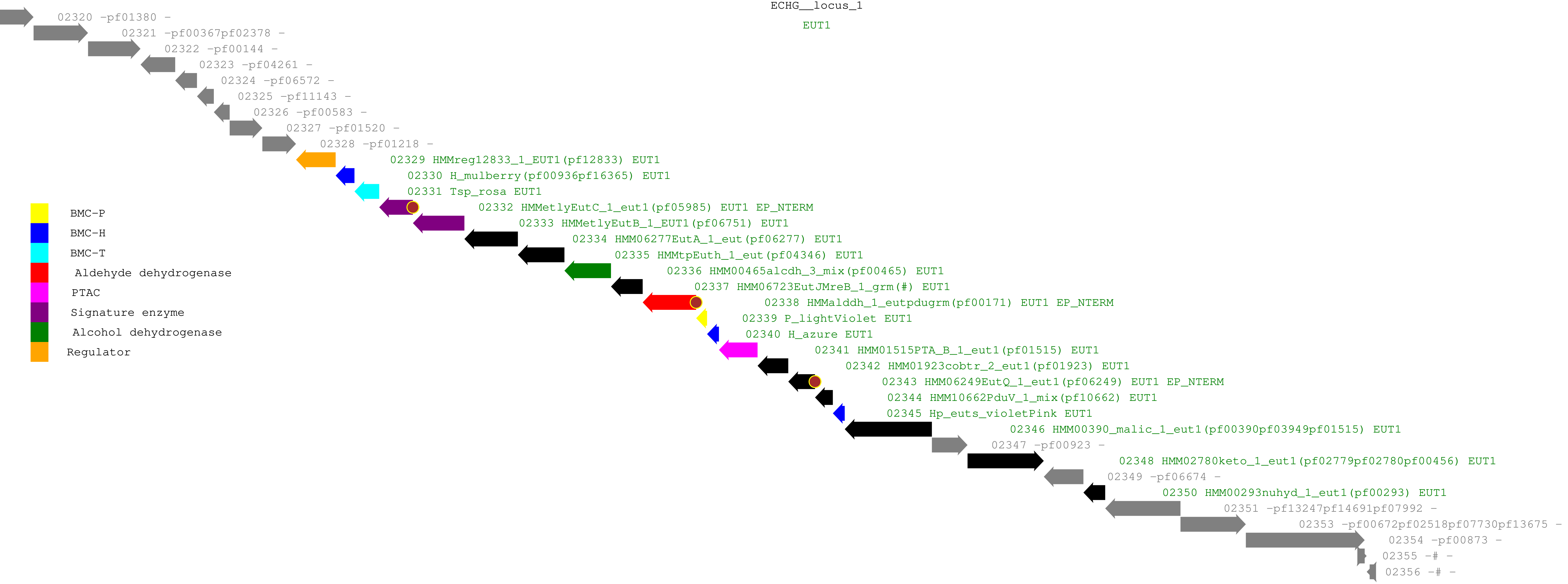

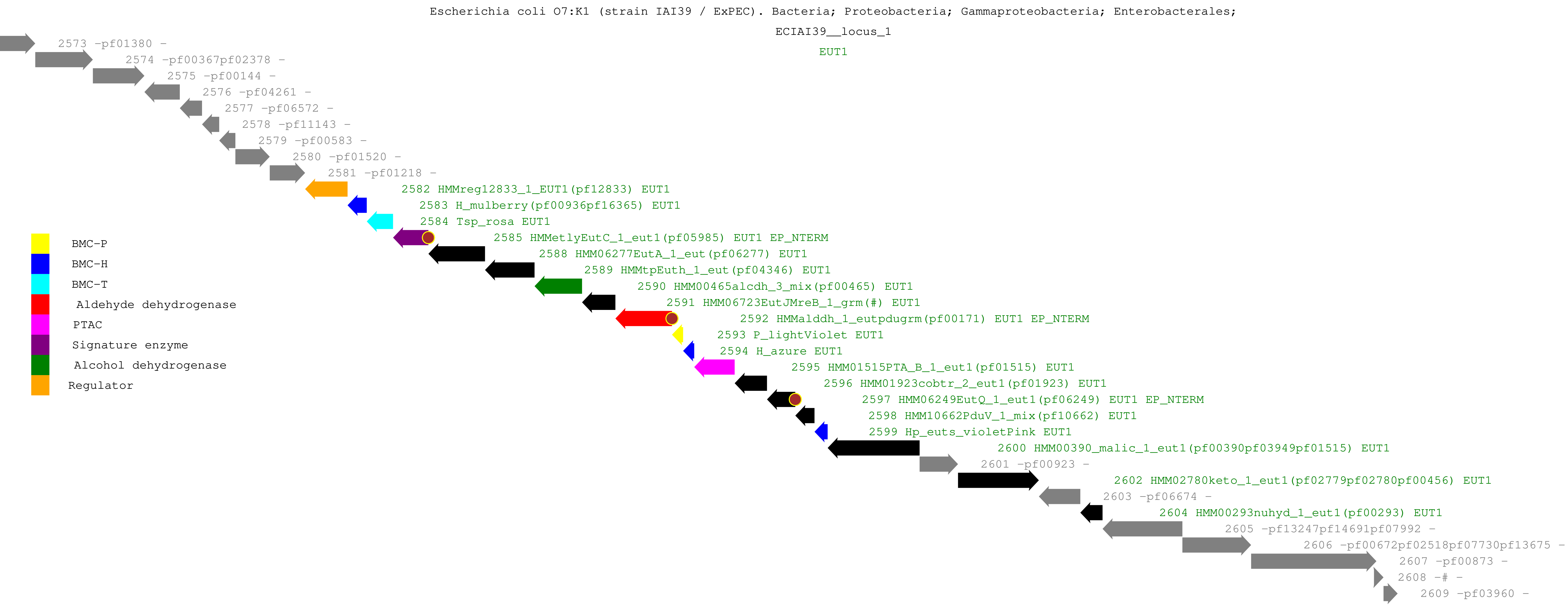
Escherichia coli O7:K1 (strain IAI39 / ExPEC). Bacteria; Proteobacteria; Gammaproteobacteria; Enterobacterales;
ECIAI39__locus_1
EUT1
2573 -pf01380 -
2574 -pf00367pf02378 -
2575 -pf00144 -
2576 -pf04261 -
2577 -pf06572 -
2578 -pf11143 -
2579 -pf00583 -
2580 -pf01520 -
2581 -pf01218 -
2582 HMMreg12833_1_EUT1(pf12833) EUT1
2583 H_mulberry(pf00936pf16365) EUT1
2584 Tsp_rosa EUT1
2585 HMMetlyEutC_1_eut1(pf05985) EUT1 EP_NTERM
2588 HMM06277EutA_1_eut(pf06277) EUT1
2589 HMMtpEuth_1_eut(pf04346) EUT1
2590 HMM00465alcdh_3_mix(pf00465) EUT1
2591 HMM06723EutJMreB_1_grm(#) EUT1
2592 HMMalddh_1_eutpdugrm(pf00171) EUT1 EP_NTERM
2593 P_lightViolet EUT1
2594 H_azure EUT1
2595 HMM01515PTA_B_1_eut1(pf01515) EUT1
2596 HMM01923cobtr_2_eut1(pf01923) EUT1
2597 HMM06249EutQ_1_eut1(pf06249) EUT1 EP_NTERM
2598 HMM10662PduV_1_mix(pf10662) EUT1
2599 Hp_euts_violetPink EUT1
2600 HMM00390_malic_1_eut1(pf00390pf03949pf01515) EUT1
2601 -pf00923 -
2602 HMM02780keto_1_eut1(pf02779pf02780pf00456) EUT1
2603 -pf06674 -
2604 HMM00293nuhyd_1_eut1(pf00293) EUT1
2605 -pf13247pf14691pf07992 -
2606 -pf00672pf02518pf07730pf13675 -
2607 -pf00873 -
2608 -# -
2609 -pf03960 -
BMC-P
BMC-H
BMC-T
Aldehyde dehydrogenase
PTAC
Signature enzyme
Alcohol dehydrogenase
Regulator

Escherichia coli M605. Bacteria; Proteobacteria; Gammaproteobacteria; Enterobacterales;

ECIG__locus_1

EUT1

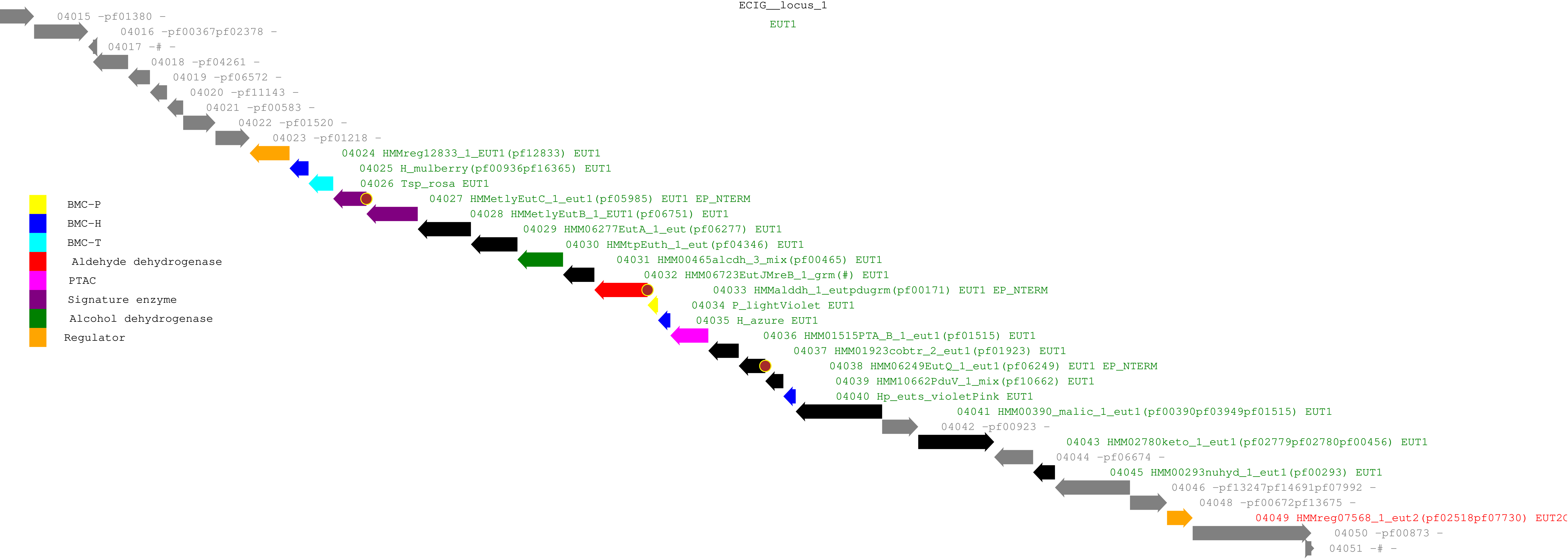

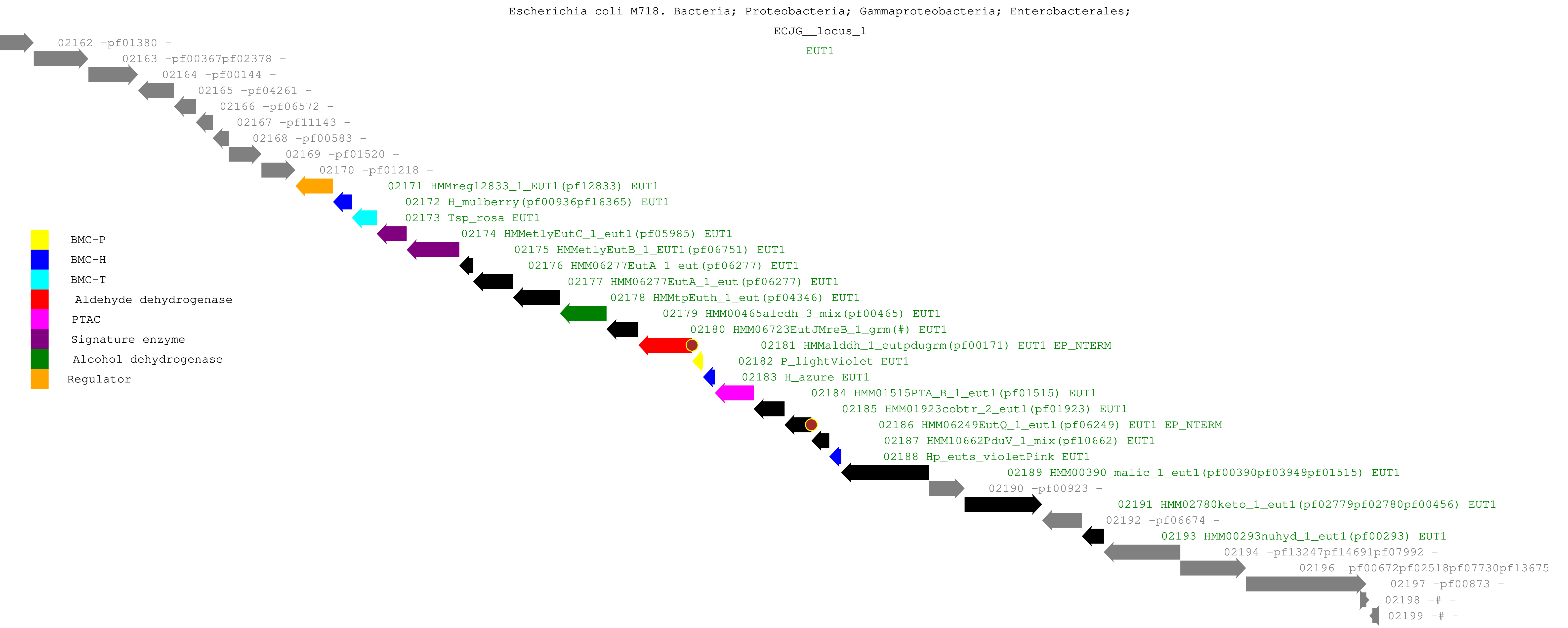
Escherichia coli M718. Bacteria; Proteobacteria; Gammaproteobacteria; Enterobacterales;
ECJG__locus_1
EUT1
02162 -pf01380 -
02163 -pf00367pf02378 -
02164 -pf00144 -
02165 -pf04261 -
02166 -pf06572 -
02167 -pf11143 -
02168 -pf00583 -
02169 -pf01520 -
02170 -pf01218 -
02171 HMMreg12833_1_EUT1(pf12833) EUT1
02172 H_mulberry(pf00936pf16365) EUT1
02173 Tsp_rosa EUT1
02174 HMMetlyEutC_1_eut1(pf05985) EUT1
02175 HMMetlyEutB_1_EUT1(pf06751) EUT1
02176 HMM06277EutA_1_eut(pf06277) EUT1
02177 HMM06277EutA_1_eut(pf06277) EUT1
02178 HMMtpEuth_1_eut(pf04346) EUT1
02179 HMM00465alcdh_3_mix(pf00465) EUT1
02180 HMM06723EutJMreB_1_grm(#) EUT1
02181 HMMalddh_1_eutpdugrm(pf00171) EUT1 EP_NTERM
02182 P_lightViolet EUT1
02183 H_azure EUT1
02184 HMM01515PTA_B_1_eut1(pf01515) EUT1
02185 HMM01923cobtr_2_eut1(pf01923) EUT1
02186 HMM06249EutQ_1_eut1(pf06249) EUT1 EP_NTERM
02187 HMM10662PduV_1_mix(pf10662) EUT1
02188 Hp_euts_violetPink EUT1
02189 HMM00390_malic_1_eut1(pf00390pf03949pf01515) EUT1
02190 -pf00923 -
02191 HMM02780keto_1_eut1(pf02779pf02780pf00456) EUT1
02192 -pf06674 -
02193 HMM00293nuhyd_1_eut1(pf00293) EUT1
02194 -pf13247pf14691pf07992 -
02196 -pf00672pf02518pf07730pf13675 -
02197 -pf00873 -
02198 -# -
02199 -# -
BMC-P
BMC-H
BMC-T
Aldehyde dehydrogenase
PTAC
Signature enzyme
Alcohol dehydrogenase
Regulator

Escherichia coli MP021561.2. Bacteria; Proteobacteria; Gammaproteobacteria; Enterobacterales;

ECMP0215612__locus_1

EUT1

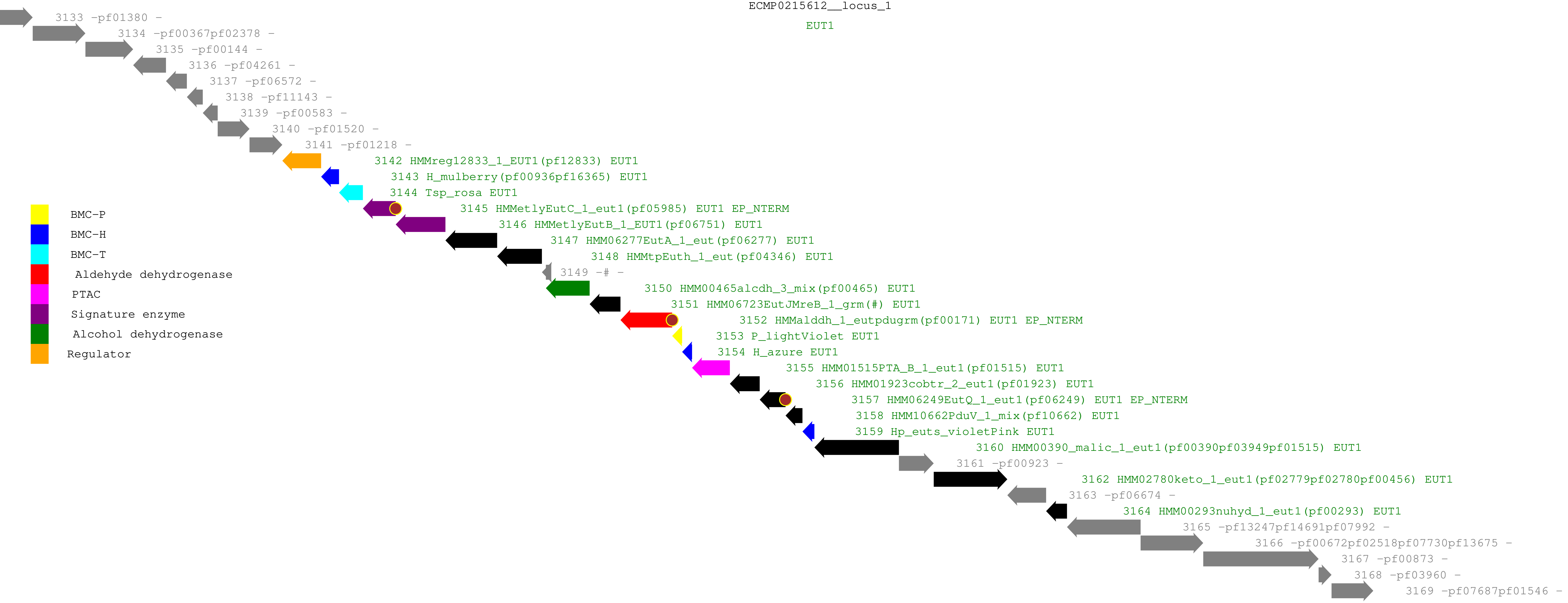

Escherichia coli O103:H2 (strain 12009 / EHEC). Bacteria; Proteobacteria; Gammaproteobacteria; Enterobacterales;

ECO103__locus_1

EUT1

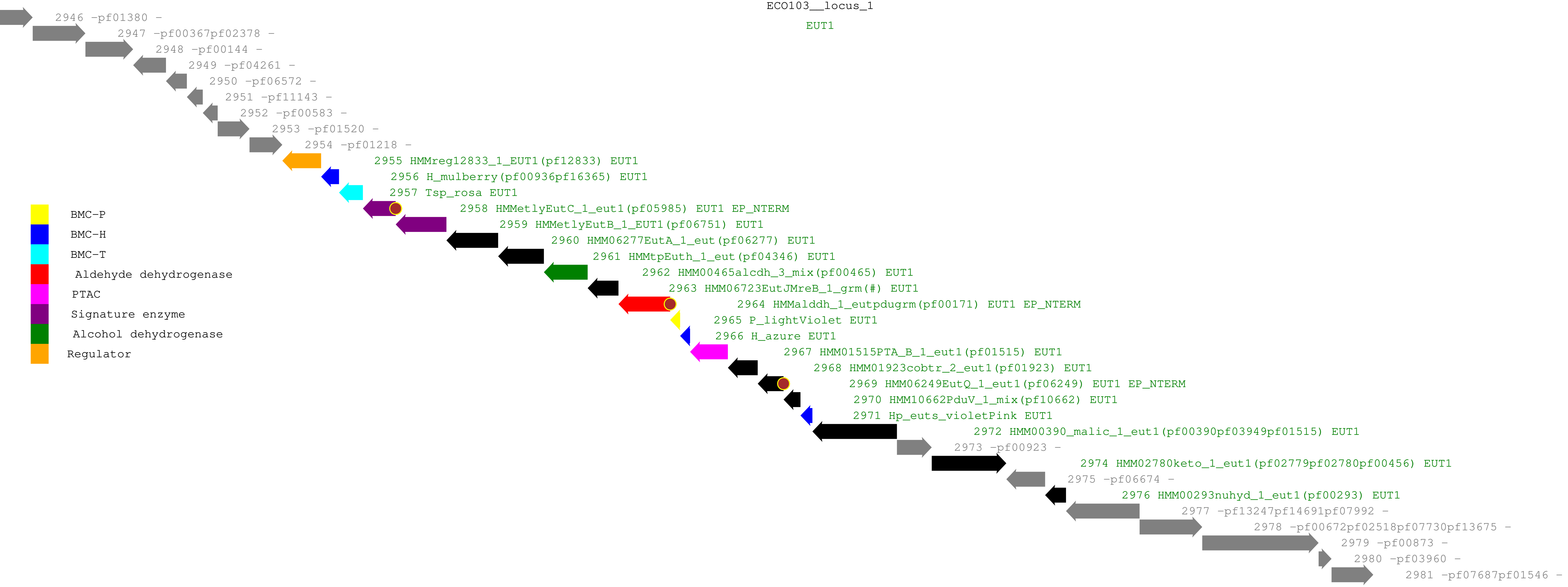

Escherichia coli O111:H- (strain 11128 / EHEC). Bacteria; Proteobacteria; Gammaproteobacteria; Enterobacterales;

ECO111__locus_1

EUT1

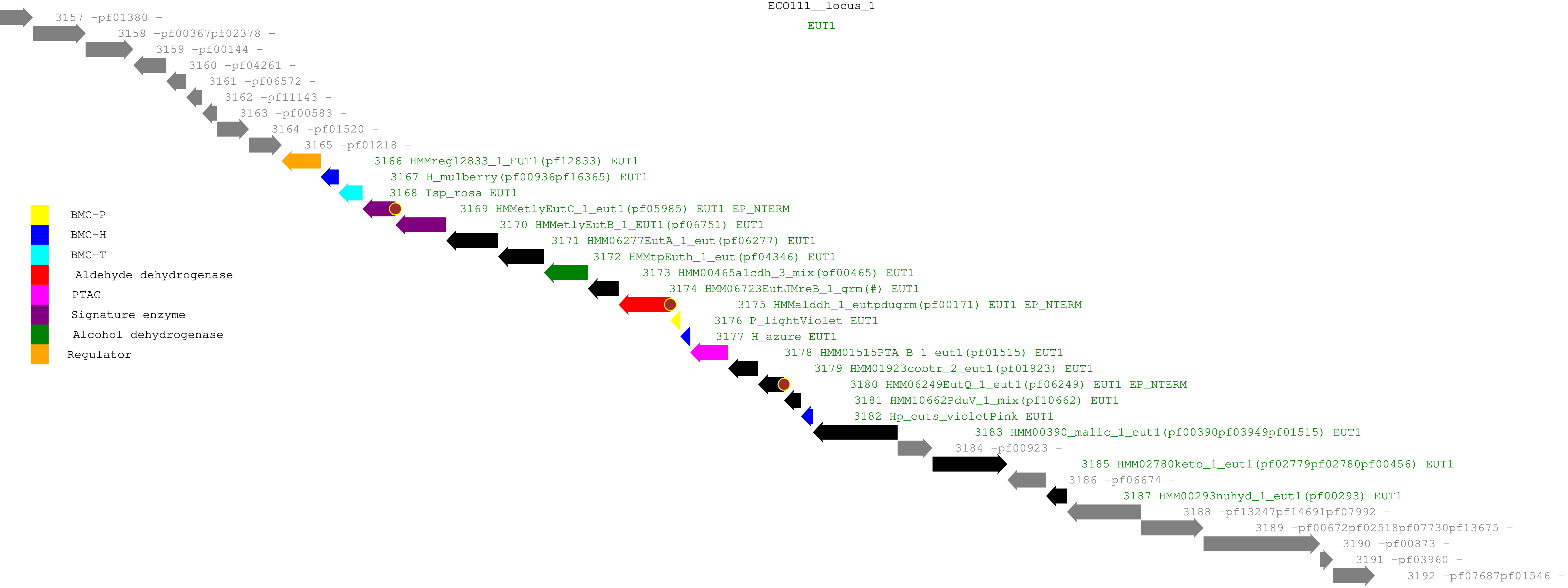

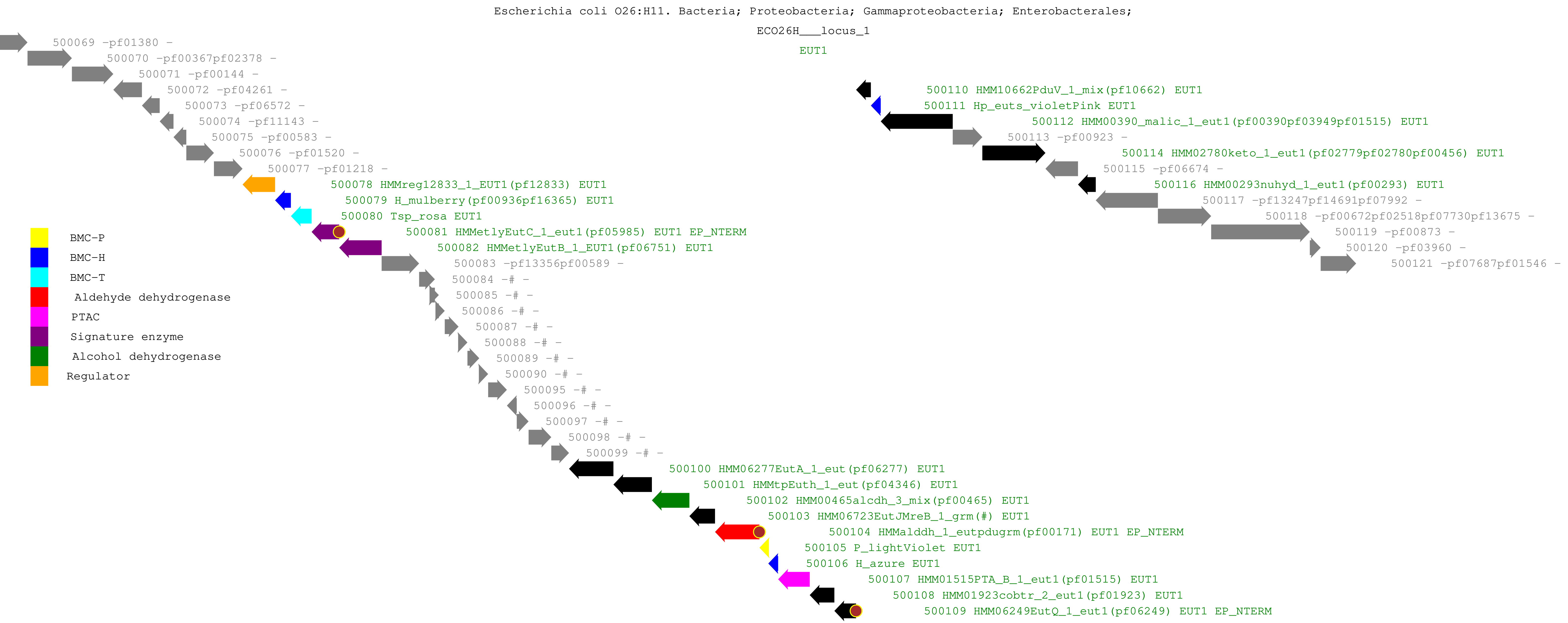
Escherichia coli O26:H11. Bacteria; Proteobacteria; Gammaproteobacteria; Enterobacterales;
ECO26H___locus_1
EUT1
500069 -pf01380 -
500070 -pf00367pf02378 -
500071 -pf00144 -
500072 -pf04261 -
500073 -pf06572 -
500074 -pf11143 -
500075 -pf00583 -
500076 -pf01520 -
500077 -pf01218 -
500078 HMMreg12833_1_EUT1(pf12833) EUT1
500079 H_mulberry(pf00936pf16365) EUT1
500080 Tsp_rosa EUT1
500081 HMMetlyEutC_1_eut1(pf05985) EUT1 EP_NTERM
500082 HMMetlyEutB_1_EUT1(pf06751) EUT1
500083 -pf13356pf00589 -
500084 -# -
500085 -# -
500086 -# -
500087 -# -
500088 -# -
500089 -# -
500090 -# -
500095 -# -
500096 -# -
500097 -# -
500098 -# -
500099 -# -
500100 HMM06277EutA_1_eut(pf06277) EUT1
500101 HMMtpEuth_1_eut(pf04346) EUT1
500102 HMM00465alcdh_3_mix(pf00465) EUT1
500103 HMM06723EutJMreB_1_grm(#) EUT1
500104 HMMaldh_1_eutpdugrm(pf00171) EUT1 EP_NTERM
500105 P_lightViolet EUT1
500106 H_azure EUT1
500107 HMM01515PTA_B_1_eut1(pf01515) EUT1
500108 HMM01923cobtr_2_eut1(pf01923) EUT1
500109 HMM06249EutQ_1_eut1(pf06249) EUT1 EP_NTERM
500110 HMM10662PduV_1_mix(pf10662) EUT1
500111 Hp_euts_violetPink EUT1
500112 HMM00390_malic_1_eut1(pf00390pf03949pf01515) EUT1
500113 -pf00923 -
500114 HMM02780keto_1_eut1(pf02779pf02780pf00456) EUT1
500115 -pf06674 -
500116 HMM00293nuhyd_1_eut1(pf00293) EUT1
500117 -pf13247pf14691pf07992 -
500118 -pf00672pf02518pf07730pf13675 -
500119 -pf00873 -
500120 -pf03960 -
500121 -pf07687pf01546 -
BMC-P
BMC-H
BMC-T
Aldehyde dehydrogenase
PTAC
Signature enzyme
Alcohol dehydrogenase
Regulator

Escherichia coli O111:H11 str. CVM9455. Bacteria; Proteobacteria; Gammaproteobacteria; Enterobacterales;

ECO9455__locus_1

EUT1

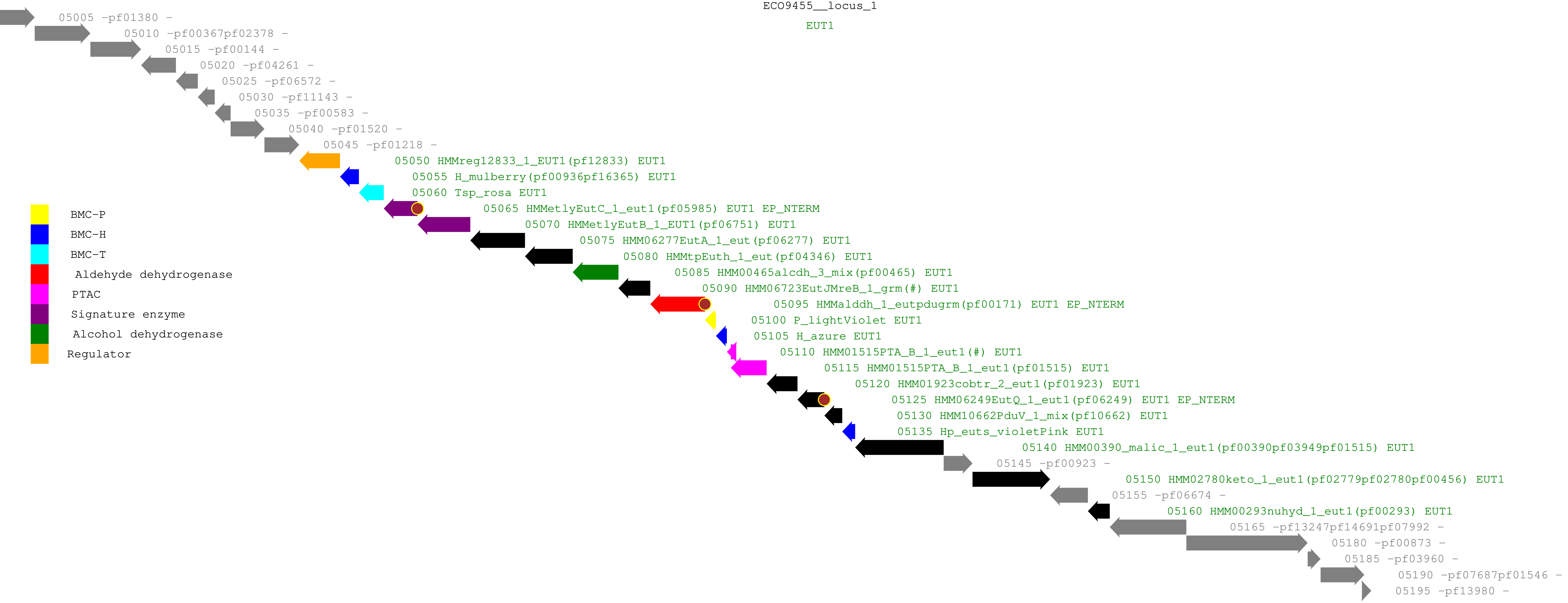

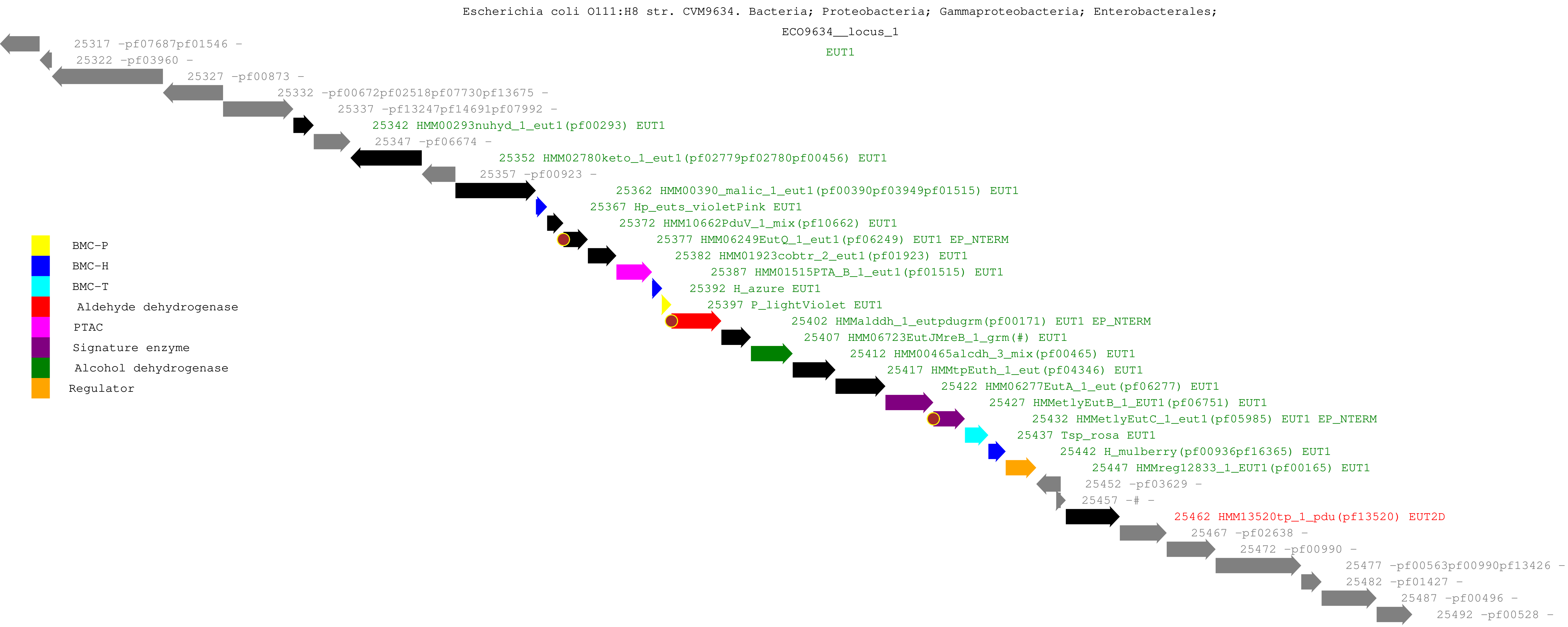
Escherichia coli O111:H8 str. CVM9634. Bacteria; Proteobacteria; Gammaproteobacteria; Enterobacterales;
ECO9634__locus_1
EUT1
25317 -pf07687pf01546 -
25322 -pf03960 -
25327 -pf00873 -
25332 -pf00672pf02518pf07730pf13675 -
25337 -pf13247pf14691pf07992 -
25342 HMM00293nuhyd_1_eut1(pf00293) EUT1
25347 -pf06674 -
25352 HMM02780keto_1_eut1(pf02779pf02780pf00456) EUT1
25357 -pf00923 -
25362 HMM00390_malic_1_eut1(pf00390pf03949pf01515) EUT1
25367 Hp_euts_violetPink EUT1
25372 HMM10662PduV_1_mix(pf10662) EUT1
25377 HMM06249EutQ_1_eut1(pf06249) EUT1 EP_NTERM
25382 HMM01923cobtr_2_eut1(pf01923) EUT1
25387 HMM01515PTA_B_1_eut1(pf01515) EUT1
25392 H_azure EUT1
25397 P_lightViolet EUT1
25402 HMMalddh_1_eutpdugrm(pf00171) EUT1 EP_NTERM
25407 HMM06723EutJMreB_1_grm(#) EUT1
25412 HMM00465alcdh_3_mix(pf00465) EUT1
25417 HMMtpEuth_1_eut(pf04346) EUT1
25422 HMM06277EutA_1_eut(pf06277) EUT1
25427 HMMetlyEutB_1_EUT1(pf06751) EUT1
25432 HMMetlyEutC_1_eut1(pf05985) EUT1 EP_NTERM
25437 Tsp_rosa EUT1
25442 H_mulberry(pf00936pf16365) EUT1
25447 HMMreg12833_1_EUT1(pf00165) EUT1
25452 -pf03629 -
25457 -# -
25462 HMM13520tp_1_pdu(pf13520) EUT2D
25467 -pf02638 -
25472 -pf00990 -
25477 -pf00563pf00990pf13426 -
25482 -pf01427 -
25487 -pf00496 -
25492 -pf00528 -
BMC-P
BMC-H
BMC-T
Aldehyde dehydrogenase
PTAC
Signature enzyme
Alcohol dehydrogenase
Regulator

Escherichia coli P0301867.11. Bacteria; Proteobacteria; Gammaproteobacteria; Enterobacterales;

ECP030186711__locus_1

EUT1

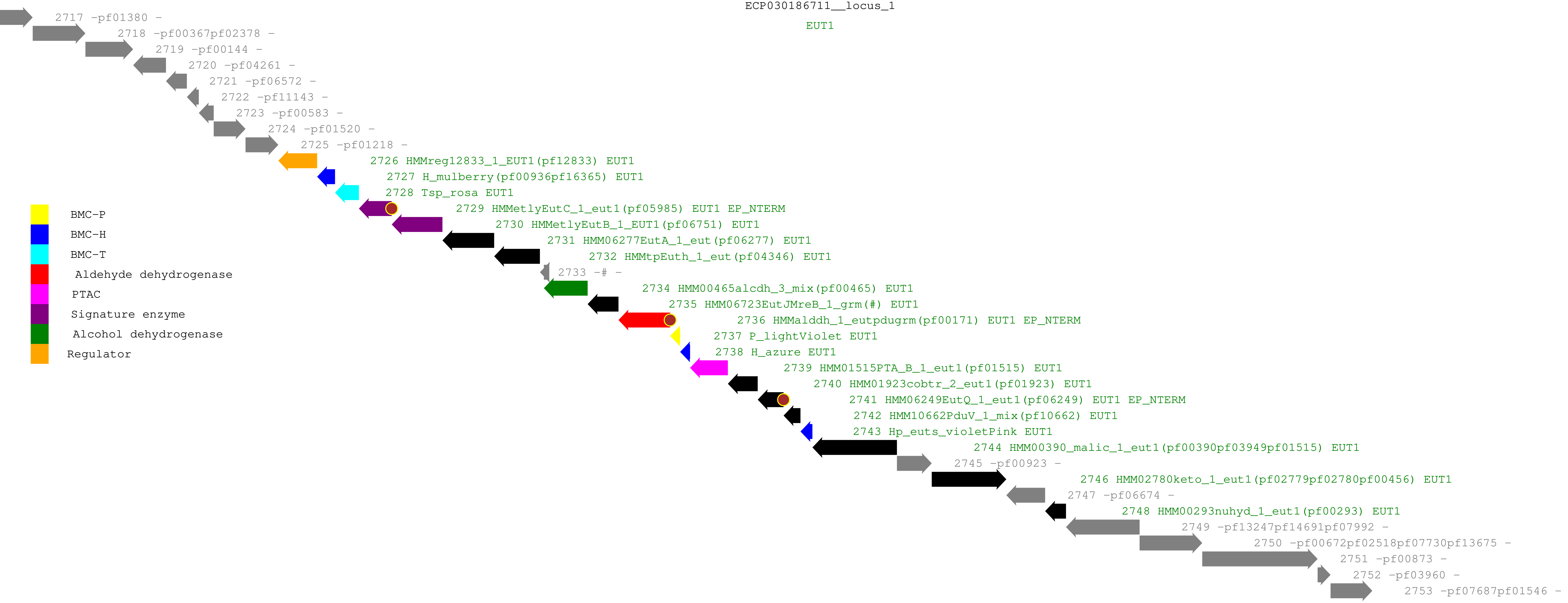

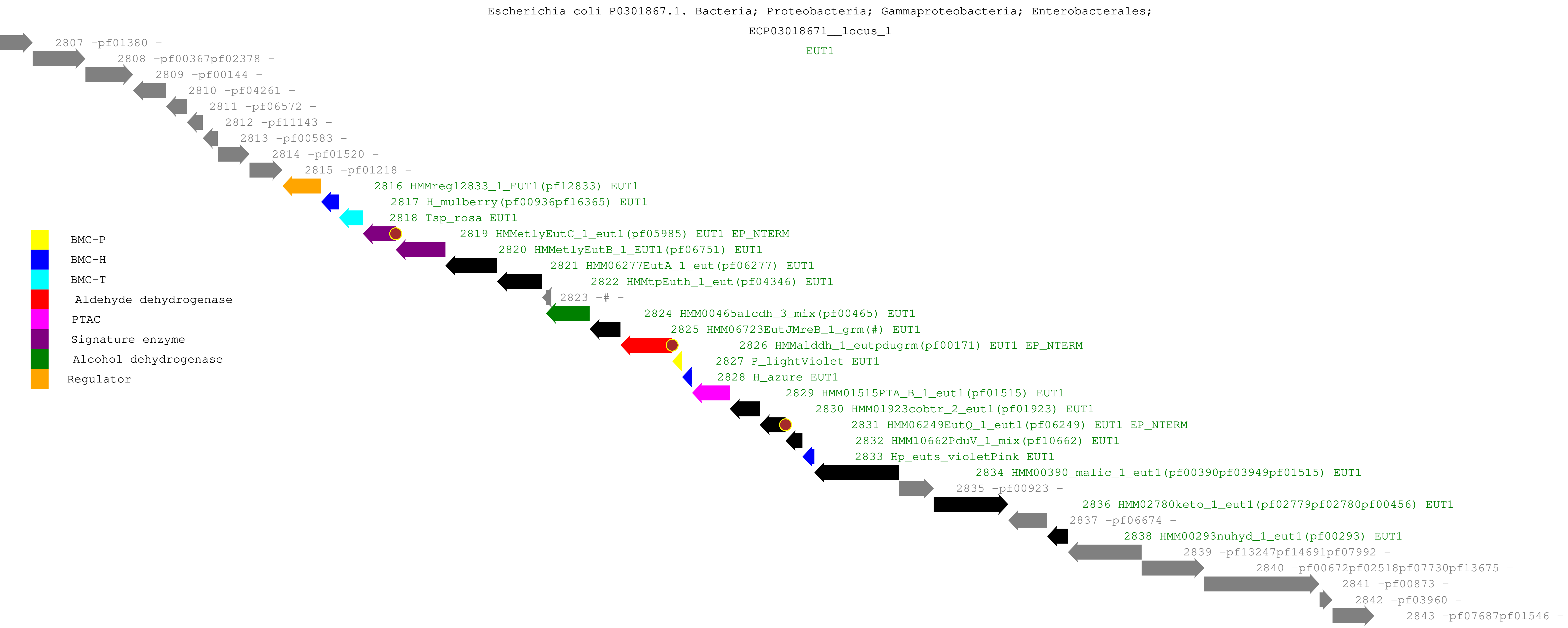
Escherichia coli P0301867.1. Bacteria; Proteobacteria; Gammaproteobacteria; Enterobacterales;
ECP03018671__locus_1
EUT1
2807 -pf01380 -
2808 -pf00367pf02378 -
2809 -pf00144 -
2810 -pf04261 -
2811 -pf06572 -
2812 -pf11143 -
2813 -pf00583 -
2814 -pf01520 -
2815 -pf01218 -
2816 HMMreg12833_1_EUT1(pf12833) EUT1
2817 H_mulberry(pf00936pf16365) EUT1
2818 Tsp_rosa EUT1
2819 HMMetlyEutC_1_eut1(pf05985) EUT1 EP_NTERM
2820 HMMetlyEutB_1_EUT1(pf06751) EUT1
2821 HMM06277EutA_1_eut(pf06277) EUT1
2822 HMMtpEuth_1_eut(pf04346) EUT1
2823 -# -
2824 HMM00465alcdh_3_mix(pf00465) EUT1
2825 HMM06723EutJMreB_1_grm(#) EUT1
2826 HMMalddh_1_eutpdugrm(pf00171) EUT1 EP_NTERM
2827 P_lightViolet EUT1
2828 H_azure EUT1
2829 HMM01515PTA_B_1_eut1(pf01515) EUT1
2830 HMM01923cobtr_2_eut1(pf01923) EUT1
2831 HMM06249EutQ_1_eut1(pf06249) EUT1 EP_NTERM
2832 HMM10662PduV_1_mix(pf10662) EUT1
2833 Hp_euts_violetPink EUT1
2834 HMM00390_malic_1_eut1(pf00390pf03949pf01515) EUT1
2835 -pf00923 -
2836 HMM02780keto_1_eut1(pf02779pf02780pf00456) EUT1
2837 -pf06674 -
2838 HMM00293nuhyd_1_eut1(pf00293) EUT1
2839 -pf13247pf14691pf07992 -
2840 -pf00672pf02518pf07730pf13675 -
2841 -pf00873 -
2842 -pf03960 -
2843 -pf07687pf01546 -
BMC-P
BMC-H
BMC-T
Aldehyde dehydrogenase
PTAC
Signature enzyme
Alcohol dehydrogenase
Regulator

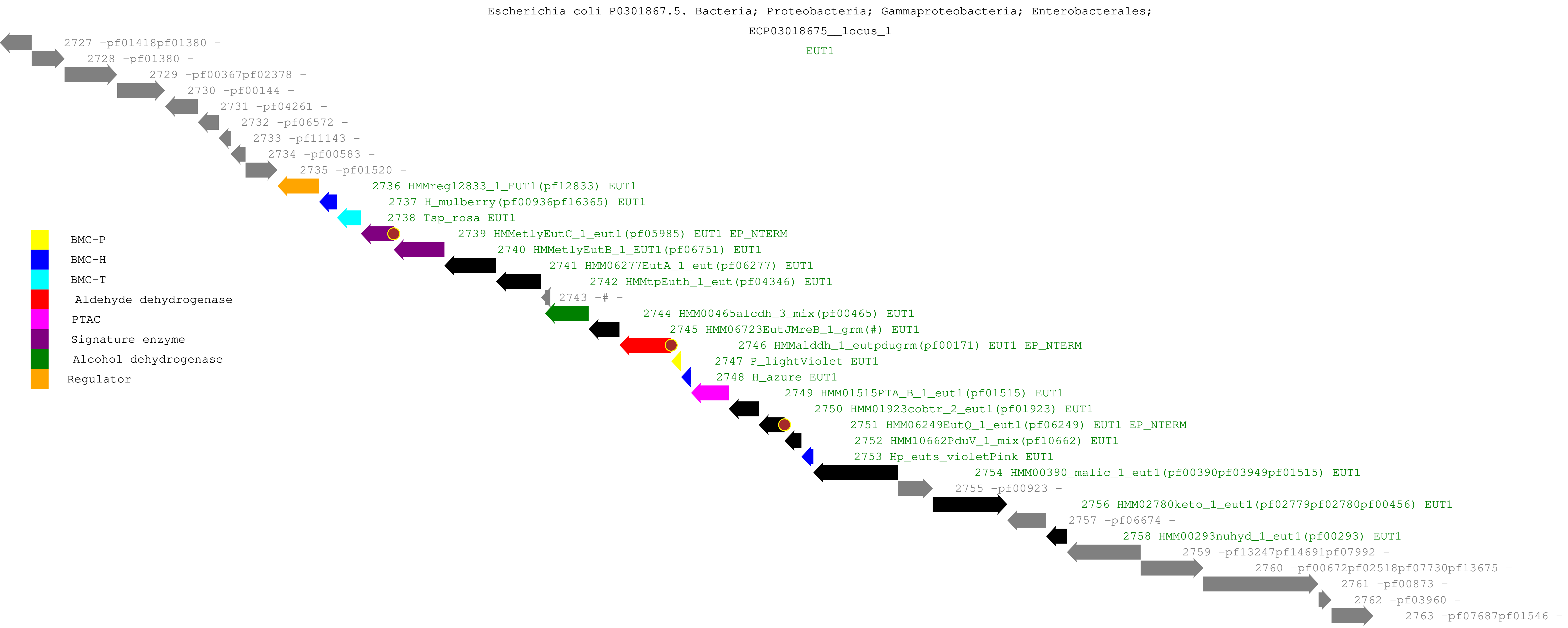
Escherichia coli P0301867.5. Bacteria; Proteobacteria; Gammaproteobacteria; Enterobacterales;
ECP03018675__locus_1
EUT1
2727 -pf01418pf01380 -
2728 -pf01380 -
2729 -pf00367pf02378 -
2730 -pf00144 -
2731 -pf04261 -
2732 -pf06572 -
2733 -pf11143 -
2734 -pf00583 -
2735 -pf01520 -
2736 HMMreg12833_1_EUT1(pf12833) EUT1
2737 H_mulberry(pf00936pf16365) EUT1
2738 Tsp_rosa EUT1
2739 HMMetlyEutC_1_eut1(pf05985) EUT1 EP_NTERM
2740 HMMetlyEutB_1_EUT1(pf06751) EUT1
2741 HMM06277EutA_1_eut(pf06277) EUT1
2742 HMMtpEuth_1_eut(pf04346) EUT1
2743 -# -
2744 HMM00465alcdh_3_mix(pf00465) EUT1
2745 HMM06723EutJMreB_1_grm(#) EUT1
2746 HMMalddh_1_eutpdugrm(pf00171) EUT1 EP_NTERM
2747 P_lightViolet EUT1
2748 H_azure EUT1
2749 HMM01515PTA_B_1_eut1(pf01515) EUT1
2750 HMM01923cobtr_2_eut1(pf01923) EUT1
2751 HMM06249EutQ_1_eut1(pf06249) EUT1 EP_NTERM
2752 HMM10662PduV_1_mix(pf10662) EUT1
2753 Hp_euts_violetPink EUT1
2754 HMM00390_malic_1_eut1(pf00390pf03949pf01515) EUT1
2755 -pf00923 -
2756 HMM02780keto_1_eut1(pf02779pf02780pf00456) EUT1
2757 -pf06674 -
2758 HMM00293nuhyd_1_eut1(pf00293) EUT1
2759 -pf13247pf14691pf07992 -
2760 -pf00672pf02518pf07730pf13675 -
2761 -pf00873 -
2762 -pf03960 -
2763 -pf07687pf01546 -
BMC-P
BMC-H
BMC-T
Aldehyde dehydrogenase
PTAC
Signature enzyme
Alcohol dehydrogenase
Regulator

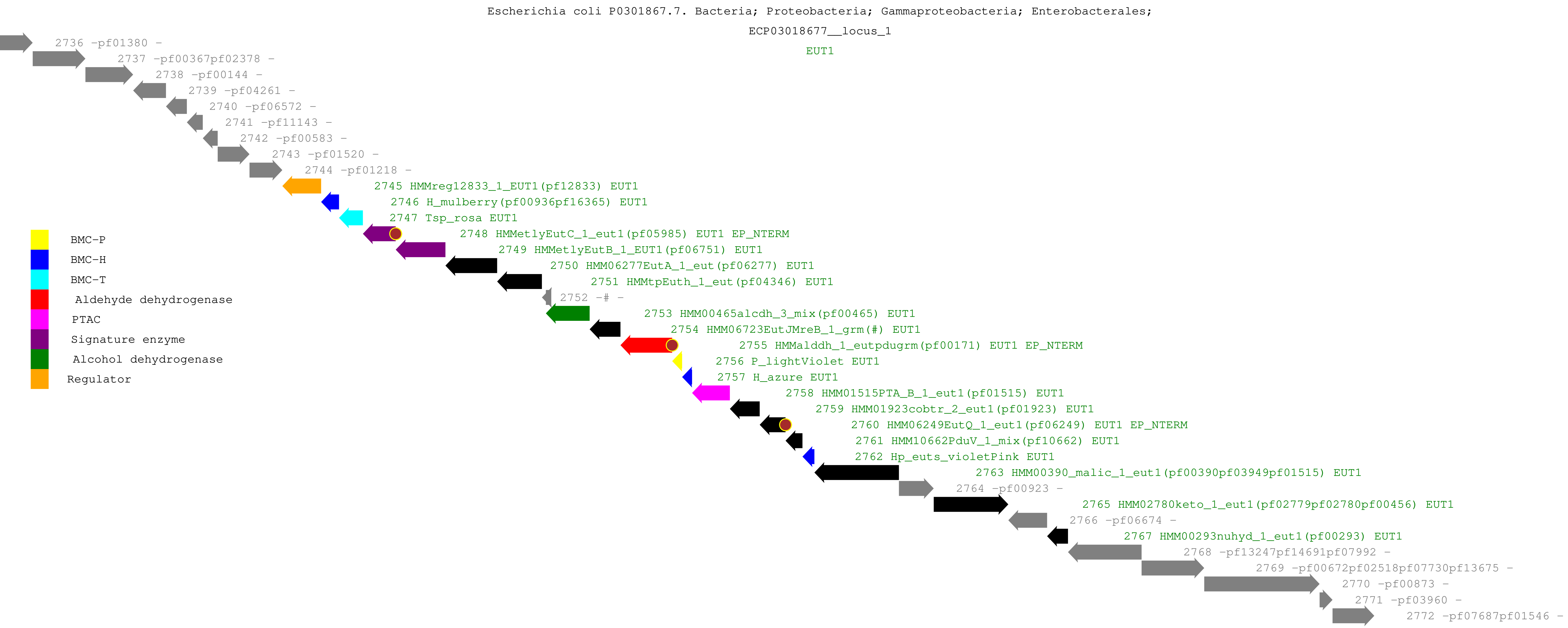
Escherichia coli P0301867.7. Bacteria; Proteobacteria; Gammaproteobacteria; Enterobacterales;
ECP03018677__locus_1
EUT1
2736 -pf01380 -
2737 -pf00367pf02378 -
2738 -pf00144 -
2739 -pf04261 -
2740 -pf06572 -
2741 -pf11143 -
2742 -pf00583 -
2743 -pf01520 -
2744 -pf01218 -
2745 HMMreg12833_1_EUT1(pf12833) EUT1
2746 H_mulberry(pf00936pf16365) EUT1
2747 Tsp_rosa EUT1
2748 HMMetlyEutC_1_eut1(pf05985) EUT1 EP_NTERM
2749 HMMetlyEutB_1_EUT1(pf06751) EUT1
2750 HMM06277EutA_1_eut(pf06277) EUT1
2751 HMMtpEuth_1_eut(pf04346) EUT1
2752 -# -
2753 HMM00465alcdh_3_mix(pf00465) EUT1
2754 HMM06723EutJMreB_1_grm(#) EUT1
2755 HMMaldh_1_eutpdugrm(pf00171) EUT1 EP_NTERM
2756 P_lightViolet EUT1
2757 H_azure EUT1
2758 HMM01515PTA_B_1_eut1(pf01515) EUT1
2759 HMM01923cobtr_2_eut1(pf01923) EUT1
2760 HMM06249EutQ_1_eut1(pf06249) EUT1 EP_NTERM
2761 HMM10662PduV_1_mix(pf10662) EUT1
2762 Hp_euts_violetPink EUT1
2763 HMM00390_malic_1_eut1(pf00390pf03949pf01515) EUT1
2764 -pf00923 -
2765 HMM02780keto_1_eut1(pf02779pf02780pf00456) EUT1
2766 -pf06674 -
2767 HMM00293nuhyd_1_eut1(pf00293) EUT1
2768 -pf13247pf14691pf07992 -
2769 -pf00672pf02518pf07730pf13675 -
2770 -pf00873 -
2771 -pf03960 -
2772 -pf07687pf01546 -
BMC-P
BMC-H
BMC-T
Aldehyde dehydrogenase
PTAC
Signature enzyme
Alcohol dehydrogenase
Regulator

Escherichia coli H591. Bacteria; Proteobacteria; Gammaproteobacteria; Enterobacterales;

ECPG__locus_1

EUT1

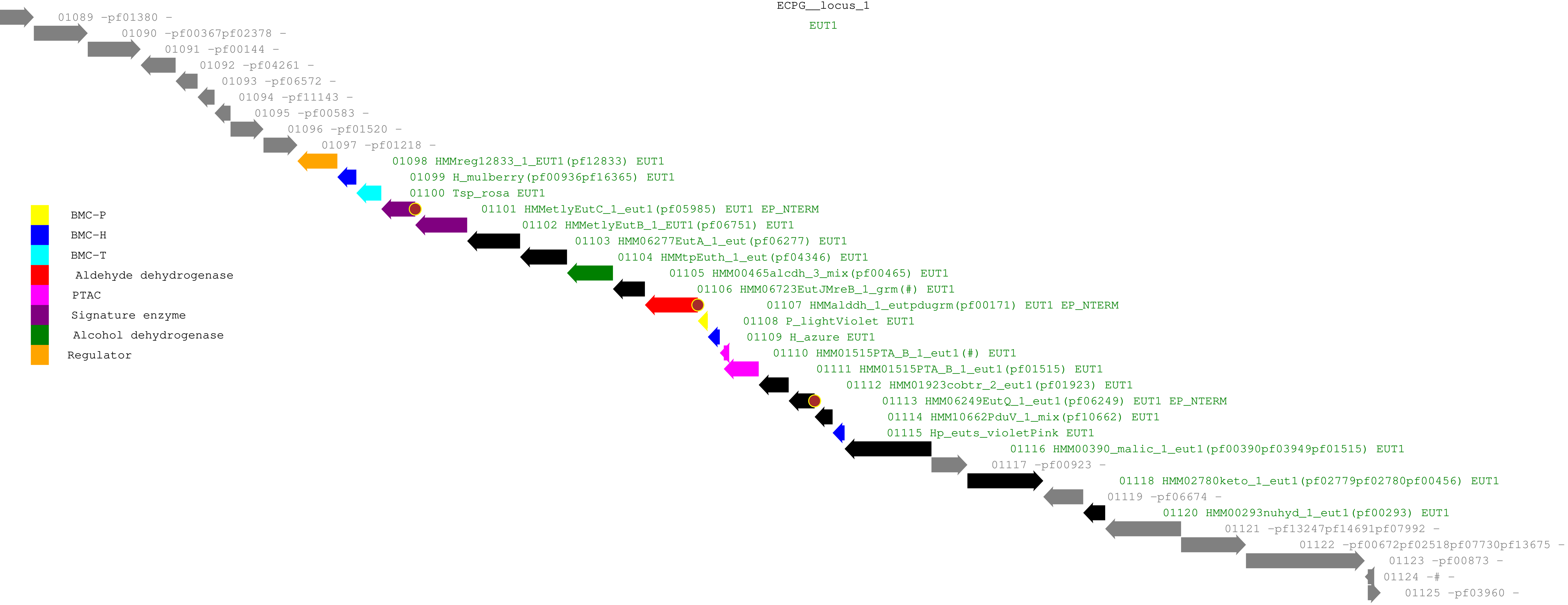

Escherichia coli O6:K15:H31 (strain 536 / UPEC). Bacteria; Proteobacteria; Gammaproteobacteria; Enterobacterales;

ECP__locus_1

EUT1

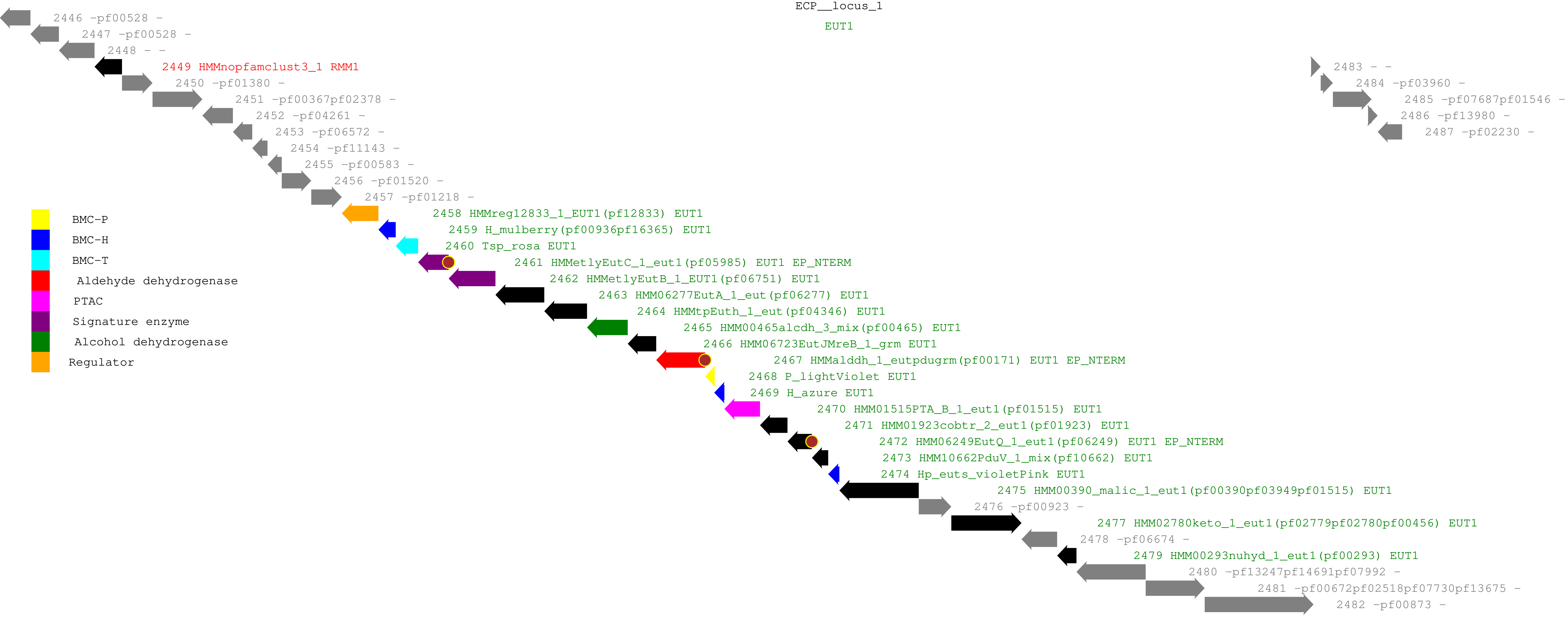

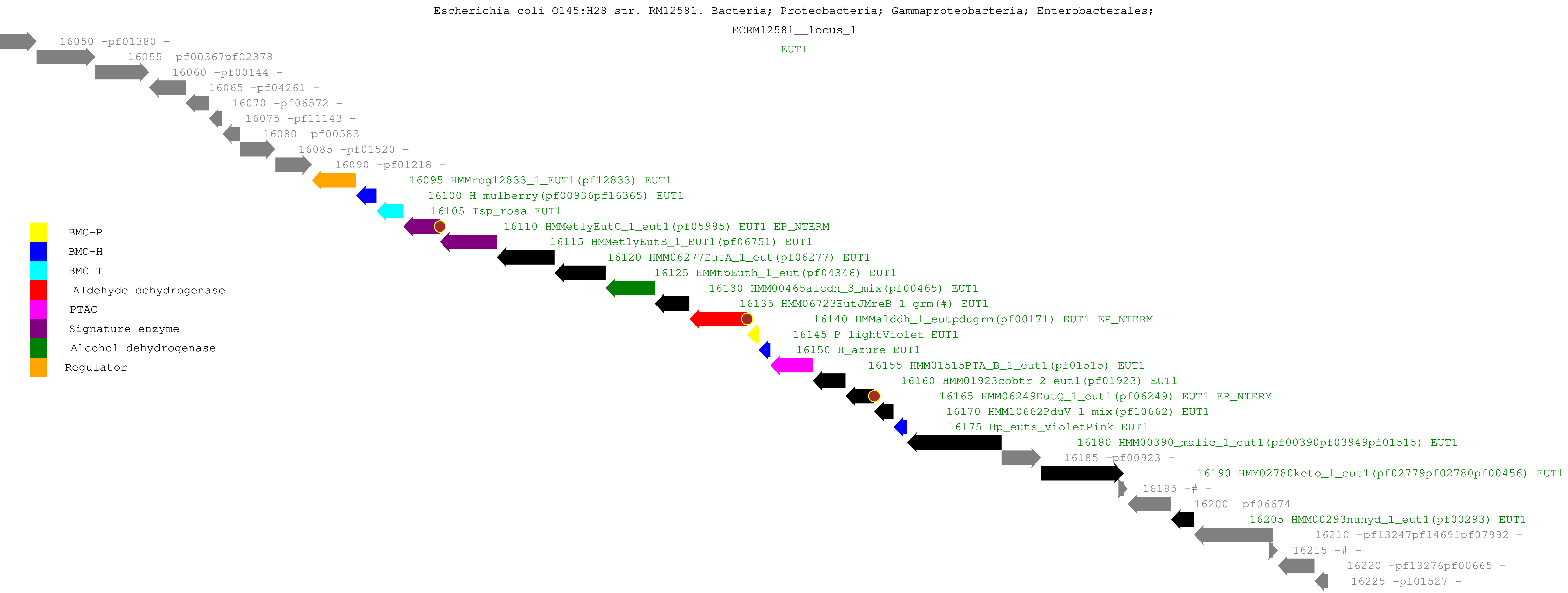
Escherichia coli O145:H28 str. RM12581. Bacteria; Proteobacteria; Gammaproteobacteria; Enterobacterales;
ECRM12581__locus_1
EUT1
16050 -pf01380 -
16055 -pf00367pf02378 -
16060 -pf00144 -
16065 -pf04261 -
16070 -pf06572 -
16075 -pf11143 -
16080 -pf00583 -
16085 -pf01520 -
16090 -pf01218 -
16095 HMMreg12833_1_EUT1(pf12833) EUT1
16100 H_mulberry(pf00936pf16365) EUT1
16105 Tsp_rosa EUT1
16110 HMMetlyEutC_1_eut1(pf05985) EUT1 EP_NTERM
16115 HMMetlyEutB_1_EUT1(pf06751) EUT1
16120 HMM06277EutA_1_eut(pf06277) EUT1
16125 HMMtpEuth_1_eut(pf04346) EUT1
16130 HMM00465alcdh_3_mix(pf00465) EUT1
16135 HMM06723EutJMreB_1_grm(#) EUT1
16140 HMMaldh_1_eutpdugrm(pf00171) EUT1 EP_NTERM
16145 P_lightViolet EUT1
16150 H_azure EUT1
16155 HMM01515PTA_B_1_eut1(pf01515) EUT1
16160 HMM01923cobtr_2_eut1(pf01923) EUT1
16165 HMM06249EutQ_1_eut1(pf06249) EUT1 EP_NTERM
16170 HMM10662PduV_1_mix(pf10662) EUT1
16175 Hp_euts_violetPink EUT1
16180 HMM00390_malic_1_eut1(pf00390pf03949pf01515) EUT1
16185 -pf00923 -
16190 HMM02780keto_1_eut1(pf02779pf02780pf00456) EUT1
16195 -# -
16200 -pf06674 -
16205 HMM00293nuhyd_1_eut1(pf00293) EUT1
16210 -pf13247pf14691pf07992 -
16215 -# -
16220 -pf13276pf00665 -
16225 -pf01527 -
BMC-P
BMC-H
BMC-T
Aldehyde dehydrogenase
PTAC
Signature enzyme
Alcohol dehydrogenase
Regulator

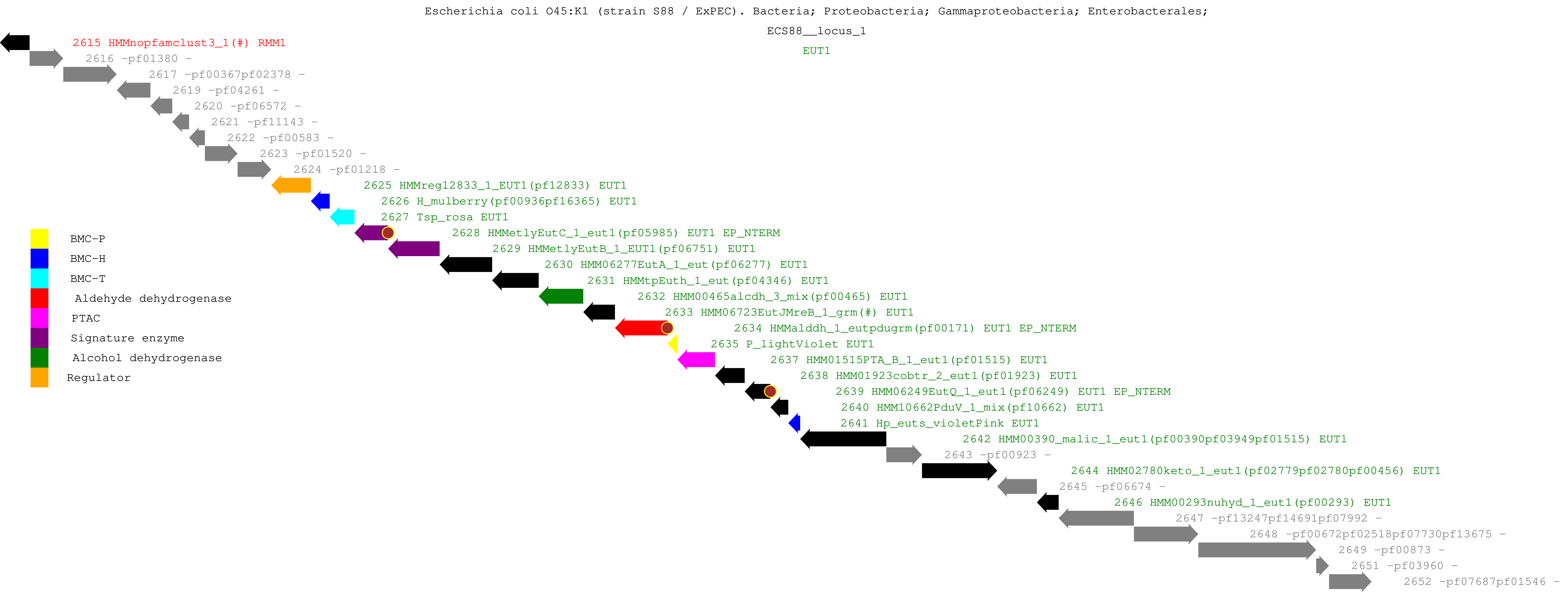
Escherichia coli O45:K1 (strain S88 / ExPEC). Bacteria; Proteobacteria; Gammaproteobacteria; Enterobacterales;
ECS88__locus_1
EUT1
2615 HMMnopfamclust3_1(#) RMM1
2616 -pf01380 -
2617 -pf00367pf02378 -
2619 -pf04261 -
2620 -pf06572 -
2621 -pf11143 -
2622 -pf00583 -
2623 -pf01520 -
2624 -pf01218 -
2625 HMMreg12833_1_EUT1(pf12833) EUT1
2626 H_mulberry(pf00936pf16365) EUT1
2627 Tsp_rosa EUT1
2628 HMMetlyEutC_1_eut1(pf05985) EUT1 EP_NTERM
2629 HMMetlyEutB_1_EUT1(pf06751) EUT1
2630 HMM06277EutA_1_eut(pf06277) EUT1
2631 HMMtpEuth_1_eut(pf04346) EUT1
2632 HMM00465alcdh_3_mix(pf00465) EUT1
2633 HMM06723EutJMreB_1_grm(#) EUT1
2634 HMMalddh_1_eutpdugrm(pf00171) EUT1 EP_NTERM
2635 P_lightViolet EUT1
2637 HMM01515PTA_B_1_eut1(pf01515) EUT1
2638 HMM01923cobtr_2_eut1(pf01923) EUT1
2639 HMM06249EutQ_1_eut1(pf06249) EUT1 EP_NTERM
2640 HMM10662PduV_1_mix(pf10662) EUT1
2641 Hp_euts_violetPink EUT1
2642 HMM00390_malic_1_eut1(pf00390pf03949pf01515) EUT1
2643 -pf00923 -
2644 HMM02780keto_1_eut1(pf02779pf02780pf00456) EUT1
2645 -pf06674 -
2646 HMM00293nuhyd_1_eut1(pf00293) EUT1
2647 -pf13247pf14691pf07992 -
2648 -pf00672pf02518pf07730pf13675 -
2649 -pf00873 -
2651 -pf03960 -
2652 -pf07687pf01546 -
BMC-P
BMC-H
BMC-T
Aldehyde dehydrogenase
PTAC
Signature enzyme
Alcohol dehydrogenase
Regulator

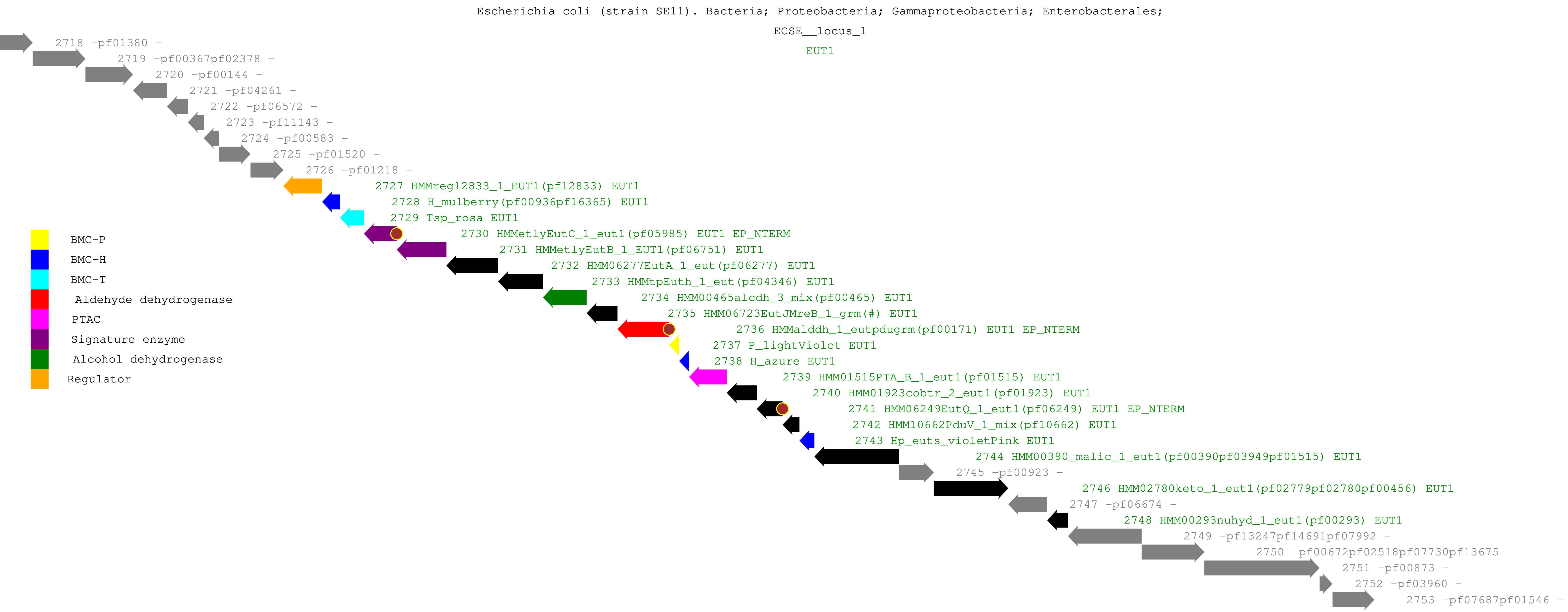

Escherichia coli (strain SE11). Bacteria; Proteobacteria; Gammaproteobacteria; Enterobacterales;
ECSE__locus_1
EUT1
2718 -pf01380 -
2719 -pf00367pf02378 -
2720 -pf00144 -
2721 -pf04261 -
2722 -pf06572 -
2723 -pf11143 -
2724 -pf00583 -
2725 -pf01520 -
2726 -pf01218 -
2727 HMMreg12833_1_EUT1(pf12833) EUT1
2728 H_mulberry(pf00936pf16365) EUT1
2729 Tsp_rosa EUT1
2730 HMMetlyEutC_1_eut1(pf05985) EUT1 EP_NTERM
2731 HMMetlyEutB_1_EUT1(pf06751) EUT1
2732 HMM06277EutA_1_eut(pf06277) EUT1
2733 HMMtpEuth_1_eut(pf04346) EUT1
2734 HMM00465alcdh_3_mix(pf00465) EUT1
2735 HMM06723EutJMreB_1_grm(#) EUT1
2736 HMMaldh_1_eutpdugrm(pf00171) EUT1 EP_NTERM
2737 P_lightViolet EUT1
2738 H_azure EUT1
2739 HMM01515PTA_B_1_eut1(pf01515) EUT1
2740 HMM01923cobtr_2_eut1(pf01923) EUT1
2741 HMM06249EutQ_1_eut1(pf06249) EUT1 EP_NTERM
2742 HMM10662PduV_1_mix(pf10662) EUT1
2743 Hp_euts_violetPink EUT1
2744 HMM00390_malic_1_eut1(pf00390pf03949pf01515) EUT1
2745 -pf00923 -
2746 HMM02780keto_1_eut1(pf02779pf02780pf00456) EUT1
2747 -pf06674 -
2748 HMM00293nuhyd_1_eut1(pf00293) EUT1
2749 -pf13247pf14691pf07992 -
2750 -pf00672pf02518pf07730pf13675 -
2751 -pf00873 -
2752 -pf03960 -
2753 -pf07687pf01546 -
BMC-P
BMC-H
BMC-T
Aldehyde dehydrogenase
PTAC
Signature enzyme
Alcohol dehydrogenase
Regulator

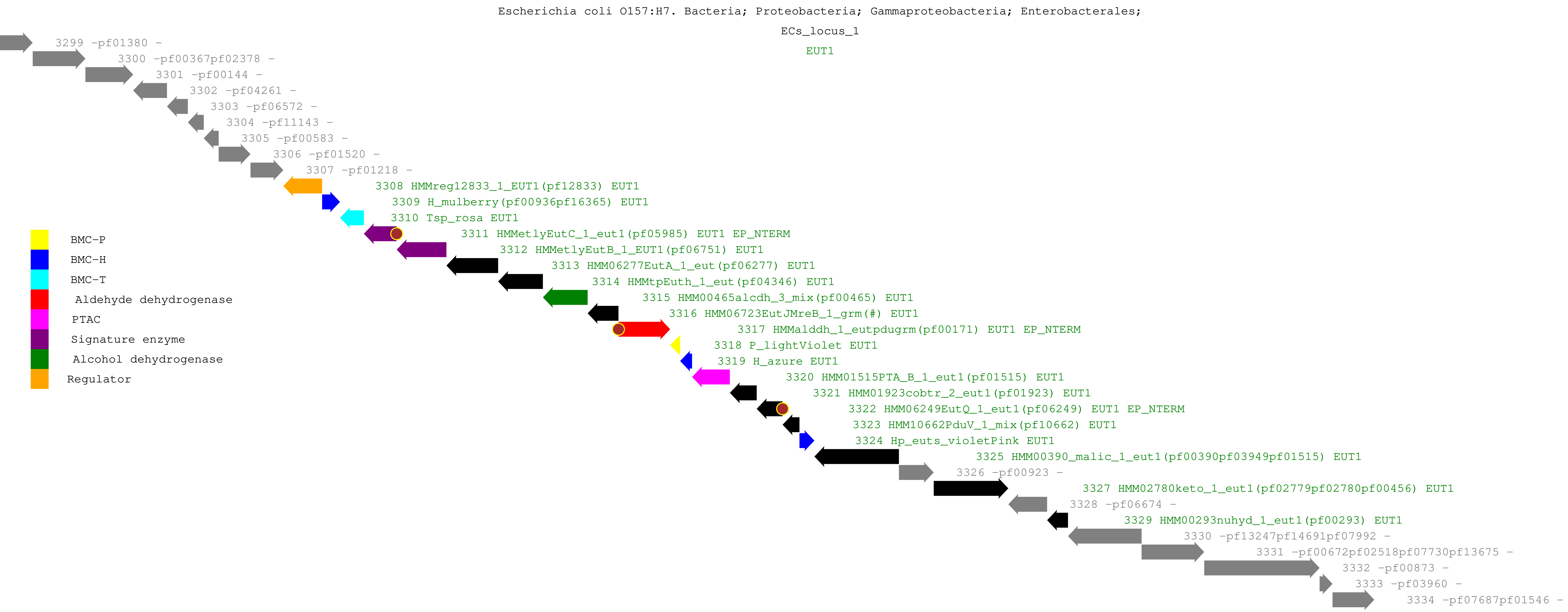

Escherichia coli O157:H7. Bacteria; Proteobacteria; Gammaproteobacteria; Enterobacterales;
ECs_locus_1
EUT1
3299 -pf01380 -
3300 -pf00367pf02378 -
3301 -pf00144 -
3302 -pf04261 -
3303 -pf06572 -
3304 -pf11143 -
3305 -pf00583 -
3306 -pf01520 -
3307 -pf01218 -
3308 HMMreg12833_1_EUT1(pf12833) EUT1
3309 H_mulberry(pf00936pf16365) EUT1
3310 Tsp_rosa EUT1
3311 HMMetlyEutC_1_eut1(pf05985) EUT1 EP_NTERM
3312 HMMetlyEutB_1_EUT1(pf06751) EUT1
3313 HMM06277EutA_1_eut(pf06277) EUT1
3314 HMMtpEuth_1_eut(pf04346) EUT1
3315 HMM00465alcdh_3_mix(pf00465) EUT1
3316 HMM06723EutJMreB_1_grm(#) EUT1
3317 HMMalddh_1_eutpdugrm(pf00171) EUT1 EP_NTERM
3318 P_lightViolet EUT1
3319 H_azure EUT1
3320 HMM01515PTA_B_1_eut1(pf01515) EUT1
3321 HMM01923cobtr_2_eut1(pf01923) EUT1
3322 HMM06249EutQ_1_eut1(pf06249) EUT1 EP_NTERM
3323 HMM10662PduV_1_mix(pf10662) EUT1
3324 Hp_euts_violetPink EUT1
3325 HMM00390_malic_1_eut1(pf00390pf03949pf01515) EUT1
3326 -pf00923 -
3327 HMM02780keto_1_eut1(pf02779pf02780pf00456) EUT1
3328 -pf06674 -
3329 HMM00293nuhyd_1_eut1(pf00293) EUT1
3330 -pf13247pf14691pf07992 -
3331 -pf00672pf02518pf07730pf13675 -
3332 -pf00873 -
3333 -pf03960 -
3334 -pf07687pf01546 -
BMC-P
BMC-H
BMC-T
Aldehyde dehydrogenase
PTAC
Signature enzyme
Alcohol dehydrogenase
Regulator

Escherichia coli (strain SMS-3-5 / SECEC). Bacteria; Proteobacteria; Gammaproteobacteria; Enterobacterales;

EcSMS35__locus_1

EUT1

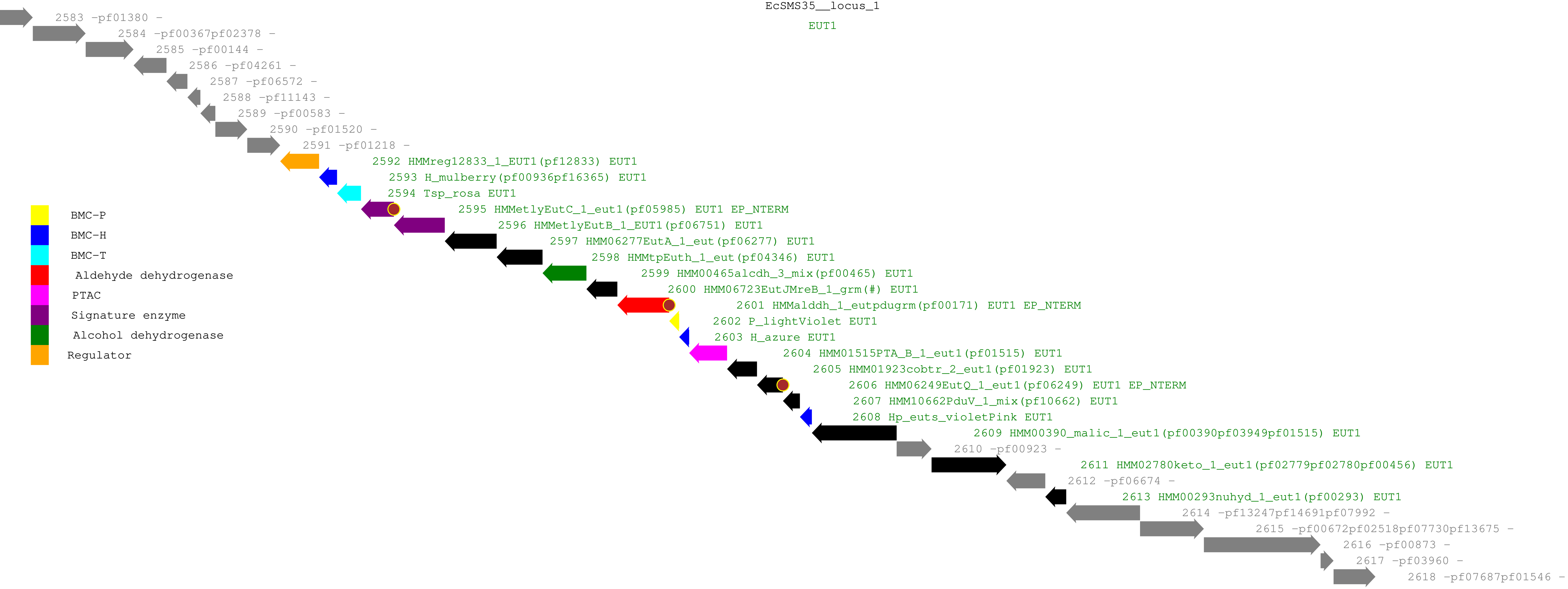

Escherichia coli O17:K52:H18 (strain UMN026 / ExPEC). Bacteria; Proteobacteria; Gammaproteobacteria; Enterobacterales;

ECUMN__locus_2

EUT1

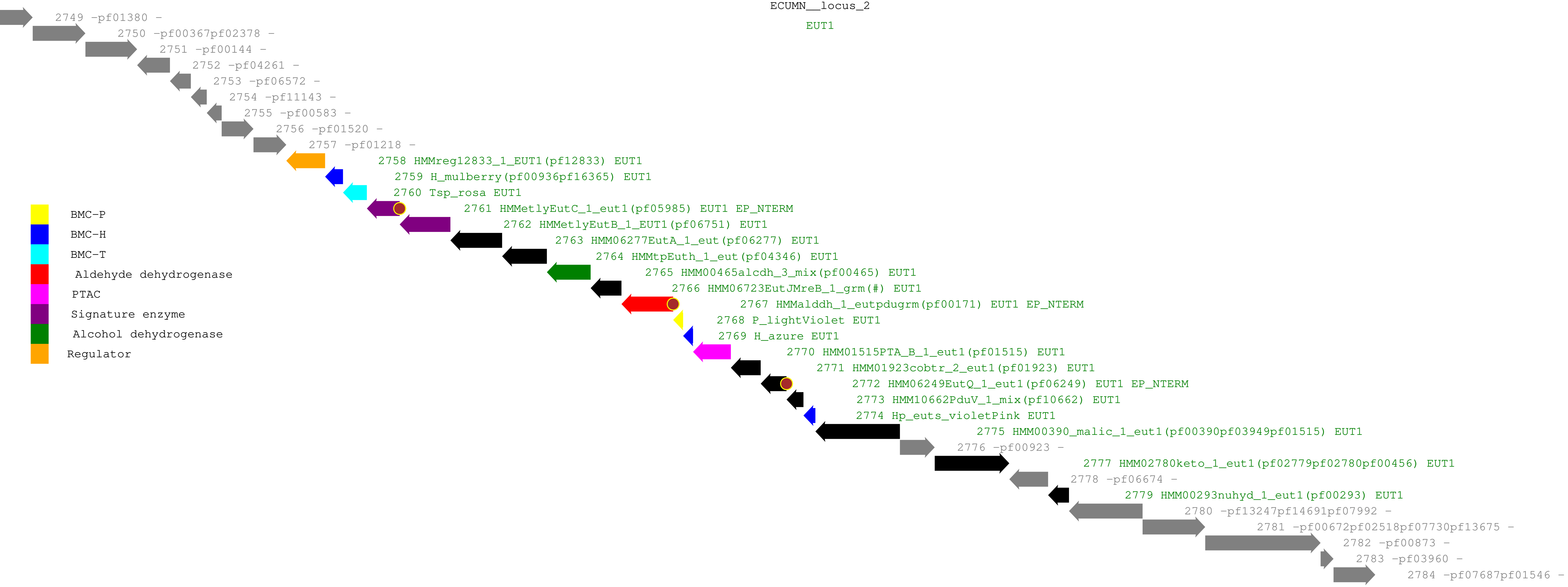

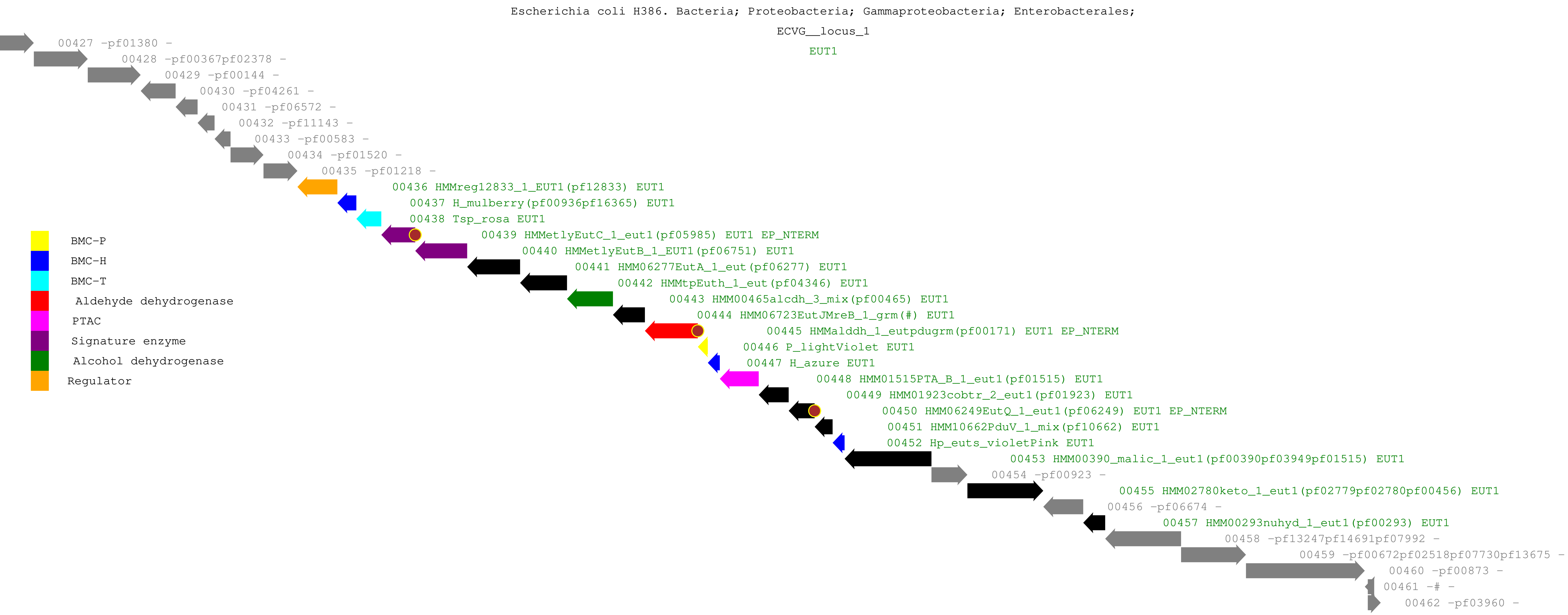

Escherichia coli H386. Bacteria; Proteobacteria; Gammaproteobacteria; Enterobacterales;
ECVG__locus_1
EUT1
00427 -pf01380 -
00428 -pf00367pf02378 -
00429 -pf00144 -
00430 -pf04261 -
00431 -pf06572 -
00432 -pf11143 -
00433 -pf00583 -
00434 -pf01520 -
00435 -pf01218 -
00436 HMMreg12833_1_EUT1(pf12833) EUT1
00437 H_mulberry(pf00936pf16365) EUT1
00438 Tsp_rosa EUT1
00439 HMMetlyEutC_1_eut1(pf05985) EUT1 EP_NTERM
00440 HMMetlyEutB_1_EUT1(pf06751) EUT1
00441 HMM06277EutA_1_eut(pf06277) EUT1
00442 HMMtpEuth_1_eut(pf04346) EUT1
00443 HMM00465alcdh_3_mix(pf00465) EUT1
00444 HMM06723EutJMreB_1_grm(#) EUT1
00445 HMMalddh_1_eutpdugrm(pf00171) EUT1 EP_NTERM
00446 P_lightViolet EUT1
00447 H_azure EUT1
00448 HMM01515PTA_B_1_eut1(pf01515) EUT1
00449 HMM01923cobtr_2_eut1(pf01923) EUT1
00450 HMM06249EutQ_1_eut1(pf06249) EUT1 EP_NTERM
00451 HMM10662PduV_1_mix(pf10662) EUT1
00452 Hp_euts_violetPink EUT1
00453 HMM00390_malic_1_eut1(pf00390pf03949pf01515) EUT1
00454 -pf00923 -
00455 HMM02780keto_1_eut1(pf02779pf02780pf00456) EUT1
00456 -pf06674 -
00457 HMM00293nuhyd_1_eut1(pf00293) EUT1
00458 -pf13247pf14691pf07992 -
00459 -pf00672pf02518pf07730pf13675 -
00460 -pf00873 -
00461 -# -
00462 -pf03960 -
BMC-P
BMC-H
BMC-T
Aldehyde dehydrogenase
PTAC
Signature enzyme
Alcohol dehydrogenase
Regulator

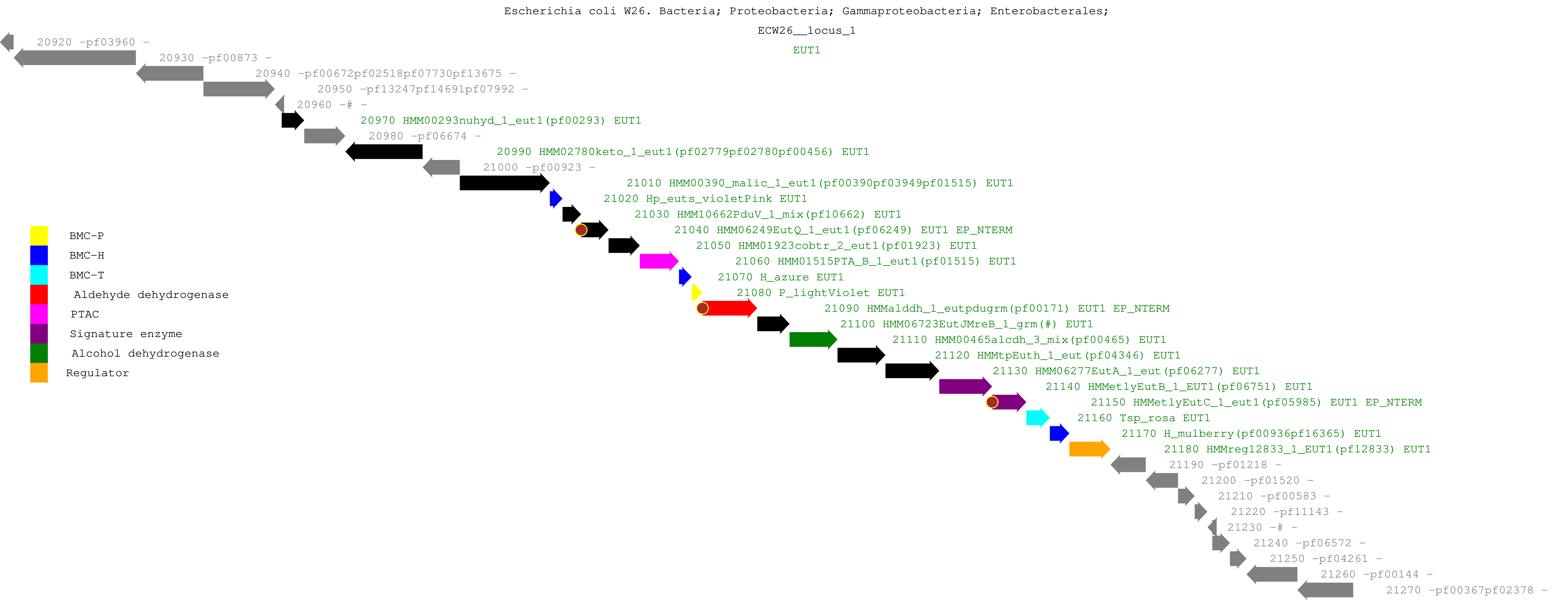

Escherichia coli W26. Bacteria; Proteobacteria; Gammaproteobacteria; Enterobacterales;
ECW26__locus_1
EUT1
20920 -pf03960 -
20930 -pf00873 -
20940 -pf00672pf02518pf07730pf13675 -
20950 -pf13247pf14691pf07992 -
20960 -# -
20970 HMM00293nuhyd_1_eut1(pf00293) EUT1
20980 -pf06674 -
20990 HMM02780keto_1_eut1(pf02779pf02780pf00456) EUT1
21000 -pf00923 -
21010 HMM00390_malic_1_eut1(pf00390pf03949pf01515) EUT1
21020 Hp_euts_violetPink EUT1
21030 HMM10662PduV_1_mix(pf10662) EUT1
21040 HMM06249EutQ_1_eut1(pf06249) EUT1 EP_NTERM
21050 HMM01923cobtr_2_eut1(pf01923) EUT1
21060 HMM01515PTA_B_1_eut1(pf01515) EUT1
21070 H_azure EUT1
21080 P_lightViolet EUT1
21090 HMMalddh_1_eutpdugrm(pf00171) EUT1 EP_NTERM
21100 HMM06723EutJMreB_1_grm(#) EUT1
21110 HMM00465alcdh_3_mix(pf00465) EUT1
21120 HMMtpEuth_1_eut(pf04346) EUT1
21130 HMM06277EutA_1_eut(pf06277) EUT1
21140 HMMetlyEutB_1_EUT1(pf06751) EUT1
21150 HMMetlyEutC_1_eut1(pf05985) EUT1 EP_NTERM
21160 Tsp_rosa EUT1
21170 H_mulberry(pf00936pf16365) EUT1
21180 HMMreg12833_1_EUT1(pf12833) EUT1
21190 -pf01218 -
21200 -pf01520 -
21210 -pf00583 -
21220 -pf11143 -
21230 -# -
21240 -pf06572 -
21250 -pf04261 -
21260 -pf00144 -
21270 -pf00367pf02378 -
BMC-P
BMC-H
BMC-T
Aldehyde dehydrogenase
PTAC
Signature enzyme
Alcohol dehydrogenase
Regulator

Escherichia coli (strain ATCC 9637 / CCM 2024 / DSM 1116 / NCIMB 8666 Bacteria; Proteobacteria; Gammaproteobacteria; Enterobacterales;

ECW_m_locus_1

EUT1

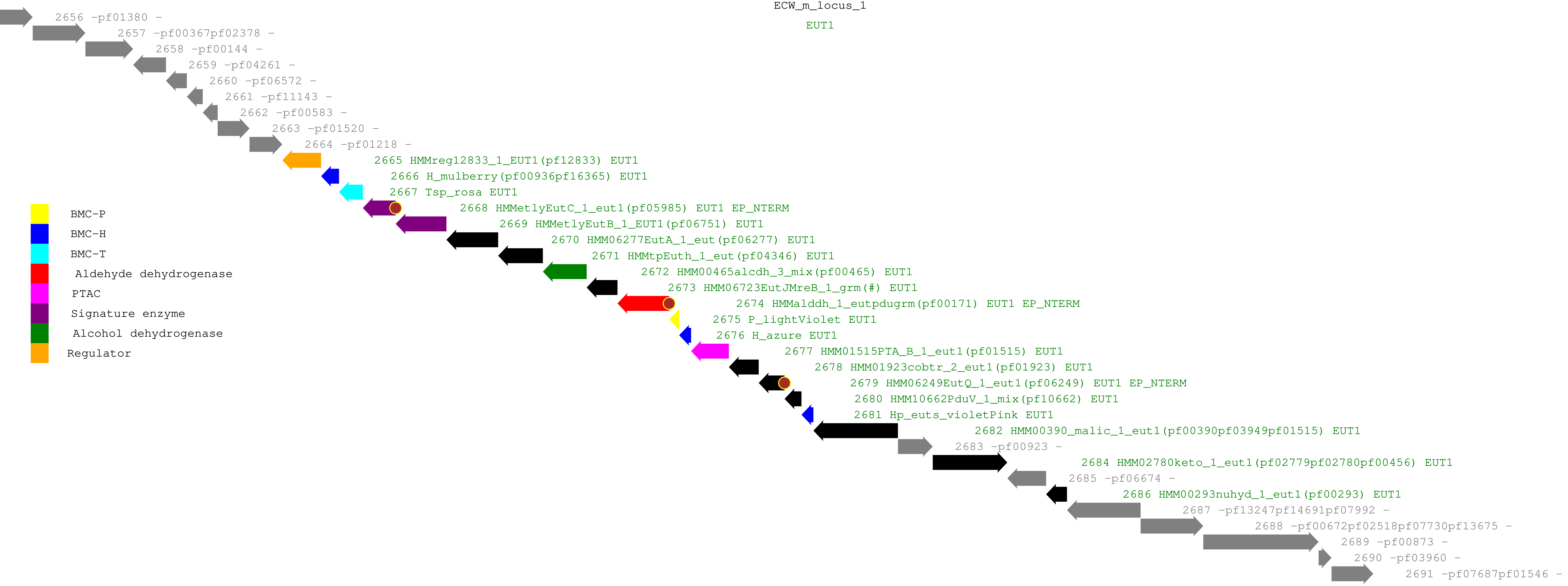

Escherichia coli TA447. Bacteria; Proteobacteria; Gammaproteobacteria; Enterobacterales;

ECXG__locus_1

EUT1

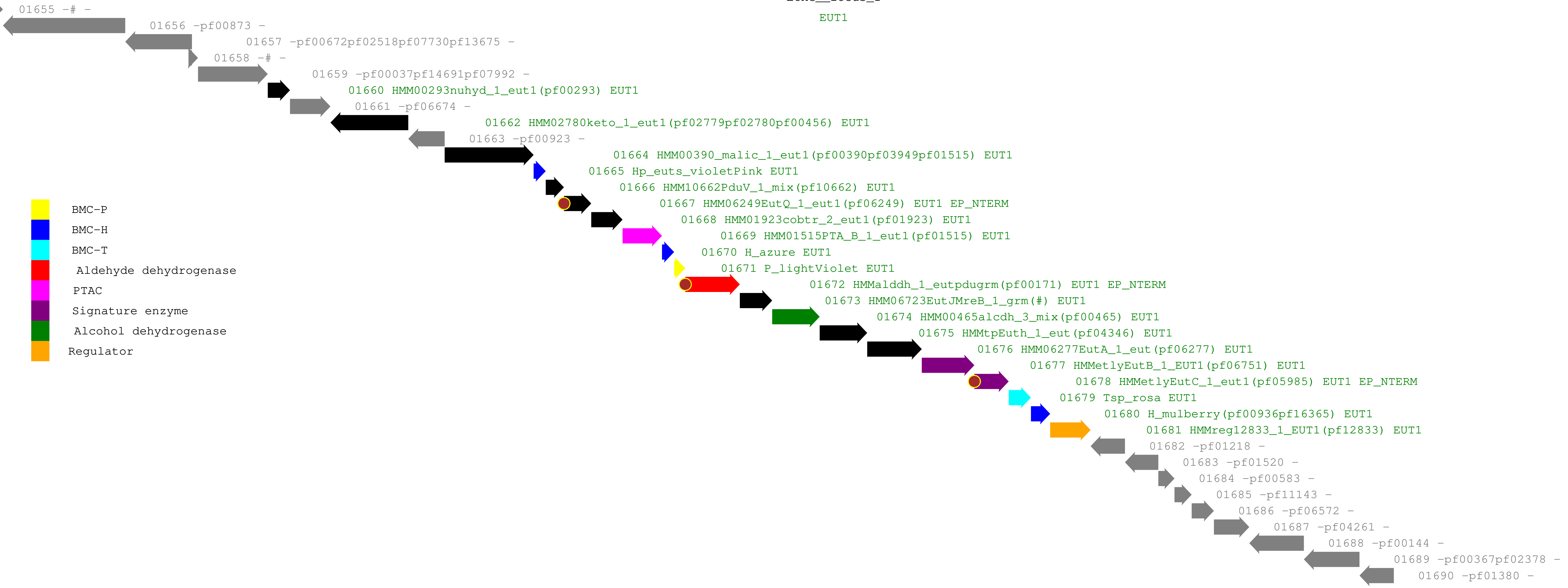

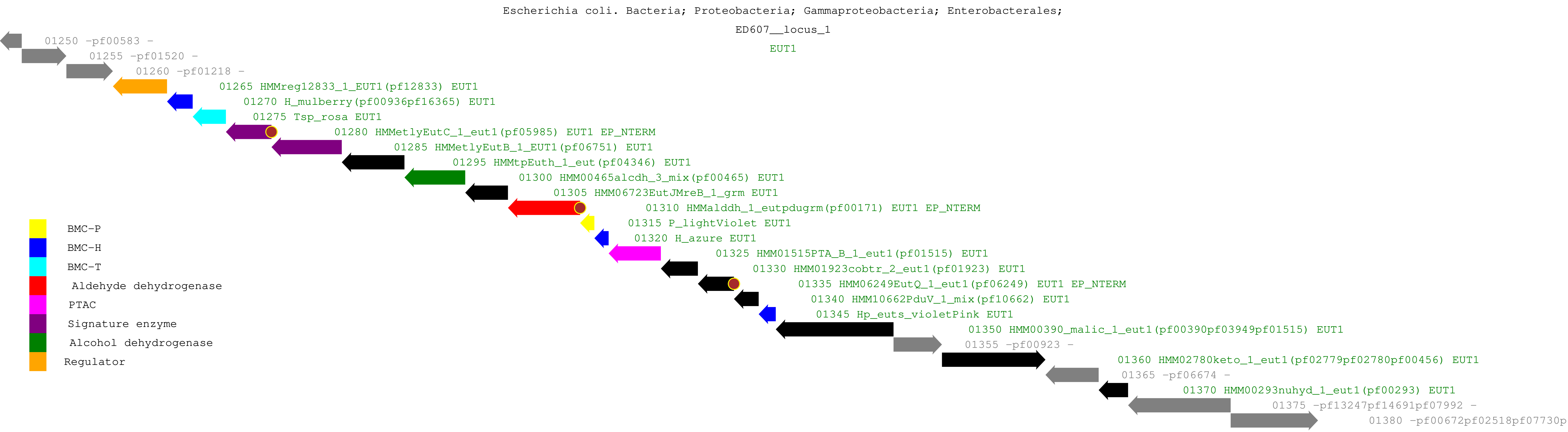
Escherichia coli. Bacteria; Proteobacteria; Gammaproteobacteria; Enterobacterales;
ED607__locus_1
EUT1
01250 -pf00583 -
01255 -pf01520 -
01260 -pf01218 -
01265 HMMreg12833_1_EUT1(pf12833) EUT1
01270 H_mulberry(pf00936pf16365) EUT1
01275 Tsp_rosa EUT1
01280 HMMetlyEutC_1_eut1(pf05985) EUT1 EP_NTERM
01285 HMMetlyEutB_1_EUT1(pf06751) EUT1
01295 HMMtpEuth_1_eut(pf04346) EUT1
01300 HMM00465alcdh_3_mix(pf00465) EUT1
01305 HMM06723EutJMreB_1_grm EUT1
01310 HMMalddh_1_eutpdugrm(pf00171) EUT1 EP_NTERM
01315 P_lightViolet EUT1
01320 H_azure EUT1
01325 HMM01515PTA_B_1_eut1(pf01515) EUT1
01330 HMM01923cobtr_2_eut1(pf01923) EUT1
01335 HMM06249EutQ_1_eut1(pf06249) EUT1 EP_NTERM
01340 HMM10662PduV_1_mix(pf10662) EUT1
01345 Hp_euts_violetPink EUT1
01350 HMM00390_malic_1_eut1(pf00390pf03949pf01515) EUT1
01355 -pf00923 -
01360 HMM02780keto_1_eut1(pf02779pf02780pf00456) EUT1
01365 -pf06674 -
01370 HMM00293nuhyd_1_eut1(pf00293) EUT1
01375 -pf13247pf14691pf07992 -
01380 -pf00672pf02518pf07730p
BMC-P
BMC-H
BMC-T
Aldehyde dehydrogenase
PTAC
Signature enzyme
Alcohol dehydrogenase
Regulator

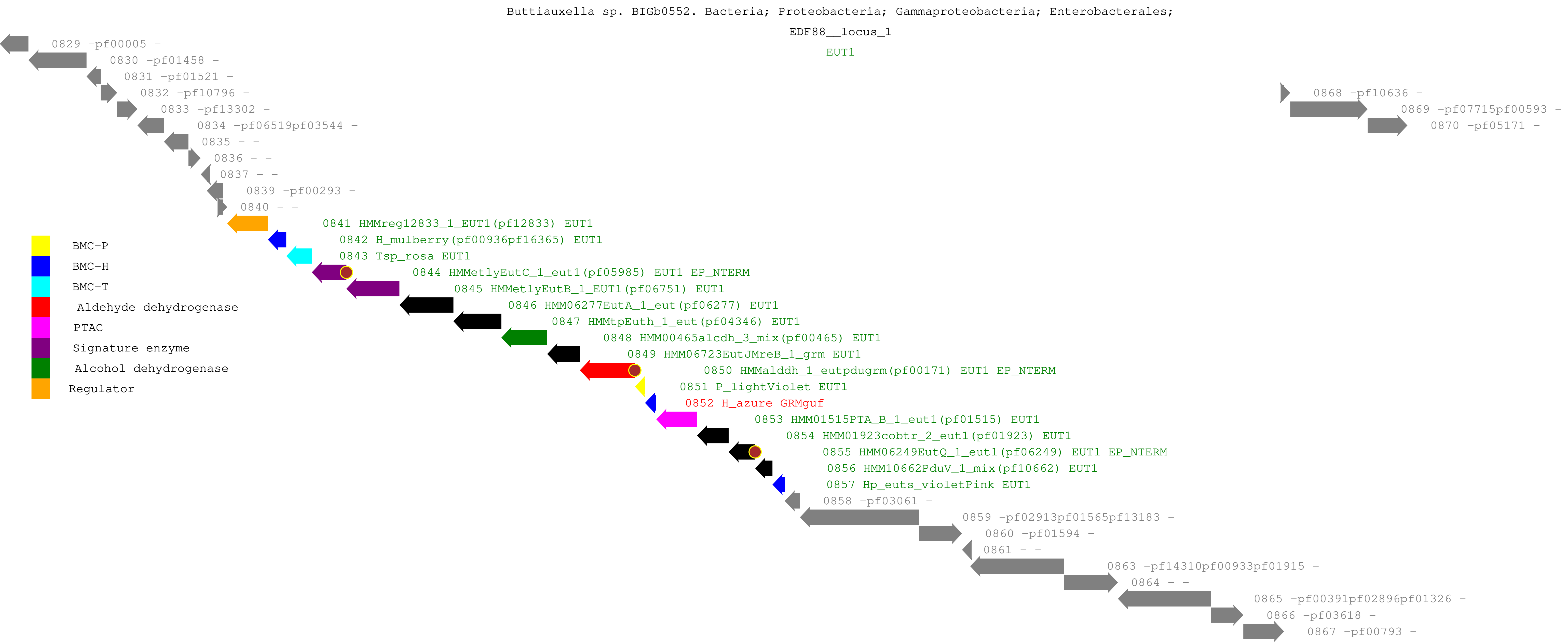
Buttiauxella sp. BIGb0552. Bacteria; Proteobacteria; Gammaproteobacteria; Enterobacterales;
EDF88__locus_1
EUT1
0829 -pf00005 -
0830 -pf01458 -
0831 -pf01521 -
0832 -pf10796 -
0833 -pf13302 -
0834 -pf06519pf03544 -
0835 - -
0836 - -
0837 - -
0839 -pf00293 -
0840 - -
0841 HMMreg12833_1_EUT1(pf12833) EUT1
0842 H_mulberry(pf00936pf16365) EUT1
0843 Tsp_rosa EUT1
0844 HMMetlyEutC_1_eut1(pf05985) EUT1 EP_NTERM
0845 HMMetlyEutB_1_EUT1(pf06751) EUT1
0846 HMM06277EutA_1_eut(pf06277) EUT1
0847 HMMtpEuth_1_eut(pf04346) EUT1
0848 HMM00465alcdh_3_mix(pf00465) EUT1
0849 HMM06723EutJMreB_1_grm EUT1
0850 HMMaldh_1_eutpdugrm(pf00171) EUT1 EP_NTERM
0851 P_lightViolet EUT1
0852 H_azure GRMguf
0853 HMM01515PTA_B_1_eut1(pf01515) EUT1
0854 HMM01923cobtr_2_eut1(pf01923) EUT1
0855 HMM06249EutQ_1_eut1(pf06249) EUT1 EP_NTERM
0856 HMM10662PduV_1_mix(pf10662) EUT1
0857 Hp_euts_violetPink EUT1
0858 -pf03061 -
0859 -pf02913pf01565pf13183 -
0860 -pf01594 -
0861 - -
0863 -pf14310pf00933pf01915 -
0864 - -
0865 -pf00391pf02896pf01326 -
0866 -pf03618 -
0867 -pf00793 -
0868 -pf10636 -
0869 -pf07715pf00593 -
0870 -pf05171 -
BMC-P
BMC-H
BMC-T
Aldehyde dehydrogenase
PTAC
Signature enzyme
Alcohol dehydrogenase
Regulator

Salmonella enterica (Salmonella choleraesuis). Bacteria; Proteobacteria; Gammaproteobacteria; Enterobacterales;

EE393__locus_1

EUT1

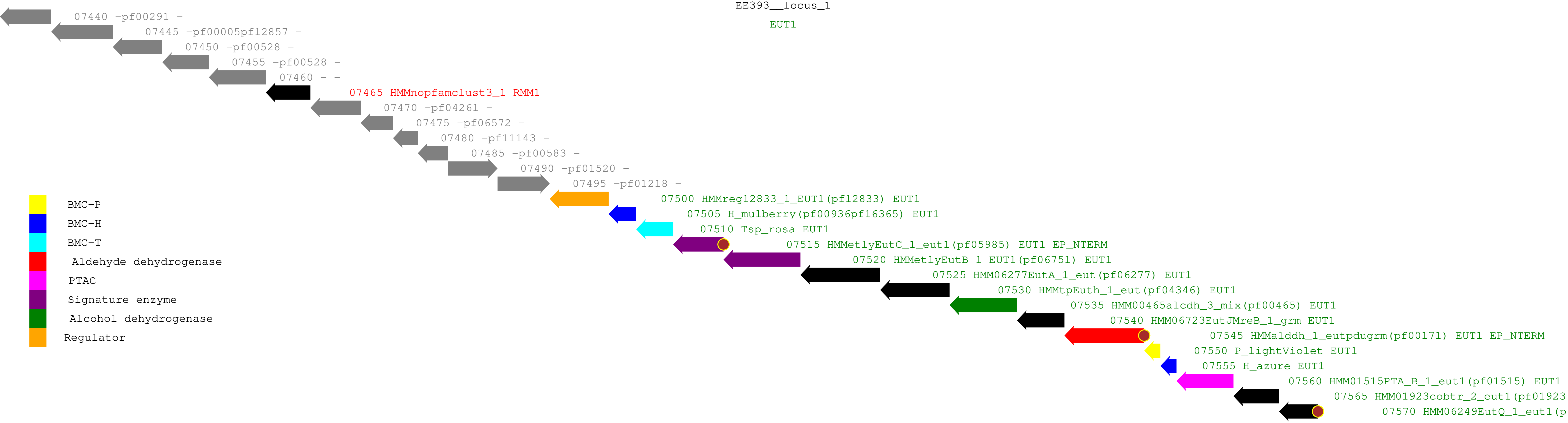

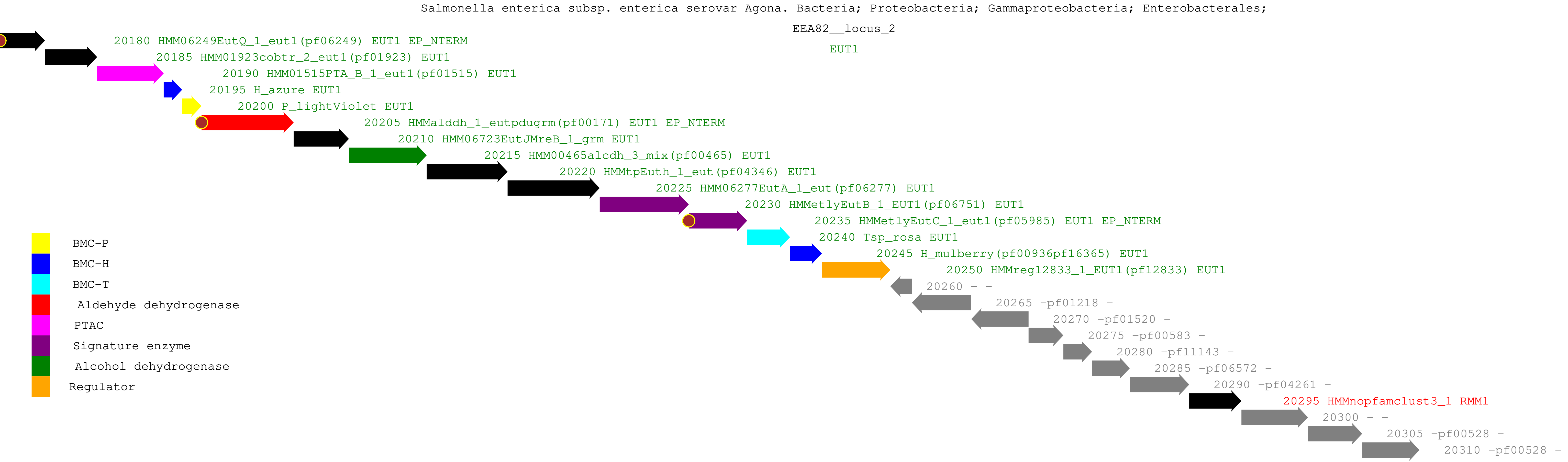
Salmonella enterica subsp. enterica serovar Agona. Bacteria; Proteobacteria; Gammaproteobacteria; Enterobacterales;
EEA82__locus_2
EUT1
20180 HMM06249EutQ_1_eut1(pf06249) EUT1 EP_NTERM
20185 HMM01923cobtr_2_eut1(pf01923) EUT1
20190 HMM01515PTA_B_1_eut1(pf01515) EUT1
20195 H_azure EUT1
20200 P_lightViolet EUT1
20205 HMMalddh_1_eutpdugrm(pf00171) EUT1 EP_NTERM
20210 HMM06723EutJMreB_1_grm EUT1
20215 HMM00465alcdh_3_mix(pf00465) EUT1
20220 HMMtpEuth_1_eut(pf04346) EUT1
20225 HMM06277EutA_1_eut(pf06277) EUT1
20230 HMMetlyEutB_1_EUT1(pf06751) EUT1
20235 HMMetlyEutC_1_eut1(pf05985) EUT1 EP_NTERM
20240 Tsp_rosa EUT1
20245 H_mulberry(pf00936pf16365) EUT1
20250 HMMreg12833_1_EUT1(pf12833) EUT1
20260 - -
20265 -pf01218 -
20270 -pf01520 -
20275 -pf00583 -
20280 -pf11143 -
20285 -pf06572 -
20290 -pf04261 -
20295 HMMnopfamclust3_1 RMM1
20300 - -
20305 -pf00528 -
20310 -pf00528 -
BMC-P
BMC-H
BMC-T
Aldehyde dehydrogenase
PTAC
Signature enzyme
Alcohol dehydrogenase
Regulator

Salmonella enterica (Salmonella choleraesuis). Bacteria; Proteobacteria; Gammaproteobacteria; Enterobacterales;

EEM01__locus_1

EUT1

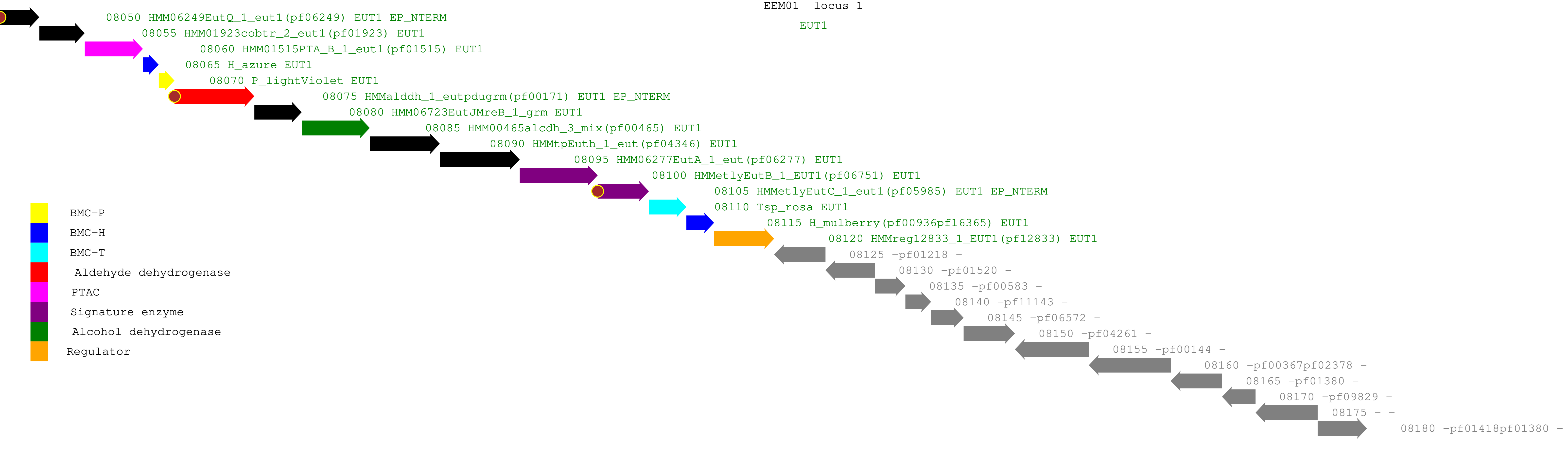

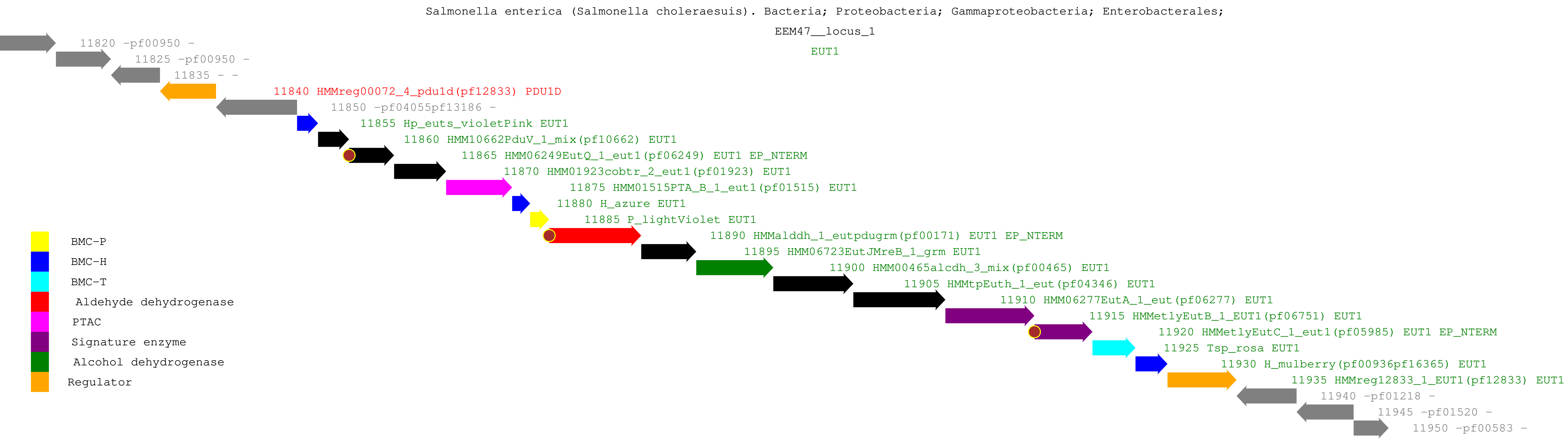
Salmonella enterica (Salmonella choleraesuis). Bacteria; Proteobacteria; Gammaproteobacteria; Enterobacterales;
EEM47__locus_1
EUT1
11820 -pf00950 -
11825 -pf00950 -
11835 - -
11840 HMMreg00072_4_pdu1d(pf12833) PDU1D
11850 -pf04055pf13186 -
11855 Hp_euts_violetPink EUT1
11860 HMM10662PduV_1_mix(pf10662) EUT1
11865 HMM06249EutQ_1_eut1(pf06249) EUT1 EP_NTERM
11870 HMM01923cobtr_2_eut1(pf01923) EUT1
11875 HMM01515PTA_B_1_eut1(pf01515) EUT1
11880 H_azure EUT1
11885 P_lightViolet EUT1
11890 HMMalddh_1_eutpdugrm(pf00171) EUT1 EP_NTERM
11895 HMM06723EutJMreB_1_grm EUT1
11900 HMM00465alcdh_3_mix(pf00465) EUT1
11905 HMMtpEuth_1_eut(pf04346) EUT1
11910 HMM06277EutA_1_eut(pf06277) EUT1
11915 HMMetlyEutB_1_EUT1(pf06751) EUT1
11920 HMMetlyEutC_1_eut1(pf05985) EUT1 EP_NTERM
11925 Tsp_rosa EUT1
11930 H_mulberry(pf00936pf16365) EUT1
11935 HMMreg12833_1_EUT1(pf12833) EUT1
11940 -pf01218 -
11945 -pf01520 -
11950 -pf00583 -
BMC-P
BMC-H
BMC-T
Aldehyde dehydrogenase
PTAC
Signature enzyme
Alcohol dehydrogenase
Regulator

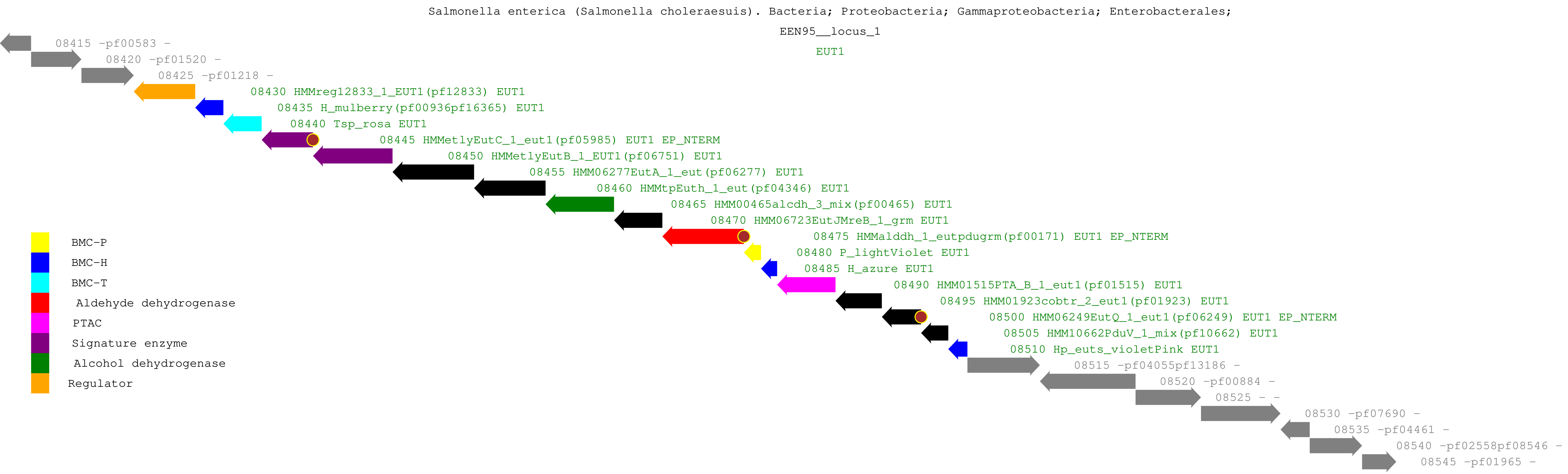
Salmonella enterica (Salmonella choleraesuis). Bacteria; Proteobacteria; Gammaproteobacteria; Enterobacterales;
EEN95__locus_1
EUT1
08415 -pf00583 -
08420 -pf01520 -
08425 -pf01218 -
08430 HMMreg12833_1_EUT1(pf12833) EUT1
08435 H_mulberry(pf00936pf16365) EUT1
08440 Tsp_rosa EUT1
08445 HMMetlyEutC_1_eut1(pf05985) EUT1 EP_NTERM
08450 HMMetlyEutB_1_EUT1(pf06751) EUT1
08455 HMM06277EutA_1_eut(pf06277) EUT1
08460 HMMtpEuth_1_eut(pf04346) EUT1
08465 HMM00465alcdh_3_mix(pf00465) EUT1
08470 HMM06723EutJMreB_1_grm EUT1
08475 HMMalddh_1_eutpdugrm(pf00171) EUT1 EP_NTERM
08480 P_lightViolet EUT1
08485 H_azure EUT1
08490 HMM01515PTA_B_1_eut1(pf01515) EUT1
08495 HMM01923cobtr_2_eut1(pf01923) EUT1
08500 HMM06249EutQ_1_eut1(pf06249) EUT1 EP_NTERM
08505 HMM10662PduV_1_mix(pf10662) EUT1
08510 Hp_euts_violetPink EUT1
08515 -pf04055pf13186 -
08520 -pf00884 -
08525 - -
08530 -pf07690 -
08535 -pf04461 -
08540 -pf02558pf08546 -
08545 -pf01965 -
BMC-P
BMC-H
BMC-T
Aldehyde dehydrogenase
PTAC
Signature enzyme
Alcohol dehydrogenase
Regulator

Salmonella enterica I. Bacteria; Proteobacteria; Gammaproteobacteria; Enterobacterales;

EER80__locus_1

EUT1

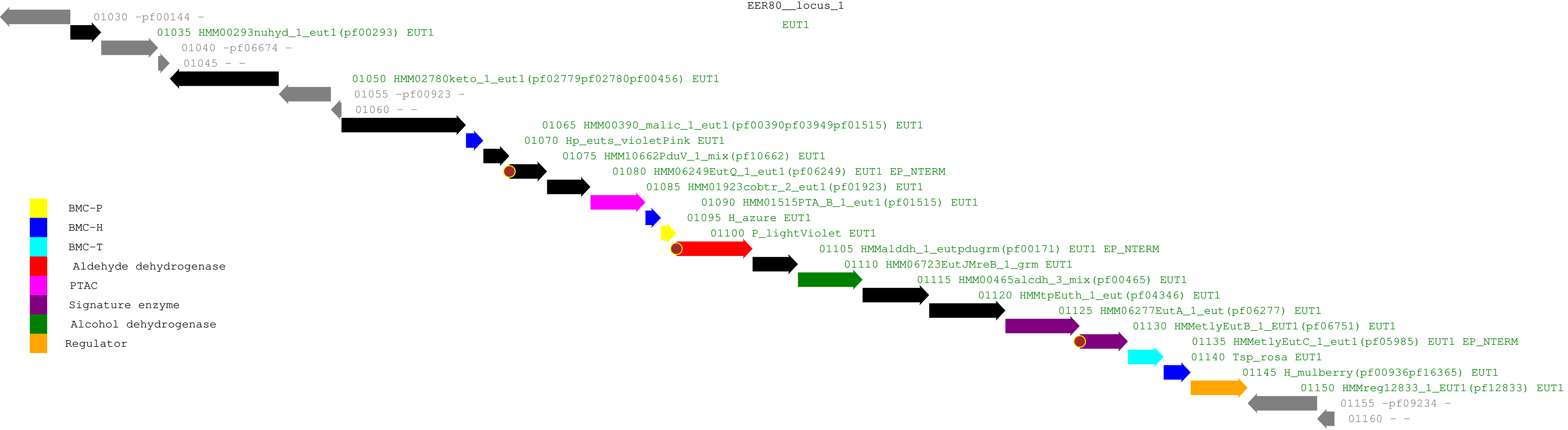

Escherichia fergusonii (strain ATCC 35469 / DSM 13698 / CDC 0568-73). Bacteria; Proteobacteria; Gammaproteobacteria; Enterobacterales;

EFER__locus_1

EUT1

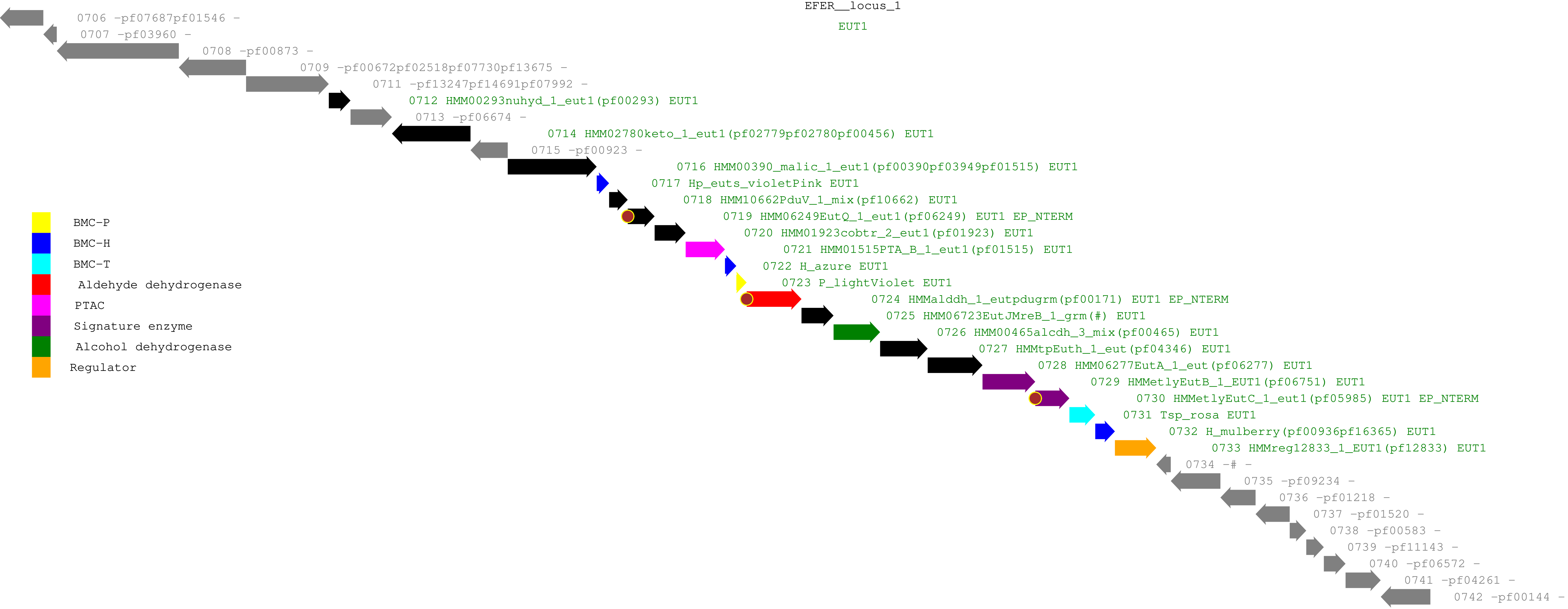

Escherichia fergusonii. Bacteria; Proteobacteria; Gammaproteobacteria; Enterobacterales;

EFSS39__locus_1

EUT1

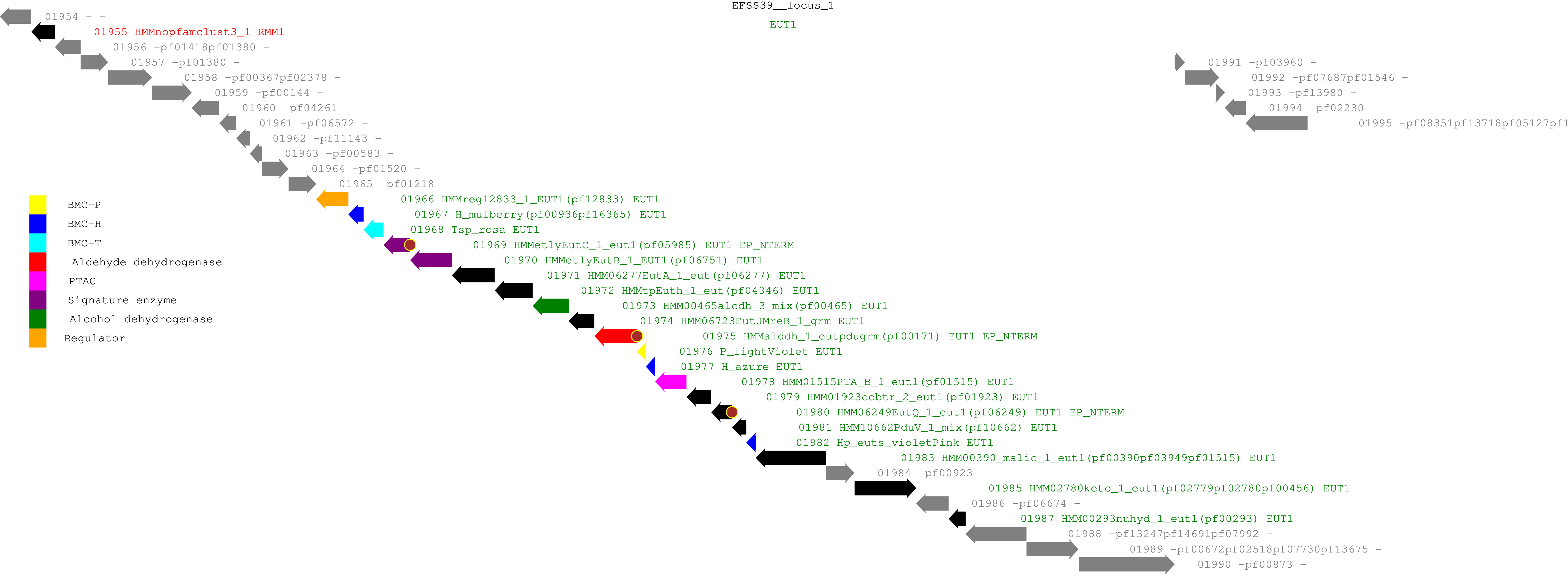

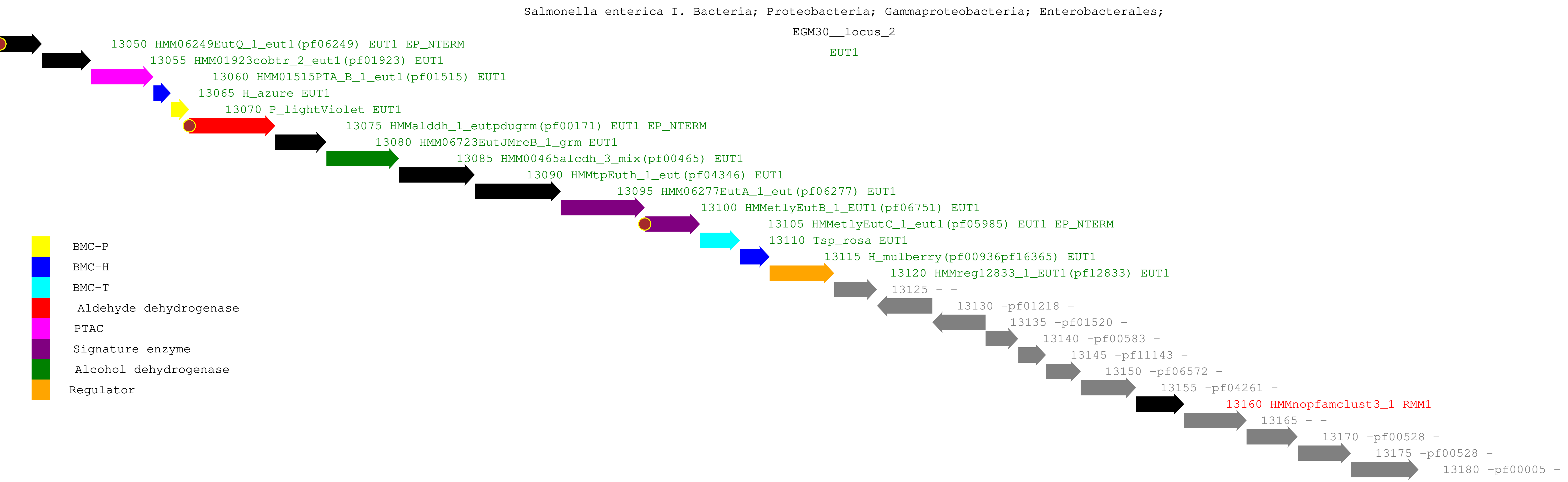
Salmonella enterica I. Bacteria; Proteobacteria; Gammaproteobacteria; Enterobacterales;
EGM30__locus_2
EUT1
13050 HMM06249EutQ_1_eut1(pf06249) EUT1 EP_NTERM
13055 HMM01923cobtr_2_eut1(pf01923) EUT1
13060 HMM01515PTA_B_1_eut1(pf01515) EUT1
13065 H_azure EUT1
13070 P_lightViolet EUT1
13075 HMMalddh_1_eutpdugrm(pf00171) EUT1 EP_NTERM
13080 HMM06723EutJMreB_1_grm EUT1
13085 HMM00465alcdh_3_mix(pf00465) EUT1
13090 HMMtpEuth_1_eut(pf04346) EUT1
13095 HMM06277EutA_1_eut(pf06277) EUT1
13100 HMMetlyEutB_1_EUT1(pf06751) EUT1
13105 HMMetlyEutC_1_eut1(pf05985) EUT1 EP_NTERM
13110 Tsp_rosa EUT1
13115 H_mulberry(pf00936pf16365) EUT1
13120 HMMreg12833_1_EUT1(pf12833) EUT1
13125 - -
13130 -pf01218 -
13135 -pf01520 -
13140 -pf00583 -
13145 -pf11143 -
13150 -pf06572 -
13155 -pf04261 -
13160 HMMnopfamclust3_1 RMM1
13165 - -
13170 -pf00528 -
13175 -pf00528 -
13180 -pf00005 -
BMC-P
BMC-H
BMC-T
Aldehyde dehydrogenase
PTAC
Signature enzyme
Alcohol dehydrogenase
Regulator

Escherichia coli. Bacteria; Proteobacteria; Gammaproteobacteria; Enterobacterales;

EIA21__locus_1

EUT1

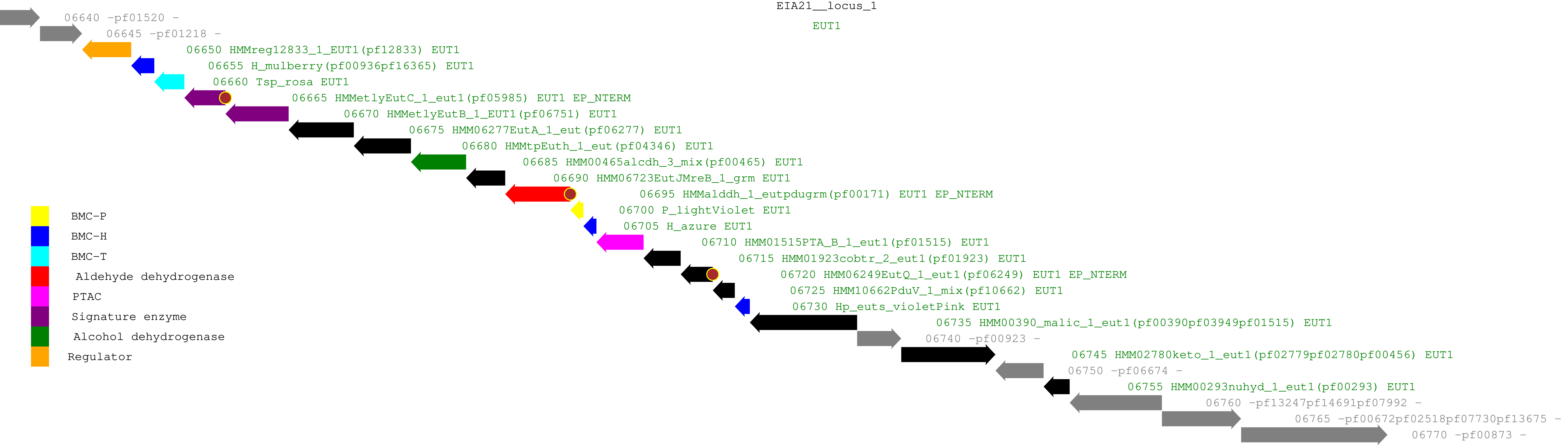

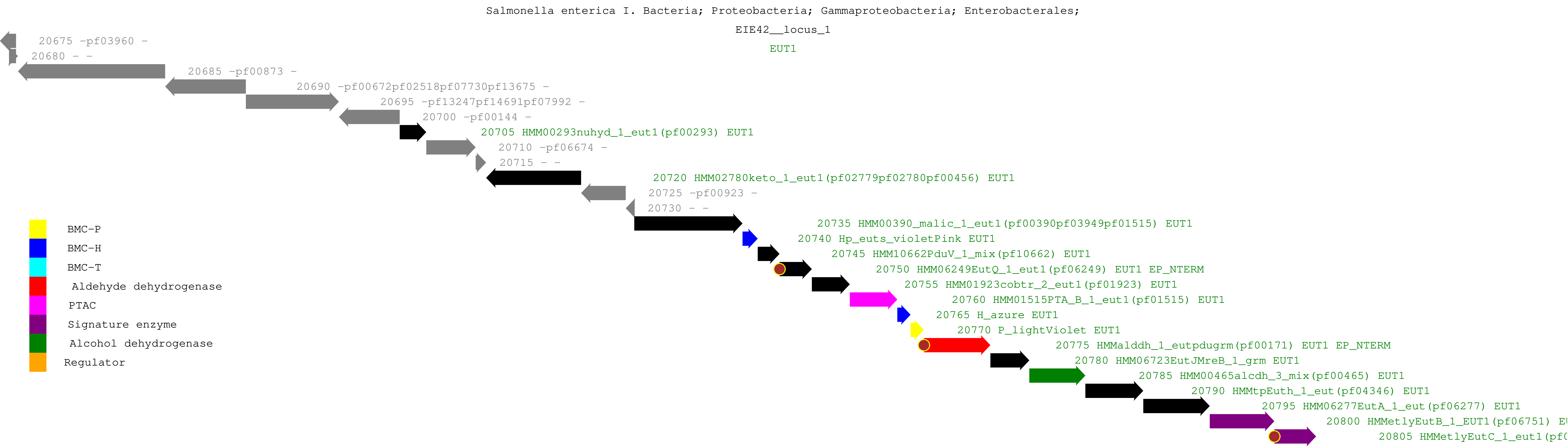
Salmonella enterica I. Bacteria; Proteobacteria; Gammaproteobacteria; Enterobacterales;
EIE42__locus_1
EUT1
20675 -pf03960 -
20680 - -
20685 -pf00873 -
20690 -pf00672pf02518pf07730pf13675 -
20695 -pf13247pf14691pf07992 -
20700 -pf00144 -
20705 HMM00293nuhyd_1_eut1(pf00293) EUT1
20710 -pf06674 -
20715 - -
20720 HMM02780keto_1_eut1(pf02779pf02780pf00456) EUT1
20725 -pf00923 -
20730 - -
20735 HMM00390_malic_1_eut1(pf00390pf03949pf01515) EUT1
20740 Hp_euts_violetPink EUT1
20745 HMM10662PduV_1_mix(pf10662) EUT1
20750 HMM06249EutQ_1_eut1(pf06249) EUT1 EP_NTERM
20755 HMM01923cobtr_2_eut1(pf01923) EUT1
20760 HMM01515PTA_B_1_eut1(pf01515) EUT1
20765 H_azure EUT1
20770 P_lightViolet EUT1
20775 HMMalddh_1_eutpdugrm(pf00171) EUT1 EP_NTERM
20780 HMM06723EutJMreB_1_grm EUT1
20785 HMM00465alcdh_3_mix(pf00465) EUT1
20790 HMMtpEuth_1_eut(pf04346) EUT1
20795 HMM06277EutA_1_eut(pf06277) EUT1
20800 HMMetlyEutB_1_EUT1(pf06751)
20805 HMMetlyEutC_1_eut1
BMC-P
BMC-H
BMC-T
Aldehyde dehydrogenase
PTAC
Signature enzyme
Alcohol dehydrogenase
Regulator

Salmonella enterica I. Bacteria; Proteobacteria; Gammaproteobacteria; Enterobacterales;

EIW58__locus_1

EUT1

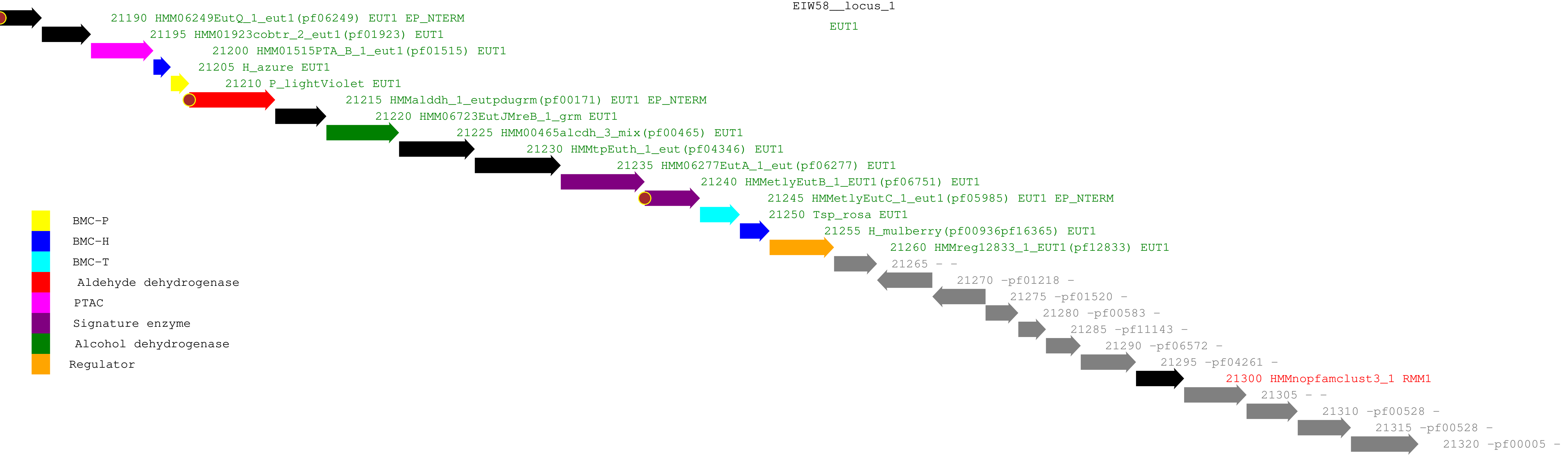

Salmonella paratyphi A. Bacteria; Proteobacteria; Gammaproteobacteria; Enterobacterales;

EJI24__locus_1

EUT1

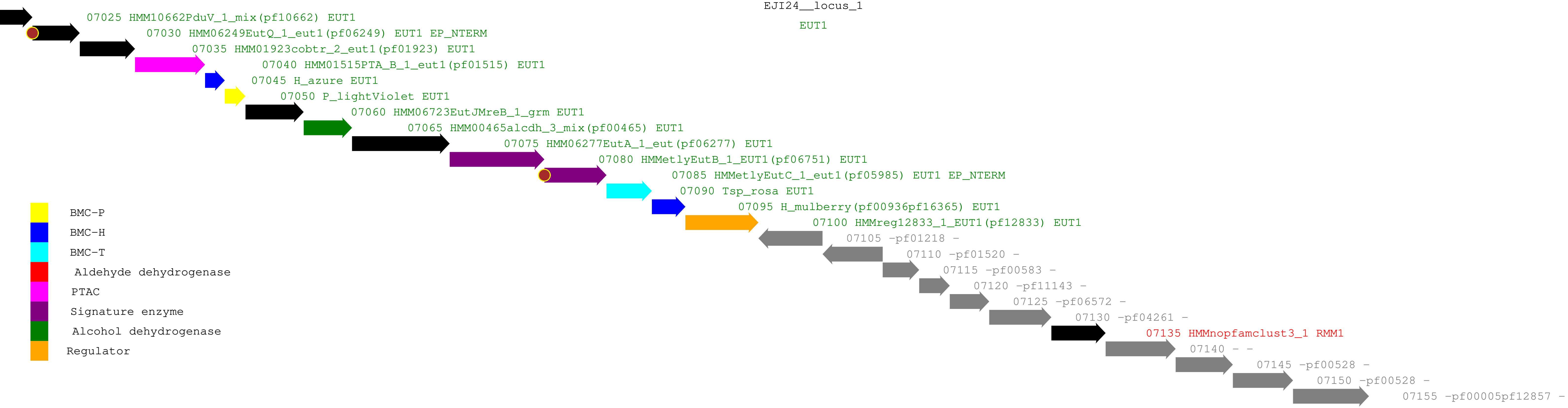

Iodobacter sp. H11R3. Bacteria; Proteobacteria; Betaproteobacteria; Neisseriales;

EJO50__locus_1

EUT1

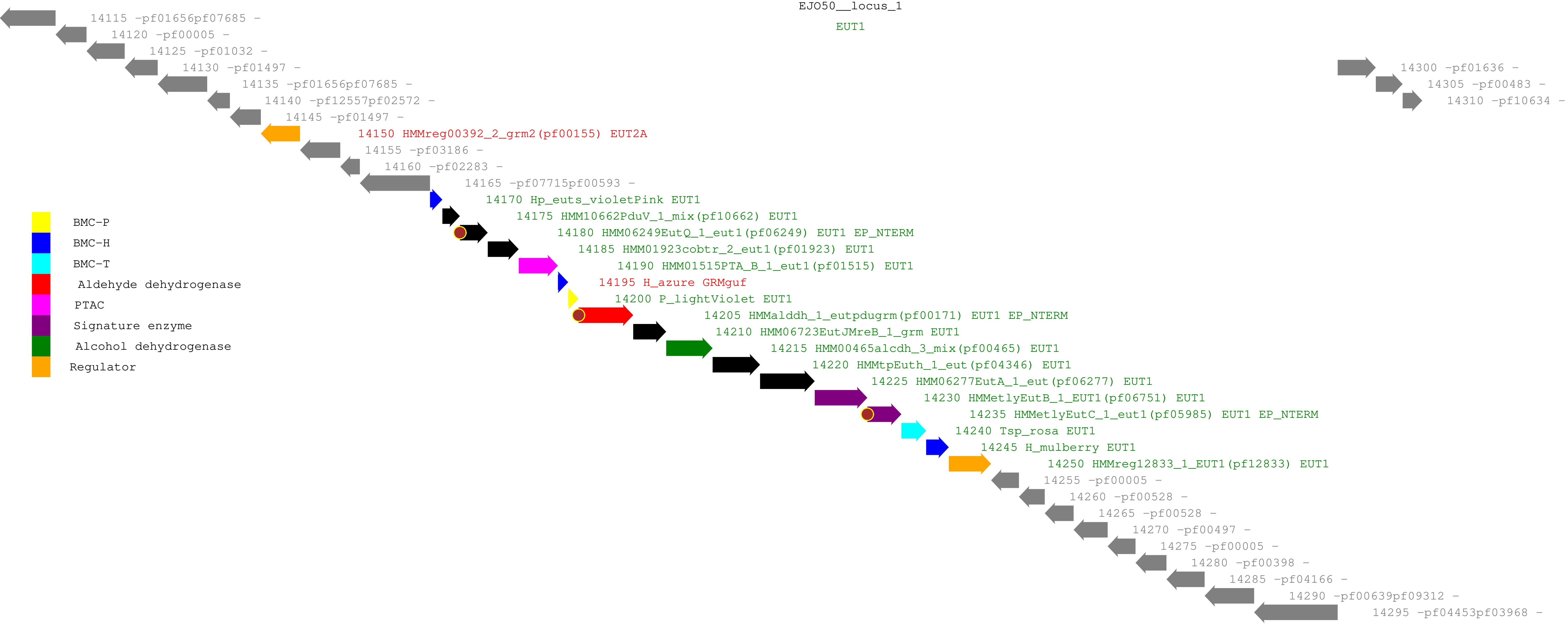

Salmonella enterica subsp. enterica serovar Karamoja. Bacteria; Proteobacteria; Gammaproteobacteria; Enterobacterales;

EL007__locus_1

EUT1

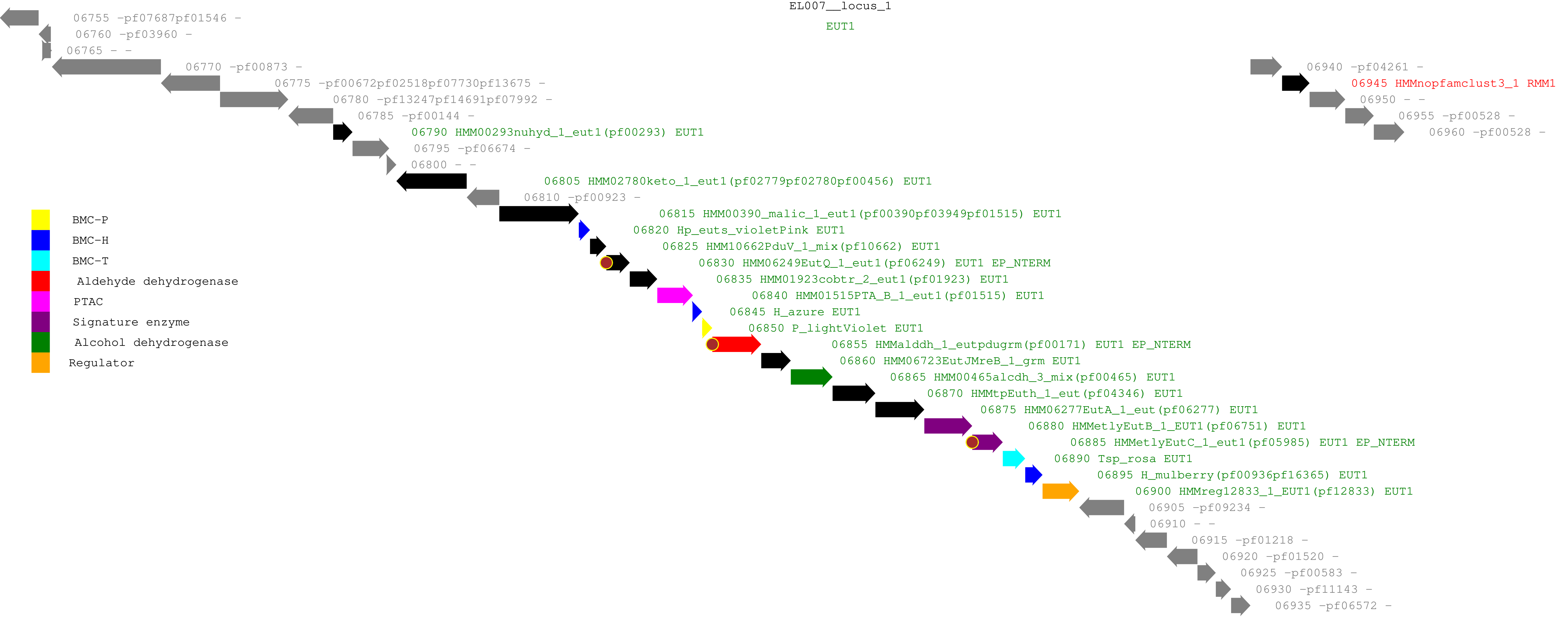

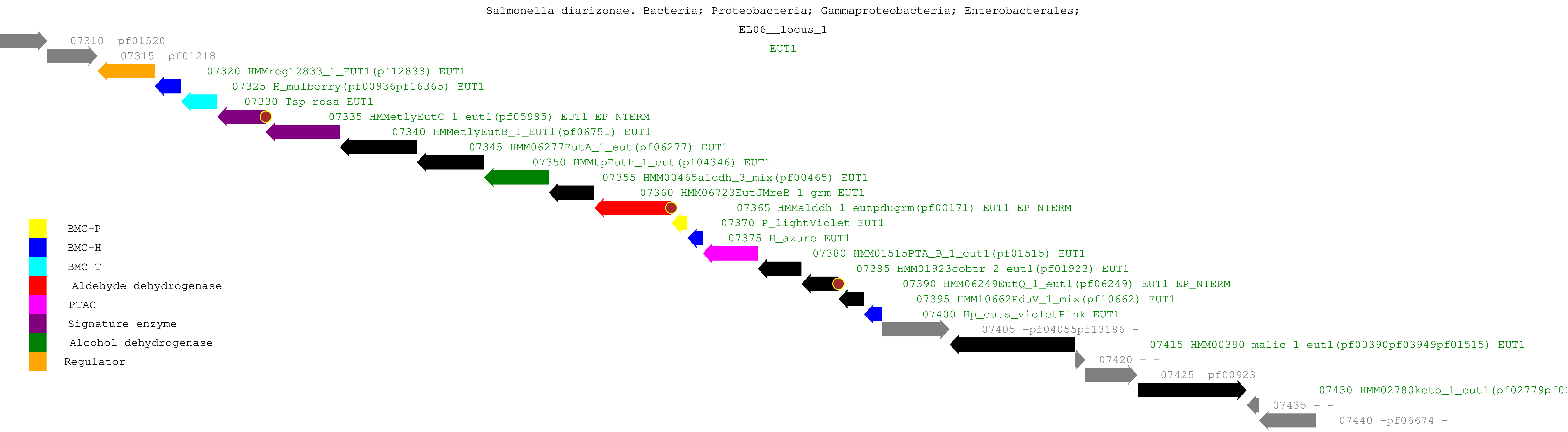
Salmonella diarizonae. Bacteria; Proteobacteria; Gammaproteobacteria; Enterobacterales;
EL06__locus_1
EUT1
07310 -pf01520 -
07315 -pf01218 -
07320 HMMreg12833_1_EUT1(pf12833) EUT1
07325 H_mulberry(pf00936pf16365) EUT1
07330 Tsp_rosa EUT1
07335 HMMetlyEutC_1_eut1(pf05985) EUT1 EP_NTERM
07340 HMMetlyEutB_1_EUT1(pf06751) EUT1
07345 HMM06277EutA_1_eut(pf06277) EUT1
07350 HMMtpEuth_1_eut(pf04346) EUT1
07355 HMM00465alcdh_3_mix(pf00465) EUT1
07360 HMM06723EutJMreB_1_grm EUT1
07365 HMMaldddh_1_eutpdugrm(pf00171) EUT1 EP_NTERM
07370 P_lightViolet EUT1
07375 H_azure EUT1
07380 HMM01515PTA_B_1_eut1(pf01515) EUT1
07385 HMM01923cobtr_2_eut1(pf01923) EUT1
07390 HMM06249EutQ_1_eut1(pf06249) EUT1 EP_NTERM
07395 HMM10662PduV_1_mix(pf10662) EUT1
07400 Hp_euts_violetPink EUT1
07405 -pf04055pf13186 -
07415 HMM00390_malic_1_eut1(pf00390pf03949pf01515) EUT1
07420 - -
07425 -pf00923 -
07430 HMM02780keto_1_eut1(pf02779pf0
07435 - -
07440 -pf06674 -
BMC-P
BMC-H
BMC-T
Aldehyde dehydrogenase
PTAC
Signature enzyme
Alcohol dehydrogenase
Regulator

Escherichia coli G3/10. Bacteria; Proteobacteria; Gammaproteobacteria; Enterobacterales;

EL76__locus_1

EUT1

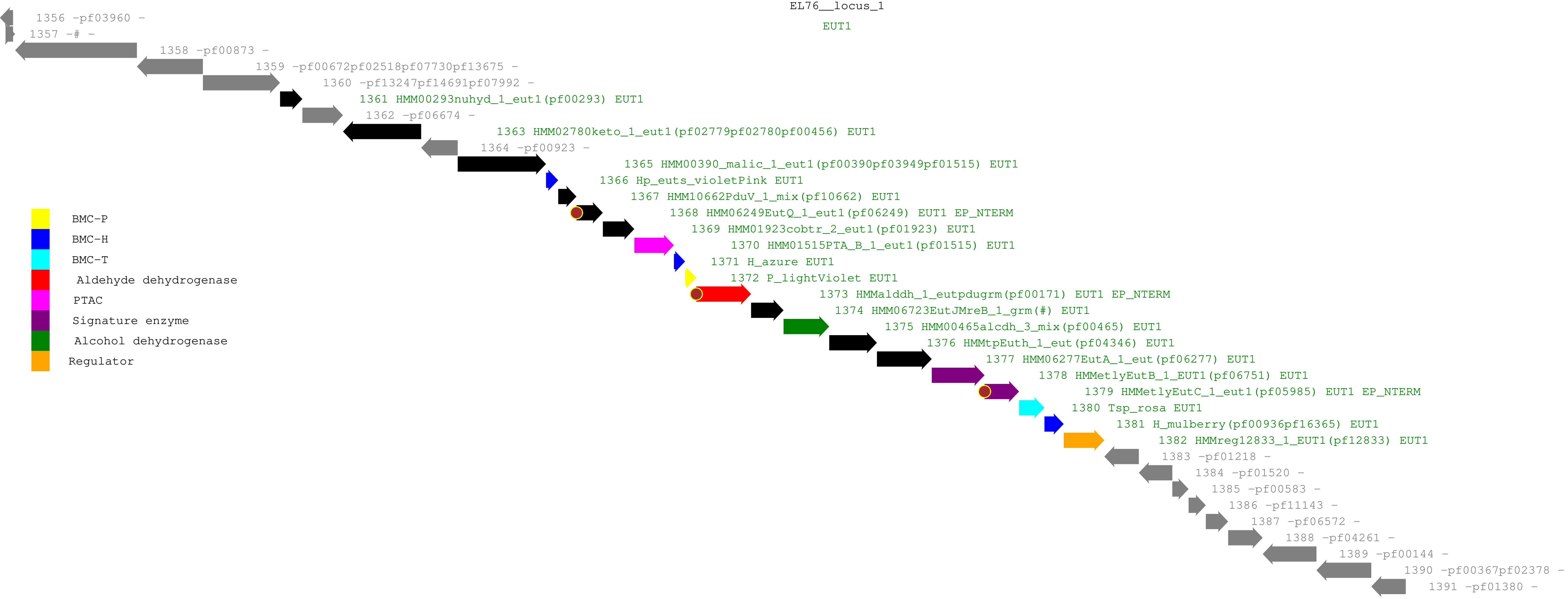

Escherichia coli. Bacteria; Proteobacteria; Gammaproteobacteria; Enterobacterales;

EL80__locus_1

EUT1

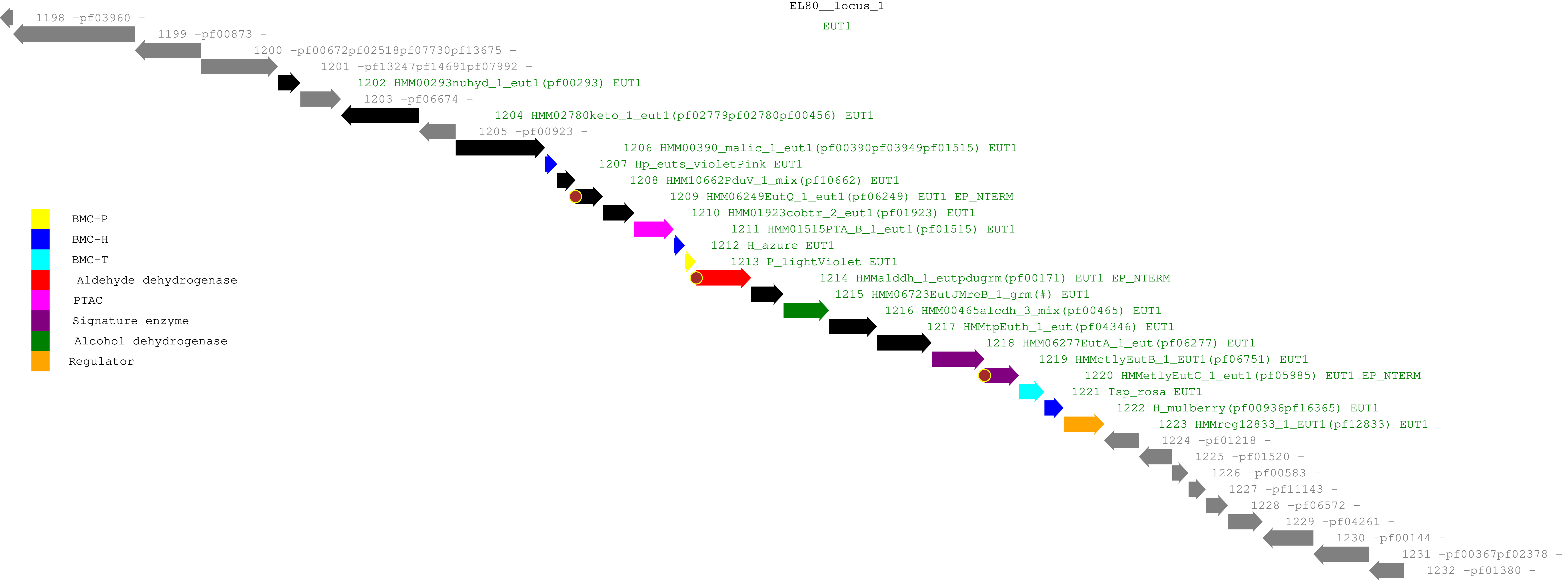

Salmonella enterica subsp. enterica serovar Moero. Bacteria; Proteobacteria; Gammaproteobacteria; Enterobacterales;

ELZ69__locus_1

EUT1

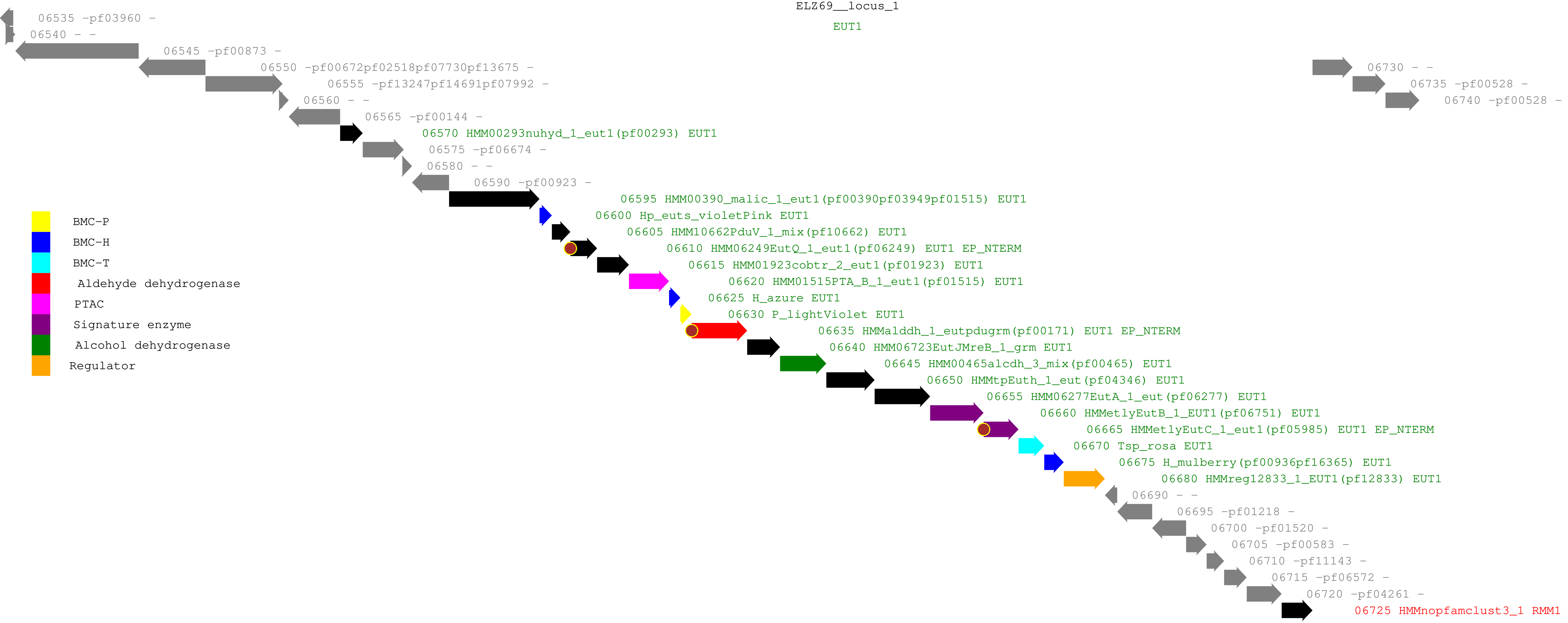

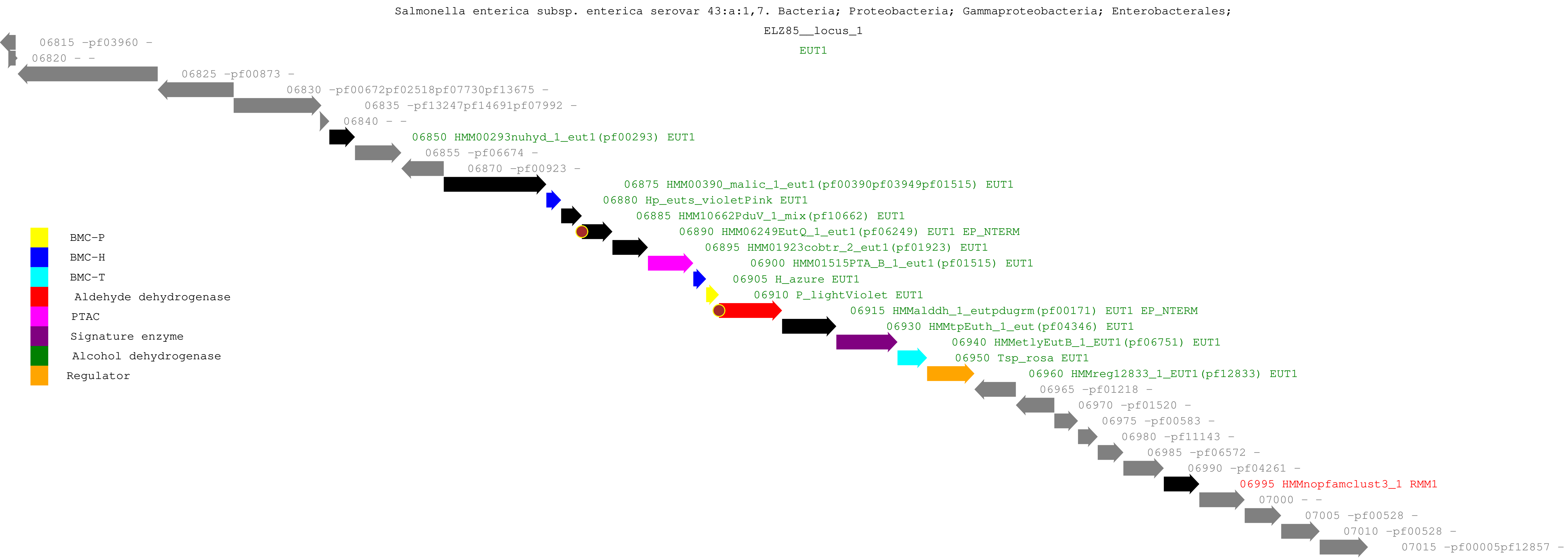
Salmonella enterica subsp. enterica serovar 43:a:1,7. Bacteria; Proteobacteria; Gammaproteobacteria; Enterobacterales;
ELZ85__locus_1
EUT1
06815 -pf03960 -
06820 - -
06825 -pf00873 -
06830 -pf00672pf02518pf07730pf13675 -
06835 -pf13247pf14691pf07992 -
06840 - -
06850 HMM00293nuhyd_1_eut1(pf00293) EUT1
06855 -pf06674 -
06870 -pf00923 -
06875 HMM00390_malic_1_eut1(pf00390pf03949pf01515) EUT1
06880 Hp_euts_violetPink EUT1
06885 HMM10662PduV_1_mix(pf10662) EUT1
06890 HMM06249EutQ_1_eut1(pf06249) EUT1 EP_NTERM
06895 HMM01923cobtr_2_eut1(pf01923) EUT1
06900 HMM01515PTA_B_1_eut1(pf01515) EUT1
06905 H_azure EUT1
06910 P_lightViolet EUT1
06915 HMMalddh_1_eutpdugrm(pf00171) EUT1 EP_NTERM
06930 HMMtpEuth_1_eut(pf04346) EUT1
06940 HMMetlyEutB_1_EUT1(pf06751) EUT1
06950 Tsp_rosa EUT1
06960 HMMreg12833_1_EUT1(pf12833) EUT1
06965 -pf01218 -
06970 -pf01520 -
06975 -pf00583 -
06980 -pf11143 -
06985 -pf06572 -
06990 -pf04261 -
06995 HMMnopfamclust3_1 RMM1
07000 - -
07005 -pf00528 -
07010 -pf00528 -
07015 -pf00005pf12857 -
BMC-P
BMC-H
BMC-T
Aldehyde dehydrogenase
PTAC
Signature enzyme
Alcohol dehydrogenase
Regulator

Salmonella enterica subsp. enterica serovar 43:a:1,7. Bacteria; Proteobacteria; Gammaproteobacteria; Enterobacterales;

ELZ87__locus_1

EUT1

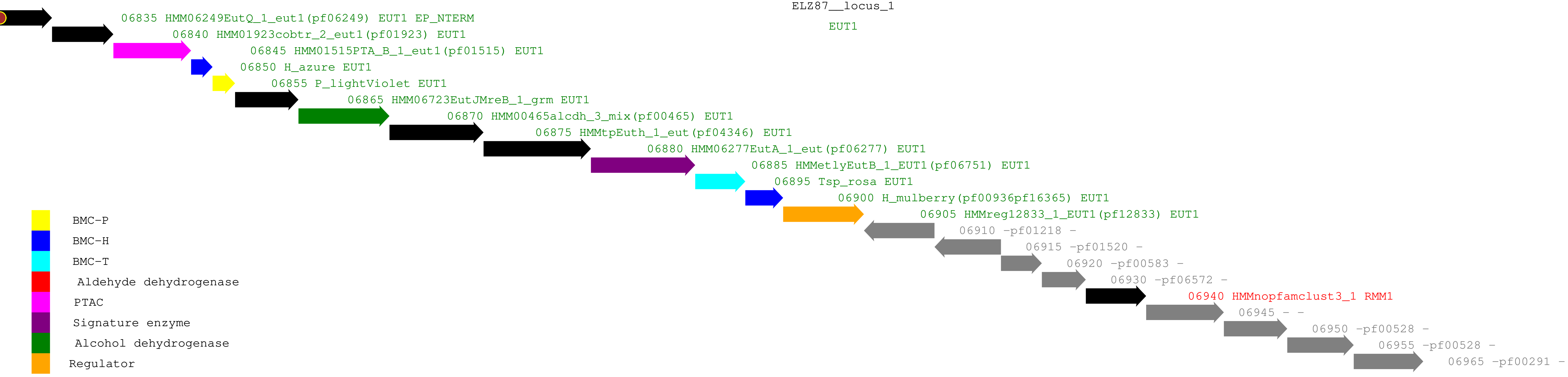

Enterobacter lignolyticus (strain SCF1). Bacteria; Proteobacteria; Gammaproteobacteria; Enterobacterales;

Entcl__locus_1

EUT1

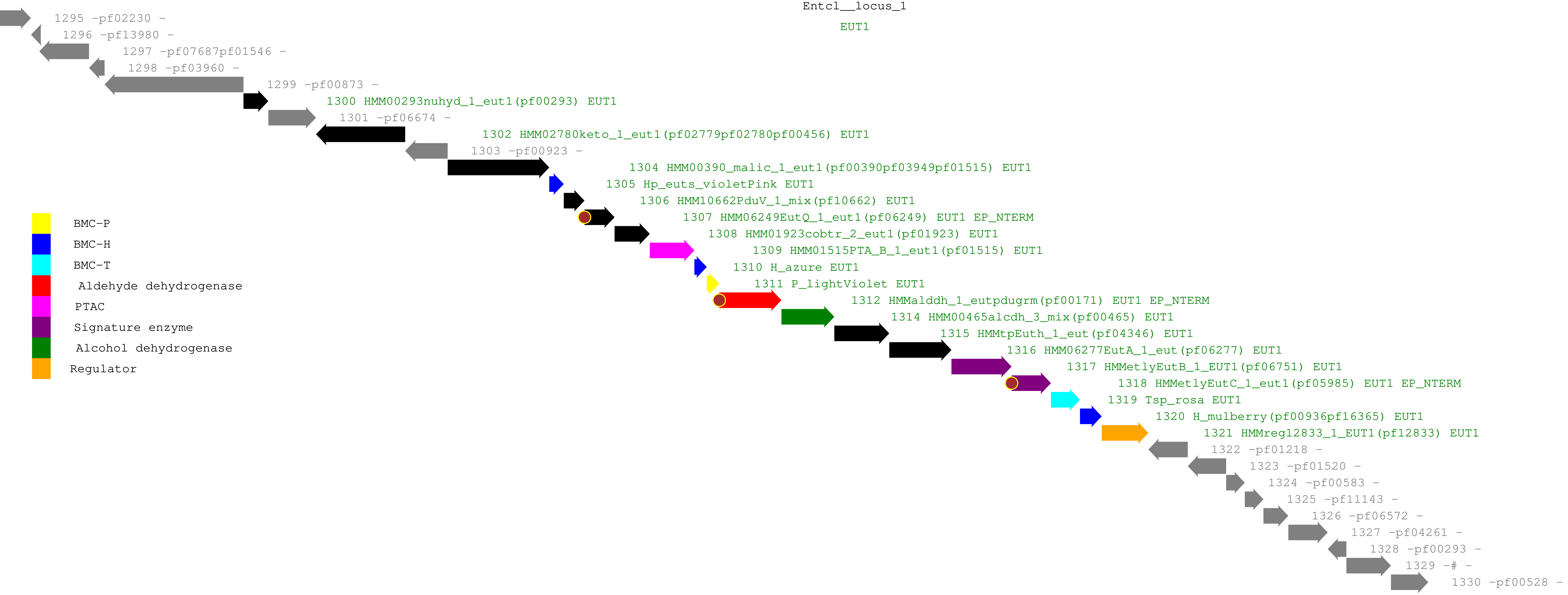

Salmonella bongori serovar 48:i:-. Bacteria; Proteobacteria; Gammaproteobacteria; Enterobacterales;

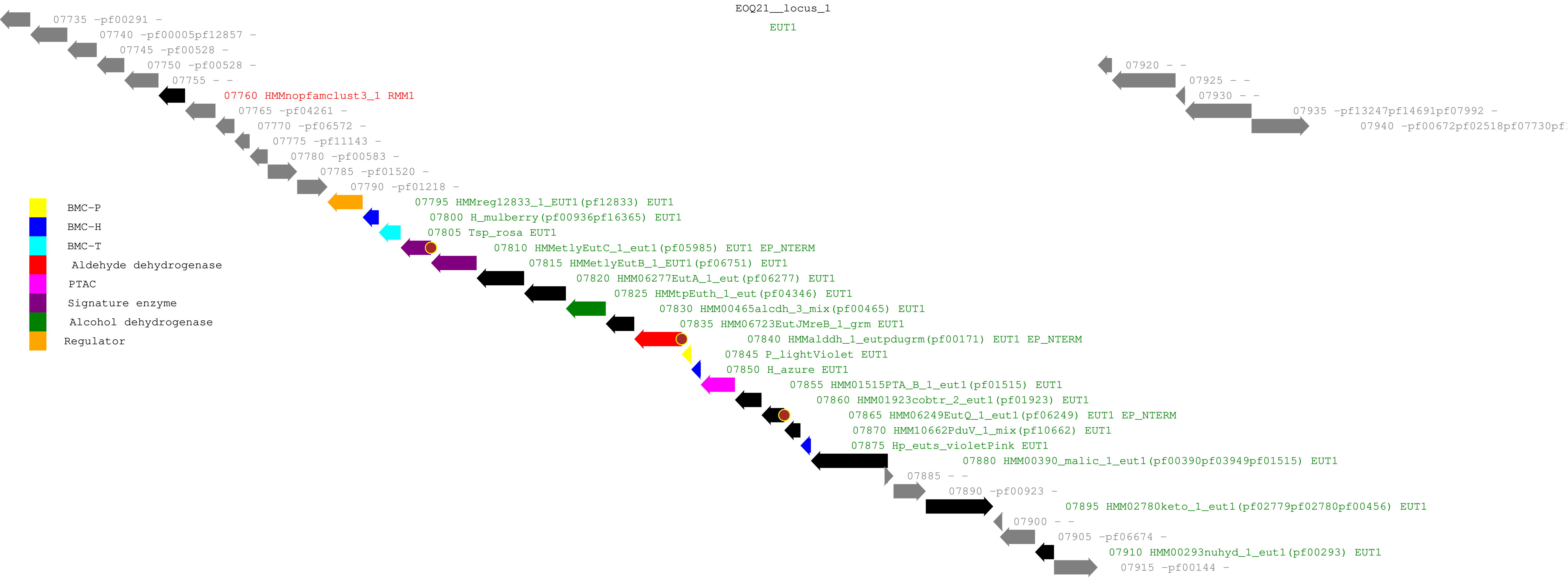

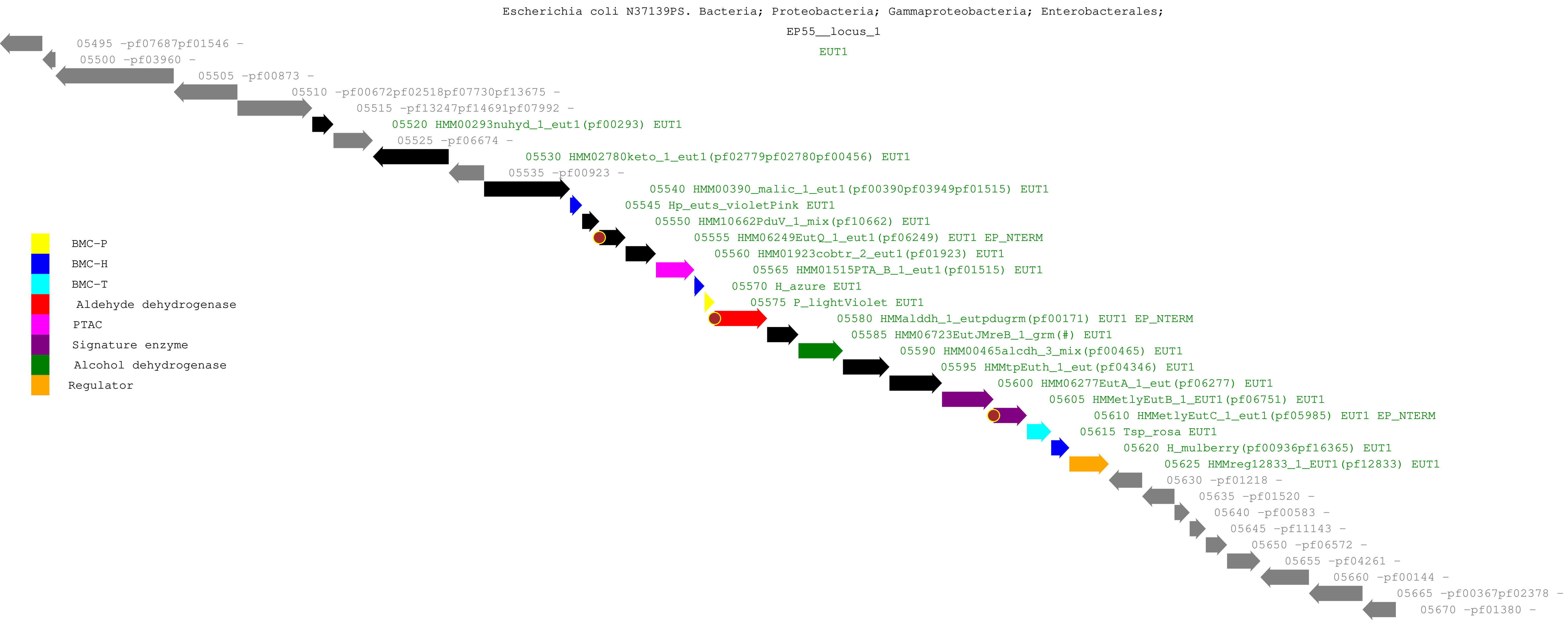
Escherichia coli N37139PS. Bacteria; Proteobacteria; Gammaproteobacteria; Enterobacterales;
EP55__locus_1
EUT1
05495 -pf07687pf01546 -
05500 -pf03960 -
05505 -pf00873 -
05510 -pf00672pf02518pf07730pf13675 -
05515 -pf13247pf14691pf07992 -
05520 HMM00293nuhyd_1_eut1(pf00293) EUT1
05525 -pf06674 -
05530 HMM02780keto_1_eut1(pf02779pf02780pf00456) EUT1
05535 -pf00923 -
05540 HMM00390_malic_1_eut1(pf00390pf03949pf01515) EUT1
05545 Hp_euts_violetPink EUT1
05550 HMM10662PduV_1_mix(pf10662) EUT1
05555 HMM06249EutQ_1_eut1(pf06249) EUT1 EP_NTERM
05560 HMM01923cobtr_2_eut1(pf01923) EUT1
05565 HMM01515PTA_B_1_eut1(pf01515) EUT1
05570 H_azure EUT1
05575 P_lightViolet EUT1
05580 HMMaldh_1_eutpdugrm(pf00171) EUT1 EP_NTERM
05585 HMM06723EutJMreB_1_grm(#) EUT1
05590 HMM00465alcdh_3_mix(pf00465) EUT1
05595 HMMtpEuth_1_eut(pf04346) EUT1
05600 HMM06277EutA_1_eut(pf06277) EUT1
05605 HMMetlyEutB_1_EUT1(pf06751) EUT1
05610 HMMetlyEutC_1_eut1(pf05985) EUT1 EP_NTERM
05615 Tsp_rosa EUT1
05620 H_mulberry(pf00936pf16365) EUT1
05625 HMMreg12833_1_EUT1(pf12833) EUT1
05630 -pf01218 -
05635 -pf01520 -
05640 -pf00583 -
05645 -pf11143 -
05650 -pf06572 -
05655 -pf04261 -
05660 -pf00144 -
05665 -pf00367pf02378 -
05670 -pf01380 -
BMC-P
BMC-H
BMC-T
Aldehyde dehydrogenase
PTAC
Signature enzyme
Alcohol dehydrogenase
Regulator

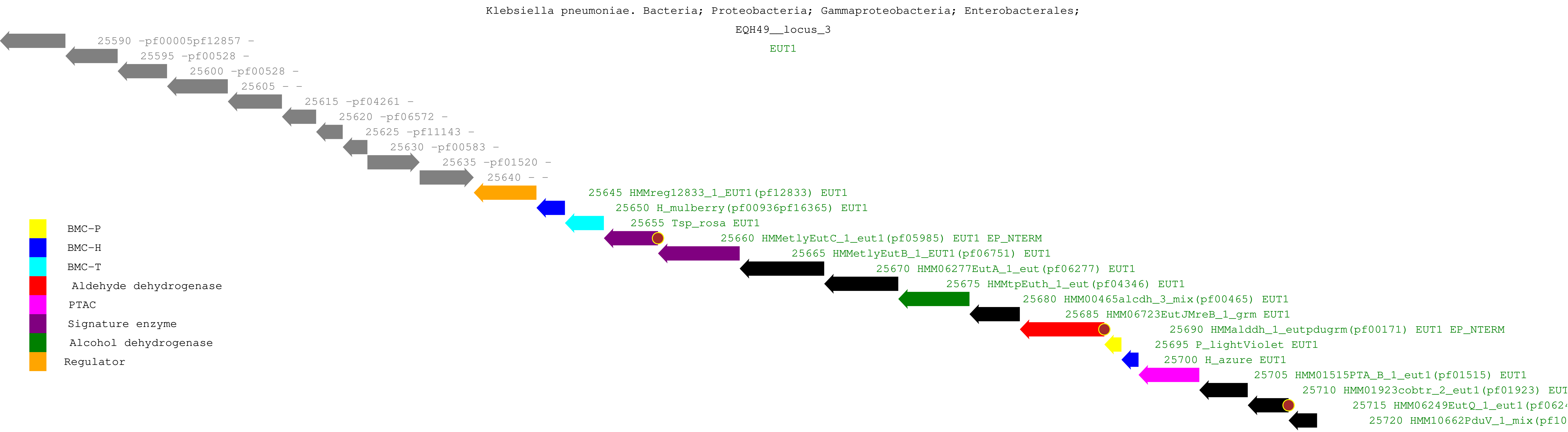
Klebsiella pneumoniae. Bacteria; Proteobacteria; Gammaproteobacteria; Enterobacterales;
EQH49__locus_3
EUT1
25590 -pf00005pf12857 -
25595 -pf00528 -
25600 -pf00528 -
25605 - -
25615 -pf04261 -
25620 -pf06572 -
25625 -pf11143 -
25630 -pf00583 -
25635 -pf01520 -
25640 - -
25645 HMMreg12833_1_EUT1(pf12833) EUT1
25650 H_mulberry(pf00936pf16365) EUT1
25655 Tsp_rosa EUT1
25660 HMMetlyEutC_1_eut1(pf05985) EUT1 EP_NTERM
25665 HMMetlyEutB_1_EUT1(pf06751) EUT1
25670 HMM06277EutA_1_eut(pf06277) EUT1
25675 HMMtpEuth_1_eut(pf04346) EUT1
25680 HMM00465alcdh_3_mix(pf00465) EUT1
25685 HMM06723EutJMreB_1_grm EUT1
25690 HMMalddh_1_eutpdugrm(pf00171) EUT1 EP_NTERM
25695 P_lightViolet EUT1
25700 H_azure EUT1
25705 HMM01515PTA_B_1_eut1(pf01515) EUT1
25710 HMM01923cobtr_2_eut1(pf01923) EUT
25715 HMM06249EutQ_1_eut1(pf062
25720 HMM10662PduV_1_mix(pf10
BMC-P
BMC-H
BMC-T
Aldehyde dehydrogenase
PTAC
Signature enzyme
Alcohol dehydrogenase
Regulator

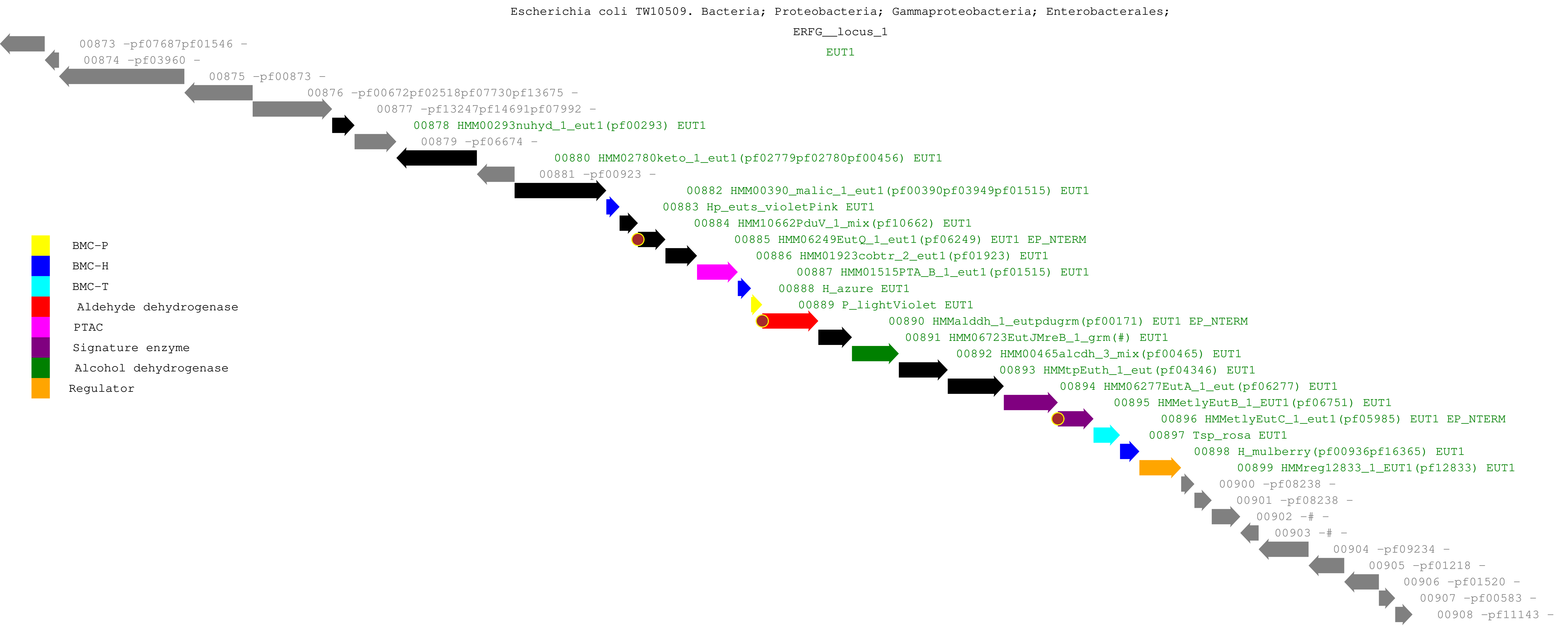
Escherichia coli TW10509. Bacteria; Proteobacteria; Gammaproteobacteria; Enterobacterales;
ERFG__locus_1
EUT1
00873 -pf07687pf01546 -
00874 -pf03960 -
00875 -pf00873 -
00876 -pf00672pf02518pf07730pf13675 -
00877 -pf13247pf14691pf07992 -
00878 HMM00293nuhyd_1_eut1(pf00293) EUT1
00879 -pf06674 -
00880 HMM02780keto_1_eut1(pf02779pf02780pf00456) EUT1
00881 -pf00923 -
00882 HMM00390_malic_1_eut1(pf00390pf03949pf01515) EUT1
00883 Hp_euts_violetPink EUT1
00884 HMM10662PduV_1_mix(pf10662) EUT1
00885 HMM06249EutQ_1_eut1(pf06249) EUT1 EP_NTERM
00886 HMM01923cobtr_2_eut1(pf01923) EUT1
00887 HMM01515PTA_B_1_eut1(pf01515) EUT1
00888 H_azure EUT1
00889 P_lightViolet EUT1
00890 HMMalddh_1_eutpdugrm(pf00171) EUT1 EP_NTERM
00891 HMM06723EutJMreB_1_grm(#) EUT1
00892 HMM00465alcdh_3_mix(pf00465) EUT1
00893 HMMtpEuth_1_eut(pf04346) EUT1
00894 HMM06277EutA_1_eut(pf06277) EUT1
00895 HMMetlyEutB_1_EUT1(pf06751) EUT1
00896 HMMetlyEutC_1_eut1(pf05985) EUT1 EP_NTERM
00897 Tsp_rosa EUT1
00898 H_mulberry(pf00936pf16365) EUT1
00899 HMMreg12833_1_EUT1(pf12833) EUT1
00900 -pf08238 -
00901 -pf08238 -
00902 -# -
00903 -# -
00904 -pf09234 -
00905 -pf01218 -
00906 -pf01520 -
00907 -pf00583 -
00908 -pf11143 -
BMC-P
BMC-H
BMC-T
Aldehyde dehydrogenase
PTAC
Signature enzyme
Alcohol dehydrogenase
Regulator

Escherichia coli M863. Bacteria; Proteobacteria; Gammaproteobacteria; Enterobacterales;

ERJG__locus_1

EUT1

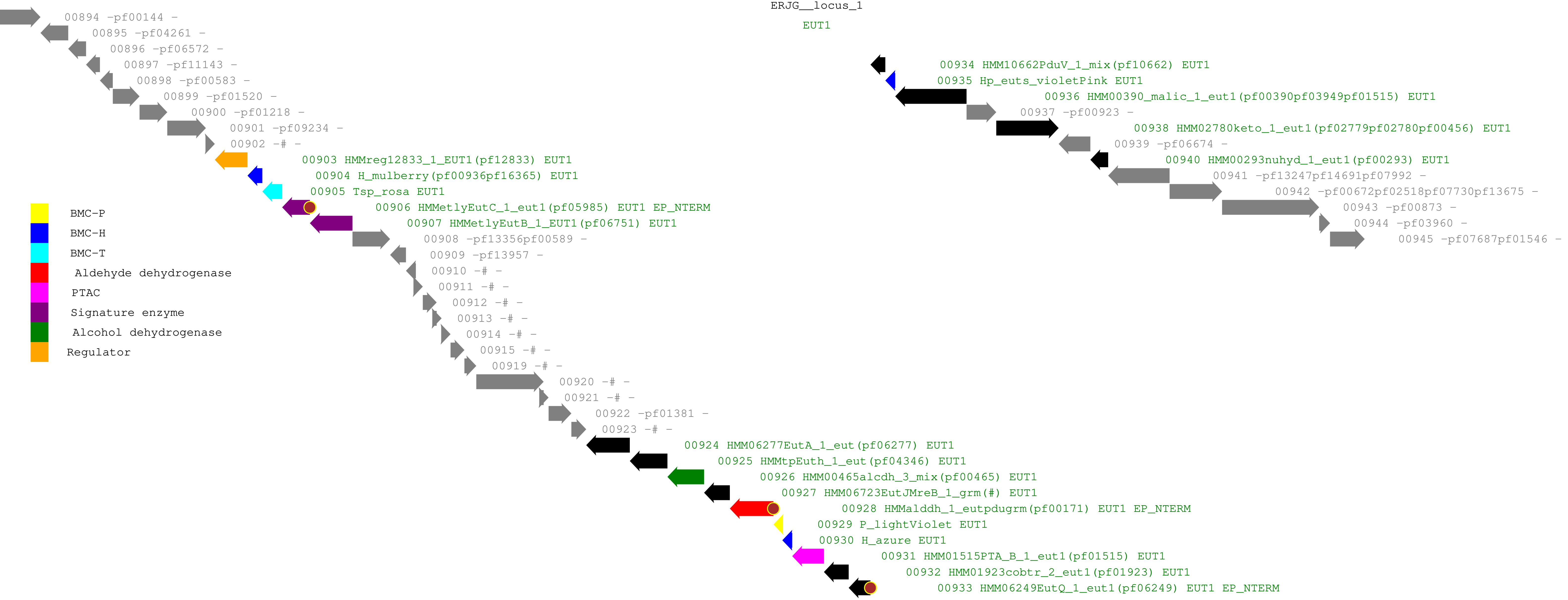

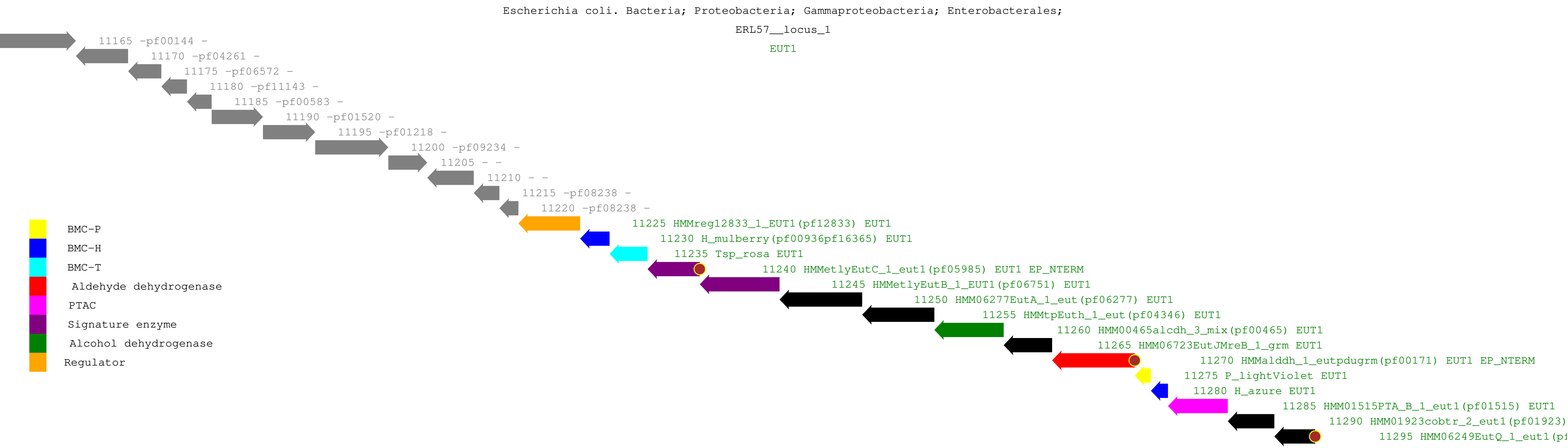

Escherichia coli. Bacteria; Proteobacteria; Gammaproteobacteria; Enterobacterales;
ERL57__locus_1
EUT1
11165 -pf00144 -
11170 -pf04261 -
11175 -pf06572 -
11180 -pf11143 -
11185 -pf00583 -
11190 -pf01520 -
11195 -pf01218 -
11200 -pf09234 -
11205 - -
11210 - -
11215 -pf08238 -
11220 -pf08238 -
11225 HMMreg12833_1_EUT1(pf12833) EUT1
11230 H_mulberry(pf00936pf16365) EUT1
11235 Tsp_rosa EUT1
11240 HMMetlyEutC_1_eut1(pf05985) EUT1 EP_NTERM
11245 HMMetlyEutB_1_EUT1(pf06751) EUT1
11250 HMM06277EutA_1_eut(pf06277) EUT1
11255 HMMtpEuth_1_eut(pf04346) EUT1
11260 HMM00465alcdh_3_mix(pf00465) EUT1
11265 HMM06723EutJMreB_1_grm EUT1
11270 HMMalddh_1_eutpdugrm(pf00171) EUT1 EP_NTERM
11275 P_lightViolet EUT1
11280 H_azure EUT1
11285 HMM01515PTA_B_1_eut1(pf01515) EUT1
11290 HMM01923cobtr_2_eut1(pf01923)
11295 HMM06249EutQ_1_eut1(pf
BMC-P
BMC-H
BMC-T
Aldehyde dehydrogenase
PTAC
Signature enzyme
Alcohol dehydrogenase
Regulator

Salmonella enterica subsp. enterica serovar Bovismorbificans. Bacteria; Proteobacteria; Gammaproteobacteria; Enterobacterales;

ERS008202__locus_2

EUT1

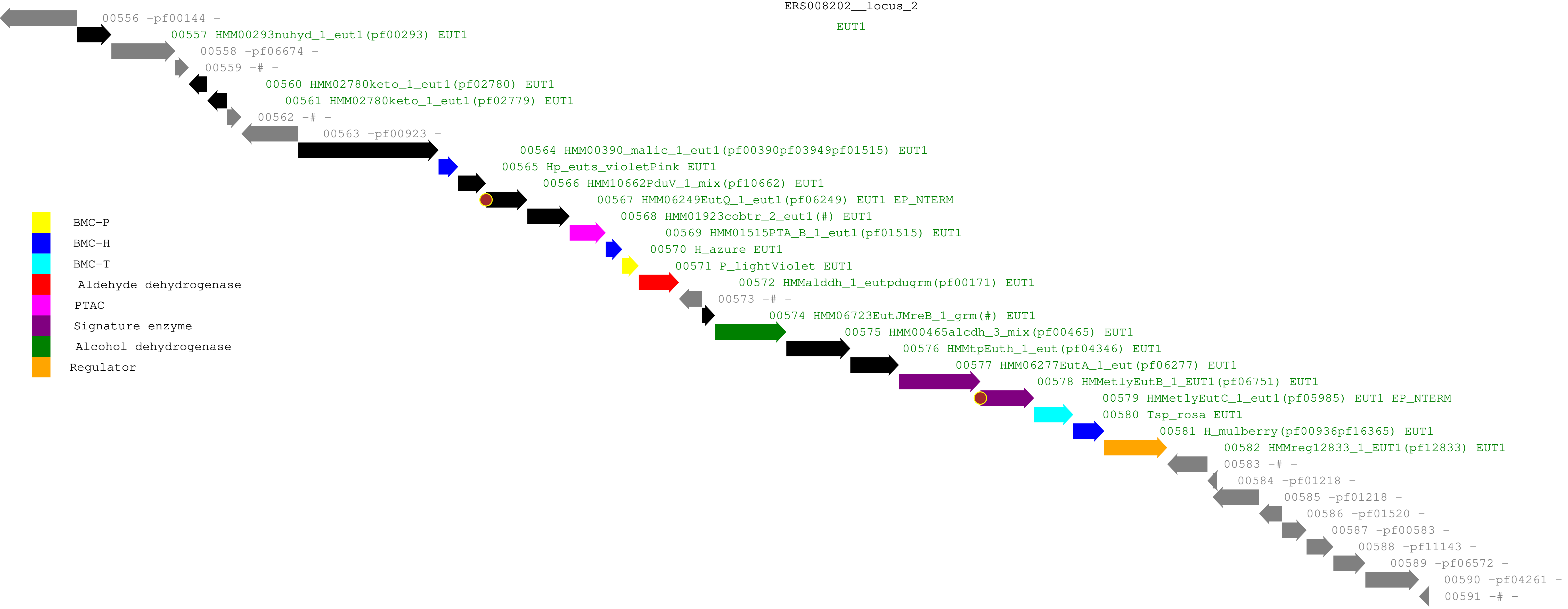

Salmonella enterica subsp. enterica serovar Bovismorbificans. Bacteria; Proteobacteria; Gammaproteobacteria; Enterobacterales;

ERS008215__locus_1

EUT1

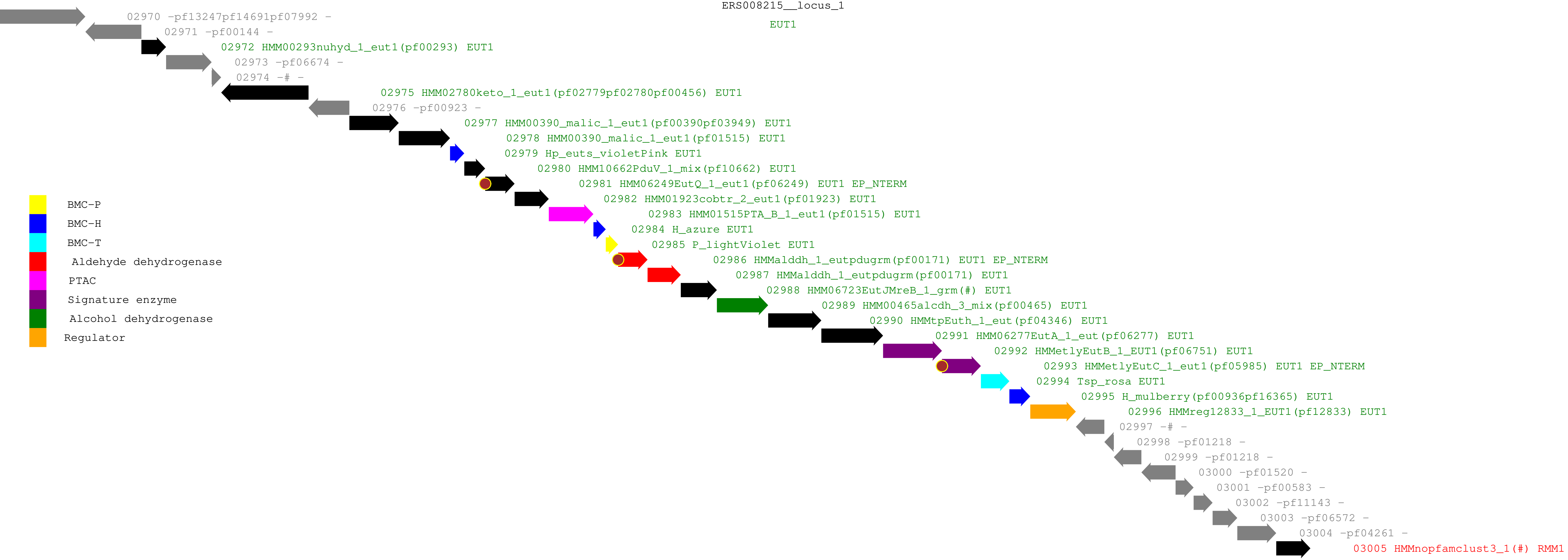

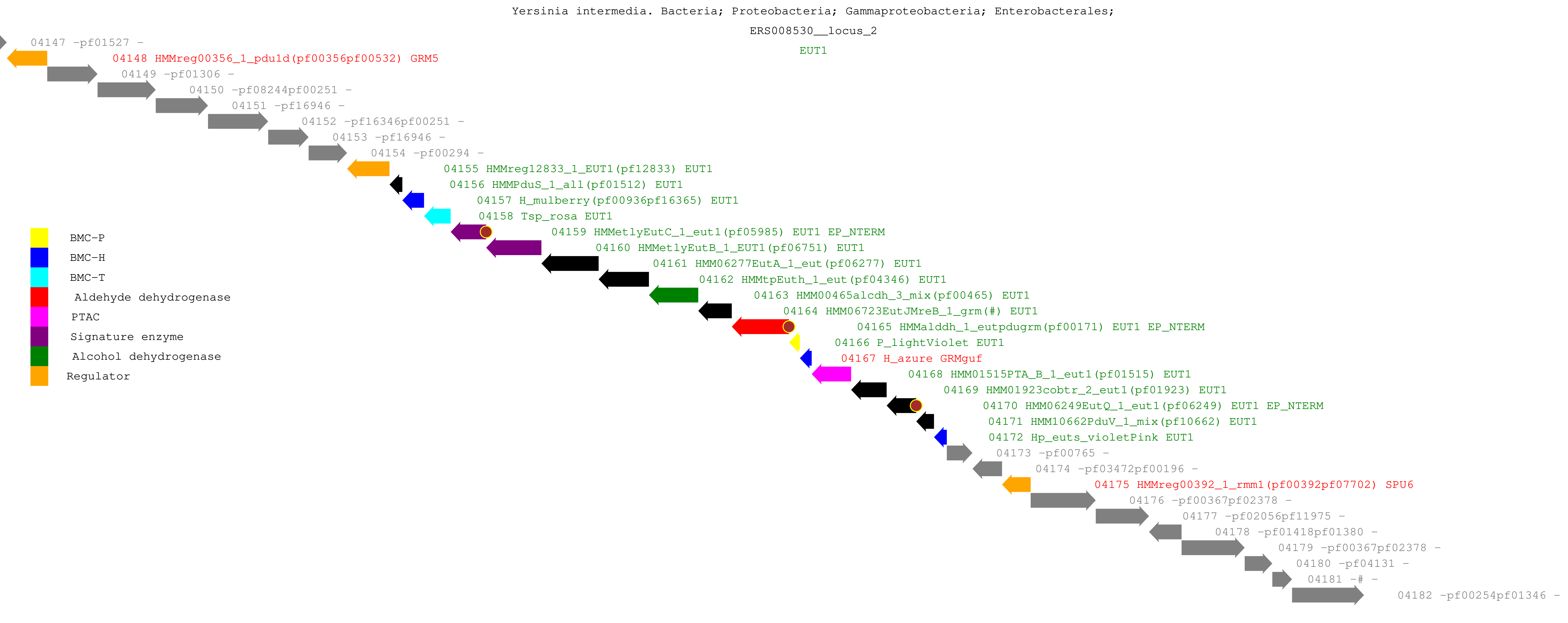
Yersinia intermedia. Bacteria; Proteobacteria; Gammaproteobacteria; Enterobacterales;
ERS008530__locus_2
EUT1
04147 -pf01527 -
04148 HMMreg00356_1_pdu1d(pf00356pf00532) GRM5
04149 -pf01306 -
04150 -pf08244pf00251 -
04151 -pf16946 -
04152 -pf16346pf00251 -
04153 -pf16946 -
04154 -pf00294 -
04155 HMMreg12833_1_EUT1(pf12833) EUT1
04156 HMMPduS_1_all(pf01512) EUT1
04157 H_mulberry(pf00936pf16365) EUT1
04158 Tsp_rosa EUT1
04159 HMMetlyEutC_1_eut1(pf05985) EUT1 EP_NTERM
04160 HMMetlyEutB_1_EUT1(pf06751) EUT1
04161 HMM06277EutA_1_eut(pf06277) EUT1
04162 HMMtpEuth_1_eut(pf04346) EUT1
04163 HMM00465alcdh_3_mix(pf00465) EUT1
04164 HMM06723EutJMreB_1_grm(#) EUT1
04165 HMMalddh_1_eutpdugrm(pf00171) EUT1 EP_NTERM
04166 P_lightViolet EUT1
04167 H_azure GRMguf
04168 HMM01515PTA_B_1_eut1(pf01515) EUT1
04169 HMM01923cobtr_2_eut1(pf01923) EUT1
04170 HMM06249EutQ_1_eut1(pf06249) EUT1 EP_NTERM
04171 HMM10662PduV_1_mix(pf10662) EUT1
04172 Hp_euts_violetPink EUT1
04173 -pf00765 -
04174 -pf03472pf00196 -
04175 HMMreg00392_1_rmm1(pf00392pf07702) SPU6
04176 -pf00367pf02378 -
04177 -pf02056pf11975 -
04178 -pf01418pf01380 -
04179 -pf00367pf02378 -
04180 -pf04131 -
04181 -# -
04182 -pf00254pf01346 -
BMC-P
BMC-H
BMC-T
Aldehyde dehydrogenase
PTAC
Signature enzyme
Alcohol dehydrogenase
Regulator

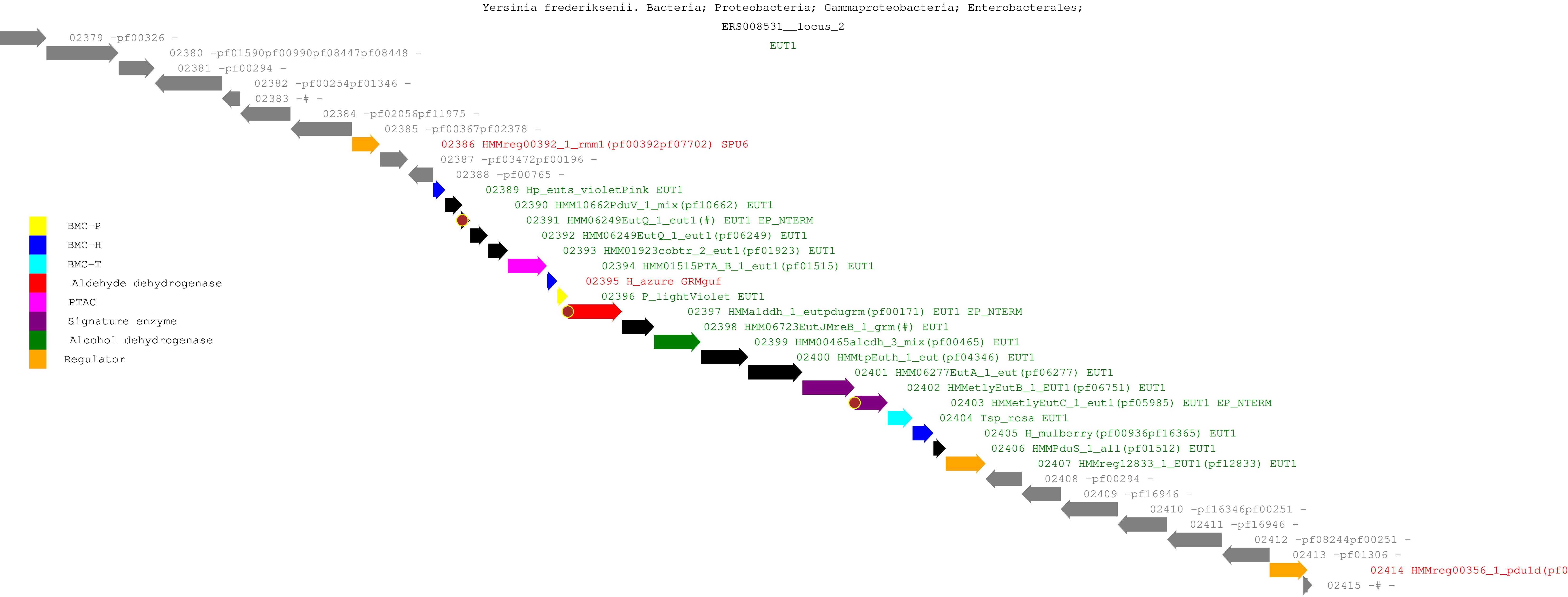
Yersinia frederiksenii. Bacteria; Proteobacteria; Gammaproteobacteria; Enterobacterales;
ERS008531__locus_2
EUT1
02379 -pf00326 -
02380 -pf01590pf00990pf08447pf08448 -
02381 -pf00294 -
02382 -pf00254pf01346 -
02383 -# -
02384 -pf02056pf11975 -
02385 -pf00367pf02378 -
02386 HMMreg00392_1_rmm1(pf00392pf07702) SPU6
02387 -pf03472pf00196 -
02388 -pf00765 -
02389 Hp_euts_violetPink EUT1
02390 HMM10662PduV_1_mix(pf10662) EUT1
02391 HMM06249EutQ_1_eut1(#) EUT1 EP_NTERM
02392 HMM06249EutQ_1_eut1(pf06249) EUT1
02393 HMM01923cobtr_2_eut1(pf01923) EUT1
02394 HMM01515PTA_B_1_eut1(pf01515) EUT1
02395 H_azure GRMguf
02396 P_lightViolet EUT1
02397 HMMaldh_1_eutpdugrm(pf00171) EUT1 EP_NTERM
02398 HMM06723EutJMreB_1_grm(#) EUT1
02399 HMM00465alcdh_3_mix(pf00465) EUT1
02400 HMMtpEuth_1_eut(pf04346) EUT1
02401 HMM06277EutA_1_eut(pf06277) EUT1
02402 HMMetlyEutB_1_EUT1(pf06751) EUT1
02403 HMMetlyEutC_1_eut1(pf05985) EUT1 EP_NTERM
02404 Tsp_rosa EUT1
02405 H_mulberry(pf00936pf16365) EUT1
02406 HMMPduS_1_all(pf01512) EUT1
02407 HMMreg12833_1_EUT1(pf12833) EUT1
02408 -pf00294 -
02409 -pf16946 -
02410 -pf16346pf00251 -
02411 -pf16946 -
02412 -pf08244pf00251 -
02413 -pf01306 -
02414 HMMreg00356_1_pdu1d(pf0
02415 -# -
BMC-P
BMC-H
BMC-T
Aldehyde dehydrogenase
PTAC
Signature enzyme
Alcohol dehydrogenase
Regulator

Shigella sonnei. Bacteria; Proteobacteria; Gammaproteobacteria; Enterobacterales;

ERS009810__locus_1_main_sat

EUT1

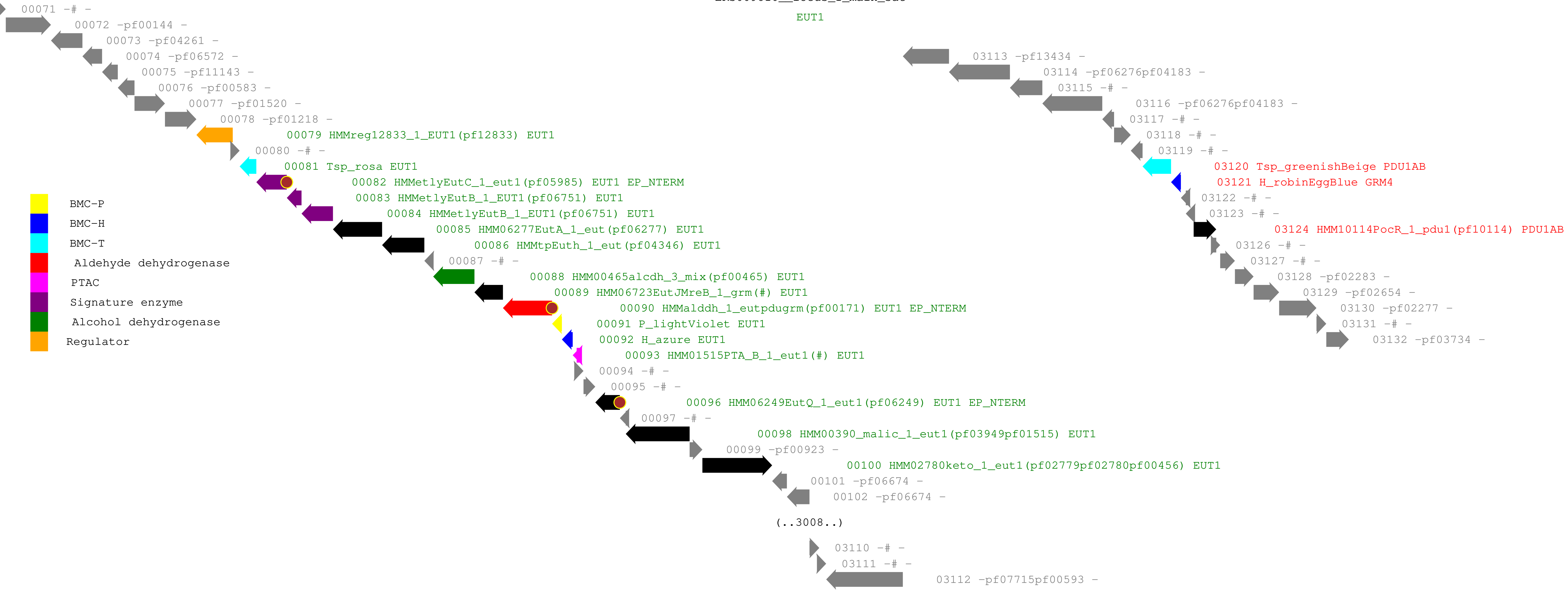

Escherichia coli. Bacteria; Proteobacteria; Gammaproteobacteria; Enterobacterales;

ERS085365__locus_1

EUT1

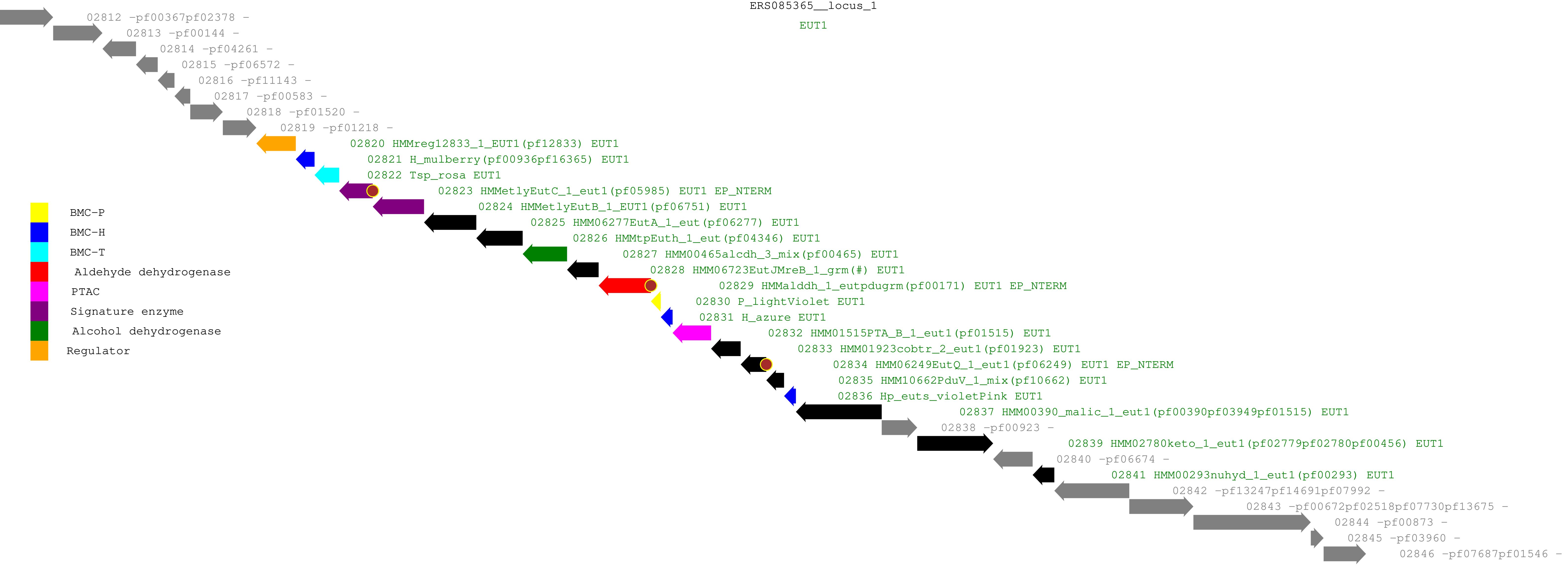

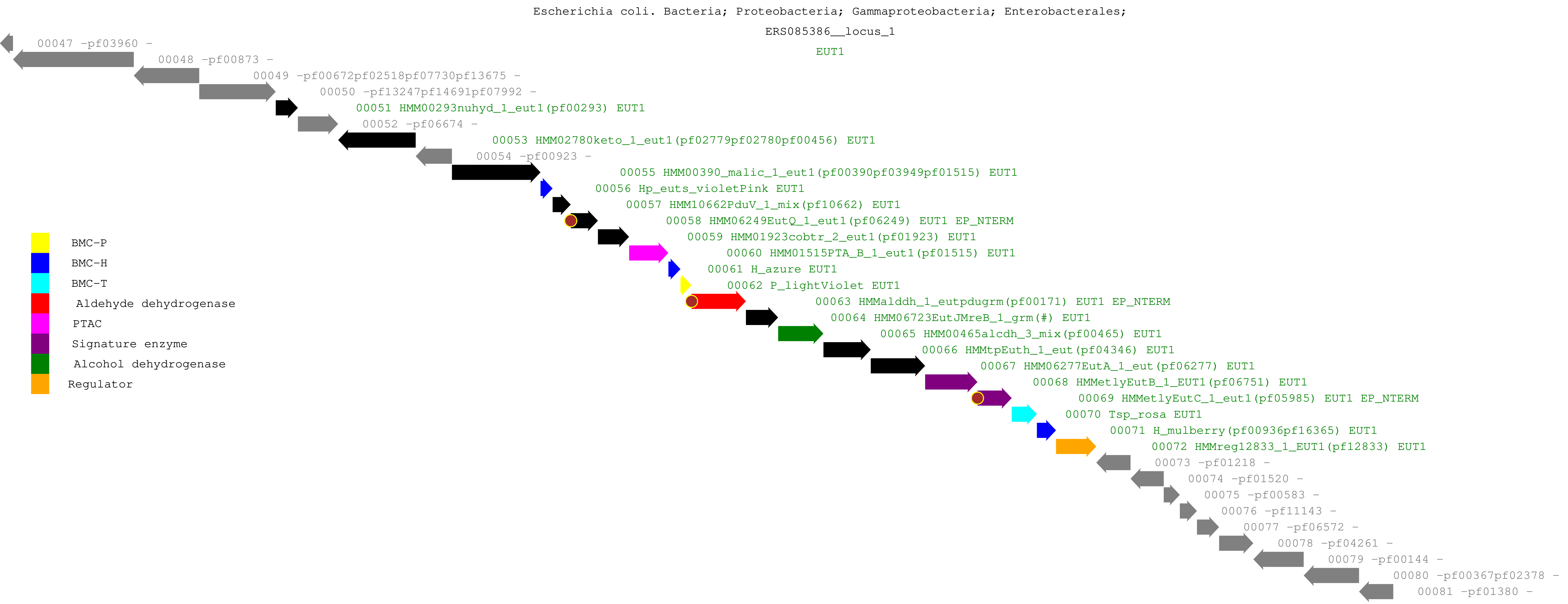

Escherichia coli. Bacteria; Proteobacteria; Gammaproteobacteria; Enterobacterales;
ERS085386__locus_1
EUT1
00047 -pf03960 -
00048 -pf00873 -
00049 -pf00672pf02518pf07730pf13675 -
00050 -pf13247pf14691pf07992 -
00051 HMM00293nuhyd_1_eut1(pf00293) EUT1
00052 -pf06674 -
00053 HMM02780keto_1_eut1(pf02779pf02780pf00456) EUT1
00054 -pf00923 -
00055 HMM00390_malic_1_eut1(pf00390pf03949pf01515) EUT1
00056 Hp_euts_violetPink EUT1
00057 HMM10662PduV_1_mix(pf10662) EUT1
00058 HMM06249EutQ_1_eut1(pf06249) EUT1 EP_NTERM
00059 HMM01923cobtr_2_eut1(pf01923) EUT1
00060 HMM01515PTA_B_1_eut1(pf01515) EUT1
00061 H_azure EUT1
00062 P_lightViolet EUT1
00063 HMMalddh_1_eutpdugrm(pf00171) EUT1 EP_NTERM
00064 HMM06723EutJMreB_1_grm(#) EUT1
00065 HMM00465alcdh_3_mix(pf00465) EUT1
00066 HMMtpEuth_1_eut(pf04346) EUT1
00067 HMM06277EutA_1_eut(pf06277) EUT1
00068 HMMetlyEutB_1_EUT1(pf06751) EUT1
00069 HMMetlyEutC_1_eut1(pf05985) EUT1 EP_NTERM
00070 Tsp_rosa EUT1
00071 H_mulberry(pf00936pf16365) EUT1
00072 HMMreg12833_1_EUT1(pf12833) EUT1
00073 -pf01218 -
00074 -pf01520 -
00075 -pf00583 -
00076 -pf11143 -
00077 -pf06572 -
00078 -pf04261 -
00079 -pf00144 -
00080 -pf00367pf02378 -
00081 -pf01380 -
BMC-P
BMC-H
BMC-T
Aldehyde dehydrogenase
PTAC
Signature enzyme
Alcohol dehydrogenase
Regulator

Escherichia coli. Bacteria; Proteobacteria; Gammaproteobacteria; Enterobacterales;

ERS085406__locus_1

EUT1

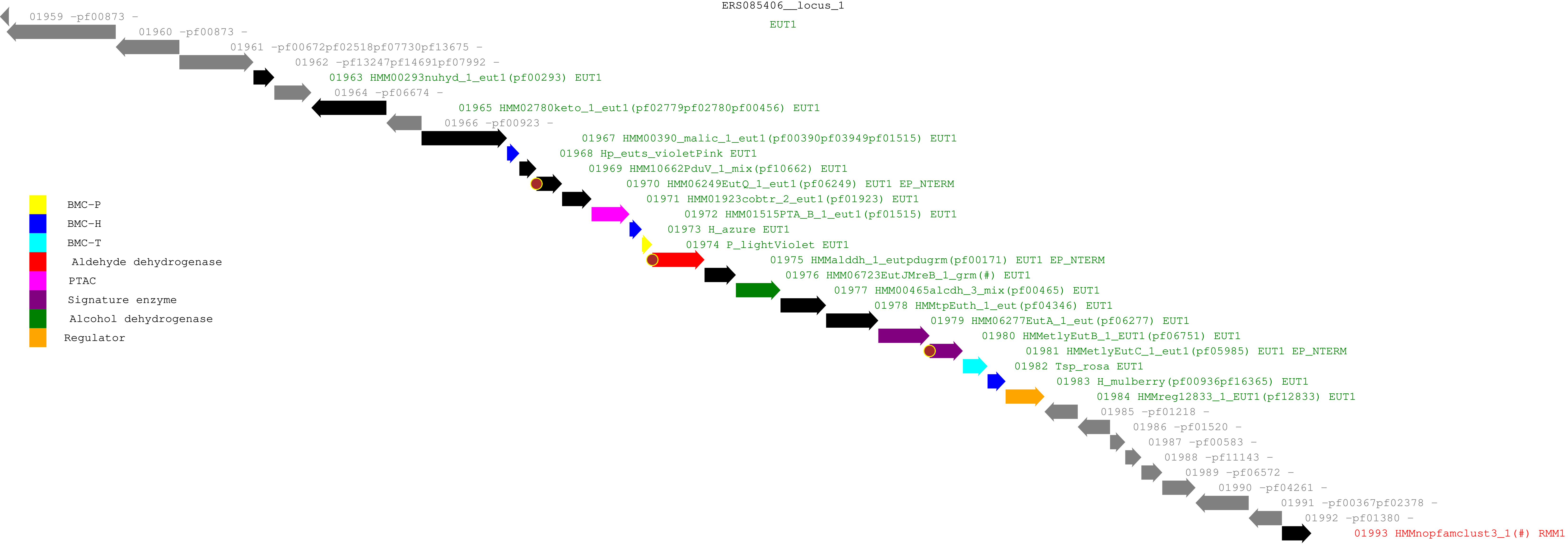

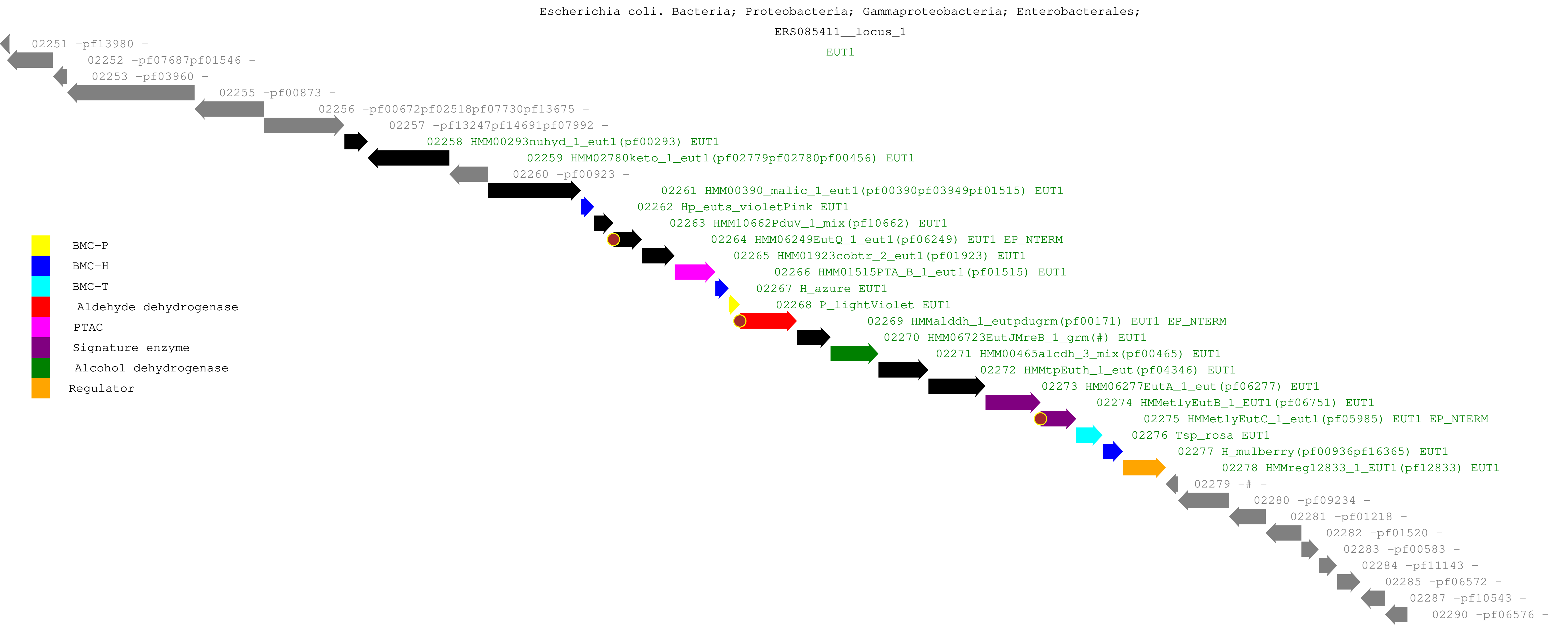

Escherichia coli. Bacteria; Proteobacteria; Gammaproteobacteria; Enterobacterales;
ERS085411__locus_1
EUT1
02251 -pf13980 -
02252 -pf07687pf01546 -
02253 -pf03960 -
02255 -pf00873 -
02256 -pf00672pf02518pf07730pf13675 -
02257 -pf13247pf14691pf07992 -
02258 HMM00293nuhyd_1_eut1(pf00293) EUT1
02259 HMM02780keto_1_eut1(pf02779pf02780pf00456) EUT1
02260 -pf00923 -
02261 HMM00390_malic_1_eut1(pf00390pf03949pf01515) EUT1
02262 Hp_euts_violetPink EUT1
02263 HMM10662PduV_1_mix(pf10662) EUT1
02264 HMM06249EutQ_1_eut1(pf06249) EUT1 EP_NTERM
02265 HMM01923cobtr_2_eut1(pf01923) EUT1
02266 HMM01515PTA_B_1_eut1(pf01515) EUT1
02267 H_azure EUT1
02268 P_lightViolet EUT1
02269 HMMalddh_1_eutpdugrm(pf00171) EUT1 EP_NTERM
02270 HMM06723EutJMreB_1_grm(#) EUT1
02271 HMM00465alcdh_3_mix(pf00465) EUT1
02272 HMMtpEuth_1_eut(pf04346) EUT1
02273 HMM06277EutA_1_eut(pf06277) EUT1
02274 HMMetlyEutB_1_EUT1(pf06751) EUT1
02275 HMMetlyEutC_1_eut1(pf05985) EUT1 EP_NTERM
02276 Tsp_rosa EUT1
02277 H_mulberry(pf00936pf16365) EUT1
02278 HMMreg12833_1_EUT1(pf12833) EUT1
02279 -# -
02280 -pf09234 -
02281 -pf01218 -
02282 -pf01520 -
02283 -pf00583 -
02284 -pf11143 -
02285 -pf06572 -
02287 -pf10543 -
02290 -pf06576 -
BMC-P
BMC-H
BMC-T
Aldehyde dehydrogenase
PTAC
Signature enzyme
Alcohol dehydrogenase
Regulator

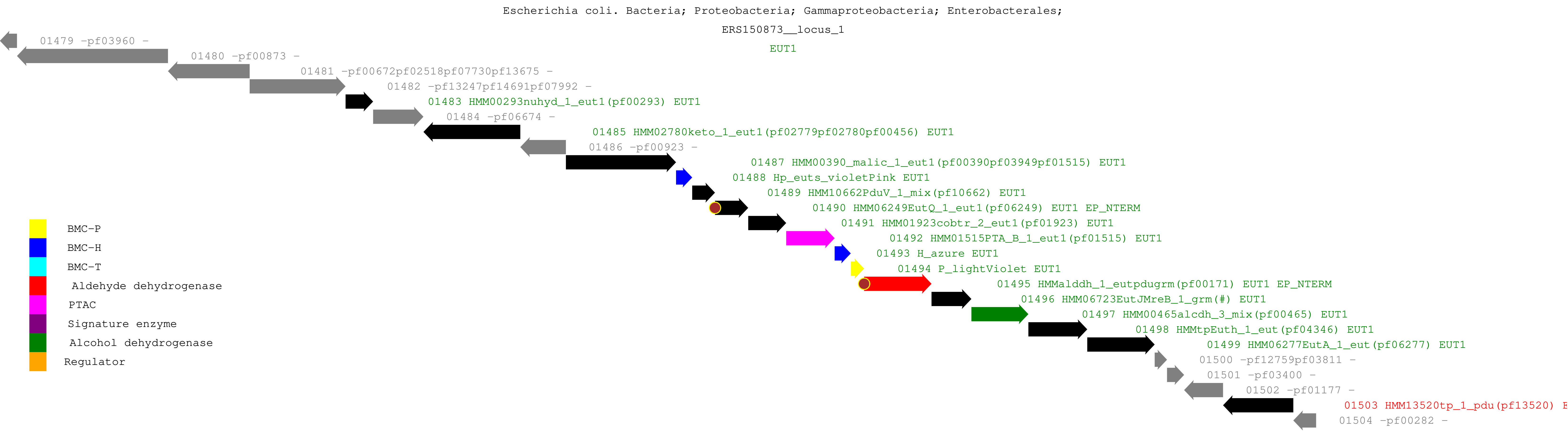
Escherichia coli. Bacteria; Proteobacteria; Gammaproteobacteria; Enterobacterales;
ERS150873__locus_1
EUT1
01479 -pf03960 -
01480 -pf00873 -
01481 -pf00672pf02518pf07730pf13675 -
01482 -pf13247pf14691pf07992 -
01483 HMM00293nuhyd_1_eut1(pf00293) EUT1
01484 -pf06674 -
01485 HMM02780keto_1_eut1(pf02779pf02780pf00456) EUT1
01486 -pf00923 -
01487 HMM00390_malic_1_eut1(pf00390pf03949pf01515) EUT1
01488 Hp_euts_violetPink EUT1
01489 HMM10662PduV_1_mix(pf10662) EUT1
01490 HMM06249EutQ_1_eut1(pf06249) EUT1 EP_NTERM
01491 HMM01923cobtr_2_eut1(pf01923) EUT1
01492 HMM01515PTA_B_1_eut1(pf01515) EUT1
01493 H_azure EUT1
01494 P_lightViolet EUT1
01495 HMMalddh_1_eutpdugrm(pf00171) EUT1 EP_NTERM
01496 HMM06723EutJMreB_1_grm(#) EUT1
01497 HMM00465alcdh_3_mix(pf00465) EUT1
01498 HMMtpEuth_1_eut(pf04346) EUT1
01499 HMM06277EutA_1_eut(pf06277) EUT1
01500 -pf12759pf03811 -
01501 -pf03400 -
01502 -pf01177 -
01503 HMM13520tp_1_pdu(pf13520) E
01504 -pf00282 -
BMC-P
BMC-H
BMC-T
Aldehyde dehydrogenase
PTAC
Signature enzyme
Alcohol dehydrogenase
Regulator

Salmonella enterica subsp. enterica serovar Typhimurium str. DT104. Bacteria; Proteobacteria; Gammaproteobacteria; Enterobacterales;

ERS157243__locus_1

EUT1

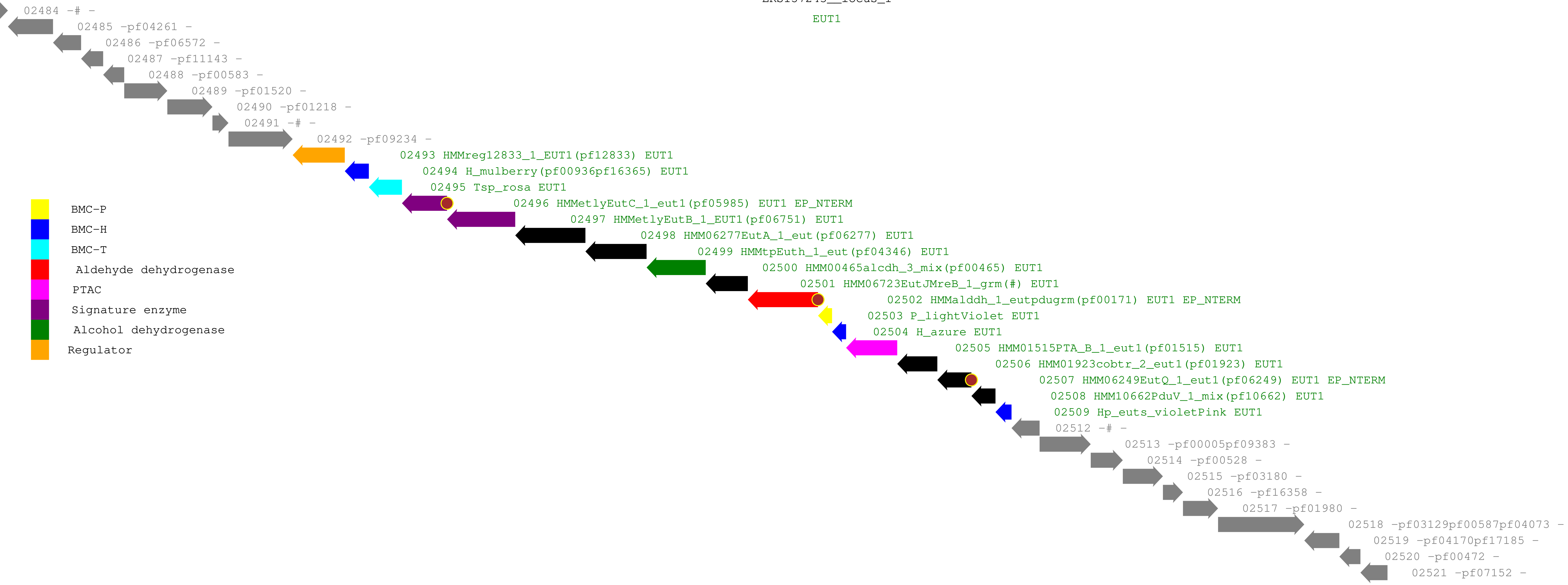

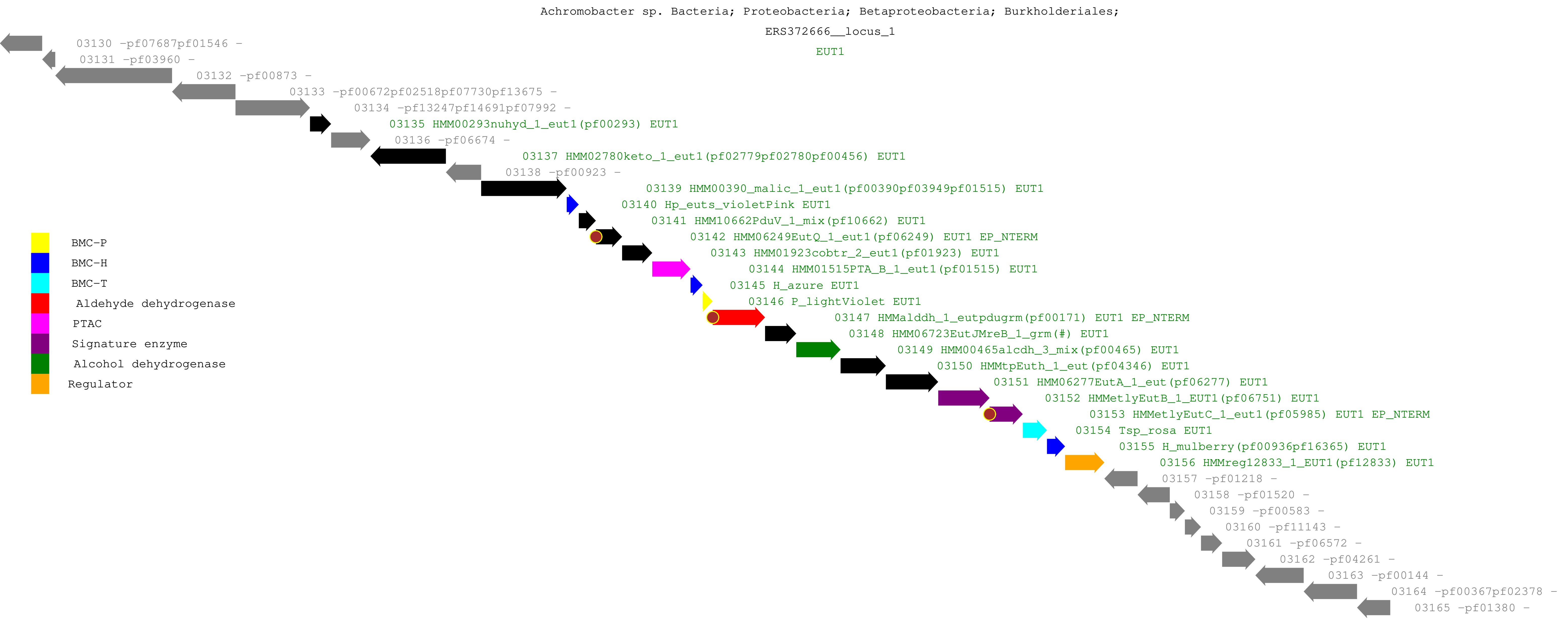

Achromobacter sp. Bacteria; Proteobacteria; Betaproteobacteria; Burkholderiales;
ERS372666__locus_1
EUT1
03130 -pf07687pf01546 -
03131 -pf03960 -
03132 -pf00873 -
03133 -pf00672pf02518pf07730pf13675 -
03134 -pf13247pf14691pf07992 -
03135 HMM00293nuhyd_1_eut1(pf00293) EUT1
03136 -pf06674 -
03137 HMM02780keto_1_eut1(pf02779pf02780pf00456) EUT1
03138 -pf00923 -
03139 HMM00390_malic_1_eut1(pf00390pf03949pf01515) EUT1
03140 Hp_euts_violetPink EUT1
03141 HMM10662PduV_1_mix(pf10662) EUT1
03142 HMM06249EutQ_1_eut1(pf06249) EUT1 EP_NTERM
03143 HMM01923cobtr_2_eut1(pf01923) EUT1
03144 HMM01515PTA_B_1_eut1(pf01515) EUT1
03145 H_azure EUT1
03146 P_lightViolet EUT1
03147 HMMalddh_1_eutpdugrm(pf00171) EUT1 EP_NTERM
03148 HMM06723EutJMreB_1_grm(#) EUT1
03149 HMM00465alcdh_3_mix(pf00465) EUT1
03150 HMMtpEuth_1_eut(pf04346) EUT1
03151 HMM06277EutA_1_eut(pf06277) EUT1
03152 HMMetlyEutB_1_EUT1(pf06751) EUT1
03153 HMMetlyEutC_1_eut1(pf05985) EUT1 EP_NTERM
03154 Tsp_rosa EUT1
03155 H_mulberry(pf00936pf16365) EUT1
03156 HMMreg12833_1_EUT1(pf12833) EUT1
03157 -pf01218 -
03158 -pf01520 -
03159 -pf00583 -
03160 -pf11143 -
03161 -pf06572 -
03162 -pf04261 -
03163 -pf00144 -
03164 -pf00367pf02378 -
03165 -pf01380 -
BMC-P
BMC-H
BMC-T
Aldehyde dehydrogenase
PTAC
Signature enzyme
Alcohol dehydrogenase
Regulator

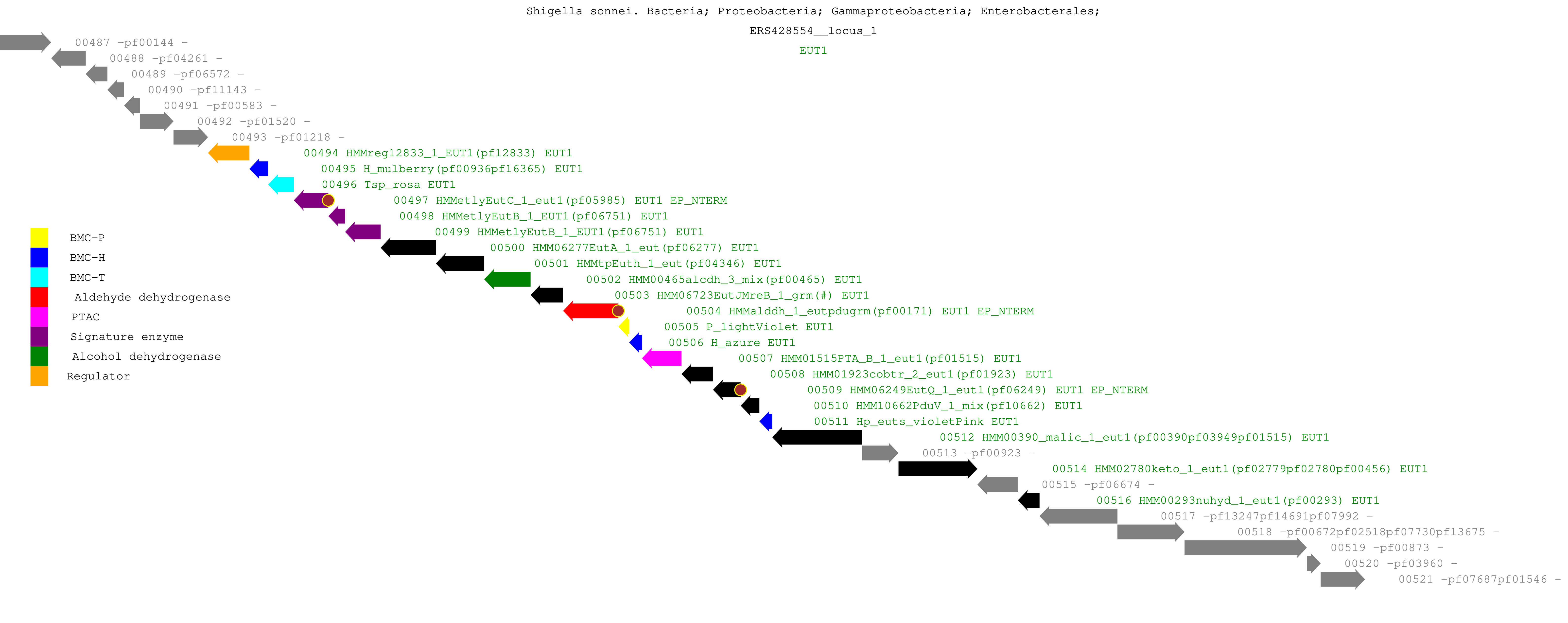
Shigella sonnei. Bacteria; Proteobacteria; Gammaproteobacteria; Enterobacterales;
ERS428554__locus_1
EUT1
00487 -pf00144 -
00488 -pf04261 -
00489 -pf06572 -
00490 -pf11143 -
00491 -pf00583 -
00492 -pf01520 -
00493 -pf01218 -
00494 HMMreg12833_1_EUT1(pf12833) EUT1
00495 H_mulberry(pf00936pf16365) EUT1
00496 Tsp_rosa EUT1
00497 HMMetlyEutC_1_eut1(pf05985) EUT1 EP_NTERM
00498 HMMetlyEutB_1_EUT1(pf06751) EUT1
00499 HMMetlyEutB_1_EUT1(pf06751) EUT1
00500 HMM06277EutA_1_eut(pf06277) EUT1
00501 HMMtpEuth_1_eut(pf04346) EUT1
00502 HMM00465alcdh_3_mix(pf00465) EUT1
00503 HMM06723EutJMreB_1_grm(#) EUT1
00504 HMMalddh_1_eutpdugrm(pf00171) EUT1 EP_NTERM
00505 P_lightViolet EUT1
00506 H_azure EUT1
00507 HMM01515PTA_B_1_eut1(pf01515) EUT1
00508 HMM01923cobtr_2_eut1(pf01923) EUT1
00509 HMM06249EutQ_1_eut1(pf06249) EUT1 EP_NTERM
00510 HMM10662PduV_1_mix(pf10662) EUT1
00511 Hp_euts_violetPink EUT1
00512 HMM00390_malic_1_eut1(pf00390pf03949pf01515) EUT1
00513 -pf00923 -
00514 HMM02780keto_1_eut1(pf02779pf02780pf00456) EUT1
00515 -pf06674 -
00516 HMM00293nuhyd_1_eut1(pf00293) EUT1
00517 -pf13247pf14691pf07992 -
00518 -pf00672pf02518pf07730pf13675 -
00519 -pf00873 -
00520 -pf03960 -
00521 -pf07687pf01546 -
BMC-P
BMC-H
BMC-T
Aldehyde dehydrogenase
PTAC
Signature enzyme
Alcohol dehydrogenase
Regulator

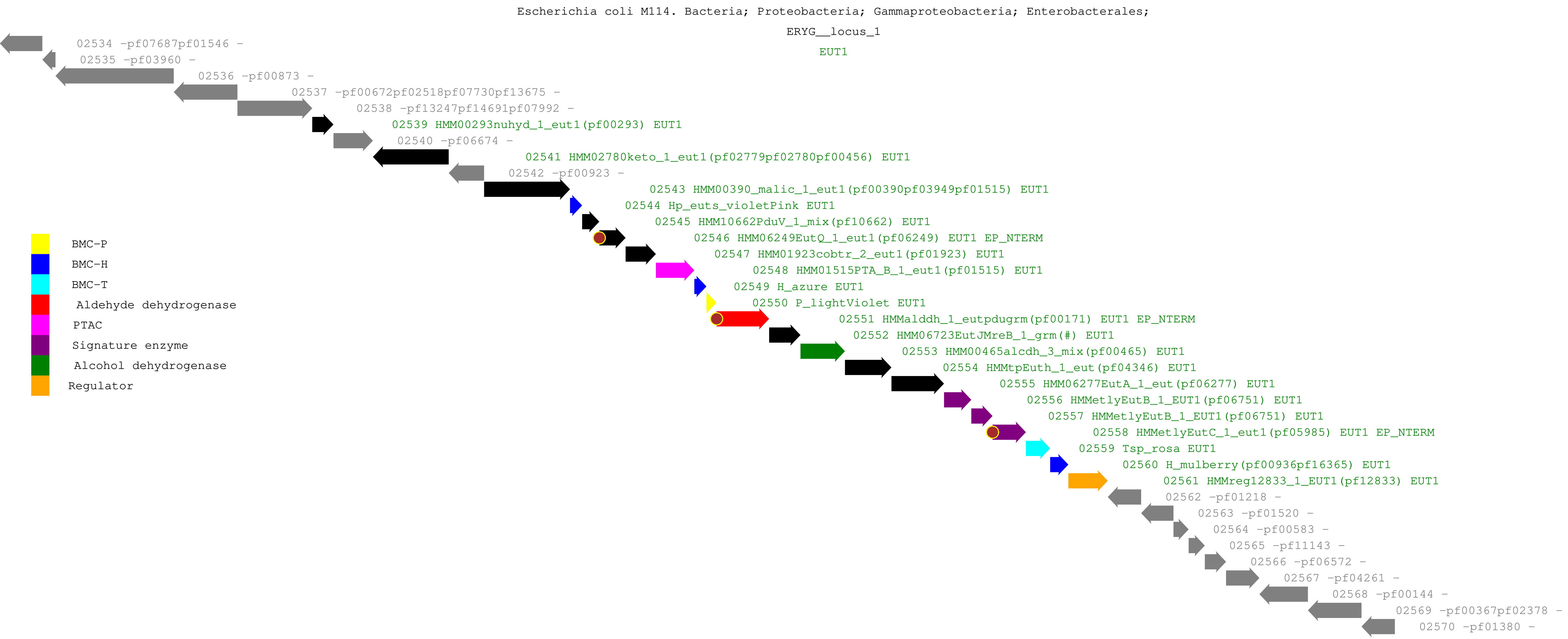
Escherichia coli M114. Bacteria; Proteobacteria; Gammaproteobacteria; Enterobacterales;
ERYG__locus_1
EUT1
02534 -pf07687pf01546 -
02535 -pf03960 -
02536 -pf00873 -
02537 -pf00672pf02518pf07730pf13675 -
02538 -pf13247pf14691pf07992 -
02539 HMM00293nuhyd_1_eut1(pf00293) EUT1
02540 -pf06674 -
02541 HMM02780keto_1_eut1(pf02779pf02780pf00456) EUT1
02542 -pf00923 -
02543 HMM00390_malic_1_eut1(pf00390pf03949pf01515) EUT1
02544 Hp_euts_violetPink EUT1
02545 HMM10662PduV_1_mix(pf10662) EUT1
02546 HMM06249EutQ_1_eut1(pf06249) EUT1 EP_NTERM
02547 HMM01923cobtr_2_eut1(pf01923) EUT1
02548 HMM01515PTA_B_1_eut1(pf01515) EUT1
02549 H_azure EUT1
02550 P_lightViolet EUT1
02551 HMMalddh_1_eutpdugrm(pf00171) EUT1 EP_NTERM
02552 HMM06723EutJMreB_1_grm(#) EUT1
02553 HMM00465alcdh_3_mix(pf00465) EUT1
02554 HMMtpEuth_1_eut(pf04346) EUT1
02555 HMM06277EutA_1_eut(pf06277) EUT1
02556 HMMetlyEutB_1_EUT1(pf06751) EUT1
02557 HMMetlyEutB_1_EUT1(pf06751) EUT1
02558 HMMetlyEutC_1_eut1(pf05985) EUT1 EP_NTERM
02559 Tsp_rosa EUT1
02560 H_mulberry(pf00936pf16365) EUT1
02561 HMMreg12833_1_EUT1(pf12833) EUT1
02562 -pf01218 -
02563 -pf01520 -
02564 -pf00583 -
02565 -pf11143 -
02566 -pf06572 -
02567 -pf04261 -
02568 -pf00144 -
02569 -pf00367pf02378 -
02570 -pf01380 -
BMC-P
BMC-H
BMC-T
Aldehyde dehydrogenase
PTAC
Signature enzyme
Alcohol dehydrogenase
Regulator

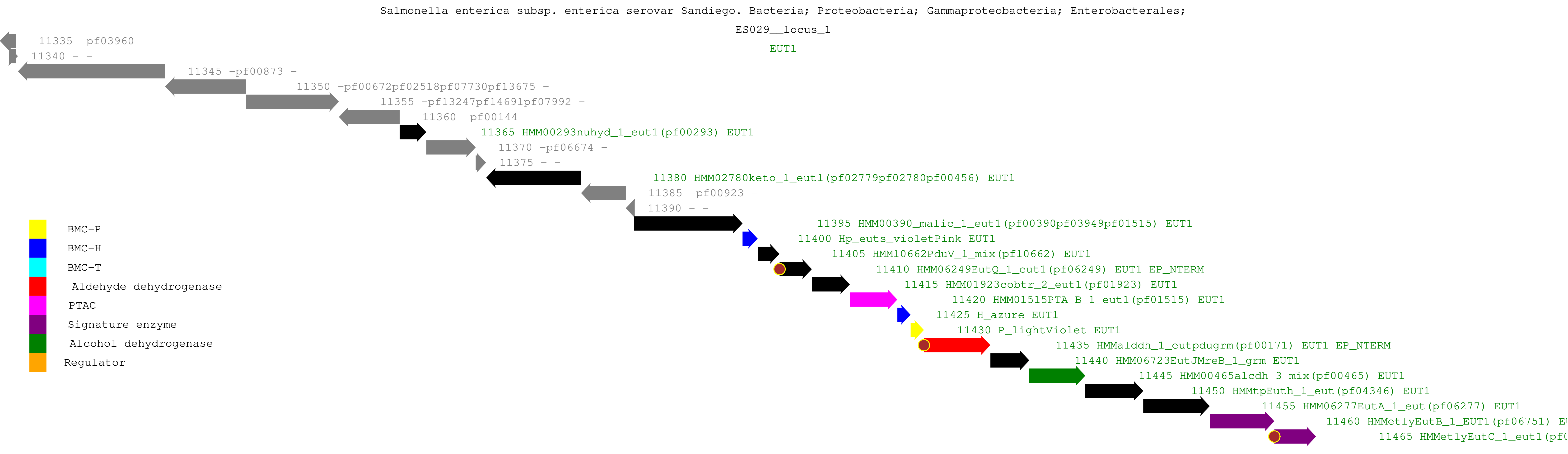
Salmonella enterica subsp. enterica serovar Sandiego. Bacteria; Proteobacteria; Gammaproteobacteria; Enterobacterales;
ES029__locus_1
EUT1
11335 -pf03960 -
11340 - -
11345 -pf00873 -
11350 -pf00672pf02518pf07730pf13675 -
11355 -pf13247pf14691pf07992 -
11360 -pf00144 -
11365 HMM00293nuhyd_1_eut1(pf00293) EUT1
11370 -pf06674 -
11375 - -
11380 HMM02780keto_1_eut1(pf02779pf02780pf00456) EUT1
11385 -pf00923 -
11390 - -
11395 HMM00390_malic_1_eut1(pf00390pf03949pf01515) EUT1
11400 Hp_euts_violetPink EUT1
11405 HMM10662PduV_1_mix(pf10662) EUT1
11410 HMM06249EutQ_1_eut1(pf06249) EUT1 EP_NTERM
11415 HMM01923cobtr_2_eut1(pf01923) EUT1
11420 HMM01515PTA_B_1_eut1(pf01515) EUT1
11425 H_azure EUT1
11430 P_lightViolet EUT1
11435 HMMalddh_1_eutpdugrm(pf00171) EUT1 EP_NTERM
11440 HMM06723EutJMreB_1_grm EUT1
11445 HMM00465alcdh_3_mix(pf00465) EUT1
11450 HMMtpEuth_1_eut(pf04346) EUT1
11455 HMM06277EutA_1_eut(pf06277) EUT1
11460 HMMetlyEutB_1_EUT1(pf06751) EU
11465 HMMetlyEutC_1_eut1(pf0
BMC-P
BMC-H
BMC-T
Aldehyde dehydrogenase
PTAC
Signature enzyme
Alcohol dehydrogenase
Regulator

Escherichia coli O78:H11 (strain H10407 / ETEC). Bacteria; Proteobacteria; Gammaproteobacteria; Enterobacterales;

ETEC__locus_1

EUT1

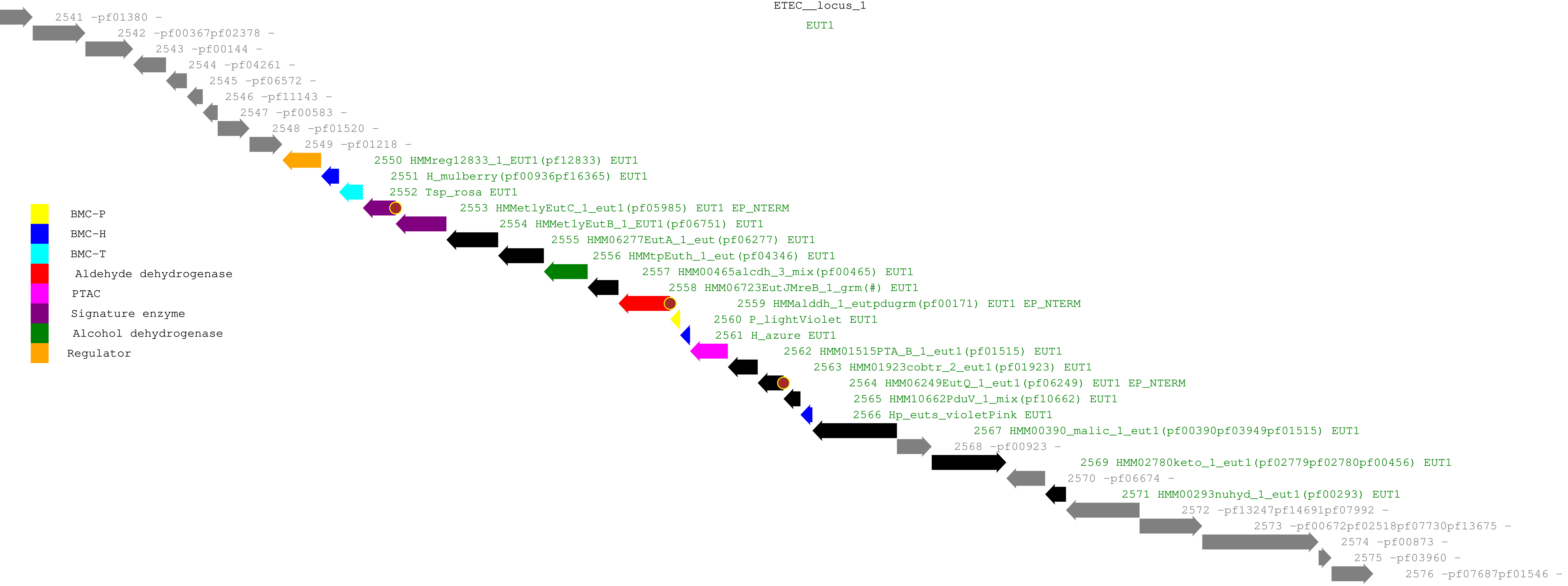

Salmonella enterica subsp. enterica serovar Bonn. Bacteria; Proteobacteria; Gammaproteobacteria; Enterobacterales;

EU363__locus_2

EUT1

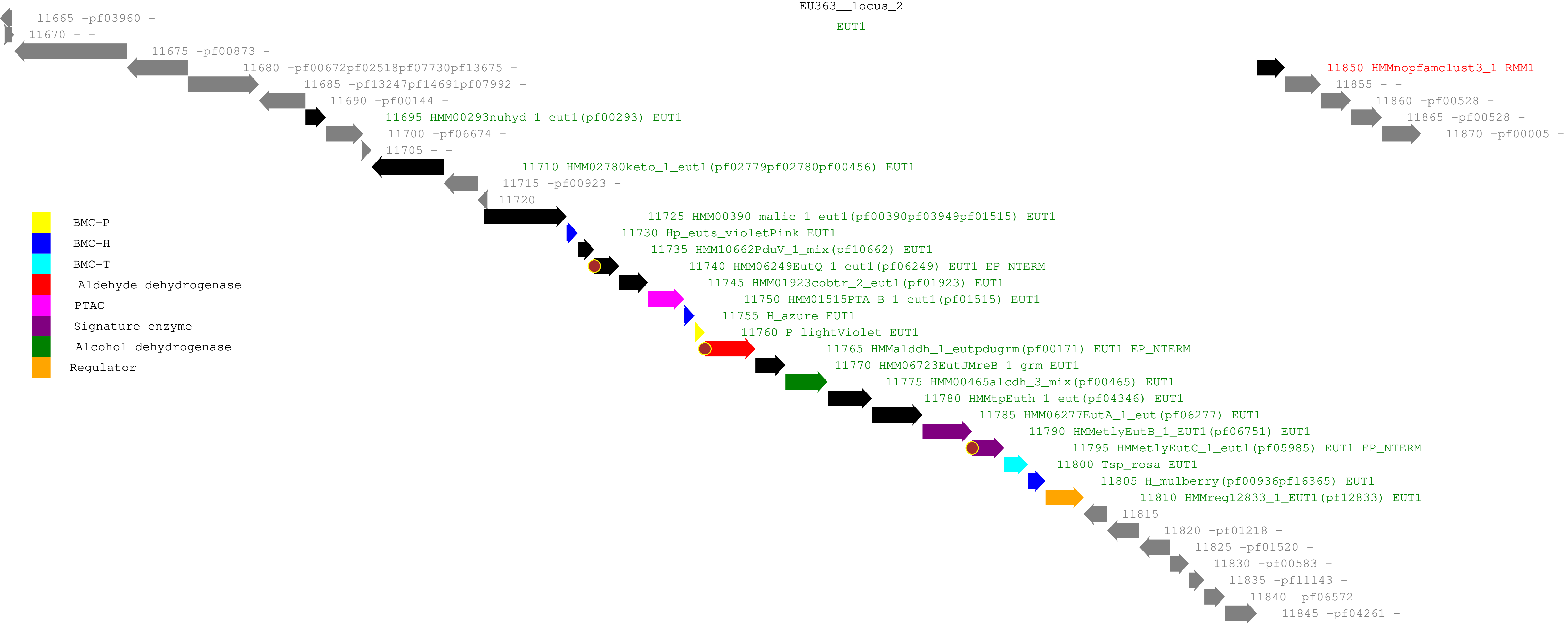

Salmonella enterica subsp. enterica serovar Neukoelln. Bacteria; Proteobacteria; Gammaproteobacteria; Enterobacterales;

EU895__locus_2

EUT1

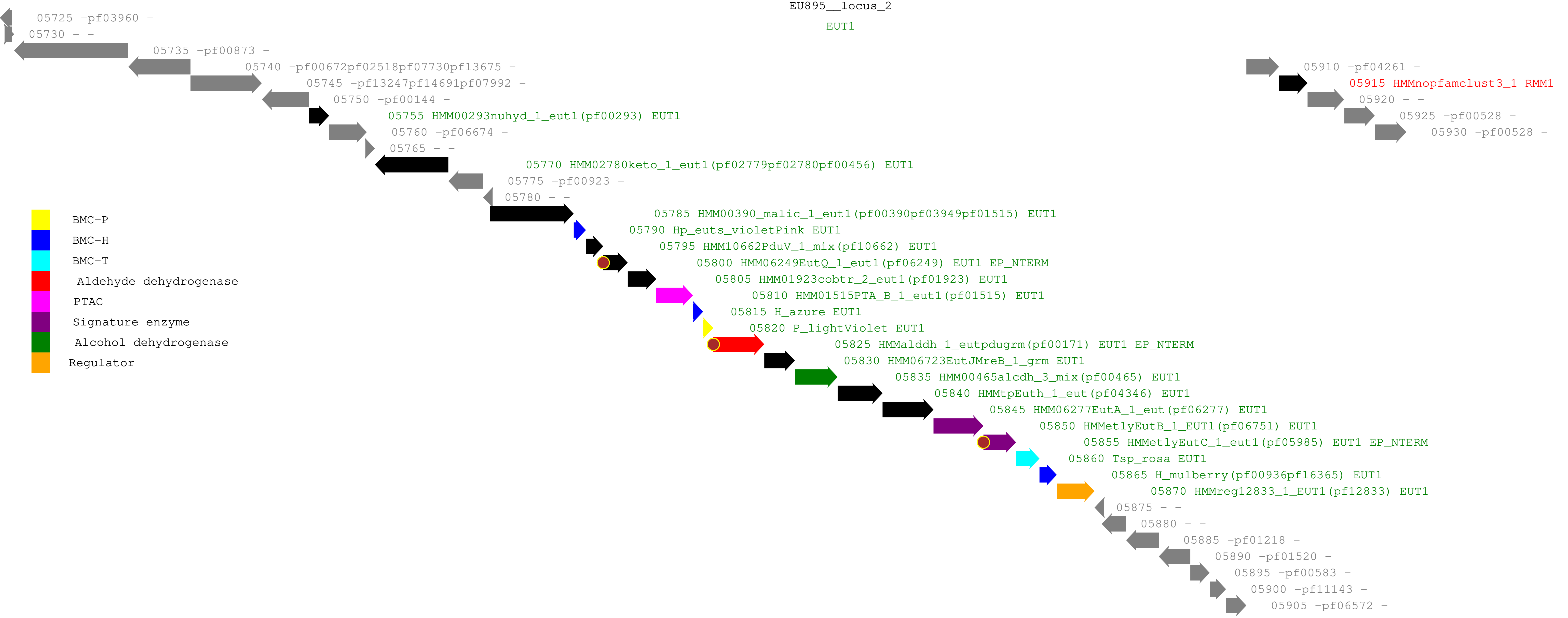

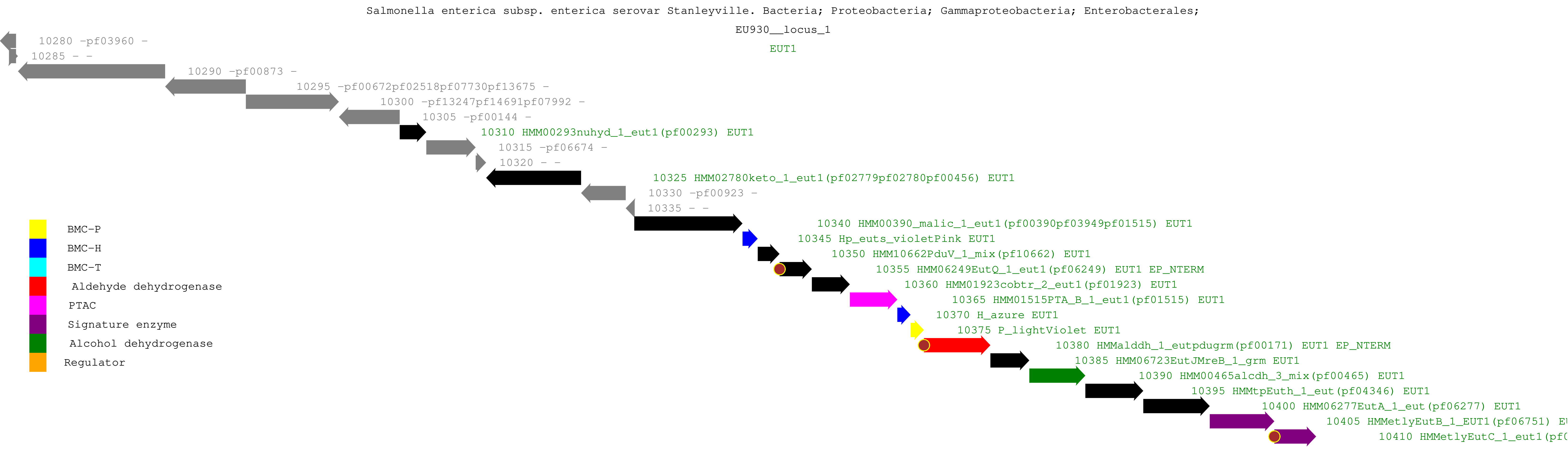
Salmonella enterica subsp. enterica serovar Stanleyville. Bacteria; Proteobacteria; Gammaproteobacteria; Enterobacterales;
EU930__locus_1
EUT1
10280 -pf03960 -
10285 - -
10290 -pf00873 -
10295 -pf00672pf02518pf07730pf13675 -
10300 -pf13247pf14691pf07992 -
10305 -pf00144 -
10310 HMM00293nuhyd_1_eut1(pf00293) EUT1
10315 -pf06674 -
10320 - -
10325 HMM02780keto_1_eut1(pf02779pf02780pf00456) EUT1
10330 -pf00923 -
10335 - -
10340 HMM00390_malic_1_eut1(pf00390pf03949pf01515) EUT1
10345 Hp_euts_violetPink EUT1
10350 HMM10662PduV_1_mix(pf10662) EUT1
10355 HMM06249EutQ_1_eut1(pf06249) EUT1 EP_NTERM
10360 HMM01923cobtr_2_eut1(pf01923) EUT1
10365 HMM01515PTA_B_1_eut1(pf01515) EUT1
10370 H_azure EUT1
10375 P_lightViolet EUT1
10380 HMMaldddh_1_eutpdugrm(pf00171) EUT1 EP_NTERM
10385 HMM06723EutJMreB_1_grm EUT1
10390 HMM00465alcdh_3_mix(pf00465) EUT1
10395 HMMtpEuth_1_eut(pf04346) EUT1
10400 HMM06277EutA_1_eut(pf06277) EUT1
10405 HMMetlyEutB_1_EUT1(pf06751)
10410 HMMetlyEutC_1_eut1(pf
BMC-P
BMC-H
BMC-T
Aldehyde dehydrogenase
PTAC
Signature enzyme
Alcohol dehydrogenase
Regulator

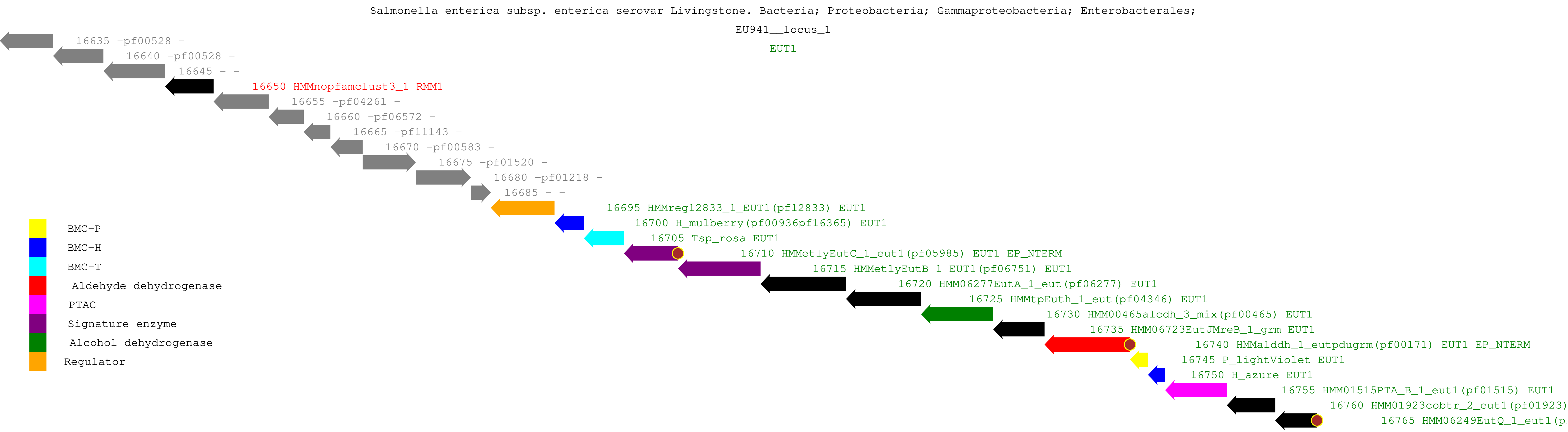

Salmonella enterica subsp. enterica serovar Livingstone. Bacteria; Proteobacteria; Gammaproteobacteria; Enterobacterales;
EU941__locus_1
EUT1
16635 -pf00528 -
16640 -pf00528 -
16645 - -
16650 HMMnopfamclust3_1 RMM1
16655 -pf04261 -
16660 -pf06572 -
16665 -pf11143 -
16670 -pf00583 -
16675 -pf01520 -
16680 -pf01218 -
16685 - -
16695 HMMreg12833_1_EUT1(pf12833) EUT1
16700 H_mulberry(pf00936pf16365) EUT1
16705 Tsp_rosa EUT1
16710 HMMetlyEutC_1_eut1(pf05985) EUT1 EP_NTERM
16715 HMMetlyEutB_1_EUT1(pf06751) EUT1
16720 HMM06277EutA_1_eut(pf06277) EUT1
16725 HMMtpEuth_1_eut(pf04346) EUT1
16730 HMM00465alcdh_3_mix(pf00465) EUT1
16735 HMM06723EutJMreB_1_grm EUT1
16740 HMMalddh_1_eutpdugrm(pf00171) EUT1 EP_NTERM
16745 P_lightViolet EUT1
16750 H_azure EUT1
16755 HMM01515PTA_B_1_eut1(pf01515) EUT1
16760 HMM01923cobtr_2_eut1(pf01923)
16765 HMM06249EutQ_1_eut1(p
BMC-P
BMC-H
BMC-T
Aldehyde dehydrogenase
PTAC
Signature enzyme
Alcohol dehydrogenase
Regulator

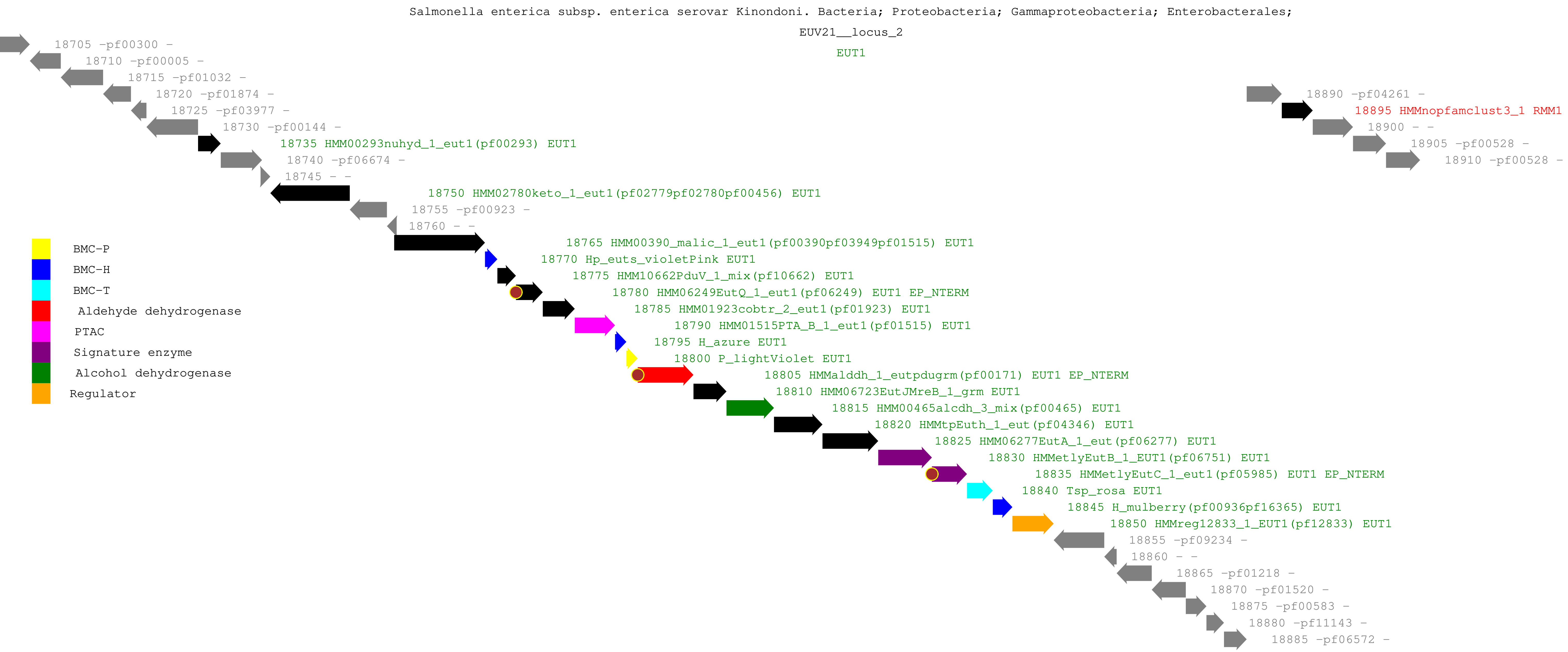
Salmonella enterica subsp. enterica serovar Kinondoni. Bacteria; Proteobacteria; Gammaproteobacteria; Enterobacterales;
EUV21__locus_2
EUT1
18705 -pf00300 -
18710 -pf00005 -
18715 -pf01032 -
18720 -pf01874 -
18725 -pf03977 -
18730 -pf00144 -
18735 HMM00293nuhyd_1_eut1(pf00293) EUT1
18740 -pf06674 -
18745 - -
18750 HMM02780keto_1_eut1(pf02779pf02780pf00456) EUT1
18755 -pf00923 -
18760 - -
18765 HMM00390_malic_1_eut1(pf00390pf03949pf01515) EUT1
18770 Hp_euts_violetPink EUT1
18775 HMM10662PduV_1_mix(pf10662) EUT1
18780 HMM06249EutQ_1_eut1(pf06249) EUT1 EP_NTERM
18785 HMM01923cobtr_2_eut1(pf01923) EUT1
18790 HMM01515PTA_B_1_eut1(pf01515) EUT1
18795 H_azure EUT1
18800 P_lightViolet EUT1
18805 HMMalddh_1_eutpdugrm(pf00171) EUT1 EP_NTERM
18810 HMM06723EutJMreB_1_grm EUT1
18815 HMM00465alcdh_3_mix(pf00465) EUT1
18820 HMMtpEuth_1_eut(pf04346) EUT1
18825 HMM06277EutA_1_eut(pf06277) EUT1
18830 HMMetlyEutB_1_EUT1(pf06751) EUT1
18835 HMMetlyEutC_1_eut1(pf05985) EUT1 EP_NTERM
18840 Tsp_rosa EUT1
18845 H_mulberry(pf00936pf16365) EUT1
18850 HMMreg12833_1_EUT1(pf12833) EUT1
18855 -pf09234 -
18860 - -
18865 -pf01218 -
18870 -pf01520 -
18875 -pf00583 -
18880 -pf11143 -
18885 -pf06572 -
18890 -pf04261 -
18895 HMMnopfamclust3_1 RMM1
18900 - -
18905 -pf00528 -
18910 -pf00528 -
BMC-P
BMC-H
BMC-T
Aldehyde dehydrogenase
PTAC
Signature enzyme
Alcohol dehydrogenase
Regulator

Salmonella enterica subsp. enterica serovar Carmel. Bacteria; Proteobacteria; Gammaproteobacteria; Enterobacterales;

EUW37__locus_2

EUT1

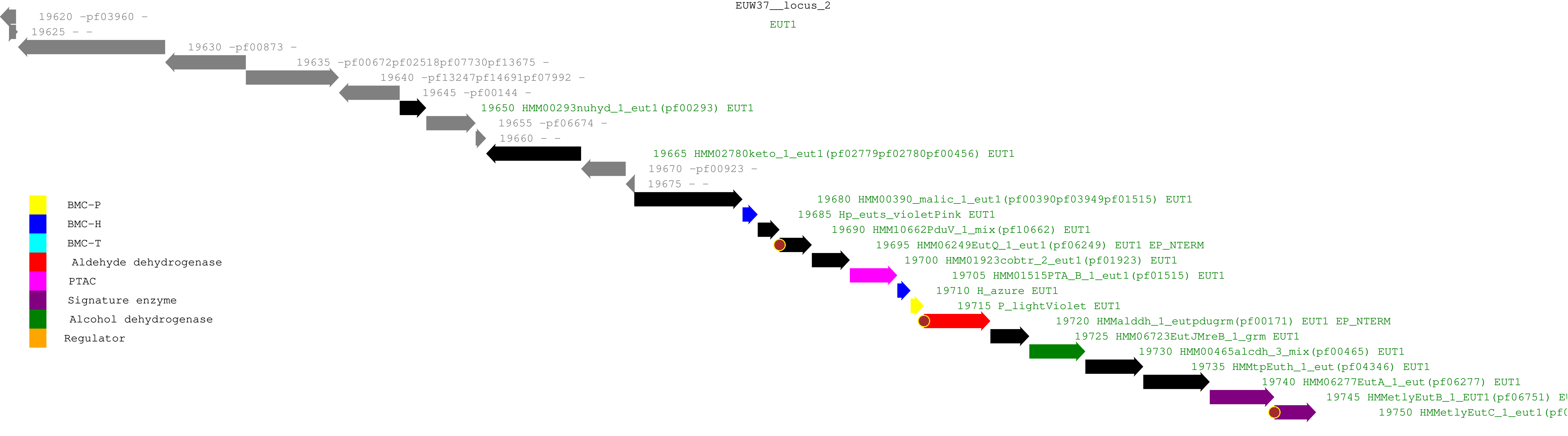

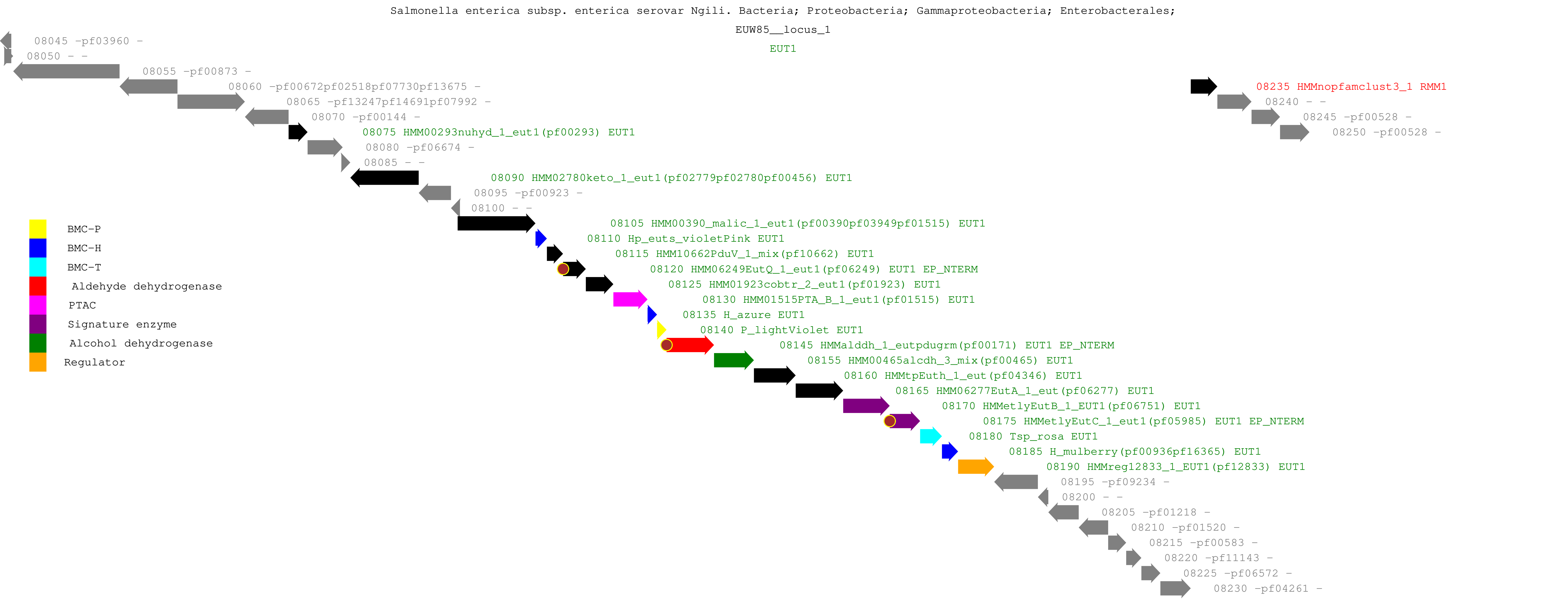

Salmonella enterica subsp. enterica serovar Ngili. Bacteria; Proteobacteria; Gammaproteobacteria; Enterobacterales;
EUW85__locus_1
EUT1
08045 -pf03960 -
08050 - -
08055 -pf00873 -
08060 -pf00672pf02518pf07730pf13675 -
08065 -pf13247pf14691pf07992 -
08070 -pf00144 -
08075 HMM00293nuhyd_1_eut1(pf00293) EUT1
08080 -pf06674 -
08085 - -
08090 HMM02780keto_1_eut1(pf02779pf02780pf00456) EUT1
08095 -pf00923 -
08100 - -
08105 HMM00390_malic_1_eut1(pf00390pf03949pf01515) EUT1
08110 Hp_euts_violetPink EUT1
08115 HMM10662PduV_1_mix(pf10662) EUT1
08120 HMM06249EutQ_1_eut1(pf06249) EUT1 EP_NTERM
08125 HMM01923cobtr_2_eut1(pf01923) EUT1
08130 HMM01515PTA_B_1_eut1(pf01515) EUT1
08135 H_azure EUT1
08140 P_lightViolet EUT1
08145 HMMalddh_1_eutpdugrm(pf00171) EUT1 EP_NTERM
08155 HMM00465alcdh_3_mix(pf00465) EUT1
08160 HMMtpEuth_1_eut(pf04346) EUT1
08165 HMM06277EutA_1_eut(pf06277) EUT1
08170 HMMetlyEutB_1_EUT1(pf06751) EUT1
08175 HMMetlyEutC_1_eut1(pf05985) EUT1 EP_NTERM
08180 Tsp_rosa EUT1
08185 H_mulberry(pf00936pf16365) EUT1
08190 HMMreg12833_1_EUT1(pf12833) EUT1
08195 -pf09234 -
08200 - -
08205 -pf01218 -
08210 -pf01520 -
08215 -pf00583 -
08220 -pf11143 -
08225 -pf06572 -
08230 -pf04261 -
08235 HMMnopfamclust3_1 RMM1
08240 - -
08245 -pf00528 -
08250 -pf00528 -
BMC-P
BMC-H
BMC-T
Aldehyde dehydrogenase
PTAC
Signature enzyme
Alcohol dehydrogenase
Regulator

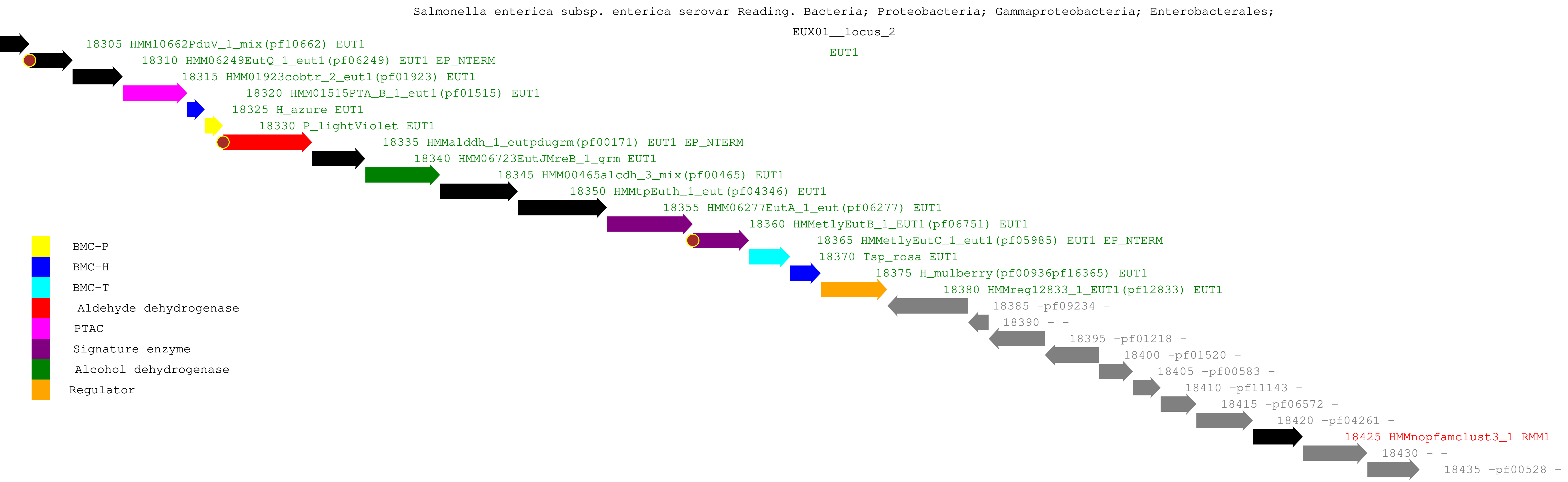
Salmonella enterica subsp. enterica serovar Reading. Bacteria; Proteobacteria; Gammaproteobacteria; Enterobacterales;
EUX01__locus_2
EUT1
18305 HMM10662PduV_1_mix(pf10662) EUT1
18310 HMM06249EutQ_1_eut1(pf06249) EUT1 EP_NTERM
18315 HMM01923cobtr_2_eut1(pf01923) EUT1
18320 HMM01515PTA_B_1_eut1(pf01515) EUT1
18325 H_azure EUT1
18330 P_lightViolet EUT1
18335 HMMalddh_1_eutpdugrm(pf00171) EUT1 EP_NTERM
18340 HMM06723EutJMreB_1_grm EUT1
18345 HMM00465alcdh_3_mix(pf00465) EUT1
18350 HMMtpEuth_1_eut(pf04346) EUT1
18355 HMM06277EutA_1_eut(pf06277) EUT1
18360 HMMetlyEutB_1_EUT1(pf06751) EUT1
18365 HMMetlyEutC_1_eut1(pf05985) EUT1 EP_NTERM
18370 Tsp_rosa EUT1
18375 H_mulberry(pf00936pf16365) EUT1
18380 HMMreg12833_1_EUT1(pf12833) EUT1
18385 -pf09234 -
18390 - -
18395 -pf01218 -
18400 -pf01520 -
18405 -pf00583 -
18410 -pf11143 -
18415 -pf06572 -
18420 -pf04261 -
18425 HMMnopfamclust3_1 RMM1
18430 - -
18435 -pf00528 -
BMC-P
BMC-H
BMC-T
Aldehyde dehydrogenase
PTAC
Signature enzyme
Alcohol dehydrogenase
Regulator

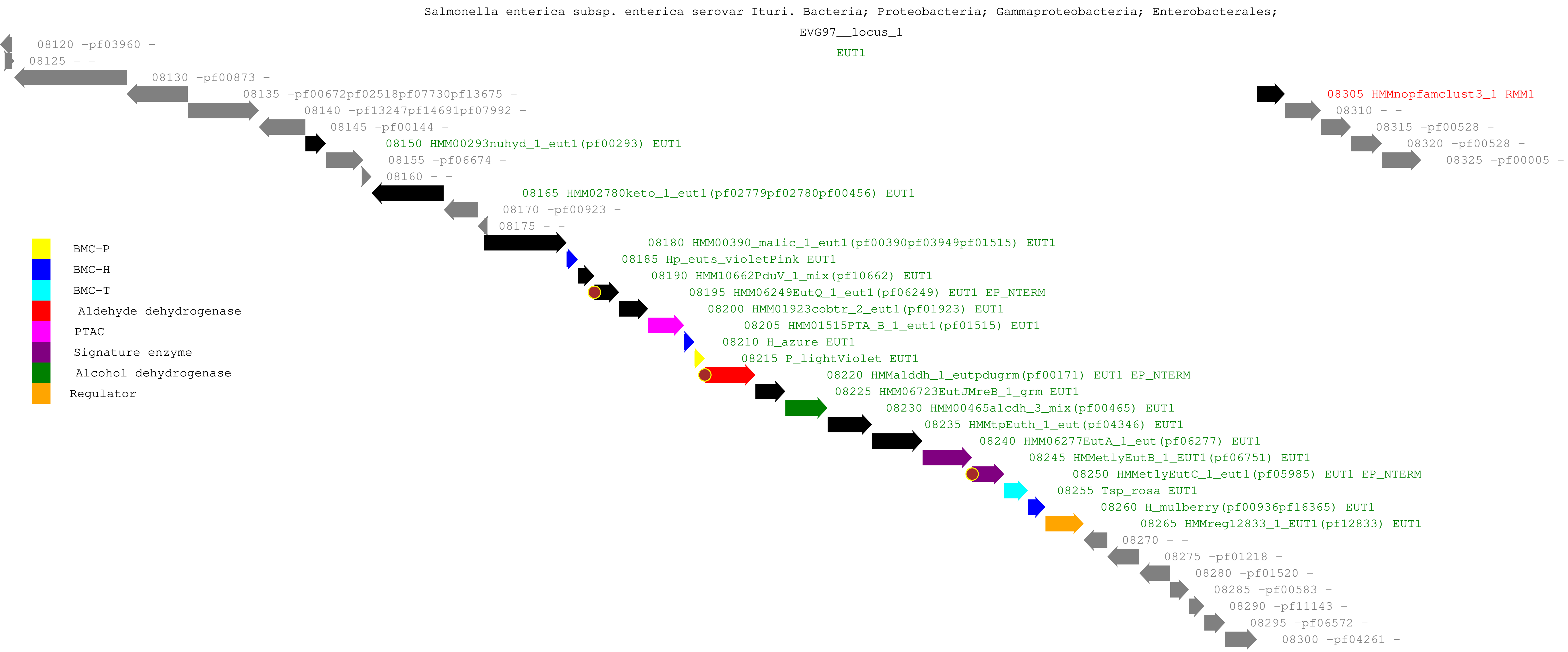
Salmonella enterica subsp. enterica serovar Ituri. Bacteria; Proteobacteria; Gammaproteobacteria; Enterobacterales;
EVG97__locus_1
EUT1
08120 -pf03960 -
08125 - -
08130 -pf00873 -
08135 -pf00672pf02518pf07730pf13675 -
08140 -pf13247pf14691pf07992 -
08145 -pf00144 -
08150 HMM00293nuhyd_1_eut1(pf00293) EUT1
08155 -pf06674 -
08160 - -
08165 HMM02780keto_1_eut1(pf02779pf02780pf00456) EUT1
08170 -pf00923 -
08175 - -
08180 HMM00390_malic_1_eut1(pf00390pf03949pf01515) EUT1
08185 Hp_euts_violetPink EUT1
08190 HMM10662PduV_1_mix(pf10662) EUT1
08195 HMM06249EutQ_1_eut1(pf06249) EUT1 EP_NTERM
08200 HMM01923cobtr_2_eut1(pf01923) EUT1
08205 HMM01515PTA_B_1_eut1(pf01515) EUT1
08210 H_azure EUT1
08215 P_lightViolet EUT1
08220 HMMalddh_1_eutpdugrm(pf00171) EUT1 EP_NTERM
08225 HMM06723EutJMreB_1_grm EUT1
08230 HMM00465alcdh_3_mix(pf00465) EUT1
08235 HMMtpEuth_1_eut(pf04346) EUT1
08240 HMM06277EutA_1_eut(pf06277) EUT1
08245 HMMetlyEutB_1_EUT1(pf06751) EUT1
08250 HMMetlyEutC_1_eut1(pf05985) EUT1 EP_NTERM
08255 Tsp_rosa EUT1
08260 H_mulberry(pf00936pf16365) EUT1
08265 HMMreg12833_1_EUT1(pf12833) EUT1
08270 - -
08275 -pf01218 -
08280 -pf01520 -
08285 -pf00583 -
08290 -pf11143 -
08295 -pf06572 -
08300 -pf04261 -
08305 HMMnopfamclust3_1 RMM1
08310 - -
08315 -pf00528 -
08320 -pf00528 -
08325 -pf00005 -
BMC-P
BMC-H
BMC-T
Aldehyde dehydrogenase
PTAC
Signature enzyme
Alcohol dehydrogenase
Regulator

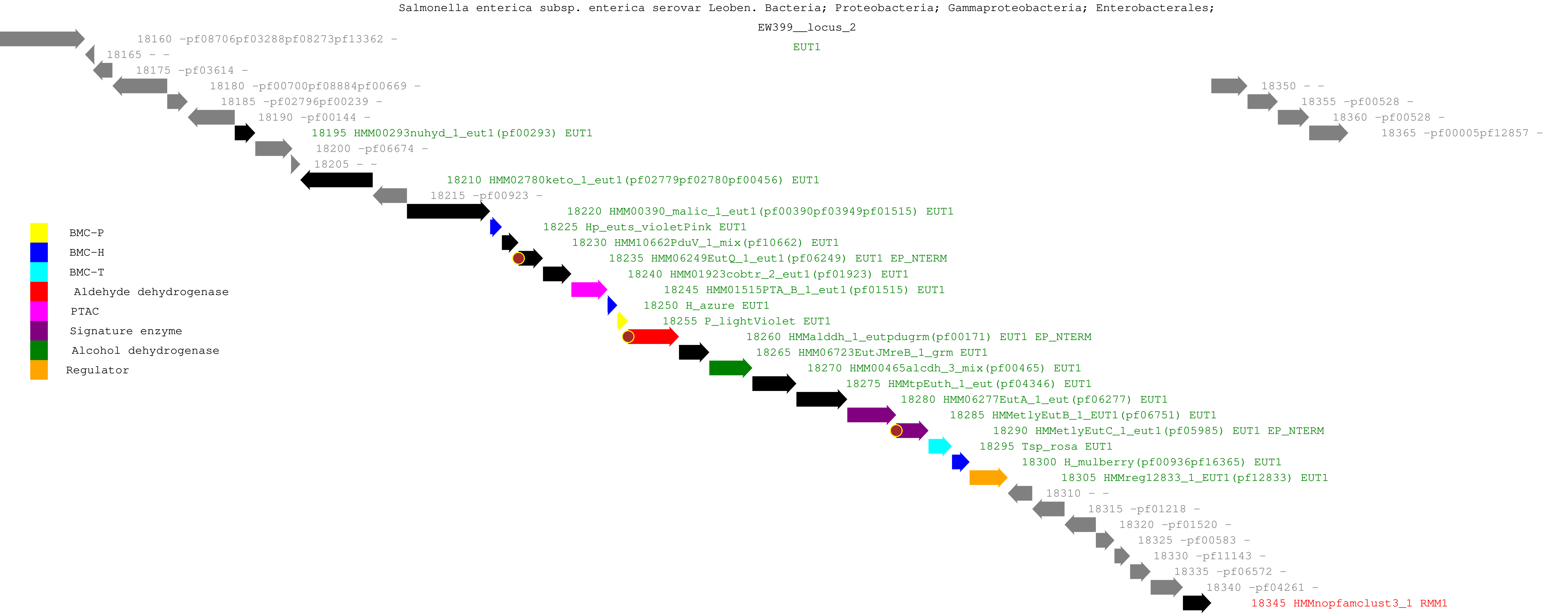
Salmonella enterica subsp. enterica serovar Leoben. Bacteria; Proteobacteria; Gammaproteobacteria; Enterobacterales;
EW399__locus_2
EUT1
18160 -pf08706pf03288pf08273pf13362 -
18165 - -
18175 -pf03614 -
18180 -pf00700pf08884pf00669 -
18185 -pf02796pf00239 -
18190 -pf00144 -
18195 HMM00293nuhyd_1_eut1(pf00293) EUT1
18200 -pf06674 -
18205 - -
18210 HMM02780keto_1_eut1(pf02779pf02780pf00456) EUT1
18215 -pf00923 -
18220 HMM00390_malic_1_eut1(pf00390pf03949pf01515) EUT1
18225 Hp_euts_violetPink EUT1
18230 HMM10662PduV_1_mix(pf10662) EUT1
18235 HMM06249EutQ_1_eut1(pf06249) EUT1 EP_NTERM
18240 HMM01923cobtr_2_eut1(pf01923) EUT1
18245 HMM01515PTA_B_1_eut1(pf01515) EUT1
18250 H_azure EUT1
18255 P_lightViolet EUT1
18260 HMMalddh_1_eutpdugrm(pf00171) EUT1 EP_NTERM
18265 HMM06723EutJMreB_1_grm EUT1
18270 HMM00465alcdh_3_mix(pf00465) EUT1
18275 HMMtpEuth_1_eut(pf04346) EUT1
18280 HMM06277EutA_1_eut(pf06277) EUT1
18285 HMMetlyEutB_1_EUT1(pf06751) EUT1
18290 HMMetlyEutC_1_eut1(pf05985) EUT1 EP_NTERM
18295 Tsp_rosa EUT1
18300 H_mulberry(pf00936pf16365) EUT1
18305 HMMreg12833_1_EUT1(pf12833) EUT1
18310 - -
18315 -pf01218 -
18320 -pf01520 -
18325 -pf00583 -
18330 -pf11143 -
18335 -pf06572 -
18340 -pf04261 -
18345 HMMnopfamclust3_1 RMM1
18350 - -
18355 -pf00528 -
18360 -pf00528 -
18365 -pf00005pf12857 -
BMC-P
BMC-H
BMC-T
Aldehyde dehydrogenase
PTAC
Signature enzyme
Alcohol dehydrogenase
Regulator

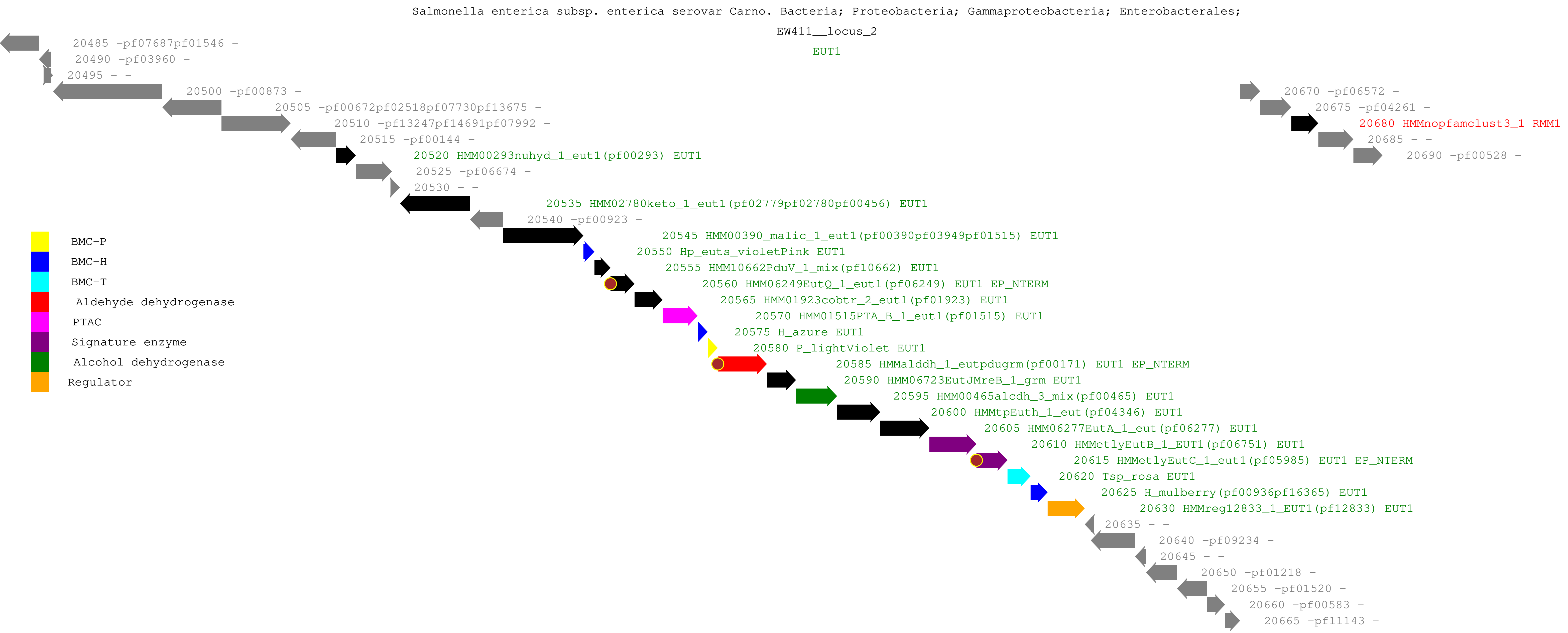
Salmonella enterica subsp. enterica serovar Carno. Bacteria; Proteobacteria; Gammaproteobacteria; Enterobacterales;
EW411__locus_2
EUT1
20485 -pf07687pf01546 -
20490 -pf03960 -
20495 - -
20500 -pf00873 -
20505 -pf00672pf02518pf07730pf13675 -
20510 -pf13247pf14691pf07992 -
20515 -pf00144 -
20520 HMM00293nuhyd_1_eut1(pf00293) EUT1
20525 -pf06674 -
20530 - -
20535 HMM02780keto_1_eut1(pf02779pf02780pf00456) EUT1
20540 -pf00923 -
20545 HMM00390_malic_1_eut1(pf00390pf03949pf01515) EUT1
20550 Hp_euts_violetPink EUT1
20555 HMM10662PduV_1_mix(pf10662) EUT1
20560 HMM06249EutQ_1_eut1(pf06249) EUT1 EP_NTERM
20565 HMM01923cobtr_2_eut1(pf01923) EUT1
20570 HMM01515PTA_B_1_eut1(pf01515) EUT1
20575 H_azure EUT1
20580 P_lightViolet EUT1
20585 HMMalddh_1_eutpdugrm(pf00171) EUT1 EP_NTERM
20590 HMM06723EutJMreB_1_grm EUT1
20595 HMM00465alcdh_3_mix(pf00465) EUT1
20600 HMMtpEuth_1_eut(pf04346) EUT1
20605 HMM06277EutA_1_eut(pf06277) EUT1
20610 HMMetlyEutB_1_EUT1(pf06751) EUT1
20615 HMMetlyEutC_1_eut1(pf05985) EUT1 EP_NTERM
20620 Tsp_rosa EUT1
20625 H_mulberry(pf00936pf16365) EUT1
20630 HMMreg12833_1_EUT1(pf12833) EUT1
20635 - -
20640 -pf09234 -
20645 - -
20650 -pf01218 -
20655 -pf01520 -
20660 -pf00583 -
20665 -pf11143 -
20670 -pf06572 -
20675 -pf04261 -
20680 HMMnopfamclust3_1 RMM1
20685 - -
20690 -pf00528 -
BMC-P
BMC-H
BMC-T
Aldehyde dehydrogenase
PTAC
Signature enzyme
Alcohol dehydrogenase
Regulator

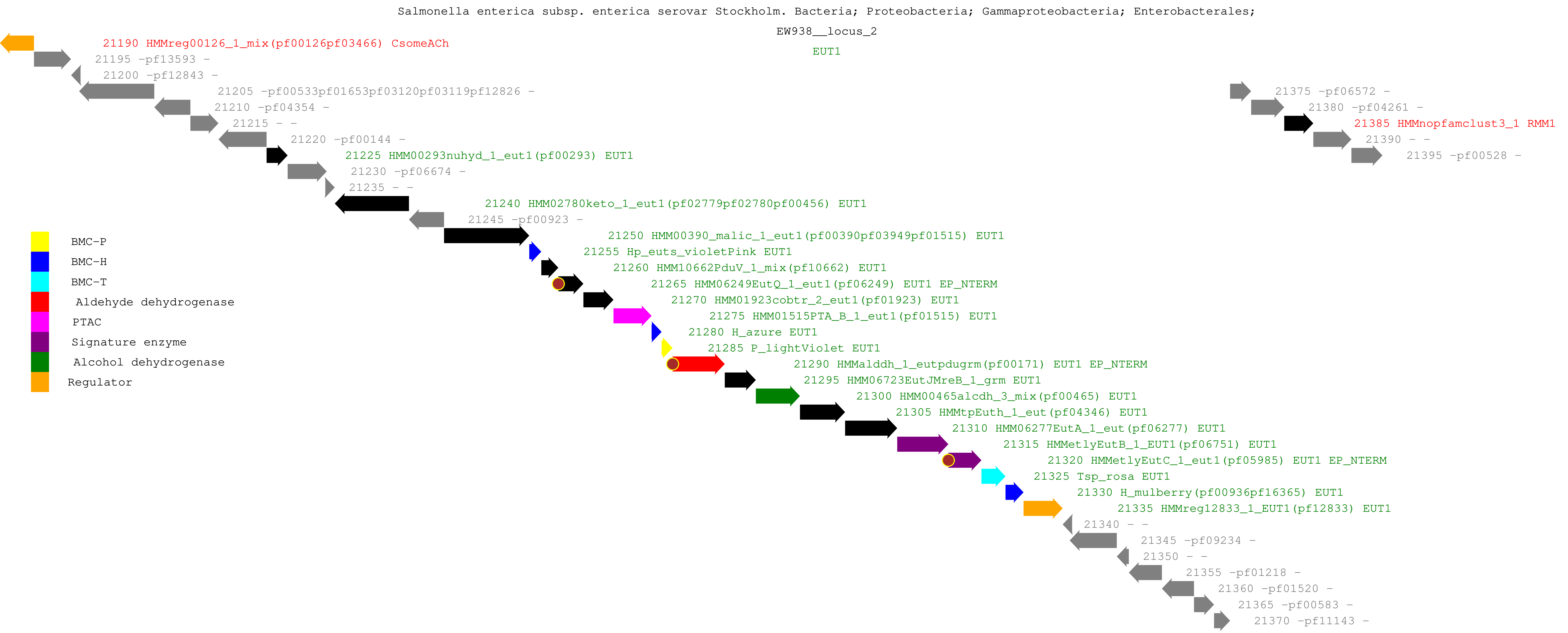

Salmonella enterica subsp. enterica serovar Stockholm. Bacteria; Proteobacteria; Gammaproteobacteria; Enterobacterales;
EW938__locus_2
EUT1
21190 HMMreg00126_1_mix(pf00126pf03466) CsomeACh
21195 -pf13593 -
21200 -pf12843 -
21205 -pf00533pf01653pf03120pf03119pf12826 -
21210 -pf04354 -
21215 - -
21220 -pf00144 -
21225 HMM00293nuhyd_1_eut1(pf00293) EUT1
21230 -pf06674 -
21235 - -
21240 HMM02780keto_1_eut1(pf02779pf02780pf00456) EUT1
21245 -pf00923 -
21250 HMM00390_malic_1_eut1(pf00390pf03949pf01515) EUT1
21255 Hp_euts_violetPink EUT1
21260 HMM10662PduV_1_mix(pf10662) EUT1
21265 HMM06249EutQ_1_eut1(pf06249) EUT1 EP_NTERM
21270 HMM01923cobtr_2_eut1(pf01923) EUT1
21275 HMM01515PTA_B_1_eut1(pf01515) EUT1
21280 H_azure EUT1
21285 P_lightViolet EUT1
21290 HMMaldd_1_eutpdugrm(pf00171) EUT1 EP_NTERM
21295 HMM06723EutJMreB_1_grm EUT1
21300 HMM00465alcdh_3_mix(pf00465) EUT1
21305 HMMtpEuth_1_eut(pf04346) EUT1
21310 HMM06277EutA_1_eut(pf06277) EUT1
21315 HMMetlyEutB_1_EUT1(pf06751) EUT1
21320 HMMetlyEutC_1_eut1(pf05985) EUT1 EP_NTERM
21325 Tsp_rosa EUT1
21330 H_mulberry(pf00936pf16365) EUT1
21335 HMMreg12833_1_EUT1(pf12833) EUT1
21340 - -
21345 -pf09234 -
21350 - -
21355 -pf01218 -
21360 -pf01520 -
21365 -pf00583 -
21370 -pf11143 -
21375 -pf06572 -
21380 -pf04261 -
21385 HMMnopfamclust3_1 RMM1
21390 - -
21395 -pf00528 -
BMC-P
BMC-H
BMC-T
Aldehyde dehydrogenase
PTAC
Signature enzyme
Alcohol dehydrogenase
Regulator

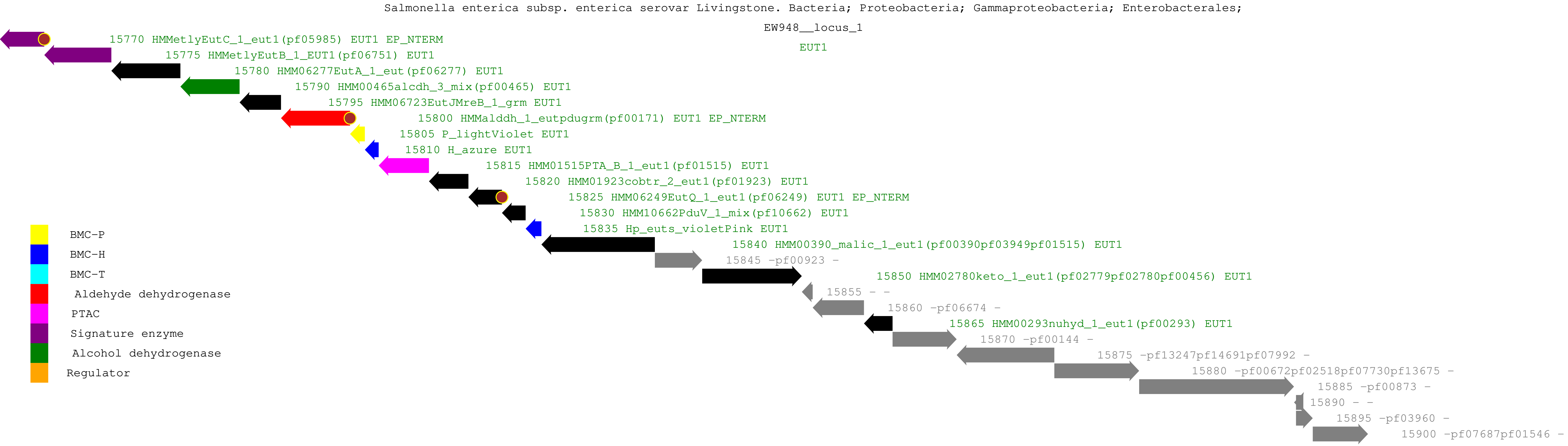
Salmonella enterica subsp. enterica serovar Livingstone. Bacteria; Proteobacteria; Gammaproteobacteria; Enterobacterales;
EW948__locus_1
EUT1
15770 HMMetlyEutC_1_eut1(pf05985) EUT1 EP_NTERM
15775 HMMetlyEutB_1_EUT1(pf06751) EUT1
15780 HMM06277EutA_1_eut(pf06277) EUT1
15790 HMM00465alcdh_3_mix(pf00465) EUT1
15795 HMM06723EutJMreB_1_grm EUT1
15800 HMMalddh_1_eutpdugrm(pf00171) EUT1 EP_NTERM
15805 P_lightViolet EUT1
15810 H_azure EUT1
15815 HMM01515PTA_B_1_eut1(pf01515) EUT1
15820 HMM01923cobtr_2_eut1(pf01923) EUT1
15825 HMM06249EutQ_1_eut1(pf06249) EUT1 EP_NTERM
15830 HMM10662PduV_1_mix(pf10662) EUT1
15835 Hp_euts_violetPink EUT1
15840 HMM00390_malic_1_eut1(pf00390pf03949pf01515) EUT1
15845 -pf00923 -
15850 HMM02780keto_1_eut1(pf02779pf02780pf00456) EUT1
15855 - -
15860 -pf06674 -
15865 HMM00293nuhyd_1_eut1(pf00293) EUT1
15870 -pf00144 -
15875 -pf13247pf14691pf07992 -
15880 -pf00672pf02518pf07730pf13675 -
15885 -pf00873 -
15890 - -
15895 -pf03960 -
15900 -pf07687pf01546 -
BMC-P
BMC-H
BMC-T
Aldehyde dehydrogenase
PTAC
Signature enzyme
Alcohol dehydrogenase
Regulator

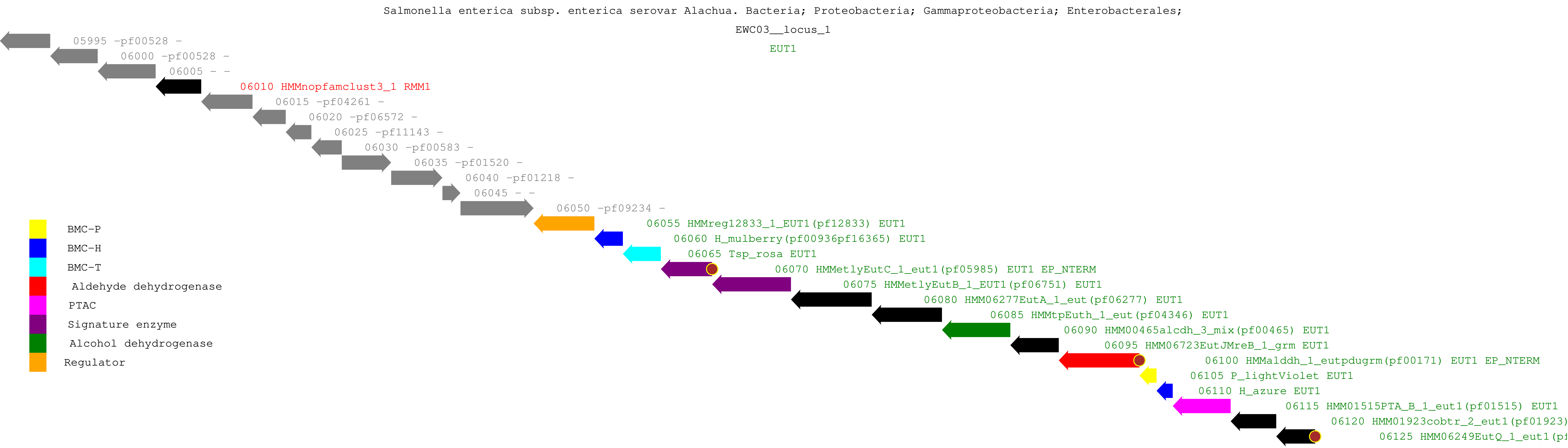

Salmonella enterica subsp. enterica serovar Alachua. Bacteria; Proteobacteria; Gammaproteobacteria; Enterobacterales;
EWC03__locus_1
EUT1
05995 -pf00528 -
06000 -pf00528 -
06005 - -
06010 HMMnopfamclust3_1 RMM1
06015 -pf04261 -
06020 -pf06572 -
06025 -pf11143 -
06030 -pf00583 -
06035 -pf01520 -
06040 -pf01218 -
06045 - -
06050 -pf09234 -
06055 HMMreg12833_1_EUT1(pf12833) EUT1
06060 H_mulberry(pf00936pf16365) EUT1
06065 Tsp_rosa EUT1
06070 HMMetlyEutC_1_eut1(pf05985) EUT1 EP_NTERM
06075 HMMetlyEutB_1_EUT1(pf06751) EUT1
06080 HMM06277EutA_1_eut(pf06277) EUT1
06085 HMMtpEuth_1_eut(pf04346) EUT1
06090 HMM00465alcdh_3_mix(pf00465) EUT1
06095 HMM06723EutJMreB_1_grm EUT1
06100 HMMalddh_1_eutpdugrm(pf00171) EUT1 EP_NTERM
06105 P_lightViolet EUT1
06110 H_azure EUT1
06115 HMM01515PTA_B_1_eut1(pf01515) EUT1
06120 HMM01923cobtr_2_eut1(pf01923)
06125 HMM06249EutQ_1_eut1(pf
BMC-P
BMC-H
BMC-T
Aldehyde dehydrogenase
PTAC
Signature enzyme
Alcohol dehydrogenase
Regulator

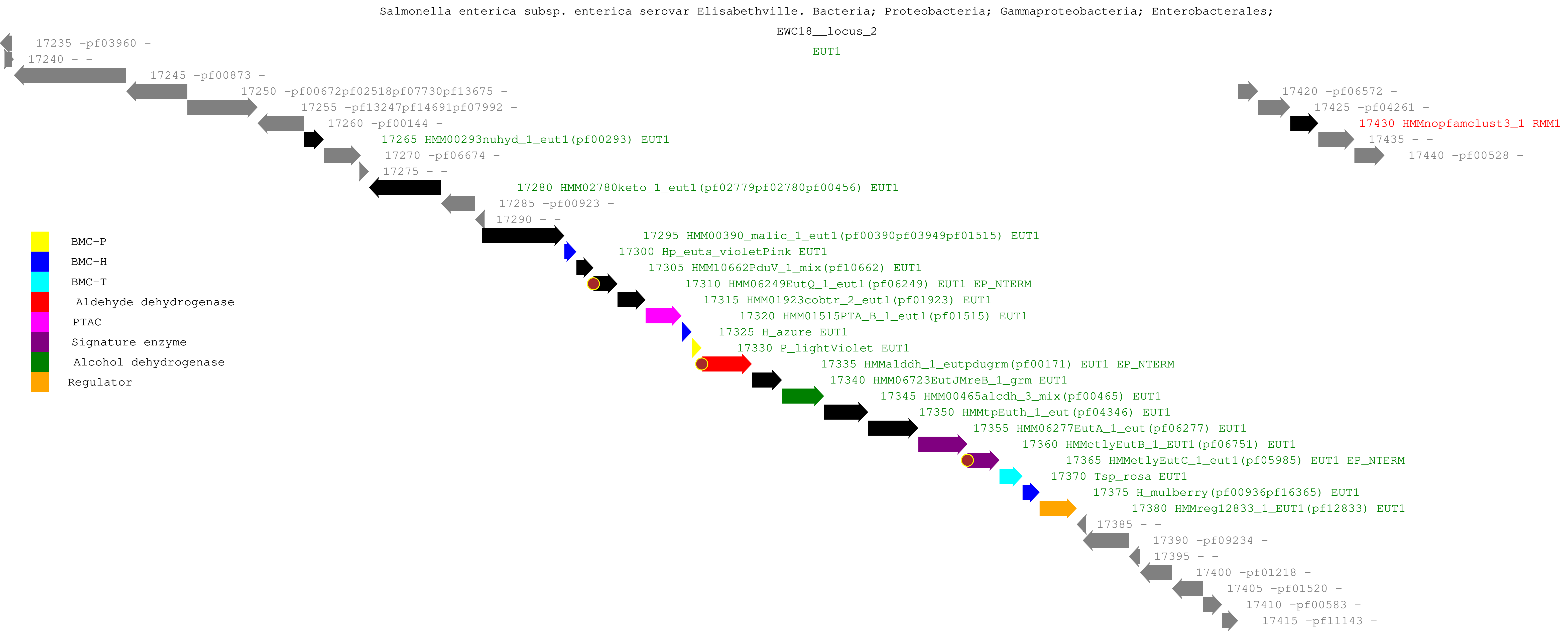

Salmonella enterica subsp. enterica serovar Elisabethville. Bacteria; Proteobacteria; Gammaproteobacteria; Enterobacterales;
EWC18__locus_2
EUT1
17235 -pf03960 -
17240 - -
17245 -pf00873 -
17250 -pf00672pf02518pf07730pf13675 -
17255 -pf13247pf14691pf07992 -
17260 -pf00144 -
17265 HMM00293nuhyd_1_eut1(pf00293) EUT1
17270 -pf06674 -
17275 - -
17280 HMM02780keto_1_eut1(pf02779pf02780pf00456) EUT1
17285 -pf00923 -
17290 - -
17295 HMM00390_malic_1_eut1(pf00390pf03949pf01515) EUT1
17300 Hp_euts_violetPink EUT1
17305 HMM10662PduV_1_mix(pf10662) EUT1
17310 HMM06249EutQ_1_eut1(pf06249) EUT1 EP_NTERM
17315 HMM01923cobtr_2_eut1(pf01923) EUT1
17320 HMM01515PTA_B_1_eut1(pf01515) EUT1
17325 H_azure EUT1
17330 P_lightViolet EUT1
17335 HMMalddh_1_eutpdugrm(pf00171) EUT1 EP_NTERM
17340 HMM06723EutJMreB_1_grm EUT1
17345 HMM00465alcdh_3_mix(pf00465) EUT1
17350 HMMtpEuth_1_eut(pf04346) EUT1
17355 HMM06277EutA_1_eut(pf06277) EUT1
17360 HMMetlyEutB_1_EUT1(pf06751) EUT1
17365 HMMetlyEutC_1_eut1(pf05985) EUT1 EP_NTERM
17370 Tsp_rosa EUT1
17375 H_mulberry(pf00936pf16365) EUT1
17380 HMMreg12833_1_EUT1(pf12833) EUT1
17385 - -
17390 -pf09234 -
17395 - -
17400 -pf01218 -
17405 -pf01520 -
17410 -pf00583 -
17415 -pf11143 -
17420 -pf06572 -
17425 -pf04261 -
17430 HMMnopfamclust3_1 RMM1
17435 - -
17440 -pf00528 -
BMC-P
BMC-H
BMC-T
Aldehyde dehydrogenase
PTAC
Signature enzyme
Alcohol dehydrogenase
Regulator

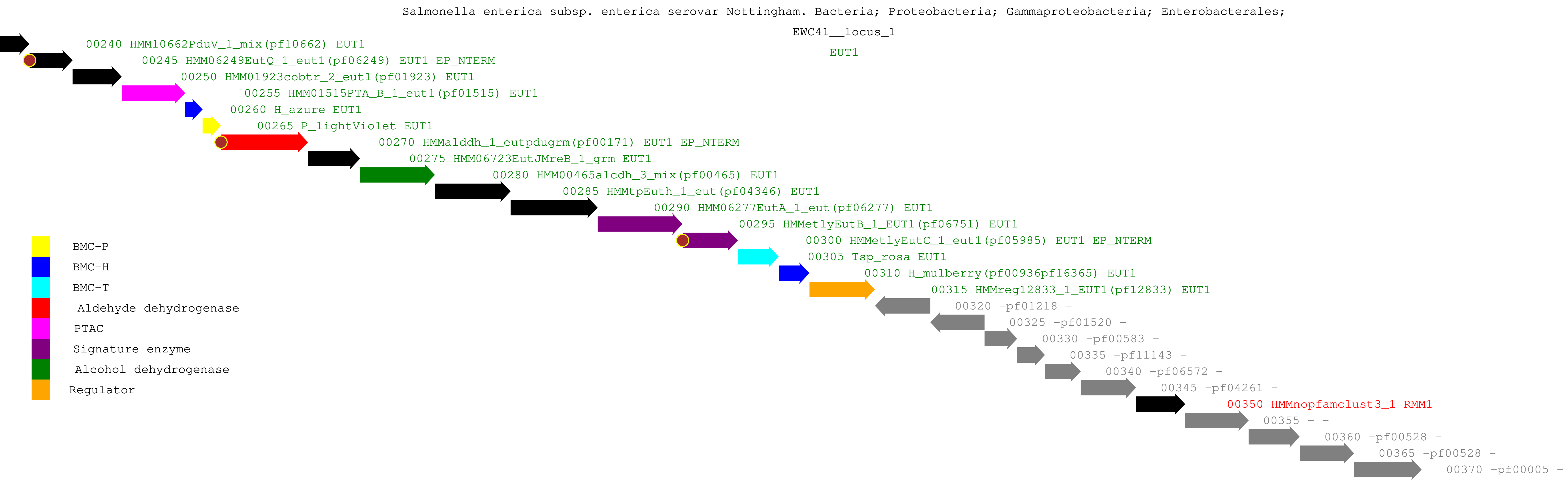
Salmonella enterica subsp. enterica serovar Nottingham. Bacteria; Proteobacteria; Gammaproteobacteria; Enterobacterales;
EWC41__locus_1
EUT1
00240 HMM10662PduV_1_mix(pf10662) EUT1
00245 HMM06249EutQ_1_eut1(pf06249) EUT1 EP_NTERM
00250 HMM01923cobtr_2_eut1(pf01923) EUT1
00255 HMM01515PTA_B_1_eut1(pf01515) EUT1
00260 H_azure EUT1
00265 P_lightViolet EUT1
00270 HMMalddh_1_eutpdugrm(pf00171) EUT1 EP_NTERM
00275 HMM06723EutJMreB_1_grm EUT1
00280 HMM00465alcdh_3_mix(pf00465) EUT1
00285 HMMtpEuth_1_eut(pf04346) EUT1
00290 HMM06277EutA_1_eut(pf06277) EUT1
00295 HMMetlyEutB_1_EUT1(pf06751) EUT1
00300 HMMetlyEutC_1_eut1(pf05985) EUT1 EP_NTERM
00305 Tsp_rosa EUT1
00310 H_mulberry(pf00936pf16365) EUT1
00315 HMMreg12833_1_EUT1(pf12833) EUT1
00320 -pf01218 -
00325 -pf01520 -
00330 -pf00583 -
00335 -pf11143 -
00340 -pf06572 -
00345 -pf04261 -
00350 HMMnopfamclust3_1 RMM1
00355 - -
00360 -pf00528 -
00365 -pf00528 -
00370 -pf00005 -
BMC-P
BMC-H
BMC-T
Aldehyde dehydrogenase
PTAC
Signature enzyme
Alcohol dehydrogenase
Regulator

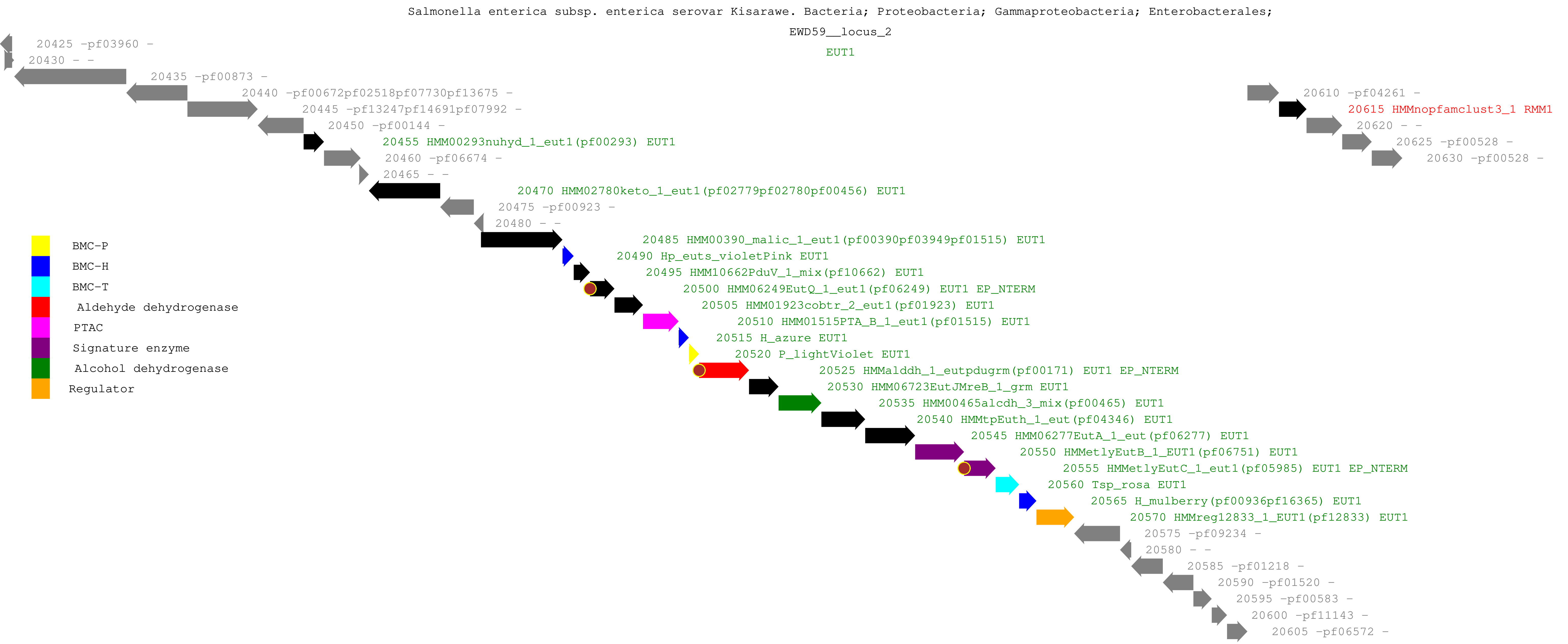
Salmonella enterica subsp. enterica serovar Kisarawe. Bacteria; Proteobacteria; Gammaproteobacteria; Enterobacterales;
EWD59__locus_2
EUT1
20425 -pf03960 -
20430 - -
20435 -pf00873 -
20440 -pf00672pf02518pf07730pf13675 -
20445 -pf13247pf14691pf07992 -
20450 -pf00144 -
20455 HMM00293nuhyd_1_eut1(pf00293) EUT1
20460 -pf06674 -
20465 - -
20470 HMM02780keto_1_eut1(pf02779pf02780pf00456) EUT1
20475 -pf00923 -
20480 - -
20485 HMM00390_malic_1_eut1(pf00390pf03949pf01515) EUT1
20490 Hp_euts_violetPink EUT1
20495 HMM10662PduV_1_mix(pf10662) EUT1
20500 HMM06249EutQ_1_eut1(pf06249) EUT1 EP_NTERM
20505 HMM01923cobtr_2_eut1(pf01923) EUT1
20510 HMM01515PTA_B_1_eut1(pf01515) EUT1
20515 H_azure EUT1
20520 P_lightViolet EUT1
20525 HMMalddh_1_eutpdugrm(pf00171) EUT1 EP_NTERM
20530 HMM06723EutJMreB_1_grm EUT1
20535 HMM00465alcdh_3_mix(pf00465) EUT1
20540 HMMtpEuth_1_eut(pf04346) EUT1
20545 HMM06277EutA_1_eut(pf06277) EUT1
20550 HMMetlyEutB_1_EUT1(pf06751) EUT1
20555 HMMetlyEutC_1_eut1(pf05985) EUT1 EP_NTERM
20560 Tsp_rosa EUT1
20565 H_mulberry(pf00936pf16365) EUT1
20570 HMMreg12833_1_EUT1(pf12833) EUT1
20575 -pf09234 -
20580 - -
20585 -pf01218 -
20590 -pf01520 -
20595 -pf00583 -
20600 -pf11143 -
20605 -pf06572 -
20610 -pf04261 -
20615 HMMnopfamclust3_1 RMM1
20620 - -
20625 -pf00528 -
20630 -pf00528 -
BMC-P
BMC-H
BMC-T
Aldehyde dehydrogenase
PTAC
Signature enzyme
Alcohol dehydrogenase
Regulator

Salmonella enterica subsp. enterica serovar Praha. Bacteria; Proteobacteria; Gammaproteobacteria; Enterobacterales;

EWD62__locus_2

EUT1

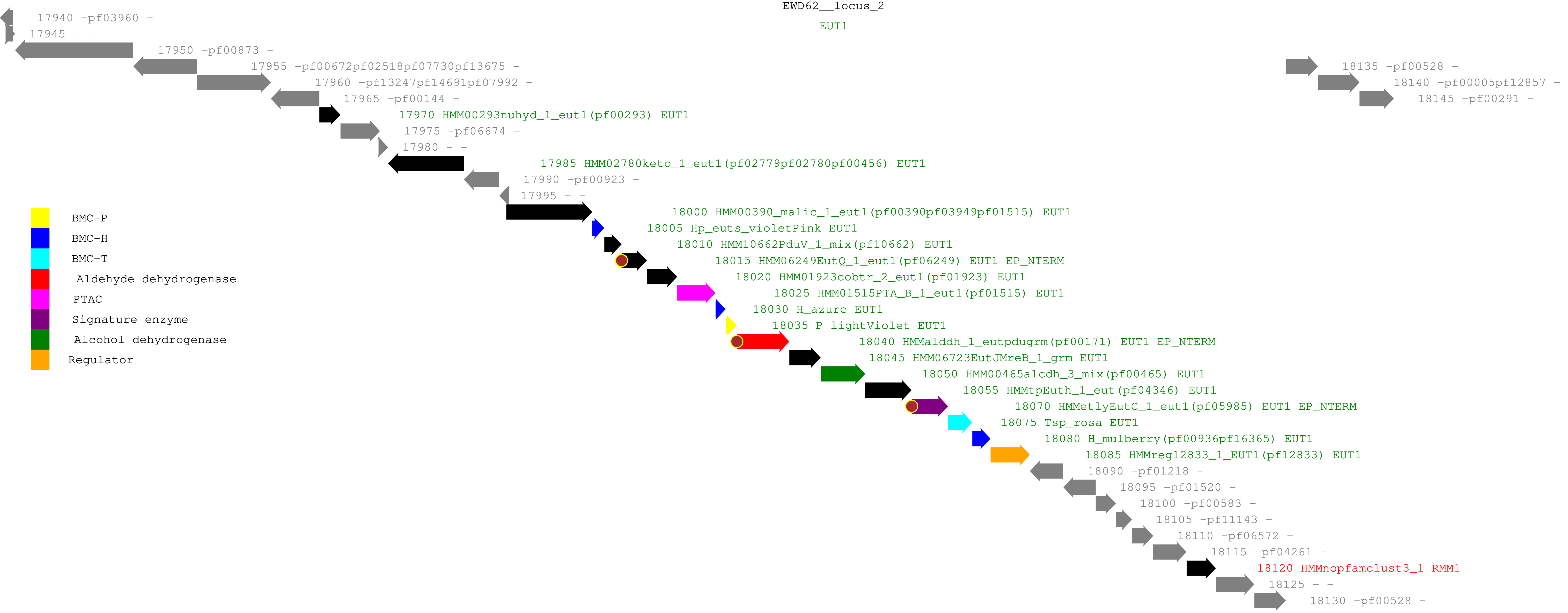

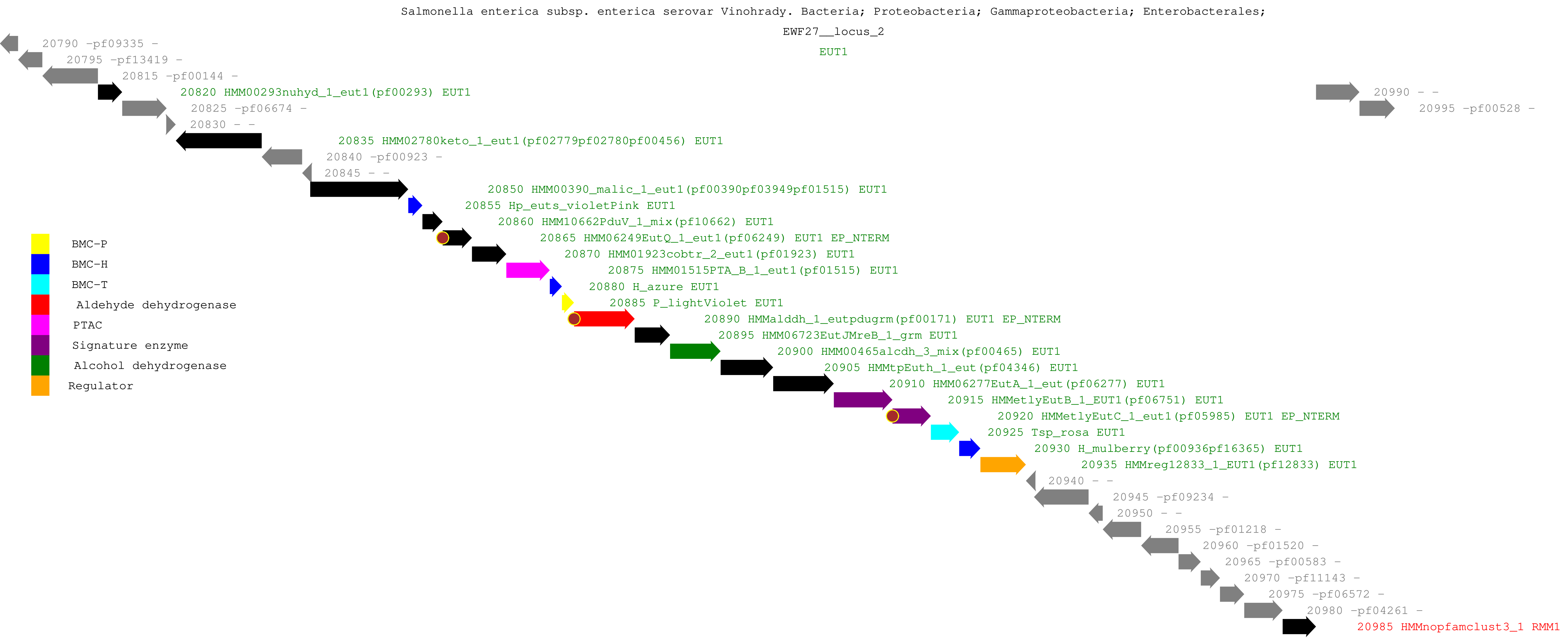

Salmonella enterica subsp. enterica serovar Vinohrady. Bacteria; Proteobacteria; Gammaproteobacteria; Enterobacterales;
EWF27__locus_2
EUT1
20790 -pf09335 -
20795 -pf13419 -
20815 -pf00144 -
20820 HMM00293nuhyd_1_eut1(pf00293) EUT1
20825 -pf06674 -
20830 - -
20835 HMM02780keto_1_eut1(pf02779pf02780pf00456) EUT1
20840 -pf00923 -
20845 - -
20850 HMM00390_malic_1_eut1(pf00390pf03949pf01515) EUT1
20855 Hp_euts_violetPink EUT1
20860 HMM10662PduV_1_mix(pf10662) EUT1
20865 HMM06249EutQ_1_eut1(pf06249) EUT1 EP_NTERM
20870 HMM01923cobtr_2_eut1(pf01923) EUT1
20875 HMM01515PTA_B_1_eut1(pf01515) EUT1
20880 H_azure EUT1
20885 P_lightViolet EUT1
20890 HMMalddh_1_eutpdugrm(pf00171) EUT1 EP_NTERM
20895 HMM06723EutJMreB_1_grm EUT1
20900 HMM00465alcdh_3_mix(pf00465) EUT1
20905 HMMtpEuth_1_eut(pf04346) EUT1
20910 HMM06277EutA_1_eut(pf06277) EUT1
20915 HMMetlyEutB_1_EUT1(pf06751) EUT1
20920 HMMetlyEutC_1_eut1(pf05985) EUT1 EP_NTERM
20925 Tsp_rosa EUT1
20930 H_mulberry(pf00936pf16365) EUT1
20935 HMMreg12833_1_EUT1(pf12833) EUT1
20940 - -
20945 -pf09234 -
20950 - -
20955 -pf01218 -
20960 -pf01520 -
20965 -pf00583 -
20970 -pf11143 -
20975 -pf06572 -
20980 -pf04261 -
20985 HMMnopfamclust3_1 RMM1
20990 - -
20995 -pf00528 -
BMC-P
BMC-H
BMC-T
Aldehyde dehydrogenase
PTAC
Signature enzyme
Alcohol dehydrogenase
Regulator

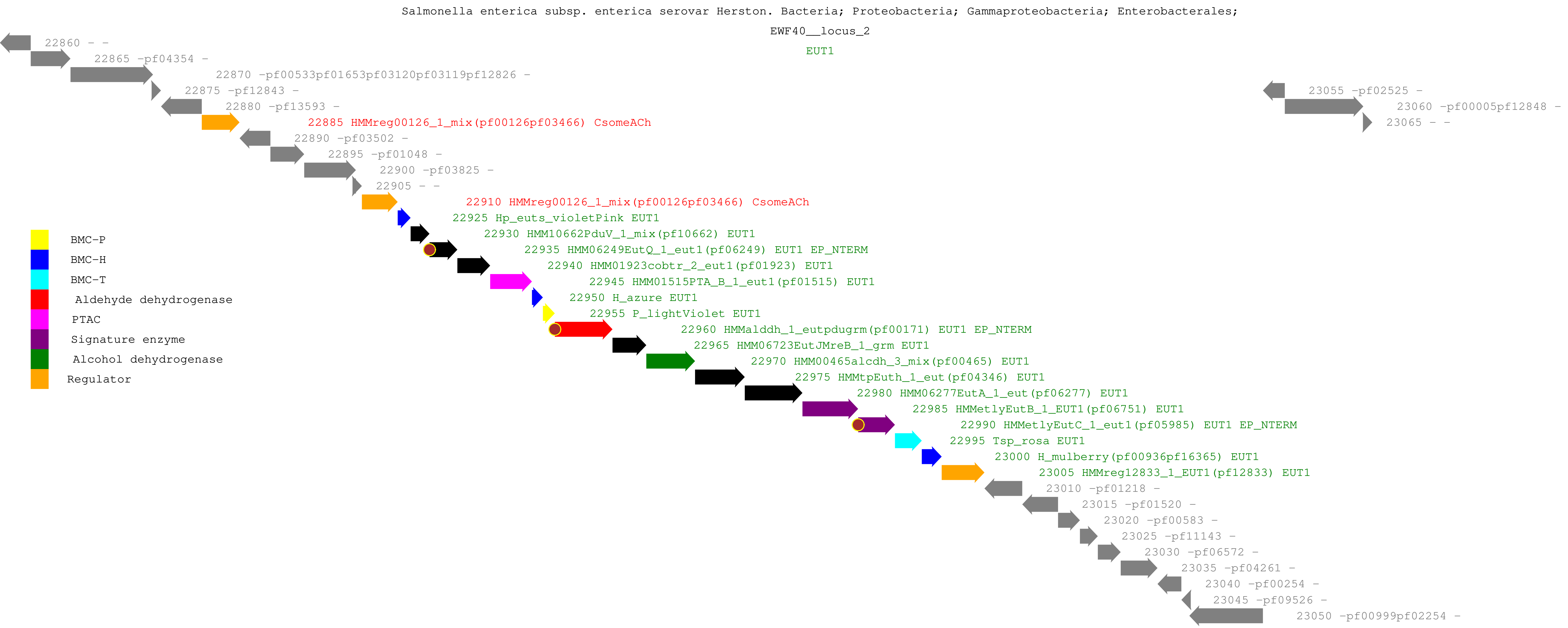

Salmonella enterica subsp. enterica serovar Herston. Bacteria; Proteobacteria; Gammaproteobacteria; Enterobacterales;
EWF40__locus_2
EUT1
22860 - -
22865 -pf04354 -
22870 -pf00533pf01653pf03120pf03119pf12826 -
22875 -pf12843 -
22880 -pf13593 -
22885 HMMreg00126_1_mix(pf00126pf03466) CsomeACh
22890 -pf03502 -
22895 -pf01048 -
22900 -pf03825 -
22905 - -
22910 HMMreg00126_1_mix(pf00126pf03466) CsomeACh
22925 Hp_euts_violetPink EUT1
22930 HMM10662PduV_1_mix(pf10662) EUT1
22935 HMM06249EutQ_1_eut1(pf06249) EUT1 EP_NTERM
22940 HMM01923cobtr_2_eut1(pf01923) EUT1
22945 HMM01515PTA_B_1_eut1(pf01515) EUT1
22950 H_azure EUT1
22955 P_lightViolet EUT1
22960 HMMalddh_1_eutpdugrm(pf00171) EUT1 EP_NTERM
22965 HMM06723EutJMreB_1_grm EUT1
22970 HMM00465alcdh_3_mix(pf00465) EUT1
22975 HMMtpEuth_1_eut(pf04346) EUT1
22980 HMM06277EutA_1_eut(pf06277) EUT1
22985 HMMetlyEutB_1_EUT1(pf06751) EUT1
22990 HMMetlyEutC_1_eut1(pf05985) EUT1 EP_NTERM
22995 Tsp_rosa EUT1
23000 H_mulberry(pf00936pf16365) EUT1
23005 HMMreg12833_1_EUT1(pf12833) EUT1
23010 -pf01218 -
23015 -pf01520 -
23020 -pf00583 -
23025 -pf11143 -
23030 -pf06572 -
23035 -pf04261 -
23040 -pf00254 -
23045 -pf09526 -
23050 -pf00999pf02254 -
23055 -pf02525 -
23060 -pf00005pf12848 -
23065 - -
BMC-P
BMC-H
BMC-T
Aldehyde dehydrogenase
PTAC
Signature enzyme
Alcohol dehydrogenase
Regulator

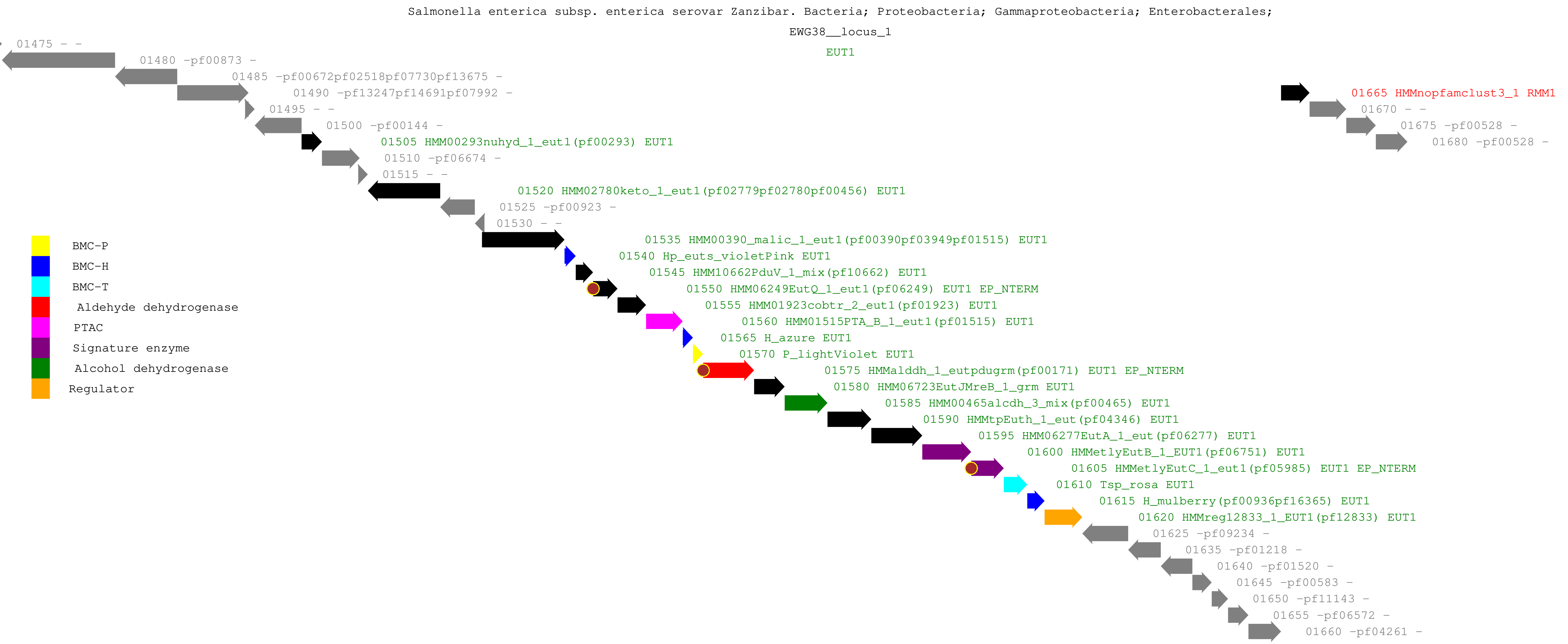
Salmonella enterica subsp. enterica serovar Zanzibar. Bacteria; Proteobacteria; Gammaproteobacteria; Enterobacterales;
EWG38__locus_1
EUT1
01475 - -
01480 -pf00873 -
01485 -pf00672pf02518pf07730pf13675 -
01490 -pf13247pf14691pf07992 -
01495 - -
01500 -pf00144 -
01505 HMM00293nuhyd_1_eut1(pf00293) EUT1
01510 -pf06674 -
01515 - -
01520 HMM02780keto_1_eut1(pf02779pf02780pf00456) EUT1
01525 -pf00923 -
01530 - -
01535 HMM00390_malic_1_eut1(pf00390pf03949pf01515) EUT1
01540 Hp_euts_violetPink EUT1
01545 HMM10662PduV_1_mix(pf10662) EUT1
01550 HMM06249EutQ_1_eut1(pf06249) EUT1 EP_NTERM
01555 HMM01923cobtr_2_eut1(pf01923) EUT1
01560 HMM01515PTA_B_1_eut1(pf01515) EUT1
01565 H_azure EUT1
01570 P_lightViolet EUT1
01575 HMMaldh_1_eutpdugrm(pf00171) EUT1 EP_NTERM
01580 HMM06723EutJMreB_1_grm EUT1
01585 HMM00465alcdh_3_mix(pf00465) EUT1
01590 HMMtpEuth_1_eut(pf04346) EUT1
01595 HMM06277EutA_1_eut(pf06277) EUT1
01600 HMMetlyEutB_1_EUT1(pf06751) EUT1
01605 HMMetlyEutC_1_eut1(pf05985) EUT1 EP_NTERM
01610 Tsp_rosa EUT1
01615 H_mulberry(pf00936pf16365) EUT1
01620 HMMreg12833_1_EUT1(pf12833) EUT1
01625 -pf09234 -
01635 -pf01218 -
01640 -pf01520 -
01645 -pf00583 -
01650 -pf11143 -
01655 -pf06572 -
01660 -pf04261 -
01665 HMMnopfamclust3_1 RMM1
01670 - -
01675 -pf00528 -
01680 -pf00528 -
BMC-P
BMC-H
BMC-T
Aldehyde dehydrogenase
PTAC
Signature enzyme
Alcohol dehydrogenase
Regulator

Salmonella enterica subsp. enterica serovar Koketime. Bacteria; Proteobacteria; Gammaproteobacteria; Enterobacterales;

EWG69__locus_2

EUT1

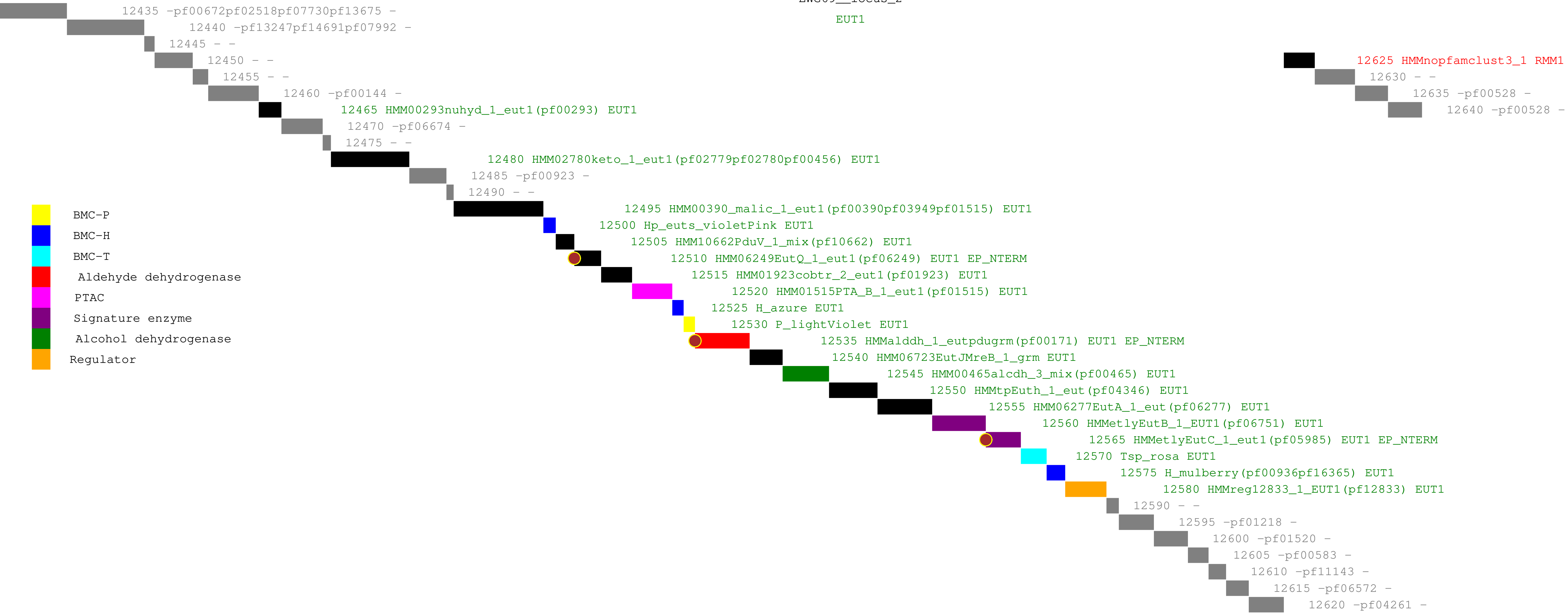

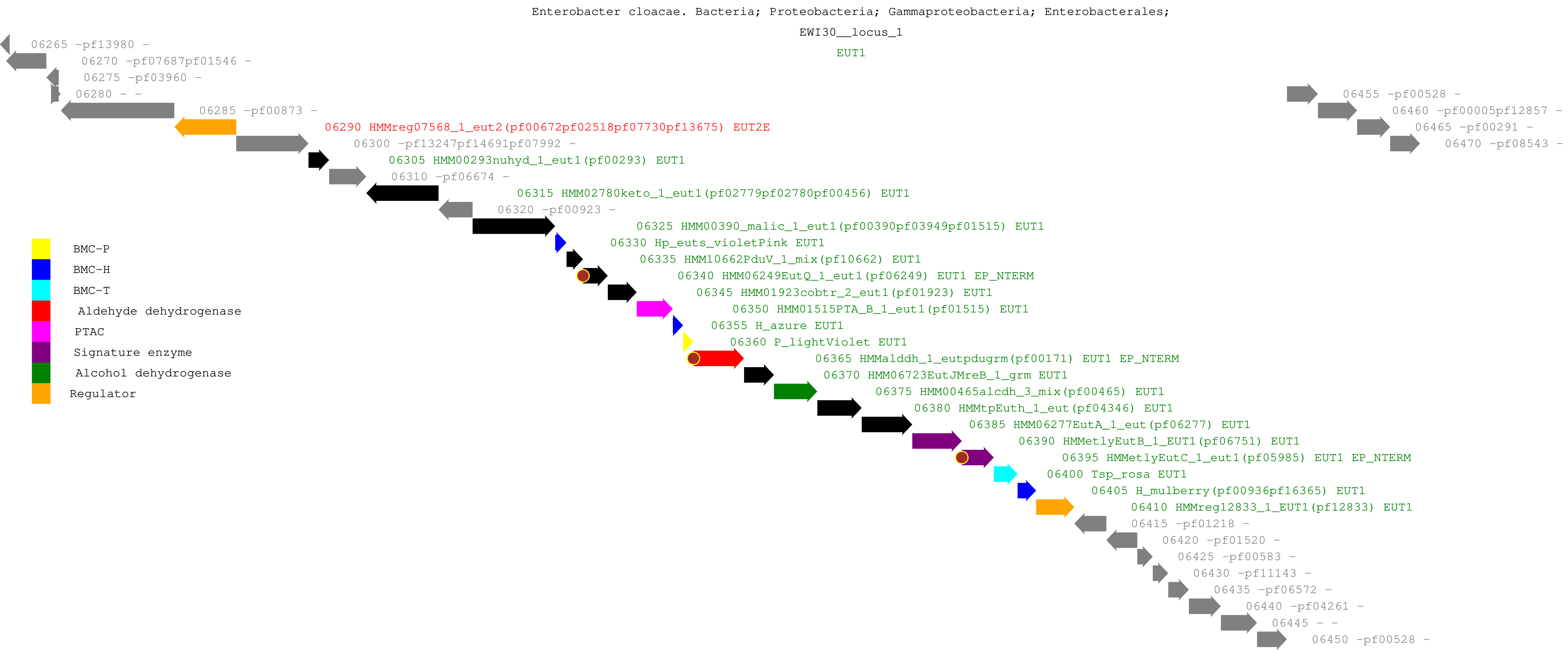

Enterobacter cloacae. Bacteria; Proteobacteria; Gammaproteobacteria; Enterobacterales;
EWI30__locus_1
EUT1
06265 -pf13980 -
06270 -pf07687pf01546 -
06275 -pf03960 -
06280 - -
06285 -pf00873 -
06290 HMMreg07568_1_eut2(pf00672pf02518pf07730pf13675) EUT2E
06300 -pf13247pf14691pf07992 -
06305 HMM00293nuhyd_1_eut1(pf00293) EUT1
06310 -pf06674 -
06315 HMM02780keto_1_eut1(pf02779pf02780pf00456) EUT1
06320 -pf00923 -
06325 HMM00390_malic_1_eut1(pf00390pf03949pf01515) EUT1
06330 Hp_euts_violetPink EUT1
06335 HMM10662PduV_1_mix(pf10662) EUT1
06340 HMM06249EutQ_1_eut1(pf06249) EUT1 EP_NTERM
06345 HMM01923cobtr_2_eut1(pf01923) EUT1
06350 HMM01515PTA_B_1_eut1(pf01515) EUT1
06355 H_azure EUT1
06360 P_lightViolet EUT1
06365 HMMaldh_1_eutpdugrm(pf00171) EUT1 EP_NTERM
06370 HMM06723EutJMreB_1_grm EUT1
06375 HMM00465alcdh_3_mix(pf00465) EUT1
06380 HMMtpEuth_1_eut(pf04346) EUT1
06385 HMM06277EutA_1_eut(pf06277) EUT1
06390 HMMetlyEutB_1_EUT1(pf06751) EUT1
06395 HMMetlyEutC_1_eut1(pf05985) EUT1 EP_NTERM
06400 Tsp_rosa EUT1
06405 H_mulberry(pf00936pf16365) EUT1
06410 HMMreg12833_1_EUT1(pf12833) EUT1
06415 -pf01218 -
06420 -pf01520 -
06425 -pf00583 -
06430 -pf11143 -
06435 -pf06572 -
06440 -pf04261 -
06445 - -
06450 -pf00528 -
06455 -pf00528 -
06460 -pf00005pf12857 -
06465 -pf00291 -
06470 -pf08543 -
BMC-P
BMC-H
BMC-T
Aldehyde dehydrogenase
PTAC
Signature enzyme
Alcohol dehydrogenase
Regulator

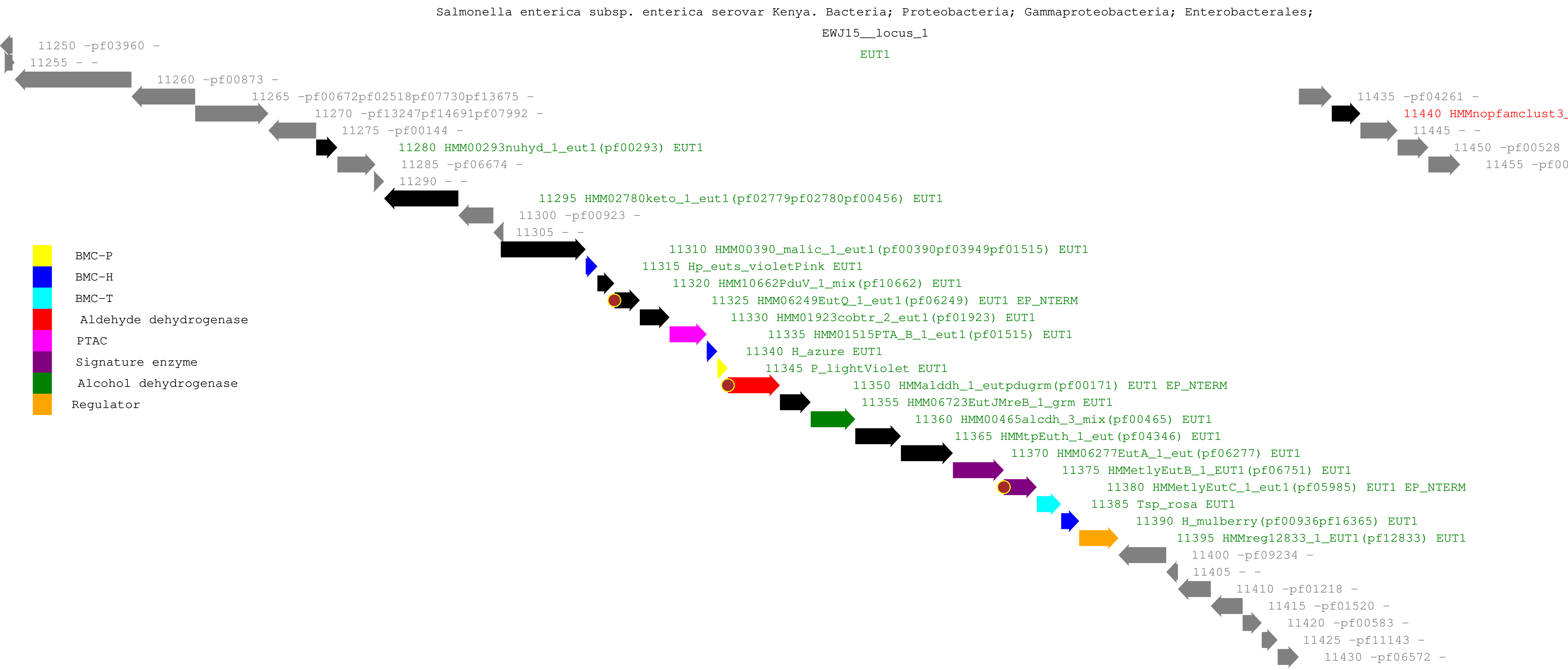

Salmonella enterica subsp. enterica serovar Kenya. Bacteria; Proteobacteria; Gammaproteobacteria; Enterobacterales;
EWJ15__locus_1
EUT1
11250 -pf03960 -
11255 - -
11260 -pf00873 -
11265 -pf00672pf02518pf07730pf13675 -
11270 -pf13247pf14691pf07992 -
11275 -pf00144 -
11280 HMM00293nuhyd_1_eut1(pf00293) EUT1
11285 -pf06674 -
11290 - -
11295 HMM02780keto_1_eut1(pf02779pf02780pf00456) EUT1
11300 -pf00923 -
11305 - -
11310 HMM00390_malic_1_eut1(pf00390pf03949pf01515) EUT1
11315 Hp_euts_violetPink EUT1
11320 HMM10662PduV_1_mix(pf10662) EUT1
11325 HMM06249EutQ_1_eut1(pf06249) EUT1 EP_NTERM
11330 HMM01923cobtr_2_eut1(pf01923) EUT1
11335 HMM01515PTA_B_1_eut1(pf01515) EUT1
11340 H_azure EUT1
11345 P_lightViolet EUT1
11350 HMMaldh_1_eutpdugrm(pf00171) EUT1 EP_NTERM
11355 HMM06723EutJMreB_1_grm EUT1
11360 HMM00465alcdh_3_mix(pf00465) EUT1
11365 HMMtpEuth_1_eut(pf04346) EUT1
11370 HMM06277EutA_1_eut(pf06277) EUT1
11375 HMMetlyEutB_1_EUT1(pf06751) EUT1
11380 HMMetlyEutC_1_eut1(pf05985) EUT1 EP_NTERM
11385 Tsp_rosa EUT1
11390 H_mulberry(pf00936pf16365) EUT1
11395 HMMreg12833_1_EUT1(pf12833) EUT1
11400 -pf09234 -
11405 - -
11410 -pf01218 -
11415 -pf01520 -
11420 -pf00583 -
11425 -pf11143 -
11430 -pf06572 -
11435 -pf04261 -
11440 HMMnopfamclust3_1 RMM1
11445 - -
11450 -pf00528 -
11455 -pf00528 -
BMC-P
BMC-H
BMC-T
Aldehyde dehydrogenase
PTAC
Signature enzyme
Alcohol dehydrogenase
Regulator

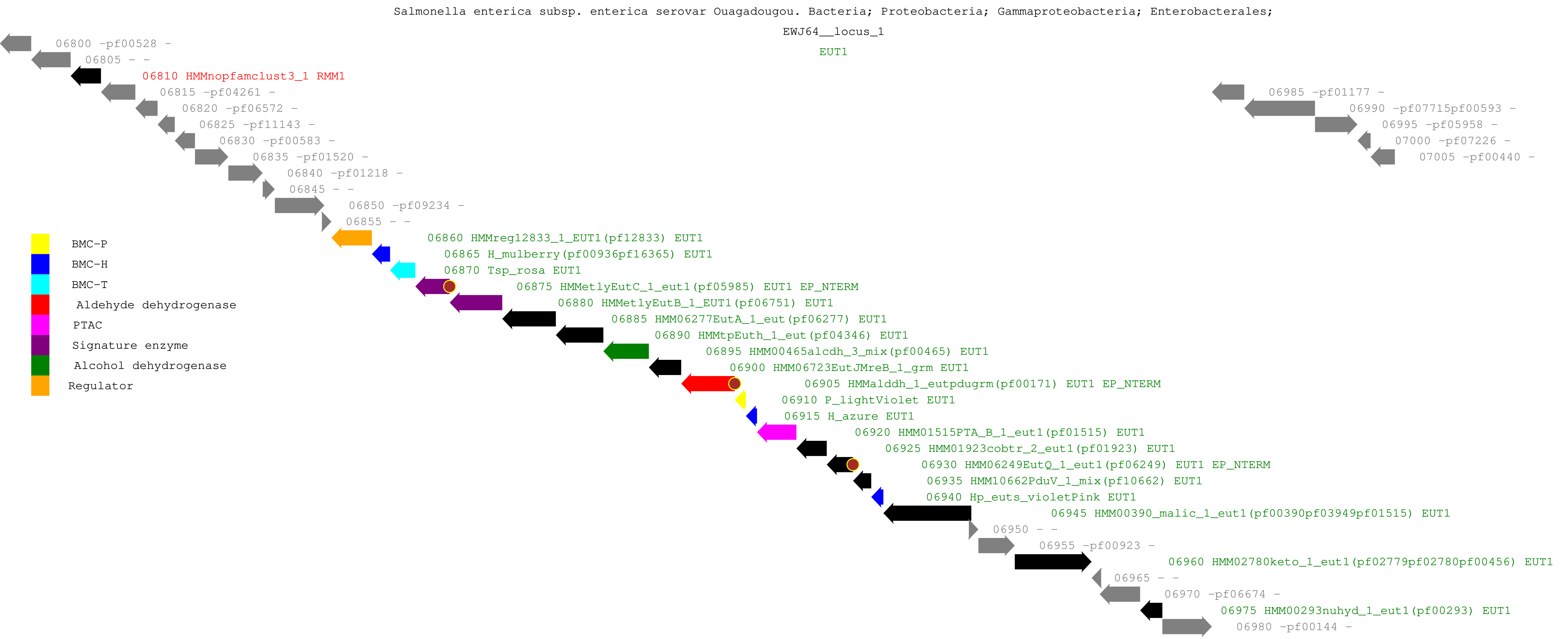

Salmonella enterica subsp. enterica serovar Ouagadougou. Bacteria; Proteobacteria; Gammaproteobacteria; Enterobacterales;
EWJ64__locus_1
EUT1
06800 -pf00528 -
06805 - -
06810 HMMnopfamclust3_1 RMM1
06815 -pf04261 -
06820 -pf06572 -
06825 -pf11143 -
06830 -pf00583 -
06835 -pf01520 -
06840 -pf01218 -
06845 - -
06850 -pf09234 -
06855 - -
06860 HMMreg12833_1_EUT1(pf12833) EUT1
06865 H_mulberry(pf00936pf16365) EUT1
06870 Tsp_rosa EUT1
06875 HMMetlyEutC_1_eut1(pf05985) EUT1 EP_NTERM
06880 HMMetlyEutB_1_EUT1(pf06751) EUT1
06885 HMM06277EutA_1_eut(pf06277) EUT1
06890 HMMtpEuth_1_eut(pf04346) EUT1
06895 HMM00465alcdh_3_mix(pf00465) EUT1
06900 HMM06723EutJMreB_1_grm EUT1
06905 HMMalddh_1_eutpdugrm(pf00171) EUT1 EP_NTERM
06910 P_lightViolet EUT1
06915 H_azure EUT1
06920 HMM01515PTA_B_1_eut1(pf01515) EUT1
06925 HMM01923cobtr_2_eut1(pf01923) EUT1
06930 HMM06249EutQ_1_eut1(pf06249) EUT1 EP_NTERM
06935 HMM10662PduV_1_mix(pf10662) EUT1
06940 Hp_euts_violetPink EUT1
06945 HMM00390_malic_1_eut1(pf00390pf03949pf01515) EUT1
06950 - -
06955 -pf00923 -
06960 HMM02780keto_1_eut1(pf02779pf02780pf00456) EUT1
06965 - -
06970 -pf06674 -
06975 HMM00293nuhyd_1_eut1(pf00293) EUT1
06980 -pf00144 -
06985 -pf01177 -
06990 -pf07715pf00593 -
06995 -pf05958 -
07000 -pf07226 -
07005 -pf00440 -
BMC-P
BMC-H
BMC-T
Aldehyde dehydrogenase
PTAC
Signature enzyme
Alcohol dehydrogenase
Regulator

Salmonella enterica subsp. enterica serovar Weybridge. Bacteria; Proteobacteria; Gammaproteobacteria; Enterobacterales;

EWJ82__locus_2

EUT1

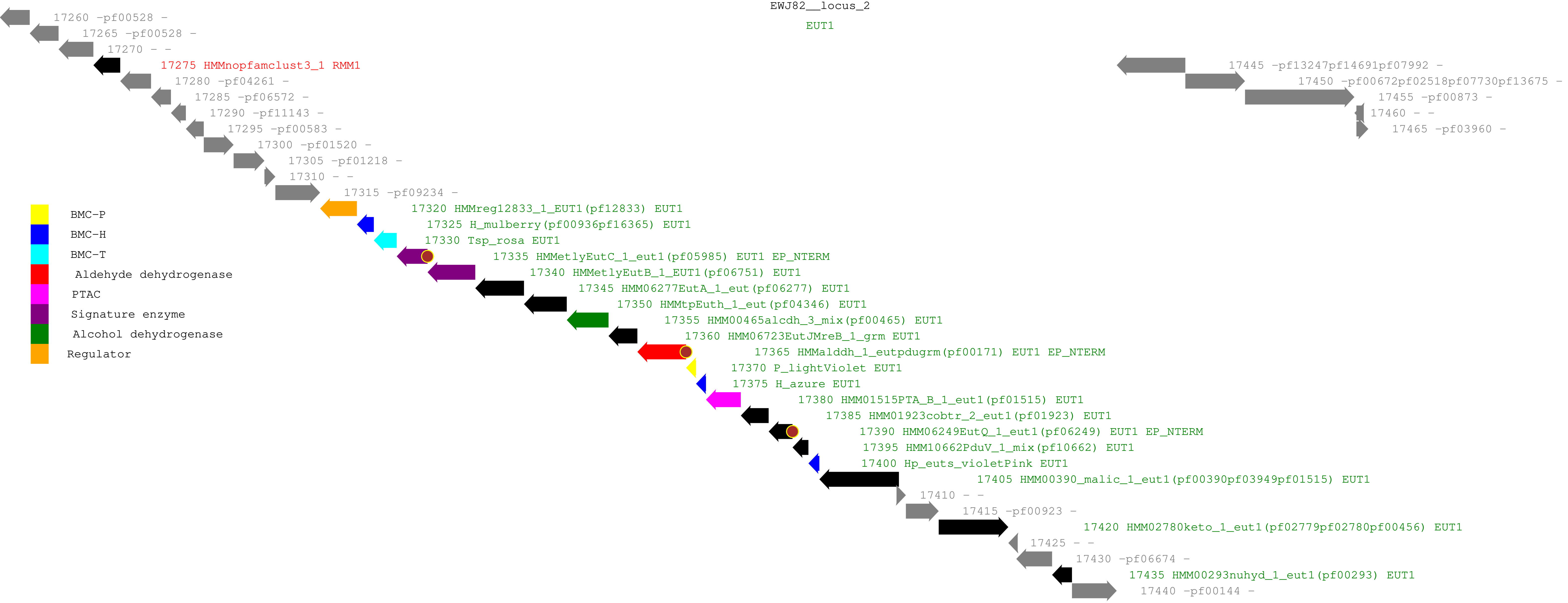

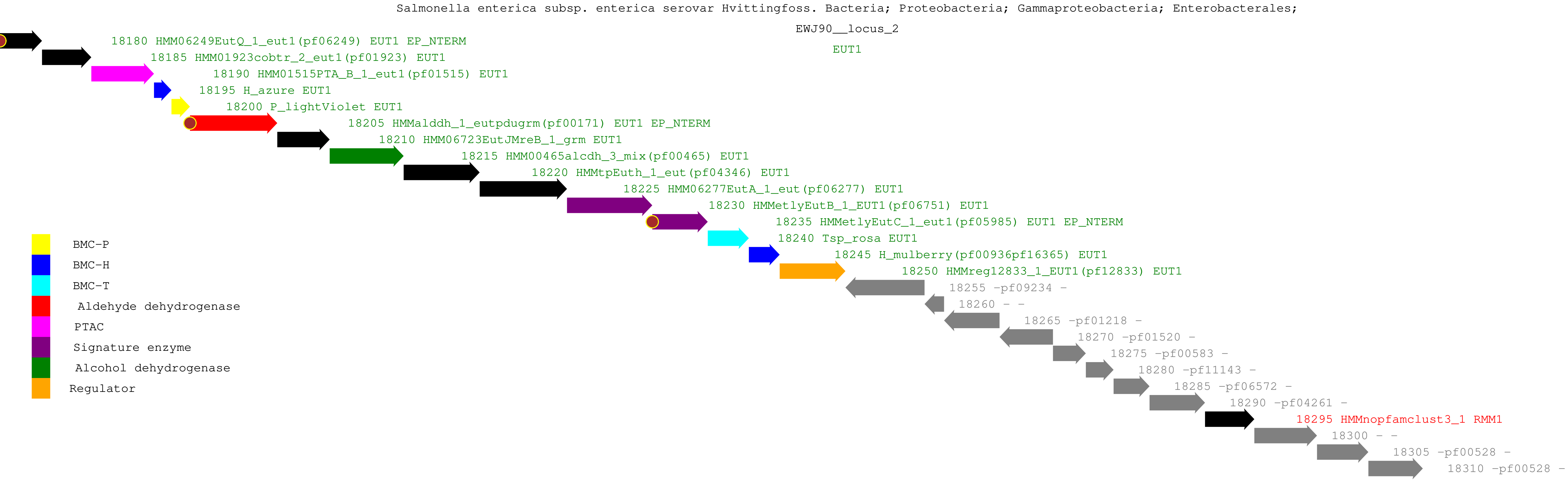

Salmonella enterica subsp. enterica serovar Hvittingfoss. Bacteria; Proteobacteria; Gammaproteobacteria; Enterobacterales;
EWJ90__locus_2
EUT1
18180 HMM06249EutQ_1_eut1(pf06249) EUT1 EP_NTERM
18185 HMM01923cobtr_2_eut1(pf01923) EUT1
18190 HMM01515PTA_B_1_eut1(pf01515) EUT1
18195 H_azure EUT1
18200 P_lightViolet EUT1
18205 HMMalddh_1_eutpdugrm(pf00171) EUT1 EP_NTERM
18210 HMM06723EutJMreB_1_grm EUT1
18215 HMM00465alcdh_3_mix(pf00465) EUT1
18220 HMMtpEuth_1_eut(pf04346) EUT1
18225 HMM06277EutA_1_eut(pf06277) EUT1
18230 HMMetlyEutB_1_EUT1(pf06751) EUT1
18235 HMMetlyEutC_1_eut1(pf05985) EUT1 EP_NTERM
18240 Tsp_rosa EUT1
18245 H_mulberry(pf00936pf16365) EUT1
18250 HMMreg12833_1_EUT1(pf12833) EUT1
18255 -pf09234 -
18260 - -
18265 -pf01218 -
18270 -pf01520 -
18275 -pf00583 -
18280 -pf11143 -
18285 -pf06572 -
18290 -pf04261 -
18295 HMMnopfamclust3_1 RMM1
18300 - -
18305 -pf00528 -
18310 -pf00528 -
BMC-P
BMC-H
BMC-T
Aldehyde dehydrogenase
PTAC
Signature enzyme
Alcohol dehydrogenase
Regulator

Salmonella enterica subsp. enterica serovar Offa. Bacteria; Proteobacteria; Gammaproteobacteria; Enterobacterales;

EWZ14__locus_2

EUT1

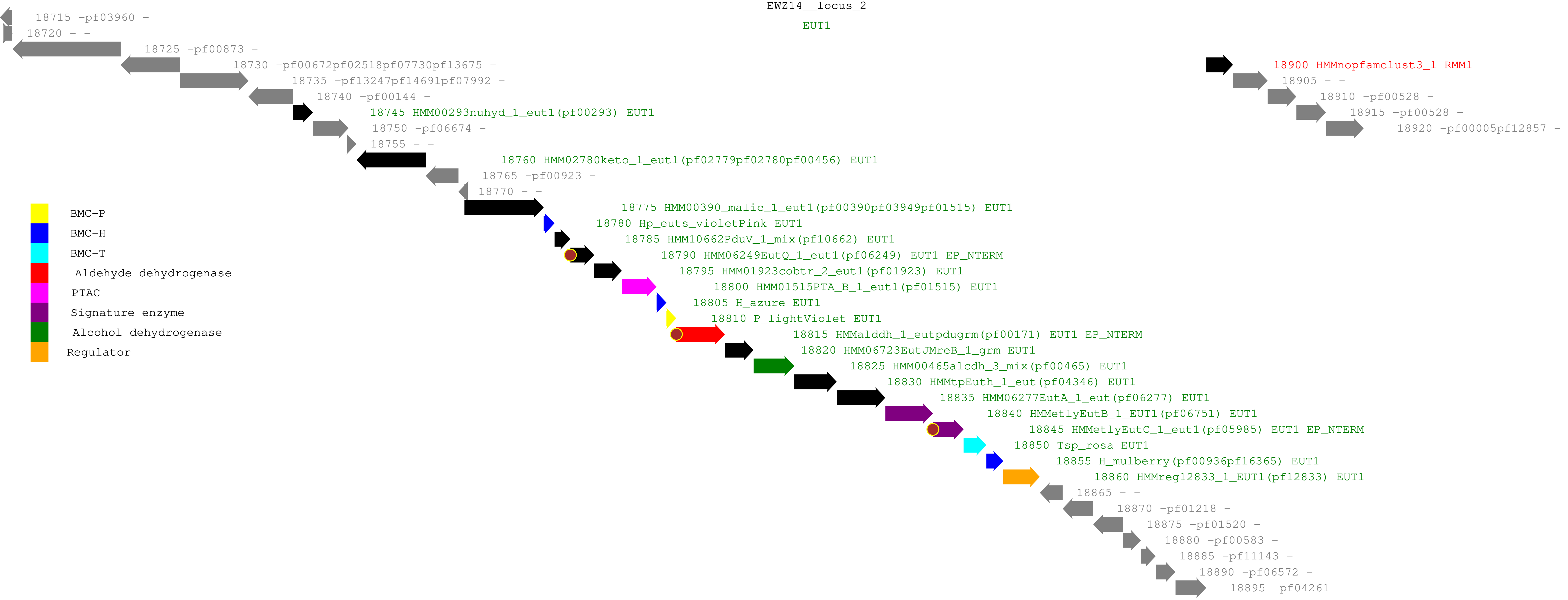

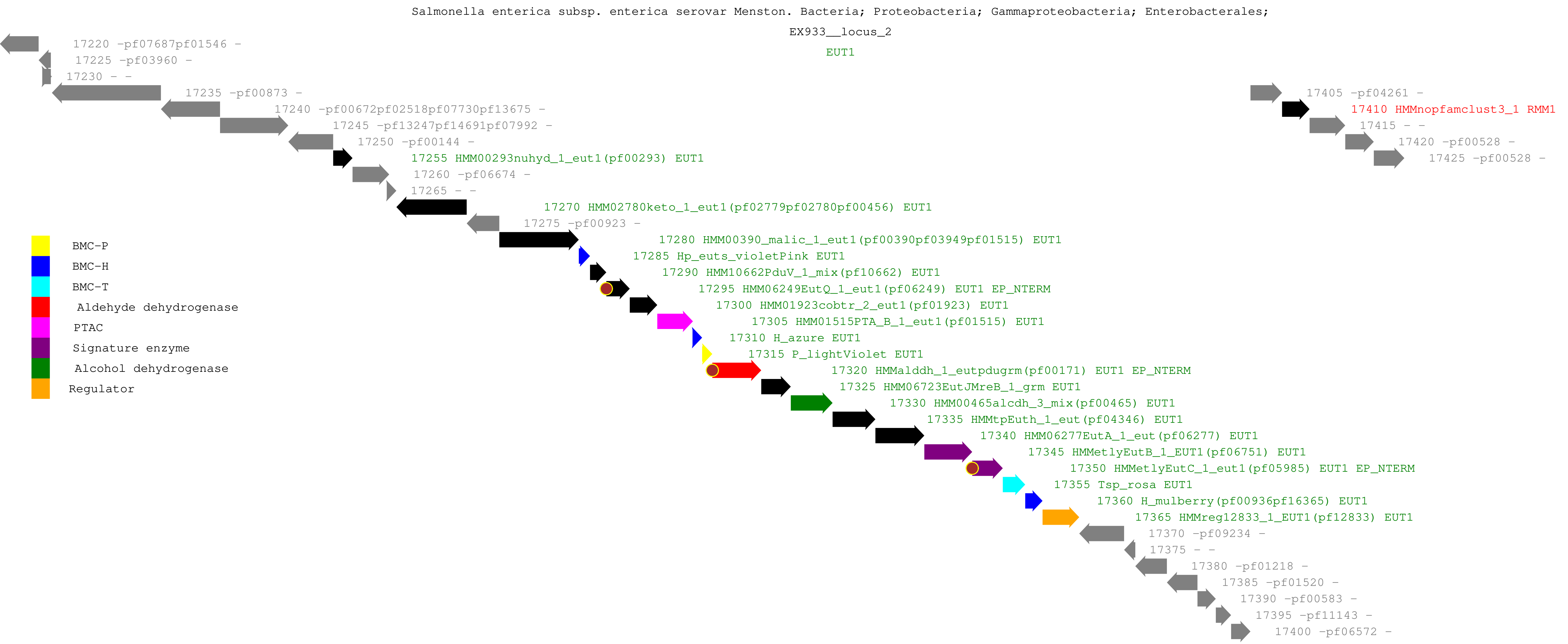
Salmonella enterica subsp. enterica serovar Menston. Bacteria; Proteobacteria; Gammaproteobacteria; Enterobacterales;
EX933__locus_2
EUT1
17220 -pf07687pf01546 -
17225 -pf03960 -
17230 - -
17235 -pf00873 -
17240 -pf00672pf02518pf07730pf13675 -
17245 -pf13247pf14691pf07992 -
17250 -pf00144 -
17255 HMM00293nuhyd_1_eut1(pf00293) EUT1
17260 -pf06674 -
17265 - -
17270 HMM02780keto_1_eut1(pf02779pf02780pf00456) EUT1
17275 -pf00923 -
17280 HMM00390_malic_1_eut1(pf00390pf03949pf01515) EUT1
17285 Hp_euts_violetPink EUT1
17290 HMM10662PduV_1_mix(pf10662) EUT1
17295 HMM06249EutQ_1_eut1(pf06249) EUT1 EP_NTERM
17300 HMM01923cobtr_2_eut1(pf01923) EUT1
17305 HMM01515PTA_B_1_eut1(pf01515) EUT1
17310 H_azure EUT1
17315 P_lightViolet EUT1
17320 HMMaldh_1_eutpdugrm(pf00171) EUT1 EP_NTERM
17325 HMM06723EutJMreB_1_grm EUT1
17330 HMM00465alcdh_3_mix(pf00465) EUT1
17335 HMMtpEuth_1_eut(pf04346) EUT1
17340 HMM06277EutA_1_eut(pf06277) EUT1
17345 HMMetlyEutB_1_EUT1(pf06751) EUT1
17350 HMMetlyEutC_1_eut1(pf05985) EUT1 EP_NTERM
17355 Tsp_rosa EUT1
17360 H_mulberry(pf00936pf16365) EUT1
17365 HMMreg12833_1_EUT1(pf12833) EUT1
17370 -pf09234 -
17375 - -
17380 -pf01218 -
17385 -pf01520 -
17390 -pf00583 -
17395 -pf11143 -
17400 -pf06572 -
17405 -pf04261 -
17410 HMMnopfamclust3_1 RMM1
17415 - -
17420 -pf00528 -
17425 -pf00528 -
BMC-P
BMC-H
BMC-T
Aldehyde dehydrogenase
PTAC
Signature enzyme
Alcohol dehydrogenase
Regulator

Salmonella enterica subsp. enterica serovar Teshie. Bacteria; Proteobacteria; Gammaproteobacteria; Enterobacterales;

EXA25__locus_2

EUT1

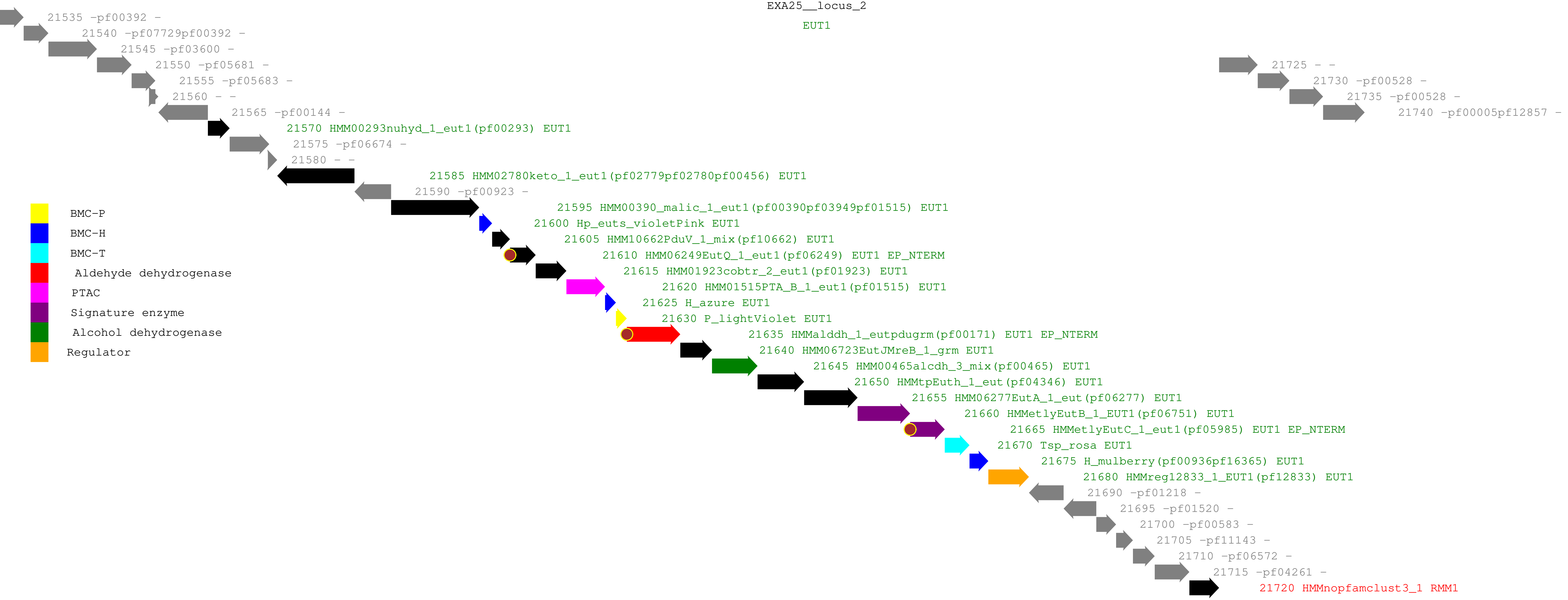

Salmonella enterica subsp. enterica serovar Korlebu. Bacteria; Proteobacteria; Gammaproteobacteria; Enterobacterales;

EXA36__locus_2

EUT1

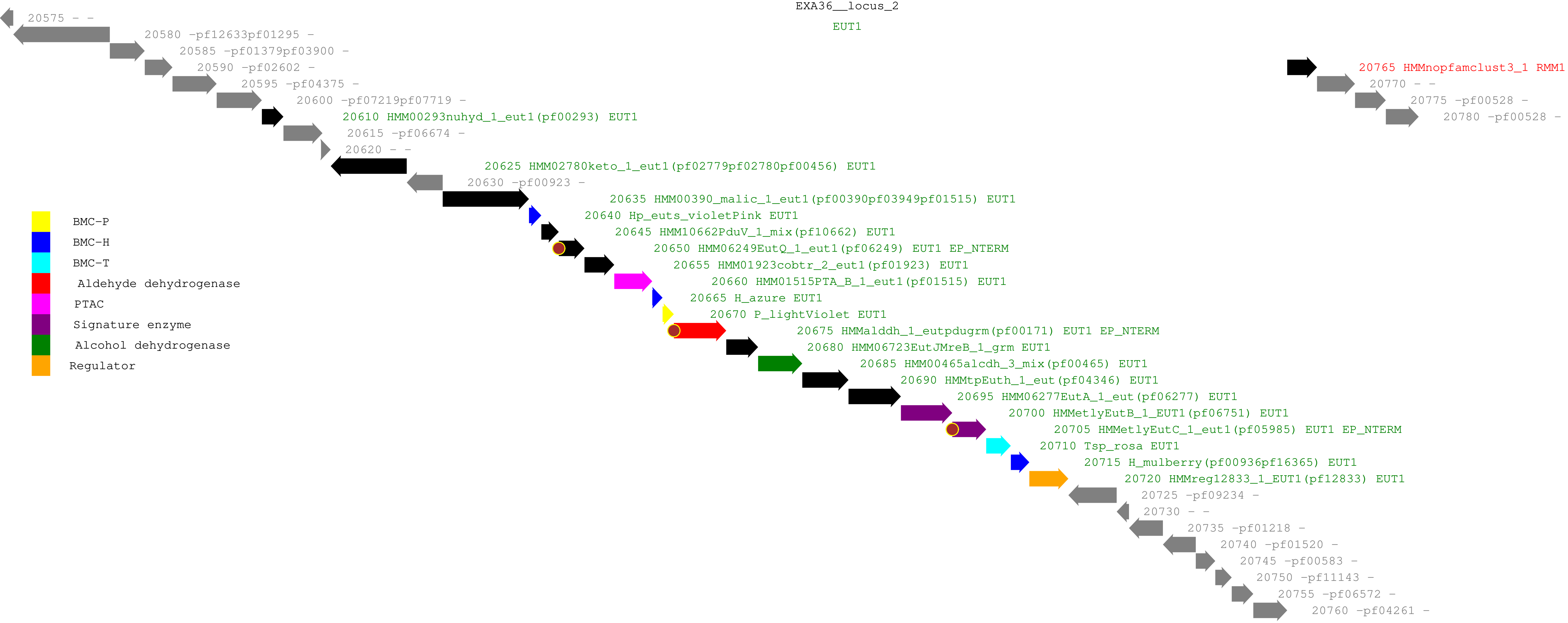

Salmonella virchow. Bacteria; Proteobacteria; Gammaproteobacteria; Enterobacterales;

EXA61__locus_1

EUT1

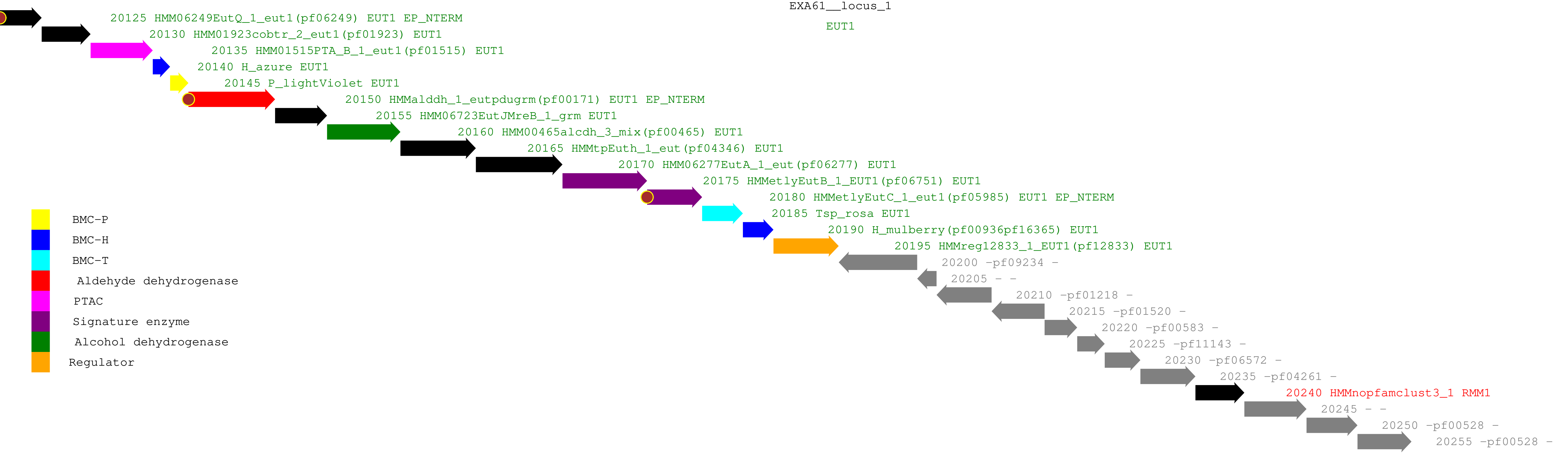

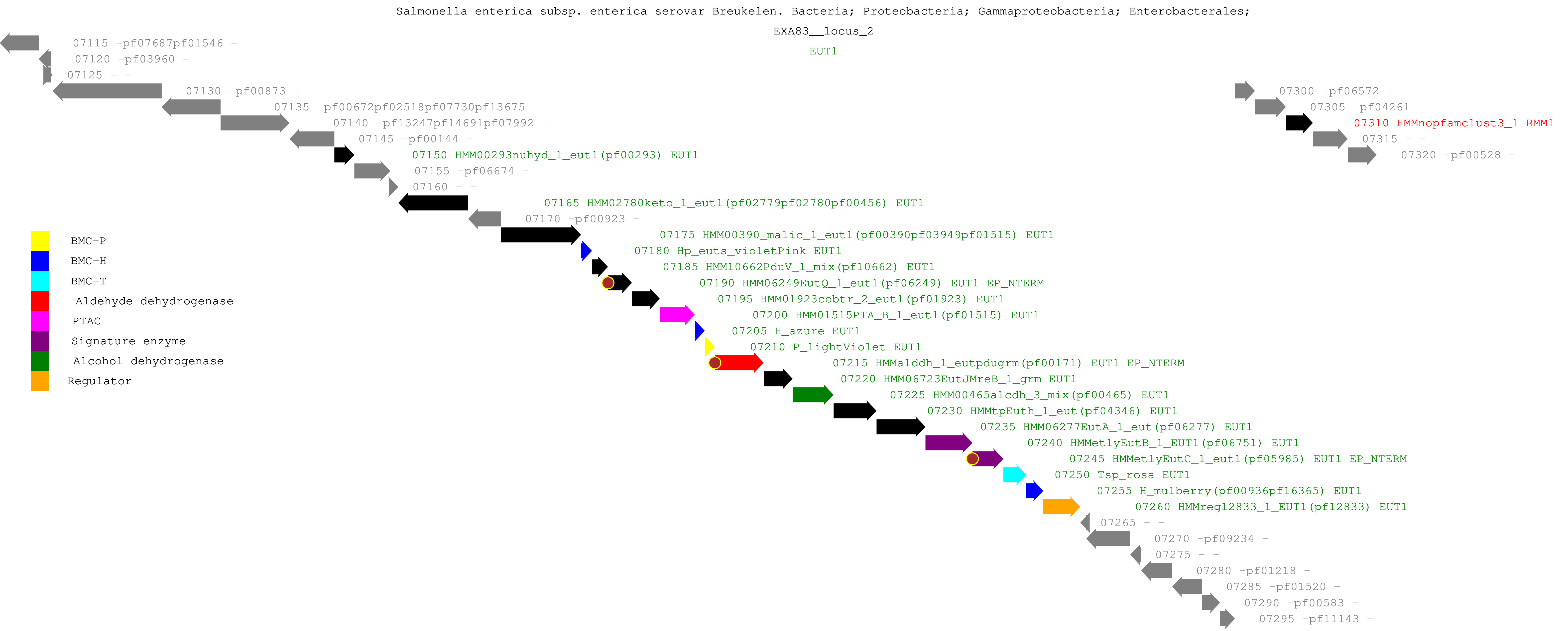
Salmonella enterica subsp. enterica serovar Breukelen. Bacteria; Proteobacteria; Gammaproteobacteria; Enterobacterales;
EXA83__locus_2
EUT1
07115 -pf07687pf01546 -
07120 -pf03960 -
07125 - -
07130 -pf00873 -
07135 -pf00672pf02518pf07730pf13675 -
07140 -pf13247pf14691pf07992 -
07145 -pf00144 -
07150 HMM00293nuhyd_1_eut1(pf00293) EUT1
07155 -pf06674 -
07160 - -
07165 HMM02780keto_1_eut1(pf02779pf02780pf00456) EUT1
07170 -pf00923 -
07175 HMM00390_malic_1_eut1(pf00390pf03949pf01515) EUT1
07180 Hp_euts_violetPink EUT1
07185 HMM10662PduV_1_mix(pf10662) EUT1
07190 HMM06249EutQ_1_eut1(pf06249) EUT1 EP_NTERM
07195 HMM01923cobtr_2_eut1(pf01923) EUT1
07200 HMM01515PTA_B_1_eut1(pf01515) EUT1
07205 H_azure EUT1
07210 P_lightViolet EUT1
07215 HMMalddh_1_eutpdugrm(pf00171) EUT1 EP_NTERM
07220 HMM06723EutJMreB_1_grm EUT1
07225 HMM00465alcdh_3_mix(pf00465) EUT1
07230 HMMtpEuth_1_eut(pf04346) EUT1
07235 HMM06277EutA_1_eut(pf06277) EUT1
07240 HMMetlyEutB_1_EUT1(pf06751) EUT1
07245 HMMetlyEutC_1_eut1(pf05985) EUT1 EP_NTERM
07250 Tsp_rosa EUT1
07255 H_mulberry(pf00936pf16365) EUT1
07260 HMMreg12833_1_EUT1(pf12833) EUT1
07265 - -
07270 -pf09234 -
07275 - -
07280 -pf01218 -
07285 -pf01520 -
07290 -pf00583 -
07295 -pf11143 -
07300 -pf06572 -
07305 -pf04261 -
07310 HMMnopfamclust3_1 RMM1
07315 - -
07320 -pf00528 -
BMC-P
BMC-H
BMC-T
Aldehyde dehydrogenase
PTAC
Signature enzyme
Alcohol dehydrogenase
Regulator

Salmonella enterica subsp. enterica serovar Aberdeen. Bacteria; Proteobacteria; Gammaproteobacteria; Enterobacterales;

EXA90__locus_1

EUT1

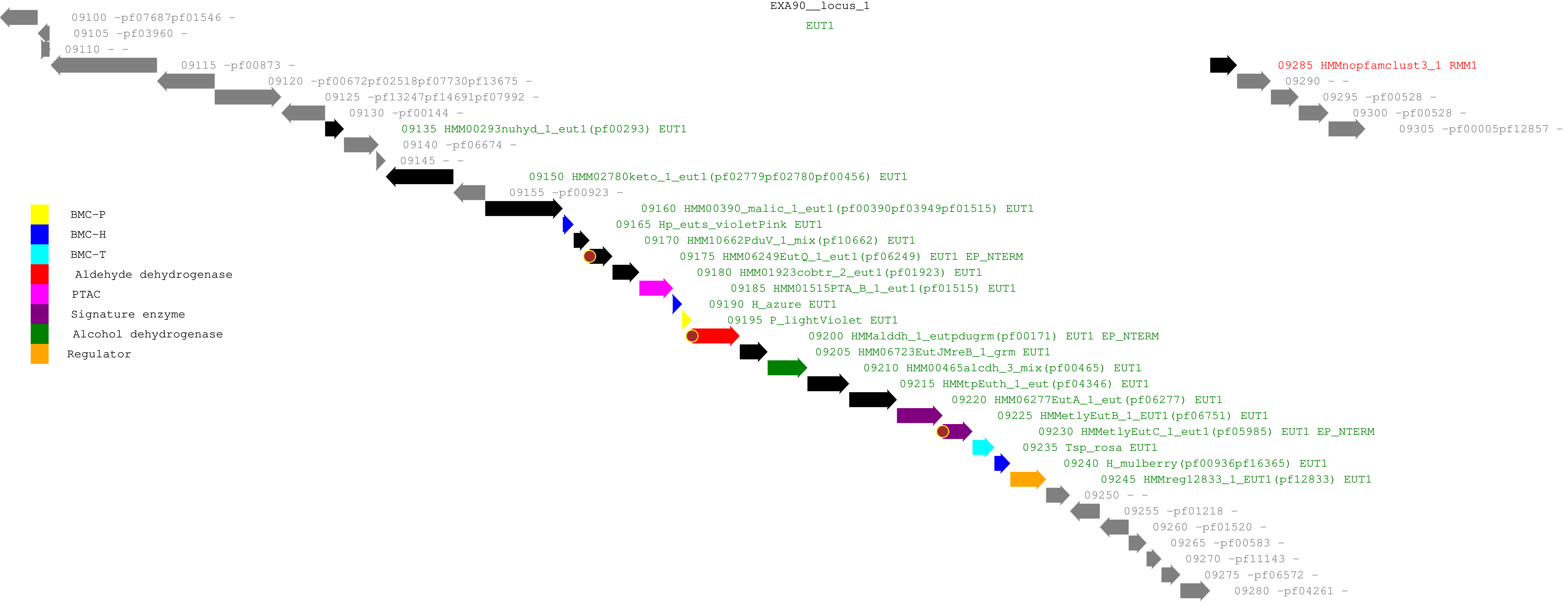

Salmonella enterica subsp. enterica serovar Fresno. Bacteria; Proteobacteria; Gammaproteobacteria; Enterobacterales;

EXB02__locus_2

EUT1

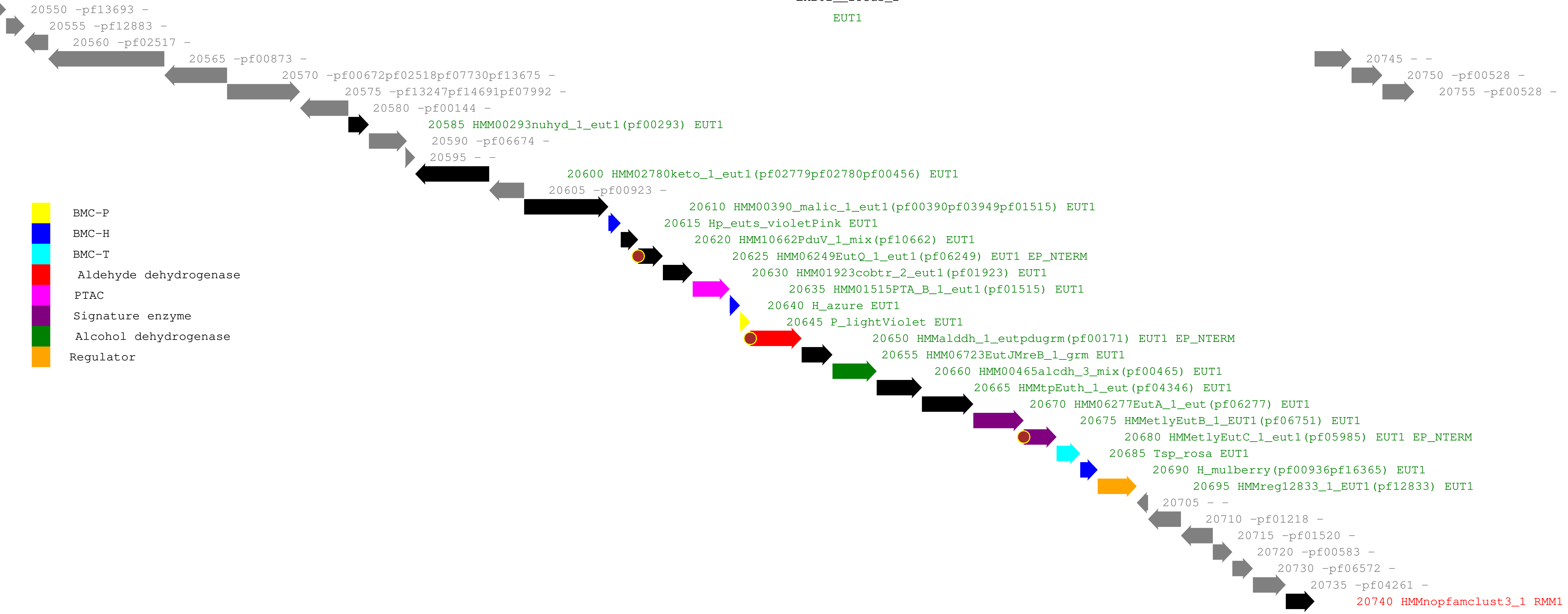

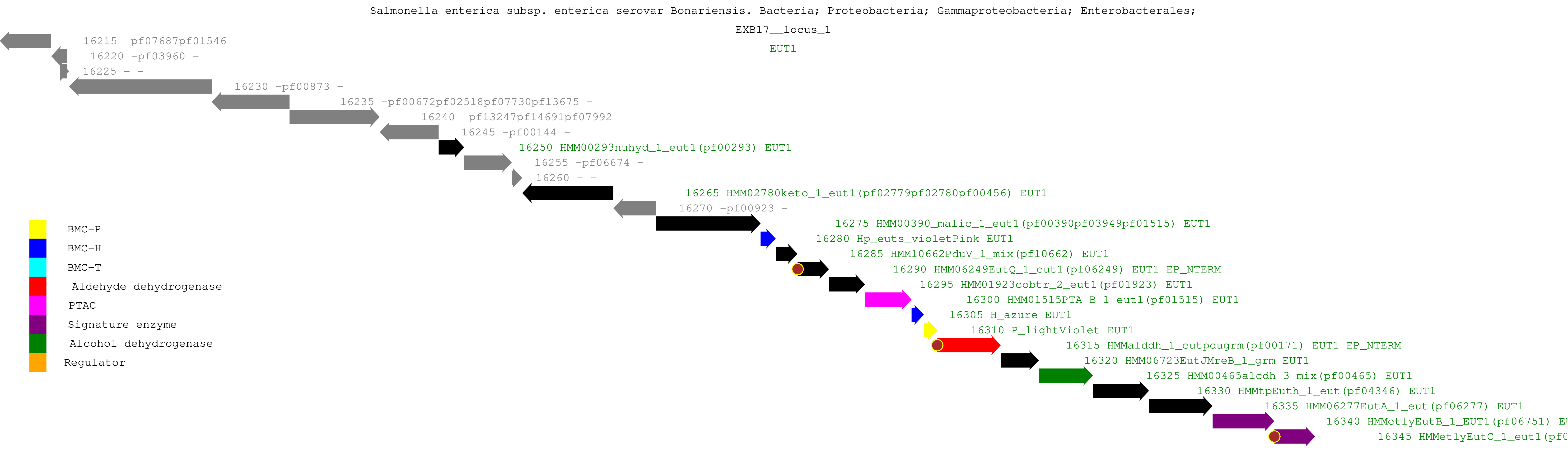

Salmonella enterica subsp. enterica serovar Bonariensis. Bacteria; Proteobacteria; Gammaproteobacteria; Enterobacterales;
EXB17__locus_1
EUT1
16215 -pf07687pf01546 -
16220 -pf03960 -
16225 - -
16230 -pf00873 -
16235 -pf00672pf02518pf07730pf13675 -
16240 -pf13247pf14691pf07992 -
16245 -pf00144 -
16250 HMM00293nuhyd_1_eut1(pf00293) EUT1
16255 -pf06674 -
16260 - -
16265 HMM02780keto_1_eut1(pf02779pf02780pf00456) EUT1
16270 -pf00923 -
16275 HMM00390_malic_1_eut1(pf00390pf03949pf01515) EUT1
16280 Hp_euts_violetPink EUT1
16285 HMM10662PduV_1_mix(pf10662) EUT1
16290 HMM06249EutQ_1_eut1(pf06249) EUT1 EP_NTERM
16295 HMM01923cobtr_2_eut1(pf01923) EUT1
16300 HMM01515PTA_B_1_eut1(pf01515) EUT1
16305 H_azure EUT1
16310 P_lightViolet EUT1
16315 HMMaldh_1_eutpdugrm(pf00171) EUT1 EP_NTERM
16320 HMM06723EutJMreB_1_grm EUT1
16325 HMM00465alcdh_3_mix(pf00465) EUT1
16330 HMMtpEuth_1_eut(pf04346) EUT1
16335 HMM06277EutA_1_eut(pf06277) EUT1
16340 HMMetlyEutB_1_EUT1(pf06751) E
16345 HMMetlyEutC_1_eut1(pf
BMC-P
BMC-H
BMC-T
Aldehyde dehydrogenase
PTAC
Signature enzyme
Alcohol dehydrogenase
Regulator

Salmonella enterica subsp. enterica serovar Napoli. Bacteria; Proteobacteria; Gammaproteobacteria; Enterobacterales;

EXB97__locus_1

EUT1

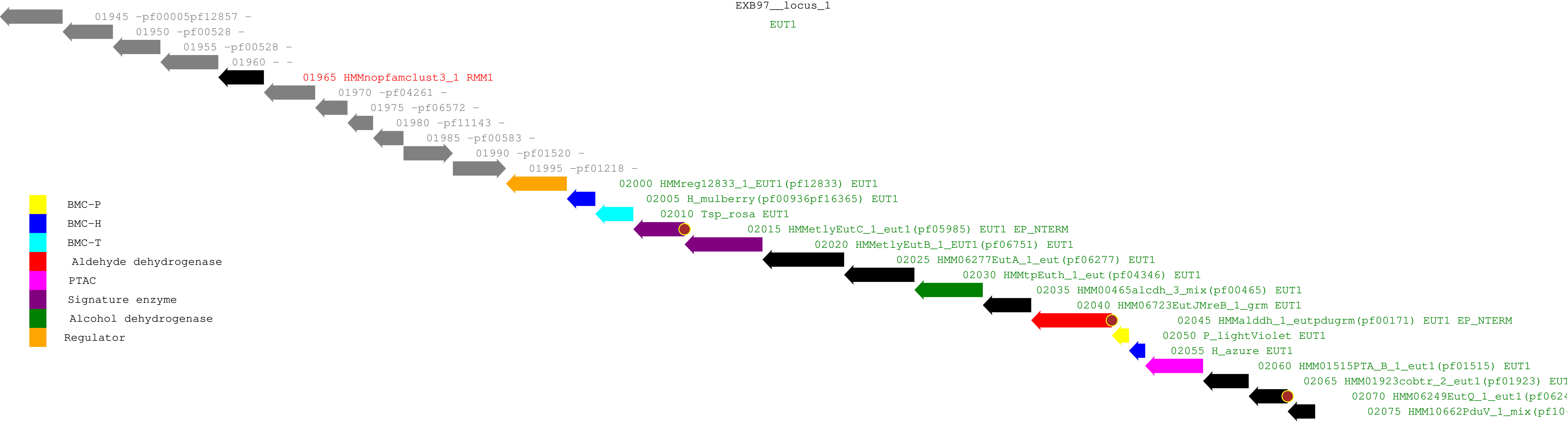

Salmonella enterica subsp. enterica serovar Agbeni. Bacteria; Proteobacteria; Gammaproteobacteria; Enterobacterales;

EXP27__locus_1

EUT1

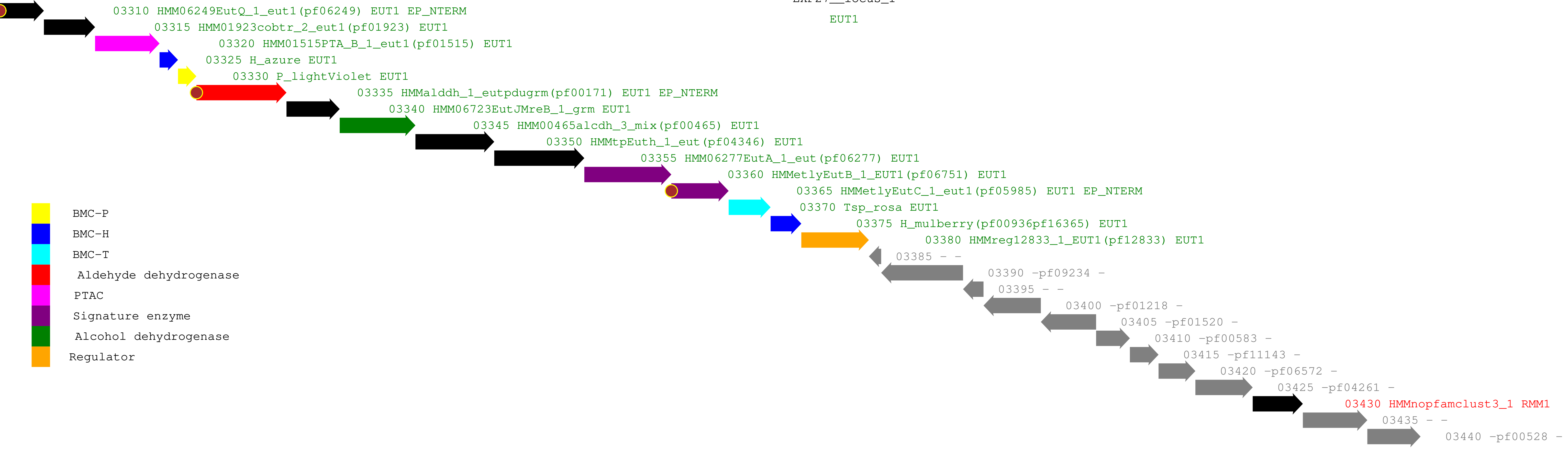

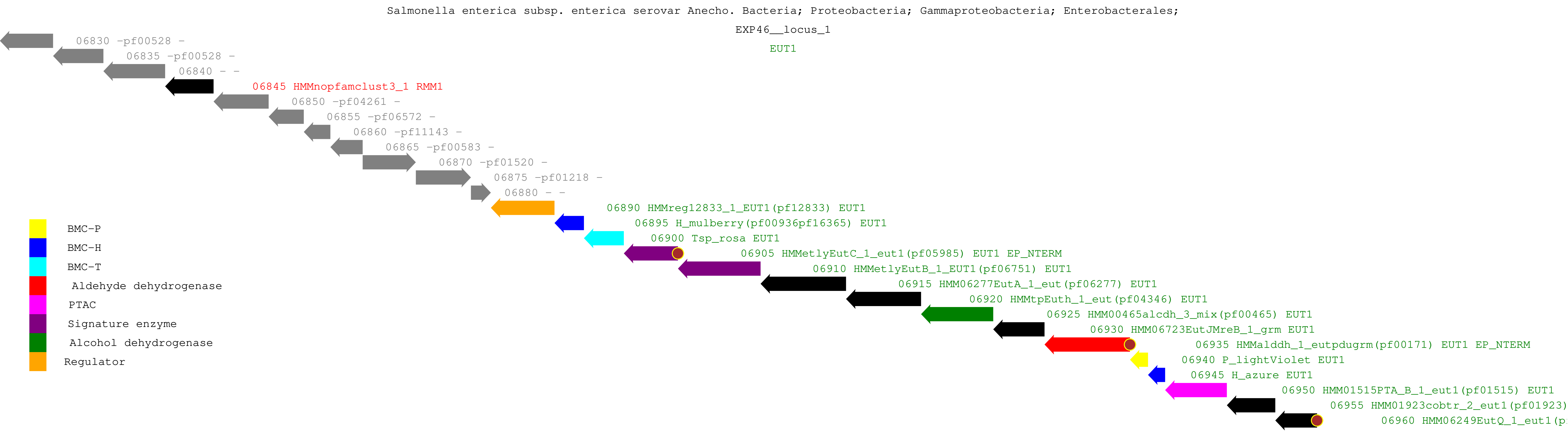

Salmonella enterica subsp. enterica serovar Anecho. Bacteria; Proteobacteria; Gammaproteobacteria; Enterobacterales;
EXP46__locus_1
EUT1
06830 -pf00528 -
06835 -pf00528 -
06840 - -
06845 HMMnopfamclust3_1 RMM1
06850 -pf04261 -
06855 -pf06572 -
06860 -pf11143 -
06865 -pf00583 -
06870 -pf01520 -
06875 -pf01218 -
06880 - -
06890 HMMreg12833_1_EUT1(pf12833) EUT1
06895 H_mulberry(pf00936pf16365) EUT1
06900 Tsp_rosa EUT1
06905 HMMetlyEutC_1_eut1(pf05985) EUT1 EP_NTERM
06910 HMMetlyEutB_1_EUT1(pf06751) EUT1
06915 HMM06277EutA_1_eut(pf06277) EUT1
06920 HMMtpEuth_1_eut(pf04346) EUT1
06925 HMM00465alcdh_3_mix(pf00465) EUT1
06930 HMM06723EutJMreB_1_grm EUT1
06935 HMMalddh_1_eutpdugrm(pf00171) EUT1 EP_NTERM
06940 P_lightViolet EUT1
06945 H_azure EUT1
06950 HMM01515PTA_B_1_eut1(pf01515) EUT1
06955 HMM01923cobtr_2_eut1(pf01923)
06960 HMM06249EutQ_1_eut1(p
BMC-P
BMC-H
BMC-T
Aldehyde dehydrogenase
PTAC
Signature enzyme
Alcohol dehydrogenase
Regulator

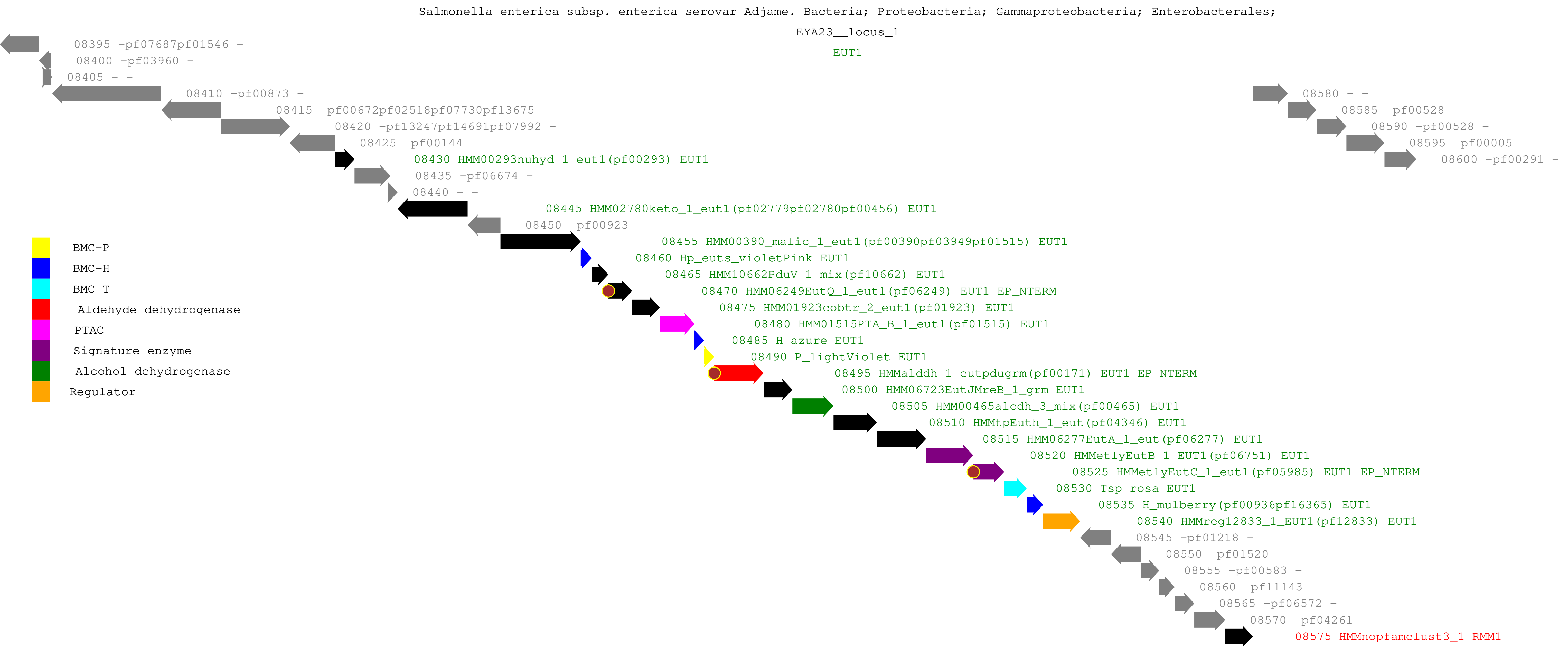

Salmonella enterica subsp. enterica serovar Adjame. Bacteria; Proteobacteria; Gammaproteobacteria; Enterobacterales;
EYA23__locus_1
EUT1
08395 -pf07687pf01546 -
08400 -pf03960 -
08405 - -
08410 -pf00873 -
08415 -pf00672pf02518pf07730pf13675 -
08420 -pf13247pf14691pf07992 -
08425 -pf00144 -
08430 HMM00293nuhyd_1_eut1(pf00293) EUT1
08435 -pf06674 -
08440 - -
08445 HMM02780keto_1_eut1(pf02779pf02780pf00456) EUT1
08450 -pf00923 -
08455 HMM00390_malic_1_eut1(pf00390pf03949pf01515) EUT1
08460 Hp_euts_violetPink EUT1
08465 HMM10662PduV_1_mix(pf10662) EUT1
08470 HMM06249EutQ_1_eut1(pf06249) EUT1 EP_NTERM
08475 HMM01923cobtr_2_eut1(pf01923) EUT1
08480 HMM01515PTA_B_1_eut1(pf01515) EUT1
08485 H_azure EUT1
08490 P_lightViolet EUT1
08495 HMMalddh_1_eutpdugrm(pf00171) EUT1 EP_NTERM
08500 HMM06723EutJMreB_1_grm EUT1
08505 HMM00465alcdh_3_mix(pf00465) EUT1
08510 HMMtpEuth_1_eut(pf04346) EUT1
08515 HMM06277EutA_1_eut(pf06277) EUT1
08520 HMMetlyEutB_1_EUT1(pf06751) EUT1
08525 HMMetlyEutC_1_eut1(pf05985) EUT1 EP_NTERM
08530 Tsp_rosa EUT1
08535 H_mulberry(pf00936pf16365) EUT1
08540 HMMreg12833_1_EUT1(pf12833) EUT1
08545 -pf01218 -
08550 -pf01520 -
08555 -pf00583 -
08560 -pf11143 -
08565 -pf06572 -
08570 -pf04261 -
08575 HMMnopfamclust3_1 RMM1
08580 - -
08585 -pf00528 -
08590 -pf00528 -
08595 -pf00005 -
08600 -pf00291 -
BMC-P
BMC-H
BMC-T
Aldehyde dehydrogenase
PTAC
Signature enzyme
Alcohol dehydrogenase
Regulator

Escherichia albertii. Bacteria; Proteobacteria; Gammaproteobacteria; Enterobacterales;

EYS06__locus_1

EUT1

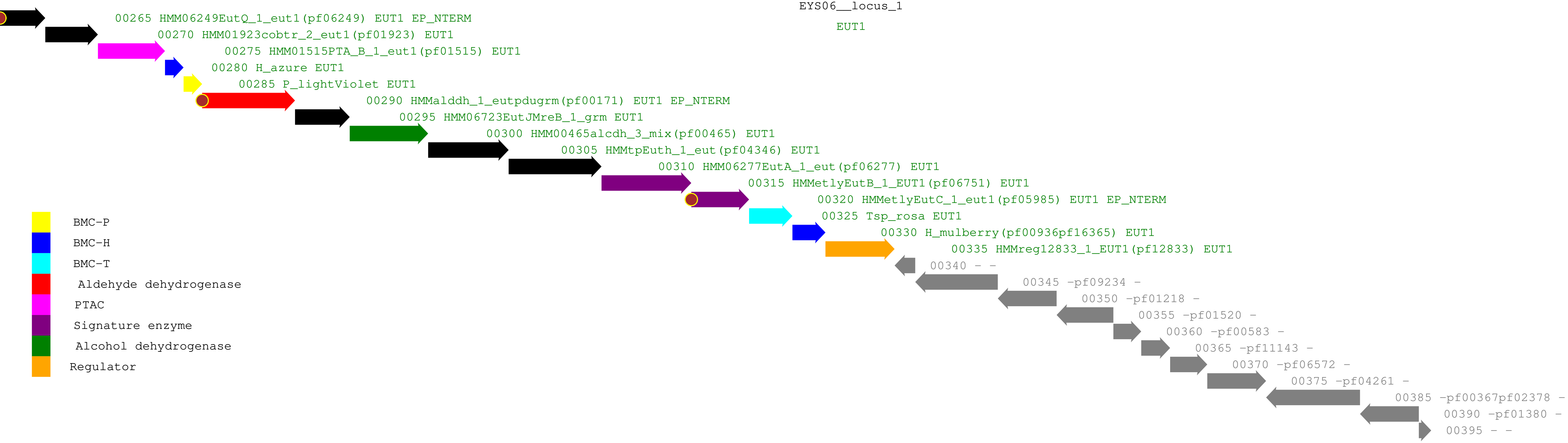

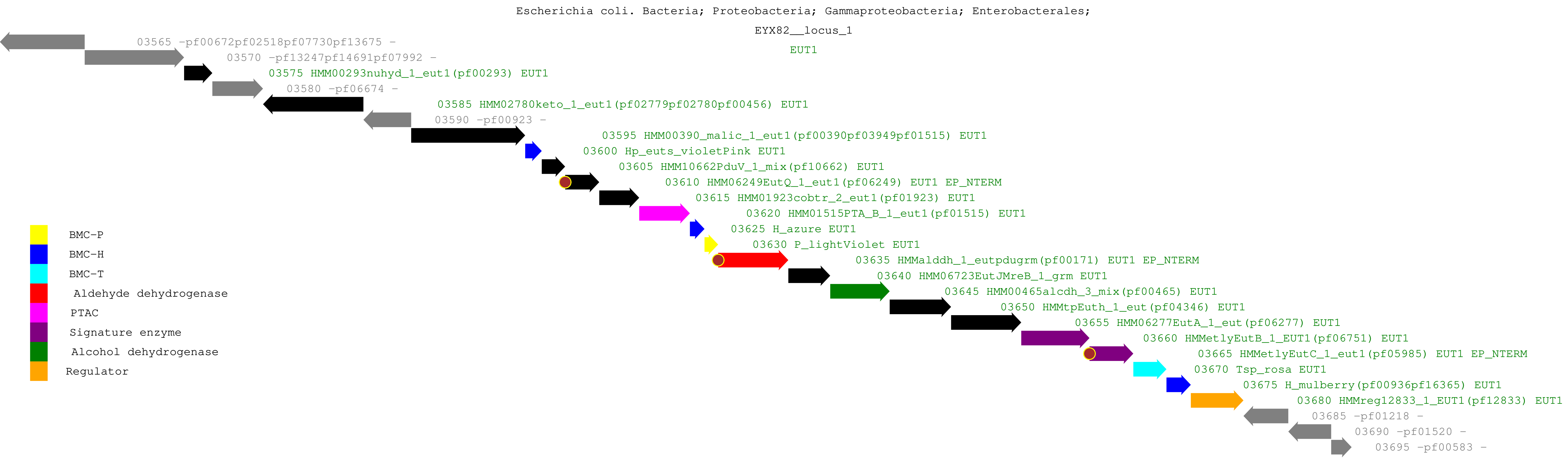

Escherichia coli. Bacteria; Proteobacteria; Gammaproteobacteria; Enterobacterales;
EYX82__locus_1
EUT1
03565 -pf00672pf02518pf07730pf13675 -
03570 -pf13247pf14691pf07992 -
03575 HMM00293nuhyd_1_eut1(pf00293) EUT1
03580 -pf06674 -
03585 HMM02780keto_1_eut1(pf02779pf02780pf00456) EUT1
03590 -pf00923 -
03595 HMM00390_malic_1_eut1(pf00390pf03949pf01515) EUT1
03600 Hp_euts_violetPink EUT1
03605 HMM10662PduV_1_mix(pf10662) EUT1
03610 HMM06249EutQ_1_eut1(pf06249) EUT1 EP_NTERM
03615 HMM01923cobtr_2_eut1(pf01923) EUT1
03620 HMM01515PTA_B_1_eut1(pf01515) EUT1
03625 H_azure EUT1
03630 P_lightViolet EUT1
03635 HMMalddh_1_eutpdugrm(pf00171) EUT1 EP_NTERM
03640 HMM06723EutJMreB_1_grm EUT1
03645 HMM00465alcdh_3_mix(pf00465) EUT1
03650 HMMtpEuth_1_eut(pf04346) EUT1
03655 HMM06277EutA_1_eut(pf06277) EUT1
03660 HMMetlyEutB_1_EUT1(pf06751) EUT1
03665 HMMetlyEutC_1_eut1(pf05985) EUT1 EP_NTERM
03670 Tsp_rosa EUT1
03675 H_mulberry(pf00936pf16365) EUT1
03680 HMMreg12833_1_EUT1(pf12833) EUT1
03685 -pf01218 -
03690 -pf01520 -
03695 -pf00583 -
BMC-P
BMC-H
BMC-T
Aldehyde dehydrogenase
PTAC
Signature enzyme
Alcohol dehydrogenase
Regulator

Obesumbacterium proteus. Bacteria; Proteobacteria; Gammaproteobacteria; Enterobacterales;

EYY98__locus_1

EUT1

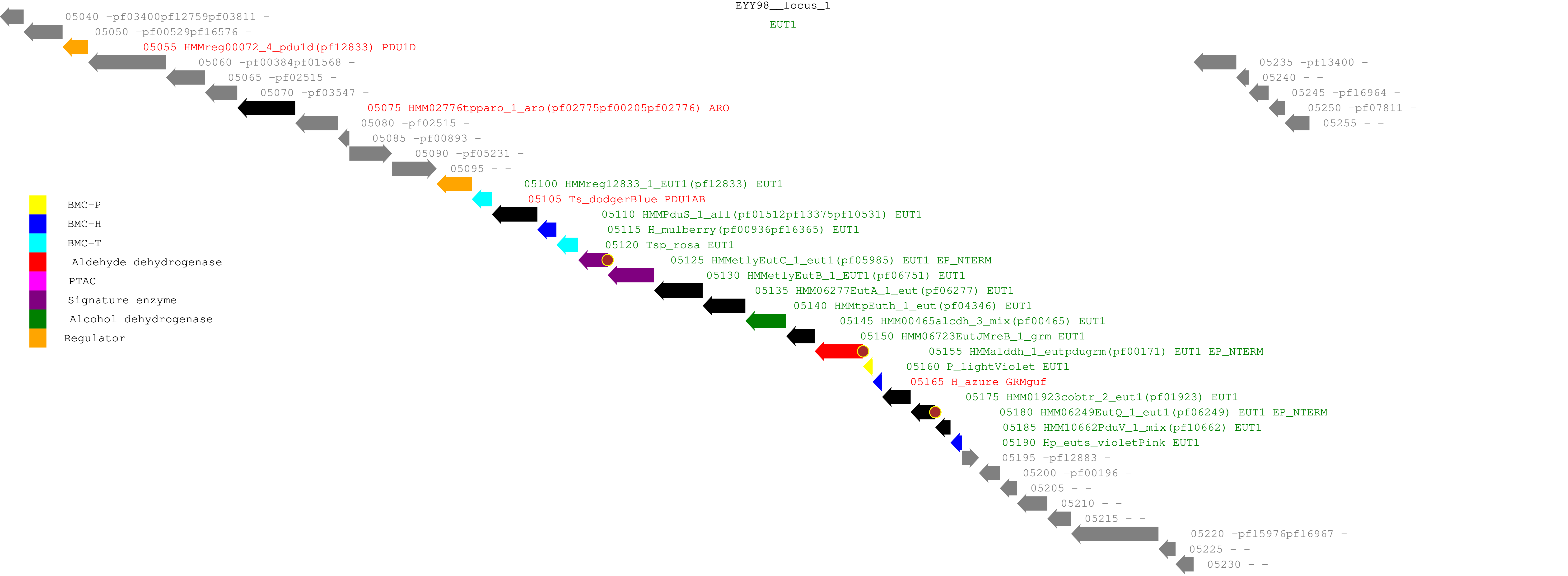

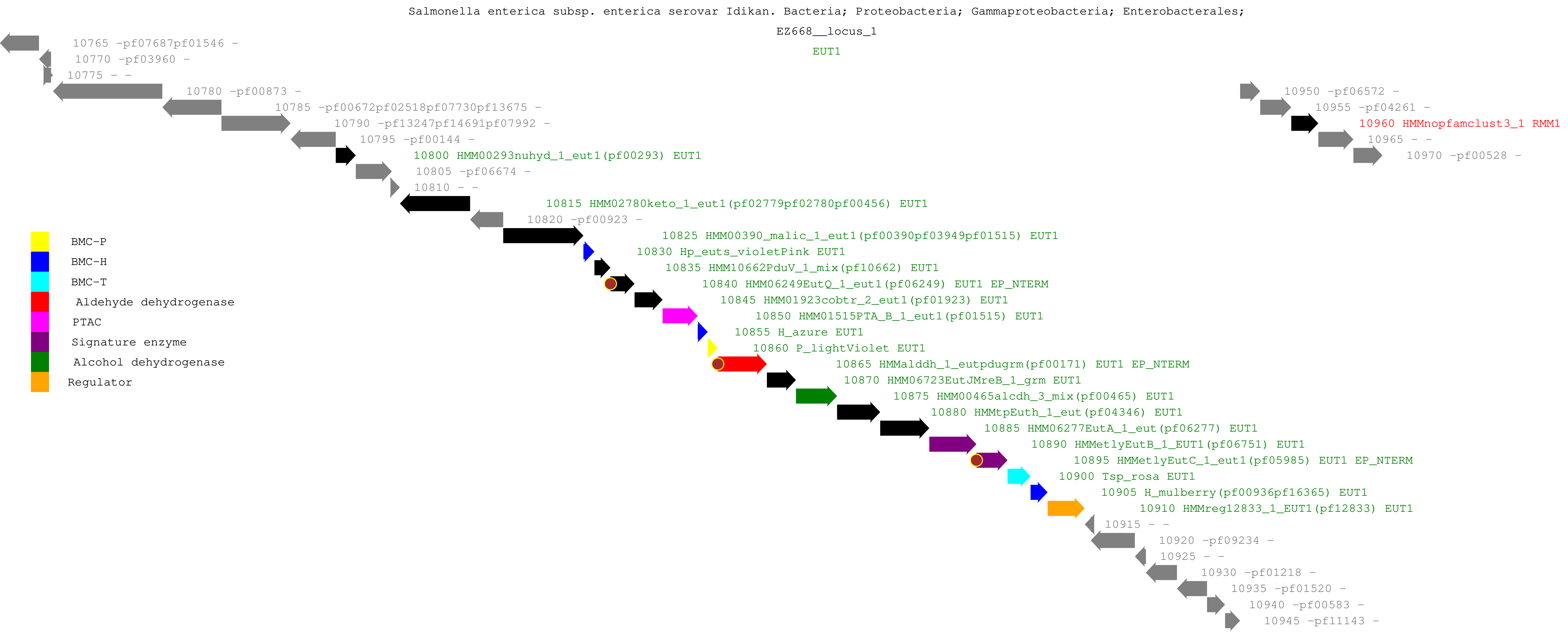
Salmonella enterica subsp. enterica serovar Idikan. Bacteria; Proteobacteria; Gammaproteobacteria; Enterobacterales;
EZ668__locus_1
EUT1
10765 -pf07687pf01546 -
10770 -pf03960 -
10775 - -
10780 -pf00873 -
10785 -pf00672pf02518pf07730pf13675 -
10790 -pf13247pf14691pf07992 -
10795 -pf00144 -
10800 HMM00293nuhyd_1_eut1(pf00293) EUT1
10805 -pf06674 -
10810 - -
10815 HMM02780keto_1_eut1(pf02779pf02780pf00456) EUT1
10820 -pf00923 -
10825 HMM00390_malic_1_eut1(pf00390pf03949pf01515) EUT1
10830 Hp_euts_violetPink EUT1
10835 HMM10662PduV_1_mix(pf10662) EUT1
10840 HMM06249EutQ_1_eut1(pf06249) EUT1 EP_NTERM
10845 HMM01923cobtr_2_eut1(pf01923) EUT1
10850 HMM01515PTA_B_1_eut1(pf01515) EUT1
10855 H_azure EUT1
10860 P_lightViolet EUT1
10865 HMMalddh_1_eutpdugrm(pf00171) EUT1 EP_NTERM
10870 HMM06723EutJMreB_1_grm EUT1
10875 HMM00465alcdh_3_mix(pf00465) EUT1
10880 HMMtpEuth_1_eut(pf04346) EUT1
10885 HMM06277EutA_1_eut(pf06277) EUT1
10890 HMMetlyEutB_1_EUT1(pf06751) EUT1
10895 HMMetlyEutC_1_eut1(pf05985) EUT1 EP_NTERM
10900 Tsp_rosa EUT1
10905 H_mulberry(pf00936pf16365) EUT1
10910 HMMreg12833_1_EUT1(pf12833) EUT1
10915 - -
10920 -pf09234 -
10925 - -
10930 -pf01218 -
10935 -pf01520 -
10940 -pf00583 -
10945 -pf11143 -
10950 -pf06572 -
10955 -pf04261 -
10960 HMMnopfamclust3_1 RMM1
10965 - -
10970 -pf00528 -
BMC-P
BMC-H
BMC-T
Aldehyde dehydrogenase
PTAC
Signature enzyme
Alcohol dehydrogenase
Regulator

Salmonella enterica subsp. enterica serovar Fulica. Bacteria; Proteobacteria; Gammaproteobacteria; Enterobacterales;

EZ771__locus_1

EUT1

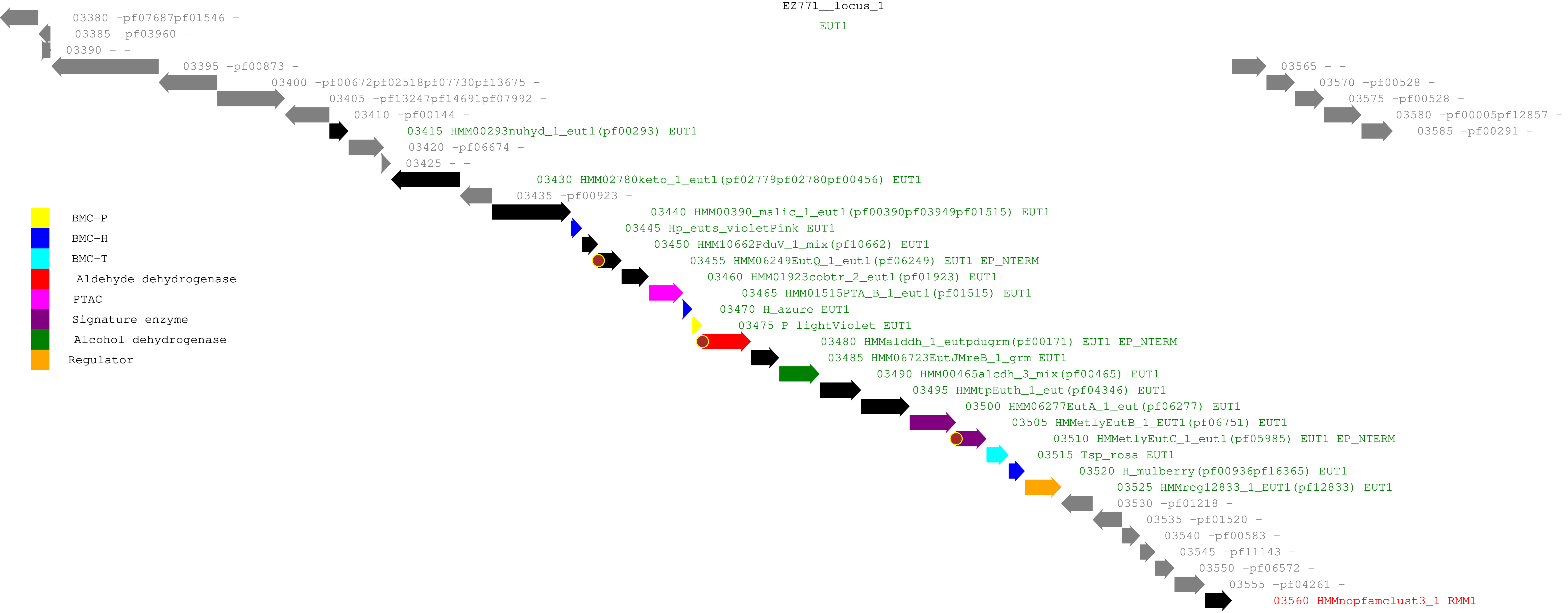

Salmonella enterica subsp. enterica serovar Vejle. Bacteria; Proteobacteria; Gammaproteobacteria; Enterobacterales;

EZ810__locus_2

EUT1

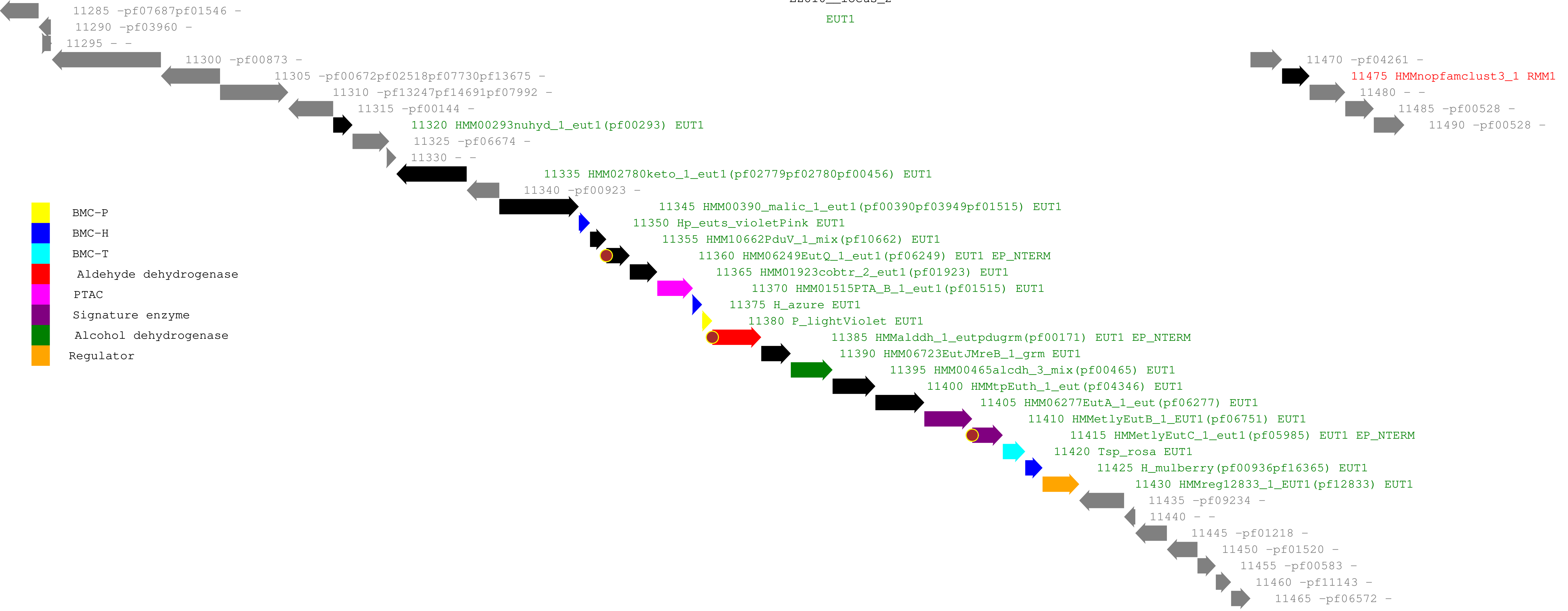

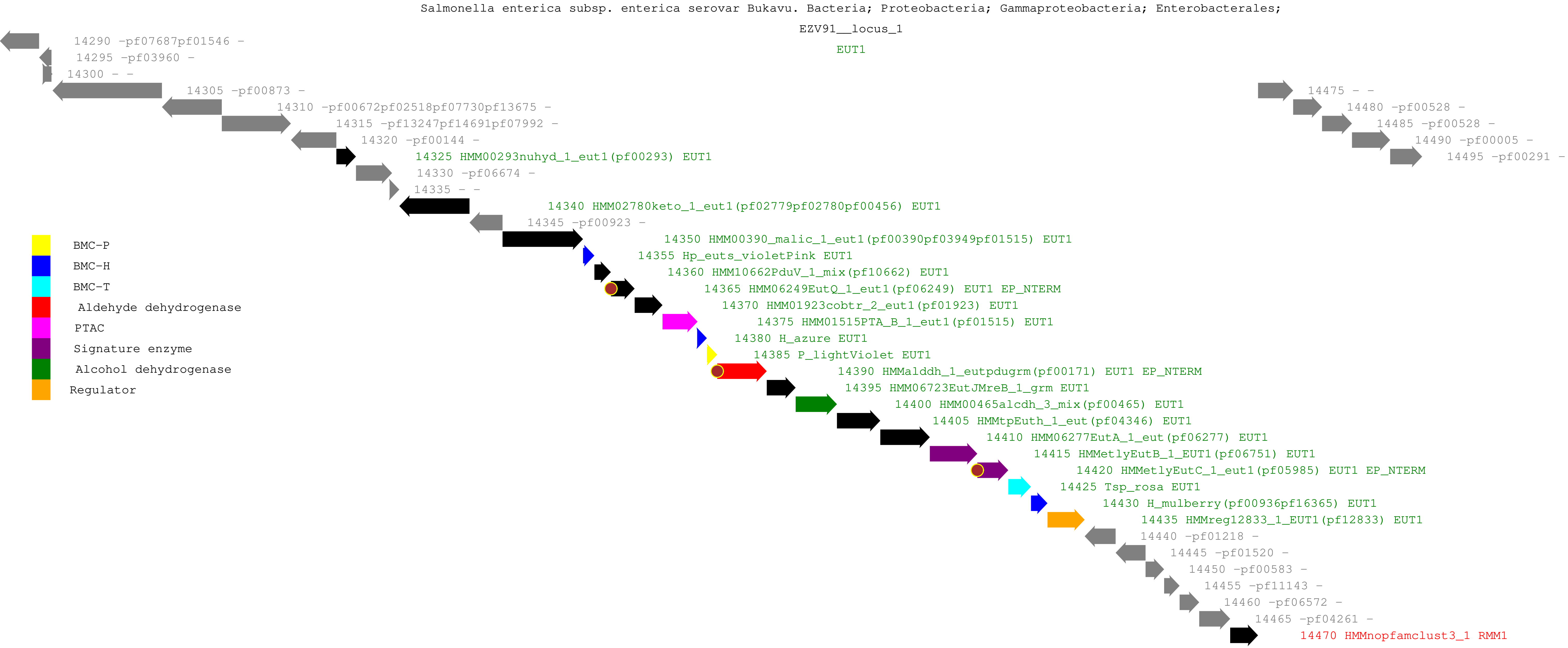
Salmonella enterica subsp. enterica serovar Bukavu. Bacteria; Proteobacteria; Gammaproteobacteria; Enterobacterales;
EZV91__locus_1
EUT1
14290 -pf07687pf01546 -
14295 -pf03960 -
14300 - -
14305 -pf00873 -
14310 -pf00672pf02518pf07730pf13675 -
14315 -pf13247pf14691pf07992 -
14320 -pf00144 -
14325 HMM00293nuhyd_1_eut1(pf00293) EUT1
14330 -pf06674 -
14335 - -
14340 HMM02780keto_1_eut1(pf02779pf02780pf00456) EUT1
14345 -pf00923 -
14350 HMM00390_malic_1_eut1(pf00390pf03949pf01515) EUT1
14355 Hp_euts_violetPink EUT1
14360 HMM10662PduV_1_mix(pf10662) EUT1
14365 HMM06249EutQ_1_eut1(pf06249) EUT1 EP_NTERM
14370 HMM01923cobtr_2_eut1(pf01923) EUT1
14375 HMM01515PTA_B_1_eut1(pf01515) EUT1
14380 H_azure EUT1
14385 P_lightViolet EUT1
14390 HMMalddh_1_eutpdugrm(pf00171) EUT1 EP_NTERM
14395 HMM06723EutJMreB_1_grm EUT1
14400 HMM00465alcdh_3_mix(pf00465) EUT1
14405 HMMtpEuth_1_eut(pf04346) EUT1
14410 HMM06277EutA_1_eut(pf06277) EUT1
14415 HMMetlyEutB_1_EUT1(pf06751) EUT1
14420 HMMetlyEutC_1_eut1(pf05985) EUT1 EP_NTERM
14425 Tsp_rosa EUT1
14430 H_mulberry(pf00936pf16365) EUT1
14435 HMMreg12833_1_EUT1(pf12833) EUT1
14440 -pf01218 -
14445 -pf01520 -
14450 -pf00583 -
14455 -pf11143 -
14460 -pf06572 -
14465 -pf04261 -
14470 HMMnopfamclust3_1 RMM1
14475 - -
14480 -pf00528 -
14485 -pf00528 -
14490 -pf00005 -
14495 -pf00291 -
BMC-P
BMC-H
BMC-T
Aldehyde dehydrogenase
PTAC
Signature enzyme
Alcohol dehydrogenase
Regulator

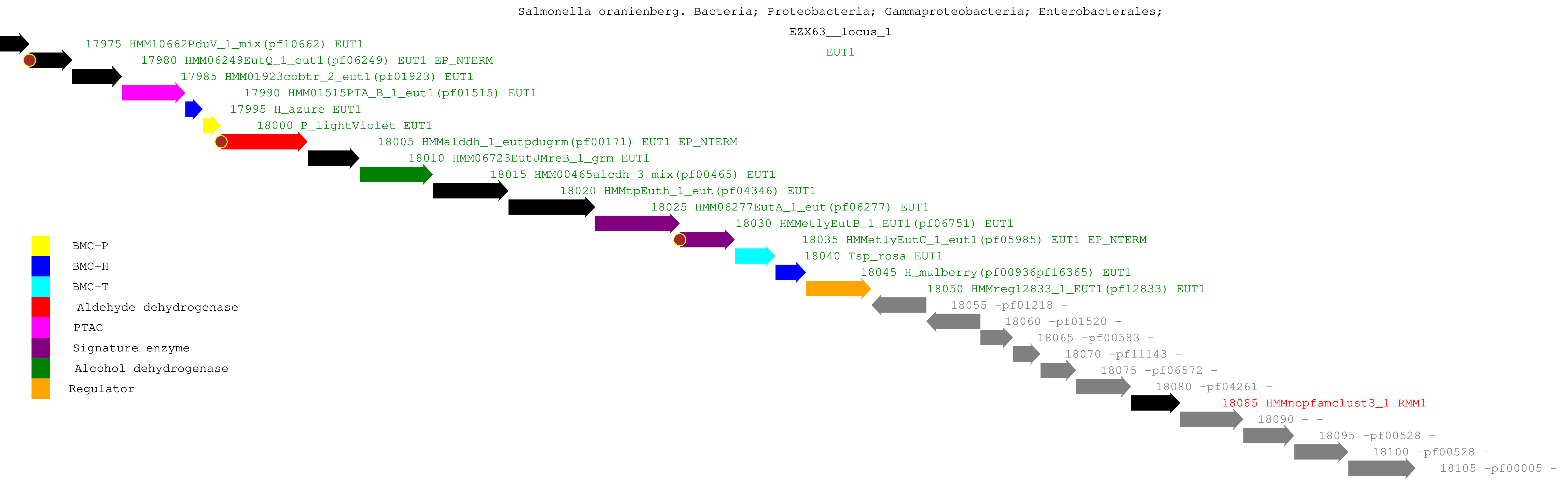

Salmonella oranienberg. Bacteria; Proteobacteria; Gammaproteobacteria; Enterobacterales;
EZX63__locus_1
EUT1
17975 HMM10662PduV_1_mix(pf10662) EUT1
17980 HMM06249EutQ_1_eut1(pf06249) EUT1 EP_NTERM
17985 HMM01923cobtr_2_eut1(pf01923) EUT1
17990 HMM01515PTA_B_1_eut1(pf01515) EUT1
17995 H_azure EUT1
18000 P_lightViolet EUT1
18005 HMMalddh_1_eutpdugrm(pf00171) EUT1 EP_NTERM
18010 HMM06723EutJMreB_1_grm EUT1
18015 HMM00465alcdh_3_mix(pf00465) EUT1
18020 HMMtpEuth_1_eut(pf04346) EUT1
18025 HMM06277EutA_1_eut(pf06277) EUT1
18030 HMMetlyEutB_1_EUT1(pf06751) EUT1
18035 HMMetlyEutC_1_eut1(pf05985) EUT1 EP_NTERM
18040 Tsp_rosa EUT1
18045 H_mulberry(pf00936pf16365) EUT1
18050 HMMreg12833_1_EUT1(pf12833) EUT1
18055 -pf01218 -
18060 -pf01520 -
18065 -pf00583 -
18070 -pf11143 -
18075 -pf06572 -
18080 -pf04261 -
18085 HMMnopfamclust3_1 RMM1
18090 - -
18095 -pf00528 -
18100 -pf00528 -
18105 -pf00005 -
BMC-P
BMC-H
BMC-T
Aldehyde dehydrogenase
PTAC
Signature enzyme
Alcohol dehydrogenase
Regulator

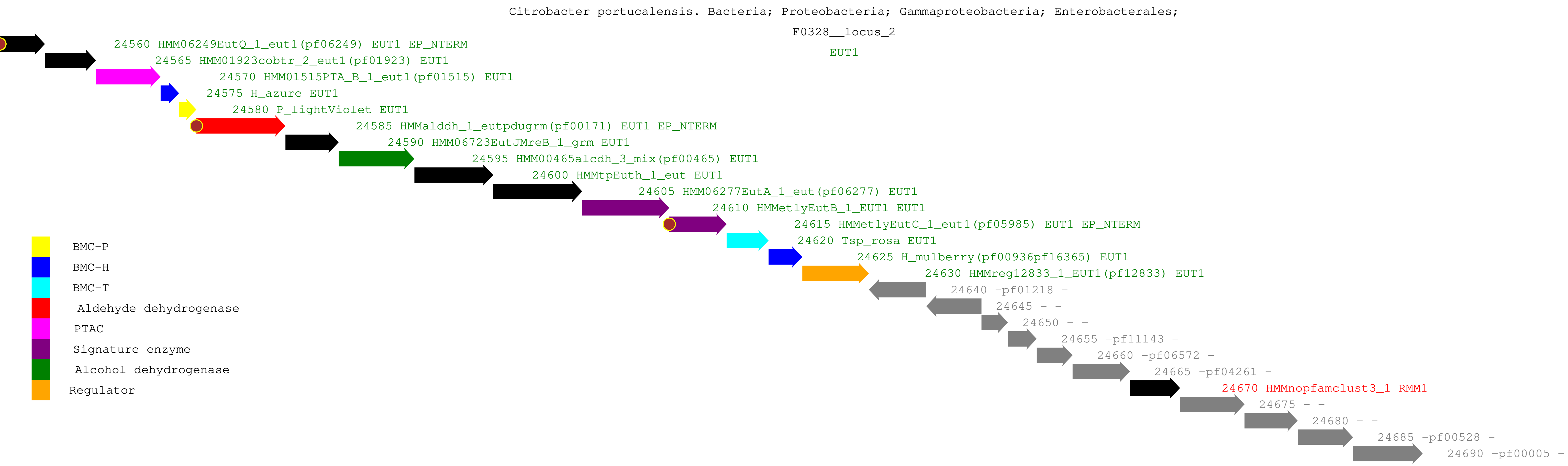
Citrobacter portucalensis. Bacteria; Proteobacteria; Gammaproteobacteria; Enterobacterales;
F0328__locus_2
EUT1
24560 HMM06249EutQ_1_eut1(pf06249) EUT1 EP_NTERM
24565 HMM01923cobtr_2_eut1(pf01923) EUT1
24570 HMM01515PTA_B_1_eut1(pf01515) EUT1
24575 H_azure EUT1
24580 P_lightViolet EUT1
24585 HMMalddh_1_eutpdugrm(pf00171) EUT1 EP_NTERM
24590 HMM06723EutJMreB_1_grm EUT1
24595 HMM00465alcdh_3_mix(pf00465) EUT1
24600 HMMtpEuth_1_eut EUT1
24605 HMM06277EutA_1_eut(pf06277) EUT1
24610 HMMetlyEutB_1_EUT1 EUT1
24615 HMMetlyEutC_1_eut1(pf05985) EUT1 EP_NTERM
24620 Tsp_rosa EUT1
24625 H_mulberry(pf00936pf16365) EUT1
24630 HMMreg12833_1_EUT1(pf12833) EUT1
24640 -pf01218 -
24645 - -
24650 - -
24655 -pf11143 -
24660 -pf06572 -
24665 -pf04261 -
24670 HMMnopfamclust3_1 RMM1
24675 - -
24680 - -
24685 -pf00528 -
24690 -pf00005 -
BMC-P
BMC-H
BMC-T
Aldehyde dehydrogenase
PTAC
Signature enzyme
Alcohol dehydrogenase
Regulator

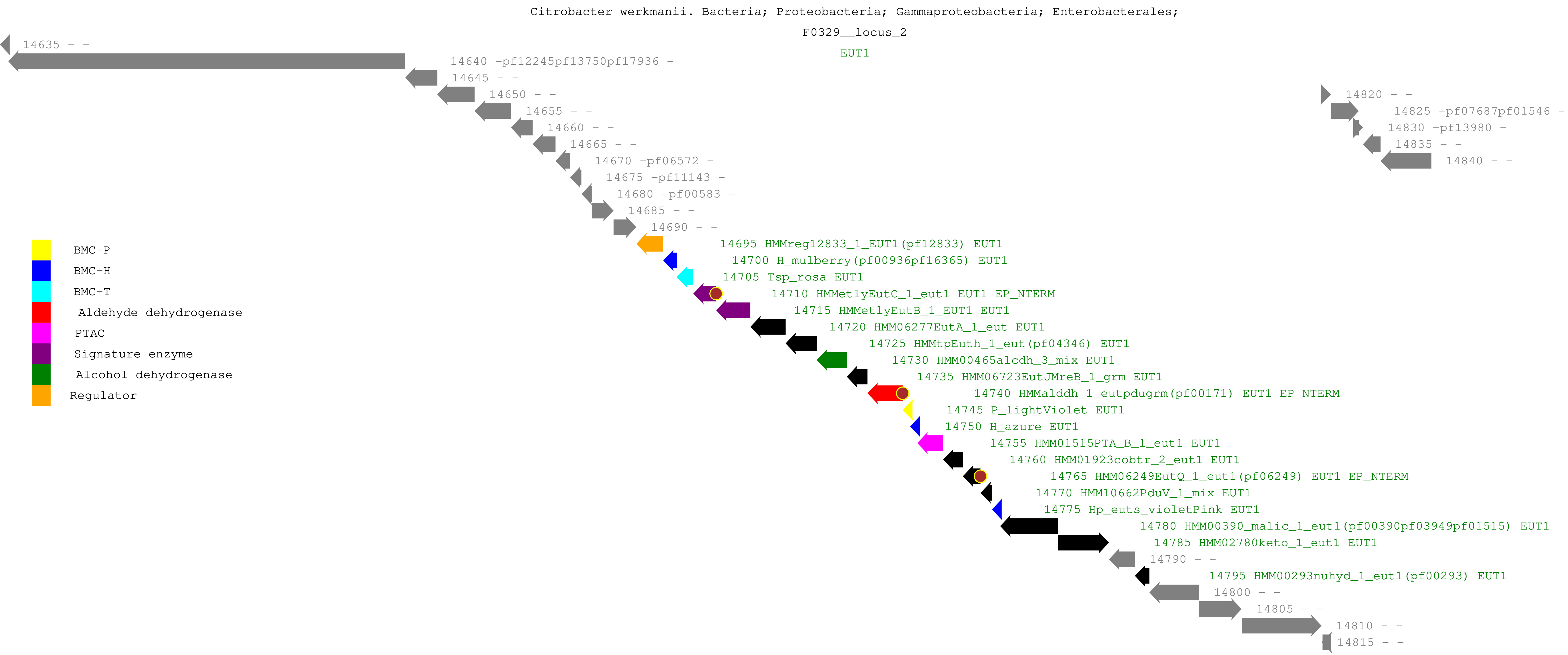
Citrobacter werkmanii. Bacteria; Proteobacteria; Gammaproteobacteria; Enterobacterales;
F0329__locus_2
EUT1
14635 - -
14640 -pf12245pf13750pf17936 -
14645 - -
14650 - -
14655 - -
14660 - -
14665 - -
14670 -pf06572 -
14675 -pf11143 -
14680 -pf00583 -
14685 - -
14690 - -
14695 HMMreg12833_1_EUT1(pf12833) EUT1
14700 H_mulberry(pf00936pf16365) EUT1
14705 Tsp_rosa EUT1
14710 HMMetlyEutC_1_eut1 EUT1 EP_NTERM
14715 HMMetlyEutB_1_EUT1 EUT1
14720 HMM06277EutA_1_eut EUT1
14725 HMMtpEuth_1_eut(pf04346) EUT1
14730 HMM00465alcdh_3_mix EUT1
14735 HMM06723EutJMreB_1_grm EUT1
14740 HMMalddh_1_eutpdugrm(pf00171) EUT1 EP_NTERM
14745 P_lightViolet EUT1
14750 H_azure EUT1
14755 HMM01515PTA_B_1_eut1 EUT1
14760 HMM01923cobtr_2_eut1 EUT1
14765 HMM06249EutQ_1_eut1(pf06249) EUT1 EP_NTERM
14770 HMM10662PduV_1_mix EUT1
14775 Hp_euts_violetPink EUT1
14780 HMM00390_malic_1_eut1(pf00390pf03949pf01515) EUT1
14785 HMM02780keto_1_eut1 EUT1
14790 - -
14795 HMM00293nuhyd_1_eut1(pf00293) EUT1
14800 - -
14805 - -
14810 - -
14815 - -
14820 - -
14825 -pf07687pf01546 -
14830 -pf13980 -
14835 - -
14840 - -
BMC-P
BMC-H
BMC-T
Aldehyde dehydrogenase
PTAC
Signature enzyme
Alcohol dehydrogenase
Regulator

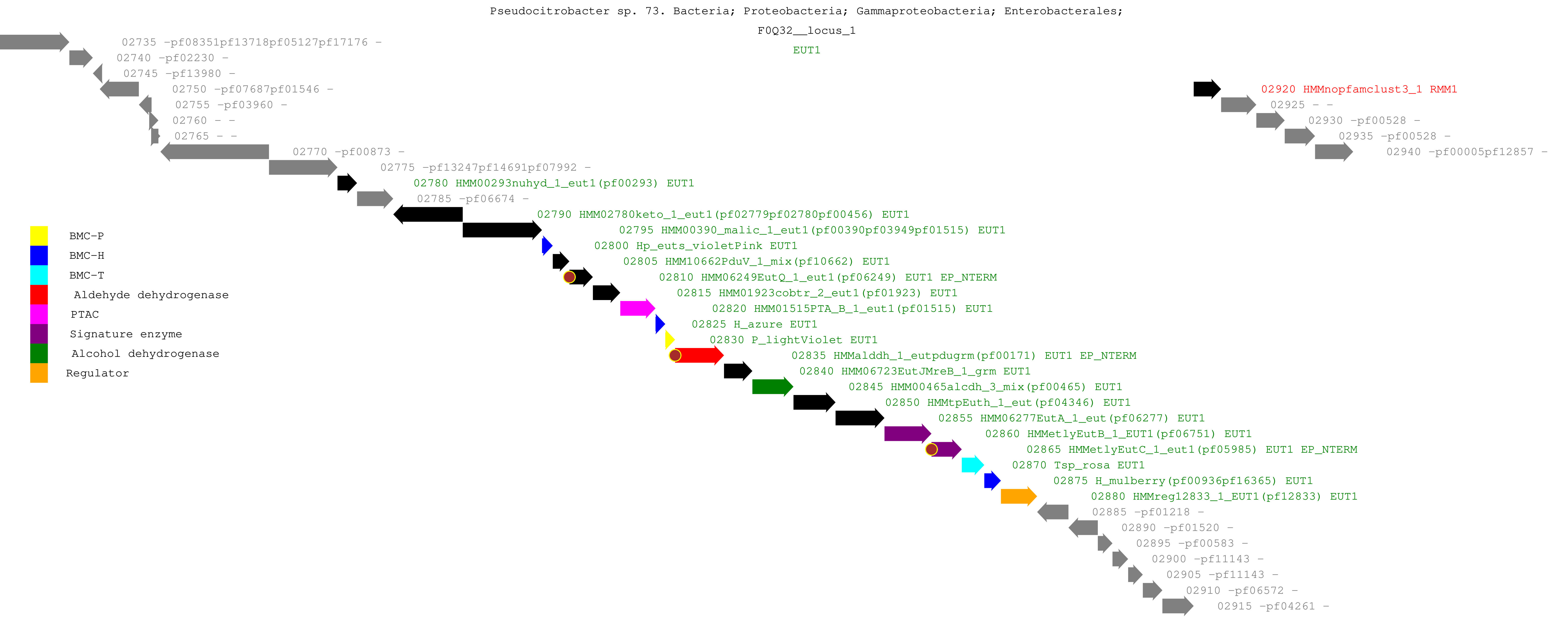
Pseudocitrobacter sp. 73. Bacteria; Proteobacteria; Gammaproteobacteria; Enterobacterales;
F0Q32__locus_1
EUT1
02735 -pf08351pf13718pf05127pf17176 -
02740 -pf02230 -
02745 -pf13980 -
02750 -pf07687pf01546 -
02755 -pf03960 -
02760 - -
02765 - -
02770 -pf00873 -
02775 -pf13247pf14691pf07992 -
02780 HMM00293nuhyd_1_eut1(pf00293) EUT1
02785 -pf06674 -
02790 HMM02780keto_1_eut1(pf02779pf02780pf00456) EUT1
02795 HMM00390_malic_1_eut1(pf00390pf03949pf01515) EUT1
02800 Hp_euts_violetPink EUT1
02805 HMM10662PduV_1_mix(pf10662) EUT1
02810 HMM06249EutQ_1_eut1(pf06249) EUT1 EP_NTERM
02815 HMM01923cobtr_2_eut1(pf01923) EUT1
02820 HMM01515PTA_B_1_eut1(pf01515) EUT1
02825 H_azure EUT1
02830 P_lightViolet EUT1
02835 HMMalddh_1_eutpdugrm(pf00171) EUT1 EP_NTERM
02840 HMM06723EutJMreB_1_grm EUT1
02845 HMM00465alcdh_3_mix(pf00465) EUT1
02850 HMMtpEuth_1_eut(pf04346) EUT1
02855 HMM06277EutA_1_eut(pf06277) EUT1
02860 HMMetlyEutB_1_EUT1(pf06751) EUT1
02865 HMMetlyEutC_1_eut1(pf05985) EUT1 EP_NTERM
02870 Tsp_rosa EUT1
02875 H_mulberry(pf00936pf16365) EUT1
02880 HMMreg12833_1_EUT1(pf12833) EUT1
02885 -pf01218 -
02890 -pf01520 -
02895 -pf00583 -
02900 -pf11143 -
02905 -pf11143 -
02910 -pf06572 -
02915 -pf04261 -
02920 HMMnopfamclust3_1 RMM1
02925 - -
02930 -pf00528 -
02935 -pf00528 -
02940 -pf00005pf12857 -
BMC-P
BMC-H
BMC-T
Aldehyde dehydrogenase
PTAC
Signature enzyme
Alcohol dehydrogenase
Regulator

Escherichia coli. Bacteria; Proteobacteria; Gammaproteobacteria; Enterobacterales;

F1E03__locus_1

EUT1

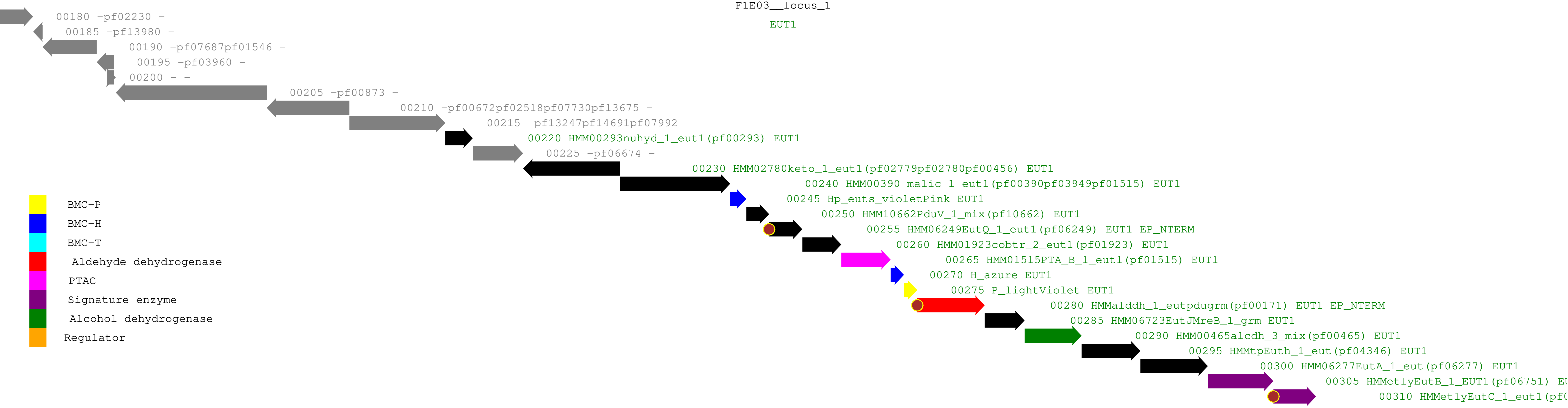

Citrobacter amalonaticus Y19. Bacteria; Proteobacteria; Gammaproteobacteria; Enterobacterales;

F384__locus_2

EUT1

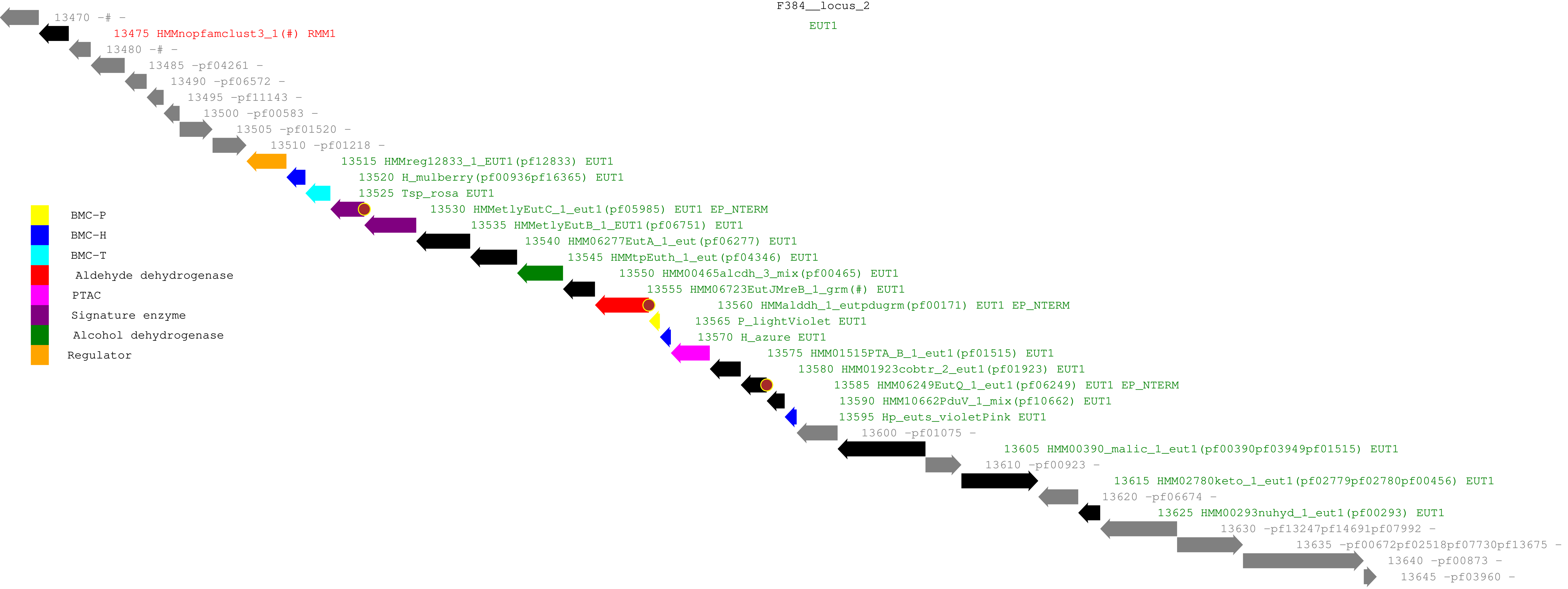

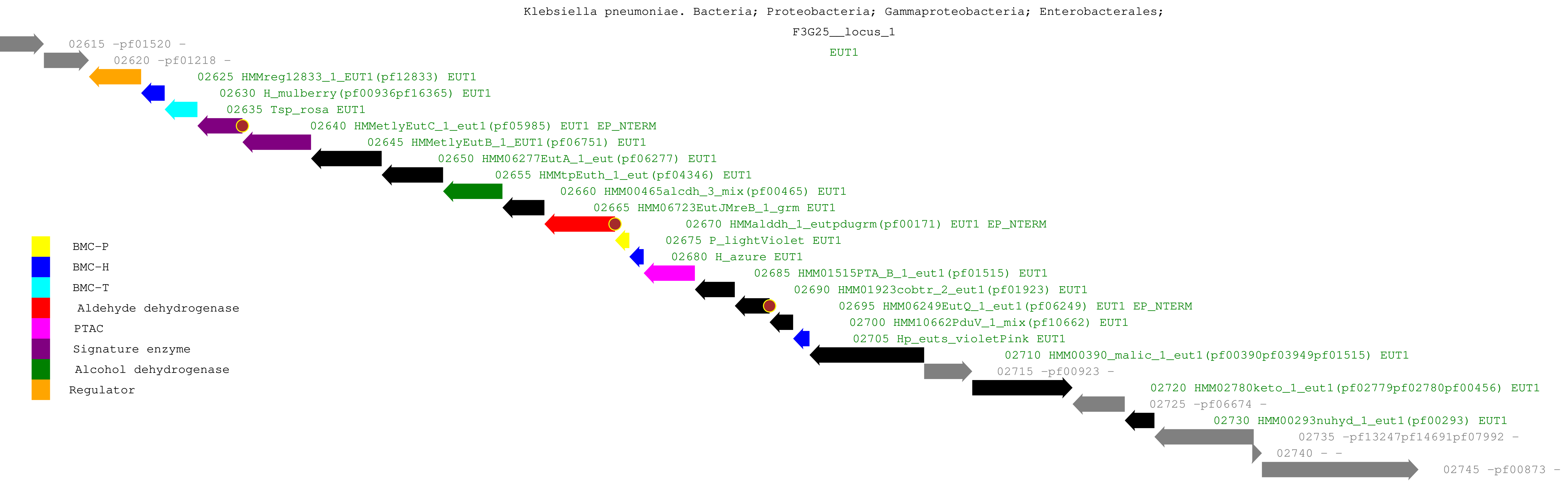
Klebsiella pneumoniae. Bacteria; Proteobacteria; Gammaproteobacteria; Enterobacterales;
F3G25__locus_1
EUT1
02615 -pf01520 -
02620 -pf01218 -
02625 HMMreg12833_1_EUT1(pf12833) EUT1
02630 H_mulberry(pf00936pf16365) EUT1
02635 Tsp_rosa EUT1
02640 HMMetlyEutC_1_eut1(pf05985) EUT1 EP_NTERM
02645 HMMetlyEutB_1_EUT1(pf06751) EUT1
02650 HMM06277EutA_1_eut(pf06277) EUT1
02655 HMMtpEuth_1_eut(pf04346) EUT1
02660 HMM00465alcdh_3_mix(pf00465) EUT1
02665 HMM06723EutJMreB_1_grm EUT1
02670 HMMalddh_1_eutpdugrm(pf00171) EUT1 EP_NTERM
02675 P_lightViolet EUT1
02680 H_azure EUT1
02685 HMM01515PTA_B_1_eut1(pf01515) EUT1
02690 HMM01923cobtr_2_eut1(pf01923) EUT1
02695 HMM06249EutQ_1_eut1(pf06249) EUT1 EP_NTERM
02700 HMM10662PduV_1_mix(pf10662) EUT1
02705 Hp_euts_violetPink EUT1
02710 HMM00390_malic_1_eut1(pf00390pf03949pf01515) EUT1
02715 -pf00923 -
02720 HMM02780keto_1_eut1(pf02779pf02780pf00456) EUT1
02725 -pf06674 -
02730 HMM00293nuhyd_1_eut1(pf00293) EUT1
02735 -pf13247pf14691pf07992 -
02740 - -
02745 -pf00873 -
BMC-P
BMC-H
BMC-T
Aldehyde dehydrogenase
PTAC
Signature enzyme
Alcohol dehydrogenase
Regulator

Salmonella enterica subsp. enterica serovar Eastbourne. Bacteria; Proteobacteria; Gammaproteobacteria; Enterobacterales;

F7754__locus_2

EUT1

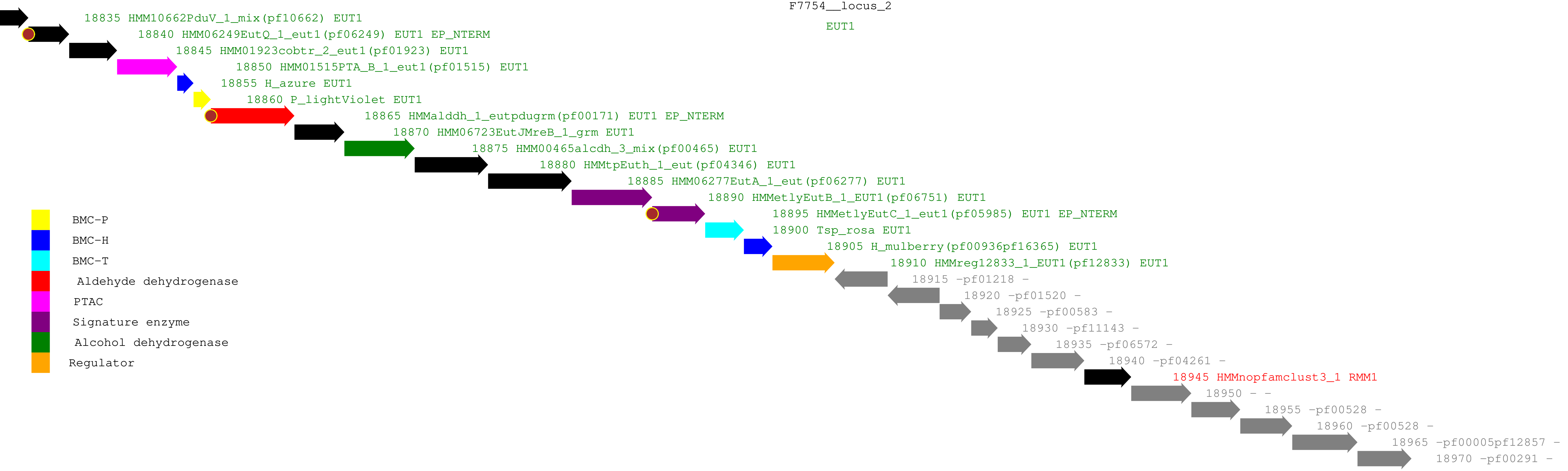

Salmonella enterica subsp. enterica serovar Nchanga. Bacteria; Proteobacteria; Gammaproteobacteria; Enterobacterales;

FA839__locus_2

EUT1

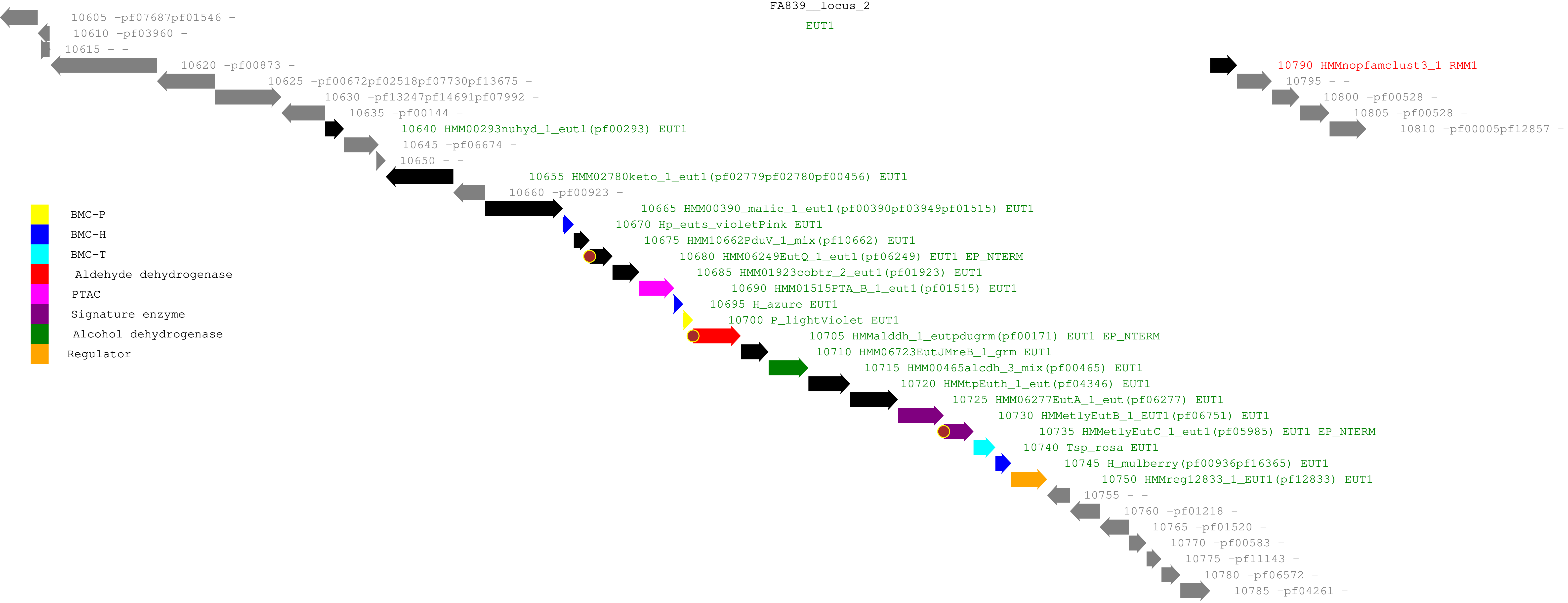

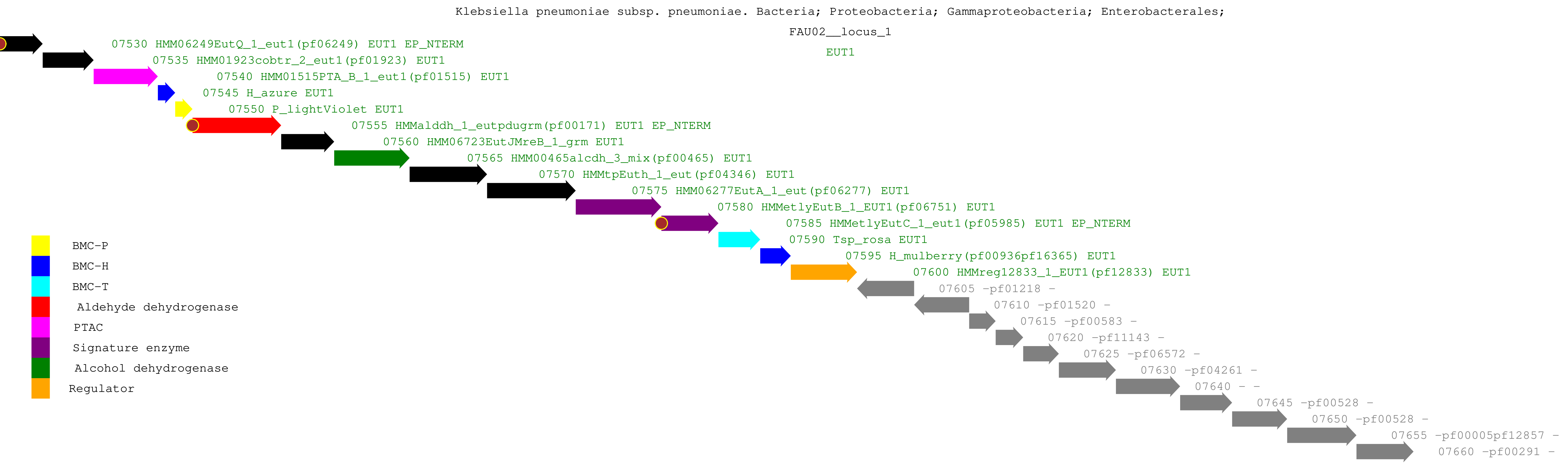
Klebsiella pneumoniae subsp. pneumoniae. Bacteria; Proteobacteria; Gammaproteobacteria; Enterobacterales;
FAU02__locus_1
EUT1
07530 HMM06249EutQ_1_eut1(pf06249) EUT1 EP_NTERM
07535 HMM01923cobtr_2_eut1(pf01923) EUT1
07540 HMM01515PTA_B_1_eut1(pf01515) EUT1
07545 H_azure EUT1
07550 P_lightViolet EUT1
07555 HMMalddh_1_eutpdugrm(pf00171) EUT1 EP_NTERM
07560 HMM06723EutJMreB_1_grm EUT1
07565 HMM00465alcdh_3_mix(pf00465) EUT1
07570 HMMtpEuth_1_eut(pf04346) EUT1
07575 HMM06277EutA_1_eut(pf06277) EUT1
07580 HMMetlyEutB_1_EUT1(pf06751) EUT1
07585 HMMetlyEutC_1_eut1(pf05985) EUT1 EP_NTERM
07590 Tsp_rosa EUT1
07595 H_mulberry(pf00936pf16365) EUT1
07600 HMMreg12833_1_EUT1(pf12833) EUT1
07605 -pf01218 -
07610 -pf01520 -
07615 -pf00583 -
07620 -pf11143 -
07625 -pf06572 -
07630 -pf04261 -
07640 - -
07645 -pf00528 -
07650 -pf00528 -
07655 -pf00005pf12857 -
07660 -pf00291 -
BMC-P
BMC-H
BMC-T
Aldehyde dehydrogenase
PTAC
Signature enzyme
Alcohol dehydrogenase
Regulator

Escherichia coli (strain K12). Bacteria; Proteobacteria; Gammaproteobacteria; Enterobacterales;

FAZ83__locus_1

EUT1

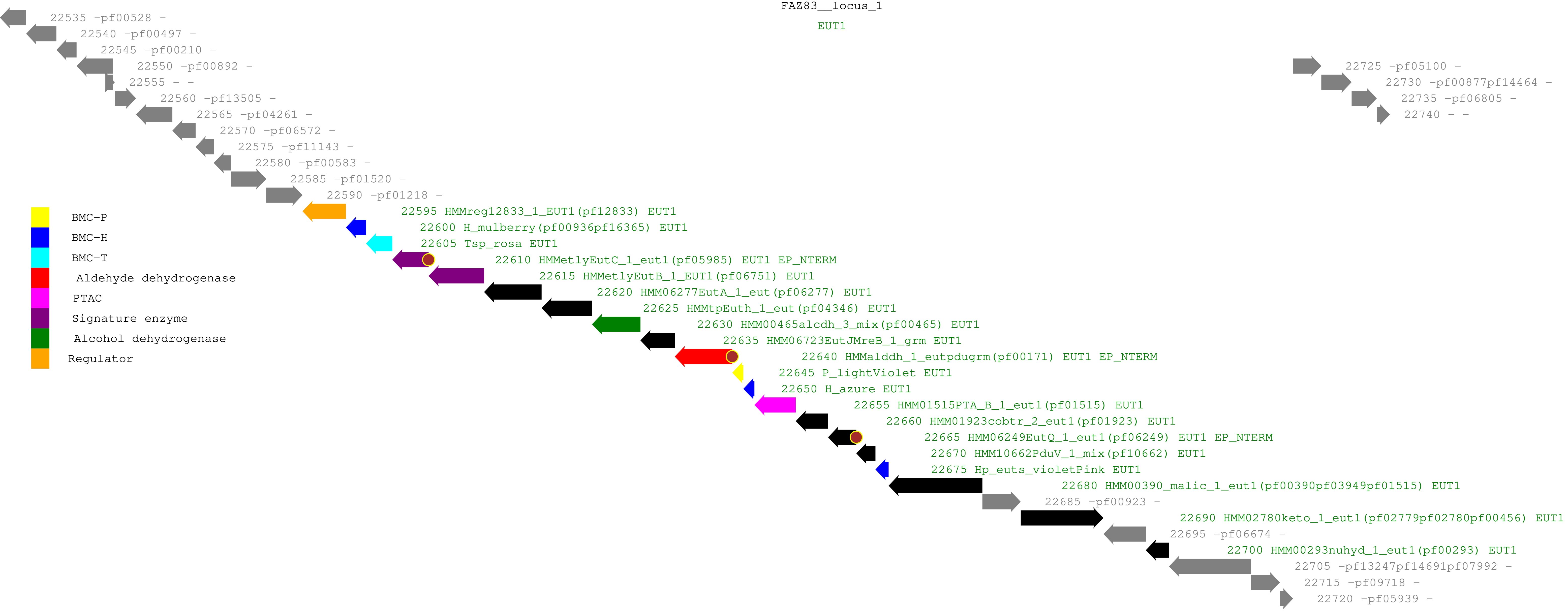

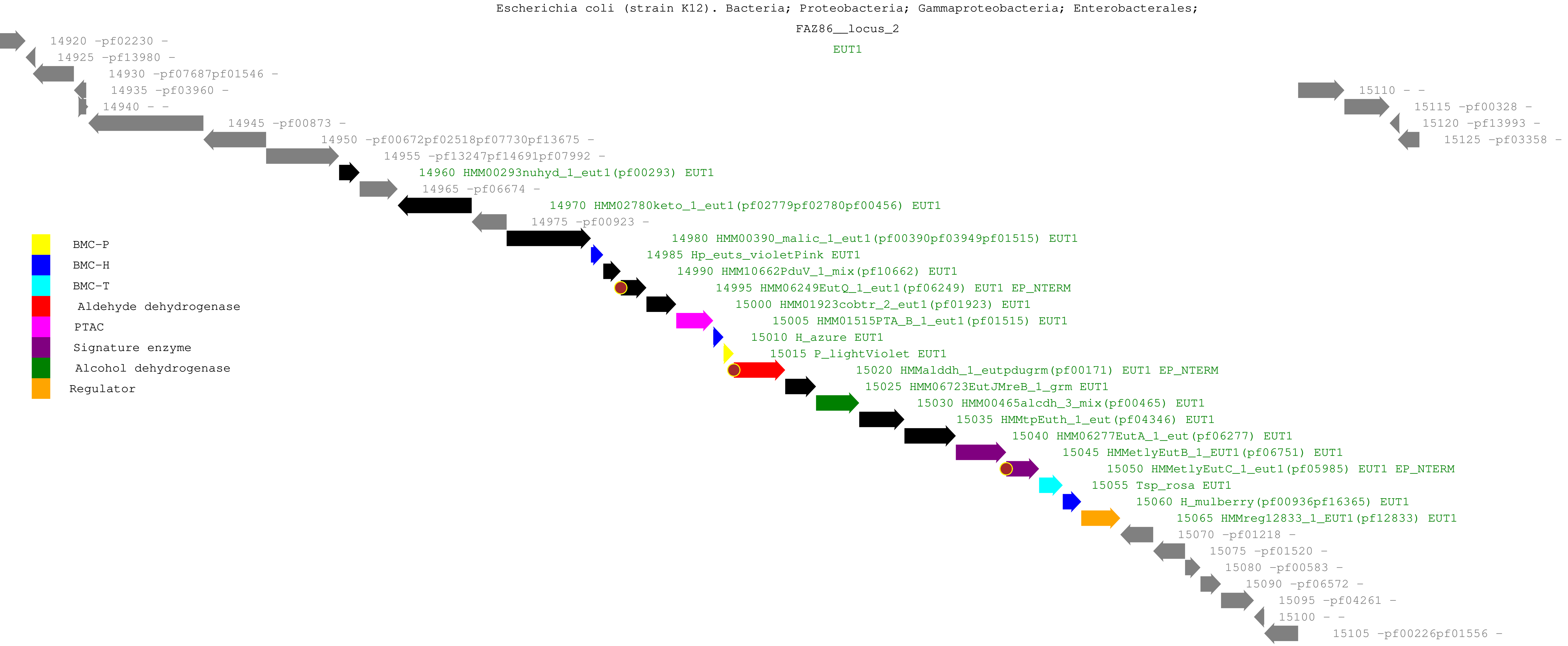
Escherichia coli (strain K12). Bacteria; Proteobacteria; Gammaproteobacteria; Enterobacterales;
FAZ86__locus_2
EUT1
14920 -pf02230 -
14925 -pf13980 -
14930 -pf07687pf01546 -
14935 -pf03960 -
14940 - -
14945 -pf00873 -
14950 -pf00672pf02518pf07730pf13675 -
14955 -pf13247pf14691pf07992 -
14960 HMM00293nuhyd_1_eut1(pf00293) EUT1
14965 -pf06674 -
14970 HMM02780keto_1_eut1(pf02779pf02780pf00456) EUT1
14975 -pf00923 -
14980 HMM00390_malic_1_eut1(pf00390pf03949pf01515) EUT1
14985 Hp_euts_violetPink EUT1
14990 HMM10662PduV_1_mix(pf10662) EUT1
14995 HMM06249EutQ_1_eut1(pf06249) EUT1 EP_NTERM
15000 HMM01923cobtr_2_eut1(pf01923) EUT1
15005 HMM01515PTA_B_1_eut1(pf01515) EUT1
15010 H_azure EUT1
15015 P_lightViolet EUT1
15020 HMMalddh_1_eutpdugrm(pf00171) EUT1 EP_NTERM
15025 HMM06723EutJMreB_1_grm EUT1
15030 HMM00465alcdh_3_mix(pf00465) EUT1
15035 HMMtpEuth_1_eut(pf04346) EUT1
15040 HMM06277EutA_1_eut(pf06277) EUT1
15045 HMMetlyEutB_1_EUT1(pf06751) EUT1
15050 HMMetlyEutC_1_eut1(pf05985) EUT1 EP_NTERM
15055 Tsp_rosa EUT1
15060 H_mulberry(pf00936pf16365) EUT1
15065 HMMreg12833_1_EUT1(pf12833) EUT1
15070 -pf01218 -
15075 -pf01520 -
15080 -pf00583 -
15090 -pf06572 -
15095 -pf04261 -
15100 - -
15105 -pf00226pf01556 -
15110 - -
15115 -pf00328 -
15120 -pf13993 -
15125 -pf03358 -
BMC-P
BMC-H
BMC-T
Aldehyde dehydrogenase
PTAC
Signature enzyme
Alcohol dehydrogenase
Regulator

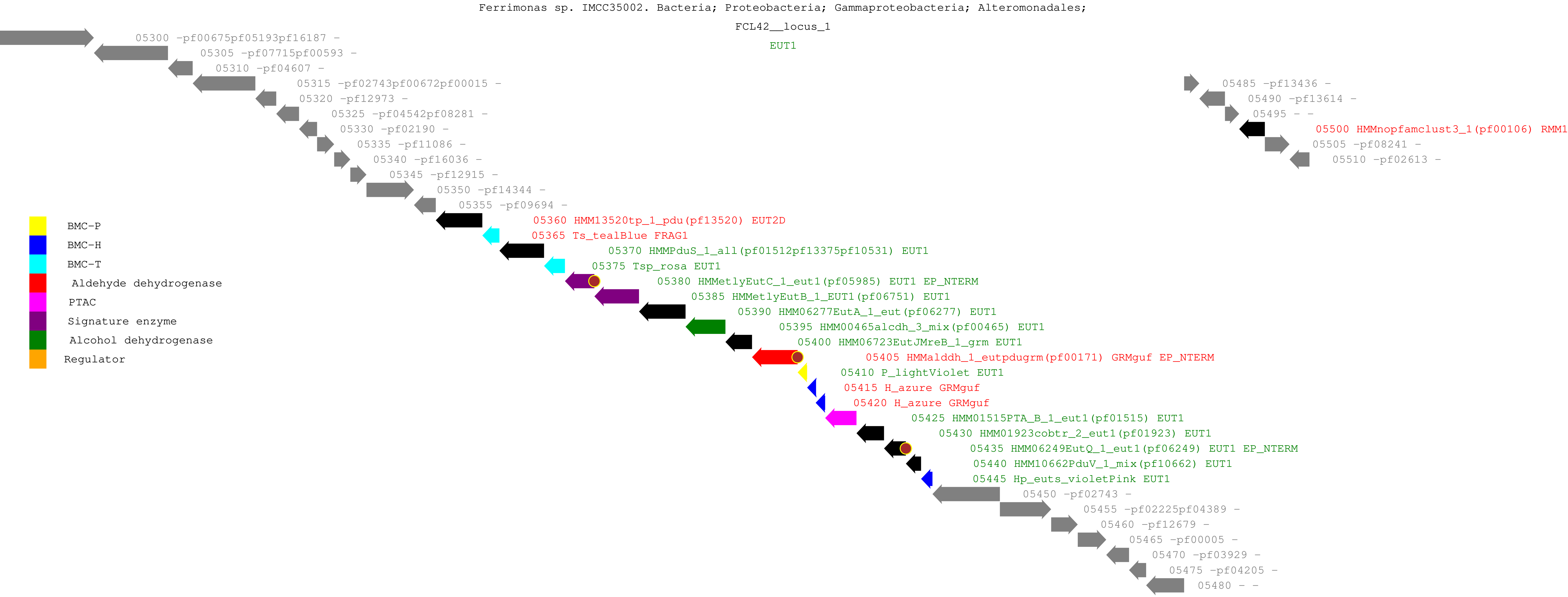
Ferrimonas sp. IMCC35002. Bacteria; Proteobacteria; Gammaproteobacteria; Alteromonadales;
FCL42__locus_1
EUT1
05300 -pf00675pf05193pf16187 -
05305 -pf07715pf00593 -
05310 -pf04607 -
05315 -pf02743pf00672pf00015 -
05320 -pf12973 -
05325 -pf04542pf08281 -
05330 -pf02190 -
05335 -pf11086 -
05340 -pf16036 -
05345 -pf12915 -
05350 -pf14344 -
05355 -pf09694 -
05360 HMM13520tp_1_pdu(pf13520) EUT2D
05365 Ts_tealBlue FRAG1
05370 HMMPduS_1_all(pf01512pf13375pf10531) EUT1
05375 Tsp_rosa EUT1
05380 HMMetlyEutC_1_eut1(pf05985) EUT1 EP_NTERM
05385 HMMetlyEutB_1_EUT1(pf06751) EUT1
05390 HMM06277EutA_1_eut(pf06277) EUT1
05395 HMM00465alcdh_3_mix(pf00465) EUT1
05400 HMM06723EutJMreB_1_grm EUT1
05405 HMMalddh_1_eutpdugrm(pf00171) GRMguf EP_NTERM
05410 P_lightViolet EUT1
05415 H_azure GRMguf
05420 H_azure GRMguf
05425 HMM01515PTA_B_1_eut1(pf01515) EUT1
05430 HMM01923cobtr_2_eut1(pf01923) EUT1
05435 HMM06249EutQ_1_eut1(pf06249) EUT1 EP_NTERM
05440 HMM10662PduV_1_mix(pf10662) EUT1
05445 Hp_euts_violetPink EUT1
05450 -pf02743 -
05455 -pf02225pf04389 -
05460 -pf12679 -
05465 -pf00005 -
05470 -pf03929 -
05475 -pf04205 -
05480 - -
05485 -pf13436 -
05490 -pf13614 -
05495 - -
05500 HMMnopfamclust3_1(pf00106) RMM1
05505 -pf08241 -
05510 -pf02613 -
BMC-P
BMC-H
BMC-T
Aldehyde dehydrogenase
PTAC
Signature enzyme
Alcohol dehydrogenase
Regulator

Comamonadaceae bacterium. Bacteria; Proteobacteria; Betaproteobacteria; Burkholderiales;

FD135__locus_1

EUT1

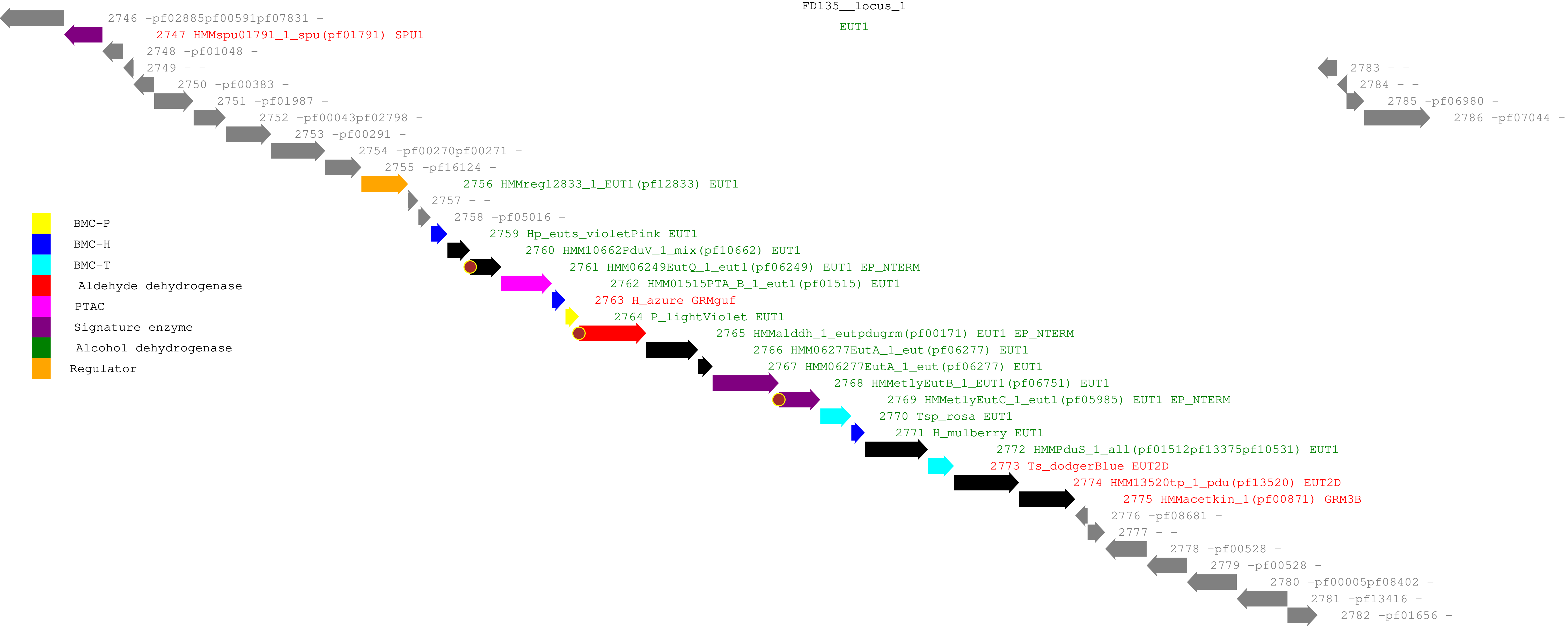

Citrobacter sp. wls715. Bacteria; Proteobacteria; Gammaproteobacteria; Enterobacterales;

FDX05__locus_2

EUT1

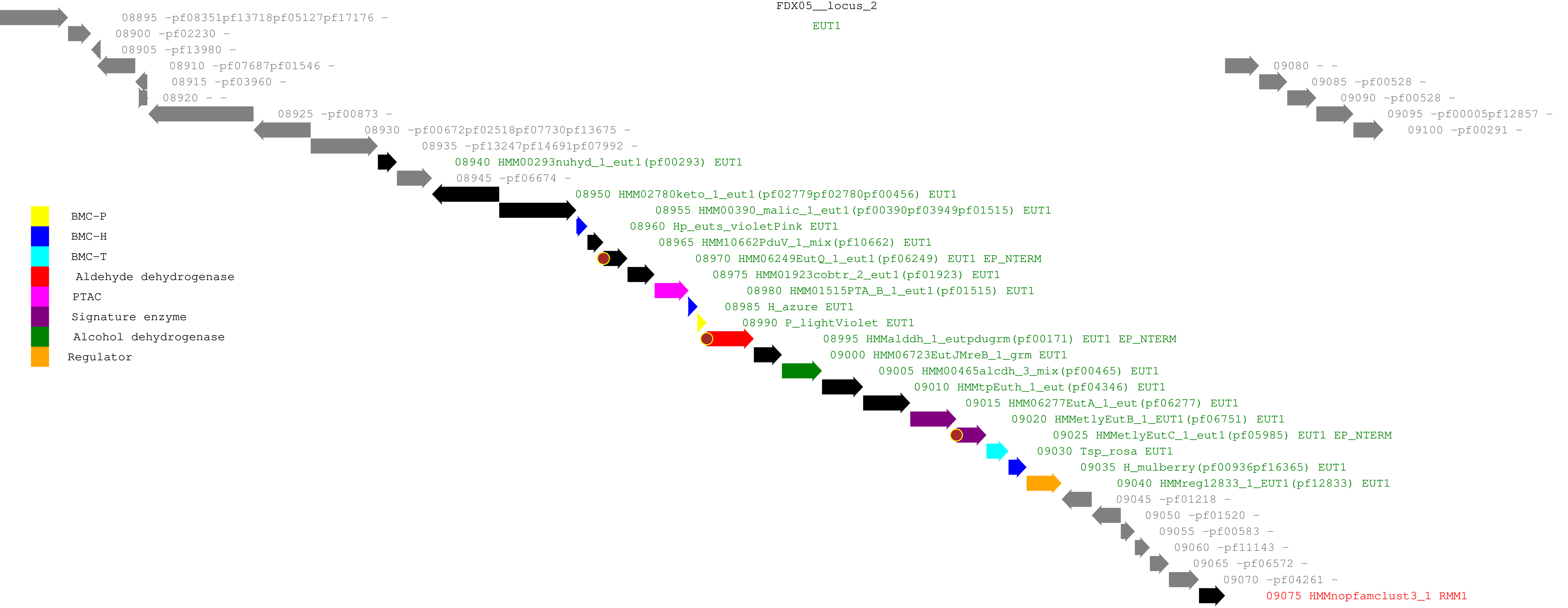

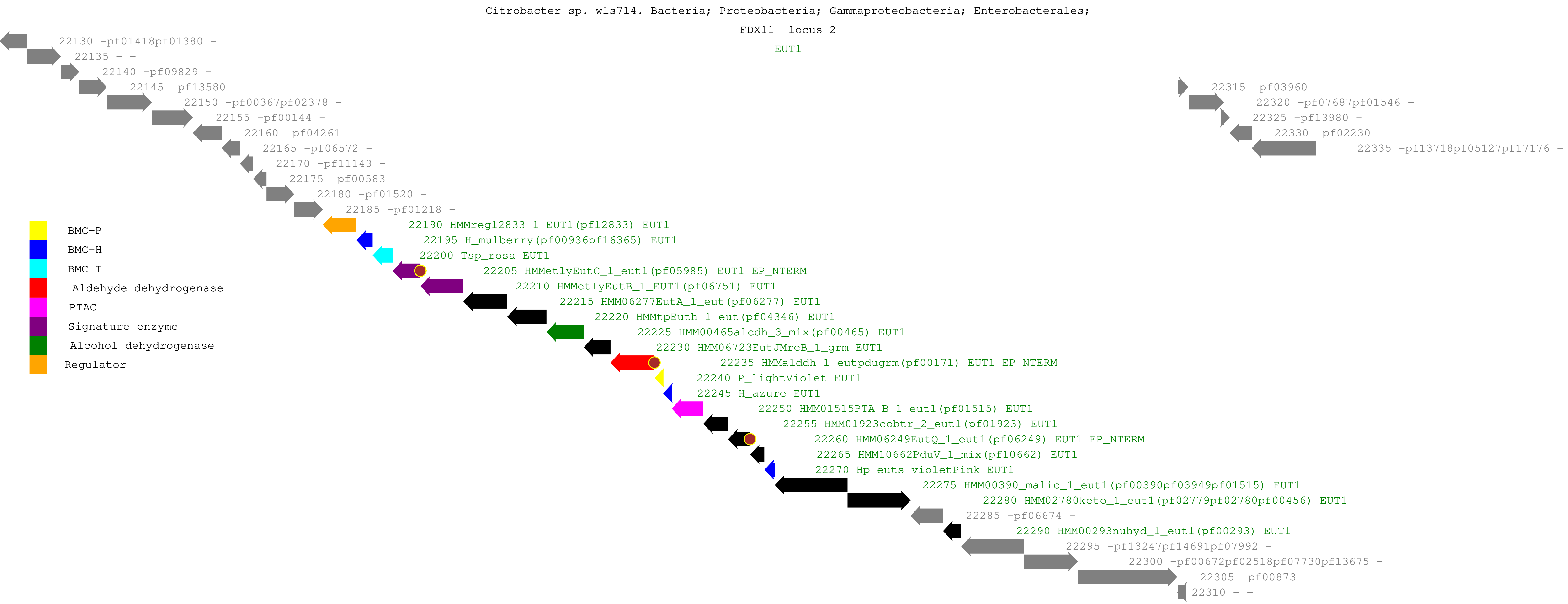

Citrobacter sp. wls714. Bacteria; Proteobacteria; Gammaproteobacteria; Enterobacterales;
FDX11__locus_2
EUT1
22130 -pf01418pf01380 -
22135 - -
22140 -pf09829 -
22145 -pf13580 -
22150 -pf00367pf02378 -
22155 -pf00144 -
22160 -pf04261 -
22165 -pf06572 -
22170 -pf11143 -
22175 -pf00583 -
22180 -pf01520 -
22185 -pf01218 -
22190 HMMreg12833_1_EUT1(pf12833) EUT1
22195 H_mulberry(pf00936pf16365) EUT1
22200 Tsp_rosa EUT1
22205 HMMetlyEutC_1_eut1(pf05985) EUT1 EP_NTERM
22210 HMMetlyEutB_1_EUT1(pf06751) EUT1
22215 HMM06277EutA_1_eut(pf06277) EUT1
22220 HMMtpEuth_1_eut(pf04346) EUT1
22225 HMM00465alcdh_3_mix(pf00465) EUT1
22230 HMM06723EutJMreB_1_grm EUT1
22235 HMMalddh_1_eutpdugrm(pf00171) EUT1 EP_NTERM
22240 P_lightViolet EUT1
22245 H_azure EUT1
22250 HMM01515PTA_B_1_eut1(pf01515) EUT1
22255 HMM01923cobtr_2_eut1(pf01923) EUT1
22260 HMM06249EutQ_1_eut1(pf06249) EUT1 EP_NTERM
22265 HMM10662PduV_1_mix(pf10662) EUT1
22270 Hp_euts_violetPink EUT1
22275 HMM00390_malic_1_eut1(pf00390pf03949pf01515) EUT1
22280 HMM02780keto_1_eut1(pf02779pf02780pf00456) EUT1
22285 -pf06674 -
22290 HMM00293nuhyd_1_eut1(pf00293) EUT1
22295 -pf13247pf14691pf07992 -
22300 -pf00672pf02518pf07730pf13675 -
22305 -pf00873 -
22310 - -
22315 -pf03960 -
22320 -pf07687pf01546 -
22325 -pf13980 -
22330 -pf02230 -
22335 -pf13718pf05127pf17176 -
BMC-P
BMC-H
BMC-T
Aldehyde dehydrogenase
PTAC
Signature enzyme
Alcohol dehydrogenase
Regulator

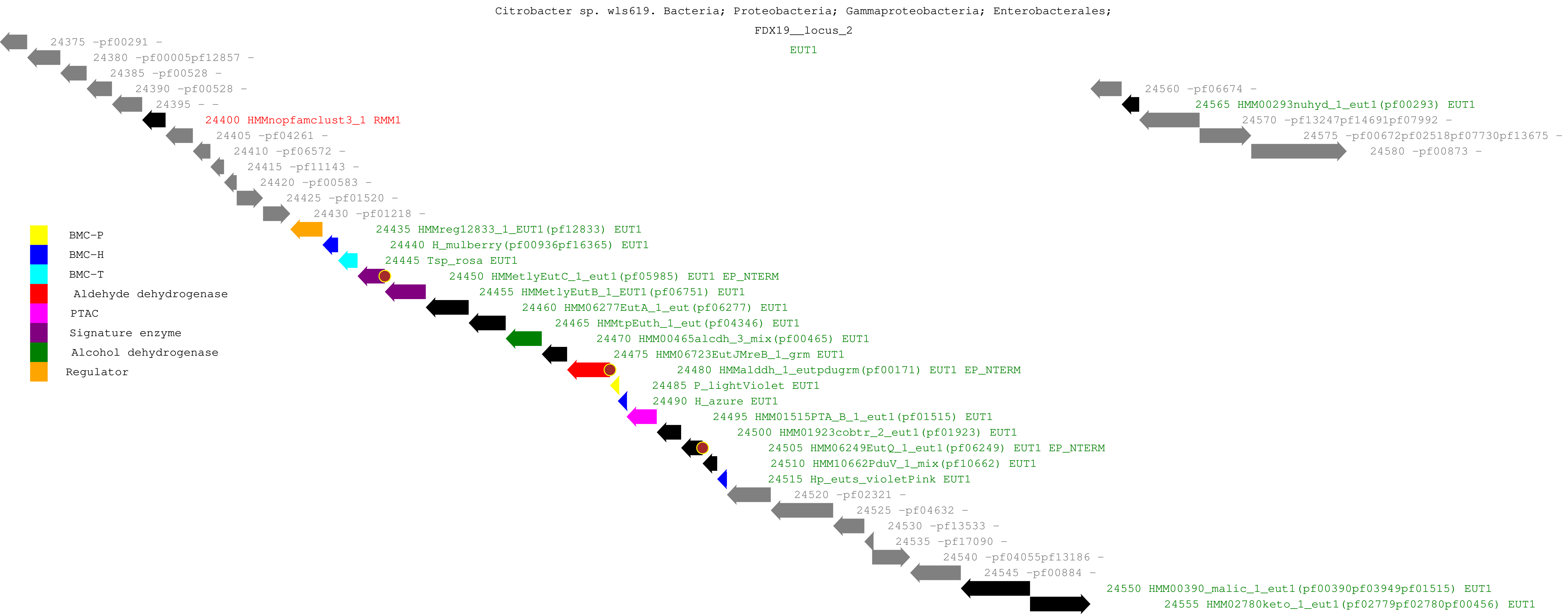
Citrobacter sp. wls619. Bacteria; Proteobacteria; Gammaproteobacteria; Enterobacterales;
FDX19__locus_2
EUT1
24375 -pf00291 -
24380 -pf00005pf12857 -
24385 -pf00528 -
24390 -pf00528 -
24395 - -
24400 HMMnopfamclust3_1 RMM1
24405 -pf04261 -
24410 -pf06572 -
24415 -pf11143 -
24420 -pf00583 -
24425 -pf01520 -
24430 -pf01218 -
24435 HMMreg12833_1_EUT1(pf12833) EUT1
24440 H_mulberry(pf00936pf16365) EUT1
24445 Tsp_rosa EUT1
24450 HMMetlyEutC_1_eut1(pf05985) EUT1 EP_NTERM
24455 HMMetlyEutB_1_EUT1(pf06751) EUT1
24460 HMM06277EutA_1_eut(pf06277) EUT1
24465 HMMtpEuth_1_eut(pf04346) EUT1
24470 HMM00465alcdh_3_mix(pf00465) EUT1
24475 HMM06723EutJMreB_1_grm EUT1
24480 HMMalddh_1_eutpdugrm(pf00171) EUT1 EP_NTERM
24485 P_lightViolet EUT1
24490 H_azure EUT1
24495 HMM01515PTA_B_1_eut1(pf01515) EUT1
24500 HMM01923cobtr_2_eut1(pf01923) EUT1
24505 HMM06249EutQ_1_eut1(pf06249) EUT1 EP_NTERM
24510 HMM10662PduV_1_mix(pf10662) EUT1
24515 Hp_euts_violetPink EUT1
24520 -pf02321 -
24525 -pf04632 -
24530 -pf13533 -
24535 -pf17090 -
24540 -pf04055pf13186 -
24545 -pf00884 -
24550 HMM00390_malic_1_eut1(pf00390pf03949pf01515) EUT1
24555 HMM02780keto_1_eut1(pf02779pf02780pf00456) EUT1
24560 -pf06674 -
24565 HMM00293nuhyd_1_eut1(pf00293) EUT1
24570 -pf13247pf14691pf07992 -
24575 -pf00672pf02518pf07730pf13675 -
24580 -pf00873 -
BMC-P
BMC-H
BMC-T
Aldehyde dehydrogenase
PTAC
Signature enzyme
Alcohol dehydrogenase
Regulator

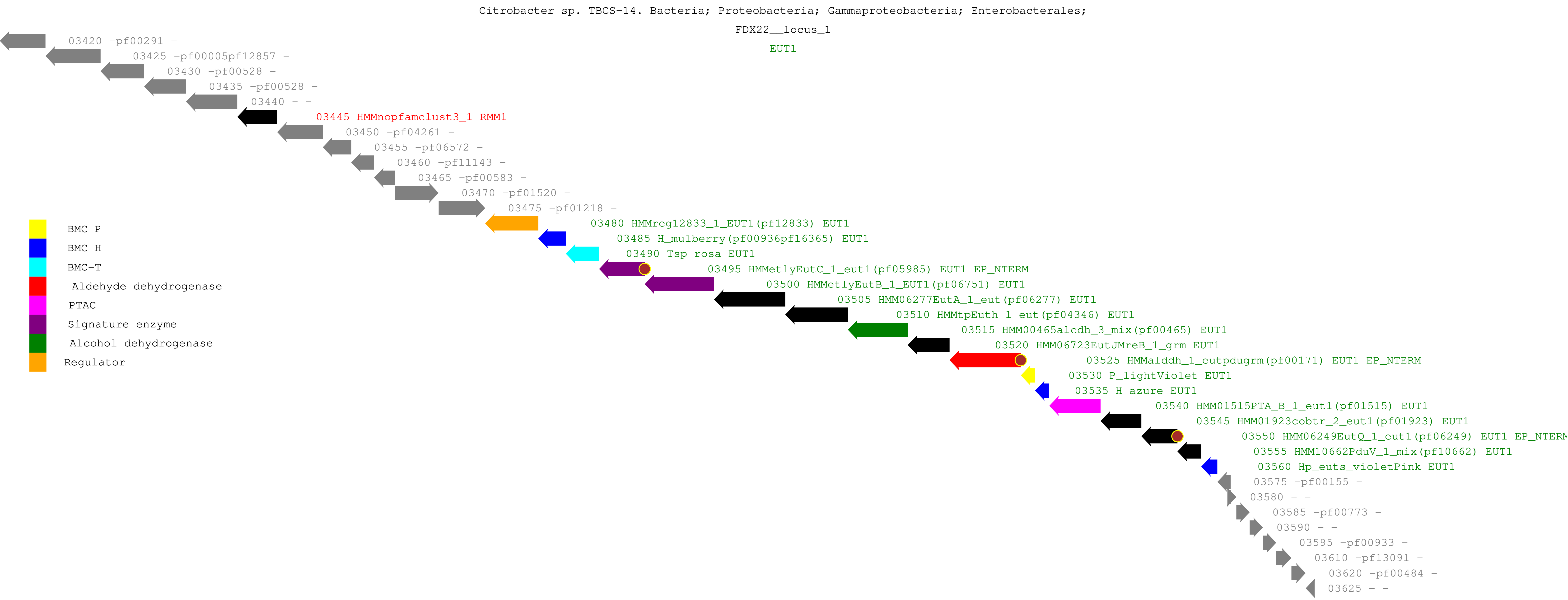

Citrobacter sp. TBCS-14. Bacteria; Proteobacteria; Gammaproteobacteria; Enterobacterales;
FDX22__locus_1
EUT1
03420 -pf00291 -
03425 -pf00005pf12857 -
03430 -pf00528 -
03435 -pf00528 -
03440 - -
03445 HMMnopfamclust3_1 RMM1
03450 -pf04261 -
03455 -pf06572 -
03460 -pf11143 -
03465 -pf00583 -
03470 -pf01520 -
03475 -pf01218 -
03480 HMMreg12833_1_EUT1(pf12833) EUT1
03485 H_mulberry(pf00936pf16365) EUT1
03490 Tsp_rosa EUT1
03495 HMMetlyEutC_1_eut1(pf05985) EUT1 EP_NTERM
03500 HMMetlyEutB_1_EUT1(pf06751) EUT1
03505 HMM06277EutA_1_eut(pf06277) EUT1
03510 HMMtpEuth_1_eut(pf04346) EUT1
03515 HMM00465alcdh_3_mix(pf00465) EUT1
03520 HMM06723EutJMreB_1_grm EUT1
03525 HMMalddh_1_eutpdugrm(pf00171) EUT1 EP_NTERM
03530 P_lightViolet EUT1
03535 H_azure EUT1
03540 HMM01515PTA_B_1_eut1(pf01515) EUT1
03545 HMM01923cobtr_2_eut1(pf01923) EUT1
03550 HMM06249EutQ_1_eut1(pf06249) EUT1 EP_NTERM
03555 HMM10662PduV_1_mix(pf10662) EUT1
03560 Hp_euts_violetPink EUT1
03575 -pf00155 -
03580 - -
03585 -pf00773 -
03590 - -
03595 -pf00933 -
03610 -pf13091 -
03620 -pf00484 -
03625 - -
BMC-P
BMC-H
BMC-T
Aldehyde dehydrogenase
PTAC
Signature enzyme
Alcohol dehydrogenase
Regulator

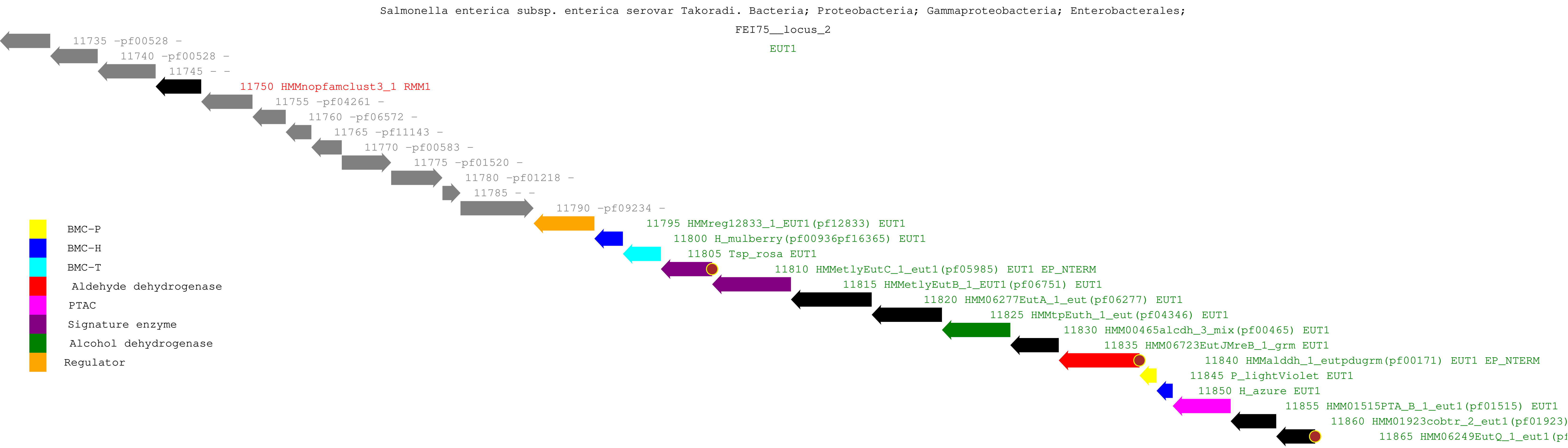

Salmonella enterica subsp. enterica serovar Takoradi. Bacteria; Proteobacteria; Gammaproteobacteria; Enterobacterales;
FEI75__locus_2
EUT1
11735 -pf00528 -
11740 -pf00528 -
11745 - -
11750 HMMnopfamclust3_1 RMM1
11755 -pf04261 -
11760 -pf06572 -
11765 -pf11143 -
11770 -pf00583 -
11775 -pf01520 -
11780 -pf01218 -
11785 - -
11790 -pf09234 -
11795 HMMreg12833_1_EUT1(pf12833) EUT1
11800 H_mulberry(pf00936pf16365) EUT1
11805 Tsp_rosa EUT1
11810 HMMetlyEutC_1_eut1(pf05985) EUT1 EP_NTERM
11815 HMMetlyEutB_1_EUT1(pf06751) EUT1
11820 HMM06277EutA_1_eut(pf06277) EUT1
11825 HMMtpEuth_1_eut(pf04346) EUT1
11830 HMM00465alcdh_3_mix(pf00465) EUT1
11835 HMM06723EutJMreB_1_grm EUT1
11840 HMMalddh_1_eutpdugrm(pf00171) EUT1 EP_NTERM
11845 P_lightViolet EUT1
11850 H_azure EUT1
11855 HMM01515PTA_B_1_eut1(pf01515) EUT1
11860 HMM01923cobtr_2_eut1(pf01923)
11865 HMM06249EutQ_1_eut1(pf
BMC-P
BMC-H
BMC-T
Aldehyde dehydrogenase
PTAC
Signature enzyme
Alcohol dehydrogenase
Regulator

Citrobacter amalonaticus. Bacteria; Proteobacteria; Gammaproteobacteria; Enterobacterales;

FEO47__locus_2

EUT1

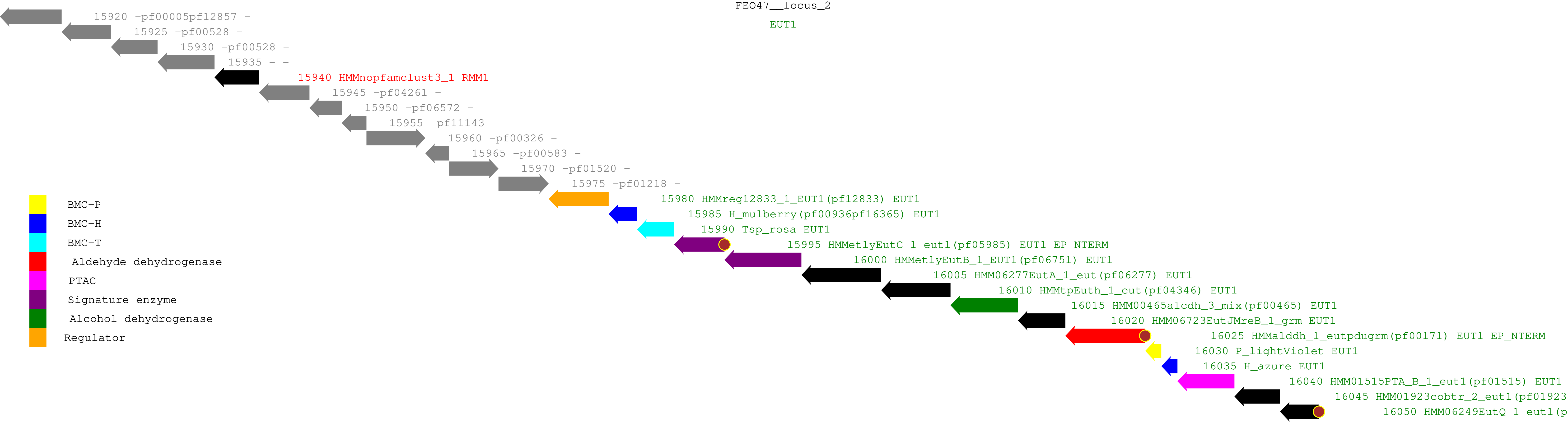

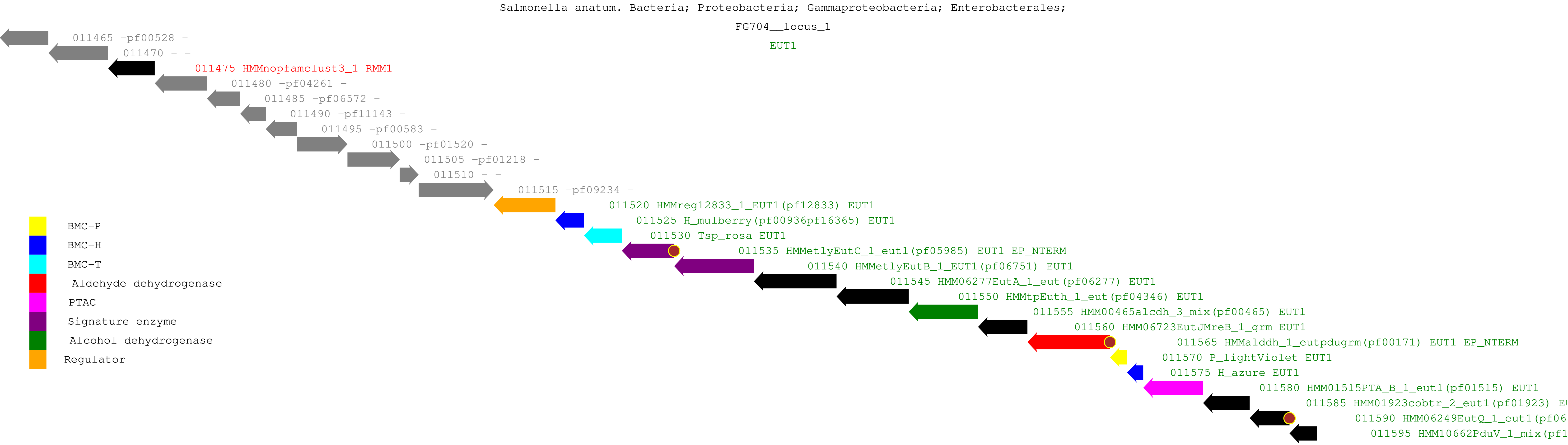
Salmonella anatum. Bacteria; Proteobacteria; Gammaproteobacteria; Enterobacterales;
FG704__locus_1
EUT1
011465 -pf00528 -
011470 - -
011475 HMMnopfamclust3_1 RMM1
011480 -pf04261 -
011485 -pf06572 -
011490 -pf11143 -
011495 -pf00583 -
011500 -pf01520 -
011505 -pf01218 -
011510 - -
011515 -pf09234 -
011520 HMMreg12833_1_EUT1(pf12833) EUT1
011525 H_mulberry(pf00936pf16365) EUT1
011530 Tsp_rosa EUT1
011535 HMMetlyEutC_1_eut1(pf05985) EUT1 EP_NTERM
011540 HMMetlyEutB_1_EUT1(pf06751) EUT1
011545 HMM06277EutA_1_eut(pf06277) EUT1
011550 HMMtpEuth_1_eut(pf04346) EUT1
011555 HMM00465alcdh_3_mix(pf00465) EUT1
011560 HMM06723EutJMreB_1_grm EUT1
011565 HMMalddh_1_eutpdugrm(pf00171) EUT1 EP_NTERM
011570 P_lightViolet EUT1
011575 H_azure EUT1
011580 HMM01515PTA_B_1_eut1(pf01515) EUT1
011585 HMM01923cobtr_2_eut1(pf01923) EU
011590 HMM06249EutQ_1_eut1(pf06
011595 HMM10662PduV_1_mix(pf1
BMC-P
BMC-H
BMC-T
Aldehyde dehydrogenase
PTAC
Signature enzyme
Alcohol dehydrogenase
Regulator

Pseudomonas sp. AAC. Bacteria; Proteobacteria; Gammaproteobacteria; Pseudomonadales;

FG99__locus_1

EUT1

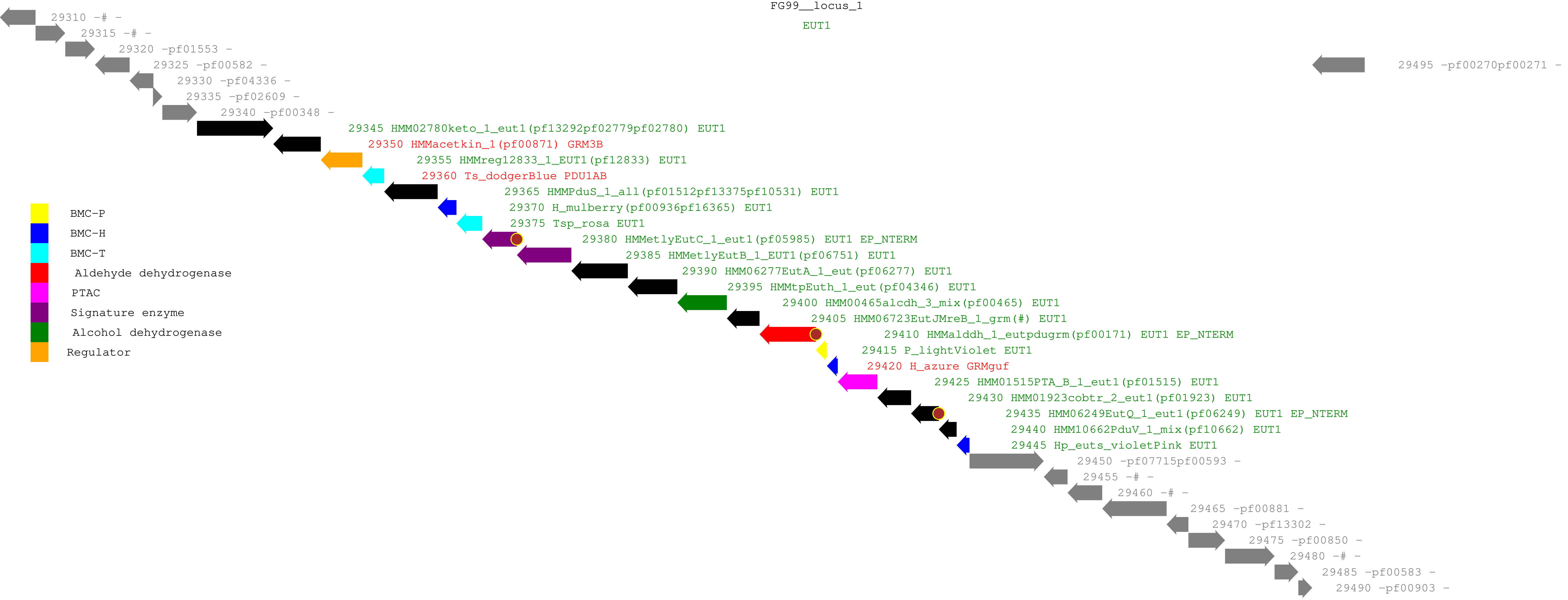

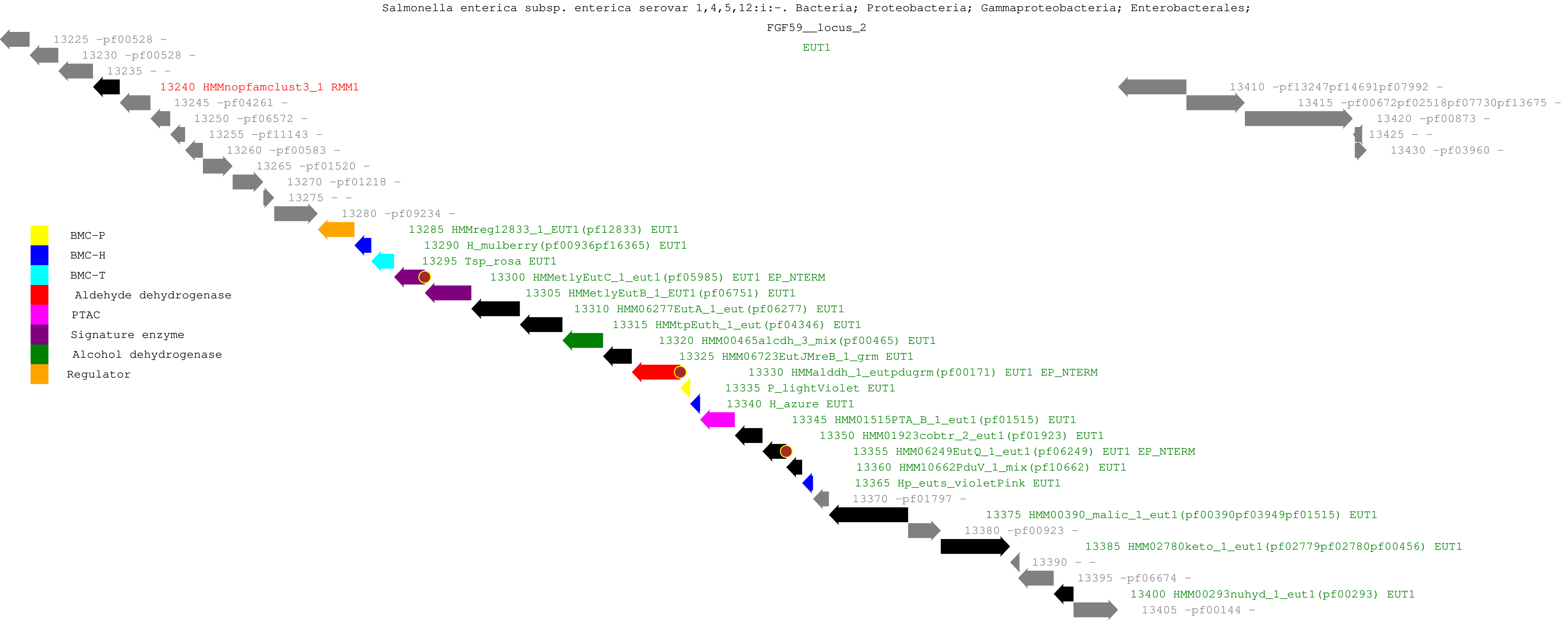

Salmonella enterica subsp. enterica serovar 1,4,5,12:i:-. Bacteria; Proteobacteria; Gammaproteobacteria; Enterobacterales;
FGF59__locus_2
EUT1
13225 -pf00528 -
13230 -pf00528 -
13235 - -
13240 HMMnopfamclust3_1 RMM1
13245 -pf04261 -
13250 -pf06572 -
13255 -pf11143 -
13260 -pf00583 -
13265 -pf01520 -
13270 -pf01218 -
13275 - -
13280 -pf09234 -
13285 HMMreg12833_1_EUT1(pf12833) EUT1
13290 H_mulberry(pf00936pf16365) EUT1
13295 Tsp_rosa EUT1
13300 HMMetlyEutC_1_eut1(pf05985) EUT1 EP_NTERM
13305 HMMetlyEutB_1_EUT1(pf06751) EUT1
13310 HMM06277EutA_1_eut(pf06277) EUT1
13315 HMMtpEuth_1_eut(pf04346) EUT1
13320 HMM00465alcdh_3_mix(pf00465) EUT1
13325 HMM06723EutJMreB_1_grm EUT1
13330 HMMalddh_1_eutpdugrm(pf00171) EUT1 EP_NTERM
13335 P_lightViolet EUT1
13340 H_azure EUT1
13345 HMM01515PTA_B_1_eut1(pf01515) EUT1
13350 HMM01923cobtr_2_eut1(pf01923) EUT1
13355 HMM06249EutQ_1_eut1(pf06249) EUT1 EP_NTERM
13360 HMM10662PduV_1_mix(pf10662) EUT1
13365 Hp_euts_violetPink EUT1
13370 -pf01797 -
13375 HMM00390_malic_1_eut1(pf00390pf03949pf01515) EUT1
13380 -pf00923 -
13385 HMM02780keto_1_eut1(pf02779pf02780pf00456) EUT1
13390 - -
13395 -pf06674 -
13400 HMM00293nuhyd_1_eut1(pf00293) EUT1
13405 -pf00144 -
13410 -pf13247pf14691pf07992 -
13415 -pf00672pf02518pf07730pf13675 -
13420 -pf00873 -
13425 - -
13430 -pf03960 -
BMC-P
BMC-H
BMC-T
Aldehyde dehydrogenase
PTAC
Signature enzyme
Alcohol dehydrogenase
Regulator

Salmonella enterica subsp. enterica serovar Eko. Bacteria; Proteobacteria; Gammaproteobacteria; Enterobacterales;

FGJ70__locus_1

EUT1

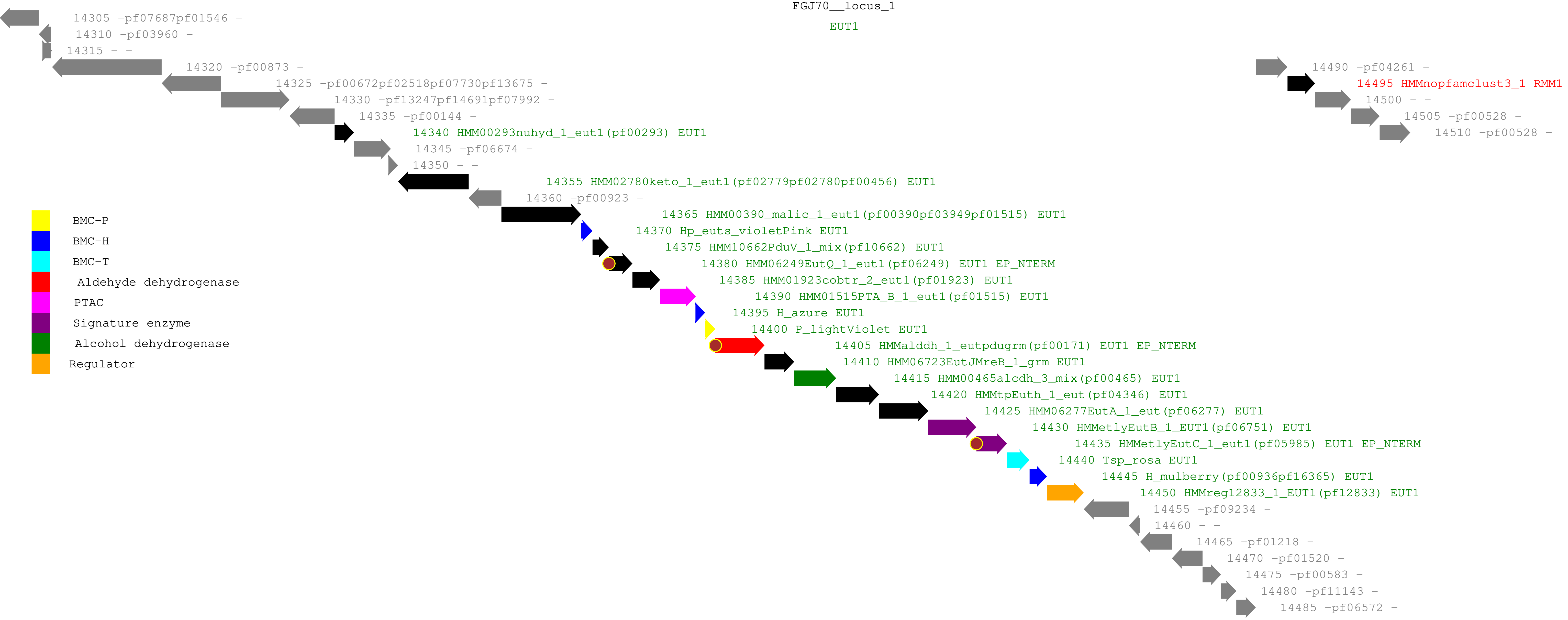

Buttiauxella sp. B2. Bacteria; Proteobacteria; Gammaproteobacteria; Enterobacterales;

FH968__locus_1

EUT1

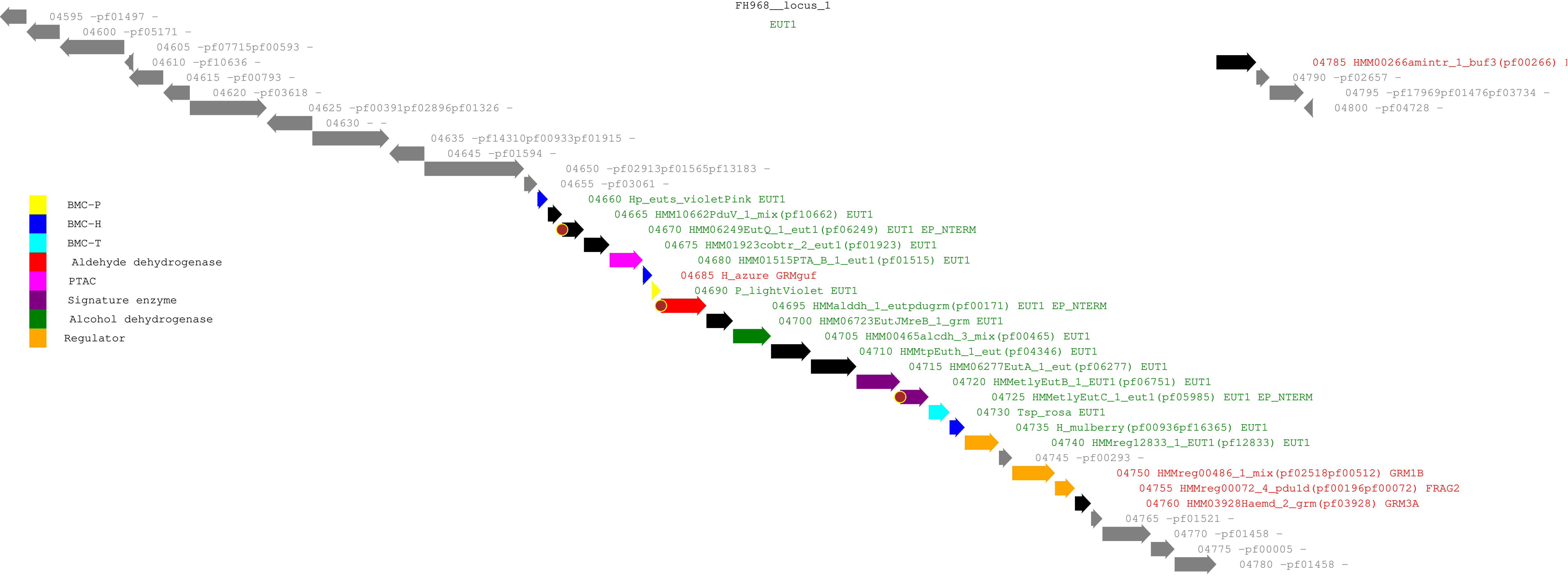

Marinobacter vinifirmus. Bacteria; Proteobacteria; Gammaproteobacteria; Alteromonadales;

FHK81__locus_1

EUT1

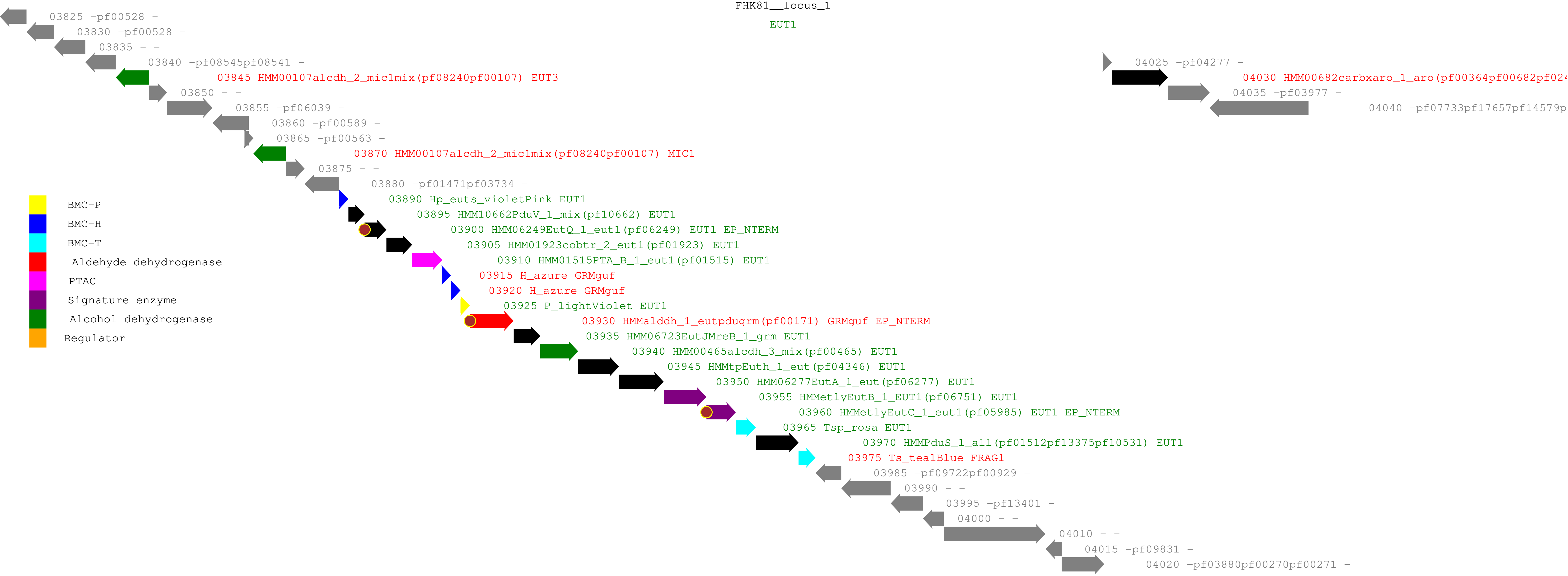

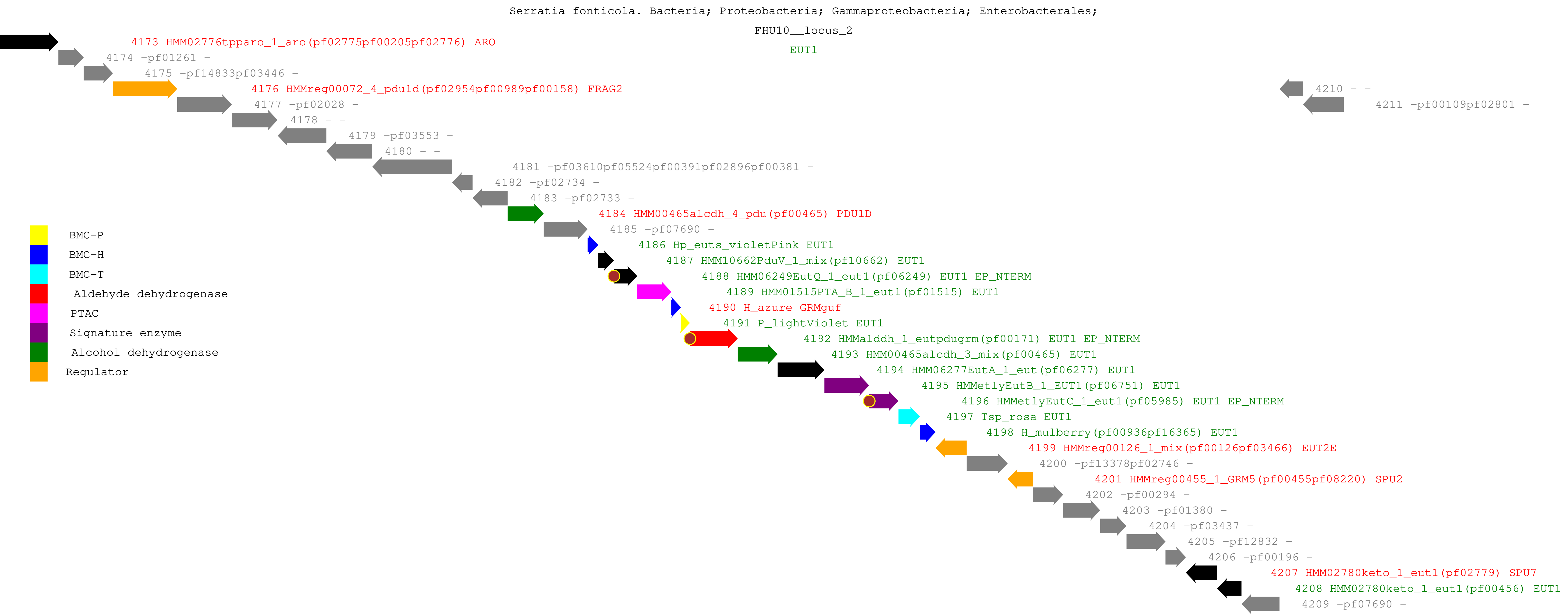
Serratia fonticola. Bacteria; Proteobacteria; Gammaproteobacteria; Enterobacterales;
FHU10__locus_2
EUT1
4173 HMM02776tpparo_1_aro(pf02775pf00205pf02776) ARO
4174 -pf01261 -
4175 -pf14833pf03446 -
4176 HMMreg00072_4_pdu1d(pf02954pf00989pf00158) FRAG2
4177 -pf02028 -
4178 - -
4179 -pf03553 -
4180 - -
4181 -pf03610pf05524pf00391pf02896pf00381 -
4182 -pf02734 -
4183 -pf02733 -
4184 HMM00465alcdh_4_pdu(pf00465) PDU1D
4185 -pf07690 -
4186 Hp_euts_violetPink EUT1
4187 HMM10662PduV_1_mix(pf10662) EUT1
4188 HMM06249EutQ_1_eut1(pf06249) EUT1 EP_NTERM
4189 HMM01515PTA_B_1_eut1(pf01515) EUT1
4190 H_azure GRMguf
4191 P_lightViolet EUT1
4192 HMMalddh_1_eutpdugrm(pf00171) EUT1 EP_NTERM
4193 HMM00465alcdh_3_mix(pf00465) EUT1
4194 HMM06277EutA_1_eut(pf06277) EUT1
4195 HMMetlyEutB_1_EUT1(pf06751) EUT1
4196 HMMetlyEutC_1_eut1(pf05985) EUT1 EP_NTERM
4197 Tsp_rosa EUT1
4198 H_mulberry(pf00936pf16365) EUT1
4199 HMMreg00126_1_mix(pf00126pf03466) EUT2E
4200 -pf13378pf02746 -
4201 HMMreg00455_1_GRM5(pf00455pf08220) SPU2
4202 -pf00294 -
4203 -pf01380 -
4204 -pf03437 -
4205 -pf12832 -
4206 -pf00196 -
4207 HMM02780keto_1_eut1(pf02779) SPU7
4208 HMM02780keto_1_eut1(pf00456) EUT1
4209 -pf07690 -
4210 - -
4211 -pf00109pf02801 -
BMC-P
BMC-H
BMC-T
Aldehyde dehydrogenase
PTAC
Signature enzyme
Alcohol dehydrogenase
Regulator

Salmonella enterica subsp. enterica serovar Nima. Bacteria; Proteobacteria; Gammaproteobacteria; Enterobacterales;

FK264__locus_1

EUT1

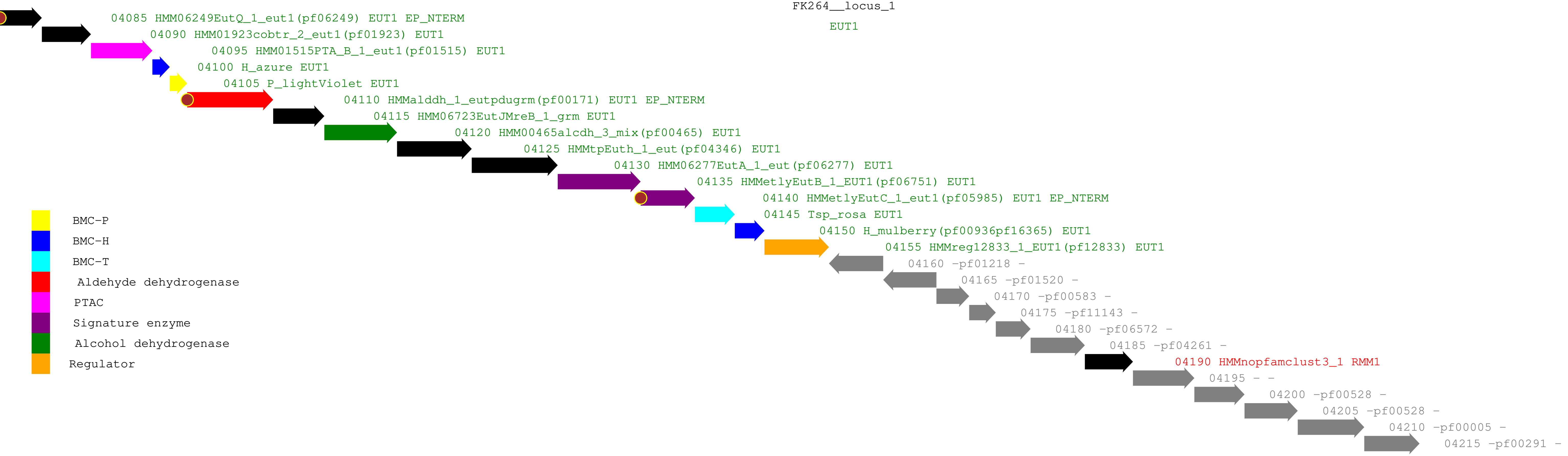

Salmonella enterica subsp. enterica serovar Bahati. Bacteria; Proteobacteria; Gammaproteobacteria; Enterobacterales;

FK693__locus_1

EUT1

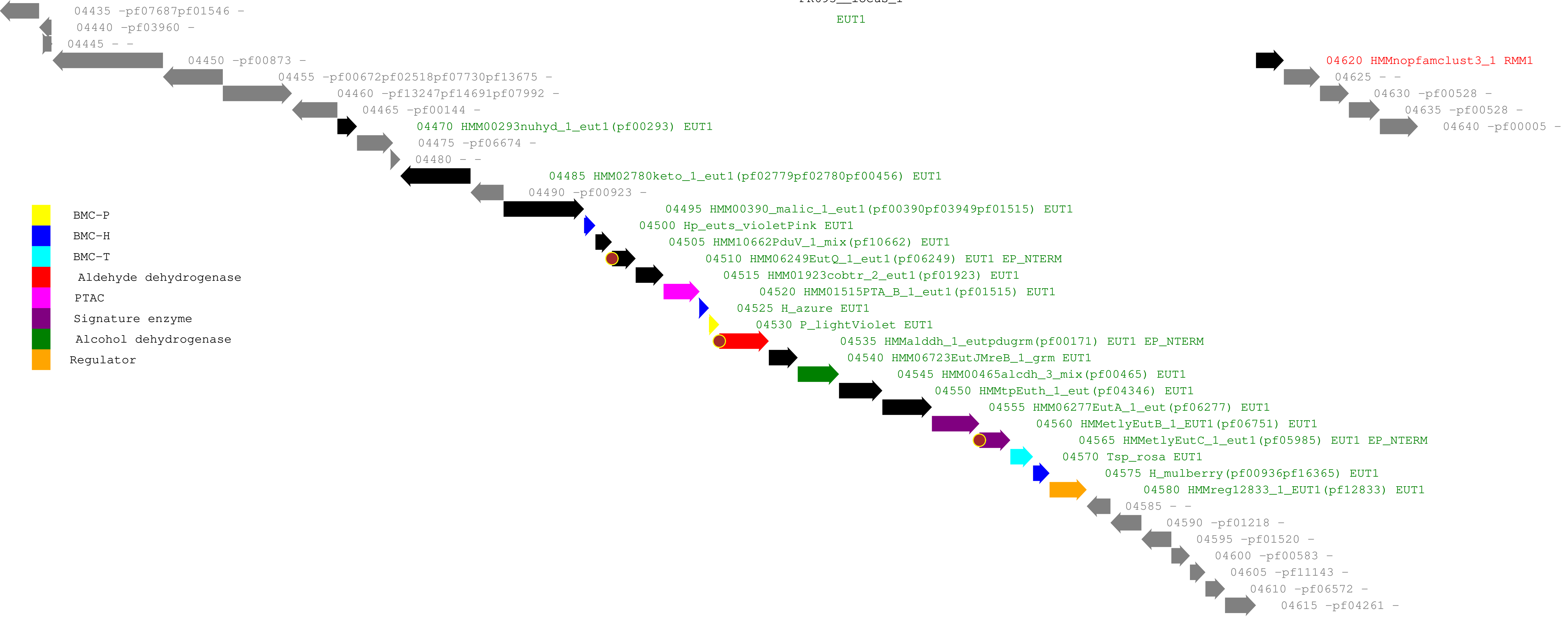

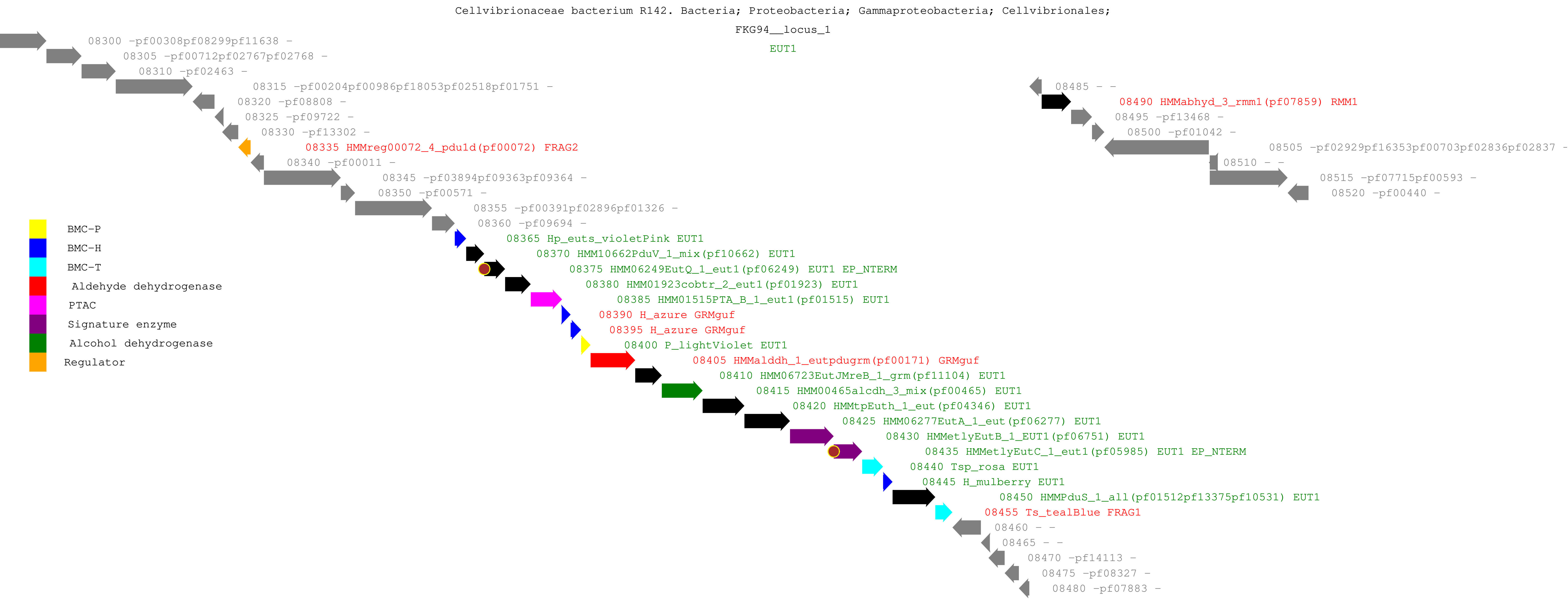
Cellvibrionaceae bacterium R142. Bacteria; Proteobacteria; Gammaproteobacteria; Cellvibrionales;
FKG94__locus_1
EUT1
08300 -pf00308pf08299pf11638 -
08305 -pf00712pf02767pf02768 -
08310 -pf02463 -
08315 -pf00204pf00986pf18053pf02518pf01751 -
08320 -pf08808 -
08325 -pf09722 -
08330 -pf13302 -
08335 HMMreg00072_4_pdu1d(pf00072) FRAG2
08340 -pf00011 -
08345 -pf03894pf09363pf09364 -
08350 -pf00571 -
08355 -pf00391pf02896pf01326 -
08360 -pf09694 -
08365 Hp_euts_violetPink EUT1
08370 HMM10662PduV_1_mix(pf10662) EUT1
08375 HMM06249EutQ_1_eut1(pf06249) EUT1 EP_NTERM
08380 HMM01923cobtr_2_eut1(pf01923) EUT1
08385 HMM01515PTA_B_1_eut1(pf01515) EUT1
08390 H_azure GRMguf
08395 H_azure GRMguf
08400 P_lightViolet EUT1
08405 HMMaldh_1_eutpdugrm(pf00171) GRMguf
08410 HMM06723EutJMreB_1_grm(pf11104) EUT1
08415 HMM00465alcdh_3_mix(pf00465) EUT1
08420 HMMtpEuth_1_eut(pf04346) EUT1
08425 HMM06277EutA_1_eut(pf06277) EUT1
08430 HMMetlyEutB_1_EUT1(pf06751) EUT1
08435 HMMetlyEutC_1_eut1(pf05985) EUT1 EP_NTERM
08440 Tsp_rosa EUT1
08445 H_mulberry EUT1
08450 HMMPduS_1_all(pf01512pf13375pf10531) EUT1
08455 Ts_tealBlue FRAG1
08460 - -
08465 - -
08470 -pf14113 -
08475 -pf08327 -
08480 -pf07883 -
08485 - -
08490 HMMabhyd_3_rmm1(pf07859) RMM1
08495 -pf13468 -
08500 -pf01042 -
08505 -pf02929pf16353pf00703pf02836pf02837 -
08510 - -
08515 -pf07715pf00593 -
08520 -pf00440 -
BMC-P
BMC-H
BMC-T
Aldehyde dehydrogenase
PTAC
Signature enzyme
Alcohol dehydrogenase
Regulator

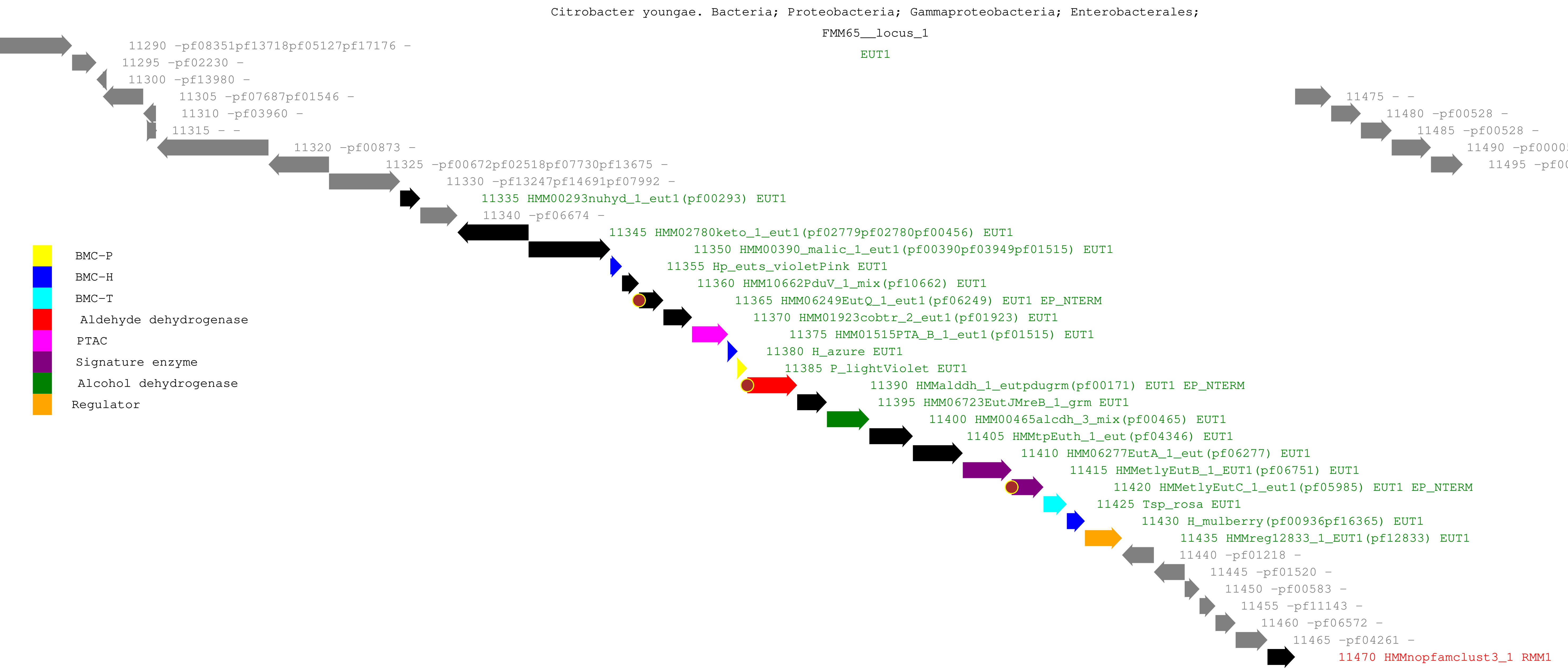
Citrobacter youngae. Bacteria; Proteobacteria; Gammaproteobacteria; Enterobacterales;
FMM65__locus_1
EUT1
11290 -pf08351pf13718pf05127pf17176 -
11295 -pf02230 -
11300 -pf13980 -
11305 -pf07687pf01546 -
11310 -pf03960 -
11315 - -
11320 -pf00873 -
11325 -pf00672pf02518pf07730pf13675 -
11330 -pf13247pf14691pf07992 -
11335 HMM00293nuhyd_1_eut1(pf00293) EUT1
11340 -pf06674 -
11345 HMM02780keto_1_eut1(pf02779pf02780pf00456) EUT1
11350 HMM00390_malic_1_eut1(pf00390pf03949pf01515) EUT1
11355 Hp_euts_violetPink EUT1
11360 HMM10662PduV_1_mix(pf10662) EUT1
11365 HMM06249EutQ_1_eut1(pf06249) EUT1 EP_NTERM
11370 HMM01923cobtr_2_eut1(pf01923) EUT1
11375 HMM01515PTA_B_1_eut1(pf01515) EUT1
11380 H_azure EUT1
11385 P_lightViolet EUT1
11390 HMMalddh_1_eutpdugrm(pf00171) EUT1 EP_NTERM
11395 HMM06723EutJMreB_1_grm EUT1
11400 HMM00465alcdh_3_mix(pf00465) EUT1
11405 HMMtpEuth_1_eut(pf04346) EUT1
11410 HMM06277EutA_1_eut(pf06277) EUT1
11415 HMMetlyEutB_1_EUT1(pf06751) EUT1
11420 HMMetlyEutC_1_eut1(pf05985) EUT1 EP_NTERM
11425 Tsp_rosa EUT1
11430 H_mulberry(pf00936pf16365) EUT1
11435 HMMreg12833_1_EUT1(pf12833) EUT1
11440 -pf01218 -
11445 -pf01520 -
11450 -pf00583 -
11455 -pf11143 -
11460 -pf06572 -
11465 -pf04261 -
11470 HMMnopfamclust3_1 RMM1
11475 - -
11480 -pf00528 -
11485 -pf00528 -
11490 -pf00005pf12857 -
11495 -pf00291 -
BMC-P
BMC-H
BMC-T
Aldehyde dehydrogenase
PTAC
Signature enzyme
Alcohol dehydrogenase
Regulator

Morganella morganii (Proteus morganii). Bacteria; Proteobacteria; Gammaproteobacteria; Enterobacterales;

FNZ58__locus_1

EUT1

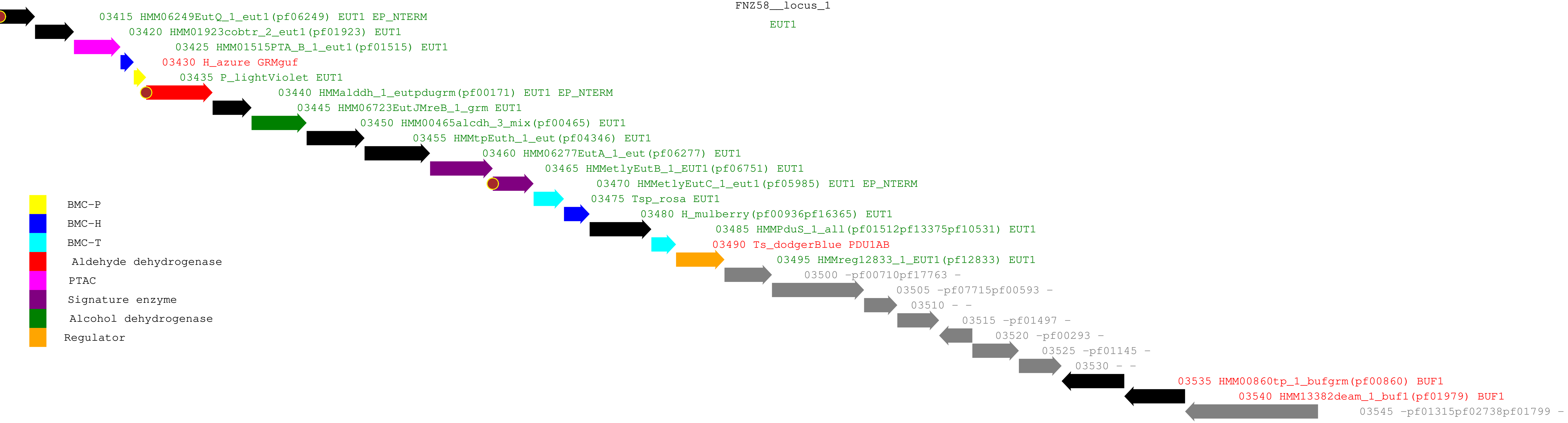

Klebsiella pneumoniae. Bacteria; Proteobacteria; Gammaproteobacteria; Enterobacterales;

FOB37__locus_1

EUT1

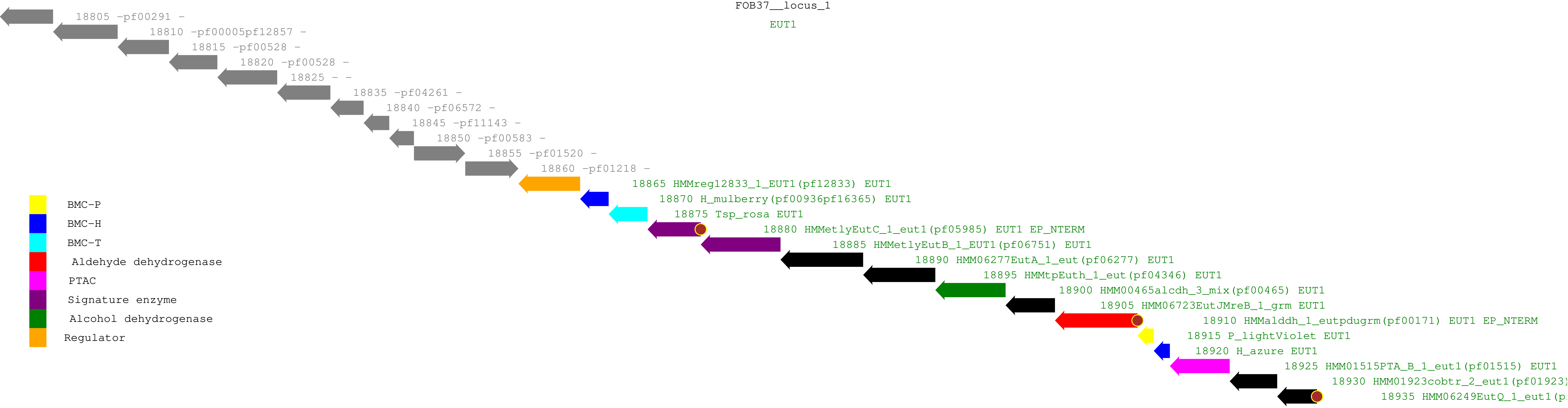

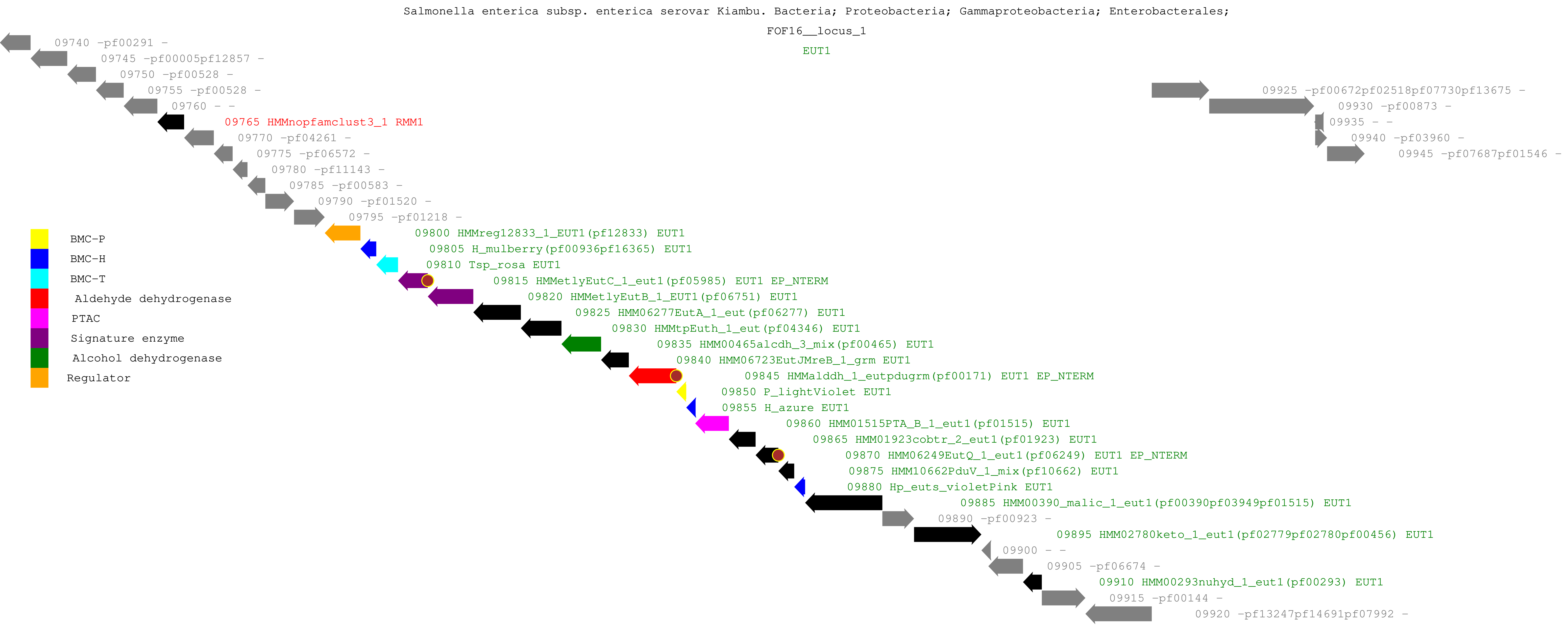
Salmonella enterica subsp. enterica serovar Kiambu. Bacteria; Proteobacteria; Gammaproteobacteria; Enterobacterales;
FOF16__locus_1
EUT1
09740 -pf00291 -
09745 -pf00005pf12857 -
09750 -pf00528 -
09755 -pf00528 -
09760 - -
09765 HMMnopfamclust3_1 RMM1
09770 -pf04261 -
09775 -pf06572 -
09780 -pf11143 -
09785 -pf00583 -
09790 -pf01520 -
09795 -pf01218 -
09800 HMMreg12833_1_EUT1(pf12833) EUT1
09805 H_mulberry(pf00936pf16365) EUT1
09810 Tsp_rosa EUT1
09815 HMMetlyEutC_1_eut1(pf05985) EUT1 EP_NTERM
09820 HMMetlyEutB_1_EUT1(pf06751) EUT1
09825 HMM06277EutA_1_eut(pf06277) EUT1
09830 HMMtpEuth_1_eut(pf04346) EUT1
09835 HMM00465alcdh_3_mix(pf00465) EUT1
09840 HMM06723EutJMreB_1_grm EUT1
09845 HMMalddh_1_eutpdugrm(pf00171) EUT1 EP_NTERM
09850 P_lightViolet EUT1
09855 H_azure EUT1
09860 HMM01515PTA_B_1_eut1(pf01515) EUT1
09865 HMM01923cobtr_2_eut1(pf01923) EUT1
09870 HMM06249EutQ_1_eut1(pf06249) EUT1 EP_NTERM
09875 HMM10662PduV_1_mix(pf10662) EUT1
09880 Hp_euts_violetPink EUT1
09885 HMM00390_malic_1_eut1(pf00390pf03949pf01515) EUT1
09890 -pf00923 -
09895 HMM02780keto_1_eut1(pf02779pf02780pf00456) EUT1
09900 - -
09905 -pf06674 -
09910 HMM00293nuhyd_1_eut1(pf00293) EUT1
09915 -pf00144 -
09920 -pf13247pf14691pf07992 -
09925 -pf00672pf02518pf07730pf13675 -
09930 -pf00873 -
09935 - -
09940 -pf03960 -
09945 -pf07687pf01546 -
BMC-P
BMC-H
BMC-T
Aldehyde dehydrogenase
PTAC
Signature enzyme
Alcohol dehydrogenase
Regulator

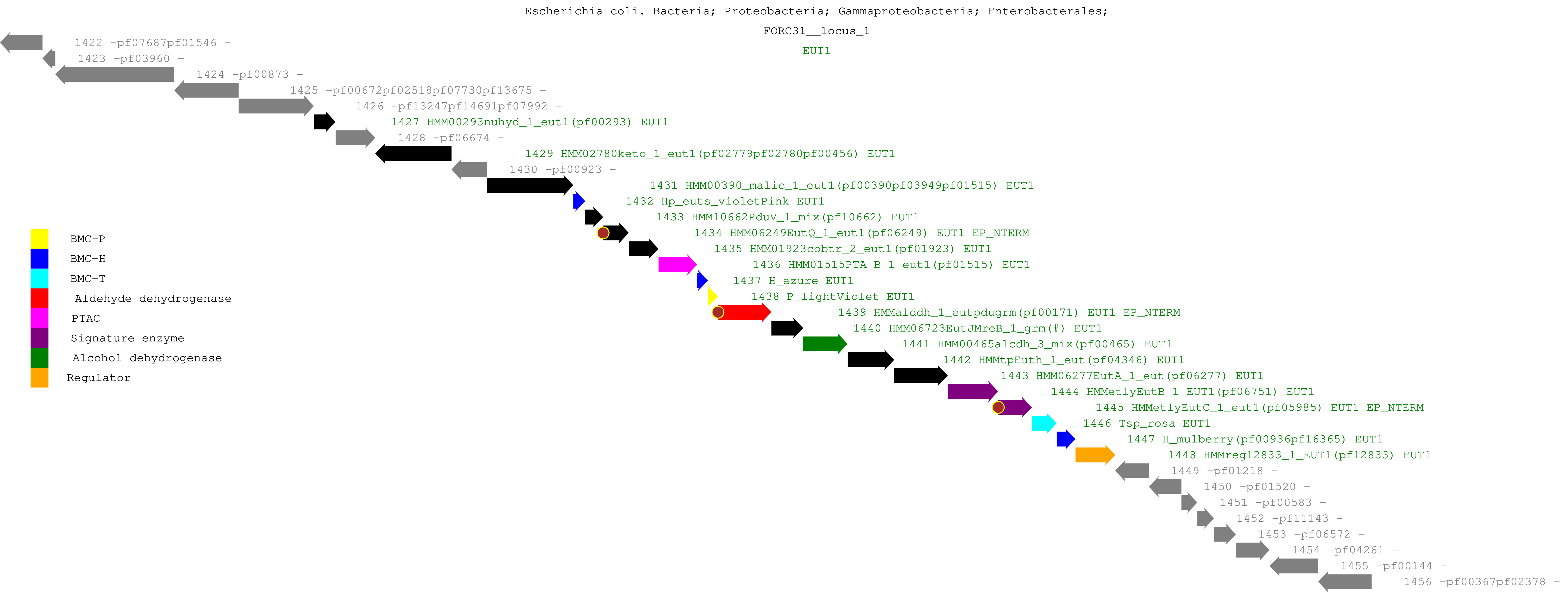

Escherichia coli. Bacteria; Proteobacteria; Gammaproteobacteria; Enterobacterales;
FORC31__locus_1
EUT1
1422 -pf07687pf01546 -
1423 -pf03960 -
1424 -pf00873 -
1425 -pf00672pf02518pf07730pf13675 -
1426 -pf13247pf14691pf07992 -
1427 HMM00293nuhyd_1_eut1(pf00293) EUT1
1428 -pf06674 -
1429 HMM02780keto_1_eut1(pf02779pf02780pf00456) EUT1
1430 -pf00923 -
1431 HMM00390_malic_1_eut1(pf00390pf03949pf01515) EUT1
1432 Hp_euts_violetPink EUT1
1433 HMM10662PduV_1_mix(pf10662) EUT1
1434 HMM06249EutQ_1_eut1(pf06249) EUT1 EP_NTERM
1435 HMM01923cobtr_2_eut1(pf01923) EUT1
1436 HMM01515PTA_B_1_eut1(pf01515) EUT1
1437 H_azure EUT1
1438 P_lightViolet EUT1
1439 HMMalddh_1_eutpdugrm(pf00171) EUT1 EP_NTERM
1440 HMM06723EutJMreB_1_grm(#) EUT1
1441 HMM00465alcdh_3_mix(pf00465) EUT1
1442 HMMtpEuth_1_eut(pf04346) EUT1
1443 HMM06277EutA_1_eut(pf06277) EUT1
1444 HMMetlyEutB_1_EUT1(pf06751) EUT1
1445 HMMetlyEutC_1_eut1(pf05985) EUT1 EP_NTERM
1446 Tsp_rosa EUT1
1447 H_mulberry(pf00936pf16365) EUT1
1448 HMMreg12833_1_EUT1(pf12833) EUT1
1449 -pf01218 -
1450 -pf01520 -
1451 -pf00583 -
1452 -pf11143 -
1453 -pf06572 -
1454 -pf04261 -
1455 -pf00144 -
1456 -pf00367pf02378 -
BMC-P
BMC-H
BMC-T
Aldehyde dehydrogenase
PTAC
Signature enzyme
Alcohol dehydrogenase
Regulator

Escherichia coli. Bacteria; Proteobacteria; Gammaproteobacteria; Enterobacterales;

FORC82__locus_1

EUT1

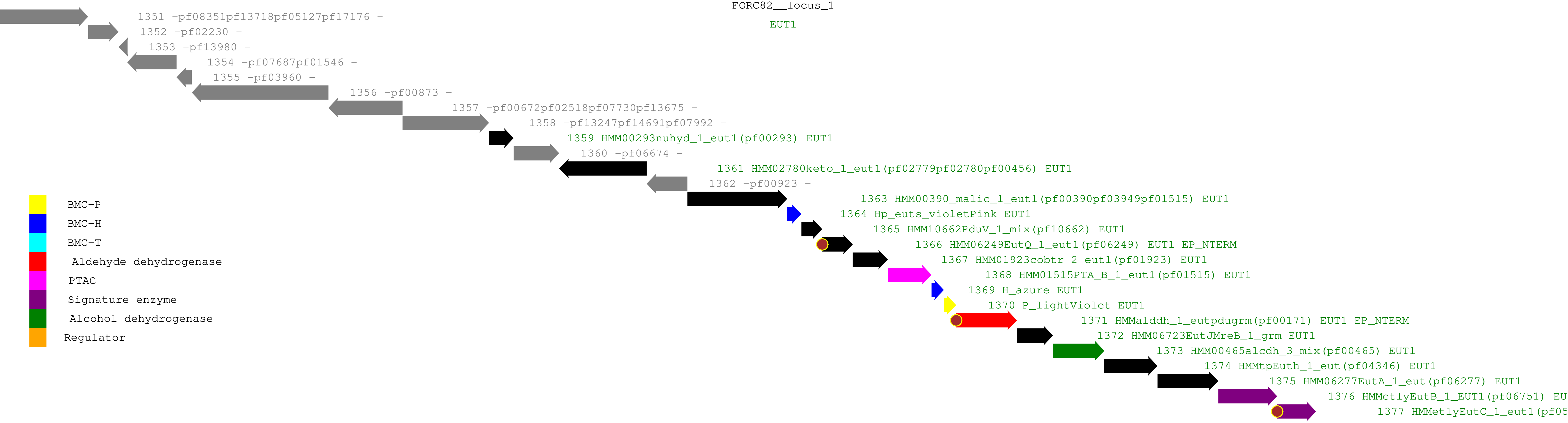

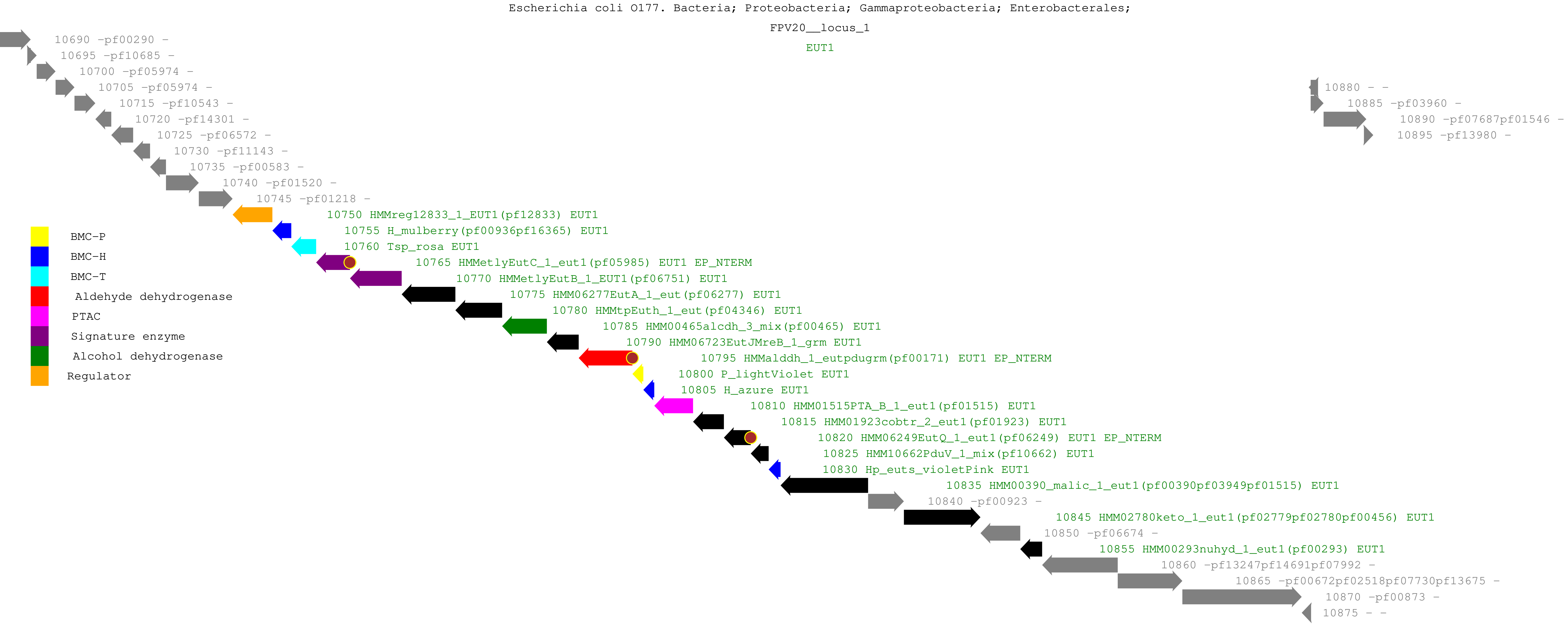

Escherichia coli O177. Bacteria; Proteobacteria; Gammaproteobacteria; Enterobacterales;
FPV20__locus_1
EUT1
10690 -pf00290 -
10695 -pf10685 -
10700 -pf05974 -
10705 -pf05974 -
10715 -pf10543 -
10720 -pf14301 -
10725 -pf06572 -
10730 -pf11143 -
10735 -pf00583 -
10740 -pf01520 -
10745 -pf01218 -
10750 HMMreg12833_1_EUT1(pf12833) EUT1
10755 H_mulberry(pf00936pf16365) EUT1
10760 Tsp_rosa EUT1
10765 HMMetlyEutC_1_eut1(pf05985) EUT1 EP_NTERM
10770 HMMetlyEutB_1_EUT1(pf06751) EUT1
10775 HMM06277EutA_1_eut(pf06277) EUT1
10780 HMMtpEuth_1_eut(pf04346) EUT1
10785 HMM00465alcdh_3_mix(pf00465) EUT1
10790 HMM06723EutJMreB_1_grm EUT1
10795 HMMalddh_1_eutpdugrm(pf00171) EUT1 EP_NTERM
10800 P_lightViolet EUT1
10805 H_azure EUT1
10810 HMM01515PTA_B_1_eut1(pf01515) EUT1
10815 HMM01923cobtr_2_eut1(pf01923) EUT1
10820 HMM06249EutQ_1_eut1(pf06249) EUT1 EP_NTERM
10825 HMM10662PduV_1_mix(pf10662) EUT1
10830 Hp_euts_violetPink EUT1
10835 HMM00390_malic_1_eut1(pf00390pf03949pf01515) EUT1
10840 -pf00923 -
10845 HMM02780keto_1_eut1(pf02779pf02780pf00456) EUT1
10850 -pf06674 -
10855 HMM00293nuhyd_1_eut1(pf00293) EUT1
10860 -pf13247pf14691pf07992 -
10865 -pf00672pf02518pf07730pf13675 -
10870 -pf00873 -
10875 - -
10880 - -
10885 -pf03960 -
10890 -pf07687pf01546 -
10895 -pf13980 -
BMC-P
BMC-H
BMC-T
Aldehyde dehydrogenase
PTAC
Signature enzyme
Alcohol dehydrogenase
Regulator

Escherichia coli. Bacteria; Proteobacteria; Gammaproteobacteria; Enterobacterales;

FTV90__locus_1

EUT1

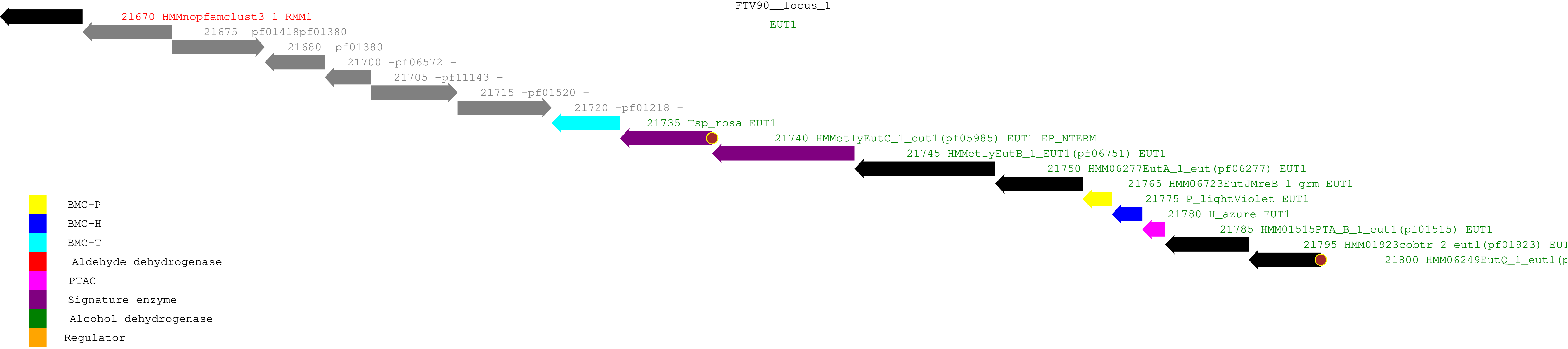

Escherichia coli. Bacteria; Proteobacteria; Gammaproteobacteria; Enterobacterales;

FTV92__locus_1

EUT1

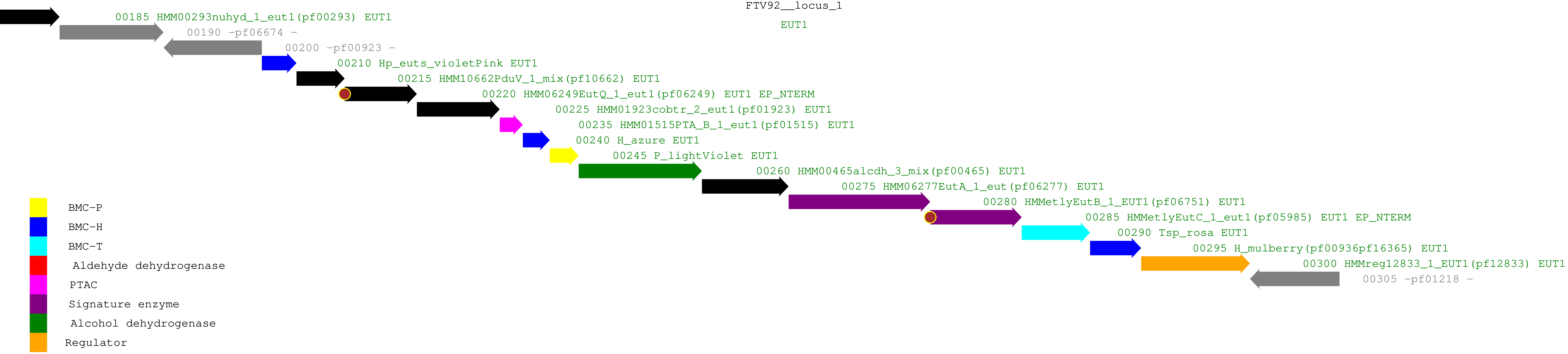

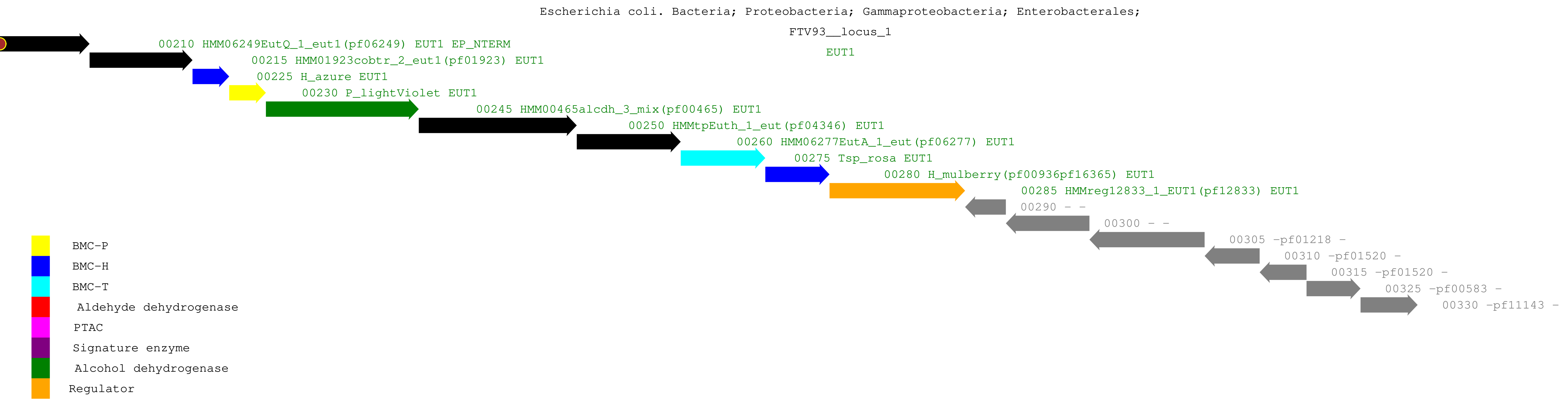
Escherichia coli. Bacteria; Proteobacteria; Gammaproteobacteria; Enterobacterales;
FTV93__locus_1
EUT1
00210 HMM06249EutQ_1_eut1(pf06249) EUT1 EP_NTERM
00215 HMM01923cobtr_2_eut1(pf01923) EUT1
00225 H_azure EUT1
00230 P_lightViolet EUT1
00245 HMM00465alcdh_3_mix(pf00465) EUT1
00250 HMMtpEuth_1_eut(pf04346) EUT1
00260 HMM06277EutA_1_eut(pf06277) EUT1
00275 Tsp_rosa EUT1
00280 H_mulberry(pf00936pf16365) EUT1
00285 HMMreg12833_1_EUT1(pf12833) EUT1
00290 - -
00300 - -
00305 -pf01218 -
00310 -pf01520 -
00315 -pf01520 -
00325 -pf00583 -
00330 -pf11143 -
BMC-P
BMC-H
BMC-T
Aldehyde dehydrogenase
PTAC
Signature enzyme
Alcohol dehydrogenase
Regulator

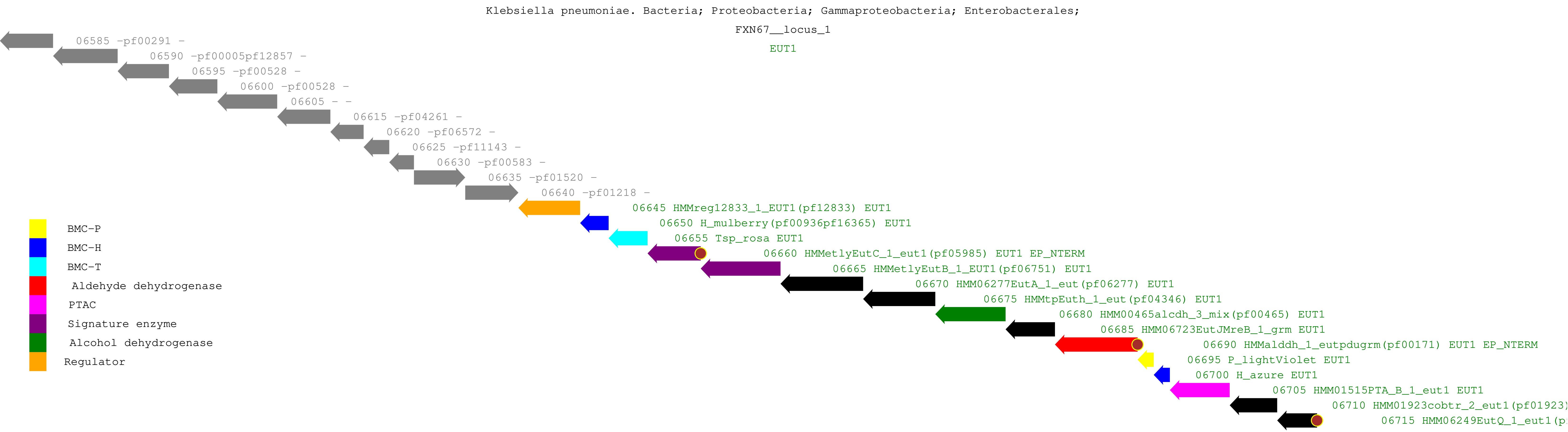

Klebsiella pneumoniae. Bacteria; Proteobacteria; Gammaproteobacteria; Enterobacterales;
FXN67__locus_1
EUT1
06585 -pf00291 -
06590 -pf00005pf12857 -
06595 -pf00528 -
06600 -pf00528 -
06605 - -
06615 -pf04261 -
06620 -pf06572 -
06625 -pf11143 -
06630 -pf00583 -
06635 -pf01520 -
06640 -pf01218 -
06645 HMMreg12833_1_EUT1(pf12833) EUT1
06650 H_mulberry(pf00936pf16365) EUT1
06655 Tsp_rosa EUT1
06660 HMMetlyEutC_1_eut1(pf05985) EUT1 EP_NTERM
06665 HMMetlyEutB_1_EUT1(pf06751) EUT1
06670 HMM06277EutA_1_eut(pf06277) EUT1
06675 HMMtpEuth_1_eut(pf04346) EUT1
06680 HMM00465alcdh_3_mix(pf00465) EUT1
06685 HMM06723EutJMreB_1_grm EUT1
06690 HMMalddh_1_eutpdugrm(pf00171) EUT1 EP_NTERM
06695 P_lightViolet EUT1
06700 H_azure EUT1
06705 HMM01515PTA_B_1_eut1 EUT1
06710 HMM01923cobtr_2_eut1(pf01923)
06715 HMM06249EutQ_1_eut1(p
BMC-P
BMC-H
BMC-T
Aldehyde dehydrogenase
PTAC
Signature enzyme
Alcohol dehydrogenase
Regulator

Escherichia coli O80:H26. Bacteria; Proteobacteria; Gammaproteobacteria; Enterobacterales;

FYA02__locus_1

EUT1

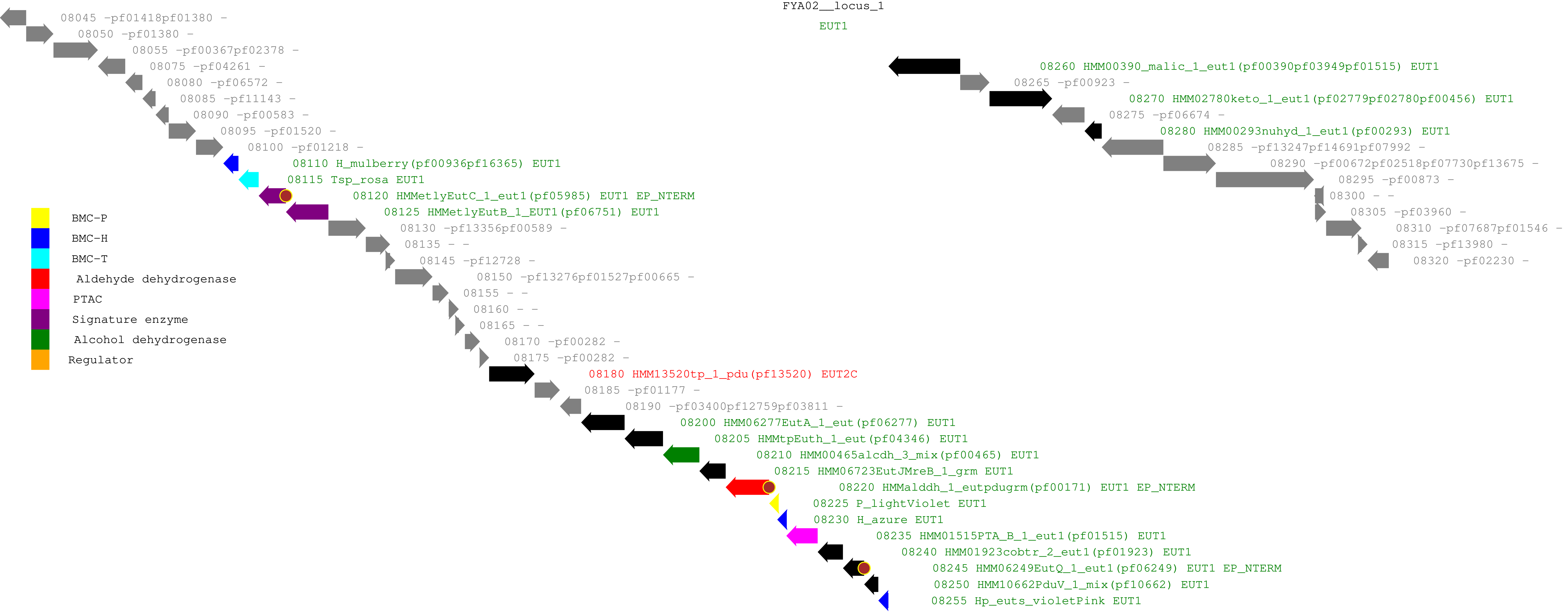

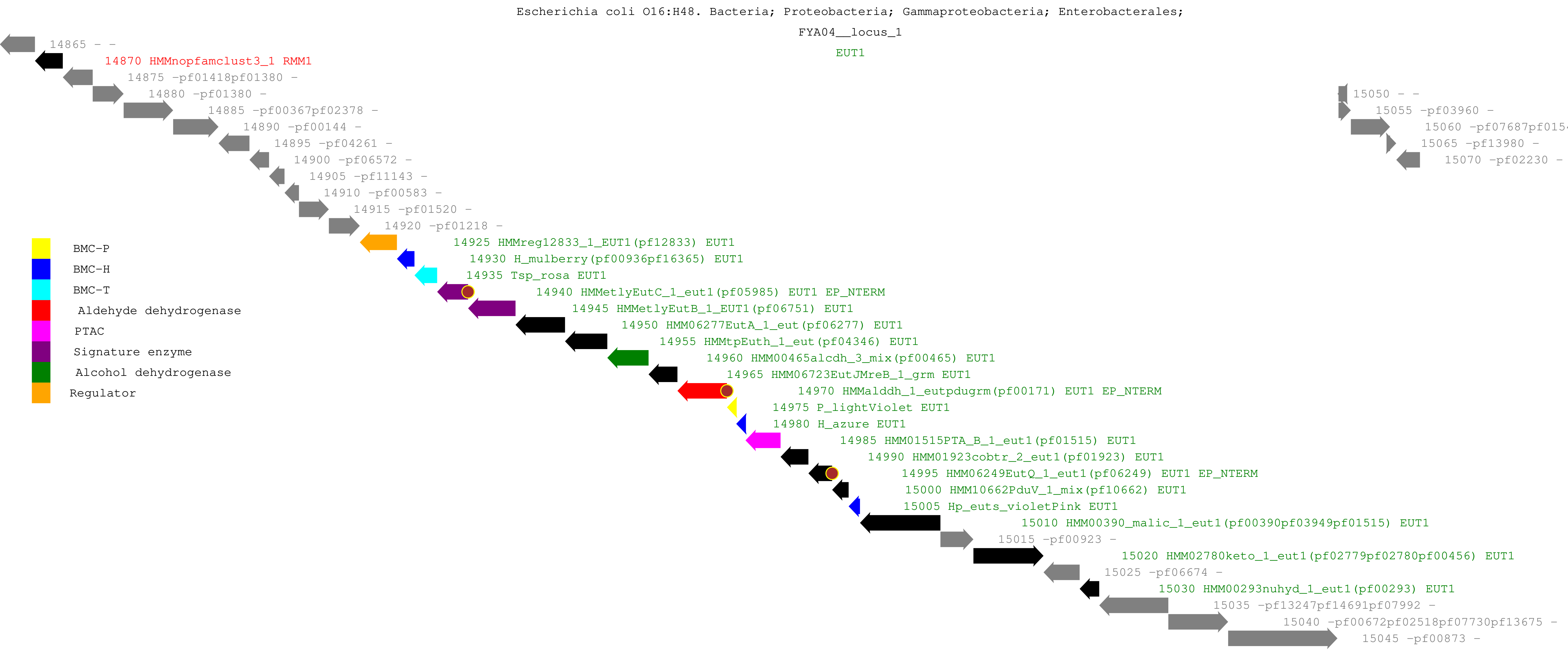
Escherichia coli O16:H48. Bacteria; Proteobacteria; Gammaproteobacteria; Enterobacterales;
FYA04__locus_1
EUT1
14865 - -
14870 HMMnopfamclust3_1 RMM1
14875 -pf01418pf01380 -
14880 -pf01380 -
14885 -pf00367pf02378 -
14890 -pf00144 -
14895 -pf04261 -
14900 -pf06572 -
14905 -pf11143 -
14910 -pf00583 -
14915 -pf01520 -
14920 -pf01218 -
14925 HMMreg12833_1_EUT1(pf12833) EUT1
14930 H_mulberry(pf00936pf16365) EUT1
14935 Tsp_rosa EUT1
14940 HMMetlyEutC_1_eut1(pf05985) EUT1 EP_NTERM
14945 HMMetlyEutB_1_EUT1(pf06751) EUT1
14950 HMM06277EutA_1_eut(pf06277) EUT1
14955 HMMtpEuth_1_eut(pf04346) EUT1
14960 HMM00465alcdh_3_mix(pf00465) EUT1
14965 HMM06723EutJMreB_1_grm EUT1
14970 HMMalddh_1_eutpdugrm(pf00171) EUT1 EP_NTERM
14975 P_lightViolet EUT1
14980 H_azure EUT1
14985 HMM01515PTA_B_1_eut1(pf01515) EUT1
14990 HMM01923cobtr_2_eut1(pf01923) EUT1
14995 HMM06249EutQ_1_eut1(pf06249) EUT1 EP_NTERM
15000 HMM10662PduV_1_mix(pf10662) EUT1
15005 Hp_euts_violetPink EUT1
15010 HMM00390_malic_1_eut1(pf00390pf03949pf01515) EUT1
15015 -pf00923 -
15020 HMM02780keto_1_eut1(pf02779pf02780pf00456) EUT1
15025 -pf06674 -
15030 HMM00293nuhyd_1_eut1(pf00293) EUT1
15035 -pf13247pf14691pf07992 -
15040 -pf00672pf02518pf07730pf13675 -
15045 -pf00873 -
15050 - -
15055 -pf03960 -
15060 -pf07687pf01546 -
15065 -pf13980 -
15070 -pf02230 -
BMC-P
BMC-H
BMC-T
Aldehyde dehydrogenase
PTAC
Signature enzyme
Alcohol dehydrogenase
Regulator

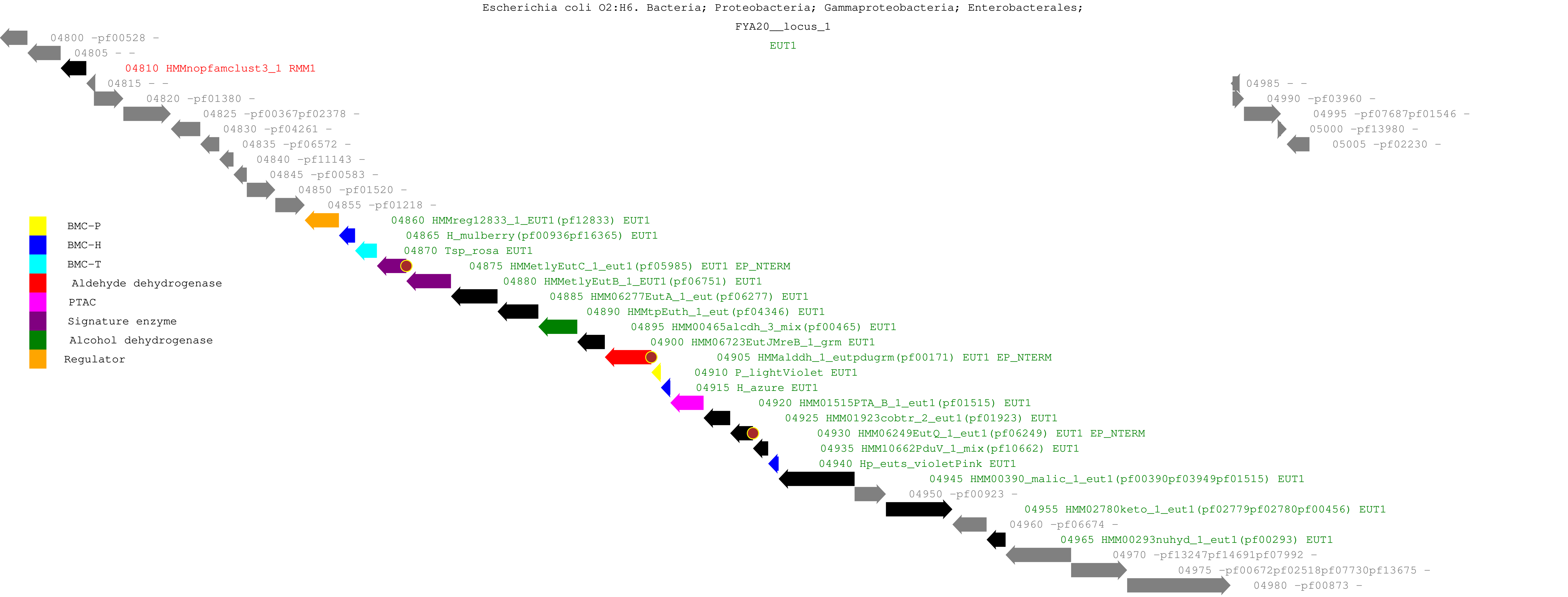
Escherichia coli O2:H6. Bacteria; Proteobacteria; Gammaproteobacteria; Enterobacterales;
FYA20__locus_1
EUT1
04800 -pf00528 -
04805 - -
04810 HMMnopfamclust3_1 RMM1
04815 - -
04820 -pf01380 -
04825 -pf00367pf02378 -
04830 -pf04261 -
04835 -pf06572 -
04840 -pf11143 -
04845 -pf00583 -
04850 -pf01520 -
04855 -pf01218 -
04860 HMMreg12833_1_EUT1(pf12833) EUT1
04865 H_mulberry(pf00936pf16365) EUT1
04870 Tsp_rosa EUT1
04875 HMMetlyEutC_1_eut1(pf05985) EUT1 EP_NTERM
04880 HMMetlyEutB_1_EUT1(pf06751) EUT1
04885 HMM06277EutA_1_eut(pf06277) EUT1
04890 HMMtpEuth_1_eut(pf04346) EUT1
04895 HMM00465alcdh_3_mix(pf00465) EUT1
04900 HMM06723EutJMreB_1_grm EUT1
04905 HMMalddh_1_eutpdugrm(pf00171) EUT1 EP_NTERM
04910 P_lightViolet EUT1
04915 H_azure EUT1
04920 HMM01515PTA_B_1_eut1(pf01515) EUT1
04925 HMM01923cobtr_2_eut1(pf01923) EUT1
04930 HMM06249EutQ_1_eut1(pf06249) EUT1 EP_NTERM
04935 HMM10662PduV_1_mix(pf10662) EUT1
04940 Hp_euts_violetPink EUT1
04945 HMM00390_malic_1_eut1(pf00390pf03949pf01515) EUT1
04950 -pf00923 -
04955 HMM02780keto_1_eut1(pf02779pf02780pf00456) EUT1
04960 -pf06674 -
04965 HMM00293nuhyd_1_eut1(pf00293) EUT1
04970 -pf13247pf14691pf07992 -
04975 -pf00672pf02518pf07730pf13675 -
04980 -pf00873 -
04985 - -
04990 -pf03960 -
04995 -pf07687pf01546 -
05000 -pf13980 -
05005 -pf02230 -
BMC-P
BMC-H
BMC-T
Aldehyde dehydrogenase
PTAC
Signature enzyme
Alcohol dehydrogenase
Regulator

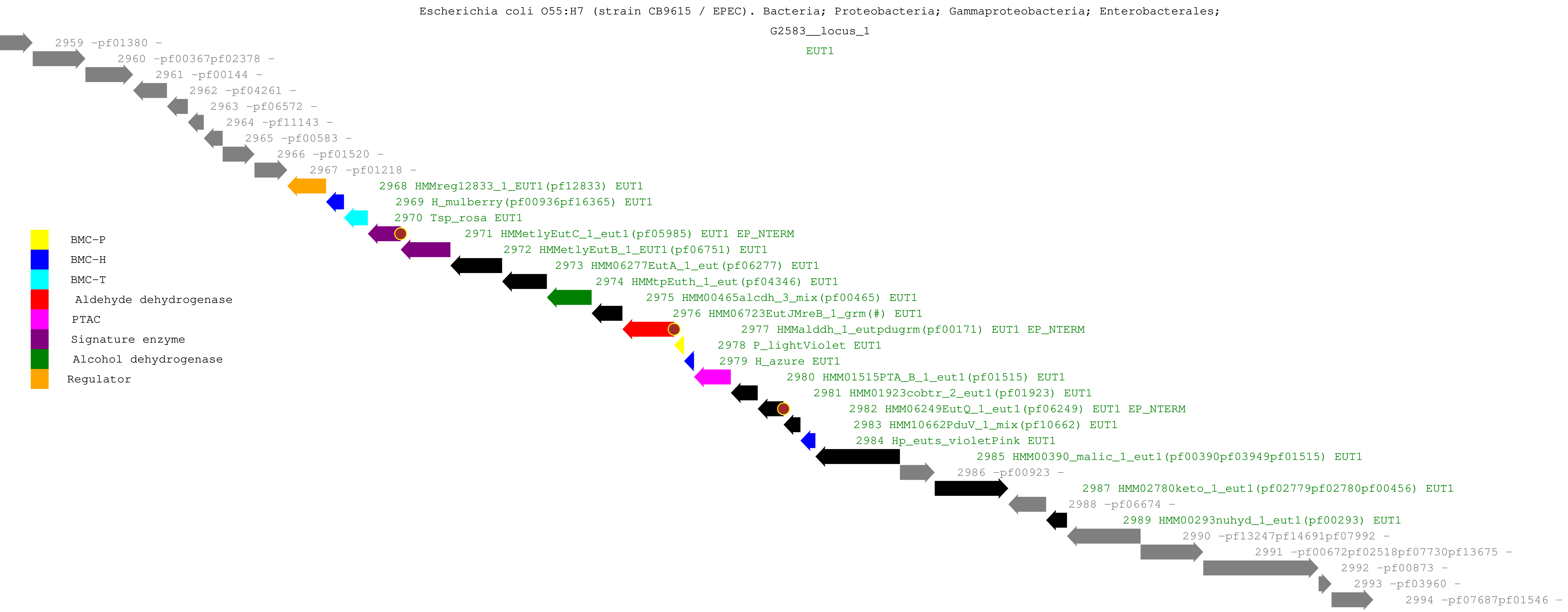
Escherichia coli O55:H7 (strain CB9615 / EPEC). Bacteria; Proteobacteria; Gammaproteobacteria; Enterobacterales;
G2583__locus_1
EUT1
2959 -pf01380 -
2960 -pf00367pf02378 -
2961 -pf00144 -
2962 -pf04261 -
2963 -pf06572 -
2964 -pf11143 -
2965 -pf00583 -
2966 -pf01520 -
2967 -pf01218 -
2968 HMMreg12833_1_EUT1(pf12833) EUT1
2969 H_mulberry(pf00936pf16365) EUT1
2970 Tsp_rosa EUT1
2971 HMMetlyEutC_1_eut1(pf05985) EUT1 EP_NTERM
2972 HMMetlyEutB_1_EUT1(pf06751) EUT1
2973 HMM06277EutA_1_eut(pf06277) EUT1
2974 HMMtpEuth_1_eut(pf04346) EUT1
2975 HMM00465alcdh_3_mix(pf00465) EUT1
2976 HMM06723EutJMreB_1_grm(#) EUT1
2977 HMMalddh_1_eutpdugrm(pf00171) EUT1 EP_NTERM
2978 P_lightViolet EUT1
2979 H_azure EUT1
2980 HMM01515PTA_B_1_eut1(pf01515) EUT1
2981 HMM01923cobtr_2_eut1(pf01923) EUT1
2982 HMM06249EutQ_1_eut1(pf06249) EUT1 EP_NTERM
2983 HMM10662PduV_1_mix(pf10662) EUT1
2984 Hp_euts_violetPink EUT1
2985 HMM00390_malic_1_eut1(pf00390pf03949pf01515) EUT1
2986 -pf00923 -
2987 HMM02780keto_1_eut1(pf02779pf02780pf00456) EUT1
2988 -pf06674 -
2989 HMM00293nuhyd_1_eut1(pf00293) EUT1
2990 -pf13247pf14691pf07992 -
2991 -pf00672pf02518pf07730pf13675 -
2992 -pf00873 -
2993 -pf03960 -
2994 -pf07687pf01546 -
BMC-P
BMC-H
BMC-T
Aldehyde dehydrogenase
PTAC
Signature enzyme
Alcohol dehydrogenase
Regulator

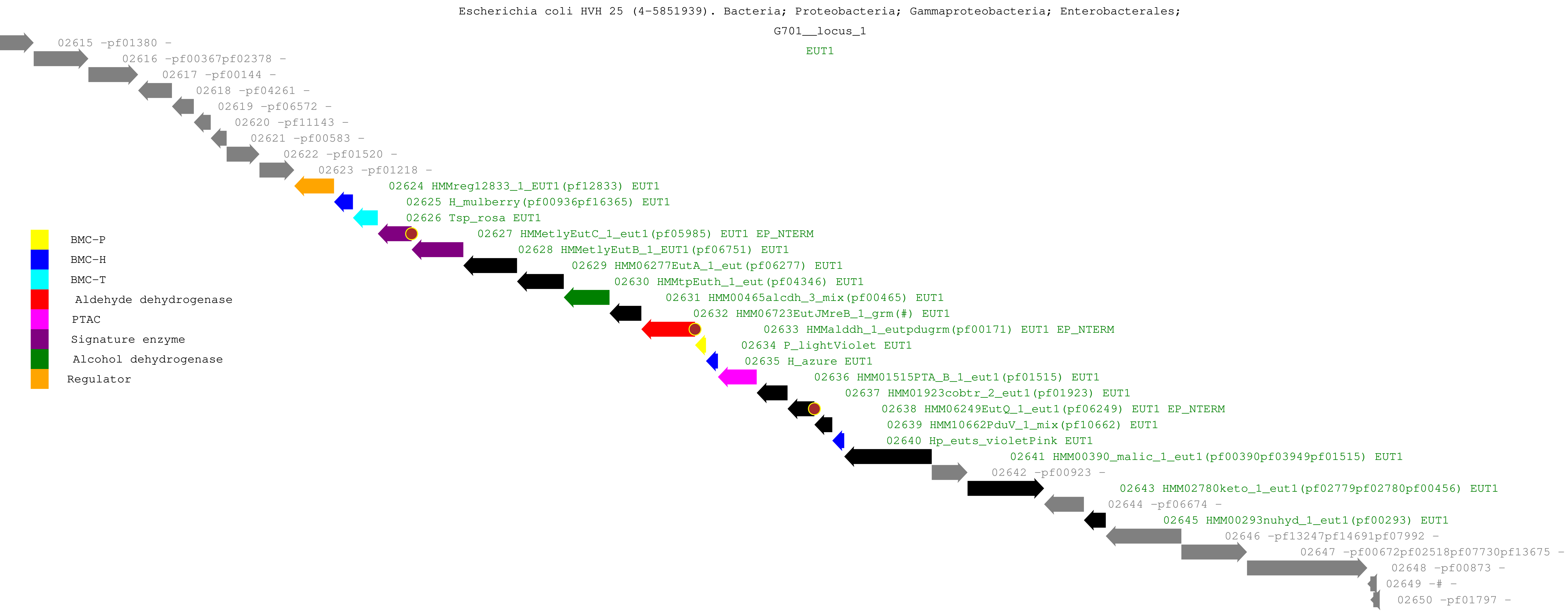

Escherichia coli HVH 25 (4-5851939). Bacteria; Proteobacteria; Gammaproteobacteria; Enterobacterales;
G701__locus_1
EUT1
02615 -pf01380 -
02616 -pf00367pf02378 -
02617 -pf00144 -
02618 -pf04261 -
02619 -pf06572 -
02620 -pf11143 -
02621 -pf00583 -
02622 -pf01520 -
02623 -pf01218 -
02624 HMMreg12833_1_EUT1(pf12833) EUT1
02625 H_mulberry(pf00936pf16365) EUT1
02626 Tsp_rosa EUT1
02627 HMMetlyEutC_1_eut1(pf05985) EUT1 EP_NTERM
02628 HMMetlyEutB_1_EUT1(pf06751) EUT1
02629 HMM06277EutA_1_eut(pf06277) EUT1
02630 HMMtpEuth_1_eut(pf04346) EUT1
02631 HMM00465alcdh_3_mix(pf00465) EUT1
02632 HMM06723EutJMreB_1_grm(#) EUT1
02633 HMMalddh_1_eutpdugrm(pf00171) EUT1 EP_NTERM
02634 P_lightViolet EUT1
02635 H_azure EUT1
02636 HMM01515PTA_B_1_eut1(pf01515) EUT1
02637 HMM01923cobtr_2_eut1(pf01923) EUT1
02638 HMM06249EutQ_1_eut1(pf06249) EUT1 EP_NTERM
02639 HMM10662PduV_1_mix(pf10662) EUT1
02640 Hp_euts_violetPink EUT1
02641 HMM00390_malic_1_eut1(pf00390pf03949pf01515) EUT1
02642 -pf00923 -
02643 HMM02780keto_1_eut1(pf02779pf02780pf00456) EUT1
02644 -pf06674 -
02645 HMM00293nuhyd_1_eut1(pf00293) EUT1
02646 -pf13247pf14691pf07992 -
02647 -pf00672pf02518pf07730pf13675 -
02648 -pf00873 -
02649 -# -
02650 -pf01797 -
BMC-P
BMC-H
BMC-T
Aldehyde dehydrogenase
PTAC
Signature enzyme
Alcohol dehydrogenase
Regulator

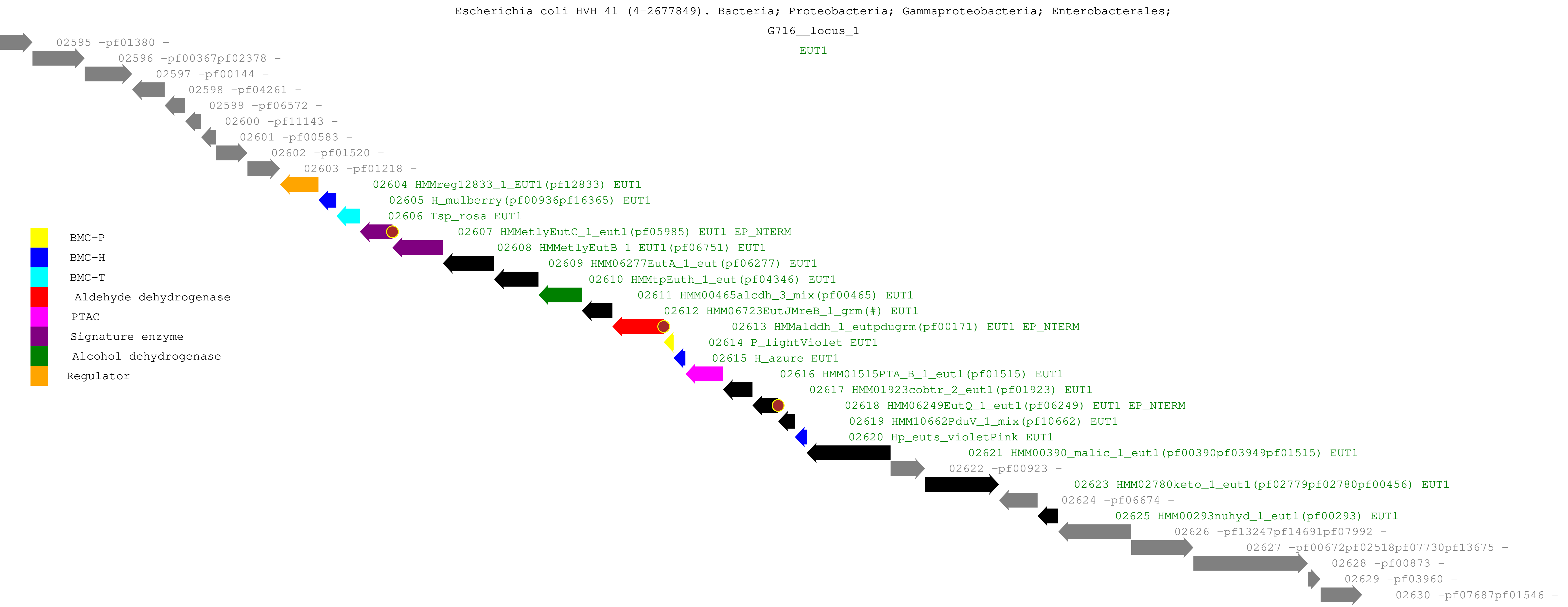
Escherichia coli HVH 41 (4-2677849). Bacteria; Proteobacteria; Gammaproteobacteria; Enterobacterales;
G716__locus_1
EUT1
02595 -pf01380 -
02596 -pf00367pf02378 -
02597 -pf00144 -
02598 -pf04261 -
02599 -pf06572 -
02600 -pf11143 -
02601 -pf00583 -
02602 -pf01520 -
02603 -pf01218 -
02604 HMMreg12833_1_EUT1(pf12833) EUT1
02605 H_mulberry(pf00936pf16365) EUT1
02606 Tsp_rosa EUT1
02607 HMMetlyEutC_1_eut1(pf05985) EUT1 EP_NTERM
02608 HMMetlyEutB_1_EUT1(pf06751) EUT1
02609 HMM06277EutA_1_eut(pf06277) EUT1
02610 HMMtpEuth_1_eut(pf04346) EUT1
02611 HMM00465alcdh_3_mix(pf00465) EUT1
02612 HMM06723EutJMreB_1_grm(#) EUT1
02613 HMMalddh_1_eutpdugrm(pf00171) EUT1 EP_NTERM
02614 P_lightViolet EUT1
02615 H_azure EUT1
02616 HMM01515PTA_B_1_eut1(pf01515) EUT1
02617 HMM01923cobtr_2_eut1(pf01923) EUT1
02618 HMM06249EutQ_1_eut1(pf06249) EUT1 EP_NTERM
02619 HMM10662PduV_1_mix(pf10662) EUT1
02620 Hp_euts_violetPink EUT1
02621 HMM00390_malic_1_eut1(pf00390pf03949pf01515) EUT1
02622 -pf00923 -
02623 HMM02780keto_1_eut1(pf02779pf02780pf00456) EUT1
02624 -pf06674 -
02625 HMM00293nuhyd_1_eut1(pf00293) EUT1
02626 -pf13247pf14691pf07992 -
02627 -pf00672pf02518pf07730pf13675 -
02628 -pf00873 -
02629 -pf03960 -
02630 -pf07687pf01546 -
BMC-P
BMC-H
BMC-T
Aldehyde dehydrogenase
PTAC
Signature enzyme
Alcohol dehydrogenase
Regulator

Escherichia coli HVH 50 (4-2593475). Bacteria; Proteobacteria; Gammaproteobacteria; Enterobacterales;

G723__locus_1

EUT1

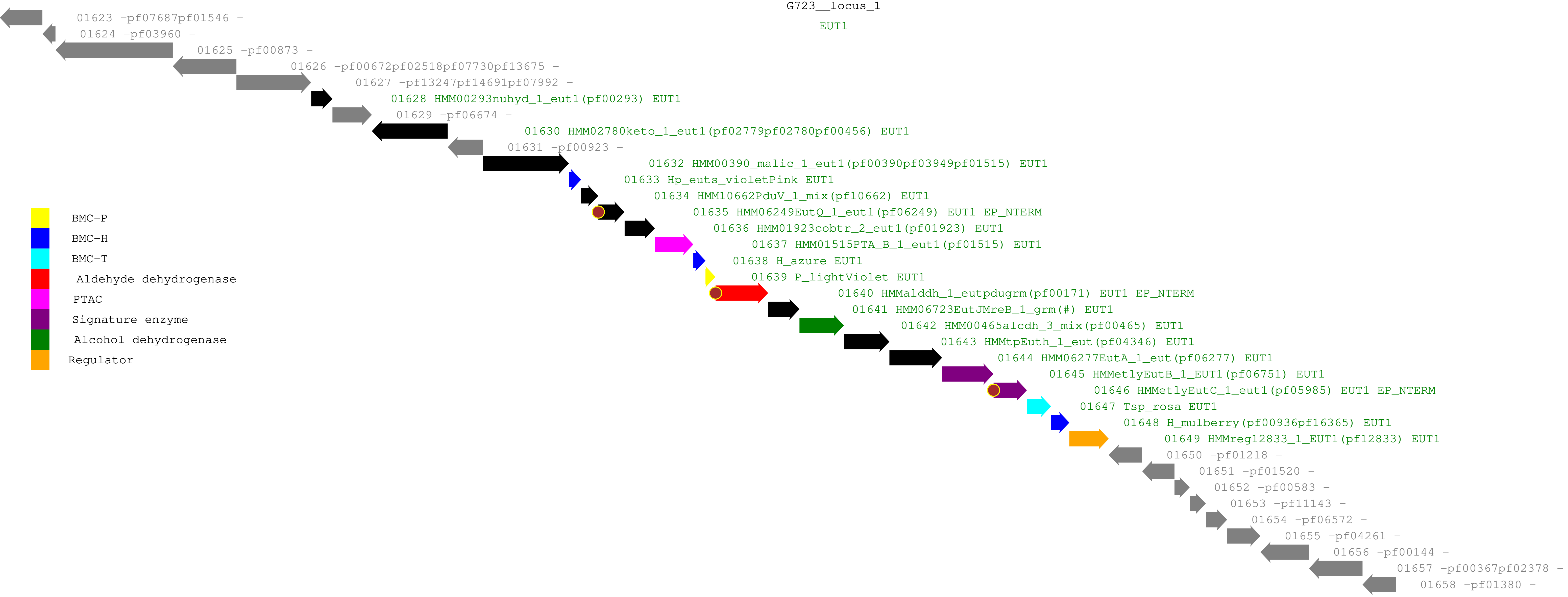

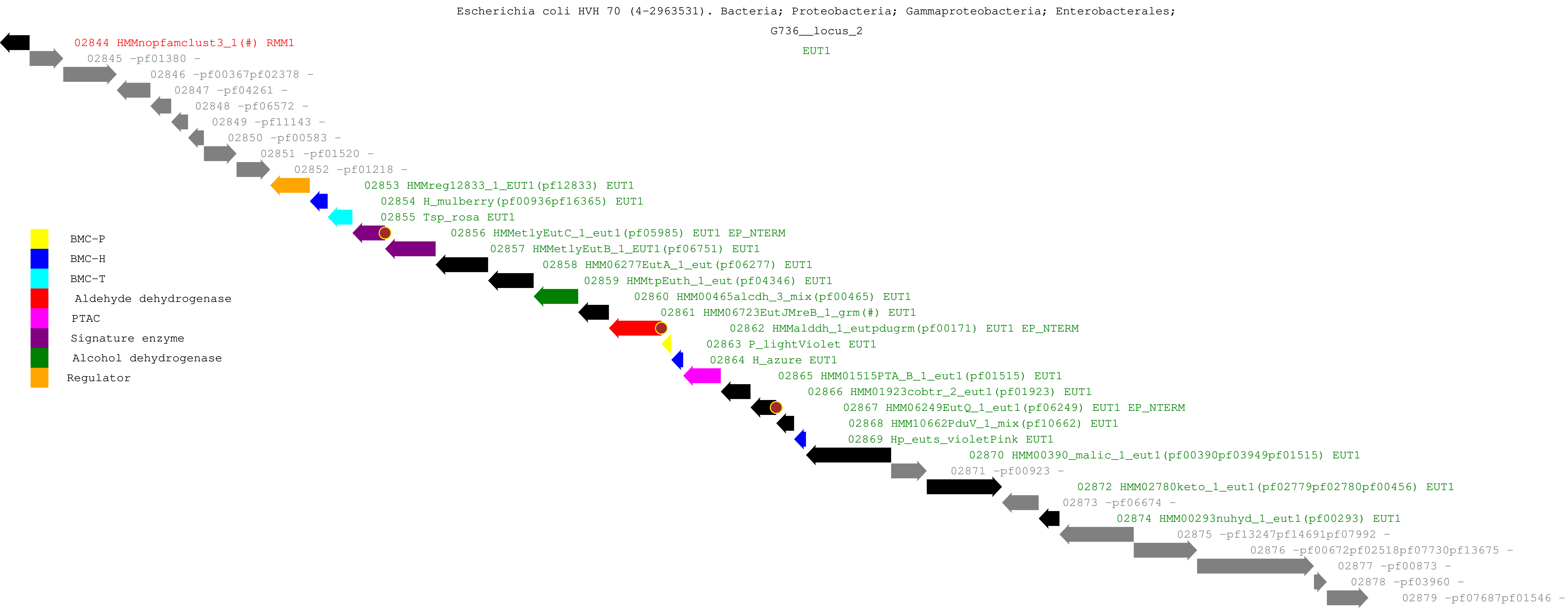

Escherichia coli HVH 70 (4-2963531). Bacteria; Proteobacteria; Gammaproteobacteria; Enterobacterales;
G736__locus_2
EUT1
02844 HMMnopfamclust3_1(#) RMM1
02845 -pf01380 -
02846 -pf00367pf02378 -
02847 -pf04261 -
02848 -pf06572 -
02849 -pf11143 -
02850 -pf00583 -
02851 -pf01520 -
02852 -pf01218 -
02853 HMMreg12833_1_EUT1(pf12833) EUT1
02854 H_mulberry(pf00936pf16365) EUT1
02855 Tsp_rosa EUT1
02856 HMMetlyEutC_1_eut1(pf05985) EUT1 EP_NTERM
02857 HMMetlyEutB_1_EUT1(pf06751) EUT1
02858 HMM06277EutA_1_eut(pf06277) EUT1
02859 HMMtpEuth_1_eut(pf04346) EUT1
02860 HMM00465alcdh_3_mix(pf00465) EUT1
02861 HMM06723EutJMreB_1_grm(#) EUT1
02862 HMMalddh_1_eutpdugrm(pf00171) EUT1 EP_NTERM
02863 P_lightViolet EUT1
02864 H_azure EUT1
02865 HMM01515PTA_B_1_eut1(pf01515) EUT1
02866 HMM01923cobtr_2_eut1(pf01923) EUT1
02867 HMM06249EutQ_1_eut1(pf06249) EUT1 EP_NTERM
02868 HMM10662PduV_1_mix(pf10662) EUT1
02869 Hp_euts_violetPink EUT1
02870 HMM00390_malic_1_eut1(pf00390pf03949pf01515) EUT1
02871 -pf00923 -
02872 HMM02780keto_1_eut1(pf02779pf02780pf00456) EUT1
02873 -pf06674 -
02874 HMM00293nuhyd_1_eut1(pf00293) EUT1
02875 -pf13247pf14691pf07992 -
02876 -pf00672pf02518pf07730pf13675 -
02877 -pf00873 -
02878 -pf03960 -
02879 -pf07687pf01546 -
BMC-P
BMC-H
BMC-T
Aldehyde dehydrogenase
PTAC
Signature enzyme
Alcohol dehydrogenase
Regulator

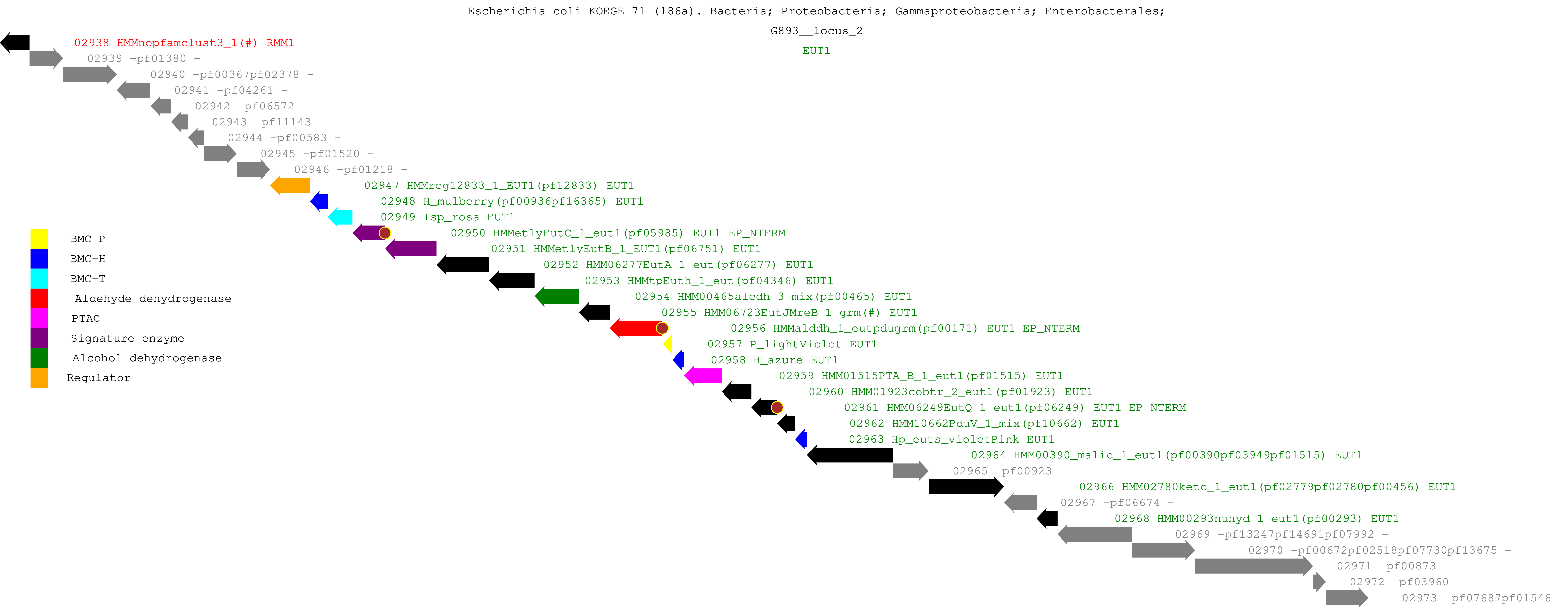
Escherichia coli KOEGE 71 (186a). Bacteria; Proteobacteria; Gammaproteobacteria; Enterobacterales;
G893__locus_2
EUT1
02938 HMMnopfamclust3_1(#) RMM1
02939 -pf01380 -
02940 -pf00367pf02378 -
02941 -pf04261 -
02942 -pf06572 -
02943 -pf11143 -
02944 -pf00583 -
02945 -pf01520 -
02946 -pf01218 -
02947 HMMreg12833_1_EUT1(pf12833) EUT1
02948 H_mulberry(pf00936pf16365) EUT1
02949 Tsp_rosa EUT1
02950 HMMetlyEutC_1_eut1(pf05985) EUT1 EP_NTERM
02951 HMMetlyEutB_1_EUT1(pf06751) EUT1
02952 HMM06277EutA_1_eut(pf06277) EUT1
02953 HMMtpEuth_1_eut(pf04346) EUT1
02954 HMM00465alcdh_3_mix(pf00465) EUT1
02955 HMM06723EutJMreB_1_grm(#) EUT1
02956 HMMalddh_1_eutpdugrm(pf00171) EUT1 EP_NTERM
02957 P_lightViolet EUT1
02958 H_azure EUT1
02959 HMM01515PTA_B_1_eut1(pf01515) EUT1
02960 HMM01923cobtr_2_eut1(pf01923) EUT1
02961 HMM06249EutQ_1_eut1(pf06249) EUT1 EP_NTERM
02962 HMM10662PduV_1_mix(pf10662) EUT1
02963 Hp_euts_violetPink EUT1
02964 HMM00390_malic_1_eut1(pf00390pf03949pf01515) EUT1
02965 -pf00923 -
02966 HMM02780keto_1_eut1(pf02779pf02780pf00456) EUT1
02967 -pf06674 -
02968 HMM00293nuhyd_1_eut1(pf00293) EUT1
02969 -pf13247pf14691pf07992 -
02970 -pf00672pf02518pf07730pf13675 -
02971 -pf00873 -
02972 -pf03960 -
02973 -pf07687pf01546 -
BMC-P
BMC-H
BMC-T
Aldehyde dehydrogenase
PTAC
Signature enzyme
Alcohol dehydrogenase
Regulator

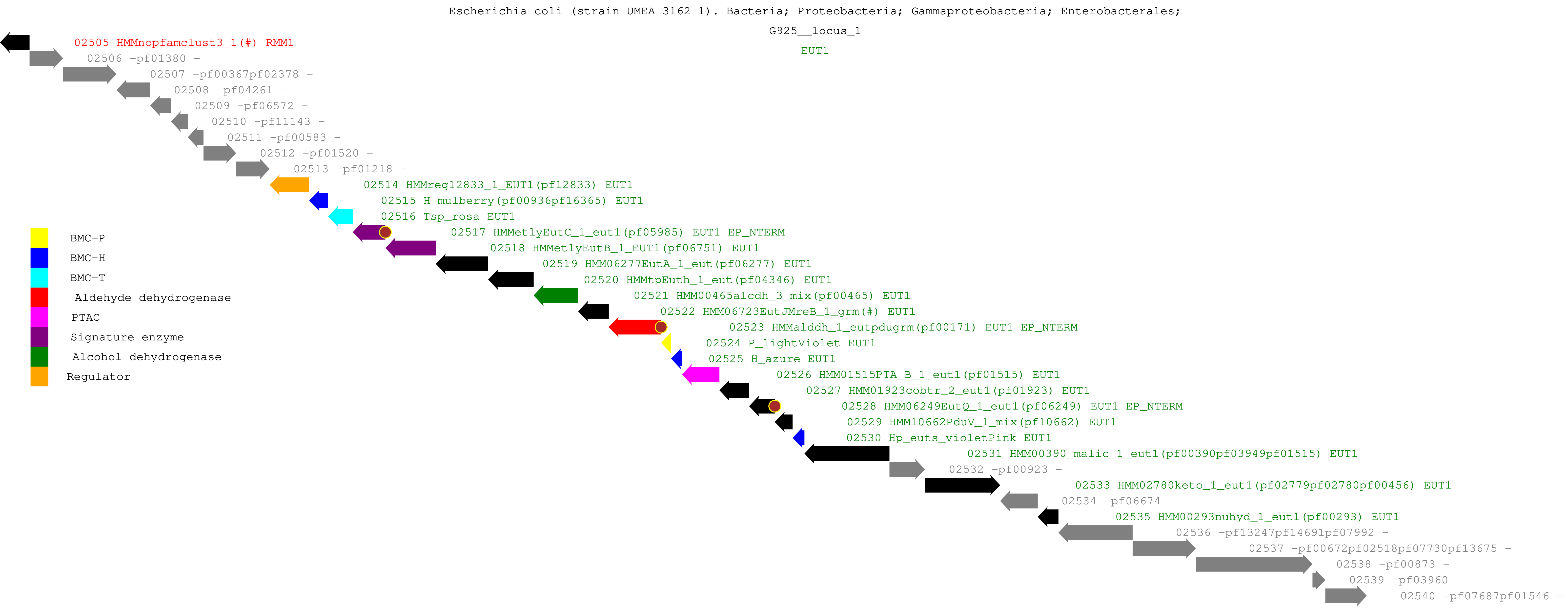
Escherichia coli (strain UMEA 3162-1). Bacteria; Proteobacteria; Gammaproteobacteria; Enterobacterales;
G925__locus_1
EUT1
02505 HMMnopfamclust3_1(#) RMM1
02506 -pf01380 -
02507 -pf00367pf02378 -
02508 -pf04261 -
02509 -pf06572 -
02510 -pf11143 -
02511 -pf00583 -
02512 -pf01520 -
02513 -pf01218 -
02514 HMMreg12833_1_EUT1(pf12833) EUT1
02515 H_mulberry(pf00936pf16365) EUT1
02516 Tsp_rosa EUT1
02517 HMMetlyEutC_1_eut1(pf05985) EUT1 EP_NTERM
02518 HMMetlyEutB_1_EUT1(pf06751) EUT1
02519 HMM06277EutA_1_eut(pf06277) EUT1
02520 HMMtpEuth_1_eut(pf04346) EUT1
02521 HMM00465alcdh_3_mix(pf00465) EUT1
02522 HMM06723EutJMreB_1_grm(#) EUT1
02523 HMMalddh_1_eutpdugrm(pf00171) EUT1 EP_NTERM
02524 P_lightViolet EUT1
02525 H_azure EUT1
02526 HMM01515PTA_B_1_eut1(pf01515) EUT1
02527 HMM01923cobtr_2_eut1(pf01923) EUT1
02528 HMM06249EutQ_1_eut1(pf06249) EUT1 EP_NTERM
02529 HMM10662PduV_1_mix(pf10662) EUT1
02530 Hp_euts_violetPink EUT1
02531 HMM00390_malic_1_eut1(pf00390pf03949pf01515) EUT1
02532 -pf00923 -
02533 HMM02780keto_1_eut1(pf02779pf02780pf00456) EUT1
02534 -pf06674 -
02535 HMM00293nuhyd_1_eut1(pf00293) EUT1
02536 -pf13247pf14691pf07992 -
02537 -pf00672pf02518pf07730pf13675 -
02538 -pf00873 -
02539 -pf03960 -
02540 -pf07687pf01546 -
BMC-P
BMC-H
BMC-T
Aldehyde dehydrogenase
PTAC
Signature enzyme
Alcohol dehydrogenase
Regulator

Escherichia coli UMEA 3200-1. Bacteria; Proteobacteria; Gammaproteobacteria; Enterobacterales;

G938__locus_2

EUT1

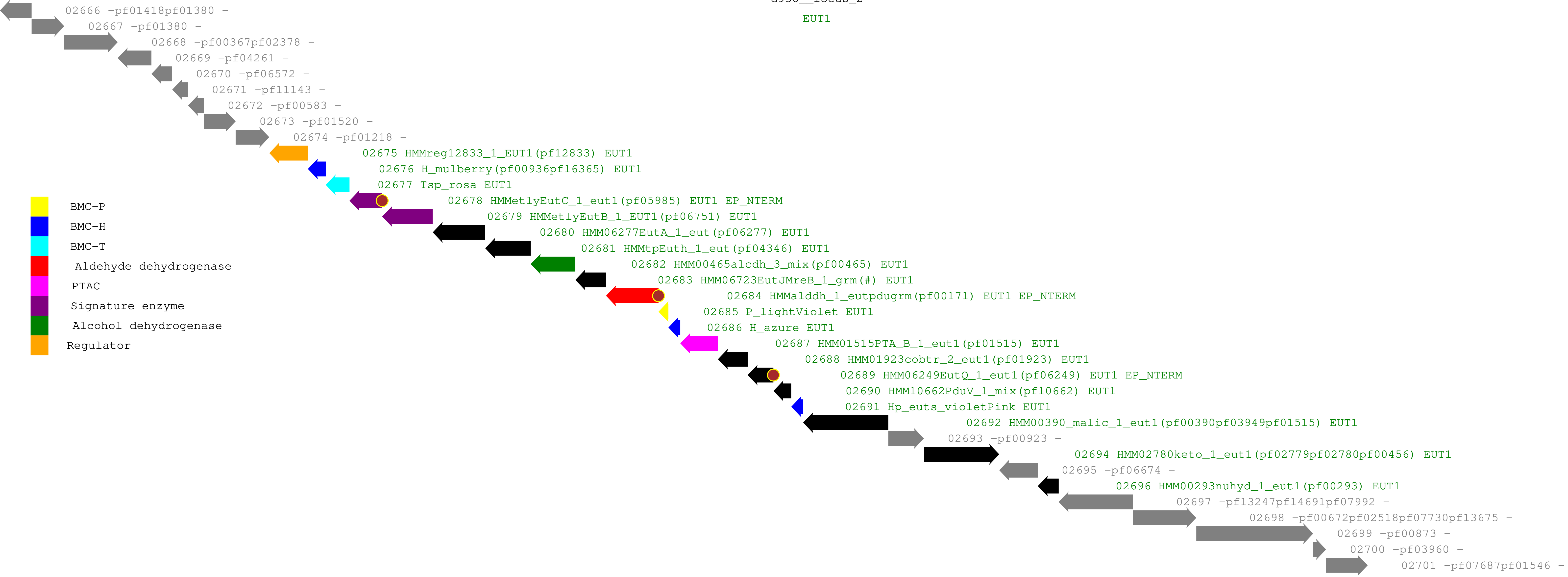

Escherichia coli UMEA 3212-1. Bacteria; Proteobacteria; Gammaproteobacteria; Enterobacterales;

G943__locus_1_main_sat

EUT1

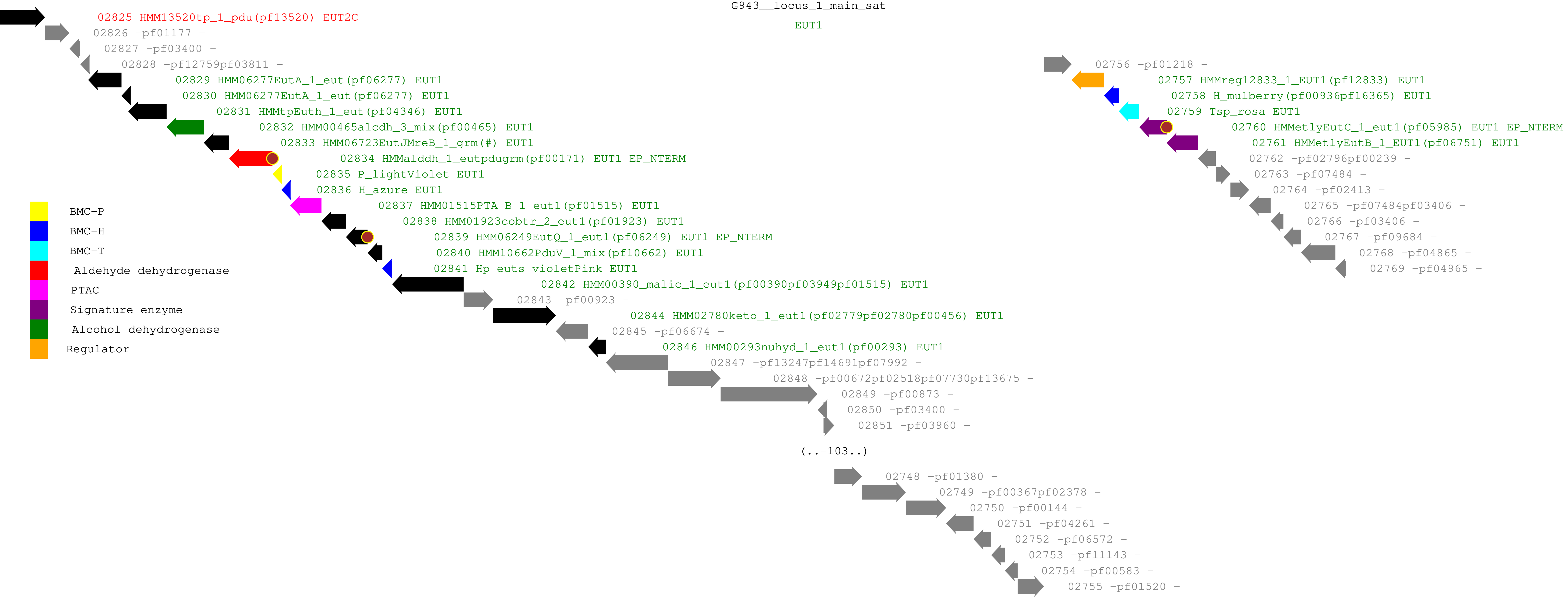

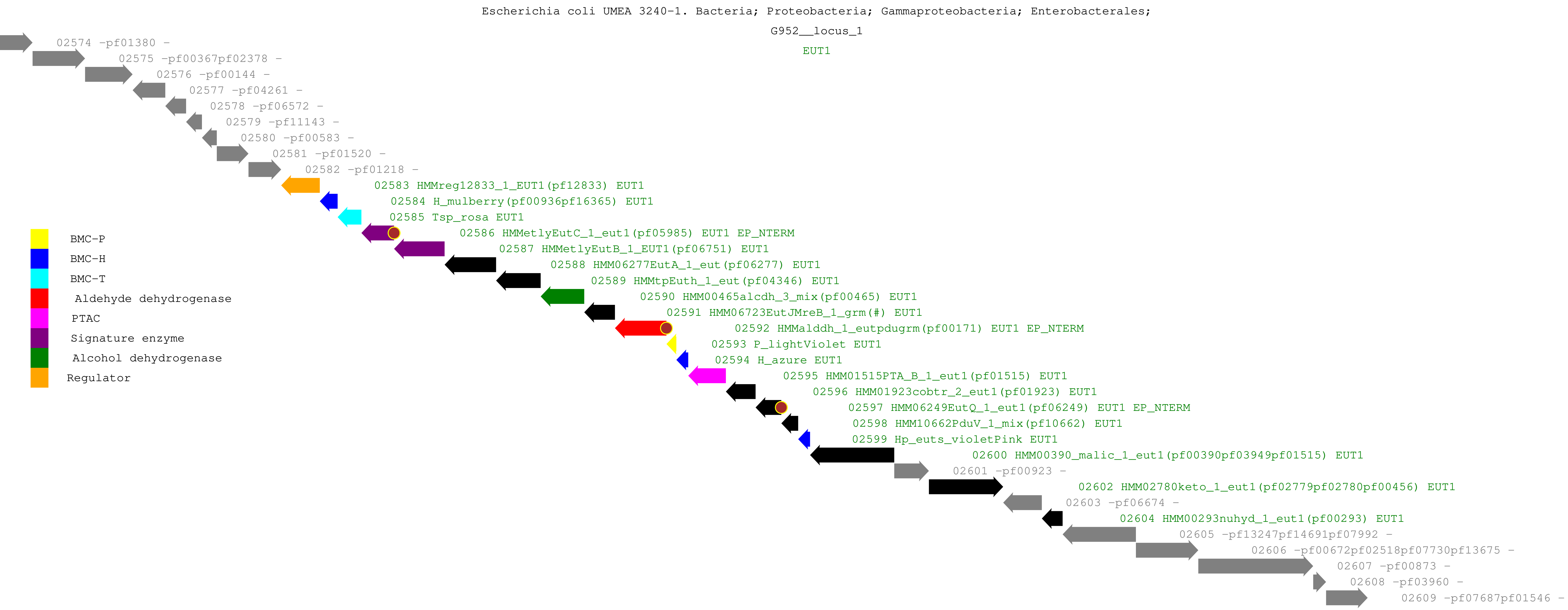

Escherichia coli UMEA 3240-1. Bacteria; Proteobacteria; Gammaproteobacteria; Enterobacterales;
G952__locus_1
EUT1
02574 -pf01380 -
02575 -pf00367pf02378 -
02576 -pf00144 -
02577 -pf04261 -
02578 -pf06572 -
02579 -pf11143 -
02580 -pf00583 -
02581 -pf01520 -
02582 -pf01218 -
02583 HMMreg12833_1_EUT1(pf12833) EUT1
02584 H_mulberry(pf00936pf16365) EUT1
02585 Tsp_rosa EUT1
02586 HMMetlyEutC_1_eut1(pf05985) EUT1 EP_NTERM
02587 HMMetlyEutB_1_EUT1(pf06751) EUT1
02588 HMM06277EutA_1_eut(pf06277) EUT1
02589 HMMtpEuth_1_eut(pf04346) EUT1
02590 HMM00465alcdh_3_mix(pf00465) EUT1
02591 HMM06723EutJMreB_1_grm(#) EUT1
02592 HMMalddh_1_eutpdugrm(pf00171) EUT1 EP_NTERM
02593 P_lightViolet EUT1
02594 H_azure EUT1
02595 HMM01515PTA_B_1_eut1(pf01515) EUT1
02596 HMM01923cobtr_2_eut1(pf01923) EUT1
02597 HMM06249EutQ_1_eut1(pf06249) EUT1 EP_NTERM
02598 HMM10662PduV_1_mix(pf10662) EUT1
02599 Hp_euts_violetPink EUT1
02600 HMM00390_malic_1_eut1(pf00390pf03949pf01515) EUT1
02601 -pf00923 -
02602 HMM02780keto_1_eut1(pf02779pf02780pf00456) EUT1
02603 -pf06674 -
02604 HMM00293nuhyd_1_eut1(pf00293) EUT1
02605 -pf13247pf14691pf07992 -
02606 -pf00672pf02518pf07730pf13675 -
02607 -pf00873 -
02608 -pf03960 -
02609 -pf07687pf01546 -
BMC-P
BMC-H
BMC-T
Aldehyde dehydrogenase
PTAC
Signature enzyme
Alcohol dehydrogenase
Regulator

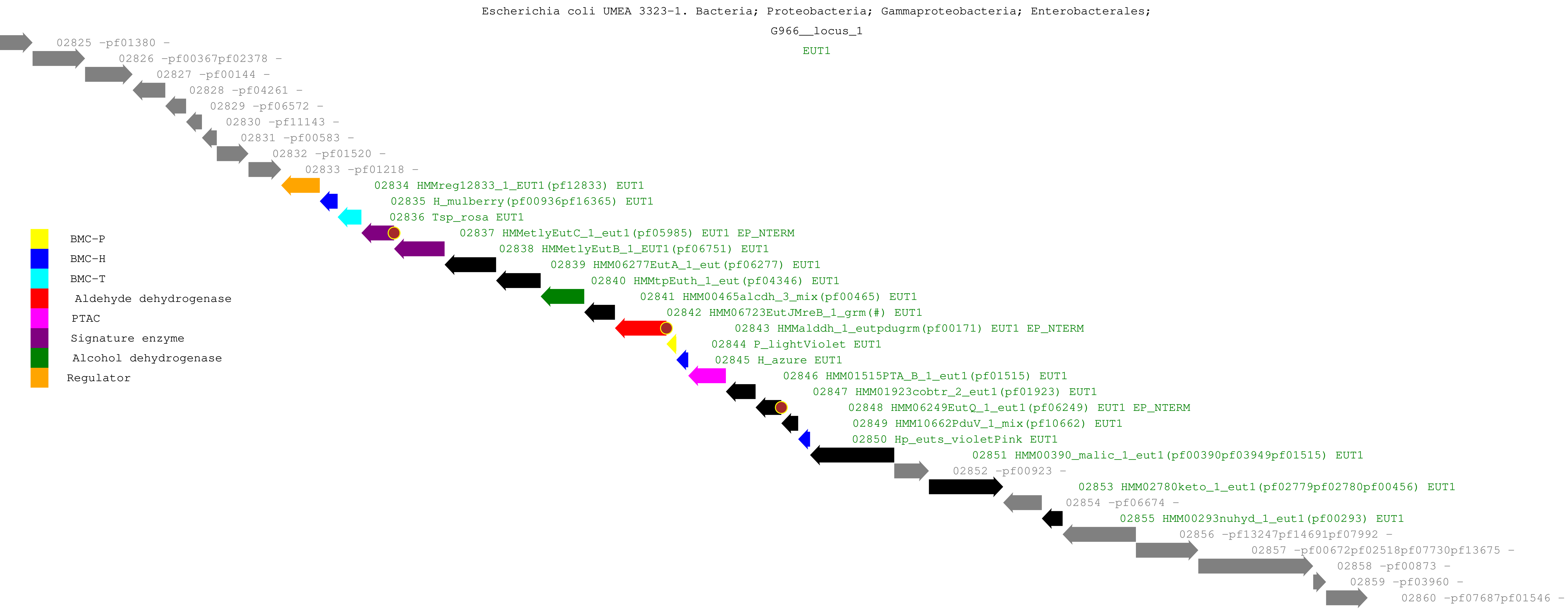
Escherichia coli UMEA 3323-1. Bacteria; Proteobacteria; Gammaproteobacteria; Enterobacterales;
G966__locus_1
EUT1
02825 -pf01380 -
02826 -pf00367pf02378 -
02827 -pf00144 -
02828 -pf04261 -
02829 -pf06572 -
02830 -pf11143 -
02831 -pf00583 -
02832 -pf01520 -
02833 -pf01218 -
02834 HMMreg12833_1_EUT1(pf12833) EUT1
02835 H_mulberry(pf00936pf16365) EUT1
02836 Tsp_rosa EUT1
02837 HMMetlyEutC_1_eut1(pf05985) EUT1 EP_NTERM
02838 HMMetlyEutB_1_EUT1(pf06751) EUT1
02839 HMM06277EutA_1_eut(pf06277) EUT1
02840 HMMtpEuth_1_eut(pf04346) EUT1
02841 HMM00465alcdh_3_mix(pf00465) EUT1
02842 HMM06723EutJMreB_1_grm(#) EUT1
02843 HMMaldh_1_eutpdugrm(pf00171) EUT1 EP_NTERM
02844 P_lightViolet EUT1
02845 H_azure EUT1
02846 HMM01515PTA_B_1_eut1(pf01515) EUT1
02847 HMM01923cobtr_2_eut1(pf01923) EUT1
02848 HMM06249EutQ_1_eut1(pf06249) EUT1 EP_NTERM
02849 HMM10662PduV_1_mix(pf10662) EUT1
02850 Hp_euts_violetPink EUT1
02851 HMM00390_malic_1_eut1(pf00390pf03949pf01515) EUT1
02852 -pf00923 -
02853 HMM02780keto_1_eut1(pf02779pf02780pf00456) EUT1
02854 -pf06674 -
02855 HMM00293nuhyd_1_eut1(pf00293) EUT1
02856 -pf13247pf14691pf07992 -
02857 -pf00672pf02518pf07730pf13675 -
02858 -pf00873 -
02859 -pf03960 -
02860 -pf07687pf01546 -
BMC-P
BMC-H
BMC-T
Aldehyde dehydrogenase
PTAC
Signature enzyme
Alcohol dehydrogenase
Regulator

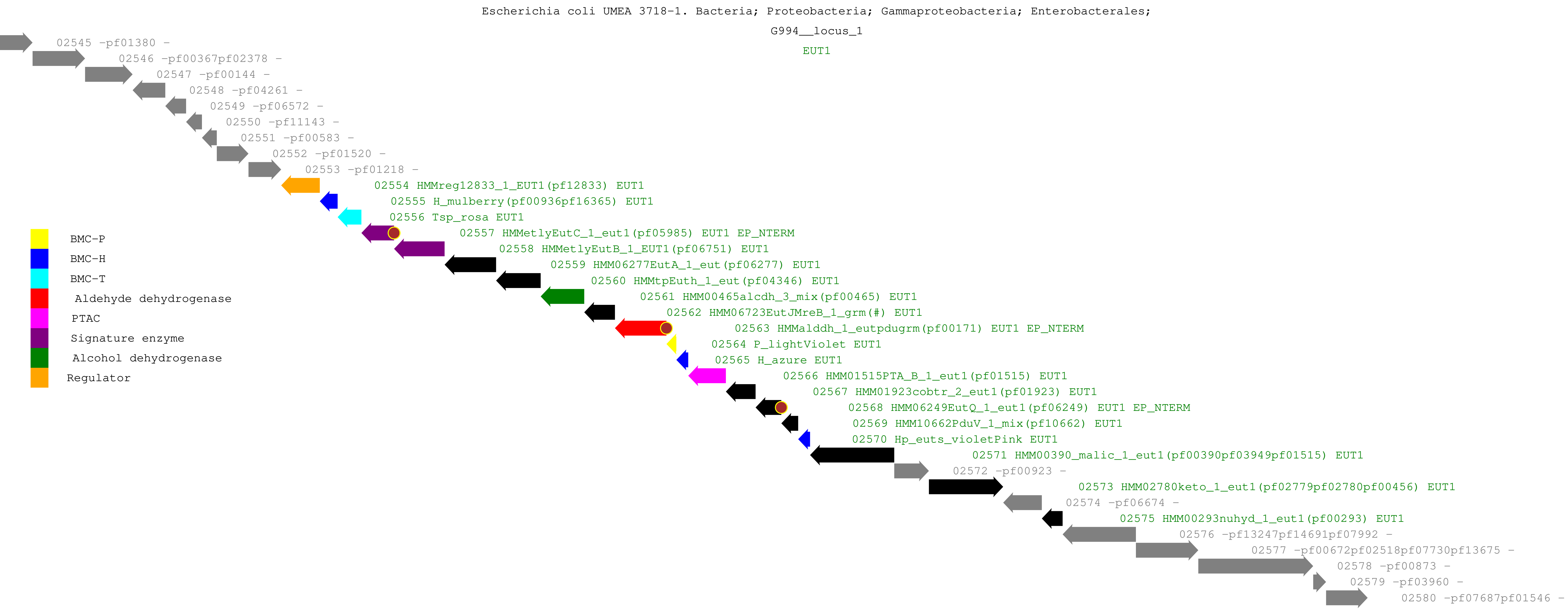
Escherichia coli UMEA 3718-1. Bacteria; Proteobacteria; Gammaproteobacteria; Enterobacterales;
G994__locus_1
EUT1
02545 -pf01380 -
02546 -pf00367pf02378 -
02547 -pf00144 -
02548 -pf04261 -
02549 -pf06572 -
02550 -pf11143 -
02551 -pf00583 -
02552 -pf01520 -
02553 -pf01218 -
02554 HMMreg12833_1_EUT1(pf12833) EUT1
02555 H_mulberry(pf00936pf16365) EUT1
02556 Tsp_rosa EUT1
02557 HMMetlyEutC_1_eut1(pf05985) EUT1 EP_NTERM
02558 HMMetlyEutB_1_EUT1(pf06751) EUT1
02559 HMM06277EutA_1_eut(pf06277) EUT1
02560 HMMtpEuth_1_eut(pf04346) EUT1
02561 HMM00465alcdh_3_mix(pf00465) EUT1
02562 HMM06723EutJMreB_1_grm(#) EUT1
02563 HMMaldh_1_eutpdugrm(pf00171) EUT1 EP_NTERM
02564 P_lightViolet EUT1
02565 H_azure EUT1
02566 HMM01515PTA_B_1_eut1(pf01515) EUT1
02567 HMM01923cobtr_2_eut1(pf01923) EUT1
02568 HMM06249EutQ_1_eut1(pf06249) EUT1 EP_NTERM
02569 HMM10662PduV_1_mix(pf10662) EUT1
02570 Hp_euts_violetPink EUT1
02571 HMM00390_malic_1_eut1(pf00390pf03949pf01515) EUT1
02572 -pf00923 -
02573 HMM02780keto_1_eut1(pf02779pf02780pf00456) EUT1
02574 -pf06674 -
02575 HMM00293nuhyd_1_eut1(pf00293) EUT1
02576 -pf13247pf14691pf07992 -
02577 -pf00672pf02518pf07730pf13675 -
02578 -pf00873 -
02579 -pf03960 -
02580 -pf07687pf01546 -
BMC-P
BMC-H
BMC-T
Aldehyde dehydrogenase
PTAC
Signature enzyme
Alcohol dehydrogenase
Regulator

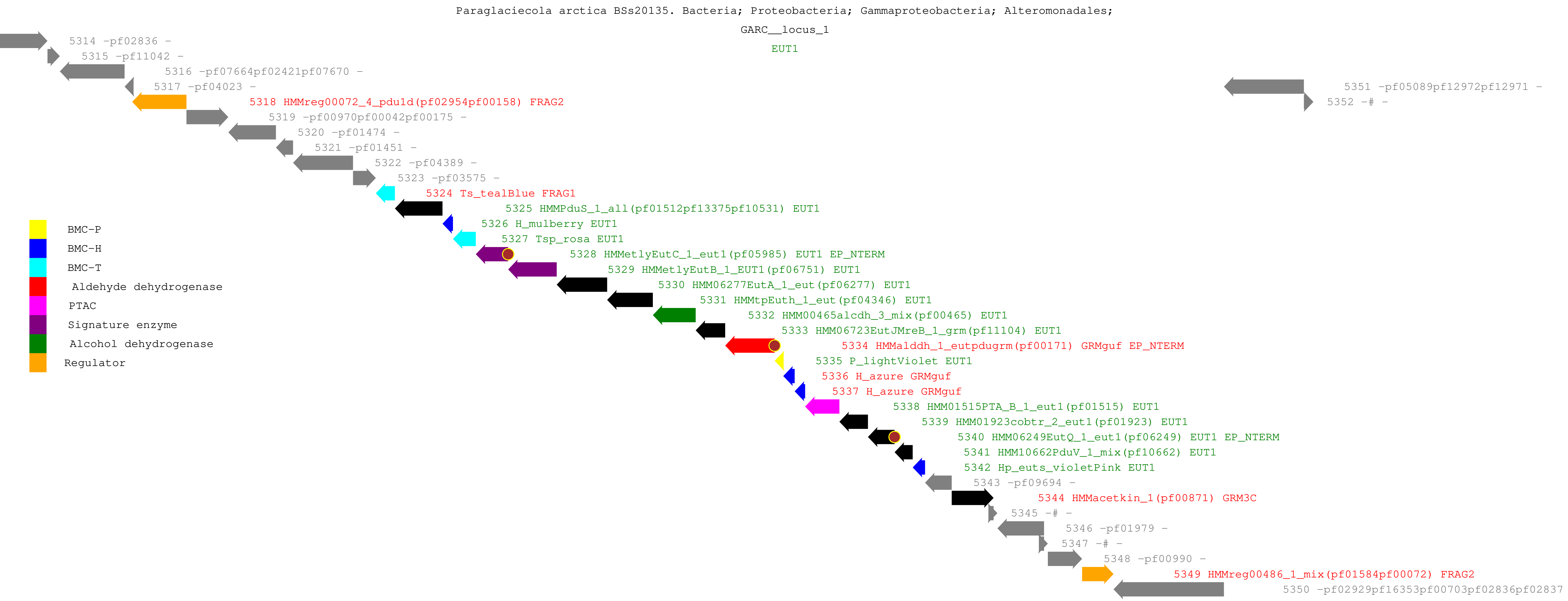
Paraglaciecola arctica BSs20135. Bacteria; Proteobacteria; Gammaproteobacteria; Alteromonadales;
GARC__locus_1
EUT1
5314 -pf02836 -
5315 -pf11042 -
5316 -pf07664pf02421pf07670 -
5317 -pf04023 -
5318 HMMreg00072_4_pdu1d(pf02954pf00158) FRAG2
5319 -pf00970pf00042pf00175 -
5320 -pf01474 -
5321 -pf01451 -
5322 -pf04389 -
5323 -pf03575 -
5324 Ts_tealBlue FRAG1
5325 HMMPduS_1_all(pf01512pf13375pf10531) EUT1
5326 H_mulberry EUT1
5327 Tsp_rosa EUT1
5328 HMMetlyEutC_1_eut1(pf05985) EUT1 EP_NTERM
5329 HMMetlyEutB_1_EUT1(pf06751) EUT1
5330 HMM06277EutA_1_eut(pf06277) EUT1
5331 HMMtpEuth_1_eut(pf04346) EUT1
5332 HMM00465alcdh_3_mix(pf00465) EUT1
5333 HMM06723EutJMreB_1_grm(pf11104) EUT1
5334 HMMalddh_1_eutpdugrm(pf00171) GRMguf EP_NTERM
5335 P_lightViolet EUT1
5336 H_azure GRMguf
5337 H_azure GRMguf
5338 HMM01515PTA_B_1_eut1(pf01515) EUT1
5339 HMM01923cobtr_2_eut1(pf01923) EUT1
5340 HMM06249EutQ_1_eut1(pf06249) EUT1 EP_NTERM
5341 HMM10662PduV_1_mix(pf10662) EUT1
5342 Hp_euts_violetPink EUT1
5343 -pf09694 -
5344 HMMacetkin_1(pf00871) GRM3C
5345 -# -
5346 -pf01979 -
5347 -# -
5348 -pf00990 -
5349 HMMreg00486_1_mix(pf01584pf00072) FRAG2
5350 -pf02929pf16353pf00703pf02836pf02837
5351 -pf05089pf12972pf12971 -
5352 -# -
BMC-P
BMC-H
BMC-T
Aldehyde dehydrogenase
PTAC
Signature enzyme
Alcohol dehydrogenase
Regulator

Hafnia alvei ATCC 13337. Bacteria; Proteobacteria; Gammaproteobacteria; Enterobacterales;

GHAL__locus_1_main_sat

EUT1

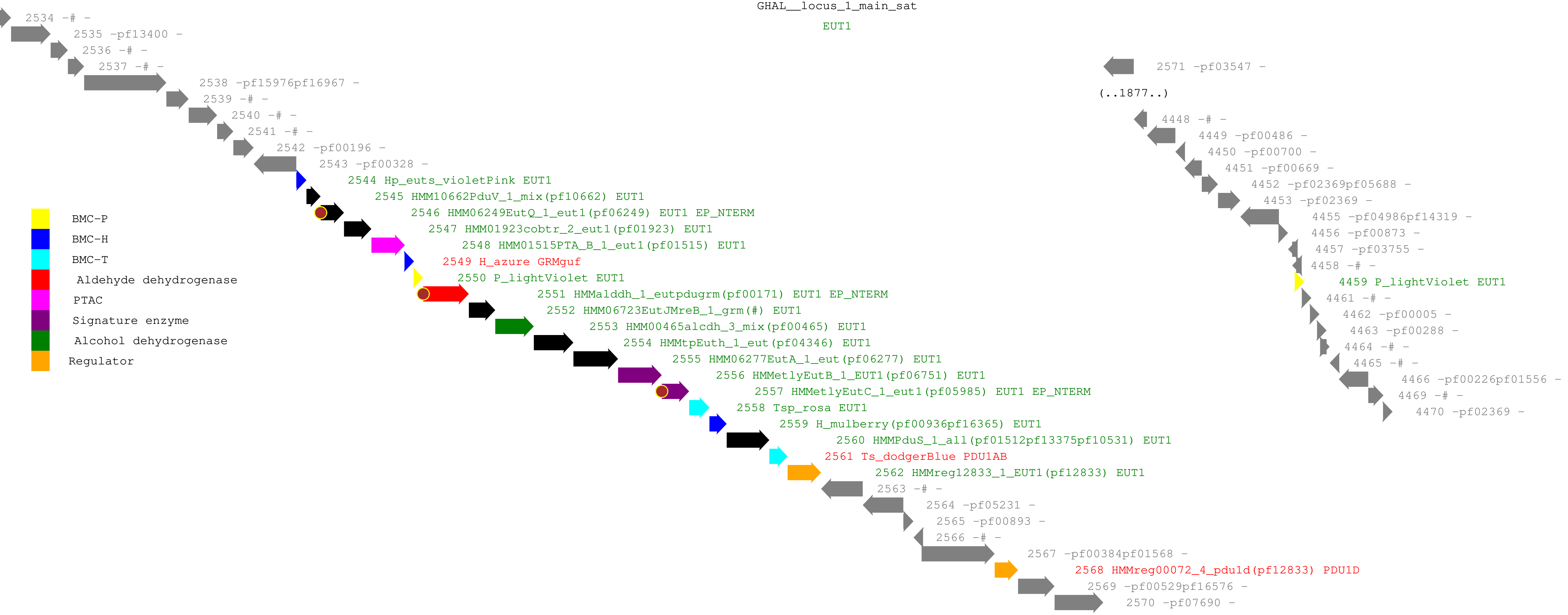

Salmonella enterica I. Bacteria; Proteobacteria; Gammaproteobacteria; Enterobacterales;

GX90__locus_1

EUT1

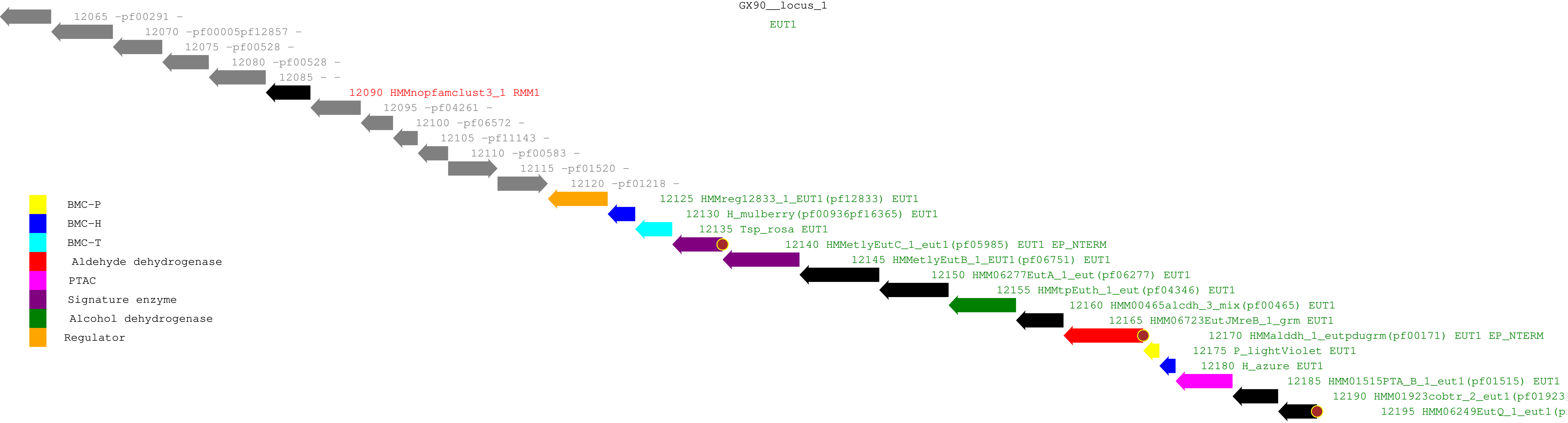

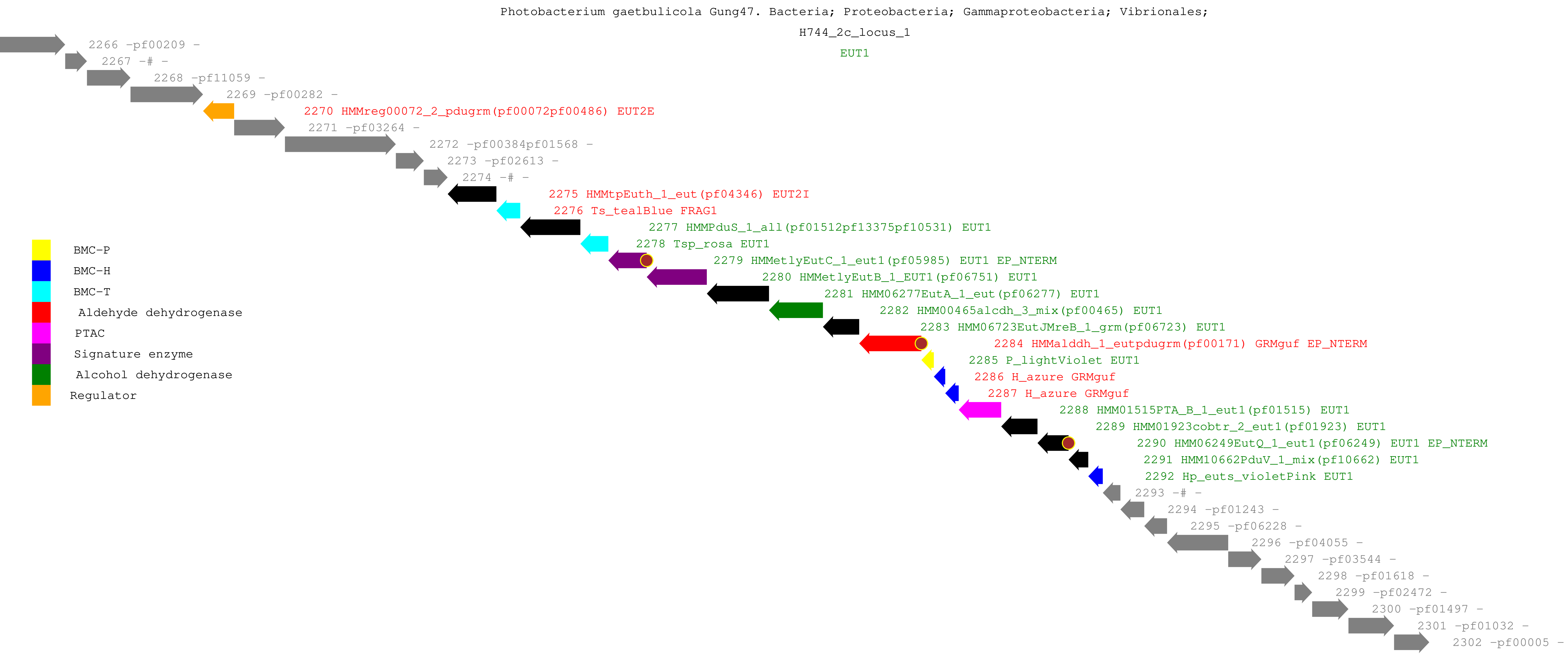
Photobacterium gaetbulicola Gung47. Bacteria; Proteobacteria; Gammaproteobacteria; Vibrionales;
H744_2c_locus_1
EUT1
2266 -pf00209 -
2267 -# -
2268 -pf11059 -
2269 -pf00282 -
2270 HMMreg00072_2_pdugrm(pf00072pf00486) EUT2E
2271 -pf03264 -
2272 -pf00384pf01568 -
2273 -pf02613 -
2274 -# -
2275 HMMtpEuth_1_eut(pf04346) EUT2I
2276 Ts_tealBlue FRAG1
2277 HMMPduS_1_all(pf01512pf13375pf10531) EUT1
2278 Tsp_rosa EUT1
2279 HMMetlyEutC_1_eut1(pf05985) EUT1 EP_NTERM
2280 HMMetlyEutB_1_EUT1(pf06751) EUT1
2281 HMM06277EutA_1_eut(pf06277) EUT1
2282 HMM00465alcdh_3_mix(pf00465) EUT1
2283 HMM06723EutJMreB_1_grm(pf06723) EUT1
2284 HMMalddh_1_eutpdugrm(pf00171) GRMguf EP_NTERM
2285 P_lightViolet EUT1
2286 H_azure GRMguf
2287 H_azure GRMguf
2288 HMM01515PTA_B_1_eut1(pf01515) EUT1
2289 HMM01923cobtr_2_eut1(pf01923) EUT1
2290 HMM06249EutQ_1_eut1(pf06249) EUT1 EP_NTERM
2291 HMM10662PduV_1_mix(pf10662) EUT1
2292 Hp_euts_violetPink EUT1
2293 -# -
2294 -pf01243 -
2295 -pf06228 -
2296 -pf04055 -
2297 -pf03544 -
2298 -pf01618 -
2299 -pf02472 -
2300 -pf01497 -
2301 -pf01032 -
2302 -pf00005 -
BMC-P
BMC-H
BMC-T
Aldehyde dehydrogenase
PTAC
Signature enzyme
Alcohol dehydrogenase
Regulator

Escherichia coli. Bacteria; Proteobacteria; Gammaproteobacteria; Enterobacterales;

HmCms184__locus_1

EUT1

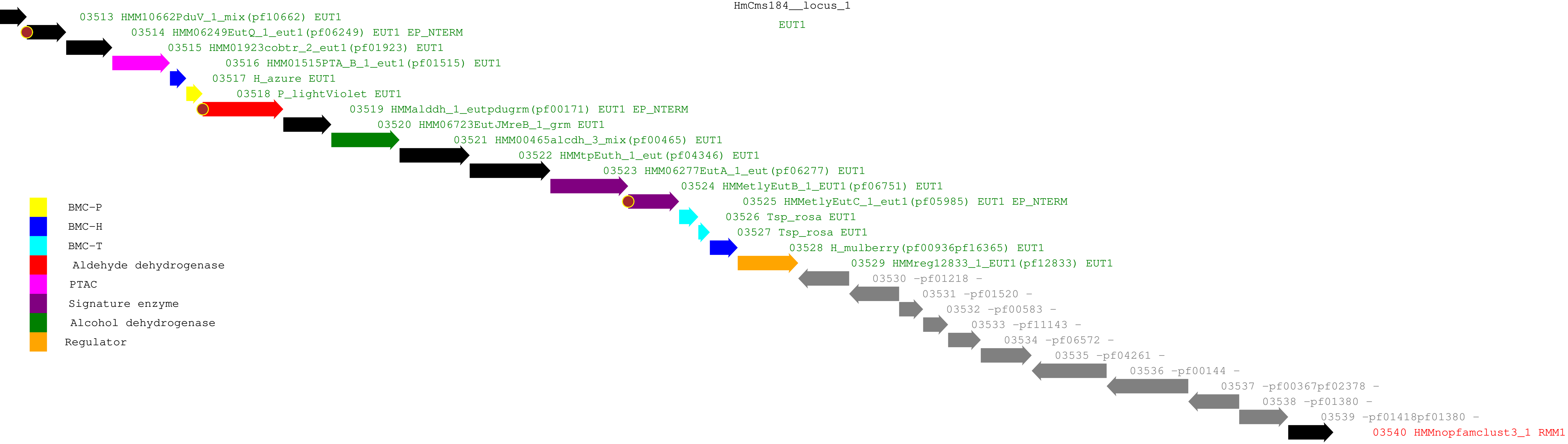

Klebsiella pneumoniae subsp. rhinoscleromatis ATCC 13884. Bacteria; Proteobacteria; Gammaproteobacteria; Enterobacterales;

HMPREF0484__locus_2

EUT1

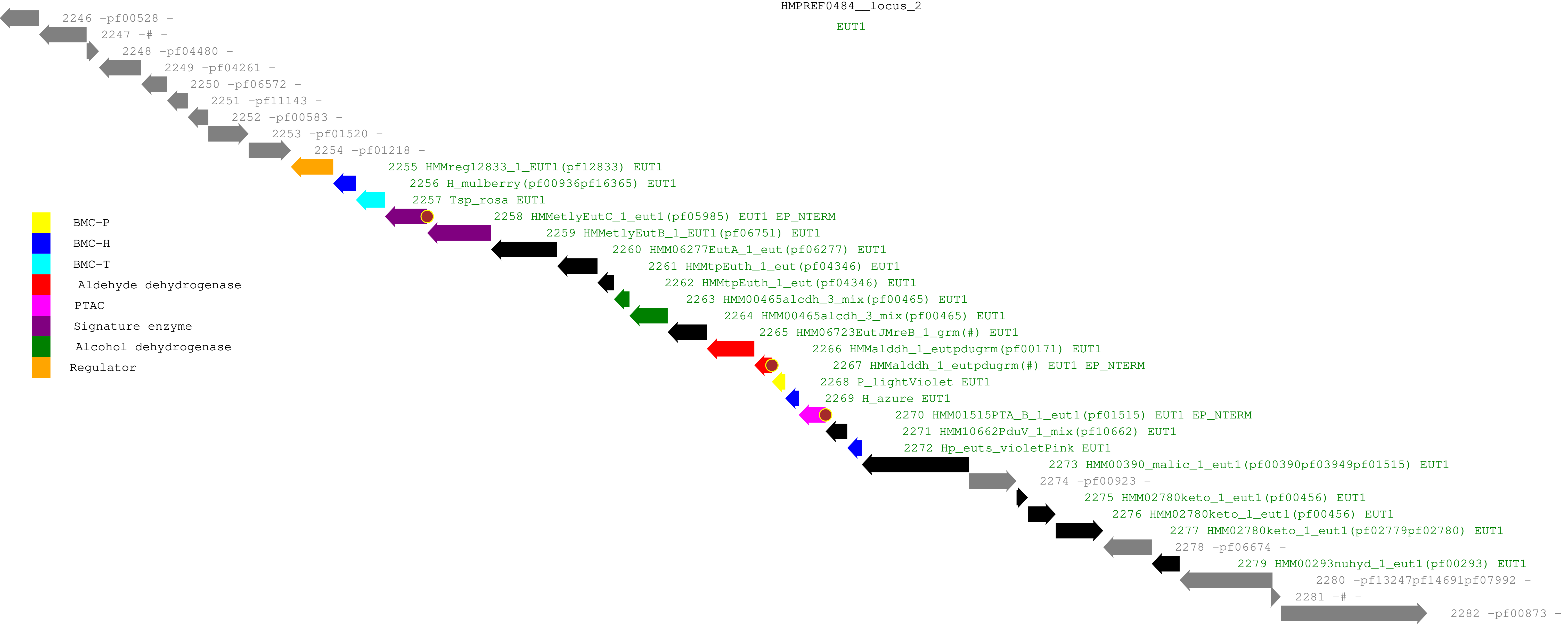

Escherichia coli 4_1_47FAA. Bacteria; Proteobacteria; Gammaproteobacteria; Enterobacterales;

HMPREF0986__locus_1

EUT1

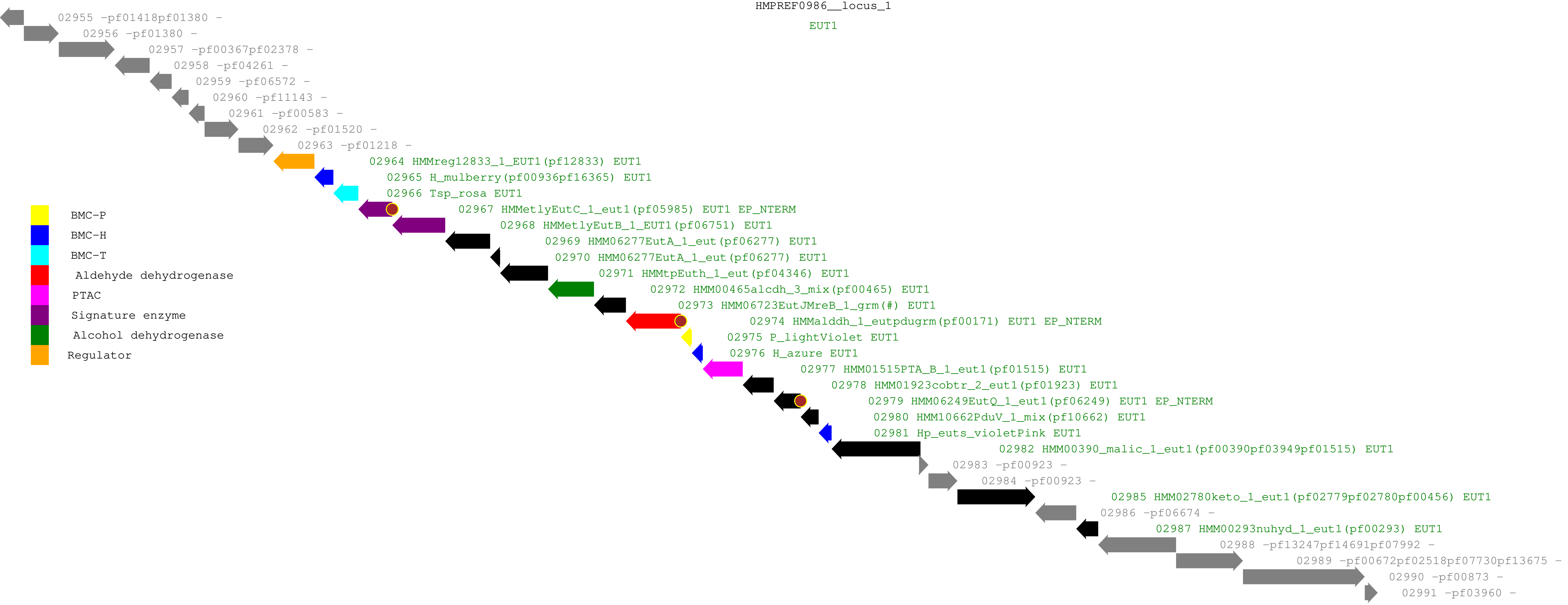

Klebsiella sp. OBRC7. Bacteria; Proteobacteria; Gammaproteobacteria; Enterobacterales;

HMPREF1144__locus_3

EUT1

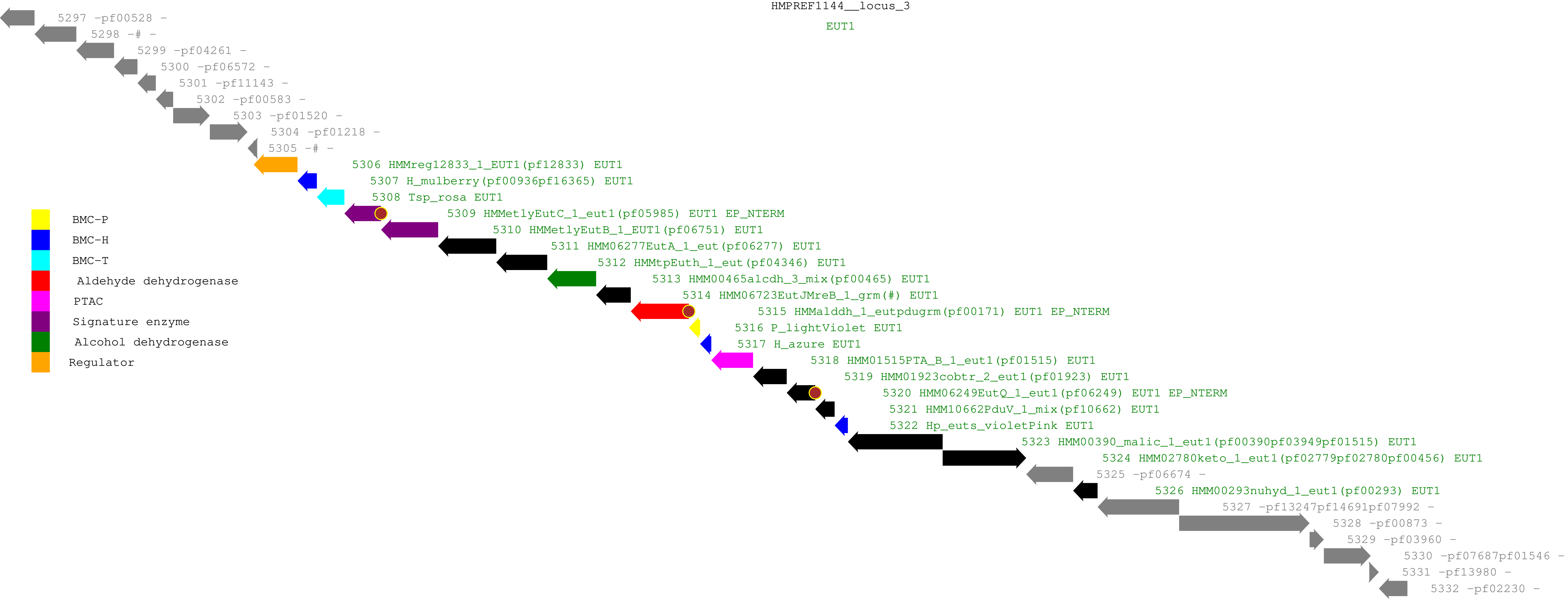

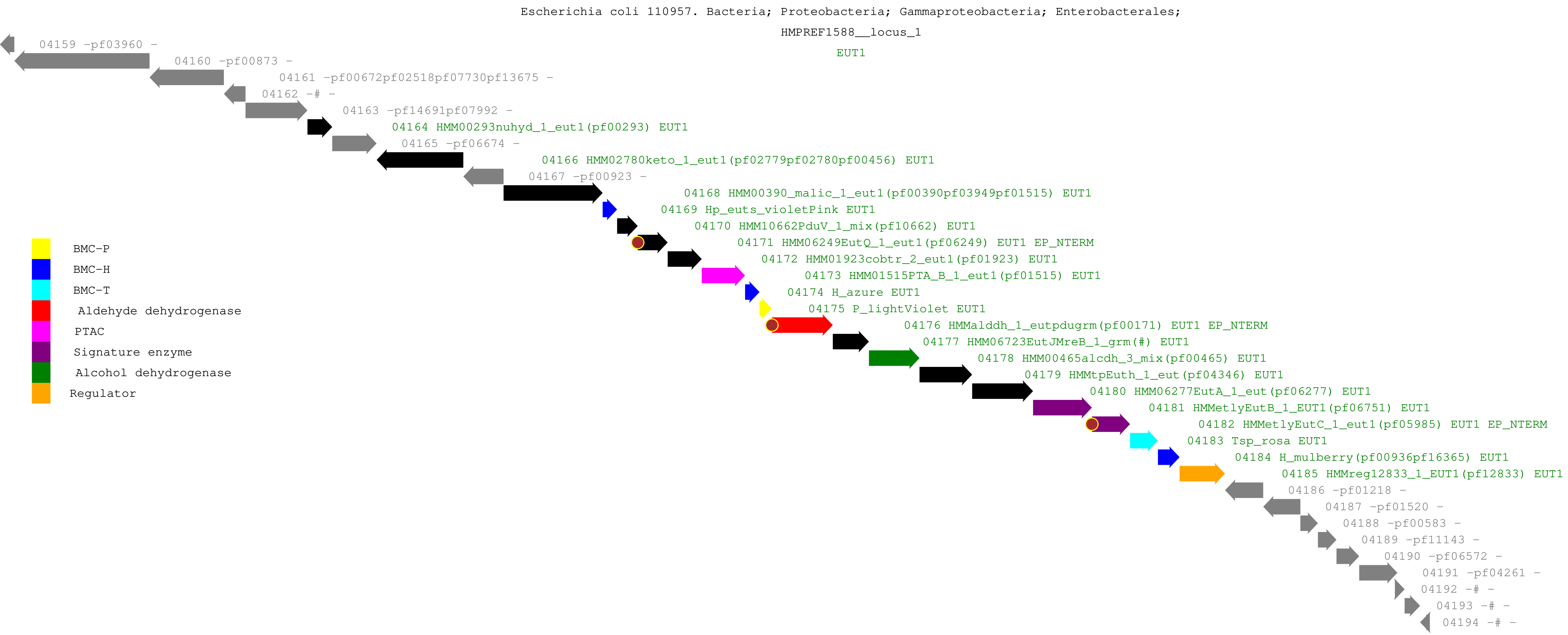
Escherichia coli 110957. Bacteria; Proteobacteria; Gammaproteobacteria; Enterobacterales;
HMPREF1588__locus_1
EUT1
04159 -pf03960 -
04160 -pf00873 -
04161 -pf00672pf02518pf07730pf13675 -
04162 -# -
04163 -pf14691pf07992 -
04164 HMM00293nuhyd_1_eut1(pf00293) EUT1
04165 -pf06674 -
04166 HMM02780keto_1_eut1(pf02779pf02780pf00456) EUT1
04167 -pf00923 -
04168 HMM00390_malic_1_eut1(pf00390pf03949pf01515) EUT1
04169 Hp_euts_violetPink EUT1
04170 HMM10662PduV_1_mix(pf10662) EUT1
04171 HMM06249EutQ_1_eut1(pf06249) EUT1 EP_NTERM
04172 HMM01923cobtr_2_eut1(pf01923) EUT1
04173 HMM01515PTA_B_1_eut1(pf01515) EUT1
04174 H_azure EUT1
04175 P_lightViolet EUT1
04176 HMMalddh_1_eutpdugrm(pf00171) EUT1 EP_NTERM
04177 HMM06723EutJMreB_1_grm(#) EUT1
04178 HMM00465alcdh_3_mix(pf00465) EUT1
04179 HMMtpEuth_1_eut(pf04346) EUT1
04180 HMM06277EutA_1_eut(pf06277) EUT1
04181 HMMetlyEutB_1_EUT1(pf06751) EUT1
04182 HMMetlyEutC_1_eut1(pf05985) EUT1 EP_NTERM
04183 Tsp_rosa EUT1
04184 H_mulberry(pf00936pf16365) EUT1
04185 HMMreg12833_1_EUT1(pf12833) EUT1
04186 -pf01218 -
04187 -pf01520 -
04188 -pf00583 -
04189 -pf11143 -
04190 -pf06572 -
04191 -pf04261 -
04192 -# -
04193 -# -
04194 -# -
BMC-P
BMC-H
BMC-T
Aldehyde dehydrogenase
PTAC
Signature enzyme
Alcohol dehydrogenase
Regulator

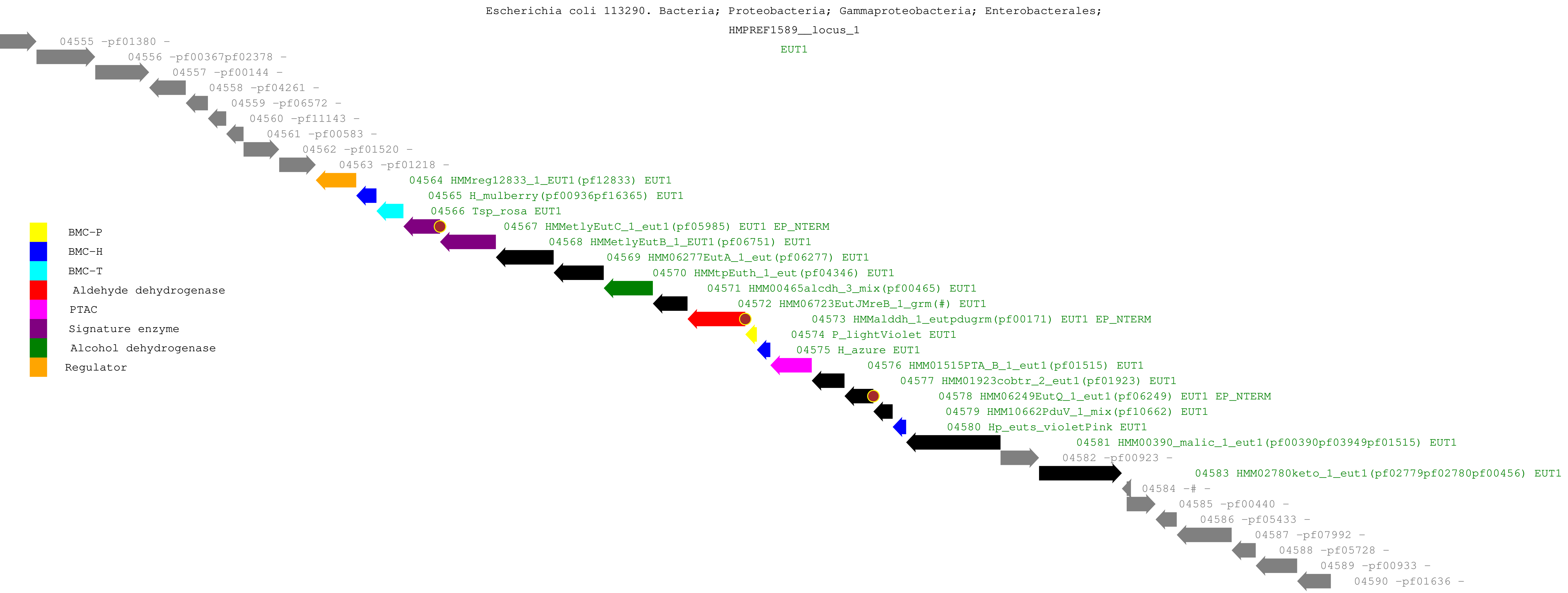
Escherichia coli 113290. Bacteria; Proteobacteria; Gammaproteobacteria; Enterobacterales;
HMPREF1589__locus_1
EUT1
04555 -pf01380 -
04556 -pf00367pf02378 -
04557 -pf00144 -
04558 -pf04261 -
04559 -pf06572 -
04560 -pf11143 -
04561 -pf00583 -
04562 -pf01520 -
04563 -pf01218 -
04564 HMMreg12833_1_EUT1(pf12833) EUT1
04565 H_mulberry(pf00936pf16365) EUT1
04566 Tsp_rosa EUT1
04567 HMMetlyEutC_1_eut1(pf05985) EUT1 EP_NTERM
04568 HMMetlyEutB_1_EUT1(pf06751) EUT1
04569 HMM06277EutA_1_eut(pf06277) EUT1
04570 HMMtpEuth_1_eut(pf04346) EUT1
04571 HMM00465alcdh_3_mix(pf00465) EUT1
04572 HMM06723EutJMreB_1_grm(#) EUT1
04573 HMMalddh_1_eutpdugrm(pf00171) EUT1 EP_NTERM
04574 P_lightViolet EUT1
04575 H_azure EUT1
04576 HMM01515PTA_B_1_eut1(pf01515) EUT1
04577 HMM01923cobtr_2_eut1(pf01923) EUT1
04578 HMM06249EutQ_1_eut1(pf06249) EUT1 EP_NTERM
04579 HMM10662PduV_1_mix(pf10662) EUT1
04580 Hp_euts_violetPink EUT1
04581 HMM00390_malic_1_eut1(pf00390pf03949pf01515) EUT1
04582 -pf00923 -
04583 HMM02780keto_1_eut1(pf02779pf02780pf00456) EUT1
04584 -# -
04585 -pf00440 -
04586 -pf05433 -
04587 -pf07992 -
04588 -pf05728 -
04589 -pf00933 -
04590 -pf01636 -
BMC-P
BMC-H
BMC-T
Aldehyde dehydrogenase
PTAC
Signature enzyme
Alcohol dehydrogenase
Regulator

Escherichia coli 113302. Bacteria; Proteobacteria; Gammaproteobacteria; Enterobacterales;

HMPREF1590__locus_1

EUT1

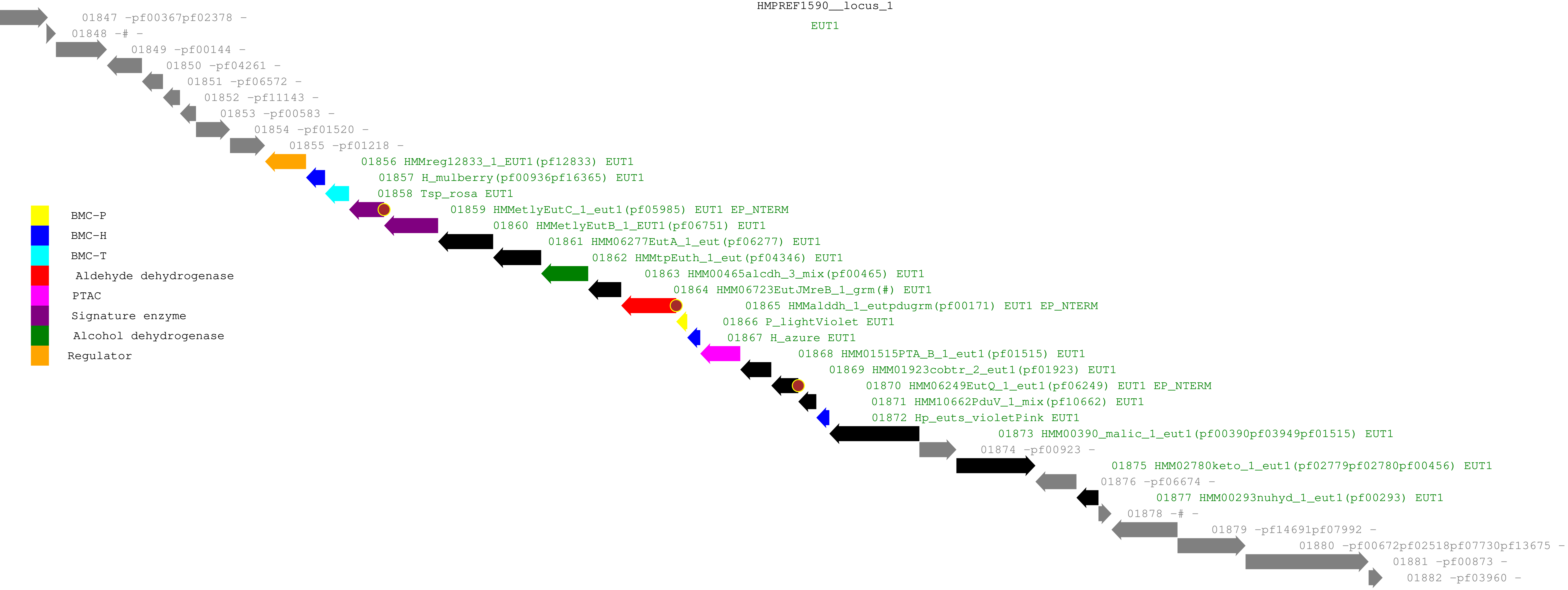

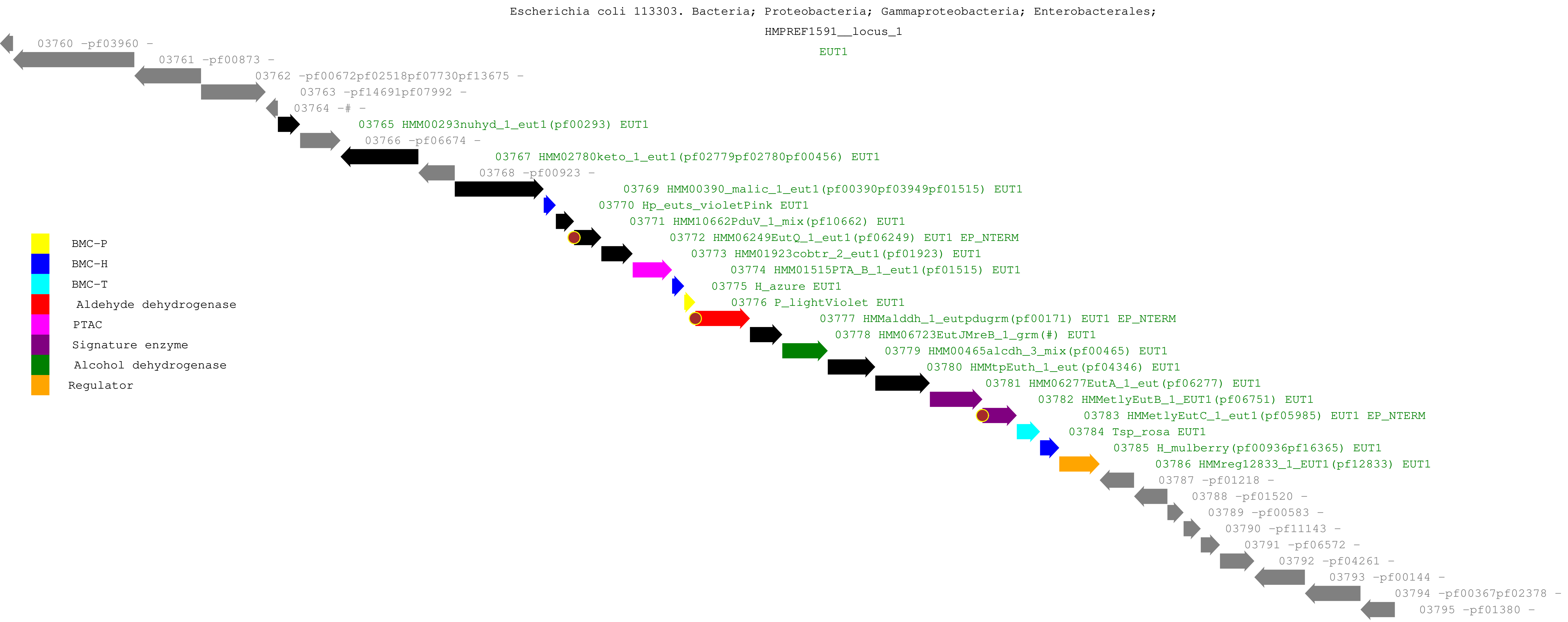
Escherichia coli 113303. Bacteria; Proteobacteria; Gammaproteobacteria; Enterobacterales;
HMPREF1591__locus_1
EUT1
03760 -pf03960 -
03761 -pf00873 -
03762 -pf00672pf02518pf07730pf13675 -
03763 -pf14691pf07992 -
03764 -# -
03765 HMM00293nuhyd_1_eut1(pf00293) EUT1
03766 -pf06674 -
03767 HMM02780keto_1_eut1(pf02779pf02780pf00456) EUT1
03768 -pf00923 -
03769 HMM00390_malic_1_eut1(pf00390pf03949pf01515) EUT1
03770 Hp_euts_violetPink EUT1
03771 HMM10662PduV_1_mix(pf10662) EUT1
03772 HMM06249EutQ_1_eut1(pf06249) EUT1 EP_NTERM
03773 HMM01923cobtr_2_eut1(pf01923) EUT1
03774 HMM01515PTA_B_1_eut1(pf01515) EUT1
03775 H_azure EUT1
03776 P_lightViolet EUT1
03777 HMMalddh_1_eutpdugrm(pf00171) EUT1 EP_NTERM
03778 HMM06723EutJMreB_1_grm(#) EUT1
03779 HMM00465alcdh_3_mix(pf00465) EUT1
03780 HMMtpEuth_1_eut(pf04346) EUT1
03781 HMM06277EutA_1_eut(pf06277) EUT1
03782 HMMetlyEutB_1_EUT1(pf06751) EUT1
03783 HMMetlyEutC_1_eut1(pf05985) EUT1 EP_NTERM
03784 Tsp_rosa EUT1
03785 H_mulberry(pf00936pf16365) EUT1
03786 HMMreg12833_1_EUT1(pf12833) EUT1
03787 -pf01218 -
03788 -pf01520 -
03789 -pf00583 -
03790 -pf11143 -
03791 -pf06572 -
03792 -pf04261 -
03793 -pf00144 -
03794 -pf00367pf02378 -
03795 -pf01380 -
BMC-P
BMC-H
BMC-T
Aldehyde dehydrogenase
PTAC
Signature enzyme
Alcohol dehydrogenase
Regulator

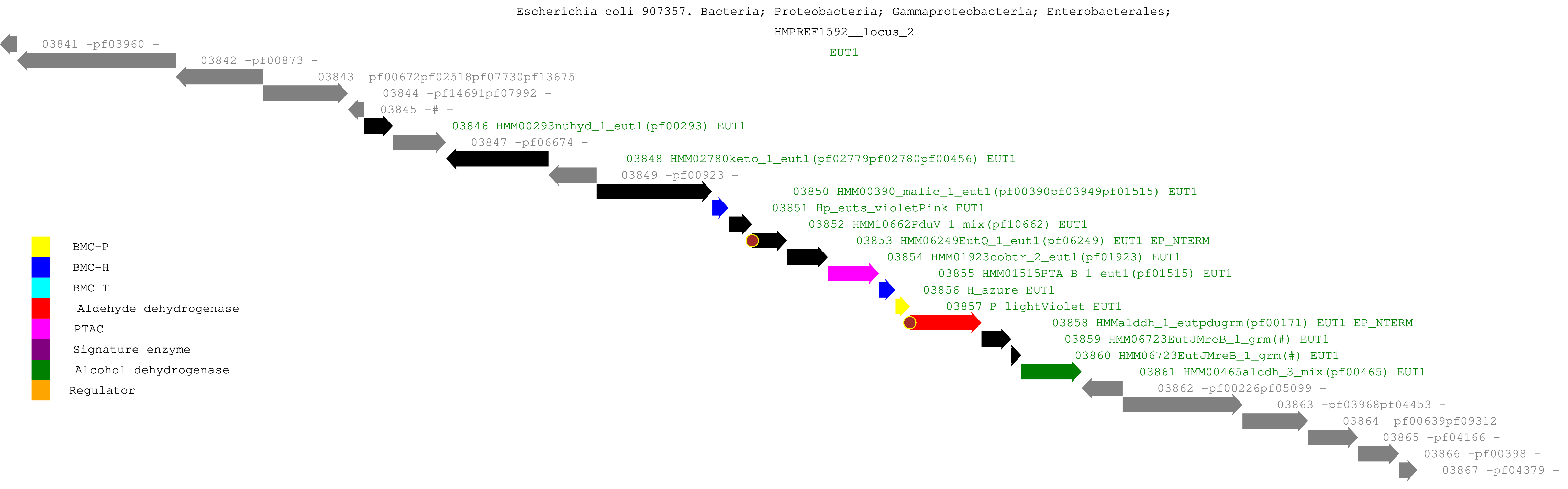

Escherichia coli 907357. Bacteria; Proteobacteria; Gammaproteobacteria; Enterobacterales;
HMPREF1592__locus_2
EUT1
03841 -pf03960 -
03842 -pf00873 -
03843 -pf00672pf02518pf07730pf13675 -
03844 -pf14691pf07992 -
03845 -# -
03846 HMM00293nuhyd_1_eut1(pf00293) EUT1
03847 -pf06674 -
03848 HMM02780keto_1_eut1(pf02779pf02780pf00456) EUT1
03849 -pf00923 -
03850 HMM00390_malic_1_eut1(pf00390pf03949pf01515) EUT1
03851 Hp_euts_violetPink EUT1
03852 HMM10662PduV_1_mix(pf10662) EUT1
03853 HMM06249EutQ_1_eut1(pf06249) EUT1 EP_NTERM
03854 HMM01923cobtr_2_eut1(pf01923) EUT1
03855 HMM01515PTA_B_1_eut1(pf01515) EUT1
03856 H_azure EUT1
03857 P_lightViolet EUT1
03858 HMMalddh_1_eutpdugrm(pf00171) EUT1 EP_NTERM
03859 HMM06723EutJMreB_1_grm(#) EUT1
03860 HMM06723EutJMreB_1_grm(#) EUT1
03861 HMM00465alcdh_3_mix(pf00465) EUT1
03862 -pf00226pf05099 -
03863 -pf03968pf04453 -
03864 -pf00639pf09312 -
03865 -pf04166 -
03866 -pf00398 -
03867 -pf04379 -
BMC-P
BMC-H
BMC-T
Aldehyde dehydrogenase
PTAC
Signature enzyme
Alcohol dehydrogenase
Regulator

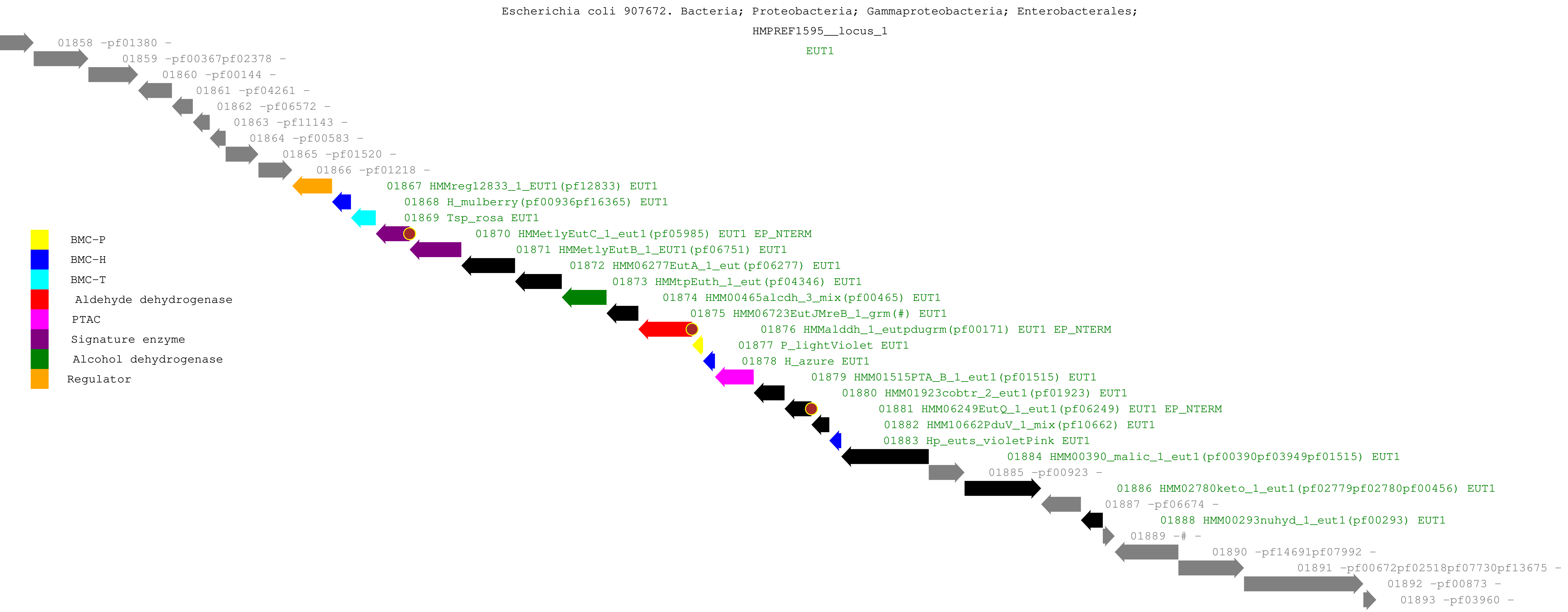
Escherichia coli 907672. Bacteria; Proteobacteria; Gammaproteobacteria; Enterobacterales;
HMPREF1595__locus_1
EUT1
01858 -pf01380 -
01859 -pf00367pf02378 -
01860 -pf00144 -
01861 -pf04261 -
01862 -pf06572 -
01863 -pf11143 -
01864 -pf00583 -
01865 -pf01520 -
01866 -pf01218 -
01867 HMMreg12833_1_EUT1(pf12833) EUT1
01868 H_mulberry(pf00936pf16365) EUT1
01869 Tsp_rosa EUT1
01870 HMMetlyEutC_1_eut1(pf05985) EUT1 EP_NTERM
01871 HMMetlyEutB_1_EUT1(pf06751) EUT1
01872 HMM06277EutA_1_eut(pf06277) EUT1
01873 HMMtpEuth_1_eut(pf04346) EUT1
01874 HMM00465alcdh_3_mix(pf00465) EUT1
01875 HMM06723EutJMreB_1_grm(#) EUT1
01876 HMMalddh_1_eutpdugrm(pf00171) EUT1 EP_NTERM
01877 P_lightViolet EUT1
01878 H_azure EUT1
01879 HMM01515PTA_B_1_eut1(pf01515) EUT1
01880 HMM01923cobtr_2_eut1(pf01923) EUT1
01881 HMM06249EutQ_1_eut1(pf06249) EUT1 EP_NTERM
01882 HMM10662PduV_1_mix(pf10662) EUT1
01883 Hp_euts_violetPink EUT1
01884 HMM00390_malic_1_eut1(pf00390pf03949pf01515) EUT1
01885 -pf00923 -
01886 HMM02780keto_1_eut1(pf02779pf02780pf00456) EUT1
01887 -pf06674 -
01888 HMM00293nuhyd_1_eut1(pf00293) EUT1
01889 -# -
01890 -pf14691pf07992 -
01891 -pf00672pf02518pf07730pf13675 -
01892 -pf00873 -
01893 -pf03960 -
BMC-P
BMC-H
BMC-T
Aldehyde dehydrogenase
PTAC
Signature enzyme
Alcohol dehydrogenase
Regulator

Escherichia coli 907710. Bacteria; Proteobacteria; Gammaproteobacteria; Enterobacterales;

HMPREF1598__locus_1

EUT1

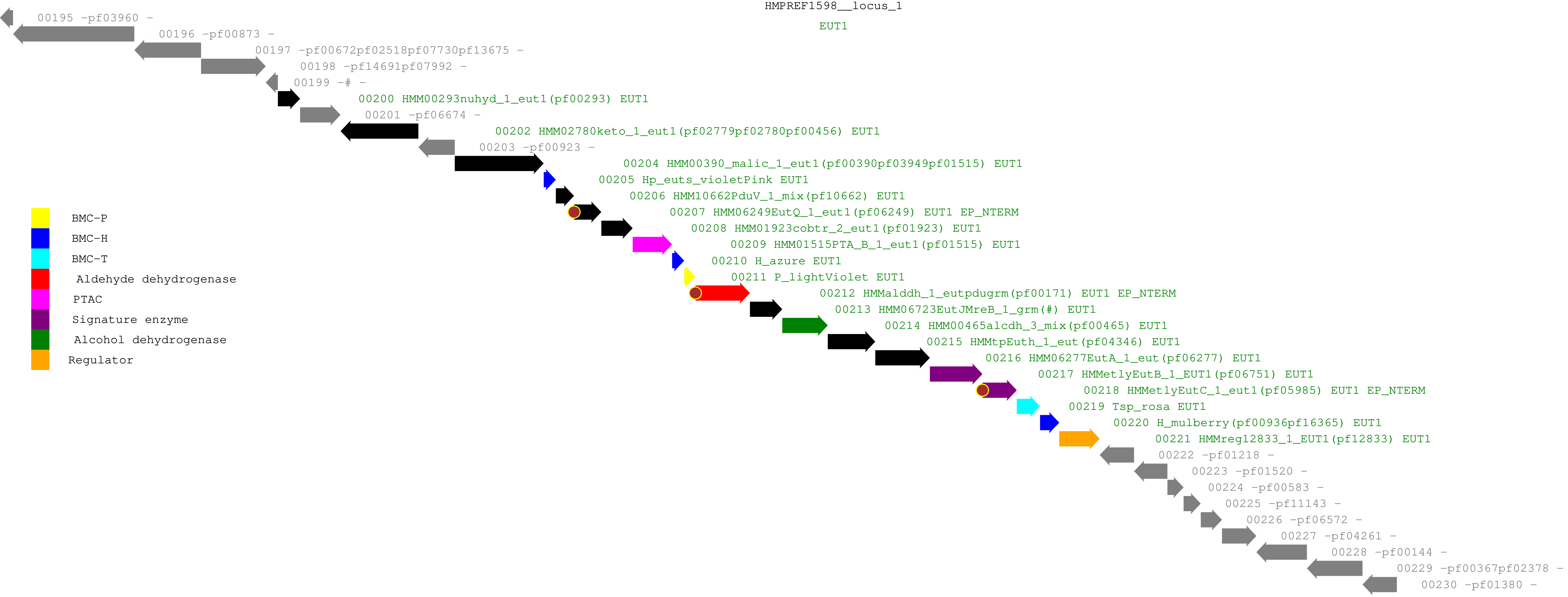

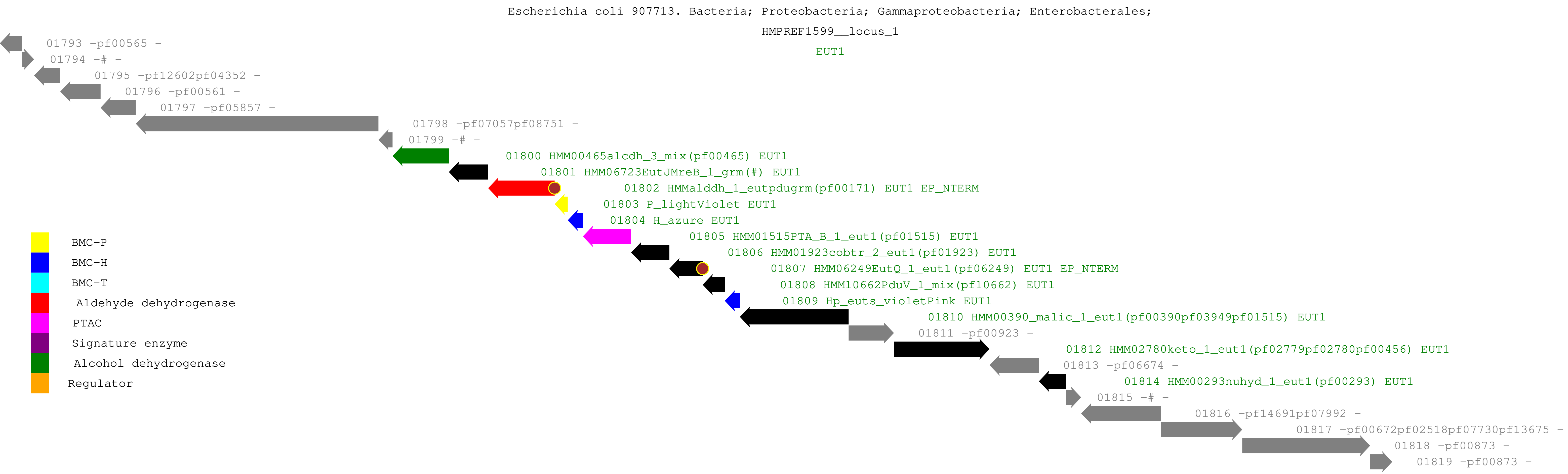
Escherichia coli 907713. Bacteria; Proteobacteria; Gammaproteobacteria; Enterobacterales;
HMPREF1599__locus_1
EUT1
01793 -pf00565 -
01794 -# -
01795 -pf12602pf04352 -
01796 -pf00561 -
01797 -pf05857 -
01798 -pf07057pf08751 -
01799 -# -
01800 HMM00465alcdh_3_mix(pf00465) EUT1
01801 HMM06723EutJMreB_1_grm(#) EUT1
01802 HMMalddh_1_eutpdugrm(pf00171) EUT1 EP_NTERM
01803 P_lightViolet EUT1
01804 H_azure EUT1
01805 HMM01515PTA_B_1_eut1(pf01515) EUT1
01806 HMM01923cobtr_2_eut1(pf01923) EUT1
01807 HMM06249EutQ_1_eut1(pf06249) EUT1 EP_NTERM
01808 HMM10662PduV_1_mix(pf10662) EUT1
01809 Hp_euts_violetPink EUT1
01810 HMM00390_malic_1_eut1(pf00390pf03949pf01515) EUT1
01811 -pf00923 -
01812 HMM02780keto_1_eut1(pf02779pf02780pf00456) EUT1
01813 -pf06674 -
01814 HMM00293nuhyd_1_eut1(pf00293) EUT1
01815 -# -
01816 -pf14691pf07992 -
01817 -pf00672pf02518pf07730pf13675 -
01818 -pf00873 -
01819 -pf00873 -
BMC-P
BMC-H
BMC-T
Aldehyde dehydrogenase
PTAC
Signature enzyme
Alcohol dehydrogenase
Regulator

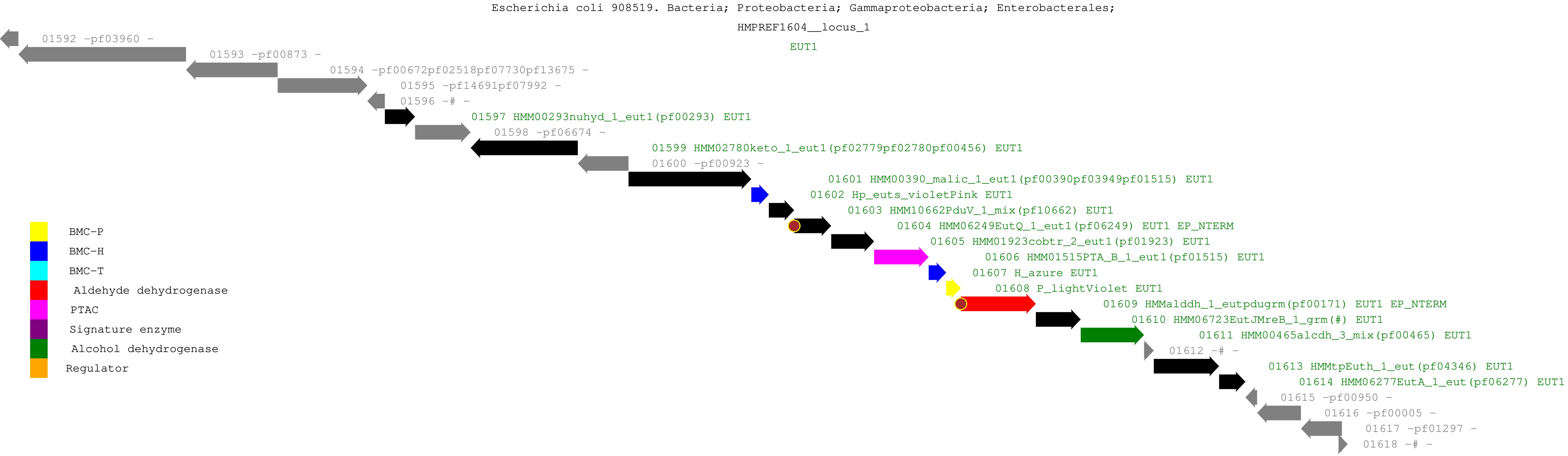

Escherichia coli 908519. Bacteria; Proteobacteria; Gammaproteobacteria; Enterobacterales;
HMPREF1604__locus_1
EUT1
01592 -pf03960 -
01593 -pf00873 -
01594 -pf00672pf02518pf07730pf13675 -
01595 -pf14691pf07992 -
01596 -# -
01597 HMM00293nuhyd_1_eut1(pf00293) EUT1
01598 -pf06674 -
01599 HMM02780keto_1_eut1(pf02779pf02780pf00456) EUT1
01600 -pf00923 -
01601 HMM00390_malic_1_eut1(pf00390pf03949pf01515) EUT1
01602 Hp_euts_violetPink EUT1
01603 HMM10662PduV_1_mix(pf10662) EUT1
01604 HMM06249EutQ_1_eut1(pf06249) EUT1 EP_NTERM
01605 HMM01923cobtr_2_eut1(pf01923) EUT1
01606 HMM01515PTA_B_1_eut1(pf01515) EUT1
01607 H_azure EUT1
01608 P_lightViolet EUT1
01609 HMMalddh_1_eutpdugrm(pf00171) EUT1 EP_NTERM
01610 HMM06723EutJMreB_1_grm(#) EUT1
01611 HMM00465alcdh_3_mix(pf00465) EUT1
01612 -# -
01613 HMMtpEuth_1_eut(pf04346) EUT1
01614 HMM06277EutA_1_eut(pf06277) EUT1
01615 -pf00950 -
01616 -pf00005 -
01617 -pf01297 -
01618 -# -
BMC-P
BMC-H
BMC-T
Aldehyde dehydrogenase
PTAC
Signature enzyme
Alcohol dehydrogenase
Regulator

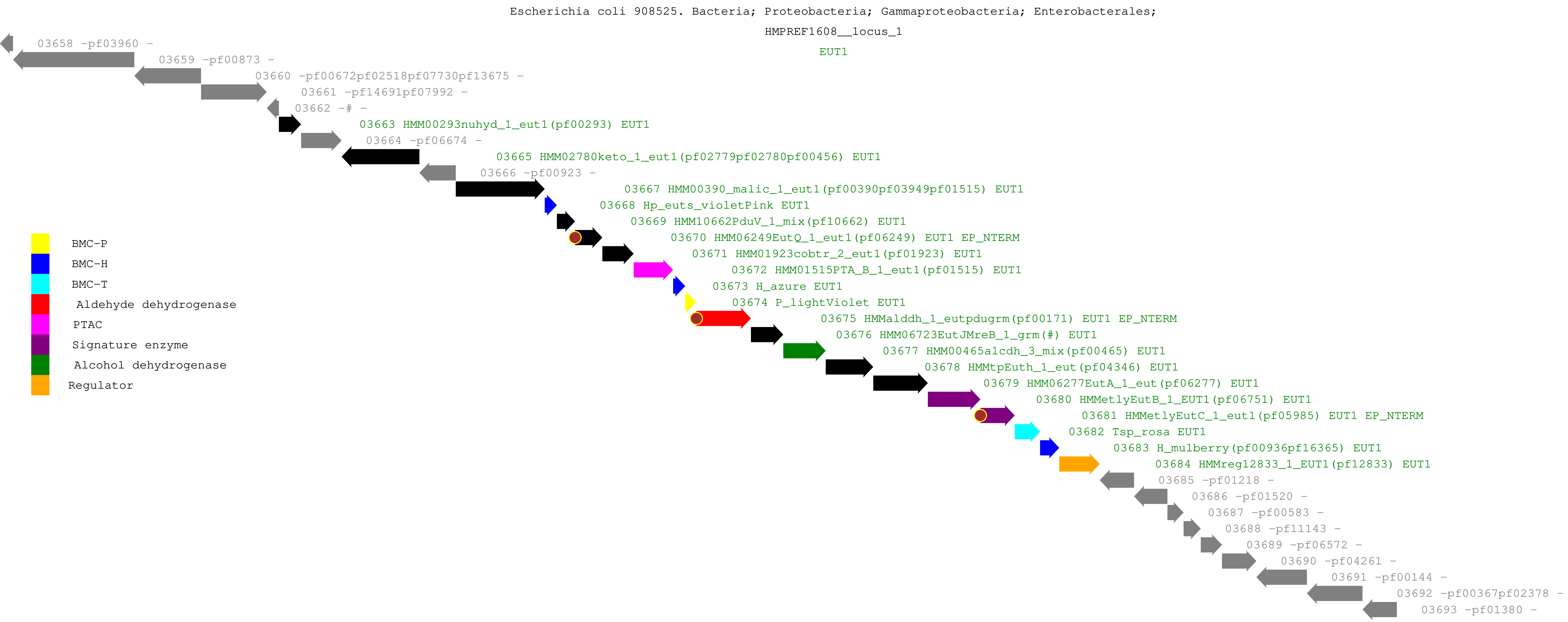
Escherichia coli 908525. Bacteria; Proteobacteria; Gammaproteobacteria; Enterobacterales;
HMPREF1608__locus_1
EUT1
03658 -pf03960 -
03659 -pf00873 -
03660 -pf00672pf02518pf07730pf13675 -
03661 -pf14691pf07992 -
03662 -# -
03663 HMM00293nuhyd_1_eut1(pf00293) EUT1
03664 -pf06674 -
03665 HMM02780keto_1_eut1(pf02779pf02780pf00456) EUT1
03666 -pf00923 -
03667 HMM00390_malic_1_eut1(pf00390pf03949pf01515) EUT1
03668 Hp_euts_violetPink EUT1
03669 HMM10662PduV_1_mix(pf10662) EUT1
03670 HMM06249EutQ_1_eut1(pf06249) EUT1 EP_NTERM
03671 HMM01923cobtr_2_eut1(pf01923) EUT1
03672 HMM01515PTA_B_1_eut1(pf01515) EUT1
03673 H_azure EUT1
03674 P_lightViolet EUT1
03675 HMMaldh_1_eutpdugrm(pf00171) EUT1 EP_NTERM
03676 HMM06723EutJMreB_1_grm(#) EUT1
03677 HMM00465alcdh_3_mix(pf00465) EUT1
03678 HMMtpEuth_1_eut(pf04346) EUT1
03679 HMM06277EutA_1_eut(pf06277) EUT1
03680 HMMetlyEutB_1_EUT1(pf06751) EUT1
03681 HMMetlyEutC_1_eut1(pf05985) EUT1 EP_NTERM
03682 Tsp_rosa EUT1
03683 H_mulberry(pf00936pf16365) EUT1
03684 HMMreg12833_1_EUT1(pf12833) EUT1
03685 -pf01218 -
03686 -pf01520 -
03687 -pf00583 -
03688 -pf11143 -
03689 -pf06572 -
03690 -pf04261 -
03691 -pf00144 -
03692 -pf00367pf02378 -
03693 -pf01380 -
BMC-P
BMC-H
BMC-T
Aldehyde dehydrogenase
PTAC
Signature enzyme
Alcohol dehydrogenase
Regulator

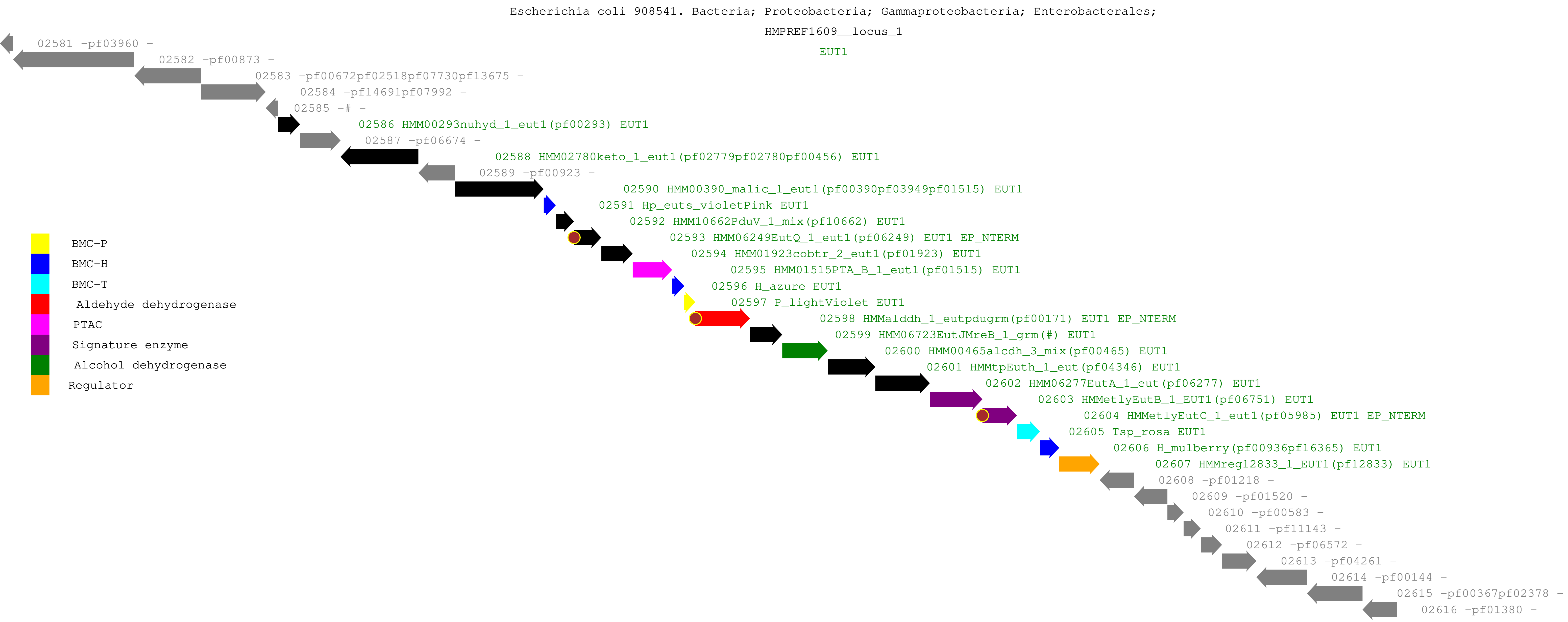

Escherichia coli 908541. Bacteria; Proteobacteria; Gammaproteobacteria; Enterobacterales;
HMPREF1609__locus_1
EUT1
02581 -pf03960 -
02582 -pf00873 -
02583 -pf00672pf02518pf07730pf13675 -
02584 -pf14691pf07992 -
02585 -# -
02586 HMM00293nuhyd_1_eut1(pf00293) EUT1
02587 -pf06674 -
02588 HMM02780keto_1_eut1(pf02779pf02780pf00456) EUT1
02589 -pf00923 -
02590 HMM00390_malic_1_eut1(pf00390pf03949pf01515) EUT1
02591 Hp_euts_violetPink EUT1
02592 HMM10662PduV_1_mix(pf10662) EUT1
02593 HMM06249EutQ_1_eut1(pf06249) EUT1 EP_NTERM
02594 HMM01923cobtr_2_eut1(pf01923) EUT1
02595 HMM01515PTA_B_1_eut1(pf01515) EUT1
02596 H_azure EUT1
02597 P_lightViolet EUT1
02598 HMMalddh_1_eutpdugrm(pf00171) EUT1 EP_NTERM
02599 HMM06723EutJMreB_1_grm(#) EUT1
02600 HMM00465alcdh_3_mix(pf00465) EUT1
02601 HMMtpEuth_1_eut(pf04346) EUT1
02602 HMM06277EutA_1_eut(pf06277) EUT1
02603 HMMetlyEutB_1_EUT1(pf06751) EUT1
02604 HMMetlyEutC_1_eut1(pf05985) EUT1 EP_NTERM
02605 Tsp_rosa EUT1
02606 H_mulberry(pf00936pf16365) EUT1
02607 HMMreg12833_1_EUT1(pf12833) EUT1
02608 -pf01218 -
02609 -pf01520 -
02610 -pf00583 -
02611 -pf11143 -
02612 -pf06572 -
02613 -pf04261 -
02614 -pf00144 -
02615 -pf00367pf02378 -
02616 -pf01380 -
BMC-P
BMC-H
BMC-T
Aldehyde dehydrogenase
PTAC
Signature enzyme
Alcohol dehydrogenase
Regulator

Escherichia coli 908573. Bacteria; Proteobacteria; Gammaproteobacteria; Enterobacterales;

HMPREF1611__locus_1

EUT1

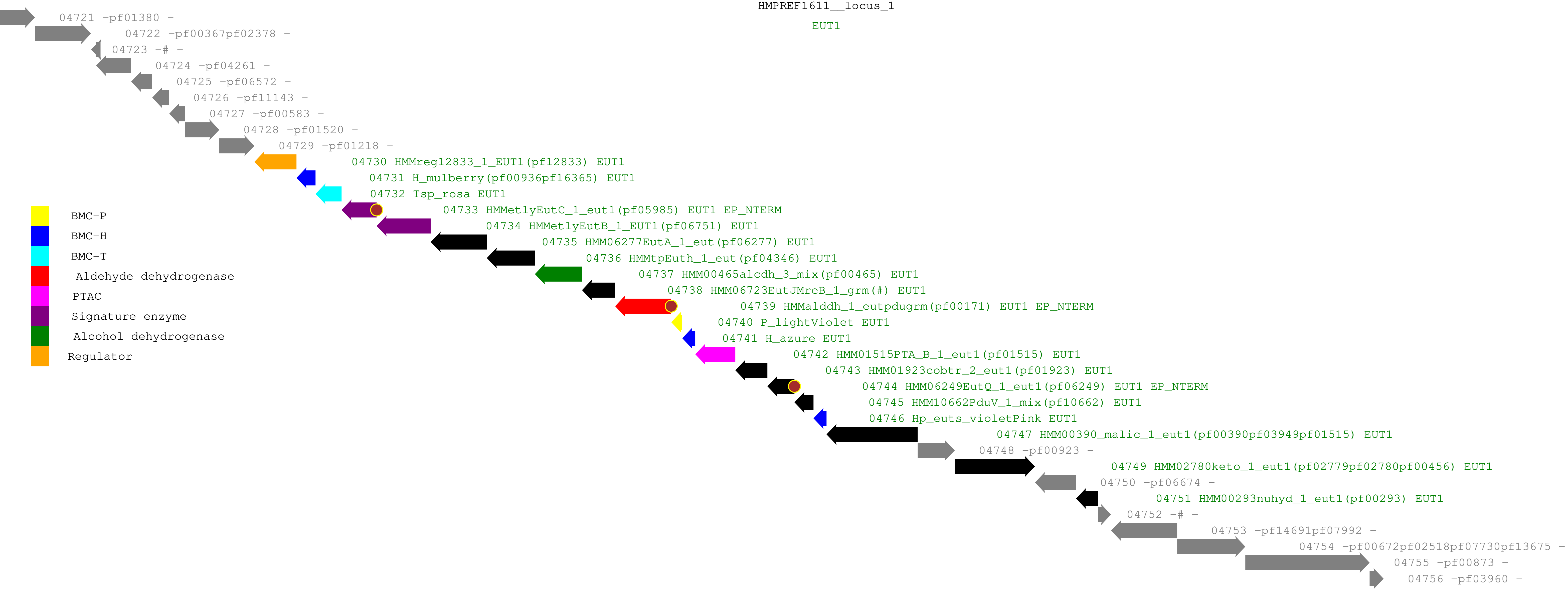

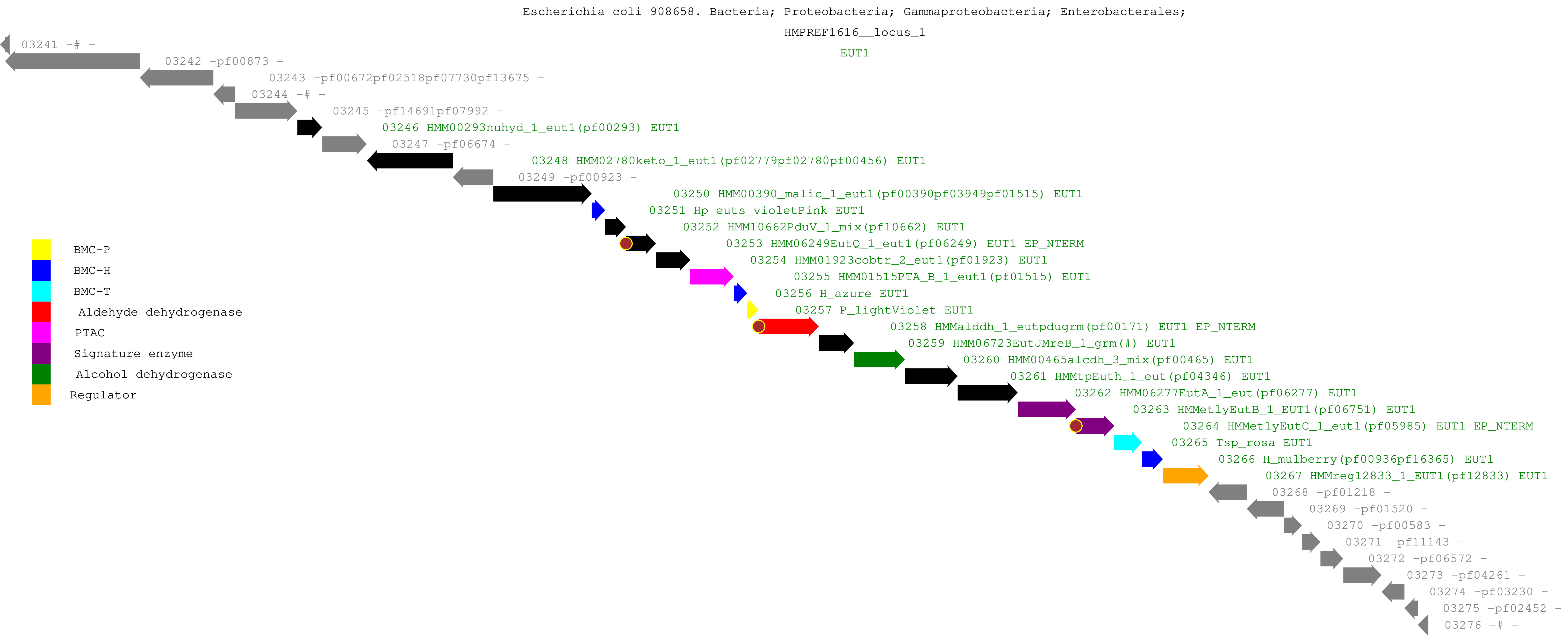
Escherichia coli 908658. Bacteria; Proteobacteria; Gammaproteobacteria; Enterobacterales;
HMPREF1616__locus_1
EUT1
03241 -# -
03242 -pf00873 -
03243 -pf00672pf02518pf07730pf13675 -
03244 -# -
03245 -pf14691pf07992 -
03246 HMM00293nuhyd_1_eut1(pf00293) EUT1
03247 -pf06674 -
03248 HMM02780keto_1_eut1(pf02779pf02780pf00456) EUT1
03249 -pf00923 -
03250 HMM00390_malic_1_eut1(pf00390pf03949pf01515) EUT1
03251 Hp_euts_violetPink EUT1
03252 HMM10662PduV_1_mix(pf10662) EUT1
03253 HMM06249EutQ_1_eut1(pf06249) EUT1 EP_NTERM
03254 HMM01923cobtr_2_eut1(pf01923) EUT1
03255 HMM01515PTA_B_1_eut1(pf01515) EUT1
03256 H_azure EUT1
03257 P_lightViolet EUT1
03258 HMMaldddh_1_eutpdugrm(pf00171) EUT1 EP_NTERM
03259 HMM06723EutJMreB_1_grm(#) EUT1
03260 HMM00465alcdh_3_mix(pf00465) EUT1
03261 HMMtpEuth_1_eut(pf04346) EUT1
03262 HMM06277EutA_1_eut(pf06277) EUT1
03263 HMMetlyEutB_1_EUT1(pf06751) EUT1
03264 HMMetlyEutC_1_eut1(pf05985) EUT1 EP_NTERM
03265 Tsp_rosa EUT1
03266 H_mulberry(pf00936pf16365) EUT1
03267 HMMreg12833_1_EUT1(pf12833) EUT1
03268 -pf01218 -
03269 -pf01520 -
03270 -pf00583 -
03271 -pf11143 -
03272 -pf06572 -
03273 -pf04261 -
03274 -pf03230 -
03275 -pf02452 -
03276 -# -
BMC-P
BMC-H
BMC-T
Aldehyde dehydrogenase
PTAC
Signature enzyme
Alcohol dehydrogenase
Regulator

Klebsiella pneumoniae 909957. Bacteria; Proteobacteria; Gammaproteobacteria; Enterobacterales;

HMPREF1619__locus_1

EUT1

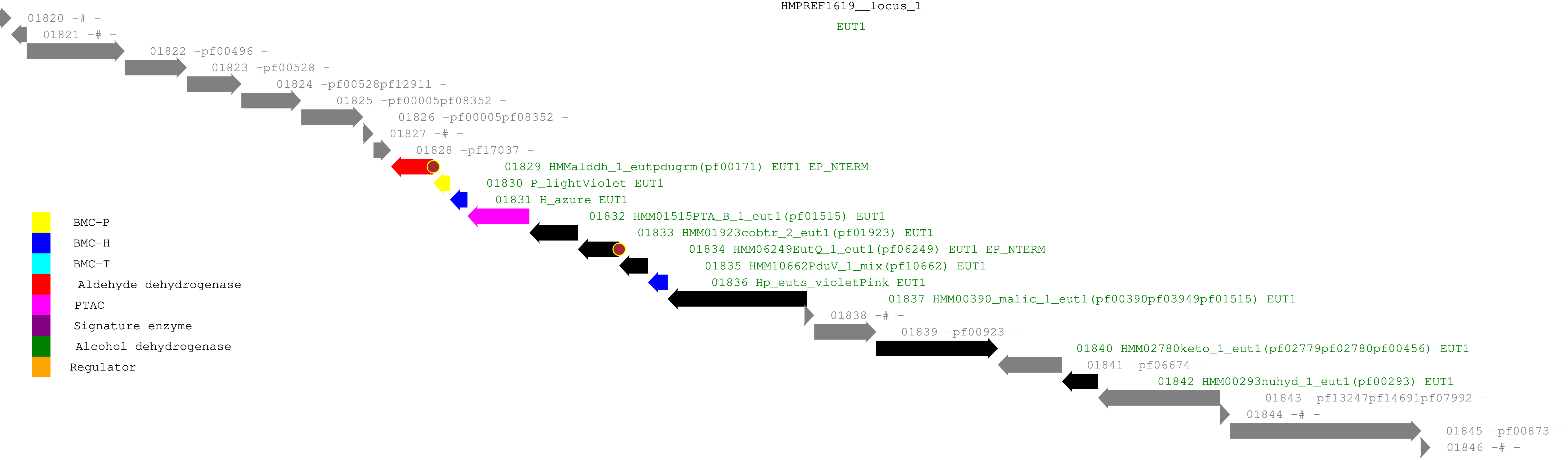

Escherichia coli 909945-2. Bacteria; Proteobacteria; Gammaproteobacteria; Enterobacterales;

HMPREF1620__locus_1

EUT1

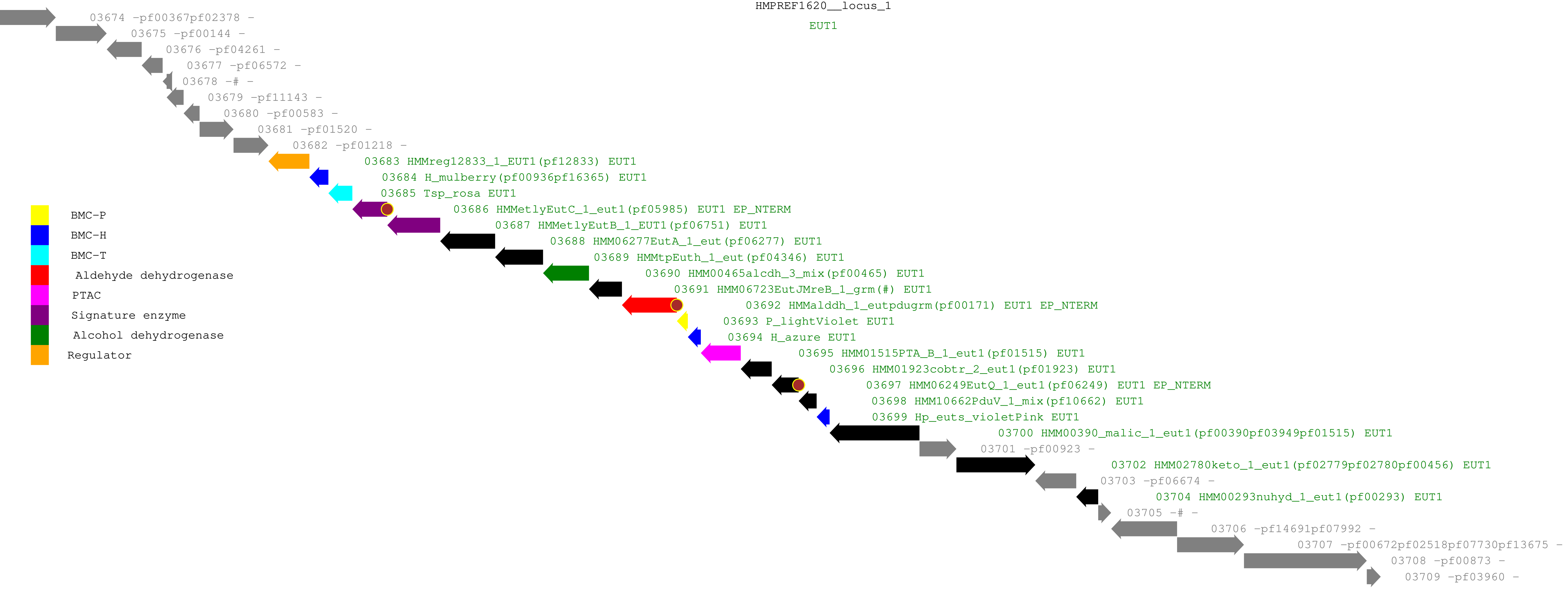

Enterobacter sp. HMSC055A11. Bacteria; Proteobacteria; Gammaproteobacteria; Enterobacterales;

HMPREF2540__locus_2

EUT1

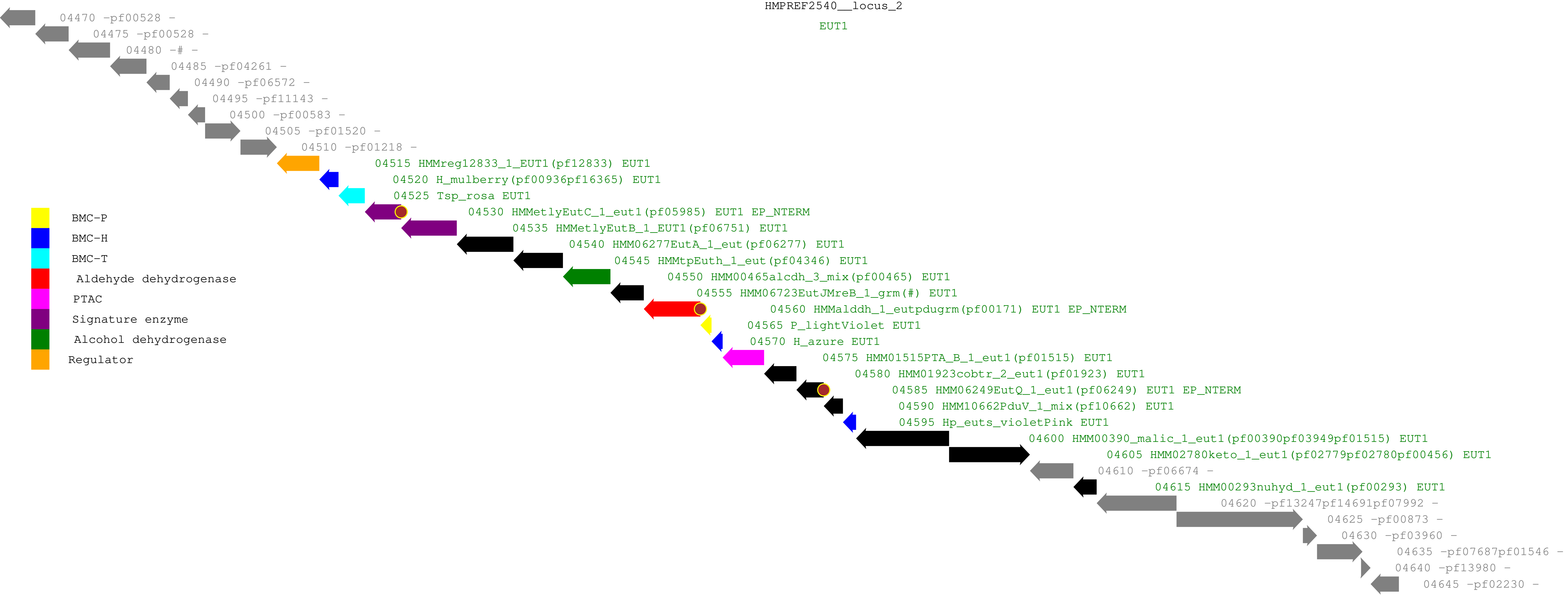

Escherichia coli. Bacteria; Proteobacteria; Gammaproteobacteria; Enterobacterales;

HMPREF3040__locus_2

EUT1

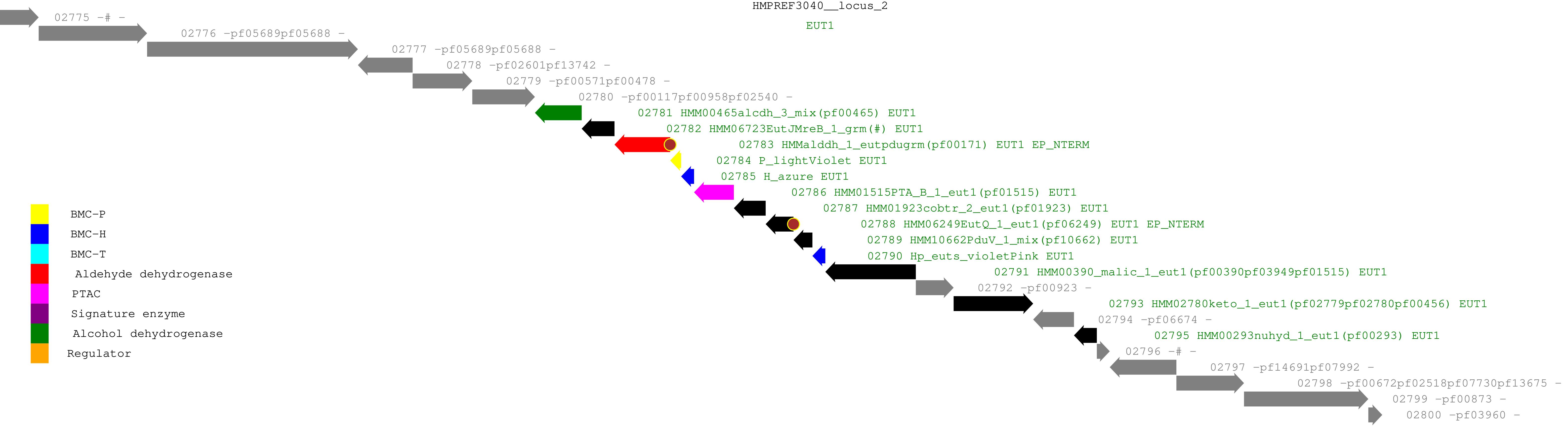

Proteus sp. HMSC10D02. Bacteria; Proteobacteria; Gammaproteobacteria; Enterobacterales;

HMPREF3111__locus_2

EUT1

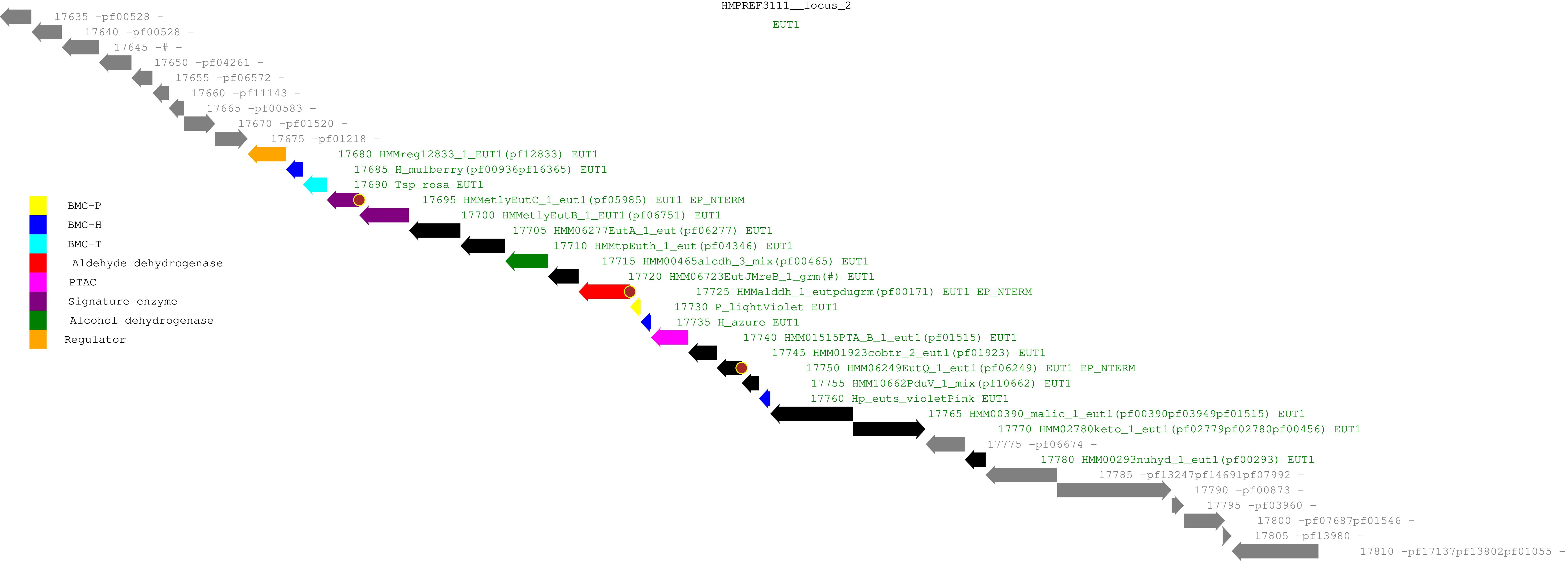

Salmonella sp. HMSC13B08. Bacteria; Proteobacteria; Gammaproteobacteria; Enterobacterales;

HMPREF3126__locus_2

EUT1

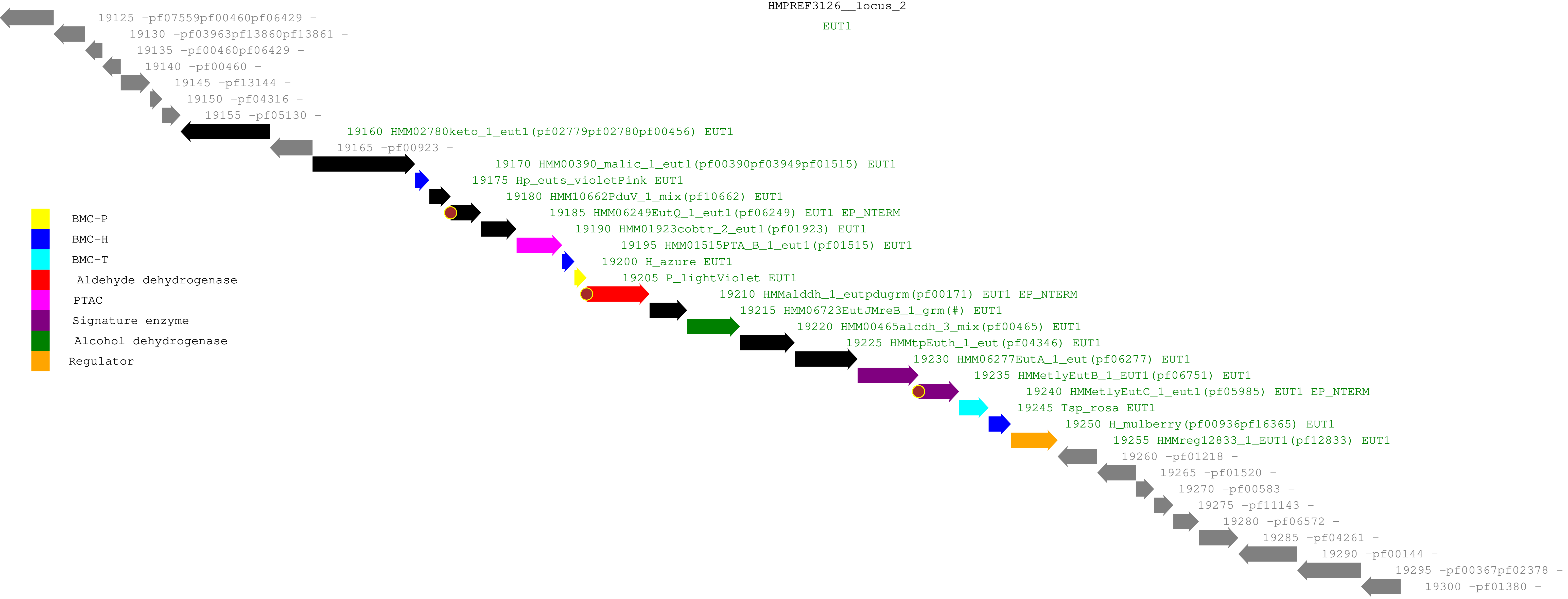

Klebsiella sp. HMSC16C06. Bacteria; Proteobacteria; Gammaproteobacteria; Enterobacterales;

HMPREF3142__locus_3

EUT1

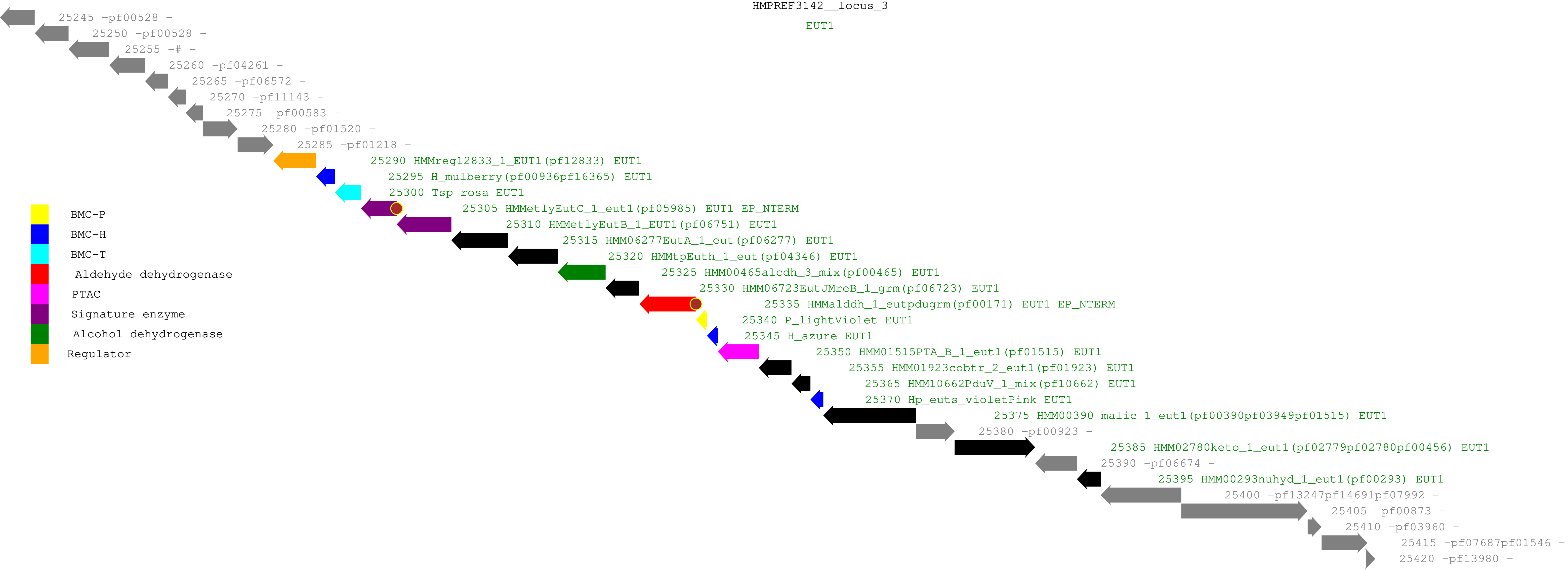

Citrobacter freundii. Bacteria; Proteobacteria; Gammaproteobacteria; Enterobacterales;

HMPREF3212__locus_2

EUT1

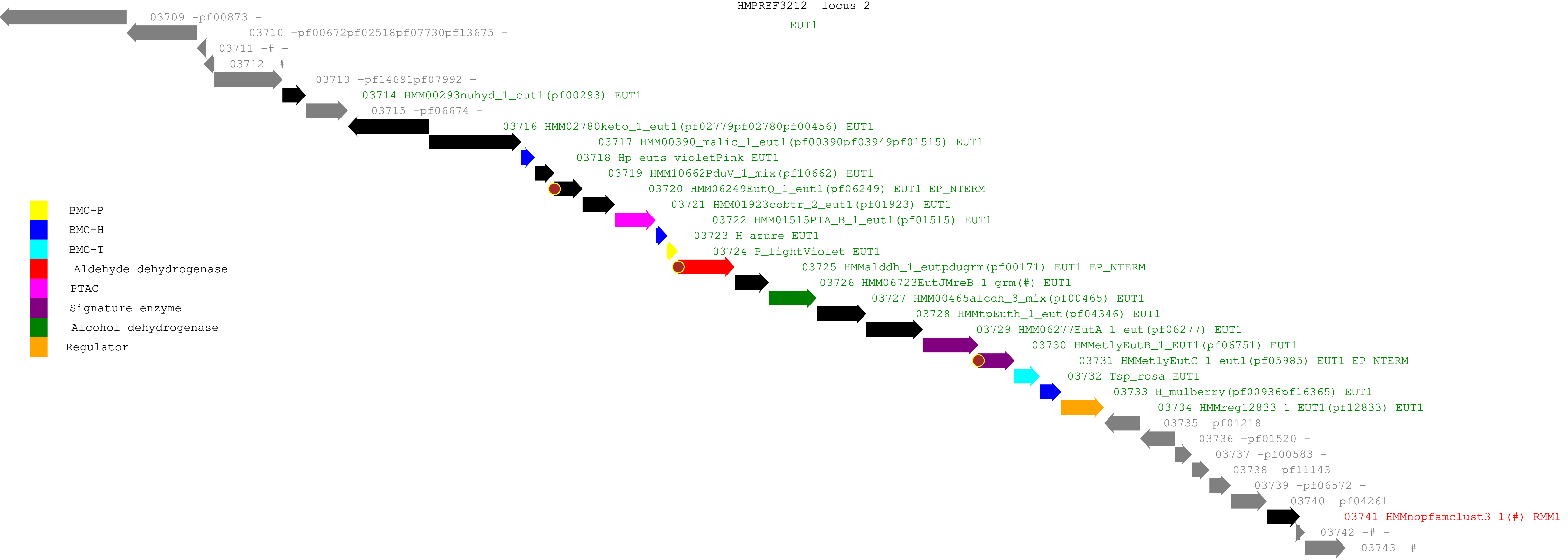

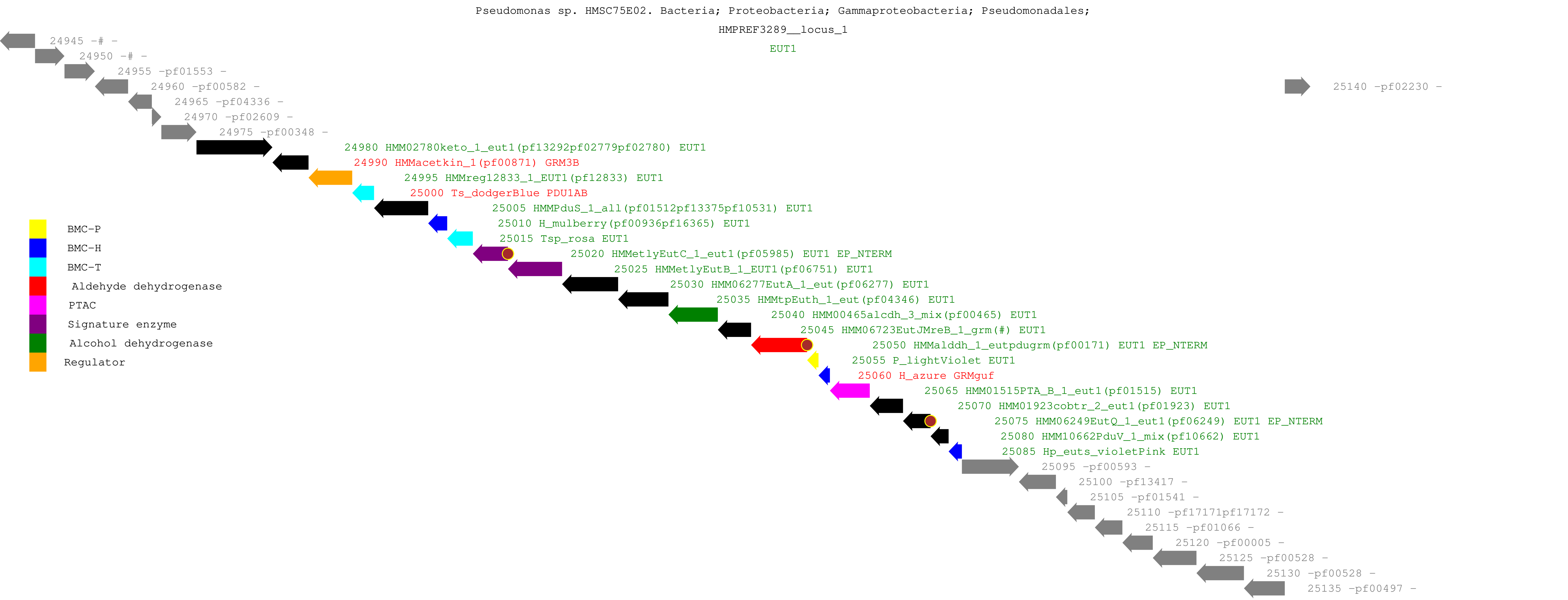
Pseudomonas sp. HMSC75E02. Bacteria; Proteobacteria; Gammaproteobacteria; Pseudomonadales;
HMPREF3289__locus_1
EUT1
24945 -# -
24950 -# -
24955 -pf01553 -
24960 -pf00582 -
24965 -pf04336 -
24970 -pf02609 -
24975 -pf00348 -
24980 HMM02780keto_1_eut1(pf13292pf02779pf02780) EUT1
24990 HMMacetkin_1(pf00871) GRM3B
24995 HMMreg12833_1_EUT1(pf12833) EUT1
25000 Ts_dodgerBlue PDU1AB
25005 HMMPduS_1_all(pf01512pf13375pf10531) EUT1
25010 H_mulberry(pf00936pf16365) EUT1
25015 Tsp_rosa EUT1
25020 HMMetlyEutC_1_eut1(pf05985) EUT1 EP_NTERM
25025 HMMetlyEutB_1_EUT1(pf06751) EUT1
25030 HMM06277EutA_1_eut(pf06277) EUT1
25035 HMMtpEuth_1_eut(pf04346) EUT1
25040 HMM00465alcdh_3_mix(pf00465) EUT1
25045 HMM06723EutJMreB_1_grm(#) EUT1
25050 HMMalddh_1_eutpdugrm(pf00171) EUT1 EP_NTERM
25055 P_lightViolet EUT1
25060 H_azure GRMguf
25065 HMM01515PTA_B_1_eut1(pf01515) EUT1
25070 HMM01923cobtr_2_eut1(pf01923) EUT1
25075 HMM06249EutQ_1_eut1(pf06249) EUT1 EP_NTERM
25080 HMM10662PduV_1_mix(pf10662) EUT1
25085 Hp_euts_violetPink EUT1
25095 -pf00593 -
25100 -pf13417 -
25105 -pf01541 -
25110 -pf17171pf17172 -
25115 -pf01066 -
25120 -pf00005 -
25125 -pf00528 -
25130 -pf00528 -
25135 -pf00497 -
25140 -pf02230 -
BMC-P
BMC-H
BMC-T
Aldehyde dehydrogenase
PTAC
Signature enzyme
Alcohol dehydrogenase
Regulator

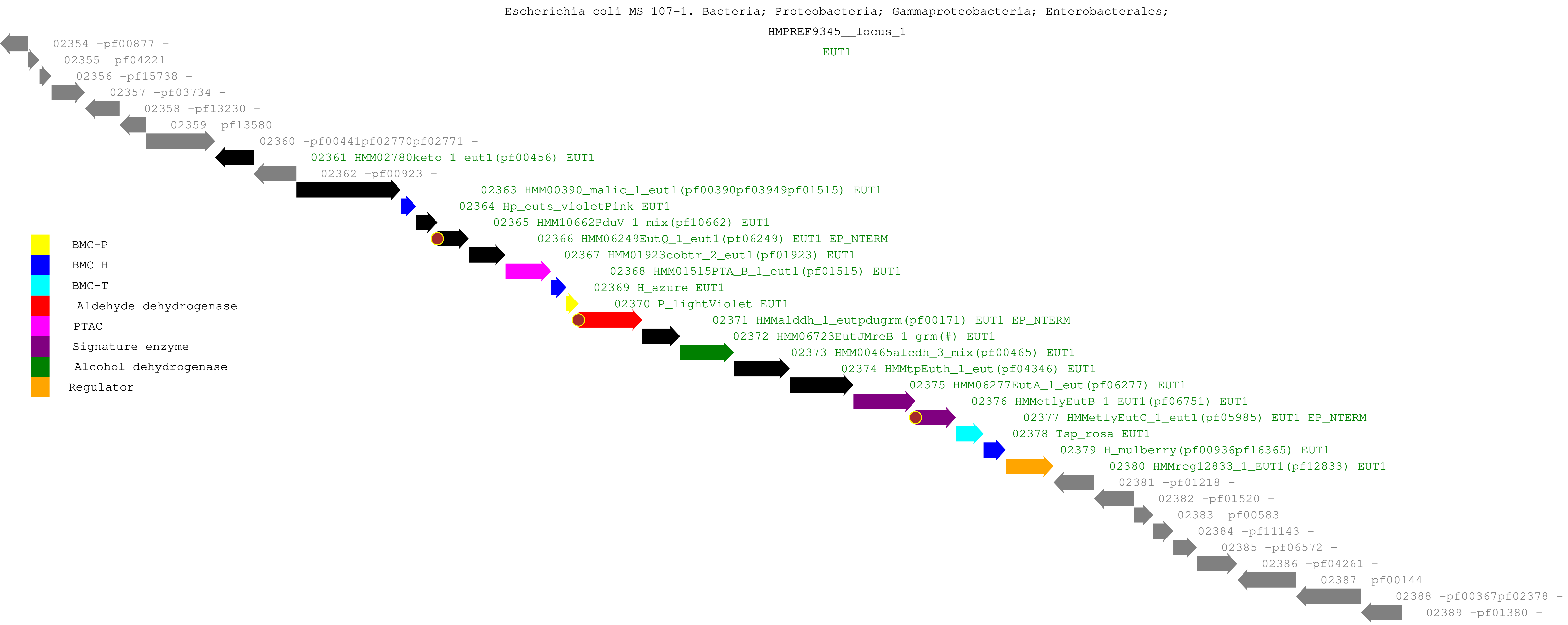
Escherichia coli MS 107-1. Bacteria; Proteobacteria; Gammaproteobacteria; Enterobacterales;
HMPREF9345__locus_1
EUT1
02354 -pf00877 -
02355 -pf04221 -
02356 -pf15738 -
02357 -pf03734 -
02358 -pf13230 -
02359 -pf13580 -
02360 -pf00441pf02770pf02771 -
02361 HMM02780keto_1_eut1(pf00456) EUT1
02362 -pf00923 -
02363 HMM00390_malic_1_eut1(pf00390pf03949pf01515) EUT1
02364 Hp_euts_violetPink EUT1
02365 HMM10662PduV_1_mix(pf10662) EUT1
02366 HMM06249EutQ_1_eut1(pf06249) EUT1 EP_NTERM
02367 HMM01923cobtr_2_eut1(pf01923) EUT1
02368 HMM01515PTA_B_1_eut1(pf01515) EUT1
02369 H_azure EUT1
02370 P_lightViolet EUT1
02371 HMMaldh_1_eutpdugrm(pf00171) EUT1 EP_NTERM
02372 HMM06723EutJMreB_1_grm(#) EUT1
02373 HMM00465alcdh_3_mix(pf00465) EUT1
02374 HMMtpEuth_1_eut(pf04346) EUT1
02375 HMM06277EutA_1_eut(pf06277) EUT1
02376 HMMetlyEutB_1_EUT1(pf06751) EUT1
02377 HMMetlyEutC_1_eut1(pf05985) EUT1 EP_NTERM
02378 Tsp_rosa EUT1
02379 H_mulberry(pf00936pf16365) EUT1
02380 HMMreg12833_1_EUT1(pf12833) EUT1
02381 -pf01218 -
02382 -pf01520 -
02383 -pf00583 -
02384 -pf11143 -
02385 -pf06572 -
02386 -pf04261 -
02387 -pf00144 -
02388 -pf00367pf02378 -
02389 -pf01380 -
BMC-P
BMC-H
BMC-T
Aldehyde dehydrogenase
PTAC
Signature enzyme
Alcohol dehydrogenase
Regulator

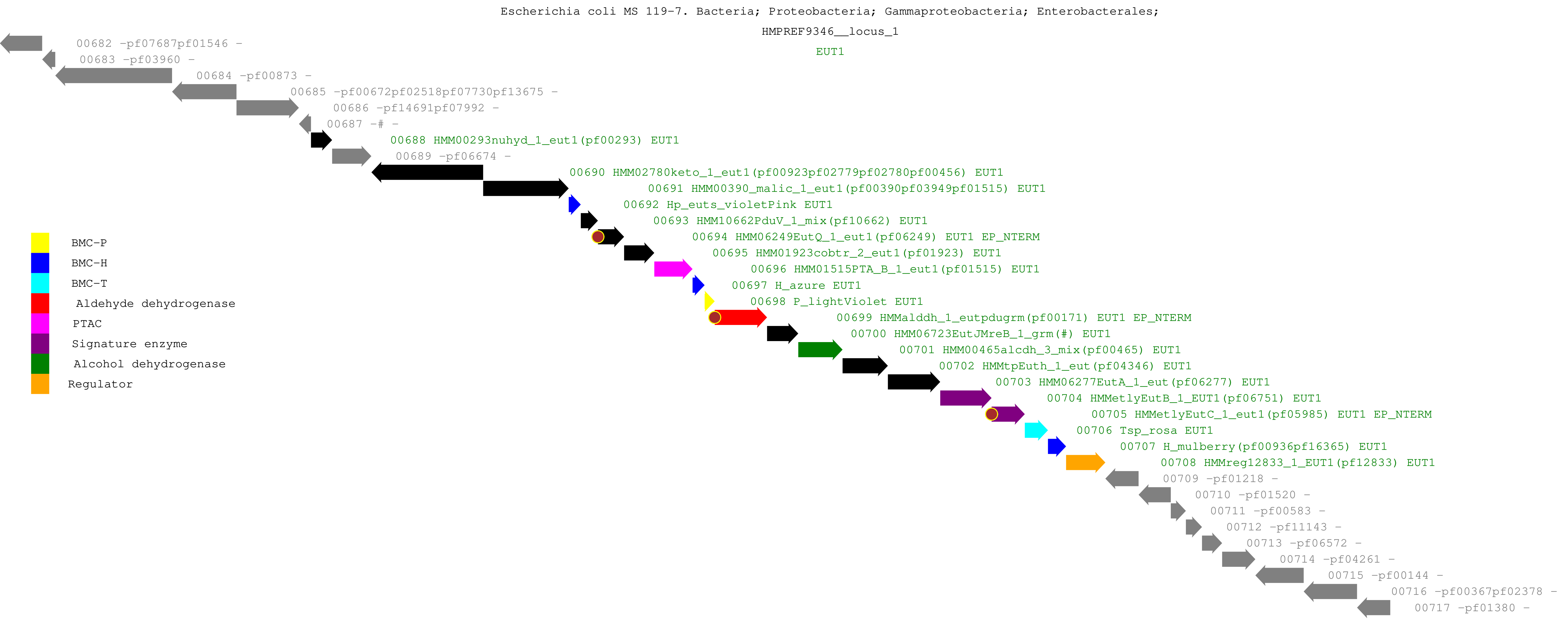
Escherichia coli MS 119-7. Bacteria; Proteobacteria; Gammaproteobacteria; Enterobacterales;
HMPREF9346__locus_1
EUT1
00682 -pf07687pf01546 -
00683 -pf03960 -
00684 -pf00873 -
00685 -pf00672pf02518pf07730pf13675 -
00686 -pf14691pf07992 -
00687 -# -
00688 HMM00293nuhyd_1_eut1(pf00293) EUT1
00689 -pf06674 -
00690 HMM02780keto_1_eut1(pf00923pf02779pf02780pf00456) EUT1
00691 HMM00390_malic_1_eut1(pf00390pf03949pf01515) EUT1
00692 Hp_euts_violetPink EUT1
00693 HMM10662PduV_1_mix(pf10662) EUT1
00694 HMM06249EutQ_1_eut1(pf06249) EUT1 EP_NTERM
00695 HMM01923cobtr_2_eut1(pf01923) EUT1
00696 HMM01515PTA_B_1_eut1(pf01515) EUT1
00697 H_azure EUT1
00698 P_lightViolet EUT1
00699 HMMalddh_1_eutpdugrm(pf00171) EUT1 EP_NTERM
00700 HMM06723EutJMreB_1_grm(#) EUT1
00701 HMM00465alcdh_3_mix(pf00465) EUT1
00702 HMMtpEuth_1_eut(pf04346) EUT1
00703 HMM06277EutA_1_eut(pf06277) EUT1
00704 HMMetlyEutB_1_EUT1(pf06751) EUT1
00705 HMMetlyEutC_1_eut1(pf05985) EUT1 EP_NTERM
00706 Tsp_rosa EUT1
00707 H_mulberry(pf00936pf16365) EUT1
00708 HMMreg12833_1_EUT1(pf12833) EUT1
00709 -pf01218 -
00710 -pf01520 -
00711 -pf00583 -
00712 -pf11143 -
00713 -pf06572 -
00714 -pf04261 -
00715 -pf00144 -
00716 -pf00367pf02378 -
00717 -pf01380 -
BMC-P
BMC-H
BMC-T
Aldehyde dehydrogenase
PTAC
Signature enzyme
Alcohol dehydrogenase
Regulator

Escherichia coli MS 124-1. Bacteria; Proteobacteria; Gammaproteobacteria; Enterobacterales;

HMPREF9347__locus_1

EUT1

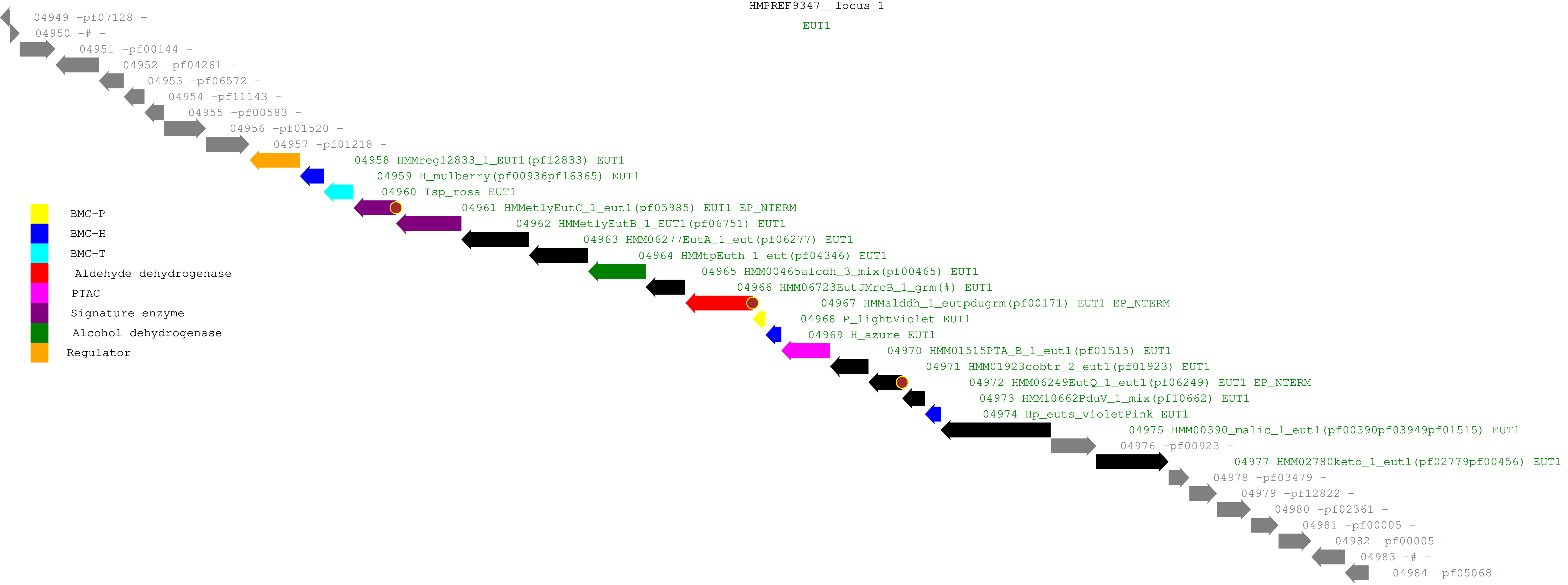

Escherichia coli MS 145-7. Bacteria; Proteobacteria; Gammaproteobacteria; Enterobacterales;

HMPREF9348__locus_1

EUT1

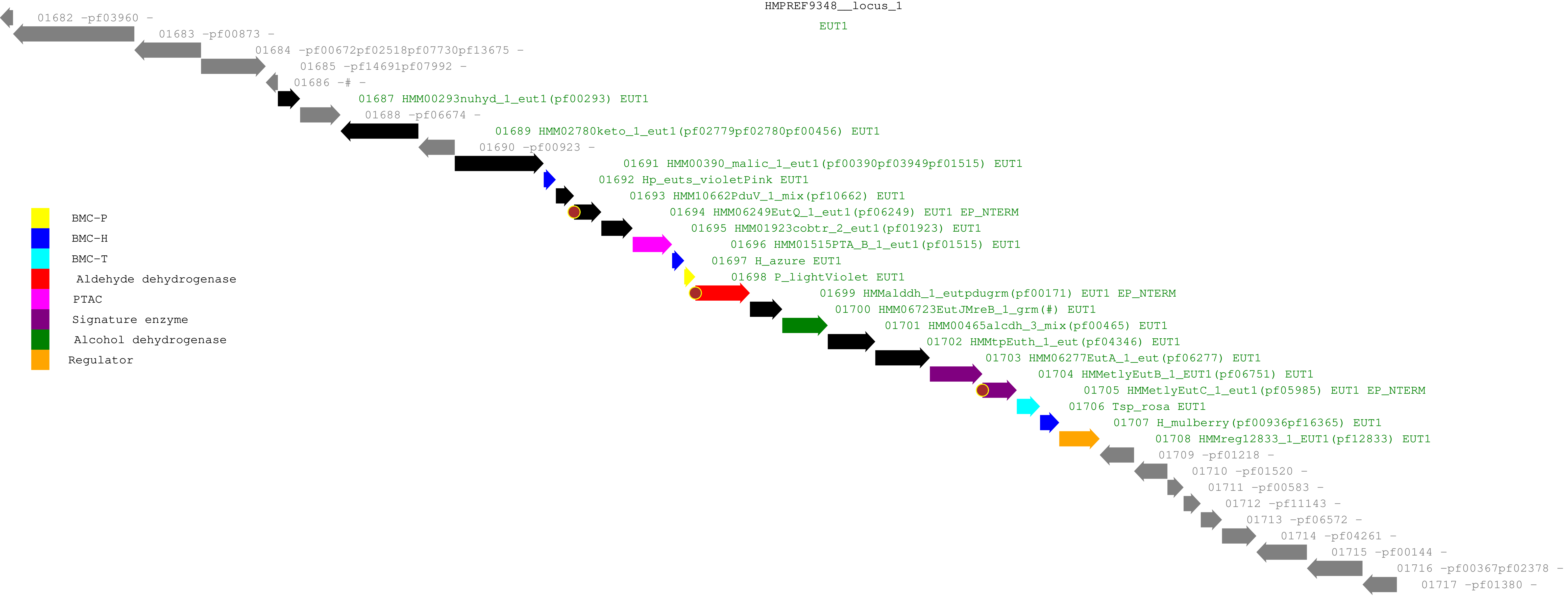

Escherichia coli MS 79-10. Bacteria; Proteobacteria; Gammaproteobacteria; Enterobacterales;

HMPREF9349__locus_1

EUT1

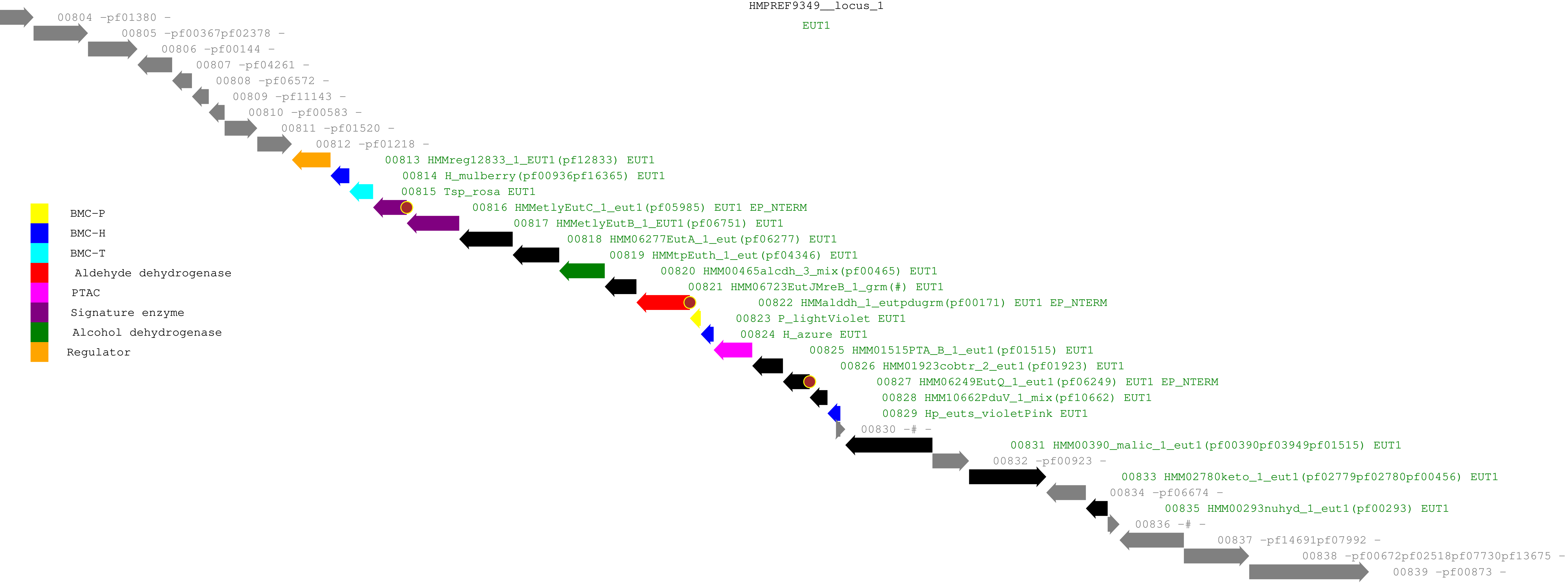

Escherichia coli MS 85-1. Bacteria; Proteobacteria; Gammaproteobacteria; Enterobacterales;

HMPREF9350__locus_1

EUT1

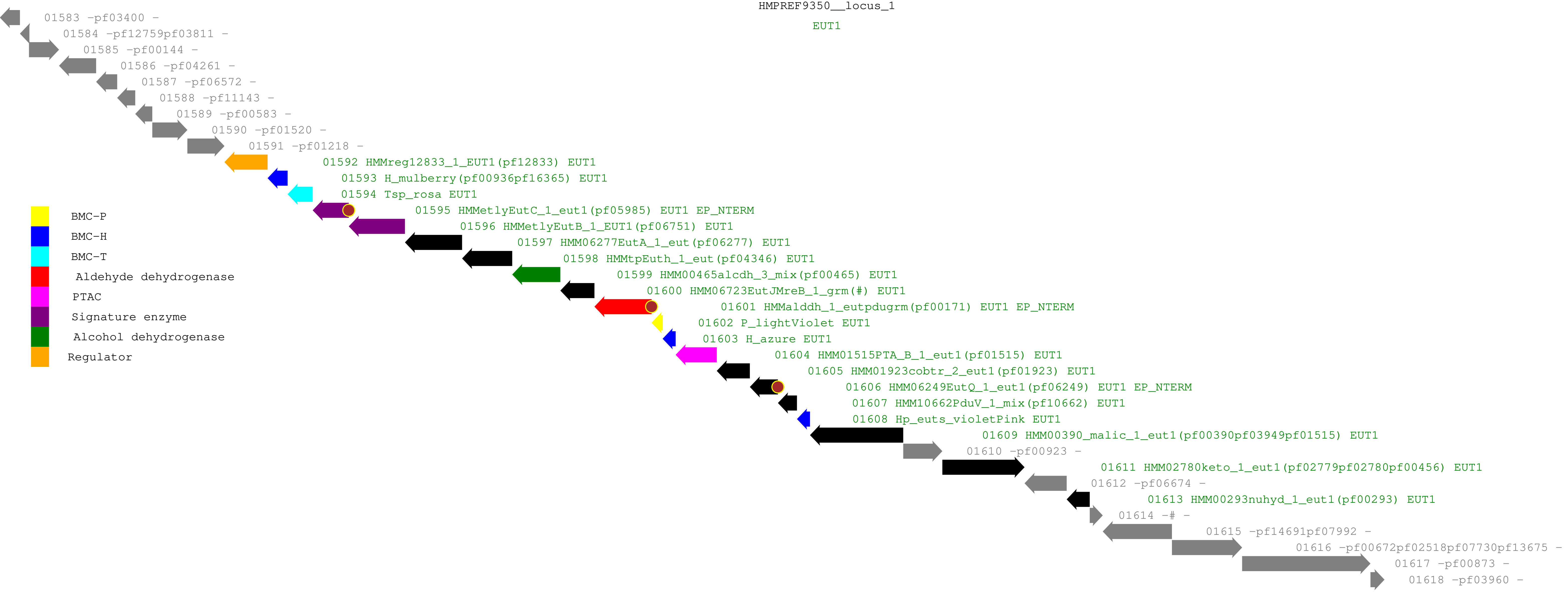

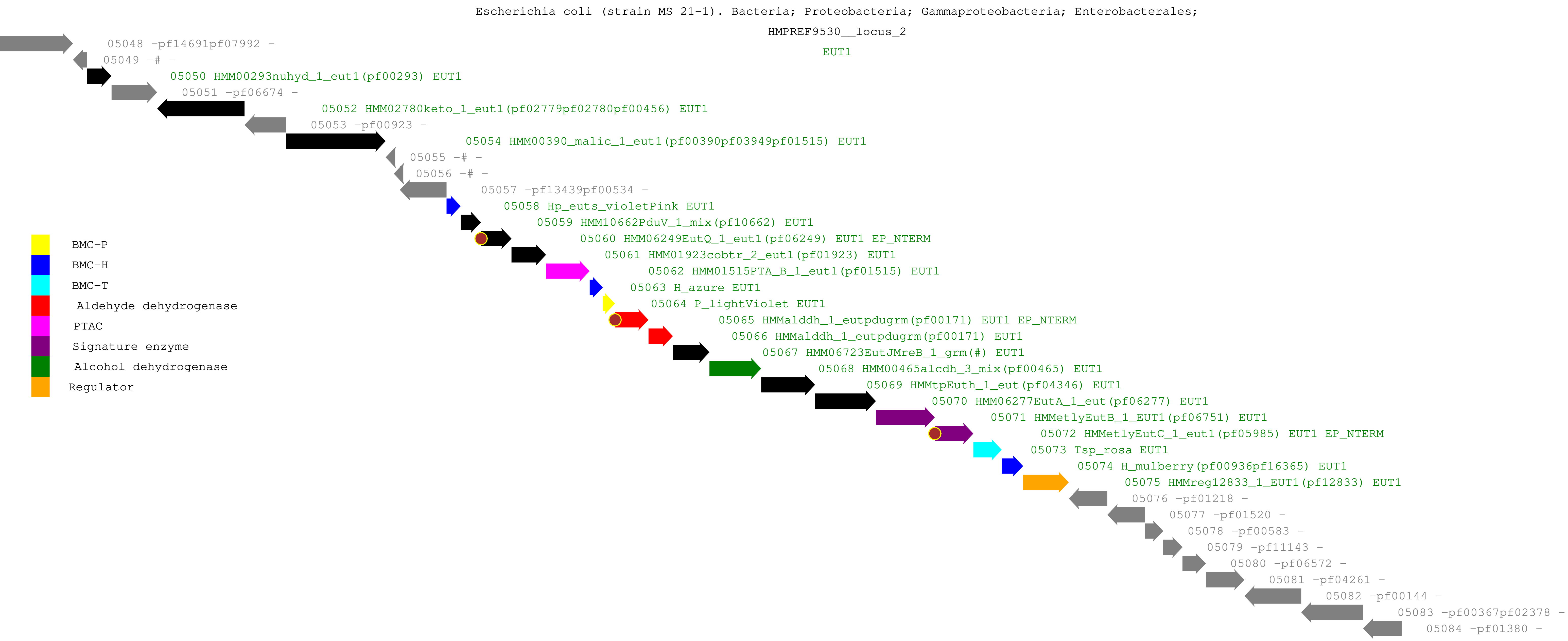
Escherichia coli (strain MS 21-1). Bacteria; Proteobacteria; Gammaproteobacteria; Enterobacterales;
HMPREF9530__locus_2
EUT1
05048 -pf14691pf07992 -
05049 -# -
05050 HMM00293nuhyd_1_eut1(pf00293) EUT1
05051 -pf06674 -
05052 HMM02780keto_1_eut1(pf02779pf02780pf00456) EUT1
05053 -pf00923 -
05054 HMM00390_malic_1_eut1(pf00390pf03949pf01515) EUT1
05055 -# -
05056 -# -
05057 -pf13439pf00534 -
05058 Hp_euts_violetPink EUT1
05059 HMM10662PduV_1_mix(pf10662) EUT1
05060 HMM06249EutQ_1_eut1(pf06249) EUT1 EP_NTERM
05061 HMM01923cobtr_2_eut1(pf01923) EUT1
05062 HMM01515PTA_B_1_eut1(pf01515) EUT1
05063 H_azure EUT1
05064 P_lightViolet EUT1
05065 HMMalddh_1_eutpdugrm(pf00171) EUT1 EP_NTERM
05066 HMMalddh_1_eutpdugrm(pf00171) EUT1
05067 HMM06723EutJMreB_1_grm(#) EUT1
05068 HMM00465alcdh_3_mix(pf00465) EUT1
05069 HMMtpEuth_1_eut(pf04346) EUT1
05070 HMM06277EutA_1_eut(pf06277) EUT1
05071 HMMetlyEutB_1_EUT1(pf06751) EUT1
05072 HMMetlyEutC_1_eut1(pf05985) EUT1 EP_NTERM
05073 Tsp_rosa EUT1
05074 H_mulberry(pf00936pf16365) EUT1
05075 HMMreg12833_1_EUT1(pf12833) EUT1
05076 -pf01218 -
05077 -pf01520 -
05078 -pf00583 -
05079 -pf11143 -
05080 -pf06572 -
05081 -pf04261 -
05082 -pf00144 -
05083 -pf00367pf02378 -
05084 -pf01380 -
BMC-P
BMC-H
BMC-T
Aldehyde dehydrogenase
PTAC
Signature enzyme
Alcohol dehydrogenase
Regulator

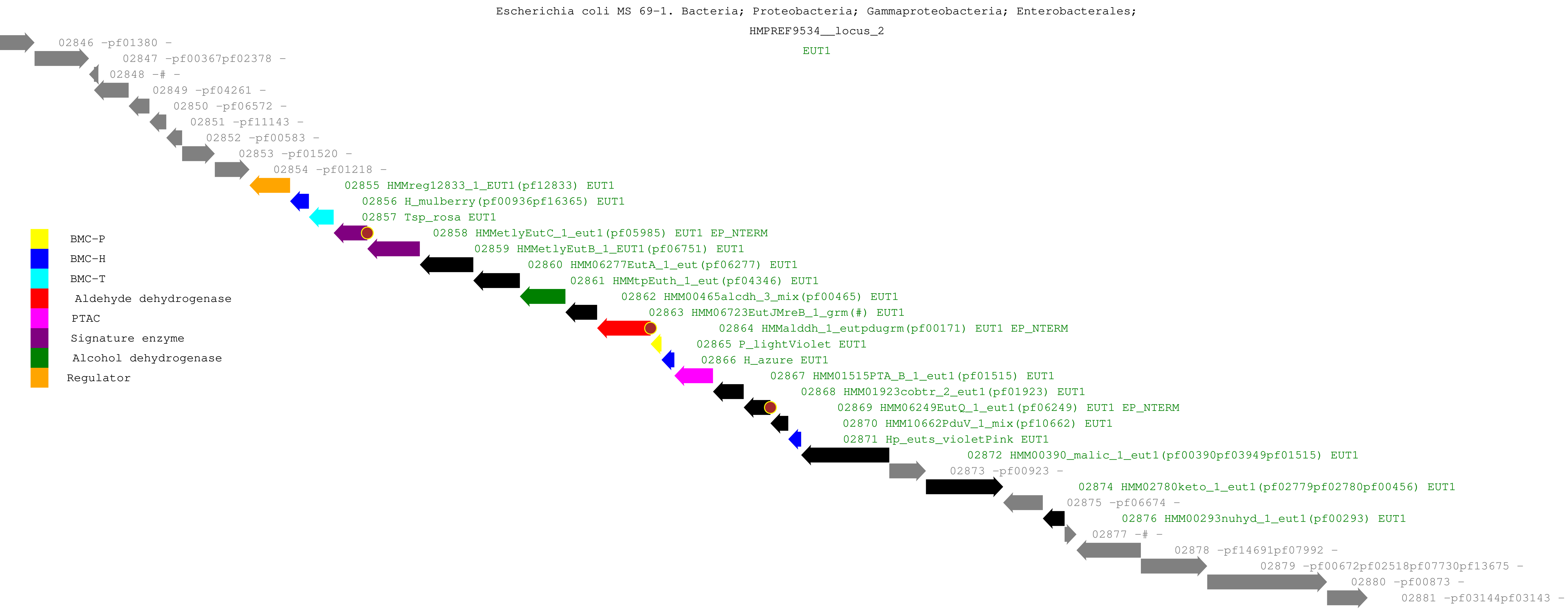
Escherichia coli MS 69-1. Bacteria; Proteobacteria; Gammaproteobacteria; Enterobacterales;
HMPREF9534__locus_2
EUT1
02846 -pf01380 -
02847 -pf00367pf02378 -
02848 -# -
02849 -pf04261 -
02850 -pf06572 -
02851 -pf11143 -
02852 -pf00583 -
02853 -pf01520 -
02854 -pf01218 -
02855 HMMreg12833_1_EUT1(pf12833) EUT1
02856 H_mulberry(pf00936pf16365) EUT1
02857 Tsp_rosa EUT1
02858 HMMetlyEutC_1_eut1(pf05985) EUT1 EP_NTERM
02859 HMMetlyEutB_1_EUT1(pf06751) EUT1
02860 HMM06277EutA_1_eut(pf06277) EUT1
02861 HMMtpEuth_1_eut(pf04346) EUT1
02862 HMM00465alcdh_3_mix(pf00465) EUT1
02863 HMM06723EutJMreB_1_grm(#) EUT1
02864 HMMalddh_1_eutpdugrm(pf00171) EUT1 EP_NTERM
02865 P_lightViolet EUT1
02866 H_azure EUT1
02867 HMM01515PTA_B_1_eut1(pf01515) EUT1
02868 HMM01923cobtr_2_eut1(pf01923) EUT1
02869 HMM06249EutQ_1_eut1(pf06249) EUT1 EP_NTERM
02870 HMM10662PduV_1_mix(pf10662) EUT1
02871 Hp_euts_violetPink EUT1
02872 HMM00390_malic_1_eut1(pf00390pf03949pf01515) EUT1
02873 -pf00923 -
02874 HMM02780keto_1_eut1(pf02779pf02780pf00456) EUT1
02875 -pf06674 -
02876 HMM00293nuhyd_1_eut1(pf00293) EUT1
02877 -# -
02878 -pf14691pf07992 -
02879 -pf00672pf02518pf07730pf13675 -
02880 -pf00873 -
02881 -pf03144pf03143 -
BMC-P
BMC-H
BMC-T
Aldehyde dehydrogenase
PTAC
Signature enzyme
Alcohol dehydrogenase
Regulator

Escherichia coli MS 78-1. Bacteria; Proteobacteria; Gammaproteobacteria; Enterobacterales;

HMPREF9535__locus_1

EUT1

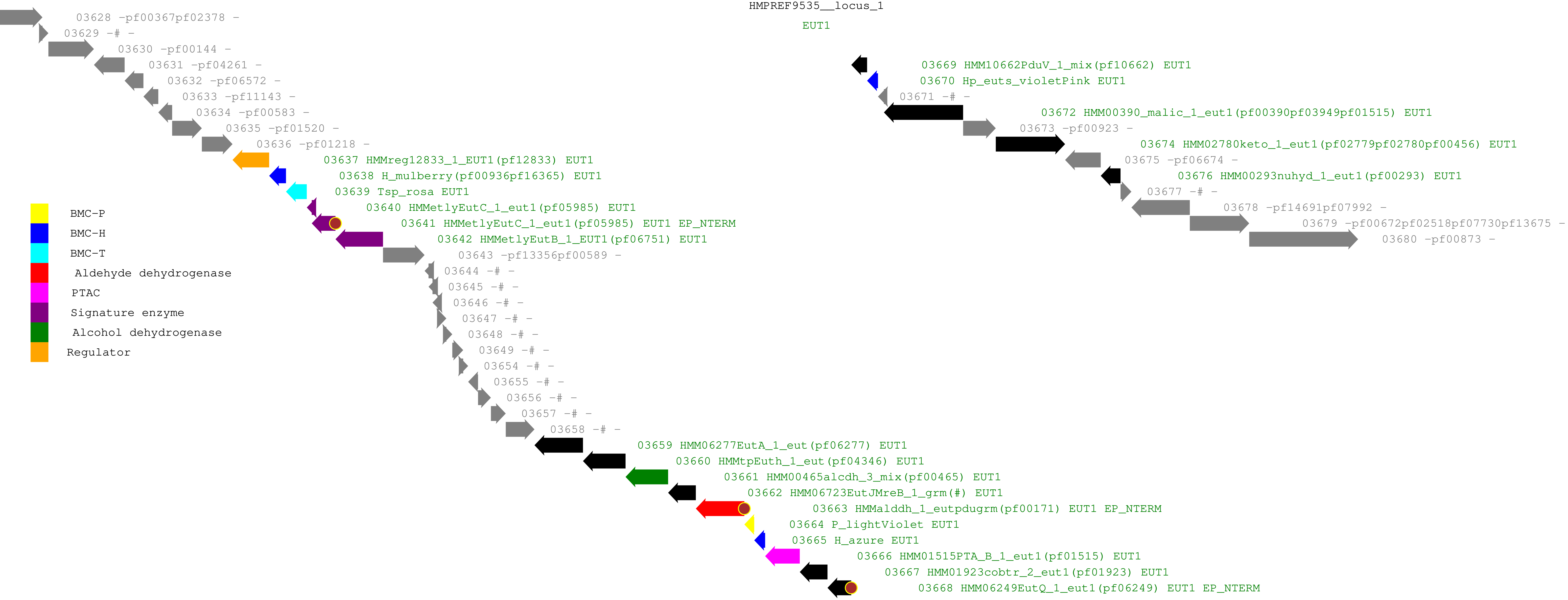

Escherichia coli MS 84-1. Bacteria; Proteobacteria; Gammaproteobacteria; Enterobacterales;

HMPREF9536__locus_1

EUT1

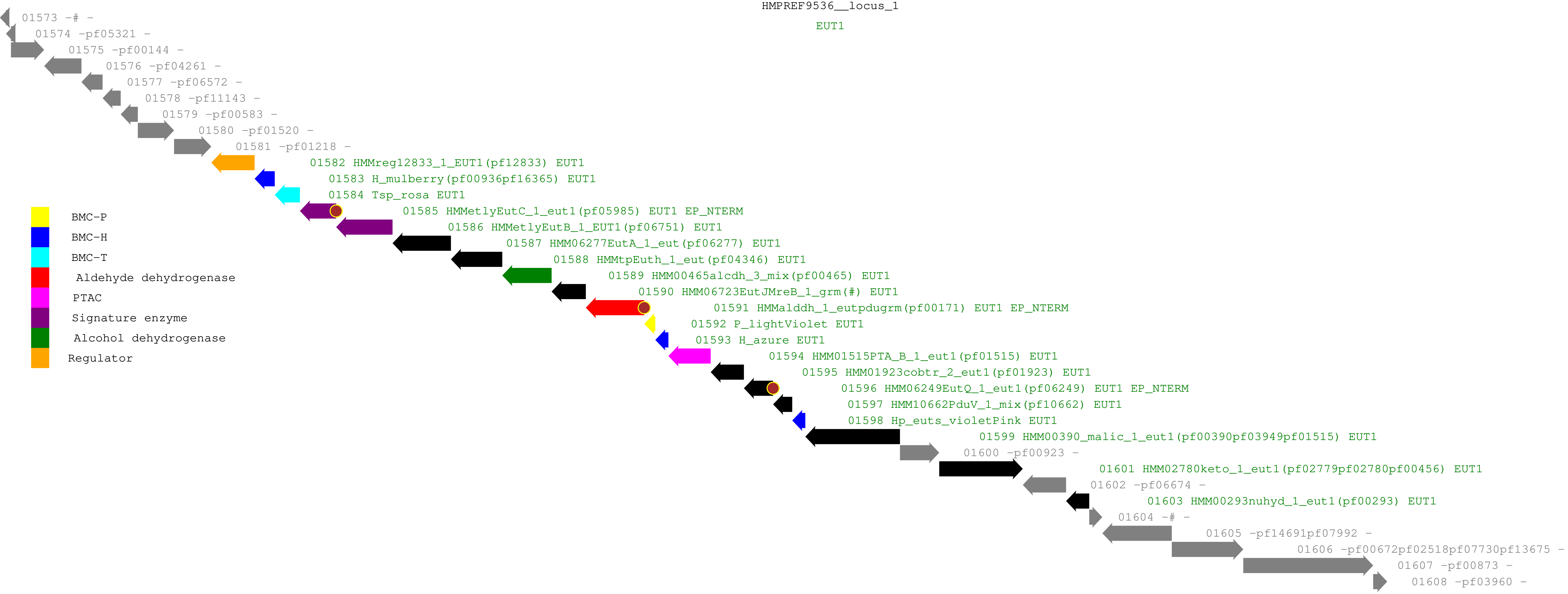

Escherichia coli MS 115-1. Bacteria; Proteobacteria; Gammaproteobacteria; Enterobacterales;

HMPREF9540__locus_2

EUT1

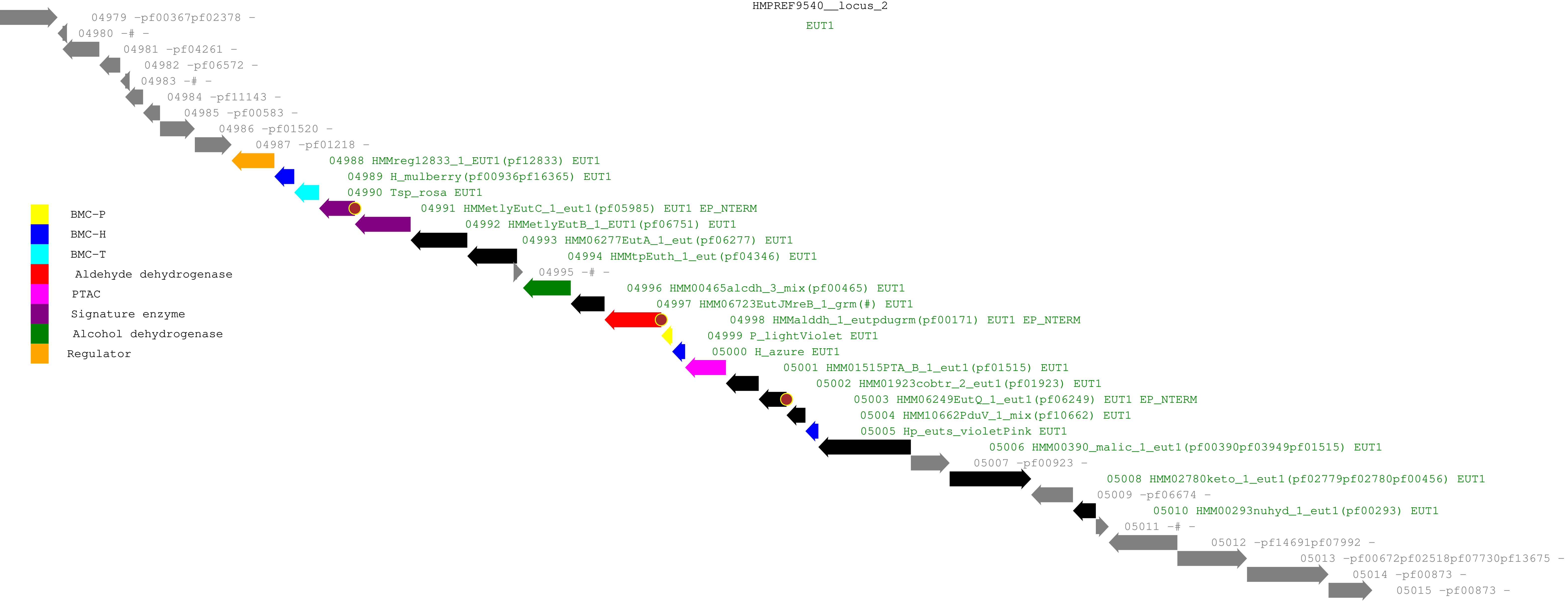

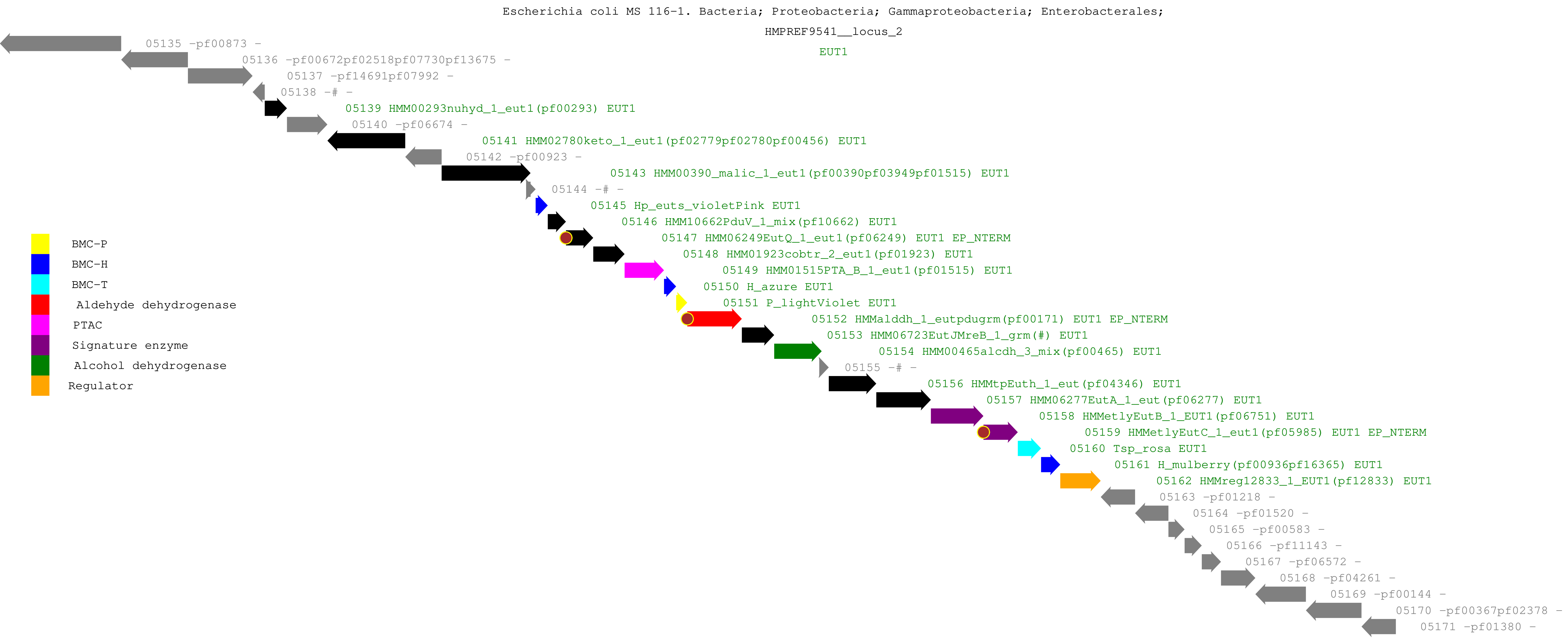
Escherichia coli MS 116-1. Bacteria; Proteobacteria; Gammaproteobacteria; Enterobacterales;
HMPREF9541__locus_2
EUT1
05135 -pf00873 -
05136 -pf00672pf02518pf07730pf13675 -
05137 -pf14691pf07992 -
05138 -# -
05139 HMM00293nuhyd_1_eut1(pf00293) EUT1
05140 -pf06674 -
05141 HMM02780keto_1_eut1(pf02779pf02780pf00456) EUT1
05142 -pf00923 -
05143 HMM00390_malic_1_eut1(pf00390pf03949pf01515) EUT1
05144 -# -
05145 Hp_euts_violetPink EUT1
05146 HMM10662PduV_1_mix(pf10662) EUT1
05147 HMM06249EutQ_1_eut1(pf06249) EUT1 EP_NTERM
05148 HMM01923cobtr_2_eut1(pf01923) EUT1
05149 HMM01515PTA_B_1_eut1(pf01515) EUT1
05150 H_azure EUT1
05151 P_lightViolet EUT1
05152 HMMalddh_1_eutpdugrm(pf00171) EUT1 EP_NTERM
05153 HMM06723EutJMreB_1_grm(#) EUT1
05154 HMM00465alcdh_3_mix(pf00465) EUT1
05155 -# -
05156 HMMtpEuth_1_eut(pf04346) EUT1
05157 HMM06277EutA_1_eut(pf06277) EUT1
05158 HMMetlyEutB_1_EUT1(pf06751) EUT1
05159 HMMetlyEutC_1_eut1(pf05985) EUT1 EP_NTERM
05160 Tsp_rosa EUT1
05161 H_mulberry(pf00936pf16365) EUT1
05162 HMMreg12833_1_EUT1(pf12833) EUT1
05163 -pf01218 -
05164 -pf01520 -
05165 -pf00583 -
05166 -pf11143 -
05167 -pf06572 -
05168 -pf04261 -
05169 -pf00144 -
05170 -pf00367pf02378 -
05171 -pf01380 -
BMC-P
BMC-H
BMC-T
Aldehyde dehydrogenase
PTAC
Signature enzyme
Alcohol dehydrogenase
Regulator

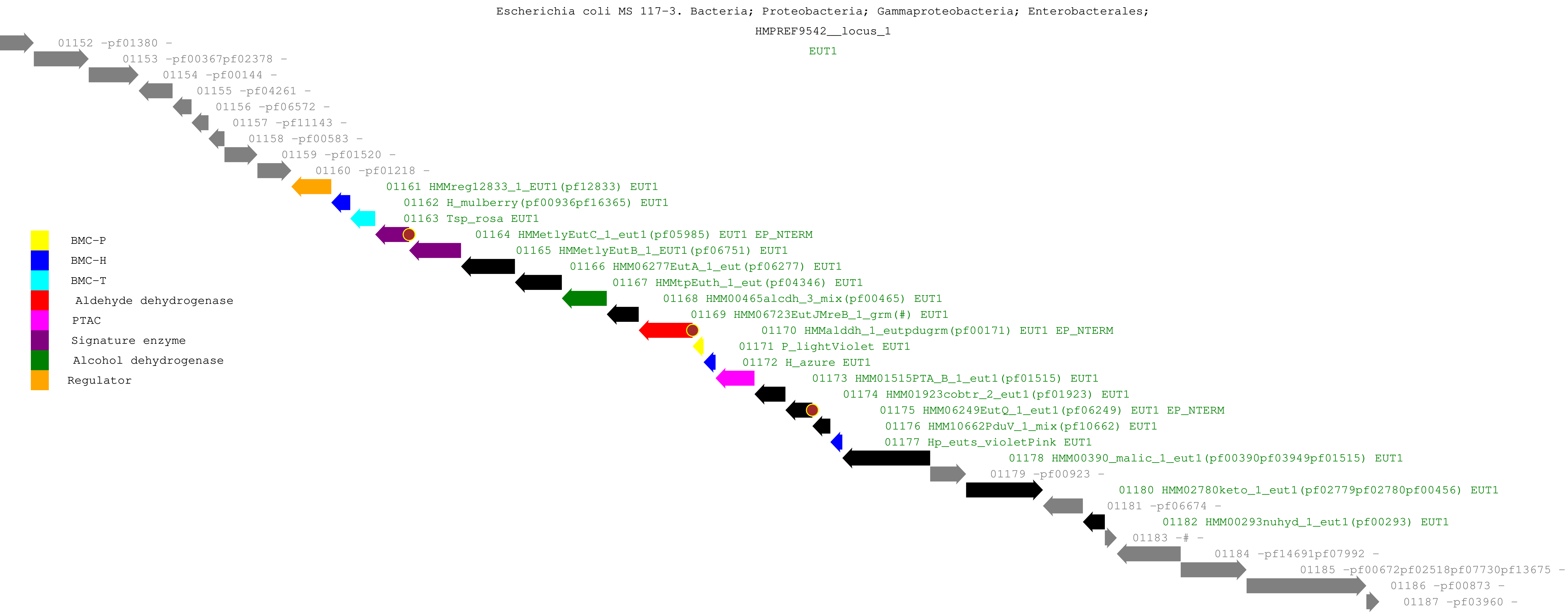
Escherichia coli MS 117-3. Bacteria; Proteobacteria; Gammaproteobacteria; Enterobacterales;
HMPREF9542__locus_1
EUT1
01152 -pf01380 -
01153 -pf00367pf02378 -
01154 -pf00144 -
01155 -pf04261 -
01156 -pf06572 -
01157 -pf11143 -
01158 -pf00583 -
01159 -pf01520 -
01160 -pf01218 -
01161 HMMreg12833_1_EUT1(pf12833) EUT1
01162 H_mulberry(pf00936pf16365) EUT1
01163 Tsp_rosa EUT1
01164 HMMetlyEutC_1_eut1(pf05985) EUT1 EP_NTERM
01165 HMMetlyEutB_1_EUT1(pf06751) EUT1
01166 HMM06277EutA_1_eut(pf06277) EUT1
01167 HMMtpEuth_1_eut(pf04346) EUT1
01168 HMM00465alcdh_3_mix(pf00465) EUT1
01169 HMM06723EutJMreB_1_grm(#) EUT1
01170 HMMalddh_1_eutpdugrm(pf00171) EUT1 EP_NTERM
01171 P_lightViolet EUT1
01172 H_azure EUT1
01173 HMM01515PTA_B_1_eut1(pf01515) EUT1
01174 HMM01923cobtr_2_eut1(pf01923) EUT1
01175 HMM06249EutQ_1_eut1(pf06249) EUT1 EP_NTERM
01176 HMM10662PduV_1_mix(pf10662) EUT1
01177 Hp_euts_violetPink EUT1
01178 HMM00390_malic_1_eut1(pf00390pf03949pf01515) EUT1
01179 -pf00923 -
01180 HMM02780keto_1_eut1(pf02779pf02780pf00456) EUT1
01181 -pf06674 -
01182 HMM00293nuhyd_1_eut1(pf00293) EUT1
01183 -# -
01184 -pf14691pf07992 -
01185 -pf00672pf02518pf07730pf13675 -
01186 -pf00873 -
01187 -pf03960 -
BMC-P
BMC-H
BMC-T
Aldehyde dehydrogenase
PTAC
Signature enzyme
Alcohol dehydrogenase
Regulator

Escherichia coli MS 146-1. Bacteria; Proteobacteria; Gammaproteobacteria; Enterobacterales;

HMPREF9543__locus_1_main_sat

EUT1

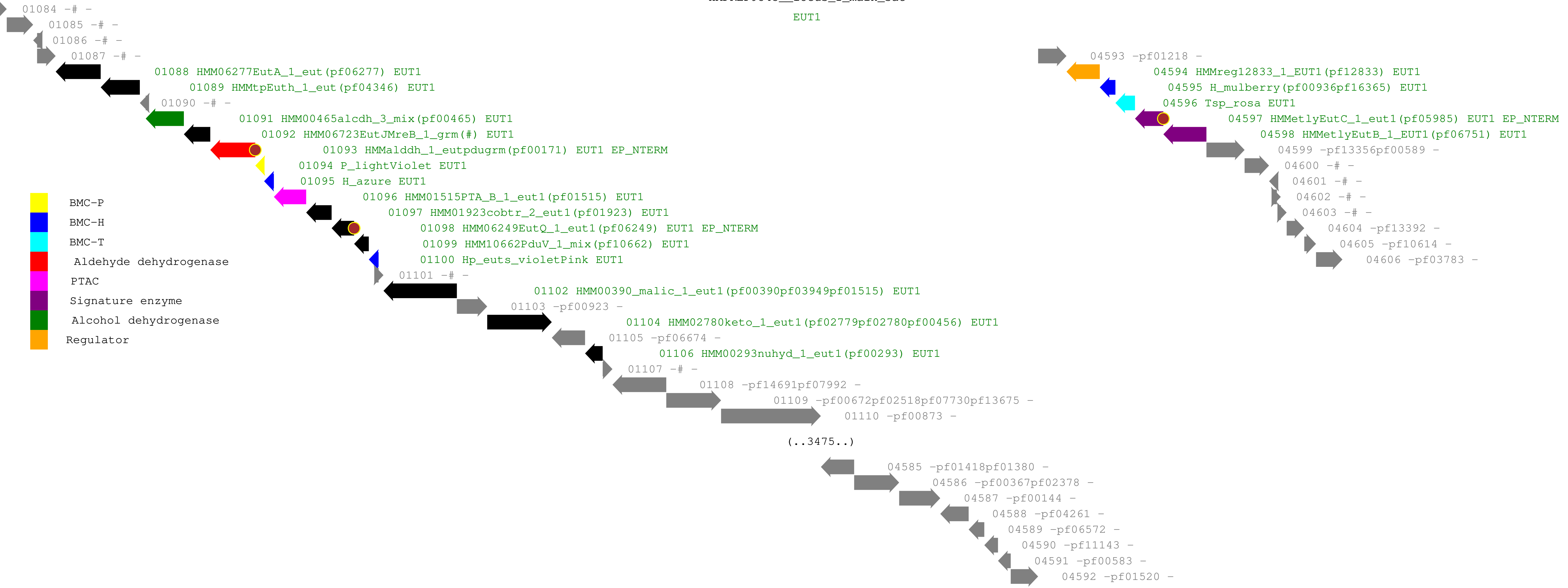

Escherichia coli MS 175-1. Bacteria; Proteobacteria; Gammaproteobacteria; Enterobacterales;

HMPREF9547__locus_1

EUT1

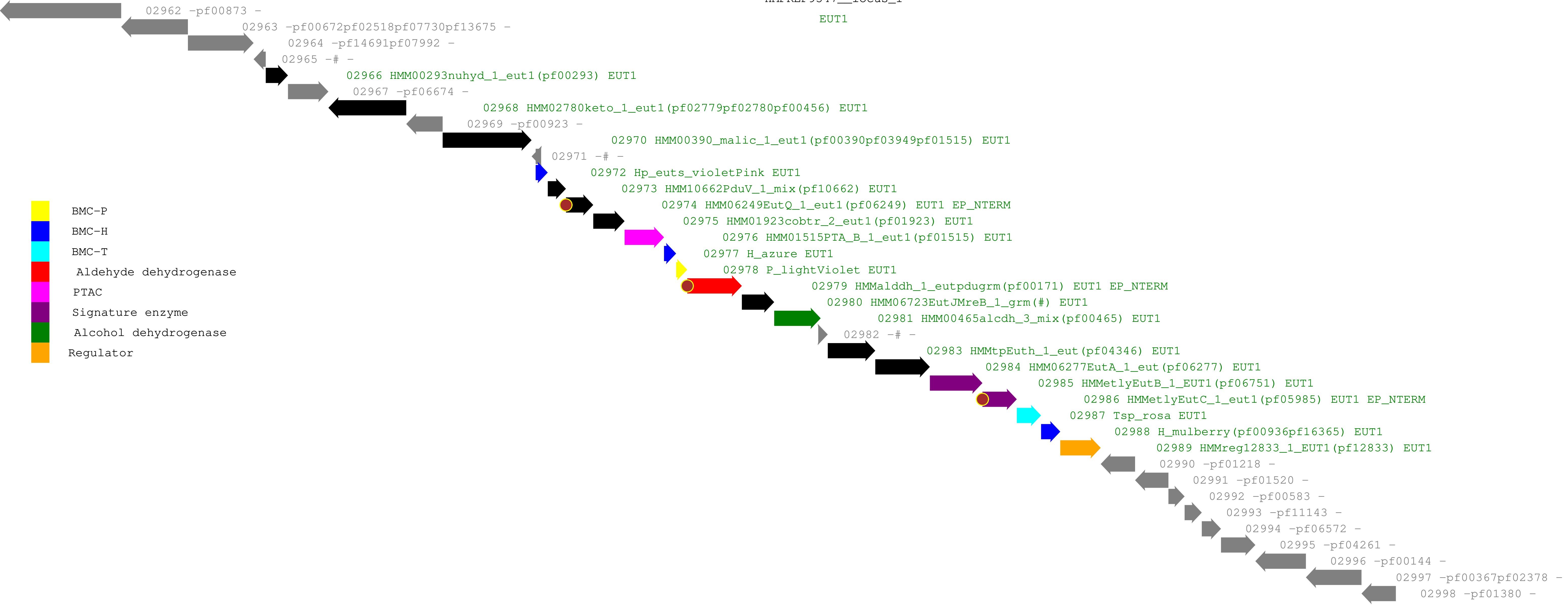

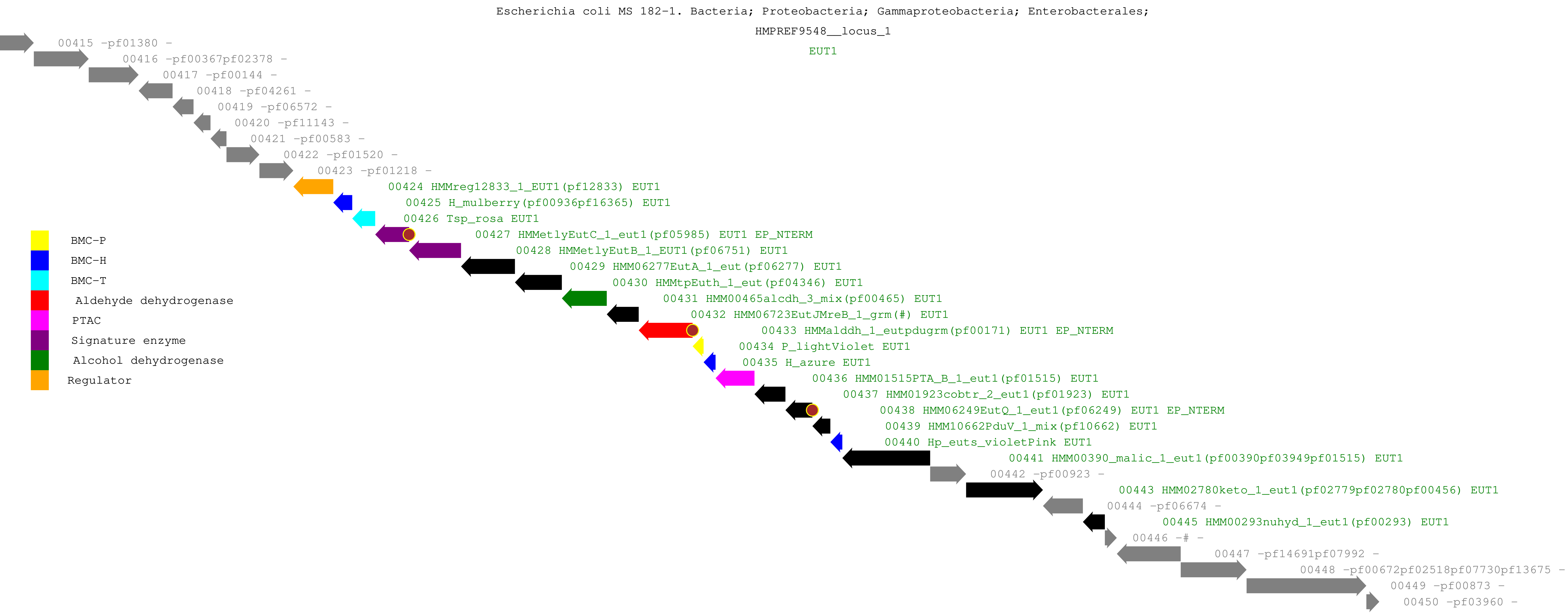
Escherichia coli MS 182-1. Bacteria; Proteobacteria; Gammaproteobacteria; Enterobacterales;
HMPREF9548__locus_1
EUT1
00415 -pf01380 -
00416 -pf00367pf02378 -
00417 -pf00144 -
00418 -pf04261 -
00419 -pf06572 -
00420 -pf11143 -
00421 -pf00583 -
00422 -pf01520 -
00423 -pf01218 -
00424 HMMreg12833_1_EUT1(pf12833) EUT1
00425 H_mulberry(pf00936pf16365) EUT1
00426 Tsp_rosa EUT1
00427 HMMetlyEutC_1_eut1(pf05985) EUT1 EP_NTERM
00428 HMMetlyEutB_1_EUT1(pf06751) EUT1
00429 HMM06277EutA_1_eut(pf06277) EUT1
00430 HMMtpEuth_1_eut(pf04346) EUT1
00431 HMM00465alcdh_3_mix(pf00465) EUT1
00432 HMM06723EutJMreB_1_grm(#) EUT1
00433 HMMalddh_1_eutpdugrm(pf00171) EUT1 EP_NTERM
00434 P_lightViolet EUT1
00435 H_azure EUT1
00436 HMM01515PTA_B_1_eut1(pf01515) EUT1
00437 HMM01923cobtr_2_eut1(pf01923) EUT1
00438 HMM06249EutQ_1_eut1(pf06249) EUT1 EP_NTERM
00439 HMM10662PduV_1_mix(pf10662) EUT1
00440 Hp_euts_violetPink EUT1
00441 HMM00390_malic_1_eut1(pf00390pf03949pf01515) EUT1
00442 -pf00923 -
00443 HMM02780keto_1_eut1(pf02779pf02780pf00456) EUT1
00444 -pf06674 -
00445 HMM00293nuhyd_1_eut1(pf00293) EUT1
00446 -# -
00447 -pf14691pf07992 -
00448 -pf00672pf02518pf07730pf13675 -
00449 -pf00873 -
00450 -pf03960 -
BMC-P
BMC-H
BMC-T
Aldehyde dehydrogenase
PTAC
Signature enzyme
Alcohol dehydrogenase
Regulator

Escherichia coli MS 196-1. Bacteria; Proteobacteria; Gammaproteobacteria; Enterobacterales;

HMPREF9551__locus_1

EUT1

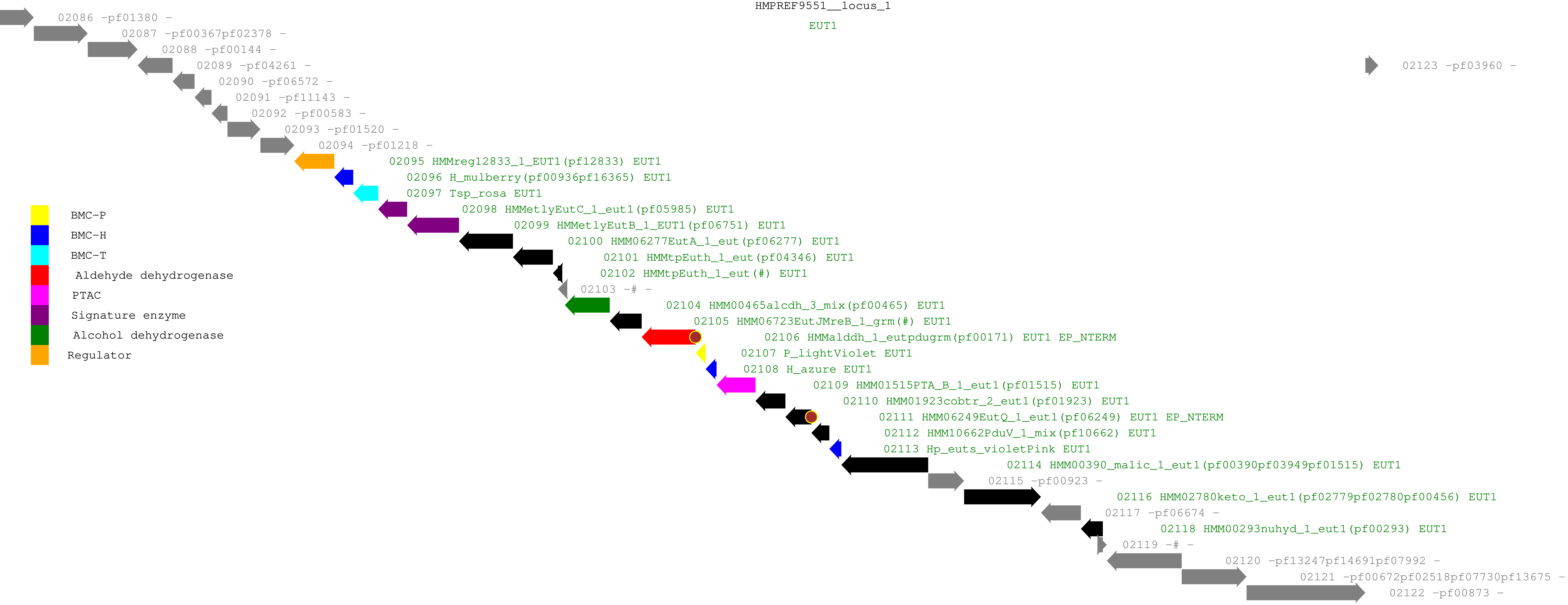

Escherichia coli MS 198-1. Bacteria; Proteobacteria; Gammaproteobacteria; Enterobacterales;

HMPREF9552__locus_1_main_sat

EUT1

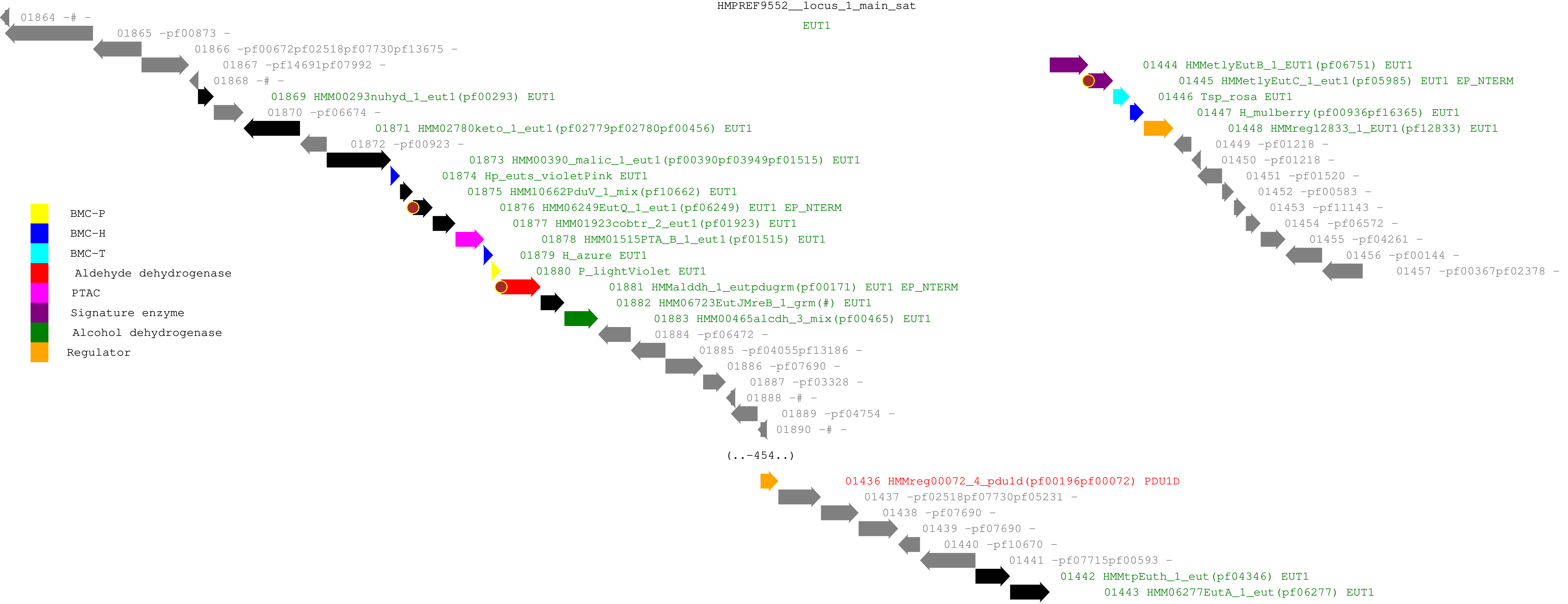

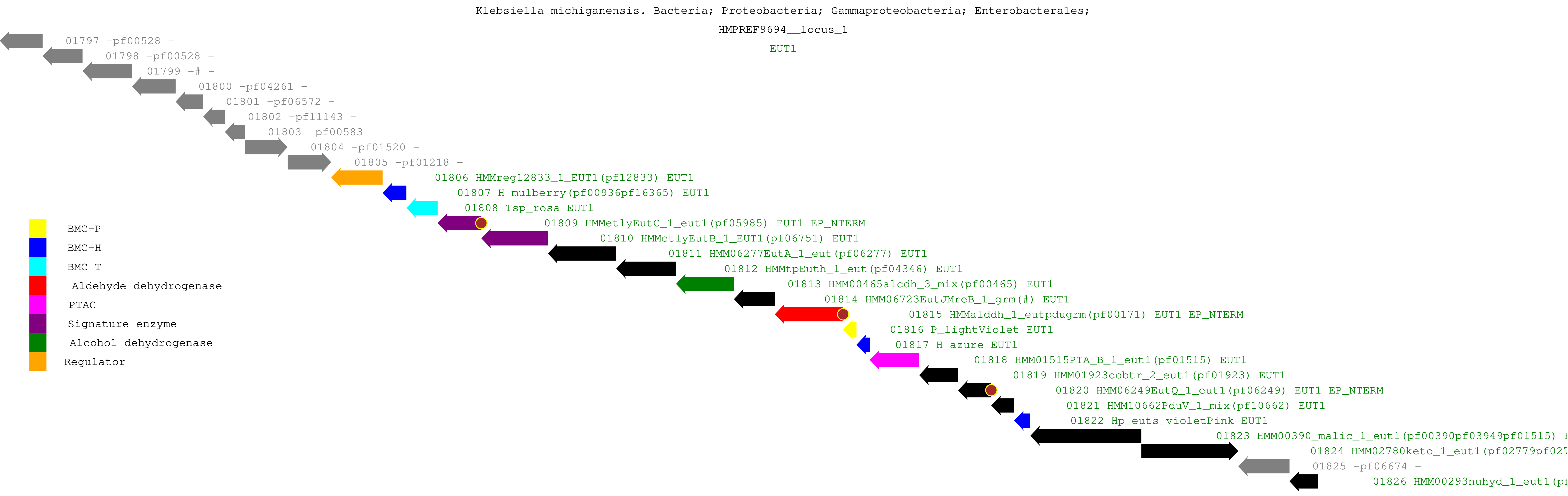
Klebsiella michiganensis. Bacteria; Proteobacteria; Gammaproteobacteria; Enterobacterales;
HMPREF9694__locus_1
EUT1
01797 -pf00528 -
01798 -pf00528 -
01799 -# -
01800 -pf04261 -
01801 -pf06572 -
01802 -pf11143 -
01803 -pf00583 -
01804 -pf01520 -
01805 -pf01218 -
01806 HMMreg12833_1_EUT1(pf12833) EUT1
01807 H_mulberry(pf00936pf16365) EUT1
01808 Tsp_rosa EUT1
01809 HMMetlyEutC_1_eut1(pf05985) EUT1 EP_NTERM
01810 HMMetlyEutB_1_EUT1(pf06751) EUT1
01811 HMM06277EutA_1_eut(pf06277) EUT1
01812 HMMtpEuth_1_eut(pf04346) EUT1
01813 HMM00465alcdh_3_mix(pf00465) EUT1
01814 HMM06723EutJMreB_1_grm(#) EUT1
01815 HMMalddh_1_eutpdugrm(pf00171) EUT1 EP_NTERM
01816 P_lightViolet EUT1
01817 H_azure EUT1
01818 HMM01515PTA_B_1_eut1(pf01515) EUT1
01819 HMM01923cobtr_2_eut1(pf01923) EUT1
01820 HMM06249EutQ_1_eut1(pf06249) EUT1 EP_NTERM
01821 HMM10662PduV_1_mix(pf10662) EUT1
01822 Hp_euts_violetPink EUT1
01823 HMM00390_malic_1_eut1(pf00390pf03949pf01515)
01824 HMM02780keto_1_eut1(pf02779pf027
01825 -pf06674 -
01826 HMM00293nuhyd_1_eut1(p
BMC-P
BMC-H
BMC-T
Aldehyde dehydrogenase
PTAC
Signature enzyme
Alcohol dehydrogenase
Regulator

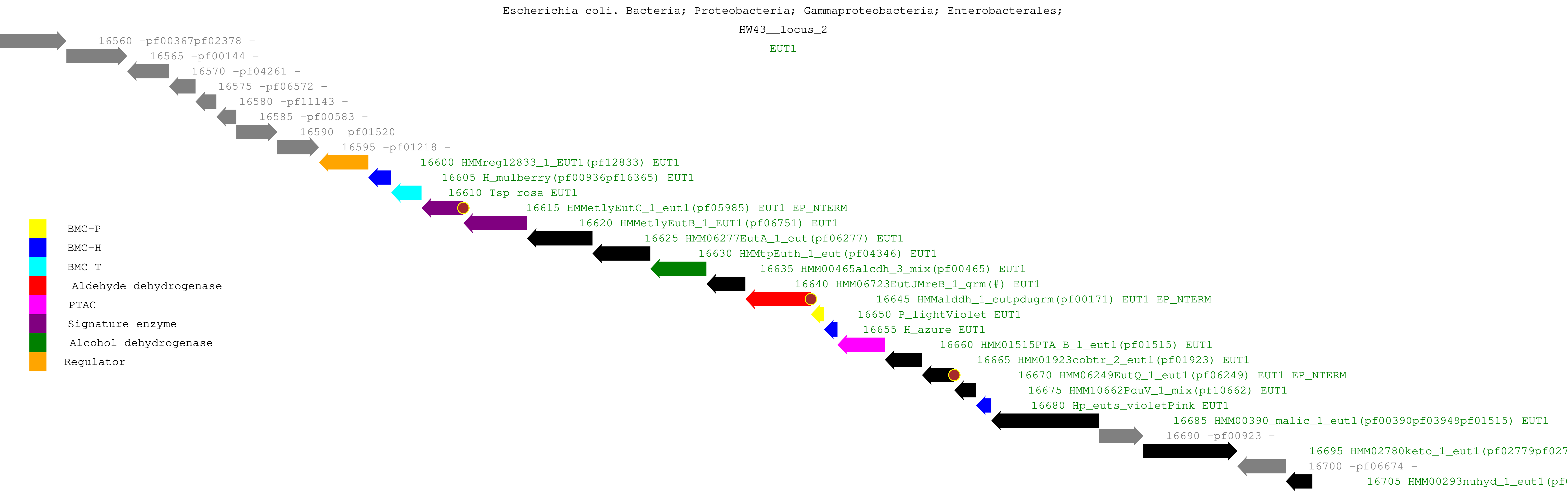
Escherichia coli. Bacteria; Proteobacteria; Gammaproteobacteria; Enterobacterales;
HW43__locus_2
EUT1
16560 -pf00367pf02378 -
16565 -pf00144 -
16570 -pf04261 -
16575 -pf06572 -
16580 -pf11143 -
16585 -pf00583 -
16590 -pf01520 -
16595 -pf01218 -
16600 HMMreg12833_1_EUT1(pf12833) EUT1
16605 H_mulberry(pf00936pf16365) EUT1
16610 Tsp_rosa EUT1
16615 HMMetlyEutC_1_eut1(pf05985) EUT1 EP_NTERM
16620 HMMetlyEutB_1_EUT1(pf06751) EUT1
16625 HMM06277EutA_1_eut(pf06277) EUT1
16630 HMMtpEuth_1_eut(pf04346) EUT1
16635 HMM00465alcdh_3_mix(pf00465) EUT1
16640 HMM06723EutJMreB_1_grm(#) EUT1
16645 HMMalddh_1_eutpdugrm(pf00171) EUT1 EP_NTERM
16650 P_lightViolet EUT1
16655 H_azure EUT1
16660 HMM01515PTA_B_1_eut1(pf01515) EUT1
16665 HMM01923cobtr_2_eut1(pf01923) EUT1
16670 HMM06249EutQ_1_eut1(pf06249) EUT1 EP_NTERM
16675 HMM10662PduV_1_mix(pf10662) EUT1
16680 Hp_euts_violetPink EUT1
16685 HMM00390_malic_1_eut1(pf00390pf03949pf01515) EUT1
16690 -pf00923 -
16695 HMM02780keto_1_eut1(pf02779pf027
16700 -pf06674 -
16705 HMM00293nuhyd_1_eut1(pf
BMC-P
BMC-H
BMC-T
Aldehyde dehydrogenase
PTAC
Signature enzyme
Alcohol dehydrogenase
Regulator

Escherichia coli O26 str. RM10386. Bacteria; Proteobacteria; Gammaproteobacteria; Enterobacterales;

I3U__locus_1

EUT1

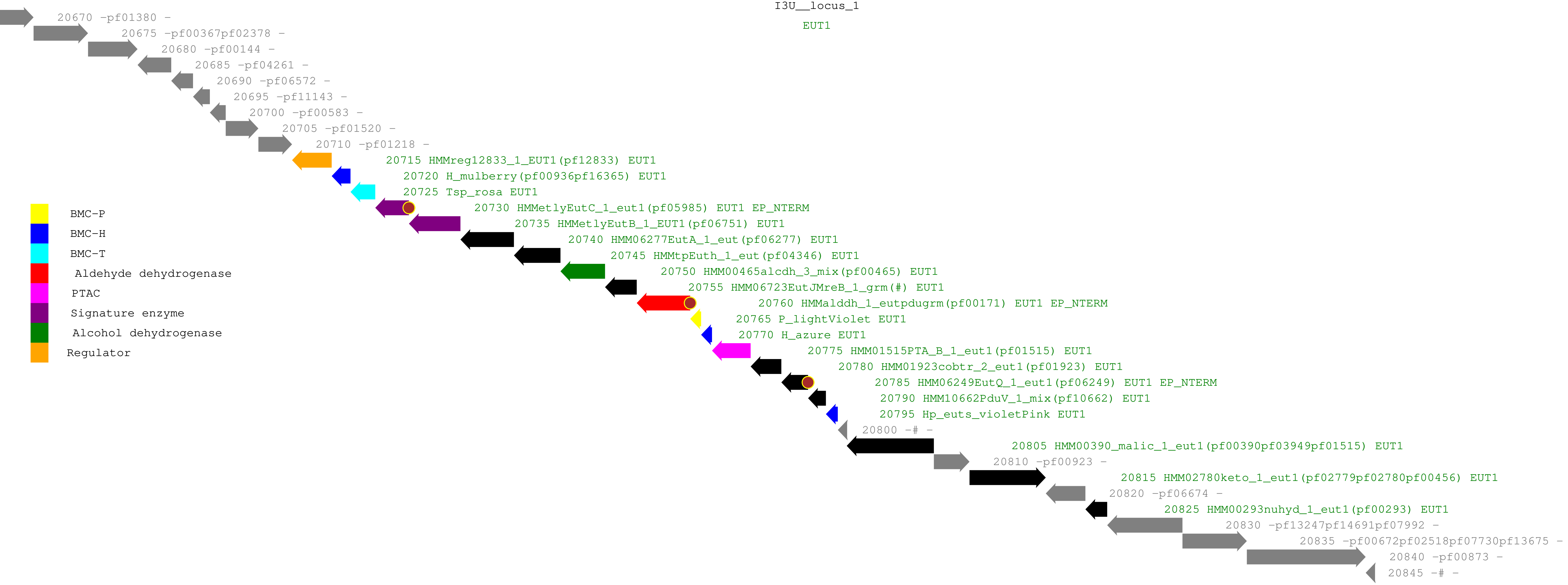

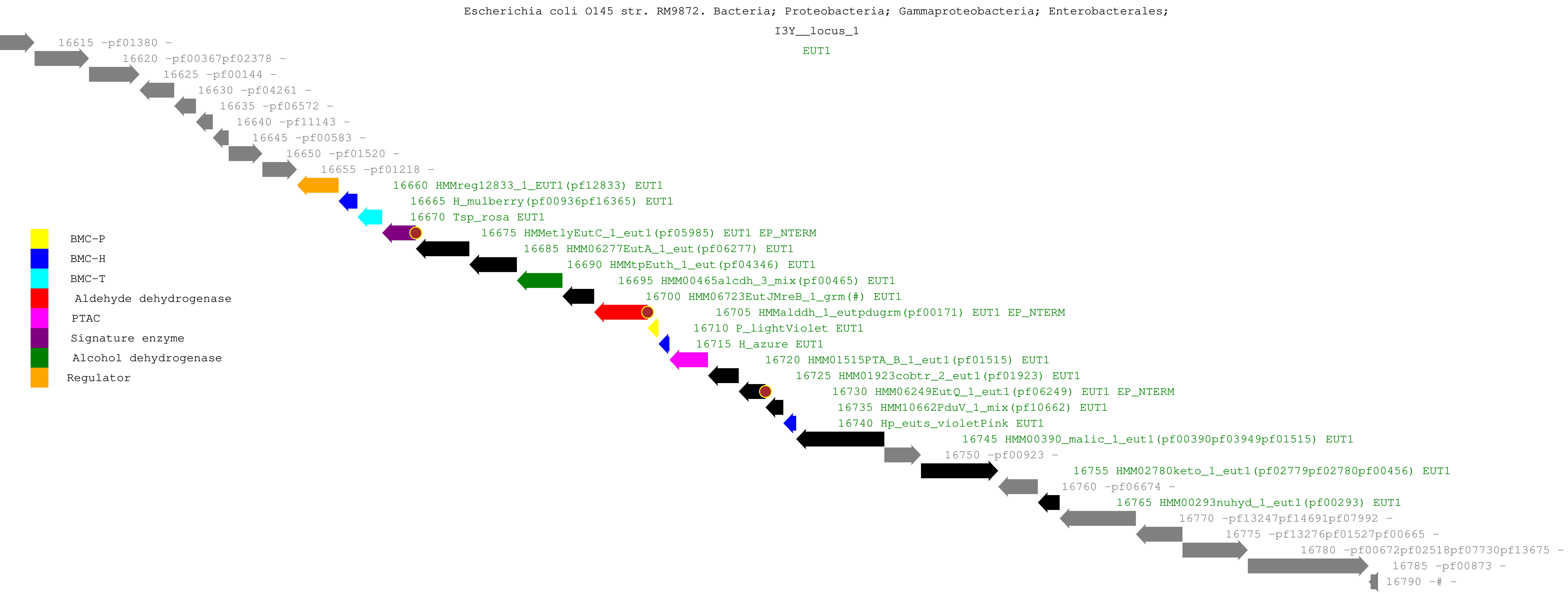

Escherichia coli O145 str. RM9872. Bacteria; Proteobacteria; Gammaproteobacteria; Enterobacterales;
I3Y__locus_1
EUT1
16615 -pf01380 -
16620 -pf00367pf02378 -
16625 -pf00144 -
16630 -pf04261 -
16635 -pf06572 -
16640 -pf11143 -
16645 -pf00583 -
16650 -pf01520 -
16655 -pf01218 -
16660 HMMreg12833_1_EUT1(pf12833) EUT1
16665 H_mulberry(pf00936pf16365) EUT1
16670 Tsp_rosa EUT1
16675 HMMetlyEutC_1_eut1(pf05985) EUT1 EP_NTERM
16685 HMM06277EutA_1_eut(pf06277) EUT1
16690 HMMtpEuth_1_eut(pf04346) EUT1
16695 HMM00465alcdh_3_mix(pf00465) EUT1
16700 HMM06723EutJMreB_1_grm(#) EUT1
16705 HMMalddh_1_eutpdugrm(pf00171) EUT1 EP_NTERM
16710 P_lightViolet EUT1
16715 H_azure EUT1
16720 HMM01515PTA_B_1_eut1(pf01515) EUT1
16725 HMM01923cobtr_2_eut1(pf01923) EUT1
16730 HMM06249EutQ_1_eut1(pf06249) EUT1 EP_NTERM
16735 HMM10662PduV_1_mix(pf10662) EUT1
16740 Hp_euts_violetPink EUT1
16745 HMM00390_malic_1_eut1(pf00390pf03949pf01515) EUT1
16750 -pf00923 -
16755 HMM02780keto_1_eut1(pf02779pf02780pf00456) EUT1
16760 -pf06674 -
16765 HMM00293nuhyd_1_eut1(pf00293) EUT1
16770 -pf13247pf14691pf07992 -
16775 -pf13276pf01527pf00665 -
16780 -pf00672pf02518pf07730pf13675 -
16785 -pf00873 -
16790 -# -
BMC-P
BMC-H
BMC-T
Aldehyde dehydrogenase
PTAC
Signature enzyme
Alcohol dehydrogenase
Regulator

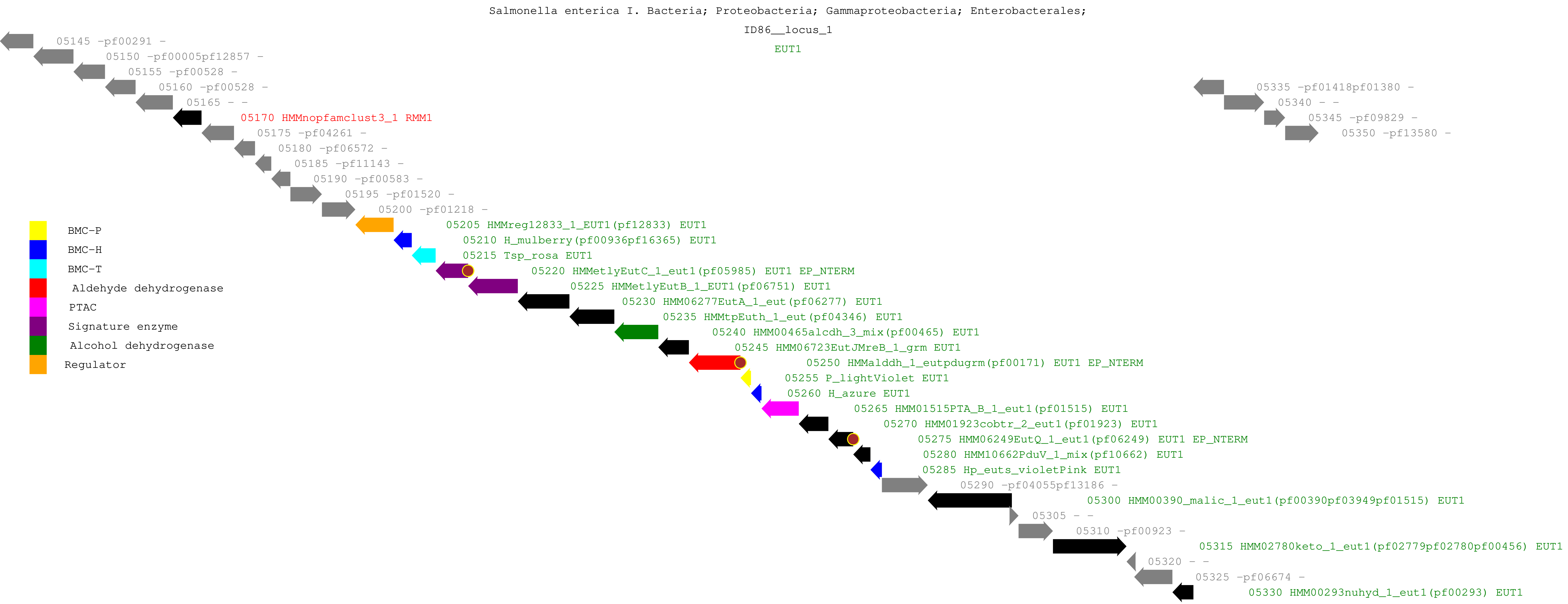

Salmonella enterica I. Bacteria; Proteobacteria; Gammaproteobacteria; Enterobacterales;
ID86__locus_1
EUT1
05145 -pf00291 -
05150 -pf00005pf12857 -
05155 -pf00528 -
05160 -pf00528 -
05165 - -
05170 HMMnopfamclust3_1 RMM1
05175 -pf04261 -
05180 -pf06572 -
05185 -pf11143 -
05190 -pf00583 -
05195 -pf01520 -
05200 -pf01218 -
05205 HMMreg12833_1_EUT1(pf12833) EUT1
05210 H_mulberry(pf00936pf16365) EUT1
05215 Tsp_rosa EUT1
05220 HMMetlyEutC_1_eut1(pf05985) EUT1 EP_NTERM
05225 HMMetlyEutB_1_EUT1(pf06751) EUT1
05230 HMM06277EutA_1_eut(pf06277) EUT1
05235 HMMtpEuth_1_eut(pf04346) EUT1
05240 HMM00465alcdh_3_mix(pf00465) EUT1
05245 HMM06723EutJMreB_1_grm EUT1
05250 HMMalddh_1_eutpdugrm(pf00171) EUT1 EP_NTERM
05255 P_lightViolet EUT1
05260 H_azure EUT1
05265 HMM01515PTA_B_1_eut1(pf01515) EUT1
05270 HMM01923cobtr_2_eut1(pf01923) EUT1
05275 HMM06249EutQ_1_eut1(pf06249) EUT1 EP_NTERM
05280 HMM10662PduV_1_mix(pf10662) EUT1
05285 Hp_euts_violetPink EUT1
05290 -pf04055pf13186 -
05300 HMM00390_malic_1_eut1(pf00390pf03949pf01515) EUT1
05305 - -
05310 -pf00923 -
05315 HMM02780keto_1_eut1(pf02779pf02780pf00456) EUT1
05320 - -
05325 -pf06674 -
05330 HMM00293nuhyd_1_eut1(pf00293) EUT1
05335 -pf01418pf01380 -
05340 - -
05345 -pf09829 -
05350 -pf13580 -
BMC-P
BMC-H
BMC-T
Aldehyde dehydrogenase
PTAC
Signature enzyme
Alcohol dehydrogenase
Regulator

Escherichia coli (strain K12). Bacteria; Proteobacteria; Gammaproteobacteria; Enterobacterales;

JW_locus_1

EUT1

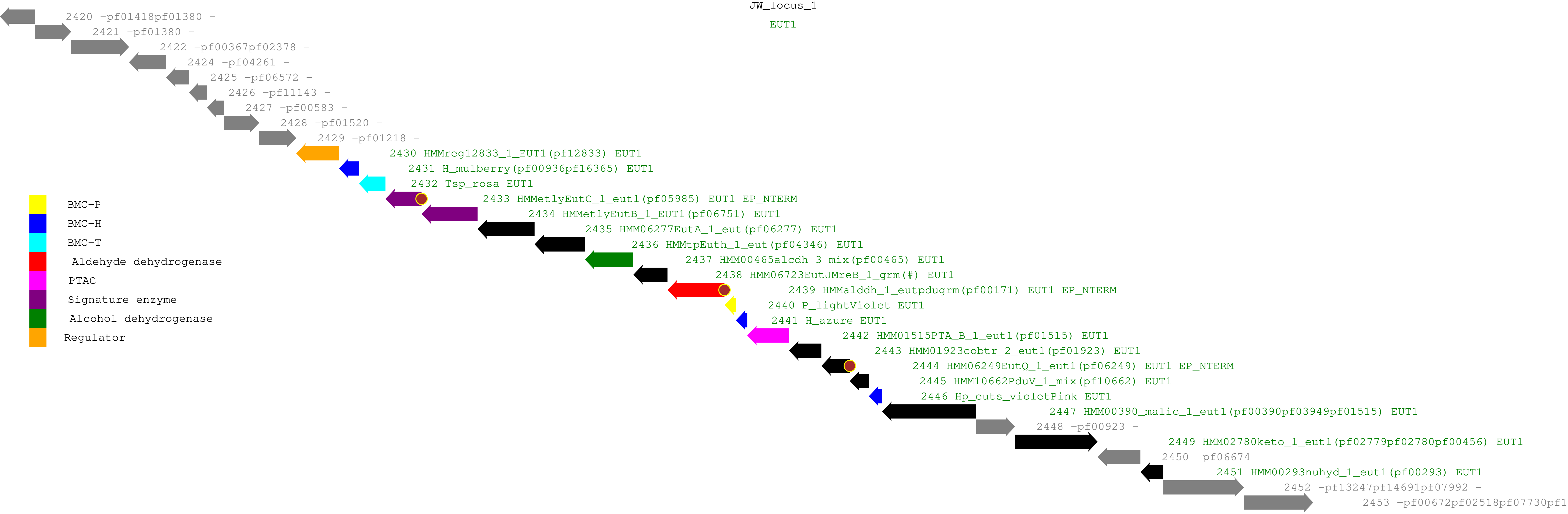

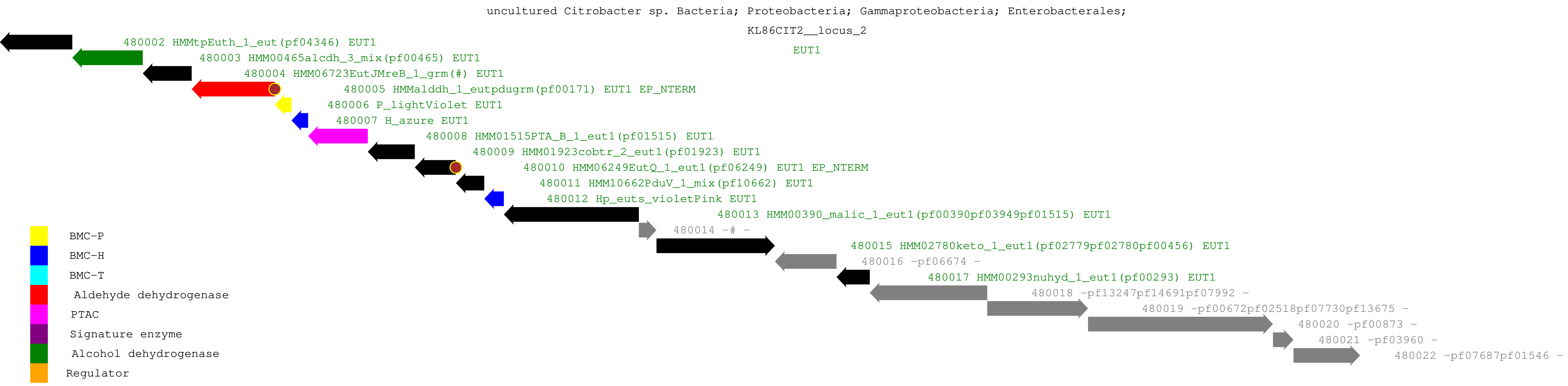
uncultured Citrobacter sp. Bacteria; Proteobacteria; Gammaproteobacteria; Enterobacterales;
KL86CIT2__locus_2
EUT1
480002 HMMtpEuth_1_eut(pf04346) EUT1
480003 HMM00465alcdh_3_mix(pf00465) EUT1
480004 HMM06723EutJMreB_1_grm(#) EUT1
480005 HMMalddh_1_eutpdugrm(pf00171) EUT1 EP_NTERM
480006 P_lightViolet EUT1
480007 H_azure EUT1
480008 HMM01515PTA_B_1_eut1(pf01515) EUT1
480009 HMM01923cobtr_2_eut1(pf01923) EUT1
480010 HMM06249EutQ_1_eut1(pf06249) EUT1 EP_NTERM
480011 HMM10662PduV_1_mix(pf10662) EUT1
480012 Hp_euts_violetPink EUT1
480013 HMM00390_malic_1_eut1(pf00390pf03949pf01515) EUT1
480014 -# -
480015 HMM02780keto_1_eut1(pf02779pf02780pf00456) EUT1
480016 -pf06674 -
480017 HMM00293nuhyd_1_eut1(pf00293) EUT1
480018 -pf13247pf14691pf07992 -
480019 -pf00672pf02518pf07730pf13675 -
480020 -pf00873 -
480021 -pf03960 -
480022 -pf07687pf01546 -
BMC-P
BMC-H
BMC-T
Aldehyde dehydrogenase
PTAC
Signature enzyme
Alcohol dehydrogenase
Regulator

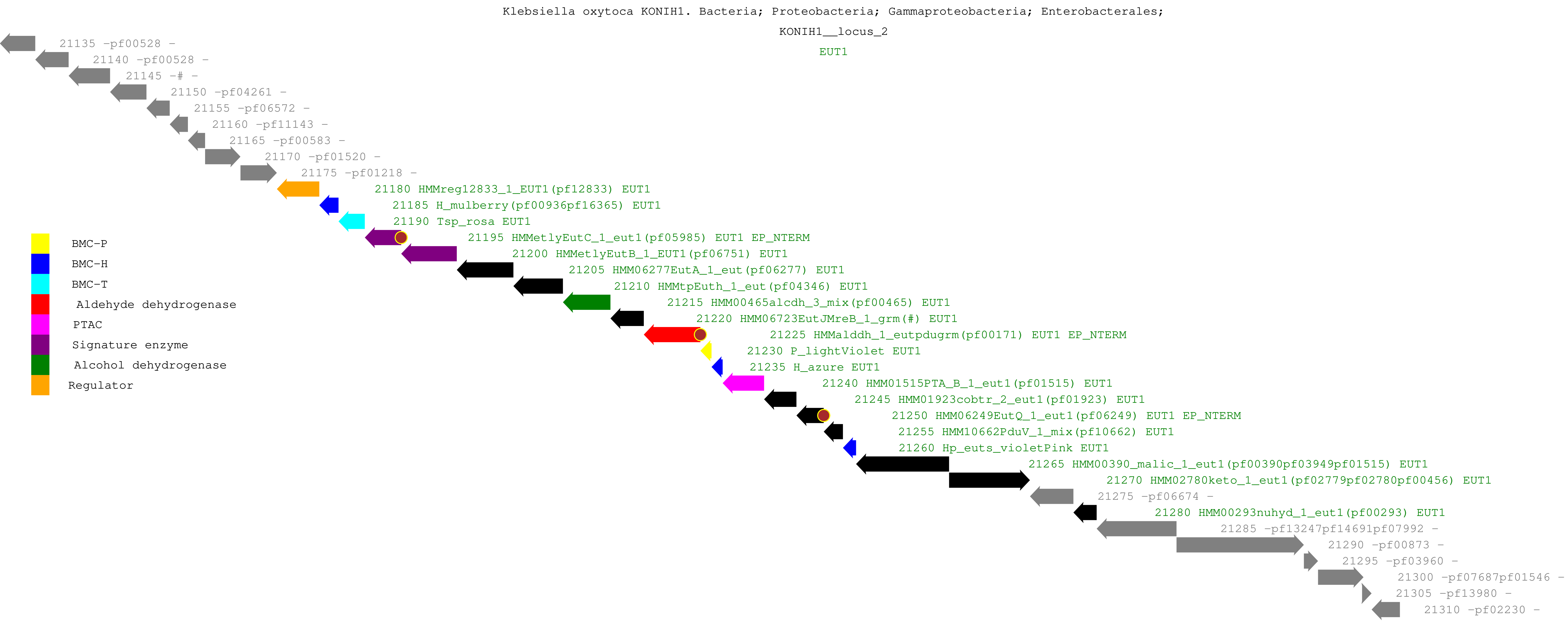
Klebsiella oxytoca KONIH1. Bacteria; Proteobacteria; Gammaproteobacteria; Enterobacterales;
KONIH1__locus_2
EUT1
21135 -pf00528 -
21140 -pf00528 -
21145 -# -
21150 -pf04261 -
21155 -pf06572 -
21160 -pf11143 -
21165 -pf00583 -
21170 -pf01520 -
21175 -pf01218 -
21180 HMMreg12833_1_EUT1(pf12833) EUT1
21185 H_mulberry(pf00936pf16365) EUT1
21190 Tsp_rosa EUT1
21195 HMMetlyEutC_1_eut1(pf05985) EUT1 EP_NTERM
21200 HMMetlyEutB_1_EUT1(pf06751) EUT1
21205 HMM06277EutA_1_eut(pf06277) EUT1
21210 HMMtpEuth_1_eut(pf04346) EUT1
21215 HMM00465alcdh_3_mix(pf00465) EUT1
21220 HMM06723EutJMreB_1_grm(#) EUT1
21225 HMMalddh_1_eutpdugrm(pf00171) EUT1 EP_NTERM
21230 P_lightViolet EUT1
21235 H_azure EUT1
21240 HMM01515PTA_B_1_eut1(pf01515) EUT1
21245 HMM01923cobtr_2_eut1(pf01923) EUT1
21250 HMM06249EutQ_1_eut1(pf06249) EUT1 EP_NTERM
21255 HMM10662PduV_1_mix(pf10662) EUT1
21260 Hp_euts_violetPink EUT1
21265 HMM00390_malic_1_eut1(pf00390pf03949pf01515) EUT1
21270 HMM02780keto_1_eut1(pf02779pf02780pf00456) EUT1
21275 -pf06674 -
21280 HMM00293nuhyd_1_eut1(pf00293) EUT1
21285 -pf13247pf14691pf07992 -
21290 -pf00873 -
21295 -pf03960 -
21300 -pf07687pf01546 -
21305 -pf13980 -
21310 -pf02230 -
BMC-P
BMC-H
BMC-T
Aldehyde dehydrogenase
PTAC
Signature enzyme
Alcohol dehydrogenase
Regulator

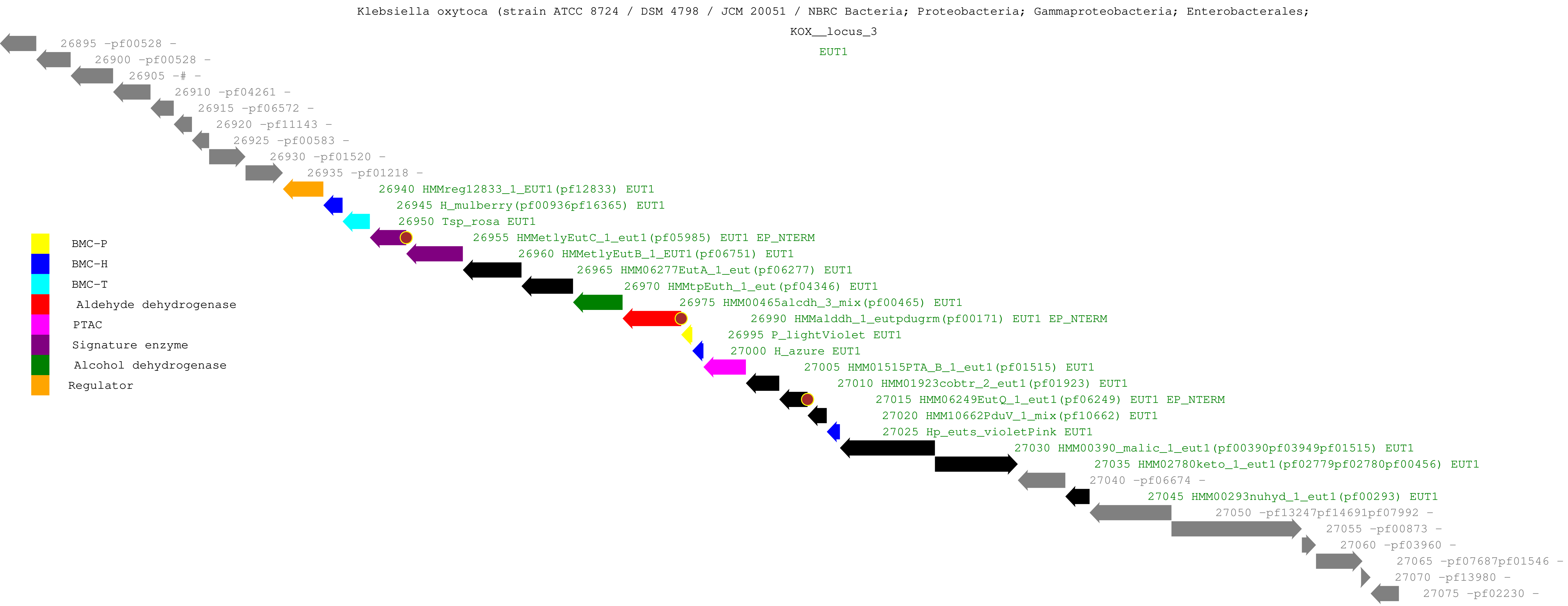
Klebsiella oxytoca (strain ATCC 8724 / DSM 4798 / JCM 20051 / NBRC Bacteria; Proteobacteria; Gammaproteobacteria; Enterobacterales;
KOX__locus_3
EUT1
26895 -pf00528 -
26900 -pf00528 -
26905 -# -
26910 -pf04261 -
26915 -pf06572 -
26920 -pf11143 -
26925 -pf00583 -
26930 -pf01520 -
26935 -pf01218 -
26940 HMMreg12833_1_EUT1(pf12833) EUT1
26945 H_mulberry(pf00936pf16365) EUT1
26950 Tsp_rosa EUT1
26955 HMMetlyEutC_1_eut1(pf05985) EUT1 EP_NTERM
26960 HMMetlyEutB_1_EUT1(pf06751) EUT1
26965 HMM06277EutA_1_eut(pf06277) EUT1
26970 HMMtpEuth_1_eut(pf04346) EUT1
26975 HMM00465alcdh_3_mix(pf00465) EUT1
26990 HMMalddh_1_eutpdugrm(pf00171) EUT1 EP_NTERM
26995 P_lightViolet EUT1
27000 H_azure EUT1
27005 HMM01515PTA_B_1_eut1(pf01515) EUT1
27010 HMM01923cobtr_2_eut1(pf01923) EUT1
27015 HMM06249EutQ_1_eut1(pf06249) EUT1 EP_NTERM
27020 HMM10662PduV_1_mix(pf10662) EUT1
27025 Hp_euts_violetPink EUT1
27030 HMM00390_malic_1_eut1(pf00390pf03949pf01515) EUT1
27035 HMM02780keto_1_eut1(pf02779pf02780pf00456) EUT1
27040 -pf06674 -
27045 HMM00293nuhyd_1_eut1(pf00293) EUT1
27050 -pf13247pf14691pf07992 -
27055 -pf00873 -
27060 -pf03960 -
27065 -pf07687pf01546 -
27070 -pf13980 -
27075 -pf02230 -
BMC-P
BMC-H
BMC-T
Aldehyde dehydrogenase
PTAC
Signature enzyme
Alcohol dehydrogenase
Regulator

Klebsiella pneumoniae subsp. pneumoniae (strain HS11286). Bacteria; Proteobacteria; Gammaproteobacteria; Enterobacterales;

KPHS__locus_1

EUT1

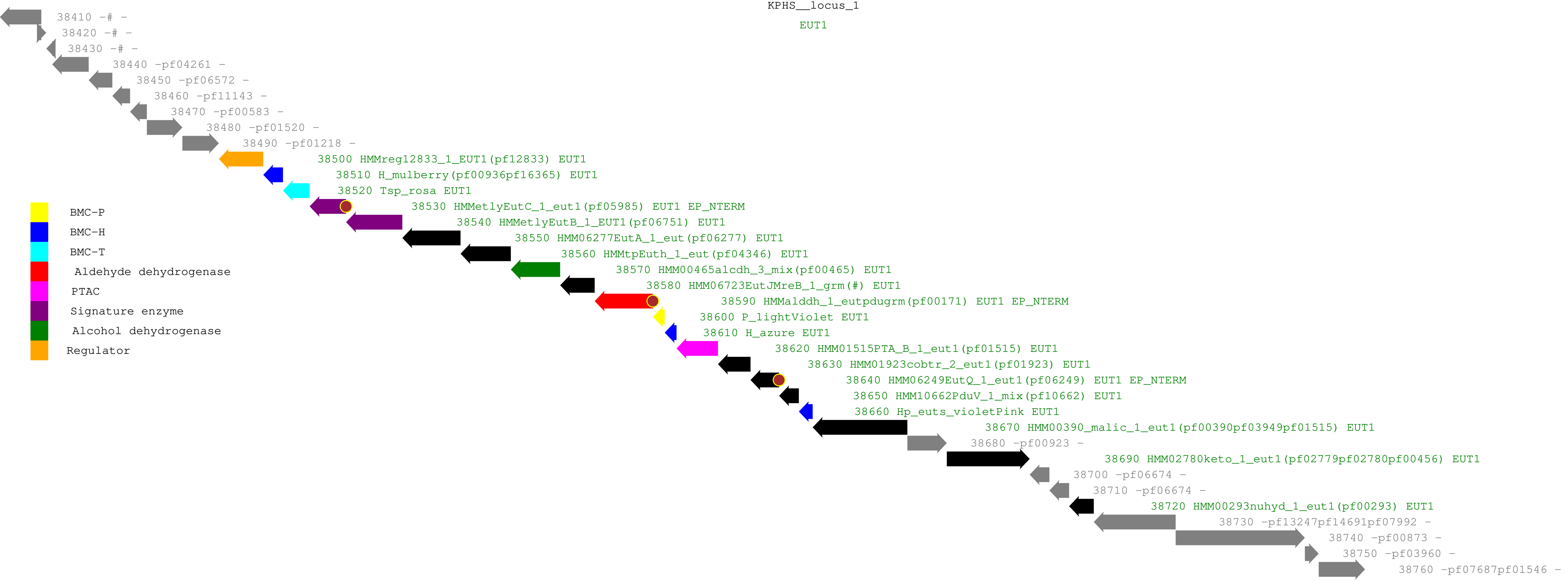

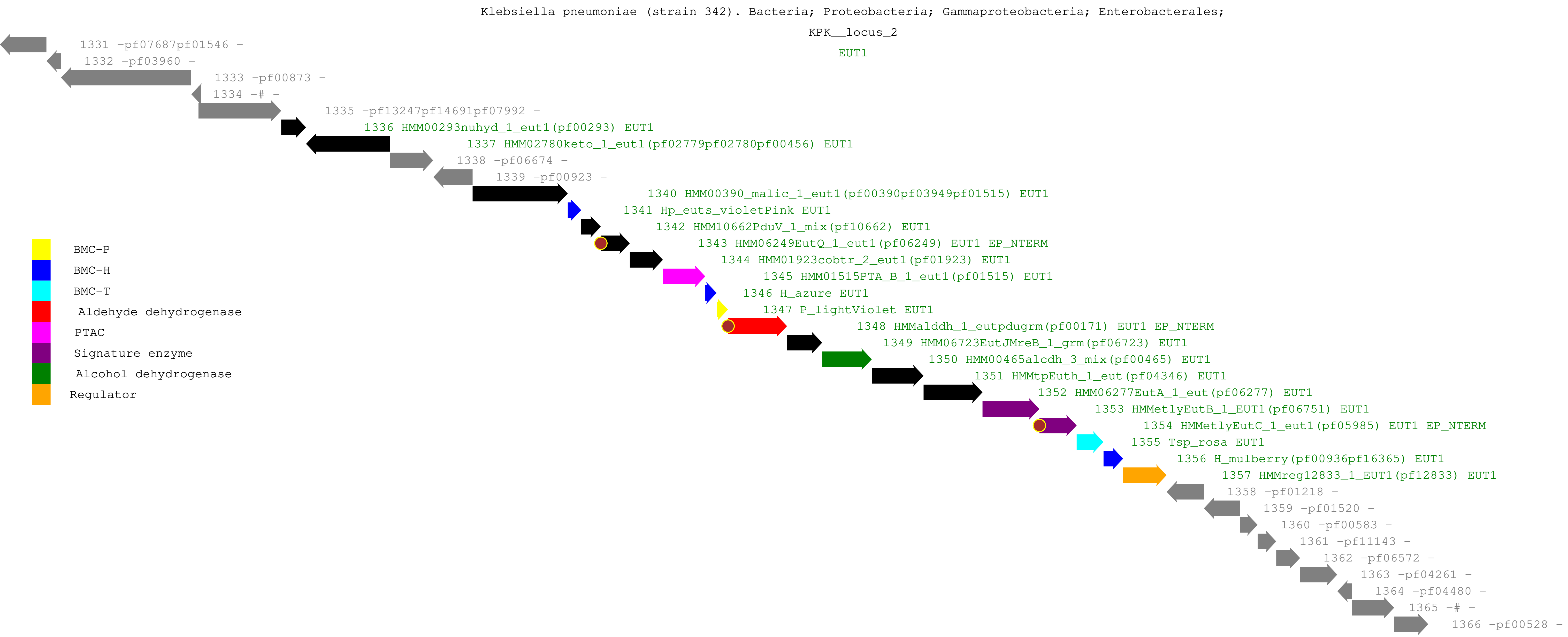
Klebsiella pneumoniae (strain 342). Bacteria; Proteobacteria; Gammaproteobacteria; Enterobacterales;
KPK__locus_2
EUT1
1331 -pf07687pf01546 -
1332 -pf03960 -
1333 -pf00873 -
1334 -# -
1335 -pf13247pf14691pf07992 -
1336 HMM00293nuhyd_1_eut1(pf00293) EUT1
1337 HMM02780keto_1_eut1(pf02779pf02780pf00456) EUT1
1338 -pf06674 -
1339 -pf00923 -
1340 HMM00390_malic_1_eut1(pf00390pf03949pf01515) EUT1
1341 Hp_euts_violetPink EUT1
1342 HMM10662PduV_1_mix(pf10662) EUT1
1343 HMM06249EutQ_1_eut1(pf06249) EUT1 EP_NTERM
1344 HMM01923cobtr_2_eut1(pf01923) EUT1
1345 HMM01515PTA_B_1_eut1(pf01515) EUT1
1346 H_azure EUT1
1347 P_lightViolet EUT1
1348 HMMalddh_1_eutpdugrm(pf00171) EUT1 EP_NTERM
1349 HMM06723EutJMreB_1_grm(pf06723) EUT1
1350 HMM00465alcdh_3_mix(pf00465) EUT1
1351 HMMtpEuth_1_eut(pf04346) EUT1
1352 HMM06277EutA_1_eut(pf06277) EUT1
1353 HMMetlyEutB_1_EUT1(pf06751) EUT1
1354 HMMetlyEutC_1_eut1(pf05985) EUT1 EP_NTERM
1355 Tsp_rosa EUT1
1356 H_mulberry(pf00936pf16365) EUT1
1357 HMMreg12833_1_EUT1(pf12833) EUT1
1358 -pf01218 -
1359 -pf01520 -
1360 -pf00583 -
1361 -pf11143 -
1362 -pf06572 -
1363 -pf04261 -
1364 -pf04480 -
1365 -# -
1366 -pf00528 -
BMC-P
BMC-H
BMC-T
Aldehyde dehydrogenase
PTAC
Signature enzyme
Alcohol dehydrogenase
Regulator

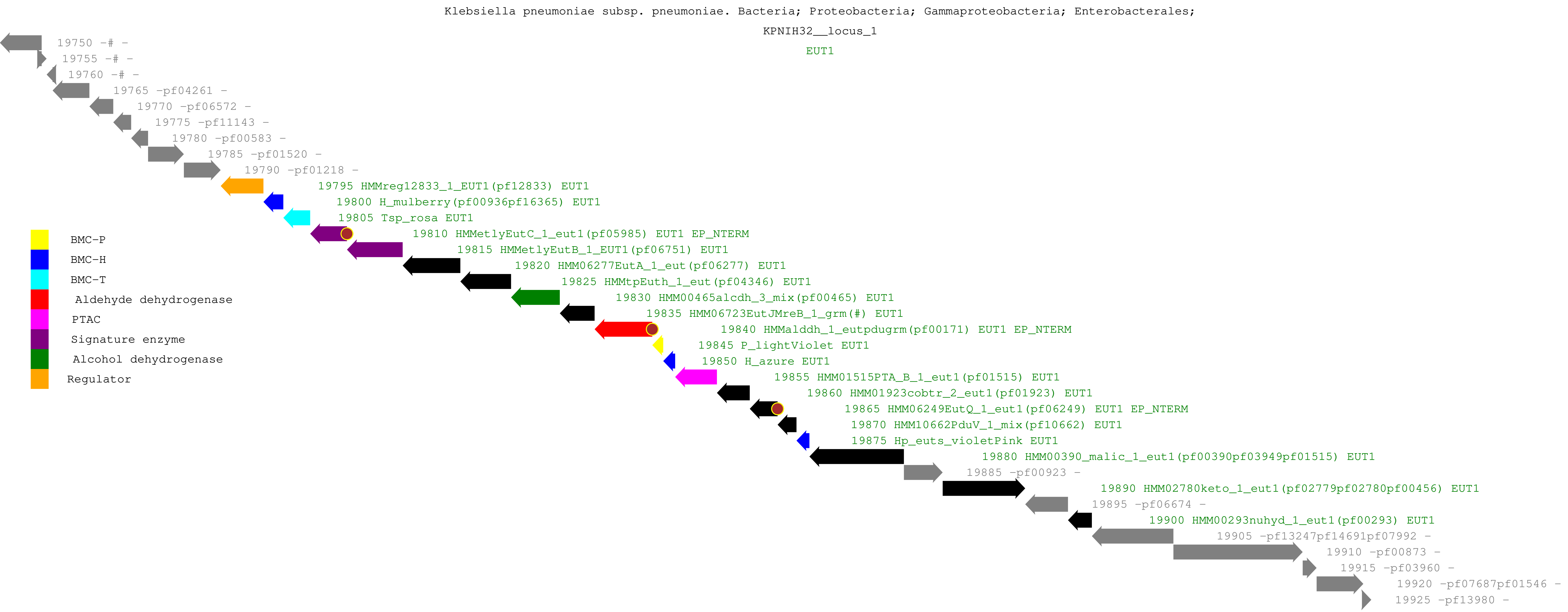
Klebsiella pneumoniae subsp. pneumoniae. Bacteria; Proteobacteria; Gammaproteobacteria; Enterobacterales;
KPNIH32__locus_1
EUT1
19750 -# -
19755 -# -
19760 -# -
19765 -pf04261 -
19770 -pf06572 -
19775 -pf11143 -
19780 -pf00583 -
19785 -pf01520 -
19790 -pf01218 -
19795 HMMreg12833_1_EUT1(pf12833) EUT1
19800 H_mulberry(pf00936pf16365) EUT1
19805 Tsp_rosa EUT1
19810 HMMetlyEutC_1_eut1(pf05985) EUT1 EP_NTERM
19815 HMMetlyEutB_1_EUT1(pf06751) EUT1
19820 HMM06277EutA_1_eut(pf06277) EUT1
19825 HMMtpEuth_1_eut(pf04346) EUT1
19830 HMM00465alcdh_3_mix(pf00465) EUT1
19835 HMM06723EutJMreB_1_grm(#) EUT1
19840 HMMalddh_1_eutpdugrm(pf00171) EUT1 EP_NTERM
19845 P_lightViolet EUT1
19850 H_azure EUT1
19855 HMM01515PTA_B_1_eut1(pf01515) EUT1
19860 HMM01923cobtr_2_eut1(pf01923) EUT1
19865 HMM06249EutQ_1_eut1(pf06249) EUT1 EP_NTERM
19870 HMM10662PduV_1_mix(pf10662) EUT1
19875 Hp_euts_violetPink EUT1
19880 HMM00390_malic_1_eut1(pf00390pf03949pf01515) EUT1
19885 -pf00923 -
19890 HMM02780keto_1_eut1(pf02779pf02780pf00456) EUT1
19895 -pf06674 -
19900 HMM00293nuhyd_1_eut1(pf00293) EUT1
19905 -pf13247pf14691pf07992 -
19910 -pf00873 -
19915 -pf03960 -
19920 -pf07687pf01546 -
19925 -pf13980 -
BMC-P
BMC-H
BMC-T
Aldehyde dehydrogenase
PTAC
Signature enzyme
Alcohol dehydrogenase
Regulator

Klebsiella pneumoniae 30660/NJST258_1. Bacteria; Proteobacteria; Gammaproteobacteria; Enterobacterales;

KPNJ1__locus_2

EUT1

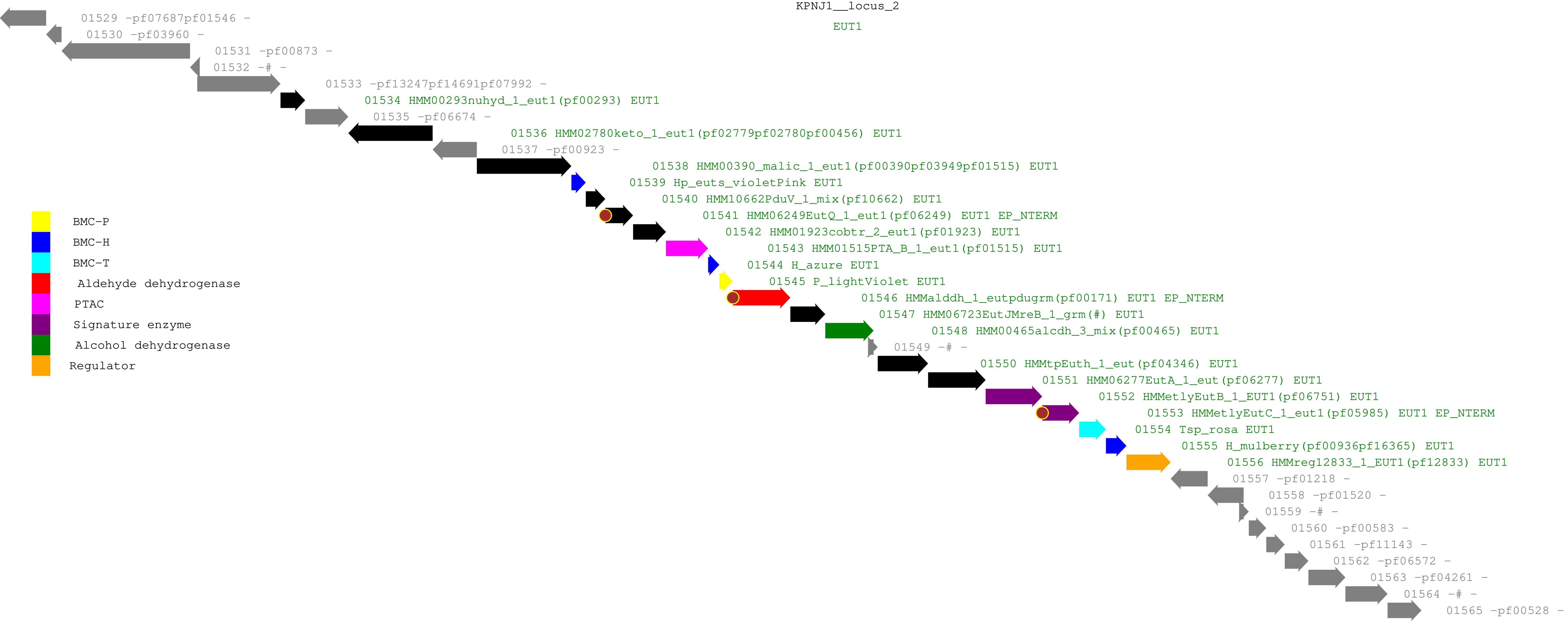

Klebsiella pneumoniae 30684/NJST258_2. Bacteria; Proteobacteria; Gammaproteobacteria; Enterobacterales;

KPNJ2__locus_2

EUT1

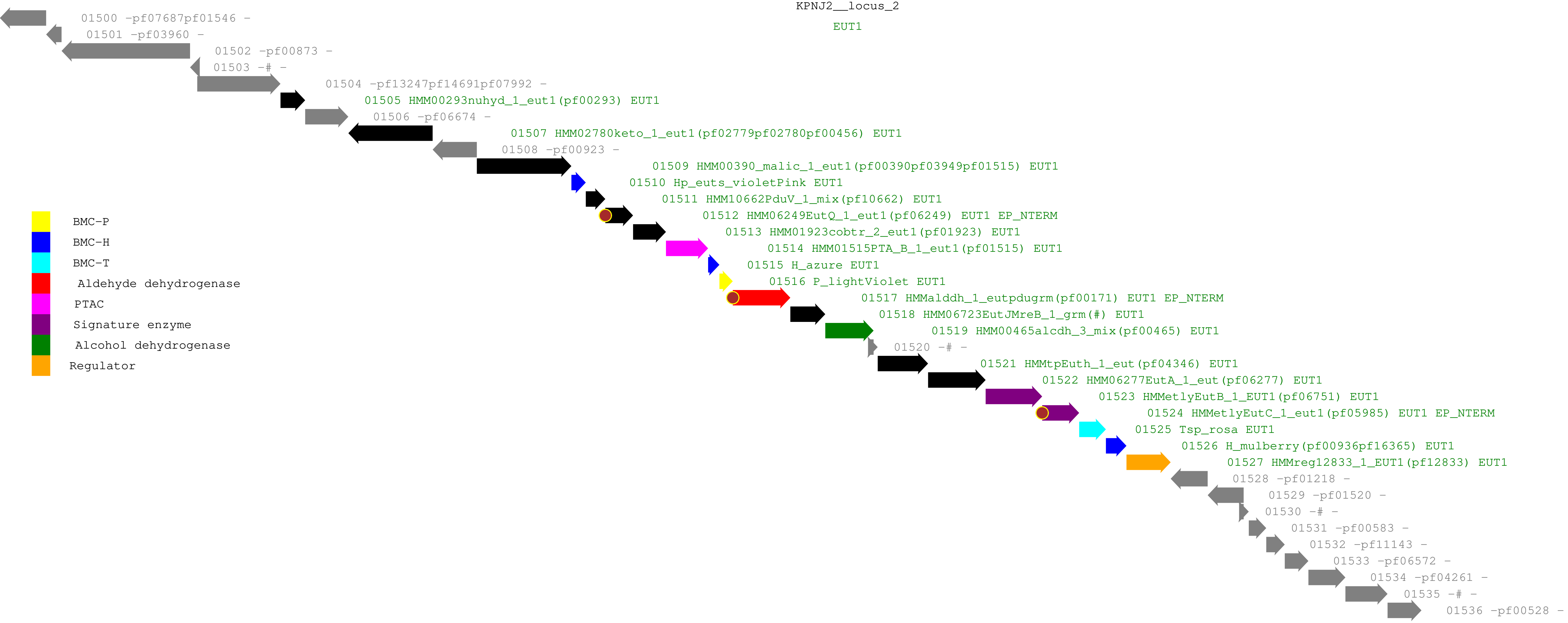

Klebsiella pneumoniae subsp. pneumoniae (strain ATCC 700721 / MGH Bacteria; Proteobacteria; Gammaproteobacteria; Enterobacterales;

KPN__locus_1

EUT1

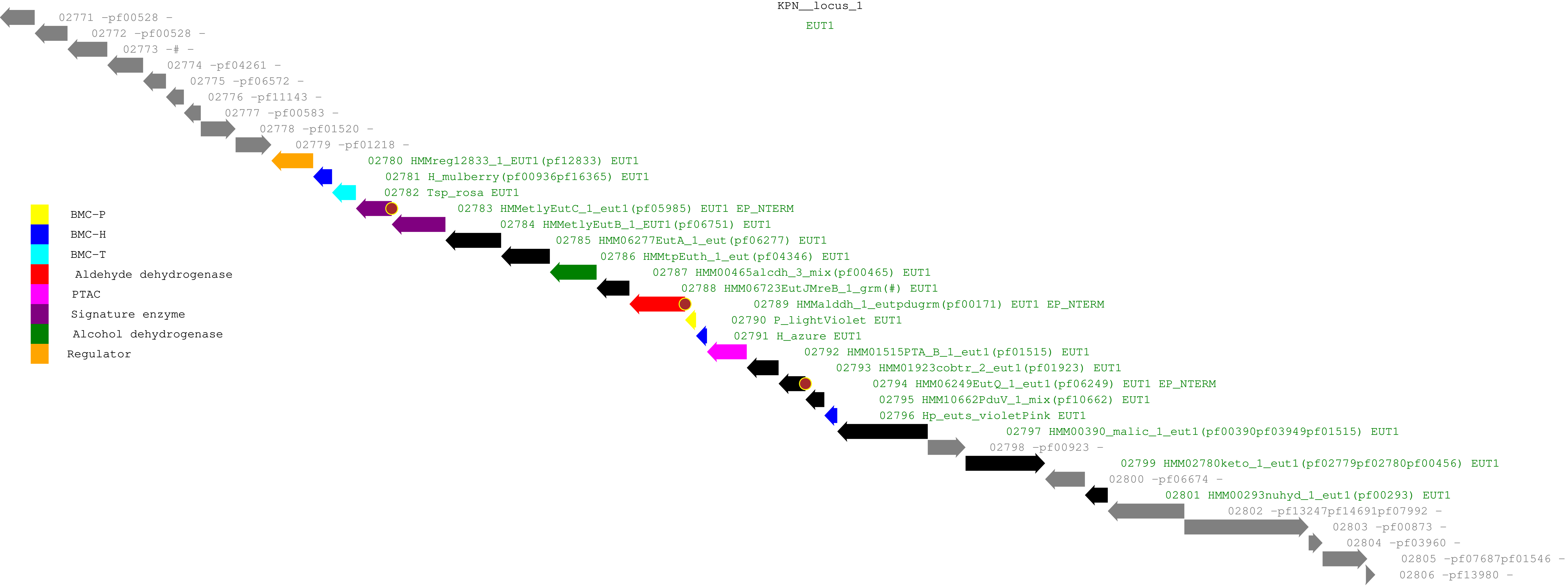

Klebsiella pneumoniae subsp. rhinoscleromatis SB3432. Bacteria; Proteobacteria; Gammaproteobacteria; Enterobacterales;

KPR__locus_3

EUT1

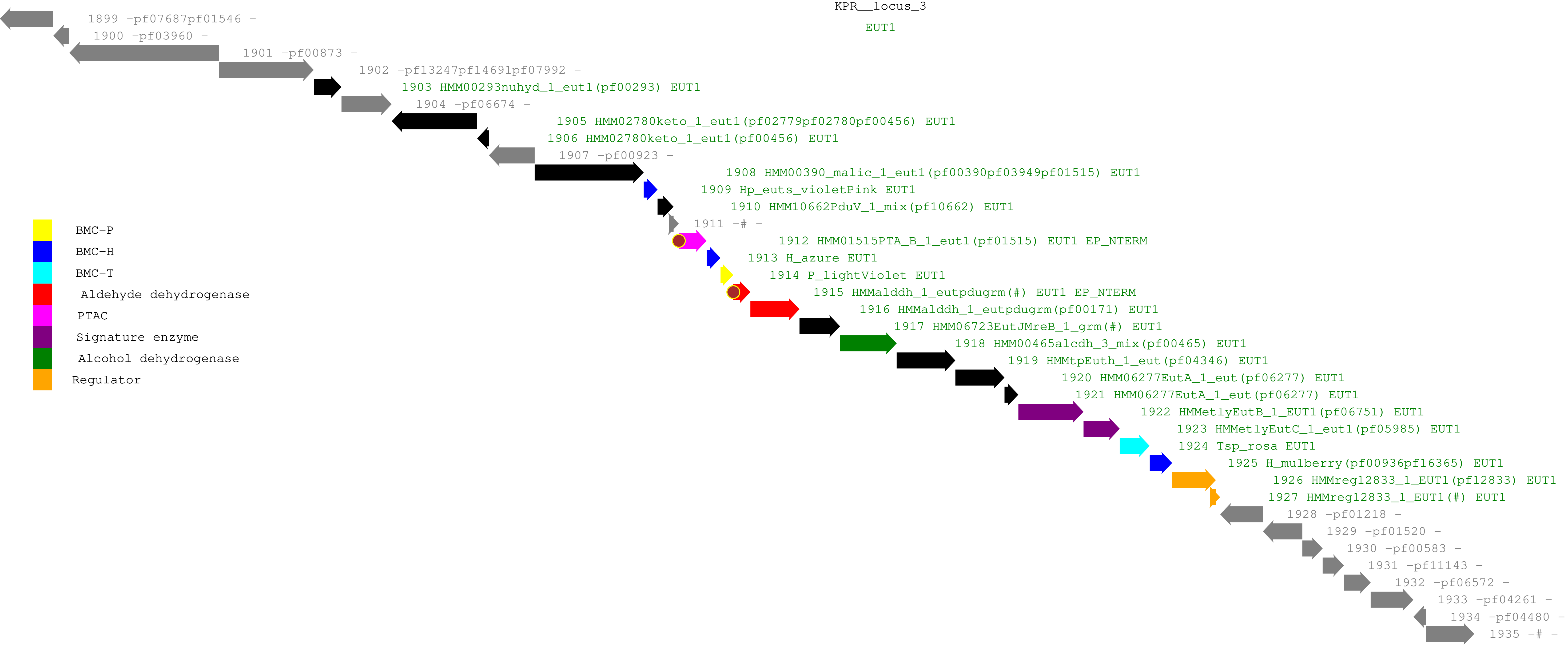

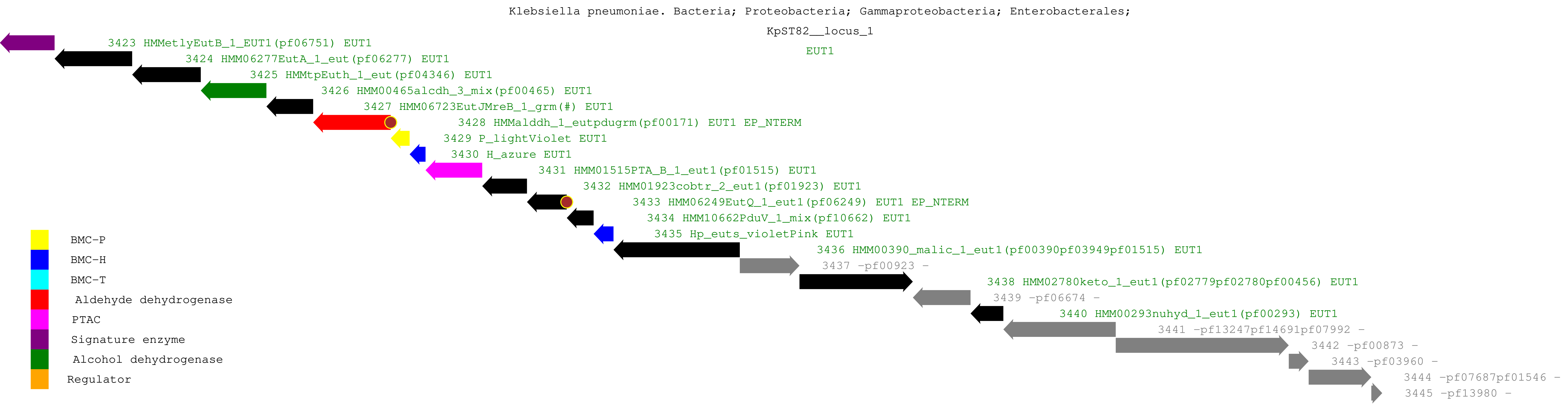
Klebsiella pneumoniae. Bacteria; Proteobacteria; Gammaproteobacteria; Enterobacterales;
KpST82__locus_1
EUT1
3423 HMMetlyEutB_1_EUT1(pf06751) EUT1
3424 HMM06277EutA_1_eut(pf06277) EUT1
3425 HMMtpEuth_1_eut(pf04346) EUT1
3426 HMM00465alcdh_3_mix(pf00465) EUT1
3427 HMM06723EutJMreB_1_grm(#) EUT1
3428 HMMalddh_1_eutpdugrm(pf00171) EUT1 EP_NTERM
3429 P_lightViolet EUT1
3430 H_azure EUT1
3431 HMM01515PTA_B_1_eut1(pf01515) EUT1
3432 HMM01923cobtr_2_eut1(pf01923) EUT1
3433 HMM06249EutQ_1_eut1(pf06249) EUT1 EP_NTERM
3434 HMM10662PduV_1_mix(pf10662) EUT1
3435 Hp_euts_violetPink EUT1
3436 HMM00390_malic_1_eut1(pf00390pf03949pf01515) EUT1
3437 -pf00923 -
3438 HMM02780keto_1_eut1(pf02779pf02780pf00456) EUT1
3439 -pf06674 -
3440 HMM00293nuhyd_1_eut1(pf00293) EUT1
3441 -pf13247pf14691pf07992 -
3442 -pf00873 -
3443 -pf03960 -
3444 -pf07687pf01546 -
3445 -pf13980 -
BMC-P
BMC-H
BMC-T
Aldehyde dehydrogenase
PTAC
Signature enzyme
Alcohol dehydrogenase
Regulator

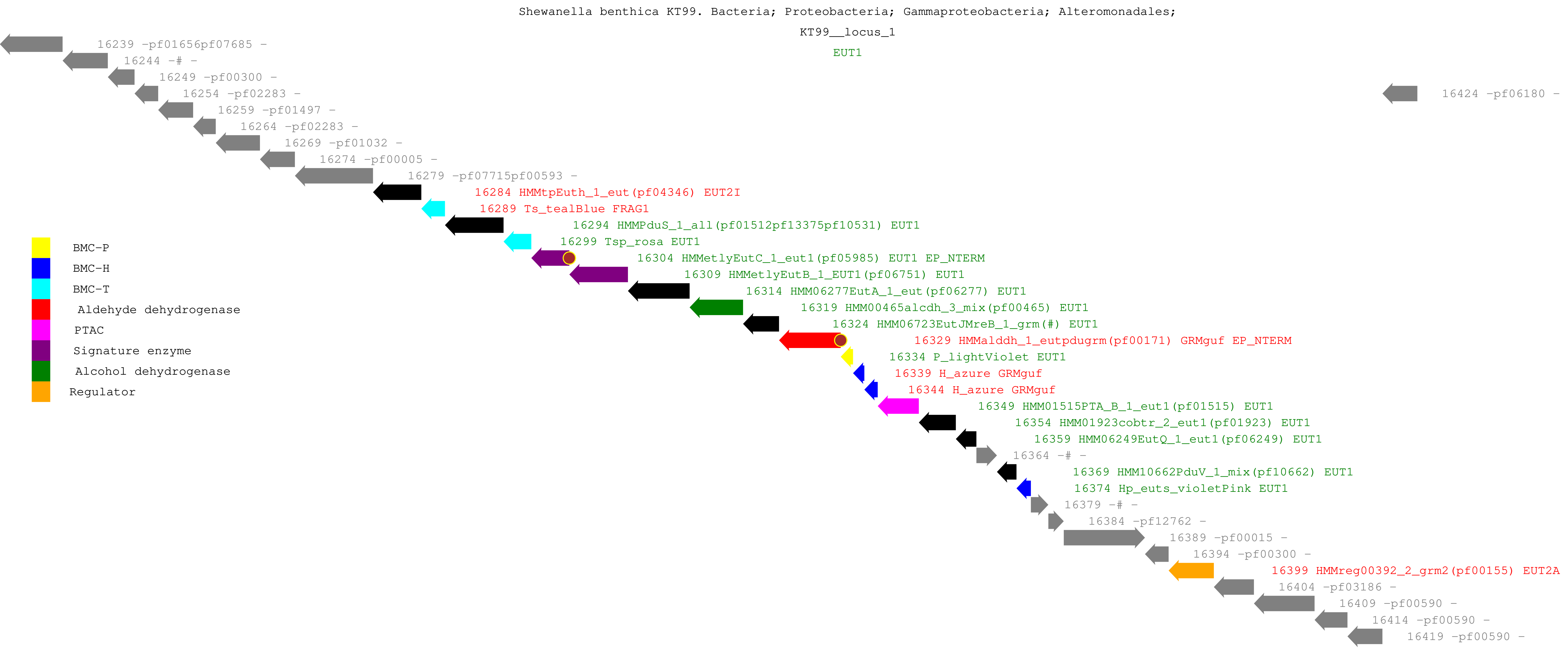

Shewanella benthica KT99. Bacteria; Proteobacteria; Gammaproteobacteria; Alteromonadales;
KT99__locus_1
EUT1
16239 -pf01656pf07685 -
16244 -# -
16249 -pf00300 -
16254 -pf02283 -
16259 -pf01497 -
16264 -pf02283 -
16269 -pf01032 -
16274 -pf00005 -
16279 -pf07715pf00593 -
16284 HMMtpEuth_1_eut(pf04346) EUT2I
16289 Ts_tealBlue FRAG1
16294 HMMPduS_1_all(pf01512pf13375pf10531) EUT1
16299 Tsp_rosa EUT1
16304 HMMetlyEutC_1_eut1(pf05985) EUT1 EP_NTERM
16309 HMMetlyEutB_1_EUT1(pf06751) EUT1
16314 HMM06277EutA_1_eut(pf06277) EUT1
16319 HMM00465alcdh_3_mix(pf00465) EUT1
16324 HMM06723EutJMreB_1_grm(#) EUT1
16329 HMMalddh_1_eutpdugrm(pf00171) GRMguf EP_NTERM
16334 P_lightViolet EUT1
16339 H_azure GRMguf
16344 H_azure GRMguf
16349 HMM01515PTA_B_1_eut1(pf01515) EUT1
16354 HMM01923cobtr_2_eut1(pf01923) EUT1
16359 HMM06249EutQ_1_eut1(pf06249) EUT1
16364 -# -
16369 HMM10662PduV_1_mix(pf10662) EUT1
16374 Hp_euts_violetPink EUT1
16379 -# -
16384 -pf12762 -
16389 -pf00015 -
16394 -pf00300 -
16399 HMMreg00392_2_grm2(pf00155) EUT2A
16404 -pf03186 -
16409 -pf00590 -
16414 -pf00590 -
16419 -pf00590 -
16424 -pf06180 -
BMC-P
BMC-H
BMC-T
Aldehyde dehydrogenase
PTAC
Signature enzyme
Alcohol dehydrogenase
Regulator

Klebsiella variicola. Bacteria; Proteobacteria; Gammaproteobacteria; Enterobacterales;

KVR801__locus_2

EUT1

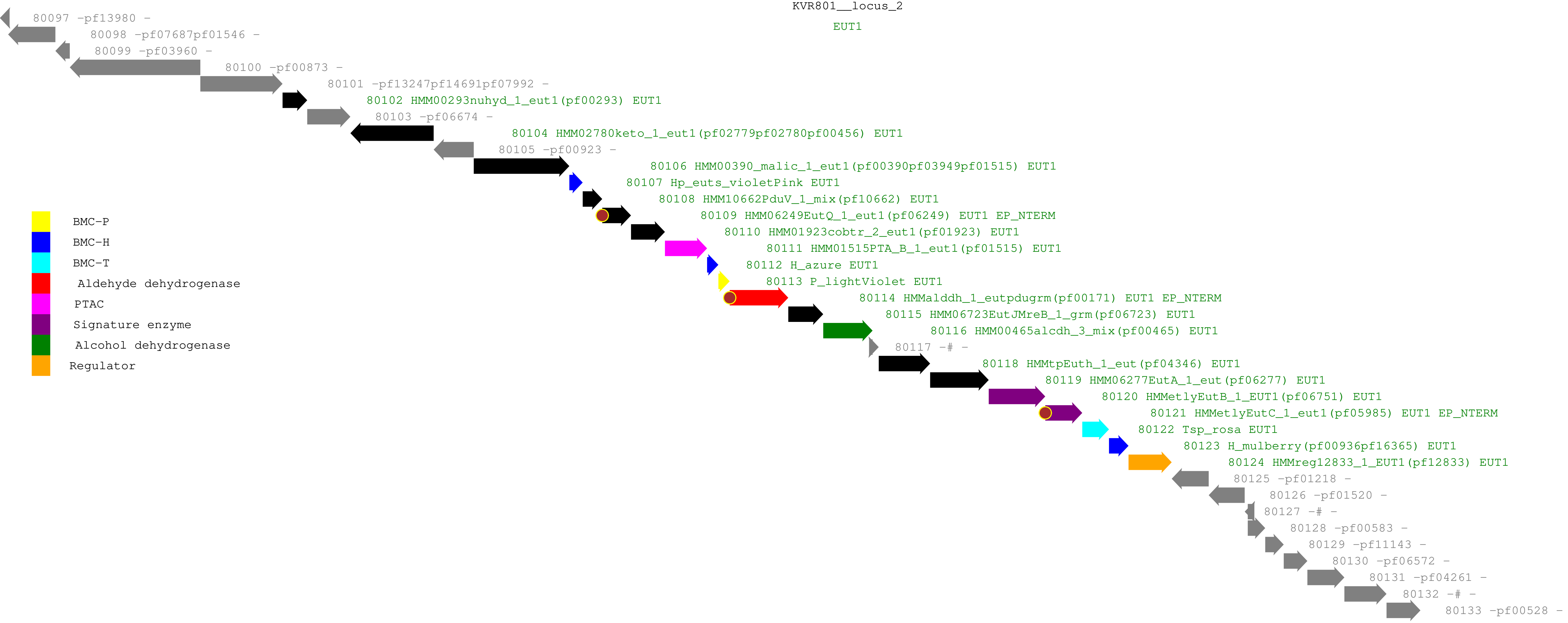

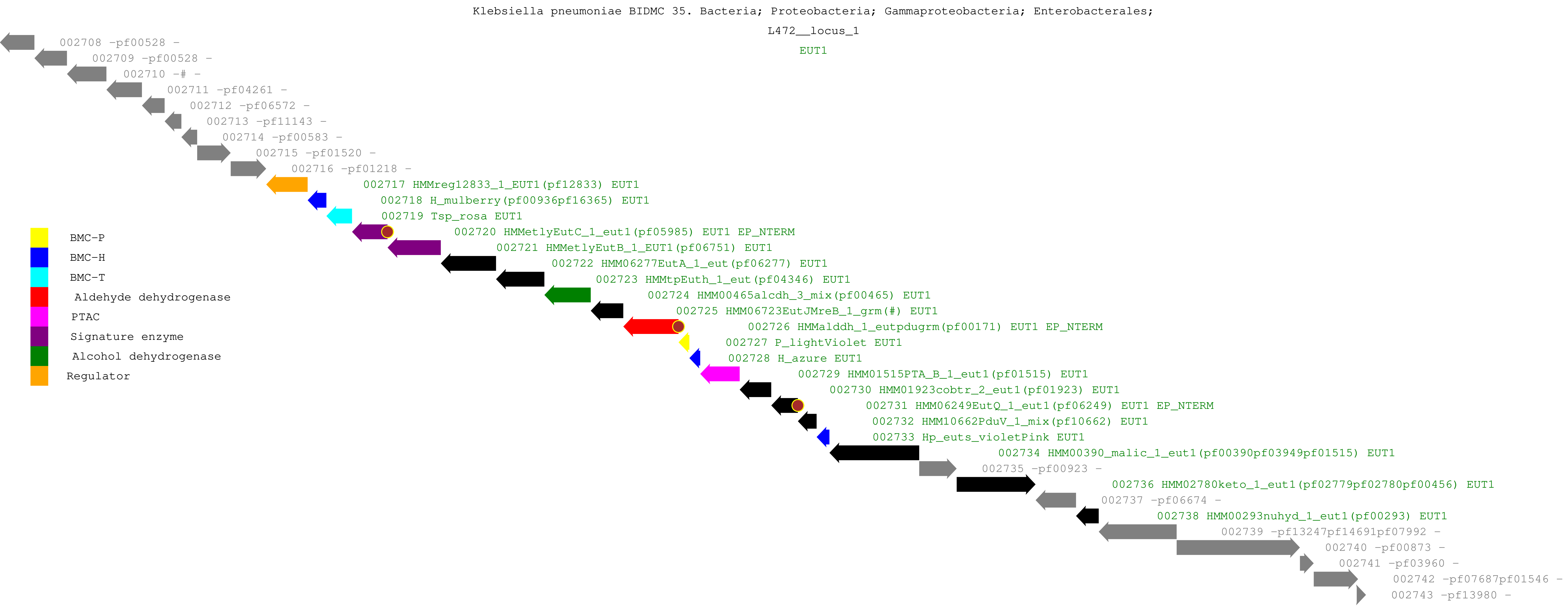
Klebsiella pneumoniae BIDMC 35. Bacteria; Proteobacteria; Gammaproteobacteria; Enterobacterales;
L472__locus_1
EUT1
002708 -pf00528 -
002709 -pf00528 -
002710 -# -
002711 -pf04261 -
002712 -pf06572 -
002713 -pf11143 -
002714 -pf00583 -
002715 -pf01520 -
002716 -pf01218 -
002717 HMMreg12833_1_EUT1(pf12833) EUT1
002718 H_mulberry(pf00936pf16365) EUT1
002719 Tsp_rosa EUT1
002720 HMMetlyEutC_1_eut1(pf05985) EUT1 EP_NTERM
002721 HMMetlyEutB_1_EUT1(pf06751) EUT1
002722 HMM06277EutA_1_eut(pf06277) EUT1
002723 HMMtpEuth_1_eut(pf04346) EUT1
002724 HMM00465alcdh_3_mix(pf00465) EUT1
002725 HMM06723EutJMreB_1_grm(#) EUT1
002726 HMMalddh_1_eutpdugrm(pf00171) EUT1 EP_NTERM
002727 P_lightViolet EUT1
002728 H_azure EUT1
002729 HMM01515PTA_B_1_eut1(pf01515) EUT1
002730 HMM01923cobtr_2_eut1(pf01923) EUT1
002731 HMM06249EutQ_1_eut1(pf06249) EUT1 EP_NTERM
002732 HMM10662PduV_1_mix(pf10662) EUT1
002733 Hp_euts_violetPink EUT1
002734 HMM00390_malic_1_eut1(pf00390pf03949pf01515) EUT1
002735 -pf00923 -
002736 HMM02780keto_1_eut1(pf02779pf02780pf00456) EUT1
002737 -pf06674 -
002738 HMM00293nuhyd_1_eut1(pf00293) EUT1
002739 -pf13247pf14691pf07992 -
002740 -pf00873 -
002741 -pf03960 -
002742 -pf07687pf01546 -
002743 -pf13980 -
BMC-P
BMC-H
BMC-T
Aldehyde dehydrogenase
PTAC
Signature enzyme
Alcohol dehydrogenase
Regulator

Salmonella enterica subsp. enterica serovar Gaminara str. SA20063285. Bacteria; Proteobacteria; Gammaproteobacteria; Enterobacterales;

LFZ12__locus_1

EUT1

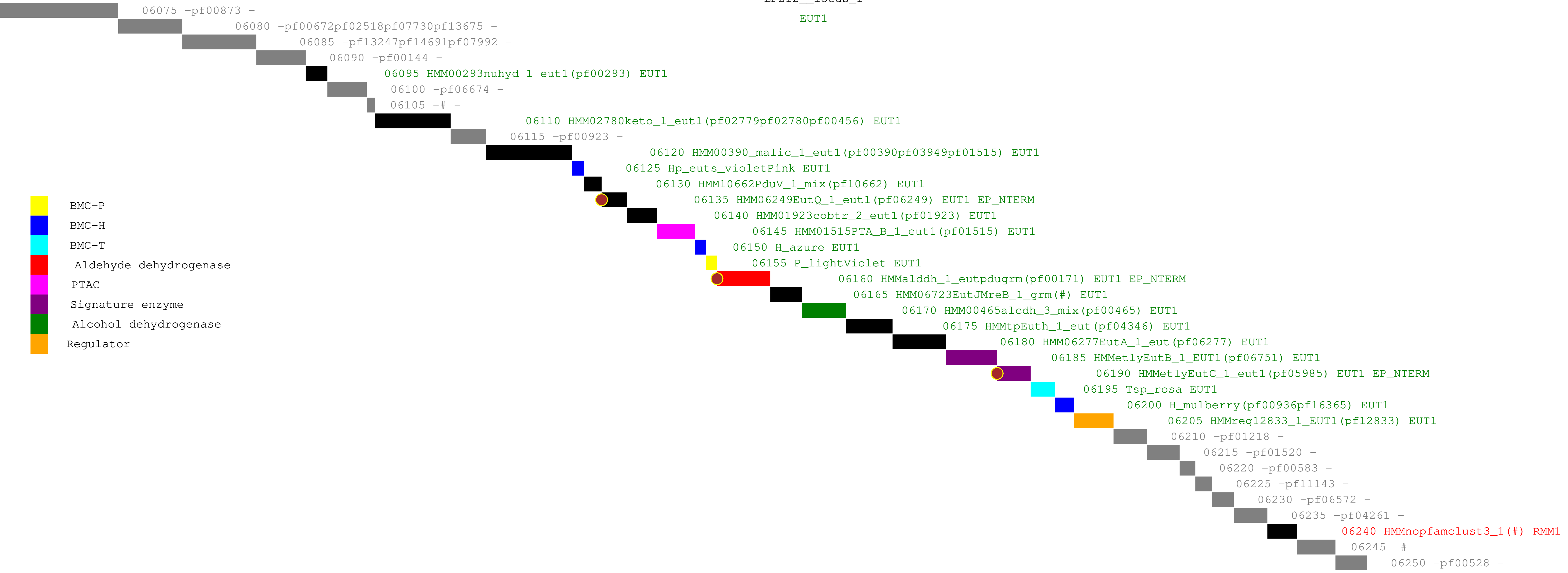

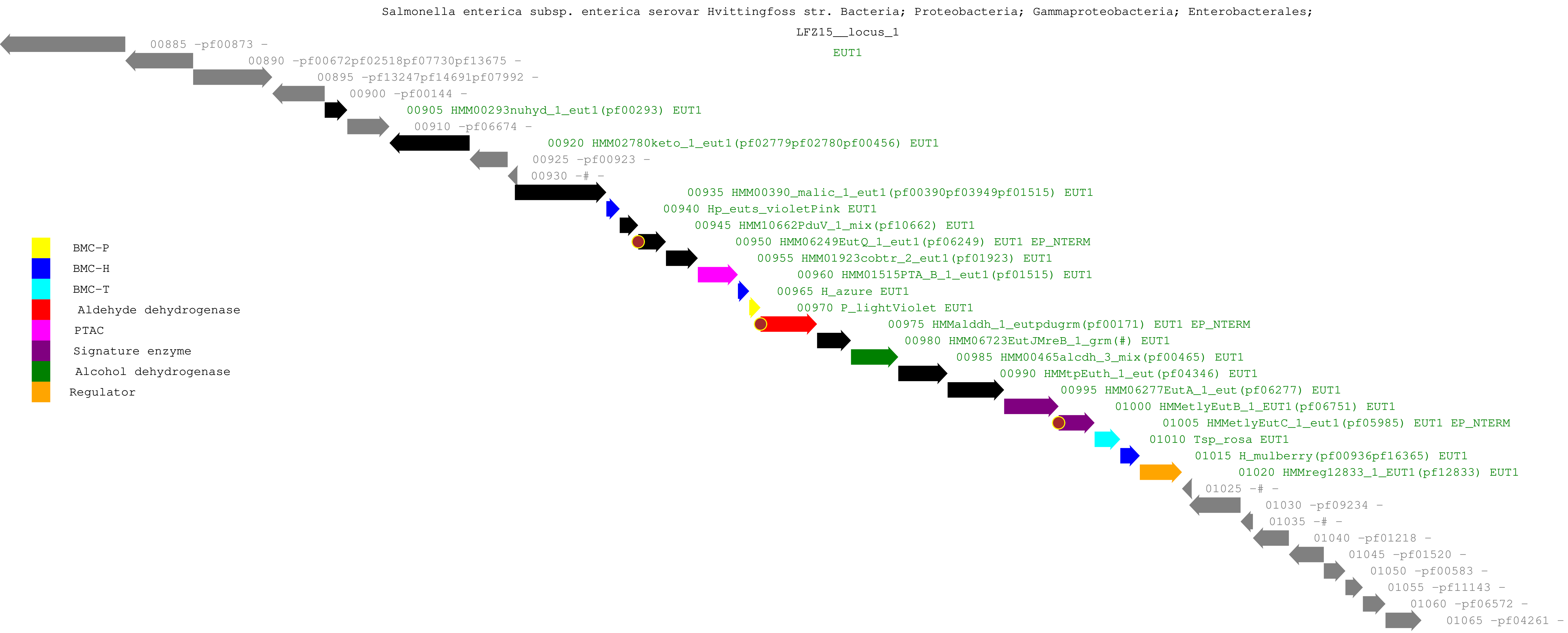
Salmonella enterica subsp. enterica serovar Hvittingfoss str. Bacteria; Proteobacteria; Gammaproteobacteria; Enterobacterales;
LFZ15__locus_1
EUT1
00885 -pf00873 -
00890 -pf00672pf02518pf07730pf13675 -
00895 -pf13247pf14691pf07992 -
00900 -pf00144 -
00905 HMM00293nuhyd_1_eut1(pf00293) EUT1
00910 -pf06674 -
00920 HMM02780keto_1_eut1(pf02779pf02780pf00456) EUT1
00925 -pf00923 -
00930 -# -
00935 HMM00390_malic_1_eut1(pf00390pf03949pf01515) EUT1
00940 Hp_euts_violetPink EUT1
00945 HMM10662PduV_1_mix(pf10662) EUT1
00950 HMM06249EutQ_1_eut1(pf06249) EUT1 EP_NTERM
00955 HMM01923cobtr_2_eut1(pf01923) EUT1
00960 HMM01515PTA_B_1_eut1(pf01515) EUT1
00965 H_azure EUT1
00970 P_lightViolet EUT1
00975 HMMaldh_1_eutpdugrm(pf00171) EUT1 EP_NTERM
00980 HMM06723EutJMreB_1_grm(#) EUT1
00985 HMM00465alcdh_3_mix(pf00465) EUT1
00990 HMMtpEuth_1_eut(pf04346) EUT1
00995 HMM06277EutA_1_eut(pf06277) EUT1
01000 HMMetlyEutB_1_EUT1(pf06751) EUT1
01005 HMMetlyEutC_1_eut1(pf05985) EUT1 EP_NTERM
01010 Tsp_rosa EUT1
01015 H_mulberry(pf00936pf16365) EUT1
01020 HMMreg12833_1_EUT1(pf12833) EUT1
01025 -# -
01030 -pf09234 -
01035 -# -
01040 -pf01218 -
01045 -pf01520 -
01050 -pf00583 -
01055 -pf11143 -
01060 -pf06572 -
01065 -pf04261 -
BMC-P
BMC-H
BMC-T
Aldehyde dehydrogenase
PTAC
Signature enzyme
Alcohol dehydrogenase
Regulator

Salmonella enterica subsp. enterica serovar India str. SA20085604. Bacteria; Proteobacteria; Gammaproteobacteria; Enterobacterales;

LFZ16__locus_1

EUT1

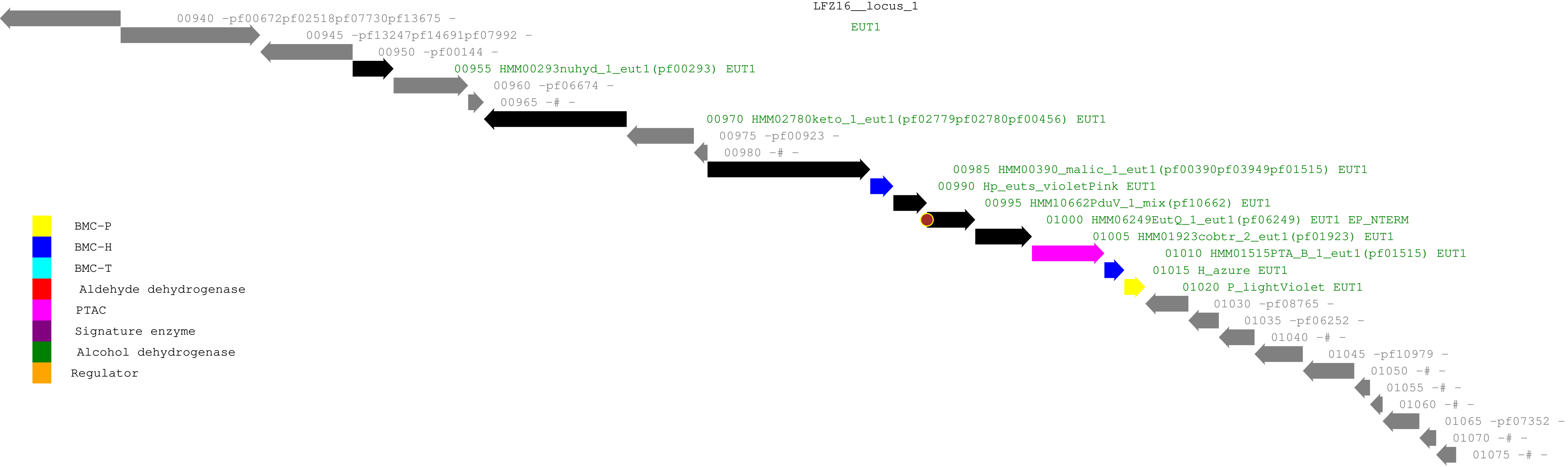

Salmonella enterica subsp. enterica serovar Kentucky str. SA20030505. Bacteria; Proteobacteria; Gammaproteobacteria; Enterobacterales;

LFZ21__locus_2

EUT1

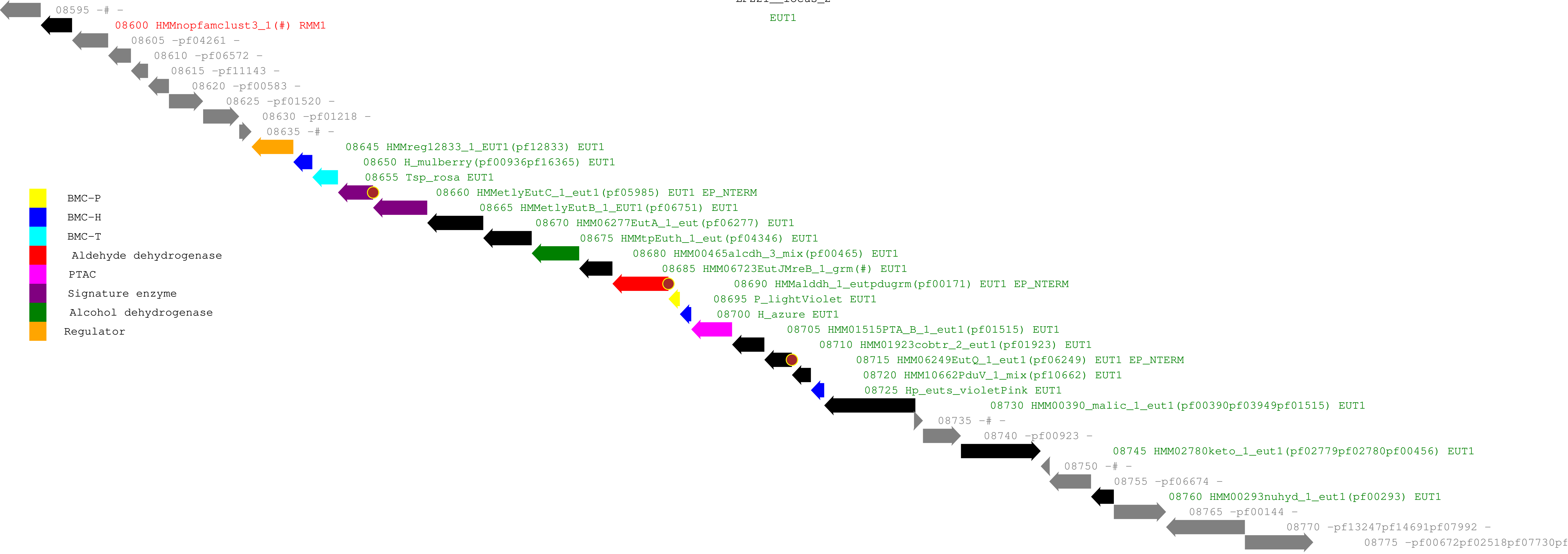

Salmonella enterica subsp. enterica serovar Krefeld str. SA20030536. Bacteria; Proteobacteria; Gammaproteobacteria; Enterobacterales;

LFZ24__locus_2

EUT1

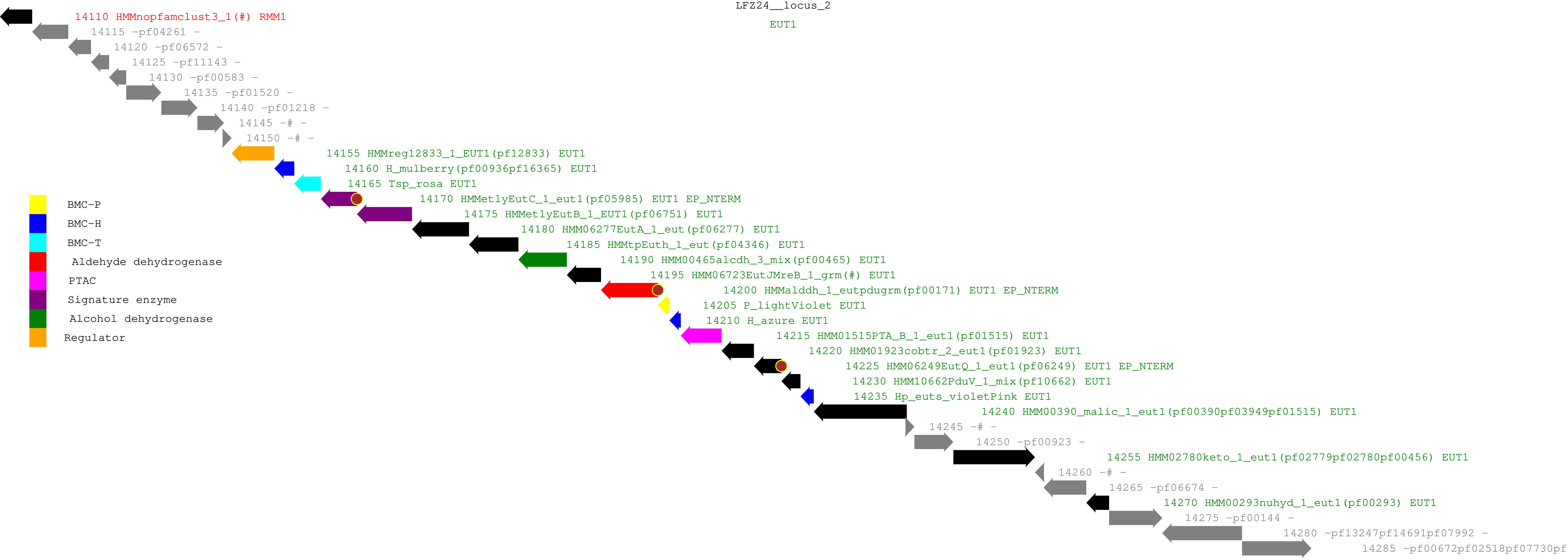

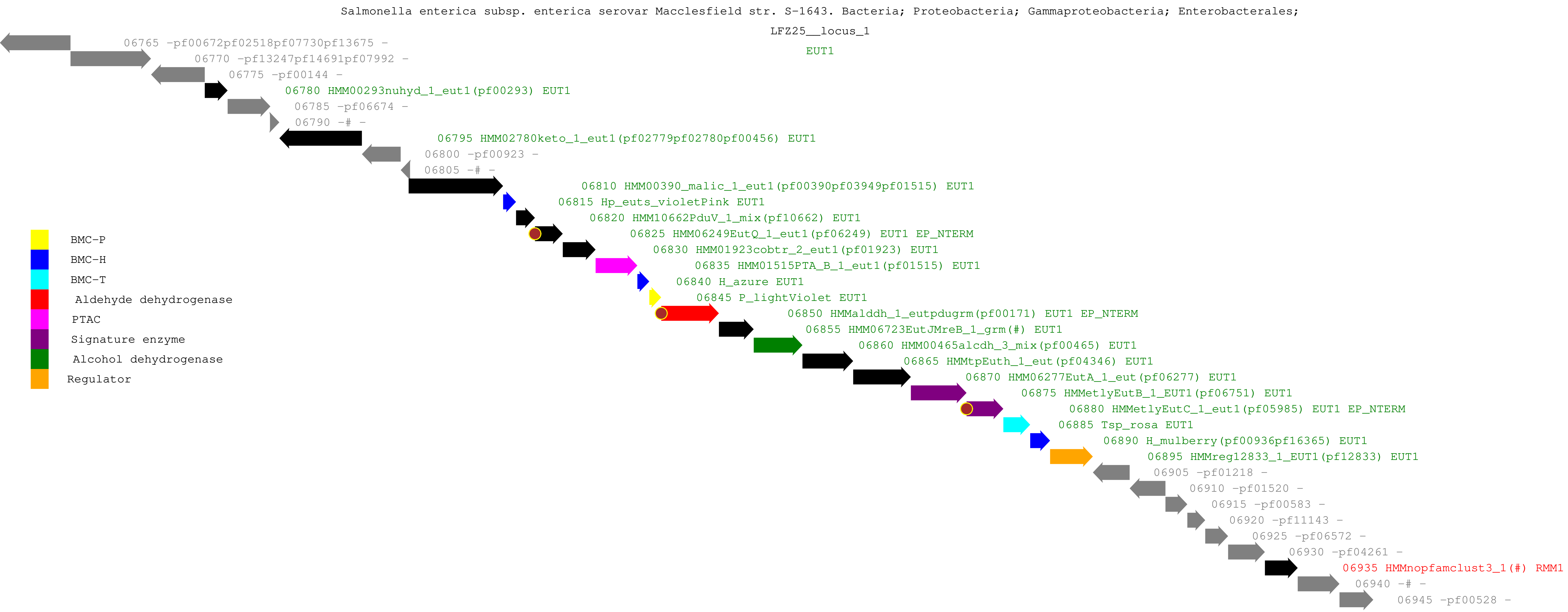
Salmonella enterica subsp. enterica serovar Macclesfield str. S-1643. Bacteria; Proteobacteria; Gammaproteobacteria; Enterobacterales;
LFZ25__locus_1
EUT1
06765 -pf00672pf02518pf07730pf13675 -
06770 -pf13247pf14691pf07992 -
06775 -pf00144 -
06780 HMM00293nuhyd_1_eut1(pf00293) EUT1
06785 -pf06674 -
06790 -# -
06795 HMM02780keto_1_eut1(pf02779pf02780pf00456) EUT1
06800 -pf00923 -
06805 -# -
06810 HMM00390_malic_1_eut1(pf00390pf03949pf01515) EUT1
06815 Hp_euts_violetPink EUT1
06820 HMM10662PduV_1_mix(pf10662) EUT1
06825 HMM06249EutQ_1_eut1(pf06249) EUT1 EP_NTERM
06830 HMM01923cobtr_2_eut1(pf01923) EUT1
06835 HMM01515PTA_B_1_eut1(pf01515) EUT1
06840 H_azure EUT1
06845 P_lightViolet EUT1
06850 HMMalddh_1_eutpdugrm(pf00171) EUT1 EP_NTERM
06855 HMM06723EutJMreB_1_grm(#) EUT1
06860 HMM00465alcdh_3_mix(pf00465) EUT1
06865 HMMtpEuth_1_eut(pf04346) EUT1
06870 HMM06277EutA_1_eut(pf06277) EUT1
06875 HMMetlyEutB_1_EUT1(pf06751) EUT1
06880 HMMetlyEutC_1_eut1(pf05985) EUT1 EP_NTERM
06885 Tsp_rosa EUT1
06890 H_mulberry(pf00936pf16365) EUT1
06895 HMMreg12833_1_EUT1(pf12833) EUT1
06905 -pf01218 -
06910 -pf01520 -
06915 -pf00583 -
06920 -pf11143 -
06925 -pf06572 -
06930 -pf04261 -
06935 HMMnopfamclust3_1(#) RMM1
06940 -# -
06945 -pf00528 -
BMC-P
BMC-H
BMC-T
Aldehyde dehydrogenase
PTAC
Signature enzyme
Alcohol dehydrogenase
Regulator

Salmonella enterica subsp. enterica serovar Wandsworth str. Bacteria; Proteobacteria; Gammaproteobacteria; Enterobacterales;

LFZ43__locus_2

EUT1

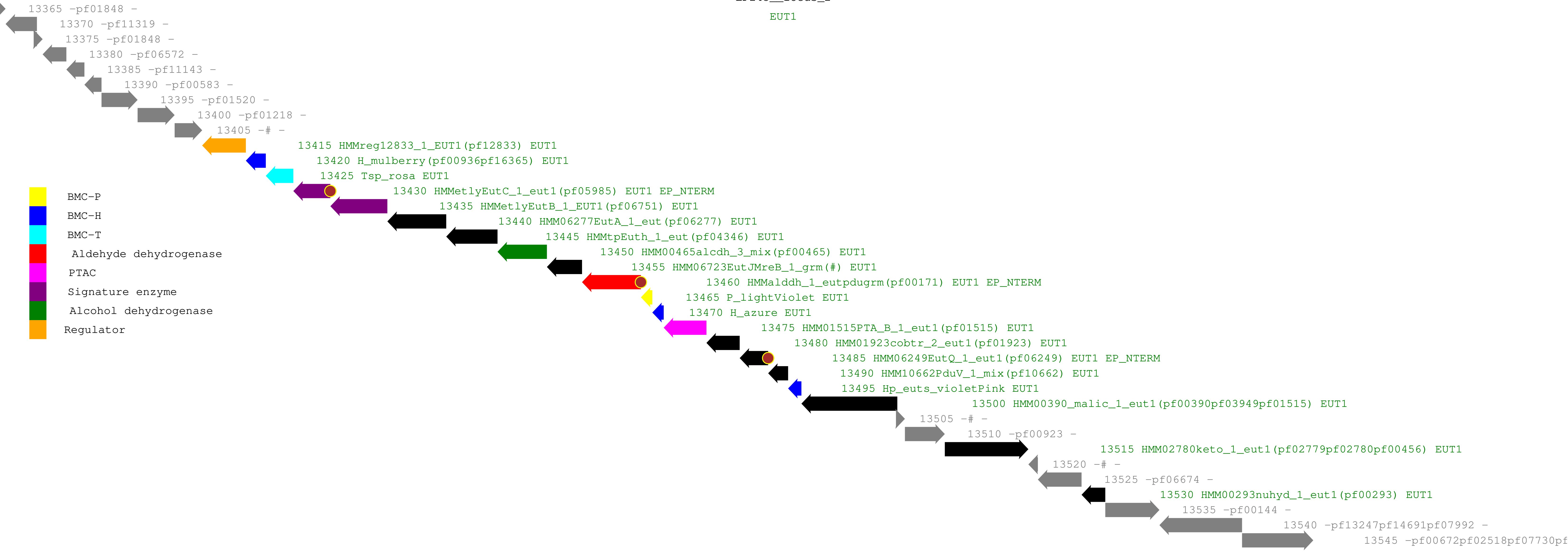

Salmonella enterica subsp. salamae serovar 55:k:z39 str. 1315K. Bacteria; Proteobacteria; Gammaproteobacteria; Enterobacterales;

LFZ47__locus_1

EUT1

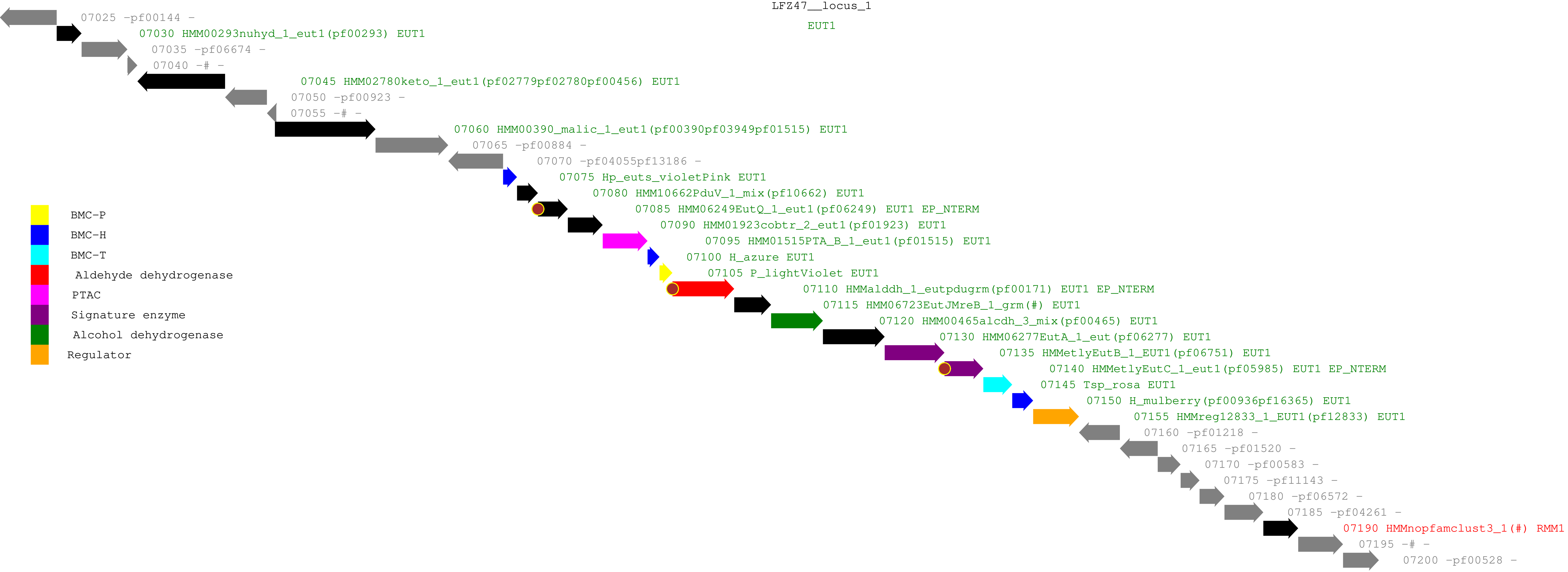

Salmonella enterica subsp. salamae serovar 56:z10:e,n,x str. 1369-73. Bacteria; Proteobacteria; Gammaproteobacteria; Enterobacterales;

LFZ48__locus_2

EUT1

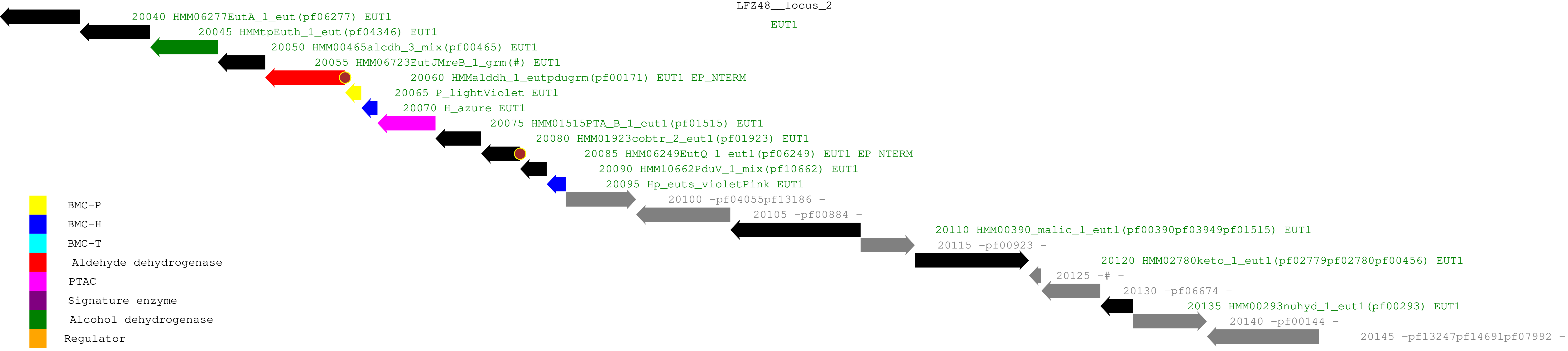

Salmonella enterica I. Bacteria; Proteobacteria; Gammaproteobacteria; Enterobacterales;

LFZ50__locus_1

EUT1

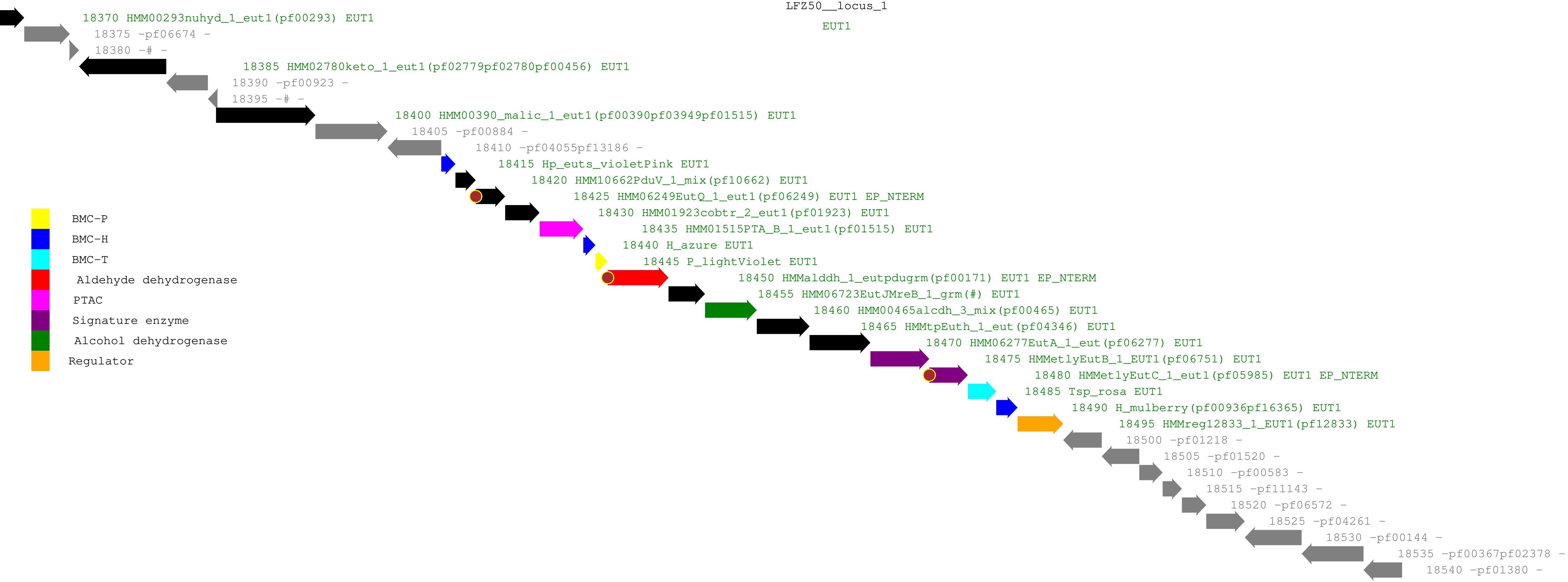

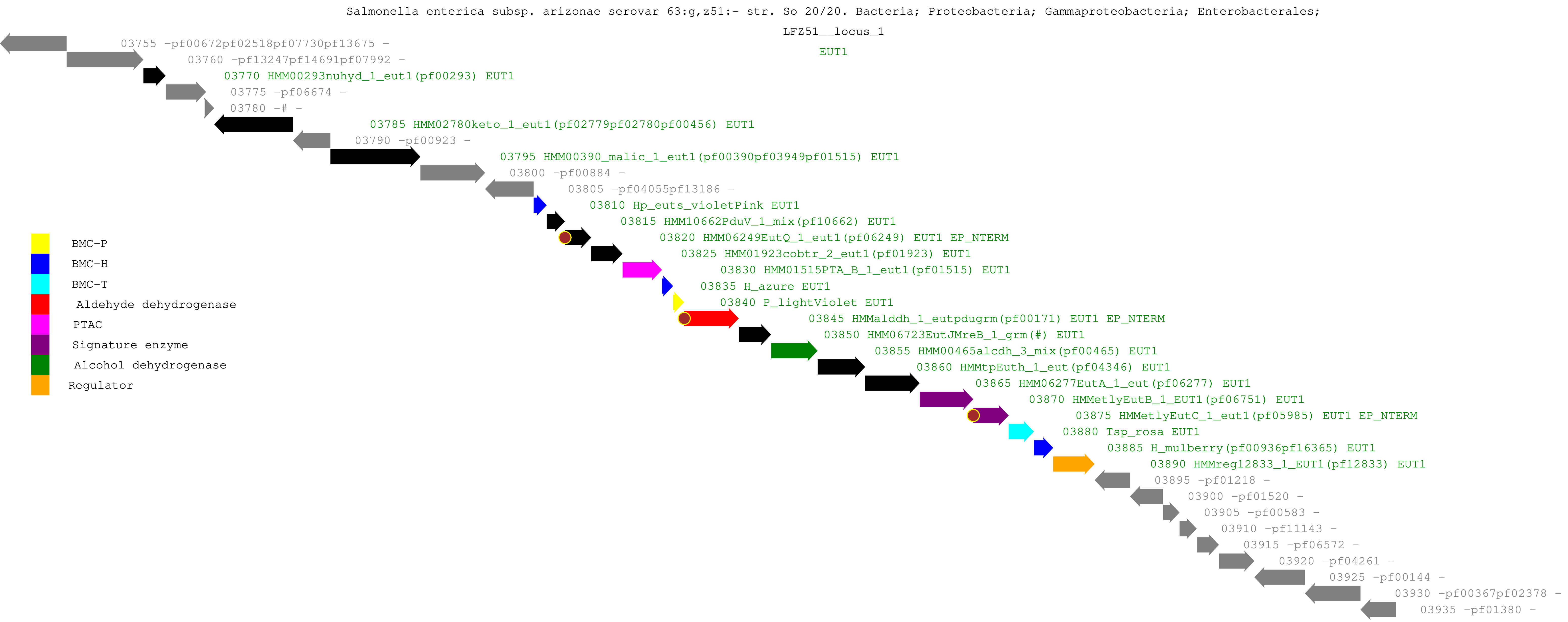
Salmonella enterica subsp. arizonae serovar 63:g,z51:- str. So 20/20. Bacteria; Proteobacteria; Gammaproteobacteria; Enterobacterales;
LFZ51__locus_1
EUT1
03755 -pf00672pf02518pf07730pf13675 -
03760 -pf13247pf14691pf07992 -
03770 HMM00293nuhyd_1_eut1(pf00293) EUT1
03775 -pf06674 -
03780 -# -
03785 HMM02780keto_1_eut1(pf02779pf02780pf00456) EUT1
03790 -pf00923 -
03795 HMM00390_malic_1_eut1(pf00390pf03949pf01515) EUT1
03800 -pf00884 -
03805 -pf04055pf13186 -
03810 Hp_euts_violetPink EUT1
03815 HMM10662PduV_1_mix(pf10662) EUT1
03820 HMM06249EutQ_1_eut1(pf06249) EUT1 EP_NTERM
03825 HMM01923cobtr_2_eut1(pf01923) EUT1
03830 HMM01515PTA_B_1_eut1(pf01515) EUT1
03835 H_azure EUT1
03840 P_lightViolet EUT1
03845 HMMalddh_1_eutpdugrm(pf00171) EUT1 EP_NTERM
03850 HMM06723EutJMreB_1_grm(#) EUT1
03855 HMM00465alcdh_3_mix(pf00465) EUT1
03860 HMMtpEuth_1_eut(pf04346) EUT1
03865 HMM06277EutA_1_eut(pf06277) EUT1
03870 HMMetlyEutB_1_EUT1(pf06751) EUT1
03875 HMMetlyEutC_1_eut1(pf05985) EUT1 EP_NTERM
03880 Tsp_rosa EUT1
03885 H_mulberry(pf00936pf16365) EUT1
03890 HMMreg12833_1_EUT1(pf12833) EUT1
03895 -pf01218 -
03900 -pf01520 -
03905 -pf00583 -
03910 -pf11143 -
03915 -pf06572 -
03920 -pf04261 -
03925 -pf00144 -
03930 -pf00367pf02378 -
03935 -pf01380 -
BMC-P
BMC-H
BMC-T
Aldehyde dehydrogenase
PTAC
Signature enzyme
Alcohol dehydrogenase
Regulator

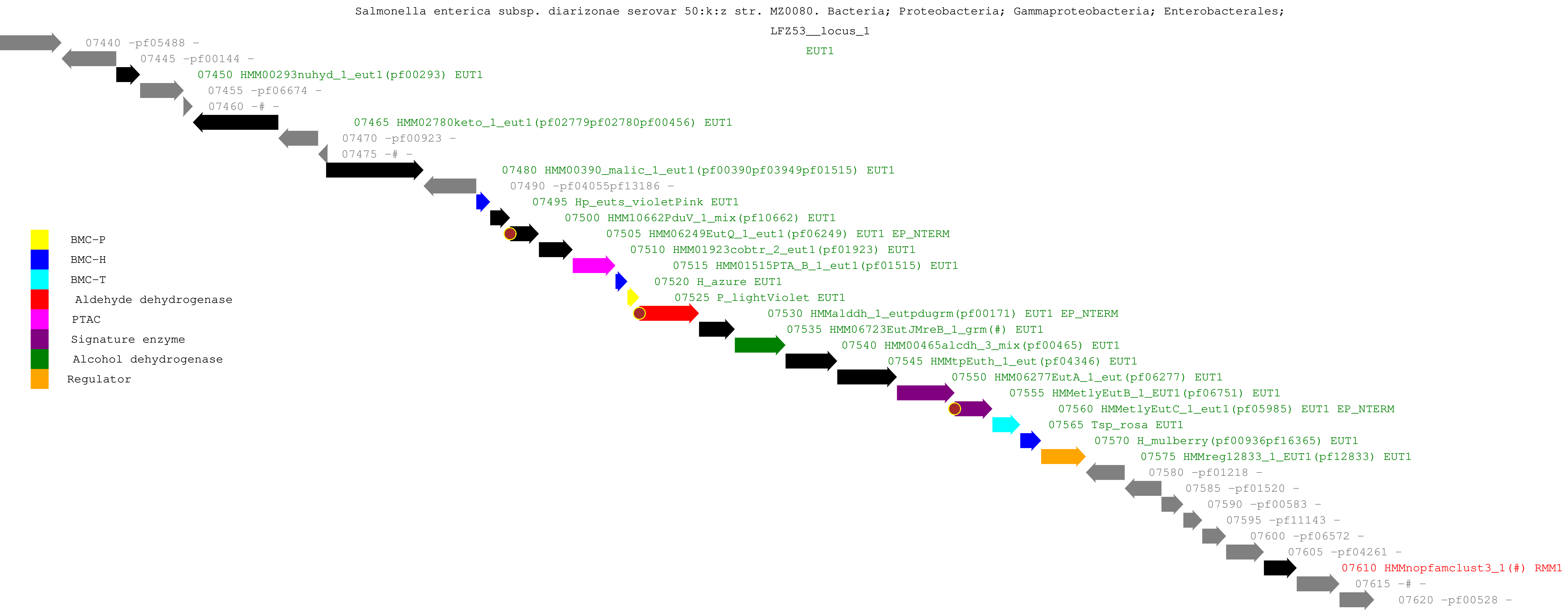

Salmonella enterica subsp. diarizonae serovar 50:k:z str. MZ0080. Bacteria; Proteobacteria; Gammaproteobacteria; Enterobacterales;
LFZ53__locus_1
EUT1
07440 -pf05488 -
07445 -pf00144 -
07450 HMM00293nuhyd_1_eut1(pf00293) EUT1
07455 -pf06674 -
07460 -# -
07465 HMM02780keto_1_eut1(pf02779pf02780pf00456) EUT1
07470 -pf00923 -
07475 -# -
07480 HMM00390_malic_1_eut1(pf00390pf03949pf01515) EUT1
07490 -pf04055pf13186 -
07495 Hp_euts_violetPink EUT1
07500 HMM10662PduV_1_mix(pf10662) EUT1
07505 HMM06249EutQ_1_eut1(pf06249) EUT1 EP_NTERM
07510 HMM01923cobtr_2_eut1(pf01923) EUT1
07515 HMM01515PTA_B_1_eut1(pf01515) EUT1
07520 H_azure EUT1
07525 P_lightViolet EUT1
07530 HMMaldDh_1_eutpdugrm(pf00171) EUT1 EP_NTERM
07535 HMM06723EutJMreB_1_grm(#) EUT1
07540 HMM00465alcdh_3_mix(pf00465) EUT1
07545 HMMtpEuth_1_eut(pf04346) EUT1
07550 HMM06277EutA_1_eut(pf06277) EUT1
07555 HMMetlyEutB_1_EUT1(pf06751) EUT1
07560 HMMetlyEutC_1_eut1(pf05985) EUT1 EP_NTERM
07565 Tsp_rosa EUT1
07570 H_mulberry(pf00936pf16365) EUT1
07575 HMMreg12833_1_EUT1(pf12833) EUT1
07580 -pf01218 -
07585 -pf01520 -
07590 -pf00583 -
07595 -pf11143 -
07600 -pf06572 -
07605 -pf04261 -
07610 HMMnopfamclust3_1(#) RMM1
07615 -# -
07620 -pf00528 -
BMC-P
BMC-H
BMC-T
Aldehyde dehydrogenase
PTAC
Signature enzyme
Alcohol dehydrogenase
Regulator

Salmonella enterica subsp. diarizonae serovar 65:c:z str. SA20044251. Bacteria; Proteobacteria; Gammaproteobacteria; Enterobacterales;

LFZ55__locus_1

EUT1

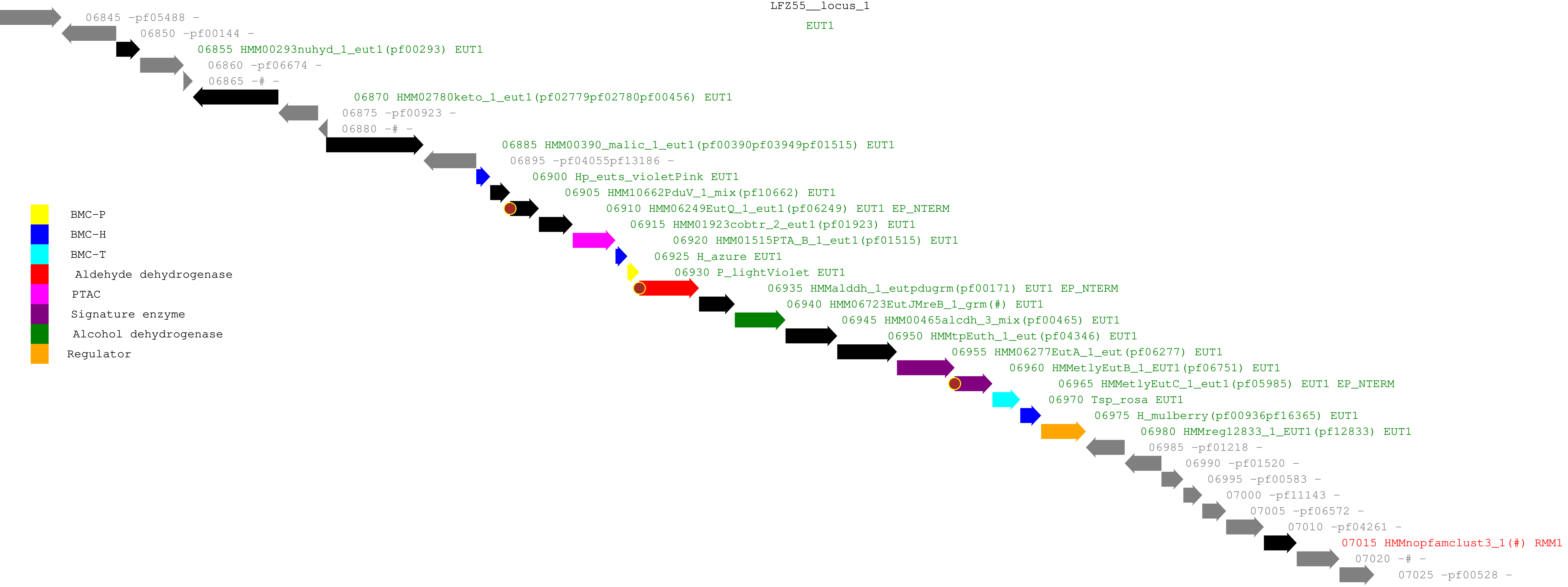

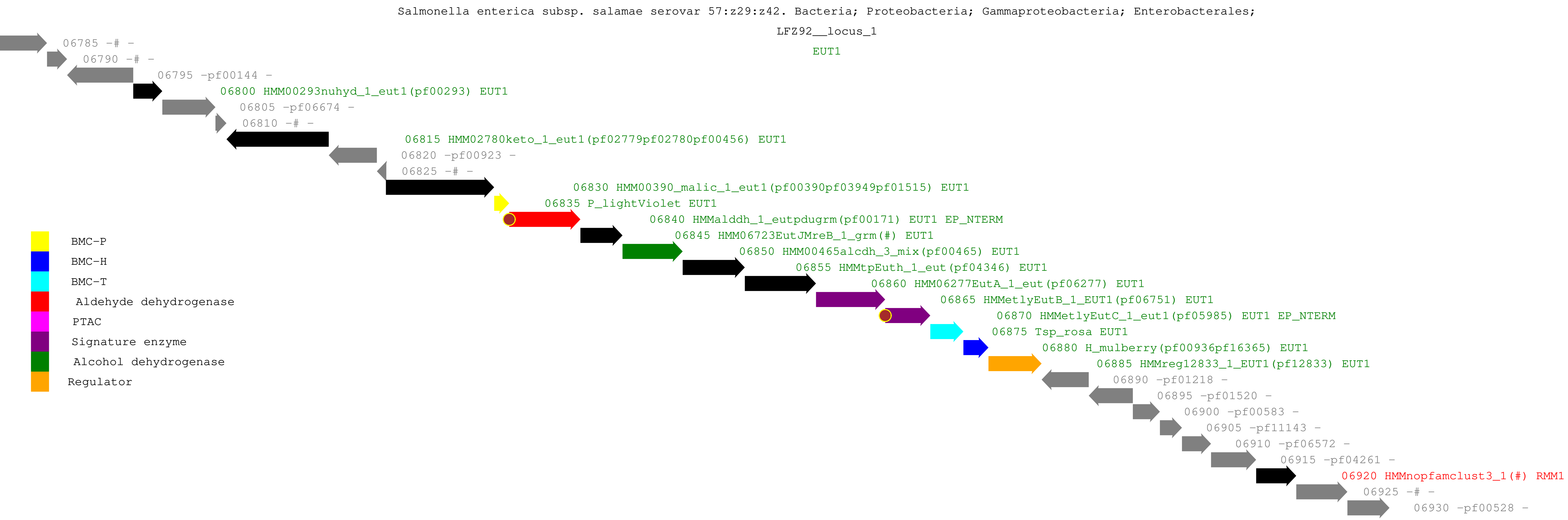
Salmonella enterica subsp. salamae serovar 57:z29:z42. Bacteria; Proteobacteria; Gammaproteobacteria; Enterobacterales;
LFZ92__locus_1
EUT1
06785 -# -
06790 -# -
06795 -pf00144 -
06800 HMM00293nuhyd_1_eut1(pf00293) EUT1
06805 -pf06674 -
06810 -# -
06815 HMM02780keto_1_eut1(pf02779pf02780pf00456) EUT1
06820 -pf00923 -
06825 -# -
06830 HMM00390_malic_1_eut1(pf00390pf03949pf01515) EUT1
06835 P_lightViolet EUT1
06840 HMMaldh_1_eutpdugrm(pf00171) EUT1 EP_NTERM
06845 HMM06723EutJMreB_1_grm(#) EUT1
06850 HMM00465alcdh_3_mix(pf00465) EUT1
06855 HMMtpEuth_1_eut(pf04346) EUT1
06860 HMM06277EutA_1_eut(pf06277) EUT1
06865 HMMetlyEutB_1_EUT1(pf06751) EUT1
06870 HMMetlyEutC_1_eut1(pf05985) EUT1 EP_NTERM
06875 Tsp_rosa EUT1
06880 H_mulberry(pf00936pf16365) EUT1
06885 HMMreg12833_1_EUT1(pf12833) EUT1
06890 -pf01218 -
06895 -pf01520 -
06900 -pf00583 -
06905 -pf11143 -
06910 -pf06572 -
06915 -pf04261 -
06920 HMMnopfamclust3_1(#) RMM1
06925 -# -
06930 -pf00528 -
BMC-P
BMC-H
BMC-T
Aldehyde dehydrogenase
PTAC
Signature enzyme
Alcohol dehydrogenase
Regulator

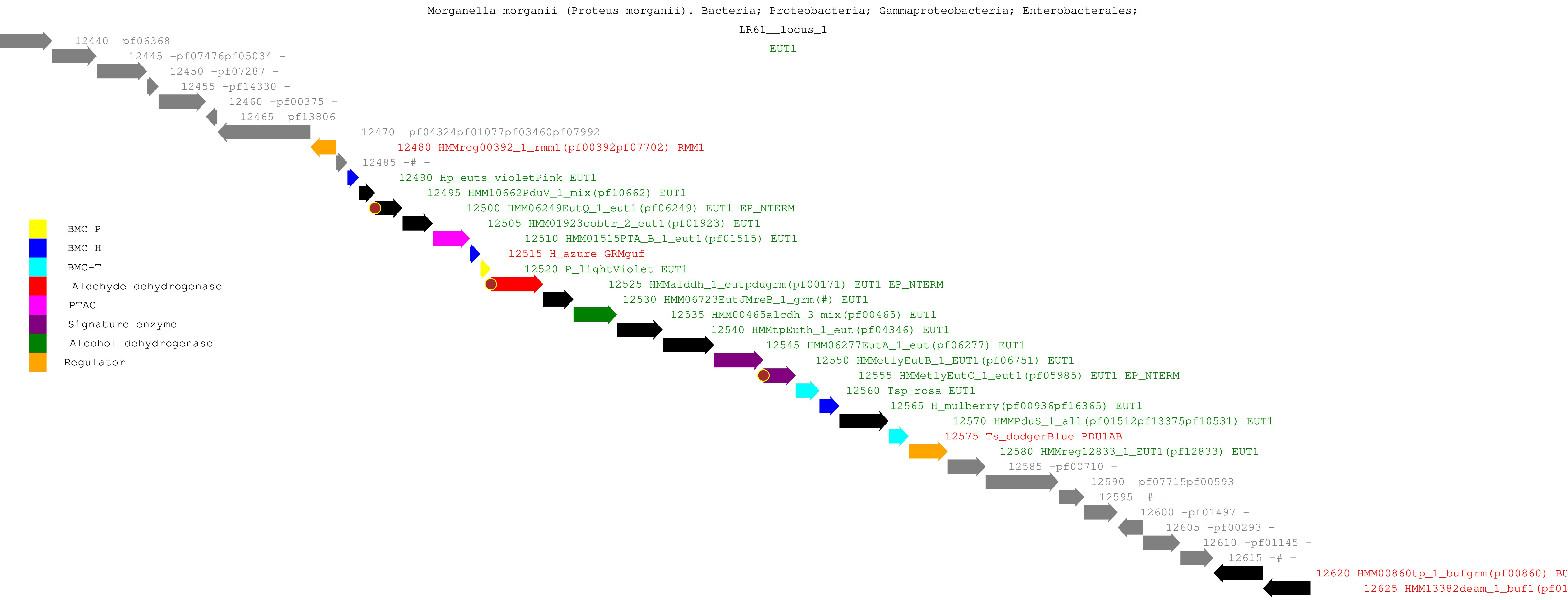
Morganella morganii (Proteus morganii). Bacteria; Proteobacteria; Gammaproteobacteria; Enterobacterales;
LR61__locus_1
EUT1
12440 -pf06368 -
12445 -pf07476pf05034 -
12450 -pf07287 -
12455 -pf14330 -
12460 -pf00375 -
12465 -pf13806 -
12470 -pf04324pf01077pf03460pf07992 -
12480 HMMreg00392_1_rmm1(pf00392pf07702) RMM1
12485 -# -
12490 Hp_euts_violetPink EUT1
12495 HMM10662PduV_1_mix(pf10662) EUT1
12500 HMM06249EutQ_1_eut1(pf06249) EUT1 EP_NTERM
12505 HMM01923cobtr_2_eut1(pf01923) EUT1
12510 HMM01515PTA_B_1_eut1(pf01515) EUT1
12515 H_azure GRMguf
12520 P_lightViolet EUT1
12525 HMMalddh_1_eutpdugrm(pf00171) EUT1 EP_NTERM
12530 HMM06723EutJMreB_1_grm(#) EUT1
12535 HMM00465alcdh_3_mix(pf00465) EUT1
12540 HMMtpEuth_1_eut(pf04346) EUT1
12545 HMM06277EutA_1_eut(pf06277) EUT1
12550 HMMetlyEutB_1_EUT1(pf06751) EUT1
12555 HMMetlyEutC_1_eut1(pf05985) EUT1 EP_NTERM
12560 Tsp_rosa EUT1
12565 H_mulberry(pf00936pf16365) EUT1
12570 HMMPduS_1_all(pf01512pf13375pf10531) EUT1
12575 Ts_dodgerBlue PDU1AB
12580 HMMreg12833_1_EUT1(pf12833) EUT1
12585 -pf00710 -
12590 -pf07715pf00593 -
12595 -# -
12600 -pf01497 -
12605 -pf00293 -
12610 -pf01145 -
12615 -# -
12620 HMM00860tp_1_bufgrm(pf00860) BU
12625 HMM13382deam_1_buf1(pf01
BMC-P
BMC-H
BMC-T
Aldehyde dehydrogenase
PTAC
Signature enzyme
Alcohol dehydrogenase
Regulator

Salmonella enterica subsp. enterica serovar Adelaide str. A4-669. Bacteria; Proteobacteria; Gammaproteobacteria; Enterobacterales;

LTSEADE__locus_2

EUT1

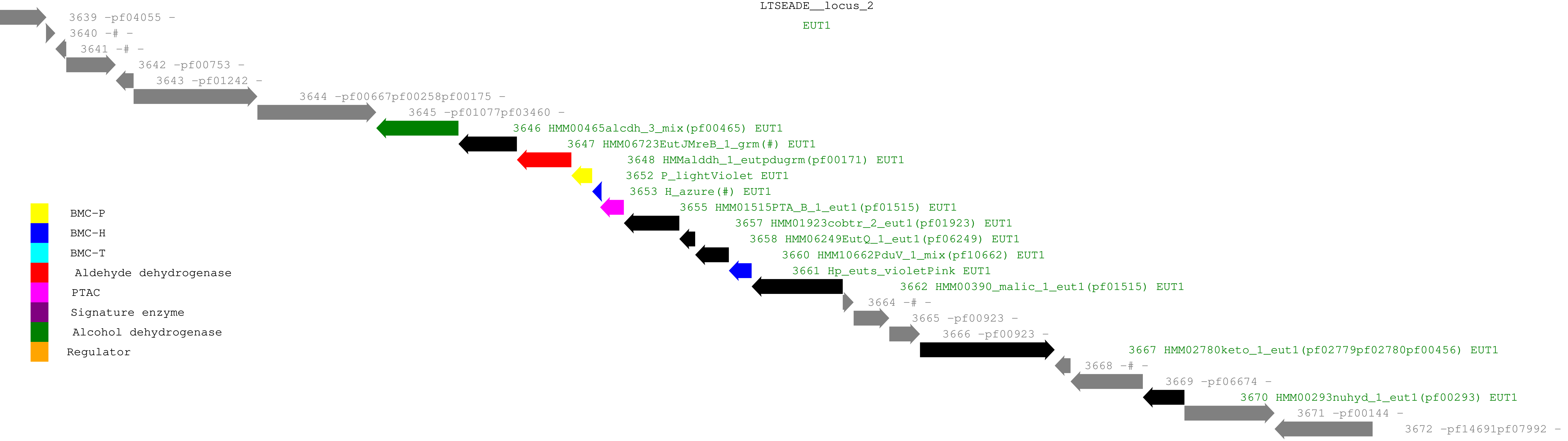

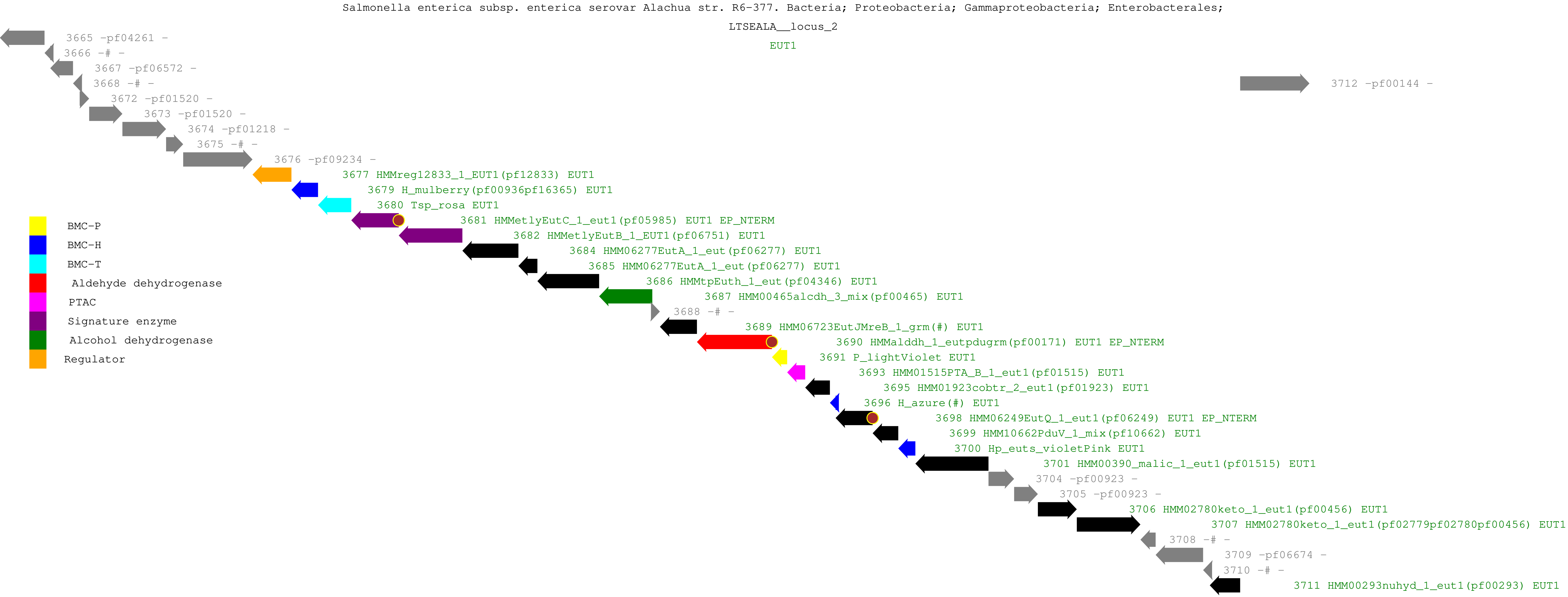
Salmonella enterica subsp. enterica serovar Alachua str. R6-377. Bacteria; Proteobacteria; Gammaproteobacteria; Enterobacterales;
LTSEALA__locus_2
EUT1
3665 -pf04261 -
3666 -# -
3667 -pf06572 -
3668 -# -
3672 -pf01520 -
3673 -pf01520 -
3674 -pf01218 -
3675 -# -
3676 -pf09234 -
3677 HMMreg12833_1_EUT1(pf12833) EUT1
3679 H_mulberry(pf00936pf16365) EUT1
3680 Tsp_rosa EUT1
3681 HMMetlyEutC_1_eut1(pf05985) EUT1 EP_NTERM
3682 HMMetlyEutB_1_EUT1(pf06751) EUT1
3684 HMM06277EutA_1_eut(pf06277) EUT1
3685 HMM06277EutA_1_eut(pf06277) EUT1
3686 HMMtpEuth_1_eut(pf04346) EUT1
3687 HMM00465alcdh_3_mix(pf00465) EUT1
3688 -# -
3689 HMM06723EutJMreB_1_grm(#) EUT1
3690 HMMaldddh_1_eutpdugrm(pf00171) EUT1 EP_NTERM
3691 P_lightViolet EUT1
3693 HMM01515PTA_B_1_eut1(pf01515) EUT1
3695 HMM01923cobtr_2_eut1(pf01923) EUT1
3696 H_azure(#) EUT1
3698 HMM06249EutQ_1_eut1(pf06249) EUT1 EP_NTERM
3699 HMM10662PduV_1_mix(pf10662) EUT1
3700 Hp_euts_violetPink EUT1
3701 HMM00390_malic_1_eut1(pf01515) EUT1
3704 -pf00923 -
3705 -pf00923 -
3706 HMM02780keto_1_eut1(pf00456) EUT1
3707 HMM02780keto_1_eut1(pf02779pf02780pf00456) EUT1
3708 -# -
3709 -pf06674 -
3710 -# -
3711 HMM00293nuhyd_1_eut1(pf00293) EUT1
3712 -pf00144 -
BMC-P
BMC-H
BMC-T
Aldehyde dehydrogenase
PTAC
Signature enzyme
Alcohol dehydrogenase
Regulator

Salmonella enterica subsp. enterica serovar Baildon str. R6-199. Bacteria; Proteobacteria; Gammaproteobacteria; Enterobacterales;

LTSEBAI__locus_2

EUT1

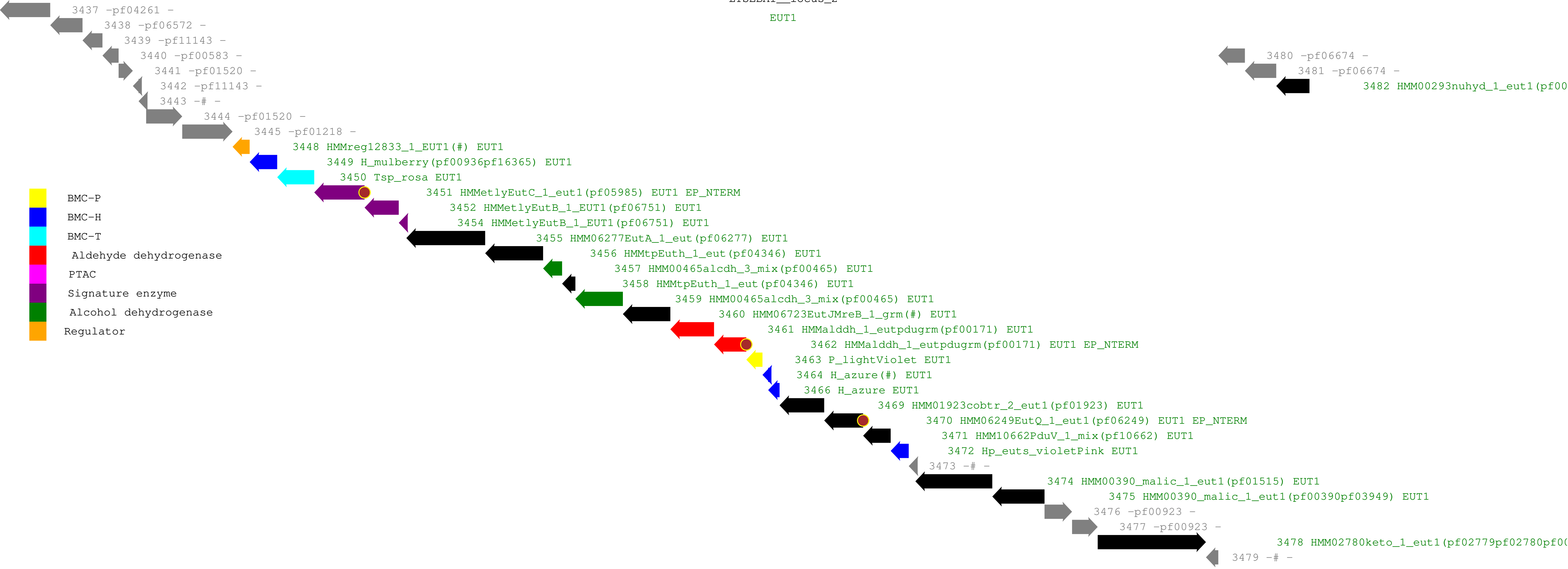

Salmonella enterica subsp. enterica serovar Give str. S5-487. Bacteria; Proteobacteria; Gammaproteobacteria; Enterobacterales;

LTSEGIV__locus_1_main_sat

EUT1

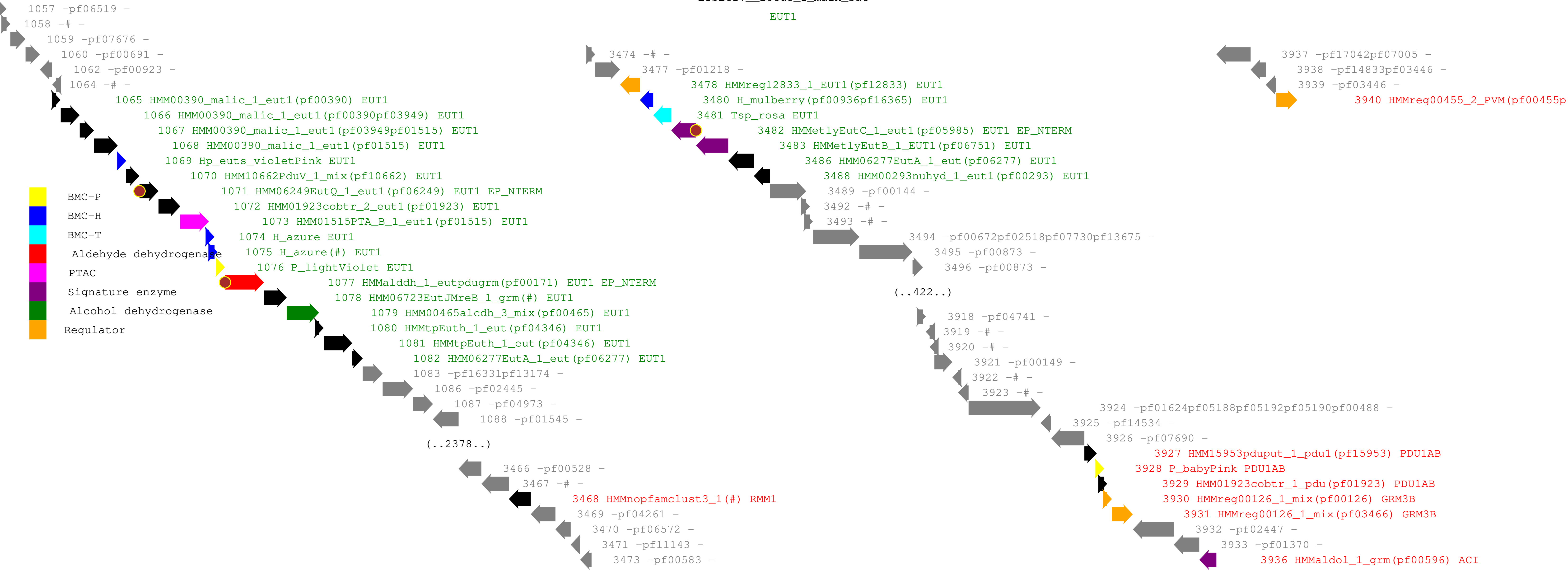

Salmonella enterica subsp. enterica serovar Inverness str. R8-3668. Bacteria; Proteobacteria; Gammaproteobacteria; Enterobacterales;

LTSEINV__locus_2

EUT1

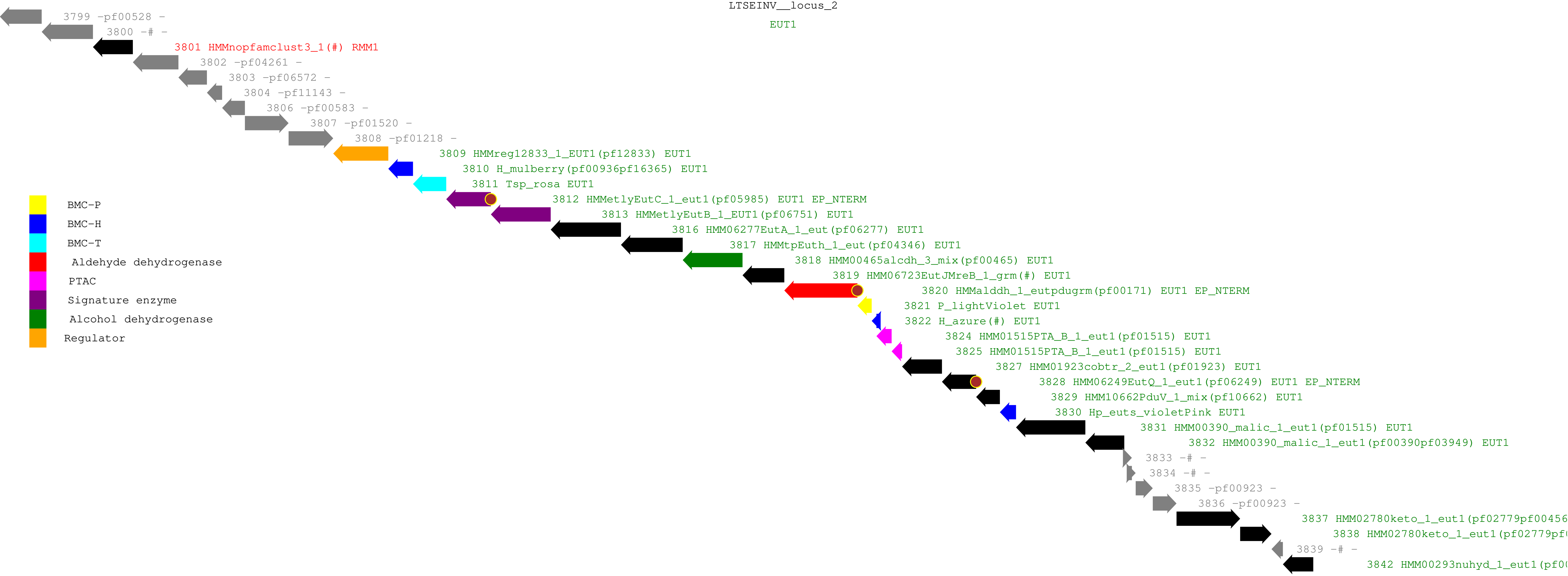

Salmonella enterica subsp. enterica serovar Minnesota str. A4-603. Bacteria; Proteobacteria; Gammaproteobacteria; Enterobacterales;

LTSEMIN__locus_2

EUT1

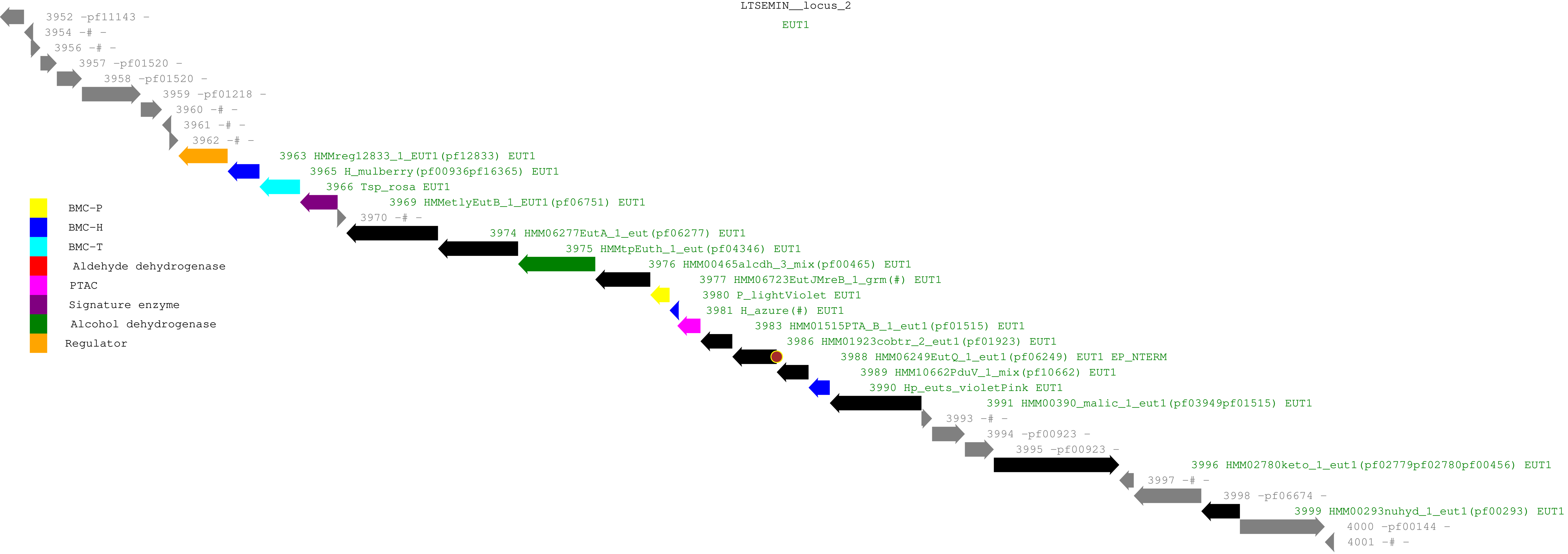

Salmonella enterica subsp. enterica serovar Montevideo str. S5-403. Bacteria; Proteobacteria; Gammaproteobacteria; Enterobacterales;

LTSEMON__locus_2

EUT1

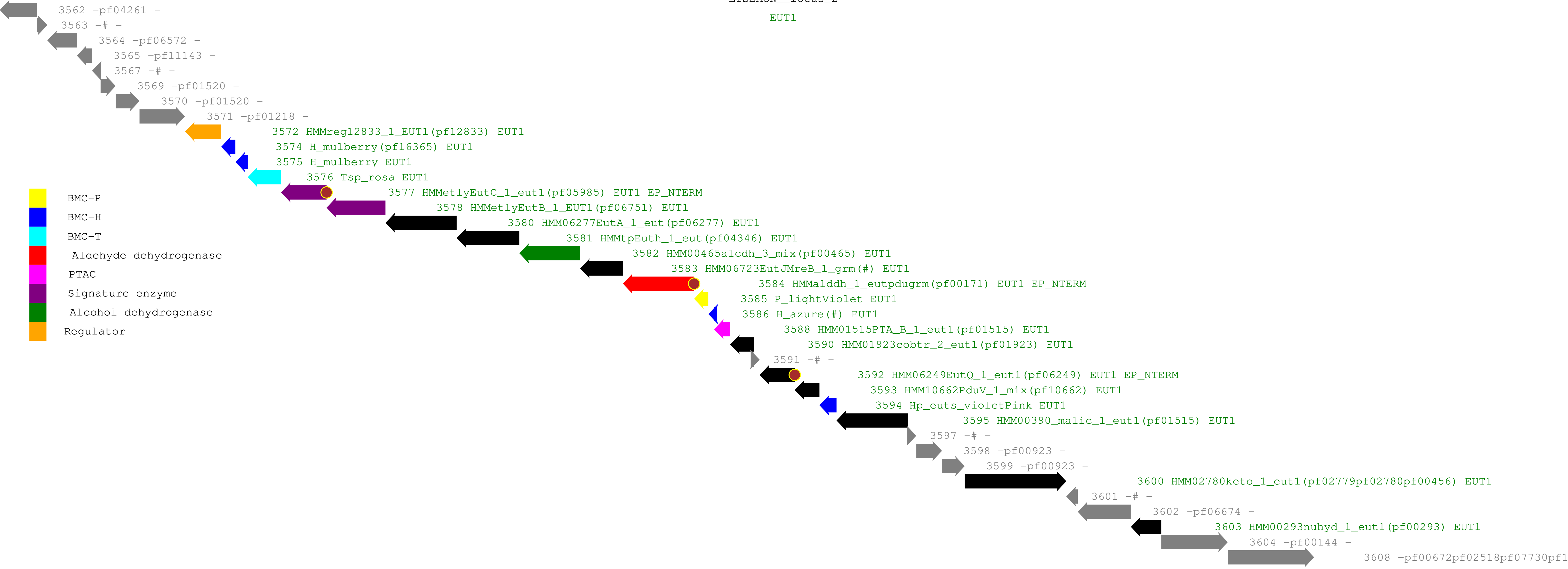

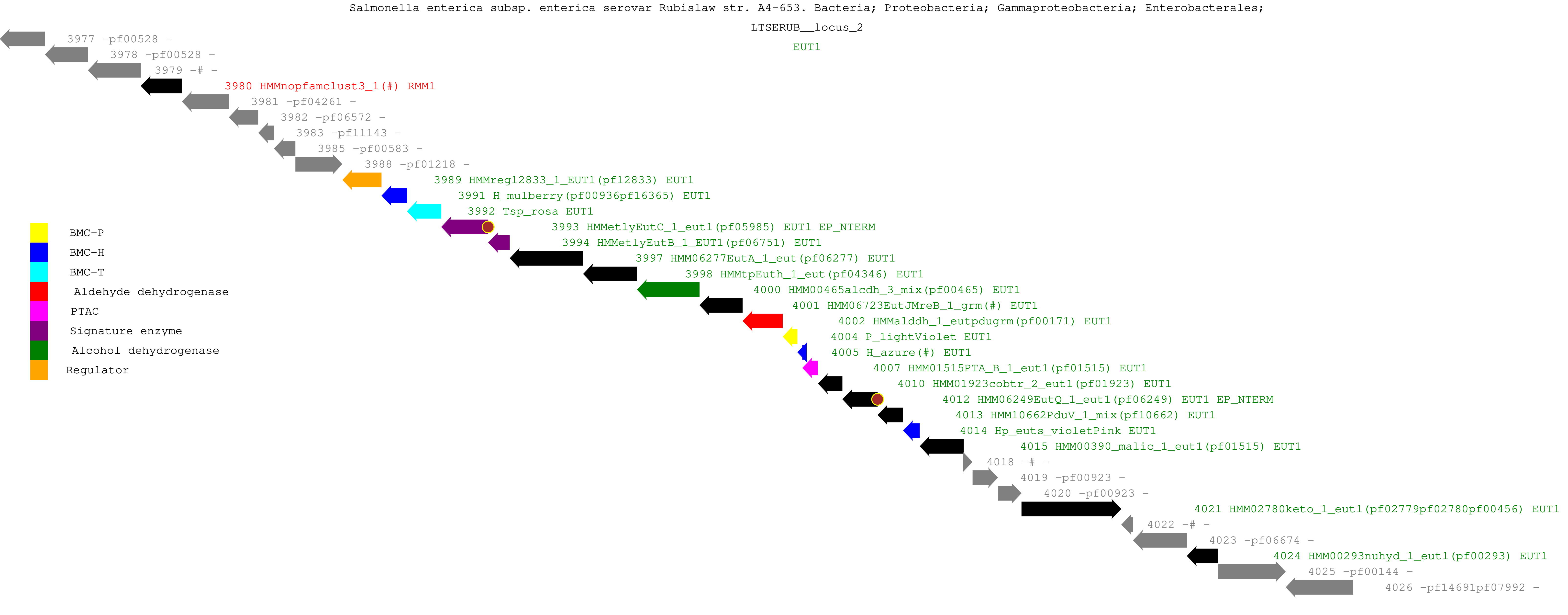
Salmonella enterica subsp. enterica serovar Rubislaw str. A4-653. Bacteria; Proteobacteria; Gammaproteobacteria; Enterobacterales;
LTSERUB__locus_2
EUT1
3977 -pf00528 -
3978 -pf00528 -
3979 -# -
3980 HMMnopfamclust3_1(#) RMM1
3981 -pf04261 -
3982 -pf06572 -
3983 -pf11143 -
3985 -pf00583 -
3988 -pf01218 -
3989 HMMreg12833_1_EUT1(pf12833) EUT1
3991 H_mulberry(pf00936pf16365) EUT1
3992 Tsp_rosa EUT1
3993 HMMetlyEutC_1_eut1(pf05985) EUT1 EP_NTERM
3994 HMMetlyEutB_1_EUT1(pf06751) EUT1
3997 HMM06277EutA_1_eut(pf06277) EUT1
3998 HMMtpEuth_1_eut(pf04346) EUT1
4000 HMM00465alcdh_3_mix(pf00465) EUT1
4001 HMM06723EutJMreB_1_grm(#) EUT1
4002 HMMalddh_1_eutpdugrm(pf00171) EUT1
4004 P_lightViolet EUT1
4005 H_azure(#) EUT1
4007 HMM01515PTA_B_1_eut1(pf01515) EUT1
4010 HMM01923cobtr_2_eut1(pf01923) EUT1
4012 HMM06249EutQ_1_eut1(pf06249) EUT1 EP_NTERM
4013 HMM10662PduV_1_mix(pf10662) EUT1
4014 Hp_euts_violetPink EUT1
4015 HMM00390_malic_1_eut1(pf01515) EUT1
4018 -# -
4019 -pf00923 -
4020 -pf00923 -
4021 HMM02780keto_1_eut1(pf02779pf02780pf00456) EUT1
4022 -# -
4023 -pf06674 -
4024 HMM00293nuhyd_1_eut1(pf00293) EUT1
4025 -pf00144 -
4026 -pf14691pf07992 -
BMC-P
BMC-H
BMC-T
Aldehyde dehydrogenase
PTAC
Signature enzyme
Alcohol dehydrogenase
Regulator

Salmonella enterica subsp. enterica serovar Senftenberg str. A4-543. Bacteria; Proteobacteria; Gammaproteobacteria; Enterobacterales;

LTSESEN__locus_2

EUT1

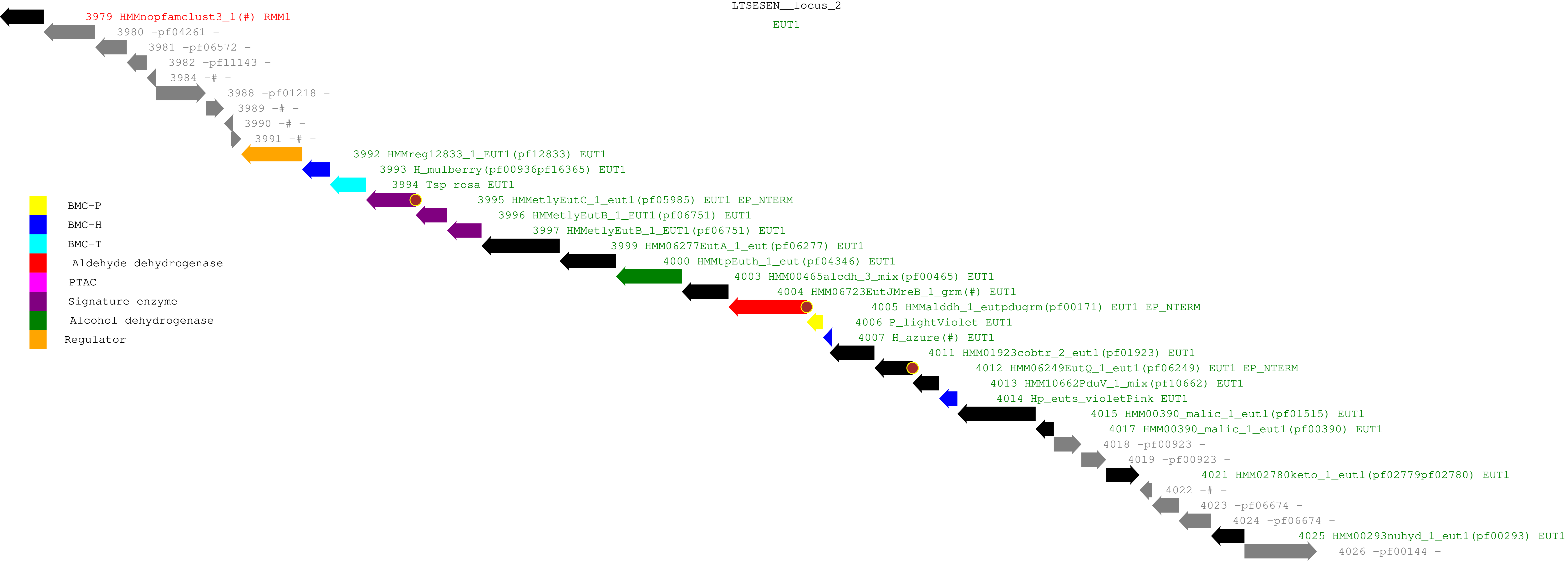

Salmonella enterica subsp. enterica serovar Uganda str. R8-3404. Bacteria; Proteobacteria; Gammaproteobacteria; Enterobacterales;

LTSEUGA__locus_2

EUT1

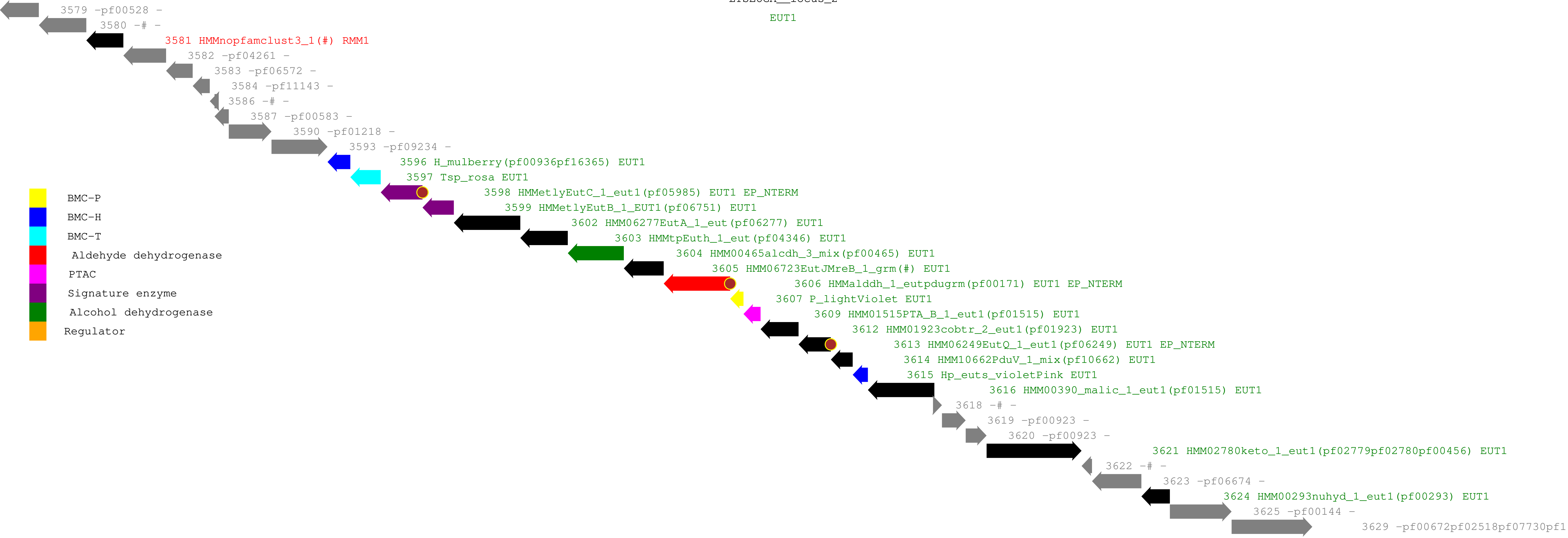

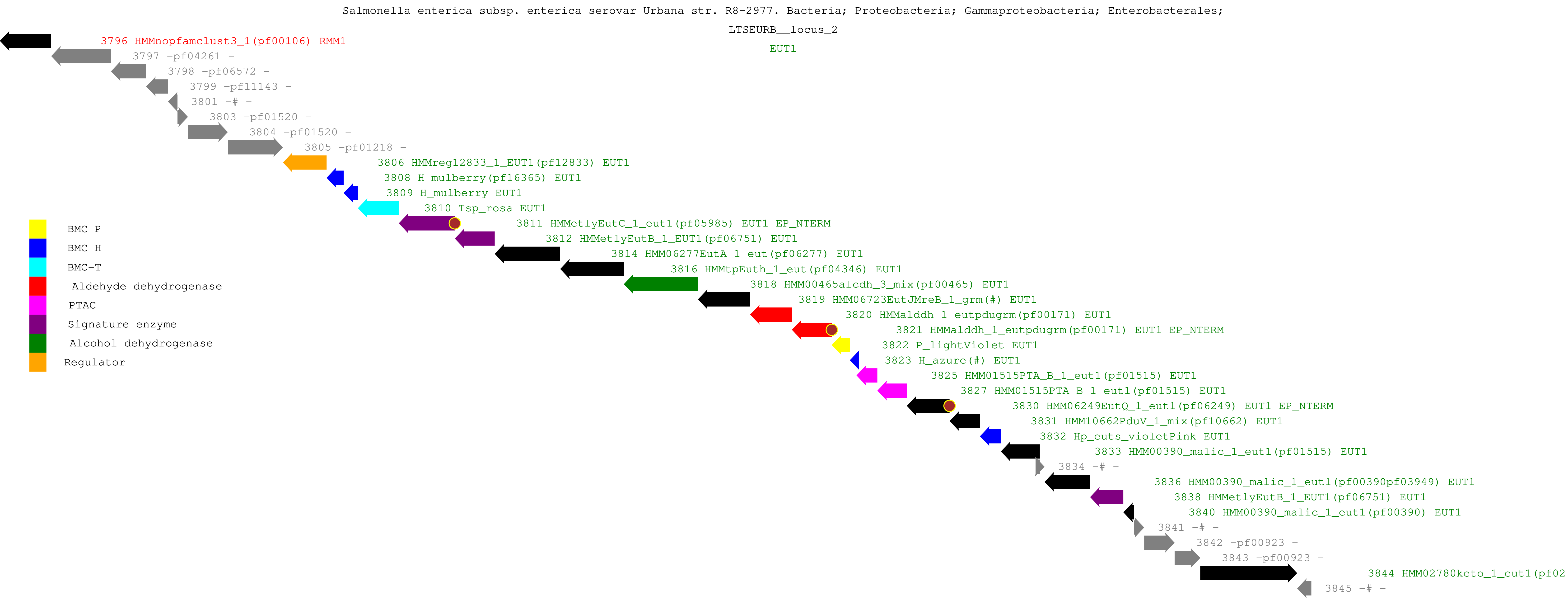
Salmonella enterica subsp. enterica serovar Urbana str. R8-2977. Bacteria; Proteobacteria; Gammaproteobacteria; Enterobacterales;
LTSEURB__locus_2
EUT1
3796 HMMnopfamclust3_1(pf00106) RMM1
3797 -pf04261 -
3798 -pf06572 -
3799 -pf11143 -
3801 -# -
3803 -pf01520 -
3804 -pf01520 -
3805 -pf01218 -
3806 HMMreg12833_1_EUT1(pf12833) EUT1
3808 H_mulberry(pf16365) EUT1
3809 H_mulberry EUT1
3810 Tsp_rosa EUT1
3811 HMMetlyEutC_1_eut1(pf05985) EUT1 EP_NTERM
3812 HMMetlyEutB_1_EUT1(pf06751) EUT1
3814 HMM06277EutA_1_eut(pf06277) EUT1
3816 HMMtpEuth_1_eut(pf04346) EUT1
3818 HMM00465alcdh_3_mix(pf00465) EUT1
3819 HMM06723EutJMreB_1_grm(#) EUT1
3820 HMMalddh_1_eutpdugrm(pf00171) EUT1
3821 HMMalddh_1_eutpdugrm(pf00171) EUT1 EP_NTERM
3822 P_lightViolet EUT1
3823 H_azure(#) EUT1
3825 HMM01515PTA_B_1_eut1(pf01515) EUT1
3827 HMM01515PTA_B_1_eut1(pf01515) EUT1
3830 HMM06249EutQ_1_eut1(pf06249) EUT1 EP_NTERM
3831 HMM10662PduV_1_mix(pf10662) EUT1
3832 Hp_euts_violetPink EUT1
3833 HMM00390_malic_1_eut1(pf01515) EUT1
3834 -# -
3836 HMM00390_malic_1_eut1(pf00390pf03949) EUT1
3838 HMMetlyEutB_1_EUT1(pf06751) EUT1
3840 HMM00390_malic_1_eut1(pf00390) EUT1
3841 -# -
3842 -pf00923 -
3843 -pf00923 -
3844 HMM02780keto_1_eut1(pf02
3845 -# -
BMC-P
BMC-H
BMC-T
Aldehyde dehydrogenase
PTAC
Signature enzyme
Alcohol dehydrogenase
Regulator

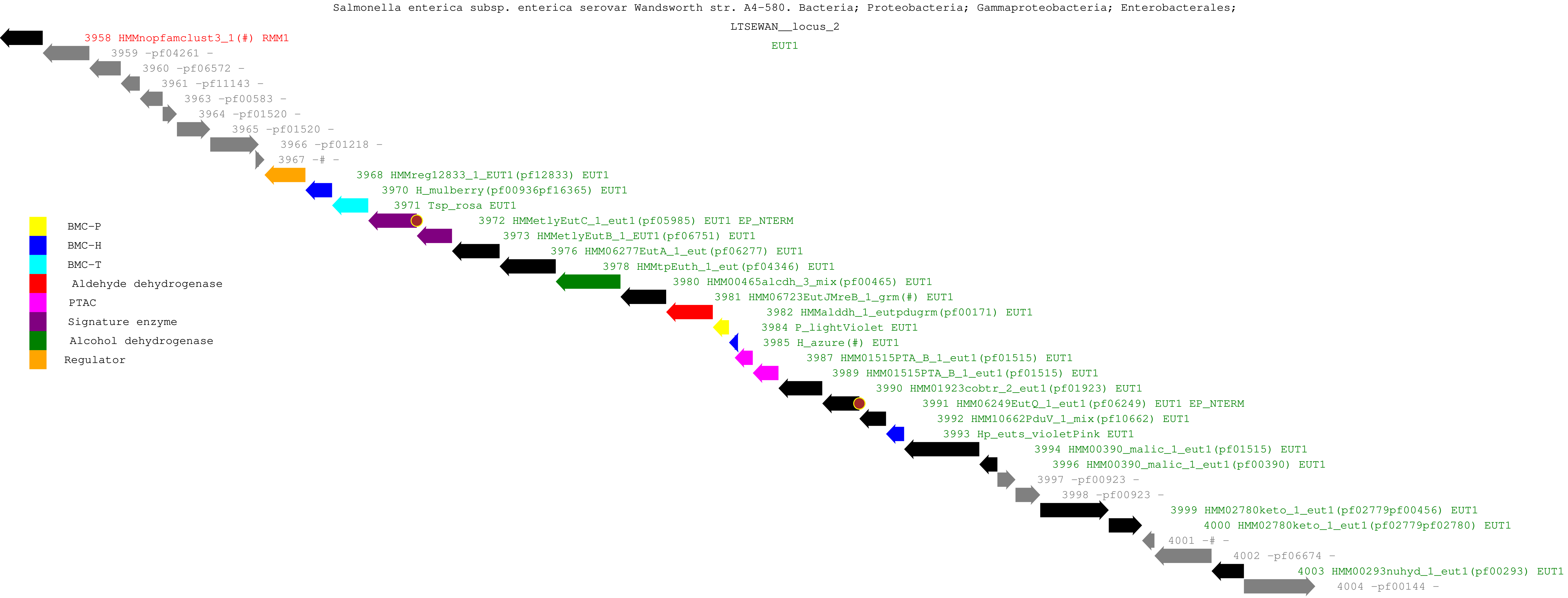
Salmonella enterica subsp. enterica serovar Wandsworth str. A4-580. Bacteria; Proteobacteria; Gammaproteobacteria; Enterobacterales;
LTSEWAN__locus_2
EUT1
3958 HMMnopfamclust3_1(#) RMM1
3959 -pf04261 -
3960 -pf06572 -
3961 -pf11143 -
3963 -pf00583 -
3964 -pf01520 -
3965 -pf01520 -
3966 -pf01218 -
3967 -# -
3968 HMMreg12833_1_EUT1(pf12833) EUT1
3970 H_mulberry(pf00936pf16365) EUT1
3971 Tsp_rosa EUT1
3972 HMMetlyEutC_1_eut1(pf05985) EUT1 EP_NTERM
3973 HMMetlyEutB_1_EUT1(pf06751) EUT1
3976 HMM06277EutA_1_eut(pf06277) EUT1
3978 HMMtpEuth_1_eut(pf04346) EUT1
3980 HMM00465alcdh_3_mix(pf00465) EUT1
3981 HMM06723EutJMreB_1_grm(#) EUT1
3982 HMMalddh_1_eutpdugrm(pf00171) EUT1
3984 P_lightViolet EUT1
3985 H_azure(#) EUT1
3987 HMM01515PTA_B_1_eut1(pf01515) EUT1
3989 HMM01515PTA_B_1_eut1(pf01515) EUT1
3990 HMM01923cobtr_2_eut1(pf01923) EUT1
3991 HMM06249EutQ_1_eut1(pf06249) EUT1 EP_NTERM
3992 HMM10662PduV_1_mix(pf10662) EUT1
3993 Hp_euts_violetPink EUT1
3994 HMM00390_malic_1_eut1(pf01515) EUT1
3996 HMM00390_malic_1_eut1(pf00390) EUT1
3997 -pf00923 -
3998 -pf00923 -
3999 HMM02780keto_1_eut1(pf02779pf00456) EUT1
4000 HMM02780keto_1_eut1(pf02779pf02780) EUT1
4001 -# -
4002 -pf06674 -
4003 HMM00293nuhyd_1_eut1(pf00293) EUT1
4004 -pf00144 -
BMC-P
BMC-H
BMC-T
Aldehyde dehydrogenase
PTAC
Signature enzyme
Alcohol dehydrogenase
Regulator

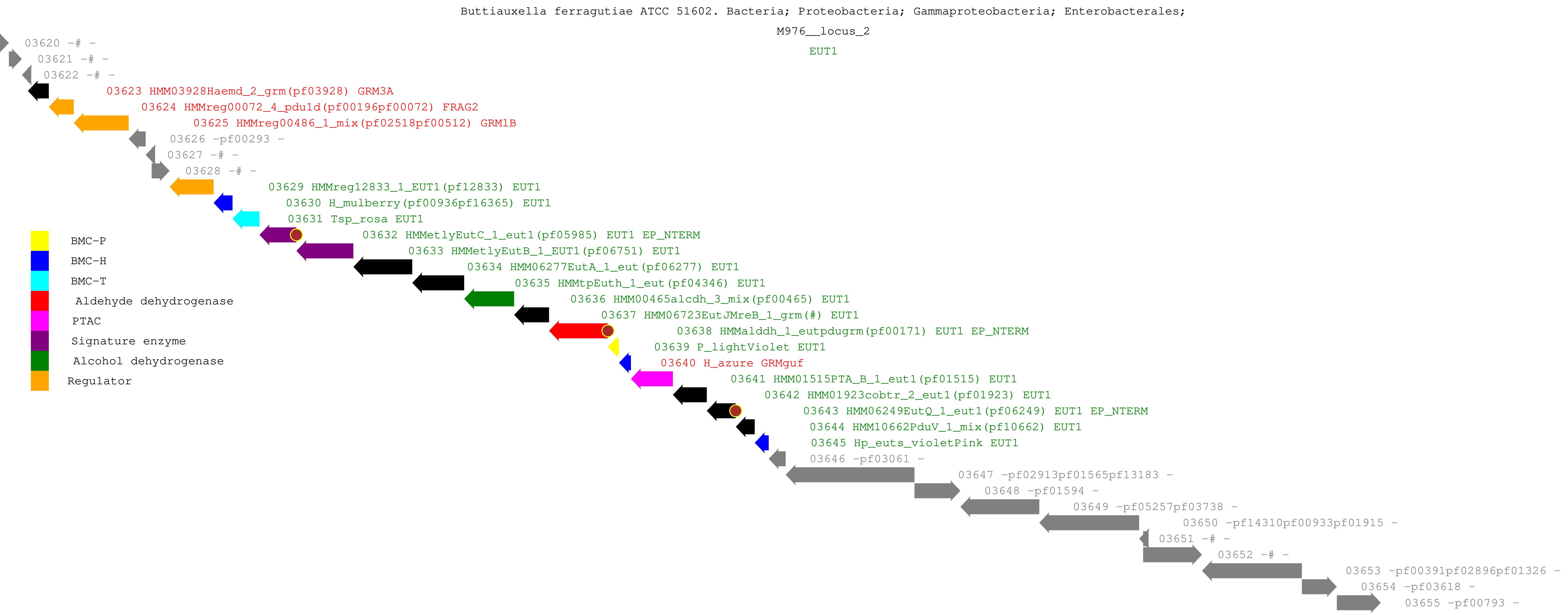
Buttiauxella ferragutiae ATCC 51602. Bacteria; Proteobacteria; Gammaproteobacteria; Enterobacterales;
M976__locus_2
EUT1
03620 -# -
03621 -# -
03622 -# -
03623 HMM03928Haemd_2_grm(pf03928) GRM3A
03624 HMMreg00072_4_pduld(pf00196pf00072) FRAG2
03625 HMMreg00486_1_mix(pf02518pf00512) GRM1B
03626 -pf00293 -
03627 -# -
03628 -# -
03629 HMMreg12833_1_EUT1(pf12833) EUT1
03630 H_mulberry(pf00936pf16365) EUT1
03631 Tsp_rosa EUT1
03632 HMMetlyEutC_1_eut1(pf05985) EUT1 EP_NTERM
03633 HMMetlyEutB_1_EUT1(pf06751) EUT1
03634 HMM06277EutA_1_eut(pf06277) EUT1
03635 HMMtpEuth_1_eut(pf04346) EUT1
03636 HMM00465alcdh_3_mix(pf00465) EUT1
03637 HMM06723EutJMreB_1_grm(#) EUT1
03638 HMMaldh_1_eutpdugrm(pf00171) EUT1 EP_NTERM
03639 P_lightViolet EUT1
03640 H_azure GRMguf
03641 HMM01515PTA_B_1_eut1(pf01515) EUT1
03642 HMM01923cobtr_2_eut1(pf01923) EUT1
03643 HMM06249EutQ_1_eut1(pf06249) EUT1 EP_NTERM
03644 HMM10662PduV_1_mix(pf10662) EUT1
03645 Hp_euts_violetPink EUT1
03646 -pf03061 -
03647 -pf02913pf01565pf13183 -
03648 -pf01594 -
03649 -pf05257pf03738 -
03650 -pf14310pf00933pf01915 -
03651 -# -
03652 -# -
03653 -pf00391pf02896pf01326 -
03654 -pf03618 -
03655 -pf00793 -
BMC-P
BMC-H
BMC-T
Aldehyde dehydrogenase
PTAC
Signature enzyme
Alcohol dehydrogenase
Regulator

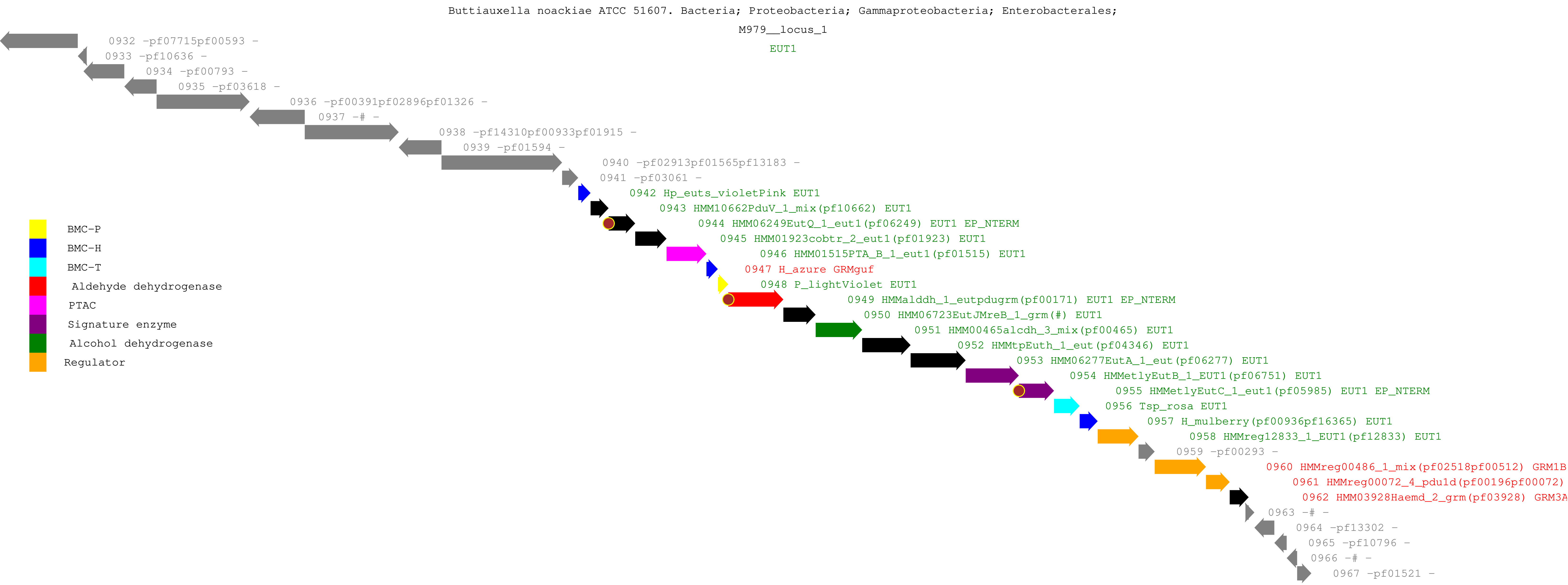
Buttiauxella noackiae ATCC 51607. Bacteria; Proteobacteria; Gammaproteobacteria; Enterobacterales;
M979__locus_1
EUT1
0932 -pf07715pf00593 -
0933 -pf10636 -
0934 -pf00793 -
0935 -pf03618 -
0936 -pf00391pf02896pf01326 -
0937 -# -
0938 -pf14310pf00933pf01915 -
0939 -pf01594 -
0940 -pf02913pf01565pf13183 -
0941 -pf03061 -
0942 Hp_euts_violetPink EUT1
0943 HMM10662PduV_1_mix(pf10662) EUT1
0944 HMM06249EutQ_1_eut1(pf06249) EUT1 EP_NTERM
0945 HMM01923cobtr_2_eut1(pf01923) EUT1
0946 HMM01515PTA_B_1_eut1(pf01515) EUT1
0947 H_azure GRMguf
0948 P_lightViolet EUT1
0949 HMMalddh_1_eutpdugrm(pf00171) EUT1 EP_NTERM
0950 HMM06723EutJMreB_1_grm(#) EUT1
0951 HMM00465alcdh_3_mix(pf00465) EUT1
0952 HMMtpEuth_1_eut(pf04346) EUT1
0953 HMM06277EutA_1_eut(pf06277) EUT1
0954 HMMetlyEutB_1_EUT1(pf06751) EUT1
0955 HMMetlyEutC_1_eut1(pf05985) EUT1 EP_NTERM
0956 Tsp_rosa EUT1
0957 H_mulberry(pf00936pf16365) EUT1
0958 HMMreg12833_1_EUT1(pf12833) EUT1
0959 -pf00293 -
0960 HMMreg00486_1_mix(pf02518pf00512) GRM1B
0961 HMMreg00072_4_pdu1d(pf00196pf00072)
0962 HMM03928Haemd_2_grm(pf03928) GRM3A
0963 -# -
0964 -pf13302 -
0965 -pf10796 -
0966 -# -
0967 -pf01521 -
BMC-P
BMC-H
BMC-T
Aldehyde dehydrogenase
PTAC
Signature enzyme
Alcohol dehydrogenase
Regulator

Obesumbacterium proteus ATCC 12841. Bacteria; Proteobacteria; Gammaproteobacteria; Enterobacterales;

M993__locus_1

EUT1

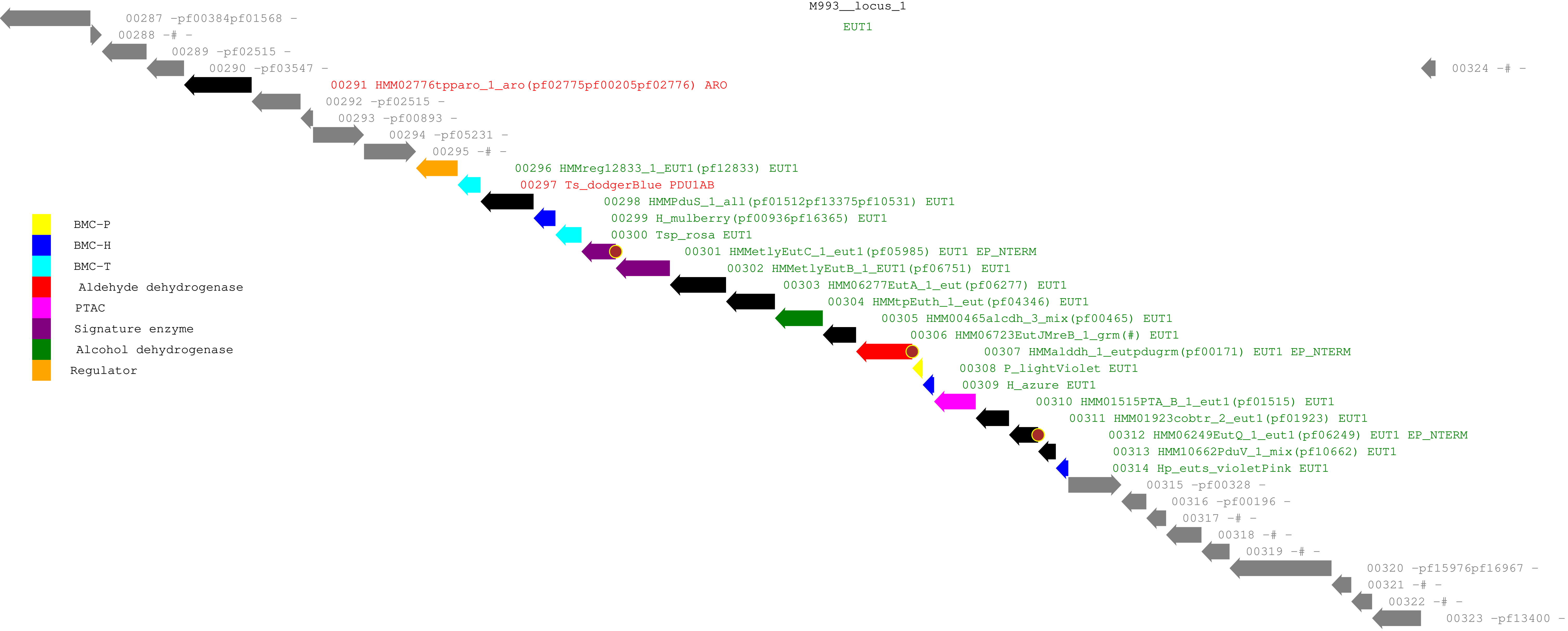

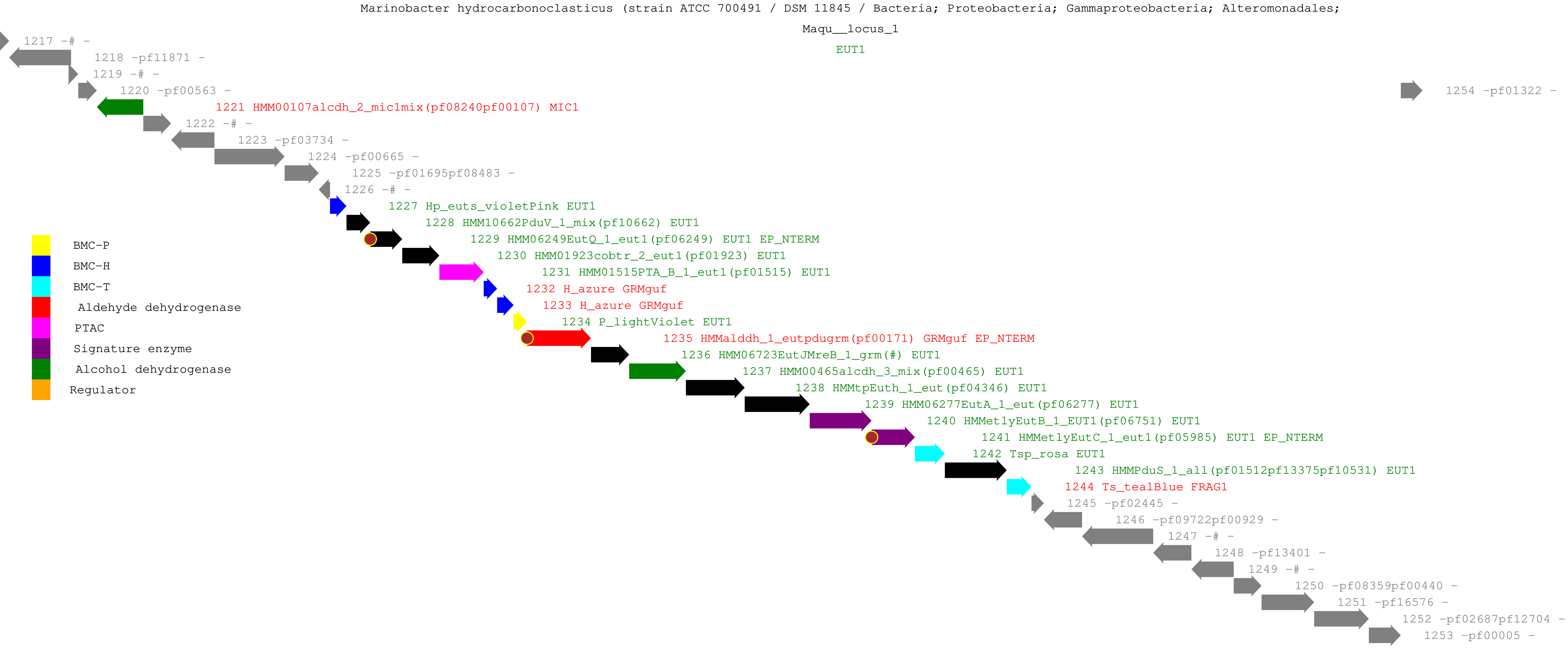
Marinobacter hydrocarbonoclasticus (strain ATCC 700491 / DSM 11845 / Bacteria; Proteobacteria; Gammaproteobacteria; Alteromonadales;
Maqu__locus_1
EUT1
1217 -# -
1218 -pf11871 -
1219 -# -
1220 -pf00563 -
1221 HMM00107alcdh_2_mic1mix(pf08240pf00107) MIC1
1222 -# -
1223 -pf03734 -
1224 -pf00665 -
1225 -pf01695pf08483 -
1226 -# -
1227 Hp_euts_violetPink EUT1
1228 HMM10662PduV_1_mix(pf10662) EUT1
1229 HMM06249EutQ_1_eut1(pf06249) EUT1 EP_NTERM
1230 HMM01923cobtr_2_eut1(pf01923) EUT1
1231 HMM01515PTA_B_1_eut1(pf01515) EUT1
1232 H_azure GRMguf
1233 H_azure GRMguf
1234 P_lightViolet EUT1
1235 HMMalddh_1_eutpdugrm(pf00171) GRMguf EP_NTERM
1236 HMM06723EutJMreB_1_grm(#) EUT1
1237 HMM00465alcdh_3_mix(pf00465) EUT1
1238 HMMtpEuth_1_eut(pf04346) EUT1
1239 HMM06277EutA_1_eut(pf06277) EUT1
1240 HMMetlyEutB_1_EUT1(pf06751) EUT1
1241 HMMetlyEutC_1_eut1(pf05985) EUT1 EP_NTERM
1242 Tsp_rosa EUT1
1243 HMMPduS_1_all(pf01512pf13375pf10531) EUT1
1244 Ts_tealBlue FRAG1
1245 -pf02445 -
1246 -pf09722pf00929 -
1247 -# -
1248 -pf13401 -
1249 -# -
1250 -pf08359pf00440 -
1251 -pf16576 -
1252 -pf02687pf12704 -
1253 -pf00005 -
1254 -pf01322 -
BMC-P
BMC-H
BMC-T
Aldehyde dehydrogenase
PTAC
Signature enzyme
Alcohol dehydrogenase
Regulator

Escherichia coli N36254PS. Bacteria; Proteobacteria; Gammaproteobacteria; Enterobacterales;

MP31__locus_2

EUT1

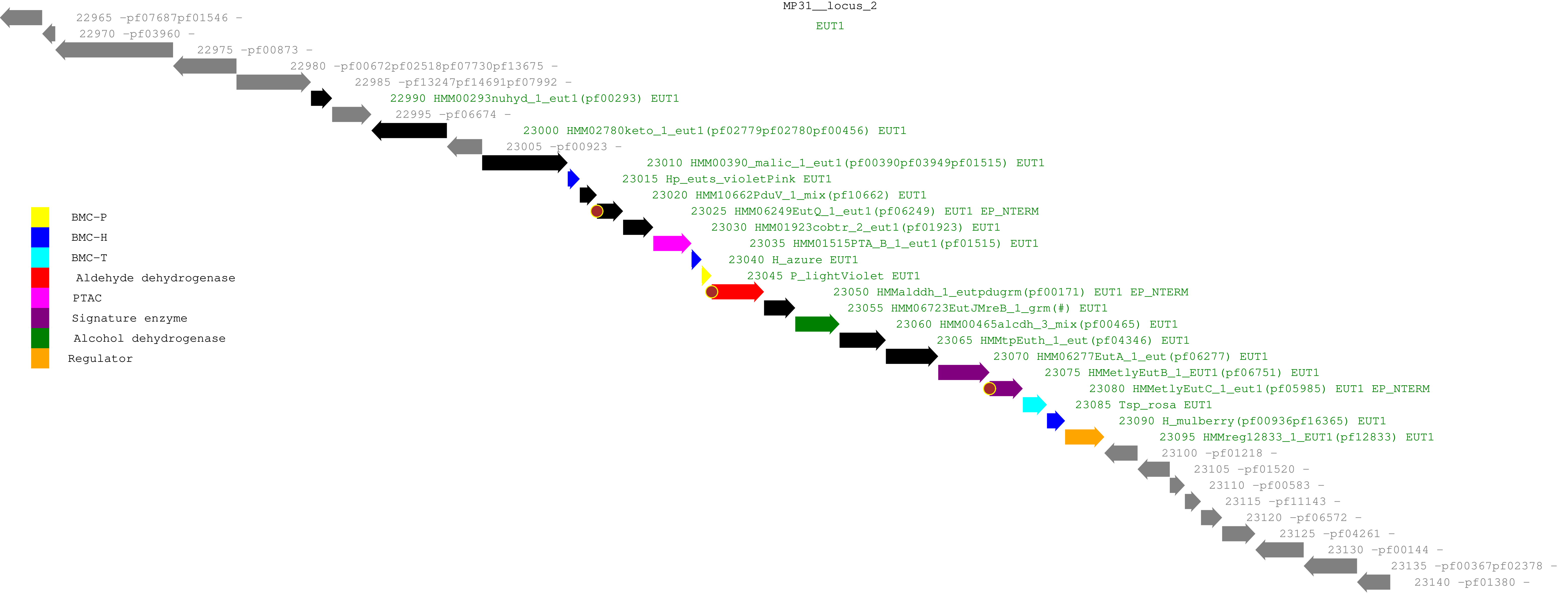

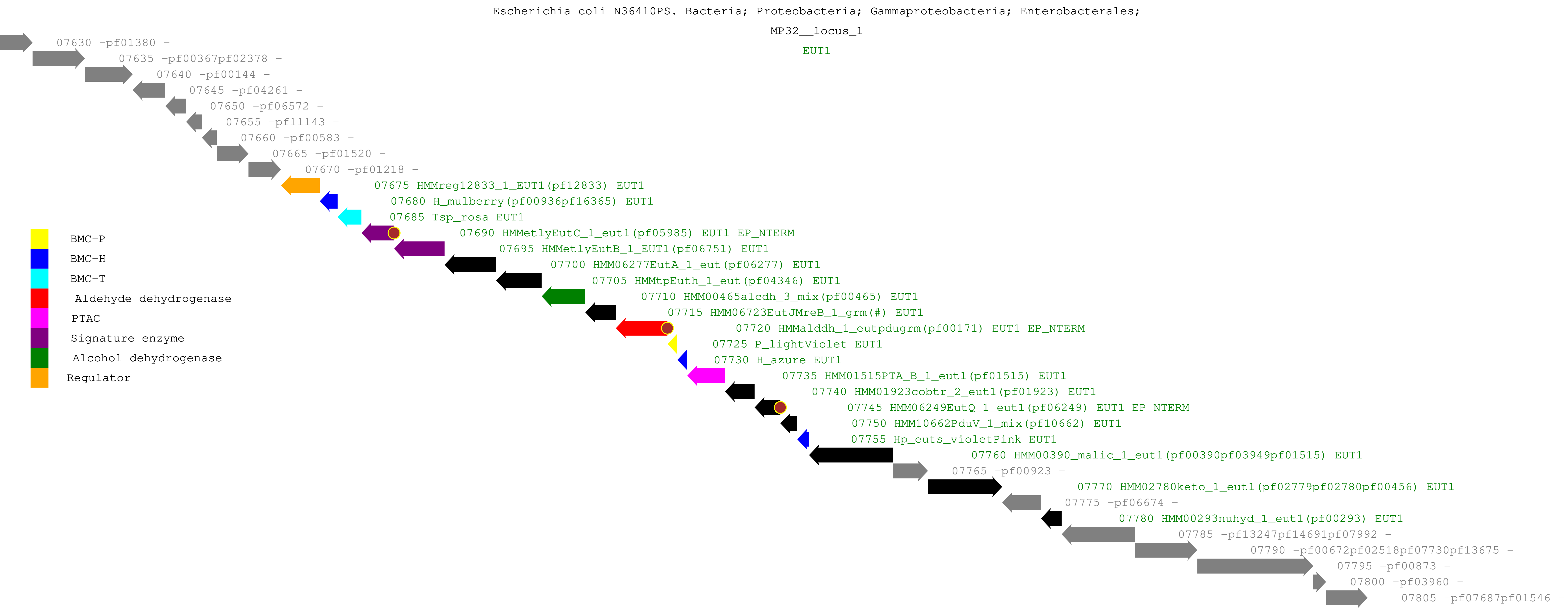
Escherichia coli N36410PS. Bacteria; Proteobacteria; Gammaproteobacteria; Enterobacterales;
MP32__locus_1
EUT1
07630 -pf01380 -
07635 -pf00367pf02378 -
07640 -pf00144 -
07645 -pf04261 -
07650 -pf06572 -
07655 -pf11143 -
07660 -pf00583 -
07665 -pf01520 -
07670 -pf01218 -
07675 HMMreg12833_1_EUT1(pf12833) EUT1
07680 H_mulberry(pf00936pf16365) EUT1
07685 Tsp_rosa EUT1
07690 HMMetlyEutC_1_eut1(pf05985) EUT1 EP_NTERM
07695 HMMetlyEutB_1_EUT1(pf06751) EUT1
07700 HMM06277EutA_1_eut(pf06277) EUT1
07705 HMMtpEuth_1_eut(pf04346) EUT1
07710 HMM00465alcdh_3_mix(pf00465) EUT1
07715 HMM06723EutJMreB_1_grm(#) EUT1
07720 HMMalddh_1_eutpdugrm(pf00171) EUT1 EP_NTERM
07725 P_lightViolet EUT1
07730 H_azure EUT1
07735 HMM01515PTA_B_1_eut1(pf01515) EUT1
07740 HMM01923cobtr_2_eut1(pf01923) EUT1
07745 HMM06249EutQ_1_eut1(pf06249) EUT1 EP_NTERM
07750 HMM10662PduV_1_mix(pf10662) EUT1
07755 Hp_euts_violetPink EUT1
07760 HMM00390_malic_1_eut1(pf00390pf03949pf01515) EUT1
07765 -pf00923 -
07770 HMM02780keto_1_eut1(pf02779pf02780pf00456) EUT1
07775 -pf06674 -
07780 HMM00293nuhyd_1_eut1(pf00293) EUT1
07785 -pf13247pf14691pf07992 -
07790 -pf00672pf02518pf07730pf13675 -
07795 -pf00873 -
07800 -pf03960 -
07805 -pf07687pf01546 -
BMC-P
BMC-H
BMC-T
Aldehyde dehydrogenase
PTAC
Signature enzyme
Alcohol dehydrogenase
Regulator

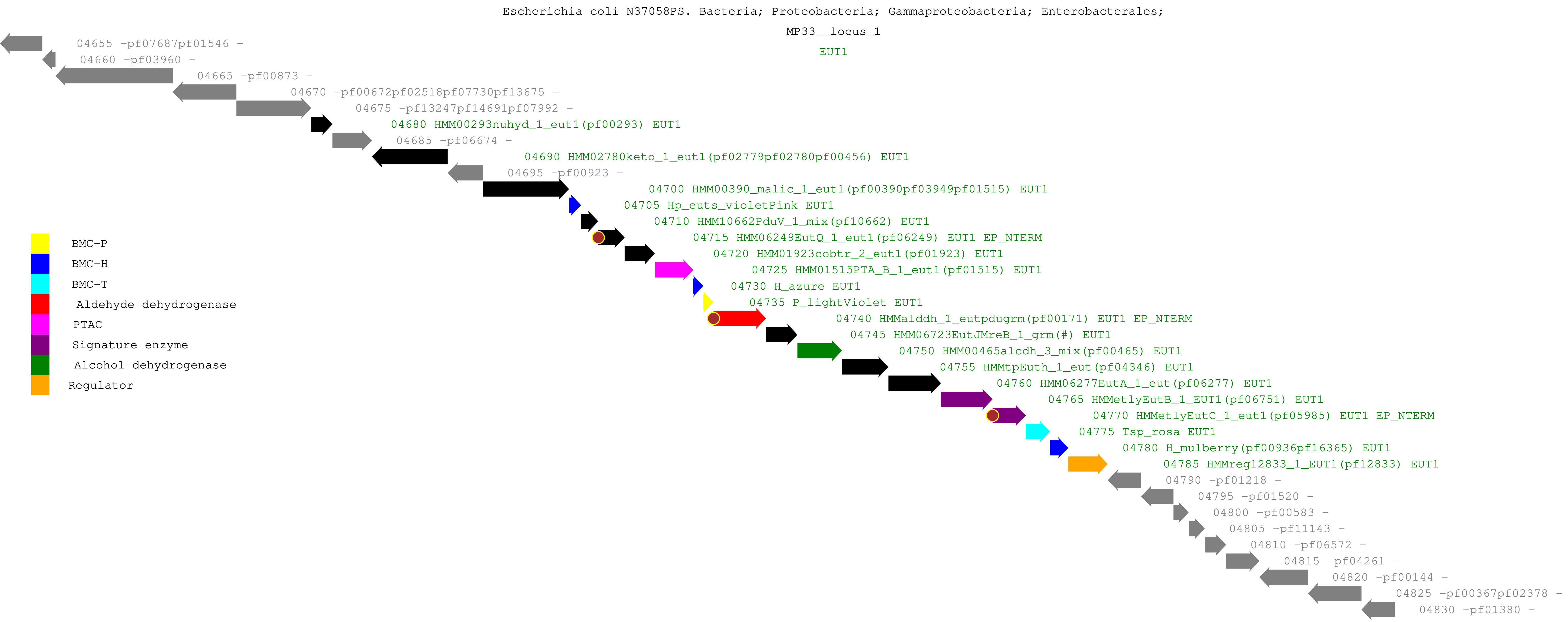
Escherichia coli N37058PS. Bacteria; Proteobacteria; Gammaproteobacteria; Enterobacterales;
MP33__locus_1
EUT1
04655 -pf07687pf01546 -
04660 -pf03960 -
04665 -pf00873 -
04670 -pf00672pf02518pf07730pf13675 -
04675 -pf13247pf14691pf07992 -
04680 HMM00293nuhyd_1_eut1(pf00293) EUT1
04685 -pf06674 -
04690 HMM02780keto_1_eut1(pf02779pf02780pf00456) EUT1
04695 -pf00923 -
04700 HMM00390_malic_1_eut1(pf00390pf03949pf01515) EUT1
04705 Hp_euts_violetPink EUT1
04710 HMM10662PduV_1_mix(pf10662) EUT1
04715 HMM06249EutQ_1_eut1(pf06249) EUT1 EP_NTERM
04720 HMM01923cobtr_2_eut1(pf01923) EUT1
04725 HMM01515PTA_B_1_eut1(pf01515) EUT1
04730 H_azure EUT1
04735 P_lightViolet EUT1
04740 HMMalddh_1_eutpdugrm(pf00171) EUT1 EP_NTERM
04745 HMM06723EutJMreB_1_grm(#) EUT1
04750 HMM00465alcdh_3_mix(pf00465) EUT1
04755 HMMtpEuth_1_eut(pf04346) EUT1
04760 HMM06277EutA_1_eut(pf06277) EUT1
04765 HMMetlyEutB_1_EUT1(pf06751) EUT1
04770 HMMetlyEutC_1_eut1(pf05985) EUT1 EP_NTERM
04775 Tsp_rosa EUT1
04780 H_mulberry(pf00936pf16365) EUT1
04785 HMMreg12833_1_EUT1(pf12833) EUT1
04790 -pf01218 -
04795 -pf01520 -
04800 -pf00583 -
04805 -pf11143 -
04810 -pf06572 -
04815 -pf04261 -
04820 -pf00144 -
04825 -pf00367pf02378 -
04830 -pf01380 -
BMC-P
BMC-H
BMC-T
Aldehyde dehydrogenase
PTAC
Signature enzyme
Alcohol dehydrogenase
Regulator

Escherichia coli N37122PS. Bacteria; Proteobacteria; Gammaproteobacteria; Enterobacterales;

MP34__locus_1

EUT1

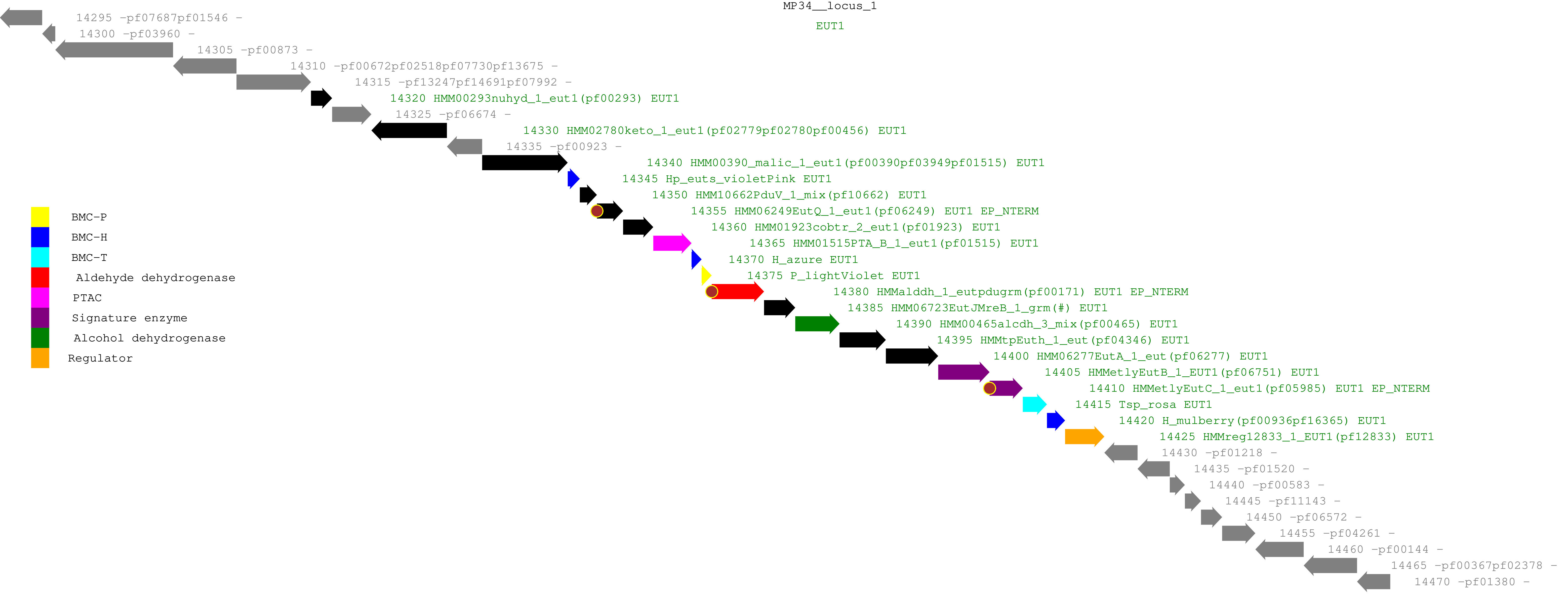

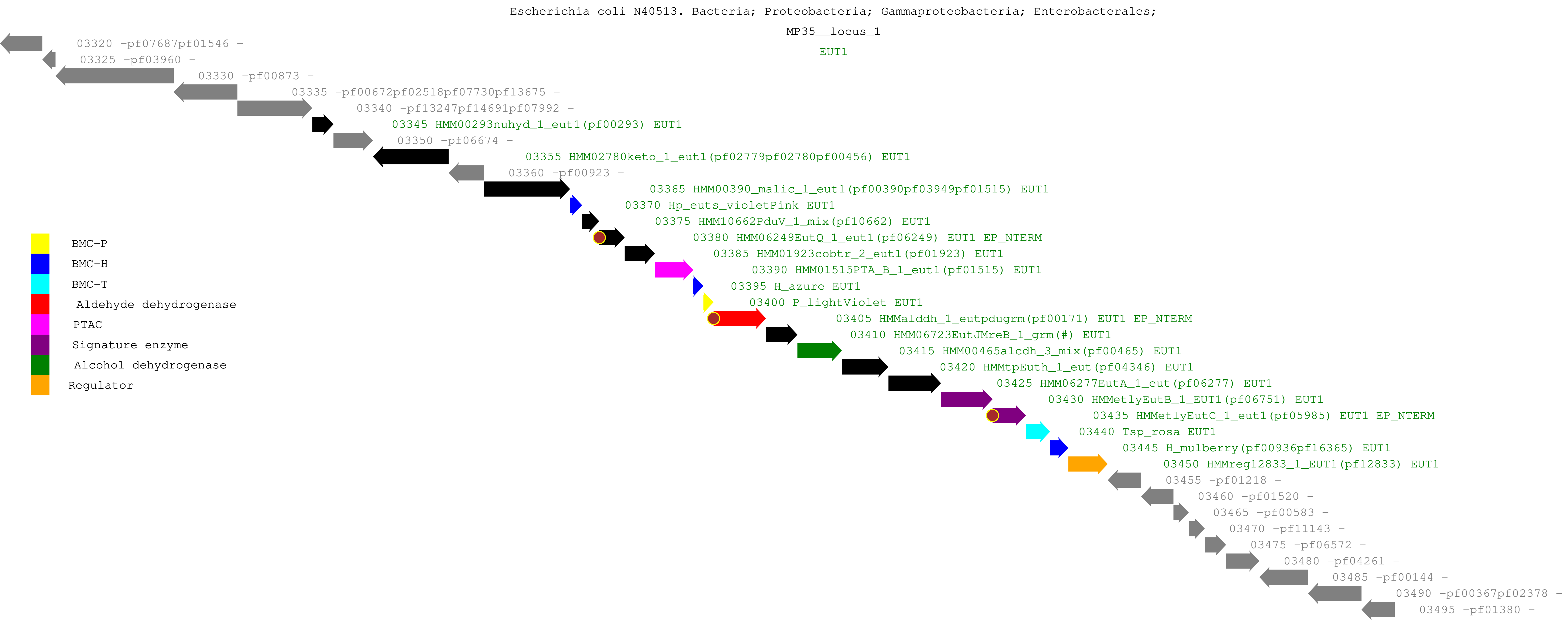
Escherichia coli N40513. Bacteria; Proteobacteria; Gammaproteobacteria; Enterobacterales;
MP35__locus_1
EUT1
03320 -pf07687pf01546 -
03325 -pf03960 -
03330 -pf00873 -
03335 -pf00672pf02518pf07730pf13675 -
03340 -pf13247pf14691pf07992 -
03345 HMM00293nuhyd_1_eut1(pf00293) EUT1
03350 -pf06674 -
03355 HMM02780keto_1_eut1(pf02779pf02780pf00456) EUT1
03360 -pf00923 -
03365 HMM00390_malic_1_eut1(pf00390pf03949pf01515) EUT1
03370 Hp_euts_violetPink EUT1
03375 HMM10662PduV_1_mix(pf10662) EUT1
03380 HMM06249EutQ_1_eut1(pf06249) EUT1 EP_NTERM
03385 HMM01923cobtr_2_eut1(pf01923) EUT1
03390 HMM01515PTA_B_1_eut1(pf01515) EUT1
03395 H_azure EUT1
03400 P_lightViolet EUT1
03405 HMMalddh_1_eutpdugrm(pf00171) EUT1 EP_NTERM
03410 HMM06723EutJMreB_1_grm(#) EUT1
03415 HMM00465alcdh_3_mix(pf00465) EUT1
03420 HMMtpEuth_1_eut(pf04346) EUT1
03425 HMM06277EutA_1_eut(pf06277) EUT1
03430 HMMetlyEutB_1_EUT1(pf06751) EUT1
03435 HMMetlyEutC_1_eut1(pf05985) EUT1 EP_NTERM
03440 Tsp_rosa EUT1
03445 H_mulberry(pf00936pf16365) EUT1
03450 HMMreg12833_1_EUT1(pf12833) EUT1
03455 -pf01218 -
03460 -pf01520 -
03465 -pf00583 -
03470 -pf11143 -
03475 -pf06572 -
03480 -pf04261 -
03485 -pf00144 -
03490 -pf00367pf02378 -
03495 -pf01380 -
BMC-P
BMC-H
BMC-T
Aldehyde dehydrogenase
PTAC
Signature enzyme
Alcohol dehydrogenase
Regulator

Morganella morganii subsp. morganii KT. Bacteria; Proteobacteria; Gammaproteobacteria; Enterobacterales;

MU9__locus_1

EUT1

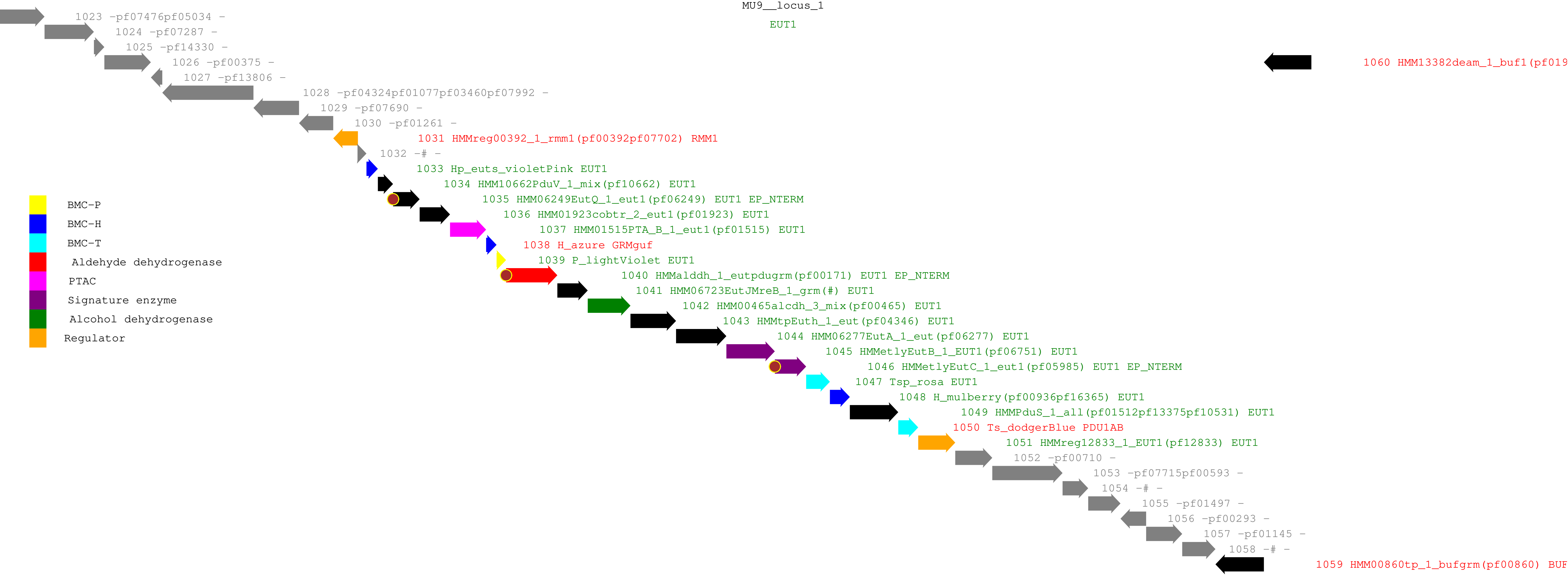

Salmonella enterica subsp. arizonae serovar 62:z36:- str. RKS2983. Bacteria; Proteobacteria; Gammaproteobacteria; Enterobacterales;

N898__locus_1

EUT1

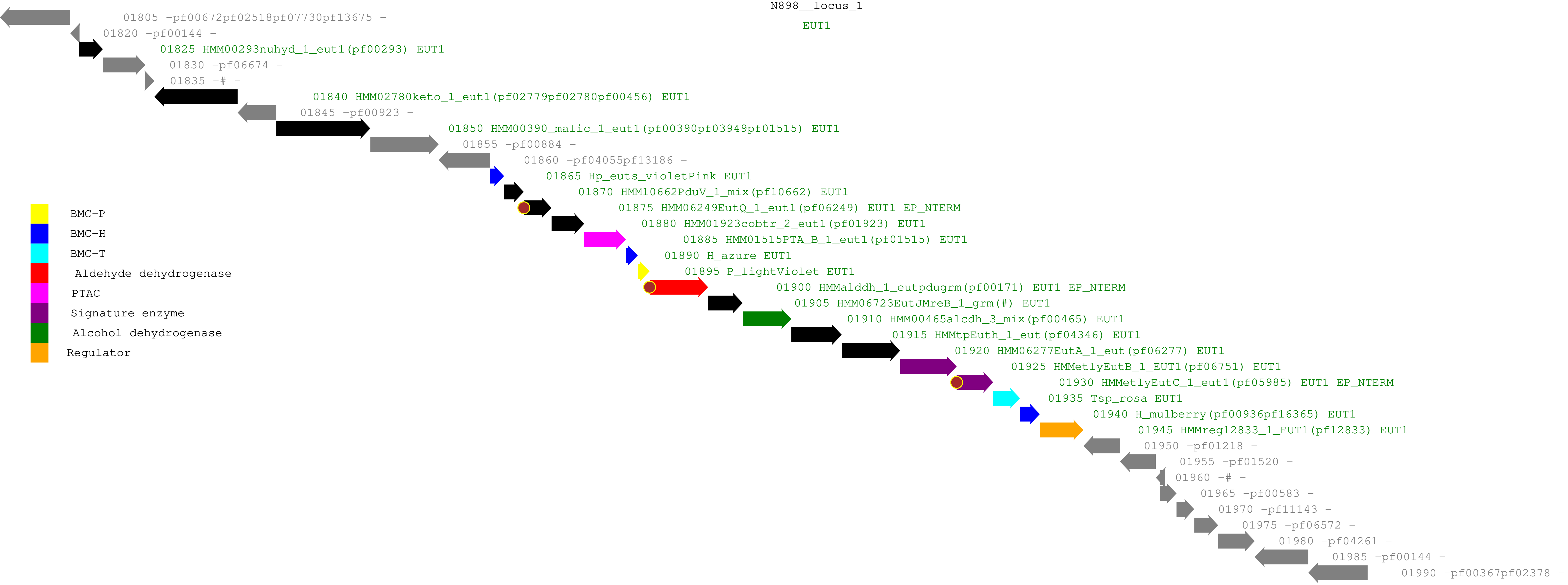

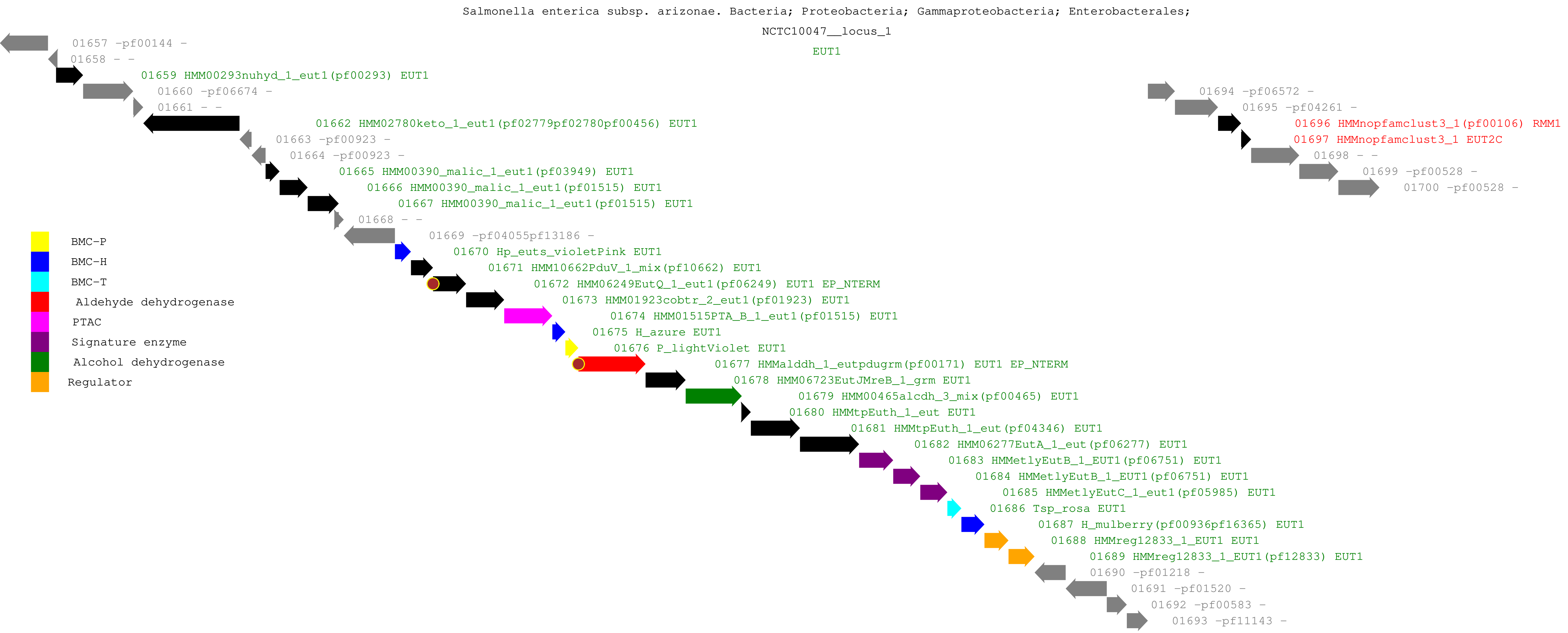
Salmonella enterica subsp. arizonae. Bacteria; Proteobacteria; Gammaproteobacteria; Enterobacterales;
NCTC10047__locus_1
EUT1
01657 -pf00144 -
01658 - -
01659 HMM00293nuhyd_1_eut1(pf00293) EUT1
01660 -pf06674 -
01661 - -
01662 HMM02780keto_1_eut1(pf02779pf02780pf00456) EUT1
01663 -pf00923 -
01664 -pf00923 -
01665 HMM00390_malic_1_eut1(pf03949) EUT1
01666 HMM00390_malic_1_eut1(pf01515) EUT1
01667 HMM00390_malic_1_eut1(pf01515) EUT1
01668 - -
01669 -pf04055pf13186 -
01670 Hp_euts_violetPink EUT1
01671 HMM10662PduV_1_mix(pf10662) EUT1
01672 HMM06249EutQ_1_eut1(pf06249) EUT1 EP_NTERM
01673 HMM01923cobtr_2_eut1(pf01923) EUT1
01674 HMM01515PTA_B_1_eut1(pf01515) EUT1
01675 H_azure EUT1
01676 P_lightViolet EUT1
01677 HMMalddh_1_eutpdugrm(pf00171) EUT1 EP_NTERM
01678 HMM06723EutJMreB_1_grm EUT1
01679 HMM00465alcdh_3_mix(pf00465) EUT1
01680 HMMtpEuth_1_eut EUT1
01681 HMMtpEuth_1_eut(pf04346) EUT1
01682 HMM06277EutA_1_eut(pf06277) EUT1
01683 HMMetlyEutB_1_EUT1(pf06751) EUT1
01684 HMMetlyEutB_1_EUT1(pf06751) EUT1
01685 HMMetlyEutC_1_eut1(pf05985) EUT1
01686 Tsp_rosa EUT1
01687 H_mulberry(pf00936pf16365) EUT1
01688 HMMreg12833_1_EUT1 EUT1
01689 HMMreg12833_1_EUT1(pf12833) EUT1
01690 -pf01218 -
01691 -pf01520 -
01692 -pf00583 -
01693 -pf11143 -
01694 -pf06572 -
01695 -pf04261 -
01696 HMMnopfamclust3_1(pf00106) RMM1
01697 HMMnopfamclust3_1 EUT2C
01698 - -
01699 -pf00528 -
01700 -pf00528 -
BMC-P
BMC-H
BMC-T
Aldehyde dehydrogenase
PTAC
Signature enzyme
Alcohol dehydrogenase
Regulator

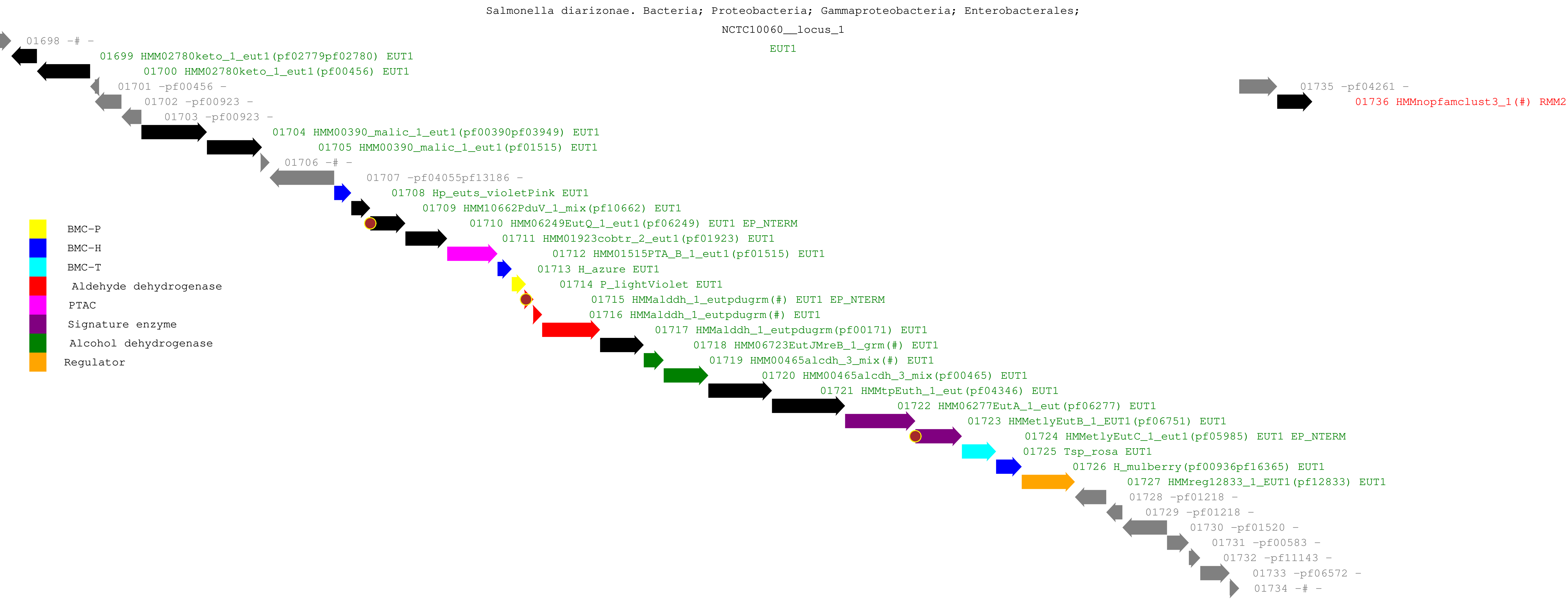
Salmonella diarizonae. Bacteria; Proteobacteria; Gammaproteobacteria; Enterobacterales;
NCTC10060__locus_1
EUT1
01698 -# -
01699 HMM02780keto_1_eut1(pf02779pf02780) EUT1
01700 HMM02780keto_1_eut1(pf00456) EUT1
01701 -pf00456 -
01702 -pf00923 -
01703 -pf00923 -
01704 HMM00390_malic_1_eut1(pf00390pf03949) EUT1
01705 HMM00390_malic_1_eut1(pf01515) EUT1
01706 -# -
01707 -pf04055pf13186 -
01708 Hp_euts_violetPink EUT1
01709 HMM10662PduV_1_mix(pf10662) EUT1
01710 HMM06249EutQ_1_eut1(pf06249) EUT1 EP_NTERM
01711 HMM01923cobtr_2_eut1(pf01923) EUT1
01712 HMM01515PTA_B_1_eut1(pf01515) EUT1
01713 H_azure EUT1
01714 P_lightViolet EUT1
01715 HMMaldh_1_eutpdugrm(#) EUT1 EP_NTERM
01716 HMMaldh_1_eutpdugrm(#) EUT1
01717 HMMaldh_1_eutpdugrm(pf00171) EUT1
01718 HMM06723EutJMreB_1_grm(#) EUT1
01719 HMM00465alcdh_3_mix(#) EUT1
01720 HMM00465alcdh_3_mix(pf00465) EUT1
01721 HMMtpEuth_1_eut(pf04346) EUT1
01722 HMM06277EutA_1_eut(pf06277) EUT1
01723 HMMetlyEutB_1_EUT1(pf06751) EUT1
01724 HMMetlyEutC_1_eut1(pf05985) EUT1 EP_NTERM
01725 Tsp_rosa EUT1
01726 H_mulberry(pf00936pf16365) EUT1
01727 HMMreg12833_1_EUT1(pf12833) EUT1
01728 -pf01218 -
01729 -pf01218 -
01730 -pf01520 -
01731 -pf00583 -
01732 -pf11143 -
01733 -pf06572 -
01734 -# -
01735 -pf04261 -
01736 HMMnopfamclust3_1(#) RMM2
BMC-P
BMC-H
BMC-T
Aldehyde dehydrogenase
PTAC
Signature enzyme
Alcohol dehydrogenase
Regulator

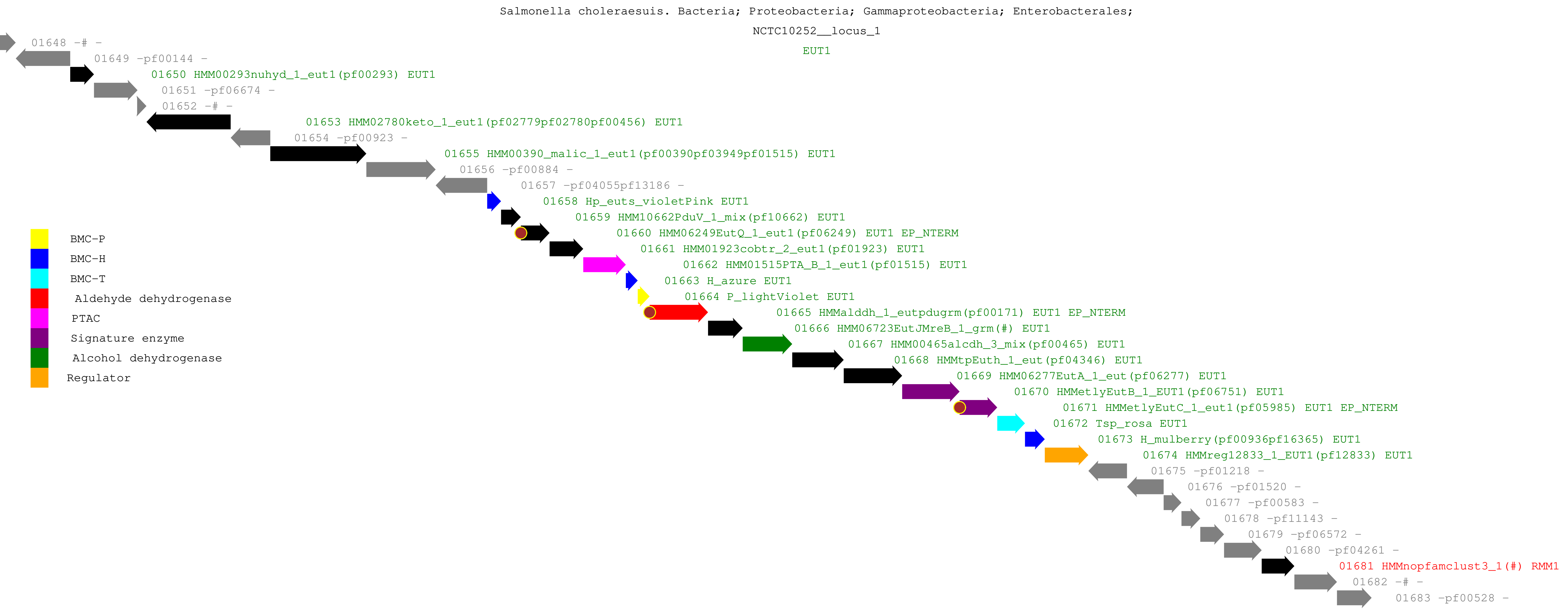
Salmonella choleraesuis. Bacteria; Proteobacteria; Gammaproteobacteria; Enterobacterales;
NCTC10252__locus_1
EUT1
01648 -# -
01649 -pf00144 -
01650 HMM00293nuhyd_1_eut1(pf00293) EUT1
01651 -pf06674 -
01652 -# -
01653 HMM02780keto_1_eut1(pf02779pf02780pf00456) EUT1
01654 -pf00923 -
01655 HMM00390_malic_1_eut1(pf00390pf03949pf01515) EUT1
01656 -pf00884 -
01657 -pf04055pf13186 -
01658 Hp_euts_violetPink EUT1
01659 HMM10662PduV_1_mix(pf10662) EUT1
01660 HMM06249EutQ_1_eut1(pf06249) EUT1 EP_NTERM
01661 HMM01923cobtr_2_eut1(pf01923) EUT1
01662 HMM01515PTA_B_1_eut1(pf01515) EUT1
01663 H_azure EUT1
01664 P_lightViolet EUT1
01665 HMMalddh_1_eutpdugrm(pf00171) EUT1 EP_NTERM
01666 HMM06723EutJMreB_1_grm(#) EUT1
01667 HMM00465alcdh_3_mix(pf00465) EUT1
01668 HMMtpEuth_1_eut(pf04346) EUT1
01669 HMM06277EutA_1_eut(pf06277) EUT1
01670 HMMetlyEutB_1_EUT1(pf06751) EUT1
01671 HMMetlyEutC_1_eut1(pf05985) EUT1 EP_NTERM
01672 Tsp_rosa EUT1
01673 H_mulberry(pf00936pf16365) EUT1
01674 HMMreg12833_1_EUT1(pf12833) EUT1
01675 -pf01218 -
01676 -pf01520 -
01677 -pf00583 -
01678 -pf11143 -
01679 -pf06572 -
01680 -pf04261 -
01681 HMMnopfamclust3_1(#) RMM1
01682 -# -
01683 -pf00528 -
BMC-P
BMC-H
BMC-T
Aldehyde dehydrogenase
PTAC
Signature enzyme
Alcohol dehydrogenase
Regulator

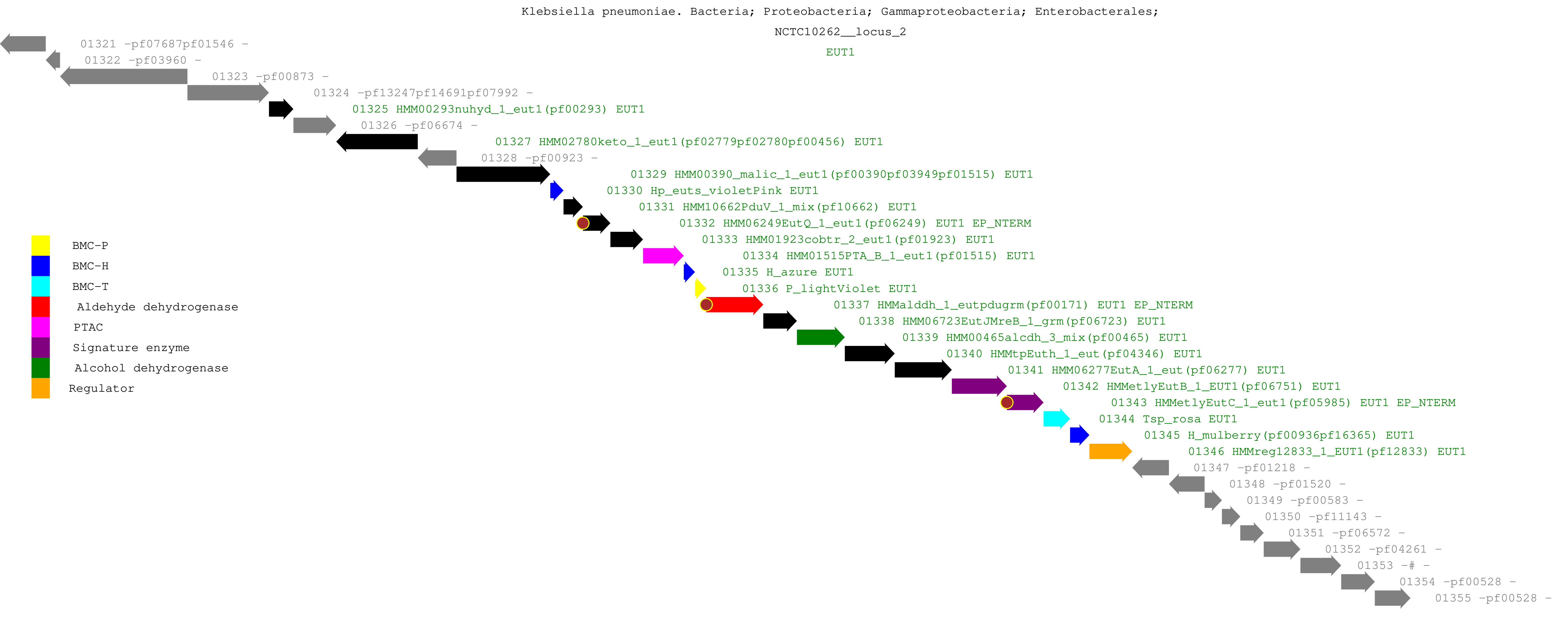
Klebsiella pneumoniae. Bacteria; Proteobacteria; Gammaproteobacteria; Enterobacterales;
NCTC10262__locus_2
EUT1
01321 -pf07687pf01546 -
01322 -pf03960 -
01323 -pf00873 -
01324 -pf13247pf14691pf07992 -
01325 HMM00293nuhyd_1_eut1(pf00293) EUT1
01326 -pf06674 -
01327 HMM02780keto_1_eut1(pf02779pf02780pf00456) EUT1
01328 -pf00923 -
01329 HMM00390_malic_1_eut1(pf00390pf03949pf01515) EUT1
01330 Hp_euts_violetPink EUT1
01331 HMM10662PduV_1_mix(pf10662) EUT1
01332 HMM06249EutQ_1_eut1(pf06249) EUT1 EP_NTERM
01333 HMM01923cobtr_2_eut1(pf01923) EUT1
01334 HMM01515PTA_B_1_eut1(pf01515) EUT1
01335 H_azure EUT1
01336 P_lightViolet EUT1
01337 HMMalddh_1_eutpdugrm(pf00171) EUT1 EP_NTERM
01338 HMM06723EutJMreB_1_grm(pf06723) EUT1
01339 HMM00465alcdh_3_mix(pf00465) EUT1
01340 HMMtpEuth_1_eut(pf04346) EUT1
01341 HMM06277EutA_1_eut(pf06277) EUT1
01342 HMMetlyEutB_1_EUT1(pf06751) EUT1
01343 HMMetlyEutC_1_eut1(pf05985) EUT1 EP_NTERM
01344 Tsp_rosa EUT1
01345 H_mulberry(pf00936pf16365) EUT1
01346 HMMreg12833_1_EUT1(pf12833) EUT1
01347 -pf01218 -
01348 -pf01520 -
01349 -pf00583 -
01350 -pf11143 -
01351 -pf06572 -
01352 -pf04261 -
01353 -# -
01354 -pf00528 -
01355 -pf00528 -
BMC-P
BMC-H
BMC-T
Aldehyde dehydrogenase
PTAC
Signature enzyme
Alcohol dehydrogenase
Regulator

Escherichia coli. Bacteria; Proteobacteria; Gammaproteobacteria; Enterobacterales;

NCTC10418__locus_1

EUT1

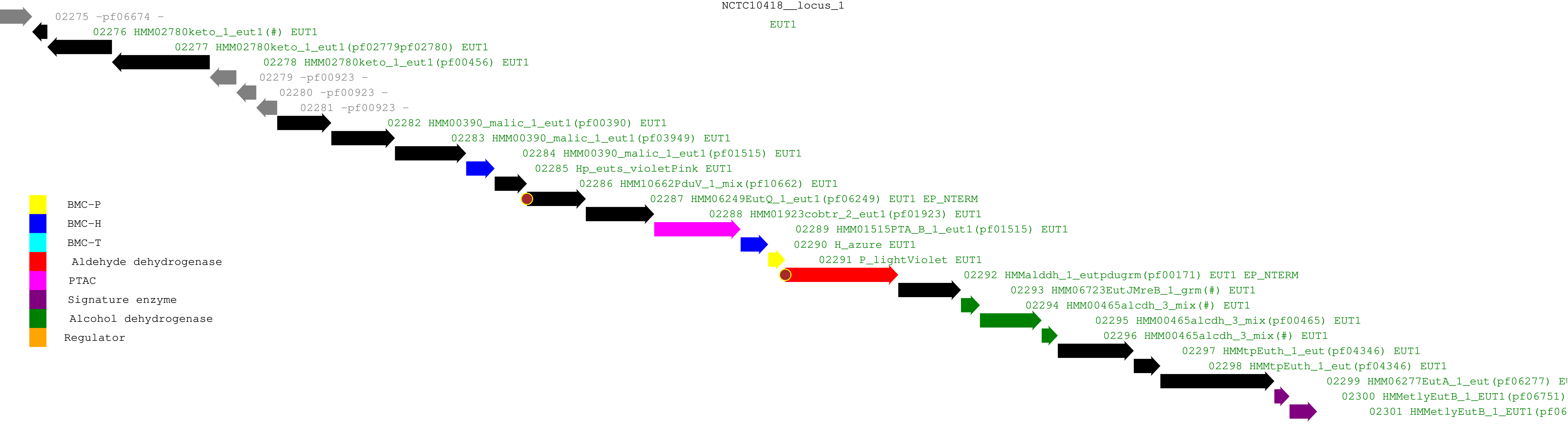

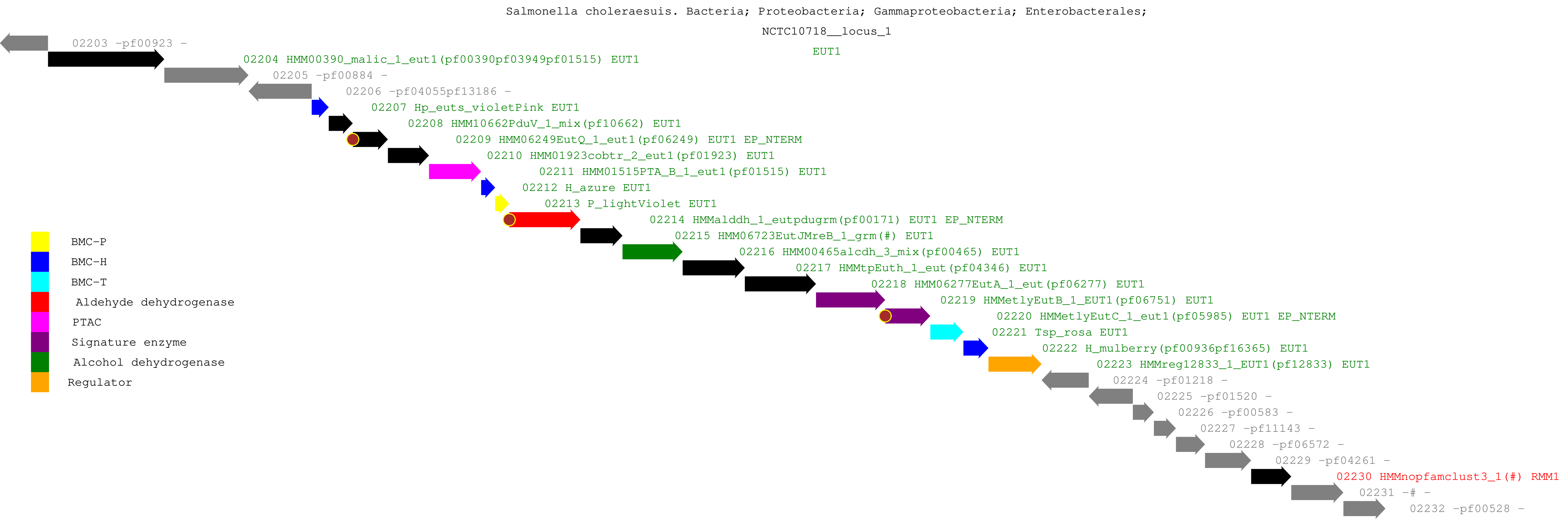
Salmonella choleraesuis. Bacteria; Proteobacteria; Gammaproteobacteria; Enterobacterales;
NCTC10718__locus_1
EUT1
02203 -pf00923 -
02204 HMM00390_malic_1_eut1(pf00390pf03949pf01515) EUT1
02205 -pf00884 -
02206 -pf04055pf13186 -
02207 Hp_euts_violetPink EUT1
02208 HMM10662PduV_1_mix(pf10662) EUT1
02209 HMM06249EutQ_1_eut1(pf06249) EUT1 EP_NTERM
02210 HMM01923cobtr_2_eut1(pf01923) EUT1
02211 HMM01515PTA_B_1_eut1(pf01515) EUT1
02212 H_azure EUT1
02213 P_lightViolet EUT1
02214 HMMalddh_1_eutpdugrm(pf00171) EUT1 EP_NTERM
02215 HMM06723EutJMreB_1_grm(#) EUT1
02216 HMM00465alcdh_3_mix(pf00465) EUT1
02217 HMMtpEuth_1_eut(pf04346) EUT1
02218 HMM06277EutA_1_eut(pf06277) EUT1
02219 HMMetlyEutB_1_EUT1(pf06751) EUT1
02220 HMMetlyEutC_1_eut1(pf05985) EUT1 EP_NTERM
02221 Tsp_rosa EUT1
02222 H_mulberry(pf00936pf16365) EUT1
02223 HMMreg12833_1_EUT1(pf12833) EUT1
02224 -pf01218 -
02225 -pf01520 -
02226 -pf00583 -
02227 -pf11143 -
02228 -pf06572 -
02229 -pf04261 -
02230 HMMnopfamclust3_1(#) RMM1
02231 -# -
02232 -pf00528 -
BMC-P
BMC-H
BMC-T
Aldehyde dehydrogenase
PTAC
Signature enzyme
Alcohol dehydrogenase
Regulator

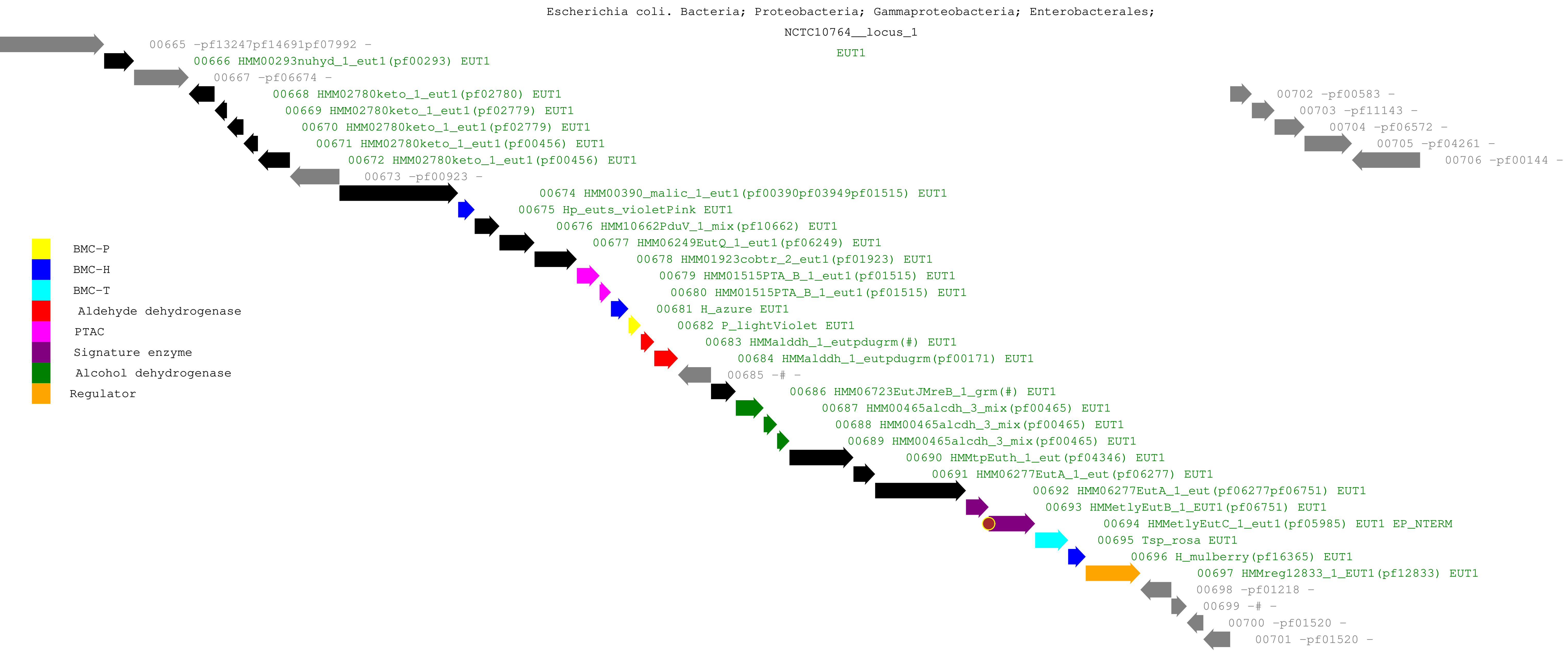
Escherichia coli. Bacteria; Proteobacteria; Gammaproteobacteria; Enterobacterales;
NCTC10764__locus_1
EUT1
00665 -pf13247pf14691pf07992 -
00666 HMM00293nuhyd_1_eut1(pf00293) EUT1
00667 -pf06674 -
00668 HMM02780keto_1_eut1(pf02780) EUT1
00669 HMM02780keto_1_eut1(pf02779) EUT1
00670 HMM02780keto_1_eut1(pf02779) EUT1
00671 HMM02780keto_1_eut1(pf00456) EUT1
00672 HMM02780keto_1_eut1(pf00456) EUT1
00673 -pf00923 -
00674 HMM00390_malic_1_eut1(pf00390pf03949pf01515) EUT1
00675 Hp_euts_violetPink EUT1
00676 HMM10662PduV_1_mix(pf10662) EUT1
00677 HMM06249EutQ_1_eut1(pf06249) EUT1
00678 HMM01923cobtr_2_eut1(pf01923) EUT1
00679 HMM01515PTA_B_1_eut1(pf01515) EUT1
00680 HMM01515PTA_B_1_eut1(pf01515) EUT1
00681 H_azure EUT1
00682 P_lightViolet EUT1
00683 HMMalddh_1_eutpdugrm(#) EUT1
00684 HMMalddh_1_eutpdugrm(pf00171) EUT1
00685 -# -
00686 HMM06723EutJMreB_1_grm(#) EUT1
00687 HMM00465alcdh_3_mix(pf00465) EUT1
00688 HMM00465alcdh_3_mix(pf00465) EUT1
00689 HMM00465alcdh_3_mix(pf00465) EUT1
00690 HMMtpEuth_1_eut(pf04346) EUT1
00691 HMM06277EutA_1_eut(pf06277) EUT1
00692 HMM06277EutA_1_eut(pf06277pf06751) EUT1
00693 HMMetlyEutB_1_EUT1(pf06751) EUT1
00694 HMMetlyEutC_1_eut1(pf05985) EUT1 EP_NTERM
00695 Tsp_rosa EUT1
00696 H_mulberry(pf16365) EUT1
00697 HMMreg12833_1_EUT1(pf12833) EUT1
00698 -pf01218 -
00699 -# -
00700 -pf01520 -
00701 -pf01520 -
00702 -pf00583 -
00703 -pf11143 -
00704 -pf06572 -
00705 -pf04261 -
00706 -pf00144 -
BMC-P
BMC-H
BMC-T
Aldehyde dehydrogenase
PTAC
Signature enzyme
Alcohol dehydrogenase
Regulator

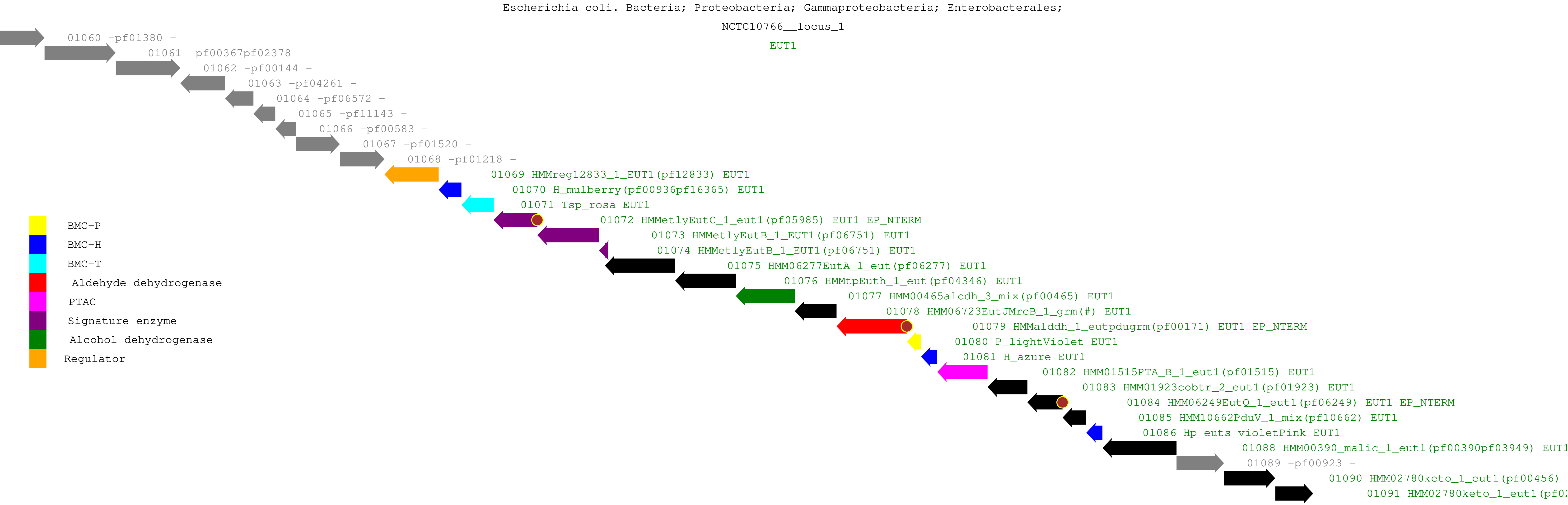

Escherichia coli. Bacteria; Proteobacteria; Gammaproteobacteria; Enterobacterales;
NCTC10766__locus_1
EUT1
01060 -pf01380 -
01061 -pf00367pf02378 -
01062 -pf00144 -
01063 -pf04261 -
01064 -pf06572 -
01065 -pf11143 -
01066 -pf00583 -
01067 -pf01520 -
01068 -pf01218 -
01069 HMMreg12833_1_EUT1(pf12833) EUT1
01070 H_mulberry(pf00936pf16365) EUT1
01071 Tsp_rosa EUT1
01072 HMMetlyEutC_1_eut1(pf05985) EUT1 EP_NTERM
01073 HMMetlyEutB_1_EUT1(pf06751) EUT1
01074 HMMetlyEutB_1_EUT1(pf06751) EUT1
01075 HMM06277EutA_1_eut(pf06277) EUT1
01076 HMMtpEuth_1_eut(pf04346) EUT1
01077 HMM00465alcdh_3_mix(pf00465) EUT1
01078 HMM06723EutJMreB_1_grm(#) EUT1
01079 HMMaldh_1_eutpdugrm(pf00171) EUT1 EP_NTERM
01080 P_lightViolet EUT1
01081 H_azure EUT1
01082 HMM01515PTA_B_1_eut1(pf01515) EUT1
01083 HMM01923cobtr_2_eut1(pf01923) EUT1
01084 HMM06249EutQ_1_eut1(pf06249) EUT1 EP_NTERM
01085 HMM10662PduV_1_mix(pf10662) EUT1
01086 Hp_euts_violetPink EUT1
01088 HMM00390_malic_1_eut1(pf00390pf03949) EUT1
01089 -pf00923 -
01090 HMM02780keto_1_eut1(pf00456)
01091 HMM02780keto_1_eut1(pf0
BMC-P
BMC-H
BMC-T
Aldehyde dehydrogenase
PTAC
Signature enzyme
Alcohol dehydrogenase
Regulator

Citrobacter koseri (Citrobacter diversus). Bacteria; Proteobacteria; Gammaproteobacteria; Enterobacterales;

NCTC10770__locus_1

EUT1

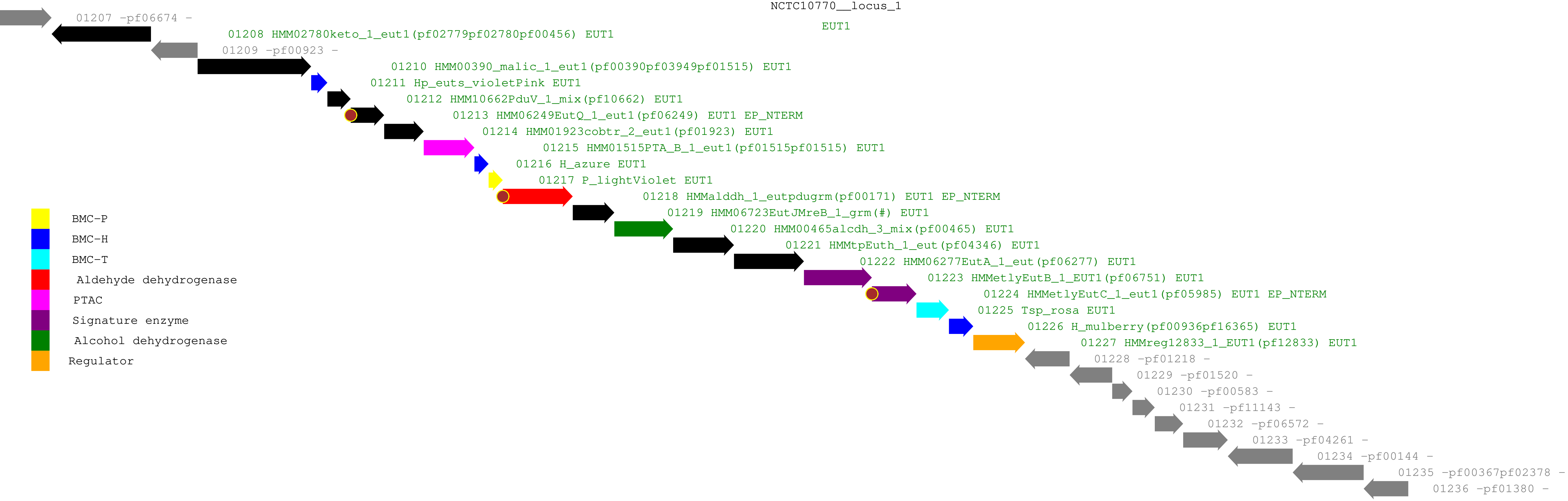

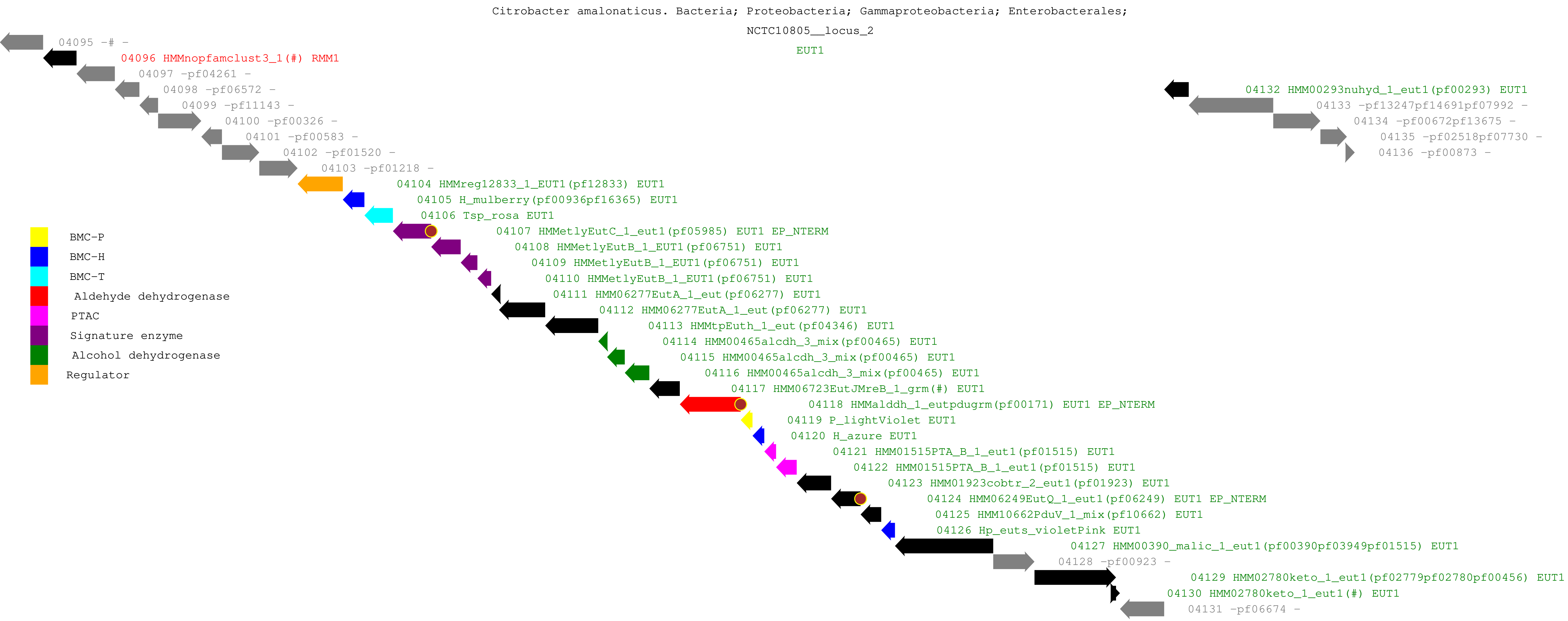

Citrobacter amalonaticus. Bacteria; Proteobacteria; Gammaproteobacteria; Enterobacterales;
NCTC10805__locus_2
EUT1
04095 -# -
04096 HMMnopfamclust3_1(#) RMM1
04097 -pf04261 -
04098 -pf06572 -
04099 -pf11143 -
04100 -pf00326 -
04101 -pf00583 -
04102 -pf01520 -
04103 -pf01218 -
04104 HMMreg12833_1_EUT1(pf12833) EUT1
04105 H_mulberry(pf00936pf16365) EUT1
04106 Tsp_rosa EUT1
04107 HMMetlyEutC_1_eut1(pf05985) EUT1 EP_NTERM
04108 HMMetlyEutB_1_EUT1(pf06751) EUT1
04109 HMMetlyEutB_1_EUT1(pf06751) EUT1
04110 HMMetlyEutB_1_EUT1(pf06751) EUT1
04111 HMM06277EutA_1_eut(pf06277) EUT1
04112 HMM06277EutA_1_eut(pf06277) EUT1
04113 HMMtpEuth_1_eut(pf04346) EUT1
04114 HMM00465alcdh_3_mix(pf00465) EUT1
04115 HMM00465alcdh_3_mix(pf00465) EUT1
04116 HMM00465alcdh_3_mix(pf00465) EUT1
04117 HMM06723EutJMreB_1_grm(#) EUT1
04118 HMMalddh_1_eutpdugrm(pf00171) EUT1 EP_NTERM
04119 P_lightViolet EUT1
04120 H_azure EUT1
04121 HMM01515PTA_B_1_eut1(pf01515) EUT1
04122 HMM01515PTA_B_1_eut1(pf01515) EUT1
04123 HMM01923cobtr_2_eut1(pf01923) EUT1
04124 HMM06249EutQ_1_eut1(pf06249) EUT1 EP_NTERM
04125 HMM10662PduV_1_mix(pf10662) EUT1
04126 Hp_euts_violetPink EUT1
04127 HMM00390_malic_1_eut1(pf00390pf03949pf01515) EUT1
04128 -pf00923 -
04129 HMM02780keto_1_eut1(pf02779pf02780pf00456) EUT1
04130 HMM02780keto_1_eut1(#) EUT1
04131 -pf06674 -
04132 HMM00293nuhyd_1_eut1(pf00293) EUT1
04133 -pf13247pf14691pf07992 -
04134 -pf00672pf13675 -
04135 -pf02518pf07730 -
04136 -pf00873 -
BMC-P
BMC-H
BMC-T
Aldehyde dehydrogenase
PTAC
Signature enzyme
Alcohol dehydrogenase
Regulator

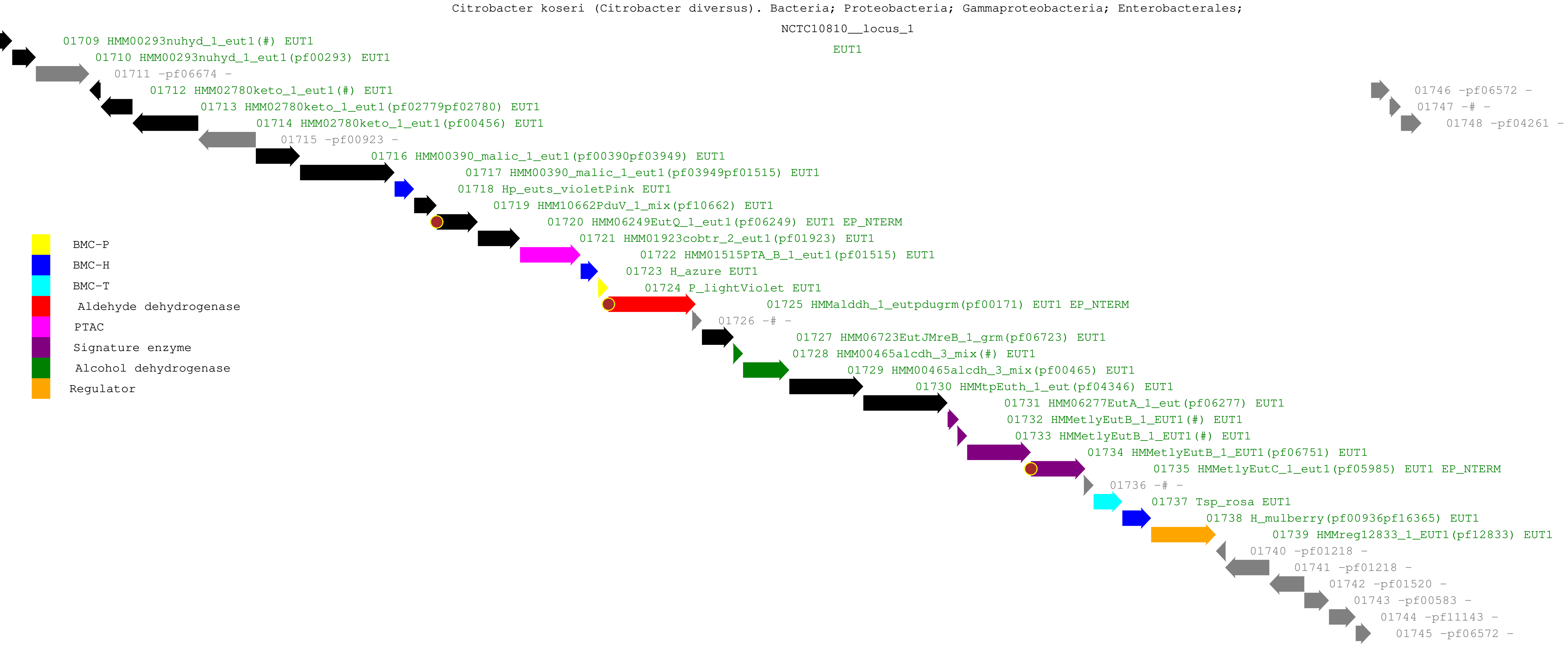
Citrobacter koseri (Citrobacter diversus). Bacteria; Proteobacteria; Gammaproteobacteria; Enterobacterales;
NCTC10810__locus_1
EUT1
01709 HMM00293nuhyd_1_eut1(#) EUT1
01710 HMM00293nuhyd_1_eut1(pf00293) EUT1
01711 -pf06674 -
01712 HMM02780keto_1_eut1(#) EUT1
01713 HMM02780keto_1_eut1(pf02779pf02780) EUT1
01714 HMM02780keto_1_eut1(pf00456) EUT1
01715 -pf00923 -
01716 HMM00390_malic_1_eut1(pf00390pf03949) EUT1
01717 HMM00390_malic_1_eut1(pf03949pf01515) EUT1
01718 Hp_euts_violetPink EUT1
01719 HMM10662PduV_1_mix(pf10662) EUT1
01720 HMM06249EutQ_1_eut1(pf06249) EUT1 EP_NTERM
01721 HMM01923cobtr_2_eut1(pf01923) EUT1
01722 HMM01515PTA_B_1_eut1(pf01515) EUT1
01723 H_azure EUT1
01724 P_lightViolet EUT1
01725 HMMaldh_1_eutpdugrm(pf00171) EUT1 EP_NTERM
01726 -# -
01727 HMM06723EutJMreB_1_grm(pf06723) EUT1
01728 HMM00465alcdh_3_mix(#) EUT1
01729 HMM00465alcdh_3_mix(pf00465) EUT1
01730 HMMtpEuth_1_eut(pf04346) EUT1
01731 HMM06277EutA_1_eut(pf06277) EUT1
01732 HMMetlyEutB_1_EUT1(#) EUT1
01733 HMMetlyEutB_1_EUT1(#) EUT1
01734 HMMetlyEutB_1_EUT1(pf06751) EUT1
01735 HMMetlyEutC_1_eut1(pf05985) EUT1 EP_NTERM
01736 -# -
01737 Tsp_rosa EUT1
01738 H_mulberry(pf00936pf16365) EUT1
01739 HMMreg12833_1_EUT1(pf12833) EUT1
01740 -pf01218 -
01741 -pf01218 -
01742 -pf01520 -
01743 -pf00583 -
01744 -pf11143 -
01745 -pf06572 -
01746 -pf06572 -
01747 -# -
01748 -pf04261 -
BMC-P
BMC-H
BMC-T
Aldehyde dehydrogenase
PTAC
Signature enzyme
Alcohol dehydrogenase
Regulator

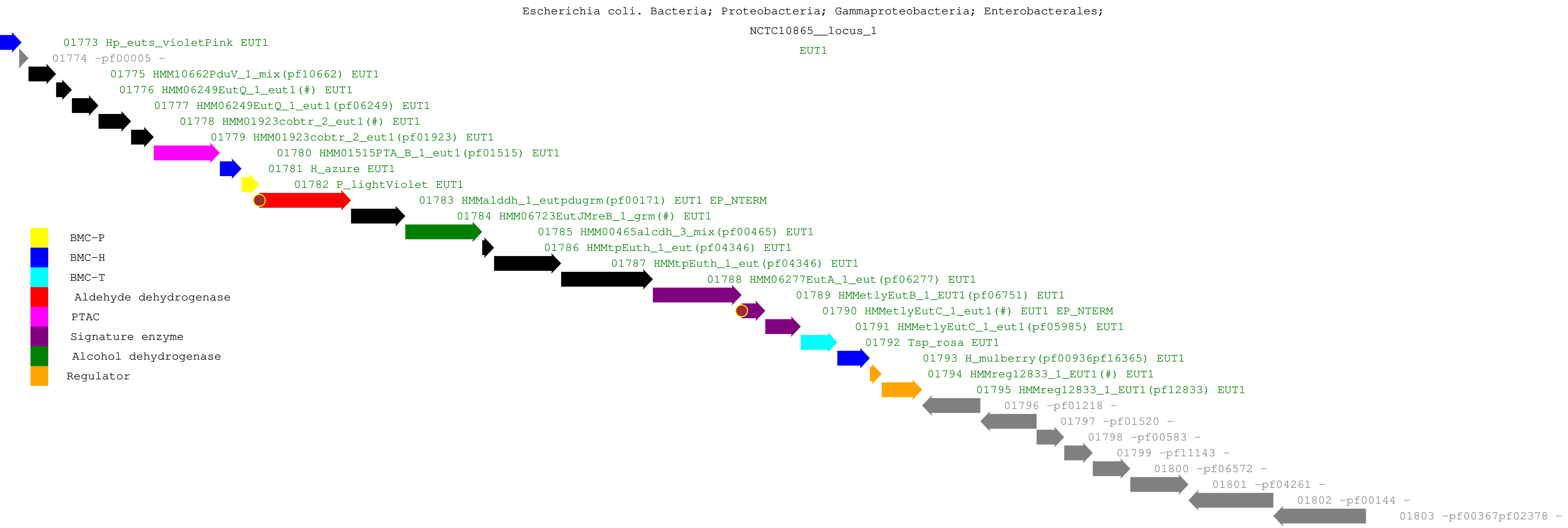
Escherichia coli. Bacteria; Proteobacteria; Gammaproteobacteria; Enterobacterales;
NCTC10865__locus_1
EUT1
01773 Hp_euts_violetPink EUT1
01774 -pf00005 -
01775 HMM10662PduV_1_mix(pf10662) EUT1
01776 HMM06249EutQ_1_eut1(#) EUT1
01777 HMM06249EutQ_1_eut1(pf06249) EUT1
01778 HMM01923cobtr_2_eut1(#) EUT1
01779 HMM01923cobtr_2_eut1(pf01923) EUT1
01780 HMM01515PTA_B_1_eut1(pf01515) EUT1
01781 H_azure EUT1
01782 P_lightViolet EUT1
01783 HMMalddh_1_eutpdugrm(pf00171) EUT1 EP_NTERM
01784 HMM06723EutJMreB_1_grm(#) EUT1
01785 HMM00465alcdh_3_mix(pf00465) EUT1
01786 HMMtpEuth_1_eut(pf04346) EUT1
01787 HMMtpEuth_1_eut(pf04346) EUT1
01788 HMM06277EutA_1_eut(pf06277) EUT1
01789 HMMetlyEutB_1_EUT1(pf06751) EUT1
01790 HMMetlyEutC_1_eut1(#) EUT1 EP_NTERM
01791 HMMetlyEutC_1_eut1(pf05985) EUT1
01792 Tsp_rosa EUT1
01793 H_mulberry(pf00936pf16365) EUT1
01794 HMMreg12833_1_EUT1(#) EUT1
01795 HMMreg12833_1_EUT1(pf12833) EUT1
01796 -pf01218 -
01797 -pf01520 -
01798 -pf00583 -
01799 -pf11143 -
01800 -pf06572 -
01801 -pf04261 -
01802 -pf00144 -
01803 -pf00367pf02378 -
BMC-P
BMC-H
BMC-T
Aldehyde dehydrogenase
PTAC
Signature enzyme
Alcohol dehydrogenase
Regulator

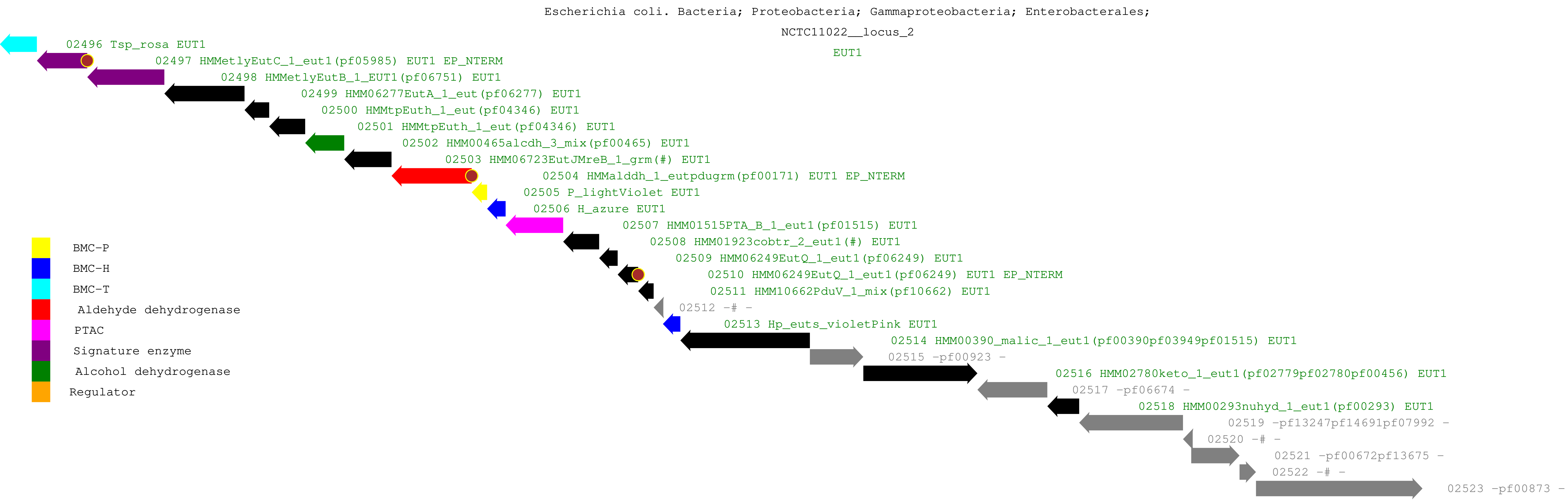
Escherichia coli. Bacteria; Proteobacteria; Gammaproteobacteria; Enterobacterales;
NCTC11022__locus_2
EUT1
02496 Tsp_rosa EUT1
02497 HMMetlyEutC_1_eut1(pf05985) EUT1 EP_NTERM
02498 HMMetlyEutB_1_EUT1(pf06751) EUT1
02499 HMM06277EutA_1_eut(pf06277) EUT1
02500 HMMtpEuth_1_eut(pf04346) EUT1
02501 HMMtpEuth_1_eut(pf04346) EUT1
02502 HMM00465alcdh_3_mix(pf00465) EUT1
02503 HMM06723EutJMreB_1_grm(#) EUT1
02504 HMMalddh_1_eutpdugrm(pf00171) EUT1 EP_NTERM
02505 P_lightViolet EUT1
02506 H_azure EUT1
02507 HMM01515PTA_B_1_eut1(pf01515) EUT1
02508 HMM01923cobtr_2_eut1(#) EUT1
02509 HMM06249EutQ_1_eut1(pf06249) EUT1
02510 HMM06249EutQ_1_eut1(pf06249) EUT1 EP_NTERM
02511 HMM10662PduV_1_mix(pf10662) EUT1
02512 -# -
02513 Hp_euts_violetPink EUT1
02514 HMM00390_malic_1_eut1(pf00390pf03949pf01515) EUT1
02515 -pf00923 -
02516 HMM02780keto_1_eut1(pf02779pf02780pf00456) EUT1
02517 -pf06674 -
02518 HMM00293nuhyd_1_eut1(pf00293) EUT1
02519 -pf13247pf14691pf07992 -
02520 -# -
02521 -pf00672pf13675 -
02522 -# -
02523 -pf00873 -
BMC-P
BMC-H
BMC-T
Aldehyde dehydrogenase
PTAC
Signature enzyme
Alcohol dehydrogenase
Regulator

Citrobacter koseri (Citrobacter diversus). Bacteria; Proteobacteria; Gammaproteobacteria; Enterobacterales;

NCTC11075__locus_2

EUT1

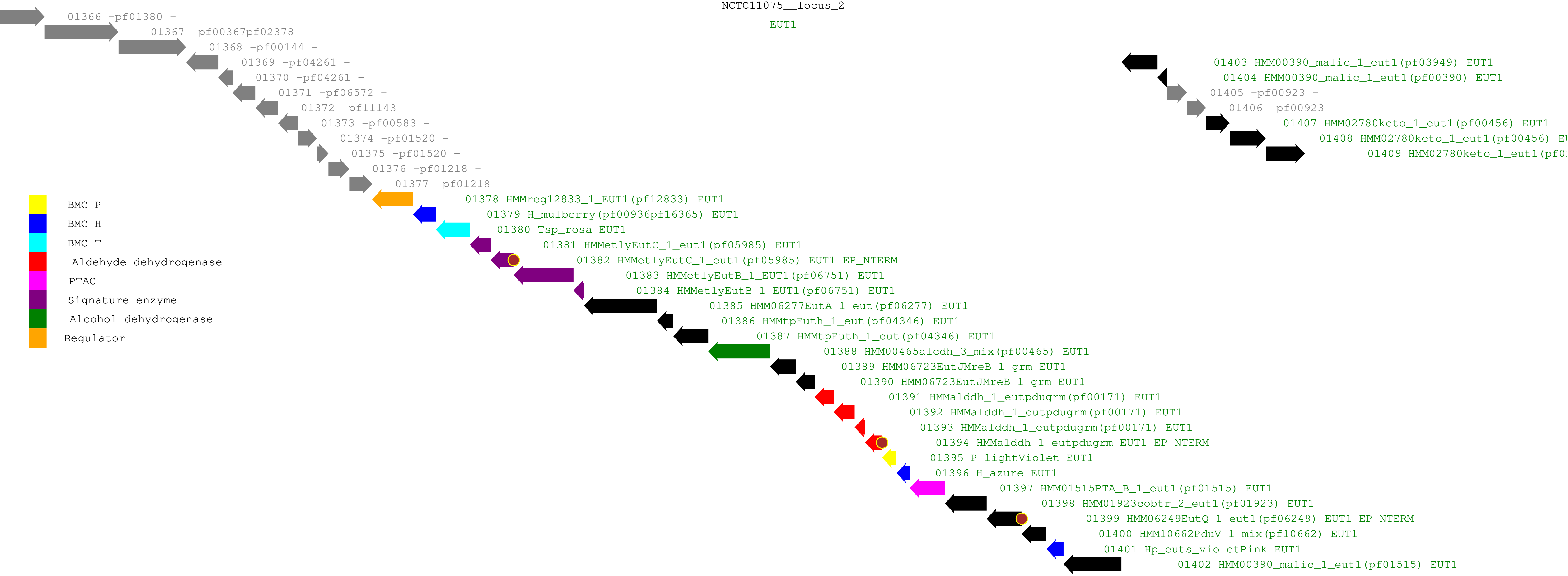

Escherichia coli. Bacteria; Proteobacteria; Gammaproteobacteria; Enterobacterales;

NCTC11112__locus_1

EUT1

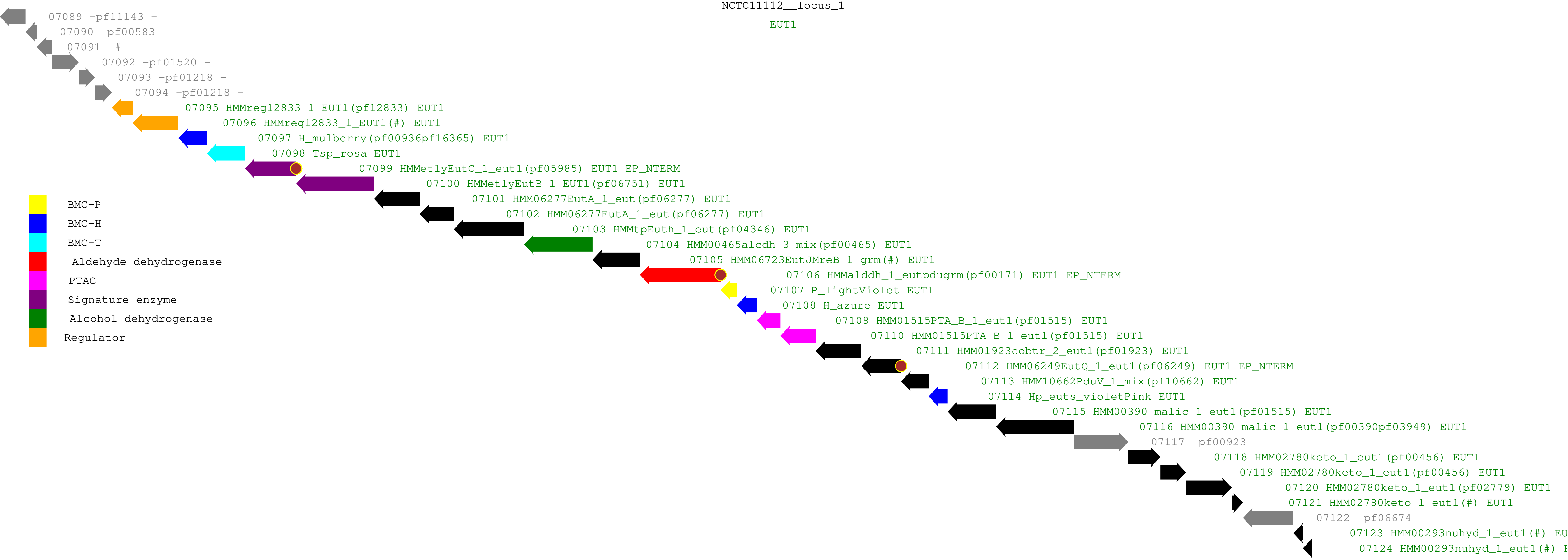

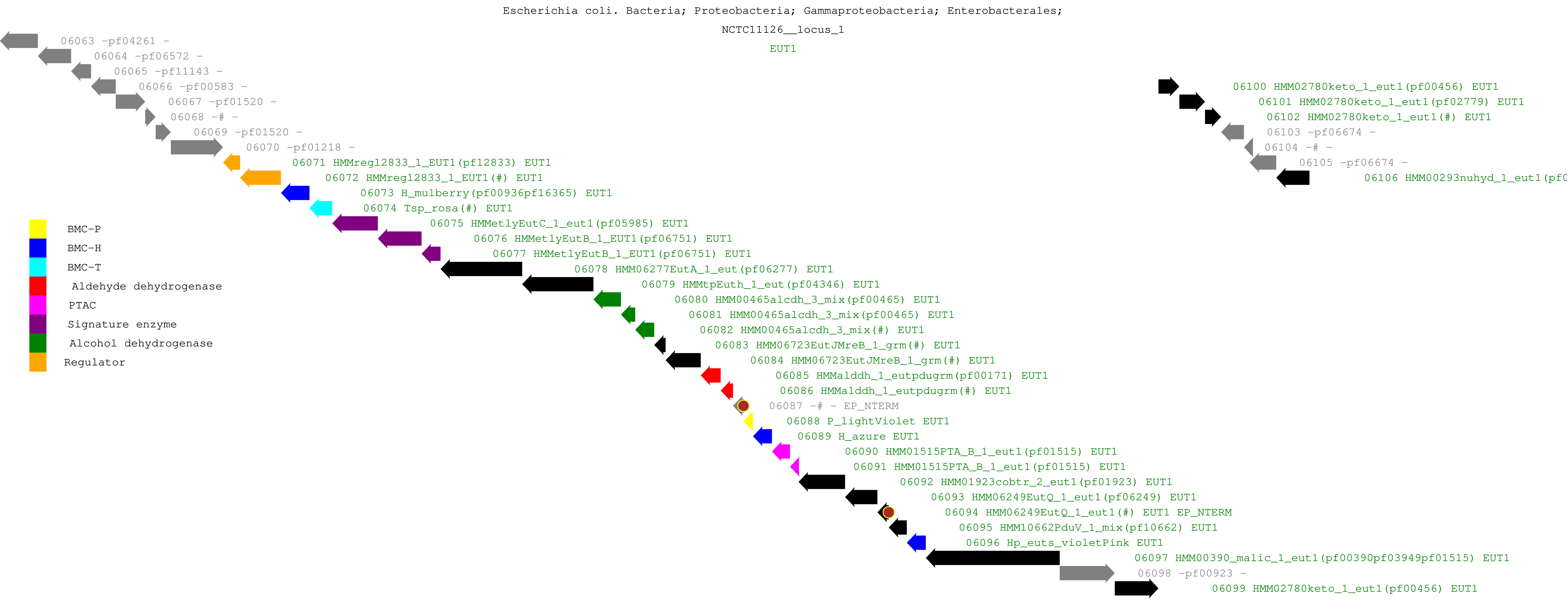

Escherichia coli. Bacteria; Proteobacteria; Gammaproteobacteria; Enterobacterales;
NCTC11126__locus_1
EUT1
06063 -pf04261 -
06064 -pf06572 -
06065 -pf11143 -
06066 -pf00583 -
06067 -pf01520 -
06068 -# -
06069 -pf01520 -
06070 -pf01218 -
06071 HMMreg12833_1_EUT1(pf12833) EUT1
06072 HMMreg12833_1_EUT1(#) EUT1
06073 H_mulberry(pf00936pf16365) EUT1
06074 Tsp_rosa(#) EUT1
06075 HMMetlyEutC_1_eut1(pf05985) EUT1
06076 HMMetlyEutB_1_EUT1(pf06751) EUT1
06077 HMMetlyEutB_1_EUT1(pf06751) EUT1
06078 HMM06277EutA_1_eut(pf06277) EUT1
06079 HMMtpEuth_1_eut(pf04346) EUT1
06080 HMM00465alcdh_3_mix(pf00465) EUT1
06081 HMM00465alcdh_3_mix(pf00465) EUT1
06082 HMM00465alcdh_3_mix(#) EUT1
06083 HMM06723EutJMreB_1_grm(#) EUT1
06084 HMM06723EutJMreB_1_grm(#) EUT1
06085 HMMaldh_1_eutpdugrm(pf00171) EUT1
06086 HMMaldh_1_eutpdugrm(#) EUT1
06087 -# - EP_NTERM
06088 P_lightViolet EUT1
06089 H_azure EUT1
06090 HMM01515PTA_B_1_eut1(pf01515) EUT1
06091 HMM01515PTA_B_1_eut1(pf01515) EUT1
06092 HMM01923cobtr_2_eut1(pf01923) EUT1
06093 HMM06249EutQ_1_eut1(pf06249) EUT1
06094 HMM06249EutQ_1_eut1(#) EUT1 EP_NTERM
06095 HMM10662PduV_1_mix(pf10662) EUT1
06096 Hp_euts_violetPink EUT1
06097 HMM00390_malic_1_eut1(pf00390pf03949pf01515) EUT1
06098 -pf00923 -
06099 HMM02780keto_1_eut1(pf00456) EUT1
06100 HMM02780keto_1_eut1(pf00456) EUT1
06101 HMM02780keto_1_eut1(pf02779) EUT1
06102 HMM02780keto_1_eut1(#) EUT1
06103 -pf06674 -
06104 -# -
06105 -pf06674 -
06106 HMM00293nuhyd_1_eut1(pf0
BMC-P
BMC-H
BMC-T
Aldehyde dehydrogenase
PTAC
Signature enzyme
Alcohol dehydrogenase
Regulator

Iodobacter fluviatilis. Bacteria; Proteobacteria; Betaproteobacteria; Neisseriales;

NCTC11159__locus_1

EUT1

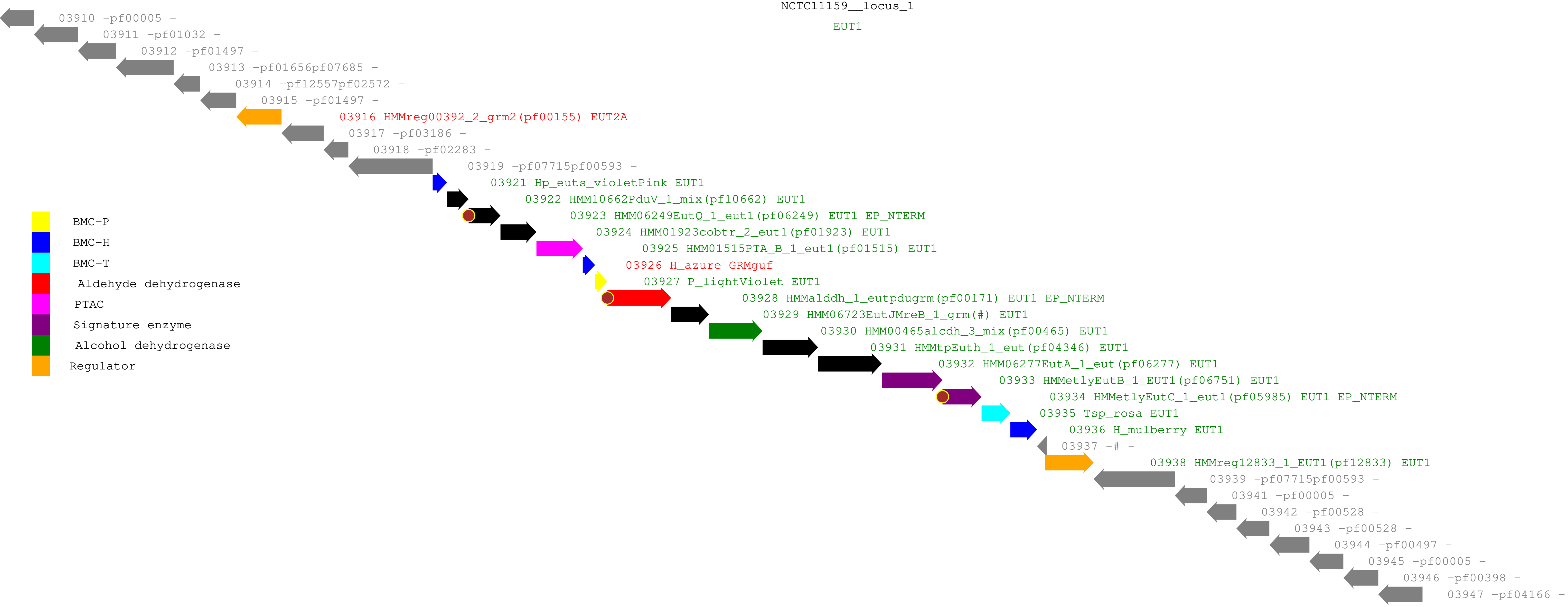

Escherichia coli. Bacteria; Proteobacteria; Gammaproteobacteria; Enterobacterales;

NCTC11181__locus_1

EUT1

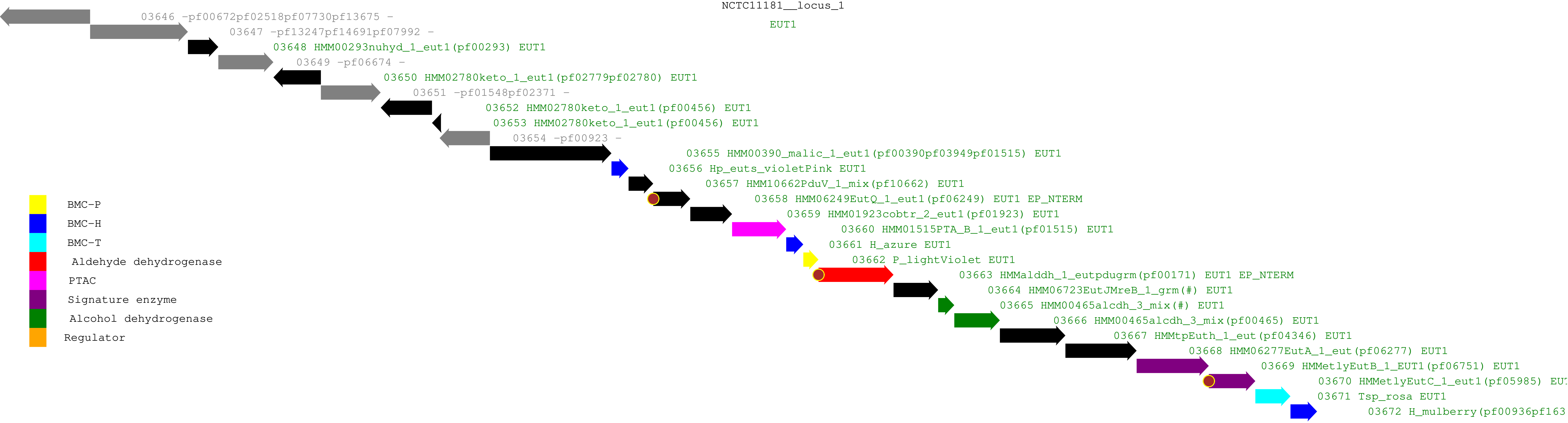

Klebsiella oxytoca. Bacteria; Proteobacteria; Gammaproteobacteria; Enterobacterales;

NCTC11355__locus_1

EUT1

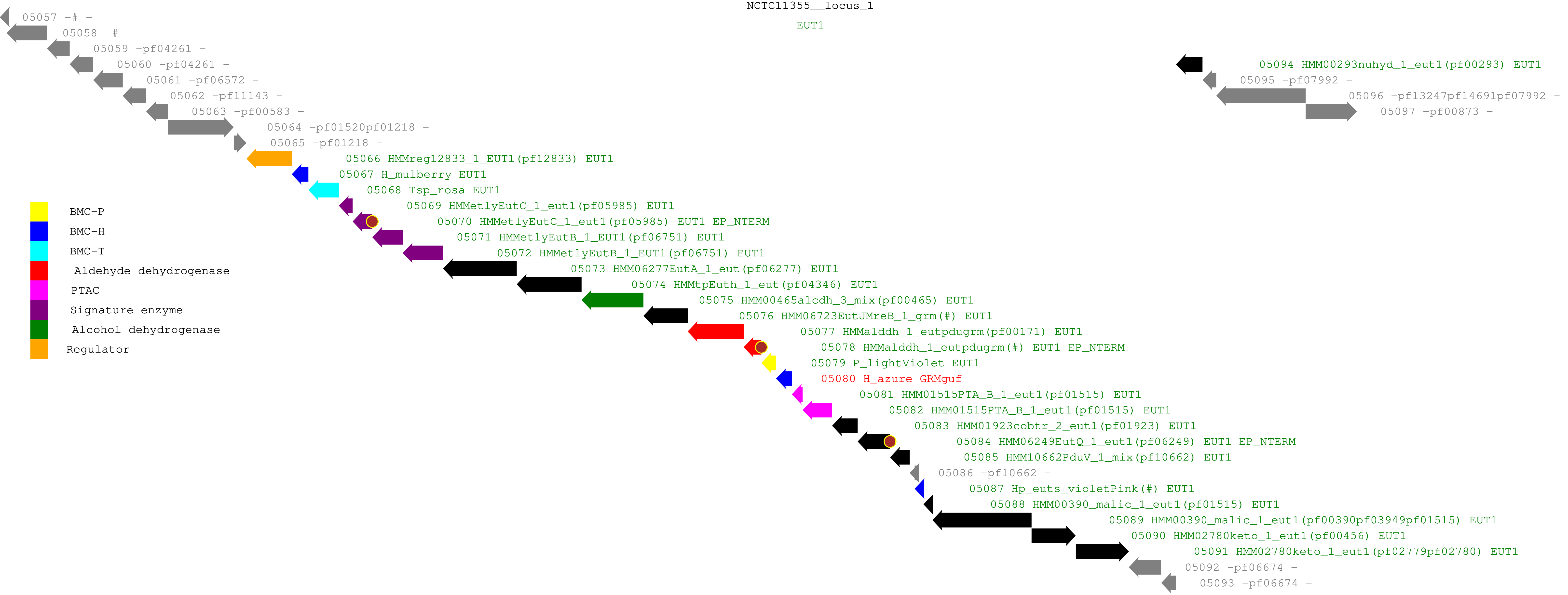

Pluralibacter gergoviae (Enterobacter gergoviae). Bacteria; Proteobacteria; Gammaproteobacteria; Enterobacterales;

NCTC11434__locus_1

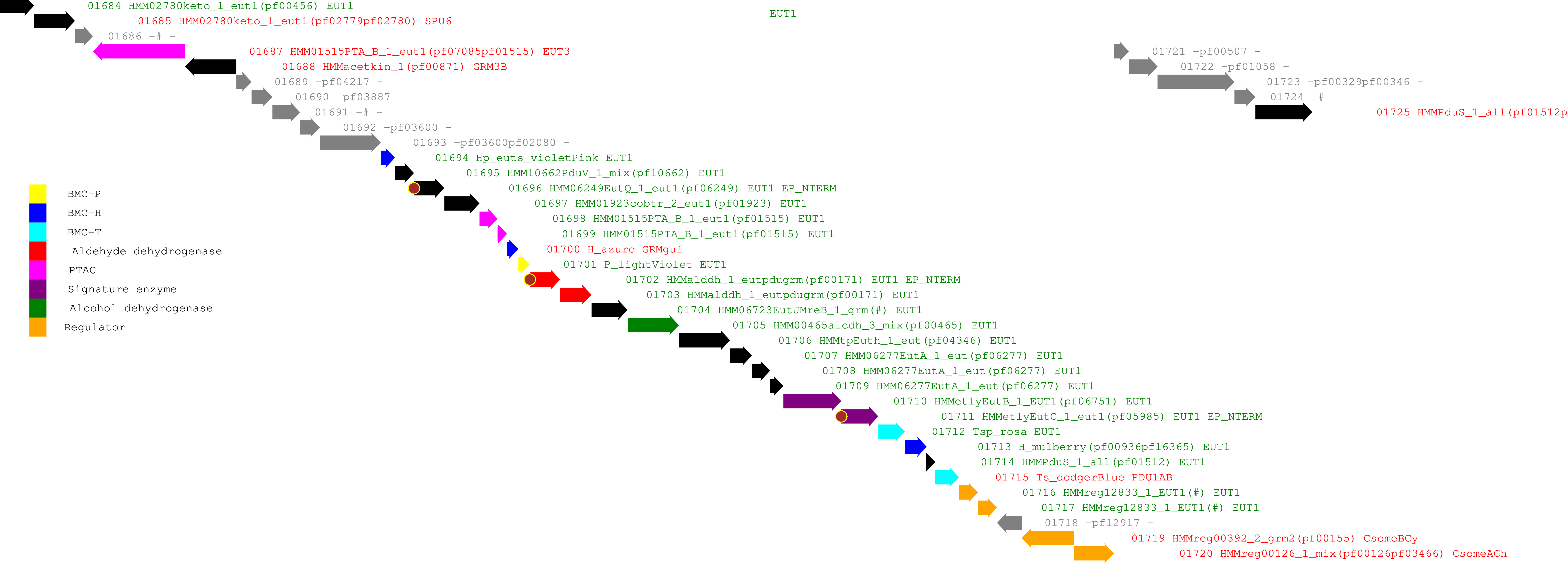

Escherichia coli. Bacteria; Proteobacteria; Gammaproteobacteria; Enterobacterales;

NCTC11560__locus_1

EUT1

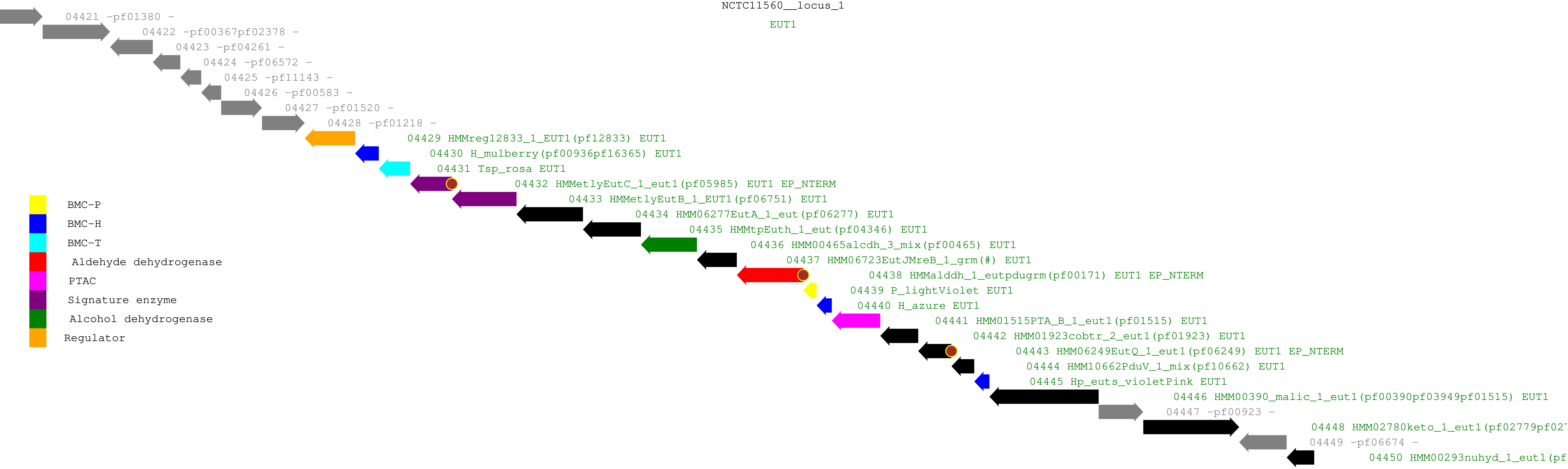

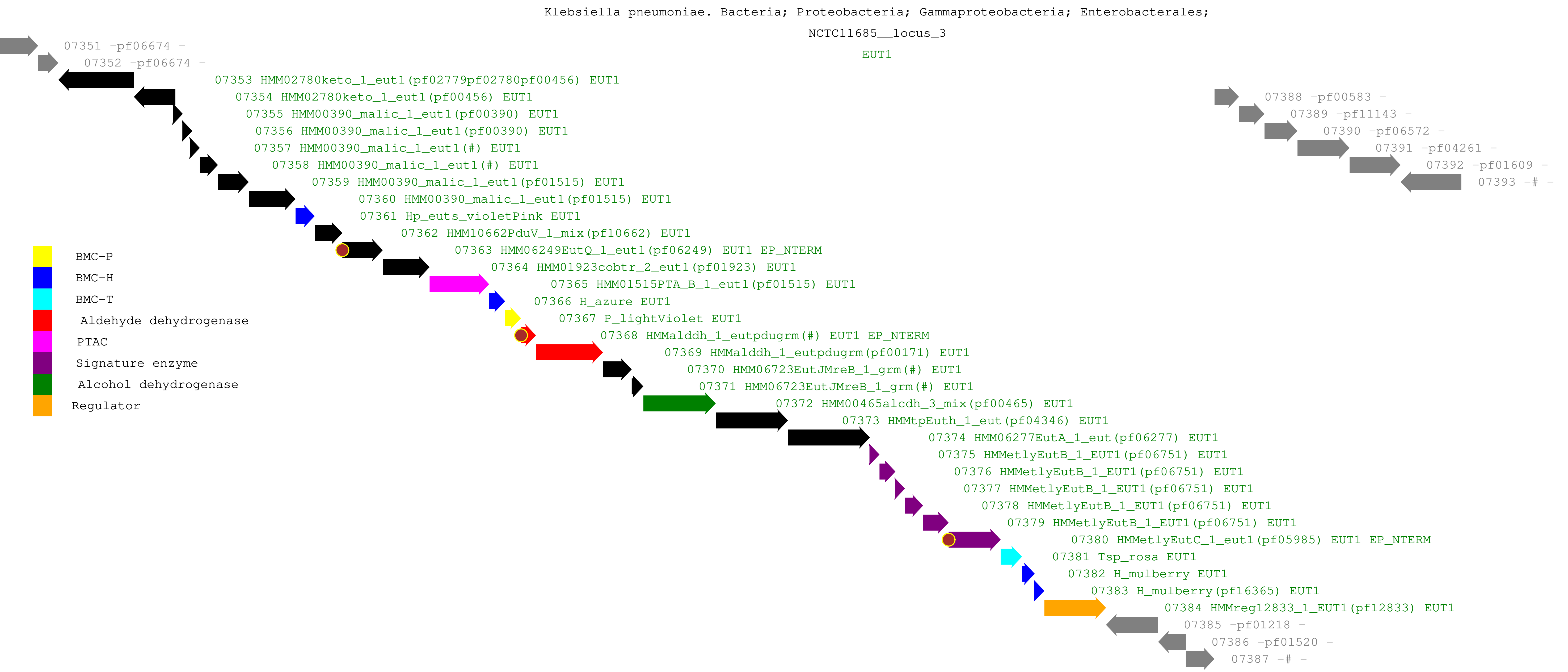
Klebsiella pneumoniae. Bacteria; Proteobacteria; Gammaproteobacteria; Enterobacterales;
NCTC11685__locus_3
EUT1
07351 -pf06674 -
07352 -pf06674 -
07353 HMM02780keto_1_eut1(pf02779pf02780pf00456) EUT1
07354 HMM02780keto_1_eut1(pf00456) EUT1
07355 HMM00390_malic_1_eut1(pf00390) EUT1
07356 HMM00390_malic_1_eut1(pf00390) EUT1
07357 HMM00390_malic_1_eut1(#) EUT1
07358 HMM00390_malic_1_eut1(#) EUT1
07359 HMM00390_malic_1_eut1(pf01515) EUT1
07360 HMM00390_malic_1_eut1(pf01515) EUT1
07361 Hp_euts_violetPink EUT1
07362 HMM10662PduV_1_mix(pf10662) EUT1
07363 HMM06249EutQ_1_eut1(pf06249) EUT1 EP_NTERM
07364 HMM01923cobtr_2_eut1(pf01923) EUT1
07365 HMM01515PTA_B_1_eut1(pf01515) EUT1
07366 H_azure EUT1
07367 P_lightViolet EUT1
07368 HMMalddh_1_eutpdugrm(#) EUT1 EP_NTERM
07369 HMMalddh_1_eutpdugrm(pf00171) EUT1
07370 HMM06723EutJMreB_1_grm(#) EUT1
07371 HMM06723EutJMreB_1_grm(#) EUT1
07372 HMM00465alcdh_3_mix(pf00465) EUT1
07373 HMMtpEuth_1_eut(pf04346) EUT1
07374 HMM06277EutA_1_eut(pf06277) EUT1
07375 HMMetlyEutB_1_EUT1(pf06751) EUT1
07376 HMMetlyEutB_1_EUT1(pf06751) EUT1
07377 HMMetlyEutB_1_EUT1(pf06751) EUT1
07378 HMMetlyEutB_1_EUT1(pf06751) EUT1
07379 HMMetlyEutB_1_EUT1(pf06751) EUT1
07380 HMMetlyEutC_1_eut1(pf05985) EUT1 EP_NTERM
07381 Tsp_rosa EUT1
07382 H_mulberry EUT1
07383 H_mulberry(pf16365) EUT1
07384 HMMreg12833_1_EUT1(pf12833) EUT1
07385 -pf01218 -
07386 -pf01520 -
07387 -# -
07388 -pf00583 -
07389 -pf11143 -
07390 -pf06572 -
07391 -pf04261 -
07392 -pf01609 -
07393 -# -
BMC-P
BMC-H
BMC-T
Aldehyde dehydrogenase
PTAC
Signature enzyme
Alcohol dehydrogenase
Regulator

Klebsiella oxytoca. Bacteria; Proteobacteria; Gammaproteobacteria; Enterobacterales;

NCTC11694__locus_2

EUT1

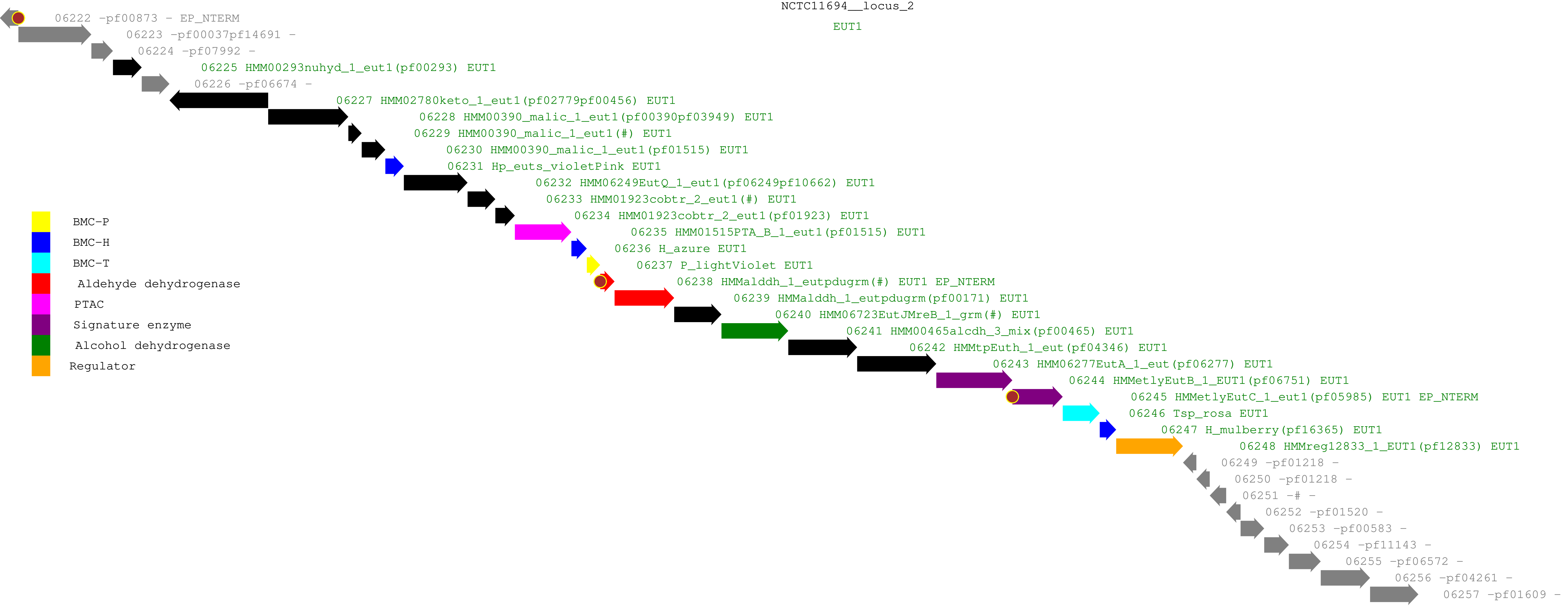

Shigella dysenteriae. Bacteria; Proteobacteria; Gammaproteobacteria; Enterobacterales;

NCTC11868__locus_1

EUT1

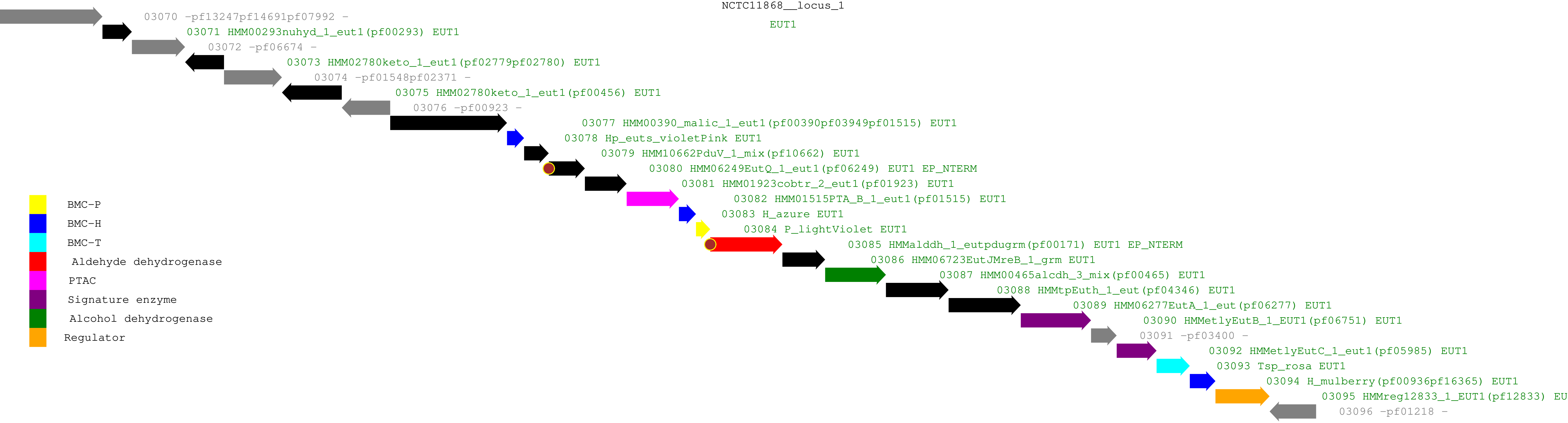

Salmonella sp. NCTC 11881. Bacteria; Proteobacteria; Gammaproteobacteria; Enterobacterales;

NCTC11881__locus_1

EUT1

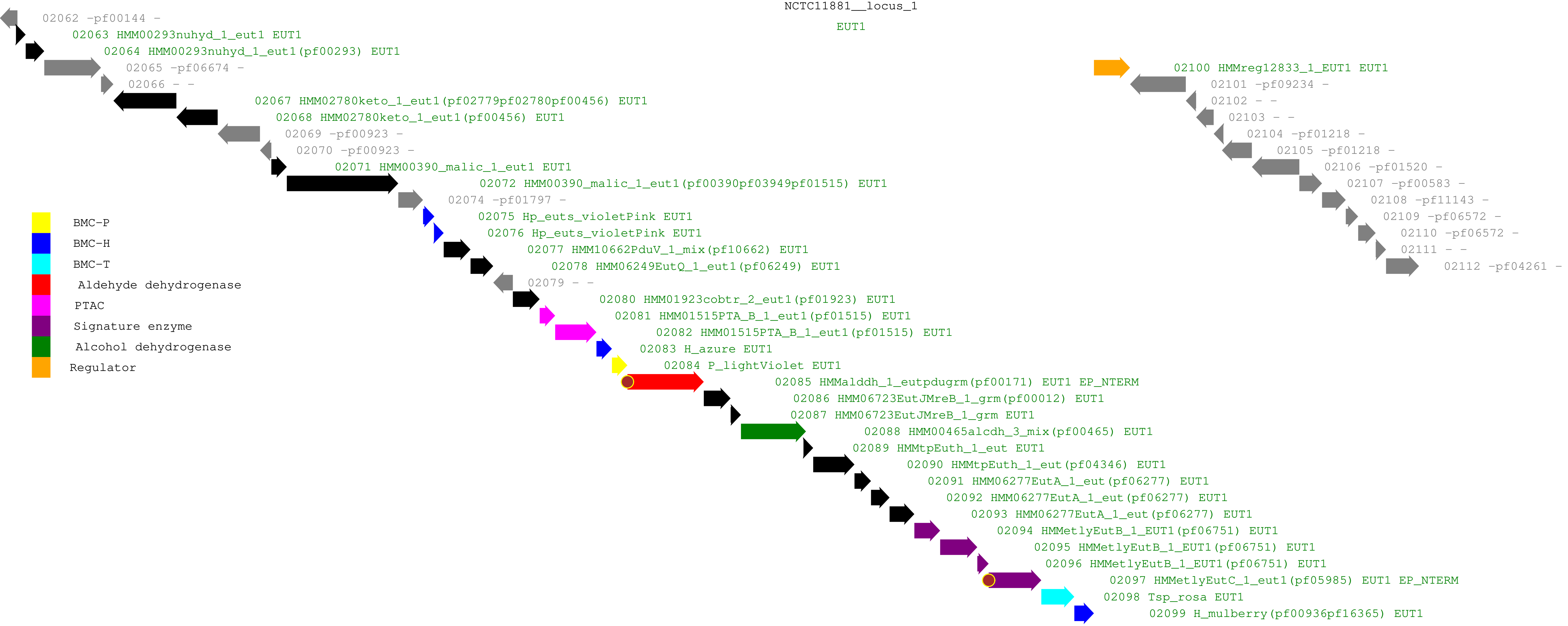

Morganella morganii (Proteus morganii). Bacteria; Proteobacteria; Gammaproteobacteria; Enterobacterales;

NCTC12358__locus_1

EUT1

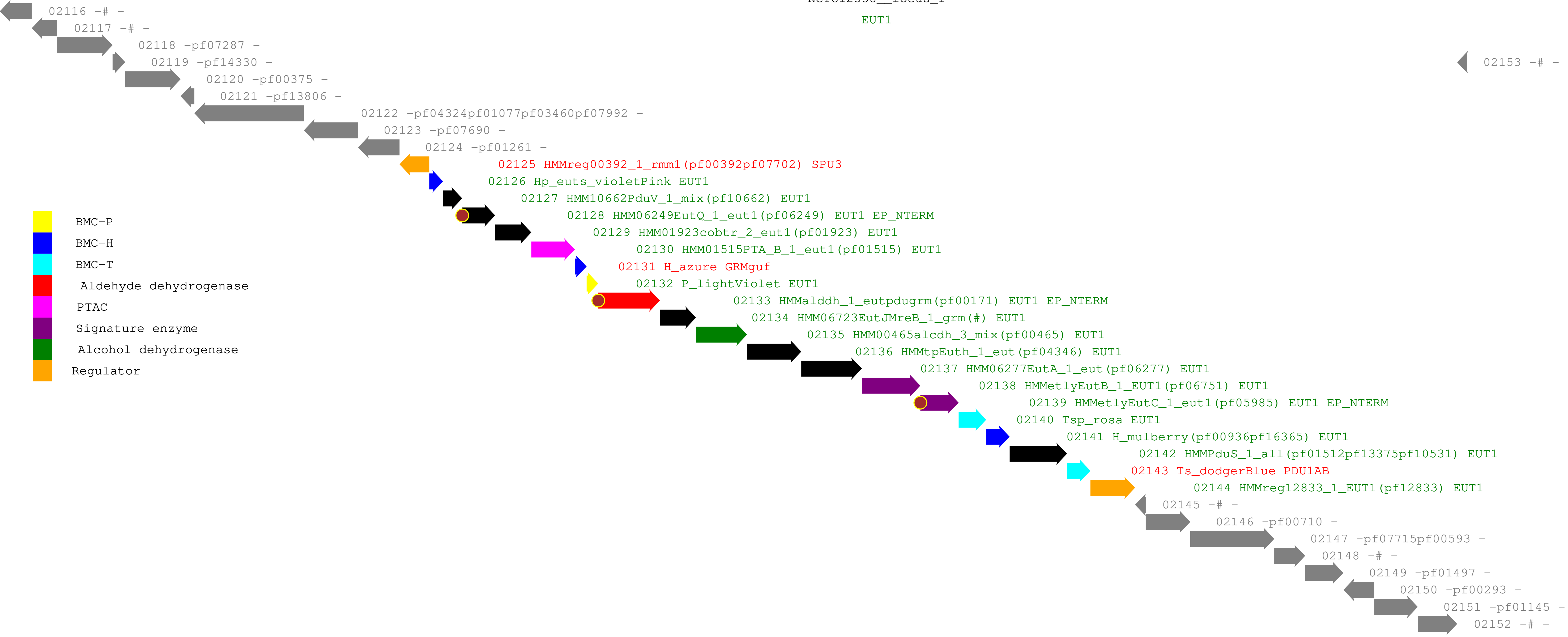

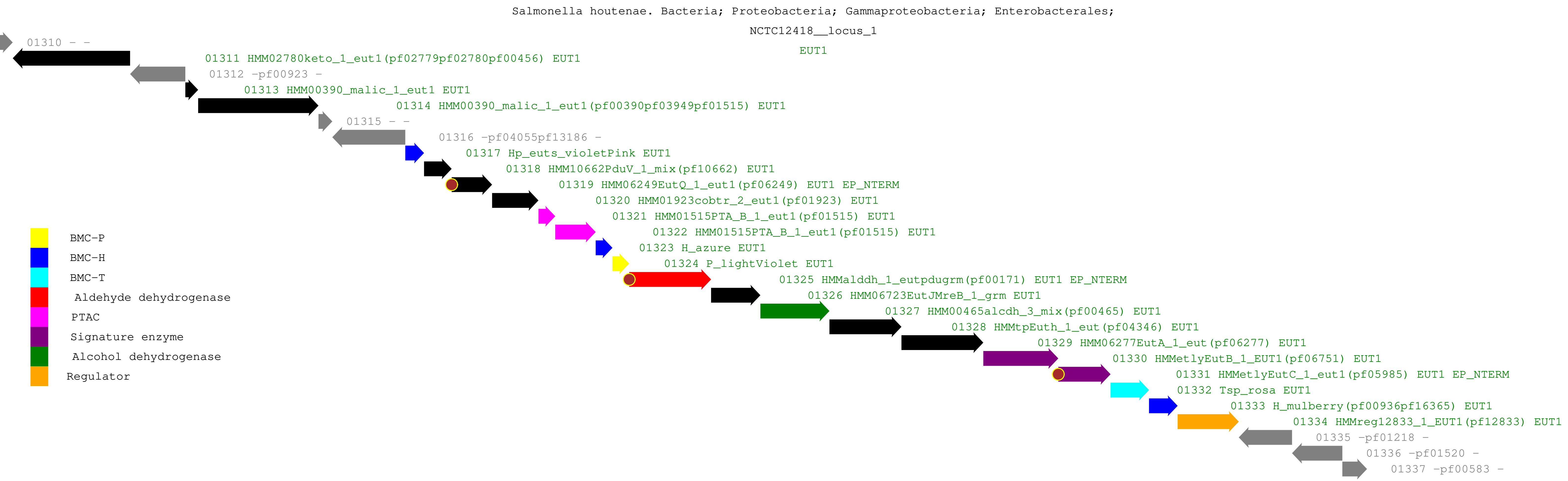
Salmonella houtenae. Bacteria; Proteobacteria; Gammaproteobacteria; Enterobacterales;
NCTC12418__locus_1
EUT1
01310 - -
01311 HMM02780keto_1_eut1(pf02779pf02780pf00456) EUT1
01312 -pf00923 -
01313 HMM00390_malic_1_eut1 EUT1
01314 HMM00390_malic_1_eut1(pf00390pf03949pf01515) EUT1
01315 - -
01316 -pf04055pf13186 -
01317 Hp_euts_violetPink EUT1
01318 HMM10662PduV_1_mix(pf10662) EUT1
01319 HMM06249EutQ_1_eut1(pf06249) EUT1 EP_NTERM
01320 HMM01923cobtr_2_eut1(pf01923) EUT1
01321 HMM01515PTA_B_1_eut1(pf01515) EUT1
01322 HMM01515PTA_B_1_eut1(pf01515) EUT1
01323 H_azure EUT1
01324 P_lightViolet EUT1
01325 HMMalddh_1_eutpdugrm(pf00171) EUT1 EP_NTERM
01326 HMM06723EutJMreB_1_grm EUT1
01327 HMM00465alcdh_3_mix(pf00465) EUT1
01328 HMMtpEuth_1_eut(pf04346) EUT1
01329 HMM06277EutA_1_eut(pf06277) EUT1
01330 HMMetlyEutB_1_EUT1(pf06751) EUT1
01331 HMMetlyEutC_1_eut1(pf05985) EUT1 EP_NTERM
01332 Tsp_rosa EUT1
01333 H_mulberry(pf00936pf16365) EUT1
01334 HMMreg12833_1_EUT1(pf12833) EUT1
01335 -pf01218 -
01336 -pf01520 -
01337 -pf00583 -
BMC-P
BMC-H
BMC-T
Aldehyde dehydrogenase
PTAC
Signature enzyme
Alcohol dehydrogenase
Regulator

Salmonella enterica subsp. indica. Bacteria; Proteobacteria; Gammaproteobacteria; Enterobacterales;

NCTC12420__locus_1

EUT1

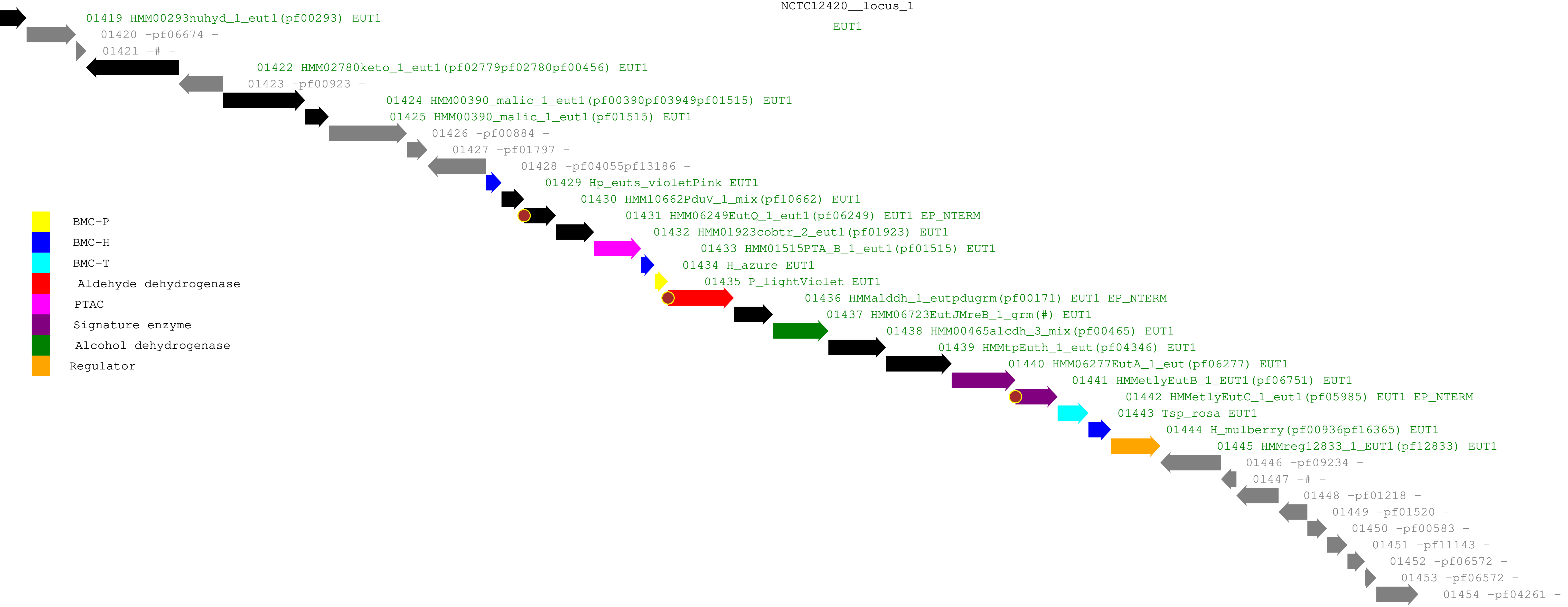

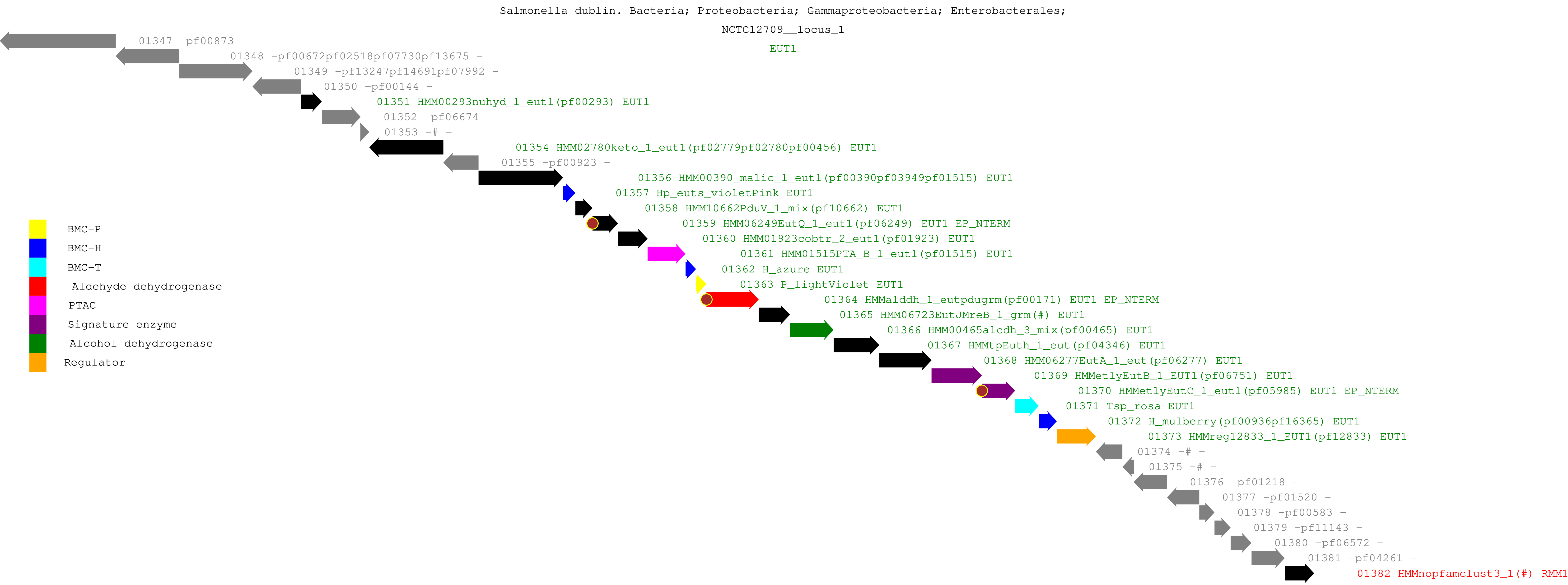

Salmonella dublin. Bacteria; Proteobacteria; Gammaproteobacteria; Enterobacterales;
NCTC12709__locus_1
EUT1
01347 -pf00873 -
01348 -pf00672pf02518pf07730pf13675 -
01349 -pf13247pf14691pf07992 -
01350 -pf00144 -
01351 HMM00293nuhyd_1_eut1(pf00293) EUT1
01352 -pf06674 -
01353 -# -
01354 HMM02780keto_1_eut1(pf02779pf02780pf00456) EUT1
01355 -pf00923 -
01356 HMM00390_malic_1_eut1(pf00390pf03949pf01515) EUT1
01357 Hp_euts_violetPink EUT1
01358 HMM10662PduV_1_mix(pf10662) EUT1
01359 HMM06249EutQ_1_eut1(pf06249) EUT1 EP_NTERM
01360 HMM01923cobtr_2_eut1(pf01923) EUT1
01361 HMM01515PTA_B_1_eut1(pf01515) EUT1
01362 H_azure EUT1
01363 P_lightViolet EUT1
01364 HMMalddh_1_eutpdugrm(pf00171) EUT1 EP_NTERM
01365 HMM06723EutJMreB_1_grm(#) EUT1
01366 HMM00465alcdh_3_mix(pf00465) EUT1
01367 HMMtpEuth_1_eut(pf04346) EUT1
01368 HMM06277EutA_1_eut(pf06277) EUT1
01369 HMMetlyEutB_1_EUT1(pf06751) EUT1
01370 HMMetlyEutC_1_eut1(pf05985) EUT1 EP_NTERM
01371 Tsp_rosa EUT1
01372 H_mulberry(pf00936pf16365) EUT1
01373 HMMreg12833_1_EUT1(pf12833) EUT1
01374 -# -
01375 -# -
01376 -pf01218 -
01377 -pf01520 -
01378 -pf00583 -
01379 -pf11143 -
01380 -pf06572 -
01381 -pf04261 -
01382 HMMnopfamclust3_1(#) RMM1
BMC-P
BMC-H
BMC-T
Aldehyde dehydrogenase
PTAC
Signature enzyme
Alcohol dehydrogenase
Regulator

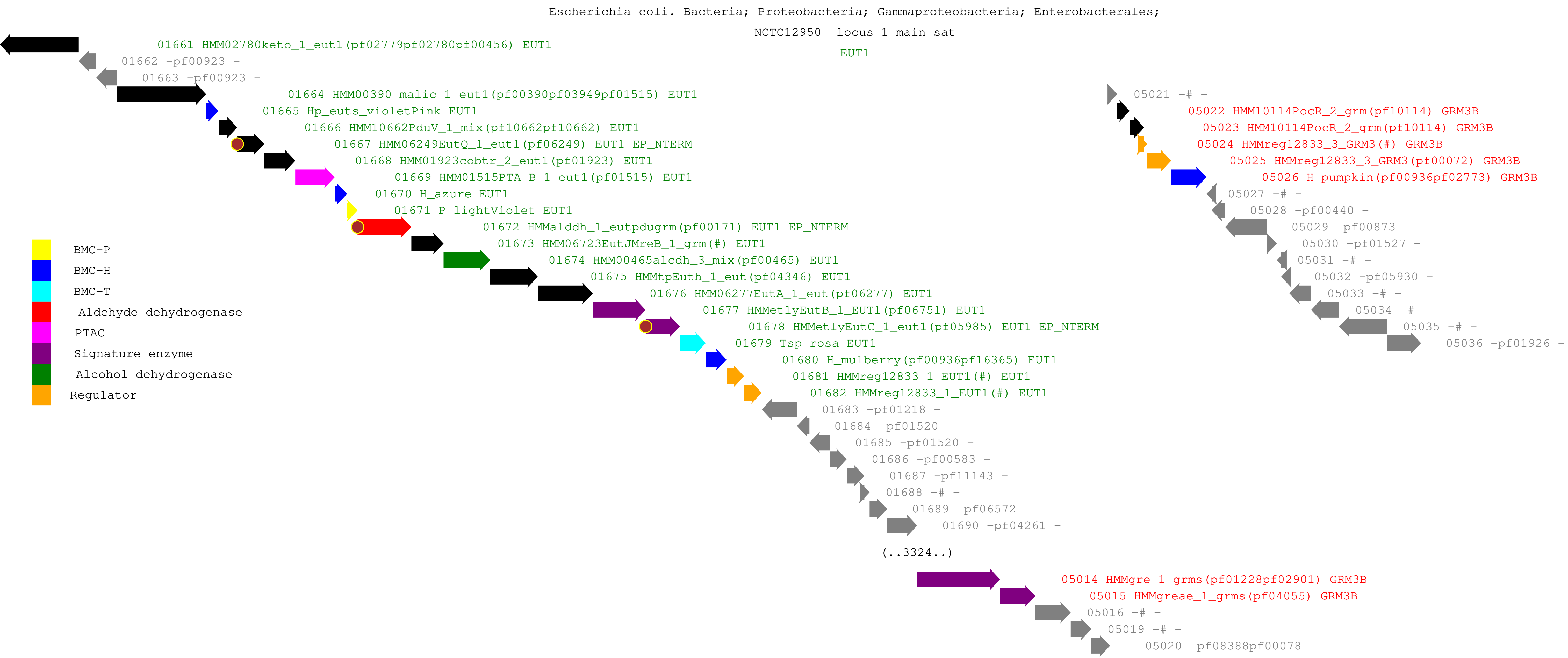
Escherichia coli. Bacteria; Proteobacteria; Gammaproteobacteria; Enterobacterales;
NCTC12950__locus_1_main_sat
EUT1
01661 HMM02780keto_1_eut1(pf02779pf02780pf00456) EUT1
01662 -pf00923 -
01663 -pf00923 -
01664 HMM00390_malic_1_eut1(pf00390pf03949pf01515) EUT1
01665 Hp_euts_violetPink EUT1
01666 HMM10662PduV_1_mix(pf10662pf10662) EUT1
01667 HMM06249EutQ_1_eut1(pf06249) EUT1 EP_NTERM
01668 HMM01923cobtr_2_eut1(pf01923) EUT1
01669 HMM01515PTA_B_1_eut1(pf01515) EUT1
01670 H_azure EUT1
01671 P_lightViolet EUT1
01672 HMMalddh_1_eutpdugrm(pf00171) EUT1 EP_NTERM
01673 HMM06723EutJMreB_1_grm(#) EUT1
01674 HMM00465alcdh_3_mix(pf00465) EUT1
01675 HMMtpEuth_1_eut(pf04346) EUT1
01676 HMM06277EutA_1_eut(pf06277) EUT1
01677 HMMetlyEutB_1_EUT1(pf06751) EUT1
01678 HMMetlyEutC_1_eut1(pf05985) EUT1 EP_NTERM
01679 Tsp_rosa EUT1
01680 H_mulberry(pf00936pf16365) EUT1
01681 HMMreg12833_1_EUT1(#) EUT1
01682 HMMreg12833_1_EUT1(#) EUT1
01683 -pf01218 -
01684 -pf01520 -
01685 -pf01520 -
01686 -pf00583 -
01687 -pf11143 -
01688 -# -
01689 -pf06572 -
01690 -pf04261 -
(..3324..)
05014 HMMgre_1_grms(pf01228pf02901) GRM3B
05015 HMMgreae_1_grms(pf04055) GRM3B
05016 -# -
05019 -# -
05020 -pf08388pf00078 -
05021 -# -
05022 HMM10114PocR_2_grm(pf10114) GRM3B
05023 HMM10114PocR_2_grm(pf10114) GRM3B
05024 HMMreg12833_3_GRM3(#) GRM3B
05025 HMMreg12833_3_GRM3(pf00072) GRM3B
05026 H_pumpkin(pf00936pf02773) GRM3B
05027 -# -
05028 -pf00440 -
05029 -pf00873 -
05030 -pf01527 -
05031 -# -
05032 -pf05930 -
05033 -# -
05034 -# -
05035 -# -
05036 -pf01926 -
BMC-P
BMC-H
BMC-T
Aldehyde dehydrogenase
PTAC
Signature enzyme
Alcohol dehydrogenase
Regulator

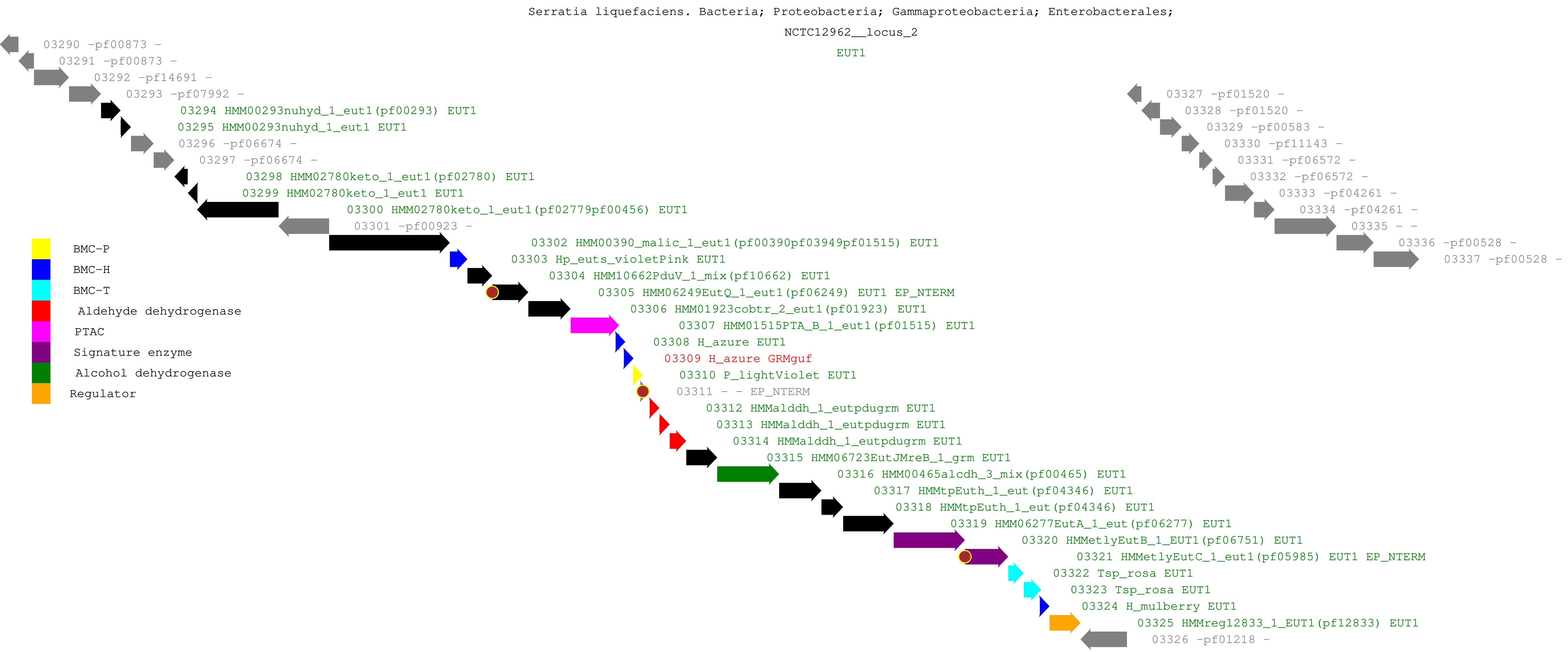

Serratia liquefaciens. Bacteria; Proteobacteria; Gammaproteobacteria; Enterobacterales;
NCTC12962__locus_2
EUT1
03290 -pf00873 -
03291 -pf00873 -
03292 -pf14691 -
03293 -pf07992 -
03294 HMM00293nuhyd_1_eut1(pf00293) EUT1
03295 HMM00293nuhyd_1_eut1 EUT1
03296 -pf06674 -
03297 -pf06674 -
03298 HMM02780keto_1_eut1(pf02780) EUT1
03299 HMM02780keto_1_eut1 EUT1
03300 HMM02780keto_1_eut1(pf02779pf00456) EUT1
03301 -pf00923 -
03302 HMM00390_malic_1_eut1(pf00390pf03949pf01515) EUT1
03303 Hp_euts_violetPink EUT1
03304 HMM10662PduV_1_mix(pf10662) EUT1
03305 HMM06249EutQ_1_eut1(pf06249) EUT1 EP_NTERM
03306 HMM01923cobtr_2_eut1(pf01923) EUT1
03307 HMM01515PTA_B_1_eut1(pf01515) EUT1
03308 H_azure EUT1
03309 H_azure GRMguf
03310 P_lightViolet EUT1
03311 - - EP_NTERM
03312 HMMalddh_1_eutpdugrm EUT1
03313 HMMalddh_1_eutpdugrm EUT1
03314 HMMalddh_1_eutpdugrm EUT1
03315 HMM06723EutJMreB_1_grm EUT1
03316 HMM00465alcdh_3_mix(pf00465) EUT1
03317 HMMtpEuth_1_eut(pf04346) EUT1
03318 HMMtpEuth_1_eut(pf04346) EUT1
03319 HMM06277EutA_1_eut(pf06277) EUT1
03320 HMMetlyEutB_1_EUT1(pf06751) EUT1
03321 HMMetlyEutC_1_eut1(pf05985) EUT1 EP_NTERM
03322 Tsp_rosa EUT1
03323 Tsp_rosa EUT1
03324 H_mulberry EUT1
03325 HMMreg12833_1_EUT1(pf12833) EUT1
03326 -pf01218 -
03327 -pf01520 -
03328 -pf01520 -
03329 -pf00583 -
03330 -pf11143 -
03331 -pf06572 -
03332 -pf06572 -
03333 -pf04261 -
03334 -pf04261 -
03335 - -
03336 -pf00528 -
03337 -pf00528 -
BMC-P
BMC-H
BMC-T
Aldehyde dehydrogenase
PTAC
Signature enzyme
Alcohol dehydrogenase
Regulator

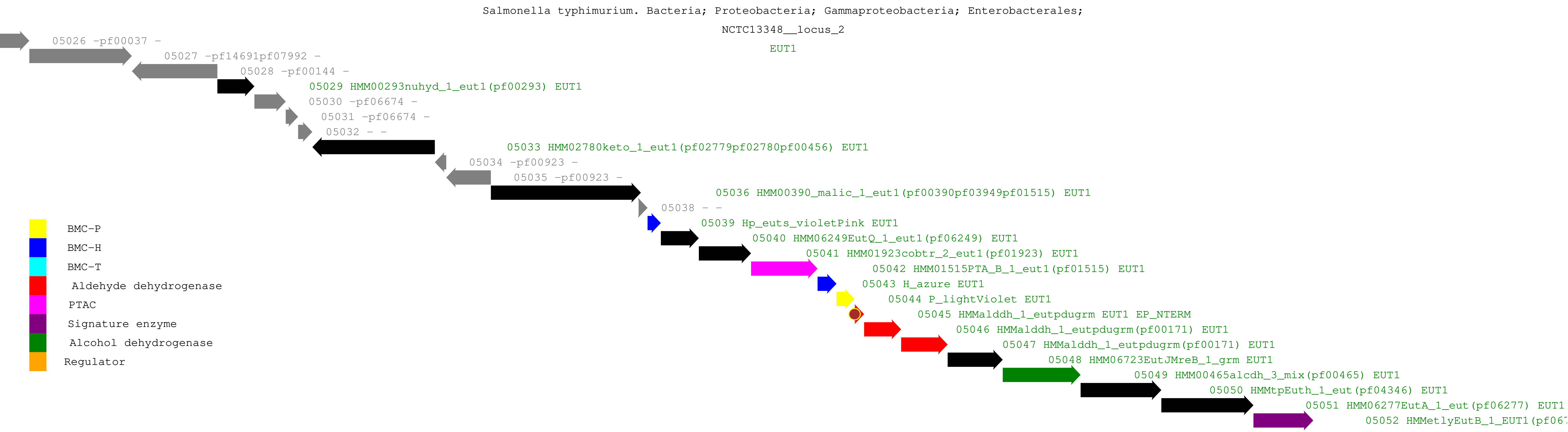

Salmonella typhimurium. Bacteria; Proteobacteria; Gammaproteobacteria; Enterobacterales;
NCTC13348__locus_2
EUT1
05026 -pf00037 -
05027 -pf14691pf07992 -
05028 -pf00144 -
05029 HMM00293nuhyd_1_eut1(pf00293) EUT1
05030 -pf06674 -
05031 -pf06674 -
05032 - -
05033 HMM02780keto_1_eut1(pf02779pf02780pf00456) EUT1
05034 -pf00923 -
05035 -pf00923 -
05036 HMM00390_malic_1_eut1(pf00390pf03949pf01515) EUT1
05038 - -
05039 Hp_euts_violetPink EUT1
05040 HMM06249EutQ_1_eut1(pf06249) EUT1
05041 HMM01923cobtr_2_eut1(pf01923) EUT1
05042 HMM01515PTA_B_1_eut1(pf01515) EUT1
05043 H_azure EUT1
05044 P_lightViolet EUT1
05045 HMMalddh_1_eutpdugrm EUT1 EP_NTERM
05046 HMMalddh_1_eutpdugrm(pf00171) EUT1
05047 HMMalddh_1_eutpdugrm(pf00171) EUT1
05048 HMM06723EutJMreB_1_grm EUT1
05049 HMM00465alcdh_3_mix(pf00465) EUT1
05050 HMMtpEuth_1_eut(pf04346) EUT1
05051 HMM06277EutA_1_eut(pf06277) EUT1
05052 HMMetlyEutB_1_EUT1(pf06
BMC-P
BMC-H
BMC-T
Aldehyde dehydrogenase
PTAC
Signature enzyme
Alcohol dehydrogenase
Regulator

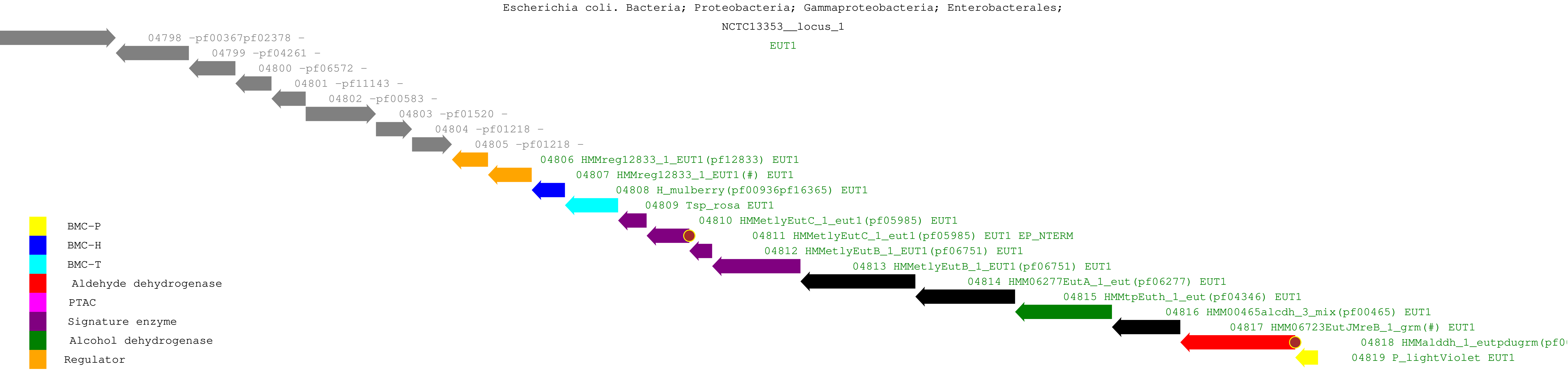

Escherichia coli. Bacteria; Proteobacteria; Gammaproteobacteria; Enterobacterales;
NCTC13353__locus_1
EUT1
04798 -pf00367pf02378 -
04799 -pf04261 -
04800 -pf06572 -
04801 -pf11143 -
04802 -pf00583 -
04803 -pf01520 -
04804 -pf01218 -
04805 -pf01218 -
04806 HMMreg12833_1_EUT1(pf12833) EUT1
04807 HMMreg12833_1_EUT1(#) EUT1
04808 H_mulberry(pf00936pf16365) EUT1
04809 Tsp_rosa EUT1
04810 HMMetlyEutC_1_eut1(pf05985) EUT1
04811 HMMetlyEutC_1_eut1(pf05985) EUT1 EP_NTERM
04812 HMMetlyEutB_1_EUT1(pf06751) EUT1
04813 HMMetlyEutB_1_EUT1(pf06751) EUT1
04814 HMM06277EutA_1_eut(pf06277) EUT1
04815 HMMtpEuth_1_eut(pf04346) EUT1
04816 HMM00465alcdh_3_mix(pf00465) EUT1
04817 HMM06723EutJMreB_1_grm(#) EUT1
04819 P_lightViolet EUT1
BMC-P
BMC-H
BMC-T
Aldehyde dehydrogenase
PTAC
Signature enzyme
Alcohol dehydrogenase
Regulator

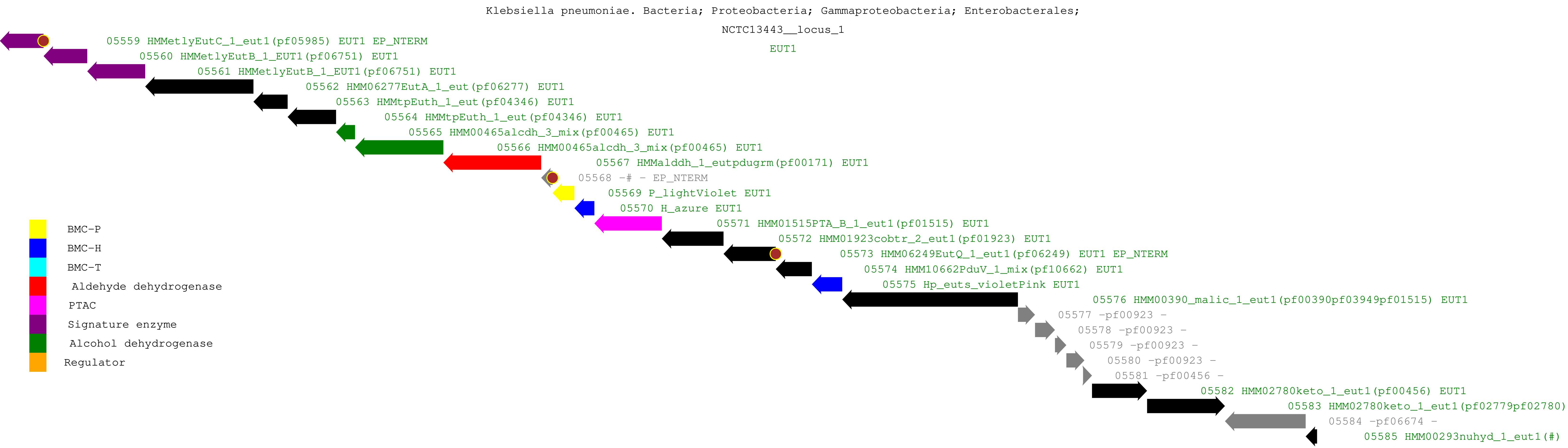
Klebsiella pneumoniae. Bacteria; Proteobacteria; Gammaproteobacteria; Enterobacterales;
NCTC13443__locus_1
EUT1
05559 HMMetlyEutC_1_eut1(pf05985) EUT1 EP_NTERM
05560 HMMetlyEutB_1_EUT1(pf06751) EUT1
05561 HMMetlyEutB_1_EUT1(pf06751) EUT1
05562 HMM06277EutA_1_eut(pf06277) EUT1
05563 HMMtpEuth_1_eut(pf04346) EUT1
05564 HMMtpEuth_1_eut(pf04346) EUT1
05565 HMM00465alcdh_3_mix(pf00465) EUT1
05566 HMM00465alcdh_3_mix(pf00465) EUT1
05567 HMMalddh_1_eutpdugrm(pf00171) EUT1
05568 -# - EP_NTERM
05569 P_lightViolet EUT1
05570 H_azure EUT1
05571 HMM01515PTA_B_1_eut1(pf01515) EUT1
05572 HMM01923cobtr_2_eut1(pf01923) EUT1
05573 HMM06249EutQ_1_eut1(pf06249) EUT1 EP_NTERM
05574 HMM10662PduV_1_mix(pf10662) EUT1
05575 Hp_euts_violetPink EUT1
05576 HMM00390_malic_1_eut1(pf00390pf03949pf01515) EUT1
05577 -pf00923 -
05578 -pf00923 -
05579 -pf00923 -
05580 -pf00923 -
05581 -pf00456 -
05582 HMM02780keto_1_eut1(pf00456) EUT1
05583 HMM02780keto_1_eut1(pf02779pf02780)
05584 -pf06674 -
05585 HMM00293nuhyd_1_eut1(#)
BMC-P
BMC-H
BMC-T
Aldehyde dehydrogenase
PTAC
Signature enzyme
Alcohol dehydrogenase
Regulator

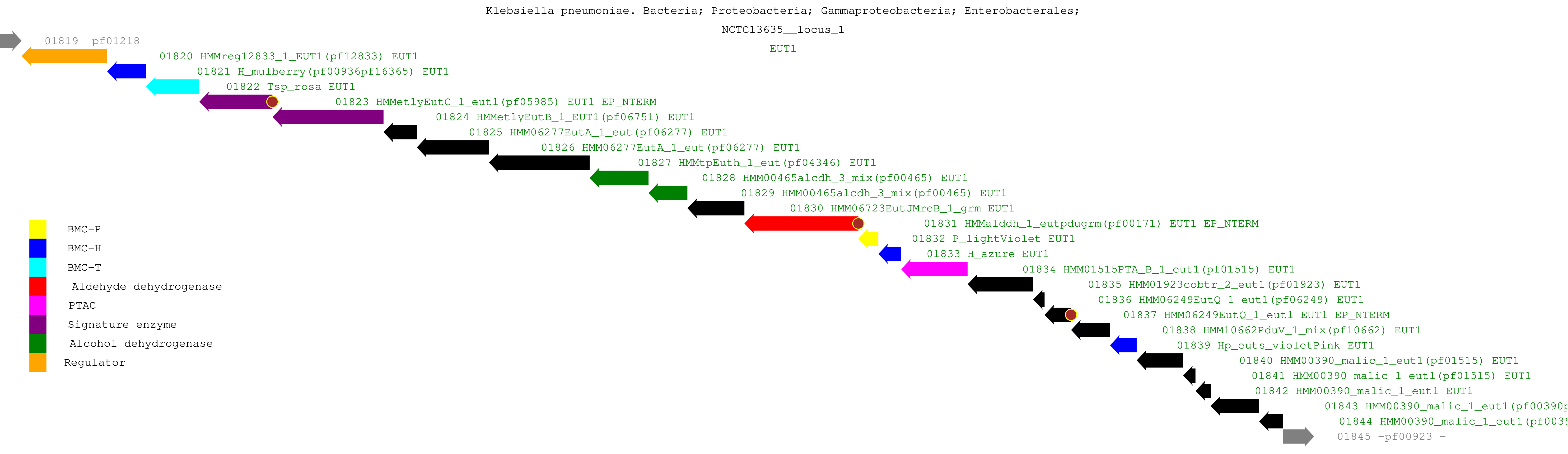
Klebsiella pneumoniae. Bacteria; Proteobacteria; Gammaproteobacteria; Enterobacterales;
NCTC13635__locus_1
EUT1
01819 -pf01218 -
01820 HMMreg12833_1_EUT1(pf12833) EUT1
01821 H_mulberry(pf00936pf16365) EUT1
01822 Tsp_rosa EUT1
01823 HMMetlyEutC_1_eut1(pf05985) EUT1 EP_NTERM
01824 HMMetlyEutB_1_EUT1(pf06751) EUT1
01825 HMM06277EutA_1_eut(pf06277) EUT1
01826 HMM06277EutA_1_eut(pf06277) EUT1
01827 HMMtpEuth_1_eut(pf04346) EUT1
01828 HMM00465alcdh_3_mix(pf00465) EUT1
01829 HMM00465alcdh_3_mix(pf00465) EUT1
01830 HMM06723EutJMreB_1_grm EUT1
01831 HMMalddh_1_eutpdugrm(pf00171) EUT1 EP_NTERM
01832 P_lightViolet EUT1
01833 H_azure EUT1
01834 HMM01515PTA_B_1_eut1(pf01515) EUT1
01835 HMM01923cobtr_2_eut1(pf01923) EUT1
01836 HMM06249EutQ_1_eut1(pf06249) EUT1
01837 HMM06249EutQ_1_eut1 EUT1 EP_NTERM
01838 HMM10662PduV_1_mix(pf10662) EUT1
01839 Hp_euts_violetPink EUT1
01840 HMM00390_malic_1_eut1(pf01515) EUT1
01841 HMM00390_malic_1_eut1(pf01515) EUT1
01842 HMM00390_malic_1_eut1 EUT1
01843 HMM00390_malic_1_eut1(pf00390
01844 HMM00390_malic_1_eut1(pf003
01845 -pf00923 -
BMC-P
BMC-H
BMC-T
Aldehyde dehydrogenase
PTAC
Signature enzyme
Alcohol dehydrogenase
Regulator

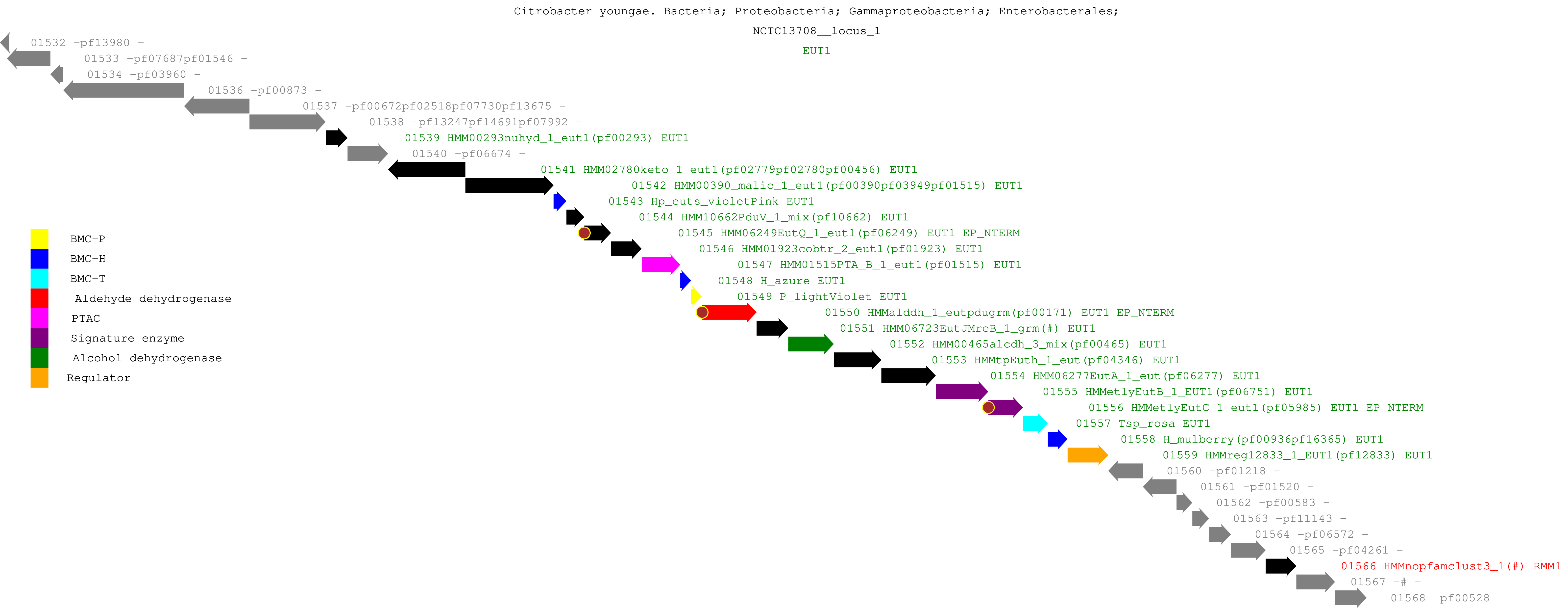
Citrobacter youngae. Bacteria; Proteobacteria; Gammaproteobacteria; Enterobacterales;
NCTC13708__locus_1
EUT1
01532 -pf13980 -
01533 -pf07687pf01546 -
01534 -pf03960 -
01536 -pf00873 -
01537 -pf00672pf02518pf07730pf13675 -
01538 -pf13247pf14691pf07992 -
01539 HMM00293nuhyd_1_eut1(pf00293) EUT1
01540 -pf06674 -
01541 HMM02780keto_1_eut1(pf02779pf02780pf00456) EUT1
01542 HMM00390_malic_1_eut1(pf00390pf03949pf01515) EUT1
01543 Hp_euts_violetPink EUT1
01544 HMM10662PduV_1_mix(pf10662) EUT1
01545 HMM06249EutQ_1_eut1(pf06249) EUT1 EP_NTERM
01546 HMM01923cobtr_2_eut1(pf01923) EUT1
01547 HMM01515PTA_B_1_eut1(pf01515) EUT1
01548 H_azure EUT1
01549 P_lightViolet EUT1
01550 HMMalddh_1_eutpdugrm(pf00171) EUT1 EP_NTERM
01551 HMM06723EutJMreB_1_grm(#) EUT1
01552 HMM00465alcdh_3_mix(pf00465) EUT1
01553 HMMtpEuth_1_eut(pf04346) EUT1
01554 HMM06277EutA_1_eut(pf06277) EUT1
01555 HMMetlyEutB_1_EUT1(pf06751) EUT1
01556 HMMetlyEutC_1_eut1(pf05985) EUT1 EP_NTERM
01557 Tsp_rosa EUT1
01558 H_mulberry(pf00936pf16365) EUT1
01559 HMMreg12833_1_EUT1(pf12833) EUT1
01560 -pf01218 -
01561 -pf01520 -
01562 -pf00583 -
01563 -pf11143 -
01564 -pf06572 -
01565 -pf04261 -
01566 HMMnopfamclust3_1(#) RMM1
01567 -# -
01568 -pf00528 -
BMC-P
BMC-H
BMC-T
Aldehyde dehydrogenase
PTAC
Signature enzyme
Alcohol dehydrogenase
Regulator

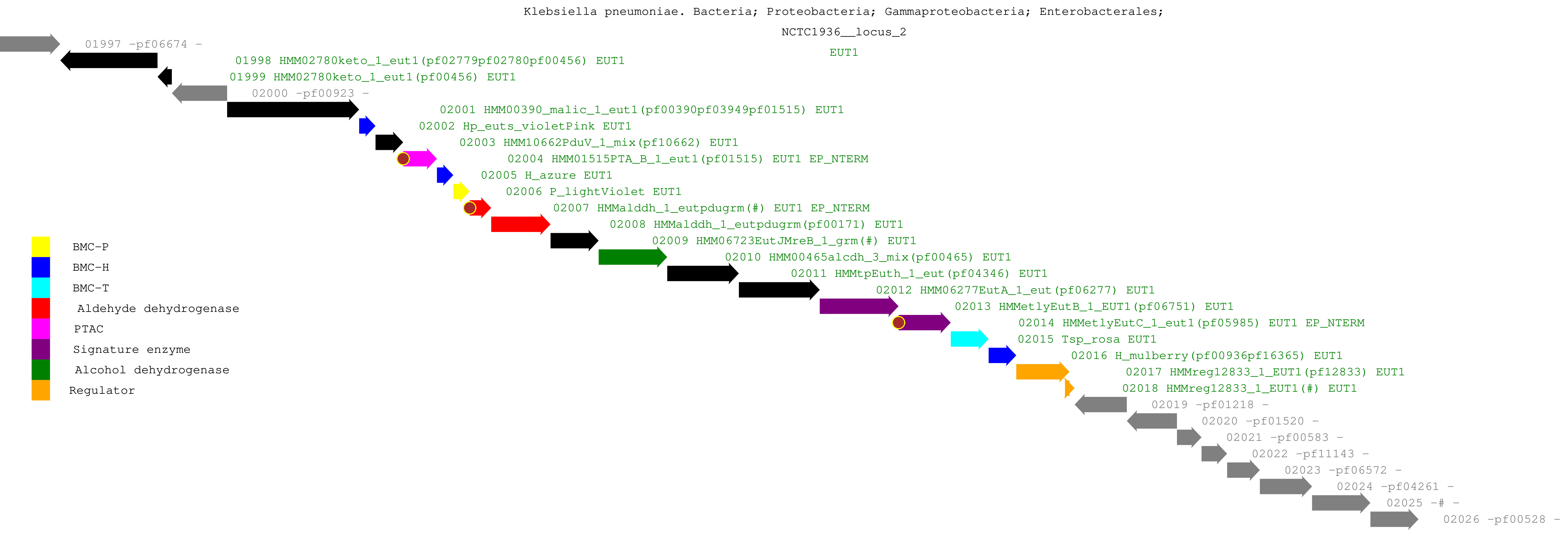
Klebsiella pneumoniae. Bacteria; Proteobacteria; Gammaproteobacteria; Enterobacterales;
NCTC1936__locus_2
EUT1
01997 -pf06674 -
01998 HMM02780keto_1_eut1(pf02779pf02780pf00456) EUT1
01999 HMM02780keto_1_eut1(pf00456) EUT1
02000 -pf00923 -
02001 HMM00390_malic_1_eut1(pf00390pf03949pf01515) EUT1
02002 Hp_euts_violetPink EUT1
02003 HMM10662PduV_1_mix(pf10662) EUT1
02004 HMM01515PTA_B_1_eut1(pf01515) EUT1 EP_NTERM
02005 H_azure EUT1
02006 P_lightViolet EUT1
02007 HMMalddh_1_eutpdugrm(#) EUT1 EP_NTERM
02008 HMMalddh_1_eutpdugrm(pf00171) EUT1
02009 HMM06723EutJMreB_1_grm(#) EUT1
02010 HMM00465alcdh_3_mix(pf00465) EUT1
02011 HMMtpEuth_1_eut(pf04346) EUT1
02012 HMM06277EutA_1_eut(pf06277) EUT1
02013 HMMetlyEutB_1_EUT1(pf06751) EUT1
02014 HMMetlyEutC_1_eut1(pf05985) EUT1 EP_NTERM
02015 Tsp_rosa EUT1
02016 H_mulberry(pf00936pf16365) EUT1
02017 HMMreg12833_1_EUT1(pf12833) EUT1
02018 HMMreg12833_1_EUT1(#) EUT1
02019 -pf01218 -
02020 -pf01520 -
02021 -pf00583 -
02022 -pf11143 -
02023 -pf06572 -
02024 -pf04261 -
02025 -# -
02026 -pf00528 -
BMC-P
BMC-H
BMC-T
Aldehyde dehydrogenase
PTAC
Signature enzyme
Alcohol dehydrogenase
Regulator

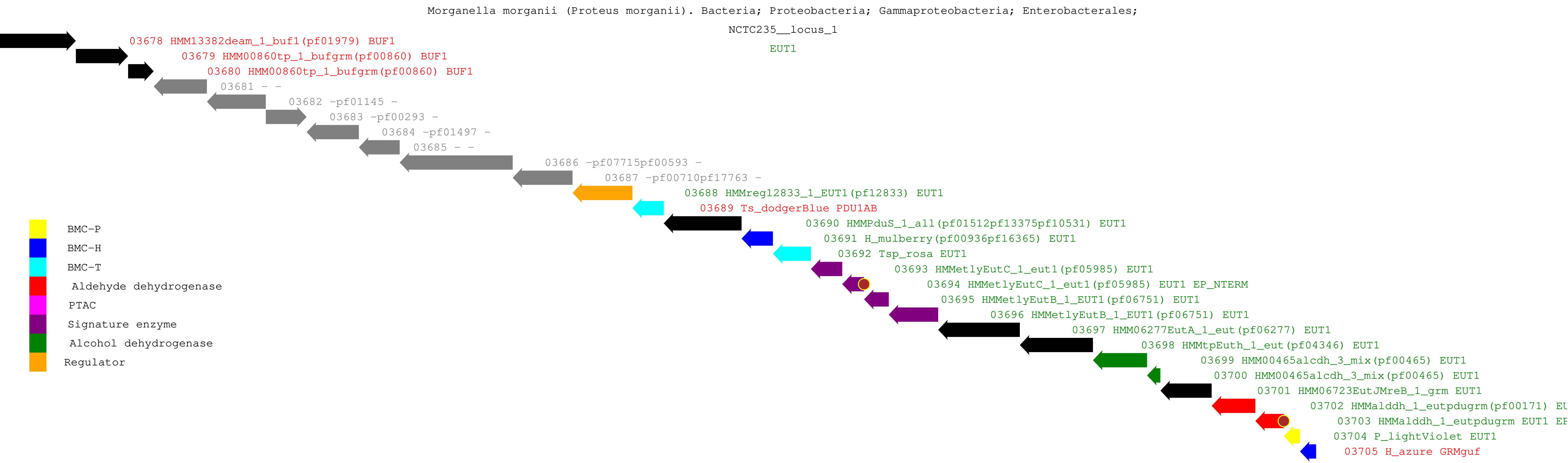
Morganella morganii (Proteus morganii). Bacteria; Proteobacteria; Gammaproteobacteria; Enterobacterales;
NCTC235__locus_1
EUT1
03678 HMM13382deam_1_buf1(pf01979) BUF1
03679 HMM00860tp_1_bufgrm(pf00860) BUF1
03680 HMM00860tp_1_bufgrm(pf00860) BUF1
03681 - -
03682 -pf01145 -
03683 -pf00293 -
03684 -pf01497 -
03685 - -
03686 -pf07715pf00593 -
03687 -pf00710pf17763 -
03688 HMMreg12833_1_EUT1(pf12833) EUT1
03689 Ts_dodgerBlue PDU1AB
03690 HMMPduS_1_all(pf01512pf13375pf10531) EUT1
03691 H_mulberry(pf00936pf16365) EUT1
03692 Tsp_rosa EUT1
03693 HMMetlyEutC_1_eut1(pf05985) EUT1
03694 HMMetlyEutC_1_eut1(pf05985) EUT1 EP_NTERM
03695 HMMetlyEutB_1_EUT1(pf06751) EUT1
03696 HMMetlyEutB_1_EUT1(pf06751) EUT1
03697 HMM06277EutA_1_eut(pf06277) EUT1
03698 HMMtpEuth_1_eut(pf04346) EUT1
03699 HMM00465alcdh_3_mix(pf00465) EUT1
03700 HMM00465alcdh_3_mix(pf00465) EUT1
03701 HMM06723EutJMreB_1_grm EUT1
03702 HMMaldh_1_eutpdugrm(pf00171) EU
03703 HMMaldh_1_eutpdugrm EUT1 EP
03704 P_lightViolet EUT1
03705 H_azure GRMguf
BMC-P
BMC-H
BMC-T
Aldehyde dehydrogenase
PTAC
Signature enzyme
Alcohol dehydrogenase
Regulator

Klebsiella aerogenes (Enterobacter aerogenes). Bacteria; Proteobacteria; Gammaproteobacteria; Enterobacterales;

NCTC243__locus_1

EUT1

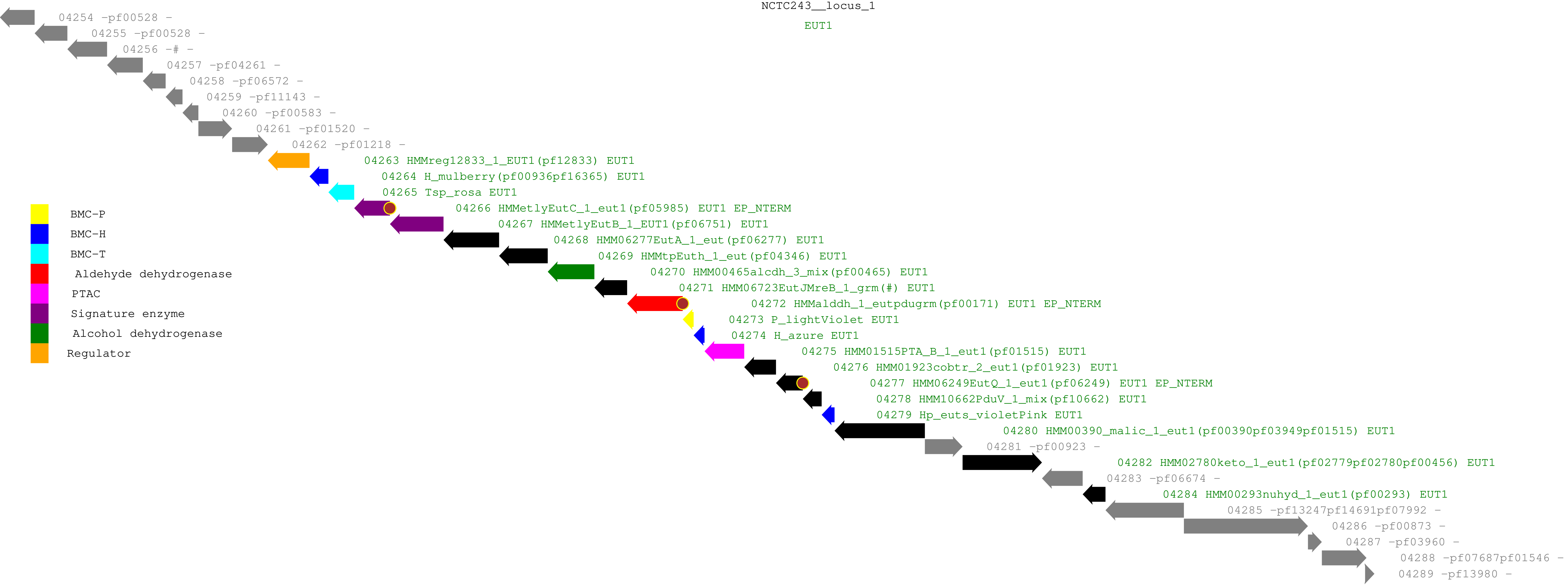

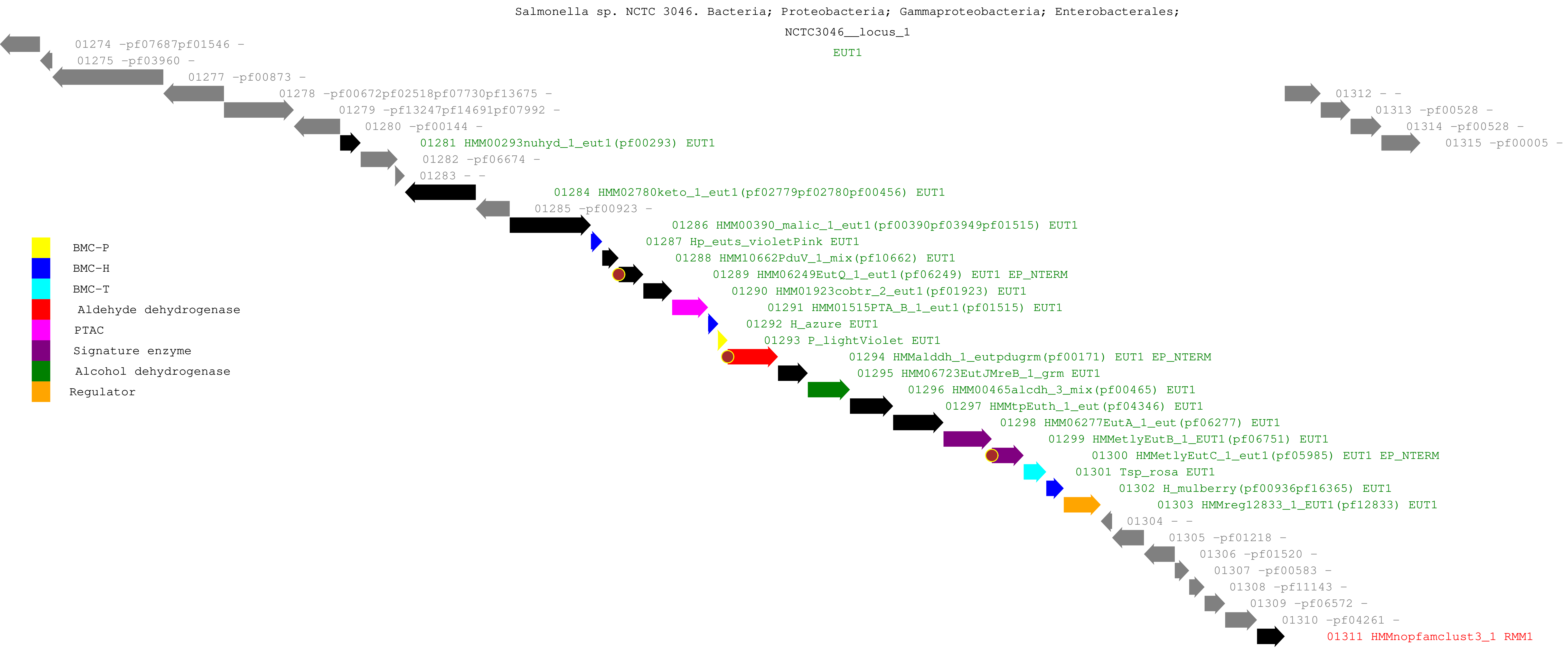
Salmonella sp. NCTC 3046. Bacteria; Proteobacteria; Gammaproteobacteria; Enterobacterales;
NCTC3046__locus_1
EUT1
01274 -pf07687pf01546 -
01275 -pf03960 -
01277 -pf00873 -
01278 -pf00672pf02518pf07730pf13675 -
01279 -pf13247pf14691pf07992 -
01280 -pf00144 -
01281 HMM00293nuhyd_1_eut1(pf00293) EUT1
01282 -pf06674 -
01283 - -
01284 HMM02780keto_1_eut1(pf02779pf02780pf00456) EUT1
01285 -pf00923 -
01286 HMM00390_malic_1_eut1(pf00390pf03949pf01515) EUT1
01287 Hp_euts_violetPink EUT1
01288 HMM10662PduV_1_mix(pf10662) EUT1
01289 HMM06249EutQ_1_eut1(pf06249) EUT1 EP_NTERM
01290 HMM01923cobtr_2_eut1(pf01923) EUT1
01291 HMM01515PTA_B_1_eut1(pf01515) EUT1
01292 H_azure EUT1
01293 P_lightViolet EUT1
01294 HMMaldh_1_eutpdugrm(pf00171) EUT1 EP_NTERM
01295 HMM06723EutJMreB_1_grm EUT1
01296 HMM00465alcdh_3_mix(pf00465) EUT1
01297 HMMtpEuth_1_eut(pf04346) EUT1
01298 HMM06277EutA_1_eut(pf06277) EUT1
01299 HMMetlyEutB_1_EUT1(pf06751) EUT1
01300 HMMetlyEutC_1_eut1(pf05985) EUT1 EP_NTERM
01301 Tsp_rosa EUT1
01302 H_mulberry(pf00936pf16365) EUT1
01303 HMMreg12833_1_EUT1(pf12833) EUT1
01304 - -
01305 -pf01218 -
01306 -pf01520 -
01307 -pf00583 -
01308 -pf11143 -
01309 -pf06572 -
01310 -pf04261 -
01311 HMMnopfamclust3_1 RMM1
01312 - -
01313 -pf00528 -
01314 -pf00528 -
01315 -pf00005 -
BMC-P
BMC-H
BMC-T
Aldehyde dehydrogenase
PTAC
Signature enzyme
Alcohol dehydrogenase
Regulator

Klebsiella pneumoniae subsp. rhinoscleromatis. Bacteria; Proteobacteria; Gammaproteobacteria; Enterobacterales;

NCTC5046__locus_3

EUT1

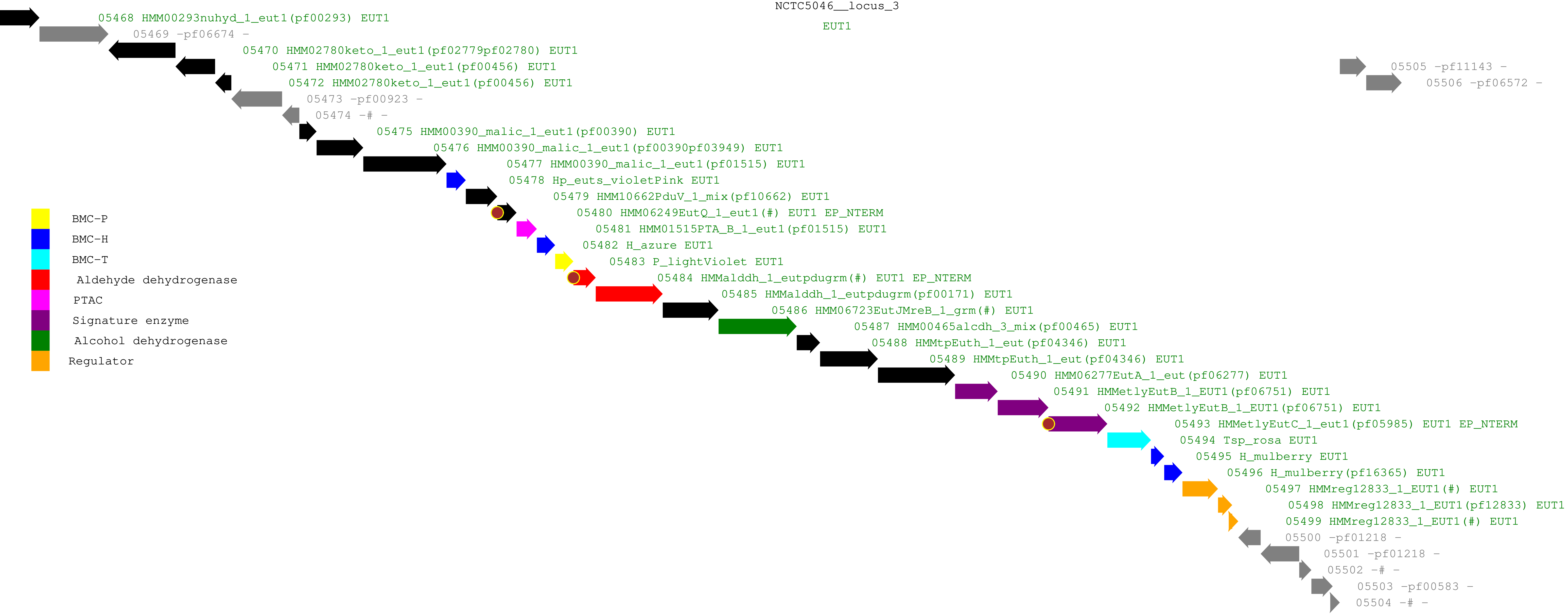

Klebsiella pneumoniae. Bacteria; Proteobacteria; Gammaproteobacteria; Enterobacterales;

NCTC5047__locus_2

EUT1

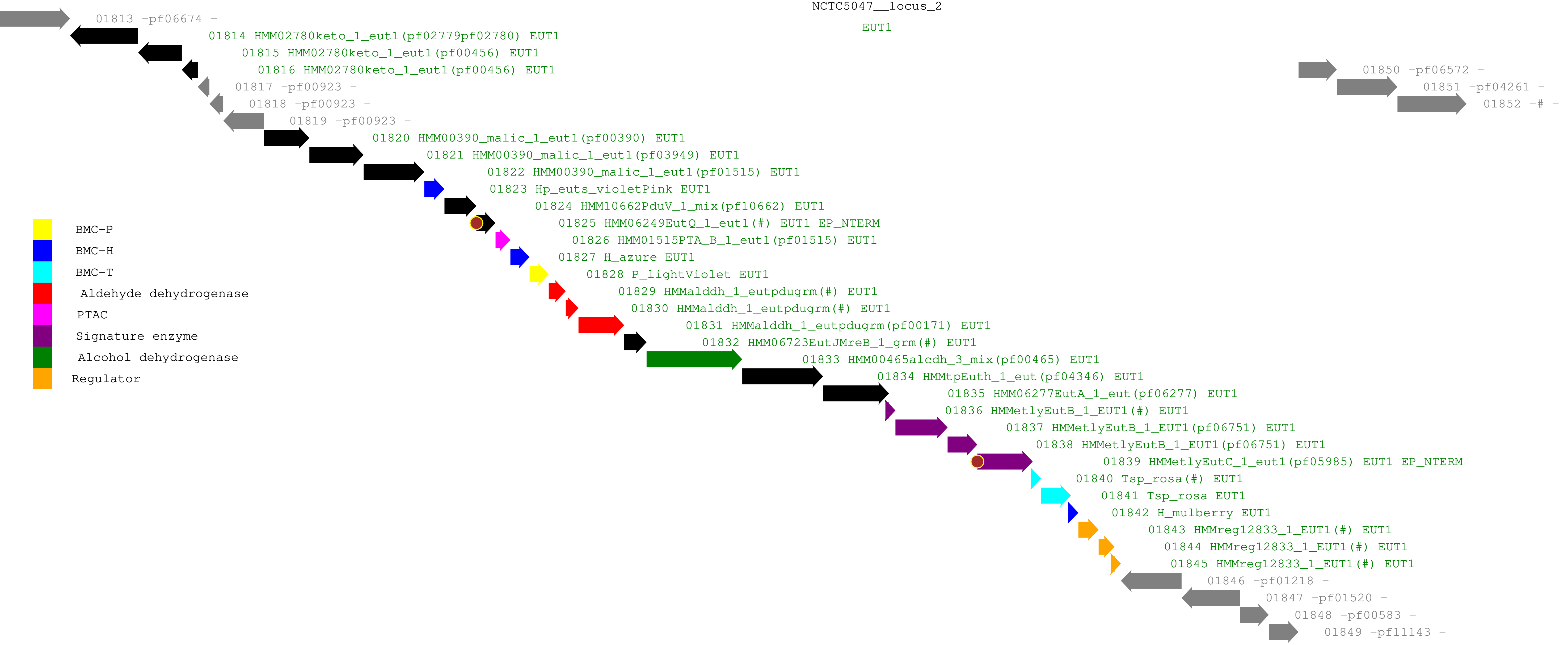

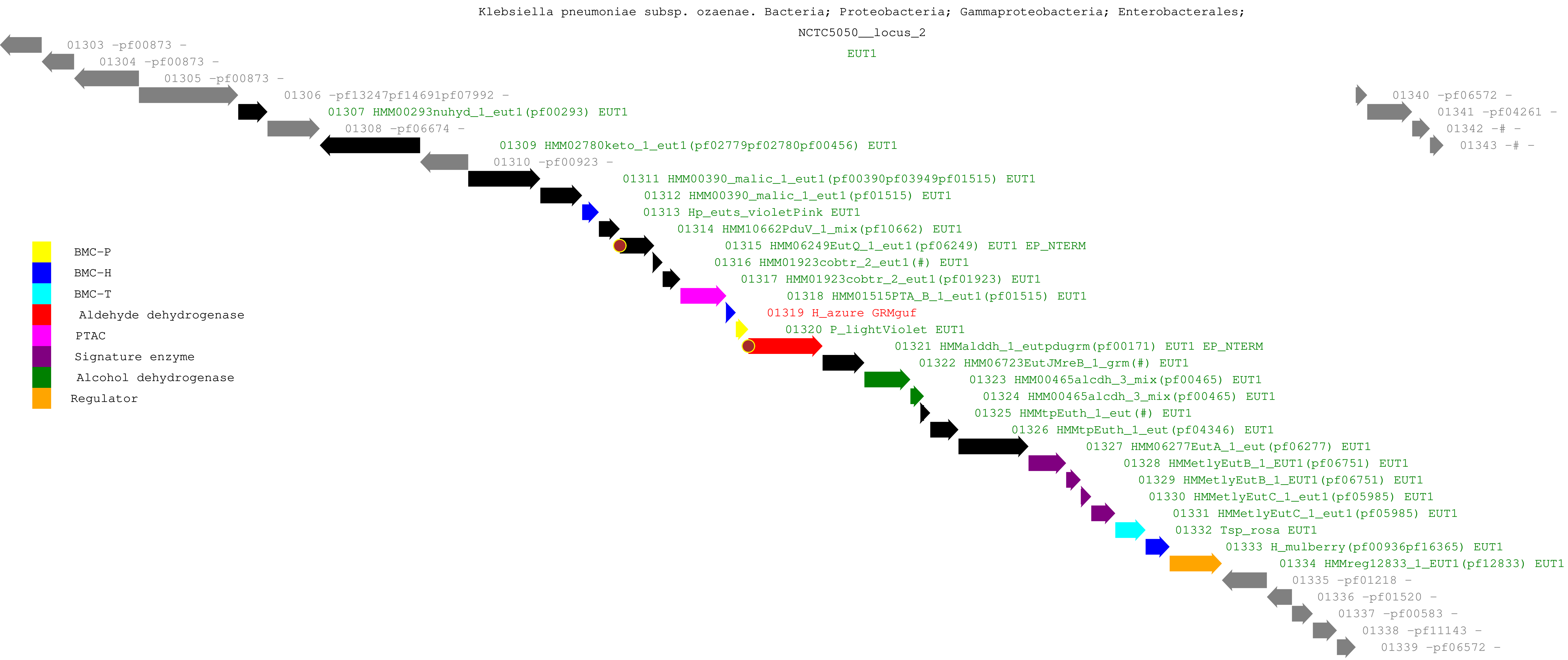

Klebsiella pneumoniae subsp. ozaenae. Bacteria; Proteobacteria; Gammaproteobacteria; Enterobacterales;
NCTC5050__locus_2
EUT1
01303 -pf00873 -
01304 -pf00873 -
01305 -pf00873 -
01306 -pf13247pf14691pf07992 -
01307 HMM00293nuhyd_1_eut1(pf00293) EUT1
01308 -pf06674 -
01309 HMM02780keto_1_eut1(pf02779pf02780pf00456) EUT1
01310 -pf00923 -
01311 HMM00390_malic_1_eut1(pf00390pf03949pf01515) EUT1
01312 HMM00390_malic_1_eut1(pf01515) EUT1
01313 Hp_euts_violetPink EUT1
01314 HMM10662PduV_1_mix(pf10662) EUT1
01315 HMM06249EutQ_1_eut1(pf06249) EUT1 EP_NTERM
01316 HMM01923cobtr_2_eut1(#) EUT1
01317 HMM01923cobtr_2_eut1(pf01923) EUT1
01318 HMM01515PTA_B_1_eut1(pf01515) EUT1
01319 H_azure GRMguf
01320 P_lightViolet EUT1
01321 HMMalddh_1_eutpdugrm(pf00171) EUT1 EP_NTERM
01322 HMM06723EutJMreB_1_grm(#) EUT1
01323 HMM00465alcdh_3_mix(pf00465) EUT1
01324 HMM00465alcdh_3_mix(pf00465) EUT1
01325 HMMtpEuth_1_eut(#) EUT1
01326 HMMtpEuth_1_eut(pf04346) EUT1
01327 HMM06277EutA_1_eut(pf06277) EUT1
01328 HMMetlyEutB_1_EUT1(pf06751) EUT1
01329 HMMetlyEutB_1_EUT1(pf06751) EUT1
01330 HMMetlyEutC_1_eut1(pf05985) EUT1
01331 HMMetlyEutC_1_eut1(pf05985) EUT1
01332 Tsp_rosa EUT1
01333 H_mulberry(pf00936pf16365) EUT1
01334 HMMreg12833_1_EUT1(pf12833) EUT1
01335 -pf01218 -
01336 -pf01520 -
01337 -pf00583 -
01338 -pf11143 -
01339 -pf06572 -
01340 -pf06572 -
01341 -pf04261 -
01342 -# -
01343 -# -
BMC-P
BMC-H
BMC-T
Aldehyde dehydrogenase
PTAC
Signature enzyme
Alcohol dehydrogenase
Regulator

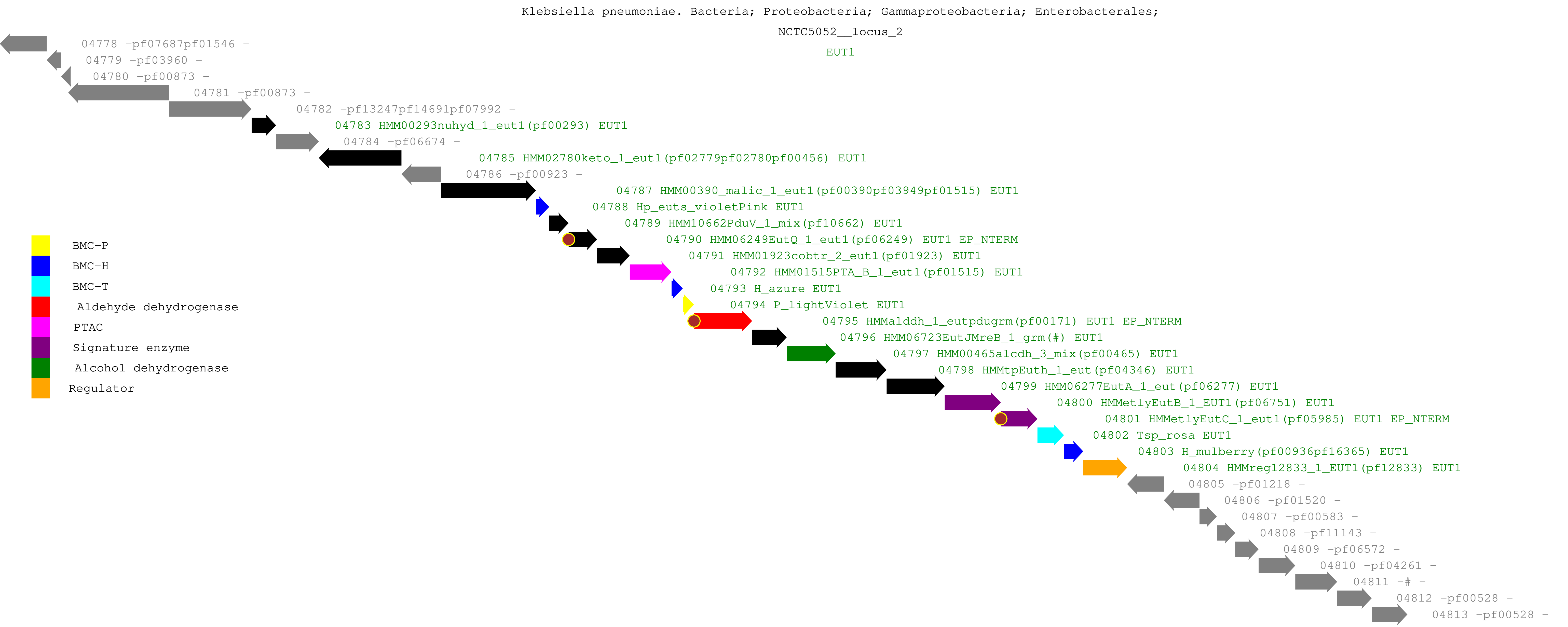
Klebsiella pneumoniae. Bacteria; Proteobacteria; Gammaproteobacteria; Enterobacterales;
NCTC5052__locus_2
EUT1
04778 -pf07687pf01546 -
04779 -pf03960 -
04780 -pf00873 -
04781 -pf00873 -
04782 -pf13247pf14691pf07992 -
04783 HMM00293nuhyd_1_eut1(pf00293) EUT1
04784 -pf06674 -
04785 HMM02780keto_1_eut1(pf02779pf02780pf00456) EUT1
04786 -pf00923 -
04787 HMM00390_malic_1_eut1(pf00390pf03949pf01515) EUT1
04788 Hp_euts_violetPink EUT1
04789 HMM10662PduV_1_mix(pf10662) EUT1
04790 HMM06249EutQ_1_eut1(pf06249) EUT1 EP_NTERM
04791 HMM01923cobtr_2_eut1(pf01923) EUT1
04792 HMM01515PTA_B_1_eut1(pf01515) EUT1
04793 H_azure EUT1
04794 P_lightViolet EUT1
04795 HMMalddh_1_eutpdugrm(pf00171) EUT1 EP_NTERM
04796 HMM06723EutJMreB_1_grm(#) EUT1
04797 HMM00465alcdh_3_mix(pf00465) EUT1
04798 HMMtpEuth_1_eut(pf04346) EUT1
04799 HMM06277EutA_1_eut(pf06277) EUT1
04800 HMMetlyEutB_1_EUT1(pf06751) EUT1
04801 HMMetlyEutC_1_eut1(pf05985) EUT1 EP_NTERM
04802 Tsp_rosa EUT1
04803 H_mulberry(pf00936pf16365) EUT1
04804 HMMreg12833_1_EUT1(pf12833) EUT1
04805 -pf01218 -
04806 -pf01520 -
04807 -pf00583 -
04808 -pf11143 -
04809 -pf06572 -
04810 -pf04261 -
04811 -# -
04812 -pf00528 -
04813 -pf00528 -
BMC-P
BMC-H
BMC-T
Aldehyde dehydrogenase
PTAC
Signature enzyme
Alcohol dehydrogenase
Regulator

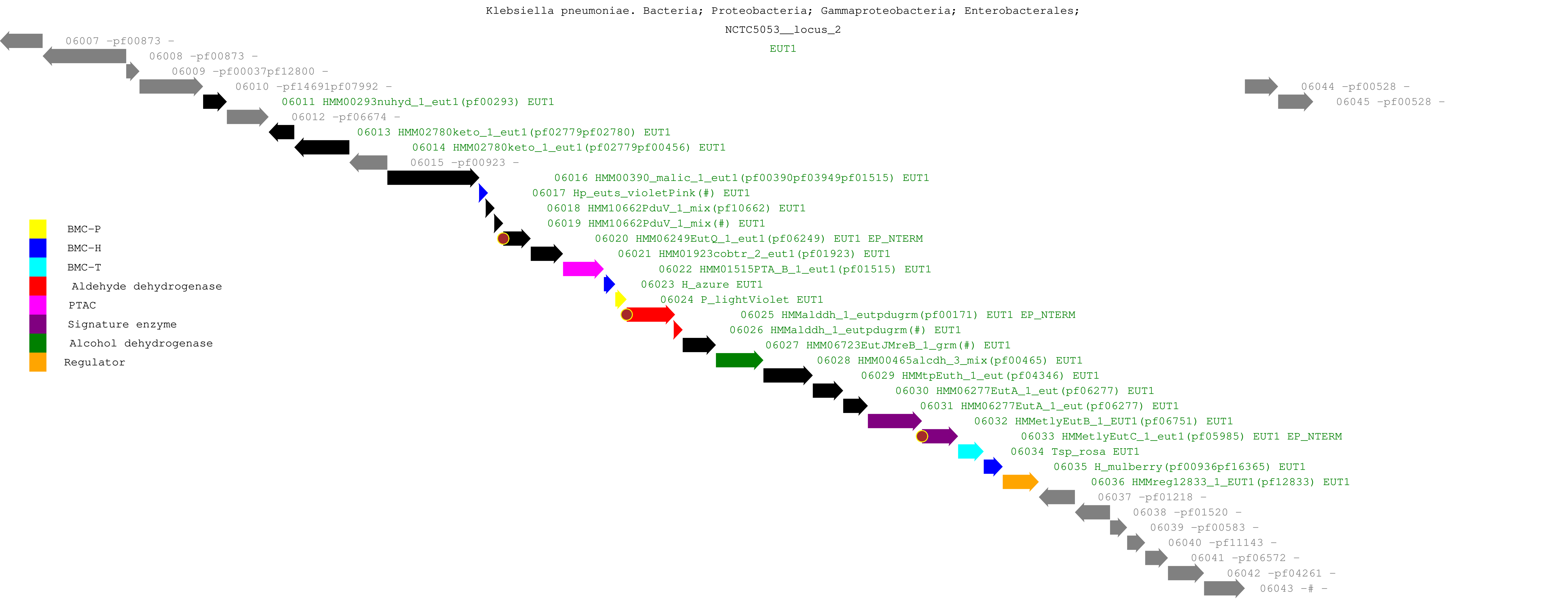
Klebsiella pneumoniae. Bacteria; Proteobacteria; Gammaproteobacteria; Enterobacterales;
NCTC5053__locus_2
EUT1
06007 -pf00873 -
06008 -pf00873 -
06009 -pf00037pf12800 -
06010 -pf14691pf07992 -
06011 HMM00293nuhyd_1_eut1(pf00293) EUT1
06012 -pf06674 -
06013 HMM02780keto_1_eut1(pf02779pf02780) EUT1
06014 HMM02780keto_1_eut1(pf02779pf00456) EUT1
06015 -pf00923 -
06016 HMM00390_malic_1_eut1(pf00390pf03949pf01515) EUT1
06017 Hp_euts_violetPink(#) EUT1
06018 HMM10662PduV_1_mix(pf10662) EUT1
06019 HMM10662PduV_1_mix(#) EUT1
06020 HMM06249EutQ_1_eut1(pf06249) EUT1 EP_NTERM
06021 HMM01923cobtr_2_eut1(pf01923) EUT1
06022 HMM01515PTA_B_1_eut1(pf01515) EUT1
06023 H_azure EUT1
06024 P_lightViolet EUT1
06025 HMMaldh_1_eutpdugrm(pf00171) EUT1 EP_NTERM
06026 HMMaldh_1_eutpdugrm(#) EUT1
06027 HMM06723EutJMreB_1_grm(#) EUT1
06028 HMM00465alcdh_3_mix(pf00465) EUT1
06029 HMMtpEuth_1_eut(pf04346) EUT1
06030 HMM06277EutA_1_eut(pf06277) EUT1
06031 HMM06277EutA_1_eut(pf06277) EUT1
06032 HMMetlyEutB_1_EUT1(pf06751) EUT1
06033 HMMetlyEutC_1_eut1(pf05985) EUT1 EP_NTERM
06034 Tsp_rosa EUT1
06035 H_mulberry(pf00936pf16365) EUT1
06036 HMMreg12833_1_EUT1(pf12833) EUT1
06037 -pf01218 -
06038 -pf01520 -
06039 -pf00583 -
06040 -pf11143 -
06041 -pf06572 -
06042 -pf04261 -
06043 -# -
06044 -pf00528 -
06045 -pf00528 -
BMC-P
BMC-H
BMC-T
Aldehyde dehydrogenase
PTAC
Signature enzyme
Alcohol dehydrogenase
Regulator

Klebsiella pneumoniae. Bacteria; Proteobacteria; Gammaproteobacteria; Enterobacterales;

NCTC5055__locus_1

EUT1

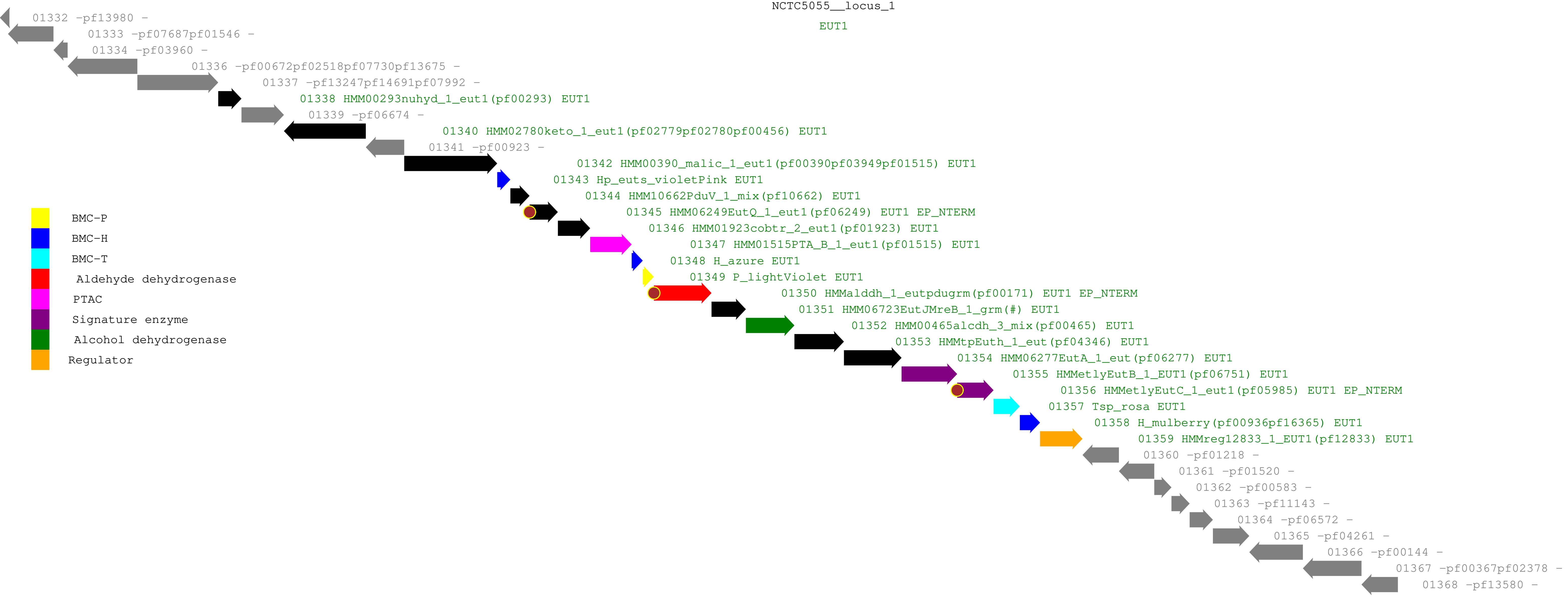

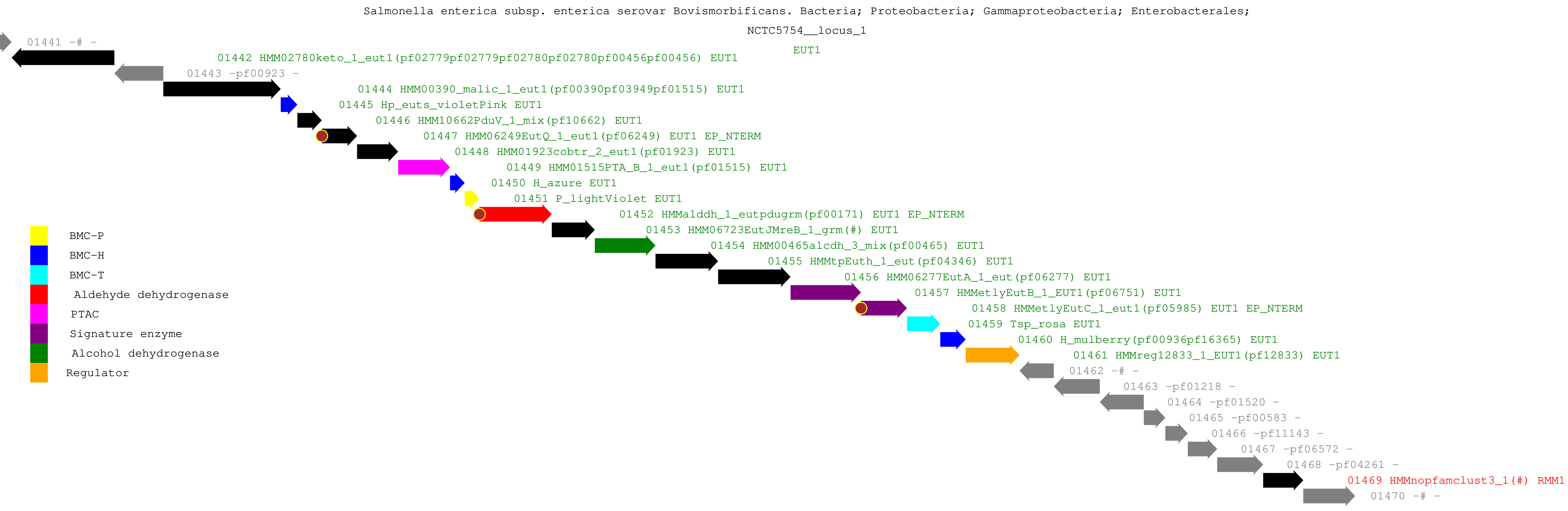

Salmonella enterica subsp. enterica serovar Bovismorbificans. Bacteria; Proteobacteria; Gammaproteobacteria; Enterobacterales;
NCTC5754__locus_1
EUT1
01441 -# -
01442 HMM02780keto_1_eut1(pf02779pf02779pf02780pf02780pf00456pf00456) EUT1
01443 -pf00923 -
01444 HMM00390_malic_1_eut1(pf00390pf03949pf01515) EUT1
01445 Hp_euts_violetPink EUT1
01446 HMM10662PduV_1_mix(pf10662) EUT1
01447 HMM06249EutQ_1_eut1(pf06249) EUT1 EP_NTERM
01448 HMM01923cobtr_2_eut1(pf01923) EUT1
01449 HMM01515PTA_B_1_eut1(pf01515) EUT1
01450 H_azure EUT1
01451 P_lightViolet EUT1
01452 HMMalddh_1_eutpdugrm(pf00171) EUT1 EP_NTERM
01453 HMM06723EutJMreB_1_grm(#) EUT1
01454 HMM00465alcdh_3_mix(pf00465) EUT1
01455 HMMtpEuth_1_eut(pf04346) EUT1
01456 HMM06277EutA_1_eut(pf06277) EUT1
01457 HMMetlyEutB_1_EUT1(pf06751) EUT1
01458 HMMetlyEutC_1_eut1(pf05985) EUT1 EP_NTERM
01459 Tsp_rosa EUT1
01460 H_mulberry(pf00936pf16365) EUT1
01461 HMMreg12833_1_EUT1(pf12833) EUT1
01462 -# -
01463 -pf01218 -
01464 -pf01520 -
01465 -pf00583 -
01466 -pf11143 -
01467 -pf06572 -
01468 -pf04261 -
01469 HMMnopfamclust3_1(#) RMM1
01470 -# -
BMC-P
BMC-H
BMC-T
Aldehyde dehydrogenase
PTAC
Signature enzyme
Alcohol dehydrogenase
Regulator

Salmonella enterica subsp. salamae. Bacteria; Proteobacteria; Gammaproteobacteria; Enterobacterales;

NCTC5773__locus_1

EUT1

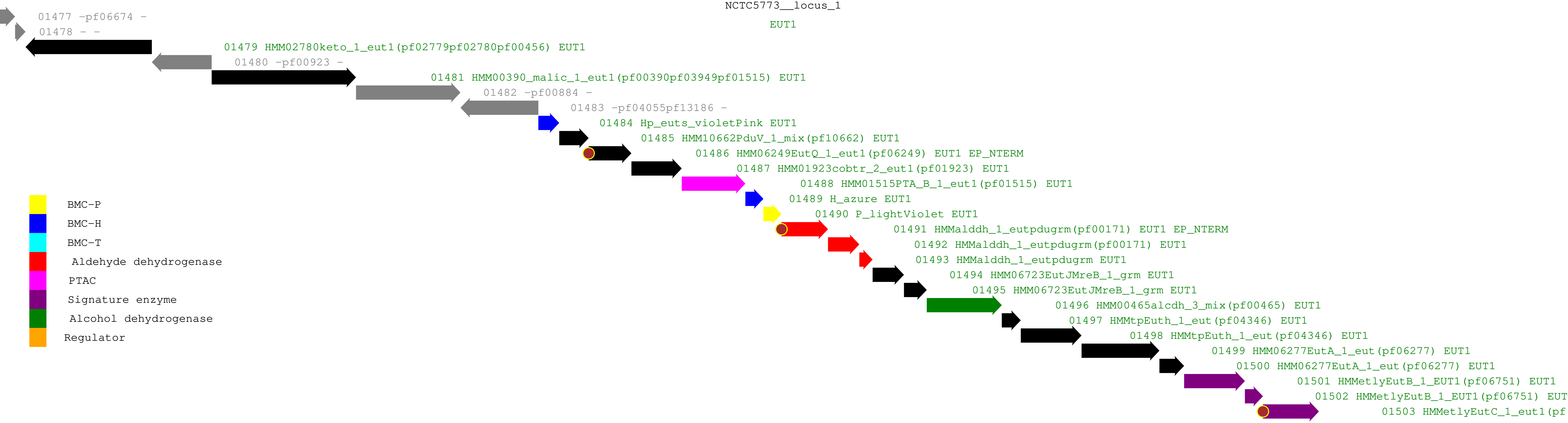

Citrobacter freundii. Bacteria; Proteobacteria; Gammaproteobacteria; Enterobacterales;

NCTC6266__locus_2

EUT1

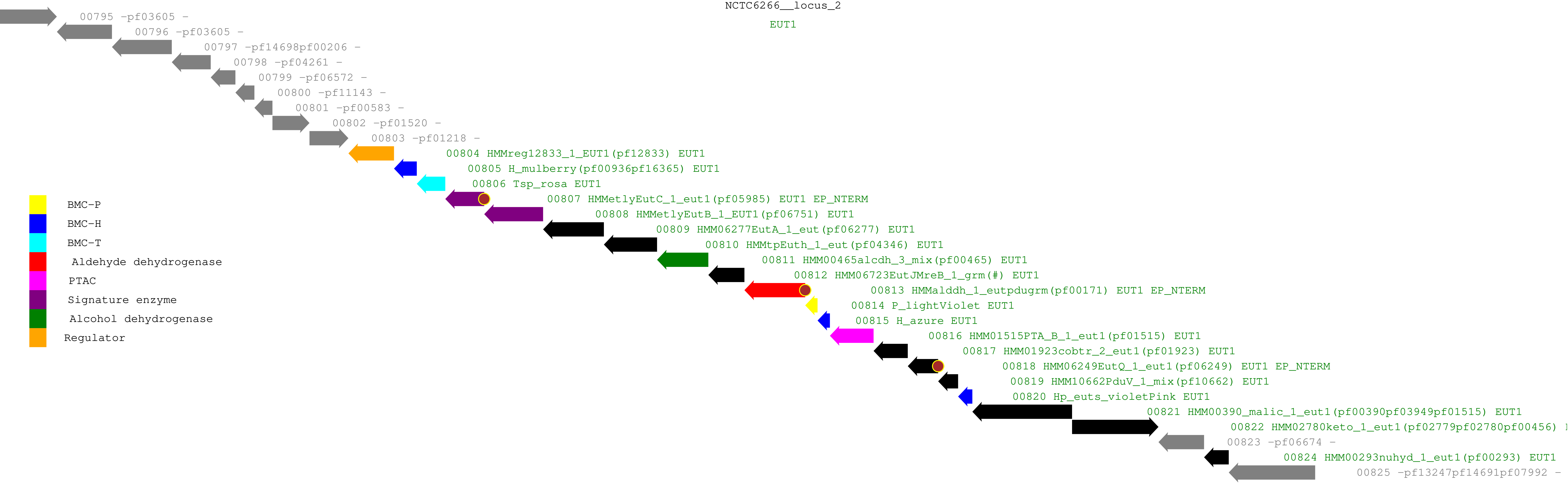

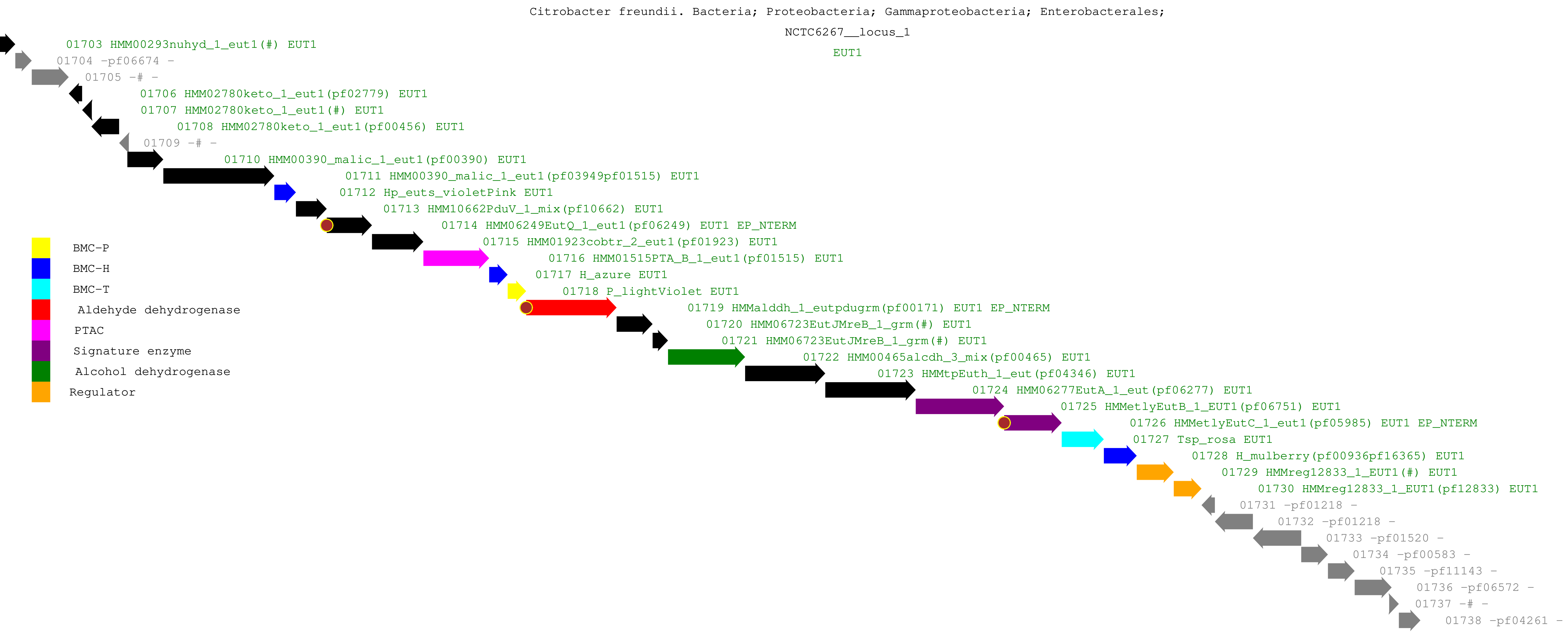
Citrobacter freundii. Bacteria; Proteobacteria; Gammaproteobacteria; Enterobacterales;
NCTC6267__locus_1
EUT1
01703 HMM00293nuhyd_1_eut1(#) EUT1
01704 -pf06674 -
01705 -# -
01706 HMM02780keto_1_eut1(pf02779) EUT1
01707 HMM02780keto_1_eut1(#) EUT1
01708 HMM02780keto_1_eut1(pf00456) EUT1
01709 -# -
01710 HMM00390_malic_1_eut1(pf00390) EUT1
01711 HMM00390_malic_1_eut1(pf03949pf01515) EUT1
01712 Hp_euts_violetPink EUT1
01713 HMM10662PduV_1_mix(pf10662) EUT1
01714 HMM06249EutQ_1_eut1(pf06249) EUT1 EP_NTERM
01715 HMM01923cobtr_2_eut1(pf01923) EUT1
01716 HMM01515PTA_B_1_eut1(pf01515) EUT1
01717 H_azure EUT1
01718 P_lightViolet EUT1
01719 HMMalddh_1_eutpdugrm(pf00171) EUT1 EP_NTERM
01720 HMM06723EutJMreB_1_grm(#) EUT1
01721 HMM06723EutJMreB_1_grm(#) EUT1
01722 HMM00465alcdh_3_mix(pf00465) EUT1
01723 HMMtpEuth_1_eut(pf04346) EUT1
01724 HMM06277EutA_1_eut(pf06277) EUT1
01725 HMMetlyEutB_1_EUT1(pf06751) EUT1
01726 HMMetlyEutC_1_eut1(pf05985) EUT1 EP_NTERM
01727 Tsp_rosa EUT1
01728 H_mulberry(pf00936pf16365) EUT1
01729 HMMreg12833_1_EUT1(#) EUT1
01730 HMMreg12833_1_EUT1(pf12833) EUT1
01731 -pf01218 -
01732 -pf01218 -
01733 -pf01520 -
01734 -pf00583 -
01735 -pf11143 -
01736 -pf06572 -
01737 -# -
01738 -pf04261 -
BMC-P
BMC-H
BMC-T
Aldehyde dehydrogenase
PTAC
Signature enzyme
Alcohol dehydrogenase
Regulator

Salmonella enterica I. Bacteria; Proteobacteria; Gammaproteobacteria; Enterobacterales;

NCTC6754__locus_1

EUT1

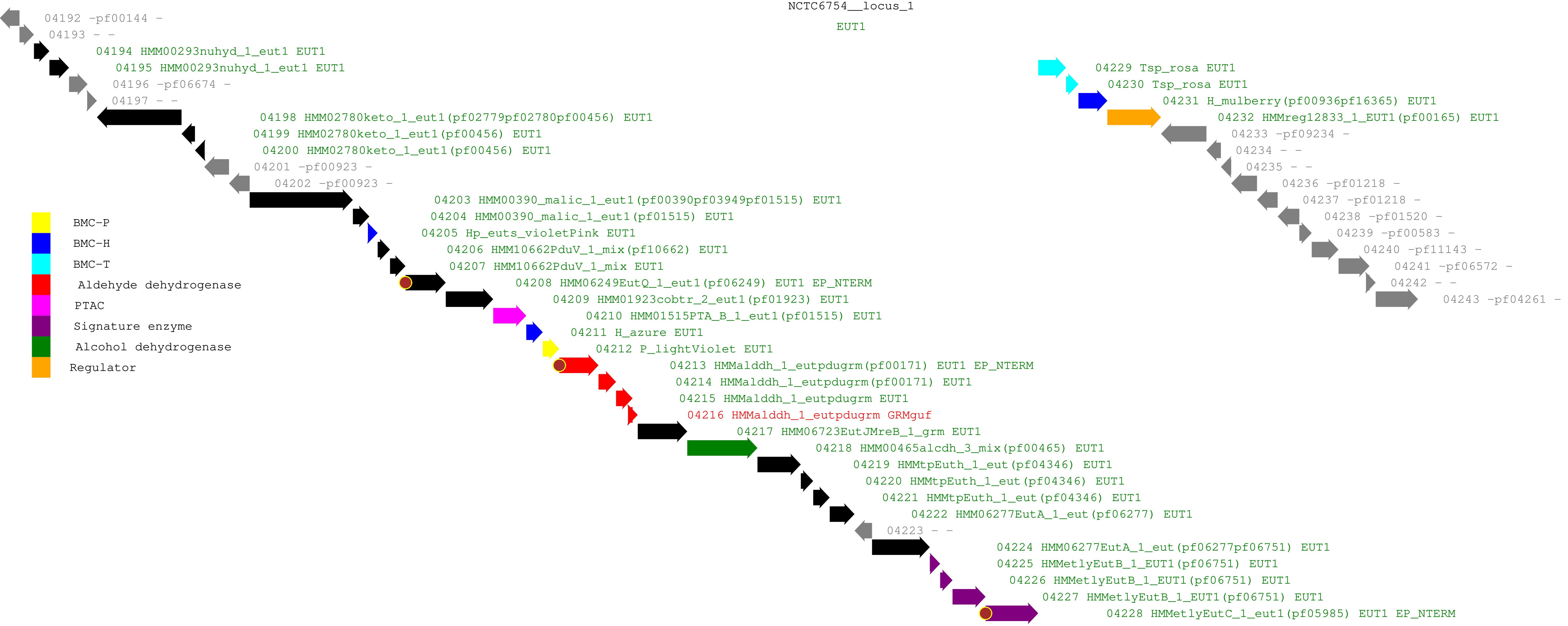

Salmonella sp. NCTC 6947. Bacteria; Proteobacteria; Gammaproteobacteria; Enterobacterales;

NCTC6947__locus_1

EUT1

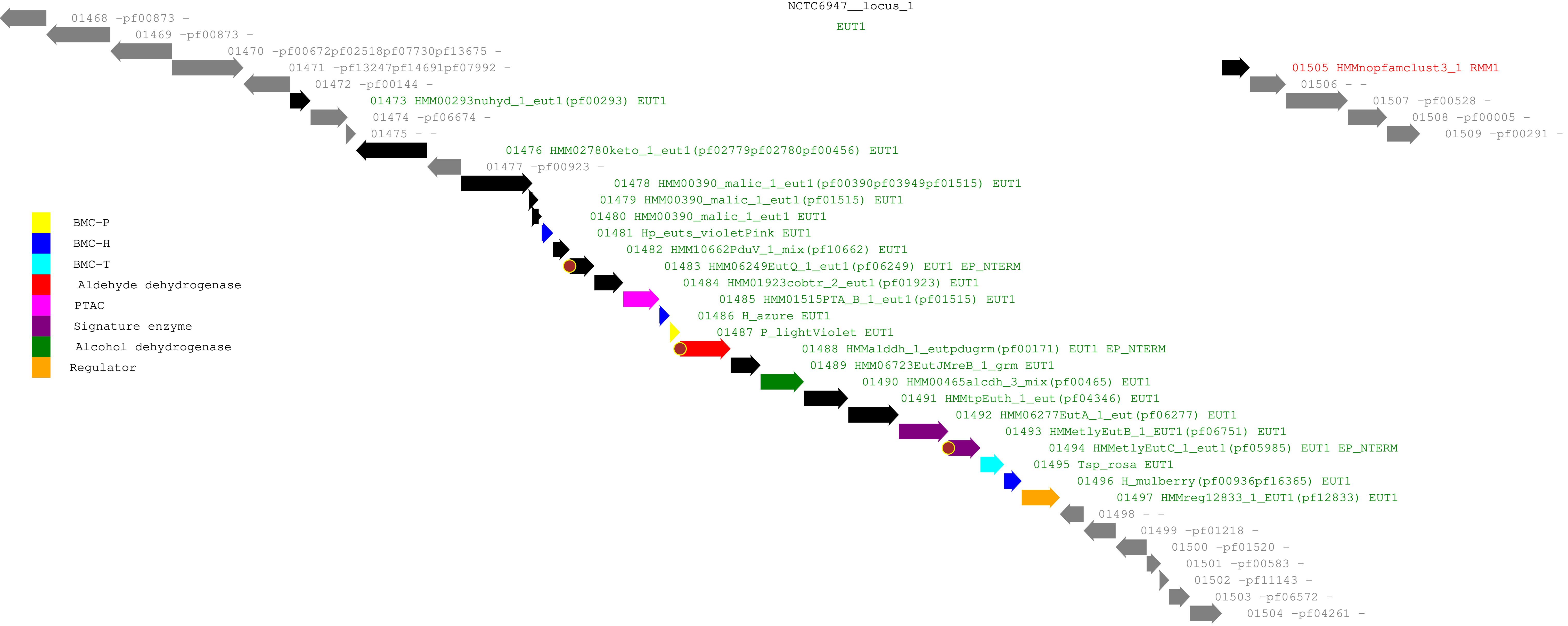

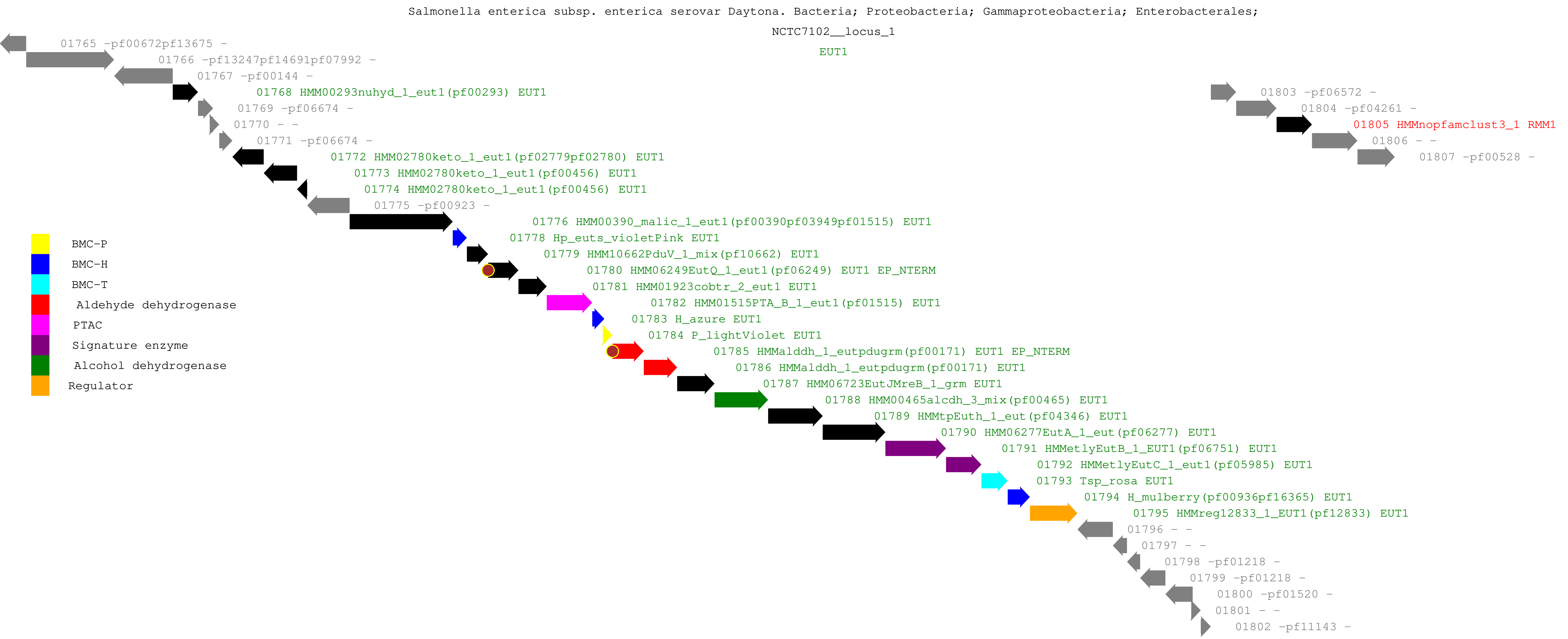

Salmonella enterica subsp. enterica serovar Daytona. Bacteria; Proteobacteria; Gammaproteobacteria; Enterobacterales;
NCTC7102__locus_1
EUT1
01765 -pf00672pf13675 -
01766 -pf13247pf14691pf07992 -
01767 -pf00144 -
01768 HMM00293nuhyd_1_eut1(pf00293) EUT1
01769 -pf06674 -
01770 - -
01771 -pf06674 -
01772 HMM02780keto_1_eut1(pf02779pf02780) EUT1
01773 HMM02780keto_1_eut1(pf00456) EUT1
01774 HMM02780keto_1_eut1(pf00456) EUT1
01775 -pf00923 -
01776 HMM00390_malic_1_eut1(pf00390pf03949pf01515) EUT1
01778 Hp_euts_violetPink EUT1
01779 HMM10662PduV_1_mix(pf10662) EUT1
01780 HMM06249EutQ_1_eut1(pf06249) EUT1 EP_NTERM
01781 HMM01923cobtr_2_eut1 EUT1
01782 HMM01515PTA_B_1_eut1(pf01515) EUT1
01783 H_azure EUT1
01784 P_lightViolet EUT1
01785 HMMalddh_1_eutpdugrm(pf00171) EUT1 EP_NTERM
01786 HMMalddh_1_eutpdugrm(pf00171) EUT1
01787 HMM06723EutJMreB_1_grm EUT1
01788 HMM00465alcdh_3_mix(pf00465) EUT1
01789 HMMtpEuth_1_eut(pf04346) EUT1
01790 HMM06277EutA_1_eut(pf06277) EUT1
01791 HMMetlyEutB_1_EUT1(pf06751) EUT1
01792 HMMetlyEutC_1_eut1(pf05985) EUT1
01793 Tsp_rosa EUT1
01794 H_mulberry(pf00936pf16365) EUT1
01795 HMMreg12833_1_EUT1(pf12833) EUT1
01796 - -
01797 - -
01798 -pf01218 -
01799 -pf01218 -
01800 -pf01520 -
01801 - -
01802 -pf11143 -
01803 -pf06572 -
01804 -pf04261 -
01805 HMMnopfamclust3_1 RMM1
01806 - -
01807 -pf00528 -
BMC-P
BMC-H
BMC-T
Aldehyde dehydrogenase
PTAC
Signature enzyme
Alcohol dehydrogenase
Regulator

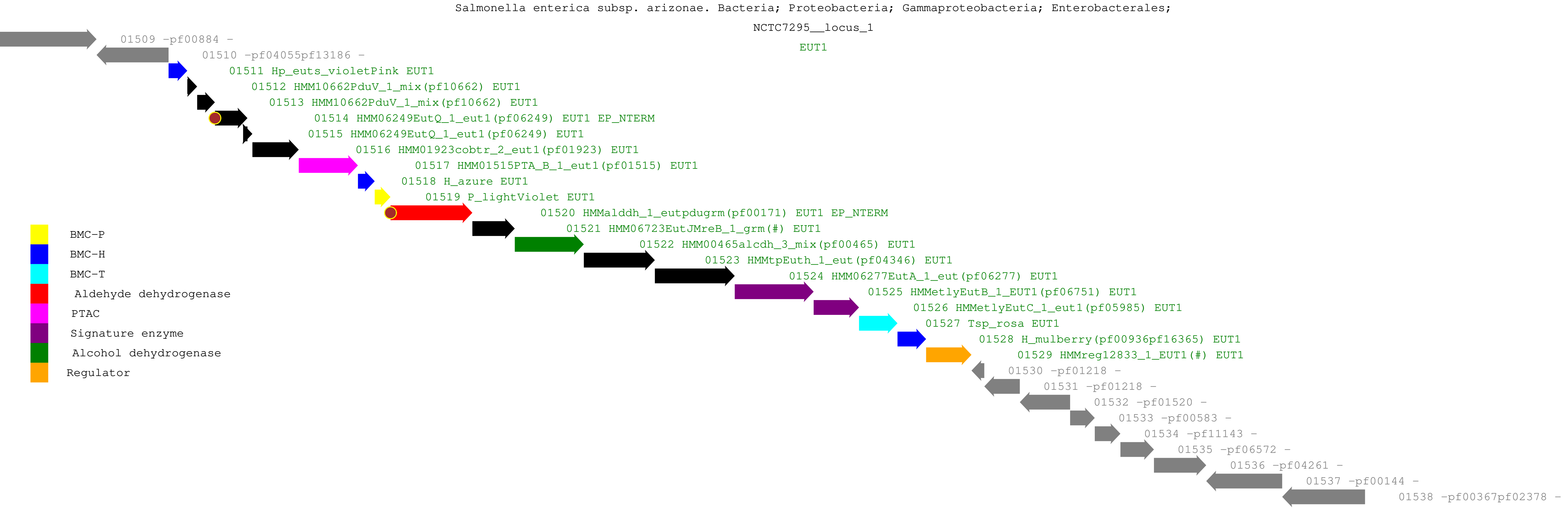
Salmonella enterica subsp. arizonae. Bacteria; Proteobacteria; Gammaproteobacteria; Enterobacterales;
NCTC7295__locus_1
EUT1
01509 -pf00884 -
01510 -pf04055pf13186 -
01511 Hp_euts_violetPink EUT1
01512 HMM10662PduV_1_mix(pf10662) EUT1
01513 HMM10662PduV_1_mix(pf10662) EUT1
01514 HMM06249EutQ_1_eut1(pf06249) EUT1 EP_NTERM
01515 HMM06249EutQ_1_eut1(pf06249) EUT1
01516 HMM01923cobtr_2_eut1(pf01923) EUT1
01517 HMM01515PTA_B_1_eut1(pf01515) EUT1
01518 H_azure EUT1
01519 P_lightViolet EUT1
01520 HMMaldh_1_eutpdugrm(pf00171) EUT1 EP_NTERM
01521 HMM06723EutJMreB_1_grm(#) EUT1
01522 HMM00465alcdh_3_mix(pf00465) EUT1
01523 HMMtpEuth_1_eut(pf04346) EUT1
01524 HMM06277EutA_1_eut(pf06277) EUT1
01525 HMMetlyEutB_1_EUT1(pf06751) EUT1
01526 HMMetlyEutC_1_eut1(pf05985) EUT1
01527 Tsp_rosa EUT1
01528 H_mulberry(pf00936pf16365) EUT1
01529 HMMreg12833_1_EUT1(#) EUT1
01530 -pf01218 -
01531 -pf01218 -
01532 -pf01520 -
01533 -pf00583 -
01534 -pf11143 -
01535 -pf06572 -
01536 -pf04261 -
01537 -pf00144 -
01538 -pf00367pf02378 -
BMC-P
BMC-H
BMC-T
Aldehyde dehydrogenase
PTAC
Signature enzyme
Alcohol dehydrogenase
Regulator

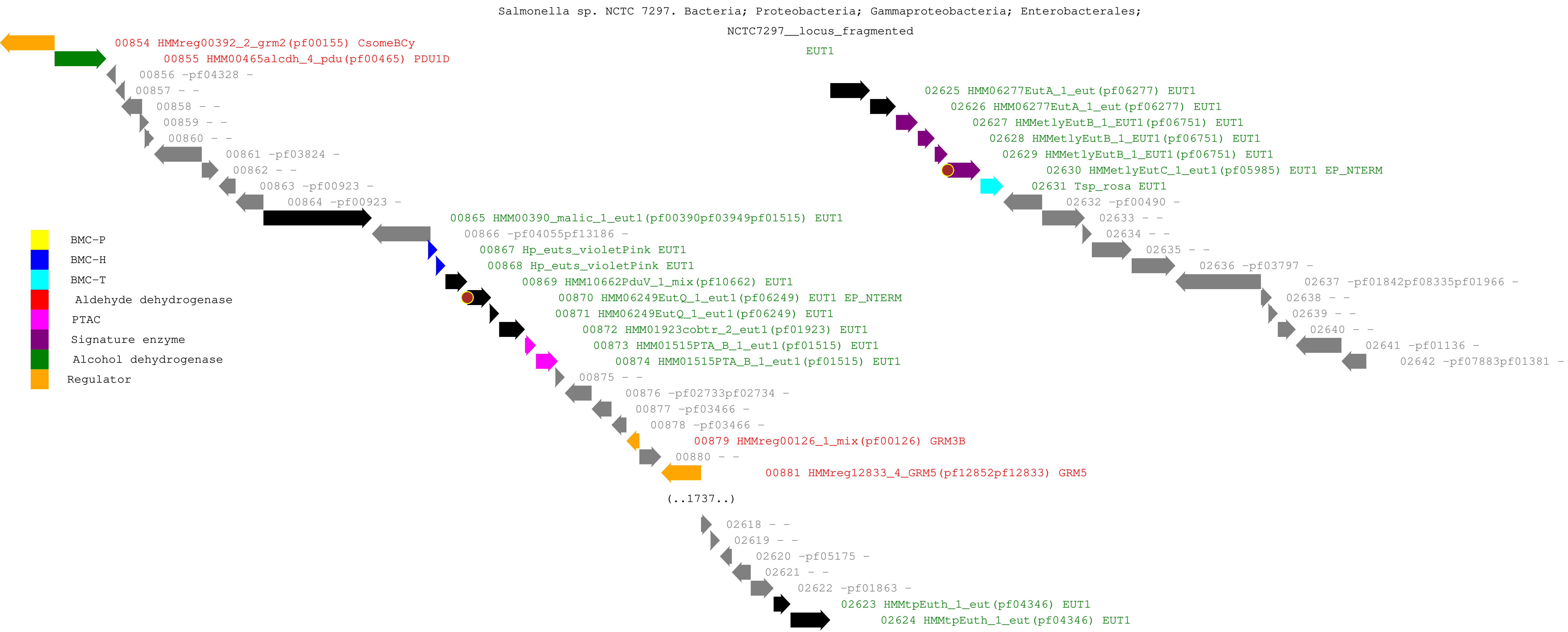

Salmonella sp. NCTC 7297. Bacteria; Proteobacteria; Gammaproteobacteria; Enterobacterales;
NCTC7297__locus_fragmented
EUT1
00854 HMMreg00392_2_grm2(pf00155) CsomeBCy
00855 HMM00465alcdh_4_pdu(pf00465) PDU1D
00856 -pf04328 -
00857 - -
00858 - -
00859 - -
00860 - -
00861 -pf03824 -
00862 - -
00863 -pf00923 -
00864 -pf00923 -
00865 HMM00390_malic_1_eut1(pf00390pf03949pf01515) EUT1
00866 -pf04055pf13186 -
00867 Hp_euts_violetPink EUT1
00868 Hp_euts_violetPink EUT1
00869 HMM10662PduV_1_mix(pf10662) EUT1
00870 HMM06249EutQ_1_eut1(pf06249) EUT1 EP_NTERM
00871 HMM06249EutQ_1_eut1(pf06249) EUT1
00872 HMM01923cobtr_2_eut1(pf01923) EUT1
00873 HMM01515PTA_B_1_eut1(pf01515) EUT1
00874 HMM01515PTA_B_1_eut1(pf01515) EUT1
00875 - -
00876 -pf02733pf02734 -
00877 -pf03466 -
00878 -pf03466 -
00879 HMMreg00126_1_mix(pf00126) GRM3B
00880 - -
00881 HMMreg12833_4_GRM5(pf12852pf12833) GRM5
(..1737..)
02618 - -
02619 - -
02620 -pf05175 -
02621 - -
02622 -pf01863 -
02623 HMMtpEuth_1_eut(pf04346) EUT1
02624 HMMtpEuth_1_eut(pf04346) EUT1
02625 HMM06277EutA_1_eut(pf06277) EUT1
02626 HMM06277EutA_1_eut(pf06277) EUT1
02627 HMMetlyEutB_1_EUT1(pf06751) EUT1
02628 HMMetlyEutB_1_EUT1(pf06751) EUT1
02629 HMMetlyEutB_1_EUT1(pf06751) EUT1
02630 HMMetlyEutC_1_eut1(pf05985) EUT1 EP_NTERM
02631 Tsp_rosa EUT1
02632 -pf00490 -
02633 - -
02634 - -
02635 - -
02636 -pf03797 -
02637 -pf01842pf08335pf01966 -
02638 - -
02639 - -
02640 - -
02641 -pf01136 -
02642 -pf07883pf01381 -
BMC-P
BMC-H
BMC-T
Aldehyde dehydrogenase
PTAC
Signature enzyme
Alcohol dehydrogenase
Regulator

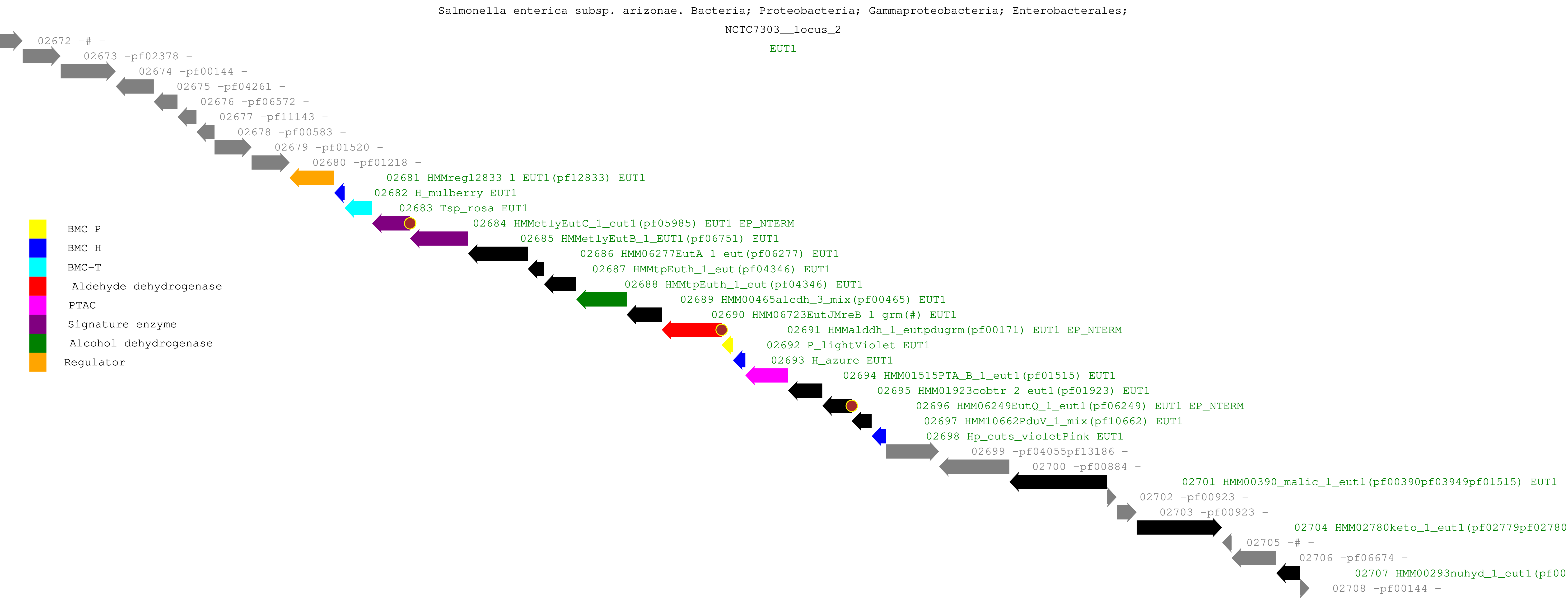

Salmonella enterica subsp. arizonae. Bacteria; Proteobacteria; Gammaproteobacteria; Enterobacterales;
NCTC7303__locus_2
EUT1
02672 -# -
02673 -pf02378 -
02674 -pf00144 -
02675 -pf04261 -
02676 -pf06572 -
02677 -pf11143 -
02678 -pf00583 -
02679 -pf01520 -
02680 -pf01218 -
02681 HMMreg12833_1_EUT1(pf12833) EUT1
02682 H_mulberry EUT1
02683 Tsp_rosa EUT1
02684 HMMetlyEutC_1_eut1(pf05985) EUT1 EP_NTERM
02685 HMMetlyEutB_1_EUT1(pf06751) EUT1
02686 HMM06277EutA_1_eut(pf06277) EUT1
02687 HMMtpEuth_1_eut(pf04346) EUT1
02688 HMMtpEuth_1_eut(pf04346) EUT1
02689 HMM00465alcdh_3_mix(pf00465) EUT1
02690 HMM06723EutJMreB_1_grm(#) EUT1
02691 HMMaldh_1_eutpdugrm(pf00171) EUT1 EP_NTERM
02692 P_lightViolet EUT1
02693 H_azure EUT1
02694 HMM01515PTA_B_1_eut1(pf01515) EUT1
02695 HMM01923cobtr_2_eut1(pf01923) EUT1
02696 HMM06249EutQ_1_eut1(pf06249) EUT1 EP_NTERM
02697 HMM10662PduV_1_mix(pf10662) EUT1
02698 Hp_euts_violetPink EUT1
02699 -pf04055pf13186 -
02700 -pf00884 -
02701 HMM00390_malic_1_eut1(pf00390pf03949pf01515) EUT1
02702 -pf00923 -
02703 -pf00923 -
02704 HMM02780keto_1_eut1(pf02779pf02780
02705 -# -
02706 -pf06674 -
02707 HMM00293nuhyd_1_eut1(pf00
02708 -pf00144 -
BMC-P
BMC-H
BMC-T
Aldehyde dehydrogenase
PTAC
Signature enzyme
Alcohol dehydrogenase
Regulator

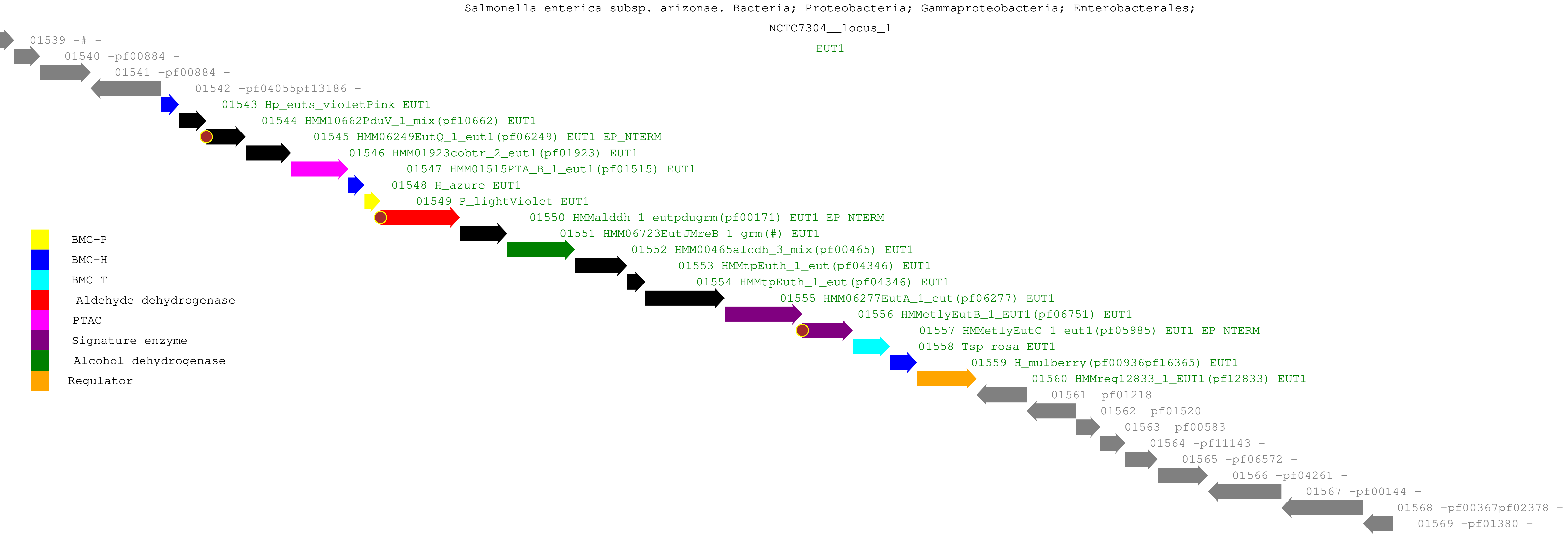

Salmonella enterica subsp. arizonae. Bacteria; Proteobacteria; Gammaproteobacteria; Enterobacterales;
NCTC7304__locus_1
EUT1
01539 -# -
01540 -pf00884 -
01541 -pf00884 -
01542 -pf04055pf13186 -
01543 Hp_euts_violetPink EUT1
01544 HMM10662PduV_1_mix(pf10662) EUT1
01545 HMM06249EutQ_1_eut1(pf06249) EUT1 EP_NTERM
01546 HMM01923cobtr_2_eut1(pf01923) EUT1
01547 HMM01515PTA_B_1_eut1(pf01515) EUT1
01548 H_azure EUT1
01549 P_lightViolet EUT1
01550 HMMaldh_1_eutpdugrm(pf00171) EUT1 EP_NTERM
01551 HMM06723EutJMreB_1_grm(#) EUT1
01552 HMM00465alcdh_3_mix(pf00465) EUT1
01553 HMMtpEuth_1_eut(pf04346) EUT1
01554 HMMtpEuth_1_eut(pf04346) EUT1
01555 HMM06277EutA_1_eut(pf06277) EUT1
01556 HMMetlyEutB_1_EUT1(pf06751) EUT1
01557 HMMetlyEutC_1_eut1(pf05985) EUT1 EP_NTERM
01558 Tsp_rosa EUT1
01559 H_mulberry(pf00936pf16365) EUT1
01560 HMMreg12833_1_EUT1(pf12833) EUT1
01561 -pf01218 -
01562 -pf01520 -
01563 -pf00583 -
01564 -pf11143 -
01565 -pf06572 -
01566 -pf04261 -
01567 -pf00144 -
01568 -pf00367pf02378 -
01569 -pf01380 -
BMC-P
BMC-H
BMC-T
Aldehyde dehydrogenase
PTAC
Signature enzyme
Alcohol dehydrogenase
Regulator

Salmonella enterica subsp. arizonae. Bacteria; Proteobacteria; Gammaproteobacteria; Enterobacterales;

NCTC7307__locus_1

EUT1

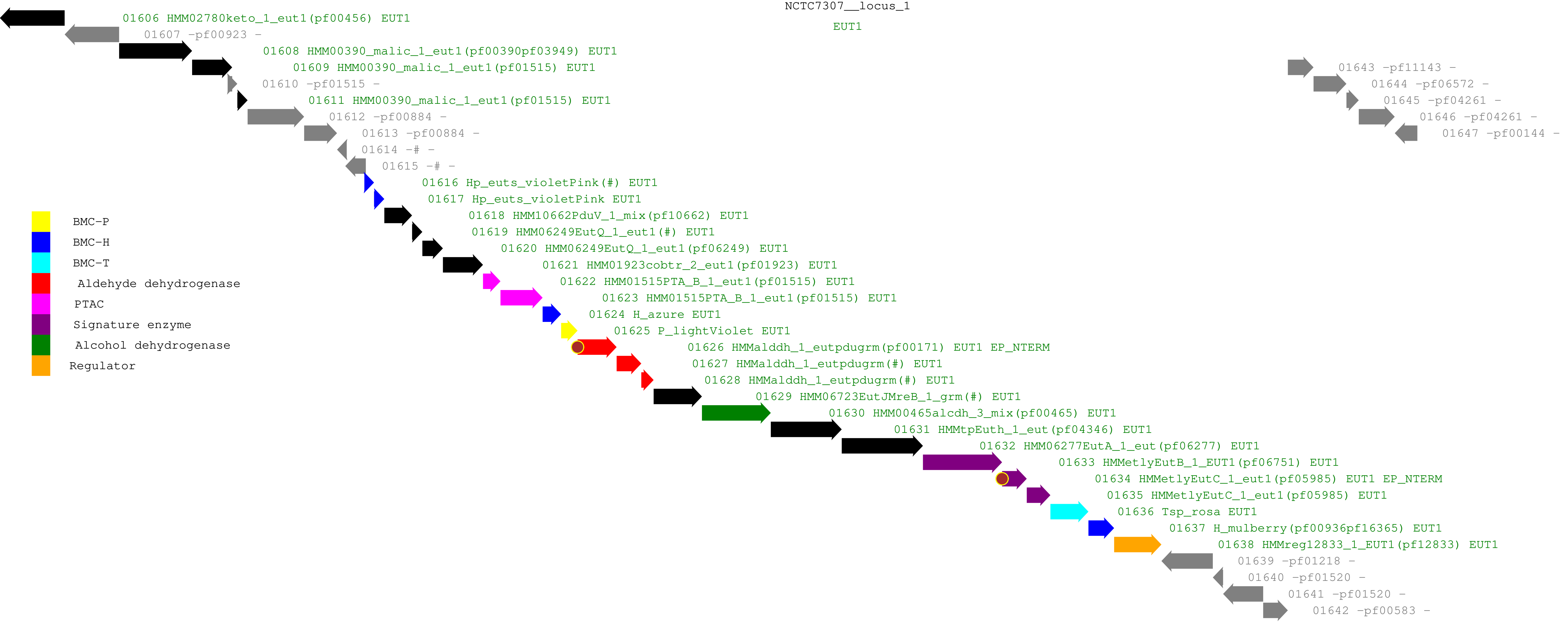

Salmonella choleraesuis. Bacteria; Proteobacteria; Gammaproteobacteria; Enterobacterales;

NCTC7346__locus_1

EUT1

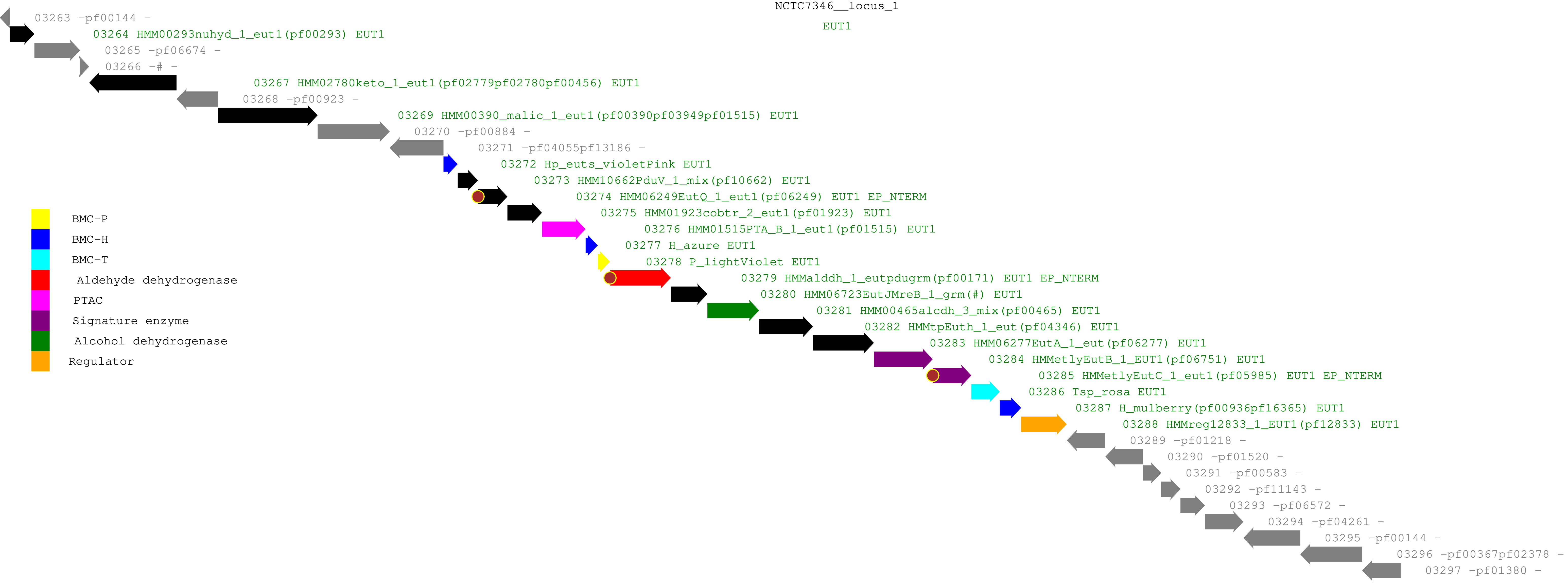

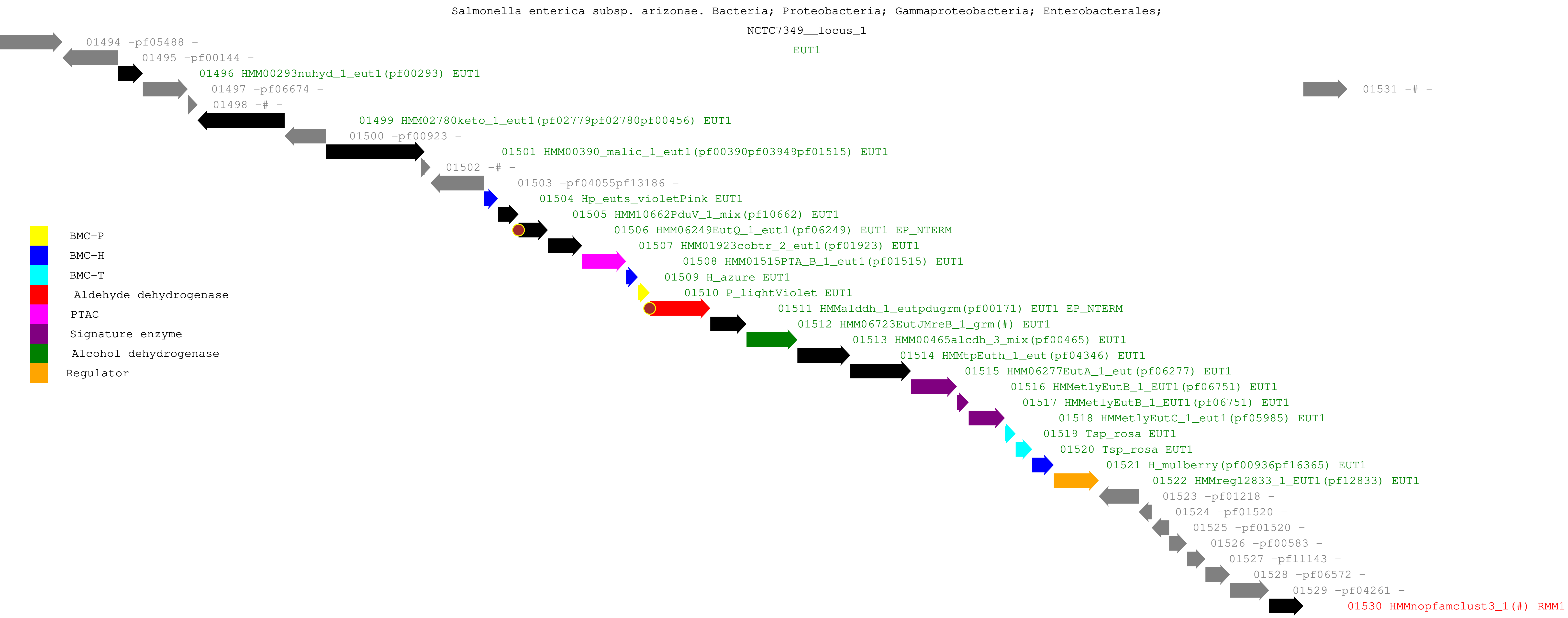

Salmonella enterica subsp. arizonae. Bacteria; Proteobacteria; Gammaproteobacteria; Enterobacterales;
NCTC7349__locus_1
EUT1
01494 -pf05488 -
01495 -pf00144 -
01496 HMM00293nuhyd_1_eut1(pf00293) EUT1
01497 -pf06674 -
01498 -# -
01499 HMM02780keto_1_eut1(pf02779pf02780pf00456) EUT1
01500 -pf00923 -
01501 HMM00390_malic_1_eut1(pf00390pf03949pf01515) EUT1
01502 -# -
01503 -pf04055pf13186 -
01504 Hp_euts_violetPink EUT1
01505 HMM10662PduV_1_mix(pf10662) EUT1
01506 HMM06249EutQ_1_eut1(pf06249) EUT1 EP_NTERM
01507 HMM01923cobtr_2_eut1(pf01923) EUT1
01508 HMM01515PTA_B_1_eut1(pf01515) EUT1
01509 H_azure EUT1
01510 P_lightViolet EUT1
01511 HMMalddh_1_eutpdugrm(pf00171) EUT1 EP_NTERM
01512 HMM06723EutJMreB_1_grm(#) EUT1
01513 HMM00465alcdh_3_mix(pf00465) EUT1
01514 HMMtpEuth_1_eut(pf04346) EUT1
01515 HMM06277EutA_1_eut(pf06277) EUT1
01516 HMMetlyEutB_1_EUT1(pf06751) EUT1
01517 HMMetlyEutB_1_EUT1(pf06751) EUT1
01518 HMMetlyEutC_1_eut1(pf05985) EUT1
01519 Tsp_rosa EUT1
01520 Tsp_rosa EUT1
01521 H_mulberry(pf00936pf16365) EUT1
01522 HMMreg12833_1_EUT1(pf12833) EUT1
01523 -pf01218 -
01524 -pf01520 -
01525 -pf01520 -
01526 -pf00583 -
01527 -pf11143 -
01528 -pf06572 -
01529 -pf04261 -
01530 HMMnopfamclust3_1(#) RMM1
01531 -# -
BMC-P
BMC-H
BMC-T
Aldehyde dehydrogenase
PTAC
Signature enzyme
Alcohol dehydrogenase
Regulator

Salmonella enterica subsp. enterica serovar Sanjuan. Bacteria; Proteobacteria; Gammaproteobacteria; Enterobacterales;

NCTC7406__locus_1

EUT1

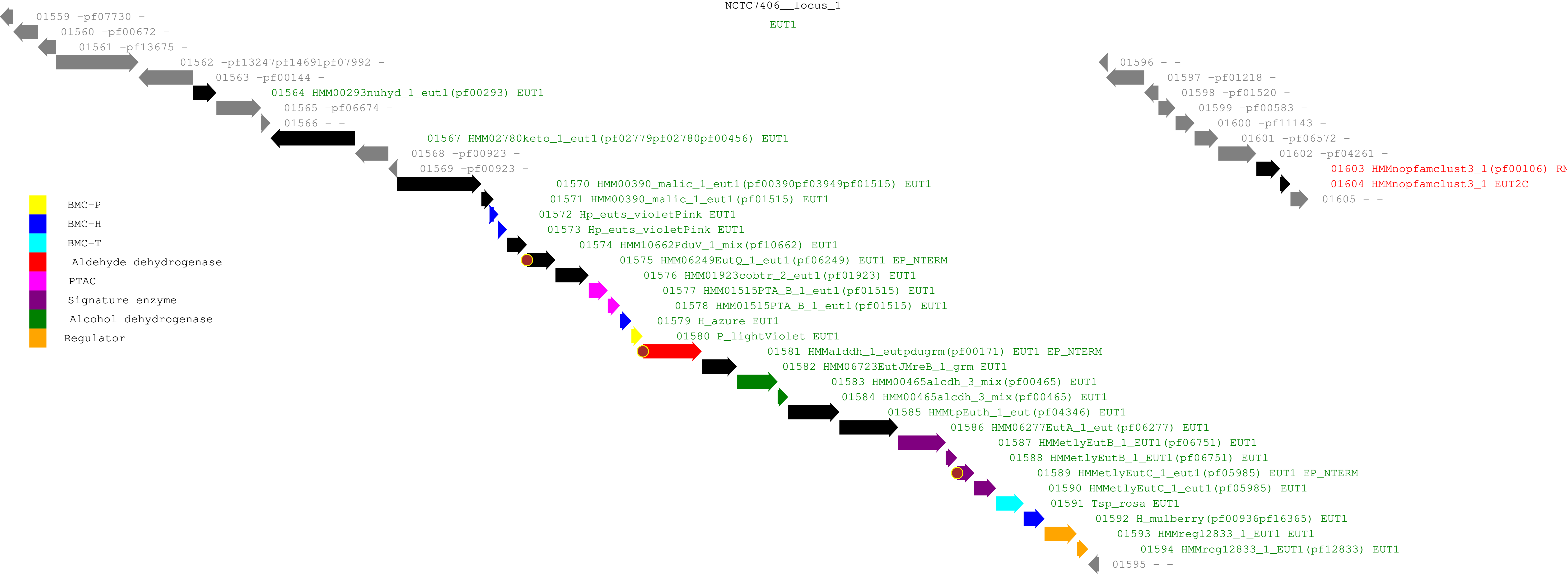

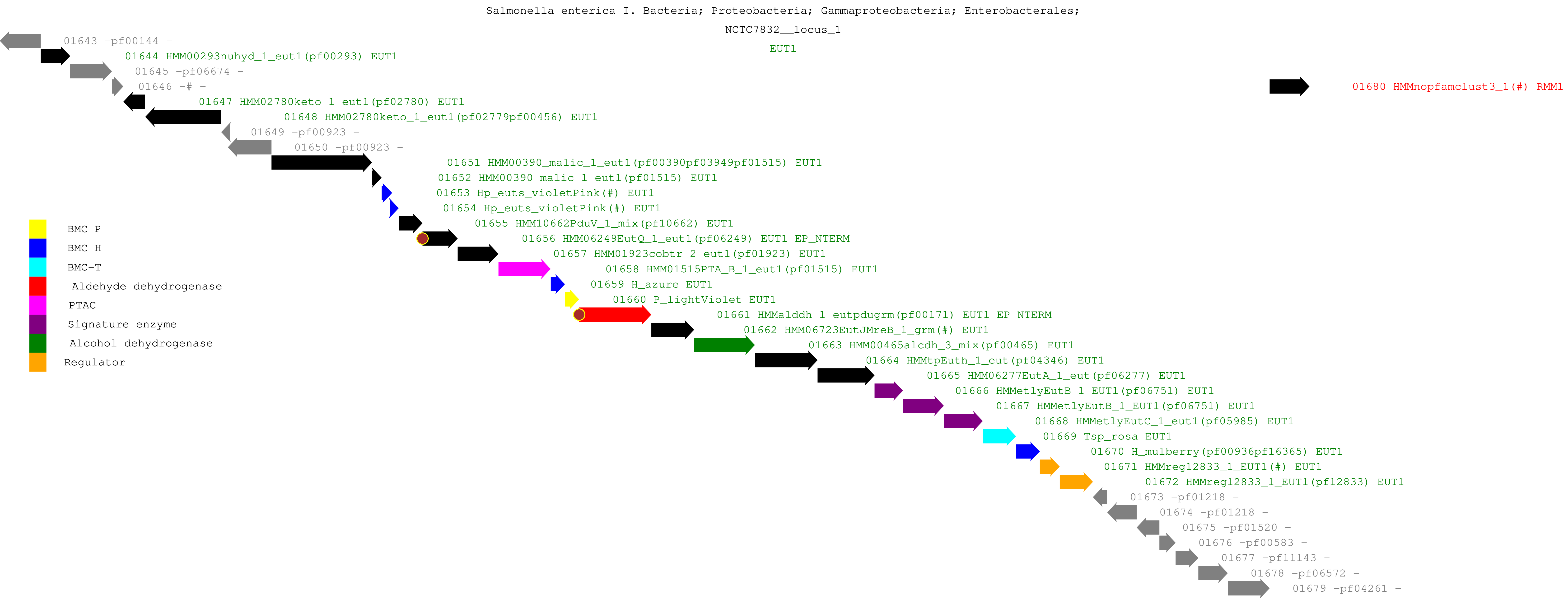
Salmonella enterica I. Bacteria; Proteobacteria; Gammaproteobacteria; Enterobacterales;
NCTC7832__locus_1
EUT1
01643 -pf00144 -
01644 HMM00293nuhyd_1_eut1(pf00293) EUT1
01645 -pf06674 -
01646 -# -
01647 HMM02780keto_1_eut1(pf02780) EUT1
01648 HMM02780keto_1_eut1(pf02779pf00456) EUT1
01649 -pf00923 -
01650 -pf00923 -
01651 HMM00390_malic_1_eut1(pf00390pf03949pf01515) EUT1
01652 HMM00390_malic_1_eut1(pf01515) EUT1
01653 Hp_euts_violetPink(#) EUT1
01654 Hp_euts_violetPink(#) EUT1
01655 HMM10662PduV_1_mix(pf10662) EUT1
01656 HMM06249EutQ_1_eut1(pf06249) EUT1 EP_NTERM
01657 HMM01923cobtr_2_eut1(pf01923) EUT1
01658 HMM01515PTA_B_1_eut1(pf01515) EUT1
01659 H_azure EUT1
01660 P_lightViolet EUT1
01661 HMMalddh_1_eutpdugrm(pf00171) EUT1 EP_NTERM
01662 HMM06723EutJMreB_1_grm(#) EUT1
01663 HMM00465alcdh_3_mix(pf00465) EUT1
01664 HMMtpEuth_1_eut(pf04346) EUT1
01665 HMM06277EutA_1_eut(pf06277) EUT1
01666 HMMetlyEutB_1_EUT1(pf06751) EUT1
01667 HMMetlyEutB_1_EUT1(pf06751) EUT1
01668 HMMetlyEutC_1_eut1(pf05985) EUT1
01669 Tsp_rosa EUT1
01670 H_mulberry(pf00936pf16365) EUT1
01671 HMMreg12833_1_EUT1(#) EUT1
01672 HMMreg12833_1_EUT1(pf12833) EUT1
01673 -pf01218 -
01674 -pf01218 -
01675 -pf01520 -
01676 -pf00583 -
01677 -pf11143 -
01678 -pf06572 -
01679 -pf04261 -
01680 HMMnopfamclust3_1(#) RMM1
BMC-P
BMC-H
BMC-T
Aldehyde dehydrogenase
PTAC
Signature enzyme
Alcohol dehydrogenase
Regulator

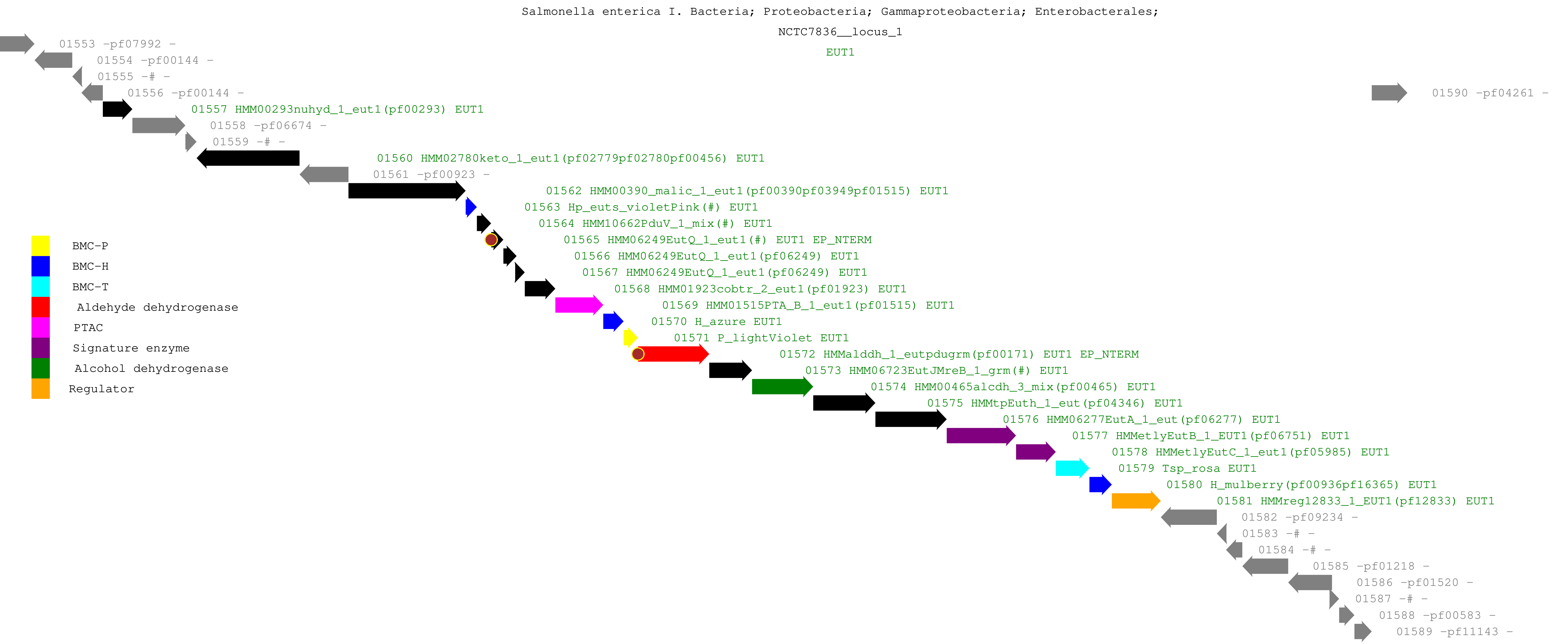

Salmonella enterica I. Bacteria; Proteobacteria; Gammaproteobacteria; Enterobacterales;
NCTC7836__locus_1
EUT1
01553 -pf07992 -
01554 -pf00144 -
01555 -# -
01556 -pf00144 -
01557 HMM00293nuhyd_1_eut1(pf00293) EUT1
01558 -pf06674 -
01559 -# -
01560 HMM02780keto_1_eut1(pf02779pf02780pf00456) EUT1
01561 -pf00923 -
01562 HMM00390_malic_1_eut1(pf00390pf03949pf01515) EUT1
01563 Hp_euts_violetPink(#) EUT1
01564 HMM10662PduV_1_mix(#) EUT1
01565 HMM06249EutQ_1_eut1(#) EUT1 EP_NTERM
01566 HMM06249EutQ_1_eut1(pf06249) EUT1
01567 HMM06249EutQ_1_eut1(pf06249) EUT1
01568 HMM01923cobtr_2_eut1(pf01923) EUT1
01569 HMM01515PTA_B_1_eut1(pf01515) EUT1
01570 H_azure EUT1
01571 P_lightViolet EUT1
01572 HMMalddh_1_eutpdugrm(pf00171) EUT1 EP_NTERM
01573 HMM06723EutJMreB_1_grm(#) EUT1
01574 HMM00465alcdh_3_mix(pf00465) EUT1
01575 HMMtpEuth_1_eut(pf04346) EUT1
01576 HMM06277EutA_1_eut(pf06277) EUT1
01577 HMMetlyEutB_1_EUT1(pf06751) EUT1
01578 HMMetlyEutC_1_eut1(pf05985) EUT1
01579 Tsp_rosa EUT1
01580 H_mulberry(pf00936pf16365) EUT1
01581 HMMreg12833_1_EUT1(pf12833) EUT1
01582 -pf09234 -
01583 -# -
01584 -# -
01585 -pf01218 -
01586 -pf01520 -
01587 -# -
01588 -pf00583 -
01589 -pf11143 -
01590 -pf04261 -
BMC-P
BMC-H
BMC-T
Aldehyde dehydrogenase
PTAC
Signature enzyme
Alcohol dehydrogenase
Regulator

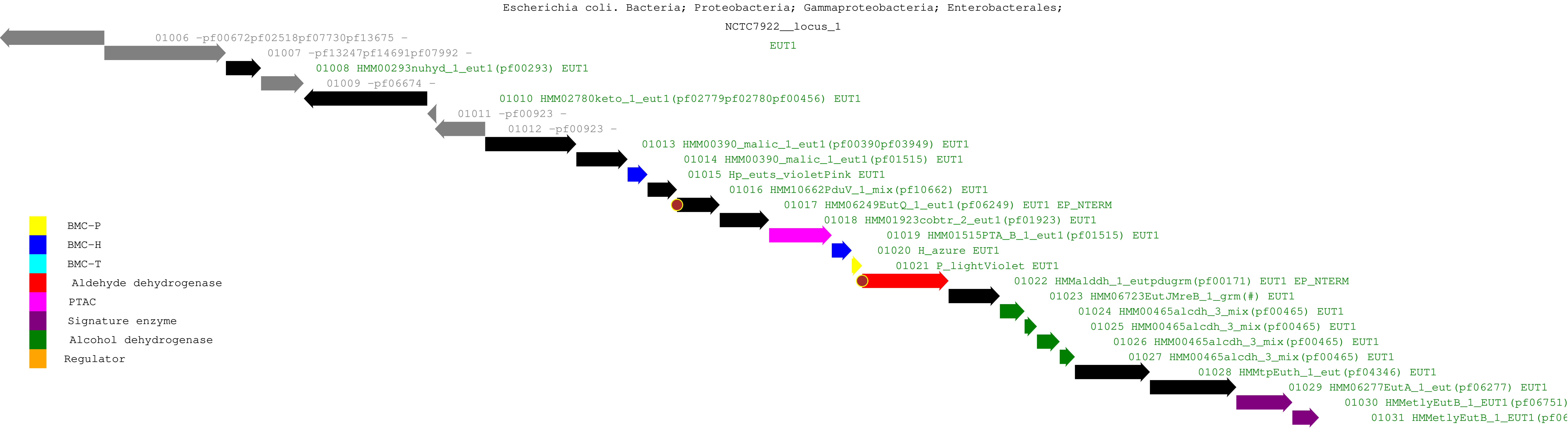
Escherichia coli. Bacteria; Proteobacteria; Gammaproteobacteria; Enterobacterales;
NCTC7922__locus_1
EUT1
01006 -pf00672pf02518pf07730pf13675 -
01007 -pf13247pf14691pf07992 -
01008 HMM00293nuhyd_1_eut1(pf00293) EUT1
01009 -pf06674 -
01010 HMM02780keto_1_eut1(pf02779pf02780pf00456) EUT1
01011 -pf00923 -
01012 -pf00923 -
01013 HMM00390_malic_1_eut1(pf00390pf03949) EUT1
01014 HMM00390_malic_1_eut1(pf01515) EUT1
01015 Hp_euts_violetPink EUT1
01016 HMM10662PduV_1_mix(pf10662) EUT1
01017 HMM06249EutQ_1_eut1(pf06249) EUT1 EP_NTERM
01018 HMM01923cobtr_2_eut1(pf01923) EUT1
01019 HMM01515PTA_B_1_eut1(pf01515) EUT1
01020 H_azure EUT1
01021 P_lightViolet EUT1
01022 HMMalddh_1_eutpdugrm(pf00171) EUT1 EP_NTERM
01023 HMM06723EutJMreB_1_grm(#) EUT1
01024 HMM00465alcdh_3_mix(pf00465) EUT1
01025 HMM00465alcdh_3_mix(pf00465) EUT1
01026 HMM00465alcdh_3_mix(pf00465) EUT1
01027 HMM00465alcdh_3_mix(pf00465) EUT1
01028 HMMtpEuth_1_eut(pf04346) EUT1
01029 HMM06277EutA_1_eut(pf06277) EUT1
01030 HMMetlyEutB_1_EUT1(pf06751)
01031 HMMetlyEutB_1_EUT1(pf06
BMC-P
BMC-H
BMC-T
Aldehyde dehydrogenase
PTAC
Signature enzyme
Alcohol dehydrogenase
Regulator

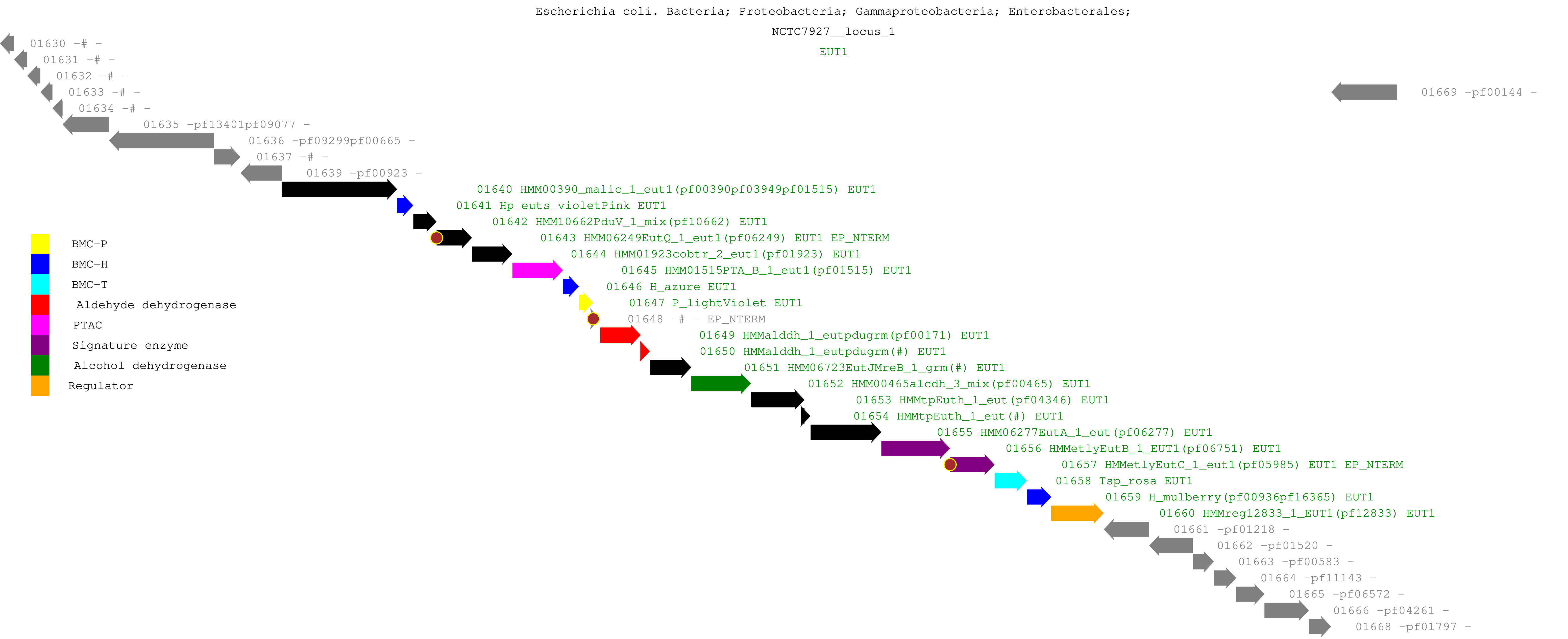
Escherichia coli. Bacteria; Proteobacteria; Gammaproteobacteria; Enterobacterales;
NCTC7927__locus_1
EUT1
01630 -# -
01631 -# -
01632 -# -
01633 -# -
01634 -# -
01635 -pf13401pf09077 -
01636 -pf09299pf00665 -
01637 -# -
01639 -pf00923 -
01640 HMM00390_malic_1_eut1(pf00390pf03949pf01515) EUT1
01641 Hp_euts_violetPink EUT1
01642 HMM10662PduV_1_mix(pf10662) EUT1
01643 HMM06249EutQ_1_eut1(pf06249) EUT1 EP_NTERM
01644 HMM01923cobtr_2_eut1(pf01923) EUT1
01645 HMM01515PTA_B_1_eut1(pf01515) EUT1
01646 H_azure EUT1
01647 P_lightViolet EUT1
01648 -# - EP_NTERM
01649 HMMalddh_1_eutpdugrm(pf00171) EUT1
01650 HMMalddh_1_eutpdugrm(#) EUT1
01651 HMM06723EutJMreB_1_grm(#) EUT1
01652 HMM00465alcdh_3_mix(pf00465) EUT1
01653 HMMtpEuth_1_eut(pf04346) EUT1
01654 HMMtpEuth_1_eut(#) EUT1
01655 HMM06277EutA_1_eut(pf06277) EUT1
01656 HMMetlyEutB_1_EUT1(pf06751) EUT1
01657 HMMetlyEutC_1_eut1(pf05985) EUT1 EP_NTERM
01658 Tsp_rosa EUT1
01659 H_mulberry(pf00936pf16365) EUT1
01660 HMMreg12833_1_EUT1(pf12833) EUT1
01661 -pf01218 -
01662 -pf01520 -
01663 -pf00583 -
01664 -pf11143 -
01665 -pf06572 -
01666 -pf04261 -
01668 -pf01797 -
01669 -pf00144 -
BMC-P
BMC-H
BMC-T
Aldehyde dehydrogenase
PTAC
Signature enzyme
Alcohol dehydrogenase
Regulator

Escherichia coli. Bacteria; Proteobacteria; Gammaproteobacteria; Enterobacterales;

NCTC7928__locus_1

EUT1

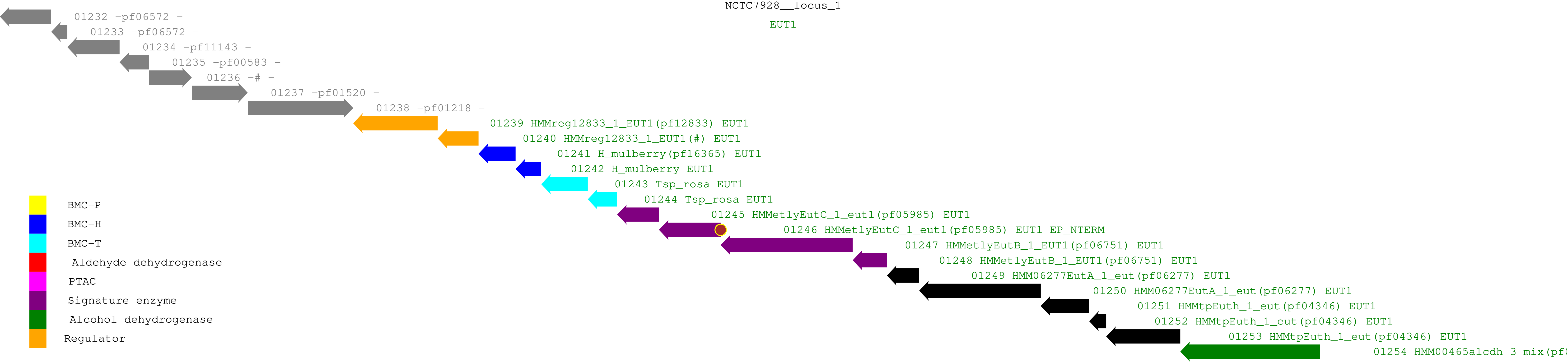

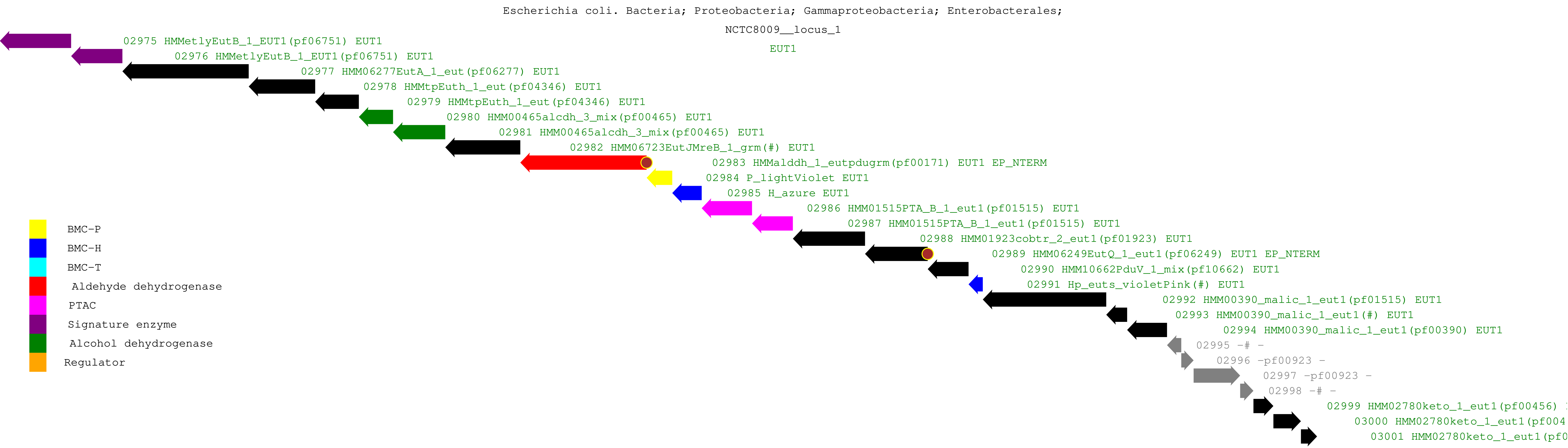
Escherichia coli. Bacteria; Proteobacteria; Gammaproteobacteria; Enterobacterales;
NCTC8009__locus_1
EUT1
02975 HMMetlyEutB_1_EUT1(pf06751) EUT1
02976 HMMetlyEutB_1_EUT1(pf06751) EUT1
02977 HMM06277EutA_1_eut(pf06277) EUT1
02978 HMMtpEuth_1_eut(pf04346) EUT1
02979 HMMtpEuth_1_eut(pf04346) EUT1
02980 HMM00465alcdh_3_mix(pf00465) EUT1
02981 HMM00465alcdh_3_mix(pf00465) EUT1
02982 HMM06723EutJMreB_1_grm(#) EUT1
02983 HMMalddh_1_eutpdugrm(pf00171) EUT1 EP_NTERM
02984 P_lightViolet EUT1
02985 H_azure EUT1
02986 HMM01515PTA_B_1_eut1(pf01515) EUT1
02987 HMM01515PTA_B_1_eut1(pf01515) EUT1
02988 HMM01923cobtr_2_eut1(pf01923) EUT1
02989 HMM06249EutQ_1_eut1(pf06249) EUT1 EP_NTERM
02990 HMM10662PduV_1_mix(pf10662) EUT1
02991 Hp_euts_violetPink(#) EUT1
02992 HMM00390_malic_1_eut1(pf01515) EUT1
02993 HMM00390_malic_1_eut1(#) EUT1
02994 HMM00390_malic_1_eut1(pf00390) EUT1
02995 -# -
02996 -pf00923 -
02997 -pf00923 -
02998 -# -
02999 HMM02780keto_1_eut1(pf00456)
03000 HMM02780keto_1_eut1(pf004
03001 HMM02780keto_1_eut1(pf0
BMC-P
BMC-H
BMC-T
Aldehyde dehydrogenase
PTAC
Signature enzyme
Alcohol dehydrogenase
Regulator

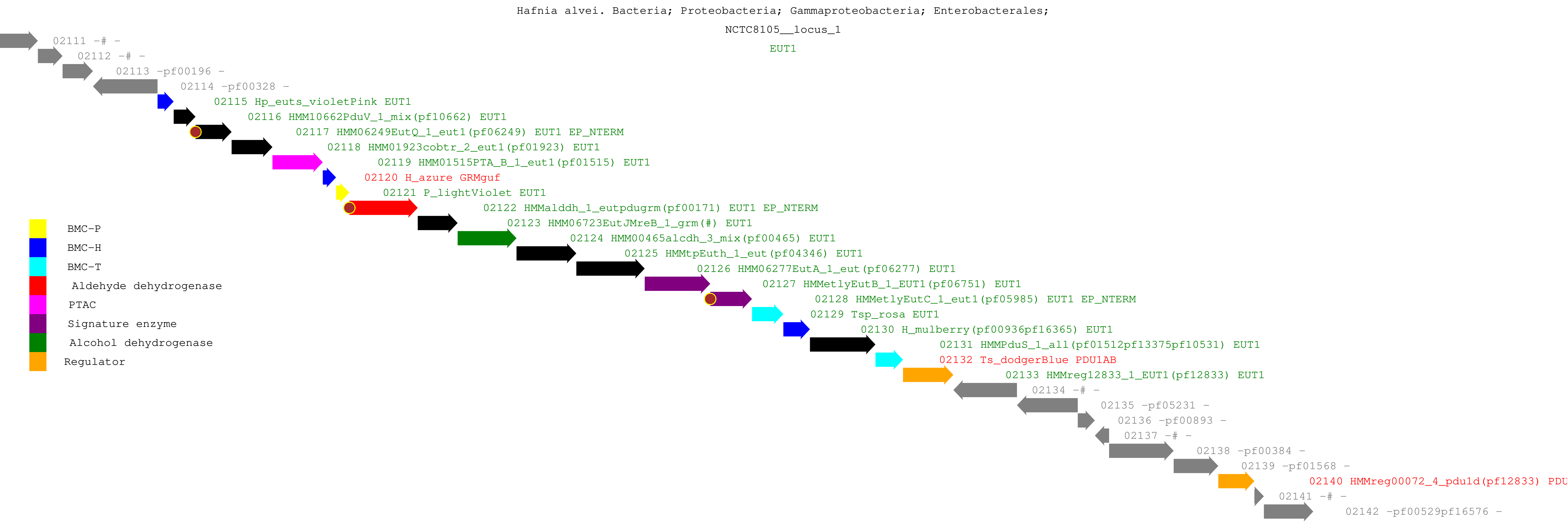
Hafnia alvei. Bacteria; Proteobacteria; Gammaproteobacteria; Enterobacterales;
NCTC8105__locus_1
EUT1
02111 -# -
02112 -# -
02113 -pf00196 -
02114 -pf00328 -
02115 Hp_euts_violetPink EUT1
02116 HMM10662PduV_1_mix(pf10662) EUT1
02117 HMM06249EutQ_1_eut1(pf06249) EUT1 EP_NTERM
02118 HMM01923cobtr_2_eut1(pf01923) EUT1
02119 HMM01515PTA_B_1_eut1(pf01515) EUT1
02120 H_azure GRMguf
02121 P_lightViolet EUT1
02122 HMMalddh_1_eutpdugrm(pf00171) EUT1 EP_NTERM
02123 HMM06723EutJMreB_1_grm(#) EUT1
02124 HMM00465alcdh_3_mix(pf00465) EUT1
02125 HMMtpEuth_1_eut(pf04346) EUT1
02126 HMM06277EutA_1_eut(pf06277) EUT1
02127 HMMetlyEutB_1_EUT1(pf06751) EUT1
02128 HMMetlyEutC_1_eut1(pf05985) EUT1 EP_NTERM
02129 Tsp_rosa EUT1
02130 H_mulberry(pf00936pf16365) EUT1
02131 HMMPduS_1_all(pf01512pf13375pf10531) EUT1
02132 Ts_dodgerBlue PDU1AB
02133 HMMreg12833_1_EUT1(pf12833) EUT1
02134 -# -
02135 -pf05231 -
02136 -pf00893 -
02137 -# -
02138 -pf00384 -
02139 -pf01568 -
02140 HMMreg00072_4_pdu1d(pf12833) PDU
02141 -# -
02142 -pf00529pf16576 -
BMC-P
BMC-H
BMC-T
Aldehyde dehydrogenase
PTAC
Signature enzyme
Alcohol dehydrogenase
Regulator

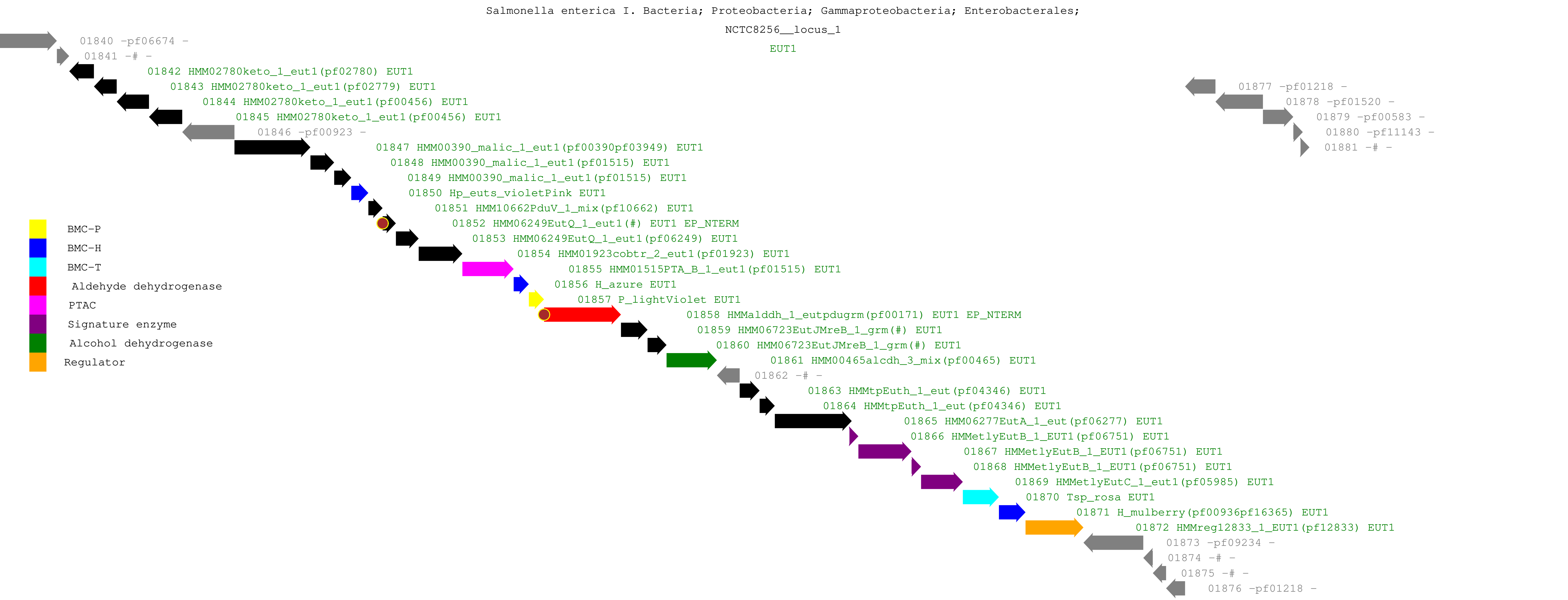
Salmonella enterica I. Bacteria; Proteobacteria; Gammaproteobacteria; Enterobacterales;
NCTC8256__locus_1
EUT1
01840 -pf06674 -
01841 -# -
01842 HMM02780keto_1_eut1(pf02780) EUT1
01843 HMM02780keto_1_eut1(pf02779) EUT1
01844 HMM02780keto_1_eut1(pf00456) EUT1
01845 HMM02780keto_1_eut1(pf00456) EUT1
01846 -pf00923 -
01847 HMM00390_malic_1_eut1(pf00390pf03949) EUT1
01848 HMM00390_malic_1_eut1(pf01515) EUT1
01849 HMM00390_malic_1_eut1(pf01515) EUT1
01850 Hp_euts_violetPink EUT1
01851 HMM10662PduV_1_mix(pf10662) EUT1
01852 HMM06249EutQ_1_eut1(#) EUT1 EP_NTERM
01853 HMM06249EutQ_1_eut1(pf06249) EUT1
01854 HMM01923cobtr_2_eut1(pf01923) EUT1
01855 HMM01515PTA_B_1_eut1(pf01515) EUT1
01856 H_azure EUT1
01857 P_lightViolet EUT1
01858 HMMalddh_1_eutpdugrm(pf00171) EUT1 EP_NTERM
01859 HMM06723EutJMreB_1_grm(#) EUT1
01860 HMM06723EutJMreB_1_grm(#) EUT1
01861 HMM00465alcdh_3_mix(pf00465) EUT1
01862 -# -
01863 HMMtpEuth_1_eut(pf04346) EUT1
01864 HMMtpEuth_1_eut(pf04346) EUT1
01865 HMM06277EutA_1_eut(pf06277) EUT1
01866 HMMetlyEutB_1_EUT1(pf06751) EUT1
01867 HMMetlyEutB_1_EUT1(pf06751) EUT1
01868 HMMetlyEutB_1_EUT1(pf06751) EUT1
01869 HMMetlyEutC_1_eut1(pf05985) EUT1
01870 Tsp_rosa EUT1
01871 H_mulberry(pf00936pf16365) EUT1
01872 HMMreg12833_1_EUT1(pf12833) EUT1
01873 -pf09234 -
01874 -# -
01875 -# -
01876 -pf01218 -
01877 -pf01218 -
01878 -pf01520 -
01879 -pf00583 -
01880 -pf11143 -
01881 -# -
BMC-P
BMC-H
BMC-T
Aldehyde dehydrogenase
PTAC
Signature enzyme
Alcohol dehydrogenase
Regulator

Salmonella enterica I. Bacteria; Proteobacteria; Gammaproteobacteria; Enterobacterales;

NCTC8261__locus_1

EUT1

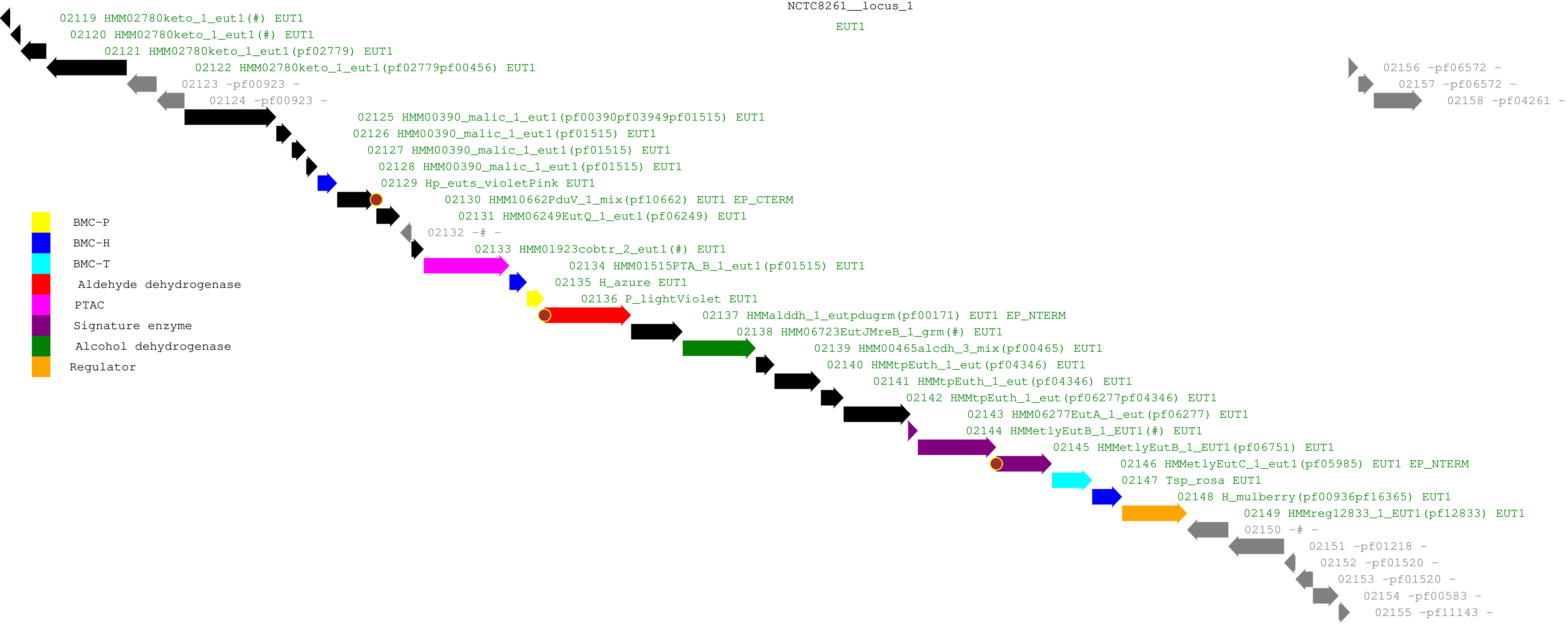

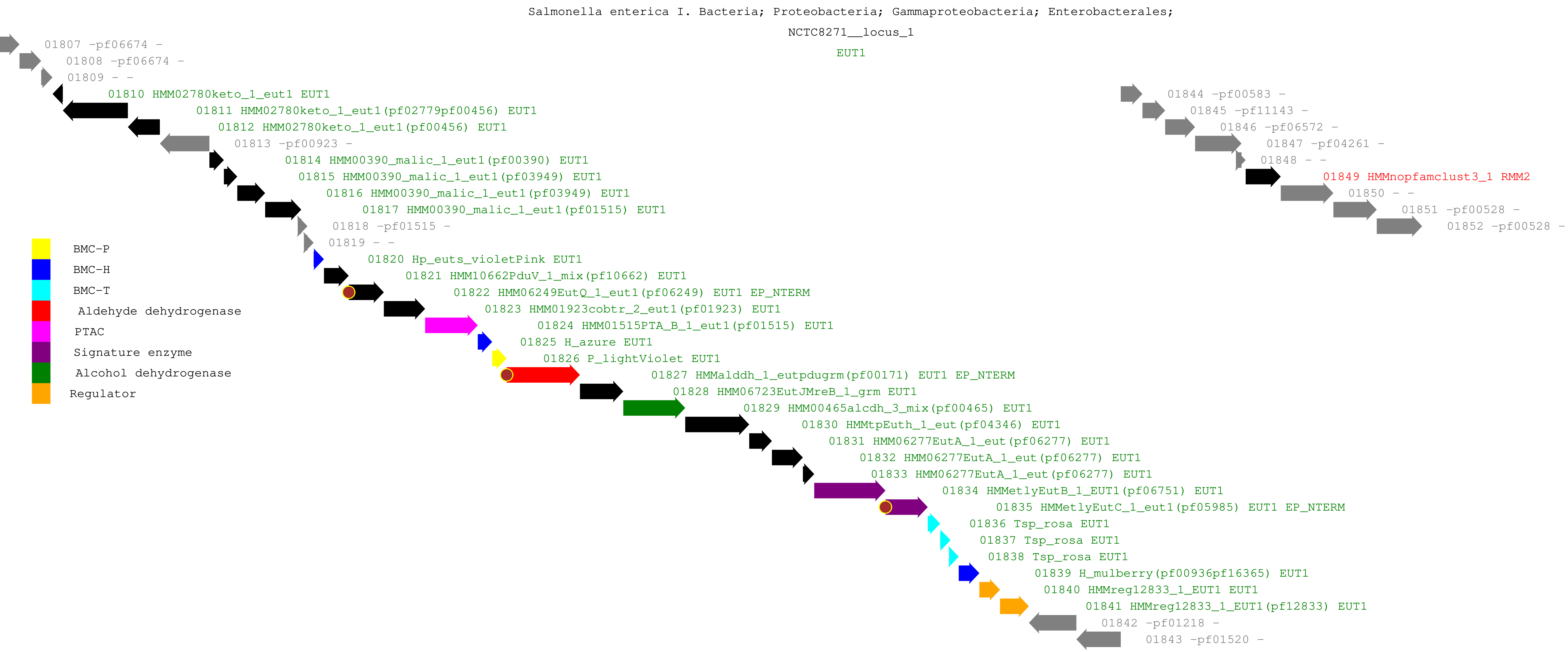
Salmonella enterica I. Bacteria; Proteobacteria; Gammaproteobacteria; Enterobacterales;
NCTC8271__locus_1
EUT1
01807 -pf06674 -
01808 -pf06674 -
01809 - -
01810 HMM02780keto_1_eut1 EUT1
01811 HMM02780keto_1_eut1(pf02779pf00456) EUT1
01812 HMM02780keto_1_eut1(pf00456) EUT1
01813 -pf00923 -
01814 HMM00390_malic_1_eut1(pf00390) EUT1
01815 HMM00390_malic_1_eut1(pf03949) EUT1
01816 HMM00390_malic_1_eut1(pf03949) EUT1
01817 HMM00390_malic_1_eut1(pf01515) EUT1
01818 -pf01515 -
01819 - -
01820 Hp_euts_violetPink EUT1
01821 HMM10662PduV_1_mix(pf10662) EUT1
01822 HMM06249EutQ_1_eut1(pf06249) EUT1 EP_NTERM
01823 HMM01923cobtr_2_eut1(pf01923) EUT1
01824 HMM01515PTA_B_1_eut1(pf01515) EUT1
01825 H_azure EUT1
01826 P_lightViolet EUT1
01827 HMMalddh_1_eutpdugrm(pf00171) EUT1 EP_NTERM
01828 HMM06723EutJMreB_1_grm EUT1
01829 HMM00465alcdh_3_mix(pf00465) EUT1
01830 HMMtpEuth_1_eut(pf04346) EUT1
01831 HMM06277EutA_1_eut(pf06277) EUT1
01832 HMM06277EutA_1_eut(pf06277) EUT1
01833 HMM06277EutA_1_eut(pf06277) EUT1
01834 HMMetlyEutB_1_EUT1(pf06751) EUT1
01835 HMMetlyEutC_1_eut1(pf05985) EUT1 EP_NTERM
01836 Tsp_rosa EUT1
01837 Tsp_rosa EUT1
01838 Tsp_rosa EUT1
01839 H_mulberry(pf00936pf16365) EUT1
01840 HMMreg12833_1_EUT1 EUT1
01841 HMMreg12833_1_EUT1(pf12833) EUT1
01842 -pf01218 -
01843 -pf01520 -
01844 -pf00583 -
01845 -pf11143 -
01846 -pf06572 -
01847 -pf04261 -
01848 - -
01849 HMMnopfamclust3_1 RMM2
01850 - -
01851 -pf00528 -
01852 -pf00528 -
BMC-P
BMC-H
BMC-T
Aldehyde dehydrogenase
PTAC
Signature enzyme
Alcohol dehydrogenase
Regulator

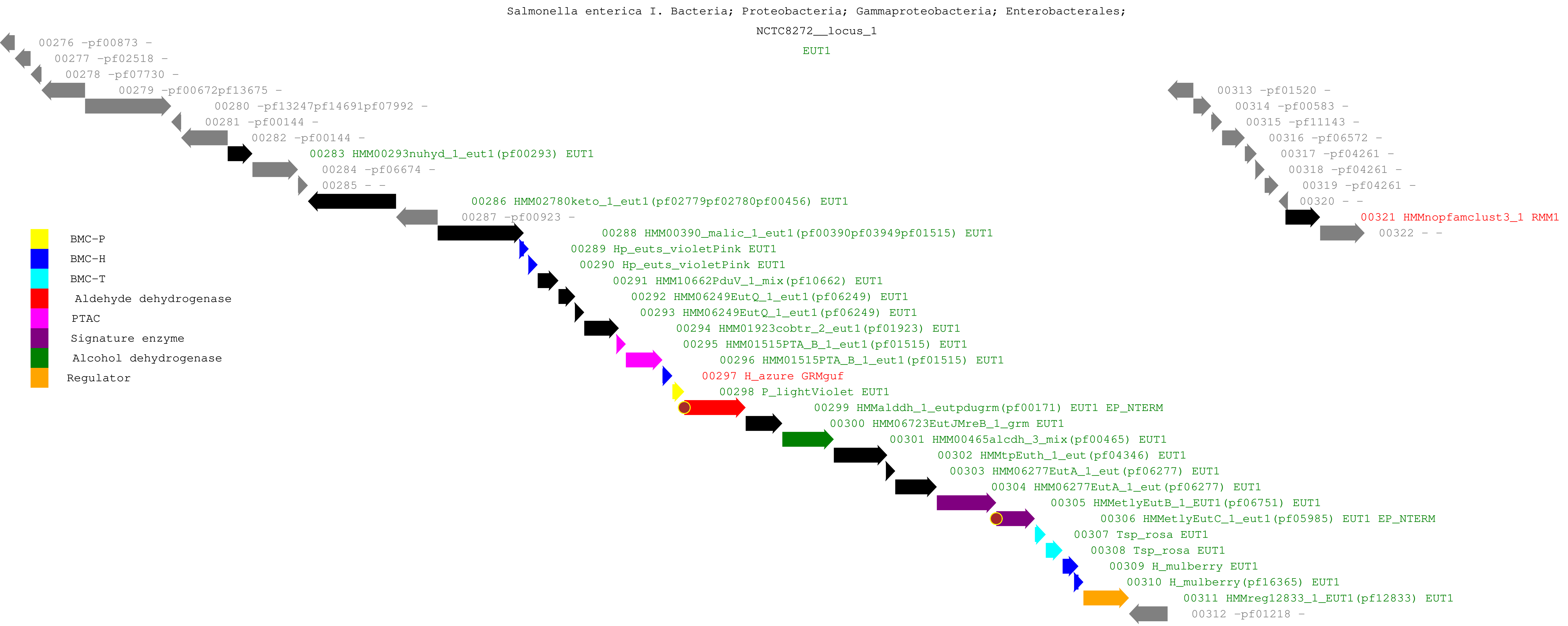
Salmonella enterica I. Bacteria; Proteobacteria; Gammaproteobacteria; Enterobacterales;
NCTC8272__locus_1
EUT1
00276 -pf00873 -
00277 -pf02518 -
00278 -pf07730 -
00279 -pf00672pf13675 -
00280 -pf13247pf14691pf07992 -
00281 -pf00144 -
00282 -pf00144 -
00283 HMM00293nuhyd_1_eut1(pf00293) EUT1
00284 -pf06674 -
00285 - -
00286 HMM02780keto_1_eut1(pf02779pf02780pf00456) EUT1
00287 -pf00923 -
00288 HMM00390_malic_1_eut1(pf00390pf03949pf01515) EUT1
00289 Hp_euts_violetPink EUT1
00290 Hp_euts_violetPink EUT1
00291 HMM10662PduV_1_mix(pf10662) EUT1
00292 HMM06249EutQ_1_eut1(pf06249) EUT1
00293 HMM06249EutQ_1_eut1(pf06249) EUT1
00294 HMM01923cobtr_2_eut1(pf01923) EUT1
00295 HMM01515PTA_B_1_eut1(pf01515) EUT1
00296 HMM01515PTA_B_1_eut1(pf01515) EUT1
00297 H_azure GRMguf
00298 P_lightViolet EUT1
00299 HMMalddh_1_eutpdugrm(pf00171) EUT1 EP_NTERM
00300 HMM06723EutJMreB_1_grm EUT1
00301 HMM00465alcdh_3_mix(pf00465) EUT1
00302 HMMtpEuth_1_eut(pf04346) EUT1
00303 HMM06277EutA_1_eut(pf06277) EUT1
00304 HMM06277EutA_1_eut(pf06277) EUT1
00305 HMMetlyEutB_1_EUT1(pf06751) EUT1
00306 HMMetlyEutC_1_eut1(pf05985) EUT1 EP_NTERM
00307 Tsp_rosa EUT1
00308 Tsp_rosa EUT1
00309 H_mulberry EUT1
00310 H_mulberry(pf16365) EUT1
00311 HMMreg12833_1_EUT1(pf12833) EUT1
00312 -pf01218 -
00313 -pf01520 -
00314 -pf00583 -
00315 -pf11143 -
00316 -pf06572 -
00317 -pf04261 -
00318 -pf04261 -
00319 -pf04261 -
00320 - -
00321 HMMnopfamclust3_1 RMM1
00322 - -
BMC-P
BMC-H
BMC-T
Aldehyde dehydrogenase
PTAC
Signature enzyme
Alcohol dehydrogenase
Regulator

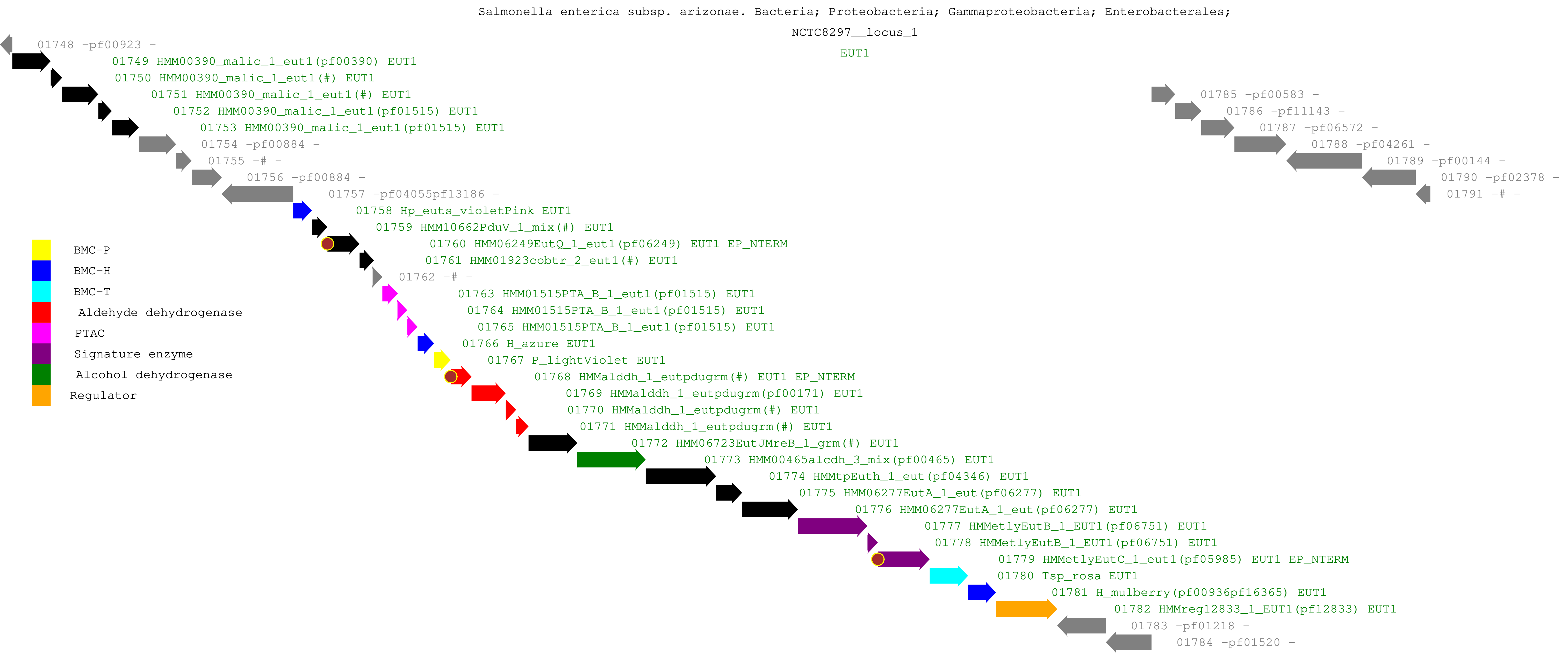
Salmonella enterica subsp. arizonae. Bacteria; Proteobacteria; Gammaproteobacteria; Enterobacterales;
NCTC8297__locus_1
EUT1
01748 -pf00923 -
01749 HMM00390_malic_1_eut1(pf00390) EUT1
01750 HMM00390_malic_1_eut1(#) EUT1
01751 HMM00390_malic_1_eut1(#) EUT1
01752 HMM00390_malic_1_eut1(pf01515) EUT1
01753 HMM00390_malic_1_eut1(pf01515) EUT1
01754 -pf00884 -
01755 -# -
01756 -pf00884 -
01757 -pf04055pf13186 -
01758 Hp_euts_violetPink EUT1
01759 HMM10662PduV_1_mix(#) EUT1
01760 HMM06249EutQ_1_eut1(pf06249) EUT1 EP_NTERM
01761 HMM01923cobtr_2_eut1(#) EUT1
01762 -# -
01763 HMM01515PTA_B_1_eut1(pf01515) EUT1
01764 HMM01515PTA_B_1_eut1(pf01515) EUT1
01765 HMM01515PTA_B_1_eut1(pf01515) EUT1
01766 H_azure EUT1
01767 P_lightViolet EUT1
01768 HMMalddh_1_eutpdugrm(#) EUT1 EP_NTERM
01769 HMMalddh_1_eutpdugrm(pf00171) EUT1
01770 HMMalddh_1_eutpdugrm(#) EUT1
01771 HMMalddh_1_eutpdugrm(#) EUT1
01772 HMM06723EutJMreB_1_grm(#) EUT1
01773 HMM00465alcdh_3_mix(pf00465) EUT1
01774 HMMtpEuth_1_eut(pf04346) EUT1
01775 HMM06277EutA_1_eut(pf06277) EUT1
01776 HMM06277EutA_1_eut(pf06277) EUT1
01777 HMMetlyEutB_1_EUT1(pf06751) EUT1
01778 HMMetlyEutB_1_EUT1(pf06751) EUT1
01779 HMMetlyEutC_1_eut1(pf05985) EUT1 EP_NTERM
01780 Tsp_rosa EUT1
01781 H_mulberry(pf00936pf16365) EUT1
01782 HMMreg12833_1_EUT1(pf12833) EUT1
01783 -pf01218 -
01784 -pf01520 -
01785 -pf00583 -
01786 -pf11143 -
01787 -pf06572 -
01788 -pf04261 -
01789 -pf00144 -
01790 -pf02378 -
01791 -# -
BMC-P
BMC-H
BMC-T
Aldehyde dehydrogenase
PTAC
Signature enzyme
Alcohol dehydrogenase
Regulator

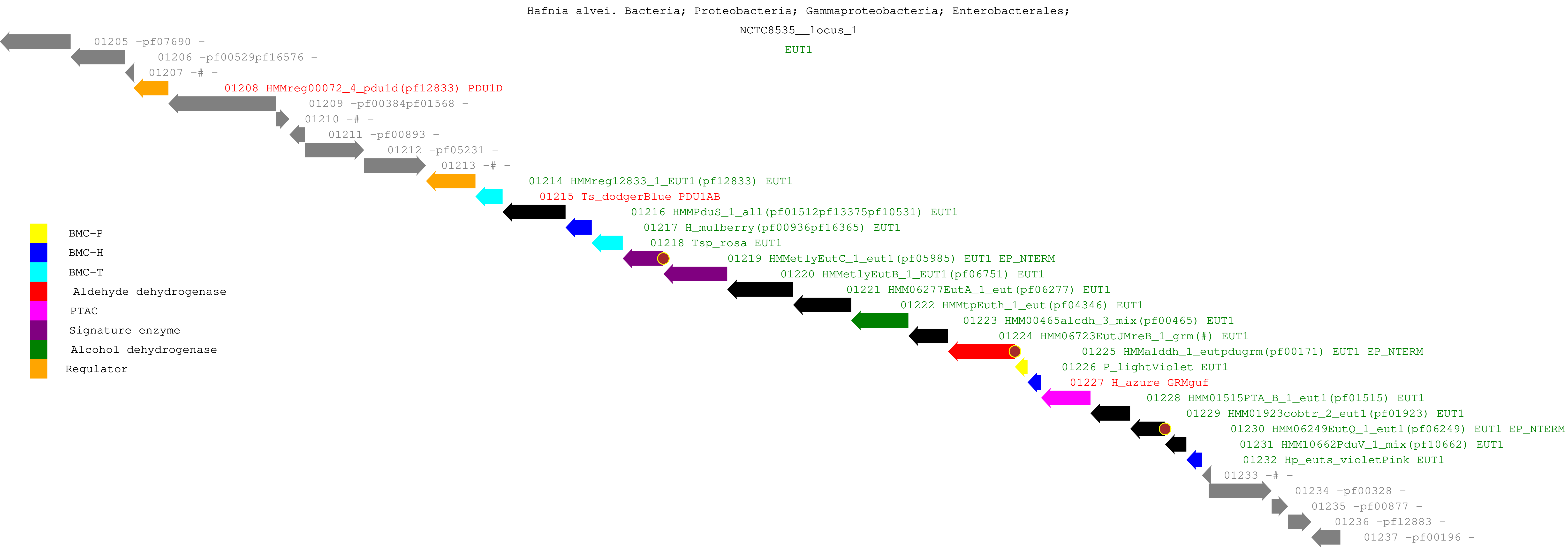
Hafnia alvei. Bacteria; Proteobacteria; Gammaproteobacteria; Enterobacterales;
NCTC8535__locus_1
EUT1
01205 -pf07690 -
01206 -pf00529pf16576 -
01207 -# -
01208 HMMreg00072_4_pdu1d(pf12833) PDU1D
01209 -pf00384pf01568 -
01210 -# -
01211 -pf00893 -
01212 -pf05231 -
01213 -# -
01214 HMMreg12833_1_EUT1(pf12833) EUT1
01215 Ts_dodgerBlue PDU1AB
01216 HMMPduS_1_all(pf01512pf13375pf10531) EUT1
01217 H_mulberry(pf00936pf16365) EUT1
01218 Tsp_rosa EUT1
01219 HMMetlyEutC_1_eut1(pf05985) EUT1 EP_NTERM
01220 HMMetlyEutB_1_EUT1(pf06751) EUT1
01221 HMM06277EutA_1_eut(pf06277) EUT1
01222 HMMtpEuth_1_eut(pf04346) EUT1
01223 HMM00465alcdh_3_mix(pf00465) EUT1
01224 HMM06723EutJMreB_1_grm(#) EUT1
01225 HMMaldh_1_eutpdugrm(pf00171) EUT1 EP_NTERM
01226 P_lightViolet EUT1
01227 H_azure GRMguf
01228 HMM01515PTA_B_1_eut1(pf01515) EUT1
01229 HMM01923cobtr_2_eut1(pf01923) EUT1
01230 HMM06249EutQ_1_eut1(pf06249) EUT1 EP_NTERM
01231 HMM10662PduV_1_mix(pf10662) EUT1
01232 Hp_euts_violetPink EUT1
01233 -# -
01234 -pf00328 -
01235 -pf00877 -
01236 -pf12883 -
01237 -pf00196 -
BMC-P
BMC-H
BMC-T
Aldehyde dehydrogenase
PTAC
Signature enzyme
Alcohol dehydrogenase
Regulator

Shigella boydii. Bacteria; Proteobacteria; Gammaproteobacteria; Enterobacterales;

NCTC8576__locus_1

EUT1

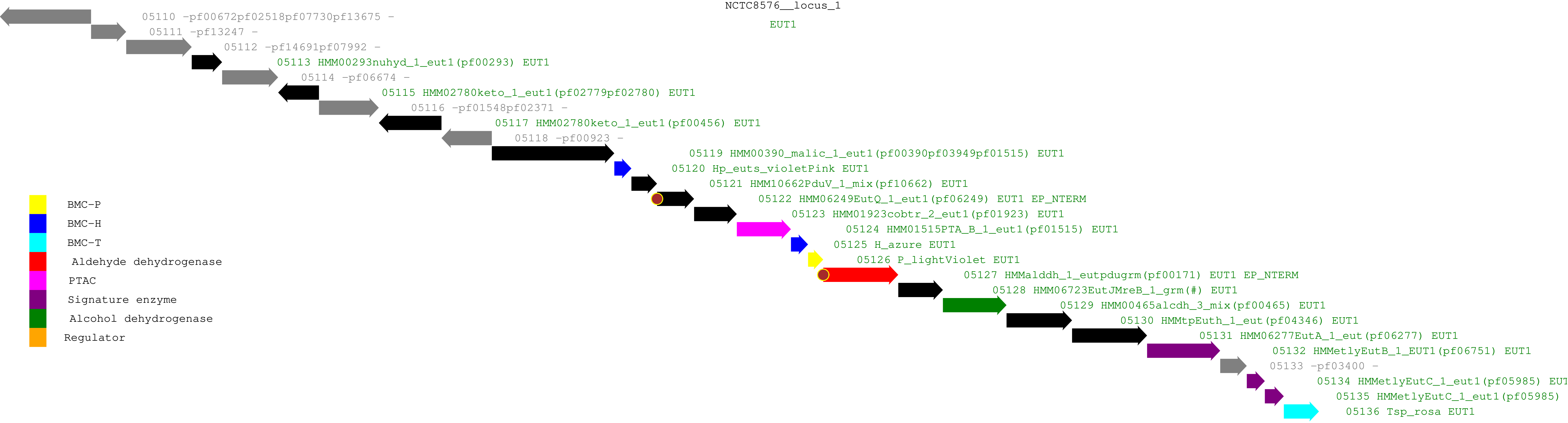

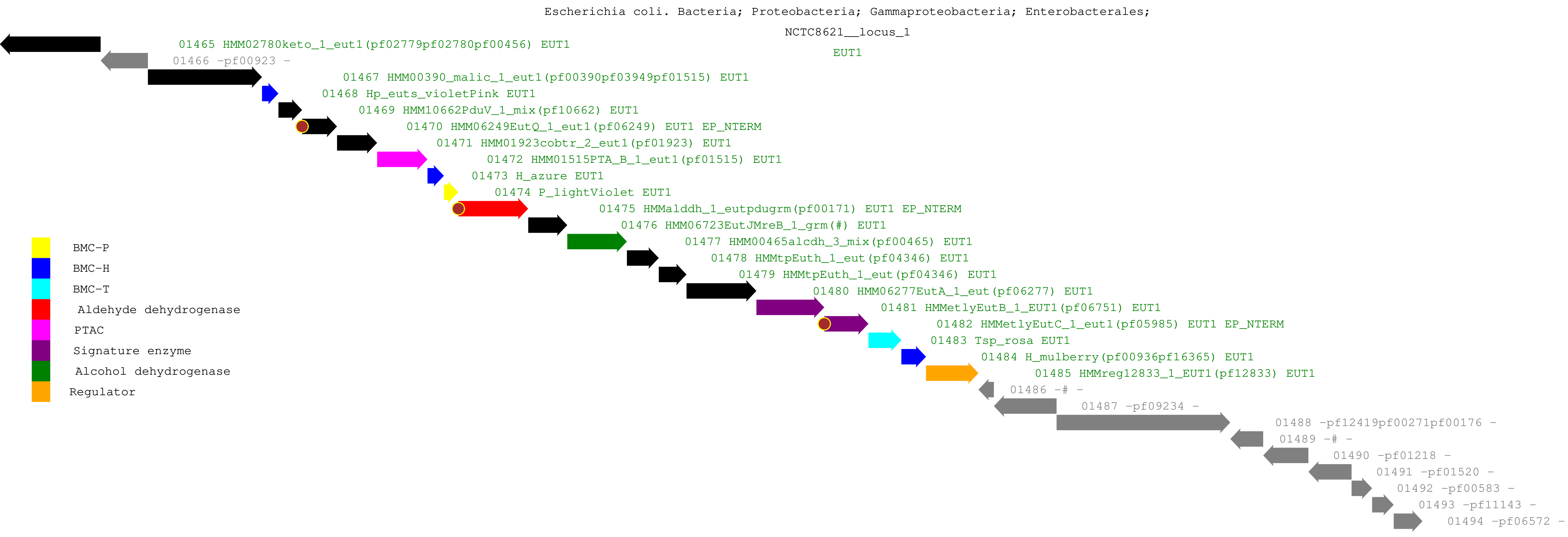
Escherichia coli. Bacteria; Proteobacteria; Gammaproteobacteria; Enterobacterales;
NCTC8621__locus_1
EUT1
01465 HMM02780keto_1_eut1(pf02779pf02780pf00456) EUT1
01466 -pf00923 -
01467 HMM00390_malic_1_eut1(pf00390pf03949pf01515) EUT1
01468 Hp_euts_violetPink EUT1
01469 HMM10662PduV_1_mix(pf10662) EUT1
01470 HMM06249EutQ_1_eut1(pf06249) EUT1 EP_NTERM
01471 HMM01923cobtr_2_eut1(pf01923) EUT1
01472 HMM01515PTA_B_1_eut1(pf01515) EUT1
01473 H_azure EUT1
01474 P_lightViolet EUT1
01475 HMMalddh_1_eutpdugrm(pf00171) EUT1 EP_NTERM
01476 HMM06723EutJMreB_1_grm(#) EUT1
01477 HMM00465alcdh_3_mix(pf00465) EUT1
01478 HMMtpEuth_1_eut(pf04346) EUT1
01479 HMMtpEuth_1_eut(pf04346) EUT1
01480 HMM06277EutA_1_eut(pf06277) EUT1
01481 HMMetlyEutB_1_EUT1(pf06751) EUT1
01482 HMMetlyEutC_1_eut1(pf05985) EUT1 EP_NTERM
01483 Tsp_rosa EUT1
01484 H_mulberry(pf00936pf16365) EUT1
01485 HMMreg12833_1_EUT1(pf12833) EUT1
01486 -# -
01487 -pf09234 -
01488 -pf12419pf00271pf00176 -
01489 -# -
01490 -pf01218 -
01491 -pf01520 -
01492 -pf00583 -
01493 -pf11143 -
01494 -pf06572 -
BMC-P
BMC-H
BMC-T
Aldehyde dehydrogenase
PTAC
Signature enzyme
Alcohol dehydrogenase
Regulator

Escherichia coli. Bacteria; Proteobacteria; Gammaproteobacteria; Enterobacterales;

NCTC8985__locus_1

EUT1

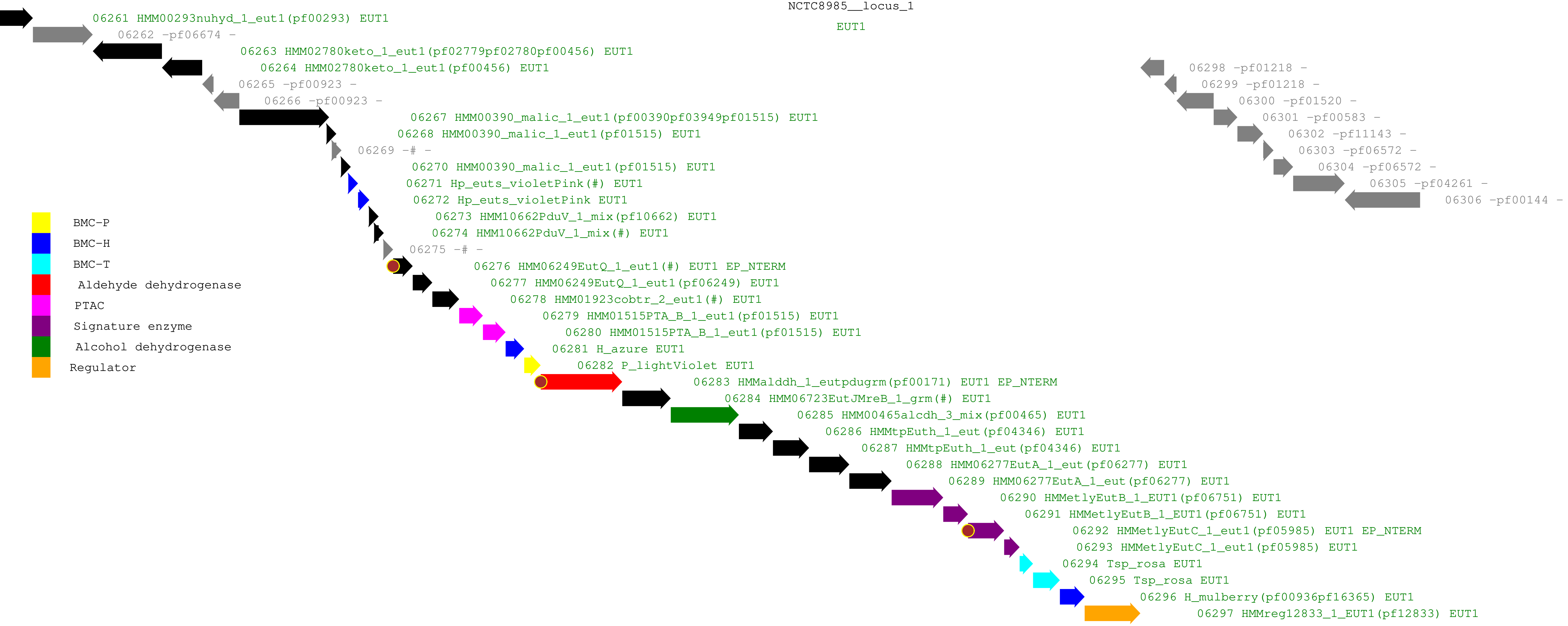

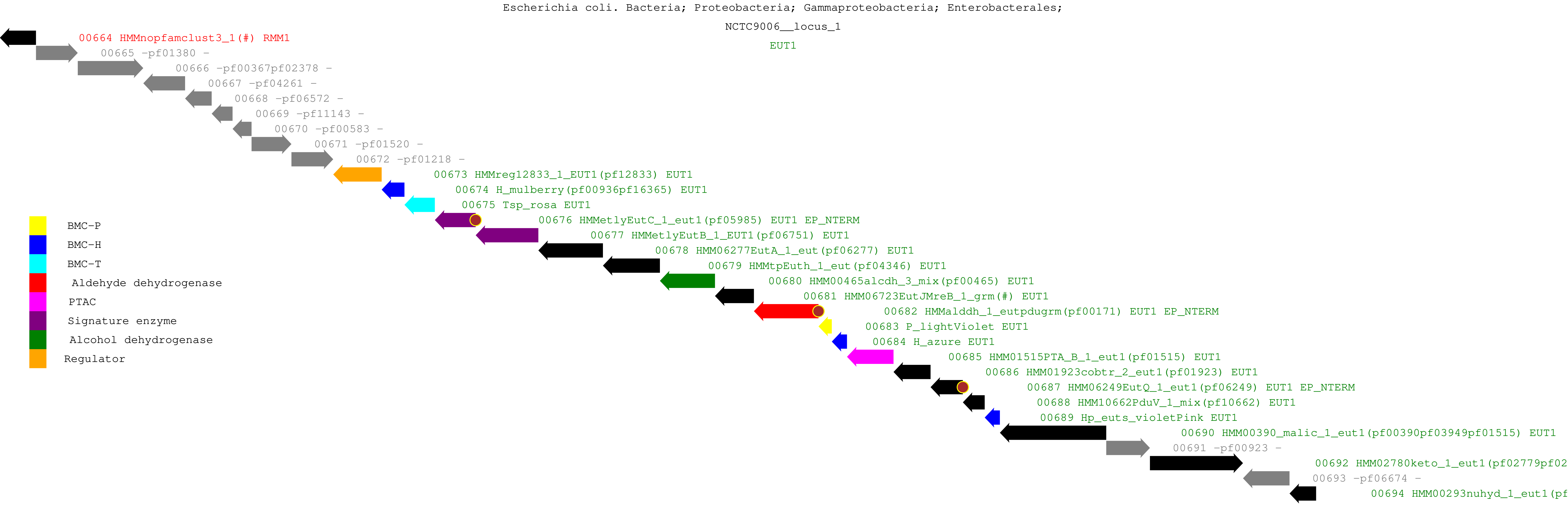
Escherichia coli. Bacteria; Proteobacteria; Gammaproteobacteria; Enterobacterales;
NCTC9006__locus_1
EUT1
00664 HMMnopfamclust3_1(#) RMM1
00665 -pf01380 -
00666 -pf00367pf02378 -
00667 -pf04261 -
00668 -pf06572 -
00669 -pf11143 -
00670 -pf00583 -
00671 -pf01520 -
00672 -pf01218 -
00673 HMMreg12833_1_EUT1(pf12833) EUT1
00674 H_mulberry(pf00936pf16365) EUT1
00675 Tsp_rosa EUT1
00676 HMMetlyEutC_1_eut1(pf05985) EUT1 EP_NTERM
00677 HMMetlyEutB_1_EUT1(pf06751) EUT1
00678 HMM06277EutA_1_eut(pf06277) EUT1
00679 HMMtpEuth_1_eut(pf04346) EUT1
00680 HMM00465alcdh_3_mix(pf00465) EUT1
00681 HMM06723EutJMreB_1_grm(#) EUT1
00682 HMMalddh_1_eutpdugrm(pf00171) EUT1 EP_NTERM
00683 P_lightViolet EUT1
00684 H_azure EUT1
00685 HMM01515PTA_B_1_eut1(pf01515) EUT1
00686 HMM01923cobtr_2_eut1(pf01923) EUT1
00687 HMM06249EutQ_1_eut1(pf06249) EUT1 EP_NTERM
00688 HMM10662PduV_1_mix(pf10662) EUT1
00689 Hp_euts_violetPink EUT1
00690 HMM00390_malic_1_eut1(pf00390pf03949pf01515) EUT1
00691 -pf00923 -
00692 HMM02780keto_1_eut1(pf02779pf02
00693 -pf06674 -
00694 HMM00293nuhyd_1_eut1(pf
BMC-P
BMC-H
BMC-T
Aldehyde dehydrogenase
PTAC
Signature enzyme
Alcohol dehydrogenase
Regulator

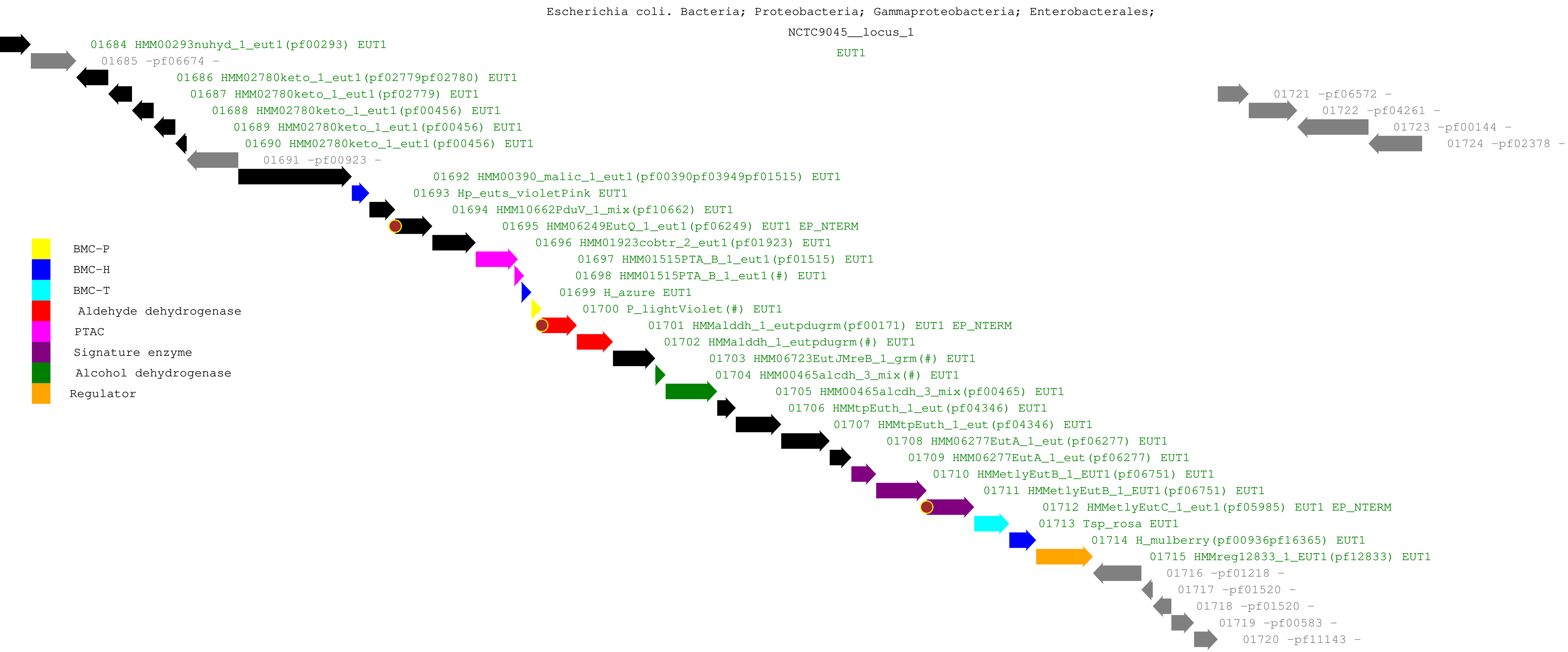

Escherichia coli. Bacteria; Proteobacteria; Gammaproteobacteria; Enterobacterales;
NCTC9045__locus_1
EUT1
01684 HMM00293nuhyd_1_eut1(pf00293) EUT1
01685 -pf06674 -
01686 HMM02780keto_1_eut1(pf02779pf02780) EUT1
01687 HMM02780keto_1_eut1(pf02779) EUT1
01688 HMM02780keto_1_eut1(pf00456) EUT1
01689 HMM02780keto_1_eut1(pf00456) EUT1
01690 HMM02780keto_1_eut1(pf00456) EUT1
01691 -pf00923 -
01692 HMM00390_malic_1_eut1(pf00390pf03949pf01515) EUT1
01693 Hp_euts_violetPink EUT1
01694 HMM10662PduV_1_mix(pf10662) EUT1
01695 HMM06249EutQ_1_eut1(pf06249) EUT1 EP_NTERM
01696 HMM01923cobtr_2_eut1(pf01923) EUT1
01697 HMM01515PTA_B_1_eut1(pf01515) EUT1
01698 HMM01515PTA_B_1_eut1(#) EUT1
01699 H_azure EUT1
01700 P_lightViolet(#) EUT1
01701 HMMalddh_1_eutpdugrm(pf00171) EUT1 EP_NTERM
01702 HMMalddh_1_eutpdugrm(#) EUT1
01703 HMM06723EutJMreB_1_grm(#) EUT1
01704 HMM00465alcdh_3_mix(#) EUT1
01705 HMM00465alcdh_3_mix(pf00465) EUT1
01706 HMMtpEuth_1_eut(pf04346) EUT1
01707 HMMtpEuth_1_eut(pf04346) EUT1
01708 HMM06277EutA_1_eut(pf06277) EUT1
01709 HMM06277EutA_1_eut(pf06277) EUT1
01710 HMMetlyEutB_1_EUT1(pf06751) EUT1
01711 HMMetlyEutB_1_EUT1(pf06751) EUT1
01712 HMMetlyEutC_1_eut1(pf05985) EUT1 EP_NTERM
01713 Tsp_rosa EUT1
01714 H_mulberry(pf00936pf16365) EUT1
01715 HMMreg12833_1_EUT1(pf12833) EUT1
01716 -pf01218 -
01717 -pf01520 -
01718 -pf01520 -
01719 -pf00583 -
01720 -pf11143 -
01721 -pf06572 -
01722 -pf04261 -
01723 -pf00144 -
01724 -pf02378 -
BMC-P
BMC-H
BMC-T
Aldehyde dehydrogenase
PTAC
Signature enzyme
Alcohol dehydrogenase
Regulator

Escherichia coli. Bacteria; Proteobacteria; Gammaproteobacteria; Enterobacterales;

NCTC9060__locus_1

EUT1

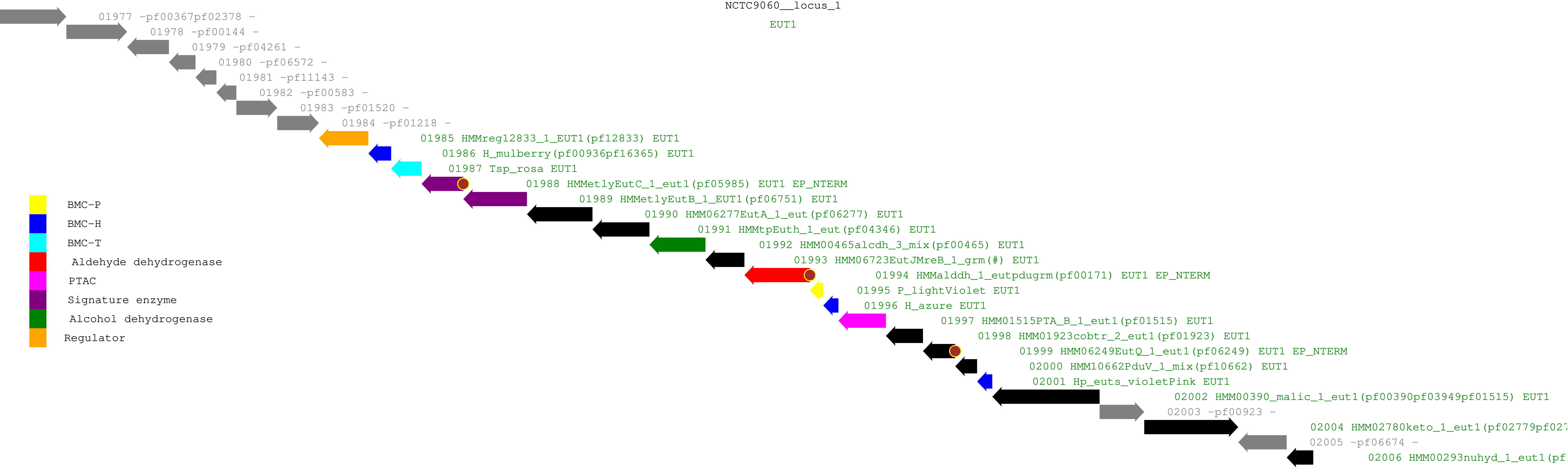

Escherichia coli. Bacteria; Proteobacteria; Gammaproteobacteria; Enterobacterales;

NCTC9067__locus_1

EUT1

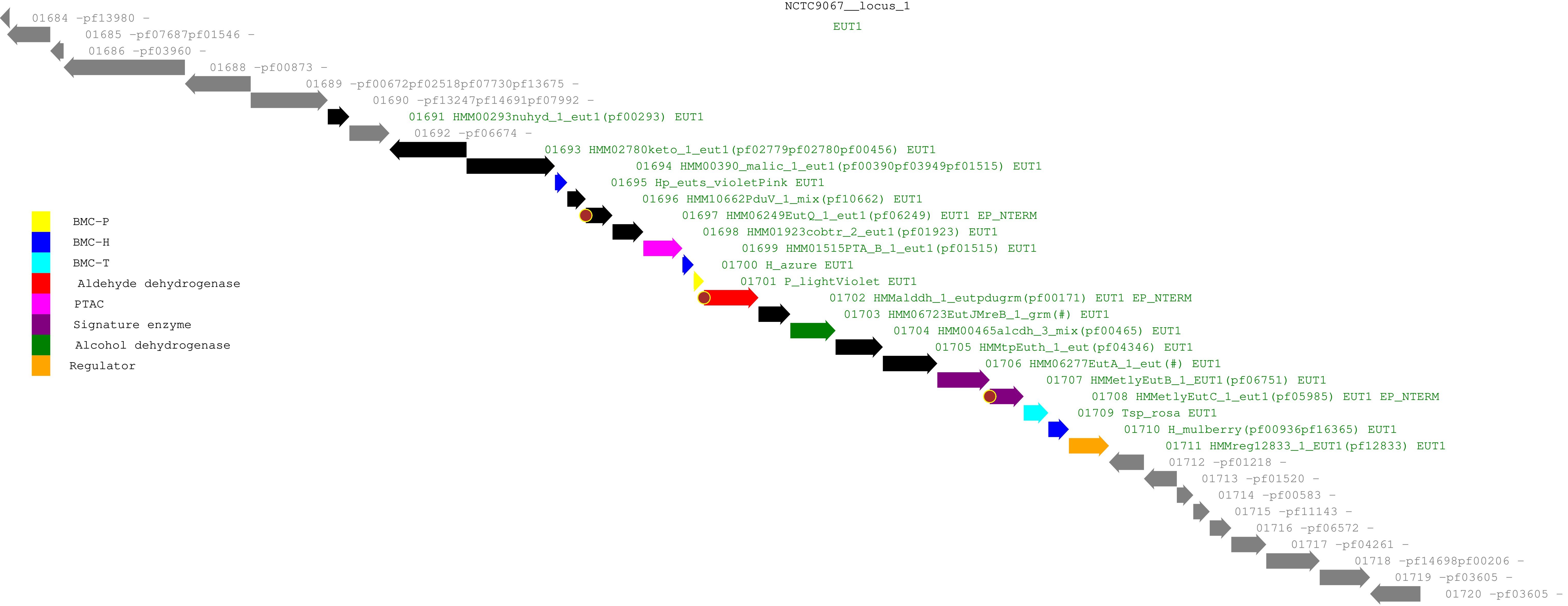

Escherichia coli. Bacteria; Proteobacteria; Gammaproteobacteria; Enterobacterales;

NCTC9094__locus_1

EUT1

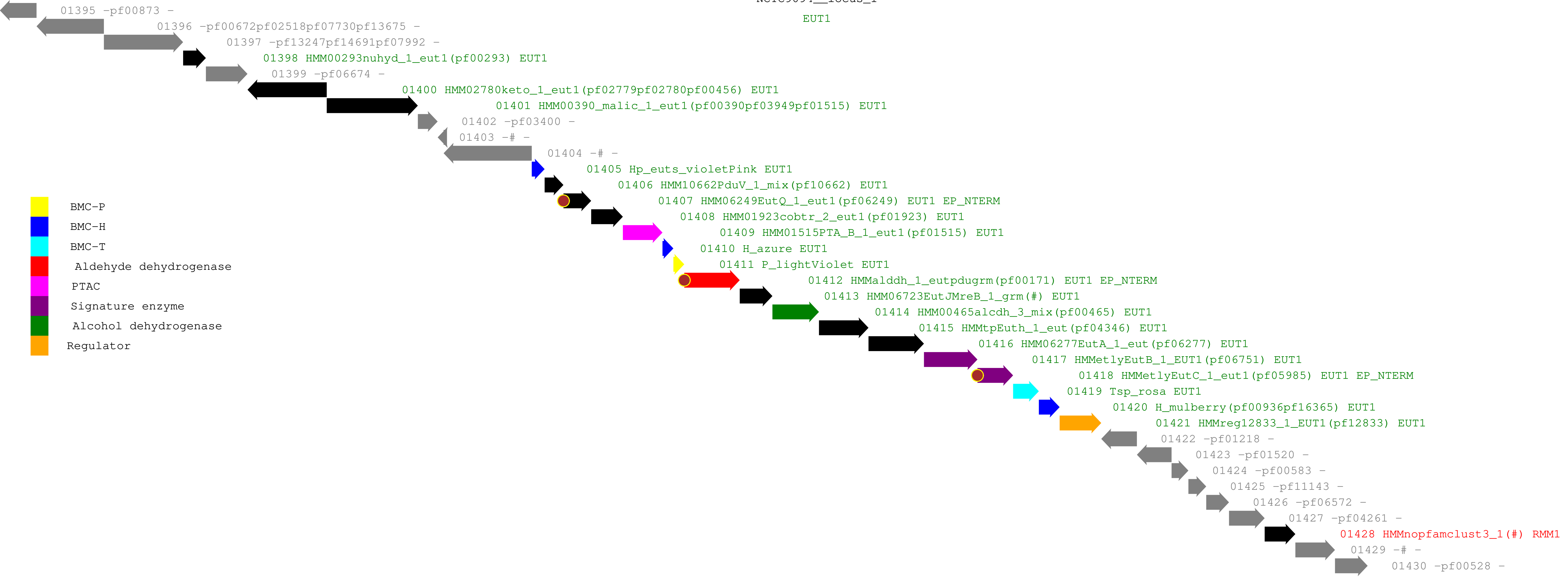

Escherichia coli. Bacteria; Proteobacteria; Gammaproteobacteria; Enterobacterales;

NCTC9110__locus_1

EUT1

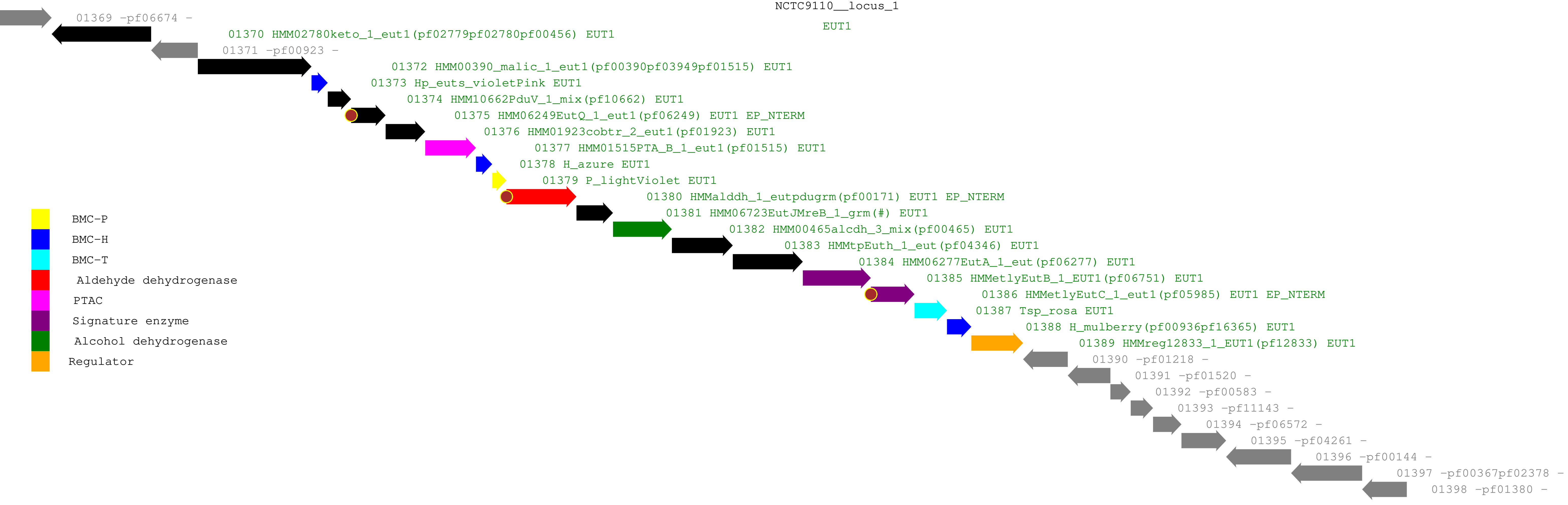

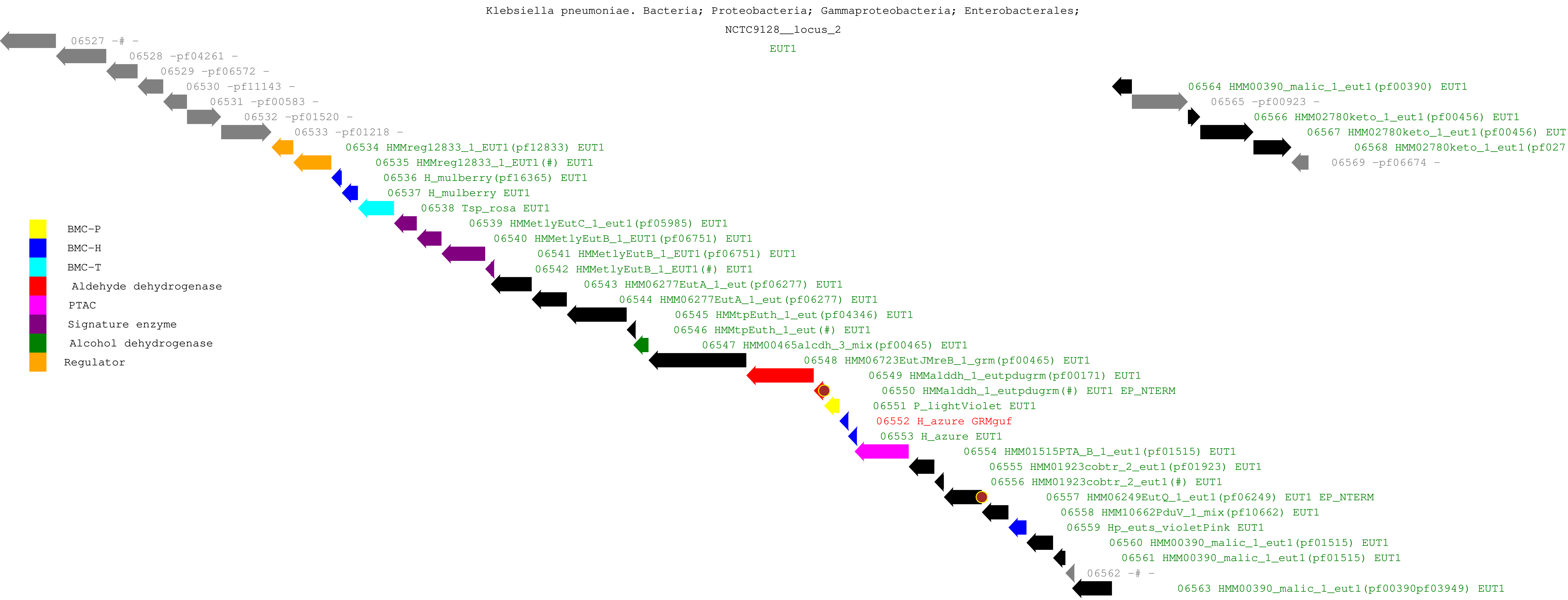
Klebsiella pneumoniae. Bacteria; Proteobacteria; Gammaproteobacteria; Enterobacterales;
NCTC9128__locus_2
EUT1
06527 -# -
06528 -pf04261 -
06529 -pf06572 -
06530 -pf11143 -
06531 -pf00583 -
06532 -pf01520 -
06533 -pf01218 -
06534 HMMreg12833_1_EUT1(pf12833) EUT1
06535 HMMreg12833_1_EUT1(#) EUT1
06536 H_mulberry(pf16365) EUT1
06537 H_mulberry EUT1
06538 Tsp_rosa EUT1
06539 HMMetlyEutC_1_eut1(pf05985) EUT1
06540 HMMetlyEutB_1_EUT1(pf06751) EUT1
06541 HMMetlyEutB_1_EUT1(pf06751) EUT1
06542 HMMetlyEutB_1_EUT1(#) EUT1
06543 HMM06277EutA_1_eut(pf06277) EUT1
06544 HMM06277EutA_1_eut(pf06277) EUT1
06545 HMMtpEuth_1_eut(pf04346) EUT1
06546 HMMtpEuth_1_eut(#) EUT1
06547 HMM00465alcdh_3_mix(pf00465) EUT1
06548 HMM06723EutJMreB_1_grm(pf00465) EUT1
06549 HMMalddh_1_eutpdugrm(pf00171) EUT1
06550 HMMaldddh_1_eutpdugrm(#) EUT1 EP_NTERM
06551 P_lightViolet EUT1
06552 H_azure GRMguf
06553 H_azure EUT1
06554 HMM01515PTA_B_1_eut1(pf01515) EUT1
06555 HMM01923cobtr_2_eut1(pf01923) EUT1
06556 HMM01923cobtr_2_eut1(#) EUT1
06557 HMM06249EutQ_1_eut1(pf06249) EUT1 EP_NTERM
06558 HMM10662PduV_1_mix(pf10662) EUT1
06559 Hp_euts_violetPink EUT1
06560 HMM00390_malic_1_eut1(pf01515) EUT1
06561 HMM00390_malic_1_eut1(pf01515) EUT1
06562 -# -
06563 HMM00390_malic_1_eut1(pf00390pf03949) EUT1
06564 HMM00390_malic_1_eut1(pf00390) EUT1
06565 -pf00923 -
06566 HMM02780keto_1_eut1(pf00456) EUT1
06567 HMM02780keto_1_eut1(pf00456) EUT
06568 HMM02780keto_1_eut1(pf027
06569 -pf06674 -
BMC-P
BMC-H
BMC-T
Aldehyde dehydrogenase
PTAC
Signature enzyme
Alcohol dehydrogenase
Regulator

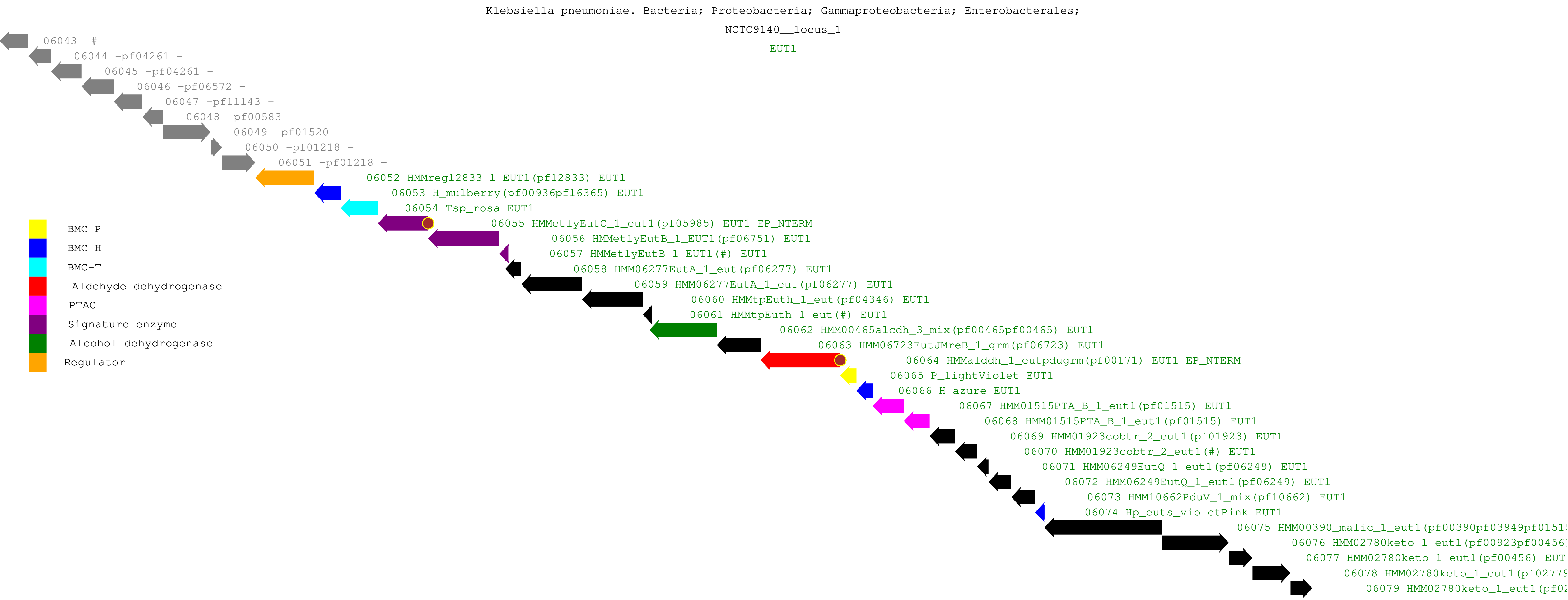

Klebsiella pneumoniae. Bacteria; Proteobacteria; Gammaproteobacteria; Enterobacterales;
NCTC9140__locus_1
EUT1
06043 -# -
06044 -pf04261 -
06045 -pf04261 -
06046 -pf06572 -
06047 -pf11143 -
06048 -pf00583 -
06049 -pf01520 -
06050 -pf01218 -
06051 -pf01218 -
06052 HMMreg12833_1_EUT1(pf12833) EUT1
06053 H_mulberry(pf00936pf16365) EUT1
06054 Tsp_rosa EUT1
06055 HMMetlyEutC_1_eut1(pf05985) EUT1 EP_NTERM
06056 HMMetlyEutB_1_EUT1(pf06751) EUT1
06057 HMMetlyEutB_1_EUT1(#) EUT1
06058 HMM06277EutA_1_eut(pf06277) EUT1
06059 HMM06277EutA_1_eut(pf06277) EUT1
06060 HMMtpEuth_1_eut(pf04346) EUT1
06061 HMMtpEuth_1_eut(#) EUT1
06062 HMM00465alcdh_3_mix(pf00465pf00465) EUT1
06063 HMM06723EutJMreB_1_grm(pf06723) EUT1
06064 HMMaldh_1_eutpdugrm(pf00171) EUT1 EP_NTERM
06065 P_lightViolet EUT1
06066 H_azure EUT1
06067 HMM01515PTA_B_1_eut1(pf01515) EUT1
06068 HMM01515PTA_B_1_eut1(pf01515) EUT1
06069 HMM01923cobtr_2_eut1(pf01923) EUT1
06070 HMM01923cobtr_2_eut1(#) EUT1
06071 HMM06249EutQ_1_eut1(pf06249) EUT1
06072 HMM06249EutQ_1_eut1(pf06249) EUT1
06073 HMM10662PduV_1_mix(pf10662) EUT1
06074 Hp_euts_violetPink EUT1
06075 HMM00390_malic_1_eut1(pf00390pf03949pf0151
06076 HMM02780keto_1_eut1(pf00923pf00456
06077 HMM02780keto_1_eut1(pf00456) EUT
06078 HMM02780keto_1_eut1(pf0277
06079 HMM02780keto_1_eut1(pf02
BMC-P
BMC-H
BMC-T
Aldehyde dehydrogenase
PTAC
Signature enzyme
Alcohol dehydrogenase
Regulator

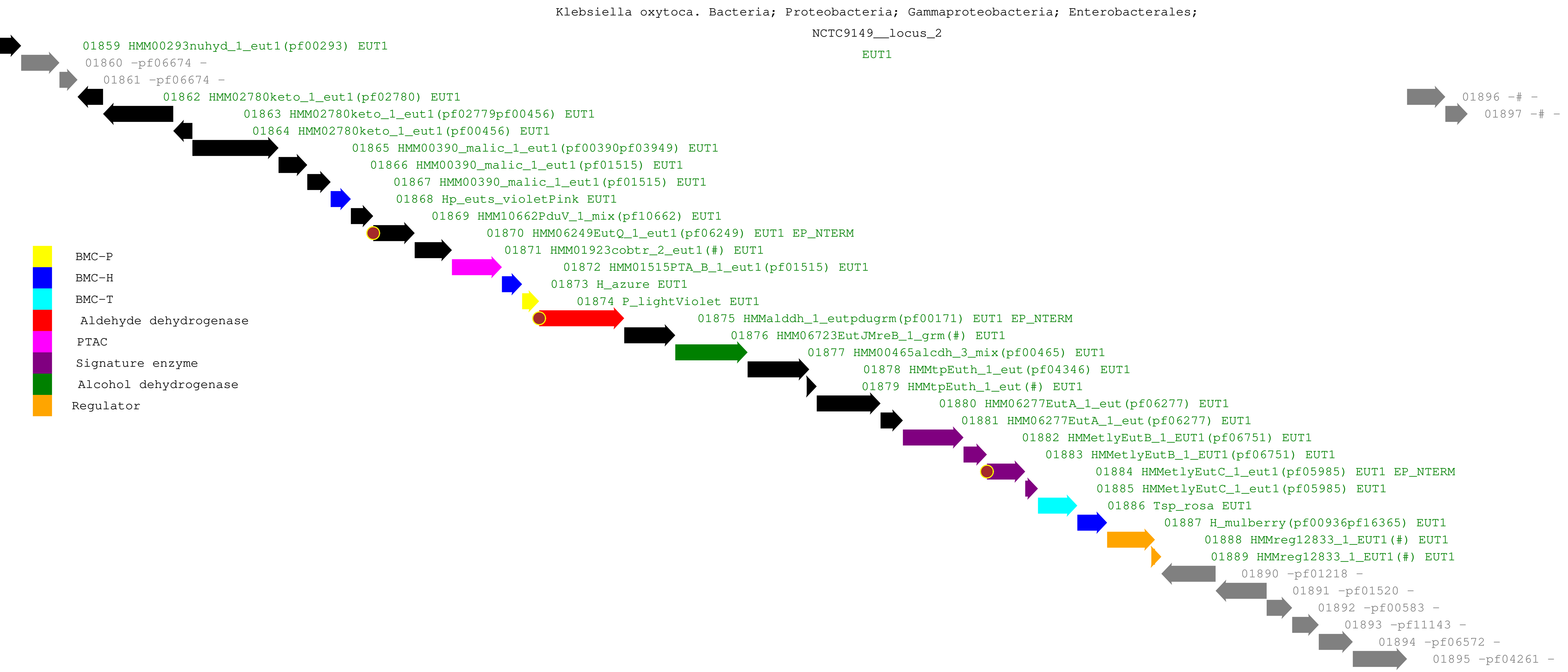
Klebsiella oxytoca. Bacteria; Proteobacteria; Gammaproteobacteria; Enterobacterales;
NCTC9149__locus_2
EUT1
01859 HMM00293nuhyd_1_eut1(pf00293) EUT1
01860 -pf06674 -
01861 -pf06674 -
01862 HMM02780keto_1_eut1(pf02780) EUT1
01863 HMM02780keto_1_eut1(pf02779pf00456) EUT1
01864 HMM02780keto_1_eut1(pf00456) EUT1
01865 HMM00390_malic_1_eut1(pf00390pf03949) EUT1
01866 HMM00390_malic_1_eut1(pf01515) EUT1
01867 HMM00390_malic_1_eut1(pf01515) EUT1
01868 Hp_euts_violetPink EUT1
01869 HMM10662PduV_1_mix(pf10662) EUT1
01870 HMM06249EutQ_1_eut1(pf06249) EUT1 EP_NTERM
01871 HMM01923cobtr_2_eut1(#) EUT1
01872 HMM01515PTA_B_1_eut1(pf01515) EUT1
01873 H_azure EUT1
01874 P_lightViolet EUT1
01875 HMMalddh_1_eutpdugrm(pf00171) EUT1 EP_NTERM
01876 HMM06723EutJMreB_1_grm(#) EUT1
01877 HMM00465alcdh_3_mix(pf00465) EUT1
01878 HMMtpEuth_1_eut(pf04346) EUT1
01879 HMMtpEuth_1_eut(#) EUT1
01880 HMM06277EutA_1_eut(pf06277) EUT1
01881 HMM06277EutA_1_eut(pf06277) EUT1
01882 HMMetlyEutB_1_EUT1(pf06751) EUT1
01883 HMMetlyEutB_1_EUT1(pf06751) EUT1
01884 HMMetlyEutC_1_eut1(pf05985) EUT1 EP_NTERM
01885 HMMetlyEutC_1_eut1(pf05985) EUT1
01886 Tsp_rosa EUT1
01887 H_mulberry(pf00936pf16365) EUT1
01888 HMMreg12833_1_EUT1(#) EUT1
01889 HMMreg12833_1_EUT1(#) EUT1
01890 -pf01218 -
01891 -pf01520 -
01892 -pf00583 -
01893 -pf11143 -
01894 -pf06572 -
01895 -pf04261 -
01896 -# -
01897 -# -
BMC-P
BMC-H
BMC-T
Aldehyde dehydrogenase
PTAC
Signature enzyme
Alcohol dehydrogenase
Regulator

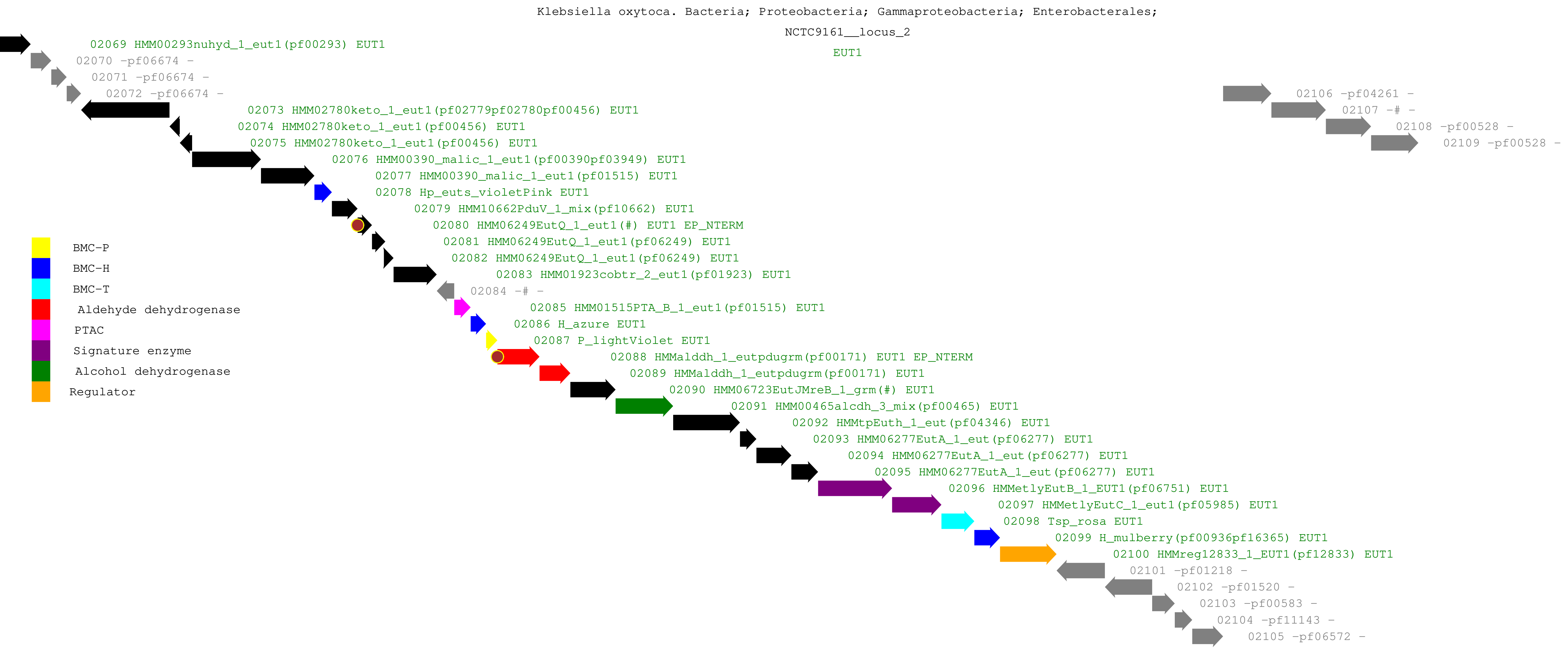
Klebsiella oxytoca. Bacteria; Proteobacteria; Gammaproteobacteria; Enterobacterales;
NCTC9161__locus_2
EUT1
02069 HMM00293nuhyd_1_eut1(pf00293) EUT1
02070 -pf06674 -
02071 -pf06674 -
02072 -pf06674 -
02073 HMM02780keto_1_eut1(pf02779pf02780pf00456) EUT1
02074 HMM02780keto_1_eut1(pf00456) EUT1
02075 HMM02780keto_1_eut1(pf00456) EUT1
02076 HMM00390_malic_1_eut1(pf00390pf03949) EUT1
02077 HMM00390_malic_1_eut1(pf01515) EUT1
02078 Hp_euts_violetPink EUT1
02079 HMM10662PduV_1_mix(pf10662) EUT1
02080 HMM06249EutQ_1_eut1(#) EUT1 EP_NTERM
02081 HMM06249EutQ_1_eut1(pf06249) EUT1
02082 HMM06249EutQ_1_eut1(pf06249) EUT1
02083 HMM01923cobtr_2_eut1(pf01923) EUT1
02084 -# -
02085 HMM01515PTA_B_1_eut1(pf01515) EUT1
02086 H_azure EUT1
02087 P_lightViolet EUT1
02088 HMMalddh_1_eutpdugrm(pf00171) EUT1 EP_NTERM
02089 HMMalddh_1_eutpdugrm(pf00171) EUT1
02090 HMM06723EutJMreB_1_grm(#) EUT1
02091 HMM00465alcdh_3_mix(pf00465) EUT1
02092 HMMtpEuth_1_eut(pf04346) EUT1
02093 HMM06277EutA_1_eut(pf06277) EUT1
02094 HMM06277EutA_1_eut(pf06277) EUT1
02095 HMM06277EutA_1_eut(pf06277) EUT1
02096 HMMetlyEutB_1_EUT1(pf06751) EUT1
02097 HMMetlyEutC_1_eut1(pf05985) EUT1
02098 Tsp_rosa EUT1
02099 H_mulberry(pf00936pf16365) EUT1
02100 HMMreg12833_1_EUT1(pf12833) EUT1
02101 -pf01218 -
02102 -pf01520 -
02103 -pf00583 -
02104 -pf11143 -
02105 -pf06572 -
02106 -pf04261 -
02107 -# -
02108 -pf00528 -
02109 -pf00528 -
BMC-P
BMC-H
BMC-T
Aldehyde dehydrogenase
PTAC
Signature enzyme
Alcohol dehydrogenase
Regulator

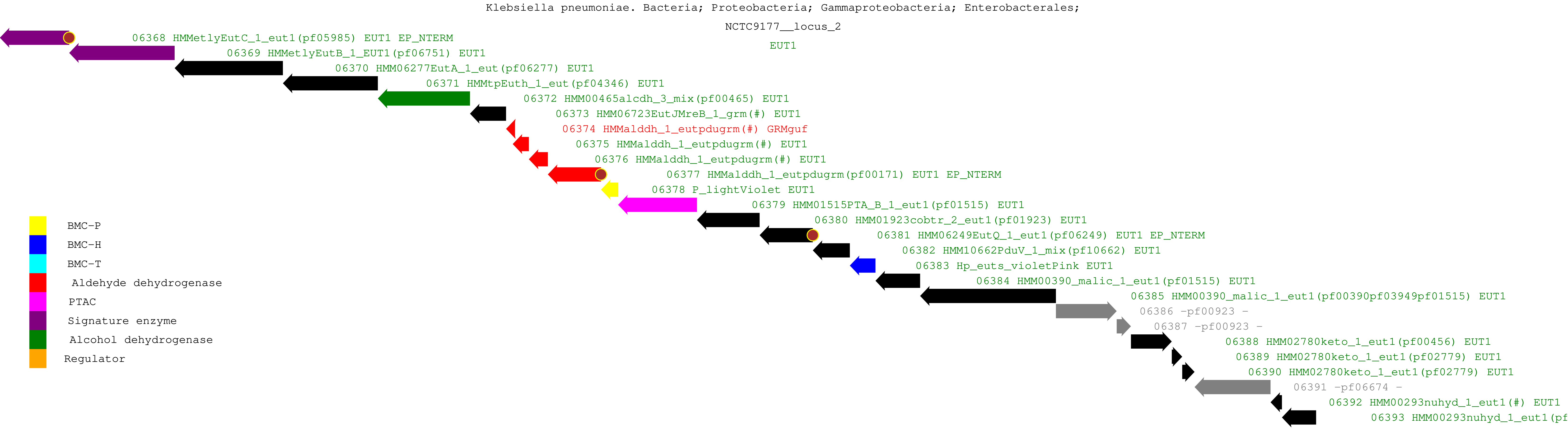
Klebsiella pneumoniae. Bacteria; Proteobacteria; Gammaproteobacteria; Enterobacterales;
NCTC9177__locus_2
EUT1
06368 HMMetlyEutC_1_eut1(pf05985) EUT1 EP_NTERM
06369 HMMetlyEutB_1_EUT1(pf06751) EUT1
06370 HMM06277EutA_1_eut(pf06277) EUT1
06371 HMMtpEuth_1_eut(pf04346) EUT1
06372 HMM00465alcdh_3_mix(pf00465) EUT1
06373 HMM06723EutJMreB_1_grm(#) EUT1
06374 HMMalddh_1_eutpdugrm(#) GRMguf
06375 HMMalddh_1_eutpdugrm(#) EUT1
06376 HMMalddh_1_eutpdugrm(#) EUT1
06377 HMMalddh_1_eutpdugrm(pf00171) EUT1 EP_NTERM
06378 P_lightViolet EUT1
06379 HMM01515PTA_B_1_eut1(pf01515) EUT1
06380 HMM01923cobtr_2_eut1(pf01923) EUT1
06381 HMM06249EutQ_1_eut1(pf06249) EUT1 EP_NTERM
06382 HMM10662PduV_1_mix(pf10662) EUT1
06383 Hp_euts_violetPink EUT1
06384 HMM00390_malic_1_eut1(pf01515) EUT1
06385 HMM00390_malic_1_eut1(pf00390pf03949pf01515) EUT1
06386 -pf00923 -
06387 -pf00923 -
06388 HMM02780keto_1_eut1(pf00456) EUT1
06389 HMM02780keto_1_eut1(pf02779) EUT1
06390 HMM02780keto_1_eut1(pf02779) EUT1
06391 -pf06674 -
06392 HMM00293nuhyd_1_eut1(#) EUT1
06393 HMM00293nuhyd_1_eut1(pf
BMC-P
BMC-H
BMC-T
Aldehyde dehydrogenase
PTAC
Signature enzyme
Alcohol dehydrogenase
Regulator

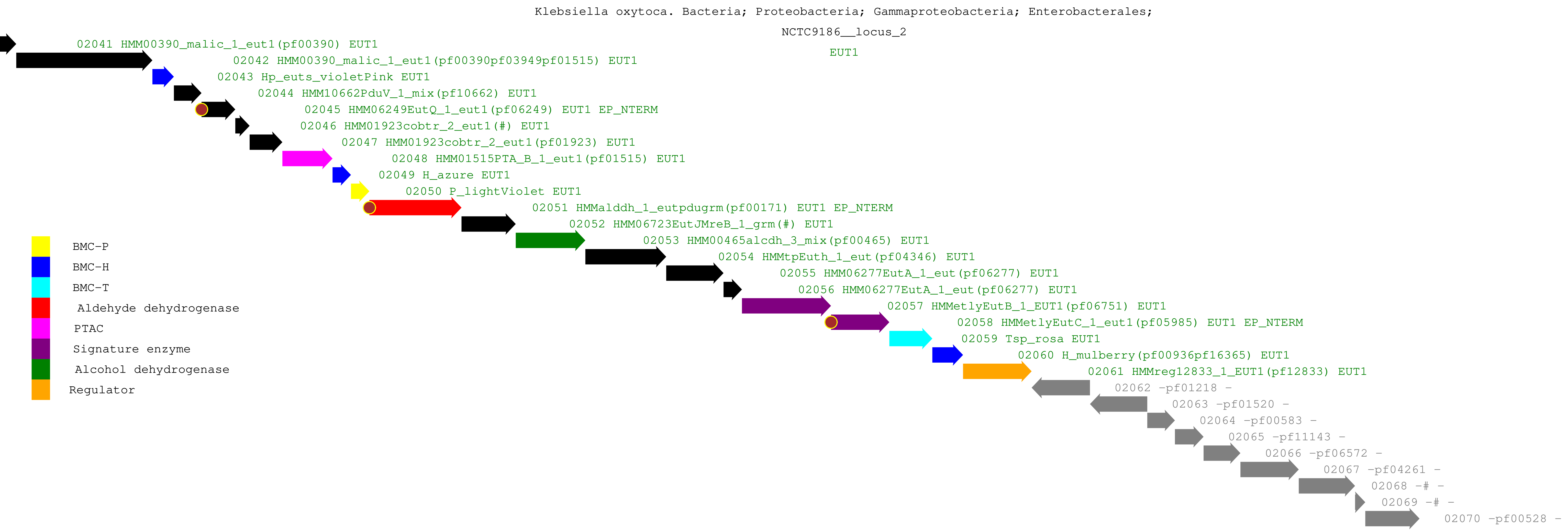
Klebsiella oxytoca. Bacteria; Proteobacteria; Gammaproteobacteria; Enterobacterales;
NCTC9186__locus_2
EUT1
02041 HMM00390_malic_1_eut1(pf00390) EUT1
02042 HMM00390_malic_1_eut1(pf00390pf03949pf01515) EUT1
02043 Hp_euts_violetPink EUT1
02044 HMM10662PduV_1_mix(pf10662) EUT1
02045 HMM06249EutQ_1_eut1(pf06249) EUT1 EP_NTERM
02046 HMM01923cobtr_2_eut1(#) EUT1
02047 HMM01923cobtr_2_eut1(pf01923) EUT1
02048 HMM01515PTA_B_1_eut1(pf01515) EUT1
02049 H_azure EUT1
02050 P_lightViolet EUT1
02051 HMMalddh_1_eutpdugrm(pf00171) EUT1 EP_NTERM
02052 HMM06723EutJMreB_1_grm(#) EUT1
02053 HMM00465alcdh_3_mix(pf00465) EUT1
02054 HMMtpEuth_1_eut(pf04346) EUT1
02055 HMM06277EutA_1_eut(pf06277) EUT1
02056 HMM06277EutA_1_eut(pf06277) EUT1
02057 HMMetlyEutB_1_EUT1(pf06751) EUT1
02058 HMMetlyEutC_1_eut1(pf05985) EUT1 EP_NTERM
02059 Tsp_rosa EUT1
02060 H_mulberry(pf00936pf16365) EUT1
02061 HMMreg12833_1_EUT1(pf12833) EUT1
02062 -pf01218 -
02063 -pf01520 -
02064 -pf00583 -
02065 -pf11143 -
02066 -pf06572 -
02067 -pf04261 -
02068 -# -
02069 -# -
02070 -pf00528 -
BMC-P
BMC-H
BMC-T
Aldehyde dehydrogenase
PTAC
Signature enzyme
Alcohol dehydrogenase
Regulator

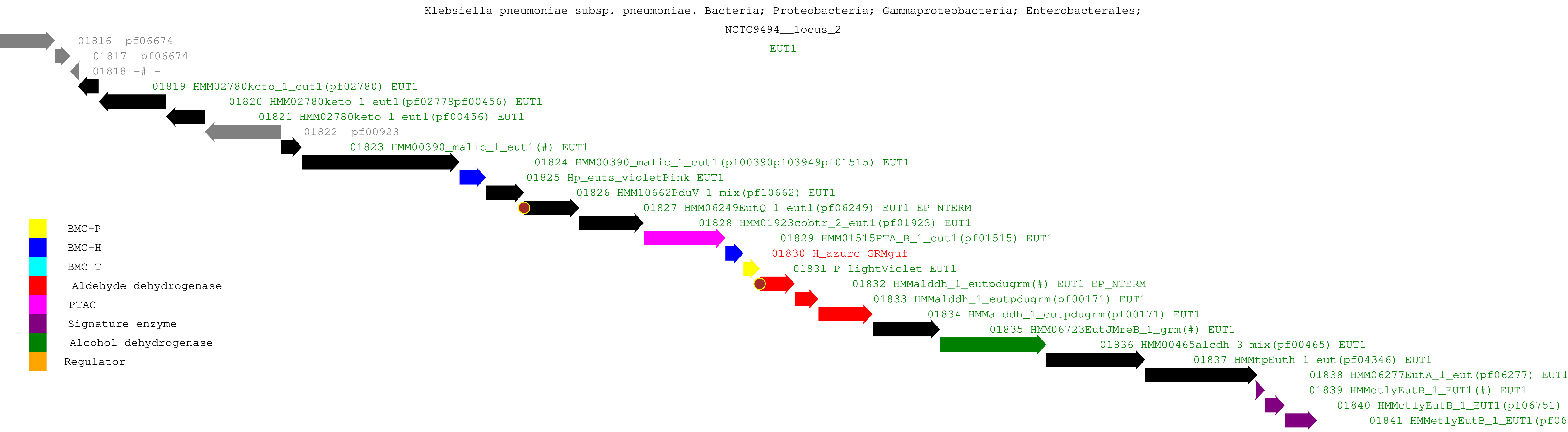
Klebsiella pneumoniae subsp. pneumoniae. Bacteria; Proteobacteria; Gammaproteobacteria; Enterobacterales;
NCTC9494__locus_2
EUT1
01816 -pf06674 -
01817 -pf06674 -
01818 -# -
01819 HMM02780keto_1_eut1(pf02780) EUT1
01820 HMM02780keto_1_eut1(pf02779pf00456) EUT1
01821 HMM02780keto_1_eut1(pf00456) EUT1
01822 -pf00923 -
01823 HMM00390_malic_1_eut1(#) EUT1
01824 HMM00390_malic_1_eut1(pf00390pf03949pf01515) EUT1
01825 Hp_euts_violetPink EUT1
01826 HMM10662PduV_1_mix(pf10662) EUT1
01827 HMM06249EutQ_1_eut1(pf06249) EUT1 EP_NTERM
01828 HMM01923cobtr_2_eut1(pf01923) EUT1
01829 HMM01515PTA_B_1_eut1(pf01515) EUT1
01830 H_azure GRMguf
01831 P_lightViolet EUT1
01832 HMMalddh_1_eutpdugrm(#) EUT1 EP_NTERM
01833 HMMalddh_1_eutpdugrm(pf00171) EUT1
01834 HMMalddh_1_eutpdugrm(pf00171) EUT1
01835 HMM06723EutJMreB_1_grm(#) EUT1
01836 HMM00465alcdh_3_mix(pf00465) EUT1
01837 HMMtpEuth_1_eut(pf04346) EUT1
01838 HMM06277EutA_1_eut(pf06277) EUT1
01839 HMMetlyEutB_1_EUT1(#) EUT1
01840 HMMetlyEutB_1_EUT1(pf06751)
01841 HMMetlyEutB_1_EUT1(pf06
BMC-P
BMC-H
BMC-T
Aldehyde dehydrogenase
PTAC
Signature enzyme
Alcohol dehydrogenase
Regulator

Klebsiella pneumoniae subsp. pneumoniae. Bacteria; Proteobacteria; Gammaproteobacteria; Enterobacterales;

NCTC9504__locus_2

EUT1

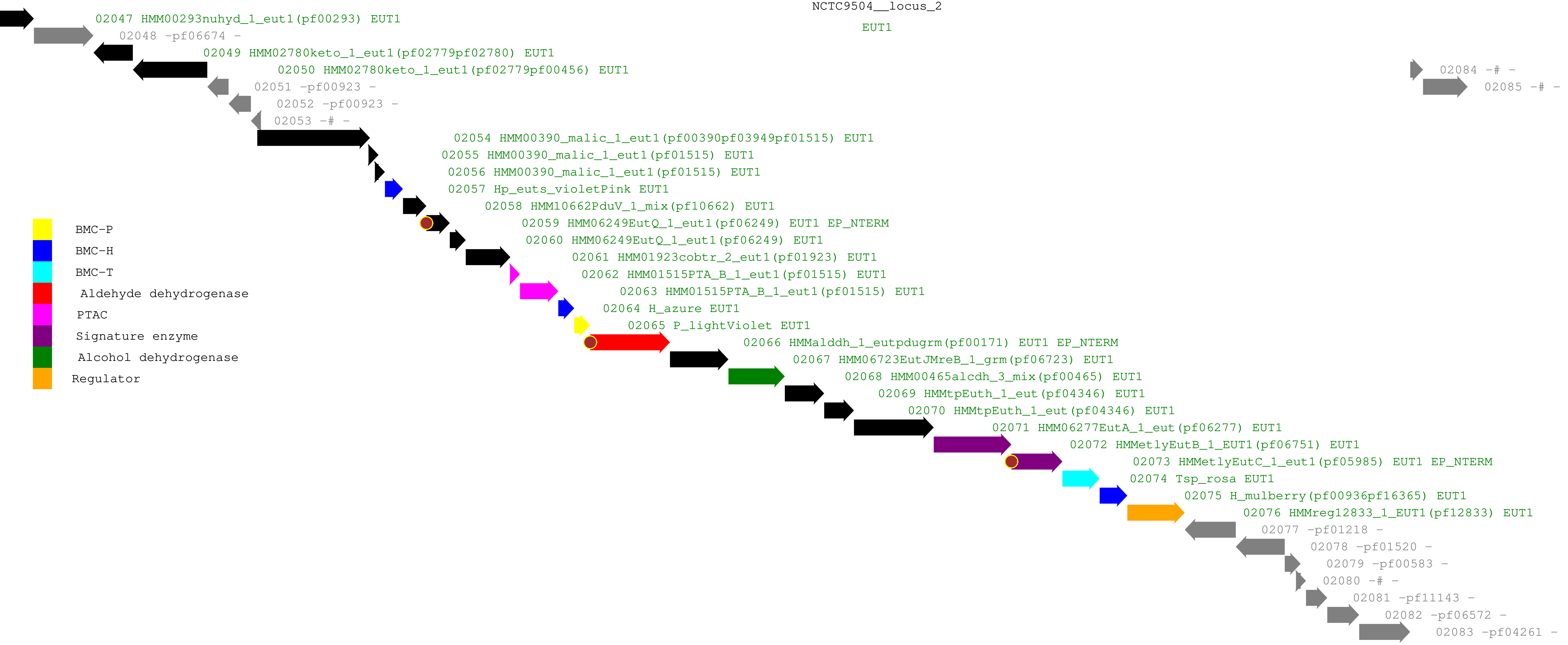

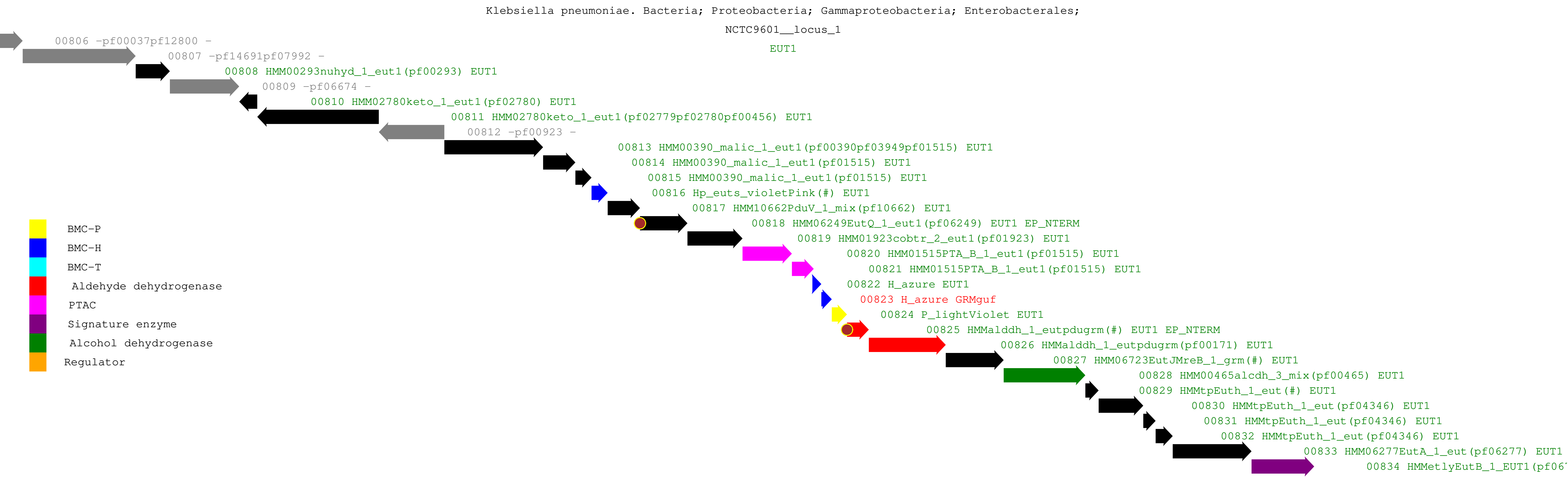
Klebsiella pneumoniae. Bacteria; Proteobacteria; Gammaproteobacteria; Enterobacterales;
NCTC9601__locus_1
EUT1
00806 -pf00037pf12800 -
00807 -pf14691pf07992 -
00808 HMM00293nuhyd_1_eut1(pf00293) EUT1
00809 -pf06674 -
00810 HMM02780keto_1_eut1(pf02780) EUT1
00811 HMM02780keto_1_eut1(pf02779pf02780pf00456) EUT1
00812 -pf00923 -
00813 HMM00390_malic_1_eut1(pf00390pf03949pf01515) EUT1
00814 HMM00390_malic_1_eut1(pf01515) EUT1
00815 HMM00390_malic_1_eut1(pf01515) EUT1
00816 Hp_euts_violetPink(#) EUT1
00817 HMM10662PduV_1_mix(pf10662) EUT1
00818 HMM06249EutQ_1_eut1(pf06249) EUT1 EP_NTERM
00819 HMM01923cobtr_2_eut1(pf01923) EUT1
00820 HMM01515PTA_B_1_eut1(pf01515) EUT1
00821 HMM01515PTA_B_1_eut1(pf01515) EUT1
00822 H_azure EUT1
00823 H_azure GRMguf
00824 P_lightViolet EUT1
00825 HMMalddh_1_eutpdugrm(#) EUT1 EP_NTERM
00826 HMMalddh_1_eutpdugrm(pf00171) EUT1
00827 HMM06723EutJMreB_1_grm(#) EUT1
00828 HMM00465alcdh_3_mix(pf00465) EUT1
00829 HMMtpEuth_1_eut(#) EUT1
00830 HMMtpEuth_1_eut(pf04346) EUT1
00831 HMMtpEuth_1_eut(pf04346) EUT1
00832 HMMtpEuth_1_eut(pf04346) EUT1
00833 HMM06277EutA_1_eut(pf06277) EUT1
00834 HMMetlyEutB_1_EUT1(pf06
BMC-P
BMC-H
BMC-T
Aldehyde dehydrogenase
PTAC
Signature enzyme
Alcohol dehydrogenase
Regulator

Klebsiella pneumoniae. Bacteria; Proteobacteria; Gammaproteobacteria; Enterobacterales;

NCTC9645__locus_1

EUT1

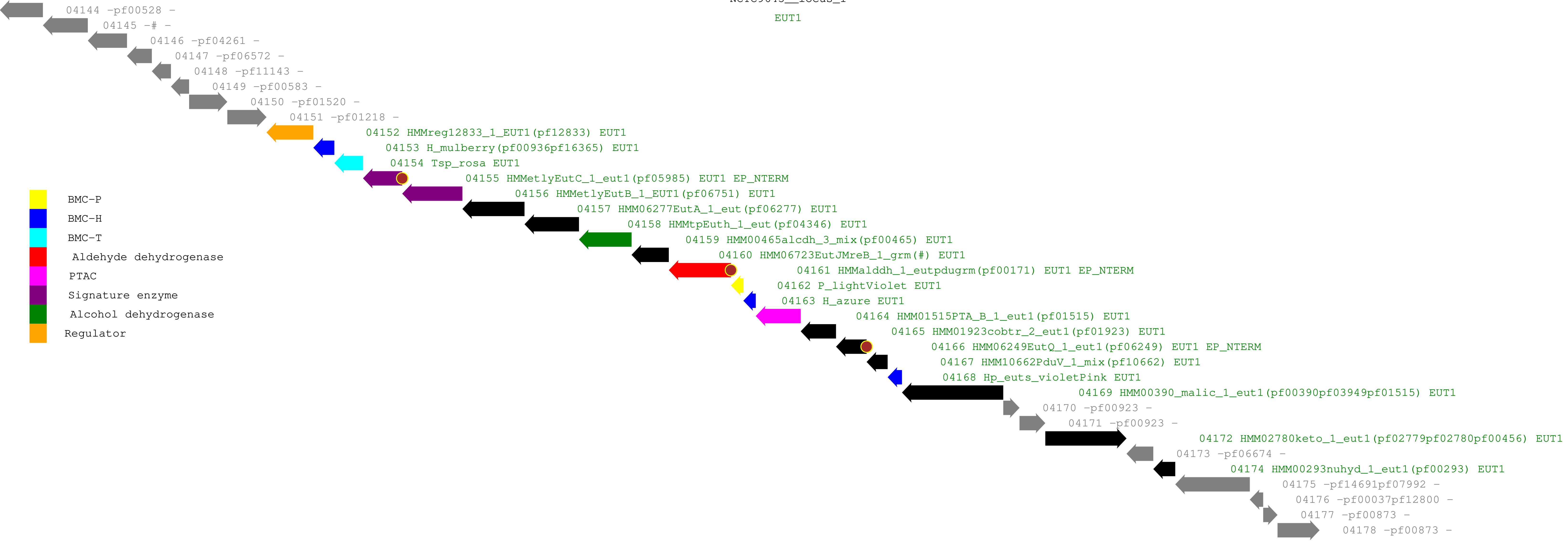

Klebsiella aerogenes (Enterobacter aerogenes). Bacteria; Proteobacteria; Gammaproteobacteria; Enterobacterales;

NCTC9652__locus_2

EUT1

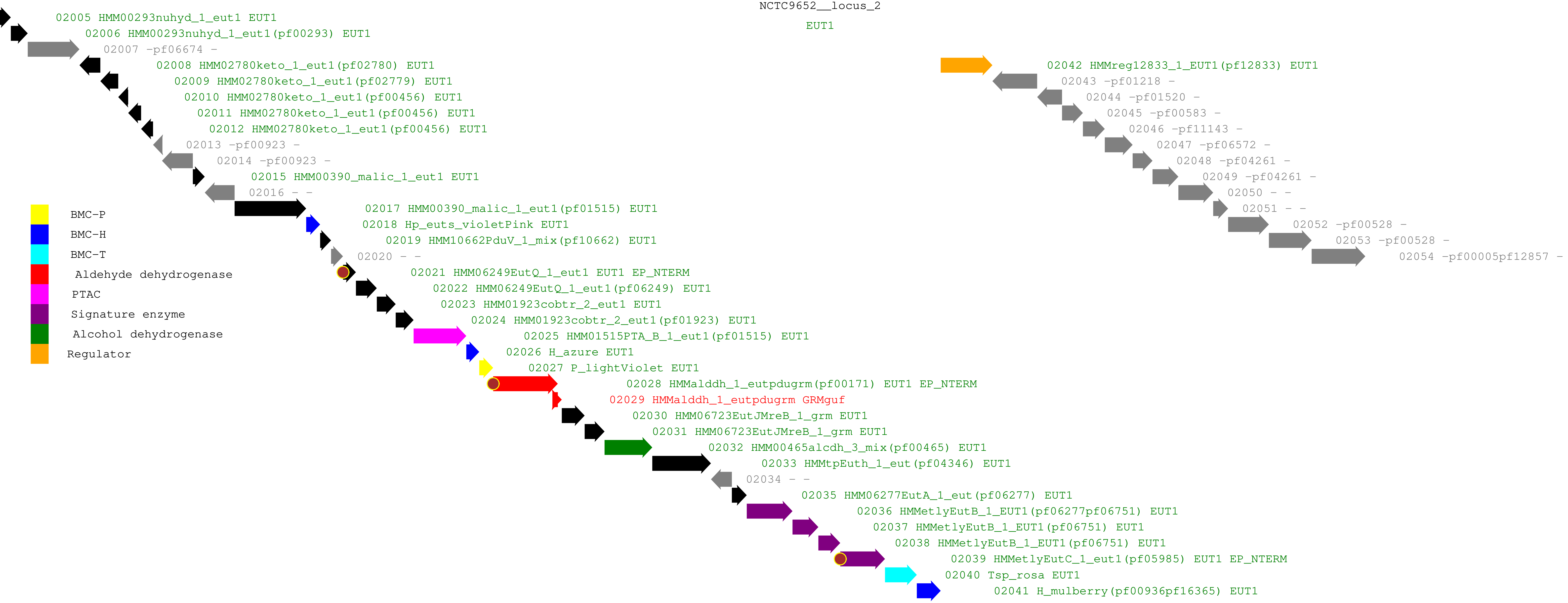

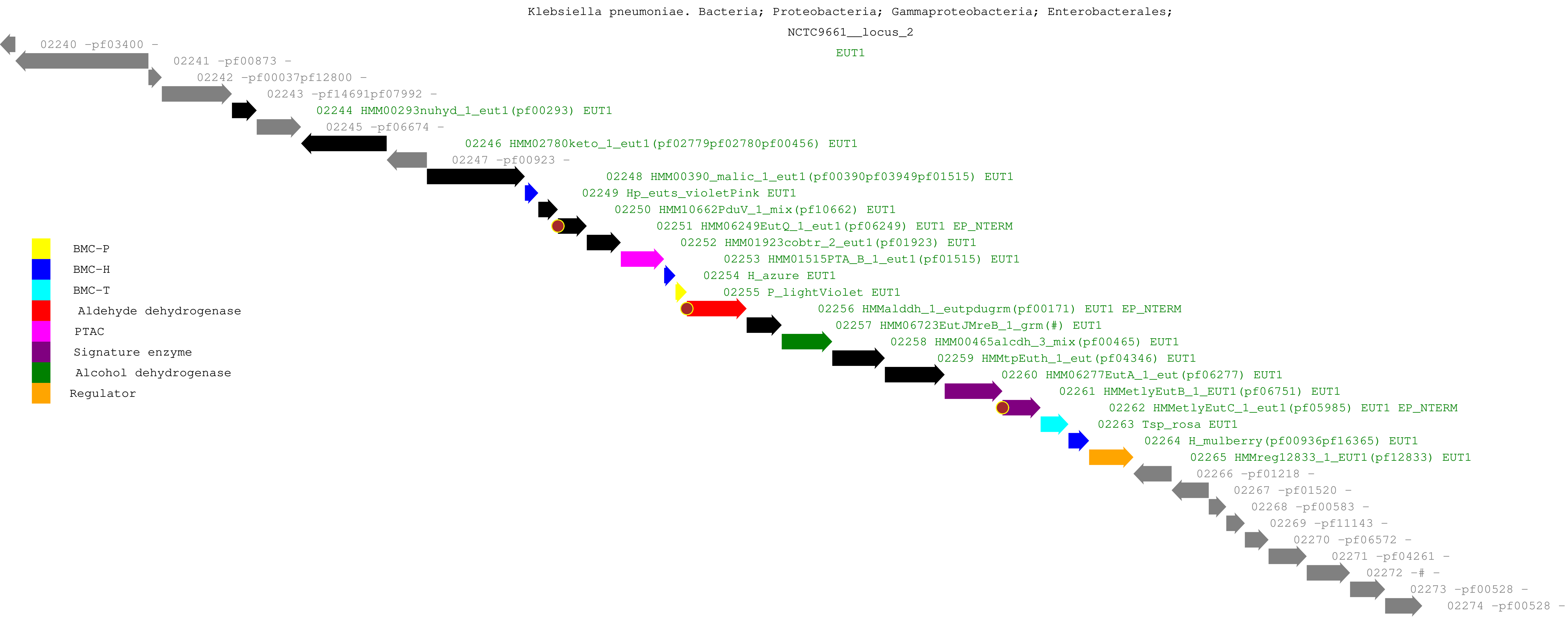
Klebsiella pneumoniae. Bacteria; Proteobacteria; Gammaproteobacteria; Enterobacterales;
NCTC9661__locus_2
EUT1
02240 -pf03400 -
02241 -pf00873 -
02242 -pf00037pf12800 -
02243 -pf14691pf07992 -
02244 HMM00293nuhyd_1_eut1(pf00293) EUT1
02245 -pf06674 -
02246 HMM02780keto_1_eut1(pf02779pf02780pf00456) EUT1
02247 -pf00923 -
02248 HMM00390_malic_1_eut1(pf00390pf03949pf01515) EUT1
02249 Hp_euts_violetPink EUT1
02250 HMM10662PduV_1_mix(pf10662) EUT1
02251 HMM06249EutQ_1_eut1(pf06249) EUT1 EP_NTERM
02252 HMM01923cobtr_2_eut1(pf01923) EUT1
02253 HMM01515PTA_B_1_eut1(pf01515) EUT1
02254 H_azure EUT1
02255 P_lightViolet EUT1
02256 HMMalddh_1_eutpdugrm(pf00171) EUT1 EP_NTERM
02257 HMM06723EutJMreB_1_grm(#) EUT1
02258 HMM00465alcdh_3_mix(pf00465) EUT1
02259 HMMtpEuth_1_eut(pf04346) EUT1
02260 HMM06277EutA_1_eut(pf06277) EUT1
02261 HMMetlyEutB_1_EUT1(pf06751) EUT1
02262 HMMetlyEutC_1_eut1(pf05985) EUT1 EP_NTERM
02263 Tsp_rosa EUT1
02264 H_mulberry(pf00936pf16365) EUT1
02265 HMMreg12833_1_EUT1(pf12833) EUT1
02266 -pf01218 -
02267 -pf01520 -
02268 -pf00583 -
02269 -pf11143 -
02270 -pf06572 -
02271 -pf04261 -
02272 -# -
02273 -pf00528 -
02274 -pf00528 -
BMC-P
BMC-H
BMC-T
Aldehyde dehydrogenase
PTAC
Signature enzyme
Alcohol dehydrogenase
Regulator

Escherichia coli. Bacteria; Proteobacteria; Gammaproteobacteria; Enterobacterales;

NCTC9775__locus_2

EUT1

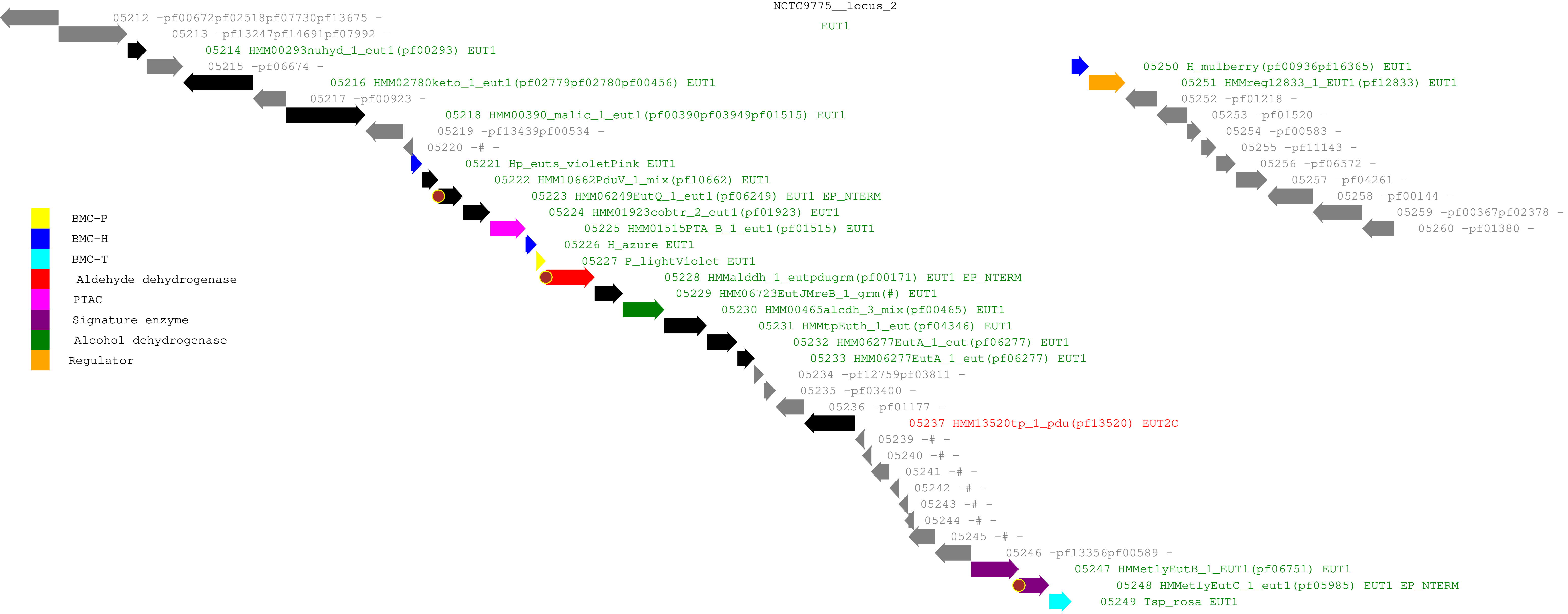

Escherichia coli. Bacteria; Proteobacteria; Gammaproteobacteria; Enterobacterales;

NCTC9777__locus_1

EUT1

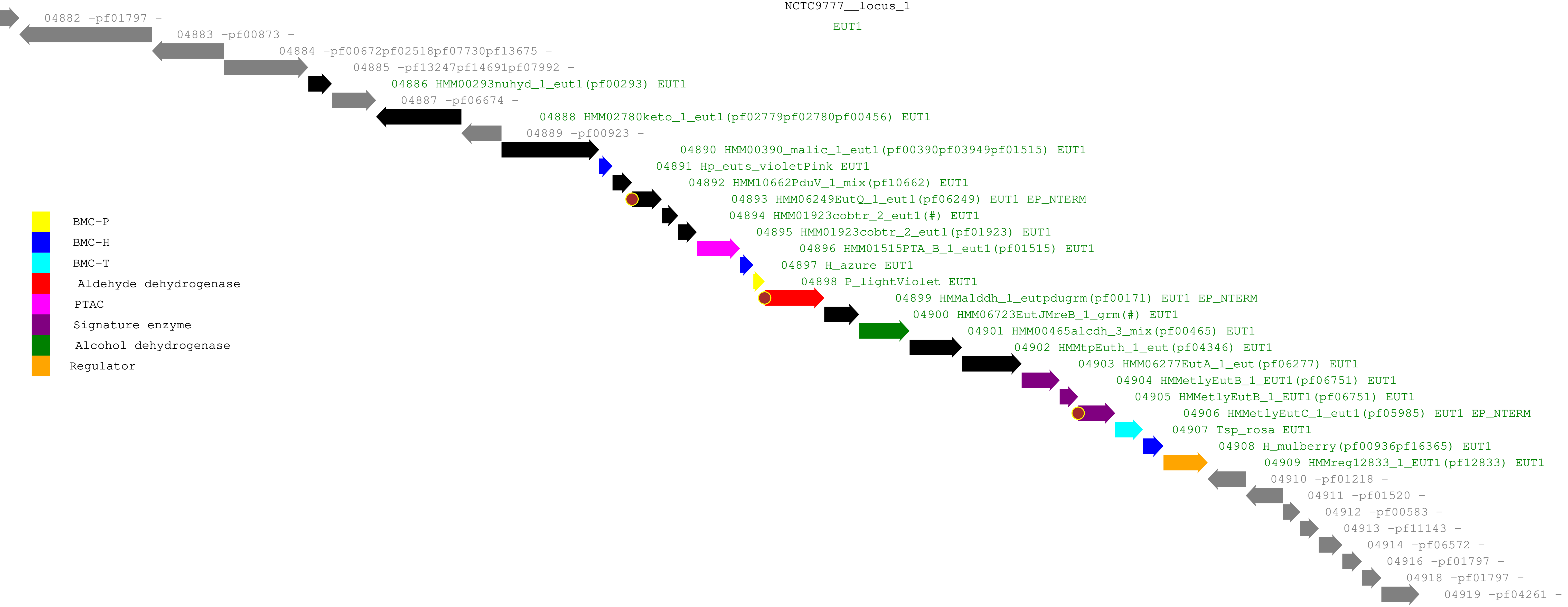

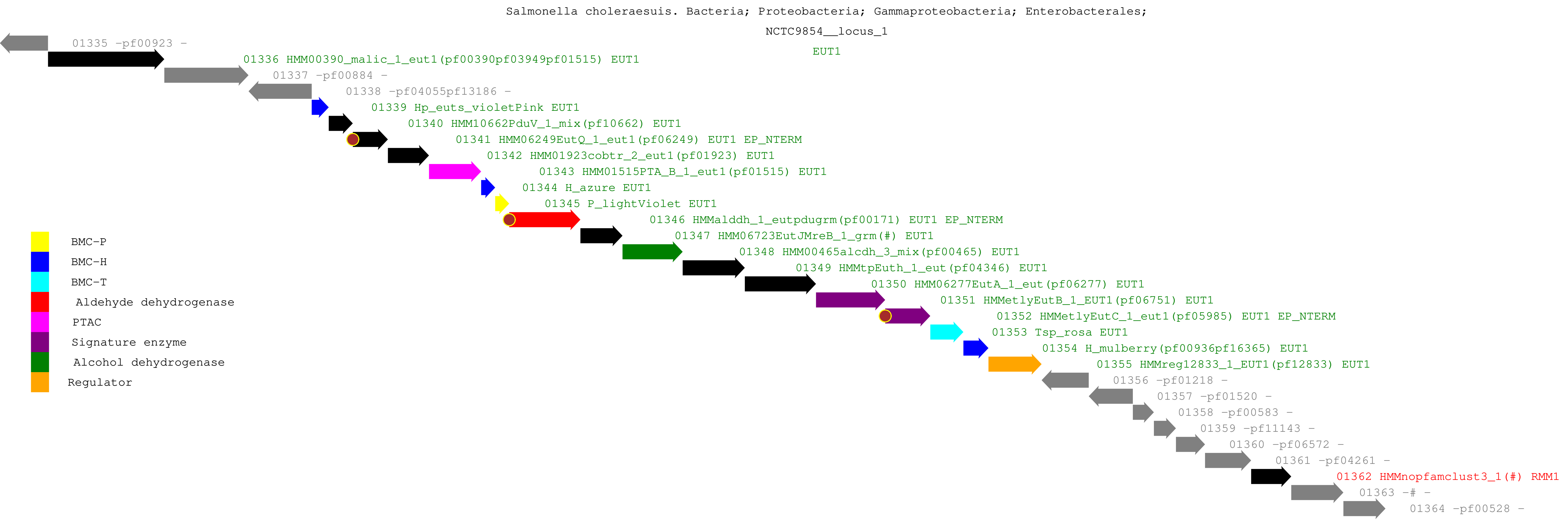

Salmonella choleraesuis. Bacteria; Proteobacteria; Gammaproteobacteria; Enterobacterales;
NCTC9854__locus_1
EUT1
01335 -pf00923 -
01336 HMM00390_malic_1_eut1(pf00390pf03949pf01515) EUT1
01337 -pf00884 -
01338 -pf04055pf13186 -
01339 Hp_euts_violetPink EUT1
01340 HMM10662PduV_1_mix(pf10662) EUT1
01341 HMM06249EutQ_1_eut1(pf06249) EUT1 EP_NTERM
01342 HMM01923cobtr_2_eut1(pf01923) EUT1
01343 HMM01515PTA_B_1_eut1(pf01515) EUT1
01344 H_azure EUT1
01345 P_lightViolet EUT1
01346 HMMalddh_1_eutpdugrm(pf00171) EUT1 EP_NTERM
01347 HMM06723EutJMreB_1_grm(#) EUT1
01348 HMM00465alcdh_3_mix(pf00465) EUT1
01349 HMMtpEuth_1_eut(pf04346) EUT1
01350 HMM06277EutA_1_eut(pf06277) EUT1
01351 HMMetlyEutB_1_EUT1(pf06751) EUT1
01352 HMMetlyEutC_1_eut1(pf05985) EUT1 EP_NTERM
01353 Tsp_rosa EUT1
01354 H_mulberry(pf00936pf16365) EUT1
01355 HMMreg12833_1_EUT1(pf12833) EUT1
01356 -pf01218 -
01357 -pf01520 -
01358 -pf00583 -
01359 -pf11143 -
01360 -pf06572 -
01361 -pf04261 -
01362 HMMnopfamclust3_1(#) RMM1
01363 -# -
01364 -pf00528 -
BMC-P
BMC-H
BMC-T
Aldehyde dehydrogenase
PTAC
Signature enzyme
Alcohol dehydrogenase
Regulator

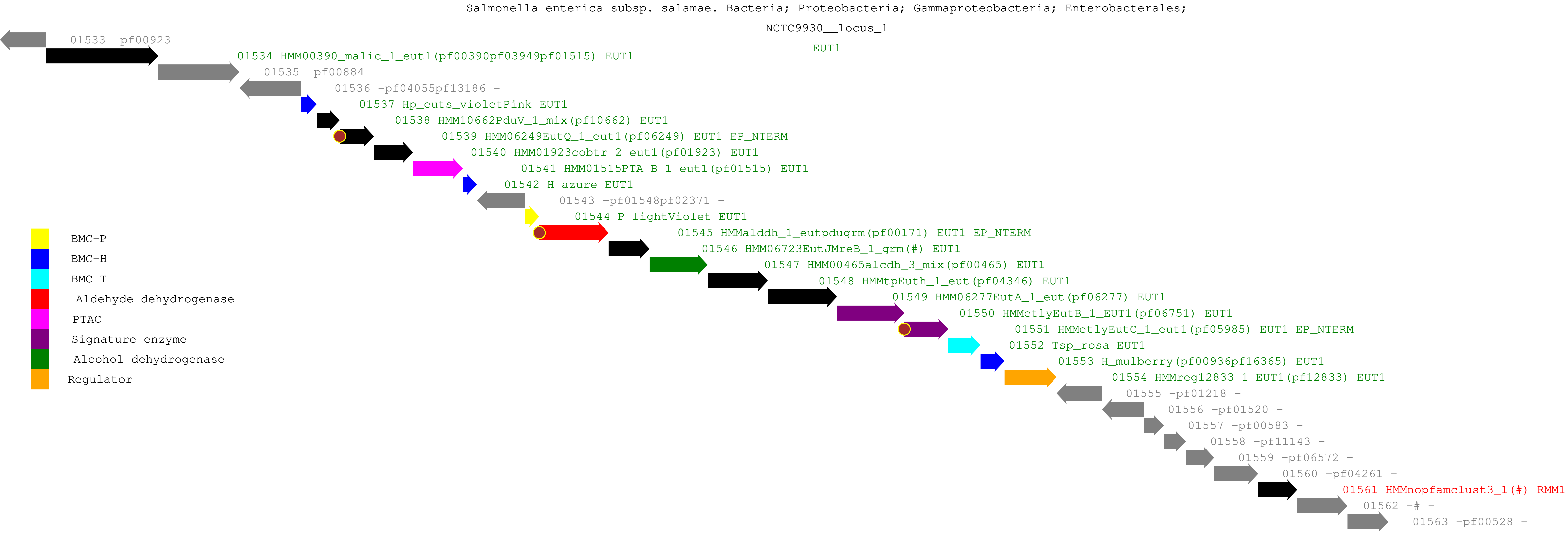
Salmonella enterica subsp. salamae. Bacteria; Proteobacteria; Gammaproteobacteria; Enterobacterales;
NCTC9930__locus_1
EUT1
01533 -pf00923 -
01534 HMM00390_malic_1_eut1(pf00390pf03949pf01515) EUT1
01535 -pf00884 -
01536 -pf04055pf13186 -
01537 Hp_euts_violetPink EUT1
01538 HMM10662PduV_1_mix(pf10662) EUT1
01539 HMM06249EutQ_1_eut1(pf06249) EUT1 EP_NTERM
01540 HMM01923cobtr_2_eut1(pf01923) EUT1
01541 HMM01515PTA_B_1_eut1(pf01515) EUT1
01542 H_azure EUT1
01543 -pf01548pf02371 -
01544 P_lightViolet EUT1
01545 HMMalddh_1_eutpdugrm(pf00171) EUT1 EP_NTERM
01546 HMM06723EutJMreB_1_grm(#) EUT1
01547 HMM00465alcdh_3_mix(pf00465) EUT1
01548 HMMtpEuth_1_eut(pf04346) EUT1
01549 HMM06277EutA_1_eut(pf06277) EUT1
01550 HMMetlyEutB_1_EUT1(pf06751) EUT1
01551 HMMetlyEutC_1_eut1(pf05985) EUT1 EP_NTERM
01552 Tsp_rosa EUT1
01553 H_mulberry(pf00936pf16365) EUT1
01554 HMMreg12833_1_EUT1(pf12833) EUT1
01555 -pf01218 -
01556 -pf01520 -
01557 -pf00583 -
01558 -pf11143 -
01559 -pf06572 -
01560 -pf04261 -
01561 HMMnopfamclust3_1(#) RMM1
01562 -# -
01563 -pf00528 -
BMC-P
BMC-H
BMC-T
Aldehyde dehydrogenase
PTAC
Signature enzyme
Alcohol dehydrogenase
Regulator

Escherichia coli. Bacteria; Proteobacteria; Gammaproteobacteria; Enterobacterales;

NCTC9962__locus_1

EUT1

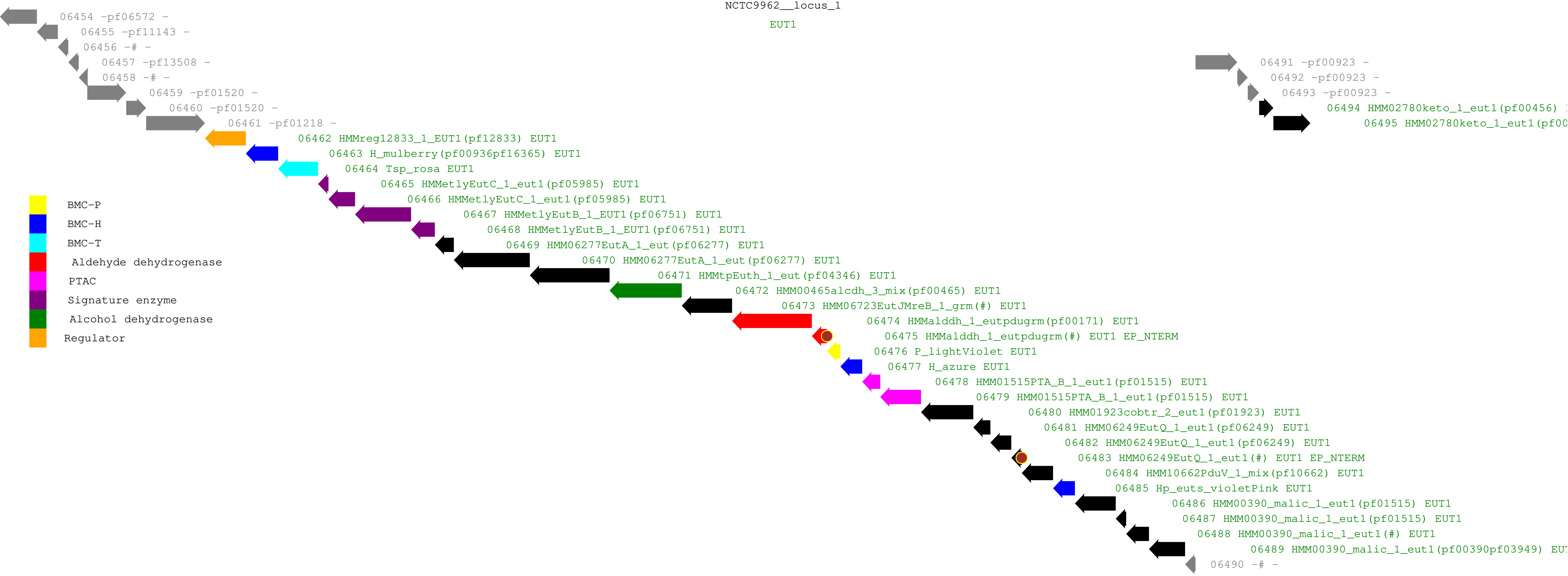

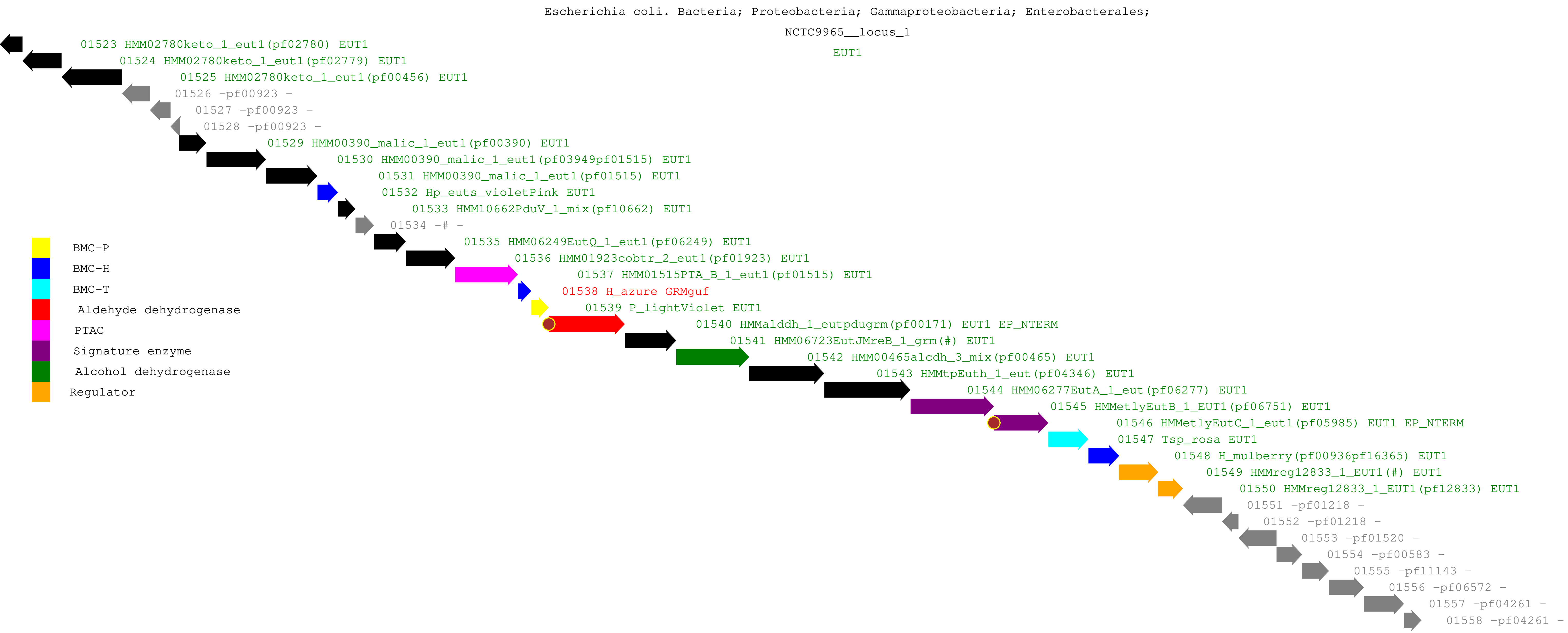

Escherichia coli. Bacteria; Proteobacteria; Gammaproteobacteria; Enterobacterales;
NCTC9965__locus_1
EUT1
01523 HMM02780keto_1_eut1(pf02780) EUT1
01524 HMM02780keto_1_eut1(pf02779) EUT1
01525 HMM02780keto_1_eut1(pf00456) EUT1
01526 -pf00923 -
01527 -pf00923 -
01528 -pf00923 -
01529 HMM00390_malic_1_eut1(pf00390) EUT1
01530 HMM00390_malic_1_eut1(pf03949pf01515) EUT1
01531 HMM00390_malic_1_eut1(pf01515) EUT1
01532 Hp_euts_violetPink EUT1
01533 HMM10662PduV_1_mix(pf10662) EUT1
01534 -# -
01535 HMM06249EutQ_1_eut1(pf06249) EUT1
01536 HMM01923cobtr_2_eut1(pf01923) EUT1
01537 HMM01515PTA_B_1_eut1(pf01515) EUT1
01538 H_azure GRMguf
01539 P_lightViolet EUT1
01540 HMMalddh_1_eutpdugrm(pf00171) EUT1 EP_NTERM
01541 HMM06723EutJMreB_1_grm(#) EUT1
01542 HMM00465alcdh_3_mix(pf00465) EUT1
01543 HMMtpEuth_1_eut(pf04346) EUT1
01544 HMM06277EutA_1_eut(pf06277) EUT1
01545 HMMetlyEutB_1_EUT1(pf06751) EUT1
01546 HMMetlyEutC_1_eut1(pf05985) EUT1 EP_NTERM
01547 Tsp_rosa EUT1
01548 H_mulberry(pf00936pf16365) EUT1
01549 HMMreg12833_1_EUT1(#) EUT1
01550 HMMreg12833_1_EUT1(pf12833) EUT1
01551 -pf01218 -
01552 -pf01218 -
01553 -pf01520 -
01554 -pf00583 -
01555 -pf11143 -
01556 -pf06572 -
01557 -pf04261 -
01558 -pf04261 -
BMC-P
BMC-H
BMC-T
Aldehyde dehydrogenase
PTAC
Signature enzyme
Alcohol dehydrogenase
Regulator

Salmonella choleraesuis. Bacteria; Proteobacteria; Gammaproteobacteria; Enterobacterales;

NGUA18__locus_1

EUT1

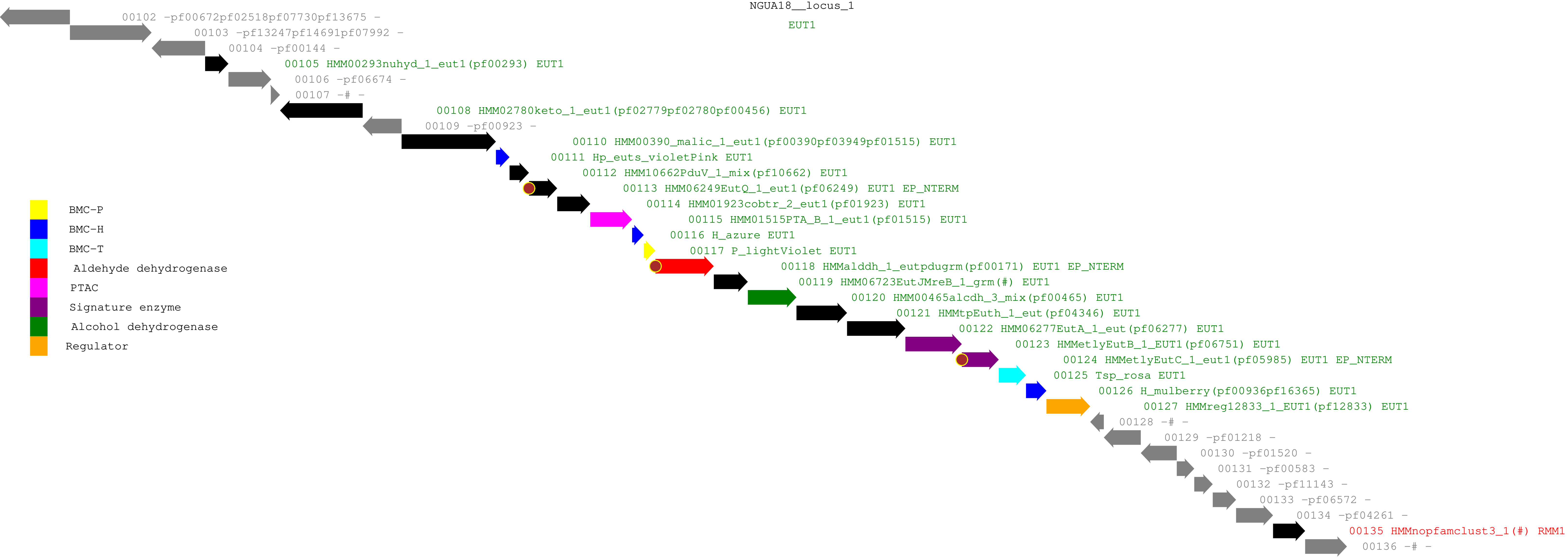

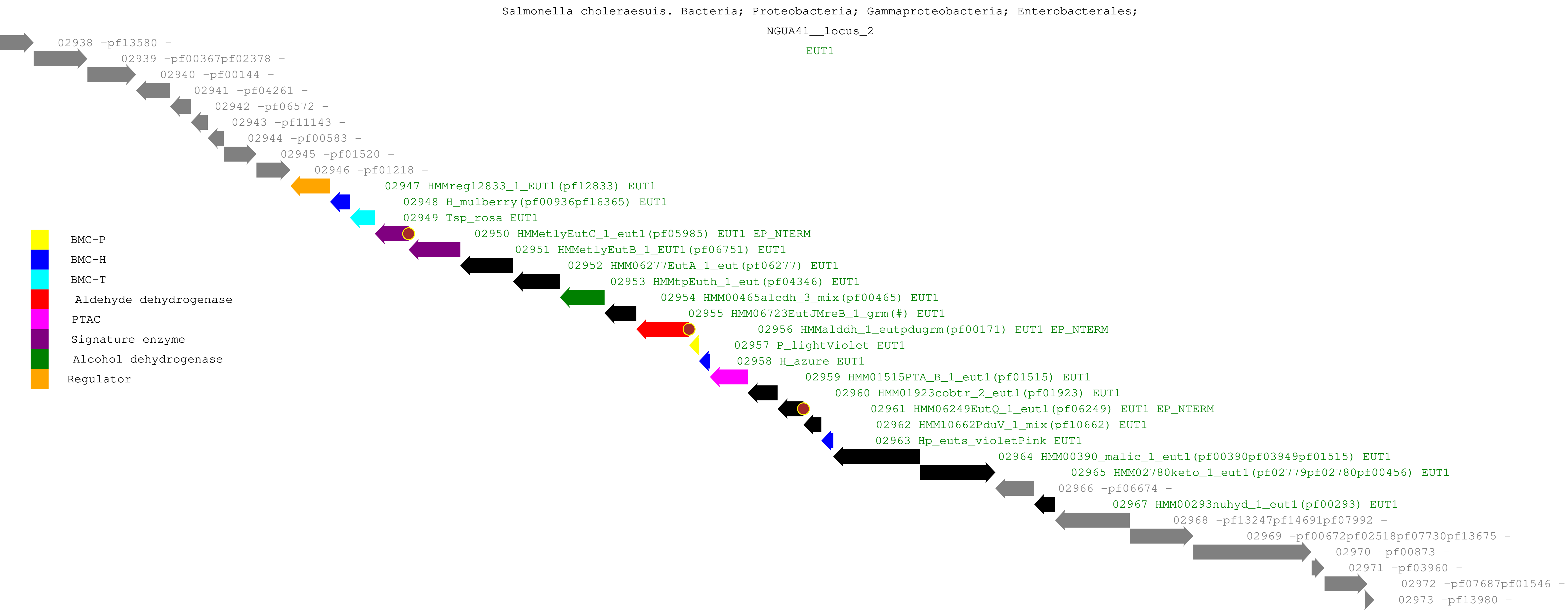
Salmonella choleraesuis. Bacteria; Proteobacteria; Gammaproteobacteria; Enterobacterales;
NGUA41__locus_2
EUT1
02938 -pf13580 -
02939 -pf00367pf02378 -
02940 -pf00144 -
02941 -pf04261 -
02942 -pf06572 -
02943 -pf11143 -
02944 -pf00583 -
02945 -pf01520 -
02946 -pf01218 -
02947 HMMreg12833_1_EUT1(pf12833) EUT1
02948 H_mulberry(pf00936pf16365) EUT1
02949 Tsp_rosa EUT1
02950 HMMetlyEutC_1_eut1(pf05985) EUT1 EP_NTERM
02951 HMMetlyEutB_1_EUT1(pf06751) EUT1
02952 HMM06277EutA_1_eut(pf06277) EUT1
02953 HMMtpEuth_1_eut(pf04346) EUT1
02954 HMM00465alcdh_3_mix(pf00465) EUT1
02955 HMM06723EutJMreB_1_grm(#) EUT1
02956 HMMalddh_1_eutpdugrm(pf00171) EUT1 EP_NTERM
02957 P_lightViolet EUT1
02958 H_azure EUT1
02959 HMM01515PTA_B_1_eut1(pf01515) EUT1
02960 HMM01923cobtr_2_eut1(pf01923) EUT1
02961 HMM06249EutQ_1_eut1(pf06249) EUT1 EP_NTERM
02962 HMM10662PduV_1_mix(pf10662) EUT1
02963 Hp_euts_violetPink EUT1
02964 HMM00390_malic_1_eut1(pf00390pf03949pf01515) EUT1
02965 HMM02780keto_1_eut1(pf02779pf02780pf00456) EUT1
02966 -pf06674 -
02967 HMM00293nuhyd_1_eut1(pf00293) EUT1
02968 -pf13247pf14691pf07992 -
02969 -pf00672pf02518pf07730pf13675 -
02970 -pf00873 -
02971 -pf03960 -
02972 -pf07687pf01546 -
02973 -pf13980 -
BMC-P
BMC-H
BMC-T
Aldehyde dehydrogenase
PTAC
Signature enzyme
Alcohol dehydrogenase
Regulator

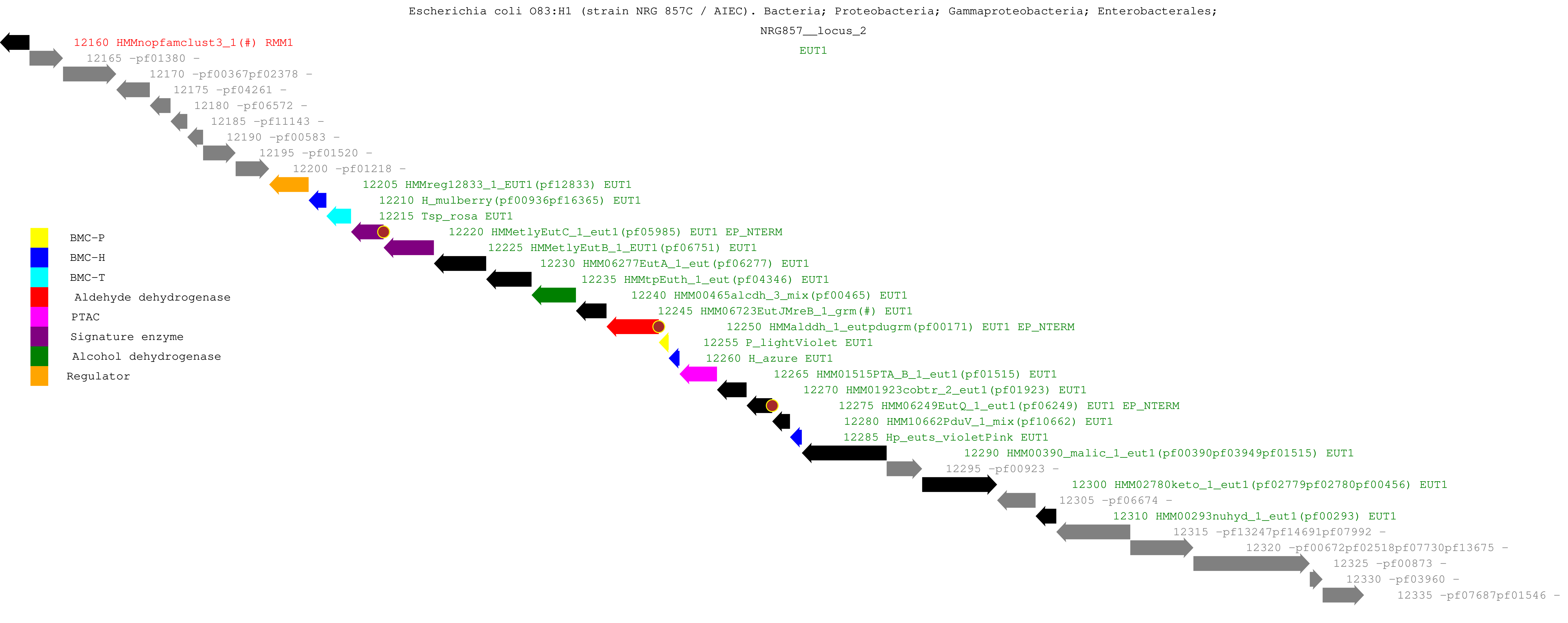
Escherichia coli O83:H1 (strain NRG 857C / AIEC). Bacteria; Proteobacteria; Gammaproteobacteria; Enterobacterales;
NRG857__locus_2
EUT1
12160 HMMnopfamclust3_1(#) RMM1
12165 -pf01380 -
12170 -pf00367pf02378 -
12175 -pf04261 -
12180 -pf06572 -
12185 -pf11143 -
12190 -pf00583 -
12195 -pf01520 -
12200 -pf01218 -
12205 HMMreg12833_1_EUT1(pf12833) EUT1
12210 H_mulberry(pf00936pf16365) EUT1
12215 Tsp_rosa EUT1
12220 HMMetlyEutC_1_eut1(pf05985) EUT1 EP_NTERM
12225 HMMetlyEutB_1_EUT1(pf06751) EUT1
12230 HMM06277EutA_1_eut(pf06277) EUT1
12235 HMMtpEuth_1_eut(pf04346) EUT1
12240 HMM00465alcdh_3_mix(pf00465) EUT1
12245 HMM06723EutJMreB_1_grm(#) EUT1
12250 HMMalddh_1_eutpdugrm(pf00171) EUT1 EP_NTERM
12255 P_lightViolet EUT1
12260 H_azure EUT1
12265 HMM01515PTA_B_1_eut1(pf01515) EUT1
12270 HMM01923cobtr_2_eut1(pf01923) EUT1
12275 HMM06249EutQ_1_eut1(pf06249) EUT1 EP_NTERM
12280 HMM10662PduV_1_mix(pf10662) EUT1
12285 Hp_euts_violetPink EUT1
12290 HMM00390_malic_1_eut1(pf00390pf03949pf01515) EUT1
12295 -pf00923 -
12300 HMM02780keto_1_eut1(pf02779pf02780pf00456) EUT1
12305 -pf06674 -
12310 HMM00293nuhyd_1_eut1(pf00293) EUT1
12315 -pf13247pf14691pf07992 -
12320 -pf00672pf02518pf07730pf13675 -
12325 -pf00873 -
12330 -pf03960 -
12335 -pf07687pf01546 -
BMC-P
BMC-H
BMC-T
Aldehyde dehydrogenase
PTAC
Signature enzyme
Alcohol dehydrogenase
Regulator

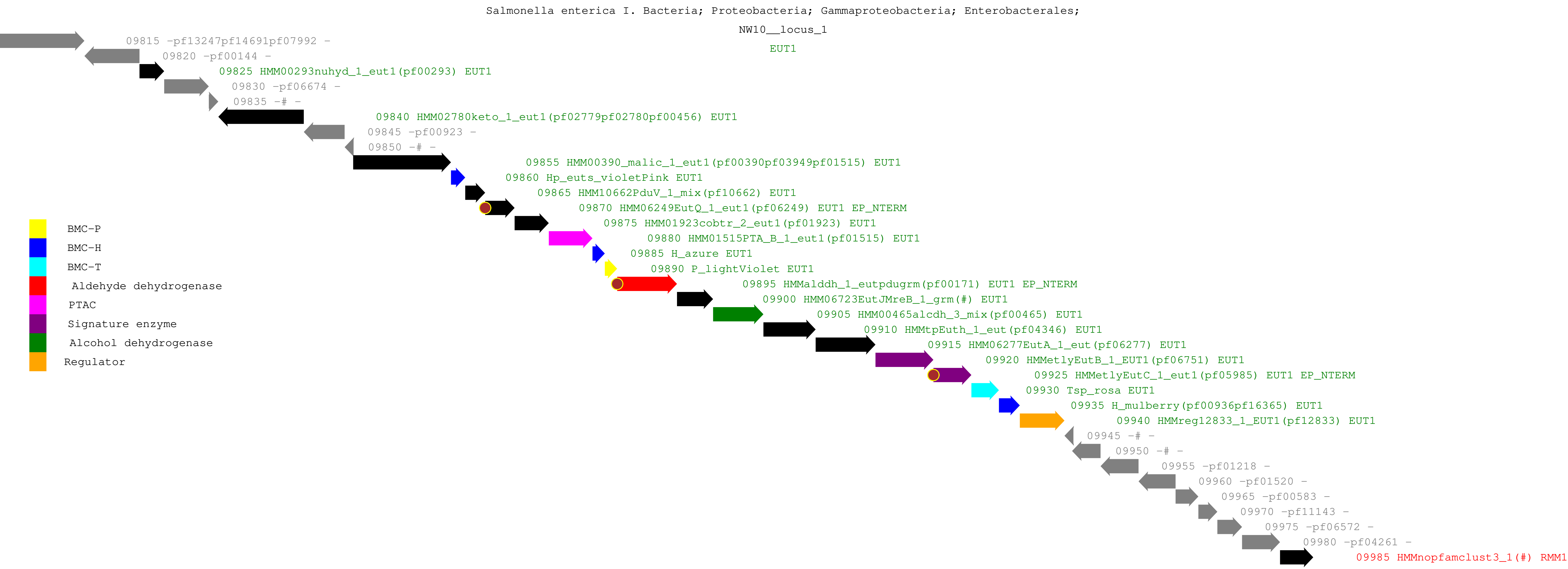

Salmonella enterica I. Bacteria; Proteobacteria; Gammaproteobacteria; Enterobacterales;
NW10__locus_1
EUT1
09815 -pf13247pf14691pf07992 -
09820 -pf00144 -
09825 HMM00293nuhyd_1_eut1(pf00293) EUT1
09830 -pf06674 -
09835 -# -
09840 HMM02780keto_1_eut1(pf02779pf02780pf00456) EUT1
09845 -pf00923 -
09850 -# -
09855 HMM00390_malic_1_eut1(pf00390pf03949pf01515) EUT1
09860 Hp_euts_violetPink EUT1
09865 HMM10662PduV_1_mix(pf10662) EUT1
09870 HMM06249EutQ_1_eut1(pf06249) EUT1 EP_NTERM
09875 HMM01923cobtr_2_eut1(pf01923) EUT1
09880 HMM01515PTA_B_1_eut1(pf01515) EUT1
09885 H_azure EUT1
09890 P_lightViolet EUT1
09895 HMMalddh_1_eutpdugrm(pf00171) EUT1 EP_NTERM
09900 HMM06723EutJMreB_1_grm(#) EUT1
09905 HMM00465alcdh_3_mix(pf00465) EUT1
09910 HMMtpEuth_1_eut(pf04346) EUT1
09915 HMM06277EutA_1_eut(pf06277) EUT1
09920 HMMetlyEutB_1_EUT1(pf06751) EUT1
09925 HMMetlyEutC_1_eut1(pf05985) EUT1 EP_NTERM
09930 Tsp_rosa EUT1
09935 H_mulberry(pf00936pf16365) EUT1
09940 HMMreg12833_1_EUT1(pf12833) EUT1
09945 -# -
09950 -# -
09955 -pf01218 -
09960 -pf01520 -
09965 -pf00583 -
09970 -pf11143 -
09975 -pf06572 -
09980 -pf04261 -
09985 HMMnopfamclust3_1(#) RMM1
BMC-P
BMC-H
BMC-T
Aldehyde dehydrogenase
PTAC
Signature enzyme
Alcohol dehydrogenase
Regulator

Escherichia coli O104:H4 (strain 2011C-3493). Bacteria; Proteobacteria; Gammaproteobacteria; Enterobacterales;

O3K__locus_1

EUT1

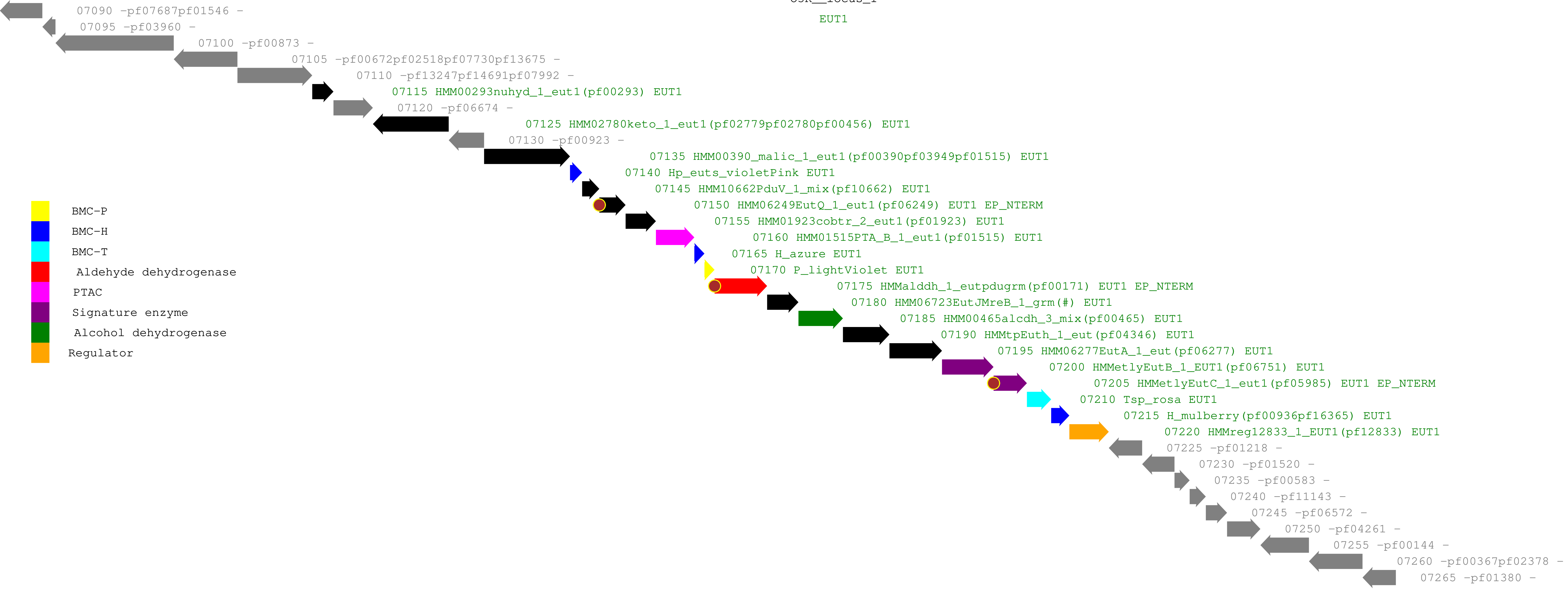

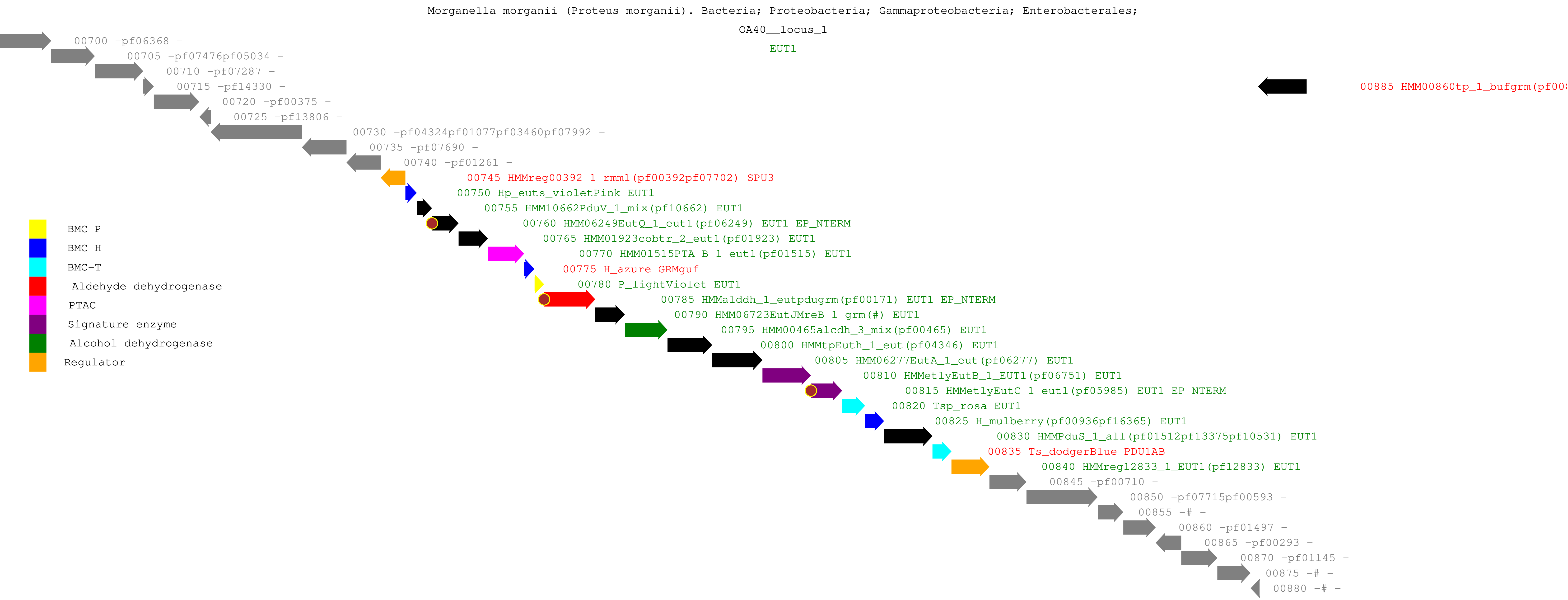
Morganella morganii (Proteus morganii). Bacteria; Proteobacteria; Gammaproteobacteria; Enterobacterales;
OA40__locus_1
EUT1
00700 -pf06368 -
00705 -pf07476pf05034 -
00710 -pf07287 -
00715 -pf14330 -
00720 -pf00375 -
00725 -pf13806 -
00730 -pf04324pf01077pf03460pf07992 -
00735 -pf07690 -
00740 -pf01261 -
00745 HMMreg00392_1_rmm1(pf00392pf07702) SPU3
00750 Hp_euts_violetPink EUT1
00755 HMM10662PduV_1_mix(pf10662) EUT1
00760 HMM06249EutQ_1_eut1(pf06249) EUT1 EP_NTERM
00765 HMM01923cobtr_2_eut1(pf01923) EUT1
00770 HMM01515PTA_B_1_eut1(pf01515) EUT1
00775 H_azure GRMguf
00780 P_lightViolet EUT1
00785 HMMalddh_1_eutpdugrm(pf00171) EUT1 EP_NTERM
00790 HMM06723EutJMreB_1_grm(#) EUT1
00795 HMM00465alcdh_3_mix(pf00465) EUT1
00800 HMMtpEuth_1_eut(pf04346) EUT1
00805 HMM06277EutA_1_eut(pf06277) EUT1
00810 HMMetlyEutB_1_EUT1(pf06751) EUT1
00815 HMMetlyEutC_1_eut1(pf05985) EUT1 EP_NTERM
00820 Tsp_rosa EUT1
00825 H_mulberry(pf00936pf16365) EUT1
00830 HMMPduS_1_all(pf01512pf13375pf10531) EUT1
00835 Ts_dodgerBlue PDU1AB
00840 HMMreg12833_1_EUT1(pf12833) EUT1
00845 -pf00710 -
00850 -pf07715pf00593 -
00855 -# -
00860 -pf01497 -
00865 -pf00293 -
00870 -pf01145 -
00875 -# -
00880 -# -
00885 HMM00860tp_1_bufgrm(pf00
BMC-P
BMC-H
BMC-T
Aldehyde dehydrogenase
PTAC
Signature enzyme
Alcohol dehydrogenase
Regulator

Salmonella enterica (Salmonella choleraesuis). Bacteria; Proteobacteria; Gammaproteobacteria; Enterobacterales;

OC65__locus_1

EUT1

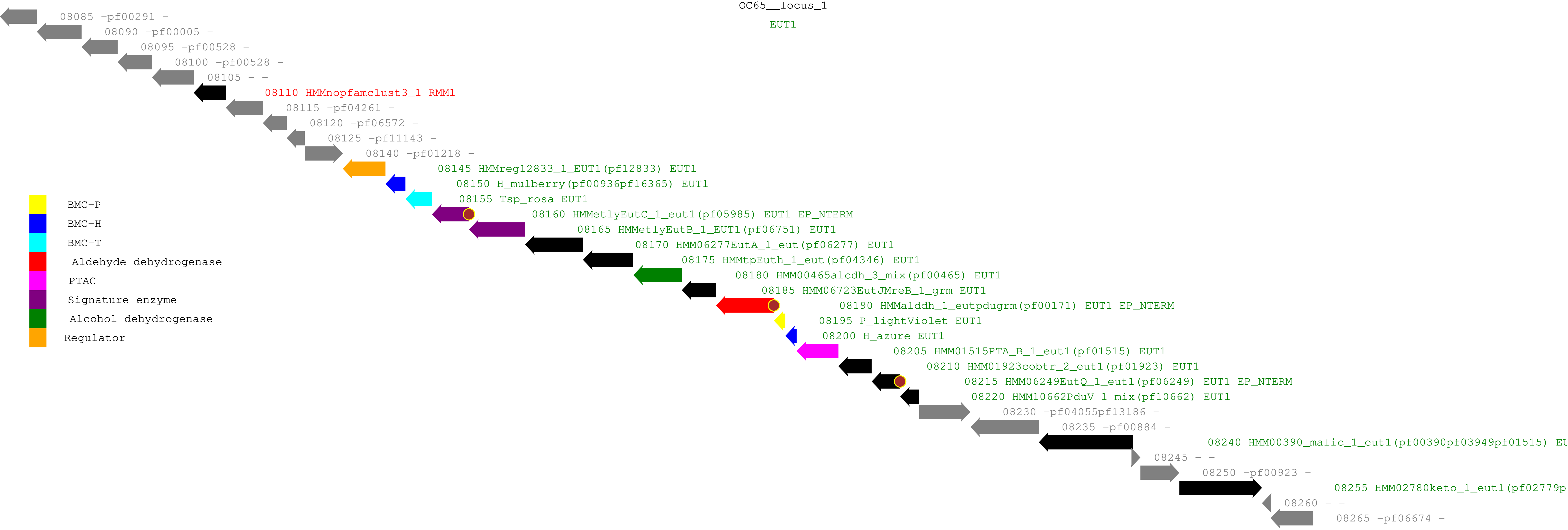

Oleispira antarctica RB-8. Bacteria; Proteobacteria; Gammaproteobacteria; Oceanospirillales;

OLEAN_C_locus_1

EUT1

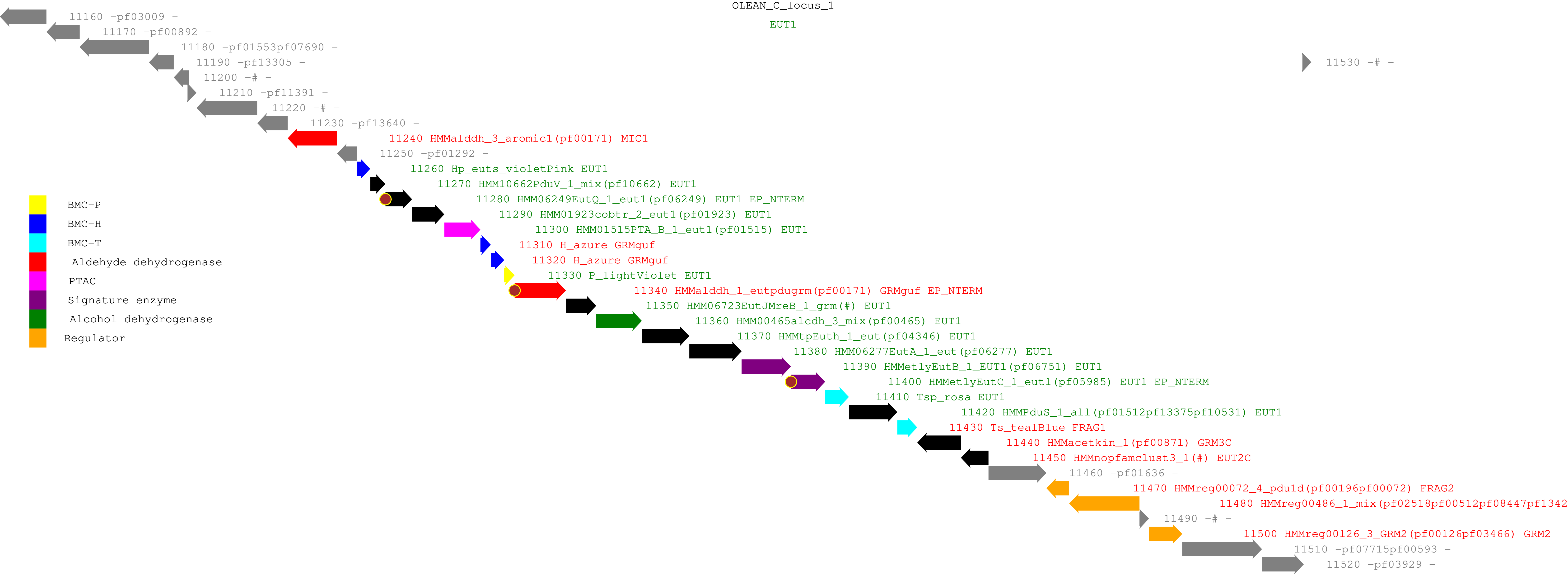

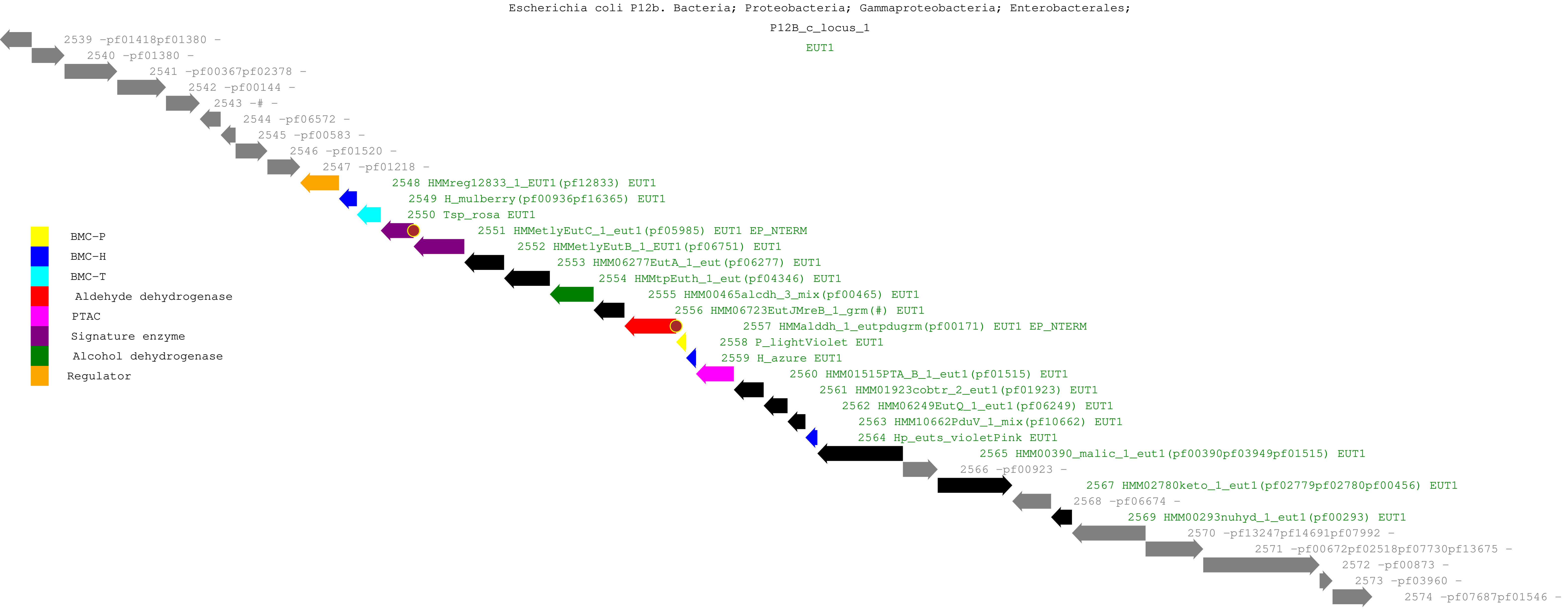
Escherichia coli P12b. Bacteria; Proteobacteria; Gammaproteobacteria; Enterobacterales;
P12B_c_locus_1
EUT1
2539 -pf01418pf01380 -
2540 -pf01380 -
2541 -pf00367pf02378 -
2542 -pf00144 -
2543 -# -
2544 -pf06572 -
2545 -pf00583 -
2546 -pf01520 -
2547 -pf01218 -
2548 HMMreg12833_1_EUT1(pf12833) EUT1
2549 H_mulberry(pf00936pf16365) EUT1
2550 Tsp_rosa EUT1
2551 HMMetlyEutC_1_eut1(pf05985) EUT1 EP_NTERM
2552 HMMetlyEutB_1_EUT1(pf06751) EUT1
2553 HMM06277EutA_1_eut(pf06277) EUT1
2554 HMMtpEuth_1_eut(pf04346) EUT1
2555 HMM00465alcdh_3_mix(pf00465) EUT1
2556 HMM06723EutJMreB_1_grm(#) EUT1
2557 HMMalddh_1_eutpdugrm(pf00171) EUT1 EP_NTERM
2558 P_lightViolet EUT1
2559 H_azure EUT1
2560 HMM01515PTA_B_1_eut1(pf01515) EUT1
2561 HMM01923cobtr_2_eut1(pf01923) EUT1
2562 HMM06249EutQ_1_eut1(pf06249) EUT1
2563 HMM10662PduV_1_mix(pf10662) EUT1
2564 Hp_euts_violetPink EUT1
2565 HMM00390_malic_1_eut1(pf00390pf03949pf01515) EUT1
2566 -pf00923 -
2567 HMM02780keto_1_eut1(pf02779pf02780pf00456) EUT1
2568 -pf06674 -
2569 HMM00293nuhyd_1_eut1(pf00293) EUT1
2570 -pf13247pf14691pf07992 -
2571 -pf00672pf02518pf07730pf13675 -
2572 -pf00873 -
2573 -pf03960 -
2574 -pf07687pf01546 -
BMC-P
BMC-H
BMC-T
Aldehyde dehydrogenase
PTAC
Signature enzyme
Alcohol dehydrogenase
Regulator

Salmonella enterica subsp. arizonae serovar 18:z4,z23:- str. CVM Bacteria; Proteobacteria; Gammaproteobacteria; Enterobacterales;

P287__locus_2

EUT1

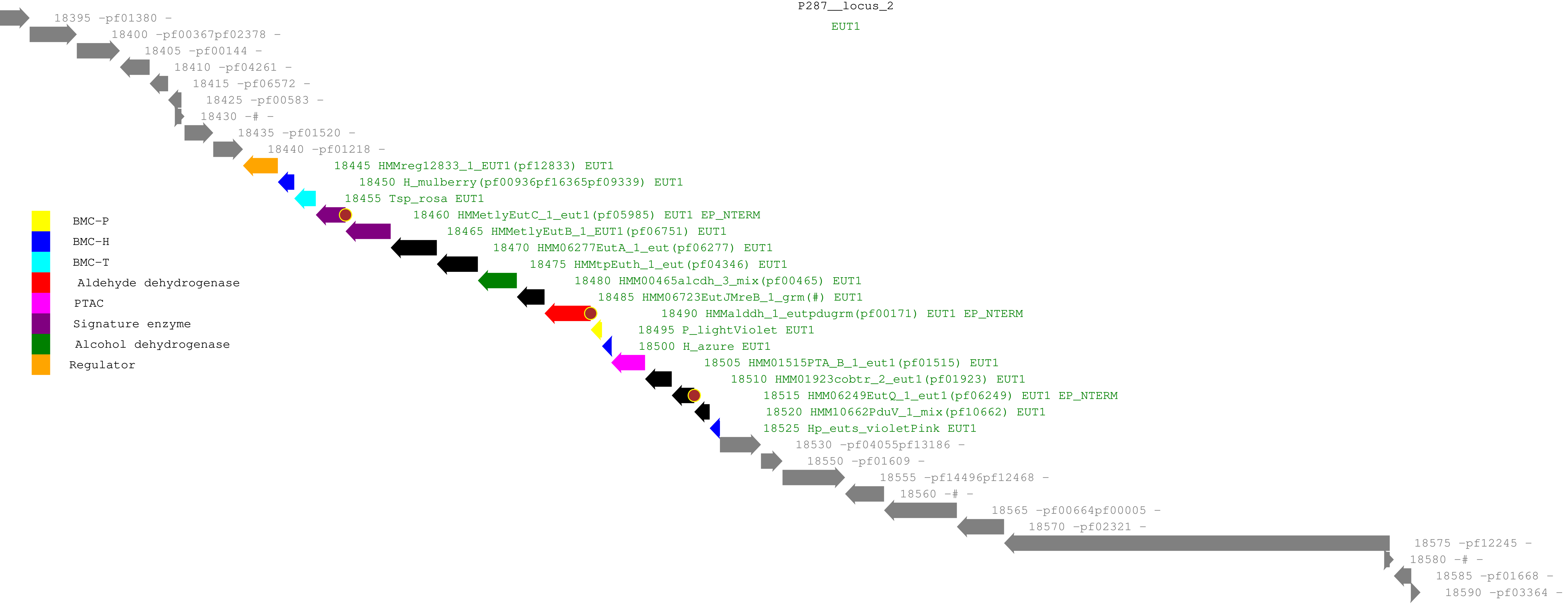

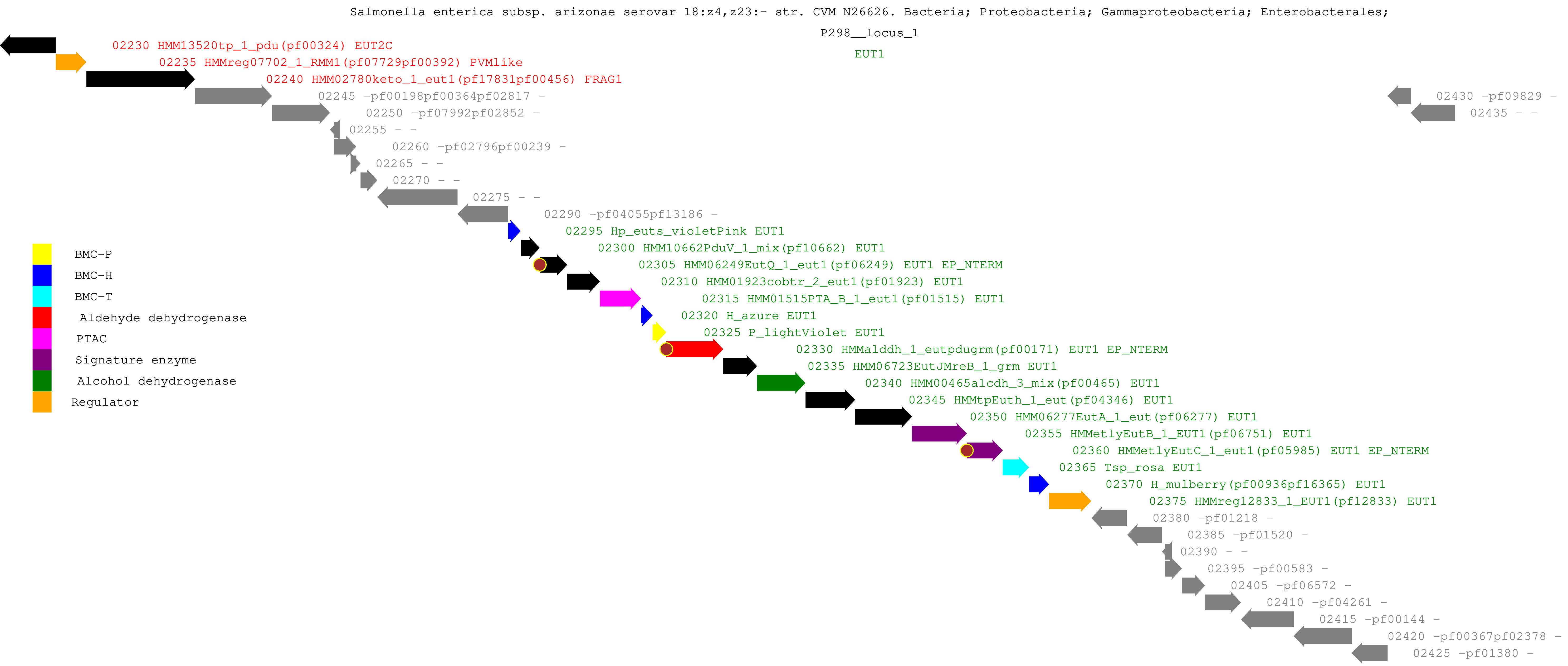

Salmonella enterica subsp. arizonae serovar 18:z4,z23:- str. CVM N26626. Bacteria; Proteobacteria; Gammaproteobacteria; Enterobacterales;
P298__locus_1
EUT1
02230 HMM13520tp_1_pdu(pf00324) EUT2C
02235 HMMreg07702_1_RMM1(pf07729pf00392) PVMlike
02240 HMM02780keto_1_eut1(pf17831pf00456) FRAG1
02245 -pf00198pf00364pf02817 -
02250 -pf07992pf02852 -
02255 - -
02260 -pf02796pf00239 -
02265 - -
02270 - -
02275 - -
02290 -pf04055pf13186 -
02295 Hp_euts_violetPink EUT1
02300 HMM10662PduV_1_mix(pf10662) EUT1
02305 HMM06249EutQ_1_eut1(pf06249) EUT1 EP_NTERM
02310 HMM01923cobtr_2_eut1(pf01923) EUT1
02315 HMM01515PTA_B_1_eut1(pf01515) EUT1
02320 H_azure EUT1
02325 P_lightViolet EUT1
02330 HMMalddh_1_eutpdugrm(pf00171) EUT1 EP_NTERM
02335 HMM06723EutJMreB_1_grm EUT1
02340 HMM00465alcdh_3_mix(pf00465) EUT1
02345 HMMtpEuth_1_eut(pf04346) EUT1
02350 HMM06277EutA_1_eut(pf06277) EUT1
02355 HMMetlyEutB_1_EUT1(pf06751) EUT1
02360 HMMetlyEutC_1_eut1(pf05985) EUT1 EP_NTERM
02365 Tsp_rosa EUT1
02370 H_mulberry(pf00936pf16365) EUT1
02375 HMMreg12833_1_EUT1(pf12833) EUT1
02380 -pf01218 -
02385 -pf01520 -
02390 - -
02395 -pf00583 -
02405 -pf06572 -
02410 -pf04261 -
02415 -pf00144 -
02420 -pf00367pf02378 -
02425 -pf01380 -
02430 -pf09829 -
02435 - -
BMC-P
BMC-H
BMC-T
Aldehyde dehydrogenase
PTAC
Signature enzyme
Alcohol dehydrogenase
Regulator

Enterobacter sp. DC4. Bacteria; Proteobacteria; Gammaproteobacteria; Enterobacterales;

P349__locus_1

EUT1

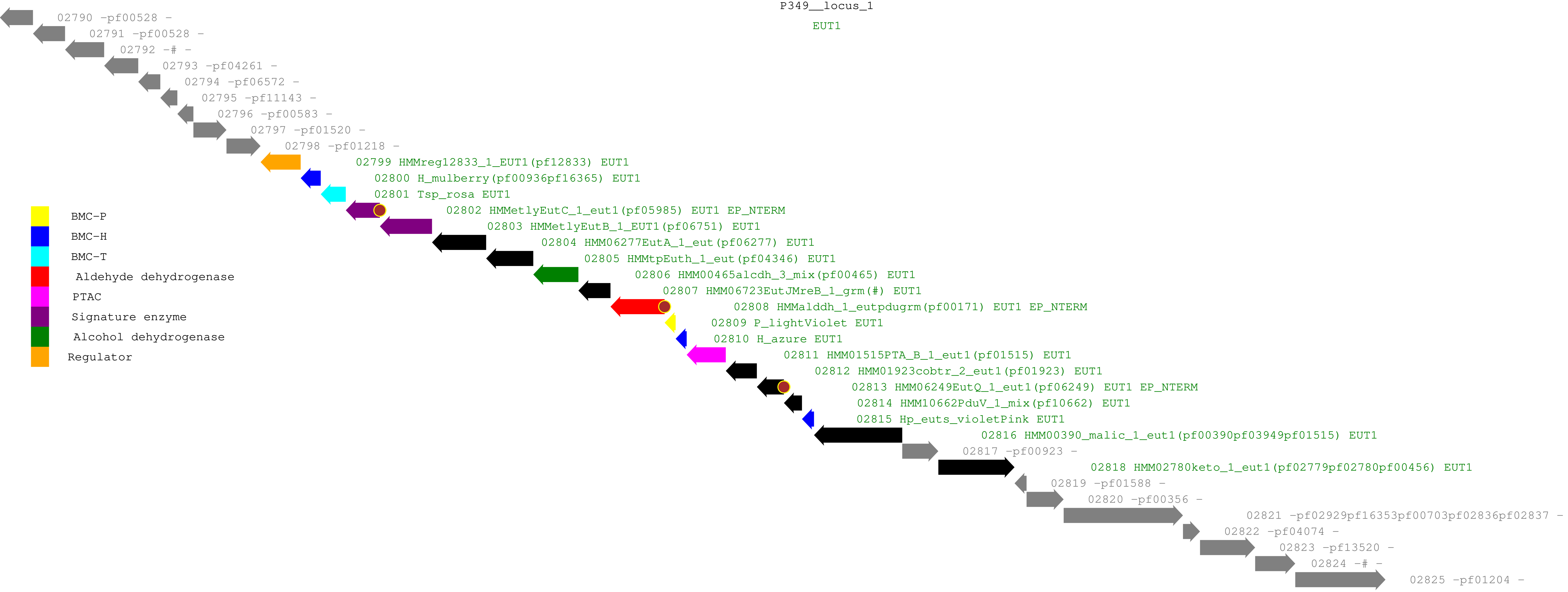

Photobacterium profundum 3TCK. Bacteria; Proteobacteria; Gammaproteobacteria; Vibrionales;

P3TCK__locus_1

EUT1

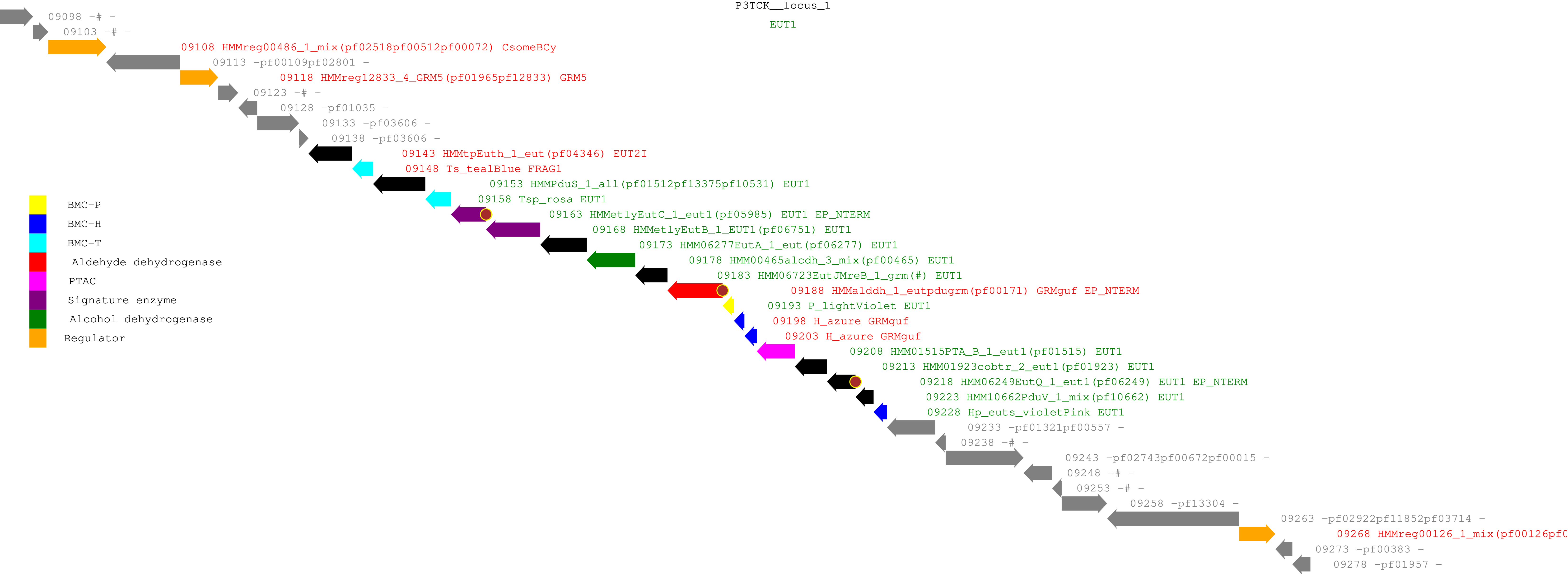

Salmonella enterica subsp. enterica serovar Newport str. CVM80_2288. Bacteria; Proteobacteria; Gammaproteobacteria; Enterobacterales;

P701__locus_2

EUT1

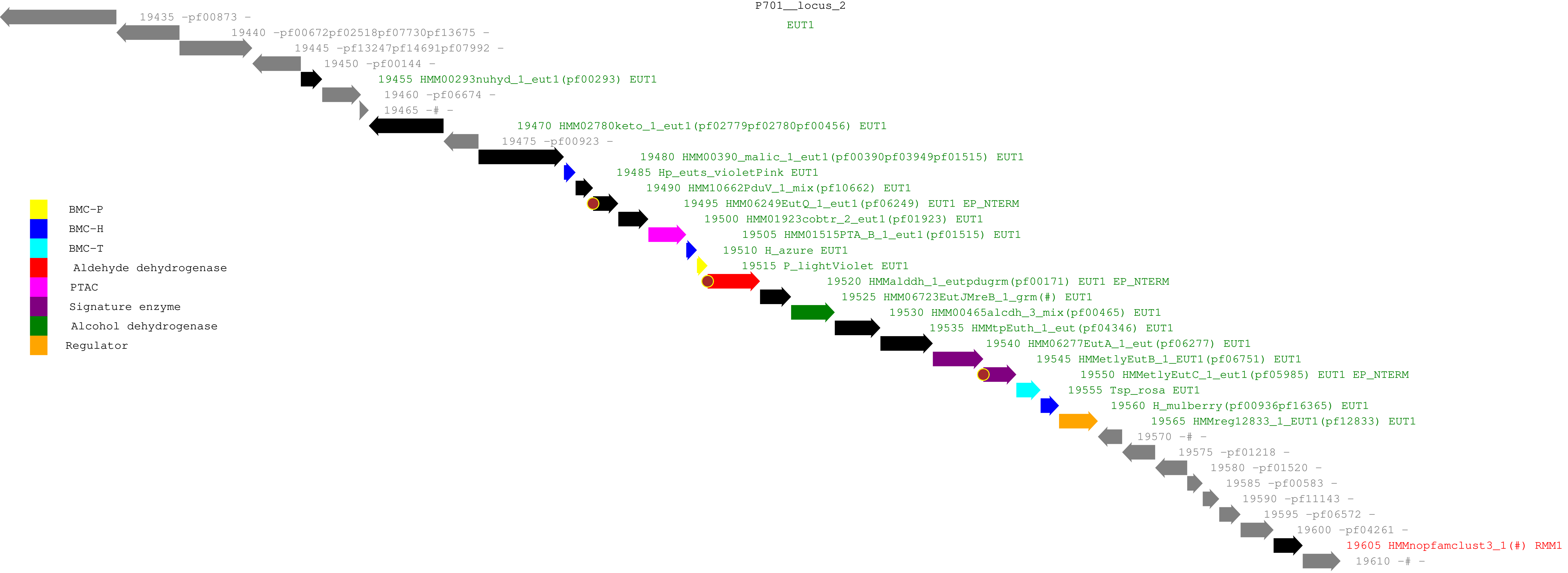

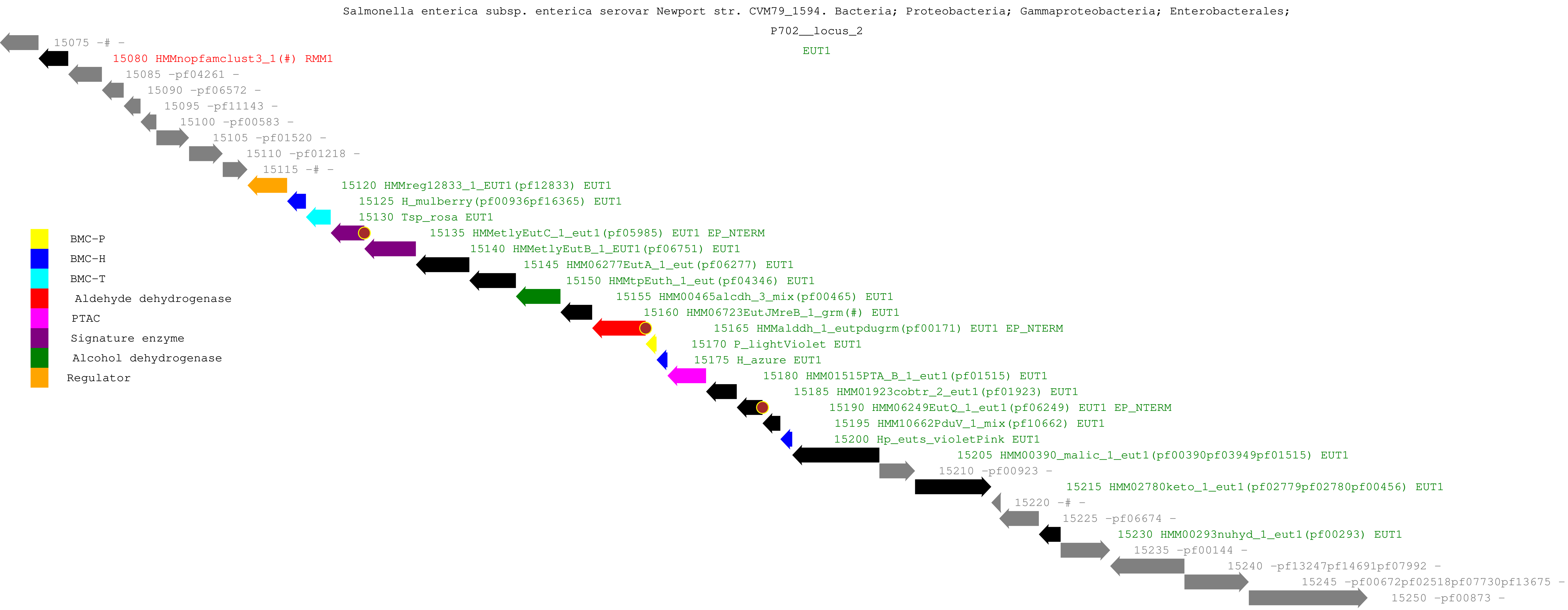
Salmonella enterica subsp. enterica serovar Newport str. CVM79_1594. Bacteria; Proteobacteria; Gammaproteobacteria; Enterobacterales;
P702__locus_2
EUT1
15075 -# -
15080 HMMnopfamclust3_1(#) RMM1
15085 -pf04261 -
15090 -pf06572 -
15095 -pf11143 -
15100 -pf00583 -
15105 -pf01520 -
15110 -pf01218 -
15115 -# -
15120 HMMreg12833_1_EUT1(pf12833) EUT1
15125 H_mulberry(pf00936pf16365) EUT1
15130 Tsp_rosa EUT1
15135 HMMetlyEutC_1_eut1(pf05985) EUT1 EP_NTERM
15140 HMMetlyEutB_1_EUT1(pf06751) EUT1
15145 HMM06277EutA_1_eut(pf06277) EUT1
15150 HMMtpEuth_1_eut(pf04346) EUT1
15155 HMM00465alcdh_3_mix(pf00465) EUT1
15160 HMM06723EutJMreB_1_grm(#) EUT1
15165 HMMalddh_1_eutpdugrm(pf00171) EUT1 EP_NTERM
15170 P_lightViolet EUT1
15175 H_azure EUT1
15180 HMM01515PTA_B_1_eut1(pf01515) EUT1
15185 HMM01923cobtr_2_eut1(pf01923) EUT1
15190 HMM06249EutQ_1_eut1(pf06249) EUT1 EP_NTERM
15195 HMM10662PduV_1_mix(pf10662) EUT1
15200 Hp_euts_violetPink EUT1
15205 HMM00390_malic_1_eut1(pf00390pf03949pf01515) EUT1
15210 -pf00923 -
15215 HMM02780keto_1_eut1(pf02779pf02780pf00456) EUT1
15220 -# -
15225 -pf06674 -
15230 HMM00293nuhyd_1_eut1(pf00293) EUT1
15235 -pf00144 -
15240 -pf13247pf14691pf07992 -
15245 -pf00672pf02518pf07730pf13675 -
15250 -pf00873 -
BMC-P
BMC-H
BMC-T
Aldehyde dehydrogenase
PTAC
Signature enzyme
Alcohol dehydrogenase
Regulator

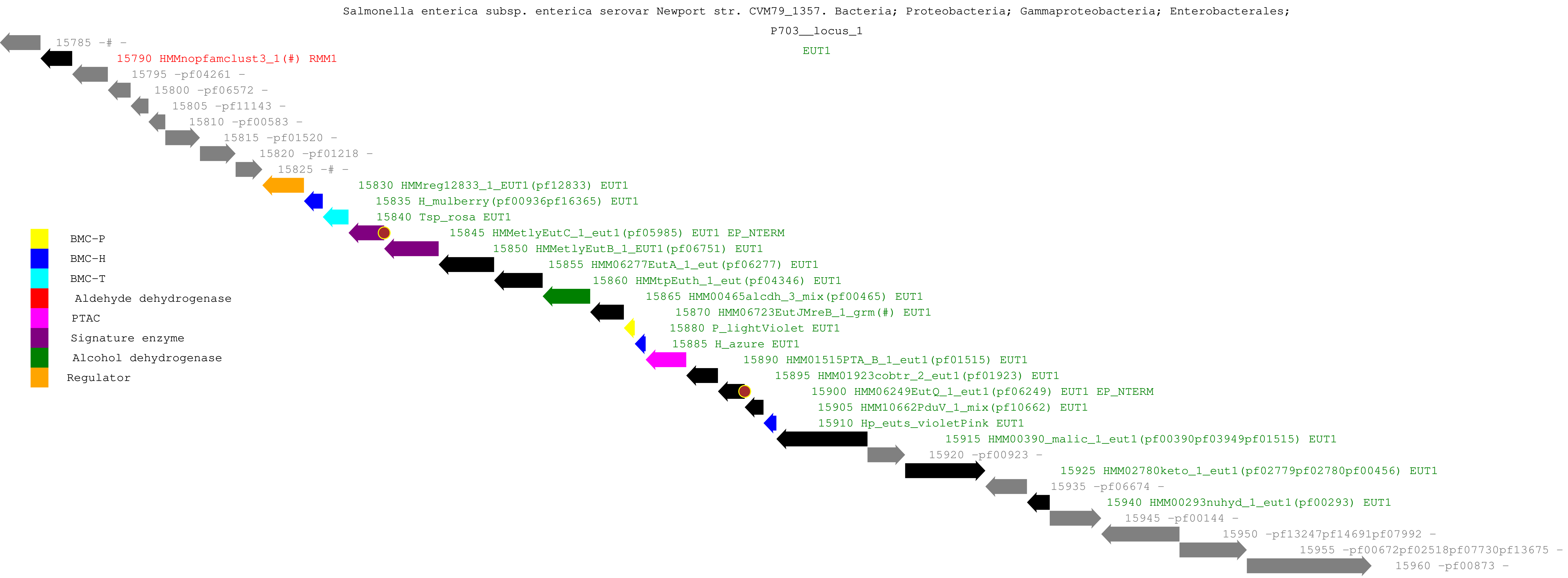
Salmonella enterica subsp. enterica serovar Newport str. CVM79_1357. Bacteria; Proteobacteria; Gammaproteobacteria; Enterobacterales;
P703__locus_1
EUT1
15785 -# -
15790 HMMnopfamclust3_1(#) RMM1
15795 -pf04261 -
15800 -pf06572 -
15805 -pf11143 -
15810 -pf00583 -
15815 -pf01520 -
15820 -pf01218 -
15825 -# -
15830 HMMreg12833_1_EUT1(pf12833) EUT1
15835 H_mulberry(pf00936pf16365) EUT1
15840 Tsp_rosa EUT1
15845 HMMetlyEutC_1_eut1(pf05985) EUT1 EP_NTERM
15850 HMMetlyEutB_1_EUT1(pf06751) EUT1
15855 HMM06277EutA_1_eut(pf06277) EUT1
15860 HMMtpEuth_1_eut(pf04346) EUT1
15865 HMM00465alcdh_3_mix(pf00465) EUT1
15870 HMM06723EutJMreB_1_grm(#) EUT1
15880 P_lightViolet EUT1
15885 H_azure EUT1
15890 HMM01515PTA_B_1_eut1(pf01515) EUT1
15895 HMM01923cobtr_2_eut1(pf01923) EUT1
15900 HMM06249EutQ_1_eut1(pf06249) EUT1 EP_NTERM
15905 HMM10662PduV_1_mix(pf10662) EUT1
15910 Hp_euts_violetPink EUT1
15915 HMM00390_malic_1_eut1(pf00390pf03949pf01515) EUT1
15920 -pf00923 -
15925 HMM02780keto_1_eut1(pf02779pf02780pf00456) EUT1
15935 -pf06674 -
15940 HMM00293nuhyd_1_eut1(pf00293) EUT1
15945 -pf00144 -
15950 -pf13247pf14691pf07992 -
15955 -pf00672pf02518pf07730pf13675 -
15960 -pf00873 -
BMC-P
BMC-H
BMC-T
Aldehyde dehydrogenase
PTAC
Signature enzyme
Alcohol dehydrogenase
Regulator

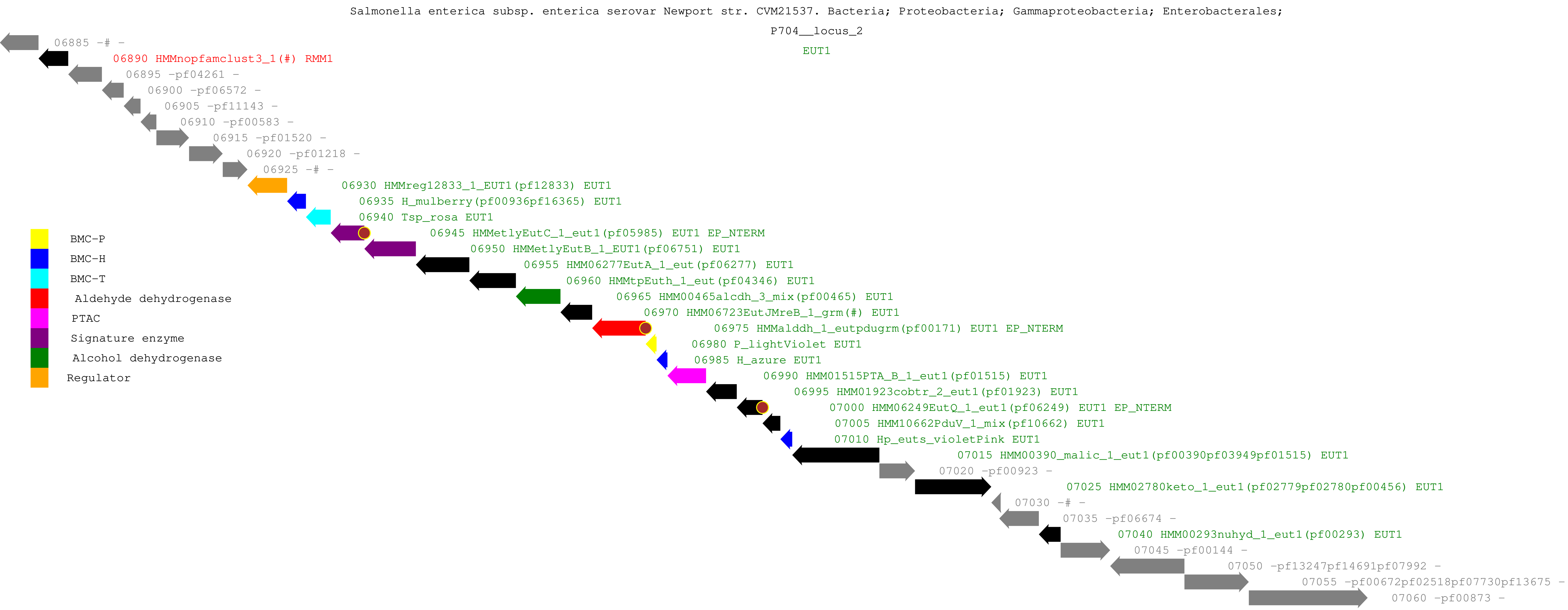
Salmonella enterica subsp. enterica serovar Newport str. CVM21537. Bacteria; Proteobacteria; Gammaproteobacteria; Enterobacterales;
P704__locus_2
EUT1
06885 -# -
06890 HMMnopfamclust3_1(#) RMM1
06895 -pf04261 -
06900 -pf06572 -
06905 -pf11143 -
06910 -pf00583 -
06915 -pf01520 -
06920 -pf01218 -
06925 -# -
06930 HMMreg12833_1_EUT1(pf12833) EUT1
06935 H_mulberry(pf00936pf16365) EUT1
06940 Tsp_rosa EUT1
06945 HMMetlyEutC_1_eut1(pf05985) EUT1 EP_NTERM
06950 HMMetlyEutB_1_EUT1(pf06751) EUT1
06955 HMM06277EutA_1_eut(pf06277) EUT1
06960 HMMtpEuth_1_eut(pf04346) EUT1
06965 HMM00465alcdh_3_mix(pf00465) EUT1
06970 HMM06723EutJMreB_1_grm(#) EUT1
06975 HMMalddh_1_eutpdugrm(pf00171) EUT1 EP_NTERM
06980 P_lightViolet EUT1
06985 H_azure EUT1
06990 HMM01515PTA_B_1_eut1(pf01515) EUT1
06995 HMM01923cobtr_2_eut1(pf01923) EUT1
07000 HMM06249EutQ_1_eut1(pf06249) EUT1 EP_NTERM
07005 HMM10662PduV_1_mix(pf10662) EUT1
07010 Hp_euts_violetPink EUT1
07015 HMM00390_malic_1_eut1(pf00390pf03949pf01515) EUT1
07020 -pf00923 -
07025 HMM02780keto_1_eut1(pf02779pf02780pf00456) EUT1
07030 -# -
07035 -pf06674 -
07040 HMM00293nuhyd_1_eut1(pf00293) EUT1
07045 -pf00144 -
07050 -pf13247pf14691pf07992 -
07055 -pf00672pf02518pf07730pf13675 -
07060 -pf00873 -
BMC-P
BMC-H
BMC-T
Aldehyde dehydrogenase
PTAC
Signature enzyme
Alcohol dehydrogenase
Regulator

Salmonella enterica subsp. enterica serovar Newport str. CVM18000. Bacteria; Proteobacteria; Gammaproteobacteria; Enterobacterales;

P705__locus_1

EUT1

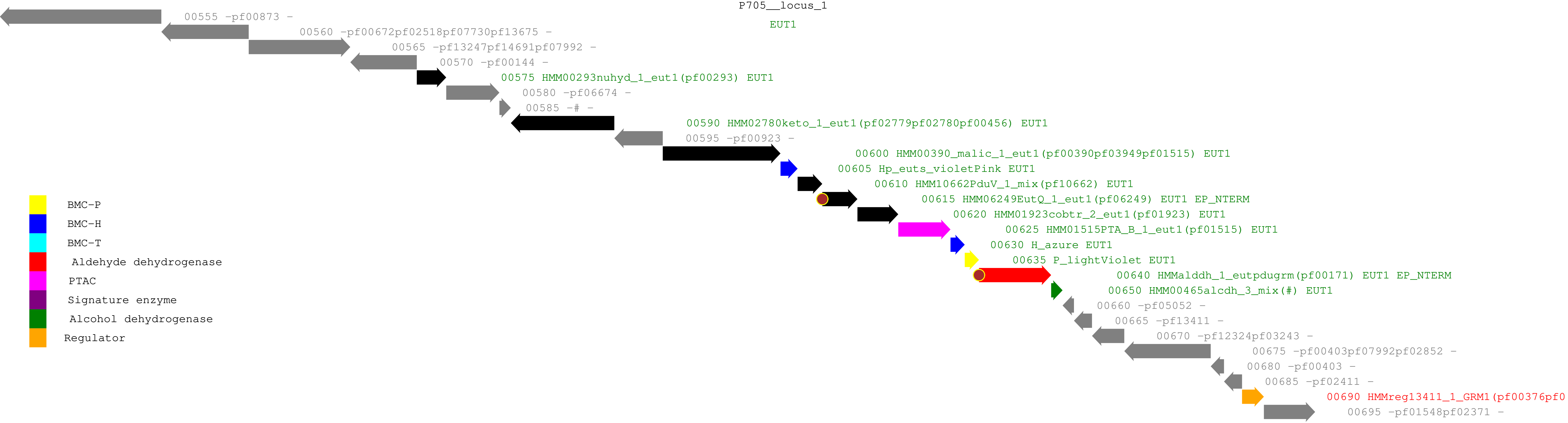

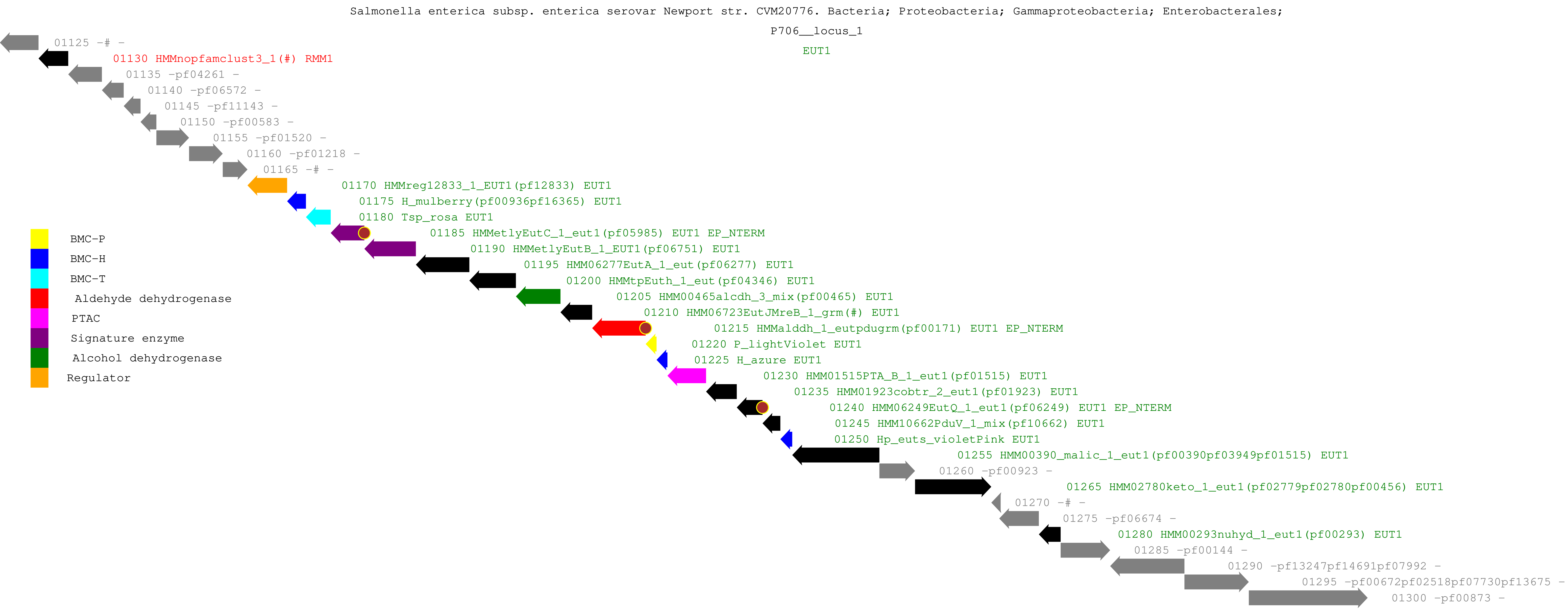
Salmonella enterica subsp. enterica serovar Newport str. CVM20776. Bacteria; Proteobacteria; Gammaproteobacteria; Enterobacterales;
P706__locus_1
EUT1
01125 -# -
01130 HMMnopfamclust3_1(#) RMM1
01135 -pf04261 -
01140 -pf06572 -
01145 -pf11143 -
01150 -pf00583 -
01155 -pf01520 -
01160 -pf01218 -
01165 -# -
01170 HMMreg12833_1_EUT1(pf12833) EUT1
01175 H_mulberry(pf00936pf16365) EUT1
01180 Tsp_rosa EUT1
01185 HMMetlyEutC_1_eut1(pf05985) EUT1 EP_NTERM
01190 HMMetlyEutB_1_EUT1(pf06751) EUT1
01195 HMM06277EutA_1_eut(pf06277) EUT1
01200 HMMtpEuth_1_eut(pf04346) EUT1
01205 HMM00465alcdh_3_mix(pf00465) EUT1
01210 HMM06723EutJMreB_1_grm(#) EUT1
01215 HMMalddh_1_eutpdugrm(pf00171) EUT1 EP_NTERM
01220 P_lightViolet EUT1
01225 H_azure EUT1
01230 HMM01515PTA_B_1_eut1(pf01515) EUT1
01235 HMM01923cobtr_2_eut1(pf01923) EUT1
01240 HMM06249EutQ_1_eut1(pf06249) EUT1 EP_NTERM
01245 HMM10662PduV_1_mix(pf10662) EUT1
01250 Hp_euts_violetPink EUT1
01255 HMM00390_malic_1_eut1(pf00390pf03949pf01515) EUT1
01260 -pf00923 -
01265 HMM02780keto_1_eut1(pf02779pf02780pf00456) EUT1
01270 -# -
01275 -pf06674 -
01280 HMM00293nuhyd_1_eut1(pf00293) EUT1
01285 -pf00144 -
01290 -pf13247pf14691pf07992 -
01295 -pf00672pf02518pf07730pf13675 -
01300 -pf00873 -
BMC-P
BMC-H
BMC-T
Aldehyde dehydrogenase
PTAC
Signature enzyme
Alcohol dehydrogenase
Regulator

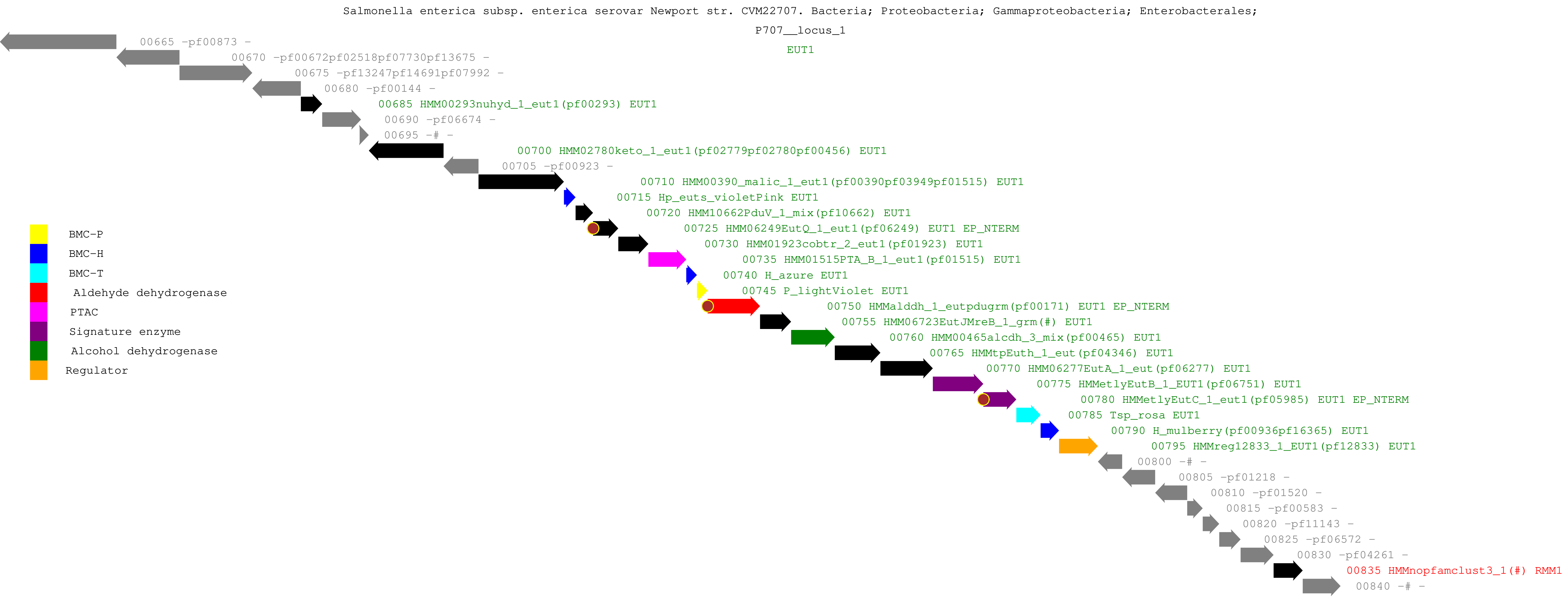
Salmonella enterica subsp. enterica serovar Newport str. CVM22707. Bacteria; Proteobacteria; Gammaproteobacteria; Enterobacterales;
P707__locus_1
EUT1
00665 -pf00873 -
00670 -pf00672pf02518pf07730pf13675 -
00675 -pf13247pf14691pf07992 -
00680 -pf00144 -
00685 HMM00293nuhyd_1_eut1(pf00293) EUT1
00690 -pf06674 -
00695 -# -
00700 HMM02780keto_1_eut1(pf02779pf02780pf00456) EUT1
00705 -pf00923 -
00710 HMM00390_malic_1_eut1(pf00390pf03949pf01515) EUT1
00715 Hp_euts_violetPink EUT1
00720 HMM10662PduV_1_mix(pf10662) EUT1
00725 HMM06249EutQ_1_eut1(pf06249) EUT1 EP_NTERM
00730 HMM01923cobtr_2_eut1(pf01923) EUT1
00735 HMM01515PTA_B_1_eut1(pf01515) EUT1
00740 H_azure EUT1
00745 P_lightViolet EUT1
00750 HMMalddh_1_eutpdugrm(pf00171) EUT1 EP_NTERM
00755 HMM06723EutJMreB_1_grm(#) EUT1
00760 HMM00465alcdh_3_mix(pf00465) EUT1
00765 HMMtpEuth_1_eut(pf04346) EUT1
00770 HMM06277EutA_1_eut(pf06277) EUT1
00775 HMMetlyEutB_1_EUT1(pf06751) EUT1
00780 HMMetlyEutC_1_eut1(pf05985) EUT1 EP_NTERM
00785 Tsp_rosa EUT1
00790 H_mulberry(pf00936pf16365) EUT1
00795 HMMreg12833_1_EUT1(pf12833) EUT1
00800 -# -
00805 -pf01218 -
00810 -pf01520 -
00815 -pf00583 -
00820 -pf11143 -
00825 -pf06572 -
00830 -pf04261 -
00835 HMMnopfamclust3_1(#) RMM1
00840 -# -
BMC-P
BMC-H
BMC-T
Aldehyde dehydrogenase
PTAC
Signature enzyme
Alcohol dehydrogenase
Regulator

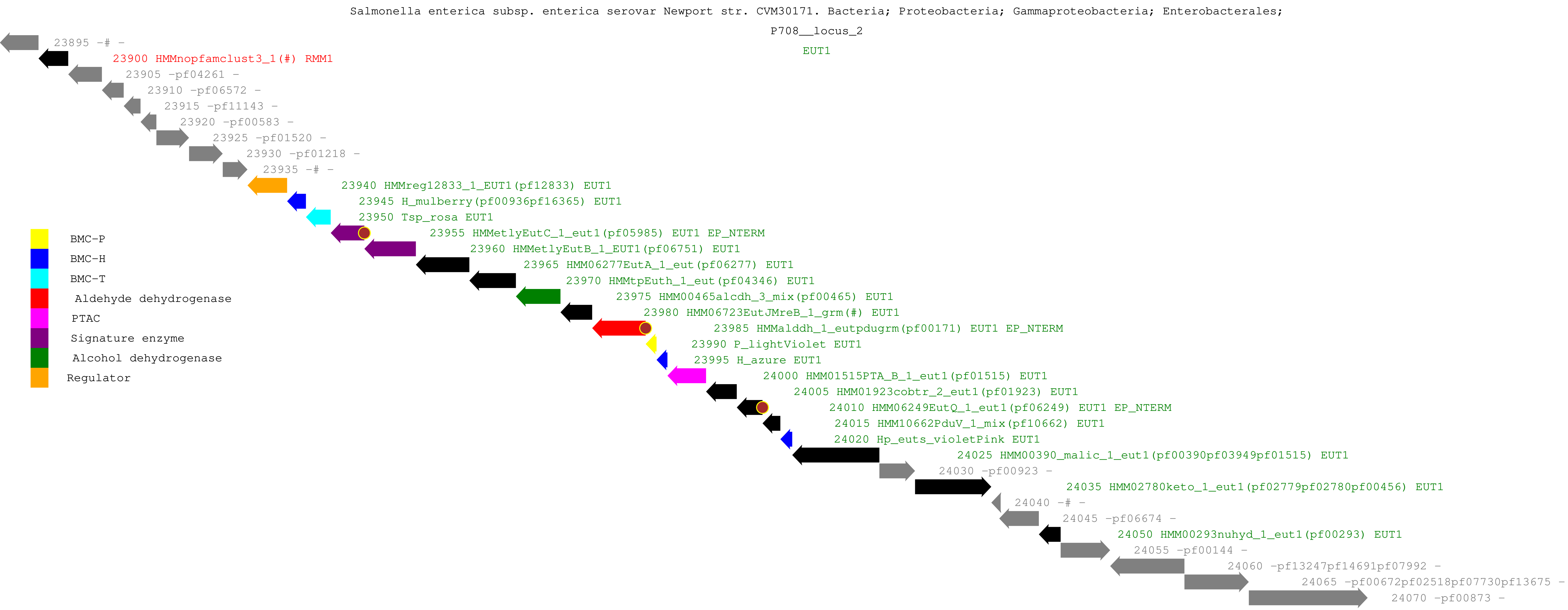
Salmonella enterica subsp. enterica serovar Newport str. CVM30171. Bacteria; Proteobacteria; Gammaproteobacteria; Enterobacterales;
P708__locus_2
EUT1
23895 -# -
23900 HMMnopfamclust3_1(#) RMM1
23905 -pf04261 -
23910 -pf06572 -
23915 -pf11143 -
23920 -pf00583 -
23925 -pf01520 -
23930 -pf01218 -
23935 -# -
23940 HMMreg12833_1_EUT1(pf12833) EUT1
23945 H_mulberry(pf00936pf16365) EUT1
23950 Tsp_rosa EUT1
23955 HMMetlyEutC_1_eut1(pf05985) EUT1 EP_NTERM
23960 HMMetlyEutB_1_EUT1(pf06751) EUT1
23965 HMM06277EutA_1_eut(pf06277) EUT1
23970 HMMtpEuth_1_eut(pf04346) EUT1
23975 HMM00465alcdh_3_mix(pf00465) EUT1
23980 HMM06723EutJMreB_1_grm(#) EUT1
23985 HMMaldh_1_eutpdugrm(pf00171) EUT1 EP_NTERM
23990 P_lightViolet EUT1
23995 H_azure EUT1
24000 HMM01515PTA_B_1_eut1(pf01515) EUT1
24005 HMM01923cobtr_2_eut1(pf01923) EUT1
24010 HMM06249EutQ_1_eut1(pf06249) EUT1 EP_NTERM
24015 HMM10662PduV_1_mix(pf10662) EUT1
24020 Hp_euts_violetPink EUT1
24025 HMM00390_malic_1_eut1(pf00390pf03949pf01515) EUT1
24030 -pf00923 -
24035 HMM02780keto_1_eut1(pf02779pf02780pf00456) EUT1
24040 -# -
24045 -pf06674 -
24050 HMM00293nuhyd_1_eut1(pf00293) EUT1
24055 -pf00144 -
24060 -pf13247pf14691pf07992 -
24065 -pf00672pf02518pf07730pf13675 -
24070 -pf00873 -
BMC-P
BMC-H
BMC-T
Aldehyde dehydrogenase
PTAC
Signature enzyme
Alcohol dehydrogenase
Regulator

Salmonella enterica subsp. enterica serovar Newport str. CVM17619. Bacteria; Proteobacteria; Gammaproteobacteria; Enterobacterales;

P709__locus_2

EUT1

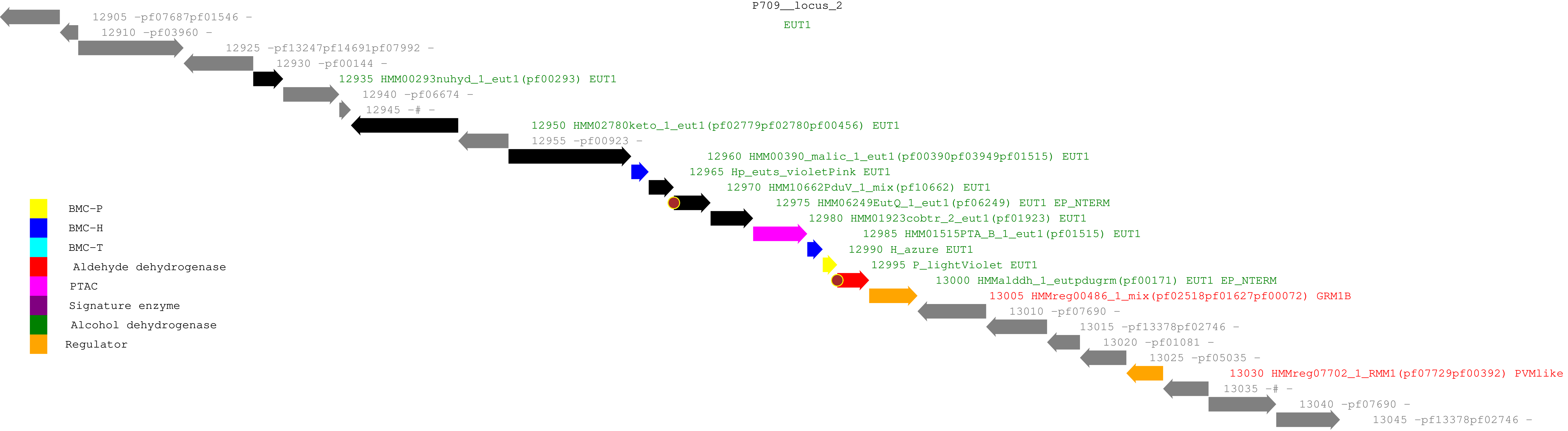

Salmonella enterica subsp. enterica serovar Newport str. CVM20775. Bacteria; Proteobacteria; Gammaproteobacteria; Enterobacterales;

P710__locus_2

EUT1

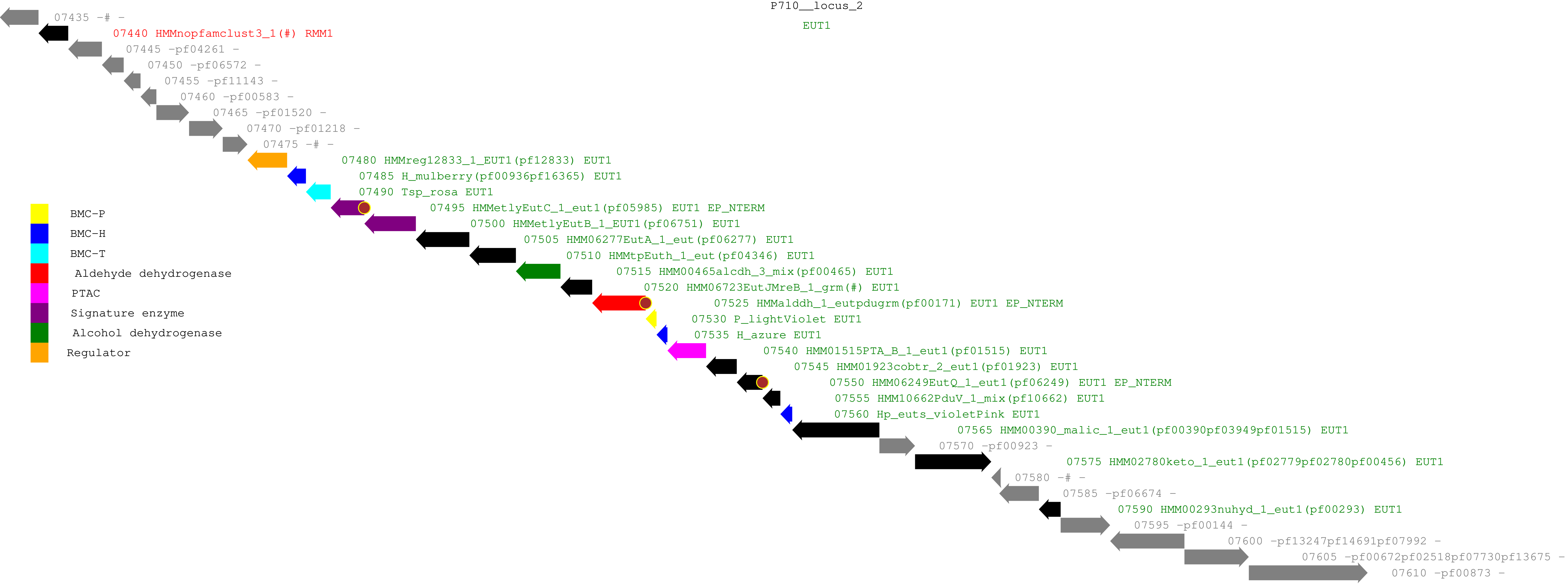

Salmonella enterica subsp. enterica serovar Newport str. CVM22697. Bacteria; Proteobacteria; Gammaproteobacteria; Enterobacterales;

P711__locus_2

EUT1

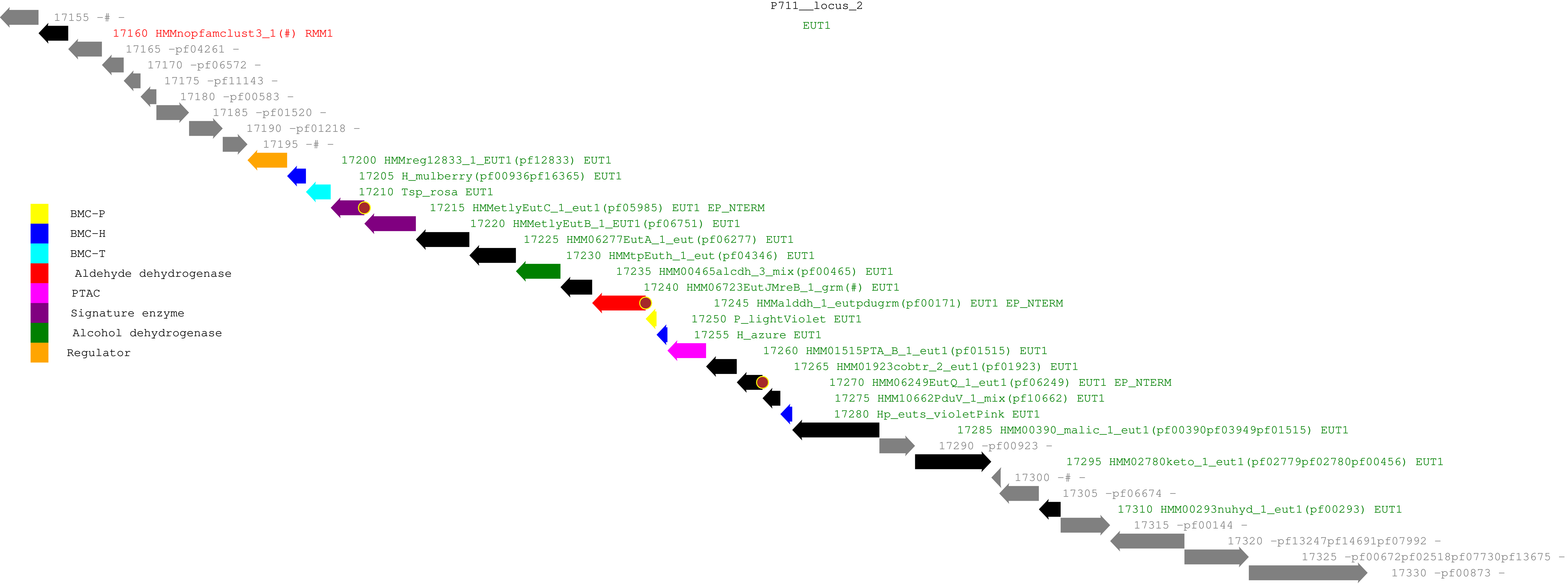

Salmonella enterica subsp. enterica serovar Newport str. CVM22699. Bacteria; Proteobacteria; Gammaproteobacteria; Enterobacterales;

P712__locus_1

EUT1

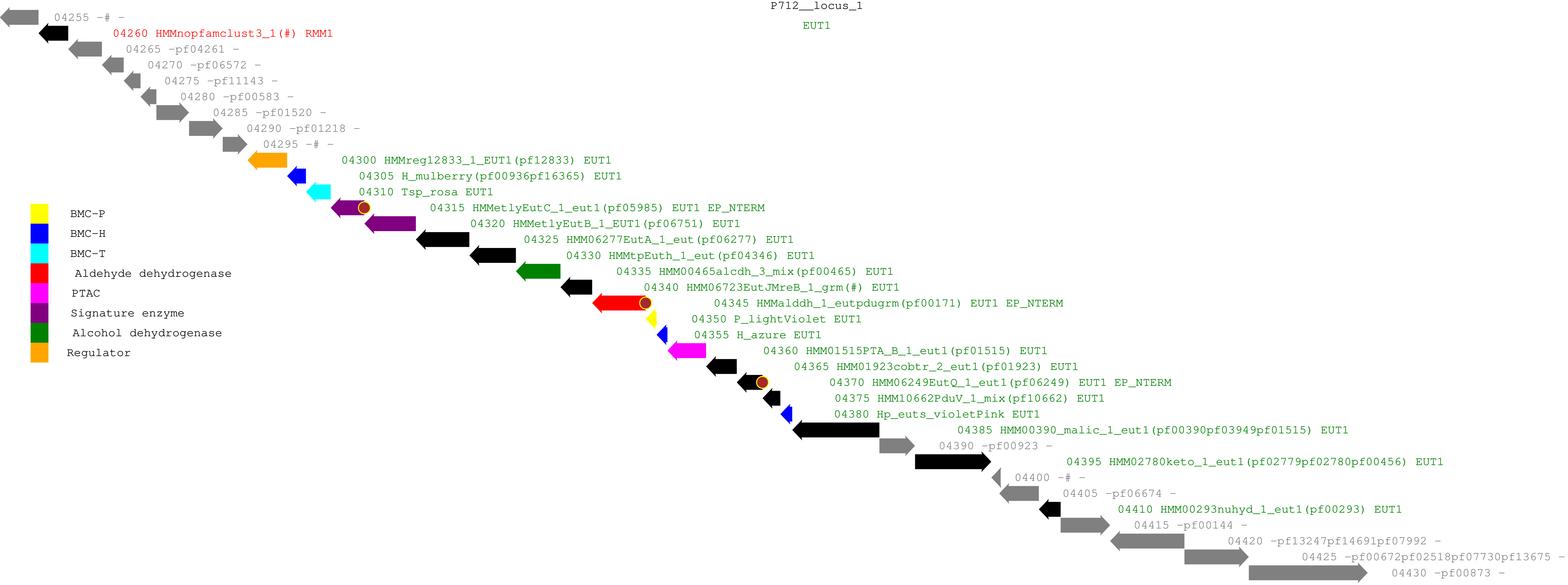

Salmonella enterica subsp. enterica serovar Newport str. CVM22443. Bacteria; Proteobacteria; Gammaproteobacteria; Enterobacterales;

P713__locus_1

EUT1

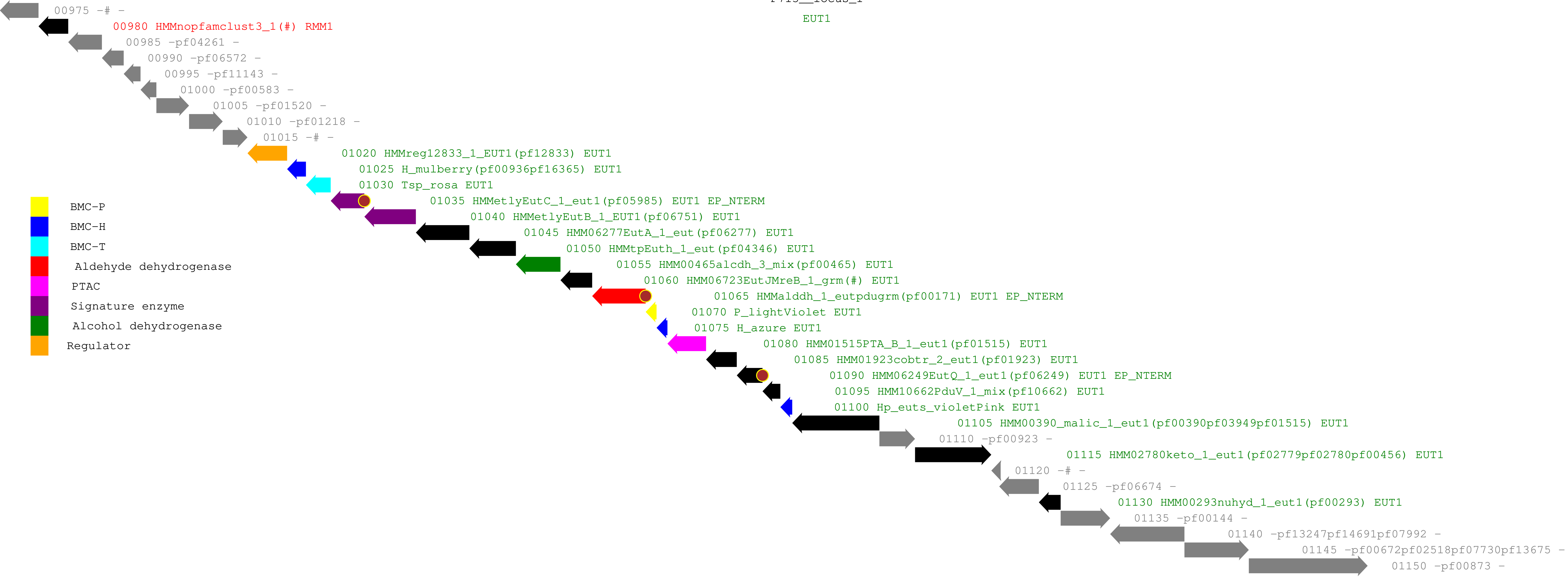

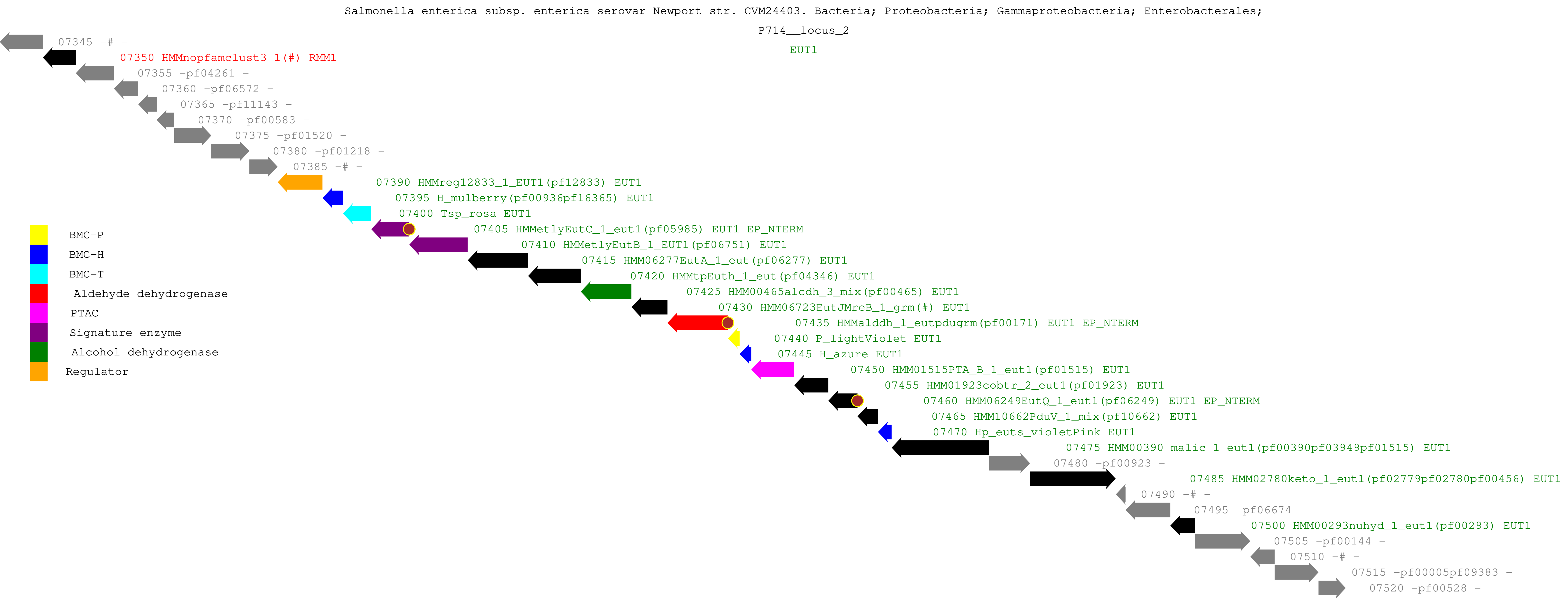
Salmonella enterica subsp. enterica serovar Newport str. CVM24403. Bacteria; Proteobacteria; Gammaproteobacteria; Enterobacterales;
P714__locus_2
EUT1
07345 -# -
07350 HMMnopfamclust3_1(#) RMM1
07355 -pf04261 -
07360 -pf06572 -
07365 -pf11143 -
07370 -pf00583 -
07375 -pf01520 -
07380 -pf01218 -
07385 -# -
07390 HMMreg12833_1_EUT1(pf12833) EUT1
07395 H_mulberry(pf00936pf16365) EUT1
07400 Tsp_rosa EUT1
07405 HMMetlyEutC_1_eut1(pf05985) EUT1 EP_NTERM
07410 HMMetlyEutB_1_EUT1(pf06751) EUT1
07415 HMM06277EutA_1_eut(pf06277) EUT1
07420 HMMtpEuth_1_eut(pf04346) EUT1
07425 HMM00465alcdh_3_mix(pf00465) EUT1
07430 HMM06723EutJMreB_1_grm(#) EUT1
07435 HMMalddh_1_eutpdugrm(pf00171) EUT1 EP_NTERM
07440 P_lightViolet EUT1
07445 H_azure EUT1
07450 HMM01515PTA_B_1_eut1(pf01515) EUT1
07455 HMM01923cobtr_2_eut1(pf01923) EUT1
07460 HMM06249EutQ_1_eut1(pf06249) EUT1 EP_NTERM
07465 HMM10662PduV_1_mix(pf10662) EUT1
07470 Hp_euts_violetPink EUT1
07475 HMM00390_malic_1_eut1(pf00390pf03949pf01515) EUT1
07480 -pf00923 -
07485 HMM02780keto_1_eut1(pf02779pf02780pf00456) EUT1
07490 -# -
07495 -pf06674 -
07500 HMM00293nuhyd_1_eut1(pf00293) EUT1
07505 -pf00144 -
07510 -# -
07515 -pf00005pf09383 -
07520 -pf00528 -
BMC-P
BMC-H
BMC-T
Aldehyde dehydrogenase
PTAC
Signature enzyme
Alcohol dehydrogenase
Regulator

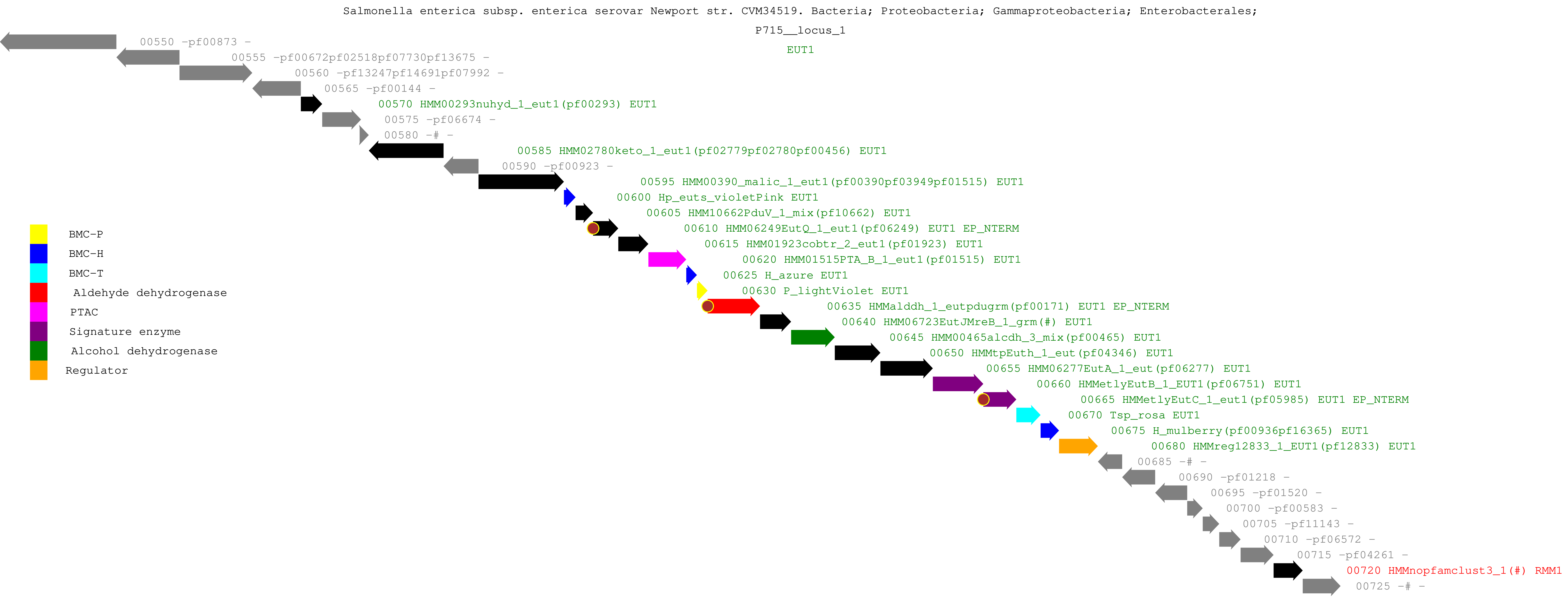

Salmonella enterica subsp. enterica serovar Newport str. CVM34519. Bacteria; Proteobacteria; Gammaproteobacteria; Enterobacterales;
P715__locus_1
EUT1
00550 -pf00873 -
00555 -pf00672pf02518pf07730pf13675 -
00560 -pf13247pf14691pf07992 -
00565 -pf00144 -
00570 HMM00293nuhyd_1_eut1(pf00293) EUT1
00575 -pf06674 -
00580 -# -
00585 HMM02780keto_1_eut1(pf02779pf02780pf00456) EUT1
00590 -pf00923 -
00595 HMM00390_malic_1_eut1(pf00390pf03949pf01515) EUT1
00600 Hp_euts_violetPink EUT1
00605 HMM10662PduV_1_mix(pf10662) EUT1
00610 HMM06249EutQ_1_eut1(pf06249) EUT1 EP_NTERM
00615 HMM01923cobtr_2_eut1(pf01923) EUT1
00620 HMM01515PTA_B_1_eut1(pf01515) EUT1
00625 H_azure EUT1
00630 P_lightViolet EUT1
00635 HMMalddh_1_eutpdugrm(pf00171) EUT1 EP_NTERM
00640 HMM06723EutJMreB_1_grm(#) EUT1
00645 HMM00465alcdh_3_mix(pf00465) EUT1
00650 HMMtpEuth_1_eut(pf04346) EUT1
00655 HMM06277EutA_1_eut(pf06277) EUT1
00660 HMMetlyEutB_1_EUT1(pf06751) EUT1
00665 HMMetlyEutC_1_eut1(pf05985) EUT1 EP_NTERM
00670 Tsp_rosa EUT1
00675 H_mulberry(pf00936pf16365) EUT1
00680 HMMreg12833_1_EUT1(pf12833) EUT1
00685 -# -
00690 -pf01218 -
00695 -pf01520 -
00700 -pf00583 -
00705 -pf11143 -
00710 -pf06572 -
00715 -pf04261 -
00720 HMMnopfamclust3_1(#) RMM1
00725 -# -
BMC-P
BMC-H
BMC-T
Aldehyde dehydrogenase
PTAC
Signature enzyme
Alcohol dehydrogenase
Regulator

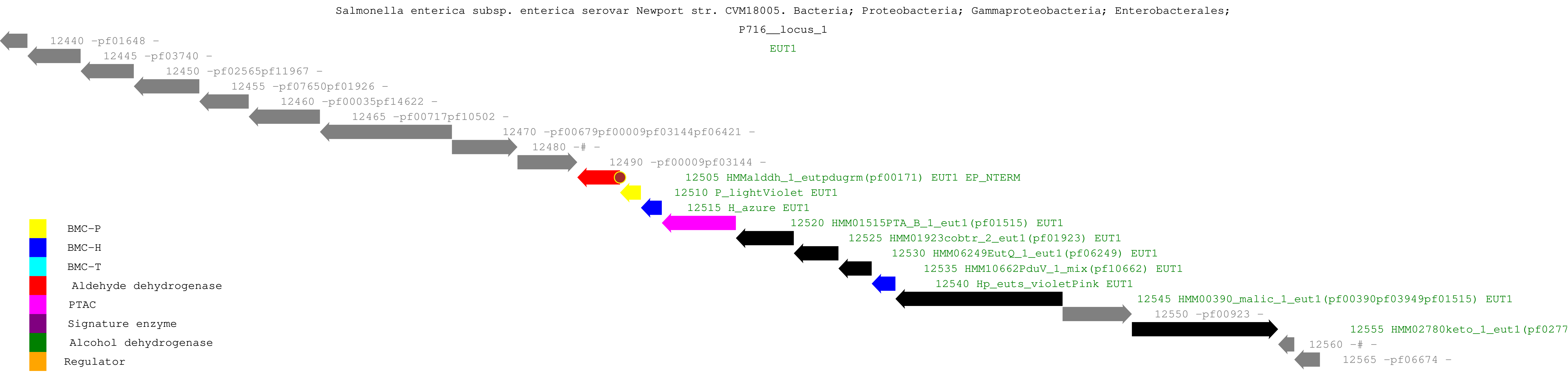
Salmonella enterica subsp. enterica serovar Newport str. CVM18005. Bacteria; Proteobacteria; Gammaproteobacteria; Enterobacterales;
P716__locus_1
EUT1
12440 -pf01648 -
12445 -pf03740 -
12450 -pf02565pf11967 -
12455 -pf07650pf01926 -
12460 -pf00035pf14622 -
12465 -pf00717pf10502 -
12470 -pf00679pf00009pf03144pf06421 -
12480 -# -
12490 -pf00009pf03144 -
12505 HMMaldh_1_eutpdugrm(pf00171) EUT1 EP_NTERM
12510 P_lightViolet EUT1
12515 H_azure EUT1
12520 HMM01515PTA_B_1_eut1(pf01515) EUT1
12525 HMM01923cobtr_2_eut1(pf01923) EUT1
12530 HMM06249EutQ_1_eut1(pf06249) EUT1
12535 HMM10662PduV_1_mix(pf10662) EUT1
12540 Hp_euts_violetPink EUT1
12545 HMM00390_malic_1_eut1(pf00390pf03949pf01515) EUT1
12550 -pf00923 -
12555 HMM02780keto_1_eut1(pf0277
12560 -# -
12565 -pf06674 -
BMC-P
BMC-H
BMC-T
Aldehyde dehydrogenase
PTAC
Signature enzyme
Alcohol dehydrogenase
Regulator

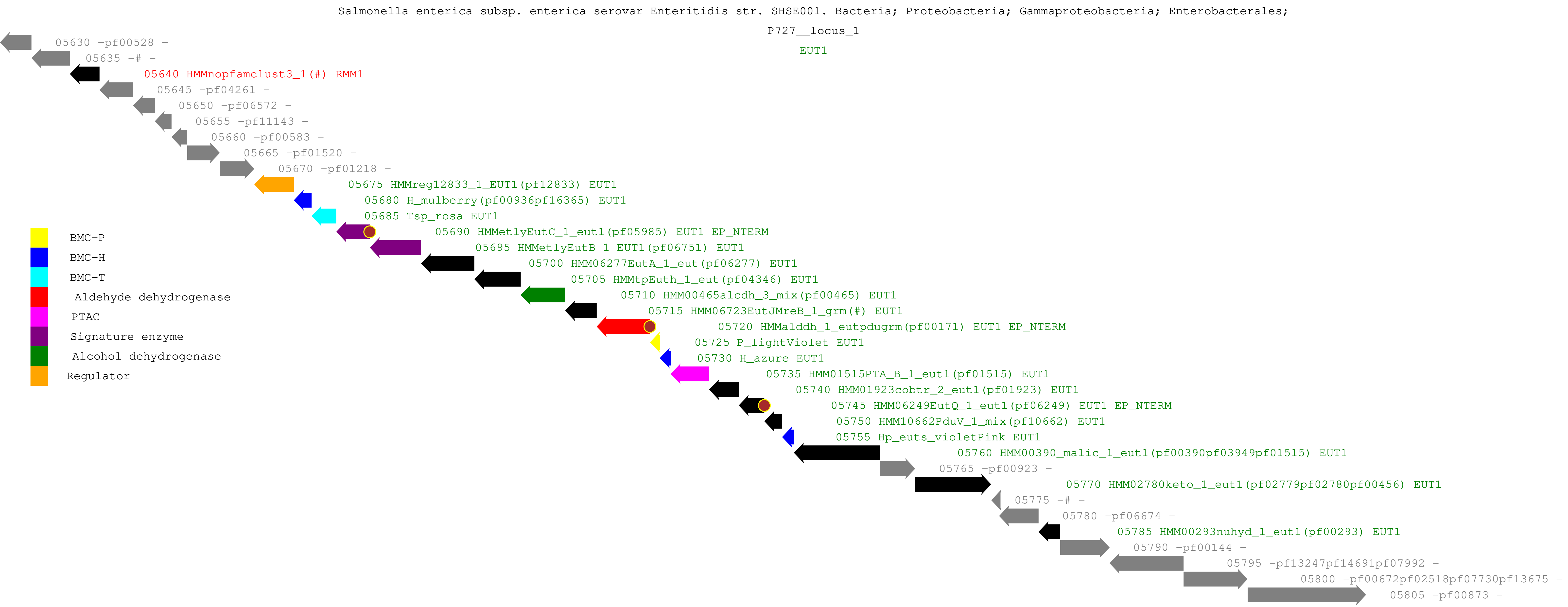
Salmonella enterica subsp. enterica serovar Enteritidis str. SHSE001. Bacteria; Proteobacteria; Gammaproteobacteria; Enterobacterales;
P727__locus_1
EUT1
05630 -pf00528 -
05635 -# -
05640 HMMnopfamclust3_1(#) RMM1
05645 -pf04261 -
05650 -pf06572 -
05655 -pf11143 -
05660 -pf00583 -
05665 -pf01520 -
05670 -pf01218 -
05675 HMMreg12833_1_EUT1(pf12833) EUT1
05680 H_mulberry(pf00936pf16365) EUT1
05685 Tsp_rosa EUT1
05690 HMMetlyEutC_1_eut1(pf05985) EUT1 EP_NTERM
05695 HMMetlyEutB_1_EUT1(pf06751) EUT1
05700 HMM06277EutA_1_eut(pf06277) EUT1
05705 HMMtpEuth_1_eut(pf04346) EUT1
05710 HMM00465alcdh_3_mix(pf00465) EUT1
05715 HMM06723EutJMreB_1_grm(#) EUT1
05720 HMMalddh_1_eutpdugrm(pf00171) EUT1 EP_NTERM
05725 P_lightViolet EUT1
05730 H_azure EUT1
05735 HMM01515PTA_B_1_eut1(pf01515) EUT1
05740 HMM01923cobtr_2_eut1(pf01923) EUT1
05745 HMM06249EutQ_1_eut1(pf06249) EUT1 EP_NTERM
05750 HMM10662PduV_1_mix(pf10662) EUT1
05755 Hp_euts_violetPink EUT1
05760 HMM00390_malic_1_eut1(pf00390pf03949pf01515) EUT1
05765 -pf00923 -
05770 HMM02780keto_1_eut1(pf02779pf02780pf00456) EUT1
05775 -# -
05780 -pf06674 -
05785 HMM00293nuhyd_1_eut1(pf00293) EUT1
05790 -pf00144 -
05795 -pf13247pf14691pf07992 -
05800 -pf00672pf02518pf07730pf13675 -
05805 -pf00873 -
BMC-P
BMC-H
BMC-T
Aldehyde dehydrogenase
PTAC
Signature enzyme
Alcohol dehydrogenase
Regulator

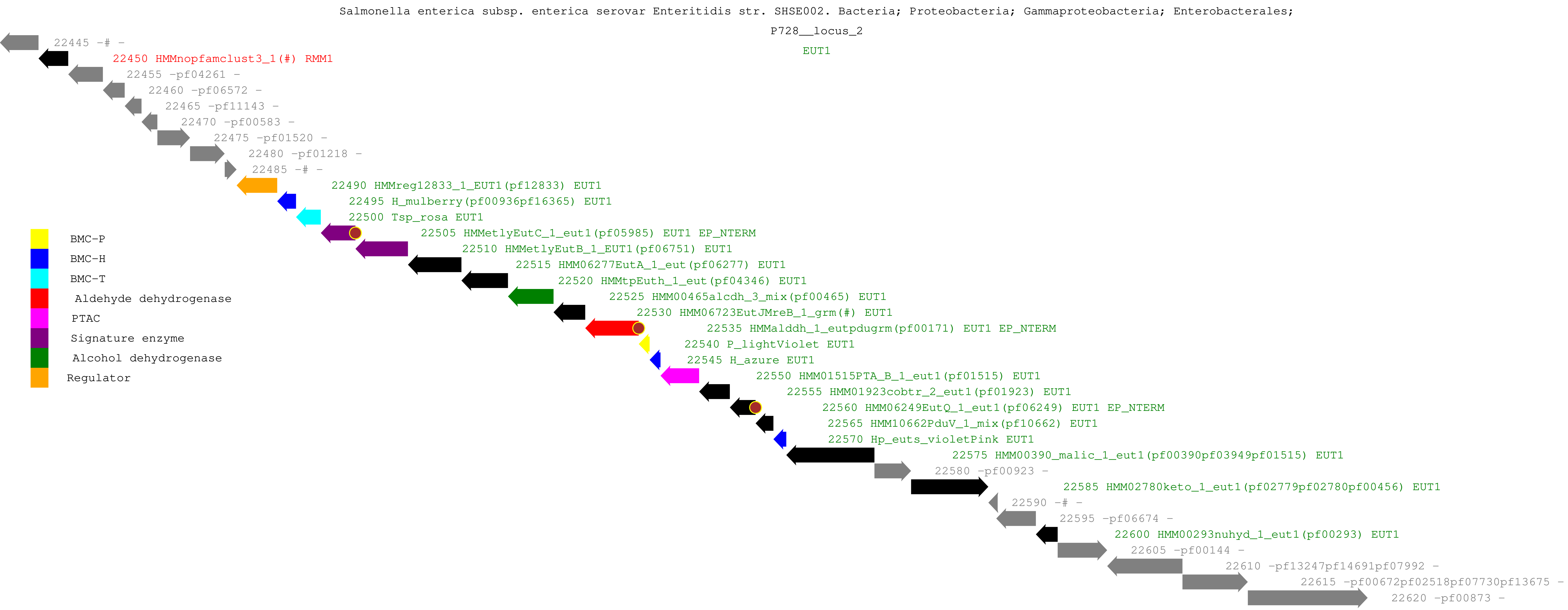
Salmonella enterica subsp. enterica serovar Enteritidis str. SHSE002. Bacteria; Proteobacteria; Gammaproteobacteria; Enterobacterales;
P728__locus_2
EUT1
22445 -# -
22450 HMMnopfamclust3_1(#) RMM1
22455 -pf04261 -
22460 -pf06572 -
22465 -pf11143 -
22470 -pf00583 -
22475 -pf01520 -
22480 -pf01218 -
22485 -# -
22490 HMMreg12833_1_EUT1(pf12833) EUT1
22495 H_mulberry(pf00936pf16365) EUT1
22500 Tsp_rosa EUT1
22505 HMMetlyEutC_1_eut1(pf05985) EUT1 EP_NTERM
22510 HMMetlyEutB_1_EUT1(pf06751) EUT1
22515 HMM06277EutA_1_eut(pf06277) EUT1
22520 HMMtpEuth_1_eut(pf04346) EUT1
22525 HMM00465alcdh_3_mix(pf00465) EUT1
22530 HMM06723EutJMreB_1_grm(#) EUT1
22535 HMMaldh_1_eutpdugrm(pf00171) EUT1 EP_NTERM
22540 P_lightViolet EUT1
22545 H_azure EUT1
22550 HMM01515PTA_B_1_eut1(pf01515) EUT1
22555 HMM01923cobtr_2_eut1(pf01923) EUT1
22560 HMM06249EutQ_1_eut1(pf06249) EUT1 EP_NTERM
22565 HMM10662PduV_1_mix(pf10662) EUT1
22570 Hp_euts_violetPink EUT1
22575 HMM00390_malic_1_eut1(pf00390pf03949pf01515) EUT1
22580 -pf00923 -
22585 HMM02780keto_1_eut1(pf02779pf02780pf00456) EUT1
22590 -# -
22595 -pf06674 -
22600 HMM00293nuhyd_1_eut1(pf00293) EUT1
22605 -pf00144 -
22610 -pf13247pf14691pf07992 -
22615 -pf00672pf02518pf07730pf13675 -
22620 -pf00873 -
BMC-P
BMC-H
BMC-T
Aldehyde dehydrogenase
PTAC
Signature enzyme
Alcohol dehydrogenase
Regulator

Salmonella enterica subsp. enterica serovar Enteritidis str. SHSE003. Bacteria; Proteobacteria; Gammaproteobacteria; Enterobacterales;

P729__locus_2

EUT1

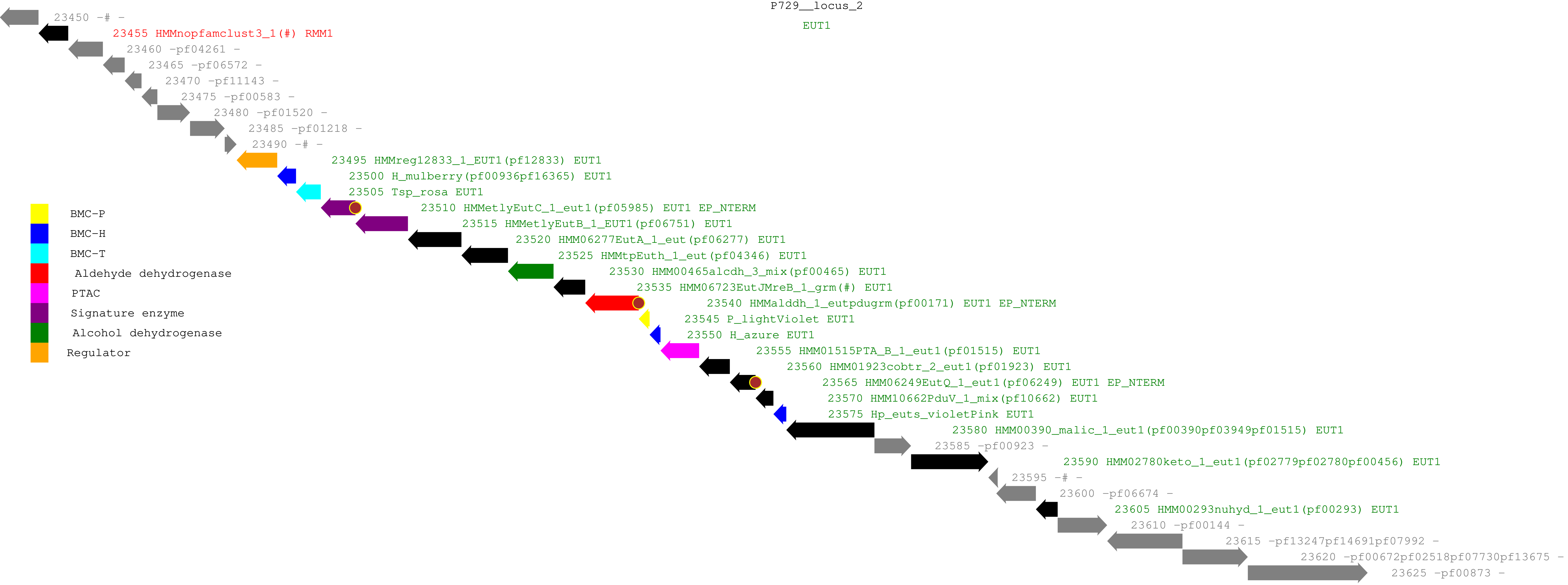

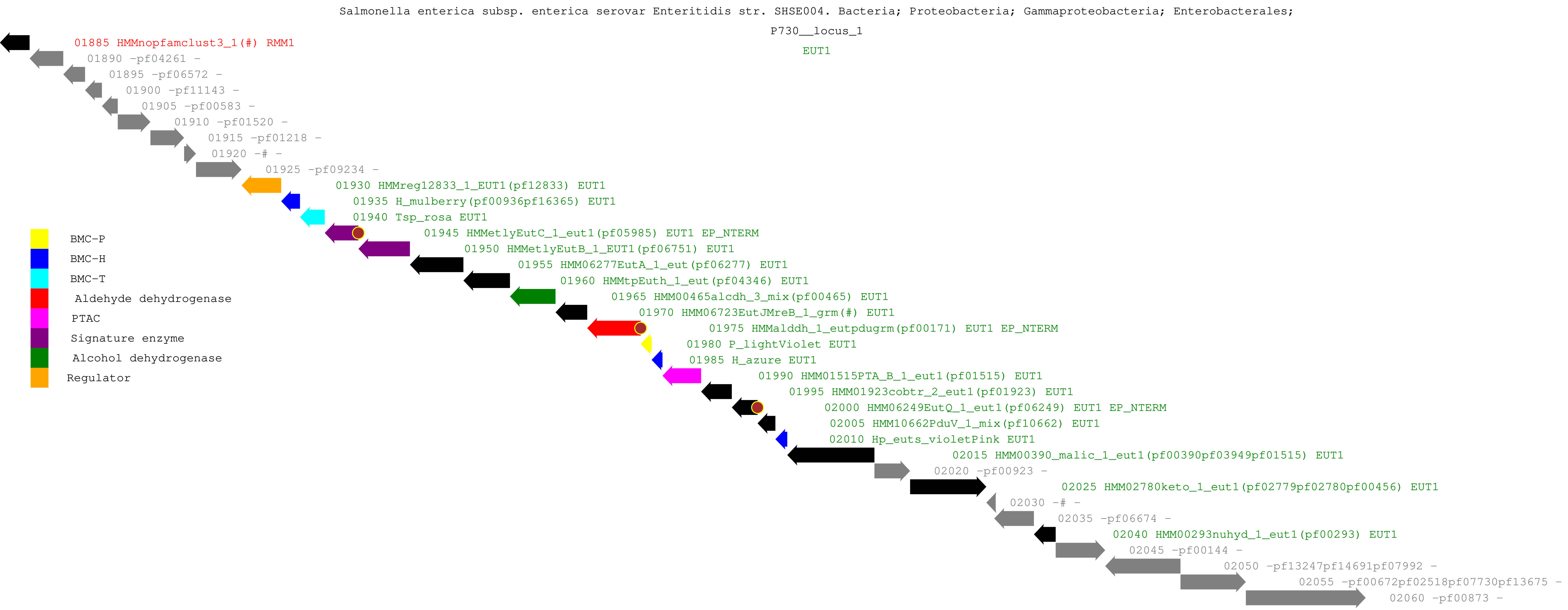
Salmonella enterica subsp. enterica serovar Enteritidis str. SHSE004. Bacteria; Proteobacteria; Gammaproteobacteria; Enterobacterales;
P730__locus_1
EUT1
01885 HMMnopfamclust3_1(#) RMM1
01890 -pf04261 -
01895 -pf06572 -
01900 -pf11143 -
01905 -pf00583 -
01910 -pf01520 -
01915 -pf01218 -
01920 -# -
01925 -pf09234 -
01930 HMMreg12833_1_EUT1(pf12833) EUT1
01935 H_mulberry(pf00936pf16365) EUT1
01940 Tsp_rosa EUT1
01945 HMMetlyEutC_1_eut1(pf05985) EUT1 EP_NTERM
01950 HMMetlyEutB_1_EUT1(pf06751) EUT1
01955 HMM06277EutA_1_eut(pf06277) EUT1
01960 HMMtpEuth_1_eut(pf04346) EUT1
01965 HMM00465alcdh_3_mix(pf00465) EUT1
01970 HMM06723EutJMreB_1_grm(#) EUT1
01975 HMMalddh_1_eutpdugrm(pf00171) EUT1 EP_NTERM
01980 P_lightViolet EUT1
01985 H_azure EUT1
01990 HMM01515PTA_B_1_eut1(pf01515) EUT1
01995 HMM01923cobtr_2_eut1(pf01923) EUT1
02000 HMM06249EutQ_1_eut1(pf06249) EUT1 EP_NTERM
02005 HMM10662PduV_1_mix(pf10662) EUT1
02010 Hp_euts_violetPink EUT1
02015 HMM00390_malic_1_eut1(pf00390pf03949pf01515) EUT1
02020 -pf00923 -
02025 HMM02780keto_1_eut1(pf02779pf02780pf00456) EUT1
02030 -# -
02035 -pf06674 -
02040 HMM00293nuhyd_1_eut1(pf00293) EUT1
02045 -pf00144 -
02050 -pf13247pf14691pf07992 -
02055 -pf00672pf02518pf07730pf13675 -
02060 -pf00873 -
BMC-P
BMC-H
BMC-T
Aldehyde dehydrogenase
PTAC
Signature enzyme
Alcohol dehydrogenase
Regulator

Pseudomonas citronellolis. Bacteria; Proteobacteria; Gammaproteobacteria; Pseudomonadales;

PcP3B5__locus_1

EUT1

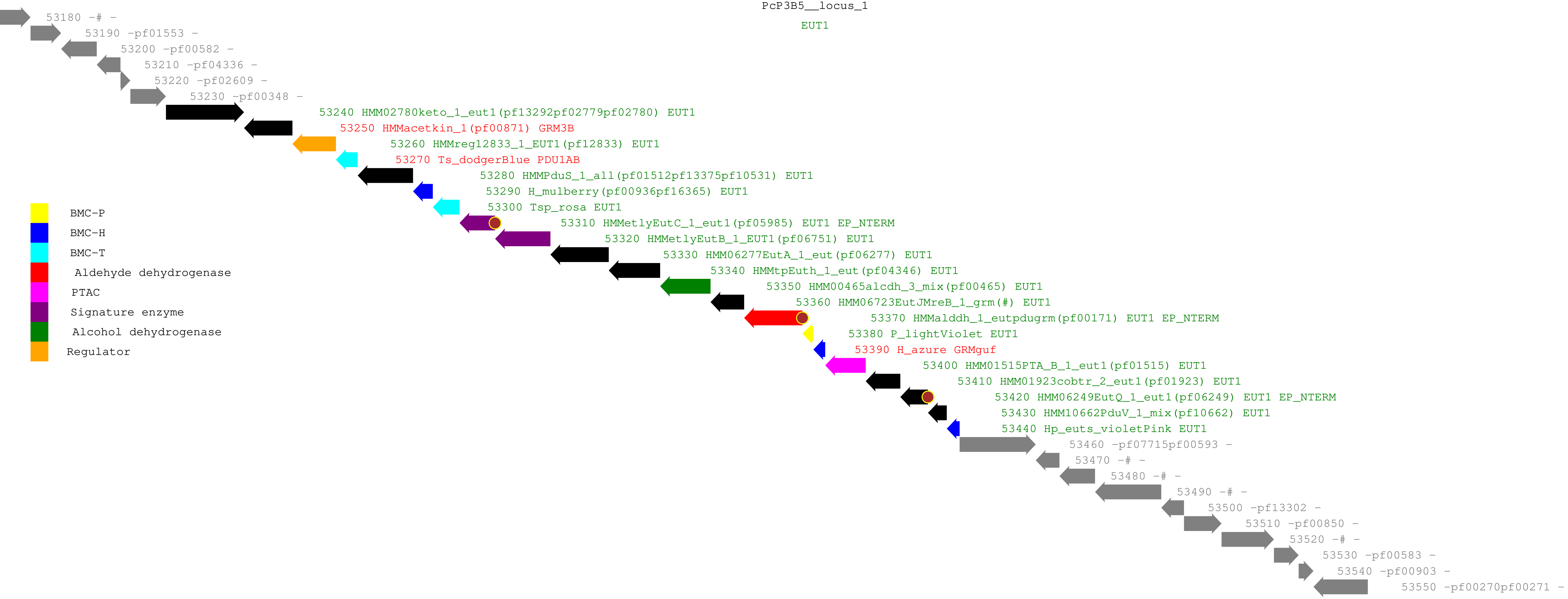

Escherichia coli D6-113.11. Bacteria; Proteobacteria; Gammaproteobacteria; Enterobacterales;

PGA__locus_1

EUT1

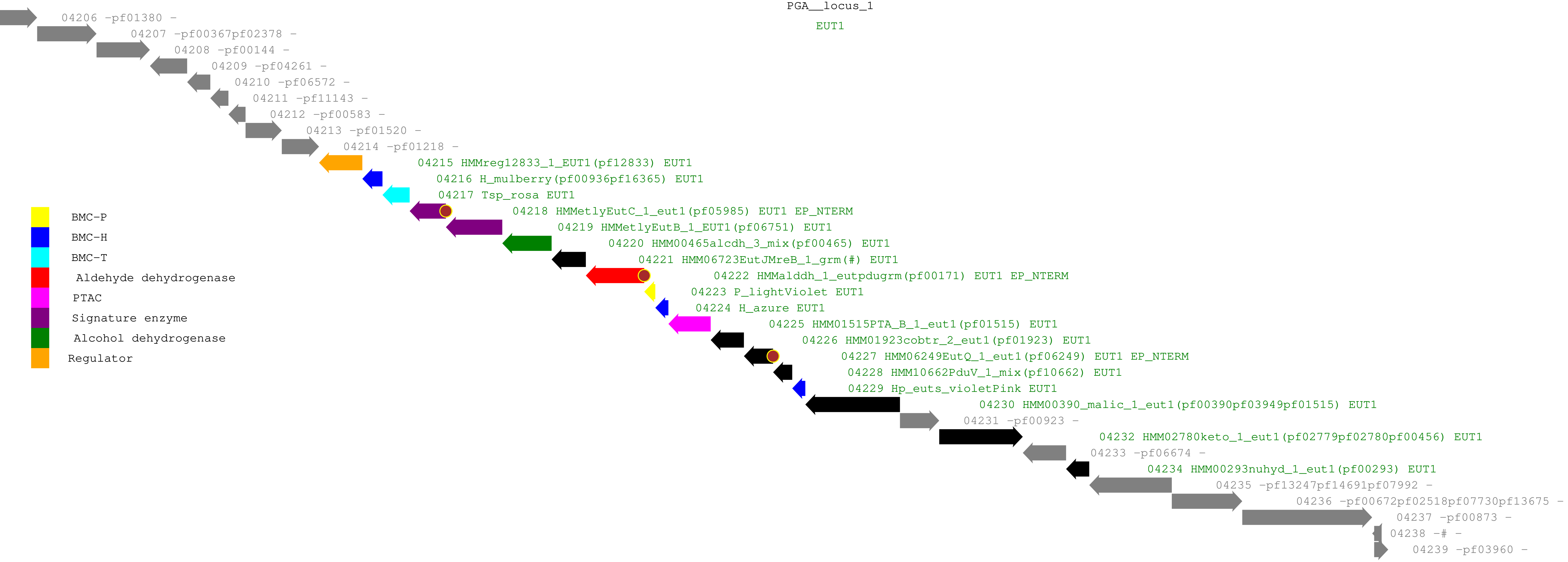

Salmonella enterica I. Bacteria; Proteobacteria; Gammaproteobacteria; Enterobacterales;

PK95__locus_2

EUT1

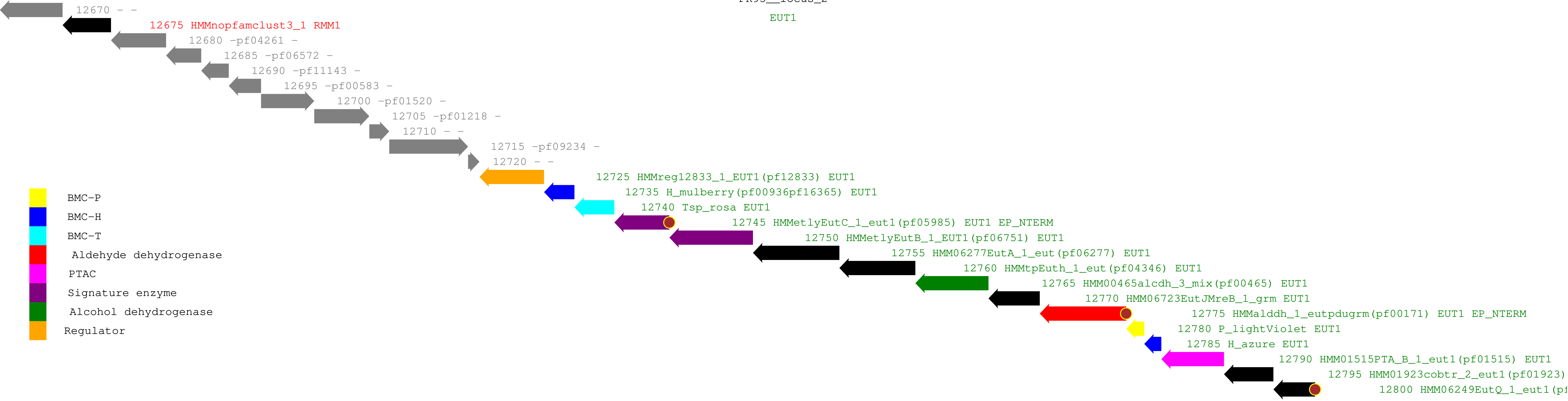

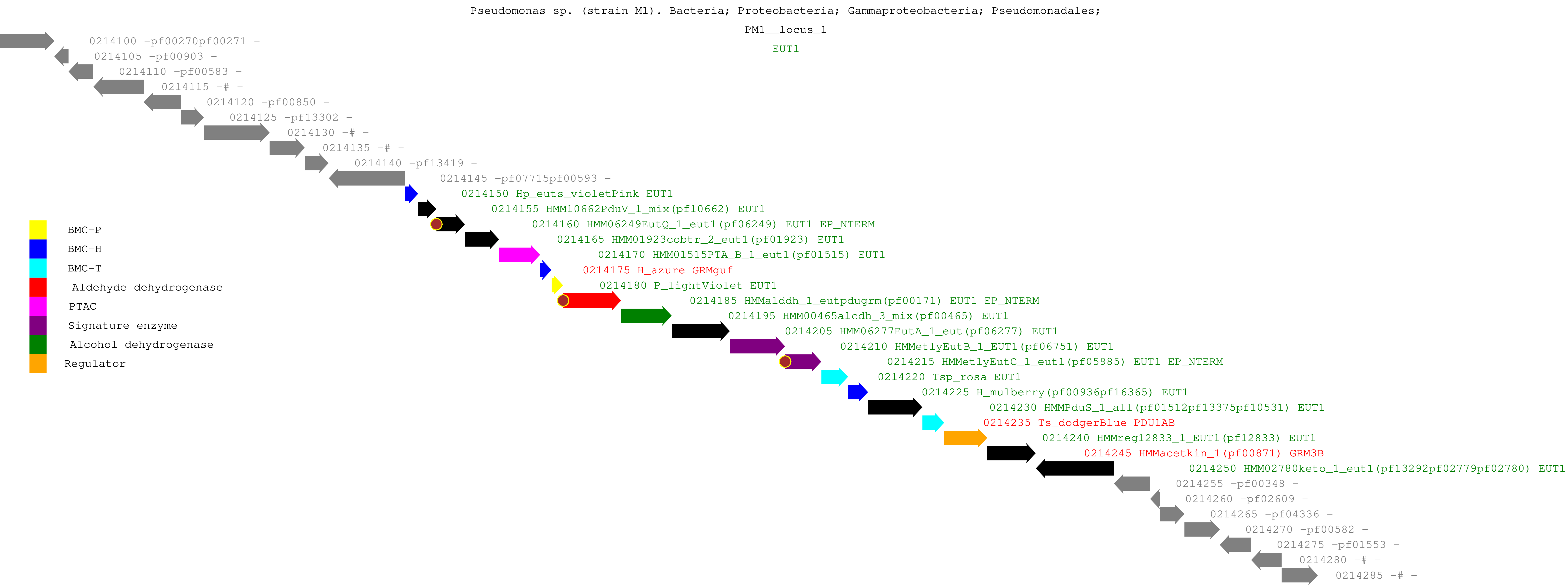
Pseudomonas sp. (strain M1). Bacteria; Proteobacteria; Gammaproteobacteria; Pseudomonadales;
PM1__locus_1
EUT1
0214100 -pf00270pf00271 -
0214105 -pf00903 -
0214110 -pf00583 -
0214115 -# -
0214120 -pf00850 -
0214125 -pf13302 -
0214130 -# -
0214135 -# -
0214140 -pf13419 -
0214145 -pf07715pf00593 -
0214150 Hp_euts_violetPink EUT1
0214155 HMM10662PduV_1_mix(pf10662) EUT1
0214160 HMM06249EutQ_1_eut1(pf06249) EUT1 EP_NTERM
0214165 HMM01923cobtr_2_eut1(pf01923) EUT1
0214170 HMM01515PTA_B_1_eut1(pf01515) EUT1
0214175 H_azure GRMguf
0214180 P_lightViolet EUT1
0214185 HMMalddh_1_eutpdugrm(pf00171) EUT1 EP_NTERM
0214195 HMM00465alcdh_3_mix(pf00465) EUT1
0214205 HMM06277EutA_1_eut(pf06277) EUT1
0214210 HMMetlyEutB_1_EUT1(pf06751) EUT1
0214215 HMMetlyEutC_1_eut1(pf05985) EUT1 EP_NTERM
0214220 Tsp_rosa EUT1
0214225 H_mulberry(pf00936pf16365) EUT1
0214230 HMMPduS_1_all(pf01512pf13375pf10531) EUT1
0214235 Ts_dodgerBlue PDU1AB
0214240 HMMreg12833_1_EUT1(pf12833) EUT1
0214245 HMMacetkin_1(pf00871) GRM3B
0214250 HMM02780keto_1_eut1(pf13292pf02779pf02780) EUT1
0214255 -pf00348 -
0214260 -pf02609 -
0214265 -pf04336 -
0214270 -pf00582 -
0214275 -pf01553 -
0214280 -# -
0214285 -# -
BMC-P
BMC-H
BMC-T
Aldehyde dehydrogenase
PTAC
Signature enzyme
Alcohol dehydrogenase
Regulator

Escherichia coli PCN033. Bacteria; Proteobacteria; Gammaproteobacteria; Enterobacterales;

PPECC33__locus_1

EUT1

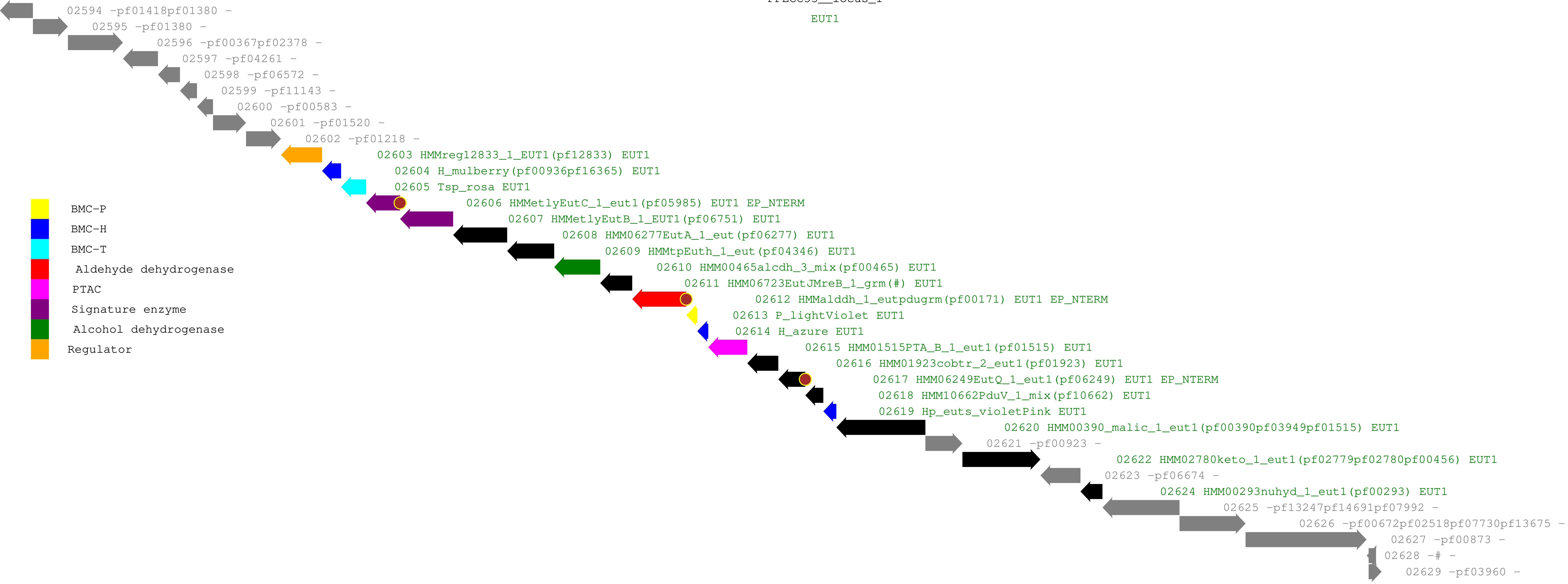

Hafnia alvei. Bacteria; Proteobacteria; Gammaproteobacteria; Enterobacterales;

PU01__locus_1

EUT1

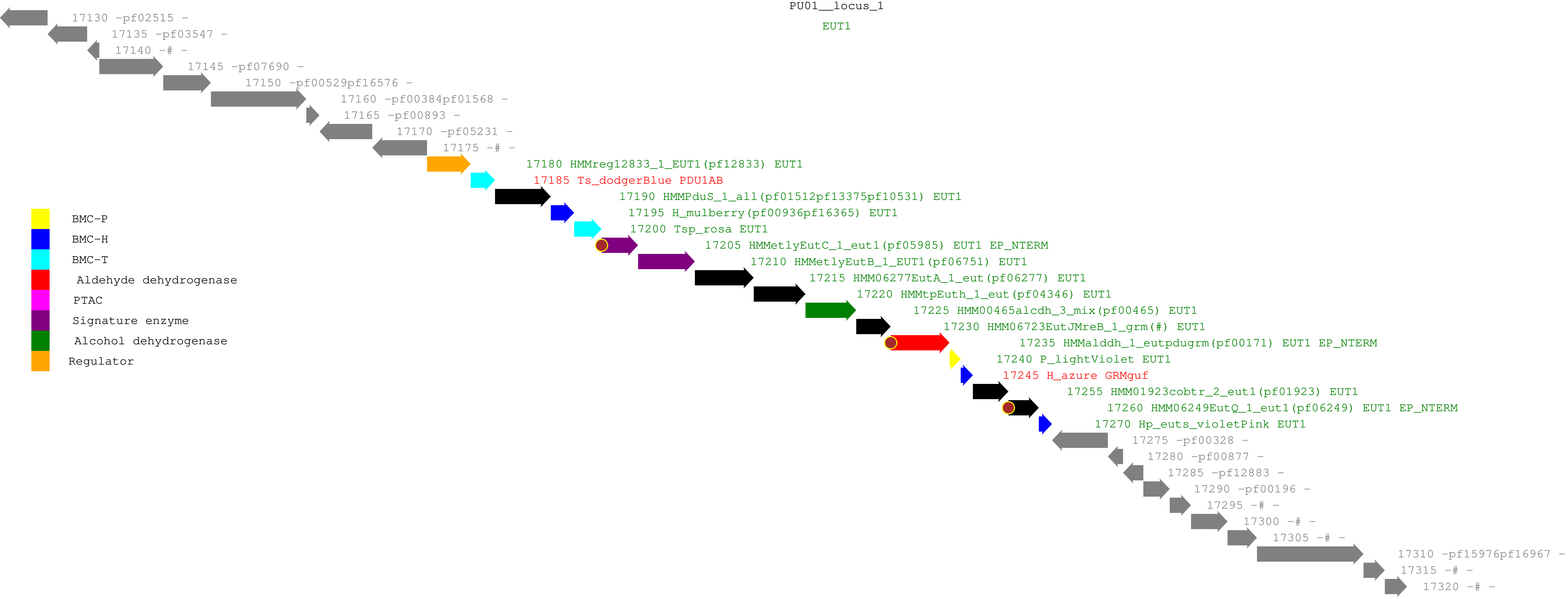

Escherichia coli ATCC BAA-2209. Bacteria; Proteobacteria; Gammaproteobacteria; Enterobacterales;

Q458__locus_1_main_sat

EUT1

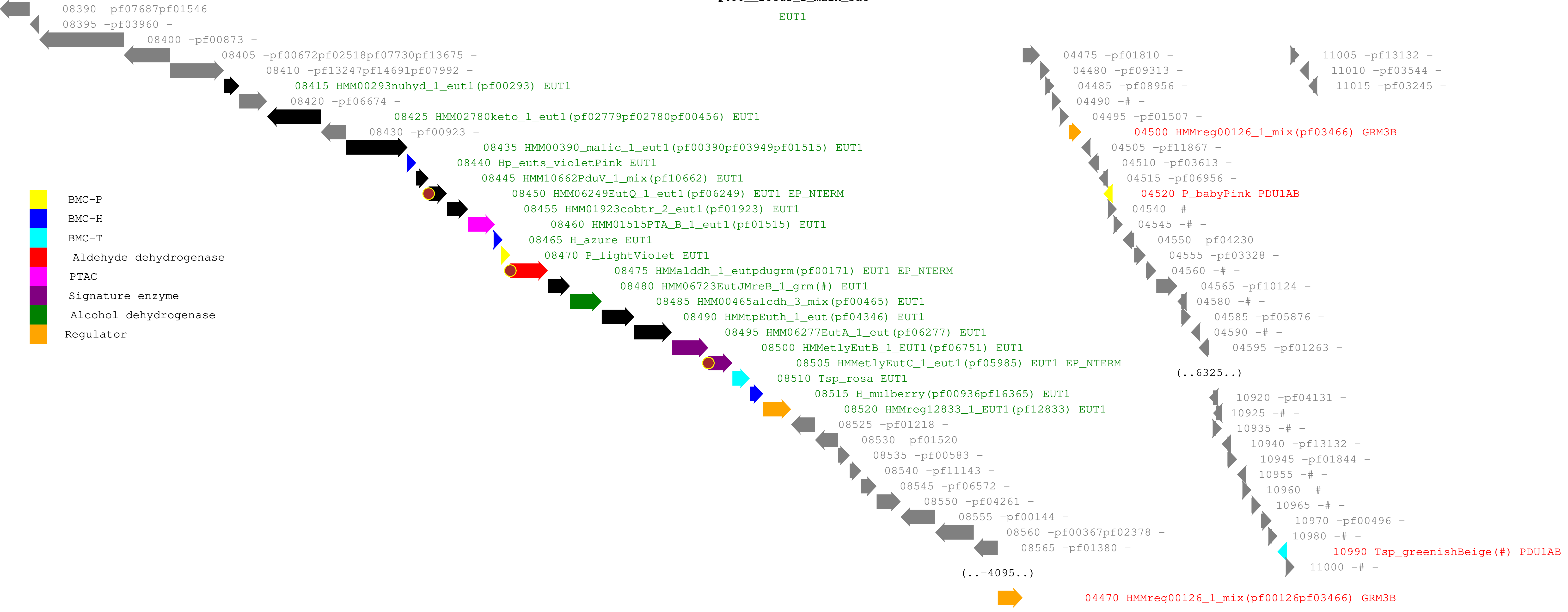

Escherichia coli N40607. Bacteria; Proteobacteria; Gammaproteobacteria; Enterobacterales;

Q676__locus_1

EUT1

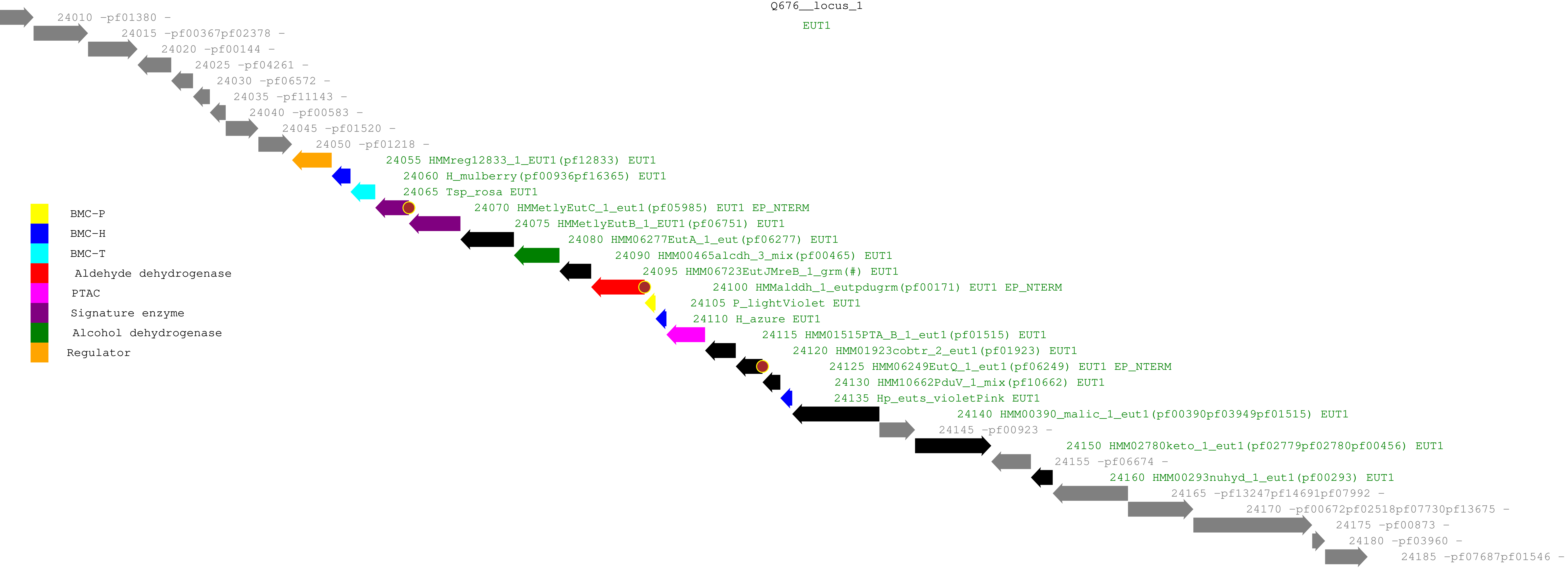

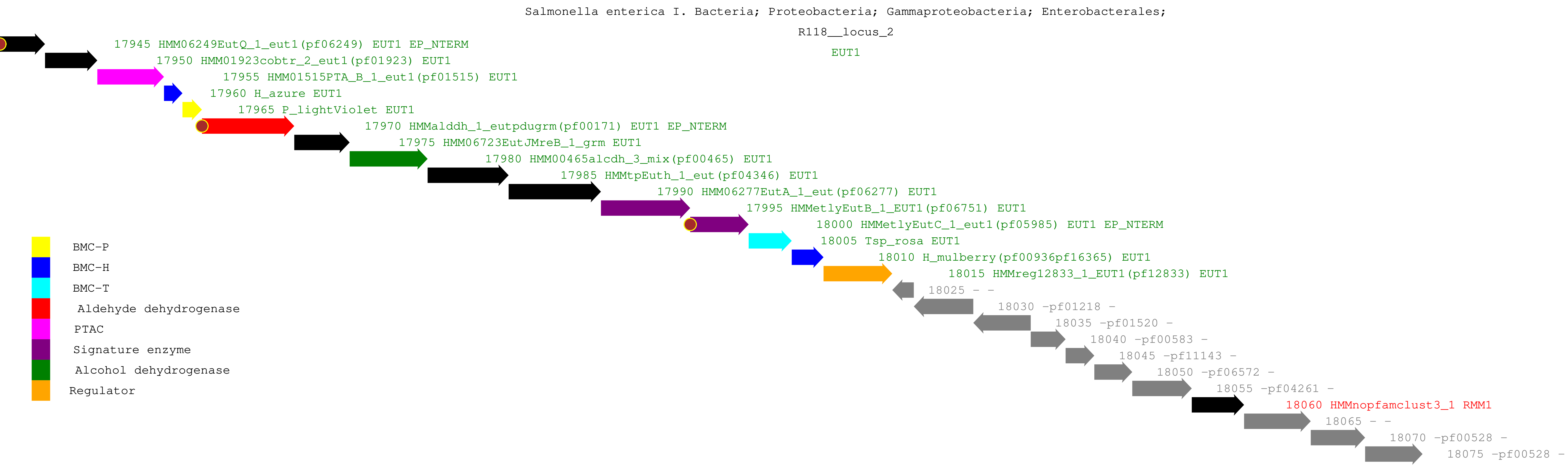
Salmonella enterica I. Bacteria; Proteobacteria; Gammaproteobacteria; Enterobacterales;
R118__locus_2
EUT1
17945 HMM06249EutQ_1_eut1(pf06249) EUT1 EP_NTERM
17950 HMM01923cobtr_2_eut1(pf01923) EUT1
17955 HMM01515PTA_B_1_eut1(pf01515) EUT1
17960 H_azure EUT1
17965 P_lightViolet EUT1
17970 HMMalddh_1_eutpdugrm(pf00171) EUT1 EP_NTERM
17975 HMM06723EutJMreB_1_grm EUT1
17980 HMM00465alcdh_3_mix(pf00465) EUT1
17985 HMMtpEuth_1_eut(pf04346) EUT1
17990 HMM06277EutA_1_eut(pf06277) EUT1
17995 HMMetlyEutB_1_EUT1(pf06751) EUT1
18000 HMMetlyEutC_1_eut1(pf05985) EUT1 EP_NTERM
18005 Tsp_rosa EUT1
18010 H_mulberry(pf00936pf16365) EUT1
18015 HMMreg12833_1_EUT1(pf12833) EUT1
18025 - -
18030 -pf01218 -
18035 -pf01520 -
18040 -pf00583 -
18045 -pf11143 -
18050 -pf06572 -
18055 -pf04261 -
18060 HMMnopfamclust3_1 RMM1
18065 - -
18070 -pf00528 -
18075 -pf00528 -
BMC-P
BMC-H
BMC-T
Aldehyde dehydrogenase
PTAC
Signature enzyme
Alcohol dehydrogenase
Regulator

Salmonella senftenberg. Bacteria; Proteobacteria; Gammaproteobacteria; Enterobacterales;

R495__locus_1

EUT1

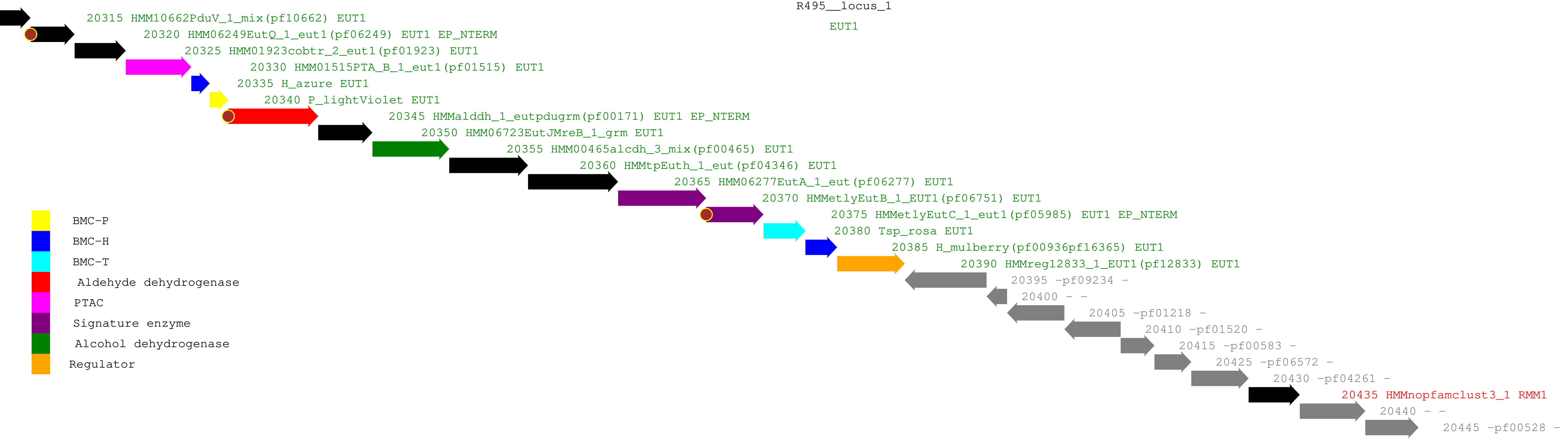

Salmonella enterica subsp. arizonae serovar 41:z4,z23:-. Bacteria; Proteobacteria; Gammaproteobacteria; Enterobacterales;

R515__locus_1

EUT1

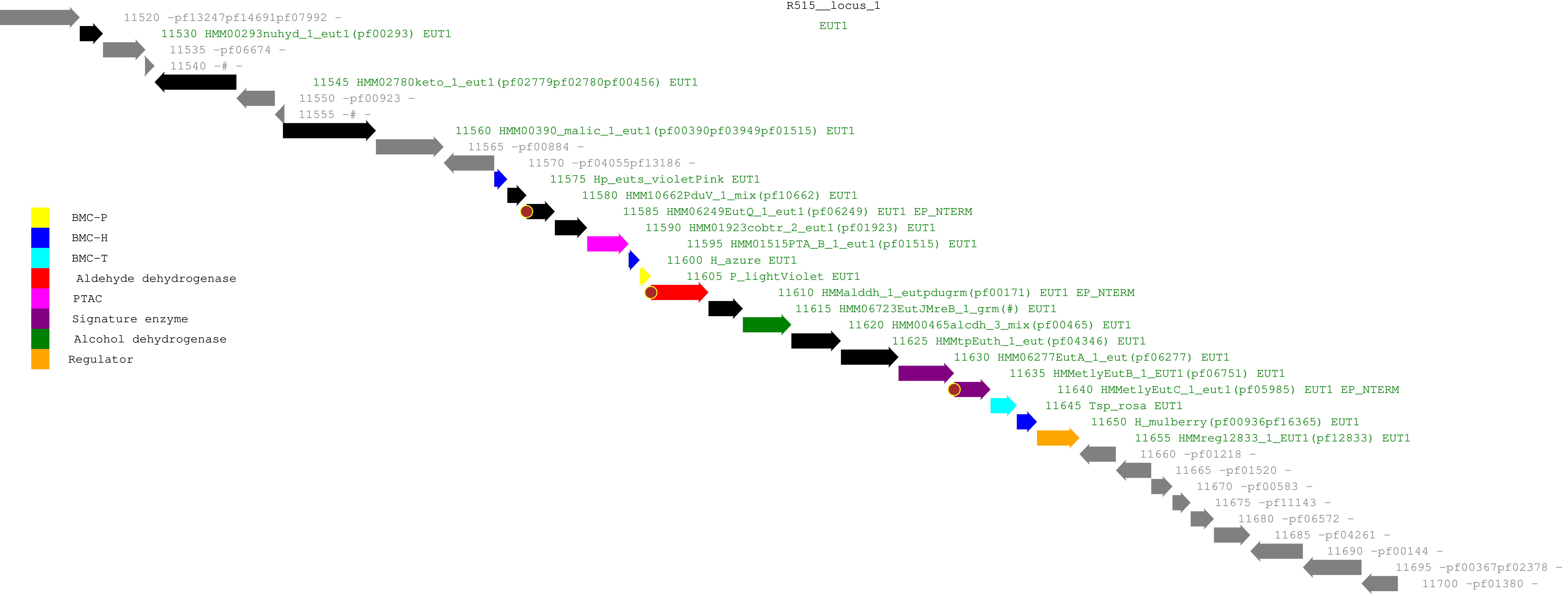

Salmonella enterica subsp. arizonae serovar 50:r:z. Bacteria; Proteobacteria; Gammaproteobacteria; Enterobacterales;

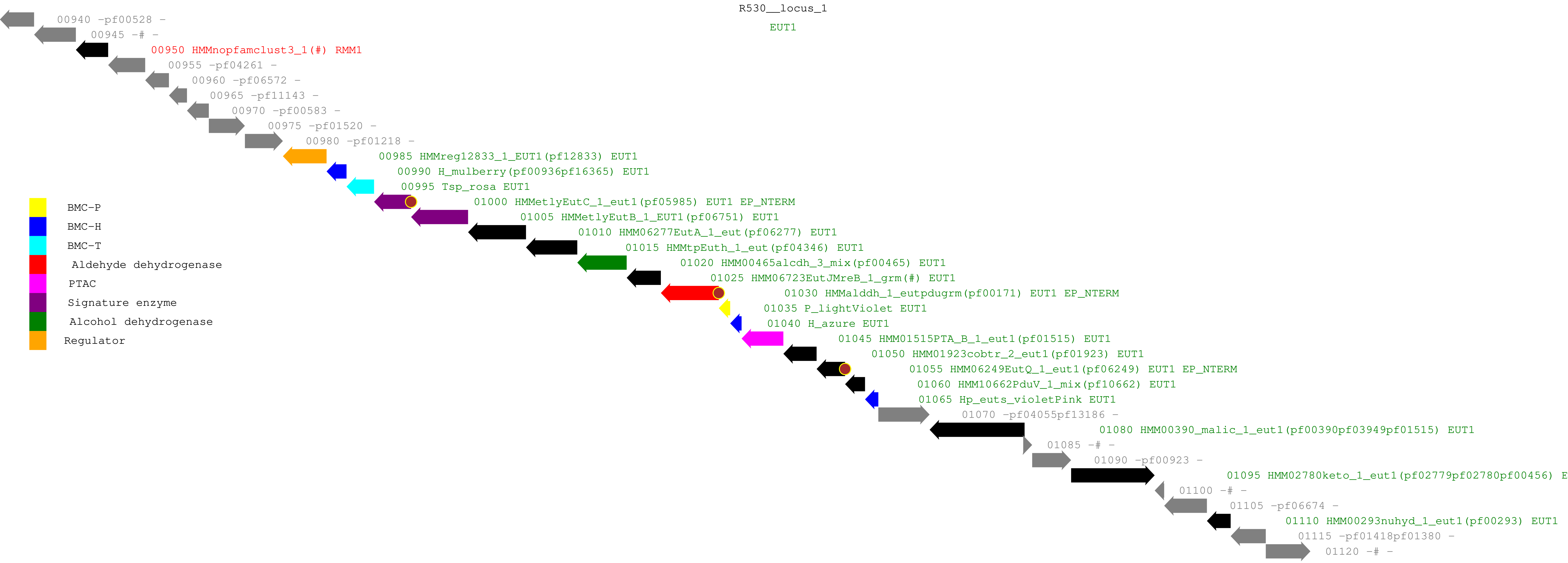

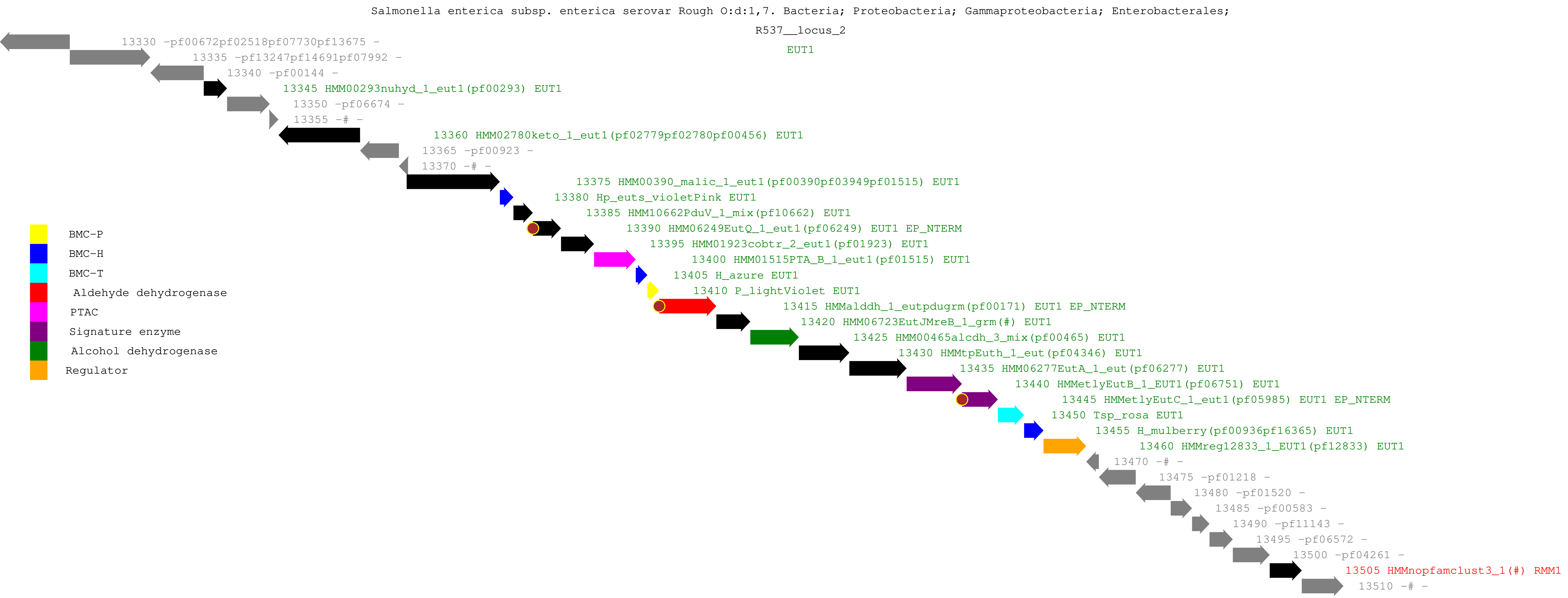
Salmonella enterica subsp. enterica serovar Rough O:d:1,7. Bacteria; Proteobacteria; Gammaproteobacteria; Enterobacterales;
R537__locus_2
EUT1
13330 -pf00672pf02518pf07730pf13675 -
13335 -pf13247pf14691pf07992 -
13340 -pf00144 -
13345 HMM00293nuhyd_1_eut1(pf00293) EUT1
13350 -pf06674 -
13355 -# -
13360 HMM02780keto_1_eut1(pf02779pf02780pf00456) EUT1
13365 -pf00923 -
13370 -# -
13375 HMM00390_malic_1_eut1(pf00390pf03949pf01515) EUT1
13380 Hp_euts_violetPink EUT1
13385 HMM10662PduV_1_mix(pf10662) EUT1
13390 HMM06249EutQ_1_eut1(pf06249) EUT1 EP_NTERM
13395 HMM01923cobtr_2_eut1(pf01923) EUT1
13400 HMM01515PTA_B_1_eut1(pf01515) EUT1
13405 H_azure EUT1
13410 P_lightViolet EUT1
13415 HMMalddh_1_eutpdugrm(pf00171) EUT1 EP_NTERM
13420 HMM06723EutJMreB_1_grm(#) EUT1
13425 HMM00465alcdh_3_mix(pf00465) EUT1
13430 HMMtpEuth_1_eut(pf04346) EUT1
13435 HMM06277EutA_1_eut(pf06277) EUT1
13440 HMMetlyEutB_1_EUT1(pf06751) EUT1
13445 HMMetlyEutC_1_eut1(pf05985) EUT1 EP_NTERM
13450 Tsp_rosa EUT1
13455 H_mulberry(pf00936pf16365) EUT1
13460 HMMreg12833_1_EUT1(pf12833) EUT1
13470 -# -
13475 -pf01218 -
13480 -pf01520 -
13485 -pf00583 -
13490 -pf11143 -
13495 -pf06572 -
13500 -pf04261 -
13505 HMMnopfamclust3_1(#) RMM1
13510 -# -
BMC-P
BMC-H
BMC-T
Aldehyde dehydrogenase
PTAC
Signature enzyme
Alcohol dehydrogenase
Regulator

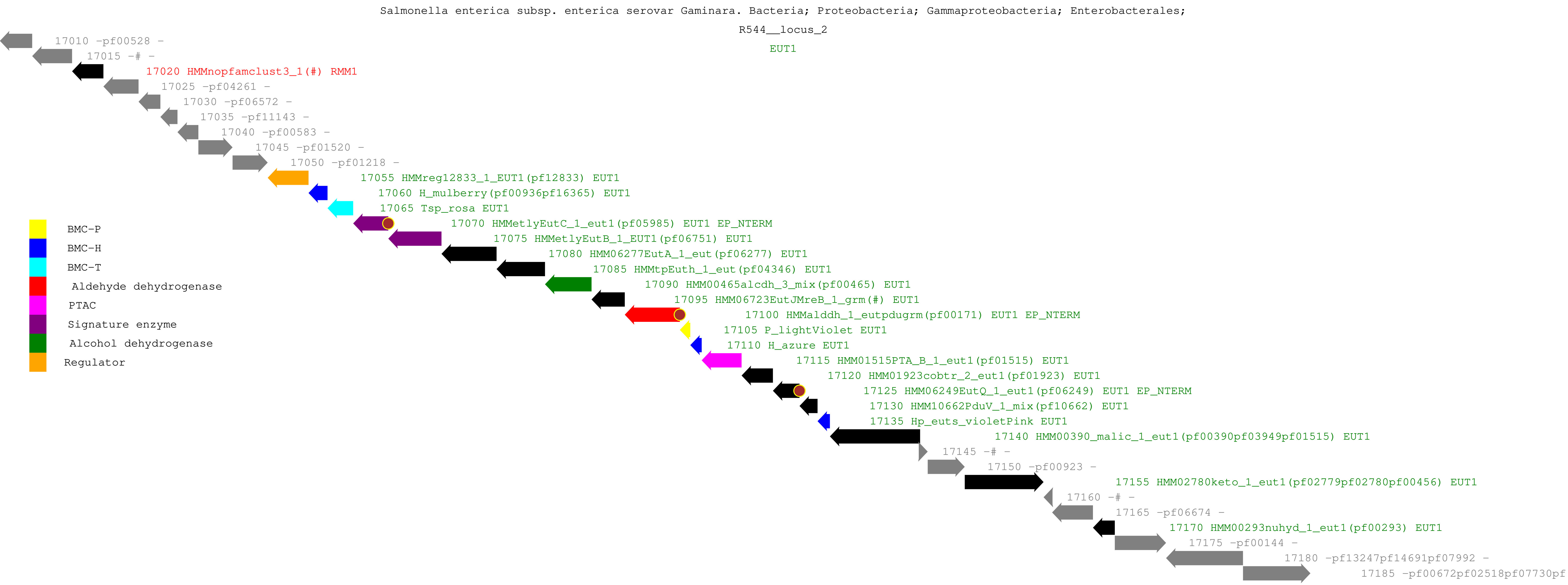
Salmonella enterica subsp. enterica serovar Gaminara. Bacteria; Proteobacteria; Gammaproteobacteria; Enterobacterales;
R544__locus_2
EUT1
17010 -pf00528 -
17015 -# -
17020 HMMnopfamclust3_1(#) RMM1
17025 -pf04261 -
17030 -pf06572 -
17035 -pf11143 -
17040 -pf00583 -
17045 -pf01520 -
17050 -pf01218 -
17055 HMMreg12833_1_EUT1(pf12833) EUT1
17060 H_mulberry(pf00936pf16365) EUT1
17065 Tsp_rosa EUT1
17070 HMMetlyEutC_1_eut1(pf05985) EUT1 EP_NTERM
17075 HMMetlyEutB_1_EUT1(pf06751) EUT1
17080 HMM06277EutA_1_eut(pf06277) EUT1
17085 HMMtpEuth_1_eut(pf04346) EUT1
17090 HMM00465alcdh_3_mix(pf00465) EUT1
17095 HMM06723EutJMreB_1_grm(#) EUT1
17100 HMMalddh_1_eutpdugrm(pf00171) EUT1 EP_NTERM
17105 P_lightViolet EUT1
17110 H_azure EUT1
17115 HMM01515PTA_B_1_eut1(pf01515) EUT1
17120 HMM01923cobtr_2_eut1(pf01923) EUT1
17125 HMM06249EutQ_1_eut1(pf06249) EUT1 EP_NTERM
17130 HMM10662PduV_1_mix(pf10662) EUT1
17135 Hp_euts_violetPink EUT1
17140 HMM00390_malic_1_eut1(pf00390pf03949pf01515) EUT1
17145 -# -
17150 -pf00923 -
17155 HMM02780keto_1_eut1(pf02779pf02780pf00456) EUT1
17160 -# -
17165 -pf06674 -
17170 HMM00293nuhyd_1_eut1(pf00293) EUT1
17175 -pf00144 -
17180 -pf13247pf14691pf07992 -
17185 -pf00672pf02518pf07730pf
BMC-P
BMC-H
BMC-T
Aldehyde dehydrogenase
PTAC
Signature enzyme
Alcohol dehydrogenase
Regulator

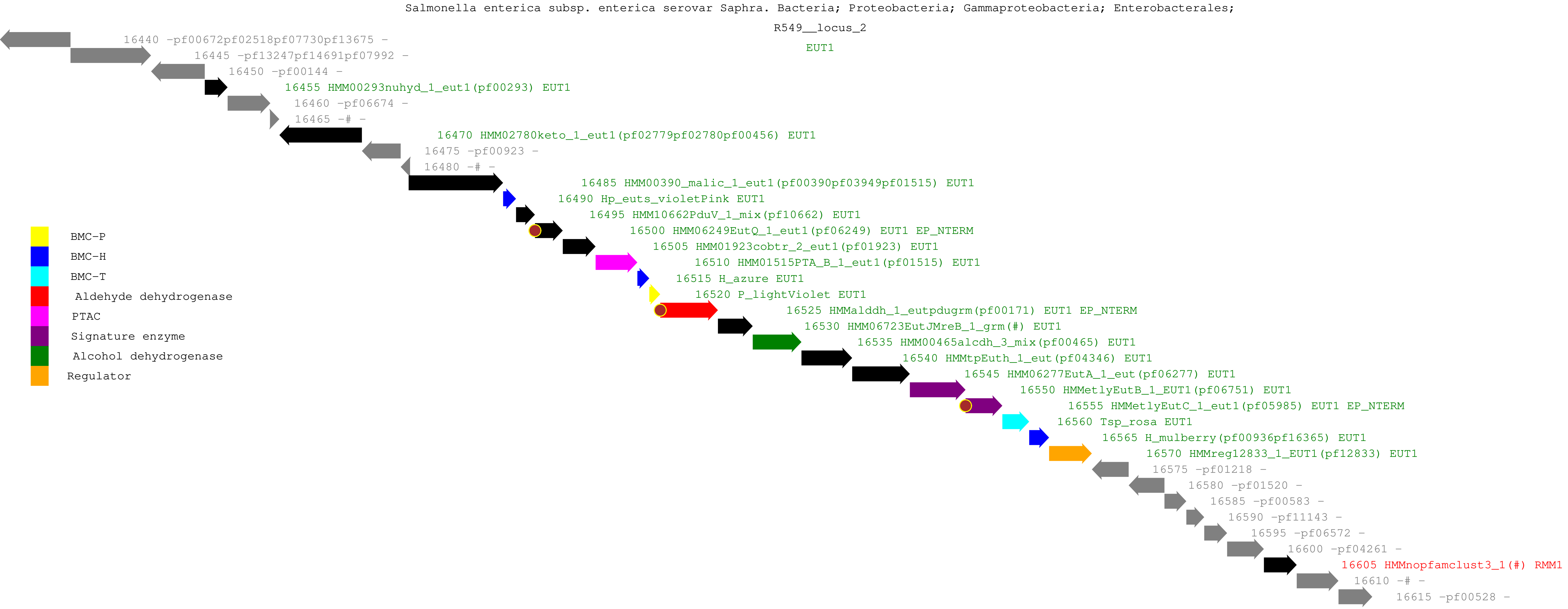
Salmonella enterica subsp. enterica serovar Saphra. Bacteria; Proteobacteria; Gammaproteobacteria; Enterobacterales;
R549__locus_2
EUT1
16440 -pf00672pf02518pf07730pf13675 -
16445 -pf13247pf14691pf07992 -
16450 -pf00144 -
16455 HMM00293nuhyd_1_eut1(pf00293) EUT1
16460 -pf06674 -
16465 -# -
16470 HMM02780keto_1_eut1(pf02779pf02780pf00456) EUT1
16475 -pf00923 -
16480 -# -
16485 HMM00390_malic_1_eut1(pf00390pf03949pf01515) EUT1
16490 Hp_euts_violetPink EUT1
16495 HMM10662PduV_1_mix(pf10662) EUT1
16500 HMM06249EutQ_1_eut1(pf06249) EUT1 EP_NTERM
16505 HMM01923cobtr_2_eut1(pf01923) EUT1
16510 HMM01515PTA_B_1_eut1(pf01515) EUT1
16515 H_azure EUT1
16520 P_lightViolet EUT1
16525 HMMalddh_1_eutpdugrm(pf00171) EUT1 EP_NTERM
16530 HMM06723EutJMreB_1_grm(#) EUT1
16535 HMM00465alcdh_3_mix(pf00465) EUT1
16540 HMMtpEuth_1_eut(pf04346) EUT1
16545 HMM06277EutA_1_eut(pf06277) EUT1
16550 HMMetlyEutB_1_EUT1(pf06751) EUT1
16555 HMMetlyEutC_1_eut1(pf05985) EUT1 EP_NTERM
16560 Tsp_rosa EUT1
16565 H_mulberry(pf00936pf16365) EUT1
16570 HMMreg12833_1_EUT1(pf12833) EUT1
16575 -pf01218 -
16580 -pf01520 -
16585 -pf00583 -
16590 -pf11143 -
16595 -pf06572 -
16600 -pf04261 -
16605 HMMnopfamclust3_1(#) RMM1
16610 -# -
16615 -pf00528 -
BMC-P
BMC-H
BMC-T
Aldehyde dehydrogenase
PTAC
Signature enzyme
Alcohol dehydrogenase
Regulator

Salmonella enterica subsp. enterica serovar Seftenburg. Bacteria; Proteobacteria; Gammaproteobacteria; Enterobacterales;

R583__locus_2

EUT1

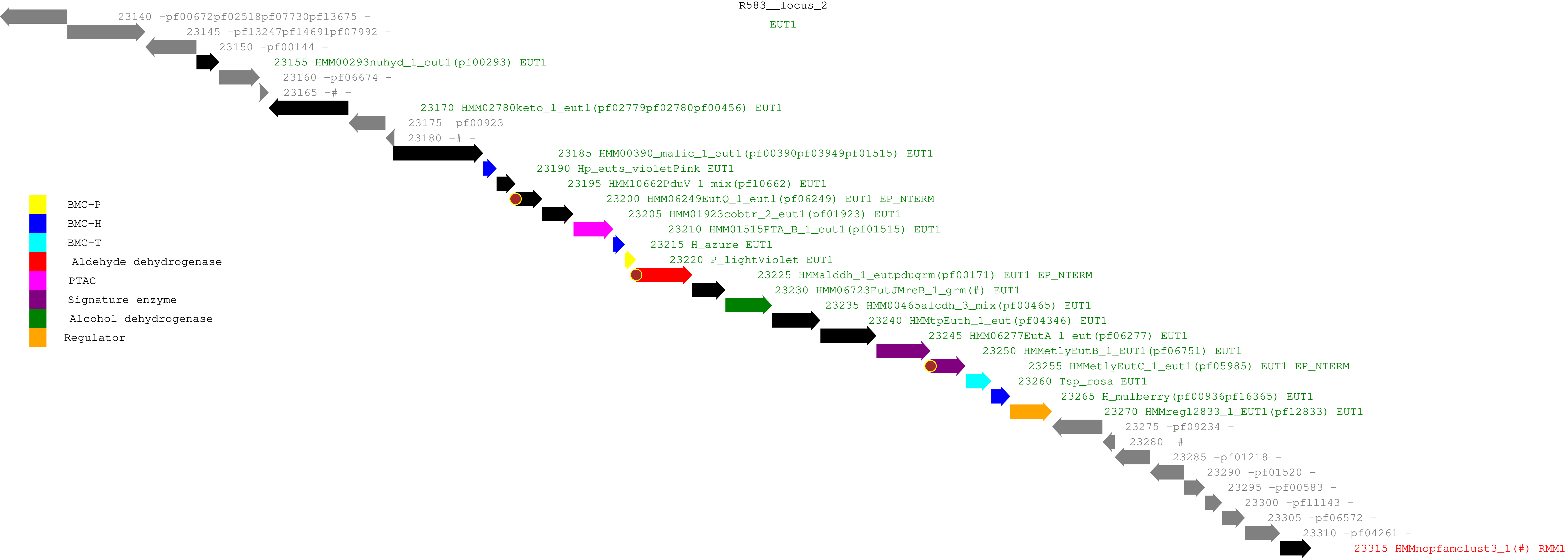

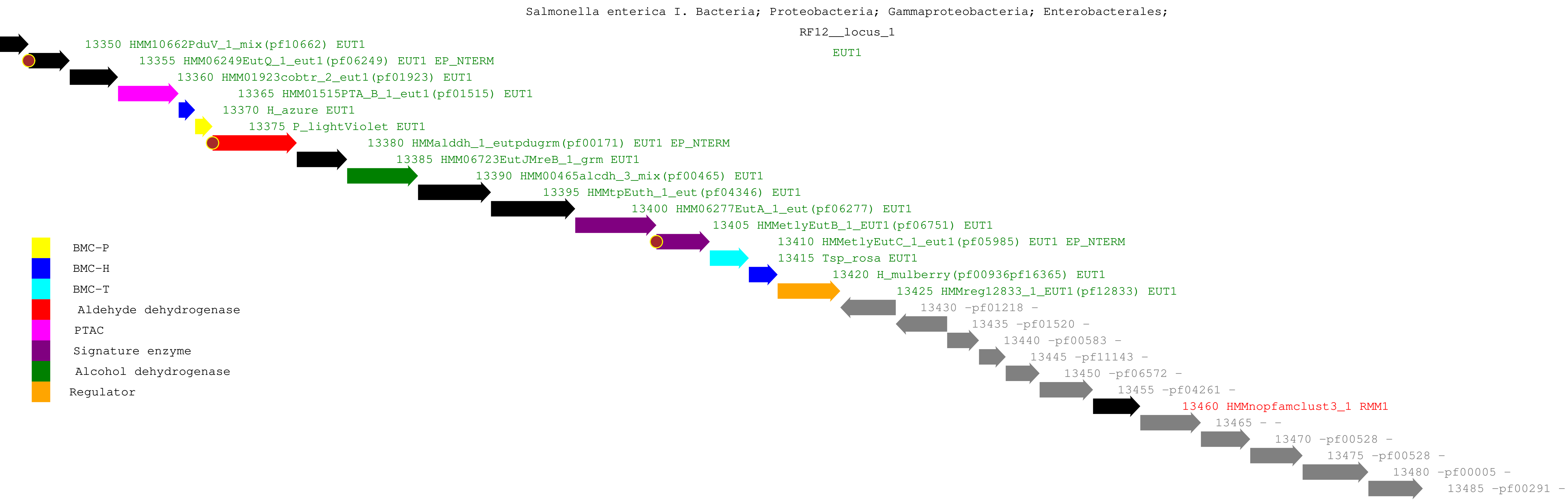
Salmonella enterica I. Bacteria; Proteobacteria; Gammaproteobacteria; Enterobacterales;
RF12__locus_1
EUT1
13350 HMM10662PduV_1_mix(pf10662) EUT1
13355 HMM06249EutQ_1_eut1(pf06249) EUT1 EP_NTERM
13360 HMM01923cobtr_2_eut1(pf01923) EUT1
13365 HMM01515PTA_B_1_eut1(pf01515) EUT1
13370 H_azure EUT1
13375 P_lightViolet EUT1
13380 HMMaldh_1_eutpdugrm(pf00171) EUT1 EP_NTERM
13385 HMM06723EutJMreB_1_grm EUT1
13390 HMM00465alcdh_3_mix(pf00465) EUT1
13395 HMMtpEuth_1_eut(pf04346) EUT1
13400 HMM06277EutA_1_eut(pf06277) EUT1
13405 HMMetlyEutB_1_EUT1(pf06751) EUT1
13410 HMMetlyEutC_1_eut1(pf05985) EUT1 EP_NTERM
13415 Tsp_rosa EUT1
13420 H_mulberry(pf00936pf16365) EUT1
13425 HMMreg12833_1_EUT1(pf12833) EUT1
13430 -pf01218 -
13435 -pf01520 -
13440 -pf00583 -
13445 -pf11143 -
13450 -pf06572 -
13455 -pf04261 -
13460 HMMnopfamclust3_1 RMM1
13465 - -
13470 -pf00528 -
13475 -pf00528 -
13480 -pf00005 -
13485 -pf00291 -
BMC-P
BMC-H
BMC-T
Aldehyde dehydrogenase
PTAC
Signature enzyme
Alcohol dehydrogenase
Regulator

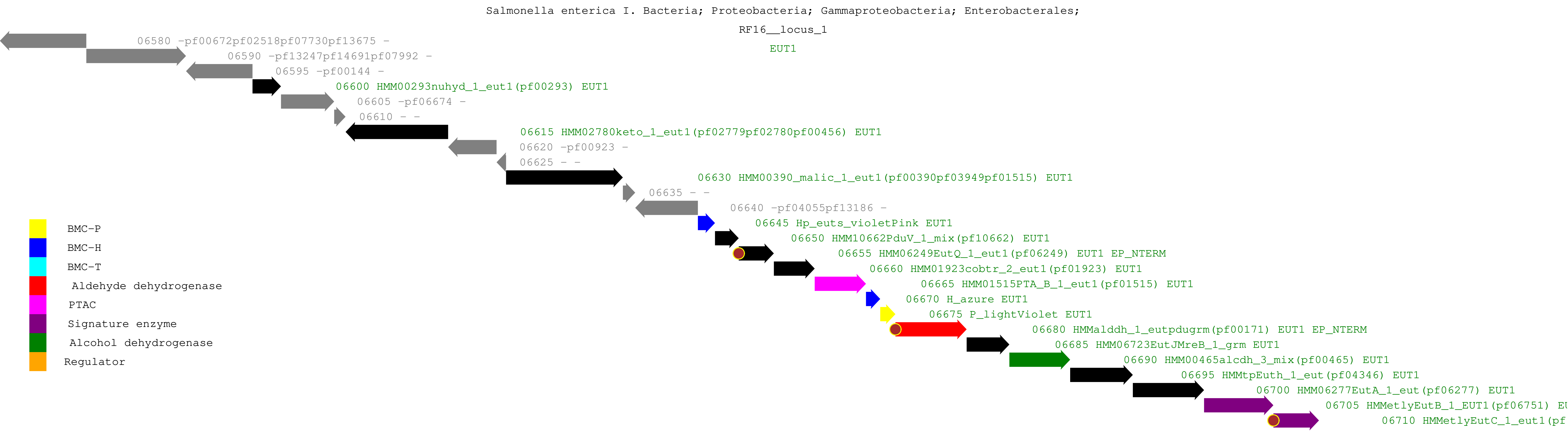
Salmonella enterica I. Bacteria; Proteobacteria; Gammaproteobacteria; Enterobacterales;
RF16__locus_1
EUT1
06580 -pf00672pf02518pf07730pf13675 -
06590 -pf13247pf14691pf07992 -
06595 -pf00144 -
06600 HMM00293nuhyd_1_eut1(pf00293) EUT1
06605 -pf06674 -
06610 - -
06615 HMM02780keto_1_eut1(pf02779pf02780pf00456) EUT1
06620 -pf00923 -
06625 - -
06630 HMM00390_malic_1_eut1(pf00390pf03949pf01515) EUT1
06635 - -
06640 -pf04055pf13186 -
06645 Hp_euts_violetPink EUT1
06650 HMM10662PduV_1_mix(pf10662) EUT1
06655 HMM06249EutQ_1_eut1(pf06249) EUT1 EP_NTERM
06660 HMM01923cobtr_2_eut1(pf01923) EUT1
06665 HMM01515PTA_B_1_eut1(pf01515) EUT1
06670 H_azure EUT1
06675 P_lightViolet EUT1
06680 HMMalddh_1_eutpdugrm(pf00171) EUT1 EP_NTERM
06685 HMM06723EutJMreB_1_grm EUT1
06690 HMM00465alcdh_3_mix(pf00465) EUT1
06695 HMMtpEuth_1_eut(pf04346) EUT1
06700 HMM06277EutA_1_eut(pf06277) EUT1
06705 HMMetlyEutB_1_EUT1(pf06751) EU
06710 HMMetlyEutC_1_eut1(pf
BMC-P
BMC-H
BMC-T
Aldehyde dehydrogenase
PTAC
Signature enzyme
Alcohol dehydrogenase
Regulator

Photobacterium gaetbulicola. Bacteria; Proteobacteria; Gammaproteobacteria; Vibrionales;

RJ45__locus_2

EUT1

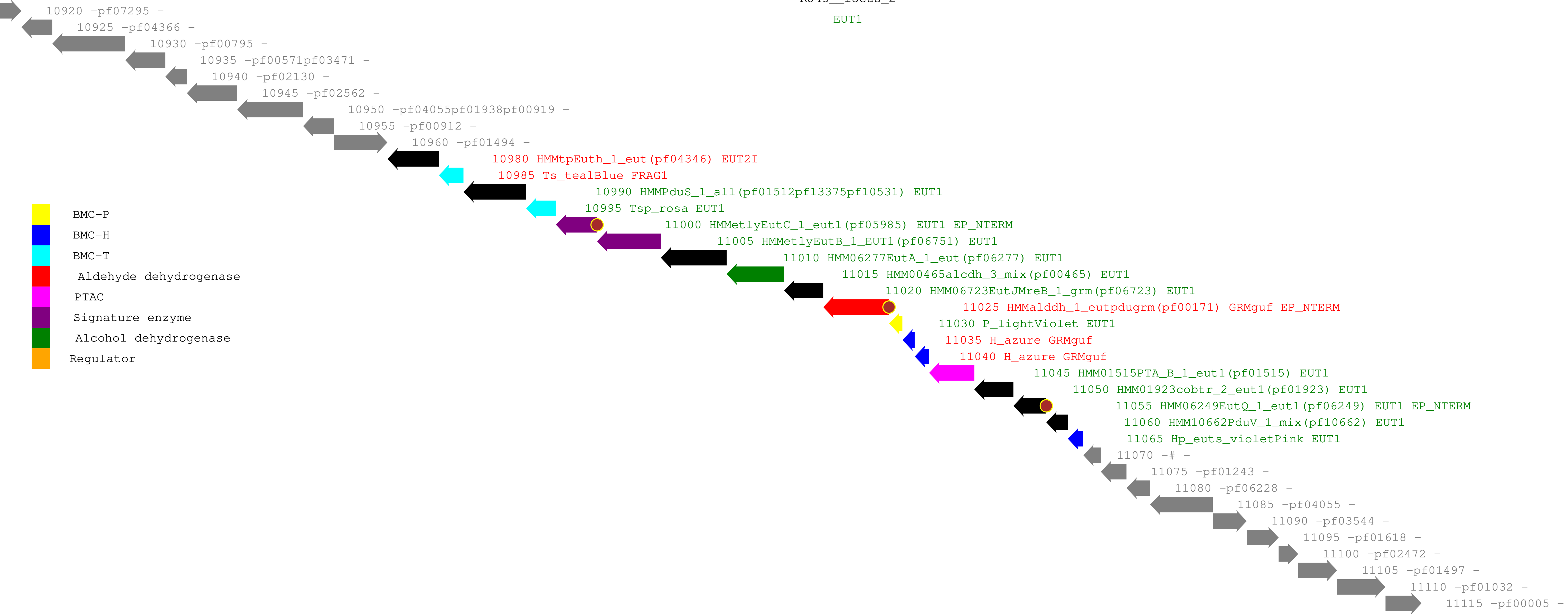

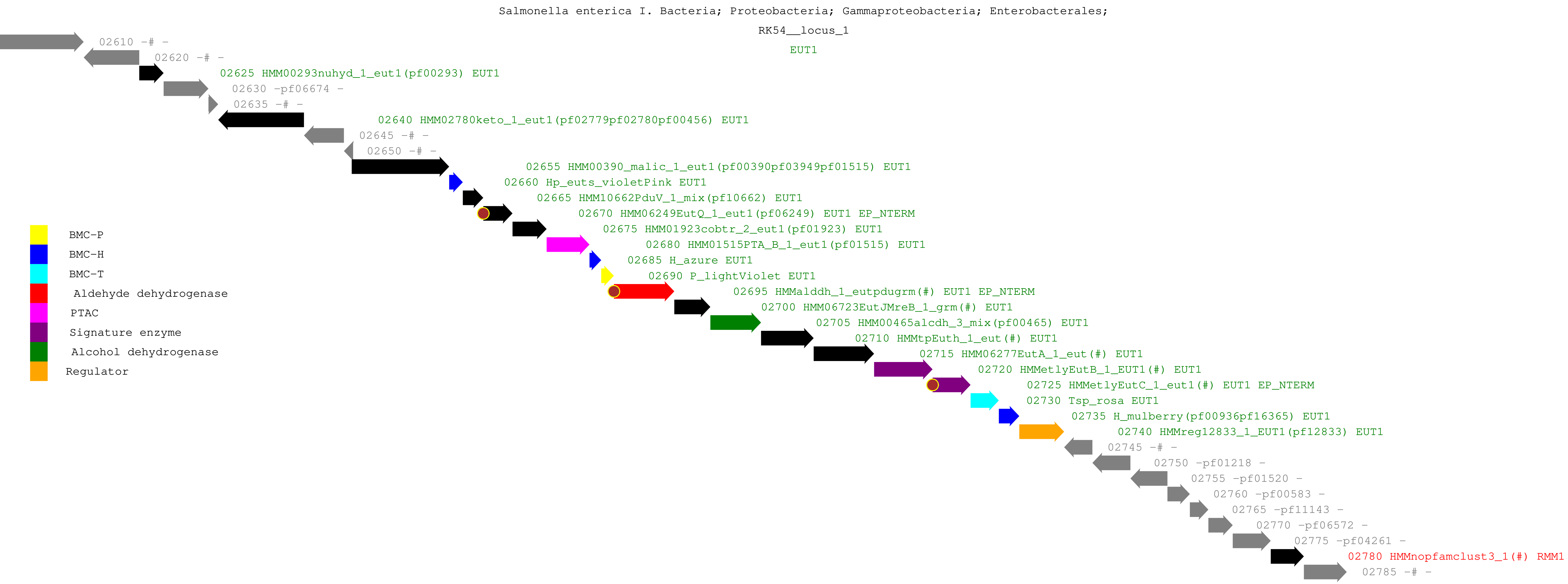
Salmonella enterica I. Bacteria; Proteobacteria; Gammaproteobacteria; Enterobacterales;
RK54__locus_1
EUT1
02610 -# -
02620 -# -
02625 HMM00293nuhyd_1_eut1(pf00293) EUT1
02630 -pf06674 -
02635 -# -
02640 HMM02780keto_1_eut1(pf02779pf02780pf00456) EUT1
02645 -# -
02650 -# -
02655 HMM00390_malic_1_eut1(pf00390pf03949pf01515) EUT1
02660 Hp_euts_violetPink EUT1
02665 HMM10662PduV_1_mix(pf10662) EUT1
02670 HMM06249EutQ_1_eut1(pf06249) EUT1 EP_NTERM
02675 HMM01923cobtr_2_eut1(pf01923) EUT1
02680 HMM01515PTA_B_1_eut1(pf01515) EUT1
02685 H_azure EUT1
02690 P_lightViolet EUT1
02695 HMMalddh_1_eutpdugrm(#) EUT1 EP_NTERM
02700 HMM06723EutJMreB_1_grm(#) EUT1
02705 HMM00465alcdh_3_mix(pf00465) EUT1
02710 HMMtpEuth_1_eut(#) EUT1
02715 HMM06277EutA_1_eut(#) EUT1
02720 HMMetlyEutB_1_EUT1(#) EUT1
02725 HMMetlyEutC_1_eut1(#) EUT1 EP_NTERM
02730 Tsp_rosa EUT1
02735 H_mulberry(pf00936pf16365) EUT1
02740 HMMreg12833_1_EUT1(pf12833) EUT1
02745 -# -
02750 -pf01218 -
02755 -pf01520 -
02760 -pf00583 -
02765 -pf11143 -
02770 -pf06572 -
02775 -pf04261 -
02780 HMMnopfamclust3_1(#) RMM1
02785 -# -
BMC-P
BMC-H
BMC-T
Aldehyde dehydrogenase
PTAC
Signature enzyme
Alcohol dehydrogenase
Regulator

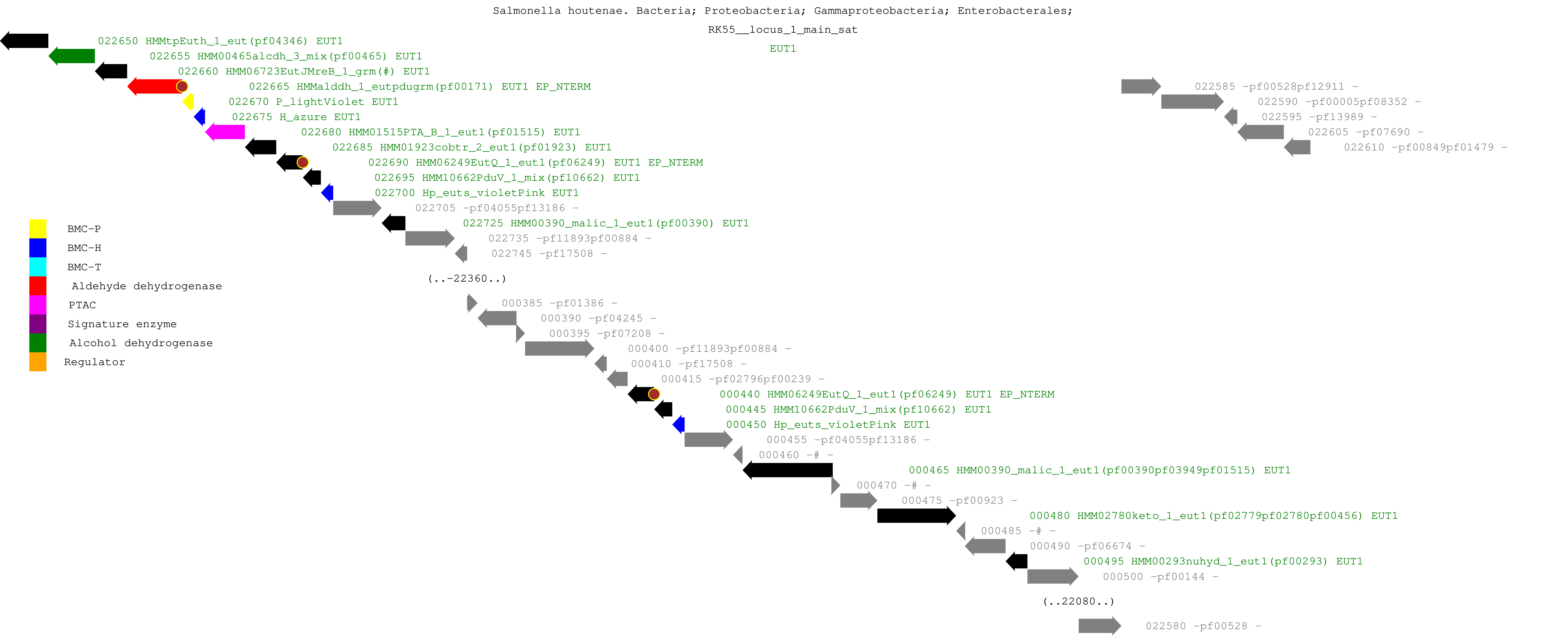

Salmonella houtenae. Bacteria; Proteobacteria; Gammaproteobacteria; Enterobacterales;
RK55__locus_1_main_sat
EUT1
022650 HMMtpEuth_1_eut(pf04346) EUT1
022655 HMM00465alcdh_3_mix(pf00465) EUT1
022660 HMM06723EutJMreB_1_grm(#) EUT1
022665 HMMalddh_1_eutpdugrm(pf00171) EUT1 EP_NTERM
022670 P_lightViolet EUT1
022675 H_azure EUT1
022680 HMM01515PTA_B_1_eut1(pf01515) EUT1
022685 HMM01923cobtr_2_eut1(pf01923) EUT1
022690 HMM06249EutQ_1_eut1(pf06249) EUT1 EP_NTERM
022695 HMM10662PduV_1_mix(pf10662) EUT1
022700 Hp_euts_violetPink EUT1
022705 -pf04055pf13186 -
022725 HMM00390_malic_1_eut1(pf00390) EUT1
022735 -pf11893pf00884 -
022745 -pf17508 -
(..-22360..)
000385 -pf01386 -
000390 -pf04245 -
000395 -pf07208 -
000400 -pf11893pf00884 -
000410 -pf17508 -
000415 -pf02796pf00239 -
000440 HMM06249EutQ_1_eut1(pf06249) EUT1 EP_NTERM
000445 HMM10662PduV_1_mix(pf10662) EUT1
000450 Hp_euts_violetPink EUT1
000455 -pf04055pf13186 -
000460 -# -
000465 HMM00390_malic_1_eut1(pf00390pf03949pf01515) EUT1
000470 -# -
000475 -pf00923 -
000480 HMM02780keto_1_eut1(pf02779pf02780pf00456) EUT1
000485 -# -
000490 -pf06674 -
000495 HMM00293nuhyd_1_eut1(pf00293) EUT1
000500 -pf00144 -
(..22080..)
022580 -pf00528 -
022585 -pf00528pf12911 -
022590 -pf00005pf08352 -
022595 -pf13989 -
022605 -pf07690 -
022610 -pf00849pf01479 -
BMC-P
BMC-H
BMC-T
Aldehyde dehydrogenase
PTAC
Signature enzyme
Alcohol dehydrogenase
Regulator

Shigella flexneri. Bacteria; Proteobacteria; Gammaproteobacteria; Enterobacterales;

RK59__locus_1

EUT1

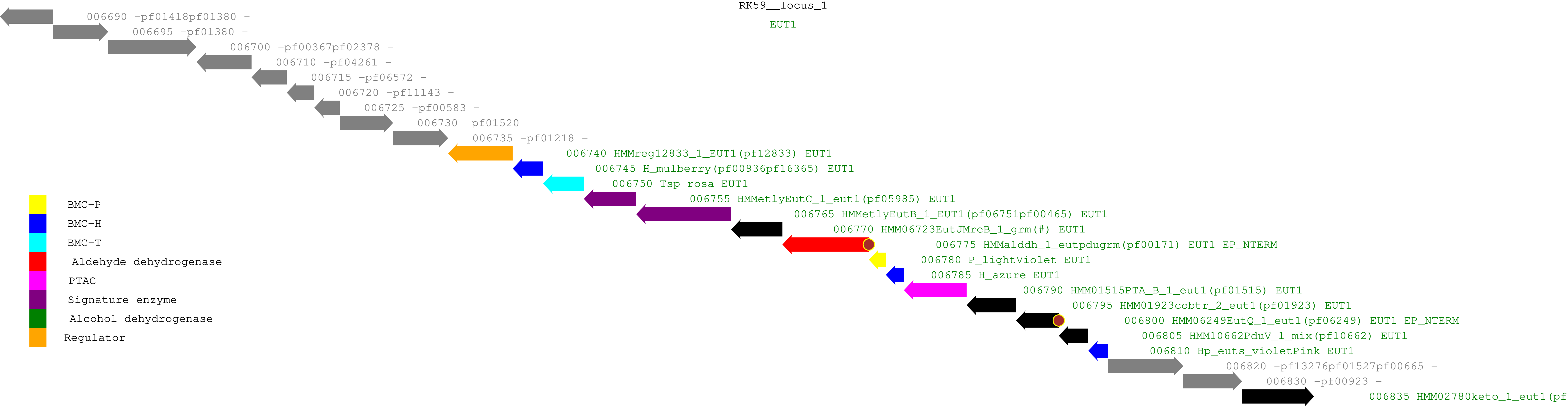

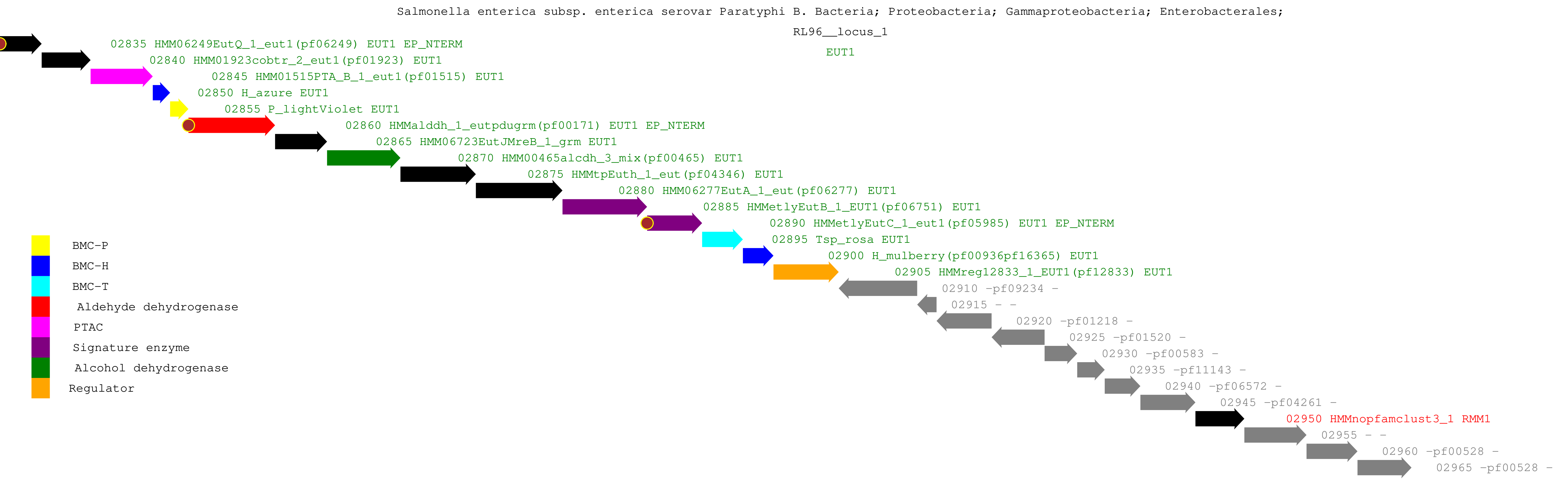
Salmonella enterica subsp. enterica serovar Paratyphi B. Bacteria; Proteobacteria; Gammaproteobacteria; Enterobacterales;
RL96__locus_1
EUT1
02835 HMM06249EutQ_1_eut1(pf06249) EUT1 EP_NTERM
02840 HMM01923cobtr_2_eut1(pf01923) EUT1
02845 HMM01515PTA_B_1_eut1(pf01515) EUT1
02850 H_azure EUT1
02855 P_lightViolet EUT1
02860 HMMalddh_1_eutpdugrm(pf00171) EUT1 EP_NTERM
02865 HMM06723EutJMreB_1_grm EUT1
02870 HMM00465alcdh_3_mix(pf00465) EUT1
02875 HMMtpEuth_1_eut(pf04346) EUT1
02880 HMM06277EutA_1_eut(pf06277) EUT1
02885 HMMetlyEutB_1_EUT1(pf06751) EUT1
02890 HMMetlyEutC_1_eut1(pf05985) EUT1 EP_NTERM
02895 Tsp_rosa EUT1
02900 H_mulberry(pf00936pf16365) EUT1
02905 HMMreg12833_1_EUT1(pf12833) EUT1
02910 -pf09234 -
02915 - -
02920 -pf01218 -
02925 -pf01520 -
02930 -pf00583 -
02935 -pf11143 -
02940 -pf06572 -
02945 -pf04261 -
02950 HMMnopfamclust3_1 RMM1
02955 - -
02960 -pf00528 -
02965 -pf00528 -
BMC-P
BMC-H
BMC-T
Aldehyde dehydrogenase
PTAC
Signature enzyme
Alcohol dehydrogenase
Regulator

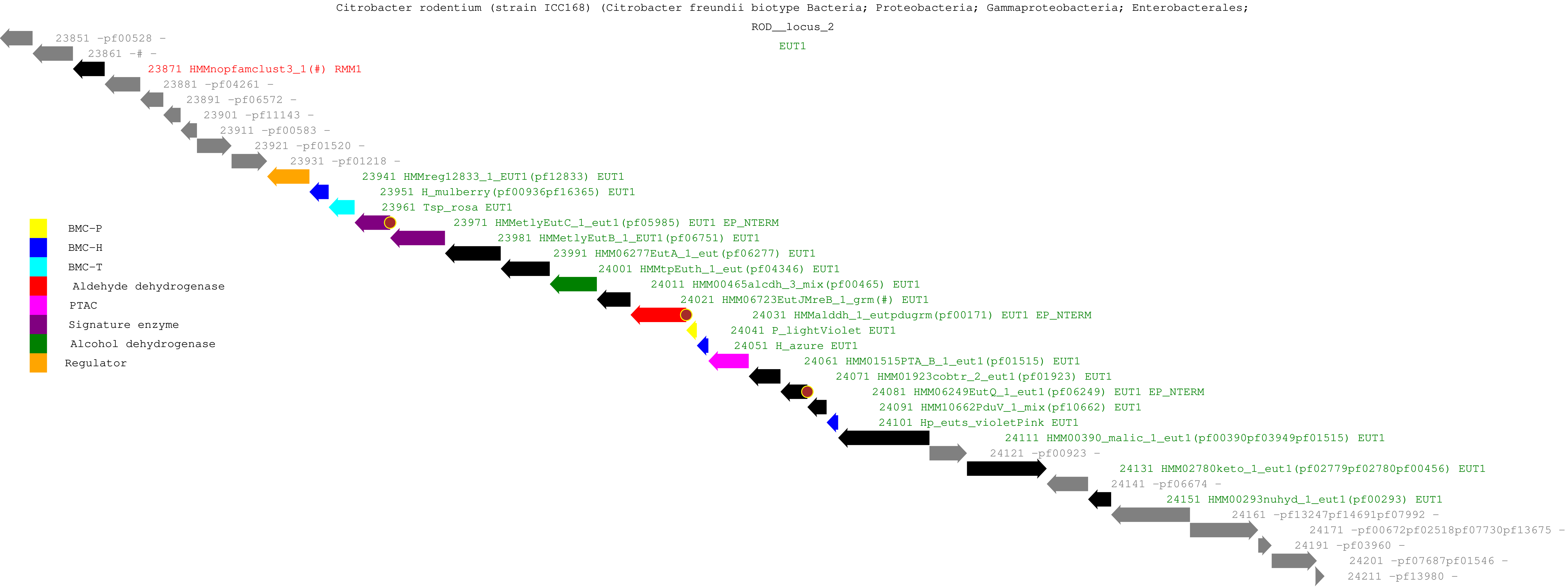
Citrobacter rodentium (strain ICC168) (Citrobacter freundii biotype Bacteria; Proteobacteria; Gammaproteobacteria; Enterobacterales;
ROD__locus_2
EUT1
23851 -pf00528 -
23861 -# -
23871 HMMnopfamclust3_1(#) RMM1
23881 -pf04261 -
23891 -pf06572 -
23901 -pf11143 -
23911 -pf00583 -
23921 -pf01520 -
23931 -pf01218 -
23941 HMMreg12833_1_EUT1(pf12833) EUT1
23951 H_mulberry(pf00936pf16365) EUT1
23961 Tsp_rosa EUT1
23971 HMMetlyEutC_1_eut1(pf05985) EUT1 EP_NTERM
23981 HMMetlyEutB_1_EUT1(pf06751) EUT1
23991 HMM06277EutA_1_eut(pf06277) EUT1
24001 HMMtpEuth_1_eut(pf04346) EUT1
24011 HMM00465alcdh_3_mix(pf00465) EUT1
24021 HMM06723EutJMreB_1_grm(#) EUT1
24031 HMMalddh_1_eutpdugrm(pf00171) EUT1 EP_NTERM
24041 P_lightViolet EUT1
24051 H_azure EUT1
24061 HMM01515PTA_B_1_eut1(pf01515) EUT1
24071 HMM01923cobtr_2_eut1(pf01923) EUT1
24081 HMM06249EutQ_1_eut1(pf06249) EUT1 EP_NTERM
24091 HMM10662PduV_1_mix(pf10662) EUT1
24101 Hp_euts_violetPink EUT1
24111 HMM00390_malic_1_eut1(pf00390pf03949pf01515) EUT1
24121 -pf00923 -
24131 HMM02780keto_1_eut1(pf02779pf02780pf00456) EUT1
24141 -pf06674 -
24151 HMM00293nuhyd_1_eut1(pf00293) EUT1
24161 -pf13247pf14691pf07992 -
24171 -pf00672pf02518pf07730pf13675 -
24191 -pf03960 -
24201 -pf07687pf01546 -
24211 -pf13980 -
BMC-P
BMC-H
BMC-T
Aldehyde dehydrogenase
PTAC
Signature enzyme
Alcohol dehydrogenase
Regulator

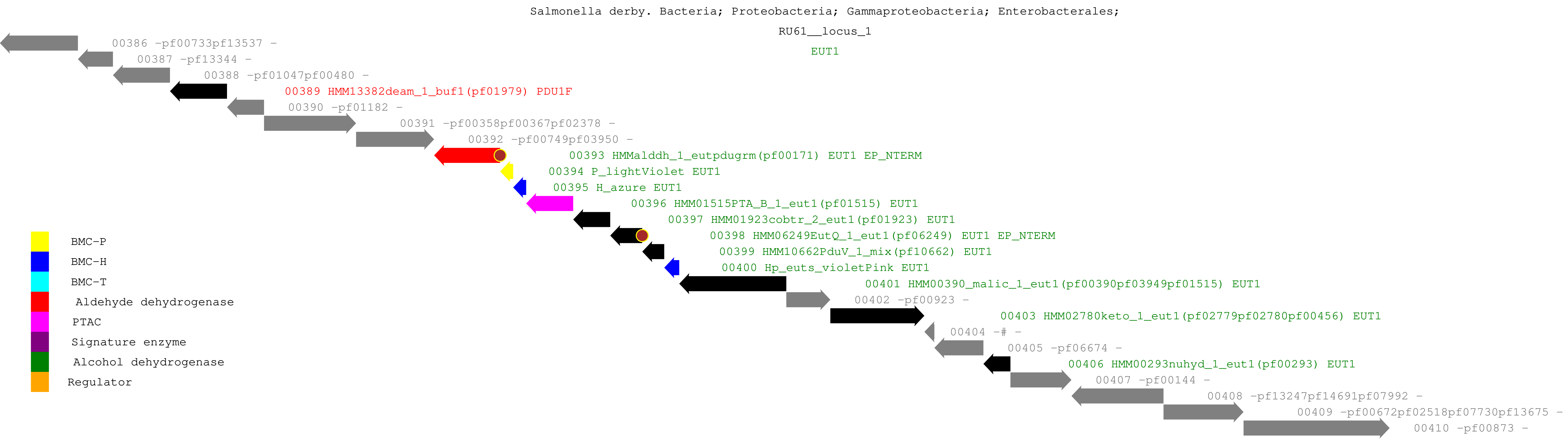

Salmonella derby. Bacteria; Proteobacteria; Gammaproteobacteria; Enterobacterales;
RU61__locus_1
EUT1
00386 -pf00733pf13537 -
00387 -pf13344 -
00388 -pf01047pf00480 -
00389 HMM13382deam_1_buf1(pf01979) PDU1F
00390 -pf01182 -
00391 -pf00358pf00367pf02378 -
00392 -pf00749pf03950 -
00393 HMMalddh_1_eutpdugrm(pf00171) EUT1 EP_NTERM
00394 P_lightViolet EUT1
00395 H_azure EUT1
00396 HMM01515PTA_B_1_eut1(pf01515) EUT1
00397 HMM01923cobtr_2_eut1(pf01923) EUT1
00398 HMM06249EutQ_1_eut1(pf06249) EUT1 EP_NTERM
00399 HMM10662PduV_1_mix(pf10662) EUT1
00400 Hp_euts_violetPink EUT1
00401 HMM00390_malic_1_eut1(pf00390pf03949pf01515) EUT1
00402 -pf00923 -
00403 HMM02780keto_1_eut1(pf02779pf02780pf00456) EUT1
00404 -# -
00405 -pf06674 -
00406 HMM00293nuhyd_1_eut1(pf00293) EUT1
00407 -pf00144 -
00408 -pf13247pf14691pf07992 -
00409 -pf00672pf02518pf07730pf13675 -
00410 -pf00873 -
BMC-P
BMC-H
BMC-T
Aldehyde dehydrogenase
PTAC
Signature enzyme
Alcohol dehydrogenase
Regulator

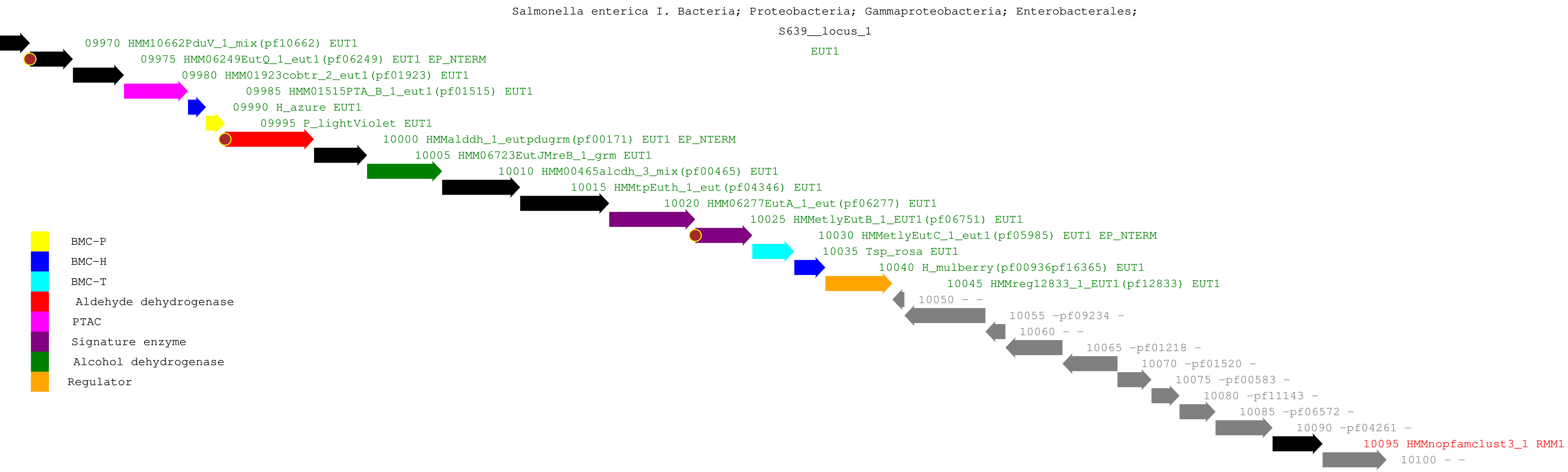

Salmonella enterica I. Bacteria; Proteobacteria; Gammaproteobacteria; Enterobacterales;
S639__locus_1
EUT1
09970 HMM10662PduV_1_mix(pf10662) EUT1
09975 HMM06249EutQ_1_eut1(pf06249) EUT1 EP_NTERM
09980 HMM01923cobtr_2_eut1(pf01923) EUT1
09985 HMM01515PTA_B_1_eut1(pf01515) EUT1
09990 H_azure EUT1
09995 P_lightViolet EUT1
10000 HMMalddh_1_eutpdugrm(pf00171) EUT1 EP_NTERM
10005 HMM06723EutJMreB_1_grm EUT1
10010 HMM00465alcdh_3_mix(pf00465) EUT1
10015 HMMtpEuth_1_eut(pf04346) EUT1
10020 HMM06277EutA_1_eut(pf06277) EUT1
10025 HMMetlyEutB_1_EUT1(pf06751) EUT1
10030 HMMetlyEutC_1_eut1(pf05985) EUT1 EP_NTERM
10035 Tsp_rosa EUT1
10040 H_mulberry(pf00936pf16365) EUT1
10045 HMMreg12833_1_EUT1(pf12833) EUT1
10050 - -
10055 -pf09234 -
10060 - -
10065 -pf01218 -
10070 -pf01520 -
10075 -pf00583 -
10080 -pf11143 -
10085 -pf06572 -
10090 -pf04261 -
10095 HMMnopfamclust3_1 RMM1
10100 - -
BMC-P
BMC-H
BMC-T
Aldehyde dehydrogenase
PTAC
Signature enzyme
Alcohol dehydrogenase
Regulator

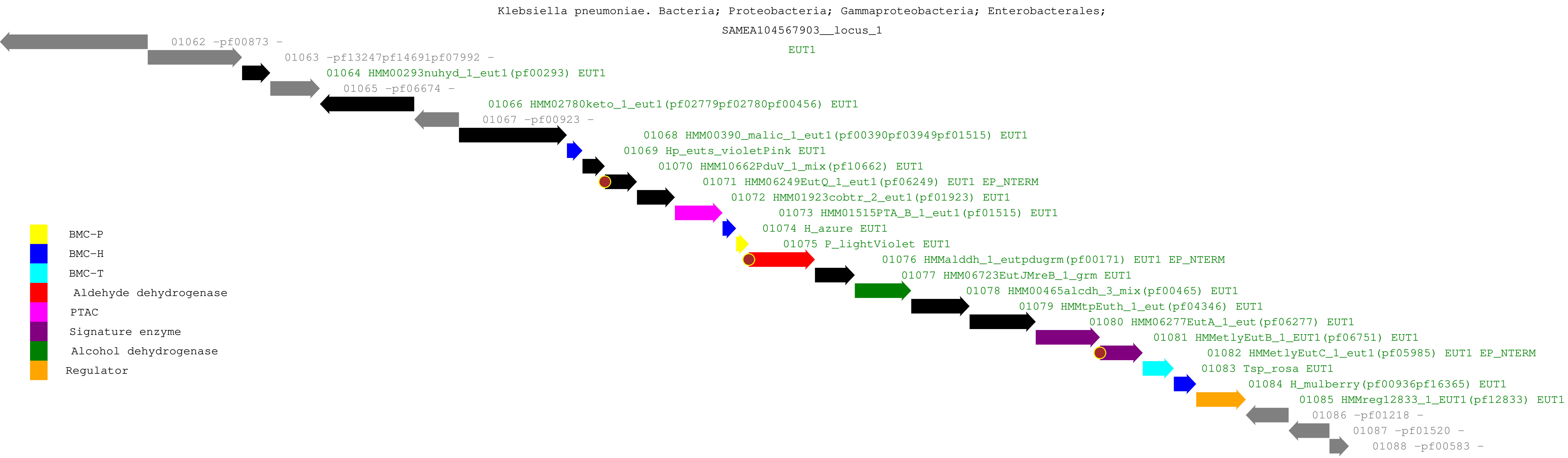
Klebsiella pneumoniae. Bacteria; Proteobacteria; Gammaproteobacteria; Enterobacterales;
SAMEA104567903__locus_1
EUT1
01062 -pf00873 -
01063 -pf13247pf14691pf07992 -
01064 HMM00293nuhyd_1_eut1(pf00293) EUT1
01065 -pf06674 -
01066 HMM02780keto_1_eut1(pf02779pf02780pf00456) EUT1
01067 -pf00923 -
01068 HMM00390_malic_1_eut1(pf00390pf03949pf01515) EUT1
01069 Hp_euts_violetPink EUT1
01070 HMM10662PduV_1_mix(pf10662) EUT1
01071 HMM06249EutQ_1_eut1(pf06249) EUT1 EP_NTERM
01072 HMM01923cobtr_2_eut1(pf01923) EUT1
01073 HMM01515PTA_B_1_eut1(pf01515) EUT1
01074 H_azure EUT1
01075 P_lightViolet EUT1
01076 HMMalddh_1_eutpdugrm(pf00171) EUT1 EP_NTERM
01077 HMM06723EutJMreB_1_grm EUT1
01078 HMM00465alcdh_3_mix(pf00465) EUT1
01079 HMMtpEuth_1_eut(pf04346) EUT1
01080 HMM06277EutA_1_eut(pf06277) EUT1
01081 HMMetlyEutB_1_EUT1(pf06751) EUT1
01082 HMMetlyEutC_1_eut1(pf05985) EUT1 EP_NTERM
01083 Tsp_rosa EUT1
01084 H_mulberry(pf00936pf16365) EUT1
01085 HMMreg12833_1_EUT1(pf12833) EUT1
01086 -pf01218 -
01087 -pf01520 -
01088 -pf00583 -
BMC-P
BMC-H
BMC-T
Aldehyde dehydrogenase
PTAC
Signature enzyme
Alcohol dehydrogenase
Regulator

Klebsiella pneumoniae. Bacteria; Proteobacteria; Gammaproteobacteria; Enterobacterales;

SAMEA1713115__locus_1

EUT1

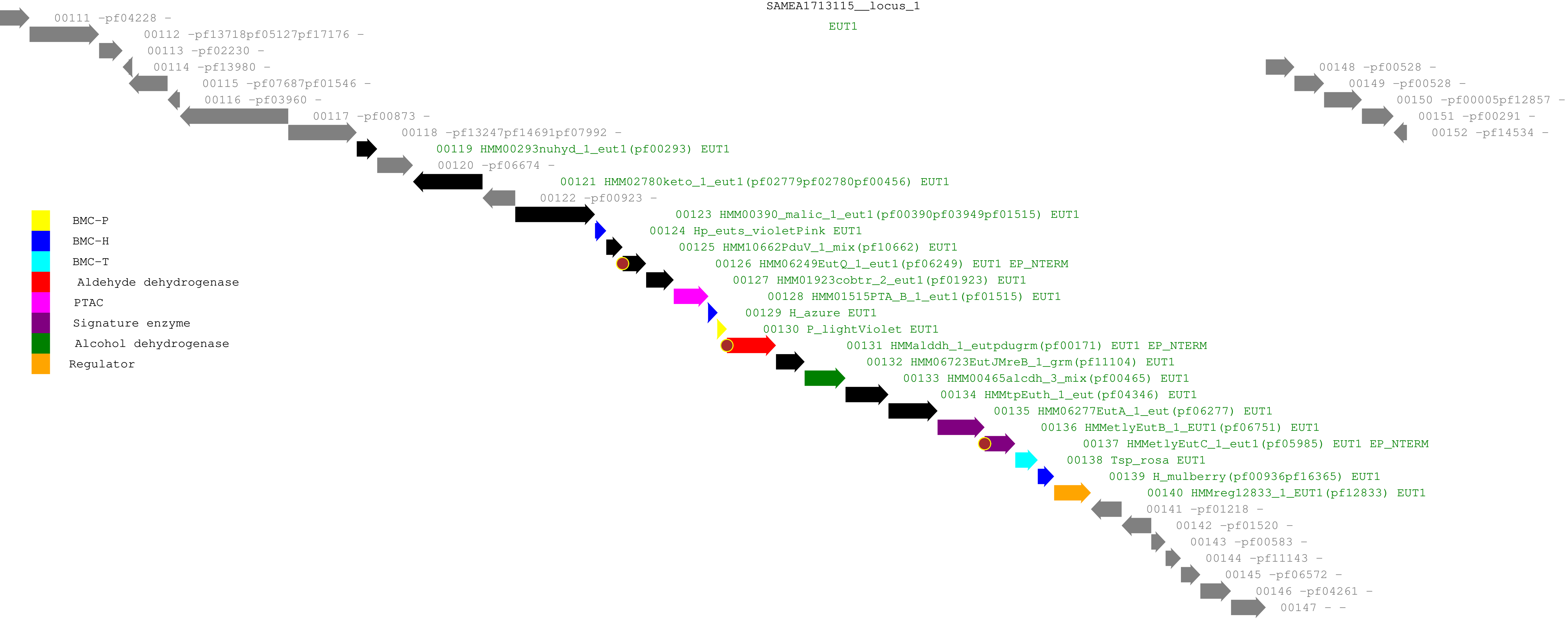

Klebsiella pneumoniae. Bacteria; Proteobacteria; Gammaproteobacteria; Enterobacterales;

SAMEA2053874__locus_2

EUT1

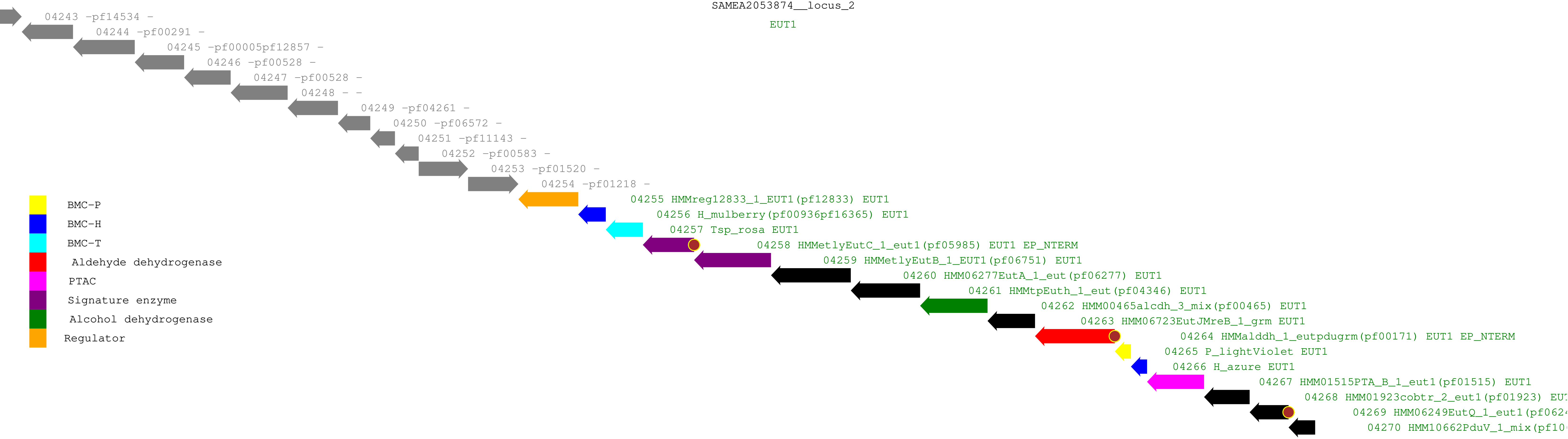

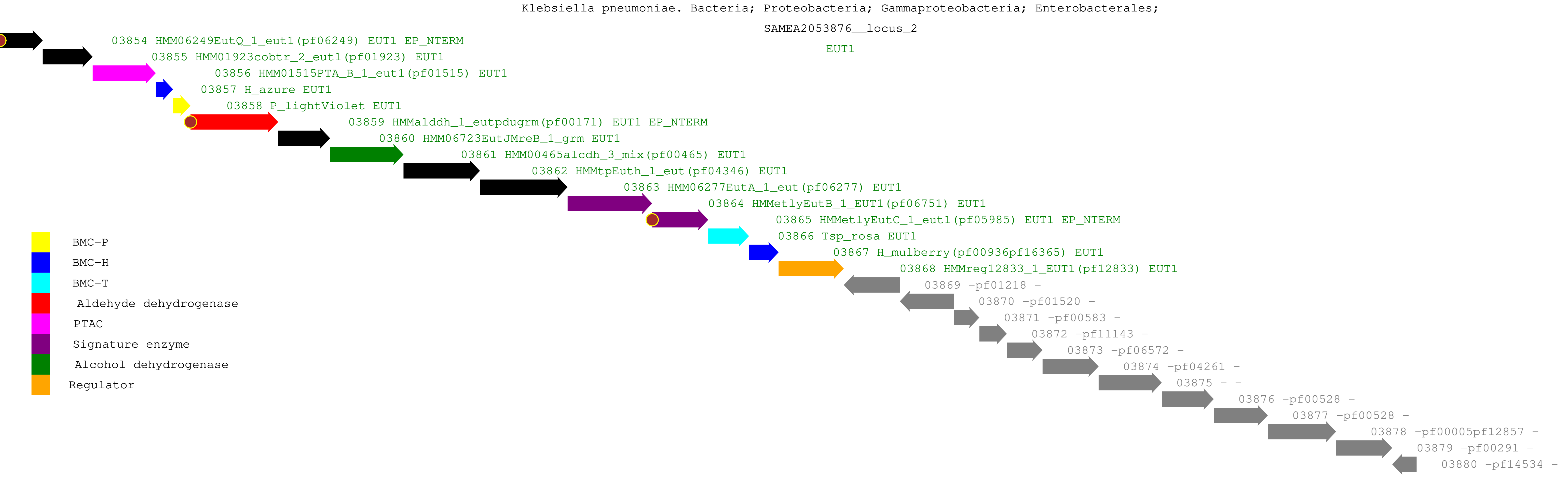

Klebsiella pneumoniae. Bacteria; Proteobacteria; Gammaproteobacteria; Enterobacterales;
SAMEA2053876__locus_2
EUT1
03854 HMM06249EutQ_1_eut1(pf06249) EUT1 EP_NTERM
03855 HMM01923cobtr_2_eut1(pf01923) EUT1
03856 HMM01515PTA_B_1_eut1(pf01515) EUT1
03857 H_azure EUT1
03858 P_lightViolet EUT1
03859 HMMalddh_1_eutpdugrm(pf00171) EUT1 EP_NTERM
03860 HMM06723EutJMreB_1_grm EUT1
03861 HMM00465alcdh_3_mix(pf00465) EUT1
03862 HMMtpEuth_1_eut(pf04346) EUT1
03863 HMM06277EutA_1_eut(pf06277) EUT1
03864 HMMetlyEutB_1_EUT1(pf06751) EUT1
03865 HMMetlyEutC_1_eut1(pf05985) EUT1 EP_NTERM
03866 Tsp_rosa EUT1
03867 H_mulberry(pf00936pf16365) EUT1
03868 HMMreg12833_1_EUT1(pf12833) EUT1
03869 -pf01218 -
03870 -pf01520 -
03871 -pf00583 -
03872 -pf11143 -
03873 -pf06572 -
03874 -pf04261 -
03875 - -
03876 -pf00528 -
03877 -pf00528 -
03878 -pf00005pf12857 -
03879 -pf00291 -
03880 -pf14534 -
BMC-P
BMC-H
BMC-T
Aldehyde dehydrogenase
PTAC
Signature enzyme
Alcohol dehydrogenase
Regulator

Klebsiella pneumoniae. Bacteria; Proteobacteria; Gammaproteobacteria; Enterobacterales;

SAMEA2053943__locus_2

EUT1

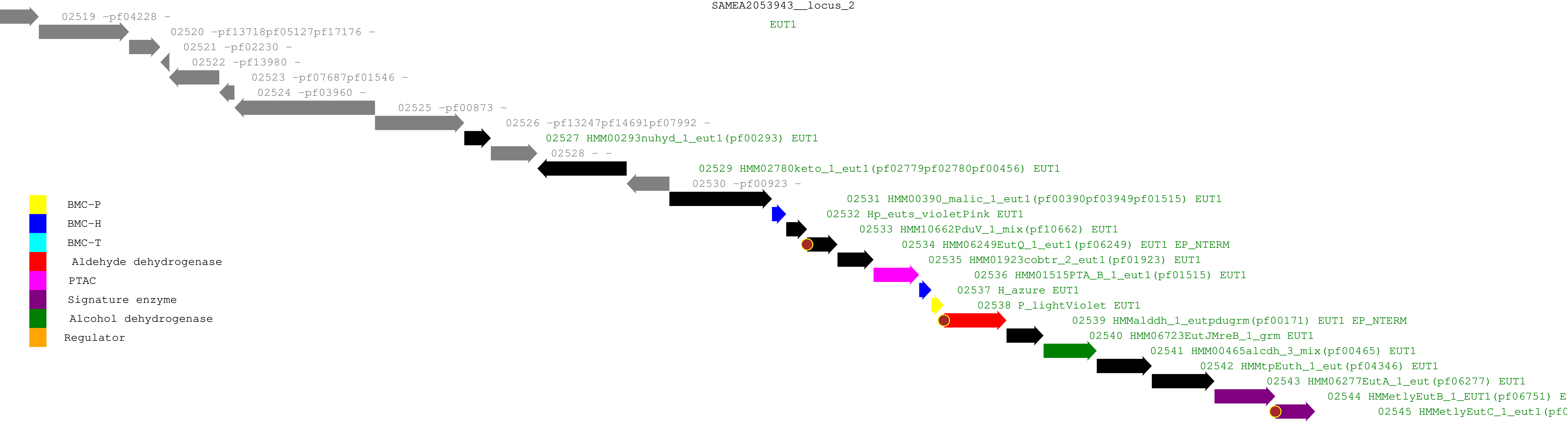

Enterobacter cloacae. Bacteria; Proteobacteria; Gammaproteobacteria; Enterobacterales;

SAMEA2273167__locus_2

EUT1

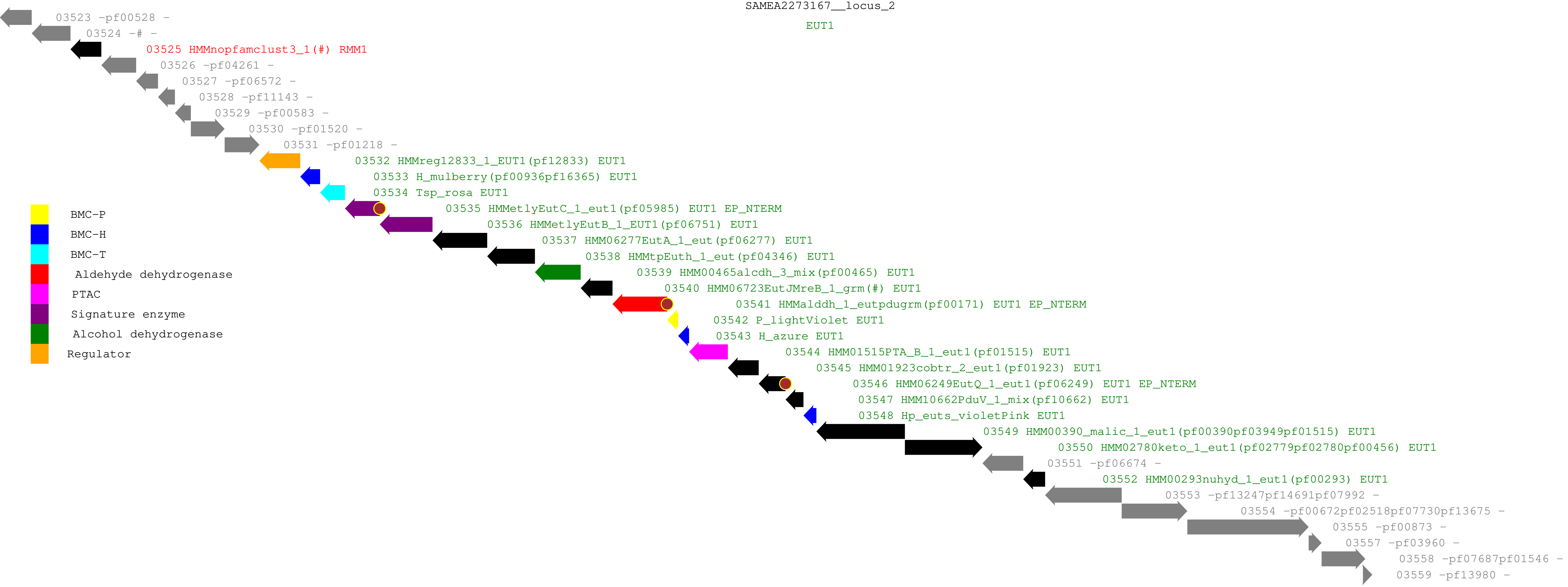

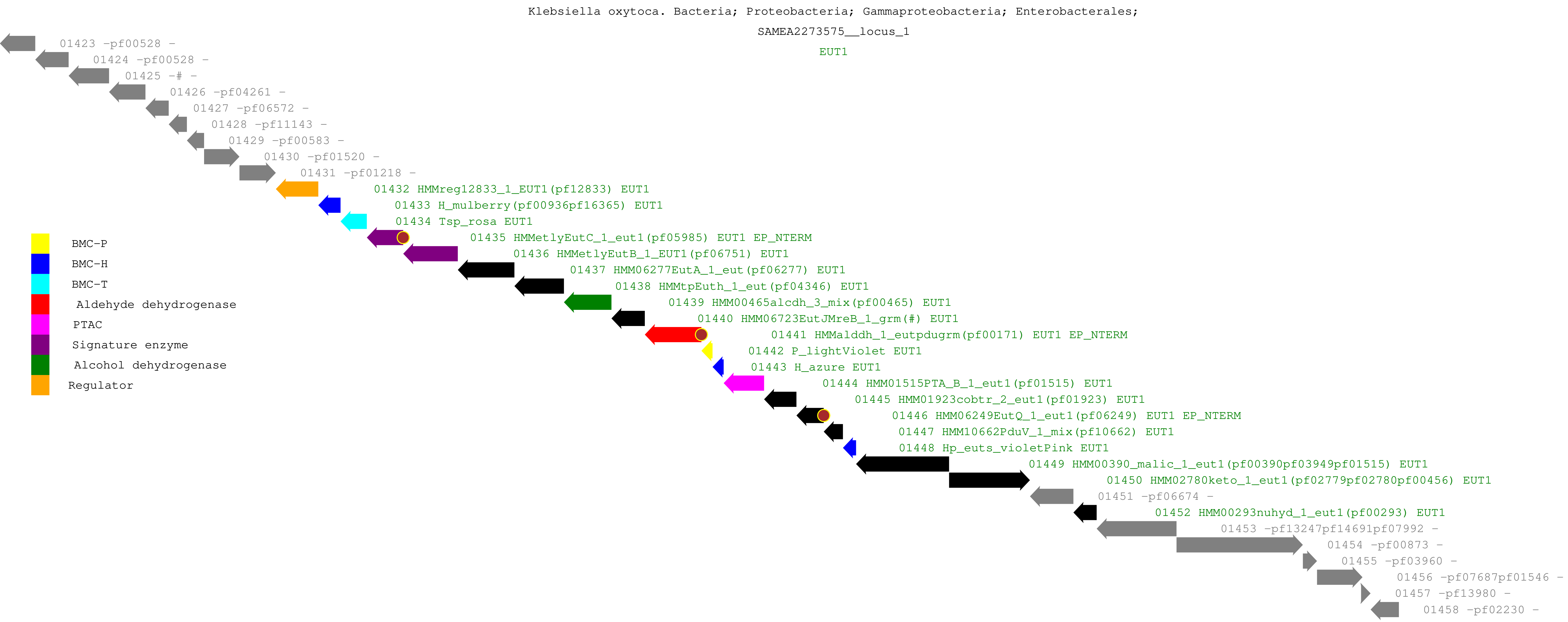

Klebsiella oxytoca. Bacteria; Proteobacteria; Gammaproteobacteria; Enterobacterales;
SAMEA2273575__locus_1
EUT1
01423 -pf00528 -
01424 -pf00528 -
01425 -# -
01426 -pf04261 -
01427 -pf06572 -
01428 -pf11143 -
01429 -pf00583 -
01430 -pf01520 -
01431 -pf01218 -
01432 HMMreg12833_1_EUT1(pf12833) EUT1
01433 H_mulberry(pf00936pf16365) EUT1
01434 Tsp_rosa EUT1
01435 HMMetlyEutC_1_eut1(pf05985) EUT1 EP_NTERM
01436 HMMetlyEutB_1_EUT1(pf06751) EUT1
01437 HMM06277EutA_1_eut(pf06277) EUT1
01438 HMMtpEuth_1_eut(pf04346) EUT1
01439 HMM00465alcdh_3_mix(pf00465) EUT1
01440 HMM06723EutJMreB_1_grm(#) EUT1
01441 HMMalddh_1_eutpdugrm(pf00171) EUT1 EP_NTERM
01442 P_lightViolet EUT1
01443 H_azure EUT1
01444 HMM01515PTA_B_1_eut1(pf01515) EUT1
01445 HMM01923cobtr_2_eut1(pf01923) EUT1
01446 HMM06249EutQ_1_eut1(pf06249) EUT1 EP_NTERM
01447 HMM10662PduV_1_mix(pf10662) EUT1
01448 Hp_euts_violetPink EUT1
01449 HMM00390_malic_1_eut1(pf00390pf03949pf01515) EUT1
01450 HMM02780keto_1_eut1(pf02779pf02780pf00456) EUT1
01451 -pf06674 -
01452 HMM00293nuhyd_1_eut1(pf00293) EUT1
01453 -pf13247pf14691pf07992 -
01454 -pf00873 -
01455 -pf03960 -
01456 -pf07687pf01546 -
01457 -pf13980 -
01458 -pf02230 -
BMC-P
BMC-H
BMC-T
Aldehyde dehydrogenase
PTAC
Signature enzyme
Alcohol dehydrogenase
Regulator

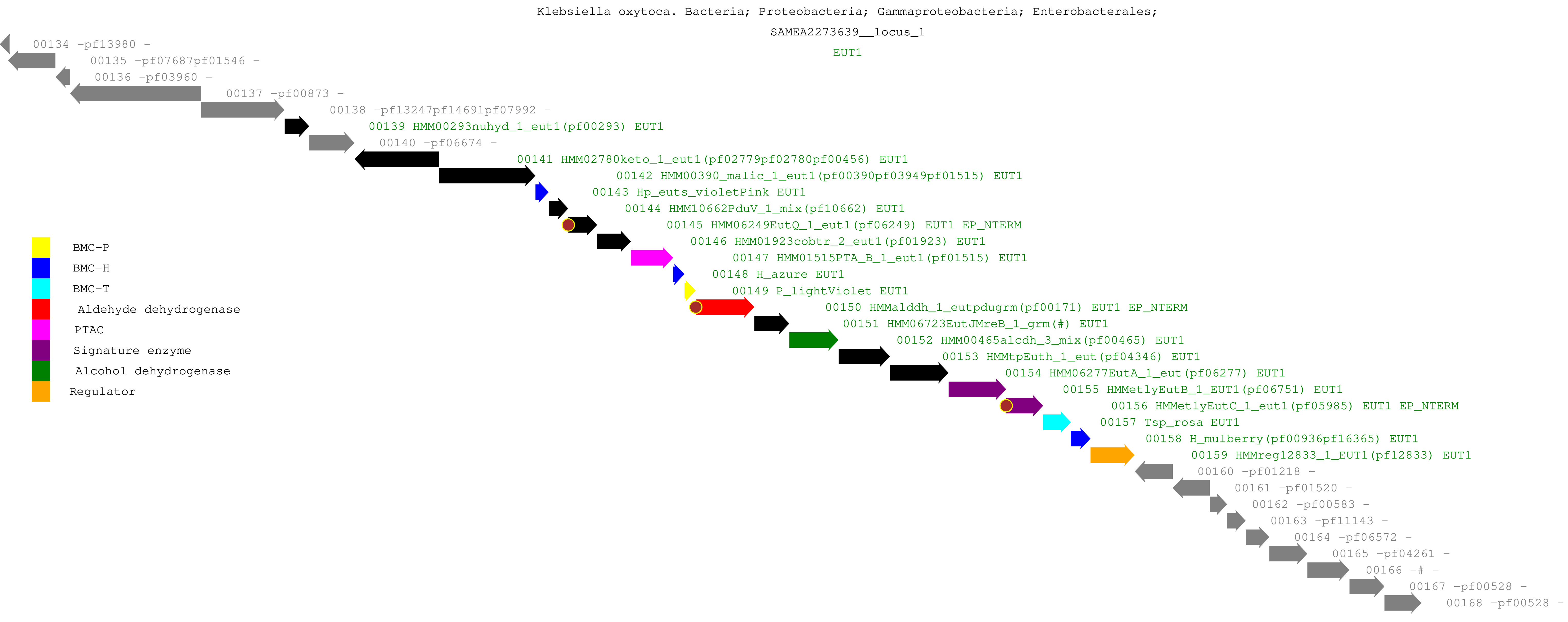
Klebsiella oxytoca. Bacteria; Proteobacteria; Gammaproteobacteria; Enterobacterales;
SAMEA2273639__locus_1
EUT1
00134 -pf13980 -
00135 -pf07687pf01546 -
00136 -pf03960 -
00137 -pf00873 -
00138 -pf13247pf14691pf07992 -
00139 HMM00293nuhyd_1_eut1(pf00293) EUT1
00140 -pf06674 -
00141 HMM02780keto_1_eut1(pf02779pf02780pf00456) EUT1
00142 HMM00390_malic_1_eut1(pf00390pf03949pf01515) EUT1
00143 Hp_euts_violetPink EUT1
00144 HMM10662PduV_1_mix(pf10662) EUT1
00145 HMM06249EutQ_1_eut1(pf06249) EUT1 EP_NTERM
00146 HMM01923cobtr_2_eut1(pf01923) EUT1
00147 HMM01515PTA_B_1_eut1(pf01515) EUT1
00148 H_azure EUT1
00149 P_lightViolet EUT1
00150 HMMalddh_1_eutpdugrm(pf00171) EUT1 EP_NTERM
00151 HMM06723EutJMreB_1_grm(#) EUT1
00152 HMM00465alcdh_3_mix(pf00465) EUT1
00153 HMMtpEuth_1_eut(pf04346) EUT1
00154 HMM06277EutA_1_eut(pf06277) EUT1
00155 HMMetlyEutB_1_EUT1(pf06751) EUT1
00156 HMMetlyEutC_1_eut1(pf05985) EUT1 EP_NTERM
00157 Tsp_rosa EUT1
00158 H_mulberry(pf00936pf16365) EUT1
00159 HMMreg12833_1_EUT1(pf12833) EUT1
00160 -pf01218 -
00161 -pf01520 -
00162 -pf00583 -
00163 -pf11143 -
00164 -pf06572 -
00165 -pf04261 -
00166 -# -
00167 -pf00528 -
00168 -pf00528 -
BMC-P
BMC-H
BMC-T
Aldehyde dehydrogenase
PTAC
Signature enzyme
Alcohol dehydrogenase
Regulator

Klebsiella oxytoca. Bacteria; Proteobacteria; Gammaproteobacteria; Enterobacterales;

SAMEA2273697__locus_1

EUT1

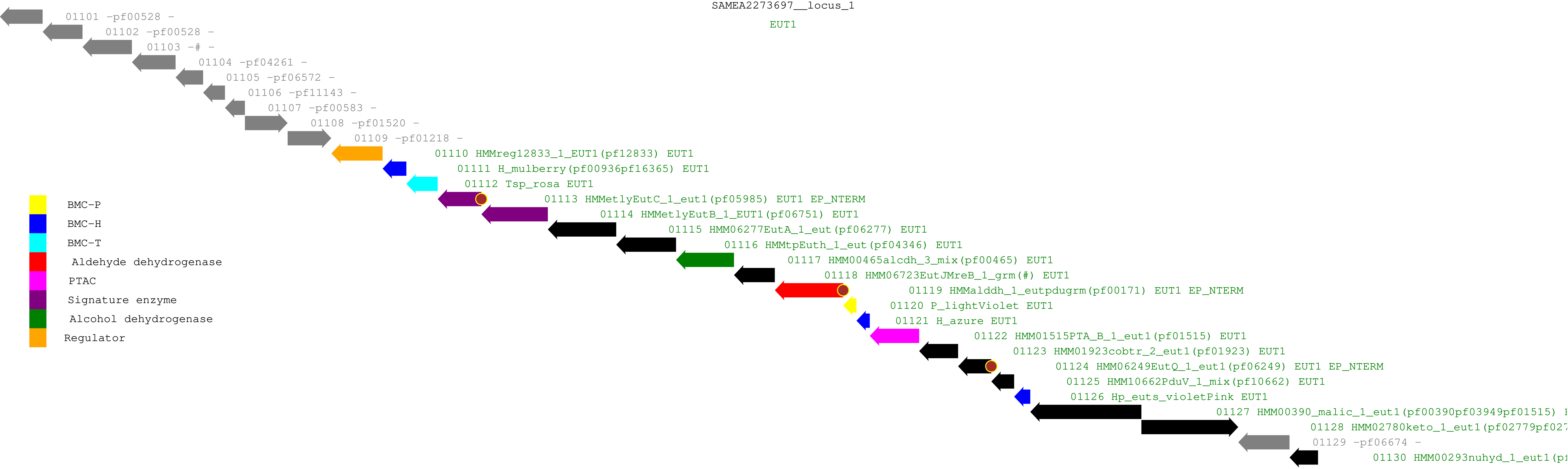

Klebsiella oxytoca. Bacteria; Proteobacteria; Gammaproteobacteria; Enterobacterales;

SAMEA2273741__locus_1

EUT1

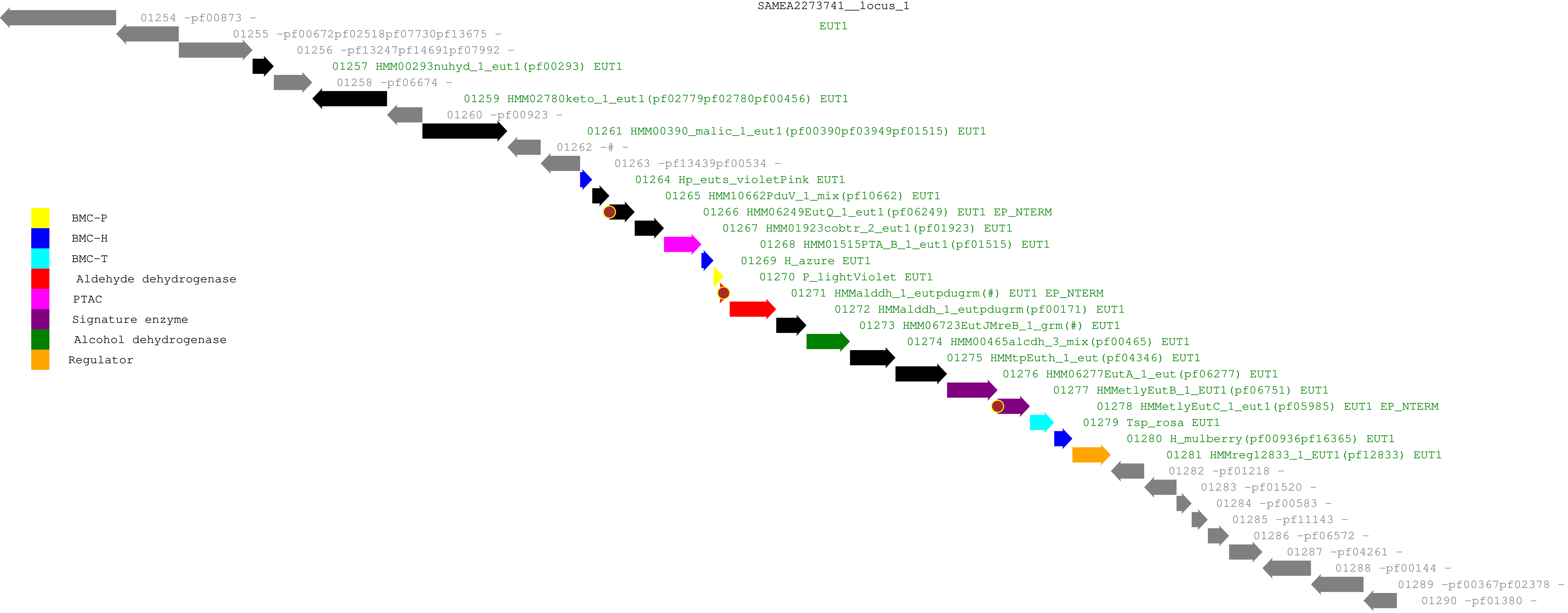

Klebsiella oxytoca. Bacteria; Proteobacteria; Gammaproteobacteria; Enterobacterales;

SAMEA2273883__locus_1

EUT1

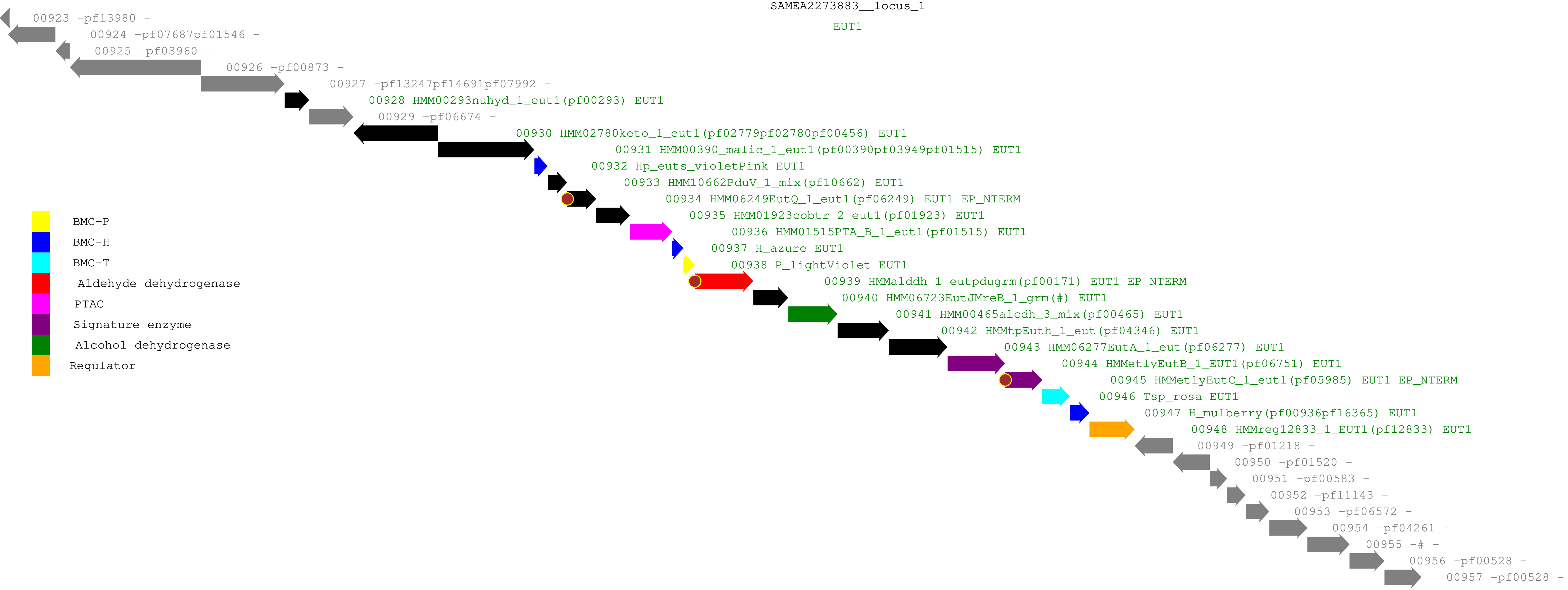

Klebsiella pneumoniae. Bacteria; Proteobacteria; Gammaproteobacteria; Enterobacterales;

SAMEA23986918__locus_1

EUT1

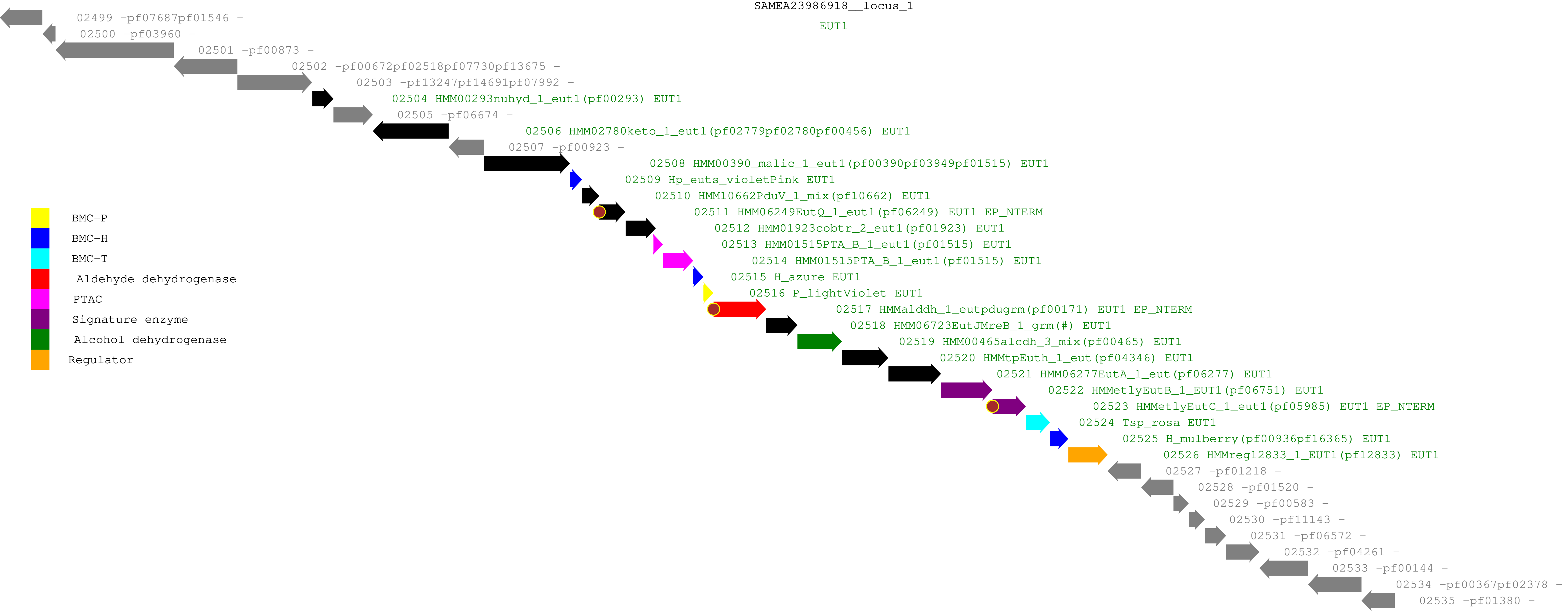

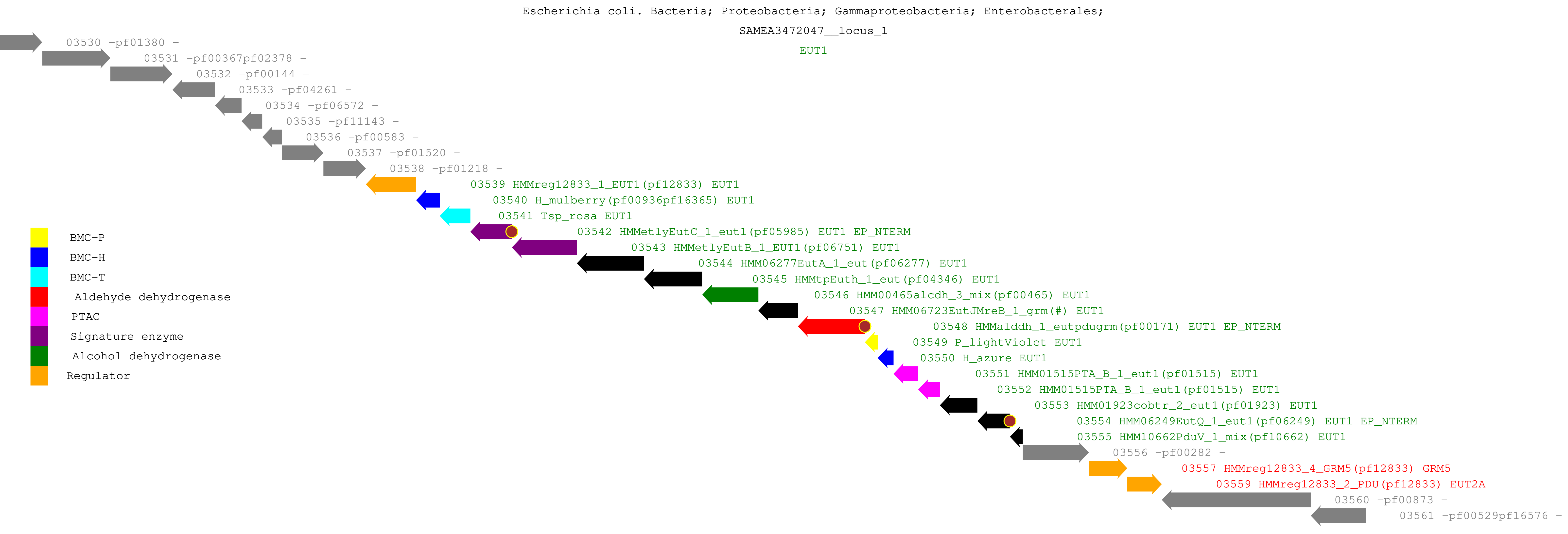
Escherichia coli. Bacteria; Proteobacteria; Gammaproteobacteria; Enterobacterales;
SAMEA3472047__locus_1
EUT1
03530 -pf01380 -
03531 -pf00367pf02378 -
03532 -pf00144 -
03533 -pf04261 -
03534 -pf06572 -
03535 -pf11143 -
03536 -pf00583 -
03537 -pf01520 -
03538 -pf01218 -
03539 HMMreg12833_1_EUT1(pf12833) EUT1
03540 H_mulberry(pf00936pf16365) EUT1
03541 Tsp_rosa EUT1
03542 HMMetlyEutC_1_eut1(pf05985) EUT1 EP_NTERM
03543 HMMetlyEutB_1_EUT1(pf06751) EUT1
03544 HMM06277EutA_1_eut(pf06277) EUT1
03545 HMMtpEuth_1_eut(pf04346) EUT1
03546 HMM00465alcdh_3_mix(pf00465) EUT1
03547 HMM06723EutJMreB_1_grm(#) EUT1
03548 HMMalddh_1_eutpdugrm(pf00171) EUT1 EP_NTERM
03549 P_lightViolet EUT1
03550 H_azure EUT1
03551 HMM01515PTA_B_1_eut1(pf01515) EUT1
03552 HMM01515PTA_B_1_eut1(pf01515) EUT1
03553 HMM01923cobtr_2_eut1(pf01923) EUT1
03554 HMM06249EutQ_1_eut1(pf06249) EUT1 EP_NTERM
03555 HMM10662PduV_1_mix(pf10662) EUT1
03556 -pf00282 -
03557 HMMreg12833_4_GRM5(pf12833) GRM5
03559 HMMreg12833_2_PDU(pf12833) EUT2A
03560 -pf00873 -
03561 -pf00529pf16576 -
BMC-P
BMC-H
BMC-T
Aldehyde dehydrogenase
PTAC
Signature enzyme
Alcohol dehydrogenase
Regulator

Escherichia coli. Bacteria; Proteobacteria; Gammaproteobacteria; Enterobacterales;

SAMEA3472115__locus_1

EUT1

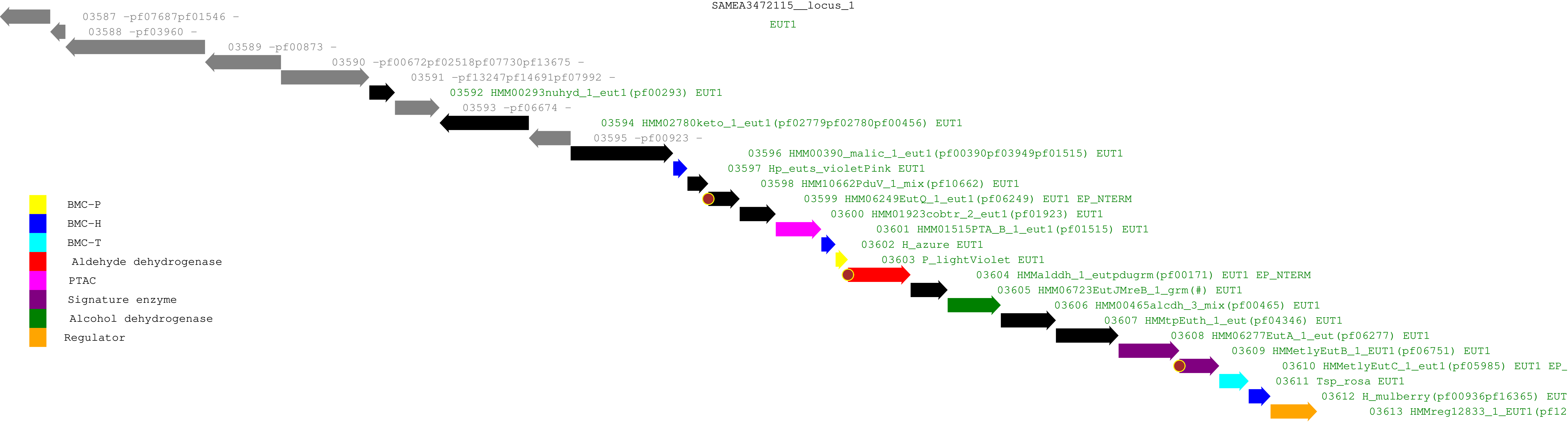

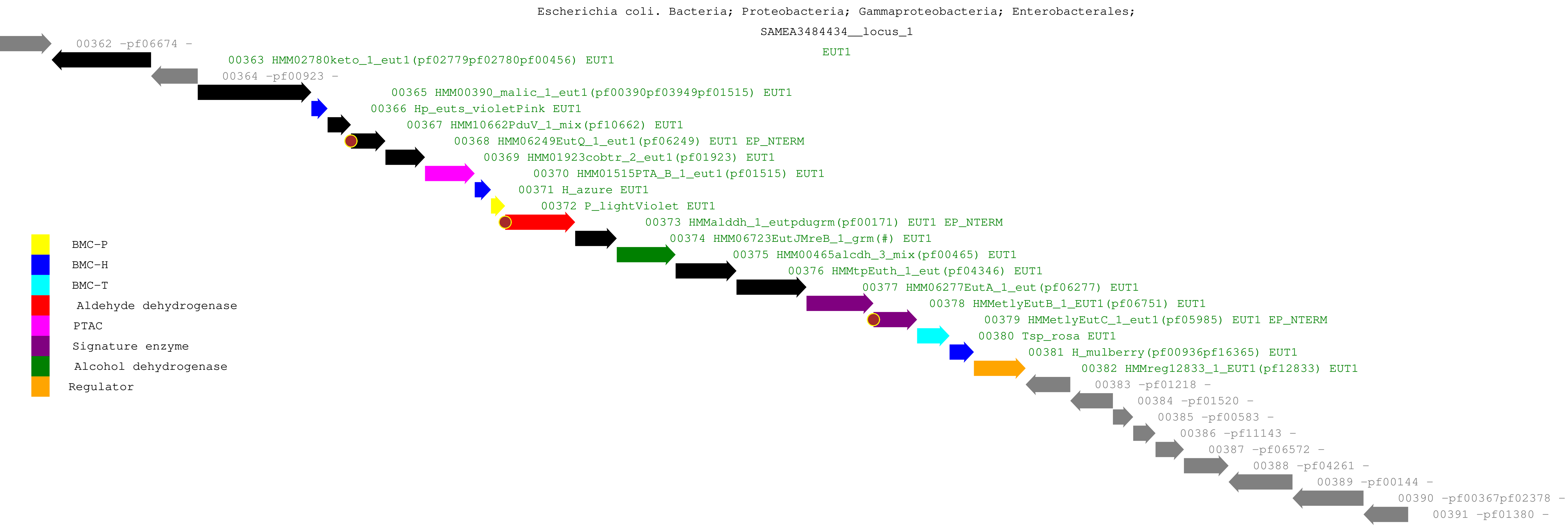

Escherichia coli. Bacteria; Proteobacteria; Gammaproteobacteria; Enterobacterales;
SAMEA3484434__locus_1
EUT1
00362 -pf06674 -
00363 HMM02780keto_1_eut1(pf02779pf02780pf00456) EUT1
00364 -pf00923 -
00365 HMM00390_malic_1_eut1(pf00390pf03949pf01515) EUT1
00366 Hp_euts_violetPink EUT1
00367 HMM10662PduV_1_mix(pf10662) EUT1
00368 HMM06249EutQ_1_eut1(pf06249) EUT1 EP_NTERM
00369 HMM01923cobtr_2_eut1(pf01923) EUT1
00370 HMM01515PTA_B_1_eut1(pf01515) EUT1
00371 H_azure EUT1
00372 P_lightViolet EUT1
00373 HMMalddh_1_eutpdugrm(pf00171) EUT1 EP_NTERM
00374 HMM06723EutJMreB_1_grm(#) EUT1
00375 HMM00465alcdh_3_mix(pf00465) EUT1
00376 HMMtpEuth_1_eut(pf04346) EUT1
00377 HMM06277EutA_1_eut(pf06277) EUT1
00378 HMMetlyEutB_1_EUT1(pf06751) EUT1
00379 HMMetlyEutC_1_eut1(pf05985) EUT1 EP_NTERM
00380 Tsp_rosa EUT1
00381 H_mulberry(pf00936pf16365) EUT1
00382 HMMreg12833_1_EUT1(pf12833) EUT1
00383 -pf01218 -
00384 -pf01520 -
00385 -pf00583 -
00386 -pf11143 -
00387 -pf06572 -
00388 -pf04261 -
00389 -pf00144 -
00390 -pf00367pf02378 -
00391 -pf01380 -
BMC-P
BMC-H
BMC-T
Aldehyde dehydrogenase
PTAC
Signature enzyme
Alcohol dehydrogenase
Regulator

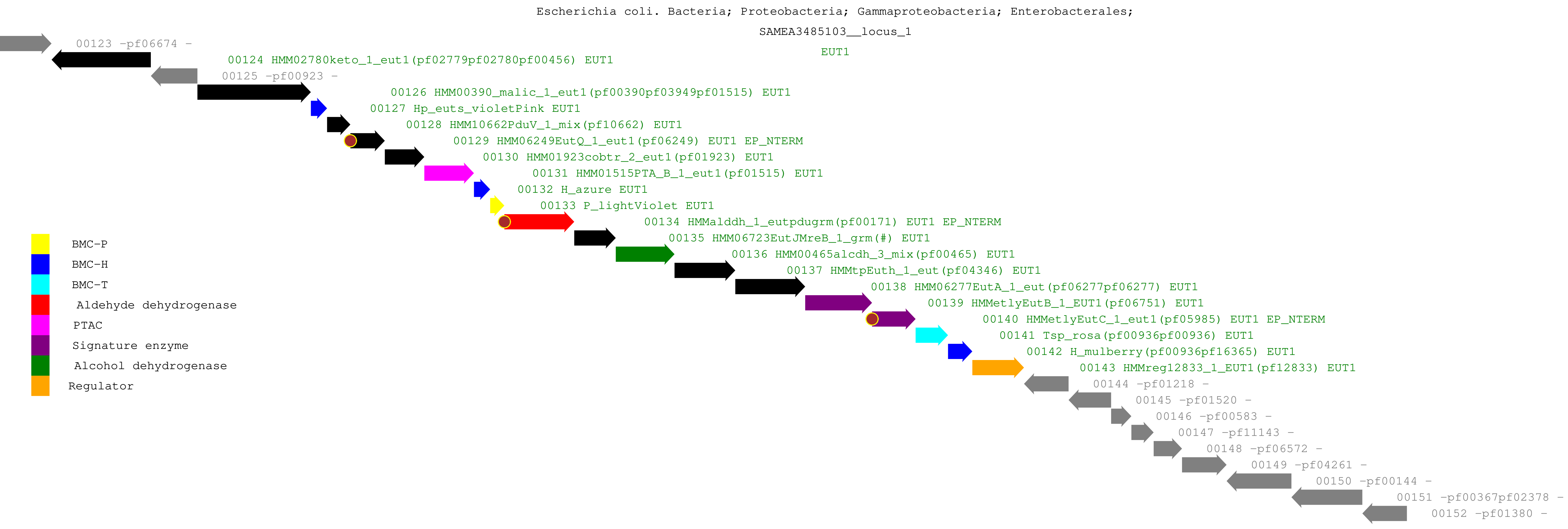

Escherichia coli. Bacteria; Proteobacteria; Gammaproteobacteria; Enterobacterales;
SAMEA3485103__locus_1
EUT1
00123 -pf06674 -
00124 HMM02780keto_1_eut1(pf02779pf02780pf00456) EUT1
00125 -pf00923 -
00126 HMM00390_malic_1_eut1(pf00390pf03949pf01515) EUT1
00127 Hp_euts_violetPink EUT1
00128 HMM10662PduV_1_mix(pf10662) EUT1
00129 HMM06249EutQ_1_eut1(pf06249) EUT1 EP_NTERM
00130 HMM01923cobtr_2_eut1(pf01923) EUT1
00131 HMM01515PTA_B_1_eut1(pf01515) EUT1
00132 H_azure EUT1
00133 P_lightViolet EUT1
00134 HMMalddh_1_eutpdugrm(pf00171) EUT1 EP_NTERM
00135 HMM06723EutJMreB_1_grm(#) EUT1
00136 HMM00465alcdh_3_mix(pf00465) EUT1
00137 HMMtpEuth_1_eut(pf04346) EUT1
00138 HMM06277EutA_1_eut(pf06277pf06277) EUT1
00139 HMMetlyEutB_1_EUT1(pf06751) EUT1
00140 HMMetlyEutC_1_eut1(pf05985) EUT1 EP_NTERM
00141 Tsp_rosa(pf00936pf00936) EUT1
00142 H_mulberry(pf00936pf16365) EUT1
00143 HMMreg12833_1_EUT1(pf12833) EUT1
00144 -pf01218 -
00145 -pf01520 -
00146 -pf00583 -
00147 -pf11143 -
00148 -pf06572 -
00149 -pf04261 -
00150 -pf00144 -
00151 -pf00367pf02378 -
00152 -pf01380 -
BMC-P
BMC-H
BMC-T
Aldehyde dehydrogenase
PTAC
Signature enzyme
Alcohol dehydrogenase
Regulator

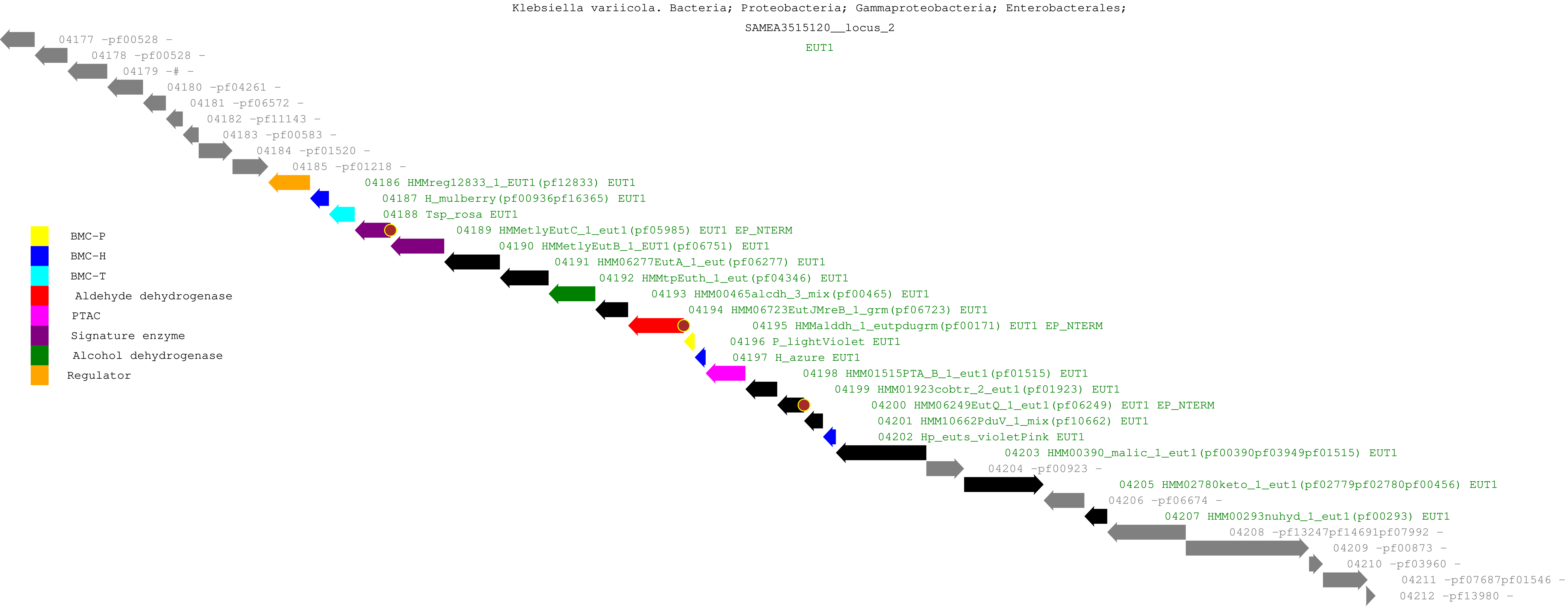
Klebsiella variicola. Bacteria; Proteobacteria; Gammaproteobacteria; Enterobacterales;
SAMEA3515120__locus_2
EUT1
04177 -pf00528 -
04178 -pf00528 -
04179 -# -
04180 -pf04261 -
04181 -pf06572 -
04182 -pf11143 -
04183 -pf00583 -
04184 -pf01520 -
04185 -pf01218 -
04186 HMMreg12833_1_EUT1(pf12833) EUT1
04187 H_mulberry(pf00936pf16365) EUT1
04188 Tsp_rosa EUT1
04189 HMMetlyEutC_1_eut1(pf05985) EUT1 EP_NTERM
04190 HMMetlyEutB_1_EUT1(pf06751) EUT1
04191 HMM06277EutA_1_eut(pf06277) EUT1
04192 HMMtpEuth_1_eut(pf04346) EUT1
04193 HMM00465alcdh_3_mix(pf00465) EUT1
04194 HMM06723EutJMreB_1_grm(pf06723) EUT1
04195 HMMaldh_1_eutpdugrm(pf00171) EUT1 EP_NTERM
04196 P_lightViolet EUT1
04197 H_azure EUT1
04198 HMM01515PTA_B_1_eut1(pf01515) EUT1
04199 HMM01923cobtr_2_eut1(pf01923) EUT1
04200 HMM06249EutQ_1_eut1(pf06249) EUT1 EP_NTERM
04201 HMM10662PduV_1_mix(pf10662) EUT1
04202 Hp_euts_violetPink EUT1
04203 HMM00390_malic_1_eut1(pf00390pf03949pf01515) EUT1
04204 -pf00923 -
04205 HMM02780keto_1_eut1(pf02779pf02780pf00456) EUT1
04206 -pf06674 -
04207 HMM00293nuhyd_1_eut1(pf00293) EUT1
04208 -pf13247pf14691pf07992 -
04209 -pf00873 -
04210 -pf03960 -
04211 -pf07687pf01546 -
04212 -pf13980 -
BMC-P
BMC-H
BMC-T
Aldehyde dehydrogenase
PTAC
Signature enzyme
Alcohol dehydrogenase
Regulator

Escherichia coli. Bacteria; Proteobacteria; Gammaproteobacteria; Enterobacterales;

SAMEA3531867__locus_1

EUT1

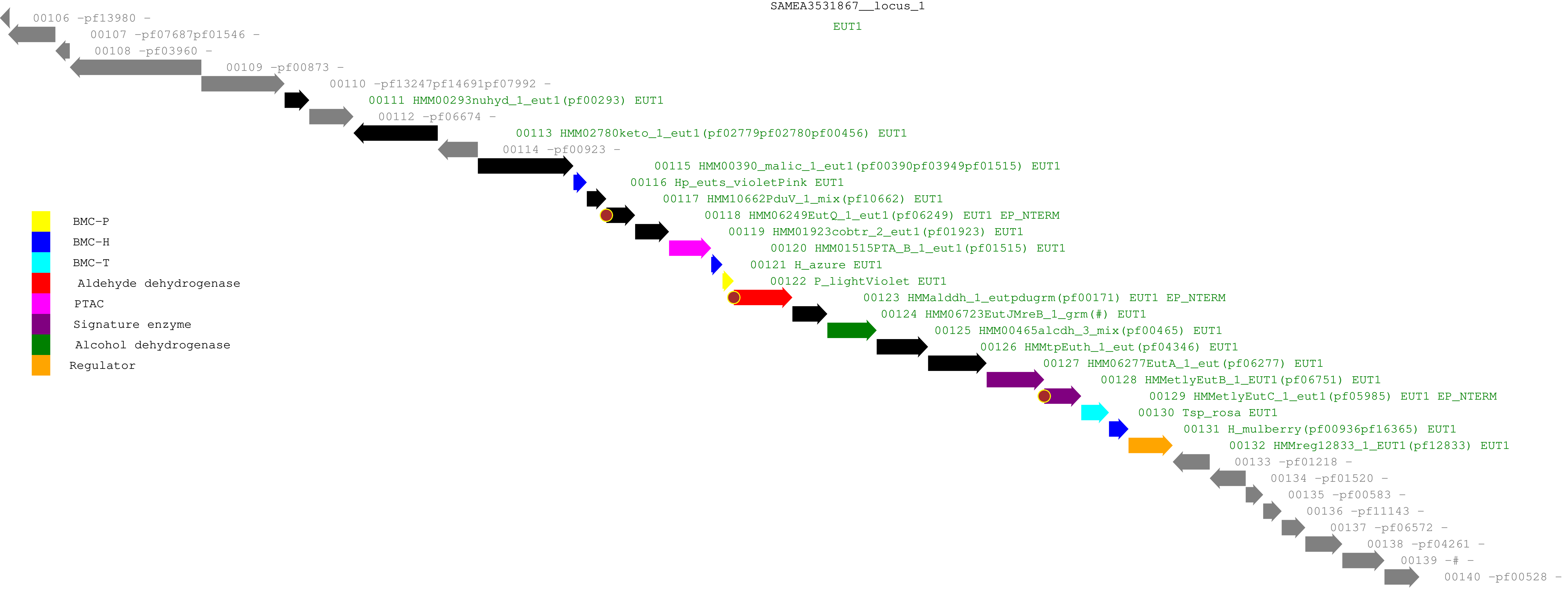

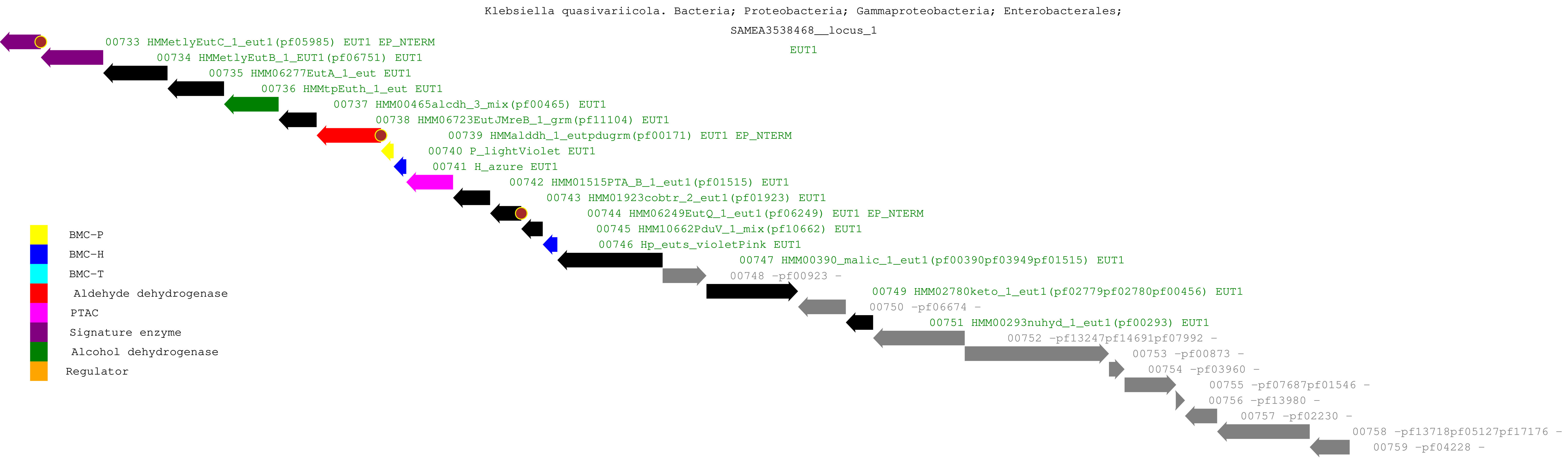
Klebsiella quasivariicola. Bacteria; Proteobacteria; Gammaproteobacteria; Enterobacterales;
SAMEA3538468__locus_1
EUT1
00733 HMMetlyEutC_1_eut1(pf05985) EUT1 EP_NTERM
00734 HMMetlyEutB_1_EUT1(pf06751) EUT1
00735 HMM06277EutA_1_eut EUT1
00736 HMMtpEuth_1_eut EUT1
00737 HMM00465alcdh_3_mix(pf00465) EUT1
00738 HMM06723EutJMreB_1_grm(pf11104) EUT1
00739 HMMalddh_1_eutpdugrm(pf00171) EUT1 EP_NTERM
00740 P_lightViolet EUT1
00741 H_azure EUT1
00742 HMM01515PTA_B_1_eut1(pf01515) EUT1
00743 HMM01923cobtr_2_eut1(pf01923) EUT1
00744 HMM06249EutQ_1_eut1(pf06249) EUT1 EP_NTERM
00745 HMM10662PduV_1_mix(pf10662) EUT1
00746 Hp_euts_violetPink EUT1
00747 HMM00390_malic_1_eut1(pf00390pf03949pf01515) EUT1
00748 -pf00923 -
00749 HMM02780keto_1_eut1(pf02779pf02780pf00456) EUT1
00750 -pf06674 -
00751 HMM00293nuhyd_1_eut1(pf00293) EUT1
00752 -pf13247pf14691pf07992 -
00753 -pf00873 -
00754 -pf03960 -
00755 -pf07687pf01546 -
00756 -pf13980 -
00757 -pf02230 -
00758 -pf13718pf05127pf17176 -
00759 -pf04228 -
BMC-P
BMC-H
BMC-T
Aldehyde dehydrogenase
PTAC
Signature enzyme
Alcohol dehydrogenase
Regulator

Klebsiella pneumoniae. Bacteria; Proteobacteria; Gammaproteobacteria; Enterobacterales;

SAMEA3649709__locus_1

EUT1

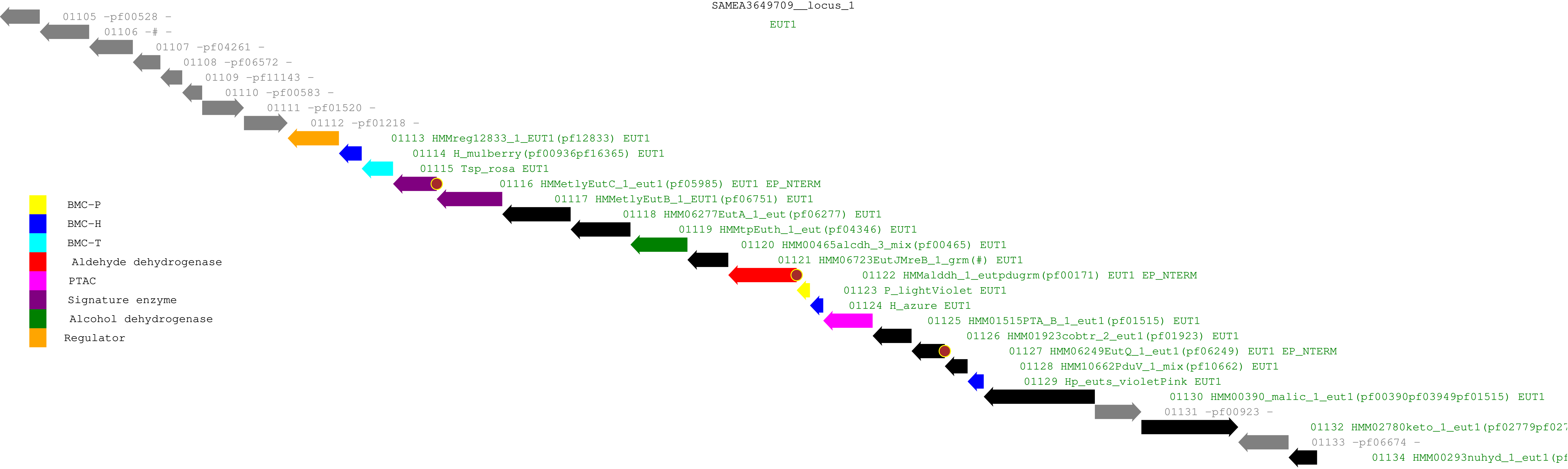

Klebsiella pneumoniae. Bacteria; Proteobacteria; Gammaproteobacteria; Enterobacterales;

SAMEA3729690__locus_1

EUT1

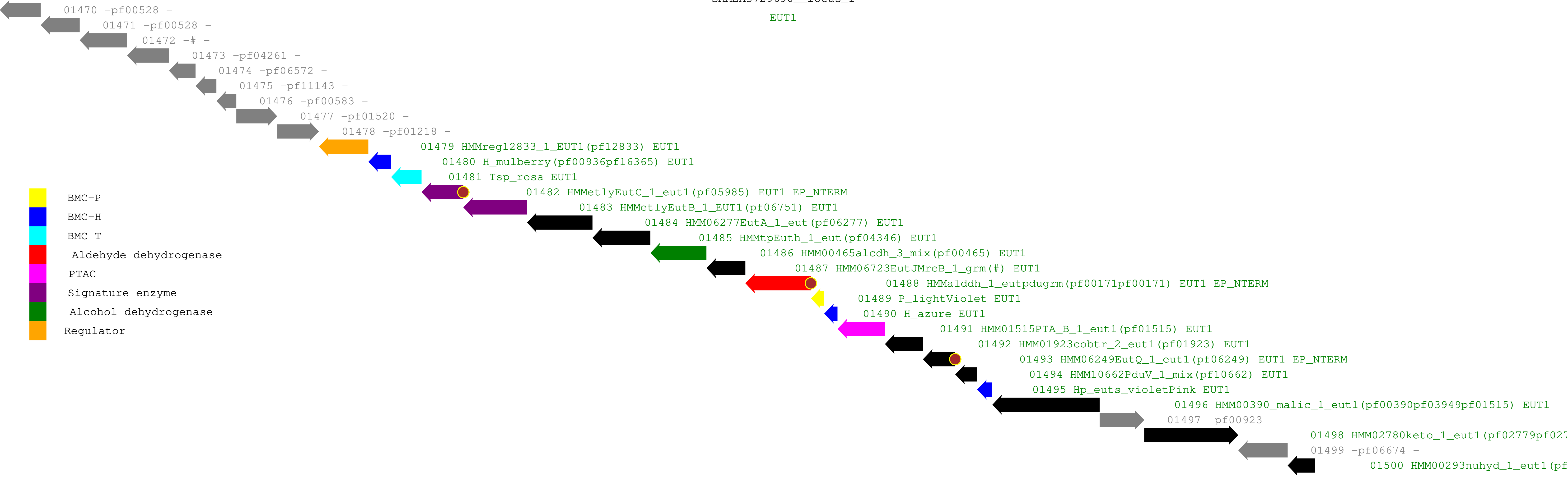

Escherichia coli. Bacteria; Proteobacteria; Gammaproteobacteria; Enterobacterales;

SAMEA3752372__locus_1

EUT1

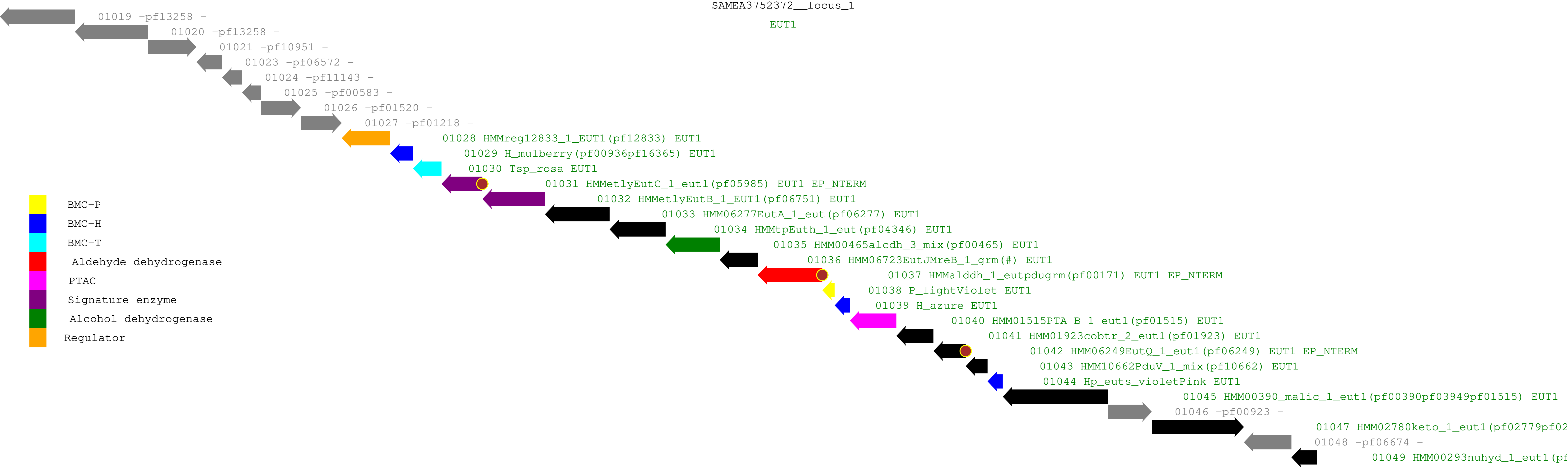

Escherichia coli. Bacteria; Proteobacteria; Gammaproteobacteria; Enterobacterales;

SAMEA3753097__locus_1

EUT1

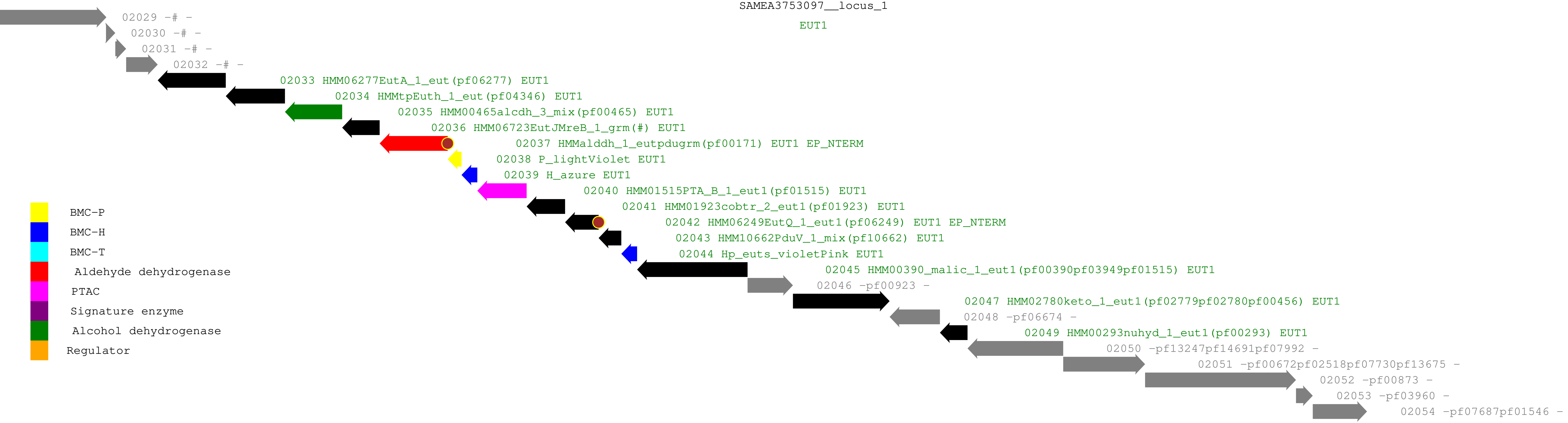

Escherichia coli. Bacteria; Proteobacteria; Gammaproteobacteria; Enterobacterales;

SAMEA3753304__locus_1

EUT1

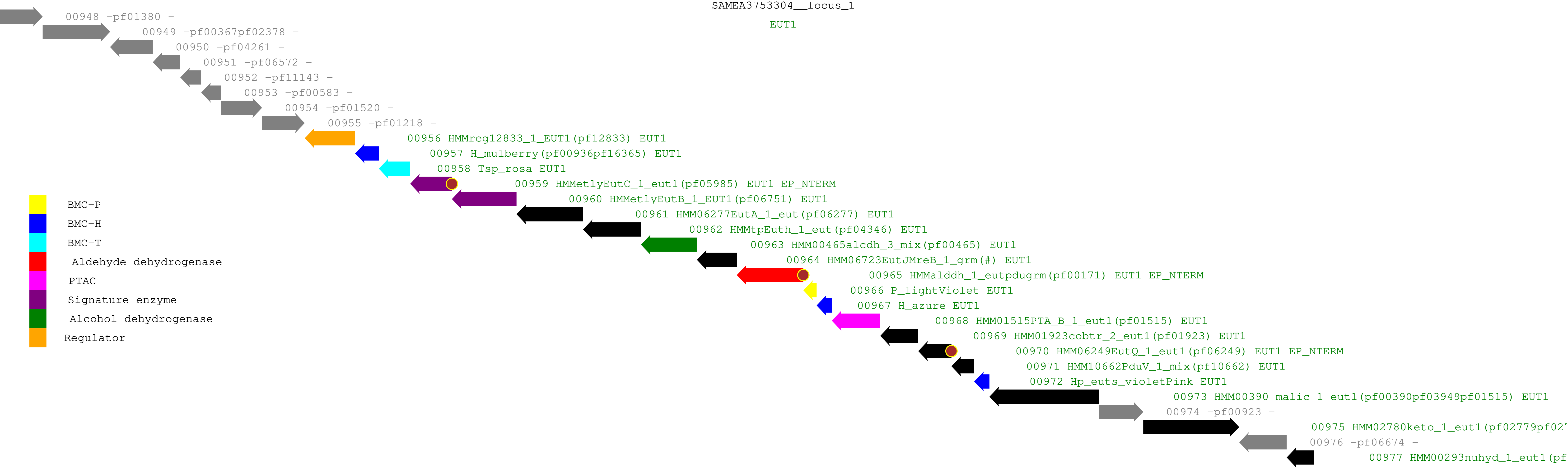

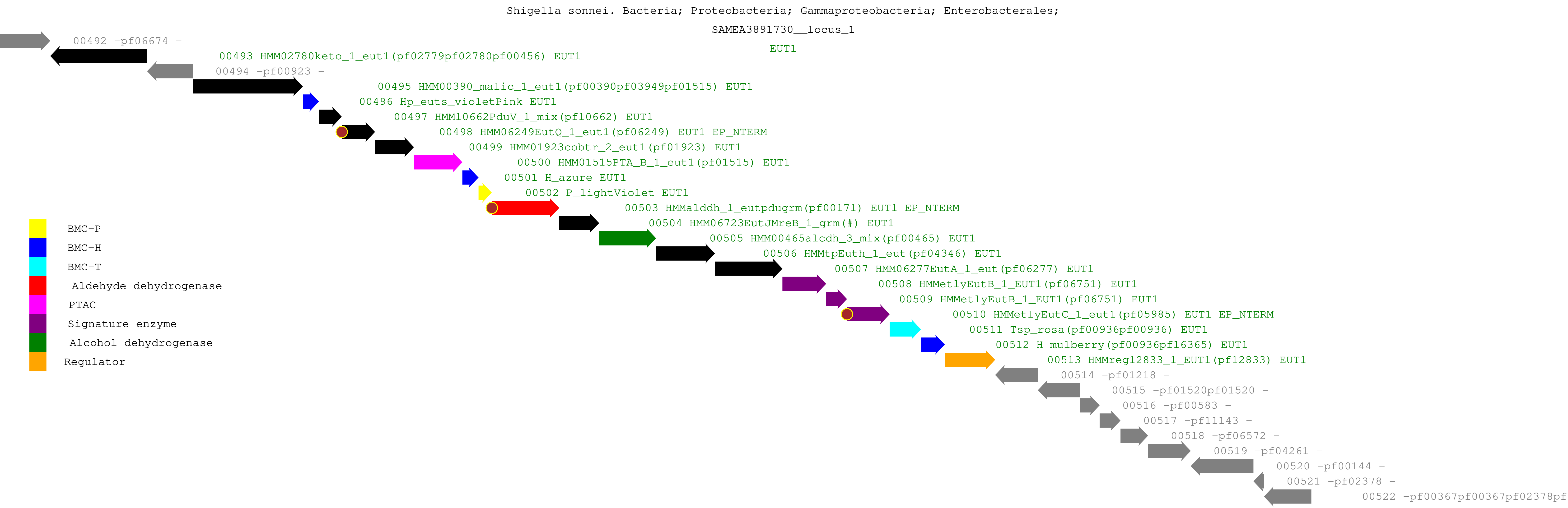

Shigella sonnei. Bacteria; Proteobacteria; Gammaproteobacteria; Enterobacterales;
SAMEA3891730__locus_1
EUT1
00492 -pf06674 -
00493 HMM02780keto_1_eut1(pf02779pf02780pf00456) EUT1
00494 -pf00923 -
00495 HMM00390_malic_1_eut1(pf00390pf03949pf01515) EUT1
00496 Hp_euts_violetPink EUT1
00497 HMM10662PduV_1_mix(pf10662) EUT1
00498 HMM06249EutQ_1_eut1(pf06249) EUT1 EP_NTERM
00499 HMM01923cobtr_2_eut1(pf01923) EUT1
00500 HMM01515PTA_B_1_eut1(pf01515) EUT1
00501 H_azure EUT1
00502 P_lightViolet EUT1
00503 HMMalddh_1_eutpdugrm(pf00171) EUT1 EP_NTERM
00504 HMM06723EutJMreB_1_grm(#) EUT1
00505 HMM00465alcdh_3_mix(pf00465) EUT1
00506 HMMtpEuth_1_eut(pf04346) EUT1
00507 HMM06277EutA_1_eut(pf06277) EUT1
00508 HMMetlyEutB_1_EUT1(pf06751) EUT1
00509 HMMetlyEutB_1_EUT1(pf06751) EUT1
00510 HMMetlyEutC_1_eut1(pf05985) EUT1 EP_NTERM
00511 Tsp_rosa(pf00936pf00936) EUT1
00512 H_mulberry(pf00936pf16365) EUT1
00513 HMMreg12833_1_EUT1(pf12833) EUT1
00514 -pf01218 -
00515 -pf01520pf01520 -
00516 -pf00583 -
00517 -pf11143 -
00518 -pf06572 -
00519 -pf04261 -
00520 -pf00144 -
00521 -pf02378 -
00522 -pf00367pf00367pf02378pf
BMC-P
BMC-H
BMC-T
Aldehyde dehydrogenase
PTAC
Signature enzyme
Alcohol dehydrogenase
Regulator

Klebsiella pneumoniae. Bacteria; Proteobacteria; Gammaproteobacteria; Enterobacterales;

SAMEA4362680__locus_1

EUT1

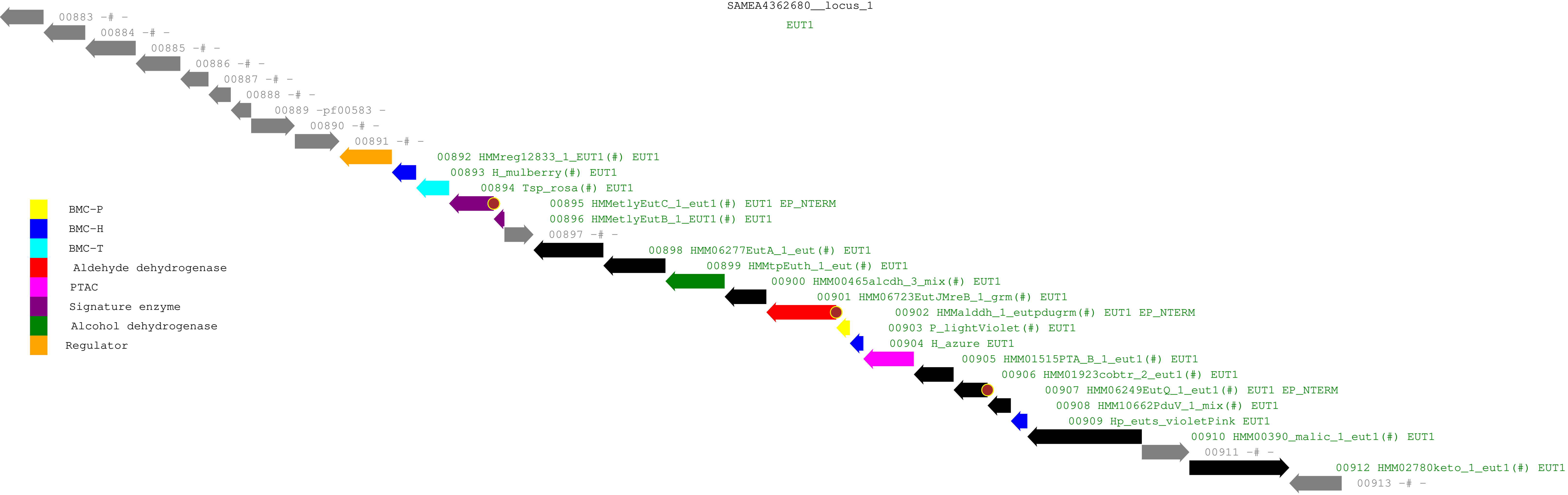

Klebsiella pneumoniae. Bacteria; Proteobacteria; Gammaproteobacteria; Enterobacterales;

SAMEA4362693__locus_2

EUT1

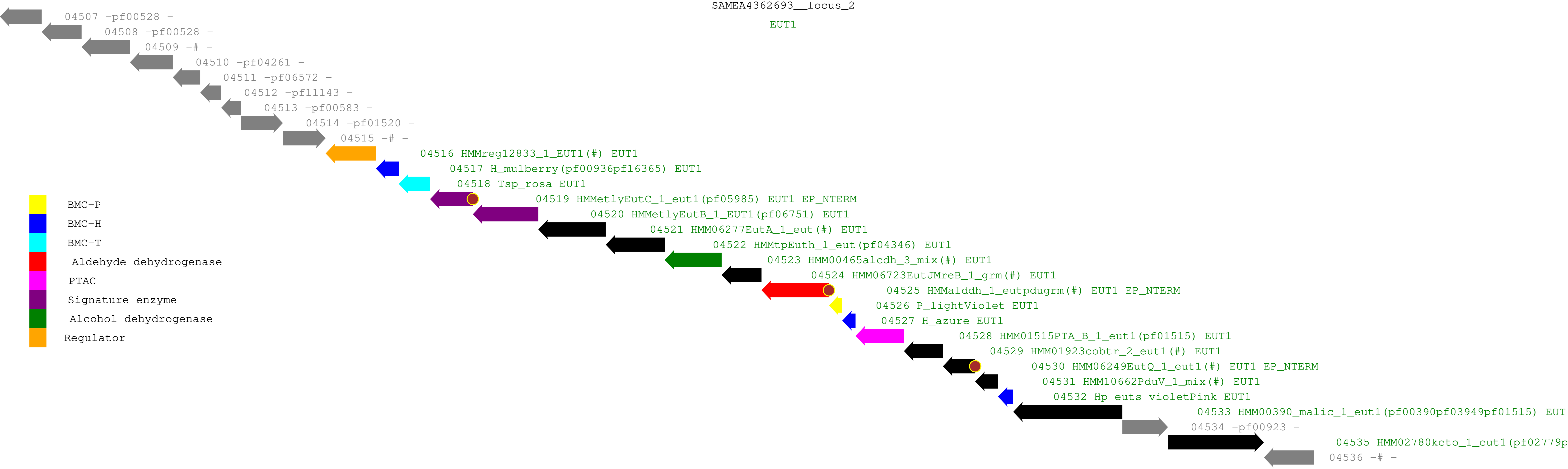

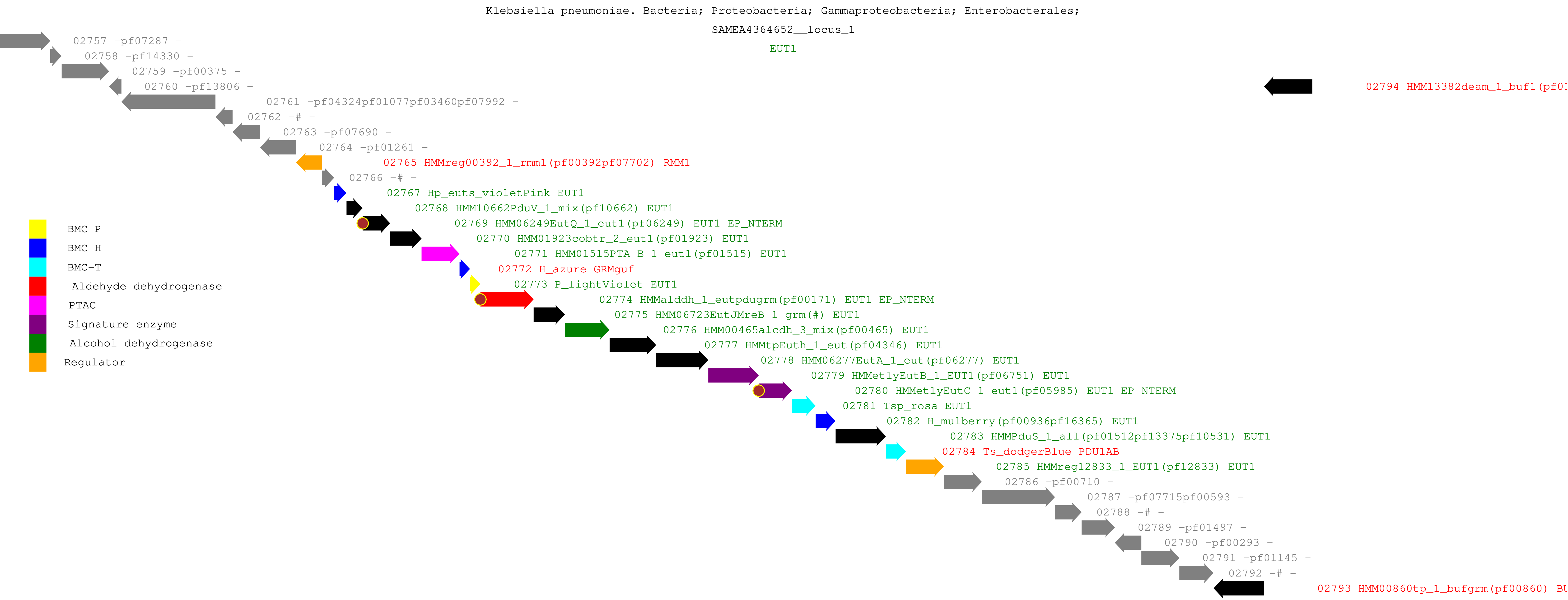
Klebsiella pneumoniae. Bacteria; Proteobacteria; Gammaproteobacteria; Enterobacterales;
SAMEA4364652__locus_1
EUT1
02757 -pf07287 -
02758 -pf14330 -
02759 -pf00375 -
02760 -pf13806 -
02761 -pf04324pf01077pf03460pf07992 -
02762 -# -
02763 -pf07690 -
02764 -pf01261 -
02765 HMMreg00392_1_rmm1(pf00392pf07702) RMM1
02766 -# -
02767 Hp_euts_violetPink EUT1
02768 HMM10662PduV_1_mix(pf10662) EUT1
02769 HMM06249EutQ_1_eut1(pf06249) EUT1 EP_NTERM
02770 HMM01923cobtr_2_eut1(pf01923) EUT1
02771 HMM01515PTA_B_1_eut1(pf01515) EUT1
02772 H_azure GRMguf
02773 P_lightViolet EUT1
02774 HMMaldh_1_eutpdugrm(pf00171) EUT1 EP_NTERM
02775 HMM06723EutJMreB_1_grm(#) EUT1
02776 HMM00465alcdh_3_mix(pf00465) EUT1
02777 HMMtpEuth_1_eut(pf04346) EUT1
02778 HMM06277EutA_1_eut(pf06277) EUT1
02779 HMMetlyEutB_1_EUT1(pf06751) EUT1
02780 HMMetlyEutC_1_eut1(pf05985) EUT1 EP_NTERM
02781 Tsp_rosa EUT1
02782 H_mulberry(pf00936pf16365) EUT1
02783 HMMPduS_1_all(pf01512pf13375pf10531) EUT1
02784 Ts_dodgerBlue PDU1AB
02785 HMMreg12833_1_EUT1(pf12833) EUT1
02786 -pf00710 -
02787 -pf07715pf00593 -
02788 -# -
02789 -pf01497 -
02790 -pf00293 -
02791 -pf01145 -
02792 -# -
02793 HMM00860tp_1_bufgrm(pf00860) BU
02794 HMM13382deam_1_buf1(pf01
BMC-P
BMC-H
BMC-T
Aldehyde dehydrogenase
PTAC
Signature enzyme
Alcohol dehydrogenase
Regulator

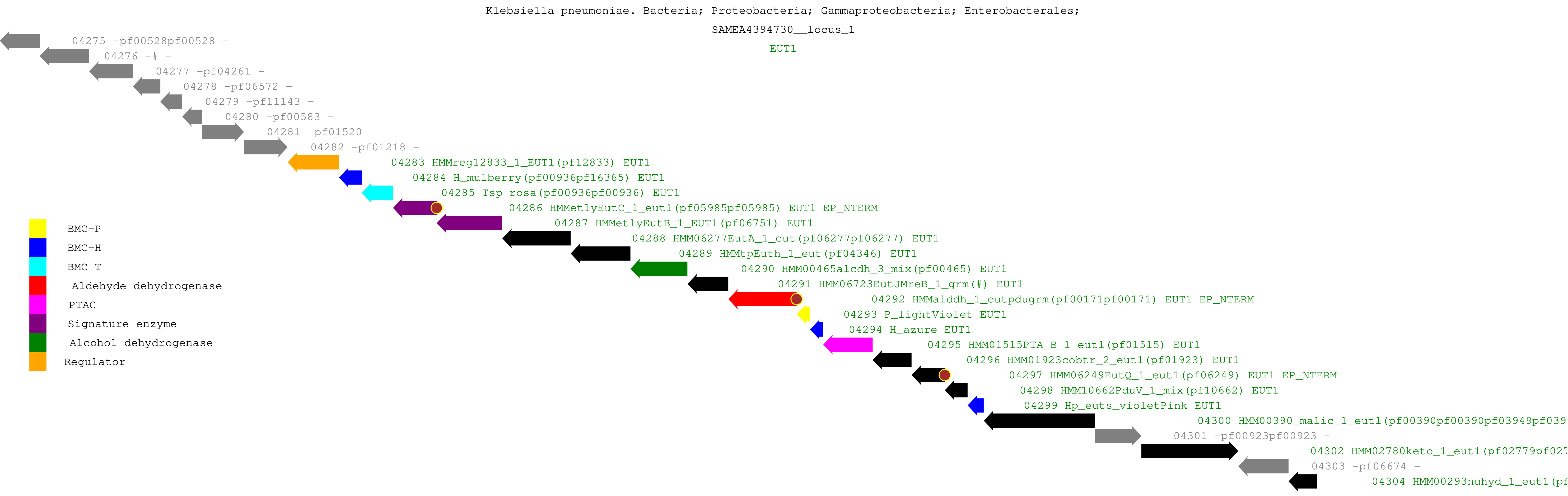
Klebsiella pneumoniae. Bacteria; Proteobacteria; Gammaproteobacteria; Enterobacterales;
SAMEA4394730__locus_1
EUT1
04275 -pf00528pf00528 -
04276 -# -
04277 -pf04261 -
04278 -pf06572 -
04279 -pf11143 -
04280 -pf00583 -
04281 -pf01520 -
04282 -pf01218 -
04283 HMMreg12833_1_EUT1(pf12833) EUT1
04284 H_mulberry(pf00936pf16365) EUT1
04285 Tsp_rosa(pf00936pf00936) EUT1
04286 HMMetlyEutC_1_eut1(pf05985pf05985) EUT1 EP_NTERM
04287 HMMetlyEutB_1_EUT1(pf06751) EUT1
04288 HMM06277EutA_1_eut(pf06277pf06277) EUT1
04289 HMMtpEuth_1_eut(pf04346) EUT1
04290 HMM00465alcdh_3_mix(pf00465) EUT1
04291 HMM06723EutJMreB_1_grm(#) EUT1
04292 HMMalddh_1_eutpdugrm(pf00171pf00171) EUT1 EP_NTERM
04293 P_lightViolet EUT1
04294 H_azure EUT1
04295 HMM01515PTA_B_1_eut1(pf01515) EUT1
04296 HMM01923cobtr_2_eut1(pf01923) EUT1
04297 HMM06249EutQ_1_eut1(pf06249) EUT1 EP_NTERM
04298 HMM10662PduV_1_mix(pf10662) EUT1
04299 Hp_euts_violetPink EUT1
04300 HMM00390_malic_1_eut1(pf00390pf00390pf03949pf039
04301 -pf00923pf00923 -
04302 HMM02780keto_1_eut1(pf02779pf027
04303 -pf06674 -
04304 HMM00293nuhyd_1_eut1(pf
BMC-P
BMC-H
BMC-T
Aldehyde dehydrogenase
PTAC
Signature enzyme
Alcohol dehydrogenase
Regulator

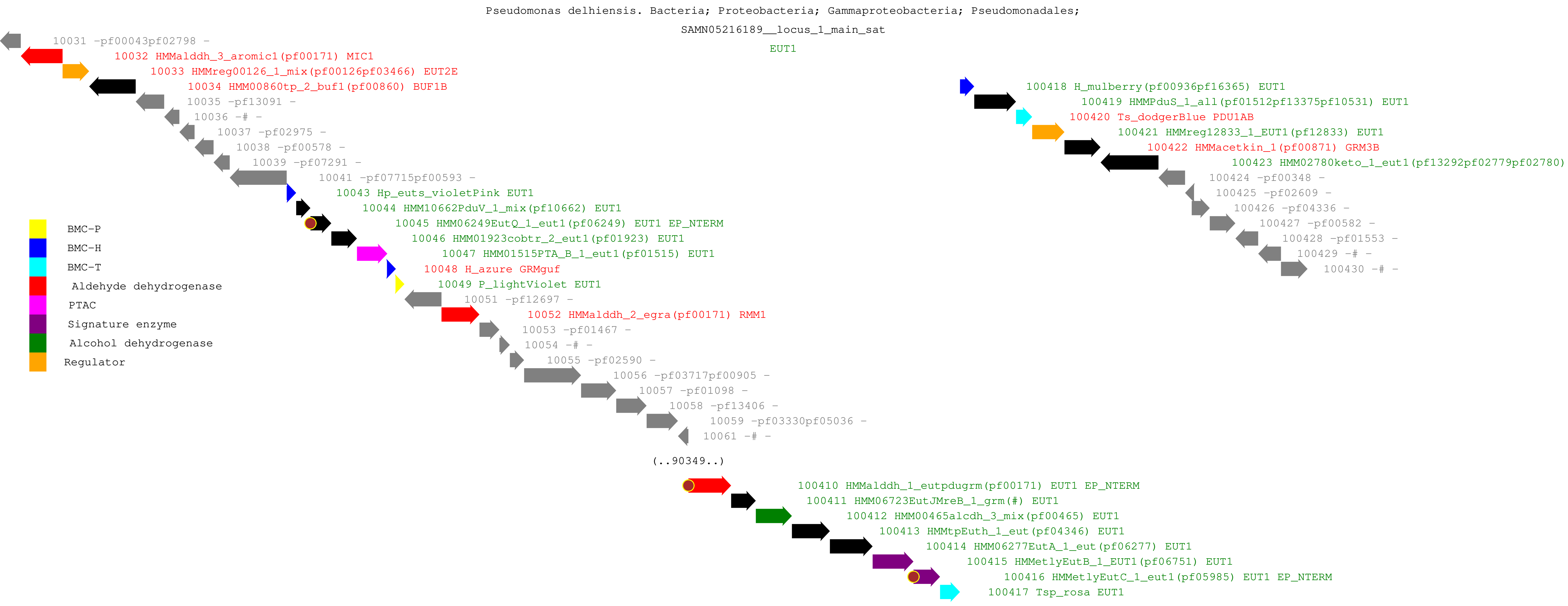
Pseudomonas delhiensis. Bacteria; Proteobacteria; Gammaproteobacteria; Pseudomonadales;
SAMN05216189__locus_1_main_sat
EUT1
10031 -pf00043pf02798 -
10032 HMMalddh_3_aromic1(pf00171) MIC1
10033 HMMreg00126_1_mix(pf00126pf03466) EUT2E
10034 HMM00860tp_2_buf1(pf00860) BUF1B
10035 -pf13091 -
10036 -# -
10037 -pf02975 -
10038 -pf00578 -
10039 -pf07291 -
10041 -pf07715pf00593 -
10043 Hp_euts_violetPink EUT1
10044 HMM10662PduV_1_mix(pf10662) EUT1
10045 HMM06249EutQ_1_eut1(pf06249) EUT1 EP_NTERM
10046 HMM01923cobtr_2_eut1(pf01923) EUT1
10047 HMM01515PTA_B_1_eut1(pf01515) EUT1
10048 H_azure GRMguf
10049 P_lightViolet EUT1
10051 -pf12697 -
10052 HMMalddh_2_egra(pf00171) RMM1
10053 -pf01467 -
10054 -# -
10055 -pf02590 -
10056 -pf03717pf00905 -
10057 -pf01098 -
10058 -pf13406 -
10059 -pf03330pf05036 -
10061 -# -
(..90349..)
100410 HMMalddh_1_eutpdugrm(pf00171) EUT1 EP_NTERM
100411 HMM06723EutJMreB_1_grm(#) EUT1
100412 HMM00465alcdh_3_mix(pf00465) EUT1
100413 HMMtpEuth_1_eut(pf04346) EUT1
100414 HMM06277EutA_1_eut(pf06277) EUT1
100415 HMMetlyEutB_1_EUT1(pf06751) EUT1
100416 HMMetlyEutC_1_eut1(pf05985) EUT1 EP_NTERM
100417 Tsp_rosa EUT1
100418 H_mulberry(pf00936pf16365) EUT1
100419 HMMPduS_1_all(pf01512pf13375pf10531) EUT1
100420 Ts_dodgerBlue PDU1AB
100421 HMMreg12833_1_EUT1(pf12833) EUT1
100422 HMMacetkin_1(pf00871) GRM3B
100423 HMM02780keto_1_eut1(pf13292pf02779pf02780)
100424 -pf00348 -
100425 -pf02609 -
100426 -pf04336 -
100427 -pf00582 -
100428 -pf01553 -
100429 -# -
100430 -# -
BMC-P
BMC-H
BMC-T
Aldehyde dehydrogenase
PTAC
Signature enzyme
Alcohol dehydrogenase
Regulator

Morganella morganii (Proteus morganii). Bacteria; Proteobacteria; Gammaproteobacteria; Enterobacterales;

SAMN05216301__locus_1

EUT1

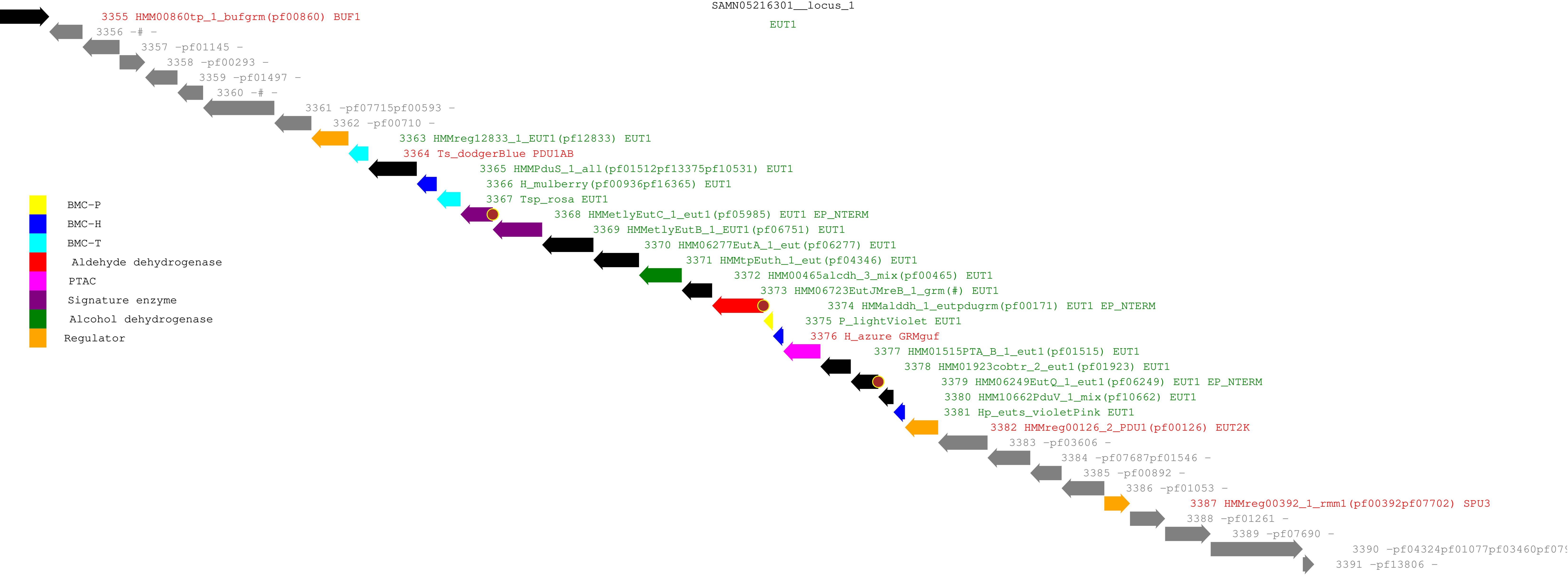

Shigella sonnei. Bacteria; Proteobacteria; Gammaproteobacteria; Enterobacterales;

SAMN05216485__locus_1

EUT1

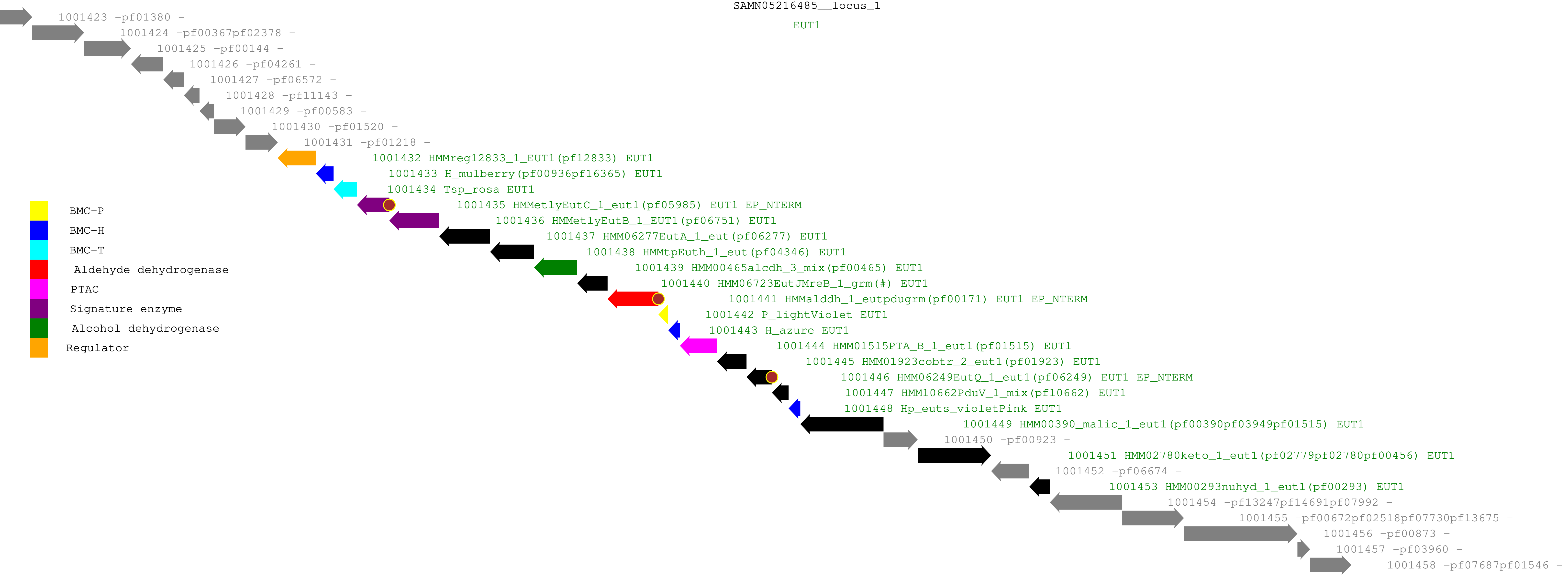

Citrobacter sp. NLAE-zl-C269. Bacteria; Proteobacteria; Gammaproteobacteria; Enterobacterales;

SAMN05216502__locus_2

EUT1

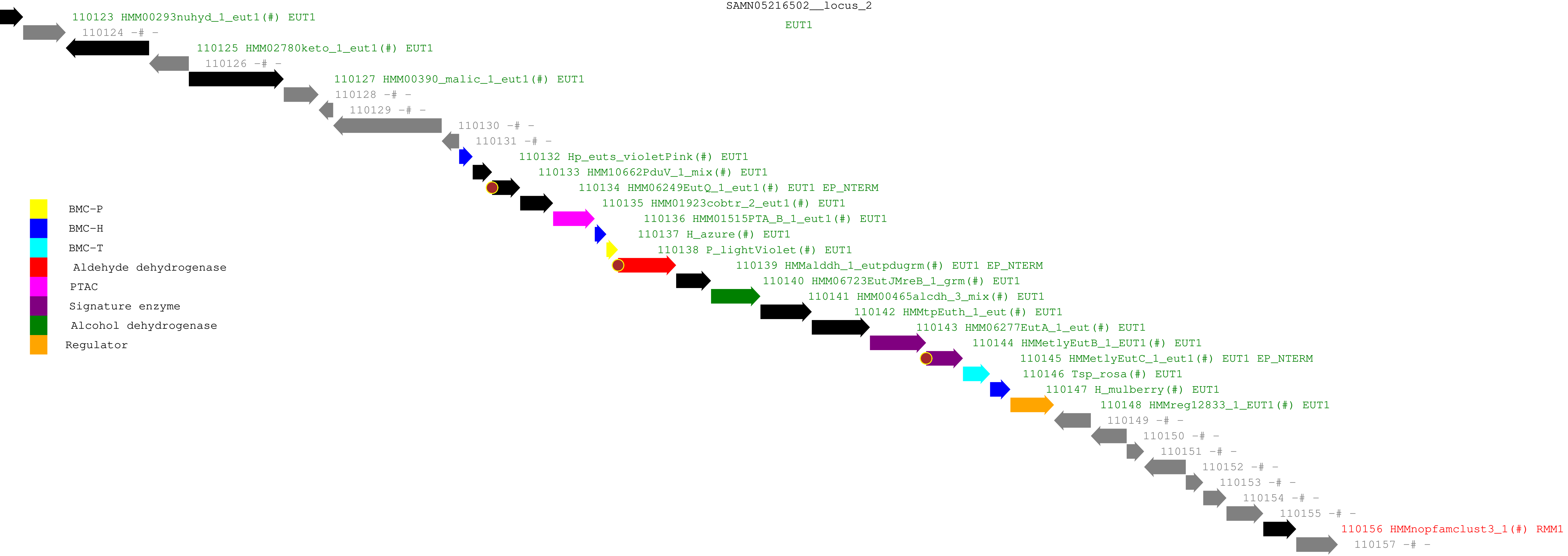

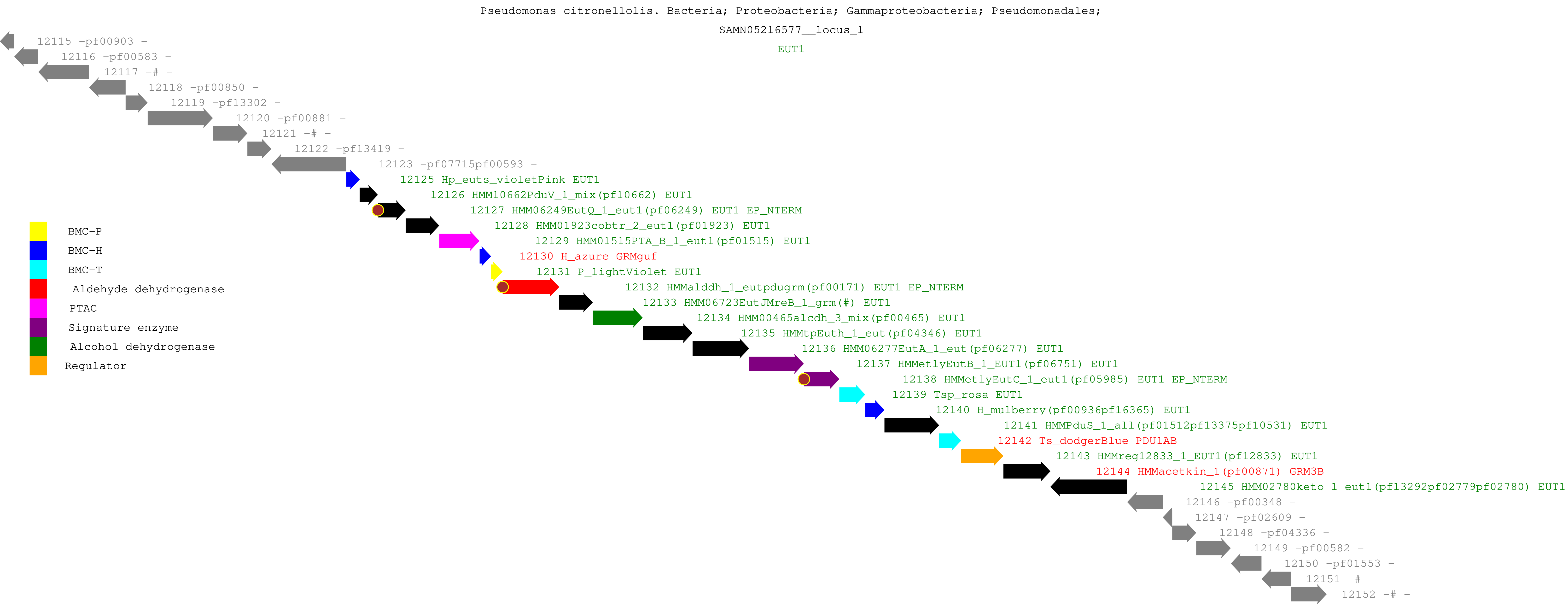
Pseudomonas citronellolis. Bacteria; Proteobacteria; Gammaproteobacteria; Pseudomonadales;
SAMN05216577__locus_1
EUT1
12115 -pf00903 -
12116 -pf00583 -
12117 -# -
12118 -pf00850 -
12119 -pf13302 -
12120 -pf00881 -
12121 -# -
12122 -pf13419 -
12123 -pf07715pf00593 -
12125 Hp_euts_violetPink EUT1
12126 HMM10662PduV_1_mix(pf10662) EUT1
12127 HMM06249EutQ_1_eut1(pf06249) EUT1 EP_NTERM
12128 HMM01923cobtr_2_eut1(pf01923) EUT1
12129 HMM01515PTA_B_1_eut1(pf01515) EUT1
12130 H_azure GRMguf
12131 P_lightViolet EUT1
12132 HMMalddh_1_eutpdugrm(pf00171) EUT1 EP_NTERM
12133 HMM06723EutJMreB_1_grm(#) EUT1
12134 HMM00465alcdh_3_mix(pf00465) EUT1
12135 HMMtpEuth_1_eut(pf04346) EUT1
12136 HMM06277EutA_1_eut(pf06277) EUT1
12137 HMMetlyEutB_1_EUT1(pf06751) EUT1
12138 HMMetlyEutC_1_eut1(pf05985) EUT1 EP_NTERM
12139 Tsp_rosa EUT1
12140 H_mulberry(pf00936pf16365) EUT1
12141 HMMPduS_1_all(pf01512pf13375pf10531) EUT1
12142 Ts_dodgerBlue PDU1AB
12143 HMMreg12833_1_EUT1(pf12833) EUT1
12144 HMMacetkin_1(pf00871) GRM3B
12145 HMM02780keto_1_eut1(pf13292pf02779pf02780) EUT1
12146 -pf00348 -
12147 -pf02609 -
12148 -pf04336 -
12149 -pf00582 -
12150 -pf01553 -
12151 -# -
12152 -# -
BMC-P
BMC-H
BMC-T
Aldehyde dehydrogenase
PTAC
Signature enzyme
Alcohol dehydrogenase
Regulator

Salmonella arizonae (strain ATCC BAA-731 / CDC346-86 / RSK2980). Bacteria; Proteobacteria; Gammaproteobacteria; Enterobacterales;

SARI__locus_1

EUT1

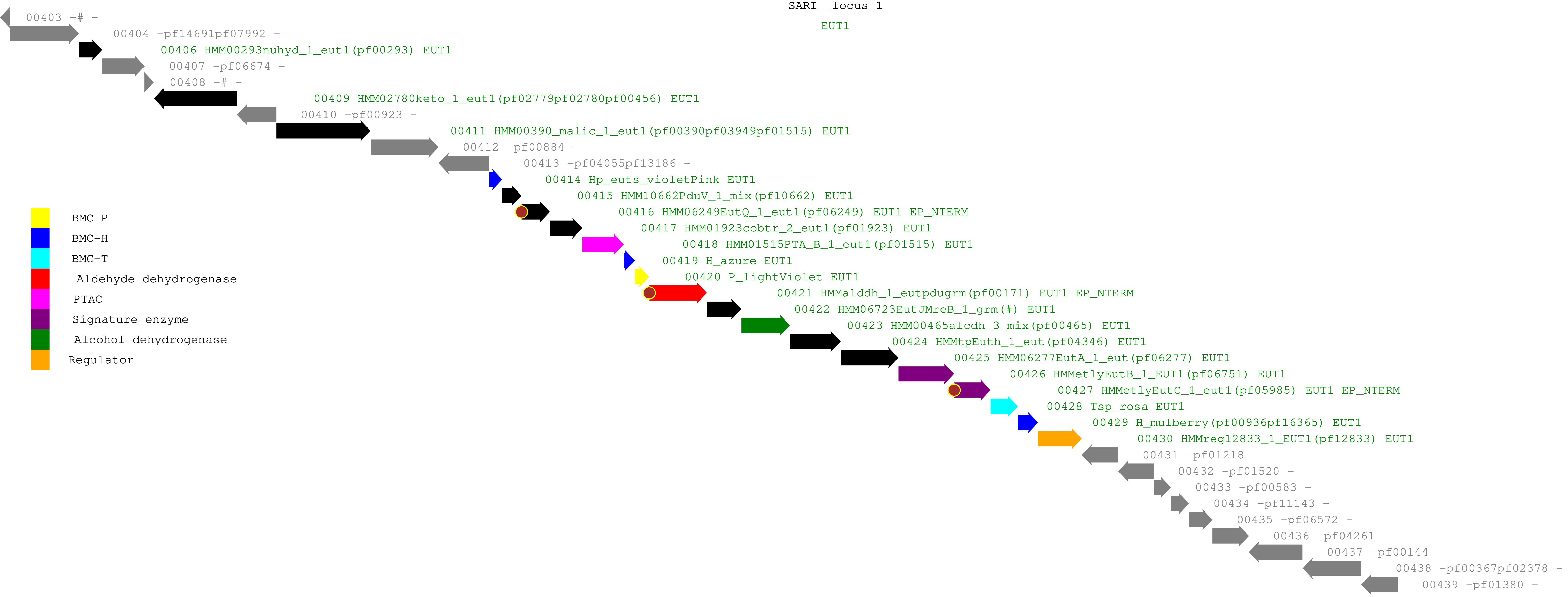

Klebsiella quasipneumoniae subsp. similipneumoniae. Bacteria; Proteobacteria; Gammaproteobacteria; Enterobacterales;

SB30__locus_2

EUT1

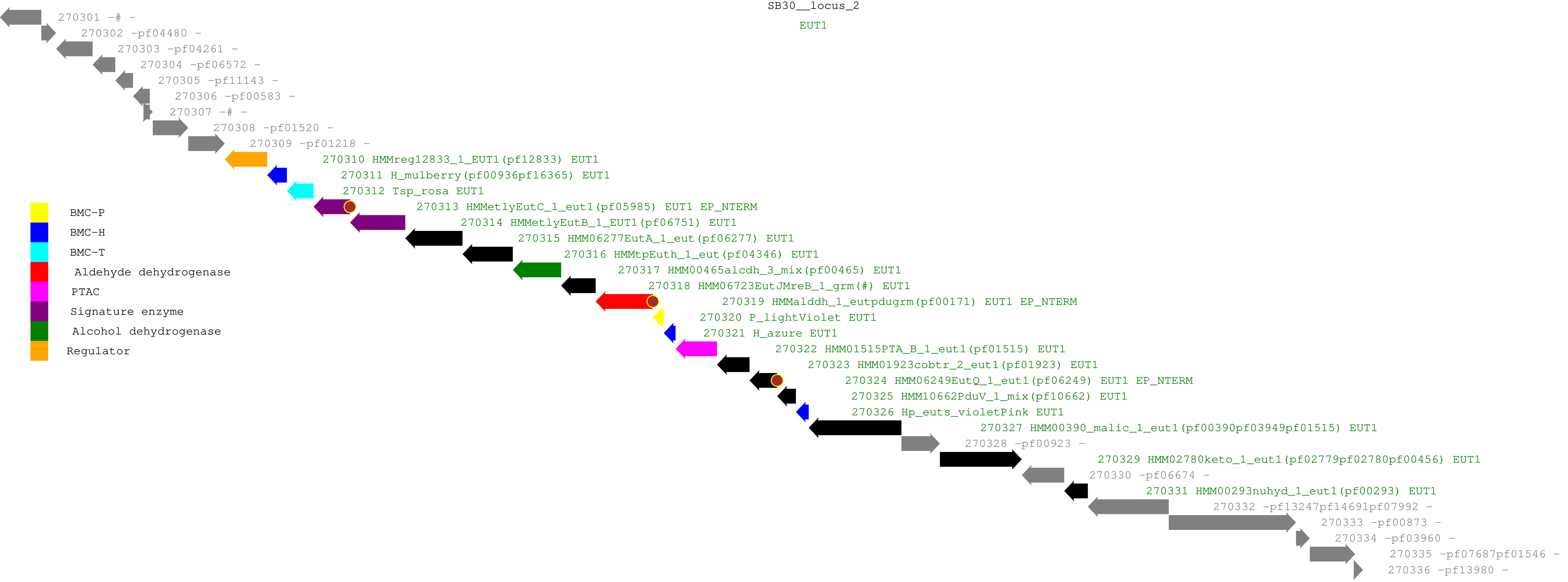

Shigella boydii 4444-74. Bacteria; Proteobacteria; Gammaproteobacteria; Enterobacterales;

SB444474__locus_1

EUT1

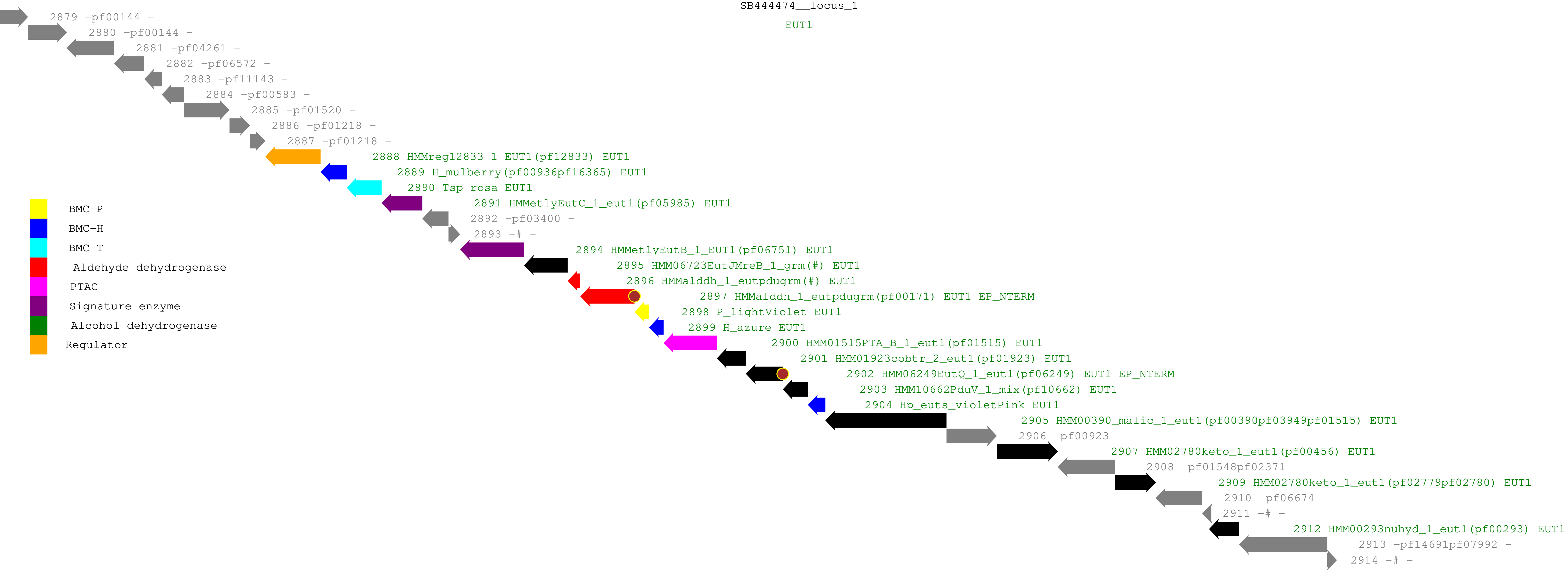

Shigella boydii 5216-82. Bacteria; Proteobacteria; Gammaproteobacteria; Enterobacterales;

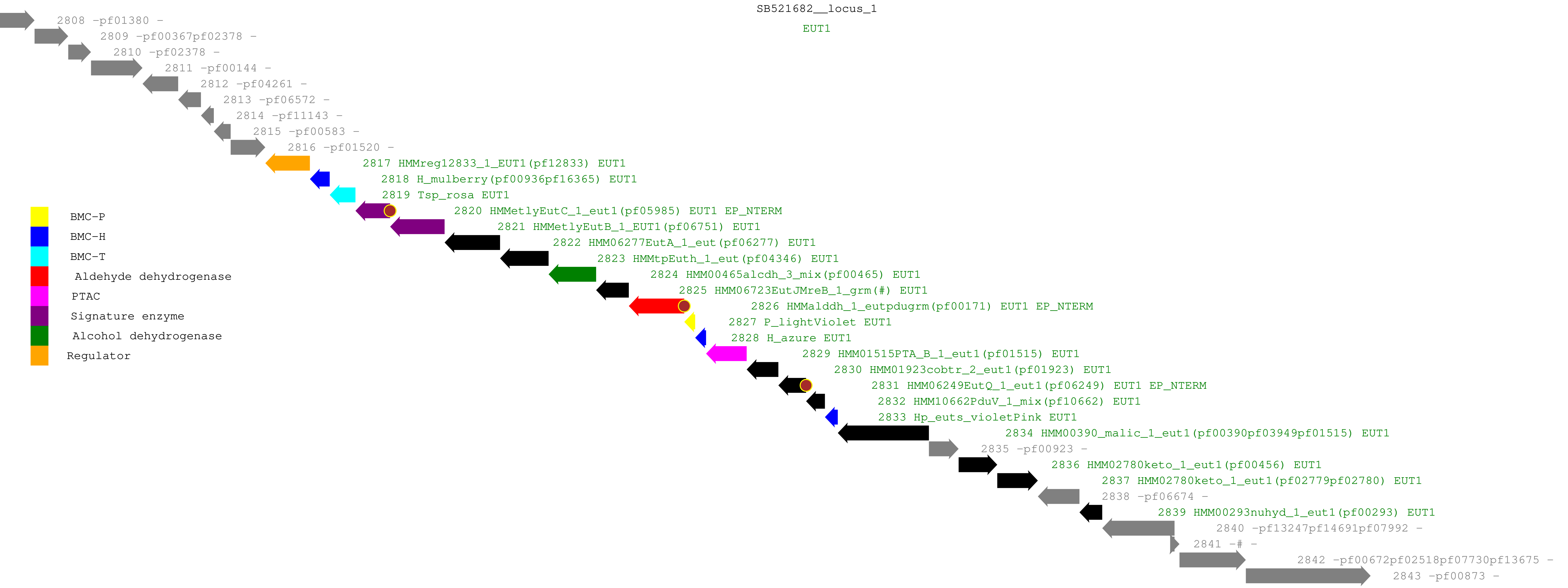

Klebsiella michiganensis. Bacteria; Proteobacteria; Gammaproteobacteria; Enterobacterales;

SB6408__locus_3

EUT1

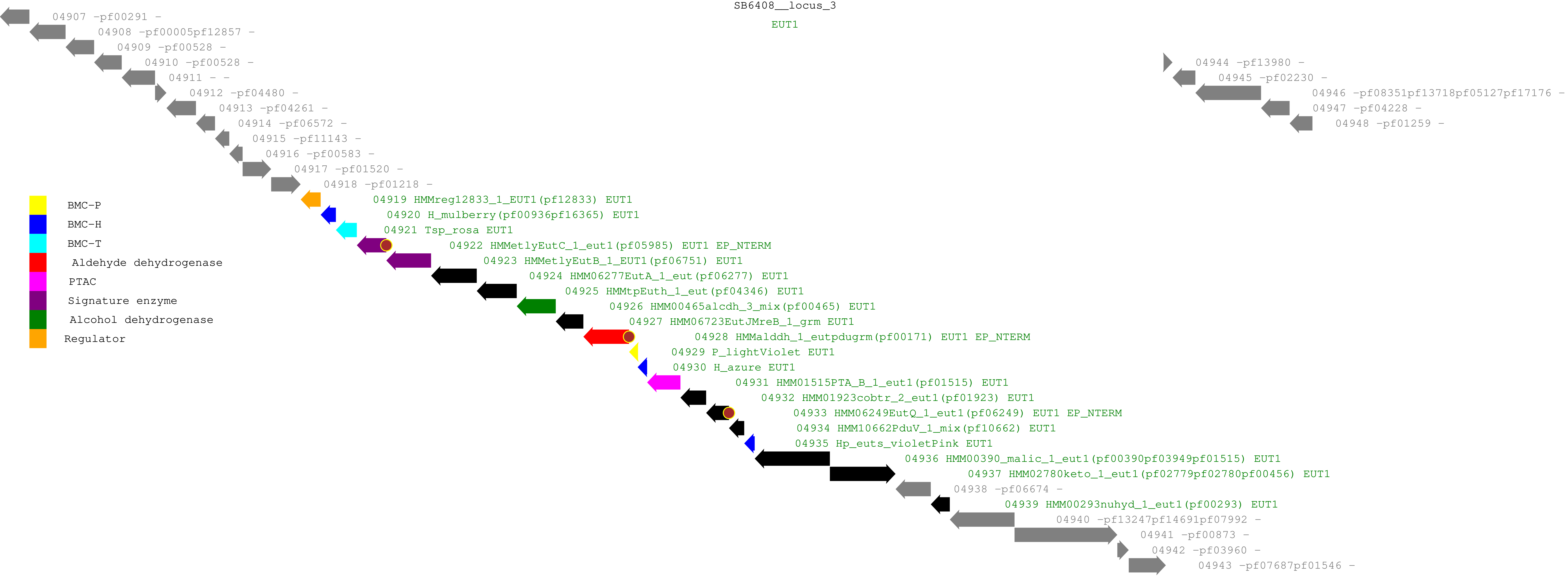

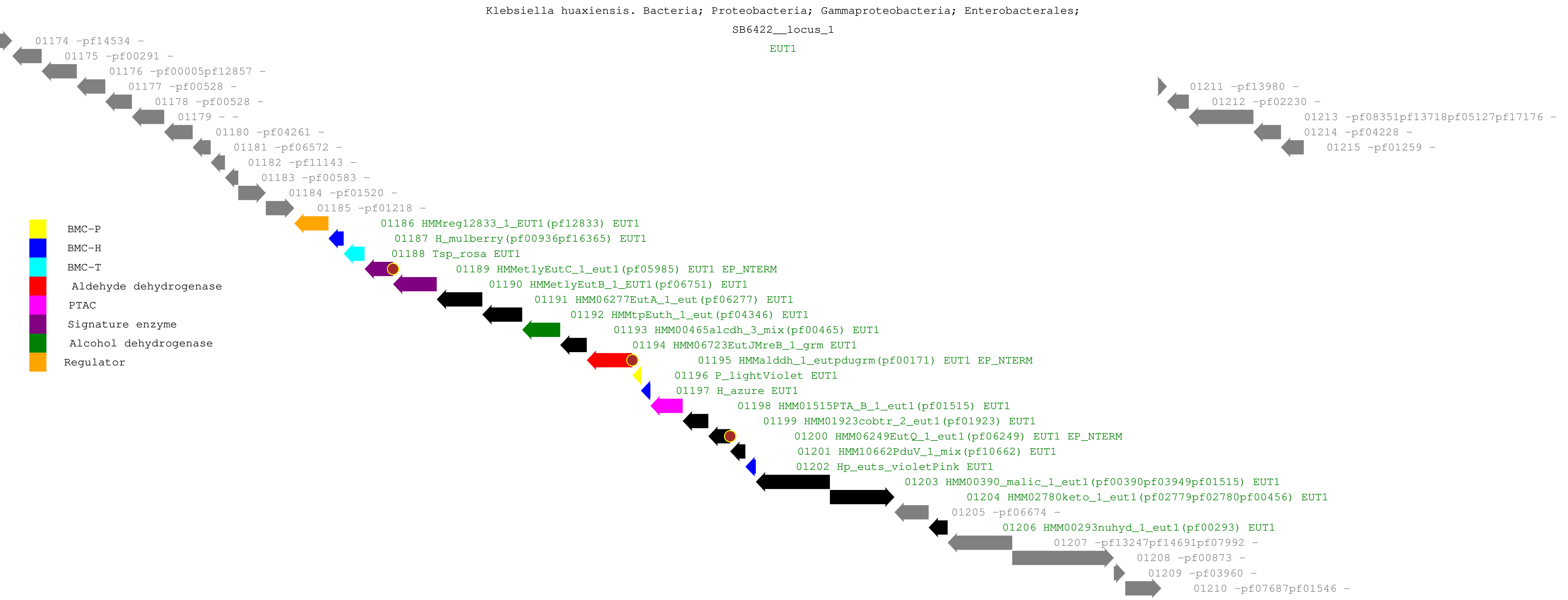
Klebsiella huaxiensis. Bacteria; Proteobacteria; Gammaproteobacteria; Enterobacterales;
SB6422__locus_1
EUT1
01174 -pf14534 -
01175 -pf00291 -
01176 -pf00005pf12857 -
01177 -pf00528 -
01178 -pf00528 -
01179 - -
01180 -pf04261 -
01181 -pf06572 -
01182 -pf11143 -
01183 -pf00583 -
01184 -pf01520 -
01185 -pf01218 -
01186 HMMreg12833_1_EUT1(pf12833) EUT1
01187 H_mulberry(pf00936pf16365) EUT1
01188 Tsp_rosa EUT1
01189 HMMetlyEutC_1_eut1(pf05985) EUT1 EP_NTERM
01190 HMMetlyEutB_1_EUT1(pf06751) EUT1
01191 HMM06277EutA_1_eut(pf06277) EUT1
01192 HMMtpEuth_1_eut(pf04346) EUT1
01193 HMM00465alcdh_3_mix(pf00465) EUT1
01194 HMM06723EutJMreB_1_grm EUT1
01195 HMMalddh_1_eutpdugrm(pf00171) EUT1 EP_NTERM
01196 P_lightViolet EUT1
01197 H_azure EUT1
01198 HMM01515PTA_B_1_eut1(pf01515) EUT1
01199 HMM01923cobtr_2_eut1(pf01923) EUT1
01200 HMM06249EutQ_1_eut1(pf06249) EUT1 EP_NTERM
01201 HMM10662PduV_1_mix(pf10662) EUT1
01202 Hp_euts_violetPink EUT1
01203 HMM00390_malic_1_eut1(pf00390pf03949pf01515) EUT1
01204 HMM02780keto_1_eut1(pf02779pf02780pf00456) EUT1
01205 -pf06674 -
01206 HMM00293nuhyd_1_eut1(pf00293) EUT1
01207 -pf13247pf14691pf07992 -
01208 -pf00873 -
01209 -pf03960 -
01210 -pf07687pf01546 -
01211 -pf13980 -
01212 -pf02230 -
01213 -pf08351pf13718pf05127pf17176 -
01214 -pf04228 -
01215 -pf01259 -
BMC-P
BMC-H
BMC-T
Aldehyde dehydrogenase
PTAC
Signature enzyme
Alcohol dehydrogenase
Regulator

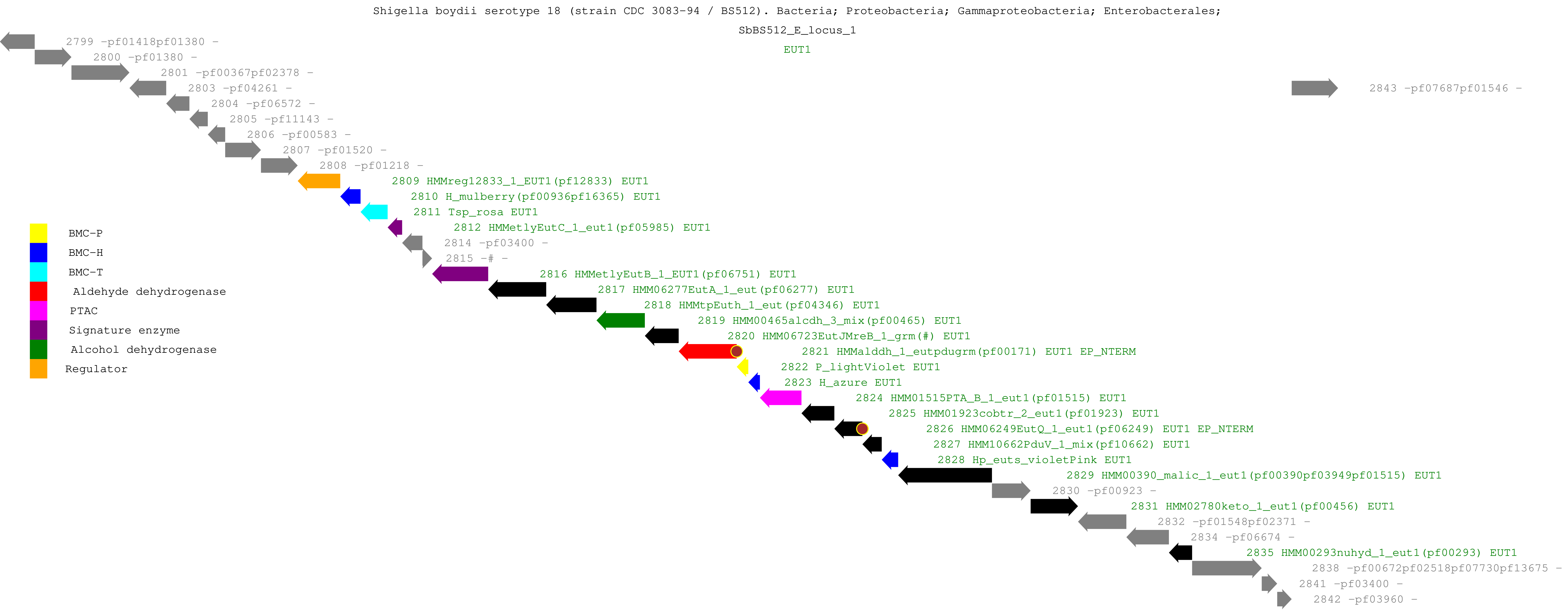
Shigella boydii serotype 18 (strain CDC 3083-94 / BS512). Bacteria; Proteobacteria; Gammaproteobacteria; Enterobacterales;
SbBS512_E_locus_1
EUT1
2799 -pf01418pf01380 -
2800 -pf01380 -
2801 -pf00367pf02378 -
2803 -pf04261 -
2804 -pf06572 -
2805 -pf11143 -
2806 -pf00583 -
2807 -pf01520 -
2808 -pf01218 -
2809 HMMreg12833_1_EUT1(pf12833) EUT1
2810 H_mulberry(pf00936pf16365) EUT1
2811 Tsp_rosa EUT1
2812 HMMetlyEutC_1_eut1(pf05985) EUT1
2814 -pf03400 -
2815 -# -
2816 HMMetlyEutB_1_EUT1(pf06751) EUT1
2817 HMM06277EutA_1_eut(pf06277) EUT1
2818 HMMtpEuth_1_eut(pf04346) EUT1
2819 HMM00465alcdh_3_mix(pf00465) EUT1
2820 HMM06723EutJMreB_1_grm(#) EUT1
2821 HMMalddh_1_eutpdugrm(pf00171) EUT1 EP_NTERM
2822 P_lightViolet EUT1
2823 H_azure EUT1
2824 HMM01515PTA_B_1_eut1(pf01515) EUT1
2825 HMM01923cobtr_2_eut1(pf01923) EUT1
2826 HMM06249EutQ_1_eut1(pf06249) EUT1 EP_NTERM
2827 HMM10662PduV_1_mix(pf10662) EUT1
2828 Hp_euts_violetPink EUT1
2829 HMM00390_malic_1_eut1(pf00390pf03949pf01515) EUT1
2830 -pf00923 -
2831 HMM02780keto_1_eut1(pf00456) EUT1
2832 -pf01548pf02371 -
2834 -pf06674 -
2835 HMM00293nuhyd_1_eut1(pf00293) EUT1
2838 -pf00672pf02518pf07730pf13675 -
2841 -pf03400 -
2842 -pf03960 -
2843 -pf07687pf01546 -
BMC-P
BMC-H
BMC-T
Aldehyde dehydrogenase
PTAC
Signature enzyme
Alcohol dehydrogenase
Regulator

Shigella boydii serotype 4 (strain Sb227). Bacteria; Proteobacteria; Gammaproteobacteria; Enterobacterales;

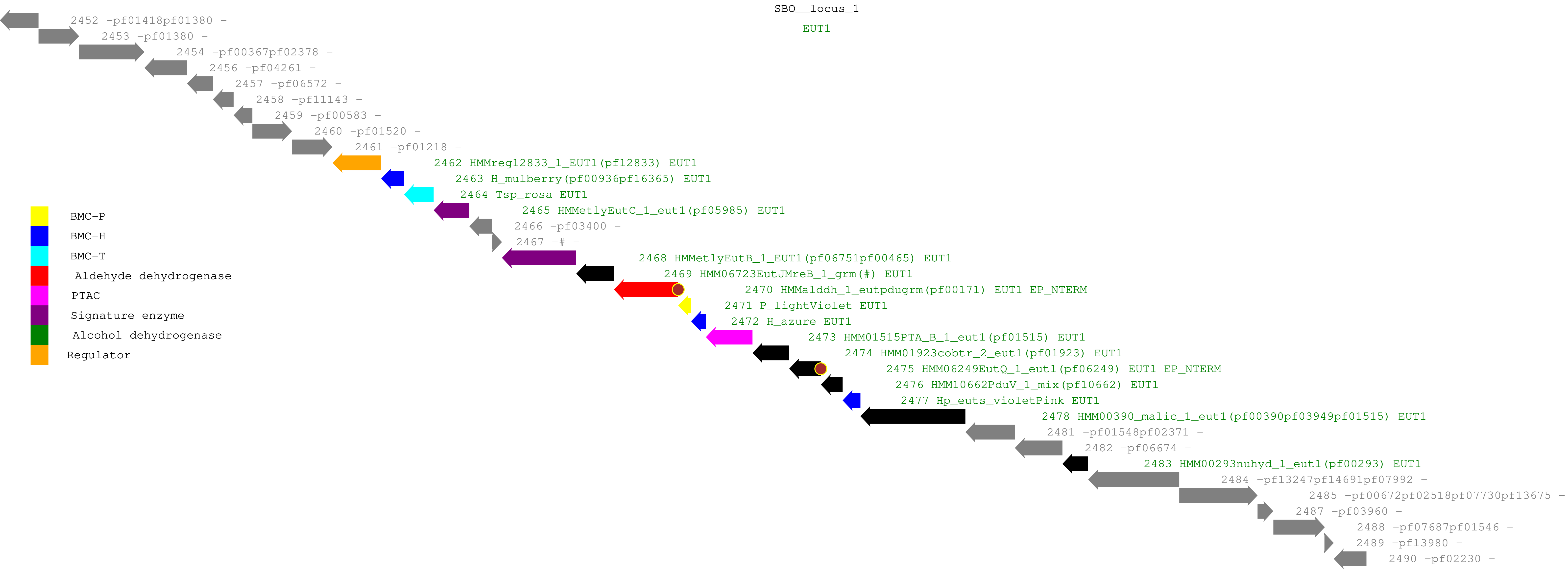

Salmonella choleraesuis (strain SC-B67). Bacteria; Proteobacteria; Gammaproteobacteria; Enterobacterales;

SCH__locus_2

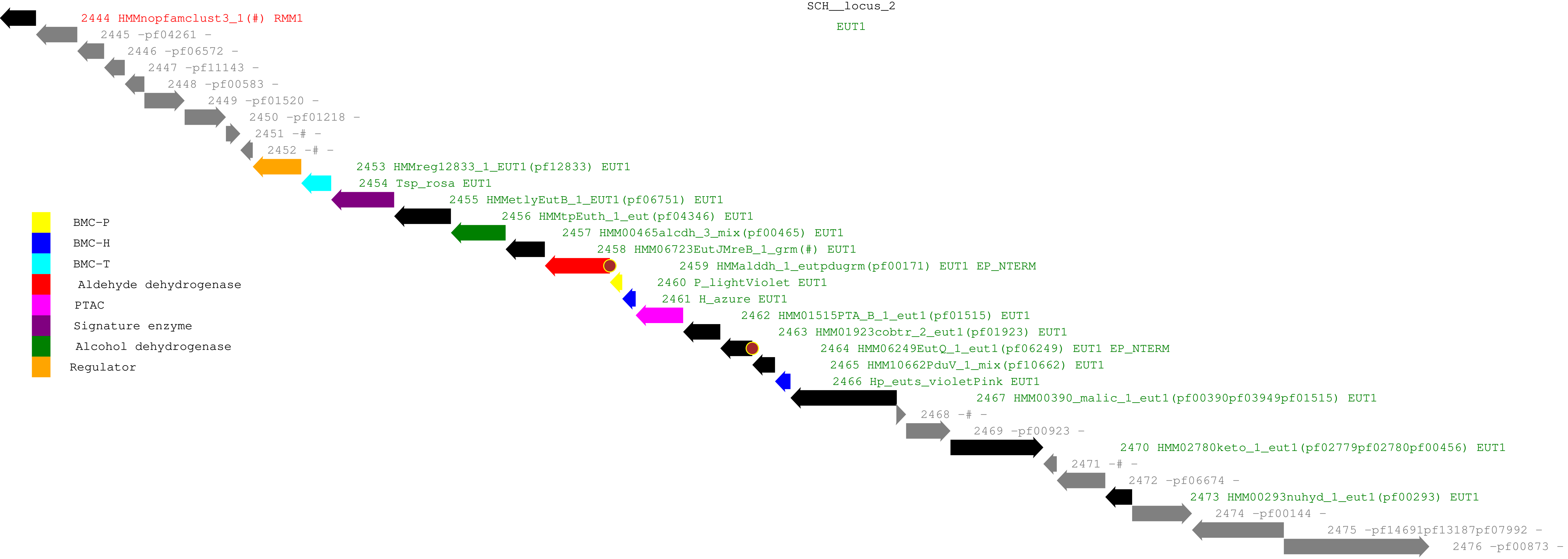

Shigella dysenteriae 1012. Bacteria; Proteobacteria; Gammaproteobacteria; Enterobacterales;

Sd1012__locus_1

EUT1

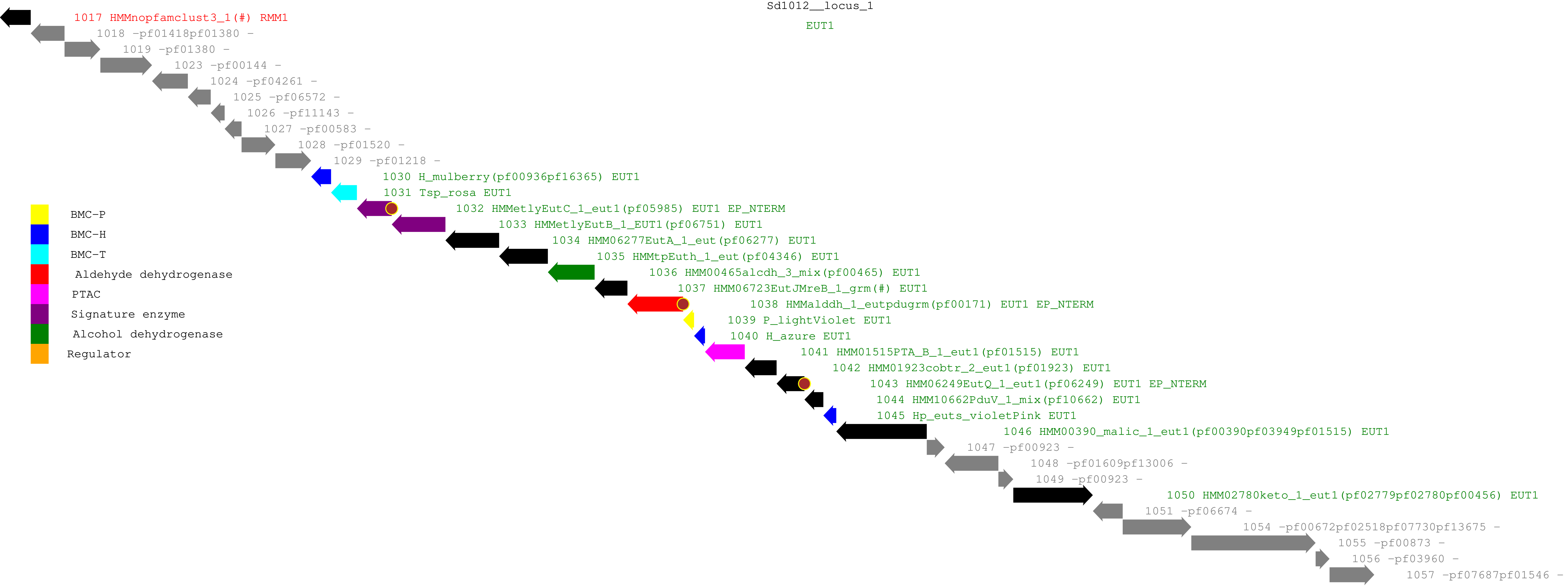

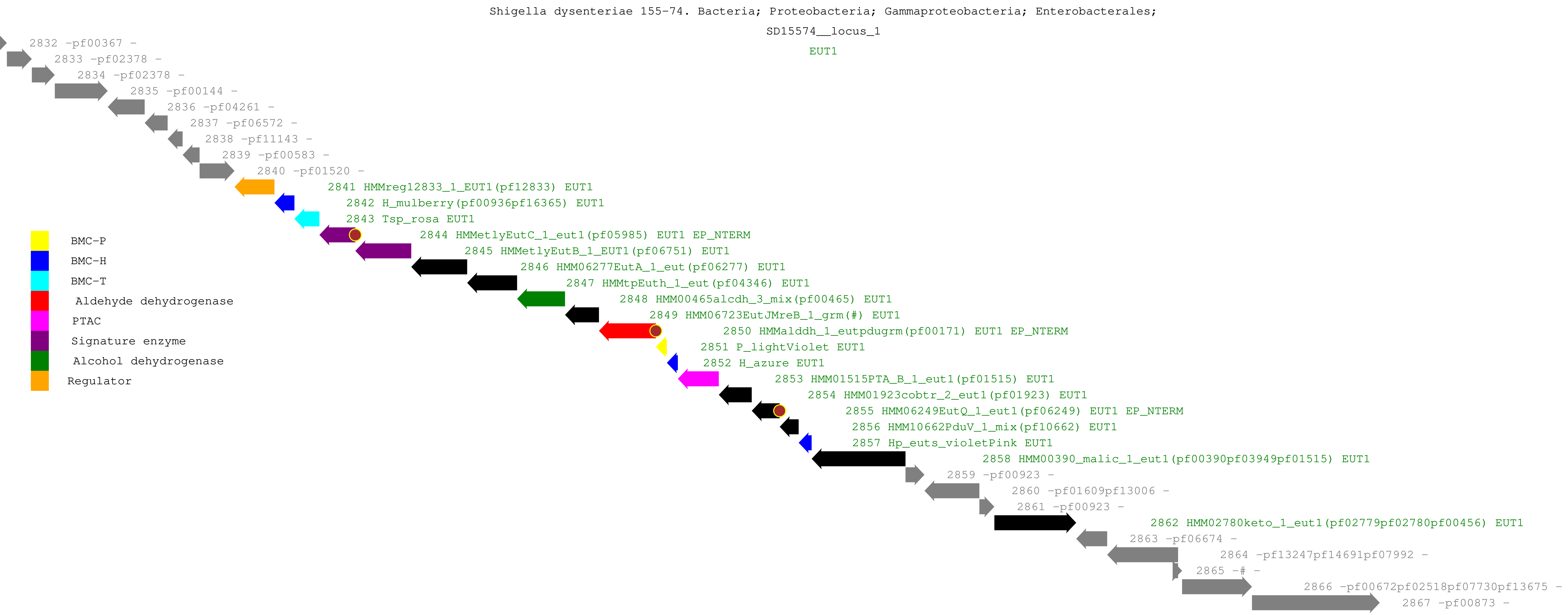

Shigella dysenteriae 155-74. Bacteria; Proteobacteria; Gammaproteobacteria; Enterobacterales;
SD15574__locus_1
EUT1
2832 -pf00367 -
2833 -pf02378 -
2834 -pf02378 -
2835 -pf00144 -
2836 -pf04261 -
2837 -pf06572 -
2838 -pf11143 -
2839 -pf00583 -
2840 -pf01520 -
2841 HMMreg12833_1_EUT1(pf12833) EUT1
2842 H_mulberry(pf00936pf16365) EUT1
2843 Tsp_rosa EUT1
2844 HMMetlyEutC_1_eut1(pf05985) EUT1 EP_NTERM
2845 HMMetlyEutB_1_EUT1(pf06751) EUT1
2846 HMM06277EutA_1_eut(pf06277) EUT1
2847 HMMtpEuth_1_eut(pf04346) EUT1
2848 HMM00465alcdh_3_mix(pf00465) EUT1
2849 HMM06723EutJMreB_1_grm(#) EUT1
2850 HMMaldh_1_eutpdugrm(pf00171) EUT1 EP_NTERM
2851 P_lightViolet EUT1
2852 H_azure EUT1
2853 HMM01515PTA_B_1_eut1(pf01515) EUT1
2854 HMM01923cobtr_2_eut1(pf01923) EUT1
2855 HMM06249EutQ_1_eut1(pf06249) EUT1 EP_NTERM
2856 HMM10662PduV_1_mix(pf10662) EUT1
2857 Hp_euts_violetPink EUT1
2858 HMM00390_malic_1_eut1(pf00390pf03949pf01515) EUT1
2859 -pf00923 -
2860 -pf01609pf13006 -
2861 -pf00923 -
2862 HMM02780keto_1_eut1(pf02779pf02780pf00456) EUT1
2863 -pf06674 -
2864 -pf13247pf14691pf07992 -
2865 -# -
2866 -pf00672pf02518pf07730pf13675 -
2867 -pf00873 -
BMC-P
BMC-H
BMC-T
Aldehyde dehydrogenase
PTAC
Signature enzyme
Alcohol dehydrogenase
Regulator

Shigella dysenteriae 225-75. Bacteria; Proteobacteria; Gammaproteobacteria; Enterobacterales;

SD22575__locus_1

EUT1

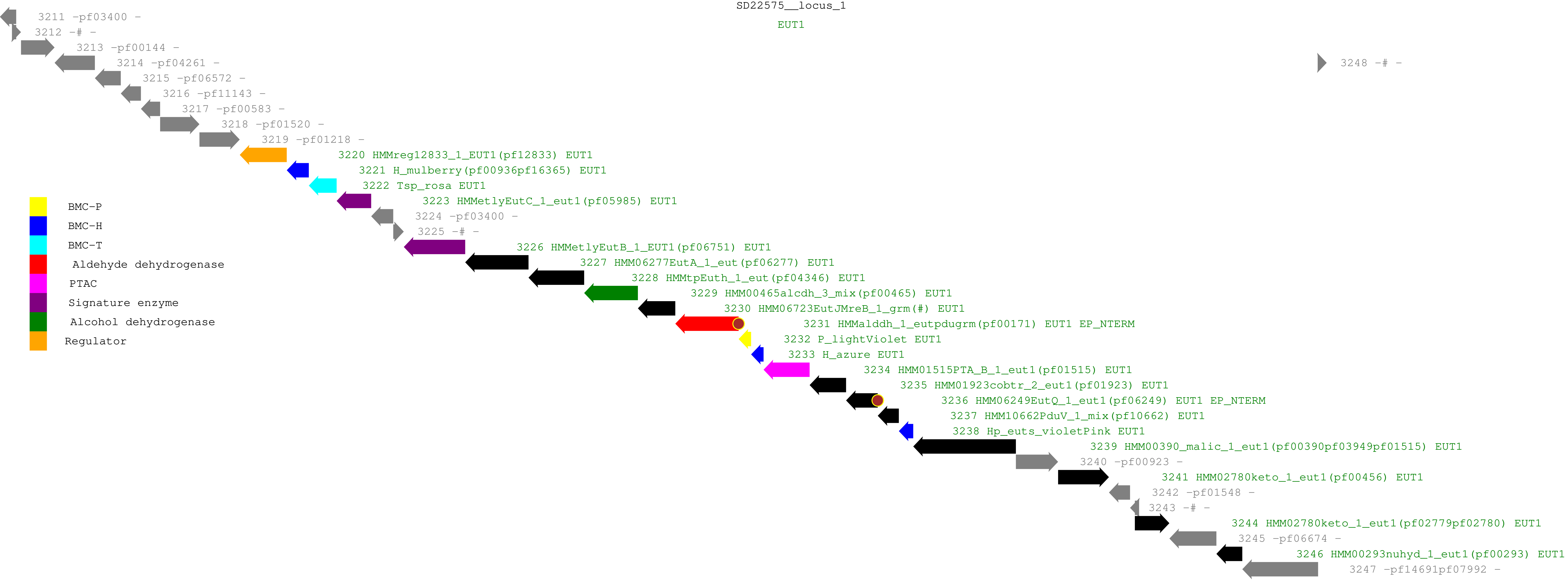

Salmonella agona (strain SL483). Bacteria; Proteobacteria; Gammaproteobacteria; Enterobacterales;

SeAg_B_locus_2

EUT1

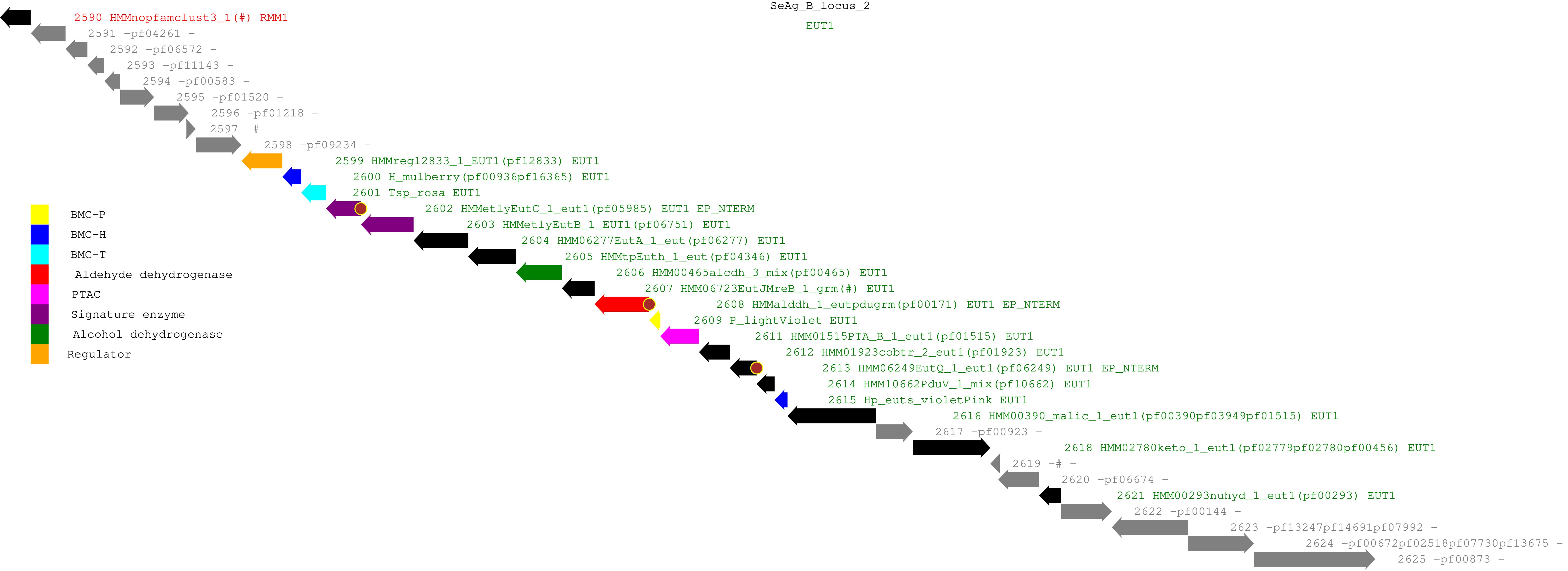

Salmonella enterica subsp. enterica serovar Chester str. ATCC 11997. Bacteria; Proteobacteria; Gammaproteobacteria; Enterobacterales;

SEECH997__locus_1

EUT1

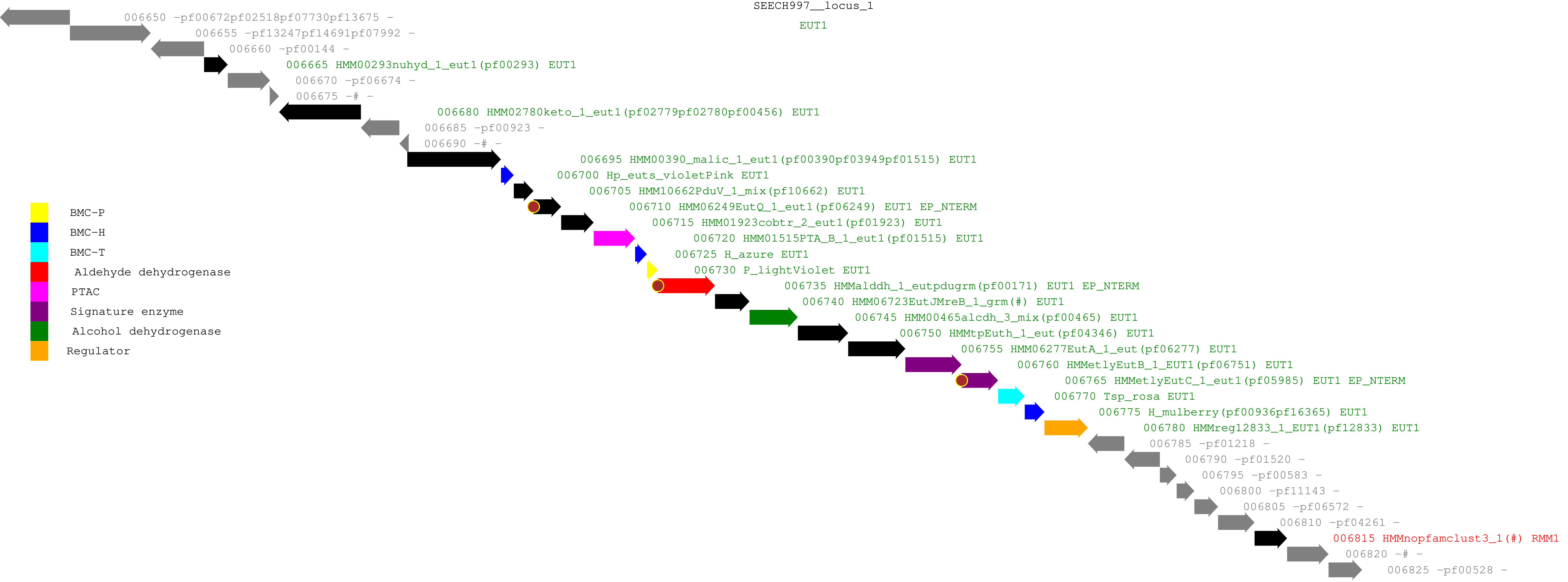

Salmonella enterica subsp. enterica serovar Muenster str. 0315. Bacteria; Proteobacteria; Gammaproteobacteria; Enterobacterales;

SEEM0315__locus_1

EUT1

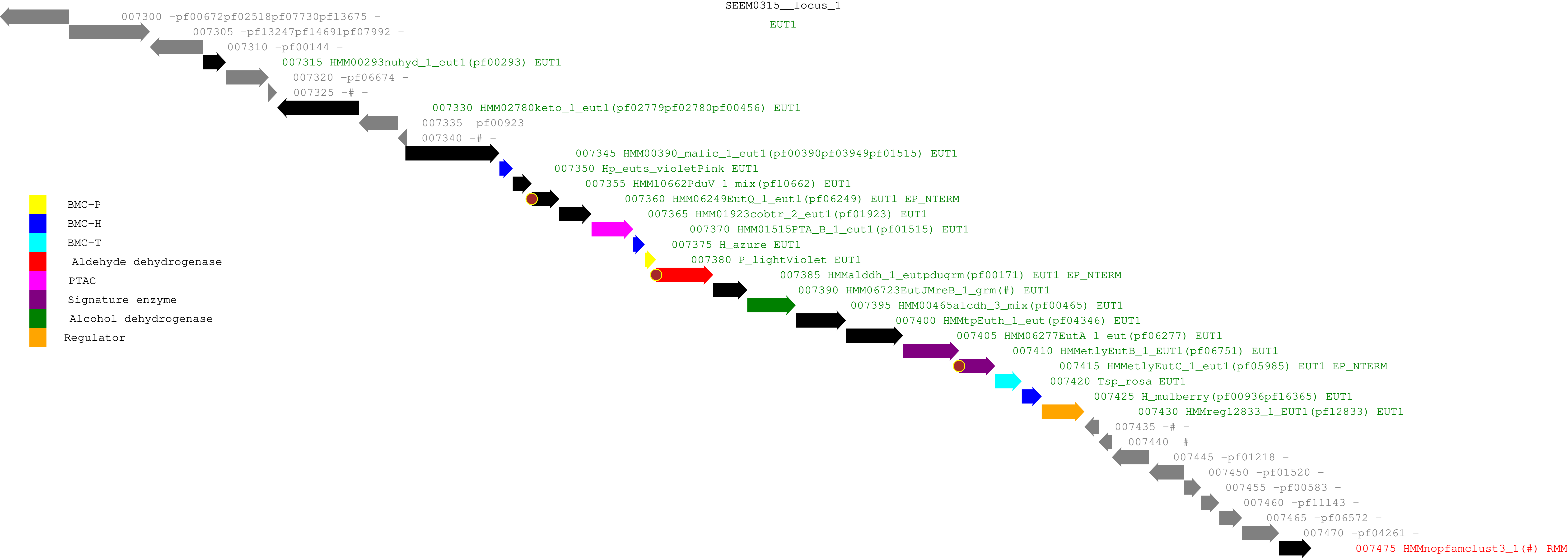

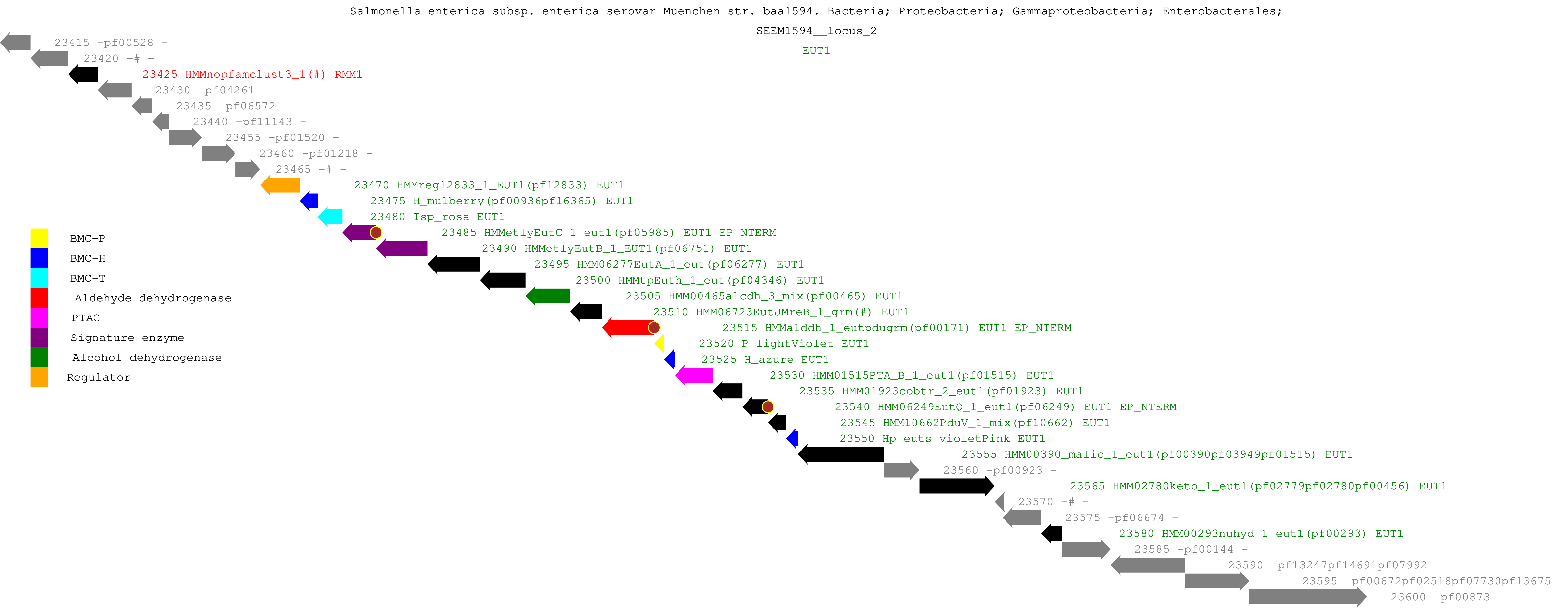
Salmonella enterica subsp. enterica serovar Muenchen str. baa1594. Bacteria; Proteobacteria; Gammaproteobacteria; Enterobacterales;
SEEM1594__locus_2
EUT1
23415 -pf00528 -
23420 -# -
23425 HMMnopfamclust3_1(#) RMM1
23430 -pf04261 -
23435 -pf06572 -
23440 -pf11143 -
23455 -pf01520 -
23460 -pf01218 -
23465 -# -
23470 HMMreg12833_1_EUT1(pf12833) EUT1
23475 H_mulberry(pf00936pf16365) EUT1
23480 Tsp_rosa EUT1
23485 HMMetlyEutC_1_eut1(pf05985) EUT1 EP_NTERM
23490 HMMetlyEutB_1_EUT1(pf06751) EUT1
23495 HMM06277EutA_1_eut(pf06277) EUT1
23500 HMMtpEuth_1_eut(pf04346) EUT1
23505 HMM00465alcdh_3_mix(pf00465) EUT1
23510 HMM06723EutJMreB_1_grm(#) EUT1
23515 HMMalddh_1_eutpdugrm(pf00171) EUT1 EP_NTERM
23520 P_lightViolet EUT1
23525 H_azure EUT1
23530 HMM01515PTA_B_1_eut1(pf01515) EUT1
23535 HMM01923cobtr_2_eut1(pf01923) EUT1
23540 HMM06249EutQ_1_eut1(pf06249) EUT1 EP_NTERM
23545 HMM10662PduV_1_mix(pf10662) EUT1
23550 Hp_euts_violetPink EUT1
23555 HMM00390_malic_1_eut1(pf00390pf03949pf01515) EUT1
23560 -pf00923 -
23565 HMM02780keto_1_eut1(pf02779pf02780pf00456) EUT1
23570 -# -
23575 -pf06674 -
23580 HMM00293nuhyd_1_eut1(pf00293) EUT1
23585 -pf00144 -
23590 -pf13247pf14691pf07992 -
23595 -pf00672pf02518pf07730pf13675 -
23600 -pf00873 -
BMC-P
BMC-H
BMC-T
Aldehyde dehydrogenase
PTAC
Signature enzyme
Alcohol dehydrogenase
Regulator

Salmonella enterica subsp. enterica serovar Mbandaka str. ATCC 51958. Bacteria; Proteobacteria; Gammaproteobacteria; Enterobacterales;

SEEM1958__locus_1

EUT1

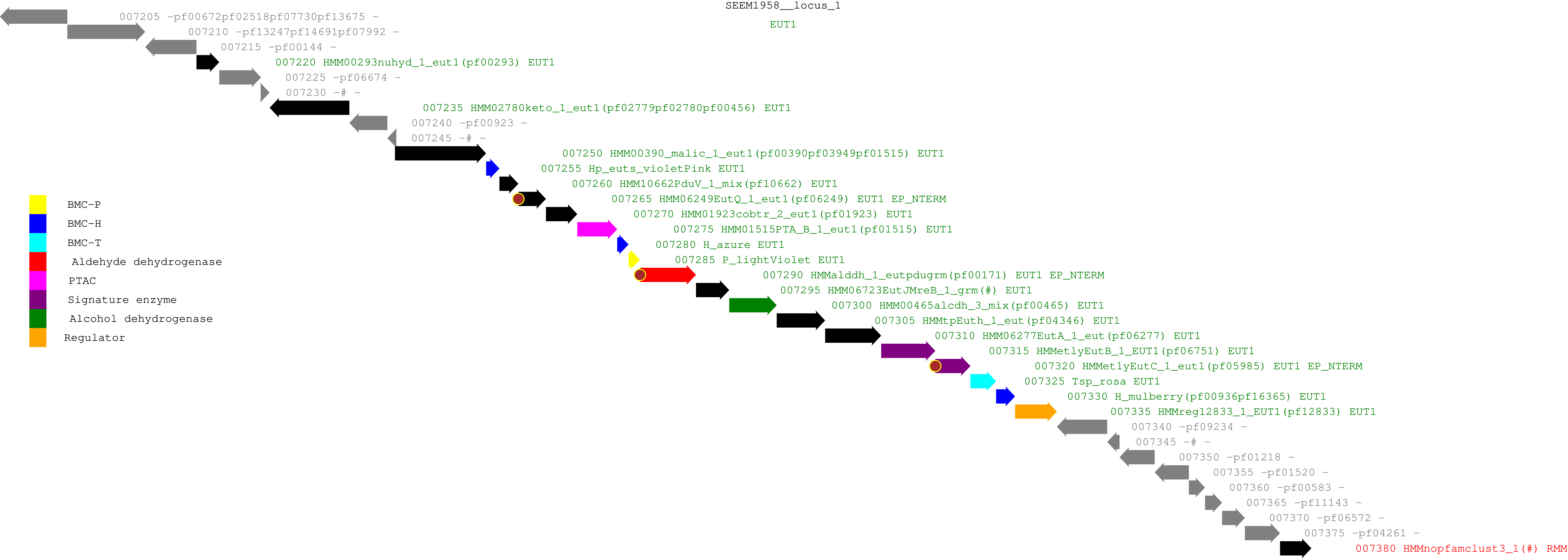

Salmonella enterica subsp. enterica serovar Montevideo str. 42N. Bacteria; Proteobacteria; Gammaproteobacteria; Enterobacterales;

SEEM42N__locus_1

EUT1

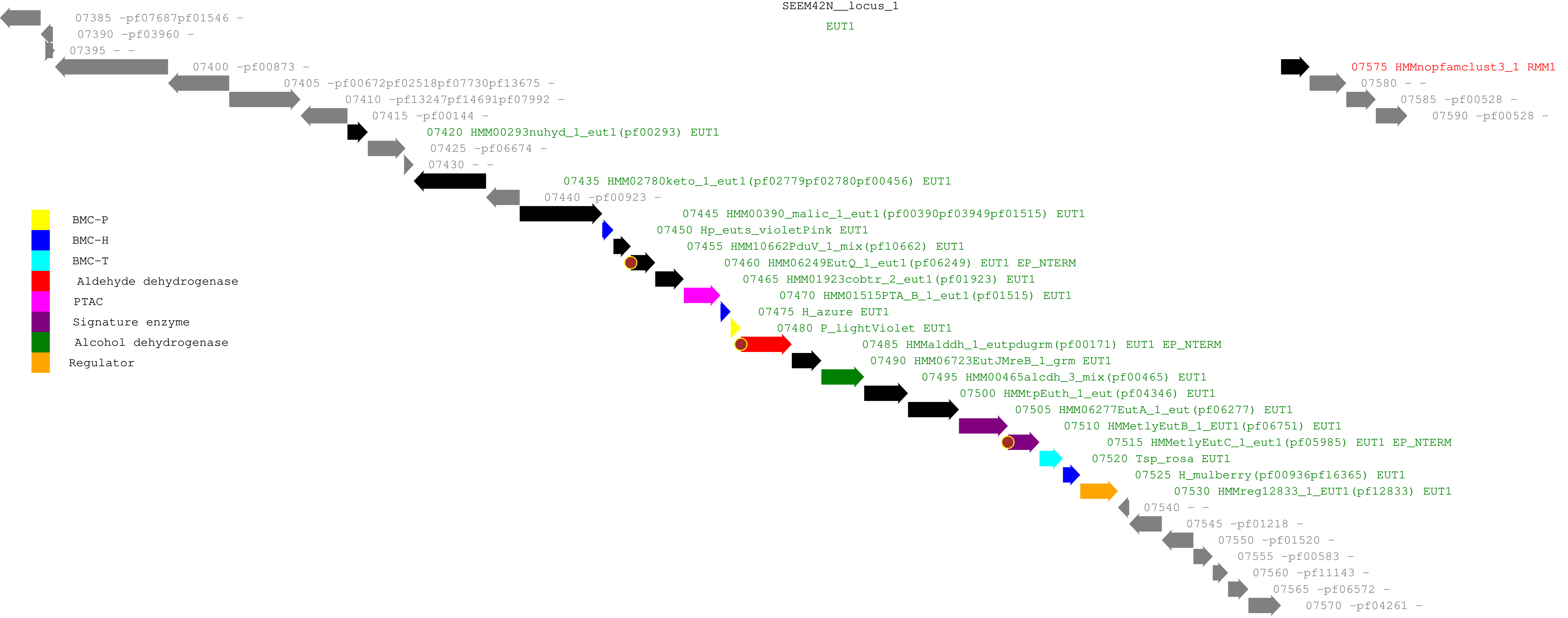

Salmonella enterica subsp. enterica serovar Minnesota str. ATCC 49284. Bacteria; Proteobacteria; Gammaproteobacteria; Enterobacterales;

SEEM9284__locus_1

EUT1

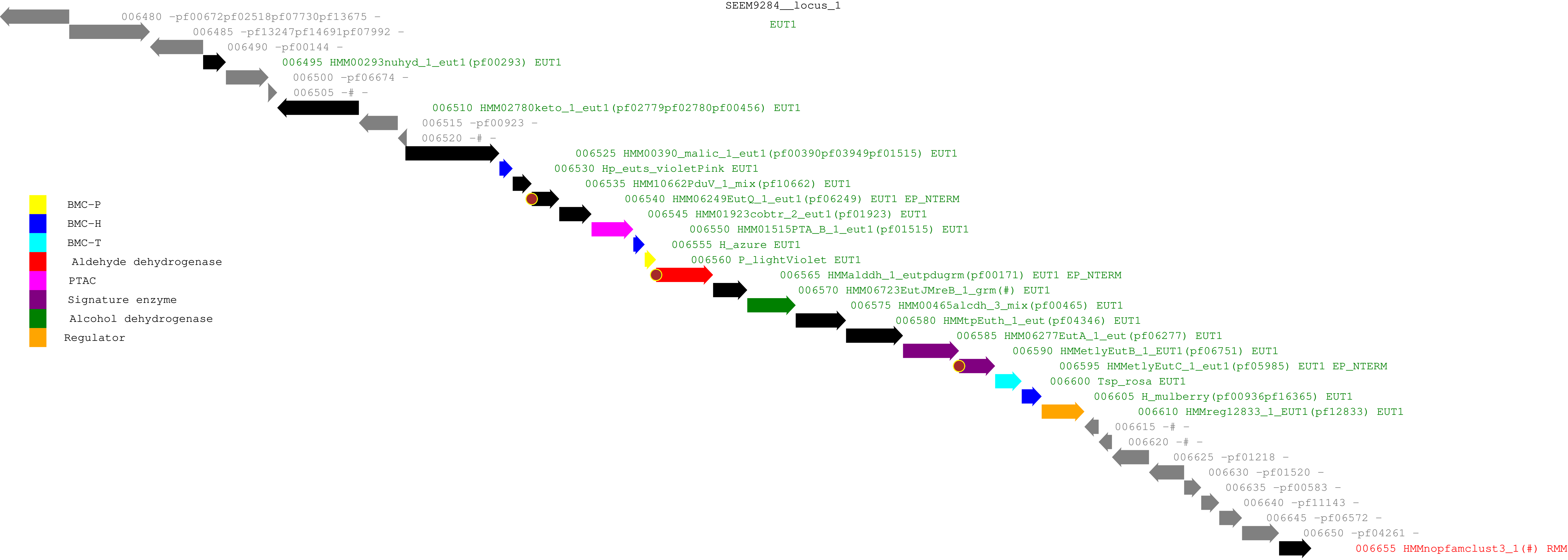

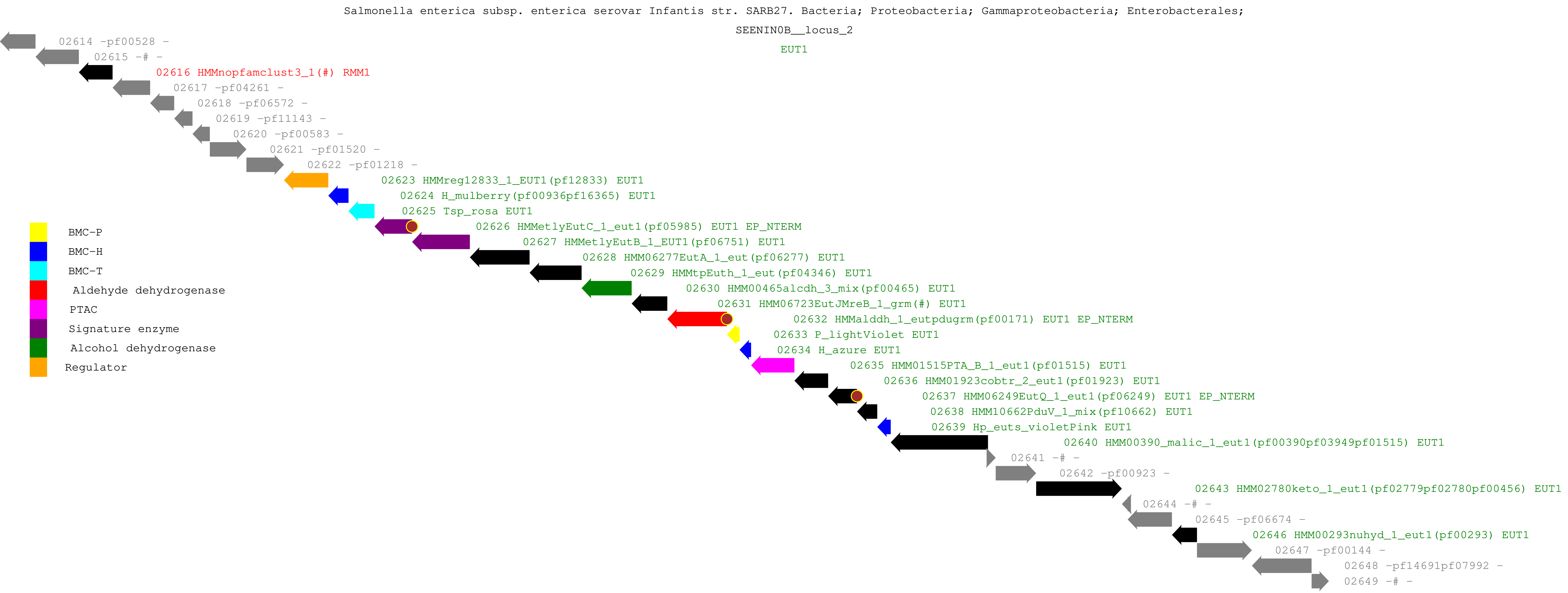
Salmonella enterica subsp. enterica serovar Infantis str. SARB27. Bacteria; Proteobacteria; Gammaproteobacteria; Enterobacterales;
SEENIN0B__locus_2
EUT1
02614 -pf00528 -
02615 -# -
02616 HMMnopfamclust3_1(#) RMM1
02617 -pf04261 -
02618 -pf06572 -
02619 -pf11143 -
02620 -pf00583 -
02621 -pf01520 -
02622 -pf01218 -
02623 HMMreg12833_1_EUT1(pf12833) EUT1
02624 H_mulberry(pf00936pf16365) EUT1
02625 Tsp_rosa EUT1
02626 HMMetlyEutC_1_eut1(pf05985) EUT1 EP_NTERM
02627 HMMetlyEutB_1_EUT1(pf06751) EUT1
02628 HMM06277EutA_1_eut(pf06277) EUT1
02629 HMMtpEuth_1_eut(pf04346) EUT1
02630 HMM00465alcdh_3_mix(pf00465) EUT1
02631 HMM06723EutJMreB_1_grm(#) EUT1
02632 HMMaldh_1_eutpdugrm(pf00171) EUT1 EP_NTERM
02633 P_lightViolet EUT1
02634 H_azure EUT1
02635 HMM01515PTA_B_1_eut1(pf01515) EUT1
02636 HMM01923cobtr_2_eut1(pf01923) EUT1
02637 HMM06249EutQ_1_eut1(pf06249) EUT1 EP_NTERM
02638 HMM10662PduV_1_mix(pf10662) EUT1
02639 Hp_euts_violetPink EUT1
02640 HMM00390_malic_1_eut1(pf00390pf03949pf01515) EUT1
02641 -# -
02642 -pf00923 -
02643 HMM02780keto_1_eut1(pf02779pf02780pf00456) EUT1
02644 -# -
02645 -pf06674 -
02646 HMM00293nuhyd_1_eut1(pf00293) EUT1
02647 -pf00144 -
02648 -pf14691pf07992 -
02649 -# -
BMC-P
BMC-H
BMC-T
Aldehyde dehydrogenase
PTAC
Signature enzyme
Alcohol dehydrogenase
Regulator

Salmonella enterica subsp. enterica serovar Rubislaw str. ATCC 10717. Bacteria; Proteobacteria; Gammaproteobacteria; Enterobacterales;

SEERU717__locus_1

EUT1

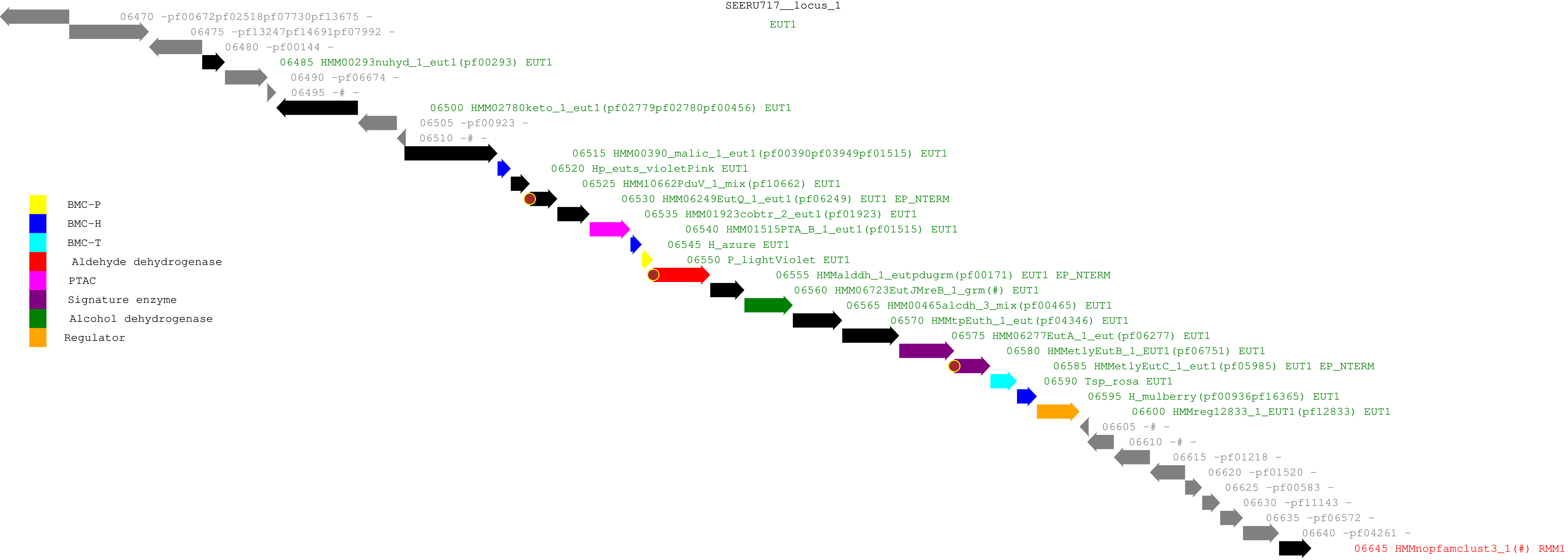

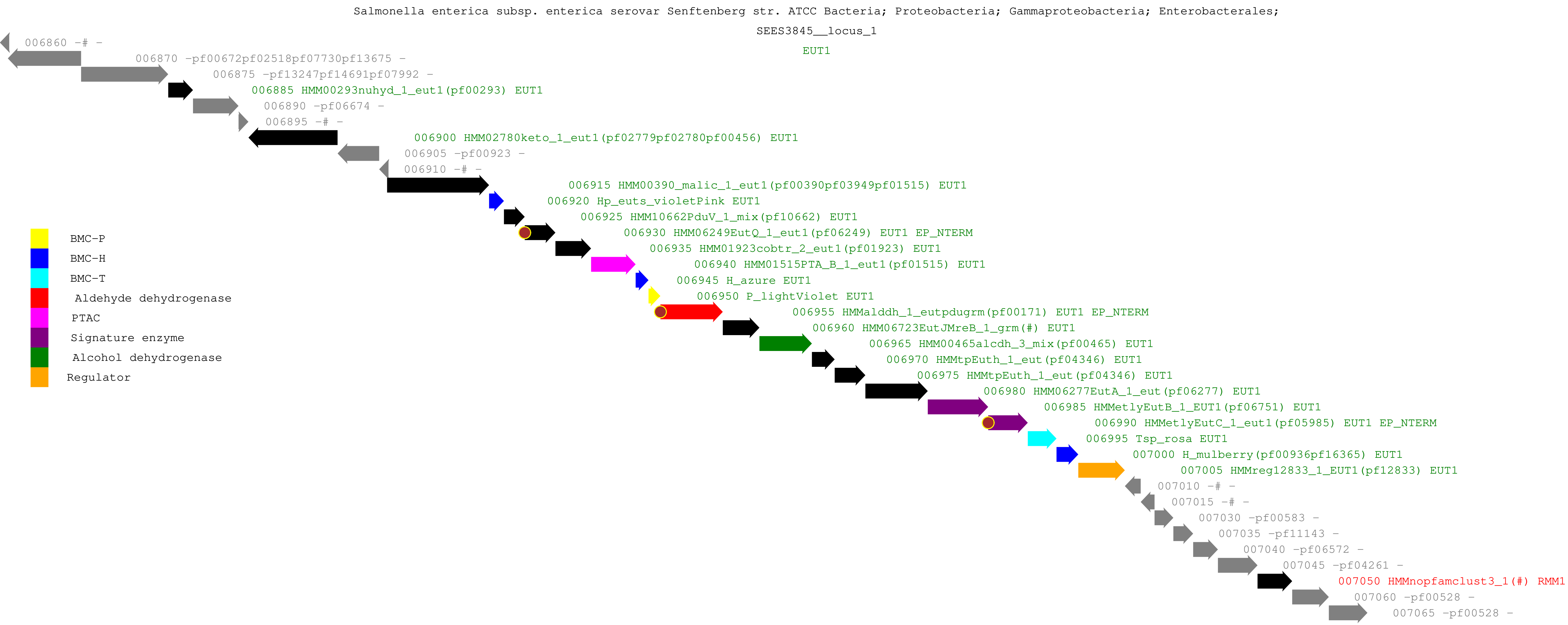
Salmonella enterica subsp. enterica serovar Senftenberg str. ATCC Bacteria; Proteobacteria; Gammaproteobacteria; Enterobacterales;
SEES3845__locus_1
EUT1
006860 -# -
006870 -pf00672pf02518pf07730pf13675 -
006875 -pf13247pf14691pf07992 -
006885 HMM00293nuhyd_1_eut1(pf00293) EUT1
006890 -pf06674 -
006895 -# -
006900 HMM02780keto_1_eut1(pf02779pf02780pf00456) EUT1
006905 -pf00923 -
006910 -# -
006915 HMM00390_malic_1_eut1(pf00390pf03949pf01515) EUT1
006920 Hp_euts_violetPink EUT1
006925 HMM10662PduV_1_mix(pf10662) EUT1
006930 HMM06249EutQ_1_eut1(pf06249) EUT1 EP_NTERM
006935 HMM01923cobtr_2_eut1(pf01923) EUT1
006940 HMM01515PTA_B_1_eut1(pf01515) EUT1
006945 H_azure EUT1
006950 P_lightViolet EUT1
006955 HMMalddh_1_eutpdugrm(pf00171) EUT1 EP_NTERM
006960 HMM06723EutJMreB_1_grm(#) EUT1
006965 HMM00465alcdh_3_mix(pf00465) EUT1
006970 HMMtpEuth_1_eut(pf04346) EUT1
006975 HMMtpEuth_1_eut(pf04346) EUT1
006980 HMM06277EutA_1_eut(pf06277) EUT1
006985 HMMetlyEutB_1_EUT1(pf06751) EUT1
006990 HMMetlyEutC_1_eut1(pf05985) EUT1 EP_NTERM
006995 Tsp_rosa EUT1
007000 H_mulberry(pf00936pf16365) EUT1
007005 HMMreg12833_1_EUT1(pf12833) EUT1
007010 -# -
007015 -# -
007030 -pf00583 -
007035 -pf11143 -
007040 -pf06572 -
007045 -pf04261 -
007050 HMMnopfamclust3_1(#) RMM1
007060 -pf00528 -
007065 -pf00528 -
BMC-P
BMC-H
BMC-T
Aldehyde dehydrogenase
PTAC
Signature enzyme
Alcohol dehydrogenase
Regulator

Salmonella enterica subsp. enterica serovar Tennessee str. Bacteria; Proteobacteria; Gammaproteobacteria; Enterobacterales;

SEET0819__locus_2

EUT1

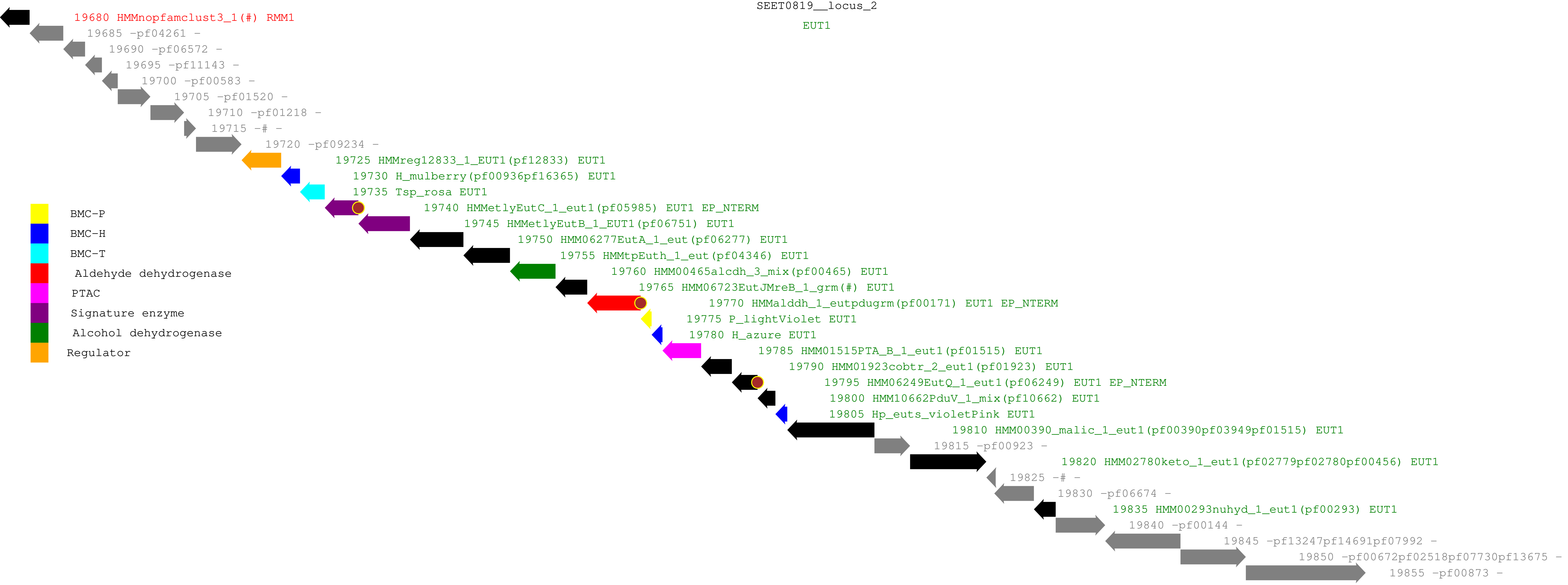

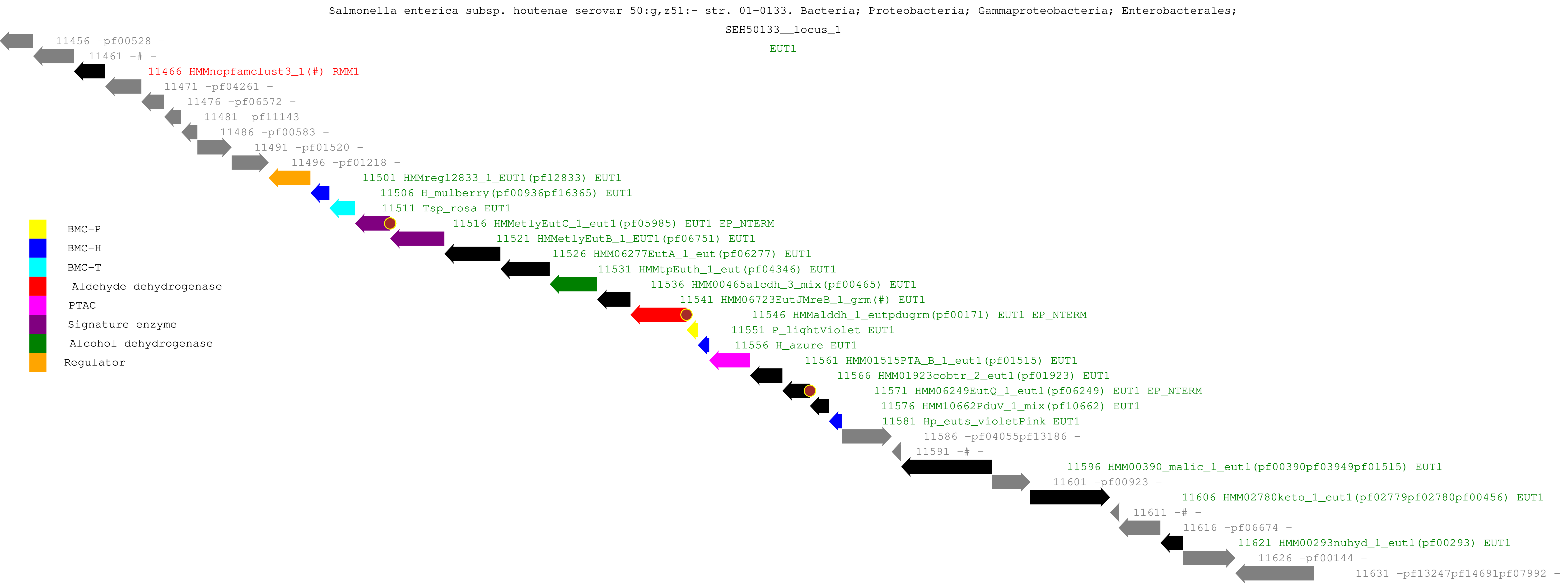

Salmonella enterica subsp. houtenae serovar 50:g,z51:- str. 01-0133. Bacteria; Proteobacteria; Gammaproteobacteria; Enterobacterales;
SEH50133__locus_1
EUT1
11456 -pf00528 -
11461 -# -
11466 HMMnopfamclust3_1(#) RMM1
11471 -pf04261 -
11476 -pf06572 -
11481 -pf11143 -
11486 -pf00583 -
11491 -pf01520 -
11496 -pf01218 -
11501 HMMreg12833_1_EUT1(pf12833) EUT1
11506 H_mulberry(pf00936pf16365) EUT1
11511 Tsp_rosa EUT1
11516 HMMetlyEutC_1_eut1(pf05985) EUT1 EP_NTERM
11521 HMMetlyEutB_1_EUT1(pf06751) EUT1
11526 HMM06277EutA_1_eut(pf06277) EUT1
11531 HMMtpEuth_1_eut(pf04346) EUT1
11536 HMM00465alcdh_3_mix(pf00465) EUT1
11541 HMM06723EutJMreB_1_grm(#) EUT1
11546 HMMaldдh_1_eutpdugrm(pf00171) EUT1 EP_NTERM
11551 P_lightViolet EUT1
11556 H_azure EUT1
11561 HMM01515PTA_B_1_eut1(pf01515) EUT1
11566 HMM01923cobtr_2_eut1(pf01923) EUT1
11571 HMM06249EutQ_1_eut1(pf06249) EUT1 EP_NTERM
11576 HMM10662PduV_1_mix(pf10662) EUT1
11581 Hp_euts_violetPink EUT1
11586 -pf04055pf13186 -
11591 -# -
11596 HMM00390_malic_1_eut1(pf00390pf03949pf01515) EUT1
11601 -pf00923 -
11606 HMM02780keto_1_eut1(pf02779pf02780pf00456) EUT1
11611 -# -
11616 -pf06674 -
11621 HMM00293nuhyd_1_eut1(pf00293) EUT1
11626 -pf00144 -
11631 -pf13247pf14691pf07992 -
BMC-P
BMC-H
BMC-T
Aldehyde dehydrogenase
PTAC
Signature enzyme
Alcohol dehydrogenase
Regulator

Salmonella enterica subsp. houtenae str. ATCC BAA-1581. Bacteria; Proteobacteria; Gammaproteobacteria; Enterobacterales;

SEHO0A__locus_1

EUT1

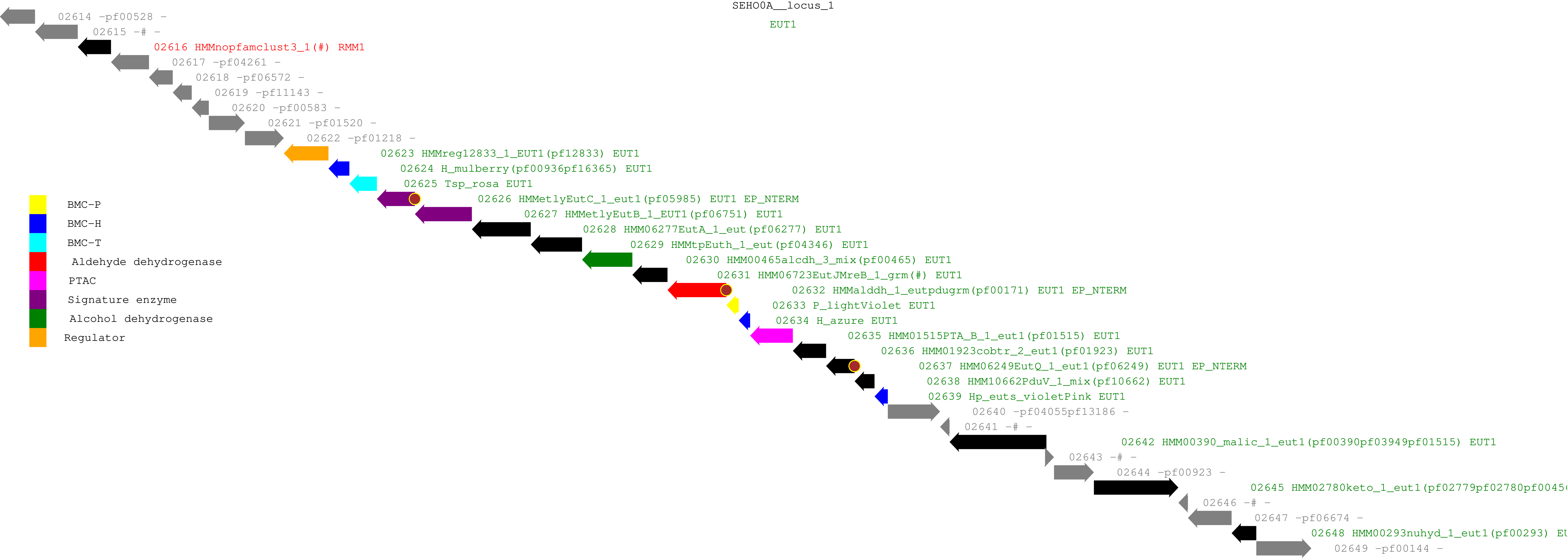

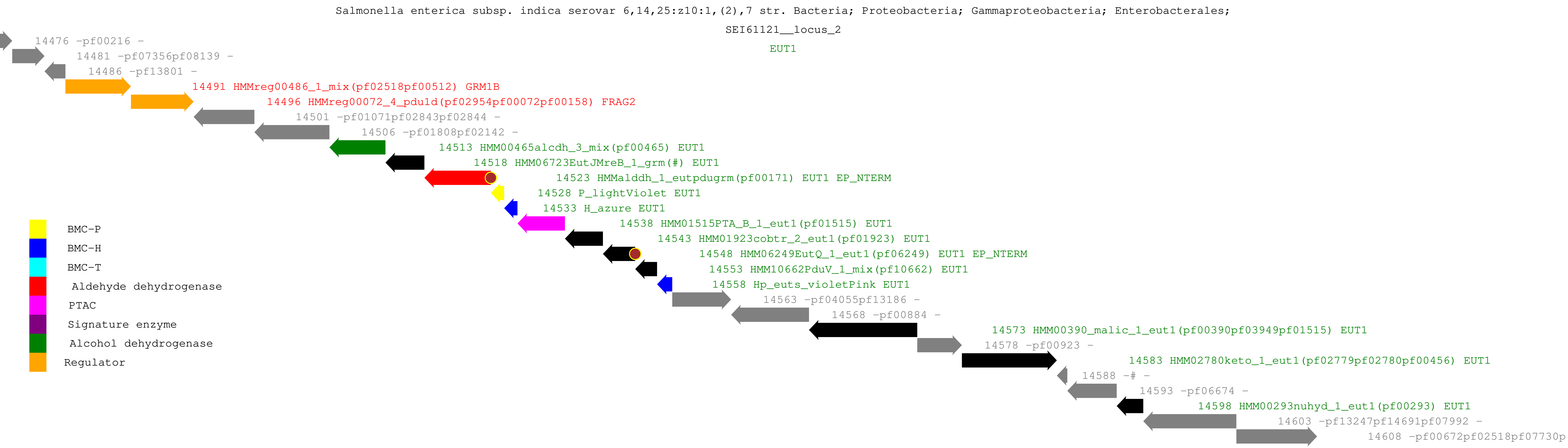
Salmonella enterica subsp. indica serovar 6,14,25:z10:1,(2),7 str. Bacteria; Proteobacteria; Gammaproteobacteria; Enterobacterales;
SEI61121__locus_2
EUT1
14476 -pf00216 -
14481 -pf07356pf08139 -
14486 -pf13801 -
14491 HMMreg00486_1_mix(pf02518pf00512) GRM1B
14496 HMMreg00072_4_pdu1d(pf02954pf00072pf00158) FRAG2
14501 -pf01071pf02843pf02844 -
14506 -pf01808pf02142 -
14513 HMM00465alcdh_3_mix(pf00465) EUT1
14518 HMM06723EutJMreB_1_grm(#) EUT1
14523 HMMalddh_1_eutpdugrm(pf00171) EUT1 EP_NTERM
14528 P_lightViolet EUT1
14533 H_azure EUT1
14538 HMM01515PTA_B_1_eut1(pf01515) EUT1
14543 HMM01923cobtr_2_eut1(pf01923) EUT1
14548 HMM06249EutQ_1_eut1(pf06249) EUT1 EP_NTERM
14553 HMM10662PduV_1_mix(pf10662) EUT1
14558 Hp_euts_violetPink EUT1
14563 -pf04055pf13186 -
14568 -pf00884 -
14573 HMM00390_malic_1_eut1(pf00390pf03949pf01515) EUT1
14578 -pf00923 -
14583 HMM02780keto_1_eut1(pf02779pf02780pf00456) EUT1
14588 -# -
14593 -pf06674 -
14598 HMM00293nuhyd_1_eut1(pf00293) EUT1
14603 -pf13247pf14691pf07992 -
14608 -pf00672pf02518pf07730p
BMC-P
BMC-H
BMC-T
Aldehyde dehydrogenase
PTAC
Signature enzyme
Alcohol dehydrogenase
Regulator

Shigella flexneri 1235-66. Bacteria; Proteobacteria; Gammaproteobacteria; Enterobacterales;

SF123566__locus_2

EUT1

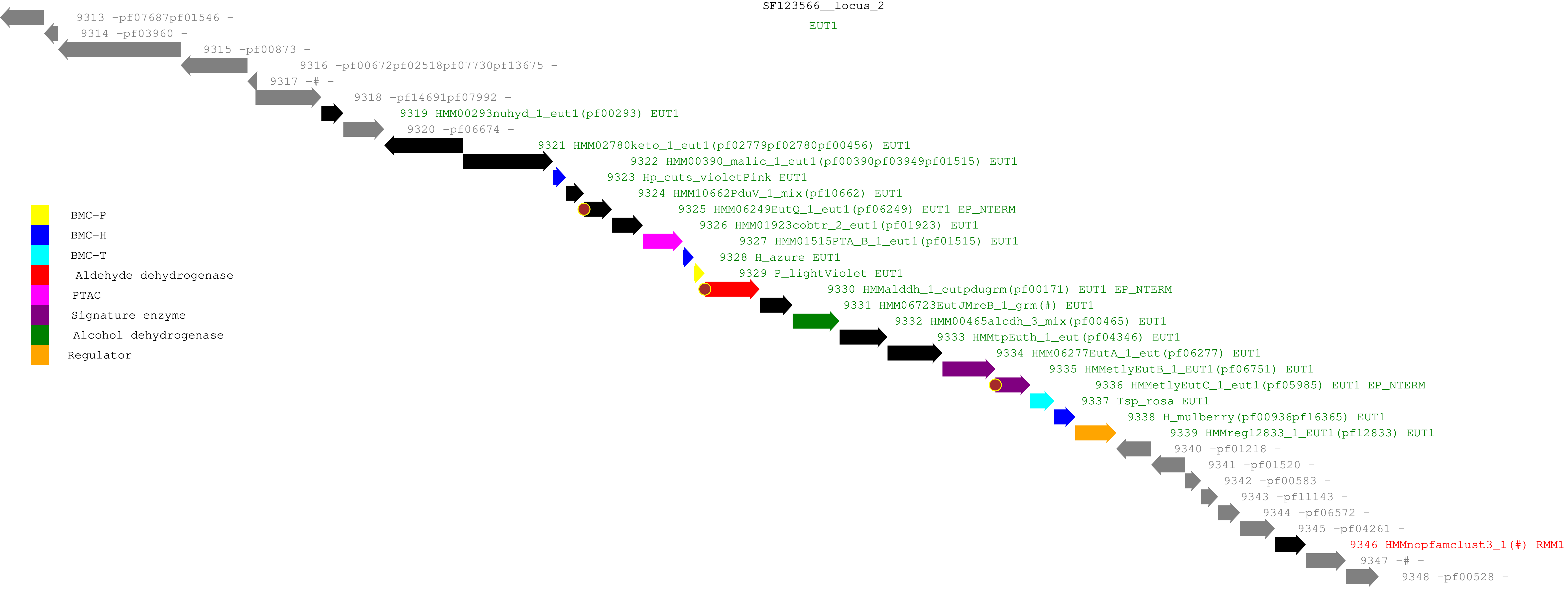

Shigella flexneri 1485-80. Bacteria; Proteobacteria; Gammaproteobacteria; Enterobacterales;

SF148580__locus_1

EUT1

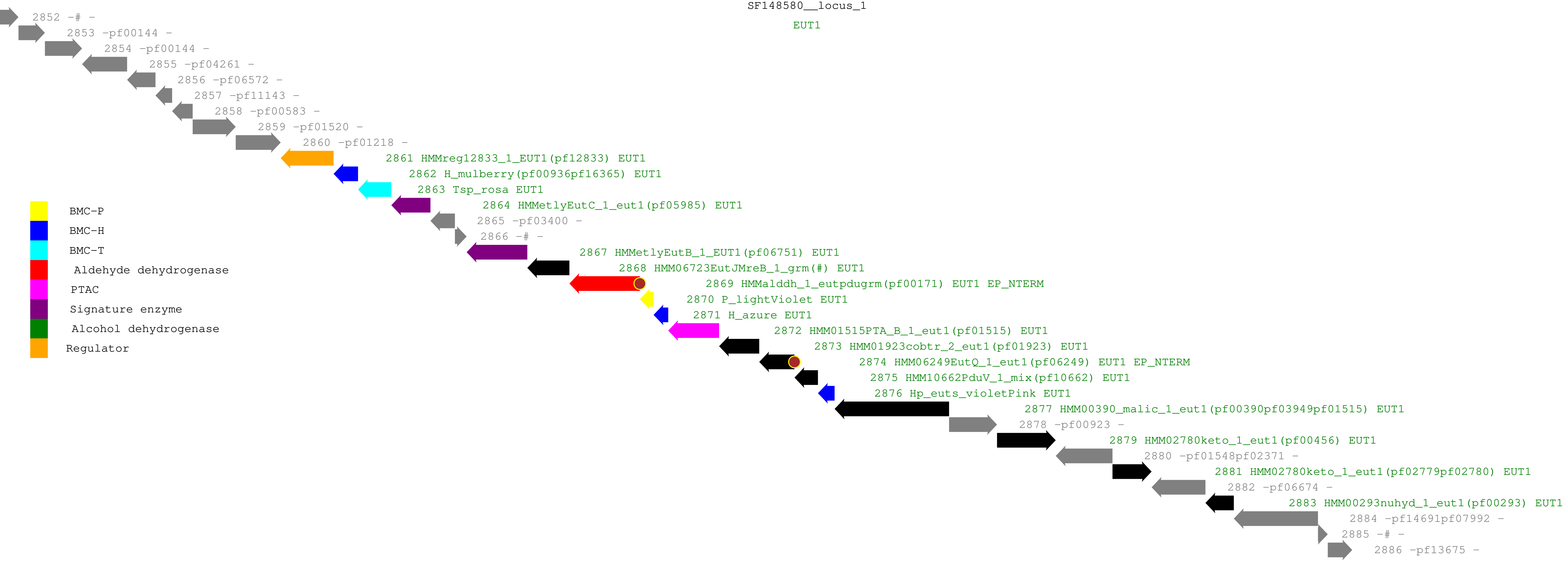

Shigella flexneri K-315. Bacteria; Proteobacteria; Gammaproteobacteria; Enterobacterales;

SFK315__locus_1

EUT1

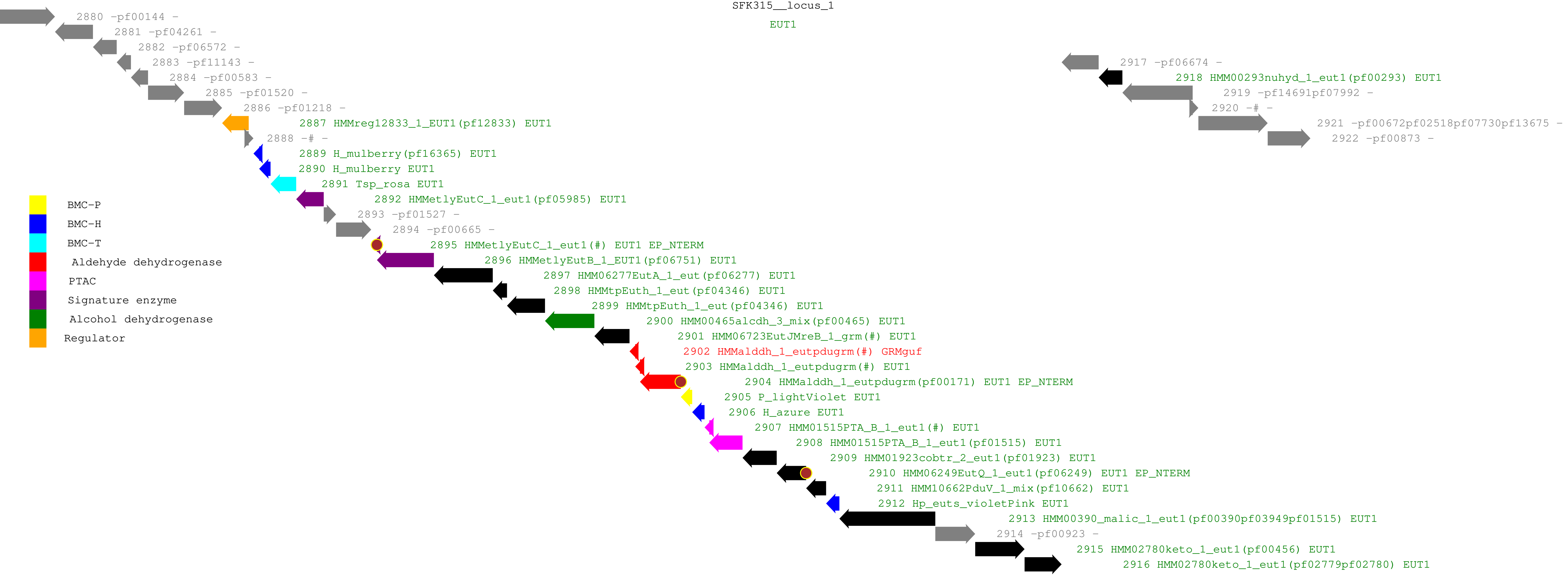

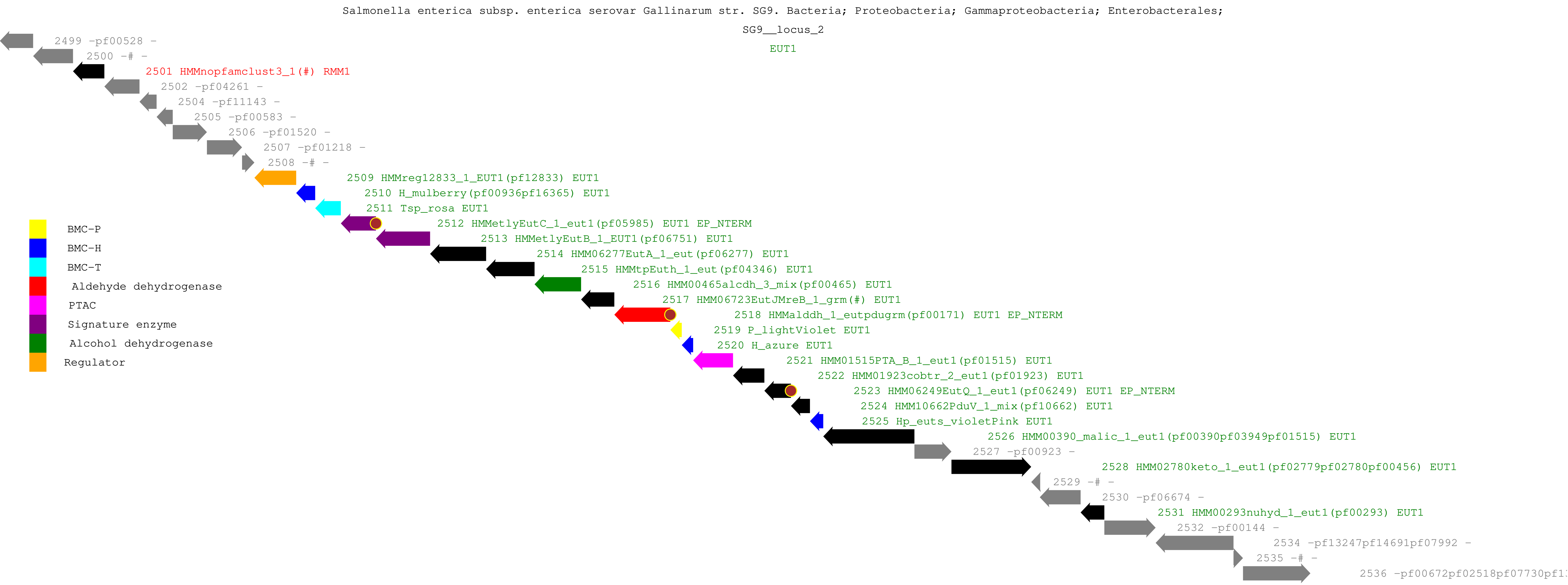
Salmonella enterica subsp. enterica serovar Gallinarum str. SG9. Bacteria; Proteobacteria; Gammaproteobacteria; Enterobacterales;
SG9__locus_2
EUT1
2499 -pf00528 -
2500 -# -
2501 HMMnopfamclust3_1(#) RMM1
2502 -pf04261 -
2504 -pf11143 -
2505 -pf00583 -
2506 -pf01520 -
2507 -pf01218 -
2508 -# -
2509 HMMreg12833_1_EUT1(pf12833) EUT1
2510 H_mulberry(pf00936pf16365) EUT1
2511 Tsp_rosa EUT1
2512 HMMetlyEutC_1_eut1(pf05985) EUT1 EP_NTERM
2513 HMMetlyEutB_1_EUT1(pf06751) EUT1
2514 HMM06277EutA_1_eut(pf06277) EUT1
2515 HMMtpEuth_1_eut(pf04346) EUT1
2516 HMM00465alcdh_3_mix(pf00465) EUT1
2517 HMM06723EutJMreB_1_grm(#) EUT1
2518 HMMalddh_1_eutpdugrm(pf00171) EUT1 EP_NTERM
2519 P_lightViolet EUT1
2520 H_azure EUT1
2521 HMM01515PTA_B_1_eut1(pf01515) EUT1
2522 HMM01923cobtr_2_eut1(pf01923) EUT1
2523 HMM06249EutQ_1_eut1(pf06249) EUT1 EP_NTERM
2524 HMM10662PduV_1_mix(pf10662) EUT1
2525 Hp_euts_violetPink EUT1
2526 HMM00390_malic_1_eut1(pf00390pf03949pf01515) EUT1
2527 -pf00923 -
2528 HMM02780keto_1_eut1(pf02779pf02780pf00456) EUT1
2529 -# -
2530 -pf06674 -
2531 HMM00293nuhyd_1_eut1(pf00293) EUT1
2532 -pf00144 -
2534 -pf13247pf14691pf07992 -
2535 -# -
2536 -pf00672pf02518pf07730pf1
BMC-P
BMC-H
BMC-T
Aldehyde dehydrogenase
PTAC
Signature enzyme
Alcohol dehydrogenase
Regulator

Shigella boydii ATCC 9905. Bacteria; Proteobacteria; Gammaproteobacteria; Enterobacterales;

SGB__locus_1

EUT1

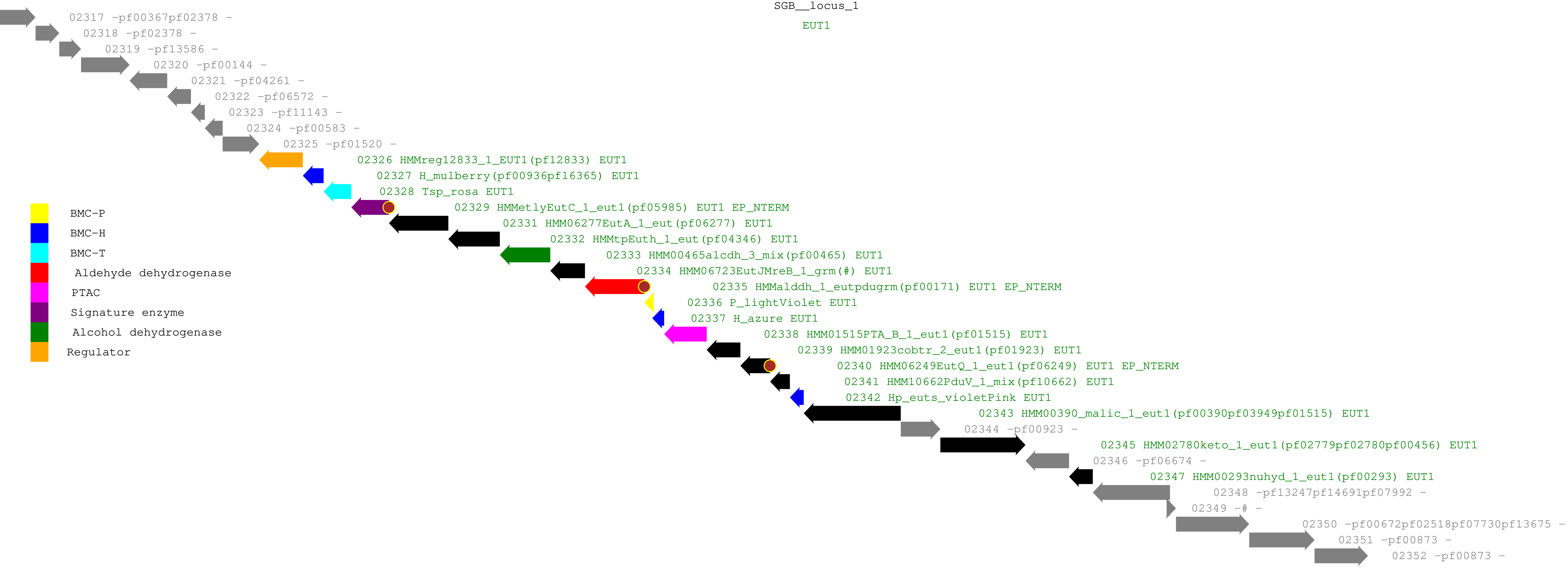

Shigella flexneri CDC 796-83. Bacteria; Proteobacteria; Gammaproteobacteria; Enterobacterales;

SGF__locus_1

EUT1

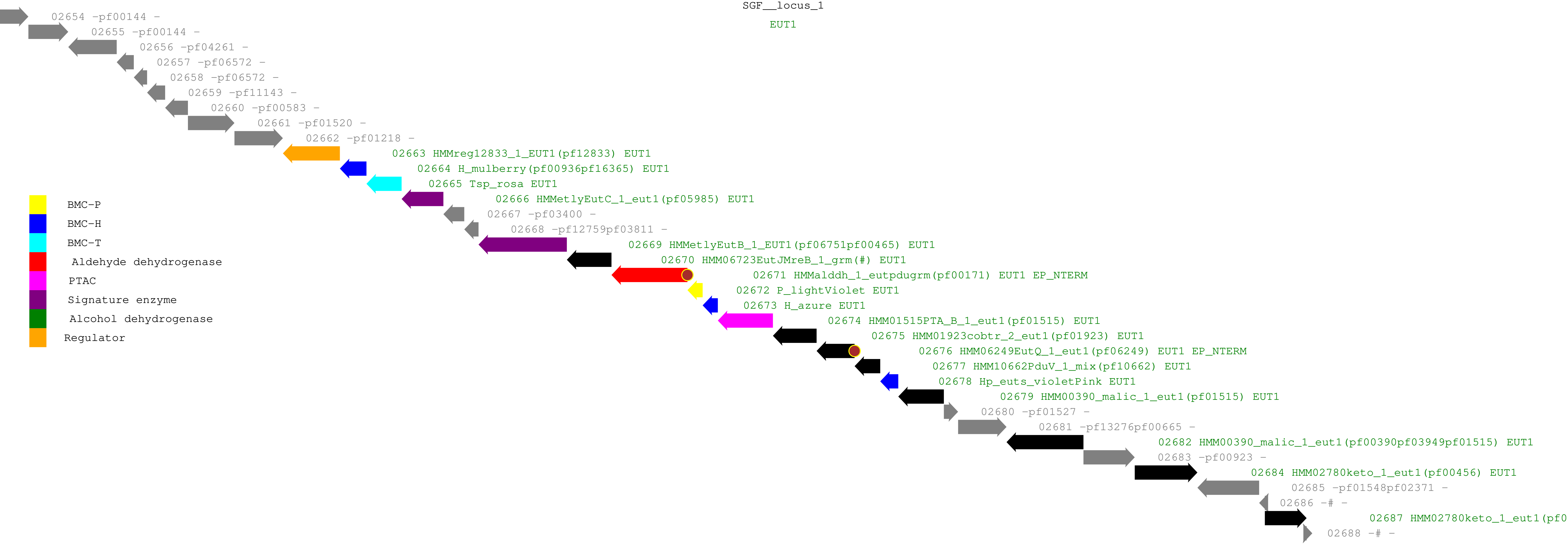

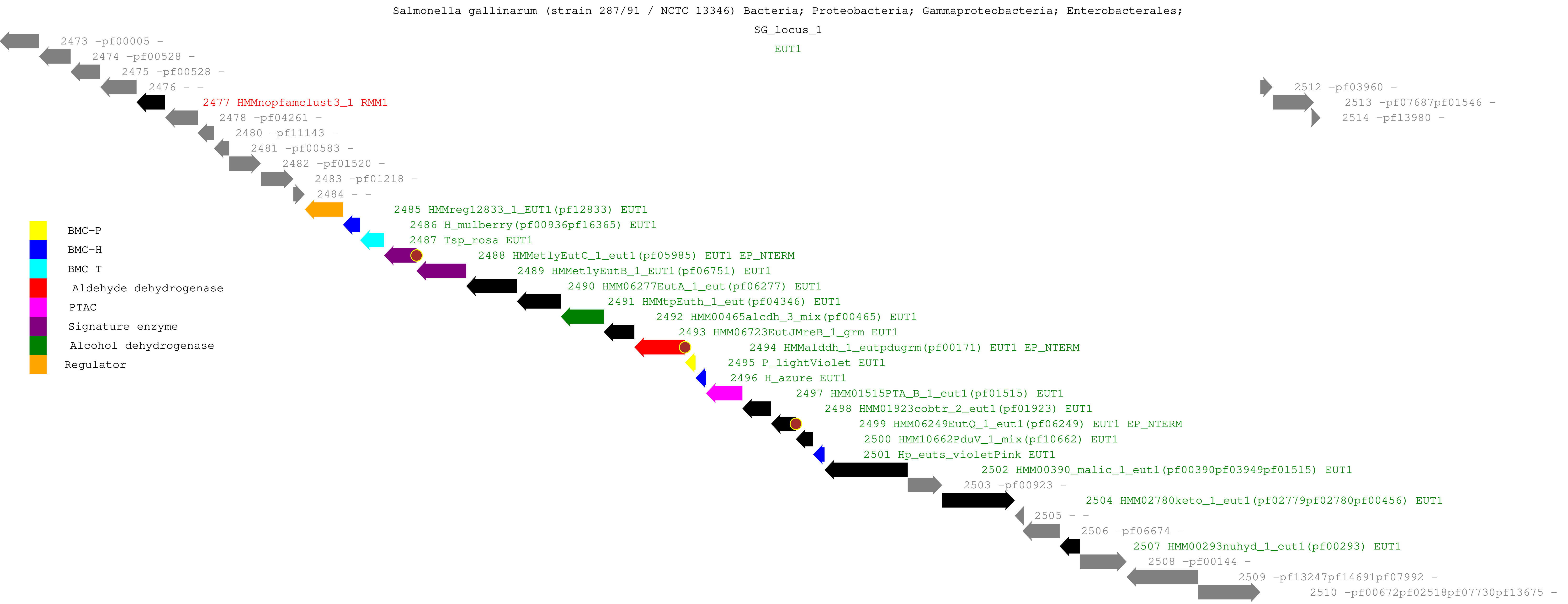

Salmonella gallinarum (strain 287/91 / NCTC 13346) Bacteria; Proteobacteria; Gammaproteobacteria; Enterobacterales;
SG_locus_1
EUT1
2473 -pf00005 -
2474 -pf00528 -
2475 -pf00528 -
2476 - -
2477 HMMnopfamclust3_1 RMM1
2478 -pf04261 -
2480 -pf11143 -
2481 -pf00583 -
2482 -pf01520 -
2483 -pf01218 -
2484 - -
2485 HMMreg12833_1_EUT1(pf12833) EUT1
2486 H_mulberry(pf00936pf16365) EUT1
2487 Tsp_rosa EUT1
2488 HMMetlyEutC_1_eut1(pf05985) EUT1 EP_NTERM
2489 HMMetlyEutB_1_EUT1(pf06751) EUT1
2490 HMM06277EutA_1_eut(pf06277) EUT1
2491 HMMtpEuth_1_eut(pf04346) EUT1
2492 HMM00465alcdh_3_mix(pf00465) EUT1
2493 HMM06723EutJMreB_1_grm EUT1
2494 HMMalddh_1_eutpdugrm(pf00171) EUT1 EP_NTERM
2495 P_lightViolet EUT1
2496 H_azure EUT1
2497 HMM01515PTA_B_1_eut1(pf01515) EUT1
2498 HMM01923cobtr_2_eut1(pf01923) EUT1
2499 HMM06249EutQ_1_eut1(pf06249) EUT1 EP_NTERM
2500 HMM10662PduV_1_mix(pf10662) EUT1
2501 Hp_euts_violetPink EUT1
2502 HMM00390_malic_1_eut1(pf00390pf03949pf01515) EUT1
2503 -pf00923 -
2504 HMM02780keto_1_eut1(pf02779pf02780pf00456) EUT1
2505 - -
2506 -pf06674 -
2507 HMM00293nuhyd_1_eut1(pf00293) EUT1
2508 -pf00144 -
2509 -pf13247pf14691pf07992 -
2510 -pf00672pf02518pf07730pf13675 -
2512 -pf03960 -
2513 -pf07687pf01546 -
2514 -pf13980 -
BMC-P
BMC-H
BMC-T
Aldehyde dehydrogenase
PTAC
Signature enzyme
Alcohol dehydrogenase
Regulator

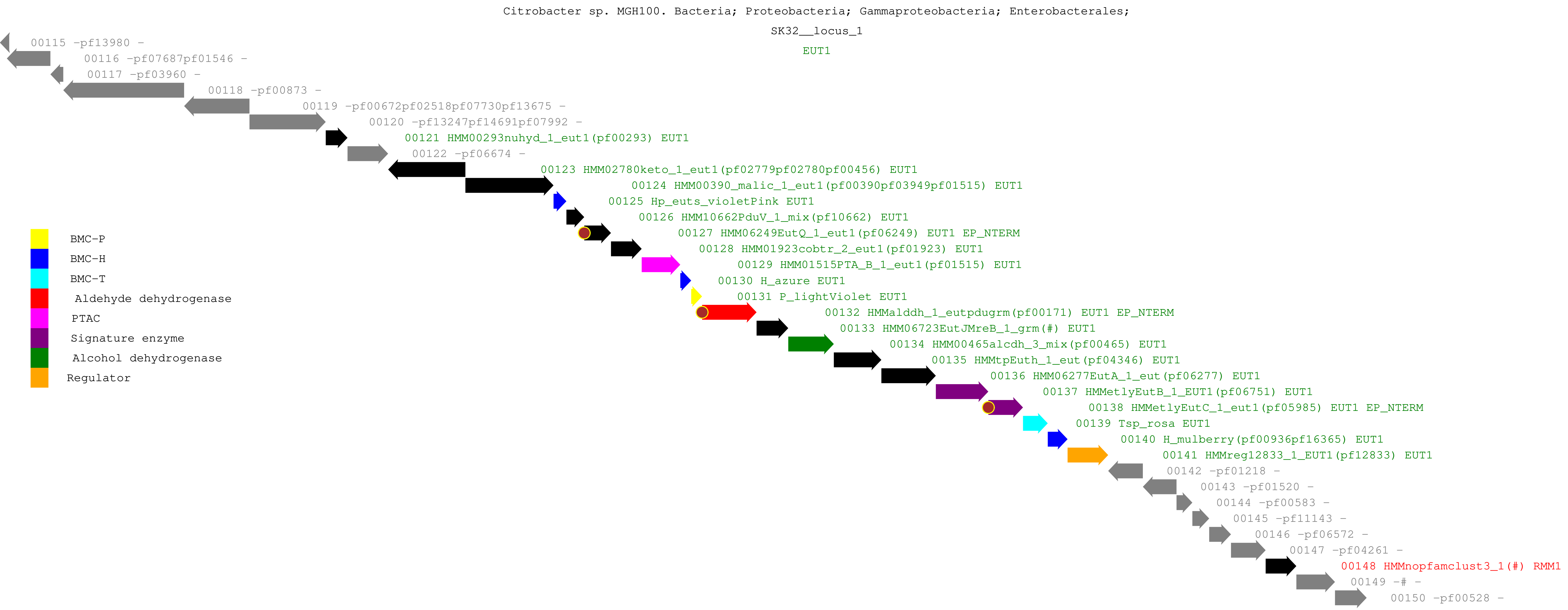
Citrobacter sp. MGH100. Bacteria; Proteobacteria; Gammaproteobacteria; Enterobacterales;
SK32__locus_1
EUT1
00115 -pf13980 -
00116 -pf07687pf01546 -
00117 -pf03960 -
00118 -pf00873 -
00119 -pf00672pf02518pf07730pf13675 -
00120 -pf13247pf14691pf07992 -
00121 HMM00293nuhyd_1_eut1(pf00293) EUT1
00122 -pf06674 -
00123 HMM02780keto_1_eut1(pf02779pf02780pf00456) EUT1
00124 HMM00390_malic_1_eut1(pf00390pf03949pf01515) EUT1
00125 Hp_euts_violetPink EUT1
00126 HMM10662PduV_1_mix(pf10662) EUT1
00127 HMM06249EutQ_1_eut1(pf06249) EUT1 EP_NTERM
00128 HMM01923cobtr_2_eut1(pf01923) EUT1
00129 HMM01515PTA_B_1_eut1(pf01515) EUT1
00130 H_azure EUT1
00131 P_lightViolet EUT1
00132 HMMalddh_1_eutpdugrm(pf00171) EUT1 EP_NTERM
00133 HMM06723EutJMreB_1_grm(#) EUT1
00134 HMM00465alcdh_3_mix(pf00465) EUT1
00135 HMMtpEuth_1_eut(pf04346) EUT1
00136 HMM06277EutA_1_eut(pf06277) EUT1
00137 HMMetlyEutB_1_EUT1(pf06751) EUT1
00138 HMMetlyEutC_1_eut1(pf05985) EUT1 EP_NTERM
00139 Tsp_rosa EUT1
00140 H_mulberry(pf00936pf16365) EUT1
00141 HMMreg12833_1_EUT1(pf12833) EUT1
00142 -pf01218 -
00143 -pf01520 -
00144 -pf00583 -
00145 -pf11143 -
00146 -pf06572 -
00147 -pf04261 -
00148 HMMnopfamclust3_1(#) RMM1
00149 -# -
00150 -pf00528 -
BMC-P
BMC-H
BMC-T
Aldehyde dehydrogenase
PTAC
Signature enzyme
Alcohol dehydrogenase
Regulator

Citrobacter sp. MGH103. Bacteria; Proteobacteria; Gammaproteobacteria; Enterobacterales;

SK33__locus_2

EUT1

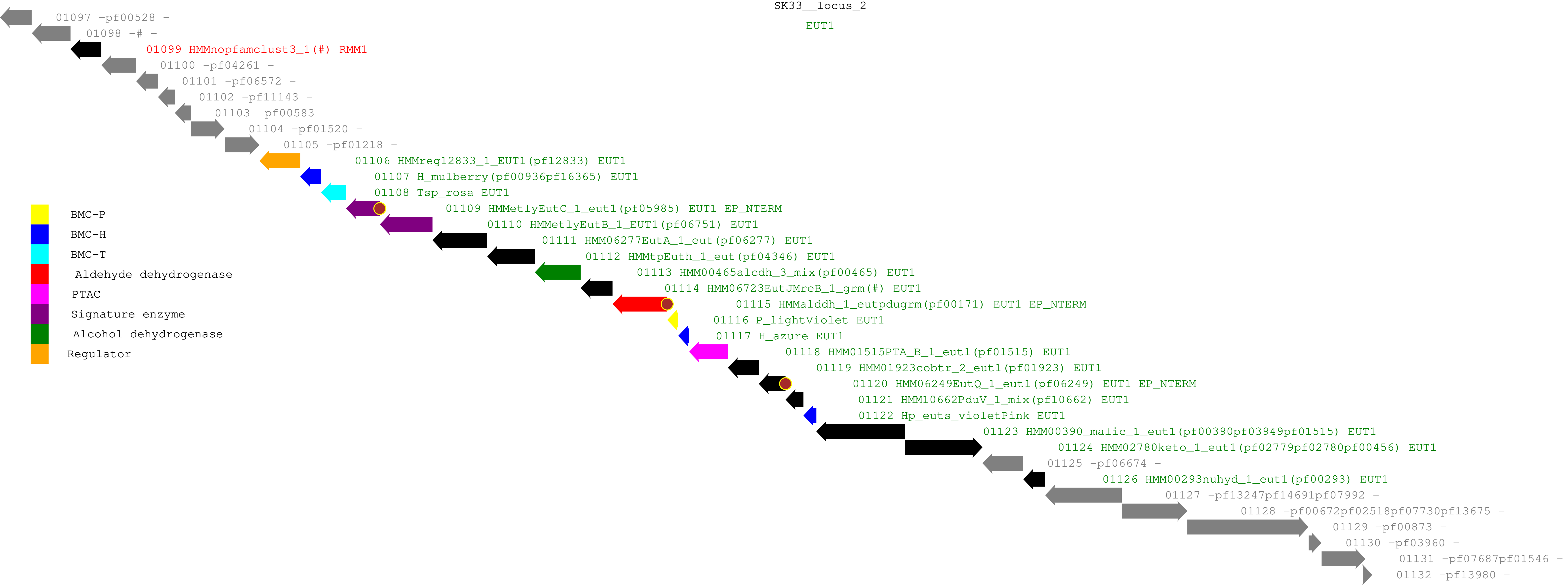

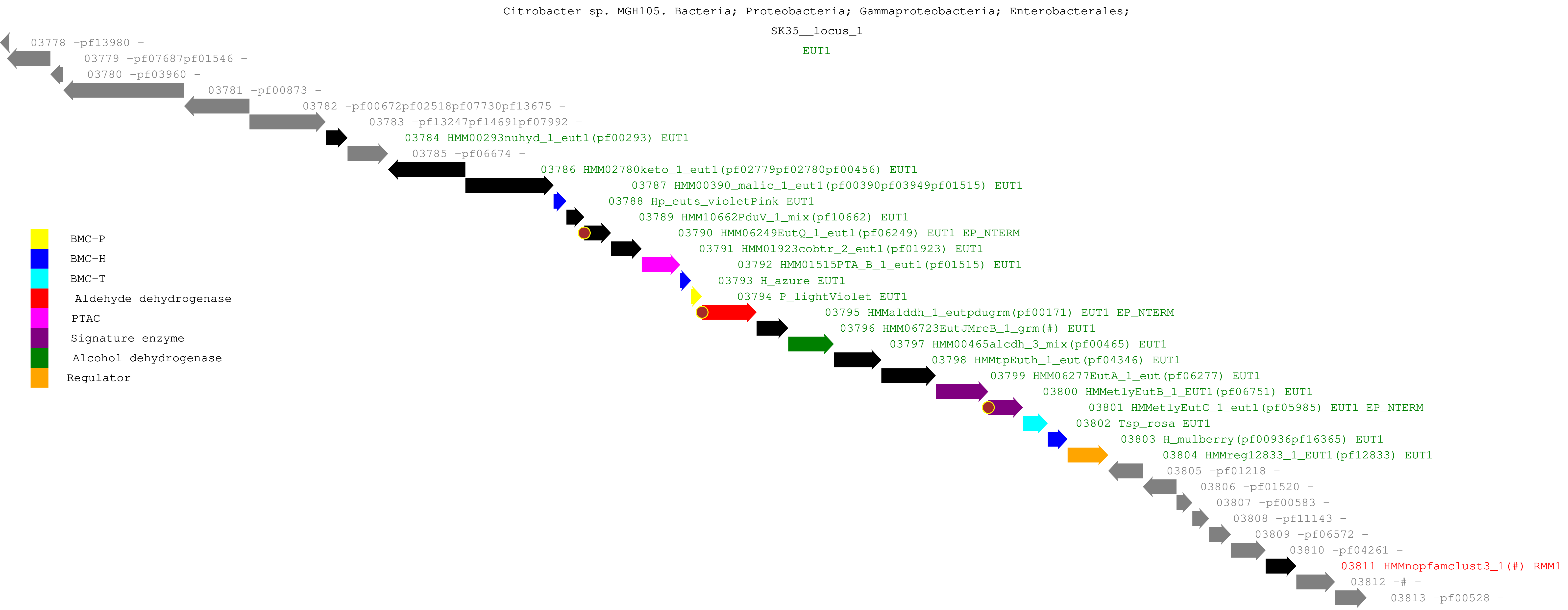
Citrobacter sp. MGH105. Bacteria; Proteobacteria; Gammaproteobacteria; Enterobacterales;
SK35__locus_1
EUT1
03778 -pf13980 -
03779 -pf07687pf01546 -
03780 -pf03960 -
03781 -pf00873 -
03782 -pf00672pf02518pf07730pf13675 -
03783 -pf13247pf14691pf07992 -
03784 HMM00293nuhyd_1_eut1(pf00293) EUT1
03785 -pf06674 -
03786 HMM02780keto_1_eut1(pf02779pf02780pf00456) EUT1
03787 HMM00390_malic_1_eut1(pf00390pf03949pf01515) EUT1
03788 Hp_euts_violetPink EUT1
03789 HMM10662PduV_1_mix(pf10662) EUT1
03790 HMM06249EutQ_1_eut1(pf06249) EUT1 EP_NTERM
03791 HMM01923cobtr_2_eut1(pf01923) EUT1
03792 HMM01515PTA_B_1_eut1(pf01515) EUT1
03793 H_azure EUT1
03794 P_lightViolet EUT1
03795 HMMalddh_1_eutpdugrm(pf00171) EUT1 EP_NTERM
03796 HMM06723EutJMreB_1_grm(#) EUT1
03797 HMM00465alcdh_3_mix(pf00465) EUT1
03798 HMMtpEuth_1_eut(pf04346) EUT1
03799 HMM06277EutA_1_eut(pf06277) EUT1
03800 HMMetlyEutB_1_EUT1(pf06751) EUT1
03801 HMMetlyEutC_1_eut1(pf05985) EUT1 EP_NTERM
03802 Tsp_rosa EUT1
03803 H_mulberry(pf00936pf16365) EUT1
03804 HMMreg12833_1_EUT1(pf12833) EUT1
03805 -pf01218 -
03806 -pf01520 -
03807 -pf00583 -
03808 -pf11143 -
03809 -pf06572 -
03810 -pf04261 -
03811 HMMnopfamclust3_1(#) RMM1
03812 -# -
03813 -pf00528 -
BMC-P
BMC-H
BMC-T
Aldehyde dehydrogenase
PTAC
Signature enzyme
Alcohol dehydrogenase
Regulator

Citrobacter sp. MGH106. Bacteria; Proteobacteria; Gammaproteobacteria; Enterobacterales;

SK36__locus_1

EUT1

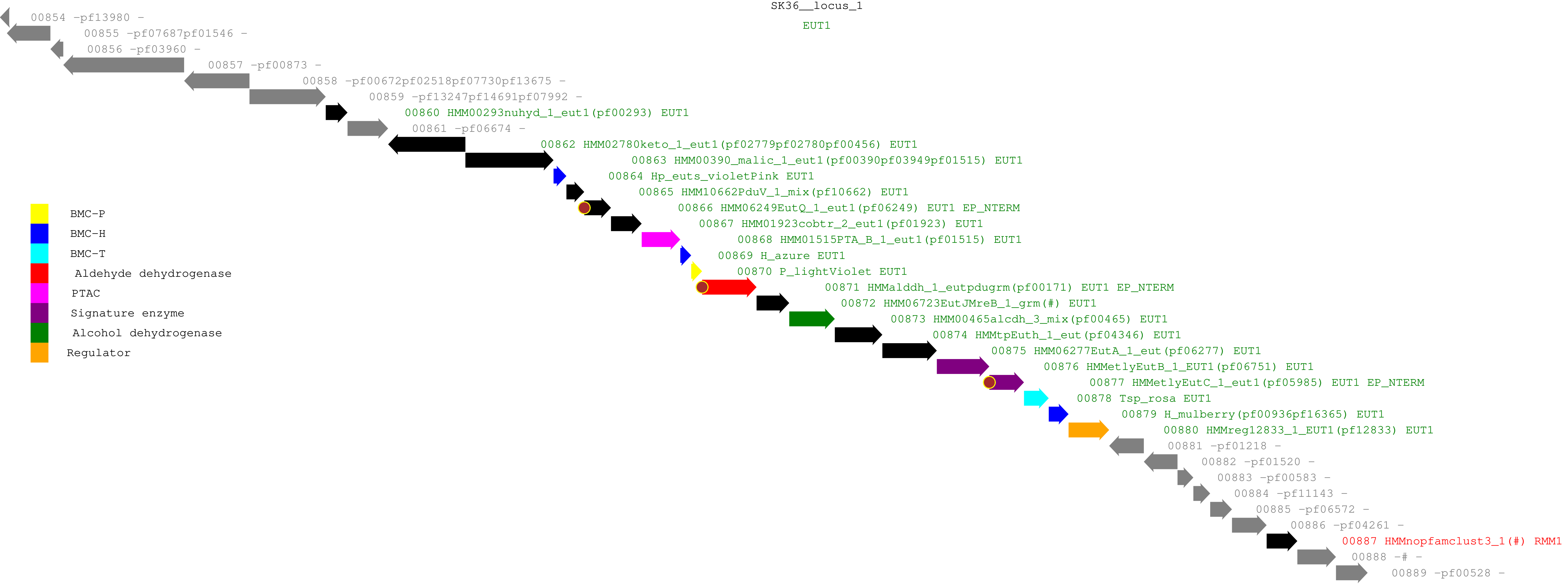

Citrobacter sp. MGH109. Bacteria; Proteobacteria; Gammaproteobacteria; Enterobacterales;

SK37__locus_1

EUT1

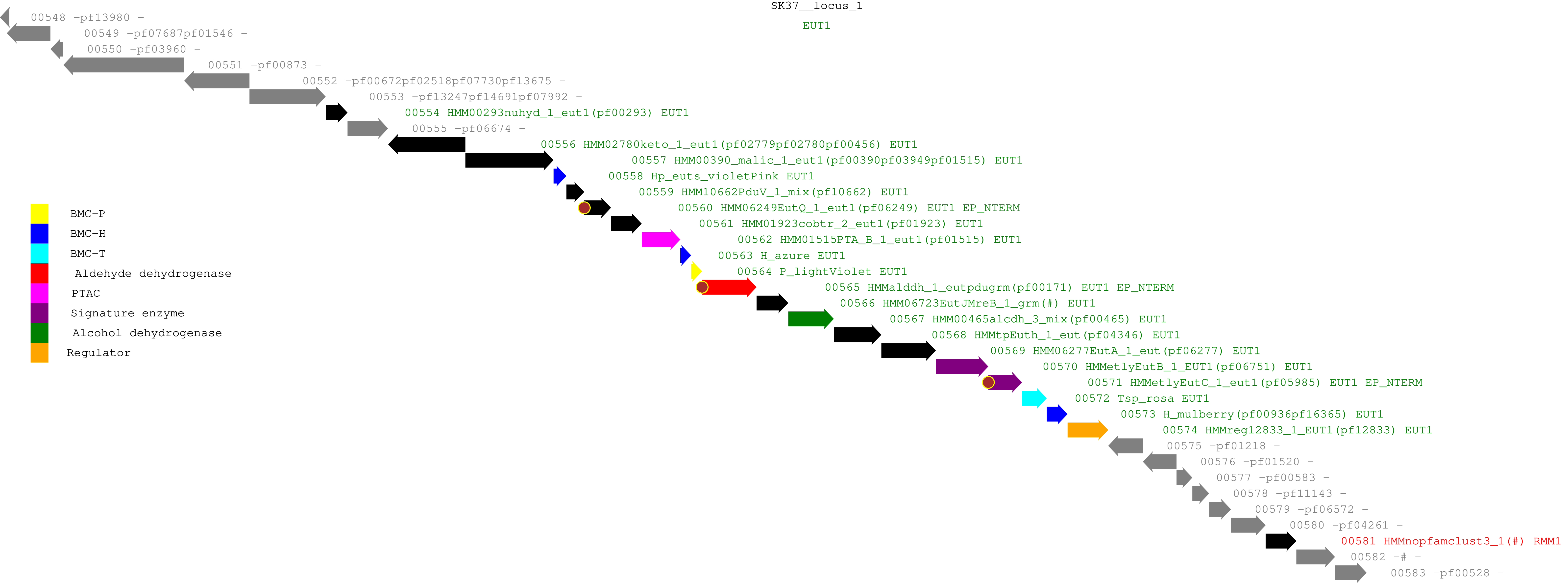

Escherichia coli. Bacteria; Proteobacteria; Gammaproteobacteria; Enterobacterales;

SK85__locus_1

EUT1

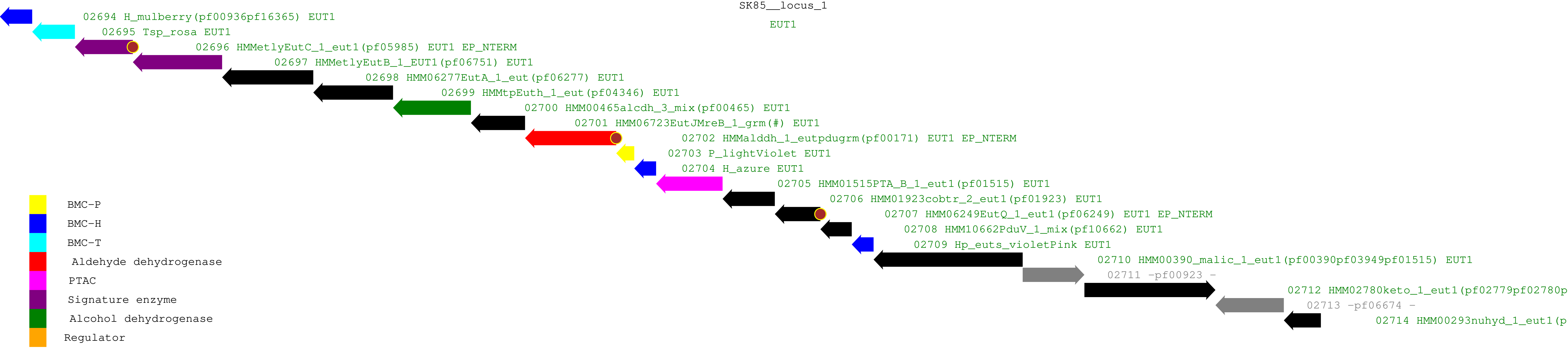

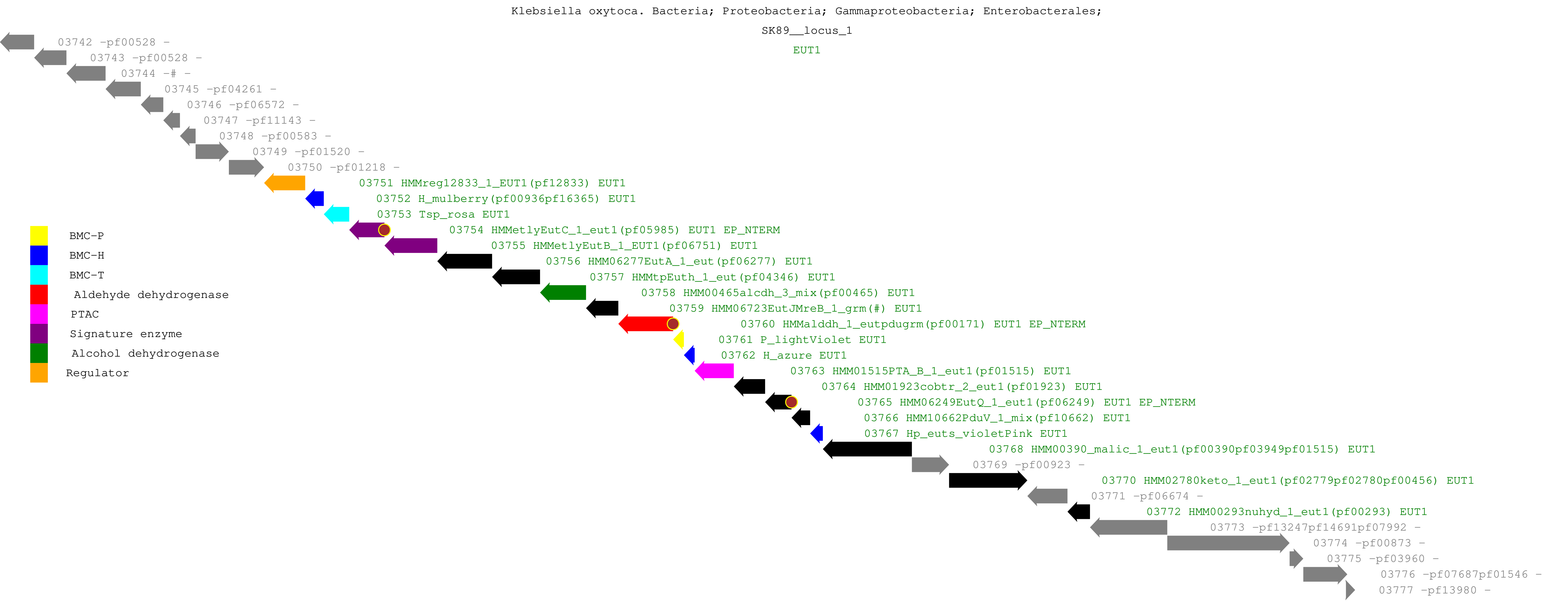
Klebsiella oxytoca. Bacteria; Proteobacteria; Gammaproteobacteria; Enterobacterales;
SK89__locus_1
EUT1
03742 -pf00528 -
03743 -pf00528 -
03744 -# -
03745 -pf04261 -
03746 -pf06572 -
03747 -pf11143 -
03748 -pf00583 -
03749 -pf01520 -
03750 -pf01218 -
03751 HMMreg12833_1_EUT1(pf12833) EUT1
03752 H_mulberry(pf00936pf16365) EUT1
03753 Tsp_rosa EUT1
03754 HMMetlyEutC_1_eut1(pf05985) EUT1 EP_NTERM
03755 HMMetlyEutB_1_EUT1(pf06751) EUT1
03756 HMM06277EutA_1_eut(pf06277) EUT1
03757 HMMtpEuth_1_eut(pf04346) EUT1
03758 HMM00465alcdh_3_mix(pf00465) EUT1
03759 HMM06723EutJMreB_1_grm(#) EUT1
03760 HMMaldh_1_eutpdugrm(pf00171) EUT1 EP_NTERM
03761 P_lightViolet EUT1
03762 H_azure EUT1
03763 HMM01515PTA_B_1_eut1(pf01515) EUT1
03764 HMM01923cobtr_2_eut1(pf01923) EUT1
03765 HMM06249EutQ_1_eut1(pf06249) EUT1 EP_NTERM
03766 HMM10662PduV_1_mix(pf10662) EUT1
03767 Hp_euts_violetPink EUT1
03768 HMM00390_malic_1_eut1(pf00390pf03949pf01515) EUT1
03769 -pf00923 -
03770 HMM02780keto_1_eut1(pf02779pf02780pf00456) EUT1
03771 -pf06674 -
03772 HMM00293nuhyd_1_eut1(pf00293) EUT1
03773 -pf13247pf14691pf07992 -
03774 -pf00873 -
03775 -pf03960 -
03776 -pf07687pf01546 -
03777 -pf13980 -
BMC-P
BMC-H
BMC-T
Aldehyde dehydrogenase
PTAC
Signature enzyme
Alcohol dehydrogenase
Regulator

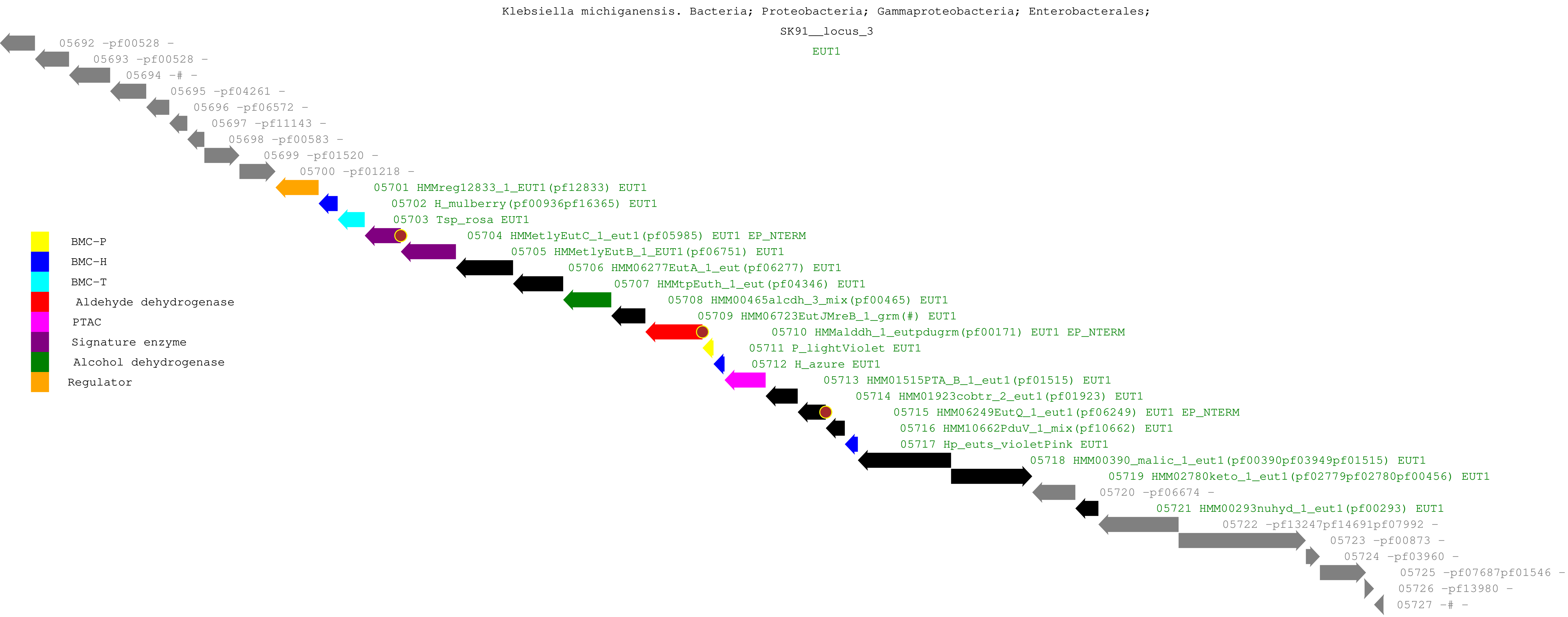

Klebsiella michiganensis. Bacteria; Proteobacteria; Gammaproteobacteria; Enterobacterales;
SK91__locus_3
EUT1
05692 -pf00528 -
05693 -pf00528 -
05694 -# -
05695 -pf04261 -
05696 -pf06572 -
05697 -pf11143 -
05698 -pf00583 -
05699 -pf01520 -
05700 -pf01218 -
05701 HMMreg12833_1_EUT1(pf12833) EUT1
05702 H_mulberry(pf00936pf16365) EUT1
05703 Tsp_rosa EUT1
05704 HMMetlyEutC_1_eut1(pf05985) EUT1 EP_NTERM
05705 HMMetlyEutB_1_EUT1(pf06751) EUT1
05706 HMM06277EutA_1_eut(pf06277) EUT1
05707 HMMtpEuth_1_eut(pf04346) EUT1
05708 HMM00465alcdh_3_mix(pf00465) EUT1
05709 HMM06723EutJMreB_1_grm(#) EUT1
05710 HMMalddh_1_eutpdugrm(pf00171) EUT1 EP_NTERM
05711 P_lightViolet EUT1
05712 H_azure EUT1
05713 HMM01515PTA_B_1_eut1(pf01515) EUT1
05714 HMM01923cobtr_2_eut1(pf01923) EUT1
05715 HMM06249EutQ_1_eut1(pf06249) EUT1 EP_NTERM
05716 HMM10662PduV_1_mix(pf10662) EUT1
05717 Hp_euts_violetPink EUT1
05718 HMM00390_malic_1_eut1(pf00390pf03949pf01515) EUT1
05719 HMM02780keto_1_eut1(pf02779pf02780pf00456) EUT1
05720 -pf06674 -
05721 HMM00293nuhyd_1_eut1(pf00293) EUT1
05722 -pf13247pf14691pf07992 -
05723 -pf00873 -
05724 -pf03960 -
05725 -pf07687pf01546 -
05726 -pf13980 -
05727 -# -
BMC-P
BMC-H
BMC-T
Aldehyde dehydrogenase
PTAC
Signature enzyme
Alcohol dehydrogenase
Regulator

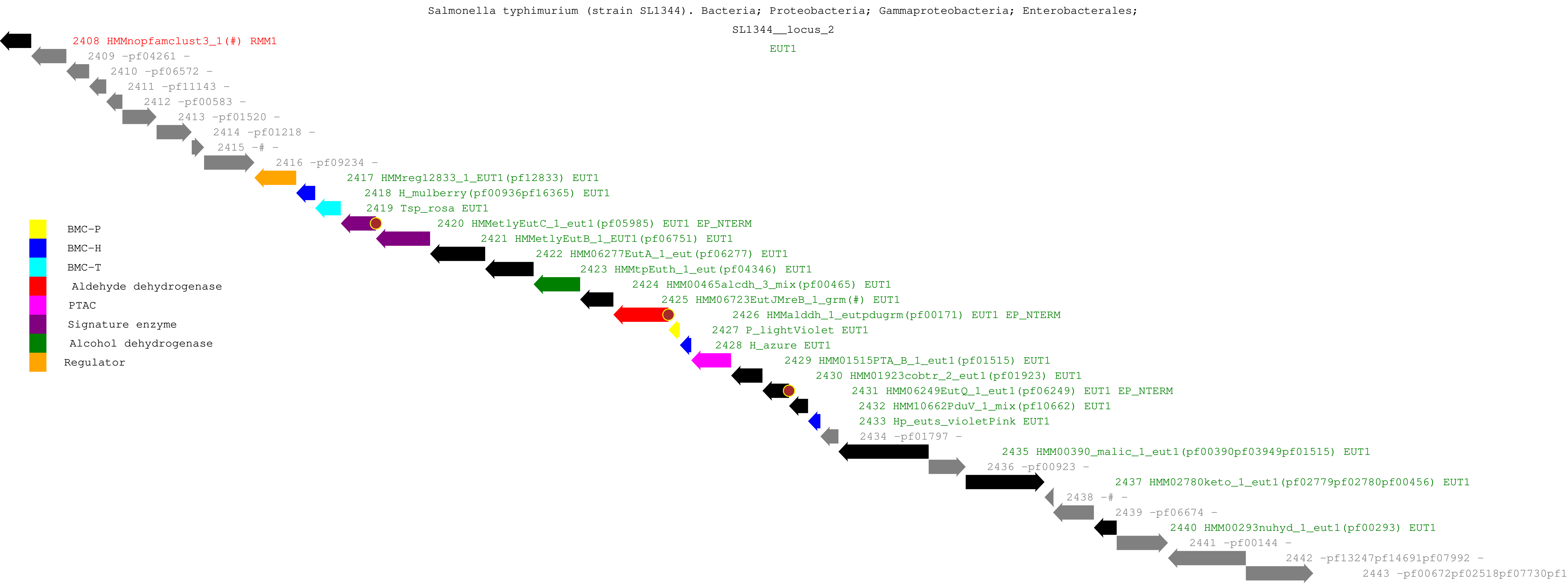

Salmonella typhimurium (strain SL1344). Bacteria; Proteobacteria; Gammaproteobacteria; Enterobacterales;
SL1344__locus_2
EUT1
2408 HMMnopfamclust3_1(#) RMM1
2409 -pf04261 -
2410 -pf06572 -
2411 -pf11143 -
2412 -pf00583 -
2413 -pf01520 -
2414 -pf01218 -
2415 -# -
2416 -pf09234 -
2417 HMMreg12833_1_EUT1(pf12833) EUT1
2418 H_mulberry(pf00936pf16365) EUT1
2419 Tsp_rosa EUT1
2420 HMMetlyEutC_1_eut1(pf05985) EUT1 EP_NTERM
2421 HMMetlyEutB_1_EUT1(pf06751) EUT1
2422 HMM06277EutA_1_eut(pf06277) EUT1
2423 HMMtpEuth_1_eut(pf04346) EUT1
2424 HMM00465alcdh_3_mix(pf00465) EUT1
2425 HMM06723EutJMreB_1_grm(#) EUT1
2426 HMMalddh_1_eutpdugrm(pf00171) EUT1 EP_NTERM
2427 P_lightViolet EUT1
2428 H_azure EUT1
2429 HMM01515PTA_B_1_eut1(pf01515) EUT1
2430 HMM01923cobtr_2_eut1(pf01923) EUT1
2431 HMM06249EutQ_1_eut1(pf06249) EUT1 EP_NTERM
2432 HMM10662PduV_1_mix(pf10662) EUT1
2433 Hp_euts_violetPink EUT1
2434 -pf01797 -
2435 HMM00390_malic_1_eut1(pf00390pf03949pf01515) EUT1
2436 -pf00923 -
2437 HMM02780keto_1_eut1(pf02779pf02780pf00456) EUT1
2438 -# -
2439 -pf06674 -
2440 HMM00293nuhyd_1_eut1(pf00293) EUT1
2441 -pf00144 -
2442 -pf13247pf14691pf07992 -
2443 -pf00672pf02518pf07730pf1
BMC-P
BMC-H
BMC-T
Aldehyde dehydrogenase
PTAC
Signature enzyme
Alcohol dehydrogenase
Regulator

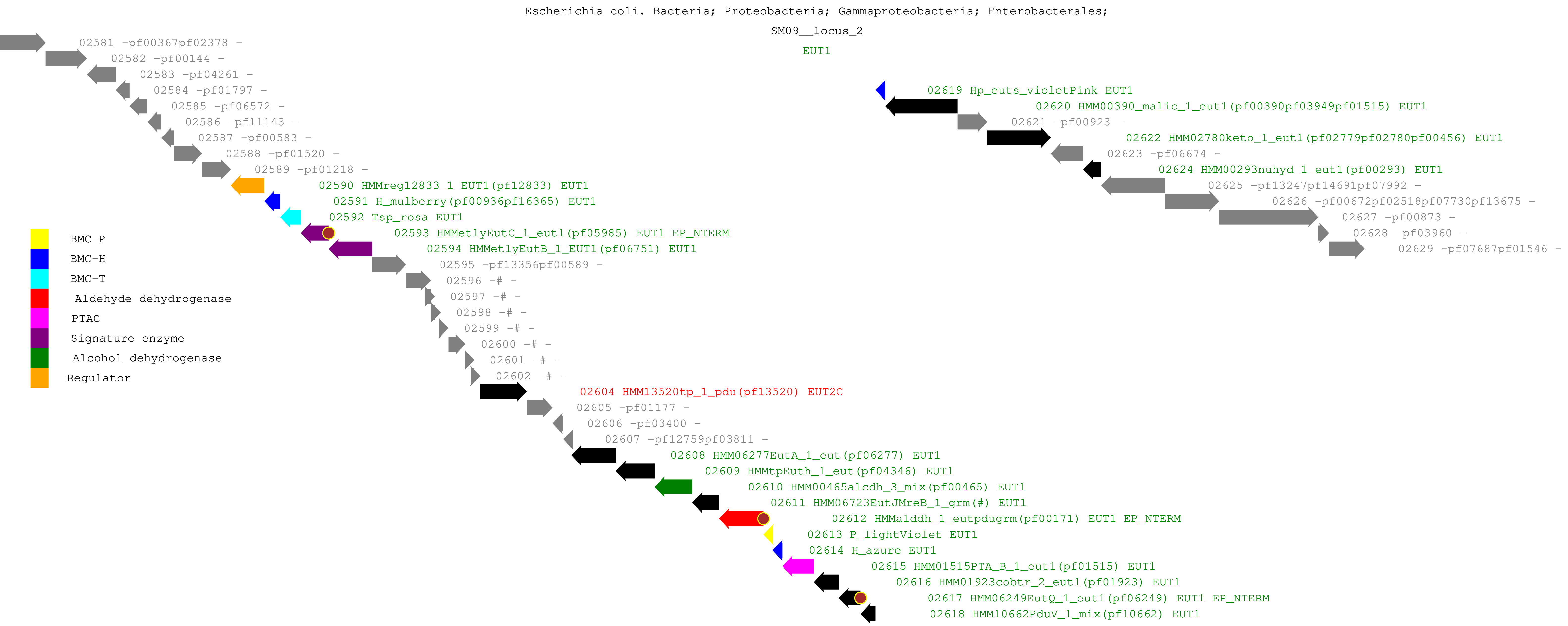
Escherichia coli. Bacteria; Proteobacteria; Gammaproteobacteria; Enterobacterales;
SM09__locus_2
EUT1
02581 -pf00367pf02378 -
02582 -pf00144 -
02583 -pf04261 -
02584 -pf01797 -
02585 -pf06572 -
02586 -pf11143 -
02587 -pf00583 -
02588 -pf01520 -
02589 -pf01218 -
02590 HMMreg12833_1_EUT1(pf12833) EUT1
02591 H_mulberry(pf00936pf16365) EUT1
02592 Tsp_rosa EUT1
02593 HMMetlyEutC_1_eut1(pf05985) EUT1 EP_NTERM
02594 HMMetlyEutB_1_EUT1(pf06751) EUT1
02595 -pf13356pf00589 -
02596 -# -
02597 -# -
02598 -# -
02599 -# -
02600 -# -
02601 -# -
02602 -# -
02604 HMM13520tp_1_pdu(pf13520) EUT2C
02605 -pf01177 -
02606 -pf03400 -
02607 -pf12759pf03811 -
02608 HMM06277EutA_1_eut(pf06277) EUT1
02609 HMMtpEuth_1_eut(pf04346) EUT1
02610 HMM00465alcdh_3_mix(pf00465) EUT1
02611 HMM06723EutJMreB_1_grm(#) EUT1
02612 HMMalddh_1_eutpdugrm(pf00171) EUT1 EP_NTERM
02613 P_lightViolet EUT1
02614 H_azure EUT1
02615 HMM01515PTA_B_1_eut1(pf01515) EUT1
02616 HMM01923cobtr_2_eut1(pf01923) EUT1
02617 HMM06249EutQ_1_eut1(pf06249) EUT1 EP_NTERM
02618 HMM10662PduV_1_mix(pf10662) EUT1
02619 Hp_euts_violetPink EUT1
02620 HMM00390_malic_1_eut1(pf00390pf03949pf01515) EUT1
02621 -pf00923 -
02622 HMM02780keto_1_eut1(pf02779pf02780pf00456) EUT1
02623 -pf06674 -
02624 HMM00293nuhyd_1_eut1(pf00293) EUT1
02625 -pf13247pf14691pf07992 -
02626 -pf00672pf02518pf07730pf13675 -
02627 -pf00873 -
02628 -pf03960 -
02629 -pf07687pf01546 -
BMC-P
BMC-H
BMC-T
Aldehyde dehydrogenase
PTAC
Signature enzyme
Alcohol dehydrogenase
Regulator

Klebsiella pneumoniae. Bacteria; Proteobacteria; Gammaproteobacteria; Enterobacterales;

SM87__locus_2

EUT1

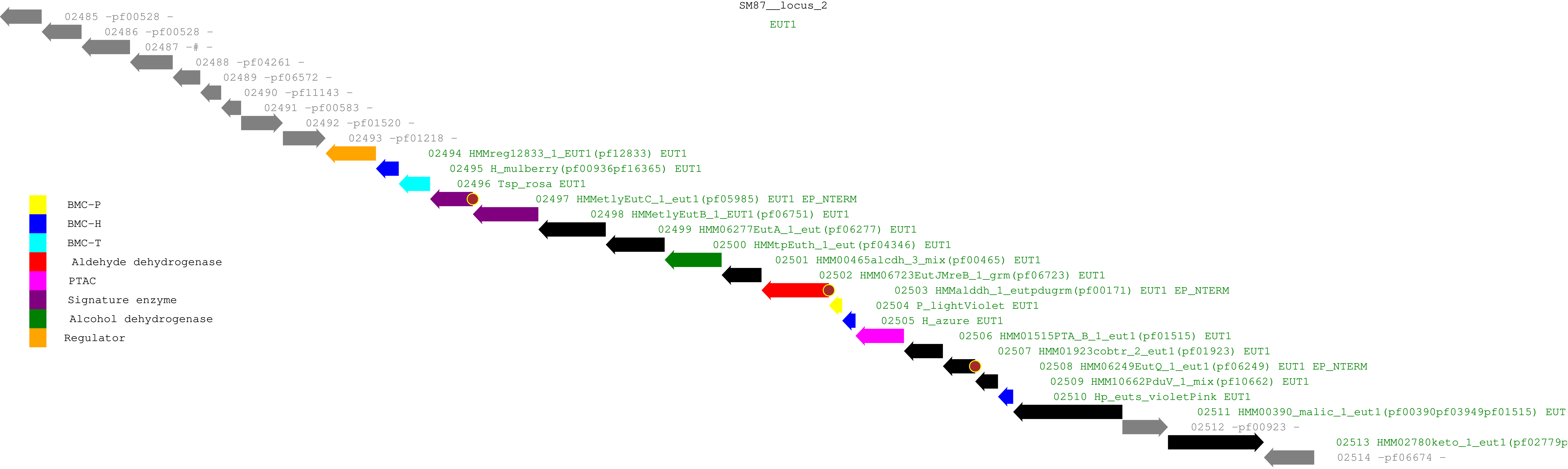

Salmonella newport (strain SL254). Bacteria; Proteobacteria; Gammaproteobacteria; Enterobacterales;

SNSL254_A_locus_2

EUT1

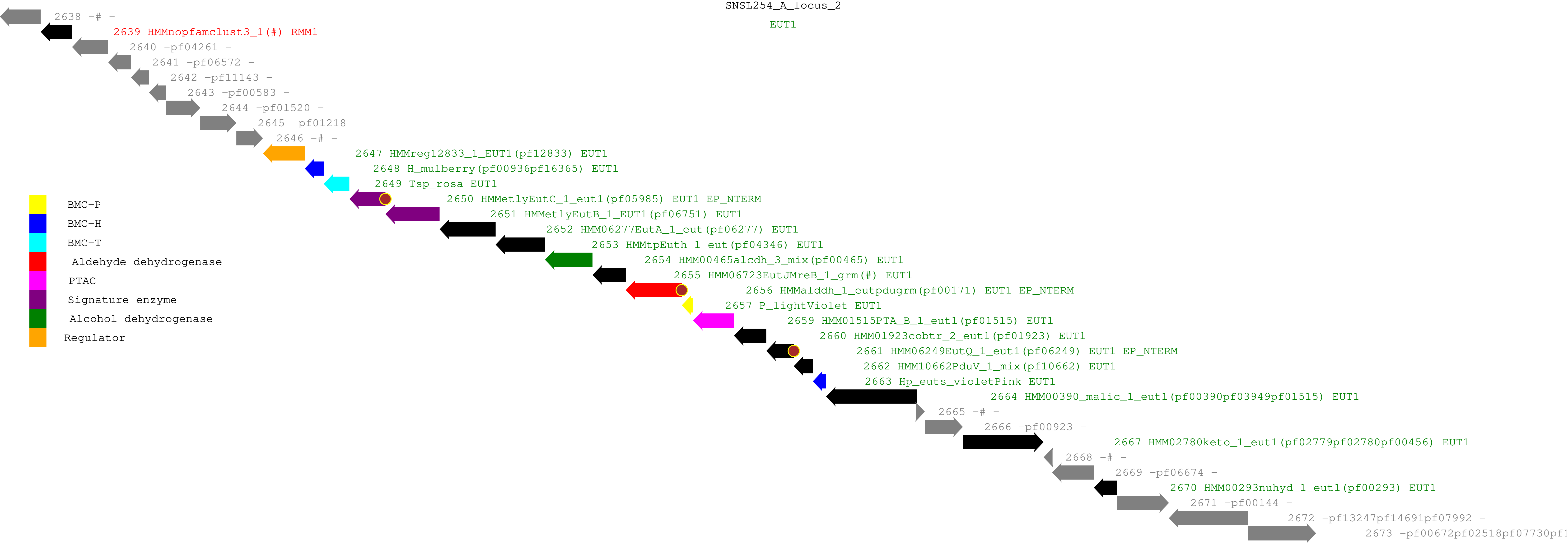

Salmonella paratyphi A (strain ATCC 9150 / SARB42). Bacteria; Proteobacteria; Gammaproteobacteria; Enterobacterales;

SPA_locus_1

EUT1

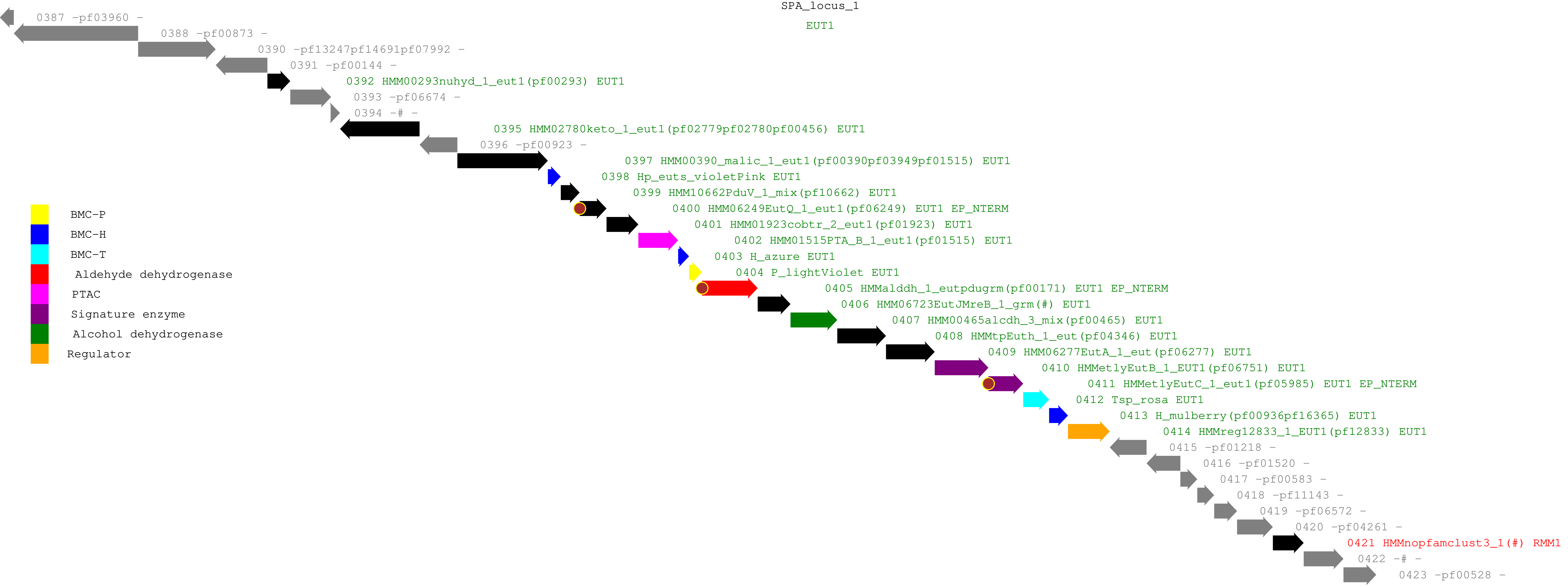

Serratia sp. DD3. Bacteria; Proteobacteria; Gammaproteobacteria; Enterobacterales;

SRDD__locus_1

EUT1

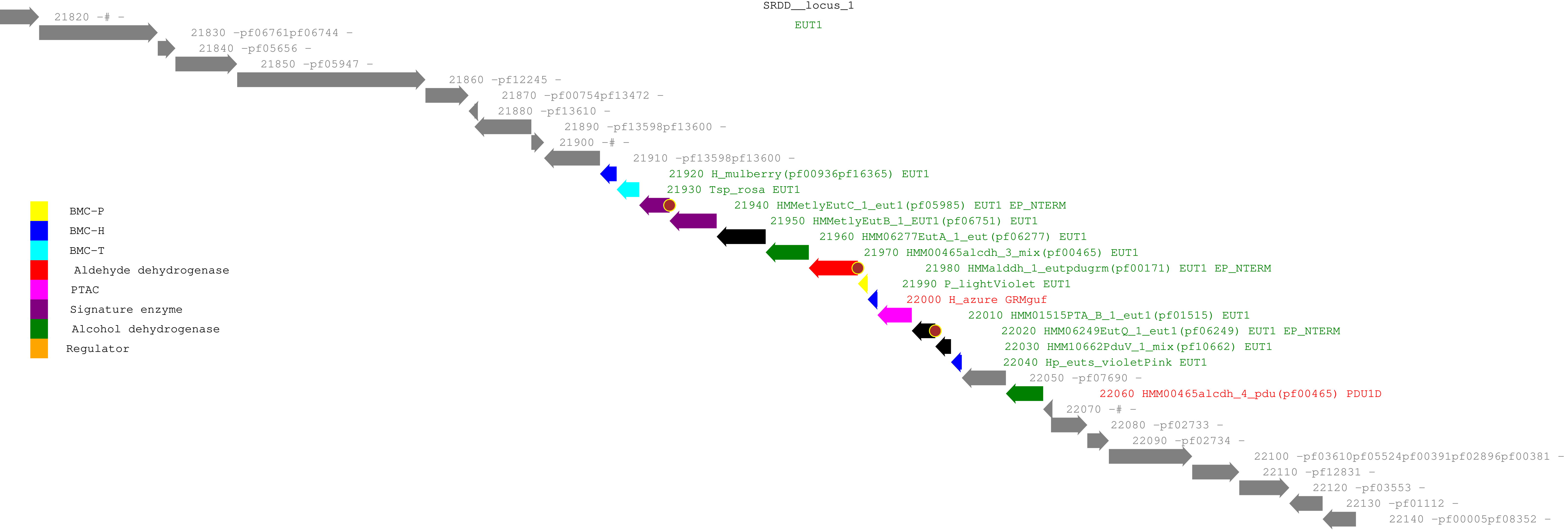

Escherichia coli O157:H7 str. SS52. Bacteria; Proteobacteria; Gammaproteobacteria; Enterobacterales;

SS52__locus_1

EUT1

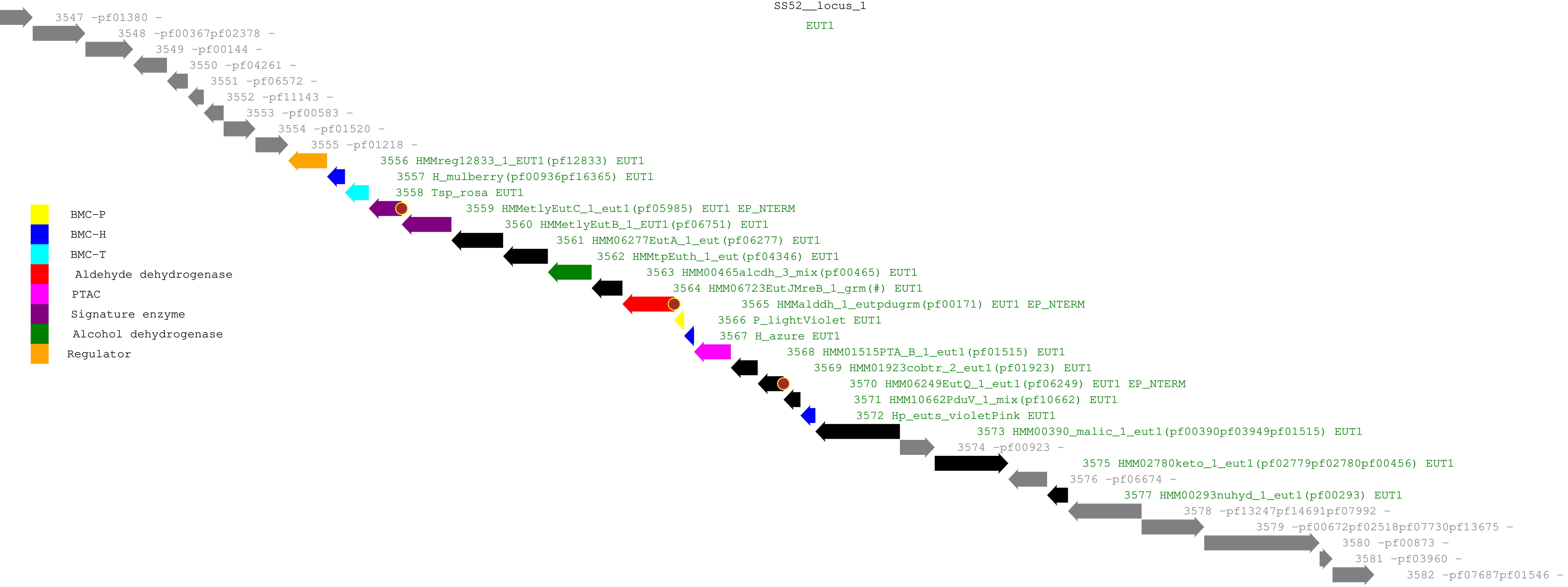

Escherichia coli D9. Bacteria; Proteobacteria; Gammaproteobacteria; Enterobacterales;

SSJG__locus_1

EUT1

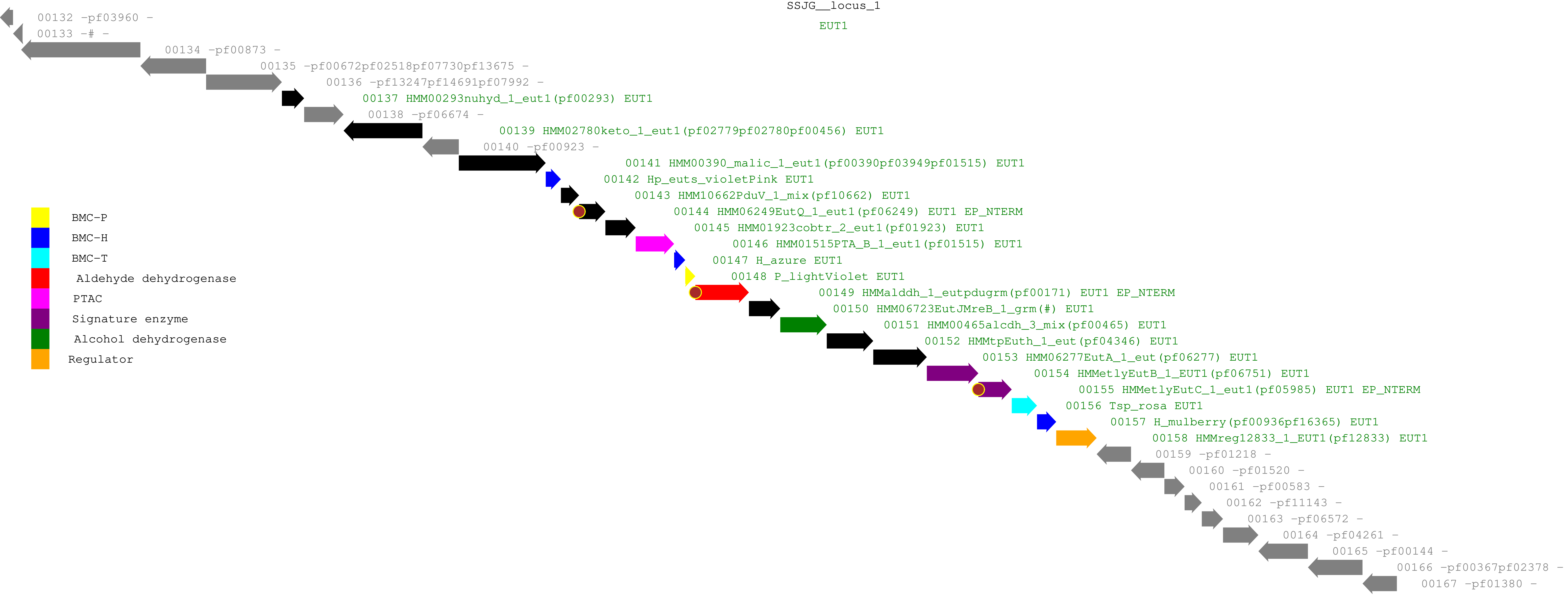

Shigella sonnei str. Moseley. Bacteria; Proteobacteria; Gammaproteobacteria; Enterobacterales;

SSMOSELEY__locus_2

EUT1

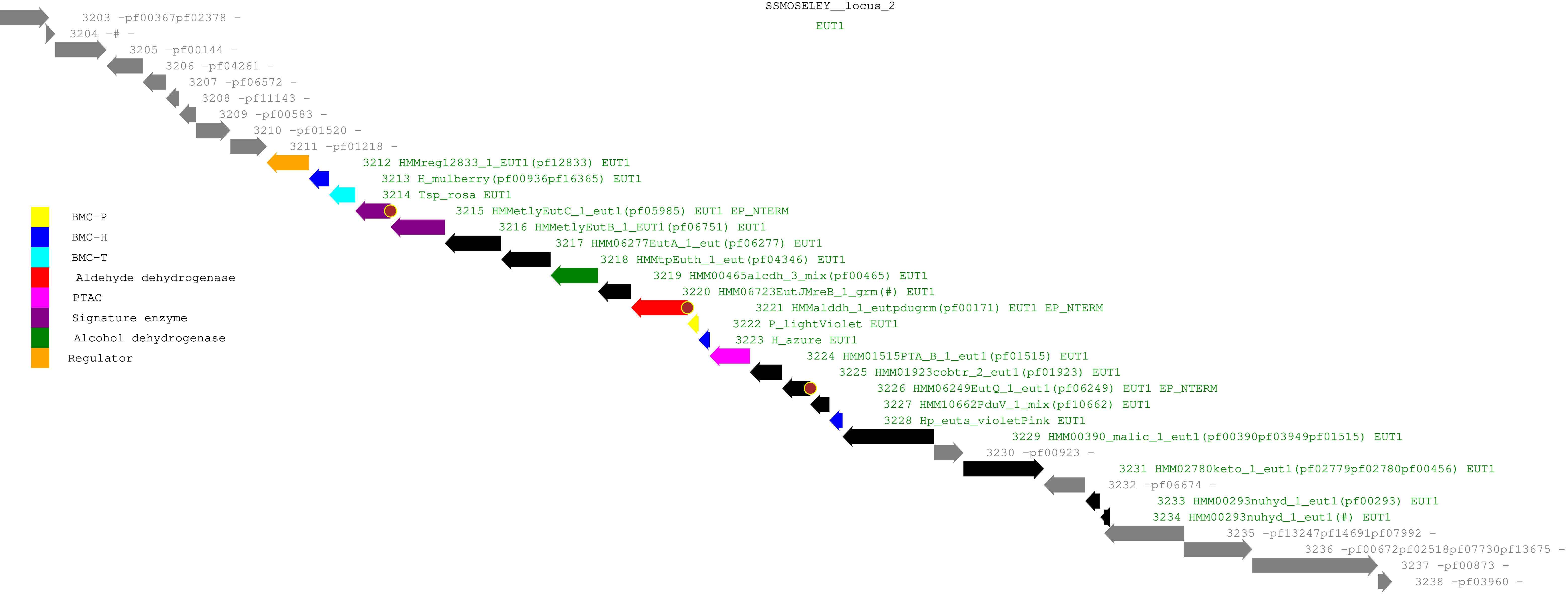

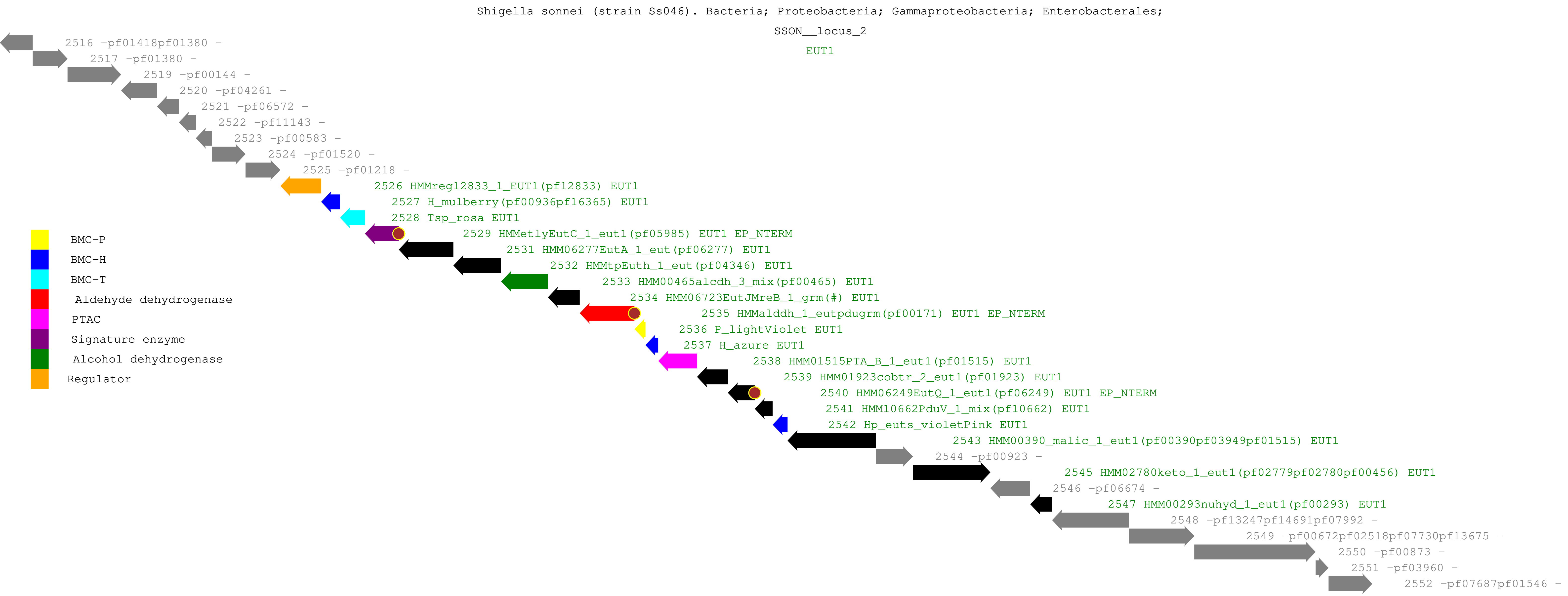
Shigella sonnei (strain Ss046). Bacteria; Proteobacteria; Gammaproteobacteria; Enterobacterales;
SSON__locus_2
EUT1
2516 -pf01418pf01380 -
2517 -pf01380 -
2519 -pf00144 -
2520 -pf04261 -
2521 -pf06572 -
2522 -pf11143 -
2523 -pf00583 -
2524 -pf01520 -
2525 -pf01218 -
2526 HMMreg12833_1_EUT1(pf12833) EUT1
2527 H_mulberry(pf00936pf16365) EUT1
2528 Tsp_rosa EUT1
2529 HMMetlyEutC_1_eut1(pf05985) EUT1 EP_NTERM
2531 HMM06277EutA_1_eut(pf06277) EUT1
2532 HMMtpEuth_1_eut(pf04346) EUT1
2533 HMM00465alcdh_3_mix(pf00465) EUT1
2534 HMM06723EutJMreB_1_grm(#) EUT1
2535 HMMalddh_1_eutpdugrm(pf00171) EUT1 EP_NTERM
2536 P_lightViolet EUT1
2537 H_azure EUT1
2538 HMM01515PTA_B_1_eut1(pf01515) EUT1
2539 HMM01923cobtr_2_eut1(pf01923) EUT1
2540 HMM06249EutQ_1_eut1(pf06249) EUT1 EP_NTERM
2541 HMM10662PduV_1_mix(pf10662) EUT1
2542 Hp_euts_violetPink EUT1
2543 HMM00390_malic_1_eut1(pf00390pf03949pf01515) EUT1
2544 -pf00923 -
2545 HMM02780keto_1_eut1(pf02779pf02780pf00456) EUT1
2546 -pf06674 -
2547 HMM00293nuhyd_1_eut1(pf00293) EUT1
2548 -pf13247pf14691pf07992 -
2549 -pf00672pf02518pf07730pf13675 -
2550 -pf00873 -
2551 -pf03960 -
2552 -pf07687pf01546 -
BMC-P
BMC-H
BMC-T
Aldehyde dehydrogenase
PTAC
Signature enzyme
Alcohol dehydrogenase
Regulator

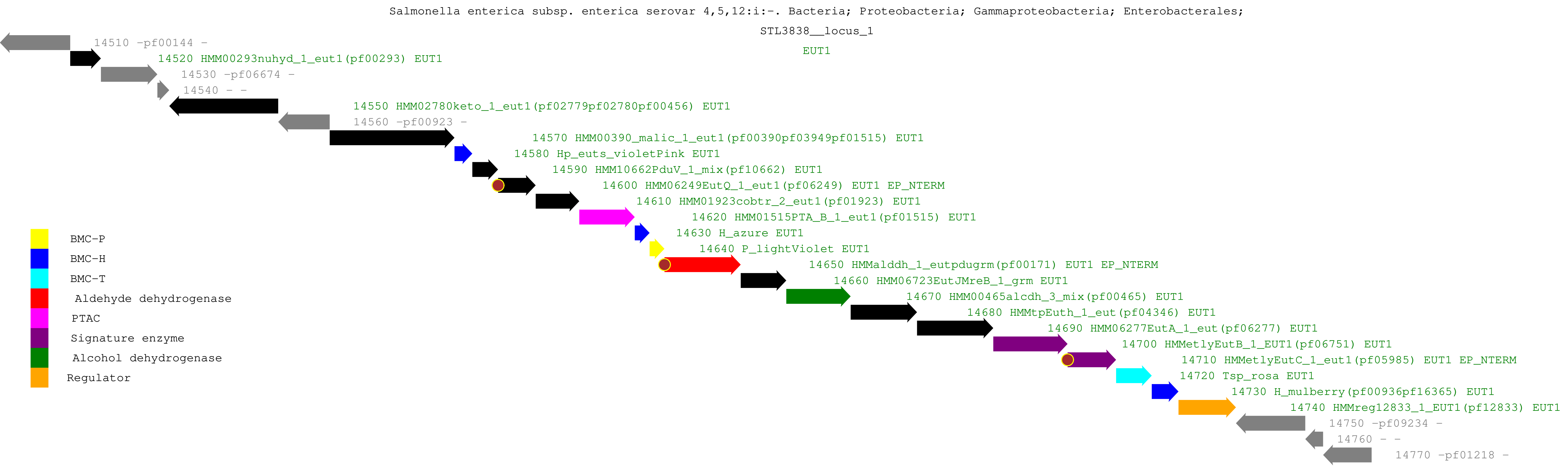
Salmonella enterica subsp. enterica serovar 4,5,12:i:-. Bacteria; Proteobacteria; Gammaproteobacteria; Enterobacterales;
STL3838__locus_1
EUT1
14510 -pf00144 -
14520 HMM00293nuhyd_1_eut1(pf00293) EUT1
14530 -pf06674 -
14540 - -
14550 HMM02780keto_1_eut1(pf02779pf02780pf00456) EUT1
14560 -pf00923 -
14570 HMM00390_malic_1_eut1(pf00390pf03949pf01515) EUT1
14580 Hp_euts_violetPink EUT1
14590 HMM10662PduV_1_mix(pf10662) EUT1
14600 HMM06249EutQ_1_eut1(pf06249) EUT1 EP_NTERM
14610 HMM01923cobtr_2_eut1(pf01923) EUT1
14620 HMM01515PTA_B_1_eut1(pf01515) EUT1
14630 H_azure EUT1
14640 P_lightViolet EUT1
14650 HMMalddh_1_eutpdugrm(pf00171) EUT1 EP_NTERM
14660 HMM06723EutJMreB_1_grm EUT1
14670 HMM00465alcdh_3_mix(pf00465) EUT1
14680 HMMtpEuth_1_eut(pf04346) EUT1
14690 HMM06277EutA_1_eut(pf06277) EUT1
14700 HMMetlyEutB_1_EUT1(pf06751) EUT1
14710 HMMetlyEutC_1_eut1(pf05985) EUT1 EP_NTERM
14720 Tsp_rosa EUT1
14730 H_mulberry(pf00936pf16365) EUT1
14740 HMMreg12833_1_EUT1(pf12833) EUT1
14750 -pf09234 -
14760 - -
14770 -pf01218 -
BMC-P
BMC-H
BMC-T
Aldehyde dehydrogenase
PTAC
Signature enzyme
Alcohol dehydrogenase
Regulator

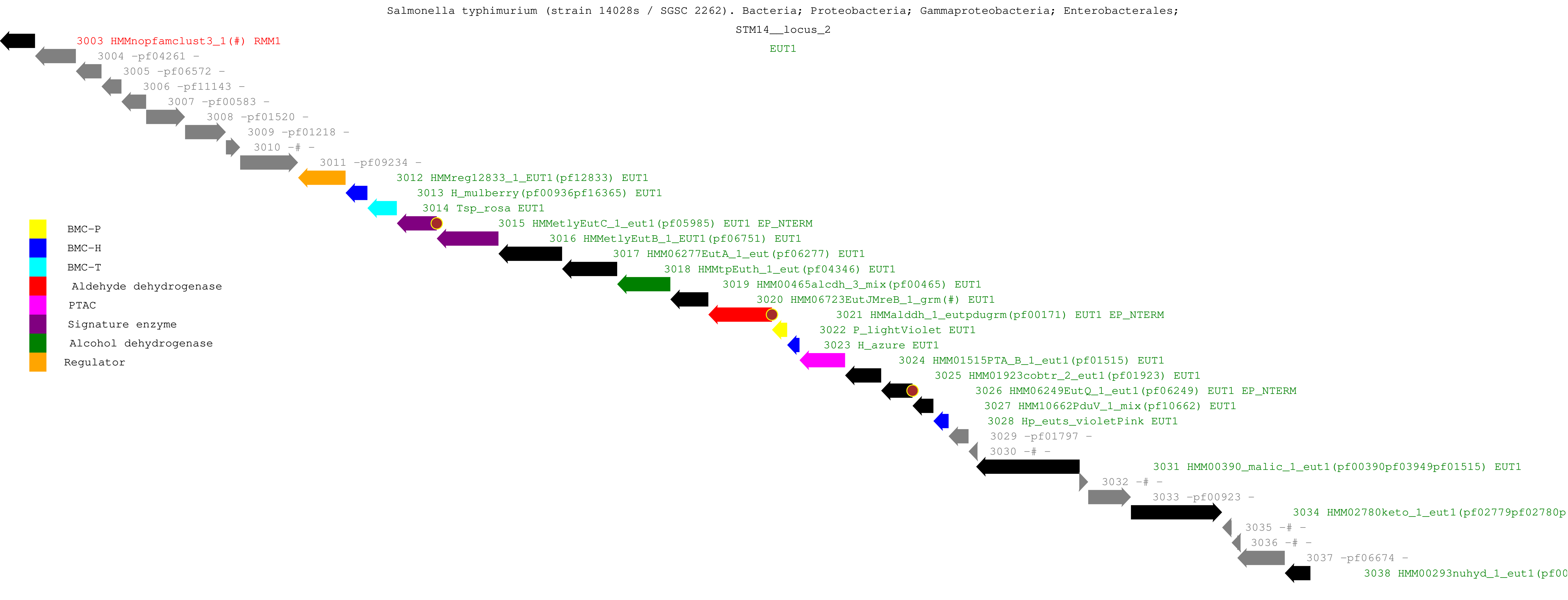

Salmonella typhimurium (strain 14028s / SGSC 2262). Bacteria; Proteobacteria; Gammaproteobacteria; Enterobacterales;
STM14__locus_2
EUT1
3003 HMMnopfamclust3_1(#) RMM1
3004 -pf04261 -
3005 -pf06572 -
3006 -pf11143 -
3007 -pf00583 -
3008 -pf01520 -
3009 -pf01218 -
3010 -# -
3011 -pf09234 -
3012 HMMreg12833_1_EUT1(pf12833) EUT1
3013 H_mulberry(pf00936pf16365) EUT1
3014 Tsp_rosa EUT1
3015 HMMetlyEutC_1_eut1(pf05985) EUT1 EP_NTERM
3016 HMMetlyEutB_1_EUT1(pf06751) EUT1
3017 HMM06277EutA_1_eut(pf06277) EUT1
3018 HMMtpEuth_1_eut(pf04346) EUT1
3019 HMM00465alcdh_3_mix(pf00465) EUT1
3020 HMM06723EutJMreB_1_grm(#) EUT1
3021 HMMalddh_1_eutpdugrm(pf00171) EUT1 EP_NTERM
3022 P_lightViolet EUT1
3023 H_azure EUT1
3024 HMM01515PTA_B_1_eut1(pf01515) EUT1
3025 HMM01923cobtr_2_eut1(pf01923) EUT1
3026 HMM06249EutQ_1_eut1(pf06249) EUT1 EP_NTERM
3027 HMM10662PduV_1_mix(pf10662) EUT1
3028 Hp_euts_violetPink EUT1
3029 -pf01797 -
3030 -# -
3031 HMM00390_malic_1_eut1(pf00390pf03949pf01515) EUT1
3032 -# -
3033 -pf00923 -
3034 HMM02780keto_1_eut1(pf02779pf02780p
3035 -# -
3036 -# -
3037 -pf06674 -
3038 HMM00293nuhyd_1_eut1(pf00
BMC-P
BMC-H
BMC-T
Aldehyde dehydrogenase
PTAC
Signature enzyme
Alcohol dehydrogenase
Regulator

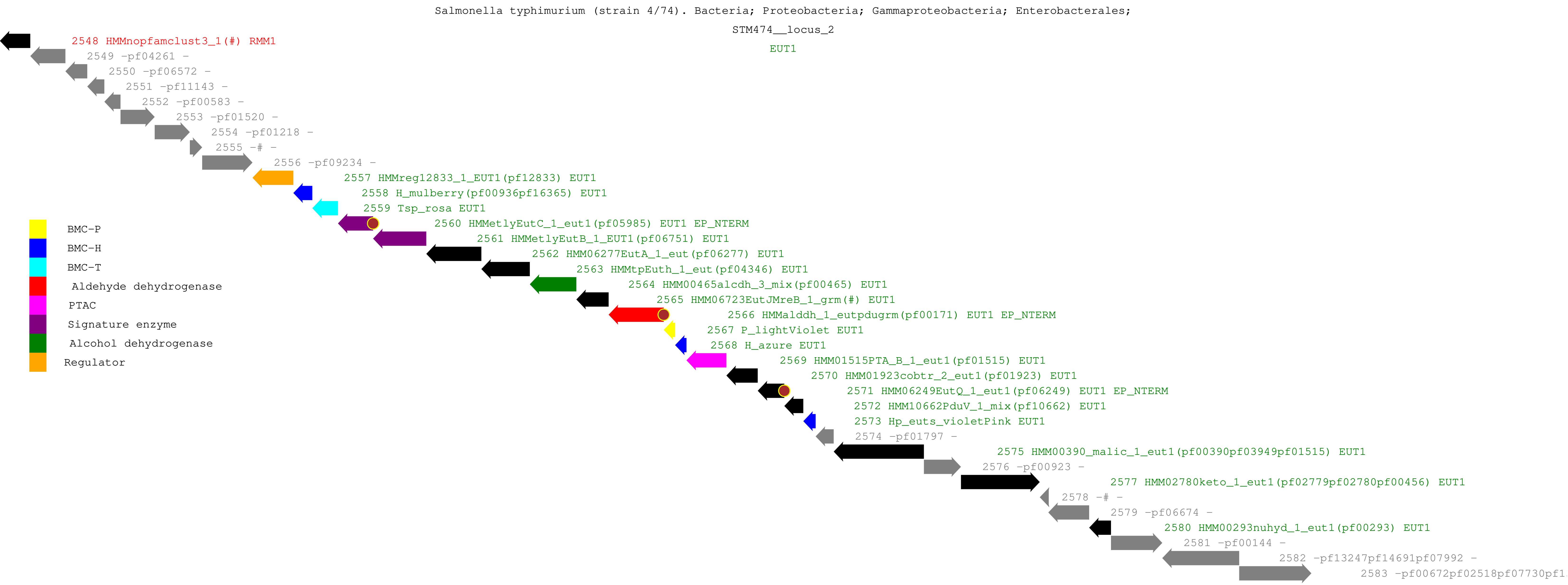

Salmonella typhimurium (strain 4/74). Bacteria; Proteobacteria; Gammaproteobacteria; Enterobacterales;
STM474__locus_2
EUT1
2548 HMMnopfamclust3_1(#) RMM1
2549 -pf04261 -
2550 -pf06572 -
2551 -pf11143 -
2552 -pf00583 -
2553 -pf01520 -
2554 -pf01218 -
2555 -# -
2556 -pf09234 -
2557 HMMreg12833_1_EUT1(pf12833) EUT1
2558 H_mulberry(pf00936pf16365) EUT1
2559 Tsp_rosa EUT1
2560 HMMetlyEutC_1_eut1(pf05985) EUT1 EP_NTERM
2561 HMMetlyEutB_1_EUT1(pf06751) EUT1
2562 HMM06277EutA_1_eut(pf06277) EUT1
2563 HMMtpEuth_1_eut(pf04346) EUT1
2564 HMM00465alcdh_3_mix(pf00465) EUT1
2565 HMM06723EutJMreB_1_grm(#) EUT1
2566 HMMalddh_1_eutpdugrm(pf00171) EUT1 EP_NTERM
2567 P_lightViolet EUT1
2568 H_azure EUT1
2569 HMM01515PTA_B_1_eut1(pf01515) EUT1
2570 HMM01923cobtr_2_eut1(pf01923) EUT1
2571 HMM06249EutQ_1_eut1(pf06249) EUT1 EP_NTERM
2572 HMM10662PduV_1_mix(pf10662) EUT1
2573 Hp_euts_violetPink EUT1
2574 -pf01797 -
2575 HMM00390_malic_1_eut1(pf00390pf03949pf01515) EUT1
2576 -pf00923 -
2577 HMM02780keto_1_eut1(pf02779pf02780pf00456) EUT1
2578 -# -
2579 -pf06674 -
2580 HMM00293nuhyd_1_eut1(pf00293) EUT1
2581 -pf00144 -
2582 -pf13247pf14691pf07992 -
2583 -pf00672pf02518pf07730pf1
BMC-P
BMC-H
BMC-T
Aldehyde dehydrogenase
PTAC
Signature enzyme
Alcohol dehydrogenase
Regulator

Salmonella typhimurium (strain LT2 / SGSC1412 / ATCC 700720). Bacteria; Proteobacteria; Gammaproteobacteria; Enterobacterales;

STM_locus_2

EUT1

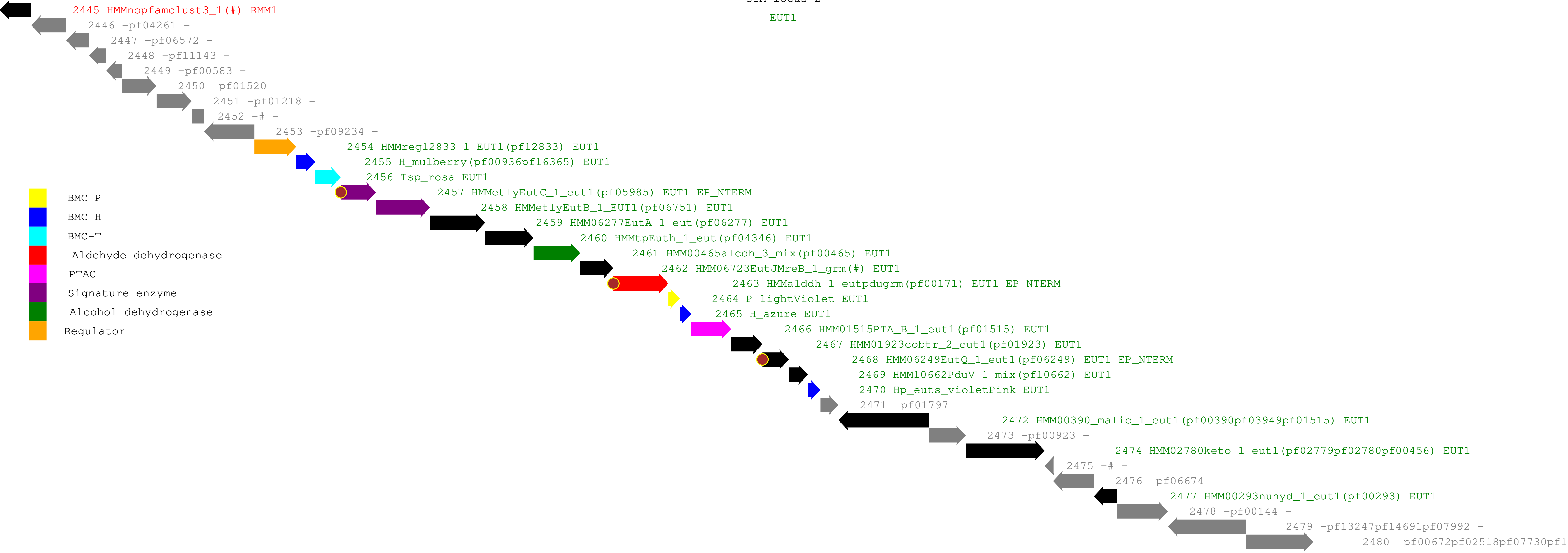

Salmonella enterica I. Bacteria; Proteobacteria; Gammaproteobacteria; Enterobacterales;

TL61__locus_2

EUT1

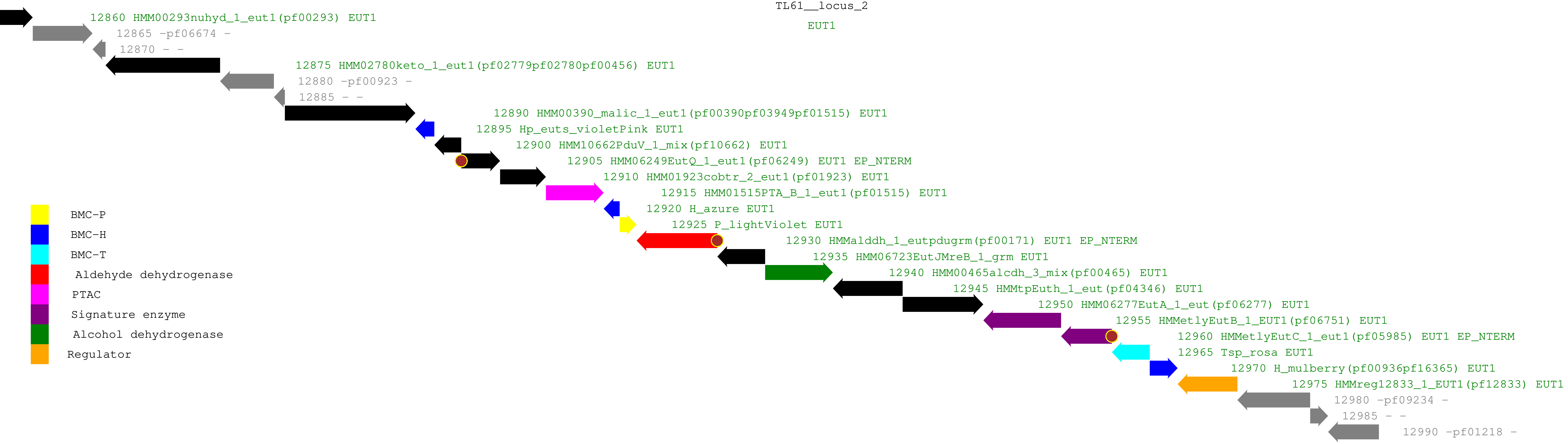

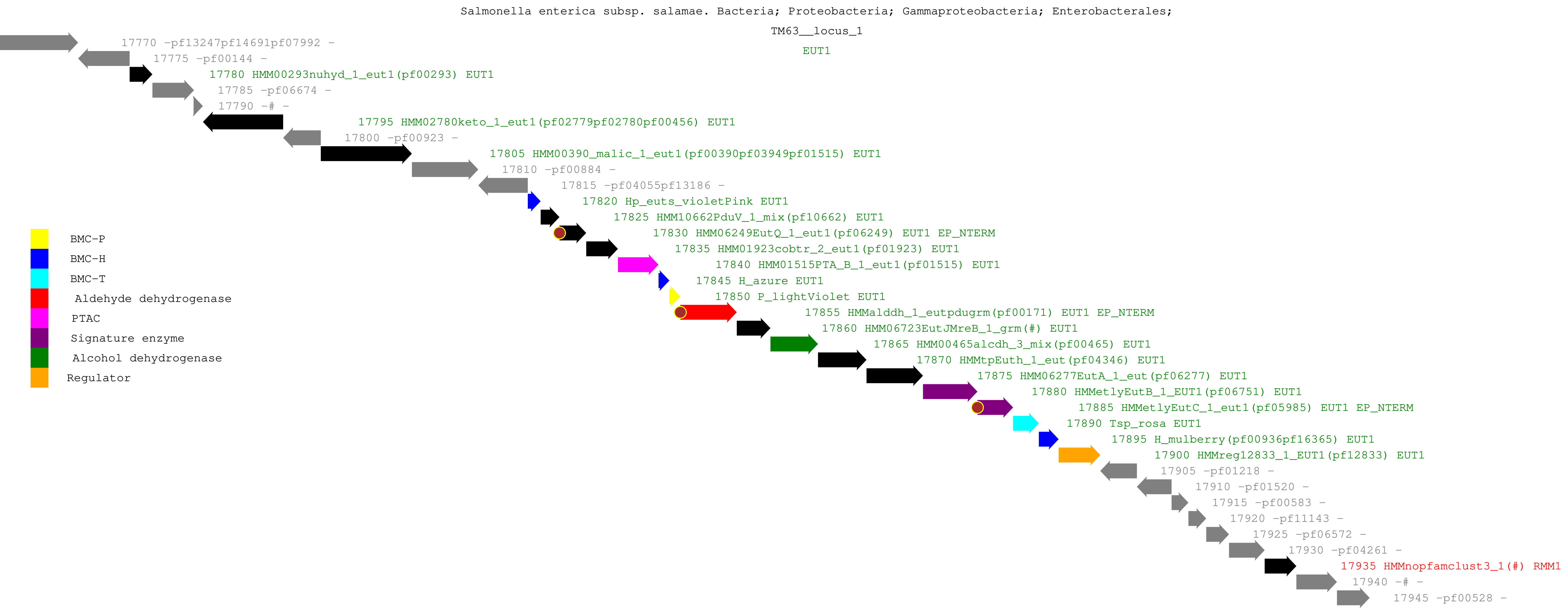
Salmonella enterica subsp. salamae. Bacteria; Proteobacteria; Gammaproteobacteria; Enterobacterales;
TM63__locus_1
EUT1
17770 -pf13247pf14691pf07992 -
17775 -pf00144 -
17780 HMM00293nuhyd_1_eut1(pf00293) EUT1
17785 -pf06674 -
17790 -# -
17795 HMM02780keto_1_eut1(pf02779pf02780pf00456) EUT1
17800 -pf00923 -
17805 HMM00390_malic_1_eut1(pf00390pf03949pf01515) EUT1
17810 -pf00884 -
17815 -pf04055pf13186 -
17820 Hp_euts_violetPink EUT1
17825 HMM10662PduV_1_mix(pf10662) EUT1
17830 HMM06249EutQ_1_eut1(pf06249) EUT1 EP_NTERM
17835 HMM01923cobtr_2_eut1(pf01923) EUT1
17840 HMM01515PTA_B_1_eut1(pf01515) EUT1
17845 H_azure EUT1
17850 P_lightViolet EUT1
17855 HMMalddh_1_eutpdugrm(pf00171) EUT1 EP_NTERM
17860 HMM06723EutJMreB_1_grm(#) EUT1
17865 HMM00465alcdh_3_mix(pf00465) EUT1
17870 HMMtpEuth_1_eut(pf04346) EUT1
17875 HMM06277EutA_1_eut(pf06277) EUT1
17880 HMMetlyEutB_1_EUT1(pf06751) EUT1
17885 HMMetlyEutC_1_eut1(pf05985) EUT1 EP_NTERM
17890 Tsp_rosa EUT1
17895 H_mulberry(pf00936pf16365) EUT1
17900 HMMreg12833_1_EUT1(pf12833) EUT1
17905 -pf01218 -
17910 -pf01520 -
17915 -pf00583 -
17920 -pf11143 -
17925 -pf06572 -
17930 -pf04261 -
17935 HMMnopfamclust3_1(#) RMM1
17940 -# -
17945 -pf00528 -
BMC-P
BMC-H
BMC-T
Aldehyde dehydrogenase
PTAC
Signature enzyme
Alcohol dehydrogenase
Regulator

Escherichia coli TW10598. Bacteria; Proteobacteria; Gammaproteobacteria; Enterobacterales;

TW10598__locus_1

EUT1

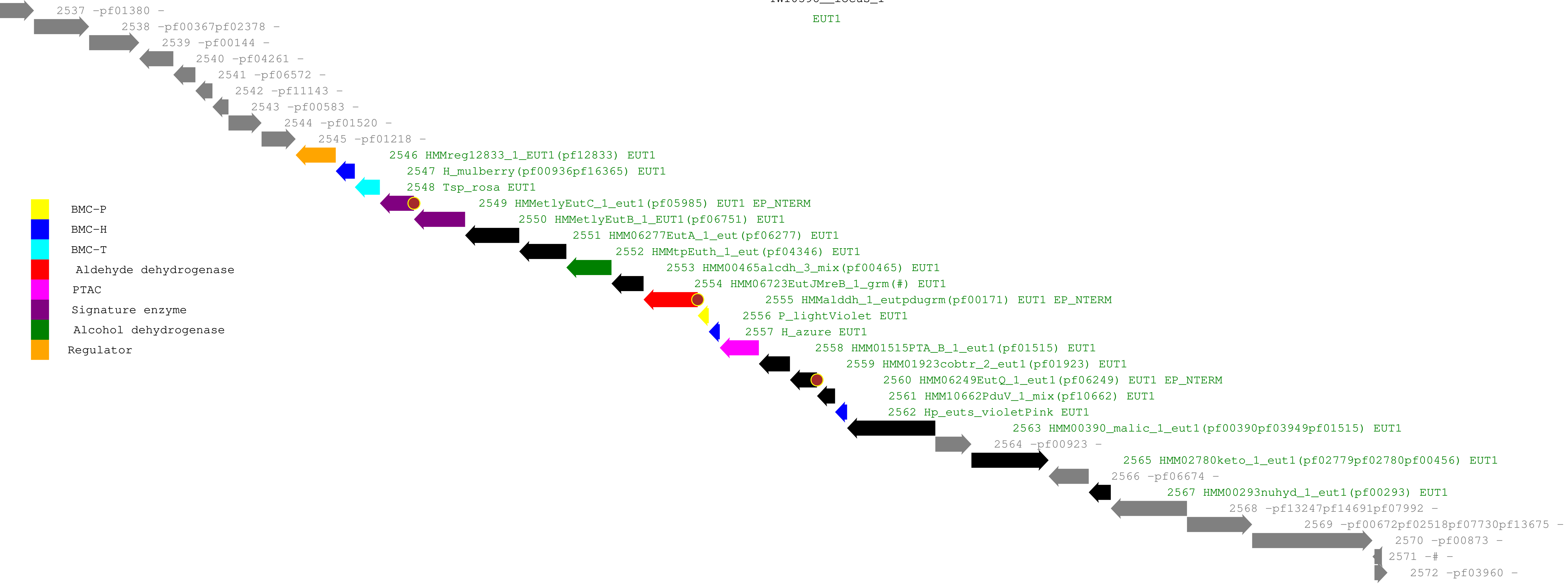

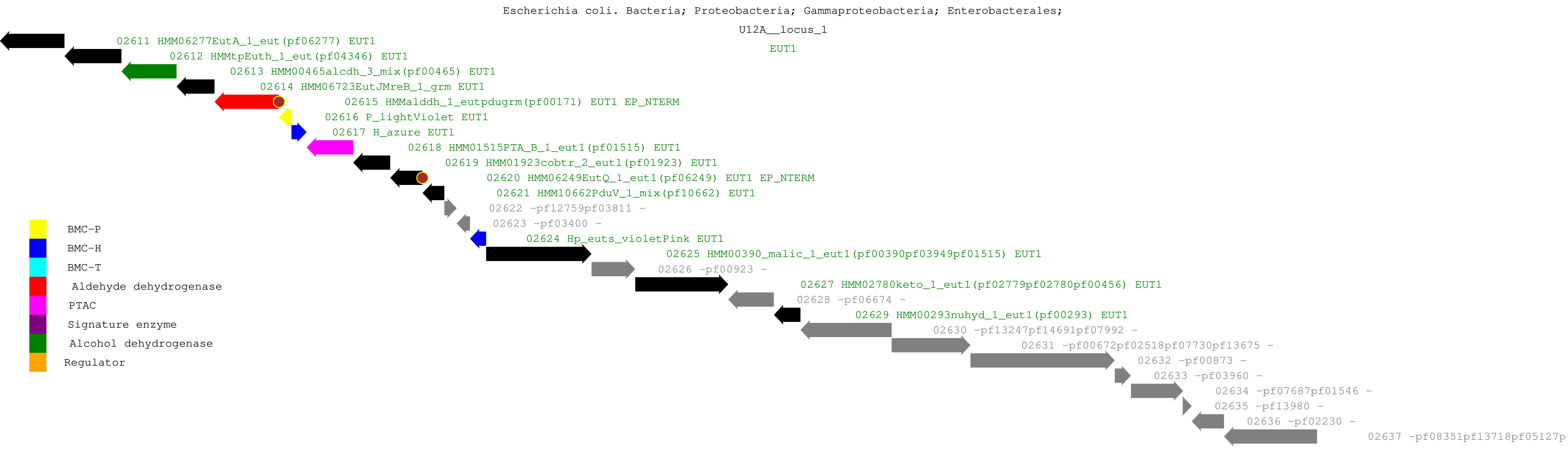
Escherichia coli. Bacteria; Proteobacteria; Gammaproteobacteria; Enterobacterales;
U12A__locus_1
EUT1
02611 HMM06277EutA_1_eut(pf06277) EUT1
02612 HMMtpEuth_1_eut(pf04346) EUT1
02613 HMM00465alcdh_3_mix(pf00465) EUT1
02614 HMM06723EutJMreB_1_grm EUT1
02615 HMMaldh_1_eutpdugrm(pf00171) EUT1 EP_NTERM
02616 P_lightViolet EUT1
02617 H_azure EUT1
02618 HMM01515PTA_B_1_eut1(pf01515) EUT1
02619 HMM01923cobtr_2_eut1(pf01923) EUT1
02620 HMM06249EutQ_1_eut1(pf06249) EUT1 EP_NTERM
02621 HMM10662PduV_1_mix(pf10662) EUT1
02622 -pf12759pf03811 -
02623 -pf03400 -
02624 Hp_euts_violetPink EUT1
02625 HMM00390_malic_1_eut1(pf00390pf03949pf01515) EUT1
02626 -pf00923 -
02627 HMM02780keto_1_eut1(pf02779pf02780pf00456) EUT1
02628 -pf06674 -
02629 HMM00293nuhyd_1_eut1(pf00293) EUT1
02630 -pf13247pf14691pf07992 -
02631 -pf00672pf02518pf07730pf13675 -
02632 -pf00873 -
02633 -pf03960 -
02634 -pf07687pf01546 -
02635 -pf13980 -
02636 -pf02230 -
02637 -pf08351pf13718pf05127p
BMC-P
BMC-H
BMC-T
Aldehyde dehydrogenase
PTAC
Signature enzyme
Alcohol dehydrogenase
Regulator

Escherichia coli O25b:H4-ST131. Bacteria; Proteobacteria; Gammaproteobacteria; Enterobacterales;

UG58__locus_1

EUT1

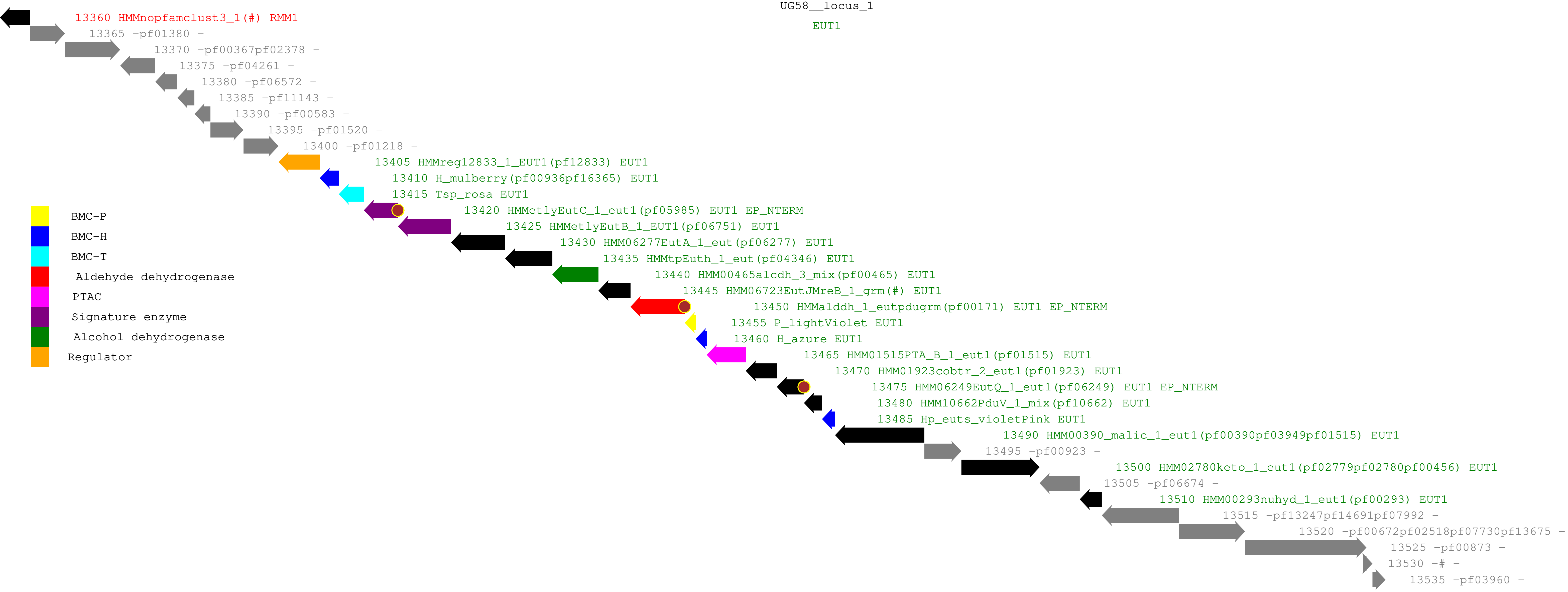

Escherichia coli UMNF18. Bacteria; Proteobacteria; Gammaproteobacteria; Enterobacterales;

UMNF18__locus_1

EUT1

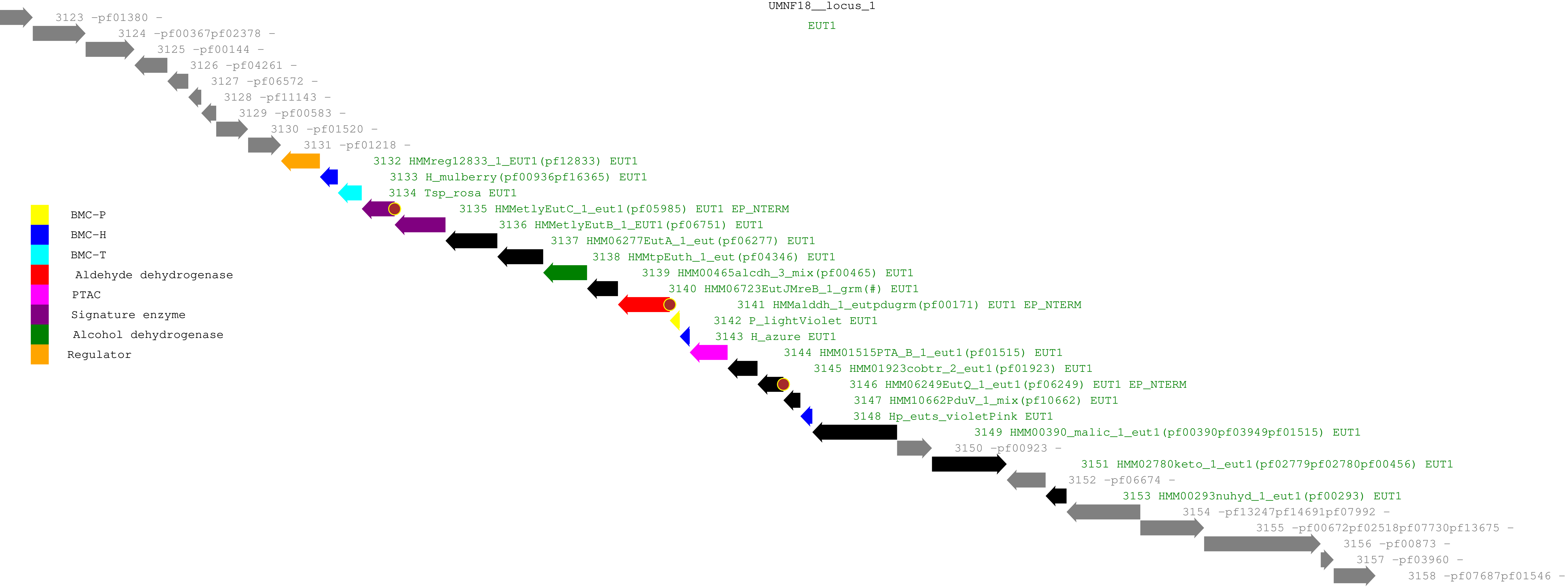

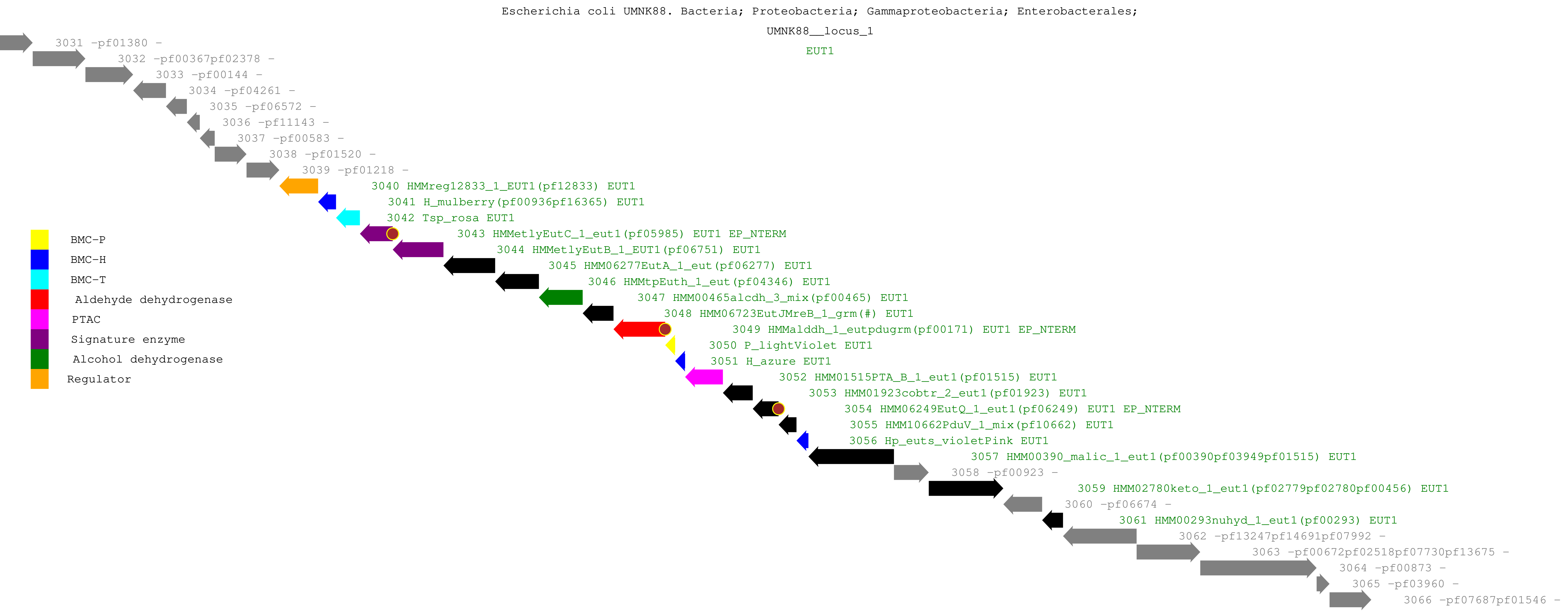
Escherichia coli UMNK88. Bacteria; Proteobacteria; Gammaproteobacteria; Enterobacterales;
UMNK88__locus_1
EUT1
3031 -pf01380 -
3032 -pf00367pf02378 -
3033 -pf00144 -
3034 -pf04261 -
3035 -pf06572 -
3036 -pf11143 -
3037 -pf00583 -
3038 -pf01520 -
3039 -pf01218 -
3040 HMMreg12833_1_EUT1(pf12833) EUT1
3041 H_mulberry(pf00936pf16365) EUT1
3042 Tsp_rosa EUT1
3043 HMMetlyEutC_1_eut1(pf05985) EUT1 EP_NTERM
3044 HMMetlyEutB_1_EUT1(pf06751) EUT1
3045 HMM06277EutA_1_eut(pf06277) EUT1
3046 HMMtpEuth_1_eut(pf04346) EUT1
3047 HMM00465alcdh_3_mix(pf00465) EUT1
3048 HMM06723EutJMreB_1_grm(#) EUT1
3049 HMMalddh_1_eutpdugrm(pf00171) EUT1 EP_NTERM
3050 P_lightViolet EUT1
3051 H_azure EUT1
3052 HMM01515PTA_B_1_eut1(pf01515) EUT1
3053 HMM01923cobtr_2_eut1(pf01923) EUT1
3054 HMM06249EutQ_1_eut1(pf06249) EUT1 EP_NTERM
3055 HMM10662PduV_1_mix(pf10662) EUT1
3056 Hp_euts_violetPink EUT1
3057 HMM00390_malic_1_eut1(pf00390pf03949pf01515) EUT1
3058 -pf00923 -
3059 HMM02780keto_1_eut1(pf02779pf02780pf00456) EUT1
3060 -pf06674 -
3061 HMM00293nuhyd_1_eut1(pf00293) EUT1
3062 -pf13247pf14691pf07992 -
3063 -pf00672pf02518pf07730pf13675 -
3064 -pf00873 -
3065 -pf03960 -
3066 -pf07687pf01546 -
BMC-P
BMC-H
BMC-T
Aldehyde dehydrogenase
PTAC
Signature enzyme
Alcohol dehydrogenase
Regulator

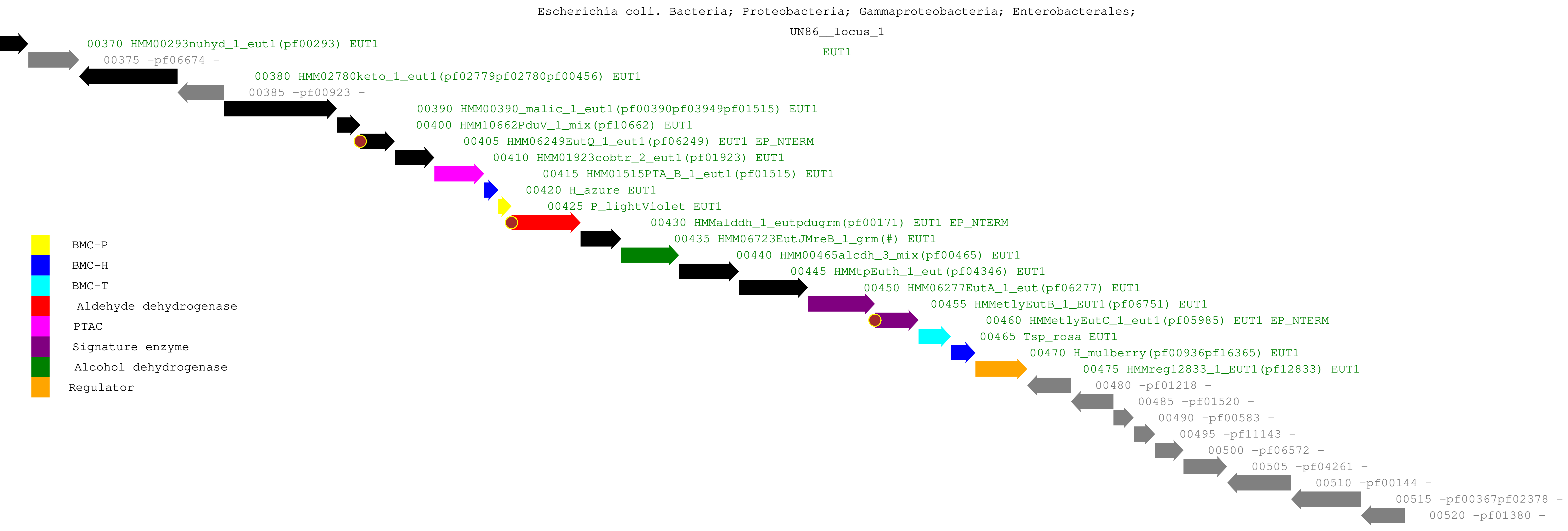
Escherichia coli. Bacteria; Proteobacteria; Gammaproteobacteria; Enterobacterales;
UN86__locus_1
EUT1
00370 HMM00293nuhyd_1_eut1(pf00293) EUT1
00375 -pf06674 -
00380 HMM02780keto_1_eut1(pf02779pf02780pf00456) EUT1
00385 -pf00923 -
00390 HMM00390_malic_1_eut1(pf00390pf03949pf01515) EUT1
00400 HMM10662PduV_1_mix(pf10662) EUT1
00405 HMM06249EutQ_1_eut1(pf06249) EUT1 EP_NTERM
00410 HMM01923cobtr_2_eut1(pf01923) EUT1
00415 HMM01515PTA_B_1_eut1(pf01515) EUT1
00420 H_azure EUT1
00425 P_lightViolet EUT1
00430 HMMalddh_1_eutpdugrm(pf00171) EUT1 EP_NTERM
00435 HMM06723EutJMreB_1_grm(#) EUT1
00440 HMM00465alcdh_3_mix(pf00465) EUT1
00445 HMMtpEuth_1_eut(pf04346) EUT1
00450 HMM06277EutA_1_eut(pf06277) EUT1
00455 HMMetlyEutB_1_EUT1(pf06751) EUT1
00460 HMMetlyEutC_1_eut1(pf05985) EUT1 EP_NTERM
00465 Tsp_rosa EUT1
00470 H_mulberry(pf00936pf16365) EUT1
00475 HMMreg12833_1_EUT1(pf12833) EUT1
00480 -pf01218 -
00485 -pf01520 -
00490 -pf00583 -
00495 -pf11143 -
00500 -pf06572 -
00505 -pf04261 -
00510 -pf00144 -
00515 -pf00367pf02378 -
00520 -pf01380 -
BMC-P
BMC-H
BMC-T
Aldehyde dehydrogenase
PTAC
Signature enzyme
Alcohol dehydrogenase
Regulator

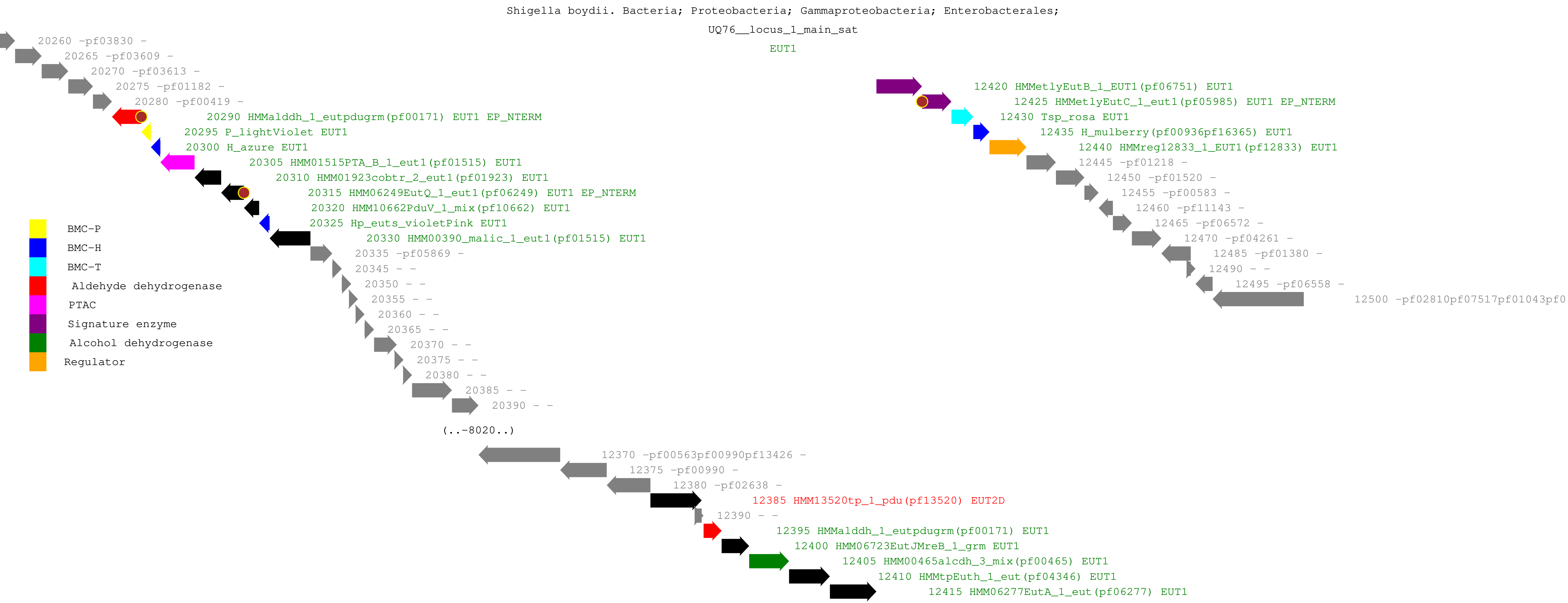

Shigella boydii. Bacteria; Proteobacteria; Gammaproteobacteria; Enterobacterales;
UQ76__locus_1_main_sat
EUT1
20260 -pf03830 -
20265 -pf03609 -
20270 -pf03613 -
20275 -pf01182 -
20280 -pf00419 -
20290 HMMalddh_1_eutpdugrm(pf00171) EUT1 EP_NTERM
20295 P_lightViolet EUT1
20300 H_azure EUT1
20305 HMM01515PTA_B_1_eut1(pf01515) EUT1
20310 HMM01923cobtr_2_eut1(pf01923) EUT1
20315 HMM06249EutQ_1_eut1(pf06249) EUT1 EP_NTERM
20320 HMM10662PduV_1_mix(pf10662) EUT1
20325 Hp_euts_violetPink EUT1
20330 HMM00390_malic_1_eut1(pf01515) EUT1
20335 -pf05869 -
20345 - -
20350 - -
20355 - -
20360 - -
20365 - -
20370 - -
20375 - -
20380 - -
20385 - -
20390 - -
(..-8020..)
12370 -pf00563pf00990pf13426 -
12375 -pf00990 -
12380 -pf02638 -
12385 HMM13520tp_1_pdu(pf13520) EUT2D
12390 - -
12395 HMMalddh_1_eutpdugrm(pf00171) EUT1
12400 HMM06723EutJMreB_1_grm EUT1
12405 HMM00465alcdh_3_mix(pf00465) EUT1
12410 HMMtpEuth_1_eut(pf04346) EUT1
12415 HMM06277EutA_1_eut(pf06277) EUT1
12420 HMMetlyEutB_1_EUT1(pf06751) EUT1
12425 HMMetlyEutC_1_eut1(pf05985) EUT1 EP_NTERM
12430 Tsp_rosa EUT1
12435 H_mulberry(pf00936pf16365) EUT1
12440 HMMreg12833_1_EUT1(pf12833) EUT1
12445 -pf01218 -
12450 -pf01520 -
12455 -pf00583 -
12460 -pf11143 -
12465 -pf06572 -
12470 -pf04261 -
12485 -pf01380 -
12490 - -
12495 -pf06558 -
12500 -pf02810pf07517pf01043pf0
BMC-P
BMC-H
BMC-T
Aldehyde dehydrogenase
PTAC
Signature enzyme
Alcohol dehydrogenase
Regulator

Escherichia coli (strain UTI89 / UPEC). Bacteria; Proteobacteria; Gammaproteobacteria; Enterobacterales;

UTI89_C_locus_1

EUT1

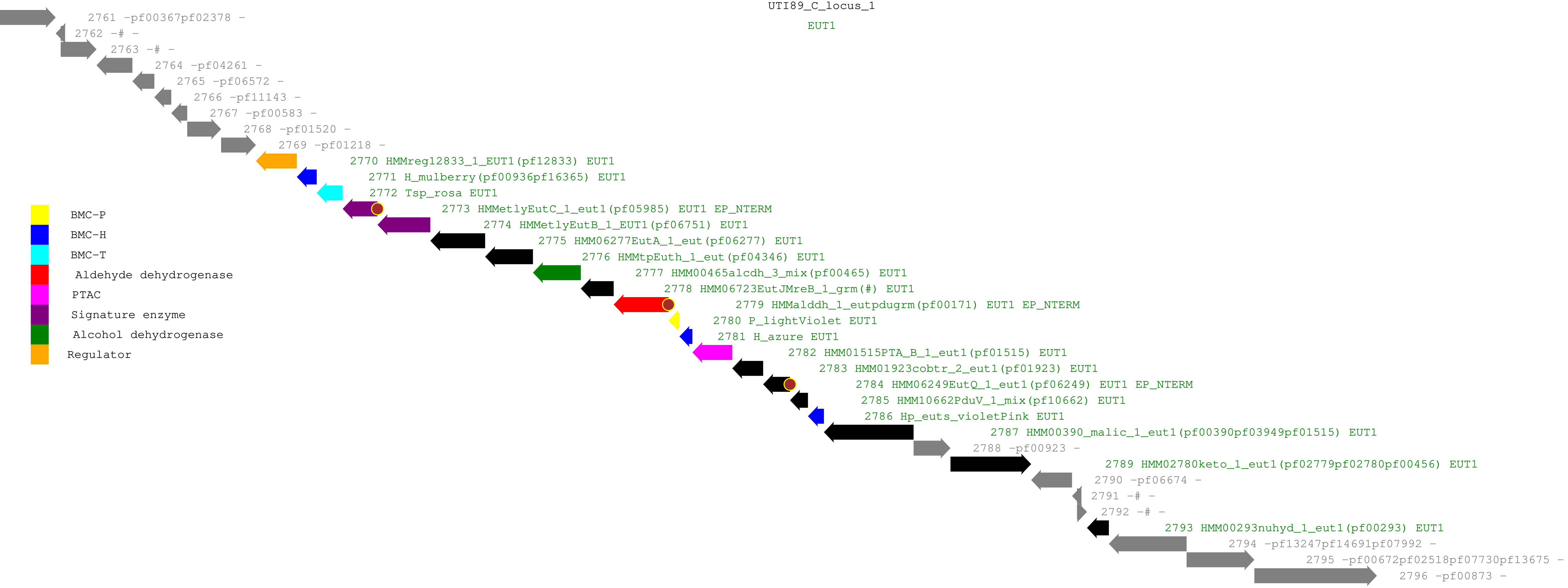

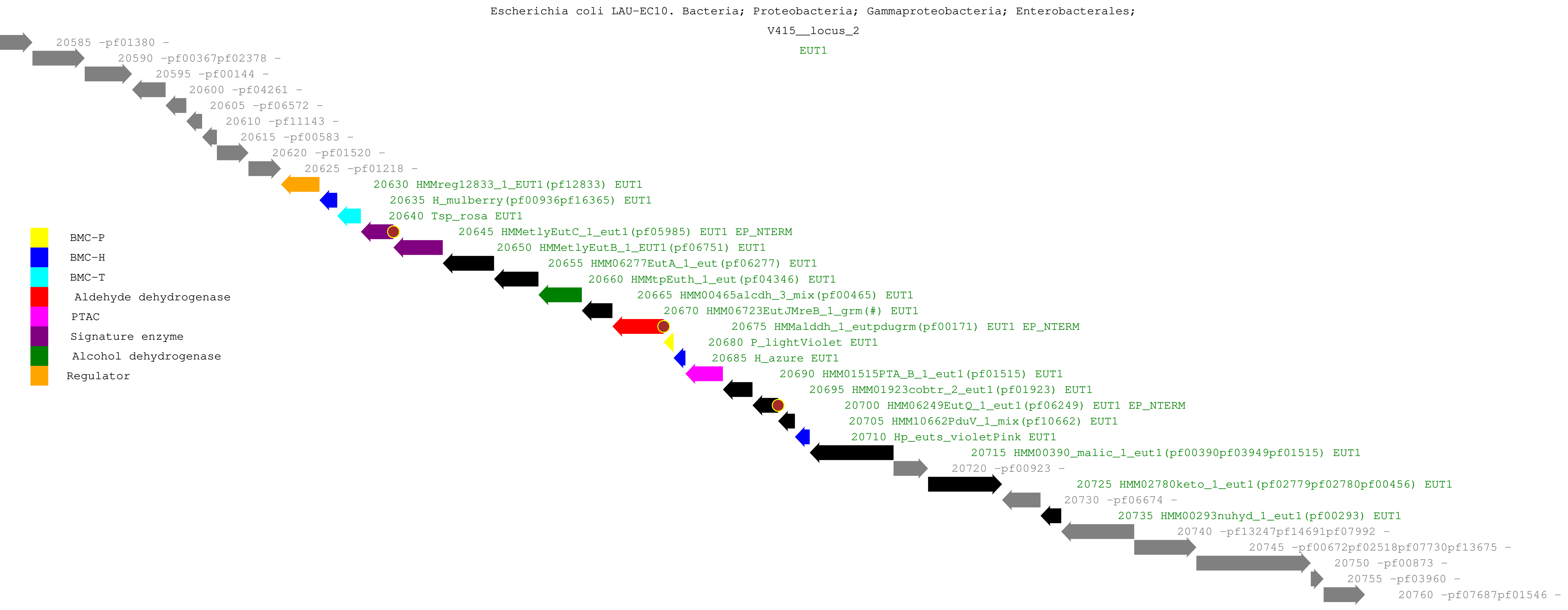
Escherichia coli LAU-EC10. Bacteria; Proteobacteria; Gammaproteobacteria; Enterobacterales;
V415__locus_2
EUT1
20585 -pf01380 -
20590 -pf00367pf02378 -
20595 -pf00144 -
20600 -pf04261 -
20605 -pf06572 -
20610 -pf11143 -
20615 -pf00583 -
20620 -pf01520 -
20625 -pf01218 -
20630 HMMreg12833_1_EUT1(pf12833) EUT1
20635 H_mulberry(pf00936pf16365) EUT1
20640 Tsp_rosa EUT1
20645 HMMetlyEutC_1_eut1(pf05985) EUT1 EP_NTERM
20650 HMMetlyEutB_1_EUT1(pf06751) EUT1
20655 HMM06277EutA_1_eut(pf06277) EUT1
20660 HMMtpEuth_1_eut(pf04346) EUT1
20665 HMM00465alcdh_3_mix(pf00465) EUT1
20670 HMM06723EutJMreB_1_grm(#) EUT1
20675 HMMalddh_1_eutpdugrm(pf00171) EUT1 EP_NTERM
20680 P_lightViolet EUT1
20685 H_azure EUT1
20690 HMM01515PTA_B_1_eut1(pf01515) EUT1
20695 HMM01923cobtr_2_eut1(pf01923) EUT1
20700 HMM06249EutQ_1_eut1(pf06249) EUT1 EP_NTERM
20705 HMM10662PduV_1_mix(pf10662) EUT1
20710 Hp_euts_violetPink EUT1
20715 HMM00390_malic_1_eut1(pf00390pf03949pf01515) EUT1
20720 -pf00923 -
20725 HMM02780keto_1_eut1(pf02779pf02780pf00456) EUT1
20730 -pf06674 -
20735 HMM00293nuhyd_1_eut1(pf00293) EUT1
20740 -pf13247pf14691pf07992 -
20745 -pf00672pf02518pf07730pf13675 -
20750 -pf00873 -
20755 -pf03960 -
20760 -pf07687pf01546 -
BMC-P
BMC-H
BMC-T
Aldehyde dehydrogenase
PTAC
Signature enzyme
Alcohol dehydrogenase
Regulator

Chromobacterium sp. LK1. Bacteria; Proteobacteria; Betaproteobacteria; Neisseriales;

VI26__locus_1

EUT1

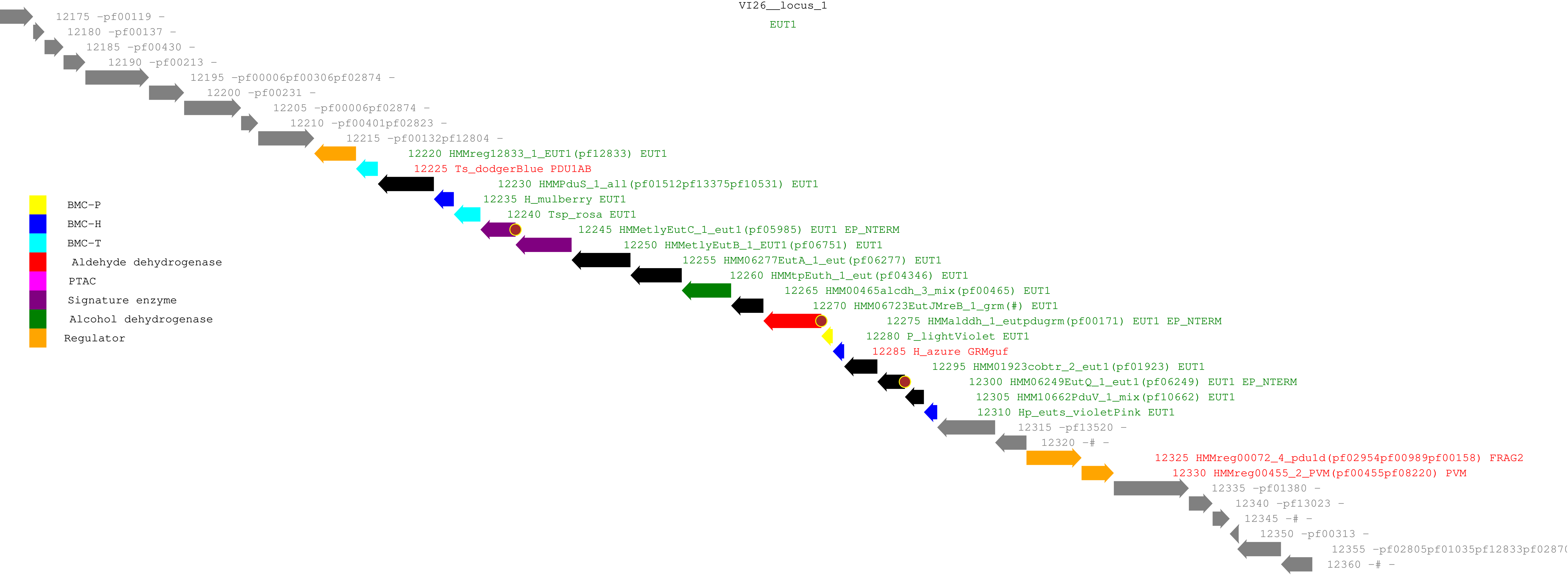

Chromobacterium sp. LK11. Bacteria; Proteobacteria; Betaproteobacteria; Neisseriales;

VK98__locus_1

EUT1

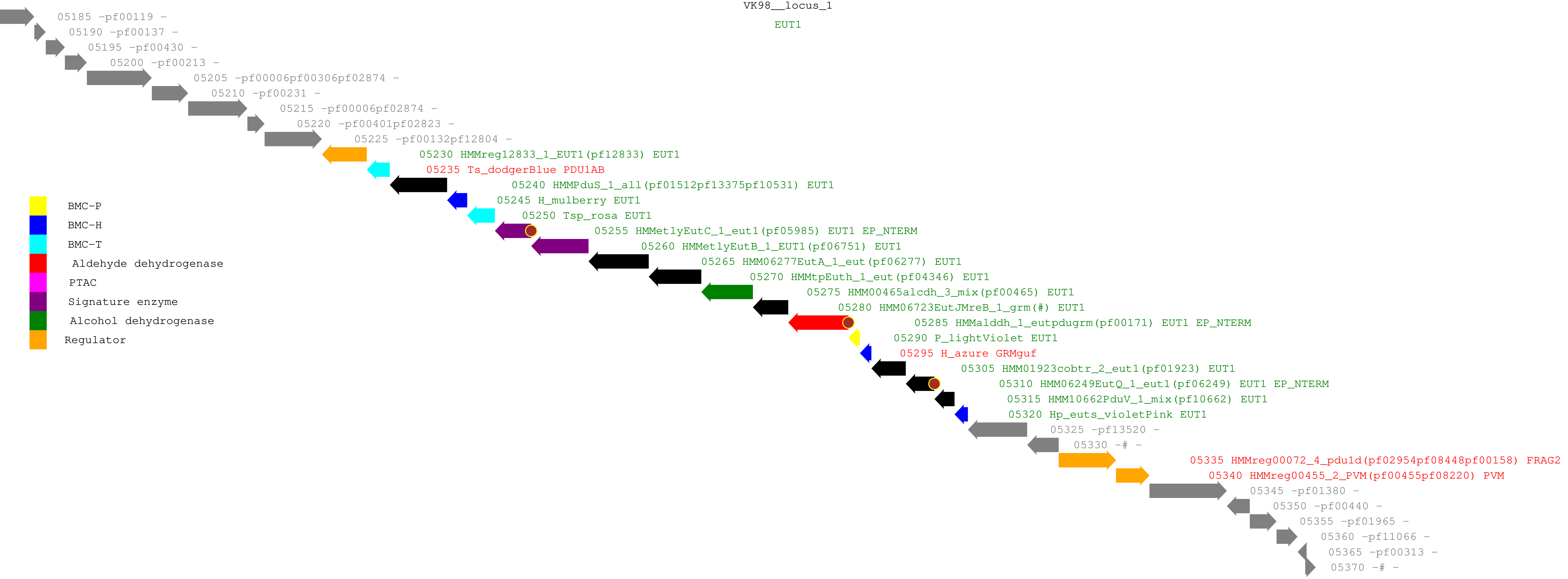

Escherichia sp. KTE114. Bacteria; Proteobacteria; Gammaproteobacteria; Enterobacterales;

WC5__locus_2

EUT1

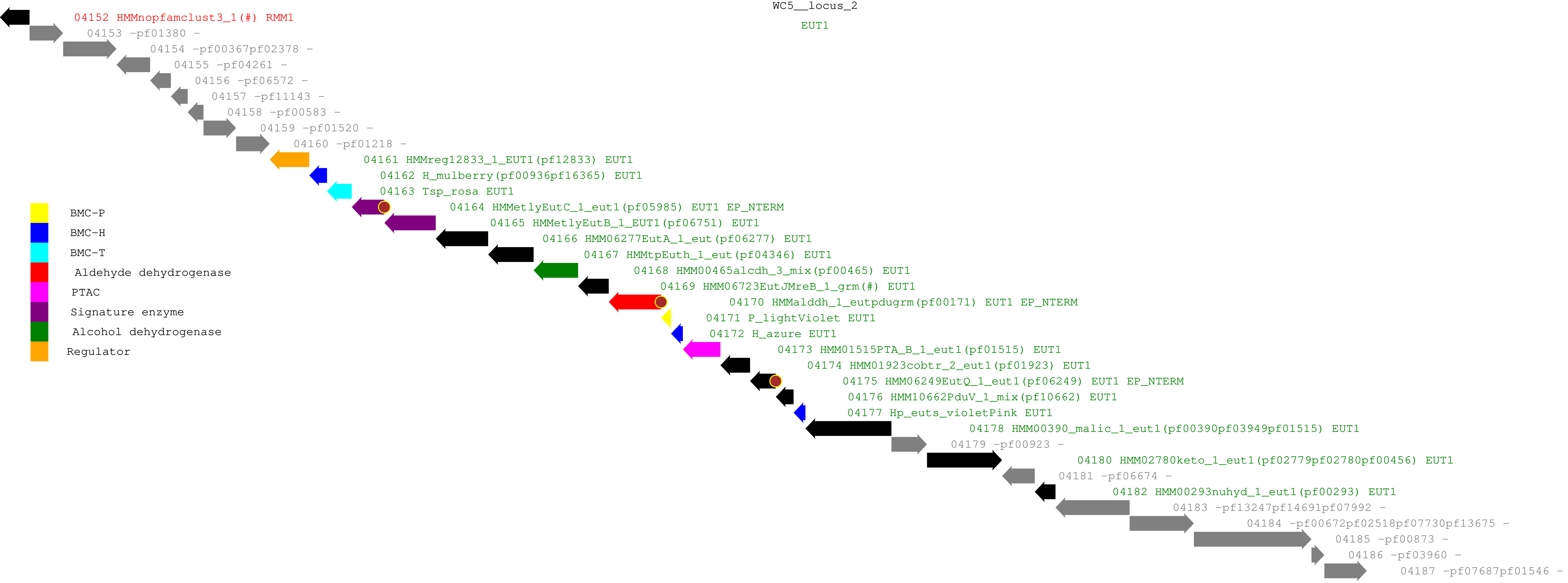

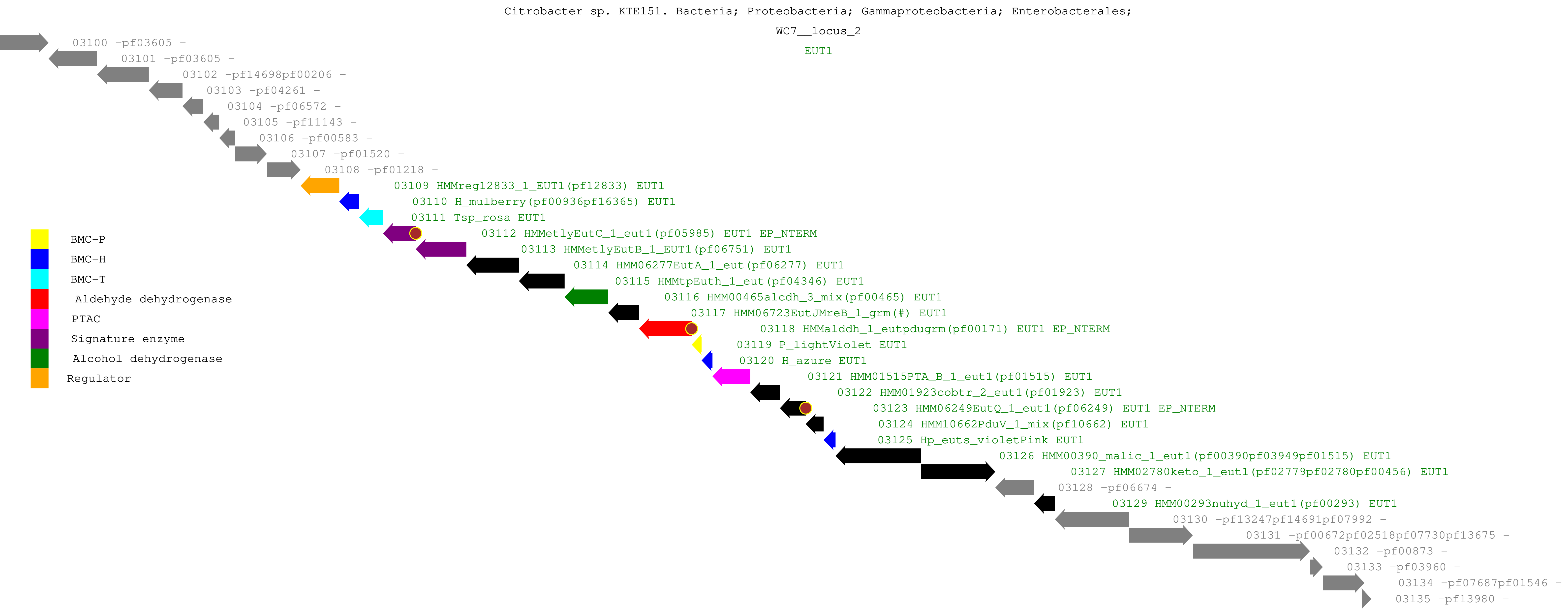
Citrobacter sp. KTE151. Bacteria; Proteobacteria; Gammaproteobacteria; Enterobacterales;
WC7__locus_2
EUT1
03100 -pf03605 -
03101 -pf03605 -
03102 -pf14698pf00206 -
03103 -pf04261 -
03104 -pf06572 -
03105 -pf11143 -
03106 -pf00583 -
03107 -pf01520 -
03108 -pf01218 -
03109 HMMreg12833_1_EUT1(pf12833) EUT1
03110 H_mulberry(pf00936pf16365) EUT1
03111 Tsp_rosa EUT1
03112 HMMetlyEutC_1_eut1(pf05985) EUT1 EP_NTERM
03113 HMMetlyEutB_1_EUT1(pf06751) EUT1
03114 HMM06277EutA_1_eut(pf06277) EUT1
03115 HMMtpEuth_1_eut(pf04346) EUT1
03116 HMM00465alcdh_3_mix(pf00465) EUT1
03117 HMM06723EutJMreB_1_grm(#) EUT1
03118 HMMalddh_1_eutpdugrm(pf00171) EUT1 EP_NTERM
03119 P_lightViolet EUT1
03120 H_azure EUT1
03121 HMM01515PTA_B_1_eut1(pf01515) EUT1
03122 HMM01923cobtr_2_eut1(pf01923) EUT1
03123 HMM06249EutQ_1_eut1(pf06249) EUT1 EP_NTERM
03124 HMM10662PduV_1_mix(pf10662) EUT1
03125 Hp_euts_violetPink EUT1
03126 HMM00390_malic_1_eut1(pf00390pf03949pf01515) EUT1
03127 HMM02780keto_1_eut1(pf02779pf02780pf00456) EUT1
03128 -pf06674 -
03129 HMM00293nuhyd_1_eut1(pf00293) EUT1
03130 -pf13247pf14691pf07992 -
03131 -pf00672pf02518pf07730pf13675 -
03132 -pf00873 -
03133 -pf03960 -
03134 -pf07687pf01546 -
03135 -pf13980 -
BMC-P
BMC-H
BMC-T
Aldehyde dehydrogenase
PTAC
Signature enzyme
Alcohol dehydrogenase
Regulator

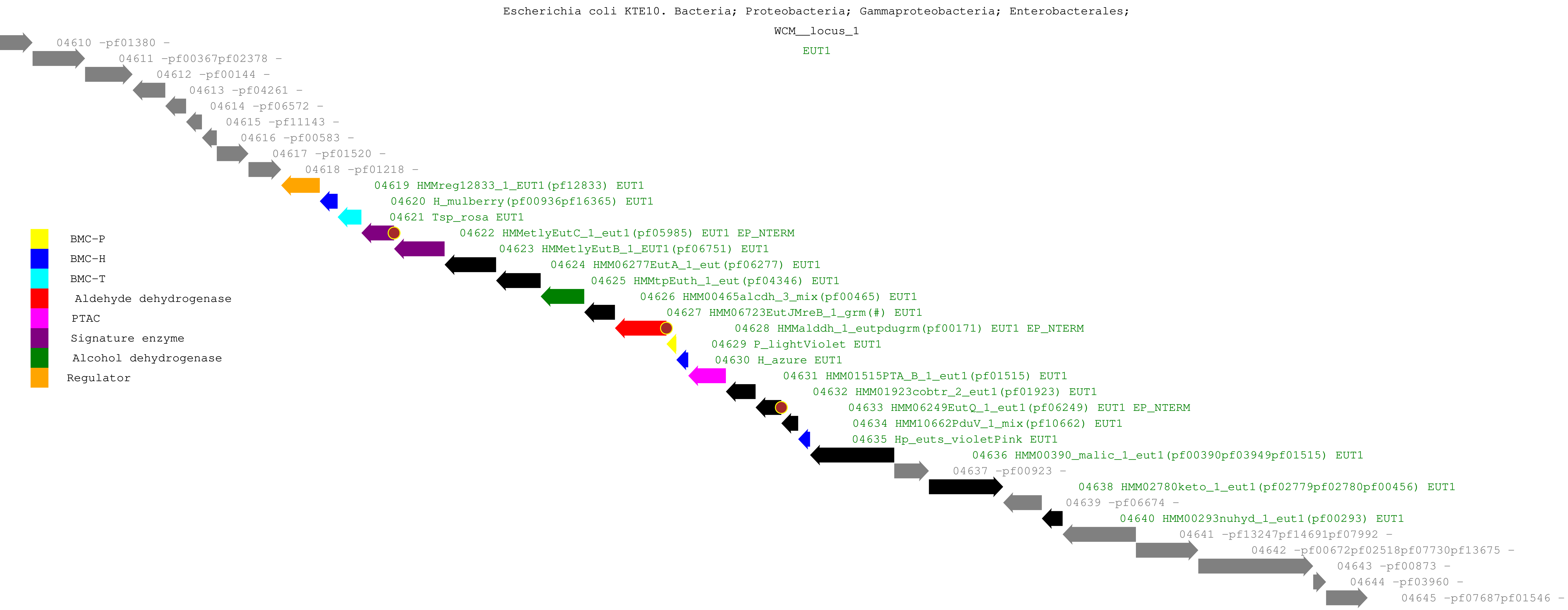

Escherichia coli KTE10. Bacteria; Proteobacteria; Gammaproteobacteria; Enterobacterales;
WCM__locus_1
EUT1
04610 -pf01380 -
04611 -pf00367pf02378 -
04612 -pf00144 -
04613 -pf04261 -
04614 -pf06572 -
04615 -pf11143 -
04616 -pf00583 -
04617 -pf01520 -
04618 -pf01218 -
04619 HMMreg12833_1_EUT1(pf12833) EUT1
04620 H_mulberry(pf00936pf16365) EUT1
04621 Tsp_rosa EUT1
04622 HMMetlyEutC_1_eut1(pf05985) EUT1 EP_NTERM
04623 HMMetlyEutB_1_EUT1(pf06751) EUT1
04624 HMM06277EutA_1_eut(pf06277) EUT1
04625 HMMtpEuth_1_eut(pf04346) EUT1
04626 HMM00465alcdh_3_mix(pf00465) EUT1
04627 HMM06723EutJMreB_1_grm(#) EUT1
04628 HMMalddh_1_eutpdugrm(pf00171) EUT1 EP_NTERM
04629 P_lightViolet EUT1
04630 H_azure EUT1
04631 HMM01515PTA_B_1_eut1(pf01515) EUT1
04632 HMM01923cobtr_2_eut1(pf01923) EUT1
04633 HMM06249EutQ_1_eut1(pf06249) EUT1 EP_NTERM
04634 HMM10662PduV_1_mix(pf10662) EUT1
04635 Hp_euts_violetPink EUT1
04636 HMM00390_malic_1_eut1(pf00390pf03949pf01515) EUT1
04637 -pf00923 -
04638 HMM02780keto_1_eut1(pf02779pf02780pf00456) EUT1
04639 -pf06674 -
04640 HMM00293nuhyd_1_eut1(pf00293) EUT1
04641 -pf13247pf14691pf07992 -
04642 -pf00672pf02518pf07730pf13675 -
04643 -pf00873 -
04644 -pf03960 -
04645 -pf07687pf01546 -
BMC-P
BMC-H
BMC-T
Aldehyde dehydrogenase
PTAC
Signature enzyme
Alcohol dehydrogenase
Regulator

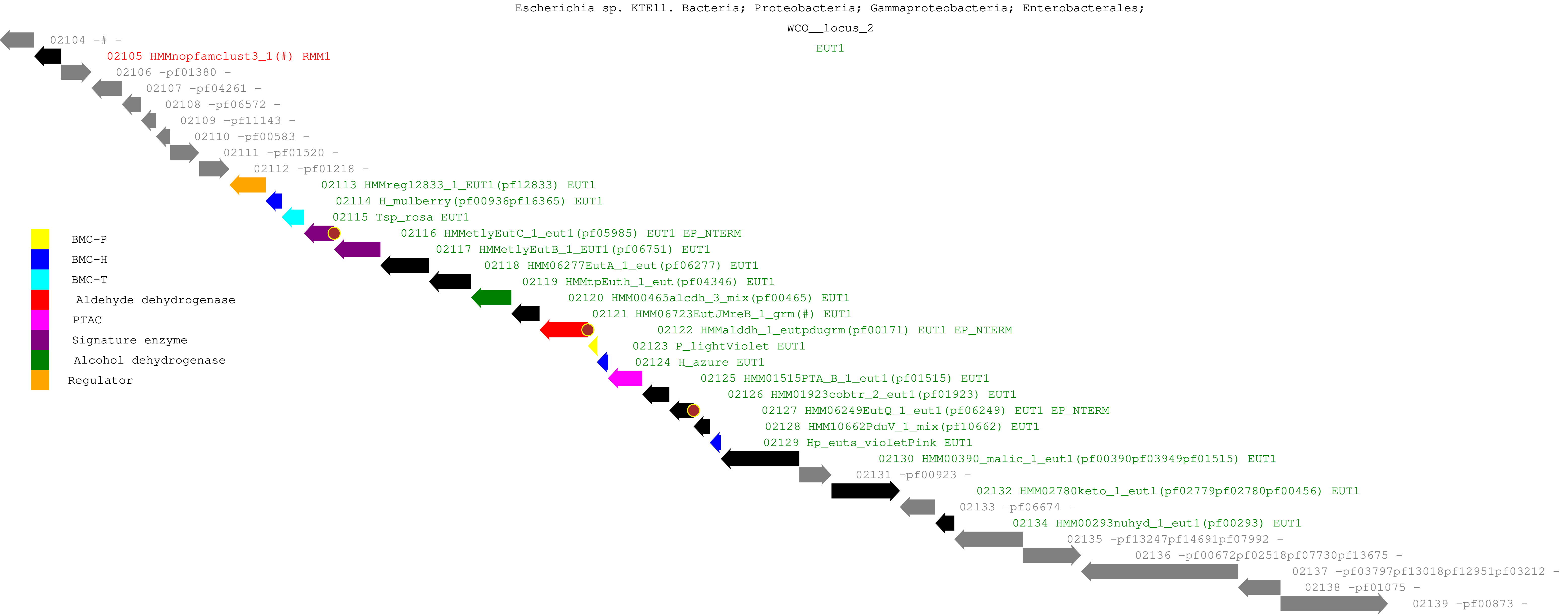
Escherichia sp. KTE11. Bacteria; Proteobacteria; Gammaproteobacteria; Enterobacterales;
WCO__locus_2
EUT1
02104 -# -
02105 HMMnopfamclust3_1(#) RMM1
02106 -pf01380 -
02107 -pf04261 -
02108 -pf06572 -
02109 -pf11143 -
02110 -pf00583 -
02111 -pf01520 -
02112 -pf01218 -
02113 HMMreg12833_1_EUT1(pf12833) EUT1
02114 H_mulberry(pf00936pf16365) EUT1
02115 Tsp_rosa EUT1
02116 HMMetlyEutC_1_eut1(pf05985) EUT1 EP_NTERM
02117 HMMetlyEutB_1_EUT1(pf06751) EUT1
02118 HMM06277EutA_1_eut(pf06277) EUT1
02119 HMMtpEuth_1_eut(pf04346) EUT1
02120 HMM00465alcdh_3_mix(pf00465) EUT1
02121 HMM06723EutJMreB_1_grm(#) EUT1
02122 HMMalddh_1_eutpdugrm(pf00171) EUT1 EP_NTERM
02123 P_lightViolet EUT1
02124 H_azure EUT1
02125 HMM01515PTA_B_1_eut1(pf01515) EUT1
02126 HMM01923cobtr_2_eut1(pf01923) EUT1
02127 HMM06249EutQ_1_eut1(pf06249) EUT1 EP_NTERM
02128 HMM10662PduV_1_mix(pf10662) EUT1
02129 Hp_euts_violetPink EUT1
02130 HMM00390_malic_1_eut1(pf00390pf03949pf01515) EUT1
02131 -pf00923 -
02132 HMM02780keto_1_eut1(pf02779pf02780pf00456) EUT1
02133 -pf06674 -
02134 HMM00293nuhyd_1_eut1(pf00293) EUT1
02135 -pf13247pf14691pf07992 -
02136 -pf00672pf02518pf07730pf13675 -
02137 -pf03797pf13018pf12951pf03212 -
02138 -pf01075 -
02139 -pf00873 -
BMC-P
BMC-H
BMC-T
Aldehyde dehydrogenase
PTAC
Signature enzyme
Alcohol dehydrogenase
Regulator

Escherichia coli KTE20. Bacteria; Proteobacteria; Gammaproteobacteria; Enterobacterales;

WE7__locus_1_main_sat

EUT1

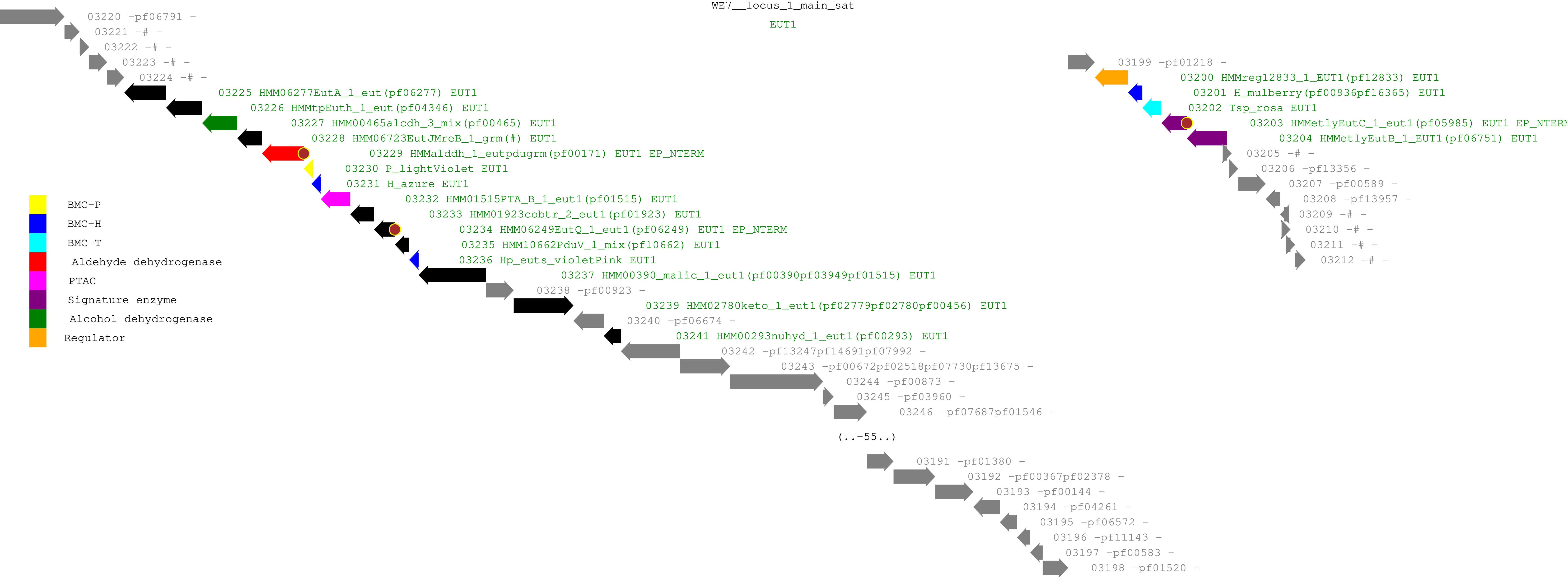

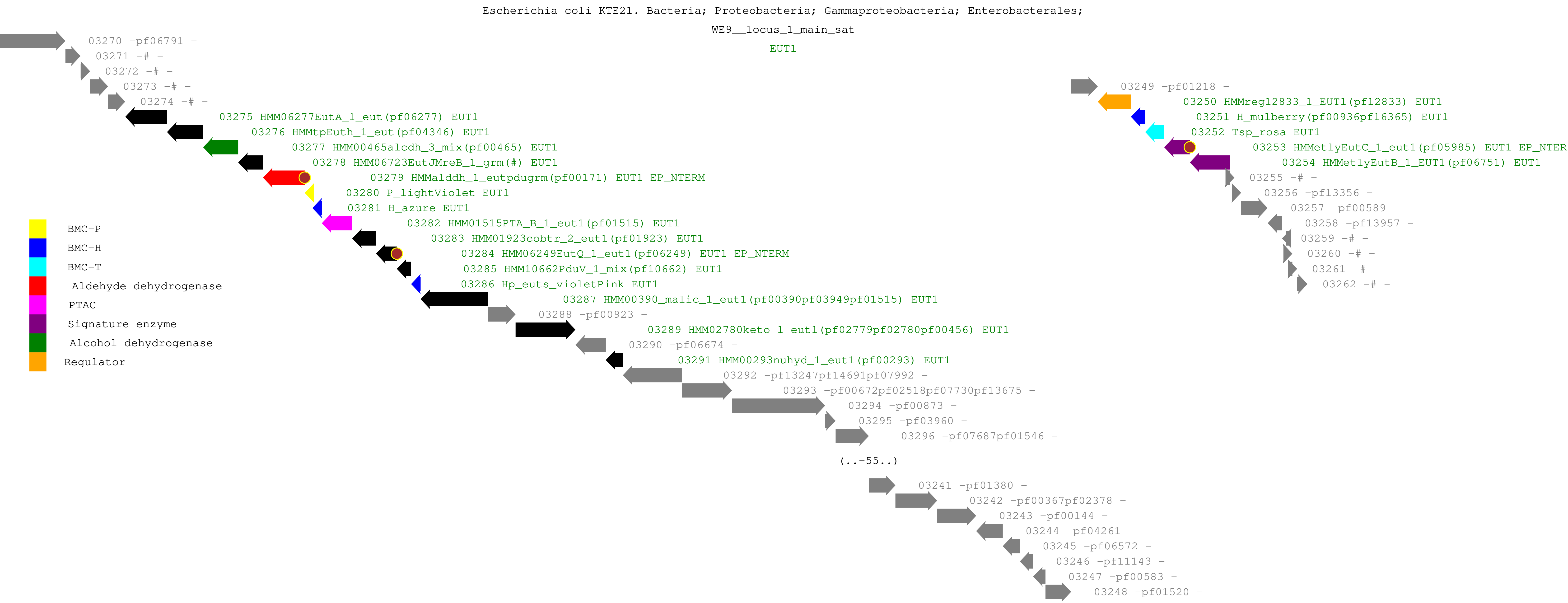
Escherichia coli KTE21. Bacteria; Proteobacteria; Gammaproteobacteria; Enterobacterales;
WE9__locus_1_main_sat
EUT1
03270 -pf06791 -
03271 -# -
03272 -# -
03273 -# -
03274 -# -
03275 HMM06277EutA_1_eut(pf06277) EUT1
03276 HMMtpEuth_1_eut(pf04346) EUT1
03277 HMM00465alcdh_3_mix(pf00465) EUT1
03278 HMM06723EutJMreB_1_grm(#) EUT1
03279 HMMalddh_1_eutpdugrm(pf00171) EUT1 EP_NTERM
03280 P_lightViolet EUT1
03281 H_azure EUT1
03282 HMM01515PTA_B_1_eut1(pf01515) EUT1
03283 HMM01923cobtr_2_eut1(pf01923) EUT1
03284 HMM06249EutQ_1_eut1(pf06249) EUT1 EP_NTERM
03285 HMM10662PduV_1_mix(pf10662) EUT1
03286 Hp_euts_violetPink EUT1
03287 HMM00390_malic_1_eut1(pf00390pf03949pf01515) EUT1
03288 -pf00923 -
03289 HMM02780keto_1_eut1(pf02779pf02780pf00456) EUT1
03290 -pf06674 -
03291 HMM00293nuhyd_1_eut1(pf00293) EUT1
03292 -pf13247pf14691pf07992 -
03293 -pf00672pf02518pf07730pf13675 -
03294 -pf00873 -
03295 -pf03960 -
03296 -pf07687pf01546 -
(..-55..)
03241 -pf01380 -
03242 -pf00367pf02378 -
03243 -pf00144 -
03244 -pf04261 -
03245 -pf06572 -
03246 -pf11143 -
03247 -pf00583 -
03248 -pf01520 -
03249 -pf01218 -
03250 HMMreg12833_1_EUT1(pf12833) EUT1
03251 H_mulberry(pf00936pf16365) EUT1
03252 Tsp_rosa EUT1
03253 HMMetlyEutC_1_eut1(pf05985) EUT1 EP_NTER
03254 HMMetlyEutB_1_EUT1(pf06751) EUT1
03255 -# -
03256 -pf13356 -
03257 -pf00589 -
03258 -pf13957 -
03259 -# -
03260 -# -
03261 -# -
03262 -# -
BMC-P
BMC-H
BMC-T
Aldehyde dehydrogenase
PTAC
Signature enzyme
Alcohol dehydrogenase
Regulator

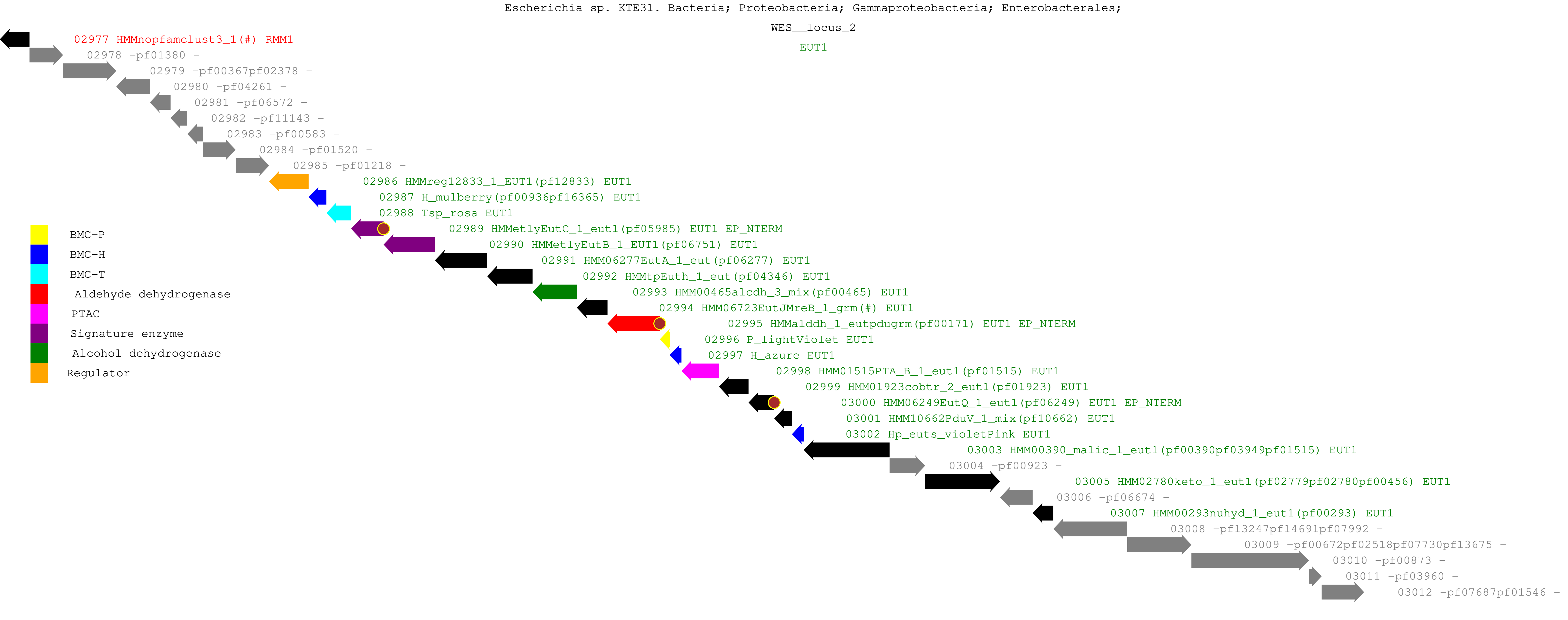
Escherichia sp. KTE31. Bacteria; Proteobacteria; Gammaproteobacteria; Enterobacterales;
WES__locus_2
EUT1
02977 HMMnopfamclust3_1(#) RMM1
02978 -pf01380 -
02979 -pf00367pf02378 -
02980 -pf04261 -
02981 -pf06572 -
02982 -pf11143 -
02983 -pf00583 -
02984 -pf01520 -
02985 -pf01218 -
02986 HMMreg12833_1_EUT1(pf12833) EUT1
02987 H_mulberry(pf00936pf16365) EUT1
02988 Tsp_rosa EUT1
02989 HMMetlyEutC_1_eut1(pf05985) EUT1 EP_NTERM
02990 HMMetlyEutB_1_EUT1(pf06751) EUT1
02991 HMM06277EutA_1_eut(pf06277) EUT1
02992 HMMtpEuth_1_eut(pf04346) EUT1
02993 HMM00465alcdh_3_mix(pf00465) EUT1
02994 HMM06723EutJMreB_1_grm(#) EUT1
02995 HMMalddh_1_eutpdugrm(pf00171) EUT1 EP_NTERM
02996 P_lightViolet EUT1
02997 H_azure EUT1
02998 HMM01515PTA_B_1_eut1(pf01515) EUT1
02999 HMM01923cobtr_2_eut1(pf01923) EUT1
03000 HMM06249EutQ_1_eut1(pf06249) EUT1 EP_NTERM
03001 HMM10662PduV_1_mix(pf10662) EUT1
03002 Hp_euts_violetPink EUT1
03003 HMM00390_malic_1_eut1(pf00390pf03949pf01515) EUT1
03004 -pf00923 -
03005 HMM02780keto_1_eut1(pf02779pf02780pf00456) EUT1
03006 -pf06674 -
03007 HMM00293nuhyd_1_eut1(pf00293) EUT1
03008 -pf13247pf14691pf07992 -
03009 -pf00672pf02518pf07730pf13675 -
03010 -pf00873 -
03011 -pf03960 -
03012 -pf07687pf01546 -
BMC-P
BMC-H
BMC-T
Aldehyde dehydrogenase
PTAC
Signature enzyme
Alcohol dehydrogenase
Regulator

Escherichia coli KTE33. Bacteria; Proteobacteria; Gammaproteobacteria; Enterobacterales;

WEW__locus_2

EUT1

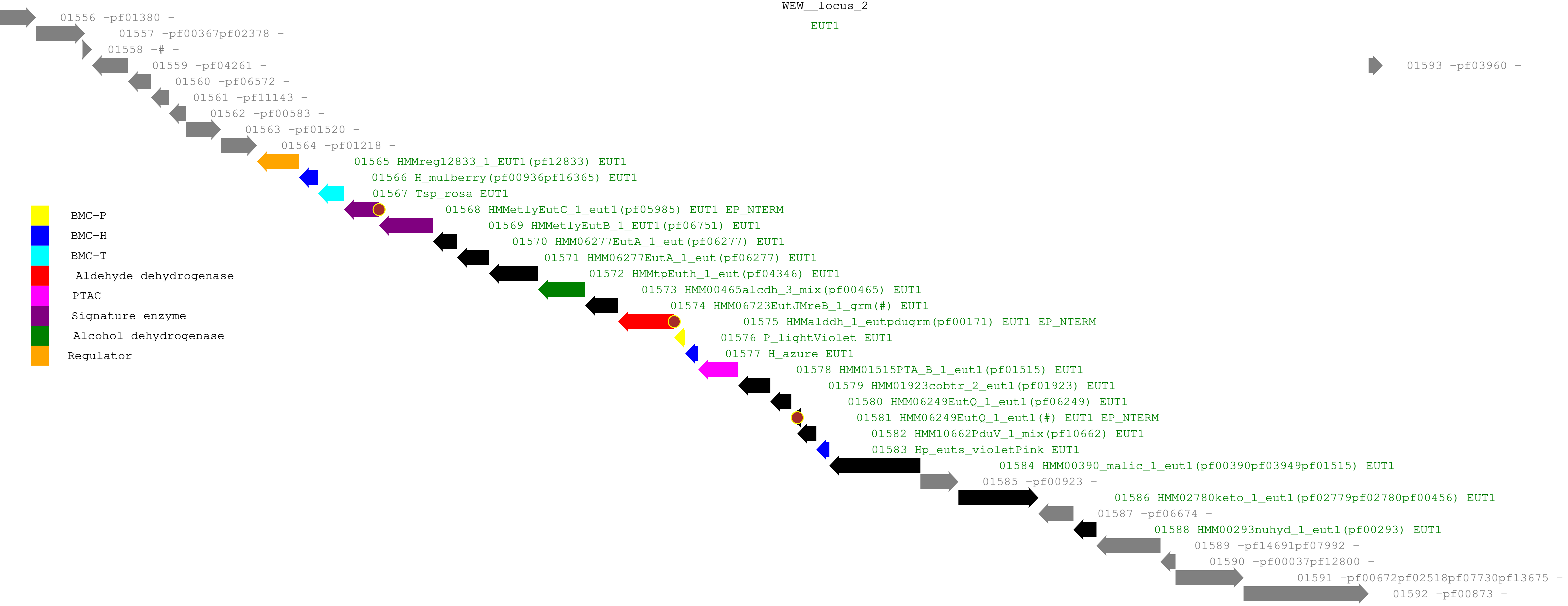

Escherichia coli KTE37. Bacteria; Proteobacteria; Gammaproteobacteria; Enterobacterales;

WG5__locus_1_main_sat

EUT1

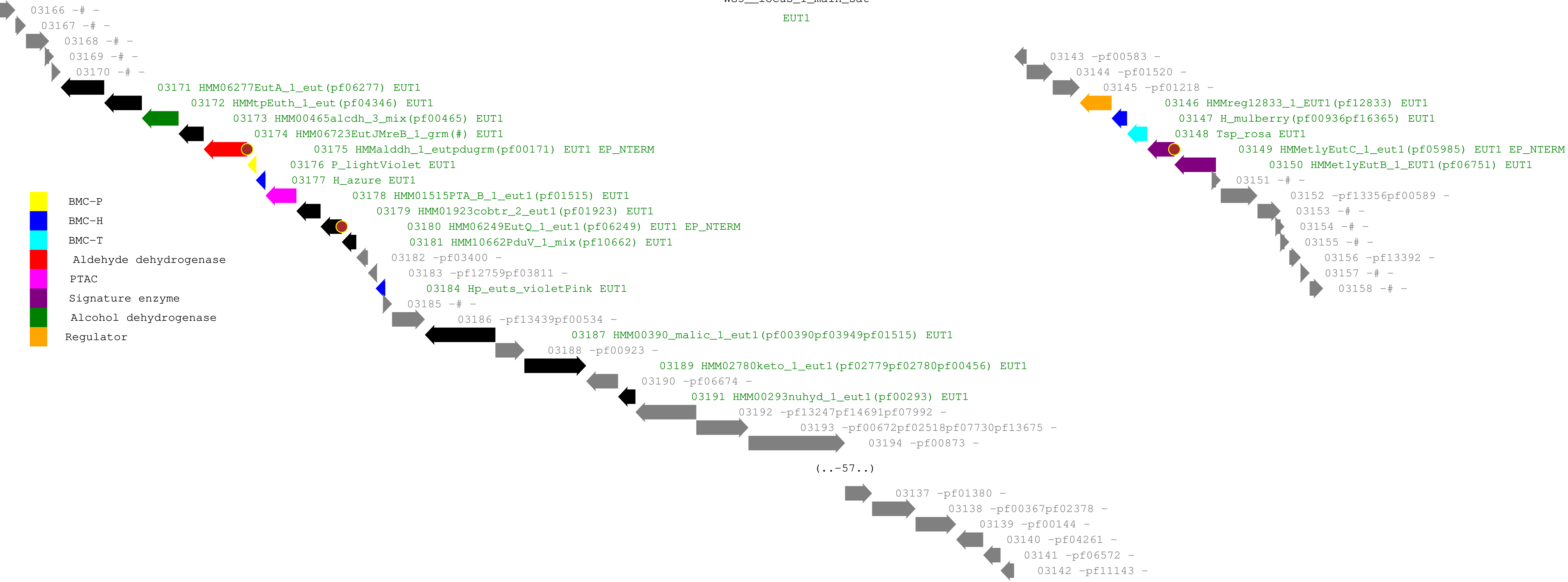

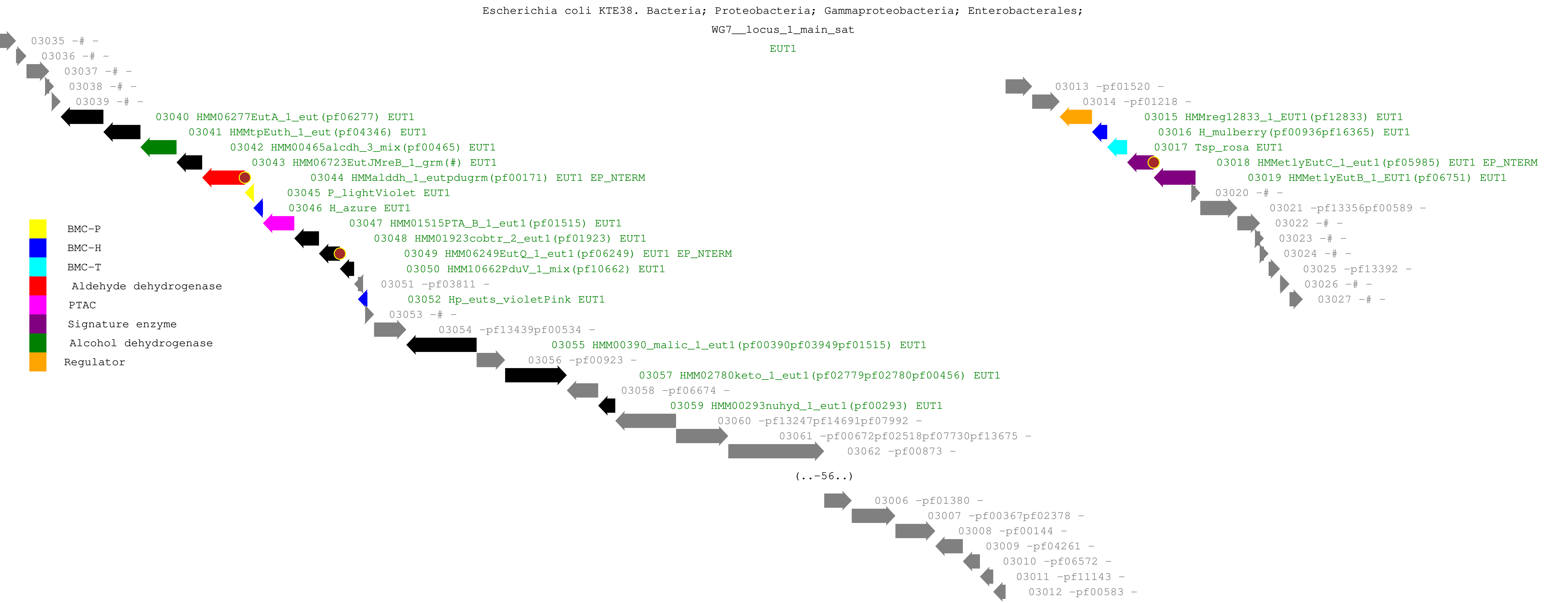
Escherichia coli KTE38. Bacteria; Proteobacteria; Gammaproteobacteria; Enterobacterales;
WG7__locus_1_main_sat
EUT1
03035 -# -
03036 -# -
03037 -# -
03038 -# -
03039 -# -
03040 HMM06277EutA_1_eut(pf06277) EUT1
03041 HMMtpEuth_1_eut(pf04346) EUT1
03042 HMM00465alcdh_3_mix(pf00465) EUT1
03043 HMM06723EutJMreB_1_grm(#) EUT1
03044 HMMalddh_1_eutpdugrm(pf00171) EUT1 EP_NTERM
03045 P_lightViolet EUT1
03046 H_azure EUT1
03047 HMM01515PTA_B_1_eut1(pf01515) EUT1
03048 HMM01923cobtr_2_eut1(pf01923) EUT1
03049 HMM06249EutQ_1_eut1(pf06249) EUT1 EP_NTERM
03050 HMM10662PduV_1_mix(pf10662) EUT1
03051 -pf03811 -
03052 Hp_euts_violetPink EUT1
03053 -# -
03054 -pf13439pf00534 -
03055 HMM00390_malic_1_eut1(pf00390pf03949pf01515) EUT1
03056 -pf00923 -
03057 HMM02780keto_1_eut1(pf02779pf02780pf00456) EUT1
03058 -pf06674 -
03059 HMM00293nuhyd_1_eut1(pf00293) EUT1
03060 -pf13247pf14691pf07992 -
03061 -pf00672pf02518pf07730pf13675 -
03062 -pf00873 -
(..-56..)
03006 -pf01380 -
03007 -pf00367pf02378 -
03008 -pf00144 -
03009 -pf04261 -
03010 -pf06572 -
03011 -pf11143 -
03012 -pf00583 -
03013 -pf01520 -
03014 -pf01218 -
03015 HMMreg12833_1_EUT1(pf12833) EUT1
03016 H_mulberry(pf00936pf16365) EUT1
03017 Tsp_rosa EUT1
03018 HMMetlyEutC_1_eut1(pf05985) EUT1 EP_NTERM
03019 HMMetlyEutB_1_EUT1(pf06751) EUT1
03020 -# -
03021 -pf13356pf00589 -
03022 -# -
03023 -# -
03024 -# -
03025 -pf13392 -
03026 -# -
03027 -# -
BMC-P
BMC-H
BMC-T
Aldehyde dehydrogenase
PTAC
Signature enzyme
Alcohol dehydrogenase
Regulator

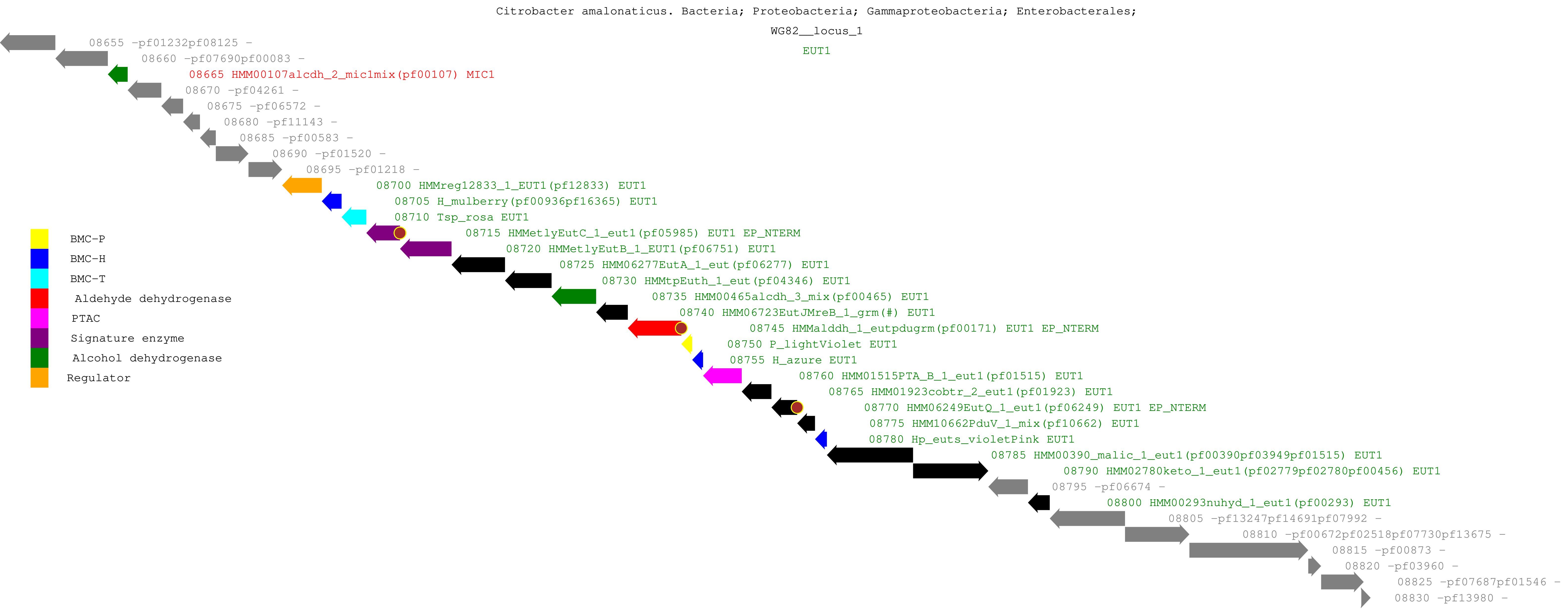

Citrobacter amalonaticus. Bacteria; Proteobacteria; Gammaproteobacteria; Enterobacterales;
WG82__locus_1
EUT1
08655 -pf01232pf08125 -
08660 -pf07690pf00083 -
08665 HMM00107alcdh_2_mic1mix(pf00107) MIC1
08670 -pf04261 -
08675 -pf06572 -
08680 -pf11143 -
08685 -pf00583 -
08690 -pf01520 -
08695 -pf01218 -
08700 HMMreg12833_1_EUT1(pf12833) EUT1
08705 H_mulberry(pf00936pf16365) EUT1
08710 Tsp_rosa EUT1
08715 HMMetlyEutC_1_eut1(pf05985) EUT1 EP_NTERM
08720 HMMetlyEutB_1_EUT1(pf06751) EUT1
08725 HMM06277EutA_1_eut(pf06277) EUT1
08730 HMMtpEuth_1_eut(pf04346) EUT1
08735 HMM00465alcdh_3_mix(pf00465) EUT1
08740 HMM06723EutJMreB_1_grm(#) EUT1
08745 HMMaldh_1_eutpdugrm(pf00171) EUT1 EP_NTERM
08750 P_lightViolet EUT1
08755 H_azure EUT1
08760 HMM01515PTA_B_1_eut1(pf01515) EUT1
08765 HMM01923cobtr_2_eut1(pf01923) EUT1
08770 HMM06249EutQ_1_eut1(pf06249) EUT1 EP_NTERM
08775 HMM10662PduV_1_mix(pf10662) EUT1
08780 Hp_euts_violetPink EUT1
08785 HMM00390_malic_1_eut1(pf00390pf03949pf01515) EUT1
08790 HMM02780keto_1_eut1(pf02779pf02780pf00456) EUT1
08795 -pf06674 -
08800 HMM00293nuhyd_1_eut1(pf00293) EUT1
08805 -pf13247pf14691pf07992 -
08810 -pf00672pf02518pf07730pf13675 -
08815 -pf00873 -
08820 -pf03960 -
08825 -pf07687pf01546 -
08830 -pf13980 -
BMC-P
BMC-H
BMC-T
Aldehyde dehydrogenase
PTAC
Signature enzyme
Alcohol dehydrogenase
Regulator

Escherichia coli KTE112. Bacteria; Proteobacteria; Gammaproteobacteria; Enterobacterales;

WIC__locus_1

EUT1

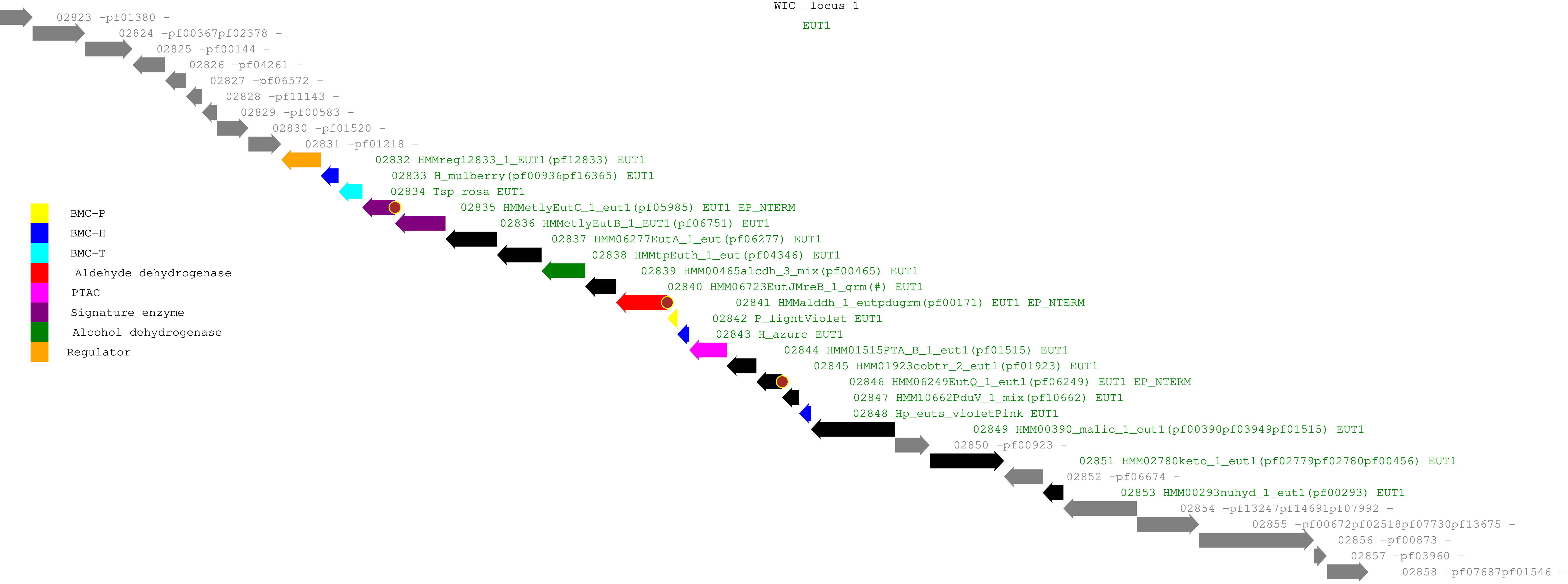

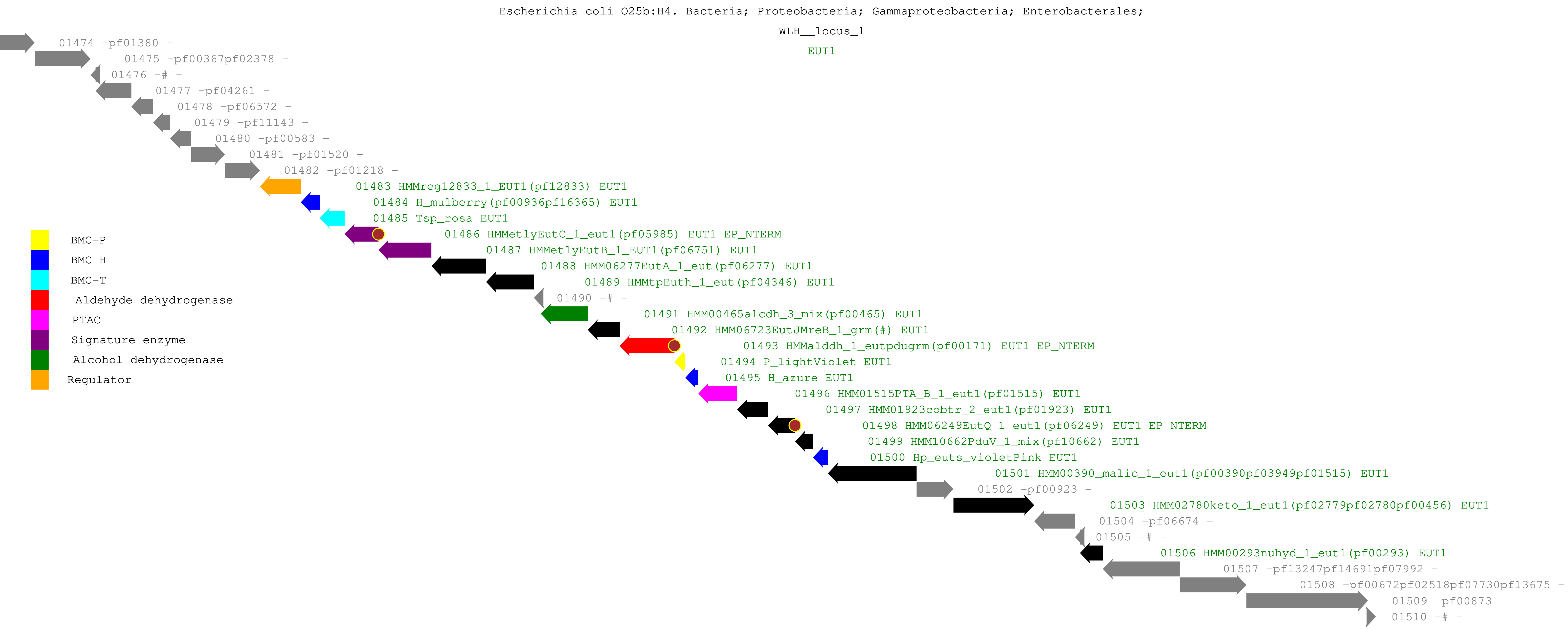
Escherichia coli O25b:H4. Bacteria; Proteobacteria; Gammaproteobacteria; Enterobacterales;
WLH__locus_1
EUT1
01474 -pf01380 -
01475 -pf00367pf02378 -
01476 -# -
01477 -pf04261 -
01478 -pf06572 -
01479 -pf11143 -
01480 -pf00583 -
01481 -pf01520 -
01482 -pf01218 -
01483 HMMreg12833_1_EUT1(pf12833) EUT1
01484 H_mulberry(pf00936pf16365) EUT1
01485 Tsp_rosa EUT1
01486 HMMetlyEutC_1_eut1(pf05985) EUT1 EP_NTERM
01487 HMMetlyEutB_1_EUT1(pf06751) EUT1
01488 HMM06277EutA_1_eut(pf06277) EUT1
01489 HMMtpEuth_1_eut(pf04346) EUT1
01490 -# -
01491 HMM00465alcdh_3_mix(pf00465) EUT1
01492 HMM06723EutJMreB_1_grm(#) EUT1
01493 HMMalddh_1_eutpdugrm(pf00171) EUT1 EP_NTERM
01494 P_lightViolet EUT1
01495 H_azure EUT1
01496 HMM01515PTA_B_1_eut1(pf01515) EUT1
01497 HMM01923cobtr_2_eut1(pf01923) EUT1
01498 HMM06249EutQ_1_eut1(pf06249) EUT1 EP_NTERM
01499 HMM10662PduV_1_mix(pf10662) EUT1
01500 Hp_euts_violetPink EUT1
01501 HMM00390_malic_1_eut1(pf00390pf03949pf01515) EUT1
01502 -pf00923 -
01503 HMM02780keto_1_eut1(pf02779pf02780pf00456) EUT1
01504 -pf06674 -
01505 -# -
01506 HMM00293nuhyd_1_eut1(pf00293) EUT1
01507 -pf13247pf14691pf07992 -
01508 -pf00672pf02518pf07730pf13675 -
01509 -pf00873 -
01510 -# -
BMC-P
BMC-H
BMC-T
Aldehyde dehydrogenase
PTAC
Signature enzyme
Alcohol dehydrogenase
Regulator

Citrobacter freundii complex sp. CFNIH2. Bacteria; Proteobacteria; Gammaproteobacteria; Enterobacterales;

WM46__locus_1

EUT1

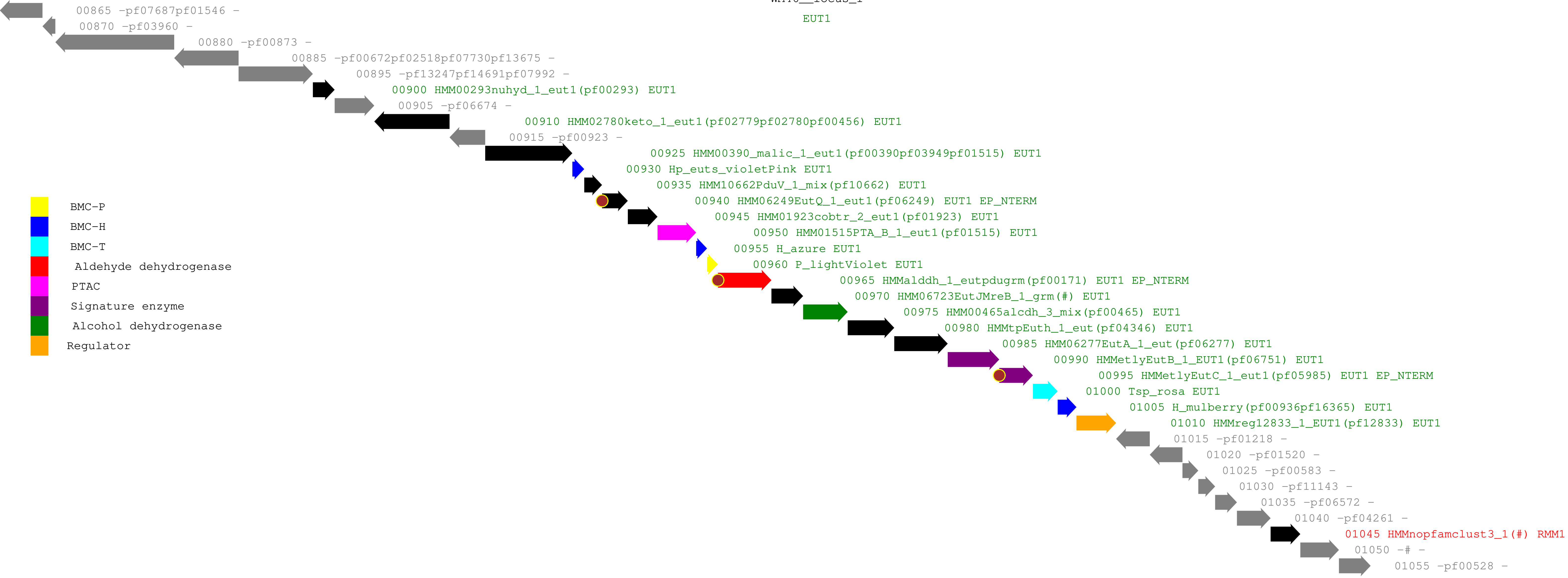

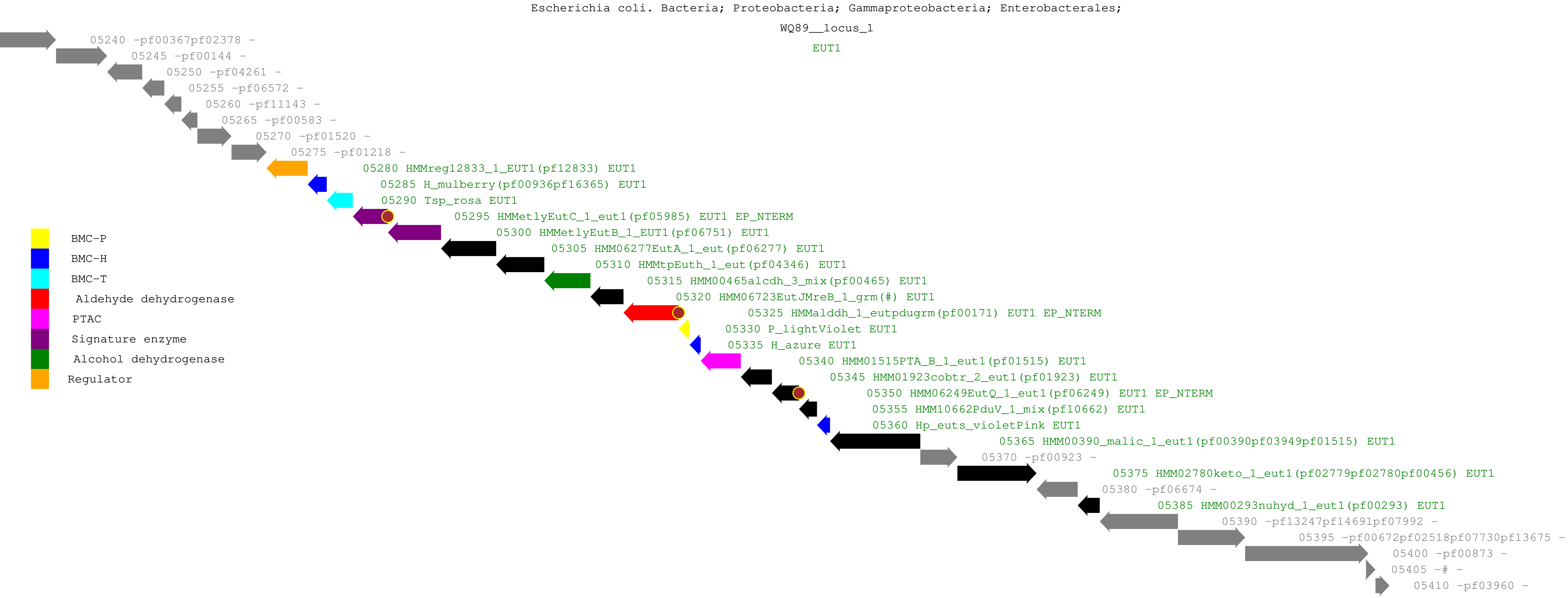

Escherichia coli. Bacteria; Proteobacteria; Gammaproteobacteria; Enterobacterales;
WQ89__locus_1
EUT1
05240 -pf00367pf02378 -
05245 -pf00144 -
05250 -pf04261 -
05255 -pf06572 -
05260 -pf11143 -
05265 -pf00583 -
05270 -pf01520 -
05275 -pf01218 -
05280 HMMreg12833_1_EUT1(pf12833) EUT1
05285 H_mulberry(pf00936pf16365) EUT1
05290 Tsp_rosa EUT1
05295 HMMetlyEutC_1_eut1(pf05985) EUT1 EP_NTERM
05300 HMMetlyEutB_1_EUT1(pf06751) EUT1
05305 HMM06277EutA_1_eut(pf06277) EUT1
05310 HMMtpEuth_1_eut(pf04346) EUT1
05315 HMM00465alcdh_3_mix(pf00465) EUT1
05320 HMM06723EutJMreB_1_grm(#) EUT1
05325 HMMalddh_1_eutpdugrm(pf00171) EUT1 EP_NTERM
05330 P_lightViolet EUT1
05335 H_azure EUT1
05340 HMM01515PTA_B_1_eut1(pf01515) EUT1
05345 HMM01923cobtr_2_eut1(pf01923) EUT1
05350 HMM06249EutQ_1_eut1(pf06249) EUT1 EP_NTERM
05355 HMM10662PduV_1_mix(pf10662) EUT1
05360 Hp_euts_violetPink EUT1
05365 HMM00390_malic_1_eut1(pf00390pf03949pf01515) EUT1
05370 -pf00923 -
05375 HMM02780keto_1_eut1(pf02779pf02780pf00456) EUT1
05380 -pf06674 -
05385 HMM00293nuhyd_1_eut1(pf00293) EUT1
05390 -pf13247pf14691pf07992 -
05395 -pf00672pf02518pf07730pf13675 -
05400 -pf00873 -
05405 -# -
05410 -pf03960 -
BMC-P
BMC-H
BMC-T
Aldehyde dehydrogenase
PTAC
Signature enzyme
Alcohol dehydrogenase
Regulator

Escherichia coli. Bacteria; Proteobacteria; Gammaproteobacteria; Enterobacterales;

WR15__locus_1

EUT1

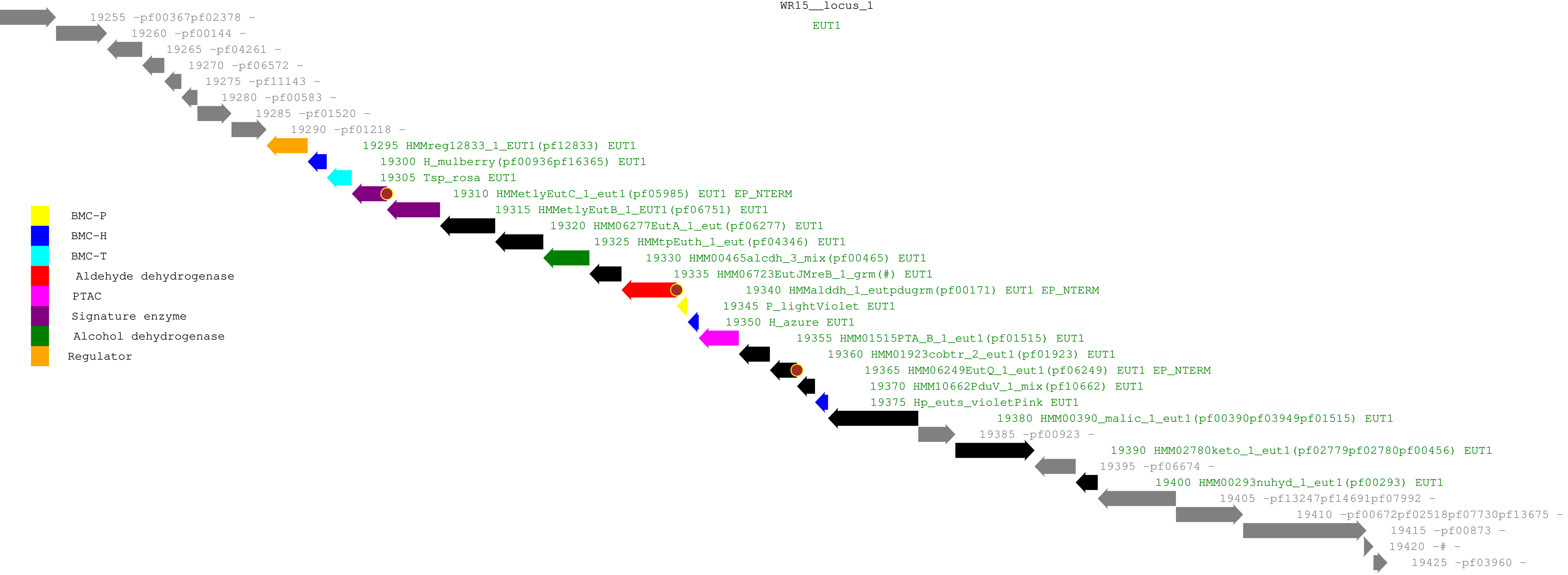

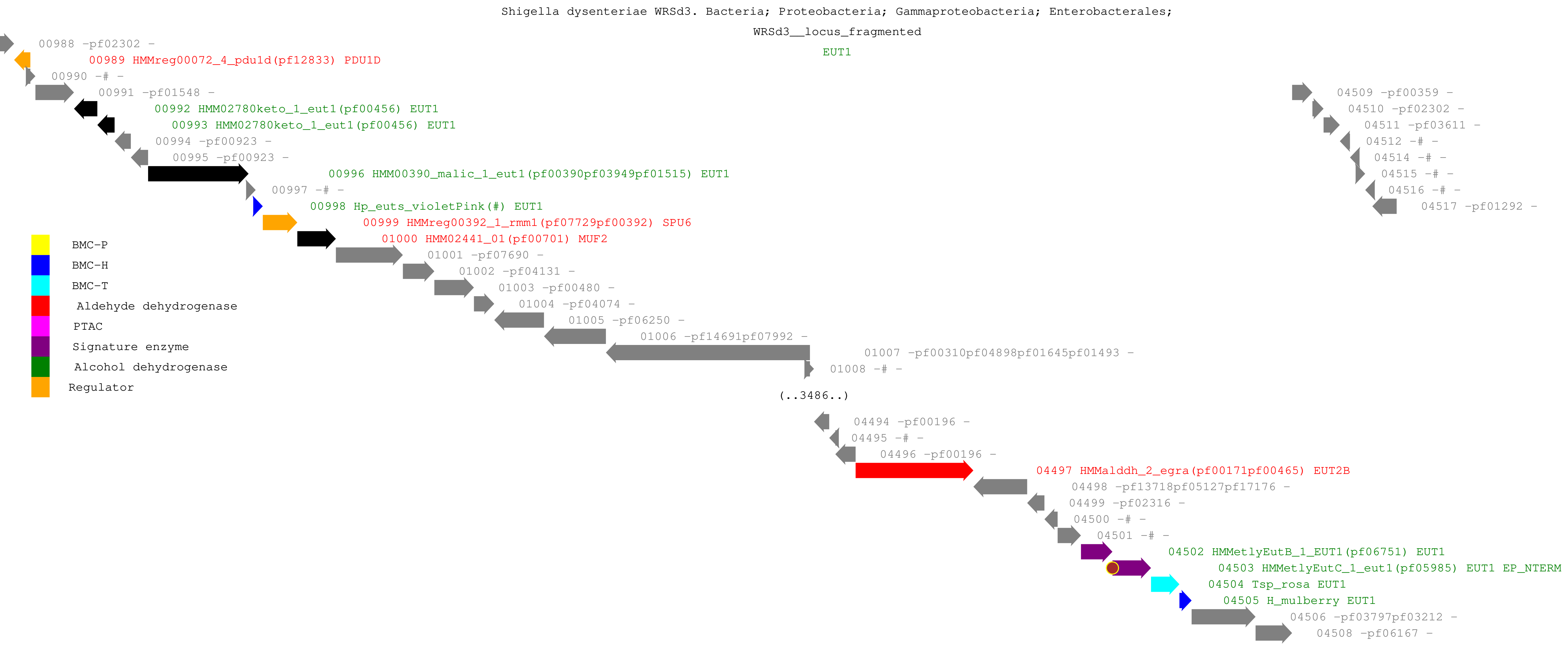

Shigella dysenteriae WRSd3. Bacteria; Proteobacteria; Gammaproteobacteria; Enterobacterales;
WRSd3__locus_fragmented
EUT1
00988 -pf02302 -
00989 HMMreg00072_4_pdu1d(pf12833) PDU1D
00990 -# -
00991 -pf01548 -
00992 HMM02780keto_1_eut1(pf00456) EUT1
00993 HMM02780keto_1_eut1(pf00456) EUT1
00994 -pf00923 -
00995 -pf00923 -
00996 HMM00390_malic_1_eut1(pf00390pf03949pf01515) EUT1
00997 -# -
00998 Hp_euts_violetPink(#) EUT1
00999 HMMreg00392_1_rmm1(pf07729pf00392) SPU6
01000 HMM02441_01(pf00701) MUF2
01001 -pf07690 -
01002 -pf04131 -
01003 -pf00480 -
01004 -pf04074 -
01005 -pf06250 -
01006 -pf14691pf07992 -
01007 -pf00310pf04898pf01645pf01493 -
01008 -# -
(..3486..)
04494 -pf00196 -
04495 -# -
04496 -pf00196 -
04497 HMMaldh_2_egra(pf00171pf00465) EUT2B
04498 -pf13718pf05127pf17176 -
04499 -pf02316 -
04500 -# -
04501 -# -
04502 HMMetlyEutB_1_EUT1(pf06751) EUT1
04503 HMMetlyEutC_1_eut1(pf05985) EUT1 EP_NTERM
04504 Tsp_rosa EUT1
04505 H_mulberry EUT1
04506 -pf03797pf03212 -
04508 -pf06167 -
04509 -pf00359 -
04510 -pf02302 -
04511 -pf03611 -
04512 -# -
04514 -# -
04515 -# -
04516 -# -
04517 -pf01292 -
BMC-P
BMC-H
BMC-T
Aldehyde dehydrogenase
PTAC
Signature enzyme
Alcohol dehydrogenase
Regulator

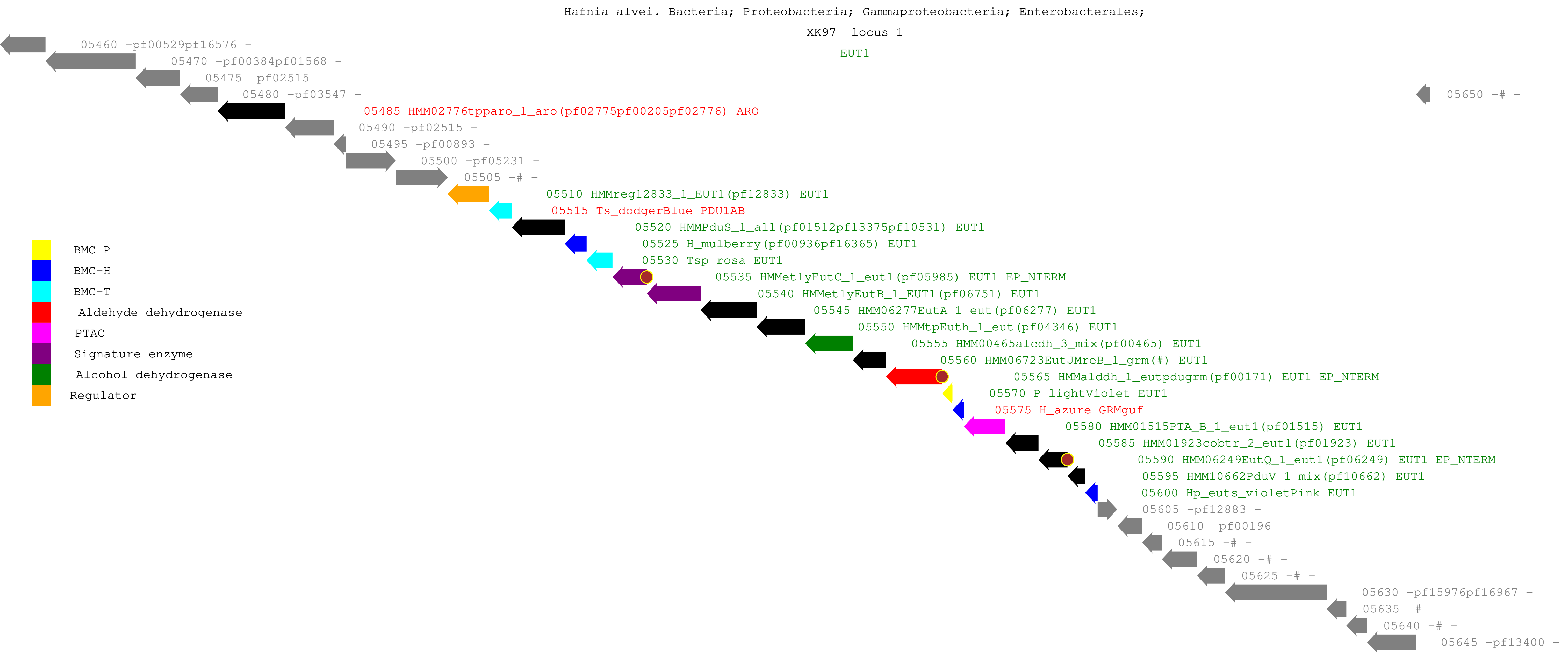
Hafnia alvei. Bacteria; Proteobacteria; Gammaproteobacteria; Enterobacterales;
XK97__locus_1
EUT1
05460 -pf00529pf16576 -
05470 -pf00384pf01568 -
05475 -pf02515 -
05480 -pf03547 -
05485 HMM02776tpparo_1_aro(pf02775pf00205pf02776) ARO
05490 -pf02515 -
05495 -pf00893 -
05500 -pf05231 -
05505 -# -
05510 HMMreg12833_1_EUT1(pf12833) EUT1
05515 Ts_dodgerBlue PDU1AB
05520 HMMPduS_1_all(pf01512pf13375pf10531) EUT1
05525 H_mulberry(pf00936pf16365) EUT1
05530 Tsp_rosa EUT1
05535 HMMetlyEutC_1_eut1(pf05985) EUT1 EP_NTERM
05540 HMMetlyEutB_1_EUT1(pf06751) EUT1
05545 HMM06277EutA_1_eut(pf06277) EUT1
05550 HMMtpEuth_1_eut(pf04346) EUT1
05555 HMM00465alcdh_3_mix(pf00465) EUT1
05560 HMM06723EutJMreB_1_grm(#) EUT1
05565 HMMaldDh_1_eutpdugrm(pf00171) EUT1 EP_NTERM
05570 P_lightViolet EUT1
05575 H_azure GRMguf
05580 HMM01515PTA_B_1_eut1(pf01515) EUT1
05585 HMM01923cobtr_2_eut1(pf01923) EUT1
05590 HMM06249EutQ_1_eut1(pf06249) EUT1 EP_NTERM
05595 HMM10662PduV_1_mix(pf10662) EUT1
05600 Hp_euts_violetPink EUT1
05605 -pf12883 -
05610 -pf00196 -
05615 -# -
05620 -# -
05625 -# -
05630 -pf15976pf16967 -
05635 -# -
05640 -# -
05645 -pf13400 -
05650 -# -
BMC-P
BMC-H
BMC-T
Aldehyde dehydrogenase
PTAC
Signature enzyme
Alcohol dehydrogenase
Regulator

Salmonella berta. Bacteria; Proteobacteria; Gammaproteobacteria; Enterobacterales;

XR79__locus_2

EUT1

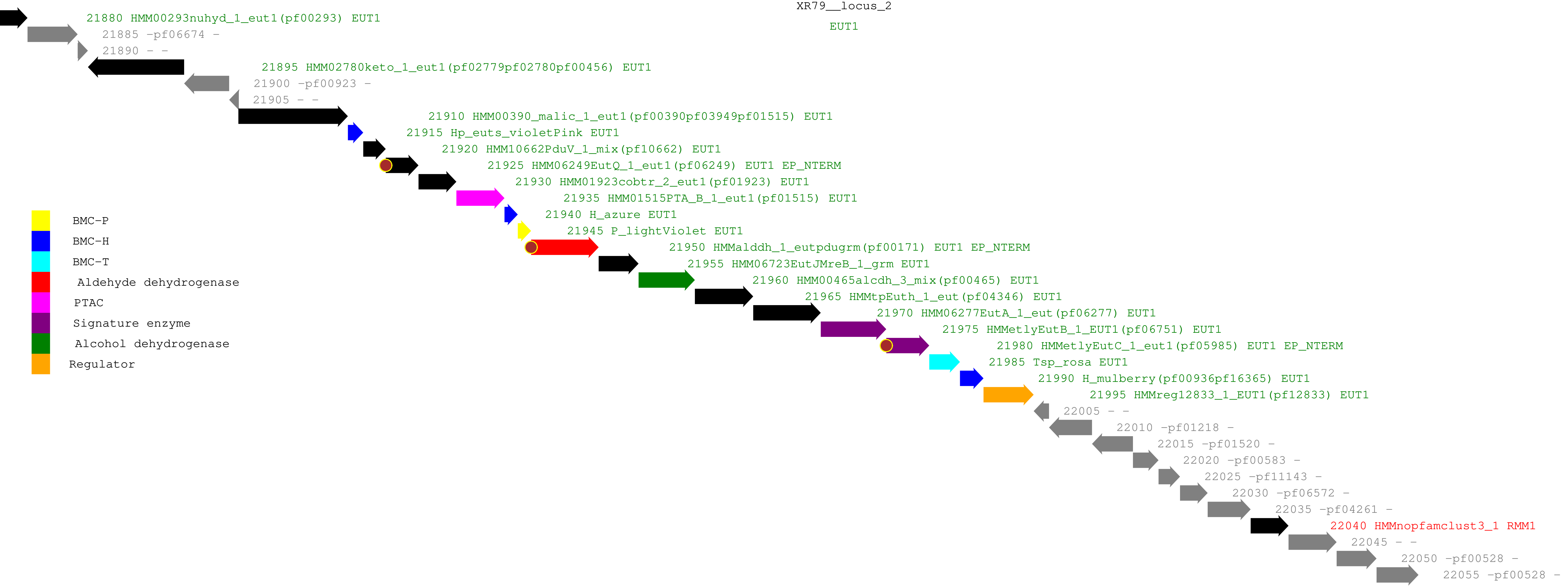

Salmonella enterica subsp. enterica serovar Infantis str. CVM Bacteria; Proteobacteria; Gammaproteobacteria; Enterobacterales;

Y082__locus_1

EUT1

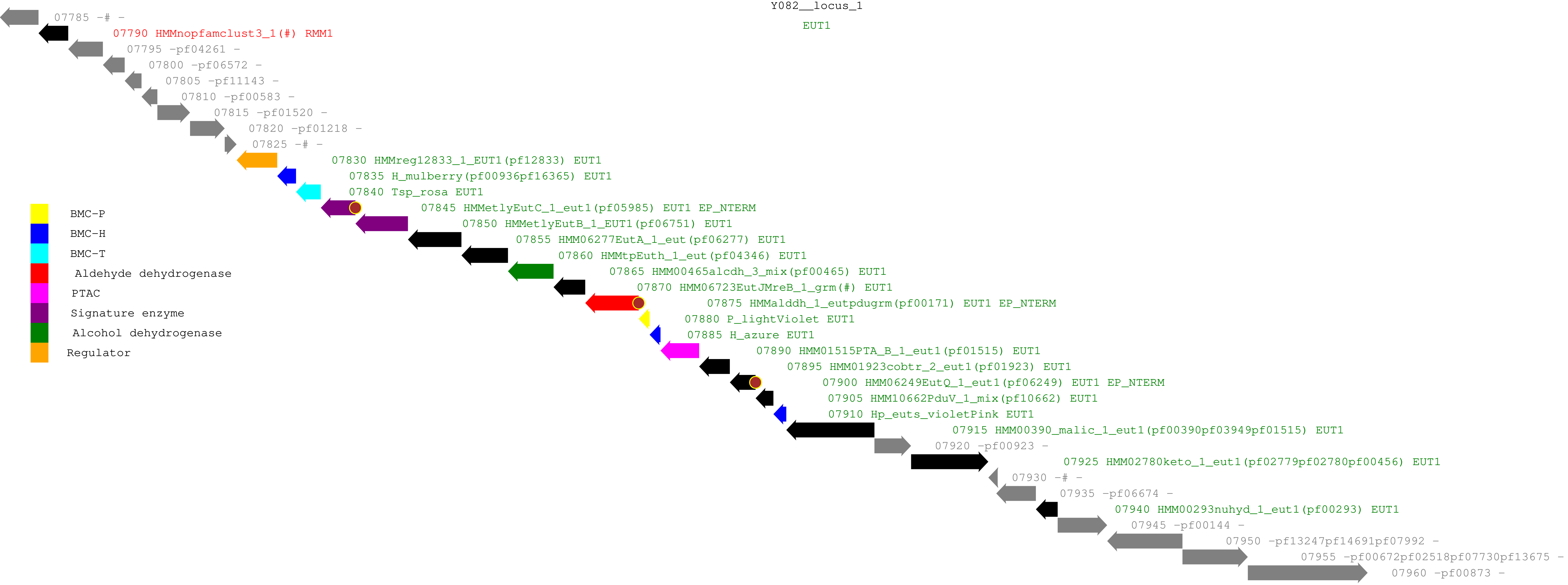

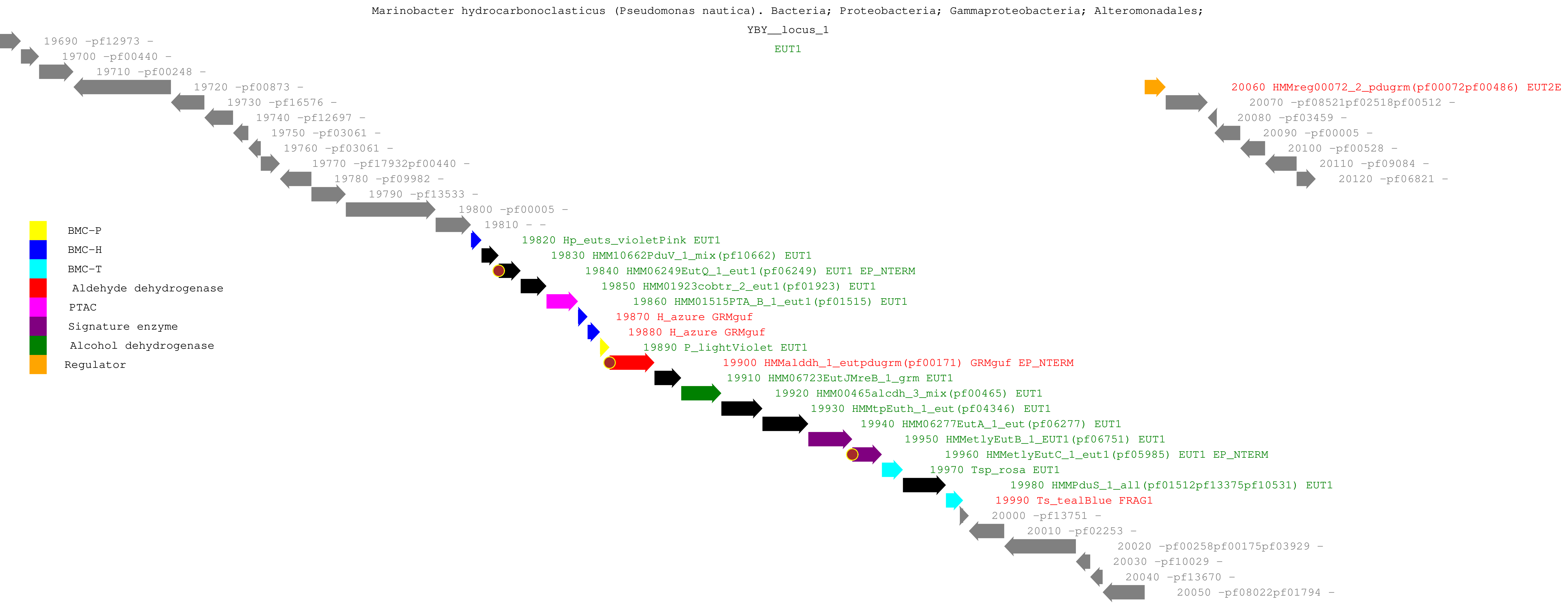
Marinobacter hydrocarbonoclasticus (Pseudomonas nautica). Bacteria; Proteobacteria; Gammaproteobacteria; Alteromonadales;
YBY__locus_1
EUT1
19690 -pf12973 -
19700 -pf00440 -
19710 -pf00248 -
19720 -pf00873 -
19730 -pf16576 -
19740 -pf12697 -
19750 -pf03061 -
19760 -pf03061 -
19770 -pf17932pf00440 -
19780 -pf09982 -
19790 -pf13533 -
19800 -pf00005 -
19810 - -
19820 Hp_euts_violetPink EUT1
19830 HMM10662PduV_1_mix(pf10662) EUT1
19840 HMM06249EutQ_1_eut1(pf06249) EUT1 EP_NTERM
19850 HMM01923cobtr_2_eut1(pf01923) EUT1
19860 HMM01515PTA_B_1_eut1(pf01515) EUT1
19870 H_azure GRMguf
19880 H_azure GRMguf
19890 P_lightViolet EUT1
19900 HMMaldh_1_eutpdugrm(pf00171) GRMguf EP_NTERM
19910 HMM06723EutJMreB_1_grm EUT1
19920 HMM00465alcdh_3_mix(pf00465) EUT1
19930 HMMtpEuth_1_eut(pf04346) EUT1
19940 HMM06277EutA_1_eut(pf06277) EUT1
19950 HMMetlyEutB_1_EUT1(pf06751) EUT1
19960 HMMetlyEutC_1_eut1(pf05985) EUT1 EP_NTERM
19970 Tsp_rosa EUT1
19980 HMMPduS_1_all(pf01512pf13375pf10531) EUT1
19990 Ts_tealBlue FRAG1
20000 -pf13751 -
20010 -pf02253 -
20020 -pf00258pf00175pf03929 -
20030 -pf10029 -
20040 -pf13670 -
20050 -pf08022pf01794 -
20060 HMMreg00072_2_pdugrm(pf00072pf00486) EUT2E
20070 -pf08521pf02518pf00512 -
20080 -pf03459 -
20090 -pf00005 -
20100 -pf00528 -
20110 -pf09084 -
20120 -pf06821 -
BMC-P
BMC-H
BMC-T
Aldehyde dehydrogenase
PTAC
Signature enzyme
Alcohol dehydrogenase
Regulator

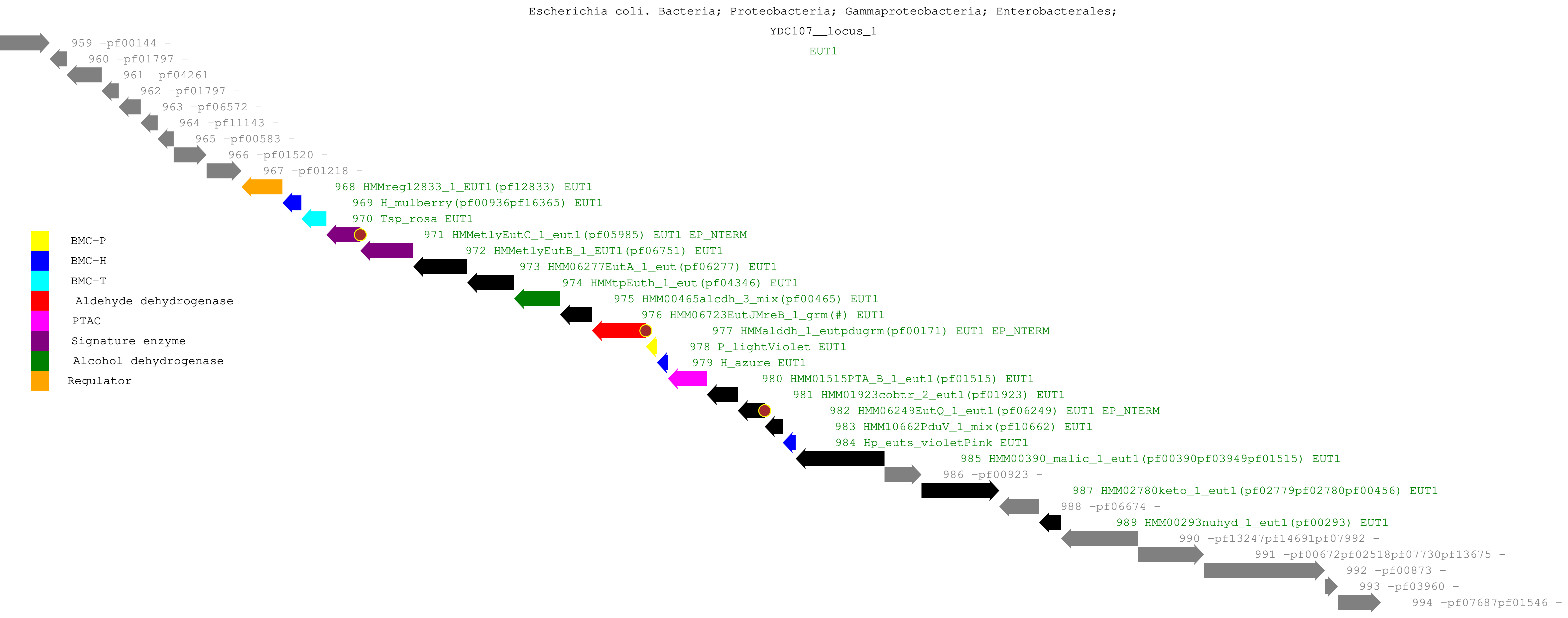

Escherichia coli. Bacteria; Proteobacteria; Gammaproteobacteria; Enterobacterales;
YDC107__locus_1
EUT1
959 -pf00144 -
960 -pf01797 -
961 -pf04261 -
962 -pf01797 -
963 -pf06572 -
964 -pf11143 -
965 -pf00583 -
966 -pf01520 -
967 -pf01218 -
968 HMMreg12833_1_EUT1(pf12833) EUT1
969 H_mulberry(pf00936pf16365) EUT1
970 Tsp_rosa EUT1
971 HMMetlyEutC_1_eut1(pf05985) EUT1 EP_NTERM
972 HMMetlyEutB_1_EUT1(pf06751) EUT1
973 HMM06277EutA_1_eut(pf06277) EUT1
974 HMMtpEuth_1_eut(pf04346) EUT1
975 HMM00465alcdh_3_mix(pf00465) EUT1
976 HMM06723EutJMreB_1_grm(#) EUT1
977 HMMalddh_1_eutpdugrm(pf00171) EUT1 EP_NTERM
978 P_lightViolet EUT1
979 H_azure EUT1
980 HMM01515PTA_B_1_eut1(pf01515) EUT1
981 HMM01923cobtr_2_eut1(pf01923) EUT1
982 HMM06249EutQ_1_eut1(pf06249) EUT1 EP_NTERM
983 HMM10662PduV_1_mix(pf10662) EUT1
984 Hp_euts_violetPink EUT1
985 HMM00390_malic_1_eut1(pf00390pf03949pf01515) EUT1
986 -pf00923 -
987 HMM02780keto_1_eut1(pf02779pf02780pf00456) EUT1
988 -pf06674 -
989 HMM00293nuhyd_1_eut1(pf00293) EUT1
990 -pf13247pf14691pf07992 -
991 -pf00672pf02518pf07730pf13675 -
992 -pf00873 -
993 -pf03960 -
994 -pf07687pf01546 -
BMC-P
BMC-H
BMC-T
Aldehyde dehydrogenase
PTAC
Signature enzyme
Alcohol dehydrogenase
Regulator

Salmonella typhimurium. Bacteria; Proteobacteria; Gammaproteobacteria; Enterobacterales;

YT65__locus_1

EUT1

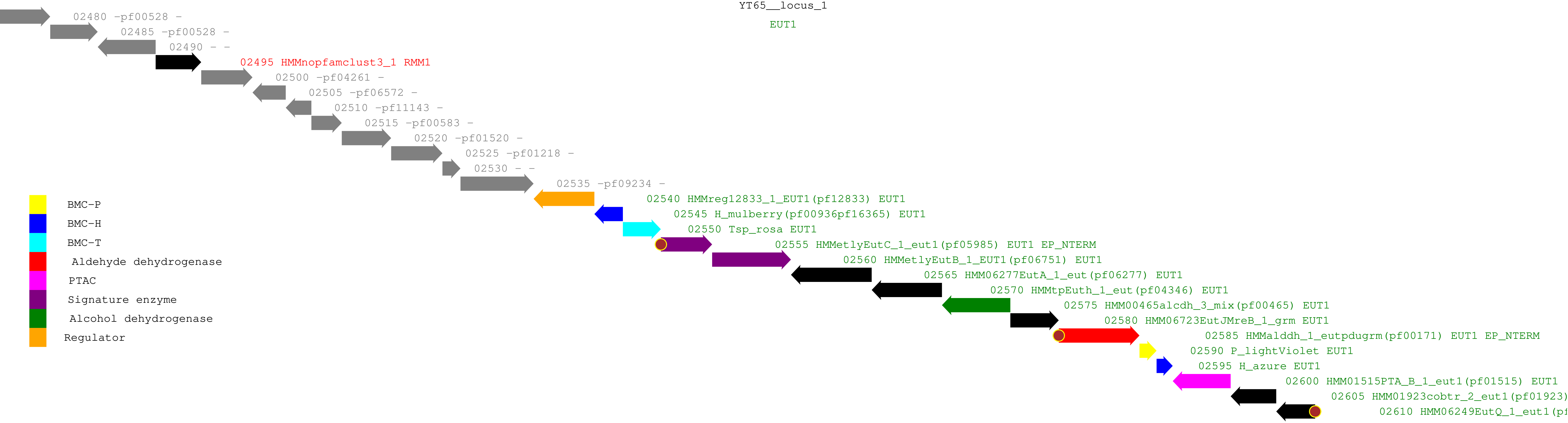

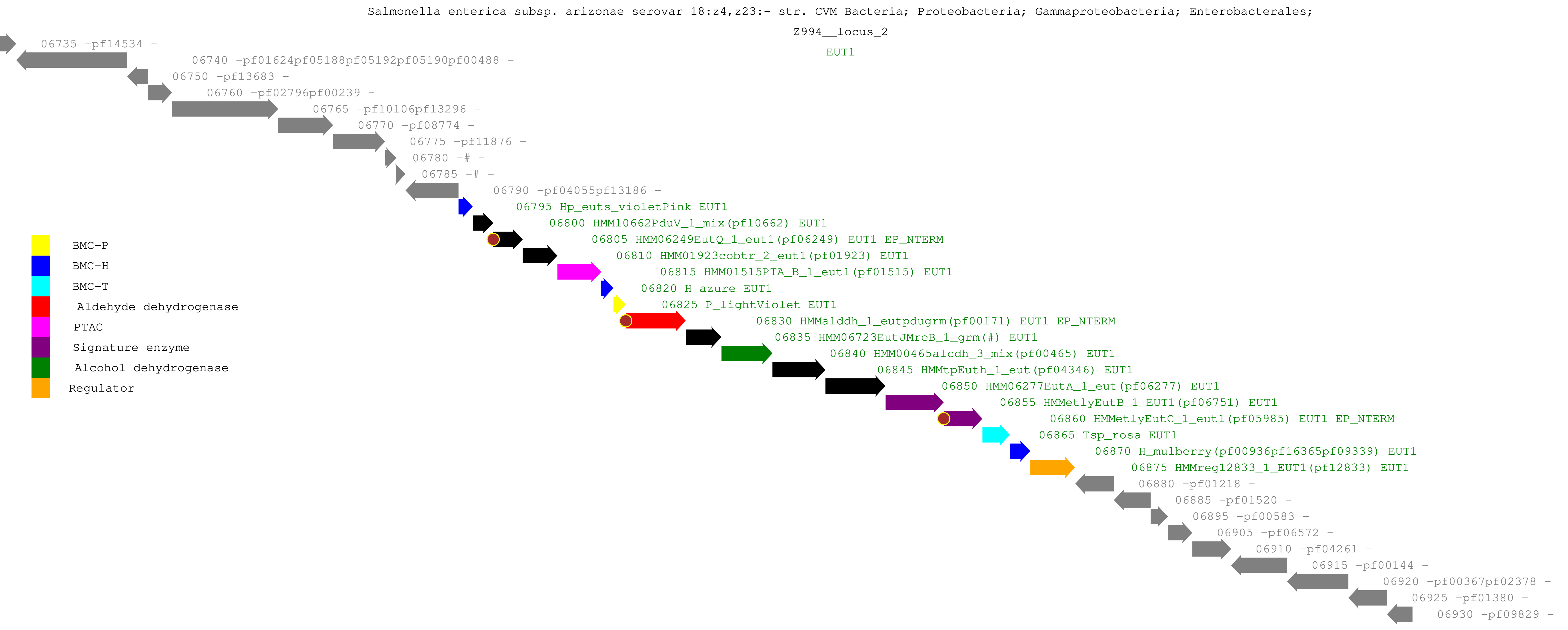
Salmonella enterica subsp. arizonae serovar 18:z4,z23:- str. CVM Bacteria; Proteobacteria; Gammaproteobacteria; Enterobacterales;
Z994__locus_2
EUT1
06735 -pf14534 -
06740 -pf01624pf05188pf05192pf05190pf00488 -
06750 -pf13683 -
06760 -pf02796pf00239 -
06765 -pf10106pf13296 -
06770 -pf08774 -
06775 -pf11876 -
06780 -# -
06785 -# -
06790 -pf04055pf13186 -
06795 Hp_euts_violetPink EUT1
06800 HMM10662PduV_1_mix(pf10662) EUT1
06805 HMM06249EutQ_1_eut1(pf06249) EUT1 EP_NTERM
06810 HMM01923cobtr_2_eut1(pf01923) EUT1
06815 HMM01515PTA_B_1_eut1(pf01515) EUT1
06820 H_azure EUT1
06825 P_lightViolet EUT1
06830 HMMalddh_1_eutpdugrm(pf00171) EUT1 EP_NTERM
06835 HMM06723EutJMreB_1_grm(#) EUT1
06840 HMM00465alcdh_3_mix(pf00465) EUT1
06845 HMMtpEuth_1_eut(pf04346) EUT1
06850 HMM06277EutA_1_eut(pf06277) EUT1
06855 HMMetlyEutB_1_EUT1(pf06751) EUT1
06860 HMMetlyEutC_1_eut1(pf05985) EUT1 EP_NTERM
06865 Tsp_rosa EUT1
06870 H_mulberry(pf00936pf16365pf09339) EUT1
06875 HMMreg12833_1_EUT1(pf12833) EUT1
06880 -pf01218 -
06885 -pf01520 -
06895 -pf00583 -
06905 -pf06572 -
06910 -pf04261 -
06915 -pf00144 -
06920 -pf00367pf02378 -
06925 -pf01380 -
06930 -pf09829 -
BMC-P
BMC-H
BMC-T
Aldehyde dehydrogenase
PTAC
Signature enzyme
Alcohol dehydrogenase
Regulator

Salmonella enterica subsp. enterica serovar Corvallis. Bacteria; Proteobacteria; Gammaproteobacteria; Enterobacterales;

ZU68__locus_2_main_sat

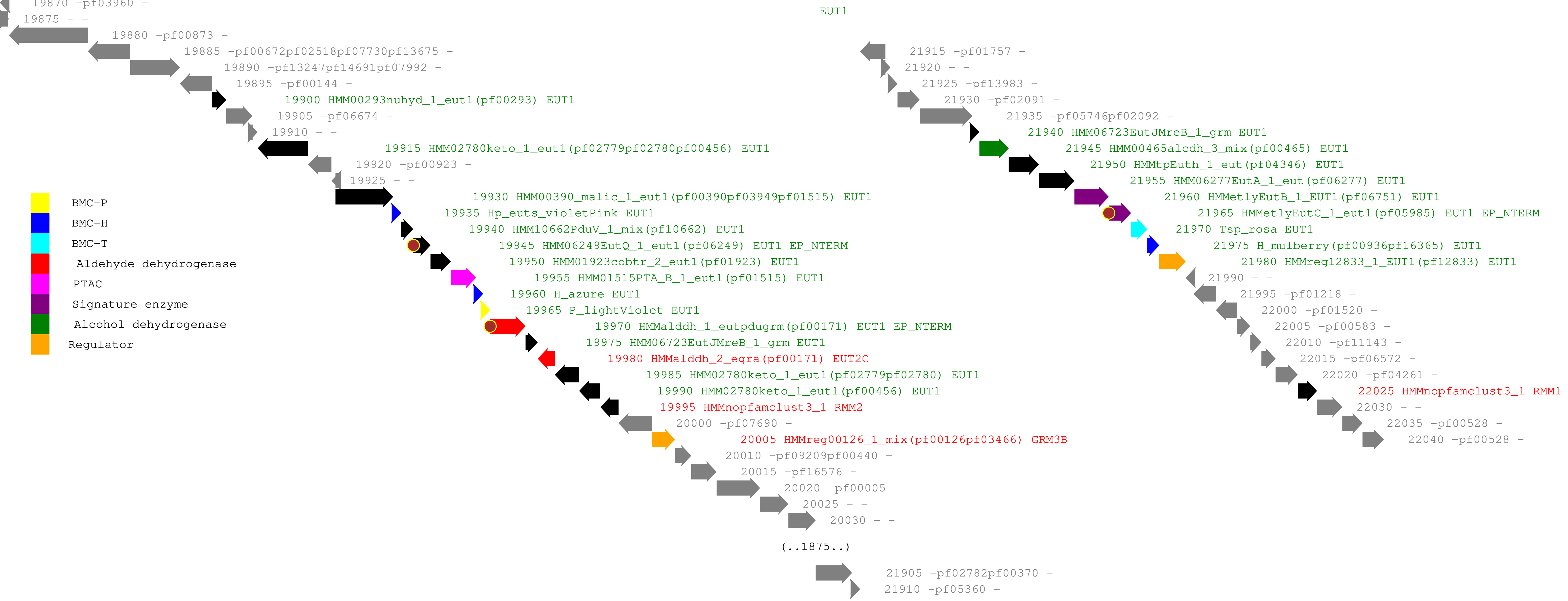

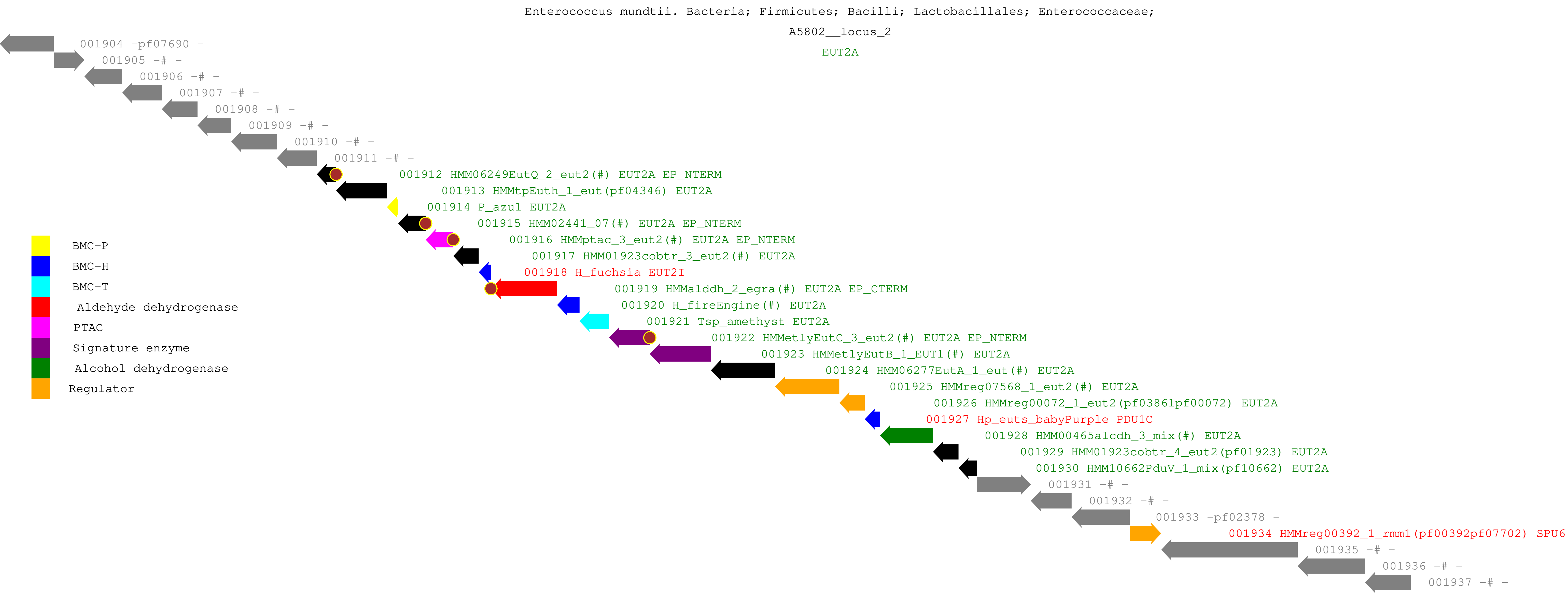

Enterococcus mundtii. Bacteria; Firmicutes; Bacilli; Lactobacillales; Enterococcaceae;
A5802__locus_2
EUT2A
001904 -pf07690 -
001905 -# -
001906 -# -
001907 -# -
001908 -# -
001909 -# -
001910 -# -
001911 -# -
001912 HMM06249EutQ_2_eut2(#) EUT2A EP_NTERM
001913 HMMtpEuth_1_eut(pf04346) EUT2A
001914 P_azul EUT2A
001915 HMM02441_07(#) EUT2A EP_NTERM
001916 HMMptac_3_eut2(#) EUT2A EP_NTERM
001917 HMM01923cobtr_3_eut2(#) EUT2A
001918 H_fuchsia EUT2I
001919 HMMalddh_2_egra(#) EUT2A EP_CTERM
001920 H_fireEngine(#) EUT2A
001921 Tsp_amethyst EUT2A
001922 HMMetlyEutC_3_eut2(#) EUT2A EP_NTERM
001923 HMMetlyEutB_1_EUT1(#) EUT2A
001924 HMM06277EutA_1_eut(#) EUT2A
001925 HMMreg07568_1_eut2(#) EUT2A
001926 HMMreg00072_1_eut2(pf03861pf00072) EUT2A
001927 Hp_euts_babyPurple PDU1C
001928 HMM00465alcdh_3_mix(#) EUT2A
001929 HMM01923cobtr_4_eut2(pf01923) EUT2A
001930 HMM10662PduV_1_mix(pf10662) EUT2A
001931 -# -
001932 -# -
001933 -pf02378 -
001934 HMMreg00392_1_rmm1(pf00392pf07702) SPU6
001935 -# -
001936 -# -
001937 -# -
BMC-P
BMC-H
BMC-T
Aldehyde dehydrogenase
PTAC
Signature enzyme
Alcohol dehydrogenase
Regulator

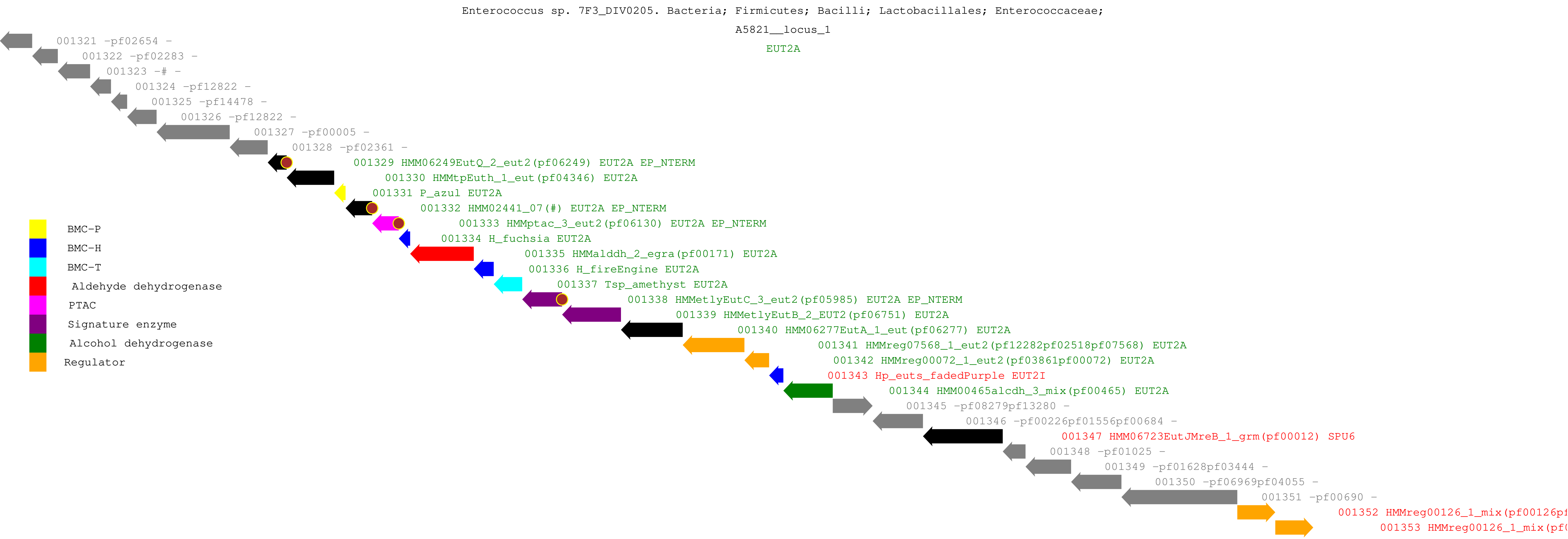

Enterococcus sp. 7F3_DIV0205. Bacteria; Firmicutes; Bacilli; Lactobacillales; Enterococcaceae;
A5821__locus_1
EUT2A
001321 -pf02654 -
001322 -pf02283 -
001323 -# -
001324 -pf12822 -
001325 -pf14478 -
001326 -pf12822 -
001327 -pf00005 -
001328 -pf02361 -
001329 HMM06249EutQ_2_eut2(pf06249) EUT2A EP_NTERM
001330 HMMtpEuth_1_eut(pf04346) EUT2A
001331 P_azul EUT2A
001332 HMM02441_07(#) EUT2A EP_NTERM
001333 HMMptac_3_eut2(pf06130) EUT2A EP_NTERM
001334 H_fuchsia EUT2A
001335 HMMalddh_2_egra(pf00171) EUT2A
001336 H_fireEngine EUT2A
001337 Tsp_amethyst EUT2A
001338 HMMetlyEutC_3_eut2(pf05985) EUT2A EP_NTERM
001339 HMMetlyEutB_2_EUT2(pf06751) EUT2A
001340 HMM06277EutA_1_eut(pf06277) EUT2A
001341 HMMreg07568_1_eut2(pf12282pf02518pf07568) EUT2A
001342 HMMreg00072_1_eut2(pf03861pf00072) EUT2A
001343 Hp_euts_fadedPurple EUT2I
001344 HMM00465alcdh_3_mix(pf00465) EUT2A
001345 -pf08279pf13280 -
001346 -pf00226pf01556pf00684 -
001347 HMM06723EutJMreB_1_grm(pf00012) SPU6
001348 -pf01025 -
001349 -pf01628pf03444 -
001350 -pf06969pf04055 -
001351 -pf00690 -
001352 HMMreg00126_1_mix(pf00126pf
001353 HMMreg00126_1_mix(pf(
BMC-P
BMC-H
BMC-T
Aldehyde dehydrogenase
PTAC
Signature enzyme
Alcohol dehydrogenase
Regulator

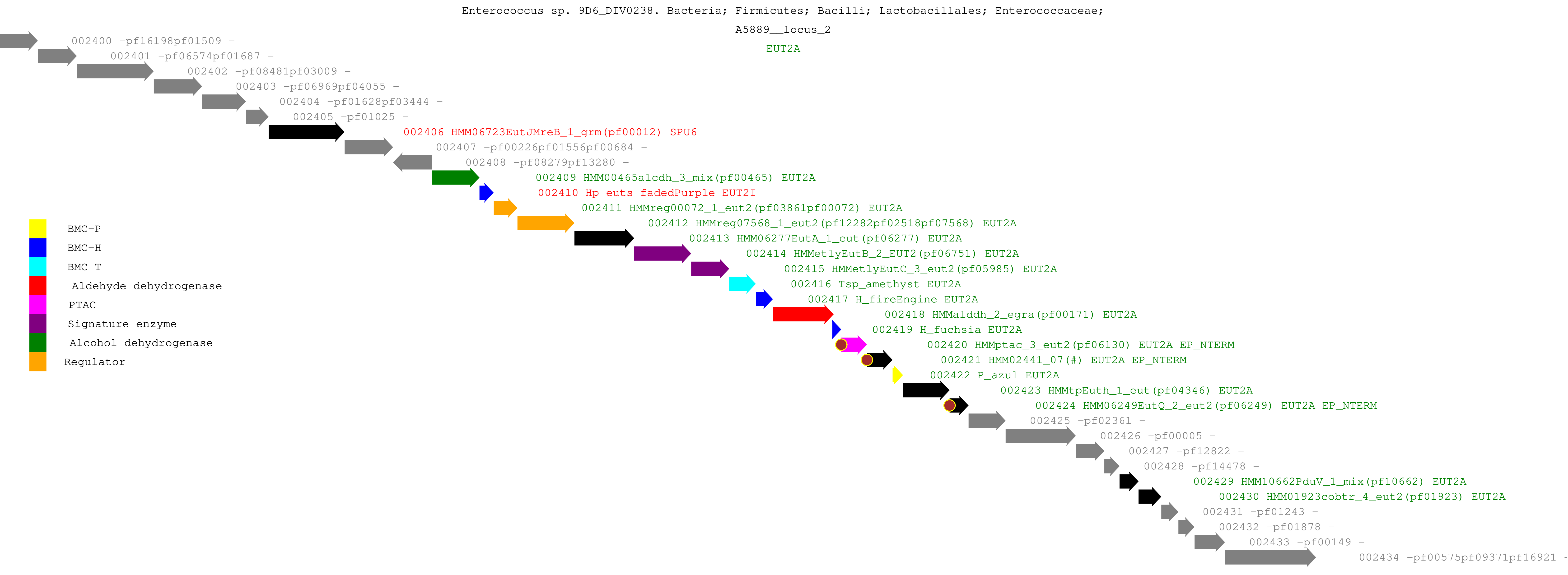
Enterococcus sp. 9D6_DIV0238. Bacteria; Firmicutes; Bacilli; Lactobacillales; Enterococcaceae;
A5889__locus_2
EUT2A
002400 -pf16198pf01509 -
002401 -pf06574pf01687 -
002402 -pf08481pf03009 -
002403 -pf06969pf04055 -
002404 -pf01628pf03444 -
002405 -pf01025 -
002406 HMM06723EutJMreB_1_grm(pf00012) SPU6
002407 -pf00226pf01556pf00684 -
002408 -pf08279pf13280 -
002409 HMM00465alcdh_3_mix(pf00465) EUT2A
002410 Hp_euts_fadedPurple EUT2I
002411 HMMreg00072_1_eut2(pf03861pf00072) EUT2A
002412 HMMreg07568_1_eut2(pf12282pf02518pf07568) EUT2A
002413 HMM06277EutA_1_eut(pf06277) EUT2A
002414 HMMetlyEutB_2_EUT2(pf06751) EUT2A
002415 HMMetlyEutC_3_eut2(pf05985) EUT2A
002416 Tsp_amethyst EUT2A
002417 H_fireEngine EUT2A
002418 HMMaldh_2_egra(pf00171) EUT2A
002419 H_fuchsia EUT2A
002420 HMMptac_3_eut2(pf06130) EUT2A EP_NTERM
002421 HMM02441_07(#) EUT2A EP_NTERM
002422 P_azul EUT2A
002423 HMMtpEuth_1_eut(pf04346) EUT2A
002424 HMM06249EutQ_2_eut2(pf06249) EUT2A EP_NTERM
002425 -pf02361 -
002426 -pf00005 -
002427 -pf12822 -
002428 -pf14478 -
002429 HMM10662PduV_1_mix(pf10662) EUT2A
002430 HMM01923cobtr_4_eut2(pf01923) EUT2A
002431 -pf01243 -
002432 -pf01878 -
002433 -pf00149 -
002434 -pf00575pf09371pf16921 -
BMC-P
BMC-H
BMC-T
Aldehyde dehydrogenase
PTAC
Signature enzyme
Alcohol dehydrogenase
Regulator

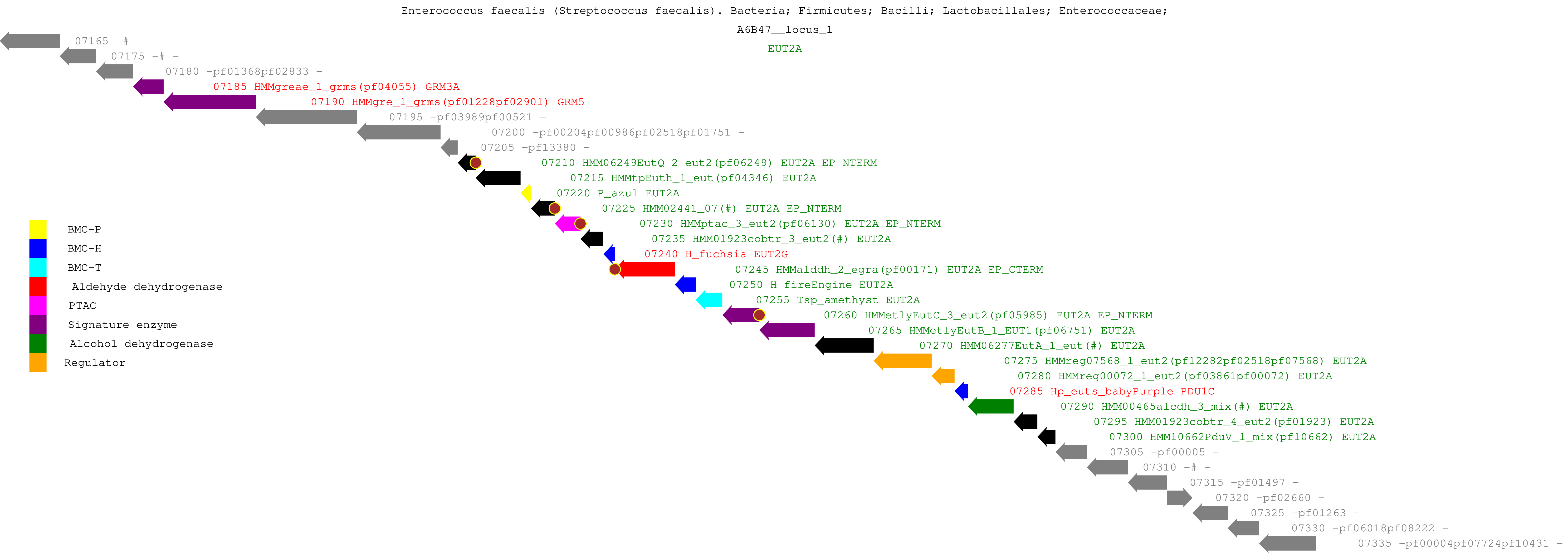

Enterococcus faecalis (Streptococcus faecalis). Bacteria; Firmicutes; Bacilli; Lactobacillales; Enterococcaceae;
A6B47__locus_1
EUT2A
07165 -# -
07175 -# -
07180 -pf01368pf02833 -
07185 HMMgreae_1_grms(pf04055) GRM3A
07190 HMMgre_1_grms(pf01228pf02901) GRM5
07195 -pf03989pf00521 -
07200 -pf00204pf00986pf02518pf01751 -
07205 -pf13380 -
07210 HMM06249EutQ_2_eut2(pf06249) EUT2A EP_NTERM
07215 HMMtpEuth_1_eut(pf04346) EUT2A
07220 P_azul EUT2A
07225 HMM02441_07(#) EUT2A EP_NTERM
07230 HMMptac_3_eut2(pf06130) EUT2A EP_NTERM
07235 HMM01923cobtr_3_eut2(#) EUT2A
07240 H_fuchsia EUT2G
07245 HMMalddh_2_egra(pf00171) EUT2A EP_CTERM
07250 H_fireEngine EUT2A
07255 Tsp_amethyst EUT2A
07260 HMMetlyEutC_3_eut2(pf05985) EUT2A EP_NTERM
07265 HMMetlyEutB_1_EUT1(pf06751) EUT2A
07270 HMM06277EutA_1_eut(#) EUT2A
07275 HMMreg07568_1_eut2(pf12282pf02518pf07568) EUT2A
07280 HMMreg00072_1_eut2(pf03861pf00072) EUT2A
07285 Hp_euts_babyPurple PDU1C
07290 HMM00465alcdh_3_mix(#) EUT2A
07295 HMM01923cobtr_4_eut2(pf01923) EUT2A
07300 HMM10662PduV_1_mix(pf10662) EUT2A
07305 -pf00005 -
07310 -# -
07315 -pf01497 -
07320 -pf02660 -
07325 -pf01263 -
07330 -pf06018pf08222 -
07335 -pf00004pf07724pf10431 -
BMC-P
BMC-H
BMC-T
Aldehyde dehydrogenase
PTAC
Signature enzyme
Alcohol dehydrogenase
Regulator

Enterococcus avium (Streptococcus avium). Bacteria; Firmicutes; Bacilli; Lactobacillales; Enterococcaceae;

AUF12__locus_1

EUT2A

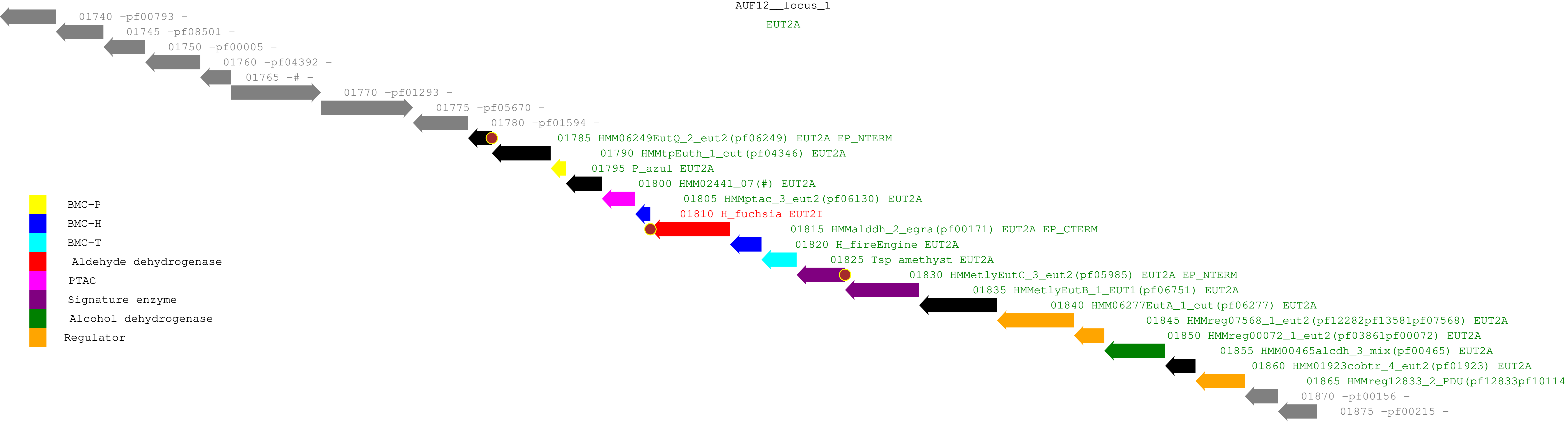

Enterococcus avium (Streptococcus avium). Bacteria; Firmicutes; Bacilli; Lactobacillales; Enterococcaceae;

AUF16__locus_1

EUT2A

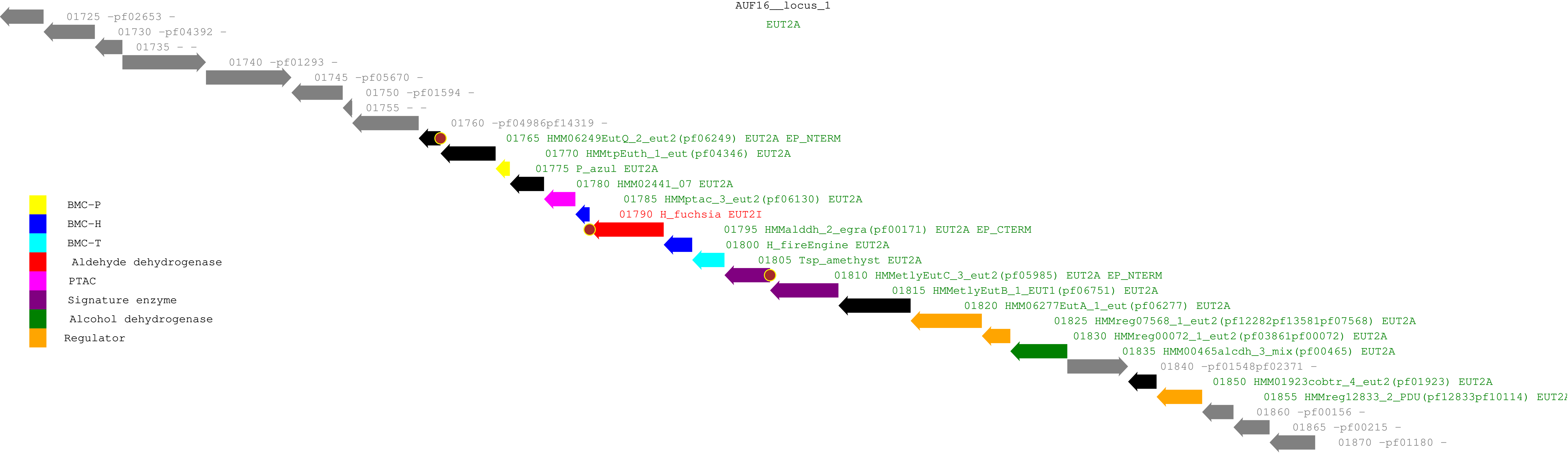

Enterococcus faecalis (Streptococcus faecalis). Bacteria; Firmicutes; Bacilli; Lactobacillales; Enterococcaceae;

B6E65__locus_1

EUT2A

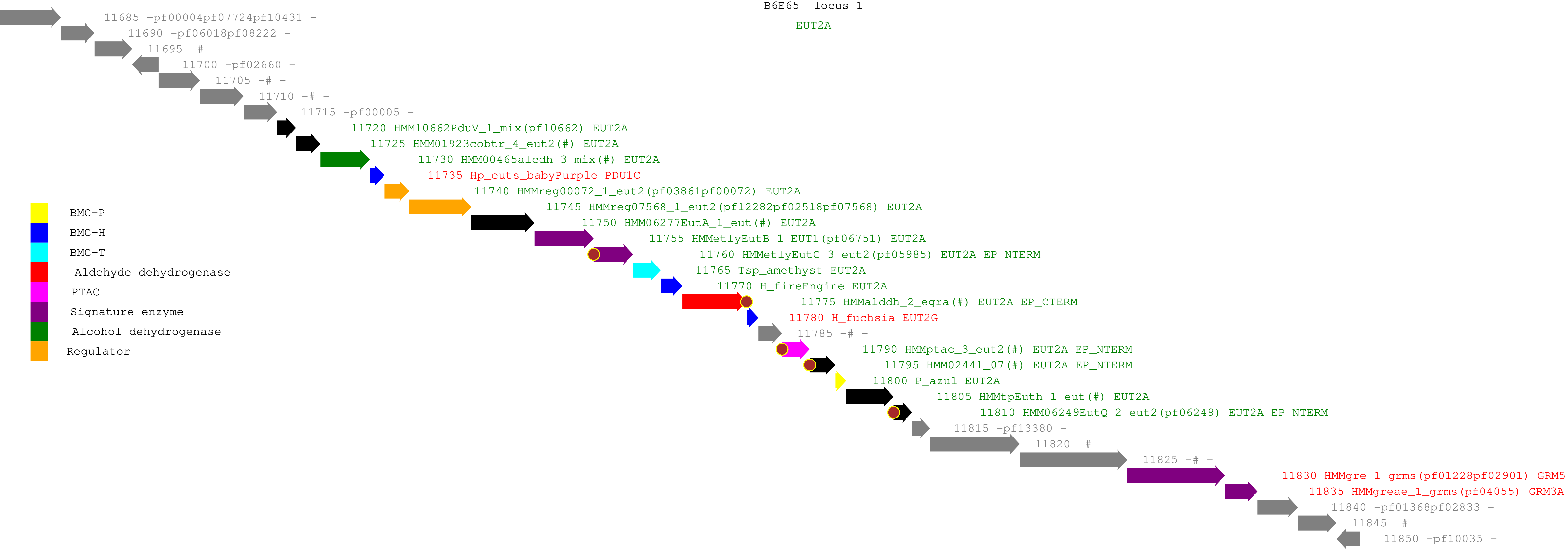

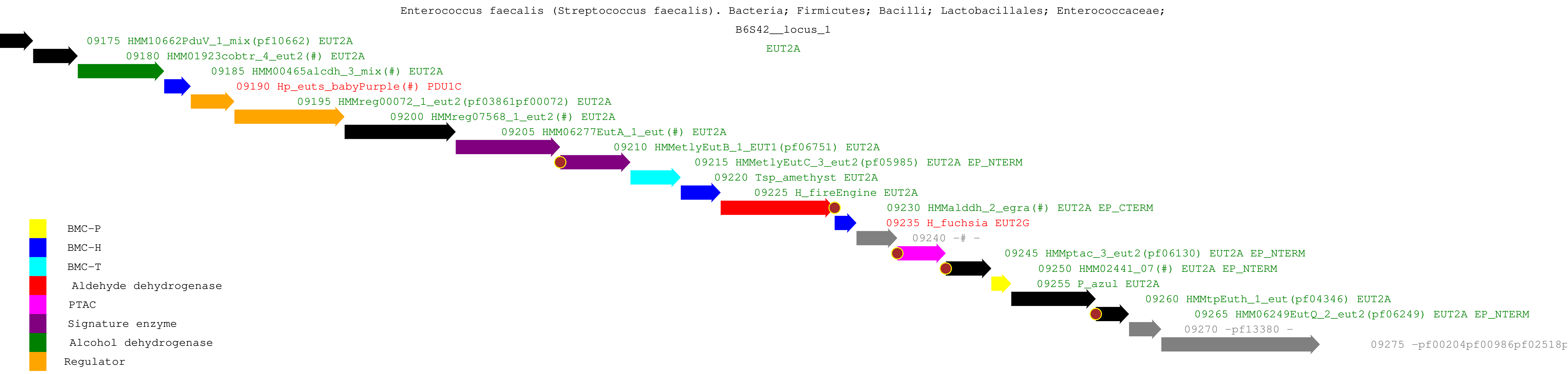
Enterococcus faecalis (Streptococcus faecalis). Bacteria; Firmicutes; Bacilli; Lactobacillales; Enterococcaceae;
B6S42__locus_1
EUT2A
09175 HMM10662PduV_1_mix(pf10662) EUT2A
09180 HMM01923cobtr_4_eut2(#) EUT2A
09185 HMM00465alcdh_3_mix(#) EUT2A
09190 Hp_euts_babyPurple(#) PDU1C
09195 HMMreg00072_1_eut2(pf03861pf00072) EUT2A
09200 HMMreg07568_1_eut2(#) EUT2A
09205 HMM06277EutA_1_eut(#) EUT2A
09210 HMMetlyEutB_1_EUT1(pf06751) EUT2A
09215 HMMetlyEutC_3_eut2(pf05985) EUT2A EP_NTERM
09220 Tsp_amethyst EUT2A
09225 H_fireEngine EUT2A
09230 HMMaldDh_2_egra(#) EUT2A EP_CTERM
09235 H_fuchsia EUT2G
09240 -# -
09245 HMMptac_3_eut2(pf06130) EUT2A EP_NTERM
09250 HMM02441_07(#) EUT2A EP_NTERM
09255 P_azul EUT2A
09260 HMMtpEuth_1_eut(pf04346) EUT2A
09265 HMM06249EutQ_2_eut2(pf06249) EUT2A EP_NTERM
09270 -pf13380 -
09275 -pf00204pf00986pf02518p
BMC-P
BMC-H
BMC-T
Aldehyde dehydrogenase
PTAC
Signature enzyme
Alcohol dehydrogenase
Regulator

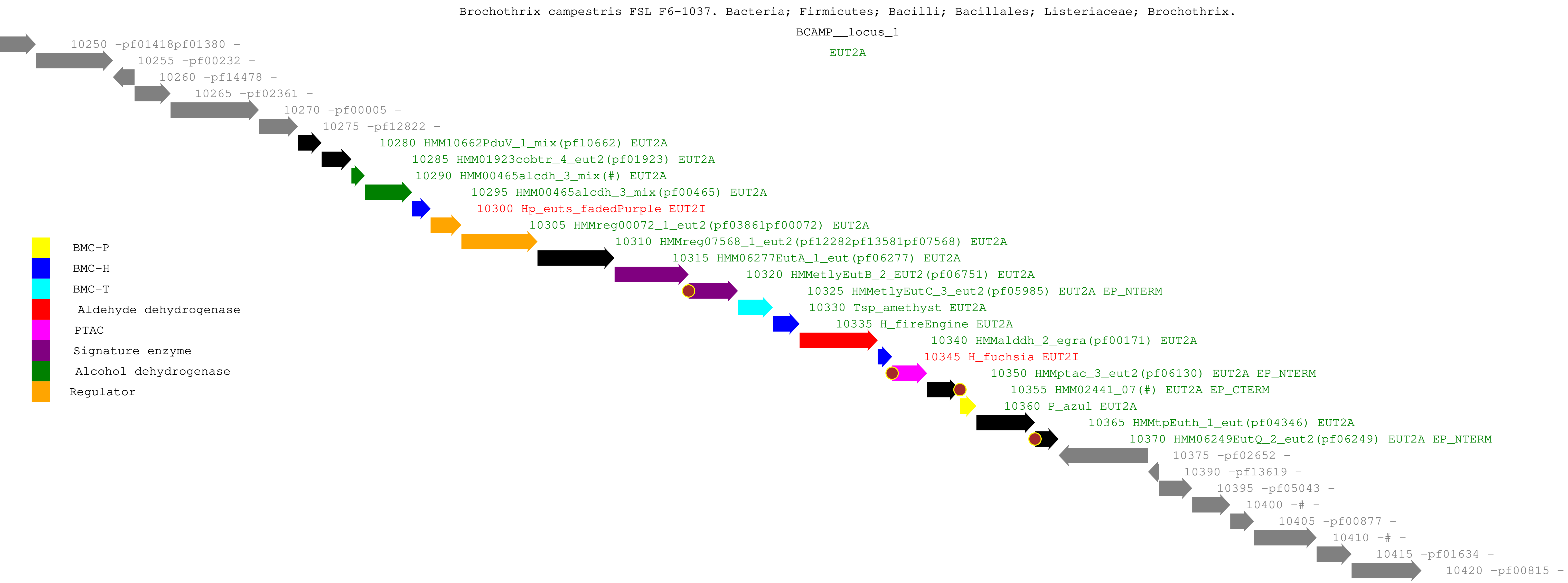
Brochothrix campestris FSL F6-1037. Bacteria; Firmicutes; Bacilli; Bacillales; Listeriaceae; Brochothrix.
BCAMP__locus_1
EUT2A
10250 -pf01418pf01380 -
10255 -pf00232 -
10260 -pf14478 -
10265 -pf02361 -
10270 -pf00005 -
10275 -pf12822 -
10280 HMM10662PduV_1_mix(pf10662) EUT2A
10285 HMM01923cobtr_4_eut2(pf01923) EUT2A
10290 HMM00465alcdh_3_mix(#) EUT2A
10295 HMM00465alcdh_3_mix(pf00465) EUT2A
10300 Hp_euts_fadedPurple EUT2I
10305 HMMreg00072_1_eut2(pf03861pf00072) EUT2A
10310 HMMreg07568_1_eut2(pf12282pf13581pf07568) EUT2A
10315 HMM06277EutA_1_eut(pf06277) EUT2A
10320 HMMetlyEutB_2_EUT2(pf06751) EUT2A
10325 HMMetlyEutC_3_eut2(pf05985) EUT2A EP_NTERM
10330 Tsp_amethyst EUT2A
10335 H_fireEngine EUT2A
10340 HMMalddh_2_egra(pf00171) EUT2A
10345 H_fuchsia EUT2I
10350 HMMptac_3_eut2(pf06130) EUT2A EP_NTERM
10355 HMM02441_07(#) EUT2A EP_CTERM
10360 P_azul EUT2A
10365 HMMtpEuth_1_eut(pf04346) EUT2A
10370 HMM06249EutQ_2_eut2(pf06249) EUT2A EP_NTERM
10375 -pf02652 -
10390 -pf13619 -
10395 -pf05043 -
10400 -# -
10405 -pf00877 -
10410 -# -
10415 -pf01634 -
10420 -pf00815 -
BMC-P
BMC-H
BMC-T
Aldehyde dehydrogenase
PTAC
Signature enzyme
Alcohol dehydrogenase
Regulator

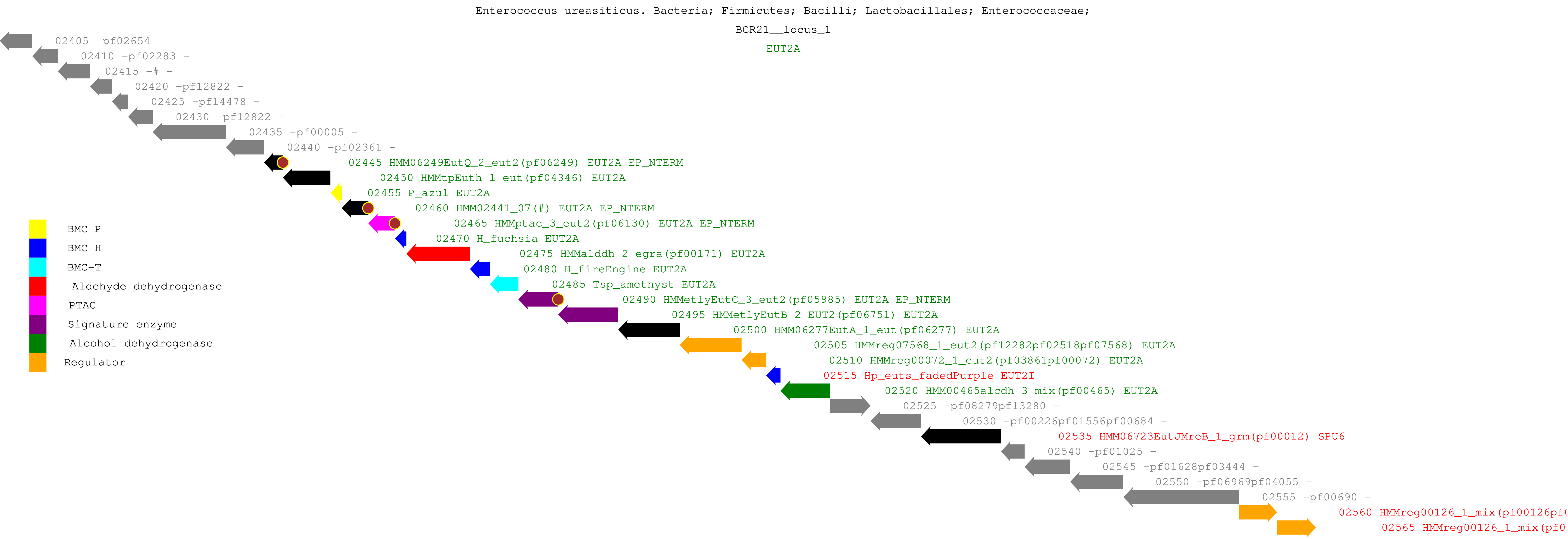
Enterococcus ureasiticus. Bacteria; Firmicutes; Bacilli; Lactobacillales; Enterococcaceae;
BCR21__locus_1
EUT2A
02405 -pf02654 -
02410 -pf02283 -
02415 -# -
02420 -pf12822 -
02425 -pf14478 -
02430 -pf12822 -
02435 -pf00005 -
02440 -pf02361 -
02445 HMM06249EutQ_2_eut2(pf06249) EUT2A EP_NTERM
02450 HMMtpEuth_1_eut(pf04346) EUT2A
02455 P_azul EUT2A
02460 HMM02441_07(#) EUT2A EP_NTERM
02465 HMMptac_3_eut2(pf06130) EUT2A EP_NTERM
02470 H_fuchsia EUT2A
02475 HMMalddh_2_egra(pf00171) EUT2A
02480 H_fireEngine EUT2A
02485 Tsp_amethyst EUT2A
02490 HMMetlyEutC_3_eut2(pf05985) EUT2A EP_NTERM
02495 HMMetlyEutB_2_EUT2(pf06751) EUT2A
02500 HMM06277EutA_1_eut(pf06277) EUT2A
02505 HMMreg07568_1_eut2(pf12282pf02518pf07568) EUT2A
02510 HMMreg00072_1_eut2(pf03861pf00072) EUT2A
02515 Hp_euts_fadedPurple EUT2I
02520 HMM00465alcdh_3_mix(pf00465) EUT2A
02525 -pf08279pf13280 -
02530 -pf00226pf01556pf00684 -
02535 HMM06723EutJMreB_1_grm(pf00012) SPU6
02540 -pf01025 -
02545 -pf01628pf03444 -
02550 -pf06969pf04055 -
02555 -pf00690 -
02560 HMMreg00126_1_mix(pf00126pf
02565 HMMreg00126_1_mix(pf0
BMC-P
BMC-H
BMC-T
Aldehyde dehydrogenase
PTAC
Signature enzyme
Alcohol dehydrogenase
Regulator

Enterococcus termitis. Bacteria; Firmicutes; Bacilli; Lactobacillales; Enterococcaceae;

BCR25__locus_1

EUT2A

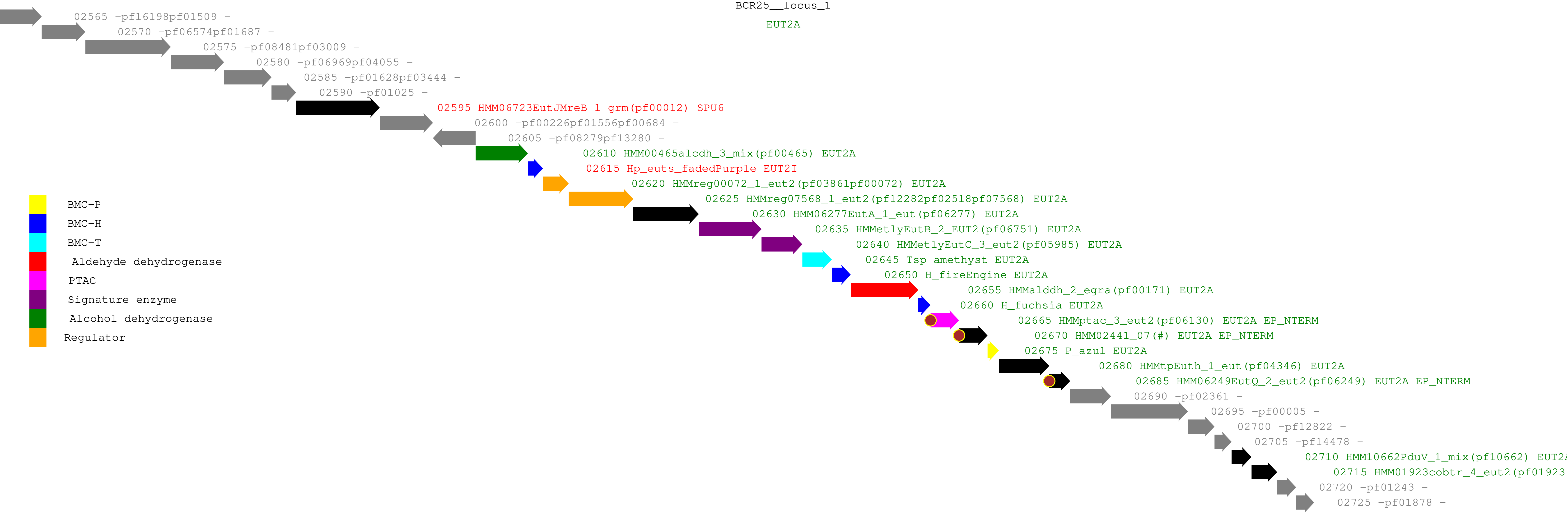

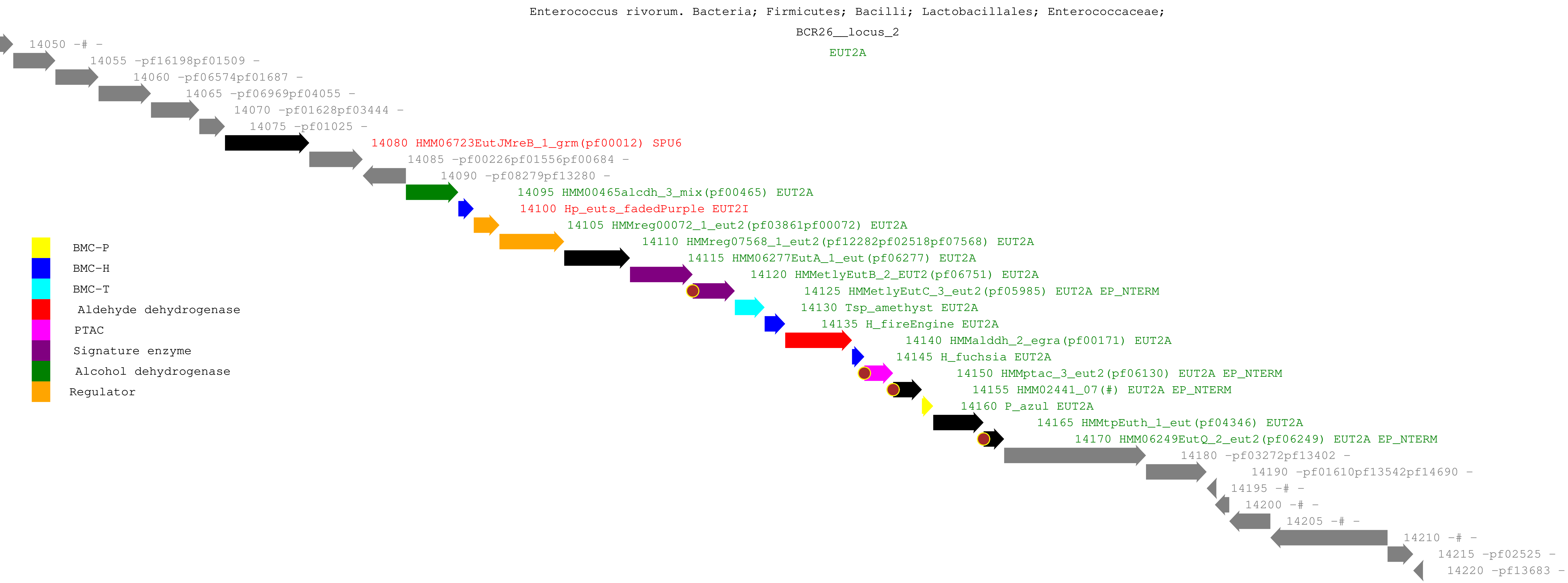
Enterococcus rivorum. Bacteria; Firmicutes; Bacilli; Lactobacillales; Enterococcaceae;
BCR26__locus_2
EUT2A
14050 -# -
14055 -pf16198pf01509 -
14060 -pf06574pf01687 -
14065 -pf06969pf04055 -
14070 -pf01628pf03444 -
14075 -pf01025 -
14080 HMM06723EutJMreB_1_grm(pf00012) SPU6
14085 -pf00226pf01556pf00684 -
14090 -pf08279pf13280 -
14095 HMM00465alcdh_3_mix(pf00465) EUT2A
14100 Hp_euts_fadedPurple EUT2I
14105 HMMreg00072_1_eut2(pf03861pf00072) EUT2A
14110 HMMreg07568_1_eut2(pf12282pf02518pf07568) EUT2A
14115 HMM06277EutA_1_eut(pf06277) EUT2A
14120 HMMetlyEutB_2_EUT2(pf06751) EUT2A
14125 HMMetlyEutC_3_eut2(pf05985) EUT2A EP_NTERM
14130 Tsp_amethyst EUT2A
14135 H_fireEngine EUT2A
14140 HMMaldh_2_egra(pf00171) EUT2A
14145 H_fuchsia EUT2A
14150 HMMptac_3_eut2(pf06130) EUT2A EP_NTERM
14155 HMM02441_07(#) EUT2A EP_NTERM
14160 P_azul EUT2A
14165 HMMtpEuth_1_eut(pf04346) EUT2A
14170 HMM06249EutQ_2_eut2(pf06249) EUT2A EP_NTERM
14180 -pf03272pf13402 -
14190 -pf01610pf13542pf14690 -
14195 -# -
14200 -# -
14205 -# -
14210 -# -
14215 -pf02525 -
14220 -pf13683 -
BMC-P
BMC-H
BMC-T
Aldehyde dehydrogenase
PTAC
Signature enzyme
Alcohol dehydrogenase
Regulator

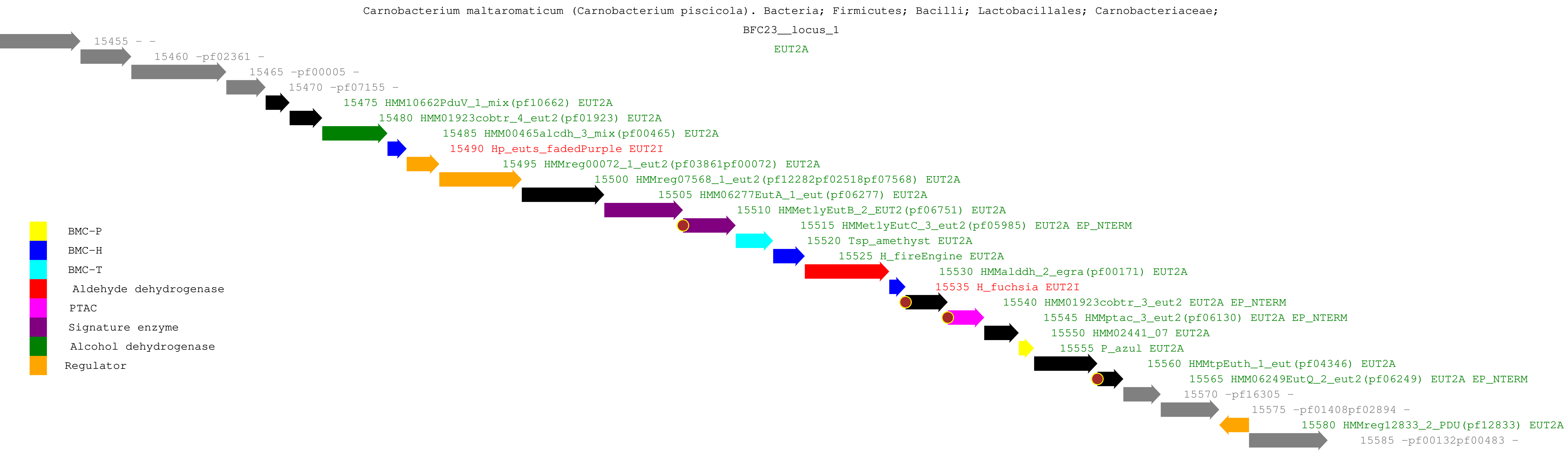
Carnobacterium maltaromaticum (Carnobacterium piscicola). Bacteria; Firmicutes; Bacilli; Lactobacillales; Carnobacteriaceae;
BFC23__locus_1
EUT2A
15455 - -
15460 -pf02361 -
15465 -pf00005 -
15470 -pf07155 -
15475 HMM10662PduV_1_mix(pf10662) EUT2A
15480 HMM01923cobtr_4_eut2(pf01923) EUT2A
15485 HMM00465alcdh_3_mix(pf00465) EUT2A
15490 Hp_euts_fadedPurple EUT2I
15495 HMMreg00072_1_eut2(pf03861pf00072) EUT2A
15500 HMMreg07568_1_eut2(pf12282pf02518pf07568) EUT2A
15505 HMM06277EutA_1_eut(pf06277) EUT2A
15510 HMMetlyEutB_2_EUT2(pf06751) EUT2A
15515 HMMetlyEutC_3_eut2(pf05985) EUT2A EP_NTERM
15520 Tsp_amethyst EUT2A
15525 H_fireEngine EUT2A
15530 HMMalddh_2_egra(pf00171) EUT2A
15535 H_fuchsia EUT2I
15540 HMM01923cobtr_3_eut2 EUT2A EP_NTERM
15545 HMMptac_3_eut2(pf06130) EUT2A EP_NTERM
15550 HMM02441_07 EUT2A
15555 P_azul EUT2A
15560 HMMtpEuth_1_eut(pf04346) EUT2A
15565 HMM06249EutQ_2_eut2(pf06249) EUT2A EP_NTERM
15570 -pf16305 -
15575 -pf01408pf02894 -
15580 HMMreg12833_2_PDU(pf12833) EUT2A
15585 -pf00132pf00483 -
BMC-P
BMC-H
BMC-T
Aldehyde dehydrogenase
PTAC
Signature enzyme
Alcohol dehydrogenase
Regulator

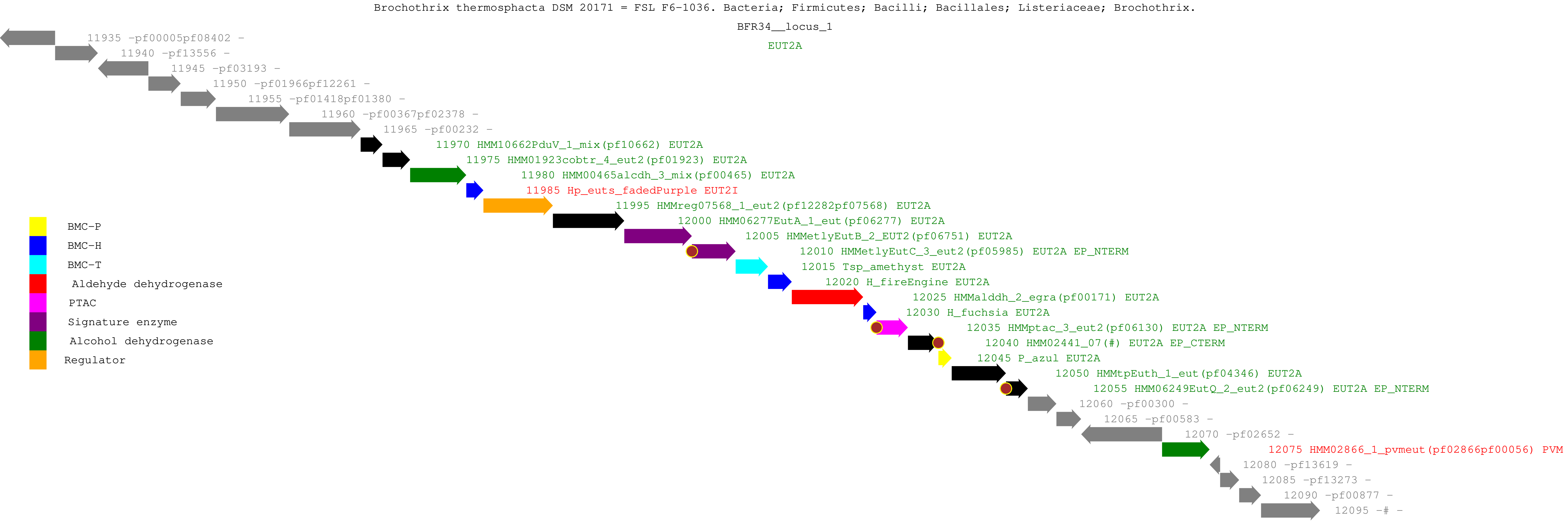

Brochothrix thermosphacta DSM 20171 = FSL F6-1036. Bacteria; Firmicutes; Bacilli; Bacillales; Listeriaceae; Brochothrix.
BFR34__locus_1
EUT2A
11935 -pf00005pf08402 -
11940 -pf13556 -
11945 -pf03193 -
11950 -pf01966pf12261 -
11955 -pf01418pf01380 -
11960 -pf00367pf02378 -
11965 -pf00232 -
11970 HMM10662PduV_1_mix(pf10662) EUT2A
11975 HMM01923cobtr_4_eut2(pf01923) EUT2A
11980 HMM00465alcdh_3_mix(pf00465) EUT2A
11985 Hp_euts_fadedPurple EUT2I
11995 HMMreg07568_1_eut2(pf12282pf07568) EUT2A
12000 HMM06277EutA_1_eut(pf06277) EUT2A
12005 HMMetlyEutB_2_EUT2(pf06751) EUT2A
12010 HMMetlyEutC_3_eut2(pf05985) EUT2A EP_NTERM
12015 Tsp_amethyst EUT2A
12020 H_fireEngine EUT2A
12025 HMMalddh_2_egra(pf00171) EUT2A
12030 H_fuchsia EUT2A
12035 HMMptac_3_eut2(pf06130) EUT2A EP_NTERM
12040 HMM02441_07(#) EUT2A EP_CTERM
12045 P_azul EUT2A
12050 HMMtpEuth_1_eut(pf04346) EUT2A
12055 HMM06249EutQ_2_eut2(pf06249) EUT2A EP_NTERM
12060 -pf00300 -
12065 -pf00583 -
12070 -pf02652 -
12075 HMM02866_1_pvmeut(pf02866pf00056) PVM
12080 -pf13619 -
12085 -pf13273 -
12090 -pf00877 -
12095 -# -
BMC-P
BMC-H
BMC-T
Aldehyde dehydrogenase
PTAC
Signature enzyme
Alcohol dehydrogenase
Regulator

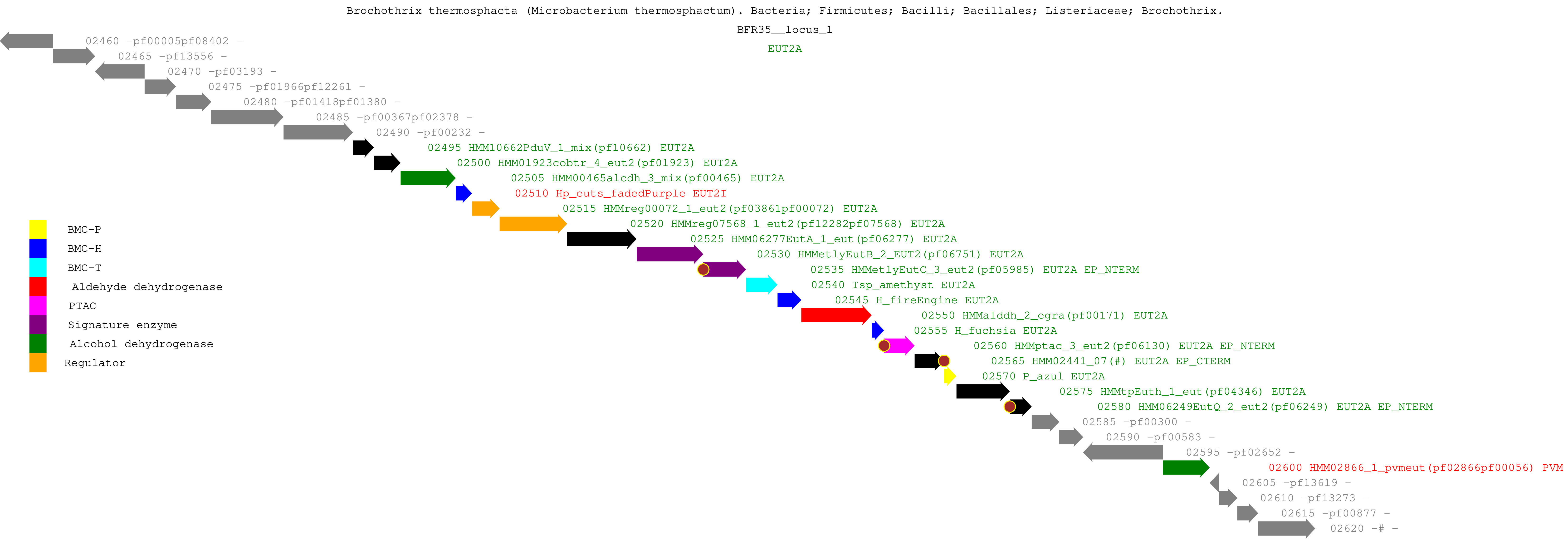

Brochothrix thermosphacta (Microbacterium thermosphactum). Bacteria; Firmicutes; Bacilli; Bacillales; Listeriaceae; Brochothrix.
BFR35__locus_1
EUT2A
02460 -pf00005pf08402 -
02465 -pf13556 -
02470 -pf03193 -
02475 -pf01966pf12261 -
02480 -pf01418pf01380 -
02485 -pf00367pf02378 -
02490 -pf00232 -
02495 HMM10662PduV_1_mix(pf10662) EUT2A
02500 HMM01923cobtr_4_eut2(pf01923) EUT2A
02505 HMM00465alcdh_3_mix(pf00465) EUT2A
02510 Hp_euts_fadedPurple EUT2I
02515 HMMreg00072_1_eut2(pf03861pf00072) EUT2A
02520 HMMreg07568_1_eut2(pf12282pf07568) EUT2A
02525 HMM06277EutA_1_eut(pf06277) EUT2A
02530 HMMetlyEutB_2_EUT2(pf06751) EUT2A
02535 HMMetlyEutC_3_eut2(pf05985) EUT2A EP_NTERM
02540 Tsp_amethyst EUT2A
02545 H_fireEngine EUT2A
02550 HMMalddh_2_egra(pf00171) EUT2A
02555 H_fuchsia EUT2A
02560 HMMptac_3_eut2(pf06130) EUT2A EP_NTERM
02565 HMM02441_07(#) EUT2A EP_CTERM
02570 P_azul EUT2A
02575 HMMtpEuth_1_eut(pf04346) EUT2A
02580 HMM06249EutQ_2_eut2(pf06249) EUT2A EP_NTERM
02585 -pf00300 -
02590 -pf00583 -
02595 -pf02652 -
02600 HMM02866_1_pvmeut(pf02866pf00056) PVM
02605 -pf13619 -
02610 -pf13273 -
02615 -pf00877 -
02620 -# -
BMC-P
BMC-H
BMC-T
Aldehyde dehydrogenase
PTAC
Signature enzyme
Alcohol dehydrogenase
Regulator

Brochothrix thermosphacta (Microbacterium thermosphactum). Bacteria; Firmicutes; Bacilli; Bacillales; Listeriaceae; Brochothrix.

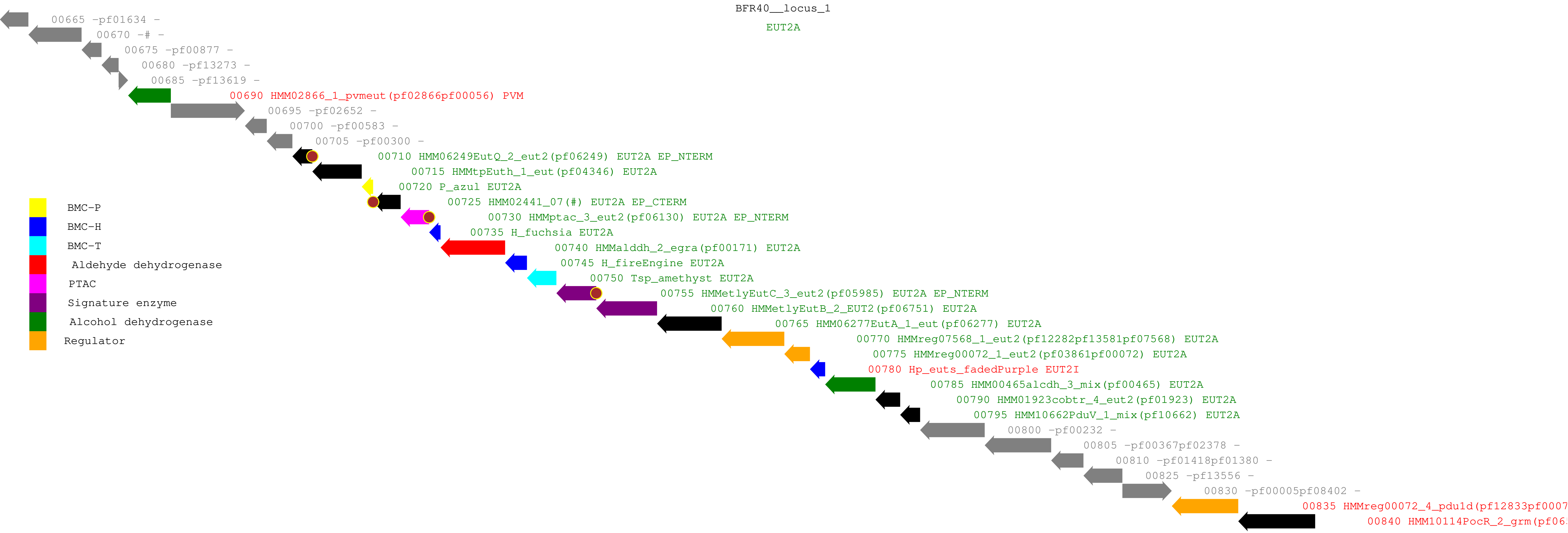

Brochothrix thermosphacta (Microbacterium thermosphactum). Bacteria; Firmicutes; Bacilli; Bacillales; Listeriaceae; Brochothrix.

BFR41__locus_1

EUT2A

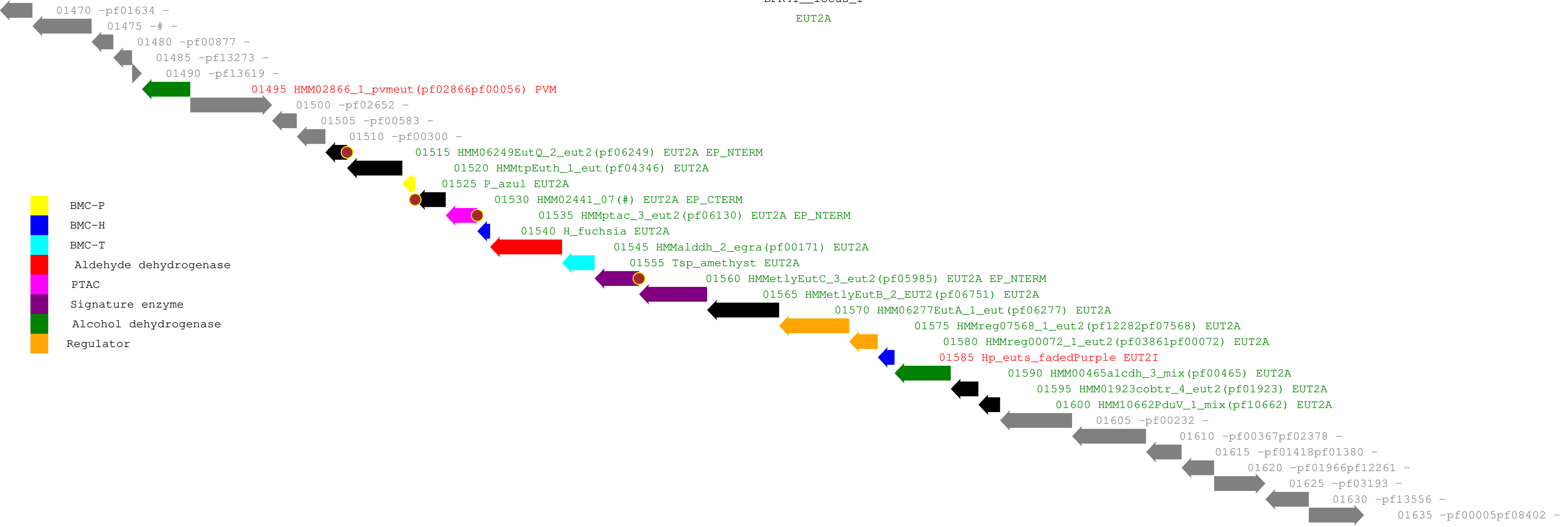

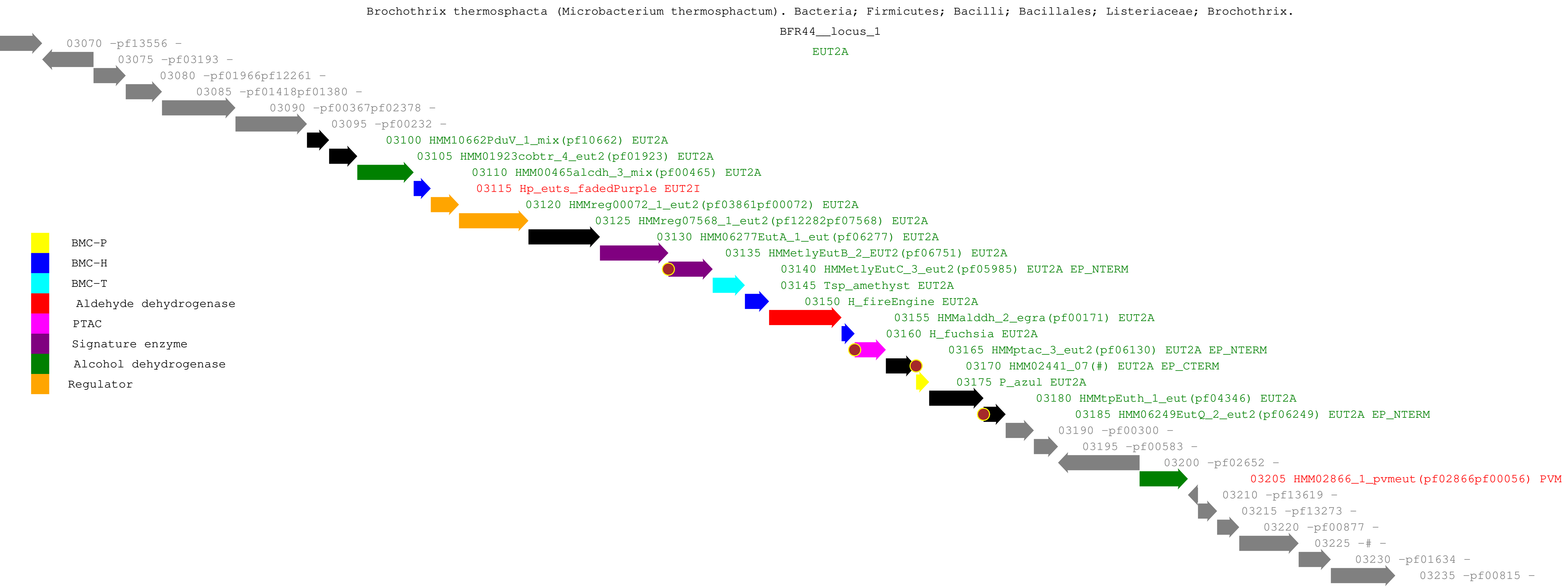
Brochothrix thermosphacta (Microbacterium thermosphactum). Bacteria; Firmicutes; Bacilli; Bacillales; Listeriaceae; Brochothrix.
BFR44__locus_1
EUT2A
03070 -pf13556 -
03075 -pf03193 -
03080 -pf01966pf12261 -
03085 -pf01418pf01380 -
03090 -pf00367pf02378 -
03095 -pf00232 -
03100 HMM10662PduV_1_mix(pf10662) EUT2A
03105 HMM01923cobtr_4_eut2(pf01923) EUT2A
03110 HMM00465alcdh_3_mix(pf00465) EUT2A
03115 Hp_euts_fadedPurple EUT2I
03120 HMMreg00072_1_eut2(pf03861pf00072) EUT2A
03125 HMMreg07568_1_eut2(pf12282pf07568) EUT2A
03130 HMM06277EutA_1_eut(pf06277) EUT2A
03135 HMMetlyEutB_2_EUT2(pf06751) EUT2A
03140 HMMetlyEutC_3_eut2(pf05985) EUT2A EP_NTERM
03145 Tsp_amethyst EUT2A
03150 H_fireEngine EUT2A
03155 HMMalddh_2_egra(pf00171) EUT2A
03160 H_fuchsia EUT2A
03165 HMMptac_3_eut2(pf06130) EUT2A EP_NTERM
03170 HMM02441_07(#) EUT2A EP_CTERM
03175 P_azul EUT2A
03180 HMMtpEuth_1_eut(pf04346) EUT2A
03185 HMM06249EutQ_2_eut2(pf06249) EUT2A EP_NTERM
03190 -pf00300 -
03195 -pf00583 -
03200 -pf02652 -
03205 HMM02866_1_pvmeut(pf02866pf00056) PVM
03210 -pf13619 -
03215 -pf13273 -
03220 -pf00877 -
03225 -# -
03230 -pf01634 -
03235 -pf00815 -
BMC-P
BMC-H
BMC-T
Aldehyde dehydrogenase
PTAC
Signature enzyme
Alcohol dehydrogenase
Regulator

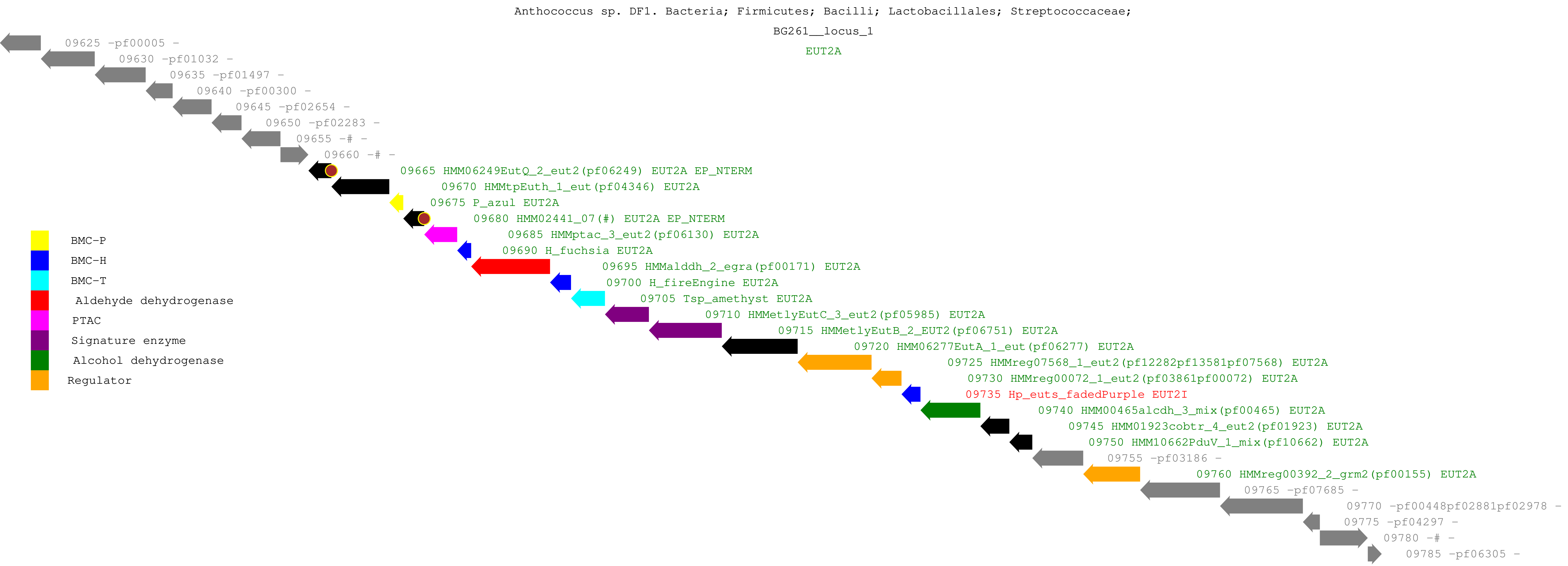

Anthococcus sp. DF1. Bacteria; Firmicutes; Bacilli; Lactobacillales; Streptococcaceae;
BG261__locus_1
EUT2A
09625 -pf00005 -
09630 -pf01032 -
09635 -pf01497 -
09640 -pf00300 -
09645 -pf02654 -
09650 -pf02283 -
09655 -# -
09660 -# -
09665 HMM06249EutQ_2_eut2(pf06249) EUT2A EP_NTERM
09670 HMMtpEuth_1_eut(pf04346) EUT2A
09675 P_azul EUT2A
09680 HMM02441_07(#) EUT2A EP_NTERM
09685 HMMptac_3_eut2(pf06130) EUT2A
09690 H_fuchsia EUT2A
09695 HMMalddh_2_egra(pf00171) EUT2A
09700 H_fireEngine EUT2A
09705 Tsp_amethyst EUT2A
09710 HMMetlyEutC_3_eut2(pf05985) EUT2A
09715 HMMetlyEutB_2_EUT2(pf06751) EUT2A
09720 HMM06277EutA_1_eut(pf06277) EUT2A
09725 HMMreg07568_1_eut2(pf12282pf13581pf07568) EUT2A
09730 HMMreg00072_1_eut2(pf03861pf00072) EUT2A
09735 Hp_euts_fadedPurple EUT2I
09740 HMM00465alcdh_3_mix(pf00465) EUT2A
09745 HMM01923cobtr_4_eut2(pf01923) EUT2A
09750 HMM10662PduV_1_mix(pf10662) EUT2A
09755 -pf03186 -
09760 HMMreg00392_2_grm2(pf00155) EUT2A
09765 -pf07685 -
09770 -pf00448pf02881pf02978 -
09775 -pf04297 -
09780 -# -
09785 -pf06305 -
BMC-P
BMC-H
BMC-T
Aldehyde dehydrogenase
PTAC
Signature enzyme
Alcohol dehydrogenase
Regulator

Enterococcus gallinarum. Bacteria; Firmicutes; Bacilli; Lactobacillales; Enterococcaceae;

BH745__locus_1

EUT2A

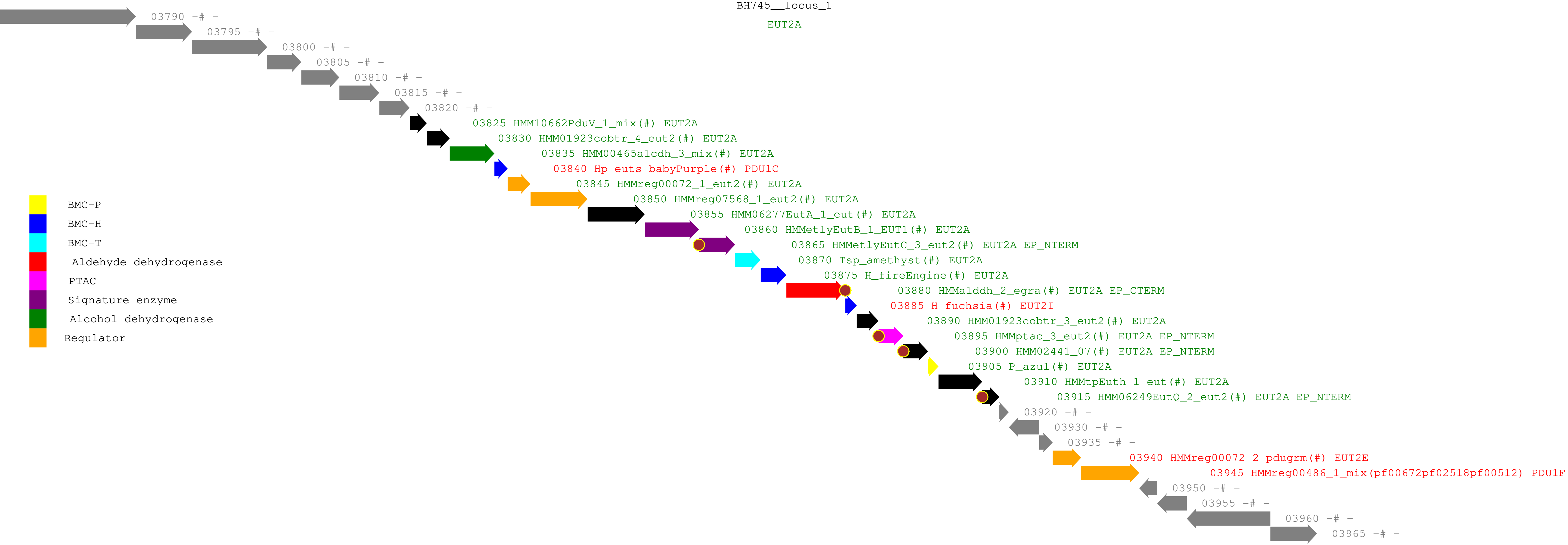

Enterococcus faecalis (Streptococcus faecalis). Bacteria; Firmicutes; Bacilli; Lactobacillales; Enterococcaceae;

BHU49__locus_1

EUT2A

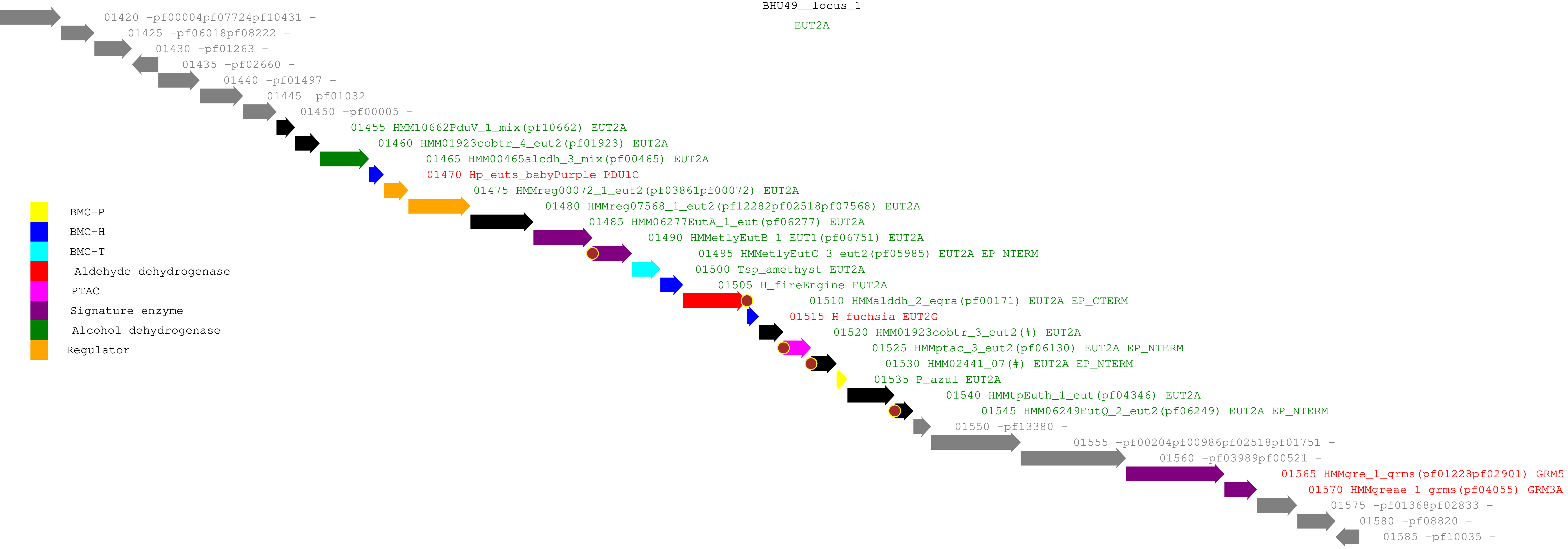

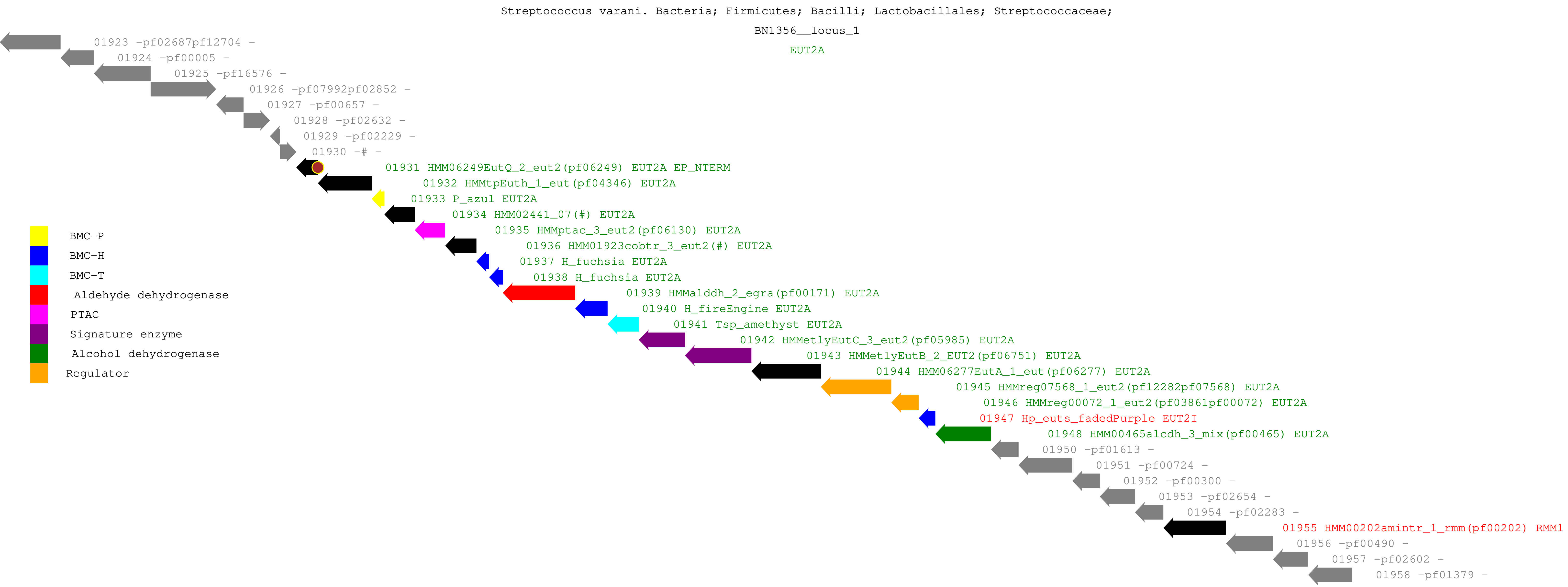

Streptococcus varani. Bacteria; Firmicutes; Bacilli; Lactobacillales; Streptococcaceae;
BN1356__locus_1
EUT2A
01923 -pf02687pf12704 -
01924 -pf00005 -
01925 -pf16576 -
01926 -pf07992pf02852 -
01927 -pf00657 -
01928 -pf02632 -
01929 -pf02229 -
01930 -# -
01931 HMM06249EutQ_2_eut2(pf06249) EUT2A EP_NTERM
01932 HMMtpEuth_1_eut(pf04346) EUT2A
01933 P_azul EUT2A
01934 HMM02441_07(#) EUT2A
01935 HMMptac_3_eut2(pf06130) EUT2A
01936 HMM01923cobtr_3_eut2(#) EUT2A
01937 H_fuchsia EUT2A
01938 H_fuchsia EUT2A
01939 HMMalddh_2_egra(pf00171) EUT2A
01940 H_fireEngine EUT2A
01941 Tsp_amethyst EUT2A
01942 HMMetlyEutC_3_eut2(pf05985) EUT2A
01943 HMMetlyEutB_2_EUT2(pf06751) EUT2A
01944 HMM06277EutA_1_eut(pf06277) EUT2A
01945 HMMreg07568_1_eut2(pf12282pf07568) EUT2A
01946 HMMreg00072_1_eut2(pf03861pf00072) EUT2A
01947 Hp_euts_fadedPurple EUT2I
01948 HMM00465alcdh_3_mix(pf00465) EUT2A
01950 -pf01613 -
01951 -pf00724 -
01952 -pf00300 -
01953 -pf02654 -
01954 -pf02283 -
01955 HMM00202amintr_1_rmm(pf00202) RMM1
01956 -pf00490 -
01957 -pf02602 -
01958 -pf01379 -
BMC-P
BMC-H
BMC-T
Aldehyde dehydrogenase
PTAC
Signature enzyme
Alcohol dehydrogenase
Regulator

Clostridium sp. C105KSO15. Bacteria; Firmicutes; Clostridia; Clostridiales; Clostridiaceae;

BN3590__locus_1

EUT2A

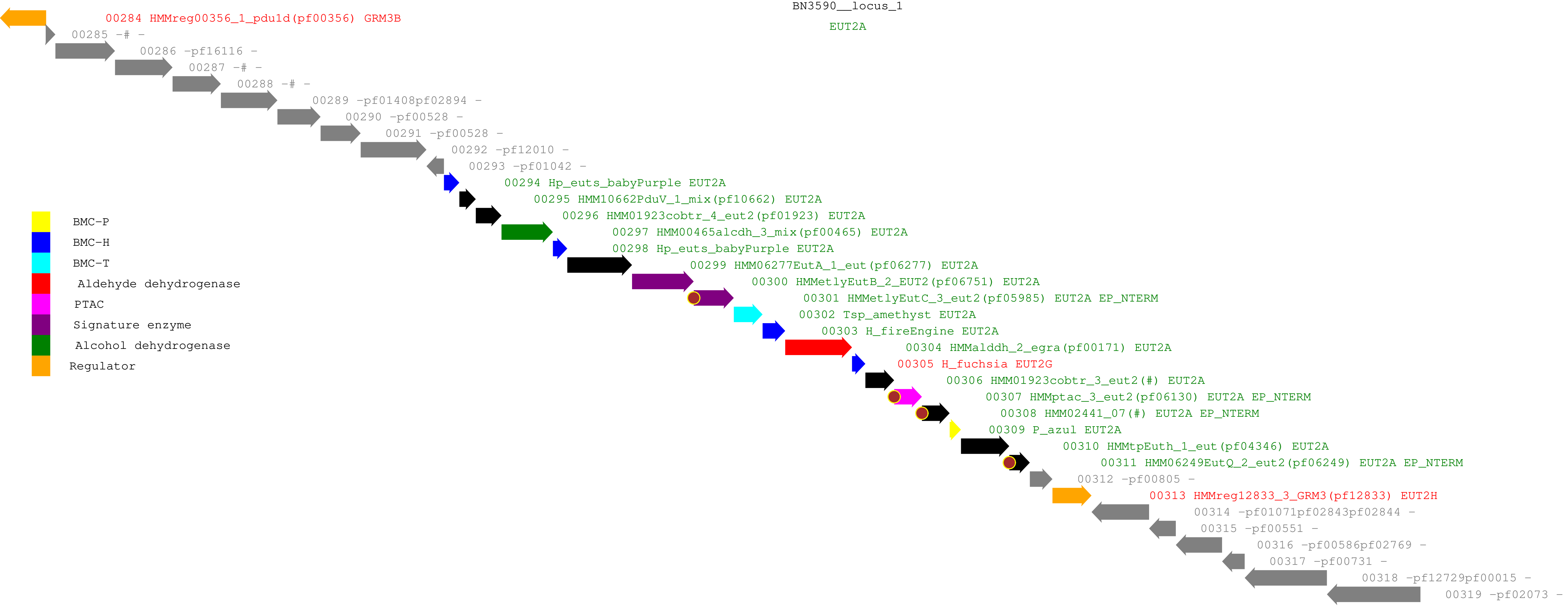

Carnobacterium maltaromaticum LMA28. Bacteria; Firmicutes; Bacilli; Lactobacillales; Carnobacteriaceae;

BN424__locus_1

EUT2A

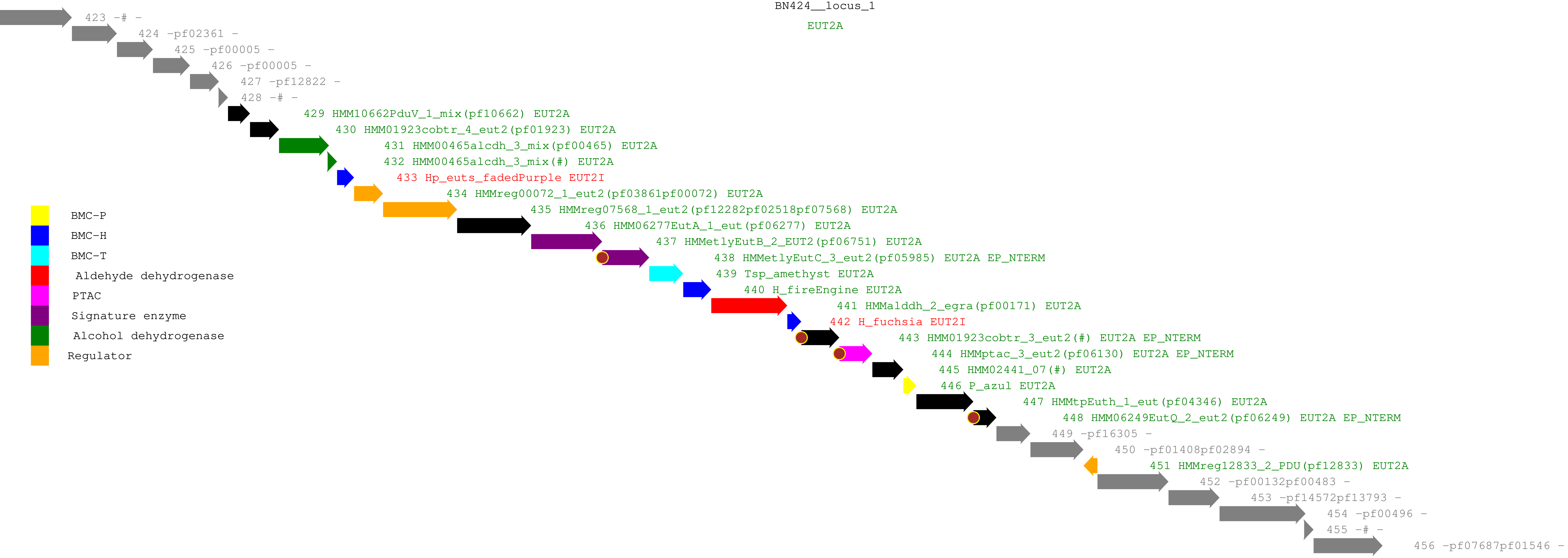

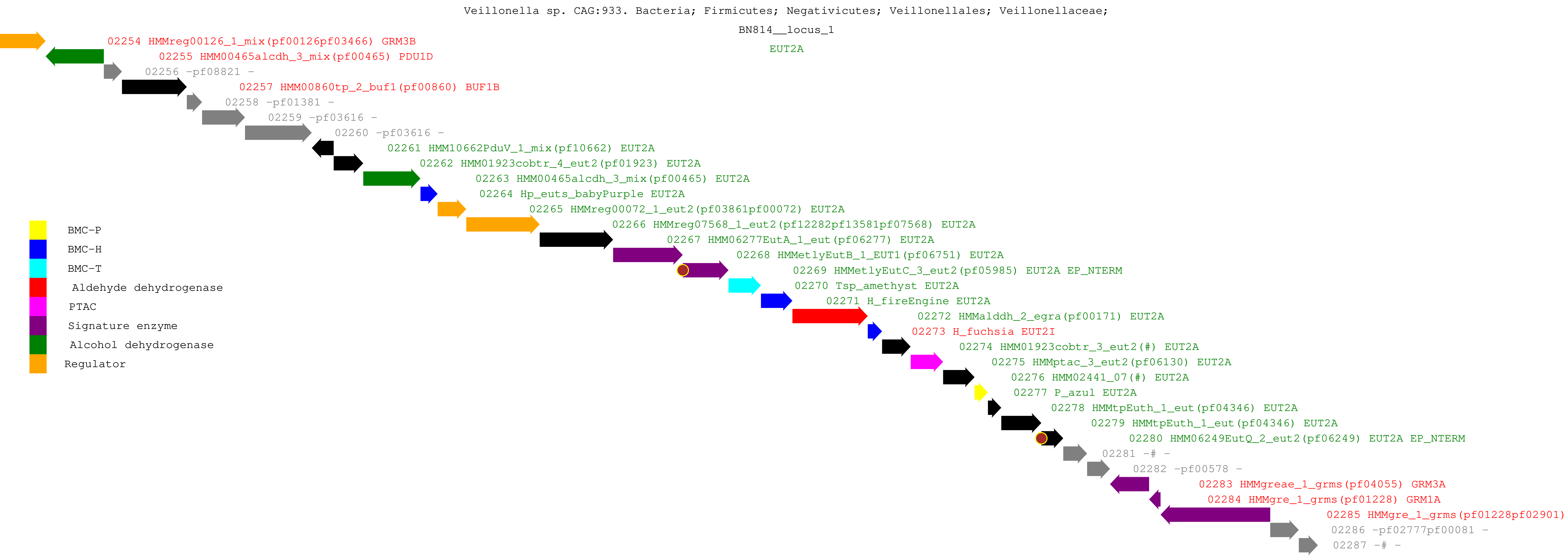

Veillonella sp. CAG:933. Bacteria; Firmicutes; Negativicutes; Veillonellales; Veillonellaceae;
BN814__locus_1
EUT2A
02254 HMMreg00126_1_mix(pf00126pf03466) GRM3B
02255 HMM00465alcdh_3_mix(pf00465) PDU1D
02256 -pf08821 -
02257 HMM00860tp_2_buf1(pf00860) BUF1B
02258 -pf01381 -
02259 -pf03616 -
02260 -pf03616 -
02261 HMM10662PduV_1_mix(pf10662) EUT2A
02262 HMM01923cobtr_4_eut2(pf01923) EUT2A
02263 HMM00465alcdh_3_mix(pf00465) EUT2A
02264 Hp_euts_babyPurple EUT2A
02265 HMMreg00072_1_eut2(pf03861pf00072) EUT2A
02266 HMMreg07568_1_eut2(pf12282pf13581pf07568) EUT2A
02267 HMM06277EutA_1_eut(pf06277) EUT2A
02268 HMMetlyEutB_1_EUT1(pf06751) EUT2A
02269 HMMetlyEutC_3_eut2(pf05985) EUT2A EP_NTERM
02270 Tsp_amethyst EUT2A
02271 H_fireEngine EUT2A
02272 HMMalddh_2_egra(pf00171) EUT2A
02273 H_fuchsia EUT2I
02274 HMM01923cobtr_3_eut2(#) EUT2A
02275 HMMptac_3_eut2(pf06130) EUT2A
02276 HMM02441_07(#) EUT2A
02277 P_azul EUT2A
02278 HMMtpEuth_1_eut(pf04346) EUT2A
02279 HMMtpEuth_1_eut(pf04346) EUT2A
02280 HMM06249EutQ_2_eut2(pf06249) EUT2A EP_NTERM
02281 -# -
02282 -pf00578 -
02283 HMMgreae_1_grms(pf04055) GRM3A
02284 HMMgre_1_grms(pf01228) GRM1A
02285 HMMgre_1_grms(pf01228pf02901)
02286 -pf02777pf00081 -
02287 -# -
BMC-P
BMC-H
BMC-T
Aldehyde dehydrogenase
PTAC
Signature enzyme
Alcohol dehydrogenase
Regulator

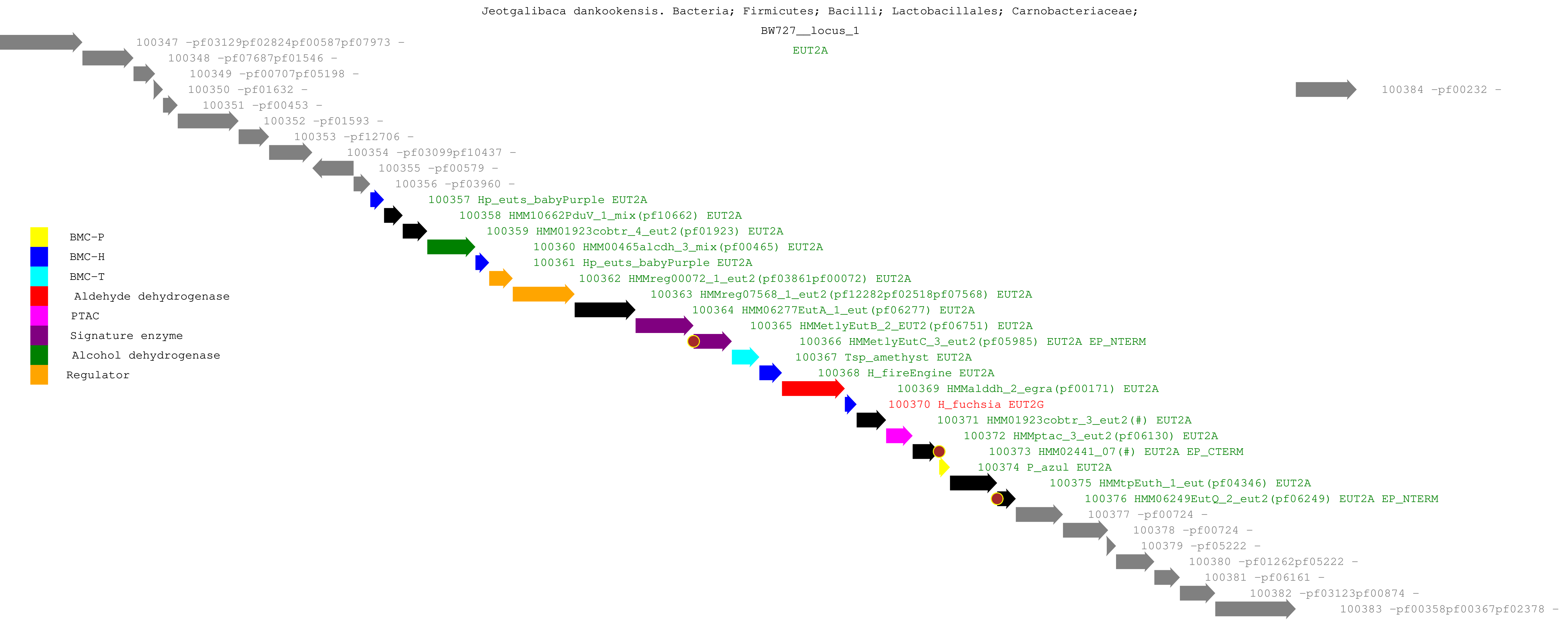
Jeotgalibaca dankookensis. Bacteria; Firmicutes; Bacilli; Lactobacillales; Carnobacteriaceae;
BW727__locus_1
EUT2A
100347 -pf03129pf02824pf00587pf07973 -
100348 -pf07687pf01546 -
100349 -pf00707pf05198 -
100350 -pf01632 -
100351 -pf00453 -
100352 -pf01593 -
100353 -pf12706 -
100354 -pf03099pf10437 -
100355 -pf00579 -
100356 -pf03960 -
100357 Hp_euts_babyPurple EUT2A
100358 HMM10662PduV_1_mix(pf10662) EUT2A
100359 HMM01923cobtr_4_eut2(pf01923) EUT2A
100360 HMM00465alcdh_3_mix(pf00465) EUT2A
100361 Hp_euts_babyPurple EUT2A
100362 HMMreg00072_1_eut2(pf03861pf00072) EUT2A
100363 HMMreg07568_1_eut2(pf12282pf02518pf07568) EUT2A
100364 HMM06277EutA_1_eut(pf06277) EUT2A
100365 HMMetlyEutB_2_EUT2(pf06751) EUT2A
100366 HMMetlyEutC_3_eut2(pf05985) EUT2A EP_NTERM
100367 Tsp_amethyst EUT2A
100368 H_fireEngine EUT2A
100369 HMMalddh_2_egra(pf00171) EUT2A
100370 H_fuchsia EUT2G
100371 HMM01923cobtr_3_eut2(#) EUT2A
100372 HMMptac_3_eut2(pf06130) EUT2A
100373 HMM02441_07(#) EUT2A EP_CTERM
100374 P_azul EUT2A
100375 HMMtpEuth_1_eut(pf04346) EUT2A
100376 HMM06249EutQ_2_eut2(pf06249) EUT2A EP_NTERM
100377 -pf00724 -
100378 -pf00724 -
100379 -pf05222 -
100380 -pf01262pf05222 -
100381 -pf06161 -
100382 -pf03123pf00874 -
100383 -pf00358pf00367pf02378 -
100384 -pf00232 -
BMC-P
BMC-H
BMC-T
Aldehyde dehydrogenase
PTAC
Signature enzyme
Alcohol dehydrogenase
Regulator

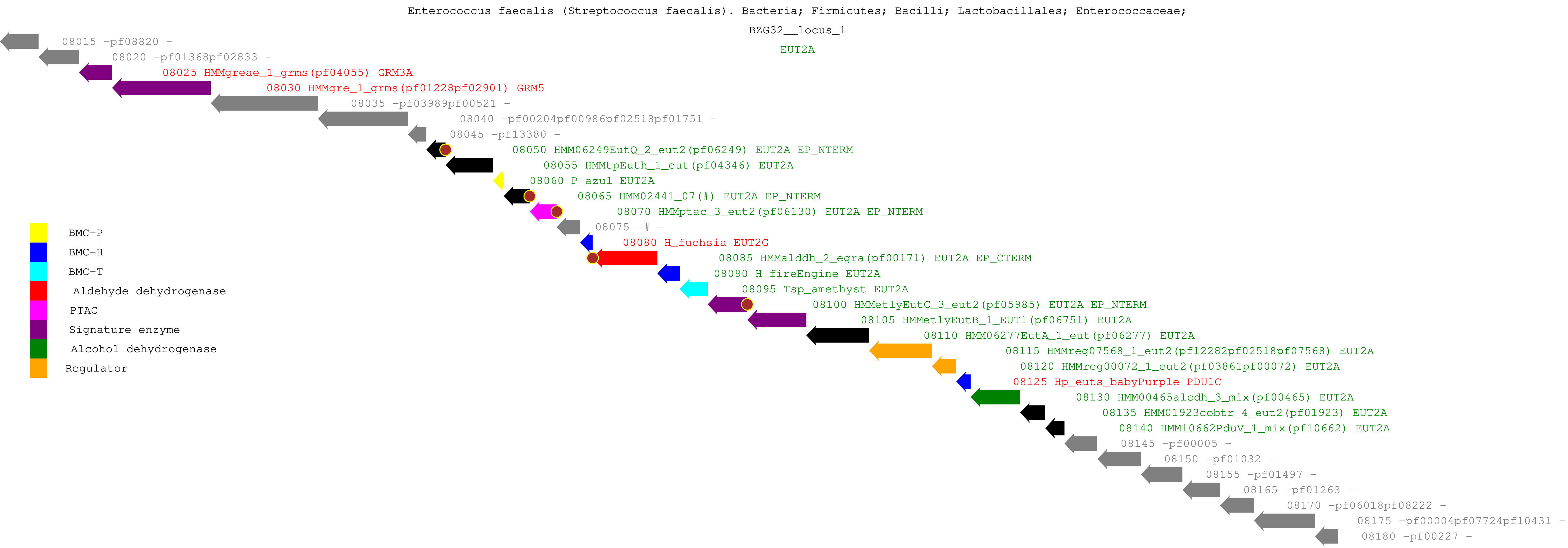

Enterococcus faecalis (Streptococcus faecalis). Bacteria; Firmicutes; Bacilli; Lactobacillales; Enterococcaceae;
BZG32__locus_1
EUT2A
08015 -pf08820 -
08020 -pf01368pf02833 -
08025 HMMgreae_1_grms(pf04055) GRM3A
08030 HMMgre_1_grms(pf01228pf02901) GRM5
08035 -pf03989pf00521 -
08040 -pf00204pf00986pf02518pf01751 -
08045 -pf13380 -
08050 HMM06249EutQ_2_eut2(pf06249) EUT2A EP_NTERM
08055 HMMtpEuth_1_eut(pf04346) EUT2A
08060 P_azul EUT2A
08065 HMM02441_07(#) EUT2A EP_NTERM
08070 HMMptac_3_eut2(pf06130) EUT2A EP_NTERM
08075 -# -
08080 H_fuchsia EUT2G
08085 HMMaldd h_2_egra(pf00171) EUT2A EP_CTERM
08090 H_fireEngine EUT2A
08095 Tsp_amethyst EUT2A
08100 HMMetlyEutC_3_eut2(pf05985) EUT2A EP_NTERM
08105 HMMetlyEutB_1_EUT1(pf06751) EUT2A
08110 HMM06277EutA_1_eut(pf06277) EUT2A
08115 HMMreg07568_1_eut2(pf12282pf02518pf07568) EUT2A
08120 HMMreg00072_1_eut2(pf03861pf00072) EUT2A
08125 Hp_euts_babyPurple PDU1C
08130 HMM00465alcdh_3_mix(pf00465) EUT2A
08135 HMM01923cobtr_4_eut2(pf01923) EUT2A
08140 HMM10662PduV_1_mix(pf10662) EUT2A
08145 -pf00005 -
08150 -pf01032 -
08155 -pf01497 -
08165 -pf01263 -
08170 -pf06018pf08222 -
08175 -pf00004pf07724pf10431 -
08180 -pf00227 -
BMC-P
BMC-H
BMC-T
Aldehyde dehydrogenase
PTAC
Signature enzyme
Alcohol dehydrogenase
Regulator

Enterococcus casseliflavus (Enterococcus flavescens). Bacteria; Firmicutes; Bacilli; Lactobacillales; Enterococcaceae;

BZK37__locus_1

EUT2A

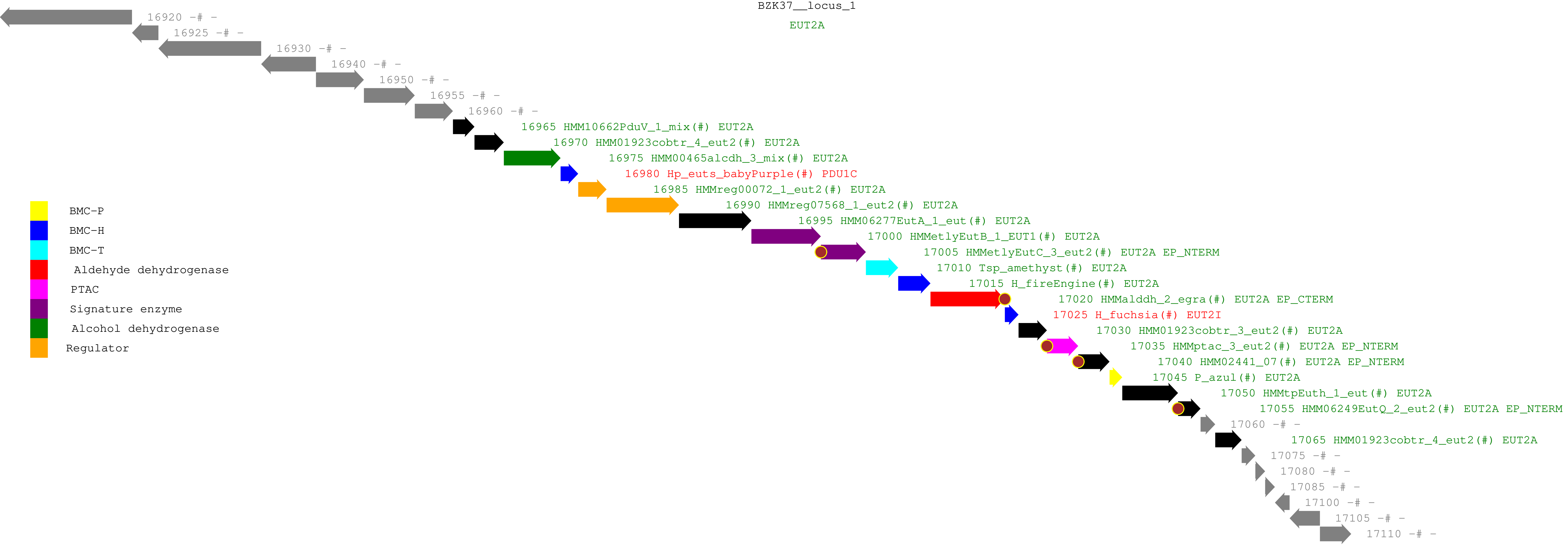

Enterococcus gallinarum. Bacteria; Firmicutes; Bacilli; Lactobacillales; Enterococcaceae;

C2L98__locus_1

EUT2A

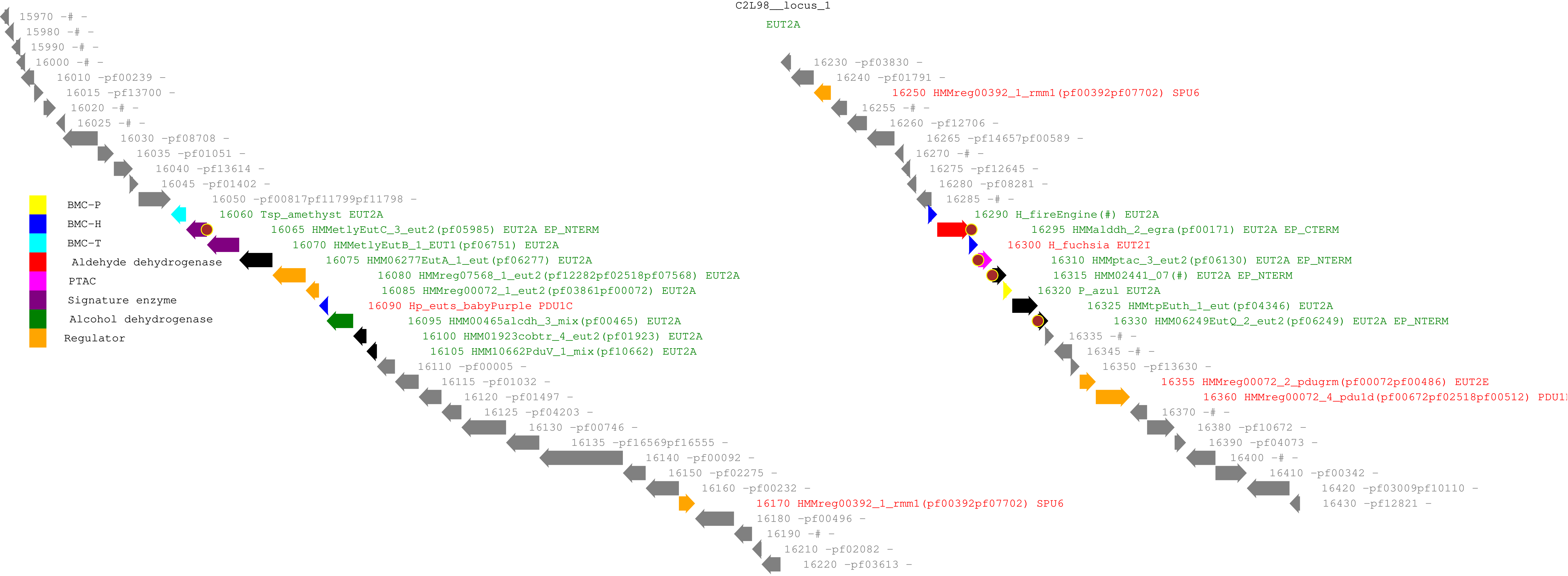

Vagococcus carniphilus. Bacteria; Firmicutes; Bacilli; Lactobacillales; Enterococcaceae;

CBF28__locus_1

EUT2A

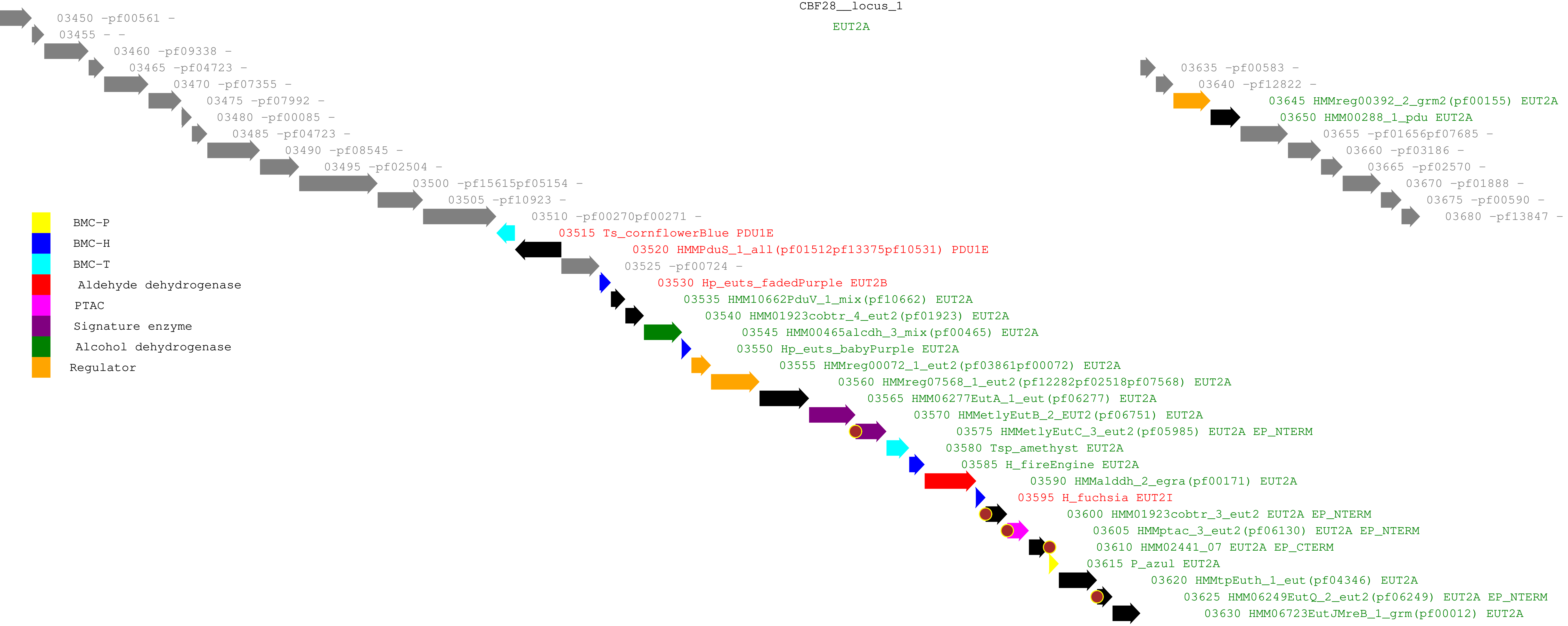

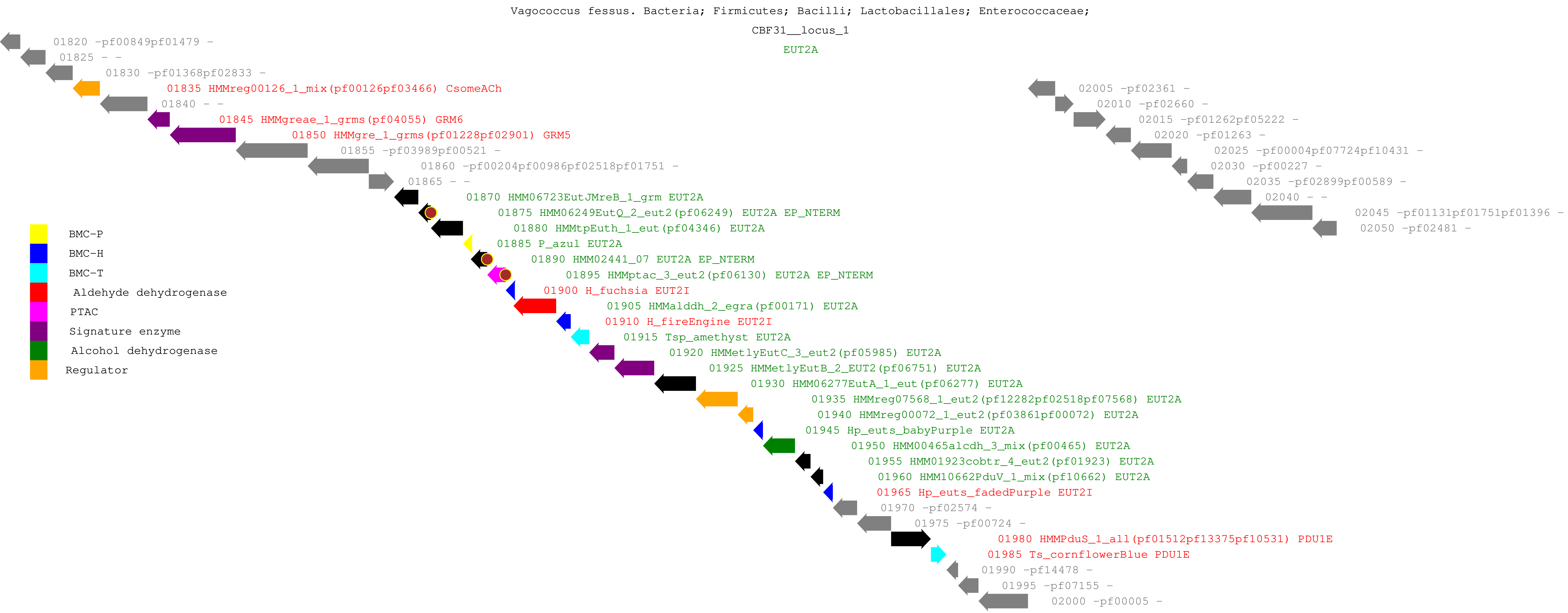

Vagococcus fessus. Bacteria; Firmicutes; Bacilli; Lactobacillales; Enterococcaceae;
CBF31__locus_1
EUT2A
01820 -pf00849pf01479 -
01825 - -
01830 -pf01368pf02833 -
01835 HMMreg00126_1_mix(pf00126pf03466) CsomeACh
01840 - -
01845 HMMgreae_1_grms(pf04055) GRM6
01850 HMMgre_1_grms(pf01228pf02901) GRM5
01855 -pf03989pf00521 -
01860 -pf00204pf00986pf02518pf01751 -
01865 - -
01870 HMM06723EutJMreB_1_grm EUT2A
01875 HMM06249EutQ_2_eut2(pf06249) EUT2A EP_NTERM
01880 HMMtpEuth_1_eut(pf04346) EUT2A
01885 P_azul EUT2A
01890 HMM02441_07 EUT2A EP_NTERM
01895 HMMptac_3_eut2(pf06130) EUT2A EP_NTERM
01900 H_fuchsia EUT2I
01905 HMMaldh_2_egra(pf00171) EUT2A
01910 H_fireEngine EUT2I
01915 Tsp_amethyst EUT2A
01920 HMMetlyEutC_3_eut2(pf05985) EUT2A
01925 HMMetlyEutB_2_EUT2(pf06751) EUT2A
01930 HMM06277EutA_1_eut(pf06277) EUT2A
01935 HMMreg07568_1_eut2(pf12282pf02518pf07568) EUT2A
01940 HMMreg00072_1_eut2(pf03861pf00072) EUT2A
01945 Hp_euts_babyPurple EUT2A
01950 HMM00465alcdh_3_mix(pf00465) EUT2A
01955 HMM01923cobtr_4_eut2(pf01923) EUT2A
01960 HMM10662PduV_1_mix(pf10662) EUT2A
01965 Hp_euts_fadedPurple EUT2I
01970 -pf02574 -
01975 -pf00724 -
01980 HMMPduS_1_all(pf01512pf13375pf10531) PDU1E
01985 Ts_cornflowerBlue PDU1E
01990 -pf14478 -
01995 -pf07155 -
02000 -pf00005 -
02005 -pf02361 -
02010 -pf02660 -
02015 -pf01262pf05222 -
02020 -pf01263 -
02025 -pf00004pf07724pf10431 -
02030 -pf00227 -
02035 -pf02899pf00589 -
02040 - -
02045 -pf01131pf01751pf01396 -
02050 -pf02481 -
BMC-P
BMC-H
BMC-T
Aldehyde dehydrogenase
PTAC
Signature enzyme
Alcohol dehydrogenase
Regulator

Vagococcus salmoninarum. Bacteria; Firmicutes; Bacilli; Lactobacillales; Enterococcaceae;

CBF35__locus_1

EUT2A

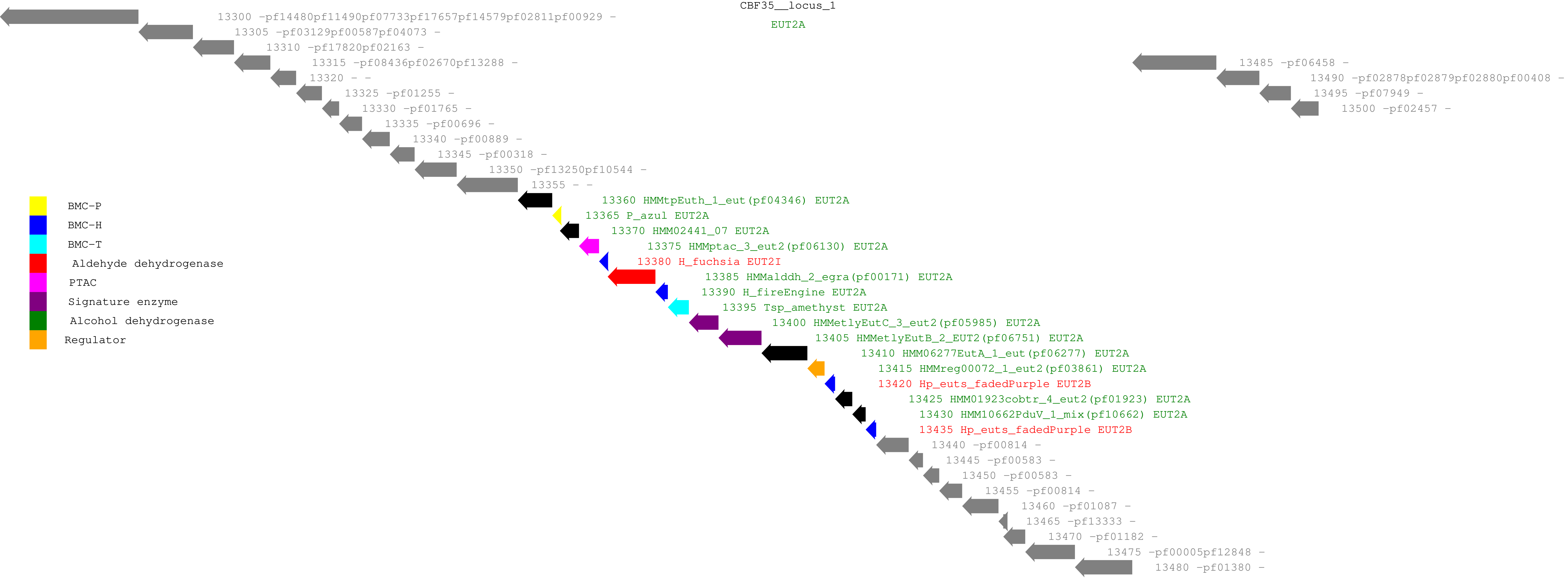

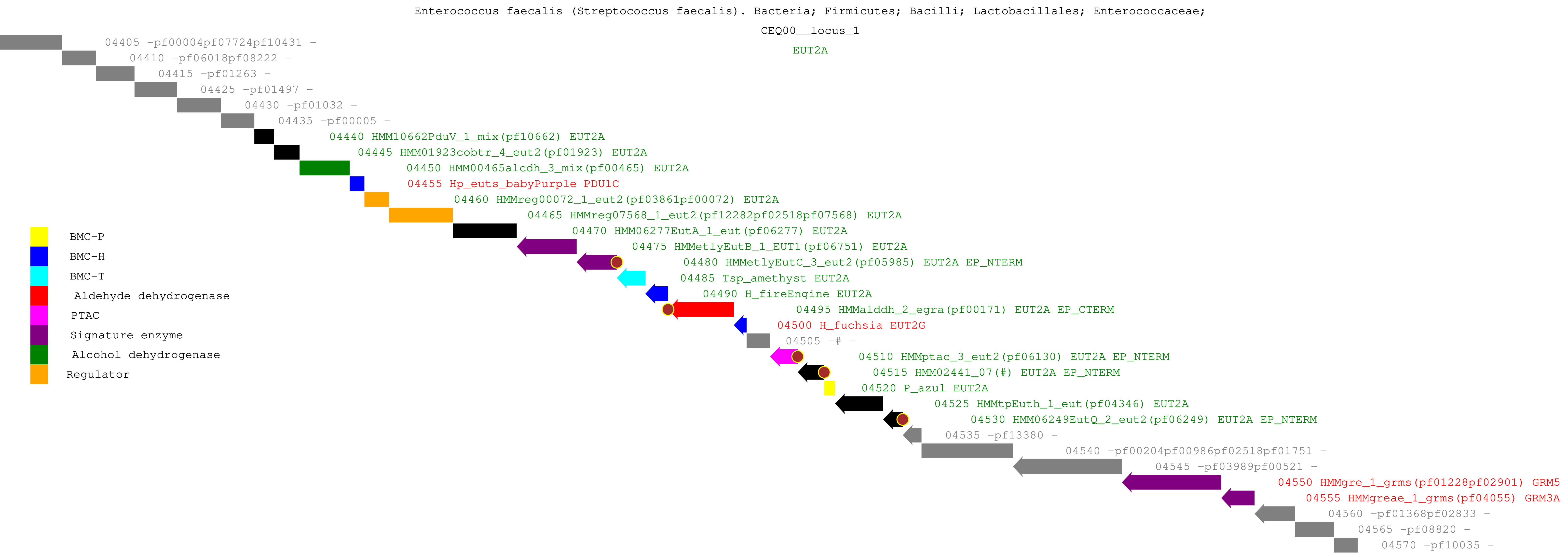
Enterococcus faecalis (Streptococcus faecalis). Bacteria; Firmicutes; Bacilli; Lactobacillales; Enterococcaceae;
CEQ00__locus_1
EUT2A
04405 -pf00004pf07724pf10431 -
04410 -pf06018pf08222 -
04415 -pf01263 -
04425 -pf01497 -
04430 -pf01032 -
04435 -pf00005 -
04440 HMM10662PduV_1_mix(pf10662) EUT2A
04445 HMM01923cobtr_4_eut2(pf01923) EUT2A
04450 HMM00465alcdh_3_mix(pf00465) EUT2A
04455 Hp_euts_babyPurple PDU1C
04460 HMMreg00072_1_eut2(pf03861pf00072) EUT2A
04465 HMMreg07568_1_eut2(pf12282pf02518pf07568) EUT2A
04470 HMM06277EutA_1_eut(pf06277) EUT2A
04475 HMMetlyEutB_1_EUT1(pf06751) EUT2A
04480 HMMetlyEutC_3_eut2(pf05985) EUT2A EP_NTERM
04485 Tsp_amethyst EUT2A
04490 H_fireEngine EUT2A
04495 HMMaldddh_2_egra(pf00171) EUT2A EP_CTERM
04500 H_fuchsia EUT2G
04505 -# -
04510 HMMptac_3_eut2(pf06130) EUT2A EP_NTERM
04515 HMM02441_07(#) EUT2A EP_NTERM
04520 P_azul EUT2A
04525 HMMtpEuth_1_eut(pf04346) EUT2A
04530 HMM06249EutQ_2_eut2(pf06249) EUT2A EP_NTERM
04535 -pf13380 -
04540 -pf00204pf00986pf02518pf01751 -
04545 -pf03989pf00521 -
04550 HMMgre_1_grms(pf01228pf02901) GRM5
04555 HMMgreae_1_grms(pf04055) GRM3A
04560 -pf01368pf02833 -
04565 -pf08820 -
04570 -pf10035 -
BMC-P
BMC-H
BMC-T
Aldehyde dehydrogenase
PTAC
Signature enzyme
Alcohol dehydrogenase
Regulator

Enterococcus faecalis (Streptococcus faecalis). Bacteria; Firmicutes; Bacilli; Lactobacillales; Enterococcaceae;

CEQ17__locus_1

EUT2A

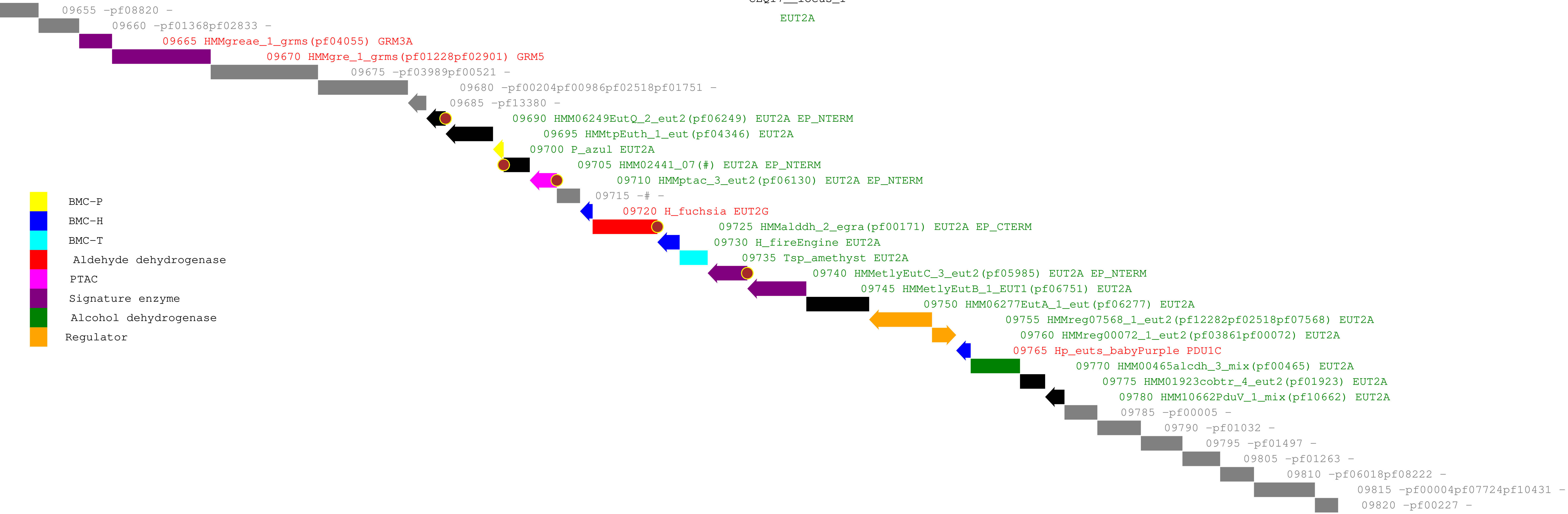

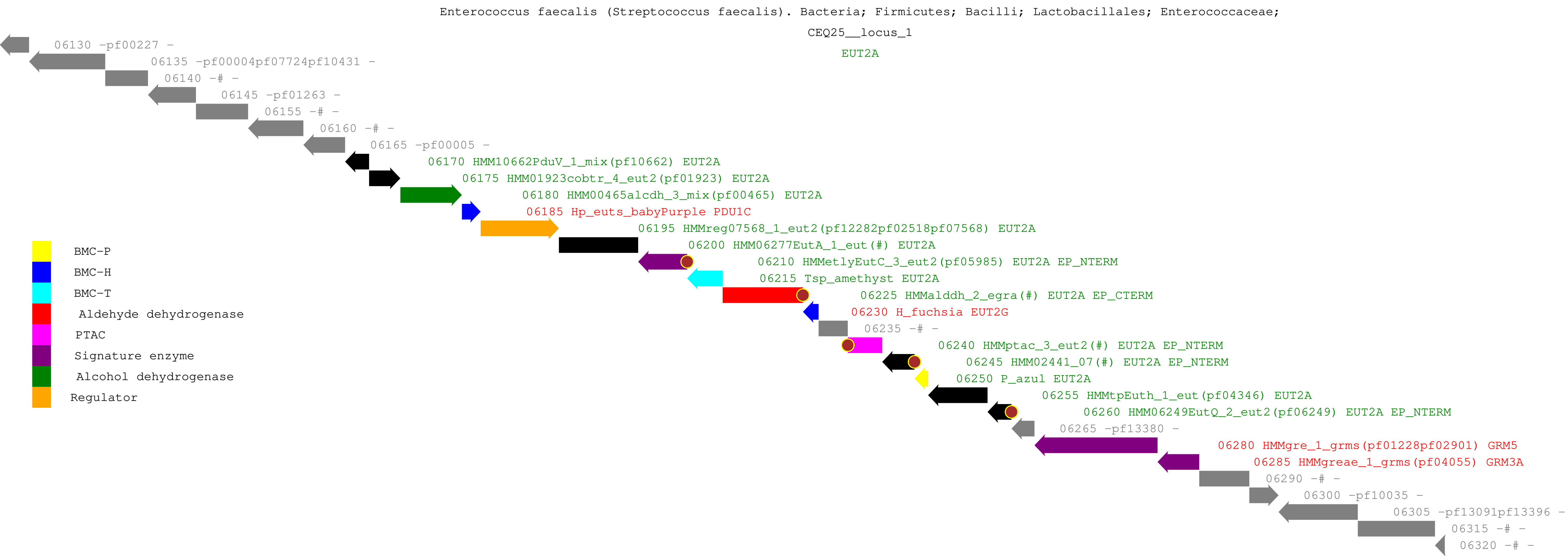

Enterococcus faecalis (Streptococcus faecalis). Bacteria; Firmicutes; Bacilli; Lactobacillales; Enterococcaceae;
CEQ25__locus_1
EUT2A
06130 -pf00227 -
06135 -pf00004pf07724pf10431 -
06140 -# -
06145 -pf01263 -
06155 -# -
06160 -# -
06165 -pf00005 -
06170 HMM10662PduV_1_mix(pf10662) EUT2A
06175 HMM01923cobtr_4_eut2(pf01923) EUT2A
06180 HMM00465alcdh_3_mix(pf00465) EUT2A
06185 Hp_euts_babyPurple PDU1C
06195 HMMreg07568_1_eut2(pf12282pf02518pf07568) EUT2A
06200 HMM06277EutA_1_eut(#) EUT2A
06210 HMMetlyEutC_3_eut2(pf05985) EUT2A EP_NTERM
06215 Tsp_amethyst EUT2A
06225 HMMaldh_2_egra(#) EUT2A EP_CTERM
06230 H_fuchsia EUT2G
06235 -# -
06240 HMMptac_3_eut2(#) EUT2A EP_NTERM
06245 HMM02441_07(#) EUT2A EP_NTERM
06250 P_azul EUT2A
06255 HMMtpEuth_1_eut(pf04346) EUT2A
06260 HMM06249EutQ_2_eut2(pf06249) EUT2A EP_NTERM
06265 -pf13380 -
06280 HMMgre_1_grms(pf01228pf02901) GRM5
06285 HMMgreae_1_grms(pf04055) GRM3A
06290 -# -
06300 -pf10035 -
06305 -pf13091pf13396 -
06315 -# -
06320 -# -
BMC-P
BMC-H
BMC-T
Aldehyde dehydrogenase
PTAC
Signature enzyme
Alcohol dehydrogenase
Regulator

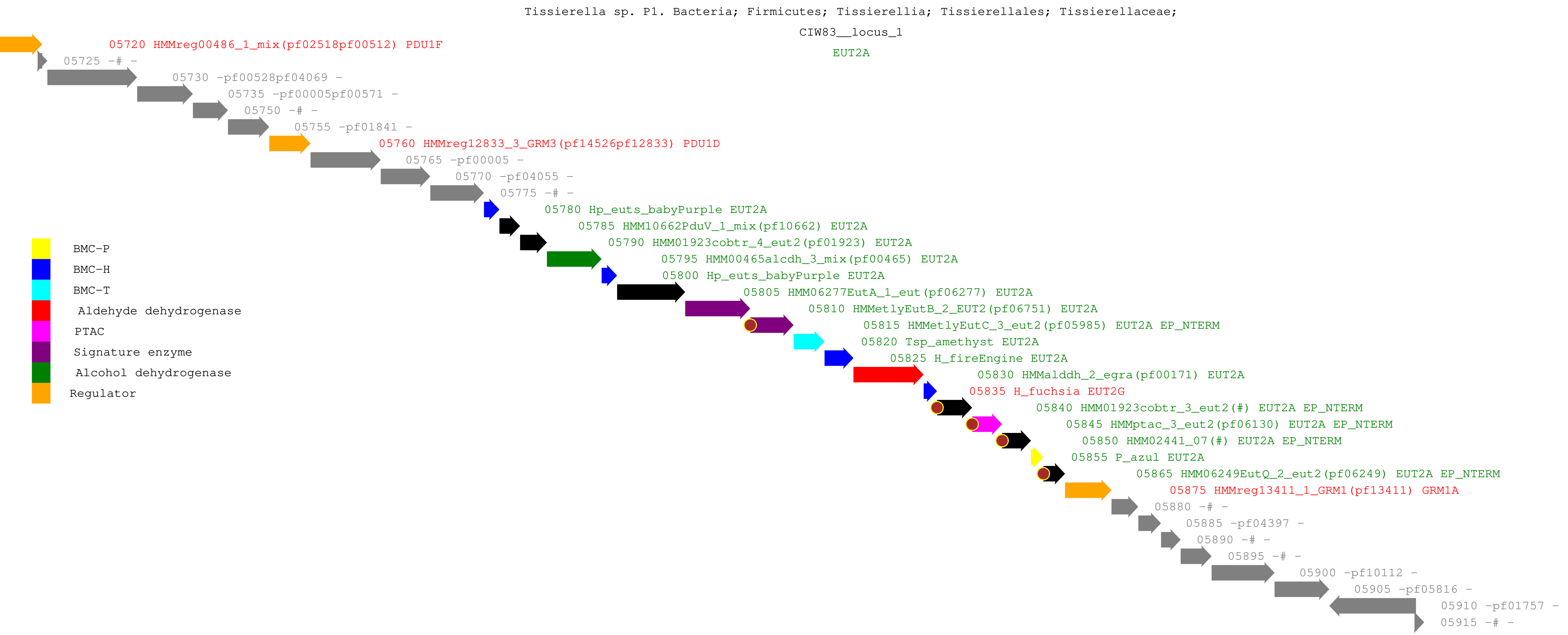

Tissierella sp. P1. Bacteria; Firmicutes; Tissierellia; Tissierellales; Tissierellaceae;
CIW83__locus_1
EUT2A
05720 HMMreg00486_1_mix(pf02518pf00512) PDU1F
05725 -# -
05730 -pf00528pf04069 -
05735 -pf00005pf00571 -
05750 -# -
05755 -pf01841 -
05760 HMMreg12833_3_GRM3(pf14526pf12833) PDU1D
05765 -pf00005 -
05770 -pf04055 -
05775 -# -
05780 Hp_euts_babyPurple EUT2A
05785 HMM10662PduV_1_mix(pf10662) EUT2A
05790 HMM01923cobtr_4_eut2(pf01923) EUT2A
05795 HMM00465alcdh_3_mix(pf00465) EUT2A
05800 Hp_euts_babyPurple EUT2A
05805 HMM06277EutA_1_eut(pf06277) EUT2A
05810 HMMetlyEutB_2_EUT2(pf06751) EUT2A
05815 HMMetlyEutC_3_eut2(pf05985) EUT2A EP_NTERM
05820 Tsp_amethyst EUT2A
05825 H_fireEngine EUT2A
05830 HMMalddh_2_egra(pf00171) EUT2A
05835 H_fuchsia EUT2G
05840 HMM01923cobtr_3_eut2(#) EUT2A EP_NTERM
05845 HMMptac_3_eut2(pf06130) EUT2A EP_NTERM
05850 HMM02441_07(#) EUT2A EP_NTERM
05855 P_azul EUT2A
05865 HMM06249EutQ_2_eut2(pf06249) EUT2A EP_NTERM
05875 HMMreg13411_1_GRM1(pf13411) GRM1A
05880 -# -
05885 -pf04397 -
05890 -# -
05895 -# -
05900 -pf10112 -
05905 -pf05816 -
05910 -pf01757 -
05915 -# -
BMC-P
BMC-H
BMC-T
Aldehyde dehydrogenase
PTAC
Signature enzyme
Alcohol dehydrogenase
Regulator

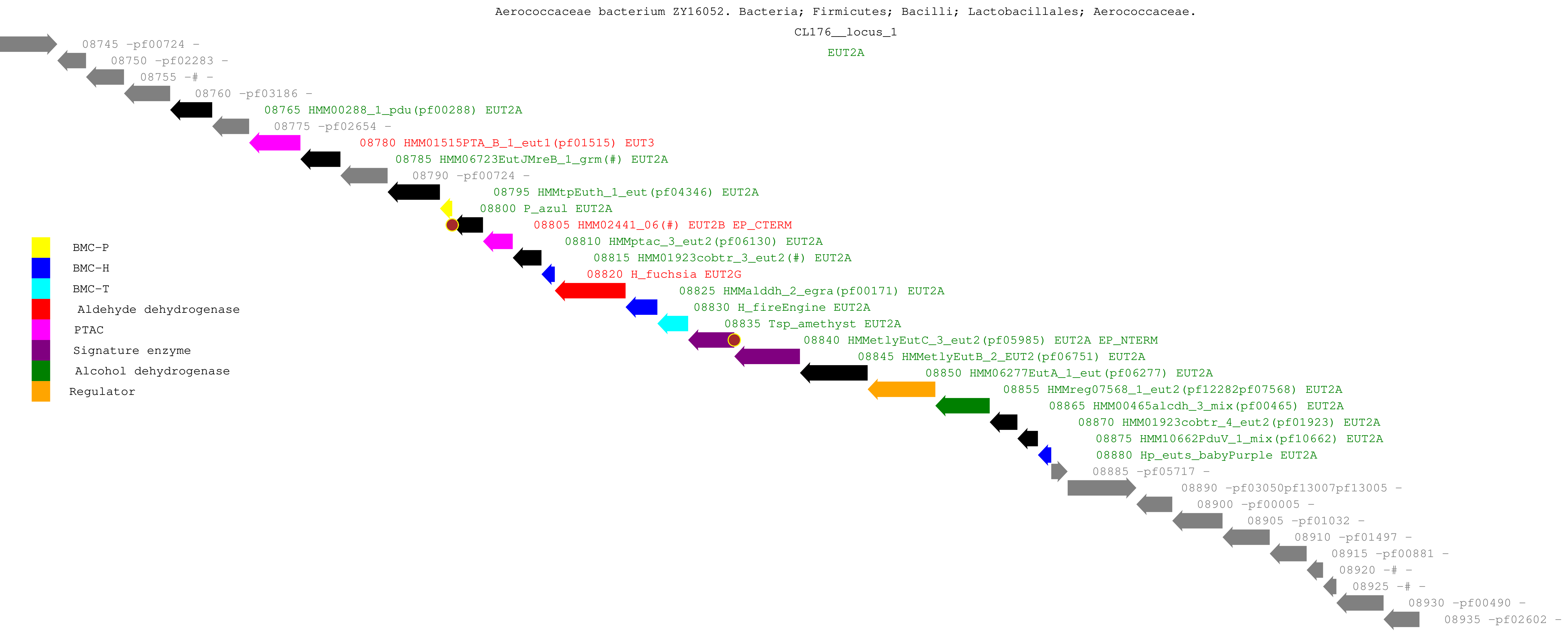
Aerococcaceae bacterium ZY16052. Bacteria; Firmicutes; Bacilli; Lactobacillales; Aerococcaceae.
CL176__locus_1
EUT2A
08745 -pf00724 -
08750 -pf02283 -
08755 -# -
08760 -pf03186 -
08765 HMM00288_1_pdu(pf00288) EUT2A
08775 -pf02654 -
08780 HMM01515PTA_B_1_eut1(pf01515) EUT3
08785 HMM06723EutJMreB_1_grm(#) EUT2A
08790 -pf00724 -
08795 HMMtpEuth_1_eut(pf04346) EUT2A
08800 P_azul EUT2A
08805 HMM02441_06(#) EUT2B EP_CTERM
08810 HMMptac_3_eut2(pf06130) EUT2A
08815 HMM01923cobtr_3_eut2(#) EUT2A
08820 H_fuchsia EUT2G
08825 HMMalddh_2_egra(pf00171) EUT2A
08830 H_fireEngine EUT2A
08835 Tsp_amethyst EUT2A
08840 HMMetlyEutC_3_eut2(pf05985) EUT2A EP_NTERM
08845 HMMetlyEutB_2_EUT2(pf06751) EUT2A
08850 HMM06277EutA_1_eut(pf06277) EUT2A
08855 HMMreg07568_1_eut2(pf12282pf07568) EUT2A
08865 HMM00465alcdh_3_mix(pf00465) EUT2A
08870 HMM01923cobtr_4_eut2(pf01923) EUT2A
08875 HMM10662PduV_1_mix(pf10662) EUT2A
08880 Hp_euts_babyPurple EUT2A
08885 -pf05717 -
08890 -pf03050pf13007pf13005 -
08900 -pf00005 -
08905 -pf01032 -
08910 -pf01497 -
08915 -pf00881 -
08920 -# -
08925 -# -
08930 -pf00490 -
08935 -pf02602 -
BMC-P
BMC-H
BMC-T
Aldehyde dehydrogenase
PTAC
Signature enzyme
Alcohol dehydrogenase
Regulator

Clostridium saccharolyticum (strain ATCC 35040 / DSM 2544 / NRCC 2533 Bacteria; Firmicutes; Clostridia; Clostridiales; Lachnospiraceae.

Closa__locus_1

EUT2A

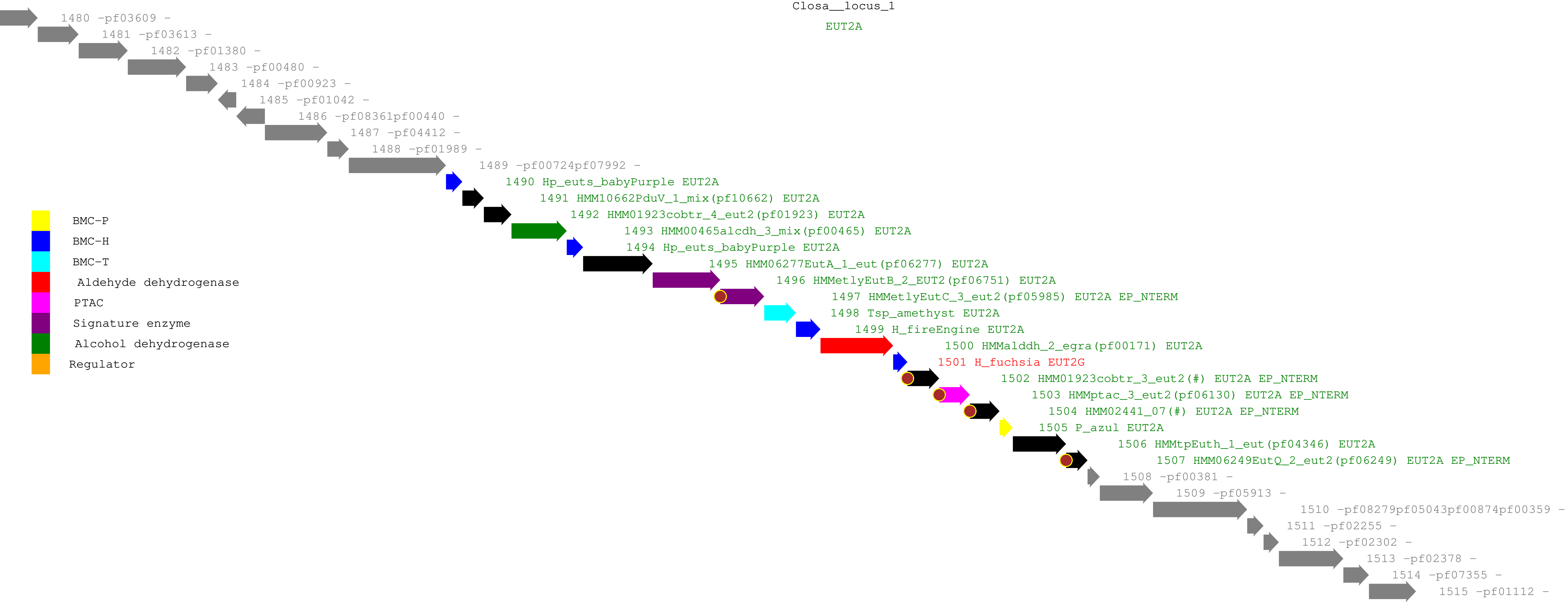

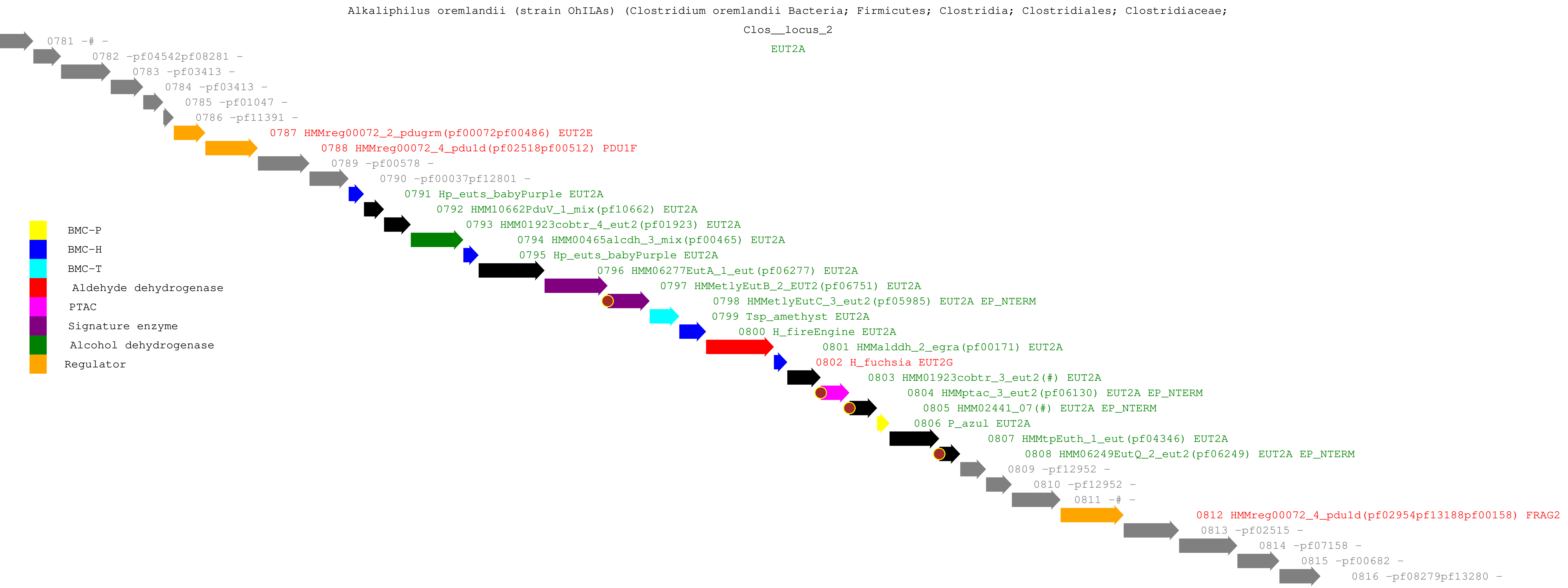
Alkaliphilus oremlandii (strain OhILAs) (Clostridium oremlandii Bacteria; Firmicutes; Clostridia; Clostridiales; Clostridiaceae;
Clos__locus_2
EUT2A
0781 -# -
0782 -pf04542pf08281 -
0783 -pf03413 -
0784 -pf03413 -
0785 -pf01047 -
0786 -pf11391 -
0787 HMMreg00072_2_pdugrm(pf00072pf00486) EUT2E
0788 HMMreg00072_4_pdu1d(pf02518pf00512) PDU1F
0789 -pf00578 -
0790 -pf00037pf12801 -
0791 Hp_euts_babyPurple EUT2A
0792 HMM10662PduV_1_mix(pf10662) EUT2A
0793 HMM01923cobtr_4_eut2(pf01923) EUT2A
0794 HMM00465alcdh_3_mix(pf00465) EUT2A
0795 Hp_euts_babyPurple EUT2A
0796 HMM06277EutA_1_eut(pf06277) EUT2A
0797 HMMetlyEutB_2_EUT2(pf06751) EUT2A
0798 HMMetlyEutC_3_eut2(pf05985) EUT2A EP_NTERM
0799 Tsp_amethyst EUT2A
0800 H_fireEngine EUT2A
0801 HMMalddh_2_egra(pf00171) EUT2A
0802 H_fuchsia EUT2G
0803 HMM01923cobtr_3_eut2(#) EUT2A
0804 HMMptac_3_eut2(pf06130) EUT2A EP_NTERM
0805 HMM02441_07(#) EUT2A EP_NTERM
0806 P_azul EUT2A
0807 HMMtpEuth_1_eut(pf04346) EUT2A
0808 HMM06249EutQ_2_eut2(pf06249) EUT2A EP_NTERM
0809 -pf12952 -
0810 -pf12952 -
0811 -# -
0812 HMMreg00072_4_pdu1d(pf02954pf13188pf00158) FRAG2
0813 -pf02515 -
0814 -pf07158 -
0815 -pf00682 -
0816 -pf08279pf13280 -
BMC-P
BMC-H
BMC-T
Aldehyde dehydrogenase
PTAC
Signature enzyme
Alcohol dehydrogenase
Regulator

Lactobacillus brevis. Bacteria; Firmicutes; Bacilli; Lactobacillales; Lactobacillaceae;

CNR29__locus_2

EUT2A

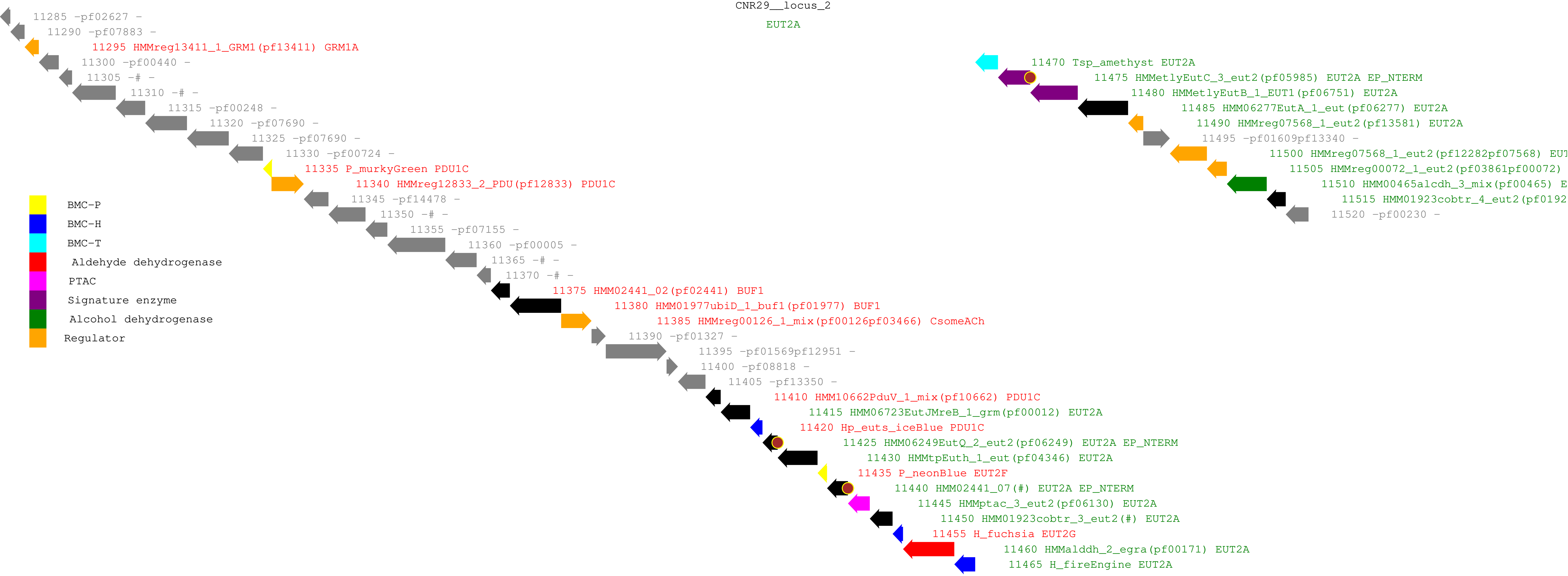

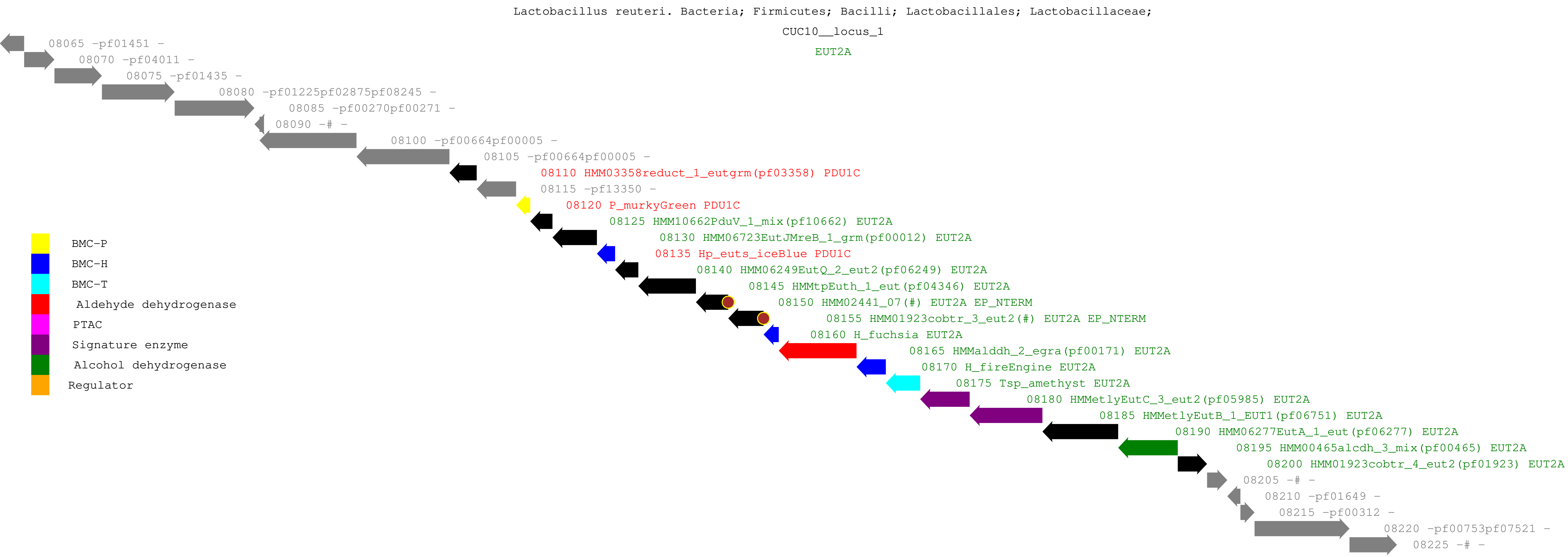

Lactobacillus reuteri. Bacteria; Firmicutes; Bacilli; Lactobacillales; Lactobacillaceae;
CUC10__locus_1
EUT2A
08065 -pf01451 -
08070 -pf04011 -
08075 -pf01435 -
08080 -pf01225pf02875pf08245 -
08085 -pf00270pf00271 -
08090 -# -
08100 -pf00664pf00005 -
08105 -pf00664pf00005 -
08110 HMM03358reduct_1_eutgrm(pf03358) PDU1C
08115 -pf13350 -
08120 P_murkyGreen PDU1C
08125 HMM10662PduV_1_mix(pf10662) EUT2A
08130 HMM06723EutJMreB_1_grm(pf00012) EUT2A
08135 Hp_euts_iceBlue PDU1C
08140 HMM06249EutQ_2_eut2(pf06249) EUT2A
08145 HMMtpEuth_1_eut(pf04346) EUT2A
08150 HMM02441_07(#) EUT2A EP_NTERM
08155 HMM01923cobtr_3_eut2(#) EUT2A EP_NTERM
08160 H_fuchsia EUT2A
08165 HMMalddh_2_egra(pf00171) EUT2A
08170 H_fireEngine EUT2A
08175 Tsp_amethyst EUT2A
08180 HMMetlyEutC_3_eut2(pf05985) EUT2A
08185 HMMetlyEutB_1_EUT1(pf06751) EUT2A
08190 HMM06277EutA_1_eut(pf06277) EUT2A
08195 HMM00465alcdh_3_mix(pf00465) EUT2A
08200 HMM01923cobtr_4_eut2(pf01923) EUT2A
08205 -# -
08210 -pf01649 -
08215 -pf00312 -
08220 -pf00753pf07521 -
08225 -# -
BMC-P
BMC-H
BMC-T
Aldehyde dehydrogenase
PTAC
Signature enzyme
Alcohol dehydrogenase
Regulator

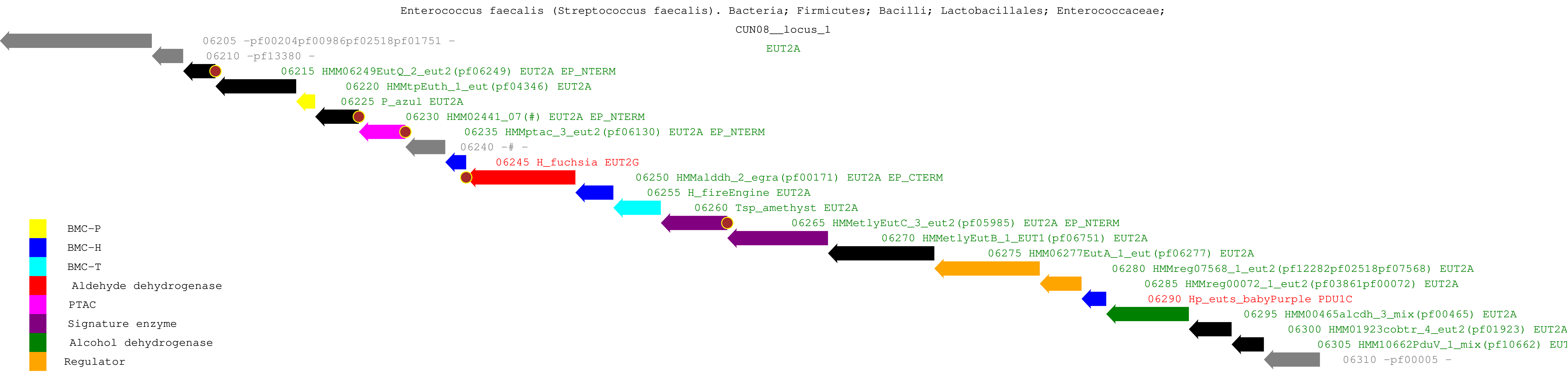
Enterococcus faecalis (Streptococcus faecalis). Bacteria; Firmicutes; Bacilli; Lactobacillales; Enterococcaceae;
CUN08__locus_1
EUT2A
06205 -pf00204pf00986pf02518pf01751 -
06210 -pf13380 -
06215 HMM06249EutQ_2_eut2(pf06249) EUT2A EP_NTERM
06220 HMMtpEuth_1_eut(pf04346) EUT2A
06225 P_azul EUT2A
06230 HMM02441_07(#) EUT2A EP_NTERM
06235 HMMptac_3_eut2(pf06130) EUT2A EP_NTERM
06240 -# -
06245 H_fuchsia EUT2G
06250 HMMalddh_2_egra(pf00171) EUT2A EP_CTERM
06255 H_fireEngine EUT2A
06260 Tsp_amethyst EUT2A
06265 HMMetlyEutC_3_eut2(pf05985) EUT2A EP_NTERM
06270 HMMetlyEutB_1_EUT1(pf06751) EUT2A
06275 HMM06277EutA_1_eut(pf06277) EUT2A
06280 HMMreg07568_1_eut2(pf12282pf02518pf07568) EUT2A
06285 HMMreg00072_1_eut2(pf03861pf00072) EUT2A
06290 Hp_euts_babyPurple PDU1C
06295 HMM00465alcdh_3_mix(pf00465) EUT2A
06300 HMM01923cobtr_4_eut2(pf01923) EUT2A
06305 HMM10662PduV_1_mix(pf10662) EUT
06310 -pf00005 -
BMC-P
BMC-H
BMC-T
Aldehyde dehydrogenase
PTAC
Signature enzyme
Alcohol dehydrogenase
Regulator

Enterococcus mundtii. Bacteria; Firmicutes; Bacilli; Lactobacillales; Enterococcaceae;

CUS89__locus_2

EUT2A

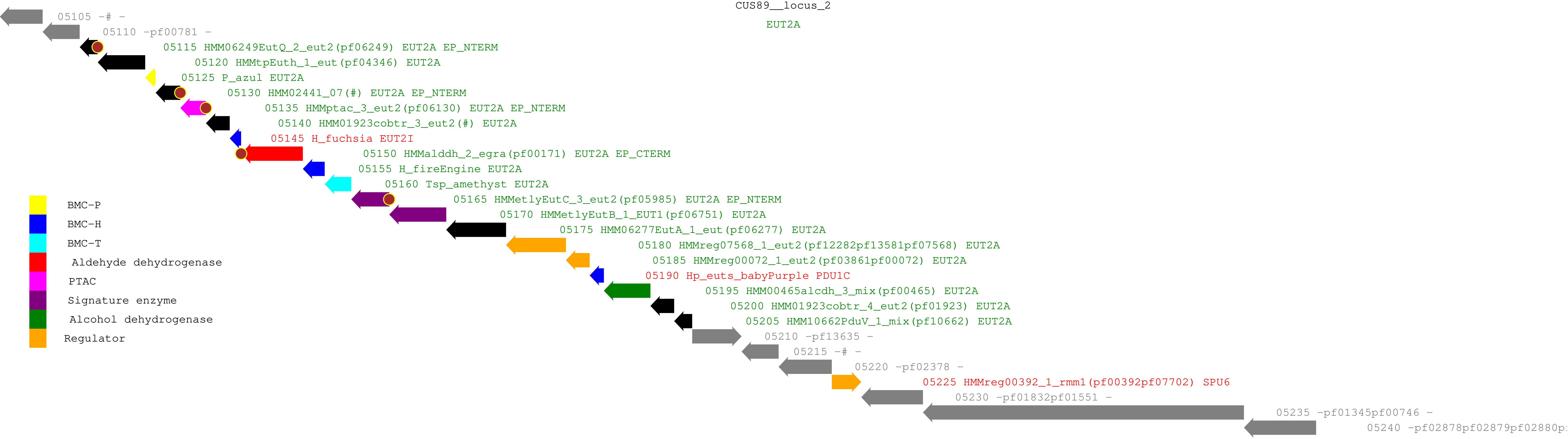

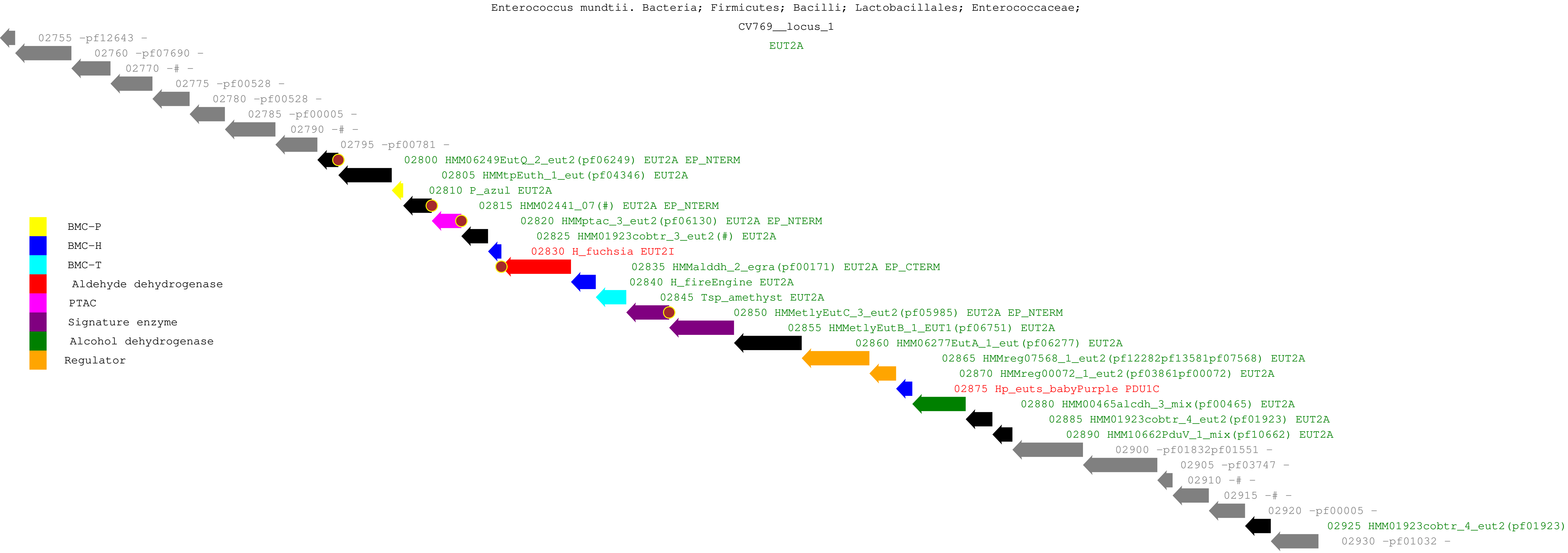
Enterococcus mundtii. Bacteria; Firmicutes; Bacilli; Lactobacillales; Enterococcaceae;
CV769__locus_1
EUT2A
02755 -pf12643 -
02760 -pf07690 -
02770 -# -
02775 -pf00528 -
02780 -pf00528 -
02785 -pf00005 -
02790 -# -
02795 -pf00781 -
02800 HMM06249EutQ_2_eut2(pf06249) EUT2A EP_NTERM
02805 HMMtpEuth_1_eut(pf04346) EUT2A
02810 P_azul EUT2A
02815 HMM02441_07(#) EUT2A EP_NTERM
02820 HMMptac_3_eut2(pf06130) EUT2A EP_NTERM
02825 HMM01923cobtr_3_eut2(#) EUT2A
02830 H_fuchsia EUT2I
02835 HMMaldh_2_egra(pf00171) EUT2A EP_CTERM
02840 H_fireEngine EUT2A
02845 Tsp_amethyst EUT2A
02850 HMMetlyEutC_3_eut2(pf05985) EUT2A EP_NTERM
02855 HMMetlyEutB_1_EUT1(pf06751) EUT2A
02860 HMM06277EutA_1_eut(pf06277) EUT2A
02865 HMMreg07568_1_eut2(pf12282pf13581pf07568) EUT2A
02870 HMMreg00072_1_eut2(pf03861pf00072) EUT2A
02875 Hp_euts_babyPurple PDU1C
02880 HMM00465alcdh_3_mix(pf00465) EUT2A
02885 HMM01923cobtr_4_eut2(pf01923) EUT2A
02890 HMM10662PduV_1_mix(pf10662) EUT2A
02900 -pf01832pf01551 -
02905 -pf03747 -
02910 -# -
02915 -# -
02920 -pf00005 -
02925 HMM01923cobtr_4_eut2(pf01923)
02930 -pf01032 -
BMC-P
BMC-H
BMC-T
Aldehyde dehydrogenase
PTAC
Signature enzyme
Alcohol dehydrogenase
Regulator

Streptococcus salivarius. Bacteria; Firmicutes; Bacilli; Lactobacillales; Streptococcaceae;

CYK23__locus_1

EUT2A

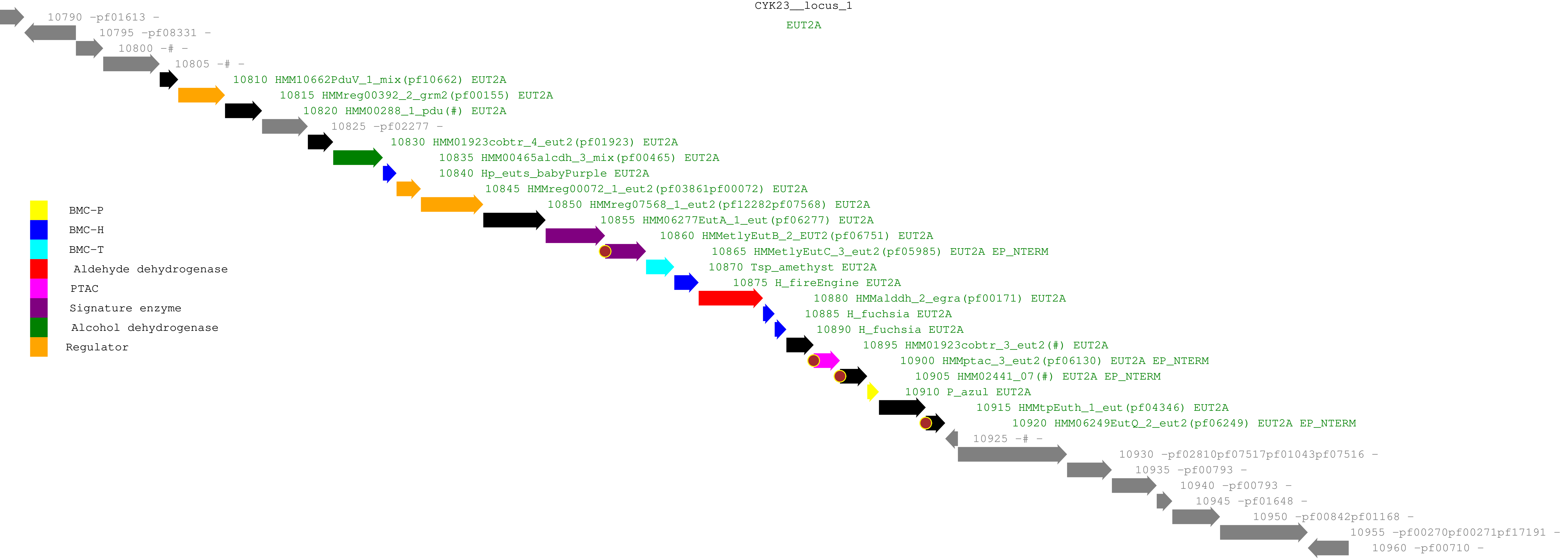

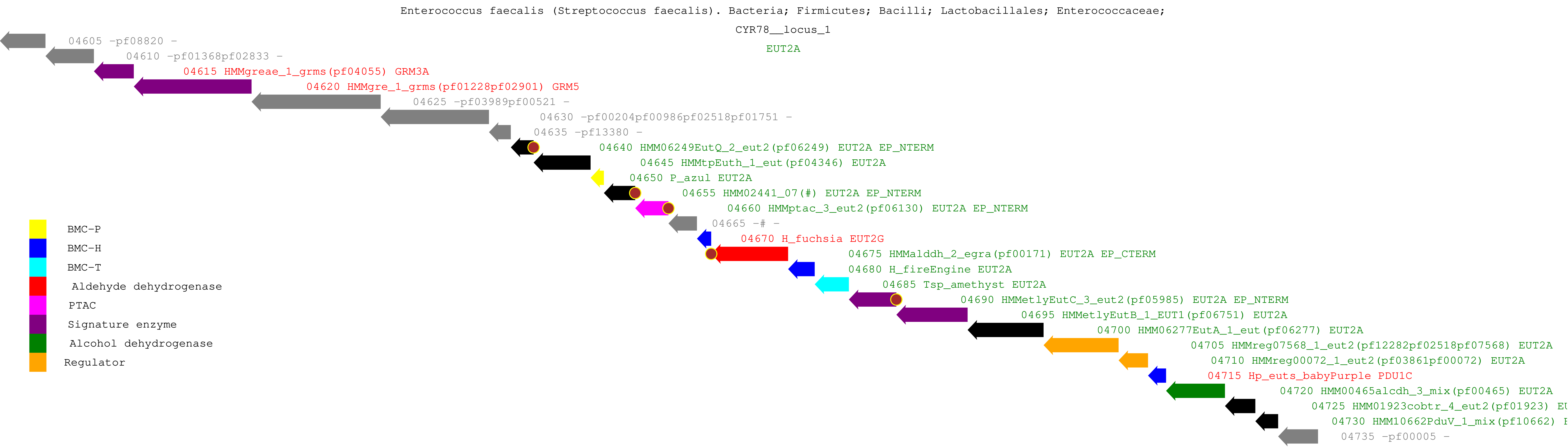
Enterococcus faecalis (Streptococcus faecalis). Bacteria; Firmicutes; Bacilli; Lactobacillales; Enterococcaceae;
CYR78__locus_1
EUT2A
04605 -pf08820 -
04610 -pf01368pf02833 -
04615 HMMgreae_1_grms(pf04055) GRM3A
04620 HMMgre_1_grms(pf01228pf02901) GRM5
04625 -pf03989pf00521 -
04630 -pf00204pf00986pf02518pf01751 -
04635 -pf13380 -
04640 HMM06249EutQ_2_eut2(pf06249) EUT2A EP_NTERM
04645 HMMtpEuth_1_eut(pf04346) EUT2A
04650 P_azul EUT2A
04655 HMM02441_07(#) EUT2A EP_NTERM
04660 HMMptac_3_eut2(pf06130) EUT2A EP_NTERM
04665 -# -
04670 H_fuchsia EUT2G
04675 HMMalddh_2_egra(pf00171) EUT2A EP_CTERM
04680 H_fireEngine EUT2A
04685 Tsp_amethyst EUT2A
04690 HMMetlyEutC_3_eut2(pf05985) EUT2A EP_NTERM
04695 HMMetlyEutB_1_EUT1(pf06751) EUT2A
04700 HMM06277EutA_1_eut(pf06277) EUT2A
04705 HMMreg07568_1_eut2(pf12282pf02518pf07568) EUT2A
04710 HMMreg00072_1_eut2(pf03861pf00072) EUT2A
04715 Hp_euts_babyPurple PDU1C
04720 HMM00465alcdh_3_mix(pf00465) EUT2A
04725 HMM01923cobtr_4_eut2(pf01923) EU
04730 HMM10662PduV_1_mix(pf10662) E
04735 -pf00005 -
BMC-P
BMC-H
BMC-T
Aldehyde dehydrogenase
PTAC
Signature enzyme
Alcohol dehydrogenase
Regulator

Carnobacterium maltaromaticum (Carnobacterium piscicola). Bacteria; Firmicutes; Bacilli; Lactobacillales; Carnobacteriaceae;

CYV28__locus_1

EUT2A

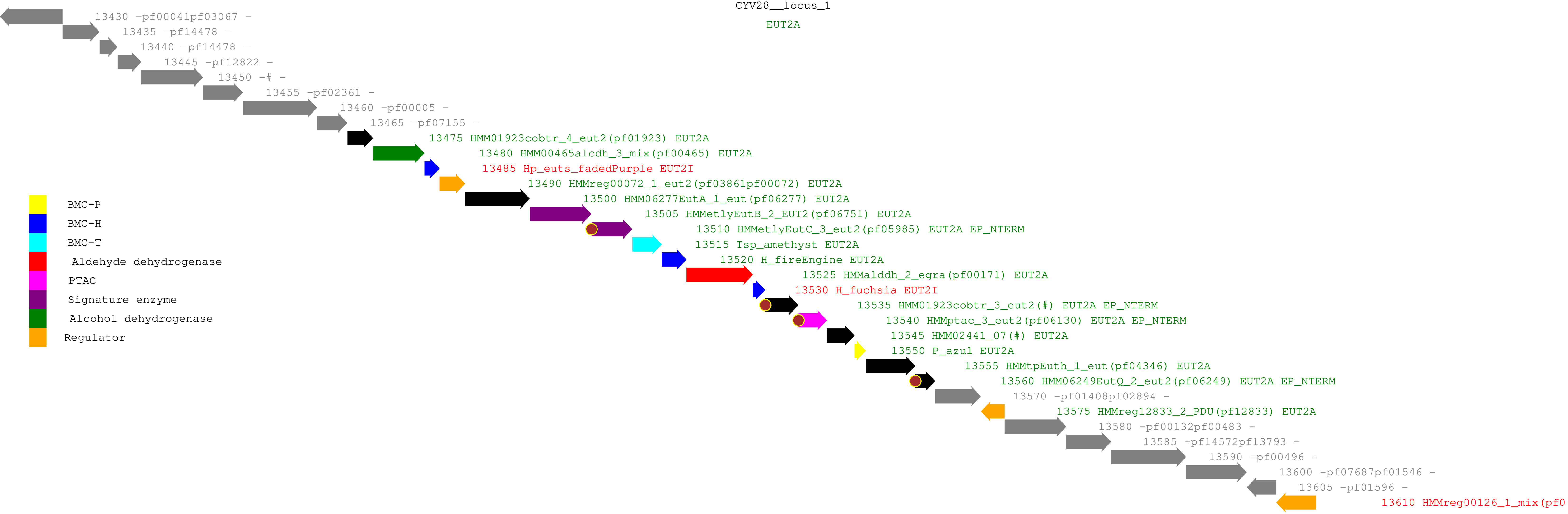

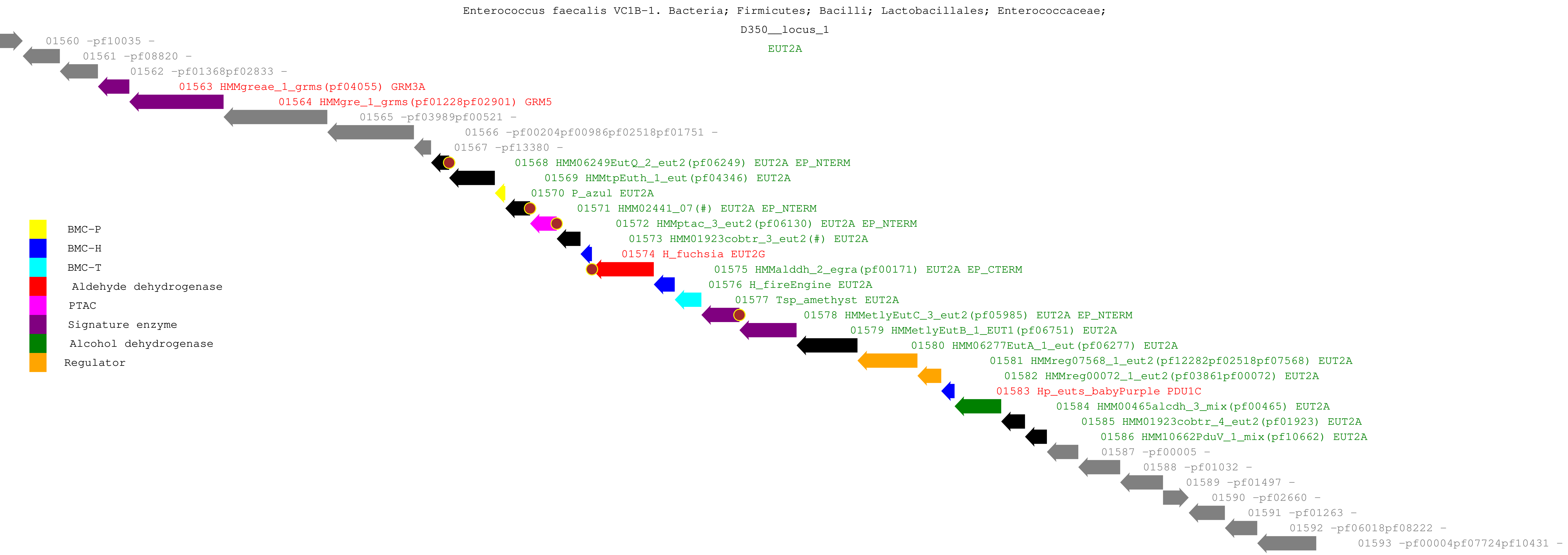
Enterococcus faecalis VC1B-1. Bacteria; Firmicutes; Bacilli; Lactobacillales; Enterococcaceae;
D350__locus_1
EUT2A
01560 -pf10035 -
01561 -pf08820 -
01562 -pf01368pf02833 -
01563 HMMgreae_1_grms(pf04055) GRM3A
01564 HMMgre_1_grms(pf01228pf02901) GRM5
01565 -pf03989pf00521 -
01566 -pf00204pf00986pf02518pf01751 -
01567 -pf13380 -
01568 HMM06249EutQ_2_eut2(pf06249) EUT2A EP_NTERM
01569 HMMtpEuth_1_eut(pf04346) EUT2A
01570 P_azul EUT2A
01571 HMM02441_07(#) EUT2A EP_NTERM
01572 HMMptac_3_eut2(pf06130) EUT2A EP_NTERM
01573 HMM01923cobtr_3_eut2(#) EUT2A
01574 H_fuchsia EUT2G
01575 HMMaldh_2_egra(pf00171) EUT2A EP_CTERM
01576 H_fireEngine EUT2A
01577 Tsp_amethyst EUT2A
01578 HMMetlyEutC_3_eut2(pf05985) EUT2A EP_NTERM
01579 HMMetlyEutB_1_EUT1(pf06751) EUT2A
01580 HMM06277EutA_1_eut(pf06277) EUT2A
01581 HMMreg07568_1_eut2(pf12282pf02518pf07568) EUT2A
01582 HMMreg00072_1_eut2(pf03861pf00072) EUT2A
01583 Hp_euts_babyPurple PDU1C
01584 HMM00465alcdh_3_mix(pf00465) EUT2A
01585 HMM01923cobtr_4_eut2(pf01923) EUT2A
01586 HMM10662PduV_1_mix(pf10662) EUT2A
01587 -pf00005 -
01588 -pf01032 -
01589 -pf01497 -
01590 -pf02660 -
01591 -pf01263 -
01592 -pf06018pf08222 -
01593 -pf00004pf07724pf10431 -
BMC-P
BMC-H
BMC-T
Aldehyde dehydrogenase
PTAC
Signature enzyme
Alcohol dehydrogenase
Regulator

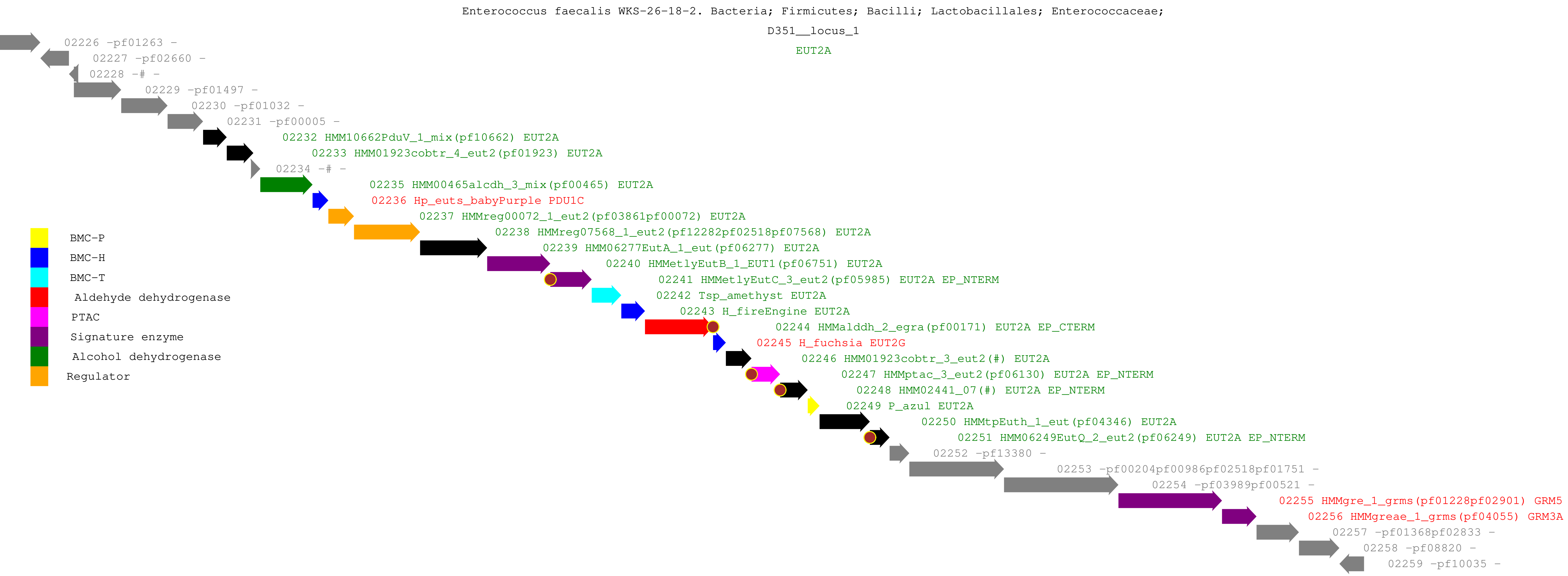

Enterococcus faecalis WKS-26-18-2. Bacteria; Firmicutes; Bacilli; Lactobacillales; Enterococcaceae;
D351__locus_1
EUT2A
02226 -pf01263 -
02227 -pf02660 -
02228 -# -
02229 -pf01497 -
02230 -pf01032 -
02231 -pf00005 -
02232 HMM10662PduV_1_mix(pf10662) EUT2A
02233 HMM01923cobtr_4_eut2(pf01923) EUT2A
02234 -# -
02235 HMM00465alcdh_3_mix(pf00465) EUT2A
02236 Hp_euts_babyPurple PDU1C
02237 HMMreg00072_1_eut2(pf03861pf00072) EUT2A
02238 HMMreg07568_1_eut2(pf12282pf02518pf07568) EUT2A
02239 HMM06277EutA_1_eut(pf06277) EUT2A
02240 HMMetlyEutB_1_EUT1(pf06751) EUT2A
02241 HMMetlyEutC_3_eut2(pf05985) EUT2A EP_NTERM
02242 Tsp_amethyst EUT2A
02243 H_fireEngine EUT2A
02244 HMMaldh_2_egra(pf00171) EUT2A EP_CTERM
02245 H_fuchsia EUT2G
02246 HMM01923cobtr_3_eut2(#) EUT2A
02247 HMMptac_3_eut2(pf06130) EUT2A EP_NTERM
02248 HMM02441_07(#) EUT2A EP_NTERM
02249 P_azul EUT2A
02250 HMMtpEuth_1_eut(pf04346) EUT2A
02251 HMM06249EutQ_2_eut2(pf06249) EUT2A EP_NTERM
02252 -pf13380 -
02253 -pf00204pf00986pf02518pf01751 -
02254 -pf03989pf00521 -
02255 HMMgre_1_grms(pf01228pf02901) GRM5
02256 HMMgreae_1_grms(pf04055) GRM3A
02257 -pf01368pf02833 -
02258 -pf08820 -
02259 -pf10035 -
BMC-P
BMC-H
BMC-T
Aldehyde dehydrogenase
PTAC
Signature enzyme
Alcohol dehydrogenase
Regulator

Enterococcus faecium SB2C-2. Bacteria; Firmicutes; Bacilli; Lactobacillales; Enterococcaceae;

D354__locus_1

EUT2A

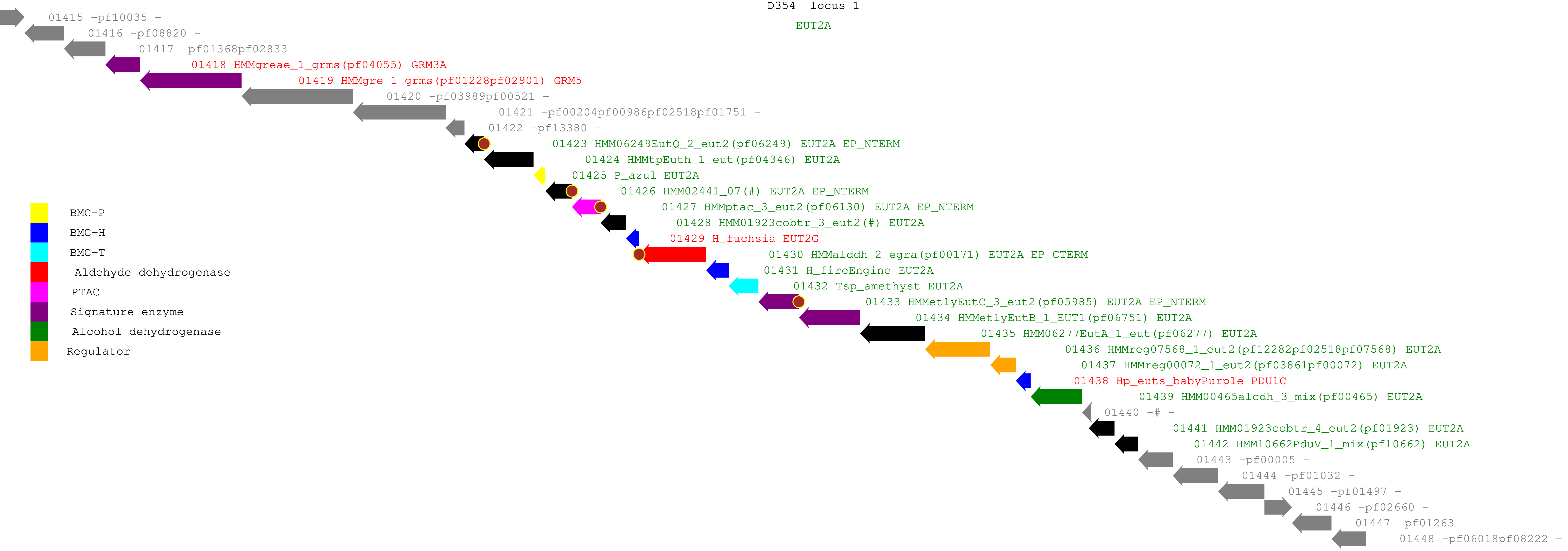

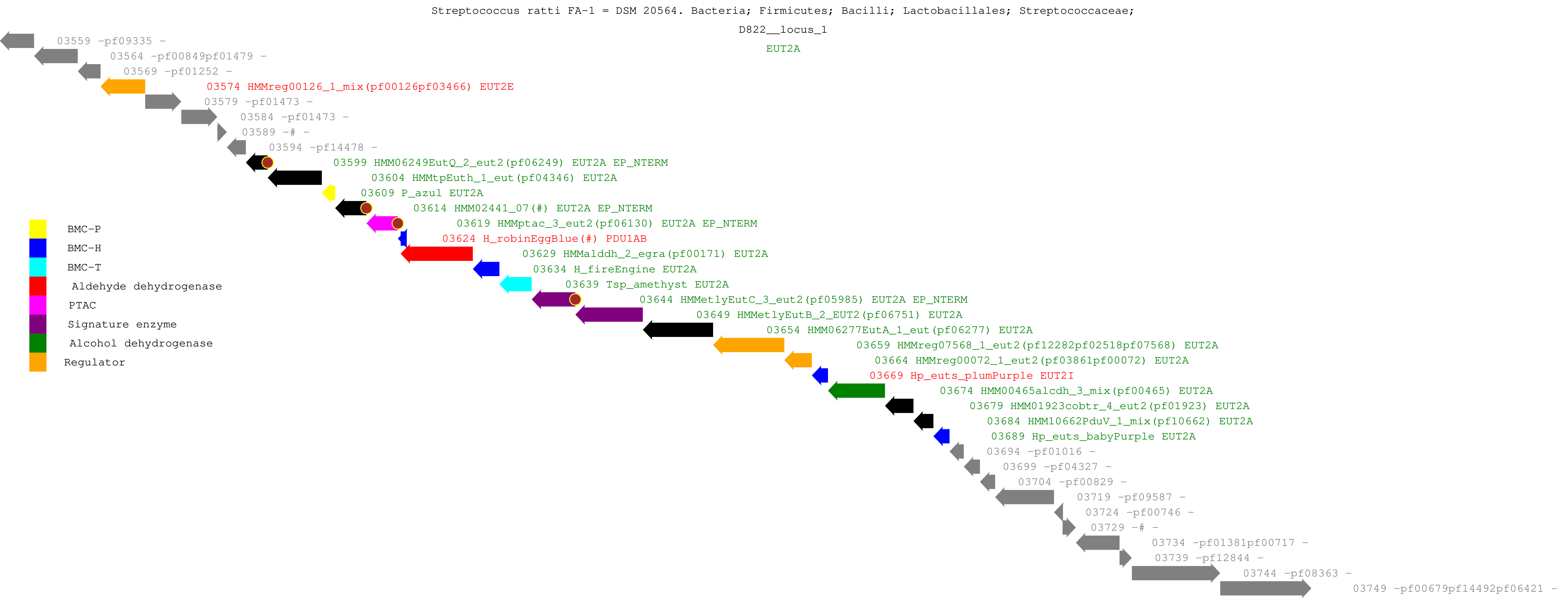

Streptococcus ratti FA-1 = DSM 20564. Bacteria; Firmicutes; Bacilli; Lactobacillales; Streptococcaceae;
D822__locus_1
EUT2A
03559 -pf09335 -
03564 -pf00849pf01479 -
03569 -pf01252 -
03574 HMMreg00126_1_mix(pf00126pf03466) EUT2E
03579 -pf01473 -
03584 -pf01473 -
03589 -# -
03594 -pf14478 -
03599 HMM06249EutQ_2_eut2(pf06249) EUT2A EP_NTERM
03604 HMMtpEuth_1_eut(pf04346) EUT2A
03609 P_azul EUT2A
03614 HMM02441_07(#) EUT2A EP_NTERM
03619 HMMptac_3_eut2(pf06130) EUT2A EP_NTERM
03624 H_robinEggBlue(#) PDU1AB
03629 HMMalddh_2_egra(pf00171) EUT2A
03634 H_fireEngine EUT2A
03639 Tsp_amethyst EUT2A
03644 HMMetlyEutC_3_eut2(pf05985) EUT2A EP_NTERM
03649 HMMetlyEutB_2_EUT2(pf06751) EUT2A
03654 HMM06277EutA_1_eut(pf06277) EUT2A
03659 HMMreg07568_1_eut2(pf12282pf02518pf07568) EUT2A
03664 HMMreg00072_1_eut2(pf03861pf00072) EUT2A
03669 Hp_euts_plumPurple EUT2I
03674 HMM00465alcdh_3_mix(pf00465) EUT2A
03679 HMM01923cobtr_4_eut2(pf01923) EUT2A
03684 HMM10662PduV_1_mix(pf10662) EUT2A
03689 Hp_euts_babyPurple EUT2A
03694 -pf01016 -
03699 -pf04327 -
03704 -pf00829 -
03719 -pf09587 -
03724 -pf00746 -
03729 -# -
03734 -pf01381pf00717 -
03739 -pf12844 -
03744 -pf08363 -
03749 -pf00679pf14492pf06421 -
BMC-P
BMC-H
BMC-T
Aldehyde dehydrogenase
PTAC
Signature enzyme
Alcohol dehydrogenase
Regulator

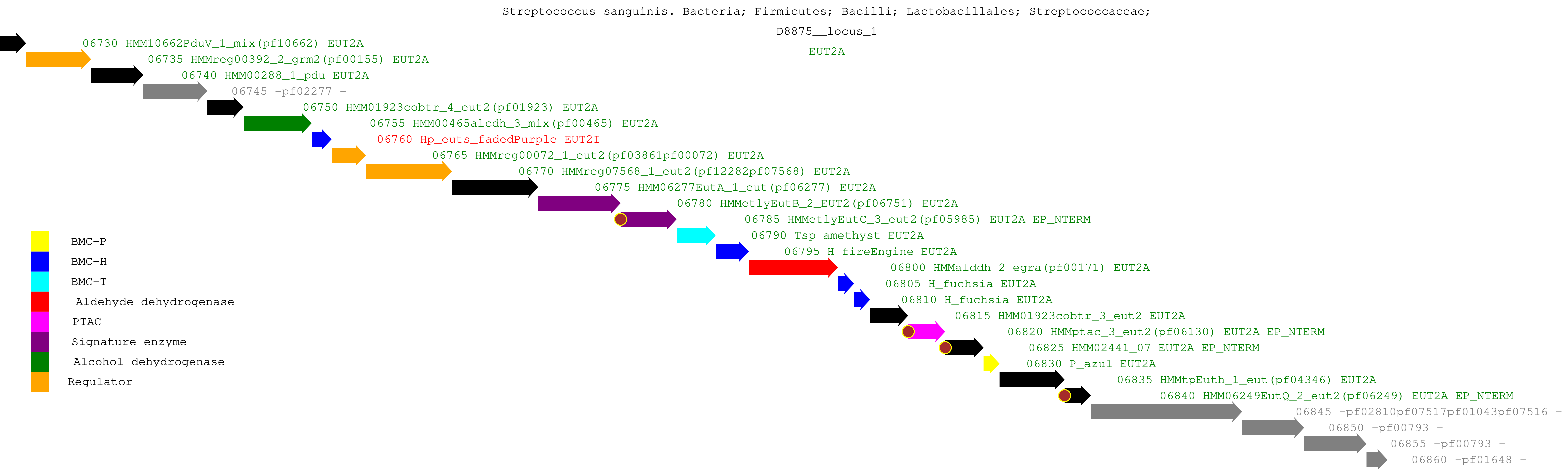
Streptococcus sanguinis. Bacteria; Firmicutes; Bacilli; Lactobacillales; Streptococcaceae;
D8875__locus_1
EUT2A
06730 HMM10662PduV_1_mix(pf10662) EUT2A
06735 HMMreg00392_2_grm2(pf00155) EUT2A
06740 HMM00288_1_pdu EUT2A
06745 -pf02277 -
06750 HMM01923cobtr_4_eut2(pf01923) EUT2A
06755 HMM00465alcdh_3_mix(pf00465) EUT2A
06760 Hp_euts_fadedPurple EUT2I
06765 HMMreg00072_1_eut2(pf03861pf00072) EUT2A
06770 HMMreg07568_1_eut2(pf12282pf07568) EUT2A
06775 HMM06277EutA_1_eut(pf06277) EUT2A
06780 HMMetlyEutB_2_EUT2(pf06751) EUT2A
06785 HMMetlyEutC_3_eut2(pf05985) EUT2A EP_NTERM
06790 Tsp_amethyst EUT2A
06795 H_fireEngine EUT2A
06800 HMMalddh_2_egra(pf00171) EUT2A
06805 H_fuchsia EUT2A
06810 H_fuchsia EUT2A
06815 HMM01923cobtr_3_eut2 EUT2A
06820 HMMptac_3_eut2(pf06130) EUT2A EP_NTERM
06825 HMM02441_07 EUT2A EP_NTERM
06830 P_azul EUT2A
06835 HMMtpEuth_1_eut(pf04346) EUT2A
06840 HMM06249EutQ_2_eut2(pf06249) EUT2A EP_NTERM
06845 -pf02810pf07517pf01043pf07516 -
06850 -pf00793 -
06855 -pf00793 -
06860 -pf01648 -
BMC-P
BMC-H
BMC-T
Aldehyde dehydrogenase
PTAC
Signature enzyme
Alcohol dehydrogenase
Regulator

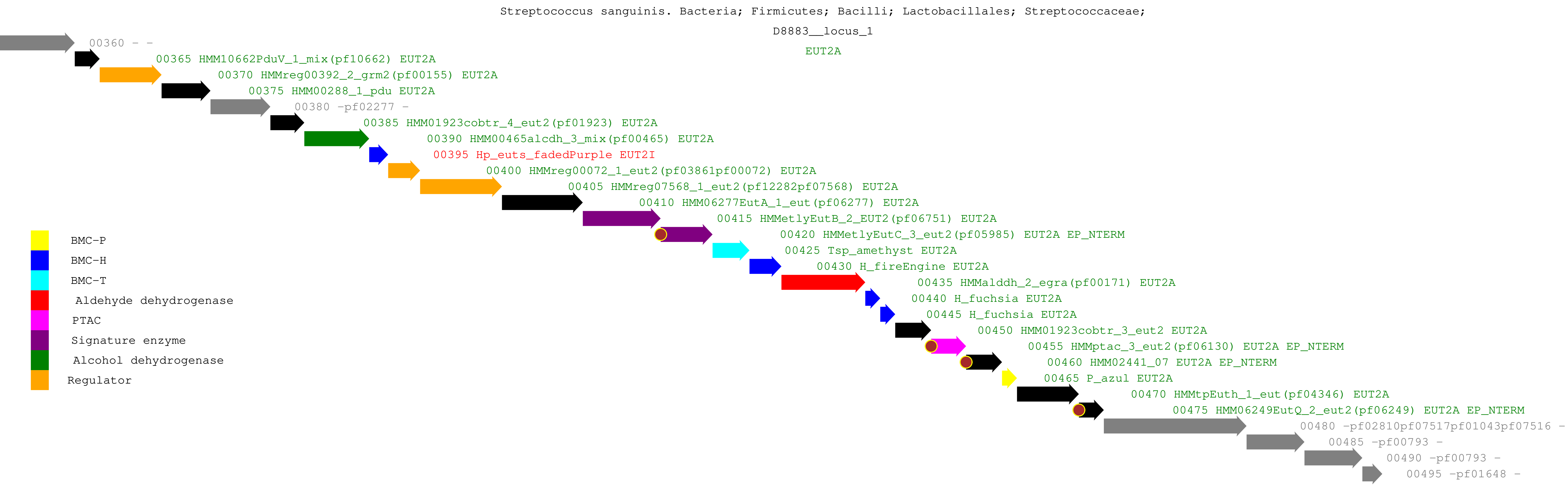

Streptococcus sanguinis. Bacteria; Firmicutes; Bacilli; Lactobacillales; Streptococcaceae;
D8883__locus_1
EUT2A
00360 - -
00365 HMM10662PduV_1_mix(pf10662) EUT2A
00370 HMMreg00392_2_grm2(pf00155) EUT2A
00375 HMM00288_1_pdu EUT2A
00380 -pf02277 -
00385 HMM01923cobtr_4_eut2(pf01923) EUT2A
00390 HMM00465alcdh_3_mix(pf00465) EUT2A
00395 Hp_euts_fadedPurple EUT2I
00400 HMMreg00072_1_eut2(pf03861pf00072) EUT2A
00405 HMMreg07568_1_eut2(pf12282pf07568) EUT2A
00410 HMM06277EutA_1_eut(pf06277) EUT2A
00415 HMMetlyEutB_2_EUT2(pf06751) EUT2A
00420 HMMetlyEutC_3_eut2(pf05985) EUT2A EP_NTERM
00425 Tsp_amethyst EUT2A
00430 H_fireEngine EUT2A
00435 HMMalddh_2_egra(pf00171) EUT2A
00440 H_fuchsia EUT2A
00445 H_fuchsia EUT2A
00450 HMM01923cobtr_3_eut2 EUT2A
00455 HMMptac_3_eut2(pf06130) EUT2A EP_NTERM
00460 HMM02441_07 EUT2A EP_NTERM
00465 P_azul EUT2A
00470 HMMtpEuth_1_eut(pf04346) EUT2A
00475 HMM06249EutQ_2_eut2(pf06249) EUT2A EP_NTERM
00480 -pf02810pf07517pf01043pf07516 -
00485 -pf00793 -
00490 -pf00793 -
00495 -pf01648 -
BMC-P
BMC-H
BMC-T
Aldehyde dehydrogenase
PTAC
Signature enzyme
Alcohol dehydrogenase
Regulator

Streptococcus sp. Bacteria; Firmicutes; Bacilli; Lactobacillales; Streptococcaceae;

D8H99__locus_1_main_sat

EUT2A

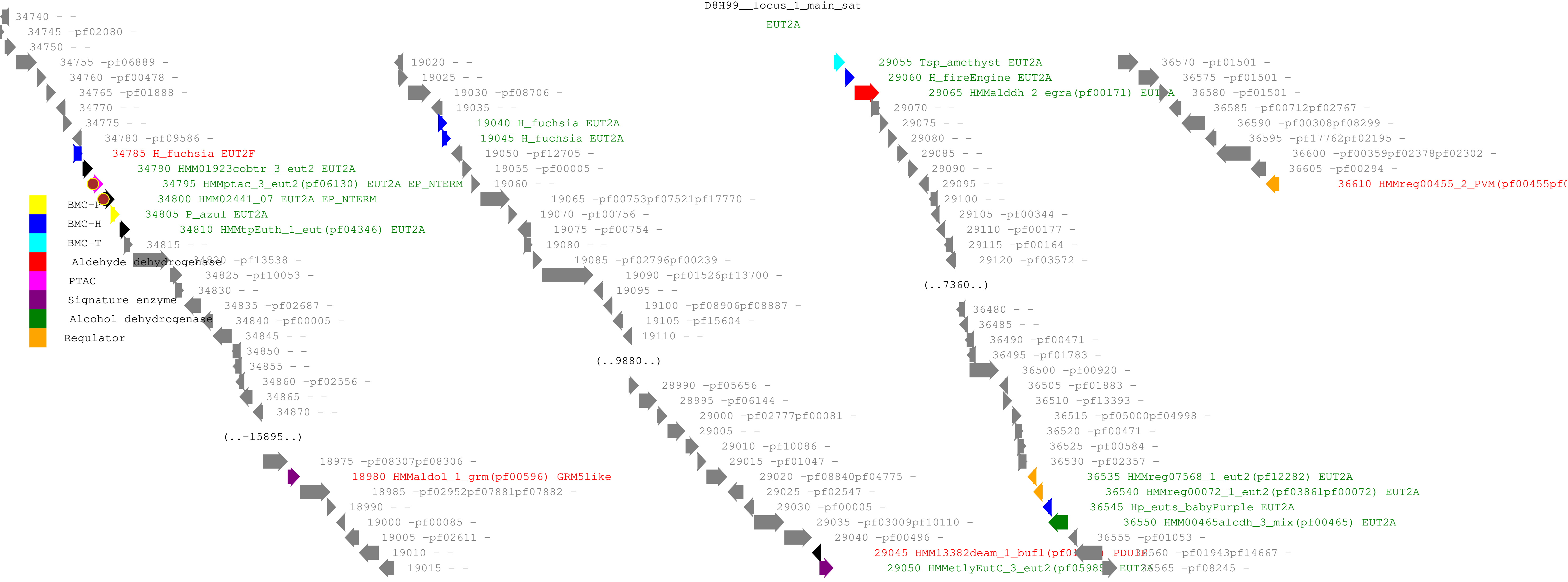

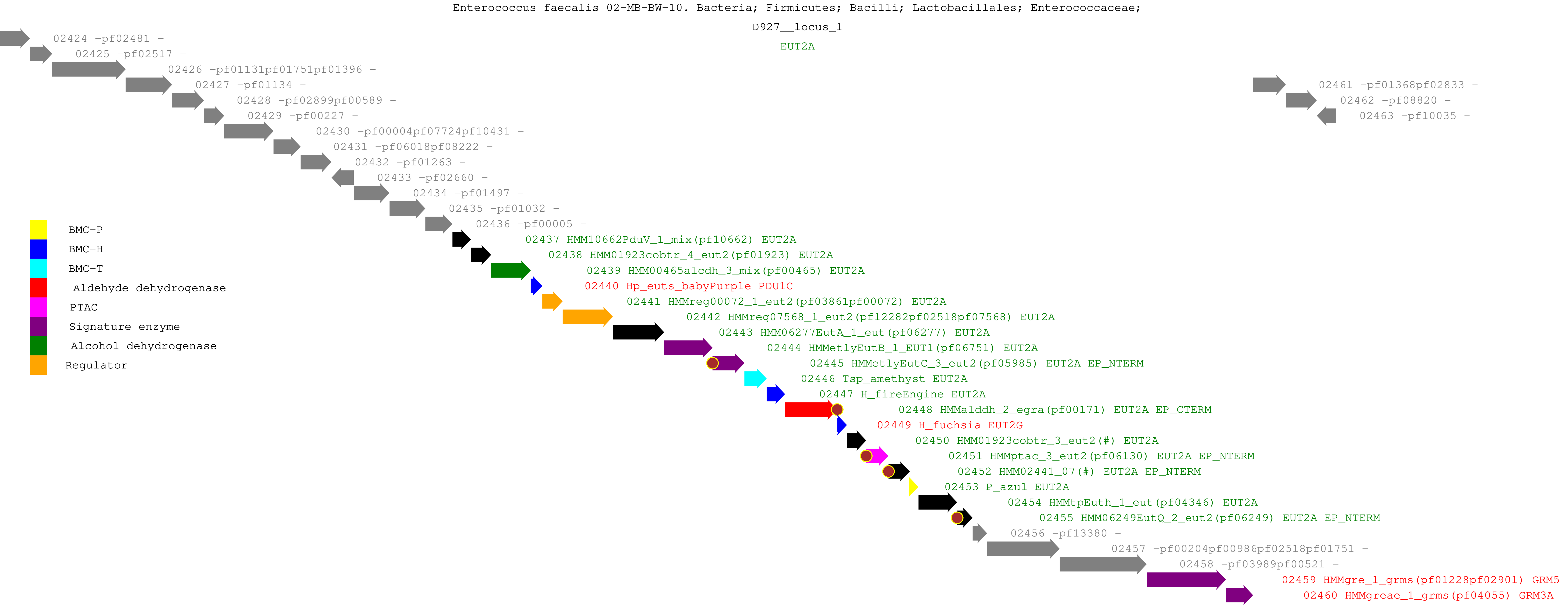
Enterococcus faecalis 02-MB-BW-10. Bacteria; Firmicutes; Bacilli; Lactobacillales; Enterococcaceae;
D927__locus_1
EUT2A
02424 -pf02481 -
02425 -pf02517 -
02426 -pf01131pf01751pf01396 -
02427 -pf01134 -
02428 -pf02899pf00589 -
02429 -pf00227 -
02430 -pf00004pf07724pf10431 -
02431 -pf06018pf08222 -
02432 -pf01263 -
02433 -pf02660 -
02434 -pf01497 -
02435 -pf01032 -
02436 -pf00005 -
02437 HMM10662PduV_1_mix(pf10662) EUT2A
02438 HMM01923cobtr_4_eut2(pf01923) EUT2A
02439 HMM00465alcdh_3_mix(pf00465) EUT2A
02440 Hp_euts_babyPurple PDU1C
02441 HMMreg00072_1_eut2(pf03861pf00072) EUT2A
02442 HMMreg07568_1_eut2(pf12282pf02518pf07568) EUT2A
02443 HMM06277EutA_1_eut(pf06277) EUT2A
02444 HMMetlyEutB_1_EUT1(pf06751) EUT2A
02445 HMMetlyEutC_3_eut2(pf05985) EUT2A EP_NTERM
02446 Tsp_amethyst EUT2A
02447 H_fireEngine EUT2A
02448 HMMalddh_2_egra(pf00171) EUT2A EP_CTERM
02449 H_fuchsia EUT2G
02450 HMM01923cobtr_3_eut2(#) EUT2A
02451 HMMptac_3_eut2(pf06130) EUT2A EP_NTERM
02452 HMM02441_07(#) EUT2A EP_NTERM
02453 P_azul EUT2A
02454 HMMtpEuth_1_eut(pf04346) EUT2A
02455 HMM06249EutQ_2_eut2(pf06249) EUT2A EP_NTERM
02456 -pf13380 -
02457 -pf00204pf00986pf02518pf01751 -
02458 -pf03989pf00521 -
02459 HMMgre_1_grms(pf01228pf02901) GRM5
02460 HMMgreae_1_grms(pf04055) GRM3A
02461 -pf01368pf02833 -
02462 -pf08820 -
02463 -pf10035 -
BMC-P
BMC-H
BMC-T
Aldehyde dehydrogenase
PTAC
Signature enzyme
Alcohol dehydrogenase
Regulator

Lactobacillus reuteri. Bacteria; Firmicutes; Bacilli; Lactobacillales; Lactobacillaceae;

DB324__locus_1

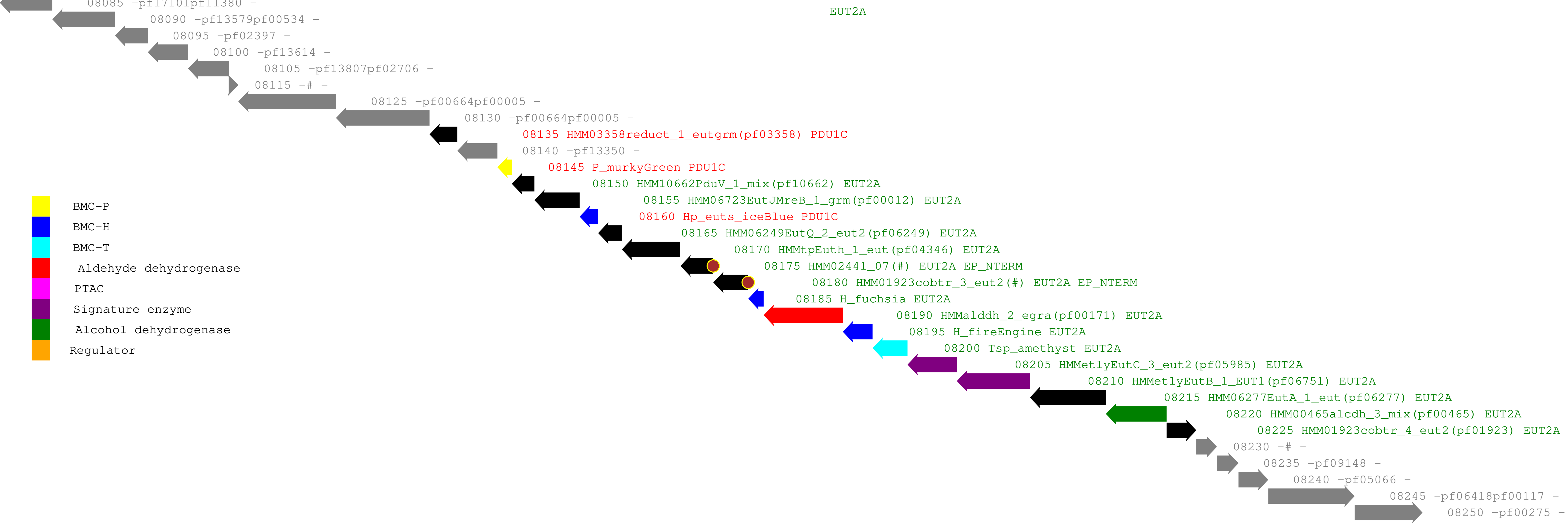

Lachnoclostridium sp. Bacteria; Firmicutes; Clostridia; Clostridiales; Lachnospiraceae.

DC053__locus_1

EUT2A

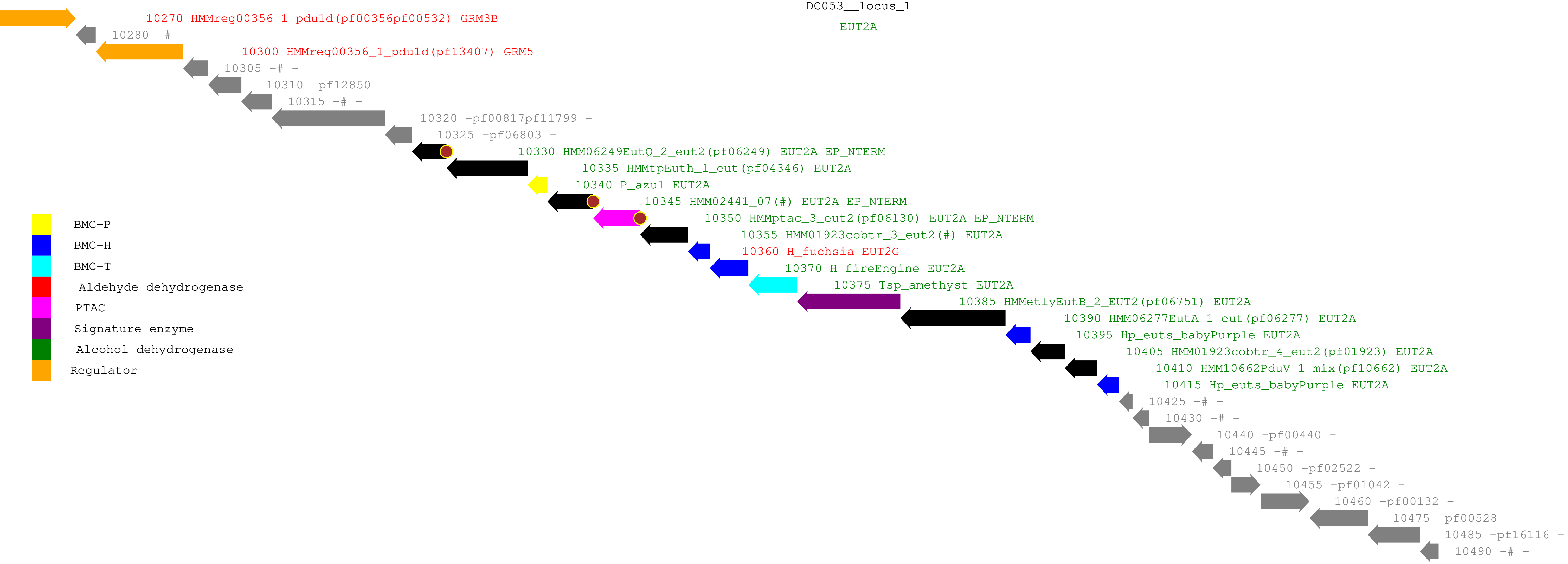

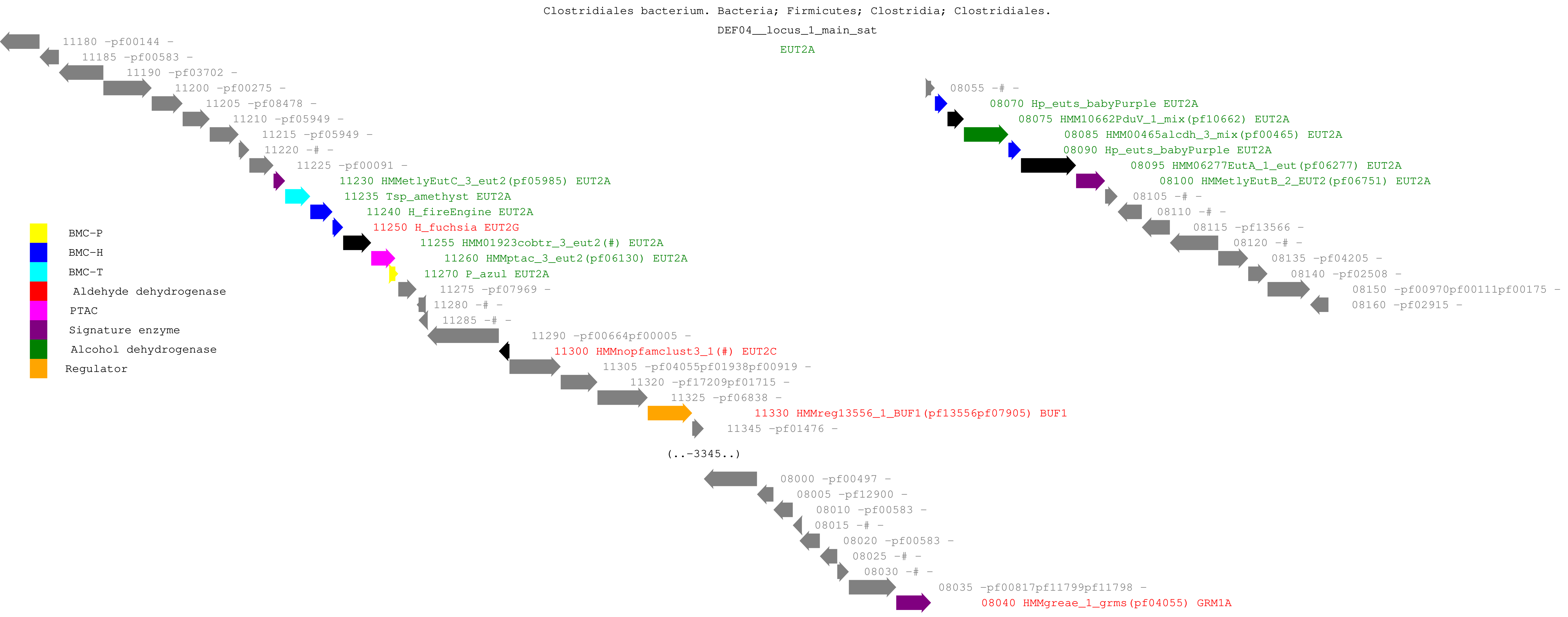

Clostridiales bacterium. Bacteria; Firmicutes; Clostridia; Clostridiales.
DEF04__locus_1_main_sat
EUT2A
11180 -pf00144 -
11185 -pf00583 -
11190 -pf03702 -
11200 -pf00275 -
11205 -pf08478 -
11210 -pf05949 -
11215 -pf05949 -
11220 -# -
11225 -pf00091 -
11230 HMMetlyEutC_3_eut2(pf05985) EUT2A
11235 Tsp_amethyst EUT2A
11240 H_fireEngine EUT2A
11250 H_fuchsia EUT2G
11255 HMM01923cobtr_3_eut2(#) EUT2A
11260 HMMptac_3_eut2(pf06130) EUT2A
11270 P_azul EUT2A
11275 -pf07969 -
11280 -# -
11285 -# -
11290 -pf00664pf00005 -
11300 HMMnopfamclust3_1(#) EUT2C
11305 -pf04055pf01938pf00919 -
11320 -pf17209pf01715 -
11325 -pf06838 -
11330 HMMreg13556_1_BUF1(pf13556pf07905) BUF1
11345 -pf01476 -
(..-3345..)
08000 -pf00497 -
08005 -pf12900 -
08010 -pf00583 -
08015 -# -
08020 -pf00583 -
08025 -# -
08030 -# -
08035 -pf00817pf11799pf11798 -
08040 HMMgreae_1_grms(pf04055) GRM1A
08055 -# -
08070 Hp_euts_babyPurple EUT2A
08075 HMM10662PduV_1_mix(pf10662) EUT2A
08085 HMM00465alcdh_3_mix(pf00465) EUT2A
08090 Hp_euts_babyPurple EUT2A
08095 HMM06277EutA_1_eut(pf06277) EUT2A
08100 HMMetlyEutB_2_EUT2(pf06751) EUT2A
08105 -# -
08110 -# -
08115 -pf13566 -
08120 -# -
08135 -pf04205 -
08140 -pf02508 -
08150 -pf00970pf00111pf00175 -
08160 -pf02915 -
BMC-P
BMC-H
BMC-T
Aldehyde dehydrogenase
PTAC
Signature enzyme
Alcohol dehydrogenase
Regulator

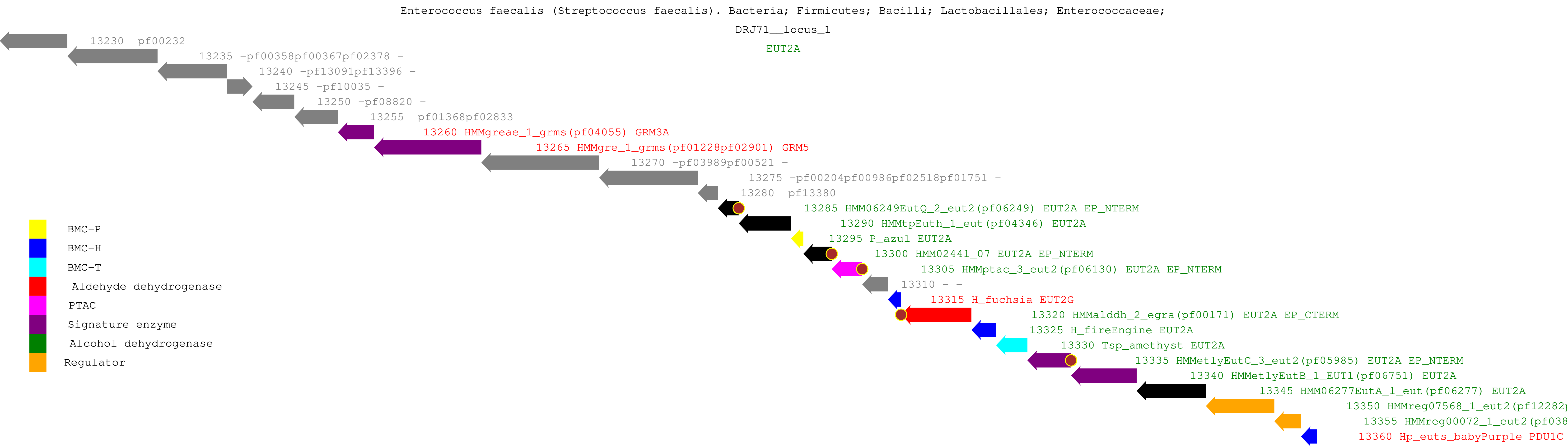
Enterococcus faecalis (Streptococcus faecalis). Bacteria; Firmicutes; Bacilli; Lactobacillales; Enterococcaceae;
DRJ71__locus_1
EUT2A
13230 -pf00232 -
13235 -pf00358pf00367pf02378 -
13240 -pf13091pf13396 -
13245 -pf10035 -
13250 -pf08820 -
13255 -pf01368pf02833 -
13260 HMMgreae_1_grms(pf04055) GRM3A
13265 HMMgre_1_grms(pf01228pf02901) GRM5
13270 -pf03989pf00521 -
13275 -pf00204pf00986pf02518pf01751 -
13280 -pf13380 -
13285 HMM06249EutQ_2_eut2(pf06249) EUT2A EP_NTERM
13290 HMMtpEuth_1_eut(pf04346) EUT2A
13295 P_azul EUT2A
13300 HMM02441_07 EUT2A EP_NTERM
13305 HMMptac_3_eut2(pf06130) EUT2A EP_NTERM
13310 - -
13315 H_fuchsia EUT2G
13320 HMMalddh_2_egra(pf00171) EUT2A EP_CTERM
13325 H_fireEngine EUT2A
13330 Tsp_amethyst EUT2A
13335 HMMetlyEutC_3_eut2(pf05985) EUT2A EP_NTERM
13340 HMMetlyEutB_1_EUT1(pf06751) EUT2A
13345 HMM06277EutA_1_eut(pf06277) EUT2A
13350 HMMreg07568_1_eut2(pf12282
13355 HMMreg00072_1_eut2(pf038
13360 Hp_euts_babyPurple PDU1C
BMC-P
BMC-H
BMC-T
Aldehyde dehydrogenase
PTAC
Signature enzyme
Alcohol dehydrogenase
Regulator

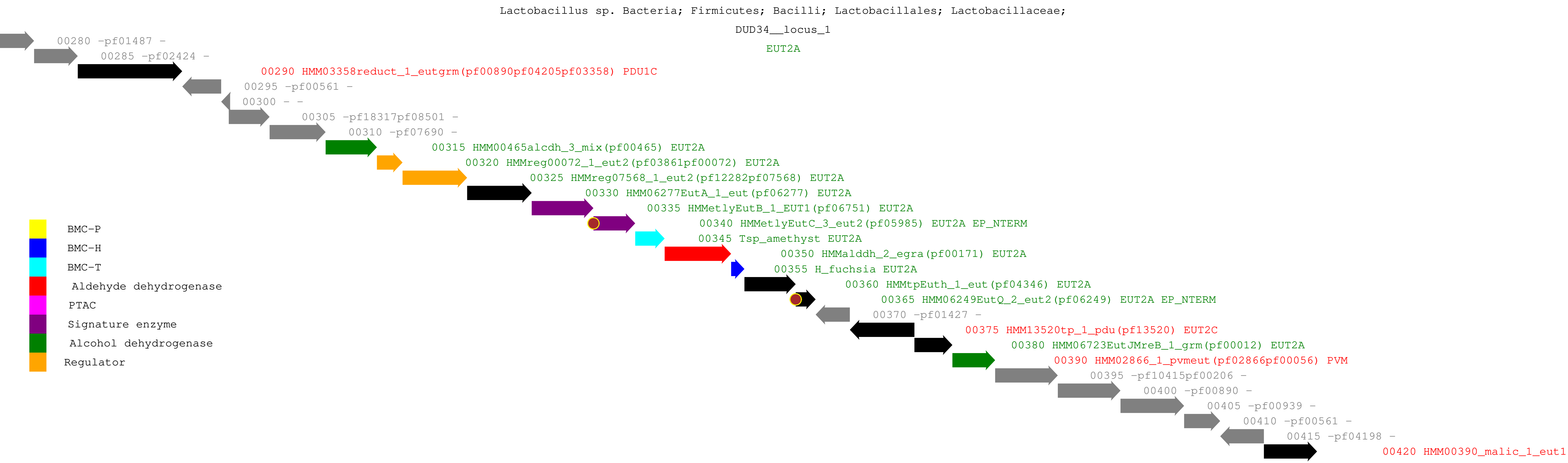
Lactobacillus sp. Bacteria; Firmicutes; Bacilli; Lactobacillales; Lactobacillaceae;
DUD34__locus_1
EUT2A
00280 -pf01487 -
00285 -pf02424 -
00290 HMM03358reduct_1_eutgrm(pf00890pf04205pf03358) PDU1C
00295 -pf00561 -
00300 - -
00305 -pf18317pf08501 -
00310 -pf07690 -
00315 HMM00465alcdh_3_mix(pf00465) EUT2A
00320 HMMreg00072_1_eut2(pf03861pf00072) EUT2A
00325 HMMreg07568_1_eut2(pf12282pf07568) EUT2A
00330 HMM06277EutA_1_eut(pf06277) EUT2A
00335 HMMetlyEutB_1_EUT1(pf06751) EUT2A
00340 HMMetlyEutC_3_eut2(pf05985) EUT2A EP_NTERM
00345 Tsp_amethyst EUT2A
00350 HMMalddh_2_egra(pf00171) EUT2A
00355 H_fuchsia EUT2A
00360 HMMtpEuth_1_eut(pf04346) EUT2A
00365 HMM06249EutQ_2_eut2(pf06249) EUT2A EP_NTERM
00370 -pf01427 -
00375 HMM13520tp_1_pdu(pf13520) EUT2C
00380 HMM06723EutJMreB_1_grm(pf00012) EUT2A
00390 HMM02866_1_pvmeut(pf02866pf00056) PVM
00395 -pf10415pf00206 -
00400 -pf00890 -
00405 -pf00939 -
00410 -pf00561 -
00415 -pf04198 -
00420 HMM00390_malic_1_eut1
BMC-P
BMC-H
BMC-T
Aldehyde dehydrogenase
PTAC
Signature enzyme
Alcohol dehydrogenase
Regulator

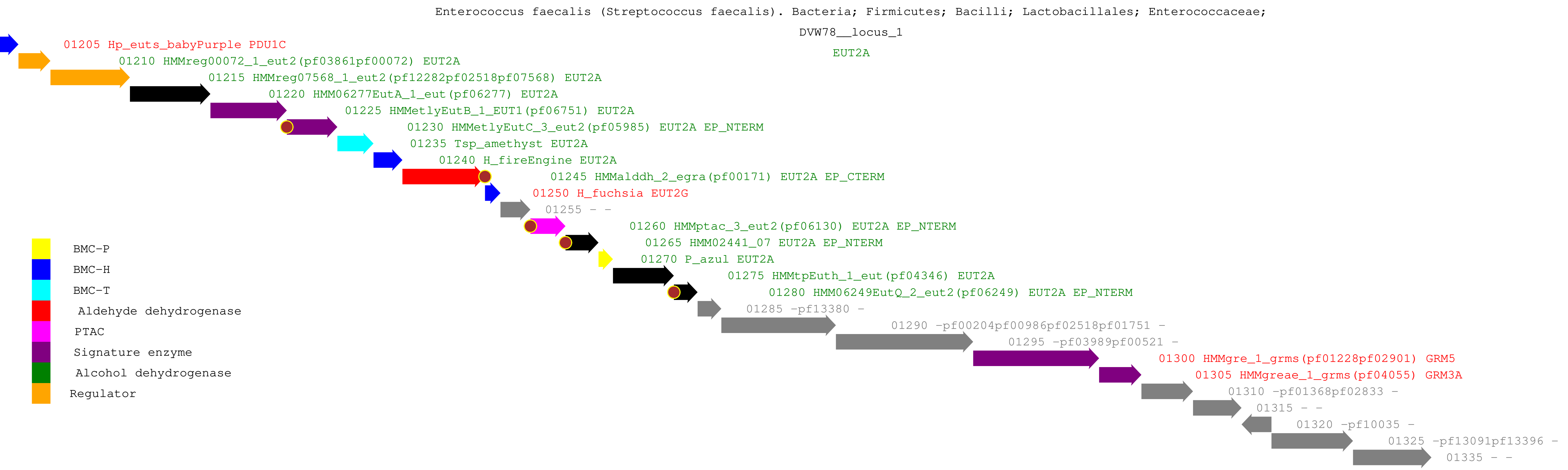
Enterococcus faecalis (Streptococcus faecalis). Bacteria; Firmicutes; Bacilli; Lactobacillales; Enterococcaceae;
DVW78__locus_1
EUT2A
01205 Hp_euts_babyPurple PDU1C
01210 HMMreg00072_1_eut2(pf03861pf00072) EUT2A
01215 HMMreg07568_1_eut2(pf12282pf02518pf07568) EUT2A
01220 HMM06277EutA_1_eut(pf06277) EUT2A
01225 HMMetlyEutB_1_EUT1(pf06751) EUT2A
01230 HMMetlyEutC_3_eut2(pf05985) EUT2A EP_NTERM
01235 Tsp_amethyst EUT2A
01240 H_fireEngine EUT2A
01245 HMMalddh_2_egra(pf00171) EUT2A EP_CTERM
01250 H_fuchsia EUT2G
01255 - -
01260 HMMptac_3_eut2(pf06130) EUT2A EP_NTERM
01265 HMM02441_07 EUT2A EP_NTERM
01270 P_azul EUT2A
01275 HMMtpEuth_1_eut(pf04346) EUT2A
01280 HMM06249EutQ_2_eut2(pf06249) EUT2A EP_NTERM
01285 -pf13380 -
01290 -pf00204pf00986pf02518pf01751 -
01295 -pf03989pf00521 -
01300 HMMgre_1_grms(pf01228pf02901) GRM5
01305 HMMgreae_1_grms(pf04055) GRM3A
01310 -pf01368pf02833 -
01315 - -
01320 -pf10035 -
01325 -pf13091pf13396 -
01335 - -
BMC-P
BMC-H
BMC-T
Aldehyde dehydrogenase
PTAC
Signature enzyme
Alcohol dehydrogenase
Regulator

Streptococcus parasanguinis. Bacteria; Firmicutes; Bacilli; Lactobacillales; Streptococcaceae;

DW820__locus_1

EUT2A

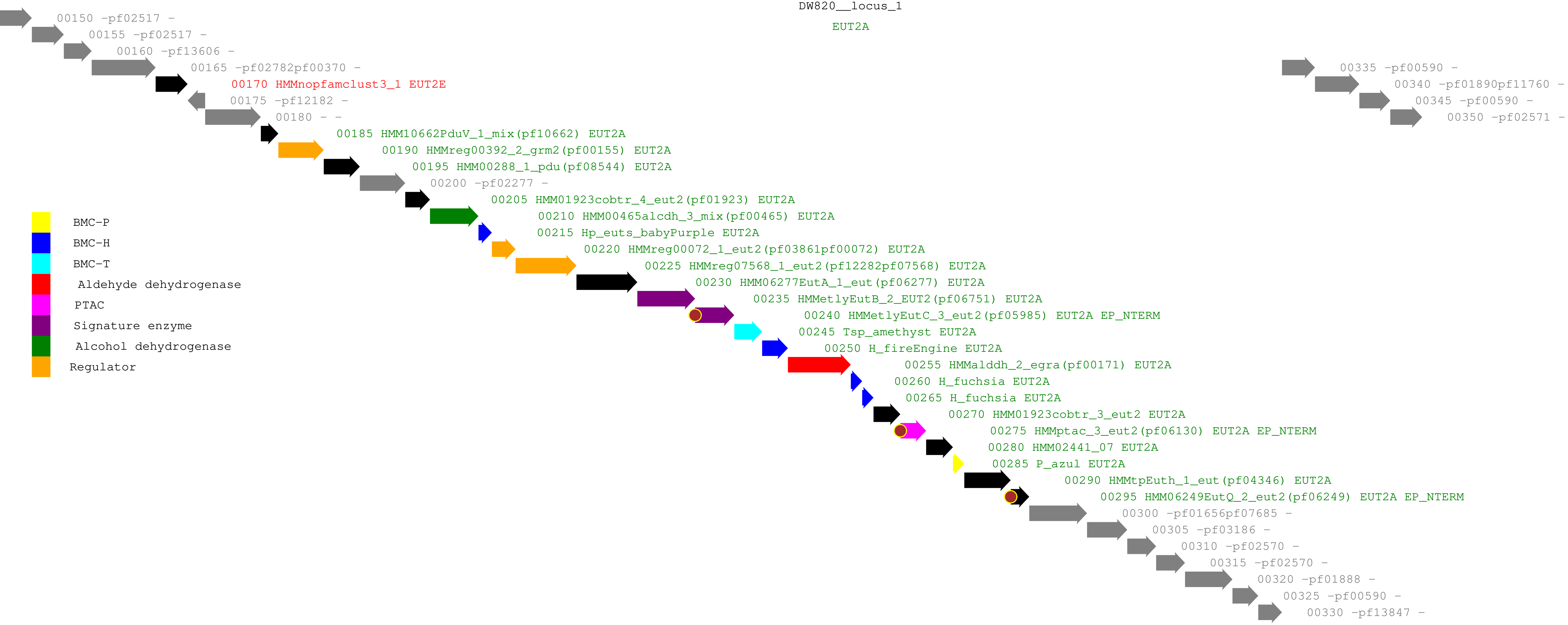

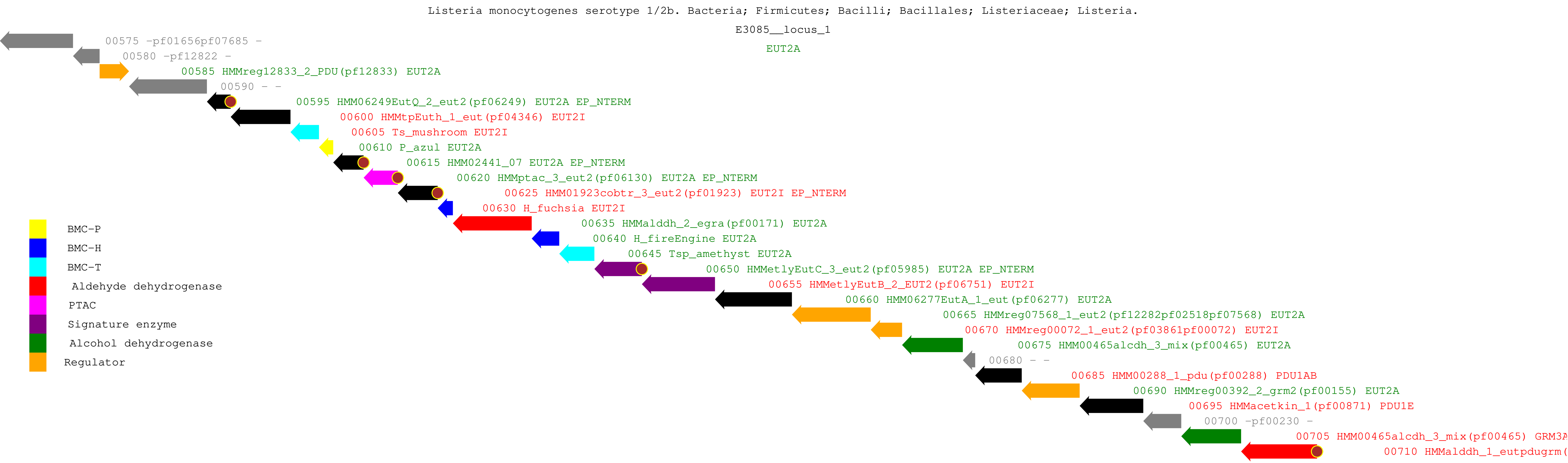
Listeria monocytogenes serotype 1/2b. Bacteria; Firmicutes; Bacilli; Bacillales; Listeriaceae; Listeria.
E3085__locus_1
EUT2A
00575 -pf01656pf07685 -
00580 -pf12822 -
00585 HMMreg12833_2_PDU(pf12833) EUT2A
00590 - -
00595 HMM06249EutQ_2_eut2(pf06249) EUT2A EP_NTERM
00600 HMMtpEuth_1_eut(pf04346) EUT2I
00605 Ts_mushroom EUT2I
00610 P_azul EUT2A
00615 HMM02441_07 EUT2A EP_NTERM
00620 HMMptac_3_eut2(pf06130) EUT2A EP_NTERM
00625 HMM01923cobtr_3_eut2(pf01923) EUT2I EP_NTERM
00630 H_fuchsia EUT2I
00635 HMMalddh_2_egra(pf00171) EUT2A
00640 H_fireEngine EUT2A
00645 Tsp_amethyst EUT2A
00650 HMMetlyEutC_3_eut2(pf05985) EUT2A EP_NTERM
00655 HMMetlyEutB_2_EUT2(pf06751) EUT2I
00660 HMM06277EutA_1_eut(pf06277) EUT2A
00665 HMMreg07568_1_eut2(pf12282pf02518pf07568) EUT2A
00670 HMMreg00072_1_eut2(pf03861pf00072) EUT2I
00675 HMM00465alcdh_3_mix(pf00465) EUT2A
00680 - -
00685 HMM00288_1_pdu(pf00288) PDU1AB
00690 HMMreg00392_2_grm2(pf00155) EUT2A
00695 HMMacetkin_1(pf00871) PDU1E
00700 -pf00230 -
00705 HMM00465alcdh_3_mix(pf00465) GRM3A
00710 HMMalddh_1_eutpdugrm(
BMC-P
BMC-H
BMC-T
Aldehyde dehydrogenase
PTAC
Signature enzyme
Alcohol dehydrogenase
Regulator

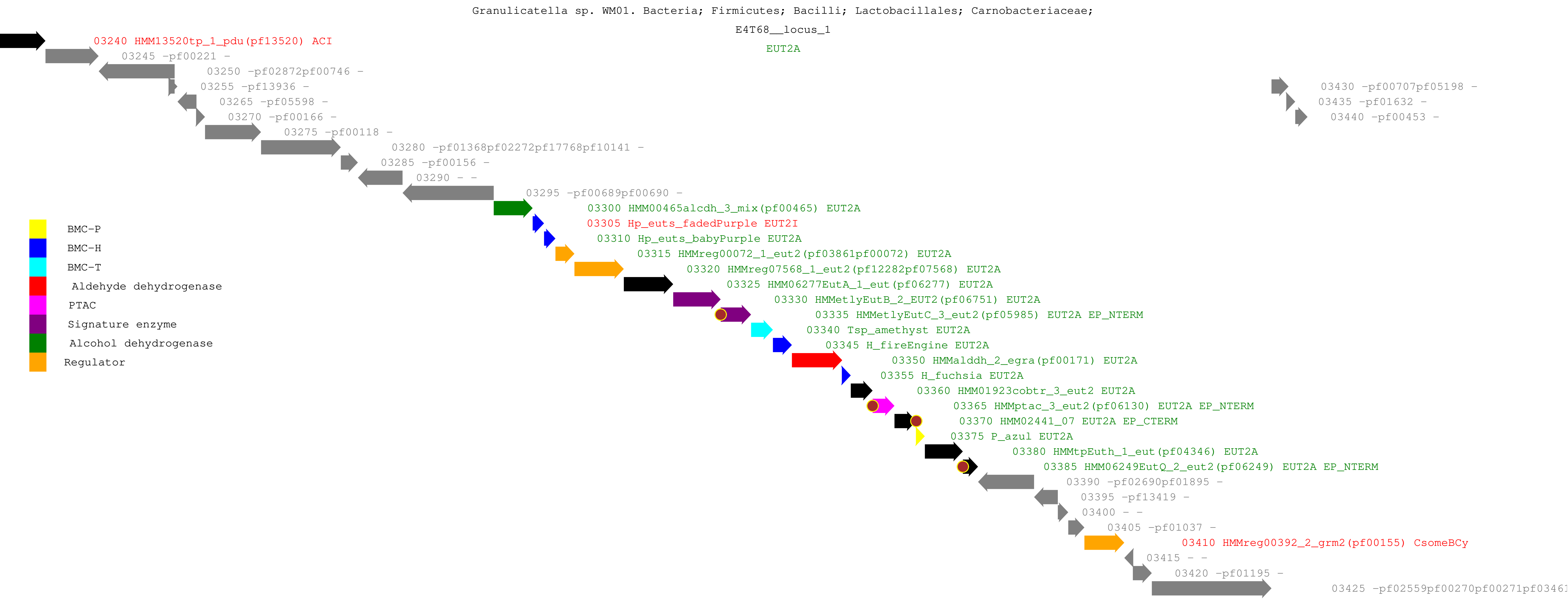
Granulicatella sp. WM01. Bacteria; Firmicutes; Bacilli; Lactobacillales; Carnobacteriaceae;
E4T68__locus_1
EUT2A
03240 HMM13520tp_1_pdu(pf13520) ACI
03245 -pf00221 -
03250 -pf02872pf00746 -
03255 -pf13936 -
03265 -pf05598 -
03270 -pf00166 -
03275 -pf00118 -
03280 -pf01368pf02272pf17768pf10141 -
03285 -pf00156 -
03290 - -
03295 -pf00689pf00690 -
03300 HMM00465alcdh_3_mix(pf00465) EUT2A
03305 Hp_euts_fadedPurple EUT2I
03310 Hp_euts_babyPurple EUT2A
03315 HMMreg00072_1_eut2(pf03861pf00072) EUT2A
03320 HMMreg07568_1_eut2(pf12282pf07568) EUT2A
03325 HMM06277EutA_1_eut(pf06277) EUT2A
03330 HMMetlyEutB_2_EUT2(pf06751) EUT2A
03335 HMMetlyEutC_3_eut2(pf05985) EUT2A EP_NTERM
03340 Tsp_amethyst EUT2A
03345 H_fireEngine EUT2A
03350 HMMalddh_2_egra(pf00171) EUT2A
03355 H_fuchsia EUT2A
03360 HMM01923cobtr_3_eut2 EUT2A
03365 HMMptac_3_eut2(pf06130) EUT2A EP_NTERM
03370 HMM02441_07 EUT2A EP_CTERM
03375 P_azul EUT2A
03380 HMMtpEuth_1_eut(pf04346) EUT2A
03385 HMM06249EutQ_2_eut2(pf06249) EUT2A EP_NTERM
03390 -pf02690pf01895 -
03395 -pf13419 -
03400 - -
03405 -pf01037 -
03410 HMMreg00392_2_grm2(pf00155) CsomeBCy
03415 - -
03420 -pf01195 -
03425 -pf02559pf00270pf00271pf0346
03430 -pf00707pf05198 -
03435 -pf01632 -
03440 -pf00453 -
BMC-P
BMC-H
BMC-T
Aldehyde dehydrogenase
PTAC
Signature enzyme
Alcohol dehydrogenase
Regulator

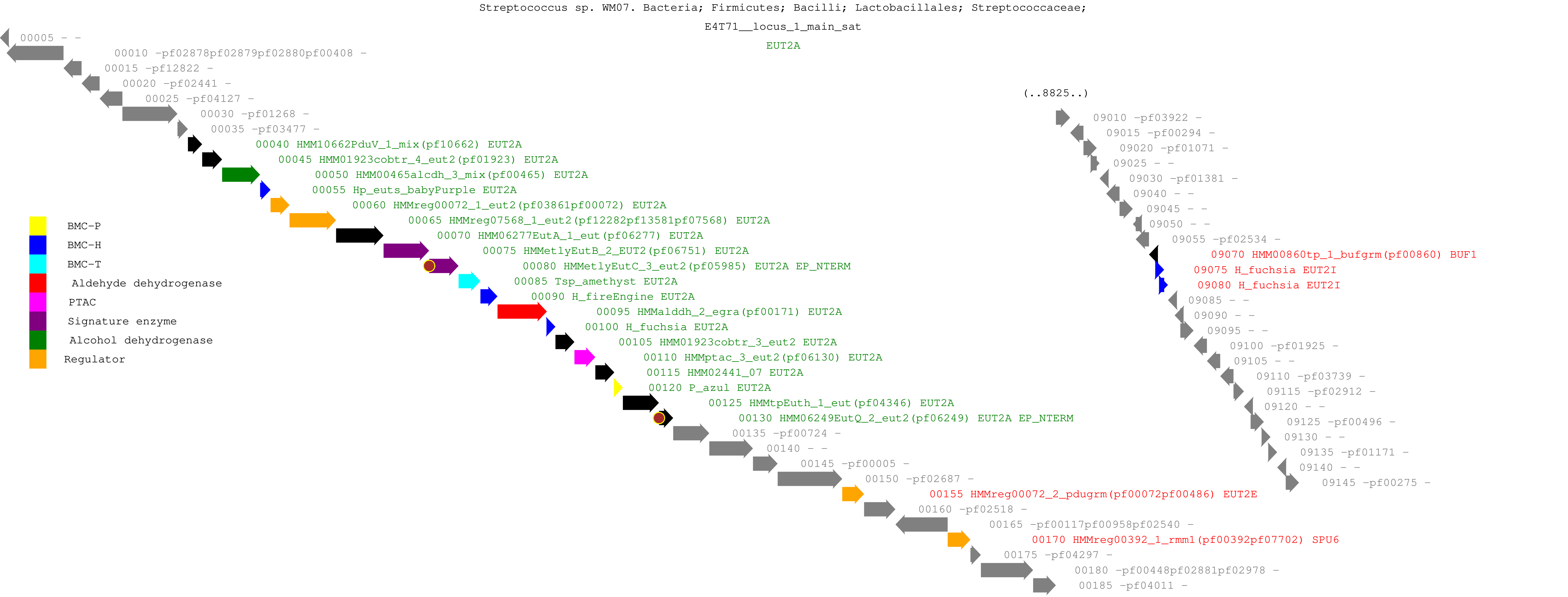
Streptococcus sp. WM07. Bacteria; Firmicutes; Bacilli; Lactobacillales; Streptococcaceae;
E4T71__locus_1_main_sat
EUT2A
00005 - -
00010 -pf02878pf02879pf02880pf00408 -
00015 -pf12822 -
00020 -pf02441 -
00025 -pf04127 -
00030 -pf01268 -
00035 -pf03477 -
00040 HMM10662PduV_1_mix(pf10662) EUT2A
00045 HMM01923cobtr_4_eut2(pf01923) EUT2A
00050 HMM00465alcdh_3_mix(pf00465) EUT2A
00055 Hp_euts_babyPurple EUT2A
00060 HMMreg00072_1_eut2(pf03861pf00072) EUT2A
00065 HMMreg07568_1_eut2(pf12282pf13581pf07568) EUT2A
00070 HMM06277EutA_1_eut(pf06277) EUT2A
00075 HMMetlyEutB_2_EUT2(pf06751) EUT2A
00080 HMMetlyEutC_3_eut2(pf05985) EUT2A EP_NTERM
00085 Tsp_amethyst EUT2A
00090 H_fireEngine EUT2A
00095 HMMalddh_2_egra(pf00171) EUT2A
00100 H_fuchsia EUT2A
00105 HMM01923cobtr_3_eut2 EUT2A
00110 HMMptac_3_eut2(pf06130) EUT2A
00115 HMM02441_07 EUT2A
00120 P_azul EUT2A
00125 HMMtpEuth_1_eut(pf04346) EUT2A
00130 HMM06249EutQ_2_eut2(pf06249) EUT2A EP_NTERM
00135 -pf00724 -
00140 - -
00145 -pf00005 -
00150 -pf02687 -
00155 HMMreg00072_2_pdugrm(pf00072pf00486) EUT2E
00160 -pf02518 -
00165 -pf00117pf00958pf02540 -
00170 HMMreg00392_1_rmm1(pf00392pf07702) SPU6
00175 -pf04297 -
00180 -pf00448pf02881pf02978 -
00185 -pf04011 -
BMC-P
BMC-H
BMC-T
Aldehyde dehydrogenase
PTAC
Signature enzyme
Alcohol dehydrogenase
Regulator
(..8825..)
09010 -pf03922 -
09015 -pf00294 -
09020 -pf01071 -
09025 - -
09030 -pf01381 -
09040 - -
09045 - -
09050 - -
09055 -pf02534 -
09070 HMM00860tp_1_bufgrm(pf00860) BUF1
09075 H_fuchsia EUT2I
09080 H_fuchsia EUT2I
09085 - -
09090 - -
09095 - -
09100 -pf01925 -
09105 - -
09110 -pf03739 -
09115 -pf02912 -
09120 - -
09125 -pf00496 -
09130 - -
09135 -pf01171 -
09140 - -
09145 -pf00275 -

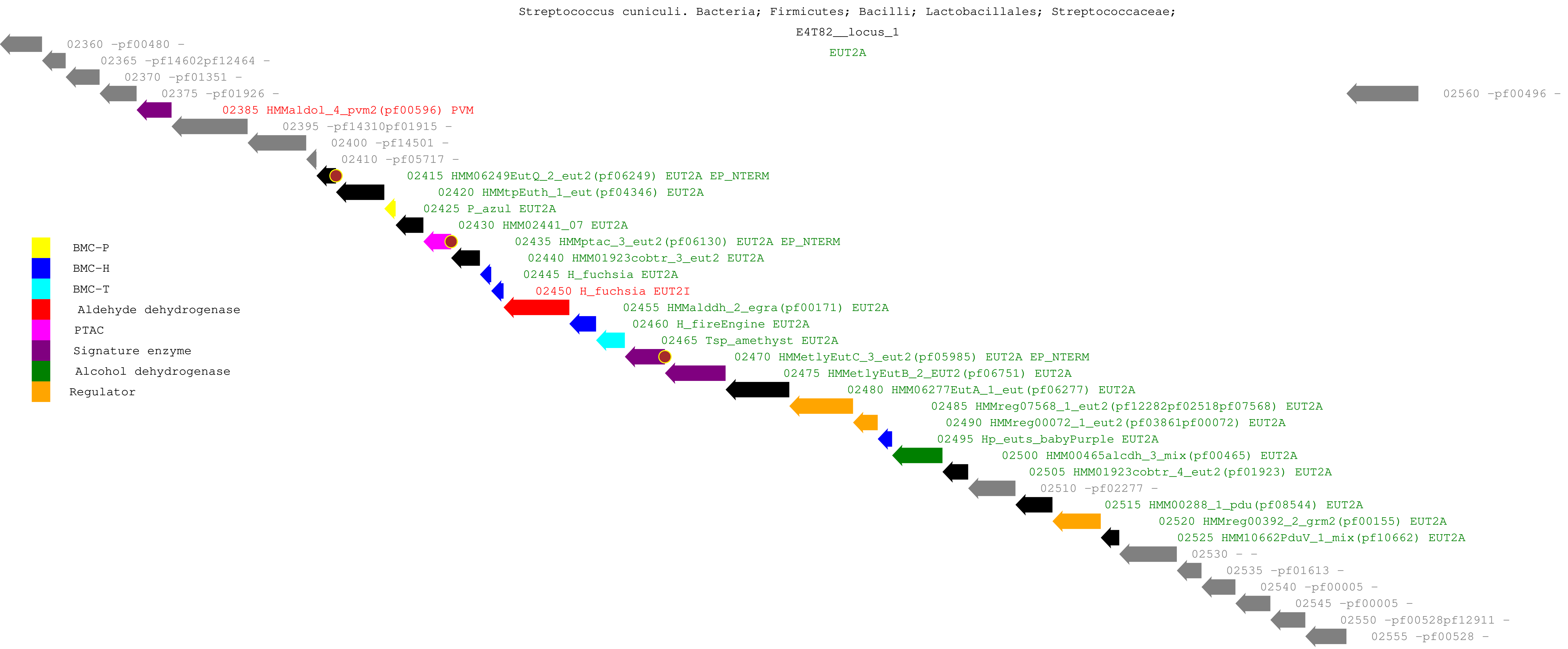
Streptococcus cuniculi. Bacteria; Firmicutes; Bacilli; Lactobacillales; Streptococcaceae;
E4T82__locus_1
EUT2A
02360 -pf00480 -
02365 -pf14602pf12464 -
02370 -pf01351 -
02375 -pf01926 -
02560 -pf00496 -
02385 HMMaldol_4_pvm2(pf00596) PVM
02395 -pf14310pf01915 -
02400 -pf14501 -
02410 -pf05717 -
02415 HMM06249EutQ_2_eut2(pf06249) EUT2A EP_NTERM
02420 HMMtpEuth_1_eut(pf04346) EUT2A
02425 P_azul EUT2A
02430 HMM02441_07 EUT2A
02435 HMMptac_3_eut2(pf06130) EUT2A EP_NTERM
02440 HMM01923cobtr_3_eut2 EUT2A
02445 H_fuchsia EUT2A
02450 H_fuchsia EUT2I
02455 HMMalddh_2_egra(pf00171) EUT2A
02460 H_fireEngine EUT2A
02465 Tsp_amethyst EUT2A
02470 HMMetlyEutC_3_eut2(pf05985) EUT2A EP_NTERM
02475 HMMetlyEutB_2_EUT2(pf06751) EUT2A
02480 HMM06277EutA_1_eut(pf06277) EUT2A
02485 HMMreg07568_1_eut2(pf12282pf02518pf07568) EUT2A
02490 HMMreg00072_1_eut2(pf03861pf00072) EUT2A
02495 Hp_euts_babyPurple EUT2A
02500 HMM00465alcdh_3_mix(pf00465) EUT2A
02505 HMM01923cobtr_4_eut2(pf01923) EUT2A
02510 -pf02277 -
02515 HMM00288_1_pdu(pf08544) EUT2A
02520 HMMreg00392_2_grm2(pf00155) EUT2A
02525 HMM10662PduV_1_mix(pf10662) EUT2A
02530 - -
02535 -pf01613 -
02540 -pf00005 -
02545 -pf00005 -
02550 -pf00528pf12911 -
02555 -pf00528 -
BMC-P
BMC-H
BMC-T
Aldehyde dehydrogenase
PTAC
Signature enzyme
Alcohol dehydrogenase
Regulator

Enterococcus avium (Streptococcus avium). Bacteria; Firmicutes; Bacilli; Lactobacillales; Enterococcaceae;

EASS39__locus_1

EUT2A

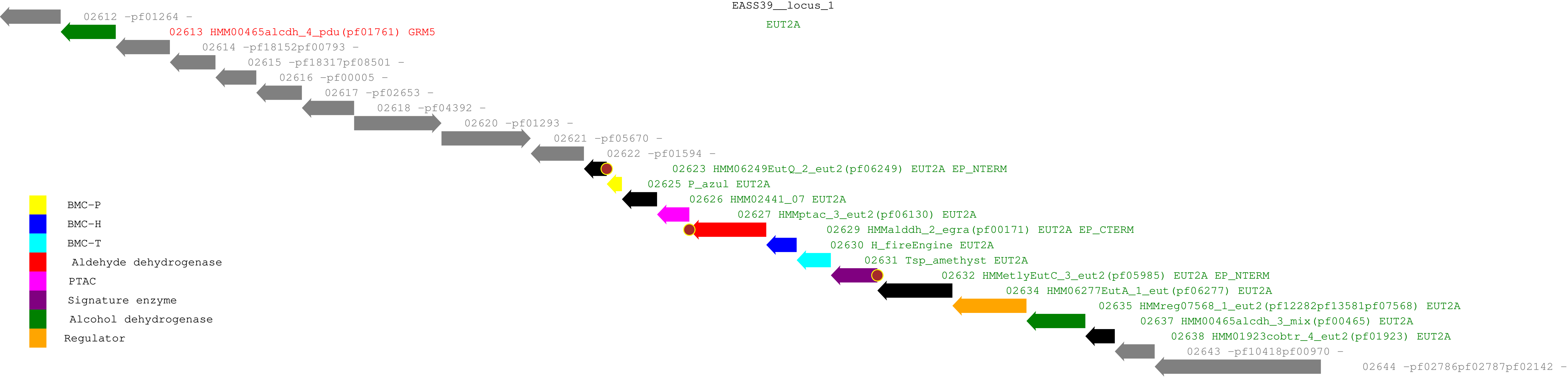

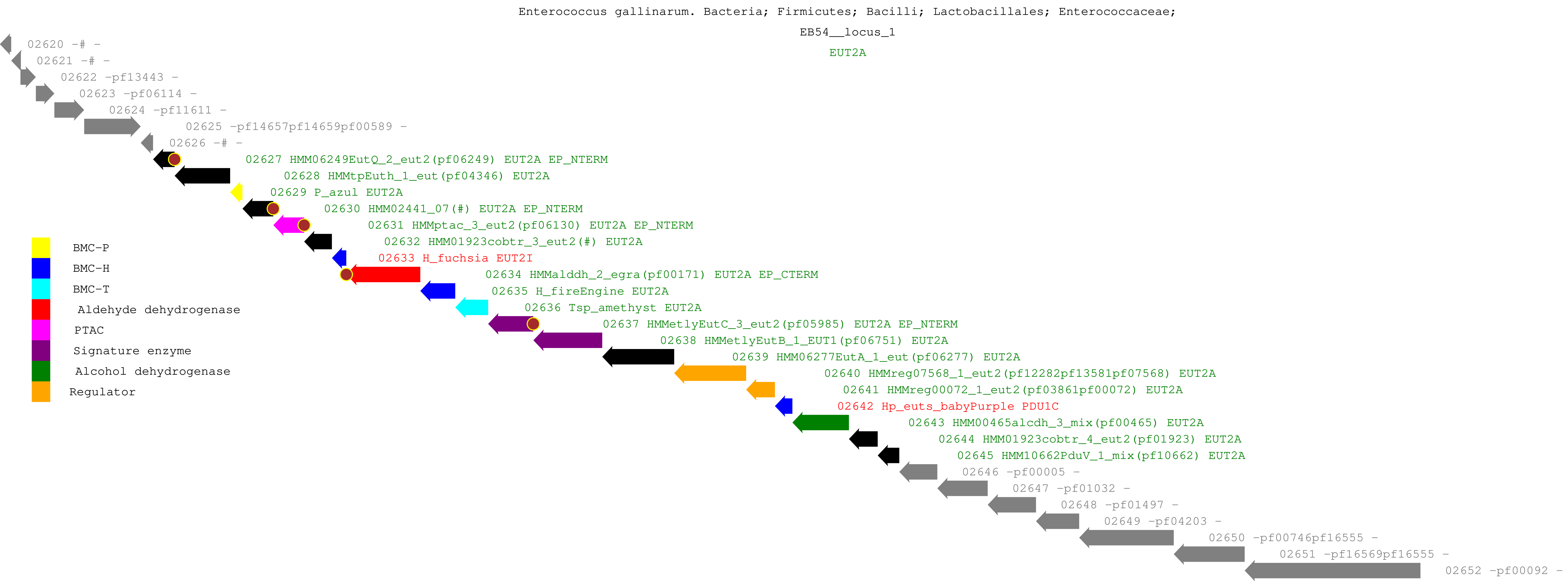
Enterococcus gallinarum. Bacteria; Firmicutes; Bacilli; Lactobacillales; Enterococcaceae;
EB54__locus_1
EUT2A
02620 -# -
02621 -# -
02622 -pf13443 -
02623 -pf06114 -
02624 -pf11611 -
02625 -pf14657pf14659pf00589 -
02626 -# -
02627 HMM06249EutQ_2_eut2(pf06249) EUT2A EP_NTERM
02628 HMMtpEuth_1_eut(pf04346) EUT2A
02629 P_azul EUT2A
02630 HMM02441_07(#) EUT2A EP_NTERM
02631 HMMptac_3_eut2(pf06130) EUT2A EP_NTERM
02632 HMM01923cobtr_3_eut2(#) EUT2A
02633 H_fuchsia EUT2I
02634 HMMalddh_2_egra(pf00171) EUT2A EP_CTERM
02635 H_fireEngine EUT2A
02636 Tsp_amethyst EUT2A
02637 HMMetlyEutC_3_eut2(pf05985) EUT2A EP_NTERM
02638 HMMetlyEutB_1_EUT1(pf06751) EUT2A
02639 HMM06277EutA_1_eut(pf06277) EUT2A
02640 HMMreg07568_1_eut2(pf12282pf13581pf07568) EUT2A
02641 HMMreg00072_1_eut2(pf03861pf00072) EUT2A
02642 Hp_euts_babyPurple PDU1C
02643 HMM00465alcdh_3_mix(pf00465) EUT2A
02644 HMM01923cobtr_4_eut2(pf01923) EUT2A
02645 HMM10662PduV_1_mix(pf10662) EUT2A
02646 -pf00005 -
02647 -pf01032 -
02648 -pf01497 -
02649 -pf04203 -
02650 -pf00746pf16555 -
02651 -pf16569pf16555 -
02652 -pf00092 -
BMC-P
BMC-H
BMC-T
Aldehyde dehydrogenase
PTAC
Signature enzyme
Alcohol dehydrogenase
Regulator

Enterococcus faecalis (Streptococcus faecalis). Bacteria; Firmicutes; Bacilli; Lactobacillales; Enterococcaceae;

EB58__locus_1

EUT2A

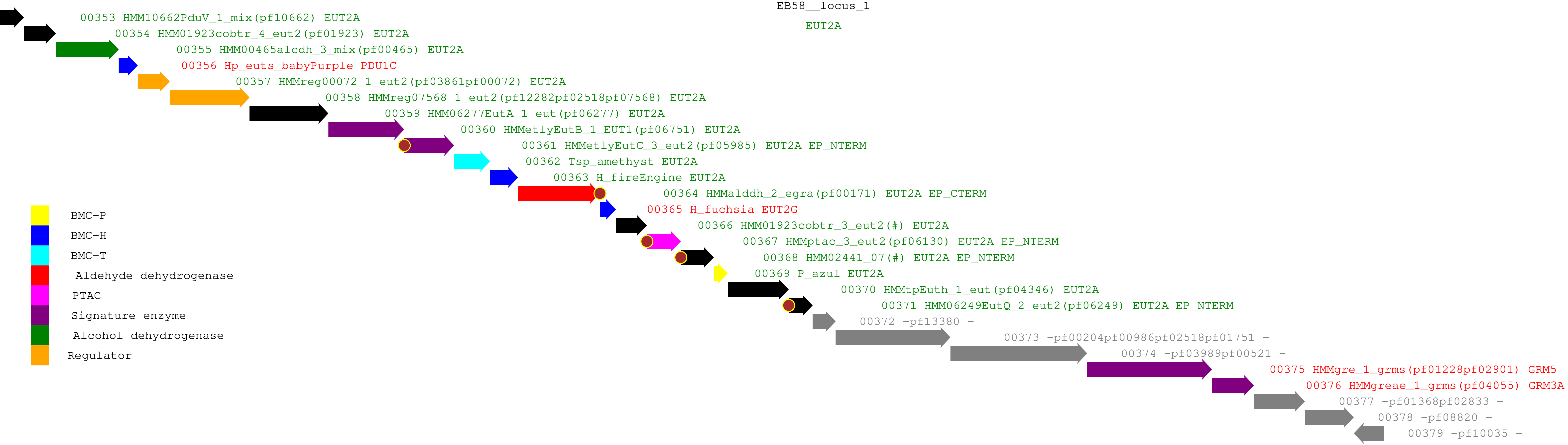

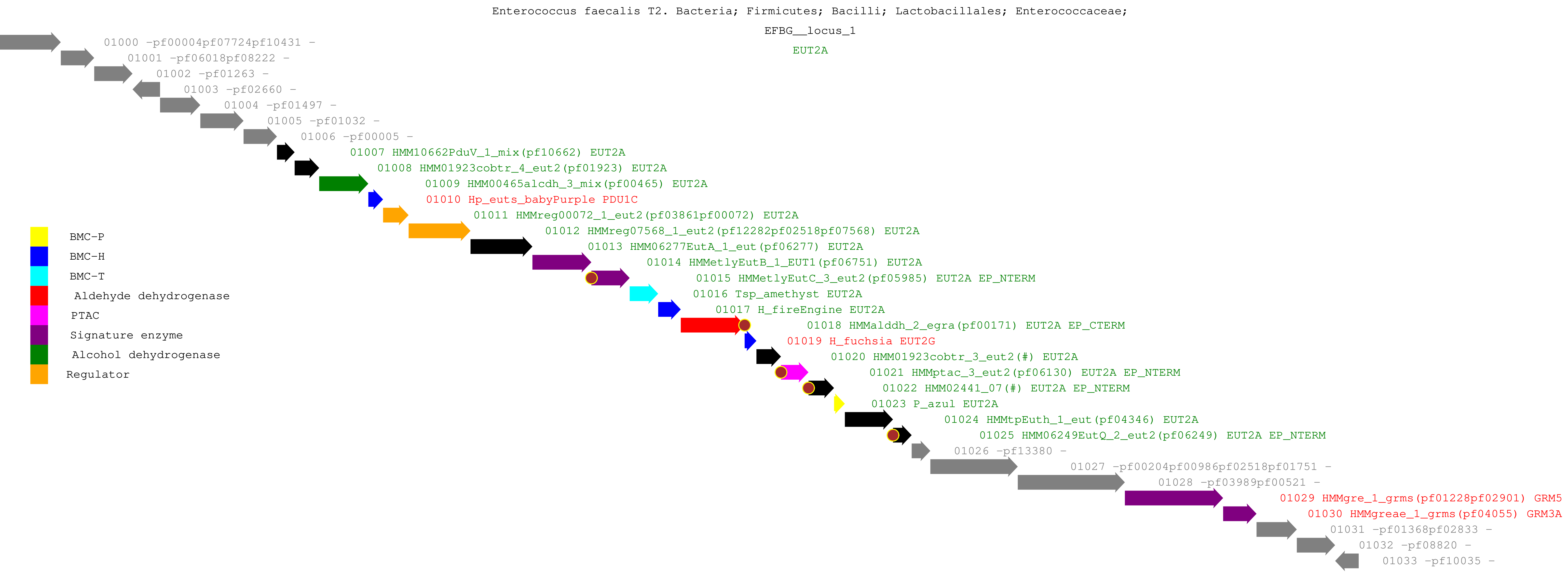
Enterococcus faecalis T2. Bacteria; Firmicutes; Bacilli; Lactobacillales; Enterococcaceae;
EFBG__locus_1
EUT2A
01000 -pf00004pf07724pf10431 -
01001 -pf06018pf08222 -
01002 -pf01263 -
01003 -pf02660 -
01004 -pf01497 -
01005 -pf01032 -
01006 -pf00005 -
01007 HMM10662PduV_1_mix(pf10662) EUT2A
01008 HMM01923cobtr_4_eut2(pf01923) EUT2A
01009 HMM00465alcdh_3_mix(pf00465) EUT2A
01010 Hp_euts_babyPurple PDU1C
01011 HMMreg00072_1_eut2(pf03861pf00072) EUT2A
01012 HMMreg07568_1_eut2(pf12282pf02518pf07568) EUT2A
01013 HMM06277EutA_1_eut(pf06277) EUT2A
01014 HMMetlyEutB_1_EUT1(pf06751) EUT2A
01015 HMMetlyEutC_3_eut2(pf05985) EUT2A EP_NTERM
01016 Tsp_amethyst EUT2A
01017 H_fireEngine EUT2A
01018 HMMalddh_2_egra(pf00171) EUT2A EP_CTERM
01019 H_fuchsia EUT2G
01020 HMM01923cobtr_3_eut2(#) EUT2A
01021 HMMptac_3_eut2(pf06130) EUT2A EP_NTERM
01022 HMM02441_07(#) EUT2A EP_NTERM
01023 P_azul EUT2A
01024 HMMtpEuth_1_eut(pf04346) EUT2A
01025 HMM06249EutQ_2_eut2(pf06249) EUT2A EP_NTERM
01026 -pf13380 -
01027 -pf00204pf00986pf02518pf01751 -
01028 -pf03989pf00521 -
01029 HMMgre_1_grms(pf01228pf02901) GRM5
01030 HMMgreae_1_grms(pf04055) GRM3A
01031 -pf01368pf02833 -
01032 -pf08820 -
01033 -pf10035 -
BMC-P
BMC-H
BMC-T
Aldehyde dehydrogenase
PTAC
Signature enzyme
Alcohol dehydrogenase
Regulator

Enterococcus faecalis Merz96. Bacteria; Firmicutes; Bacilli; Lactobacillales; Enterococcaceae;

EFGG__locus_1

EUT2A

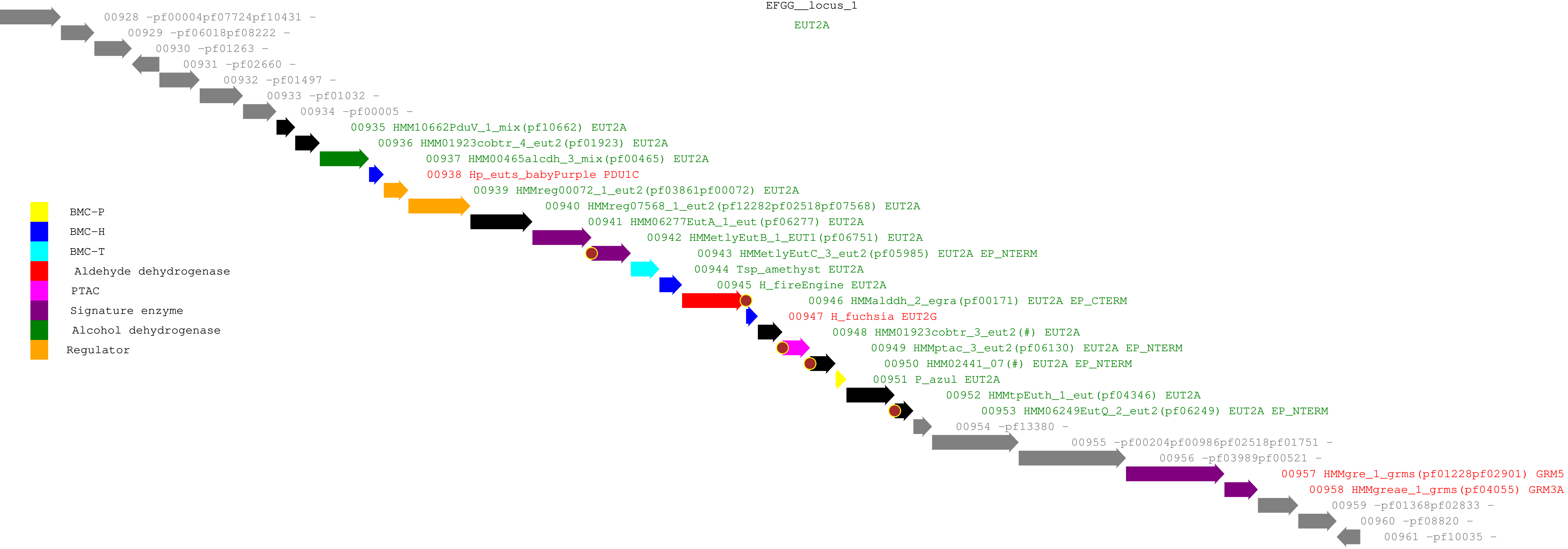

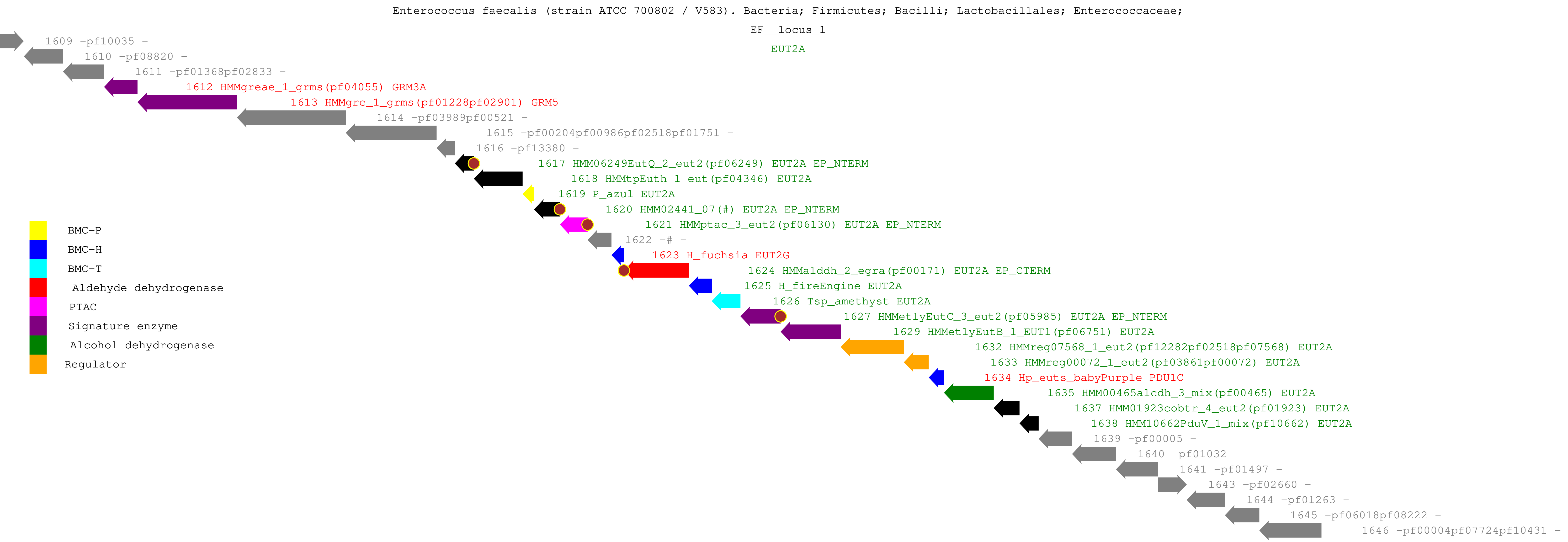
Enterococcus faecalis (strain ATCC 700802 / V583). Bacteria; Firmicutes; Bacilli; Lactobacillales; Enterococcaceae;
EF__locus_1
EUT2A
1609 -pf10035 -
1610 -pf08820 -
1611 -pf01368pf02833 -
1612 HMMgreae_1_grms(pf04055) GRM3A
1613 HMMgre_1_grms(pf01228pf02901) GRM5
1614 -pf03989pf00521 -
1615 -pf00204pf00986pf02518pf01751 -
1616 -pf13380 -
1617 HMM06249EutQ_2_eut2(pf06249) EUT2A EP_NTERM
1618 HMMtpEuth_1_eut(pf04346) EUT2A
1619 P_azul EUT2A
1620 HMM02441_07(#) EUT2A EP_NTERM
1621 HMMptac_3_eut2(pf06130) EUT2A EP_NTERM
1622 -# -
1623 H_fuchsia EUT2G
1624 HMMalddh_2_egra(pf00171) EUT2A EP_CTERM
1625 H_fireEngine EUT2A
1626 Tsp_amethyst EUT2A
1627 HMMetlyEutC_3_eut2(pf05985) EUT2A EP_NTERM
1629 HMMetlyEutB_1_EUT1(pf06751) EUT2A
1632 HMMreg07568_1_eut2(pf12282pf02518pf07568) EUT2A
1633 HMMreg00072_1_eut2(pf03861pf00072) EUT2A
1634 Hp_euts_babyPurple PDU1C
1635 HMM00465alcdh_3_mix(pf00465) EUT2A
1637 HMM01923cobtr_4_eut2(pf01923) EUT2A
1638 HMM10662PduV_1_mix(pf10662) EUT2A
1639 -pf00005 -
1640 -pf01032 -
1641 -pf01497 -
1643 -pf02660 -
1644 -pf01263 -
1645 -pf06018pf08222 -
1646 -pf00004pf07724pf10431 -
BMC-P
BMC-H
BMC-T
Aldehyde dehydrogenase
PTAC
Signature enzyme
Alcohol dehydrogenase
Regulator

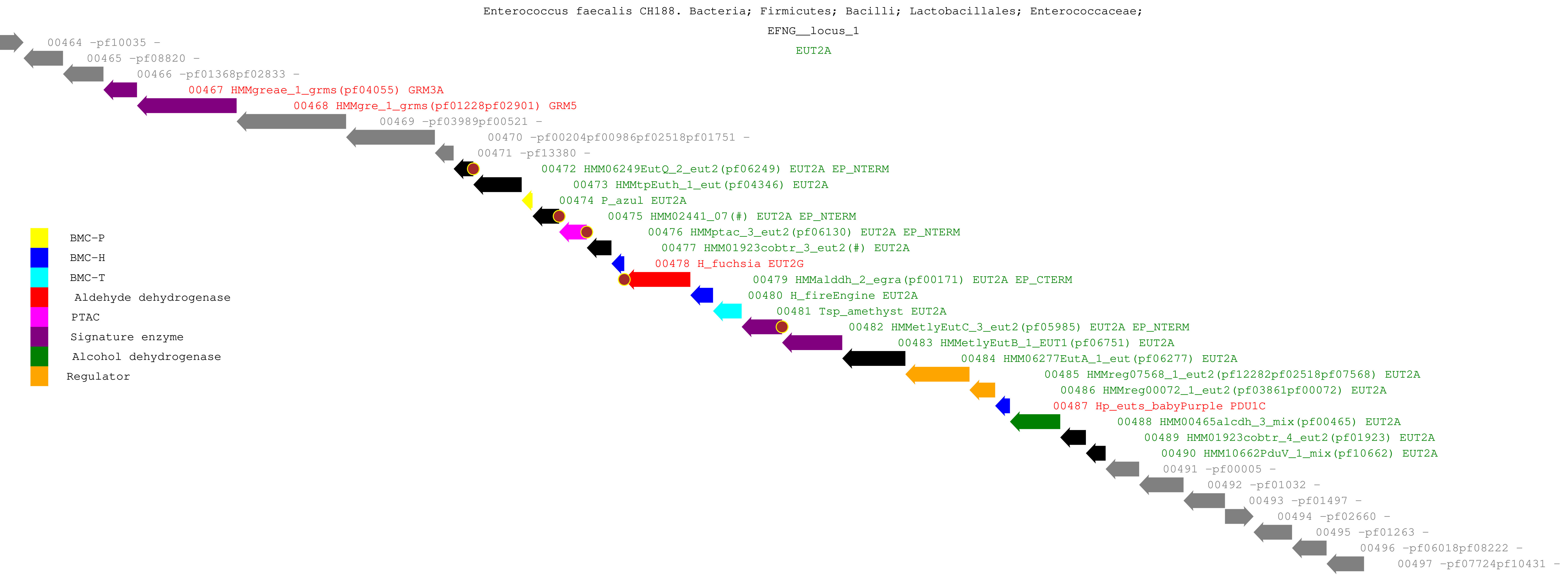
Enterococcus faecalis CH188. Bacteria; Firmicutes; Bacilli; Lactobacillales; Enterococcaceae;
EFNG__locus_1
EUT2A
00464 -pf10035 -
00465 -pf08820 -
00466 -pf01368pf02833 -
00467 HMMgreae_1_grms(pf04055) GRM3A
00468 HMMgre_1_grms(pf01228pf02901) GRM5
00469 -pf03989pf00521 -
00470 -pf00204pf00986pf02518pf01751 -
00471 -pf13380 -
00472 HMM06249EutQ_2_eut2(pf06249) EUT2A EP_NTERM
00473 HMMtpEuth_1_eut(pf04346) EUT2A
00474 P_azul EUT2A
00475 HMM02441_07(#) EUT2A EP_NTERM
00476 HMMptac_3_eut2(pf06130) EUT2A EP_NTERM
00477 HMM01923cobtr_3_eut2(#) EUT2A
00478 H_fuchsia EUT2G
00479 HMMalddh_2_egra(pf00171) EUT2A EP_CTERM
00480 H_fireEngine EUT2A
00481 Tsp_amethyst EUT2A
00482 HMMetlyEutC_3_eut2(pf05985) EUT2A EP_NTERM
00483 HMMetlyEutB_1_EUT1(pf06751) EUT2A
00484 HMM06277EutA_1_eut(pf06277) EUT2A
00485 HMMreg07568_1_eut2(pf12282pf02518pf07568) EUT2A
00486 HMMreg00072_1_eut2(pf03861pf00072) EUT2A
00487 Hp_euts_babyPurple PDU1C
00488 HMM00465alcdh_3_mix(pf00465) EUT2A
00489 HMM01923cobtr_4_eut2(pf01923) EUT2A
00490 HMM10662PduV_1_mix(pf10662) EUT2A
00491 -pf00005 -
00492 -pf01032 -
00493 -pf01497 -
00494 -pf02660 -
00495 -pf01263 -
00496 -pf06018pf08222 -
00497 -pf07724pf10431 -
BMC-P
BMC-H
BMC-T
Aldehyde dehydrogenase
PTAC
Signature enzyme
Alcohol dehydrogenase
Regulator

Enterococcus faecalis (Streptococcus faecalis). Bacteria; Firmicutes; Bacilli; Lactobacillales; Enterococcaceae;

EFW11__locus_1

EUT2A

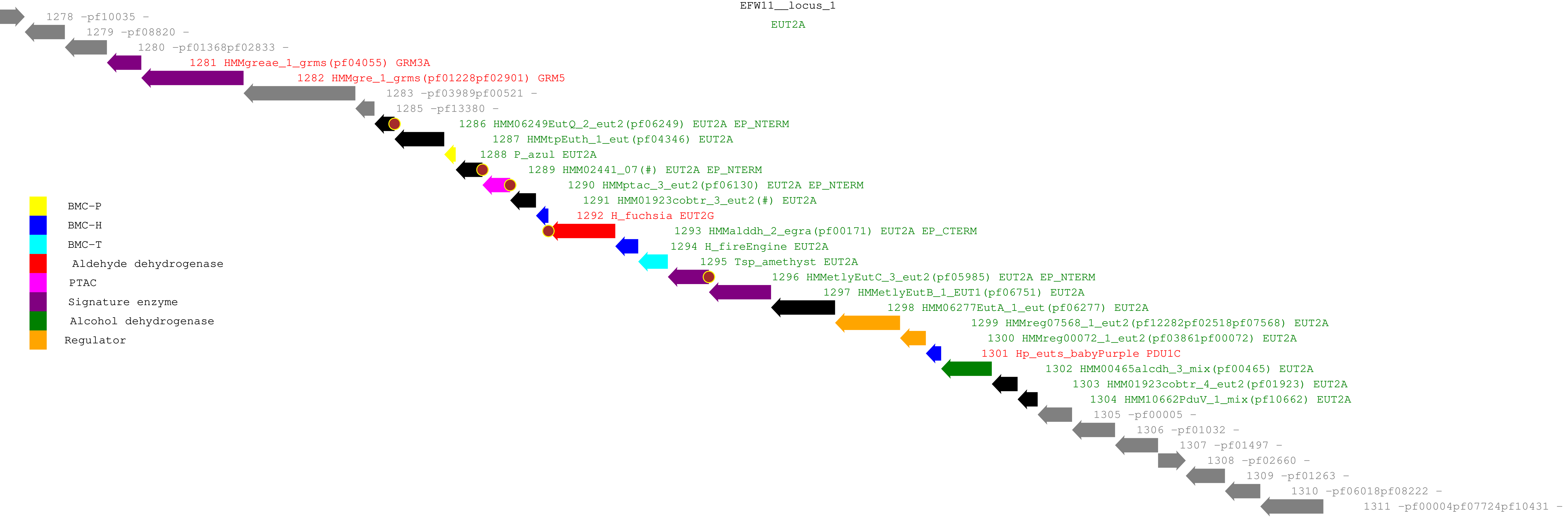

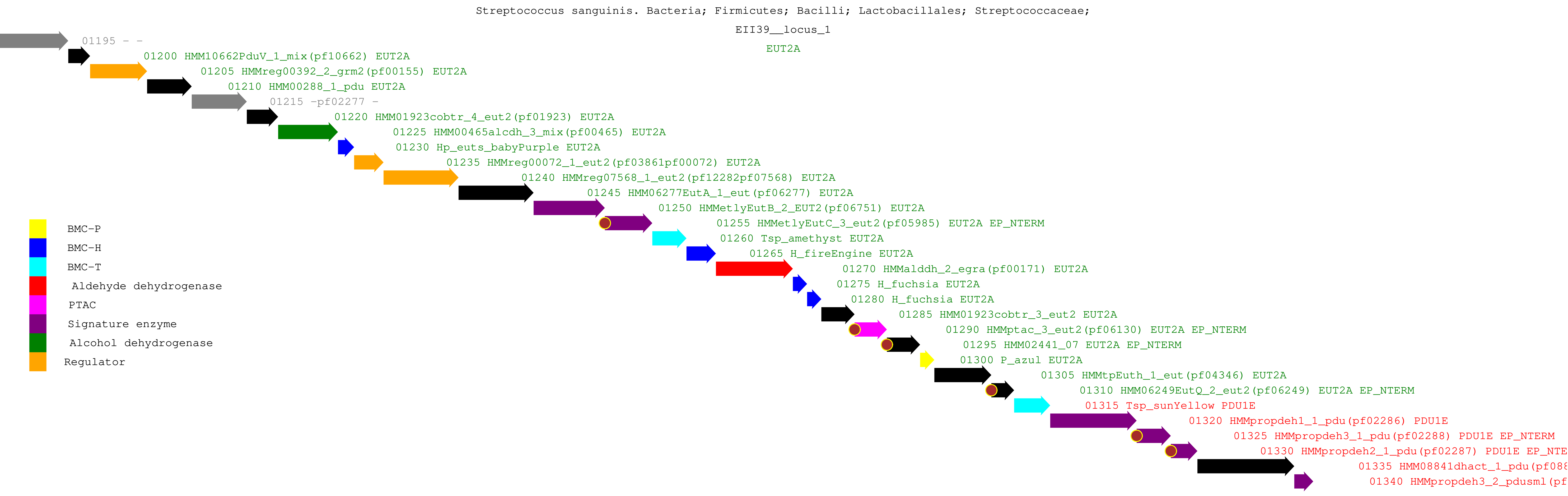
Streptococcus sanguinis. Bacteria; Firmicutes; Bacilli; Lactobacillales; Streptococcaceae;
EII39__locus_1
EUT2A
01195 - -
01200 HMM10662PduV_1_mix(pf10662) EUT2A
01205 HMMreg00392_2_grm2(pf00155) EUT2A
01210 HMM00288_1_pdu EUT2A
01215 -pf02277 -
01220 HMM01923cobtr_4_eut2(pf01923) EUT2A
01225 HMM00465alcdh_3_mix(pf00465) EUT2A
01230 Hp_euts_babyPurple EUT2A
01235 HMMreg00072_1_eut2(pf03861pf00072) EUT2A
01240 HMMreg07568_1_eut2(pf12282pf07568) EUT2A
01245 HMM06277EutA_1_eut(pf06277) EUT2A
01250 HMMetlyEutB_2_EUT2(pf06751) EUT2A
01255 HMMetlyEutC_3_eut2(pf05985) EUT2A EP_NTERM
01260 Tsp_amethyst EUT2A
01265 H_fireEngine EUT2A
01270 HMMalddh_2_egra(pf00171) EUT2A
01275 H_fuchsia EUT2A
01280 H_fuchsia EUT2A
01285 HMM01923cobtr_3_eut2 EUT2A
01290 HMMptac_3_eut2(pf06130) EUT2A EP_NTERM
01295 HMM02441_07 EUT2A EP_NTERM
01300 P_azul EUT2A
01305 HMMtpEuth_1_eut(pf04346) EUT2A
01310 HMM06249EutQ_2_eut2(pf06249) EUT2A EP_NTERM
01315 Tsp_sunYellow PDU1E
01320 HMMpropdeh1_1_pdu(pf02286) PDU1E
01325 HMMpropdeh3_1_pdu(pf02288) PDU1E EP_NTERM
01330 HMMpropdeh2_1_pdu(pf02287) PDU1E EP_NTE
01335 HMM08841dhact_1_pdu(pf088
01340 HMMpropdeh3_2_pdusml(pf
BMC-P
BMC-H
BMC-T
Aldehyde dehydrogenase
PTAC
Signature enzyme
Alcohol dehydrogenase
Regulator

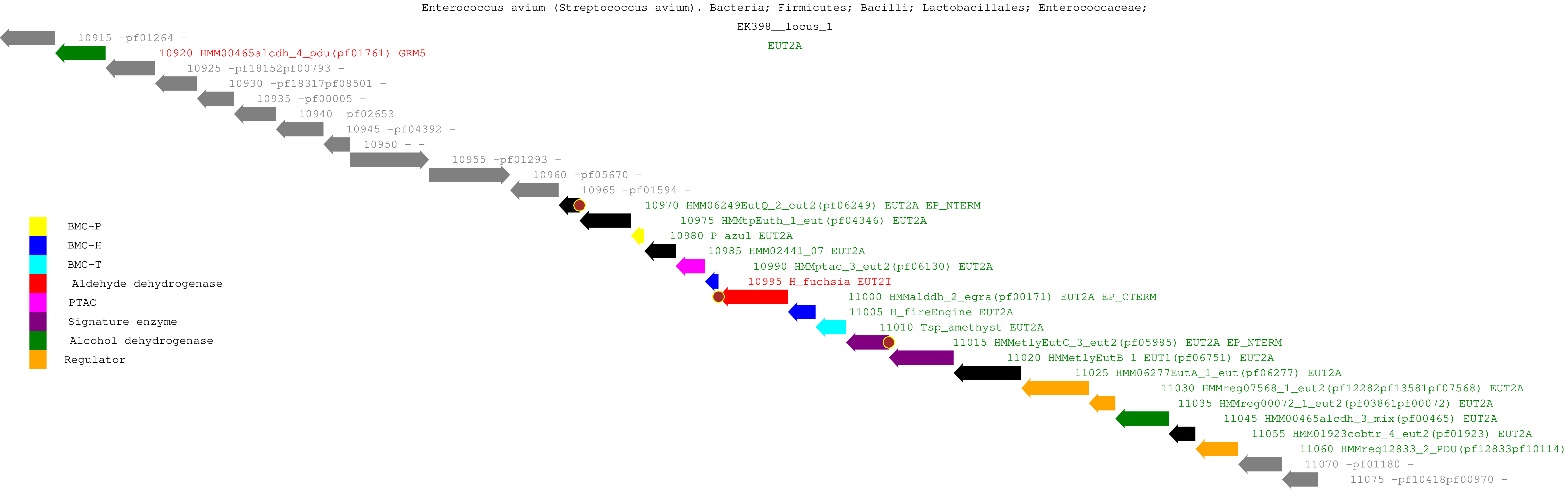
Enterococcus avium (Streptococcus avium). Bacteria; Firmicutes; Bacilli; Lactobacillales; Enterococcaceae;
EK398__locus_1
EUT2A
10915 -pf01264 -
10920 HMM00465alcdh_4_pdu(pf01761) GRM5
10925 -pf18152pf00793 -
10930 -pf18317pf08501 -
10935 -pf00005 -
10940 -pf02653 -
10945 -pf04392 -
10950 - -
10955 -pf01293 -
10960 -pf05670 -
10965 -pf01594 -
10970 HMM06249EutQ_2_eut2(pf06249) EUT2A EP_NTERM
10975 HMMtpEuth_1_eut(pf04346) EUT2A
10980 P_azul EUT2A
10985 HMM02441_07 EUT2A
10990 HMMptac_3_eut2(pf06130) EUT2A
10995 H_fuchsia EUT2I
11000 HMMaldh_2_egra(pf00171) EUT2A EP_CTERM
11005 H_fireEngine EUT2A
11010 Tsp_amethyst EUT2A
11015 HMMetlyEutC_3_eut2(pf05985) EUT2A EP_NTERM
11020 HMMetlyEutB_1_EUT1(pf06751) EUT2A
11025 HMM06277EutA_1_eut(pf06277) EUT2A
11030 HMMreg07568_1_eut2(pf12282pf13581pf07568) EUT2A
11035 HMMreg00072_1_eut2(pf03861pf00072) EUT2A
11045 HMM00465alcdh_3_mix(pf00465) EUT2A
11055 HMM01923cobtr_4_eut2(pf01923) EUT2A
11060 HMMreg12833_2_PDU(pf12833pf10114)
11070 -pf01180 -
11075 -pf10418pf00970 -
BMC-P
BMC-H
BMC-T
Aldehyde dehydrogenase
PTAC
Signature enzyme
Alcohol dehydrogenase
Regulator

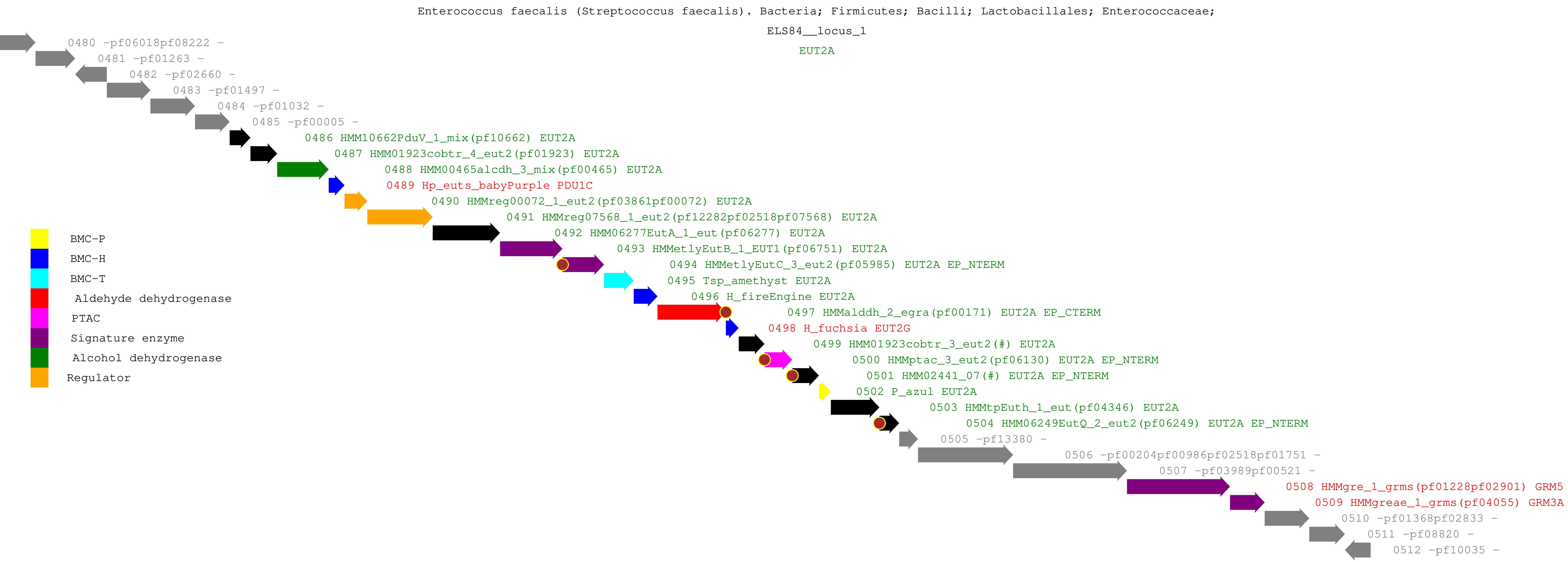

Enterococcus faecalis (Streptococcus faecalis). Bacteria; Firmicutes; Bacilli; Lactobacillales; Enterococcaceae;
ELS84__locus_1
EUT2A
0480 -pf06018pf08222 -
0481 -pf01263 -
0482 -pf02660 -
0483 -pf01497 -
0484 -pf01032 -
0485 -pf00005 -
0486 HMM10662PduV_1_mix(pf10662) EUT2A
0487 HMM01923cobtr_4_eut2(pf01923) EUT2A
0488 HMM00465alcdh_3_mix(pf00465) EUT2A
0489 Hp_euts_babyPurple PDU1C
0490 HMMreg00072_1_eut2(pf03861pf00072) EUT2A
0491 HMMreg07568_1_eut2(pf12282pf02518pf07568) EUT2A
0492 HMM06277EutA_1_eut(pf06277) EUT2A
0493 HMMetlyEutB_1_EUT1(pf06751) EUT2A
0494 HMMetlyEutC_3_eut2(pf05985) EUT2A EP_NTERM
0495 Tsp_amethyst EUT2A
0496 H_fireEngine EUT2A
0497 HMMalddh_2_egra(pf00171) EUT2A EP_CTERM
0498 H_fuchsia EUT2G
0499 HMM01923cobtr_3_eut2(#) EUT2A
0500 HMMptac_3_eut2(pf06130) EUT2A EP_NTERM
0501 HMM02441_07(#) EUT2A EP_NTERM
0502 P_azul EUT2A
0503 HMMtpEuth_1_eut(pf04346) EUT2A
0504 HMM06249EutQ_2_eut2(pf06249) EUT2A EP_NTERM
0505 -pf13380 -
0506 -pf00204pf00986pf02518pf01751 -
0507 -pf03989pf00521 -
0508 HMMgre_1_grms(pf01228pf02901) GRM5
0509 HMMgreae_1_grms(pf04055) GRM3A
0510 -pf01368pf02833 -
0511 -pf08820 -
0512 -pf10035 -
BMC-P
BMC-H
BMC-T
Aldehyde dehydrogenase
PTAC
Signature enzyme
Alcohol dehydrogenase
Regulator

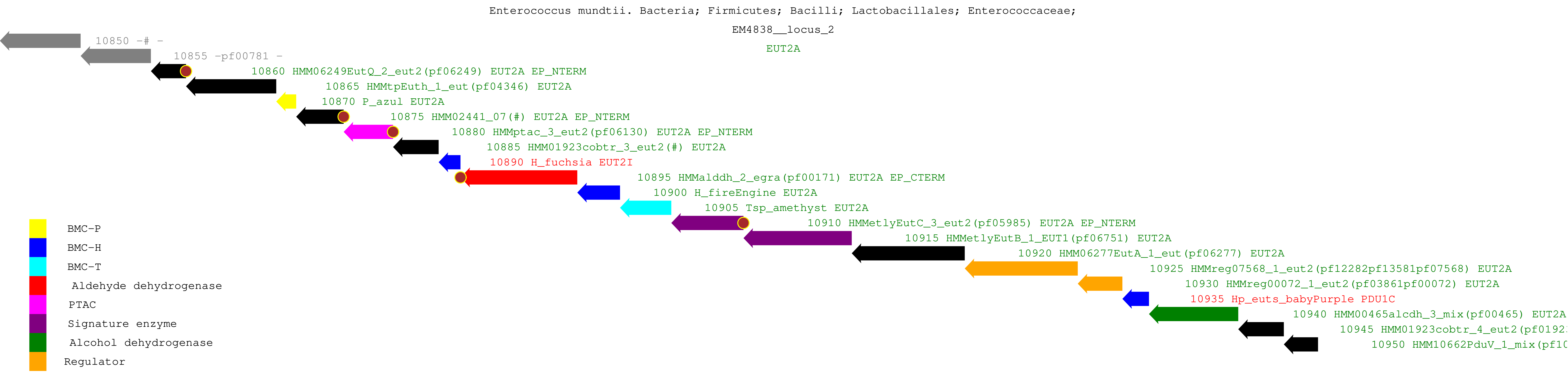
Enterococcus mundtii. Bacteria; Firmicutes; Bacilli; Lactobacillales; Enterococcaceae;
EM4838__locus_2
EUT2A
10850 -# -
10855 -pf00781 -
10860 HMM06249EutQ_2_eut2(pf06249) EUT2A EP_NTERM
10865 HMMtpEuth_1_eut(pf04346) EUT2A
10870 P_azul EUT2A
10875 HMM02441_07(#) EUT2A EP_NTERM
10880 HMMptac_3_eut2(pf06130) EUT2A EP_NTERM
10885 HMM01923cobtr_3_eut2(#) EUT2A
10890 H_fuchsia EUT2I
10895 HMMalddh_2_egra(pf00171) EUT2A EP_CTERM
10900 H_fireEngine EUT2A
10905 Tsp_amethyst EUT2A
10910 HMMetlyEutC_3_eut2(pf05985) EUT2A EP_NTERM
10915 HMMetlyEutB_1_EUT1(pf06751) EUT2A
10920 HMM06277EutA_1_eut(pf06277) EUT2A
10925 HMMreg07568_1_eut2(pf12282pf13581pf07568) EUT2A
10930 HMMreg00072_1_eut2(pf03861pf00072) EUT2A
10935 Hp_euts_babyPurple PDU1C
10940 HMM00465alcdh_3_mix(pf00465) EUT2A
10945 HMM01923cobtr_4_eut2(pf0192
10950 HMM10662PduV_1_mix(pf10
BMC-P
BMC-H
BMC-T
Aldehyde dehydrogenase
PTAC
Signature enzyme
Alcohol dehydrogenase
Regulator

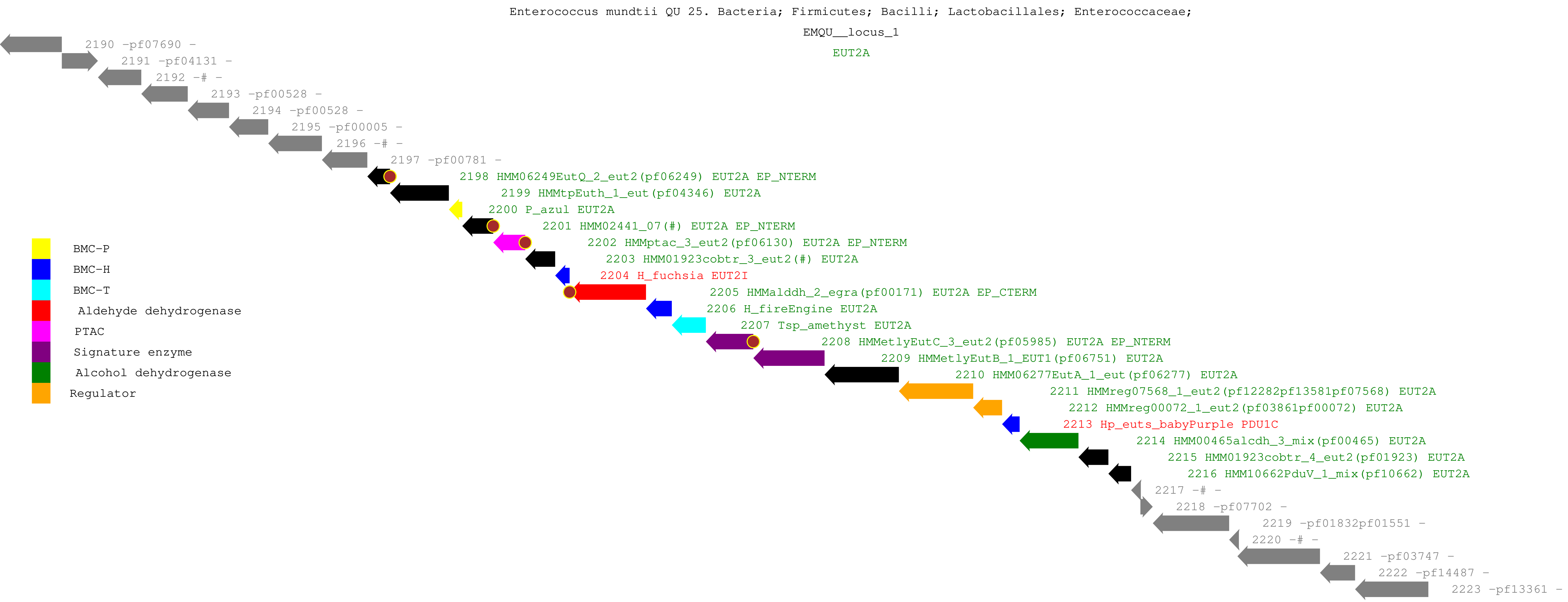
Enterococcus mundtii QU 25. Bacteria; Firmicutes; Bacilli; Lactobacillales; Enterococcaceae;
EMQU__locus_1
EUT2A
2190 -pf07690 -
2191 -pf04131 -
2192 -# -
2193 -pf00528 -
2194 -pf00528 -
2195 -pf00005 -
2196 -# -
2197 -pf00781 -
2198 HMM06249EutQ_2_eut2(pf06249) EUT2A EP_NTERM
2199 HMMtpEuth_1_eut(pf04346) EUT2A
2200 P_azul EUT2A
2201 HMM02441_07(#) EUT2A EP_NTERM
2202 HMMptac_3_eut2(pf06130) EUT2A EP_NTERM
2203 HMM01923cobtr_3_eut2(#) EUT2A
2204 H_fuchsia EUT2I
2205 HMMaldddh_2_egra(pf00171) EUT2A EP_CTERM
2206 H_fireEngine EUT2A
2207 Tsp_amethyst EUT2A
2208 HMMetlyEutC_3_eut2(pf05985) EUT2A EP_NTERM
2209 HMMetlyEutB_1_EUT1(pf06751) EUT2A
2210 HMM06277EutA_1_eut(pf06277) EUT2A
2211 HMMreg07568_1_eut2(pf12282pf13581pf07568) EUT2A
2212 HMMreg00072_1_eut2(pf03861pf00072) EUT2A
2213 Hp_euts_babyPurple PDU1C
2214 HMM00465alcdh_3_mix(pf00465) EUT2A
2215 HMM01923cobtr_4_eut2(pf01923) EUT2A
2216 HMM10662PduV_1_mix(pf10662) EUT2A
2217 -# -
2218 -pf07702 -
2219 -pf01832pf01551 -
2220 -# -
2221 -pf03747 -
2222 -pf14487 -
2223 -pf13361 -
BMC-P
BMC-H
BMC-T
Aldehyde dehydrogenase
PTAC
Signature enzyme
Alcohol dehydrogenase
Regulator

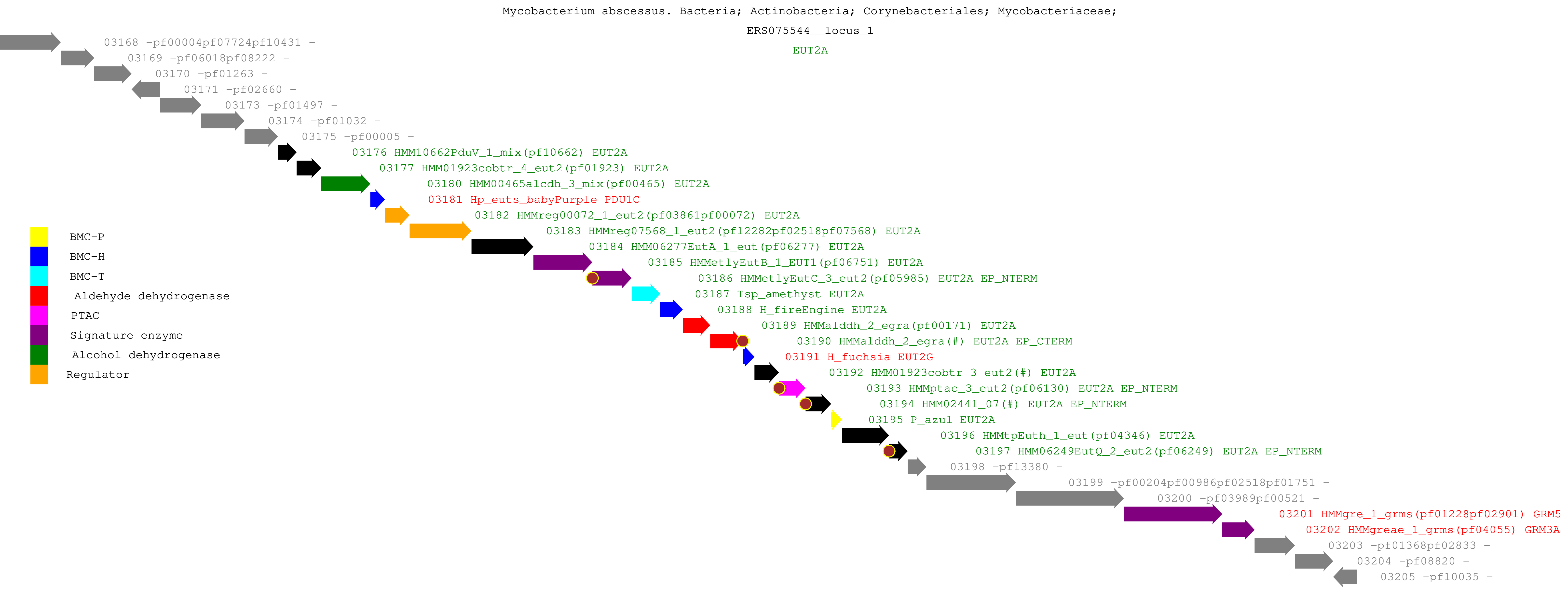

Mycobacterium abscessus. Bacteria; Actinobacteria; Corynebacteriales; Mycobacteriaceae;
ERS075544__locus_1
EUT2A
03168 -pf00004pf07724pf10431 -
03169 -pf06018pf08222 -
03170 -pf01263 -
03171 -pf02660 -
03173 -pf01497 -
03174 -pf01032 -
03175 -pf00005 -
03176 HMM10662PduV_1_mix(pf10662) EUT2A
03177 HMM01923cobtr_4_eut2(pf01923) EUT2A
03180 HMM00465alcdh_3_mix(pf00465) EUT2A
03181 Hp_euts_babyPurple PDU1C
03182 HMMreg00072_1_eut2(pf03861pf00072) EUT2A
03183 HMMreg07568_1_eut2(pf12282pf02518pf07568) EUT2A
03184 HMM06277EutA_1_eut(pf06277) EUT2A
03185 HMMetlyEutB_1_EUT1(pf06751) EUT2A
03186 HMMetlyEutC_3_eut2(pf05985) EUT2A EP_NTERM
03187 Tsp_amethyst EUT2A
03188 H_fireEngine EUT2A
03189 HMMalddh_2_egra(pf00171) EUT2A
03190 HMMalddh_2_egra(#) EUT2A EP_CTERM
03191 H_fuchsia EUT2G
03192 HMM01923cobtr_3_eut2(#) EUT2A
03193 HMMptac_3_eut2(pf06130) EUT2A EP_NTERM
03194 HMM02441_07(#) EUT2A EP_NTERM
03195 P_azul EUT2A
03196 HMMtpEuth_1_eut(pf04346) EUT2A
03197 HMM06249EutQ_2_eut2(pf06249) EUT2A EP_NTERM
03198 -pf13380 -
03199 -pf00204pf00986pf02518pf01751 -
03200 -pf03989pf00521 -
03201 HMMgre_1_grms(pf01228pf02901) GRM5
03202 HMMgreae_1_grms(pf04055) GRM3A
03203 -pf01368pf02833 -
03204 -pf08820 -
03205 -pf10035 -
BMC-P
BMC-H
BMC-T
Aldehyde dehydrogenase
PTAC
Signature enzyme
Alcohol dehydrogenase
Regulator

Chlamydia trachomatis. Bacteria; Chlamydiae; Chlamydiales; Chlamydiaceae;

ERS095036__locus_1_main_sat

EUT2A

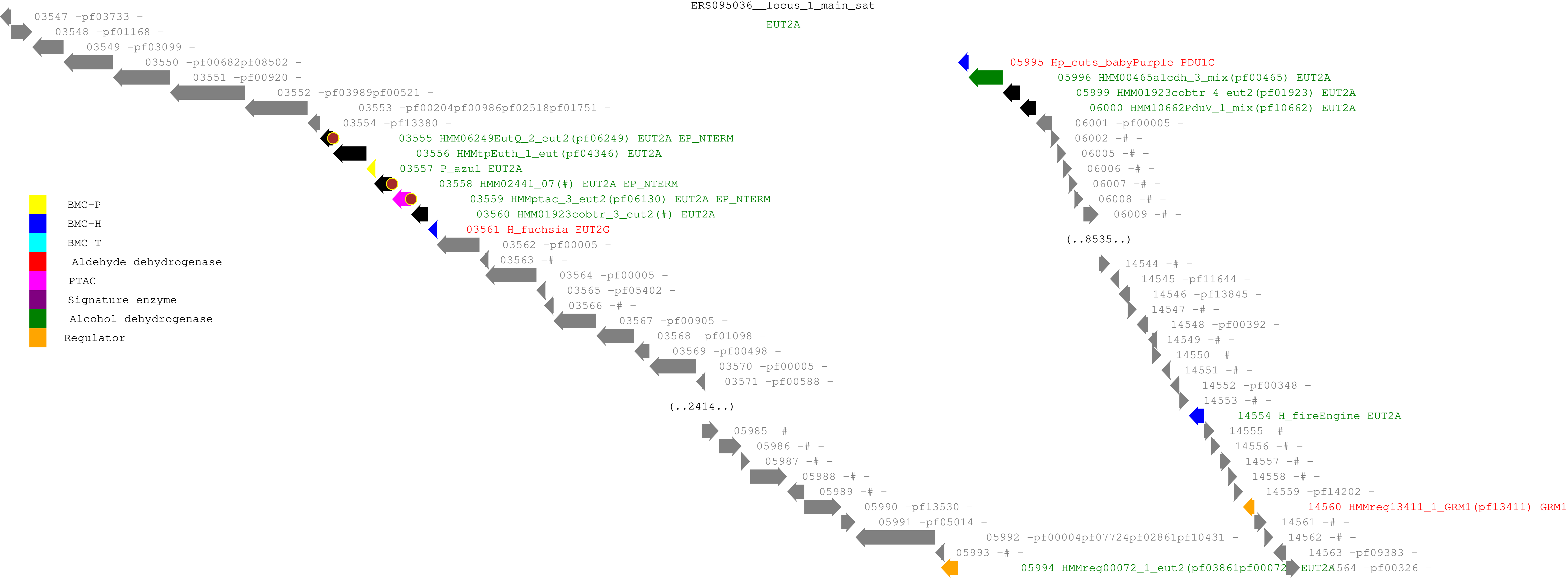

Streptococcus pneumoniae. Bacteria; Firmicutes; Bacilli; Lactobacillales; Streptococcaceae;

ERS409405__locus_1

EUT2A

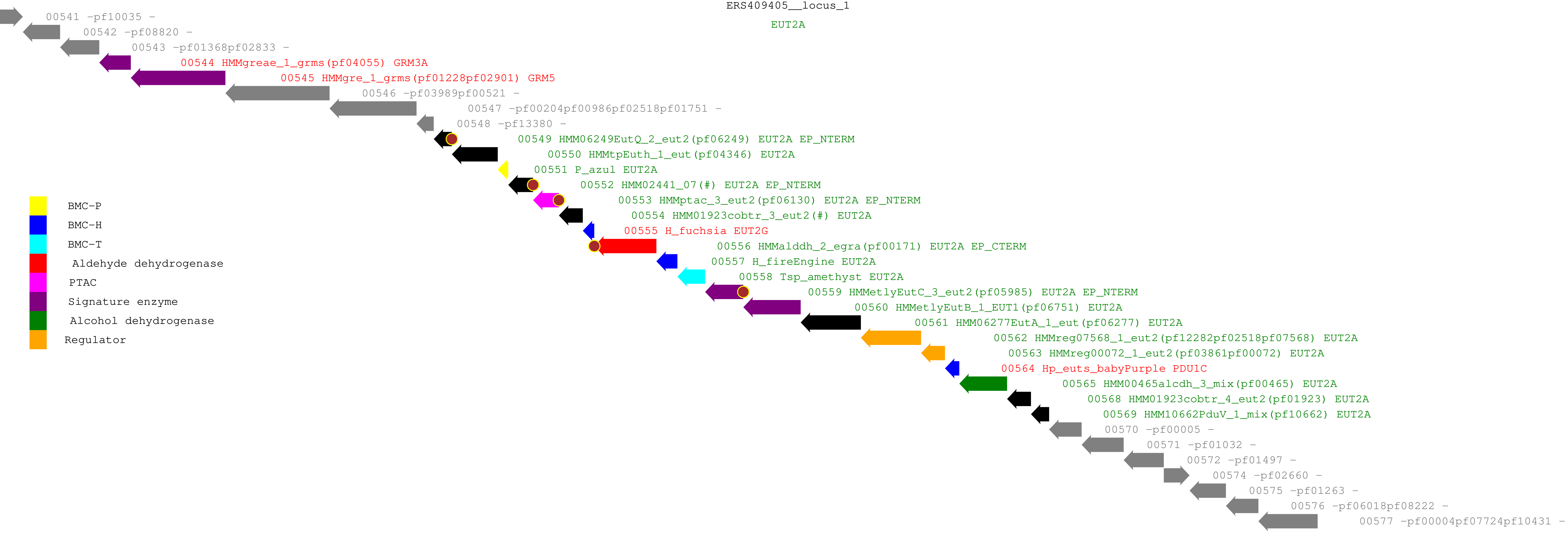

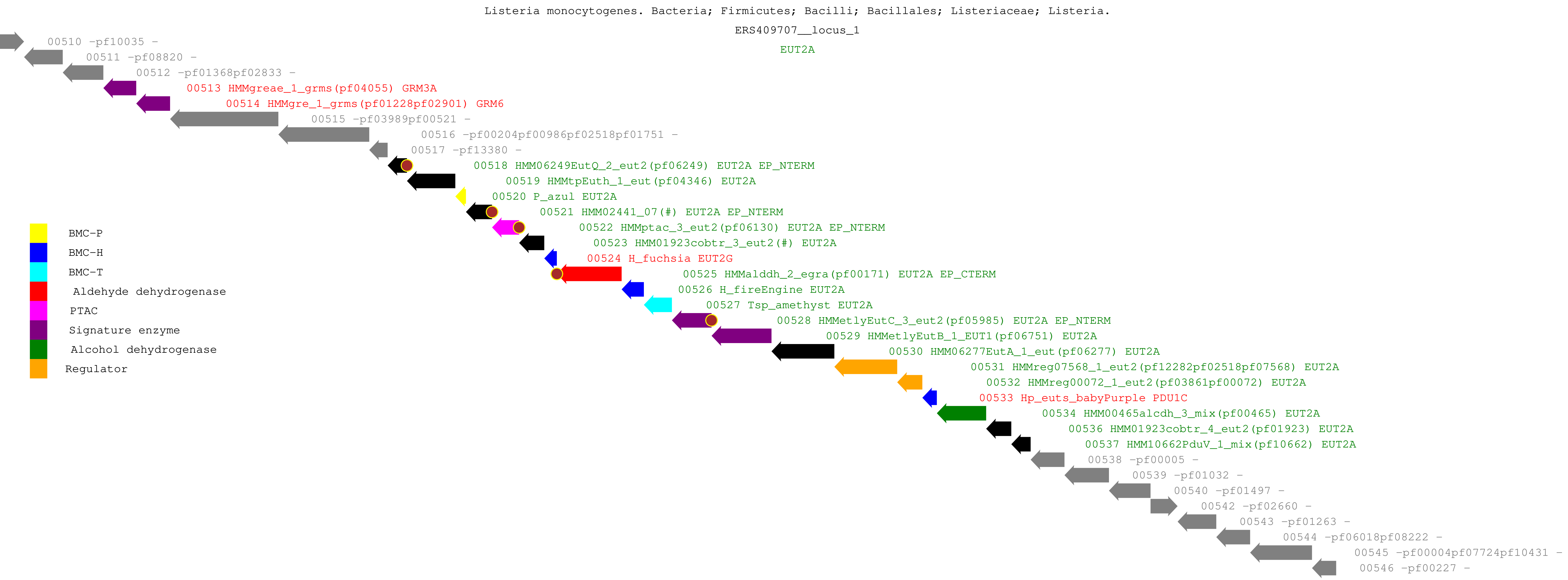

Listeria monocytogenes. Bacteria; Firmicutes; Bacilli; Bacillales; Listeriaceae; Listeria.
ERS409707__locus_1
EUT2A
00510 -pf10035 -
00511 -pf08820 -
00512 -pf01368pf02833 -
00513 HMMgreae_1_grms(pf04055) GRM3A
00514 HMMgre_1_grms(pf01228pf02901) GRM6
00515 -pf03989pf00521 -
00516 -pf00204pf00986pf02518pf01751 -
00517 -pf13380 -
00518 HMM06249EutQ_2_eut2(pf06249) EUT2A EP_NTERM
00519 HMMtpEuth_1_eut(pf04346) EUT2A
00520 P_azul EUT2A
00521 HMM02441_07(#) EUT2A EP_NTERM
00522 HMMptac_3_eut2(pf06130) EUT2A EP_NTERM
00523 HMM01923cobtr_3_eut2(#) EUT2A
00524 H_fuchsia EUT2G
00525 HMMalddh_2_egra(pf00171) EUT2A EP_CTERM
00526 H_fireEngine EUT2A
00527 Tsp_amethyst EUT2A
00528 HMMetlyEutC_3_eut2(pf05985) EUT2A EP_NTERM
00529 HMMetlyEutB_1_EUT1(pf06751) EUT2A
00530 HMM06277EutA_1_eut(pf06277) EUT2A
00531 HMMreg07568_1_eut2(pf12282pf02518pf07568) EUT2A
00532 HMMreg00072_1_eut2(pf03861pf00072) EUT2A
00533 Hp_euts_babyPurple PDU1C
00534 HMM00465alcdh_3_mix(pf00465) EUT2A
00536 HMM01923cobtr_4_eut2(pf01923) EUT2A
00537 HMM10662PduV_1_mix(pf10662) EUT2A
00538 -pf00005 -
00539 -pf01032 -
00540 -pf01497 -
00542 -pf02660 -
00543 -pf01263 -
00544 -pf06018pf08222 -
00545 -pf00004pf07724pf10431 -
00546 -pf00227 -
BMC-P
BMC-H
BMC-T
Aldehyde dehydrogenase
PTAC
Signature enzyme
Alcohol dehydrogenase
Regulator

Lactobacillus rennini DSM 20253. Bacteria; Firmicutes; Bacilli; Lactobacillales; Lactobacillaceae;

FC24_GL_locus_2

EUT2A

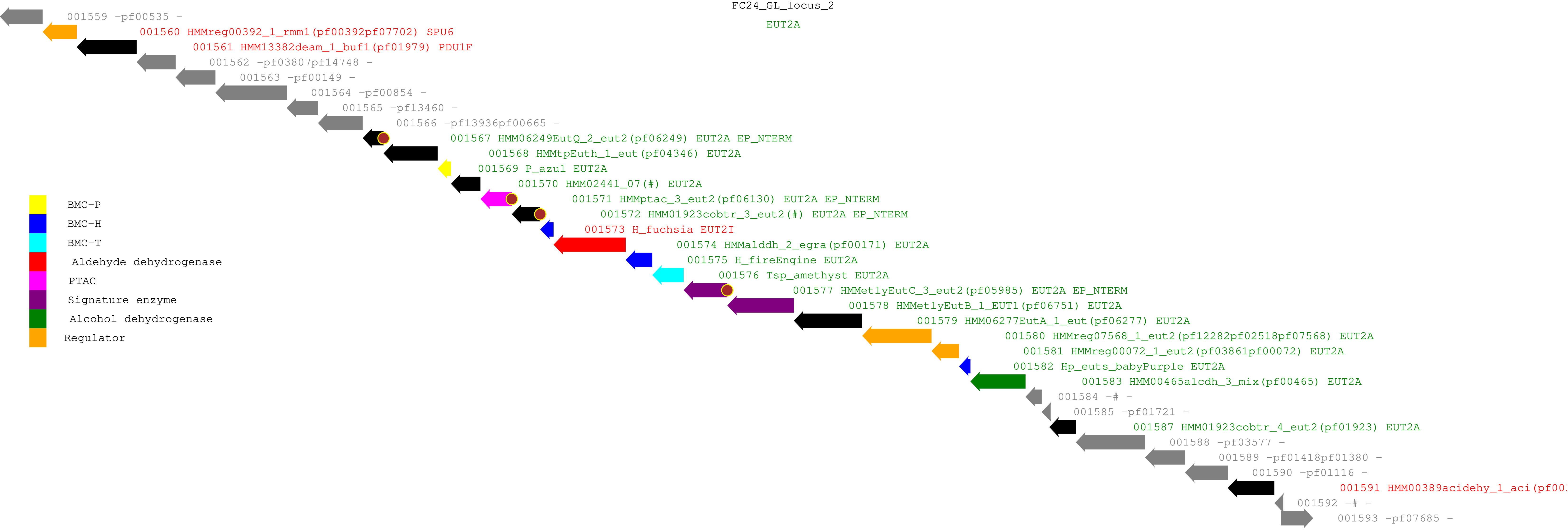

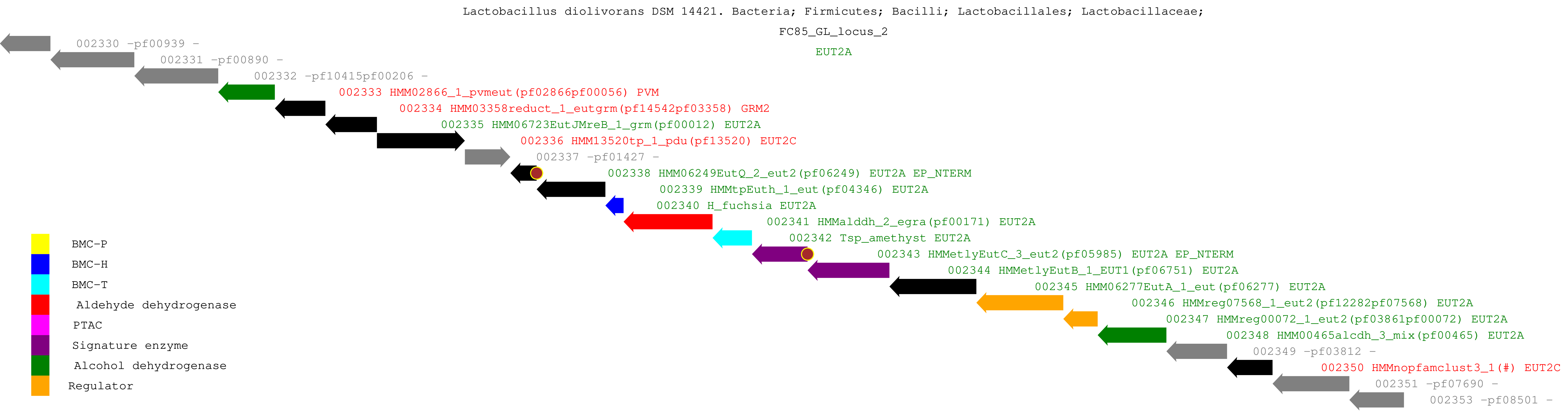
Lactobacillus diolivorans DSM 14421. Bacteria; Firmicutes; Bacilli; Lactobacillales; Lactobacillaceae;
FC85_GL_locus_2
EUT2A
002330 -pf00939 -
002331 -pf00890 -
002332 -pf10415pf00206 -
002333 HMM02866_1_pvmeut(pf02866pf00056) PVM
002334 HMM03358reduct_1_eutgrm(pf14542pf03358) GRM2
002335 HMM06723EutJMreB_1_grm(pf00012) EUT2A
002336 HMM13520tp_1_pdu(pf13520) EUT2C
002337 -pf01427 -
002338 HMM06249EutQ_2_eut2(pf06249) EUT2A EP_NTERM
002339 HMMtpEuth_1_eut(pf04346) EUT2A
002340 H_fuchsia EUT2A
002341 HMMalddh_2_egra(pf00171) EUT2A
002342 Tsp_amethyst EUT2A
002343 HMMetlyEutC_3_eut2(pf05985) EUT2A EP_NTERM
002344 HMMetlyEutB_1_EUT1(pf06751) EUT2A
002345 HMM06277EutA_1_eut(pf06277) EUT2A
002346 HMMreg07568_1_eut2(pf12282pf07568) EUT2A
002347 HMMreg00072_1_eut2(pf03861pf00072) EUT2A
002348 HMM00465alcdh_3_mix(pf00465) EUT2A
002349 -pf03812 -
002350 HMMnopfamclust3_1(#) EUT2C
002351 -pf07690 -
002353 -pf08501 -
BMC-P
BMC-H
BMC-T
Aldehyde dehydrogenase
PTAC
Signature enzyme
Alcohol dehydrogenase
Regulator

Lactobacillus similis DSM 23365 = JCM 2765. Bacteria; Firmicutes; Bacilli; Lactobacillales; Lactobacillaceae;

FD14_GL_locus_1

EUT2A

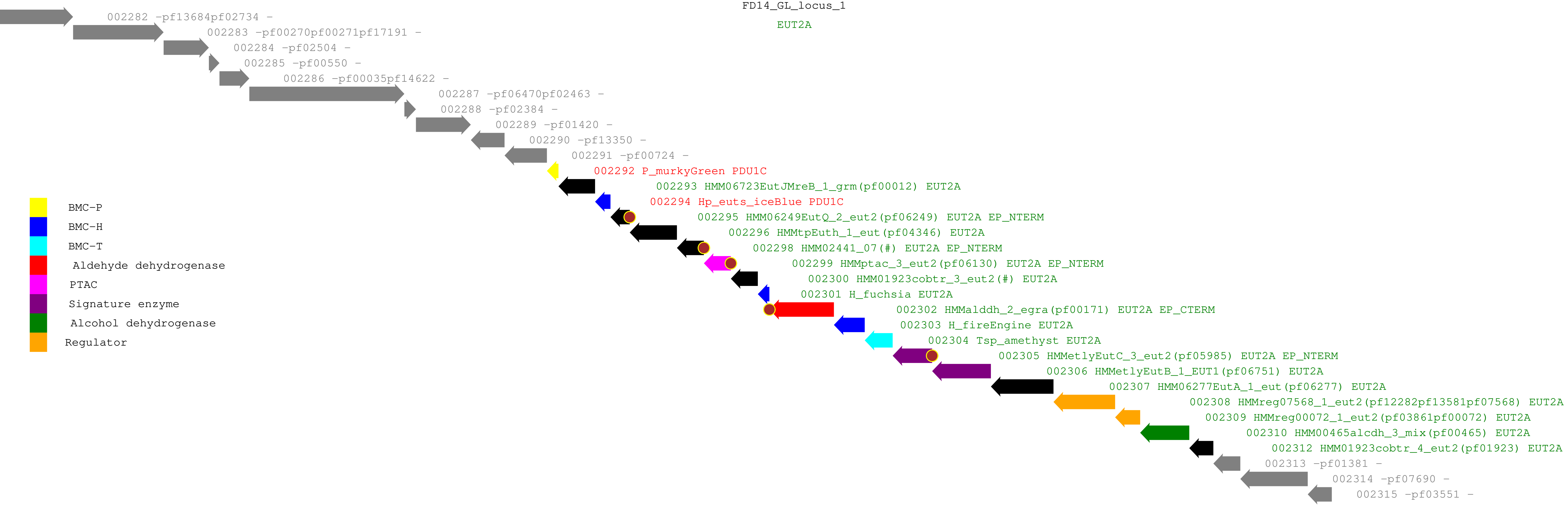

Lactobacillus rossiae DSM 15814. Bacteria; Firmicutes; Bacilli; Lactobacillales; Lactobacillaceae;

FD35_GL_locus_1

EUT2A

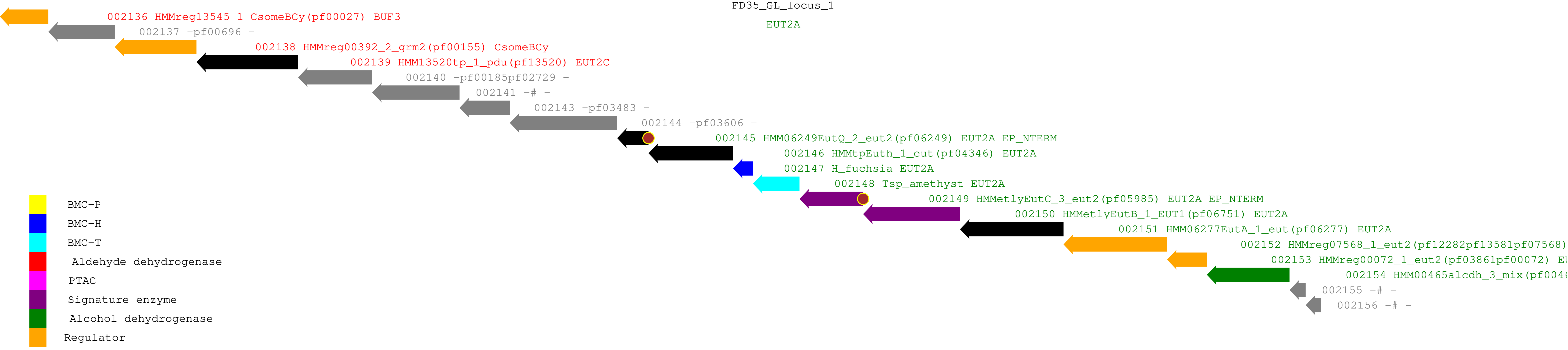

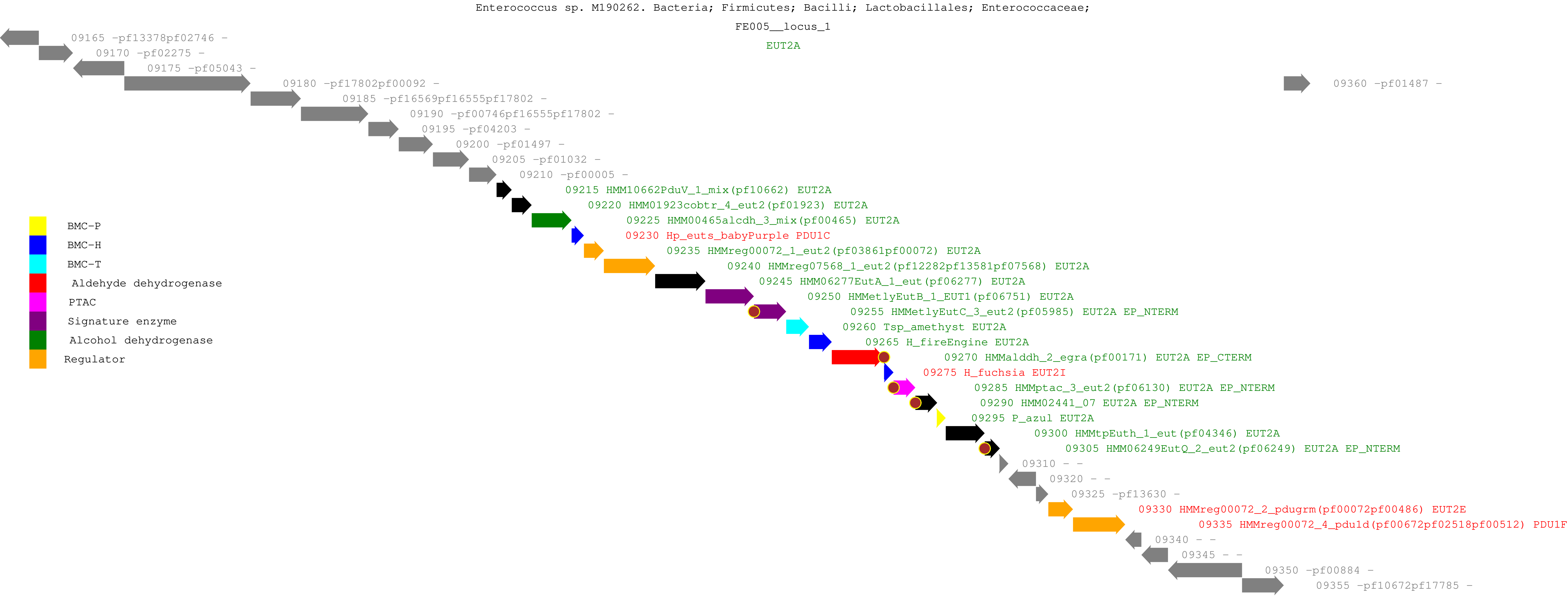
Enterococcus sp. M190262. Bacteria; Firmicutes; Bacilli; Lactobacillales; Enterococcaceae;
FE005__locus_1
EUT2A
09165 -pf13378pf02746 -
09170 -pf02275 -
09175 -pf05043 -
09180 -pf17802pf00092 -
09185 -pf16569pf16555pf17802 -
09190 -pf00746pf16555pf17802 -
09195 -pf04203 -
09200 -pf01497 -
09205 -pf01032 -
09210 -pf00005 -
09215 HMM10662PduV_1_mix(pf10662) EUT2A
09220 HMM01923cobtr_4_eut2(pf01923) EUT2A
09225 HMM00465alcdh_3_mix(pf00465) EUT2A
09230 Hp_euts_babyPurple PDU1C
09235 HMMreg00072_1_eut2(pf03861pf00072) EUT2A
09240 HMMreg07568_1_eut2(pf12282pf13581pf07568) EUT2A
09245 HMM06277EutA_1_eut(pf06277) EUT2A
09250 HMMetlyEutB_1_EUT1(pf06751) EUT2A
09255 HMMetlyEutC_3_eut2(pf05985) EUT2A EP_NTERM
09260 Tsp_amethyst EUT2A
09265 H_fireEngine EUT2A
09270 HMMalddh_2_egra(pf00171) EUT2A EP_CTERM
09275 H_fuchsia EUT2I
09285 HMMptac_3_eut2(pf06130) EUT2A EP_NTERM
09290 HMM02441_07 EUT2A EP_NTERM
09295 P_azul EUT2A
09300 HMMtpEuth_1_eut(pf04346) EUT2A
09305 HMM06249EutQ_2_eut2(pf06249) EUT2A EP_NTERM
09310 - -
09320 - -
09325 -pf13630 -
09330 HMMreg00072_2_pdugrm(pf00072pf00486) EUT2E
09335 HMMreg00072_4_pdu1d(pf00672pf02518pf00512) PDU1F
09340 - -
09345 - -
09350 -pf00884 -
09355 -pf10672pf17785 -
09360 -pf01487 -
BMC-P
BMC-H
BMC-T
Aldehyde dehydrogenase
PTAC
Signature enzyme
Alcohol dehydrogenase
Regulator

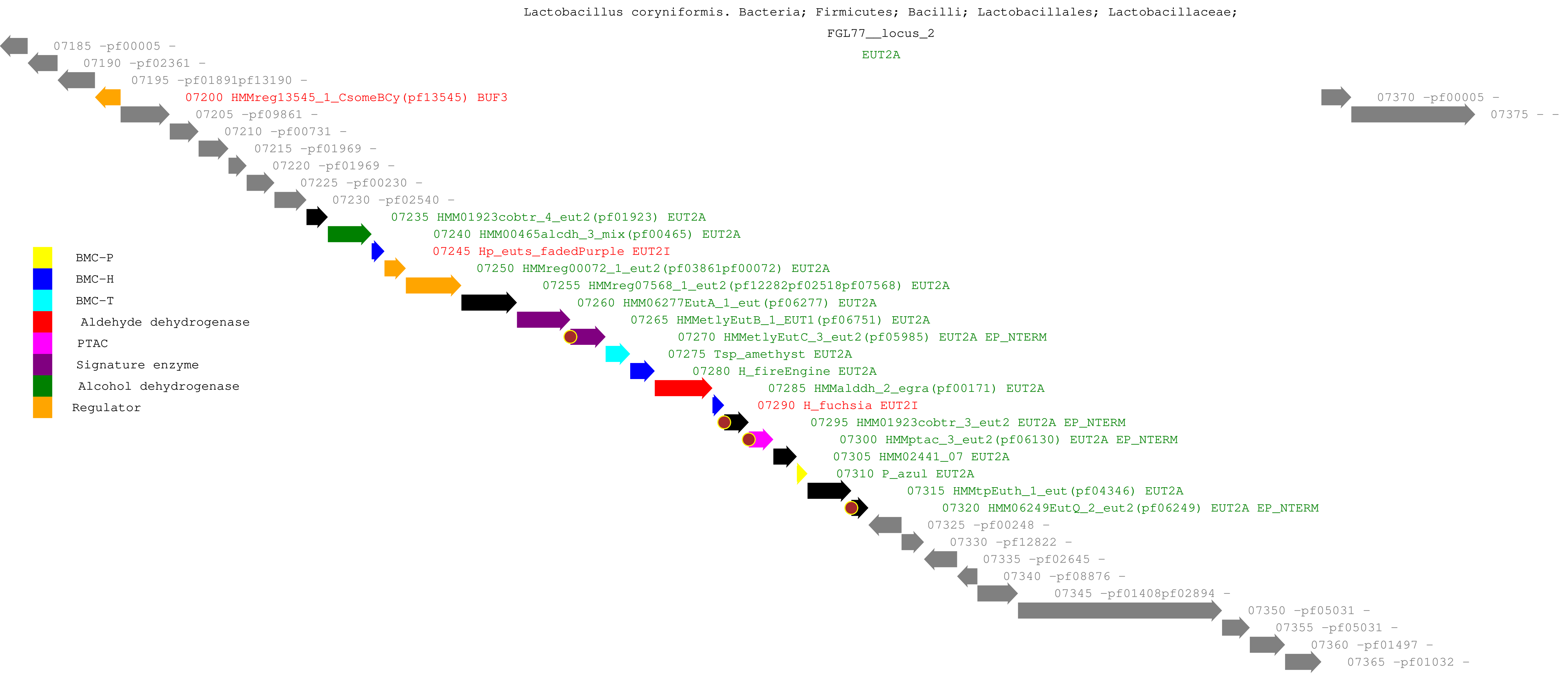
Lactobacillus coryniformis. Bacteria; Firmicutes; Bacilli; Lactobacillales; Lactobacillaceae;
FGL77__locus_2
EUT2A
07185 -pf00005 -
07190 -pf02361 -
07195 -pf01891pf13190 -
07200 HMMreg13545_1_CsomeBCy(pf13545) BUF3
07205 -pf09861 -
07210 -pf00731 -
07215 -pf01969 -
07220 -pf01969 -
07225 -pf00230 -
07230 -pf02540 -
07235 HMM01923cobtr_4_eut2(pf01923) EUT2A
07240 HMM00465alcdh_3_mix(pf00465) EUT2A
07245 Hp_euts_fadedPurple EUT2I
07250 HMMreg00072_1_eut2(pf03861pf00072) EUT2A
07255 HMMreg07568_1_eut2(pf12282pf02518pf07568) EUT2A
07260 HMM06277EutA_1_eut(pf06277) EUT2A
07265 HMMetlyEutB_1_EUT1(pf06751) EUT2A
07270 HMMetlyEutC_3_eut2(pf05985) EUT2A EP_NTERM
07275 Tsp_amethyst EUT2A
07280 H_fireEngine EUT2A
07285 HMMaldh_2_egra(pf00171) EUT2A
07290 H_fuchsia EUT2I
07295 HMM01923cobtr_3_eut2 EUT2A EP_NTERM
07300 HMMptac_3_eut2(pf06130) EUT2A EP_NTERM
07305 HMM02441_07 EUT2A
07310 P_azul EUT2A
07315 HMMtpEuth_1_eut(pf04346) EUT2A
07320 HMM06249EutQ_2_eut2(pf06249) EUT2A EP_NTERM
07325 -pf00248 -
07330 -pf12822 -
07335 -pf02645 -
07340 -pf08876 -
07345 -pf01408pf02894 -
07350 -pf05031 -
07355 -pf05031 -
07360 -pf01497 -
07365 -pf01032 -
07370 -pf00005 -
07375 - -
BMC-P
BMC-H
BMC-T
Aldehyde dehydrogenase
PTAC
Signature enzyme
Alcohol dehydrogenase
Regulator

Brachybacterium faecium. Bacteria; Actinobacteria; Micrococcales; Dermabacteraceae;

FM106__locus_1

EUT2A

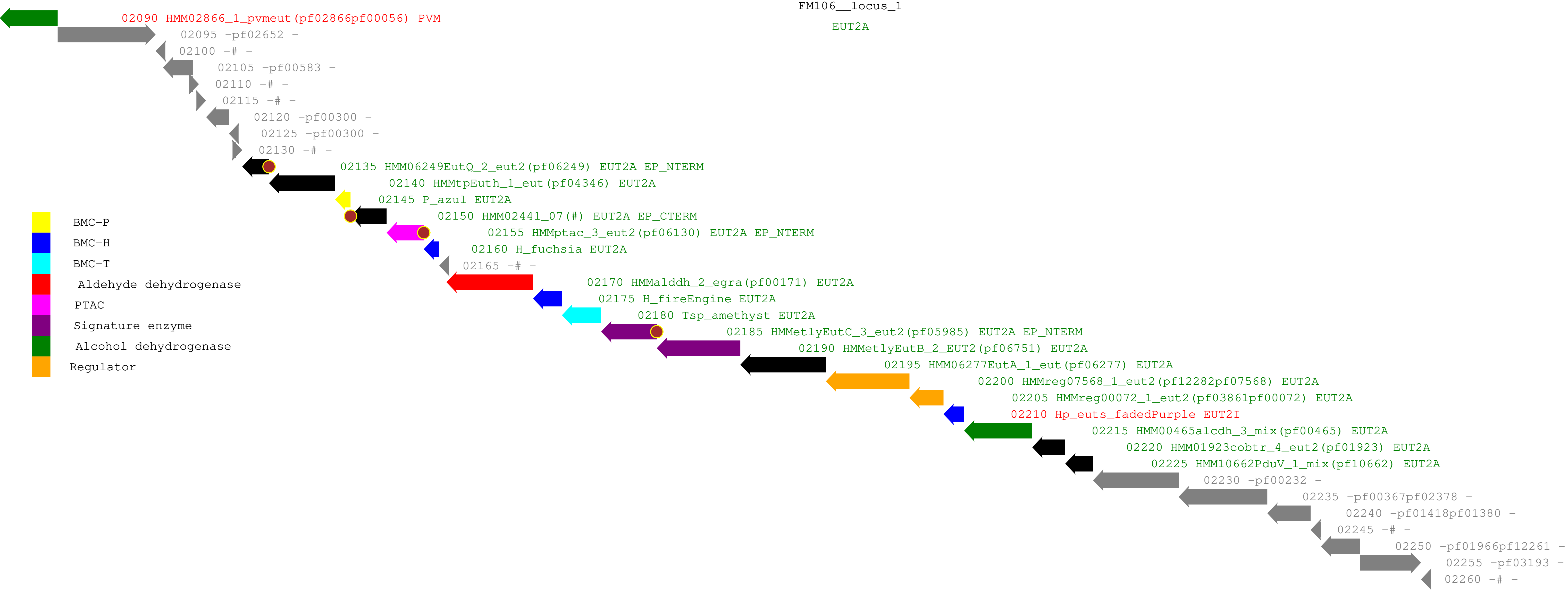

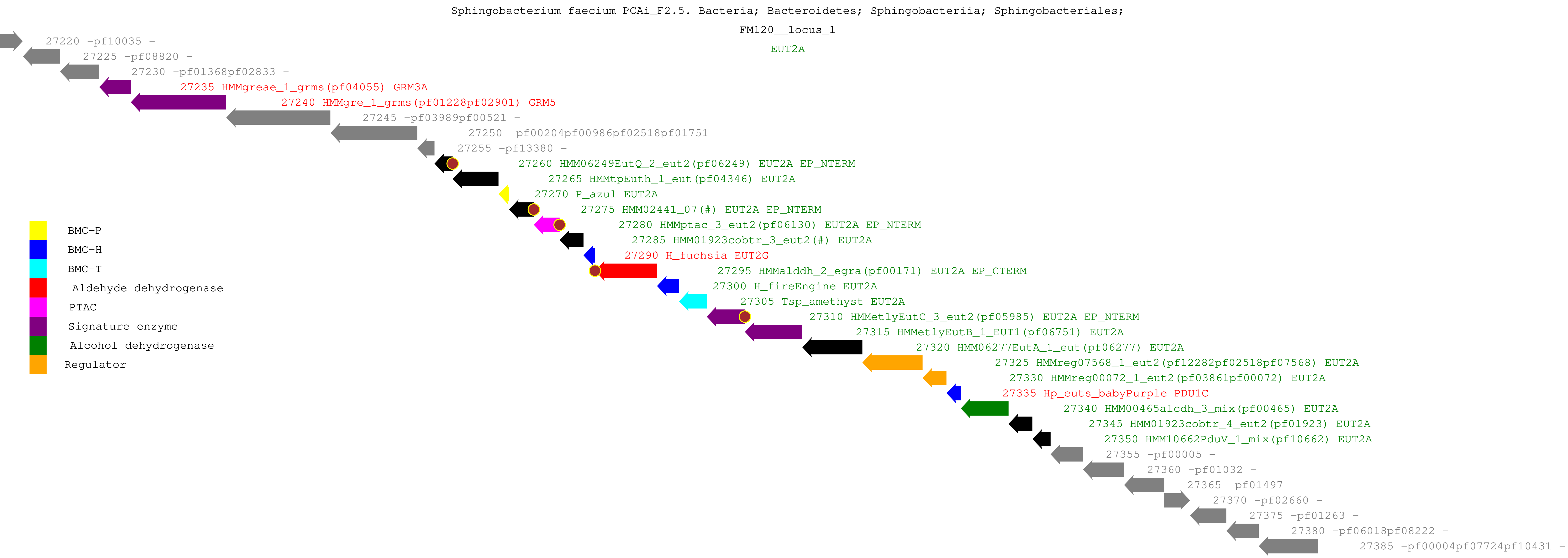

Sphingobacterium faecium PCAi_F2.5. Bacteria; Bacteroidetes; Sphingobacteriia; Sphingobacteriales;
FM120__locus_1
EUT2A
27220 -pf10035 -
27225 -pf08820 -
27230 -pf01368pf02833 -
27235 HMMgreae_1_grms(pf04055) GRM3A
27240 HMMgre_1_grms(pf01228pf02901) GRM5
27245 -pf03989pf00521 -
27250 -pf00204pf00986pf02518pf01751 -
27255 -pf13380 -
27260 HMM06249EutQ_2_eut2(pf06249) EUT2A EP_NTERM
27265 HMMtpEuth_1_eut(pf04346) EUT2A
27270 P_azul EUT2A
27275 HMM02441_07(#) EUT2A EP_NTERM
27280 HMMptac_3_eut2(pf06130) EUT2A EP_NTERM
27285 HMM01923cobtr_3_eut2(#) EUT2A
27290 H_fuchsia EUT2G
27295 HMMalddh_2_egra(pf00171) EUT2A EP_CTERM
27300 H_fireEngine EUT2A
27305 Tsp_amethyst EUT2A
27310 HMMetlyEutC_3_eut2(pf05985) EUT2A EP_NTERM
27315 HMMetlyEutB_1_EUT1(pf06751) EUT2A
27320 HMM06277EutA_1_eut(pf06277) EUT2A
27325 HMMreg07568_1_eut2(pf12282pf02518pf07568) EUT2A
27330 HMMreg00072_1_eut2(pf03861pf00072) EUT2A
27335 Hp_euts_babyPurple PDU1C
27340 HMM00465alcdh_3_mix(pf00465) EUT2A
27345 HMM01923cobtr_4_eut2(pf01923) EUT2A
27350 HMM10662PduV_1_mix(pf10662) EUT2A
27355 -pf00005 -
27360 -pf01032 -
27365 -pf01497 -
27370 -pf02660 -
27375 -pf01263 -
27380 -pf06018pf08222 -
27385 -pf00004pf07724pf10431 -
BMC-P
BMC-H
BMC-T
Aldehyde dehydrogenase
PTAC
Signature enzyme
Alcohol dehydrogenase
Regulator

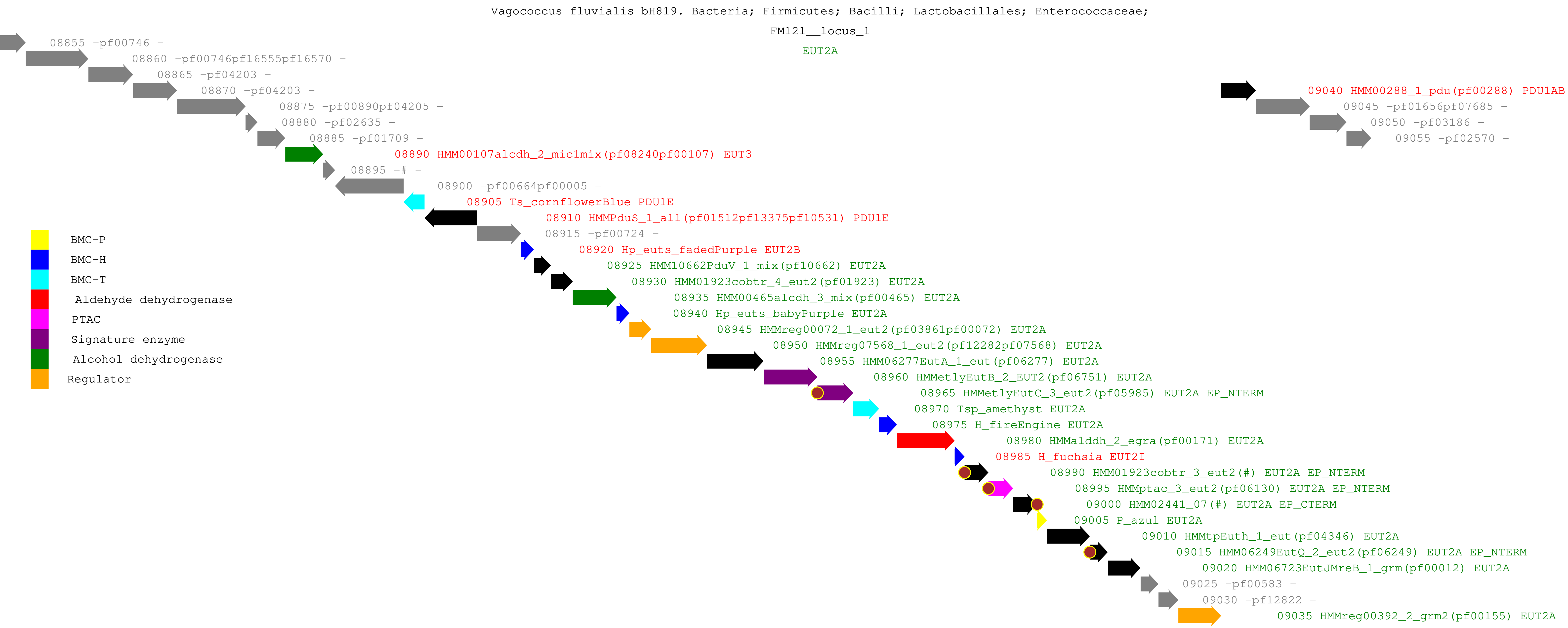

Vagococcus fluvialis bH819. Bacteria; Firmicutes; Bacilli; Lactobacillales; Enterococcaceae;
FM121__locus_1
EUT2A
08855 -pf00746 -
08860 -pf00746pf16555pf16570 -
08865 -pf04203 -
08870 -pf04203 -
08875 -pf00890pf04205 -
08880 -pf02635 -
08885 -pf01709 -
08890 HMM00107alcdh_2_mic1mix(pf08240pf00107) EUT3
08895 -# -
08900 -pf00664pf00005 -
08905 Ts_cornflowerBlue PDU1E
08910 HMMPduS_1_all(pf01512pf13375pf10531) PDU1E
08915 -pf00724 -
08920 Hp_euts_fadedPurple EUT2B
08925 HMM10662PduV_1_mix(pf10662) EUT2A
08930 HMM01923cobtr_4_eut2(pf01923) EUT2A
08935 HMM00465alcdh_3_mix(pf00465) EUT2A
08940 Hp_euts_babyPurple EUT2A
08945 HMMreg00072_1_eut2(pf03861pf00072) EUT2A
08950 HMMreg07568_1_eut2(pf12282pf07568) EUT2A
08955 HMM06277EutA_1_eut(pf06277) EUT2A
08960 HMMetlyEutB_2_EUT2(pf06751) EUT2A
08965 HMMetlyEutC_3_eut2(pf05985) EUT2A EP_NTERM
08970 Tsp_amethyst EUT2A
08975 H_fireEngine EUT2A
08980 HMMaldd h_2_egra(pf00171) EUT2A
08985 H_fuchsia EUT2I
08990 HMM01923cobtr_3_eut2(#) EUT2A EP_NTERM
08995 HMMptac_3_eut2(pf06130) EUT2A EP_NTERM
09000 HMM02441_07(#) EUT2A EP_CTERM
09005 P_azul EUT2A
09010 HMMtpEuth_1_eut(pf04346) EUT2A
09015 HMM06249EutQ_2_eut2(pf06249) EUT2A EP_NTERM
09020 HMM06723EutJMreB_1_grm(pf00012) EUT2A
09025 -pf00583 -
09030 -pf12822 -
09035 HMMreg00392_2_grm2(pf00155) EUT2A
09040 HMM00288_1_pdu(pf00288) PDU1AB
09045 -pf01656pf07685 -
09050 -pf03186 -
09055 -pf02570 -
BMC-P
BMC-H
BMC-T
Aldehyde dehydrogenase
PTAC
Signature enzyme
Alcohol dehydrogenase
Regulator

Enterococcus gallinarum. Bacteria; Firmicutes; Bacilli; Lactobacillales; Enterococcaceae;

FSZ06__locus_1_main_sat

EUT2A

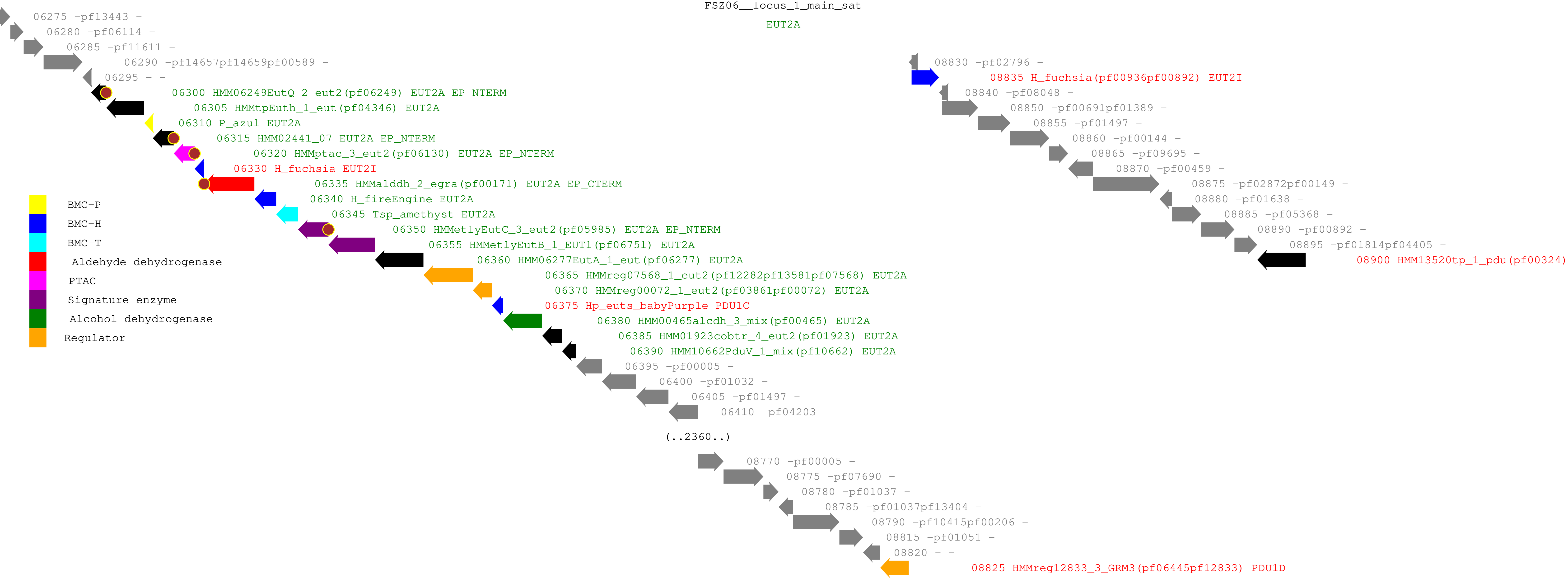

Streptococcus ratti. Bacteria; Firmicutes; Bacilli; Lactobacillales; Streptococcaceae;

FY406__locus_1

EUT2A

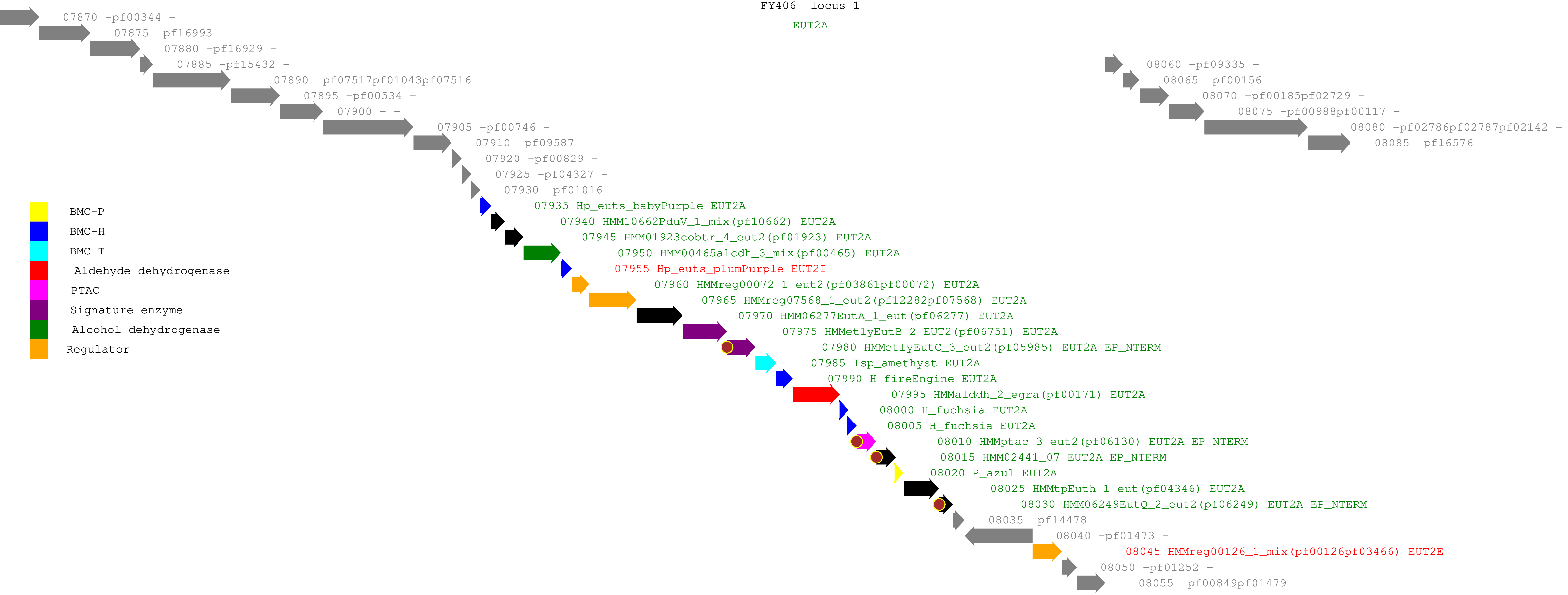

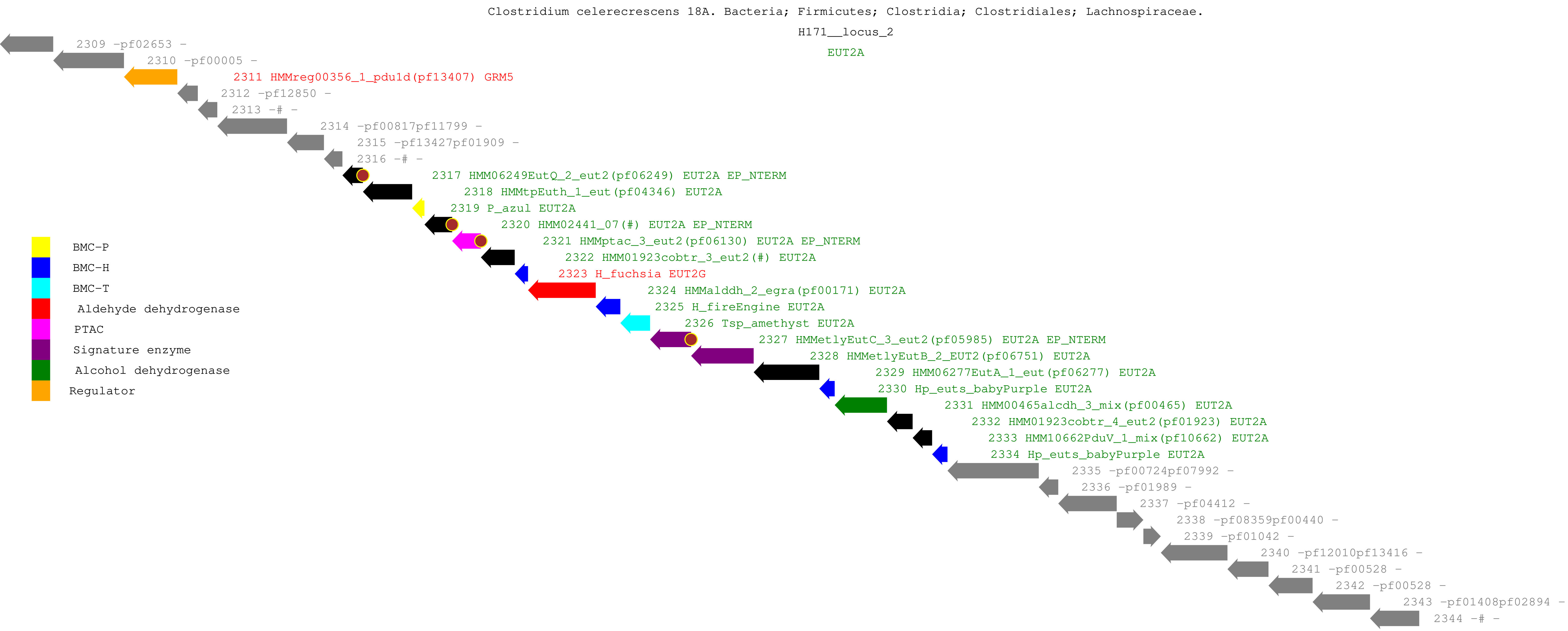
Clostridium celerecrescens 18A. Bacteria; Firmicutes; Clostridia; Clostridiales; Lachnospiraceae.
H171__locus_2
EUT2A
2309 -pf02653 -
2310 -pf00005 -
2311 HMMreg00356_1_pdu1d(pf13407) GRM5
2312 -pf12850 -
2313 -# -
2314 -pf00817pf11799 -
2315 -pf13427pf01909 -
2316 -# -
2317 HMM06249EutQ_2_eut2(pf06249) EUT2A EP_NTERM
2318 HMMtpEuth_1_eut(pf04346) EUT2A
2319 P_azul EUT2A
2320 HMM02441_07(#) EUT2A EP_NTERM
2321 HMMptac_3_eut2(pf06130) EUT2A EP_NTERM
2322 HMM01923cobtr_3_eut2(#) EUT2A
2323 H_fuchsia EUT2G
2324 HMMalddh_2_egra(pf00171) EUT2A
2325 H_fireEngine EUT2A
2326 Tsp_amethyst EUT2A
2327 HMMetlyEutC_3_eut2(pf05985) EUT2A EP_NTERM
2328 HMMetlyEutB_2_EUT2(pf06751) EUT2A
2329 HMM06277EutA_1_eut(pf06277) EUT2A
2330 Hp_euts_babyPurple EUT2A
2331 HMM00465alcdh_3_mix(pf00465) EUT2A
2332 HMM01923cobtr_4_eut2(pf01923) EUT2A
2333 HMM10662PduV_1_mix(pf10662) EUT2A
2334 Hp_euts_babyPurple EUT2A
2335 -pf00724pf07992 -
2336 -pf01989 -
2337 -pf04412 -
2338 -pf08359pf00440 -
2339 -pf01042 -
2340 -pf12010pf13416 -
2341 -pf00528 -
2342 -pf00528 -
2343 -pf01408pf02894 -
2344 -# -
BMC-P
BMC-H
BMC-T
Aldehyde dehydrogenase
PTAC
Signature enzyme
Alcohol dehydrogenase
Regulator

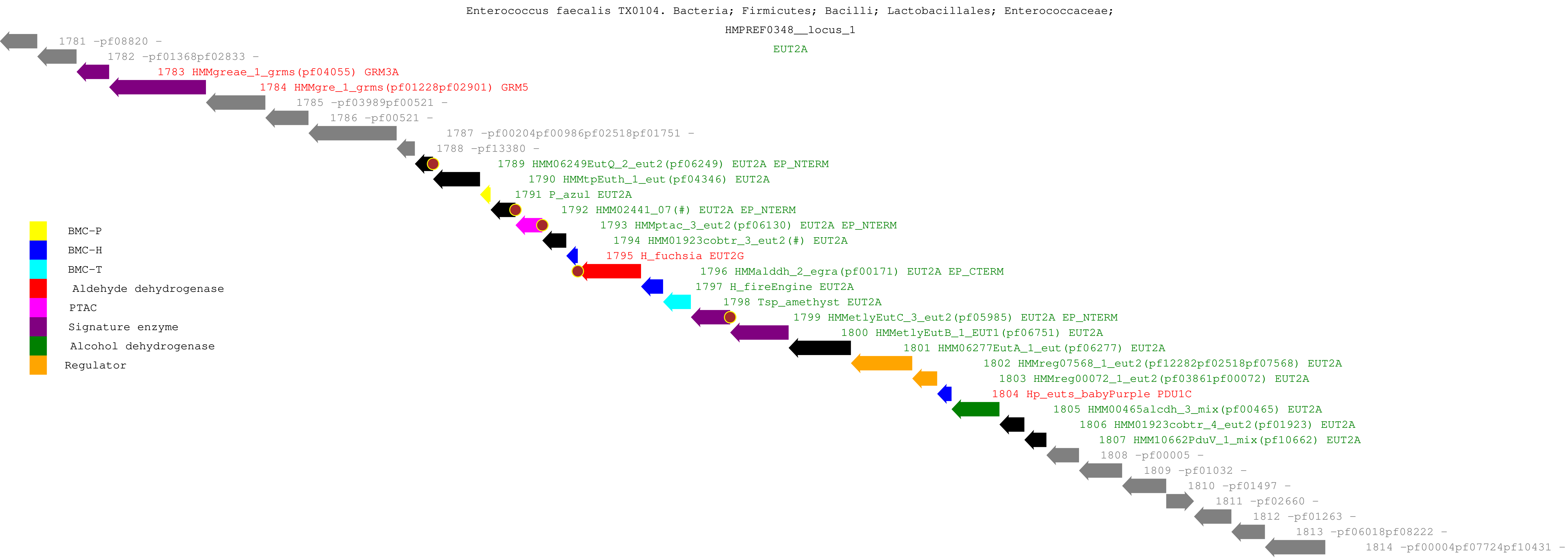
Enterococcus faecalis TX0104. Bacteria; Firmicutes; Bacilli; Lactobacillales; Enterococcaceae;
HMPREF0348__locus_1
EUT2A
1781 -pf08820 -
1782 -pf01368pf02833 -
1783 HMMgreae_1_grms(pf04055) GRM3A
1784 HMMgre_1_grms(pf01228pf02901) GRM5
1785 -pf03989pf00521 -
1786 -pf00521 -
1787 -pf00204pf00986pf02518pf01751 -
1788 -pf13380 -
1789 HMM06249EutQ_2_eut2(pf06249) EUT2A EP_NTERM
1790 HMMtpEuth_1_eut(pf04346) EUT2A
1791 P_azul EUT2A
1792 HMM02441_07(#) EUT2A EP_NTERM
1793 HMMptac_3_eut2(pf06130) EUT2A EP_NTERM
1794 HMM01923cobtr_3_eut2(#) EUT2A
1795 H_fuchsia EUT2G
1796 HMMalddh_2_egra(pf00171) EUT2A EP_CTERM
1797 H_fireEngine EUT2A
1798 Tsp_amethyst EUT2A
1799 HMMetlyEutC_3_eut2(pf05985) EUT2A EP_NTERM
1800 HMMetlyEutB_1_EUT1(pf06751) EUT2A
1801 HMM06277EutA_1_eut(pf06277) EUT2A
1802 HMMreg07568_1_eut2(pf12282pf02518pf07568) EUT2A
1803 HMMreg00072_1_eut2(pf03861pf00072) EUT2A
1804 Hp_euts_babyPurple PDU1C
1805 HMM00465alcdh_3_mix(pf00465) EUT2A
1806 HMM01923cobtr_4_eut2(pf01923) EUT2A
1807 HMM10662PduV_1_mix(pf10662) EUT2A
1808 -pf00005 -
1809 -pf01032 -
1810 -pf01497 -
1811 -pf02660 -
1812 -pf01263 -
1813 -pf06018pf08222 -
1814 -pf00004pf07724pf10431 -
BMC-P
BMC-H
BMC-T
Aldehyde dehydrogenase
PTAC
Signature enzyme
Alcohol dehydrogenase
Regulator

Lactobacillus brevis ATCC 14869 = DSM 20054. Bacteria; Firmicutes; Bacilli; Lactobacillales; Lactobacillaceae;

HMPREF0495__locus_1

EUT2A

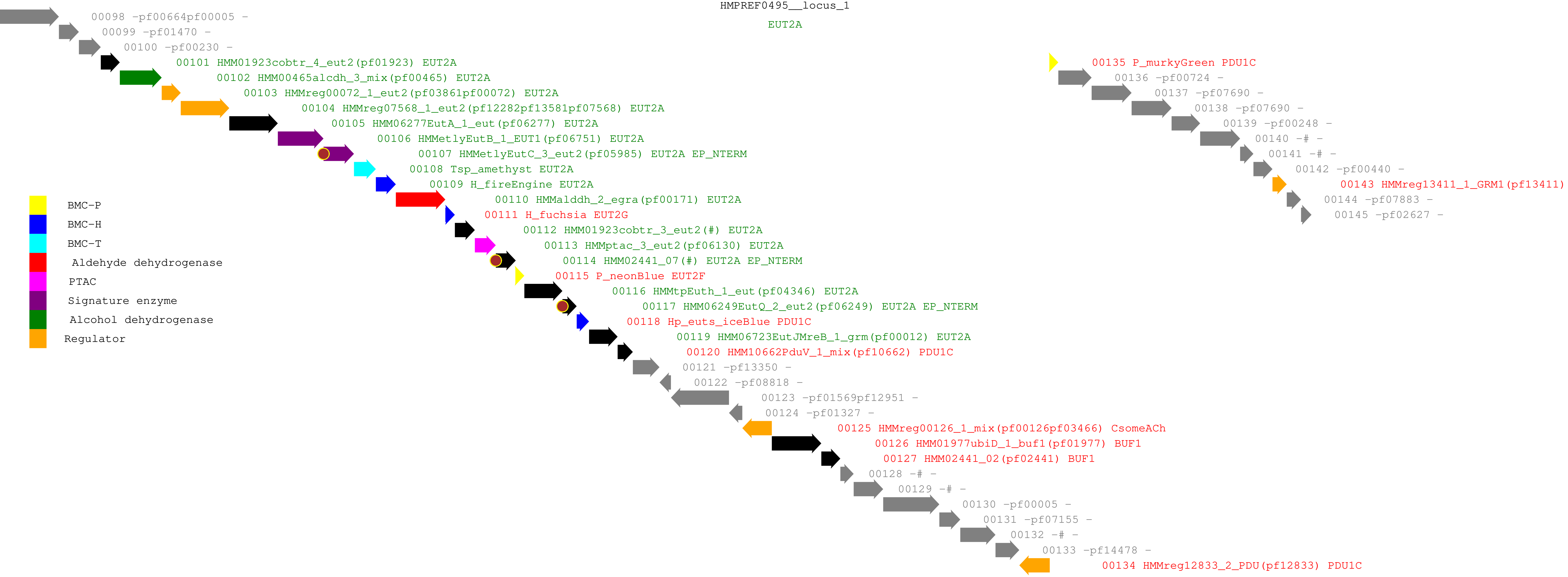

Listeria innocua ATCC 33091. Bacteria; Firmicutes; Bacilli; Bacillales; Listeriaceae; Listeria.

HMPREF0557__locus_2

EUT2A

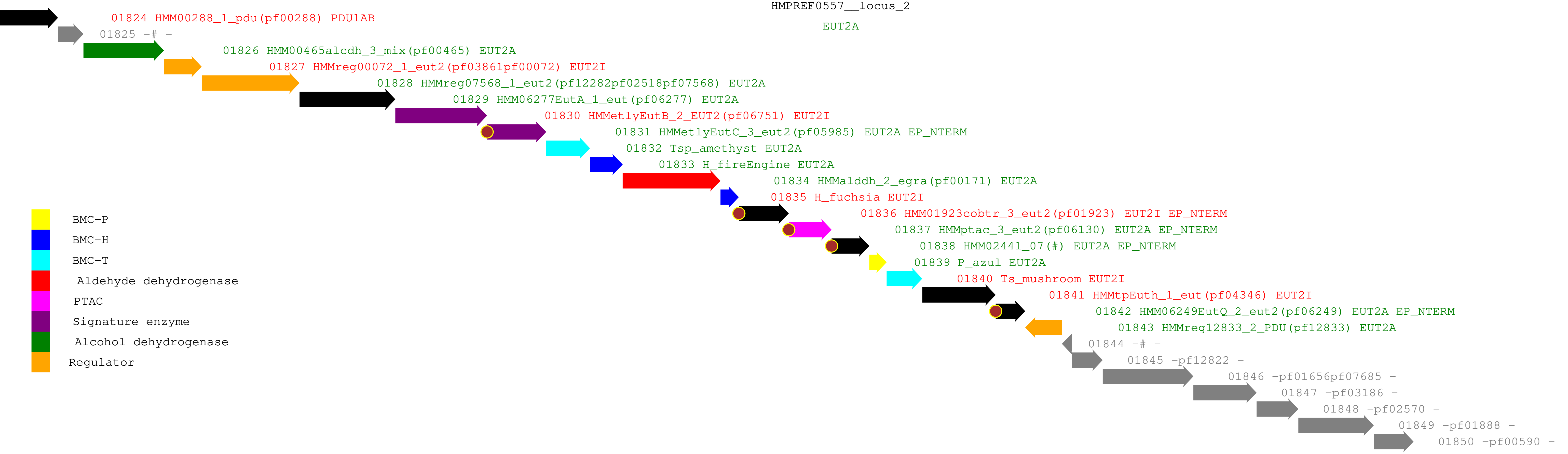

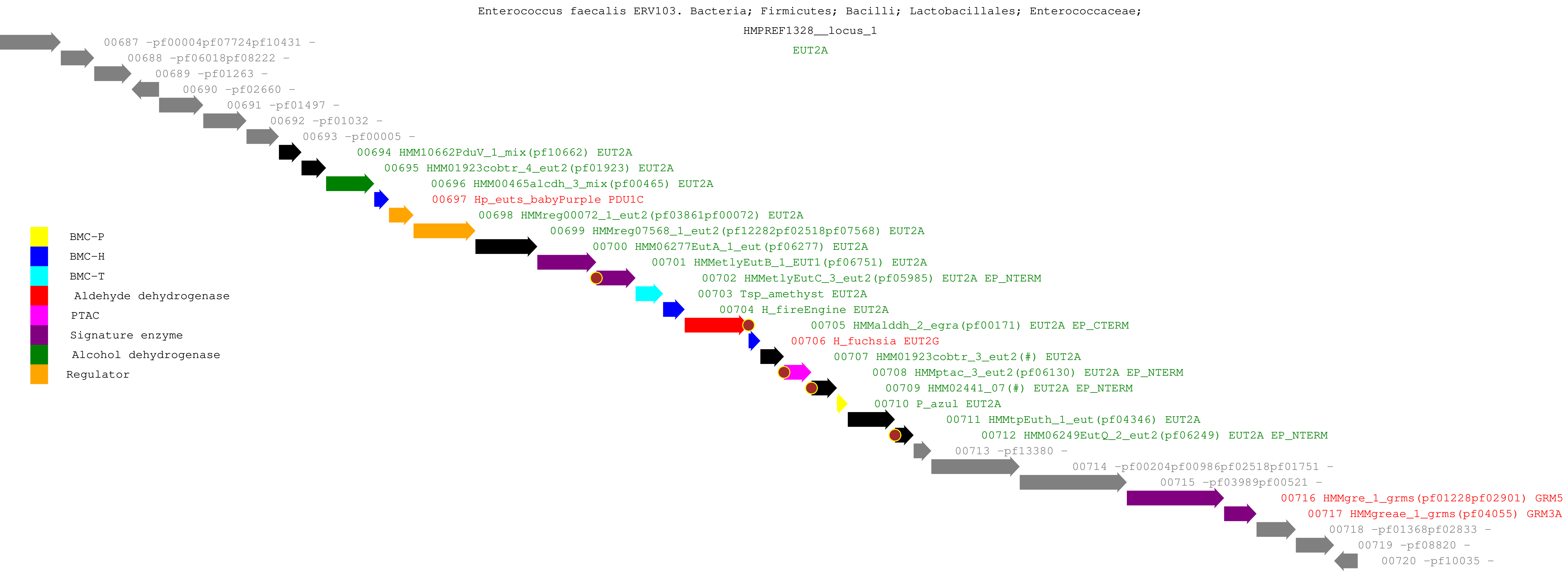
Enterococcus faecalis ERV103. Bacteria; Firmicutes; Bacilli; Lactobacillales; Enterococcaceae;
HMPREF1328__locus_1
EUT2A
00687 -pf00004pf07724pf10431 -
00688 -pf06018pf08222 -
00689 -pf01263 -
00690 -pf02660 -
00691 -pf01497 -
00692 -pf01032 -
00693 -pf00005 -
00694 HMM10662PduV_1_mix(pf10662) EUT2A
00695 HMM01923cobtr_4_eut2(pf01923) EUT2A
00696 HMM00465alcdh_3_mix(pf00465) EUT2A
00697 Hp_euts_babyPurple PDU1C
00698 HMMreg00072_1_eut2(pf03861pf00072) EUT2A
00699 HMMreg07568_1_eut2(pf12282pf02518pf07568) EUT2A
00700 HMM06277EutA_1_eut(pf06277) EUT2A
00701 HMMetlyEutB_1_EUT1(pf06751) EUT2A
00702 HMMetlyEutC_3_eut2(pf05985) EUT2A EP_NTERM
00703 Tsp_amethyst EUT2A
00704 H_fireEngine EUT2A
00705 HMMalddh_2_egra(pf00171) EUT2A EP_CTERM
00706 H_fuchsia EUT2G
00707 HMM01923cobtr_3_eut2(#) EUT2A
00708 HMMptac_3_eut2(pf06130) EUT2A EP_NTERM
00709 HMM02441_07(#) EUT2A EP_NTERM
00710 P_azul EUT2A
00711 HMMtpEuth_1_eut(pf04346) EUT2A
00712 HMM06249EutQ_2_eut2(pf06249) EUT2A EP_NTERM
00713 -pf13380 -
00714 -pf00204pf00986pf02518pf01751 -
00715 -pf03989pf00521 -
00716 HMMgre_1_grms(pf01228pf02901) GRM5
00717 HMMgreae_1_grms(pf04055) GRM3A
00718 -pf01368pf02833 -
00719 -pf08820 -
00720 -pf10035 -
BMC-P
BMC-H
BMC-T
Aldehyde dehydrogenase
PTAC
Signature enzyme
Alcohol dehydrogenase
Regulator

Enterococcus faecalis ERV116. Bacteria; Firmicutes; Bacilli; Lactobacillales; Enterococcaceae;

HMPREF1329__locus_1

EUT2A

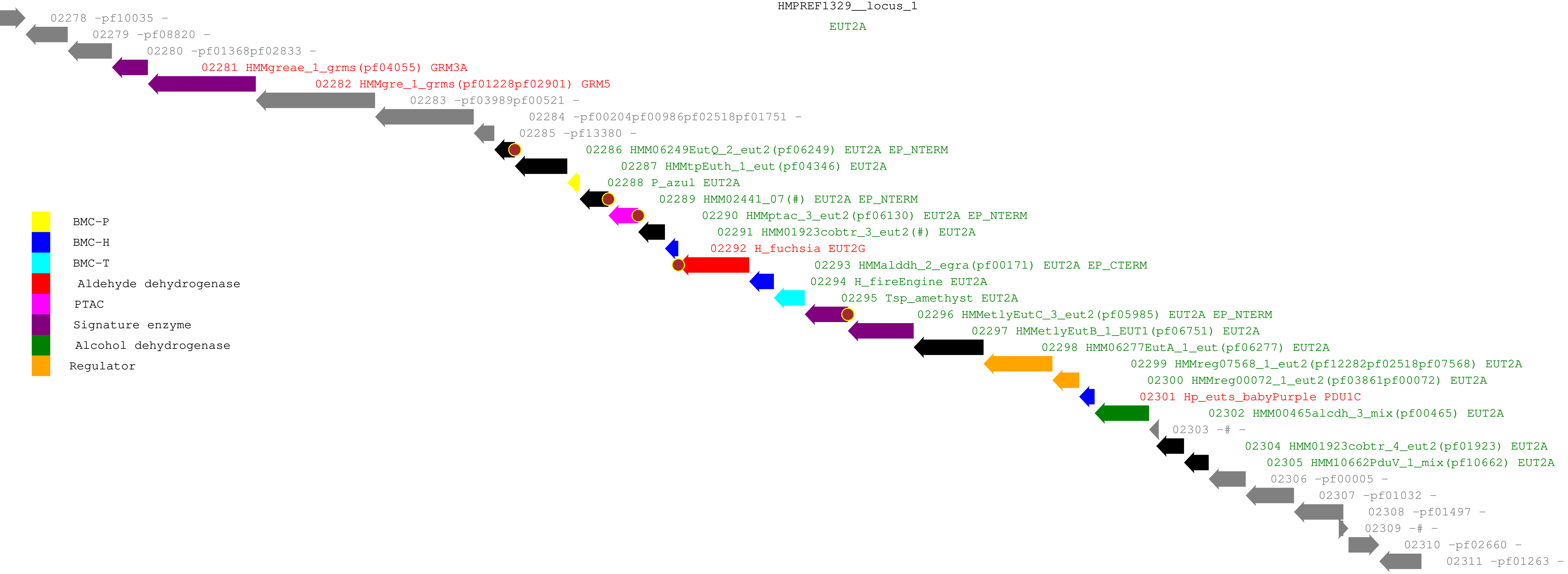

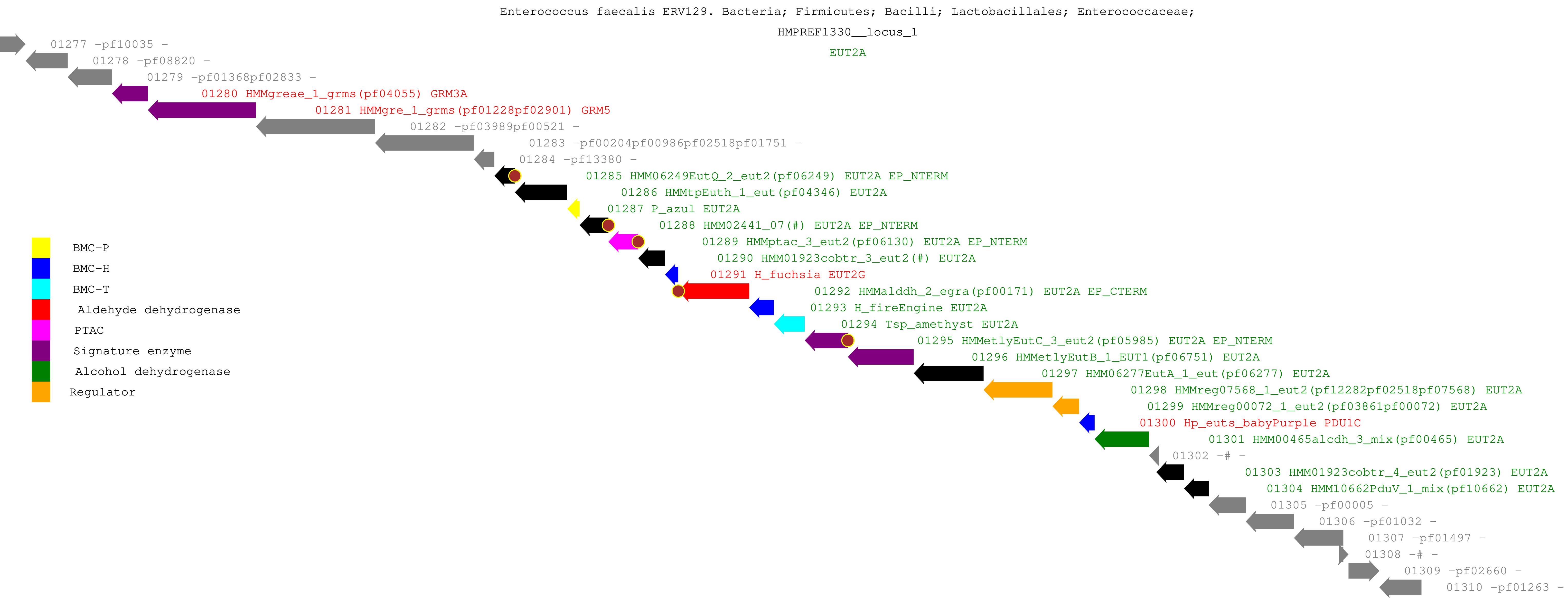
Enterococcus faecalis ERV129. Bacteria; Firmicutes; Bacilli; Lactobacillales; Enterococcaceae;
HMPREF1330__locus_1
EUT2A
01277 -pf10035 -
01278 -pf08820 -
01279 -pf01368pf02833 -
01280 HMMgreae_1_grms(pf04055) GRM3A
01281 HMMgre_1_grms(pf01228pf02901) GRM5
01282 -pf03989pf00521 -
01283 -pf00204pf00986pf02518pf01751 -
01284 -pf13380 -
01285 HMM06249EutQ_2_eut2(pf06249) EUT2A EP_NTERM
01286 HMMtpEuth_1_eut(pf04346) EUT2A
01287 P_azul EUT2A
01288 HMM02441_07(#) EUT2A EP_NTERM
01289 HMMptac_3_eut2(pf06130) EUT2A EP_NTERM
01290 HMM01923cobtr_3_eut2(#) EUT2A
01291 H_fuchsia EUT2G
01292 HMMalddh_2_egra(pf00171) EUT2A EP_CTERM
01293 H_fireEngine EUT2A
01294 Tsp_amethyst EUT2A
01295 HMMetlyEutC_3_eut2(pf05985) EUT2A EP_NTERM
01296 HMMetlyEutB_1_EUT1(pf06751) EUT2A
01297 HMM06277EutA_1_eut(pf06277) EUT2A
01298 HMMreg07568_1_eut2(pf12282pf02518pf07568) EUT2A
01299 HMMreg00072_1_eut2(pf03861pf00072) EUT2A
01300 Hp_euts_babyPurple PDU1C
01301 HMM00465alcdh_3_mix(pf00465) EUT2A
01302 -# -
01303 HMM01923cobtr_4_eut2(pf01923) EUT2A
01304 HMM10662PduV_1_mix(pf10662) EUT2A
01305 -pf00005 -
01306 -pf01032 -
01307 -pf01497 -
01308 -# -
01309 -pf02660 -
01310 -pf01263 -
BMC-P
BMC-H
BMC-T
Aldehyde dehydrogenase
PTAC
Signature enzyme
Alcohol dehydrogenase
Regulator

Enterococcus faecalis ERV25. Bacteria; Firmicutes; Bacilli; Lactobacillales; Enterococcaceae;

HMPREF1331__locus_1

EUT2A

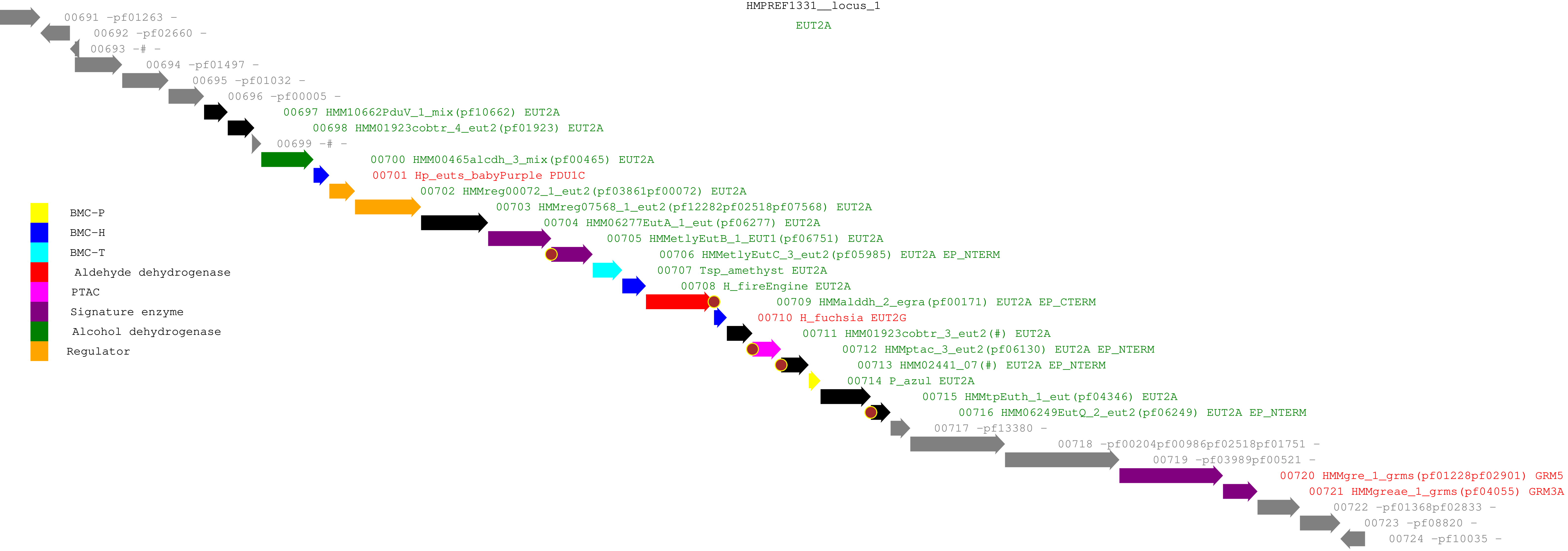

Enterococcus faecalis ERV31. Bacteria; Firmicutes; Bacilli; Lactobacillales; Enterococcaceae;

HMPREF1332__locus_1

EUT2A

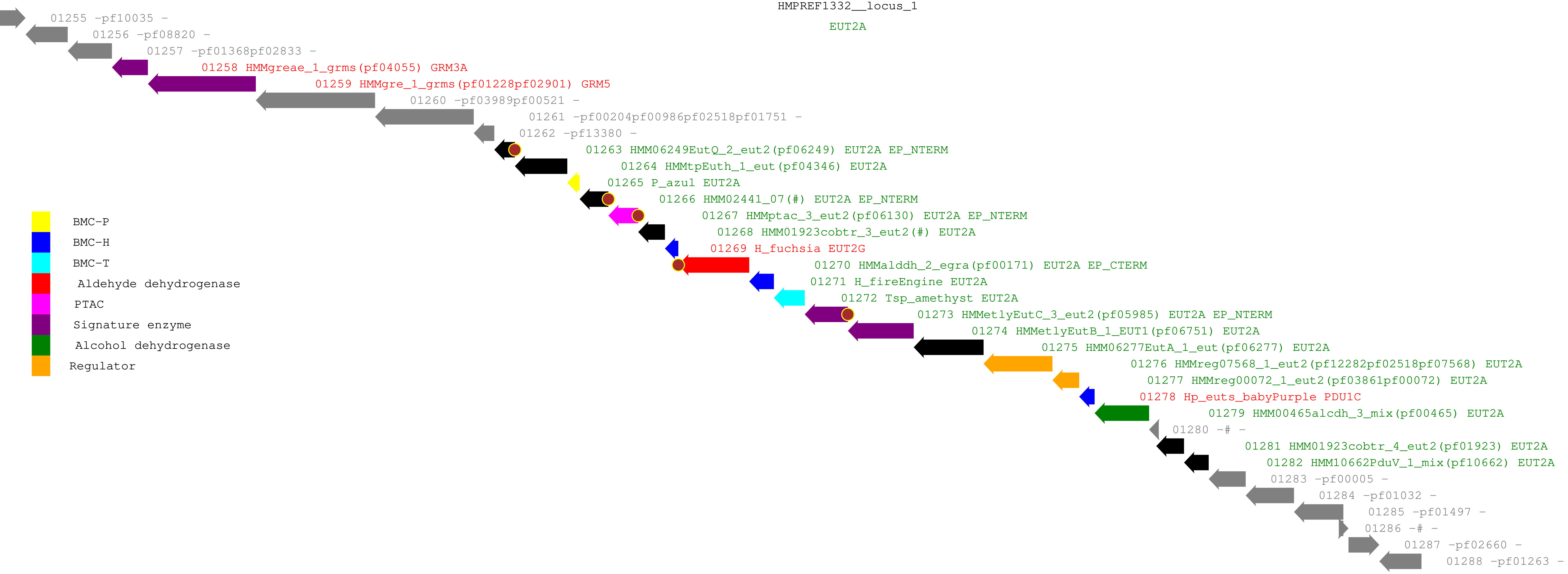

Enterococcus faecalis ERV37. Bacteria; Firmicutes; Bacilli; Lactobacillales; Enterococcaceae;

HMPREF1333__locus_1

EUT2A

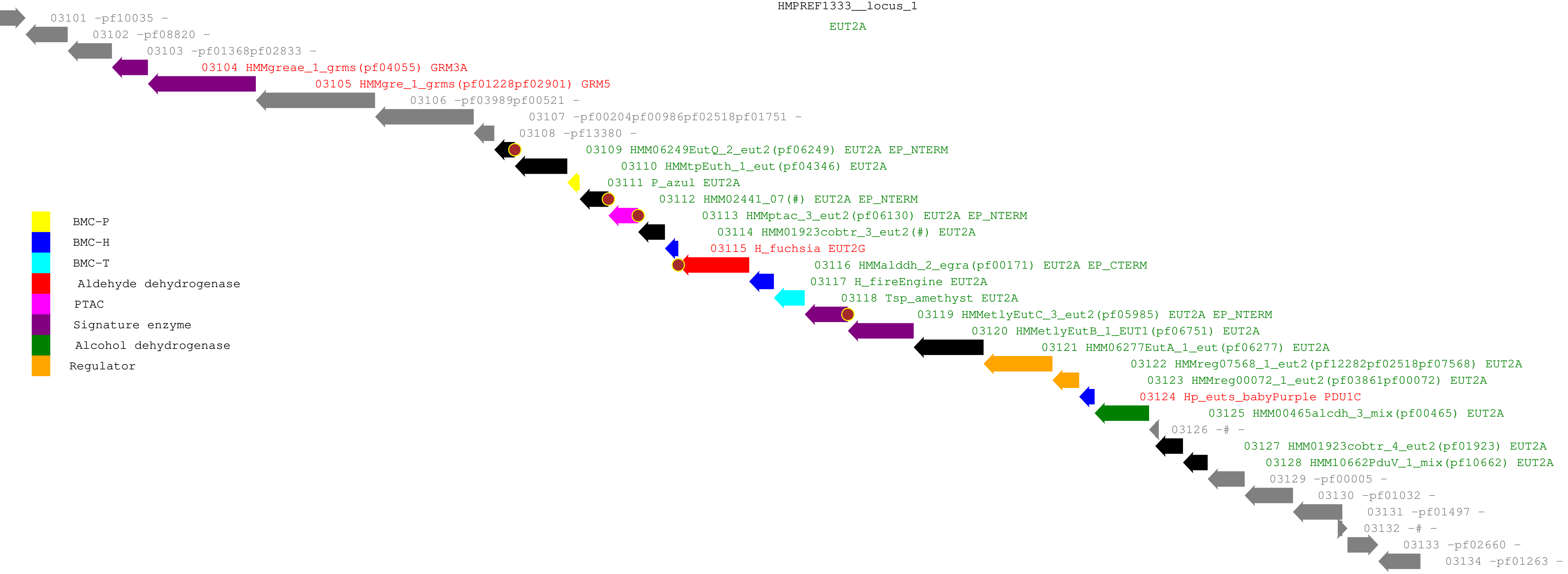

Enterococcus faecalis ERV41. Bacteria; Firmicutes; Bacilli; Lactobacillales; Enterococcaceae;

HMPREF1334__locus_1

EUT2A

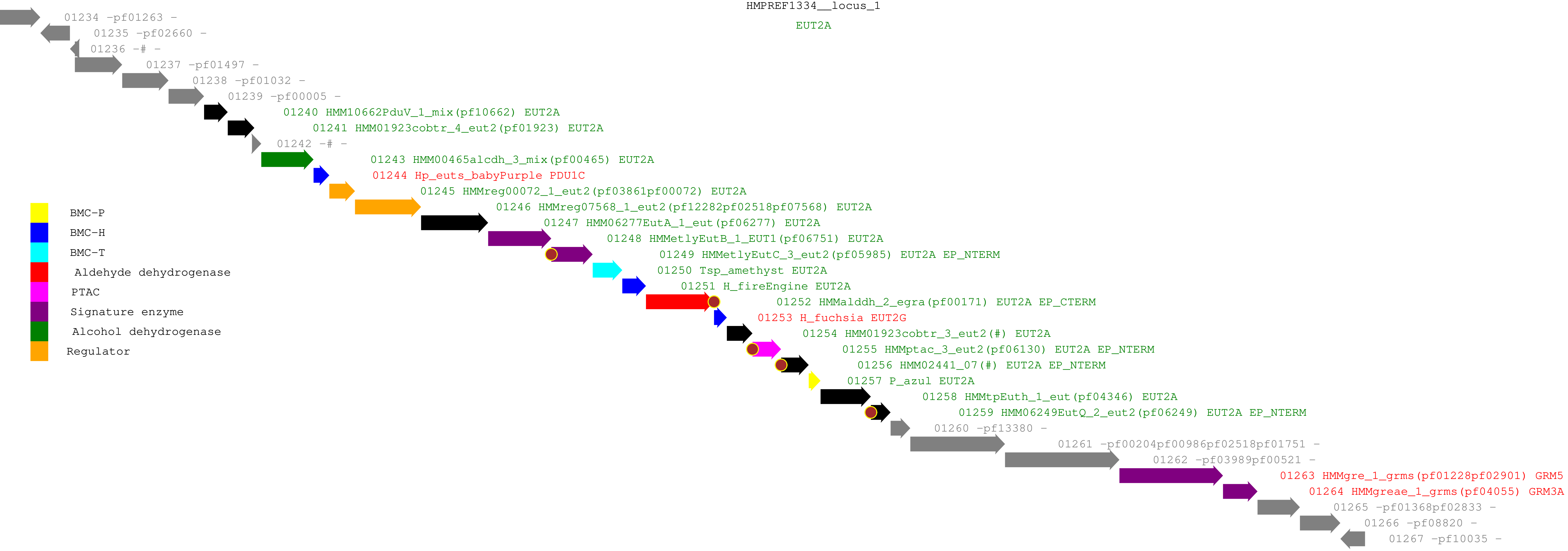

Enterococcus faecalis ERV62. Bacteria; Firmicutes; Bacilli; Lactobacillales; Enterococcaceae;

HMPREF1335__locus_1

EUT2A

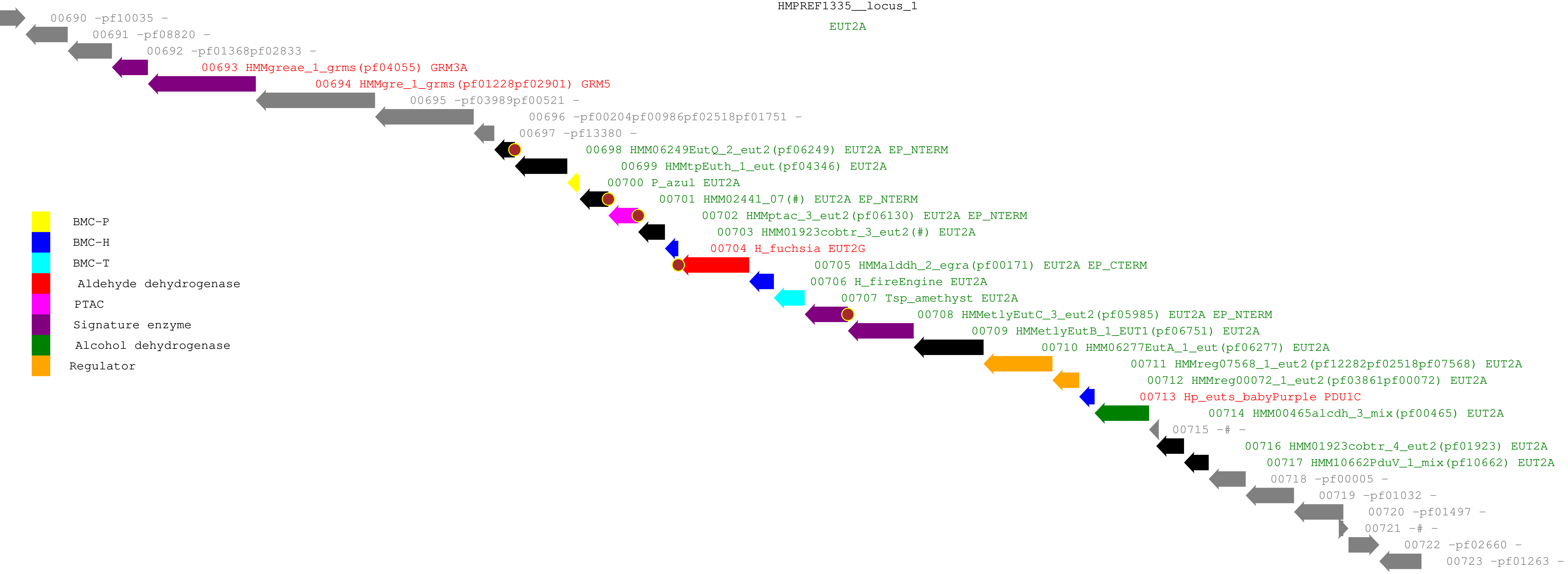

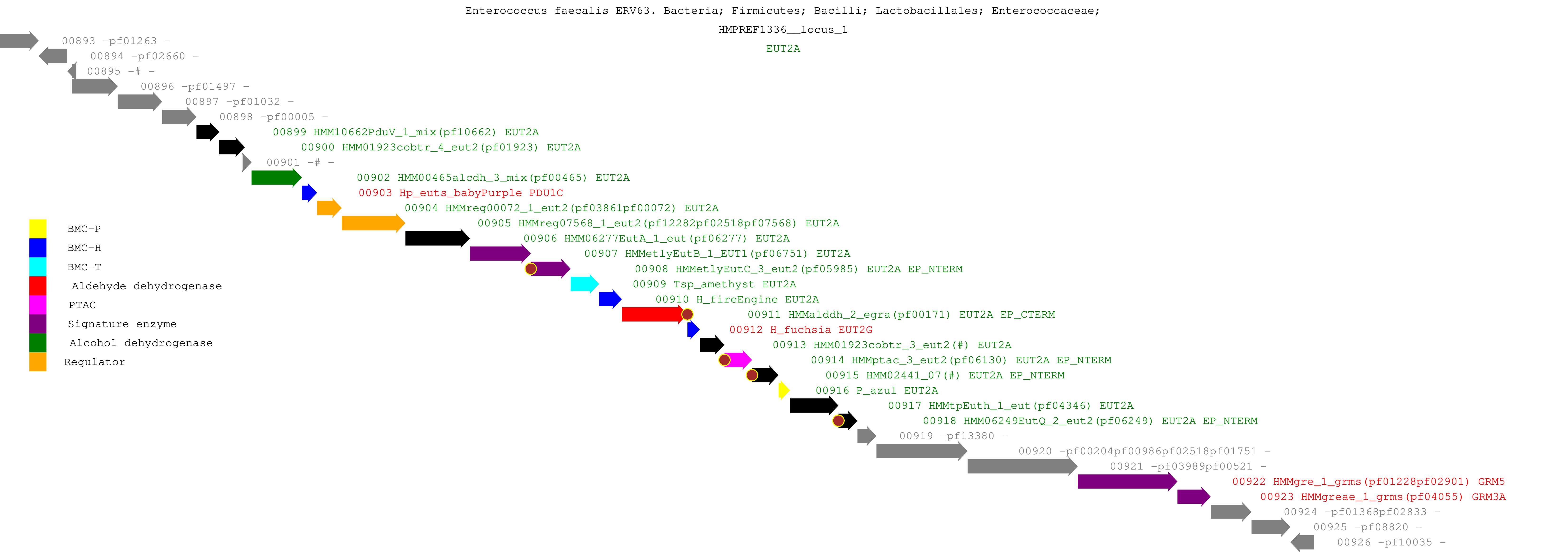
Enterococcus faecalis ERV63. Bacteria; Firmicutes; Bacilli; Lactobacillales; Enterococcaceae;
HMPREF1336__locus_1
EUT2A
00893 -pf01263 -
00894 -pf02660 -
00895 -# -
00896 -pf01497 -
00897 -pf01032 -
00898 -pf00005 -
00899 HMM10662PduV_1_mix(pf10662) EUT2A
00900 HMM01923cobtr_4_eut2(pf01923) EUT2A
00901 -# -
00902 HMM00465alcdh_3_mix(pf00465) EUT2A
00903 Hp_euts_babyPurple PDU1C
00904 HMMreg00072_1_eut2(pf03861pf00072) EUT2A
00905 HMMreg07568_1_eut2(pf12282pf02518pf07568) EUT2A
00906 HMM06277EutA_1_eut(pf06277) EUT2A
00907 HMMetlyEutB_1_EUT1(pf06751) EUT2A
00908 HMMetlyEutC_3_eut2(pf05985) EUT2A EP_NTERM
00909 Tsp_amethyst EUT2A
00910 H_fireEngine EUT2A
00911 HMMalddh_2_egra(pf00171) EUT2A EP_CTERM
00912 H_fuchsia EUT2G
00913 HMM01923cobtr_3_eut2(#) EUT2A
00914 HMMptac_3_eut2(pf06130) EUT2A EP_NTERM
00915 HMM02441_07(#) EUT2A EP_NTERM
00916 P_azul EUT2A
00917 HMMtpEuth_1_eut(pf04346) EUT2A
00918 HMM06249EutQ_2_eut2(pf06249) EUT2A EP_NTERM
00919 -pf13380 -
00920 -pf00204pf00986pf02518pf01751 -
00921 -pf03989pf00521 -
00922 HMMgre_1_grms(pf01228pf02901) GRM5
00923 HMMgreae_1_grms(pf04055) GRM3A
00924 -pf01368pf02833 -
00925 -pf08820 -
00926 -pf10035 -
BMC-P
BMC-H
BMC-T
Aldehyde dehydrogenase
PTAC
Signature enzyme
Alcohol dehydrogenase
Regulator

Enterococcus faecalis ERV65. Bacteria; Firmicutes; Bacilli; Lactobacillales; Enterococcaceae;

HMPREF1337__locus_1

EUT2A

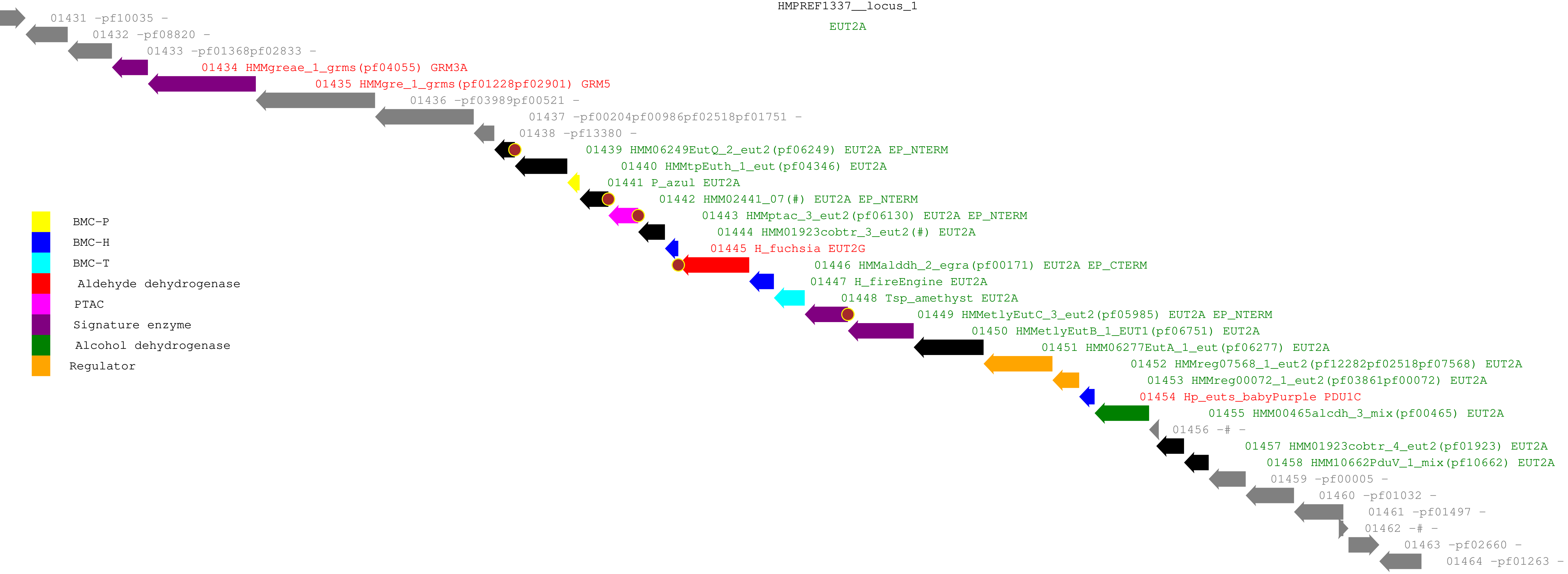

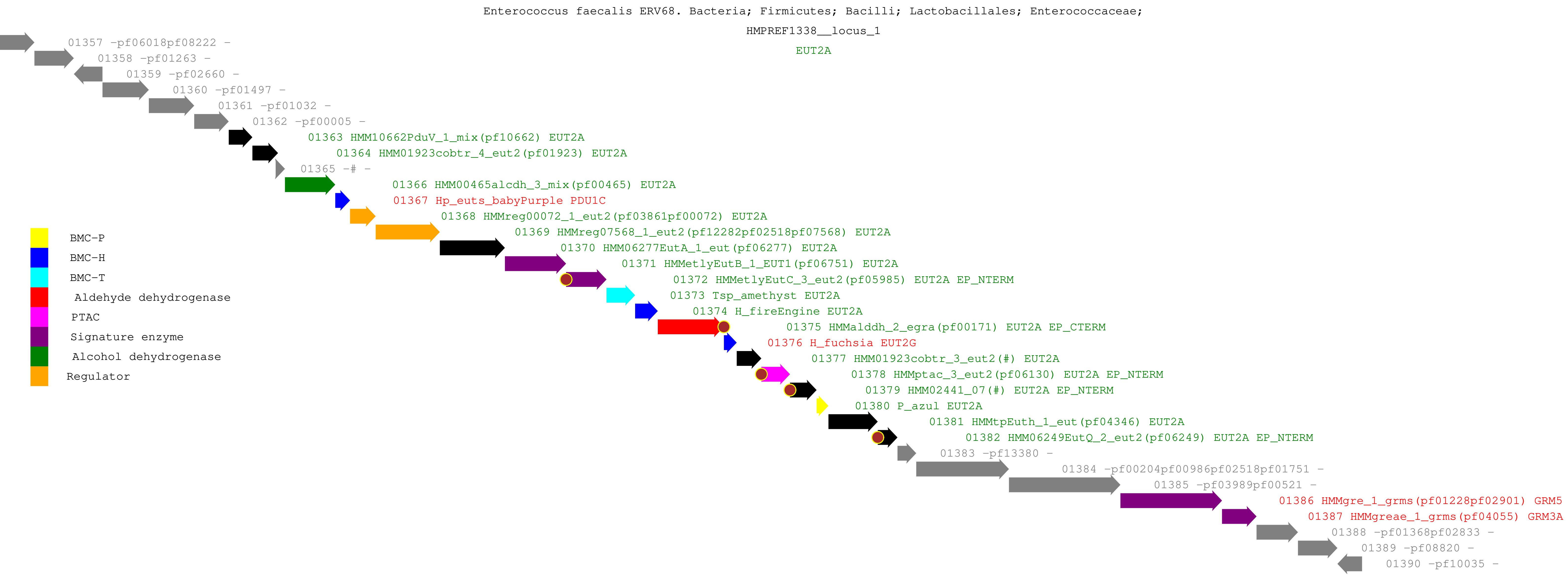
Enterococcus faecalis ERV68. Bacteria; Firmicutes; Bacilli; Lactobacillales; Enterococcaceae;
HMPREF1338__locus_1
EUT2A
01357 -pf06018pf08222 -
01358 -pf01263 -
01359 -pf02660 -
01360 -pf01497 -
01361 -pf01032 -
01362 -pf00005 -
01363 HMM10662PduV_1_mix(pf10662) EUT2A
01364 HMM01923cobtr_4_eut2(pf01923) EUT2A
01365 -# -
01366 HMM00465alcdh_3_mix(pf00465) EUT2A
01367 Hp_euts_babyPurple PDU1C
01368 HMMreg00072_1_eut2(pf03861pf00072) EUT2A
01369 HMMreg07568_1_eut2(pf12282pf02518pf07568) EUT2A
01370 HMM06277EutA_1_eut(pf06277) EUT2A
01371 HMMetlyEutB_1_EUT1(pf06751) EUT2A
01372 HMMetlyEutC_3_eut2(pf05985) EUT2A EP_NTERM
01373 Tsp_amethyst EUT2A
01374 H_fireEngine EUT2A
01375 HMMalddh_2_egra(pf00171) EUT2A EP_CTERM
01376 H_fuchsia EUT2G
01377 HMM01923cobtr_3_eut2(#) EUT2A
01378 HMMptac_3_eut2(pf06130) EUT2A EP_NTERM
01379 HMM02441_07(#) EUT2A EP_NTERM
01380 P_azul EUT2A
01381 HMMtpEuth_1_eut(pf04346) EUT2A
01382 HMM06249EutQ_2_eut2(pf06249) EUT2A EP_NTERM
01383 -pf13380 -
01384 -pf00204pf00986pf02518pf01751 -
01385 -pf03989pf00521 -
01386 HMMgre_1_grms(pf01228pf02901) GRM5
01387 HMMgreae_1_grms(pf04055) GRM3A
01388 -pf01368pf02833 -
01389 -pf08820 -
01390 -pf10035 -
BMC-P
BMC-H
BMC-T
Aldehyde dehydrogenase
PTAC
Signature enzyme
Alcohol dehydrogenase
Regulator

Enterococcus faecalis ERV72. Bacteria; Firmicutes; Bacilli; Lactobacillales; Enterococcaceae;

HMPREF1339__locus_1

EUT2A

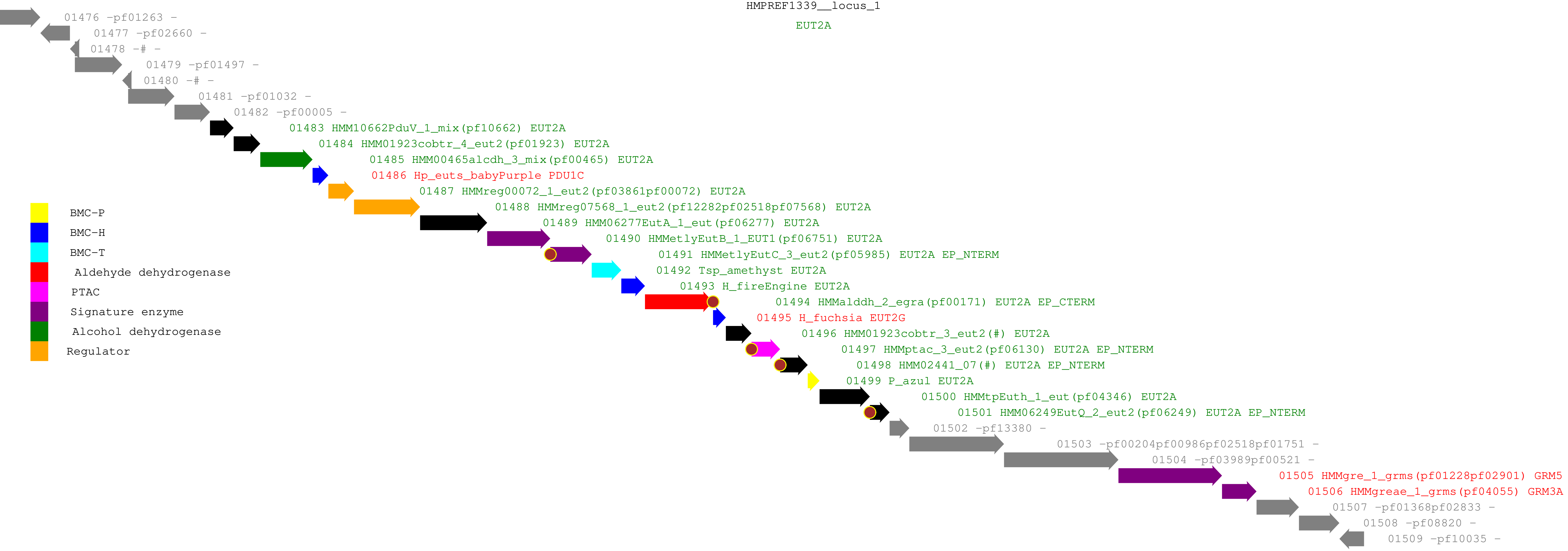

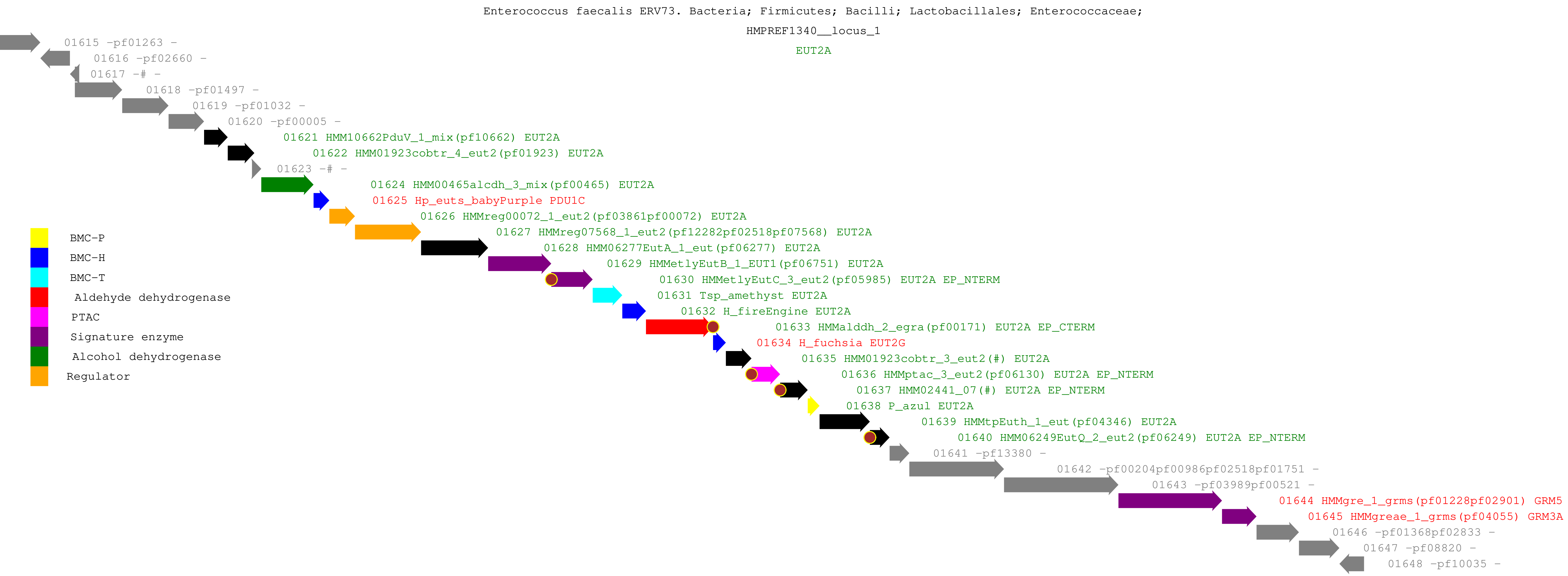
Enterococcus faecalis ERV73. Bacteria; Firmicutes; Bacilli; Lactobacillales; Enterococcaceae;
HMPREF1340__locus_1
EUT2A
01615 -pf01263 -
01616 -pf02660 -
01617 -# -
01618 -pf01497 -
01619 -pf01032 -
01620 -pf00005 -
01621 HMM10662PduV_1_mix(pf10662) EUT2A
01622 HMM01923cobtr_4_eut2(pf01923) EUT2A
01623 -# -
01624 HMM00465alcdh_3_mix(pf00465) EUT2A
01625 Hp_euts_babyPurple PDU1C
01626 HMMreg00072_1_eut2(pf03861pf00072) EUT2A
01627 HMMreg07568_1_eut2(pf12282pf02518pf07568) EUT2A
01628 HMM06277EutA_1_eut(pf06277) EUT2A
01629 HMMetlyEutB_1_EUT1(pf06751) EUT2A
01630 HMMetlyEutC_3_eut2(pf05985) EUT2A EP_NTERM
01631 Tsp_amethyst EUT2A
01632 H_fireEngine EUT2A
01633 HMMalddh_2_egra(pf00171) EUT2A EP_CTERM
01634 H_fuchsia EUT2G
01635 HMM01923cobtr_3_eut2(#) EUT2A
01636 HMMptac_3_eut2(pf06130) EUT2A EP_NTERM
01637 HMM02441_07(#) EUT2A EP_NTERM
01638 P_azul EUT2A
01639 HMMtpEuth_1_eut(pf04346) EUT2A
01640 HMM06249EutQ_2_eut2(pf06249) EUT2A EP_NTERM
01641 -pf13380 -
01642 -pf00204pf00986pf02518pf01751 -
01643 -pf03989pf00521 -
01644 HMMgre_1_grms(pf01228pf02901) GRM5
01645 HMMgreae_1_grms(pf04055) GRM3A
01646 -pf01368pf02833 -
01647 -pf08820 -
01648 -pf10035 -
BMC-P
BMC-H
BMC-T
Aldehyde dehydrogenase
PTAC
Signature enzyme
Alcohol dehydrogenase
Regulator

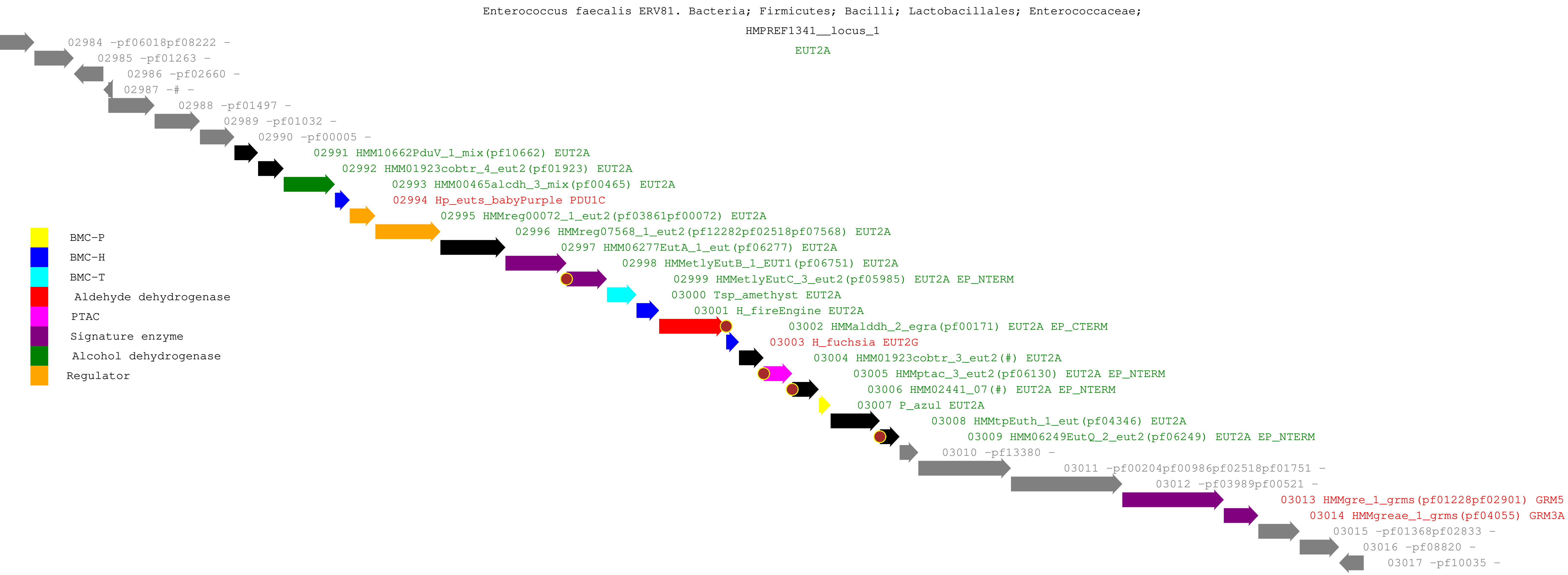
Enterococcus faecalis ERV81. Bacteria; Firmicutes; Bacilli; Lactobacillales; Enterococcaceae;
HMPREF1341__locus_1
EUT2A
02984 -pf06018pf08222 -
02985 -pf01263 -
02986 -pf02660 -
02987 -# -
02988 -pf01497 -
02989 -pf01032 -
02990 -pf00005 -
02991 HMM10662PduV_1_mix(pf10662) EUT2A
02992 HMM01923cobtr_4_eut2(pf01923) EUT2A
02993 HMM00465alcdh_3_mix(pf00465) EUT2A
02994 Hp_euts_babyPurple PDU1C
02995 HMMreg00072_1_eut2(pf03861pf00072) EUT2A
02996 HMMreg07568_1_eut2(pf12282pf02518pf07568) EUT2A
02997 HMM06277EutA_1_eut(pf06277) EUT2A
02998 HMMetlyEutB_1_EUT1(pf06751) EUT2A
02999 HMMetlyEutC_3_eut2(pf05985) EUT2A EP_NTERM
03000 Tsp_amethyst EUT2A
03001 H_fireEngine EUT2A
03002 HMMalddh_2_egra(pf00171) EUT2A EP_CTERM
03003 H_fuchsia EUT2G
03004 HMM01923cobtr_3_eut2(#) EUT2A
03005 HMMptac_3_eut2(pf06130) EUT2A EP_NTERM
03006 HMM02441_07(#) EUT2A EP_NTERM
03007 P_azul EUT2A
03008 HMMtpEuth_1_eut(pf04346) EUT2A
03009 HMM06249EutQ_2_eut2(pf06249) EUT2A EP_NTERM
03010 -pf13380 -
03011 -pf00204pf00986pf02518pf01751 -
03012 -pf03989pf00521 -
03013 HMMgre_1_grms(pf01228pf02901) GRM5
03014 HMMgreae_1_grms(pf04055) GRM3A
03015 -pf01368pf02833 -
03016 -pf08820 -
03017 -pf10035 -
BMC-P
BMC-H
BMC-T
Aldehyde dehydrogenase
PTAC
Signature enzyme
Alcohol dehydrogenase
Regulator

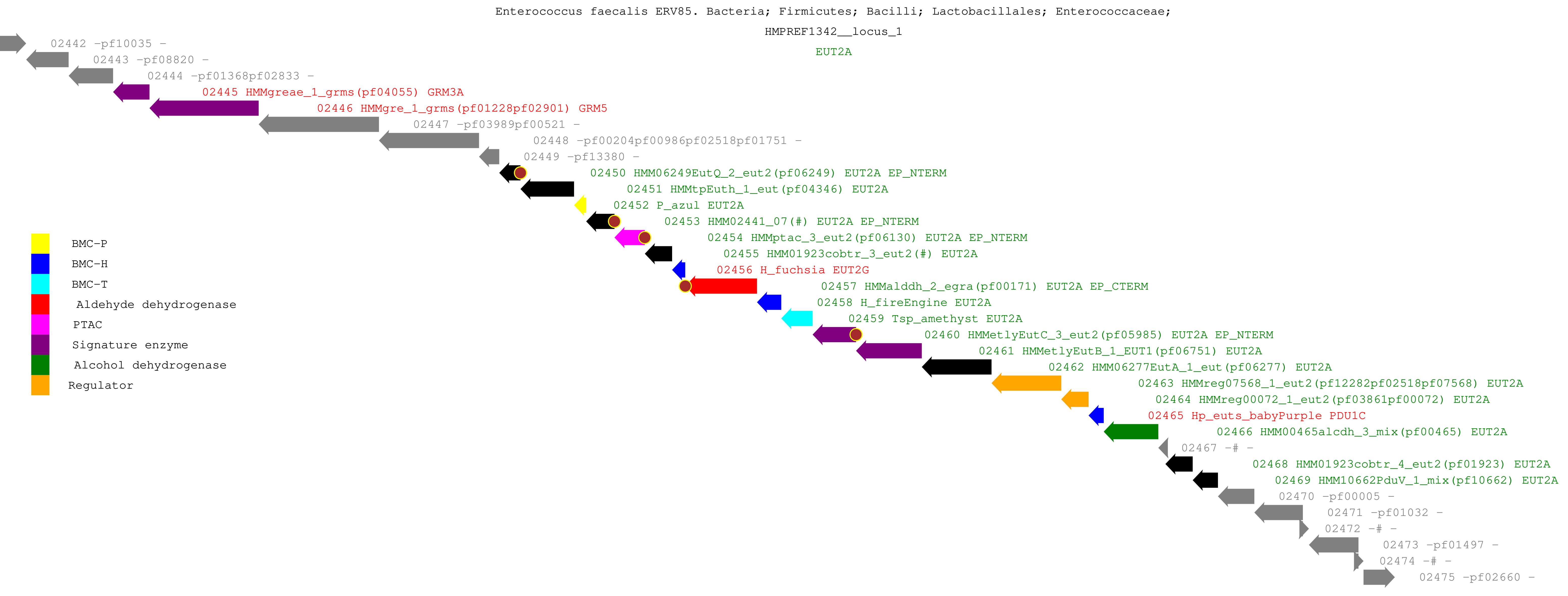

Enterococcus faecalis ERV85. Bacteria; Firmicutes; Bacilli; Lactobacillales; Enterococcaceae;
HMPREF1342__locus_1
EUT2A
02442 -pf10035 -
02443 -pf08820 -
02444 -pf01368pf02833 -
02445 HMMgreae_1_grms(pf04055) GRM3A
02446 HMMgre_1_grms(pf01228pf02901) GRM5
02447 -pf03989pf00521 -
02448 -pf00204pf00986pf02518pf01751 -
02449 -pf13380 -
02450 HMM06249EutQ_2_eut2(pf06249) EUT2A EP_NTERM
02451 HMMtpEuth_1_eut(pf04346) EUT2A
02452 P_azul EUT2A
02453 HMM02441_07(#) EUT2A EP_NTERM
02454 HMMptac_3_eut2(pf06130) EUT2A EP_NTERM
02455 HMM01923cobtr_3_eut2(#) EUT2A
02456 H_fuchsia EUT2G
02457 HMMaldh_2_egra(pf00171) EUT2A EP_CTERM
02458 H_fireEngine EUT2A
02459 Tsp_amethyst EUT2A
02460 HMMetlyEutC_3_eut2(pf05985) EUT2A EP_NTERM
02461 HMMetlyEutB_1_EUT1(pf06751) EUT2A
02462 HMM06277EutA_1_eut(pf06277) EUT2A
02463 HMMreg07568_1_eut2(pf12282pf02518pf07568) EUT2A
02464 HMMreg00072_1_eut2(pf03861pf00072) EUT2A
02465 Hp_euts_babyPurple PDU1C
02466 HMM00465alcdh_3_mix(pf00465) EUT2A
02467 -# -
02468 HMM01923cobtr_4_eut2(pf01923) EUT2A
02469 HMM10662PduV_1_mix(pf10662) EUT2A
02470 -pf00005 -
02471 -pf01032 -
02472 -# -
02473 -pf01497 -
02474 -# -
02475 -pf02660 -
BMC-P
BMC-H
BMC-T
Aldehyde dehydrogenase
PTAC
Signature enzyme
Alcohol dehydrogenase
Regulator

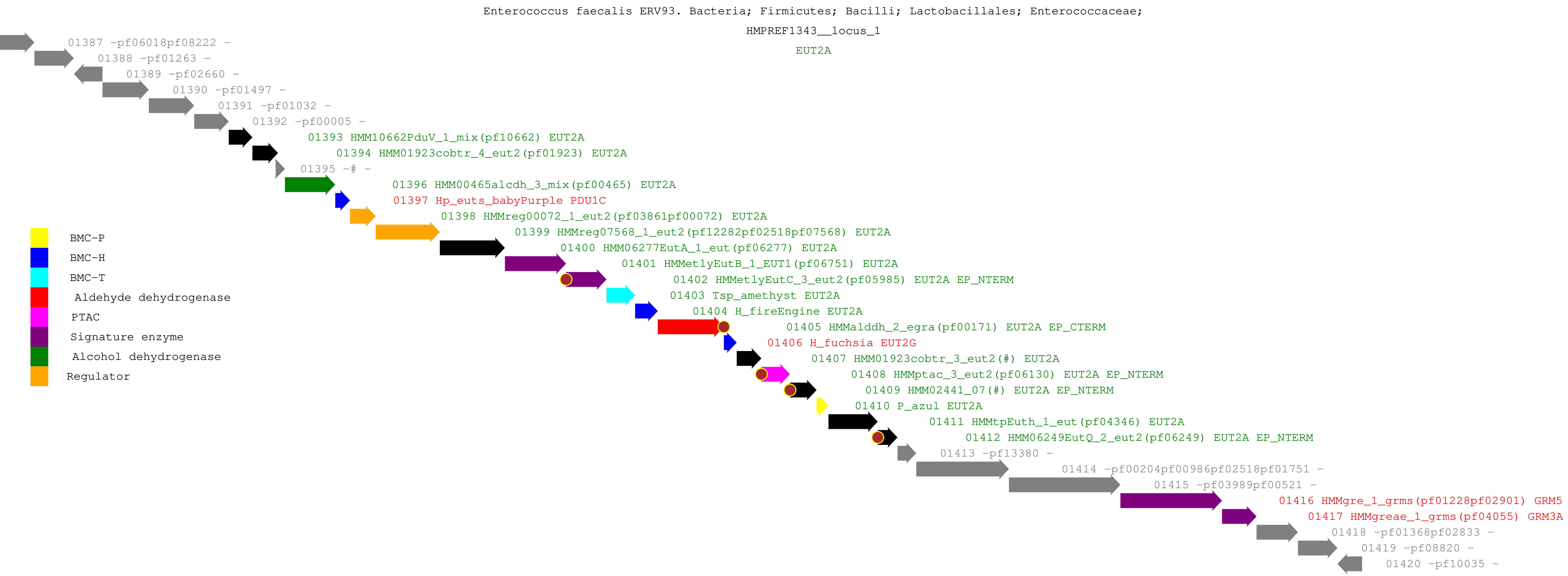
Enterococcus faecalis ERV93. Bacteria; Firmicutes; Bacilli; Lactobacillales; Enterococcaceae;
HMPREF1343__locus_1
EUT2A
01387 -pf06018pf08222 -
01388 -pf01263 -
01389 -pf02660 -
01390 -pf01497 -
01391 -pf01032 -
01392 -pf00005 -
01393 HMM10662PduV_1_mix(pf10662) EUT2A
01394 HMM01923cobtr_4_eut2(pf01923) EUT2A
01395 -# -
01396 HMM00465alcdh_3_mix(pf00465) EUT2A
01397 Hp_euts_babyPurple PDU1C
01398 HMMreg00072_1_eut2(pf03861pf00072) EUT2A
01399 HMMreg07568_1_eut2(pf12282pf02518pf07568) EUT2A
01400 HMM06277EutA_1_eut(pf06277) EUT2A
01401 HMMetlyEutB_1_EUT1(pf06751) EUT2A
01402 HMMetlyEutC_3_eut2(pf05985) EUT2A EP_NTERM
01403 Tsp_amethyst EUT2A
01404 H_fireEngine EUT2A
01405 HMMalddh_2_egra(pf00171) EUT2A EP_CTERM
01406 H_fuchsia EUT2G
01407 HMM01923cobtr_3_eut2(#) EUT2A
01408 HMMptac_3_eut2(pf06130) EUT2A EP_NTERM
01409 HMM02441_07(#) EUT2A EP_NTERM
01410 P_azul EUT2A
01411 HMMtpEuth_1_eut(pf04346) EUT2A
01412 HMM06249EutQ_2_eut2(pf06249) EUT2A EP_NTERM
01413 -pf13380 -
01414 -pf00204pf00986pf02518pf01751 -
01415 -pf03989pf00521 -
01416 HMMgre_1_grms(pf01228pf02901) GRM5
01417 HMMgreae_1_grms(pf04055) GRM3A
01418 -pf01368pf02833 -
01419 -pf08820 -
01420 -pf10035 -
BMC-P
BMC-H
BMC-T
Aldehyde dehydrogenase
PTAC
Signature enzyme
Alcohol dehydrogenase
Regulator

Enterococcus faecalis 918. Bacteria; Firmicutes; Bacilli; Lactobacillales; Enterococcaceae;

HMPREF2097__locus_1

EUT2A

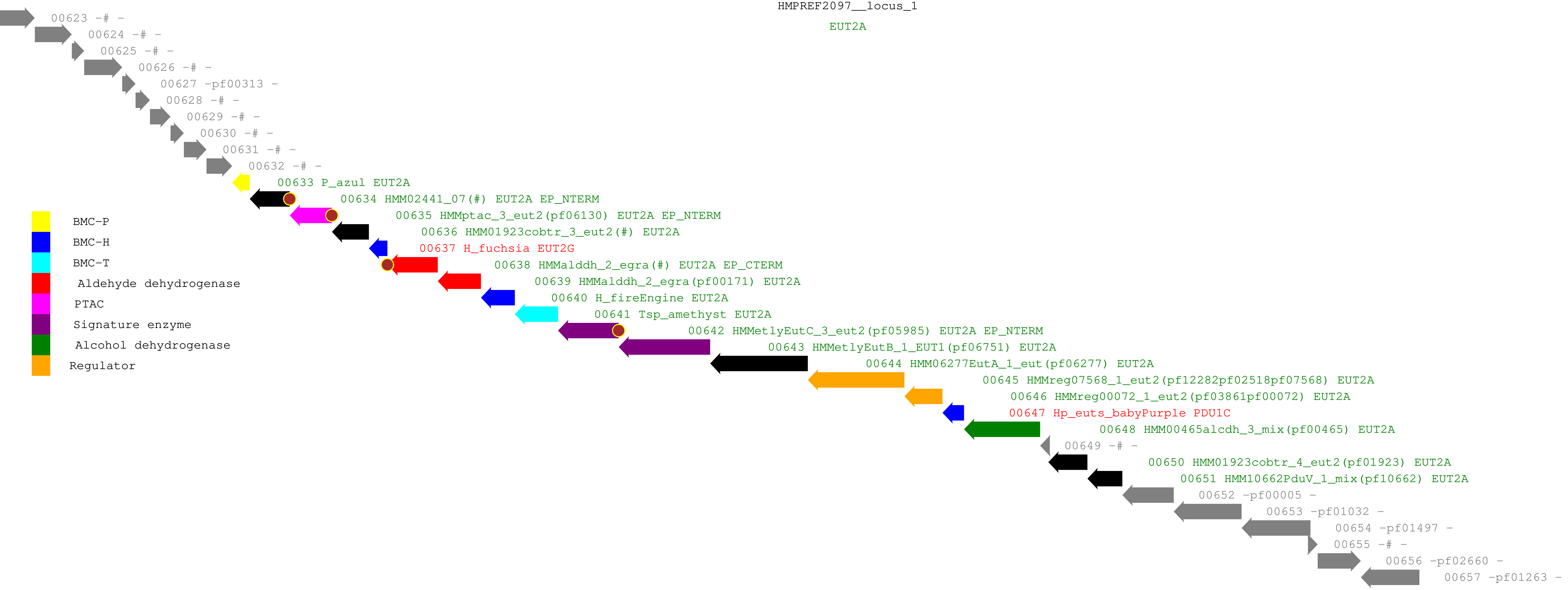

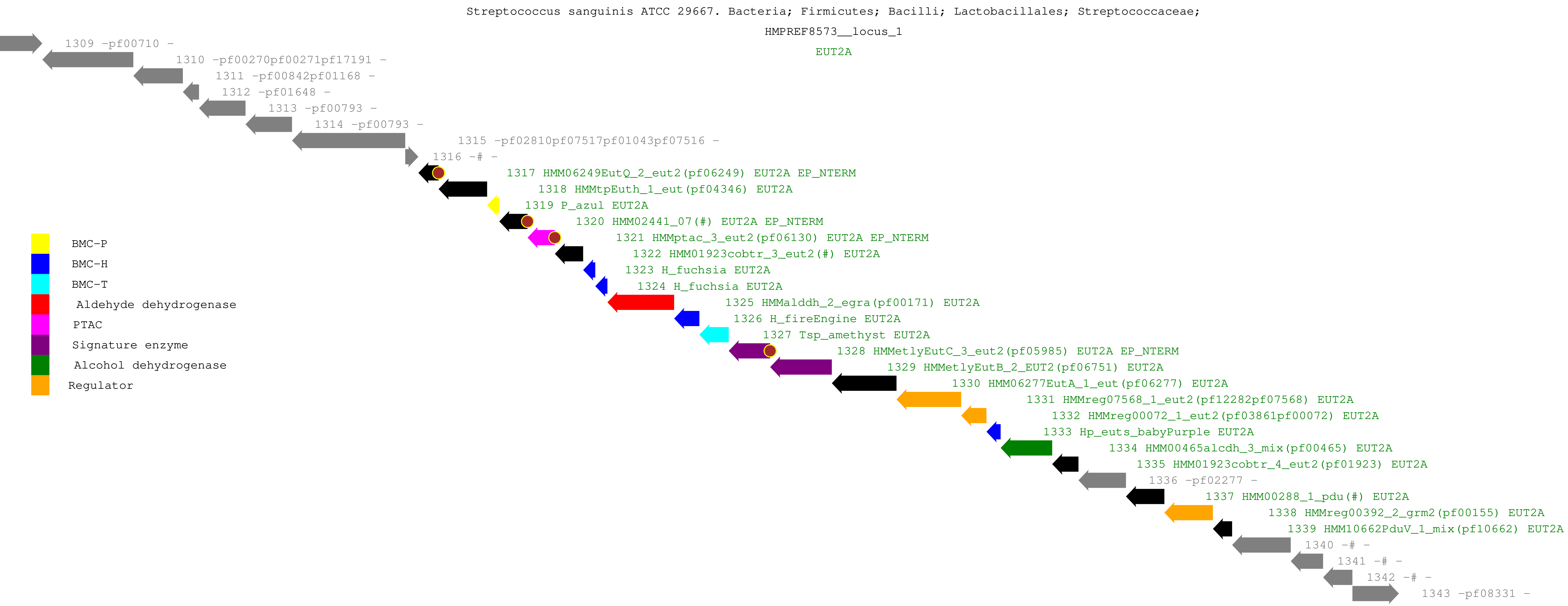
Streptococcus sanguinis ATCC 29667. Bacteria; Firmicutes; Bacilli; Lactobacillales; Streptococcaceae;
HMPREF8573__locus_1
EUT2A
1309 -pf00710 -
1310 -pf00270pf00271pf17191 -
1311 -pf00842pf01168 -
1312 -pf01648 -
1313 -pf00793 -
1314 -pf00793 -
1315 -pf02810pf07517pf01043pf07516 -
1316 -# -
1317 HMM06249EutQ_2_eut2(pf06249) EUT2A EP_NTERM
1318 HMMtpEuth_1_eut(pf04346) EUT2A
1319 P_azul EUT2A
1320 HMM02441_07(#) EUT2A EP_NTERM
1321 HMMptac_3_eut2(pf06130) EUT2A EP_NTERM
1322 HMM01923cobtr_3_eut2(#) EUT2A
1323 H_fuchsia EUT2A
1324 H_fuchsia EUT2A
1325 HMMaldh_2_egra(pf00171) EUT2A
1326 H_fireEngine EUT2A
1327 Tsp_amethyst EUT2A
1328 HMMetlyEutC_3_eut2(pf05985) EUT2A EP_NTERM
1329 HMMetlyEutB_2_EUT2(pf06751) EUT2A
1330 HMM06277EutA_1_eut(pf06277) EUT2A
1331 HMMreg07568_1_eut2(pf12282pf07568) EUT2A
1332 HMMreg00072_1_eut2(pf03861pf00072) EUT2A
1333 Hp_euts_babyPurple EUT2A
1334 HMM00465alcdh_3_mix(pf00465) EUT2A
1335 HMM01923cobtr_4_eut2(pf01923) EUT2A
1336 -pf02277 -
1337 HMM00288_1_pdu(#) EUT2A
1338 HMMreg00392_2_grm2(pf00155) EUT2A
1339 HMM10662PduV_1_mix(pf10662) EUT2A
1340 -# -
1341 -# -
1342 -# -
1343 -pf08331 -
BMC-P
BMC-H
BMC-T
Aldehyde dehydrogenase
PTAC
Signature enzyme
Alcohol dehydrogenase
Regulator

Streptococcus parasanguinis ATCC 903. Bacteria; Firmicutes; Bacilli; Lactobacillales; Streptococcaceae;

HMPREF8577__locus_1

EUT2A

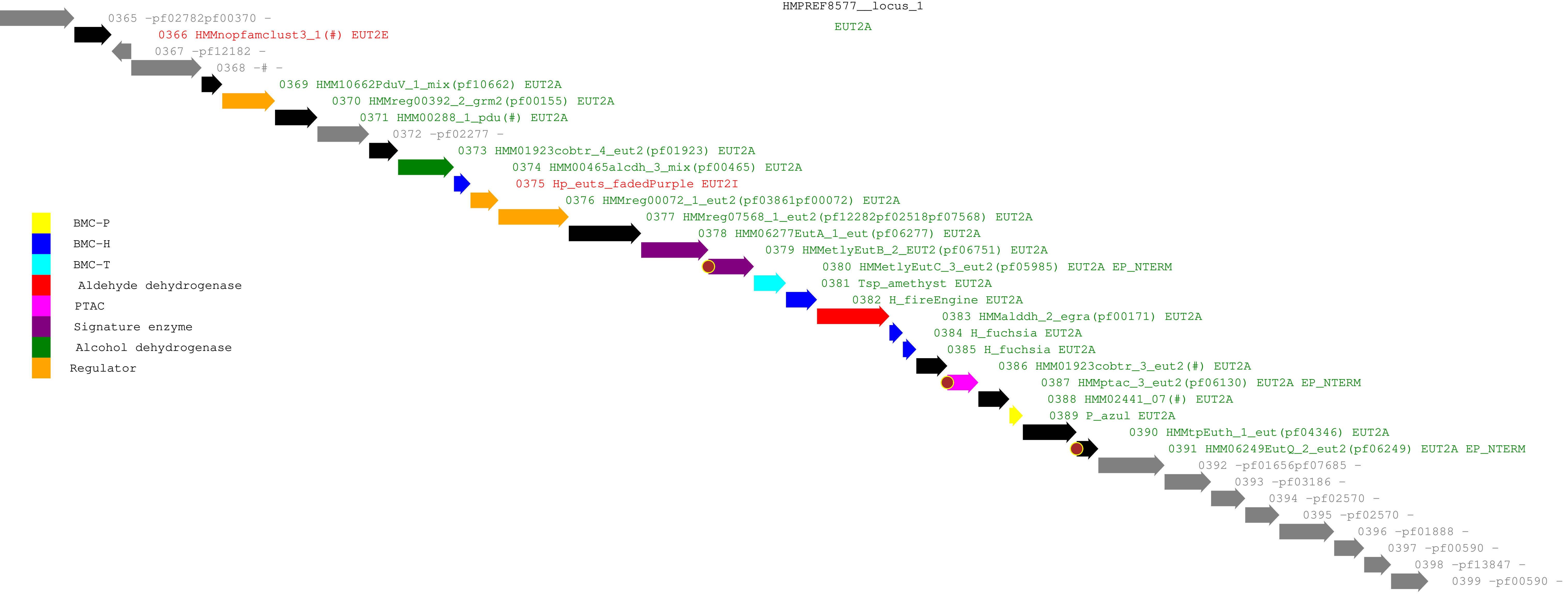

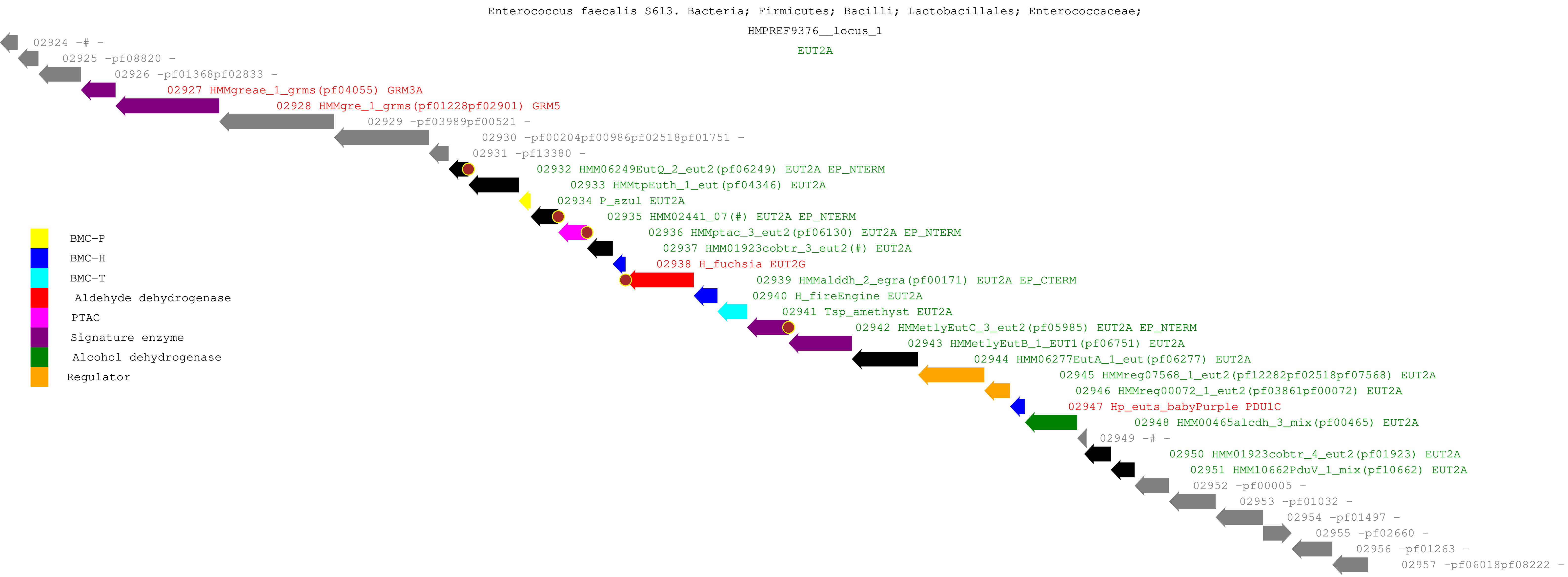
Enterococcus faecalis S613. Bacteria; Firmicutes; Bacilli; Lactobacillales; Enterococcaceae;
HMPREF9376__locus_1
EUT2A
02924 -# -
02925 -pf08820 -
02926 -pf01368pf02833 -
02927 HMMgreae_1_grms(pf04055) GRM3A
02928 HMMgre_1_grms(pf01228pf02901) GRM5
02929 -pf03989pf00521 -
02930 -pf00204pf00986pf02518pf01751 -
02931 -pf13380 -
02932 HMM06249EutQ_2_eut2(pf06249) EUT2A EP_NTERM
02933 HMMtpEuth_1_eut(pf04346) EUT2A
02934 P_azul EUT2A
02935 HMM02441_07(#) EUT2A EP_NTERM
02936 HMMptac_3_eut2(pf06130) EUT2A EP_NTERM
02937 HMM01923cobtr_3_eut2(#) EUT2A
02938 H_fuchsia EUT2G
02939 HMMalddh_2_egra(pf00171) EUT2A EP_CTERM
02940 H_fireEngine EUT2A
02941 Tsp_amethyst EUT2A
02942 HMMetlyEutC_3_eut2(pf05985) EUT2A EP_NTERM
02943 HMMetlyEutB_1_EUT1(pf06751) EUT2A
02944 HMM06277EutA_1_eut(pf06277) EUT2A
02945 HMMreg07568_1_eut2(pf12282pf02518pf07568) EUT2A
02946 HMMreg00072_1_eut2(pf03861pf00072) EUT2A
02947 Hp_euts_babyPurple PDU1C
02948 HMM00465alcdh_3_mix(pf00465) EUT2A
02949 -# -
02950 HMM01923cobtr_4_eut2(pf01923) EUT2A
02951 HMM10662PduV_1_mix(pf10662) EUT2A
02952 -pf00005 -
02953 -pf01032 -
02954 -pf01497 -
02955 -pf02660 -
02956 -pf01263 -
02957 -pf06018pf08222 -
BMC-P
BMC-H
BMC-T
Aldehyde dehydrogenase
PTAC
Signature enzyme
Alcohol dehydrogenase
Regulator

Enterococcus faecalis R712. Bacteria; Firmicutes; Bacilli; Lactobacillales; Enterococcaceae;

HMPREF9377__locus_1

EUT2A

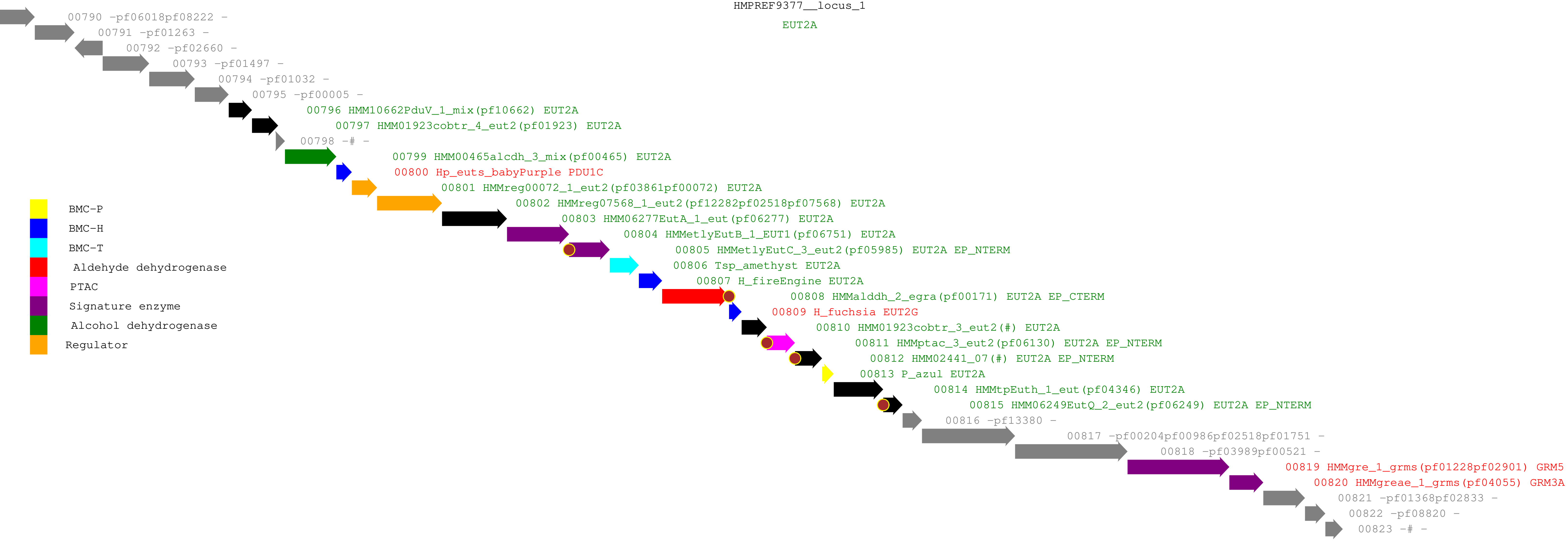

Streptococcus sanguinis SK49. Bacteria; Firmicutes; Bacilli; Lactobacillales; Streptococcaceae;

HMPREF9380__locus_1

EUT2A

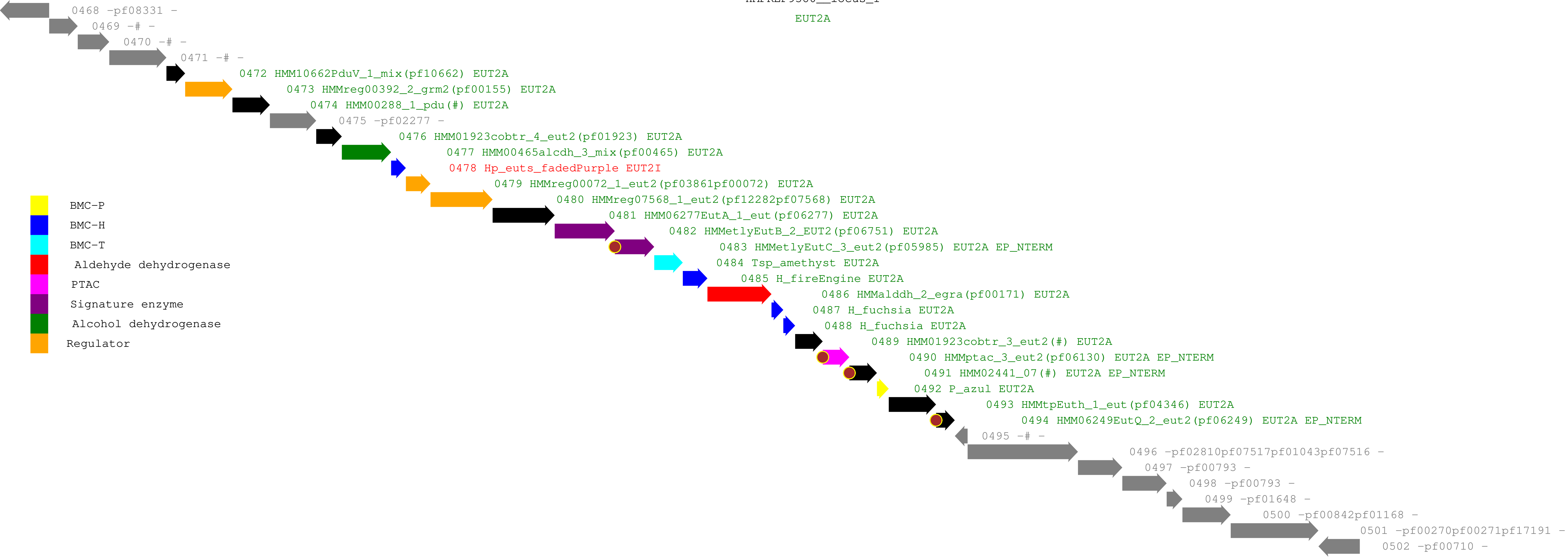

Streptococcus sanguinis SK160. Bacteria; Firmicutes; Bacilli; Lactobacillales; Streptococcaceae;

HMPREF9384__locus_1

EUT2A

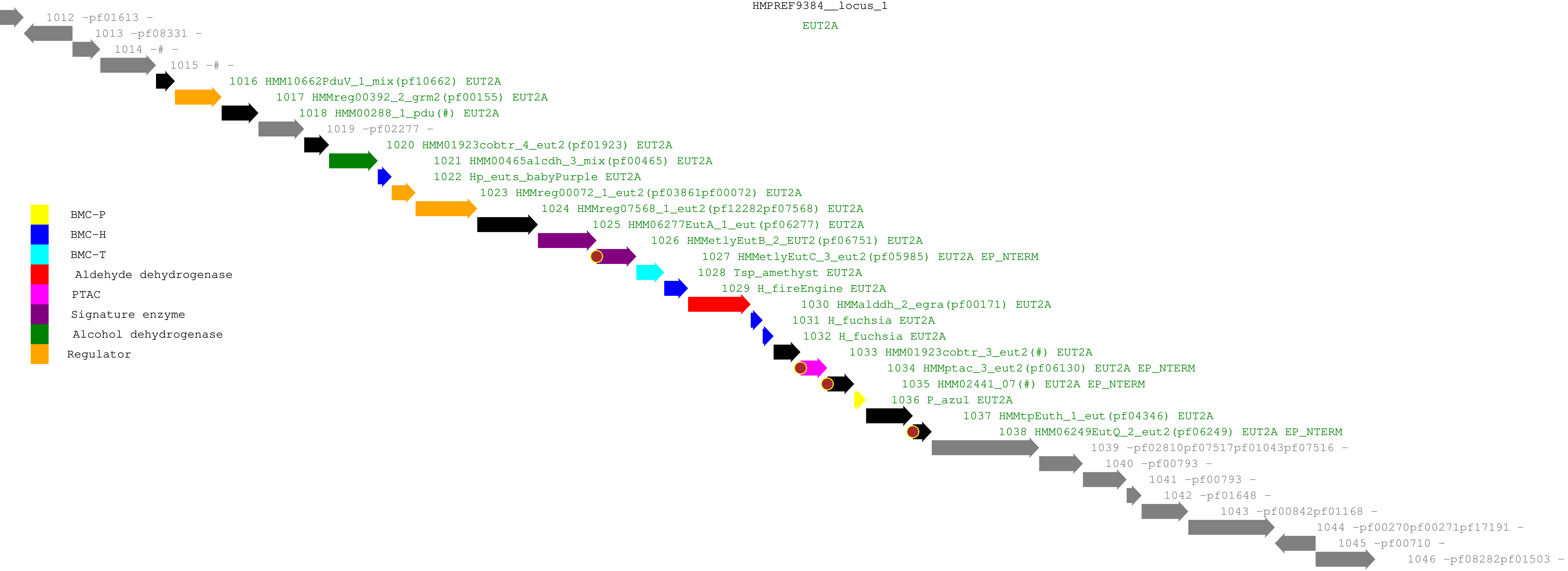

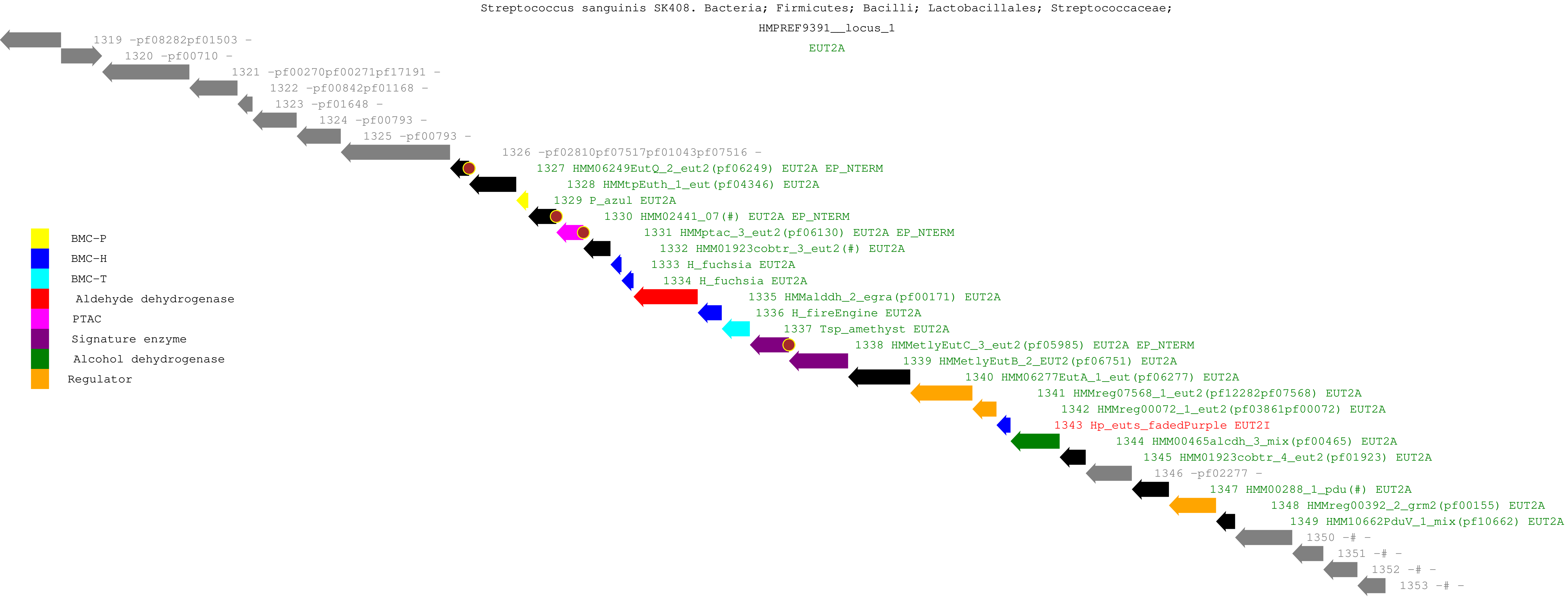
Streptococcus sanguinis SK408. Bacteria; Firmicutes; Bacilli; Lactobacillales; Streptococcaceae;
HMPREF9391__locus_1
EUT2A
1319 -pf08282pf01503 -
1320 -pf00710 -
1321 -pf00270pf00271pf17191 -
1322 -pf00842pf01168 -
1323 -pf01648 -
1324 -pf00793 -
1325 -pf00793 -
1326 -pf02810pf07517pf01043pf07516 -
1327 HMM06249EutQ_2_eut2(pf06249) EUT2A EP_NTERM
1328 HMMtpEuth_1_eut(pf04346) EUT2A
1329 P_azul EUT2A
1330 HMM02441_07(#) EUT2A EP_NTERM
1331 HMMptac_3_eut2(pf06130) EUT2A EP_NTERM
1332 HMM01923cobtr_3_eut2(#) EUT2A
1333 H_fuchsia EUT2A
1334 H_fuchsia EUT2A
1335 HMMaldh_2_egra(pf00171) EUT2A
1336 H_fireEngine EUT2A
1337 Tsp_amethyst EUT2A
1338 HMMetlyEutC_3_eut2(pf05985) EUT2A EP_NTERM
1339 HMMetlyEutB_2_EUT2(pf06751) EUT2A
1340 HMM06277EutA_1_eut(pf06277) EUT2A
1341 HMMreg07568_1_eut2(pf12282pf07568) EUT2A
1342 HMMreg00072_1_eut2(pf03861pf00072) EUT2A
1343 Hp_euts_fadedPurple EUT2I
1344 HMM00465alcdh_3_mix(pf00465) EUT2A
1345 HMM01923cobtr_4_eut2(pf01923) EUT2A
1346 -pf02277 -
1347 HMM00288_1_pdu(#) EUT2A
1348 HMMreg00392_2_grm2(pf00155) EUT2A
1349 HMM10662PduV_1_mix(pf10662) EUT2A
1350 -# -
1351 -# -
1352 -# -
1353 -# -
BMC-P
BMC-H
BMC-T
Aldehyde dehydrogenase
PTAC
Signature enzyme
Alcohol dehydrogenase
Regulator

Streptococcus sanguinis SK678. Bacteria; Firmicutes; Bacilli; Lactobacillales; Streptococcaceae;

HMPREF9392__locus_1

EUT2A

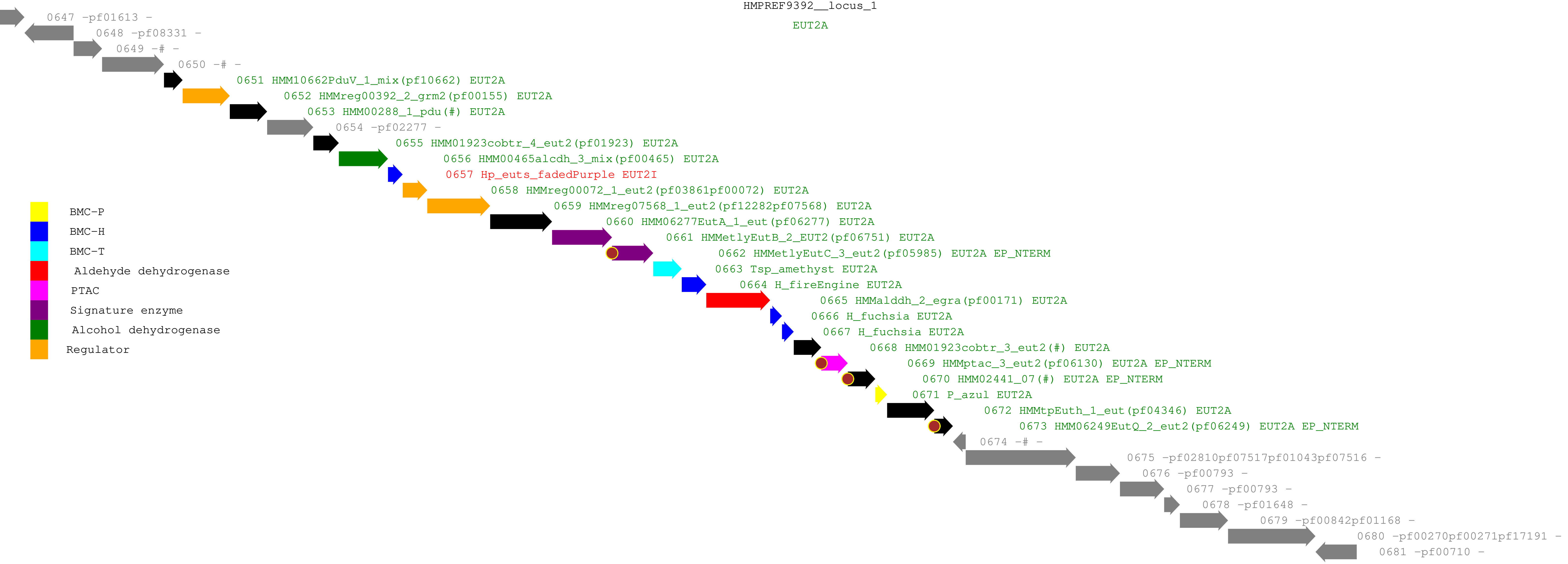

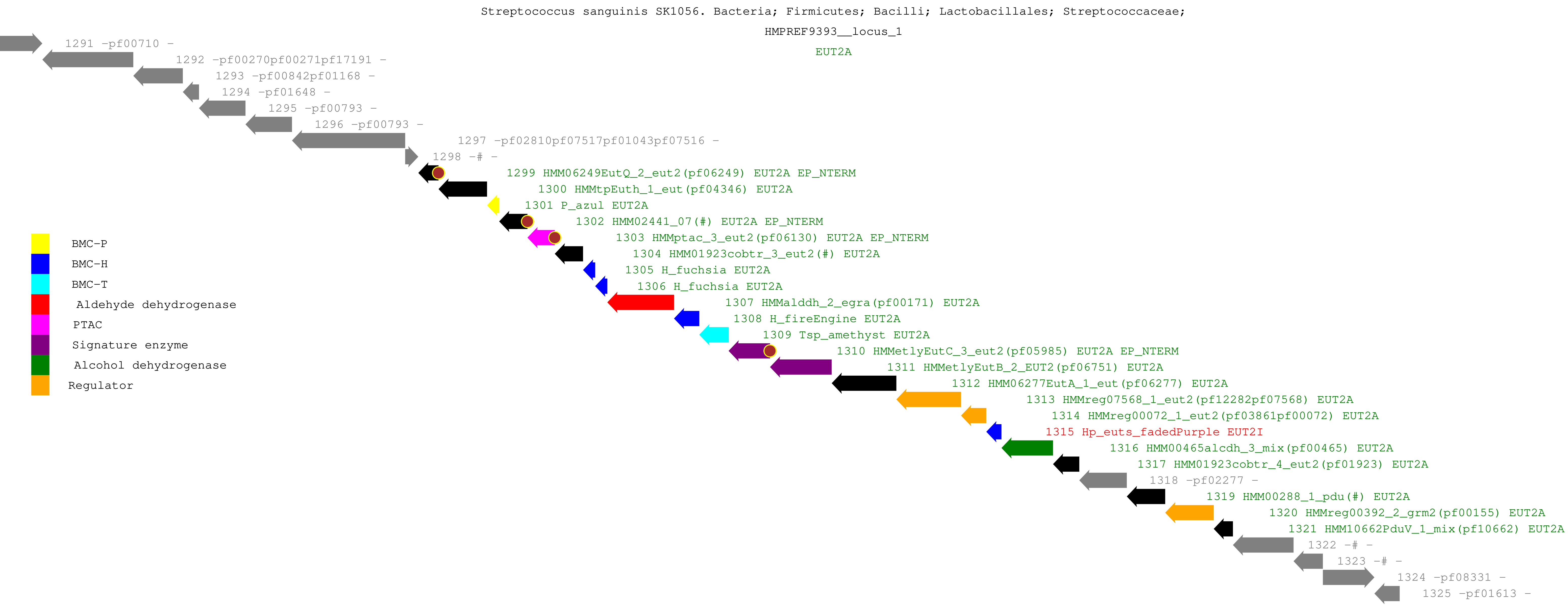
Streptococcus sanguinis SK1056. Bacteria; Firmicutes; Bacilli; Lactobacillales; Streptococcaceae;
HMPREF9393__locus_1
EUT2A
1291 -pf00710 -
1292 -pf00270pf00271pf17191 -
1293 -pf00842pf01168 -
1294 -pf01648 -
1295 -pf00793 -
1296 -pf00793 -
1297 -pf02810pf07517pf01043pf07516 -
1298 -# -
1299 HMM06249EutQ_2_eut2(pf06249) EUT2A EP_NTERM
1300 HMMtpEuth_1_eut(pf04346) EUT2A
1301 P_azul EUT2A
1302 HMM02441_07(#) EUT2A EP_NTERM
1303 HMMptac_3_eut2(pf06130) EUT2A EP_NTERM
1304 HMM01923cobtr_3_eut2(#) EUT2A
1305 H_fuchsia EUT2A
1306 H_fuchsia EUT2A
1307 HMMaldh_2_egra(pf00171) EUT2A
1308 H_fireEngine EUT2A
1309 Tsp_amethyst EUT2A
1310 HMMetlyEutC_3_eut2(pf05985) EUT2A EP_NTERM
1311 HMMetlyEutB_2_EUT2(pf06751) EUT2A
1312 HMM06277EutA_1_eut(pf06277) EUT2A
1313 HMMreg07568_1_eut2(pf12282pf07568) EUT2A
1314 HMMreg00072_1_eut2(pf03861pf00072) EUT2A
1315 Hp_euts_fadedPurple EUT2I
1316 HMM00465alcdh_3_mix(pf00465) EUT2A
1317 HMM01923cobtr_4_eut2(pf01923) EUT2A
1318 -pf02277 -
1319 HMM00288_1_pdu(#) EUT2A
1320 HMMreg00392_2_grm2(pf00155) EUT2A
1321 HMM10662PduV_1_mix(pf10662) EUT2A
1322 -# -
1323 -# -
1324 -pf08331 -
1325 -pf01613 -
BMC-P
BMC-H
BMC-T
Aldehyde dehydrogenase
PTAC
Signature enzyme
Alcohol dehydrogenase
Regulator

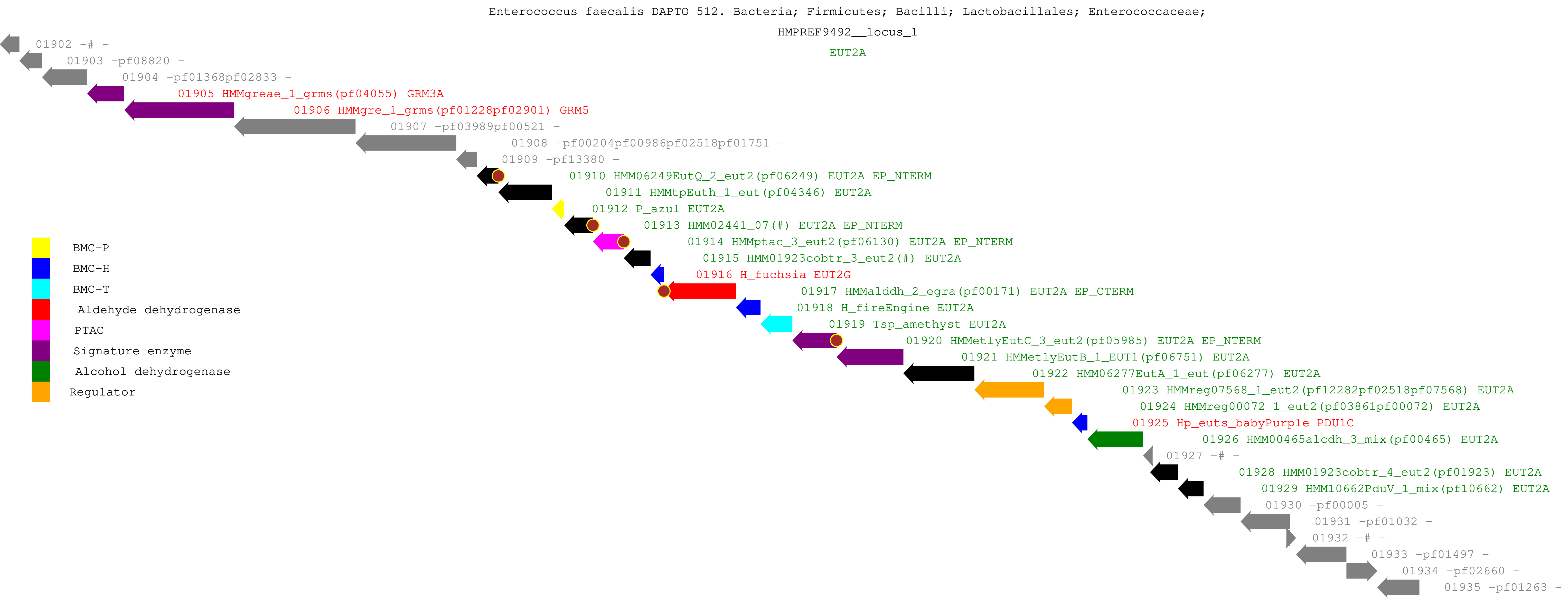
Enterococcus faecalis DAPTO 512. Bacteria; Firmicutes; Bacilli; Lactobacillales; Enterococcaceae;
HMPREF9492__locus_1
EUT2A
01902 -# -
01903 -pf08820 -
01904 -pf01368pf02833 -
01905 HMMgreae_1_grms(pf04055) GRM3A
01906 HMMgre_1_grms(pf01228pf02901) GRM5
01907 -pf03989pf00521 -
01908 -pf00204pf00986pf02518pf01751 -
01909 -pf13380 -
01910 HMM06249EutQ_2_eut2(pf06249) EUT2A EP_NTERM
01911 HMMtpEuth_1_eut(pf04346) EUT2A
01912 P_azul EUT2A
01913 HMM02441_07(#) EUT2A EP_NTERM
01914 HMMptac_3_eut2(pf06130) EUT2A EP_NTERM
01915 HMM01923cobtr_3_eut2(#) EUT2A
01916 H_fuchsia EUT2G
01917 HMMalddh_2_egra(pf00171) EUT2A EP_CTERM
01918 H_fireEngine EUT2A
01919 Tsp_amethyst EUT2A
01920 HMMetlyEutC_3_eut2(pf05985) EUT2A EP_NTERM
01921 HMMetlyEutB_1_EUT1(pf06751) EUT2A
01922 HMM06277EutA_1_eut(pf06277) EUT2A
01923 HMMreg07568_1_eut2(pf12282pf02518pf07568) EUT2A
01924 HMMreg00072_1_eut2(pf03861pf00072) EUT2A
01925 Hp_euts_babyPurple PDU1C
01926 HMM00465alcdh_3_mix(pf00465) EUT2A
01927 -# -
01928 HMM01923cobtr_4_eut2(pf01923) EUT2A
01929 HMM10662PduV_1_mix(pf10662) EUT2A
01930 -pf00005 -
01931 -pf01032 -
01932 -# -
01933 -pf01497 -
01934 -pf02660 -
01935 -pf01263 -
BMC-P
BMC-H
BMC-T
Aldehyde dehydrogenase
PTAC
Signature enzyme
Alcohol dehydrogenase
Regulator

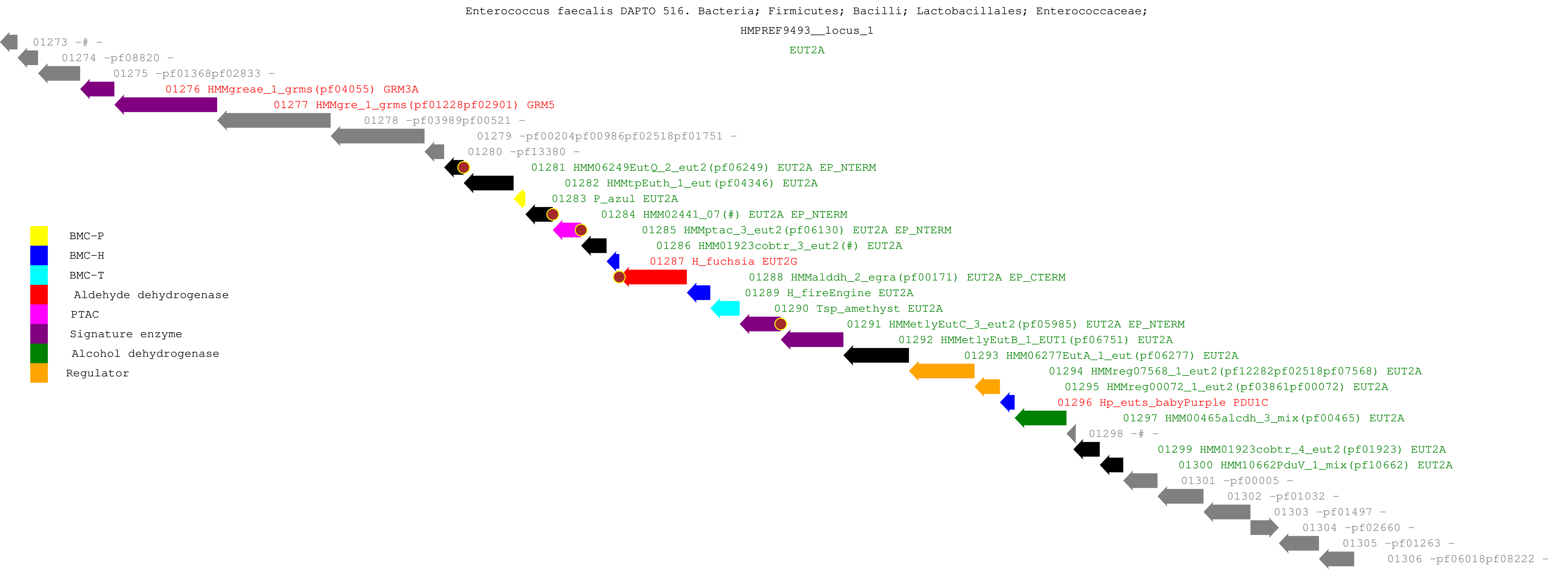
Enterococcus faecalis DAPTO 516. Bacteria; Firmicutes; Bacilli; Lactobacillales; Enterococcaceae;
HMPREF9493__locus_1
EUT2A
01273 -# -
01274 -pf08820 -
01275 -pf01368pf02833 -
01276 HMMgreae_1_grms(pf04055) GRM3A
01277 HMMgre_1_grms(pf01228pf02901) GRM5
01278 -pf03989pf00521 -
01279 -pf00204pf00986pf02518pf01751 -
01280 -pf13380 -
01281 HMM06249EutQ_2_eut2(pf06249) EUT2A EP_NTERM
01282 HMMtpEuth_1_eut(pf04346) EUT2A
01283 P_azul EUT2A
01284 HMM02441_07(#) EUT2A EP_NTERM
01285 HMMptac_3_eut2(pf06130) EUT2A EP_NTERM
01286 HMM01923cobtr_3_eut2(#) EUT2A
01287 H_fuchsia EUT2G
01288 HMMalddh_2_egra(pf00171) EUT2A EP_CTERM
01289 H_fireEngine EUT2A
01290 Tsp_amethyst EUT2A
01291 HMMetlyEutC_3_eut2(pf05985) EUT2A EP_NTERM
01292 HMMetlyEutB_1_EUT1(pf06751) EUT2A
01293 HMM06277EutA_1_eut(pf06277) EUT2A
01294 HMMreg07568_1_eut2(pf12282pf02518pf07568) EUT2A
01295 HMMreg00072_1_eut2(pf03861pf00072) EUT2A
01296 Hp_euts_babyPurple PDU1C
01297 HMM00465alcdh_3_mix(pf00465) EUT2A
01298 -# -
01299 HMM01923cobtr_4_eut2(pf01923) EUT2A
01300 HMM10662PduV_1_mix(pf10662) EUT2A
01301 -pf00005 -
01302 -pf01032 -
01303 -pf01497 -
01304 -pf02660 -
01305 -pf01263 -
01306 -pf06018pf08222 -
BMC-P
BMC-H
BMC-T
Aldehyde dehydrogenase
PTAC
Signature enzyme
Alcohol dehydrogenase
Regulator

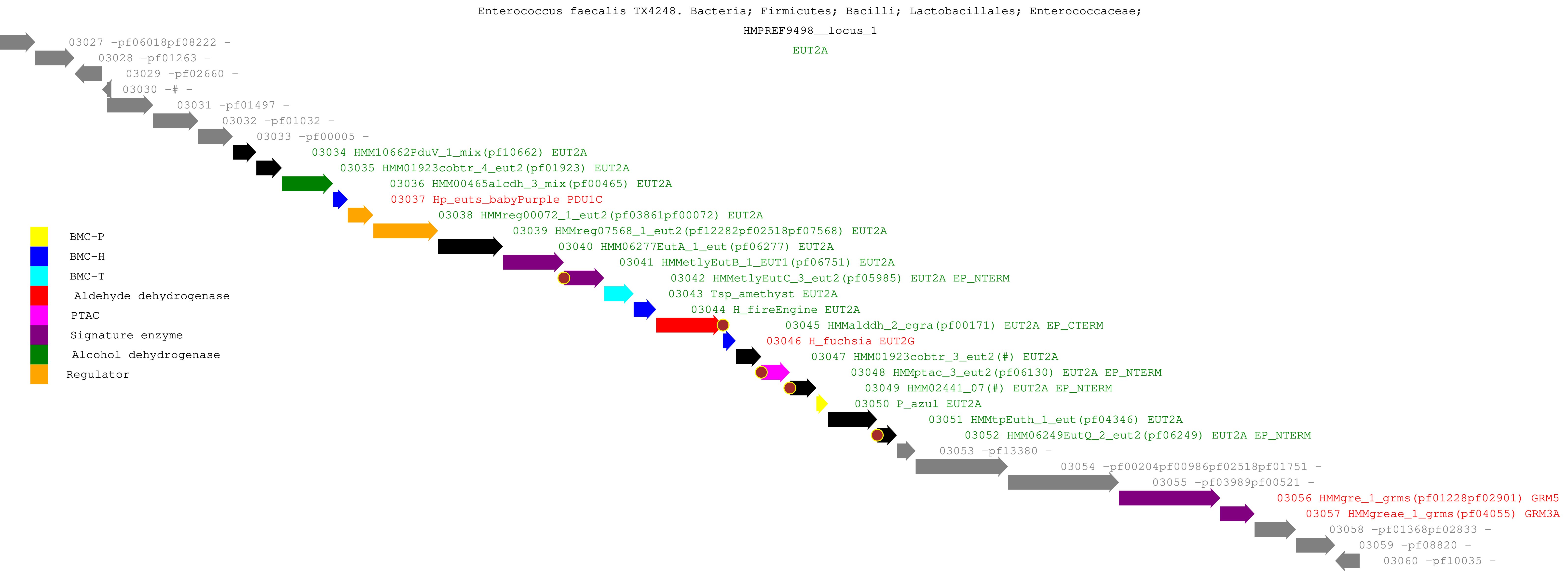
Enterococcus faecalis TX4248. Bacteria; Firmicutes; Bacilli; Lactobacillales; Enterococcaceae;
HMPREF9498__locus_1
EUT2A
03027 -pf06018pf08222 -
03028 -pf01263 -
03029 -pf02660 -
03030 -# -
03031 -pf01497 -
03032 -pf01032 -
03033 -pf00005 -
03034 HMM10662PduV_1_mix(pf10662) EUT2A
03035 HMM01923cobtr_4_eut2(pf01923) EUT2A
03036 HMM00465alcdh_3_mix(pf00465) EUT2A
03037 Hp_euts_babyPurple PDU1C
03038 HMMreg00072_1_eut2(pf03861pf00072) EUT2A
03039 HMMreg07568_1_eut2(pf12282pf02518pf07568) EUT2A
03040 HMM06277EutA_1_eut(pf06277) EUT2A
03041 HMMetlyEutB_1_EUT1(pf06751) EUT2A
03042 HMMetlyEutC_3_eut2(pf05985) EUT2A EP_NTERM
03043 Tsp_amethyst EUT2A
03044 H_fireEngine EUT2A
03045 HMMalddh_2_egra(pf00171) EUT2A EP_CTERM
03046 H_fuchsia EUT2G
03047 HMM01923cobtr_3_eut2(#) EUT2A
03048 HMMptac_3_eut2(pf06130) EUT2A EP_NTERM
03049 HMM02441_07(#) EUT2A EP_NTERM
03050 P_azul EUT2A
03051 HMMtpEuth_1_eut(pf04346) EUT2A
03052 HMM06249EutQ_2_eut2(pf06249) EUT2A EP_NTERM
03053 -pf13380 -
03054 -pf00204pf00986pf02518pf01751 -
03055 -pf03989pf00521 -
03056 HMMgre_1_grms(pf01228pf02901) GRM5
03057 HMMgreae_1_grms(pf04055) GRM3A
03058 -pf01368pf02833 -
03059 -pf08820 -
03060 -pf10035 -
BMC-P
BMC-H
BMC-T
Aldehyde dehydrogenase
PTAC
Signature enzyme
Alcohol dehydrogenase
Regulator

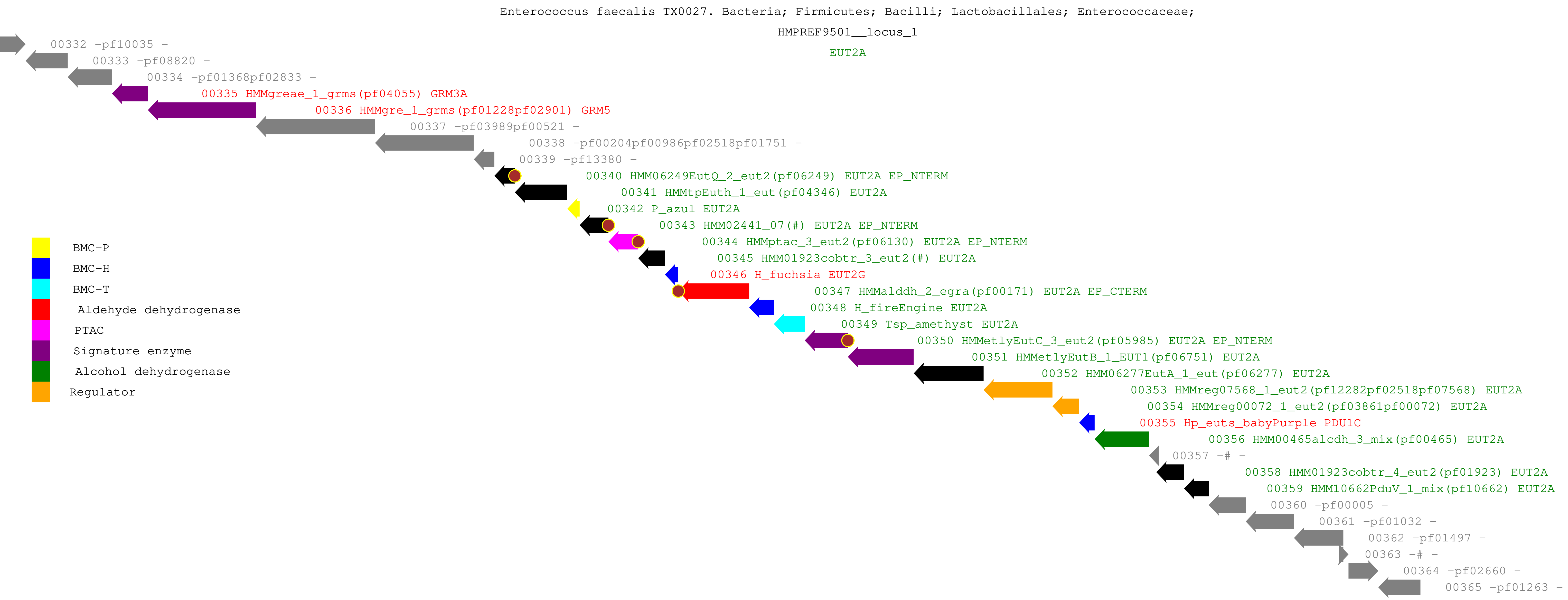

Enterococcus faecalis TX0027. Bacteria; Firmicutes; Bacilli; Lactobacillales; Enterococcaceae;
HMPREF9501__locus_1
EUT2A
00332 -pf10035 -
00333 -pf08820 -
00334 -pf01368pf02833 -
00335 HMMgreae_1_grms(pf04055) GRM3A
00336 HMMgre_1_grms(pf01228pf02901) GRM5
00337 -pf03989pf00521 -
00338 -pf00204pf00986pf02518pf01751 -
00339 -pf13380 -
00340 HMM06249EutQ_2_eut2(pf06249) EUT2A EP_NTERM
00341 HMMtpEuth_1_eut(pf04346) EUT2A
00342 P_azul EUT2A
00343 HMM02441_07(#) EUT2A EP_NTERM
00344 HMMptac_3_eut2(pf06130) EUT2A EP_NTERM
00345 HMM01923cobtr_3_eut2(#) EUT2A
00346 H_fuchsia EUT2G
00347 HMMalddh_2_egra(pf00171) EUT2A EP_CTERM
00348 H_fireEngine EUT2A
00349 Tsp_amethyst EUT2A
00350 HMMetlyEutC_3_eut2(pf05985) EUT2A EP_NTERM
00351 HMMetlyEutB_1_EUT1(pf06751) EUT2A
00352 HMM06277EutA_1_eut(pf06277) EUT2A
00353 HMMreg07568_1_eut2(pf12282pf02518pf07568) EUT2A
00354 HMMreg00072_1_eut2(pf03861pf00072) EUT2A
00355 Hp_euts_babyPurple PDU1C
00356 HMM00465alcdh_3_mix(pf00465) EUT2A
00357 -# -
00358 HMM01923cobtr_4_eut2(pf01923) EUT2A
00359 HMM10662PduV_1_mix(pf10662) EUT2A
00360 -pf00005 -
00361 -pf01032 -
00362 -pf01497 -
00363 -# -
00364 -pf02660 -
00365 -pf01263 -
BMC-P
BMC-H
BMC-T
Aldehyde dehydrogenase
PTAC
Signature enzyme
Alcohol dehydrogenase
Regulator

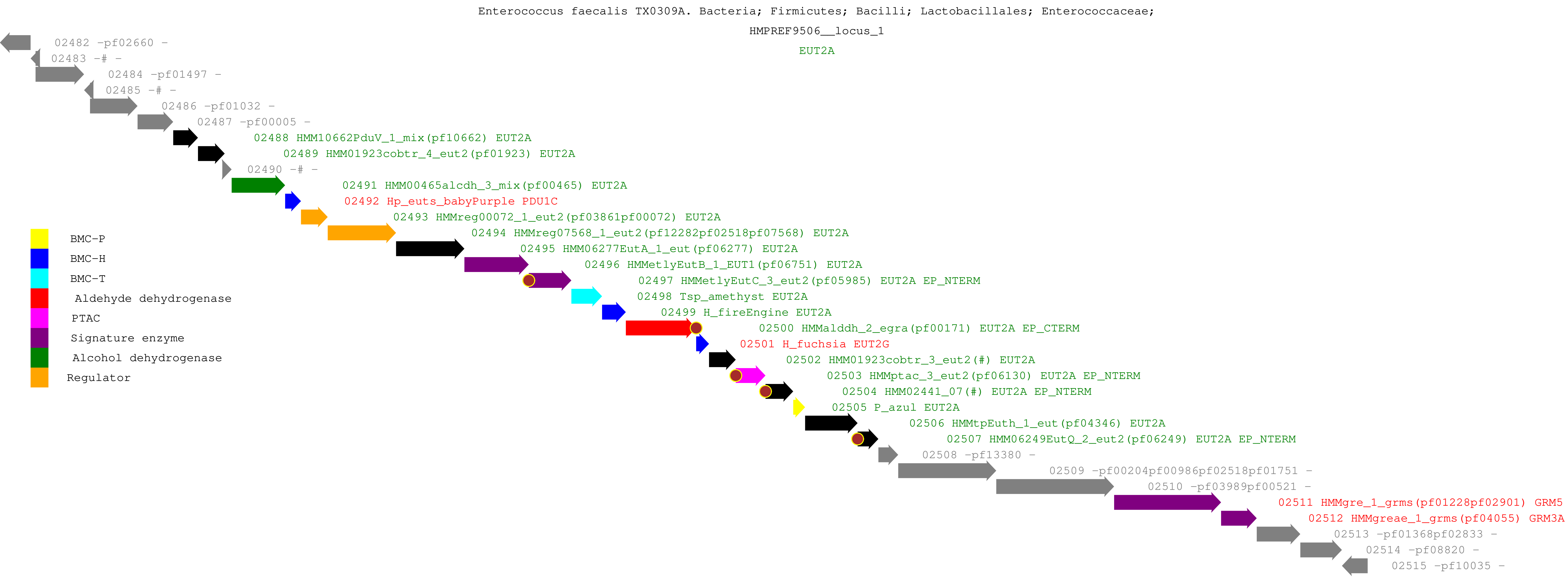

Enterococcus faecalis TX0309A. Bacteria; Firmicutes; Bacilli; Lactobacillales; Enterococcaceae;
HMPREF9506__locus_1
EUT2A
02482 -pf02660 -
02483 -# -
02484 -pf01497 -
02485 -# -
02486 -pf01032 -
02487 -pf00005 -
02488 HMM10662PduV_1_mix(pf10662) EUT2A
02489 HMM01923cobtr_4_eut2(pf01923) EUT2A
02490 -# -
02491 HMM00465alcdh_3_mix(pf00465) EUT2A
02492 Hp_euts_babyPurple PDU1C
02493 HMMreg00072_1_eut2(pf03861pf00072) EUT2A
02494 HMMreg07568_1_eut2(pf12282pf02518pf07568) EUT2A
02495 HMM06277EutA_1_eut(pf06277) EUT2A
02496 HMMetlyEutB_1_EUT1(pf06751) EUT2A
02497 HMMetlyEutC_3_eut2(pf05985) EUT2A EP_NTERM
02498 Tsp_amethyst EUT2A
02499 H_fireEngine EUT2A
02500 HMMaldh_2_egra(pf00171) EUT2A EP_CTERM
02501 H_fuchsia EUT2G
02502 HMM01923cobtr_3_eut2(#) EUT2A
02503 HMMptac_3_eut2(pf06130) EUT2A EP_NTERM
02504 HMM02441_07(#) EUT2A EP_NTERM
02505 P_azul EUT2A
02506 HMMtpEuth_1_eut(pf04346) EUT2A
02507 HMM06249EutQ_2_eut2(pf06249) EUT2A EP_NTERM
02508 -pf13380 -
02509 -pf00204pf00986pf02518pf01751 -
02510 -pf03989pf00521 -
02511 HMMgre_1_grms(pf01228pf02901) GRM5
02512 HMMgreae_1_grms(pf04055) GRM3A
02513 -pf01368pf02833 -
02514 -pf08820 -
02515 -pf10035 -
BMC-P
BMC-H
BMC-T
Aldehyde dehydrogenase
PTAC
Signature enzyme
Alcohol dehydrogenase
Regulator

Enterococcus faecalis TX0309B. Bacteria; Firmicutes; Bacilli; Lactobacillales; Enterococcaceae;

HMPREF9507__locus_1

EUT2A

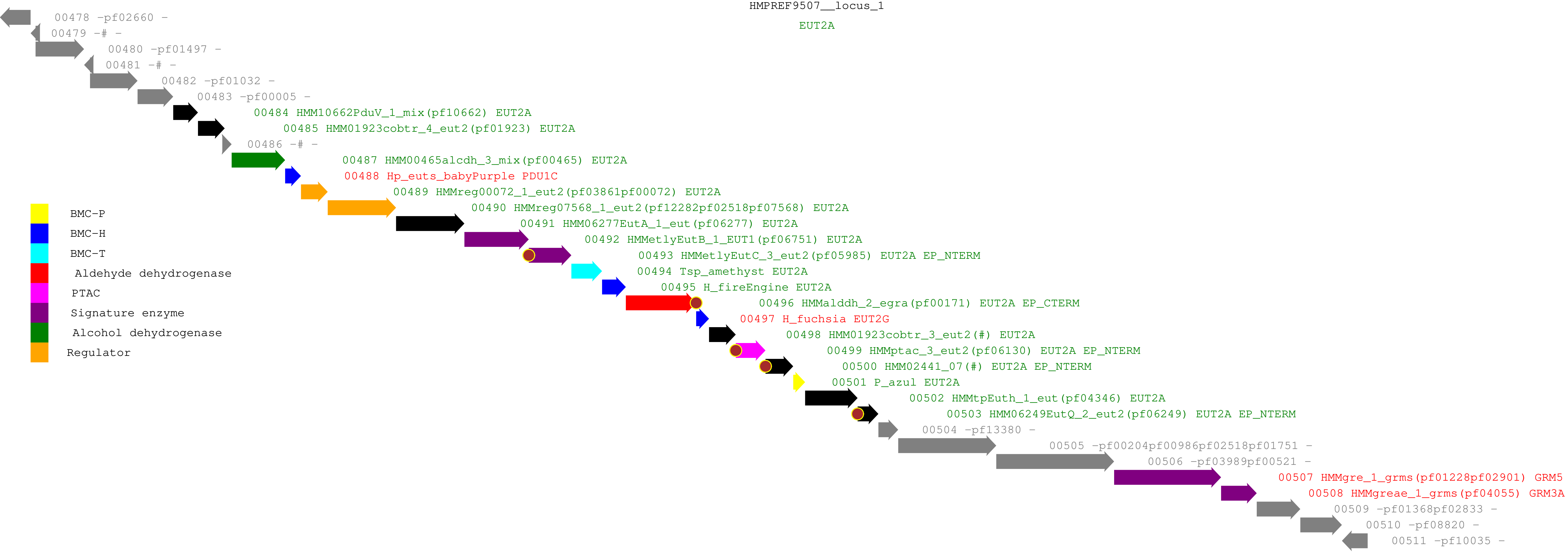

Enterococcus faecalis TX0411. Bacteria; Firmicutes; Bacilli; Lactobacillales; Enterococcaceae;

HMPREF9509__locus_1

EUT2A

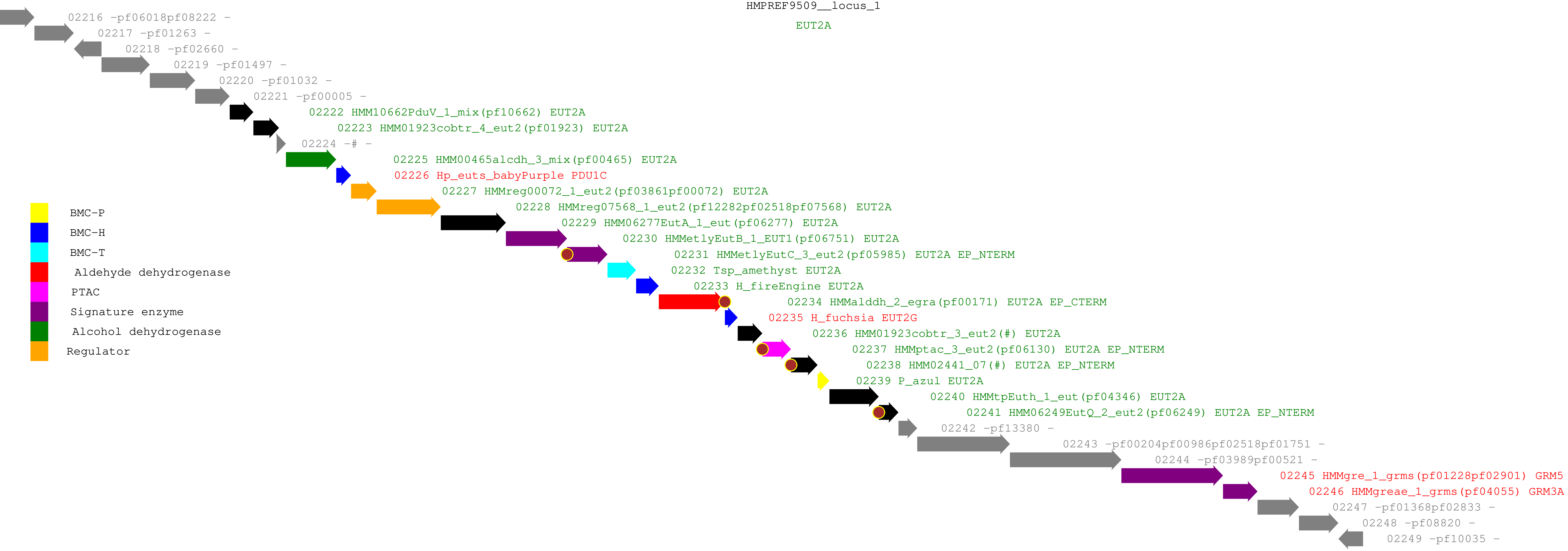

Enterococcus faecalis TX0630. Bacteria; Firmicutes; Bacilli; Lactobacillales; Enterococcaceae;

HMPREF9511__locus_1

EUT2A

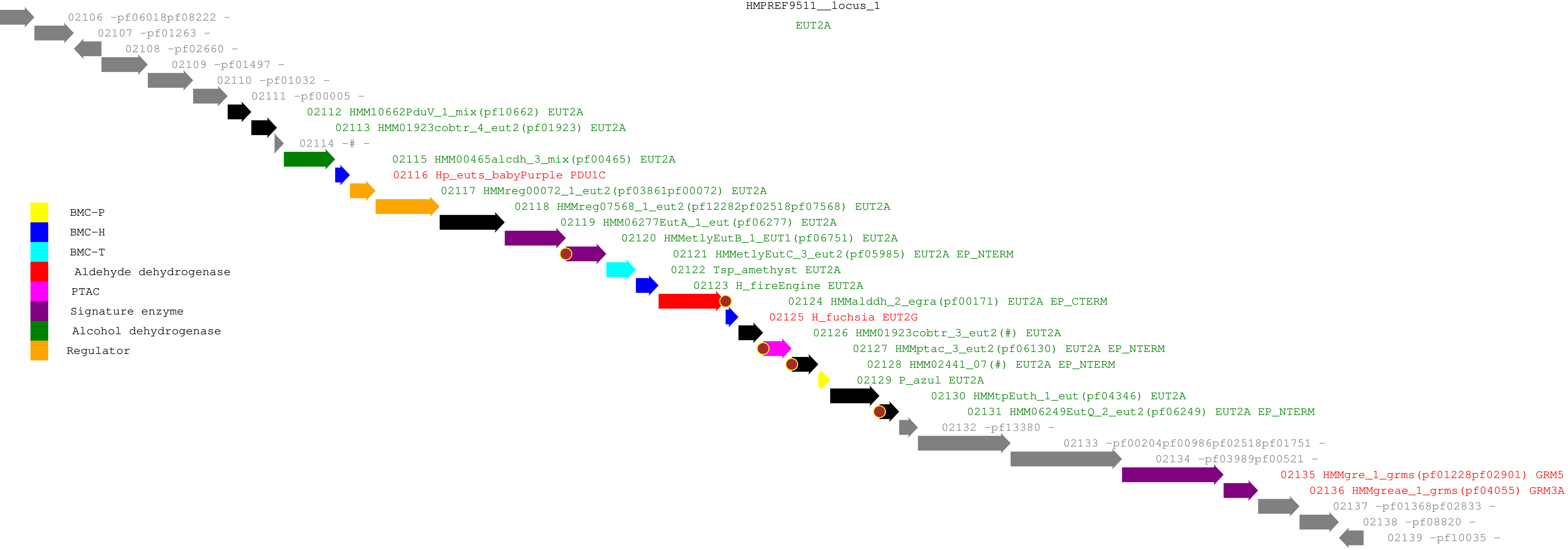

Enterococcus faecalis EnGen0311. Bacteria; Firmicutes; Bacilli; Lactobacillales; Enterococcaceae;

HMPREF9512__locus_1

EUT2A

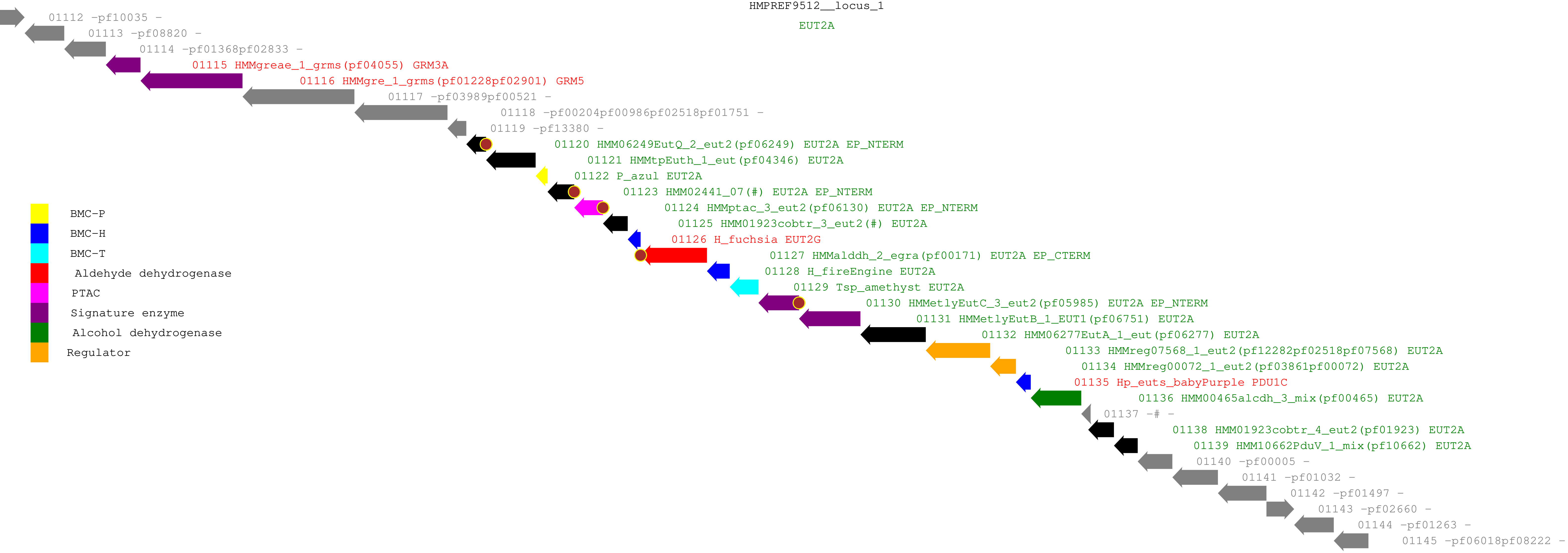

Enterococcus faecalis TX0645. Bacteria; Firmicutes; Bacilli; Lactobacillales; Enterococcaceae;

HMPREF9513__locus_1

EUT2A

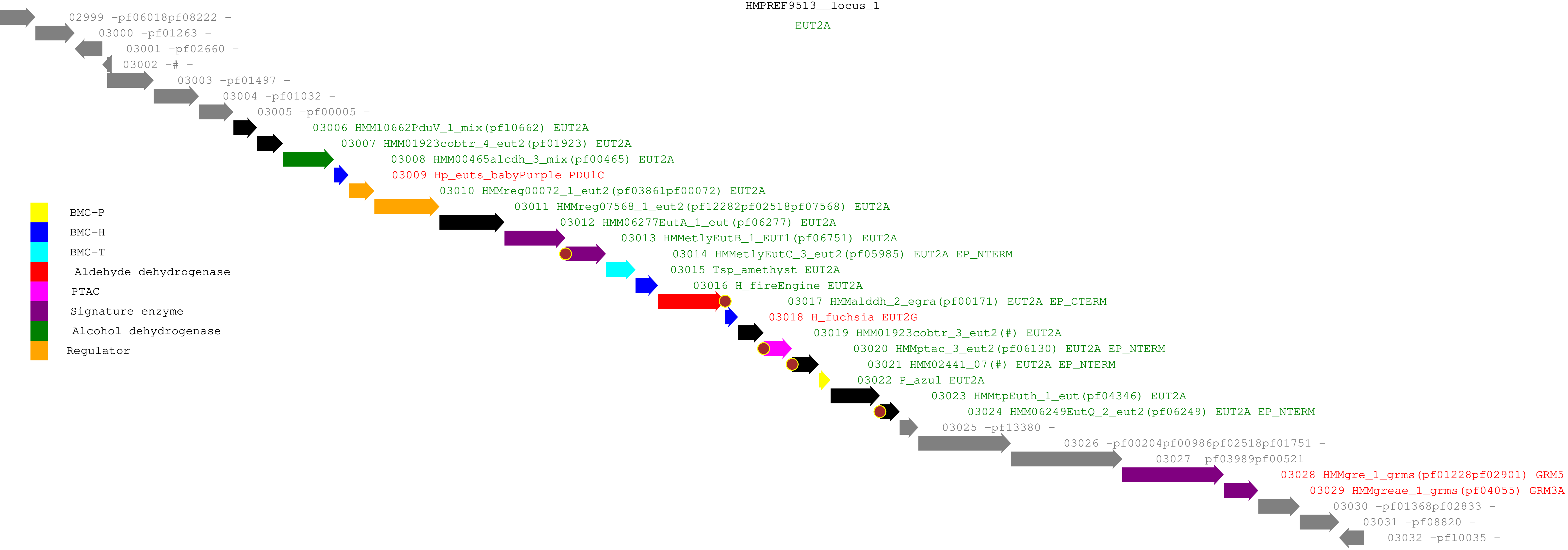

Clostridium celerecrescens. Bacteria; Firmicutes; Clostridia; Clostridiales; Lachnospiraceae.

IO98__locus_2

EUT2A

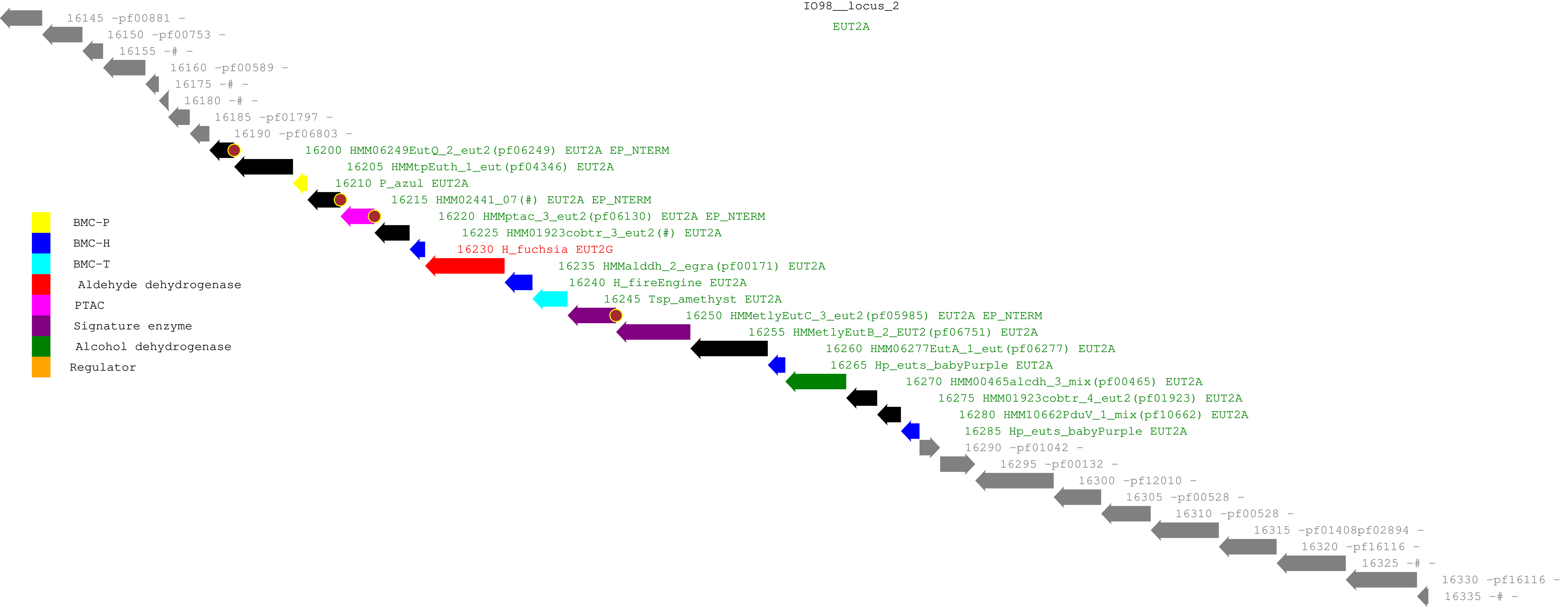

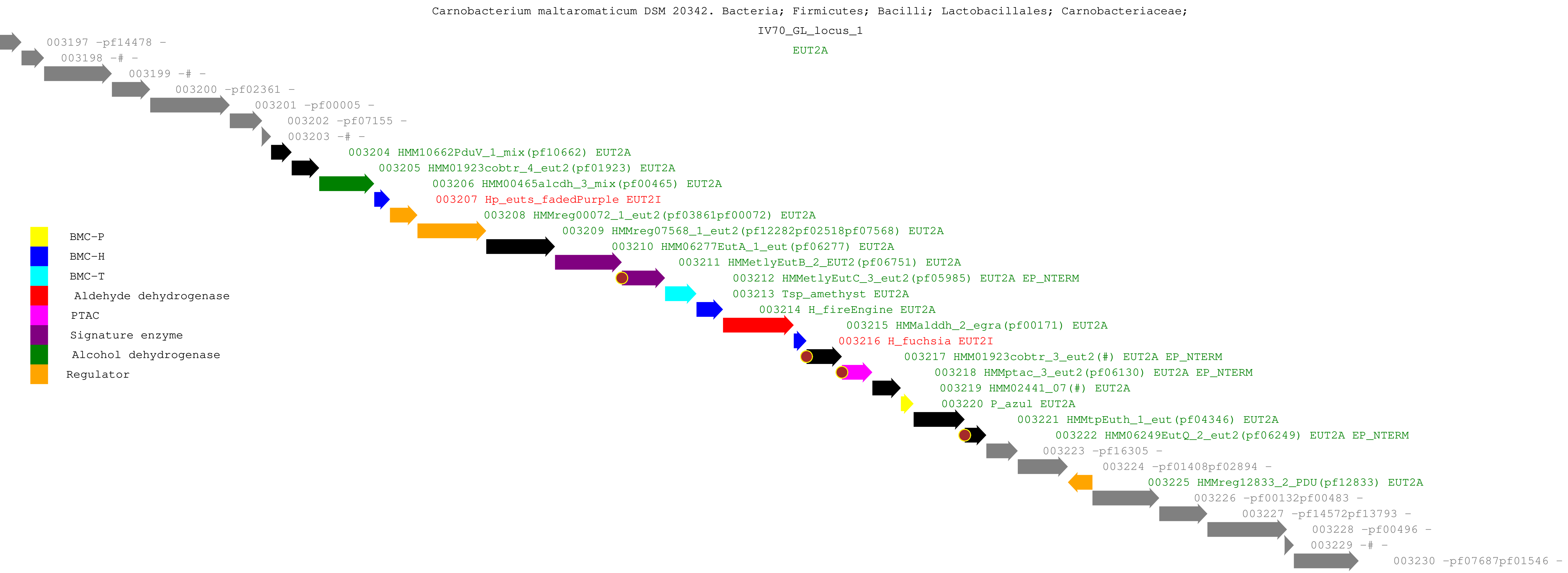
Carnobacterium maltaromaticum DSM 20342. Bacteria; Firmicutes; Bacilli; Lactobacillales; Carnobacteriaceae;
IV70_GL_locus_1
EUT2A
003197 -pf14478 -
003198 -# -
003199 -# -
003200 -pf02361 -
003201 -pf00005 -
003202 -pf07155 -
003203 -# -
003204 HMM10662PduV_1_mix(pf10662) EUT2A
003205 HMM01923cobtr_4_eut2(pf01923) EUT2A
003206 HMM00465alcdh_3_mix(pf00465) EUT2A
003207 Hp_euts_fadedPurple EUT2I
003208 HMMreg00072_1_eut2(pf03861pf00072) EUT2A
003209 HMMreg07568_1_eut2(pf12282pf02518pf07568) EUT2A
003210 HMM06277EutA_1_eut(pf06277) EUT2A
003211 HMMetlyEutB_2_EUT2(pf06751) EUT2A
003212 HMMetlyEutC_3_eut2(pf05985) EUT2A EP_NTERM
003213 Tsp_amethyst EUT2A
003214 H_fireEngine EUT2A
003215 HMMalddh_2_egra(pf00171) EUT2A
003216 H_fuchsia EUT2I
003217 HMM01923cobtr_3_eut2(#) EUT2A EP_NTERM
003218 HMMptac_3_eut2(pf06130) EUT2A EP_NTERM
003219 HMM02441_07(#) EUT2A
003220 P_azul EUT2A
003221 HMMtpEuth_1_eut(pf04346) EUT2A
003222 HMM06249EutQ_2_eut2(pf06249) EUT2A EP_NTERM
003223 -pf16305 -
003224 -pf01408pf02894 -
003225 HMMreg12833_2_PDU(pf12833) EUT2A
003226 -pf00132pf00483 -
003227 -pf14572pf13793 -
003228 -pf00496 -
003229 -# -
003230 -pf07687pf01546 -
BMC-P
BMC-H
BMC-T
Aldehyde dehydrogenase
PTAC
Signature enzyme
Alcohol dehydrogenase
Regulator

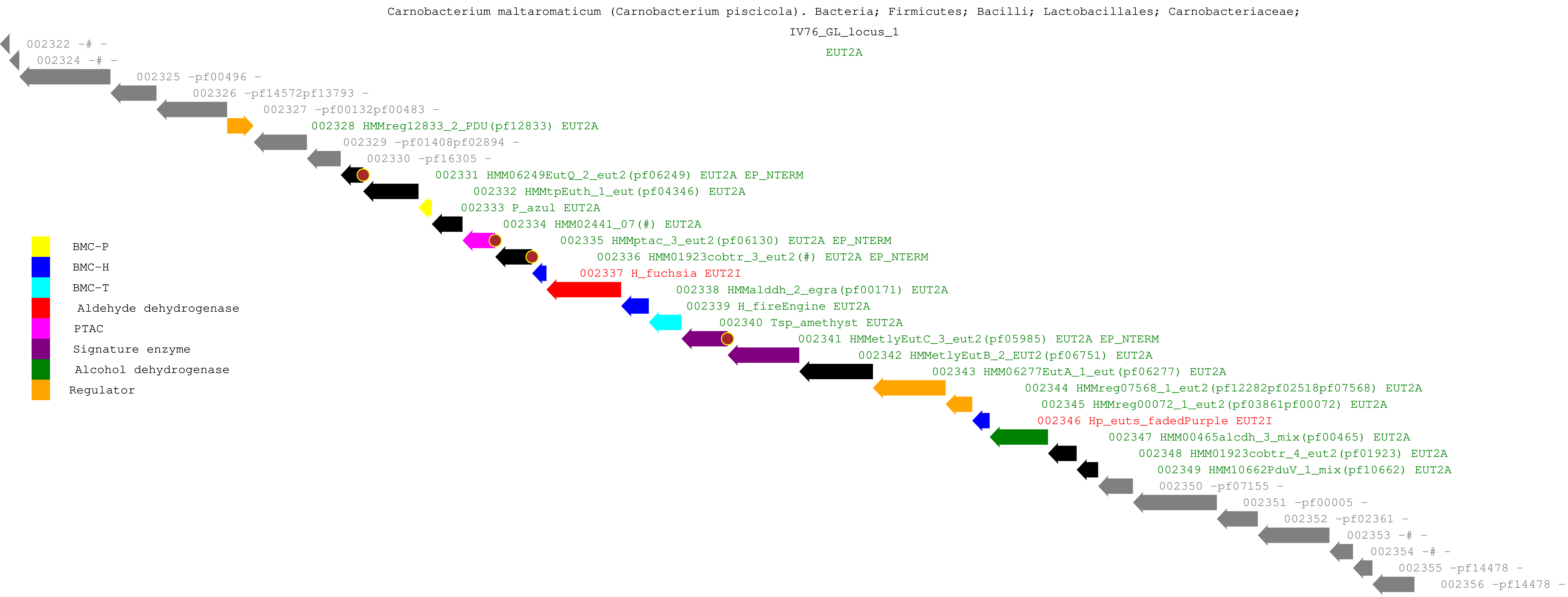

Carnobacterium maltaromaticum (Carnobacterium piscicola). Bacteria; Firmicutes; Bacilli; Lactobacillales; Carnobacteriaceae;
IV76_GL_locus_1
EUT2A
002322 -# -
002324 -# -
002325 -pf00496 -
002326 -pf14572pf13793 -
002327 -pf00132pf00483 -
002328 HMMreg12833_2_PDU(pf12833) EUT2A
002329 -pf01408pf02894 -
002330 -pf16305 -
002331 HMM06249EutQ_2_eut2(pf06249) EUT2A EP_NTERM
002332 HMMtpEuth_1_eut(pf04346) EUT2A
002333 P_azul EUT2A
002334 HMM02441_07(#) EUT2A
002335 HMMptac_3_eut2(pf06130) EUT2A EP_NTERM
002336 HMM01923cobtr_3_eut2(#) EUT2A EP_NTERM
002337 H_fuchsia EUT2I
002338 HMMalddh_2_egra(pf00171) EUT2A
002339 H_fireEngine EUT2A
002340 Tsp_amethyst EUT2A
002341 HMMetlyEutC_3_eut2(pf05985) EUT2A EP_NTERM
002342 HMMetlyEutB_2_EUT2(pf06751) EUT2A
002343 HMM06277EutA_1_eut(pf06277) EUT2A
002344 HMMreg07568_1_eut2(pf12282pf02518pf07568) EUT2A
002345 HMMreg00072_1_eut2(pf03861pf00072) EUT2A
002346 Hp_euts_fadedPurple EUT2I
002347 HMM00465alcdh_3_mix(pf00465) EUT2A
002348 HMM01923cobtr_4_eut2(pf01923) EUT2A
002349 HMM10662PduV_1_mix(pf10662) EUT2A
002350 -pf07155 -
002351 -pf00005 -
002352 -pf02361 -
002353 -# -
002354 -# -
002355 -pf14478 -
002356 -pf14478 -
BMC-P
BMC-H
BMC-T
Aldehyde dehydrogenase
PTAC
Signature enzyme
Alcohol dehydrogenase
Regulator

Lactobacillus mellis. Bacteria; Firmicutes; Bacilli; Lactobacillales; Lactobacillaceae;

JG29__locus_1

EUT2A

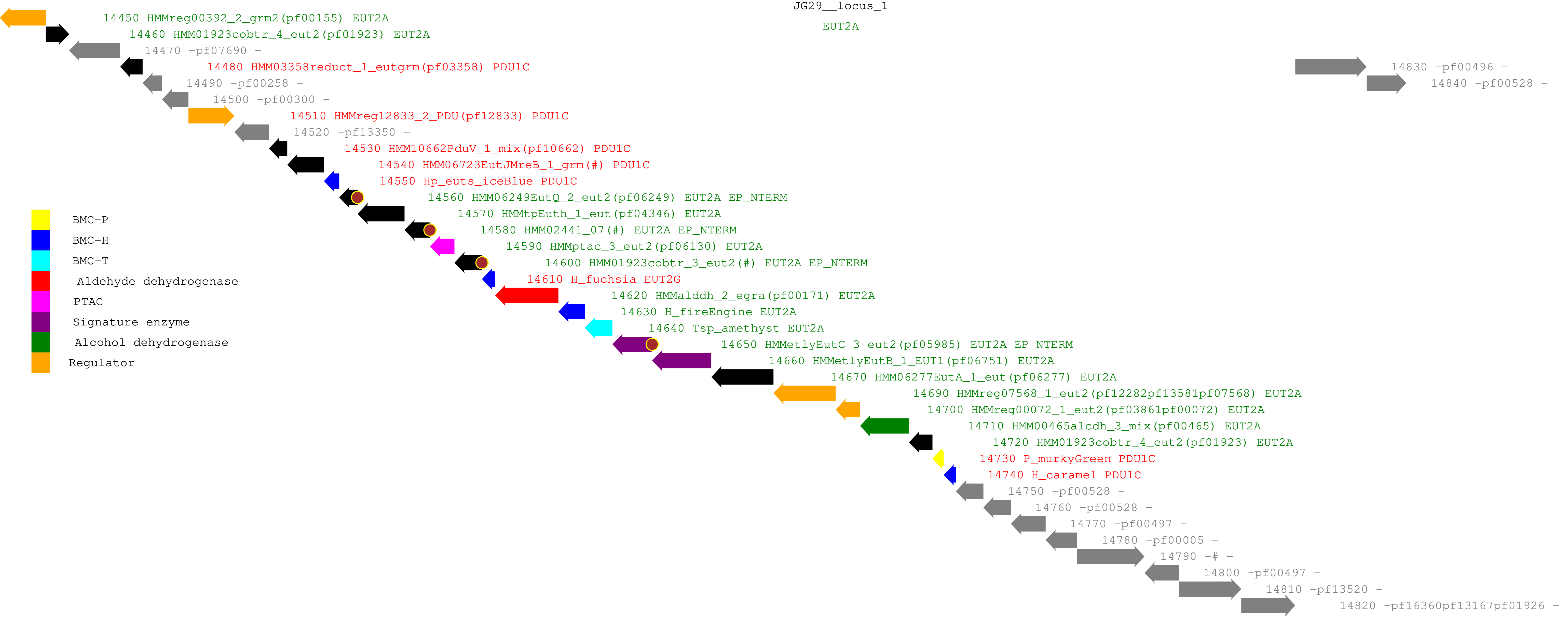

Clostridium sp. ASBs410. Bacteria; Firmicutes; Clostridia; Clostridiales; Clostridiaceae;

K413DRAFT__locus_2

EUT2A

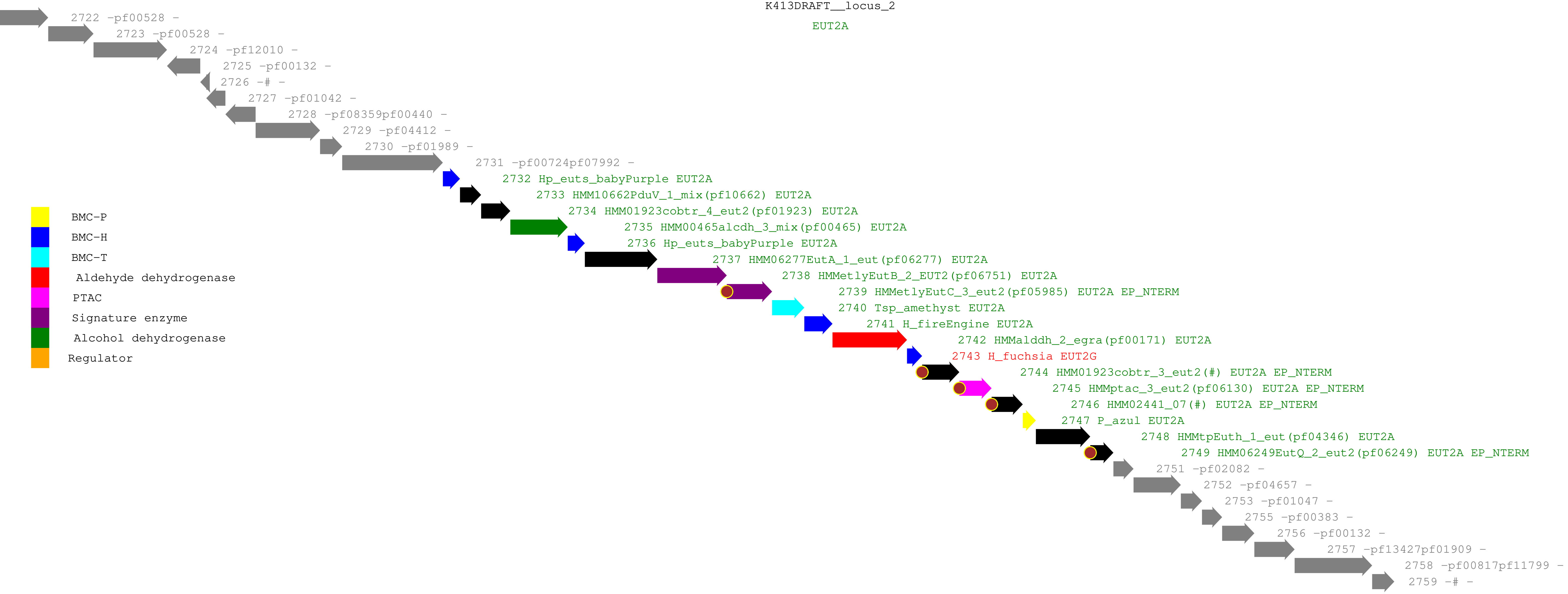

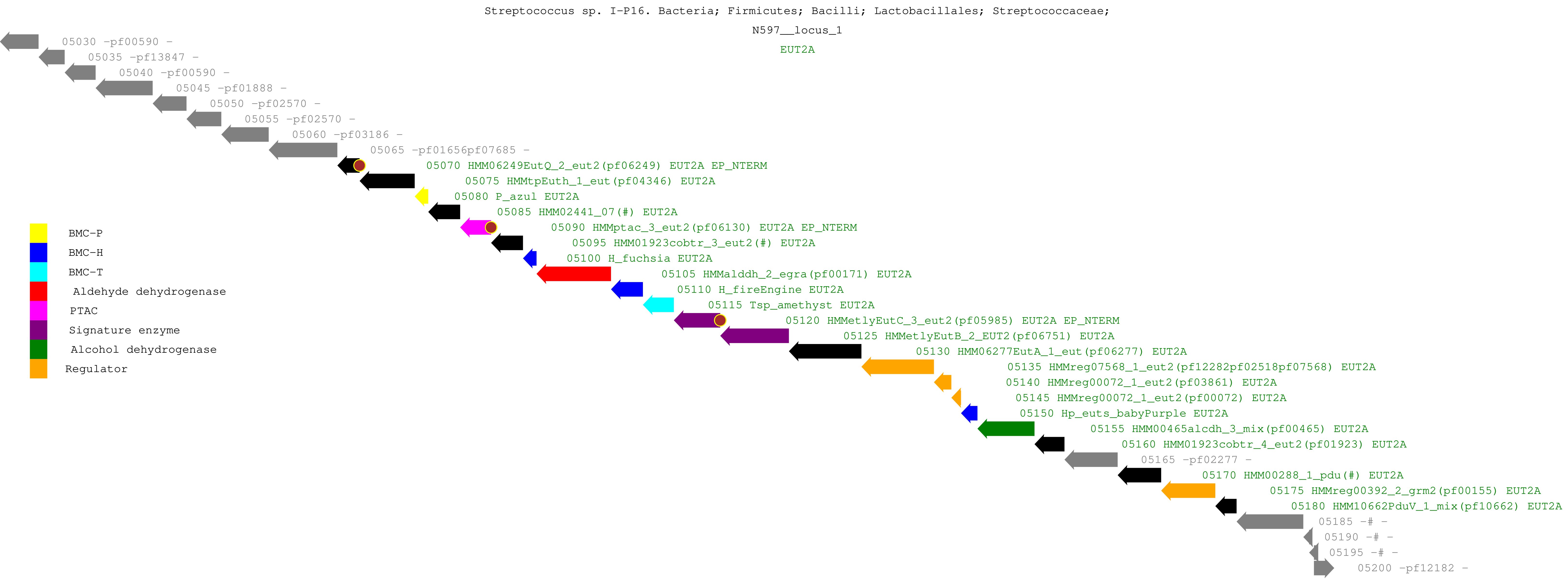
Streptococcus sp. I-P16. Bacteria; Firmicutes; Bacilli; Lactobacillales; Streptococcaceae;
N597__locus_1
EUT2A
05030 -pf00590 -
05035 -pf13847 -
05040 -pf00590 -
05045 -pf01888 -
05050 -pf02570 -
05055 -pf02570 -
05060 -pf03186 -
05065 -pf01656pf07685 -
05070 HMM06249EutQ_2_eut2(pf06249) EUT2A EP_NTERM
05075 HMMtpEuth_1_eut(pf04346) EUT2A
05080 P_azul EUT2A
05085 HMM02441_07(#) EUT2A
05090 HMMptac_3_eut2(pf06130) EUT2A EP_NTERM
05095 HMM01923cobtr_3_eut2(#) EUT2A
05100 H_fuchsia EUT2A
05105 HMMaldh_2_egra(pf00171) EUT2A
05110 H_fireEngine EUT2A
05115 Tsp_amethyst EUT2A
05120 HMMetlyEutC_3_eut2(pf05985) EUT2A EP_NTERM
05125 HMMetlyEutB_2_EUT2(pf06751) EUT2A
05130 HMM06277EutA_1_eut(pf06277) EUT2A
05135 HMMreg07568_1_eut2(pf12282pf02518pf07568) EUT2A
05140 HMMreg00072_1_eut2(pf03861) EUT2A
05145 HMMreg00072_1_eut2(pf00072) EUT2A
05150 Hp_euts_babyPurple EUT2A
05155 HMM00465alcdh_3_mix(pf00465) EUT2A
05160 HMM01923cobtr_4_eut2(pf01923) EUT2A
05165 -pf02277 -
05170 HMM00288_1_pdu(#) EUT2A
05175 HMMreg00392_2_grm2(pf00155) EUT2A
05180 HMM10662PduV_1_mix(pf10662) EUT2A
05185 -# -
05190 -# -
05195 -# -
05200 -pf12182 -
BMC-P
BMC-H
BMC-T
Aldehyde dehydrogenase
PTAC
Signature enzyme
Alcohol dehydrogenase
Regulator

Streptococcus sanguinis. Bacteria; Firmicutes; Bacilli; Lactobacillales; Streptococcaceae;

NCTC11085__locus_1

EUT2A

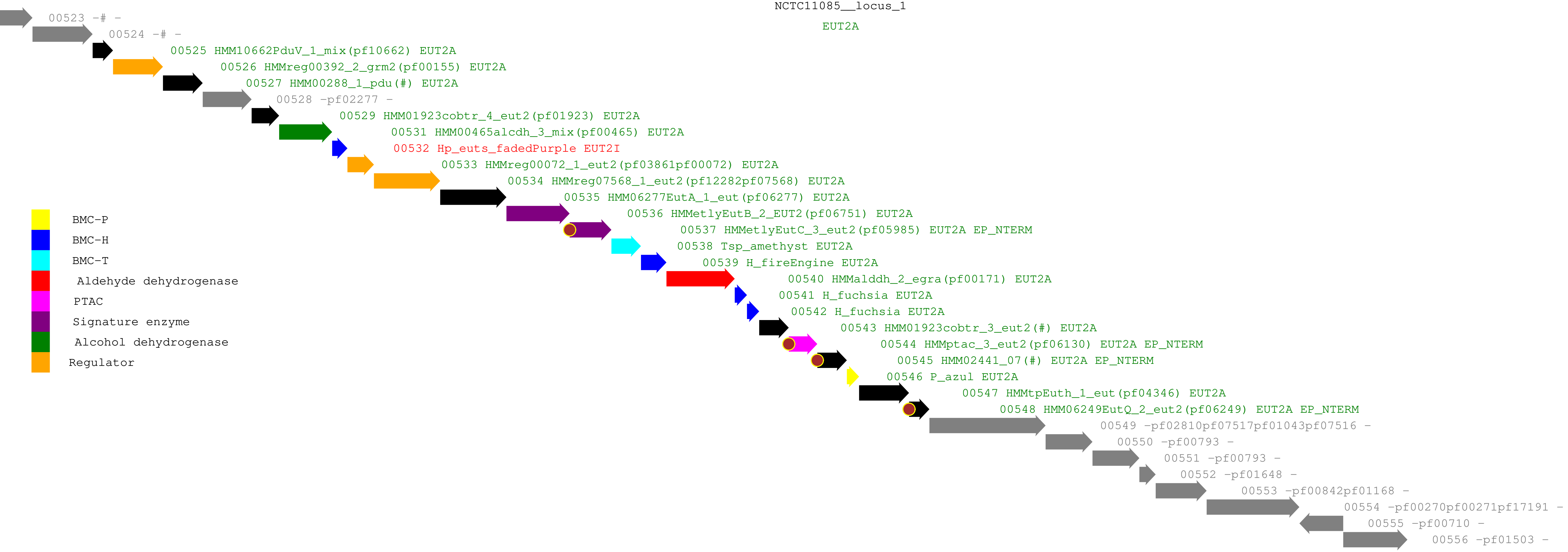

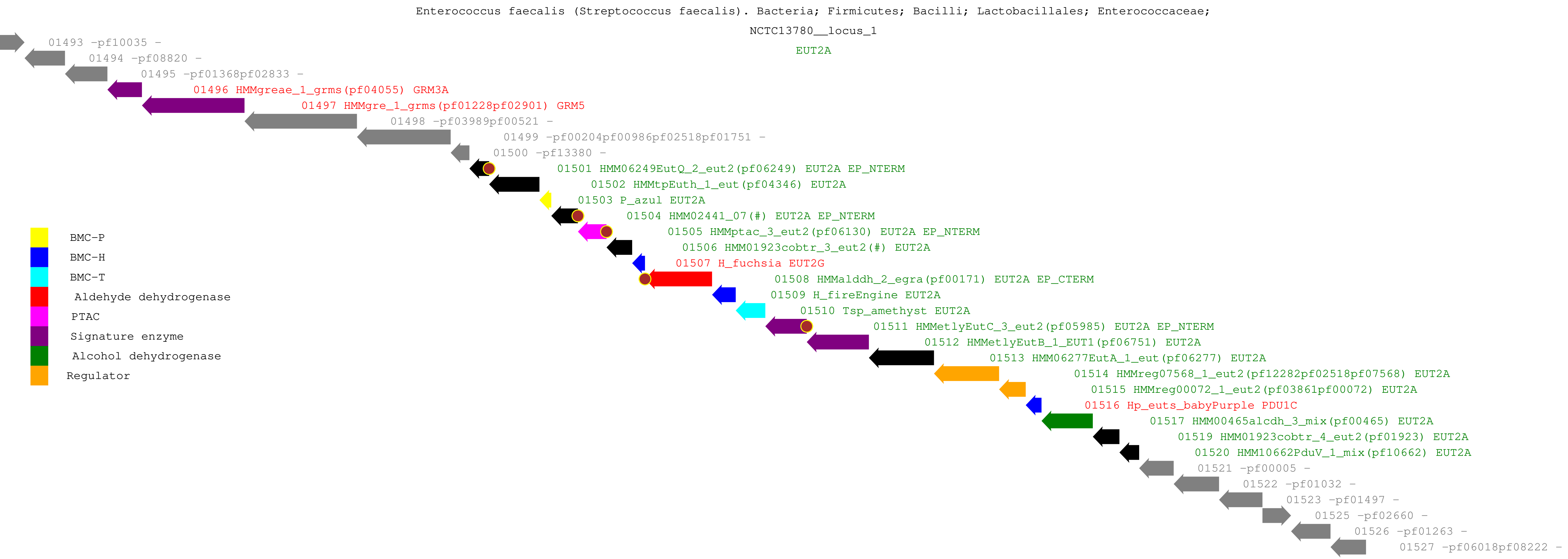

Enterococcus faecalis (Streptococcus faecalis). Bacteria; Firmicutes; Bacilli; Lactobacillales; Enterococcaceae;
NCTC13780__locus_1
EUT2A
01493 -pf10035 -
01494 -pf08820 -
01495 -pf01368pf02833 -
01496 HMMgreae_1_grms(pf04055) GRM3A
01497 HMMgre_1_grms(pf01228pf02901) GRM5
01498 -pf03989pf00521 -
01499 -pf00204pf00986pf02518pf01751 -
01500 -pf13380 -
01501 HMM06249EutQ_2_eut2(pf06249) EUT2A EP_NTERM
01502 HMMtpEuth_1_eut(pf04346) EUT2A
01503 P_azul EUT2A
01504 HMM02441_07(#) EUT2A EP_NTERM
01505 HMMptac_3_eut2(pf06130) EUT2A EP_NTERM
01506 HMM01923cobtr_3_eut2(#) EUT2A
01507 H_fuchsia EUT2G
01508 HMMalddh_2_egra(pf00171) EUT2A EP_CTERM
01509 H_fireEngine EUT2A
01510 Tsp_amethyst EUT2A
01511 HMMetlyEutC_3_eut2(pf05985) EUT2A EP_NTERM
01512 HMMetlyEutB_1_EUT1(pf06751) EUT2A
01513 HMM06277EutA_1_eut(pf06277) EUT2A
01514 HMMreg07568_1_eut2(pf12282pf02518pf07568) EUT2A
01515 HMMreg00072_1_eut2(pf03861pf00072) EUT2A
01516 Hp_euts_babyPurple PDU1C
01517 HMM00465alcdh_3_mix(pf00465) EUT2A
01519 HMM01923cobtr_4_eut2(pf01923) EUT2A
01520 HMM10662PduV_1_mix(pf10662) EUT2A
01521 -pf00005 -
01522 -pf01032 -
01523 -pf01497 -
01525 -pf02660 -
01526 -pf01263 -
01527 -pf06018pf08222 -
BMC-P
BMC-H
BMC-T
Aldehyde dehydrogenase
PTAC
Signature enzyme
Alcohol dehydrogenase
Regulator

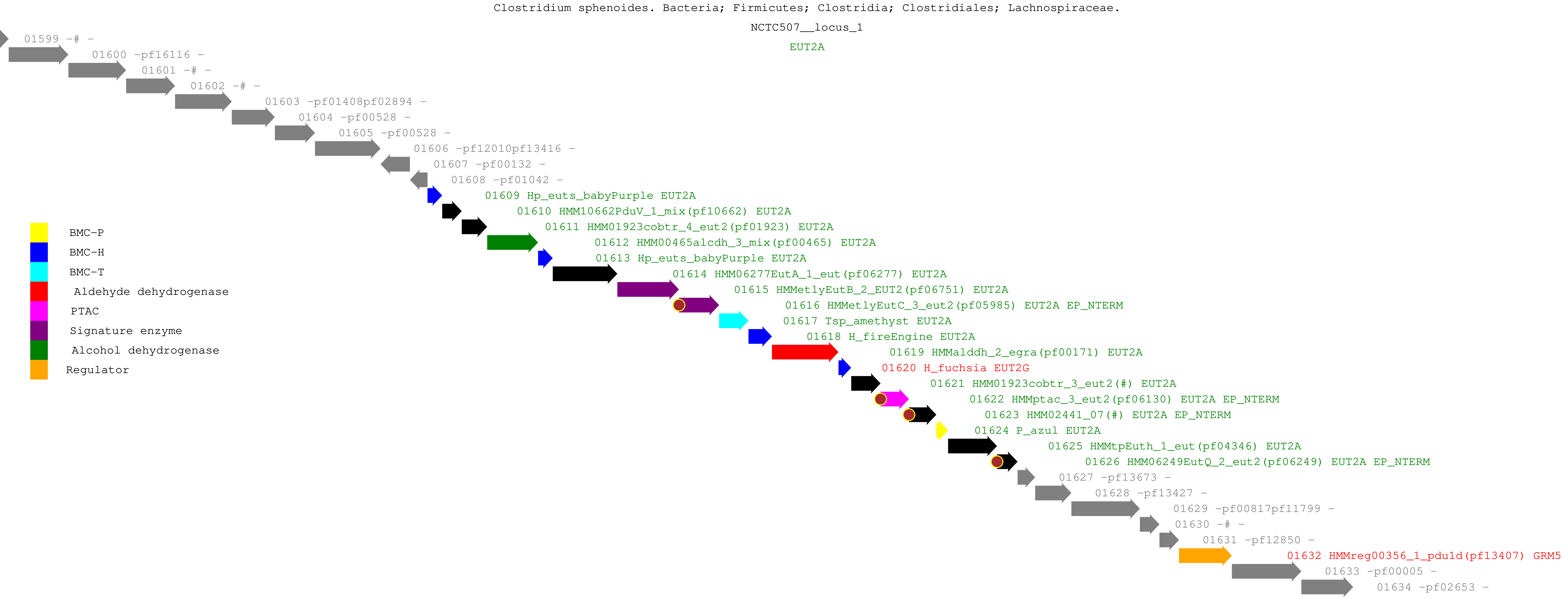
Clostridium sphenoides. Bacteria; Firmicutes; Clostridia; Clostridiales; Lachnospiraceae.
NCTC507__locus_1
EUT2A
01599 -# -
01600 -pf16116 -
01601 -# -
01602 -# -
01603 -pf01408pf02894 -
01604 -pf00528 -
01605 -pf00528 -
01606 -pf12010pf13416 -
01607 -pf00132 -
01608 -pf01042 -
01609 Hp_euts_babyPurple EUT2A
01610 HMM10662PduV_1_mix(pf10662) EUT2A
01611 HMM01923cobtr_4_eut2(pf01923) EUT2A
01612 HMM00465alcdh_3_mix(pf00465) EUT2A
01613 Hp_euts_babyPurple EUT2A
01614 HMM06277EutA_1_eut(pf06277) EUT2A
01615 HMMetlyEutB_2_EUT2(pf06751) EUT2A
01616 HMMetlyEutC_3_eut2(pf05985) EUT2A EP_NTERM
01617 Tsp_amethyst EUT2A
01618 H_fireEngine EUT2A
01619 HMMalddh_2_egra(pf00171) EUT2A
01620 H_fuchsia EUT2G
01621 HMM01923cobtr_3_eut2(#) EUT2A
01622 HMMptac_3_eut2(pf06130) EUT2A EP_NTERM
01623 HMM02441_07(#) EUT2A EP_NTERM
01624 P_azul EUT2A
01625 HMMtpEuth_1_eut(pf04346) EUT2A
01626 HMM06249EutQ_2_eut2(pf06249) EUT2A EP_NTERM
01627 -pf13673 -
01628 -pf13427 -
01629 -pf00817pf11799 -
01630 -# -
01631 -pf12850 -
01632 HMMreg00356_1_pdu1d(pf13407) GRM5
01633 -pf00005 -
01634 -pf02653 -
BMC-P
BMC-H
BMC-T
Aldehyde dehydrogenase
PTAC
Signature enzyme
Alcohol dehydrogenase
Regulator

Enterococcus avium ATCC 14025. Bacteria; Firmicutes; Bacilli; Lactobacillales; Enterococcaceae;

OMU__locus_2

EUT2A

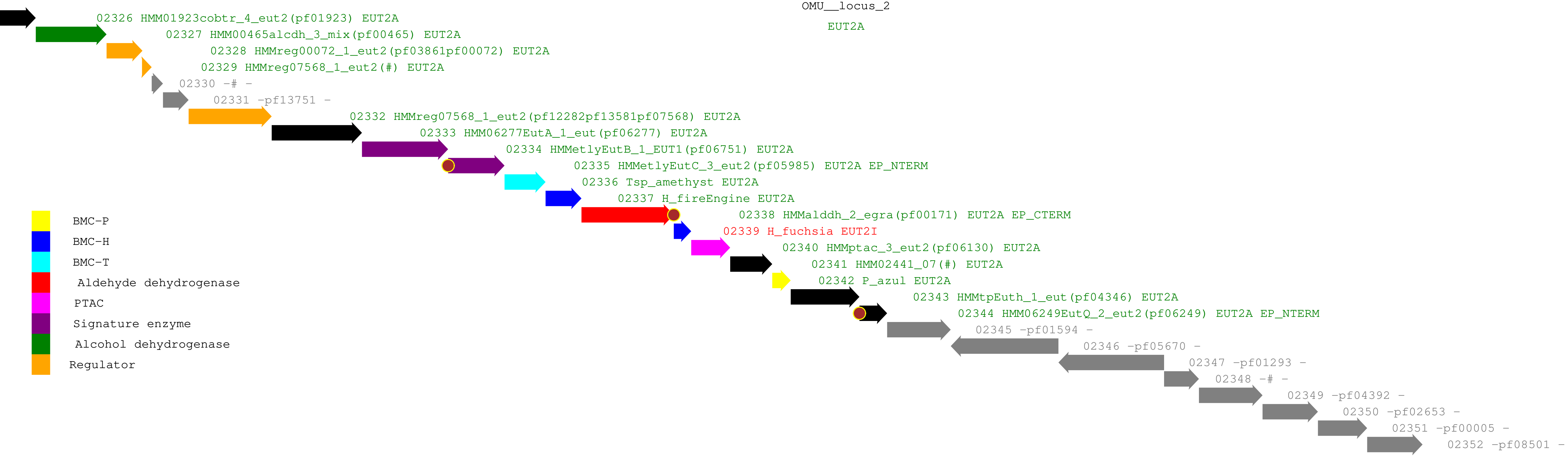

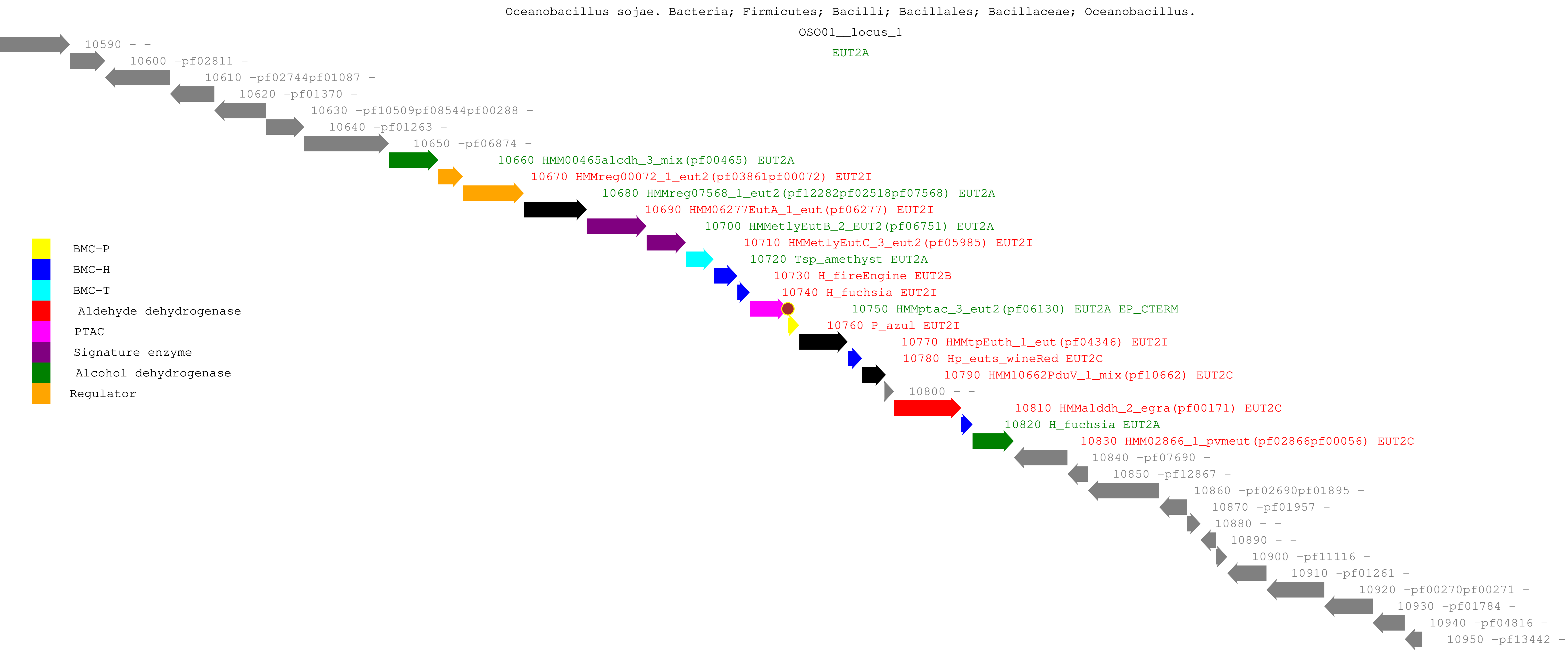

Oceanobacillus sojae. Bacteria; Firmicutes; Bacilli; Bacillales; Bacillaceae; Oceanobacillus.
OSO01__locus_1
EUT2A
10590 - -
10600 -pf02811 -
10610 -pf02744pf01087 -
10620 -pf01370 -
10630 -pf10509pf08544pf00288 -
10640 -pf01263 -
10650 -pf06874 -
10660 HMM00465alcdh_3_mix(pf00465) EUT2A
10670 HMMreg00072_1_eut2(pf03861pf00072) EUT2I
10680 HMMreg07568_1_eut2(pf12282pf02518pf07568) EUT2A
10690 HMM06277EutA_1_eut(pf06277) EUT2I
10700 HMMetlyEutB_2_EUT2(pf06751) EUT2A
10710 HMMetlyEutC_3_eut2(pf05985) EUT2I
10720 Tsp_amethyst EUT2A
10730 H_fireEngine EUT2B
10740 H_fuchsia EUT2I
10750 HMMptac_3_eut2(pf06130) EUT2A EP_CTERM
10760 P_azul EUT2I
10770 HMMtpEuth_1_eut(pf04346) EUT2I
10780 Hp_euts_wineRed EUT2C
10790 HMM10662PduV_1_mix(pf10662) EUT2C
10800 - -
10810 HMMalddh_2_egra(pf00171) EUT2C
10820 H_fuchsia EUT2A
10830 HMM02866_1_pvmeut(pf02866pf00056) EUT2C
10840 -pf07690 -
10850 -pf12867 -
10860 -pf02690pf01895 -
10870 -pf01957 -
10880 - -
10890 - -
10900 -pf11116 -
10910 -pf01261 -
10920 -pf00270pf00271 -
10930 -pf01784 -
10940 -pf04816 -
10950 -pf13442 -
BMC-P
BMC-H
BMC-T
Aldehyde dehydrogenase
PTAC
Signature enzyme
Alcohol dehydrogenase
Regulator

Enterococcus faecalis EnGen0426. Bacteria; Firmicutes; Bacilli; Lactobacillales; Enterococcaceae;

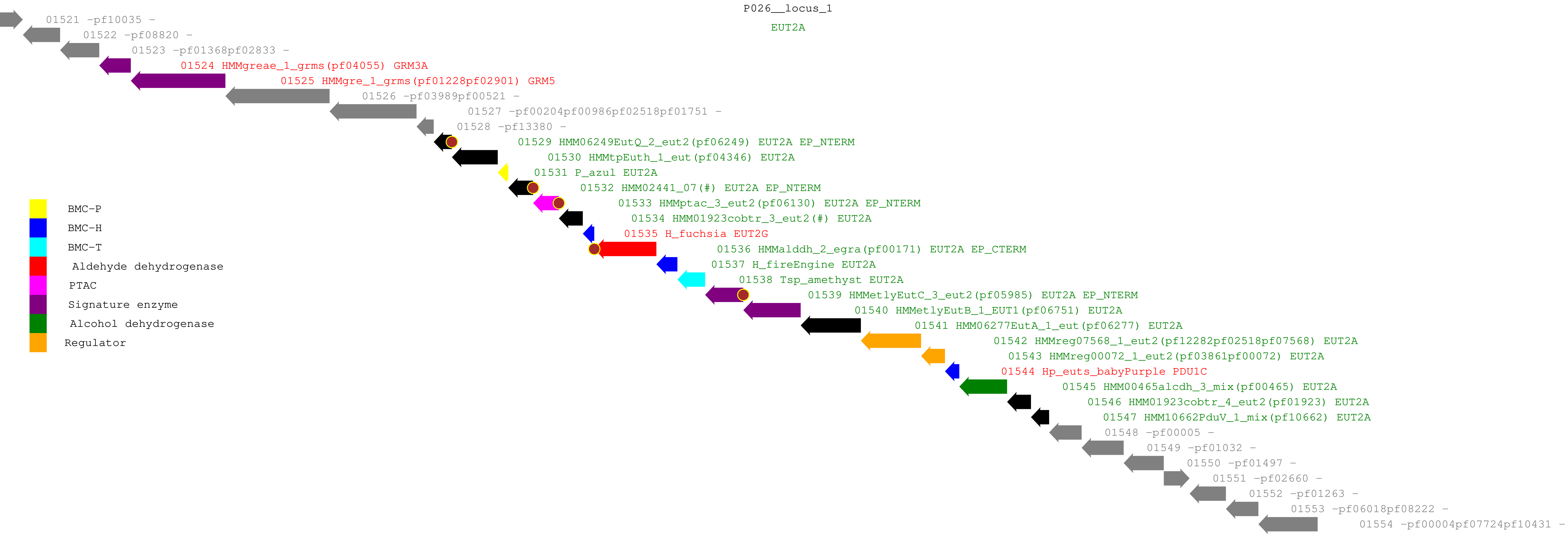

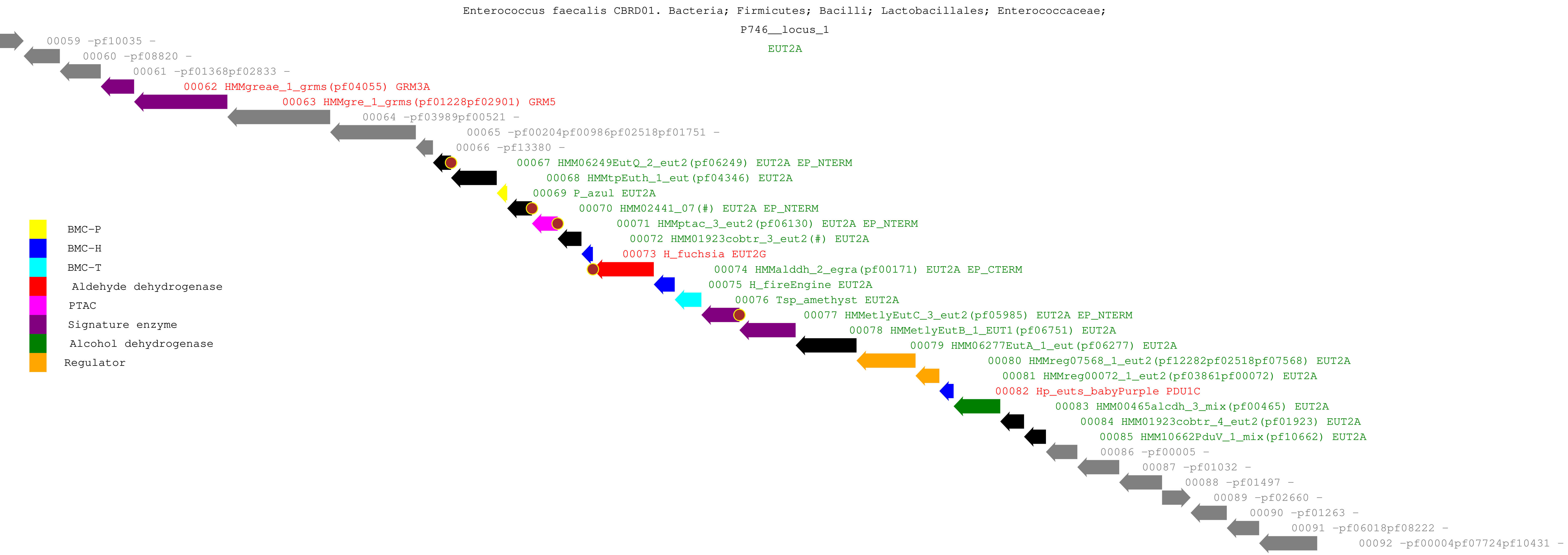
Enterococcus faecalis CBRD01. Bacteria; Firmicutes; Bacilli; Lactobacillales; Enterococcaceae;
P746__locus_1
EUT2A
00059 -pf10035 -
00060 -pf08820 -
00061 -pf01368pf02833 -
00062 HMMgreae_1_grms(pf04055) GRM3A
00063 HMMgre_1_grms(pf01228pf02901) GRM5
00064 -pf03989pf00521 -
00065 -pf00204pf00986pf02518pf01751 -
00066 -pf13380 -
00067 HMM06249EutQ_2_eut2(pf06249) EUT2A EP_NTERM
00068 HMMtpEuth_1_eut(pf04346) EUT2A
00069 P_azul EUT2A
00070 HMM02441_07(#) EUT2A EP_NTERM
00071 HMMptac_3_eut2(pf06130) EUT2A EP_NTERM
00072 HMM01923cobtr_3_eut2(#) EUT2A
00073 H_fuchsia EUT2G
00074 HMMalddh_2_egra(pf00171) EUT2A EP_CTERM
00075 H_fireEngine EUT2A
00076 Tsp_amethyst EUT2A
00077 HMMetlyEutC_3_eut2(pf05985) EUT2A EP_NTERM
00078 HMMetlyEutB_1_EUT1(pf06751) EUT2A
00079 HMM06277EutA_1_eut(pf06277) EUT2A
00080 HMMreg07568_1_eut2(pf12282pf02518pf07568) EUT2A
00081 HMMreg00072_1_eut2(pf03861pf00072) EUT2A
00082 Hp_euts_babyPurple PDU1C
00083 HMM00465alcdh_3_mix(pf00465) EUT2A
00084 HMM01923cobtr_4_eut2(pf01923) EUT2A
00085 HMM10662PduV_1_mix(pf10662) EUT2A
00086 -pf00005 -
00087 -pf01032 -
00088 -pf01497 -
00089 -pf02660 -
00090 -pf01263 -
00091 -pf06018pf08222 -
00092 -pf00004pf07724pf10431 -
BMC-P
BMC-H
BMC-T
Aldehyde dehydrogenase
PTAC
Signature enzyme
Alcohol dehydrogenase
Regulator

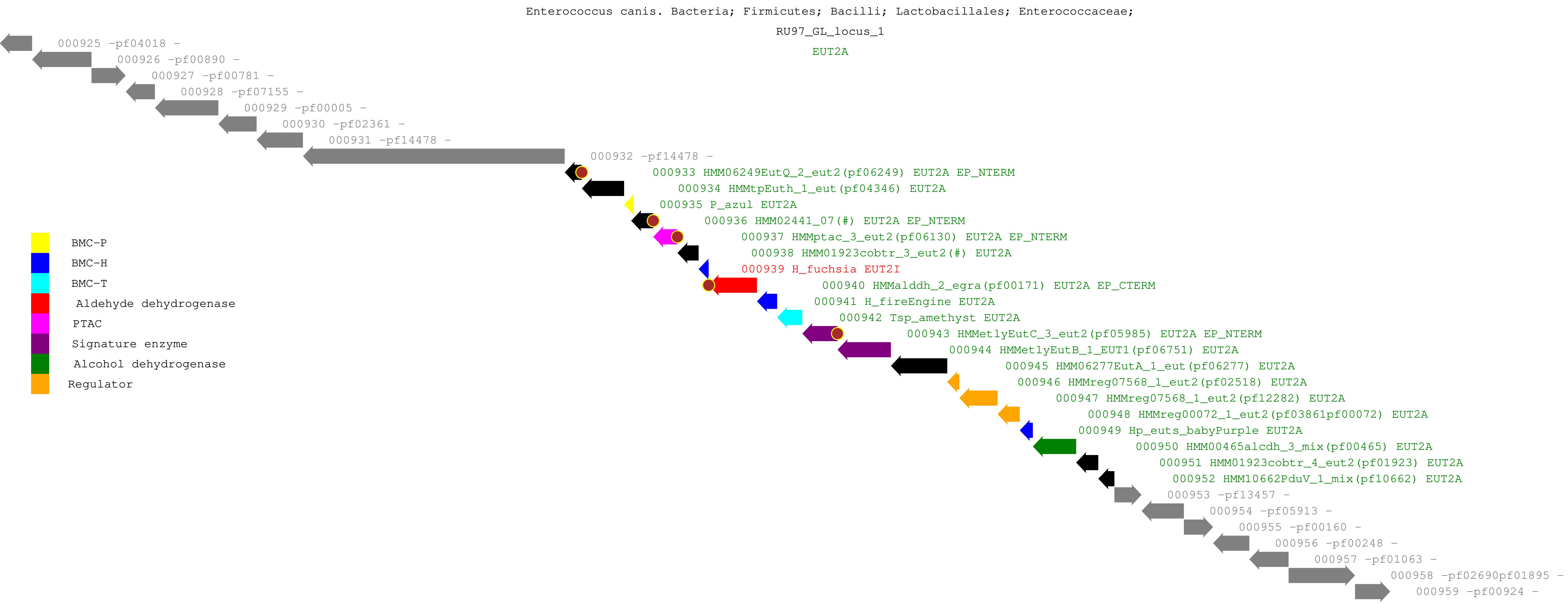
Enterococcus canis. Bacteria; Firmicutes; Bacilli; Lactobacillales; Enterococcaceae;
RU97_GL_locus_1
EUT2A
000925 -pf04018 -
000926 -pf00890 -
000927 -pf00781 -
000928 -pf07155 -
000929 -pf00005 -
000930 -pf02361 -
000931 -pf14478 -
000932 -pf14478 -
000933 HMM06249EutQ_2_eut2(pf06249) EUT2A EP_NTERM
000934 HMMtpEuth_1_eut(pf04346) EUT2A
000935 P_azul EUT2A
000936 HMM02441_07(#) EUT2A EP_NTERM
000937 HMMptac_3_eut2(pf06130) EUT2A EP_NTERM
000938 HMM01923cobtr_3_eut2(#) EUT2A
000939 H_fuchsia EUT2I
000940 HMMalddh_2_egra(pf00171) EUT2A EP_CTERM
000941 H_fireEngine EUT2A
000942 Tsp_amethyst EUT2A
000943 HMMetlyEutC_3_eut2(pf05985) EUT2A EP_NTERM
000944 HMMetlyEutB_1_EUT1(pf06751) EUT2A
000945 HMM06277EutA_1_eut(pf06277) EUT2A
000946 HMMreg07568_1_eut2(pf02518) EUT2A
000947 HMMreg07568_1_eut2(pf12282) EUT2A
000948 HMMreg00072_1_eut2(pf03861pf00072) EUT2A
000949 Hp_euts_babyPurple EUT2A
000950 HMM00465alcdh_3_mix(pf00465) EUT2A
000951 HMM01923cobtr_4_eut2(pf01923) EUT2A
000952 HMM10662PduV_1_mix(pf10662) EUT2A
000953 -pf13457 -
000954 -pf05913 -
000955 -pf00160 -
000956 -pf00248 -
000957 -pf01063 -
000958 -pf02690pf01895 -
000959 -pf00924 -
BMC-P
BMC-H
BMC-T
Aldehyde dehydrogenase
PTAC
Signature enzyme
Alcohol dehydrogenase
Regulator

Enterococcus mundtii. Bacteria; Firmicutes; Bacilli; Lactobacillales; Enterococcaceae;

RV08_GL_locus_1

EUT2A

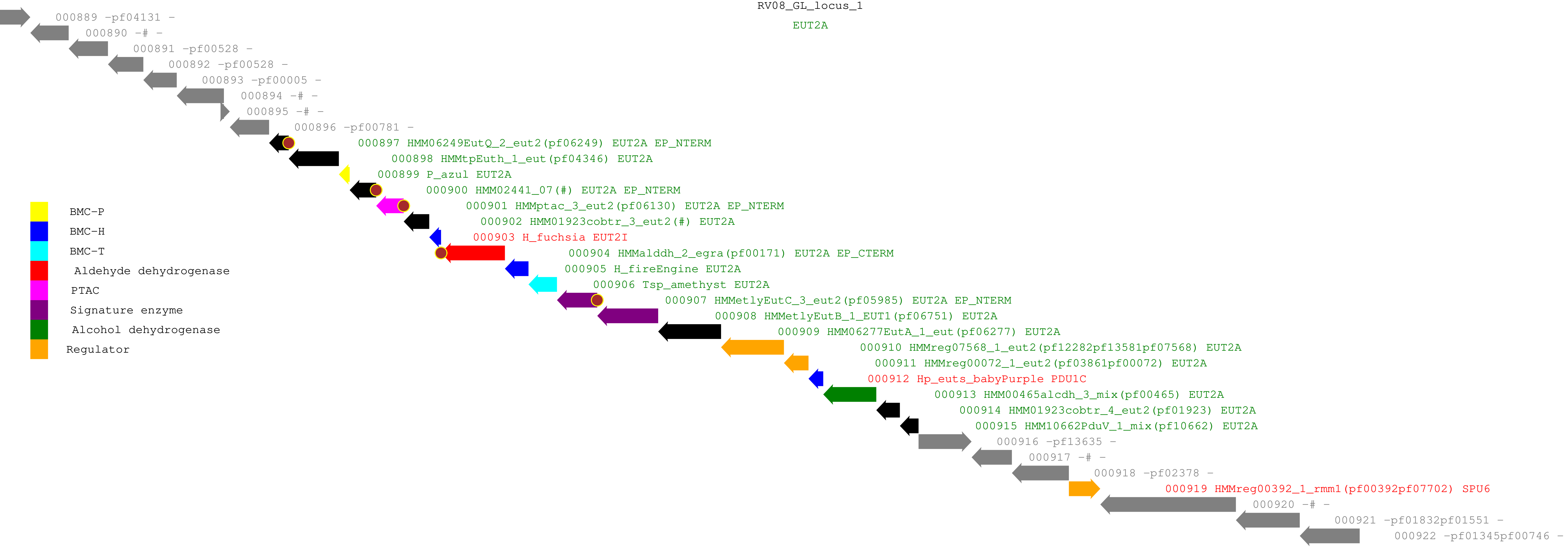

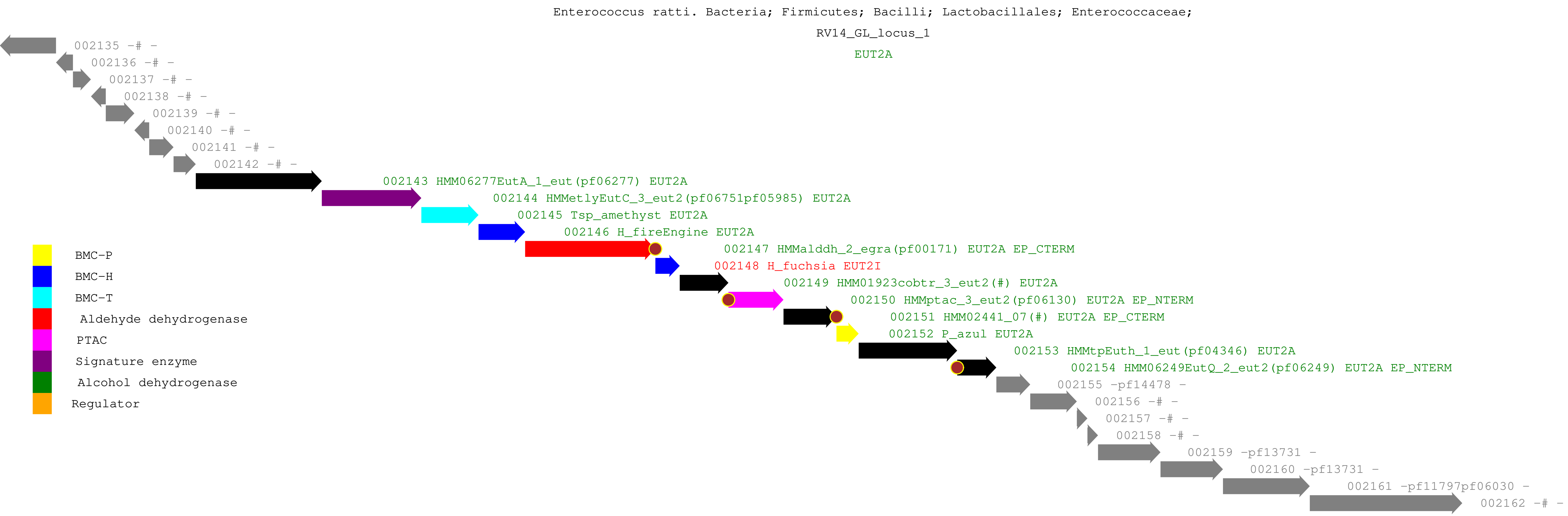
Enterococcus ratti. Bacteria; Firmicutes; Bacilli; Lactobacillales; Enterococcaceae;
RV14_GL_locus_1
EUT2A
002135 -# -
002136 -# -
002137 -# -
002138 -# -
002139 -# -
002140 -# -
002141 -# -
002142 -# -
002143 HMM06277EutA_1_eut(pf06277) EUT2A
002144 HMMetlyEutC_3_eut2(pf06751pf05985) EUT2A
002145 Tsp_amethyst EUT2A
002146 H_fireEngine EUT2A
002147 HMMalddh_2_egra(pf00171) EUT2A EP_CTERM
002148 H_fuchsia EUT2I
002149 HMM01923cobtr_3_eut2(#) EUT2A
002150 HMMptac_3_eut2(pf06130) EUT2A EP_NTERM
002151 HMM02441_07(#) EUT2A EP_CTERM
002152 P_azul EUT2A
002153 HMMtpEuth_1_eut(pf04346) EUT2A
002154 HMM06249EutQ_2_eut2(pf06249) EUT2A EP_NTERM
002155 -pf14478 -
002156 -# -
002157 -# -
002158 -# -
002159 -pf13731 -
002160 -pf13731 -
002161 -pf11797pf06030 -
002162 -# -
BMC-P
BMC-H
BMC-T
Aldehyde dehydrogenase
PTAC
Signature enzyme
Alcohol dehydrogenase
Regulator

Streptococcus merionis. Bacteria; Firmicutes; Bacilli; Lactobacillales; Streptococcaceae;

SAMEA4412692__locus_1

EUT2A

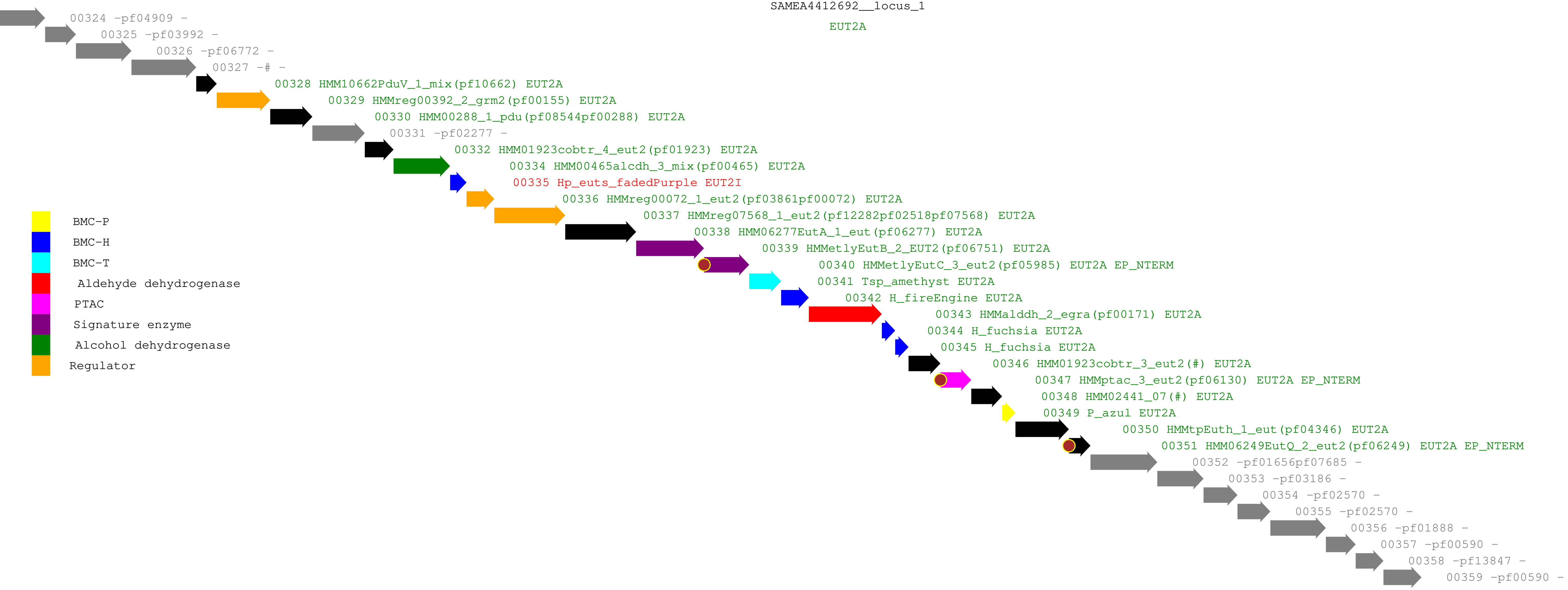

Pilibacter termitis. Bacteria; Firmicutes; Bacilli; Lactobacillales; Enterococcaceae;

SAMN02745116__locus_1

EUT2A

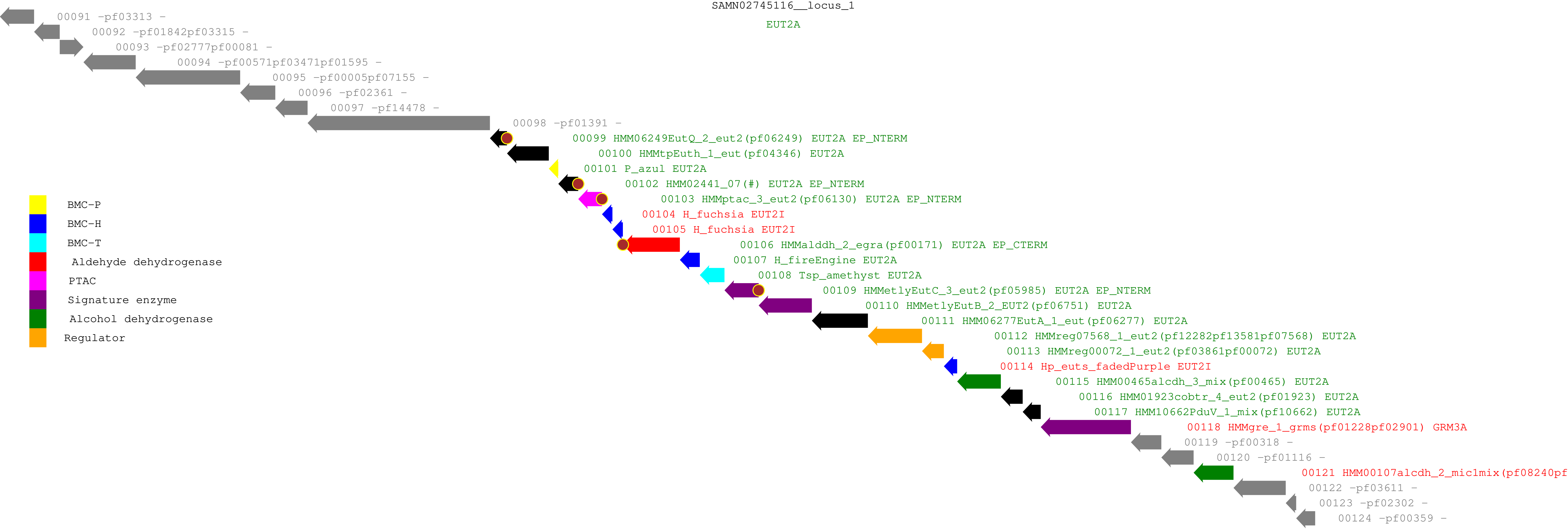

Clostridium sphenoides JCM 1415. Bacteria; Firmicutes; Clostridia; Clostridiales; Lachnospiraceae.

SAMN02745906__locus_1

EUT2A

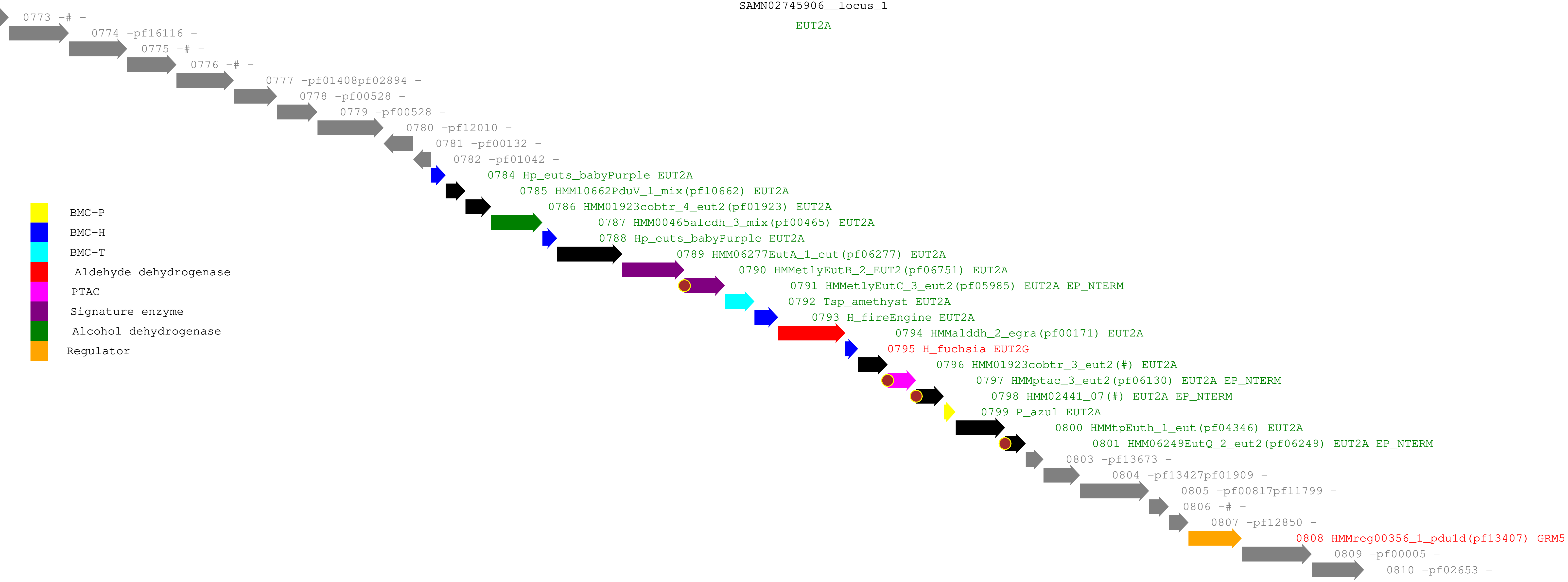

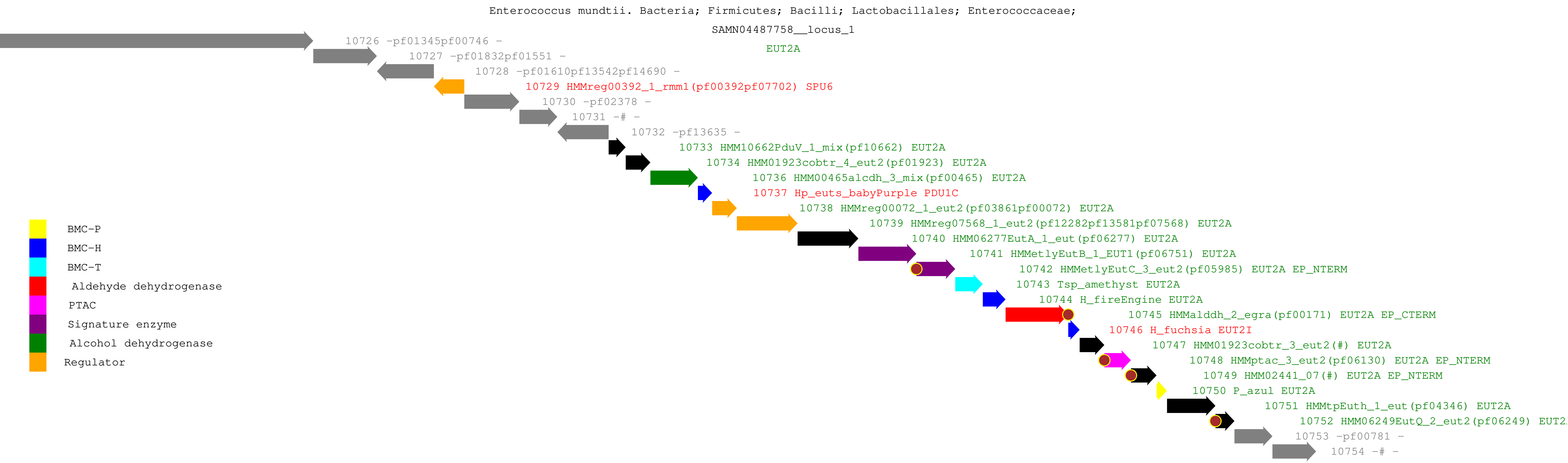

Enterococcus mundtii. Bacteria; Firmicutes; Bacilli; Lactobacillales; Enterococcaceae;
SAMN04487758__locus_1
EUT2A
10726 -pf01345pf00746 -
10727 -pf01832pf01551 -
10728 -pf01610pf13542pf14690 -
10729 HMMreg00392_1_rmm1(pf00392pf07702) SPU6
10730 -pf02378 -
10731 -# -
10732 -pf13635 -
10733 HMM10662PduV_1_mix(pf10662) EUT2A
10734 HMM01923cobtr_4_eut2(pf01923) EUT2A
10736 HMM00465alcdh_3_mix(pf00465) EUT2A
10737 Hp_euts_babyPurple PDU1C
10738 HMMreg00072_1_eut2(pf03861pf00072) EUT2A
10739 HMMreg07568_1_eut2(pf12282pf13581pf07568) EUT2A
10740 HMM06277EutA_1_eut(pf06277) EUT2A
10741 HMMetlyEutB_1_EUT1(pf06751) EUT2A
10742 HMMetlyEutC_3_eut2(pf05985) EUT2A EP_NTERM
10743 Tsp_amethyst EUT2A
10744 H_fireEngine EUT2A
10745 HMMaldh_2_egra(pf00171) EUT2A EP_CTERM
10746 H_fuchsia EUT2I
10747 HMM01923cobtr_3_eut2(#) EUT2A
10748 HMMptac_3_eut2(pf06130) EUT2A EP_NTERM
10749 HMM02441_07(#) EUT2A EP_NTERM
10750 P_azul EUT2A
10751 HMMtpEuth_1_eut(pf04346) EUT2A
10752 HMM06249EutQ_2_eut2(pf06249) EUT2
10753 -pf00781 -
10754 -# -
BMC-P
BMC-H
BMC-T
Aldehyde dehydrogenase
PTAC
Signature enzyme
Alcohol dehydrogenase
Regulator

Ignavigranum ruoffiae. Bacteria; Firmicutes; Bacilli; Lactobacillales; Aerococcaceae;

SAMN04488558__locus_1_main_sat

EUT2A

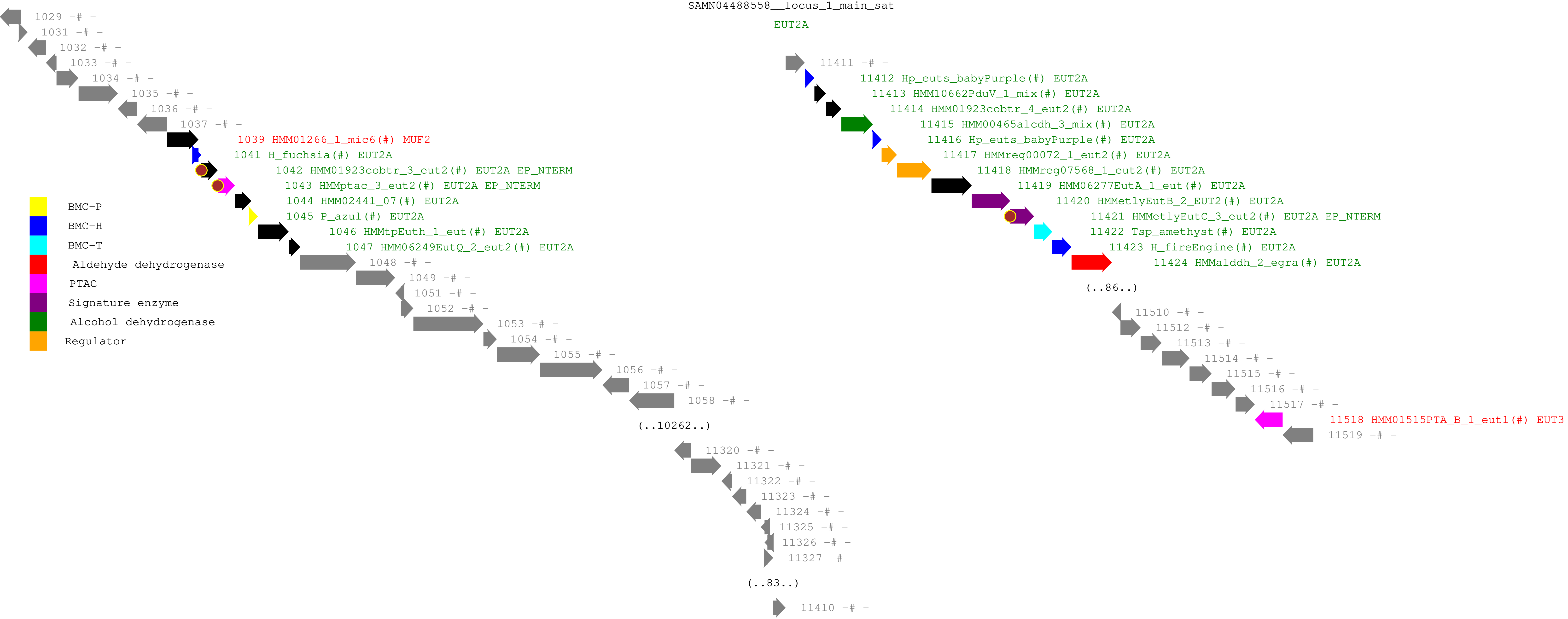

Isobaculum melis. Bacteria; Firmicutes; Bacilli; Lactobacillales; Carnobacteriaceae;

SAMN04488559__locus_1

EUT2A

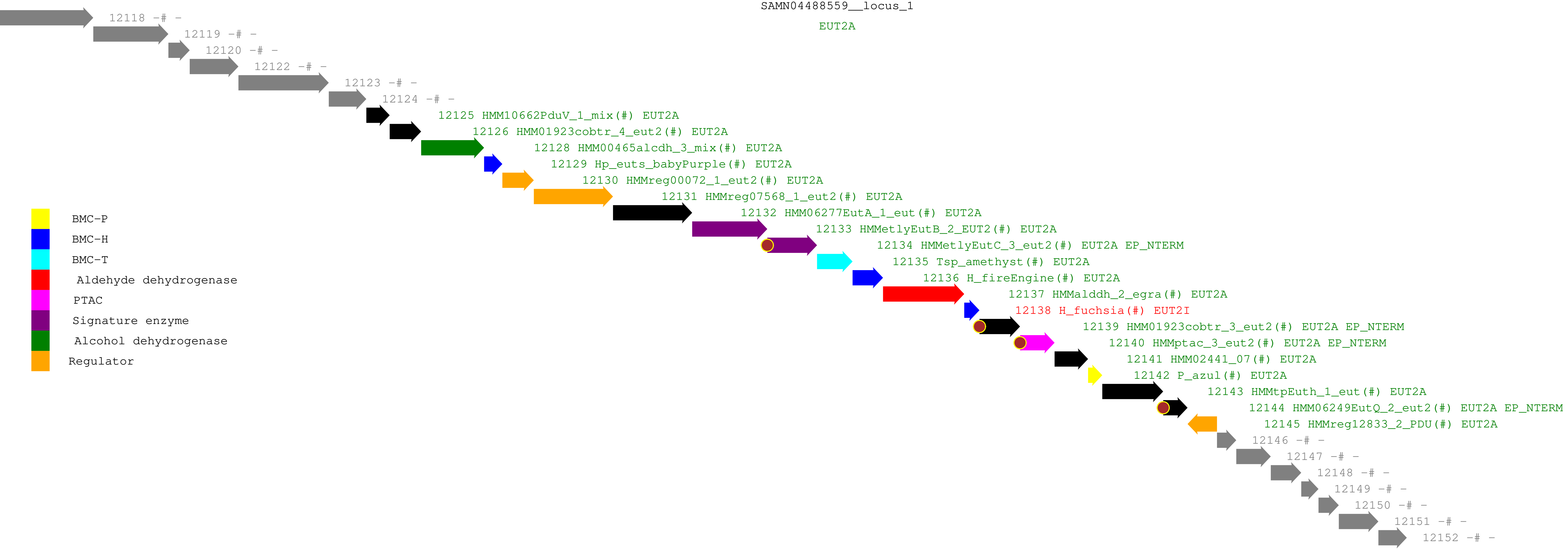

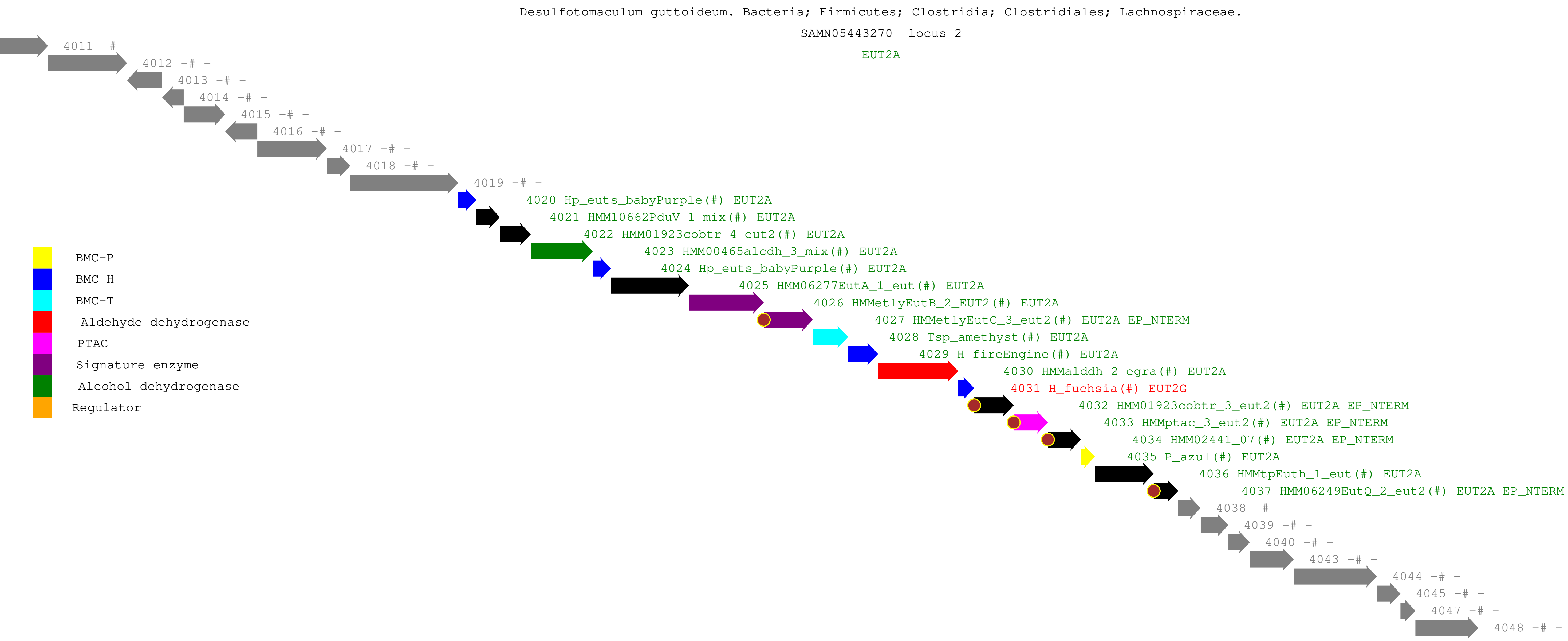
Desulfotomaculum guttoideum. Bacteria; Firmicutes; Clostridia; Clostridiales; Lachnospiraceae.
SAMN05443270__locus_2
EUT2A
4011 -# -
4012 -# -
4013 -# -
4014 -# -
4015 -# -
4016 -# -
4017 -# -
4018 -# -
4019 -# -
4020 Hp_euts_babyPurple(#) EUT2A
4021 HMM10662PduV_1_mix(#) EUT2A
4022 HMM01923cobtr_4_eut2(#) EUT2A
4023 HMM00465alcdh_3_mix(#) EUT2A
4024 Hp_euts_babyPurple(#) EUT2A
4025 HMM06277EutA_1_eut(#) EUT2A
4026 HMMetlyEutB_2_EUT2(#) EUT2A
4027 HMMetlyEutC_3_eut2(#) EUT2A EP_NTERM
4028 Tsp_amethyst(#) EUT2A
4029 H_fireEngine(#) EUT2A
4030 HMMalddh_2_egra(#) EUT2A
4031 H_fuchsia(#) EUT2G
4032 HMM01923cobtr_3_eut2(#) EUT2A EP_NTERM
4033 HMMptac_3_eut2(#) EUT2A EP_NTERM
4034 HMM02441_07(#) EUT2A EP_NTERM
4035 P_azul(#) EUT2A
4036 HMMtpEuth_1_eut(#) EUT2A
4037 HMM06249EutQ_2_eut2(#) EUT2A EP_NTERM
4038 -# -
4039 -# -
4040 -# -
4043 -# -
4044 -# -
4045 -# -
4047 -# -
4048 -# -
BMC-P
BMC-H
BMC-T
Aldehyde dehydrogenase
PTAC
Signature enzyme
Alcohol dehydrogenase
Regulator

Streptococcus pluranimalium. Bacteria; Firmicutes; Bacilli; Lactobacillales; Streptococcaceae;

Sp14A__locus_1

EUT2A

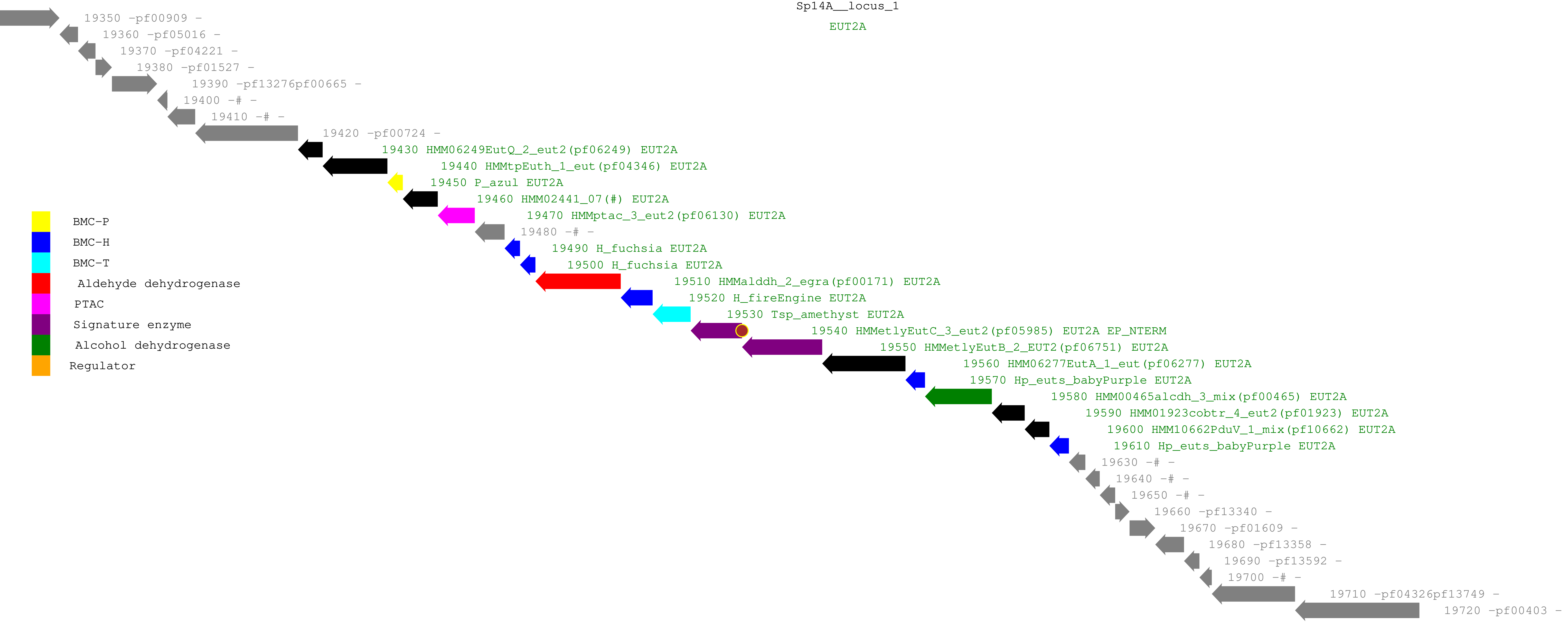

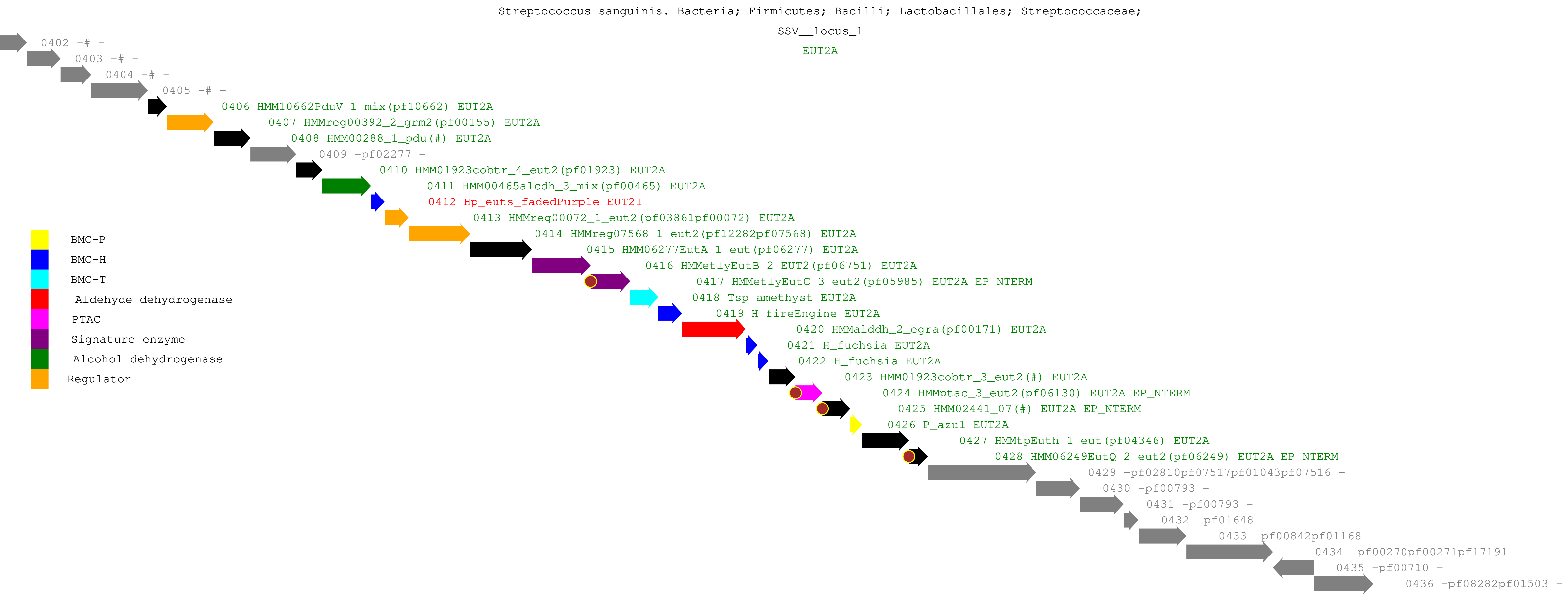

Streptococcus sanguinis. Bacteria; Firmicutes; Bacilli; Lactobacillales; Streptococcaceae;
SSV__locus_1
EUT2A
0402 -# -
0403 -# -
0404 -# -
0405 -# -
0406 HMM10662PduV_1_mix(pf10662) EUT2A
0407 HMMreg00392_2_grm2(pf00155) EUT2A
0408 HMM00288_1_pdu(#) EUT2A
0409 -pf02277 -
0410 HMM01923cobtr_4_eut2(pf01923) EUT2A
0411 HMM00465alcdh_3_mix(pf00465) EUT2A
0412 Hp_euts_fadedPurple EUT2I
0413 HMMreg00072_1_eut2(pf03861pf00072) EUT2A
0414 HMMreg07568_1_eut2(pf12282pf07568) EUT2A
0415 HMM06277EutA_1_eut(pf06277) EUT2A
0416 HMMetlyEutB_2_EUT2(pf06751) EUT2A
0417 HMMetlyEutC_3_eut2(pf05985) EUT2A EP_NTERM
0418 Tsp_amethyst EUT2A
0419 H_fireEngine EUT2A
0420 HMMaldh_2_egra(pf00171) EUT2A
0421 H_fuchsia EUT2A
0422 H_fuchsia EUT2A
0423 HMM01923cobtr_3_eut2(#) EUT2A
0424 HMMptac_3_eut2(pf06130) EUT2A EP_NTERM
0425 HMM02441_07(#) EUT2A EP_NTERM
0426 P_azul EUT2A
0427 HMMtpEuth_1_eut(pf04346) EUT2A
0428 HMM06249EutQ_2_eut2(pf06249) EUT2A EP_NTERM
0429 -pf02810pf07517pf01043pf07516 -
0430 -pf00793 -
0431 -pf00793 -
0432 -pf01648 -
0433 -pf00842pf01168 -
0434 -pf00270pf00271pf17191 -
0435 -pf00710 -
0436 -pf08282pf01503 -
BMC-P
BMC-H
BMC-T
Aldehyde dehydrogenase
PTAC
Signature enzyme
Alcohol dehydrogenase
Regulator

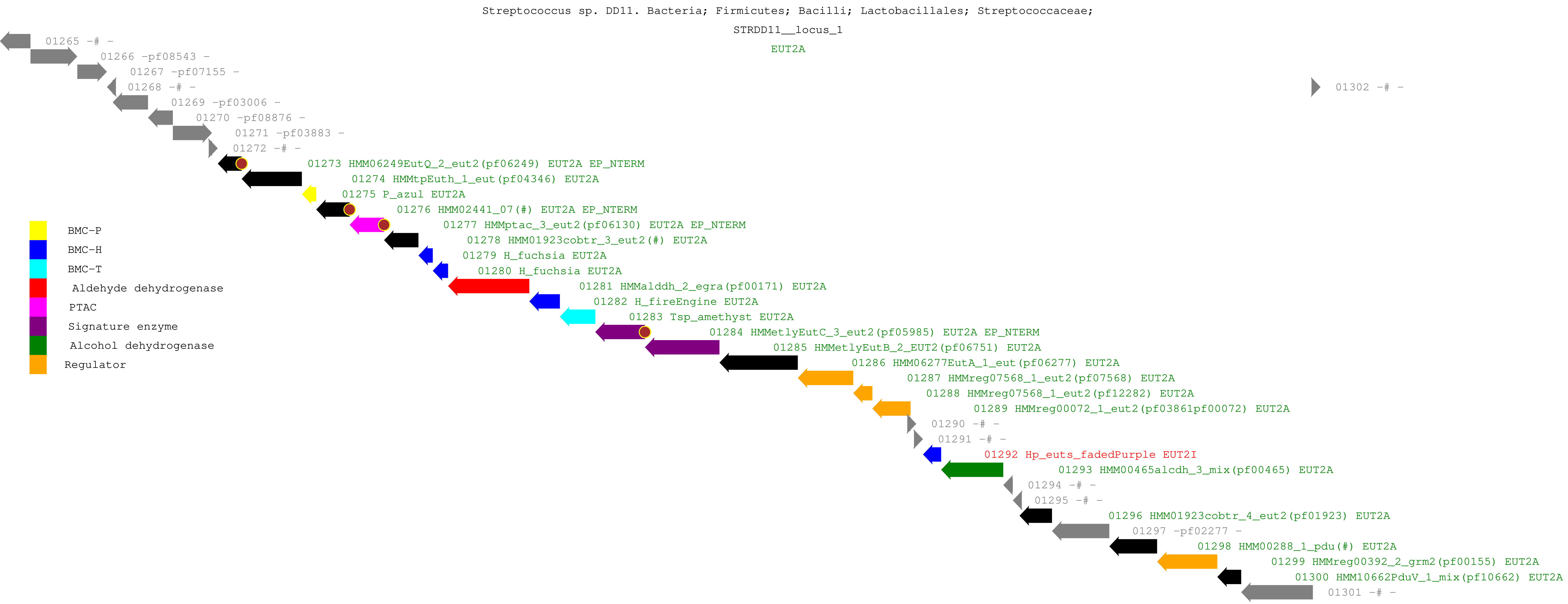
Streptococcus sp. DD11. Bacteria; Firmicutes; Bacilli; Lactobacillales; Streptococcaceae;
STRDD11__locus_1
EUT2A
01265 -# -
01266 -pf08543 -
01267 -pf07155 -
01268 -# -
01269 -pf03006 -
01270 -pf08876 -
01271 -pf03883 -
01272 -# -
01273 HMM06249EutQ_2_eut2(pf06249) EUT2A EP_NTERM
01274 HMMtpEuth_1_eut(pf04346) EUT2A
01275 P_azul EUT2A
01276 HMM02441_07(#) EUT2A EP_NTERM
01277 HMMptac_3_eut2(pf06130) EUT2A EP_NTERM
01278 HMM01923cobtr_3_eut2(#) EUT2A
01279 H_fuchsia EUT2A
01280 H_fuchsia EUT2A
01281 HMMalddh_2_egra(pf00171) EUT2A
01282 H_fireEngine EUT2A
01283 Tsp_amethyst EUT2A
01284 HMMetlyEutC_3_eut2(pf05985) EUT2A EP_NTERM
01285 HMMetlyEutB_2_EUT2(pf06751) EUT2A
01286 HMM06277EutA_1_eut(pf06277) EUT2A
01287 HMMreg07568_1_eut2(pf07568) EUT2A
01288 HMMreg07568_1_eut2(pf12282) EUT2A
01289 HMMreg00072_1_eut2(pf03861pf00072) EUT2A
01290 -# -
01291 -# -
01292 Hp_euts_fadedPurple EUT2I
01293 HMM00465alcdh_3_mix(pf00465) EUT2A
01294 -# -
01295 -# -
01296 HMM01923cobtr_4_eut2(pf01923) EUT2A
01297 -pf02277 -
01298 HMM00288_1_pdu(#) EUT2A
01299 HMMreg00392_2_grm2(pf00155) EUT2A
01300 HMM10662PduV_1_mix(pf10662) EUT2A
01301 -# -
01302 -# -
BMC-P
BMC-H
BMC-T
Aldehyde dehydrogenase
PTAC
Signature enzyme
Alcohol dehydrogenase
Regulator

Enterococcus faecalis PF3. Bacteria; Firmicutes; Bacilli; Lactobacillales; Enterococcaceae;

T481__locus_1_main_sat

EUT2A

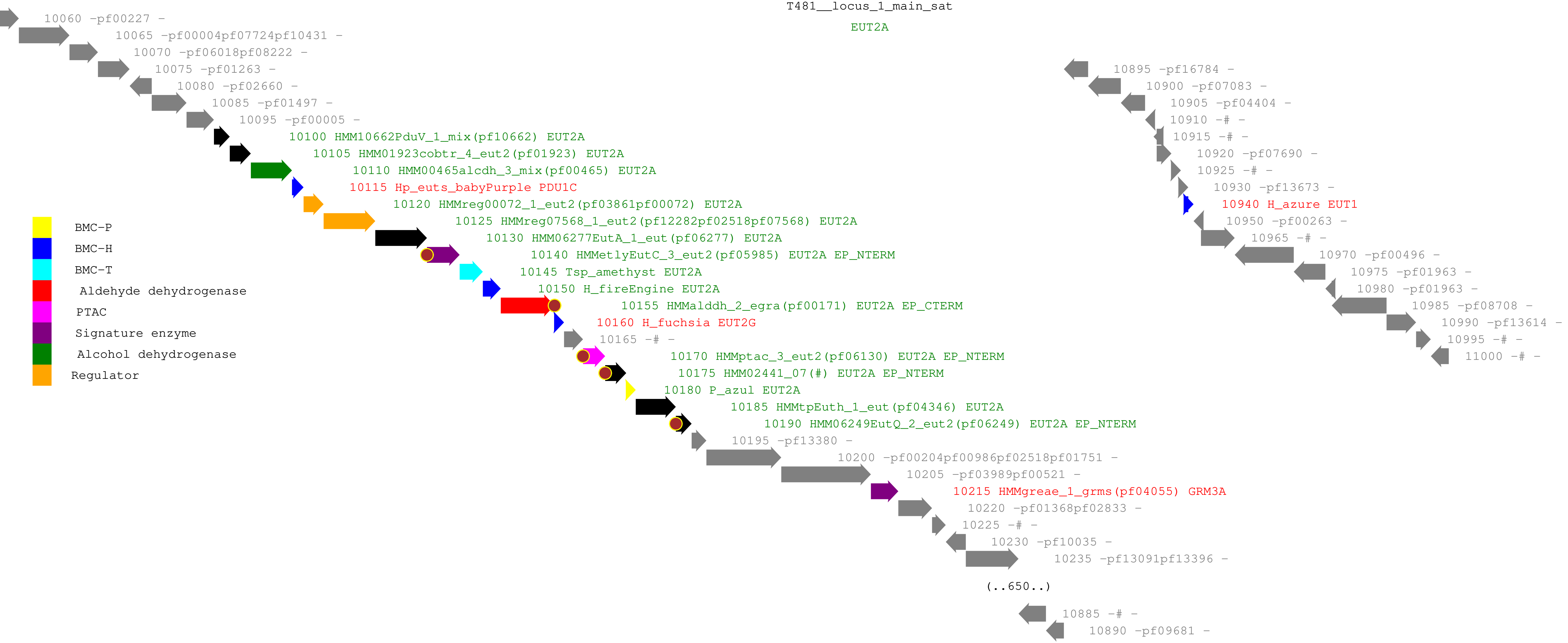

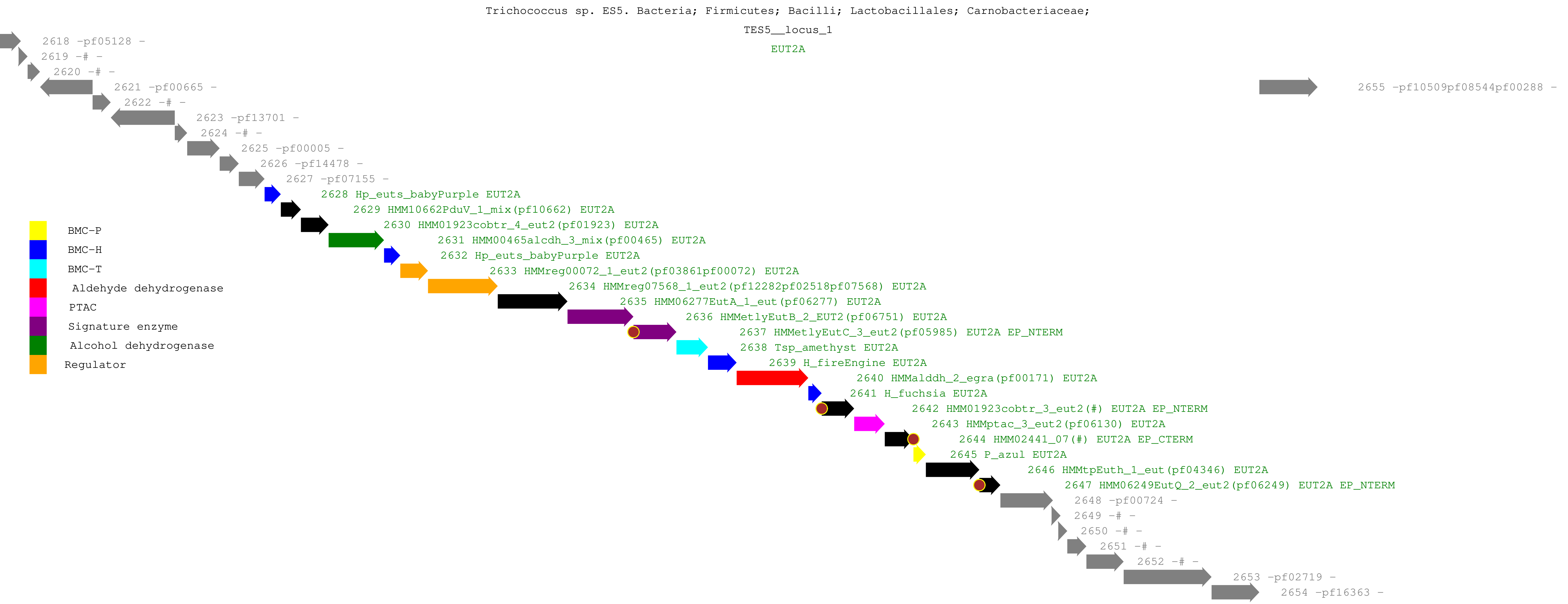

Trichococcus sp. ES5. Bacteria; Firmicutes; Bacilli; Lactobacillales; Carnobacteriaceae;
TES5__locus_1
EUT2A
2618 -pf05128 -
2619 -# -
2620 -# -
2621 -pf00665 -
2622 -# -
2623 -pf13701 -
2624 -# -
2625 -pf00005 -
2626 -pf14478 -
2627 -pf07155 -
2628 Hp_euts_babyPurple EUT2A
2629 HMM10662PduV_1_mix(pf10662) EUT2A
2630 HMM01923cobtr_4_eut2(pf01923) EUT2A
2631 HMM00465alcdh_3_mix(pf00465) EUT2A
2632 Hp_euts_babyPurple EUT2A
2633 HMMreg00072_1_eut2(pf03861pf00072) EUT2A
2634 HMMreg07568_1_eut2(pf12282pf02518pf07568) EUT2A
2635 HMM06277EutA_1_eut(pf06277) EUT2A
2636 HMMetlyEutB_2_EUT2(pf06751) EUT2A
2637 HMMetlyEutC_3_eut2(pf05985) EUT2A EP_NTERM
2638 Tsp_amethyst EUT2A
2639 H_fireEngine EUT2A
2640 HMMalddh_2_egra(pf00171) EUT2A
2641 H_fuchsia EUT2A
2642 HMM01923cobtr_3_eut2(#) EUT2A EP_NTERM
2643 HMMptac_3_eut2(pf06130) EUT2A
2644 HMM02441_07(#) EUT2A EP_CTERM
2645 P_azul EUT2A
2646 HMMtpEuth_1_eut(pf04346) EUT2A
2647 HMM06249EutQ_2_eut2(pf06249) EUT2A EP_NTERM
2648 -pf00724 -
2649 -# -
2650 -# -
2651 -# -
2652 -# -
2653 -pf02719 -
2654 -pf16363 -
2655 -pf10509pf08544pf00288 -
BMC-P
BMC-H
BMC-T
Aldehyde dehydrogenase
PTAC
Signature enzyme
Alcohol dehydrogenase
Regulator

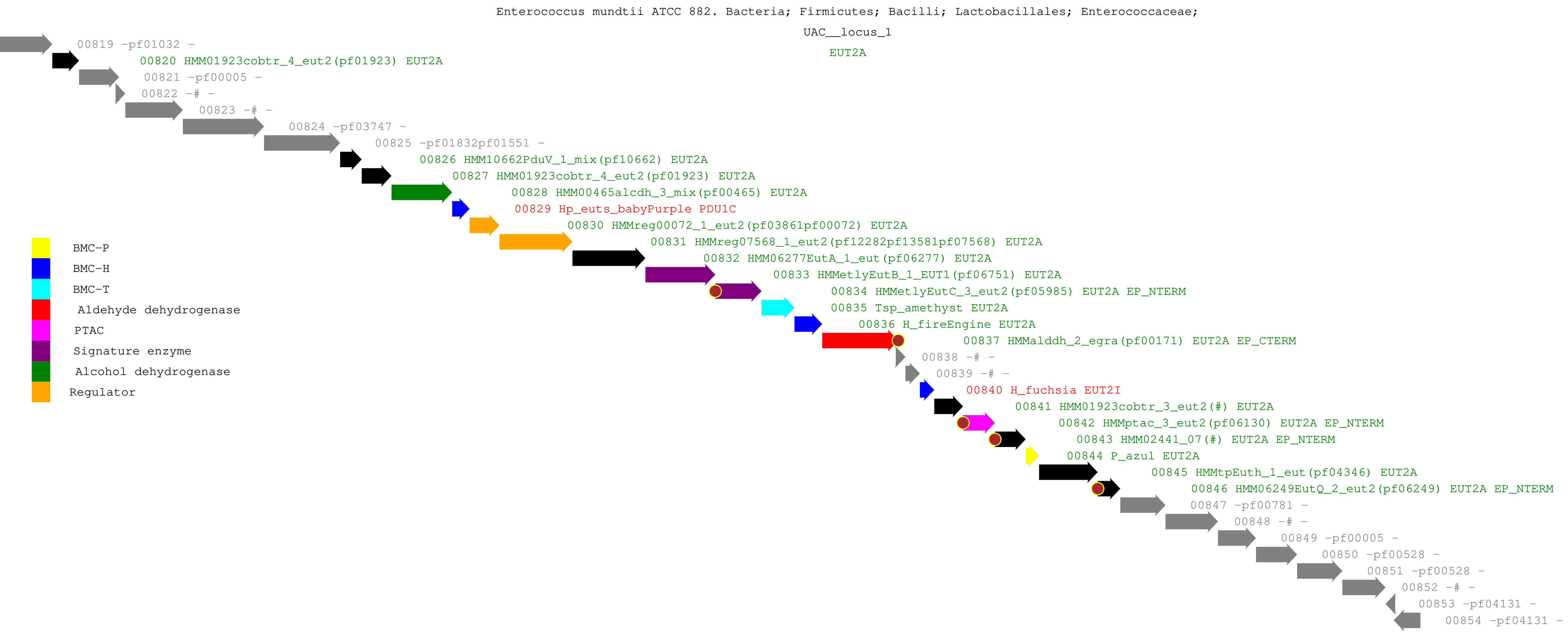
Enterococcus mundtii ATCC 882. Bacteria; Firmicutes; Bacilli; Lactobacillales; Enterococcaceae;
UAC__locus_1
EUT2A
00819 -pf01032 -
00820 HMM01923cobtr_4_eut2(pf01923) EUT2A
00821 -pf00005 -
00822 -# -
00823 -# -
00824 -pf03747 -
00825 -pf01832pf01551 -
00826 HMM10662PduV_1_mix(pf10662) EUT2A
00827 HMM01923cobtr_4_eut2(pf01923) EUT2A
00828 HMM00465alcdh_3_mix(pf00465) EUT2A
00829 Hp_euts_babyPurple PDU1C
00830 HMMreg00072_1_eut2(pf03861pf00072) EUT2A
00831 HMMreg07568_1_eut2(pf12282pf13581pf07568) EUT2A
00832 HMM06277EutA_1_eut(pf06277) EUT2A
00833 HMMetlyEutB_1_EUT1(pf06751) EUT2A
00834 HMMetlyEutC_3_eut2(pf05985) EUT2A EP_NTERM
00835 Tsp_amethyst EUT2A
00836 H_fireEngine EUT2A
00837 HMMalddh_2_egra(pf00171) EUT2A EP_CTERM
00838 -# -
00839 -# -
00840 H_fuchsia EUT2I
00841 HMM01923cobtr_3_eut2(#) EUT2A
00842 HMMptac_3_eut2(pf06130) EUT2A EP_NTERM
00843 HMM02441_07(#) EUT2A EP_NTERM
00844 P_azul EUT2A
00845 HMMtpEuth_1_eut(pf04346) EUT2A
00846 HMM06249EutQ_2_eut2(pf06249) EUT2A EP_NTERM
00847 -pf00781 -
00848 -# -
00849 -pf00005 -
00850 -pf00528 -
00851 -pf00528 -
00852 -# -
00853 -pf04131 -
00854 -pf04131 -
BMC-P
BMC-H
BMC-T
Aldehyde dehydrogenase
PTAC
Signature enzyme
Alcohol dehydrogenase
Regulator

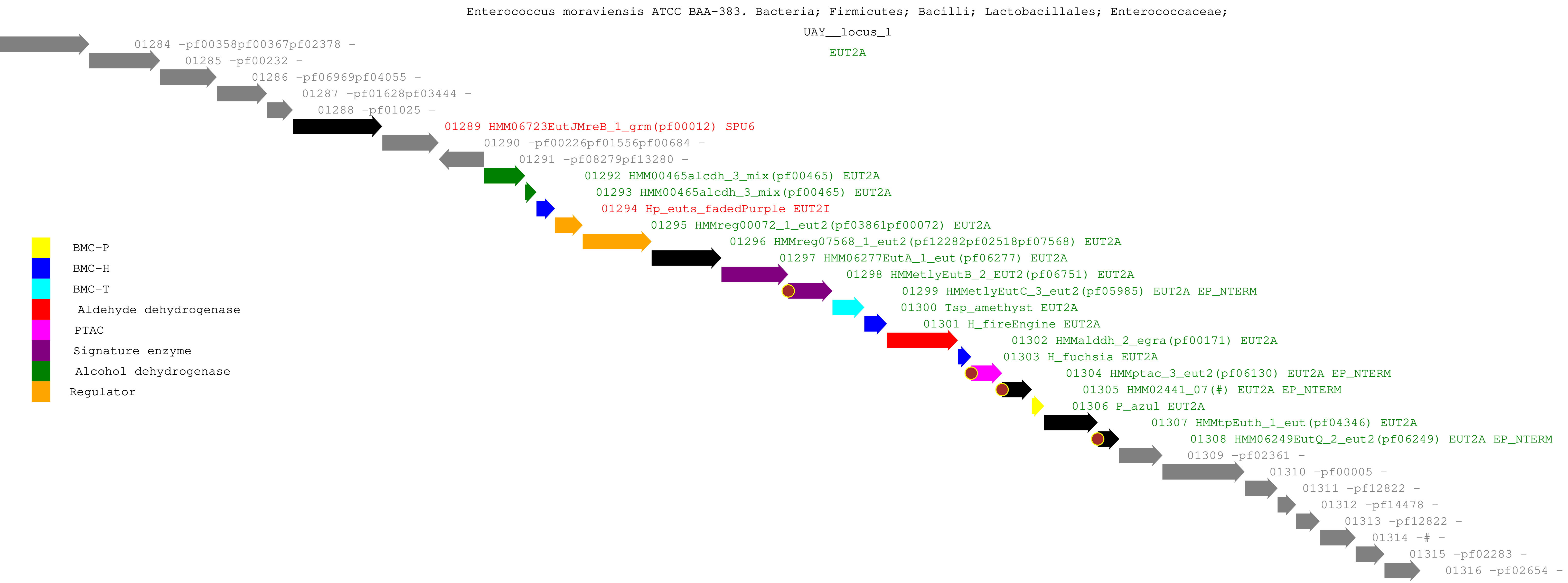
Enterococcus moraviensis ATCC BAA-383. Bacteria; Firmicutes; Bacilli; Lactobacillales; Enterococcaceae;
UAY__locus_1
EUT2A
01284 -pf00358pf00367pf02378 -
01285 -pf00232 -
01286 -pf06969pf04055 -
01287 -pf01628pf03444 -
01288 -pf01025 -
01289 HMM06723EutJMreB_1_grm(pf00012) SPU6
01290 -pf00226pf01556pf00684 -
01291 -pf08279pf13280 -
01292 HMM00465alcdh_3_mix(pf00465) EUT2A
01293 HMM00465alcdh_3_mix(pf00465) EUT2A
01294 Hp_euts_fadedPurple EUT2I
01295 HMMreg00072_1_eut2(pf03861pf00072) EUT2A
01296 HMMreg07568_1_eut2(pf12282pf02518pf07568) EUT2A
01297 HMM06277EutA_1_eut(pf06277) EUT2A
01298 HMMetlyEutB_2_EUT2(pf06751) EUT2A
01299 HMMetlyEutC_3_eut2(pf05985) EUT2A EP_NTERM
01300 Tsp_amethyst EUT2A
01301 H_fireEngine EUT2A
01302 HMMalddh_2_egra(pf00171) EUT2A
01303 H_fuchsia EUT2A
01304 HMMptac_3_eut2(pf06130) EUT2A EP_NTERM
01305 HMM02441_07(#) EUT2A EP_NTERM
01306 P_azul EUT2A
01307 HMMtpEuth_1_eut(pf04346) EUT2A
01308 HMM06249EutQ_2_eut2(pf06249) EUT2A EP_NTERM
01309 -pf02361 -
01310 -pf00005 -
01311 -pf12822 -
01312 -pf14478 -
01313 -pf12822 -
01314 -# -
01315 -pf02283 -
01316 -pf02654 -
BMC-P
BMC-H
BMC-T
Aldehyde dehydrogenase
PTAC
Signature enzyme
Alcohol dehydrogenase
Regulator

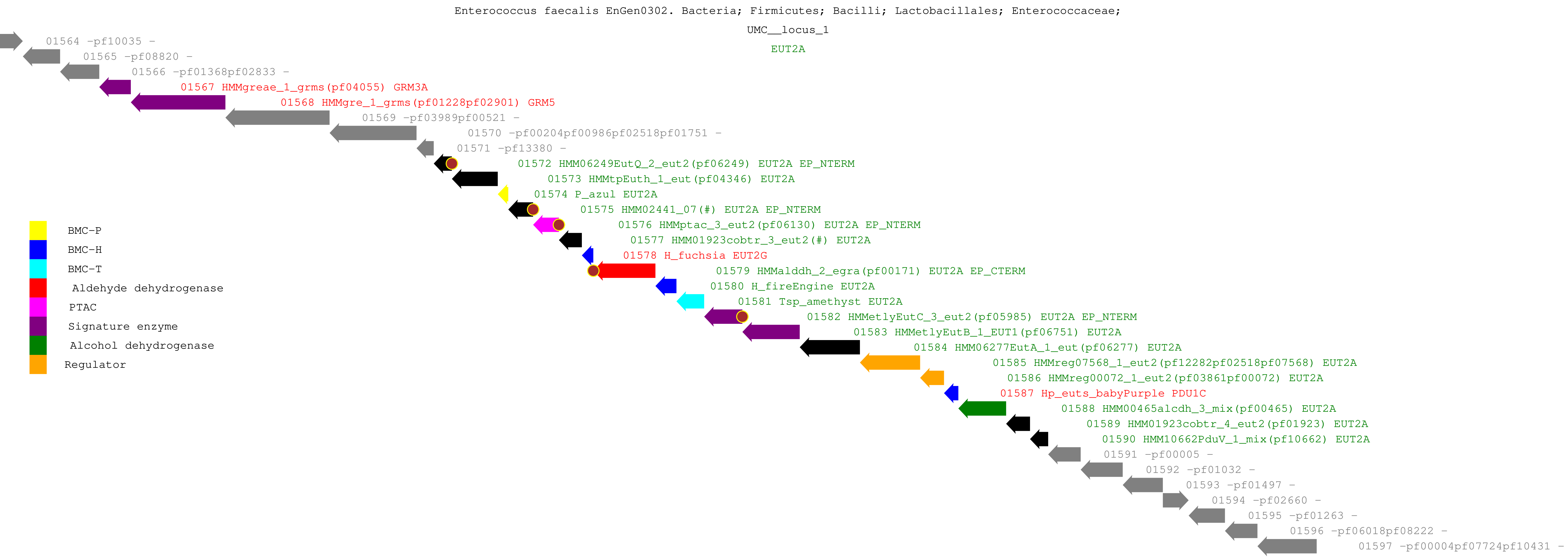

Enterococcus faecalis EnGen0302. Bacteria; Firmicutes; Bacilli; Lactobacillales; Enterococcaceae;
UMC__locus_1
EUT2A
01564 -pf10035 -
01565 -pf08820 -
01566 -pf01368pf02833 -
01567 HMMgreae_1_grms(pf04055) GRM3A
01568 HMMgre_1_grms(pf01228pf02901) GRM5
01569 -pf03989pf00521 -
01570 -pf00204pf00986pf02518pf01751 -
01571 -pf13380 -
01572 HMM06249EutQ_2_eut2(pf06249) EUT2A EP_NTERM
01573 HMMtpEuth_1_eut(pf04346) EUT2A
01574 P_azul EUT2A
01575 HMM02441_07(#) EUT2A EP_NTERM
01576 HMMptac_3_eut2(pf06130) EUT2A EP_NTERM
01577 HMM01923cobtr_3_eut2(#) EUT2A
01578 H_fuchsia EUT2G
01579 HMMalddh_2_egra(pf00171) EUT2A EP_CTERM
01580 H_fireEngine EUT2A
01581 Tsp_amethyst EUT2A
01582 HMMetlyEutC_3_eut2(pf05985) EUT2A EP_NTERM
01583 HMMetlyEutB_1_EUT1(pf06751) EUT2A
01584 HMM06277EutA_1_eut(pf06277) EUT2A
01585 HMMreg07568_1_eut2(pf12282pf02518pf07568) EUT2A
01586 HMMreg00072_1_eut2(pf03861pf00072) EUT2A
01587 Hp_euts_babyPurple PDU1C
01588 HMM00465alcdh_3_mix(pf00465) EUT2A
01589 HMM01923cobtr_4_eut2(pf01923) EUT2A
01590 HMM10662PduV_1_mix(pf10662) EUT2A
01591 -pf00005 -
01592 -pf01032 -
01593 -pf01497 -
01594 -pf02660 -
01595 -pf01263 -
01596 -pf06018pf08222 -
01597 -pf00004pf07724pf10431 -
BMC-P
BMC-H
BMC-T
Aldehyde dehydrogenase
PTAC
Signature enzyme
Alcohol dehydrogenase
Regulator

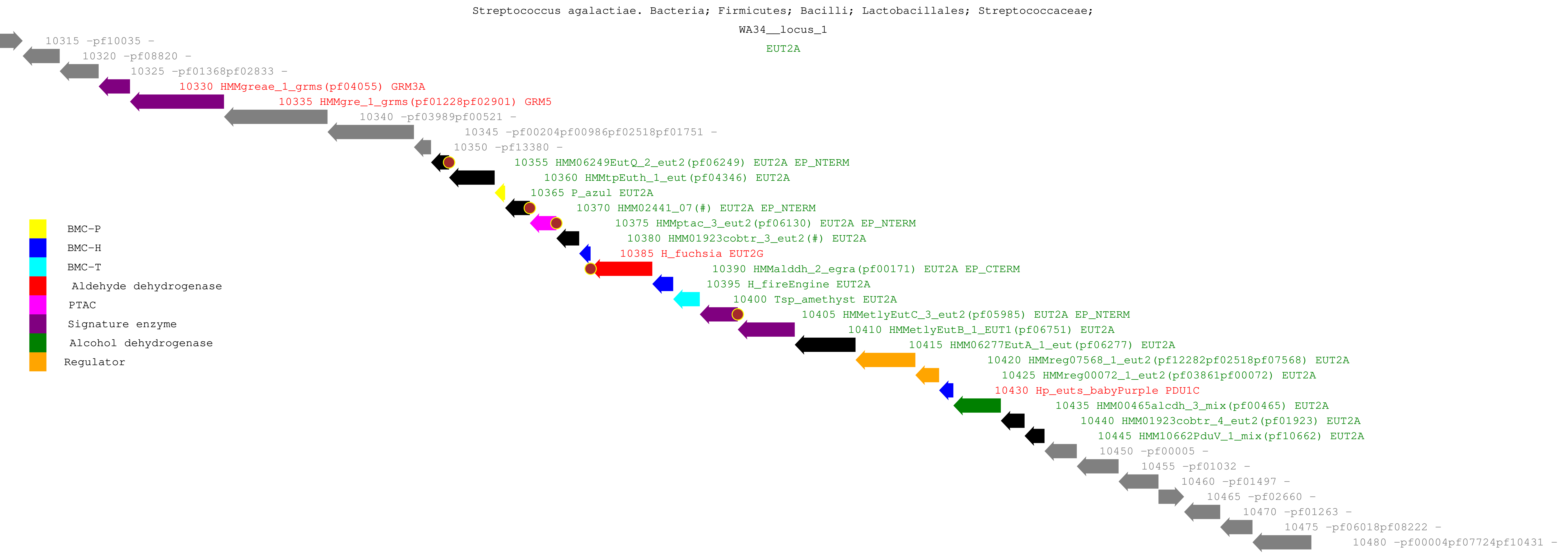
Streptococcus agalactiae. Bacteria; Firmicutes; Bacilli; Lactobacillales; Streptococcaceae;
WA34__locus_1
EUT2A
10315 -pf10035 -
10320 -pf08820 -
10325 -pf01368pf02833 -
10330 HMMgreae_1_grms(pf04055) GRM3A
10335 HMMgre_1_grms(pf01228pf02901) GRM5
10340 -pf03989pf00521 -
10345 -pf00204pf00986pf02518pf01751 -
10350 -pf13380 -
10355 HMM06249EutQ_2_eut2(pf06249) EUT2A EP_NTERM
10360 HMMtpEuth_1_eut(pf04346) EUT2A
10365 P_azul EUT2A
10370 HMM02441_07(#) EUT2A EP_NTERM
10375 HMMptac_3_eut2(pf06130) EUT2A EP_NTERM
10380 HMM01923cobtr_3_eut2(#) EUT2A
10385 H_fuchsia EUT2G
10390 HMMaldddh_2_egra(pf00171) EUT2A EP_CTERM
10395 H_fireEngine EUT2A
10400 Tsp_amethyst EUT2A
10405 HMMetlyEutC_3_eut2(pf05985) EUT2A EP_NTERM
10410 HMMetlyEutB_1_EUT1(pf06751) EUT2A
10415 HMM06277EutA_1_eut(pf06277) EUT2A
10420 HMMreg07568_1_eut2(pf12282pf02518pf07568) EUT2A
10425 HMMreg00072_1_eut2(pf03861pf00072) EUT2A
10430 Hp_euts_babyPurple PDU1C
10435 HMM00465alcdh_3_mix(pf00465) EUT2A
10440 HMM01923cobtr_4_eut2(pf01923) EUT2A
10445 HMM10662PduV_1_mix(pf10662) EUT2A
10450 -pf00005 -
10455 -pf01032 -
10460 -pf01497 -
10465 -pf02660 -
10470 -pf01263 -
10475 -pf06018pf08222 -
10480 -pf00004pf07724pf10431 -
BMC-P
BMC-H
BMC-T
Aldehyde dehydrogenase
PTAC
Signature enzyme
Alcohol dehydrogenase
Regulator

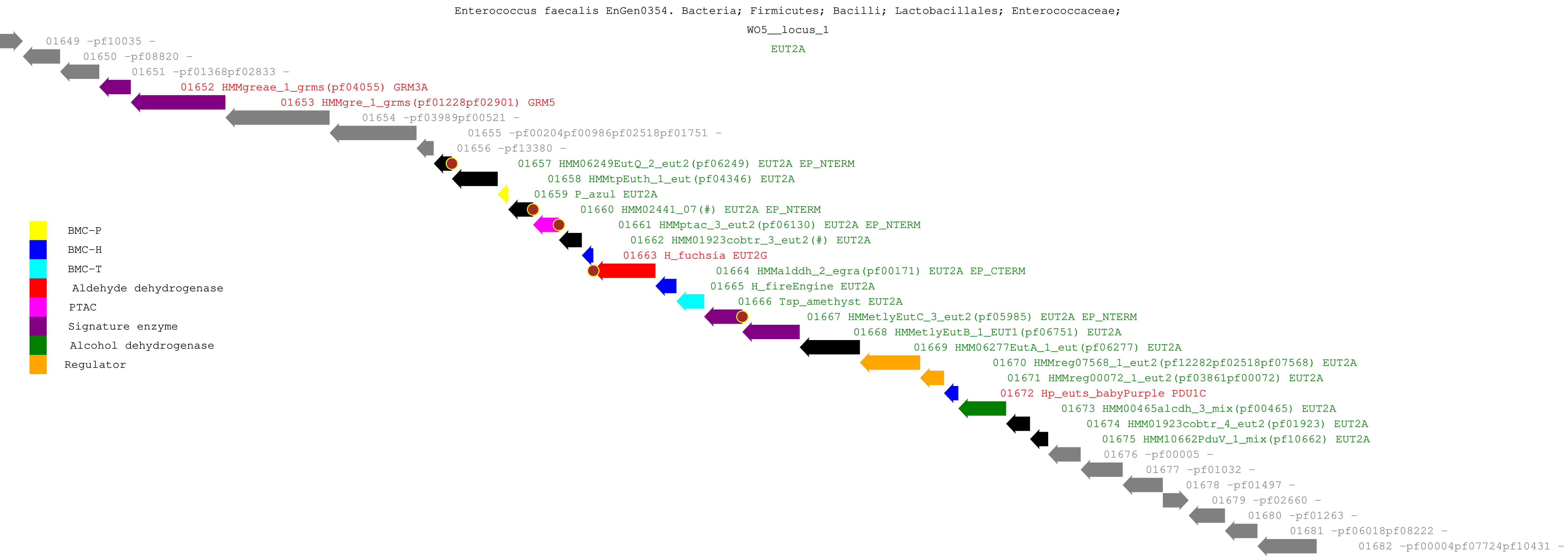

Enterococcus faecalis EnGen0354. Bacteria; Firmicutes; Bacilli; Lactobacillales; Enterococcaceae;
WO5__locus_1
EUT2A
01649 -pf10035 -
01650 -pf08820 -
01651 -pf01368pf02833 -
01652 HMMgreae_1_grms(pf04055) GRM3A
01653 HMMgre_1_grms(pf01228pf02901) GRM5
01654 -pf03989pf00521 -
01655 -pf00204pf00986pf02518pf01751 -
01656 -pf13380 -
01657 HMM06249EutQ_2_eut2(pf06249) EUT2A EP_NTERM
01658 HMMtpEuth_1_eut(pf04346) EUT2A
01659 P_azul EUT2A
01660 HMM02441_07(#) EUT2A EP_NTERM
01661 HMMptac_3_eut2(pf06130) EUT2A EP_NTERM
01662 HMM01923cobtr_3_eut2(#) EUT2A
01663 H_fuchsia EUT2G
01664 HMMalddh_2_egra(pf00171) EUT2A EP_CTERM
01665 H_fireEngine EUT2A
01666 Tsp_amethyst EUT2A
01667 HMMetlyEutC_3_eut2(pf05985) EUT2A EP_NTERM
01668 HMMetlyEutB_1_EUT1(pf06751) EUT2A
01669 HMM06277EutA_1_eut(pf06277) EUT2A
01670 HMMreg07568_1_eut2(pf12282pf02518pf07568) EUT2A
01671 HMMreg00072_1_eut2(pf03861pf00072) EUT2A
01672 Hp_euts_babyPurple PDU1C
01673 HMM00465alcdh_3_mix(pf00465) EUT2A
01674 HMM01923cobtr_4_eut2(pf01923) EUT2A
01675 HMM10662PduV_1_mix(pf10662) EUT2A
01676 -pf00005 -
01677 -pf01032 -
01678 -pf01497 -
01679 -pf02660 -
01680 -pf01263 -
01681 -pf06018pf08222 -
01682 -pf00004pf07724pf10431 -
BMC-P
BMC-H
BMC-T
Aldehyde dehydrogenase
PTAC
Signature enzyme
Alcohol dehydrogenase
Regulator

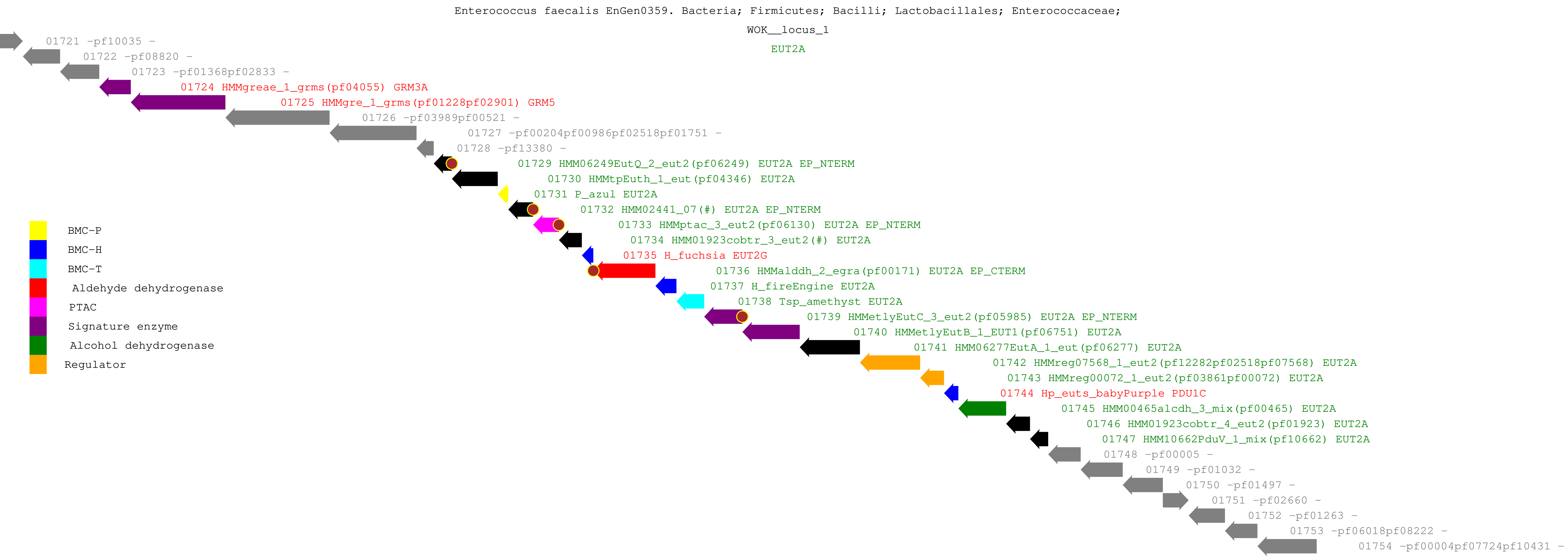

Enterococcus faecalis EnGen0359. Bacteria; Firmicutes; Bacilli; Lactobacillales; Enterococcaceae;
WOK__locus_1
EUT2A
01721 -pf10035 -
01722 -pf08820 -
01723 -pf01368pf02833 -
01724 HMMgreae_1_grms(pf04055) GRM3A
01725 HMMgre_1_grms(pf01228pf02901) GRM5
01726 -pf03989pf00521 -
01727 -pf00204pf00986pf02518pf01751 -
01728 -pf13380 -
01729 HMM06249EutQ_2_eut2(pf06249) EUT2A EP_NTERM
01730 HMMtpEuth_1_eut(pf04346) EUT2A
01731 P_azul EUT2A
01732 HMM02441_07(#) EUT2A EP_NTERM
01733 HMMptac_3_eut2(pf06130) EUT2A EP_NTERM
01734 HMM01923cobtr_3_eut2(#) EUT2A
01735 H_fuchsia EUT2G
01736 HMMalddh_2_egra(pf00171) EUT2A EP_CTERM
01737 H_fireEngine EUT2A
01738 Tsp_amethyst EUT2A
01739 HMMetlyEutC_3_eut2(pf05985) EUT2A EP_NTERM
01740 HMMetlyEutB_1_EUT1(pf06751) EUT2A
01741 HMM06277EutA_1_eut(pf06277) EUT2A
01742 HMMreg07568_1_eut2(pf12282pf02518pf07568) EUT2A
01743 HMMreg00072_1_eut2(pf03861pf00072) EUT2A
01744 Hp_euts_babyPurple PDU1C
01745 HMM00465alcdh_3_mix(pf00465) EUT2A
01746 HMM01923cobtr_4_eut2(pf01923) EUT2A
01747 HMM10662PduV_1_mix(pf10662) EUT2A
01748 -pf00005 -
01749 -pf01032 -
01750 -pf01497 -
01751 -pf02660 -
01752 -pf01263 -
01753 -pf06018pf08222 -
01754 -pf00004pf07724pf10431 -
BMC-P
BMC-H
BMC-T
Aldehyde dehydrogenase
PTAC
Signature enzyme
Alcohol dehydrogenase
Regulator

Enterococcus faecalis ATCC 6055. Bacteria; Firmicutes; Bacilli; Lactobacillales; Enterococcaceae;

WOU__locus_1

EUT2A

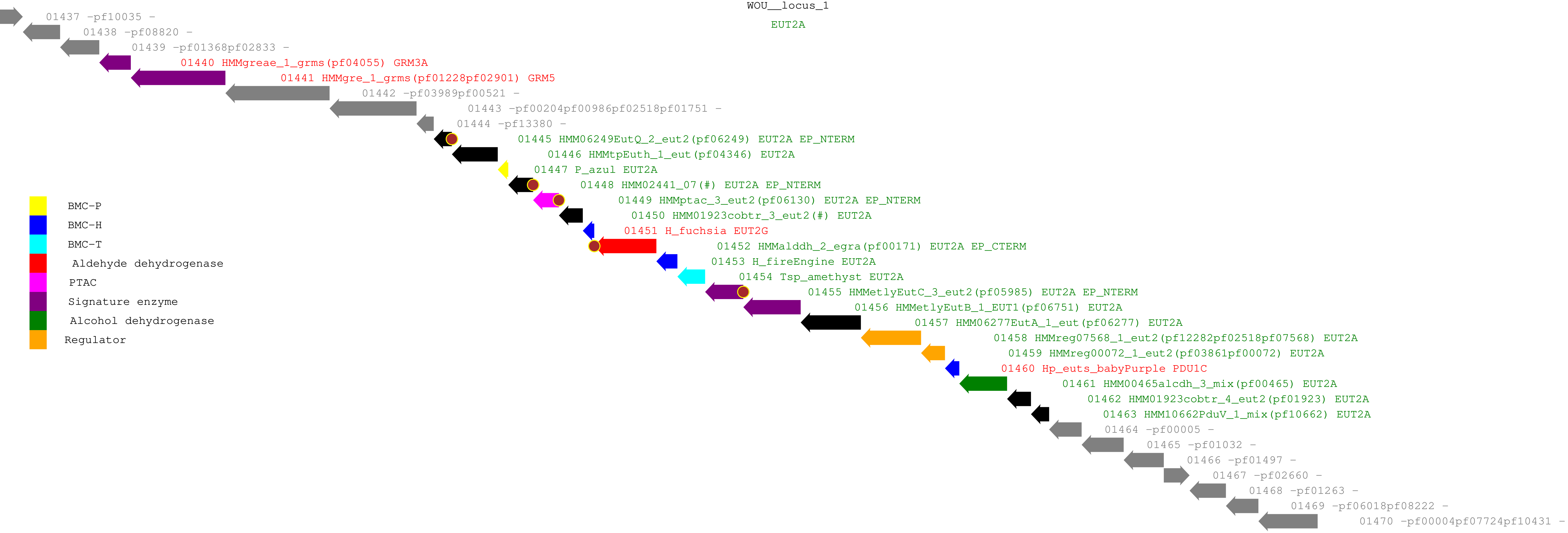

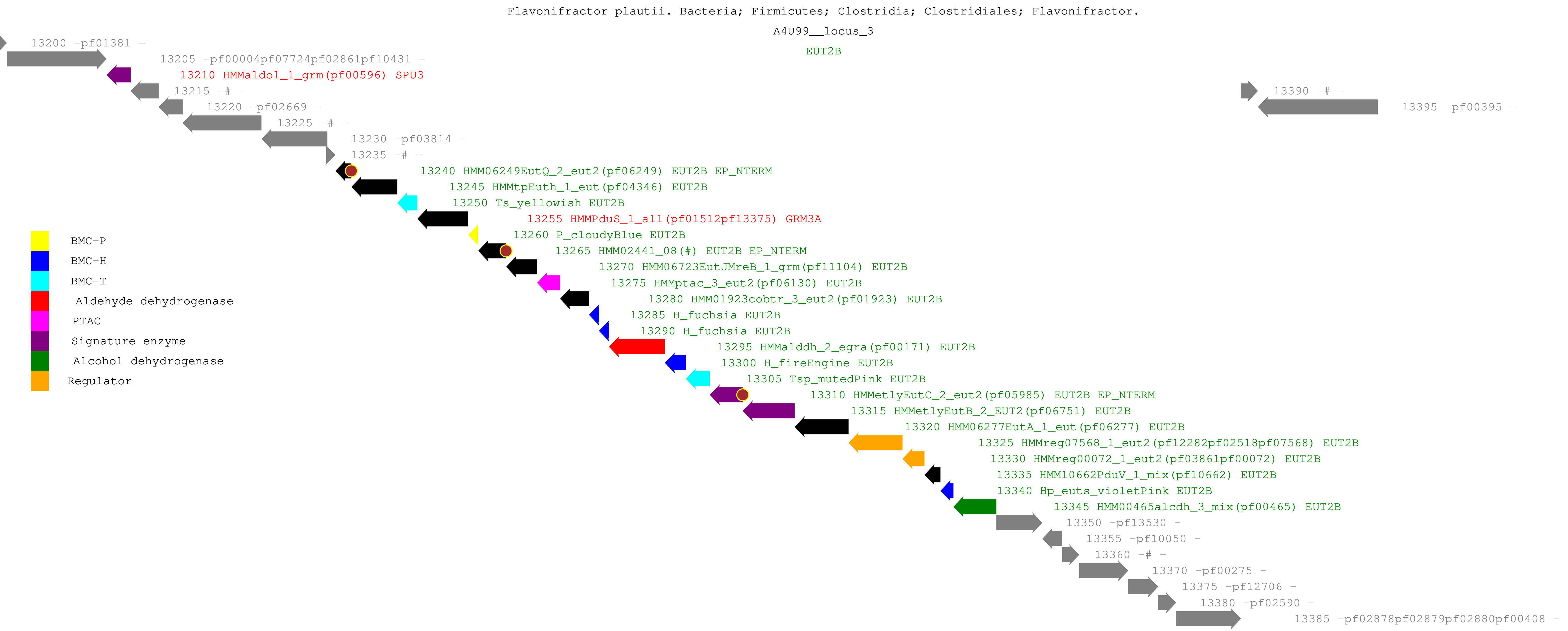
Flavonifractor plautii. Bacteria; Firmicutes; Clostridia; Clostridiales; Flavonifractor.
A4U99__locus_3
EUT2B
13200 -pf01381 -
13205 -pf00004pf07724pf02861pf10431 -
13210 HMMaldol_1_grm(pf00596) SPU3
13215 -# -
13220 -pf02669 -
13225 -# -
13230 -pf03814 -
13235 -# -
13240 HMM06249EutQ_2_eut2(pf06249) EUT2B EP_NTERM
13245 HMMtpEuth_1_eut(pf04346) EUT2B
13250 Ts_yellowish EUT2B
13255 HMMPduS_1_all(pf01512pf13375) GRM3A
13260 P_cloudyBlue EUT2B
13265 HMM02441_08(#) EUT2B EP_NTERM
13270 HMM06723EutJMreB_1_grm(pf11104) EUT2B
13275 HMMptac_3_eut2(pf06130) EUT2B
13280 HMM01923cobtr_3_eut2(pf01923) EUT2B
13285 H_fuchsia EUT2B
13290 H_fuchsia EUT2B
13295 HMMalddh_2_egra(pf00171) EUT2B
13300 H_fireEngine EUT2B
13305 Tsp_mutedPink EUT2B
13310 HMMetlyEutC_2_eut2(pf05985) EUT2B EP_NTERM
13315 HMMetlyEutB_2_EUT2(pf06751) EUT2B
13320 HMM06277EutA_1_eut(pf06277) EUT2B
13325 HMMreg07568_1_eut2(pf12282pf02518pf07568) EUT2B
13330 HMMreg00072_1_eut2(pf03861pf00072) EUT2B
13335 HMM10662PduV_1_mix(pf10662) EUT2B
13340 Hp_euts_violetPink EUT2B
13345 HMM00465alcdh_3_mix(pf00465) EUT2B
13350 -pf13530 -
13355 -pf10050 -
13360 -# -
13370 -pf00275 -
13375 -pf12706 -
13380 -pf02590 -
13385 -pf02878pf02879pf02880pf00408 -
13390 -# -
13395 -pf00395 -
BMC-P
BMC-H
BMC-T
Aldehyde dehydrogenase
PTAC
Signature enzyme
Alcohol dehydrogenase
Regulator

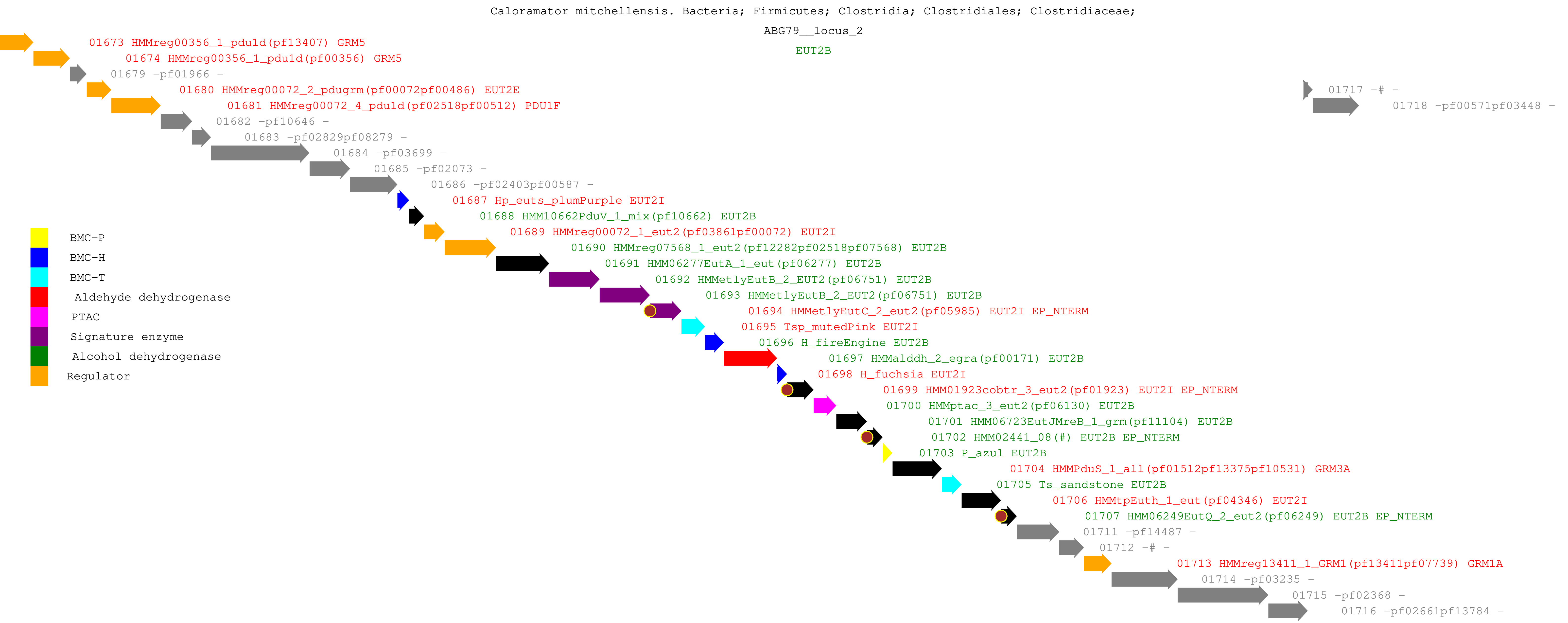
Caloramator mitchellensis. Bacteria; Firmicutes; Clostridia; Clostridiales; Clostridiaceae;
ABG79__locus_2
EUT2B
01673 HMMreg00356_1_pdu1d(pf13407) GRM5
01674 HMMreg00356_1_pdu1d(pf00356) GRM5
01679 -pf01966 -
01680 HMMreg00072_2_pdugrm(pf00072pf00486) EUT2E
01681 HMMreg00072_4_pdu1d(pf02518pf00512) PDU1F
01682 -pf10646 -
01683 -pf02829pf08279 -
01684 -pf03699 -
01685 -pf02073 -
01686 -pf02403pf00587 -
01687 Hp_euts_plumPurple EUT2I
01688 HMM10662PduV_1_mix(pf10662) EUT2B
01689 HMMreg00072_1_eut2(pf03861pf00072) EUT2I
01690 HMMreg07568_1_eut2(pf12282pf02518pf07568) EUT2B
01691 HMM06277EutA_1_eut(pf06277) EUT2B
01692 HMMetlyEutB_2_EUT2(pf06751) EUT2B
01693 HMMetlyEutB_2_EUT2(pf06751) EUT2B
01694 HMMetlyEutC_2_eut2(pf05985) EUT2I EP_NTERM
01695 Tsp_mutedPink EUT2I
01696 H_fireEngine EUT2B
01697 HMMalddh_2_egra(pf00171) EUT2B
01698 H_fuchsia EUT2I
01699 HMM01923cobtr_3_eut2(pf01923) EUT2I EP_NTERM
01700 HMMptac_3_eut2(pf06130) EUT2B
01701 HMM06723EutJMreB_1_grm(pf11104) EUT2B
01702 HMM02441_08(#) EUT2B EP_NTERM
01703 P_azul EUT2B
01704 HMMPduS_1_all(pf01512pf13375pf10531) GRM3A
01705 Ts_sandstone EUT2B
01706 HMMtpEuth_1_eut(pf04346) EUT2I
01707 HMM06249EutQ_2_eut2(pf06249) EUT2B EP_NTERM
01711 -pf14487 -
01712 -# -
01713 HMMreg13411_1_GRM1(pf13411pf07739) GRM1A
01714 -pf03235 -
01715 -pf02368 -
01716 -pf02661pf13784 -
01717 -# -
01718 -pf00571pf03448 -
BMC-P
BMC-H
BMC-T
Aldehyde dehydrogenase
PTAC
Signature enzyme
Alcohol dehydrogenase
Regulator

Acetobacterium bakii. Bacteria; Firmicutes; Clostridia; Clostridiales; Eubacteriaceae;

AKG39__locus_1

EUT2B

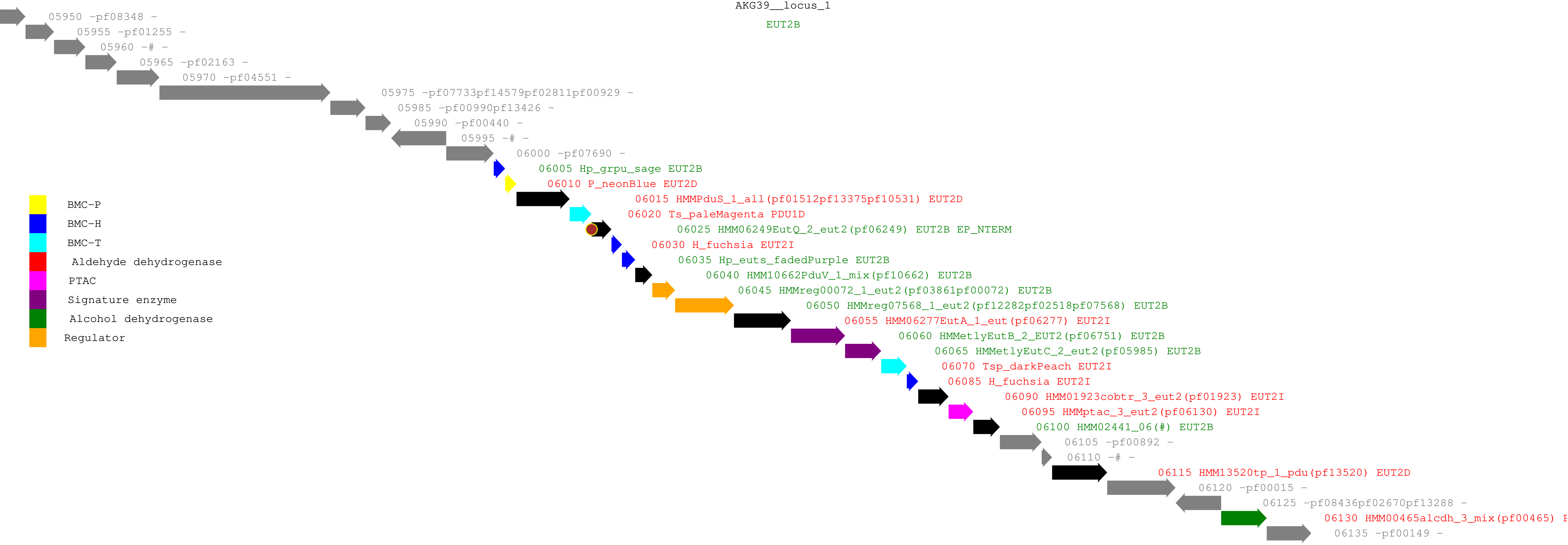

Alkaliphilus metalliredigens (strain QYMF). Bacteria; Firmicutes; Clostridia; Clostridiales; Clostridiaceae;

Amet__locus_1

EUT2B

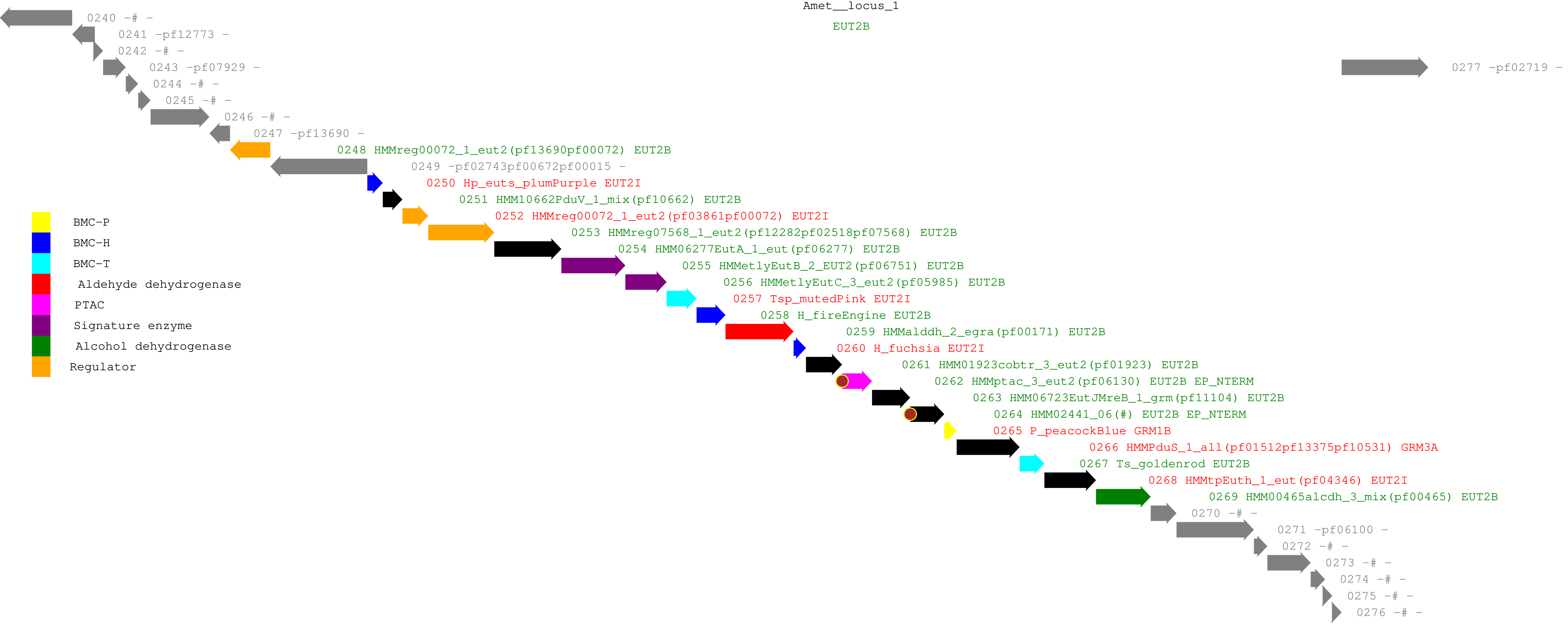

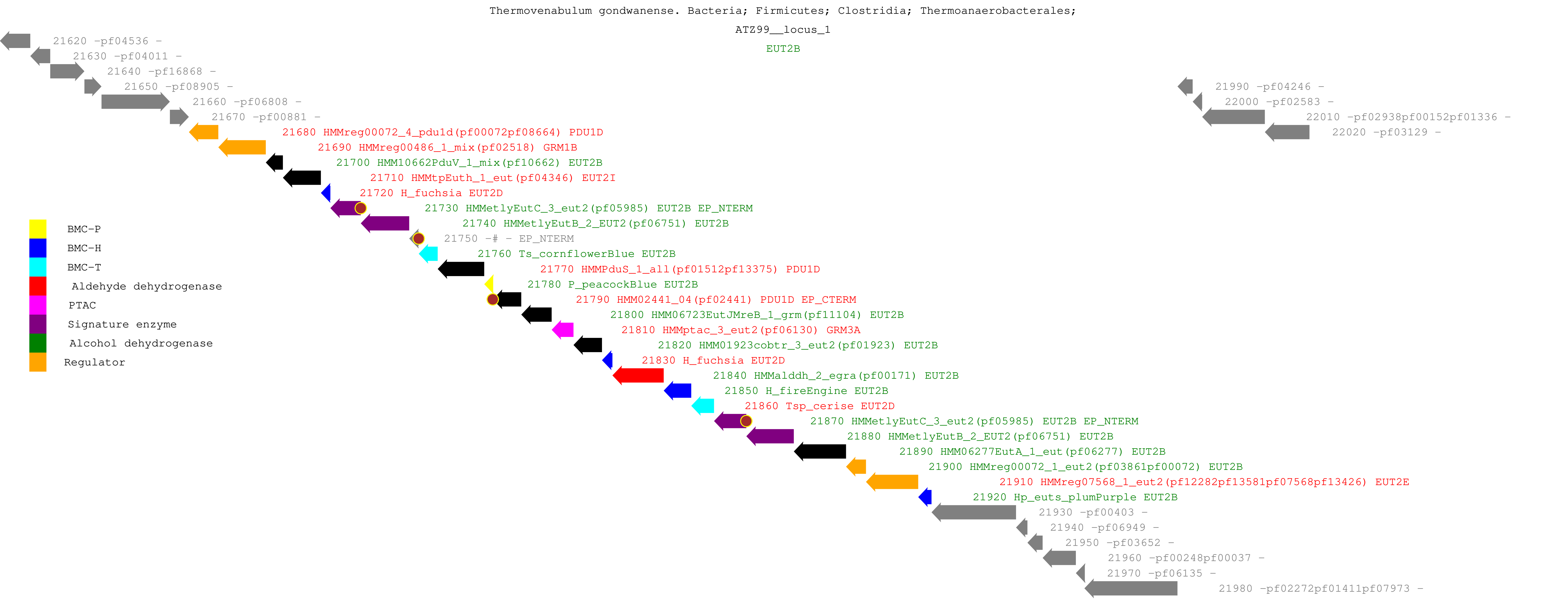
Thermovenabulum gondwanense. Bacteria; Firmicutes; Clostridia; Thermoanaerobacterales;
ATZ99__locus_1
EUT2B
21620 -pf04536 -
21630 -pf04011 -
21640 -pf16868 -
21650 -pf08905 -
21660 -pf06808 -
21670 -pf00881 -
21680 HMMreg00072_4_pdu1d(pf00072pf08664) PDU1D
21690 HMMreg00486_1_mix(pf02518) GRM1B
21700 HMM10662PduV_1_mix(pf10662) EUT2B
21710 HMMtpEuth_1_eut(pf04346) EUT2I
21720 H_fuchsia EUT2D
21730 HMMetlyEutC_3_eut2(pf05985) EUT2B EP_NTERM
21740 HMMetlyEutB_2_EUT2(pf06751) EUT2B
21750 -# - EP_NTERM
21760 Ts_cornflowerBlue EUT2B
21770 HMMPduS_1_all(pf01512pf13375) PDU1D
21780 P_peacockBlue EUT2B
21790 HMM02441_04(pf02441) PDU1D EP_CTERM
21800 HMM06723EutJMreB_1_grm(pf11104) EUT2B
21810 HMMptac_3_eut2(pf06130) GRM3A
21820 HMM01923cobtr_3_eut2(pf01923) EUT2B
21830 H_fuchsia EUT2D
21840 HMMalddh_2_egra(pf00171) EUT2B
21850 H_fireEngine EUT2B
21860 Tsp_cerise EUT2D
21870 HMMetlyEutC_3_eut2(pf05985) EUT2B EP_NTERM
21880 HMMetlyEutB_2_EUT2(pf06751) EUT2B
21890 HMM06277EutA_1_eut(pf06277) EUT2B
21900 HMMreg00072_1_eut2(pf03861pf00072) EUT2B
21910 HMMreg07568_1_eut2(pf12282pf13581pf07568pf13426) EUT2E
21920 Hp_euts_plumPurple EUT2B
21930 -pf00403 -
21940 -pf06949 -
21950 -pf03652 -
21960 -pf00248pf00037 -
21970 -pf06135 -
21980 -pf02272pf01411pf07973 -
21990 -pf04246 -
22000 -pf02583 -
22010 -pf02938pf00152pf01336 -
22020 -pf03129 -
BMC-P
BMC-H
BMC-T
Aldehyde dehydrogenase
PTAC
Signature enzyme
Alcohol dehydrogenase
Regulator

Halanaerobium sp. T82-1. Bacteria; Firmicutes; Clostridia; Halanaerobiales; Halanaerobiaceae;

AWL62__locus_1

EUT2B

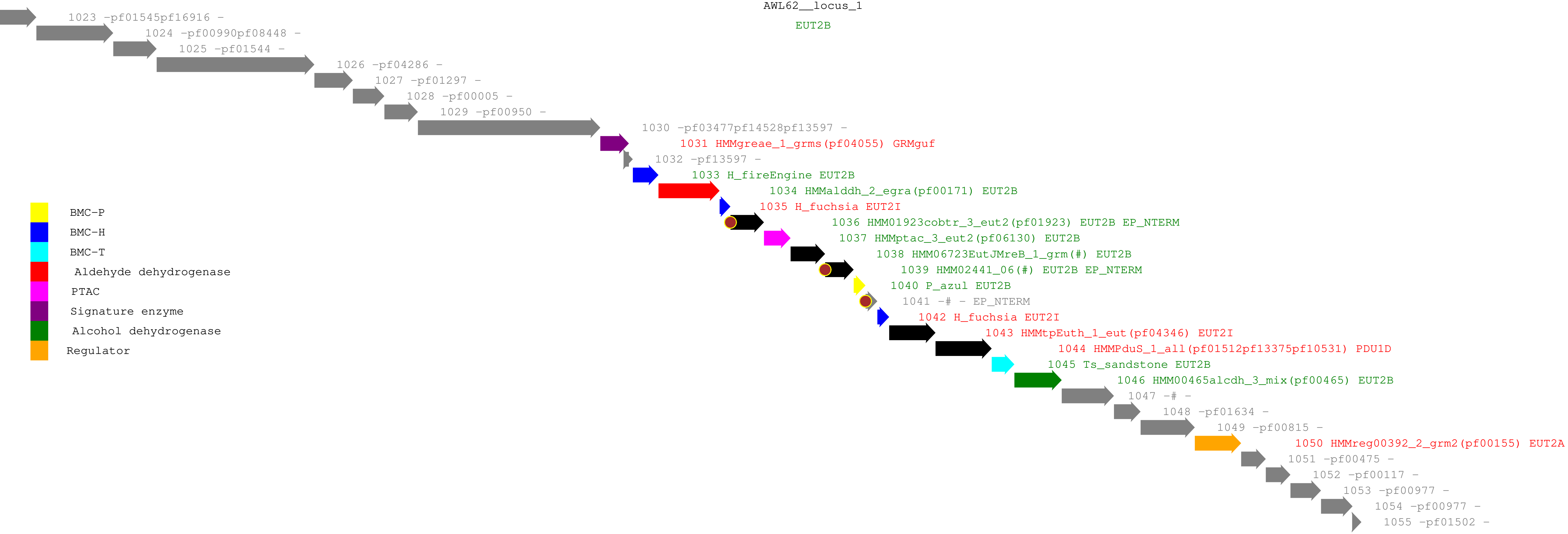

Eubacterium minutum ATCC 700079. Bacteria; Firmicutes; Clostridia; Clostridiales;

AXF21__locus_1

EUT2B

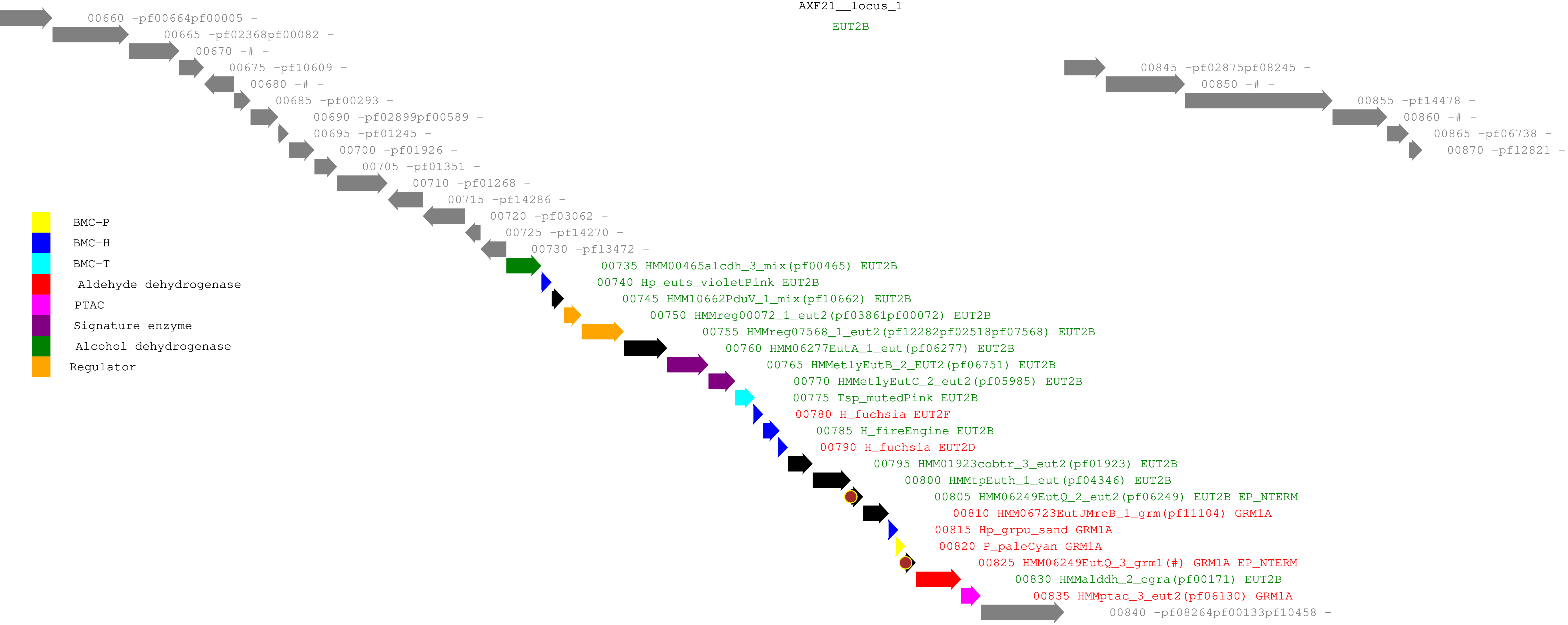

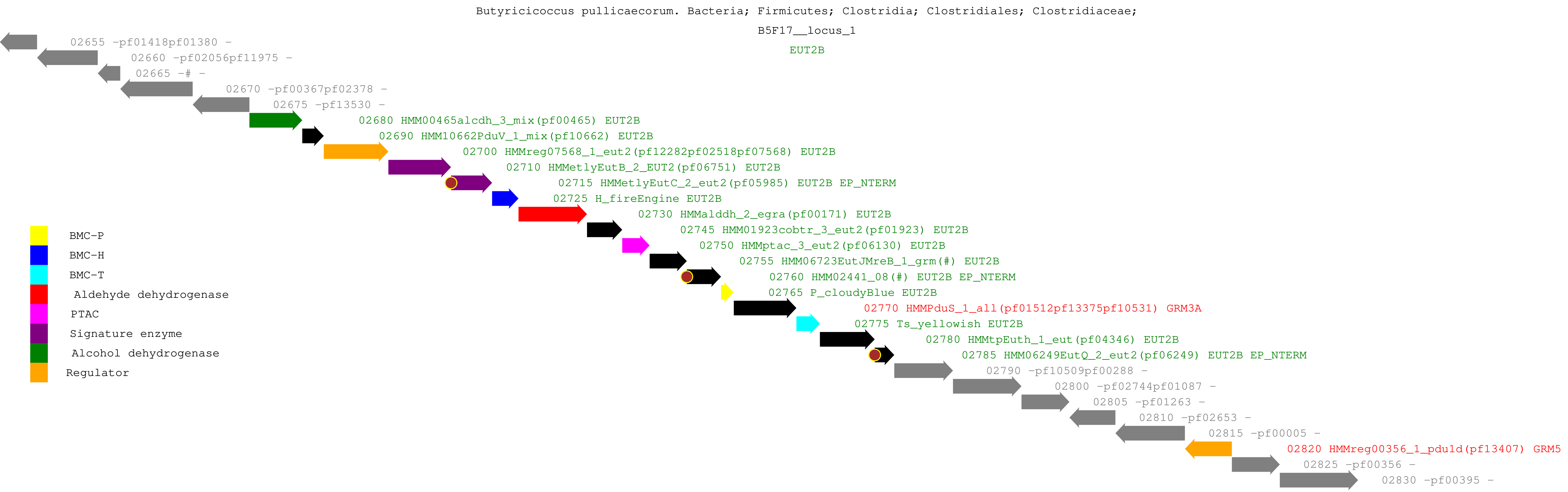

Butyricicoccus pullicaecorum. Bacteria; Firmicutes; Clostridia; Clostridiales; Clostridiaceae;
B5F17__locus_1
EUT2B
02655 -pf01418pf01380 -
02660 -pf02056pf11975 -
02665 -# -
02670 -pf00367pf02378 -
02675 -pf13530 -
02680 HMM00465alcdh_3_mix(pf00465) EUT2B
02690 HMM10662PduV_1_mix(pf10662) EUT2B
02700 HMMreg07568_1_eut2(pf12282pf02518pf07568) EUT2B
02710 HMMetlyEutB_2_EUT2(pf06751) EUT2B
02715 HMMetlyEutC_2_eut2(pf05985) EUT2B EP_NTERM
02725 H_fireEngine EUT2B
02730 HMMalddh_2_egra(pf00171) EUT2B
02745 HMM01923cobtr_3_eut2(pf01923) EUT2B
02750 HMMptac_3_eut2(pf06130) EUT2B
02755 HMM06723EutJMreB_1_grm(#) EUT2B
02760 HMM02441_08(#) EUT2B EP_NTERM
02765 P_cloudyBlue EUT2B
02770 HMMPduS_1_all(pf01512pf13375pf10531) GRM3A
02775 Ts_yellowish EUT2B
02780 HMMtpEuth_1_eut(pf04346) EUT2B
02785 HMM06249EutQ_2_eut2(pf06249) EUT2B EP_NTERM
02790 -pf10509pf00288 -
02800 -pf02744pf01087 -
02805 -pf01263 -
02810 -pf02653 -
02815 -pf00005 -
02820 HMMreg00356_1_pdu1d(pf13407) GRM5
02825 -pf00356 -
02830 -pf00395 -
BMC-P
BMC-H
BMC-T
Aldehyde dehydrogenase
PTAC
Signature enzyme
Alcohol dehydrogenase
Regulator

Lachnoclostridium sp. An196. Bacteria; Firmicutes; Clostridia; Clostridiales; Lachnospiraceae.

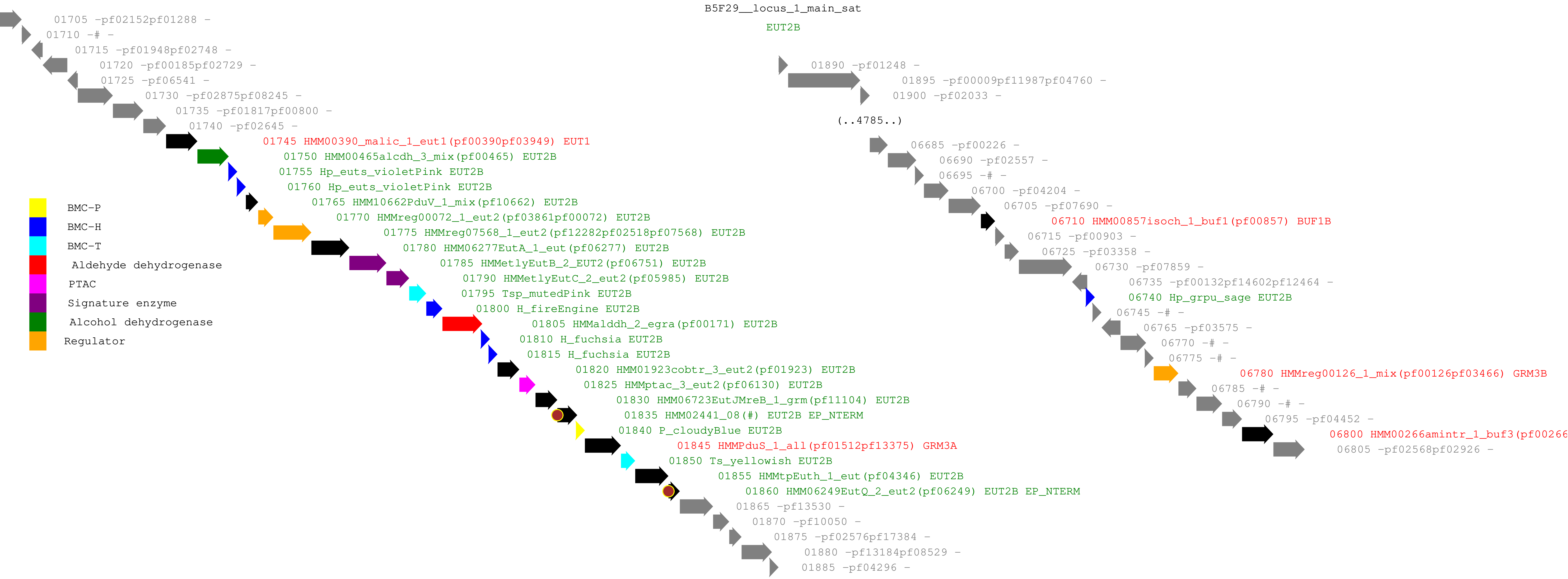

Flavonifractor plautii. Bacteria; Firmicutes; Clostridia; Clostridiales; Flavonifractor.

B5F52__locus_3

EUT2B

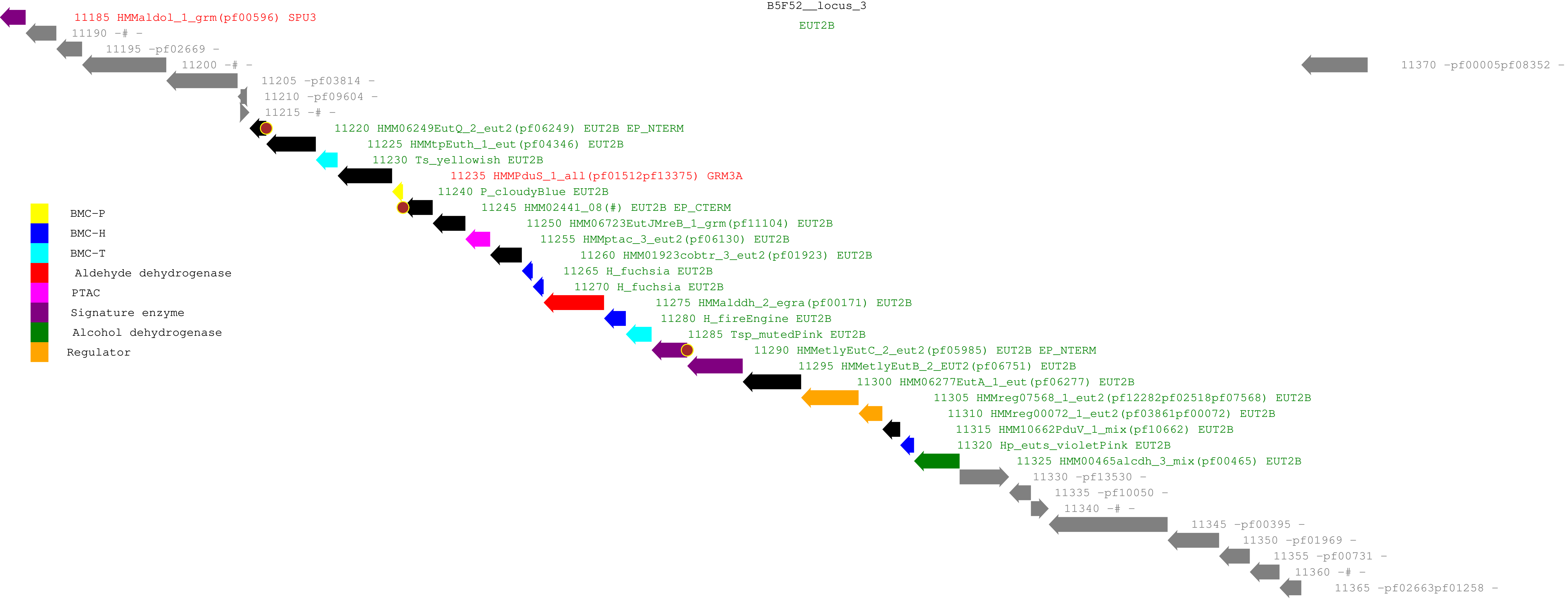

Bacteroidetes bacterium 4572_77. Bacteria; Bacteroidetes.

B7C24__locus_1

EUT2B

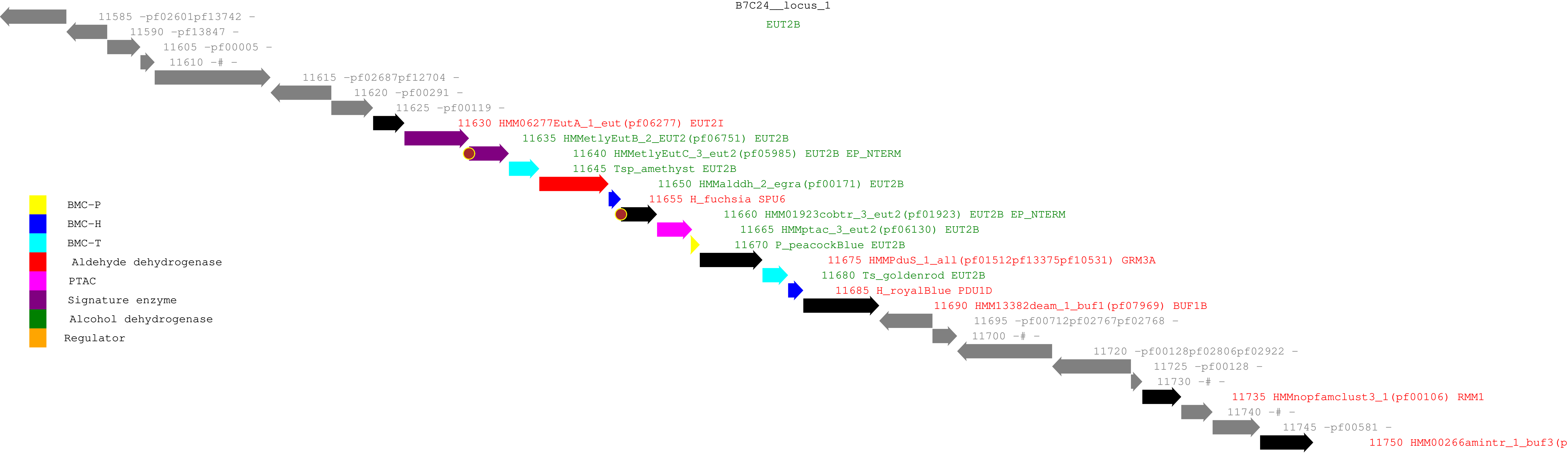

Pseudoflavonifractor capillosus ATCC 29799. Bacteria; Firmicutes; Clostridia; Clostridiales; Pseudoflavonifractor.

BACCAP__locus_2

EUT2B

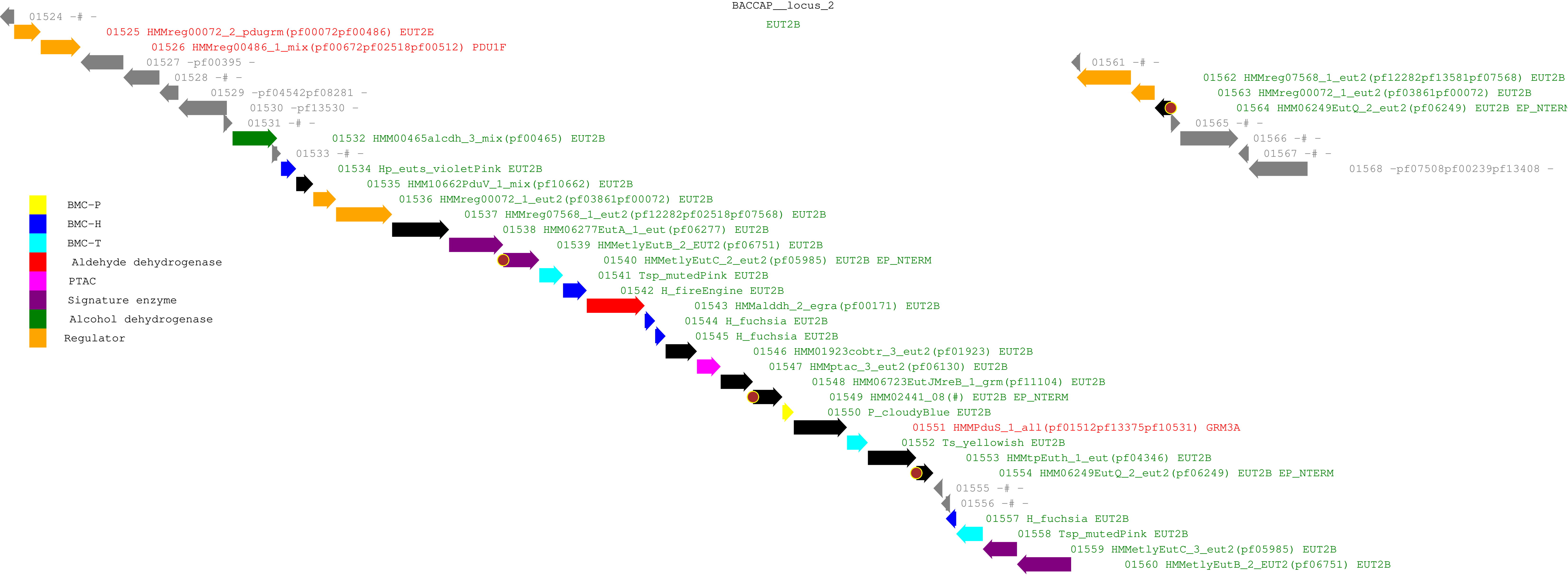

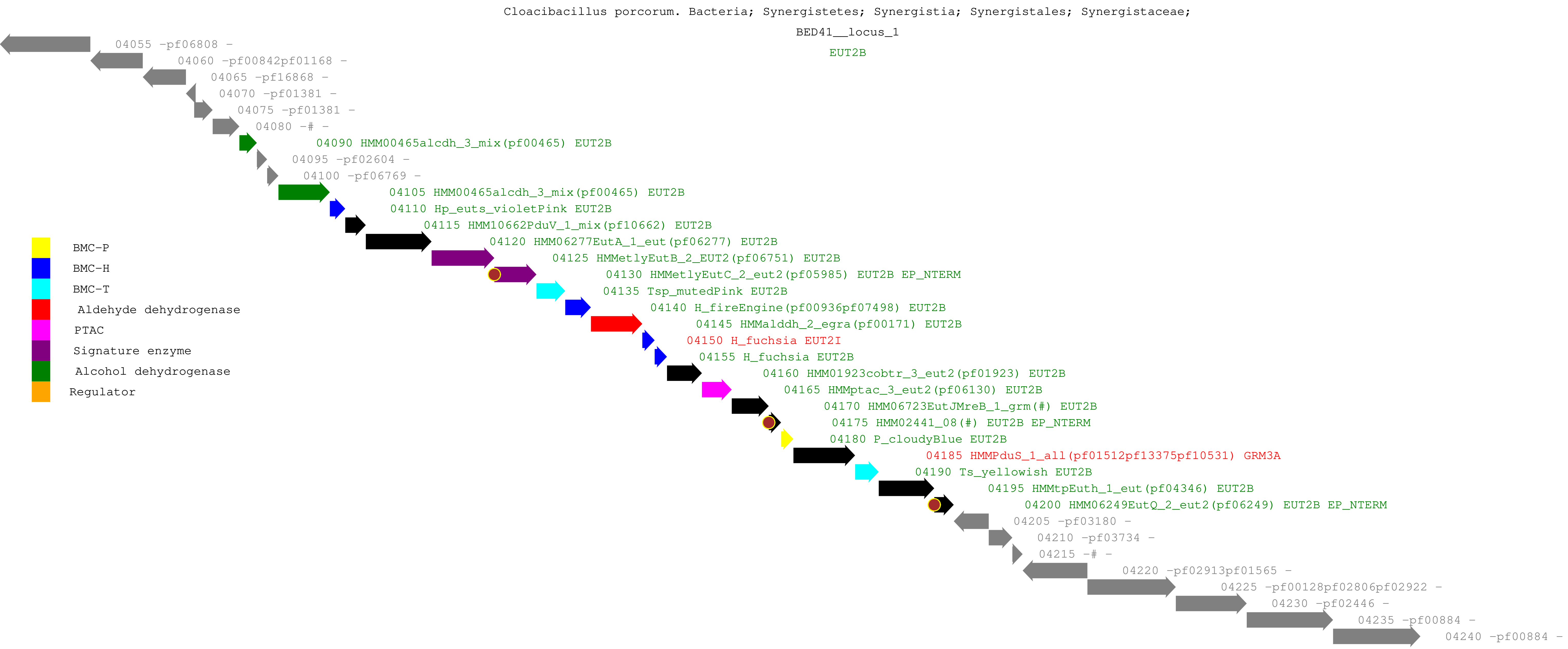
Cloacibacillus porcorum. Bacteria; Synergistetes; Synergistia; Synergistales; Synergistaceae;
BED41__locus_1
EUT2B
04055 -pf06808 -
04060 -pf00842pf01168 -
04065 -pf16868 -
04070 -pf01381 -
04075 -pf01381 -
04080 -# -
04090 HMM00465alcdh_3_mix(pf00465) EUT2B
04095 -pf02604 -
04100 -pf06769 -
04105 HMM00465alcdh_3_mix(pf00465) EUT2B
04110 Hp_euts_violetPink EUT2B
04115 HMM10662PduV_1_mix(pf10662) EUT2B
04120 HMM06277EutA_1_eut(pf06277) EUT2B
04125 HMMetlyEutB_2_EUT2(pf06751) EUT2B
04130 HMMetlyEutC_2_eut2(pf05985) EUT2B EP_NTERM
04135 Tsp_mutedPink EUT2B
04140 H_fireEngine(pf00936pf07498) EUT2B
04145 HMMalddh_2_egra(pf00171) EUT2B
04150 H_fuchsia EUT2I
04155 H_fuchsia EUT2B
04160 HMM01923cobtr_3_eut2(pf01923) EUT2B
04165 HMMptac_3_eut2(pf06130) EUT2B
04170 HMM06723EutJMreB_1_grm(#) EUT2B
04175 HMM02441_08(#) EUT2B EP_NTERM
04180 P_cloudyBlue EUT2B
04185 HMMPduS_1_all(pf01512pf13375pf10531) GRM3A
04190 Ts_yellowish EUT2B
04195 HMMtpEuth_1_eut(pf04346) EUT2B
04200 HMM06249EutQ_2_eut2(pf06249) EUT2B EP_NTERM
04205 -pf03180 -
04210 -pf03734 -
04215 -# -
04220 -pf02913pf01565 -
04225 -pf00128pf02806pf02922 -
04230 -pf02446 -
04235 -pf00884 -
04240 -pf00884 -
BMC-P
BMC-H
BMC-T
Aldehyde dehydrogenase
PTAC
Signature enzyme
Alcohol dehydrogenase
Regulator

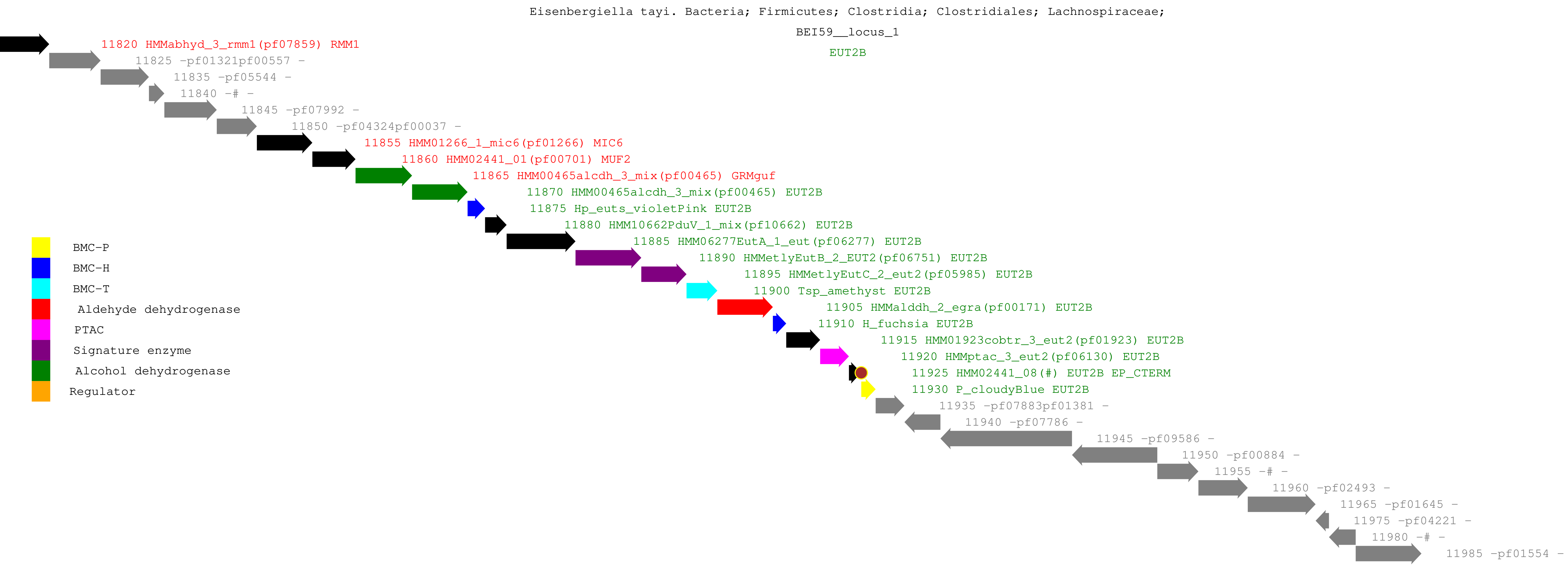

Eisenbergiella tayi. Bacteria; Firmicutes; Clostridia; Clostridiales; Lachnospiraceae;
BEI59__locus_1
EUT2B
11820 HMMabhyd_3_rmm1(pf07859) RMM1
11825 -pf01321pf00557 -
11835 -pf05544 -
11840 -# -
11845 -pf07992 -
11850 -pf04324pf00037 -
11855 HMM01266_1_mic6(pf01266) MIC6
11860 HMM02441_01(pf00701) MUF2
11865 HMM00465alcdh_3_mix(pf00465) GRMguf
11870 HMM00465alcdh_3_mix(pf00465) EUT2B
11875 Hp_euts_violetPink EUT2B
11880 HMM10662PduV_1_mix(pf10662) EUT2B
11885 HMM06277EutA_1_eut(pf06277) EUT2B
11890 HMMetlyEutB_2_EUT2(pf06751) EUT2B
11895 HMMetlyEutC_2_eut2(pf05985) EUT2B
11900 Tsp_amethyst EUT2B
11905 HMMalddh_2_egra(pf00171) EUT2B
11910 H_fuchsia EUT2B
11915 HMM01923cobtr_3_eut2(pf01923) EUT2B
11920 HMMptac_3_eut2(pf06130) EUT2B
11925 HMM02441_08(#) EUT2B EP_CTERM
11930 P_cloudyBlue EUT2B
11935 -pf07883pf01381 -
11940 -pf07786 -
11945 -pf09586 -
11950 -pf00884 -
11955 -# -
11960 -pf02493 -
11965 -pf01645 -
11975 -pf04221 -
11980 -# -
11985 -pf01554 -
BMC-P
BMC-H
BMC-T
Aldehyde dehydrogenase
PTAC
Signature enzyme
Alcohol dehydrogenase
Regulator

Halanaerobium sp. MDAL1. Bacteria; Firmicutes; Clostridia; Halanaerobiales; Halanaerobiaceae;

BHK79__locus_1

EUT2B

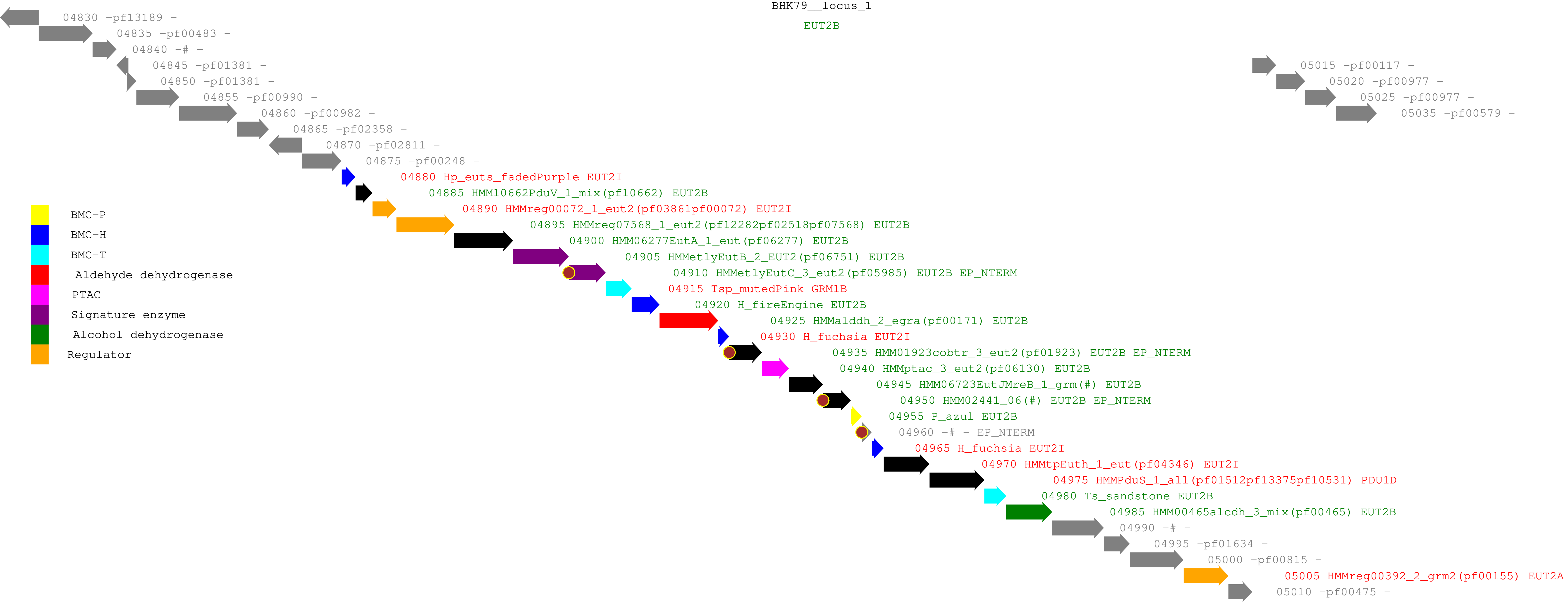

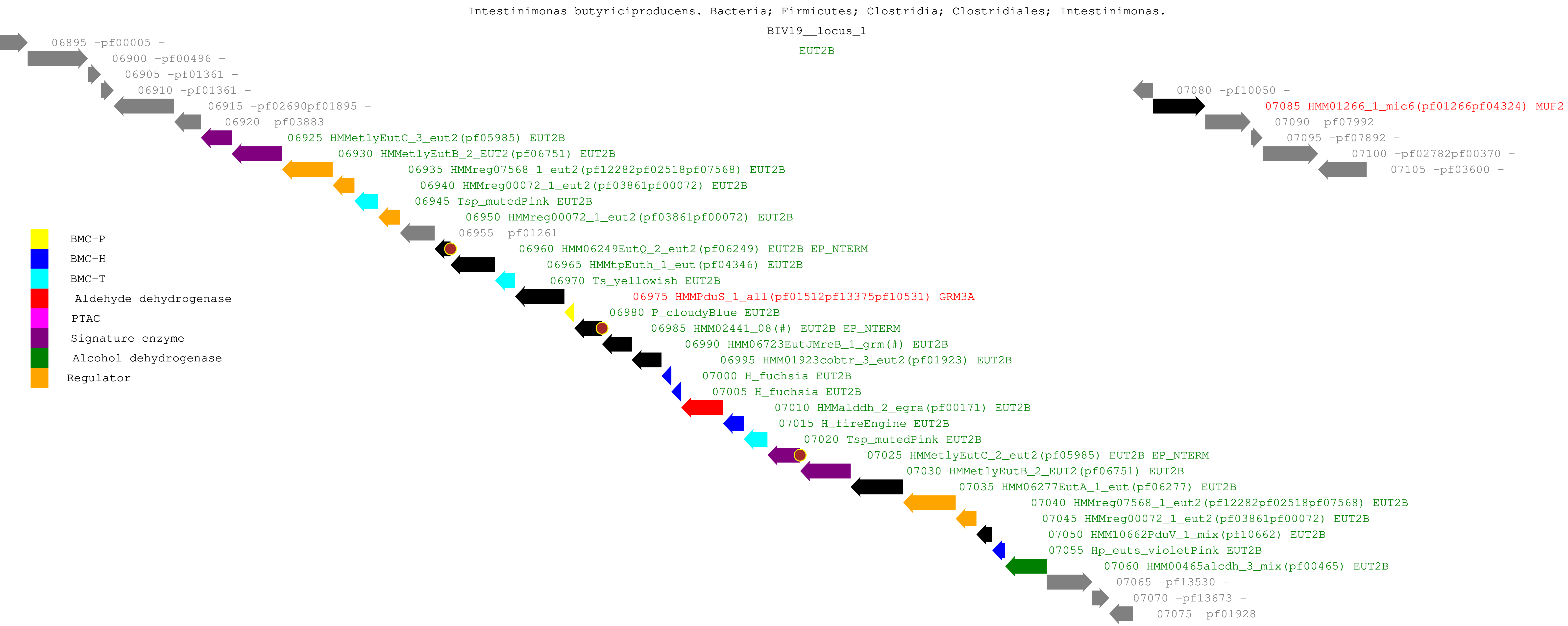

Intestinimonas butyriciproducens. Bacteria; Firmicutes; Clostridia; Clostridiales; Intestinimonas.
BIV19__locus_1
EUT2B
06895 -pf00005 -
06900 -pf00496 -
06905 -pf01361 -
06910 -pf01361 -
06915 -pf02690pf01895 -
06920 -pf03883 -
06925 HMMetlyEutC_3_eut2(pf05985) EUT2B
06930 HMMetlyEutB_2_EUT2(pf06751) EUT2B
06935 HMMreg07568_1_eut2(pf12282pf02518pf07568) EUT2B
06940 HMMreg00072_1_eut2(pf03861pf00072) EUT2B
06945 Tsp_mutedPink EUT2B
06950 HMMreg00072_1_eut2(pf03861pf00072) EUT2B
06955 -pf01261 -
06960 HMM06249EutQ_2_eut2(pf06249) EUT2B EP_NTERM
06965 HMMtpEuth_1_eut(pf04346) EUT2B
06970 Ts_yellowish EUT2B
06975 HMMPduS_1_all(pf01512pf13375pf10531) GRM3A
06980 P_cloudyBlue EUT2B
06985 HMM02441_08(#) EUT2B EP_NTERM
06990 HMM06723EutJMreB_1_grm(#) EUT2B
06995 HMM01923cobtr_3_eut2(pf01923) EUT2B
07000 H_fuchsia EUT2B
07005 H_fuchsia EUT2B
07010 HMMalddh_2_egra(pf00171) EUT2B
07015 H_fireEngine EUT2B
07020 Tsp_mutedPink EUT2B
07025 HMMetlyEutC_2_eut2(pf05985) EUT2B EP_NTERM
07030 HMMetlyEutB_2_EUT2(pf06751) EUT2B
07035 HMM06277EutA_1_eut(pf06277) EUT2B
07040 HMMreg07568_1_eut2(pf12282pf02518pf07568) EUT2B
07045 HMMreg00072_1_eut2(pf03861pf00072) EUT2B
07050 HMM10662PduV_1_mix(pf10662) EUT2B
07055 Hp_euts_violetPink EUT2B
07060 HMM00465alcdh_3_mix(pf00465) EUT2B
07065 -pf13530 -
07070 -pf13673 -
07075 -pf01928 -
07080 -pf10050 -
07085 HMM01266_1_mic6(pf01266pf04324) MUF2
07090 -pf07992 -
07095 -pf07892 -
07100 -pf02782pf00370 -
07105 -pf03600 -
BMC-P
BMC-H
BMC-T
Aldehyde dehydrogenase
PTAC
Signature enzyme
Alcohol dehydrogenase
Regulator

Clostridium formicaceticum. Bacteria; Firmicutes; Clostridia; Clostridiales; Clostridiaceae;

BJL90__locus_2

EUT2B

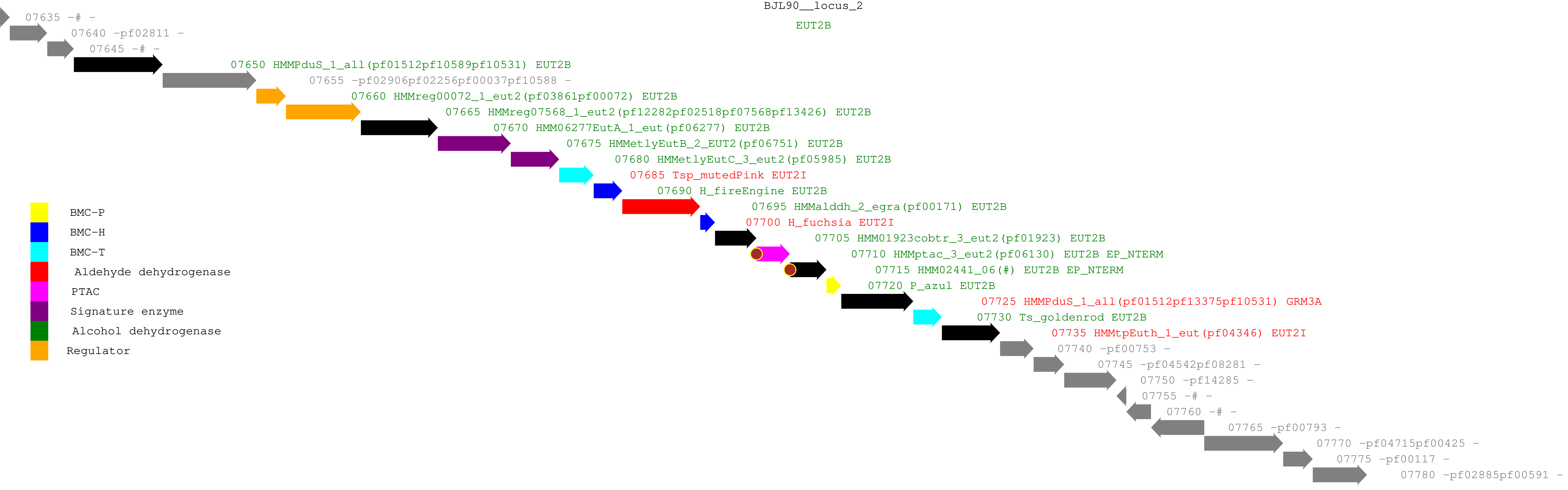

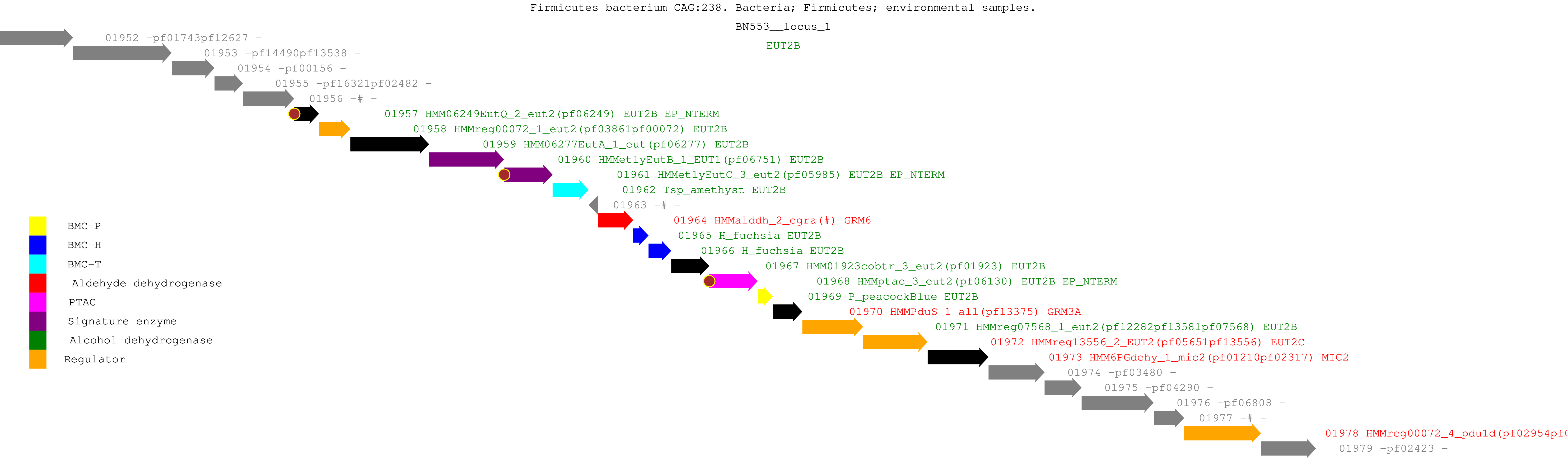
Firmicutes bacterium CAG:238. Bacteria; Firmicutes; environmental samples.
BN553__locus_1
EUT2B
01952 -pf01743pf12627 -
01953 -pf14490pf13538 -
01954 -pf00156 -
01955 -pf16321pf02482 -
01956 -# -
01957 HMM06249EutQ_2_eut2(pf06249) EUT2B EP_NTERM
01958 HMMreg00072_1_eut2(pf03861pf00072) EUT2B
01959 HMM06277EutA_1_eut(pf06277) EUT2B
01960 HMMetlyEutB_1_EUT1(pf06751) EUT2B
01961 HMMetlyEutC_3_eut2(pf05985) EUT2B EP_NTERM
01962 Tsp_amethyst EUT2B
01963 -# -
01964 HMMalddh_2_egra(#) GRM6
01965 H_fuchsia EUT2B
01966 H_fuchsia EUT2B
01967 HMM01923cobtr_3_eut2(pf01923) EUT2B
01968 HMMptac_3_eut2(pf06130) EUT2B EP_NTERM
01969 P_peacockBlue EUT2B
01970 HMMPduS_1_all(pf13375) GRM3A
01971 HMMreg07568_1_eut2(pf12282pf13581pf07568) EUT2B
01972 HMMreg13556_2_EUT2(pf05651pf13556) EUT2C
01973 HMM6PGdehy_1_mic2(pf01210pf02317) MIC2
01974 -pf03480 -
01975 -pf04290 -
01976 -pf06808 -
01977 -# -
01978 HMMreg00072_4_pdu1d(pf02954pf0
01979 -pf02423 -
BMC-P
BMC-H
BMC-T
Aldehyde dehydrogenase
PTAC
Signature enzyme
Alcohol dehydrogenase
Regulator

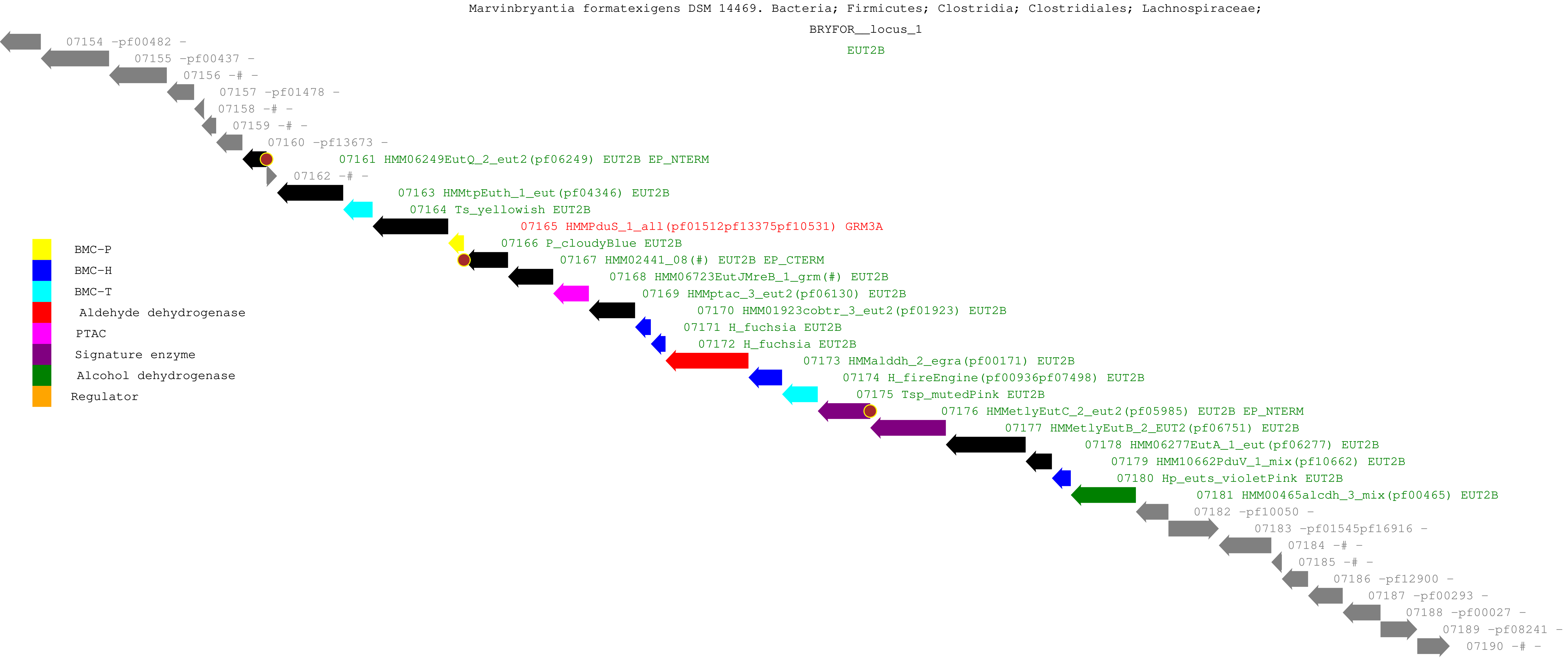
Marvinbryantia formatexigens DSM 14469. Bacteria; Firmicutes; Clostridia; Clostridiales; Lachnospiraceae;
BRYFOR__locus_1
EUT2B
07154 -pf00482 -
07155 -pf00437 -
07156 -# -
07157 -pf01478 -
07158 -# -
07159 -# -
07160 -pf13673 -
07161 HMM06249EutQ_2_eut2(pf06249) EUT2B EP_NTERM
07162 -# -
07163 HMMtpEuth_1_eut(pf04346) EUT2B
07164 Ts_yellowish EUT2B
07165 HMMPduS_1_all(pf01512pf13375pf10531) GRM3A
07166 P_cloudyBlue EUT2B
07167 HMM02441_08(#) EUT2B EP_CTERM
07168 HMM06723EutJMreB_1_grm(#) EUT2B
07169 HMMptac_3_eut2(pf06130) EUT2B
07170 HMM01923cobtr_3_eut2(pf01923) EUT2B
07171 H_fuchsia EUT2B
07172 H_fuchsia EUT2B
07173 HMMalddh_2_egra(pf00171) EUT2B
07174 H_fireEngine(pf00936pf07498) EUT2B
07175 Tsp_mutedPink EUT2B
07176 HMMetlyEutC_2_eut2(pf05985) EUT2B EP_NTERM
07177 HMMetlyEutB_2_EUT2(pf06751) EUT2B
07178 HMM06277EutA_1_eut(pf06277) EUT2B
07179 HMM10662PduV_1_mix(pf10662) EUT2B
07180 Hp_euts_violetPink EUT2B
07181 HMM00465alcdh_3_mix(pf00465) EUT2B
07182 -pf10050 -
07183 -pf01545pf16916 -
07184 -# -
07185 -# -
07186 -pf12900 -
07187 -pf00293 -
07188 -pf00027 -
07189 -pf08241 -
07190 -# -
BMC-P
BMC-H
BMC-T
Aldehyde dehydrogenase
PTAC
Signature enzyme
Alcohol dehydrogenase
Regulator

Firmicutes bacterium ADurb.Bin182. Bacteria; Firmicutes.

BWY11__locus_1

EUT2B

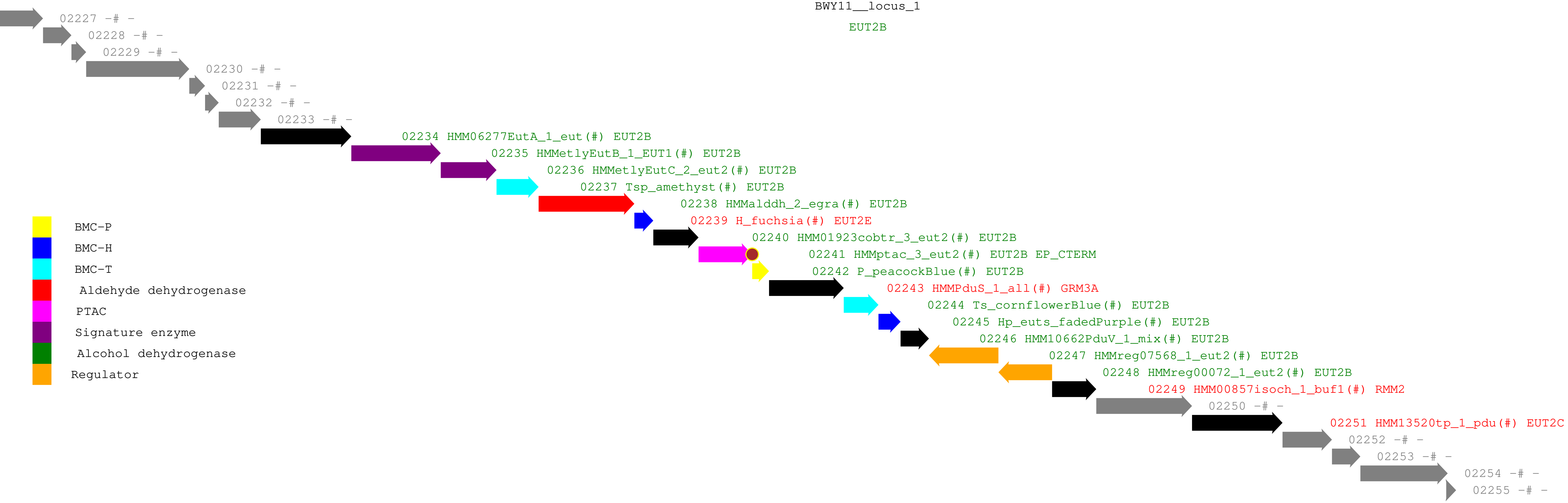

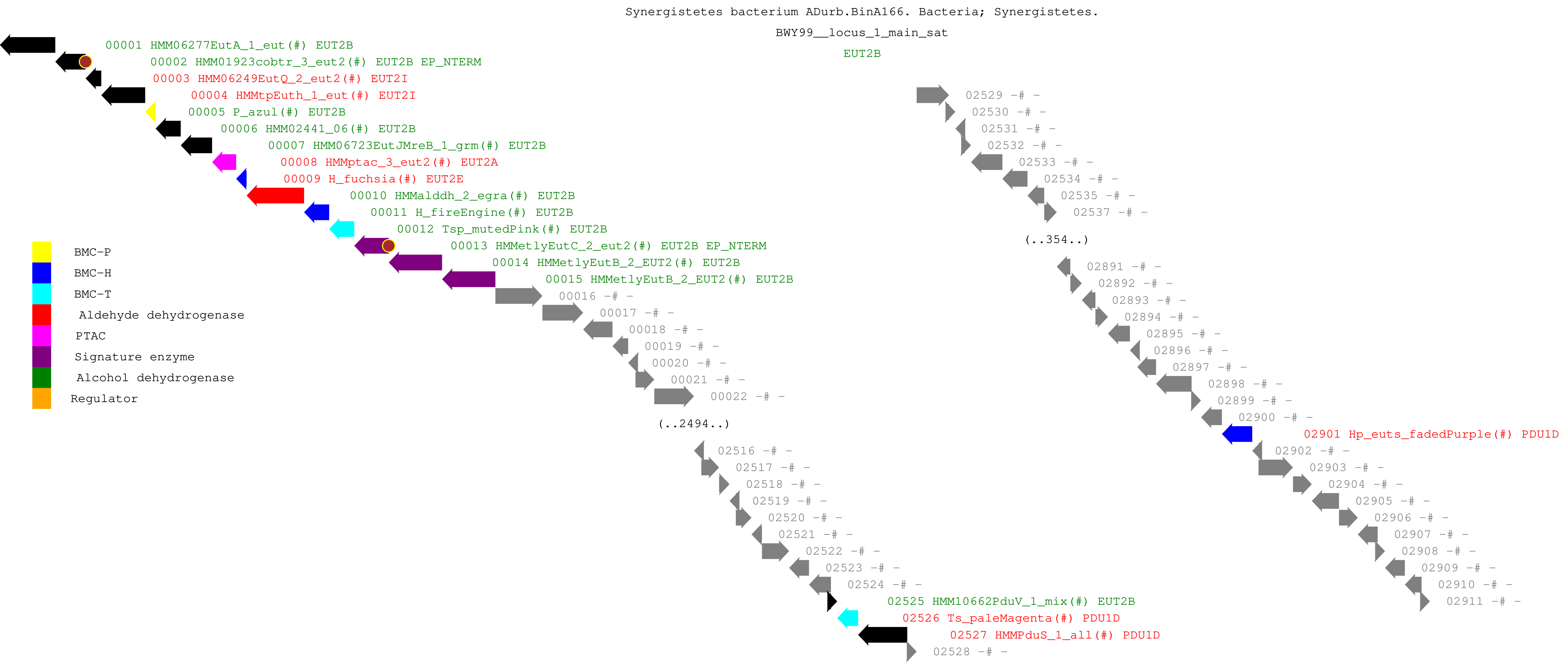
Synergistetes bacterium ADurb.BinA166. Bacteria; Synergistetes.
BWY99__locus_1_main_sat
EUT2B
00001 HMM06277EutA_1_eut(#) EUT2B
00002 HMM01923cobtr_3_eut2(#) EUT2B EP_NTERM
00003 HMM06249EutQ_2_eut2(#) EUT2I
00004 HMMtpEuth_1_eut(#) EUT2I
00005 P_azul(#) EUT2B
00006 HMM02441_06(#) EUT2B
00007 HMM06723EutJMreB_1_grm(#) EUT2B
00008 HMMptac_3_eut2(#) EUT2A
00009 H_fuchsia(#) EUT2E
00010 HMMalddh_2_egra(#) EUT2B
00011 H_fireEngine(#) EUT2B
00012 Tsp_mutedPink(#) EUT2B
00013 HMMetlyEutC_2_eut2(#) EUT2B EP_NTERM
00014 HMMetlyEutB_2_EUT2(#) EUT2B
00015 HMMetlyEutB_2_EUT2(#) EUT2B
00016 -# -
00017 -# -
00018 -# -
00019 -# -
00020 -# -
00021 -# -
00022 -# -
(..2494..)
02516 -# -
02517 -# -
02518 -# -
02519 -# -
02520 -# -
02521 -# -
02522 -# -
02523 -# -
02524 -# -
02525 HMM10662PduV_1_mix(#) EUT2B
02526 Ts_paleMagenta(#) PDU1D
02527 HMMPduS_1_all(#) PDU1D
02528 -# -
02529 -# -
02530 -# -
02531 -# -
02532 -# -
02533 -# -
02534 -# -
02535 -# -
02537 -# -
(..354..)
02891 -# -
02892 -# -
02893 -# -
02894 -# -
02895 -# -
02896 -# -
02897 -# -
02898 -# -
02899 -# -
02900 -# -
02901 Hp_euts_fadedPurple(#) PDU1D
02902 -# -
02903 -# -
02904 -# -
02905 -# -
02906 -# -
02907 -# -
02908 -# -
02909 -# -
02910 -# -
02911 -# -
BMC-P
BMC-H
BMC-T
Aldehyde dehydrogenase
PTAC
Signature enzyme
Alcohol dehydrogenase
Regulator

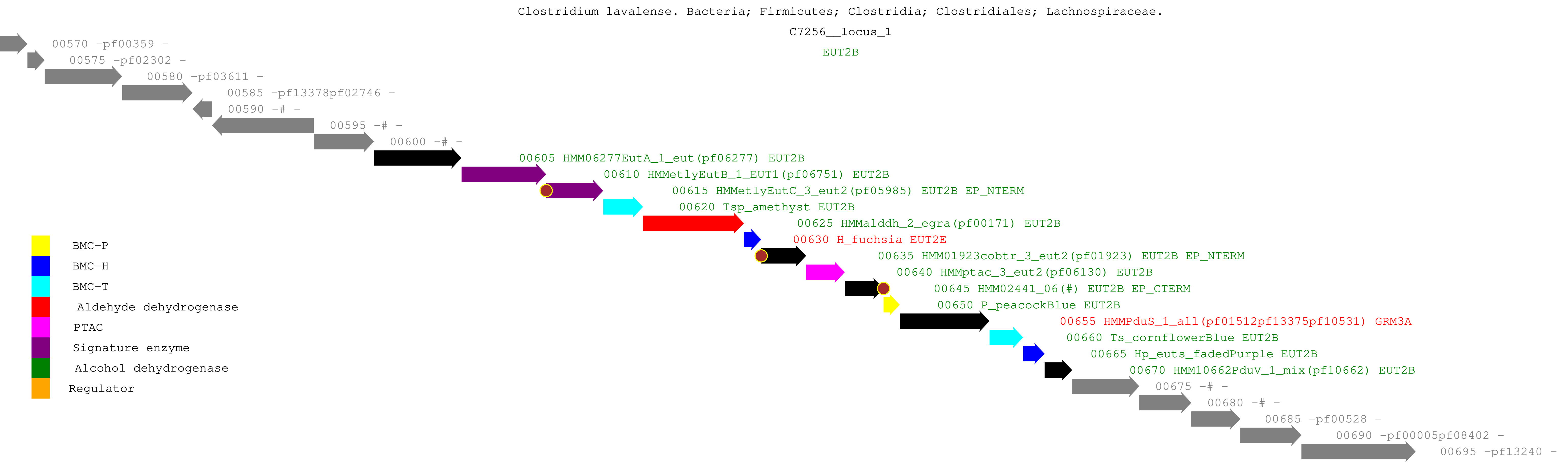

Clostridium lavalense. Bacteria; Firmicutes; Clostridia; Clostridiales; Lachnospiraceae.
C7256__locus_1
EUT2B
00570 -pf00359 -
00575 -pf02302 -
00580 -pf03611 -
00585 -pf13378pf02746 -
00590 -# -
00595 -# -
00600 -# -
00605 HMM06277EutA_1_eut(pf06277) EUT2B
00610 HMMetlyEutB_1_EUT1(pf06751) EUT2B
00615 HMMetlyEutC_3_eut2(pf05985) EUT2B EP_NTERM
00620 Tsp_amethyst EUT2B
00625 HMMaldh_2_egra(pf00171) EUT2B
00630 H_fuchsia EUT2E
00635 HMM01923cobtr_3_eut2(pf01923) EUT2B EP_NTERM
00640 HMMptac_3_eut2(pf06130) EUT2B
00645 HMM02441_06(#) EUT2B EP_CTERM
00650 P_peacockBlue EUT2B
00655 HMMPduS_1_all(pf01512pf13375pf10531) GRM3A
00660 Ts_cornflowerBlue EUT2B
00665 Hp_euts_fadedPurple EUT2B
00670 HMM10662PduV_1_mix(pf10662) EUT2B
00675 -# -
00680 -# -
00685 -pf00528 -
00690 -pf00005pf08402 -
00695 -pf13240 -
BMC-P
BMC-H
BMC-T
Aldehyde dehydrogenase
PTAC
Signature enzyme
Alcohol dehydrogenase
Regulator

Intestinimonas butyriciproducens. Bacteria; Firmicutes; Clostridia; Clostridiales; Intestinimonas.

C7373__locus_2

EUT2B

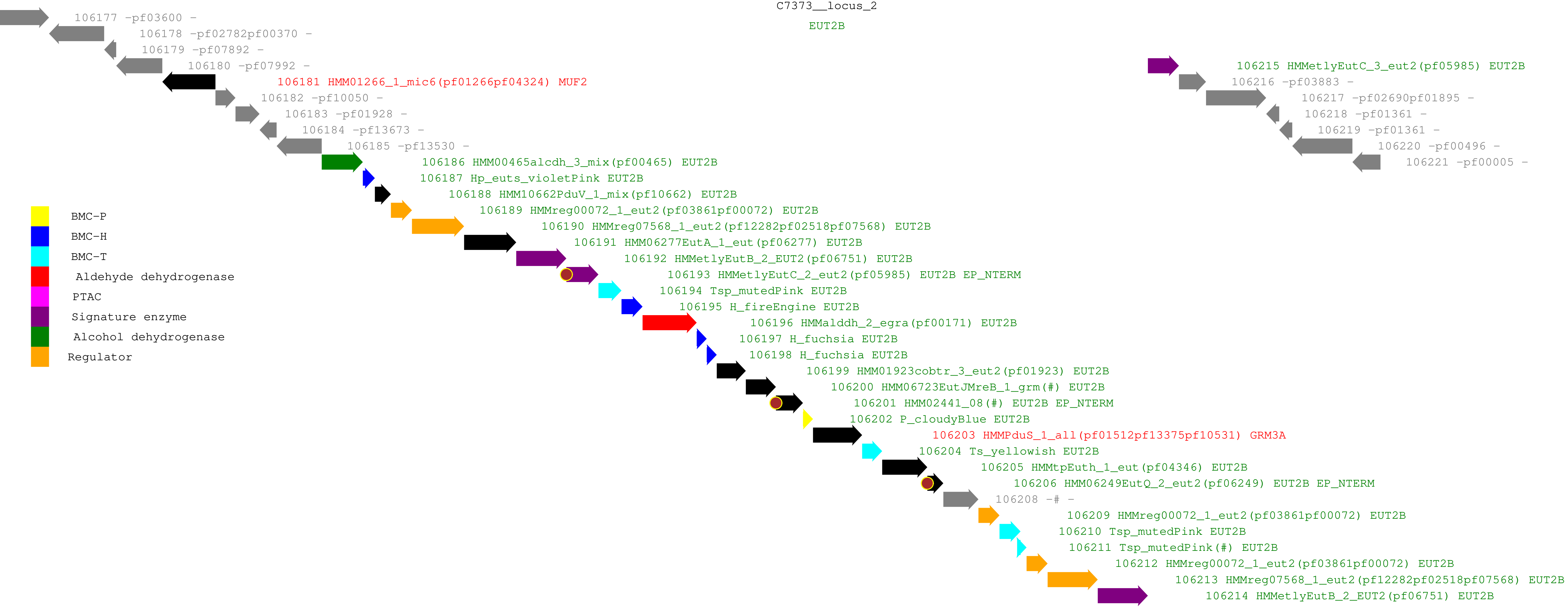

Anaerotruncus sp. G3(2012). Bacteria; Firmicutes; Clostridia; Clostridiales; Ruminococcaceae;

C814__locus_1

EUT2B

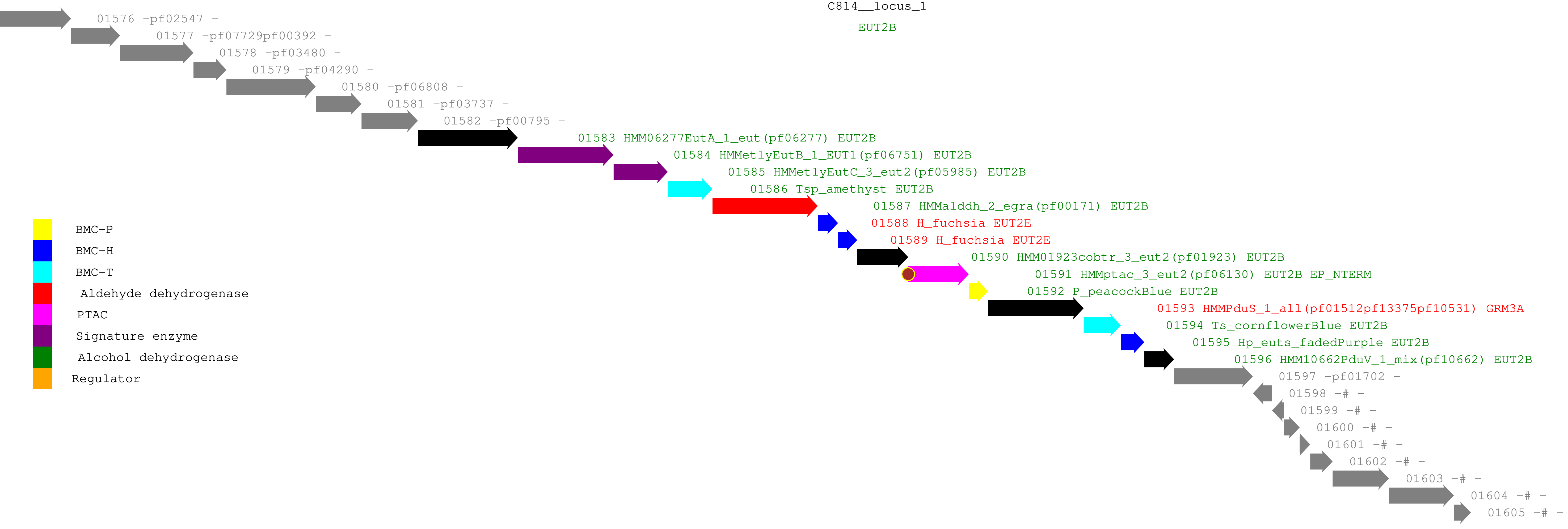

Lachnotalea glycerini. Bacteria; Firmicutes; Clostridia; Clostridiales; Lachnospiraceae;

C8E03__locus_1

EUT2B

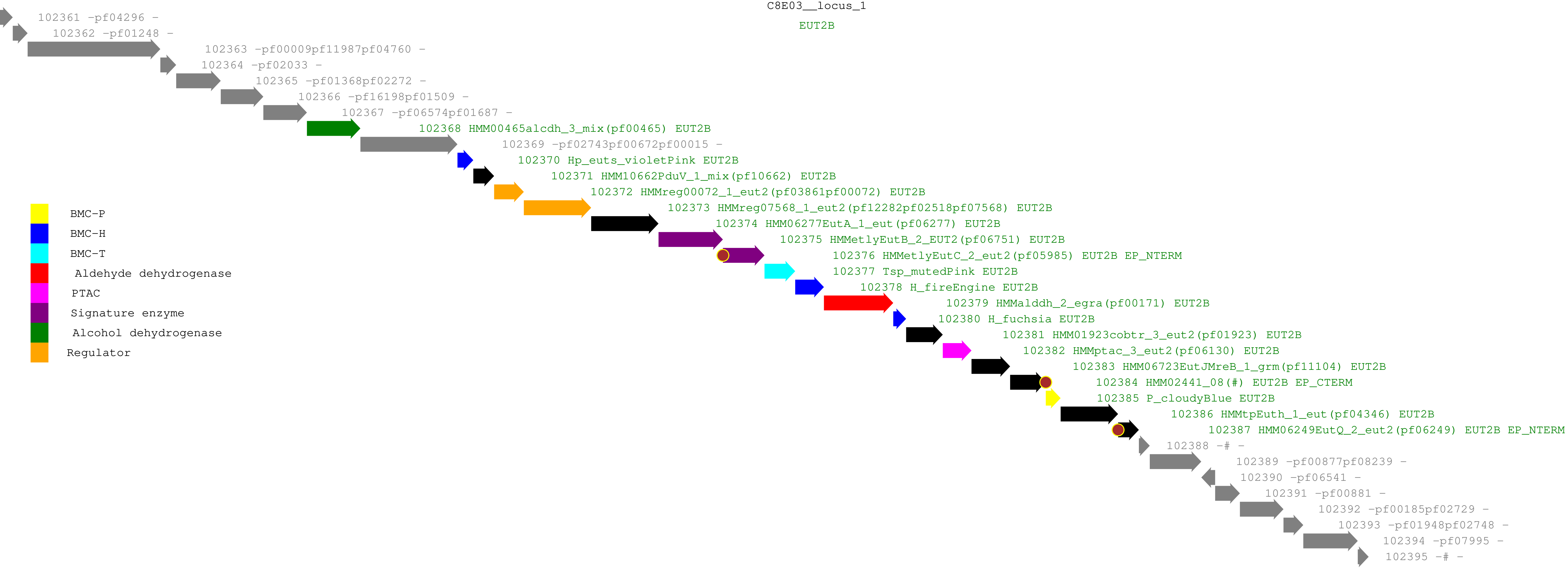

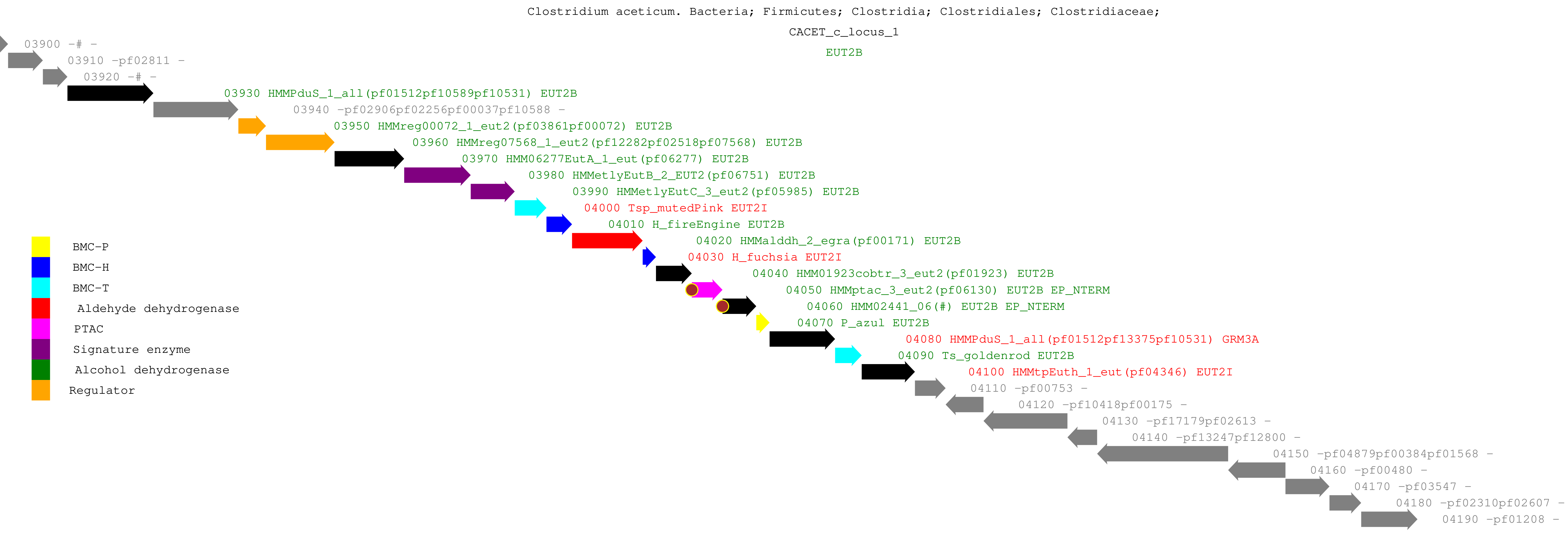

Clostridium aceticum. Bacteria; Firmicutes; Clostridia; Clostridiales; Clostridiaceae;
CACET_c_locus_1
EUT2B
03900 -# -
03910 -pf02811 -
03920 -# -
03930 HMMPduS_1_all(pf01512pf10589pf10531) EUT2B
03940 -pf02906pf02256pf00037pf10588 -
03950 HMMreg00072_1_eut2(pf03861pf00072) EUT2B
03960 HMMreg07568_1_eut2(pf12282pf02518pf07568) EUT2B
03970 HMM06277EutA_1_eut(pf06277) EUT2B
03980 HMMetlyEutB_2_EUT2(pf06751) EUT2B
03990 HMMetlyEutC_3_eut2(pf05985) EUT2B
04000 Tsp_mutedPink EUT2I
04010 H_fireEngine EUT2B
04020 HMMaldh_2_egra(pf00171) EUT2B
04030 H_fuchsia EUT2I
04040 HMM01923cobtr_3_eut2(pf01923) EUT2B
04050 HMMptac_3_eut2(pf06130) EUT2B EP_NTERM
04060 HMM02441_06(#) EUT2B EP_NTERM
04070 P_azul EUT2B
04080 HMMPduS_1_all(pf01512pf13375pf10531) GRM3A
04090 Ts_goldenrod EUT2B
04100 HMMtpEuth_1_eut(pf04346) EUT2I
04110 -pf00753 -
04120 -pf10418pf00175 -
04130 -pf17179pf02613 -
04140 -pf13247pf12800 -
04150 -pf04879pf00384pf01568 -
04160 -pf00480 -
04170 -pf03547 -
04180 -pf02310pf02607 -
04190 -pf01208 -
BMC-P
BMC-H
BMC-T
Aldehyde dehydrogenase
PTAC
Signature enzyme
Alcohol dehydrogenase
Regulator

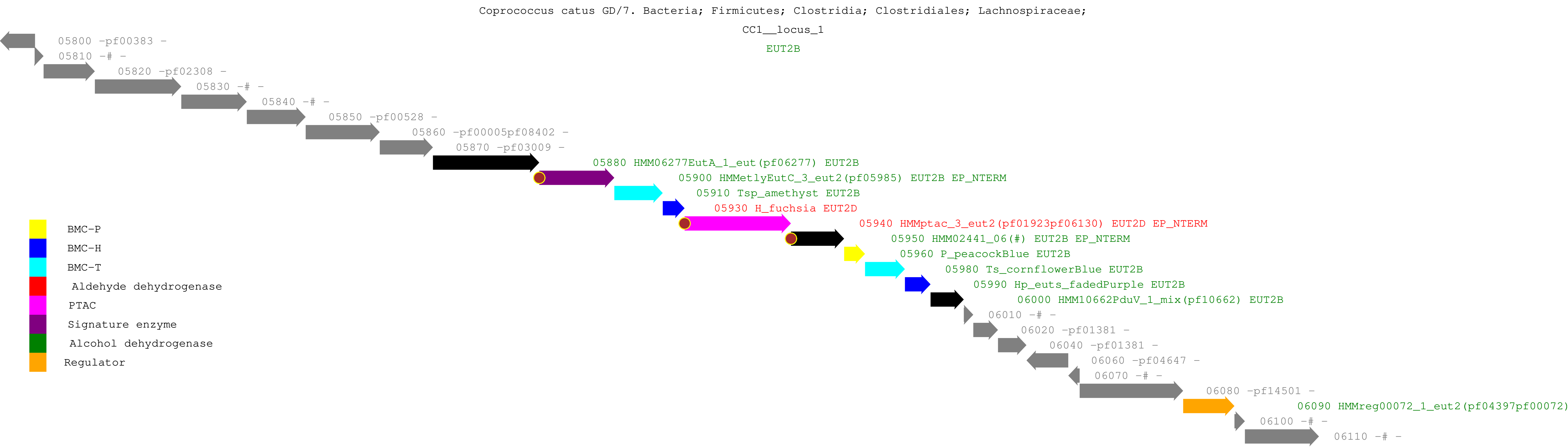
Coprococcus catus GD/7. Bacteria; Firmicutes; Clostridia; Clostridiales; Lachnospiraceae;
CC1__locus_1
EUT2B
05800 -pf00383 -
05810 -# -
05820 -pf02308 -
05830 -# -
05840 -# -
05850 -pf00528 -
05860 -pf00005pf08402 -
05870 -pf03009 -
05880 HMM06277EutA_1_eut(pf06277) EUT2B
05900 HMMetlyEutC_3_eut2(pf05985) EUT2B EP_NTERM
05910 Tsp_amethyst EUT2B
05930 H_fuchsia EUT2D
05940 HMMptac_3_eut2(pf01923pf06130) EUT2D EP_NTERM
05950 HMM02441_06(#) EUT2B EP_NTERM
05960 P_peacockBlue EUT2B
05980 Ts_cornflowerBlue EUT2B
05990 Hp_euts_fadedPurple EUT2B
06000 HMM10662PduV_1_mix(pf10662) EUT2B
06010 -# -
06020 -pf01381 -
06040 -pf01381 -
06060 -pf04647 -
06070 -# -
06080 -pf14501 -
06090 HMMreg00072_1_eut2(pf04397pf00072)
06100 -# -
06110 -# -
BMC-P
BMC-H
BMC-T
Aldehyde dehydrogenase
PTAC
Signature enzyme
Alcohol dehydrogenase
Regulator

Lachnotalea glycerini. Bacteria; Firmicutes; Clostridia; Clostridiales; Lachnospiraceae;

CG710__locus_1

EUT2B

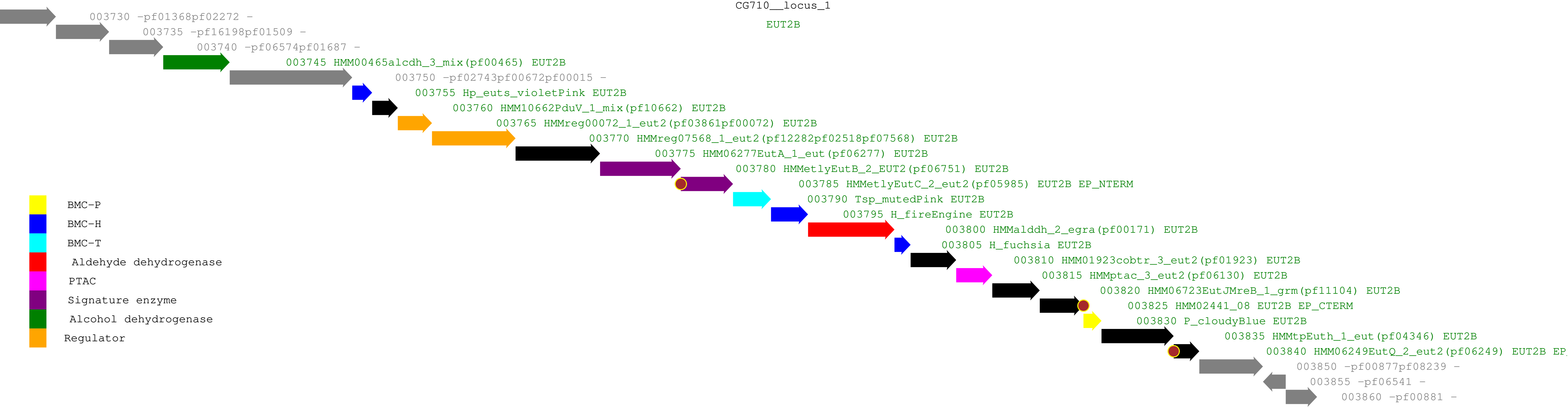

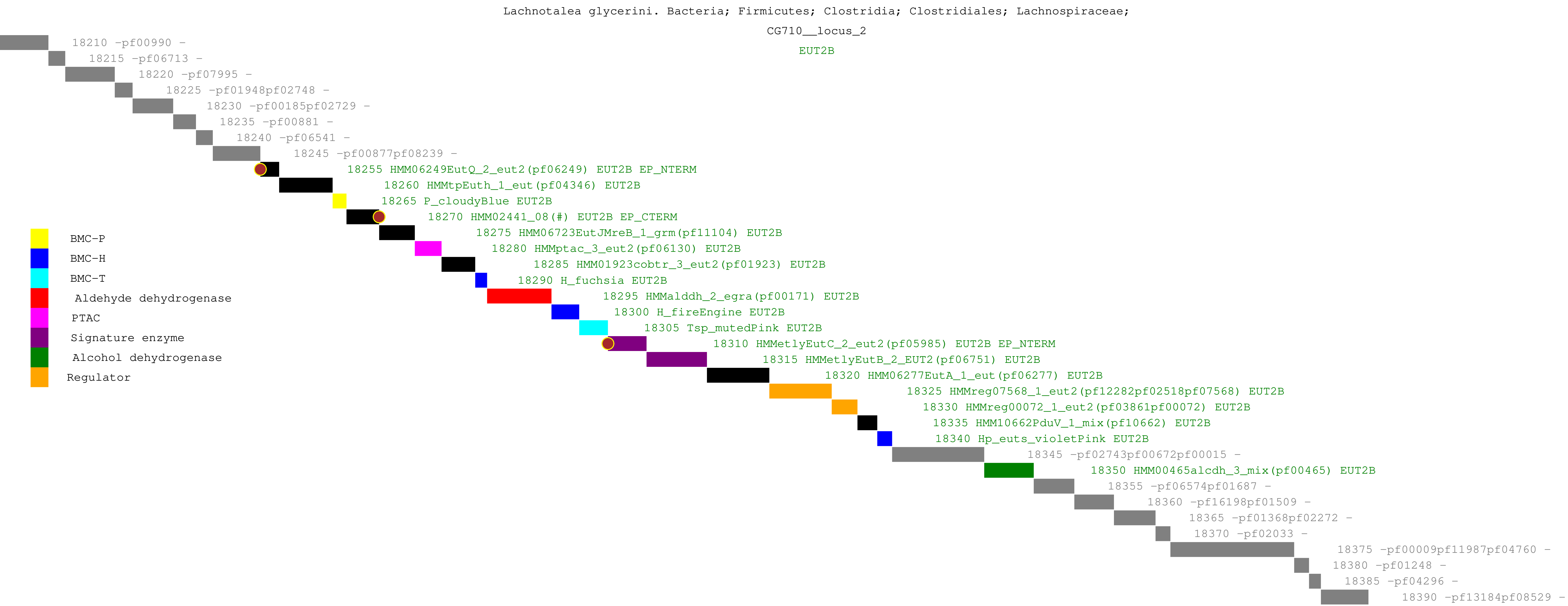

Lachnotalea glycerini. Bacteria; Firmicutes; Clostridia; Clostridiales; Lachnospiraceae;
CG710__locus_2
EUT2B
18210 -pf00990 -
18215 -pf06713 -
18220 -pf07995 -
18225 -pf01948pf02748 -
18230 -pf00185pf02729 -
18235 -pf00881 -
18240 -pf06541 -
18245 -pf00877pf08239 -
18255 HMM06249EutQ_2_eut2(pf06249) EUT2B EP_NTERM
18260 HMMtpEuth_1_eut(pf04346) EUT2B
18265 P_cloudyBlue EUT2B
18270 HMM02441_08(#) EUT2B EP_CTERM
18275 HMM06723EutJMreB_1_grm(pf11104) EUT2B
18280 HMMptac_3_eut2(pf06130) EUT2B
18285 HMM01923cobtr_3_eut2(pf01923) EUT2B
18290 H_fuchsia EUT2B
18295 HMMalddh_2_egra(pf00171) EUT2B
18300 H_fireEngine EUT2B
18305 Tsp_mutedPink EUT2B
18310 HMMetlyEutC_2_eut2(pf05985) EUT2B EP_NTERM
18315 HMMetlyEutB_2_EUT2(pf06751) EUT2B
18320 HMM06277EutA_1_eut(pf06277) EUT2B
18325 HMMreg07568_1_eut2(pf12282pf02518pf07568) EUT2B
18330 HMMreg00072_1_eut2(pf03861pf00072) EUT2B
18335 HMM10662PduV_1_mix(pf10662) EUT2B
18340 Hp_euts_violetPink EUT2B
18345 -pf02743pf00672pf00015 -
18350 HMM00465alcdh_3_mix(pf00465) EUT2B
18355 -pf06574pf01687 -
18360 -pf16198pf01509 -
18365 -pf01368pf02272 -
18370 -pf02033 -
18375 -pf00009pf11987pf04760 -
18380 -pf01248 -
18385 -pf04296 -
18390 -pf13184pf08529 -
BMC-P
BMC-H
BMC-T
Aldehyde dehydrogenase
PTAC
Signature enzyme
Alcohol dehydrogenase
Regulator

Cloacibacillus evryensis DSM 19522. Bacteria; Synergistetes; Synergistia; Synergistales; Synergistaceae;

Cloev__locus_1

EUT2B

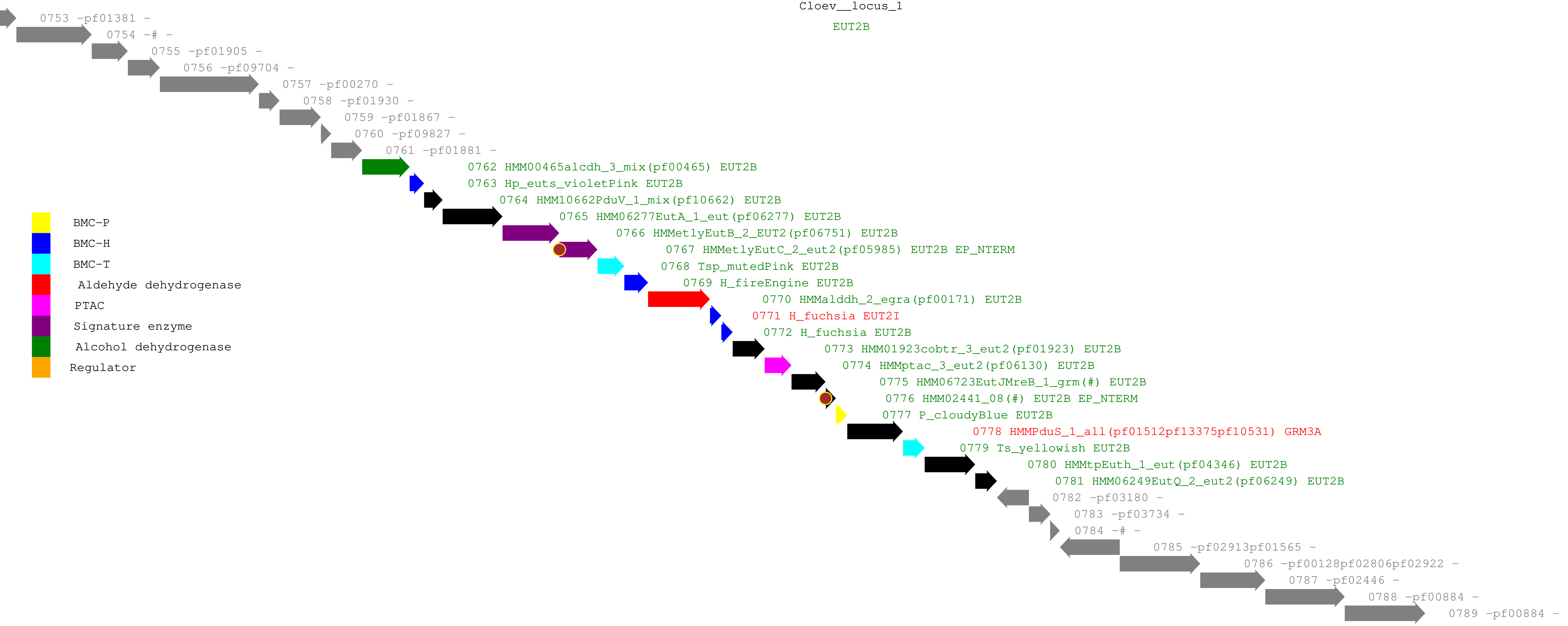

Clostridium asparagiforme DSM 15981. Bacteria; Firmicutes; Clostridia; Clostridiales; Lachnospiraceae.

CLOSTASPAR__locus_1

EUT2B

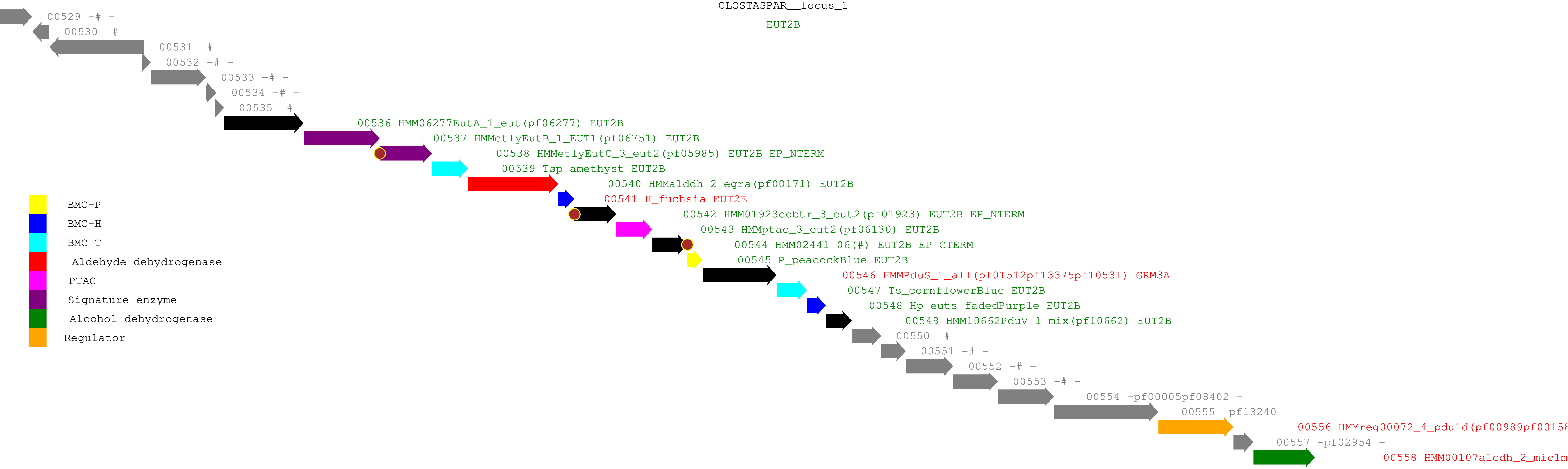

Acetoanaerobium sticklandii (strain ATCC 12662 / DSM 519 / JCM 1433 / Bacteria; Firmicutes; Clostridia; Clostridiales;

CLOST__locus_1

EUT2B

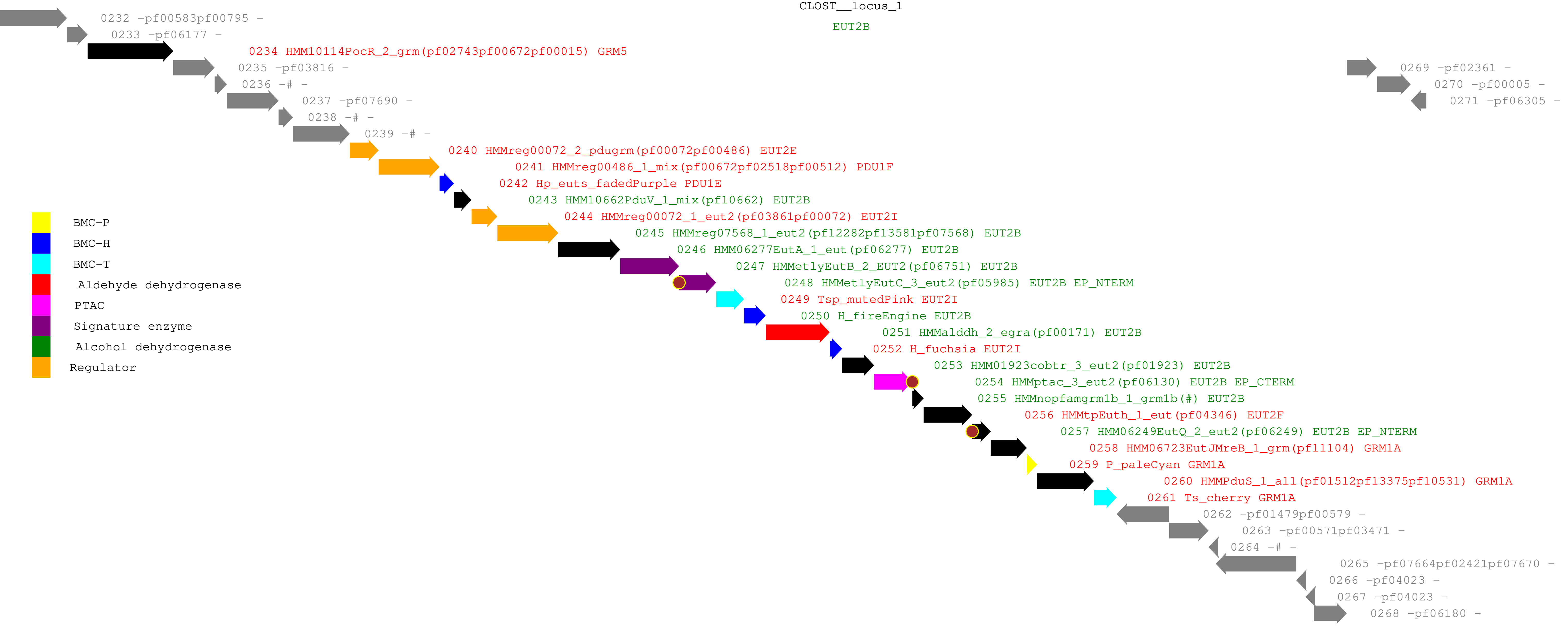

Clostridium symbiosum ATCC 14940. Bacteria; Firmicutes; Clostridia; Clostridiales; Lachnospiraceae.

CLOSYM__locus_1

EUT2B

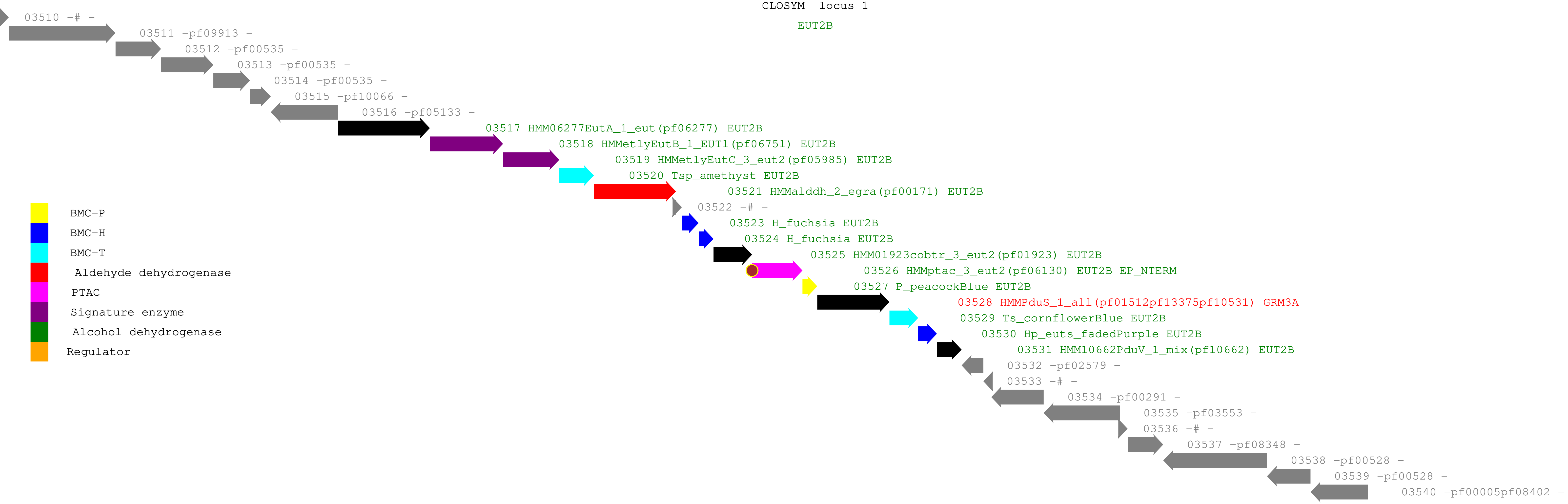

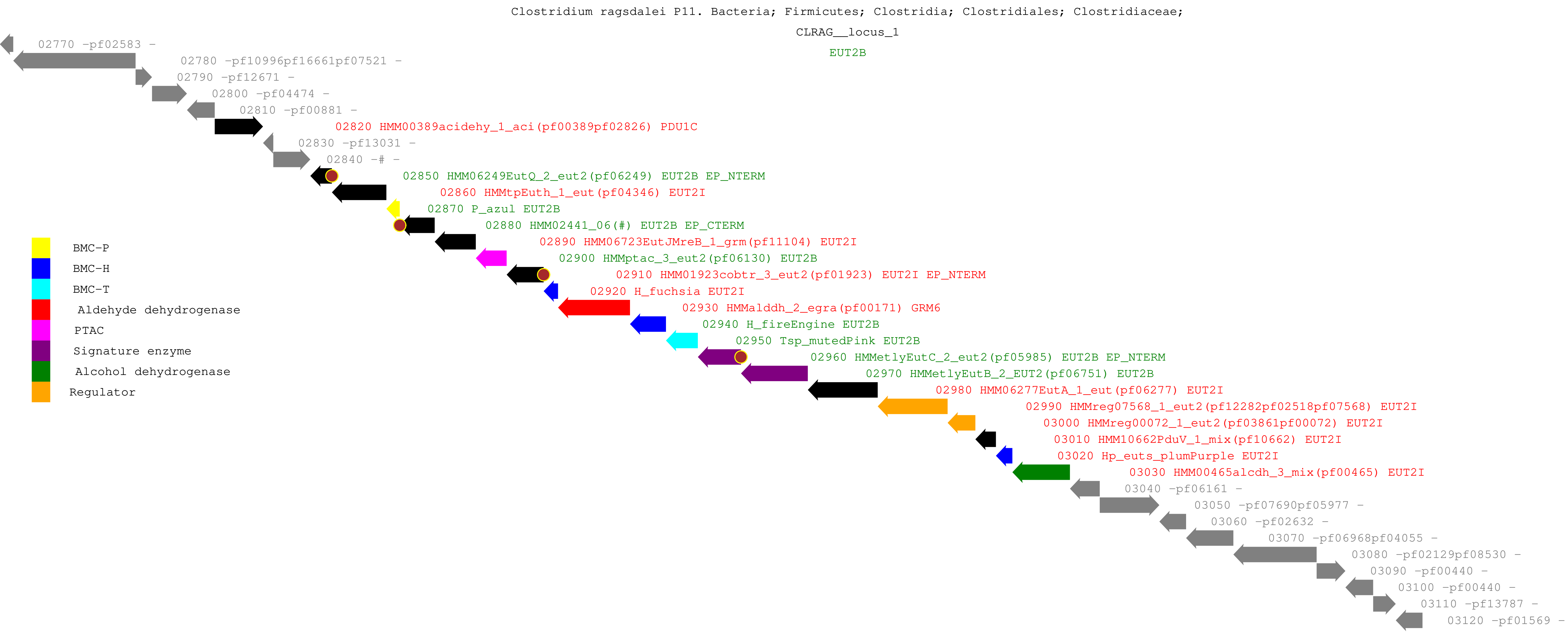
Clostridium ragsdalei P11. Bacteria; Firmicutes; Clostridia; Clostridiales; Clostridiaceae;
CLRAG__locus_1
EUT2B
02770 -pf02583 -
02780 -pf10996pf16661pf07521 -
02790 -pf12671 -
02800 -pf04474 -
02810 -pf00881 -
02820 HMM00389acidehy_1_aci(pf00389pf02826) PDU1C
02830 -pf13031 -
02840 -# -
02850 HMM06249EutQ_2_eut2(pf06249) EUT2B EP_NTERM
02860 HMMtpEuth_1_eut(pf04346) EUT2I
02870 P_azul EUT2B
02880 HMM02441_06(#) EUT2B EP_CTERM
02890 HMM06723EutJMreB_1_grm(pf11104) EUT2I
02900 HMMptac_3_eut2(pf06130) EUT2B
02910 HMM01923cobtr_3_eut2(pf01923) EUT2I EP_NTERM
02920 H_fuchsia EUT2I
02930 HMMalddh_2_egra(pf00171) GRM6
02940 H_fireEngine EUT2B
02950 Tsp_mutedPink EUT2B
02960 HMMetlyEutC_2_eut2(pf05985) EUT2B EP_NTERM
02970 HMMetlyEutB_2_EUT2(pf06751) EUT2B
02980 HMM06277EutA_1_eut(pf06277) EUT2I
02990 HMMreg07568_1_eut2(pf12282pf02518pf07568) EUT2I
03000 HMMreg00072_1_eut2(pf03861pf00072) EUT2I
03010 HMM10662PduV_1_mix(pf10662) EUT2I
03020 Hp_euts_plumPurple EUT2I
03030 HMM00465alcdh_3_mix(pf00465) EUT2I
03040 -pf06161 -
03050 -pf07690pf05977 -
03060 -pf02632 -
03070 -pf06968pf04055 -
03080 -pf02129pf08530 -
03090 -pf00440 -
03100 -pf00440 -
03110 -pf13787 -
03120 -pf01569 -
BMC-P
BMC-H
BMC-T
Aldehyde dehydrogenase
PTAC
Signature enzyme
Alcohol dehydrogenase
Regulator

Alkaliphilus sp. Bacteria; Firmicutes; Clostridia; Clostridiales; Clostridiaceae;

COA82__locus_2

EUT2B

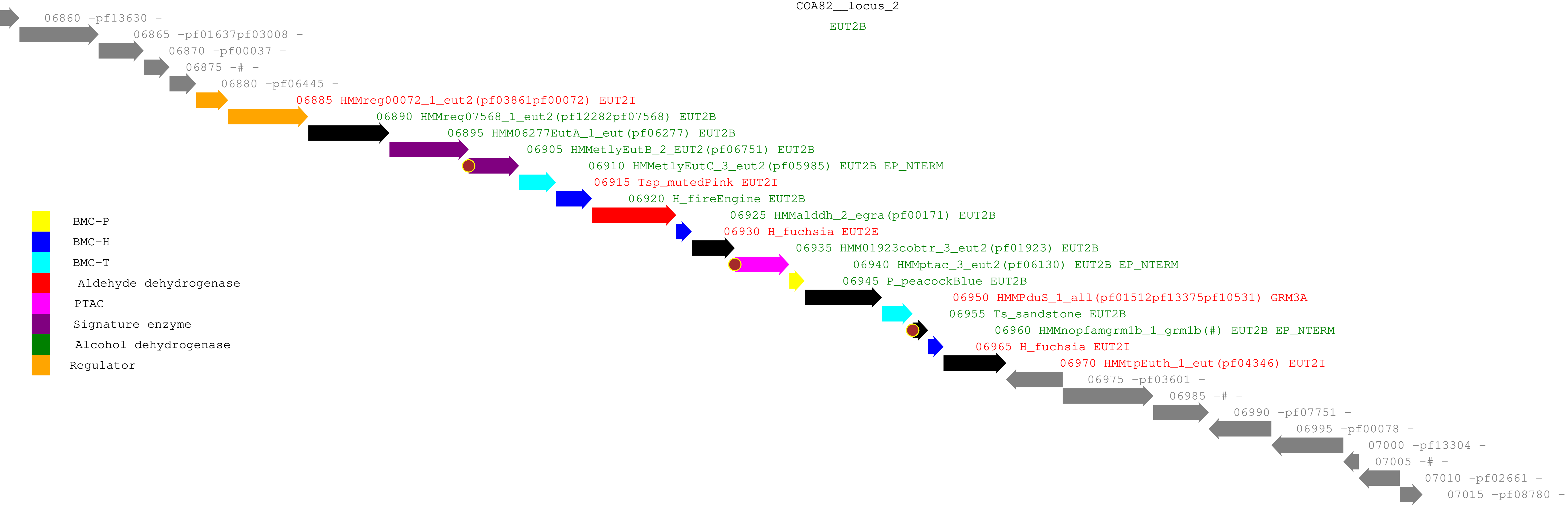

Clostridium sp. HMb25. Bacteria; Firmicutes; Clostridia; Clostridiales; Clostridiaceae;

CRH03__locus_1

EUT2B

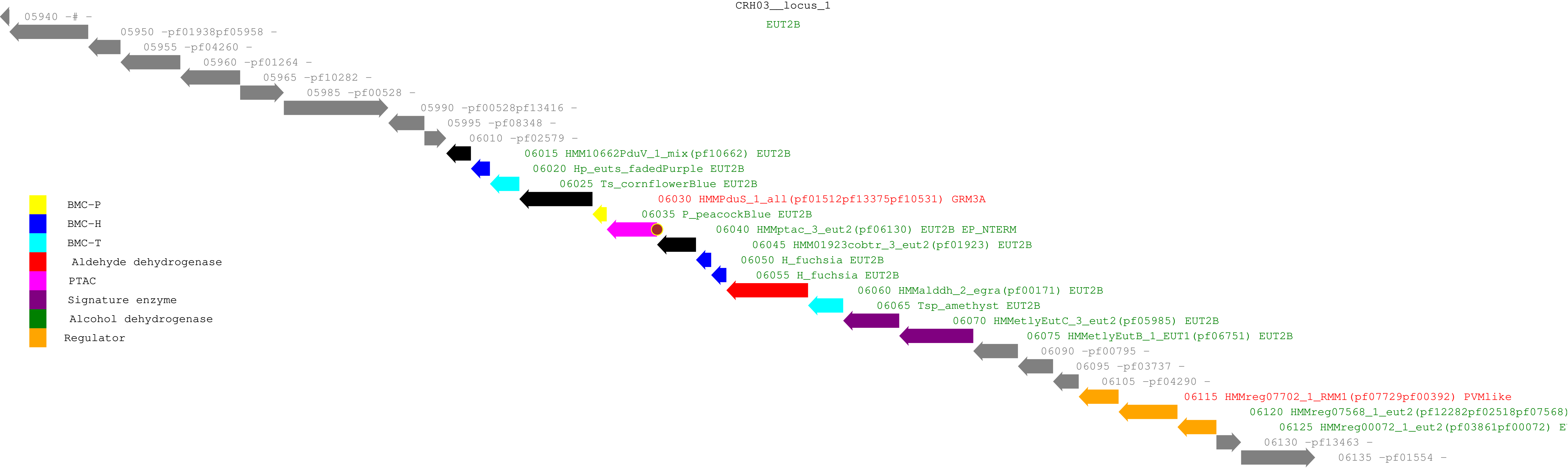

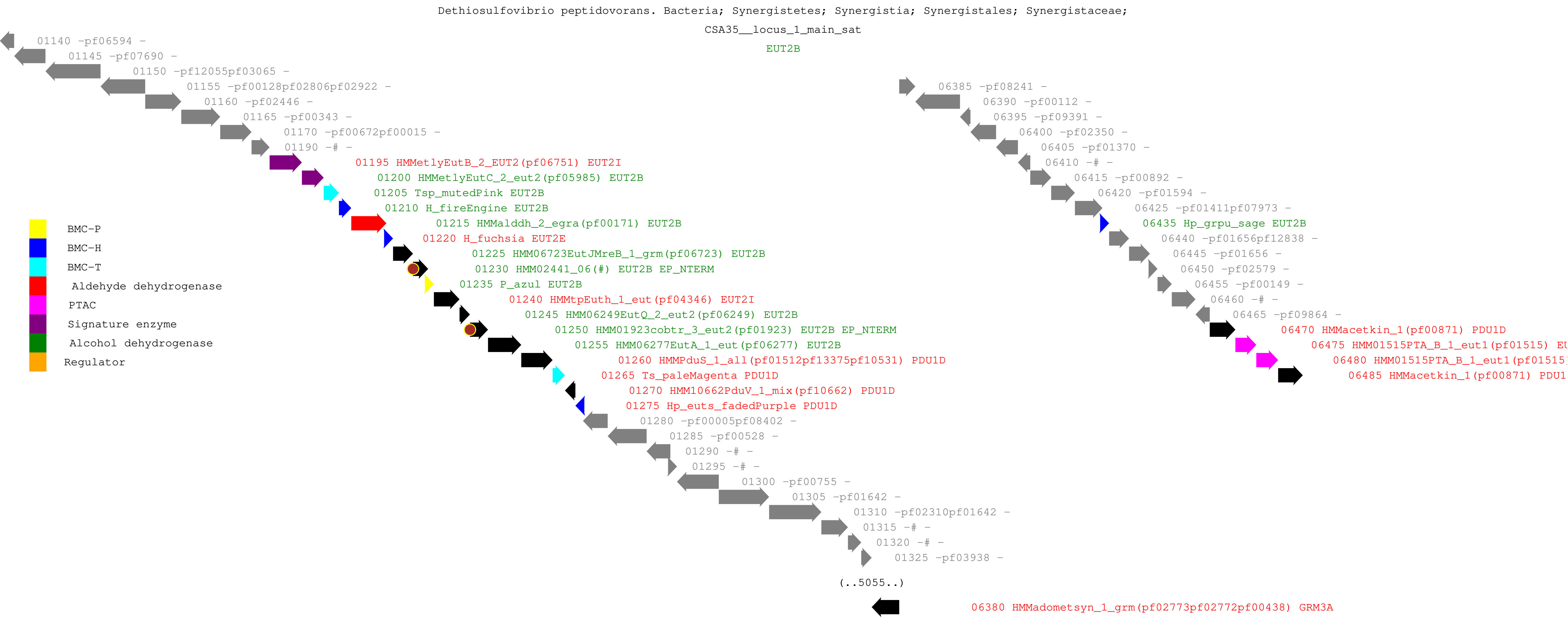

Dethiosulfovibrio peptidovorans. Bacteria; Synergistetes; Synergistia; Synergistales; Synergistaceae;
CSA35__locus_1_main_sat
EUT2B
01140 -pf06594 -
01145 -pf07690 -
01150 -pf12055pf03065 -
01155 -pf00128pf02806pf02922 -
01160 -pf02446 -
01165 -pf00343 -
01170 -pf00672pf00015 -
01190 -# -
01195 HMMetlyEutB_2_EUT2(pf06751) EUT2I
01200 HMMetlyEutC_2_eut2(pf05985) EUT2B
01205 Tsp_mutedPink EUT2B
01210 H_fireEngine EUT2B
01215 HMMaldh_2_egra(pf00171) EUT2B
01220 H_fuchsia EUT2E
01225 HMM06723EutJMreB_1_grm(pf06723) EUT2B
01230 HMM02441_06(#) EUT2B EP_NTERM
01235 P_azul EUT2B
01240 HMMtpEuth_1_eut(pf04346) EUT2I
01245 HMM06249EutQ_2_eut2(pf06249) EUT2B
01250 HMM01923cobtr_3_eut2(pf01923) EUT2B EP_NTERM
01255 HMM06277EutA_1_eut(pf06277) EUT2B
01260 HMMPduS_1_all(pf01512pf13375pf10531) PDU1D
01265 Ts_paleMagenta PDU1D
01270 HMM10662PduV_1_mix(pf10662) PDU1D
01275 Hp_euts_fadedPurple PDU1D
01280 -pf00005pf08402 -
01285 -pf00528 -
01290 -# -
01295 -# -
01300 -pf00755 -
01305 -pf01642 -
01310 -pf02310pf01642 -
01315 -# -
01320 -# -
01325 -pf03938 -
(..5055..)
06380 HMMadometsyn_1_grm(pf02773pf02772pf00438) GRM3A
06385 -pf08241 -
06390 -pf00112 -
06395 -pf09391 -
06400 -pf02350 -
06405 -pf01370 -
06410 -# -
06415 -pf00892 -
06420 -pf01594 -
06425 -pf01411pf07973 -
06435 Hp_grpu_sage EUT2B
06440 -pf01656pf12838 -
06445 -pf01656 -
06450 -pf02579 -
06455 -pf00149 -
06460 -# -
06465 -pf09864 -
06470 HMMacetkin_1(pf00871) PDU1D
06475 HMM01515PTA_B_1_eut1(pf01515) EU
06480 HMM01515PTA_B_1_eut1(pf01515
06485 HMMacetkin_1(pf00871) PDU1
BMC-P
BMC-H
BMC-T
Aldehyde dehydrogenase
PTAC
Signature enzyme
Alcohol dehydrogenase
Regulator

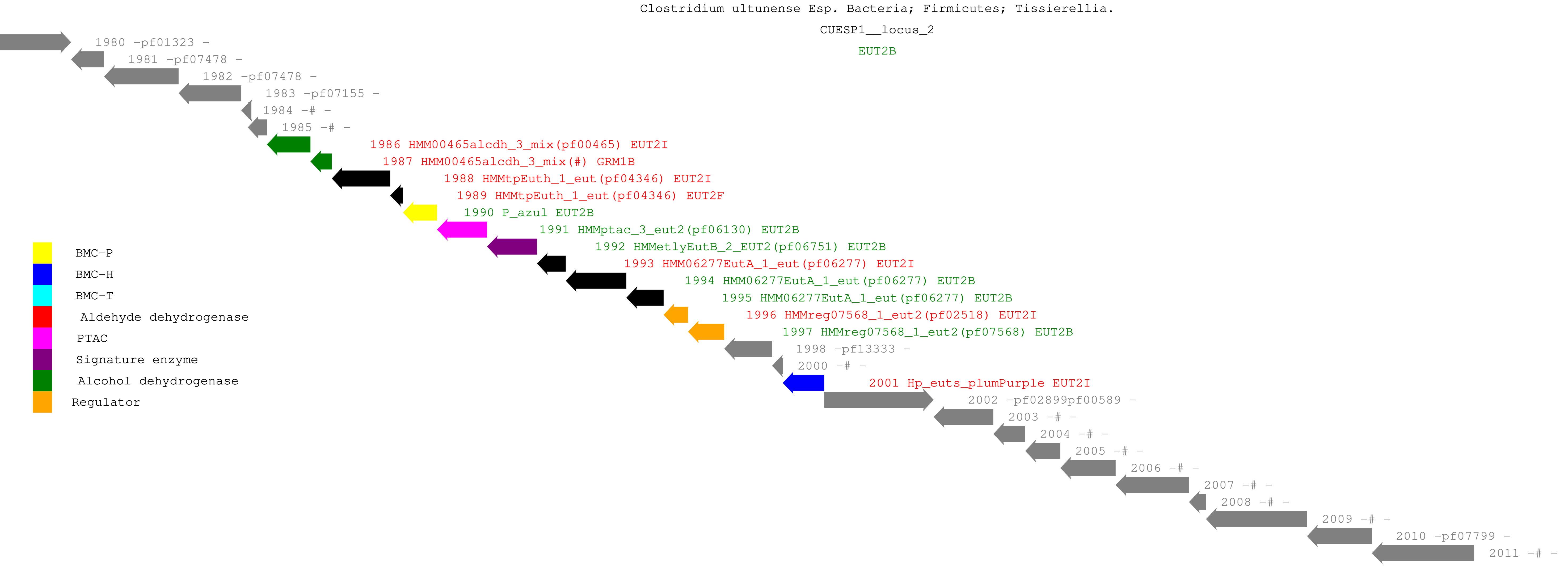
Clostridium ultunense Esp. Bacteria; Firmicutes; Tissierellia.
CUESP1__locus_2
EUT2B
1980 -pf01323 -
1981 -pf07478 -
1982 -pf07478 -
1983 -pf07155 -
1984 -# -
1985 -# -
1986 HMM00465alcdh_3_mix(pf00465) EUT2I
1987 HMM00465alcdh_3_mix(#) GRM1B
1988 HMMtpEuth_1_eut(pf04346) EUT2I
1989 HMMtpEuth_1_eut(pf04346) EUT2F
1990 P_azul EUT2B
1991 HMMptac_3_eut2(pf06130) EUT2B
1992 HMMetlyEutB_2_EUT2(pf06751) EUT2B
1993 HMM06277EutA_1_eut(pf06277) EUT2I
1994 HMM06277EutA_1_eut(pf06277) EUT2B
1995 HMM06277EutA_1_eut(pf06277) EUT2B
1996 HMMreg07568_1_eut2(pf02518) EUT2I
1997 HMMreg07568_1_eut2(pf07568) EUT2B
1998 -pf13333 -
2000 -# -
2001 Hp_euts_plumPurple EUT2I
2002 -pf02899pf00589 -
2003 -# -
2004 -# -
2005 -# -
2006 -# -
2007 -# -
2008 -# -
2009 -# -
2010 -pf07799 -
2011 -# -
BMC-P
BMC-H
BMC-T
Aldehyde dehydrogenase
PTAC
Signature enzyme
Alcohol dehydrogenase
Regulator

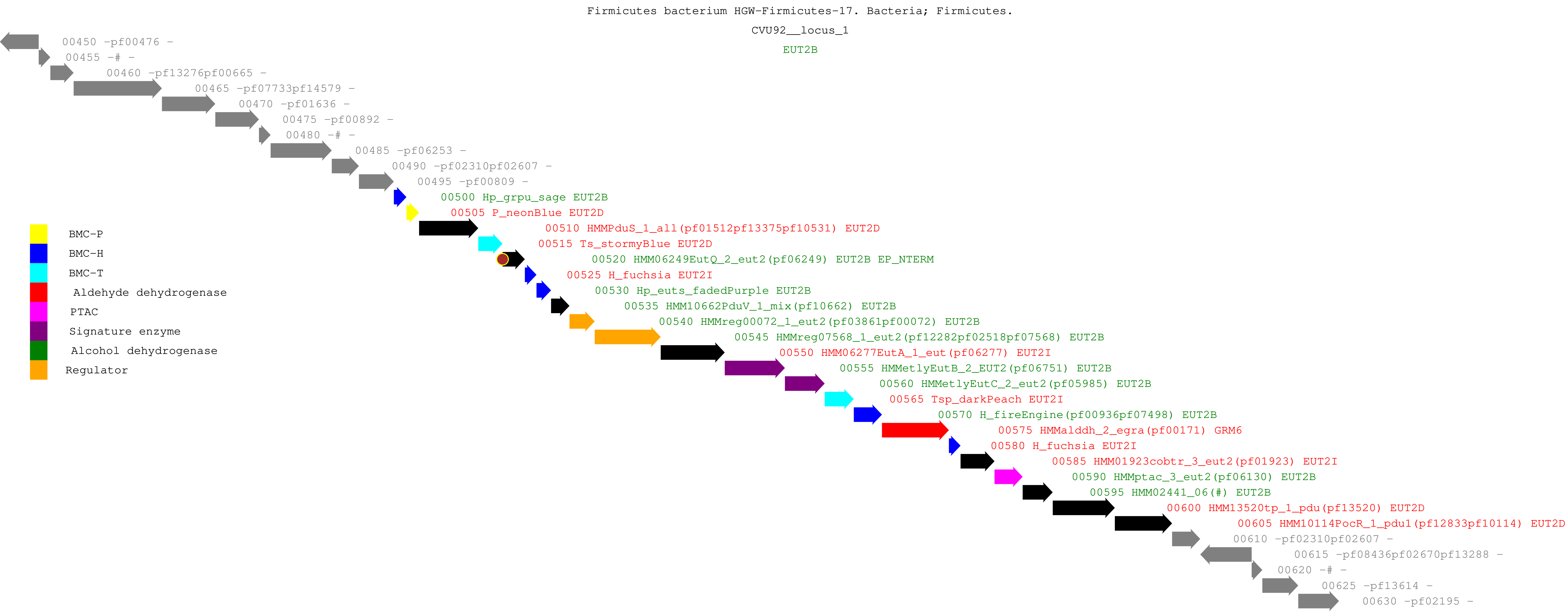

Firmicutes bacterium HGW-Firmicutes-17. Bacteria; Firmicutes.
CVU92__locus_1
EUT2B
00450 -pf00476 -
00455 -# -
00460 -pf13276pf00665 -
00465 -pf07733pf14579 -
00470 -pf01636 -
00475 -pf00892 -
00480 -# -
00485 -pf06253 -
00490 -pf02310pf02607 -
00495 -pf00809 -
00500 Hp_grpu_sage EUT2B
00505 P_neonBlue EUT2D
00510 HMMPduS_1_all(pf01512pf13375pf10531) EUT2D
00515 Ts_stormyBlue EUT2D
00520 HMM06249EutQ_2_eut2(pf06249) EUT2B EP_NTERM
00525 H_fuchsia EUT2I
00530 Hp_euts_fadedPurple EUT2B
00535 HMM10662PduV_1_mix(pf10662) EUT2B
00540 HMMreg00072_1_eut2(pf03861pf00072) EUT2B
00545 HMMreg07568_1_eut2(pf12282pf02518pf07568) EUT2B
00550 HMM06277EutA_1_eut(pf06277) EUT2I
00555 HMMetlyEutB_2_EUT2(pf06751) EUT2B
00560 HMMetlyEutC_2_eut2(pf05985) EUT2B
00565 Tsp_darkPeach EUT2I
00570 H_fireEngine(pf00936pf07498) EUT2B
00575 HMMalddh_2_egra(pf00171) GRM6
00580 H_fuchsia EUT2I
00585 HMM01923cobtr_3_eut2(pf01923) EUT2I
00590 HMMptac_3_eut2(pf06130) EUT2B
00595 HMM02441_06(#) EUT2B
00600 HMM13520tp_1_pdu(pf13520) EUT2D
00605 HMM10114PocR_1_pdu1(pf12833pf10114) EUT2D
00610 -pf02310pf02607 -
00615 -pf08436pf02670pf13288 -
00620 -# -
00625 -pf13614 -
00630 -pf02195 -
BMC-P
BMC-H
BMC-T
Aldehyde dehydrogenase
PTAC
Signature enzyme
Alcohol dehydrogenase
Regulator

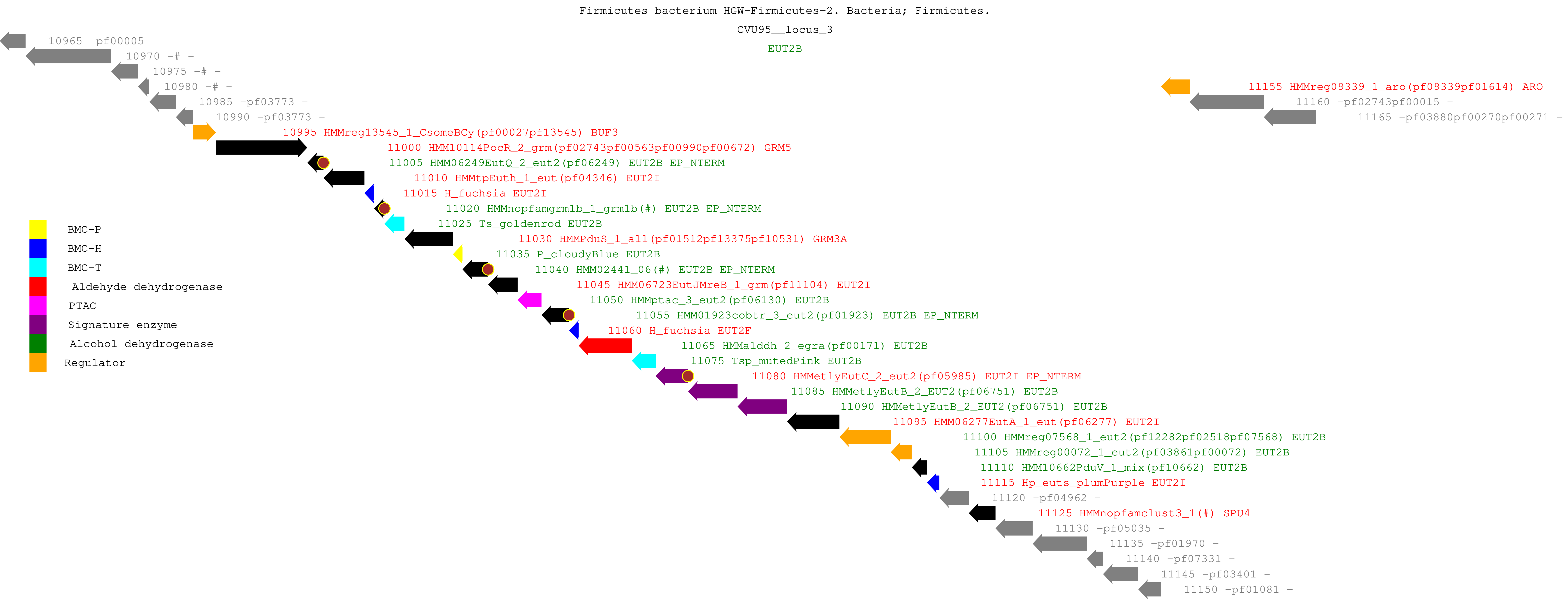

Firmicutes bacterium HGW-Firmicutes-2. Bacteria; Firmicutes.
CVU95__locus_3
EUT2B
10965 -pf00005 -
10970 -# -
10975 -# -
10980 -# -
10985 -pf03773 -
10990 -pf03773 -
10995 HMMreg13545_1_CsomeBCy(pf00027pf13545) BUF3
11000 HMM10114PocR_2_grm(pf02743pf00563pf00990pf00672) GRM5
11005 HMM06249EutQ_2_eut2(pf06249) EUT2B EP_NTERM
11010 HMMtpEuth_1_eut(pf04346) EUT2I
11015 H_fuchsia EUT2I
11020 HMMnopfamgrm1b_1_grm1b(#) EUT2B EP_NTERM
11025 Ts_goldenrod EUT2B
11030 HMMPduS_1_all(pf01512pf13375pf10531) GRM3A
11035 P_cloudyBlue EUT2B
11040 HMM02441_06(#) EUT2B EP_NTERM
11045 HMM06723EutJMreB_1_grm(pf11104) EUT2I
11050 HMMptac_3_eut2(pf06130) EUT2B
11055 HMM01923cobtr_3_eut2(pf01923) EUT2B EP_NTERM
11060 H_fuchsia EUT2F
11065 HMMalddh_2_egra(pf00171) EUT2B
11075 Tsp_mutedPink EUT2B
11080 HMMetlyEutC_2_eut2(pf05985) EUT2I EP_NTERM
11085 HMMetlyEutB_2_EUT2(pf06751) EUT2B
11090 HMMetlyEutB_2_EUT2(pf06751) EUT2B
11095 HMM06277EutA_1_eut(pf06277) EUT2I
11100 HMMreg07568_1_eut2(pf12282pf02518pf07568) EUT2B
11105 HMMreg00072_1_eut2(pf03861pf00072) EUT2B
11110 HMM10662PduV_1_mix(pf10662) EUT2B
11115 Hp_euts_plumPurple EUT2I
11120 -pf04962 -
11125 HMMnopfamclust3_1(#) SPU4
11130 -pf05035 -
11135 -pf01970 -
11140 -pf07331 -
11145 -pf03401 -
11150 -pf01081 -
11155 HMMreg09339_1_aro(pf09339pf01614) ARO
11160 -pf02743pf00015 -
11165 -pf03880pf00270pf00271 -
BMC-P
BMC-H
BMC-T
Aldehyde dehydrogenase
PTAC
Signature enzyme
Alcohol dehydrogenase
Regulator

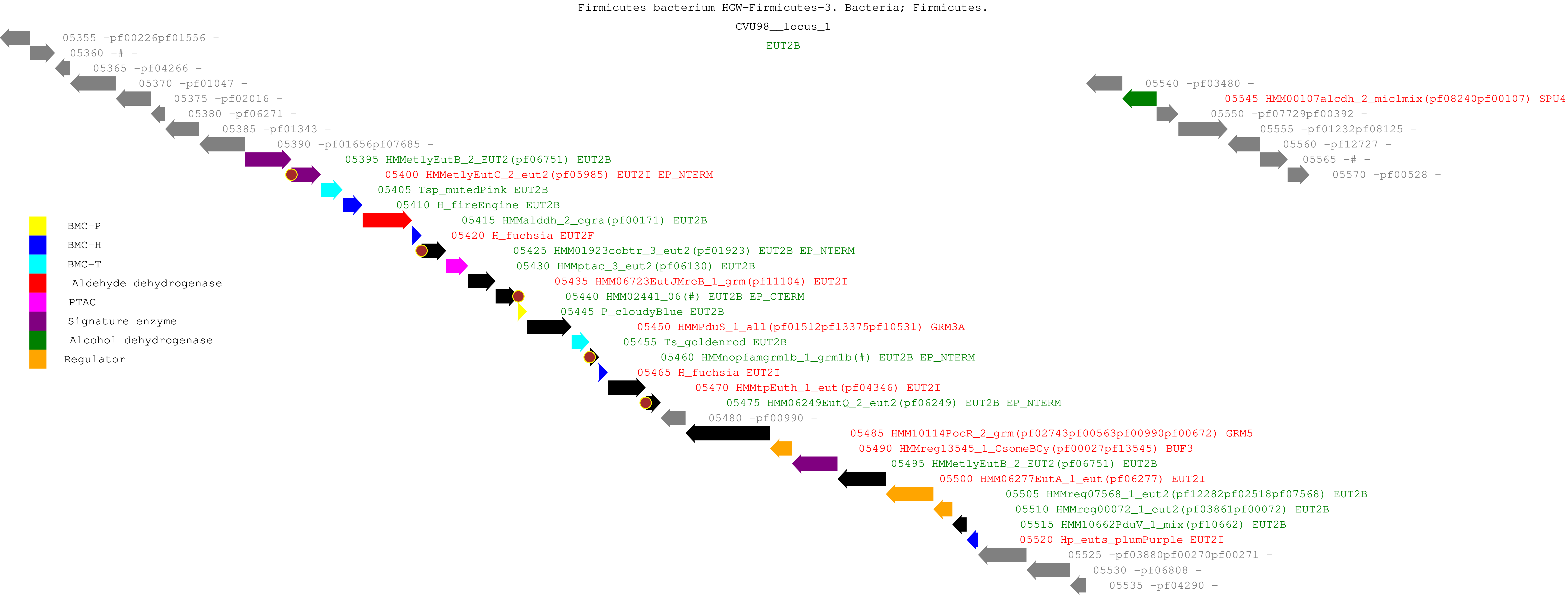
Firmicutes bacterium HGW-Firmicutes-3. Bacteria; Firmicutes.
CVU98__locus_1
EUT2B
05355 -pf00226pf01556 -
05360 -# -
05365 -pf04266 -
05370 -pf01047 -
05375 -pf02016 -
05380 -pf06271 -
05385 -pf01343 -
05390 -pf01656pf07685 -
05395 HMMetlyEutB_2_EUT2(pf06751) EUT2B
05400 HMMetlyEutC_2_eut2(pf05985) EUT2I EP_NTERM
05405 Tsp_mutedPink EUT2B
05410 H_fireEngine EUT2B
05415 HMMaldh_2_egra(pf00171) EUT2B
05420 H_fuchsia EUT2F
05425 HMM01923cobtr_3_eut2(pf01923) EUT2B EP_NTERM
05430 HMMptac_3_eut2(pf06130) EUT2B
05435 HMM06723EutJMreB_1_grm(pf11104) EUT2I
05440 HMM02441_06(#) EUT2B EP_CTERM
05445 P_cloudyBlue EUT2B
05450 HMMPduS_1_all(pf01512pf13375pf10531) GRM3A
05455 Ts_goldenrod EUT2B
05460 HMMnopfamgrm1b_1_grm1b(#) EUT2B EP_NTERM
05465 H_fuchsia EUT2I
05470 HMMtpEuth_1_eut(pf04346) EUT2I
05475 HMM06249EutQ_2_eut2(pf06249) EUT2B EP_NTERM
05480 -pf00990 -
05485 HMM10114PocR_2_grm(pf02743pf00563pf00990pf00672) GRM5
05490 HMMreg13545_1_CsomeBCy(pf00027pf13545) BUF3
05495 HMMetlyEutB_2_EUT2(pf06751) EUT2B
05500 HMM06277EutA_1_eut(pf06277) EUT2I
05505 HMMreg07568_1_eut2(pf12282pf02518pf07568) EUT2B
05510 HMMreg00072_1_eut2(pf03861pf00072) EUT2B
05515 HMM10662PduV_1_mix(pf10662) EUT2B
05520 Hp_euts_plumPurple EUT2I
05525 -pf03880pf00270pf00271 -
05530 -pf06808 -
05535 -pf04290 -
05540 -pf03480 -
05545 HMM00107alcdh_2_mic1mix(pf08240pf00107) SPU4
05550 -pf07729pf00392 -
05555 -pf01232pf08125 -
05560 -pf12727 -
05565 -# -
05570 -pf00528 -
BMC-P
BMC-H
BMC-T
Aldehyde dehydrogenase
PTAC
Signature enzyme
Alcohol dehydrogenase
Regulator

Firmicutes bacterium HGW-Firmicutes-4. Bacteria; Firmicutes.

CVU99__locus_1

EUT2B

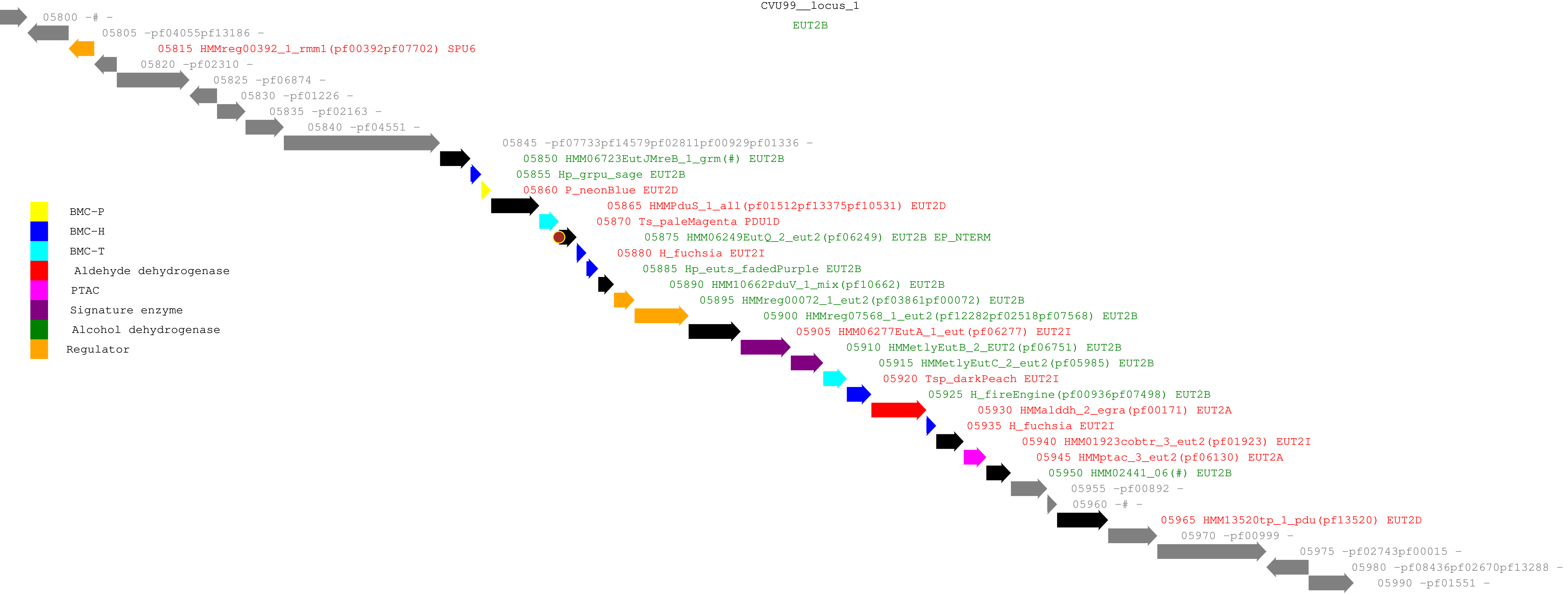

Firmicutes bacterium HGW-Firmicutes-7. Bacteria; Firmicutes.

CVV02__locus_2

EUT2B

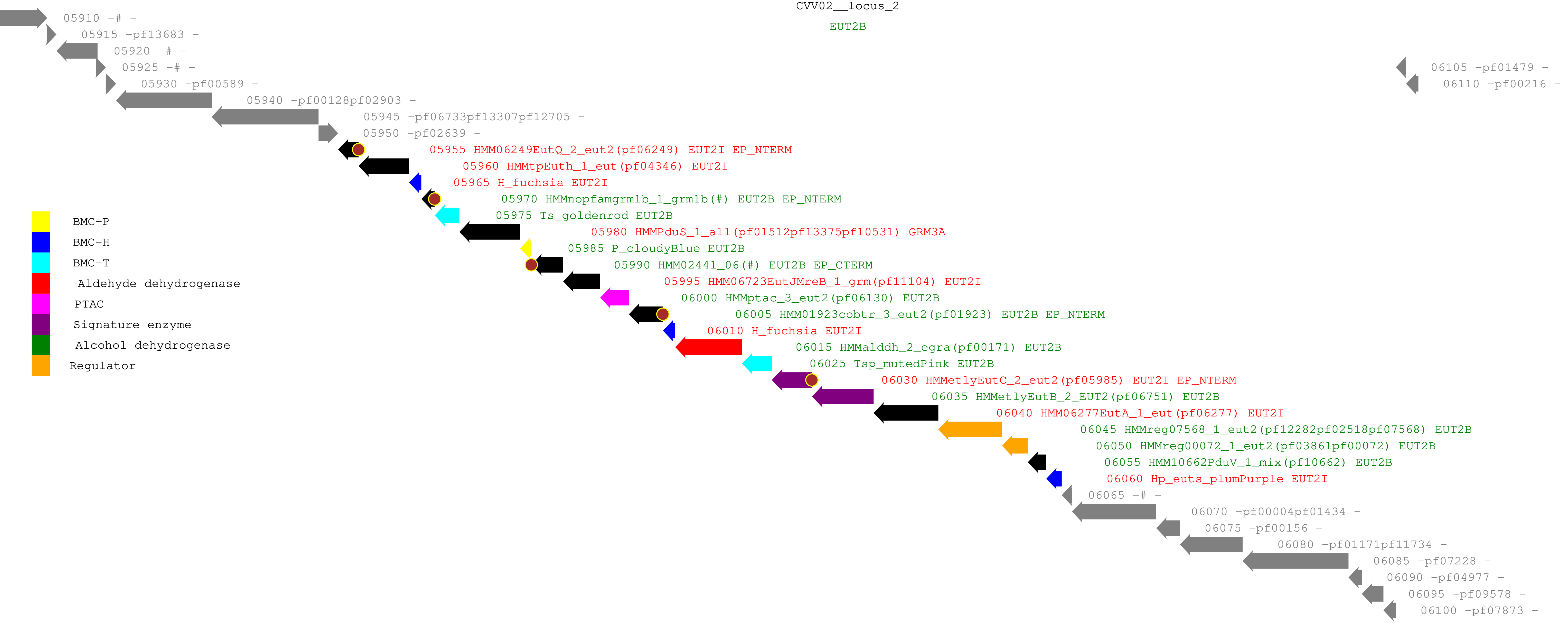

Anaerotruncus sp. 1XD22-93. Bacteria; Firmicutes; Clostridia; Clostridiales; Ruminococcaceae;

D7Y41__locus_1

EUT2B

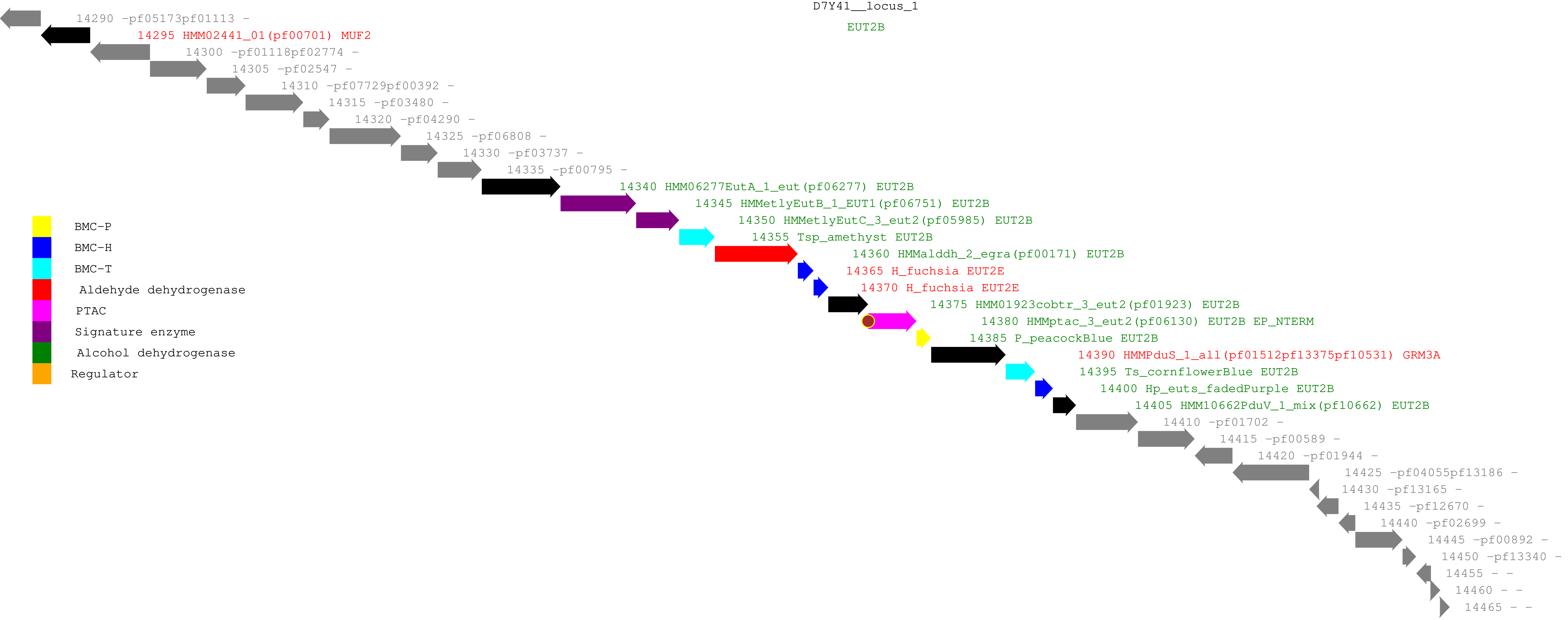

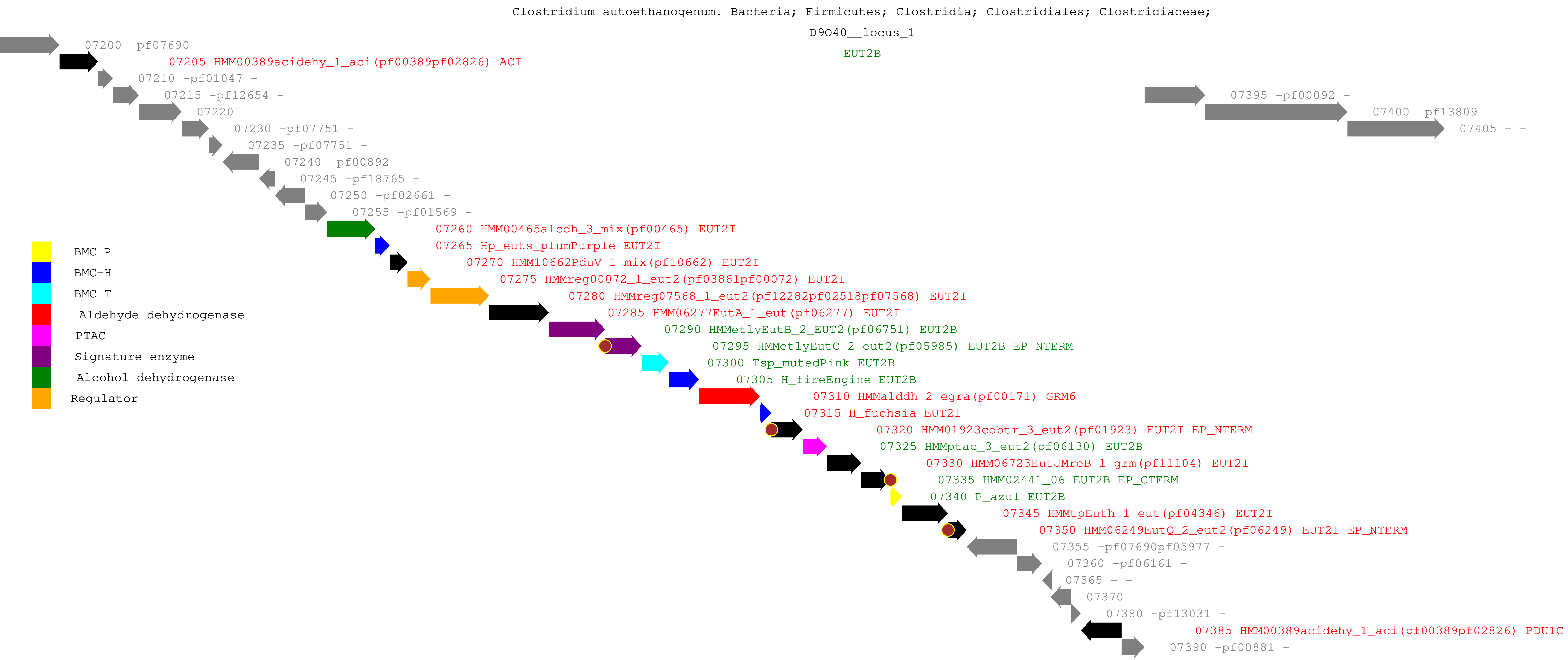

Clostridium autoethanogenum. Bacteria; Firmicutes; Clostridia; Clostridiales; Clostridiaceae;
D9040__locus_1
EUT2B
07200 -pf07690 -
07205 HMM00389acidehy_1_aci(pf00389pf02826) ACI
07210 -pf01047 -
07215 -pf12654 -
07220 - -
07230 -pf07751 -
07235 -pf07751 -
07240 -pf00892 -
07245 -pf18765 -
07250 -pf02661 -
07255 -pf01569 -
07260 HMM00465alcdh_3_mix(pf00465) EUT2I
07265 Hp_euts_plumPurple EUT2I
07270 HMM10662PduV_1_mix(pf10662) EUT2I
07275 HMMreg00072_1_eut2(pf03861pf00072) EUT2I
07280 HMMreg07568_1_eut2(pf12282pf02518pf07568) EUT2I
07285 HMM06277EutA_1_eut(pf06277) EUT2I
07290 HMMetlyEutB_2_EUT2(pf06751) EUT2B
07295 HMMetlyEutC_2_eut2(pf05985) EUT2B EP_NTERM
07300 Tsp_mutedPink EUT2B
07305 H_fireEngine EUT2B
07310 HMMalddh_2_egra(pf00171) GRM6
07315 H_fuchsia EUT2I
07320 HMM01923cobtr_3_eut2(pf01923) EUT2I EP_NTERM
07325 HMMptac_3_eut2(pf06130) EUT2B
07330 HMM06723EutJMreB_1_grm(pf11104) EUT2I
07335 HMM02441_06 EUT2B EP_CTERM
07340 P_azul EUT2B
07345 HMMtpEuth_1_eut(pf04346) EUT2I
07350 HMM06249EutQ_2_eut2(pf06249) EUT2I EP_NTERM
07355 -pf07690pf05977 -
07360 -pf06161 -
07365 - -
07370 - -
07380 -pf13031 -
07385 HMM00389acidehy_1_aci(pf00389pf02826) PDU1C
07390 -pf00881 -
07395 -pf00092 -
07400 -pf13809 -
07405 - -
BMC-P
BMC-H
BMC-T
Aldehyde dehydrogenase
PTAC
Signature enzyme
Alcohol dehydrogenase
Regulator

Clostridiales bacterium. Bacteria; Firmicutes; Clostridia; Clostridiales.

DBX66__locus_1

EUT2B

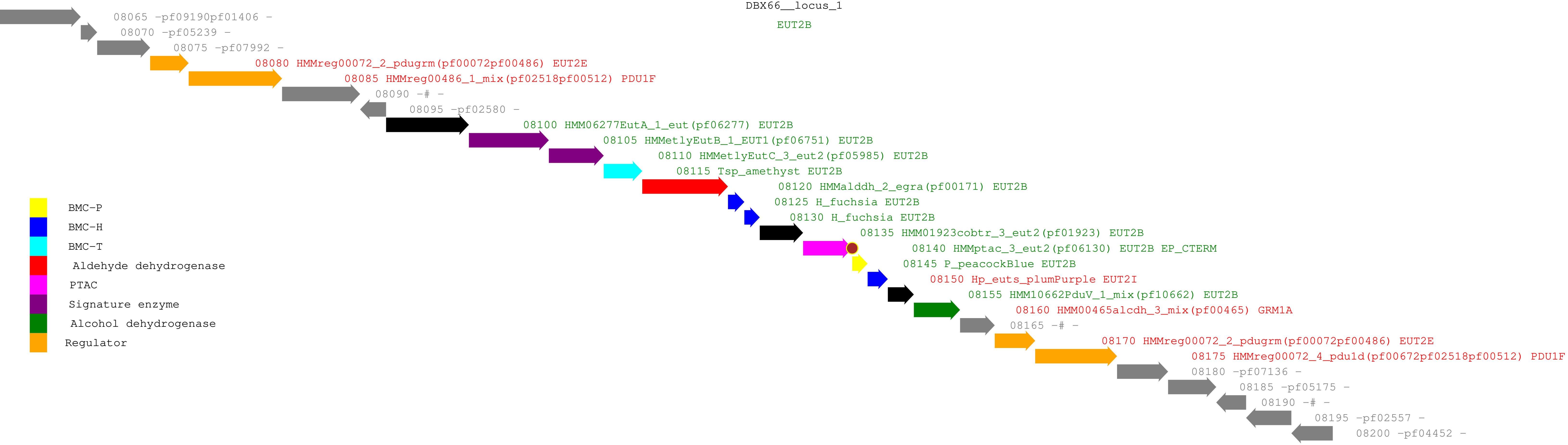

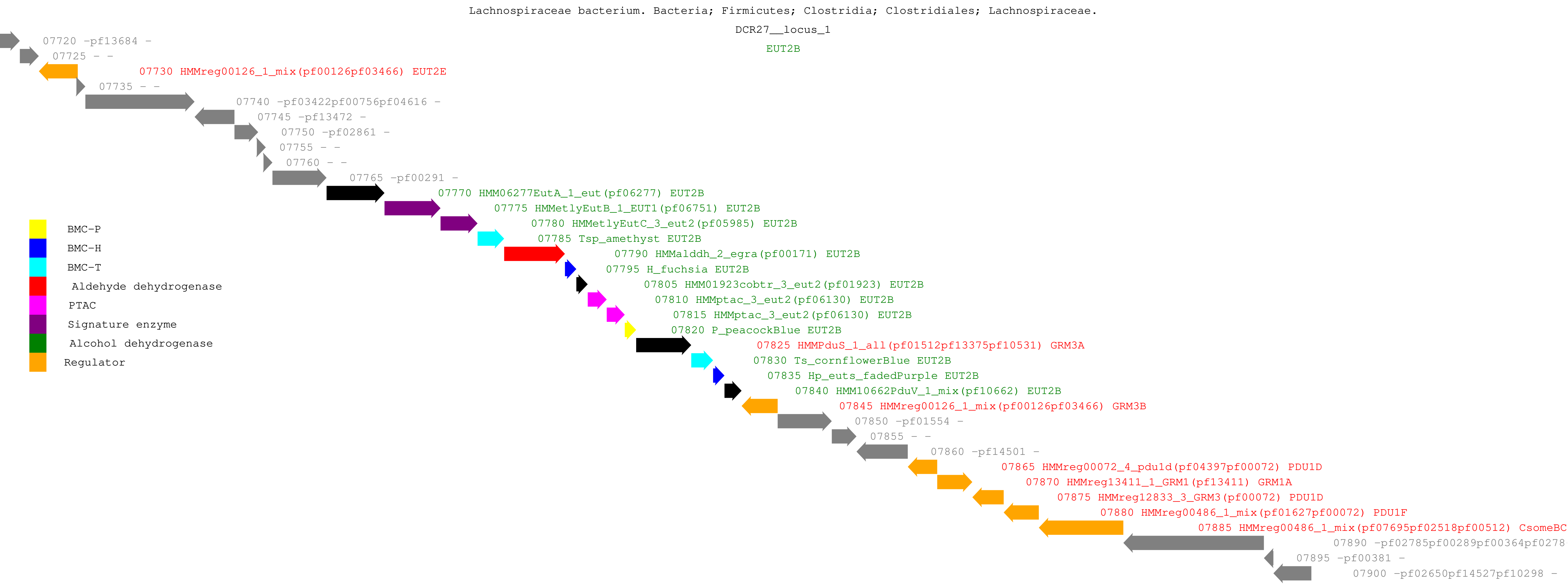
Lachnospiraceae bacterium. Bacteria; Firmicutes; Clostridia; Clostridiales; Lachnospiraceae.
DCR27__locus_1
EUT2B
07720 -pf13684 -
07725 - -
07730 HMMreg00126_1_mix(pf00126pf03466) EUT2E
07735 - -
07740 -pf03422pf00756pf04616 -
07745 -pf13472 -
07750 -pf02861 -
07755 - -
07760 - -
07765 -pf00291 -
07770 HMM06277EutA_1_eut(pf06277) EUT2B
07775 HMMetlyEutB_1_EUT1(pf06751) EUT2B
07780 HMMetlyEutC_3_eut2(pf05985) EUT2B
07785 Tsp_amethyst EUT2B
07790 HMMalddh_2_egra(pf00171) EUT2B
07795 H_fuchsia EUT2B
07805 HMM01923cobtr_3_eut2(pf01923) EUT2B
07810 HMMptac_3_eut2(pf06130) EUT2B
07815 HMMptac_3_eut2(pf06130) EUT2B
07820 P_peacockBlue EUT2B
07825 HMMPduS_1_all(pf01512pf13375pf10531) GRM3A
07830 Ts_cornflowerBlue EUT2B
07835 Hp_euts_fadedPurple EUT2B
07840 HMM10662PduV_1_mix(pf10662) EUT2B
07845 HMMreg00126_1_mix(pf00126pf03466) GRM3B
07850 -pf01554 -
07855 - -
07860 -pf14501 -
07865 HMMreg00072_4_pdu1d(pf04397pf00072) PDU1D
07870 HMMreg13411_1_GRM1(pf13411) GRM1A
07875 HMMreg12833_3_GRM3(pf00072) PDU1D
07880 HMMreg00486_1_mix(pf01627pf00072) PDU1F
07885 HMMreg00486_1_mix(pf07695pf02518pf00512) CsomeBC
07890 -pf02785pf00289pf00364pf0278
07895 -pf00381 -
07900 -pf02650pf14527pf10298 -
BMC-P
BMC-H
BMC-T
Aldehyde dehydrogenase
PTAC
Signature enzyme
Alcohol dehydrogenase
Regulator

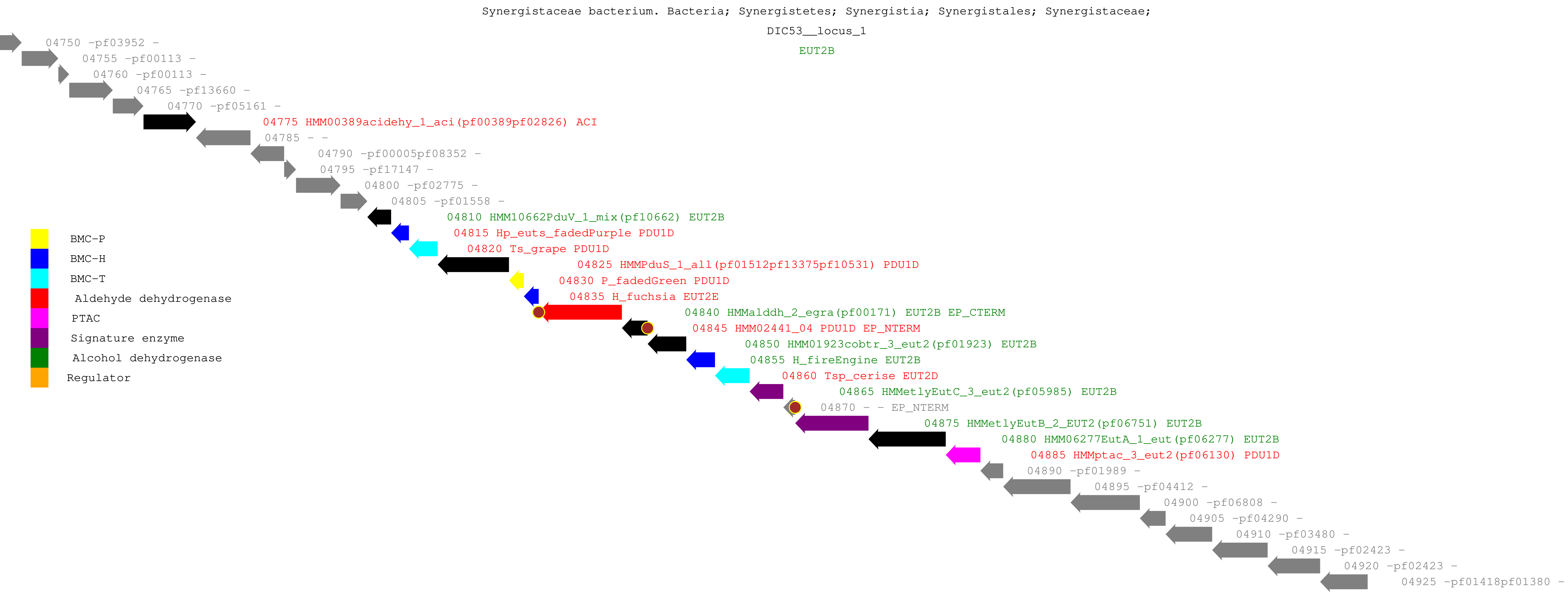
Synergistaceae bacterium. Bacteria; Synergistetes; Synergistia; Synergistales; Synergistaceae;
DIC53__locus_1
EUT2B
04750 -pf03952 -
04755 -pf00113 -
04760 -pf00113 -
04765 -pf13660 -
04770 -pf05161 -
04775 HMM00389acidehy_1_aci(pf00389pf02826) ACI
04785 - -
04790 -pf00005pf08352 -
04795 -pf17147 -
04800 -pf02775 -
04805 -pf01558 -
04810 HMM10662PduV_1_mix(pf10662) EUT2B
04815 Hp_euts_fadedPurple PDU1D
04820 Ts_grape PDU1D
04825 HMMPduS_1_all(pf01512pf13375pf10531) PDU1D
04830 P_fadedGreen PDU1D
04835 H_fuchsia EUT2E
04840 HMMalddh_2_egra(pf00171) EUT2B EP_CTERM
04845 HMM02441_04 PDU1D EP_NTERM
04850 HMM01923cobtr_3_eut2(pf01923) EUT2B
04855 H_fireEngine EUT2B
04860 Tsp_cerise EUT2D
04865 HMMetlyEutC_3_eut2(pf05985) EUT2B
04870 - - EP_NTERM
04875 HMMetlyEutB_2_EUT2(pf06751) EUT2B
04880 HMM06277EutA_1_eut(pf06277) EUT2B
04885 HMMptac_3_eut2(pf06130) PDU1D
04890 -pf01989 -
04895 -pf04412 -
04900 -pf06808 -
04905 -pf04290 -
04910 -pf03480 -
04915 -pf02423 -
04920 -pf02423 -
04925 -pf01418pf01380 -
BMC-P
BMC-H
BMC-T
Aldehyde dehydrogenase
PTAC
Signature enzyme
Alcohol dehydrogenase
Regulator

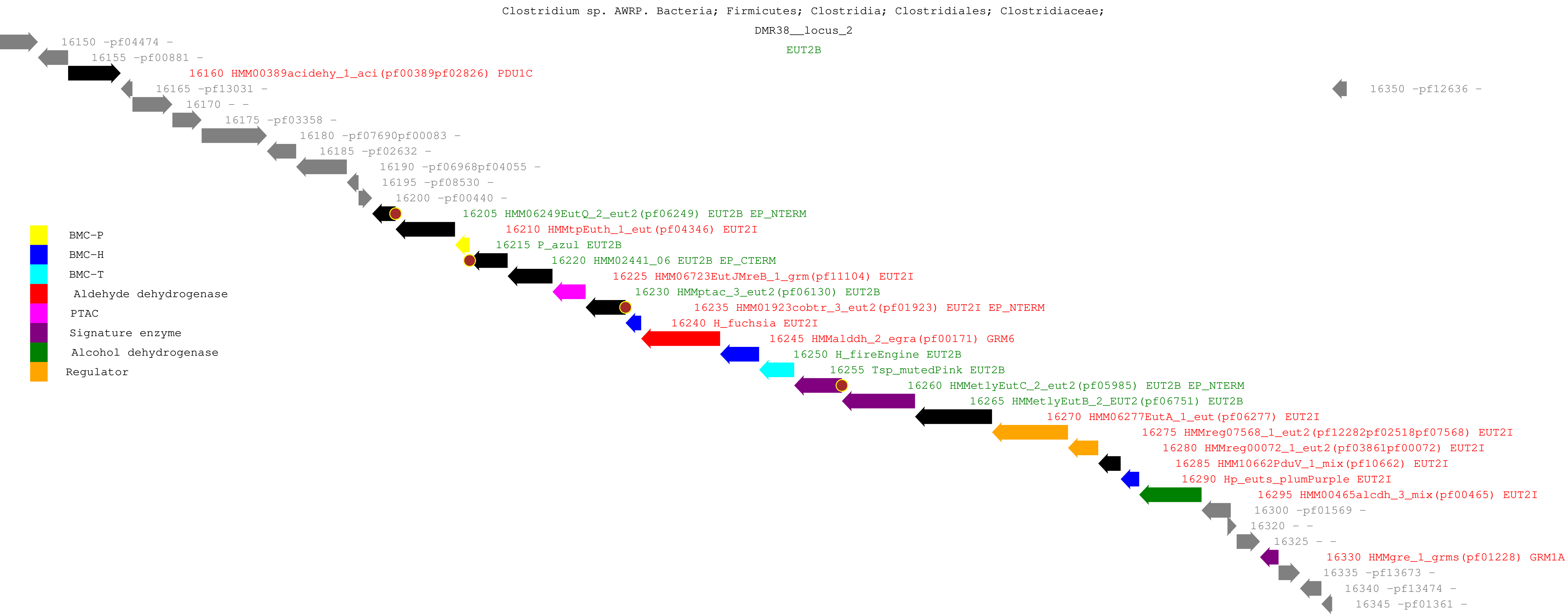
Clostridium sp. AWRP. Bacteria; Firmicutes; Clostridia; Clostridiales; Clostridiaceae;
DMR38__locus_2
EUT2B
16150 -pf04474 -
16155 -pf00881 -
16160 HMM00389acidehy_1_aci(pf00389pf02826) PDU1C
16165 -pf13031 -
16170 - -
16175 -pf03358 -
16180 -pf07690pf00083 -
16185 -pf02632 -
16190 -pf06968pf04055 -
16195 -pf08530 -
16200 -pf00440 -
16205 HMM06249EutQ_2_eut2(pf06249) EUT2B EP_NTERM
16210 HMMtpEuth_1_eut(pf04346) EUT2I
16215 P_azul EUT2B
16220 HMM02441_06 EUT2B EP_CTERM
16225 HMM06723EutJMreB_1_grm(pf11104) EUT2I
16230 HMMptac_3_eut2(pf06130) EUT2B
16235 HMM01923cobtr_3_eut2(pf01923) EUT2I EP_NTERM
16240 H_fuchsia EUT2I
16245 HMMalddh_2_egra(pf00171) GRM6
16250 H_fireEngine EUT2B
16255 Tsp_mutedPink EUT2B
16260 HMMetlyEutC_2_eut2(pf05985) EUT2B EP_NTERM
16265 HMMetlyEutB_2_EUT2(pf06751) EUT2B
16270 HMM06277EutA_1_eut(pf06277) EUT2I
16275 HMMreg07568_1_eut2(pf12282pf02518pf07568) EUT2I
16280 HMMreg00072_1_eut2(pf03861pf00072) EUT2I
16285 HMM10662PduV_1_mix(pf10662) EUT2I
16290 Hp_euts_plumPurple EUT2I
16295 HMM00465alcdh_3_mix(pf00465) EUT2I
16300 -pf01569 -
16320 - -
16325 - -
16330 HMMgre_1_grms(pf01228) GRM1A
16335 -pf13673 -
16340 -pf13474 -
16345 -pf01361 -
16350 -pf12636 -
BMC-P
BMC-H
BMC-T
Aldehyde dehydrogenase
PTAC
Signature enzyme
Alcohol dehydrogenase
Regulator

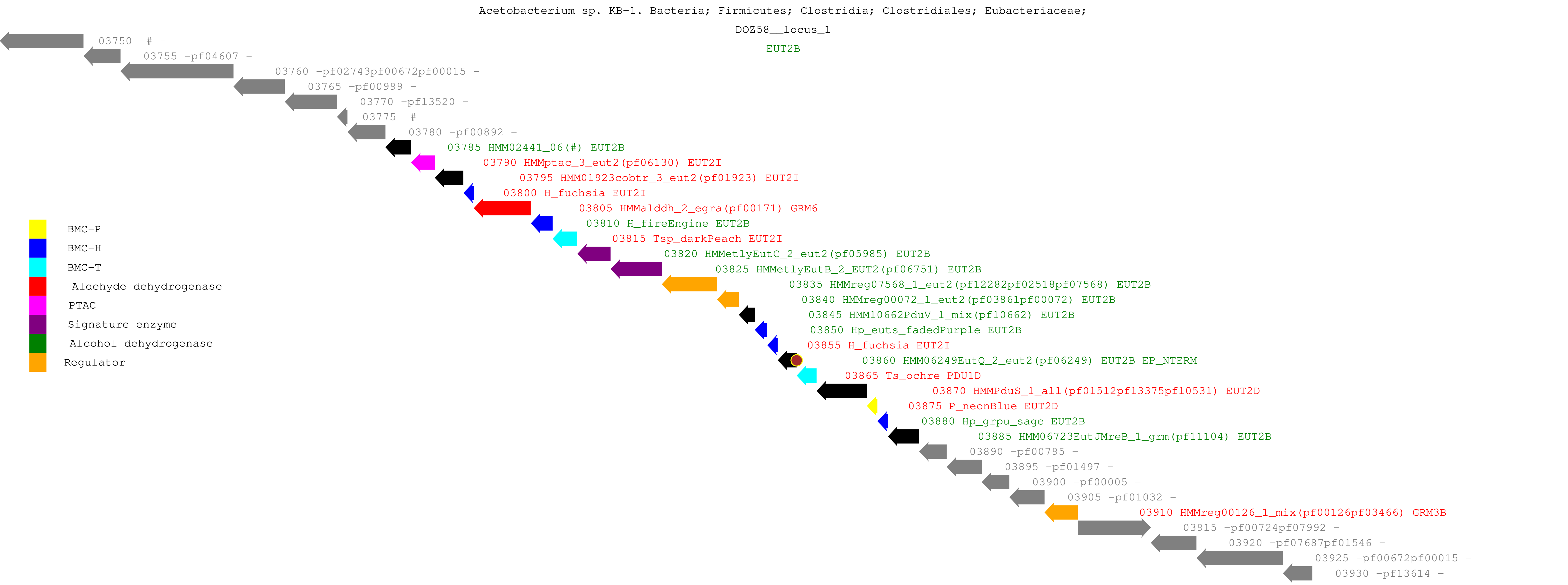
Acetobacterium sp. KB-1. Bacteria; Firmicutes; Clostridia; Clostridiales; Eubacteriaceae;
DOZ58__locus_1
EUT2B
03750 -# -
03755 -pf04607 -
03760 -pf02743pf00672pf00015 -
03765 -pf00999 -
03770 -pf13520 -
03775 -# -
03780 -pf00892 -
03785 HMM02441_06(#) EUT2B
03790 HMMptac_3_eut2(pf06130) EUT2I
03795 HMM01923cobtr_3_eut2(pf01923) EUT2I
03800 H_fuchsia EUT2I
03805 HMMalddh_2_egra(pf00171) GRM6
03810 H_fireEngine EUT2B
03815 Tsp_darkPeach EUT2I
03820 HMMetlyEutC_2_eut2(pf05985) EUT2B
03825 HMMetlyEutB_2_EUT2(pf06751) EUT2B
03835 HMMreg07568_1_eut2(pf12282pf02518pf07568) EUT2B
03840 HMMreg00072_1_eut2(pf03861pf00072) EUT2B
03845 HMM10662PduV_1_mix(pf10662) EUT2B
03850 Hp_euts_fadedPurple EUT2B
03855 H_fuchsia EUT2I
03860 HMM06249EutQ_2_eut2(pf06249) EUT2B EP_NTERM
03865 Ts_ochre PDU1D
03870 HMMPduS_1_all(pf01512pf13375pf10531) EUT2D
03875 P_neonBlue EUT2D
03880 Hp_grpu_sage EUT2B
03885 HMM06723EutJMreB_1_grm(pf11104) EUT2B
03890 -pf00795 -
03895 -pf01497 -
03900 -pf00005 -
03905 -pf01032 -
03910 HMMreg00126_1_mix(pf00126pf03466) GRM3B
03915 -pf00724pf07992 -
03920 -pf07687pf01546 -
03925 -pf00672pf00015 -
03930 -pf13614 -
BMC-P
BMC-H
BMC-T
Aldehyde dehydrogenase
PTAC
Signature enzyme
Alcohol dehydrogenase
Regulator

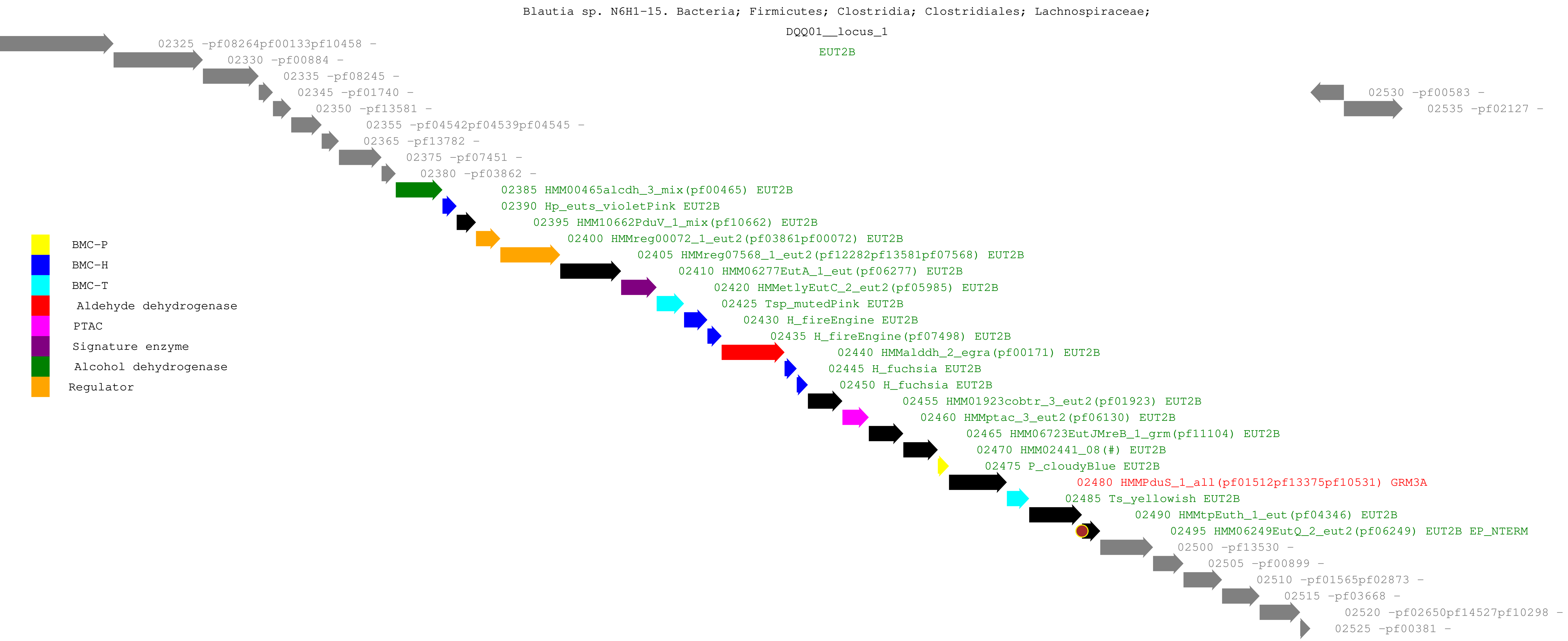
Blautia sp. N6H1-15. Bacteria; Firmicutes; Clostridia; Clostridiales; Lachnospiraceae;
DQQ01__locus_1
EUT2B
02325 -pf08264pf00133pf10458 -
02330 -pf00884 -
02335 -pf08245 -
02345 -pf01740 -
02350 -pf13581 -
02355 -pf04542pf04539pf04545 -
02365 -pf13782 -
02375 -pf07451 -
02380 -pf03862 -
02385 HMM00465alcdh_3_mix(pf00465) EUT2B
02390 Hp_euts_violetPink EUT2B
02395 HMM10662PduV_1_mix(pf10662) EUT2B
02400 HMMreg00072_1_eut2(pf03861pf00072) EUT2B
02405 HMMreg07568_1_eut2(pf12282pf13581pf07568) EUT2B
02410 HMM06277EutA_1_eut(pf06277) EUT2B
02420 HMMetlyEutC_2_eut2(pf05985) EUT2B
02425 Tsp_mutedPink EUT2B
02430 H_fireEngine EUT2B
02435 H_fireEngine(pf07498) EUT2B
02440 HMMaldh_2_egra(pf00171) EUT2B
02445 H_fuchsia EUT2B
02450 H_fuchsia EUT2B
02455 HMM01923cobtr_3_eut2(pf01923) EUT2B
02460 HMMptac_3_eut2(pf06130) EUT2B
02465 HMM06723EutJMreB_1_grm(pf11104) EUT2B
02470 HMM02441_08(#) EUT2B
02475 P_cloudyBlue EUT2B
02480 HMMPduS_1_all(pf01512pf13375pf10531) GRM3A
02485 Ts_yellowish EUT2B
02490 HMMtpEuth_1_eut(pf04346) EUT2B
02495 HMM06249EutQ_2_eut2(pf06249) EUT2B EP_NTERM
02500 -pf13530 -
02505 -pf00899 -
02510 -pf01565pf02873 -
02515 -pf03668 -
02520 -pf02650pf14527pf10298 -
02525 -pf00381 -
02530 -pf00583 -
02535 -pf02127 -
BMC-P
BMC-H
BMC-T
Aldehyde dehydrogenase
PTAC
Signature enzyme
Alcohol dehydrogenase
Regulator

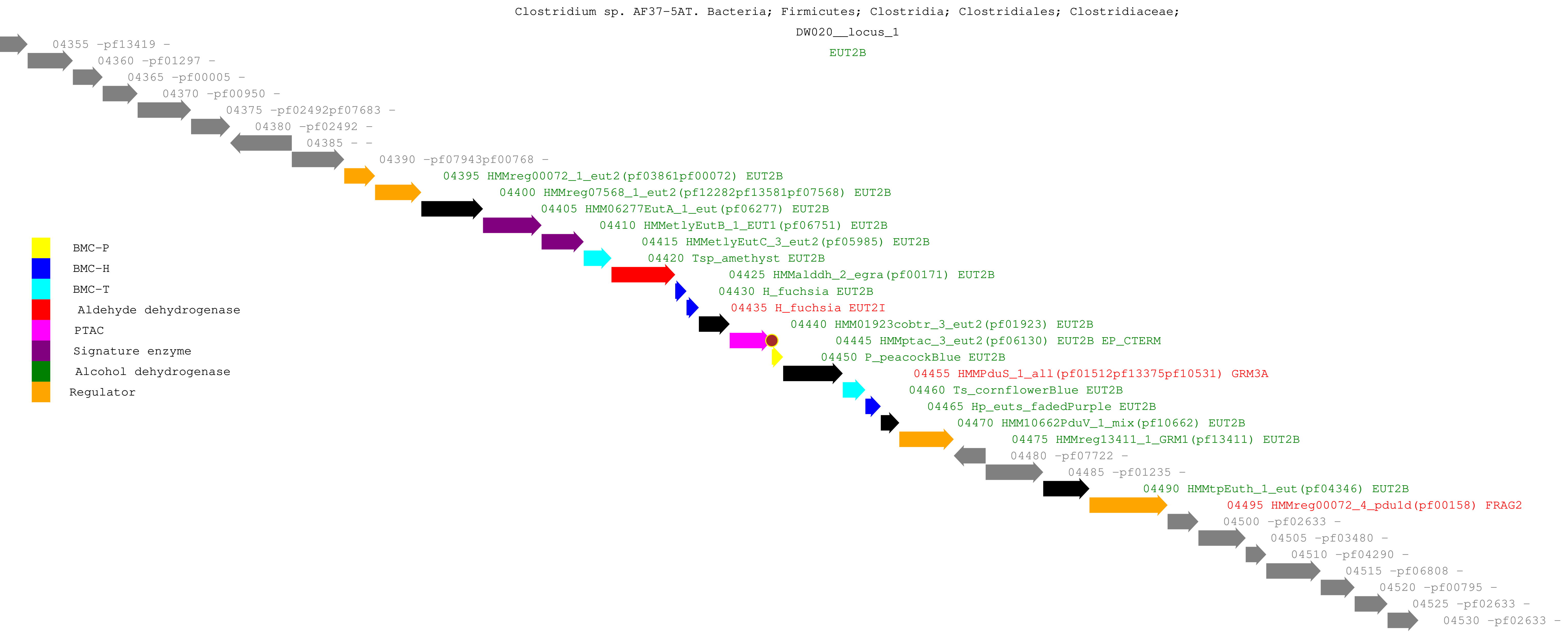

Clostridium sp. AF37-5AT. Bacteria; Firmicutes; Clostridia; Clostridiales; Clostridiaceae;
DW020__locus_1
EUT2B
04355 -pf13419 -
04360 -pf01297 -
04365 -pf00005 -
04370 -pf00950 -
04375 -pf02492pf07683 -
04380 -pf02492 -
04385 - -
04390 -pf07943pf00768 -
04395 HMMreg00072_1_eut2(pf03861pf00072) EUT2B
04400 HMMreg07568_1_eut2(pf12282pf13581pf07568) EUT2B
04405 HMM06277EutA_1_eut(pf06277) EUT2B
04410 HMMetlyEutB_1_EUT1(pf06751) EUT2B
04415 HMMetlyEutC_3_eut2(pf05985) EUT2B
04420 Tsp_amethyst EUT2B
04425 HMMalddh_2_egra(pf00171) EUT2B
04430 H_fuchsia EUT2B
04435 H_fuchsia EUT2I
04440 HMM01923cobtr_3_eut2(pf01923) EUT2B
04445 HMMptac_3_eut2(pf06130) EUT2B EP_CTERM
04450 P_peacockBlue EUT2B
04455 HMMPduS_1_all(pf01512pf13375pf10531) GRM3A
04460 Ts_cornflowerBlue EUT2B
04465 Hp_euts_fadedPurple EUT2B
04470 HMM10662PduV_1_mix(pf10662) EUT2B
04475 HMMreg13411_1_GRM1(pf13411) EUT2B
04480 -pf07722 -
04485 -pf01235 -
04490 HMMtpEuth_1_eut(pf04346) EUT2B
04495 HMMreg00072_4_pdu1d(pf00158) FRAG2
04500 -pf02633 -
04505 -pf03480 -
04510 -pf04290 -
04515 -pf06808 -
04520 -pf00795 -
04525 -pf02633 -
04530 -pf02633 -
BMC-P
BMC-H
BMC-T
Aldehyde dehydrogenase
PTAC
Signature enzyme
Alcohol dehydrogenase
Regulator

Clostridiaceae bacterium AF42-6. Bacteria; Firmicutes; Clostridia; Clostridiales; Clostridiaceae.

DW058__locus_1

EUT2B

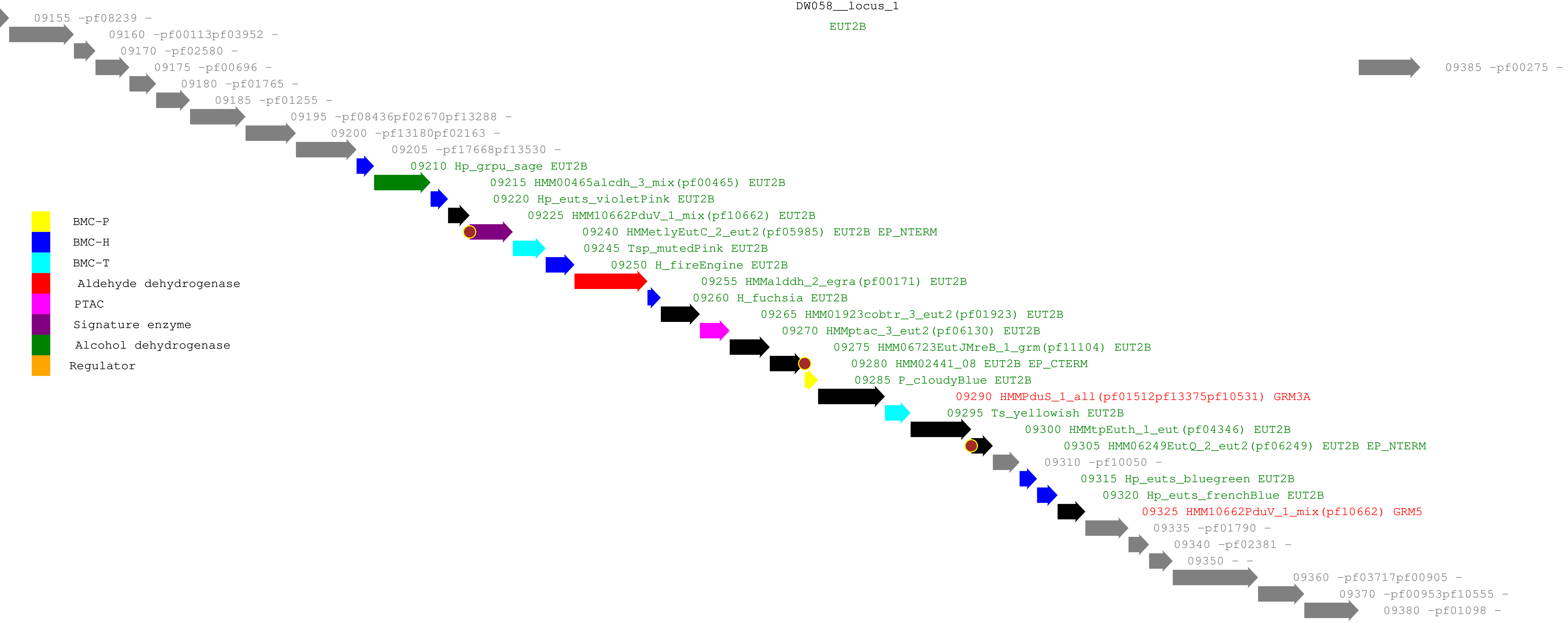

Harryflintia acetispora. Bacteria; Firmicutes; Clostridia; Clostridiales; Ruminococcaceae;

DW086__locus_1

EUT2B

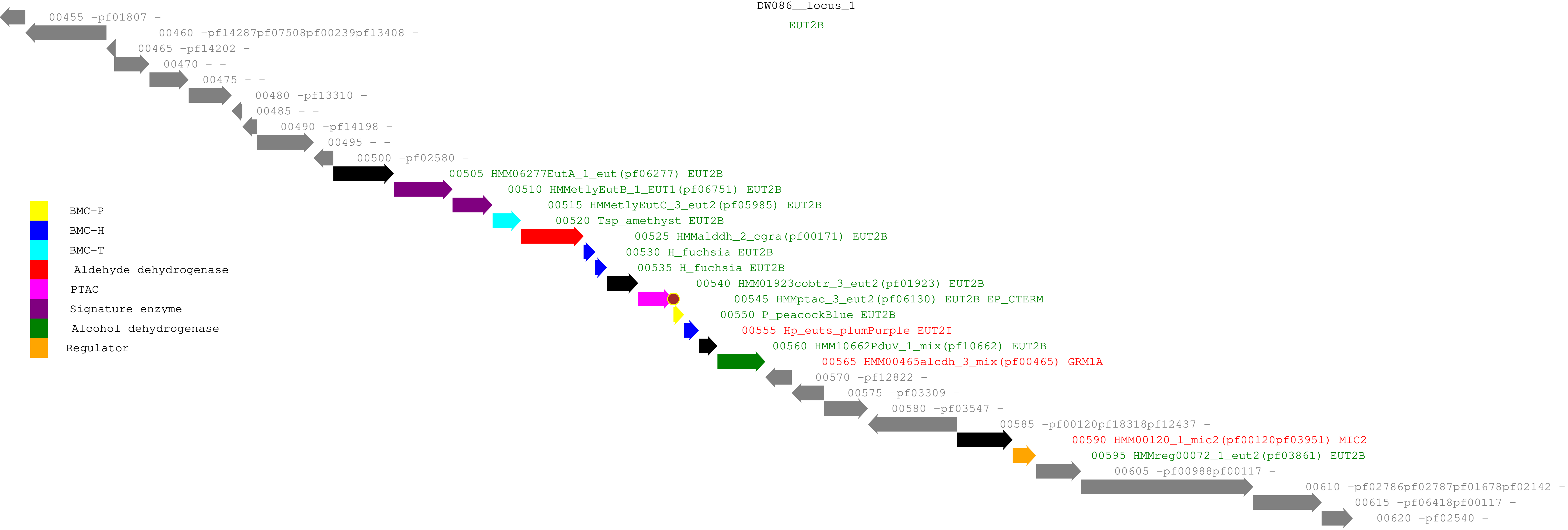

Emergencia timonensis. Bacteria; Firmicutes; Clostridia; Clostridiales;

DW099__locus_1

EUT2B

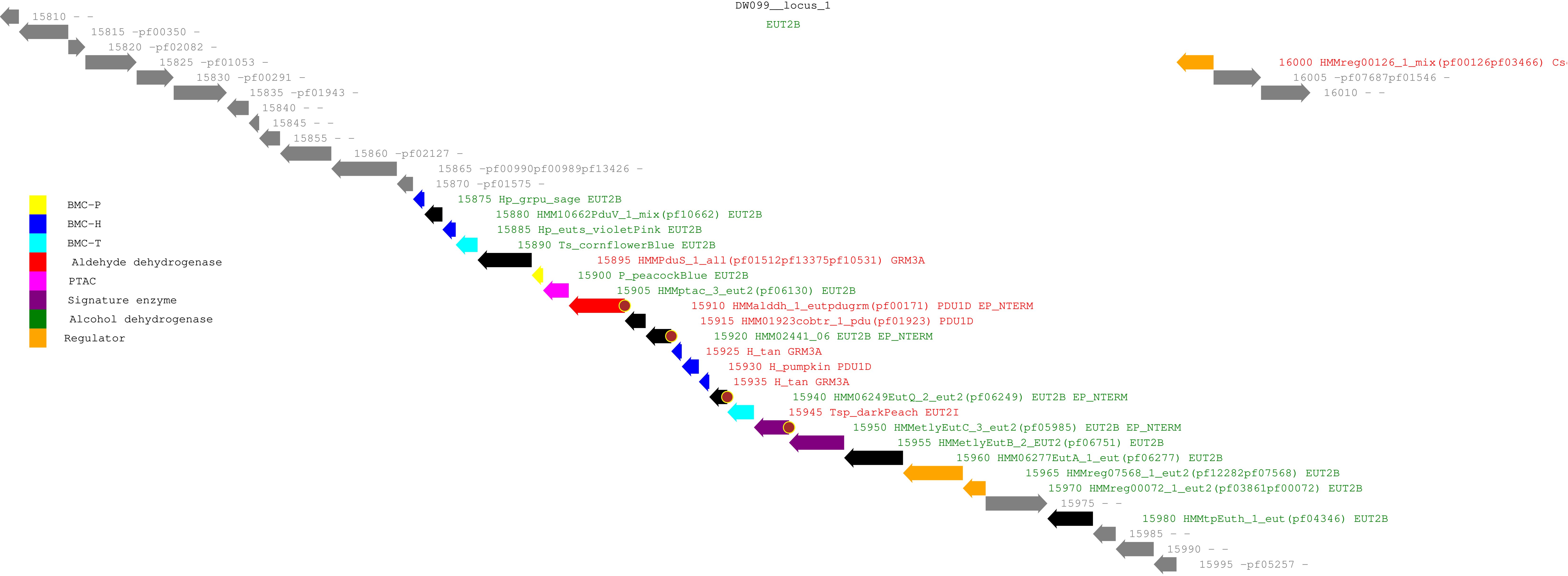

Clostridium sp. AM16-23. Bacteria; Firmicutes; Clostridia; Clostridiales; Clostridiaceae;

DW181__locus_1

EUT2B

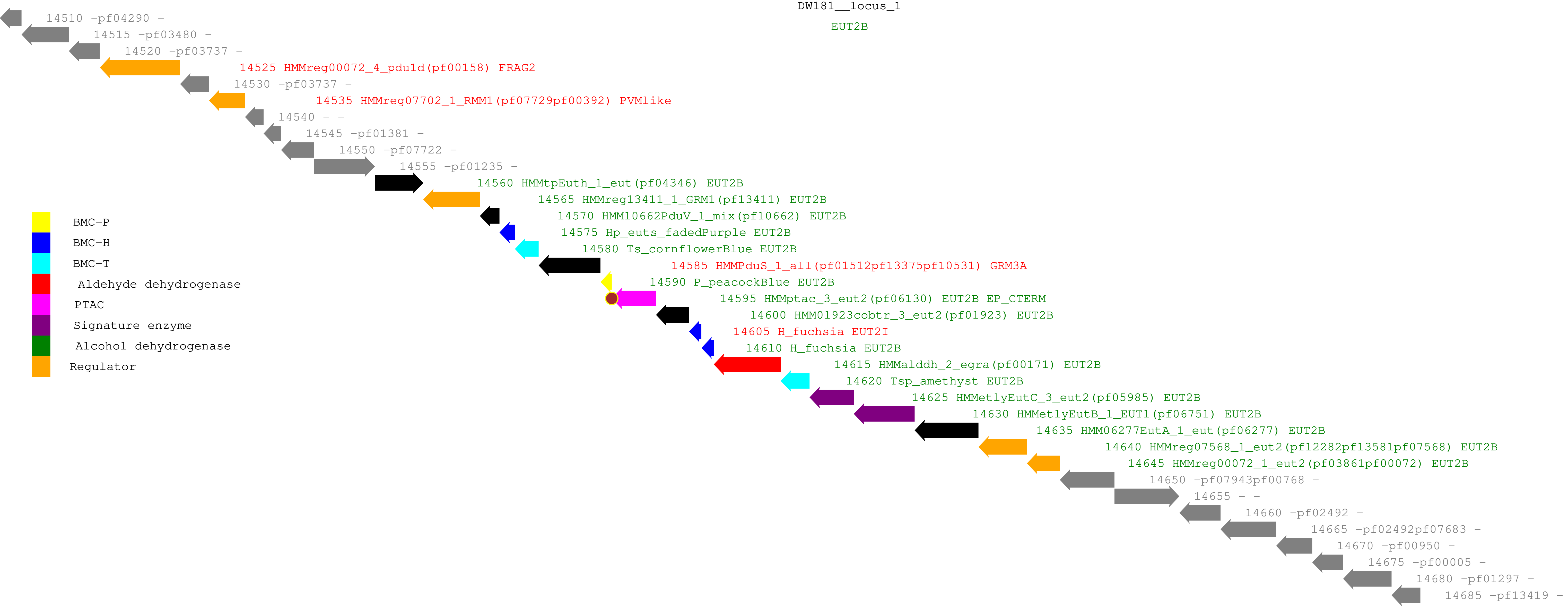

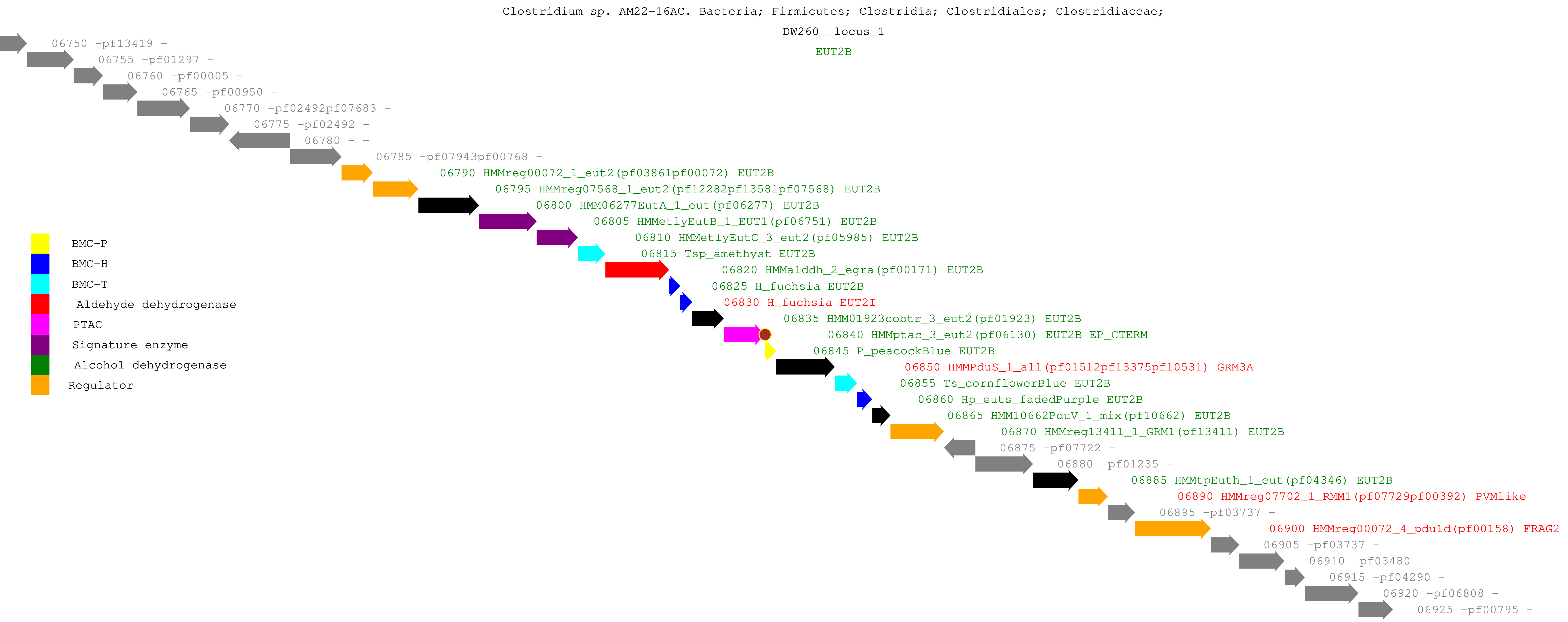
Clostridium sp. AM22-16AC. Bacteria; Firmicutes; Clostridia; Clostridiales; Clostridiaceae;
DW260__locus_1
EUT2B
06750 -pf13419 -
06755 -pf01297 -
06760 -pf00005 -
06765 -pf00950 -
06770 -pf02492pf07683 -
06775 -pf02492 -
06780 - -
06785 -pf07943pf00768 -
06790 HMMreg00072_1_eut2(pf03861pf00072) EUT2B
06795 HMMreg07568_1_eut2(pf12282pf13581pf07568) EUT2B
06800 HMM06277EutA_1_eut(pf06277) EUT2B
06805 HMMetlyEutB_1_EUT1(pf06751) EUT2B
06810 HMMetlyEutC_3_eut2(pf05985) EUT2B
06815 Tsp_amethyst EUT2B
06820 HMMalddh_2_egra(pf00171) EUT2B
06825 H_fuchsia EUT2B
06830 H_fuchsia EUT2I
06835 HMM01923cobtr_3_eut2(pf01923) EUT2B
06840 HMMptac_3_eut2(pf06130) EUT2B EP_CTERM
06845 P_peacockBlue EUT2B
06850 HMMPduS_1_all(pf01512pf13375pf10531) GRM3A
06855 Ts_cornflowerBlue EUT2B
06860 Hp_euts_fadedPurple EUT2B
06865 HMM10662PduV_1_mix(pf10662) EUT2B
06870 HMMreg13411_1_GRM1(pf13411) EUT2B
06875 -pf07722 -
06880 -pf01235 -
06885 HMMtpEuth_1_eut(pf04346) EUT2B
06890 HMMreg07702_1_RMM1(pf07729pf00392) PVMlike
06895 -pf03737 -
06900 HMMreg00072_4_pdu1d(pf00158) FRAG2
06905 -pf03737 -
06910 -pf03480 -
06915 -pf04290 -
06920 -pf06808 -
06925 -pf00795 -
BMC-P
BMC-H
BMC-T
Aldehyde dehydrogenase
PTAC
Signature enzyme
Alcohol dehydrogenase
Regulator

Clostridiales bacterium AM23-16LB. Bacteria; Firmicutes; Clostridia; Clostridiales.

DW644__locus_1

EUT2B

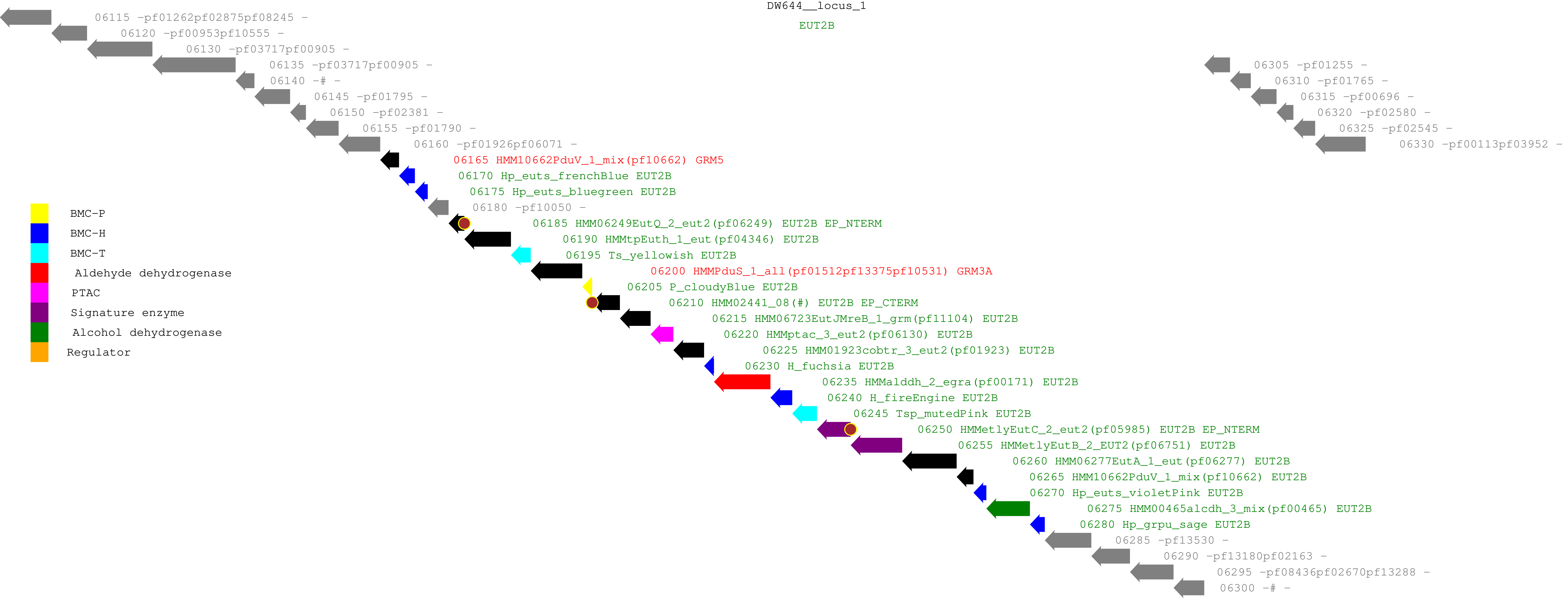

Clostridiaceae bacterium AM27-36LB. Bacteria; Firmicutes; Clostridia; Clostridiales; Clostridiaceae.

DW725__locus_1

EUT2B

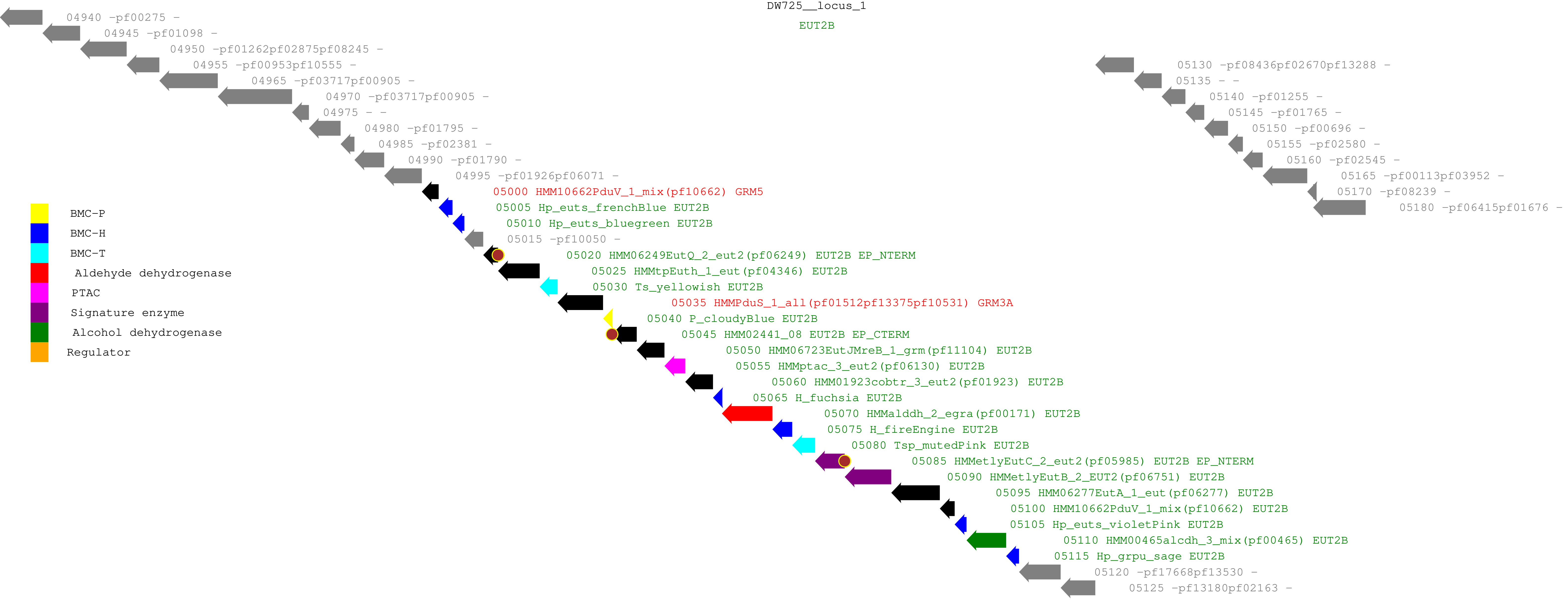

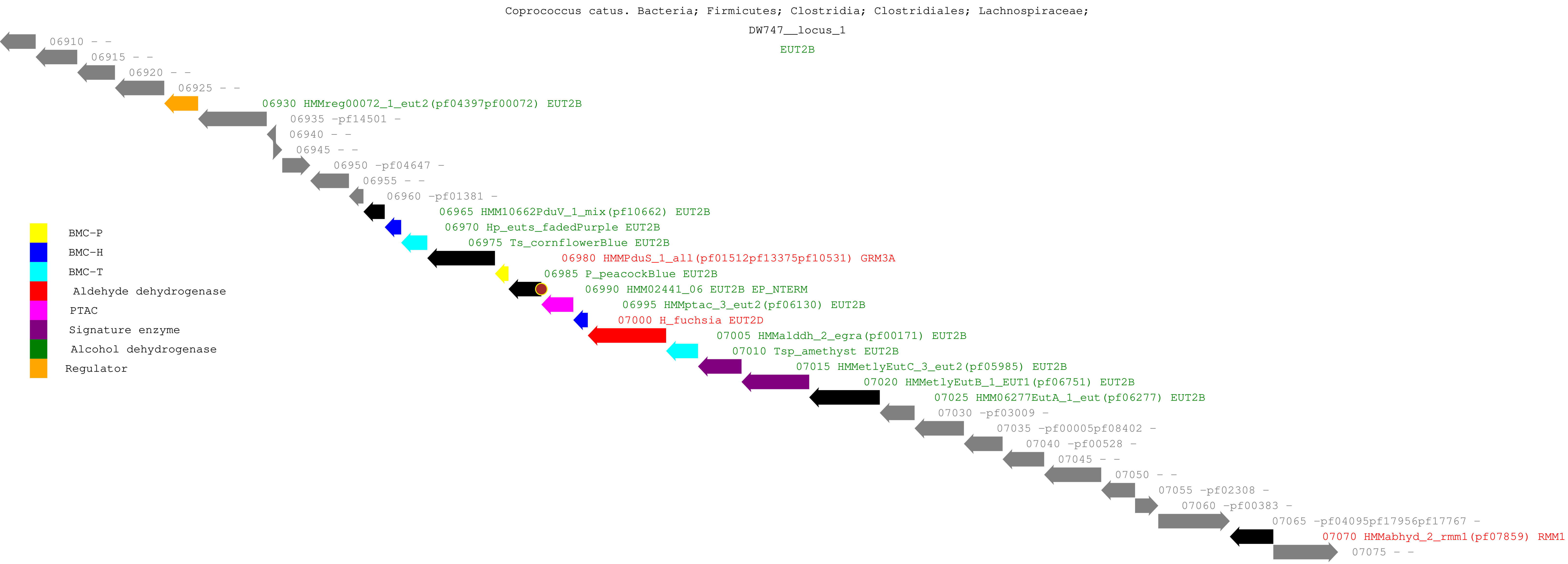
Coprococcus catus. Bacteria; Firmicutes; Clostridia; Clostridiales; Lachnospiraceae;
DW747__locus_1
EUT2B
06910 - -
06915 - -
06920 - -
06925 - -
06930 HMMreg00072_1_eut2(pf04397pf00072) EUT2B
06935 -pf14501 -
06940 - -
06945 - -
06950 -pf04647 -
06955 - -
06960 -pf01381 -
06965 HMM10662PduV_1_mix(pf10662) EUT2B
06970 Hp_euts_fadedPurple EUT2B
06975 Ts_cornflowerBlue EUT2B
06980 HMMPduS_1_all(pf01512pf13375pf10531) GRM3A
06985 P_peacockBlue EUT2B
06990 HMM02441_06 EUT2B EP_NTERM
06995 HMMptac_3_eut2(pf06130) EUT2B
07000 H_fuchsia EUT2D
07005 HMMaldh_2_egra(pf00171) EUT2B
07010 Tsp_amethyst EUT2B
07015 HMMetlyEutC_3_eut2(pf05985) EUT2B
07020 HMMetlyEutB_1_EUT1(pf06751) EUT2B
07025 HMM06277EutA_1_eut(pf06277) EUT2B
07030 -pf03009 -
07035 -pf00005pf08402 -
07040 -pf00528 -
07045 - -
07050 - -
07055 -pf02308 -
07060 -pf00383 -
07065 -pf04095pf17956pf17767 -
07070 HMMabhyd_2_rmm1(pf07859) RMM1
07075 - -
BMC-P
BMC-H
BMC-T
Aldehyde dehydrogenase
PTAC
Signature enzyme
Alcohol dehydrogenase
Regulator

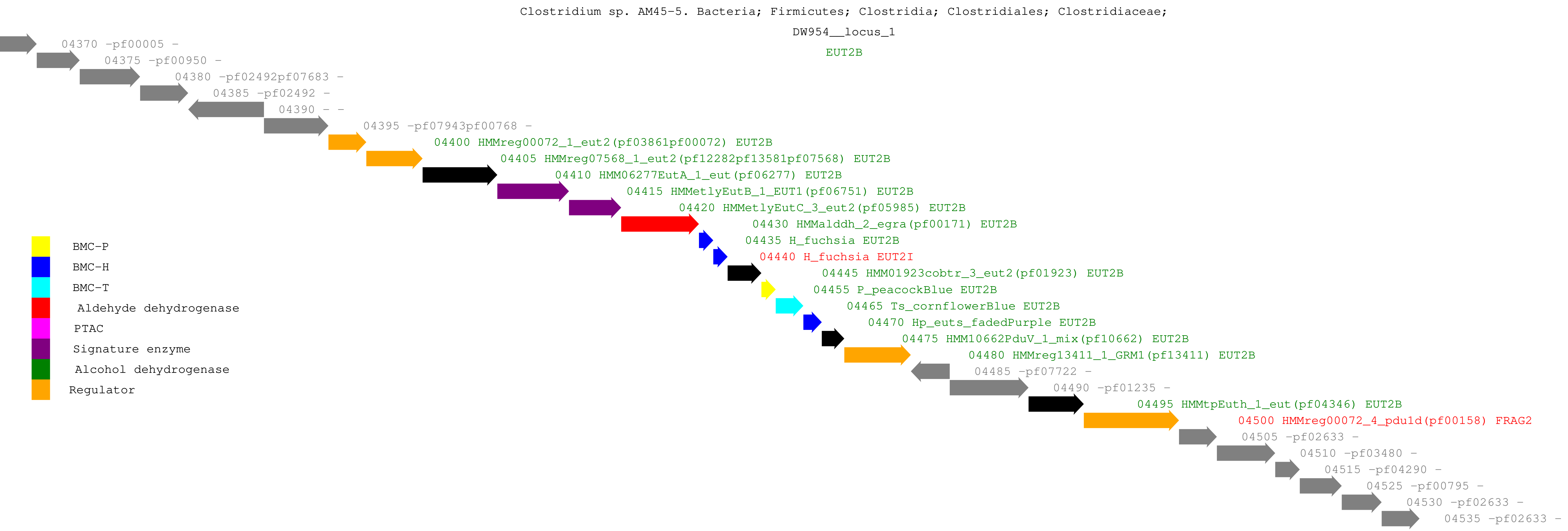
Clostridium sp. AM45-5. Bacteria; Firmicutes; Clostridia; Clostridiales; Clostridiaceae;
DW954__locus_1
EUT2B
04370 -pf00005 -
04375 -pf00950 -
04380 -pf02492pf07683 -
04385 -pf02492 -
04390 - -
04395 -pf07943pf00768 -
04400 HMMreg00072_1_eut2(pf03861pf00072) EUT2B
04405 HMMreg07568_1_eut2(pf12282pf13581pf07568) EUT2B
04410 HMM06277EutA_1_eut(pf06277) EUT2B
04415 HMMetlyEutB_1_EUT1(pf06751) EUT2B
04420 HMMetlyEutC_3_eut2(pf05985) EUT2B
04430 HMMalddh_2_egra(pf00171) EUT2B
04435 H_fuchsia EUT2B
04440 H_fuchsia EUT2I
04445 HMM01923cobtr_3_eut2(pf01923) EUT2B
04455 P_peacockBlue EUT2B
04465 Ts_cornflowerBlue EUT2B
04470 Hp_euts_fadedPurple EUT2B
04475 HMM10662PduV_1_mix(pf10662) EUT2B
04480 HMMreg13411_1_GRM1(pf13411) EUT2B
04485 -pf07722 -
04490 -pf01235 -
04495 HMMtpEuth_1_eut(pf04346) EUT2B
04500 HMMreg00072_4_pdu1d(pf00158) FRAG2
04505 -pf02633 -
04510 -pf03480 -
04515 -pf04290 -
04525 -pf00795 -
04530 -pf02633 -
04535 -pf02633 -
BMC-P
BMC-H
BMC-T
Aldehyde dehydrogenase
PTAC
Signature enzyme
Alcohol dehydrogenase
Regulator

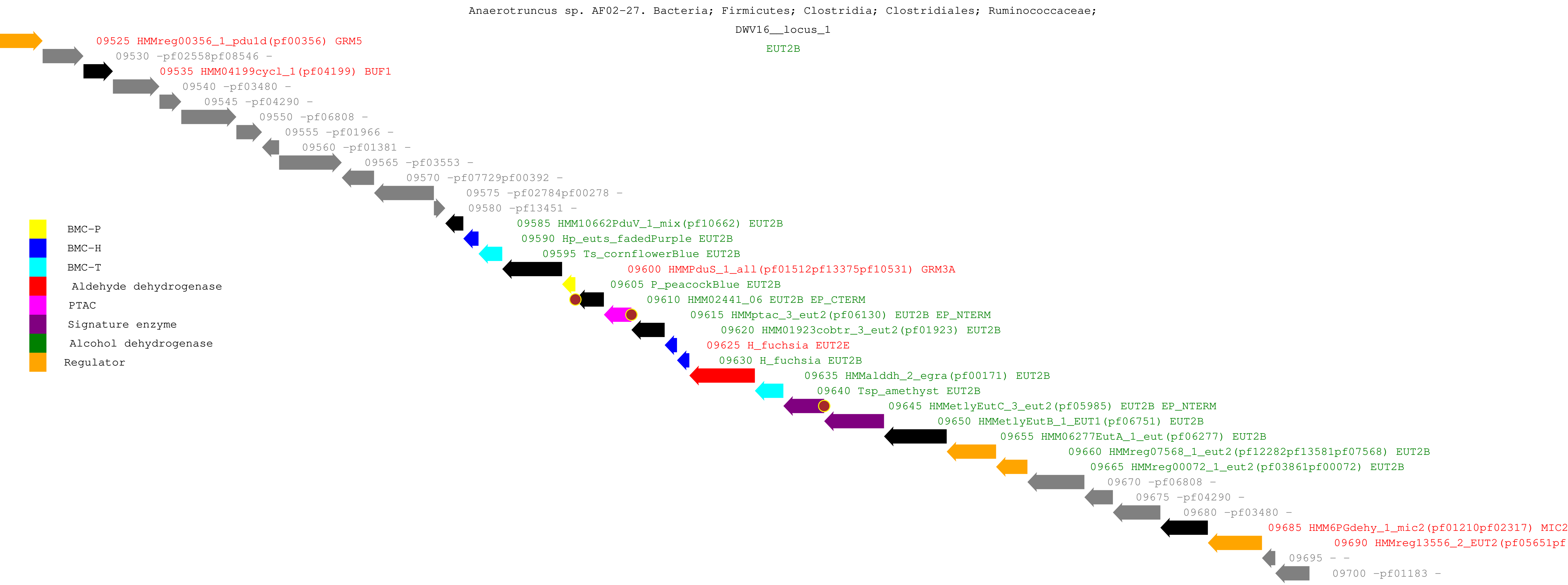
Anaerotruncus sp. AF02-27. Bacteria; Firmicutes; Clostridia; Clostridiales; Ruminococcaceae;
DWV16__locus_1
EUT2B
09525 HMMreg00356_1_pdu1d(pf00356) GRM5
09530 -pf02558pf08546 -
09535 HMM04199cycl_1(pf04199) BUF1
09540 -pf03480 -
09545 -pf04290 -
09550 -pf06808 -
09555 -pf01966 -
09560 -pf01381 -
09565 -pf03553 -
09570 -pf07729pf00392 -
09575 -pf02784pf00278 -
09580 -pf13451 -
09585 HMM10662PduV_1_mix(pf10662) EUT2B
09590 Hp_euts_fadedPurple EUT2B
09595 Ts_cornflowerBlue EUT2B
09600 HMMPduS_1_all(pf01512pf13375pf10531) GRM3A
09605 P_peacockBlue EUT2B
09610 HMM02441_06 EUT2B EP_CTERM
09615 HMMptac_3_eut2(pf06130) EUT2B EP_NTERM
09620 HMM01923cobtr_3_eut2(pf01923) EUT2B
09625 H_fuchsia EUT2E
09630 H_fuchsia EUT2B
09635 HMMalddh_2_egra(pf00171) EUT2B
09640 Tsp_amethyst EUT2B
09645 HMMetlyEutC_3_eut2(pf05985) EUT2B EP_NTERM
09650 HMMetlyEutB_1_EUT1(pf06751) EUT2B
09655 HMM06277EutA_1_eut(pf06277) EUT2B
09660 HMMreg07568_1_eut2(pf12282pf13581pf07568) EUT2B
09665 HMMreg00072_1_eut2(pf03861pf00072) EUT2B
09670 -pf06808 -
09675 -pf04290 -
09680 -pf03480 -
09685 HMM6PGdehy_1_mic2(pf01210pf02317) MIC2
09690 HMMreg13556_2_EUT2(pf05651pf
09695 - -
09700 -pf01183 -
BMC-P
BMC-H
BMC-T
Aldehyde dehydrogenase
PTAC
Signature enzyme
Alcohol dehydrogenase
Regulator

Clostridium asparagiforme. Bacteria; Firmicutes; Clostridia; Clostridiales; Lachnospiraceae.

DWV29__locus_1

EUT2B

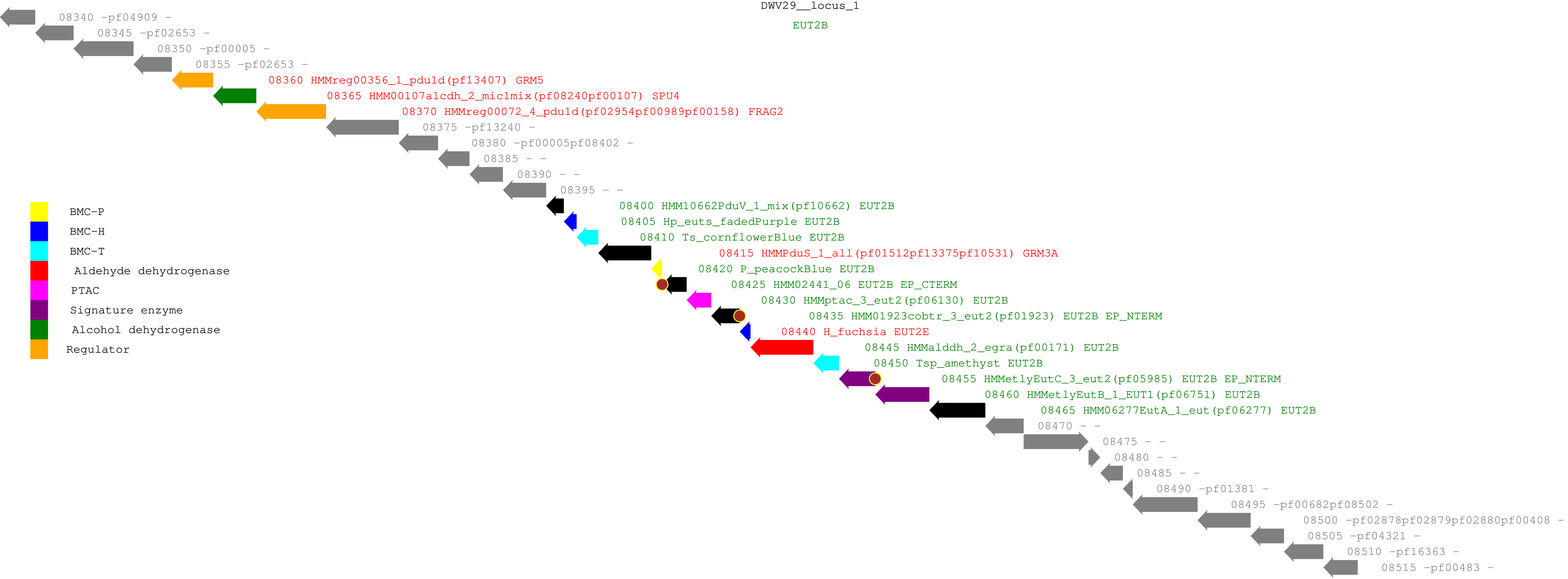

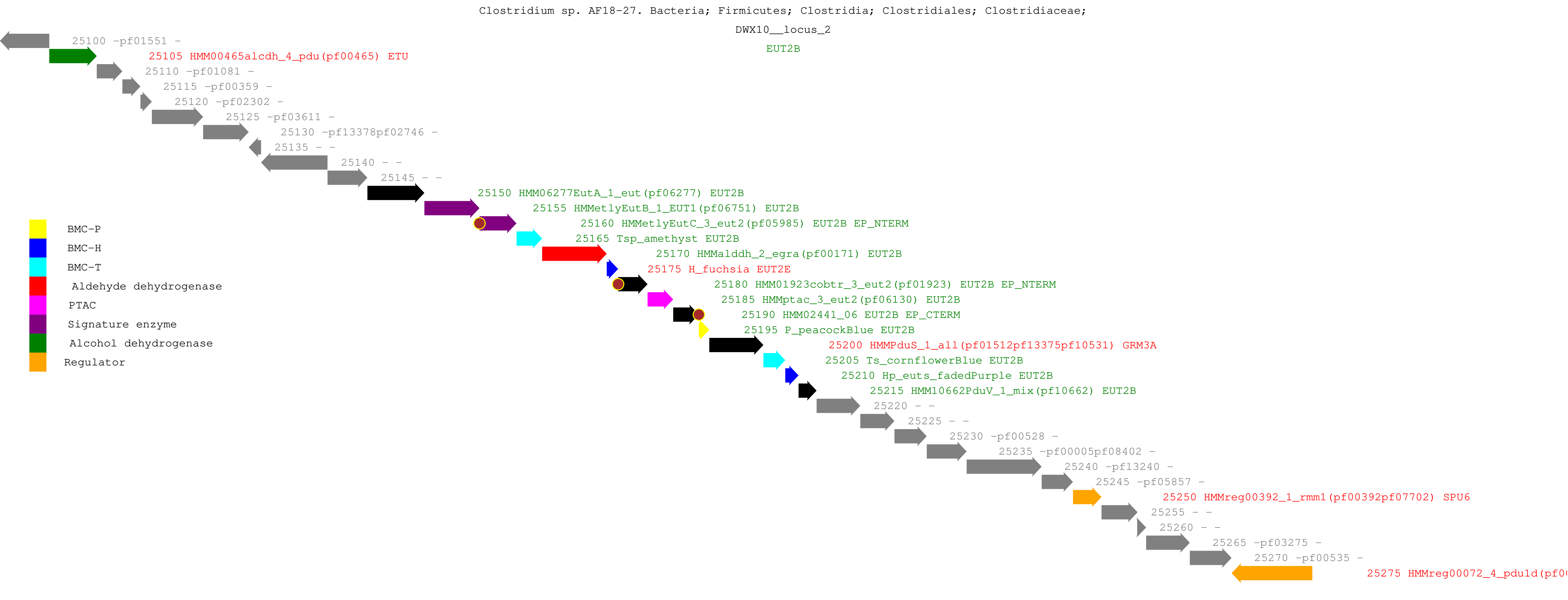

Clostridium sp. AF18-27. Bacteria; Firmicutes; Clostridia; Clostridiales; Clostridiaceae;
DWX10__locus_2
EUT2B
25100 -pf01551 -
25105 HMM00465alcdh_4_pdu(pf00465) ETU
25110 -pf01081 -
25115 -pf00359 -
25120 -pf02302 -
25125 -pf03611 -
25130 -pf13378pf02746 -
25135 - -
25140 - -
25145 - -
25150 HMM06277EutA_1_eut(pf06277) EUT2B
25155 HMMetlyEutB_1_EUT1(pf06751) EUT2B
25160 HMMetlyEutC_3_eut2(pf05985) EUT2B EP_NTERM
25165 Tsp_amethyst EUT2B
25170 HMMaldh_2_egra(pf00171) EUT2B
25175 H_fuchsia EUT2E
25180 HMM01923cobtr_3_eut2(pf01923) EUT2B EP_NTERM
25185 HMMptac_3_eut2(pf06130) EUT2B
25190 HMM02441_06 EUT2B EP_CTERM
25195 P_peacockBlue EUT2B
25200 HMMPduS_1_all(pf01512pf13375pf10531) GRM3A
25205 Ts_cornflowerBlue EUT2B
25210 Hp_euts_fadedPurple EUT2B
25215 HMM10662PduV_1_mix(pf10662) EUT2B
25220 - -
25225 - -
25230 -pf00528 -
25235 -pf00005pf08402 -
25240 -pf13240 -
25245 -pf05857 -
25250 HMMreg00392_1_rmm1(pf00392pf07702) SPU6
25255 - -
25260 - -
25265 -pf03275 -
25270 -pf00535 -
25275 HMMreg00072_4_pdu1d(pf0
BMC-P
BMC-H
BMC-T
Aldehyde dehydrogenase
PTAC
Signature enzyme
Alcohol dehydrogenase
Regulator

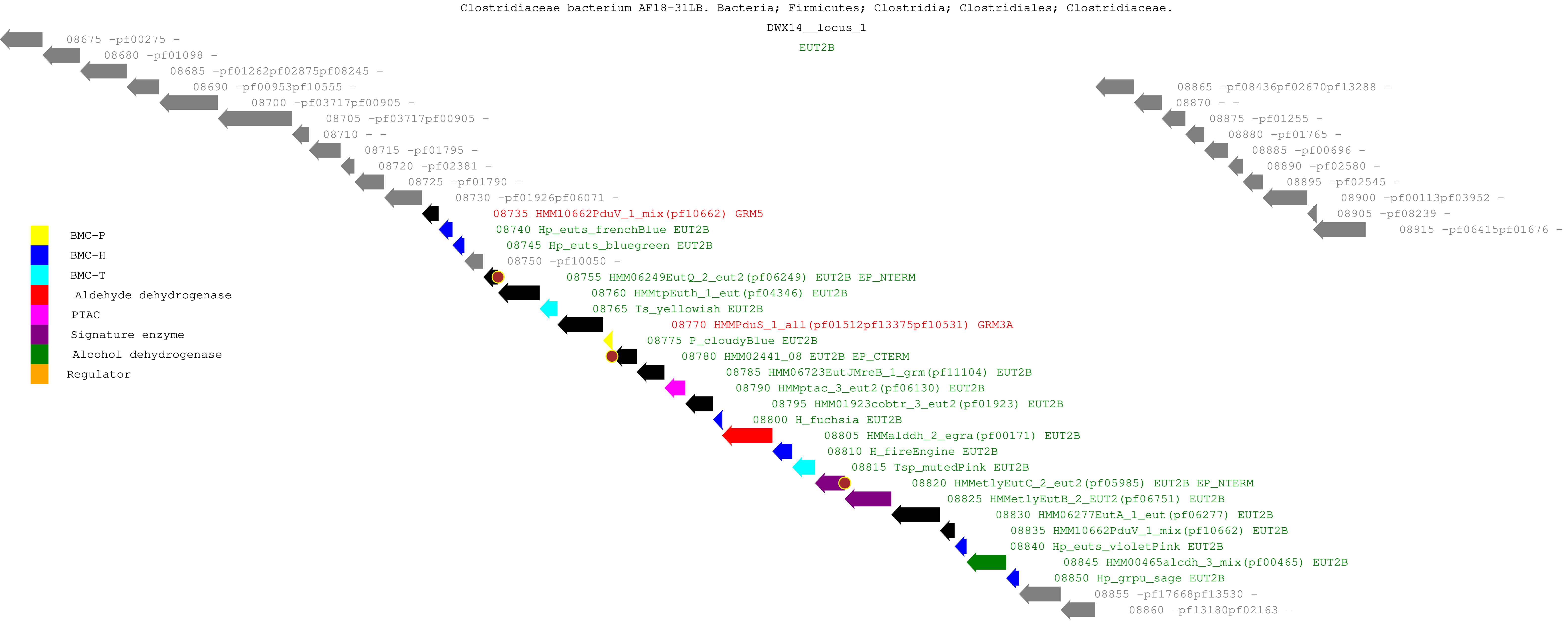
Clostridiaceae bacterium AF18-31LB. Bacteria; Firmicutes; Clostridia; Clostridiales; Clostridiaceae.
DWX14__locus_1
EUT2B
08675 -pf00275 -
08680 -pf01098 -
08685 -pf01262pf02875pf08245 -
08690 -pf00953pf10555 -
08700 -pf03717pf00905 -
08705 -pf03717pf00905 -
08710 - -
08715 -pf01795 -
08720 -pf02381 -
08725 -pf01790 -
08730 -pf01926pf06071 -
08735 HMM10662PduV_1_mix(pf10662) GRM5
08740 Hp_euts_frenchBlue EUT2B
08745 Hp_euts_bluegreen EUT2B
08750 -pf10050 -
08755 HMM06249EutQ_2_eut2(pf06249) EUT2B EP_NTERM
08760 HMMtpEuth_1_eut(pf04346) EUT2B
08765 Ts_yellowish EUT2B
08770 HMMPduS_1_all(pf01512pf13375pf10531) GRM3A
08775 P_cloudyBlue EUT2B
08780 HMM02441_08 EUT2B EP_CTERM
08785 HMM06723EutJMreB_1_grm(pf11104) EUT2B
08790 HMMptac_3_eut2(pf06130) EUT2B
08795 HMM01923cobtr_3_eut2(pf01923) EUT2B
08800 H_fuchsia EUT2B
08805 HMMalddh_2_egra(pf00171) EUT2B
08810 H_fireEngine EUT2B
08815 Tsp_mutedPink EUT2B
08820 HMMetlyEutC_2_eut2(pf05985) EUT2B EP_NTERM
08825 HMMetlyEutB_2_EUT2(pf06751) EUT2B
08830 HMM06277EutA_1_eut(pf06277) EUT2B
08835 HMM10662PduV_1_mix(pf10662) EUT2B
08840 Hp_euts_violetPink EUT2B
08845 HMM00465alcdh_3_mix(pf00465) EUT2B
08850 Hp_grpu_sage EUT2B
08855 -pf17668pf13530 -
08860 -pf13180pf02163 -
08865 -pf08436pf02670pf13288 -
08870 - -
08875 -pf01255 -
08880 -pf01765 -
08885 -pf00696 -
08890 -pf02580 -
08895 -pf02545 -
08900 -pf00113pf03952 -
08905 -pf08239 -
08915 -pf06415pf01676 -
BMC-P
BMC-H
BMC-T
Aldehyde dehydrogenase
PTAC
Signature enzyme
Alcohol dehydrogenase
Regulator

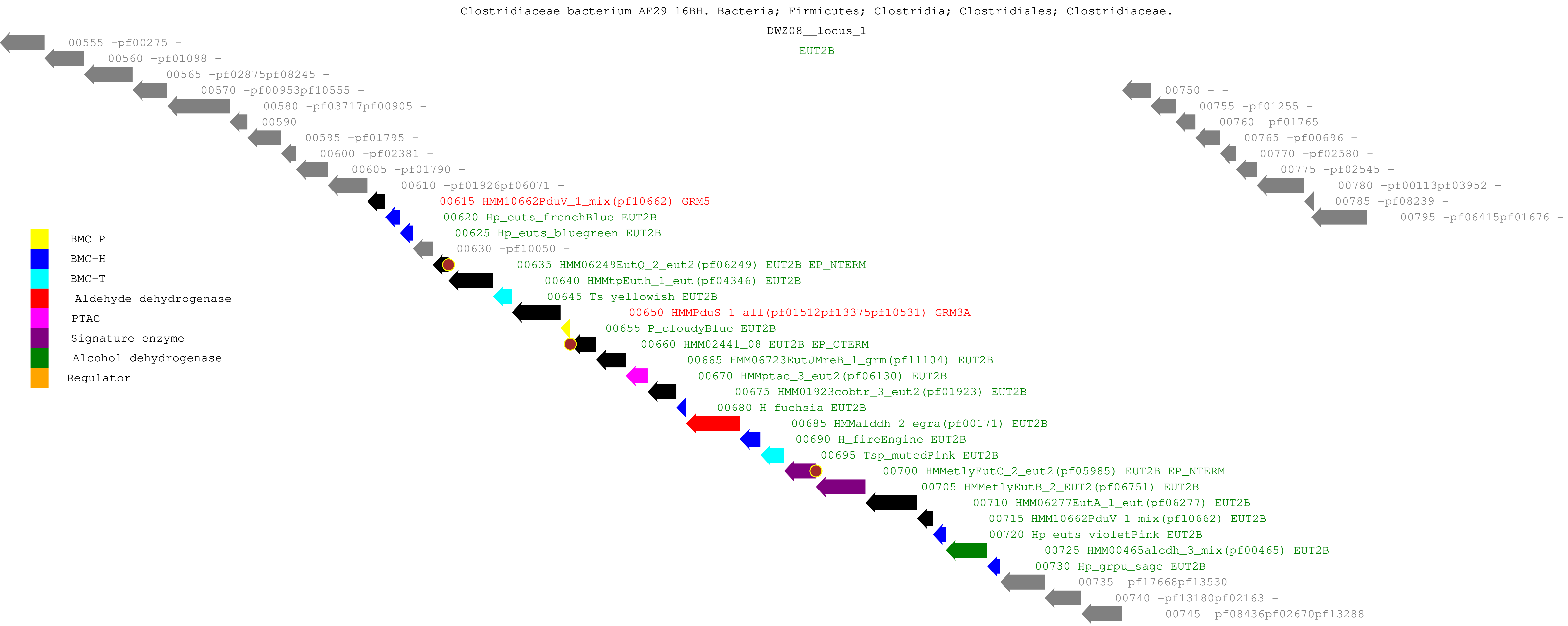
Clostridiaceae bacterium AF29-16BH. Bacteria; Firmicutes; Clostridia; Clostridiales; Clostridiaceae.
DWZ08__locus_1
EUT2B
00555 -pf00275 -
00560 -pf01098 -
00565 -pf02875pf08245 -
00570 -pf00953pf10555 -
00580 -pf03717pf00905 -
00590 - -
00595 -pf01795 -
00600 -pf02381 -
00605 -pf01790 -
00610 -pf01926pf06071 -
00615 HMM10662PduV_1_mix(pf10662) GRM5
00620 Hp_euts_frenchBlue EUT2B
00625 Hp_euts_bluegreen EUT2B
00630 -pf10050 -
00635 HMM06249EutQ_2_eut2(pf06249) EUT2B EP_NTERM
00640 HMMtpEuth_1_eut(pf04346) EUT2B
00645 Ts_yellowish EUT2B
00650 HMMPduS_1_all(pf01512pf13375pf10531) GRM3A
00655 P_cloudyBlue EUT2B
00660 HMM02441_08 EUT2B EP_CTERM
00665 HMM06723EutJMreB_1_grm(pf11104) EUT2B
00670 HMMptac_3_eut2(pf06130) EUT2B
00675 HMM01923cobtr_3_eut2(pf01923) EUT2B
00680 H_fuchsia EUT2B
00685 HMMalddh_2_egra(pf00171) EUT2B
00690 H_fireEngine EUT2B
00695 Tsp_mutedPink EUT2B
00700 HMMetlyEutC_2_eut2(pf05985) EUT2B EP_NTERM
00705 HMMetlyEutB_2_EUT2(pf06751) EUT2B
00710 HMM06277EutA_1_eut(pf06277) EUT2B
00715 HMM10662PduV_1_mix(pf10662) EUT2B
00720 Hp_euts_violetPink EUT2B
00725 HMM00465alcdh_3_mix(pf00465) EUT2B
00730 Hp_grpu_sage EUT2B
00735 -pf17668pf13530 -
00740 -pf13180pf02163 -
00745 -pf08436pf02670pf13288 -
00750 - -
00755 -pf01255 -
00760 -pf01765 -
00765 -pf00696 -
00770 -pf02580 -
00775 -pf02545 -
00780 -pf00113pf03952 -
00785 -pf08239 -
00795 -pf06415pf01676 -
BMC-P
BMC-H
BMC-T
Aldehyde dehydrogenase
PTAC
Signature enzyme
Alcohol dehydrogenase
Regulator

Clostridiaceae bacterium AF31-3BH. Bacteria; Firmicutes; Clostridia; Clostridiales; Clostridiaceae.

DWZ37__locus_1

EUT2B

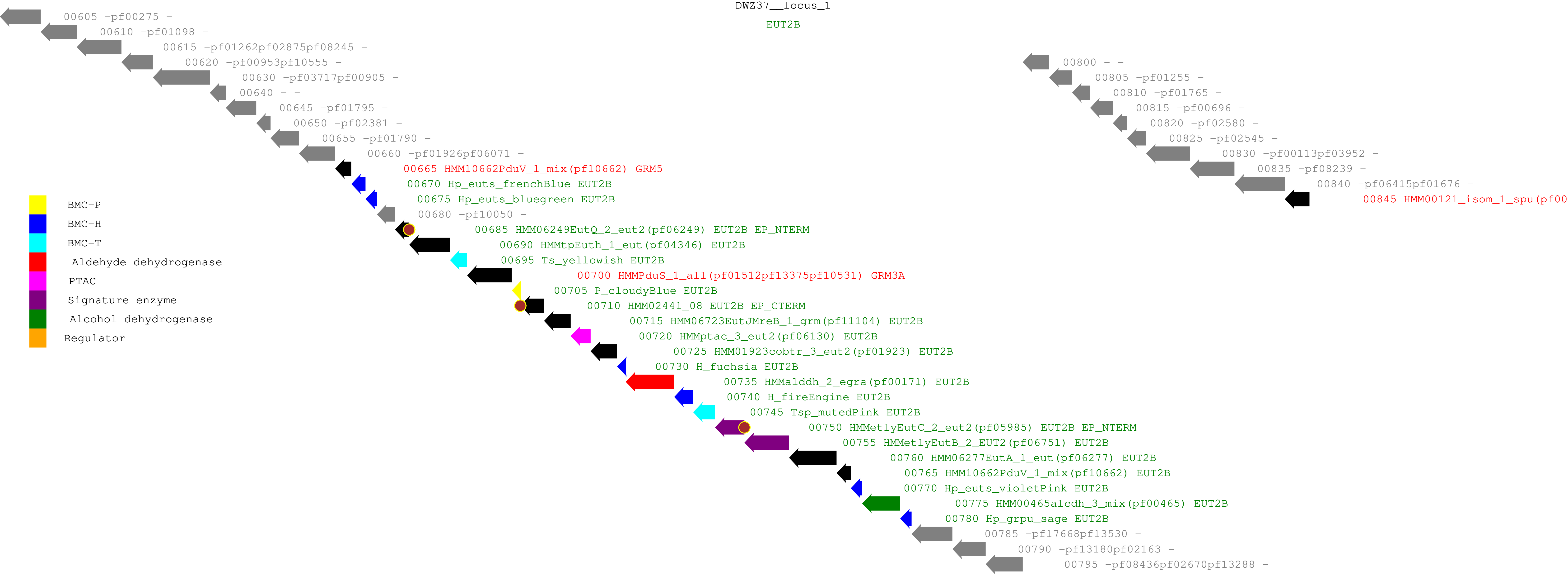

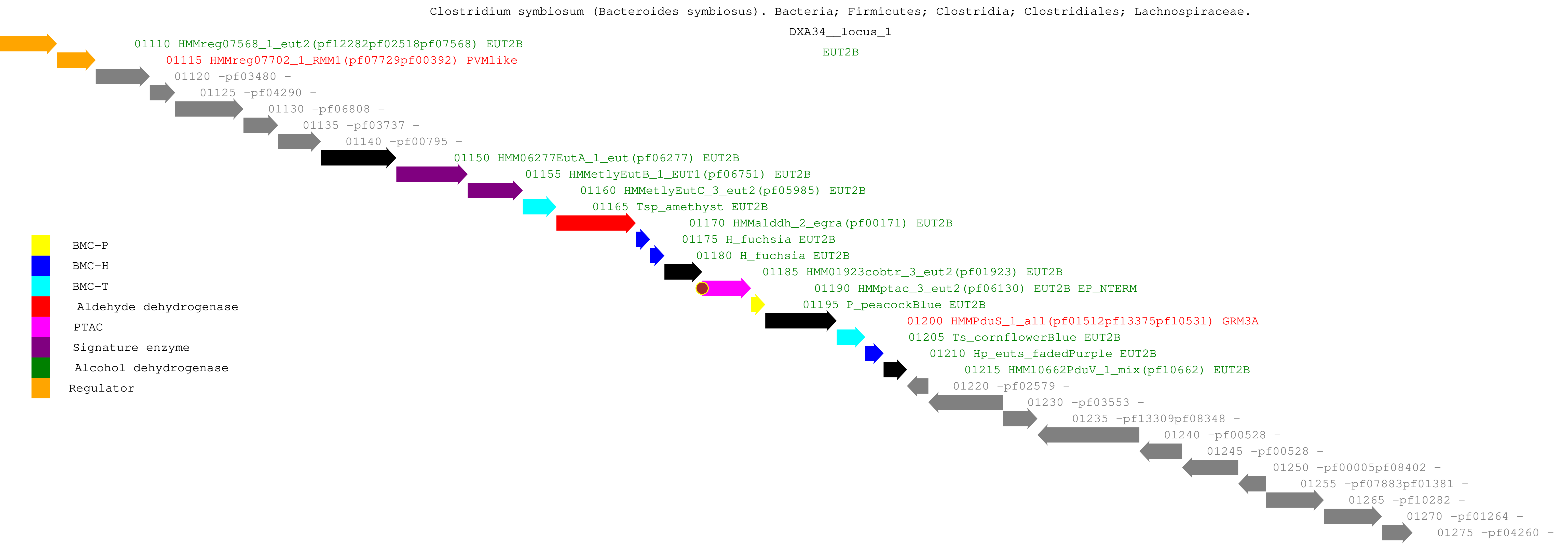
Clostridium symbiosum (Bacteroides symbiosus). Bacteria; Firmicutes; Clostridia; Clostridiales; Lachnospiraceae.
DXA34__locus_1
EUT2B
01110 HMMreg07568_1_eut2(pf12282pf02518pf07568) EUT2B
01115 HMMreg07702_1_RMM1(pf07729pf00392) PVMlike
01120 -pf03480 -
01125 -pf04290 -
01130 -pf06808 -
01135 -pf03737 -
01140 -pf00795 -
01150 HMM06277EutA_1_eut(pf06277) EUT2B
01155 HMMetlyEutB_1_EUT1(pf06751) EUT2B
01160 HMMetlyEutC_3_eut2(pf05985) EUT2B
01165 Tsp_amethyst EUT2B
01170 HMMaldd h_2_egra(pf00171) EUT2B
01175 H_fuchsia EUT2B
01180 H_fuchsia EUT2B
01185 HMM01923cobtr_3_eut2(pf01923) EUT2B
01190 HMMptac_3_eut2(pf06130) EUT2B EP_NTERM
01195 P_peacockBlue EUT2B
01200 HMMPduS_1_all(pf01512pf13375pf10531) GRM3A
01205 Ts_cornflowerBlue EUT2B
01210 Hp_euts_fadedPurple EUT2B
01215 HMM10662PduV_1_mix(pf10662) EUT2B
01220 -pf02579 -
01230 -pf03553 -
01235 -pf13309pf08348 -
01240 -pf00528 -
01245 -pf00528 -
01250 -pf00005pf08402 -
01255 -pf07883pf01381 -
01265 -pf10282 -
01270 -pf01264 -
01275 -pf04260 -
BMC-P
BMC-H
BMC-T
Aldehyde dehydrogenase
PTAC
Signature enzyme
Alcohol dehydrogenase
Regulator

Clostridium sp. OF03-18AA. Bacteria; Firmicutes; Clostridia; Clostridiales; Clostridiaceae;

DXA59__locus_1

EUT2B

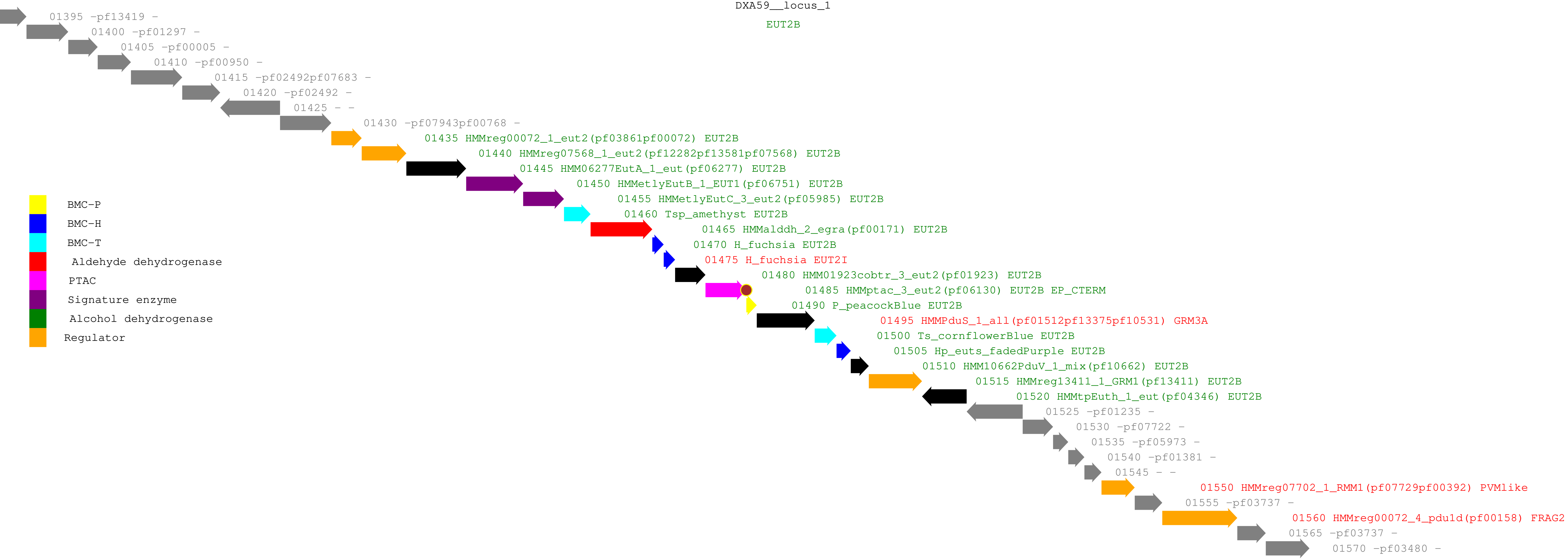

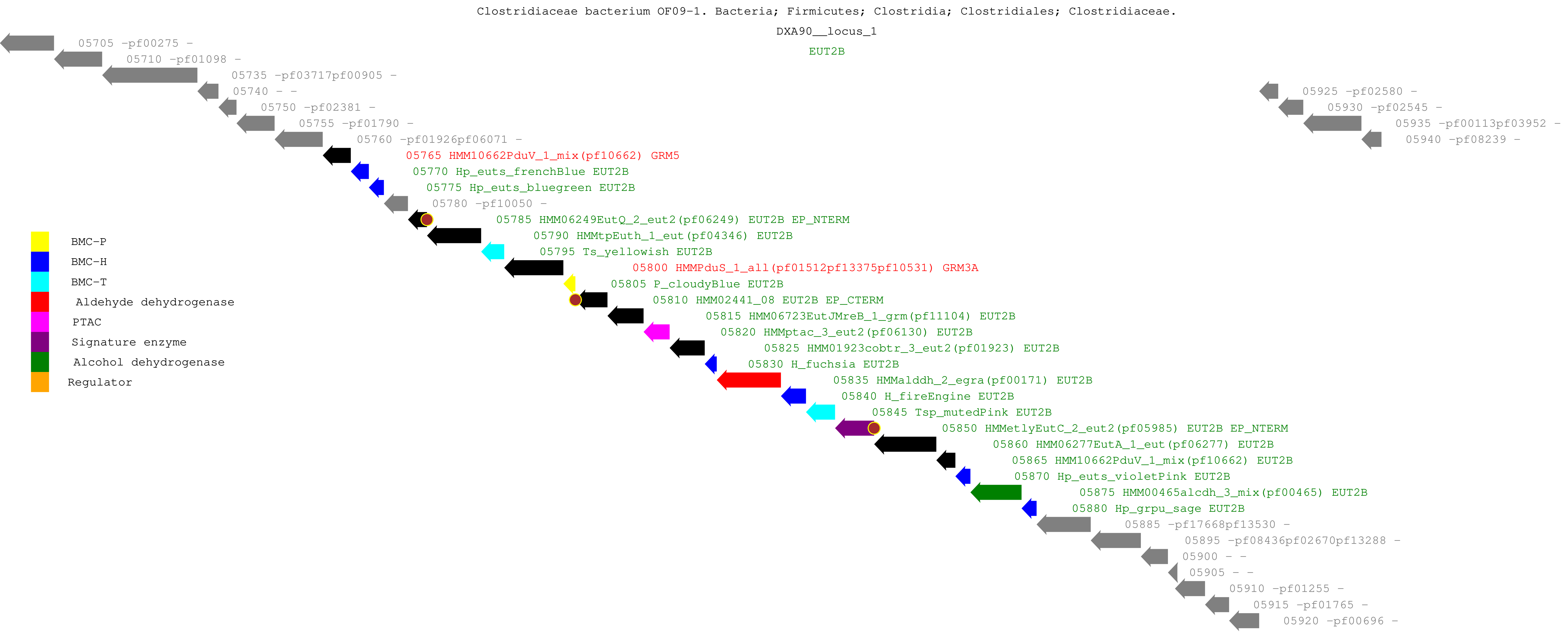
Clostridiaceae bacterium OF09-1. Bacteria; Firmicutes; Clostridia; Clostridiales; Clostridiaceae.
DXA90__locus_1
EUT2B
05705 -pf00275 -
05710 -pf01098 -
05735 -pf03717pf00905 -
05740 - -
05750 -pf02381 -
05755 -pf01790 -
05760 -pf01926pf06071 -
05765 HMM10662PduV_1_mix(pf10662) GRM5
05770 Hp_euts_frenchBlue EUT2B
05775 Hp_euts_bluegreen EUT2B
05780 -pf10050 -
05785 HMM06249EutQ_2_eut2(pf06249) EUT2B EP_NTERM
05790 HMMtpEuth_1_eut(pf04346) EUT2B
05795 Ts_yellowish EUT2B
05800 HMMPduS_1_all(pf01512pf13375pf10531) GRM3A
05805 P_cloudyBlue EUT2B
05810 HMM02441_08 EUT2B EP_CTERM
05815 HMM06723EutJMreB_1_grm(pf11104) EUT2B
05820 HMMptac_3_eut2(pf06130) EUT2B
05825 HMM01923cobtr_3_eut2(pf01923) EUT2B
05830 H_fuchsia EUT2B
05835 HMMalddh_2_egra(pf00171) EUT2B
05840 H_fireEngine EUT2B
05845 Tsp_mutedPink EUT2B
05850 HMMetlyEutC_2_eut2(pf05985) EUT2B EP_NTERM
05860 HMM06277EutA_1_eut(pf06277) EUT2B
05865 HMM10662PduV_1_mix(pf10662) EUT2B
05870 Hp_euts_violetPink EUT2B
05875 HMM00465alcdh_3_mix(pf00465) EUT2B
05880 Hp_grpu_sage EUT2B
05885 -pf17668pf13530 -
05895 -pf08436pf02670pf13288 -
05900 - -
05905 - -
05910 -pf01255 -
05915 -pf01765 -
05920 -pf00696 -
05925 -pf02580 -
05930 -pf02545 -
05935 -pf00113pf03952 -
05940 -pf08239 -
BMC-P
BMC-H
BMC-T
Aldehyde dehydrogenase
PTAC
Signature enzyme
Alcohol dehydrogenase
Regulator

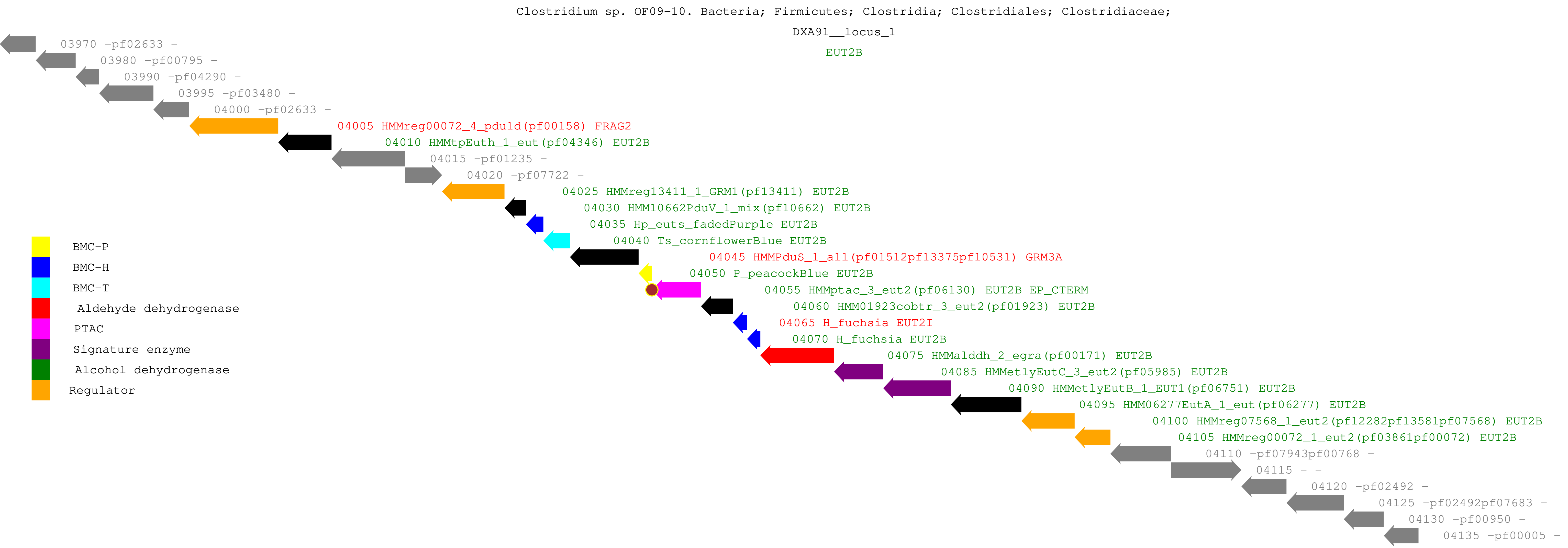
Clostridium sp. OF09-10. Bacteria; Firmicutes; Clostridia; Clostridiales; Clostridiaceae;
DXA91__locus_1
EUT2B
03970 -pf02633 -
03980 -pf00795 -
03990 -pf04290 -
03995 -pf03480 -
04000 -pf02633 -
04005 HMMreg00072_4_pdu1d(pf00158) FRAG2
04010 HMMtpEuth_1_eut(pf04346) EUT2B
04015 -pf01235 -
04020 -pf07722 -
04025 HMMreg13411_1_GRM1(pf13411) EUT2B
04030 HMM10662PduV_1_mix(pf10662) EUT2B
04035 Hp_euts_fadedPurple EUT2B
04040 Ts_cornflowerBlue EUT2B
04045 HMMPduS_1_all(pf01512pf13375pf10531) GRM3A
04050 P_peacockBlue EUT2B
04055 HMMptac_3_eut2(pf06130) EUT2B EP_CTERM
04060 HMM01923cobtr_3_eut2(pf01923) EUT2B
04065 H_fuchsia EUT2I
04070 H_fuchsia EUT2B
04075 HMMalddh_2_egra(pf00171) EUT2B
04085 HMMetlyEutC_3_eut2(pf05985) EUT2B
04090 HMMetlyEutB_1_EUT1(pf06751) EUT2B
04095 HMM06277EutA_1_eut(pf06277) EUT2B
04100 HMMreg07568_1_eut2(pf12282pf13581pf07568) EUT2B
04105 HMMreg00072_1_eut2(pf03861pf00072) EUT2B
04110 -pf07943pf00768 -
04115 - -
04120 -pf02492 -
04125 -pf02492pf07683 -
04130 -pf00950 -
04135 -pf00005 -
BMC-P
BMC-H
BMC-T
Aldehyde dehydrogenase
PTAC
Signature enzyme
Alcohol dehydrogenase
Regulator

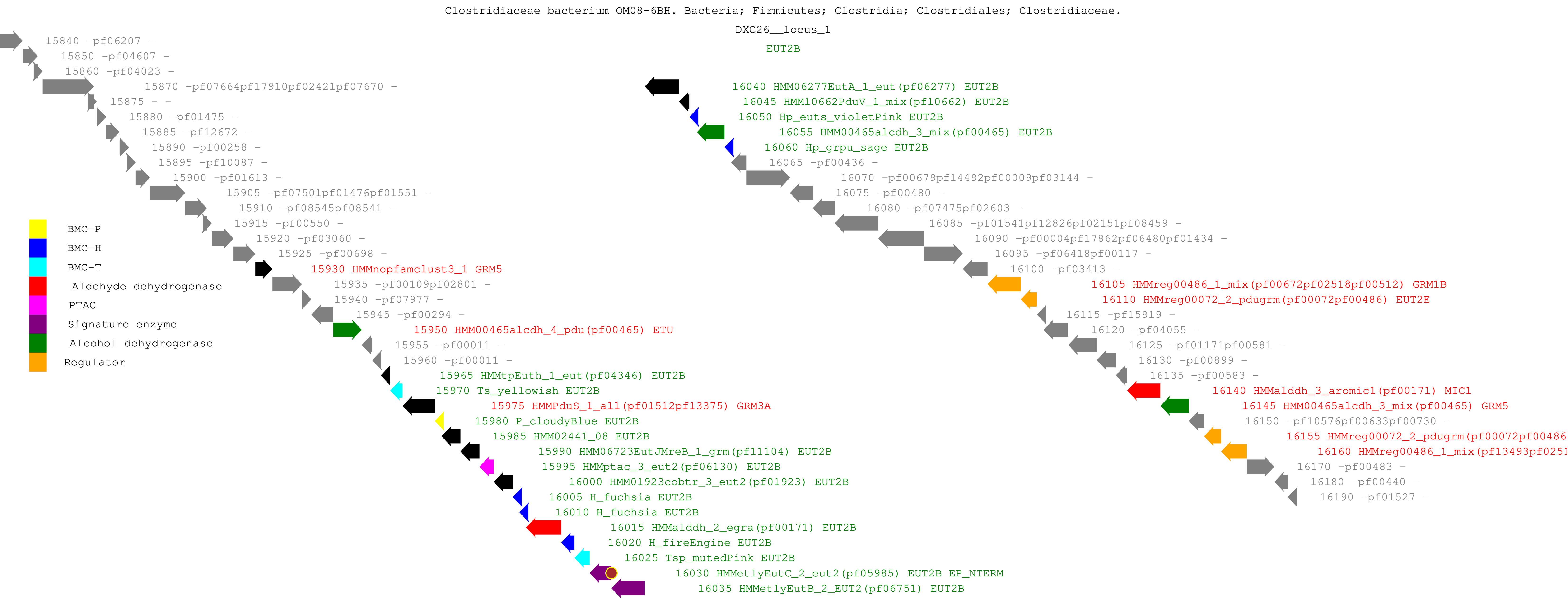

Clostridiaceae bacterium OM08-6BH. Bacteria; Firmicutes; Clostridia; Clostridiales; Clostridiaceae.
DXC26__locus_1
EUT2B
15840 -pf06207 -
15850 -pf04607 -
15860 -pf04023 -
15870 -pf07664pf17910pf02421pf07670 -
15875 - -
15880 -pf01475 -
15885 -pf12672 -
15890 -pf00258 -
15895 -pf10087 -
15900 -pf01613 -
15905 -pf07501pf01476pf01551 -
15910 -pf08545pf08541 -
15915 -pf00550 -
15920 -pf03060 -
15925 -pf00698 -
15930 HMMnopfamclust3_1 GRM5
15935 -pf00109pf02801 -
15940 -pf07977 -
15945 -pf00294 -
15950 HMM00465alcdh_4_pdu(pf00465) ETU
15955 -pf00011 -
15960 -pf00011 -
15965 HMMtpEuth_1_eut(pf04346) EUT2B
15970 Ts_yellowish EUT2B
15975 HMMPduS_1_all(pf01512pf13375) GRM3A
15980 P_cloudyBlue EUT2B
15985 HMM02441_08 EUT2B
15990 HMM06723EutJMreB_1_grm(pf11104) EUT2B
15995 HMMptac_3_eut2(pf06130) EUT2B
16000 HMM01923cobtr_3_eut2(pf01923) EUT2B
16005 H_fuchsia EUT2B
16010 H_fuchsia EUT2B
16015 HMMalddh_2_egra(pf00171) EUT2B
16020 H_fireEngine EUT2B
16025 Tsp_mutedPink EUT2B
16030 HMMetlyEutC_2_eut2(pf05985) EUT2B EP_NTERM
16035 HMMetlyEutB_2_EUT2(pf06751) EUT2B
16040 HMM06277EutA_1_eut(pf06277) EUT2B
16045 HMM10662PduV_1_mix(pf10662) EUT2B
16050 Hp_euts_violetPink EUT2B
16055 HMM00465alcdh_3_mix(pf00465) EUT2B
16060 Hp_grpu_sage EUT2B
16065 -pf00436 -
16070 -pf00679pf14492pf00009pf03144 -
16075 -pf00480 -
16080 -pf07475pf02603 -
16085 -pf01541pf12826pf02151pf08459 -
16090 -pf00004pf17862pf06480pf01434 -
16095 -pf06418pf00117 -
16100 -pf03413 -
16105 HMMreg00486_1_mix(pf00672pf02518pf00512) GRM1B
16110 HMMreg00072_2_pdugrm(pf00072pf00486) EUT2E
16115 -pf15919 -
16120 -pf04055 -
16125 -pf01171pf00581 -
16130 -pf00899 -
16135 -pf00583 -
16140 HMMalddh_3_aromic1(pf00171) MIC1
16145 HMM00465alcdh_3_mix(pf00465) GRM5
16150 -pf10576pf00633pf00730 -
16155 HMMreg00072_2_pdugrm(pf00072pf00486
16160 HMMreg00486_1_mix(pf13493pf0251
16170 -pf00483 -
16180 -pf00440 -
16190 -pf01527 -
BMC-P
BMC-H
BMC-T
Aldehyde dehydrogenase
PTAC
Signature enzyme
Alcohol dehydrogenase
Regulator

Clostridiaceae bacterium OM08-6BH. Bacteria; Firmicutes; Clostridia; Clostridiales; Clostridiaceae.

DXC26__locus_2

EUT2B

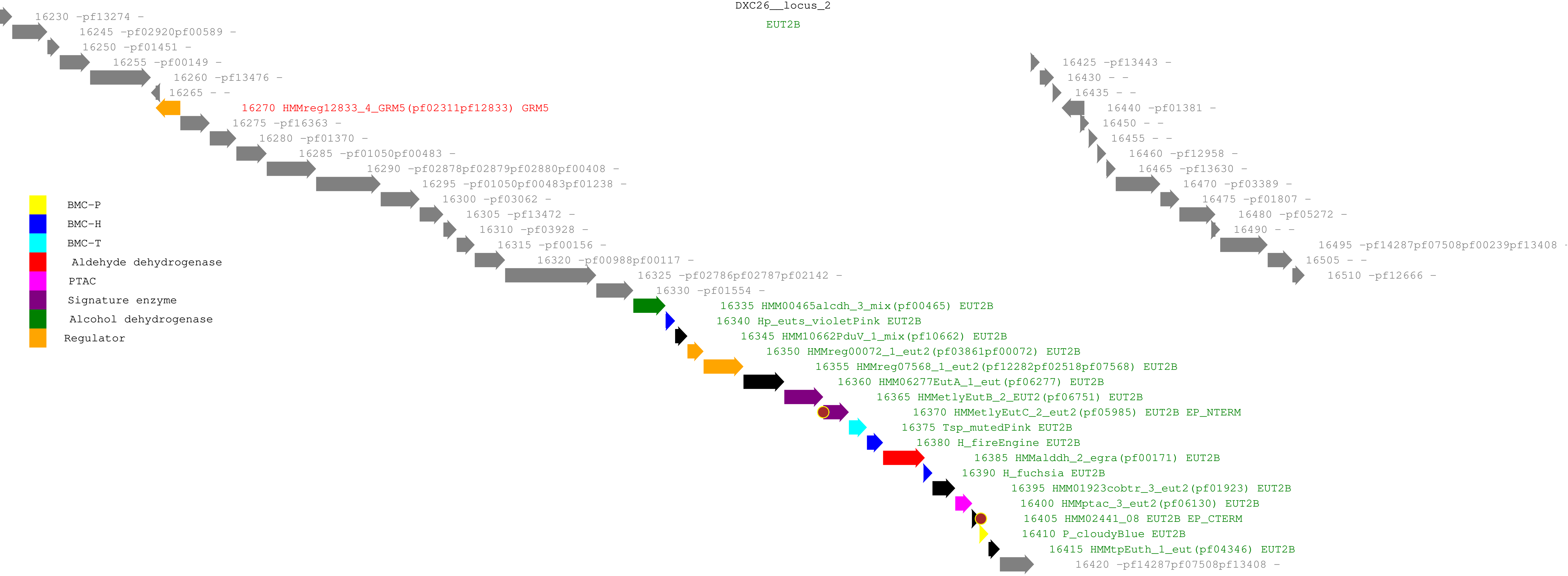

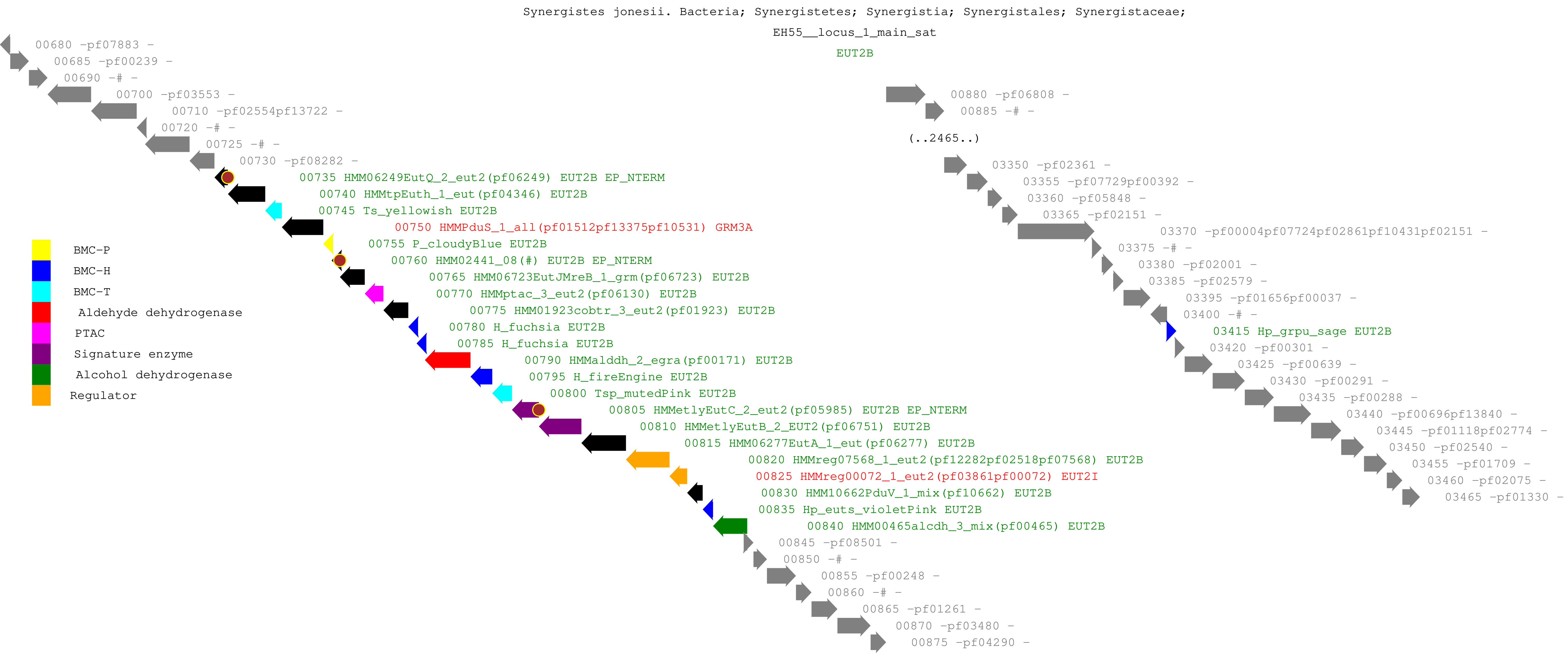

Synergistes jonesii. Bacteria; Synergistetes; Synergistia; Synergistales; Synergistaceae;
EH55__locus_1_main_sat
EUT2B
00680 -pf07883 -
00685 -pf00239 -
00690 -# -
00700 -pf03553 -
00710 -pf02554pf13722 -
00720 -# -
00725 -# -
00730 -pf08282 -
00735 HMM06249EutQ_2_eut2(pf06249) EUT2B EP_NTERM
00740 HMMtpEuth_1_eut(pf04346) EUT2B
00745 Ts_yellowish EUT2B
00750 HMMPduS_1_all(pf01512pf13375pf10531) GRM3A
00755 P_cloudyBlue EUT2B
00760 HMM02441_08(#) EUT2B EP_NTERM
00765 HMM06723EutJMreB_1_grm(pf06723) EUT2B
00770 HMMptac_3_eut2(pf06130) EUT2B
00775 HMM01923cobtr_3_eut2(pf01923) EUT2B
00780 H_fuchsia EUT2B
00785 H_fuchsia EUT2B
00790 HMMaldh_2_egra(pf00171) EUT2B
00795 H_fireEngine EUT2B
00800 Tsp_mutedPink EUT2B
00805 HMMetlyEutC_2_eut2(pf05985) EUT2B EP_NTERM
00810 HMMetlyEutB_2_EUT2(pf06751) EUT2B
00815 HMM06277EutA_1_eut(pf06277) EUT2B
00820 HMMreg07568_1_eut2(pf12282pf02518pf07568) EUT2B
00825 HMMreg00072_1_eut2(pf03861pf00072) EUT2I
00830 HMM10662PduV_1_mix(pf10662) EUT2B
00835 Hp_euts_violetPink EUT2B
00840 HMM00465alcdh_3_mix(pf00465) EUT2B
00845 -pf08501 -
00850 -# -
00855 -pf00248 -
00860 -# -
00865 -pf01261 -
00870 -pf03480 -
00875 -pf04290 -
00880 -pf06808 -
00885 -# -
(..2465..)
03350 -pf02361 -
03355 -pf07729pf00392 -
03360 -pf05848 -
03365 -pf02151 -
03370 -pf00004pf07724pf02861pf10431pf02151 -
03375 -# -
03380 -pf02001 -
03385 -pf02579 -
03395 -pf01656pf00037 -
03400 -# -
03415 Hp_grpu_sage EUT2B
03420 -pf00301 -
03425 -pf00639 -
03430 -pf00291 -
03435 -pf00288 -
03440 -pf00696pf13840 -
03445 -pf01118pf02774 -
03450 -pf02540 -
03455 -pf01709 -
03460 -pf02075 -
03465 -pf01330 -
BMC-P
BMC-H
BMC-T
Aldehyde dehydrogenase
PTAC
Signature enzyme
Alcohol dehydrogenase
Regulator

Fretibacterium sp. OH1220_COT-178. Bacteria; Synergistetes; Synergistia; Synergistales; Synergistaceae;

EII26__locus_1

EUT2B

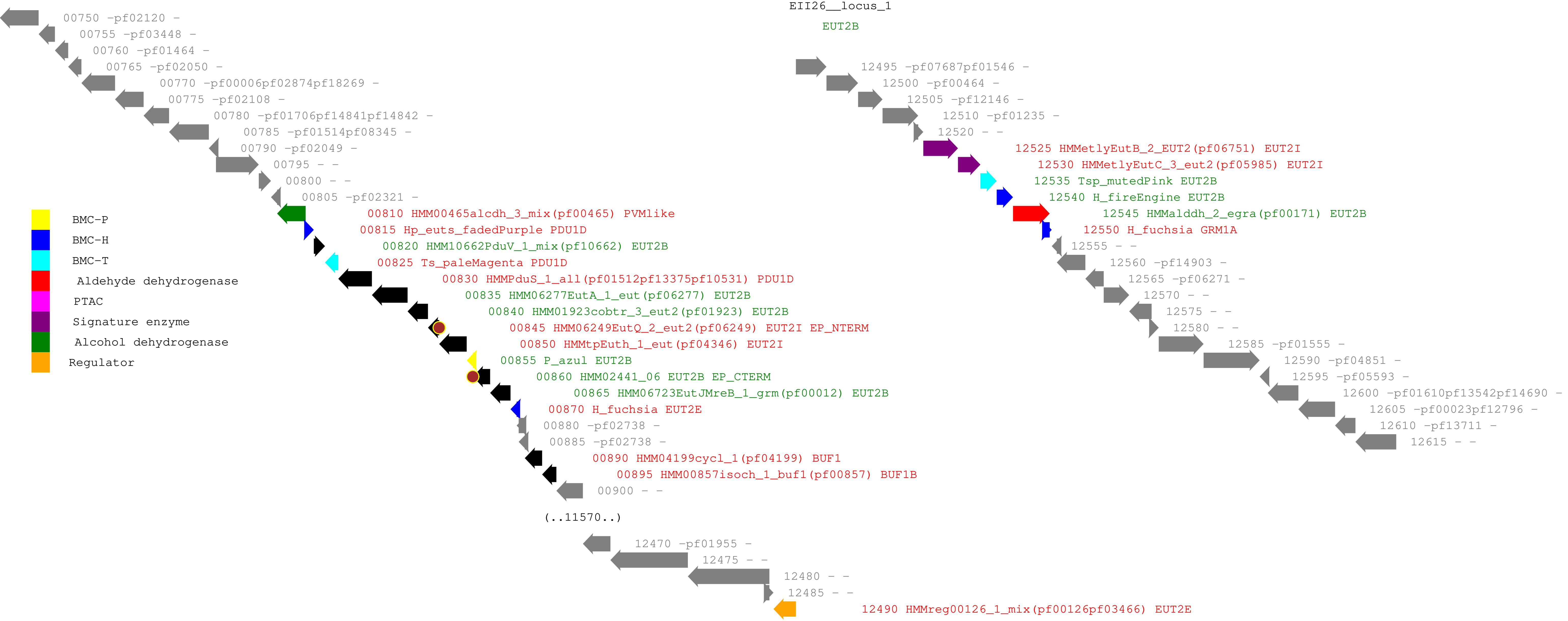

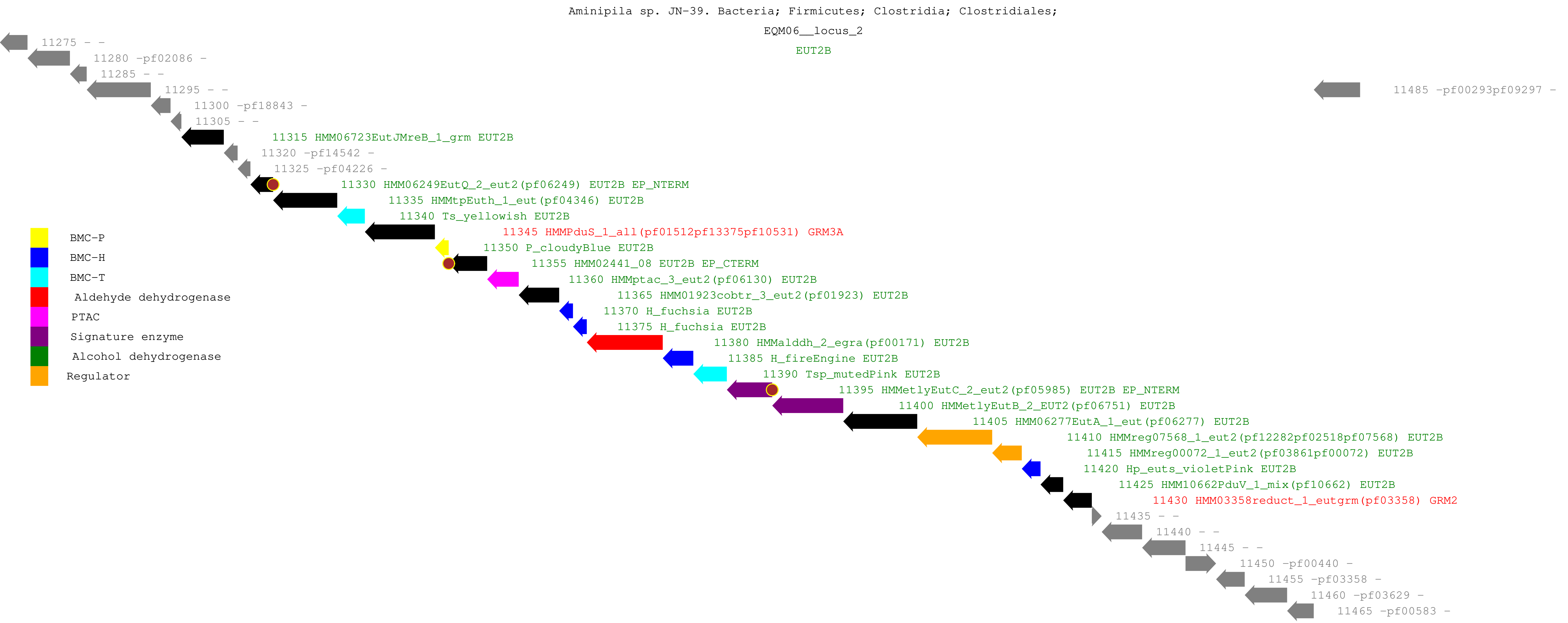
Aminipila sp. JN-39. Bacteria; Firmicutes; Clostridia; Clostridiales;
EQM06__locus_2
EUT2B
11275 - -
11280 -pf02086 -
11285 - -
11295 - -
11300 -pf18843 -
11305 - -
11315 HMM06723EutJMreB_1_grm EUT2B
11320 -pf14542 -
11325 -pf04226 -
11330 HMM06249EutQ_2_eut2(pf06249) EUT2B EP_NTERM
11335 HMMtpEuth_1_eut(pf04346) EUT2B
11340 Ts_yellowish EUT2B
11345 HMMPduS_1_all(pf01512pf13375pf10531) GRM3A
11350 P_cloudyBlue EUT2B
11355 HMM02441_08 EUT2B EP_CTERM
11360 HMMptac_3_eut2(pf06130) EUT2B
11365 HMM01923cobtr_3_eut2(pf01923) EUT2B
11370 H_fuchsia EUT2B
11375 H_fuchsia EUT2B
11380 HMMalddh_2_egra(pf00171) EUT2B
11385 H_fireEngine EUT2B
11390 Tsp_mutedPink EUT2B
11395 HMMetlyEutC_2_eut2(pf05985) EUT2B EP_NTERM
11400 HMMetlyEutB_2_EUT2(pf06751) EUT2B
11405 HMM06277EutA_1_eut(pf06277) EUT2B
11410 HMMreg07568_1_eut2(pf12282pf02518pf07568) EUT2B
11415 HMMreg00072_1_eut2(pf03861pf00072) EUT2B
11420 Hp_euts_violetPink EUT2B
11425 HMM10662PduV_1_mix(pf10662) EUT2B
11430 HMM03358reduct_1_eutgrm(pf03358) GRM2
11435 - -
11440 - -
11445 - -
11450 -pf00440 -
11455 -pf03358 -
11460 -pf03629 -
11465 -pf00583 -
11485 -pf00293pf09297 -
BMC-P
BMC-H
BMC-T
Aldehyde dehydrogenase
PTAC
Signature enzyme
Alcohol dehydrogenase
Regulator

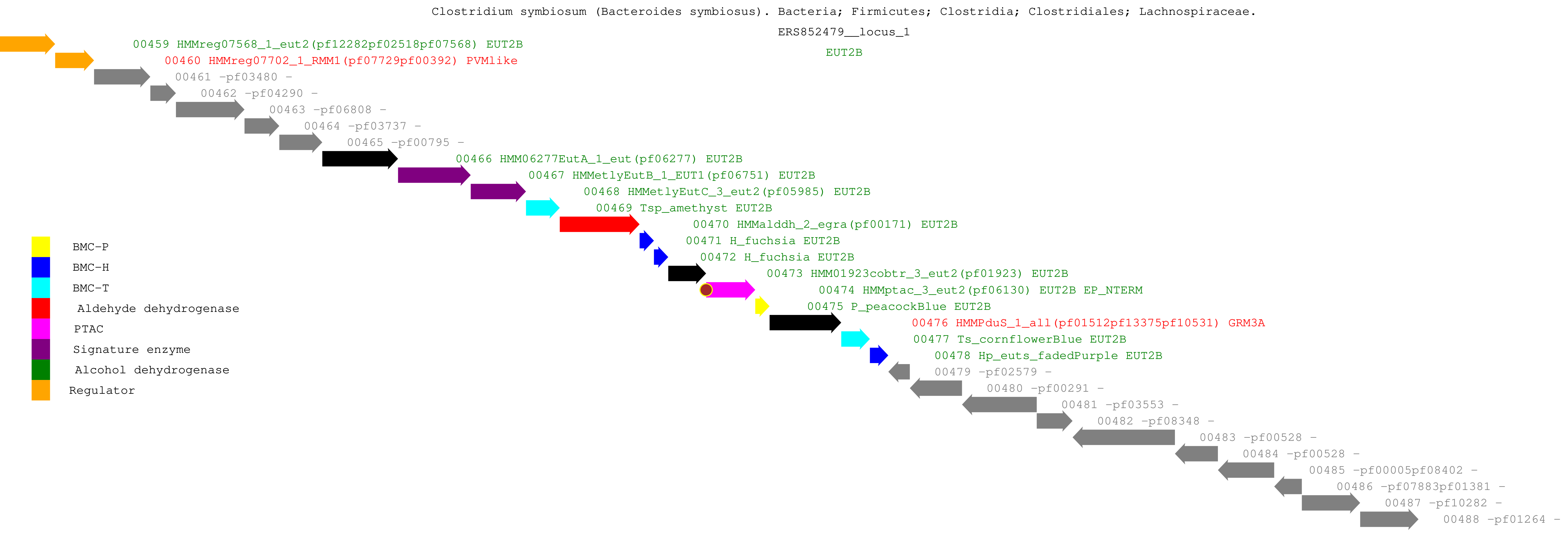

Clostridium symbiosum (Bacteroides symbiosus). Bacteria; Firmicutes; Clostridia; Clostridiales; Lachnospiraceae.
ERS852479__locus_1
EUT2B
00459 HMMreg07568_1_eut2(pf12282pf02518pf07568) EUT2B
00460 HMMreg07702_1_RMM1(pf07729pf00392) PVMlike
00461 -pf03480 -
00462 -pf04290 -
00463 -pf06808 -
00464 -pf03737 -
00465 -pf00795 -
00466 HMM06277EutA_1_eut(pf06277) EUT2B
00467 HMMetlyEutB_1_EUT1(pf06751) EUT2B
00468 HMMetlyEutC_3_eut2(pf05985) EUT2B
00469 Tsp_amethyst EUT2B
00470 HMMaldh_2_egra(pf00171) EUT2B
00471 H_fuchsia EUT2B
00472 H_fuchsia EUT2B
00473 HMM01923cobtr_3_eut2(pf01923) EUT2B
00474 HMMptac_3_eut2(pf06130) EUT2B EP_NTERM
00475 P_peacockBlue EUT2B
00476 HMMPduS_1_all(pf01512pf13375pf10531) GRM3A
00477 Ts_cornflowerBlue EUT2B
00478 Hp_euts_fadedPurple EUT2B
00479 -pf02579 -
00480 -pf00291 -
00481 -pf03553 -
00482 -pf08348 -
00483 -pf00528 -
00484 -pf00528 -
00485 -pf00005pf08402 -
00486 -pf07883pf01381 -
00487 -pf10282 -
00488 -pf01264 -
BMC-P
BMC-H
BMC-T
Aldehyde dehydrogenase
PTAC
Signature enzyme
Alcohol dehydrogenase
Regulator

Flavonifractor plautii. Bacteria; Firmicutes; Clostridia; Clostridiales; Flavonifractor.

ERS852504__locus_2

EUT2B

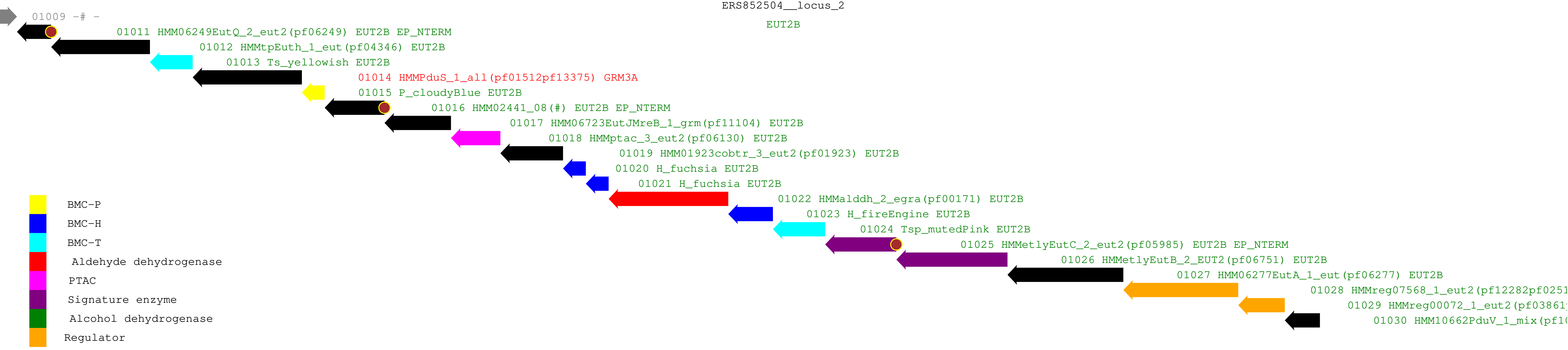

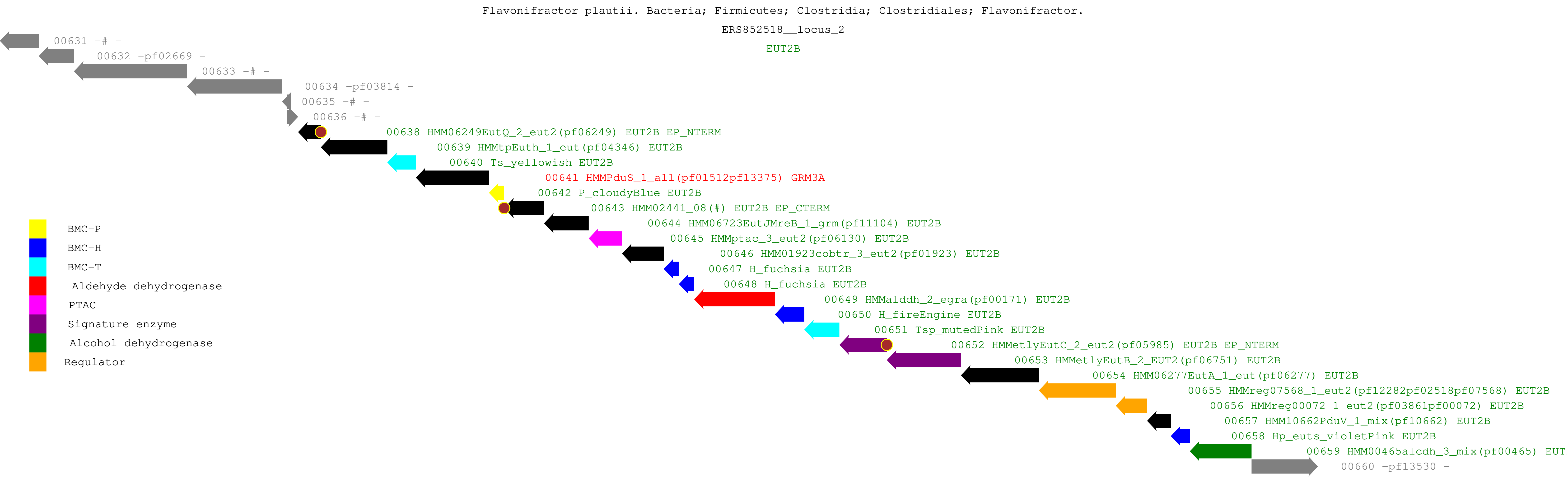
Flavonifractor plautii. Bacteria; Firmicutes; Clostridia; Clostridiales; Flavonifractor.
ERS852518__locus_2
EUT2B
00631 -# -
00632 -pf02669 -
00633 -# -
00634 -pf03814 -
00635 -# -
00636 -# -
00638 HMM06249EutQ_2_eut2(pf06249) EUT2B EP_NTERM
00639 HMMtpEuth_1_eut(pf04346) EUT2B
00640 Ts_yellowish EUT2B
00641 HMMPduS_1_all(pf01512pf13375) GRM3A
00642 P_cloudyBlue EUT2B
00643 HMM02441_08(#) EUT2B EP_CTERM
00644 HMM06723EutJMreB_1_grm(pf11104) EUT2B
00645 HMMptac_3_eut2(pf06130) EUT2B
00646 HMM01923cobtr_3_eut2(pf01923) EUT2B
00647 H_fuchsia EUT2B
00648 H_fuchsia EUT2B
00649 HMMalddh_2_egra(pf00171) EUT2B
00650 H_fireEngine EUT2B
00651 Tsp_mutedPink EUT2B
00652 HMMetlyEutC_2_eut2(pf05985) EUT2B EP_NTERM
00653 HMMetlyEutB_2_EUT2(pf06751) EUT2B
00654 HMM06277EutA_1_eut(pf06277) EUT2B
00655 HMMreg07568_1_eut2(pf12282pf02518pf07568) EUT2B
00656 HMMreg00072_1_eut2(pf03861pf00072) EUT2B
00657 HMM10662PduV_1_mix(pf10662) EUT2B
00658 Hp_euts_violetPink EUT2B
00659 HMM00465alcdh_3_mix(pf00465) EUT
00660 -pf13530 -
BMC-P
BMC-H
BMC-T
Aldehyde dehydrogenase
PTAC
Signature enzyme
Alcohol dehydrogenase
Regulator

Fusicatenibacter. Bacteria; Firmicutes; Clostridia; Clostridiales; Lachnospiraceae.

ERS852537__locus_1

EUT2B

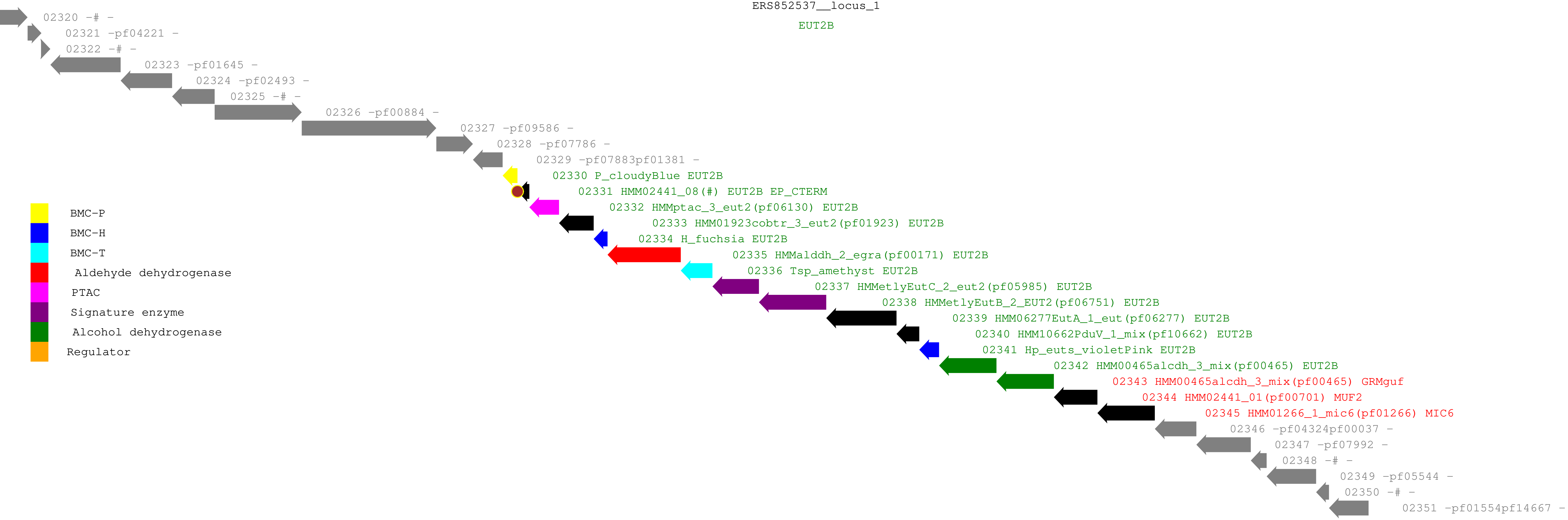

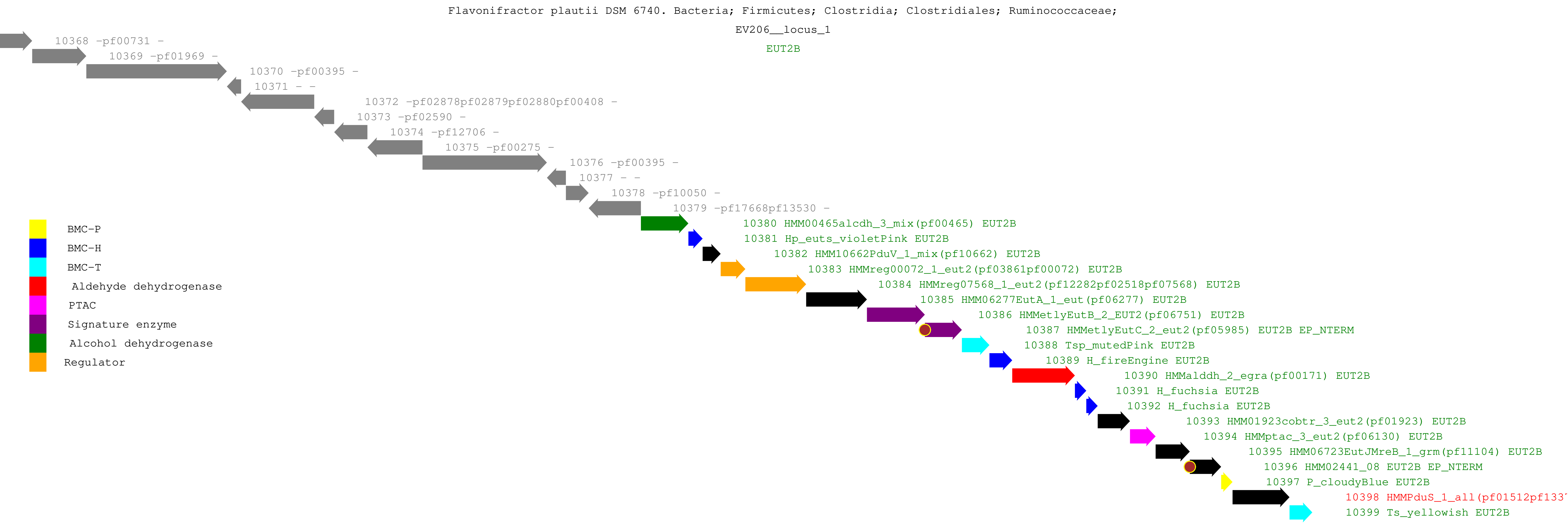

Flavonifractor plautii DSM 6740. Bacteria; Firmicutes; Clostridia; Clostridiales; Ruminococcaceae;
EV206__locus_1
EUT2B
10368 -pf00731 -
10369 -pf01969 -
10370 -pf00395 -
10371 - -
10372 -pf02878pf02879pf02880pf00408 -
10373 -pf02590 -
10374 -pf12706 -
10375 -pf00275 -
10376 -pf00395 -
10377 - -
10378 -pf10050 -
10379 -pf17668pf13530 -
10380 HMM00465alcdh_3_mix(pf00465) EUT2B
10381 Hp_euts_violetPink EUT2B
10382 HMM10662PduV_1_mix(pf10662) EUT2B
10383 HMMreg00072_1_eut2(pf03861pf00072) EUT2B
10384 HMMreg07568_1_eut2(pf12282pf02518pf07568) EUT2B
10385 HMM06277EutA_1_eut(pf06277) EUT2B
10386 HMMetlyEutB_2_EUT2(pf06751) EUT2B
10387 HMMetlyEutC_2_eut2(pf05985) EUT2B EP_NTERM
10388 Tsp_mutedPink EUT2B
10389 H_fireEngine EUT2B
10390 HMMalddh_2_egra(pf00171) EUT2B
10391 H_fuchsia EUT2B
10392 H_fuchsia EUT2B
10393 HMM01923cobtr_3_eut2(pf01923) EUT2B
10394 HMMptac_3_eut2(pf06130) EUT2B
10395 HMM06723EutJMreB_1_grm(pf11104) EUT2B
10396 HMM02441_08 EUT2B EP_NTERM
10397 P_cloudyBlue EUT2B
10398 HMMPduS_1_all(pf01512pf133
10399 Ts_yellowish EUT2B
BMC-P
BMC-H
BMC-T
Aldehyde dehydrogenase
PTAC
Signature enzyme
Alcohol dehydrogenase
Regulator

Frisingicoccus caecimuris. Bacteria; Firmicutes; Clostridia; Clostridiales; Lachnospiraceae;

EV212__locus_1

EUT2B

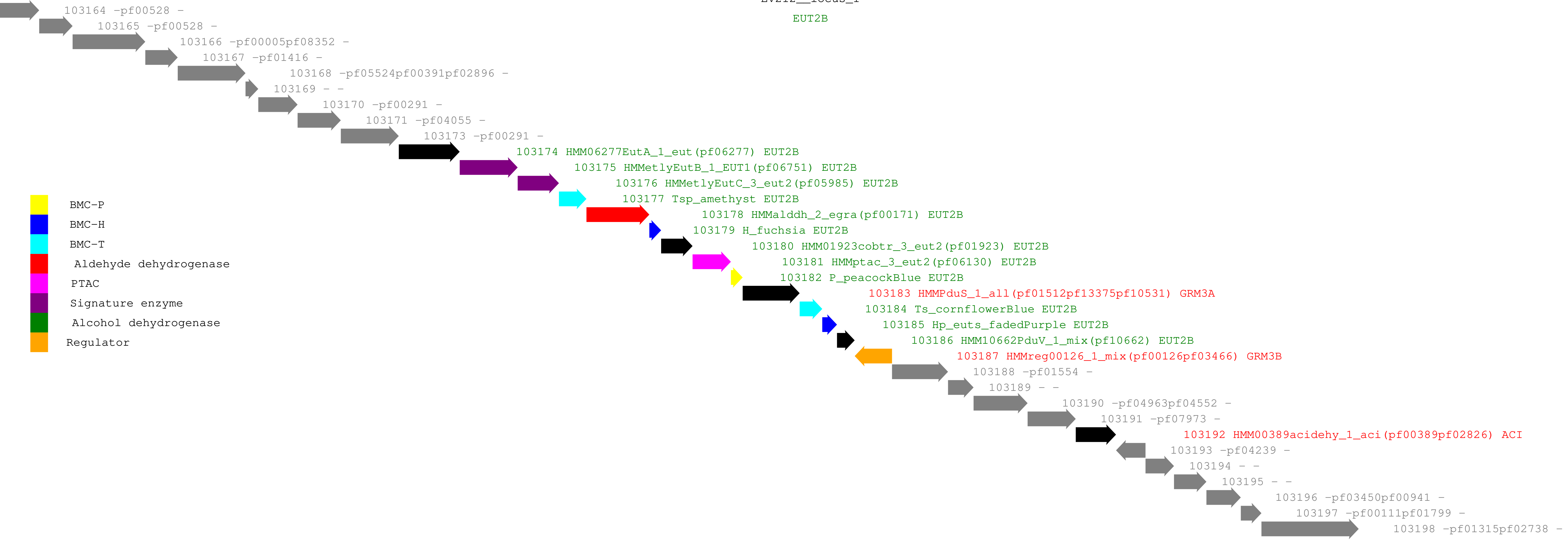

Fusibacter sp. 3D3. Bacteria; Firmicutes; Clostridia; Clostridiales;

F3D3__locus_1

EUT2B

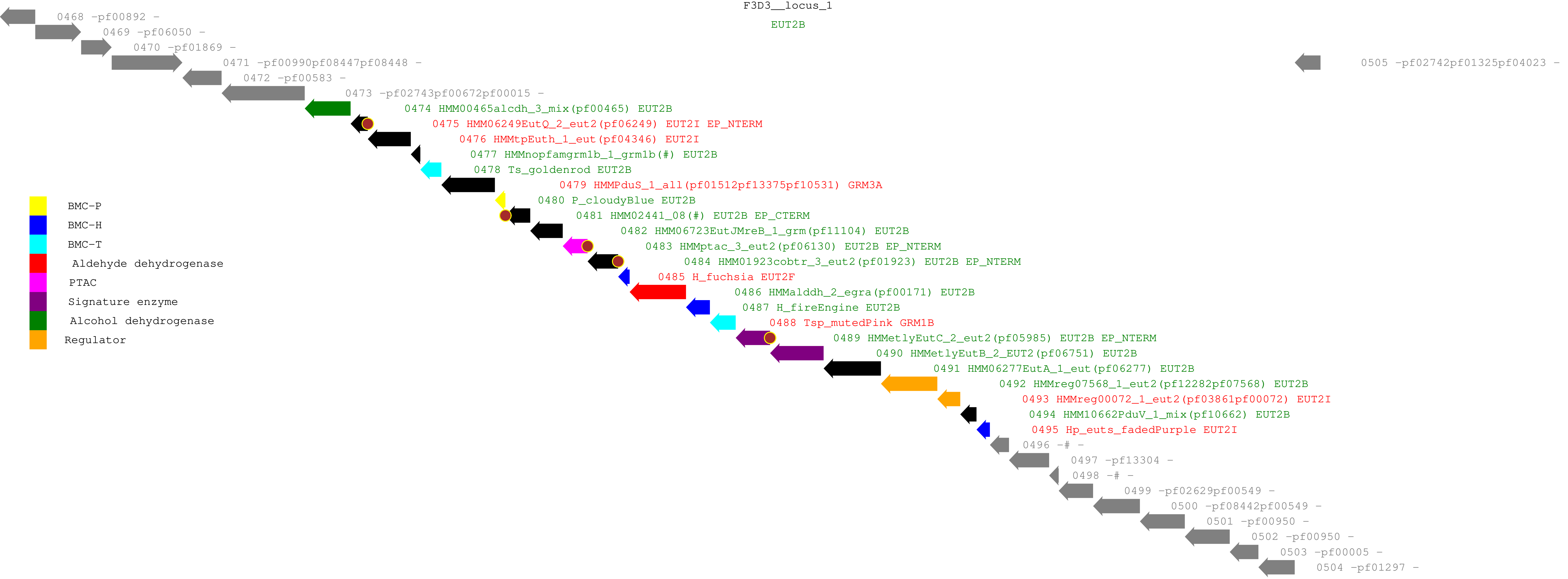

Crassaminicella sp. SY095. Bacteria; Firmicutes; Clostridia; Clostridiales; Clostridiaceae;

FQB35__locus_2

EUT2B

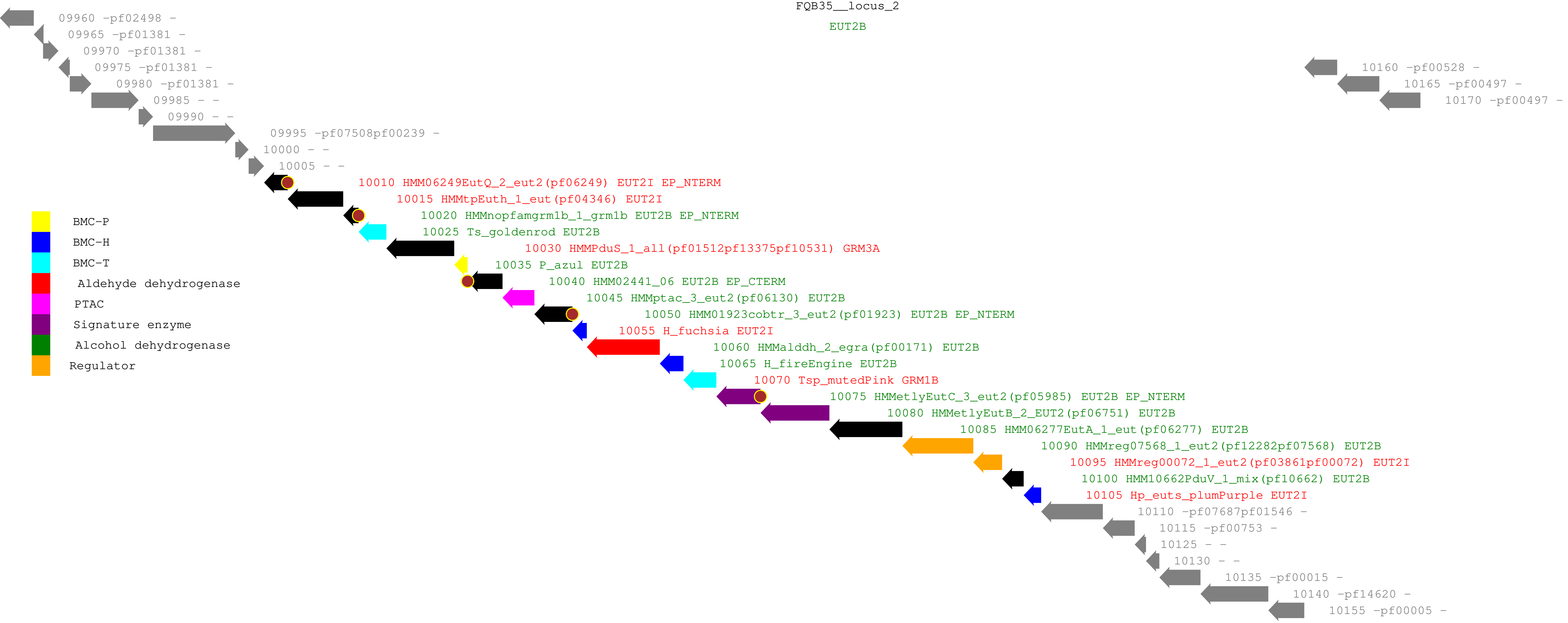

Leptotrichia hofstadii F0254. Bacteria; Fusobacteria; Fusobacteriales; Leptotrichiaceae;

GCWU000323__locus_1

EUT2B

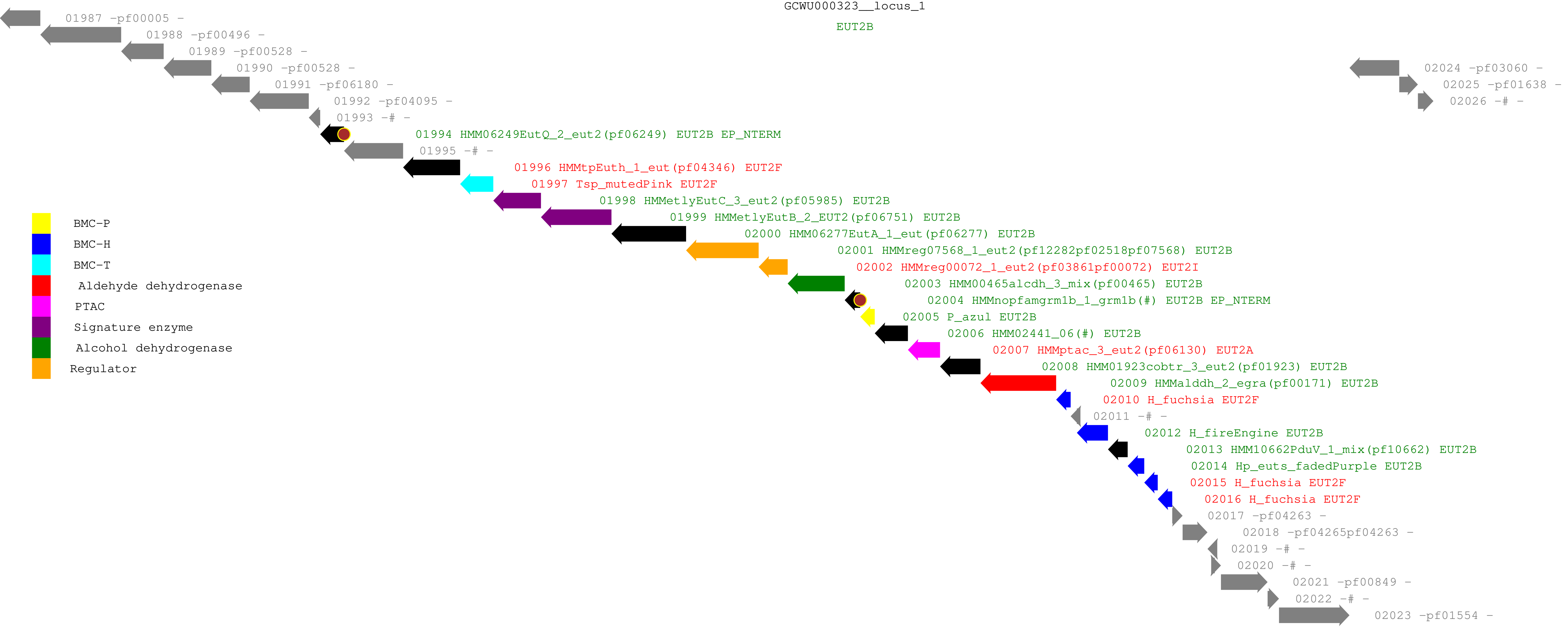

Geosporobacter ferrireducens. Bacteria; Firmicutes; Clostridia; Clostridiales; Clostridiaceae;

Gferi__locus_1

EUT2B

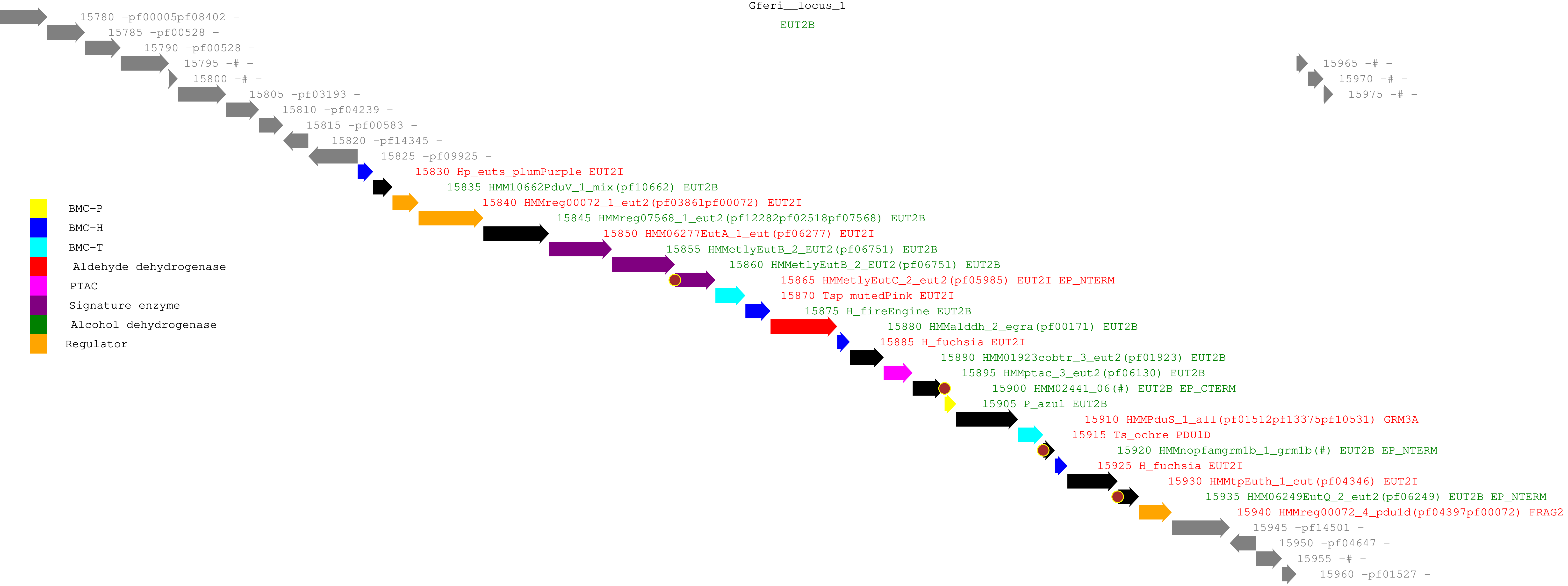

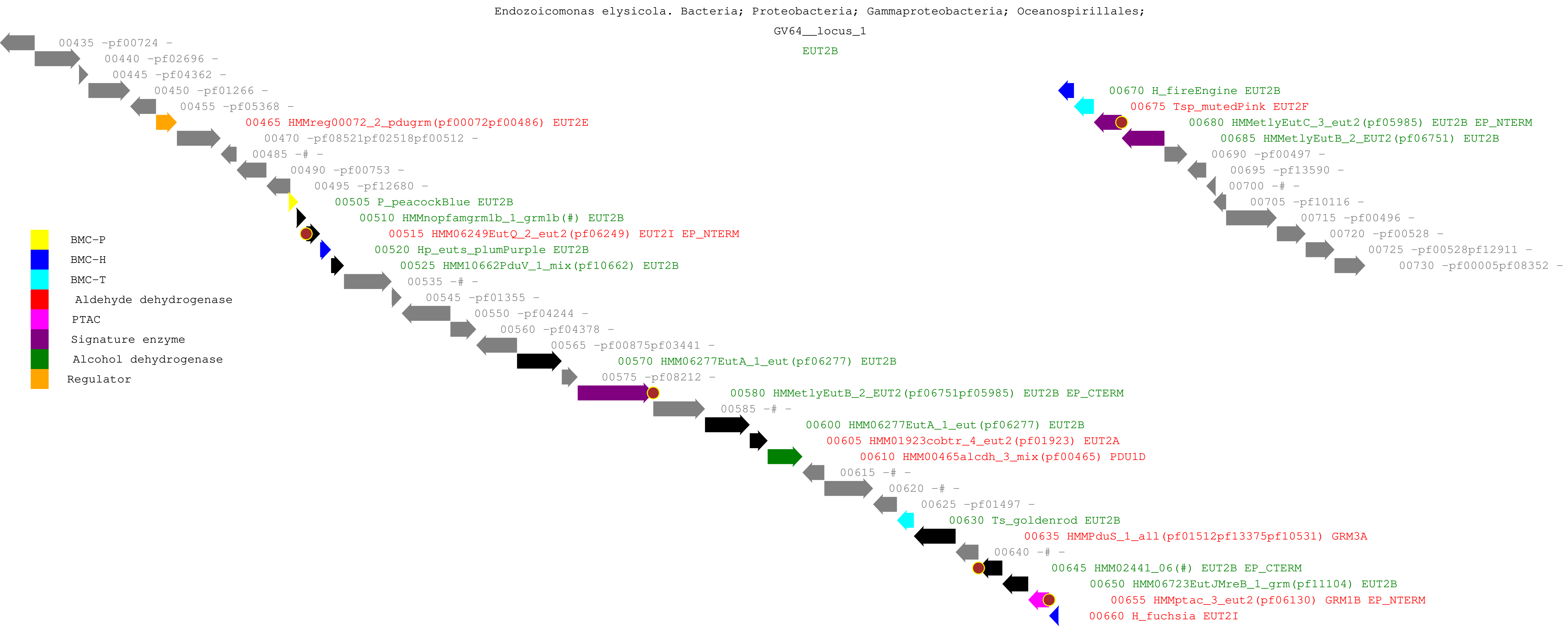
Endozoicomonas elysicola. Bacteria; Proteobacteria; Gammaproteobacteria; Oceanospirillales;
GV64__locus_1
EUT2B
00435 -pf00724 -
00440 -pf02696 -
00445 -pf04362 -
00450 -pf01266 -
00455 -pf05368 -
00465 HMMreg00072_2_pdugrm(pf00072pf00486) EUT2E
00470 -pf08521pf02518pf00512 -
00485 -# -
00490 -pf00753 -
00495 -pf12680 -
00505 P_peacockBlue EUT2B
00510 HMMnopfamgrm1b_1_grm1b(#) EUT2B
00515 HMM06249EutQ_2_eut2(pf06249) EUT2I EP_NTERM
00520 Hp_euts_plumPurple EUT2B
00525 HMM10662PduV_1_mix(pf10662) EUT2B
00535 -# -
00545 -pf01355 -
00550 -pf04244 -
00560 -pf04378 -
00565 -pf00875pf03441 -
00570 HMM06277EutA_1_eut(pf06277) EUT2B
00575 -pf08212 -
00580 HMMetlyEutB_2_EUT2(pf06751pf05985) EUT2B EP_CTERM
00585 -# -
00600 HMM06277EutA_1_eut(pf06277) EUT2B
00605 HMM01923cobtr_4_eut2(pf01923) EUT2A
00610 HMM00465alcdh_3_mix(pf00465) PDU1D
00615 -# -
00620 -# -
00625 -pf01497 -
00630 Ts_goldenrod EUT2B
00635 HMMPduS_1_all(pf01512pf13375pf10531) GRM3A
00640 -# -
00645 HMM02441_06(#) EUT2B EP_CTERM
00650 HMM06723EutJMreB_1_grm(pf11104) EUT2B
00655 HMMptac_3_eut2(pf06130) GRM1B EP_NTERM
00660 H_fuchsia EUT2I
00670 H_fireEngine EUT2B
00675 Tsp_mutedPink EUT2F
00680 HMMetlyEutC_3_eut2(pf05985) EUT2B EP_NTERM
00685 HMMetlyEutB_2_EUT2(pf06751) EUT2B
00690 -pf00497 -
00695 -pf13590 -
00700 -# -
00705 -pf10116 -
00715 -pf00496 -
00720 -pf00528 -
00725 -pf00528pf12911 -
00730 -pf00005pf08352 -
BMC-P
BMC-H
BMC-T
Aldehyde dehydrogenase
PTAC
Signature enzyme
Alcohol dehydrogenase
Regulator

Pseudoramibacter alactolyticus ATCC 23263. Bacteria; Firmicutes; Clostridia; Clostridiales; Eubacteriaceae;

HMP0721__locus_1

EUT2B

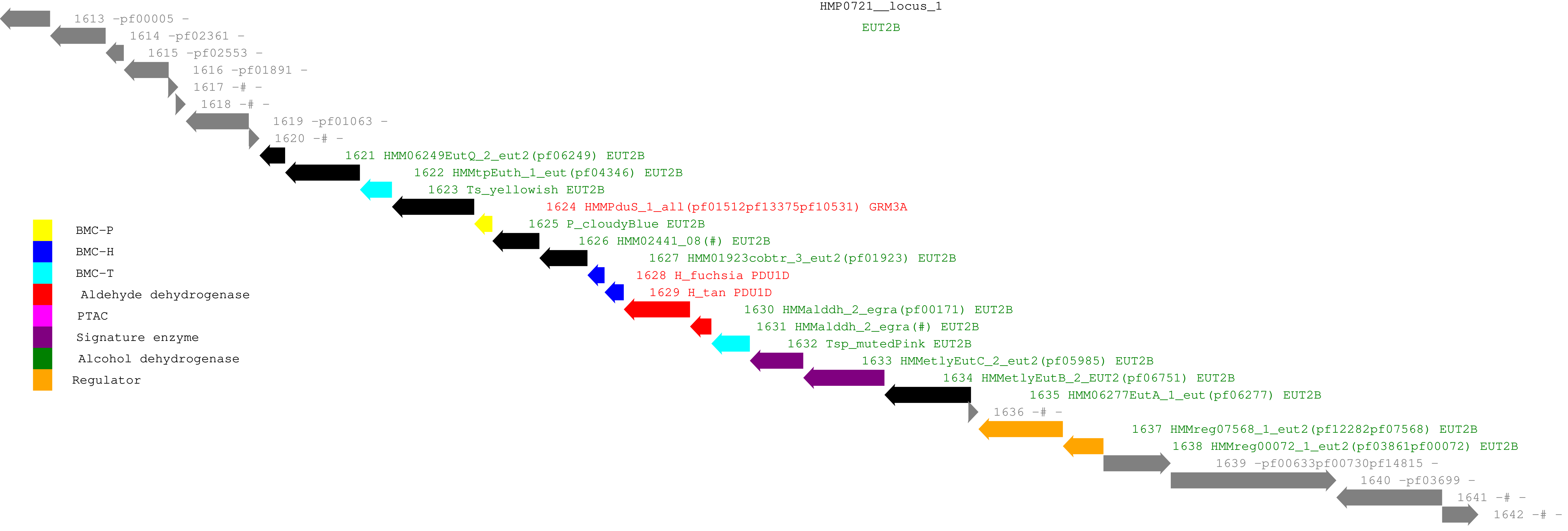

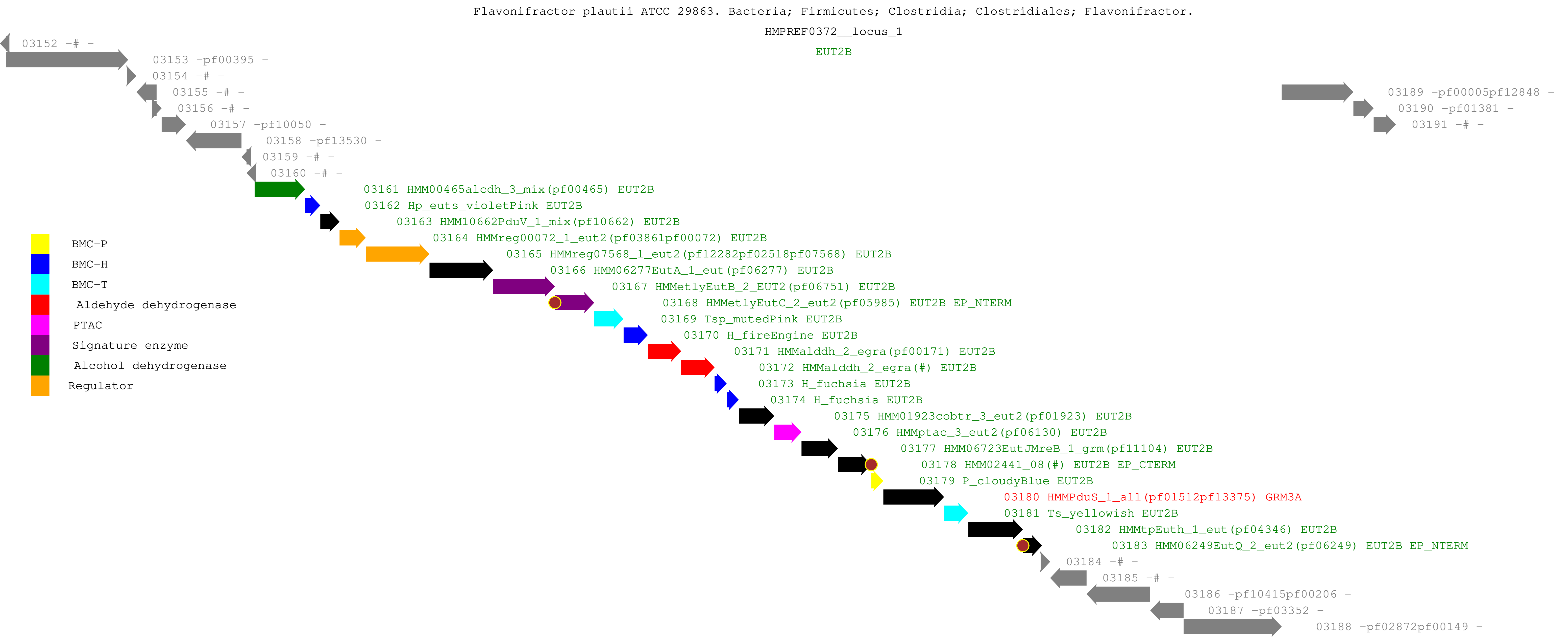

Flavonifractor plautii ATCC 29863. Bacteria; Firmicutes; Clostridia; Clostridiales; Flavonifractor.
HMPREF0372__locus_1
EUT2B
03152 -# -
03153 -pf00395 -
03154 -# -
03155 -# -
03156 -# -
03157 -pf10050 -
03158 -pf13530 -
03159 -# -
03160 -# -
03161 HMM00465alcdh_3_mix(pf00465) EUT2B
03162 Hp_euts_violetPink EUT2B
03163 HMM10662PduV_1_mix(pf10662) EUT2B
03164 HMMreg00072_1_eut2(pf03861pf00072) EUT2B
03165 HMMreg07568_1_eut2(pf12282pf02518pf07568) EUT2B
03166 HMM06277EutA_1_eut(pf06277) EUT2B
03167 HMMetlyEutB_2_EUT2(pf06751) EUT2B
03168 HMMetlyEutC_2_eut2(pf05985) EUT2B EP_NTERM
03169 Tsp_mutedPink EUT2B
03170 H_fireEngine EUT2B
03171 HMMalddh_2_egra(pf00171) EUT2B
03172 HMMalddh_2_egra(#) EUT2B
03173 H_fuchsia EUT2B
03174 H_fuchsia EUT2B
03175 HMM01923cobtr_3_eut2(pf01923) EUT2B
03176 HMMptac_3_eut2(pf06130) EUT2B
03177 HMM06723EutJMreB_1_grm(pf11104) EUT2B
03178 HMM02441_08(#) EUT2B EP_CTERM
03179 P_cloudyBlue EUT2B
03180 HMMPduS_1_all(pf01512pf13375) GRM3A
03181 Ts_yellowish EUT2B
03182 HMMtpEuth_1_eut(pf04346) EUT2B
03183 HMM06249EutQ_2_eut2(pf06249) EUT2B EP_NTERM
03184 -# -
03185 -# -
03186 -pf10415pf00206 -
03187 -pf03352 -
03188 -pf02872pf00149 -
03189 -pf00005pf12848 -
03190 -pf01381 -
03191 -# -
BMC-P
BMC-H
BMC-T
Aldehyde dehydrogenase
PTAC
Signature enzyme
Alcohol dehydrogenase
Regulator

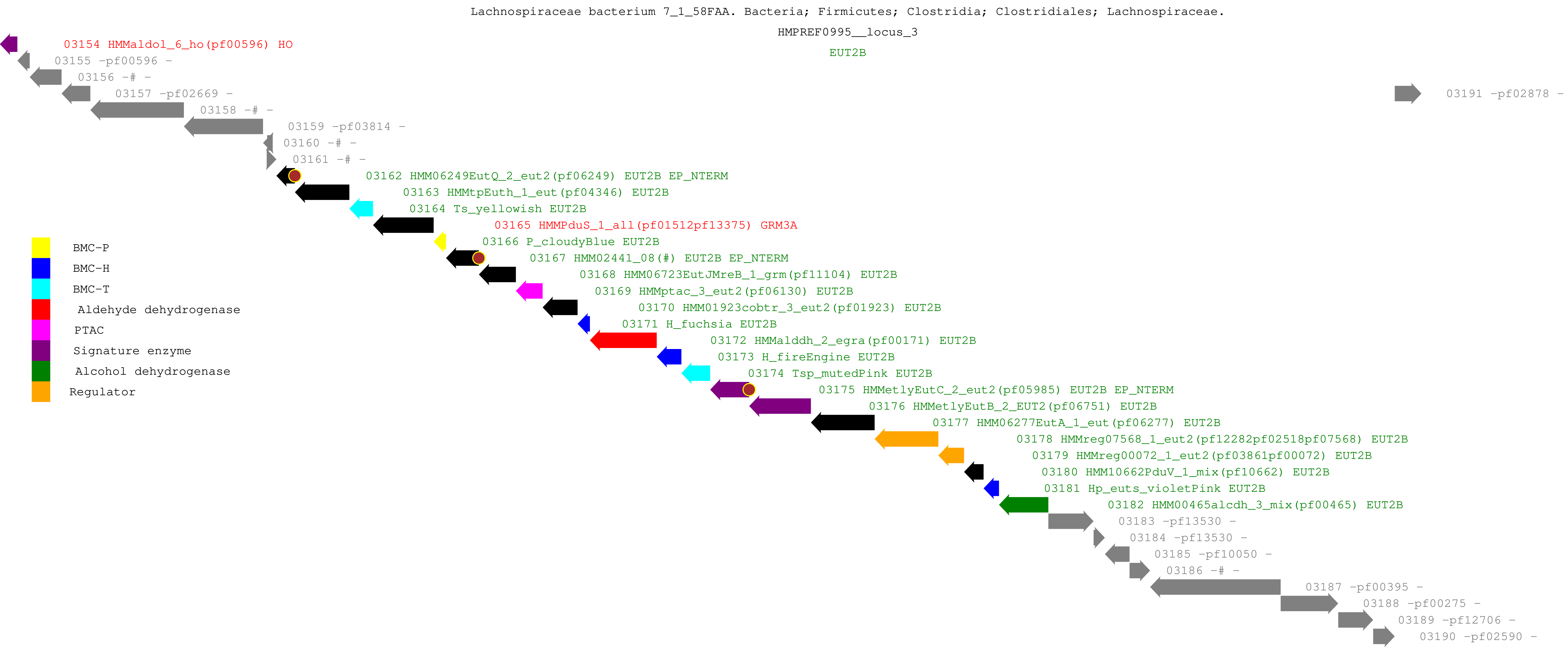
Lachnospiraceae bacterium 7_1_58FAA. Bacteria; Firmicutes; Clostridia; Clostridiales; Lachnospiraceae.
HMPREF0995__locus_3
EUT2B
03154 HMMaldol_6_ho(pf00596) HO
03155 -pf00596 -
03156 -# -
03157 -pf02669 -
03158 -# -
03159 -pf03814 -
03160 -# -
03161 -# -
03162 HMM06249EutQ_2_eut2(pf06249) EUT2B EP_NTERM
03163 HMMtpEuth_1_eut(pf04346) EUT2B
03164 Ts_yellowish EUT2B
03165 HMMPduS_1_all(pf01512pf13375) GRM3A
03166 P_cloudyBlue EUT2B
03167 HMM02441_08(#) EUT2B EP_NTERM
03168 HMM06723EutJMreB_1_grm(pf11104) EUT2B
03169 HMMptac_3_eut2(pf06130) EUT2B
03170 HMM01923cobtr_3_eut2(pf01923) EUT2B
03171 H_fuchsia EUT2B
03172 HMMaldh_2_egra(pf00171) EUT2B
03173 H_fireEngine EUT2B
03174 Tsp_mutedPink EUT2B
03175 HMMetlyEutC_2_eut2(pf05985) EUT2B EP_NTERM
03176 HMMetlyEutB_2_EUT2(pf06751) EUT2B
03177 HMM06277EutA_1_eut(pf06277) EUT2B
03178 HMMreg07568_1_eut2(pf12282pf02518pf07568) EUT2B
03179 HMMreg00072_1_eut2(pf03861pf00072) EUT2B
03180 HMM10662PduV_1_mix(pf10662) EUT2B
03181 Hp_euts_violetPink EUT2B
03182 HMM00465alcdh_3_mix(pf00465) EUT2B
03183 -pf13530 -
03184 -pf13530 -
03185 -pf10050 -
03186 -# -
03187 -pf00395 -
03188 -pf00275 -
03189 -pf12706 -
03190 -pf02590 -
03191 -pf02878 -
BMC-P
BMC-H
BMC-T
Aldehyde dehydrogenase
PTAC
Signature enzyme
Alcohol dehydrogenase
Regulator

Synergistes sp. 3_1_syn1. Bacteria; Synergistetes; Synergistia; Synergistales; Synergistaceae;

HMPREF1006__locus_1

EUT2B

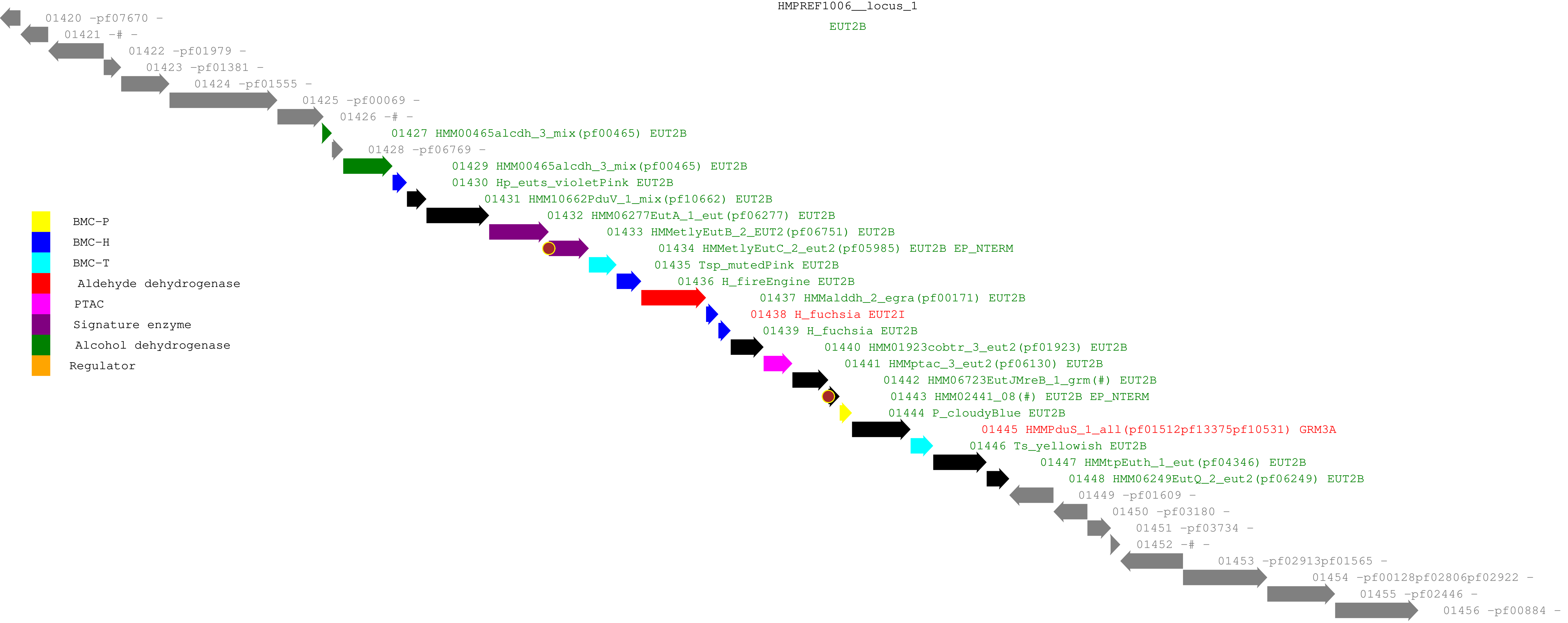

Clostridium sp. 7_3_54FAA. Bacteria; Firmicutes; Clostridia; Clostridiales; Clostridiaceae;

HMPREF1020__locus_1

EUT2B

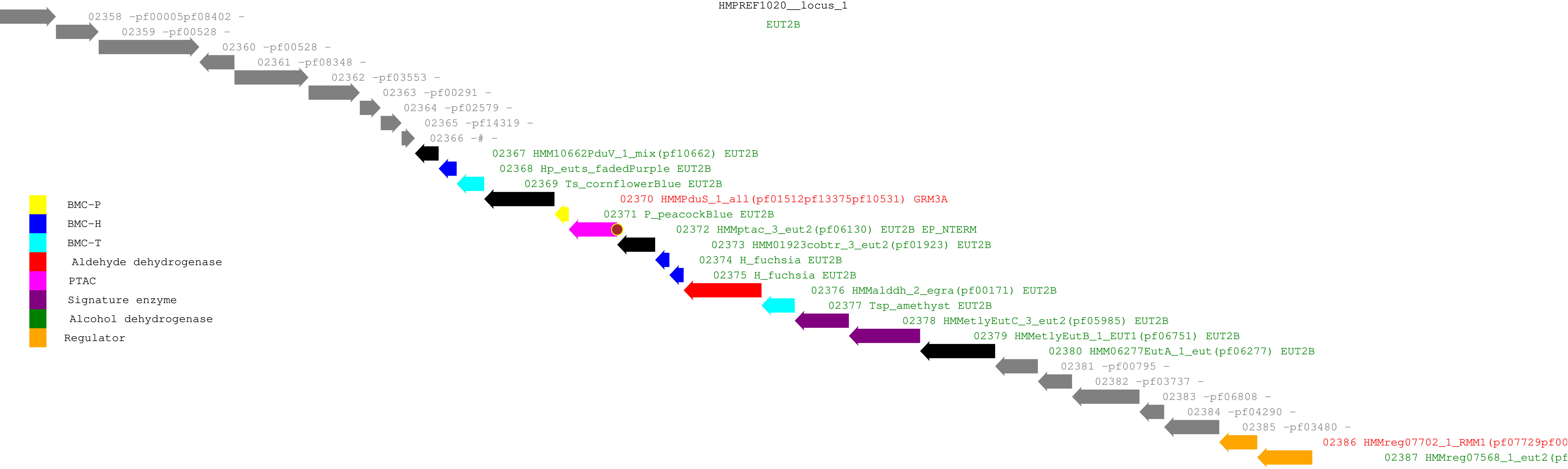

Butyricicoccus pullicaecorum 1.2. Bacteria; Firmicutes; Clostridia; Clostridiales; Clostridiaceae;

HMPREF1526__locus_1

EUT2B

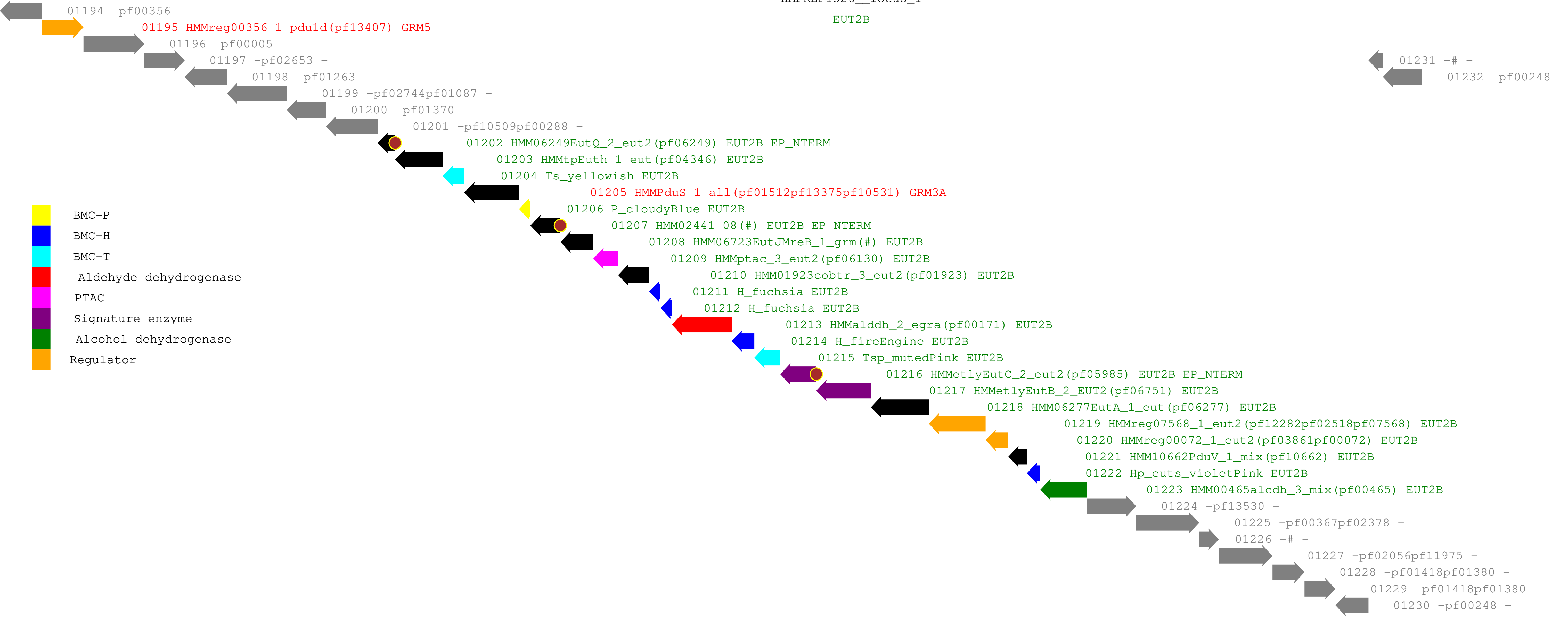

Clostridium sp. KLE 1755. Bacteria; Firmicutes; Clostridia; Clostridiales; Clostridiaceae;

HMPREF1548__locus_2

EUT2B

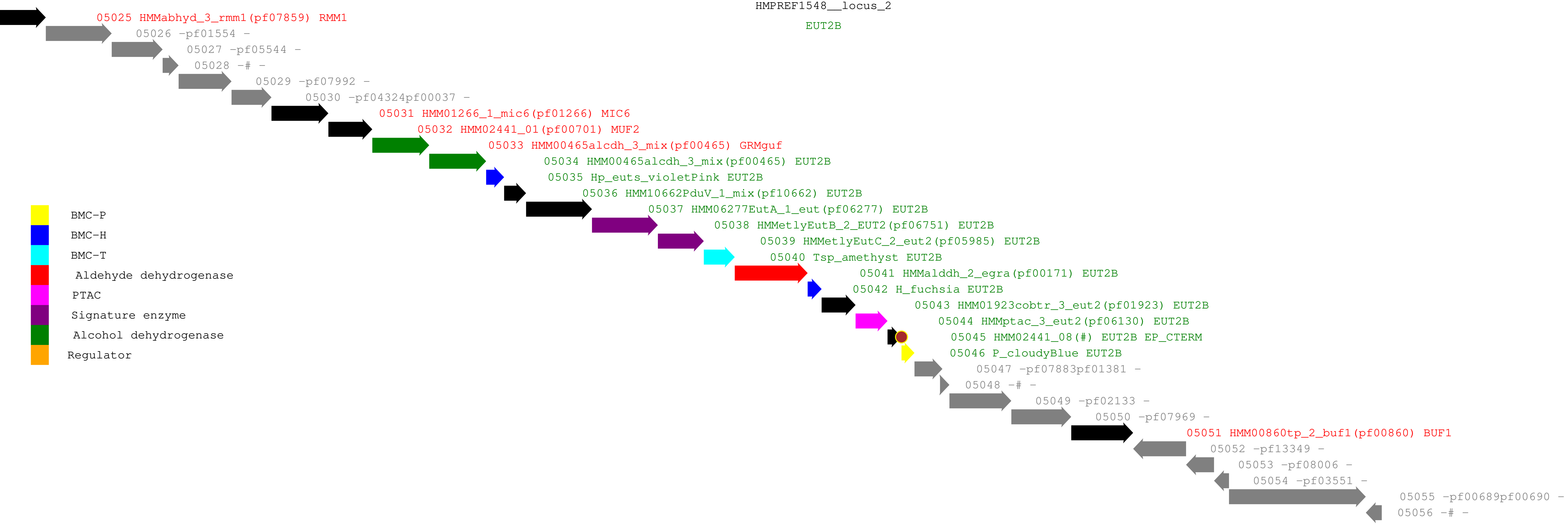

Leptotrichia sp. oral taxon 879 str. F0557. Bacteria; Fusobacteria; Fusobacteriales; Leptotrichiaceae;

HMPREF1552__locus_1

EUT2B

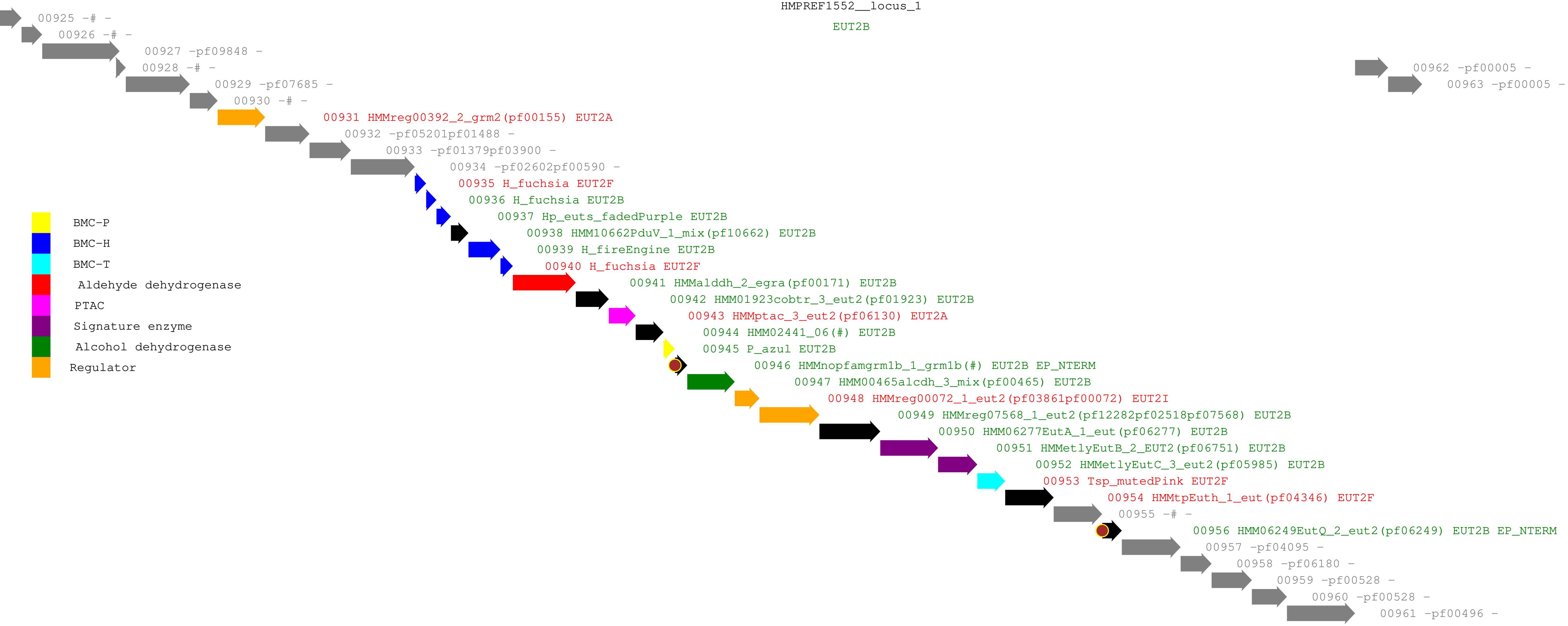

Tissierellia bacterium S5-A11. Bacteria; Firmicutes; Tissierellia.

HMPREF1633__locus_1

EUT2B

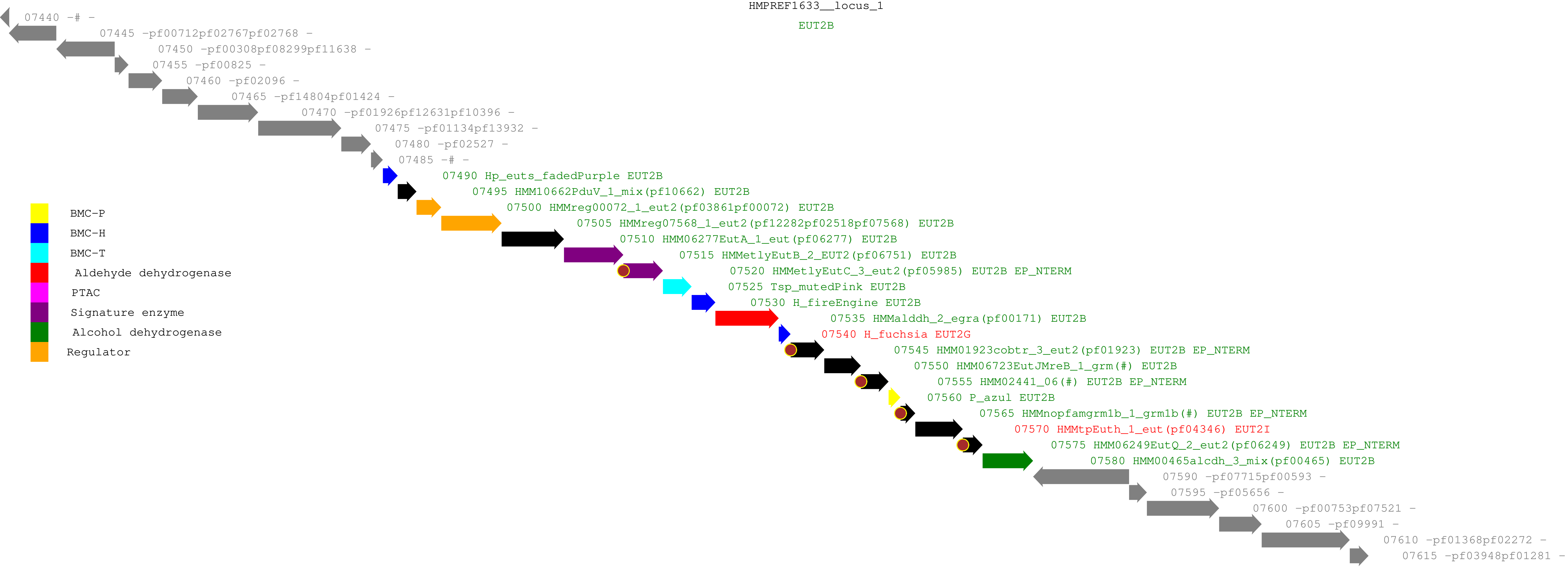

Leptotrichia sp. oral taxon 225 str. F0581. Bacteria; Fusobacteria; Fusobacteriales; Leptotrichiaceae;

HMPREF9108__locus_1_main_sat

EUT2B

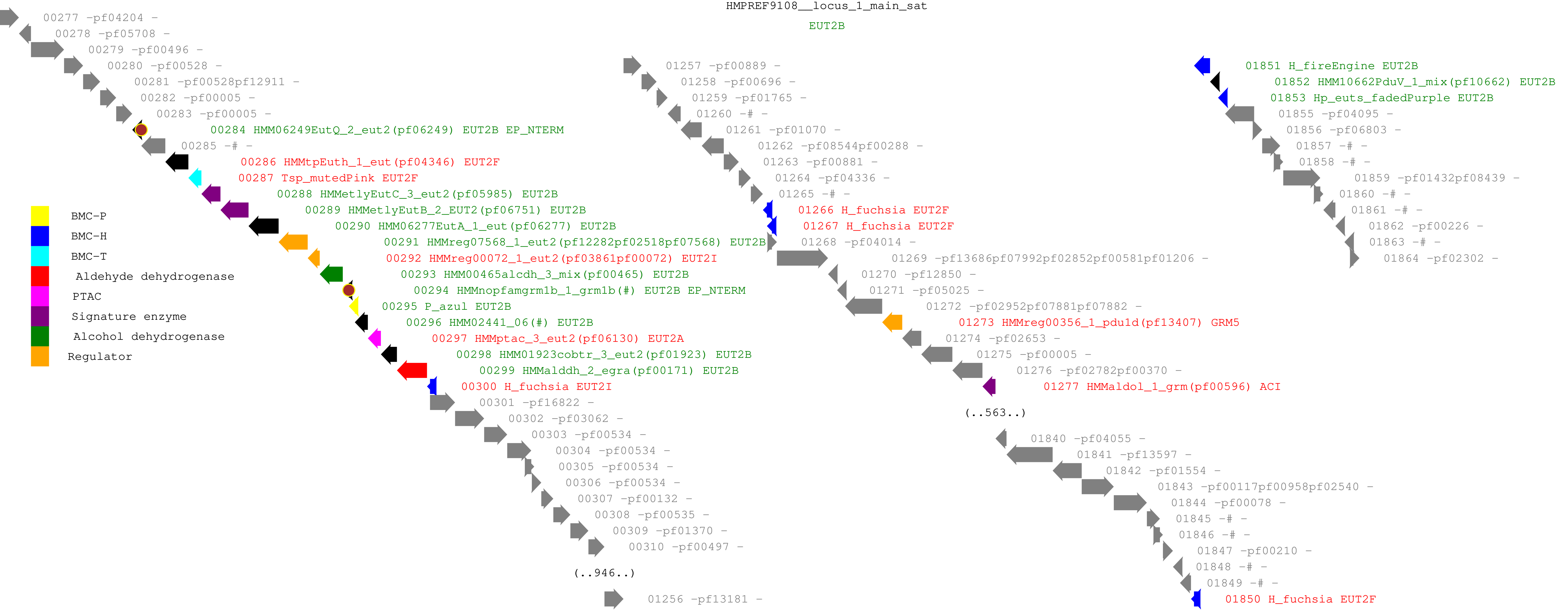

Peptoniphilus sp. oral taxon 375 str. F0436. Bacteria; Firmicutes; Tissierellia; Tissierellales; Peptoniphilaceae;

HMPREF9130__locus_1_main_sat

EUT2B

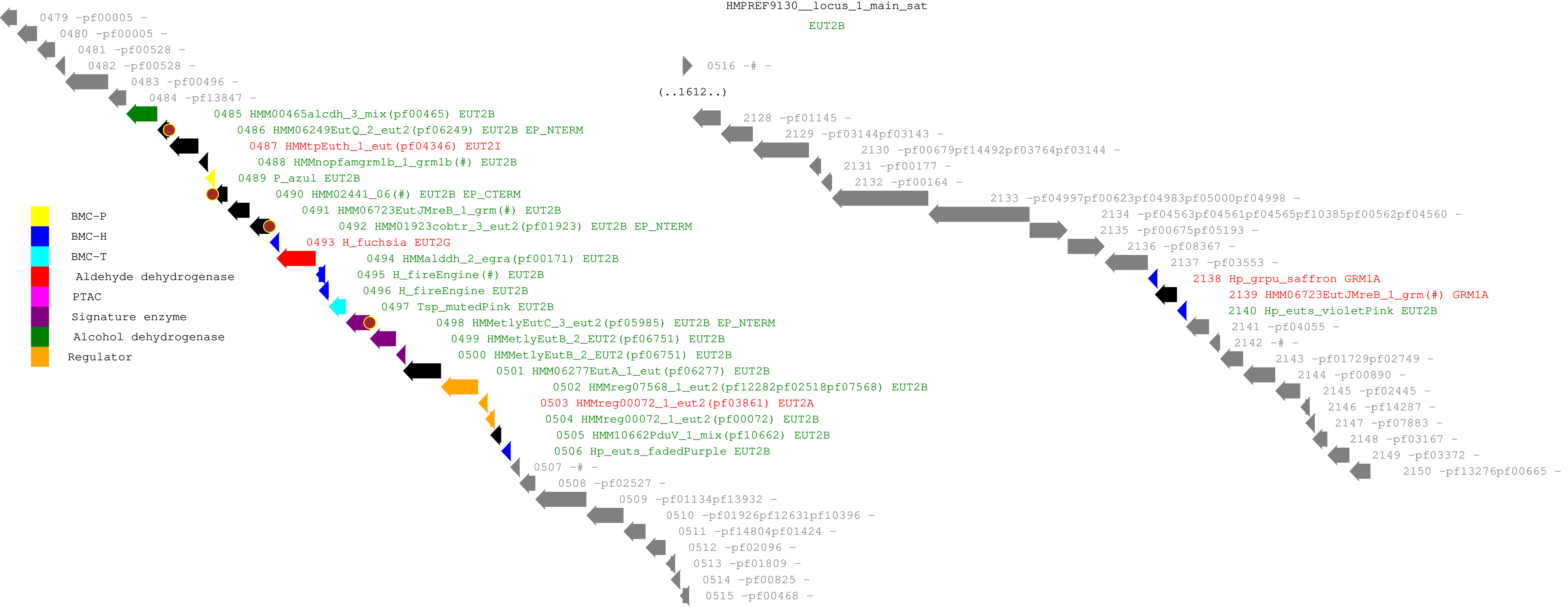

Clostridium orbiscindens 1_3_50AFAA. Bacteria; Firmicutes; Clostridia; Clostridiales; Flavonifractor.

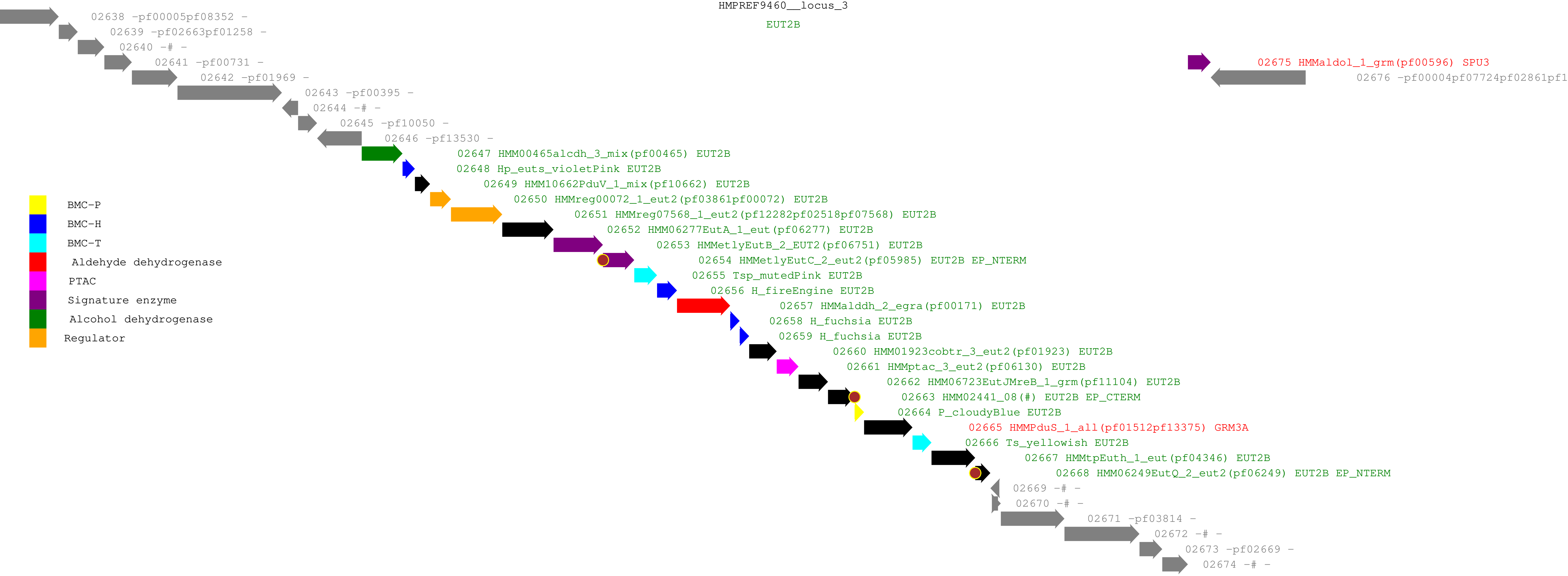

Clostridium symbiosum WAL-14163. Bacteria; Firmicutes; Clostridia; Clostridiales; Lachnospiraceae.

HMPREF9474__locus_1

EUT2B

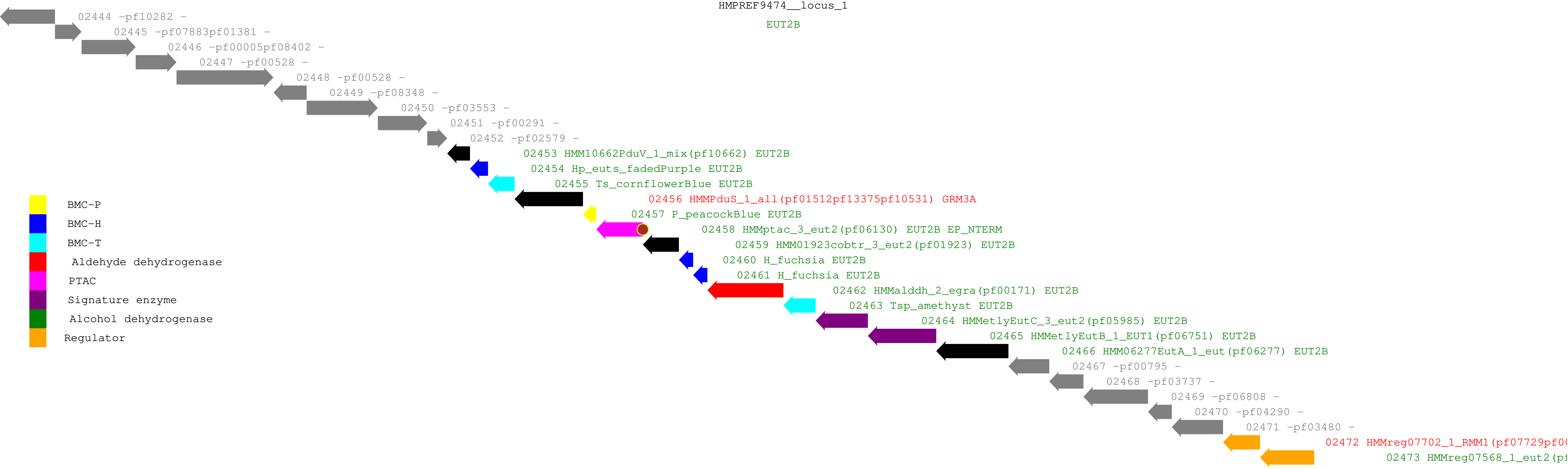

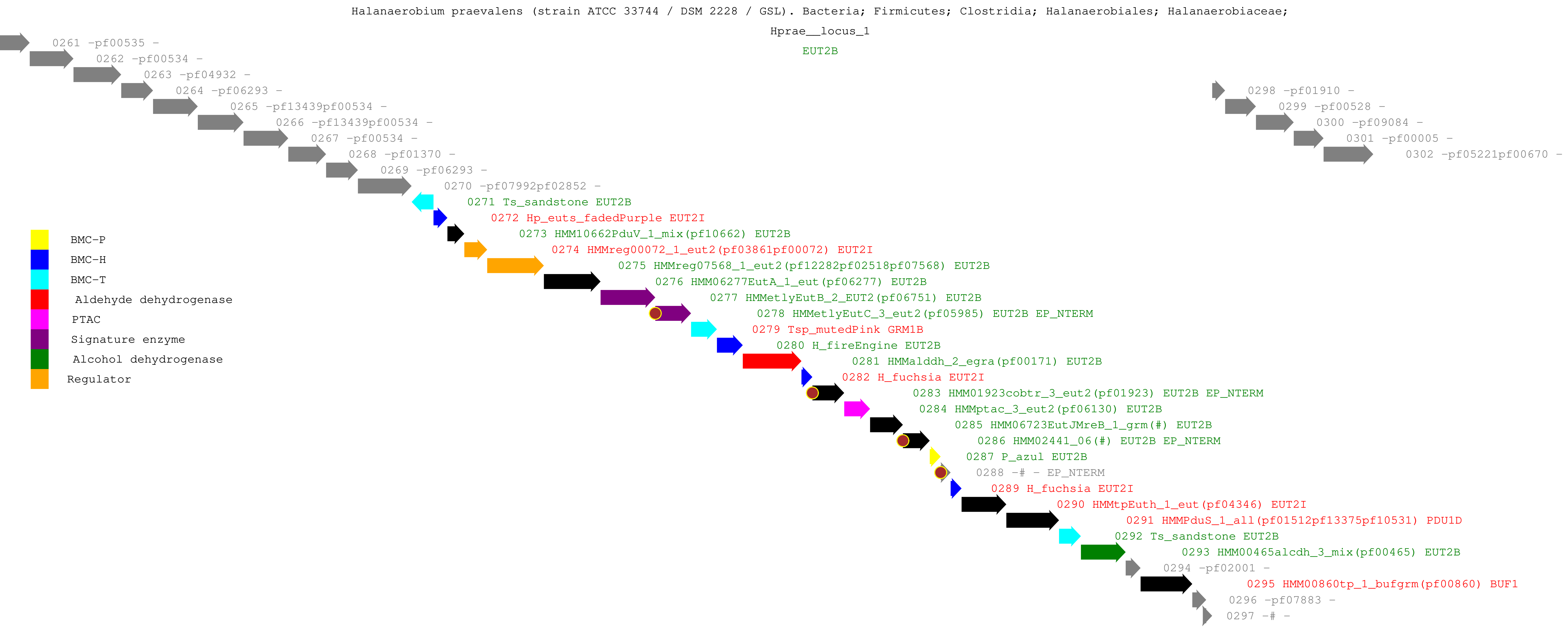
Halanaerobium praevalens (strain ATCC 33744 / DSM 2228 / GSL). Bacteria; Firmicutes; Clostridia; Halanaerobiales; Halanaerobiaceae;
Hprae__locus_1
EUT2B
0261 -pf00535 -
0262 -pf00534 -
0263 -pf04932 -
0264 -pf06293 -
0265 -pf13439pf00534 -
0266 -pf13439pf00534 -
0267 -pf00534 -
0268 -pf01370 -
0269 -pf06293 -
0270 -pf07992pf02852 -
0271 Ts_sandstone EUT2B
0272 Hp_euts_fadedPurple EUT2I
0273 HMM10662PduV_1_mix(pf10662) EUT2B
0274 HMMreg00072_1_eut2(pf03861pf00072) EUT2I
0275 HMMreg07568_1_eut2(pf12282pf02518pf07568) EUT2B
0276 HMM06277EutA_1_eut(pf06277) EUT2B
0277 HMMetlyEutB_2_EUT2(pf06751) EUT2B
0278 HMMetlyEutC_3_eut2(pf05985) EUT2B EP_NTERM
0279 Tsp_mutedPink GRM1B
0280 H_fireEngine EUT2B
0281 HMMalddh_2_egra(pf00171) EUT2B
0282 H_fuchsia EUT2I
0283 HMM01923cobtr_3_eut2(pf01923) EUT2B EP_NTERM
0284 HMMptac_3_eut2(pf06130) EUT2B
0285 HMM06723EutJMreB_1_grm(#) EUT2B
0286 HMM02441_06(#) EUT2B EP_NTERM
0287 P_azul EUT2B
0288 -# - EP_NTERM
0289 H_fuchsia EUT2I
0290 HMMtpEuth_1_eut(pf04346) EUT2I
0291 HMMPduS_1_all(pf01512pf13375pf10531) PDU1D
0292 Ts_sandstone EUT2B
0293 HMM00465alcdh_3_mix(pf00465) EUT2B
0294 -pf02001 -
0295 HMM00860tp_1_bufgrm(pf00860) BUF1
0296 -pf07883 -
0297 -# -
0298 -pf01910 -
0299 -pf00528 -
0300 -pf09084 -
0301 -pf00005 -
0302 -pf05221pf00670 -
BMC-P
BMC-H
BMC-T
Aldehyde dehydrogenase
PTAC
Signature enzyme
Alcohol dehydrogenase
Regulator

Leptotrichia hofstadii. Bacteria; Fusobacteria; Fusobacteriales; Leptotrichiaceae; Leptotrichia.

JCM16775__locus_1

EUT2B

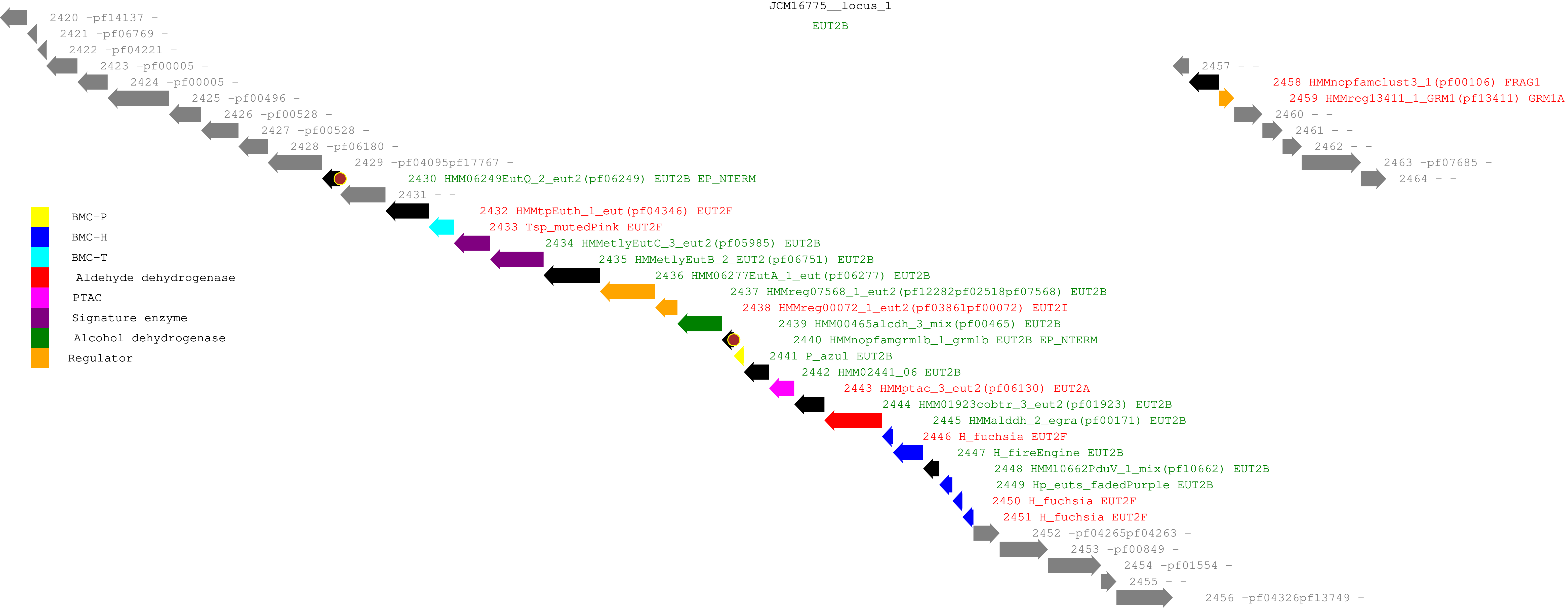

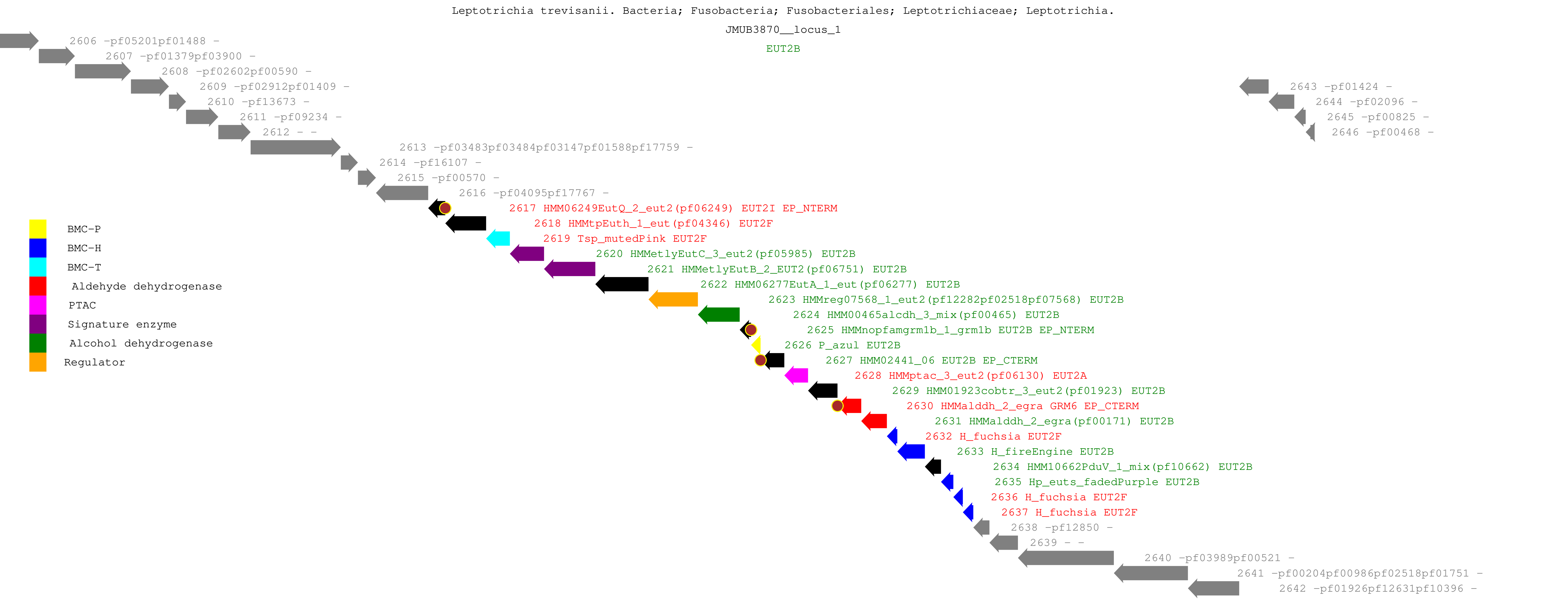
Leptotrichia trevisanii. Bacteria; Fusobacteria; Fusobacteriales; Leptotrichiaceae; Leptotrichia.
JMUB3870__locus_1
EUT2B
2606 -pf05201pf01488 -
2607 -pf01379pf03900 -
2608 -pf02602pf00590 -
2609 -pf02912pf01409 -
2610 -pf13673 -
2611 -pf09234 -
2612 - -
2613 -pf03483pf03484pf03147pf01588pf17759 -
2614 -pf16107 -
2615 -pf00570 -
2616 -pf04095pf17767 -
2617 HMM06249EutQ_2_eut2(pf06249) EUT2I EP_NTERM
2618 HMMtpEuth_1_eut(pf04346) EUT2F
2619 Tsp_mutedPink EUT2F
2620 HMMetlyEutC_3_eut2(pf05985) EUT2B
2621 HMMetlyEutB_2_EUT2(pf06751) EUT2B
2622 HMM06277EutA_1_eut(pf06277) EUT2B
2623 HMMreg07568_1_eut2(pf12282pf02518pf07568) EUT2B
2624 HMM00465alcdh_3_mix(pf00465) EUT2B
2625 HMMnopfamgrmlb_1_grmlb EUT2B EP_NTERM
2626 P_azul EUT2B
2627 HMM02441_06 EUT2B EP_CTERM
2628 HMMptac_3_eut2(pf06130) EUT2A
2629 HMM01923cobtr_3_eut2(pf01923) EUT2B
2630 HMMaldh_2_egra GRM6 EP_CTERM
2631 HMMaldh_2_egra(pf00171) EUT2B
2632 H_fuchsia EUT2F
2633 H_fireEngine EUT2B
2634 HMM10662PduV_1_mix(pf10662) EUT2B
2635 Hp_euts_fadedPurple EUT2B
2636 H_fuchsia EUT2F
2637 H_fuchsia EUT2F
2638 -pf12850 -
2639 - -
2640 -pf03989pf00521 -
2641 -pf00204pf00986pf02518pf01751 -
2642 -pf01926pf12631pf10396 -
2643 -pf01424 -
2644 -pf02096 -
2645 -pf00825 -
2646 -pf00468 -
BMC-P
BMC-H
BMC-T
Aldehyde dehydrogenase
PTAC
Signature enzyme
Alcohol dehydrogenase
Regulator

Leptotrichia hongkongensis. Bacteria; Fusobacteria; Fusobacteriales; Leptotrichiaceae; Leptotrichia.

JMUB5056__locus_1

EUT2B

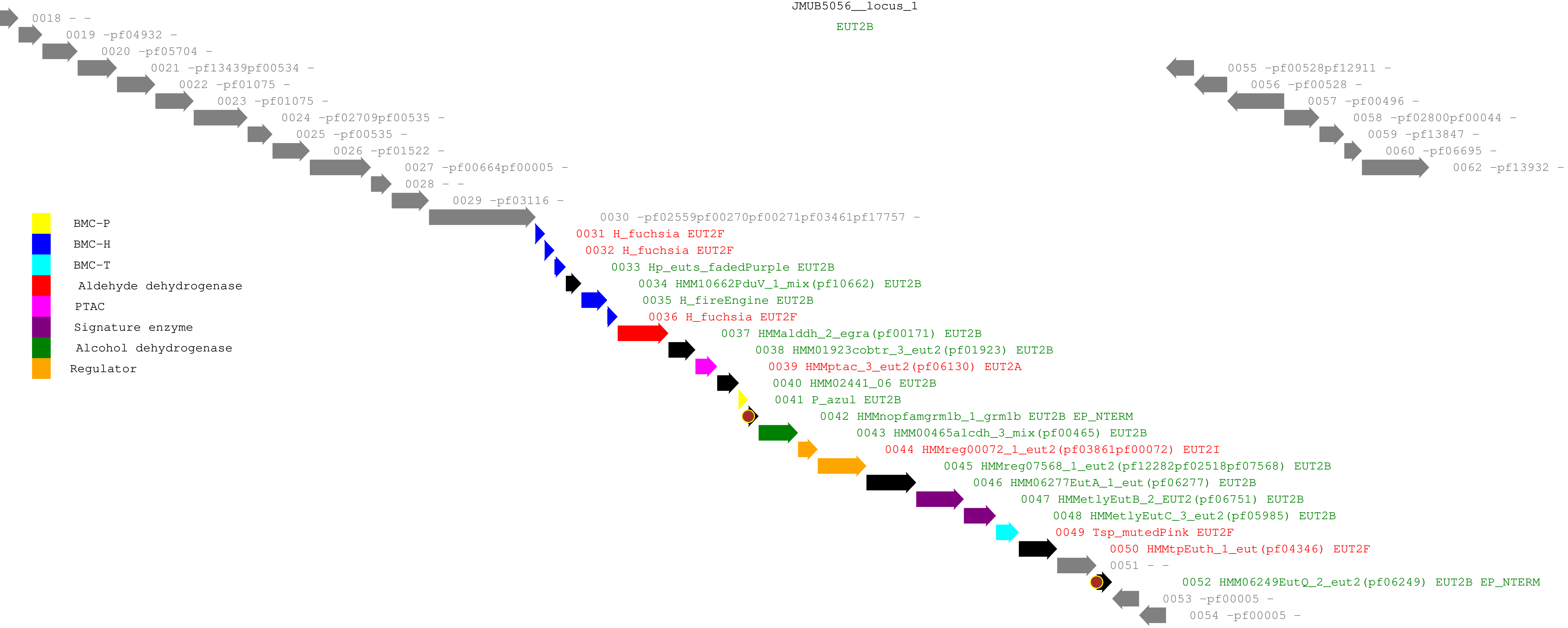

uncultured Clostridiales bacterium. Bacteria; Firmicutes; Clostridia; Clostridiales;

KL86CLO1__locus_1_main_sat

EUT2B

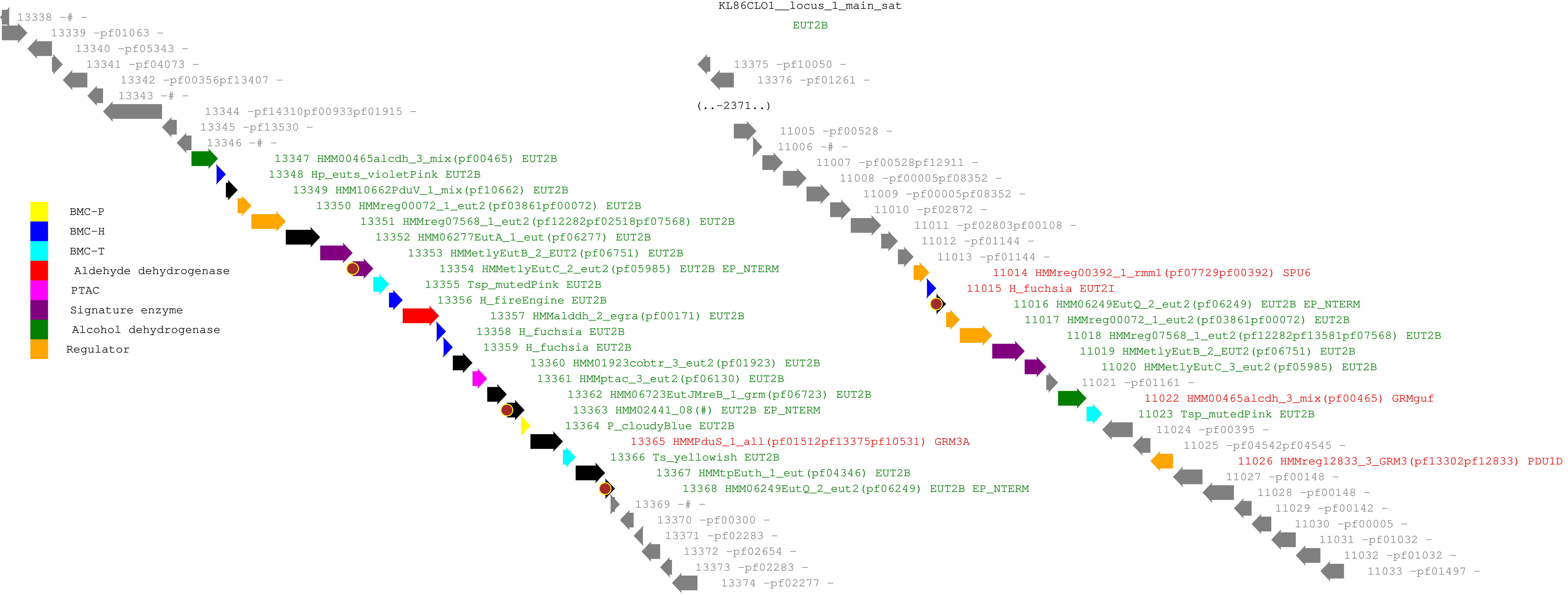

Leptotrichia buccalis (strain ATCC 14201 / DSM 1135 / JCM 12969 / NCTC Bacteria; Fusobacteria; Fusobacteriales; Leptotrichiaceae;

Lebu__locus_1

EUT2B

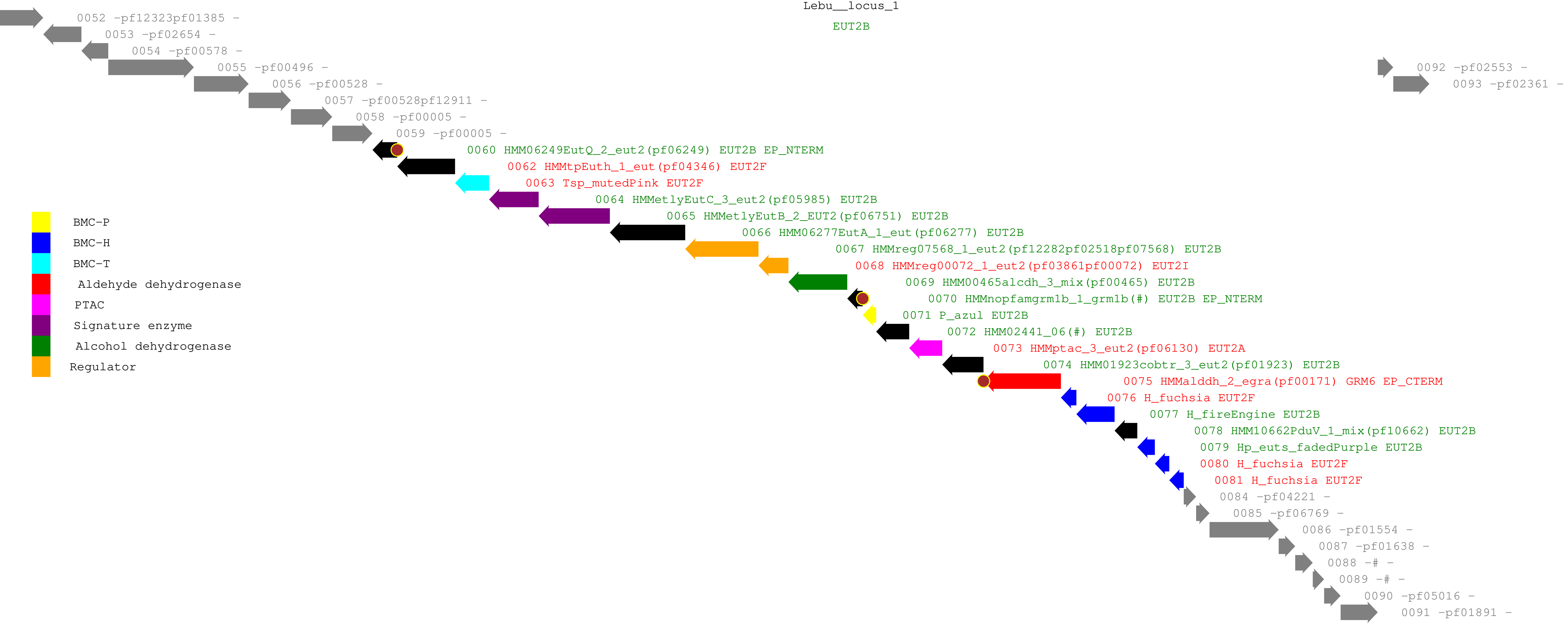

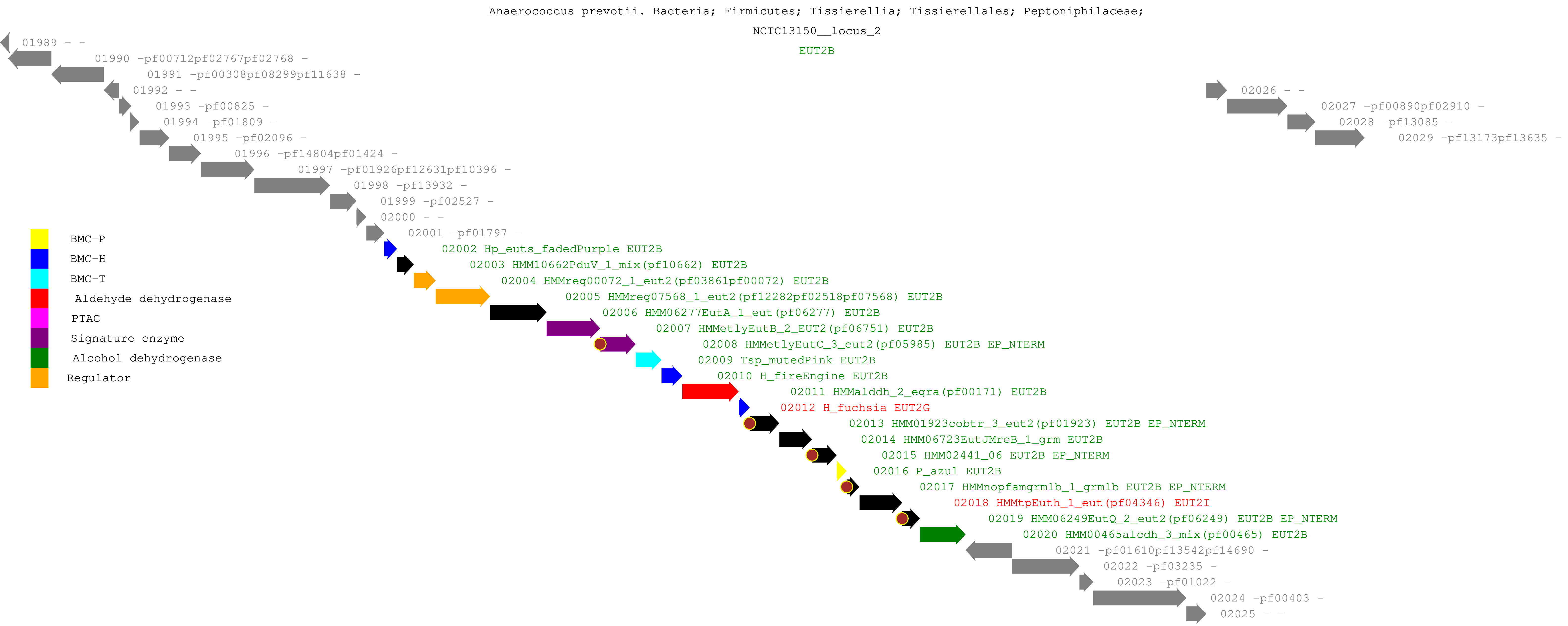
Anaerococcus prevotii. Bacteria; Firmicutes; Tissierellia; Tissierellales; Peptoniphilaceae;
NCTC13150__locus_2
EUT2B
01989 - -
01990 -pf00712pf02767pf02768 -
01991 -pf00308pf08299pf11638 -
01992 - -
01993 -pf00825 -
01994 -pf01809 -
01995 -pf02096 -
01996 -pf14804pf01424 -
01997 -pf01926pf12631pf10396 -
01998 -pf13932 -
01999 -pf02527 -
02000 - -
02001 -pf01797 -
02002 Hp_euts_fadedPurple EUT2B
02003 HMM10662PduV_1_mix(pf10662) EUT2B
02004 HMMreg00072_1_eut2(pf03861pf00072) EUT2B
02005 HMMreg07568_1_eut2(pf12282pf02518pf07568) EUT2B
02006 HMM06277EutA_1_eut(pf06277) EUT2B
02007 HMMetlyEutB_2_EUT2(pf06751) EUT2B
02008 HMMetlyEutC_3_eut2(pf05985) EUT2B EP_NTERM
02009 Tsp_mutedPink EUT2B
02010 H_fireEngine EUT2B
02011 HMMalddh_2_egra(pf00171) EUT2B
02012 H_fuchsia EUT2G
02013 HMM01923cobtr_3_eut2(pf01923) EUT2B EP_NTERM
02014 HMM06723EutJMreB_1_grm EUT2B
02015 HMM02441_06 EUT2B EP_NTERM
02016 P_azul EUT2B
02017 HMMnopfamgrm1b_1_grm1b EUT2B EP_NTERM
02018 HMMtpEuth_1_eut(pf04346) EUT2I
02019 HMM06249EutQ_2_eut2(pf06249) EUT2B EP_NTERM
02020 HMM00465alcdh_3_mix(pf00465) EUT2B
02021 -pf01610pf13542pf14690 -
02022 -pf03235 -
02023 -pf01022 -
02024 -pf00403 -
02025 - -
02026 - -
02027 -pf00890pf02910 -
02028 -pf13085 -
02029 -pf13173pf13635 -
BMC-P
BMC-H
BMC-T
Aldehyde dehydrogenase
PTAC
Signature enzyme
Alcohol dehydrogenase
Regulator

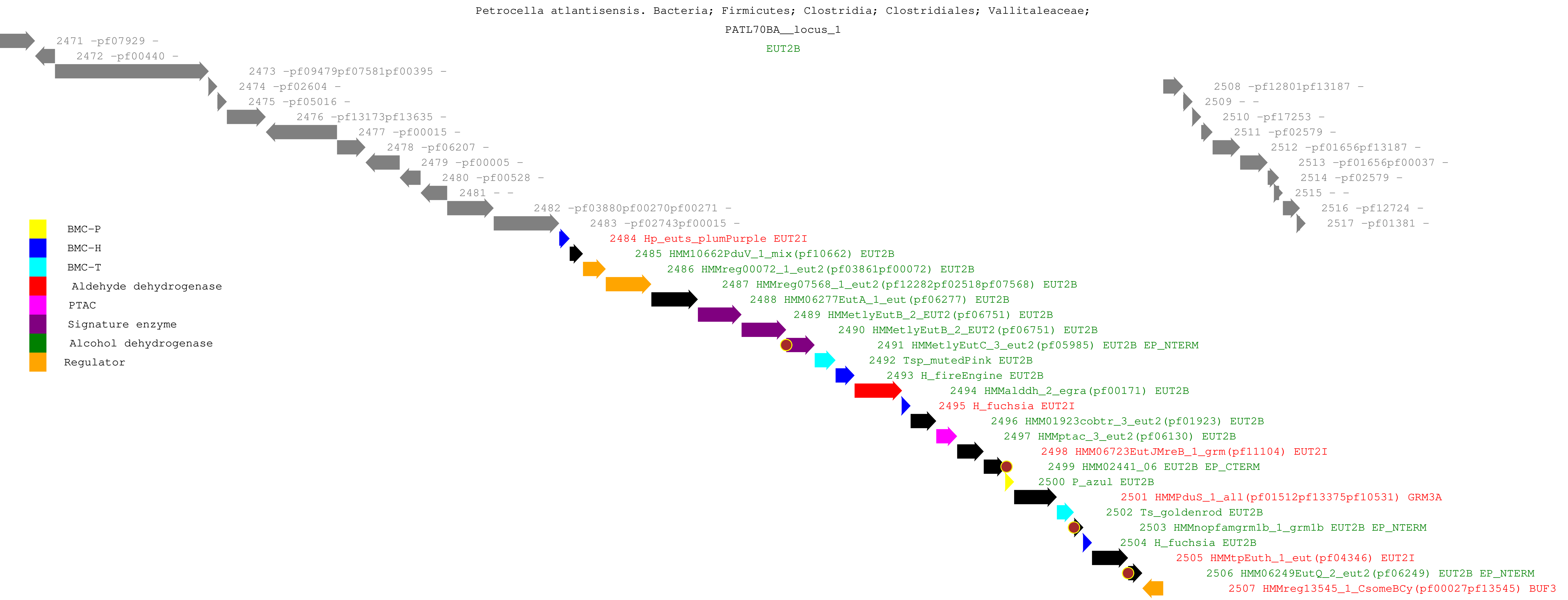
Petrocella atlantisensis. Bacteria; Firmicutes; Clostridia; Clostridiales; Vallitaleaceae;
PATL70BA__locus_1
EUT2B
2471 -pf07929 -
2472 -pf00440 -
2473 -pf09479pf07581pf00395 -
2474 -pf02604 -
2475 -pf05016 -
2476 -pf13173pf13635 -
2477 -pf00015 -
2478 -pf06207 -
2479 -pf00005 -
2480 -pf00528 -
2481 - -
2482 -pf03880pf00270pf00271 -
2483 -pf02743pf00015 -
2484 Hp_euts_plumPurple EUT2I
2485 HMM10662PduV_1_mix(pf10662) EUT2B
2486 HMMreg00072_1_eut2(pf03861pf00072) EUT2B
2487 HMMreg07568_1_eut2(pf12282pf02518pf07568) EUT2B
2488 HMM06277EutA_1_eut(pf06277) EUT2B
2489 HMMetlyEutB_2_EUT2(pf06751) EUT2B
2490 HMMetlyEutB_2_EUT2(pf06751) EUT2B
2491 HMMetlyEutC_3_eut2(pf05985) EUT2B EP_NTERM
2492 Tsp_mutedPink EUT2B
2493 H_fireEngine EUT2B
2494 HMMalddh_2_egra(pf00171) EUT2B
2495 H_fuchsia EUT2I
2496 HMM01923cobtr_3_eut2(pf01923) EUT2B
2497 HMMptac_3_eut2(pf06130) EUT2B
2498 HMM06723EutJMreB_1_grm(pf11104) EUT2I
2499 HMM02441_06 EUT2B EP_CTERM
2500 P_azul EUT2B
2501 HMMPduS_1_all(pf01512pf13375pf10531) GRM3A
2502 Ts_goldenrod EUT2B
2503 HMMnopfamgrm1b_1_grm1b EUT2B EP_NTERM
2504 H_fuchsia EUT2B
2505 HMMtpEuth_1_eut(pf04346) EUT2I
2506 HMM06249EutQ_2_eut2(pf06249) EUT2B EP_NTERM
2507 HMMreg13545_1_CsomeBCy(pf00027pf13545) BUF3
2508 -pf12801pf13187 -
2509 - -
2510 -pf17253 -
2511 -pf02579 -
2512 -pf01656pf13187 -
2513 -pf01656pf00037 -
2514 -pf02579 -
2515 - -
2516 -pf12724 -
2517 -pf01381 -
BMC-P
BMC-H
BMC-T
Aldehyde dehydrogenase
PTAC
Signature enzyme
Alcohol dehydrogenase
Regulator

uncultured Roseburia sp. Bacteria; Firmicutes; Clostridia; Clostridiales; Lachnospiraceae;

SAMEA3545351__locus_1

EUT2B

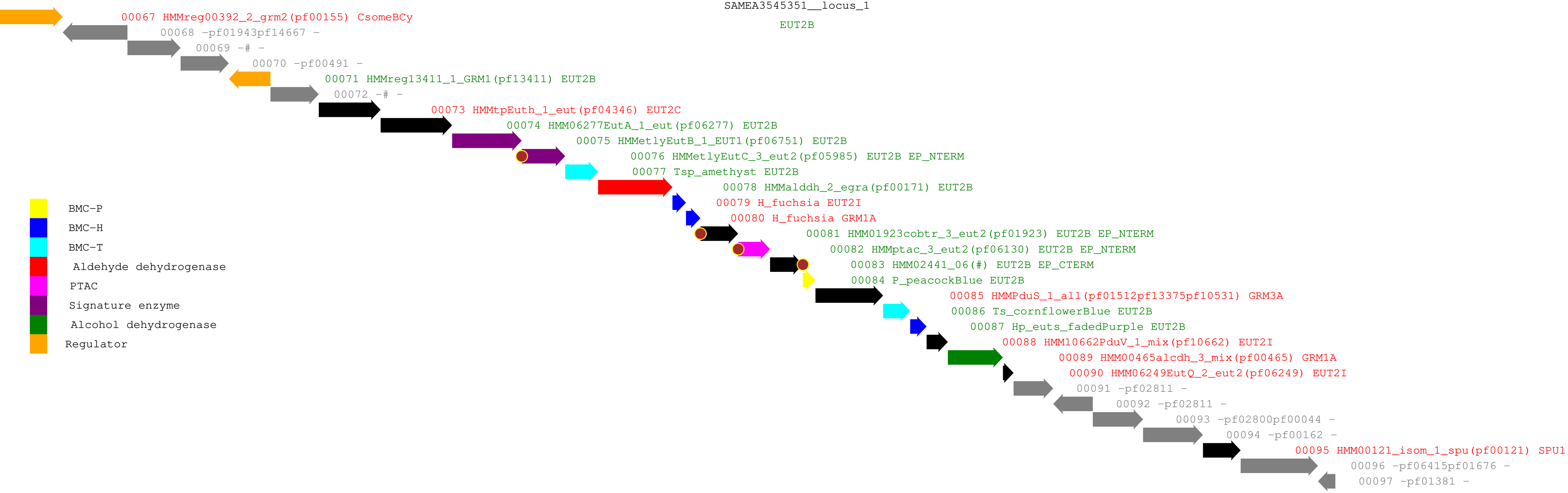

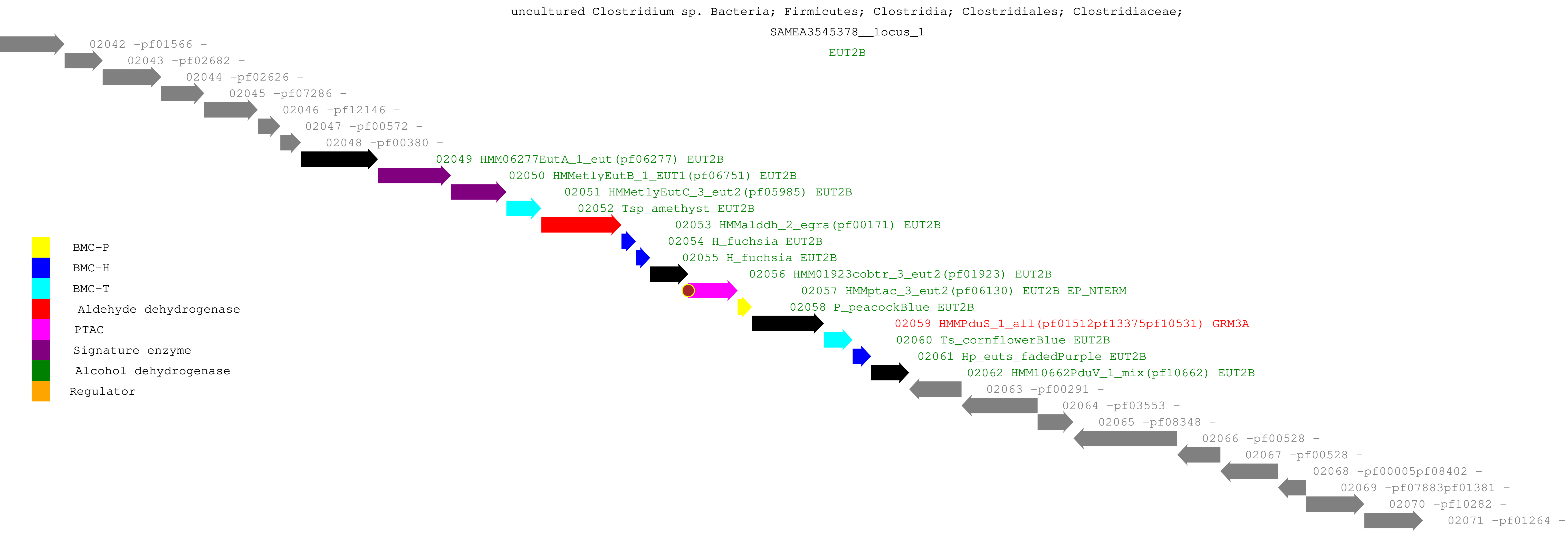
uncultured Clostridium sp. Bacteria; Firmicutes; Clostridia; Clostridiales; Clostridiaceae;
SAMEA3545378__locus_1
EUT2B
02042 -pf01566 -
02043 -pf02682 -
02044 -pf02626 -
02045 -pf07286 -
02046 -pf12146 -
02047 -pf00572 -
02048 -pf00380 -
02049 HMM06277EutA_1_eut(pf06277) EUT2B
02050 HMMetlyEutB_1_EUT1(pf06751) EUT2B
02051 HMMetlyEutC_3_eut2(pf05985) EUT2B
02052 Tsp_amethyst EUT2B
02053 HMMaldh_2_egra(pf00171) EUT2B
02054 H_fuchsia EUT2B
02055 H_fuchsia EUT2B
02056 HMM01923cobtr_3_eut2(pf01923) EUT2B
02057 HMMptac_3_eut2(pf06130) EUT2B EP_NTERM
02058 P_peacockBlue EUT2B
02059 HMMPduS_1_all(pf01512pf13375pf10531) GRM3A
02060 Ts_cornflowerBlue EUT2B
02061 Hp_euts_fadedPurple EUT2B
02062 HMM10662PduV_1_mix(pf10662) EUT2B
02063 -pf00291 -
02064 -pf03553 -
02065 -pf08348 -
02066 -pf00528 -
02067 -pf00528 -
02068 -pf00005pf08402 -
02069 -pf07883pf01381 -
02070 -pf10282 -
02071 -pf01264 -
BMC-P
BMC-H
BMC-T
Aldehyde dehydrogenase
PTAC
Signature enzyme
Alcohol dehydrogenase
Regulator

uncultured Clostridium sp. Bacteria; Firmicutes; Clostridia; Clostridiales; Clostridiaceae;

SAMEA3545387__locus_1

EUT2B

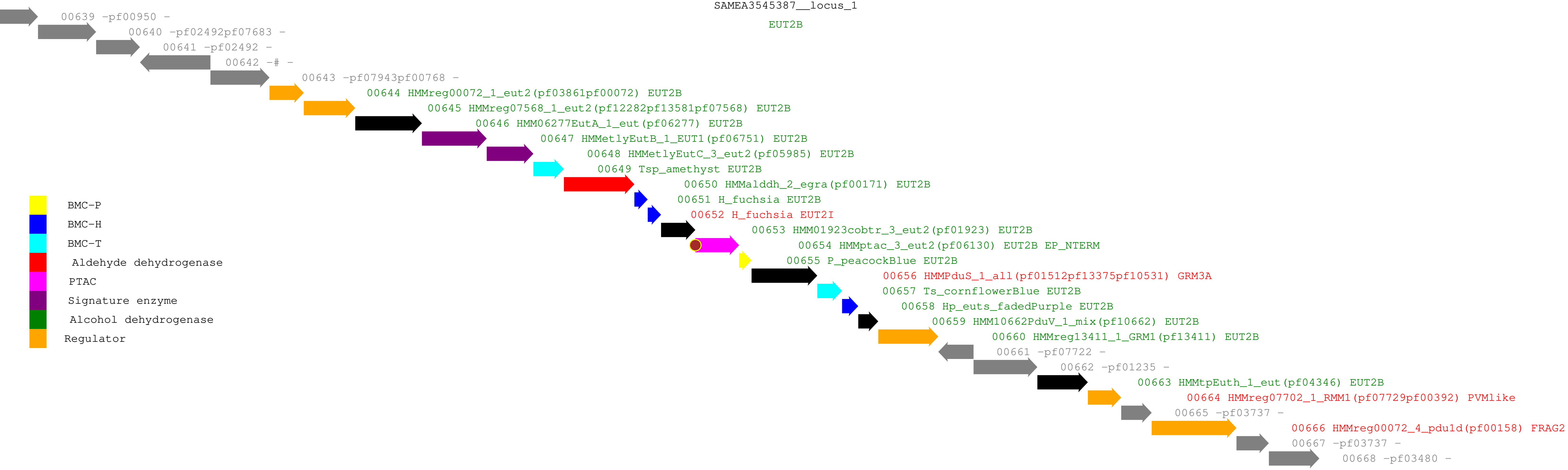

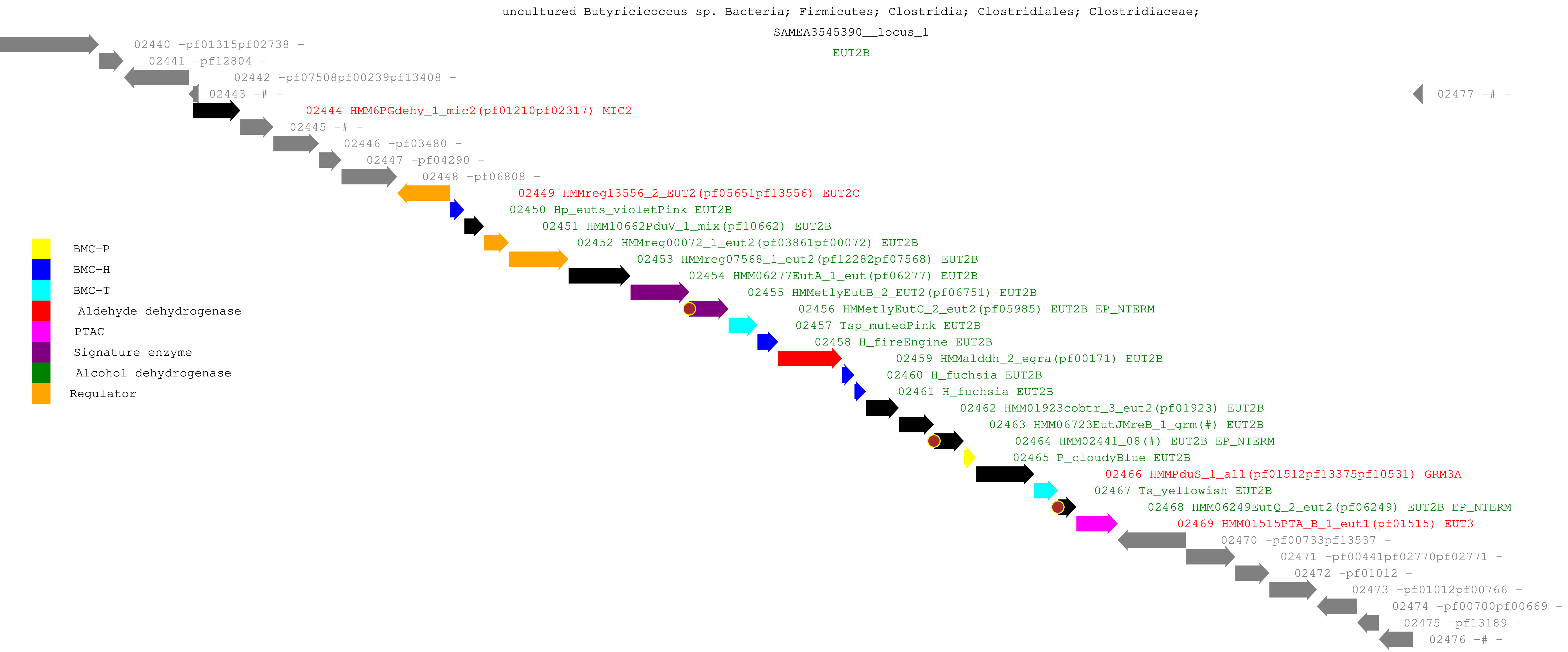
uncultured Butyricicoccus sp. Bacteria; Firmicutes; Clostridia; Clostridiales; Clostridiaceae;
SAMEA3545390__locus_1
EUT2B
02440 -pf01315pf02738 -
02441 -pf12804 -
02442 -pf07508pf00239pf13408 -
02443 -# -
02444 HMM6PGdehy_1_mic2(pf01210pf02317) MIC2
02445 -# -
02446 -pf03480 -
02447 -pf04290 -
02448 -pf06808 -
02449 HMMreg13556_2_EUT2(pf05651pf13556) EUT2C
02450 Hp_euts_violetPink EUT2B
02451 HMM10662PduV_1_mix(pf10662) EUT2B
02452 HMMreg00072_1_eut2(pf03861pf00072) EUT2B
02453 HMMreg07568_1_eut2(pf12282pf07568) EUT2B
02454 HMM06277EutA_1_eut(pf06277) EUT2B
02455 HMMetlyEutB_2_EUT2(pf06751) EUT2B
02456 HMMetlyEutC_2_eut2(pf05985) EUT2B EP_NTERM
02457 Tsp_mutedPink EUT2B
02458 H_fireEngine EUT2B
02459 HMMalddh_2_egra(pf00171) EUT2B
02460 H_fuchsia EUT2B
02461 H_fuchsia EUT2B
02462 HMM01923cobtr_3_eut2(pf01923) EUT2B
02463 HMM06723EutJMreB_1_grm(#) EUT2B
02464 HMM02441_08(#) EUT2B EP_NTERM
02465 P_cloudyBlue EUT2B
02466 HMMPduS_1_all(pf01512pf13375pf10531) GRM3A
02467 Ts_yellowish EUT2B
02468 HMM06249EutQ_2_eut2(pf06249) EUT2B EP_NTERM
02469 HMM01515PTA_B_1_eut1(pf01515) EUT3
02470 -pf00733pf13537 -
02471 -pf00441pf02770pf02771 -
02472 -pf01012 -
02473 -pf01012pf00766 -
02474 -pf00700pf00669 -
02475 -pf13189 -
02476 -# -
02477 -# -
BMC-P
BMC-H
BMC-T
Aldehyde dehydrogenase
PTAC
Signature enzyme
Alcohol dehydrogenase
Regulator

uncultured Flavonifractor sp. Bacteria; Firmicutes; Clostridia; Clostridiales; Flavonifractor;

SAMEA3545396__locus_1

EUT2B

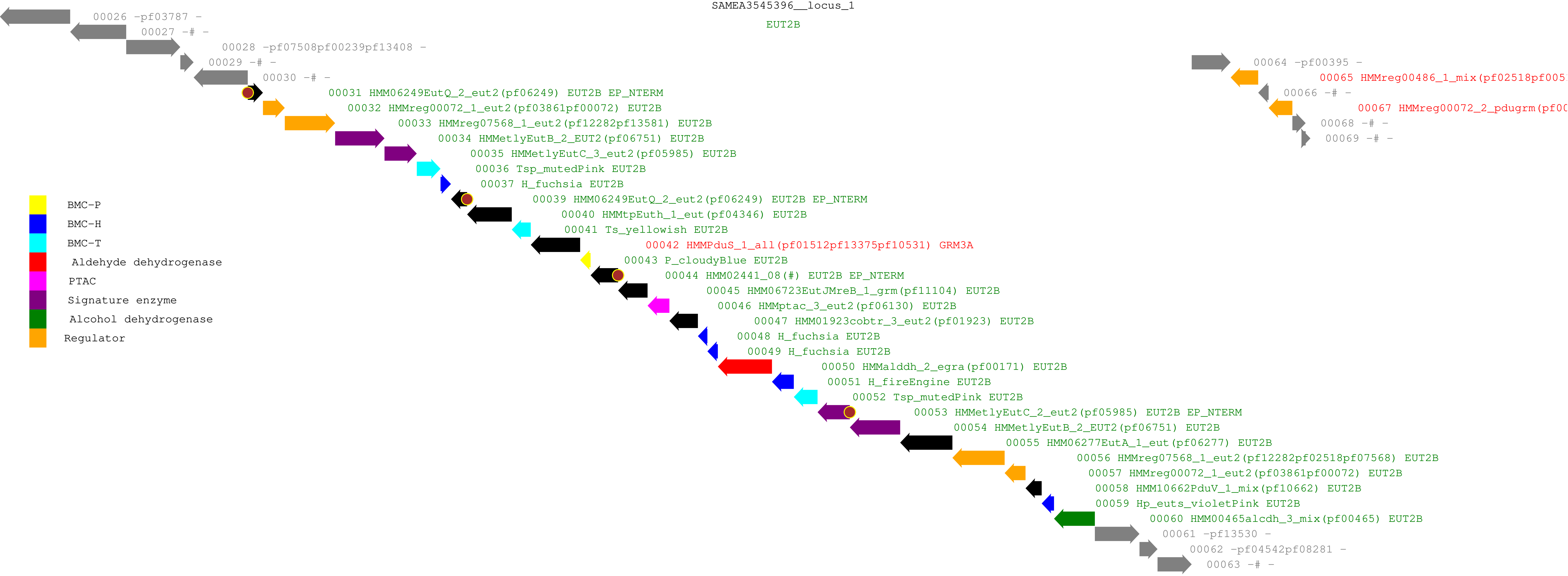

uncultured Clostridium sp. Bacteria; Firmicutes; Clostridia; Clostridiales; Clostridiaceae;

SAMEA3545399__locus_2_main_sat

EUT2B

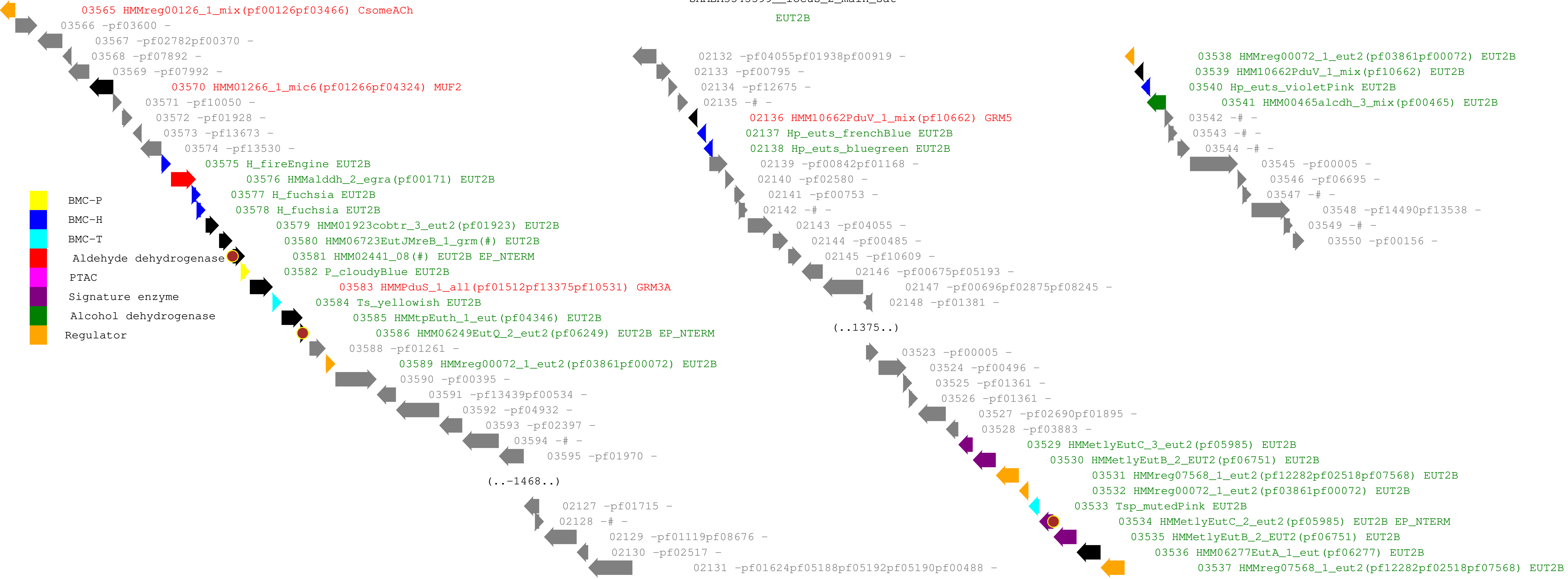

Maledivibacter halophilus. Bacteria; Firmicutes; Clostridia; Clostridiales; Clostridiaceae;

SAMN02194393__locus_2

EUT2B

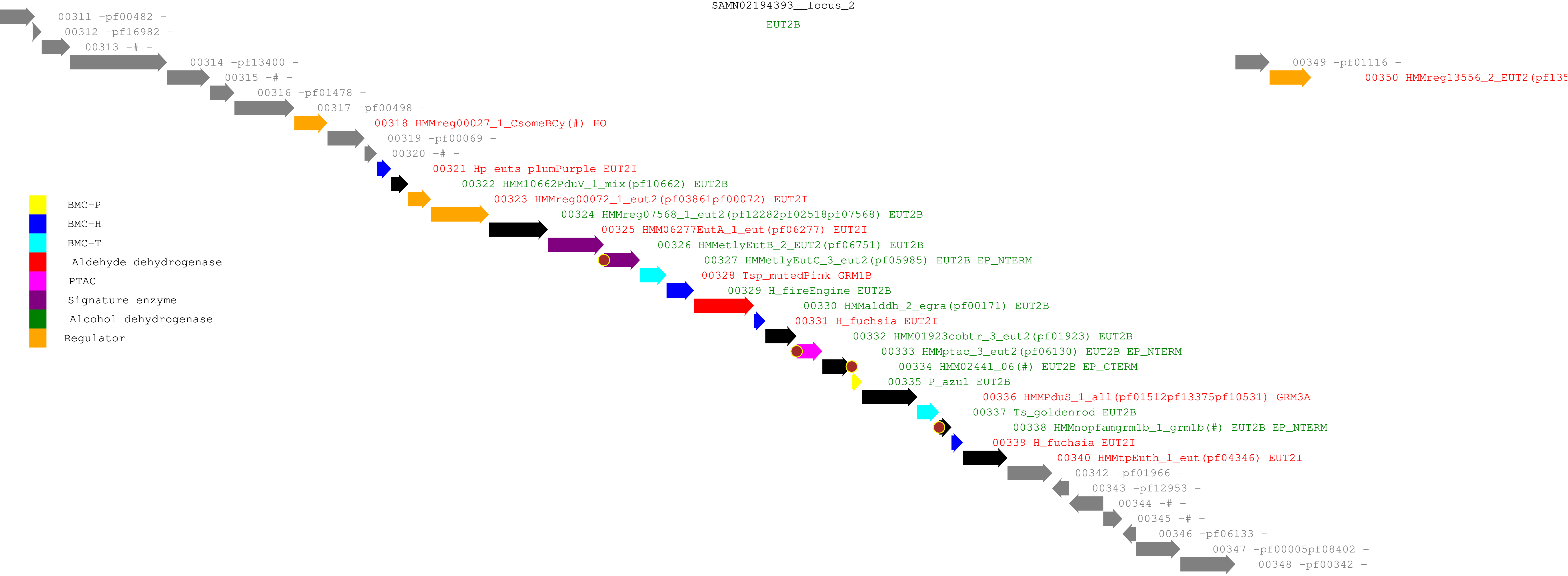

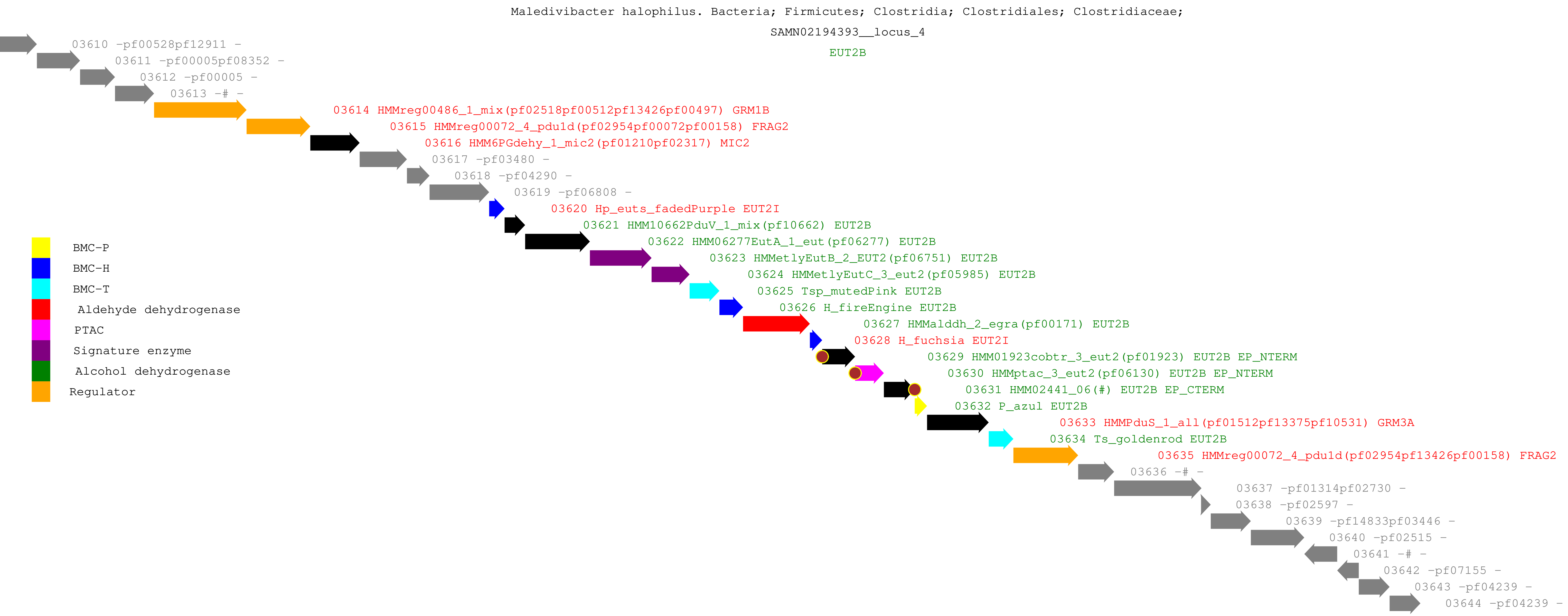
Maledivibacter halophilus. Bacteria; Firmicutes; Clostridia; Clostridiales; Clostridiaceae;
SAMN02194393__locus_4
EUT2B
03610 -pf00528pf12911 -
03611 -pf00005pf08352 -
03612 -pf00005 -
03613 -# -
03614 HMMreg00486_1_mix(pf02518pf00512pf13426pf00497) GRM1B
03615 HMMreg00072_4_pdu1d(pf02954pf00072pf00158) FRAG2
03616 HMM6PGdehy_1_mic2(pf01210pf02317) MIC2
03617 -pf03480 -
03618 -pf04290 -
03619 -pf06808 -
03620 Hp_euts_fadedPurple EUT2I
03621 HMM10662PduV_1_mix(pf10662) EUT2B
03622 HMM06277EutA_1_eut(pf06277) EUT2B
03623 HMMetlyEutB_2_EUT2(pf06751) EUT2B
03624 HMMetlyEutC_3_eut2(pf05985) EUT2B
03625 Tsp_mutedPink EUT2B
03626 H_fireEngine EUT2B
03627 HMMalddh_2_egra(pf00171) EUT2B
03628 H_fuchsia EUT2I
03629 HMM01923cobtr_3_eut2(pf01923) EUT2B EP_NTERM
03630 HMMptac_3_eut2(pf06130) EUT2B EP_NTERM
03631 HMM02441_06(#) EUT2B EP_CTERM
03632 P_azul EUT2B
03633 HMMPduS_1_all(pf01512pf13375pf10531) GRM3A
03634 Ts_goldenrod EUT2B
03635 HMMreg00072_4_pdu1d(pf02954pf13426pf00158) FRAG2
03636 -# -
03637 -pf01314pf02730 -
03638 -pf02597 -
03639 -pf14833pf03446 -
03640 -pf02515 -
03641 -# -
03642 -pf07155 -
03643 -pf04239 -
03644 -pf04239 -
BMC-P
BMC-H
BMC-T
Aldehyde dehydrogenase
PTAC
Signature enzyme
Alcohol dehydrogenase
Regulator

Acetoanaerobium noterae. Bacteria; Firmicutes; Clostridia; Clostridiales;

SAMN02745120__locus_1

EUT2B

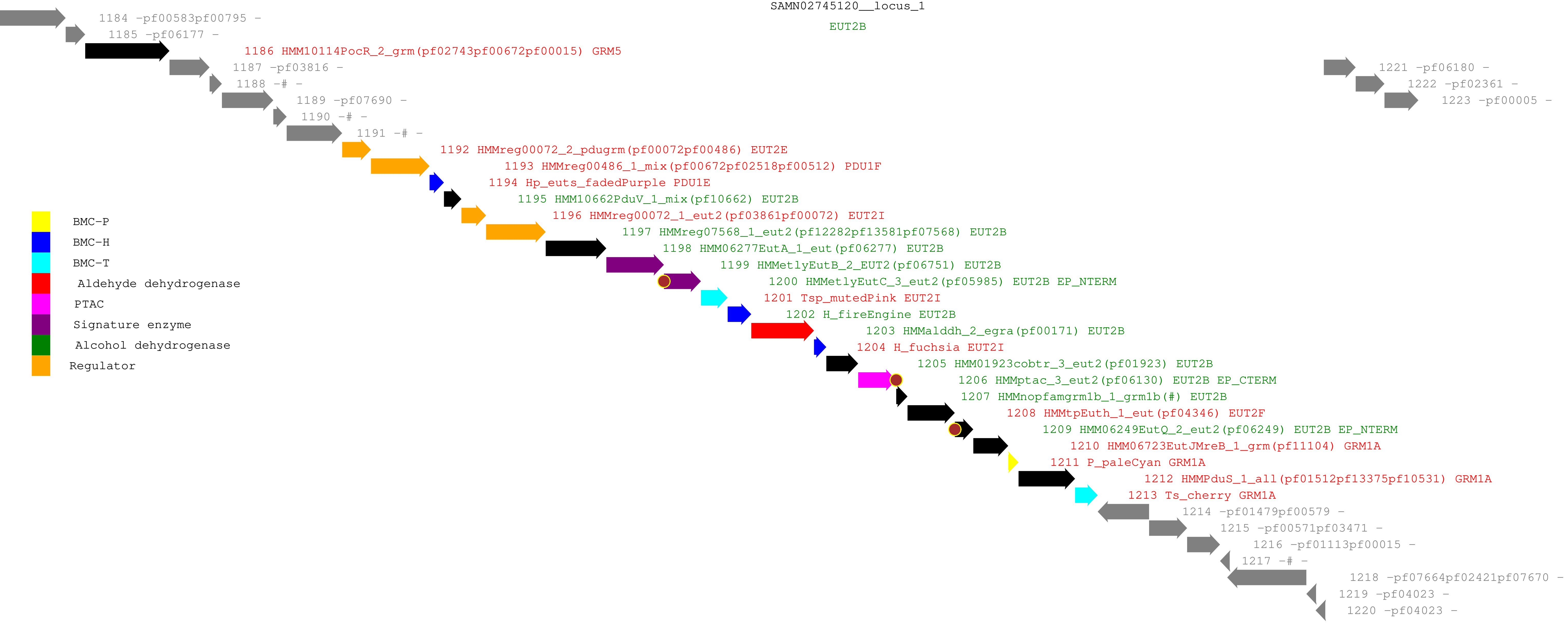

Lutispora thermophila DSM 19022. Bacteria; Firmicutes; Clostridia; Clostridiales; Clostridiaceae;

SAMN02745176__locus_1

EUT2B

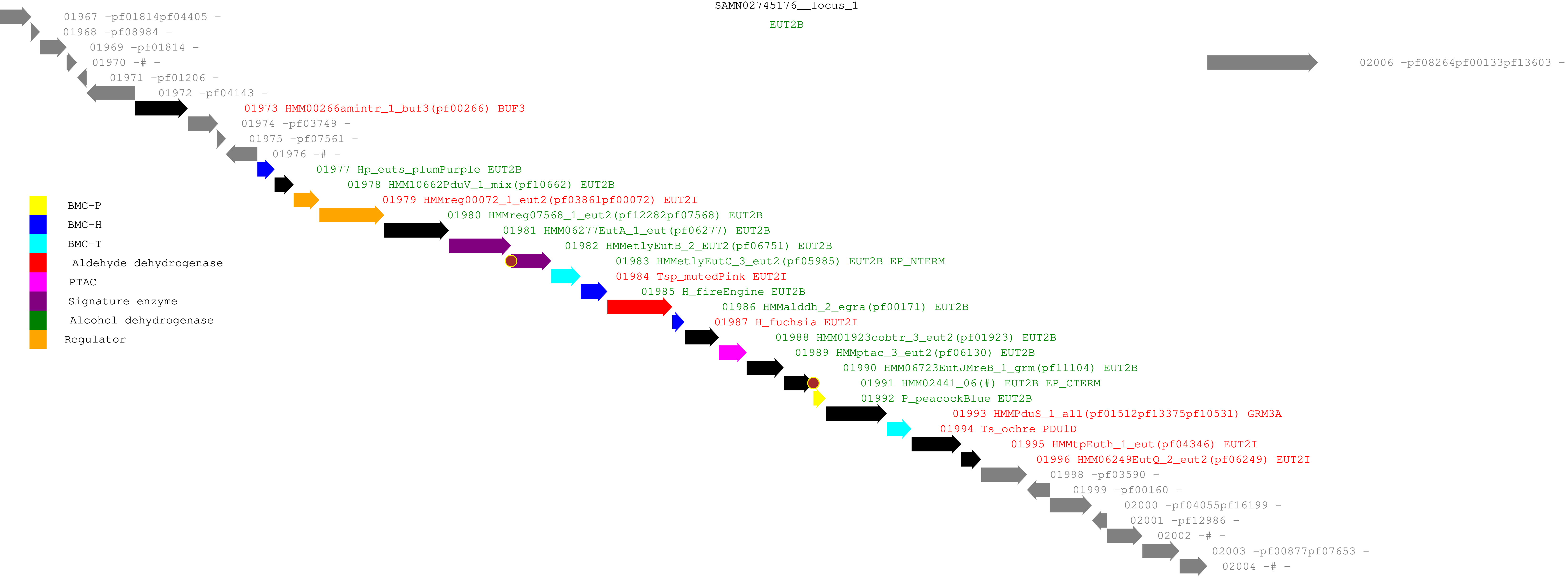

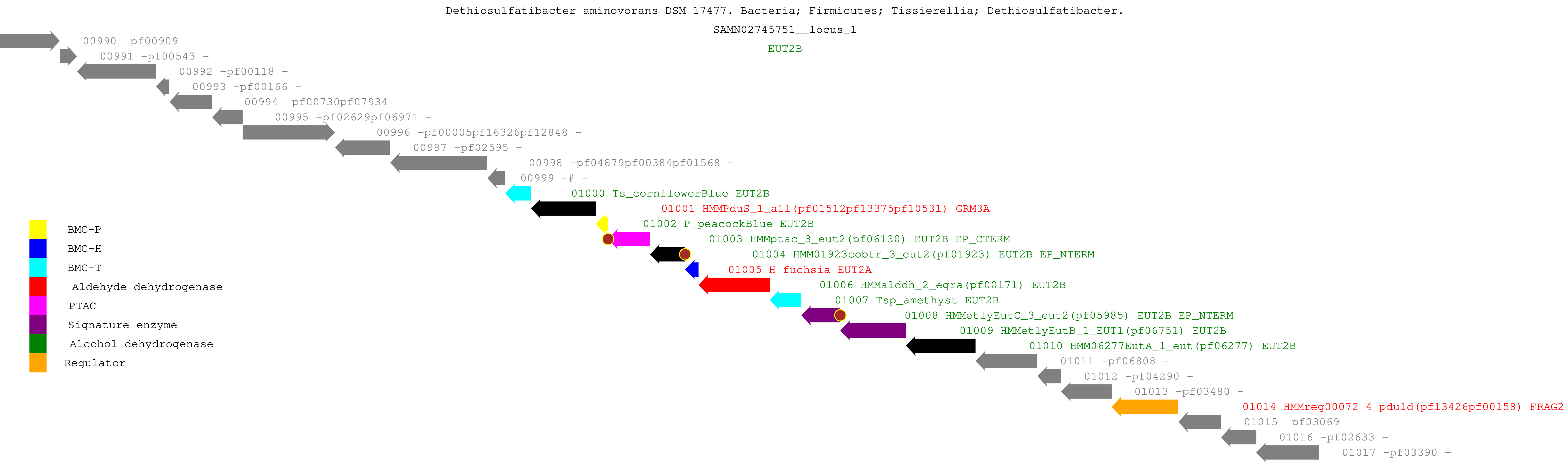
Dethiosulfatibacter aminovorans DSM 17477. Bacteria; Firmicutes; Tissierellia; Dethiosulfatibacter.
SAMN02745751__locus_1
EUT2B
00990 -pf00909 -
00991 -pf00543 -
00992 -pf00118 -
00993 -pf00166 -
00994 -pf00730pf07934 -
00995 -pf02629pf06971 -
00996 -pf00005pf16326pf12848 -
00997 -pf02595 -
00998 -pf04879pf00384pf01568 -
00999 -# -
01000 Ts_cornflowerBlue EUT2B
01001 HMMPduS_1_all(pf01512pf13375pf10531) GRM3A
01002 P_peacockBlue EUT2B
01003 HMMptac_3_eut2(pf06130) EUT2B EP_CTERM
01004 HMM01923cobtr_3_eut2(pf01923) EUT2B EP_NTERM
01005 H_fuchsia EUT2A
01006 HMMaldddh_2_egra(pf00171) EUT2B
01007 Tsp_amethyst EUT2B
01008 HMMetlyEutC_3_eut2(pf05985) EUT2B EP_NTERM
01009 HMMetlyEutB_1_EUT1(pf06751) EUT2B
01010 HMM06277EutA_1_eut(pf06277) EUT2B
01011 -pf06808 -
01012 -pf04290 -
01013 -pf03480 -
01014 HMMreg00072_4_pdu1d(pf13426pf00158) FRAG2
01015 -pf03069 -
01016 -pf02633 -
01017 -pf03390 -
BMC-P
BMC-H
BMC-T
Aldehyde dehydrogenase
PTAC
Signature enzyme
Alcohol dehydrogenase
Regulator

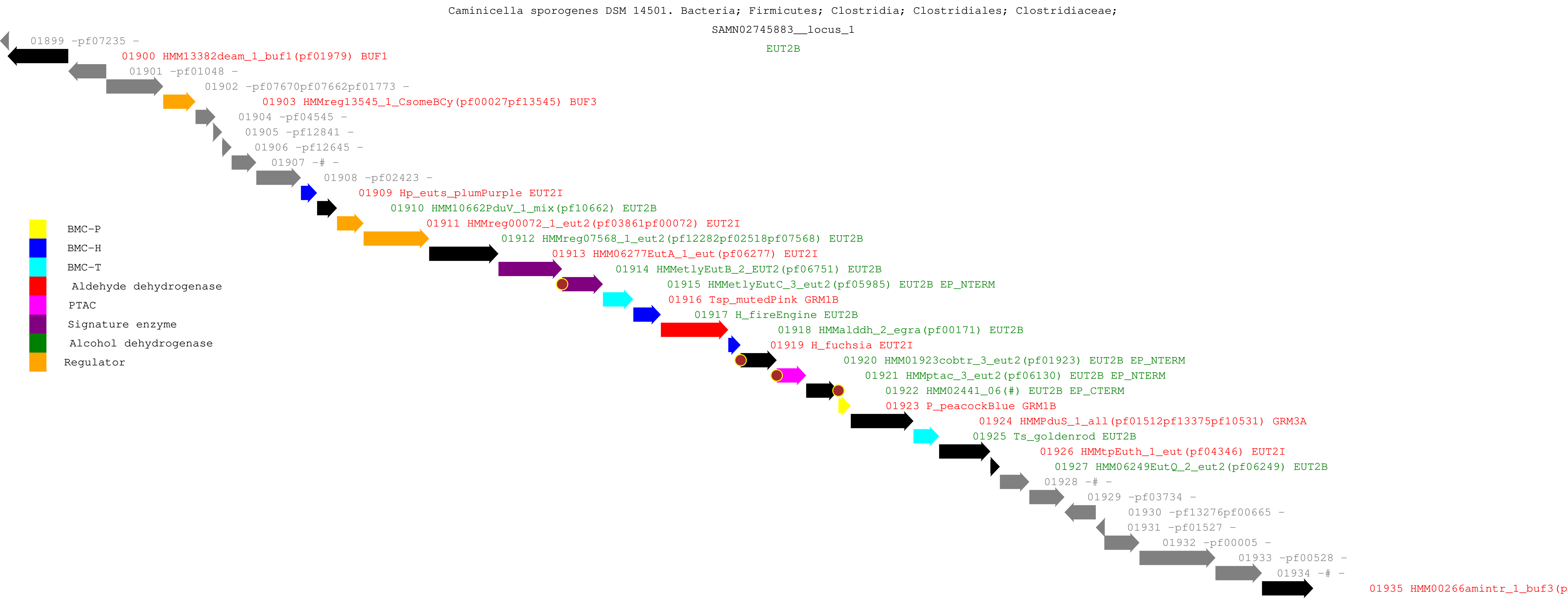
Caminicella sporogenes DSM 14501. Bacteria; Firmicutes; Clostridia; Clostridiales; Clostridiaceae;
SAMN02745883__locus_1
EUT2B
01899 -pf07235 -
01900 HMM13382deam_1_buf1(pf01979) BUF1
01901 -pf01048 -
01902 -pf07670pf07662pf01773 -
01903 HMMreg13545_1_CsomeBCy(pf00027pf13545) BUF3
01904 -pf04545 -
01905 -pf12841 -
01906 -pf12645 -
01907 -# -
01908 -pf02423 -
01909 Hp_euts_plumPurple EUT2I
01910 HMM10662PduV_1_mix(pf10662) EUT2B
01911 HMMreg00072_1_eut2(pf03861pf00072) EUT2I
01912 HMMreg07568_1_eut2(pf12282pf02518pf07568) EUT2B
01913 HMM06277EutA_1_eut(pf06277) EUT2I
01914 HMMetlyEutB_2_EUT2(pf06751) EUT2B
01915 HMMetlyEutC_3_eut2(pf05985) EUT2B EP_NTERM
01916 Tsp_mutedPink GRM1B
01917 H_fireEngine EUT2B
01918 HMMaldh_2_egra(pf00171) EUT2B
01919 H_fuchsia EUT2I
01920 HMM01923cobtr_3_eut2(pf01923) EUT2B EP_NTERM
01921 HMMptac_3_eut2(pf06130) EUT2B EP_NTERM
01922 HMM02441_06(#) EUT2B EP_CTERM
01923 P_peacockBlue GRM1B
01924 HMMPduS_1_all(pf01512pf13375pf10531) GRM3A
01925 Ts_goldenrod EUT2B
01926 HMMtpEuth_1_eut(pf04346) EUT2I
01927 HMM06249EutQ_2_eut2(pf06249) EUT2B
01928 -# -
01929 -pf03734 -
01930 -pf13276pf00665 -
01931 -pf01527 -
01932 -pf00005 -
01933 -pf00528 -
01934 -# -
01935 HMM00266amintr_1_buf3(p
BMC-P
BMC-H
BMC-T
Aldehyde dehydrogenase
PTAC
Signature enzyme
Alcohol dehydrogenase
Regulator

Clostridium caminithermale DSM 15212. Bacteria; Firmicutes; Clostridia; Clostridiales; Clostridiaceae;

SAMN02745912__locus_2

EUT2B

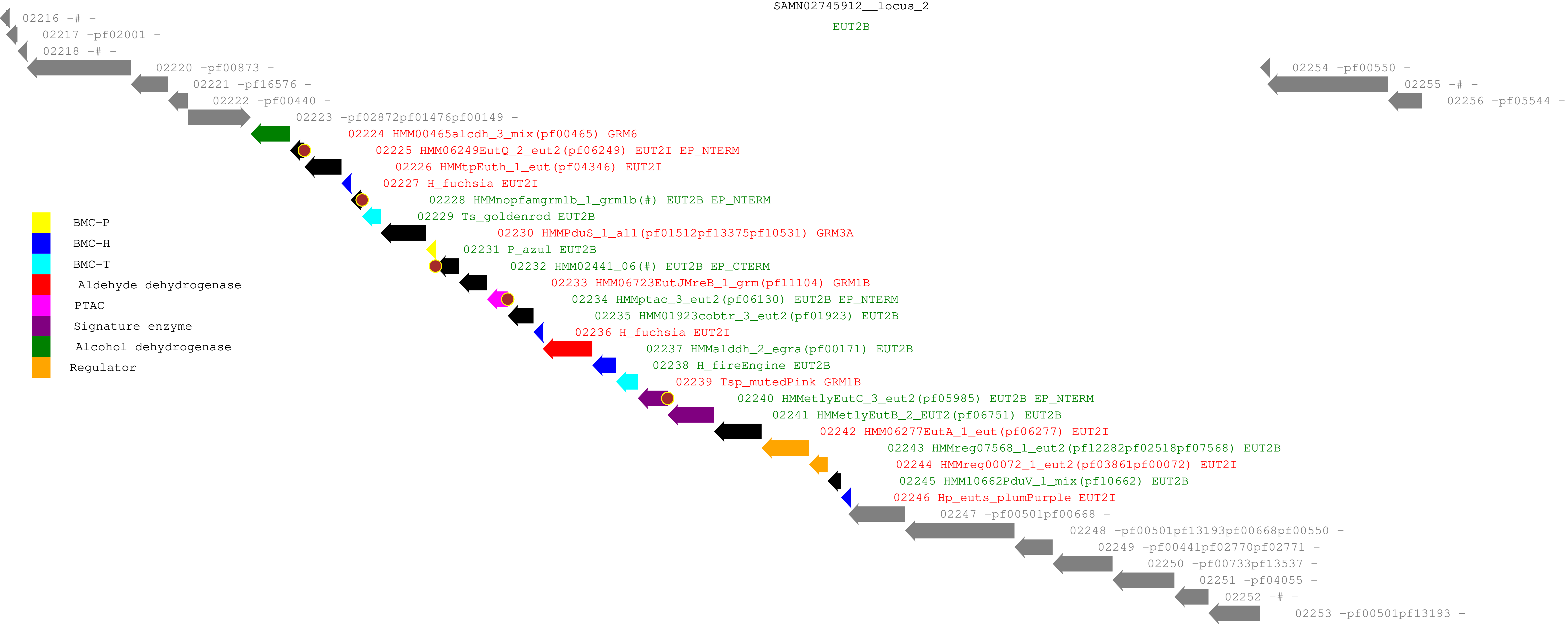

Geosporobacter subterraneus DSM 17957. Bacteria; Firmicutes; Clostridia; Clostridiales; Clostridiaceae;

SAMN02745975__locus_2

EUT2B

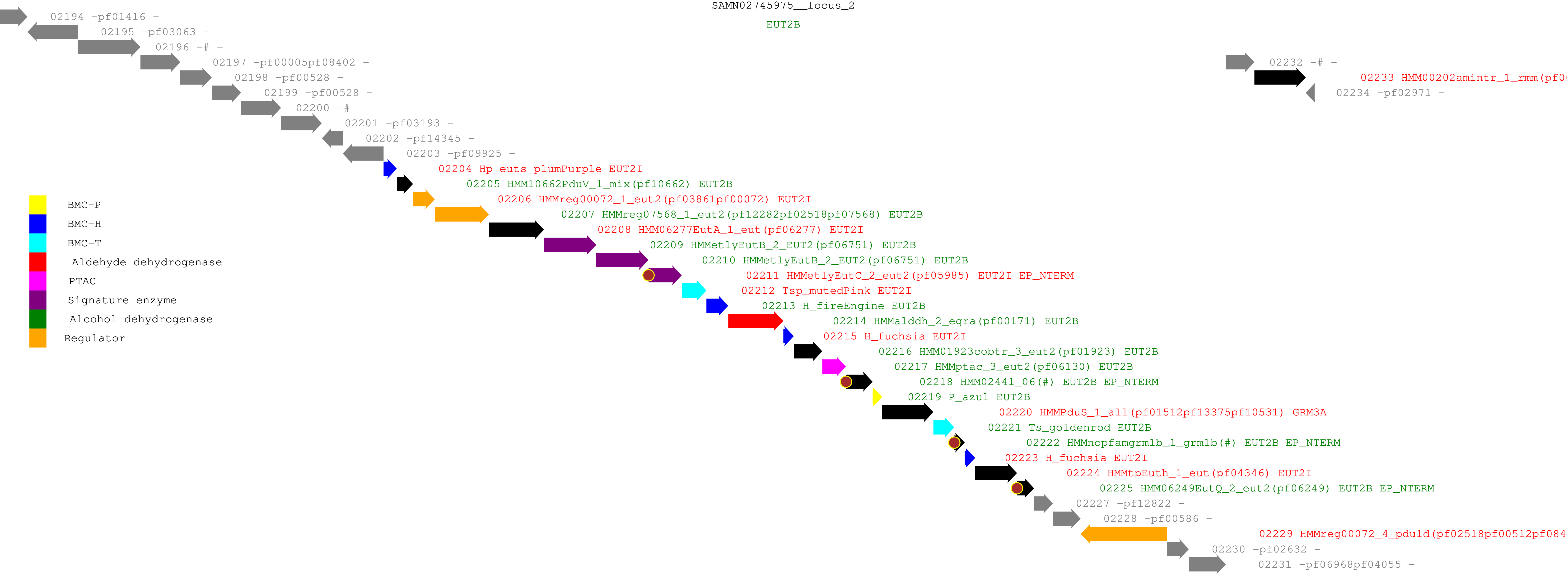

Butyricicoccus pullicaecorum DSM 23266. Bacteria; Firmicutes; Clostridia; Clostridiales; Clostridiaceae;

SAMN02745978__locus_1

EUT2B

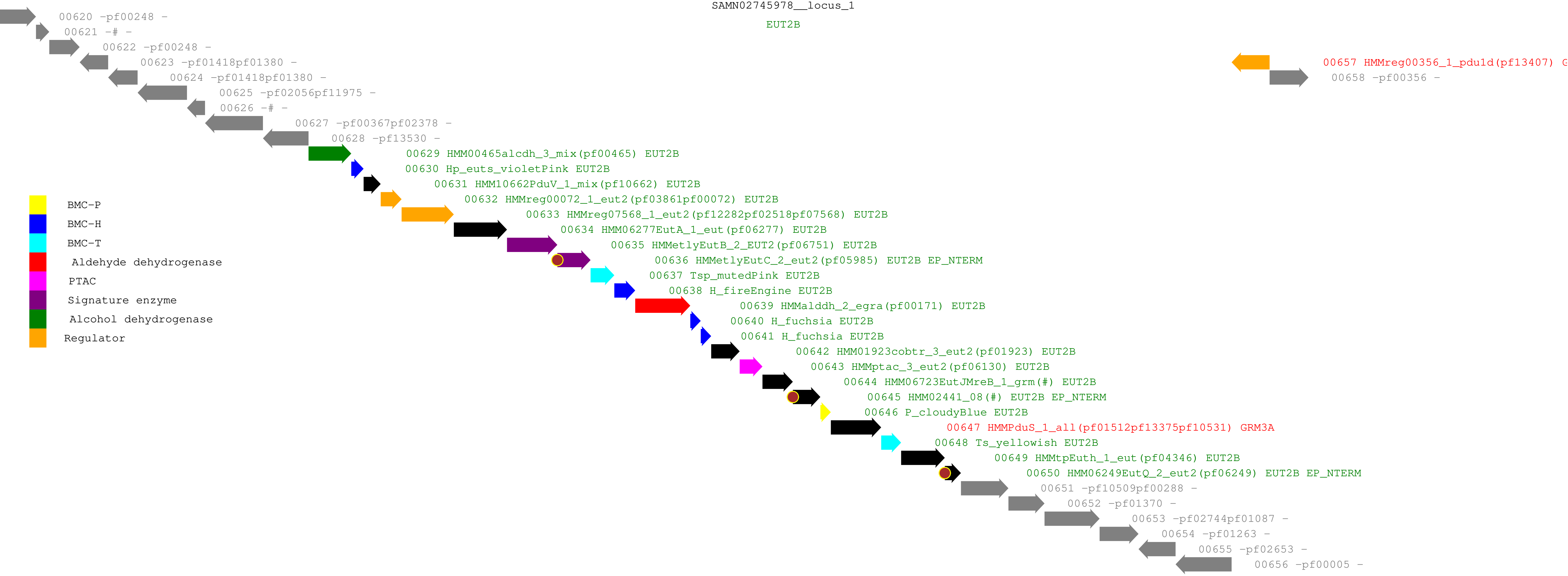

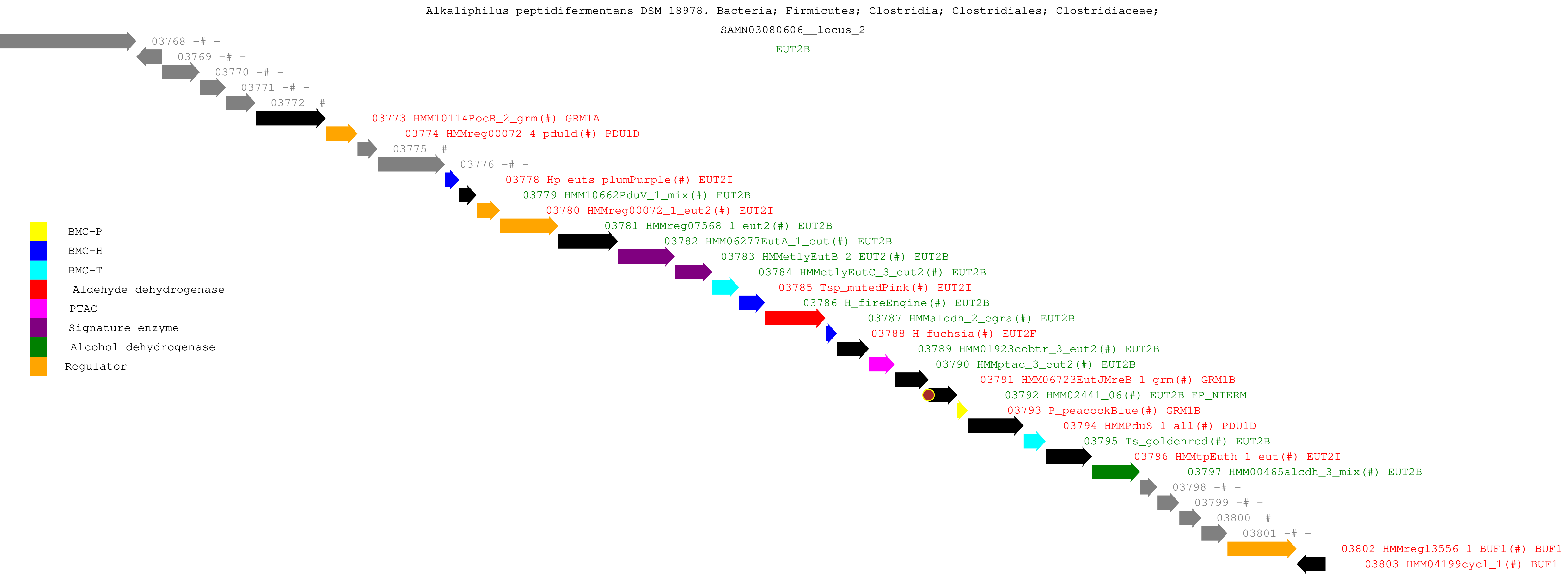
Alkaliphilus peptidifermentans DSM 18978. Bacteria; Firmicutes; Clostridia; Clostridiales; Clostridiaceae;
SAMN03080606__locus_2
EUT2B
03768 -# -
03769 -# -
03770 -# -
03771 -# -
03772 -# -
03773 HMM10114PocR_2_grm(#) GRM1A
03774 HMMreg00072_4_pdu1d(#) PDU1D
03775 -# -
03776 -# -
03778 Hp_euts_plumPurple(#) EUT2I
03779 HMM10662PduV_1_mix(#) EUT2B
03780 HMMreg00072_1_eut2(#) EUT2I
03781 HMMreg07568_1_eut2(#) EUT2B
03782 HMM06277EutA_1_eut(#) EUT2B
03783 HMMetlyEutB_2_EUT2(#) EUT2B
03784 HMMetlyEutC_3_eut2(#) EUT2B
03785 Tsp_mutedPink(#) EUT2I
03786 H_fireEngine(#) EUT2B
03787 HMMalddh_2_egra(#) EUT2B
03788 H_fuchsia(#) EUT2F
03789 HMM01923cobtr_3_eut2(#) EUT2B
03790 HMMptac_3_eut2(#) EUT2B
03791 HMM06723EutJMreB_1_grm(#) GRM1B
03792 HMM02441_06(#) EUT2B EP_NTERM
03793 P_peacockBlue(#) GRM1B
03794 HMMPduS_1_all(#) PDU1D
03795 Ts_goldenrod(#) EUT2B
03796 HMMtpEuth_1_eut(#) EUT2I
03797 HMM00465alcdh_3_mix(#) EUT2B
03798 -# -
03799 -# -
03800 -# -
03801 -# -
03802 HMMreg13556_1_BUF1(#) BUF1
03803 HMM04199cycl_1(#) BUF1
BMC-P
BMC-H
BMC-T
Aldehyde dehydrogenase
PTAC
Signature enzyme
Alcohol dehydrogenase
Regulator

Eubacterium barkeri (Clostridium barkeri). Bacteria; Firmicutes; Clostridia; Clostridiales; Eubacteriaceae;

SAMN04488579__locus_1

EUT2B

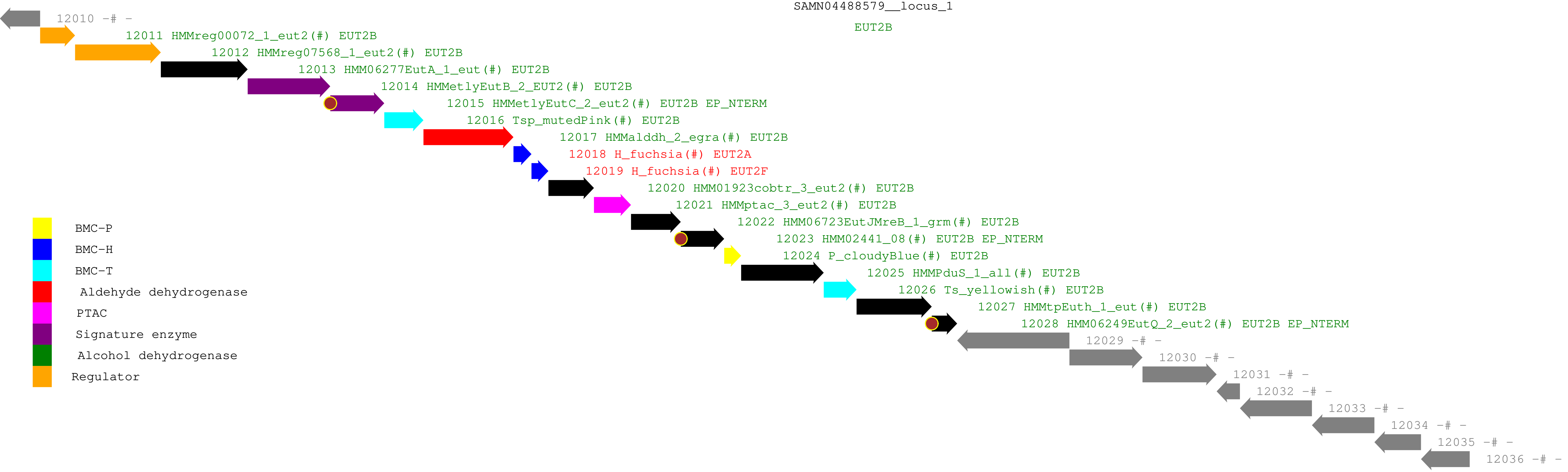

Halanaerobium congolense. Bacteria; Firmicutes; Clostridia; Halanaerobiales; Halanaerobiaceae;

SAMN04488598__locus_1

EUT2B

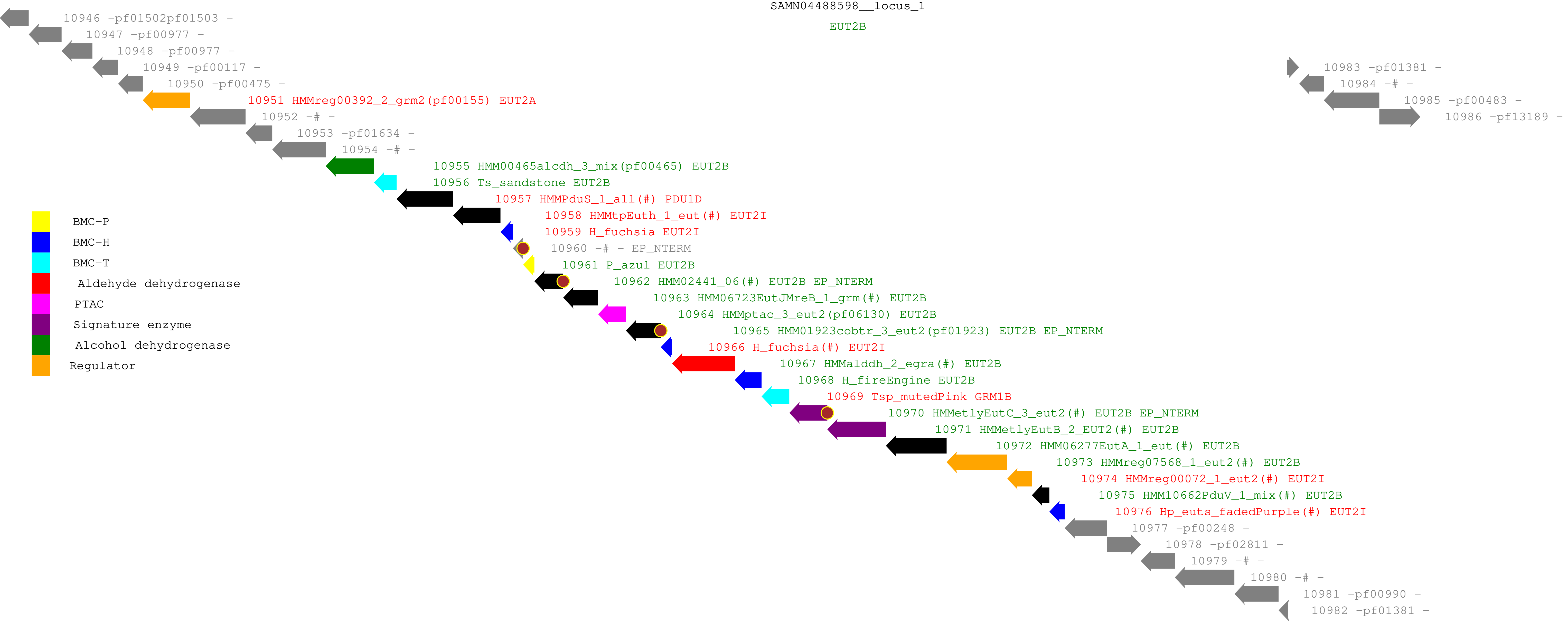

Proteiniclasticum ruminis. Bacteria; Firmicutes; Clostridia; Clostridiales; Clostridiaceae;

SAMN04488695__locus_1

EUT2B

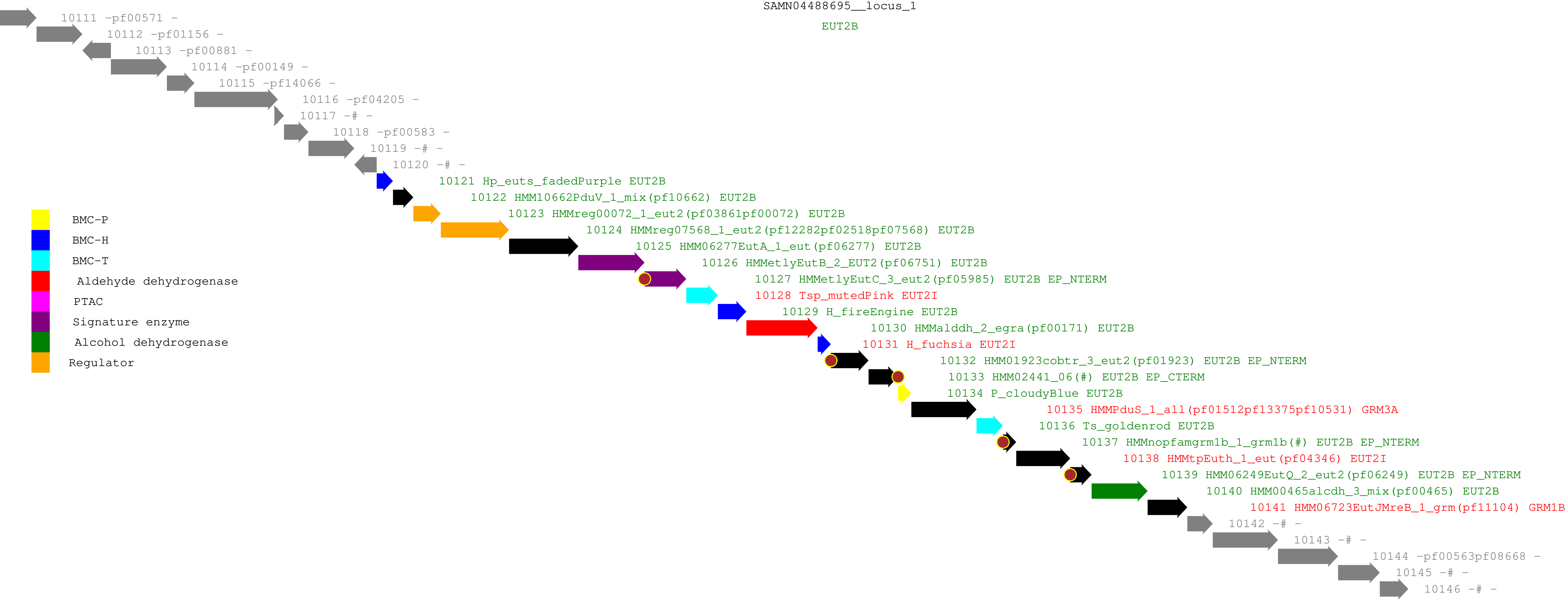

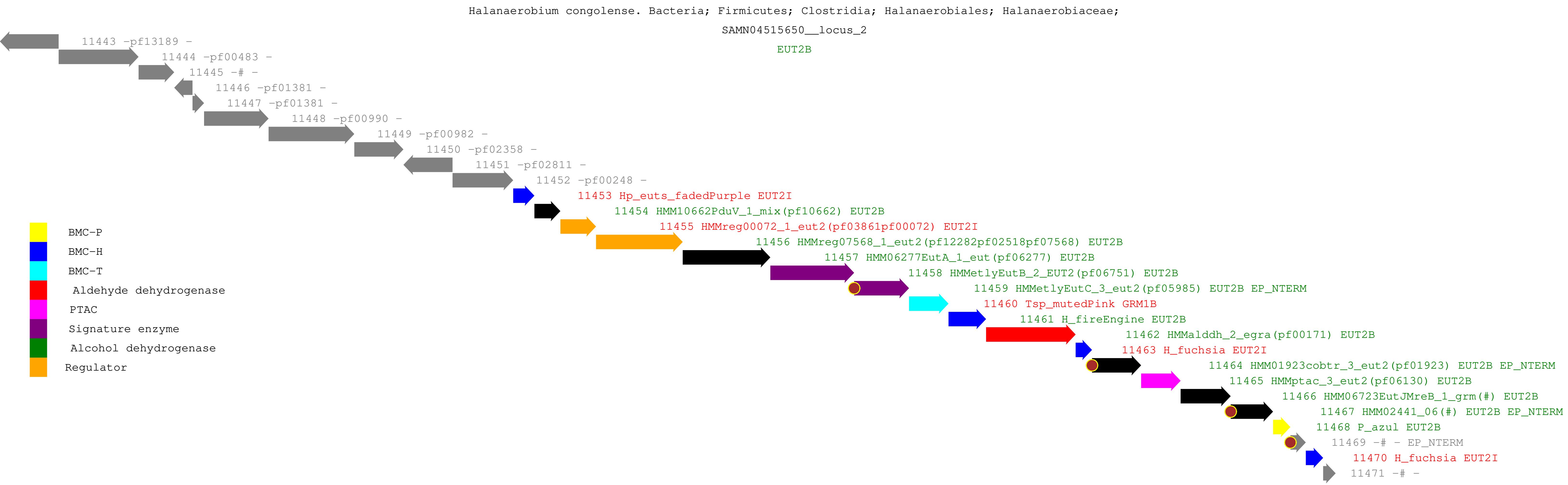
Halanaerobium congolense. Bacteria; Firmicutes; Clostridia; Halanaerobiales; Halanaerobiaceae;
SAMN04515650__locus_2
EUT2B
11443 -pf13189 -
11444 -pf00483 -
11445 -# -
11446 -pf01381 -
11447 -pf01381 -
11448 -pf00990 -
11449 -pf00982 -
11450 -pf02358 -
11451 -pf02811 -
11452 -pf00248 -
11453 Hp_euts_fadedPurple EUT2I
11454 HMM10662PduV_1_mix(pf10662) EUT2B
11455 HMMreg00072_1_eut2(pf03861pf00072) EUT2I
11456 HMMreg07568_1_eut2(pf12282pf02518pf07568) EUT2B
11457 HMM06277EutA_1_eut(pf06277) EUT2B
11458 HMMetlyEutB_2_EUT2(pf06751) EUT2B
11459 HMMetlyEutC_3_eut2(pf05985) EUT2B EP_NTERM
11460 Tsp_mutedPink GRM1B
11461 H_fireEngine EUT2B
11462 HMMalddh_2_egra(pf00171) EUT2B
11463 H_fuchsia EUT2I
11464 HMM01923cobtr_3_eut2(pf01923) EUT2B EP_NTERM
11465 HMMptac_3_eut2(pf06130) EUT2B
11466 HMM06723EutJMreB_1_grm(#) EUT2B
11467 HMM02441_06(#) EUT2B EP_NTERM
11468 P_azul EUT2B
11469 -# - EP_NTERM
11470 H_fuchsia EUT2I
11471 -# -
BMC-P
BMC-H
BMC-T
Aldehyde dehydrogenase
PTAC
Signature enzyme
Alcohol dehydrogenase
Regulator

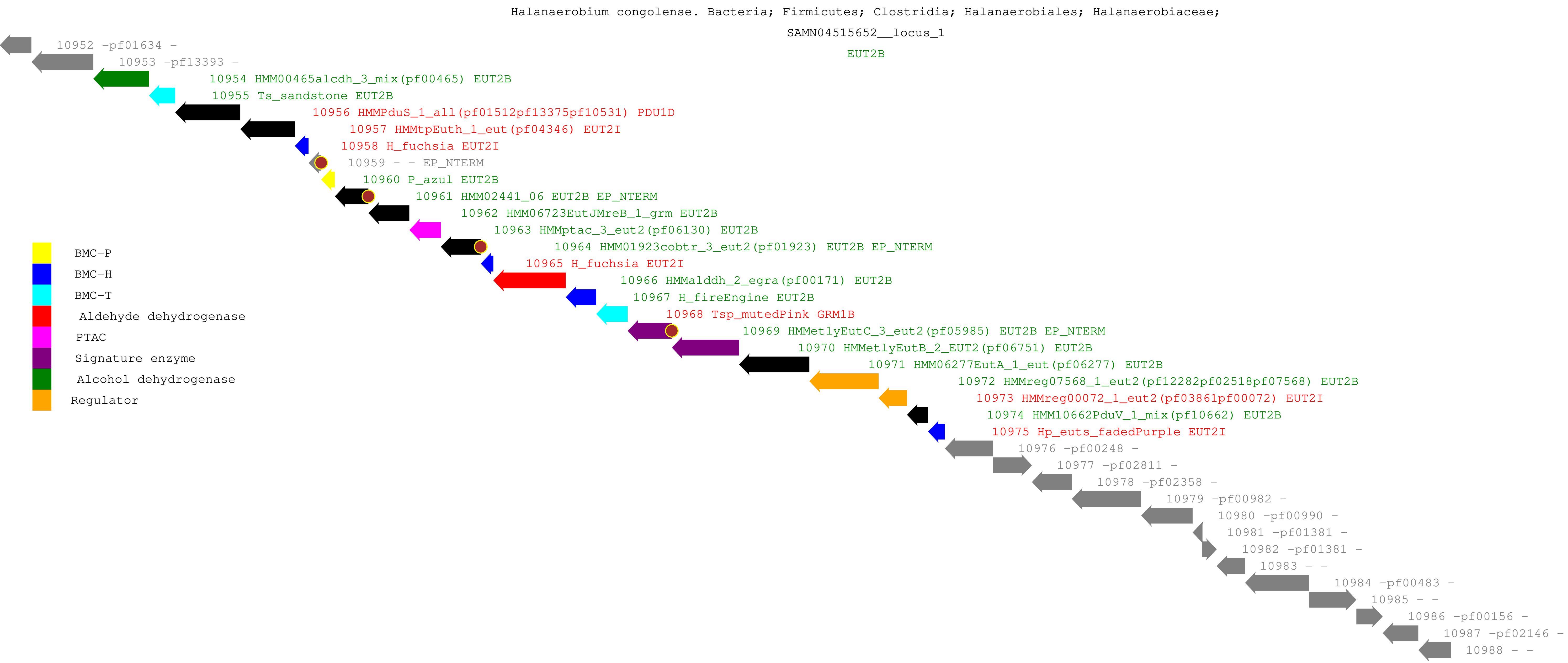
Halanaerobium congolense. Bacteria; Firmicutes; Clostridia; Halanaerobiales; Halanaerobiaceae;
SAMN04515652__locus_1
EUT2B
10952 -pf01634 -
10953 -pf13393 -
10954 HMM00465alcdh_3_mix(pf00465) EUT2B
10955 Ts_sandstone EUT2B
10956 HMMPduS_1_all(pf01512pf13375pf10531) PDU1D
10957 HMMtpEuth_1_eut(pf04346) EUT2I
10958 H_fuchsia EUT2I
10959 - - EP_NTERM
10960 P_azul EUT2B
10961 HMM02441_06 EUT2B EP_NTERM
10962 HMM06723EutJMreB_1_grm EUT2B
10963 HMMptac_3_eut2(pf06130) EUT2B
10964 HMM01923cobtr_3_eut2(pf01923) EUT2B EP_NTERM
10965 H_fuchsia EUT2I
10966 HMMalddh_2_egra(pf00171) EUT2B
10967 H_fireEngine EUT2B
10968 Tsp_mutedPink GRM1B
10969 HMMetlyEutC_3_eut2(pf05985) EUT2B EP_NTERM
10970 HMMetlyEutB_2_EUT2(pf06751) EUT2B
10971 HMM06277EutA_1_eut(pf06277) EUT2B
10972 HMMreg07568_1_eut2(pf12282pf02518pf07568) EUT2B
10973 HMMreg00072_1_eut2(pf03861pf00072) EUT2I
10974 HMM10662PduV_1_mix(pf10662) EUT2B
10975 Hp_euts_fadedPurple EUT2I
10976 -pf00248 -
10977 -pf02811 -
10978 -pf02358 -
10979 -pf00982 -
10980 -pf00990 -
10981 -pf01381 -
10982 -pf01381 -
10983 - -
10984 -pf00483 -
10985 - -
10986 -pf00156 -
10987 -pf02146 -
10988 - -
BMC-P
BMC-H
BMC-T
Aldehyde dehydrogenase
PTAC
Signature enzyme
Alcohol dehydrogenase
Regulator

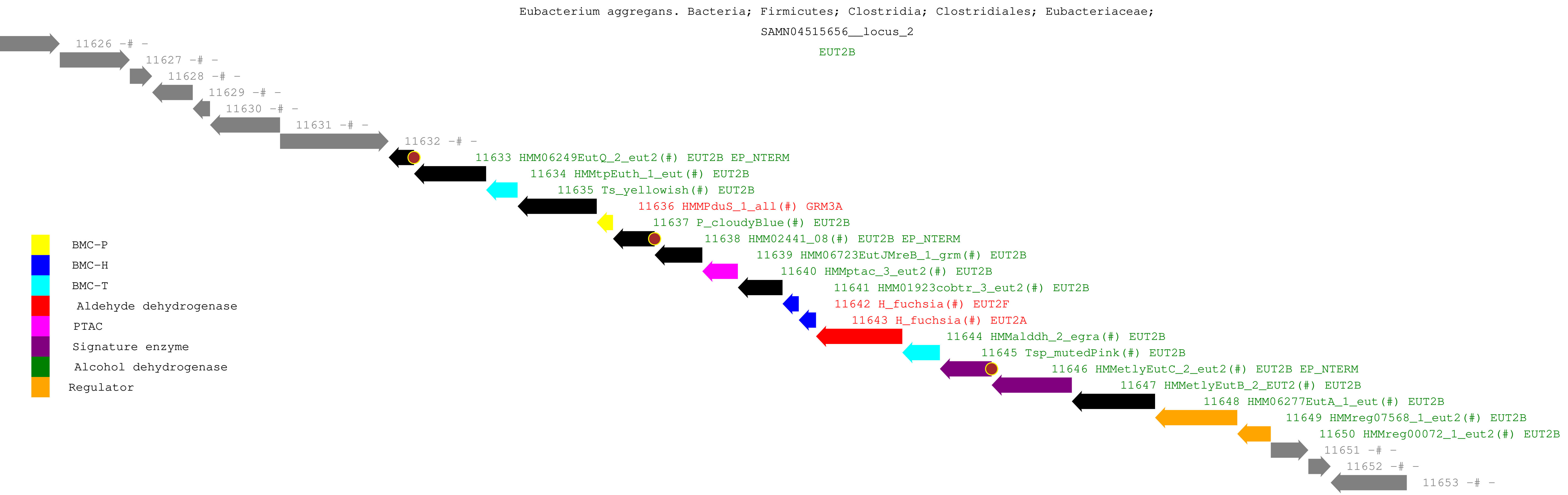
Eubacterium aggregans. Bacteria; Firmicutes; Clostridia; Clostridiales; Eubacteriaceae;
SAMN04515656__locus_2
EUT2B
11626 -# -
11627 -# -
11628 -# -
11629 -# -
11630 -# -
11631 -# -
11632 -# -
11633 HMM06249EutQ_2_eut2(#) EUT2B EP_NTERM
11634 HMMtpEuth_1_eut(#) EUT2B
11635 Ts_yellowish(#) EUT2B
11636 HMMPduS_1_all(#) GRM3A
11637 P_cloudyBlue(#) EUT2B
11638 HMM02441_08(#) EUT2B EP_NTERM
11639 HMM06723EutJMreB_1_grm(#) EUT2B
11640 HMMptac_3_eut2(#) EUT2B
11641 HMM01923cobtr_3_eut2(#) EUT2B
11642 H_fuchsia(#) EUT2F
11643 H_fuchsia(#) EUT2A
11644 HMMaldh_2_egra(#) EUT2B
11645 Tsp_mutedPink(#) EUT2B
11646 HMMetlyEutC_2_eut2(#) EUT2B EP_NTERM
11647 HMMetlyEutB_2_EUT2(#) EUT2B
11648 HMM06277EutA_1_eut(#) EUT2B
11649 HMMreg07568_1_eut2(#) EUT2B
11650 HMMreg00072_1_eut2(#) EUT2B
11651 -# -
11652 -# -
11653 -# -
BMC-P
BMC-H
BMC-T
Aldehyde dehydrogenase
PTAC
Signature enzyme
Alcohol dehydrogenase
Regulator

Tindallia californiensis. Bacteria; Firmicutes; Clostridia; Clostridiales; Clostridiaceae;

SAMN05192546__locus_1

EUT2B

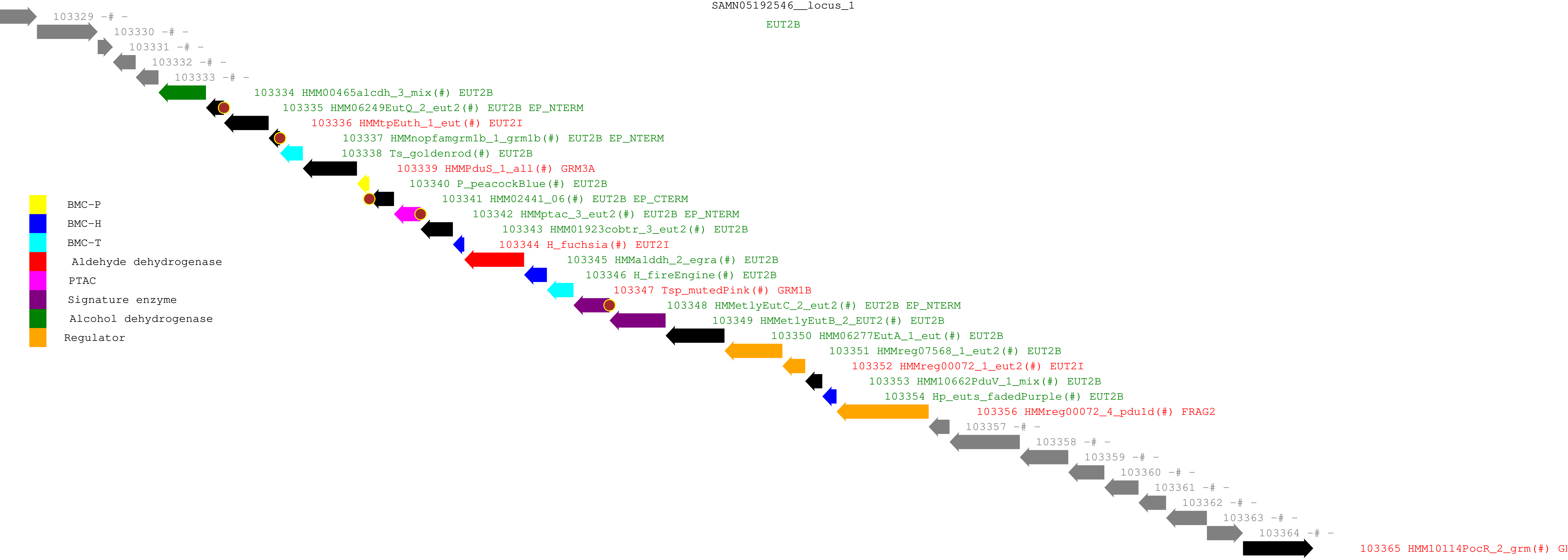

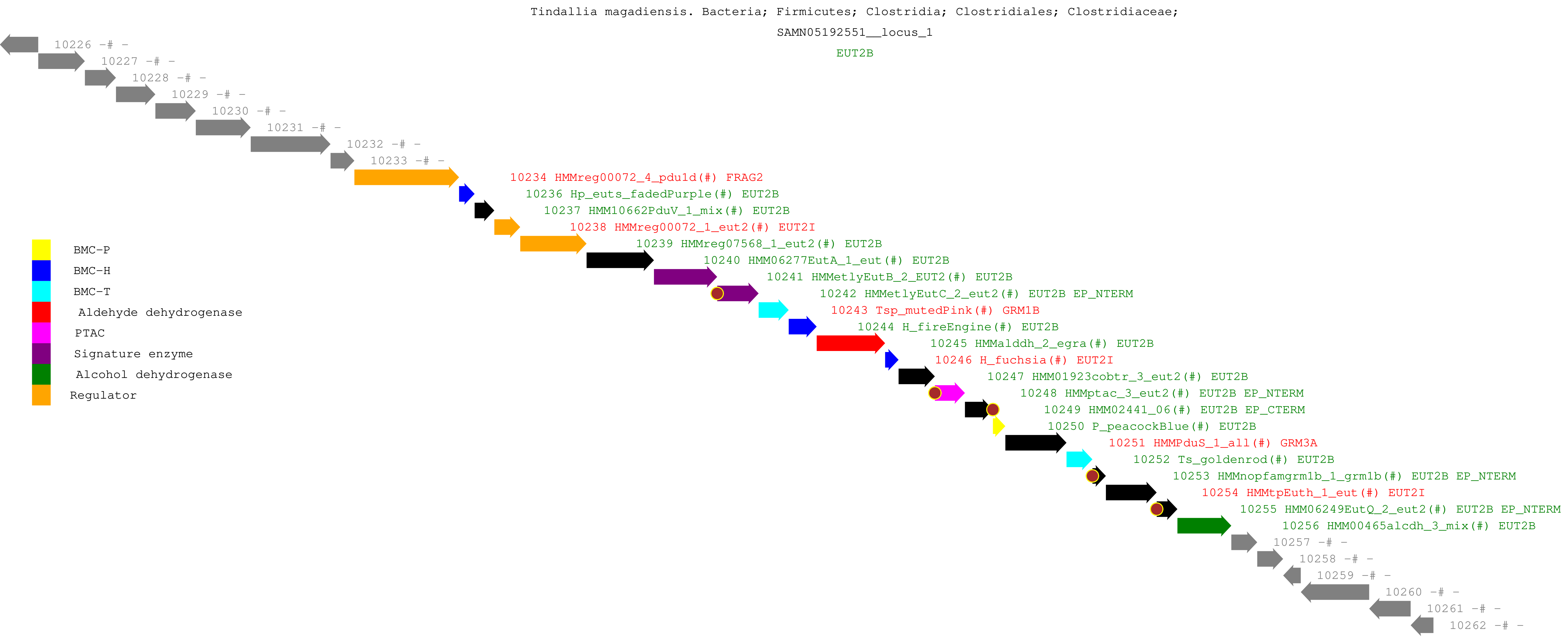
Tindallia magadiensis. Bacteria; Firmicutes; Clostridia; Clostridiales; Clostridiaceae;
SAMN05192551__locus_1
EUT2B
10226 -# -
10227 -# -
10228 -# -
10229 -# -
10230 -# -
10231 -# -
10232 -# -
10233 -# -
10234 HMMreg00072_4_pdu1d(#) FRAG2
10236 Hp_euts_fadedPurple(#) EUT2B
10237 HMM10662PduV_1_mix(#) EUT2B
10238 HMMreg00072_1_eut2(#) EUT2I
10239 HMMreg07568_1_eut2(#) EUT2B
10240 HMM06277EutA_1_eut(#) EUT2B
10241 HMMetlyEutB_2_EUT2(#) EUT2B
10242 HMMetlyEutC_2_eut2(#) EUT2B EP_NTERM
10243 Tsp_mutedPink(#) GRM1B
10244 H_fireEngine(#) EUT2B
10245 HMMalddh_2_egra(#) EUT2B
10246 H_fuchsia(#) EUT2I
10247 HMM01923cobtr_3_eut2(#) EUT2B
10248 HMMptac_3_eut2(#) EUT2B EP_NTERM
10249 HMM02441_06(#) EUT2B EP_CTERM
10250 P_peacockBlue(#) EUT2B
10251 HMMPduS_1_all(#) GRM3A
10252 Ts_goldenrod(#) EUT2B
10253 HMMnopfamgrm1b_1_grm1b(#) EUT2B EP_NTERM
10254 HMMtpEuth_1_eut(#) EUT2I
10255 HMM06249EutQ_2_eut2(#) EUT2B EP_NTERM
10256 HMM00465alcdh_3_mix(#) EUT2B
10257 -# -
10258 -# -
10259 -# -
10260 -# -
10261 -# -
10262 -# -
BMC-P
BMC-H
BMC-T
Aldehyde dehydrogenase
PTAC
Signature enzyme
Alcohol dehydrogenase
Regulator

Hydrogenoanaerobacterium saccharovorans. Bacteria; Firmicutes; Clostridia; Clostridiales; Ruminococcaceae;

SAMN05216180__locus_1

EUT2B

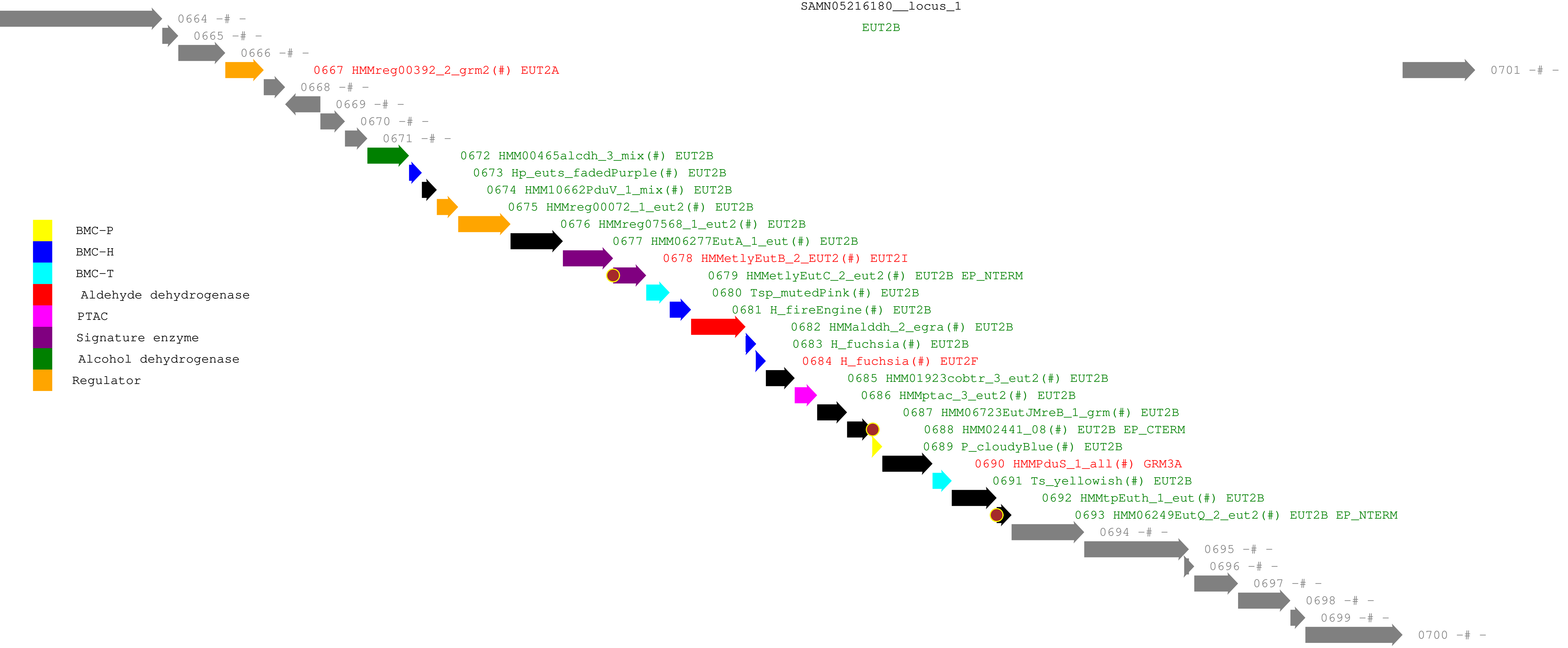

Acetitomaculum ruminis DSM 5522. Bacteria; Firmicutes; Clostridia; Clostridiales; Lachnospiraceae;

SAMN05216249__locus_1

EUT2B

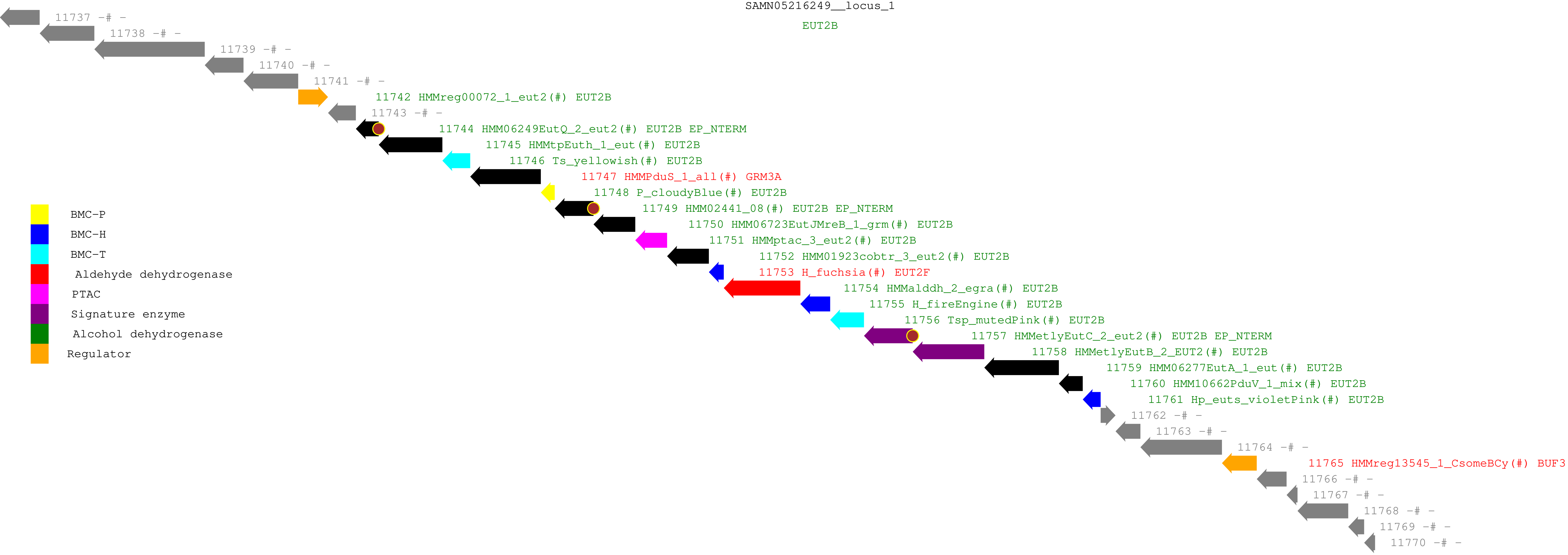

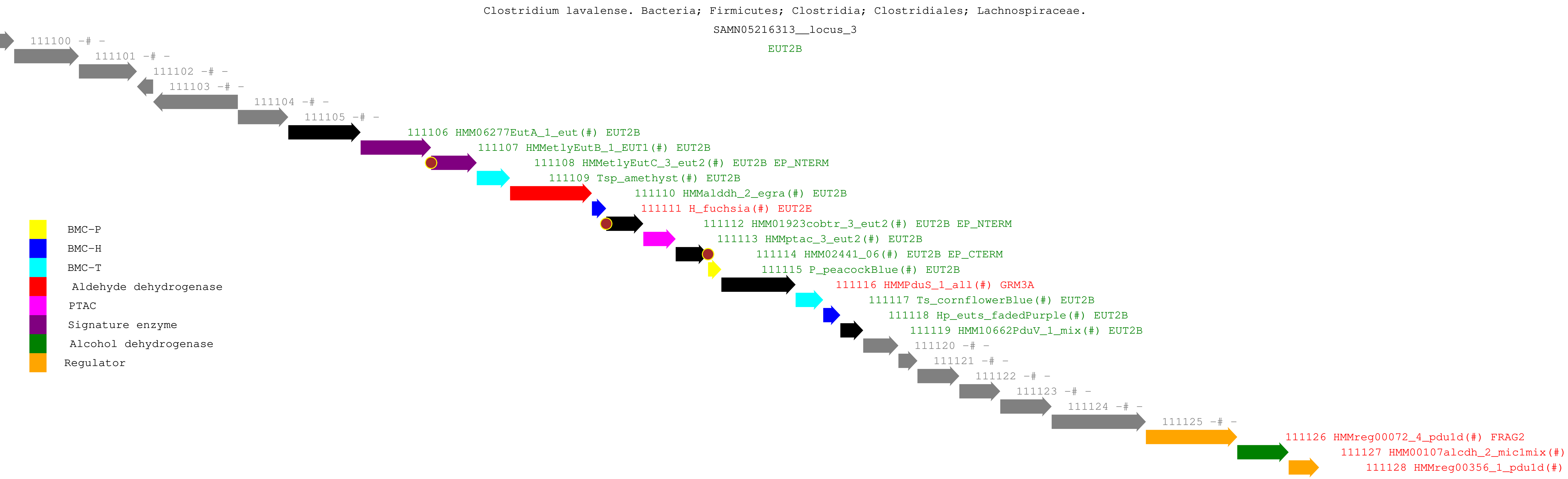

Clostridium lavalense. Bacteria; Firmicutes; Clostridia; Clostridiales; Lachnospiraceae.
SAMN05216313__locus_3
EUT2B
111100 -# -
111101 -# -
111102 -# -
111103 -# -
111104 -# -
111105 -# -
111106 HMM06277EutA_1_eut(#) EUT2B
111107 HMMetlyEutB_1_EUT1(#) EUT2B
111108 HMMetlyEutC_3_eut2(#) EUT2B EP_NTERM
111109 Tsp_amethyst(#) EUT2B
111110 HMMaldh_2_egra(#) EUT2B
111111 H_fuchsia(#) EUT2E
111112 HMM01923cobtr_3_eut2(#) EUT2B EP_NTERM
111113 HMMptac_3_eut2(#) EUT2B
111114 HMM02441_06(#) EUT2B EP_CTERM
111115 P_peacockBlue(#) EUT2B
111116 HMMPduS_1_all(#) GRM3A
111117 Ts_cornflowerBlue(#) EUT2B
111118 Hp_euts_fadedPurple(#) EUT2B
111119 HMM10662PduV_1_mix(#) EUT2B
111120 -# -
111121 -# -
111122 -# -
111123 -# -
111124 -# -
111125 -# -
111126 HMMreg00072_4_pdu1d(#) FRAG2
111127 HMM00107alcdh_2_mic1mix(#)
111128 HMMreg00356_1_pdu1d(#)
BMC-P
BMC-H
BMC-T
Aldehyde dehydrogenase
PTAC
Signature enzyme
Alcohol dehydrogenase
Regulator

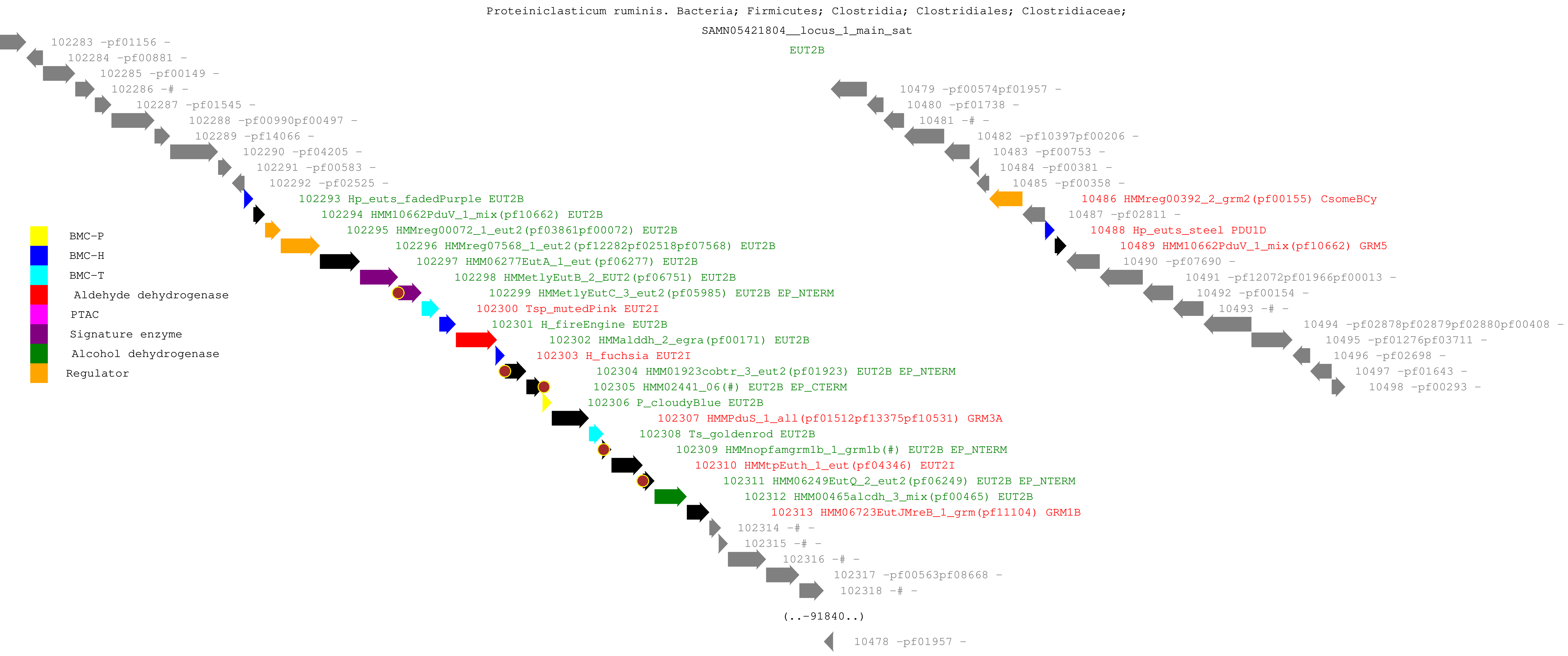
Proteiniclasticum ruminis. Bacteria; Firmicutes; Clostridia; Clostridiales; Clostridiaceae;
SAMN05421804__locus_1_main_sat
EUT2B
102283 -pf01156 -
102284 -pf00881 -
102285 -pf00149 -
102286 -# -
102287 -pf01545 -
102288 -pf00990pf00497 -
102289 -pf14066 -
102290 -pf04205 -
102291 -pf00583 -
102292 -pf02525 -
102293 Hp_euts_fadedPurple EUT2B
102294 HMM10662PduV_1_mix(pf10662) EUT2B
102295 HMMreg00072_1_eut2(pf03861pf00072) EUT2B
102296 HMMreg07568_1_eut2(pf12282pf02518pf07568) EUT2B
102297 HMM06277EutA_1_eut(pf06277) EUT2B
102298 HMMetlyEutB_2_EUT2(pf06751) EUT2B
102299 HMMetlyEutC_3_eut2(pf05985) EUT2B EP_NTERM
102300 Tsp_mutedPink EUT2I
102301 H_fireEngine EUT2B
102302 HMMaldh_2_egra(pf00171) EUT2B
102303 H_fuchsia EUT2I
102304 HMM01923cobtr_3_eut2(pf01923) EUT2B EP_NTERM
102305 HMM02441_06(#) EUT2B EP_CTERM
102306 P_cloudyBlue EUT2B
102307 HMMPduS_1_all(pf01512pf13375pf10531) GRM3A
102308 Ts_goldenrod EUT2B
102309 HMMnopfamgrm1b_1_grm1b(#) EUT2B EP_NTERM
102310 HMMtpEuth_1_eut(pf04346) EUT2I
102311 HMM06249EutQ_2_eut2(pf06249) EUT2B EP_NTERM
102312 HMM00465alcdh_3_mix(pf00465) EUT2B
102313 HMM06723EutJMreB_1_grm(pf11104) GRM1B
102314 -# -
102315 -# -
102316 -# -
102317 -pf00563pf08668 -
102318 -# -
(..-91840..)
10478 -pf01957 -
10479 -pf00574pf01957 -
10480 -pf01738 -
10481 -# -
10482 -pf10397pf00206 -
10483 -pf00753 -
10484 -pf00381 -
10485 -pf00358 -
10486 HMMreg00392_2_grm2(pf00155) CsomeBCy
10487 -pf02811 -
10488 Hp_euts_steel PDU1D
10489 HMM10662PduV_1_mix(pf10662) GRM5
10490 -pf07690 -
10491 -pf12072pf01966pf00013 -
10492 -pf00154 -
10493 -# -
10494 -pf02878pf02879pf02880pf00408 -
10495 -pf01276pf03711 -
10496 -pf02698 -
10497 -pf01643 -
10498 -pf00293 -
BMC-P
BMC-H
BMC-T
Aldehyde dehydrogenase
PTAC
Signature enzyme
Alcohol dehydrogenase
Regulator

Anaerovirgula multivorans. Bacteria; Firmicutes; Clostridia; Clostridiales; Anaerovirgula.

SAMN05446037__locus_2

EUT2B

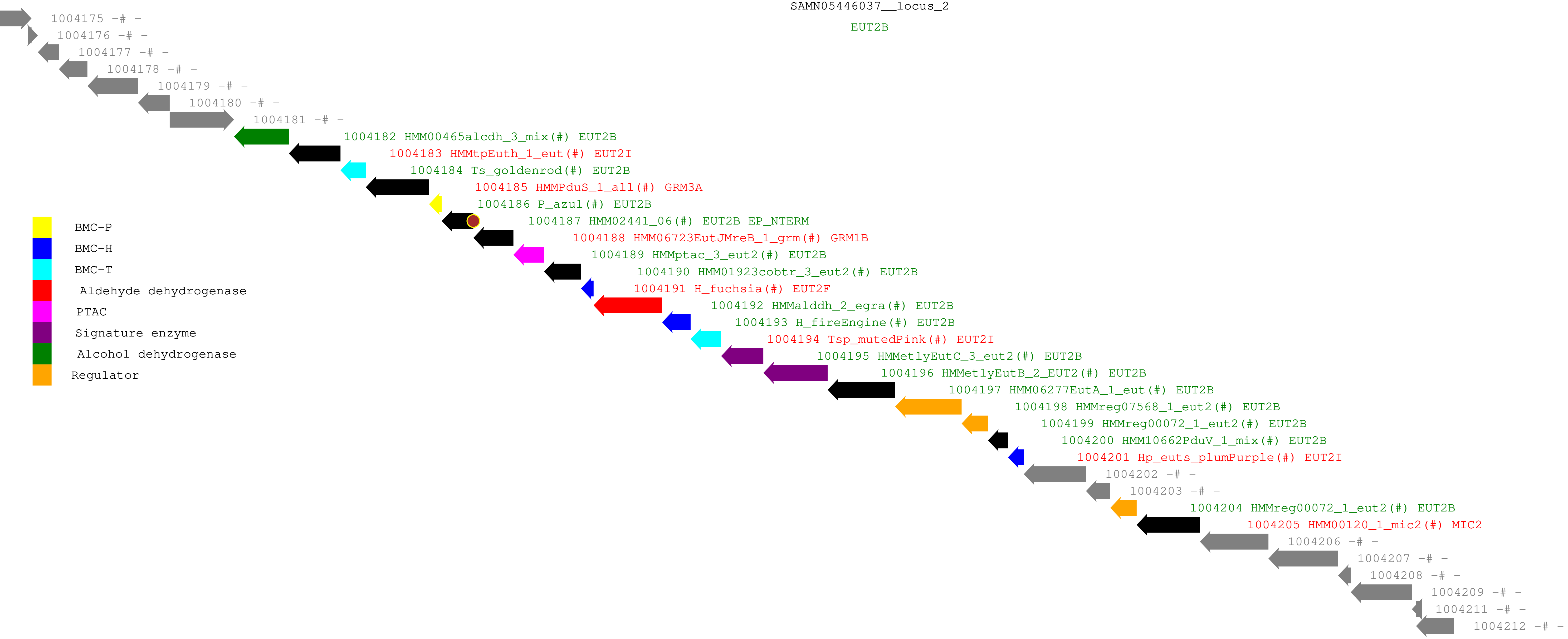

Natronincola peptidivorans. Bacteria; Firmicutes; Clostridia; Clostridiales; Clostridiaceae;

SAMN05660297__locus_3

EUT2B

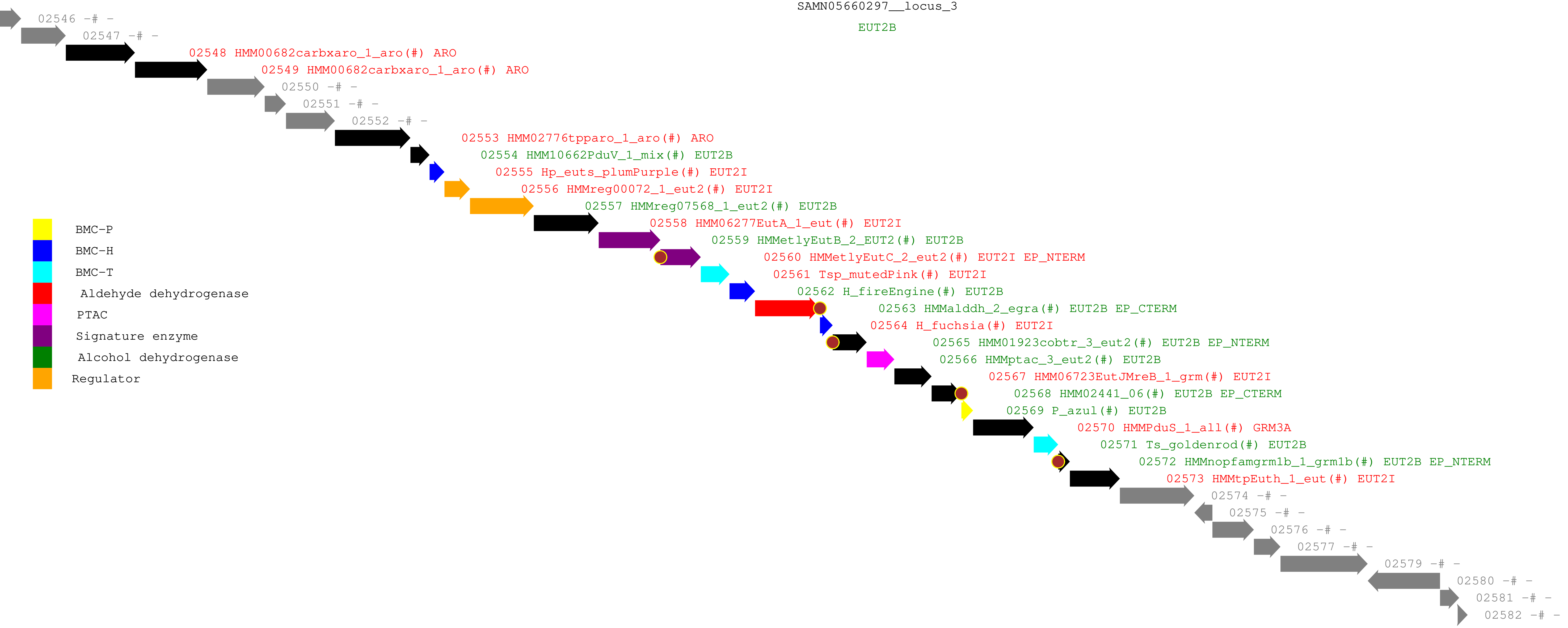

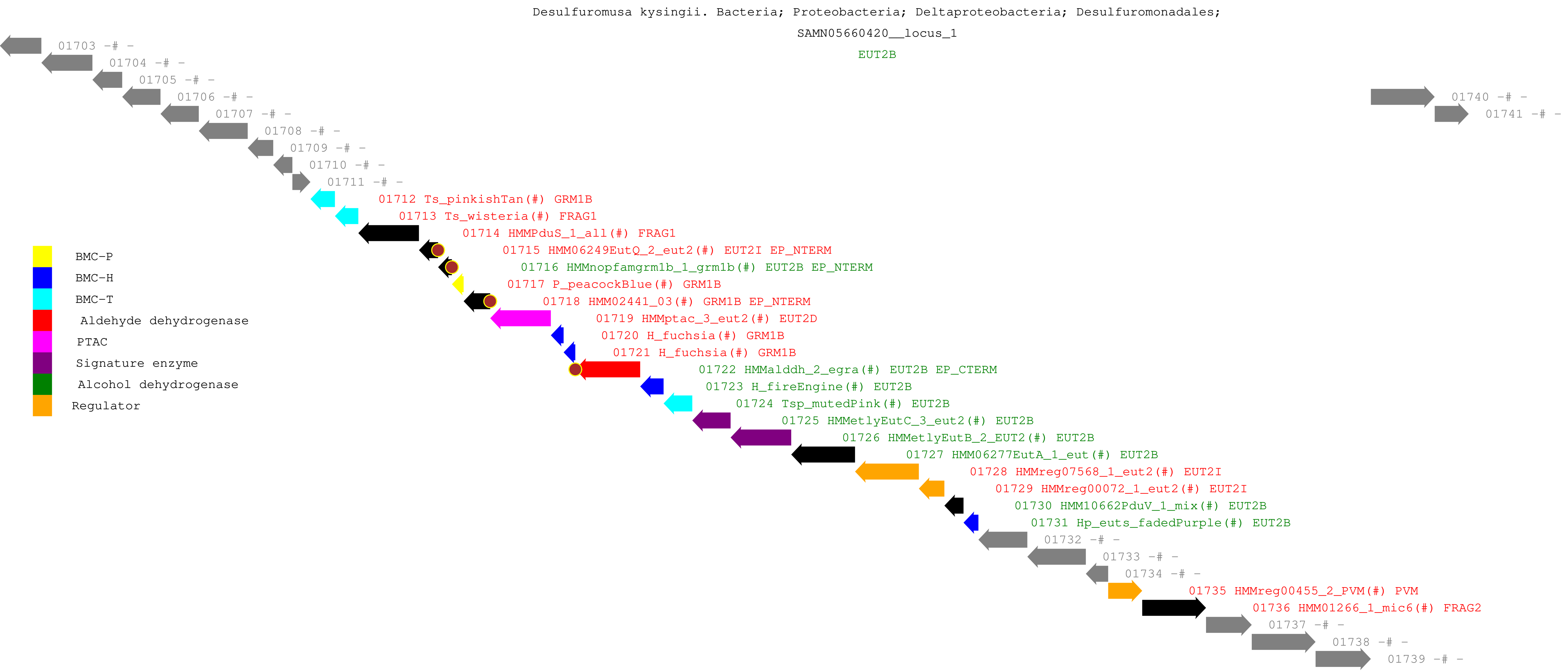

Desulfuromusa kysingii. Bacteria; Proteobacteria; Deltaproteobacteria; Desulfuromonadales;
SAMN05660420__locus_1
EUT2B
01703 -# -
01704 -# -
01705 -# -
01706 -# -
01707 -# -
01708 -# -
01709 -# -
01710 -# -
01711 -# -
01712 Ts_pinkishTan(#) GRM1B
01713 Ts_wisteria(#) FRAG1
01714 HMMPduS_1_all(#) FRAG1
01715 HMM06249EutQ_2_eut2(#) EUT2I EP_NTERM
01716 HMMnopfamgrm1b_1_grm1b(#) EUT2B EP_NTERM
01717 P_peacockBlue(#) GRM1B
01718 HMM02441_03(#) GRM1B EP_NTERM
01719 HMMptac_3_eut2(#) EUT2D
01720 H_fuchsia(#) GRM1B
01721 H_fuchsia(#) GRM1B
01722 HMMaldh_2_egra(#) EUT2B EP_CTERM
01723 H_fireEngine(#) EUT2B
01724 Tsp_mutedPink(#) EUT2B
01725 HMMetlyEutC_3_eut2(#) EUT2B
01726 HMMetlyEutB_2_EUT2(#) EUT2B
01727 HMM06277EutA_1_eut(#) EUT2B
01728 HMMreg07568_1_eut2(#) EUT2I
01729 HMMreg00072_1_eut2(#) EUT2I
01730 HMM10662PduV_1_mix(#) EUT2B
01731 Hp_euts_fadedPurple(#) EUT2B
01732 -# -
01733 -# -
01734 -# -
01735 HMMreg00455_2_PVM(#) PVM
01736 HMM01266_1_mic6(#) FRAG2
01737 -# -
01738 -# -
01739 -# -
01740 -# -
01741 -# -
BMC-P
BMC-H
BMC-T
Aldehyde dehydrogenase
PTAC
Signature enzyme
Alcohol dehydrogenase
Regulator

Thermanaerovibrio acidaminovorans (strain ATCC 49978 / DSM 6589 / Bacteria; Synergistetes; Synergistia; Synergistales; Synergistaceae;

Taci__locus_1

EUT2B

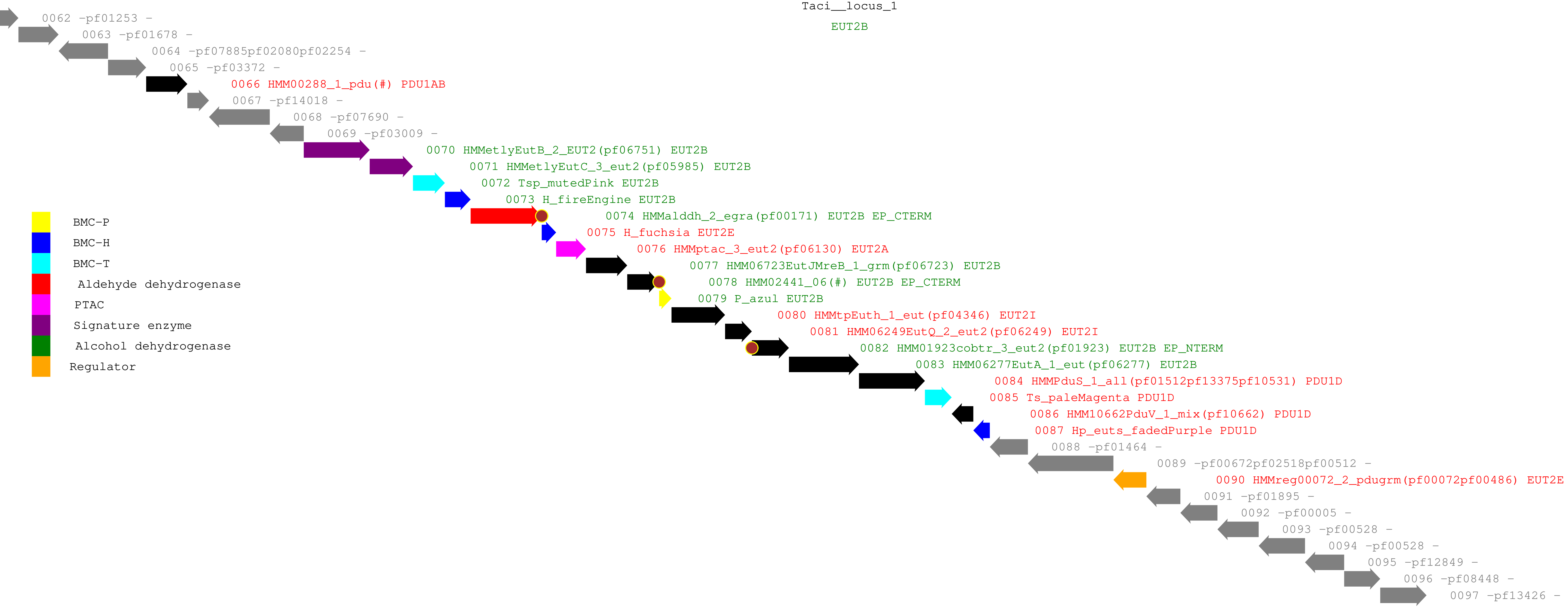

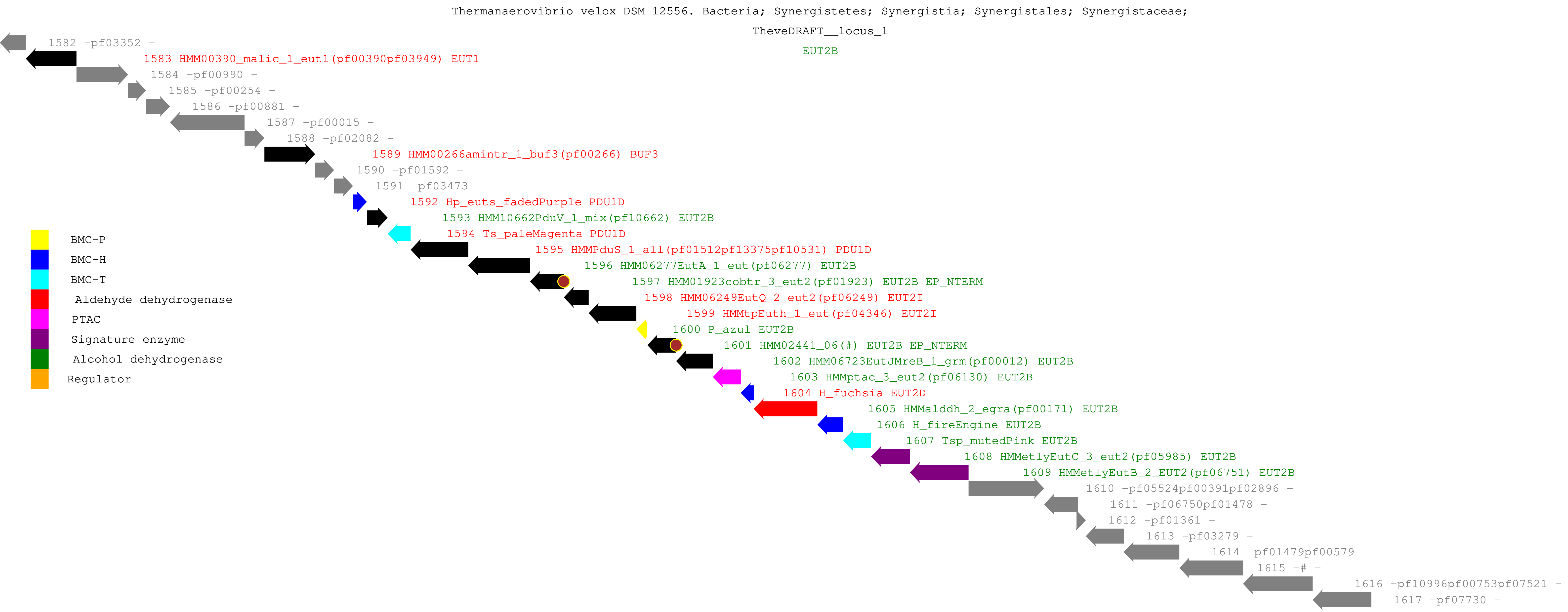
Thermanaerovibrio velox DSM 12556. Bacteria; Synergistetes; Synergistia; Synergistales; Synergistaceae;
TheveDRAFT__locus_1
EUT2B
1582 -pf03352 -
1583 HMM00390_malic_1_eut1(pf00390pf03949) EUT1
1584 -pf00990 -
1585 -pf00254 -
1586 -pf00881 -
1587 -pf00015 -
1588 -pf02082 -
1589 HMM00266amintr_1_buf3(pf00266) BUF3
1590 -pf01592 -
1591 -pf03473 -
1592 Hp_euts_fadedPurple PDU1D
1593 HMM10662PduV_1_mix(pf10662) EUT2B
1594 Ts_paleMagenta PDU1D
1595 HMMPduS_1_all(pf01512pf13375pf10531) PDU1D
1596 HMM06277EutA_1_eut(pf06277) EUT2B
1597 HMM01923cobtr_3_eut2(pf01923) EUT2B EP_NTERM
1598 HMM06249EutQ_2_eut2(pf06249) EUT2I
1599 HMMtpEuth_1_eut(pf04346) EUT2I
1600 P_azul EUT2B
1601 HMM02441_06(#) EUT2B EP_NTERM
1602 HMM06723EutJMreB_1_grm(pf00012) EUT2B
1603 HMMptac_3_eut2(pf06130) EUT2B
1604 H_fuchsia EUT2D
1605 HMMalddh_2_egra(pf00171) EUT2B
1606 H_fireEngine EUT2B
1607 Tsp_mutedPink EUT2B
1608 HMMetlyEutC_3_eut2(pf05985) EUT2B
1609 HMMetlyEutB_2_EUT2(pf06751) EUT2B
1610 -pf05524pf00391pf02896 -
1611 -pf06750pf01478 -
1612 -pf01361 -
1613 -pf03279 -
1614 -pf01479pf00579 -
1615 -# -
1616 -pf10996pf00753pf07521 -
1617 -pf07730 -
BMC-P
BMC-H
BMC-T
Aldehyde dehydrogenase
PTAC
Signature enzyme
Alcohol dehydrogenase
Regulator

Thermacetogenium phaeum (strain ATCC BAA-254 / DSM 12270 / PB). Bacteria; Firmicutes; Clostridia; Thermoanaerobacterales;

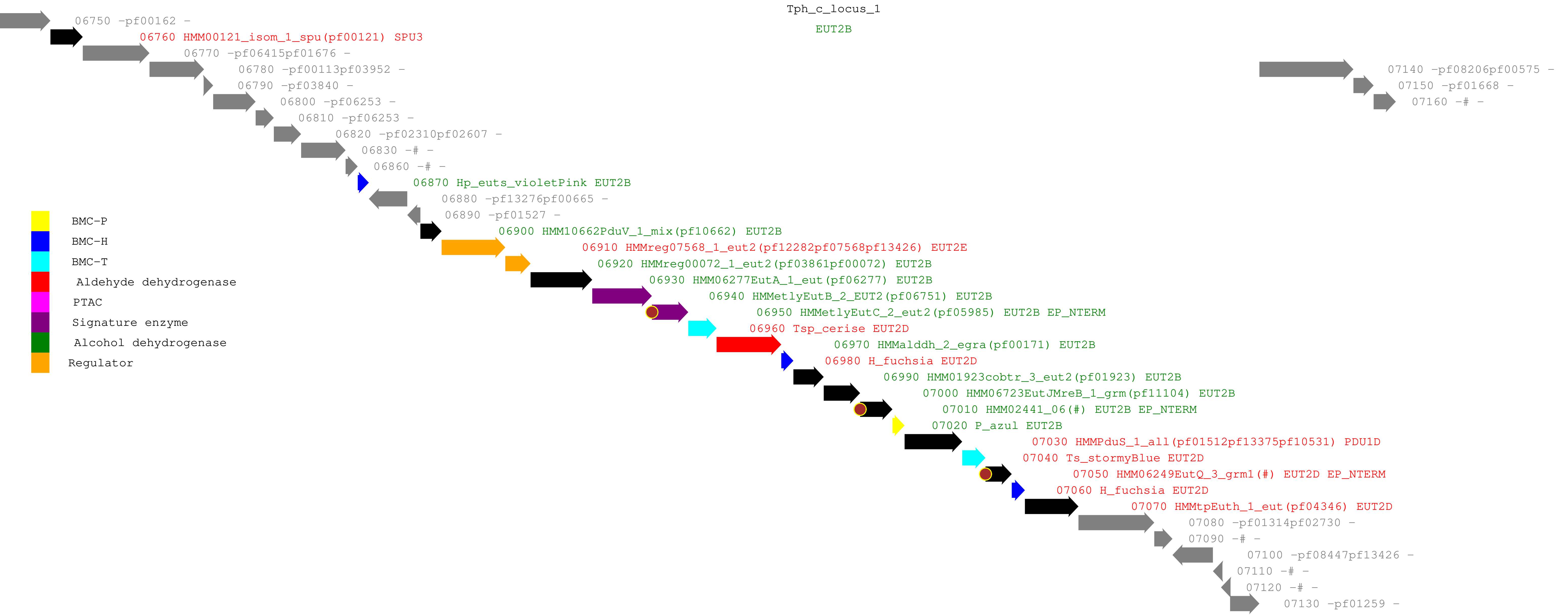

Tepidanaerobacter syntrophicus. Bacteria; Firmicutes; Clostridia; Thermoanaerobacterales;

TSYNT__locus_1

EUT2B

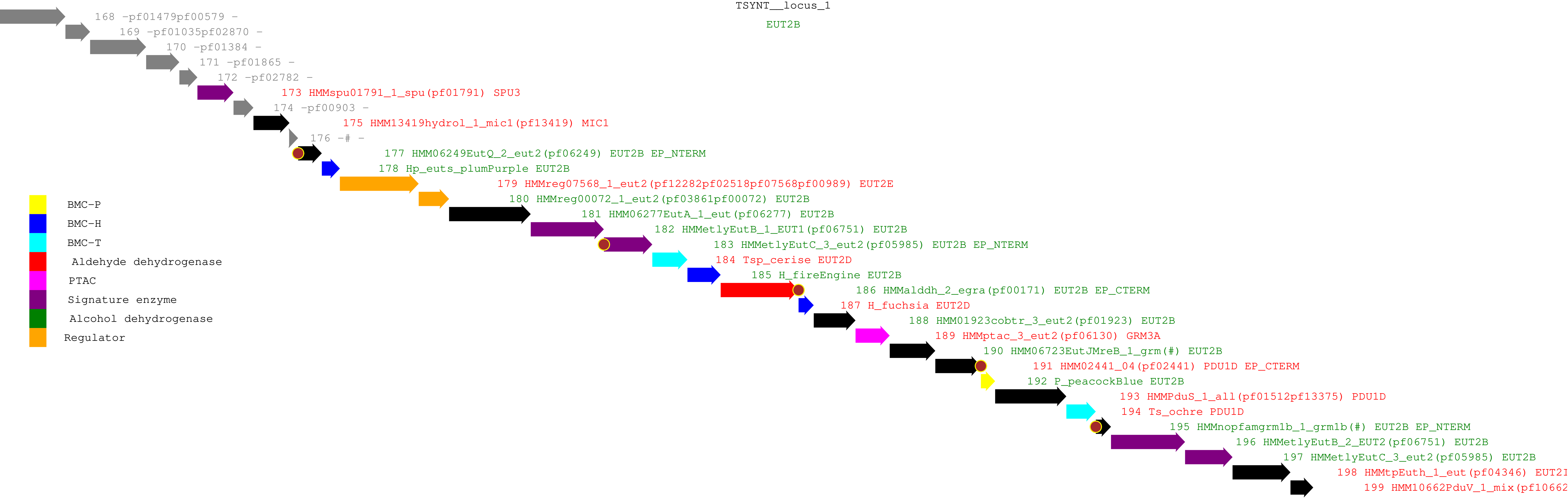

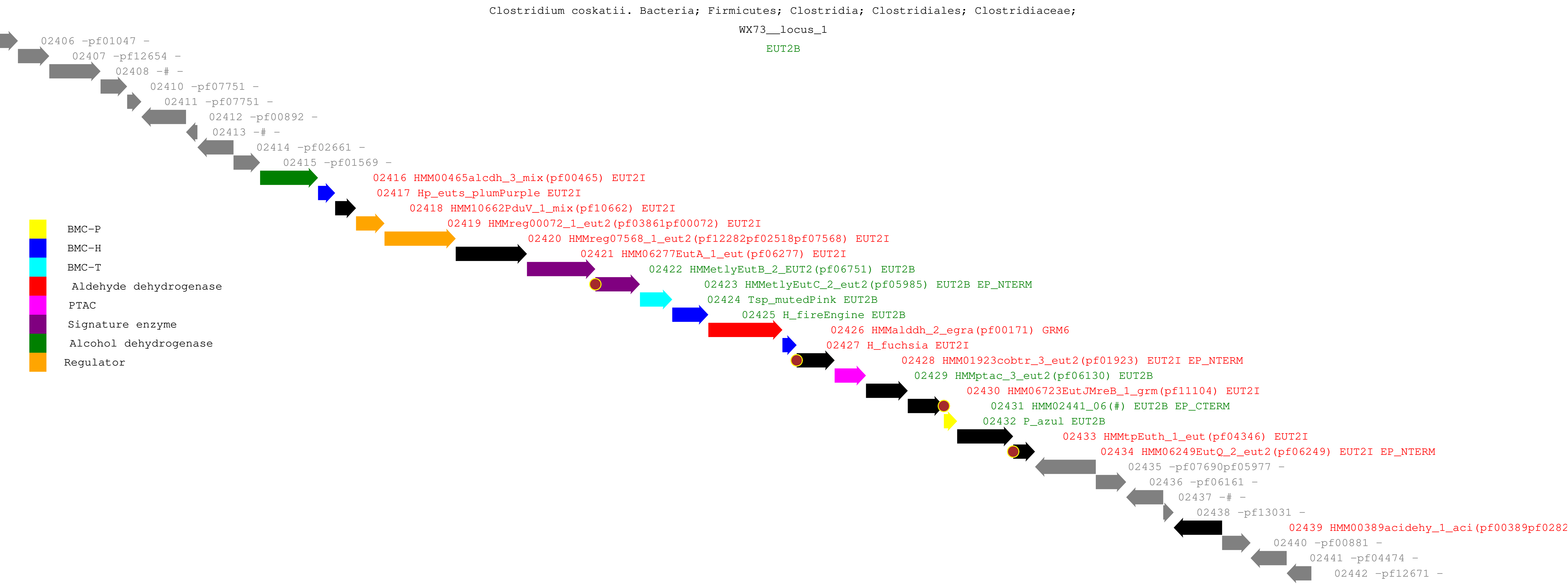
Clostridium coskatii. Bacteria; Firmicutes; Clostridia; Clostridiales; Clostridiaceae;
WX73__locus_1
EUT2B
02406 -pf01047 -
02407 -pf12654 -
02408 -# -
02410 -pf07751 -
02411 -pf07751 -
02412 -pf00892 -
02413 -# -
02414 -pf02661 -
02415 -pf01569 -
02416 HMM00465alcdh_3_mix(pf00465) EUT2I
02417 Hp_euts_plumPurple EUT2I
02418 HMM10662PduV_1_mix(pf10662) EUT2I
02419 HMMreg00072_1_eut2(pf03861pf00072) EUT2I
02420 HMMreg07568_1_eut2(pf12282pf02518pf07568) EUT2I
02421 HMM06277EutA_1_eut(pf06277) EUT2I
02422 HMMetlyEutB_2_EUT2(pf06751) EUT2B
02423 HMMetlyEutC_2_eut2(pf05985) EUT2B EP_NTERM
02424 Tsp_mutedPink EUT2B
02425 H_fireEngine EUT2B
02426 HMMalddh_2_egra(pf00171) GRM6
02427 H_fuchsia EUT2I
02428 HMM01923cobtr_3_eut2(pf01923) EUT2I EP_NTERM
02429 HMMptac_3_eut2(pf06130) EUT2B
02430 HMM06723EutJMreB_1_grm(pf11104) EUT2I
02431 HMM02441_06(#) EUT2B EP_CTERM
02432 P_azul EUT2B
02433 HMMtpEuth_1_eut(pf04346) EUT2I
02434 HMM06249EutQ_2_eut2(pf06249) EUT2I EP_NTERM
02435 -pf07690pf05977 -
02436 -pf06161 -
02437 -# -
02438 -pf13031 -
02439 HMM00389acidehy_1_aci(pf00389pf0282
02440 -pf00881 -
02441 -pf04474 -
02442 -pf12671 -
BMC-P
BMC-H
BMC-T
Aldehyde dehydrogenase
PTAC
Signature enzyme
Alcohol dehydrogenase
Regulator

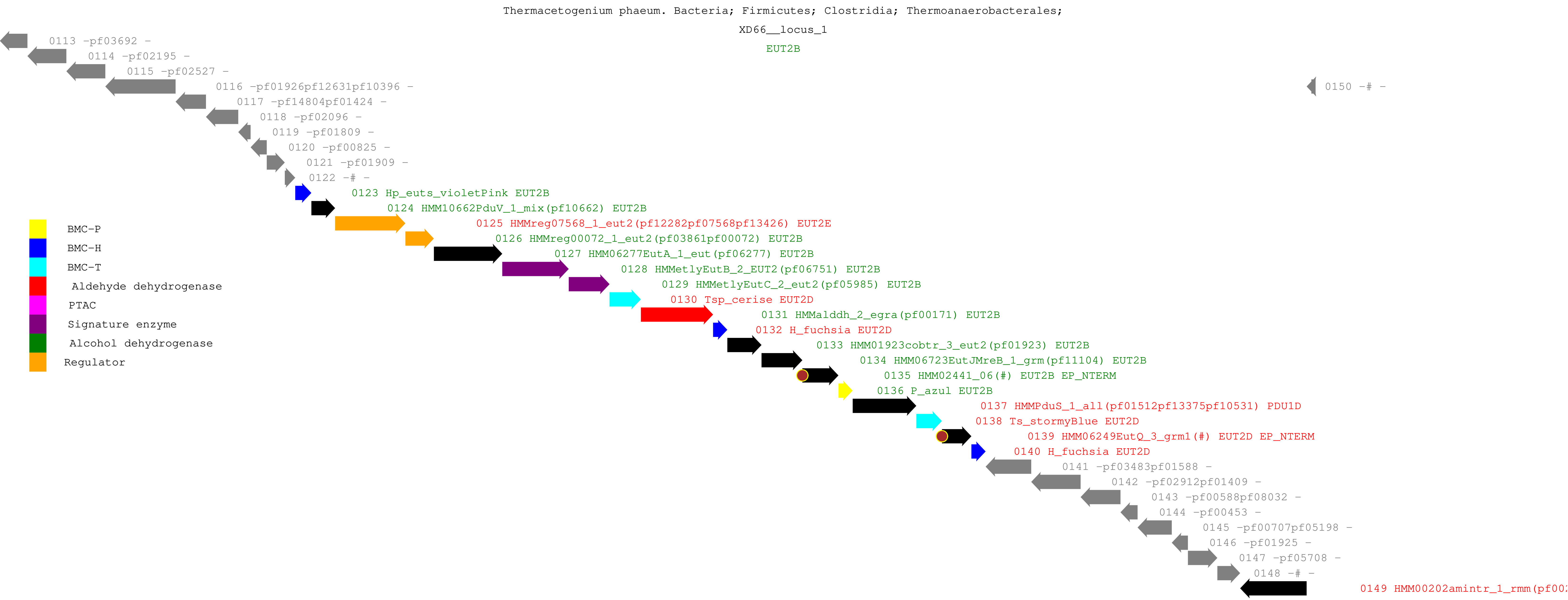
Thermacetogenium phaeum. Bacteria; Firmicutes; Clostridia; Thermoanaerobacterales;
XD66__locus_1
EUT2B
0113 -pf03692 -
0114 -pf02195 -
0115 -pf02527 -
0116 -pf01926pf12631pf10396 -
0117 -pf14804pf01424 -
0118 -pf02096 -
0119 -pf01809 -
0120 -pf00825 -
0121 -pf01909 -
0122 -# -
0123 Hp_euts_violetPink EUT2B
0124 HMM10662PduV_1_mix(pf10662) EUT2B
0125 HMMreg07568_1_eut2(pf12282pf07568pf13426) EUT2E
0126 HMMreg00072_1_eut2(pf03861pf00072) EUT2B
0127 HMM06277EutA_1_eut(pf06277) EUT2B
0128 HMMetlyEutB_2_EUT2(pf06751) EUT2B
0129 HMMetlyEutC_2_eut2(pf05985) EUT2B
0130 Tsp_cerise EUT2D
0131 HMMalddh_2_egra(pf00171) EUT2B
0132 H_fuchsia EUT2D
0133 HMM01923cobtr_3_eut2(pf01923) EUT2B
0134 HMM06723EutJMreB_1_grm(pf11104) EUT2B
0135 HMM02441_06(#) EUT2B EP_NTERM
0136 P_azul EUT2B
0137 HMMPduS_1_all(pf01512pf13375pf10531) PDU1D
0138 Ts_stormyBlue EUT2D
0139 HMM06249EutQ_3_grm1(#) EUT2D EP_NTERM
0140 H_fuchsia EUT2D
0141 -pf03483pf01588 -
0142 -pf02912pf01409 -
0143 -pf00588pf08032 -
0144 -pf00453 -
0145 -pf00707pf05198 -
0146 -pf01925 -
0147 -pf05708 -
0148 -# -
0149 HMM00202amintr_1_rmm(pf00
0150 -# -
BMC-P
BMC-H
BMC-T
Aldehyde dehydrogenase
PTAC
Signature enzyme
Alcohol dehydrogenase
Regulator

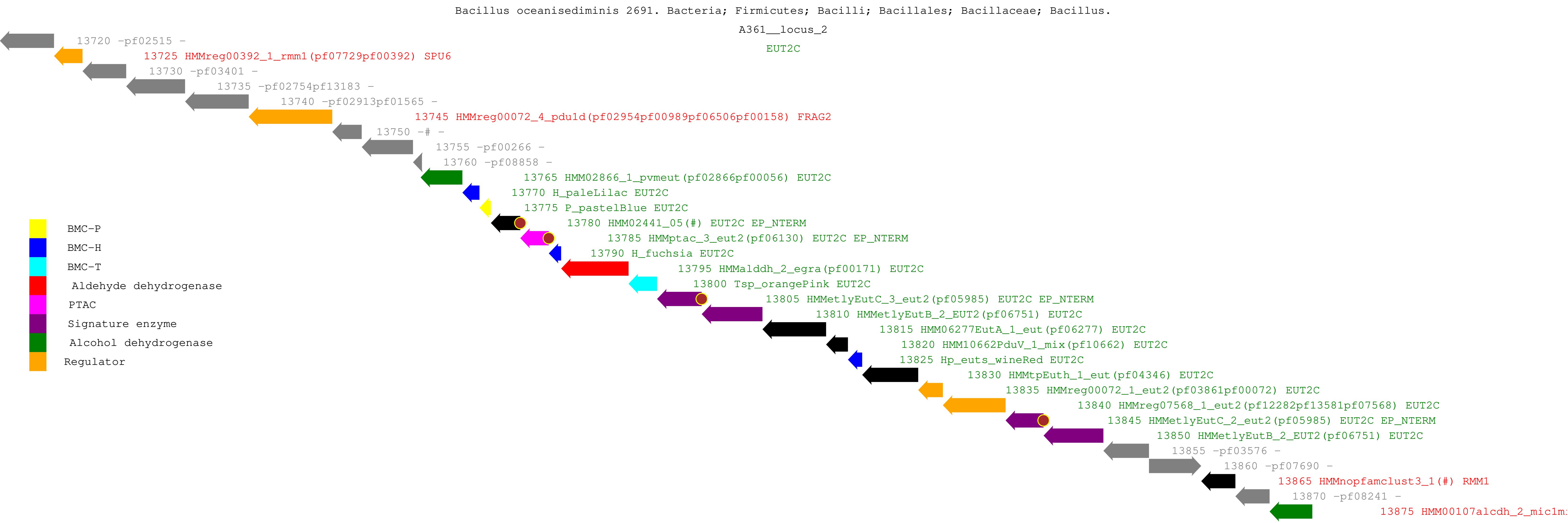
Bacillus oceanisediminis 2691. Bacteria; Firmicutes; Bacilli; Bacillales; Bacillaceae; Bacillus.
A361__locus_2
EUT2C
13720 -pf02515 -
13725 HMMreg00392_1_rmm1(pf07729pf00392) SPU6
13730 -pf03401 -
13735 -pf02754pf13183 -
13740 -pf02913pf01565 -
13745 HMMreg00072_4_pdu1d(pf02954pf00989pf06506pf00158) FRAG2
13750 -# -
13755 -pf00266 -
13760 -pf08858 -
13765 HMM02866_1_pvmeut(pf02866pf00056) EUT2C
13770 H_paleLilac EUT2C
13775 P_pastelBlue EUT2C
13780 HMM02441_05(#) EUT2C EP_NTERM
13785 HMMptac_3_eut2(pf06130) EUT2C EP_NTERM
13790 H_fuchsia EUT2C
13795 HMMalddh_2_egra(pf00171) EUT2C
13800 Tsp_orangePink EUT2C
13805 HMMetlyEutC_3_eut2(pf05985) EUT2C EP_NTERM
13810 HMMetlyEutB_2_EUT2(pf06751) EUT2C
13815 HMM06277EutA_1_eut(pf06277) EUT2C
13820 HMM10662PduV_1_mix(pf10662) EUT2C
13825 Hp_euts_wineRed EUT2C
13830 HMMtpEuth_1_eut(pf04346) EUT2C
13835 HMMreg00072_1_eut2(pf03861pf00072) EUT2C
13840 HMMreg07568_1_eut2(pf12282pf13581pf07568) EUT2C
13845 HMMetlyEutC_2_eut2(pf05985) EUT2C EP_NTERM
13850 HMMetlyEutB_2_EUT2(pf06751) EUT2C
13855 -pf03576 -
13860 -pf07690 -
13865 HMMnopfamclust3_1(#) RMM1
13870 -pf08241 -
13875 HMM00107alcdh_2_mic1m
BMC-P
BMC-H
BMC-T
Aldehyde dehydrogenase
PTAC
Signature enzyme
Alcohol dehydrogenase
Regulator

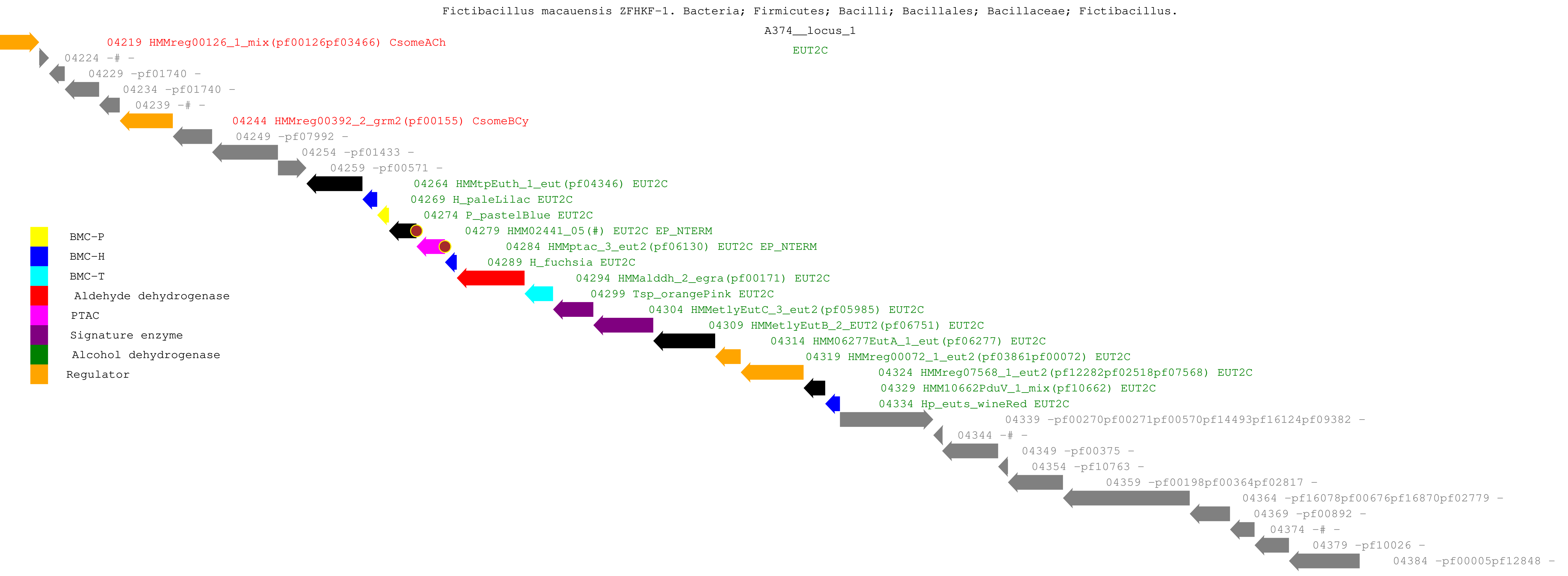
Fictibacillus macauensis ZFHKF-1. Bacteria; Firmicutes; Bacilli; Bacillales; Bacillaceae; Fictibacillus.
A374__locus_1
EUT2C
04219 HMMreg00126_1_mix(pf00126pf03466) CsomeACh
04224 -# -
04229 -pf01740 -
04234 -pf01740 -
04239 -# -
04244 HMMreg00392_2_grm2(pf00155) CsomeBCy
04249 -pf07992 -
04254 -pf01433 -
04259 -pf00571 -
04264 HMMtpEuth_1_eut(pf04346) EUT2C
04269 H_paleLilac EUT2C
04274 P_pastelBlue EUT2C
04279 HMM02441_05(#) EUT2C EP_NTERM
04284 HMMptac_3_eut2(pf06130) EUT2C EP_NTERM
04289 H_fuchsia EUT2C
04294 HMMalddh_2_egra(pf00171) EUT2C
04299 Tsp_orangePink EUT2C
04304 HMMetlyEutC_3_eut2(pf05985) EUT2C
04309 HMMetlyEutB_2_EUT2(pf06751) EUT2C
04314 HMM06277EutA_1_eut(pf06277) EUT2C
04319 HMMreg00072_1_eut2(pf03861pf00072) EUT2C
04324 HMMreg07568_1_eut2(pf12282pf02518pf07568) EUT2C
04329 HMM10662PduV_1_mix(pf10662) EUT2C
04334 Hp_euts_wineRed EUT2C
04339 -pf00270pf00271pf00570pf14493pf16124pf09382 -
04344 -# -
04349 -pf00375 -
04354 -pf10763 -
04359 -pf00198pf00364pf02817 -
04364 -pf16078pf00676pf16870pf02779 -
04369 -pf00892 -
04374 -# -
04379 -pf10026 -
04384 -pf00005pf12848 -
BMC-P
BMC-H
BMC-T
Aldehyde dehydrogenase
PTAC
Signature enzyme
Alcohol dehydrogenase
Regulator

Bacillus badius. Bacteria; Firmicutes; Bacilli; Bacillales; Bacillaceae; Bacillus.

A3781__locus_1

EUT2C

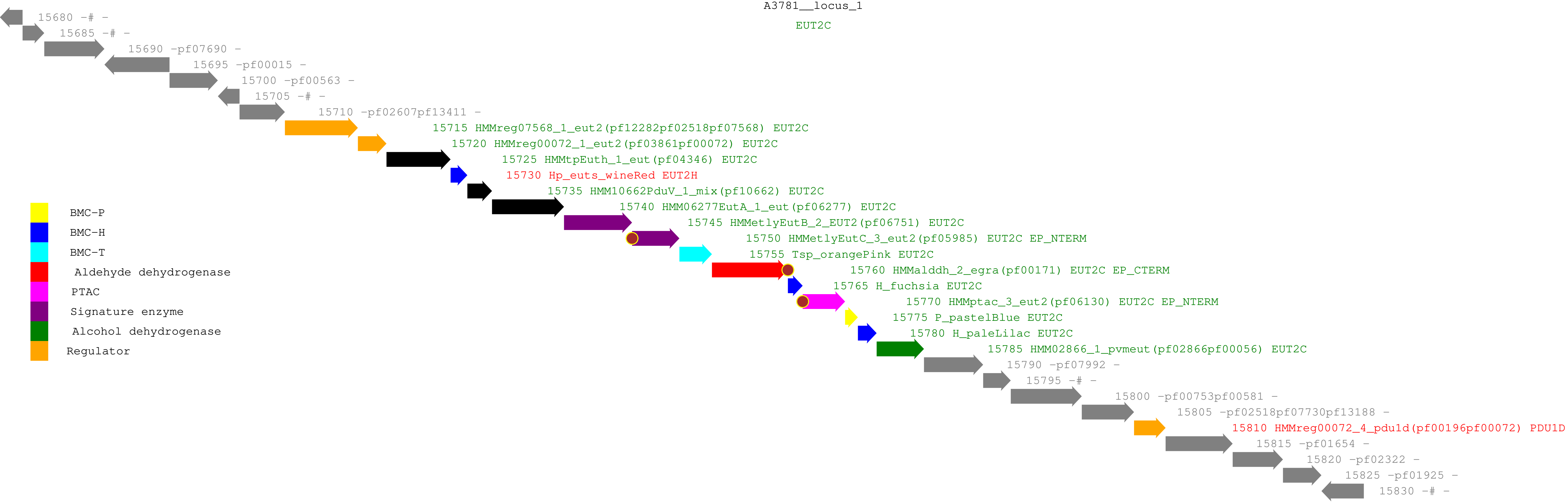

Bacillus encimensis. Bacteria; Firmicutes; Bacilli; Bacillales; Bacillaceae; Bacillus.

A4244__locus_1

EUT2C

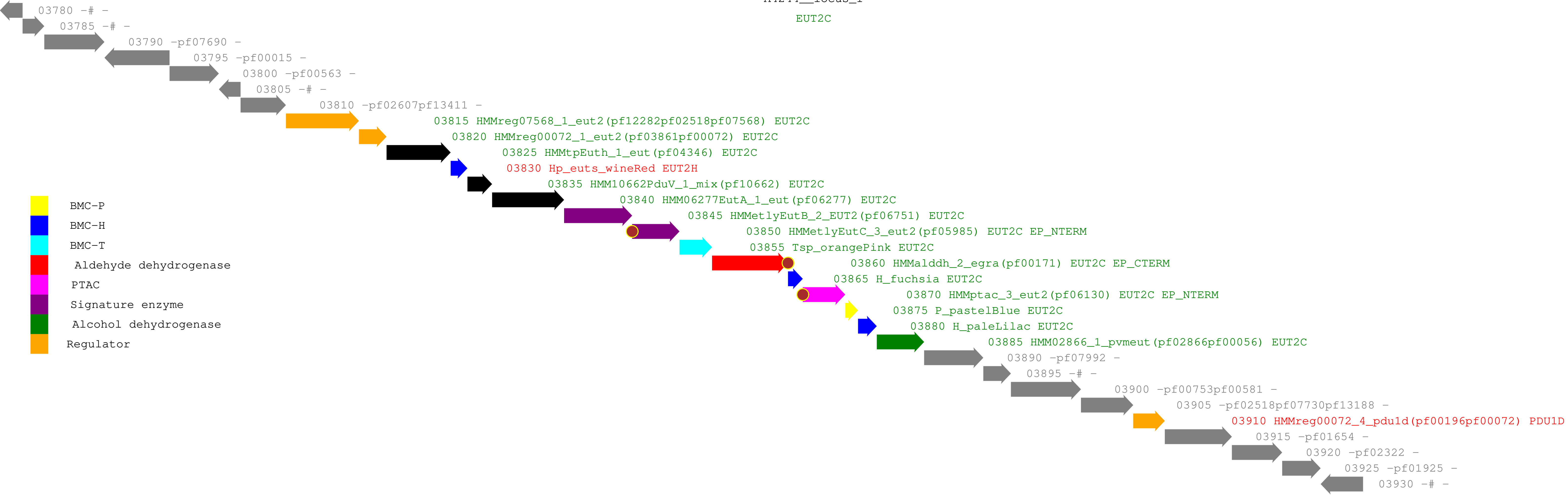

Bacillus sp. FJAT-27215. Bacteria; Firmicutes; Bacilli; Bacillales; Bacillaceae; Bacillus.

A8F95__locus_1

EUT2C

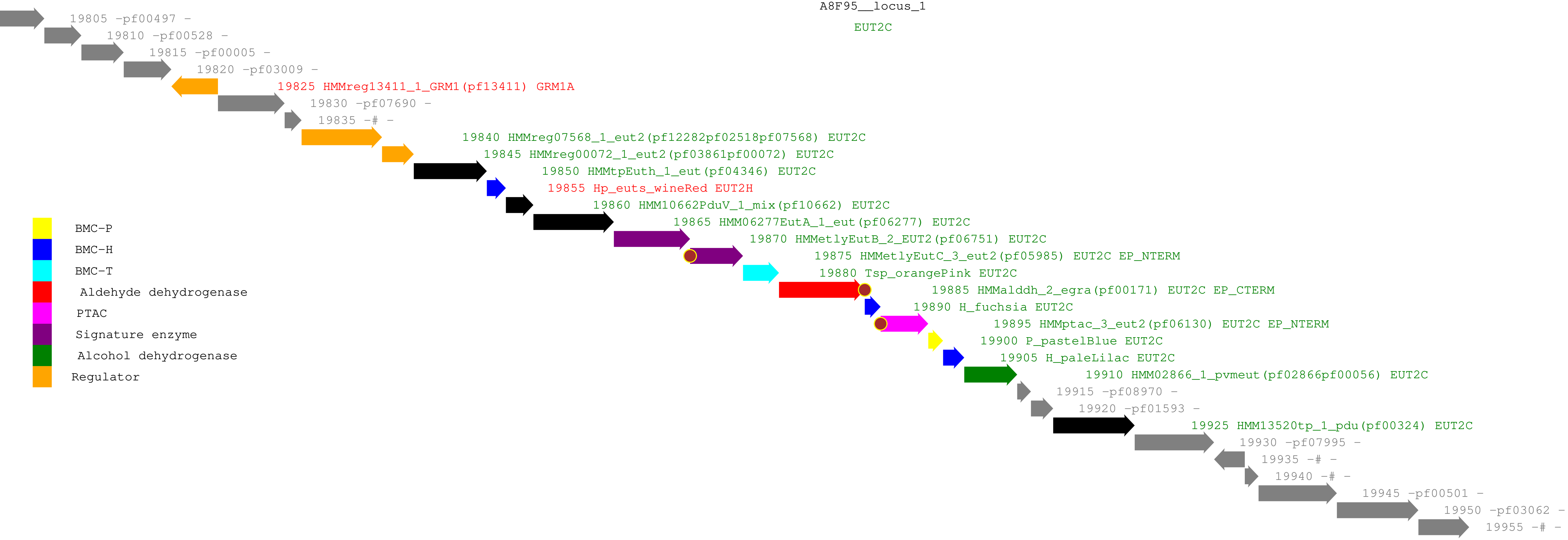

Bacillus sp. FJAT-27231. Bacteria; Firmicutes; Bacilli; Bacillales; Bacillaceae; Bacillus.

AC623__locus_1

EUT2C

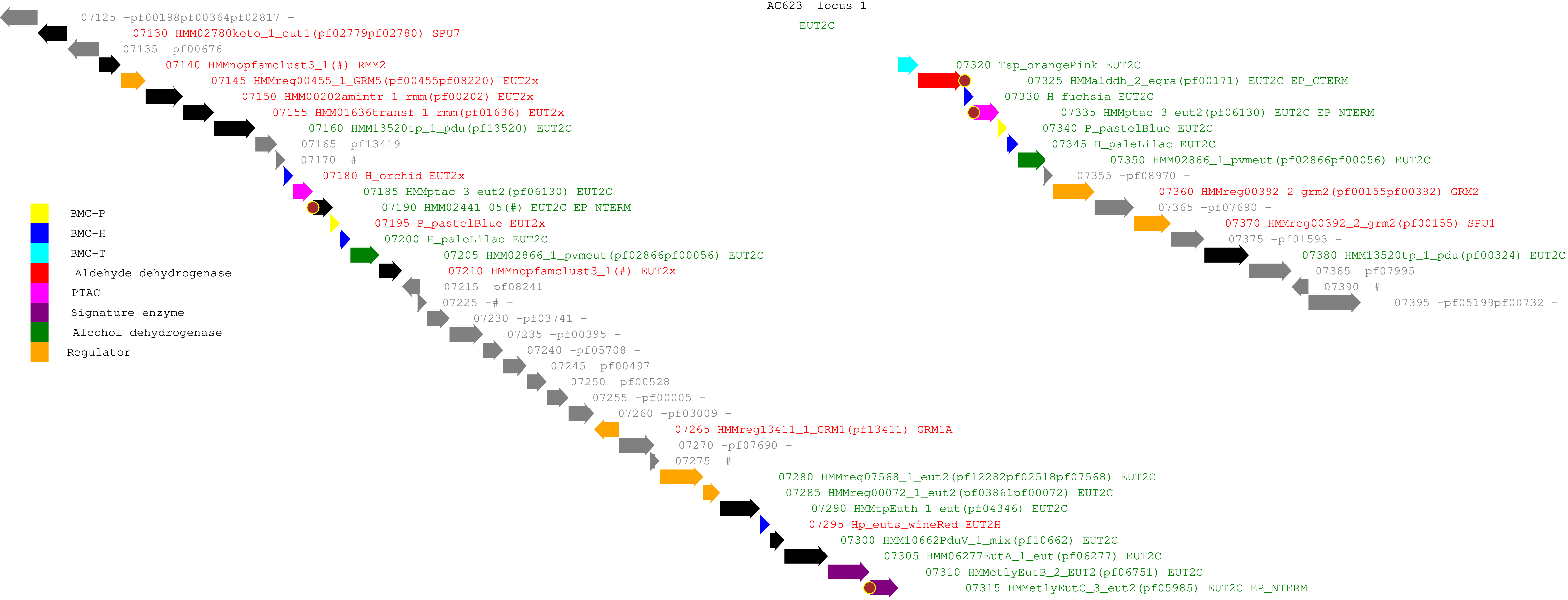

Aneurinibacillus sp. XH2. Bacteria; Firmicutes; Bacilli; Bacillales; Paenibacillaceae;

ACH33__locus_1

EUT2C

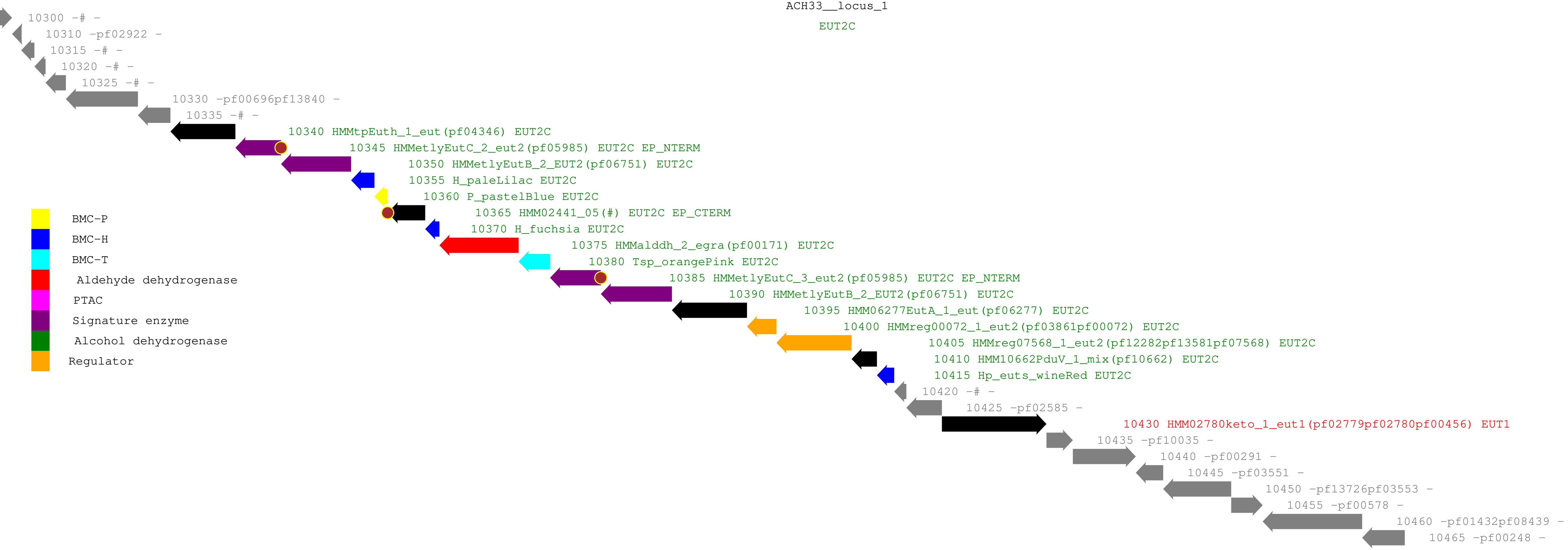

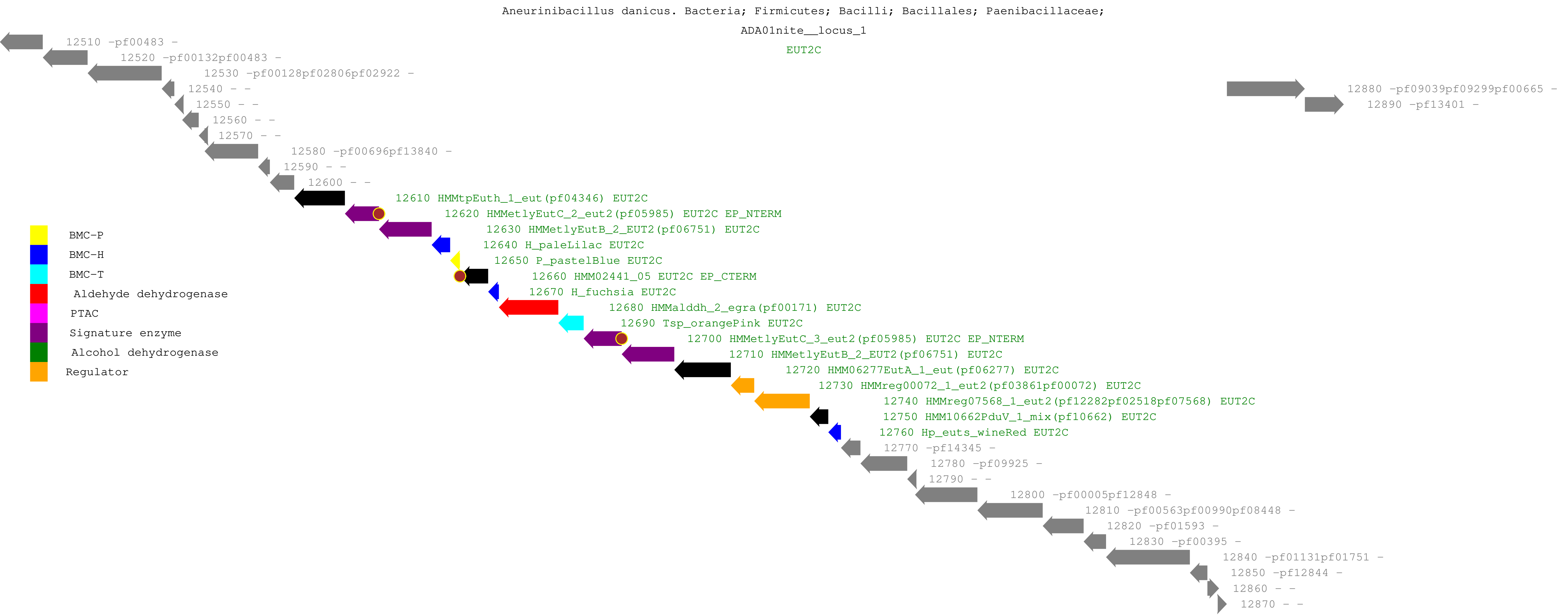
Aneurinibacillus danicus. Bacteria; Firmicutes; Bacilli; Bacillales; Paenibacillaceae;
ADA01nite__locus_1
EUT2C
12510 -pf00483 -
12520 -pf00132pf00483 -
12530 -pf00128pf02806pf02922 -
12540 - -
12550 - -
12560 - -
12570 - -
12580 -pf00696pf13840 -
12590 - -
12600 - -
12610 HMMtpEuth_1_eut(pf04346) EUT2C
12620 HMMetlyEutC_2_eut2(pf05985) EUT2C EP_NTERM
12630 HMMetlyEutB_2_EUT2(pf06751) EUT2C
12640 H_paleLilac EUT2C
12650 P_pastelBlue EUT2C
12660 HMM02441_05 EUT2C EP_CTERM
12670 H_fuchsia EUT2C
12680 HMMalddh_2_egra(pf00171) EUT2C
12690 Tsp_orangePink EUT2C
12700 HMMetlyEutC_3_eut2(pf05985) EUT2C EP_NTERM
12710 HMMetlyEutB_2_EUT2(pf06751) EUT2C
12720 HMM06277EutA_1_eut(pf06277) EUT2C
12730 HMMreg00072_1_eut2(pf03861pf00072) EUT2C
12740 HMMreg07568_1_eut2(pf12282pf02518pf07568) EUT2C
12750 HMM10662PduV_1_mix(pf10662) EUT2C
12760 Hp_euts_wineRed EUT2C
12770 -pf14345 -
12780 -pf09925 -
12790 - -
12800 -pf00005pf12848 -
12810 -pf00563pf00990pf08448 -
12820 -pf01593 -
12830 -pf00395 -
12840 -pf01131pf01751 -
12850 -pf12844 -
12860 - -
12870 - -
12880 -pf09039pf09299pf00665 -
12890 -pf13401 -
BMC-P
BMC-H
BMC-T
Aldehyde dehydrogenase
PTAC
Signature enzyme
Alcohol dehydrogenase
Regulator

Sporosarcina globispora (Bacillus globisporus). Bacteria; Firmicutes; Bacilli; Bacillales; Planococcaceae;

AF332__locus_1

EUT2C

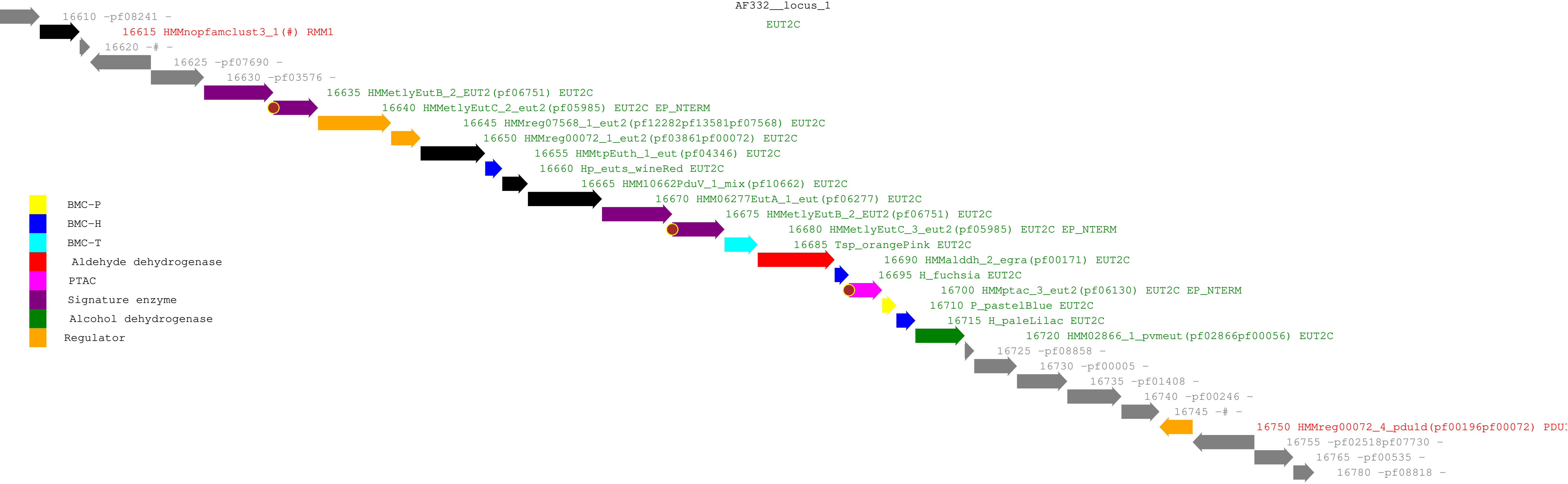

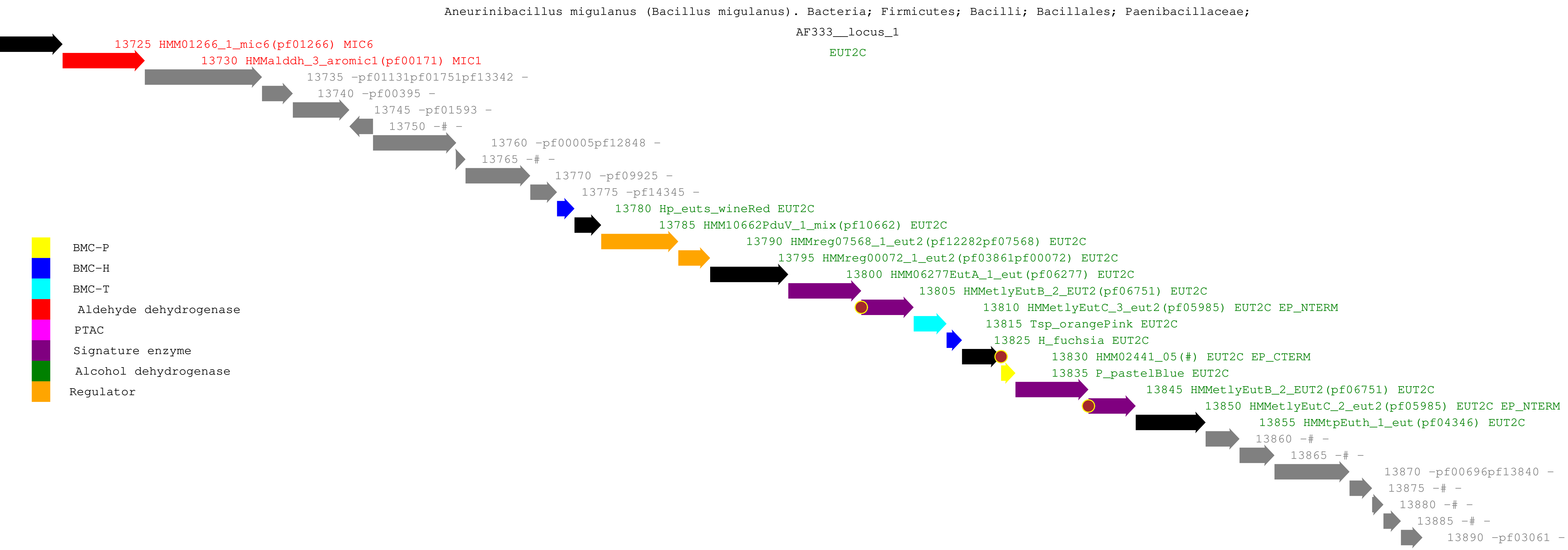
Aneurinibacillus migulanus (Bacillus migulanus). Bacteria; Firmicutes; Bacilli; Bacillales; Paenibacillaceae;
AF333__locus_1
EUT2C
13725 HMM01266_1_mic6(pf01266) MIC6
13730 HMMalddh_3_aromic1(pf00171) MIC1
13735 -pf01131pf01751pf13342 -
13740 -pf00395 -
13745 -pf01593 -
13750 -# -
13760 -pf00005pf12848 -
13765 -# -
13770 -pf09925 -
13775 -pf14345 -
13780 Hp_euts_wineRed EUT2C
13785 HMM10662PduV_1_mix(pf10662) EUT2C
13790 HMMreg07568_1_eut2(pf12282pf07568) EUT2C
13795 HMMreg00072_1_eut2(pf03861pf00072) EUT2C
13800 HMM06277EutA_1_eut(pf06277) EUT2C
13805 HMMetlyEutB_2_EUT2(pf06751) EUT2C
13810 HMMetlyEutC_3_eut2(pf05985) EUT2C EP_NTERM
13815 Tsp_orangePink EUT2C
13825 H_fuchsia EUT2C
13830 HMM02441_05(#) EUT2C EP_CTERM
13835 P_pastelBlue EUT2C
13845 HMMetlyEutB_2_EUT2(pf06751) EUT2C
13850 HMMetlyEutC_2_eut2(pf05985) EUT2C EP_NTERM
13855 HMMtpEuth_1_eut(pf04346) EUT2C
13860 -# -
13865 -# -
13870 -pf00696pf13840 -
13875 -# -
13880 -# -
13885 -# -
13890 -pf03061 -
BMC-P
BMC-H
BMC-T
Aldehyde dehydrogenase
PTAC
Signature enzyme
Alcohol dehydrogenase
Regulator

Bacillus sp. FJAT-21351. Bacteria; Firmicutes; Bacilli; Bacillales; Bacillaceae; Bacillus.

AMS61__locus_1

EUT2C

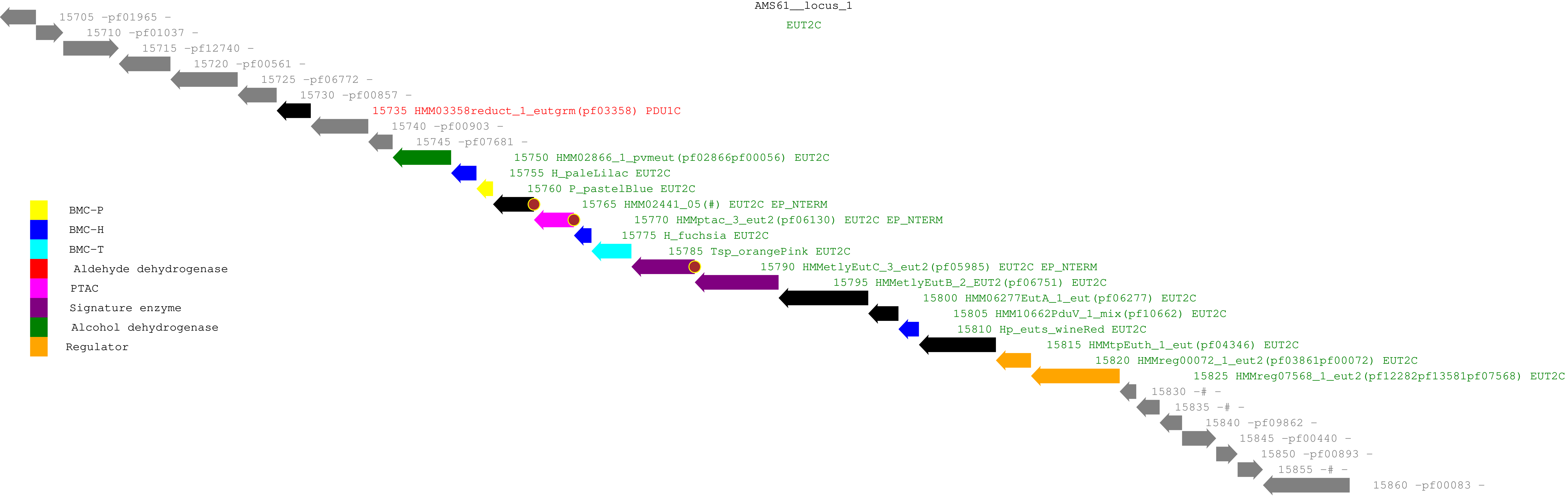

Bacillus sp. Root147. Bacteria; Firmicutes; Bacilli; Bacillales; Bacillaceae; Bacillus.

ASE51__locus_1

EUT2C

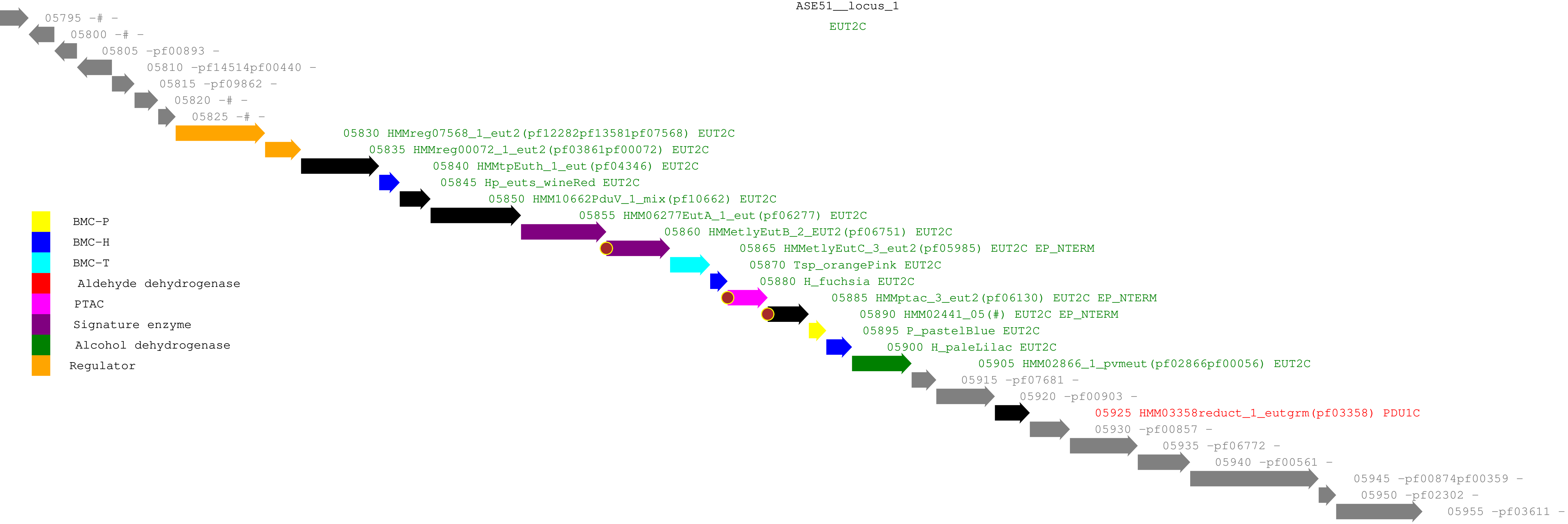

Bacillus sp. Leaf75. Bacteria; Firmicutes; Bacilli; Bacillales; Bacillaceae; Bacillus.

ASG61__locus_1

EUT2C

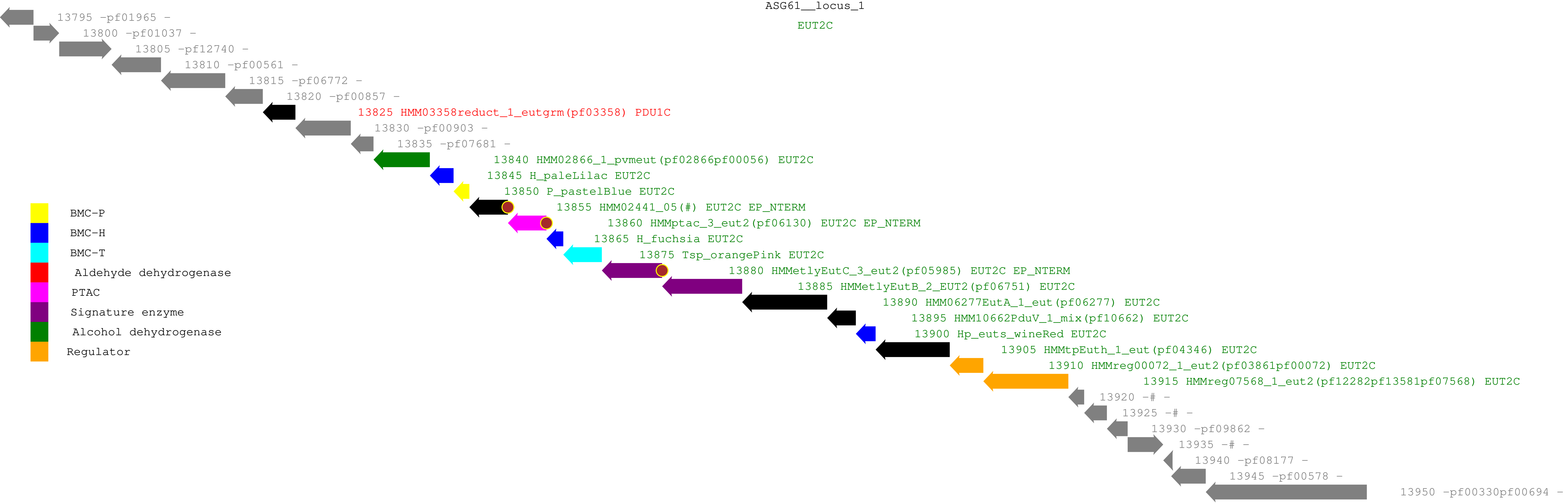

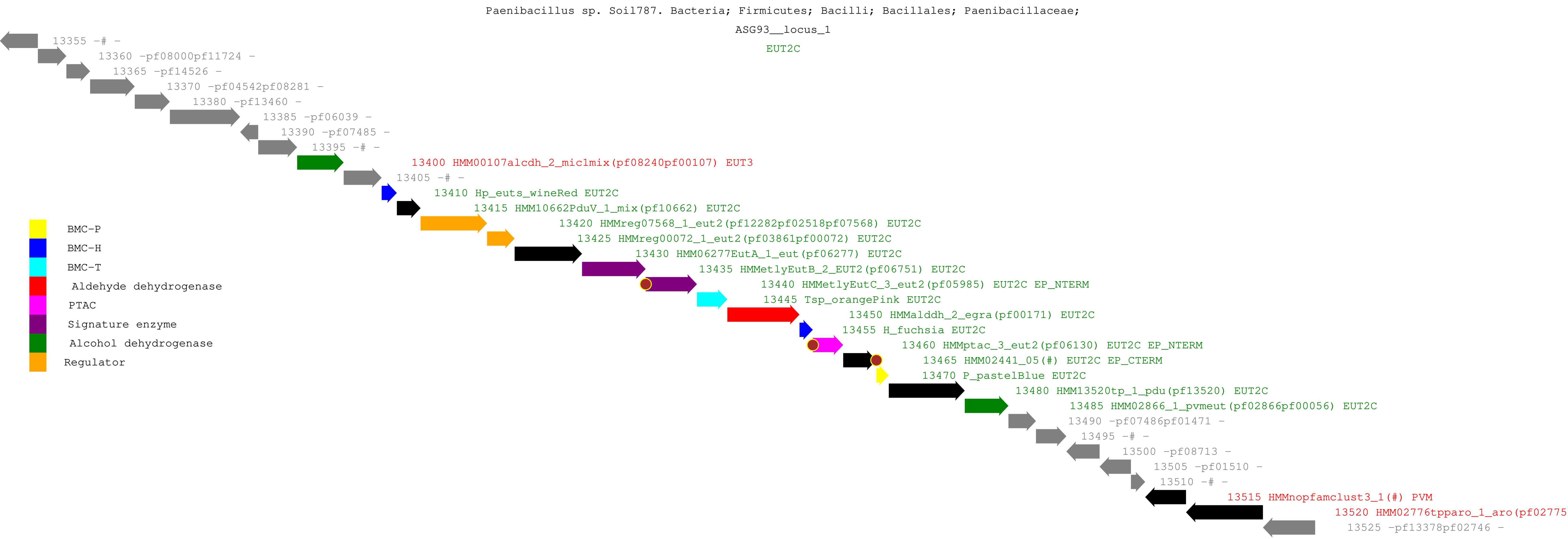
Paenibacillus sp. Soil787. Bacteria; Firmicutes; Bacilli; Bacillales; Paenibacillaceae;
ASG93__locus_1
EUT2C
13355 -# -
13360 -pf08000pf11724 -
13365 -pf14526 -
13370 -pf04542pf08281 -
13380 -pf13460 -
13385 -pf06039 -
13390 -pf07485 -
13395 -# -
13400 HMM00107alcdh_2_mic1mix(pf08240pf00107) EUT3
13405 -# -
13410 Hp_euts_wineRed EUT2C
13415 HMM10662PduV_1_mix(pf10662) EUT2C
13420 HMMreg07568_1_eut2(pf12282pf02518pf07568) EUT2C
13425 HMMreg00072_1_eut2(pf03861pf00072) EUT2C
13430 HMM06277EutA_1_eut(pf06277) EUT2C
13435 HMMetlyEutB_2_EUT2(pf06751) EUT2C
13440 HMMetlyEutC_3_eut2(pf05985) EUT2C EP_NTERM
13445 Tsp_orangePink EUT2C
13450 HMMalddh_2_egra(pf00171) EUT2C
13455 H_fuchsia EUT2C
13460 HMMptac_3_eut2(pf06130) EUT2C EP_NTERM
13465 HMM02441_05(#) EUT2C EP_CTERM
13470 P_pastelBlue EUT2C
13480 HMM13520tp_1_pdu(pf13520) EUT2C
13485 HMM02866_1_pvmeut(pf02866pf00056) EUT2C
13490 -pf07486pf01471 -
13495 -# -
13500 -pf08713 -
13505 -pf01510 -
13510 -# -
13515 HMMnopfamclust3_1(#) PVM
13520 HMM02776tpparo_1_aro(pf02775
13525 -pf13378pf02746 -
BMC-P
BMC-H
BMC-T
Aldehyde dehydrogenase
PTAC
Signature enzyme
Alcohol dehydrogenase
Regulator

Bacillus sp. Soil531. Bacteria; Firmicutes; Bacilli; Bacillales; Bacillaceae; Bacillus.

ASG98__locus_1

EUT2C

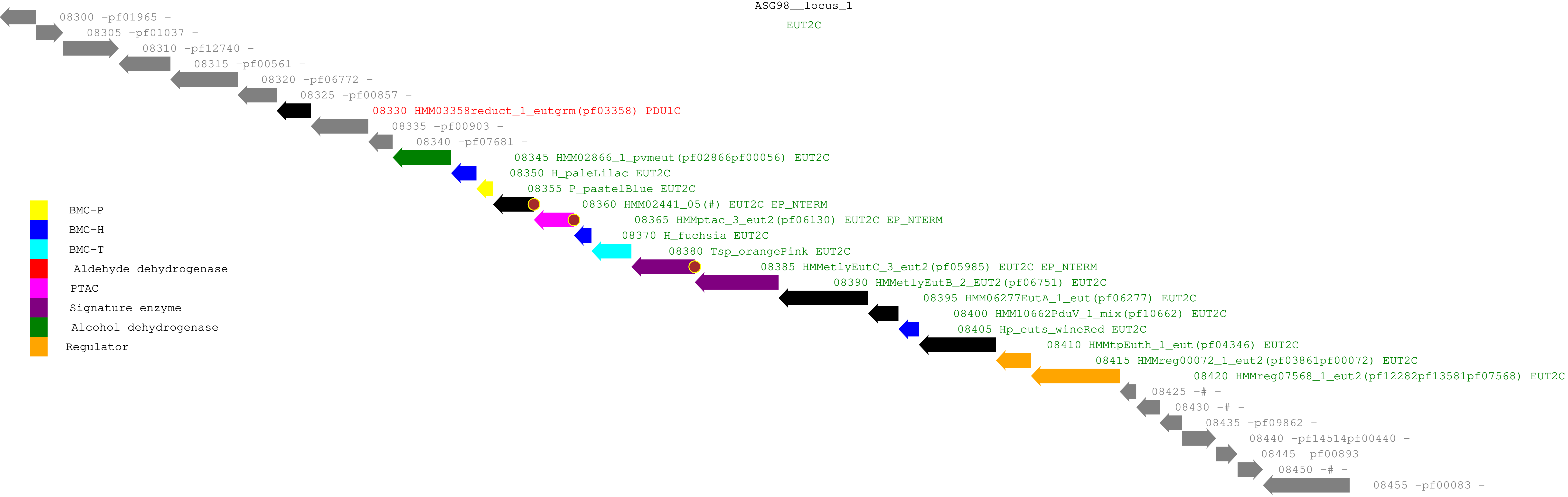

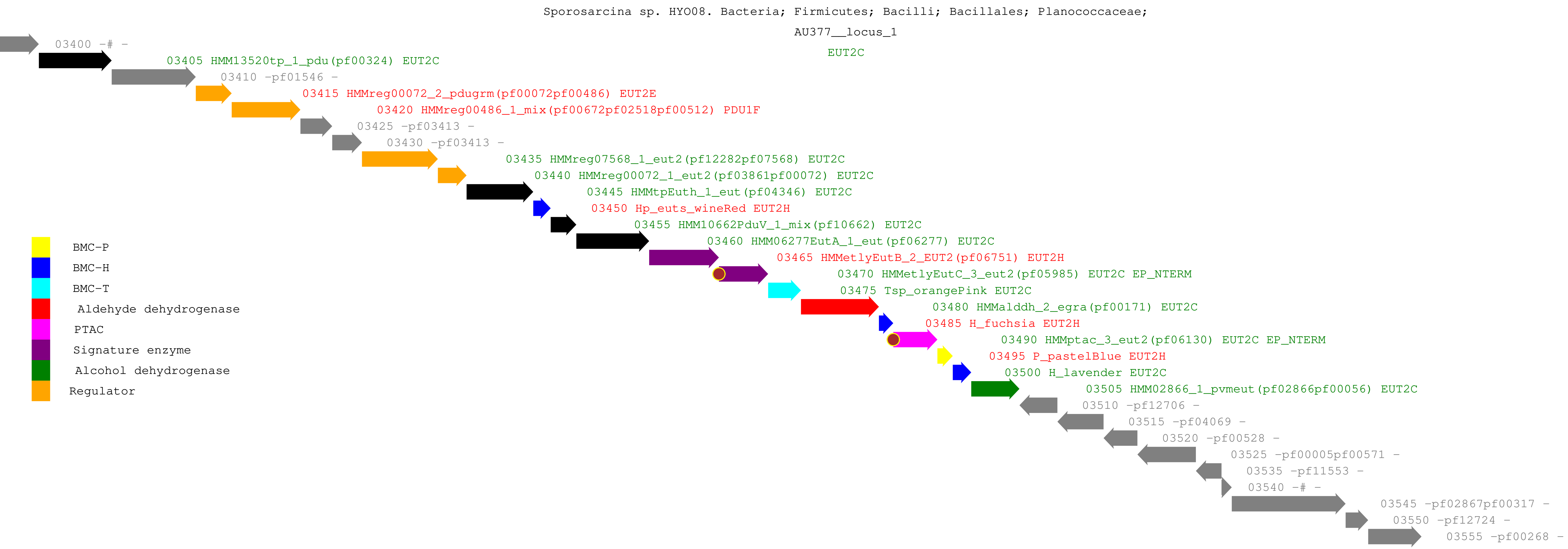
Sporosarcina sp. HYO08. Bacteria; Firmicutes; Bacilli; Bacillales; Planococcaceae;
AU377__locus_1
EUT2C
03400 -# -
03405 HMM13520tp_1_pdu(pf00324) EUT2C
03410 -pf01546 -
03415 HMMreg00072_2_pdugrm(pf00072pf00486) EUT2E
03420 HMMreg00486_1_mix(pf00672pf02518pf00512) PDU1F
03425 -pf03413 -
03430 -pf03413 -
03435 HMMreg07568_1_eut2(pf12282pf07568) EUT2C
03440 HMMreg00072_1_eut2(pf03861pf00072) EUT2C
03445 HMMtpEuth_1_eut(pf04346) EUT2C
03450 Hp_euts_wineRed EUT2H
03455 HMM10662PduV_1_mix(pf10662) EUT2C
03460 HMM06277EutA_1_eut(pf06277) EUT2C
03465 HMMetlyEutB_2_EUT2(pf06751) EUT2H
03470 HMMetlyEutC_3_eut2(pf05985) EUT2C EP_NTERM
03475 Tsp_orangePink EUT2C
03480 HMMalddh_2_egra(pf00171) EUT2C
03485 H_fuchsia EUT2H
03490 HMMptac_3_eut2(pf06130) EUT2C EP_NTERM
03495 P_pastelBlue EUT2H
03500 H_lavender EUT2C
03505 HMM02866_1_pvmeut(pf02866pf00056) EUT2C
03510 -pf12706 -
03515 -pf04069 -
03520 -pf00528 -
03525 -pf00005pf00571 -
03535 -pf11553 -
03540 -# -
03545 -pf02867pf00317 -
03550 -pf12724 -
03555 -pf00268 -
BMC-P
BMC-H
BMC-T
Aldehyde dehydrogenase
PTAC
Signature enzyme
Alcohol dehydrogenase
Regulator

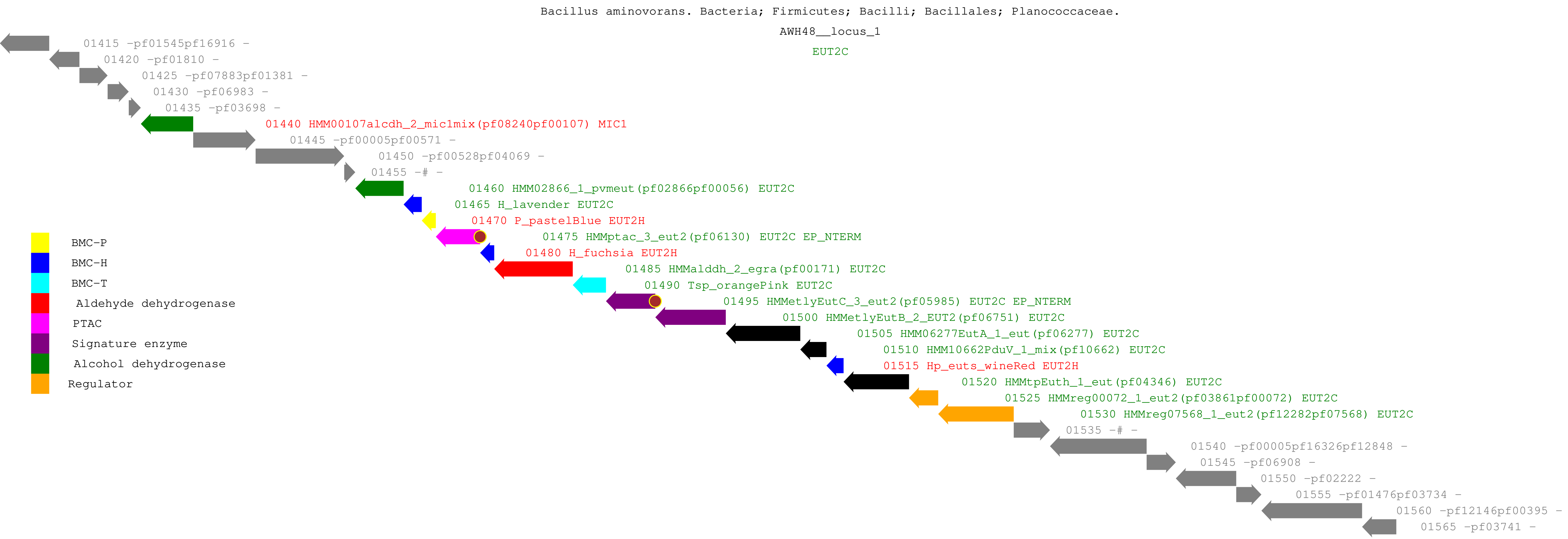

Bacillus aminovorans. Bacteria; Firmicutes; Bacilli; Bacillales; Planococcaceae.
AWH48__locus_1
EUT2C
01415 -pf01545pf16916 -
01420 -pf01810 -
01425 -pf07883pf01381 -
01430 -pf06983 -
01435 -pf03698 -
01440 HMM00107alcdh_2_mic1mix(pf08240pf00107) MIC1
01445 -pf00005pf00571 -
01450 -pf00528pf04069 -
01455 -# -
01460 HMM02866_1_pvmeut(pf02866pf00056) EUT2C
01465 H_lavender EUT2C
01470 P_pastelBlue EUT2H
01475 HMMptac_3_eut2(pf06130) EUT2C EP_NTERM
01480 H_fuchsia EUT2H
01485 HMMalddh_2_egra(pf00171) EUT2C
01490 Tsp_orangePink EUT2C
01495 HMMetlyEutC_3_eut2(pf05985) EUT2C EP_NTERM
01500 HMMetlyEutB_2_EUT2(pf06751) EUT2C
01505 HMM06277EutA_1_eut(pf06277) EUT2C
01510 HMM10662PduV_1_mix(pf10662) EUT2C
01515 Hp_euts_wineRed EUT2H
01520 HMMtpEuth_1_eut(pf04346) EUT2C
01525 HMMreg00072_1_eut2(pf03861pf00072) EUT2C
01530 HMMreg07568_1_eut2(pf12282pf07568) EUT2C
01535 -# -
01540 -pf00005pf16326pf12848 -
01545 -pf06908 -
01550 -pf02222 -
01555 -pf01476pf03734 -
01560 -pf12146pf00395 -
01565 -pf03741 -
BMC-P
BMC-H
BMC-T
Aldehyde dehydrogenase
PTAC
Signature enzyme
Alcohol dehydrogenase
Regulator

Bacillus aminovorans. Bacteria; Firmicutes; Bacilli; Bacillales; Planococcaceae.

AWH49__locus_1

EUT2C

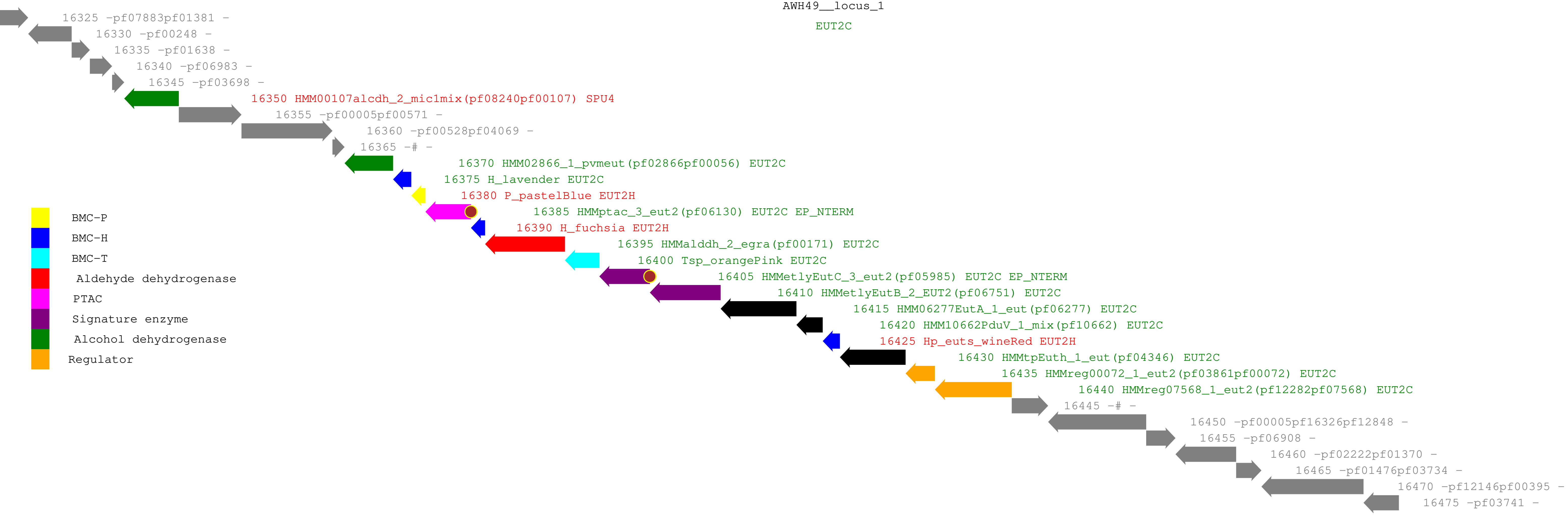

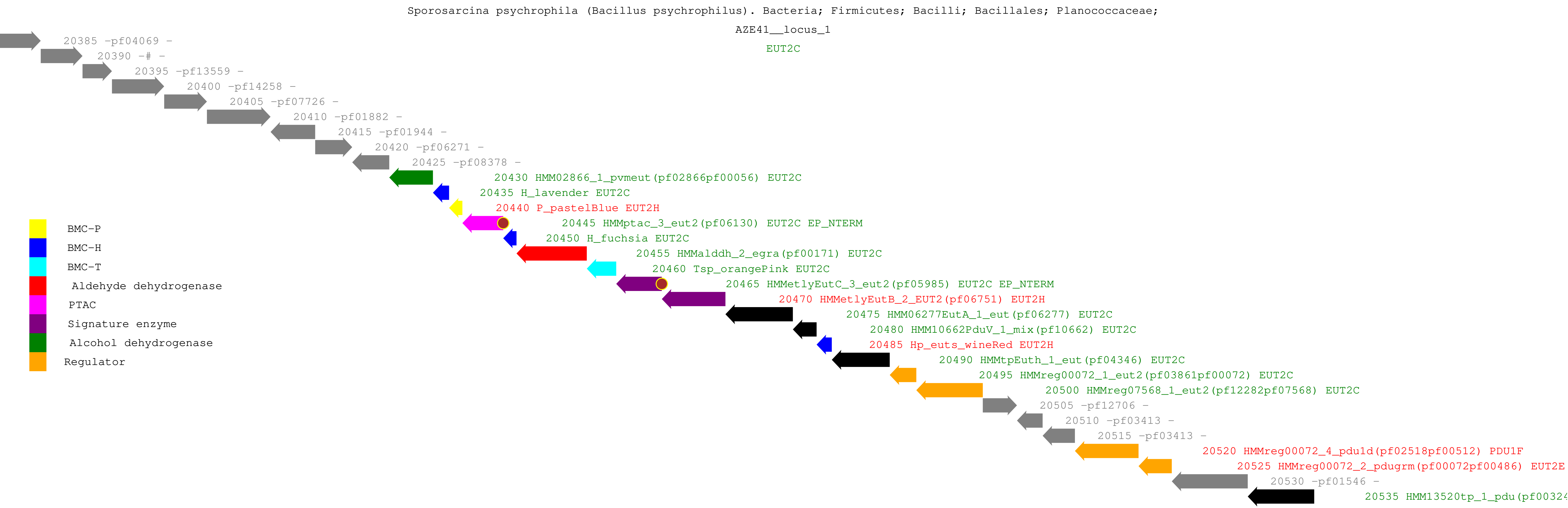

Sporosarcina psychrophila (Bacillus psychrophilus). Bacteria; Firmicutes; Bacilli; Bacillales; Planococcaceae;
AZE41__locus_1
EUT2C
20385 -pf04069 -
20390 -# -
20395 -pf13559 -
20400 -pf14258 -
20405 -pf07726 -
20410 -pf01882 -
20415 -pf01944 -
20420 -pf06271 -
20425 -pf08378 -
20430 HMM02866_1_pvmeut(pf02866pf00056) EUT2C
20435 H_lavender EUT2C
20440 P_pastelBlue EUT2H
20445 HMMptac_3_eut2(pf06130) EUT2C EP_NTERM
20450 H_fuchsia EUT2C
20455 HMMalddh_2_egra(pf00171) EUT2C
20460 Tsp_orangePink EUT2C
20465 HMMetlyEutC_3_eut2(pf05985) EUT2C EP_NTERM
20470 HMMetlyEutB_2_EUT2(pf06751) EUT2H
20475 HMM06277EutA_1_eut(pf06277) EUT2C
20480 HMM10662PduV_1_mix(pf10662) EUT2C
20485 Hp_euts_wineRed EUT2H
20490 HMMtpEuth_1_eut(pf04346) EUT2C
20495 HMMreg00072_1_eut2(pf03861pf00072) EUT2C
20500 HMMreg07568_1_eut2(pf12282pf07568) EUT2C
20505 -pf12706 -
20510 -pf03413 -
20515 -pf03413 -
20520 HMMreg00072_4_pdu1d(pf02518pf00512) PDU1F
20525 HMMreg00072_2_pdugrm(pf00072pf00486) EUT2E
20530 -pf01546 -
20535 HMM13520tp_1_pdu(pf00324
BMC-P
BMC-H
BMC-T
Aldehyde dehydrogenase
PTAC
Signature enzyme
Alcohol dehydrogenase
Regulator

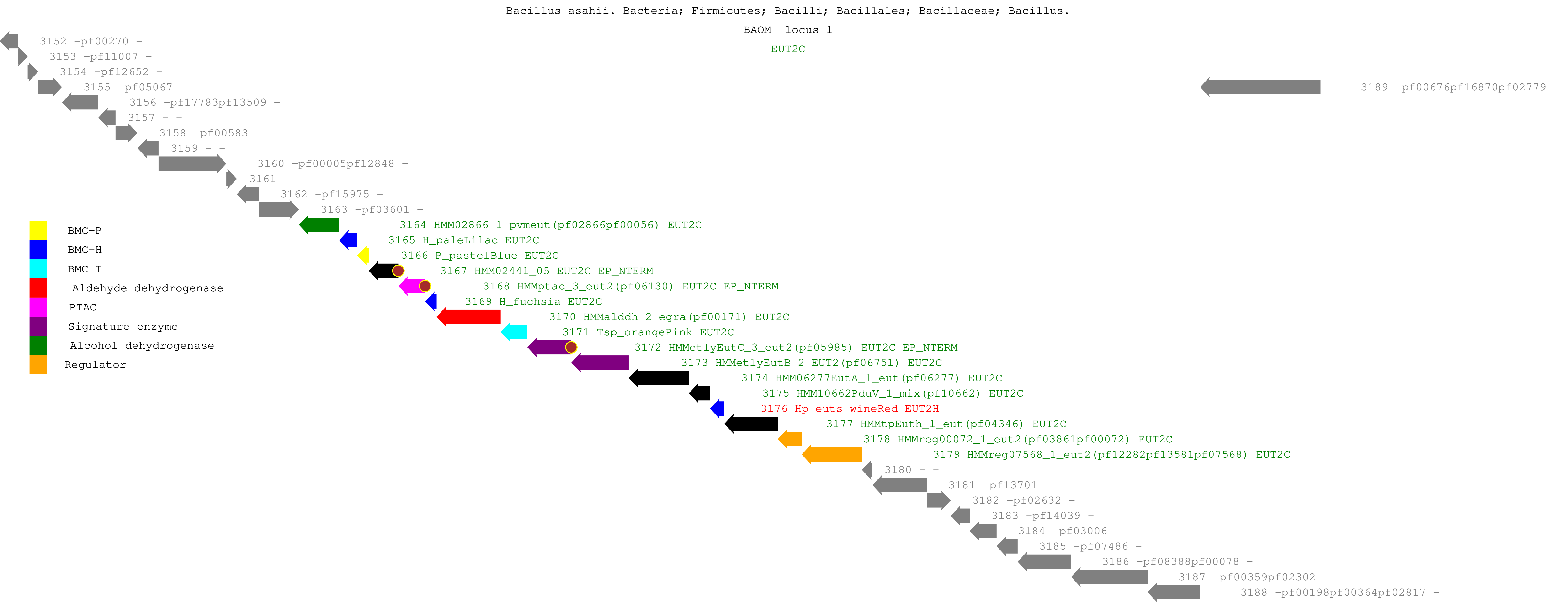

Bacillus asahii. Bacteria; Firmicutes; Bacilli; Bacillales; Bacillaceae; Bacillus.
BAOM__locus_1
EUT2C
3152 -pf00270 -
3153 -pf11007 -
3154 -pf12652 -
3155 -pf05067 -
3156 -pf17783pf13509 -
3157 - -
3158 -pf00583 -
3159 - -
3160 -pf00005pf12848 -
3161 - -
3162 -pf15975 -
3163 -pf03601 -
3164 HMM02866_1_pvmeut(pf02866pf00056) EUT2C
3165 H_paleLilac EUT2C
3166 P_pastelBlue EUT2C
3167 HMM02441_05 EUT2C EP_NTERM
3168 HMMptac_3_eut2(pf06130) EUT2C EP_NTERM
3169 H_fuchsia EUT2C
3170 HMMalddh_2_egra(pf00171) EUT2C
3171 Tsp_orangePink EUT2C
3172 HMMetlyEutC_3_eut2(pf05985) EUT2C EP_NTERM
3173 HMMetlyEutB_2_EUT2(pf06751) EUT2C
3174 HMM06277EutA_1_eut(pf06277) EUT2C
3175 HMM10662PduV_1_mix(pf10662) EUT2C
3176 Hp_euts_wineRed EUT2H
3177 HMMtpEuth_1_eut(pf04346) EUT2C
3178 HMMreg00072_1_eut2(pf03861pf00072) EUT2C
3179 HMMreg07568_1_eut2(pf12282pf13581pf07568) EUT2C
3180 - -
3181 -pf13701 -
3182 -pf02632 -
3183 -pf14039 -
3184 -pf03006 -
3185 -pf07486 -
3186 -pf08388pf00078 -
3187 -pf00359pf02302 -
3188 -pf00198pf00364pf02817 -
3189 -pf00676pf16870pf02779 -
BMC-P
BMC-H
BMC-T
Aldehyde dehydrogenase
PTAC
Signature enzyme
Alcohol dehydrogenase
Regulator

Bacillus oceanisediminis. Bacteria; Firmicutes; Bacilli; Bacillales; Bacillaceae; Bacillus.

BBV17__locus_2

EUT2C

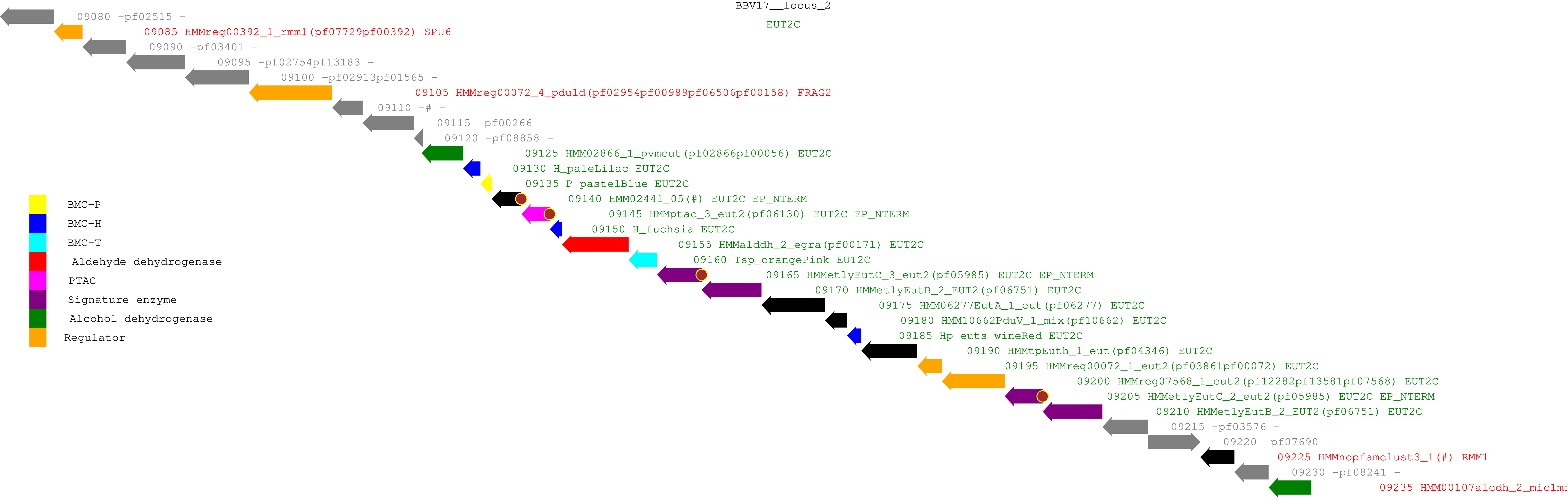

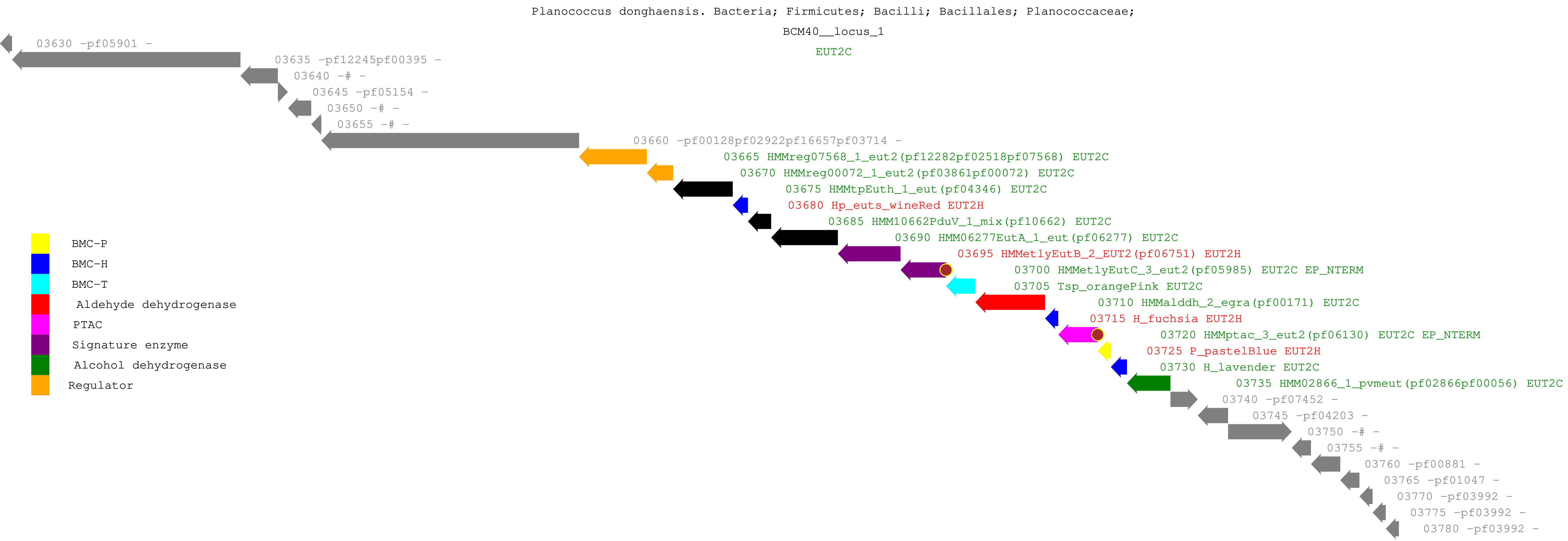

Planococcus donghaensis. Bacteria; Firmicutes; Bacilli; Bacillales; Planococcaceae;
BCM40__locus_1
EUT2C
03630 -pf05901 -
03635 -pf12245pf00395 -
03640 -# -
03645 -pf05154 -
03650 -# -
03655 -# -
03660 -pf00128pf02922pf16657pf03714 -
03665 HMMreg07568_1_eut2(pf12282pf02518pf07568) EUT2C
03670 HMMreg00072_1_eut2(pf03861pf00072) EUT2C
03675 HMMtpEuth_1_eut(pf04346) EUT2C
03680 Hp_euts_wineRed EUT2H
03685 HMM10662PduV_1_mix(pf10662) EUT2C
03690 HMM06277EutA_1_eut(pf06277) EUT2C
03695 HMMetlyEutB_2_EUT2(pf06751) EUT2H
03700 HMMetlyEutC_3_eut2(pf05985) EUT2C EP_NTERM
03705 Tsp_orangePink EUT2C
03710 HMMaldh_2_egra(pf00171) EUT2C
03715 H_fuchsia EUT2H
03720 HMMptac_3_eut2(pf06130) EUT2C EP_NTERM
03725 P_pastelBlue EUT2H
03730 H_lavender EUT2C
03735 HMM02866_1_pvmeut(pf02866pf00056) EUT2C
03740 -pf07452 -
03745 -pf04203 -
03750 -# -
03755 -# -
03760 -pf00881 -
03765 -pf01047 -
03770 -pf03992 -
03775 -pf03992 -
03780 -pf03992 -
BMC-P
BMC-H
BMC-T
Aldehyde dehydrogenase
PTAC
Signature enzyme
Alcohol dehydrogenase
Regulator

Bacillus aryabhattai. Bacteria; Firmicutes; Bacilli; Bacillales; Bacillaceae; Bacillus.

BCV52__locus_1

EUT2C

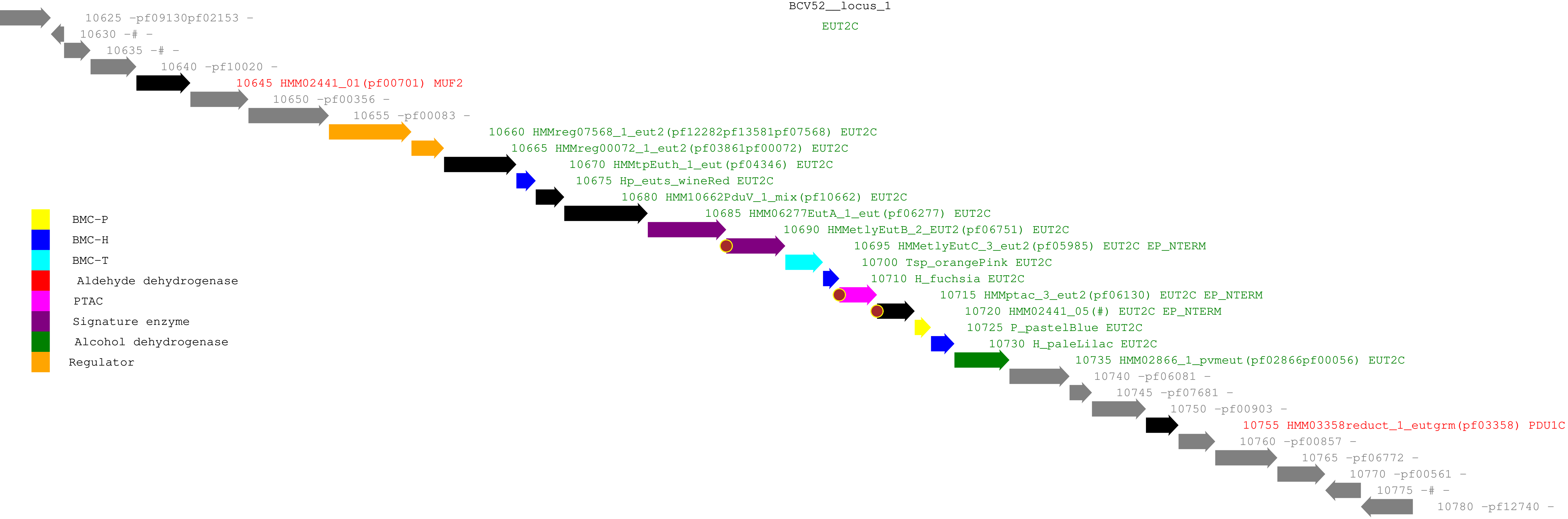

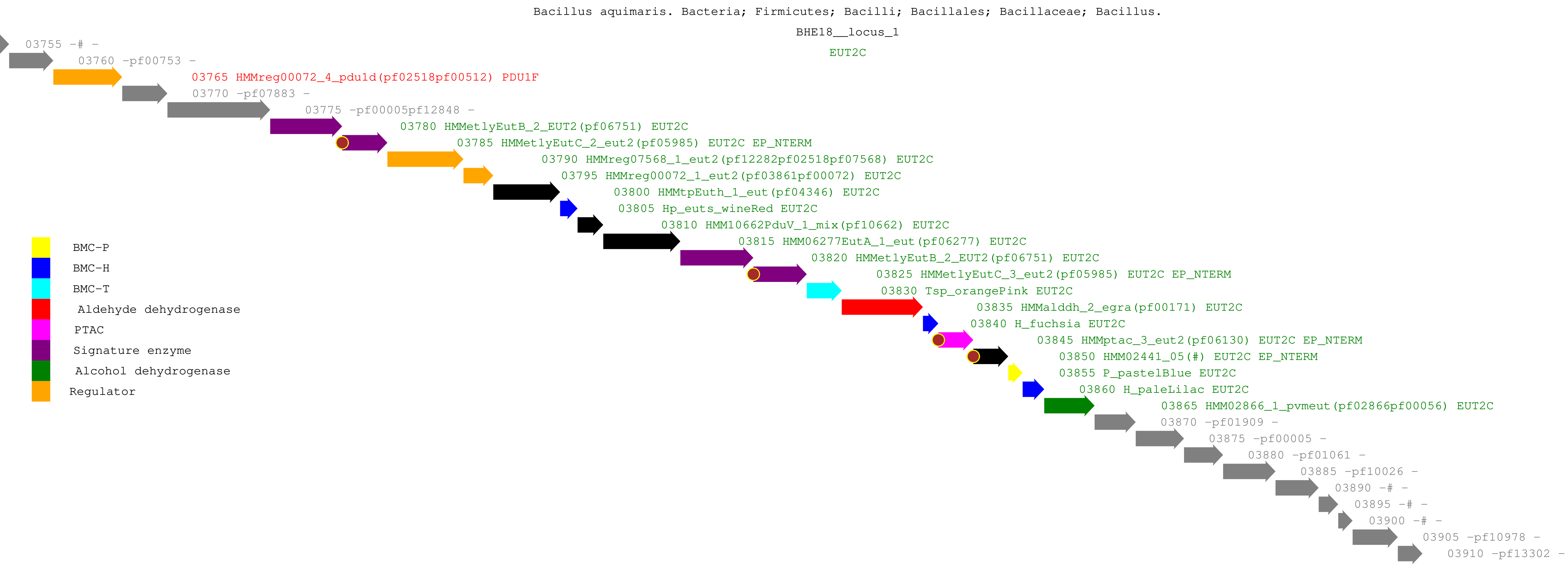
Bacillus aquimaris. Bacteria; Firmicutes; Bacilli; Bacillales; Bacillaceae; Bacillus.
BHE18__locus_1
EUT2C
03755 -# -
03760 -pf00753 -
03765 HMMreg00072_4_pdu1d(pf02518pf00512) PDU1F
03770 -pf07883 -
03775 -pf00005pf12848 -
03780 HMMetlyEutB_2_EUT2(pf06751) EUT2C
03785 HMMetlyEutC_2_eut2(pf05985) EUT2C EP_NTERM
03790 HMMreg07568_1_eut2(pf12282pf02518pf07568) EUT2C
03795 HMMreg00072_1_eut2(pf03861pf00072) EUT2C
03800 HMMtpEuth_1_eut(pf04346) EUT2C
03805 Hp_euts_wineRed EUT2C
03810 HMM10662PduV_1_mix(pf10662) EUT2C
03815 HMM06277EutA_1_eut(pf06277) EUT2C
03820 HMMetlyEutB_2_EUT2(pf06751) EUT2C
03825 HMMetlyEutC_3_eut2(pf05985) EUT2C EP_NTERM
03830 Tsp_orangePink EUT2C
03835 HMMalddh_2_egra(pf00171) EUT2C
03840 H_fuchsia EUT2C
03845 HMMptac_3_eut2(pf06130) EUT2C EP_NTERM
03850 HMM02441_05(#) EUT2C EP_NTERM
03855 P_pastelBlue EUT2C
03860 H_paleLilac EUT2C
03865 HMM02866_1_pvmeut(pf02866pf00056) EUT2C
03870 -pf01909 -
03875 -pf00005 -
03880 -pf01061 -
03885 -pf10026 -
03890 -# -
03895 -# -
03900 -# -
03905 -pf10978 -
03910 -pf13302 -
BMC-P
BMC-H
BMC-T
Aldehyde dehydrogenase
PTAC
Signature enzyme
Alcohol dehydrogenase
Regulator

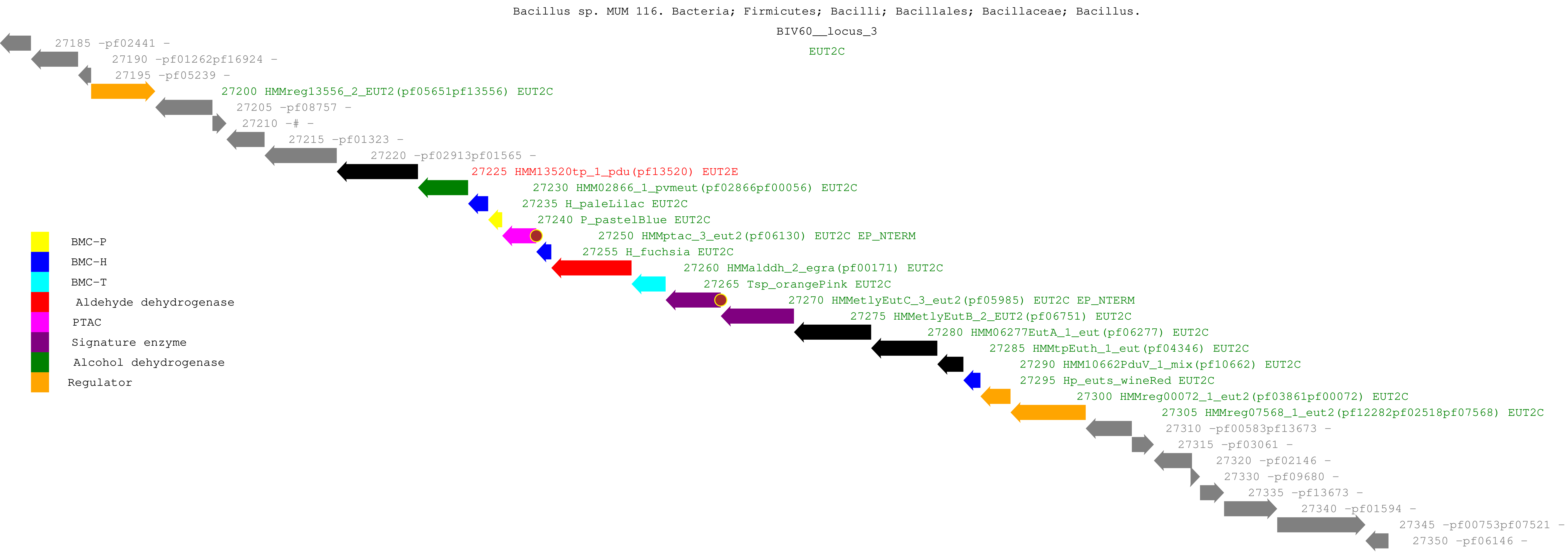
Bacillus sp. MUM 116. Bacteria; Firmicutes; Bacilli; Bacillales; Bacillaceae; Bacillus.
BIV60__locus_3
EUT2C
27185 -pf02441 -
27190 -pf01262pf16924 -
27195 -pf05239 -
27200 HMMreg13556_2_EUT2(pf05651pf13556) EUT2C
27205 -pf08757 -
27210 -# -
27215 -pf01323 -
27220 -pf02913pf01565 -
27225 HMM13520tp_1_pdu(pf13520) EUT2E
27230 HMM02866_1_pvmeut(pf02866pf00056) EUT2C
27235 H_paleLilac EUT2C
27240 P_pastelBlue EUT2C
27250 HMMptac_3_eut2(pf06130) EUT2C EP_NTERM
27255 H_fuchsia EUT2C
27260 HMMaldh_2_egra(pf00171) EUT2C
27265 Tsp_orangePink EUT2C
27270 HMMetlyEutC_3_eut2(pf05985) EUT2C EP_NTERM
27275 HMMetlyEutB_2_EUT2(pf06751) EUT2C
27280 HMM06277EutA_1_eut(pf06277) EUT2C
27285 HMMtpEuth_1_eut(pf04346) EUT2C
27290 HMM10662PduV_1_mix(pf10662) EUT2C
27295 Hp_euts_wineRed EUT2C
27300 HMMreg00072_1_eut2(pf03861pf00072) EUT2C
27305 HMMreg07568_1_eut2(pf12282pf02518pf07568) EUT2C
27310 -pf00583pf13673 -
27315 -pf03061 -
27320 -pf02146 -
27330 -pf09680 -
27335 -pf13673 -
27340 -pf01594 -
27345 -pf00753pf07521 -
27350 -pf06146 -
BMC-P
BMC-H
BMC-T
Aldehyde dehydrogenase
PTAC
Signature enzyme
Alcohol dehydrogenase
Regulator

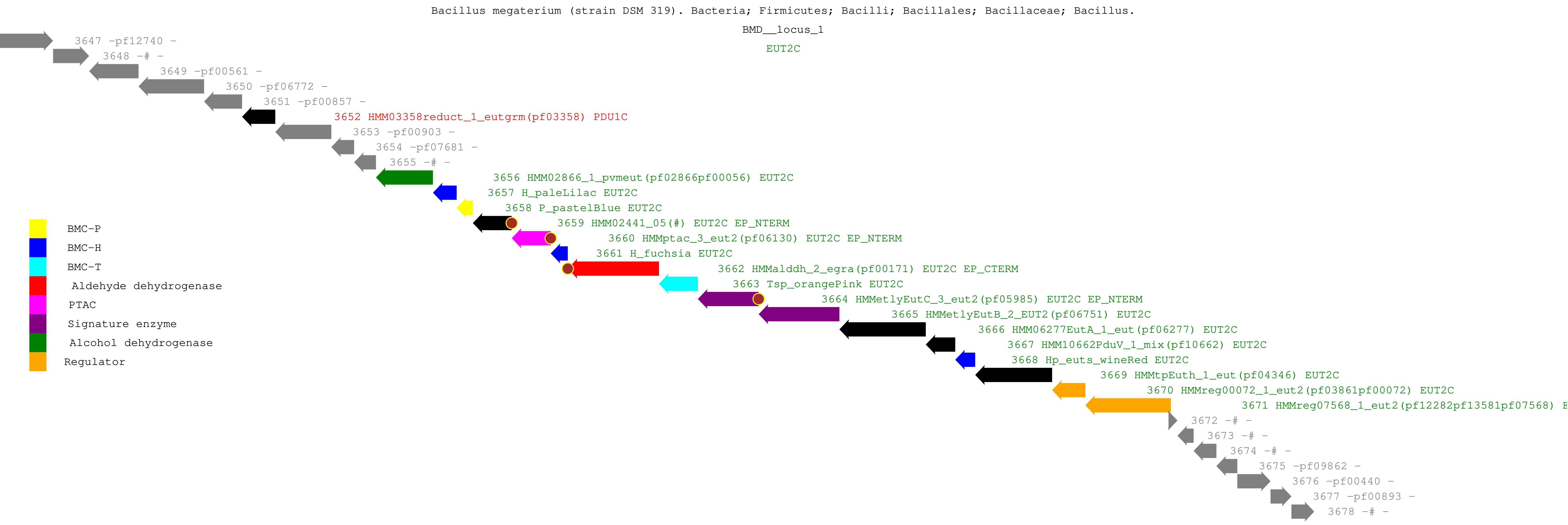
Bacillus megaterium (strain DSM 319). Bacteria; Firmicutes; Bacilli; Bacillales; Bacillaceae; Bacillus.
BMD__locus_1
EUT2C
3647 -pf12740 -
3648 -# -
3649 -pf00561 -
3650 -pf06772 -
3651 -pf00857 -
3652 HMM03358reduct_1_eutgrm(pf03358) PDU1C
3653 -pf00903 -
3654 -pf07681 -
3655 -# -
3656 HMM02866_1_pvmeut(pf02866pf00056) EUT2C
3657 H_paleLilac EUT2C
3658 P_pastelBlue EUT2C
3659 HMM02441_05(#) EUT2C EP_NTERM
3660 HMMptac_3_eut2(pf06130) EUT2C EP_NTERM
3661 H_fuchsia EUT2C
3662 HMMalddh_2_egra(pf00171) EUT2C EP_CTERM
3663 Tsp_orangePink EUT2C
3664 HMMetlyEutC_3_eut2(pf05985) EUT2C EP_NTERM
3665 HMMetlyEutB_2_EUT2(pf06751) EUT2C
3666 HMM06277EutA_1_eut(pf06277) EUT2C
3667 HMM10662PduV_1_mix(pf10662) EUT2C
3668 Hp_euts_wineRed EUT2C
3669 HMMtpEuth_1_eut(pf04346) EUT2C
3670 HMMreg00072_1_eut2(pf03861pf00072) EUT2C
3671 HMMreg07568_1_eut2(pf12282pf13581pf07568) E
3672 -# -
3673 -# -
3674 -# -
3675 -pf09862 -
3676 -pf00440 -
3677 -pf00893 -
3678 -# -
BMC-P
BMC-H
BMC-T
Aldehyde dehydrogenase
PTAC
Signature enzyme
Alcohol dehydrogenase
Regulator

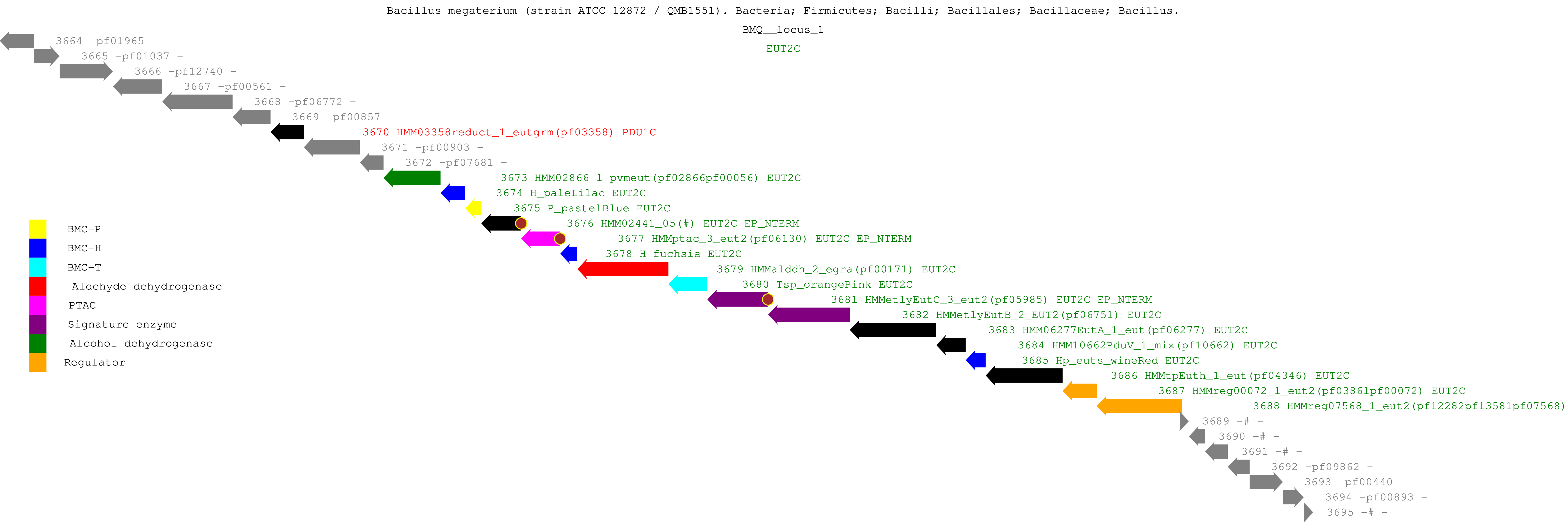

Bacillus megaterium (strain ATCC 12872 / QMB1551). Bacteria; Firmicutes; Bacilli; Bacillales; Bacillaceae; Bacillus.
BMQ__locus_1
EUT2C
3664 -pf01965 -
3665 -pf01037 -
3666 -pf12740 -
3667 -pf00561 -
3668 -pf06772 -
3669 -pf00857 -
3670 HMM03358reduct_1_eutgrm(pf03358) PDU1C
3671 -pf00903 -
3672 -pf07681 -
3673 HMM02866_1_pvmeut(pf02866pf00056) EUT2C
3674 H_paleLilac EUT2C
3675 P_pastelBlue EUT2C
3676 HMM02441_05(#) EUT2C EP_NTERM
3677 HMMptac_3_eut2(pf06130) EUT2C EP_NTERM
3678 H_fuchsia EUT2C
3679 HMMalddh_2_egra(pf00171) EUT2C
3680 Tsp_orangePink EUT2C
3681 HMMetlyEutC_3_eut2(pf05985) EUT2C EP_NTERM
3682 HMMetlyEutB_2_EUT2(pf06751) EUT2C
3683 HMM06277EutA_1_eut(pf06277) EUT2C
3684 HMM10662PduV_1_mix(pf10662) EUT2C
3685 Hp_euts_wineRed EUT2C
3686 HMMtpEuth_1_eut(pf04346) EUT2C
3687 HMMreg00072_1_eut2(pf03861pf00072) EUT2C
3688 HMMreg07568_1_eut2(pf12282pf13581pf07568)
3689 -# -
3690 -# -
3691 -# -
3692 -pf09862 -
3693 -pf00440 -
3694 -pf00893 -
3695 -# -
BMC-P
BMC-H
BMC-T
Aldehyde dehydrogenase
PTAC
Signature enzyme
Alcohol dehydrogenase
Regulator

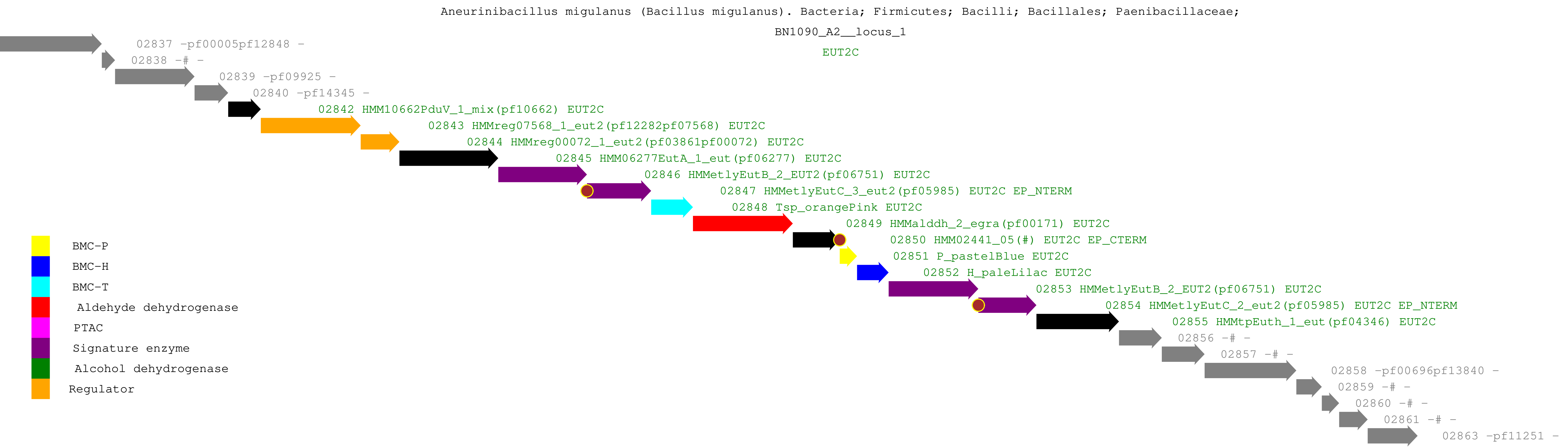
Aneurinibacillus migulanus (Bacillus migulanus). Bacteria; Firmicutes; Bacilli; Bacillales; Paenibacillaceae;
BN1090_A2__locus_1
EUT2C
02837 -pf00005pf12848 -
02838 -# -
02839 -pf09925 -
02840 -pf14345 -
02842 HMM10662PduV_1_mix(pf10662) EUT2C
02843 HMMreg07568_1_eut2(pf12282pf07568) EUT2C
02844 HMMreg00072_1_eut2(pf03861pf00072) EUT2C
02845 HMM06277EutA_1_eut(pf06277) EUT2C
02846 HMMetlyEutB_2_EUT2(pf06751) EUT2C
02847 HMMetlyEutC_3_eut2(pf05985) EUT2C EP_NTERM
02848 Tsp_orangePink EUT2C
02849 HMMalddh_2_egra(pf00171) EUT2C
02850 HMM02441_05(#) EUT2C EP_CTERM
02851 P_pastelBlue EUT2C
02852 H_paleLilac EUT2C
02853 HMMetlyEutB_2_EUT2(pf06751) EUT2C
02854 HMMetlyEutC_2_eut2(pf05985) EUT2C EP_NTERM
02855 HMMtpEuth_1_eut(pf04346) EUT2C
02856 -# -
02857 -# -
02858 -pf00696pf13840 -
02859 -# -
02860 -# -
02861 -# -
02863 -pf11251 -
BMC-P
BMC-H
BMC-T
Aldehyde dehydrogenase
PTAC
Signature enzyme
Alcohol dehydrogenase
Regulator

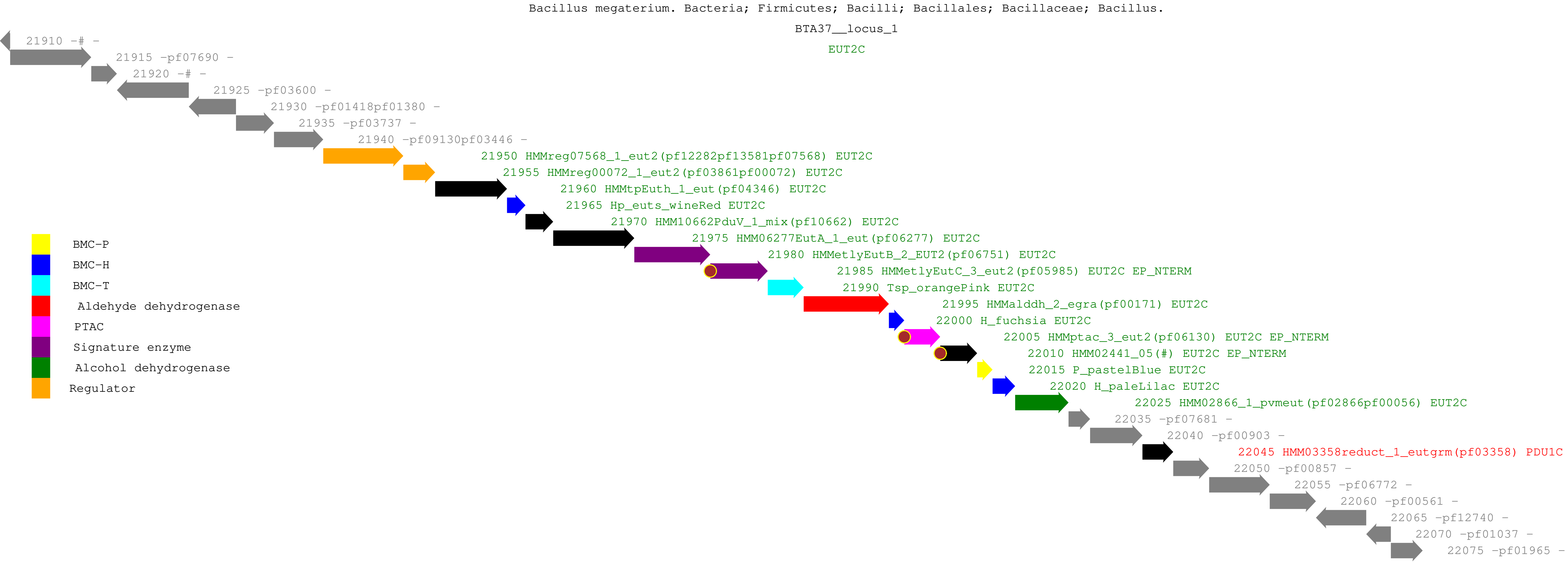
Bacillus megaterium. Bacteria; Firmicutes; Bacilli; Bacillales; Bacillaceae; Bacillus.
BTA37__locus_1
EUT2C
21910 -# -
21915 -pf07690 -
21920 -# -
21925 -pf03600 -
21930 -pf01418pf01380 -
21935 -pf03737 -
21940 -pf09130pf03446 -
21950 HMMreg07568_1_eut2(pf12282pf13581pf07568) EUT2C
21955 HMMreg00072_1_eut2(pf03861pf00072) EUT2C
21960 HMMtpEuth_1_eut(pf04346) EUT2C
21965 Hp_euts_wineRed EUT2C
21970 HMM10662PduV_1_mix(pf10662) EUT2C
21975 HMM06277EutA_1_eut(pf06277) EUT2C
21980 HMMetlyEutB_2_EUT2(pf06751) EUT2C
21985 HMMetlyEutC_3_eut2(pf05985) EUT2C EP_NTERM
21990 Tsp_orangePink EUT2C
21995 HMMalddh_2_egra(pf00171) EUT2C
22000 H_fuchsia EUT2C
22005 HMMptac_3_eut2(pf06130) EUT2C EP_NTERM
22010 HMM02441_05(#) EUT2C EP_NTERM
22015 P_pastelBlue EUT2C
22020 H_paleLilac EUT2C
22025 HMM02866_1_pvmeut(pf02866pf00056) EUT2C
22035 -pf07681 -
22040 -pf00903 -
22045 HMM03358reduct_1_eutgrm(pf03358) PDU1C
22050 -pf00857 -
22055 -pf06772 -
22060 -pf00561 -
22065 -pf12740 -
22070 -pf01037 -
22075 -pf01965 -
BMC-P
BMC-H
BMC-T
Aldehyde dehydrogenase
PTAC
Signature enzyme
Alcohol dehydrogenase
Regulator

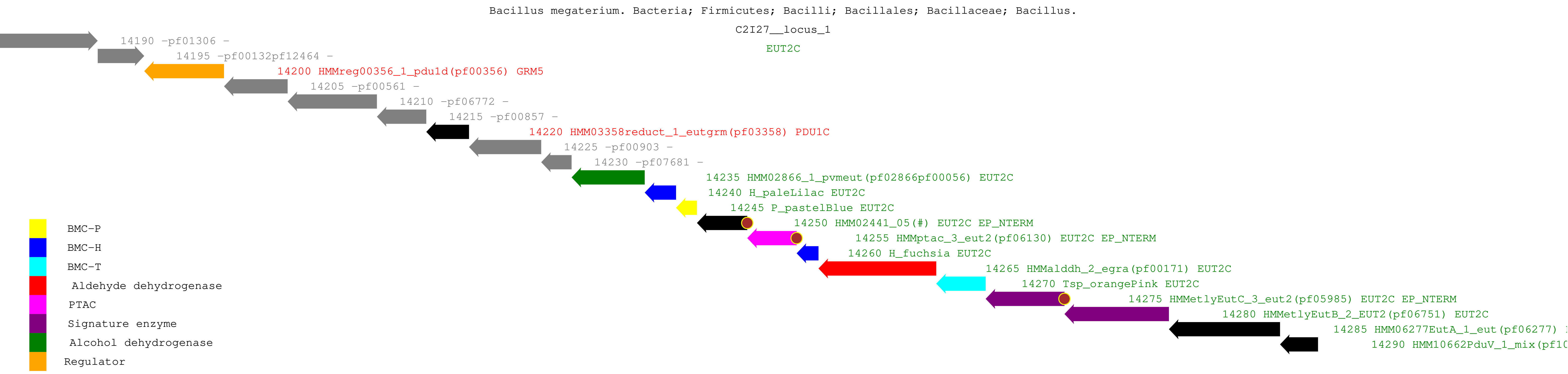

Bacillus megaterium. Bacteria; Firmicutes; Bacilli; Bacillales; Bacillaceae; Bacillus.
C2I27__locus_1
EUT2C
14190 -pf01306 -
14195 -pf00132pf12464 -
14200 HMMreg00356_1_pdu1d(pf00356) GRM5
14205 -pf00561 -
14210 -pf06772 -
14215 -pf00857 -
14220 HMM03358reduct_1_eutgrm(pf03358) PDU1C
14225 -pf00903 -
14230 -pf07681 -
14235 HMM02866_1_pvmeut(pf02866pf00056) EUT2C
14240 H_paleLilac EUT2C
14245 P_pastelBlue EUT2C
14250 HMM02441_05(#) EUT2C EP_NTERM
14255 HMMptac_3_eut2(pf06130) EUT2C EP_NTERM
14260 H_fuchsia EUT2C
14265 HMMalddh_2_egra(pf00171) EUT2C
14270 Tsp_orangePink EUT2C
14275 HMMetlyEutC_3_eut2(pf05985) EUT2C EP_NTERM
14280 HMMetlyEutB_2_EUT2(pf06751) EUT2C
14285 HMM06277EutA_1_eut(pf06277)
BMC-P
BMC-H
BMC-T
Aldehyde dehydrogenase
PTAC
Signature enzyme
Alcohol dehydrogenase
Regulator

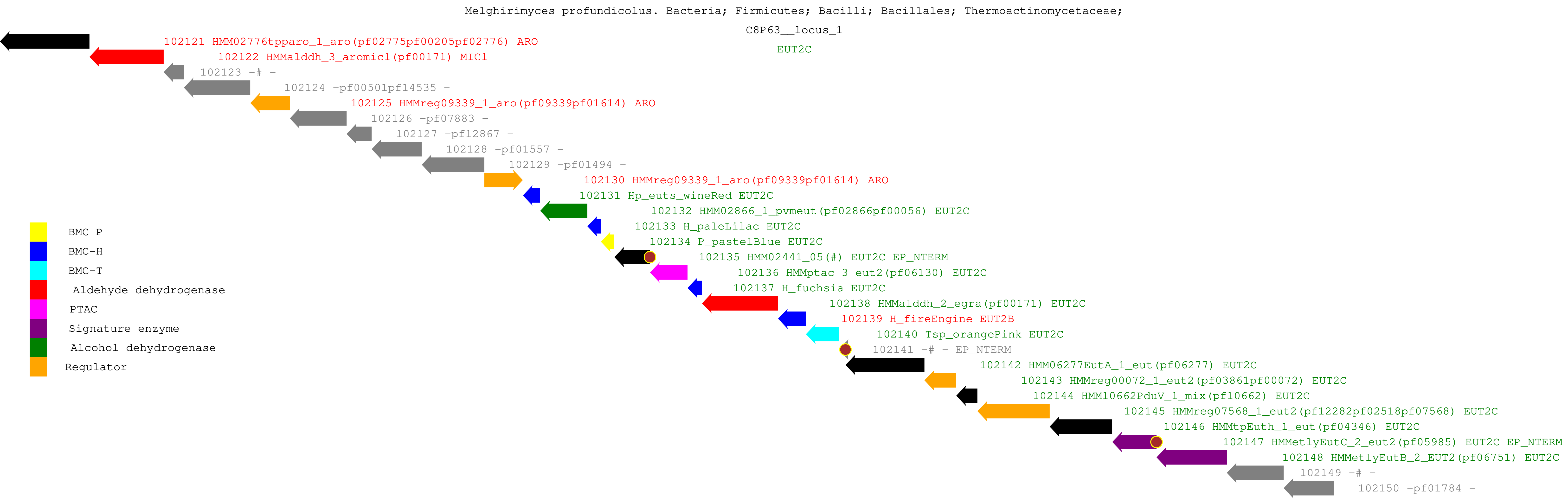
Melghirimyces profundicolus. Bacteria; Firmicutes; Bacilli; Bacillales; Thermoactinomycetaceae;
C8P63__locus_1
EUT2C
102121 HMM02776tpparo_1_aro(pf02775pf00205pf02776) ARO
102122 HMMalddh_3_aromic1(pf00171) MIC1
102123 -# -
102124 -pf00501pf14535 -
102125 HMMreg09339_1_aro(pf09339pf01614) ARO
102126 -pf07883 -
102127 -pf12867 -
102128 -pf01557 -
102129 -pf01494 -
102130 HMMreg09339_1_aro(pf09339pf01614) ARO
102131 Hp_euts_wineRed EUT2C
102132 HMM02866_1_pvmeut(pf02866pf00056) EUT2C
102133 H_paleLilac EUT2C
102134 P_pastelBlue EUT2C
102135 HMM02441_05(#) EUT2C EP_NTERM
102136 HMMptac_3_eut2(pf06130) EUT2C
102137 H_fuchsia EUT2C
102138 HMMalddh_2_egra(pf00171) EUT2C
102139 H_fireEngine EUT2B
102140 Tsp_orangePink EUT2C
102141 -# - EP_NTERM
102142 HMM06277EutA_1_eut(pf06277) EUT2C
102143 HMMreg00072_1_eut2(pf03861pf00072) EUT2C
102144 HMM10662PduV_1_mix(pf10662) EUT2C
102145 HMMreg07568_1_eut2(pf12282pf02518pf07568) EUT2C
102146 HMMtpEuth_1_eut(pf04346) EUT2C
102147 HMMetlyEutC_2_eut2(pf05985) EUT2C EP_NTERM
102148 HMMetlyEutB_2_EUT2(pf06751) EUT2C
102149 -# -
102150 -pf01784 -
BMC-P
BMC-H
BMC-T
Aldehyde dehydrogenase
PTAC
Signature enzyme
Alcohol dehydrogenase
Regulator

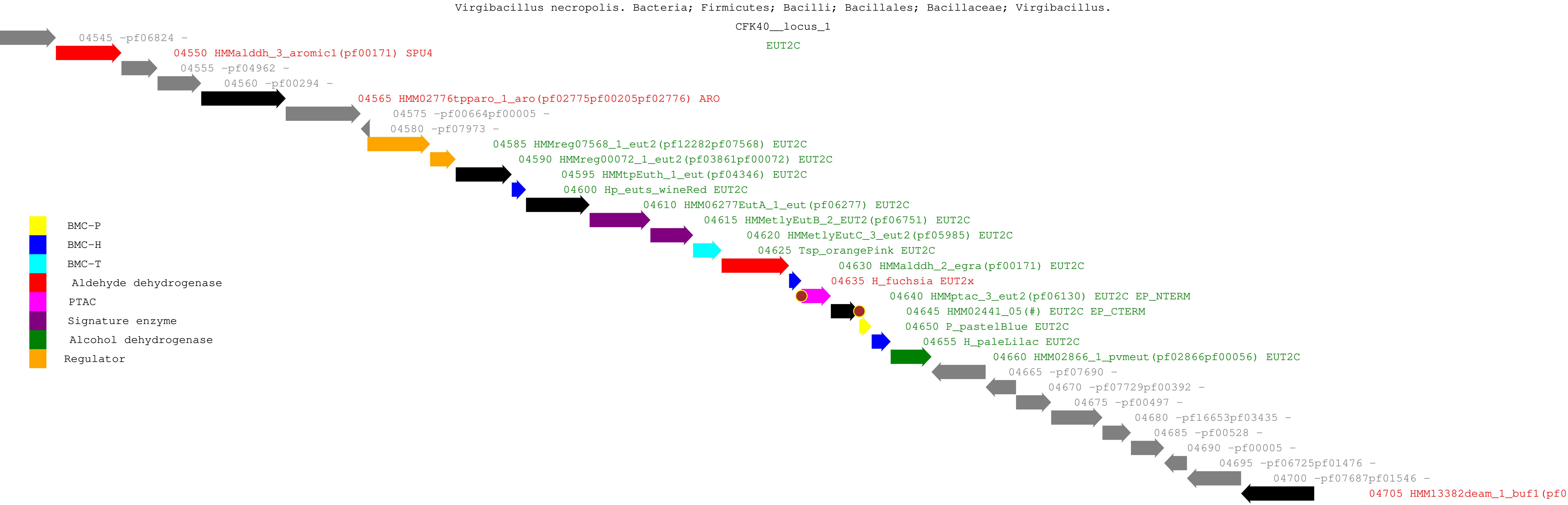
Virgibacillus necropolis. Bacteria; Firmicutes; Bacilli; Bacillales; Bacillaceae; Virgibacillus.
CFK40__locus_1
EUT2C
04545 -pf06824 -
04550 HMMalddh_3_aromic1(pf00171) SPU4
04555 -pf04962 -
04560 -pf00294 -
04565 HMM02776tpparo_1_aro(pf02775pf00205pf02776) ARO
04575 -pf00664pf00005 -
04580 -pf07973 -
04585 HMMreg07568_1_eut2(pf12282pf07568) EUT2C
04590 HMMreg00072_1_eut2(pf03861pf00072) EUT2C
04595 HMMtpEuth_1_eut(pf04346) EUT2C
04600 Hp_euts_wineRed EUT2C
04610 HMM06277EutA_1_eut(pf06277) EUT2C
04615 HMMetlyEutB_2_EUT2(pf06751) EUT2C
04620 HMMetlyEutC_3_eut2(pf05985) EUT2C
04625 Tsp_orangePink EUT2C
04630 HMMalddh_2_egra(pf00171) EUT2C
04635 H_fuchsia EUT2x
04640 HMMptac_3_eut2(pf06130) EUT2C EP_NTERM
04645 HMM02441_05(#) EUT2C EP_CTERM
04650 P_pastelBlue EUT2C
04655 H_paleLilac EUT2C
04660 HMM02866_1_pvmeut(pf02866pf00056) EUT2C
04665 -pf07690 -
04670 -pf07729pf00392 -
04675 -pf00497 -
04680 -pf16653pf03435 -
04685 -pf00528 -
04690 -pf00005 -
04695 -pf06725pf01476 -
04700 -pf07687pf01546 -
04705 HMM13382deam_1_buf1(pf0
BMC-P
BMC-H
BMC-T
Aldehyde dehydrogenase
PTAC
Signature enzyme
Alcohol dehydrogenase
Regulator

Fictibacillus sp. GDSW-R2A3. Bacteria; Firmicutes; Bacilli; Bacillales; Bacillaceae; Fictibacillus.

CGZ90__locus_1_main_sat

EUT2C

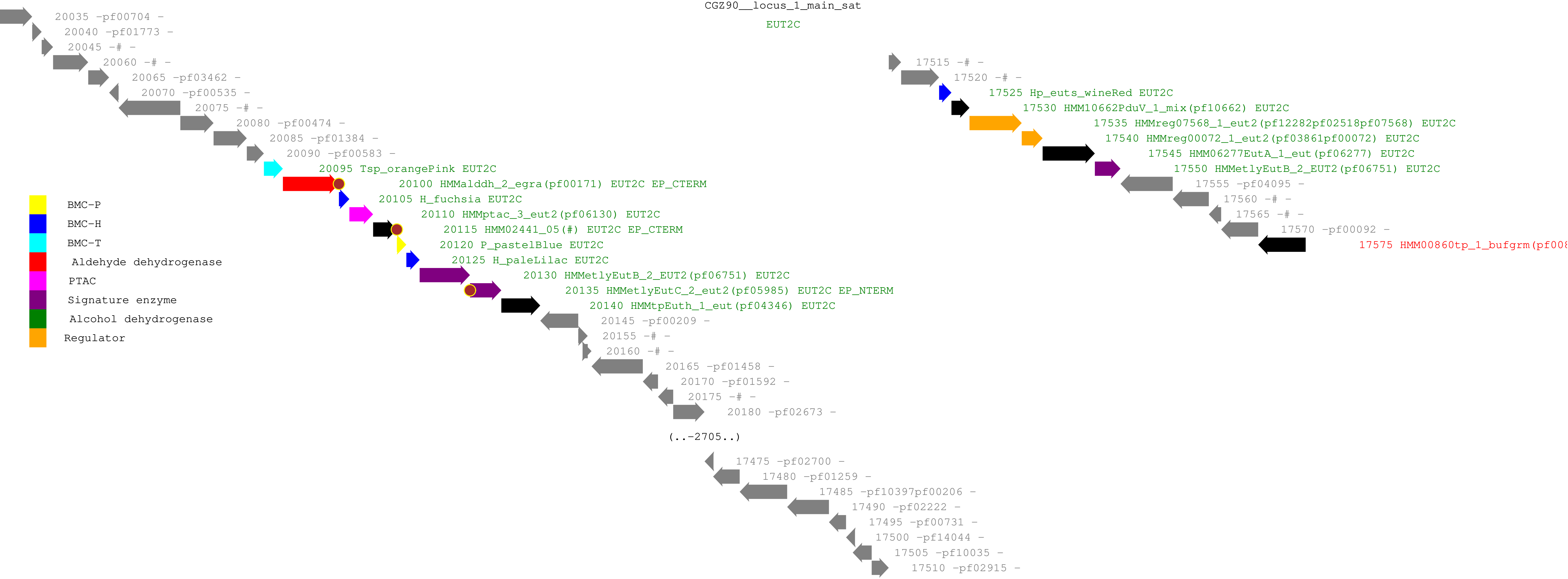

Bacillus mesonae. Bacteria; Firmicutes; Bacilli; Bacillales; Bacillaceae; Bacillus.

CHR53__locus_2

EUT2C

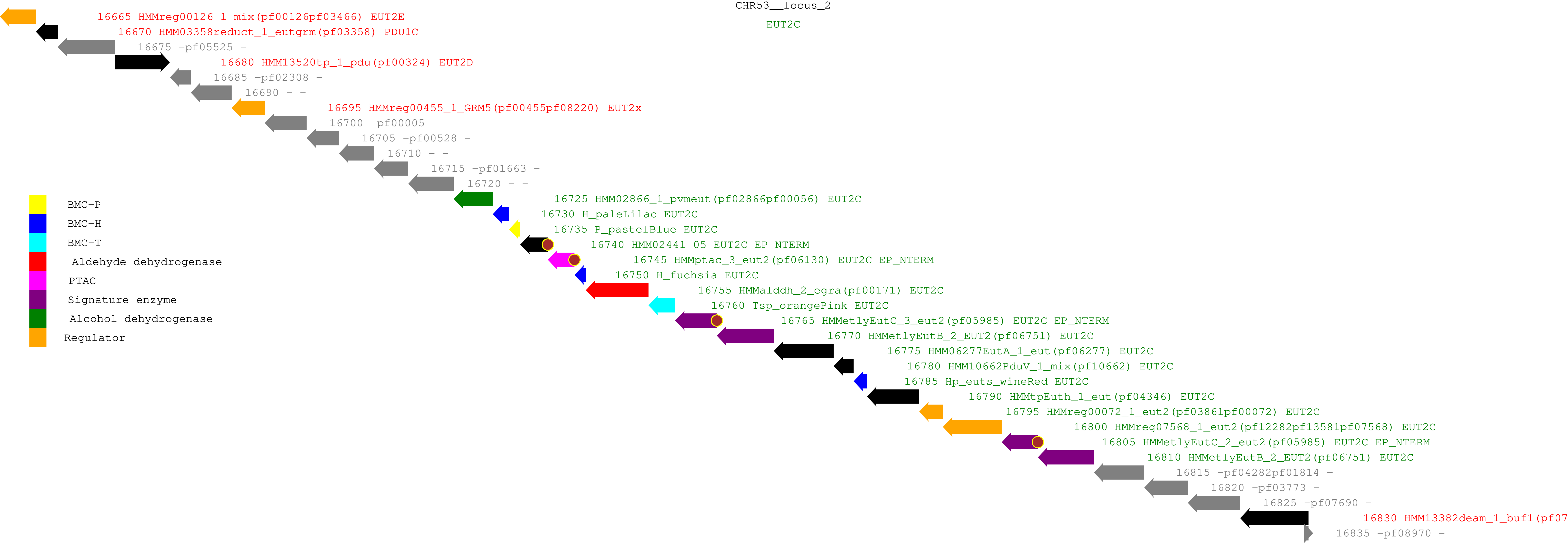

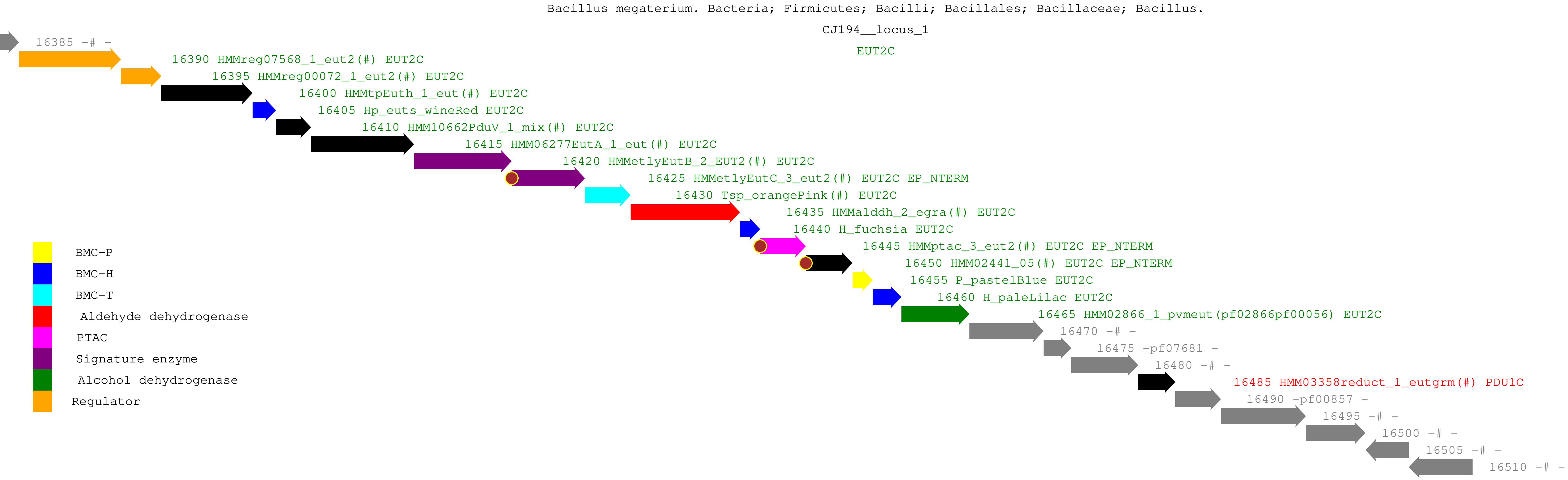

Bacillus megaterium. Bacteria; Firmicutes; Bacilli; Bacillales; Bacillaceae; Bacillus.
CJ194__locus_1
EUT2C
16385 -# -
16390 HMMreg07568_1_eut2(#) EUT2C
16395 HMMreg00072_1_eut2(#) EUT2C
16400 HMMtpEuth_1_eut(#) EUT2C
16405 Hp_euts_wineRed EUT2C
16410 HMM10662PduV_1_mix(#) EUT2C
16415 HMM06277EutA_1_eut(#) EUT2C
16420 HMMetlyEutB_2_EUT2(#) EUT2C
16425 HMMetlyEutC_3_eut2(#) EUT2C EP_NTERM
16430 Tsp_orangePink(#) EUT2C
16435 HMMaldh_2_egra(#) EUT2C
16440 H_fuchsia EUT2C
16445 HMMptac_3_eut2(#) EUT2C EP_NTERM
16450 HMM02441_05(#) EUT2C EP_NTERM
16455 P_pastelBlue EUT2C
16460 H_paleLilac EUT2C
16465 HMM02866_1_pvmeut(pf02866pf00056) EUT2C
16470 -# -
16475 -pf07681 -
16480 -# -
16485 HMM03358reduct_1_eutgrm(#) PDU1C
16490 -pf00857 -
16495 -# -
16500 -# -
16505 -# -
16510 -# -
BMC-P
BMC-H
BMC-T
Aldehyde dehydrogenase
PTAC
Signature enzyme
Alcohol dehydrogenase
Regulator

Bacillus sp. PK3_68. Bacteria; Firmicutes; Bacilli; Bacillales; Bacillaceae; Bacillus.

CJ483__locus_1

EUT2C

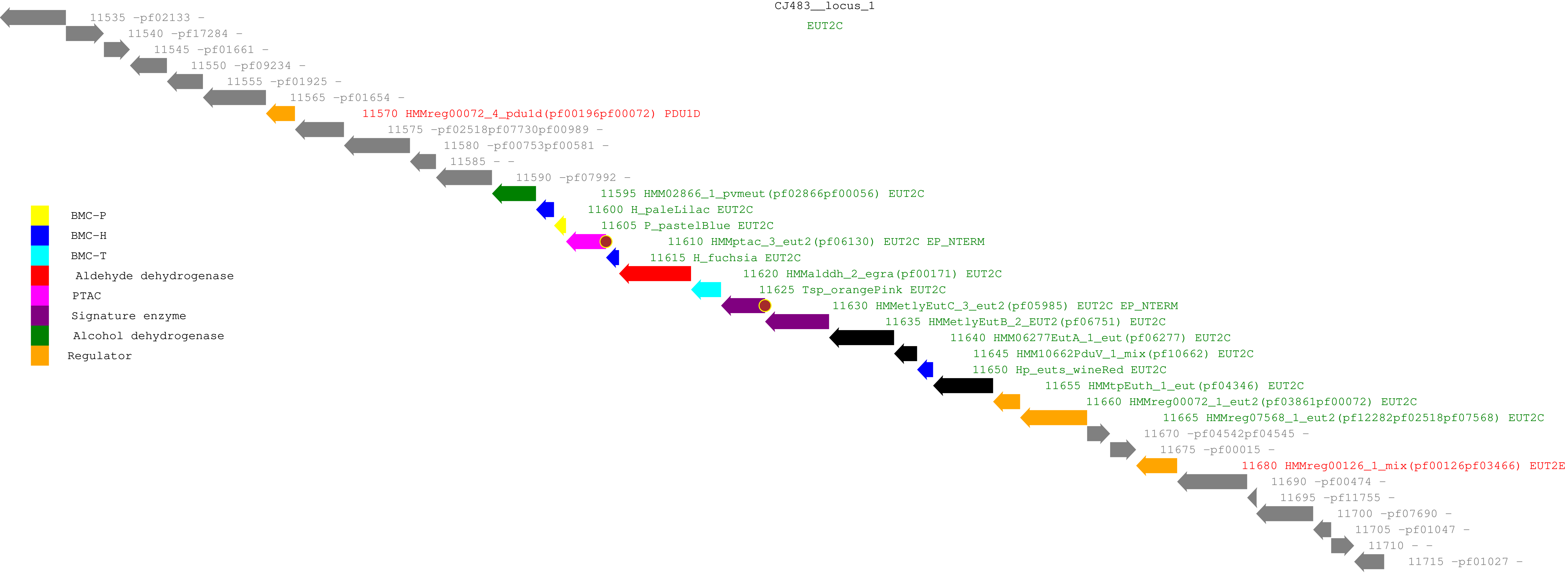

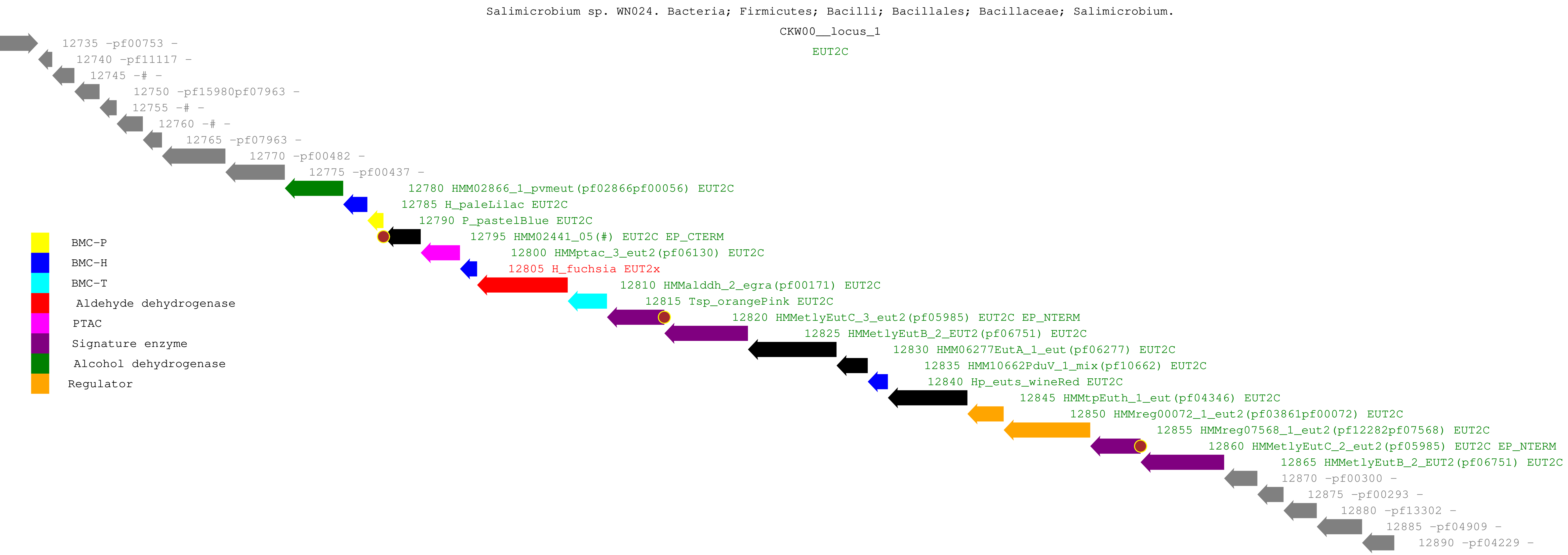
Salimicrobium sp. WN024. Bacteria; Firmicutes; Bacilli; Bacillales; Bacillaceae; Salimicrobium.
CKW00__locus_1
EUT2C
12735 -pf00753 -
12740 -pf11117 -
12745 -# -
12750 -pf15980pf07963 -
12755 -# -
12760 -# -
12765 -pf07963 -
12770 -pf00482 -
12775 -pf00437 -
12780 HMM02866_1_pvmeut(pf02866pf00056) EUT2C
12785 H_paleLilac EUT2C
12790 P_pastelBlue EUT2C
12795 HMM02441_05(#) EUT2C EP_CTERM
12800 HMMptac_3_eut2(pf06130) EUT2C
12805 H_fuchsia EUT2x
12810 HMMalddh_2_egra(pf00171) EUT2C
12815 Tsp_orangePink EUT2C
12820 HMMetlyEutC_3_eut2(pf05985) EUT2C EP_NTERM
12825 HMMetlyEutB_2_EUT2(pf06751) EUT2C
12830 HMM06277EutA_1_eut(pf06277) EUT2C
12835 HMM10662PduV_1_mix(pf10662) EUT2C
12840 Hp_euts_wineRed EUT2C
12845 HMMtpEuth_1_eut(pf04346) EUT2C
12850 HMMreg00072_1_eut2(pf03861pf00072) EUT2C
12855 HMMreg07568_1_eut2(pf12282pf07568) EUT2C
12860 HMMetlyEutC_2_eut2(pf05985) EUT2C EP_NTERM
12865 HMMetlyEutB_2_EUT2(pf06751) EUT2C
12870 -pf00300 -
12875 -pf00293 -
12880 -pf13302 -
12885 -pf04909 -
12890 -pf04229 -
BMC-P
BMC-H
BMC-T
Aldehyde dehydrogenase
PTAC
Signature enzyme
Alcohol dehydrogenase
Regulator

Bacillus megaterium. Bacteria; Firmicutes; Bacilli; Bacillales; Bacillaceae; Bacillus.

COI84__locus_1

EUT2C

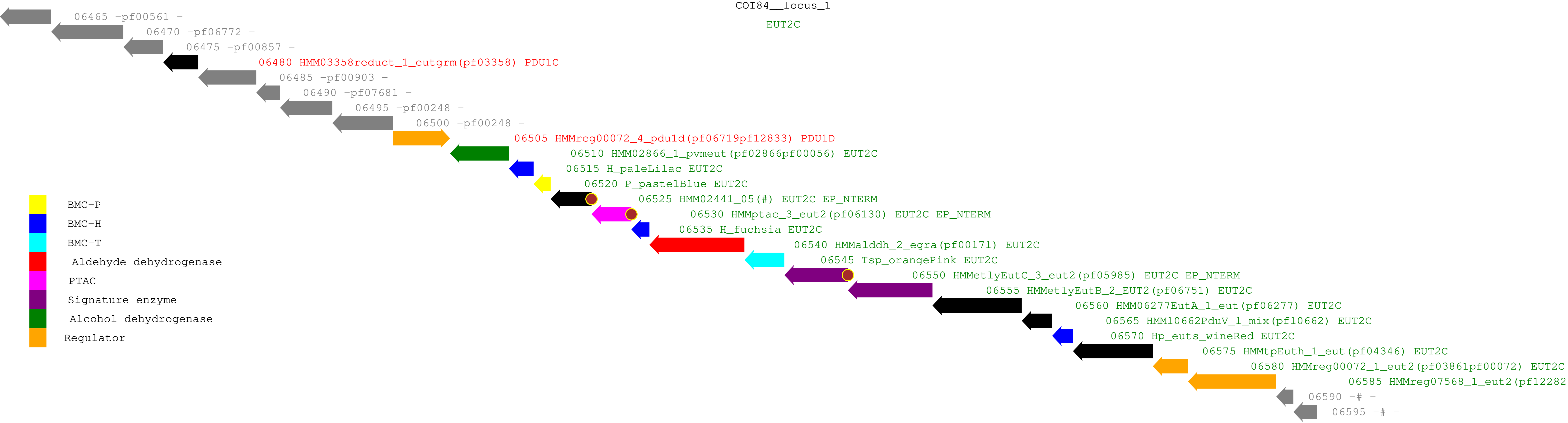

Bacillus megaterium. Bacteria; Firmicutes; Bacilli; Bacillales; Bacillaceae; Bacillus.

COJ21__locus_1

EUT2C

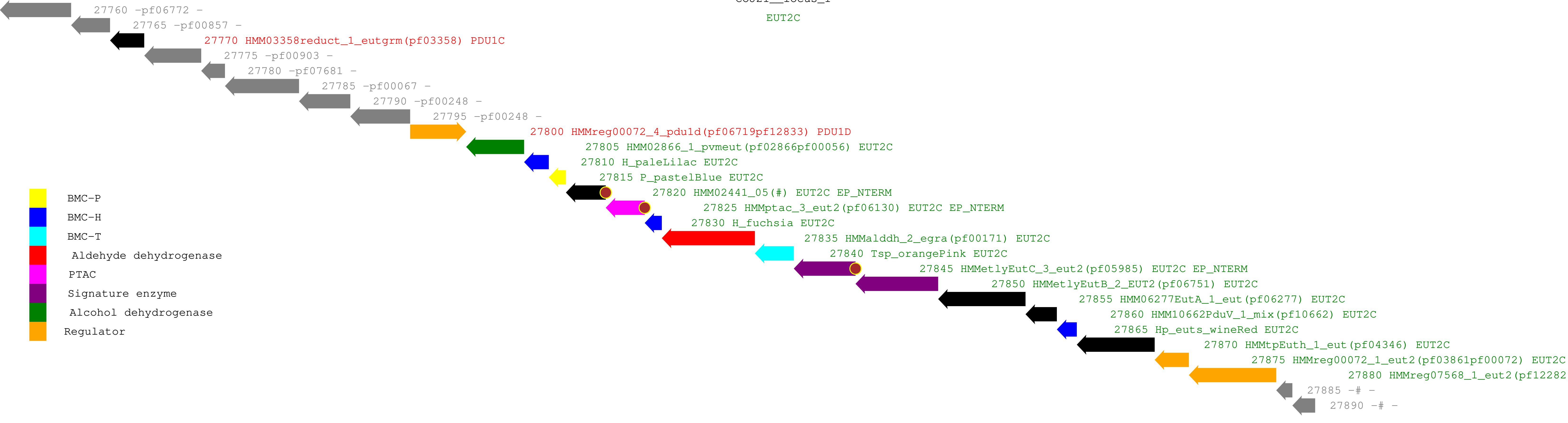

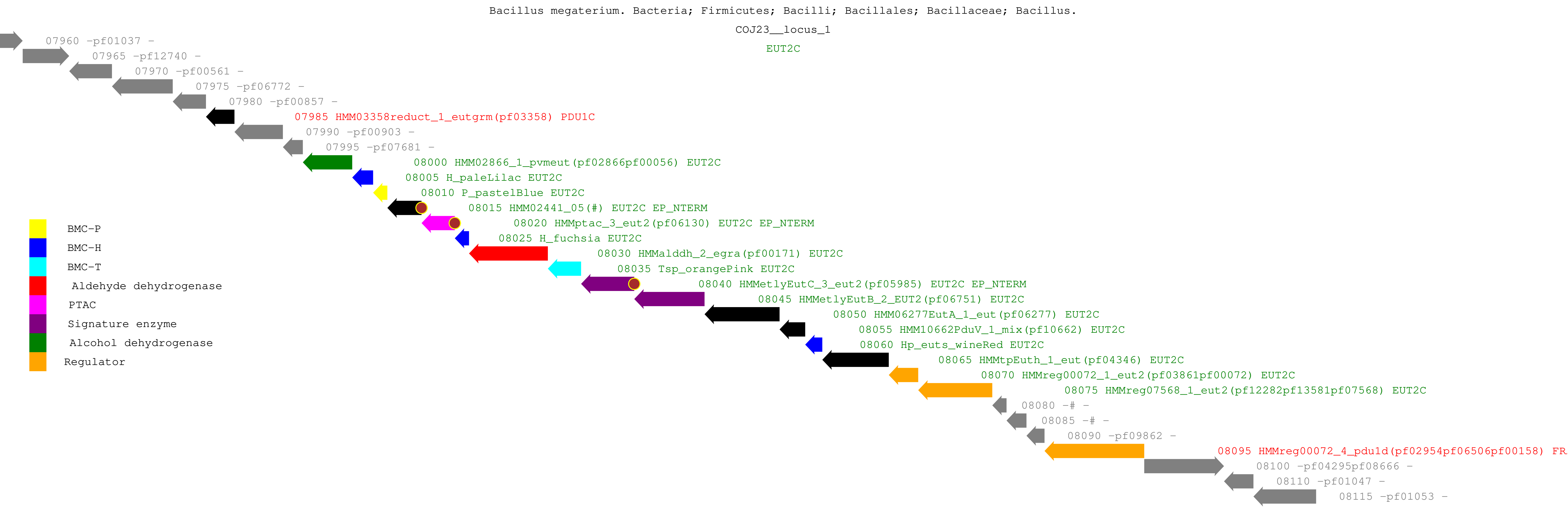
Bacillus megaterium. Bacteria; Firmicutes; Bacilli; Bacillales; Bacillaceae; Bacillus.
COJ23__locus_1
EUT2C
07960 -pf01037 -
07965 -pf12740 -
07970 -pf00561 -
07975 -pf06772 -
07980 -pf00857 -
07985 HMM03358reduct_1_eutgrm(pf03358) PDU1C
07990 -pf00903 -
07995 -pf07681 -
08000 HMM02866_1_pvmeut(pf02866pf00056) EUT2C
08005 H_paleLilac EUT2C
08010 P_pastelBlue EUT2C
08015 HMM02441_05(#) EUT2C EP_NTERM
08020 HMMptac_3_eut2(pf06130) EUT2C EP_NTERM
08025 H_fuchsia EUT2C
08030 HMMalddh_2_egra(pf00171) EUT2C
08035 Tsp_orangePink EUT2C
08040 HMMetlyEutC_3_eut2(pf05985) EUT2C EP_NTERM
08045 HMMetlyEutB_2_EUT2(pf06751) EUT2C
08050 HMM06277EutA_1_eut(pf06277) EUT2C
08055 HMM10662PduV_1_mix(pf10662) EUT2C
08060 Hp_euts_wineRed EUT2C
08065 HMMtpEuth_1_eut(pf04346) EUT2C
08070 HMMreg00072_1_eut2(pf03861pf00072) EUT2C
08075 HMMreg07568_1_eut2(pf12282pf13581pf07568) EUT2C
08080 -# -
08085 -# -
08090 -pf09862 -
08095 HMMreg00072_4_pdu1d(pf02954pf06506pf00158) FR
08100 -pf04295pf08666 -
08110 -pf01047 -
08115 -pf01053 -
BMC-P
BMC-H
BMC-T
Aldehyde dehydrogenase
PTAC
Signature enzyme
Alcohol dehydrogenase
Regulator

Bacillus megaterium. Bacteria; Firmicutes; Bacilli; Bacillales; Bacillaceae; Bacillus.

COK90__locus_1

EUT2C

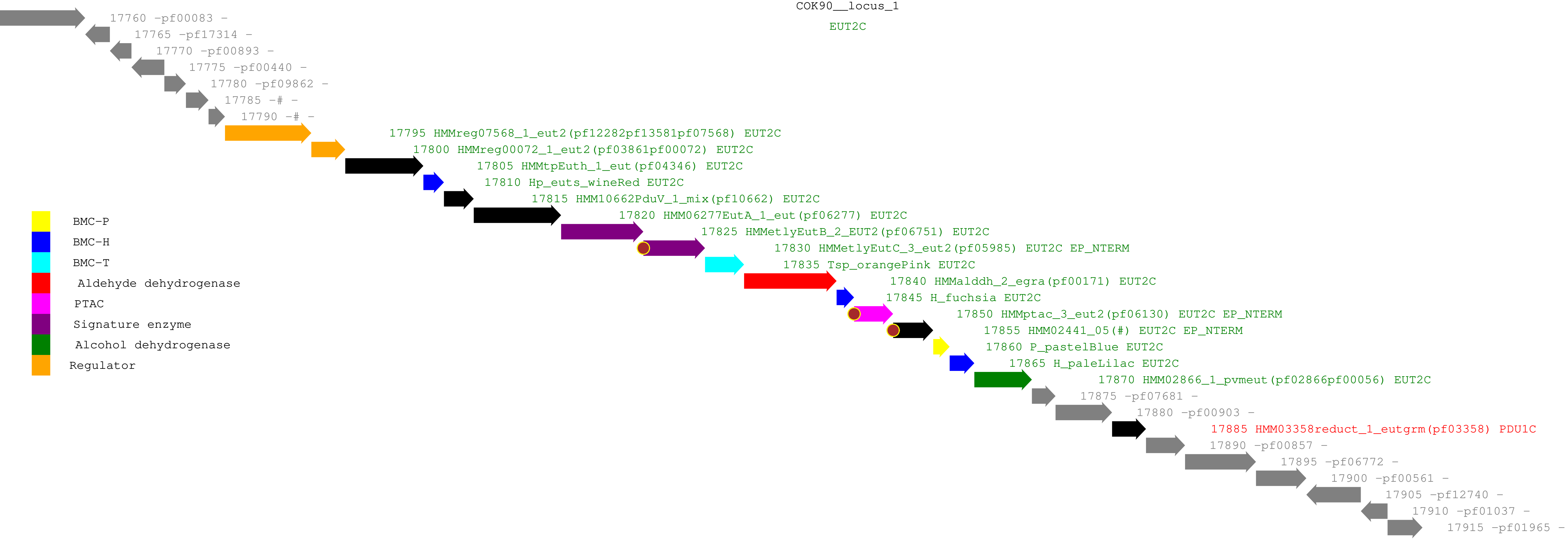

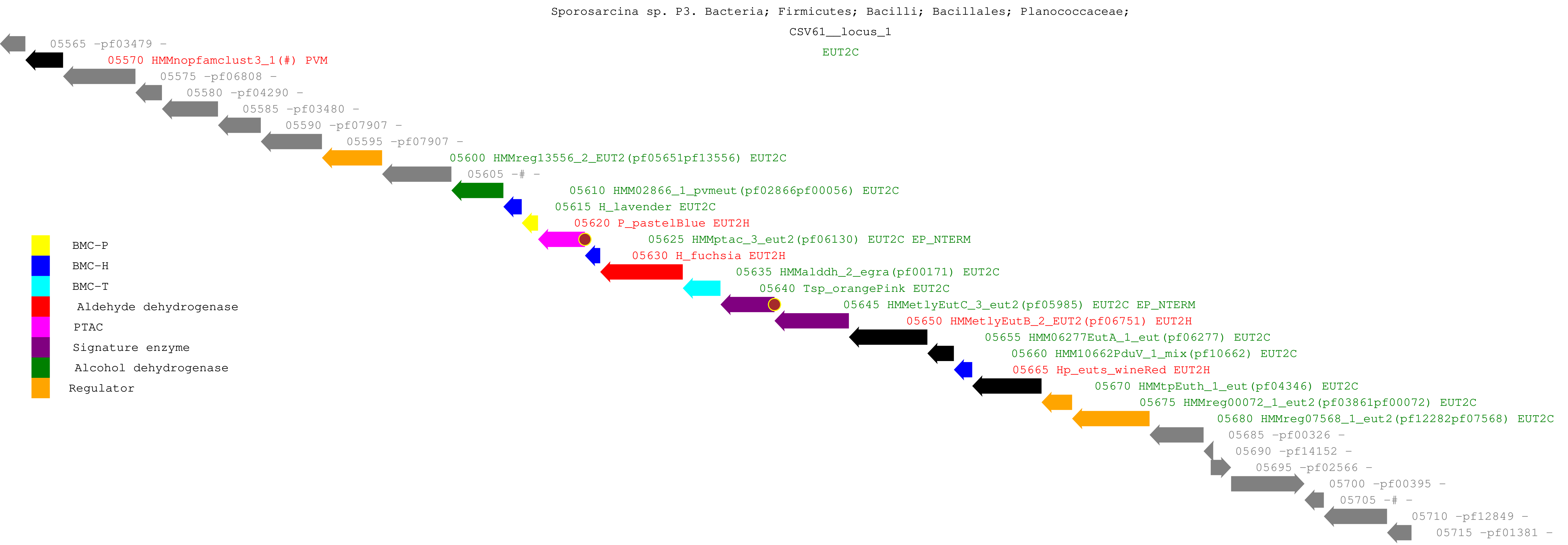
Sporosarcina sp. P3. Bacteria; Firmicutes; Bacilli; Bacillales; Planococcaceae;
CSV61__locus_1
EUT2C
05565 -pf03479 -
05570 HMMnopfamclust3_1(#) PVM
05575 -pf06808 -
05580 -pf04290 -
05585 -pf03480 -
05590 -pf07907 -
05595 -pf07907 -
05600 HMMreg13556_2_EUT2(pf05651pf13556) EUT2C
05605 -# -
05610 HMM02866_1_pvmeut(pf02866pf00056) EUT2C
05615 H_lavender EUT2C
05620 P_pastelBlue EUT2H
05625 HMMptac_3_eut2(pf06130) EUT2C EP_NTERM
05630 H_fuchsia EUT2H
05635 HMMalddh_2_egra(pf00171) EUT2C
05640 Tsp_orangePink EUT2C
05645 HMMetlyEutC_3_eut2(pf05985) EUT2C EP_NTERM
05650 HMMetlyEutB_2_EUT2(pf06751) EUT2H
05655 HMM06277EutA_1_eut(pf06277) EUT2C
05660 HMM10662PduV_1_mix(pf10662) EUT2C
05665 Hp_euts_wineRed EUT2H
05670 HMMtpEuth_1_eut(pf04346) EUT2C
05675 HMMreg00072_1_eut2(pf03861pf00072) EUT2C
05680 HMMreg07568_1_eut2(pf12282pf07568) EUT2C
05685 -pf00326 -
05690 -pf14152 -
05695 -pf02566 -
05700 -pf00395 -
05705 -# -
05710 -pf12849 -
05715 -pf01381 -
BMC-P
BMC-H
BMC-T
Aldehyde dehydrogenase
PTAC
Signature enzyme
Alcohol dehydrogenase
Regulator

Sporosarcina sp. P35. Bacteria; Firmicutes; Bacilli; Bacillales; Planococcaceae;

CSV62__locus_1

EUT2C

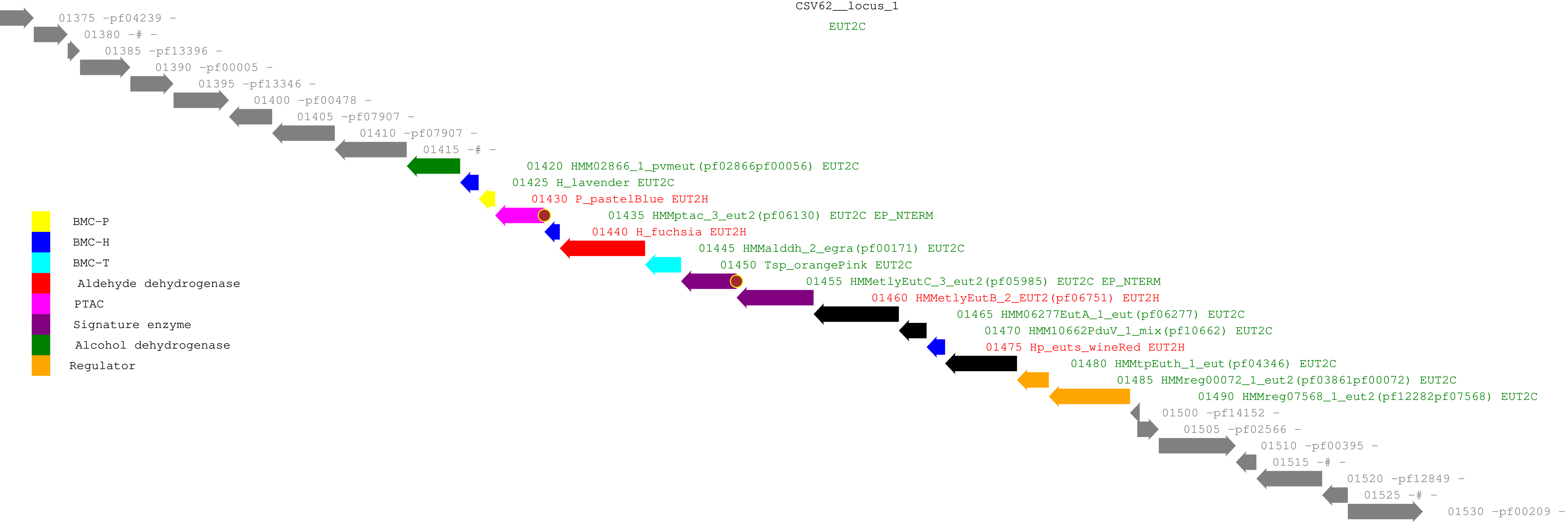

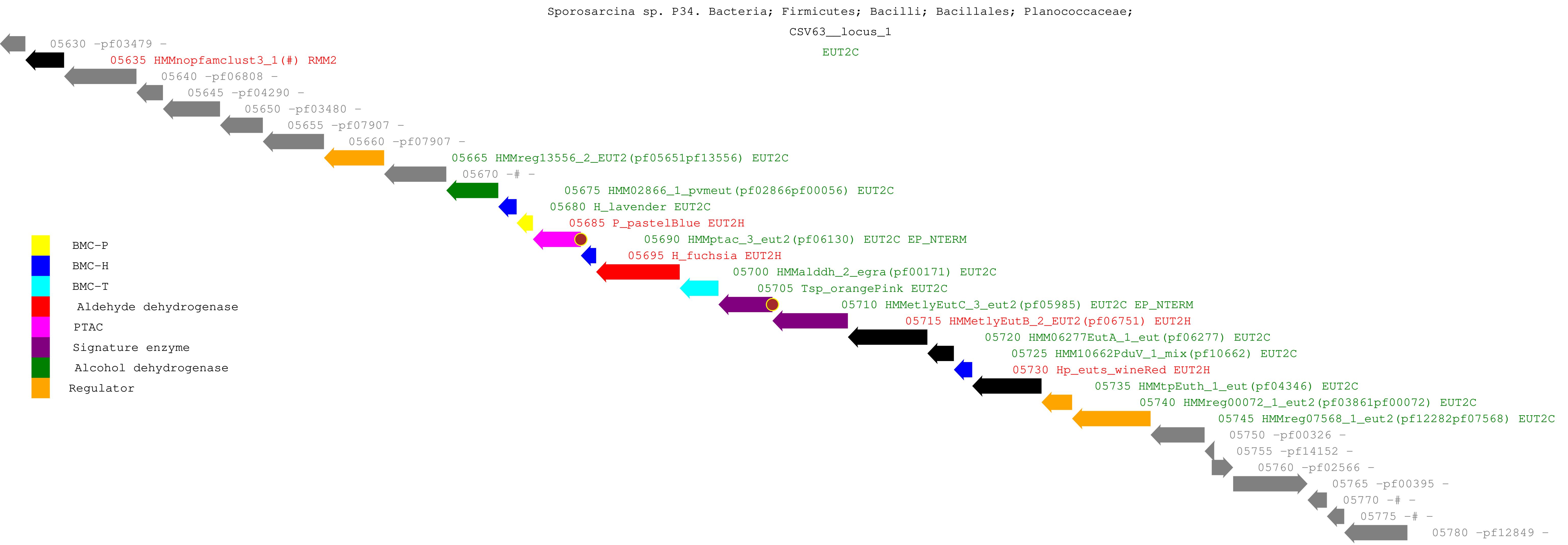
Sporosarcina sp. P34. Bacteria; Firmicutes; Bacilli; Bacillales; Planococcaceae;
CSV63__locus_1
EUT2C
05630 -pf03479 -
05635 HMMnopfamclust3_1(#) RMM2
05640 -pf06808 -
05645 -pf04290 -
05650 -pf03480 -
05655 -pf07907 -
05660 -pf07907 -
05665 HMMreg13556_2_EUT2(pf05651pf13556) EUT2C
05670 -# -
05675 HMM02866_1_pvmeut(pf02866pf00056) EUT2C
05680 H_lavender EUT2C
05685 P_pastelBlue EUT2H
05690 HMMptac_3_eut2(pf06130) EUT2C EP_NTERM
05695 H_fuchsia EUT2H
05700 HMMaldh_2_egra(pf00171) EUT2C
05705 Tsp_orangePink EUT2C
05710 HMMetlyEutC_3_eut2(pf05985) EUT2C EP_NTERM
05715 HMMetlyEutB_2_EUT2(pf06751) EUT2H
05720 HMM06277EutA_1_eut(pf06277) EUT2C
05725 HMM10662PduV_1_mix(pf10662) EUT2C
05730 Hp_euts_wineRed EUT2H
05735 HMMtpEuth_1_eut(pf04346) EUT2C
05740 HMMreg00072_1_eut2(pf03861pf00072) EUT2C
05745 HMMreg07568_1_eut2(pf12282pf07568) EUT2C
05750 -pf00326 -
05755 -pf14152 -
05760 -pf02566 -
05765 -pf00395 -
05770 -# -
05775 -# -
05780 -pf12849 -
BMC-P
BMC-H
BMC-T
Aldehyde dehydrogenase
PTAC
Signature enzyme
Alcohol dehydrogenase
Regulator

Sporosarcina sp. P29. Bacteria; Firmicutes; Bacilli; Bacillales; Planococcaceae;

CSV68__locus_2

EUT2C

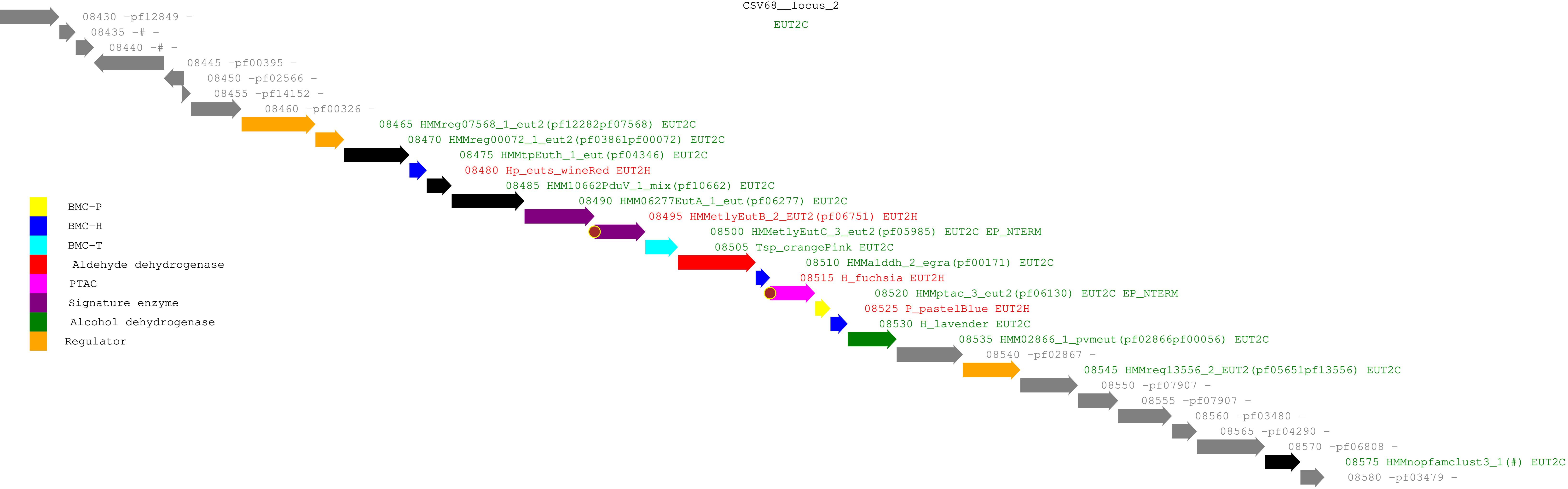

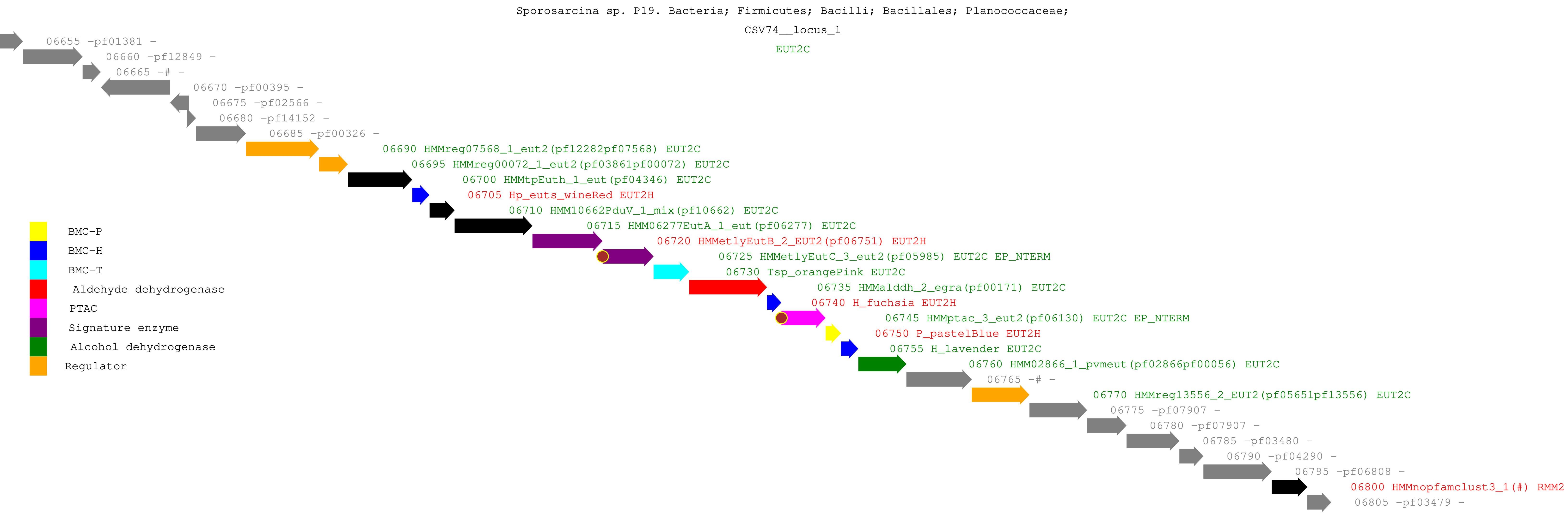
Sporosarcina sp. P19. Bacteria; Firmicutes; Bacilli; Bacillales; Planococcaceae;
CSV74__locus_1
EUT2C
06655 -pf01381 -
06660 -pf12849 -
06665 -# -
06670 -pf00395 -
06675 -pf02566 -
06680 -pf14152 -
06685 -pf00326 -
06690 HMMreg07568_1_eut2(pf12282pf07568) EUT2C
06695 HMMreg00072_1_eut2(pf03861pf00072) EUT2C
06700 HMMtpEuth_1_eut(pf04346) EUT2C
06705 Hp_euts_wineRed EUT2H
06710 HMM10662PduV_1_mix(pf10662) EUT2C
06715 HMM06277EutA_1_eut(pf06277) EUT2C
06720 HMMetlyEutB_2_EUT2(pf06751) EUT2H
06725 HMMetlyEutC_3_eut2(pf05985) EUT2C EP_NTERM
06730 Tsp_orangePink EUT2H
06735 HMMalddh_2_egra(pf00171) EUT2C
06740 H_fuchsia EUT2H
06745 HMMptac_3_eut2(pf06130) EUT2C EP_NTERM
06750 P_pastelBlue EUT2H
06755 H_lavender EUT2C
06760 HMM02866_1_pvmeut(pf02866pf00056) EUT2C
06765 -# -
06770 HMMreg13556_2_EUT2(pf05651pf13556) EUT2C
06775 -pf07907 -
06780 -pf07907 -
06785 -pf03480 -
06790 -pf04290 -
06795 -pf06808 -
06800 HMMnopfamclust3_1(#) RMM2
06805 -pf03479 -
BMC-P
BMC-H
BMC-T
Aldehyde dehydrogenase
PTAC
Signature enzyme
Alcohol dehydrogenase
Regulator

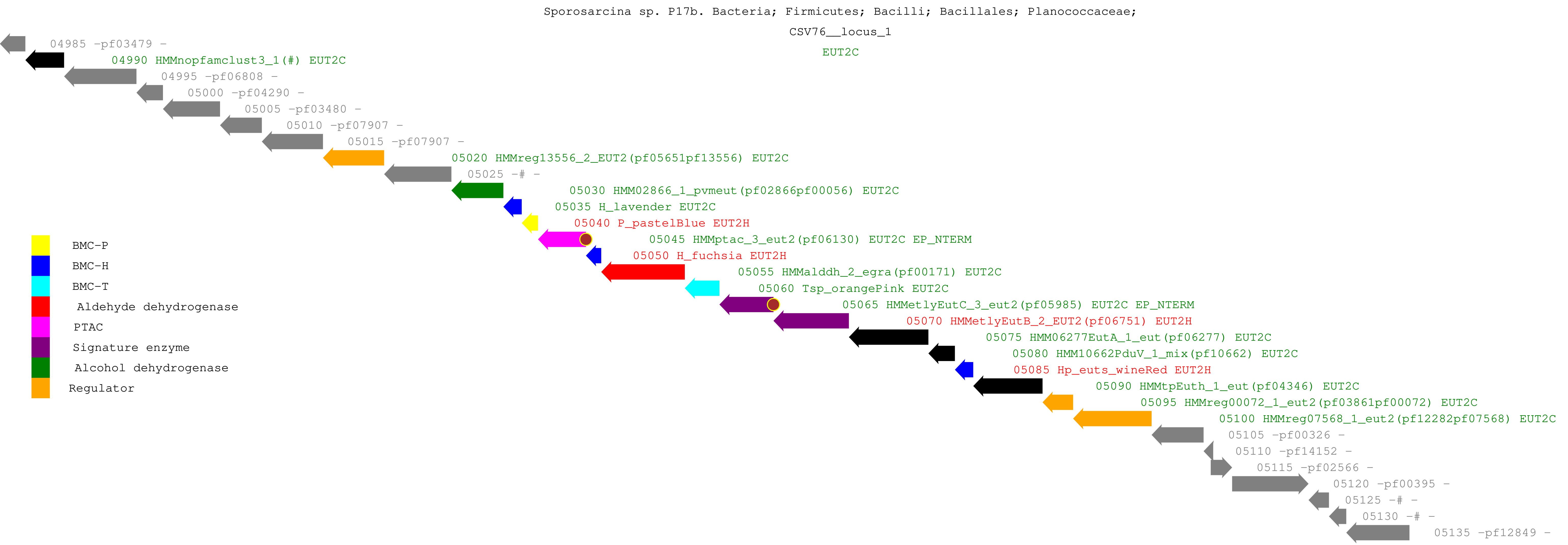
Sporosarcina sp. P17b. Bacteria; Firmicutes; Bacilli; Bacillales; Planococcaceae;
CSV76__locus_1
EUT2C
04985 -pf03479 -
04990 HMMnopfamclust3_1(#) EUT2C
04995 -pf06808 -
05000 -pf04290 -
05005 -pf03480 -
05010 -pf07907 -
05015 -pf07907 -
05020 HMMreg13556_2_EUT2(pf05651pf13556) EUT2C
05025 -# -
05030 HMM02866_1_pvmeut(pf02866pf00056) EUT2C
05035 H_lavender EUT2C
05040 P_pastelBlue EUT2H
05045 HMMptac_3_eut2(pf06130) EUT2C EP_NTERM
05050 H_fuchsia EUT2H
05055 HMMalddh_2_egra(pf00171) EUT2C
05060 Tsp_orangePink EUT2C
05065 HMMetlyEutC_3_eut2(pf05985) EUT2C EP_NTERM
05070 HMMetlyEutB_2_EUT2(pf06751) EUT2H
05075 HMM06277EutA_1_eut(pf06277) EUT2C
05080 HMM10662PduV_1_mix(pf10662) EUT2C
05085 Hp_euts_wineRed EUT2H
05090 HMMtpEuth_1_eut(pf04346) EUT2C
05095 HMMreg00072_1_eut2(pf03861pf00072) EUT2C
05100 HMMreg07568_1_eut2(pf12282pf07568) EUT2C
05105 -pf00326 -
05110 -pf14152 -
05115 -pf02566 -
05120 -pf00395 -
05125 -# -
05130 -# -
05135 -pf12849 -
BMC-P
BMC-H
BMC-T
Aldehyde dehydrogenase
PTAC
Signature enzyme
Alcohol dehydrogenase
Regulator

Sporosarcina sp. P16b. Bacteria; Firmicutes; Bacilli; Bacillales; Planococcaceae;

CSV77__locus_2

EUT2C

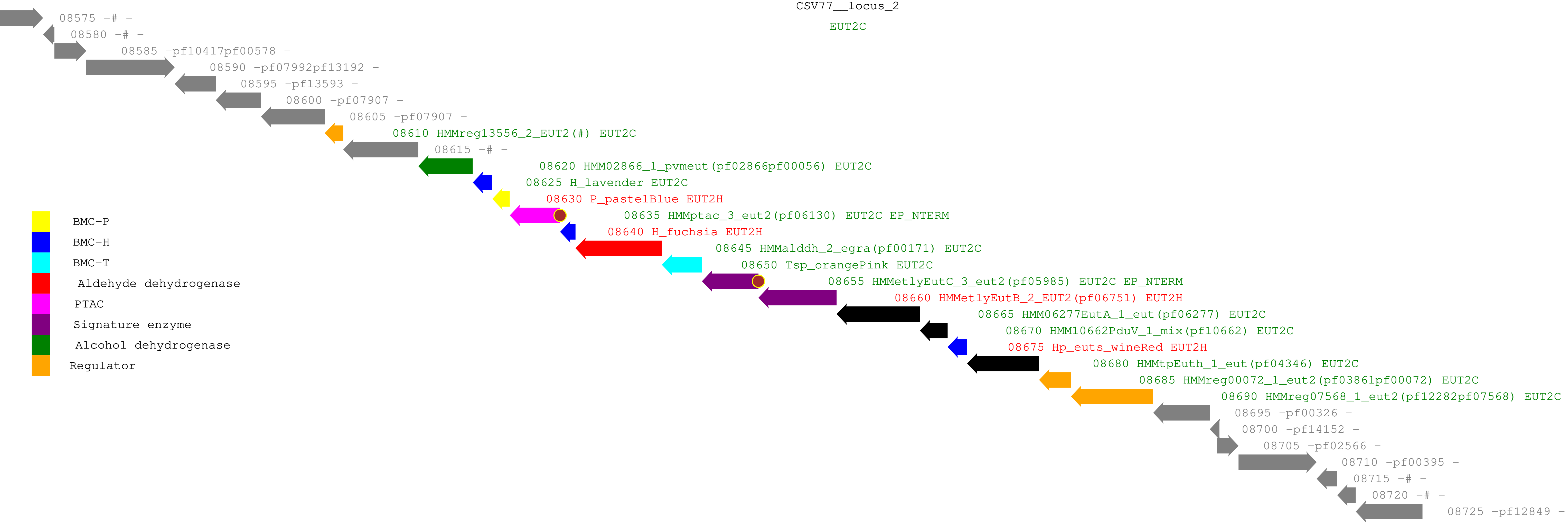

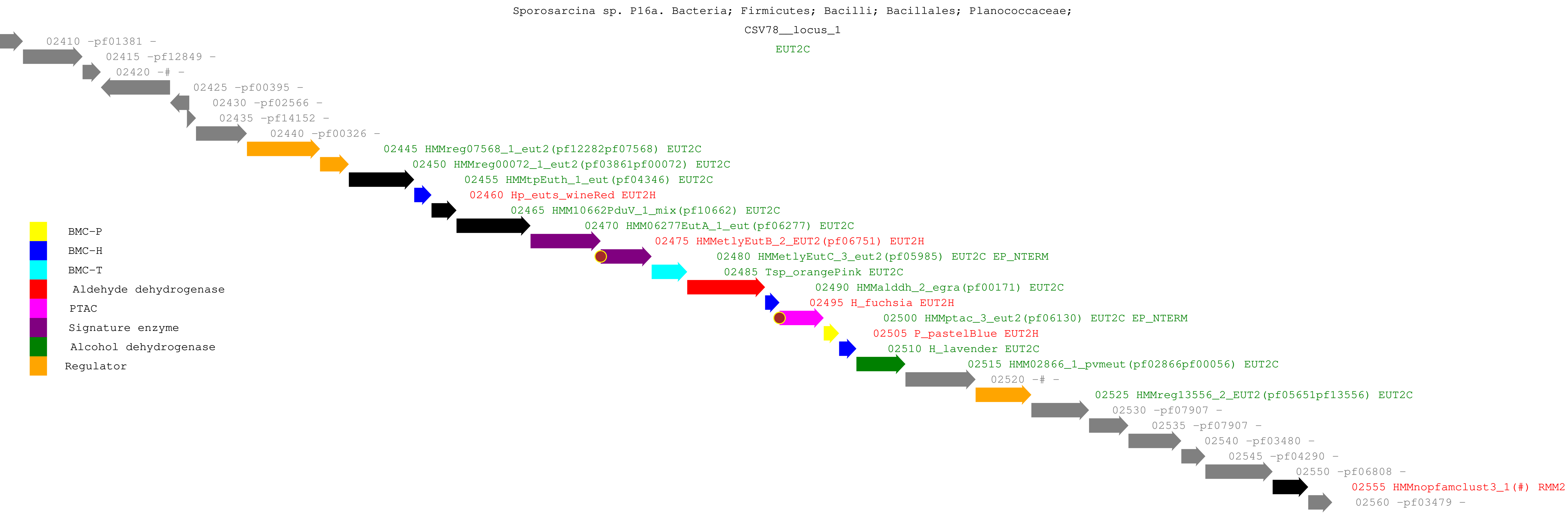
Sporosarcina sp. P16a. Bacteria; Firmicutes; Bacilli; Bacillales; Planococcaceae;
CSV78__locus_1
EUT2C
02410 -pf01381 -
02415 -pf12849 -
02420 -# -
02425 -pf00395 -
02430 -pf02566 -
02435 -pf14152 -
02440 -pf00326 -
02445 HMMreg07568_1_eut2(pf12282pf07568) EUT2C
02450 HMMreg00072_1_eut2(pf03861pf00072) EUT2C
02455 HMMtpEuth_1_eut(pf04346) EUT2C
02460 Hp_euts_wineRed EUT2H
02465 HMM10662PduV_1_mix(pf10662) EUT2C
02470 HMM06277EutA_1_eut(pf06277) EUT2C
02475 HMMetlyEutB_2_EUT2(pf06751) EUT2H
02480 HMMetlyEutC_3_eut2(pf05985) EUT2C EP_NTERM
02485 Tsp_orangePink EUT2C
02490 HMMalddh_2_egra(pf00171) EUT2C
02495 H_fuchsia EUT2H
02500 HMMptac_3_eut2(pf06130) EUT2C EP_NTERM
02505 P_pastelBlue EUT2H
02510 H_lavender EUT2C
02515 HMM02866_1_pvmeut(pf02866pf00056) EUT2C
02520 -# -
02525 HMMreg13556_2_EUT2(pf05651pf13556) EUT2C
02530 -pf07907 -
02535 -pf07907 -
02540 -pf03480 -
02545 -pf04290 -
02550 -pf06808 -
02555 HMMnopfamclust3_1(#) RMM2
02560 -pf03479 -
BMC-P
BMC-H
BMC-T
Aldehyde dehydrogenase
PTAC
Signature enzyme
Alcohol dehydrogenase
Regulator

Sporosarcina sp. P13. Bacteria; Firmicutes; Bacilli; Bacillales; Planococcaceae;

CSV79__locus_1

EUT2C

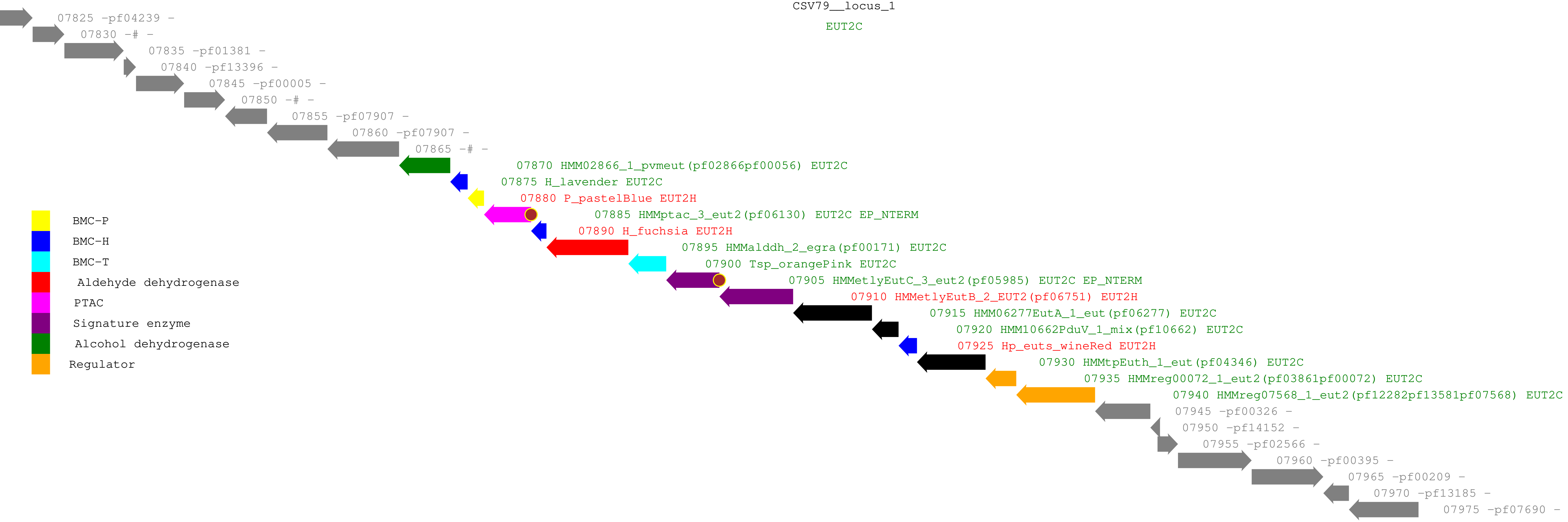

Sporosarcina sp. P10. Bacteria; Firmicutes; Bacilli; Bacillales; Planococcaceae;

CSV81__locus_1

EUT2C

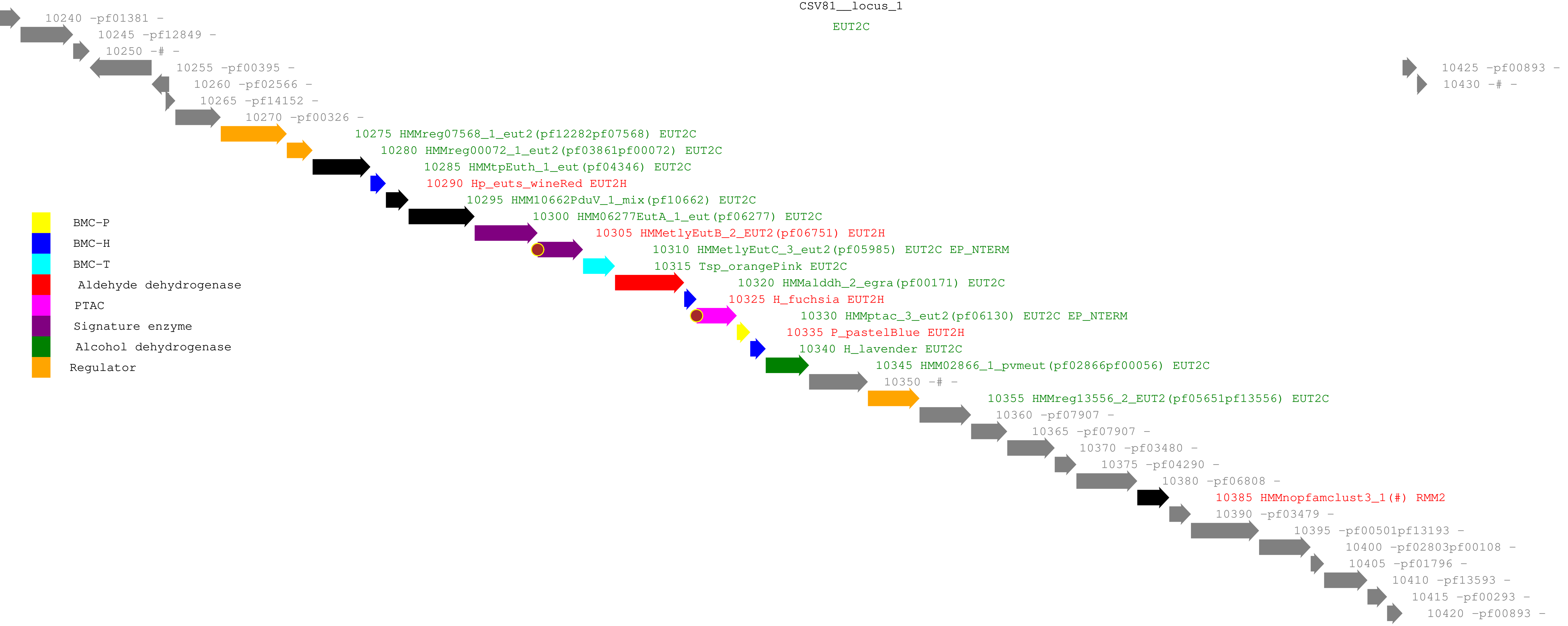

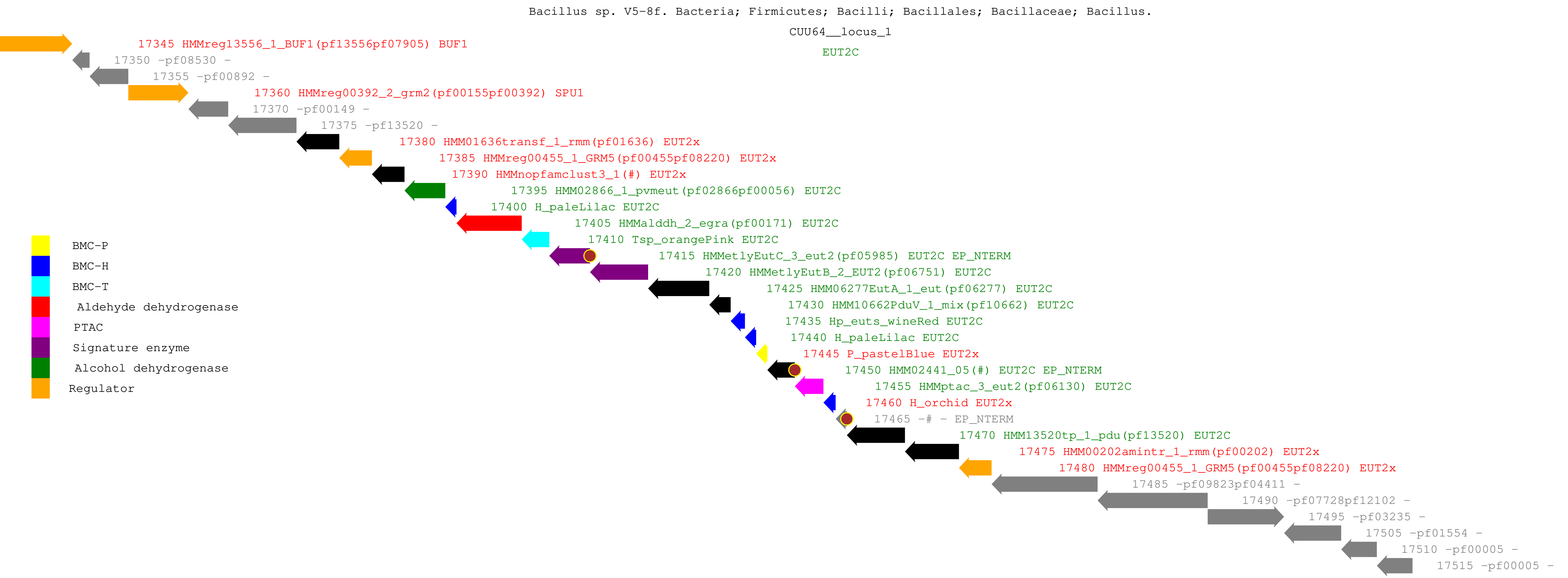
Bacillus sp. V5-8f. Bacteria; Firmicutes; Bacilli; Bacillales; Bacillaceae; Bacillus.
CUU64__locus_1
EUT2C
17345 HMMreg13556_1_BUF1(pf13556pf07905) BUF1
17350 -pf08530 -
17355 -pf00892 -
17360 HMMreg00392_2_grm2(pf00155pf00392) SPU1
17370 -pf00149 -
17375 -pf13520 -
17380 HMM01636transf_1_rmm(pf01636) EUT2x
17385 HMMreg00455_1_GRM5(pf00455pf08220) EUT2x
17390 HMMnopfamclust3_1(#) EUT2x
17395 HMM02866_1_pvmeut(pf02866pf00056) EUT2C
17400 H_paleLilac EUT2C
17405 HMMalddh_2_egra(pf00171) EUT2C
17410 Tsp_orangePink EUT2C
17415 HMMetlyEutC_3_eut2(pf05985) EUT2C EP_NTERM
17420 HMMetlyEutB_2_EUT2(pf06751) EUT2C
17425 HMM06277EutA_1_eut(pf06277) EUT2C
17430 HMM10662PduV_1_mix(pf10662) EUT2C
17435 Hp_euts_wineRed EUT2C
17440 H_paleLilac EUT2C
17445 P_pastelBlue EUT2x
17450 HMM02441_05(#) EUT2C EP_NTERM
17455 HMMptac_3_eut2(pf06130) EUT2C
17460 H_orchid EUT2x
17465 -# - EP_NTERM
17470 HMM13520tp_1_pdu(pf13520) EUT2C
17475 HMM00202amintr_1_rmm(pf00202) EUT2x
17480 HMMreg00455_1_GRM5(pf00455pf08220) EUT2x
17485 -pf09823pf04411 -
17490 -pf07728pf12102 -
17495 -pf03235 -
17505 -pf01554 -
17510 -pf00005 -
17515 -pf00005 -
BMC-P
BMC-H
BMC-T
Aldehyde dehydrogenase
PTAC
Signature enzyme
Alcohol dehydrogenase
Regulator

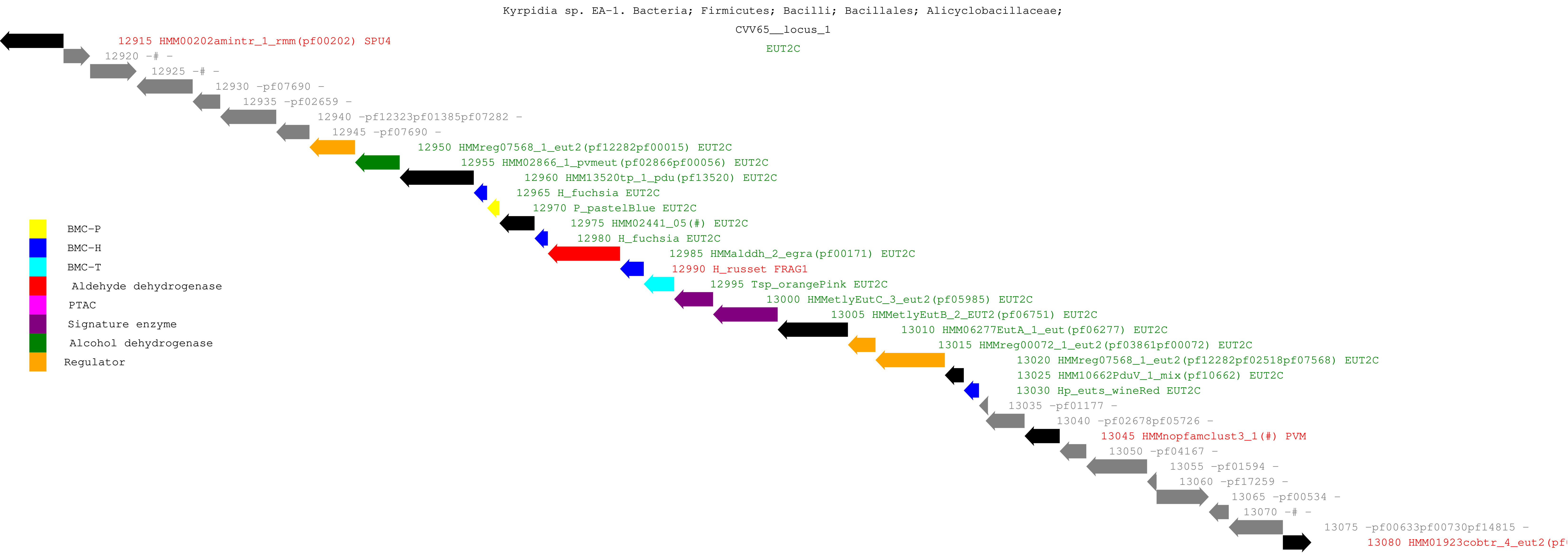
Kyrpidia sp. EA-1. Bacteria; Firmicutes; Bacilli; Bacillales; Alicyclobacillaceae;
CVV65__locus_1
EUT2C
12915 HMM00202amintr_1_rmm(pf00202) SPU4
12920 -# -
12925 -# -
12930 -pf07690 -
12935 -pf02659 -
12940 -pf12323pf01385pf07282 -
12945 -pf07690 -
12950 HMMreg07568_1_eut2(pf12282pf00015) EUT2C
12955 HMM02866_1_pvmeut(pf02866pf00056) EUT2C
12960 HMM13520tp_1_pdu(pf13520) EUT2C
12965 H_fuchsia EUT2C
12970 P_pastelBlue EUT2C
12975 HMM02441_05(#) EUT2C
12980 H_fuchsia EUT2C
12985 HMMalddh_2_egra(pf00171) EUT2C
12990 H_russet FRAG1
12995 Tsp_orangePink EUT2C
13000 HMMetlyEutC_3_eut2(pf05985) EUT2C
13005 HMMetlyEutB_2_EUT2(pf06751) EUT2C
13010 HMM06277EutA_1_eut(pf06277) EUT2C
13015 HMMreg00072_1_eut2(pf03861pf00072) EUT2C
13020 HMMreg07568_1_eut2(pf12282pf02518pf07568) EUT2C
13025 HMM10662PduV_1_mix(pf10662) EUT2C
13030 Hp_euts_wineRed EUT2C
13035 -pf01177 -
13040 -pf02678pf05726 -
13045 HMMnopfamclust3_1(#) PVM
13050 -pf04167 -
13055 -pf01594 -
13060 -pf17259 -
13065 -pf00534 -
13070 -# -
13075 -pf00633pf00730pf14815 -
13080 HMM01923cobtr_4_eut2(pf
BMC-P
BMC-H
BMC-T
Aldehyde dehydrogenase
PTAC
Signature enzyme
Alcohol dehydrogenase
Regulator

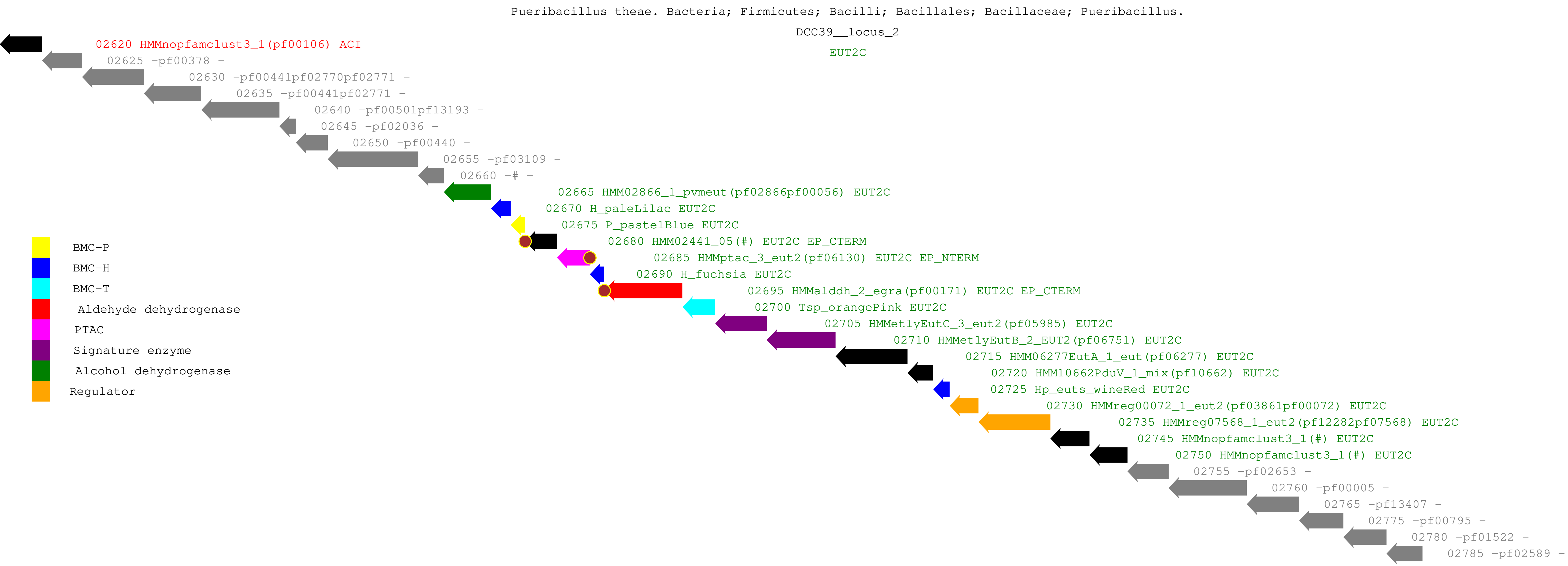

Pueribacillus theae. Bacteria; Firmicutes; Bacilli; Bacillales; Bacillaceae; Pueribacillus.
DCC39__locus_2
EUT2C
02620 HMMnopfamclust3_1(pf00106) ACI
02625 -pf00378 -
02630 -pf00441pf02770pf02771 -
02635 -pf00441pf02771 -
02640 -pf00501pf13193 -
02645 -pf02036 -
02650 -pf00440 -
02655 -pf03109 -
02660 -# -
02665 HMM02866_1_pvmeut(pf02866pf00056) EUT2C
02670 H_paleLilac EUT2C
02675 P_pastelBlue EUT2C
02680 HMM02441_05(#) EUT2C EP_CTERM
02685 HMMptac_3_eut2(pf06130) EUT2C EP_NTERM
02690 H_fuchsia EUT2C
02695 HMMaldh_2_egra(pf00171) EUT2C EP_CTERM
02700 Tsp_orangePink EUT2C
02705 HMMetlyEutC_3_eut2(pf05985) EUT2C
02710 HMMetlyEutB_2_EUT2(pf06751) EUT2C
02715 HMM06277EutA_1_eut(pf06277) EUT2C
02720 HMM10662PduV_1_mix(pf10662) EUT2C
02725 Hp_euts_wineRed EUT2C
02730 HMMreg00072_1_eut2(pf03861pf00072) EUT2C
02735 HMMreg07568_1_eut2(pf12282pf07568) EUT2C
02745 HMMnopfamclust3_1(#) EUT2C
02750 HMMnopfamclust3_1(#) EUT2C
02755 -pf02653 -
02760 -pf00005 -
02765 -pf13407 -
02775 -pf00795 -
02780 -pf01522 -
02785 -pf02589 -
BMC-P
BMC-H
BMC-T
Aldehyde dehydrogenase
PTAC
Signature enzyme
Alcohol dehydrogenase
Regulator

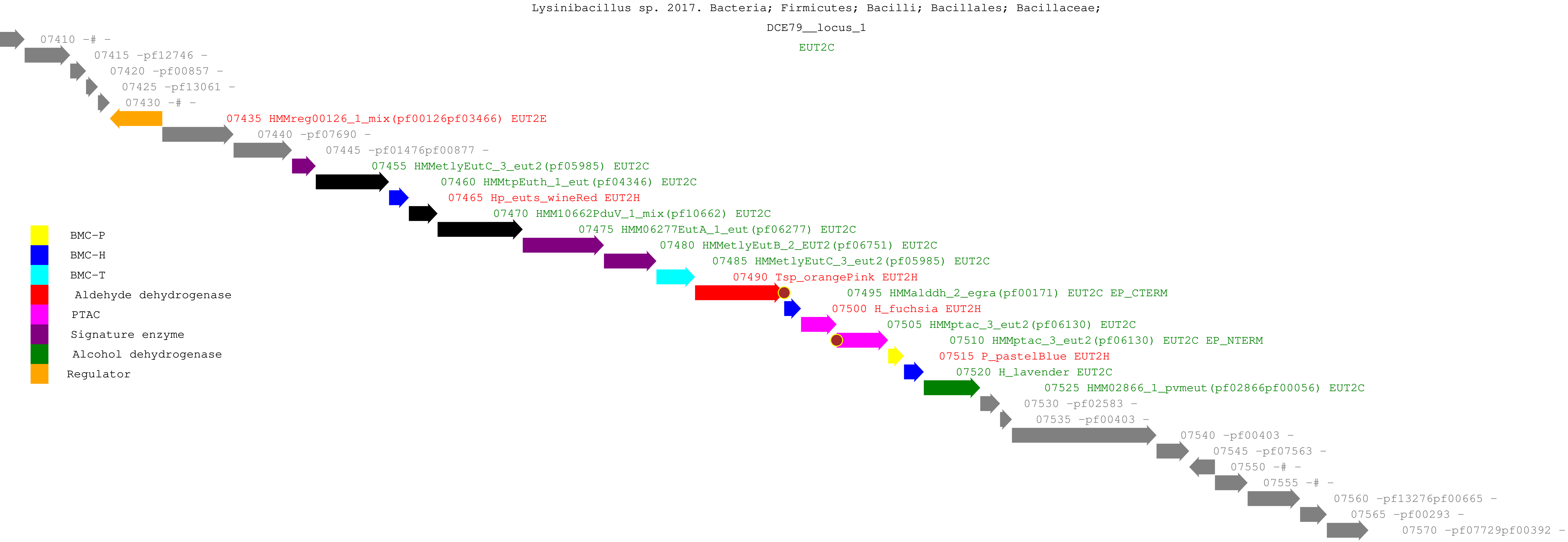

Lysinibacillus sp. 2017. Bacteria; Firmicutes; Bacilli; Bacillales; Bacillaceae;
DCE79__locus_1
EUT2C
07410 -# -
07415 -pf12746 -
07420 -pf00857 -
07425 -pf13061 -
07430 -# -
07435 HMMreg00126_1_mix(pf00126pf03466) EUT2E
07440 -pf07690 -
07445 -pf01476pf00877 -
07455 HMMetlyEutC_3_eut2(pf05985) EUT2C
07460 HMMtpEuth_1_eut(pf04346) EUT2C
07465 Hp_euts_wineRed EUT2H
07470 HMM10662PduV_1_mix(pf10662) EUT2C
07475 HMM06277EutA_1_eut(pf06277) EUT2C
07480 HMMetlyEutB_2_EUT2(pf06751) EUT2C
07485 HMMetlyEutC_3_eut2(pf05985) EUT2C
07490 Tsp_orangePink EUT2H
07495 HMMalddh_2_egra(pf00171) EUT2C EP_CTERM
07500 H_fuchsia EUT2H
07505 HMMptac_3_eut2(pf06130) EUT2C
07510 HMMptac_3_eut2(pf06130) EUT2C EP_NTERM
07515 P_pastelBlue EUT2H
07520 H_lavender EUT2C
07525 HMM02866_1_pvmeut(pf02866pf00056) EUT2C
07530 -pf02583 -
07535 -pf00403 -
07540 -pf00403 -
07545 -pf07563 -
07550 -# -
07555 -# -
07560 -pf13276pf00665 -
07565 -pf00293 -
07570 -pf07729pf00392 -
BMC-P
BMC-H
BMC-T
Aldehyde dehydrogenase
PTAC
Signature enzyme
Alcohol dehydrogenase
Regulator

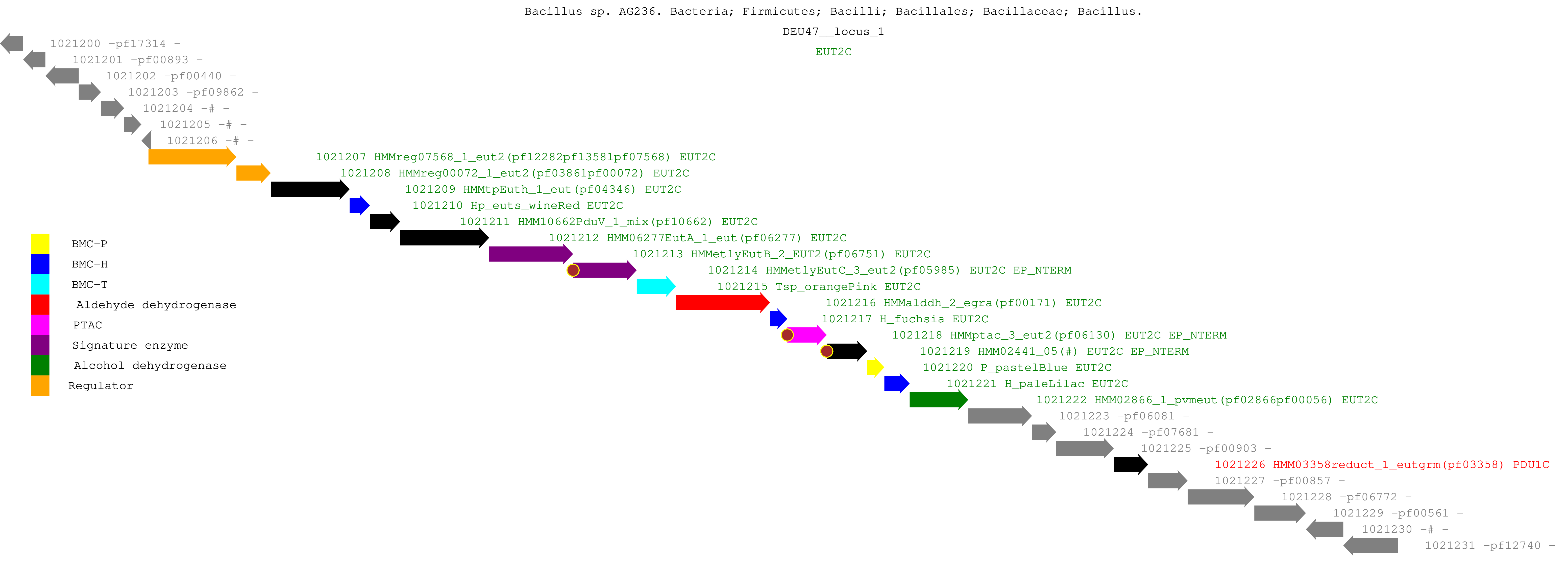

Bacillus sp. AG236. Bacteria; Firmicutes; Bacilli; Bacillales; Bacillaceae; Bacillus.
DEU47__locus_1
EUT2C
1021200 -pf17314 -
1021201 -pf00893 -
1021202 -pf00440 -
1021203 -pf09862 -
1021204 -# -
1021205 -# -
1021206 -# -
1021207 HMMreg07568_1_eut2(pf12282pf13581pf07568) EUT2C
1021208 HMMreg00072_1_eut2(pf03861pf00072) EUT2C
1021209 HMMtpEuth_1_eut(pf04346) EUT2C
1021210 Hp_euts_wineRed EUT2C
1021211 HMM10662PduV_1_mix(pf10662) EUT2C
1021212 HMM06277EutA_1_eut(pf06277) EUT2C
1021213 HMMetlyEutB_2_EUT2(pf06751) EUT2C
1021214 HMMetlyEutC_3_eut2(pf05985) EUT2C EP_NTERM
1021215 Tsp_orangePink EUT2C
1021216 HMMalddh_2_egra(pf00171) EUT2C
1021217 H_fuchsia EUT2C
1021218 HMMptac_3_eut2(pf06130) EUT2C EP_NTERM
1021219 HMM02441_05(#) EUT2C EP_NTERM
1021220 P_pastelBlue EUT2C
1021221 H_paleLilac EUT2C
1021222 HMM02866_1_pvmeut(pf02866pf00056) EUT2C
1021223 -pf06081 -
1021224 -pf07681 -
1021225 -pf00903 -
1021226 HMM03358reduct_1_eutgrm(pf03358) PDU1C
1021227 -pf00857 -
1021228 -pf06772 -
1021229 -pf00561 -
1021230 -# -
1021231 -pf12740 -
BMC-P
BMC-H
BMC-T
Aldehyde dehydrogenase
PTAC
Signature enzyme
Alcohol dehydrogenase
Regulator

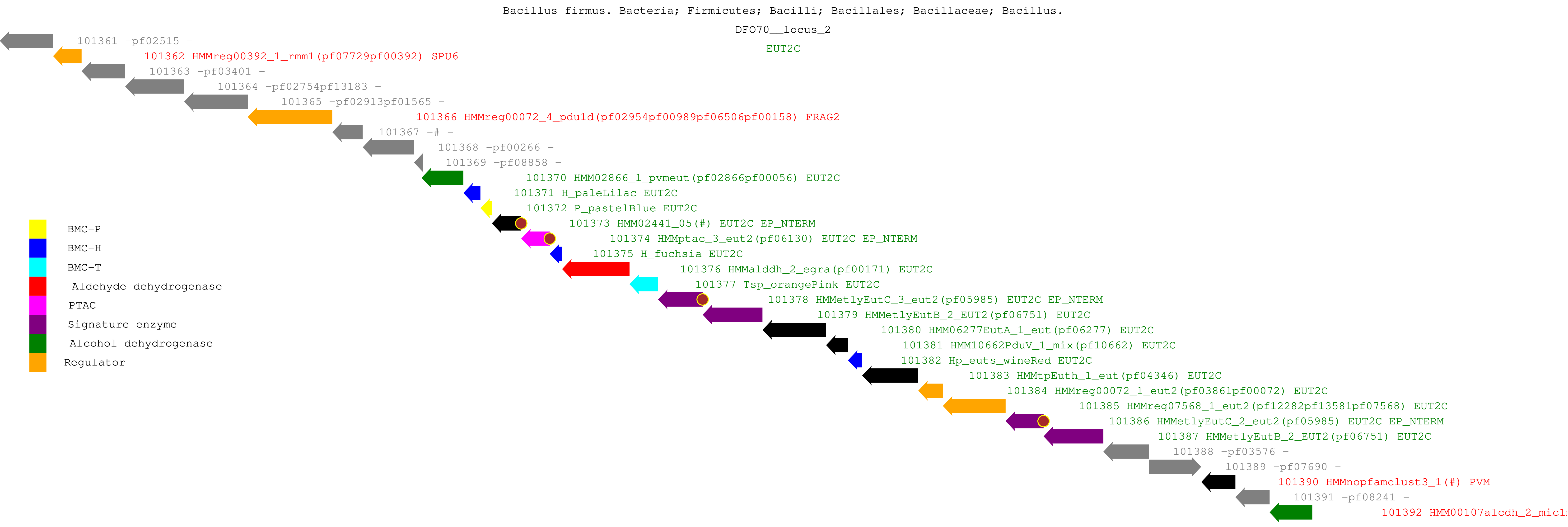
Bacillus firmus. Bacteria; Firmicutes; Bacilli; Bacillales; Bacillaceae; Bacillus.
DFO70__locus_2
EUT2C
101361 -pf02515 -
101362 HMMreg00392_1_rmm1(pf07729pf00392) SPU6
101363 -pf03401 -
101364 -pf02754pf13183 -
101365 -pf02913pf01565 -
101366 HMMreg00072_4_pdu1d(pf02954pf00989pf06506pf00158) FRAG2
101367 -# -
101368 -pf00266 -
101369 -pf08858 -
101370 HMM02866_1_pvmeut(pf02866pf00056) EUT2C
101371 H_paleLilac EUT2C
101372 P_pastelBlue EUT2C
101373 HMM02441_05(#) EUT2C EP_NTERM
101374 HMMptac_3_eut2(pf06130) EUT2C EP_NTERM
101375 H_fuchsia EUT2C
101376 HMMaldh_2_egra(pf00171) EUT2C
101377 Tsp_orangePink EUT2C
101378 HMMetlyEutC_3_eut2(pf05985) EUT2C EP_NTERM
101379 HMMetlyEutB_2_EUT2(pf06751) EUT2C
101380 HMM06277EutA_1_eut(pf06277) EUT2C
101381 HMM10662PduV_1_mix(pf10662) EUT2C
101382 Hp_euts_wineRed EUT2C
101383 HMMtpEuth_1_eut(pf04346) EUT2C
101384 HMMreg00072_1_eut2(pf03861pf00072) EUT2C
101385 HMMreg07568_1_eut2(pf12282pf13581pf07568) EUT2C
101386 HMMetlyEutC_2_eut2(pf05985) EUT2C EP_NTERM
101387 HMMetlyEutB_2_EUT2(pf06751) EUT2C
101388 -pf03576 -
101389 -pf07690 -
101390 HMMnopfamclust3_1(#) PVM
101391 -pf08241 -
101392 HMM00107alcdh_2_mic1
BMC-P
BMC-H
BMC-T
Aldehyde dehydrogenase
PTAC
Signature enzyme
Alcohol dehydrogenase
Regulator

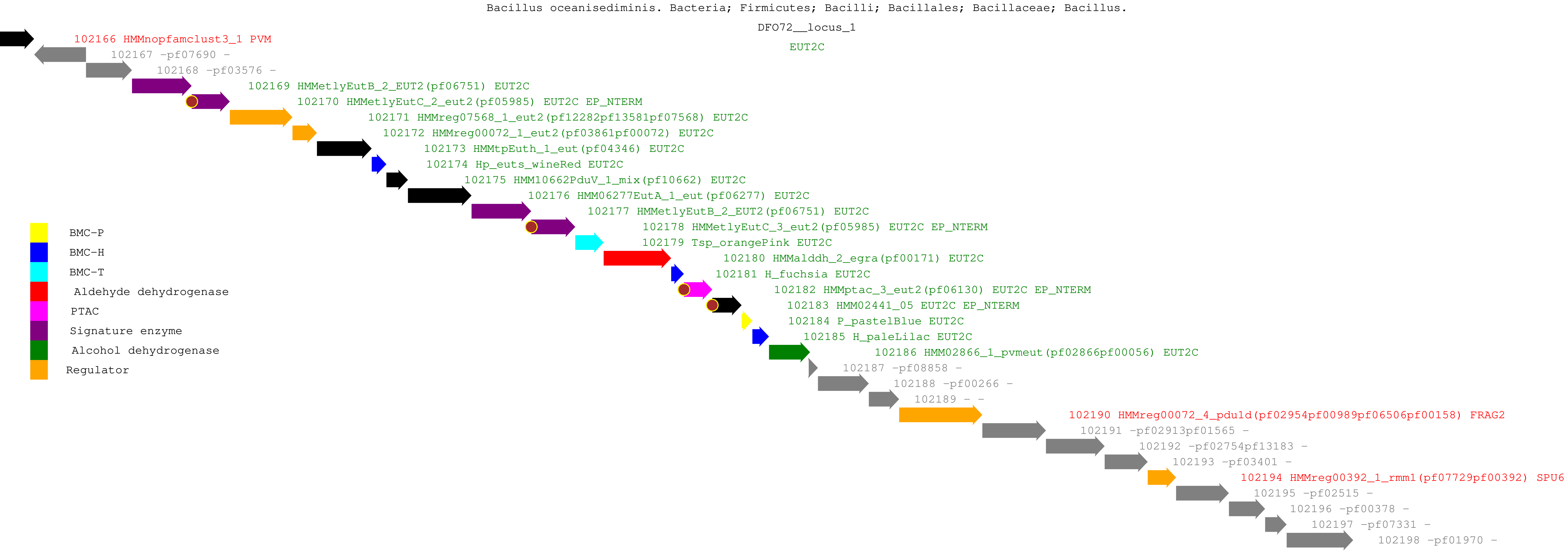
Bacillus oceanisediminis. Bacteria; Firmicutes; Bacilli; Bacillales; Bacillaceae; Bacillus.
DFO72__locus_1
EUT2C
102166 HMMnopfamclust3_1 PVM
102167 -pf07690 -
102168 -pf03576 -
102169 HMMetlyEutB_2_EUT2(pf06751) EUT2C
102170 HMMetlyEutC_2_eut2(pf05985) EUT2C EP_NTERM
102171 HMMreg07568_1_eut2(pf12282pf13581pf07568) EUT2C
102172 HMMreg00072_1_eut2(pf03861pf00072) EUT2C
102173 HMMtpEuth_1_eut(pf04346) EUT2C
102174 Hp_euts_wineRed EUT2C
102175 HMM10662PduV_1_mix(pf10662) EUT2C
102176 HMM06277EutA_1_eut(pf06277) EUT2C
102177 HMMetlyEutB_2_EUT2(pf06751) EUT2C
102178 HMMetlyEutC_3_eut2(pf05985) EUT2C EP_NTERM
102179 Tsp_orangePink EUT2C
102180 HMMalddh_2_egra(pf00171) EUT2C
102181 H_fuchsia EUT2C
102182 HMMptac_3_eut2(pf06130) EUT2C EP_NTERM
102183 HMM02441_05 EUT2C EP_NTERM
102184 P_pastelBlue EUT2C
102185 H_paleLilac EUT2C
102186 HMM02866_1_pvmeut(pf02866pf00056) EUT2C
102187 -pf08858 -
102188 -pf00266 -
102189 - -
102190 HMMreg00072_4_pdu1d(pf02954pf00989pf06506pf00158) FRAG2
102191 -pf02913pf01565 -
102192 -pf02754pf13183 -
102193 -pf03401 -
102194 HMMreg00392_1_rmm1(pf07729pf00392) SPU6
102195 -pf02515 -
102196 -pf00378 -
102197 -pf07331 -
102198 -pf01970 -
BMC-P
BMC-H
BMC-T
Aldehyde dehydrogenase
PTAC
Signature enzyme
Alcohol dehydrogenase
Regulator

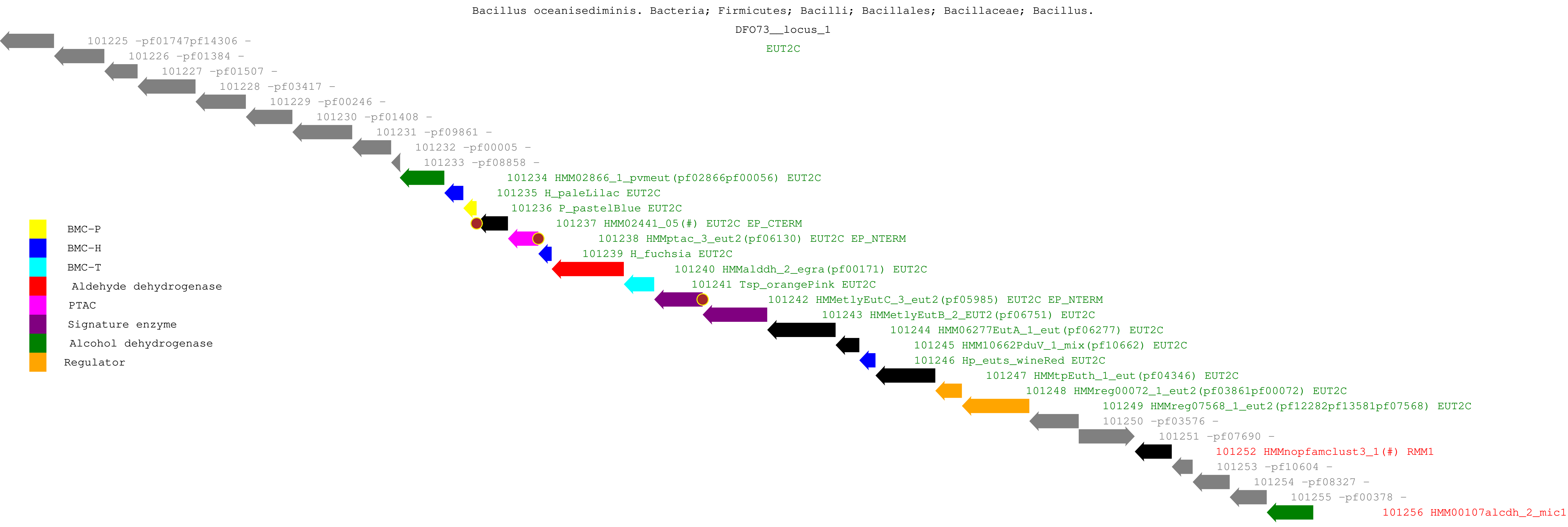

Bacillus oceanisediminis. Bacteria; Firmicutes; Bacilli; Bacillales; Bacillaceae; Bacillus.
DFO73__locus_1
EUT2C
101225 -pf01747pf14306 -
101226 -pf01384 -
101227 -pf01507 -
101228 -pf03417 -
101229 -pf00246 -
101230 -pf01408 -
101231 -pf09861 -
101232 -pf00005 -
101233 -pf08858 -
101234 HMM02866_1_pvmeut(pf02866pf00056) EUT2C
101235 H_paleLilac EUT2C
101236 P_pastelBlue EUT2C
101237 HMM02441_05(#) EUT2C EP_CTERM
101238 HMMptac_3_eut2(pf06130) EUT2C EP_NTERM
101239 H_fuchsia EUT2C
101240 HMMalddh_2_egra(pf00171) EUT2C
101241 Tsp_orangePink EUT2C
101242 HMMetlyEutC_3_eut2(pf05985) EUT2C EP_NTERM
101243 HMMetlyEutB_2_EUT2(pf06751) EUT2C
101244 HMM06277EutA_1_eut(pf06277) EUT2C
101245 HMM10662PduV_1_mix(pf10662) EUT2C
101246 Hp_euts_wineRed EUT2C
101247 HMMtpEuth_1_eut(pf04346) EUT2C
101248 HMMreg00072_1_eut2(pf03861pf00072) EUT2C
101249 HMMreg07568_1_eut2(pf12282pf13581pf07568) EUT2C
101250 -pf03576 -
101251 -pf07690 -
101252 HMMnopfamclust3_1(#) RMM1
101253 -pf10604 -
101254 -pf08327 -
101255 -pf00378 -
101256 HMM00107alcdh_2_mic1
BMC-P
BMC-H
BMC-T
Aldehyde dehydrogenase
PTAC
Signature enzyme
Alcohol dehydrogenase
Regulator

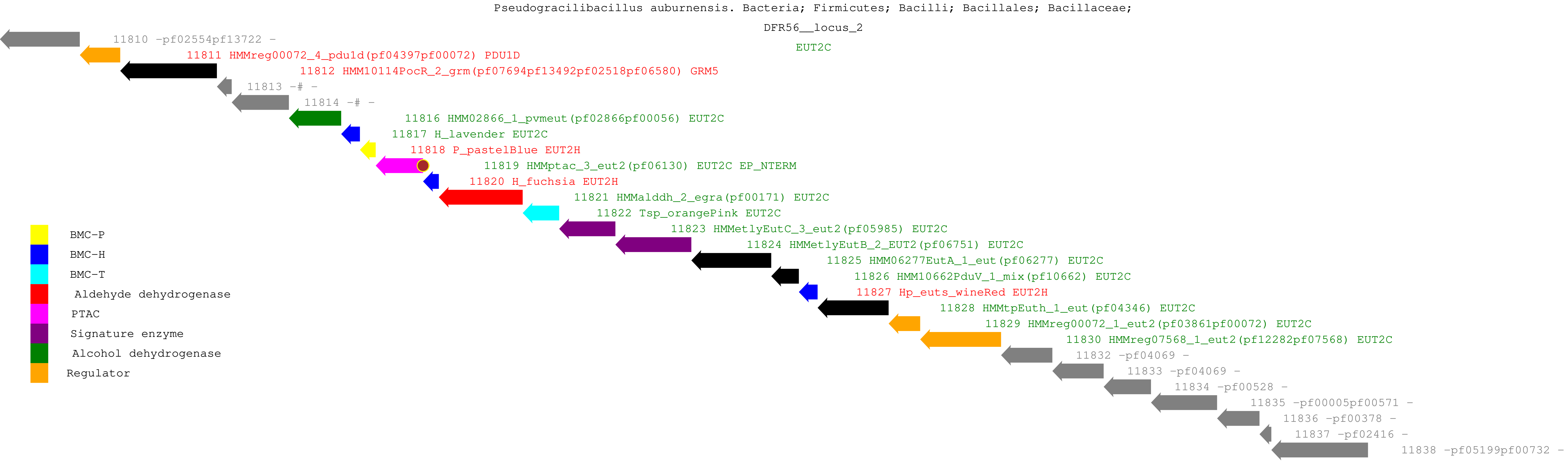
Pseudogracilibacillus auburnensis. Bacteria; Firmicutes; Bacilli; Bacillales; Bacillaceae;
DFR56__locus_2
EUT2C
11810 -pf02554pf13722 -
11811 HMMreg00072_4_pdu1d(pf04397pf00072) PDU1D
11812 HMM10114PocR_2_grm(pf07694pf13492pf02518pf06580) GRM5
11813 -# -
11814 -# -
11816 HMM02866_1_pvmeut(pf02866pf00056) EUT2C
11817 H_lavender EUT2C
11818 P_pastelBlue EUT2H
11819 HMMptac_3_eut2(pf06130) EUT2C EP_NTERM
11820 H_fuchsia EUT2H
11821 HMMalddh_2_egra(pf00171) EUT2C
11822 Tsp_orangePink EUT2C
11823 HMMetlyEutC_3_eut2(pf05985) EUT2C
11824 HMMetlyEutB_2_EUT2(pf06751) EUT2C
11825 HMM06277EutA_1_eut(pf06277) EUT2C
11826 HMM10662PduV_1_mix(pf10662) EUT2C
11827 Hp_euts_wineRed EUT2H
11828 HMMtpEuth_1_eut(pf04346) EUT2C
11829 HMMreg00072_1_eut2(pf03861pf00072) EUT2C
11830 HMMreg07568_1_eut2(pf12282pf07568) EUT2C
11832 -pf04069 -
11833 -pf04069 -
11834 -pf00528 -
11835 -pf00005pf00571 -
11836 -pf00378 -
11837 -pf02416 -
11838 -pf05199pf00732 -
BMC-P
BMC-H
BMC-T
Aldehyde dehydrogenase
PTAC
Signature enzyme
Alcohol dehydrogenase
Regulator

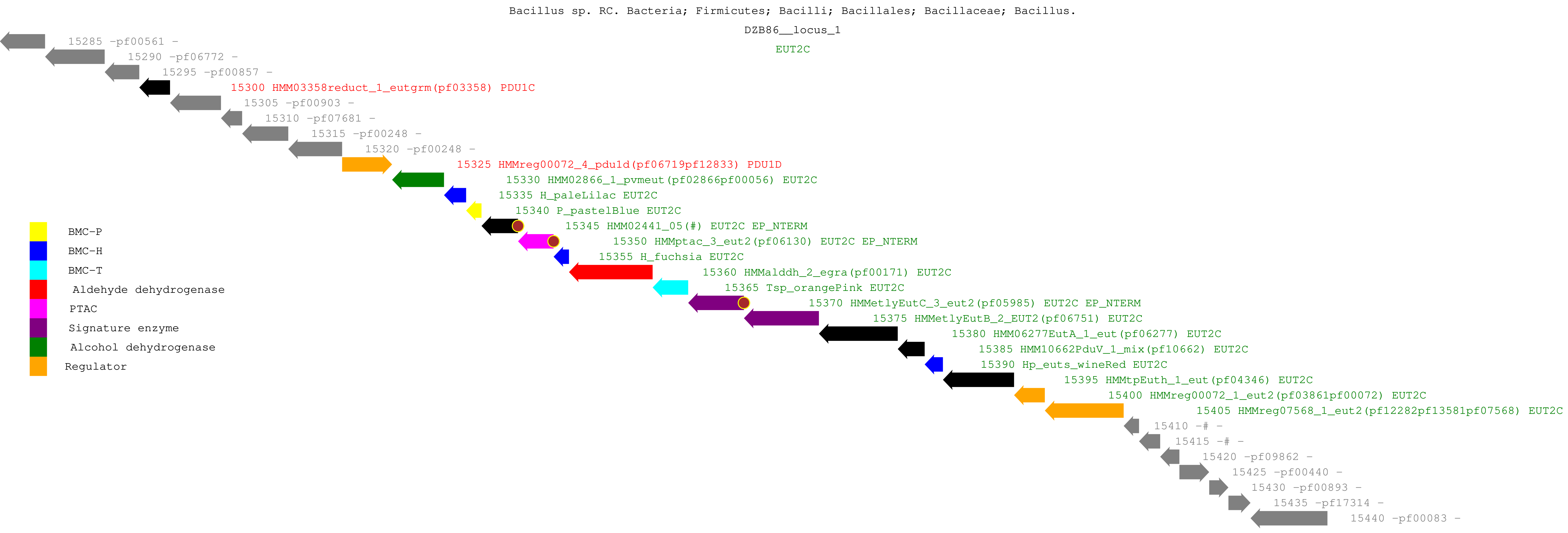

Bacillus sp. RC. Bacteria; Firmicutes; Bacilli; Bacillales; Bacillaceae; Bacillus.
DZB86__locus_1
EUT2C
15285 -pf00561 -
15290 -pf06772 -
15295 -pf00857 -
15300 HMM03358reduct_1_eutgrm(pf03358) PDU1C
15305 -pf00903 -
15310 -pf07681 -
15315 -pf00248 -
15320 -pf00248 -
15325 HMMreg00072_4_pdu1d(pf06719pf12833) PDU1D
15330 HMM02866_1_pvmeut(pf02866pf00056) EUT2C
15335 H_paleLilac EUT2C
15340 P_pastelBlue EUT2C
15345 HMM02441_05(#) EUT2C EP_NTERM
15350 HMMptac_3_eut2(pf06130) EUT2C EP_NTERM
15355 H_fuchsia EUT2C
15360 HMMaldh_2_egra(pf00171) EUT2C
15365 Tsp_orangePink EUT2C
15370 HMMetlyEutC_3_eut2(pf05985) EUT2C EP_NTERM
15375 HMMetlyEutB_2_EUT2(pf06751) EUT2C
15380 HMM06277EutA_1_eut(pf06277) EUT2C
15385 HMM10662PduV_1_mix(pf10662) EUT2C
15390 Hp_euts_wineRed EUT2C
15395 HMMtpEuth_1_eut(pf04346) EUT2C
15400 HMMreg00072_1_eut2(pf03861pf00072) EUT2C
15405 HMMreg07568_1_eut2(pf12282pf13581pf07568) EUT2C
15410 -# -
15415 -# -
15420 -pf09862 -
15425 -pf00440 -
15430 -pf00893 -
15435 -pf17314 -
15440 -pf00083 -
BMC-P
BMC-H
BMC-T
Aldehyde dehydrogenase
PTAC
Signature enzyme
Alcohol dehydrogenase
Regulator

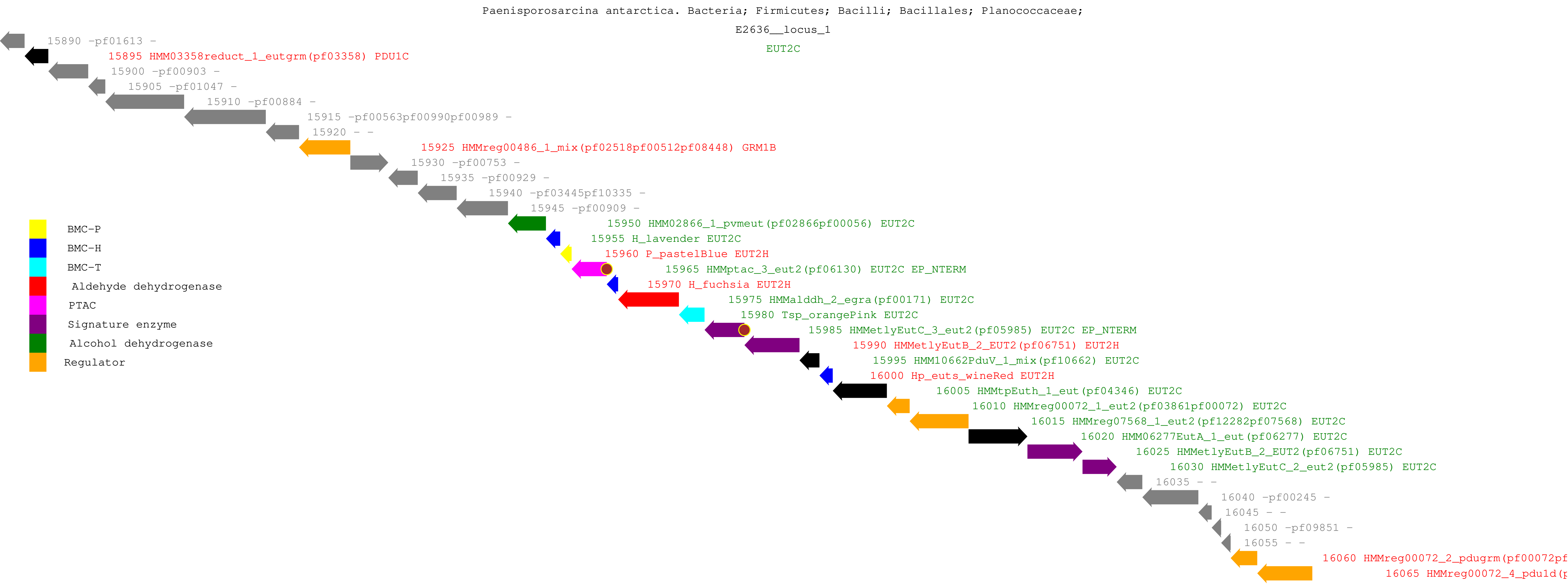

Paenisporosarcina antarctica. Bacteria; Firmicutes; Bacilli; Bacillales; Planococcaceae;
E2636__locus_1
EUT2C
15890 -pf01613 -
15895 HMM03358reduct_1_eutgrm(pf03358) PDU1C
15900 -pf00903 -
15905 -pf01047 -
15910 -pf00884 -
15915 -pf00563pf00990pf00989 -
15920 - -
15925 HMMreg00486_1_mix(pf02518pf00512pf08448) GRM1B
15930 -pf00753 -
15935 -pf00929 -
15940 -pf03445pf10335 -
15945 -pf00909 -
15950 HMM02866_1_pvmeut(pf02866pf00056) EUT2C
15955 H_lavender EUT2C
15960 P_pastelBlue EUT2H
15965 HMMptac_3_eut2(pf06130) EUT2C EP_NTERM
15970 H_fuchsia EUT2H
15975 HMMalddh_2_egra(pf00171) EUT2C
15980 Tsp_orangePink EUT2C
15985 HMMetlyEutC_3_eut2(pf05985) EUT2C EP_NTERM
15990 HMMetlyEutB_2_EUT2(pf06751) EUT2H
15995 HMM10662PduV_1_mix(pf10662) EUT2C
16000 Hp_euts_wineRed EUT2H
16005 HMMtpEuth_1_eut(pf04346) EUT2C
16010 HMMreg00072_1_eut2(pf03861pf00072) EUT2C
16015 HMMreg07568_1_eut2(pf12282pf07568) EUT2C
16020 HMM06277EutA_1_eut(pf06277) EUT2C
16025 HMMetlyEutB_2_EUT2(pf06751) EUT2C
16030 HMMetlyEutC_2_eut2(pf05985) EUT2C
16035 - -
16040 -pf00245 -
16045 - -
16050 -pf09851 -
16055 - -
16060 HMMreg00072_2_pdugrm(pf00072pf
16065 HMMreg00072_4_pdu1d(p
BMC-P
BMC-H
BMC-T
Aldehyde dehydrogenase
PTAC
Signature enzyme
Alcohol dehydrogenase
Regulator

Bacillus sp. WN066. Bacteria; Firmicutes; Bacilli; Bacillales; Bacillaceae; Bacillus.

E2K98__locus_2

EUT2C

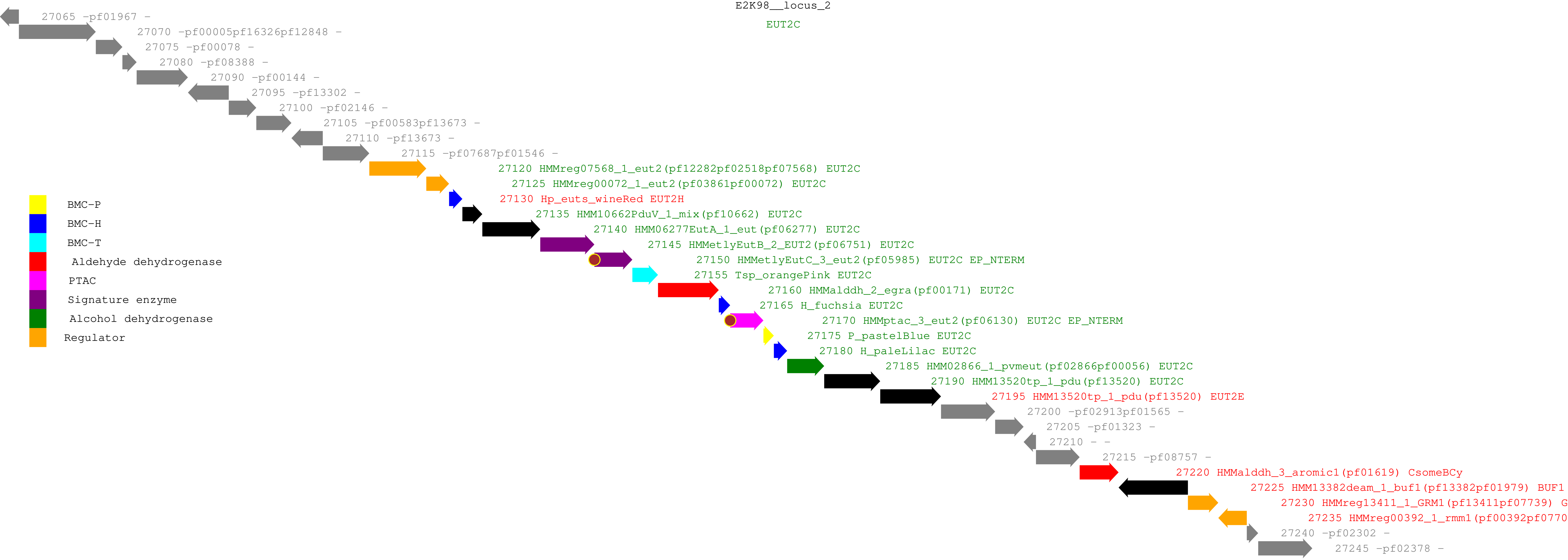

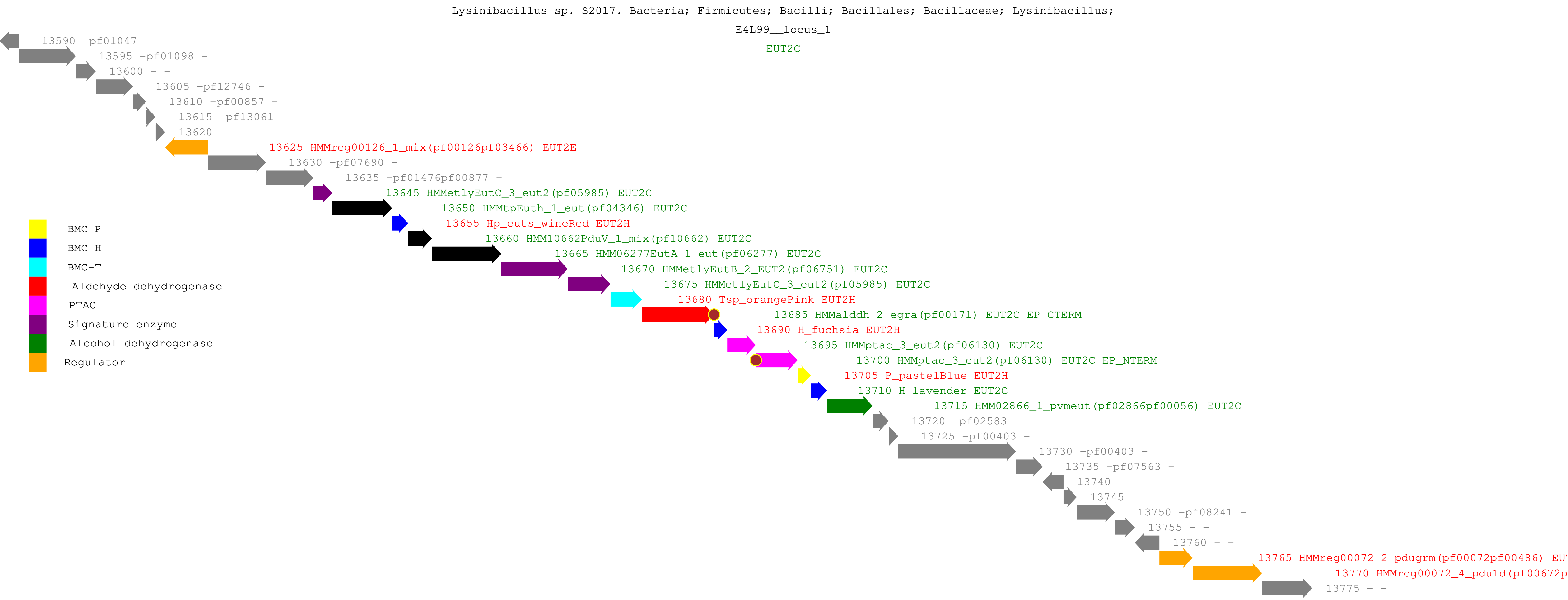
Lysinibacillus sp. S2017. Bacteria; Firmicutes; Bacilli; Bacillales; Bacillaceae; Lysinibacillus;
E4L99__locus_1
EUT2C
13590 -pf01047 -
13595 -pf01098 -
13600 - -
13605 -pf12746 -
13610 -pf00857 -
13615 -pf13061 -
13620 - -
13625 HMMreg00126_1_mix(pf00126pf03466) EUT2E
13630 -pf07690 -
13635 -pf01476pf00877 -
13645 HMMetlyEutC_3_eut2(pf05985) EUT2C
13650 HMMtpEuth_1_eut(pf04346) EUT2C
13655 Hp_euts_wineRed EUT2H
13660 HMM10662PduV_1_mix(pf10662) EUT2C
13665 HMM06277EutA_1_eut(pf06277) EUT2C
13670 HMMetlyEutB_2_EUT2(pf06751) EUT2C
13675 HMMetlyEutC_3_eut2(pf05985) EUT2C
13680 Tsp_orangePink EUT2H
13685 HMMalddh_2_egra(pf00171) EUT2C EP_CTERM
13690 H_fuchsia EUT2H
13695 HMMptac_3_eut2(pf06130) EUT2C
13700 HMMptac_3_eut2(pf06130) EUT2C EP_NTERM
13705 P_pastelBlue EUT2H
13710 H_lavender EUT2C
13715 HMM02866_1_pvmeut(pf02866pf00056) EUT2C
13720 -pf02583 -
13725 -pf00403 -
13730 -pf00403 -
13735 -pf07563 -
13740 - -
13745 - -
13750 -pf08241 -
13755 - -
13760 - -
13765 HMMreg00072_2_pdugrm(pf00072pf00486) EU
13770 HMMreg00072_4_pdu1d(pf00672p
13775 - -
BMC-P
BMC-H
BMC-T
Aldehyde dehydrogenase
PTAC
Signature enzyme
Alcohol dehydrogenase
Regulator

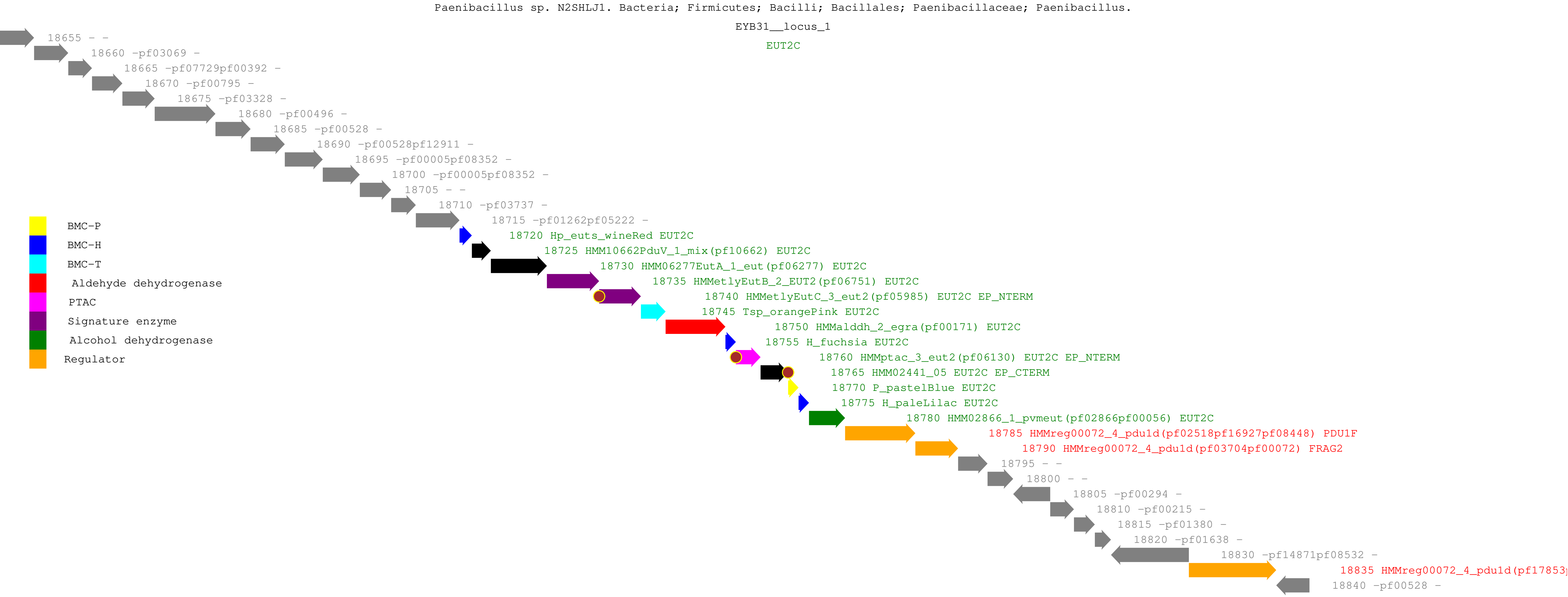
Paenibacillus sp. N2SHLJ1. Bacteria; Firmicutes; Bacilli; Bacillales; Paenibacillaceae; Paenibacillus.
EYB31__locus_1
EUT2C
18655 - -
18660 -pf03069 -
18665 -pf07729pf00392 -
18670 -pf00795 -
18675 -pf03328 -
18680 -pf00496 -
18685 -pf00528 -
18690 -pf00528pf12911 -
18695 -pf00005pf08352 -
18700 -pf00005pf08352 -
18705 - -
18710 -pf03737 -
18715 -pf01262pf05222 -
18720 Hp_euts_wineRed EUT2C
18725 HMM10662PduV_1_mix(pf10662) EUT2C
18730 HMM06277EutA_1_eut(pf06277) EUT2C
18735 HMMetlyEutB_2_EUT2(pf06751) EUT2C
18740 HMMetlyEutC_3_eut2(pf05985) EUT2C EP_NTERM
18745 Tsp_orangePink EUT2C
18750 HMMalddh_2_egra(pf00171) EUT2C
18755 H_fuchsia EUT2C
18760 HMMptac_3_eut2(pf06130) EUT2C EP_NTERM
18765 HMM02441_05 EUT2C EP_CTERM
18770 P_pastelBlue EUT2C
18775 H_paleLilac EUT2C
18780 HMM02866_1_pvmeut(pf02866pf00056) EUT2C
18785 HMMreg00072_4_pdu1d(pf02518pf16927pf08448) PDU1F
18790 HMMreg00072_4_pdu1d(pf03704pf00072) FRAG2
18795 - -
18800 - -
18805 -pf00294 -
18810 -pf00215 -
18815 -pf01380 -
18820 -pf01638 -
18830 -pf14871pf08532 -
18835 HMMreg00072_4_pdu1d(pf17853
18840 -pf00528 -
BMC-P
BMC-H
BMC-T
Aldehyde dehydrogenase
PTAC
Signature enzyme
Alcohol dehydrogenase
Regulator

Bacillus sp. S3. Bacteria; Firmicutes; Bacilli; Bacillales; Bacillaceae; Bacillus.

FAY30__locus_2

EUT2C

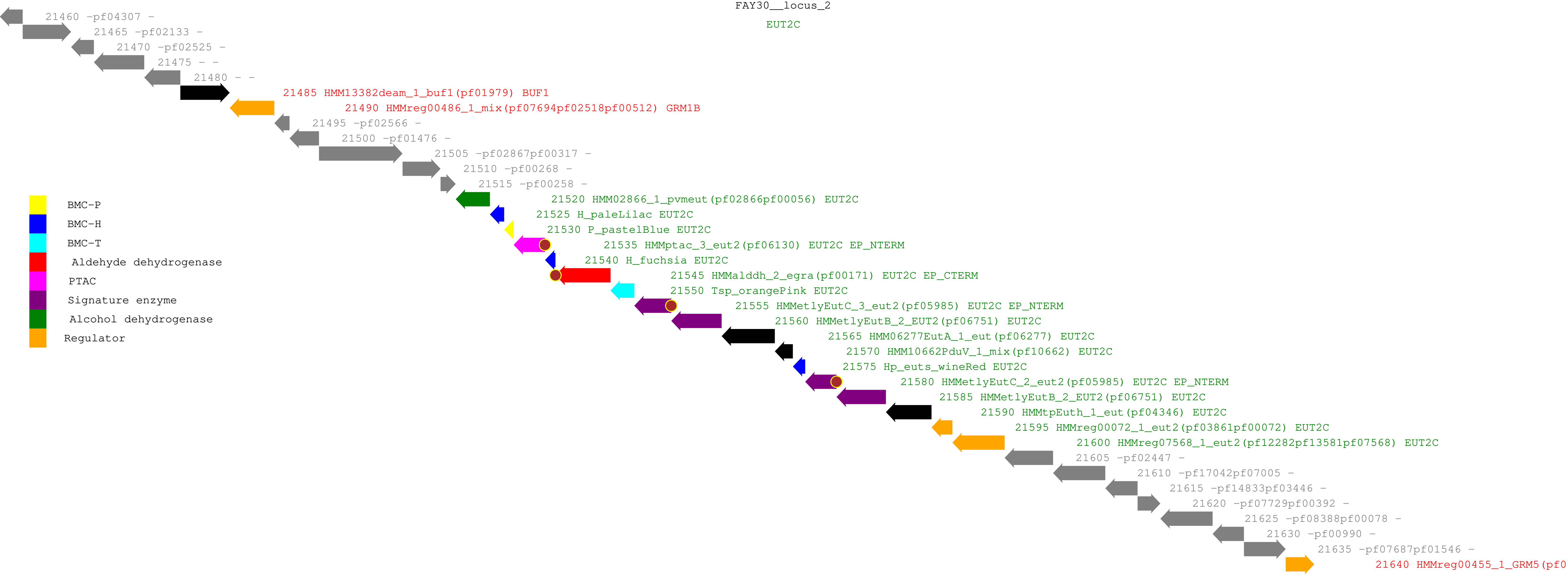

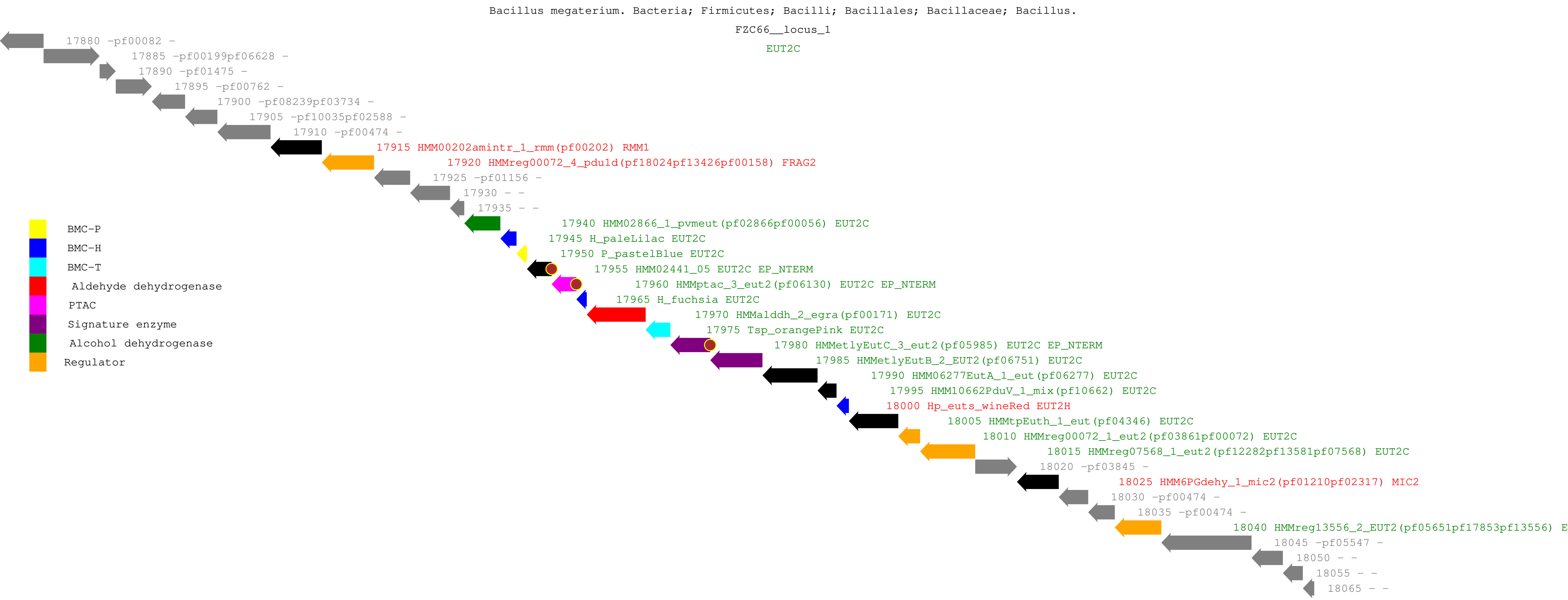
Bacillus megaterium. Bacteria; Firmicutes; Bacilli; Bacillales; Bacillaceae; Bacillus.
FZC66__locus_1
EUT2C
17880 -pf00082 -
17885 -pf00199pf06628 -
17890 -pf01475 -
17895 -pf00762 -
17900 -pf08239pf03734 -
17905 -pf10035pf02588 -
17910 -pf00474 -
17915 HMM00202amintr_1_rmm(pf00202) RMM1
17920 HMMreg00072_4_pdu1d(pf18024pf13426pf00158) FRAG2
17925 -pf01156 -
17930 - -
17935 - -
17940 HMM02866_1_pvmeut(pf02866pf00056) EUT2C
17945 H_paleLilac EUT2C
17950 P_pastelBlue EUT2C
17955 HMM02441_05 EUT2C EP_NTERM
17960 HMMptac_3_eut2(pf06130) EUT2C EP_NTERM
17965 H_fuchsia EUT2C
17970 HMMalddh_2_egra(pf00171) EUT2C
17975 Tsp_orangePink EUT2C
17980 HMMetlyEutC_3_eut2(pf05985) EUT2C EP_NTERM
17985 HMMetlyEutB_2_EUT2(pf06751) EUT2C
17990 HMM06277EutA_1_eut(pf06277) EUT2C
17995 HMM10662PduV_1_mix(pf10662) EUT2C
18000 Hp_euts_wineRed EUT2H
18005 HMMtpEuth_1_eut(pf04346) EUT2C
18010 HMMreg00072_1_eut2(pf03861pf00072) EUT2C
18015 HMMreg07568_1_eut2(pf12282pf13581pf07568) EUT2C
18020 -pf03845 -
18025 HMM6PGdehy_1_mic2(pf01210pf02317) MIC2
18030 -pf00474 -
18035 -pf00474 -
18040 HMMreg13556_2_EUT2(pf05651pf17853pf13556) E
18045 -pf05547 -
18050 - -
18055 - -
18065 - -
BMC-P
BMC-H
BMC-T
Aldehyde dehydrogenase
PTAC
Signature enzyme
Alcohol dehydrogenase
Regulator

Planococcus donghaensis MPA1U2. Bacteria; Firmicutes; Bacilli; Bacillales; Planococcaceae;

GPDM__locus_1

EUT2C

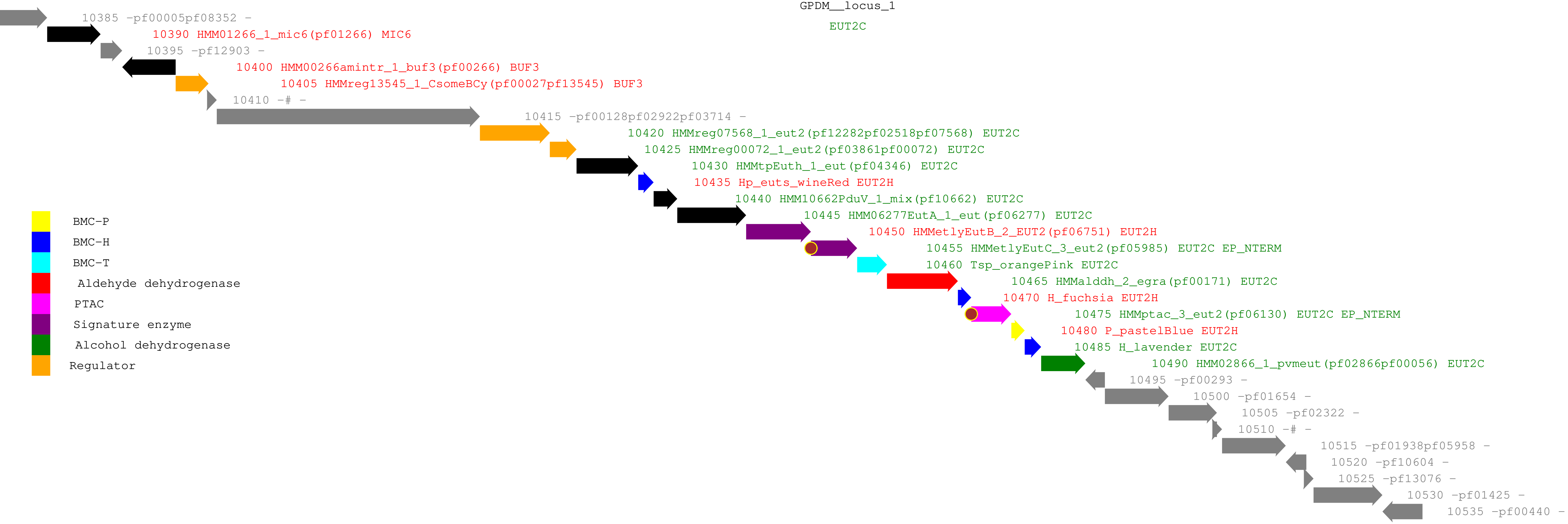

Halobacillus faecis. Bacteria; Firmicutes; Bacilli; Bacillales; Bacillaceae; Halobacillus.

HFA01__locus_1

EUT2C

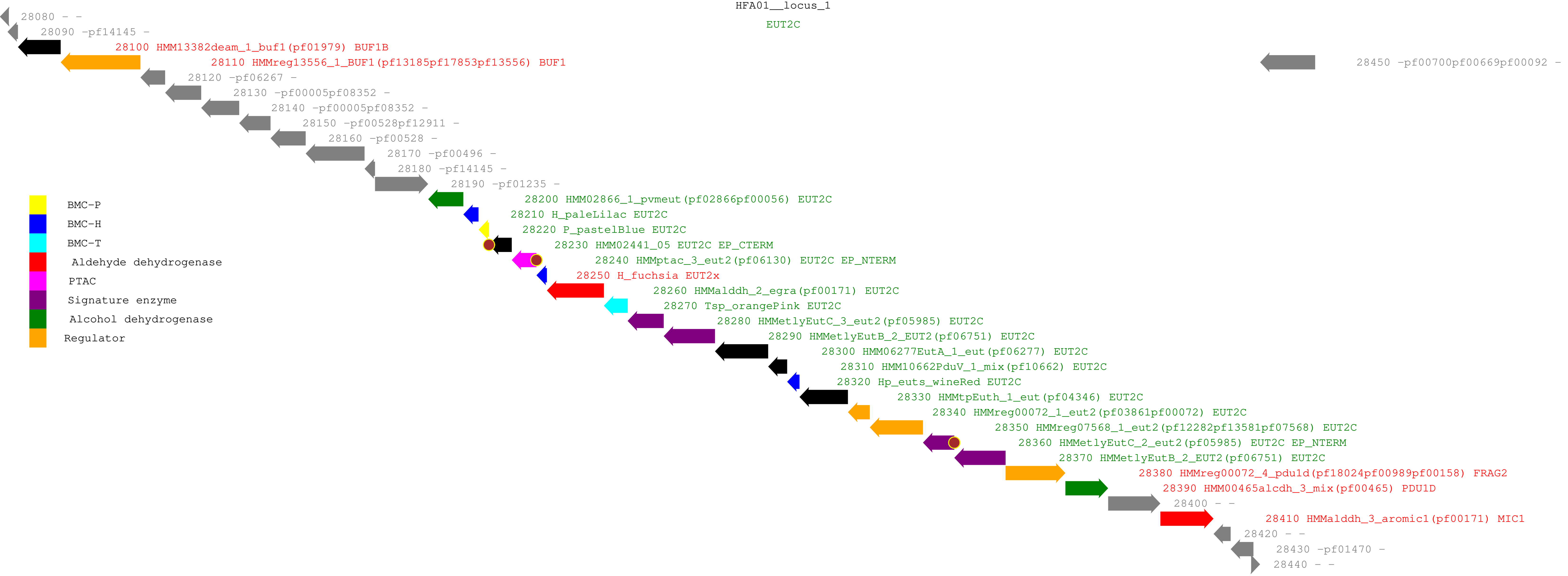

Halobacillus mangrovi. Bacteria; Firmicutes; Bacilli; Bacillales; Bacillaceae; Halobacillus.

HM131__locus_1

EUT2C

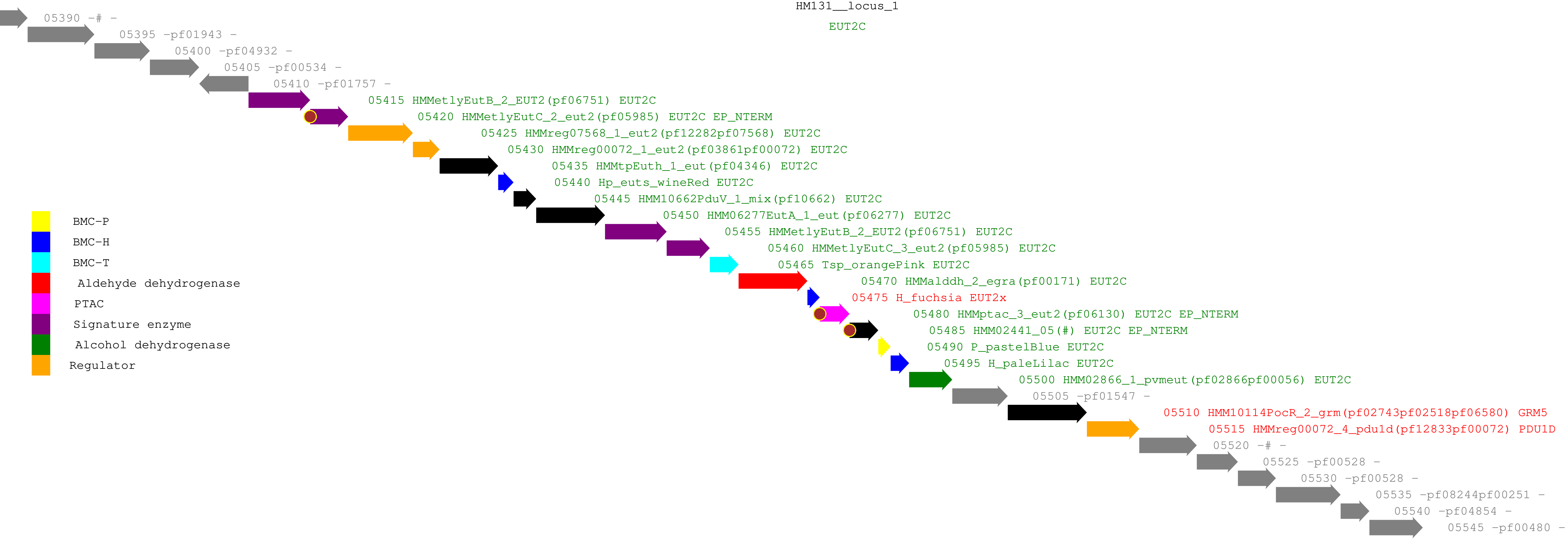

Aneurinibacillus aneurinilyticus ATCC 12856. Bacteria; Firmicutes; Bacilli; Bacillales; Paenibacillaceae;

HMPREF0083__locus_1

EUT2C

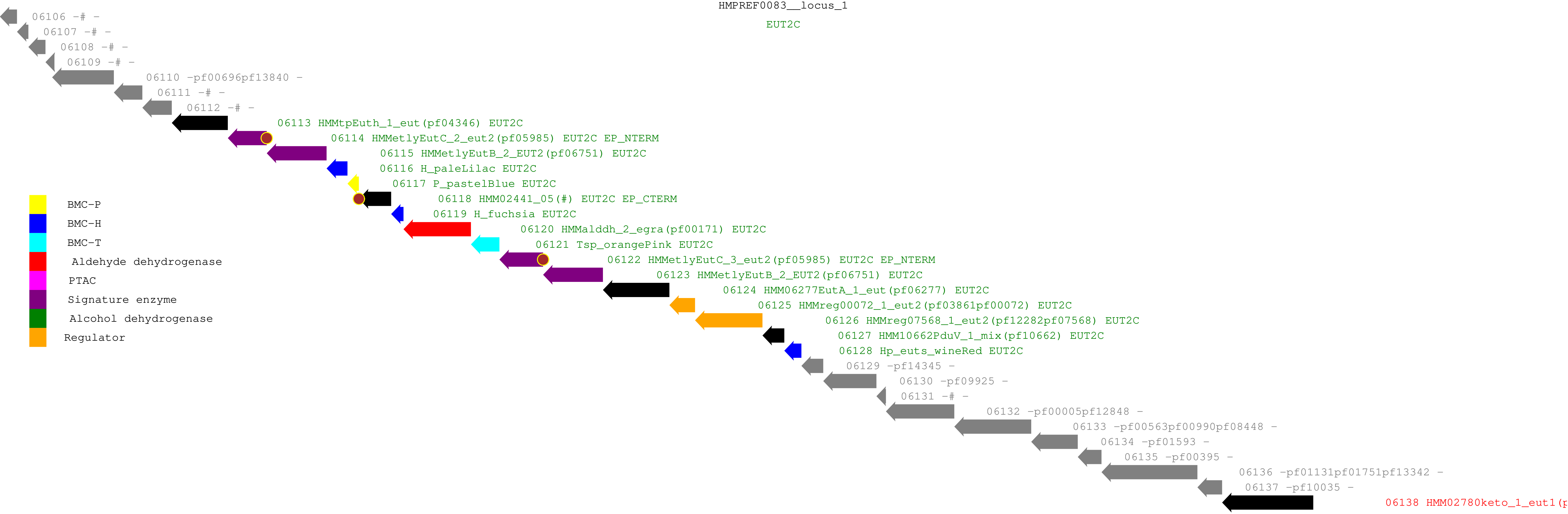

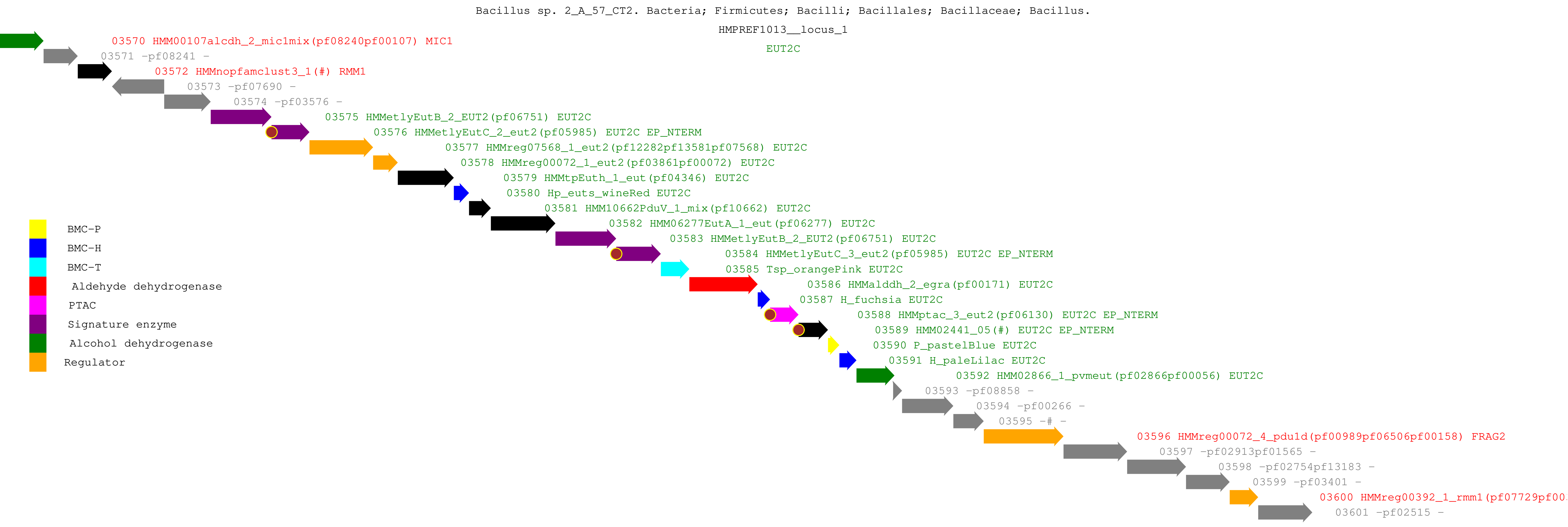
Bacillus sp. 2_A_57_CT2. Bacteria; Firmicutes; Bacilli; Bacillales; Bacillaceae; Bacillus.
HMPREF1013__locus_1
EUT2C
03570 HMM00107alcdh_2_mic1mix(pf08240pf00107) MIC1
03571 -pf08241 -
03572 HMMnopfamclust3_1(#) RMM1
03573 -pf07690 -
03574 -pf03576 -
03575 HMMetlyEutB_2_EUT2(pf06751) EUT2C
03576 HMMetlyEutC_2_eut2(pf05985) EUT2C EP_NTERM
03577 HMMreg07568_1_eut2(pf12282pf13581pf07568) EUT2C
03578 HMMreg00072_1_eut2(pf03861pf00072) EUT2C
03579 HMMtpEuth_1_eut(pf04346) EUT2C
03580 Hp_euts_wineRed EUT2C
03581 HMM10662PduV_1_mix(pf10662) EUT2C
03582 HMM06277EutA_1_eut(pf06277) EUT2C
03583 HMMetlyEutB_2_EUT2(pf06751) EUT2C
03584 HMMetlyEutC_3_eut2(pf05985) EUT2C EP_NTERM
03585 Tsp_orangePink EUT2C
03586 HMMalddh_2_egra(pf00171) EUT2C
03587 H_fuchsia EUT2C
03588 HMMptac_3_eut2(pf06130) EUT2C EP_NTERM
03589 HMM02441_05(#) EUT2C EP_NTERM
03590 P_pastelBlue EUT2C
03591 H_paleLilac EUT2C
03592 HMM02866_1_pvmeut(pf02866pf00056) EUT2C
03593 -pf08858 -
03594 -pf00266 -
03595 -# -
03596 HMMreg00072_4_pdu1d(pf00989pf06506pf00158) FRAG2
03597 -pf02913pf01565 -
03598 -pf02754pf13183 -
03599 -pf03401 -
03600 HMMreg00392_1_rmm1(pf07729pf00
03601 -pf02515 -
BMC-P
BMC-H
BMC-T
Aldehyde dehydrogenase
PTAC
Signature enzyme
Alcohol dehydrogenase
Regulator

Sporosarcina newyorkensis 2681. Bacteria; Firmicutes; Bacilli; Bacillales; Planococcaceae;

HMPREF9372__locus_1

EUT2C

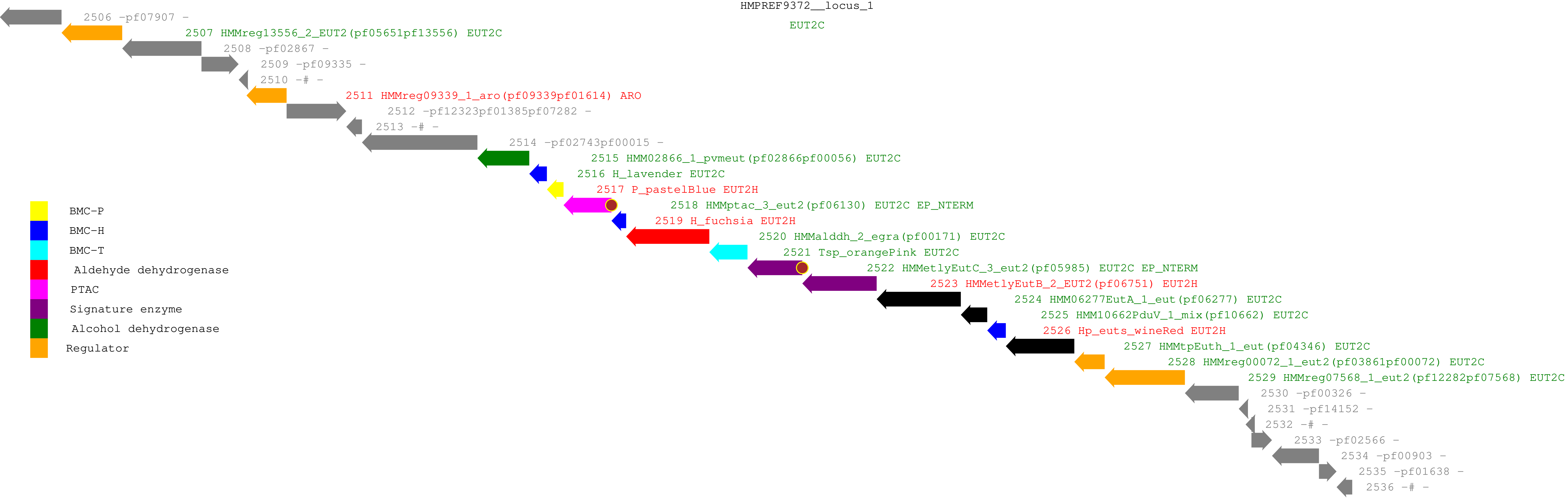

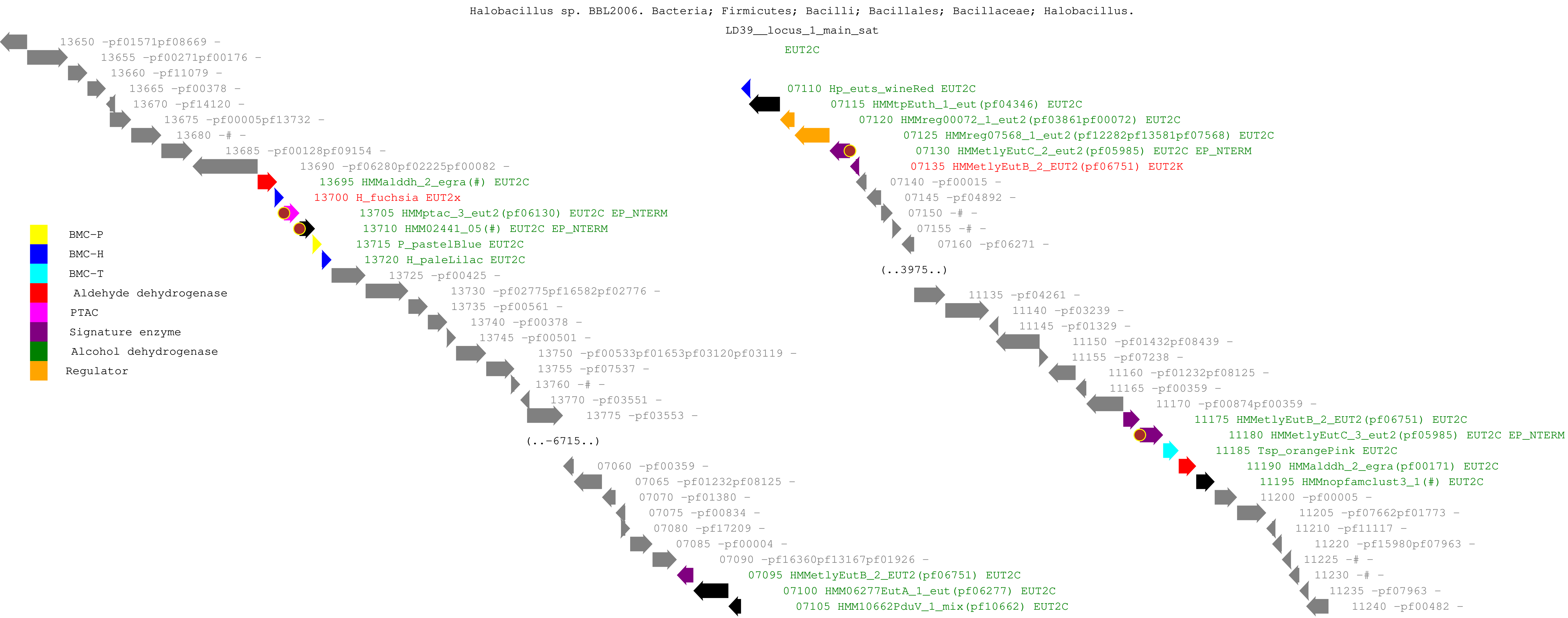
Halobacillus sp. BBL2006. Bacteria; Firmicutes; Bacilli; Bacillales; Bacillaceae; Halobacillus.
LD39__locus_1_main_sat
EUT2C
13650 -pf01571pf08669 -
13655 -pf00271pf00176 -
13660 -pf11079 -
13665 -pf00378 -
13670 -pf14120 -
13675 -pf00005pf13732 -
13680 -# -
13685 -pf00128pf09154 -
13690 -pf06280pf02225pf00082 -
13695 HMMaldh_2_egra(#) EUT2C
13700 H_fuchsia EUT2x
13705 HMMptac_3_eut2(pf06130) EUT2C EP_NTERM
13710 HMM02441_05(#) EUT2C EP_NTERM
13715 P_pastelBlue EUT2C
13720 H_paleLilac EUT2C
13725 -pf00425 -
13730 -pf02775pf16582pf02776 -
13735 -pf00561 -
13740 -pf00378 -
13745 -pf00501 -
13750 -pf00533pf01653pf03120pf03119 -
13755 -pf07537 -
13760 -# -
13770 -pf03551 -
13775 -pf03553 -
(..-6715..)
07060 -pf00359 -
07065 -pf01232pf08125 -
07070 -pf01380 -
07075 -pf00834 -
07080 -pf17209 -
07085 -pf00004 -
07090 -pf16360pf13167pf01926 -
07095 HMMetlyEutB_2_EUT2(pf06751) EUT2C
07100 HMM06277EutA_1_eut(pf06277) EUT2C
07105 HMM10662PduV_1_mix(pf10662) EUT2C
BMC-P
BMC-H
BMC-T
Aldehyde dehydrogenase
PTAC
Signature enzyme
Alcohol dehydrogenase
Regulator
07110 Hp_euts_wineRed EUT2C
07115 HMMtpEuth_1_eut(pf04346) EUT2C
07120 HMMreg00072_1_eut2(pf03861pf00072) EUT2C
07125 HMMreg07568_1_eut2(pf12282pf13581pf07568) EUT2C
07130 HMMetlyEutC_2_eut2(pf05985) EUT2C EP_NTERM
07135 HMMetlyEutB_2_EUT2(pf06751) EUT2K
07140 -pf00015 -
07145 -pf04892 -
07150 -# -
07155 -# -
07160 -pf06271 -
(..3975..)
11135 -pf04261 -
11140 -pf03239 -
11145 -pf01329 -
11150 -pf01432pf08439 -
11155 -pf07238 -
11160 -pf01232pf08125 -
11165 -pf00359 -
11170 -pf00874pf00359 -
11175 HMMetlyEutB_2_EUT2(pf06751) EUT2C
11180 HMMetlyEutC_3_eut2(pf05985) EUT2C EP_NTERM
11185 Tsp_orangePink EUT2C
11190 HMMaldh_2_egra(pf00171) EUT2C
11195 HMMnopfamclust3_1(#) EUT2C
11200 -pf00005 -
11205 -pf07662pf01773 -
11210 -pf11117 -
11220 -pf15980pf07963 -
11225 -# -
11230 -# -
11235 -pf07963 -
11240 -pf00482 -

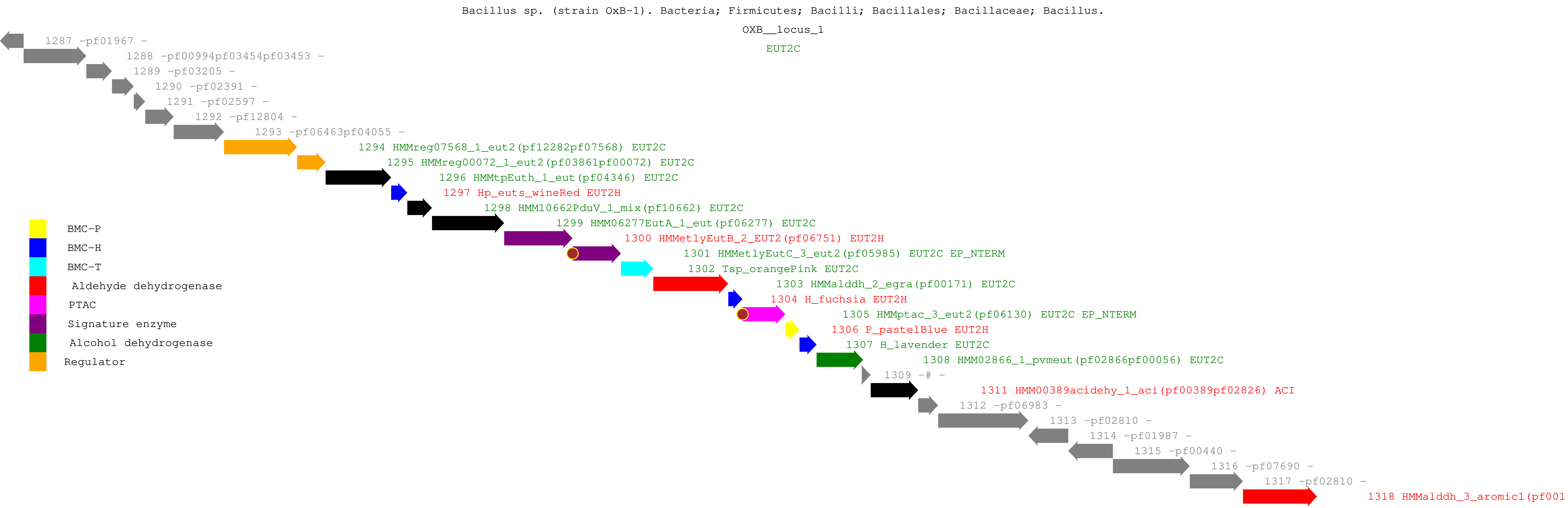
Bacillus sp. (strain OxB-1). Bacteria; Firmicutes; Bacilli; Bacillales; Bacillaceae; Bacillus.
OXB__locus_1
EUT2C
1287 -pf01967 -
1288 -pf00994pf03454pf03453 -
1289 -pf03205 -
1290 -pf02391 -
1291 -pf02597 -
1292 -pf12804 -
1293 -pf06463pf04055 -
1294 HMMreg07568_1_eut2(pf12282pf07568) EUT2C
1295 HMMreg00072_1_eut2(pf03861pf00072) EUT2C
1296 HMMtpEuth_1_eut(pf04346) EUT2C
1297 Hp_euts_wineRed EUT2H
1298 HMM10662PduV_1_mix(pf10662) EUT2C
1299 HMM06277EutA_1_eut(pf06277) EUT2C
1300 HMMetlyEutB_2_EUT2(pf06751) EUT2H
1301 HMMetlyEutC_3_eut2(pf05985) EUT2C EP_NTERM
1302 Tsp_orangePink EUT2C
1303 HMMalddh_2_egra(pf00171) EUT2C
1304 H_fuchsia EUT2H
1305 HMMptac_3_eut2(pf06130) EUT2C EP_NTERM
1306 P_pastelBlue EUT2H
1307 H_lavender EUT2C
1308 HMM02866_1_pvmeut(pf02866pf00056) EUT2C
1309 -# -
1311 HMM00389acidehy_1_aci(pf00389pf02826) ACI
1312 -pf06983 -
1313 -pf02810 -
1314 -pf01987 -
1315 -pf00440 -
1316 -pf07690 -
1317 -pf02810 -
1318 HMMalddh_3_aromic1(pf001
BMC-P
BMC-H
BMC-T
Aldehyde dehydrogenase
PTAC
Signature enzyme
Alcohol dehydrogenase
Regulator

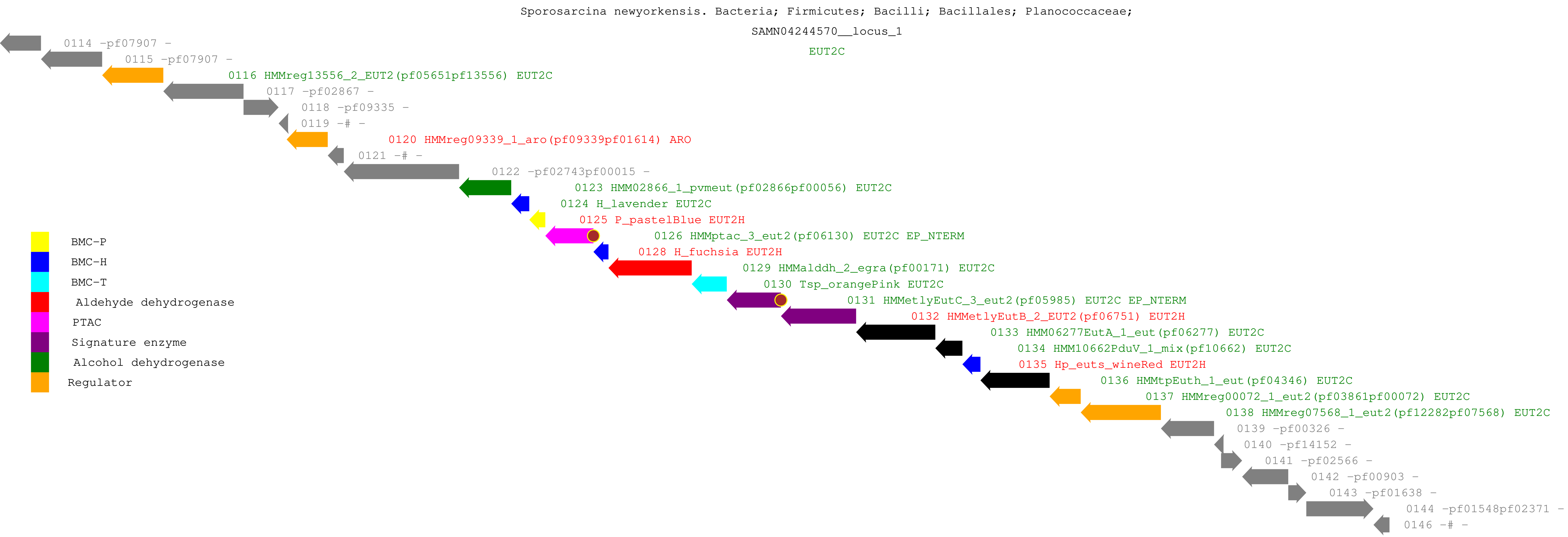

Sporosarcina newyorkensis. Bacteria; Firmicutes; Bacilli; Bacillales; Planococcaceae;
SAMN04244570__locus_1
EUT2C
0114 -pf07907 -
0115 -pf07907 -
0116 HMMreg13556_2_EUT2(pf05651pf13556) EUT2C
0117 -pf02867 -
0118 -pf09335 -
0119 -# -
0120 HMMreg09339_1_aro(pf09339pf01614) ARO
0121 -# -
0122 -pf02743pf00015 -
0123 HMM02866_1_pvmeut(pf02866pf00056) EUT2C
0124 H_lavender EUT2C
0125 P_pastelBlue EUT2H
0126 HMMptac_3_eut2(pf06130) EUT2C EP_NTERM
0128 H_fuchsia EUT2H
0129 HMMalddh_2_egra(pf00171) EUT2C
0130 Tsp_orangePink EUT2C
0131 HMMetlyEutC_3_eut2(pf05985) EUT2C EP_NTERM
0132 HMMetlyEutB_2_EUT2(pf06751) EUT2H
0133 HMM06277EutA_1_eut(pf06277) EUT2C
0134 HMM10662PduV_1_mix(pf10662) EUT2C
0135 Hp_euts_wineRed EUT2H
0136 HMMtpEuth_1_eut(pf04346) EUT2C
0137 HMMreg00072_1_eut2(pf03861pf00072) EUT2C
0138 HMMreg07568_1_eut2(pf12282pf07568) EUT2C
0139 -pf00326 -
0140 -pf14152 -
0141 -pf02566 -
0142 -pf00903 -
0143 -pf01638 -
0144 -pf01548pf02371 -
0146 -# -
BMC-P
BMC-H
BMC-T
Aldehyde dehydrogenase
PTAC
Signature enzyme
Alcohol dehydrogenase
Regulator

Bacillus megaterium. Bacteria; Firmicutes; Bacilli; Bacillales; Bacillaceae; Bacillus.

SAMN04487776__locus_1

EUT2C

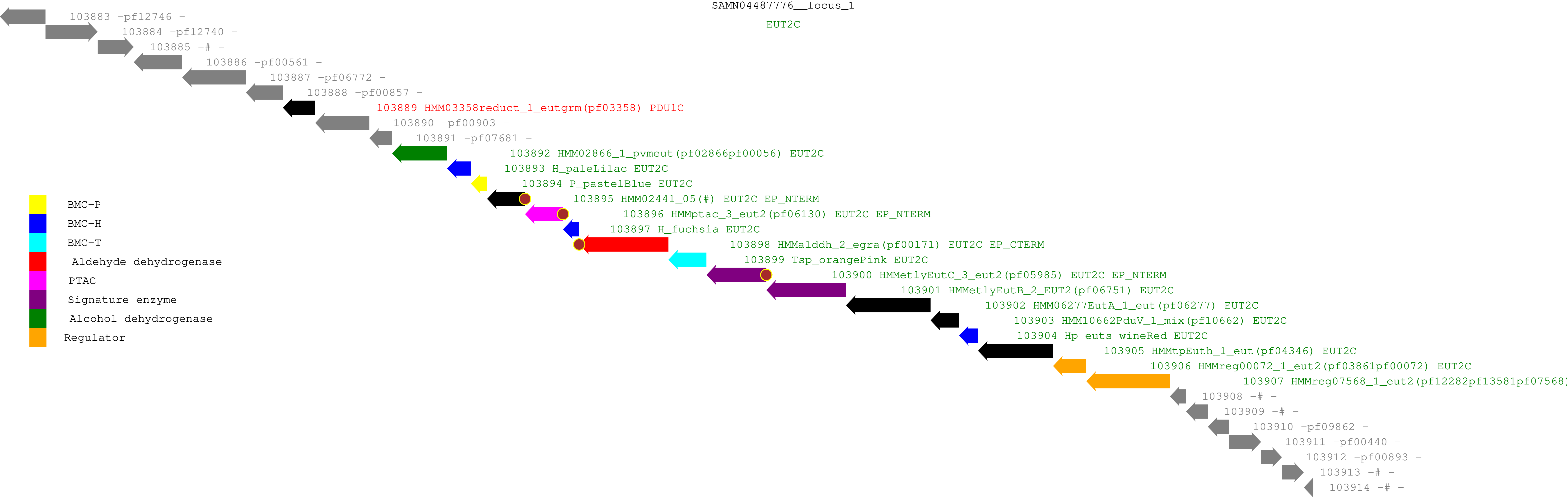

Bacillus aryabhattai B8W22. Bacteria; Firmicutes; Bacilli; Bacillales; Bacillaceae; Bacillus.

SAMN04487777__locus_1

EUT2C

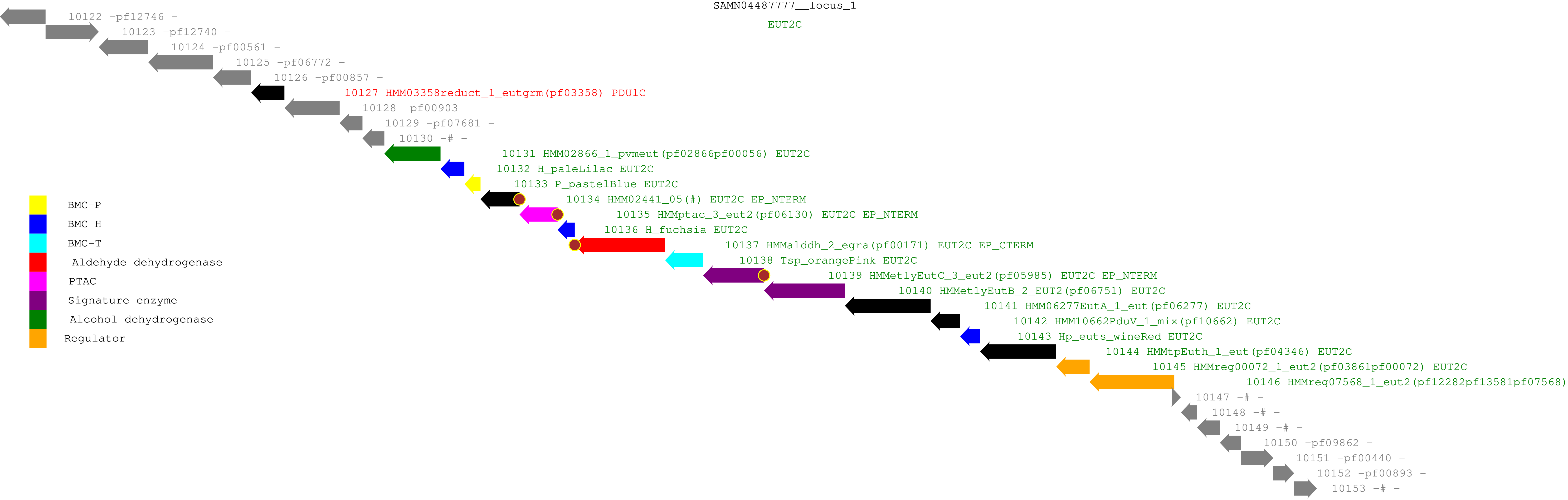

Aneurinibacillus migulanus (Bacillus migulanus). Bacteria; Firmicutes; Bacilli; Bacillales; Paenibacillaceae;

SAMN04487909__locus_1

EUT2C

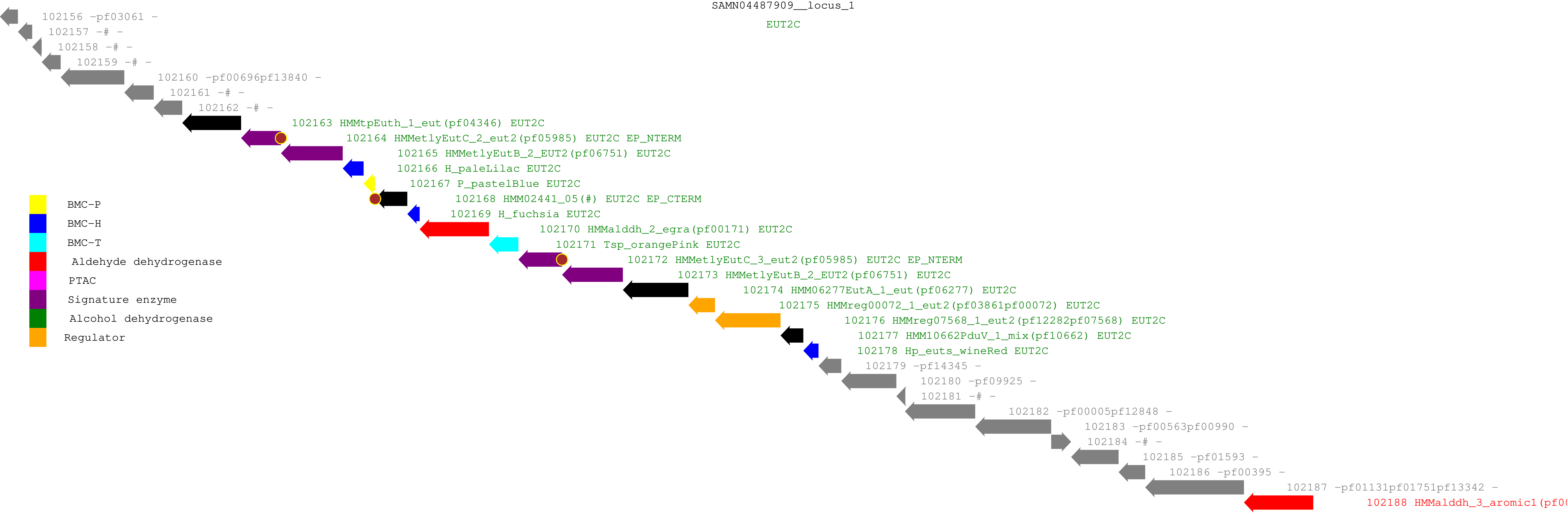

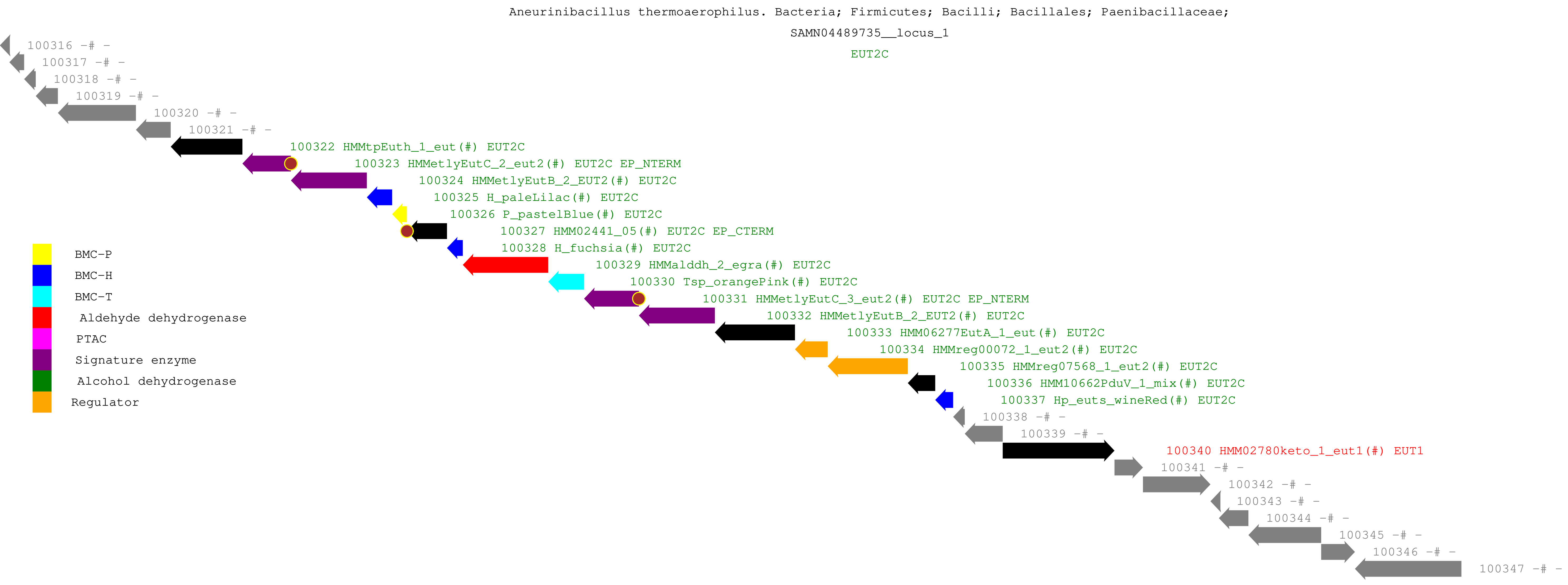
Aneurinibacillus thermoaerophilus. Bacteria; Firmicutes; Bacilli; Bacillales; Paenibacillaceae;
SAMN04489735__locus_1
EUT2C
100316 -# -
100317 -# -
100318 -# -
100319 -# -
100320 -# -
100321 -# -
100322 HMMtpEuth_1_eut(#) EUT2C
100323 HMMetlyEutC_2_eut2(#) EUT2C EP_NTERM
100324 HMMetlyEutB_2_EUT2(#) EUT2C
100325 H_paleLilac(#) EUT2C
100326 P_pastelBlue(#) EUT2C
100327 HMM02441_05(#) EUT2C EP_CTERM
100328 H_fuchsia(#) EUT2C
100329 HMMalddh_2_egra(#) EUT2C
100330 Tsp_orangePink(#) EUT2C
100331 HMMetlyEutC_3_eut2(#) EUT2C EP_NTERM
100332 HMMetlyEutB_2_EUT2(#) EUT2C
100333 HMM06277EutA_1_eut(#) EUT2C
100334 HMMreg00072_1_eut2(#) EUT2C
100335 HMMreg07568_1_eut2(#) EUT2C
100336 HMM10662PduV_1_mix(#) EUT2C
100337 Hp_euts_wineRed(#) EUT2C
100338 -# -
100339 -# -
100340 HMM02780keto_1_eut1(#) EUT1
100341 -# -
100342 -# -
100343 -# -
100344 -# -
100345 -# -
100346 -# -
100347 -# -
BMC-P
BMC-H
BMC-T
Aldehyde dehydrogenase
PTAC
Signature enzyme
Alcohol dehydrogenase
Regulator

Domibacillus enclensis. Bacteria; Firmicutes; Bacilli; Bacillales; Bacillaceae; Domibacillus.

SAMN05443094__locus_1

EUT2C

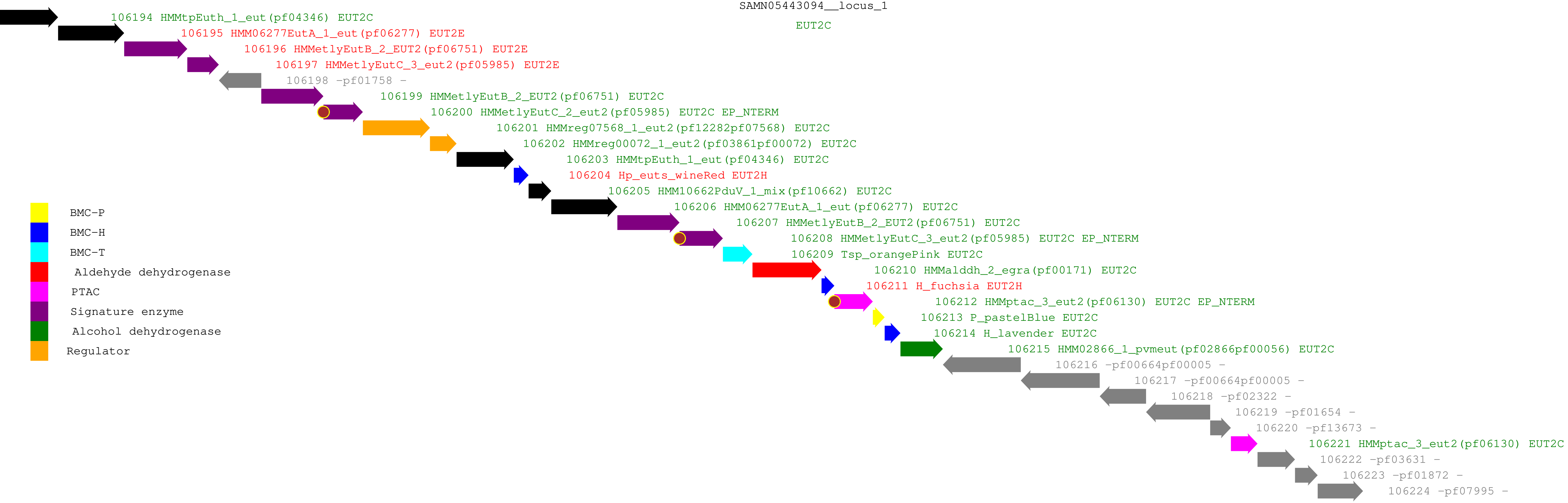

Psychrobacillus sp. OK028. Bacteria; Firmicutes; Bacilli; Bacillales; Bacillaceae;

SAMN05518871__locus_1

EUT2C

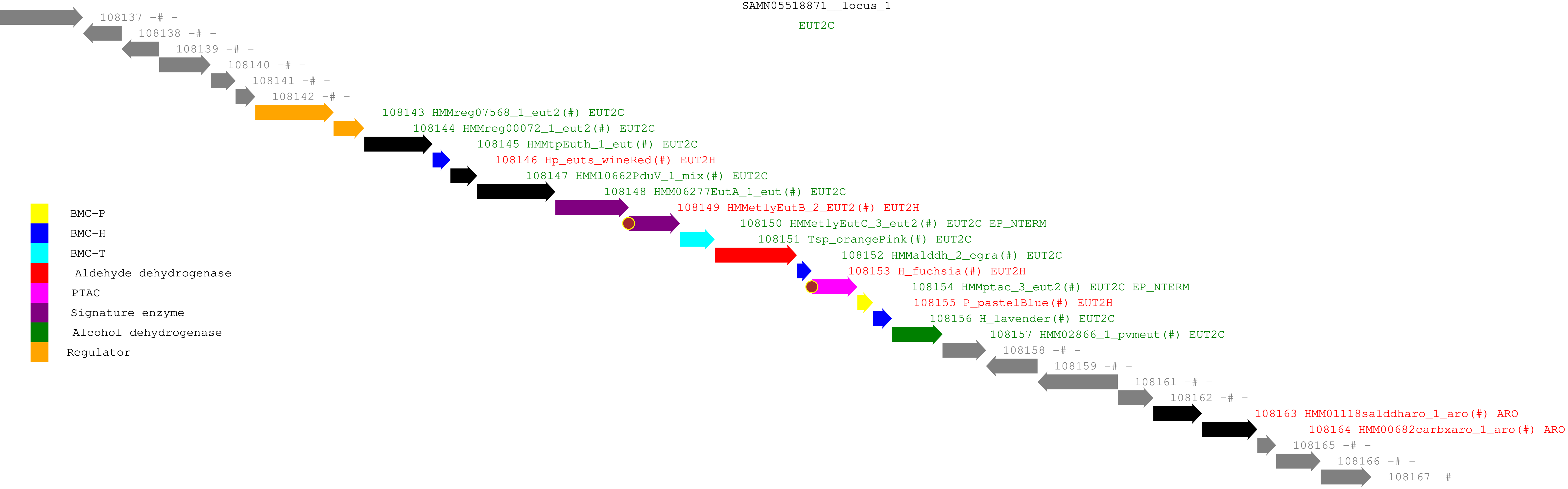

Sporosarcina ureae. Bacteria; Firmicutes; Bacilli; Bacillales; Planococcaceae;

SporoP17a__locus_1

EUT2C

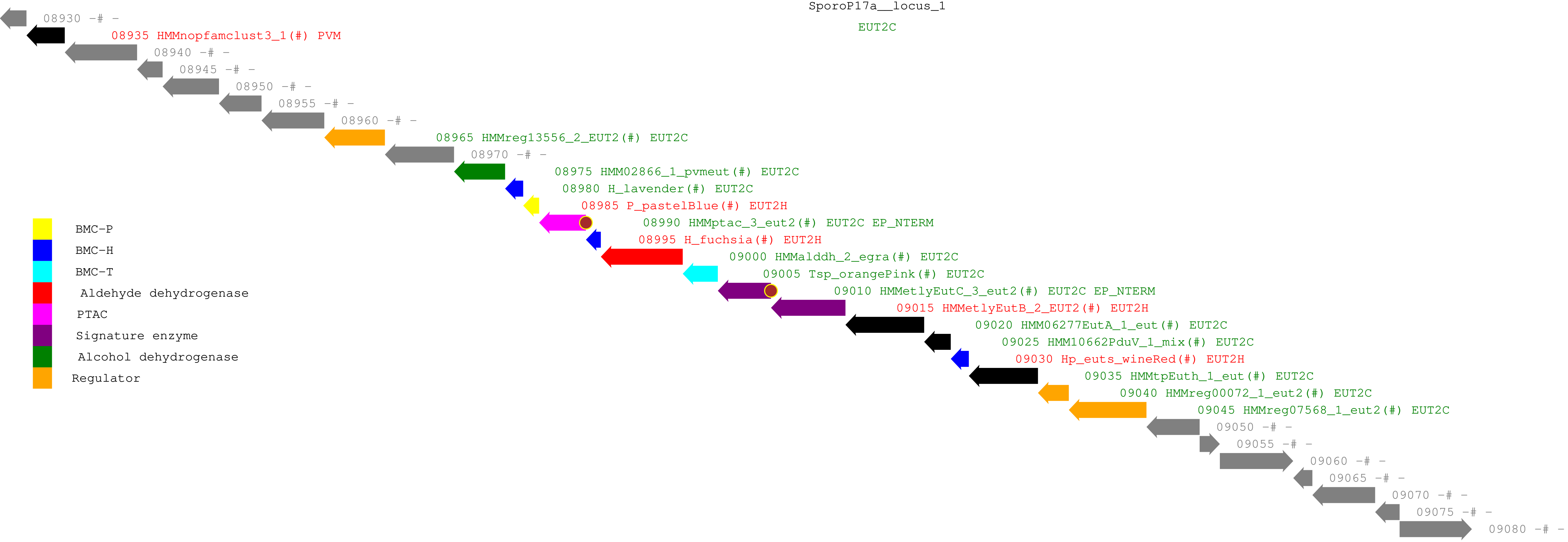

Sporosarcina ureae. Bacteria; Firmicutes; Bacilli; Bacillales; Planococcaceae;

SporoP32a__locus_1

EUT2C

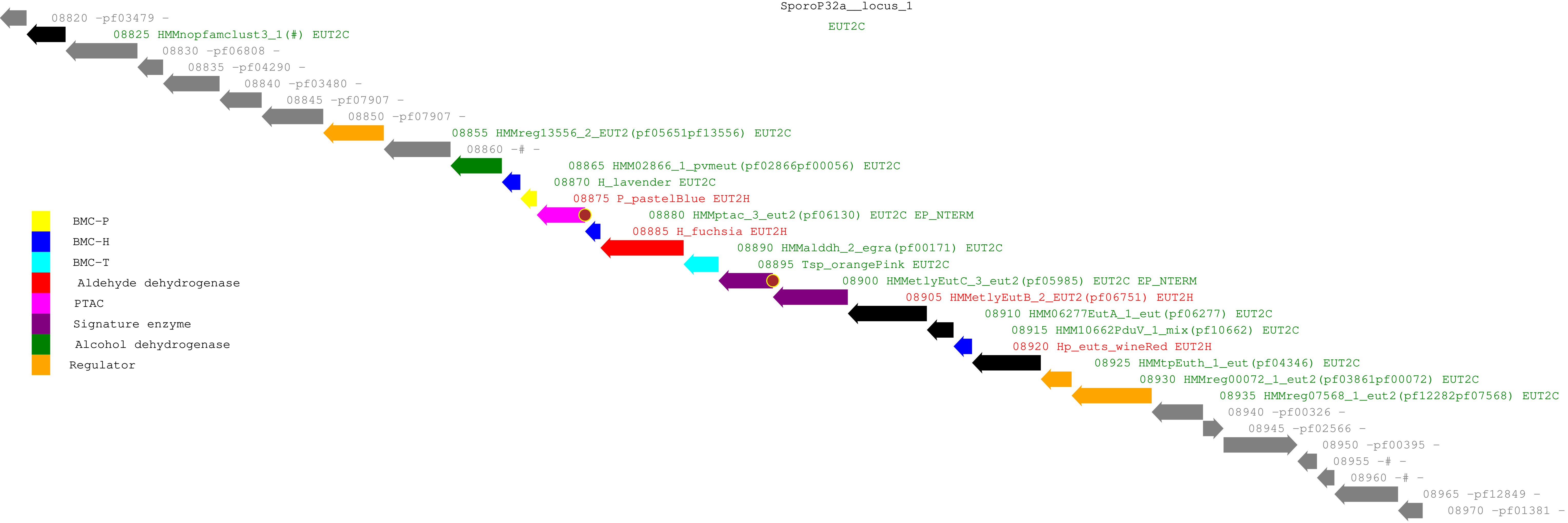

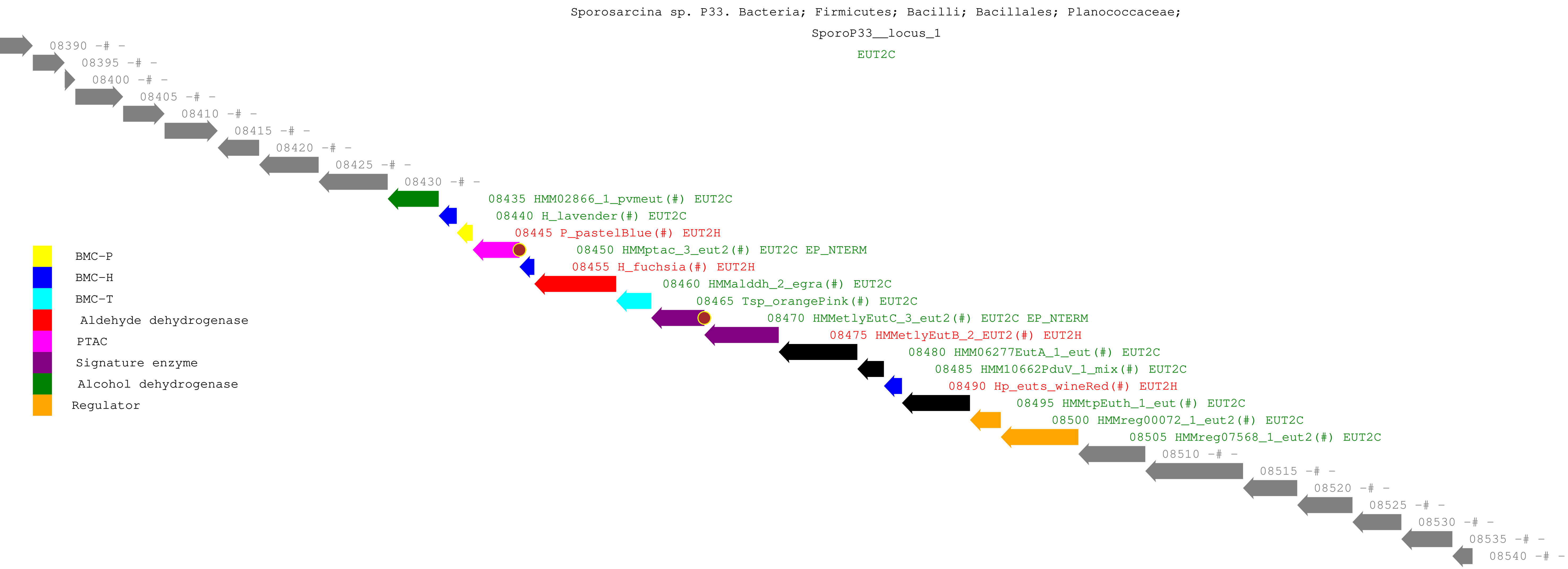
Sporosarcina sp. P33. Bacteria; Firmicutes; Bacilli; Bacillales; Planococcaceae;
SporoP33__locus_1
EUT2C
08390 -# -
08395 -# -
08400 -# -
08405 -# -
08410 -# -
08415 -# -
08420 -# -
08425 -# -
08430 -# -
08435 HMM02866_1_pvmeut(#) EUT2C
08440 H_lavender(#) EUT2C
08445 P_pastelBlue(#) EUT2H
08450 HMMptac_3_eut2(#) EUT2C EP_NTERM
08455 H_fuchsia(#) EUT2H
08460 HMMalddh_2_egra(#) EUT2C
08465 Tsp_orangePink(#) EUT2C
08470 HMMetlyEutC_3_eut2(#) EUT2C EP_NTERM
08475 HMMetlyEutB_2_EUT2(#) EUT2H
08480 HMM06277EutA_1_eut(#) EUT2C
08485 HMM10662PduV_1_mix(#) EUT2C
08490 Hp_euts_wineRed(#) EUT2H
08495 HMMtpEuth_1_eut(#) EUT2C
08500 HMMreg00072_1_eut2(#) EUT2C
08505 HMMreg07568_1_eut2(#) EUT2C
08510 -# -
08515 -# -
08520 -# -
08525 -# -
08530 -# -
08535 -# -
08540 -# -
BMC-P
BMC-H
BMC-T
Aldehyde dehydrogenase
PTAC
Signature enzyme
Alcohol dehydrogenase
Regulator

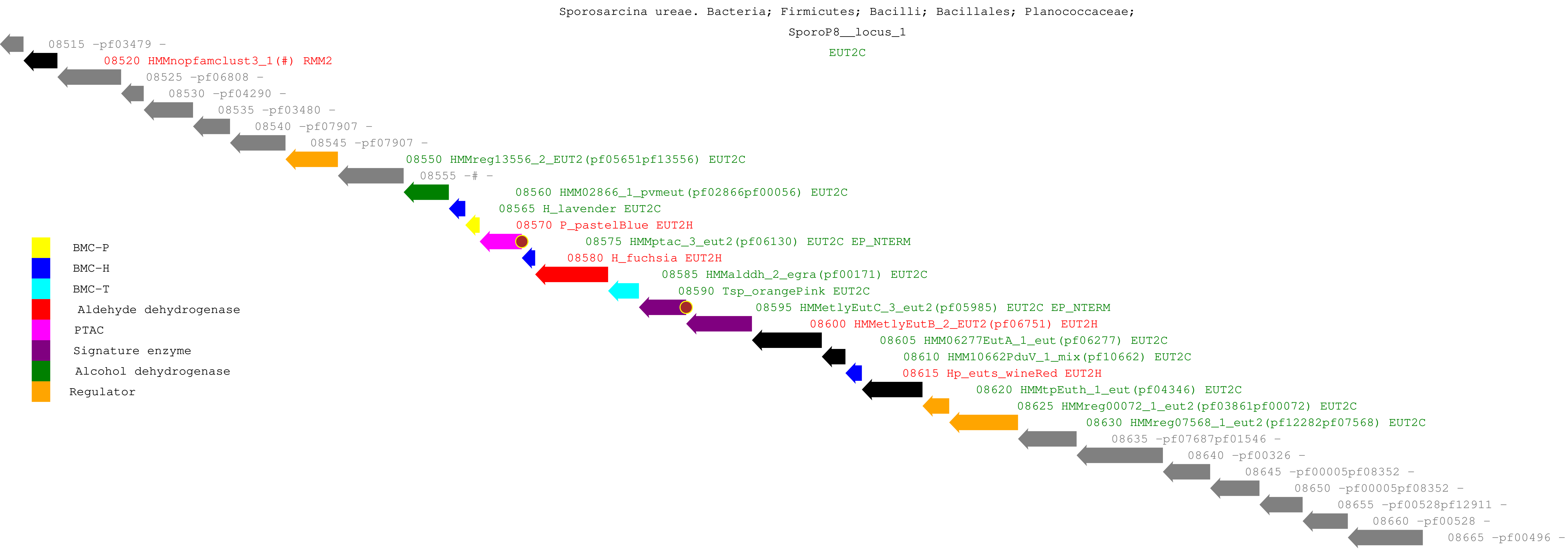
Sporosarcina ureae. Bacteria; Firmicutes; Bacilli; Bacillales; Planococcaceae;
SporoP8__locus_1
EUT2C
08515 -pf03479 -
08520 HMMnopfamclust3_1(#) RMM2
08525 -pf06808 -
08530 -pf04290 -
08535 -pf03480 -
08540 -pf07907 -
08545 -pf07907 -
08550 HMMreg13556_2_EUT2(pf05651pf13556) EUT2C
08555 -# -
08560 HMM02866_1_pvmeut(pf02866pf00056) EUT2C
08565 H_lavender EUT2C
08570 P_pastelBlue EUT2H
08575 HMMptac_3_eut2(pf06130) EUT2C EP_NTERM
08580 H_fuchsia EUT2H
08585 HMMalddh_2_egra(pf00171) EUT2C
08590 Tsp_orangePink EUT2C
08595 HMMetlyEutC_3_eut2(pf05985) EUT2C EP_NTERM
08600 HMMetlyEutB_2_EUT2(pf06751) EUT2H
08605 HMM06277EutA_1_eut(pf06277) EUT2C
08610 HMM10662PduV_1_mix(pf10662) EUT2C
08615 Hp_euts_wineRed EUT2H
08620 HMMtpEuth_1_eut(pf04346) EUT2C
08625 HMMreg00072_1_eut2(pf03861pf00072) EUT2C
08630 HMMreg07568_1_eut2(pf12282pf07568) EUT2C
08635 -pf07687pf01546 -
08640 -pf00326 -
08645 -pf00005pf08352 -
08650 -pf00005pf08352 -
08655 -pf00528pf12911 -
08660 -pf00528 -
08665 -pf00496 -
BMC-P
BMC-H
BMC-T
Aldehyde dehydrogenase
PTAC
Signature enzyme
Alcohol dehydrogenase
Regulator

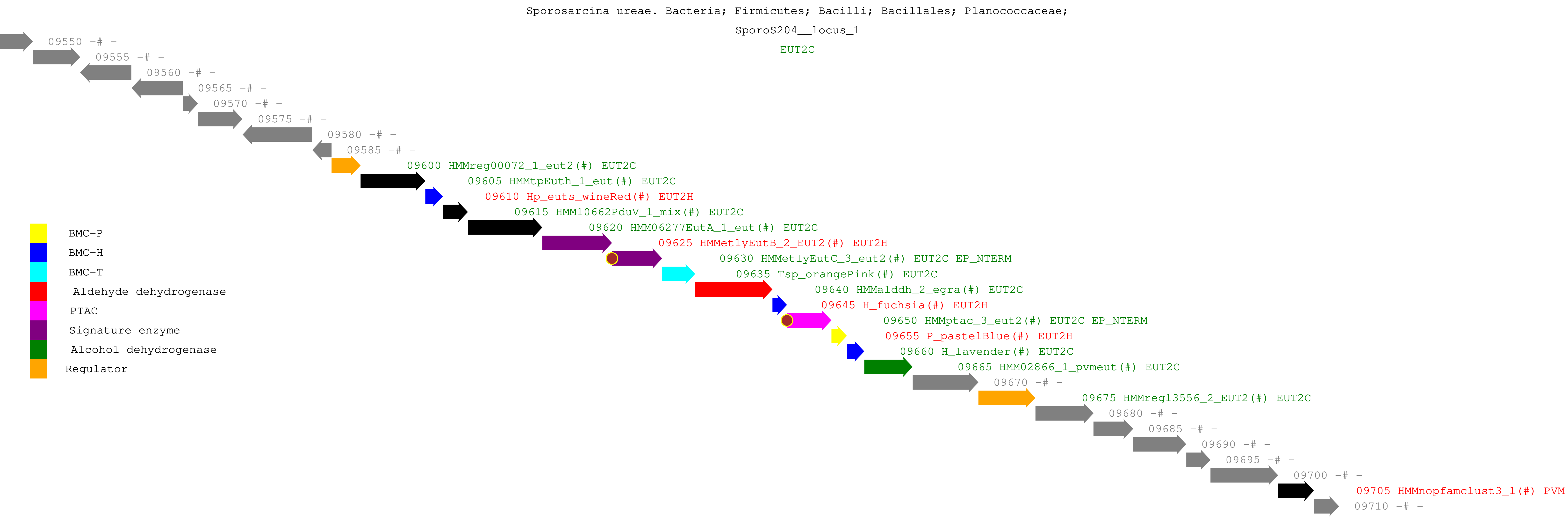

Sporosarcina ureae. Bacteria; Firmicutes; Bacilli; Bacillales; Planococcaceae;
SporoS204__locus_1
EUT2C
09550 -# -
09555 -# -
09560 -# -
09565 -# -
09570 -# -
09575 -# -
09580 -# -
09585 -# -
09600 HMMreg00072_1_eut2(#) EUT2C
09605 HMMtpEuth_1_eut(#) EUT2C
09610 Hp_euts_wineRed(#) EUT2H
09615 HMM10662PduV_1_mix(#) EUT2C
09620 HMM06277EutA_1_eut(#) EUT2C
09625 HMMetlyEutB_2_EUT2(#) EUT2H
09630 HMMetlyEutC_3_eut2(#) EUT2C EP_NTERM
09635 Tsp_orangePink(#) EUT2C
09640 HMMaldh_2_egra(#) EUT2C
09645 H_fuchsia(#) EUT2H
09650 HMMptac_3_eut2(#) EUT2C EP_NTERM
09655 P_pastelBlue(#) EUT2H
09660 H_lavender(#) EUT2C
09665 HMM02866_1_pvmeut(#) EUT2C
09670 -# -
09675 HMMreg13556_2_EUT2(#) EUT2C
09680 -# -
09685 -# -
09690 -# -
09695 -# -
09700 -# -
09705 HMMnopfamclust3_1(#) PVM
09710 -# -
BMC-P
BMC-H
BMC-T
Aldehyde dehydrogenase
PTAC
Signature enzyme
Alcohol dehydrogenase
Regulator

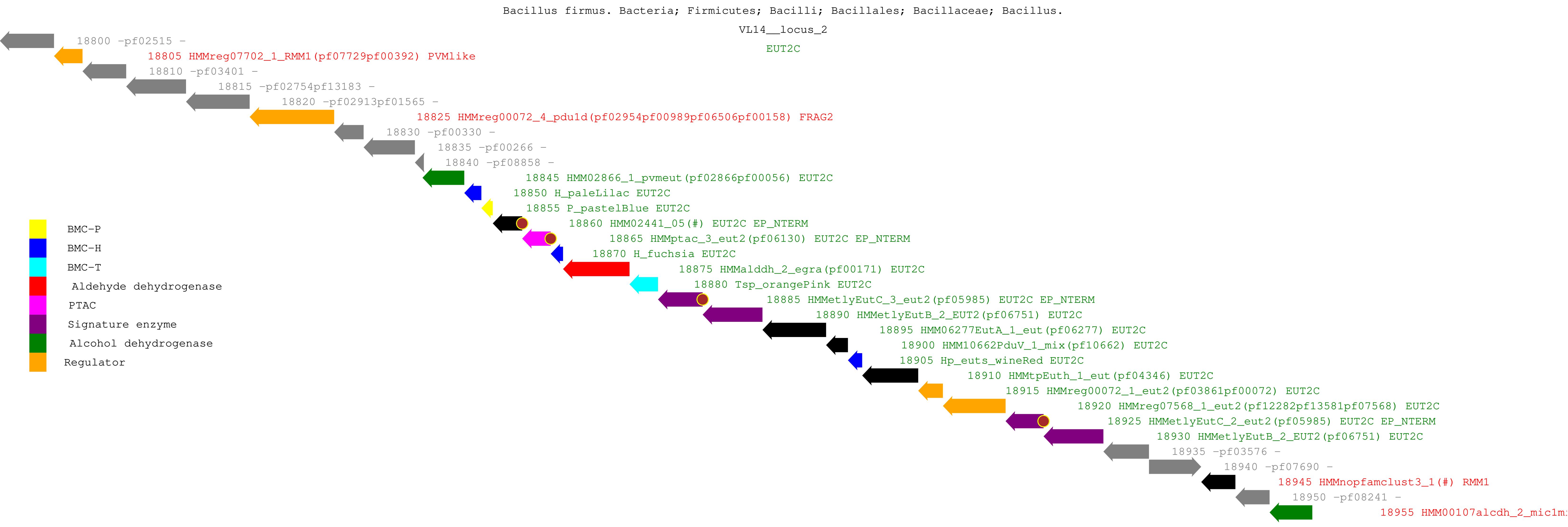
Bacillus firmus. Bacteria; Firmicutes; Bacilli; Bacillales; Bacillaceae; Bacillus.
VL14__locus_2
EUT2C
18800 -pf02515 -
18805 HMMreg07702_1_RMM1(pf07729pf00392) PVMlike
18810 -pf03401 -
18815 -pf02754pf13183 -
18820 -pf02913pf01565 -
18825 HMMreg00072_4_pdu1d(pf02954pf00989pf06506pf00158) FRAG2
18830 -pf00330 -
18835 -pf00266 -
18840 -pf08858 -
18845 HMM02866_1_pvmeut(pf02866pf00056) EUT2C
18850 H_paleLilac EUT2C
18855 P_pastelBlue EUT2C
18860 HMM02441_05(#) EUT2C EP_NTERM
18865 HMMptac_3_eut2(pf06130) EUT2C EP_NTERM
18870 H_fuchsia EUT2C
18875 HMMaldh_2_egra(pf00171) EUT2C
18880 Tsp_orangePink EUT2C
18885 HMMetlyEutC_3_eut2(pf05985) EUT2C EP_NTERM
18890 HMMetlyEutB_2_EUT2(pf06751) EUT2C
18895 HMM06277EutA_1_eut(pf06277) EUT2C
18900 HMM10662PduV_1_mix(pf10662) EUT2C
18905 Hp_euts_wineRed EUT2C
18910 HMMtpEuth_1_eut(pf04346) EUT2C
18915 HMMreg00072_1_eut2(pf03861pf00072) EUT2C
18920 HMMreg07568_1_eut2(pf12282pf13581pf07568) EUT2C
18925 HMMetlyEutC_2_eut2(pf05985) EUT2C EP_NTERM
18930 HMMetlyEutB_2_EUT2(pf06751) EUT2C
18935 -pf03576 -
18940 -pf07690 -
18945 HMMnopfamclust3_1(#) RMM1
18950 -pf08241 -
18955 HMM00107alcdh_2_mic1m
BMC-P
BMC-H
BMC-T
Aldehyde dehydrogenase
PTAC
Signature enzyme
Alcohol dehydrogenase
Regulator

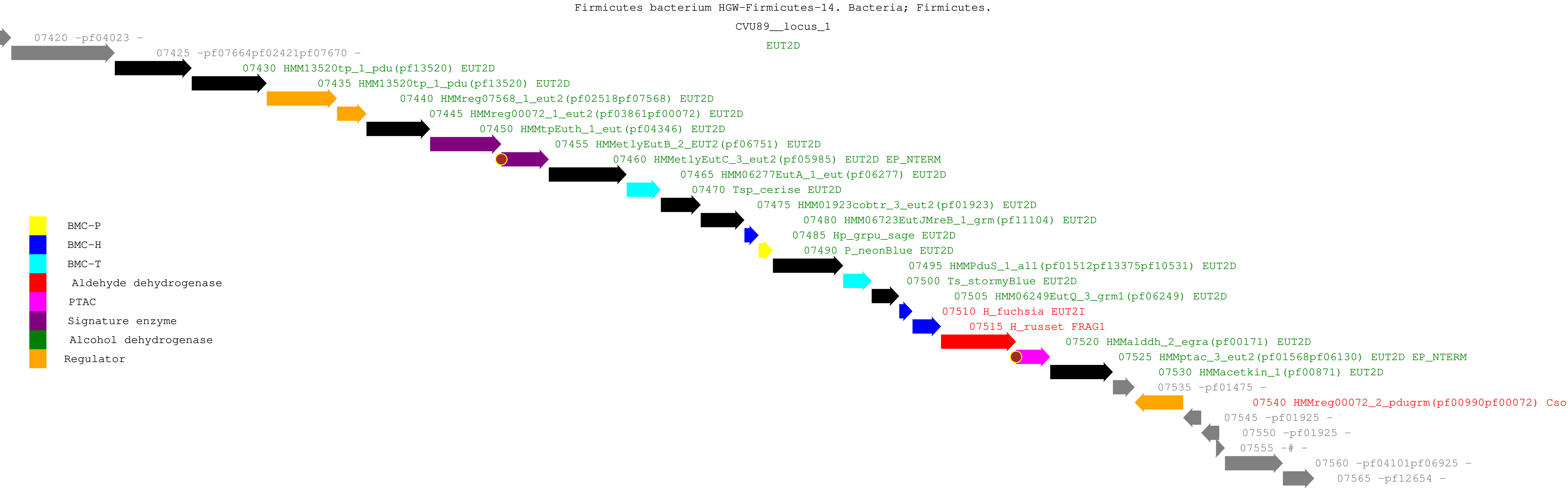
Firmicutes bacterium HGW-Firmicutes-14. Bacteria; Firmicutes.
CVU89__locus_1
EUT2D
07420 -pf04023 -
07425 -pf07664pf02421pf07670 -
07430 HMM13520tp_1_pdu(pf13520) EUT2D
07435 HMM13520tp_1_pdu(pf13520) EUT2D
07440 HMMreg07568_1_eut2(pf02518pf07568) EUT2D
07445 HMMreg00072_1_eut2(pf03861pf00072) EUT2D
07450 HMMtpEuth_1_eut(pf04346) EUT2D
07455 HMMetlyEutB_2_EUT2(pf06751) EUT2D
07460 HMMetlyEutC_3_eut2(pf05985) EUT2D EP_NTERM
07465 HMM06277EutA_1_eut(pf06277) EUT2D
07470 Tsp_cerise EUT2D
07475 HMM01923cobtr_3_eut2(pf01923) EUT2D
07480 HMM06723EutJMreB_1_grm(pf11104) EUT2D
07485 Hp_grpu_sage EUT2D
07490 P_neonBlue EUT2D
07495 HMMPduS_1_all(pf01512pf13375pf10531) EUT2D
07500 Ts_stormyBlue EUT2D
07505 HMM06249EutQ_3_grm1(pf06249) EUT2D
07510 H_fuchsia EUT2I
07515 H_russet FRAG1
07520 HMMalddh_2_egra(pf00171) EUT2D
07525 HMMptac_3_eut2(pf01568pf06130) EUT2D EP_NTERM
07530 HMMacetkin_1(pf00871) EUT2D
07535 -pf01475 -
07540 HMMreg00072_2_pdugrm(pf00990pf00072) Cso
07545 -pf01925 -
07550 -pf01925 -
07555 -# -
07560 -pf04101pf06925 -
07565 -pf12654 -
BMC-P
BMC-H
BMC-T
Aldehyde dehydrogenase
PTAC
Signature enzyme
Alcohol dehydrogenase
Regulator

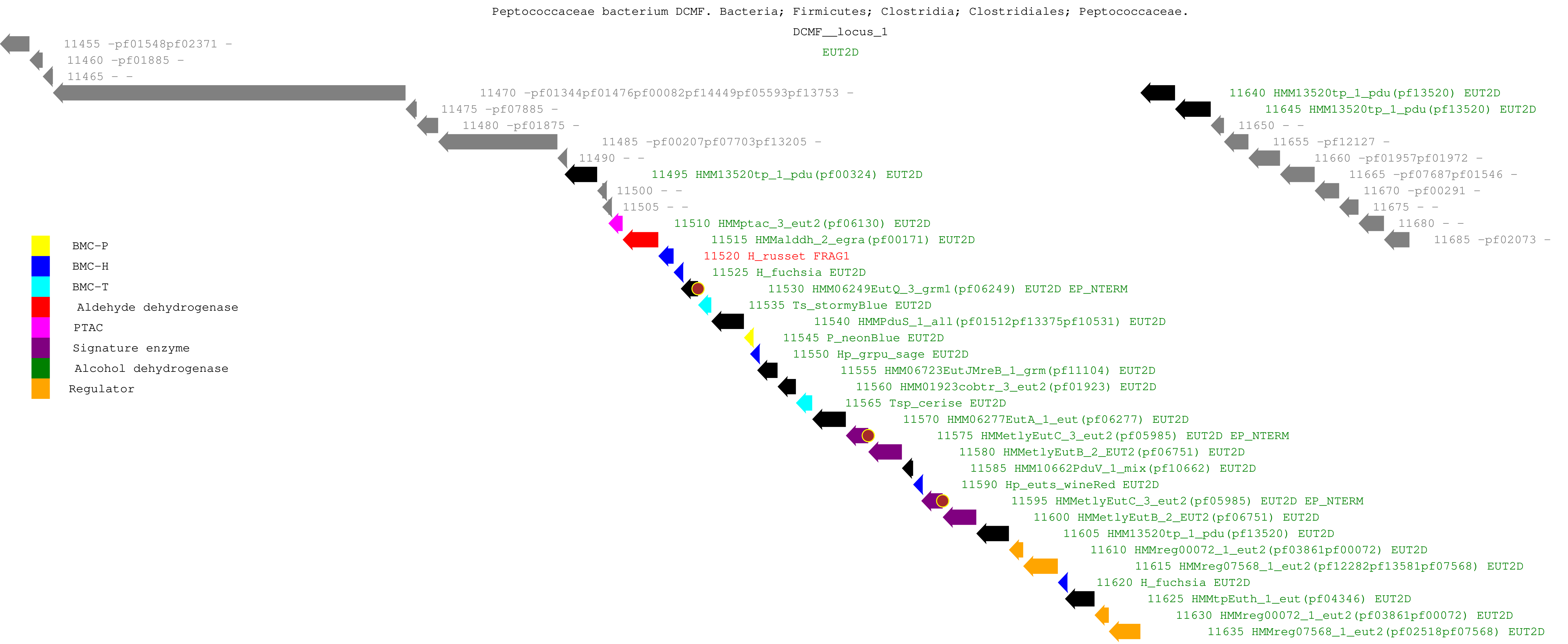
Peptococcaceae bacterium DCMF. Bacteria; Firmicutes; Clostridia; Clostridiales; Peptococcaceae.
DCMF__locus_1
EUT2D
11455 -pf01548pf02371 -
11460 -pf01885 -
11465 - -
11470 -pf01344pf01476pf00082pf14449pf05593pf13753 -
11475 -pf07885 -
11480 -pf01875 -
11485 -pf00207pf07703pf13205 -
11490 - -
11495 HMM13520tp_1_pdu(pf00324) EUT2D
11500 - -
11505 - -
11510 HMMptac_3_eut2(pf06130) EUT2D
11515 HMMalddh_2_egra(pf00171) EUT2D
11520 H_russet FRAG1
11525 H_fuchsia EUT2D
11530 HMM06249EutQ_3_grm1(pf06249) EUT2D EP_NTERM
11535 Ts_stormyBlue EUT2D
11540 HMMPduS_1_all(pf01512pf13375pf10531) EUT2D
11545 P_neonBlue EUT2D
11550 Hp_grpu_sage EUT2D
11555 HMM06723EutJMreB_1_grm(pf11104) EUT2D
11560 HMM01923cobtr_3_eut2(pf01923) EUT2D
11565 Tsp_cerise EUT2D
11570 HMM06277EutA_1_eut(pf06277) EUT2D
11575 HMMetlyEutC_3_eut2(pf05985) EUT2D EP_NTERM
11580 HMMetlyEutB_2_EUT2(pf06751) EUT2D
11585 HMM10662PduV_1_mix(pf10662) EUT2D
11590 Hp_euts_wineRed EUT2D
11595 HMMetlyEutC_3_eut2(pf05985) EUT2D EP_NTERM
11600 HMMetlyEutB_2_EUT2(pf06751) EUT2D
11605 HMM13520tp_1_pdu(pf13520) EUT2D
11610 HMMreg00072_1_eut2(pf03861pf00072) EUT2D
11615 HMMreg07568_1_eut2(pf12282pf13581pf07568) EUT2D
11620 H_fuchsia EUT2D
11625 HMMtpEuth_1_eut(pf04346) EUT2D
11630 HMMreg00072_1_eut2(pf03861pf00072) EUT2D
11635 HMMreg07568_1_eut2(pf02518pf07568) EUT2D
11640 HMM13520tp_1_pdu(pf13520) EUT2D
11645 HMM13520tp_1_pdu(pf13520) EUT2D
11650 - -
11655 -pf12127 -
11660 -pf01957pf01972 -
11665 -pf07687pf01546 -
11670 -pf00291 -
11675 - -
11680 - -
11685 -pf02073 -
BMC-P
BMC-H
BMC-T
Aldehyde dehydrogenase
PTAC
Signature enzyme
Alcohol dehydrogenase
Regulator

Desulfosporosinus sp. Bacteria; Firmicutes; Clostridia; Clostridiales; Peptococcaceae;

DEF42__locus_1

EUT2D

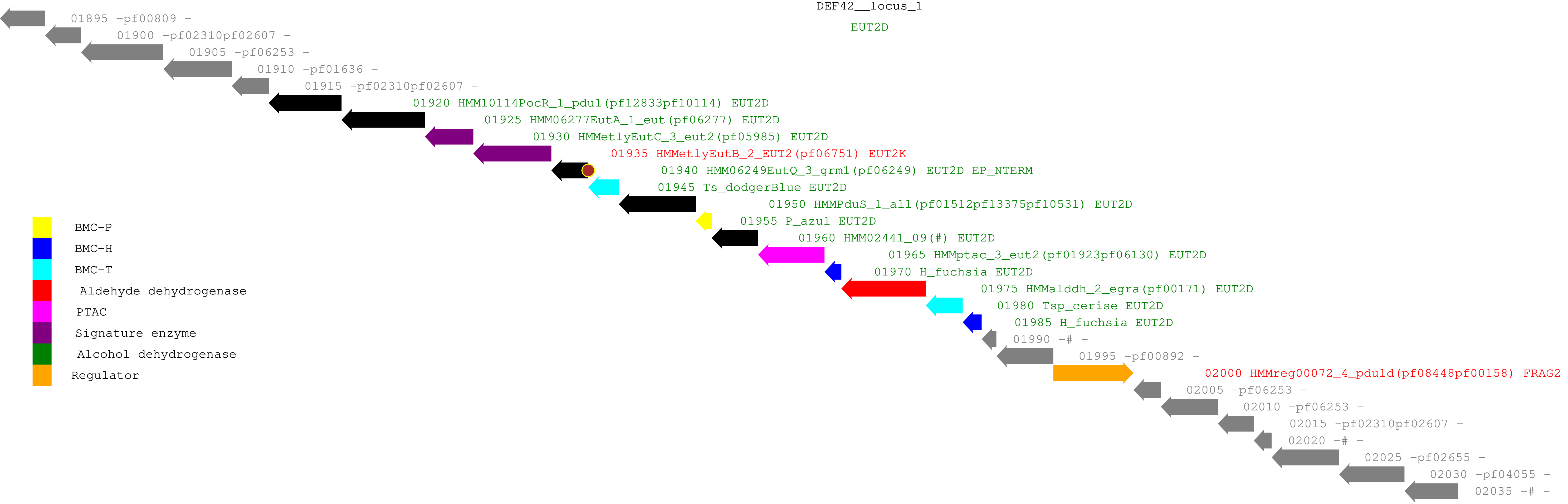

Desulfosporosinus sp. Bacteria; Firmicutes; Clostridia; Clostridiales; Peptococcaceae;

DEF42__locus_4

EUT2D

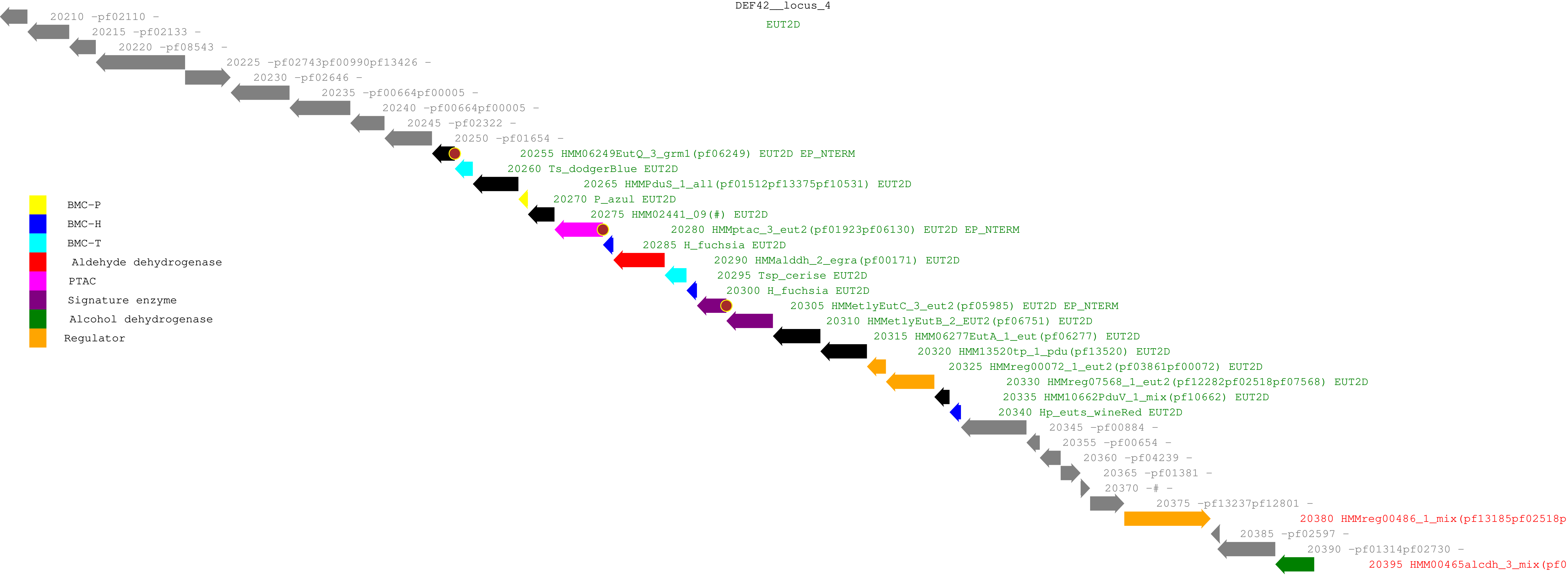

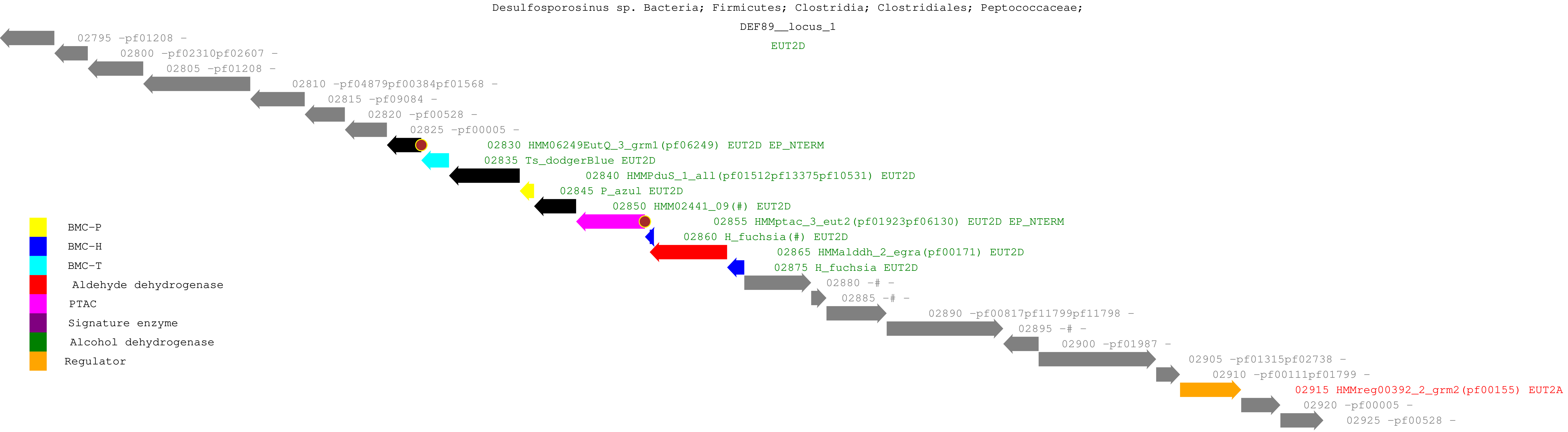
Desulfosporosinus sp. Bacteria; Firmicutes; Clostridia; Clostridiales; Peptococcaceae;
DEF89__locus_1
EUT2D
02795 -pf01208 -
02800 -pf02310pf02607 -
02805 -pf01208 -
02810 -pf04879pf00384pf01568 -
02815 -pf09084 -
02820 -pf00528 -
02825 -pf00005 -
02830 HMM06249EutQ_3_grm1(pf06249) EUT2D EP_NTERM
02835 Ts_dodgerBlue EUT2D
02840 HMMPduS_1_all(pf01512pf13375pf10531) EUT2D
02845 P_azul EUT2D
02850 HMM02441_09(#) EUT2D
02855 HMMptac_3_eut2(pf01923pf06130) EUT2D EP_NTERM
02860 H_fuchsia(#) EUT2D
02865 HMMalddh_2_egra(pf00171) EUT2D
02875 H_fuchsia EUT2D
02880 -# -
02885 -# -
02890 -pf00817pf11799pf11798 -
02895 -# -
02900 -pf01987 -
02905 -pf01315pf02738 -
02910 -pf00111pf01799 -
02915 HMMreg00392_2_grm2(pf00155) EUT2A
02920 -pf00005 -
02925 -pf00528 -
BMC-P
BMC-H
BMC-T
Aldehyde dehydrogenase
PTAC
Signature enzyme
Alcohol dehydrogenase
Regulator

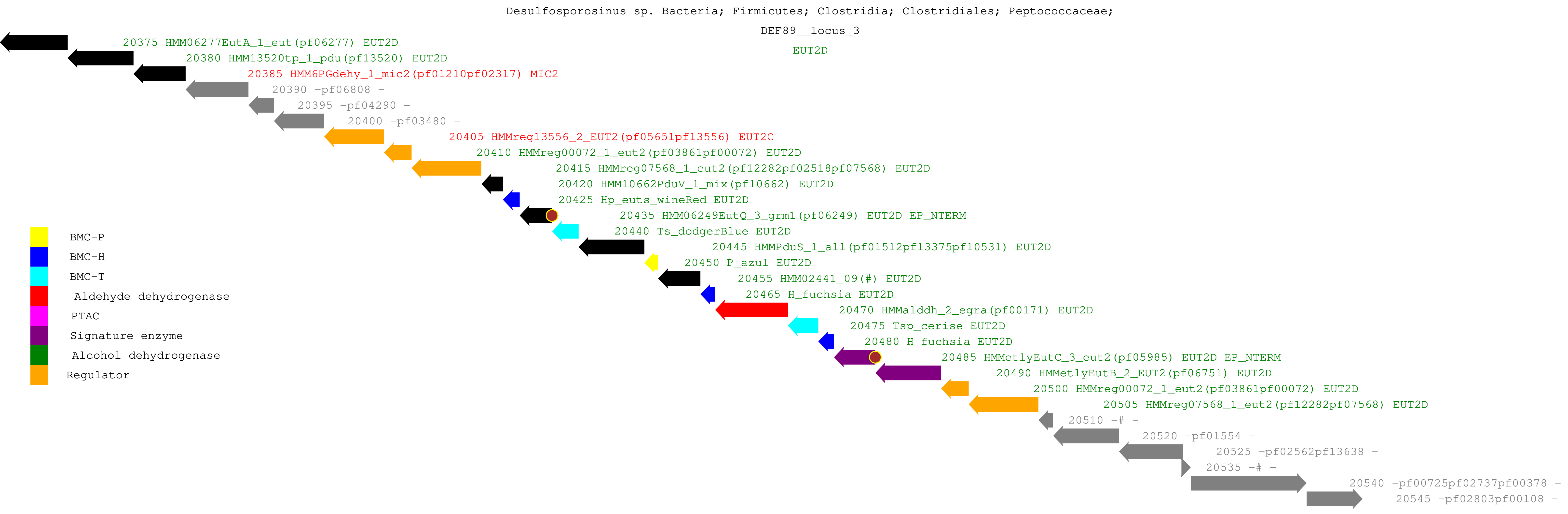

Desulfosporosinus sp. Bacteria; Firmicutes; Clostridia; Clostridiales; Peptococcaceae;
DEF89__locus_3
EUT2D
20375 HMM06277EutA_1_eut(pf06277) EUT2D
20380 HMM13520tp_1_pdu(pf13520) EUT2D
20385 HMM6PGdehy_1_mic2(pf01210pf02317) MIC2
20390 -pf06808 -
20395 -pf04290 -
20400 -pf03480 -
20405 HMMreg13556_2_EUT2(pf05651pf13556) EUT2C
20410 HMMreg00072_1_eut2(pf03861pf00072) EUT2D
20415 HMMreg07568_1_eut2(pf12282pf02518pf07568) EUT2D
20420 HMM10662PduV_1_mix(pf10662) EUT2D
20425 Hp_euts_wineRed EUT2D
20435 HMM06249EutQ_3_grm1(pf06249) EUT2D EP_NTERM
20440 Ts_dodgerBlue EUT2D
20445 HMMPduS_1_all(pf01512pf13375pf10531) EUT2D
20450 P_azul EUT2D
20455 HMM02441_09(#) EUT2D
20465 H_fuchsia EUT2D
20470 HMMaldh_2_egra(pf00171) EUT2D
20475 Tsp_cerise EUT2D
20480 H_fuchsia EUT2D
20485 HMMetlyEutC_3_eut2(pf05985) EUT2D EP_NTERM
20490 HMMetlyEutB_2_EUT2(pf06751) EUT2D
20500 HMMreg00072_1_eut2(pf03861pf00072) EUT2D
20505 HMMreg07568_1_eut2(pf12282pf07568) EUT2D
20510 -# -
20520 -pf01554 -
20525 -pf02562pf13638 -
20535 -# -
20540 -pf00725pf02737pf00378 -
20545 -pf02803pf00108 -
BMC-P
BMC-H
BMC-T
Aldehyde dehydrogenase
PTAC
Signature enzyme
Alcohol dehydrogenase
Regulator

Desulfosporosinus meridiei (strain ATCC BAA-275 / DSM 13257 / NCIMB Bacteria; Firmicutes; Clostridia; Clostridiales; Peptococcaceae;

Desmer__locus_1

EUT2D

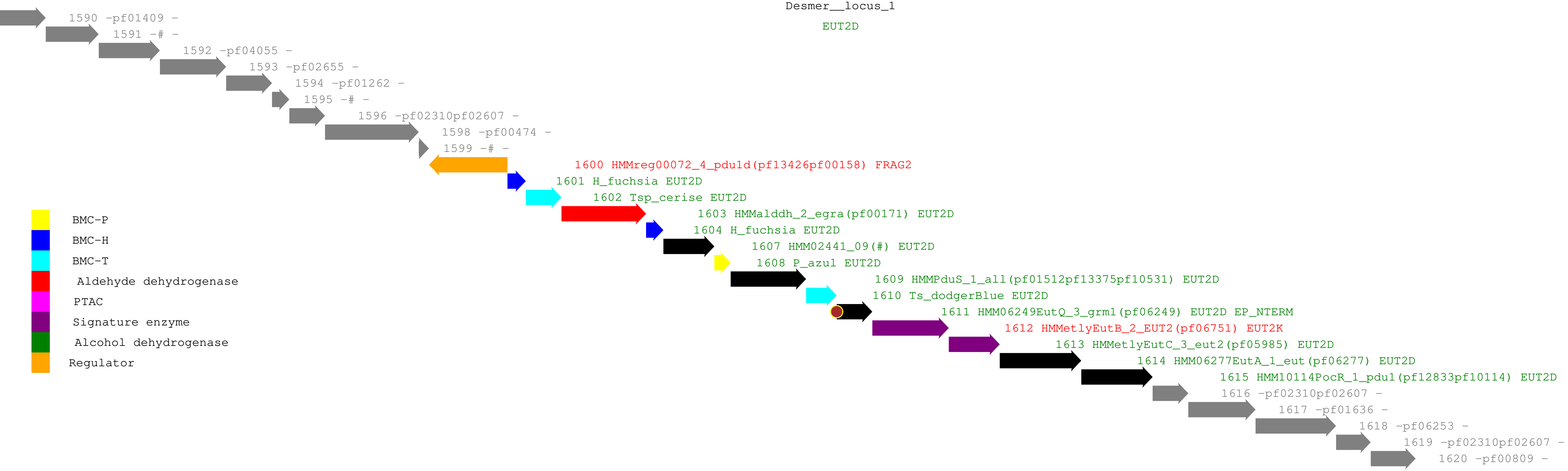

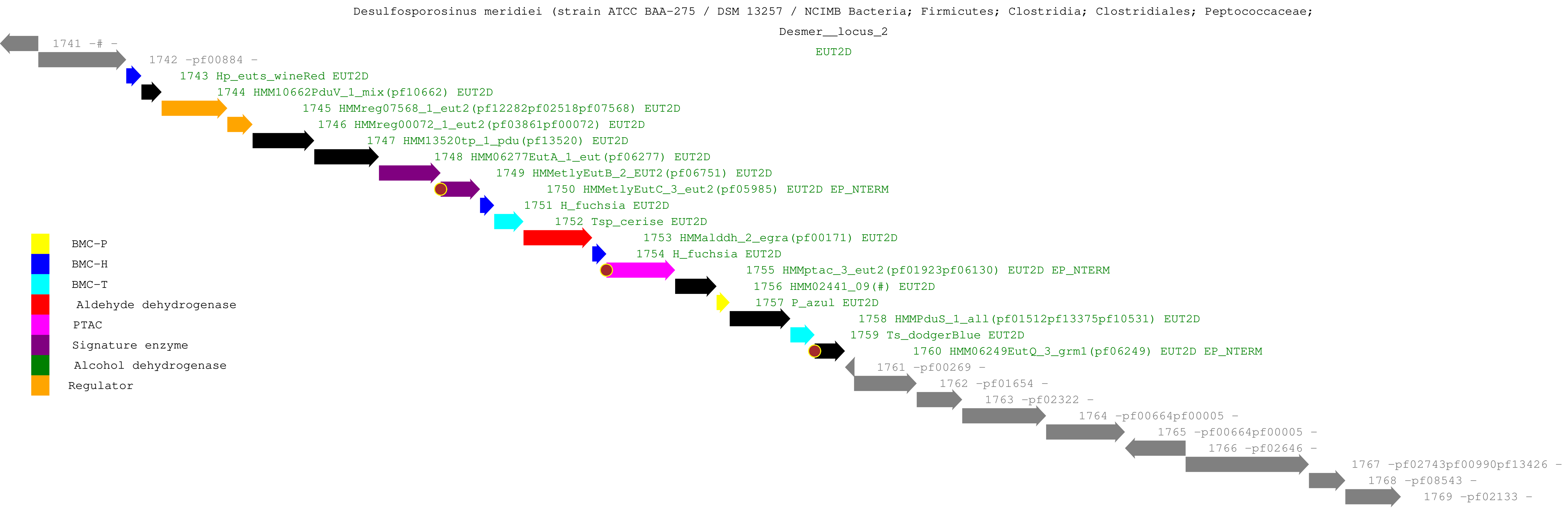
Desulfosporosinus meridiei (strain ATCC BAA-275 / DSM 13257 / NCIMB Bacteria; Firmicutes; Clostridia; Clostridiales; Peptococcaceae;
Desmer__locus_2
EUT2D
1741 -# -
1742 -pf00884 -
1743 Hp_euts_wineRed EUT2D
1744 HMM10662PduV_1_mix(pf10662) EUT2D
1745 HMMreg07568_1_eut2(pf12282pf02518pf07568) EUT2D
1746 HMMreg00072_1_eut2(pf03861pf00072) EUT2D
1747 HMM13520tp_1_pdu(pf13520) EUT2D
1748 HMM06277EutA_1_eut(pf06277) EUT2D
1749 HMMetlyEutB_2_EUT2(pf06751) EUT2D
1750 HMMetlyEutC_3_eut2(pf05985) EUT2D EP_NTERM
1751 H_fuchsia EUT2D
1752 Tsp_cerise EUT2D
1753 HMMaldh_2_egra(pf00171) EUT2D
1754 H_fuchsia EUT2D
1755 HMMptac_3_eut2(pf01923pf06130) EUT2D EP_NTERM
1756 HMM02441_09(#) EUT2D
1757 P_azul EUT2D
1758 HMMPduS_1_all(pf01512pf13375pf10531) EUT2D
1759 Ts_dodgerBlue EUT2D
1760 HMM06249EutQ_3_grm1(pf06249) EUT2D EP_NTERM
1761 -pf00269 -
1762 -pf01654 -
1763 -pf02322 -
1764 -pf00664pf00005 -
1765 -pf00664pf00005 -
1766 -pf02646 -
1767 -pf02743pf00990pf13426 -
1768 -pf08543 -
1769 -pf02133 -
BMC-P
BMC-H
BMC-T
Aldehyde dehydrogenase
PTAC
Signature enzyme
Alcohol dehydrogenase
Regulator

Desulfosporosinus orientis (strain ATCC 19365 / DSM 765 / NCIMB 8382 / Bacteria; Firmicutes; Clostridia; Clostridiales; Peptococcaceae;

Desor__locus_1

EUT2D

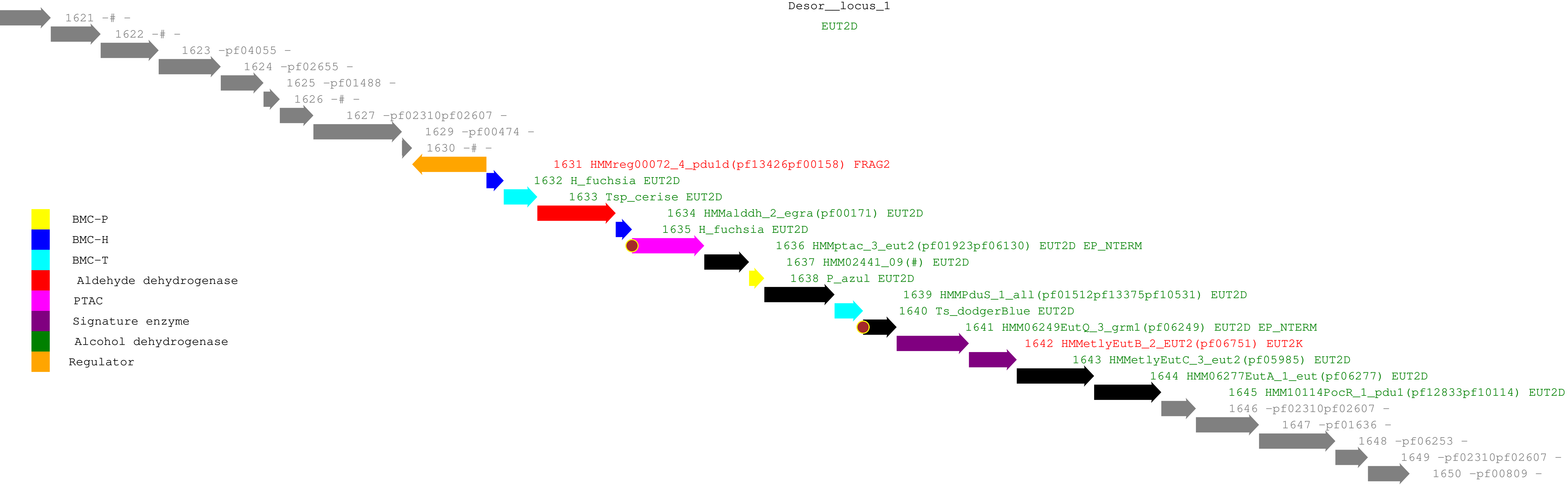

Desulfosporosinus orientis (strain ATCC 19365 / DSM 765 / NCIMB 8382 / Bacteria; Firmicutes; Clostridia; Clostridiales; Peptococcaceae;

Desor__locus_2

EUT2D

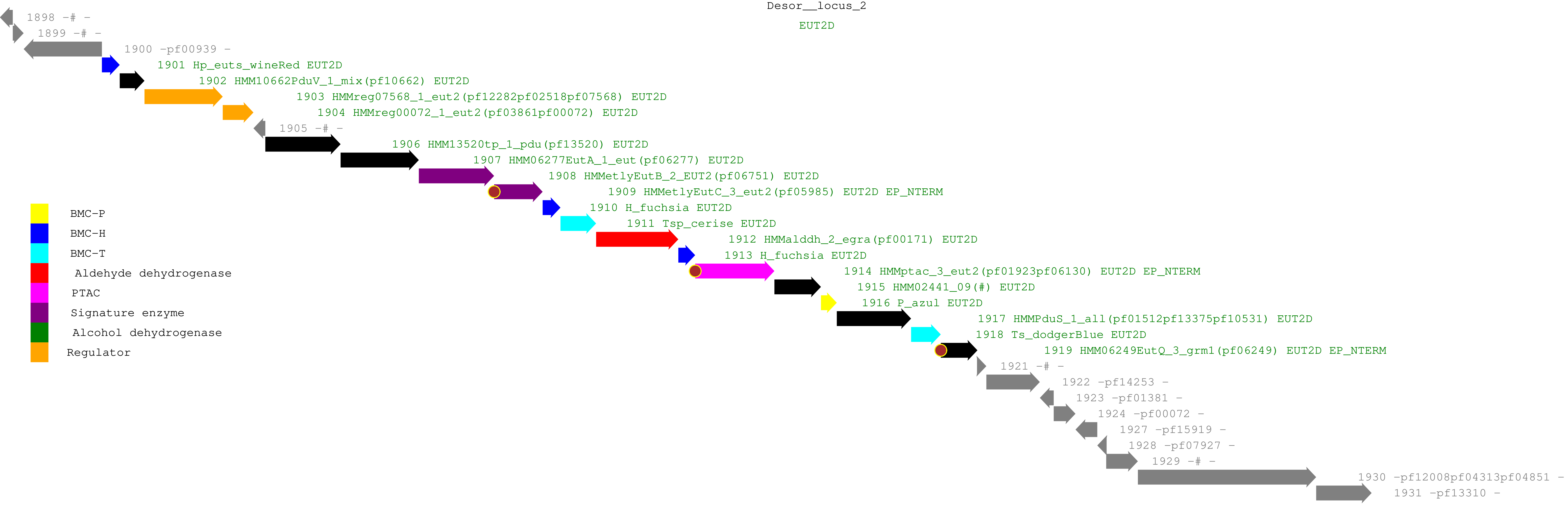

Desulfosporosinus youngiae DSM 17734. Bacteria; Firmicutes; Clostridia; Clostridiales; Peptococcaceae;

DesyoDRAFT__locus_1

EUT2D

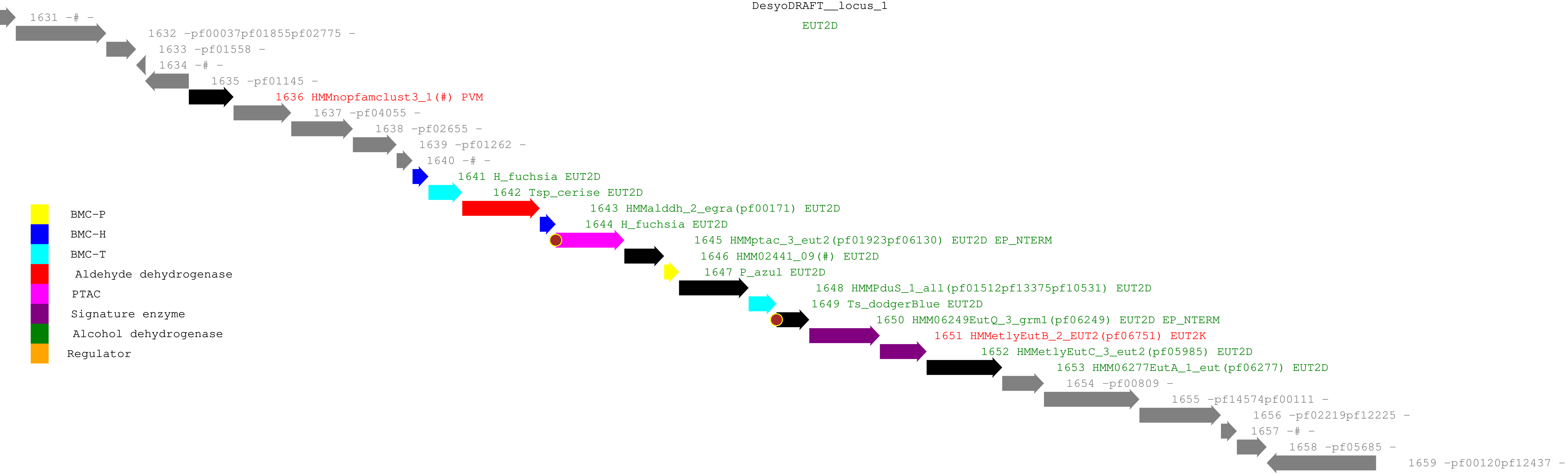

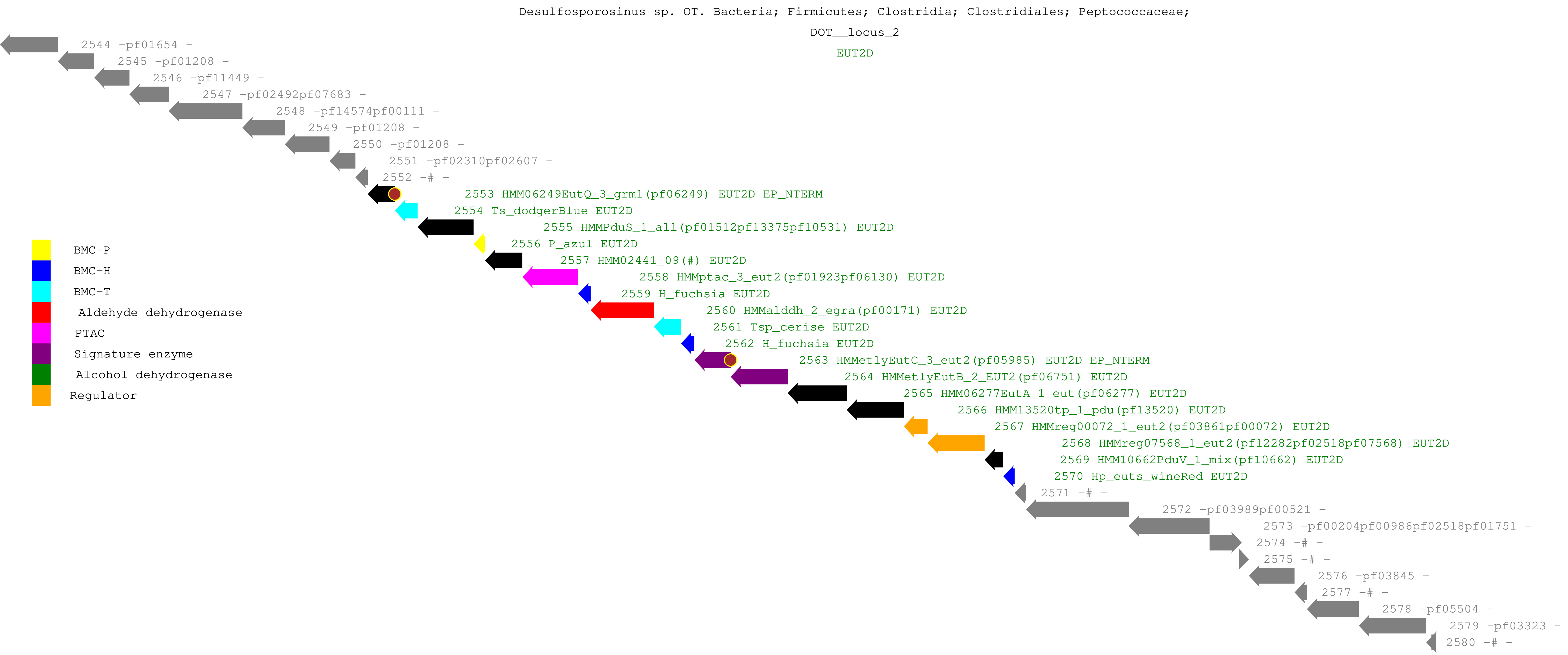
Desulfosporosinus sp. OT. Bacteria; Firmicutes; Clostridia; Clostridiales; Peptococcaceae;
DOT__locus_2
EUT2D
2544 -pf01654 -
2545 -pf01208 -
2546 -pf11449 -
2547 -pf02492pf07683 -
2548 -pf14574pf00111 -
2549 -pf01208 -
2550 -pf01208 -
2551 -pf02310pf02607 -
2552 -# -
2553 HMM06249EutQ_3_grm1(pf06249) EUT2D EP_NTERM
2554 Ts_dodgerBlue EUT2D
2555 HMMPduS_1_all(pf01512pf13375pf10531) EUT2D
2556 P_azul EUT2D
2557 HMM02441_09(#) EUT2D
2558 HMMptac_3_eut2(pf01923pf06130) EUT2D
2559 H_fuchsia EUT2D
2560 HMMalddh_2_egra(pf00171) EUT2D
2561 Tsp_cerise EUT2D
2562 H_fuchsia EUT2D
2563 HMMetlyEutC_3_eut2(pf05985) EUT2D EP_NTERM
2564 HMMetlyEutB_2_EUT2(pf06751) EUT2D
2565 HMM06277EutA_1_eut(pf06277) EUT2D
2566 HMM13520tp_1_pdu(pf13520) EUT2D
2567 HMMreg00072_1_eut2(pf03861pf00072) EUT2D
2568 HMMreg07568_1_eut2(pf12282pf02518pf07568) EUT2D
2569 HMM10662PduV_1_mix(pf10662) EUT2D
2570 Hp_euts_wineRed EUT2D
2571 -# -
2572 -pf03989pf00521 -
2573 -pf00204pf00986pf02518pf01751 -
2574 -# -
2575 -# -
2576 -pf03845 -
2577 -# -
2578 -pf05504 -
2579 -pf03323 -
2580 -# -
BMC-P
BMC-H
BMC-T
Aldehyde dehydrogenase
PTAC
Signature enzyme
Alcohol dehydrogenase
Regulator

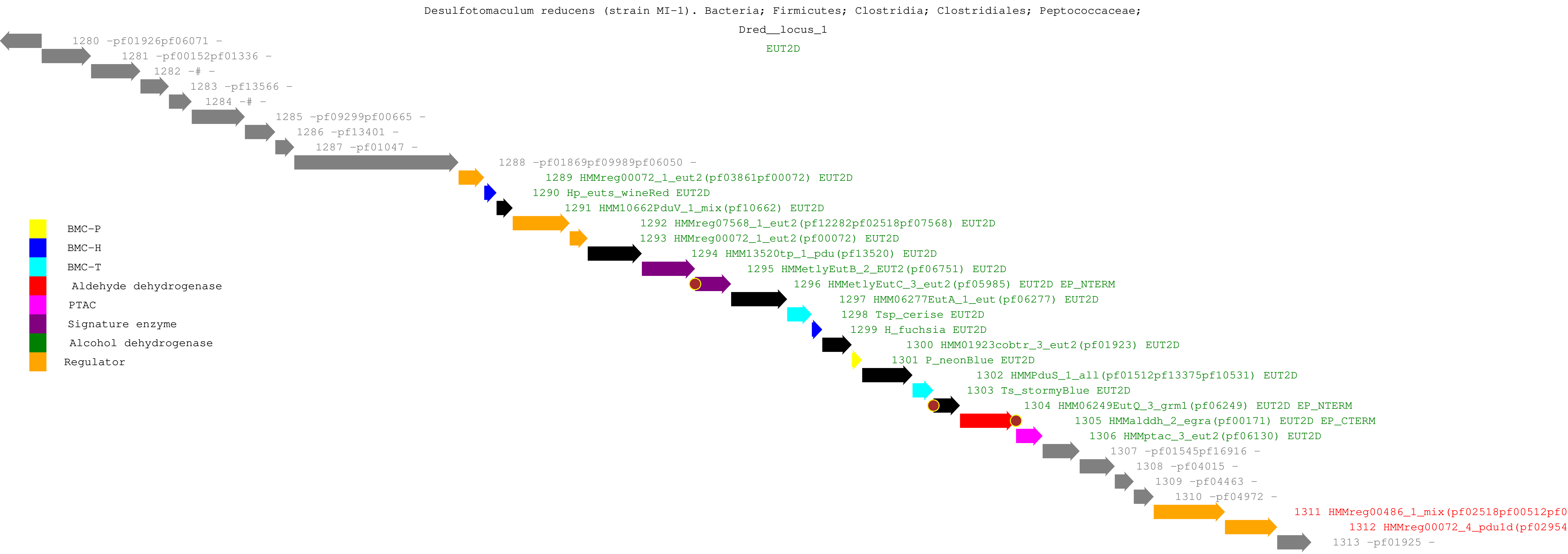

Desulfotomaculum reducens (strain MI-1). Bacteria; Firmicutes; Clostridia; Clostridiales; Peptococcaceae;
Dred__locus_1
EUT2D
1280 -pf01926pf06071 -
1281 -pf00152pf01336 -
1282 -# -
1283 -pf13566 -
1284 -# -
1285 -pf09299pf00665 -
1286 -pf13401 -
1287 -pf01047 -
1288 -pf01869pf09989pf06050 -
1289 HMMreg00072_1_eut2(pf03861pf00072) EUT2D
1290 Hp_euts_wineRed EUT2D
1291 HMM10662PduV_1_mix(pf10662) EUT2D
1292 HMMreg07568_1_eut2(pf12282pf02518pf07568) EUT2D
1293 HMMreg00072_1_eut2(pf00072) EUT2D
1294 HMM13520tp_1_pdu(pf13520) EUT2D
1295 HMMetlyEutB_2_EUT2(pf06751) EUT2D
1296 HMMetlyEutC_3_eut2(pf05985) EUT2D EP_NTERM
1297 HMM06277EutA_1_eut(pf06277) EUT2D
1298 Tsp_cerise EUT2D
1299 H_fuchsia EUT2D
1300 HMM01923cobtr_3_eut2(pf01923) EUT2D
1301 P_neonBlue EUT2D
1302 HMMPduS_1_all(pf01512pf13375pf10531) EUT2D
1303 Ts_stormyBlue EUT2D
1304 HMM06249EutQ_3_grm1(pf06249) EUT2D EP_NTERM
1305 HMMalddh_2_egra(pf00171) EUT2D EP_CTERM
1306 HMMptac_3_eut2(pf06130) EUT2D
1307 -pf01545pf16916 -
1308 -pf04015 -
1309 -pf04463 -
1310 -pf04972 -
1311 HMMreg00486_1_mix(pf02518pf00512pf0
1312 HMMreg00072_4_pdu1d(pf02954
1313 -pf01925 -
BMC-P
BMC-H
BMC-T
Aldehyde dehydrogenase
PTAC
Signature enzyme
Alcohol dehydrogenase
Regulator

Desulfosporosinus sp. OL. Bacteria; Firmicutes; Clostridia; Clostridiales; Peptococcaceae;

DSOL__locus_1_main_sat

EUT2D

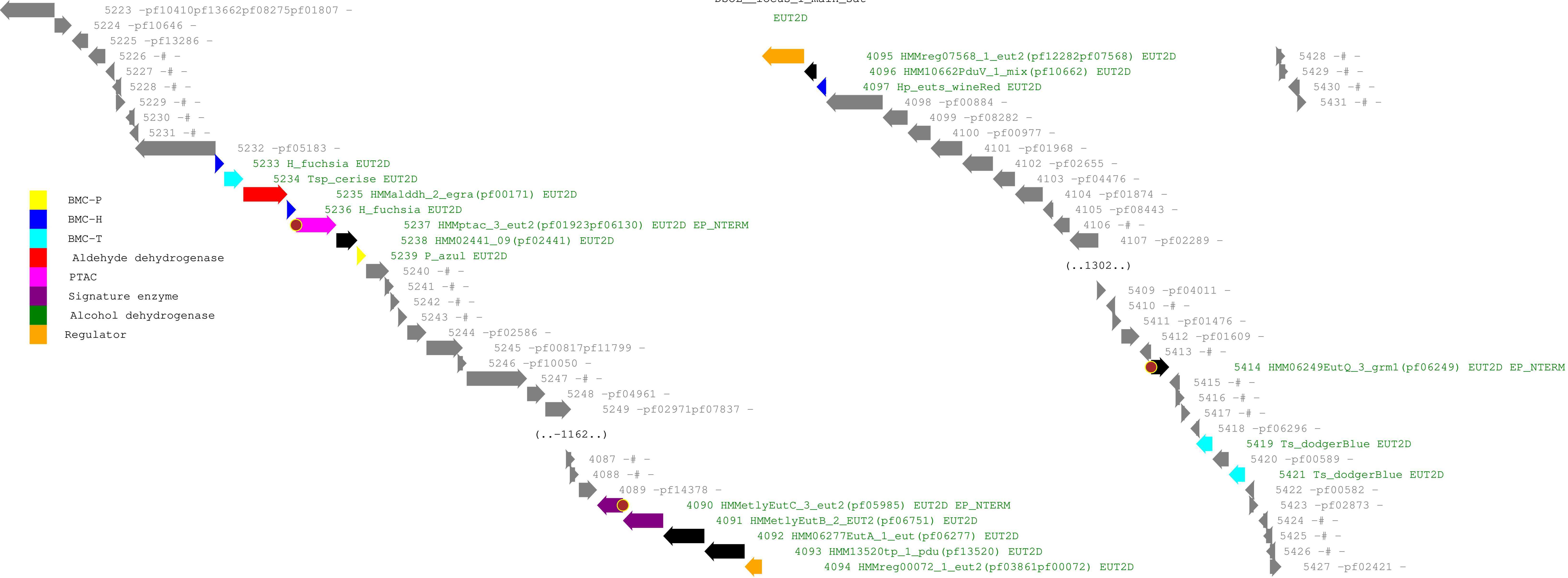

Desulfosporosinus fructosivorans. Bacteria; Firmicutes; Clostridia; Clostridiales; Peptococcaceae;

E4K67__locus_3

EUT2D

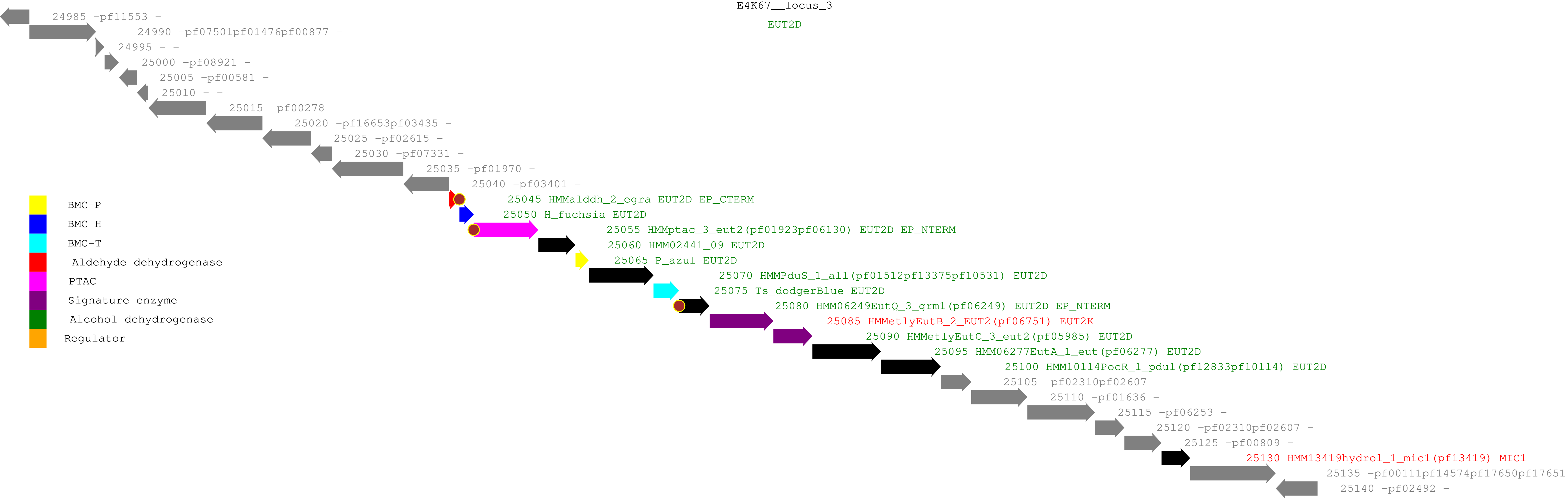

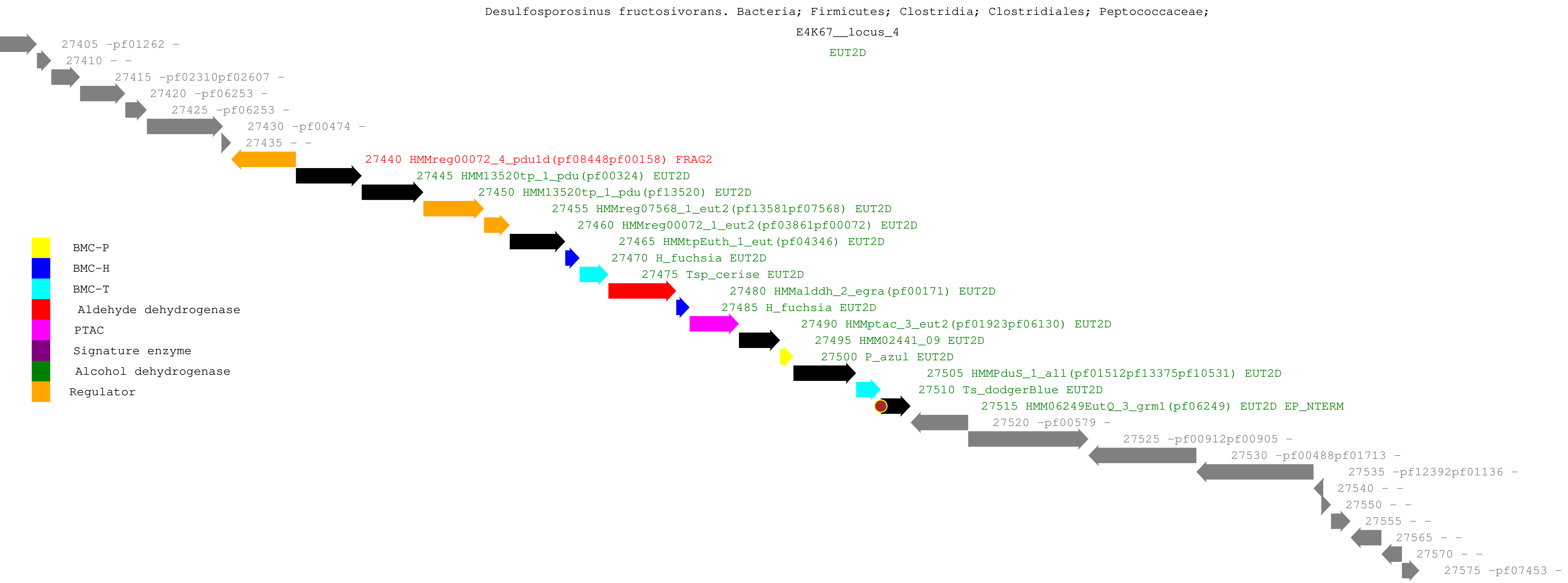

Desulfosporosinus fructosivorans. Bacteria; Firmicutes; Clostridia; Clostridiales; Peptococcaceae;
E4K67__locus_4
EUT2D
27405 -pf01262 -
27410 - -
27415 -pf02310pf02607 -
27420 -pf06253 -
27425 -pf06253 -
27430 -pf00474 -
27435 - -
27440 HMMreg00072_4_pdu1d(pf08448pf00158) FRAG2
27445 HMM13520tp_1_pdu(pf00324) EUT2D
27450 HMM13520tp_1_pdu(pf13520) EUT2D
27455 HMMreg07568_1_eut2(pf13581pf07568) EUT2D
27460 HMMreg00072_1_eut2(pf03861pf00072) EUT2D
27465 HMMtpEuth_1_eut(pf04346) EUT2D
27470 H_fuchsia EUT2D
27475 Tsp_cerise EUT2D
27480 HMMaldh_2_egra(pf00171) EUT2D
27485 H_fuchsia EUT2D
27490 HMMptac_3_eut2(pf01923pf06130) EUT2D
27495 HMM02441_09 EUT2D
27500 P_azul EUT2D
27505 HMMPduS_1_all(pf01512pf13375pf10531) EUT2D
27510 Ts_dodgerBlue EUT2D
27515 HMM06249EutQ_3_grm1(pf06249) EUT2D EP_NTERM
27520 -pf00579 -
27525 -pf00912pf00905 -
27530 -pf00488pf01713 -
27535 -pf12392pf01136 -
27540 - -
27550 - -
27555 - -
27565 - -
27570 - -
27575 -pf07453 -
BMC-P
BMC-H
BMC-T
Aldehyde dehydrogenase
PTAC
Signature enzyme
Alcohol dehydrogenase
Regulator

Clostridia bacterium k32. Bacteria; Firmicutes; Clostridia.

EYS13__locus_1_main_sat

EUT2D

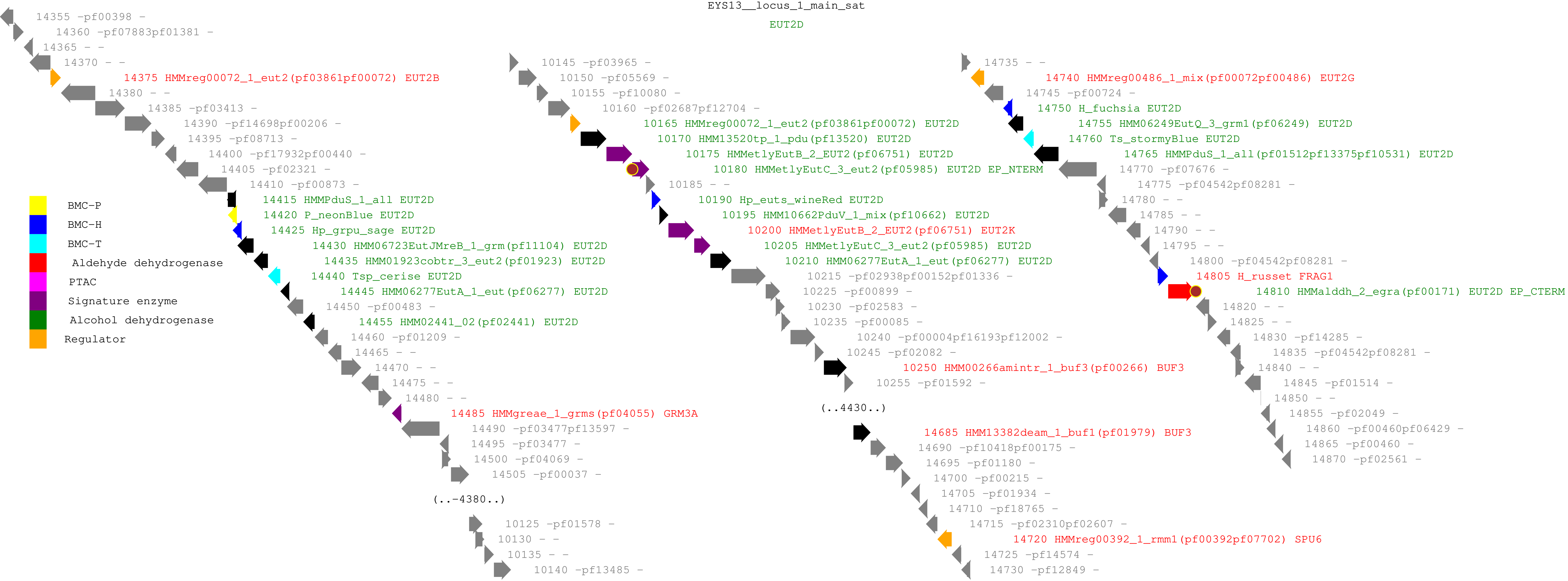

Desulfitobacterium chlororespirans DSM 11544. Bacteria; Firmicutes; Clostridia; Clostridiales; Peptococcaceae;

SAMN02745215__locus_1

EUT2D

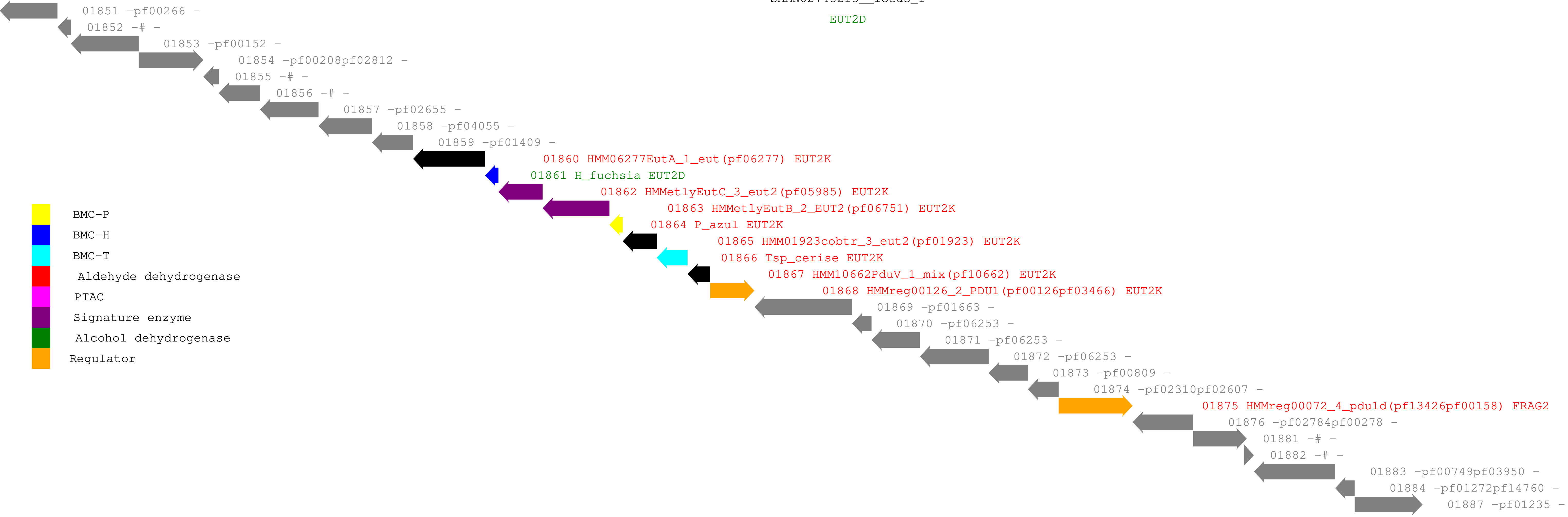

Desulfosporosinus lacus DSM 15449. Bacteria; Firmicutes; Clostridia; Clostridiales; Peptococcaceae;

SAMN02746098__locus_2_main_sat

EUT2D

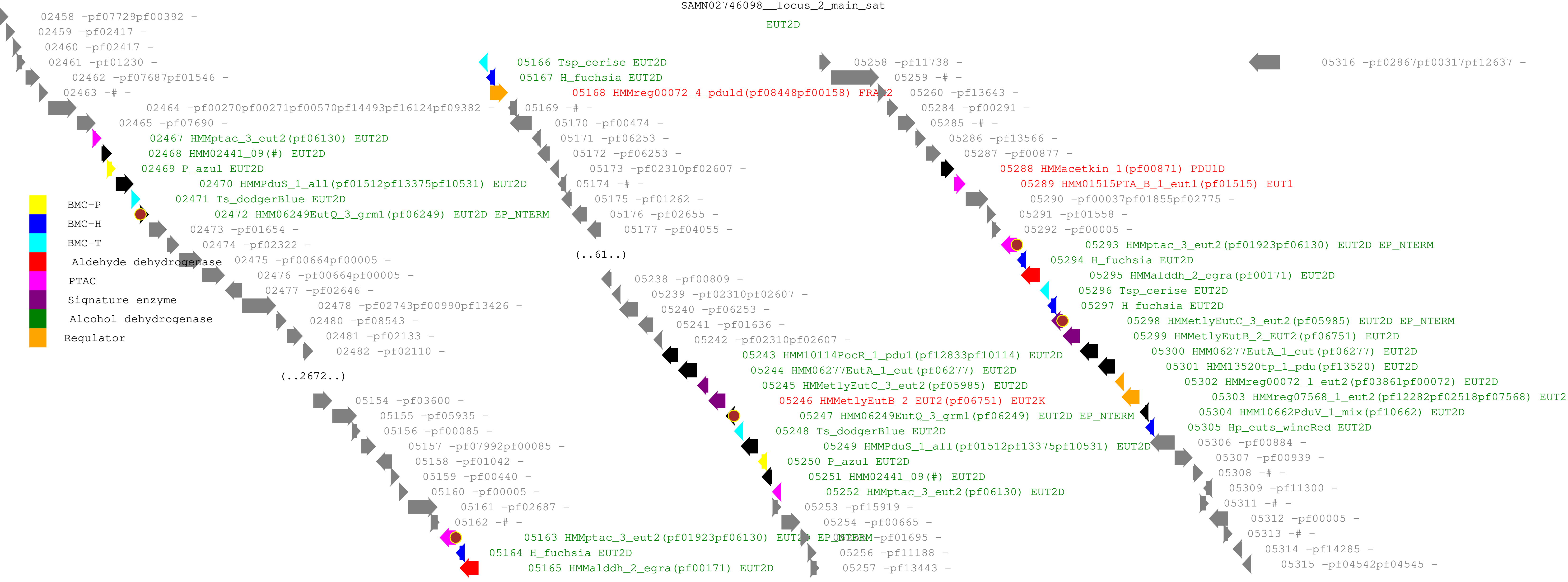

Sporomusa acidovorans. Bacteria; Firmicutes; Negativicutes; Selenomonadales; Sporomusaceae;

SAMN04488499__locus_1

EUT2D

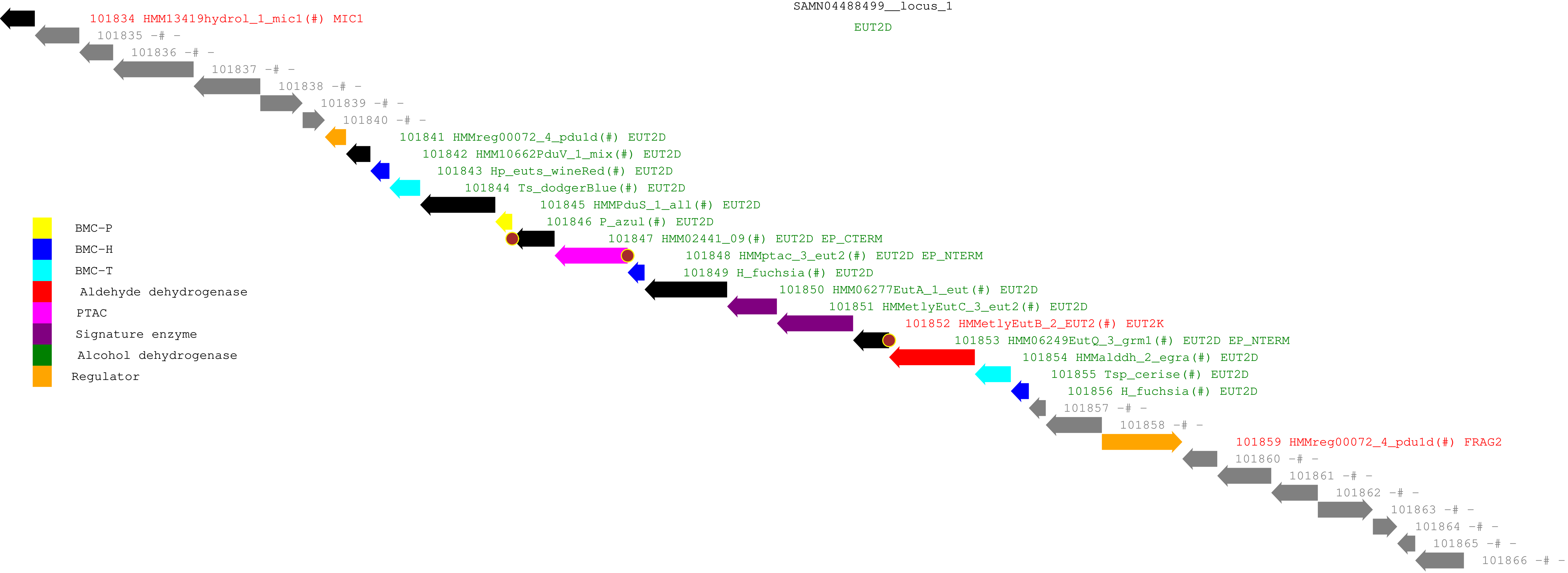

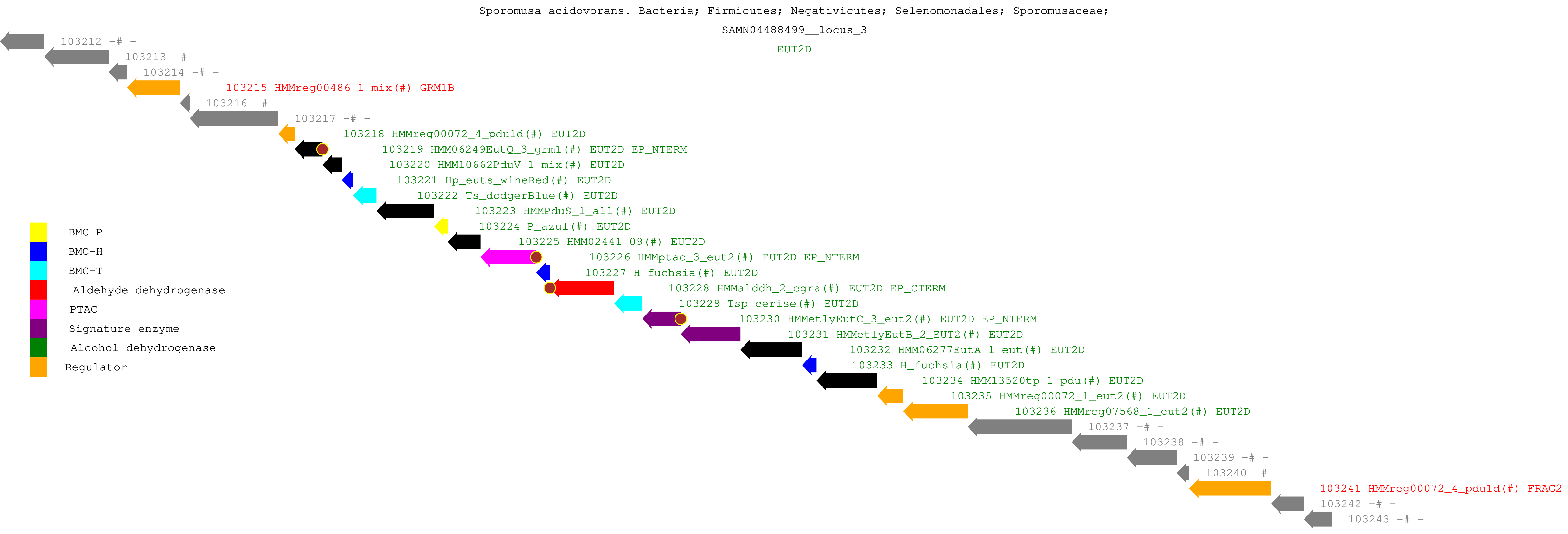

Sporomusa acidovorans. Bacteria; Firmicutes; Negativicutes; Selenomonadales; Sporomusaceae;
SAMN04488499__locus_3
EUT2D
103212 -# -
103213 -# -
103214 -# -
103215 HMMreg00486_1_mix(#) GRM1B
103216 -# -
103217 -# -
103218 HMMreg00072_4_pdu1d(#) EUT2D
103219 HMM06249EutQ_3_grm1(#) EUT2D EP_NTERM
103220 HMM10662PduV_1_mix(#) EUT2D
103221 Hp_euts_wineRed(#) EUT2D
103222 Ts_dodgerBlue(#) EUT2D
103223 HMMPduS_1_all(#) EUT2D
103224 P_azul(#) EUT2D
103225 HMM02441_09(#) EUT2D
103226 HMMptac_3_eut2(#) EUT2D EP_NTERM
103227 H_fuchsia(#) EUT2D
103228 HMMalddh_2_egra(#) EUT2D EP_CTERM
103229 Tsp_cerise(#) EUT2D
103230 HMMetlyEutC_3_eut2(#) EUT2D EP_NTERM
103231 HMMetlyEutB_2_EUT2(#) EUT2D
103232 HMM06277EutA_1_eut(#) EUT2D
103233 H_fuchsia(#) EUT2D
103234 HMM13520tp_1_pdu(#) EUT2D
103235 HMMreg00072_1_eut2(#) EUT2D
103236 HMMreg07568_1_eut2(#) EUT2D
103237 -# -
103238 -# -
103239 -# -
103240 -# -
103241 HMMreg00072_4_pdu1d(#) FRAG2
103242 -# -
103243 -# -
BMC-P
BMC-H
BMC-T
Aldehyde dehydrogenase
PTAC
Signature enzyme
Alcohol dehydrogenase
Regulator

Desulfosporosinus sp. SbF1. Bacteria; Firmicutes; Clostridia; Clostridiales; Peptococcaceae;

SBF1__locus_1

EUT2D

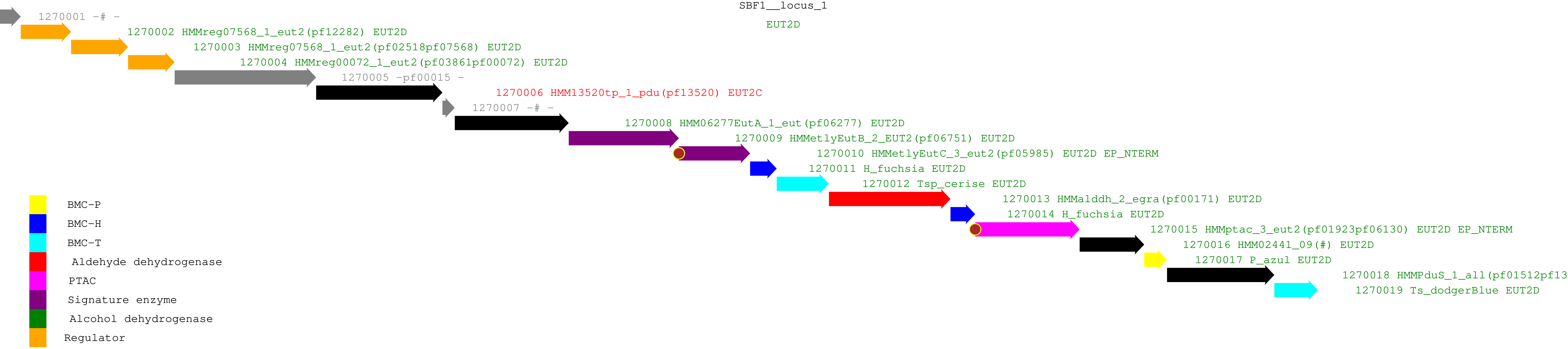

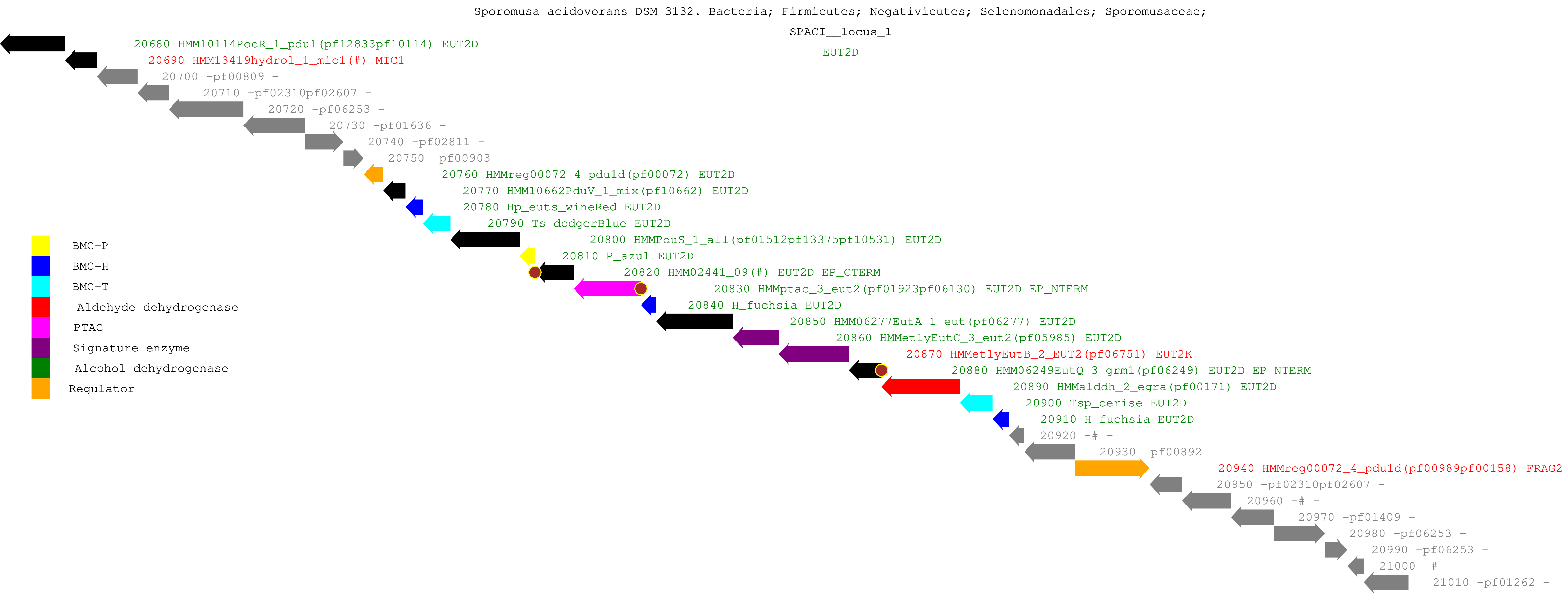

Sporomusa acidovorans DSM 3132. Bacteria; Firmicutes; Negativicutes; Selenomonadales; Sporomusaceae;
SPACI__locus_1
EUT2D
20680 HMM10114PocR_1_pdu1(pf12833pf10114) EUT2D
20690 HMM13419hydrol_1_mic1(#) MIC1
20700 -pf00809 -
20710 -pf02310pf02607 -
20720 -pf06253 -
20730 -pf01636 -
20740 -pf02811 -
20750 -pf00903 -
20760 HMMreg00072_4_pdu1d(pf00072) EUT2D
20770 HMM10662PduV_1_mix(pf10662) EUT2D
20780 Hp_euts_wineRed EUT2D
20790 Ts_dodgerBlue EUT2D
20800 HMMPduS_1_all(pf01512pf13375pf10531) EUT2D
20810 P_azul EUT2D
20820 HMM02441_09(#) EUT2D EP_CTERM
20830 HMMptac_3_eut2(pf01923pf06130) EUT2D EP_NTERM
20840 H_fuchsia EUT2D
20850 HMM06277EutA_1_eut(pf06277) EUT2D
20860 HMMetlyEutC_3_eut2(pf05985) EUT2D
20870 HMMetlyEutB_2_EUT2(pf06751) EUT2K
20880 HMM06249EutQ_3_grm1(pf06249) EUT2D EP_NTERM
20890 HMMalddh_2_egra(pf00171) EUT2D
20900 Tsp_cerise EUT2D
20910 H_fuchsia EUT2D
20920 -# -
20930 -pf00892 -
20940 HMMreg00072_4_pdu1d(pf00989pf00158) FRAG2
20950 -pf02310pf02607 -
20960 -# -
20970 -pf01409 -
20980 -pf06253 -
20990 -pf06253 -
21000 -# -
21010 -pf01262 -
BMC-P
BMC-H
BMC-T
Aldehyde dehydrogenase
PTAC
Signature enzyme
Alcohol dehydrogenase
Regulator

Sporomusa acidovorans DSM 3132. Bacteria; Firmicutes; Negativicutes; Selenomonadales; Sporomusaceae;

SPACI__locus_2

EUT2D

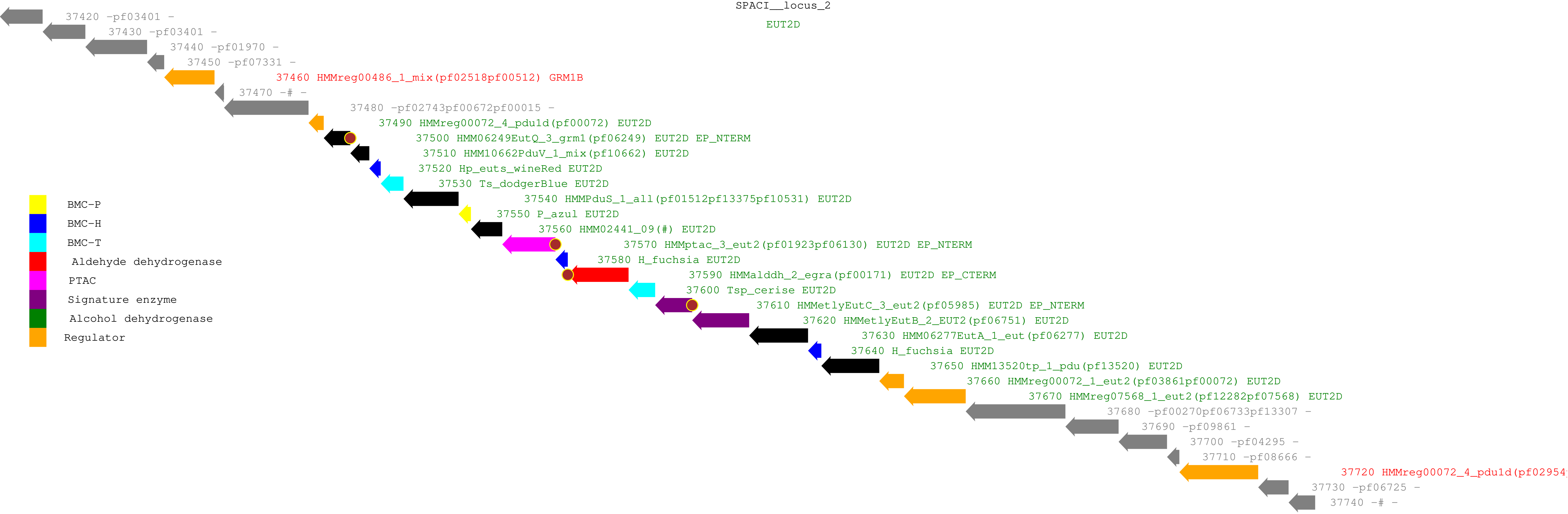

Thermincola ferriacetica. Bacteria; Firmicutes; Clostridia; Clostridiales; Peptococcaceae;

Tfer__locus_2

EUT2D

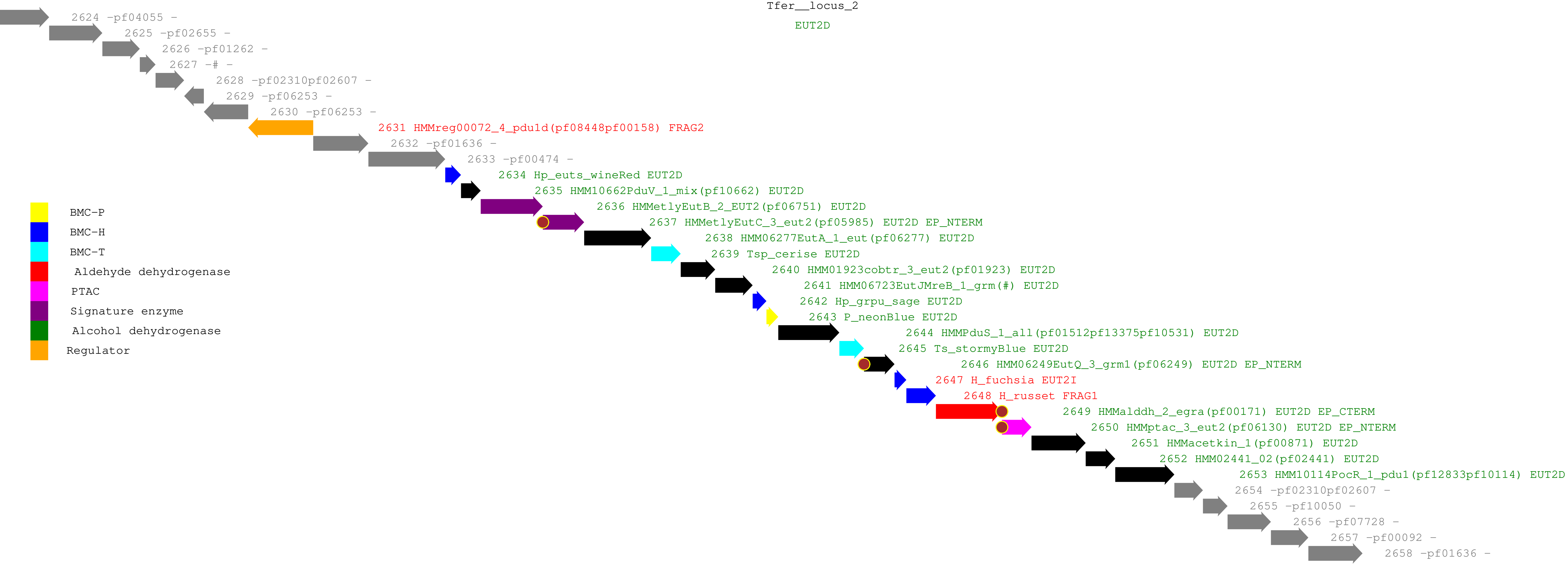

Thermincola potens (strain JR). Bacteria; Firmicutes; Clostridia; Clostridiales; Peptococcaceae;

TherJR__locus_1

EUT2D

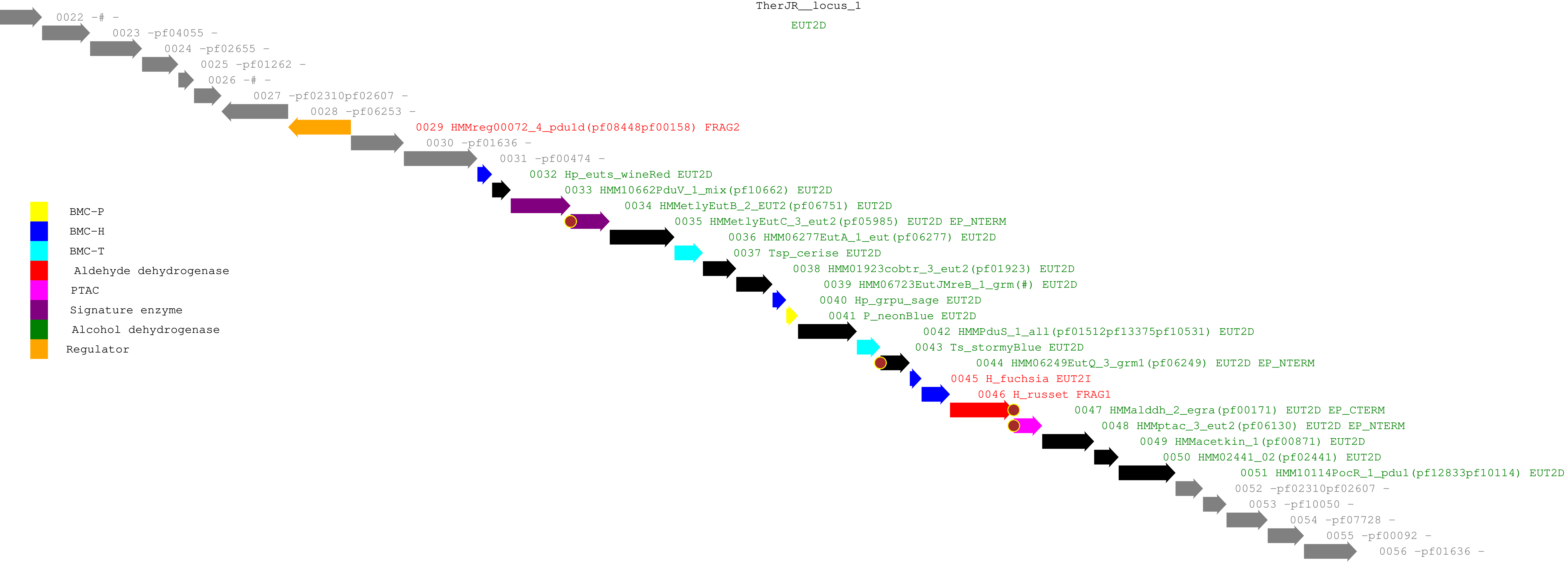

Peptococcaceae bacterium BRH_c23. Bacteria; Firmicutes; Clostridia; Clostridiales; Peptococcaceae.

VR66__locus_1

EUT2D

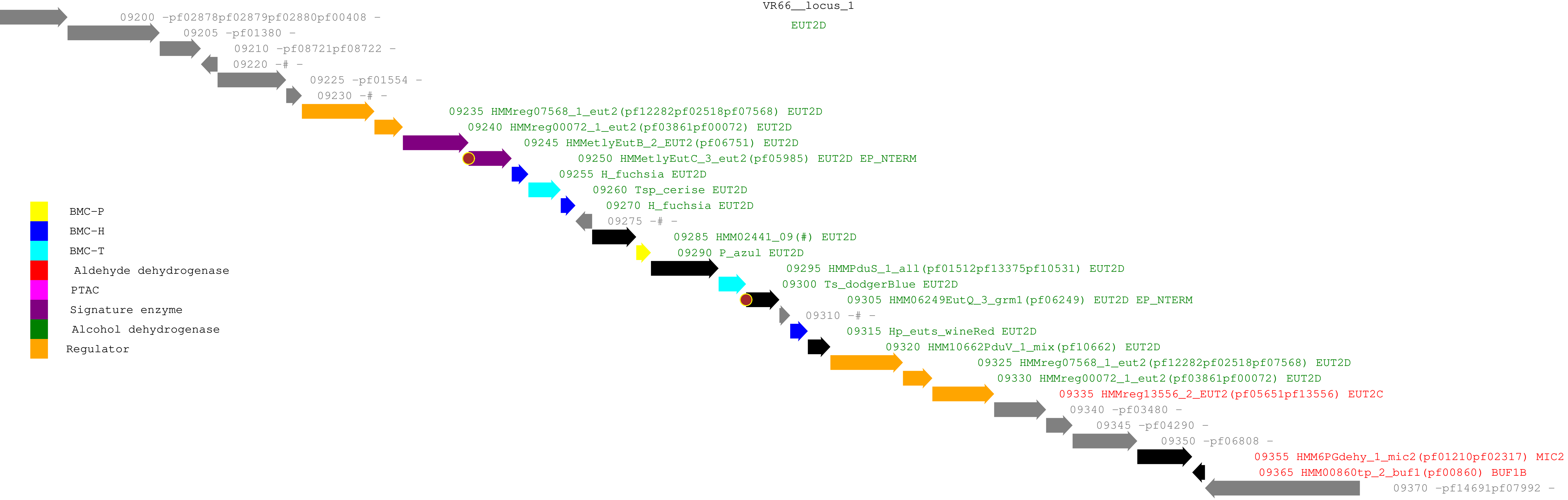

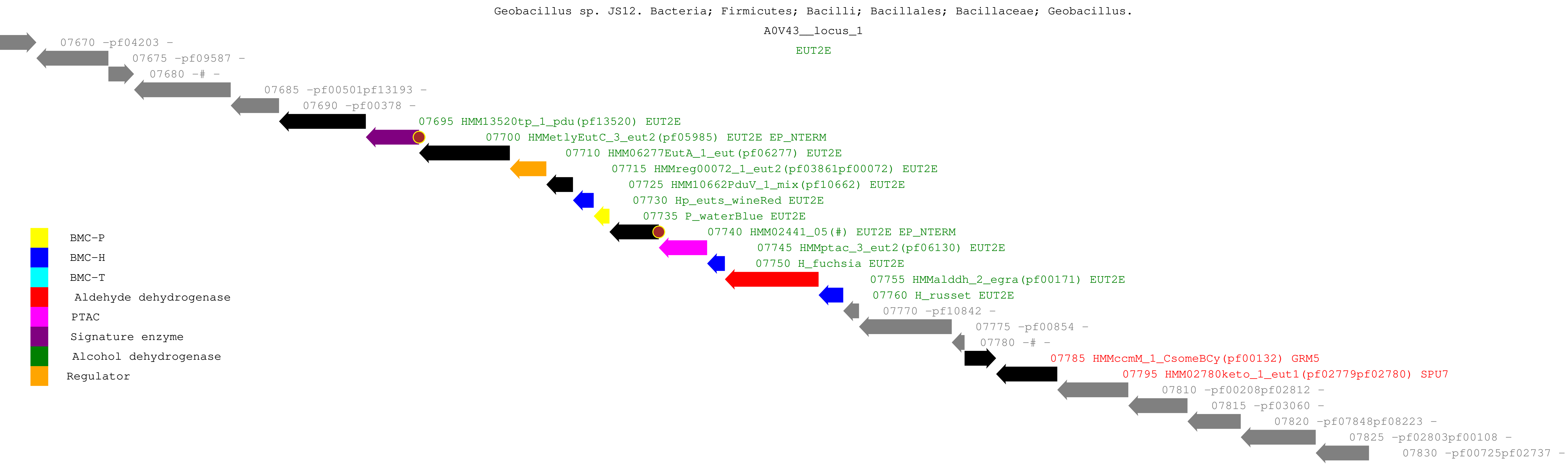
Geobacillus sp. JS12. Bacteria; Firmicutes; Bacilli; Bacillales; Bacillaceae; Geobacillus.
A0V43__locus_1
EUT2E
07670 -pf04203 -
07675 -pf09587 -
07680 -# -
07685 -pf00501pf13193 -
07690 -pf00378 -
07695 HMM13520tp_1_pdu(pf13520) EUT2E
07700 HMMetlyEutC_3_eut2(pf05985) EUT2E EP_NTERM
07710 HMM06277EutA_1_eut(pf06277) EUT2E
07715 HMMreg00072_1_eut2(pf03861pf00072) EUT2E
07725 HMM10662PduV_1_mix(pf10662) EUT2E
07730 Hp_euts_wineRed EUT2E
07735 P_waterBlue EUT2E
07740 HMM02441_05(#) EUT2E EP_NTERM
07745 HMMptac_3_eut2(pf06130) EUT2E
07750 H_fuchsia EUT2E
07755 HMMalddh_2_egra(pf00171) EUT2E
07760 H_russet EUT2E
07770 -pf10842 -
07775 -pf00854 -
07780 -# -
07785 HMMccmM_1_CsomeBCy(pf00132) GRM5
07795 HMM02780keto_1_eut1(pf02779pf02780) SPU7
07810 -pf00208pf02812 -
07815 -pf03060 -
07820 -pf07848pf08223 -
07825 -pf02803pf00108 -
07830 -pf00725pf02737 -
BMC-P
BMC-H
BMC-T
Aldehyde dehydrogenase
PTAC
Signature enzyme
Alcohol dehydrogenase
Regulator

Geobacillus thermoleovorans (Bacillus thermoleovorans). Bacteria; Firmicutes; Bacilli; Bacillales; Bacillaceae; Geobacillus;

A5N86__locus_1_main_sat

EUT2E

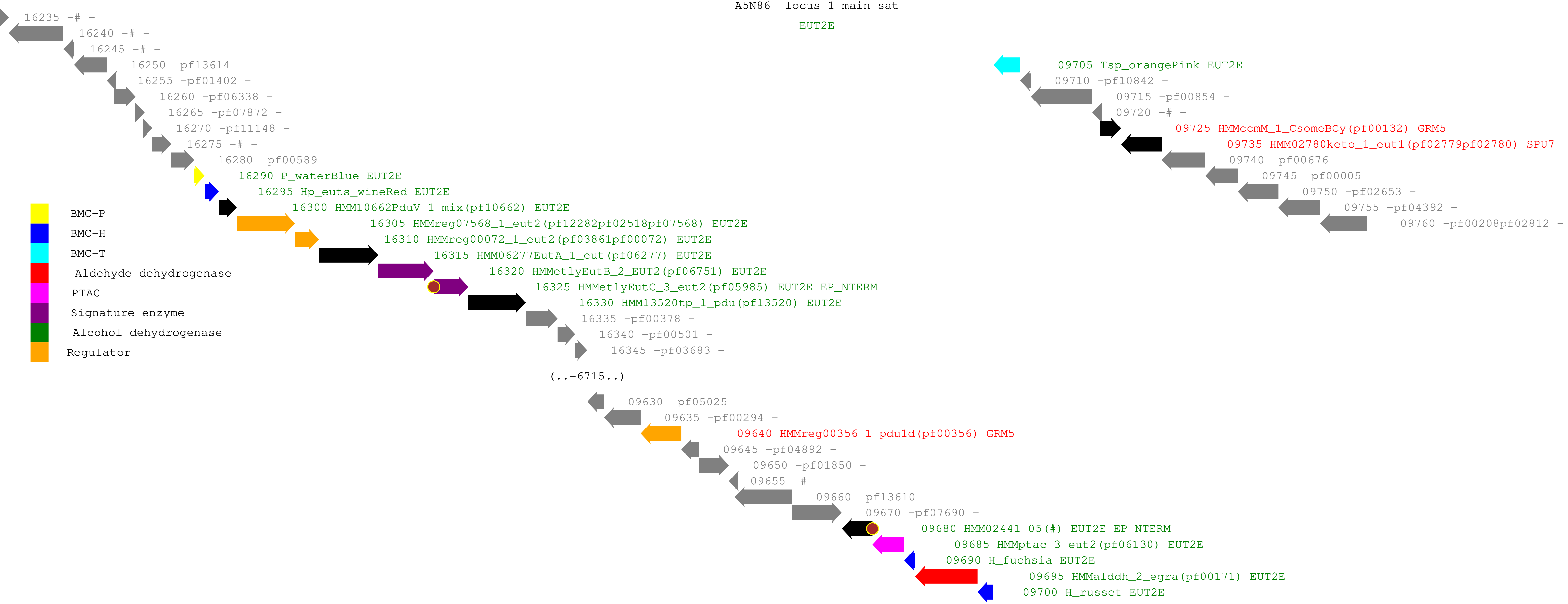

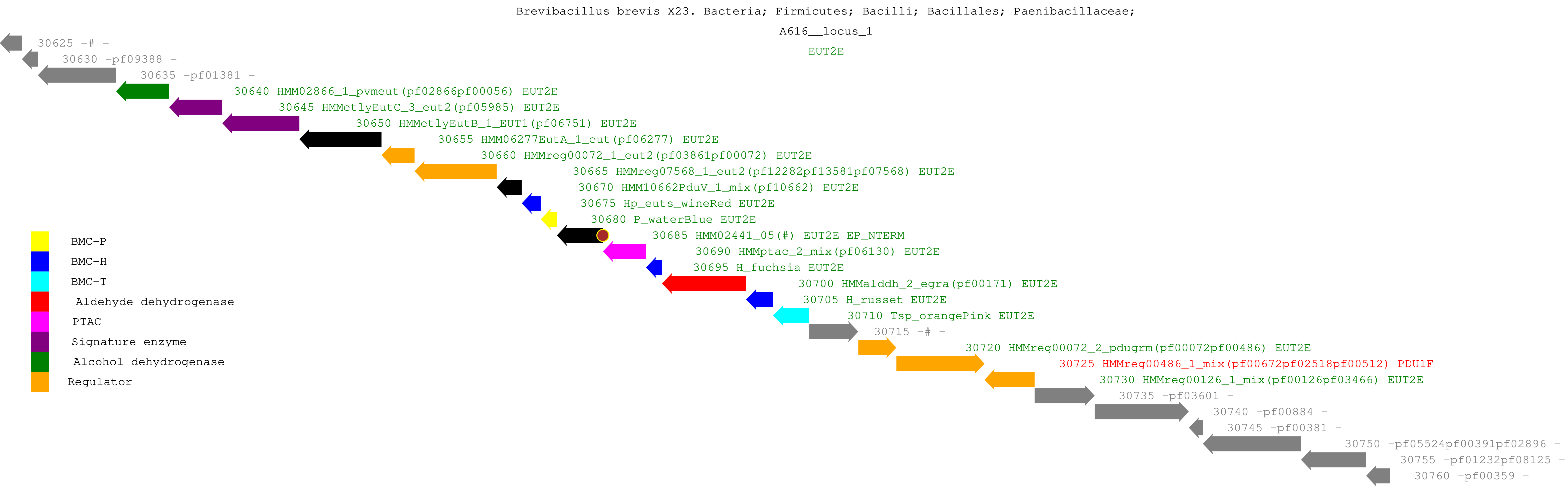

Brevibacillus brevis X23. Bacteria; Firmicutes; Bacilli; Bacillales; Paenibacillaceae;
A616__locus_1
EUT2E
30625 -# -
30630 -pf09388 -
30635 -pf01381 -
30640 HMM02866_1_pvmeut(pf02866pf00056) EUT2E
30645 HMMetlyEutC_3_eut2(pf05985) EUT2E
30650 HMMetlyEutB_1_EUT1(pf06751) EUT2E
30655 HMM06277EutA_1_eut(pf06277) EUT2E
30660 HMMreg00072_1_eut2(pf03861pf00072) EUT2E
30665 HMMreg07568_1_eut2(pf12282pf13581pf07568) EUT2E
30670 HMM10662PduV_1_mix(pf10662) EUT2E
30675 Hp_euts_wineRed EUT2E
30680 P_waterBlue EUT2E
30685 HMM02441_05(#) EUT2E EP_NTERM
30690 HMMptac_2_mix(pf06130) EUT2E
30695 H_fuchsia EUT2E
30700 HMMalddh_2_egra(pf00171) EUT2E
30705 H_russet EUT2E
30710 Tsp_orangePink EUT2E
30715 -# -
30720 HMMreg00072_2_pdugrm(pf00072pf00486) EUT2E
30725 HMMreg00486_1_mix(pf00672pf02518pf00512) PDU1F
30730 HMMreg00126_1_mix(pf00126pf03466) EUT2E
30735 -pf03601 -
30740 -pf00884 -
30745 -pf00381 -
30750 -pf05524pf00391pf02896 -
30755 -pf01232pf08125 -
30760 -pf00359 -
BMC-P
BMC-H
BMC-T
Aldehyde dehydrogenase
PTAC
Signature enzyme
Alcohol dehydrogenase
Regulator

Bacillus sp. FJAT-27215. Bacteria; Firmicutes; Bacilli; Bacillales; Bacillaceae; Bacillus.

A8F95__locus_2

EUT2E

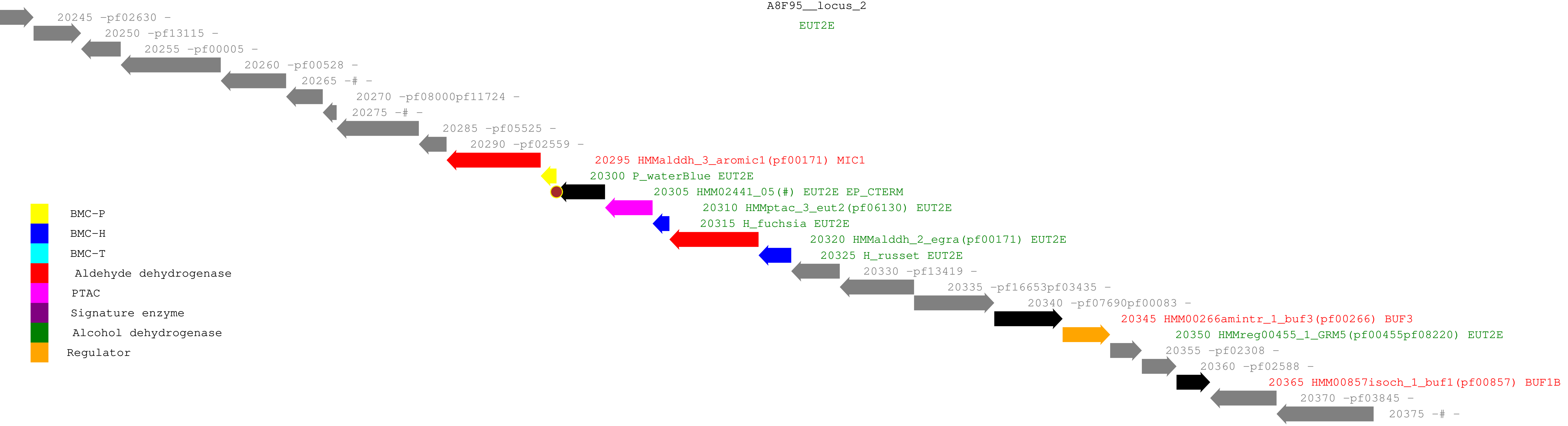

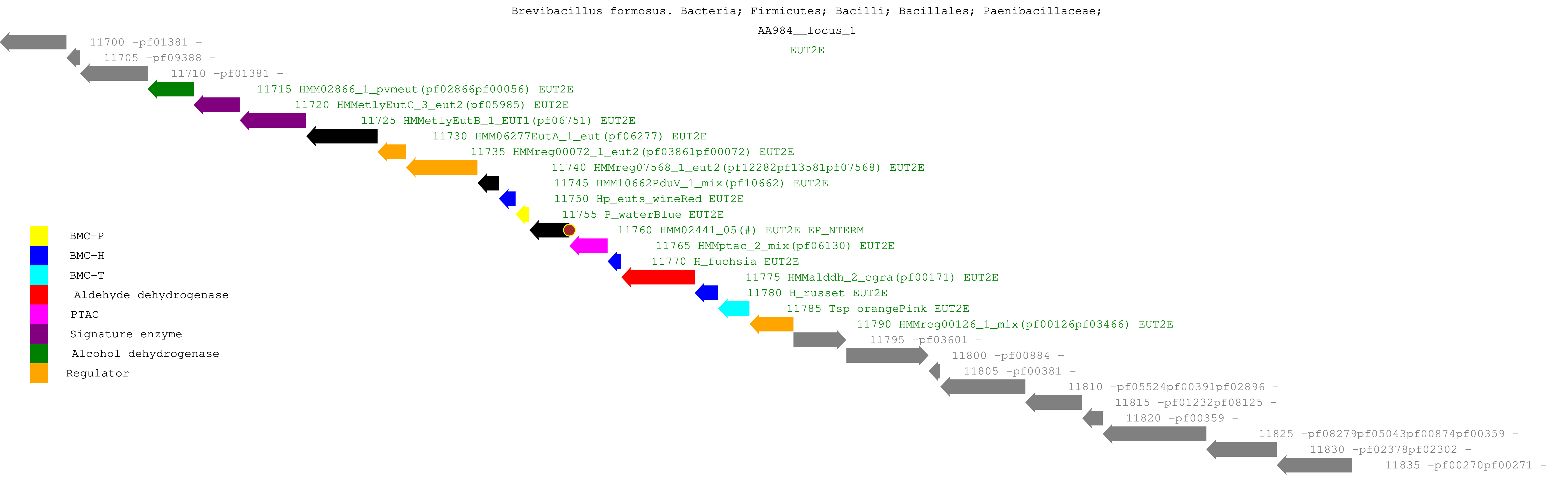

Brevibacillus formosus. Bacteria; Firmicutes; Bacilli; Bacillales; Paenibacillaceae;
AA984__locus_1
EUT2E
11700 -pf01381 -
11705 -pf09388 -
11710 -pf01381 -
11715 HMM02866_1_pvmeut(pf02866pf00056) EUT2E
11720 HMMetlyEutC_3_eut2(pf05985) EUT2E
11725 HMMetlyEutB_1_EUT1(pf06751) EUT2E
11730 HMM06277EutA_1_eut(pf06277) EUT2E
11735 HMMreg00072_1_eut2(pf03861pf00072) EUT2E
11740 HMMreg07568_1_eut2(pf12282pf13581pf07568) EUT2E
11745 HMM10662PduV_1_mix(pf10662) EUT2E
11750 Hp_euts_wineRed EUT2E
11755 P_waterBlue EUT2E
11760 HMM02441_05(#) EUT2E EP_NTERM
11765 HMMptac_2_mix(pf06130) EUT2E
11770 H_fuchsia EUT2E
11775 HMMaldh_2_egra(pf00171) EUT2E
11780 H_russet EUT2E
11785 Tsp_orangePink EUT2E
11790 HMMreg00126_1_mix(pf00126pf03466) EUT2E
11795 -pf03601 -
11800 -pf00884 -
11805 -pf00381 -
11810 -pf05524pf00391pf02896 -
11815 -pf01232pf08125 -
11820 -pf00359 -
11825 -pf08279pf05043pf00874pf00359 -
11830 -pf02378pf02302 -
11835 -pf00270pf00271 -
BMC-P
BMC-H
BMC-T
Aldehyde dehydrogenase
PTAC
Signature enzyme
Alcohol dehydrogenase
Regulator

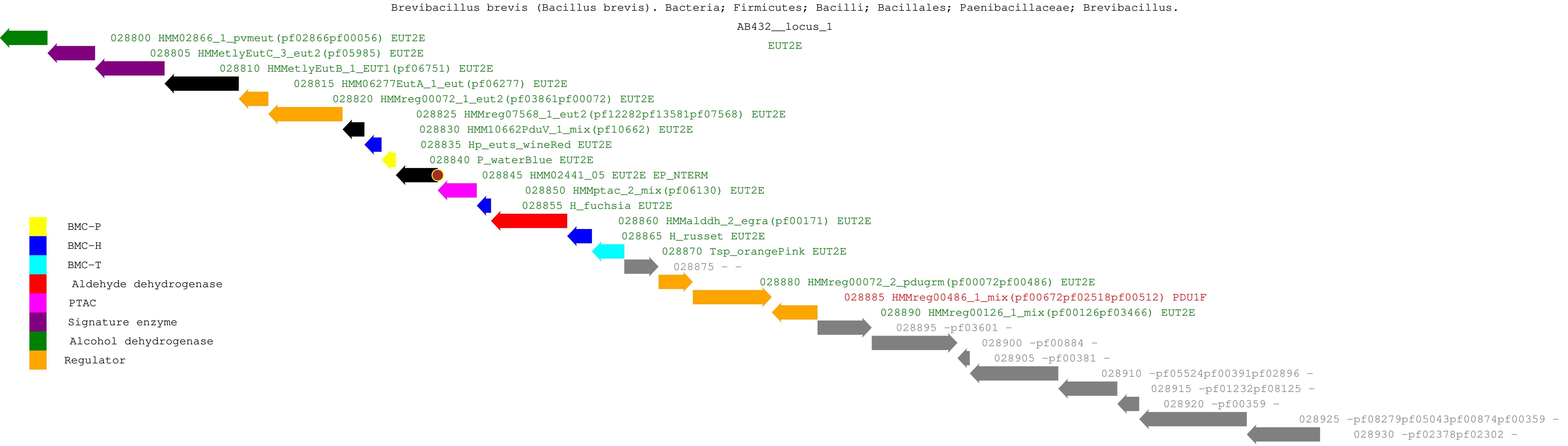
Brevibacillus brevis (Bacillus brevis). Bacteria; Firmicutes; Bacilli; Bacillales; Paenibacillaceae; Brevibacillus.
AB432__locus_1
EUT2E
028800 HMM02866_1_pvmeut(pf02866pf00056) EUT2E
028805 HMMetlyEutC_3_eut2(pf05985) EUT2E
028810 HMMetlyEutB_1_EUT1(pf06751) EUT2E
028815 HMM06277EutA_1_eut(pf06277) EUT2E
028820 HMMreg00072_1_eut2(pf03861pf00072) EUT2E
028825 HMMreg07568_1_eut2(pf12282pf13581pf07568) EUT2E
028830 HMM10662PduV_1_mix(pf10662) EUT2E
028835 Hp_euts_wineRed EUT2E
028840 P_waterBlue EUT2E
028845 HMM02441_05 EUT2E EP_NTERM
028850 HMMptac_2_mix(pf06130) EUT2E
028855 H_fuchsia EUT2E
028860 HMMalddh_2_egra(pf00171) EUT2E
028865 H_russet EUT2E
028870 Tsp_orangePink EUT2E
028875 - -
028880 HMMreg00072_2_pdugrm(pf00072pf00486) EUT2E
028885 HMMreg00486_1_mix(pf00672pf02518pf00512) PDU1F
028890 HMMreg00126_1_mix(pf00126pf03466) EUT2E
028895 -pf03601 -
028900 -pf00884 -
028905 -pf00381 -
028910 -pf05524pf00391pf02896 -
028915 -pf01232pf08125 -
028920 -pf00359 -
028925 -pf08279pf05043pf00874pf00359 -
028930 -pf02378pf02302 -
BMC-P
BMC-H
BMC-T
Aldehyde dehydrogenase
PTAC
Signature enzyme
Alcohol dehydrogenase
Regulator

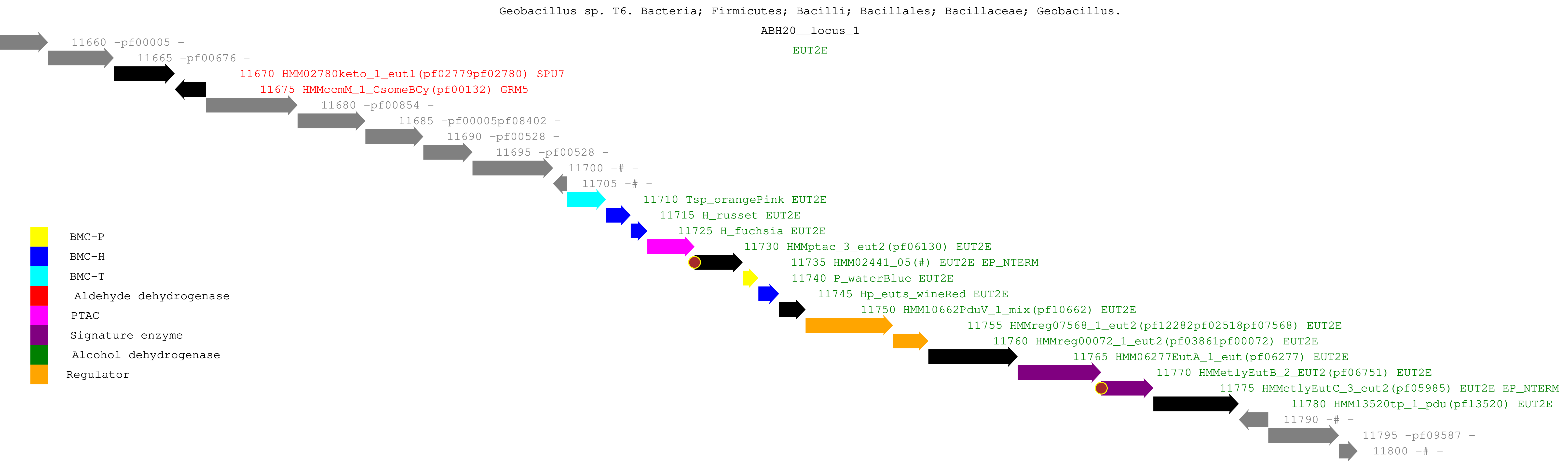

Geobacillus sp. T6. Bacteria; Firmicutes; Bacilli; Bacillales; Bacillaceae; Geobacillus.
ABH20__locus_1
EUT2E
11660 -pf00005 -
11665 -pf00676 -
11670 HMM02780keto_1_eut1(pf02779pf02780) SPU7
11675 HMMccmM_1_CsomeBCy(pf00132) GRM5
11680 -pf00854 -
11685 -pf00005pf08402 -
11690 -pf00528 -
11695 -pf00528 -
11700 -# -
11705 -# -
11710 Tsp_orangePink EUT2E
11715 H_russet EUT2E
11725 H_fuchsia EUT2E
11730 HMMptac_3_eut2(pf06130) EUT2E
11735 HMM02441_05(#) EUT2E EP_NTERM
11740 P_waterBlue EUT2E
11745 Hp_euts_wineRed EUT2E
11750 HMM10662PduV_1_mix(pf10662) EUT2E
11755 HMMreg07568_1_eut2(pf12282pf02518pf07568) EUT2E
11760 HMMreg00072_1_eut2(pf03861pf00072) EUT2E
11765 HMM06277EutA_1_eut(pf06277) EUT2E
11770 HMMetlyEutB_2_EUT2(pf06751) EUT2E
11775 HMMetlyEutC_3_eut2(pf05985) EUT2E EP_NTERM
11780 HMM13520tp_1_pdu(pf13520) EUT2E
11790 -# -
11795 -pf09587 -
11800 -# -
BMC-P
BMC-H
BMC-T
Aldehyde dehydrogenase
PTAC
Signature enzyme
Alcohol dehydrogenase
Regulator

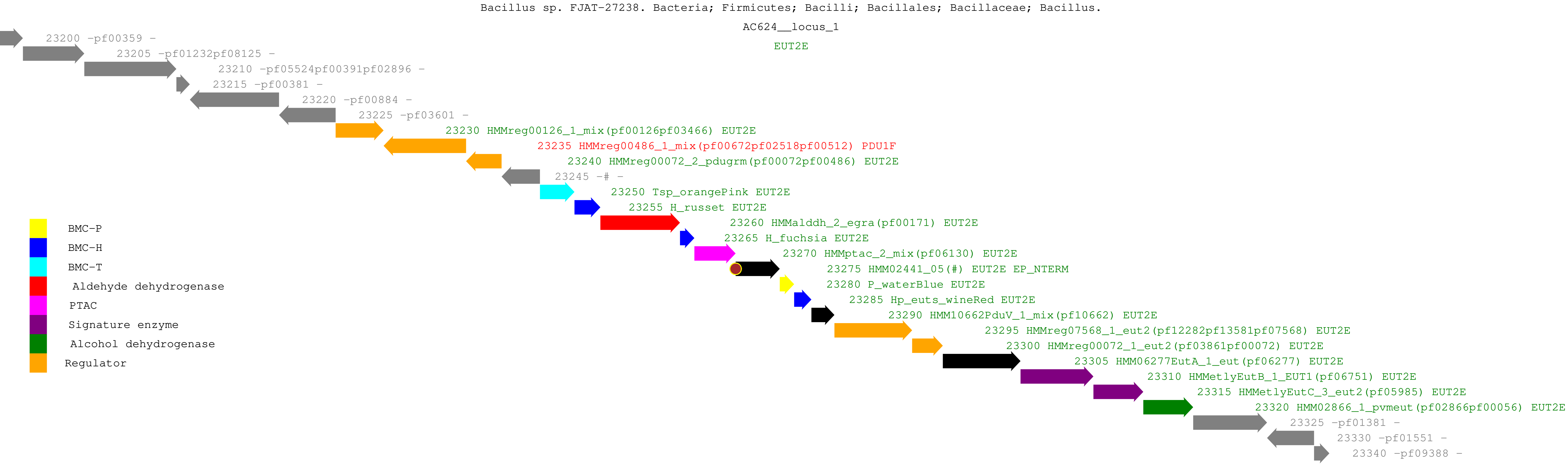
Bacillus sp. FJAT-27238. Bacteria; Firmicutes; Bacilli; Bacillales; Bacillaceae; Bacillus.
AC624__locus_1
EUT2E
23200 -pf00359 -
23205 -pf01232pf08125 -
23210 -pf05524pf00391pf02896 -
23215 -pf00381 -
23220 -pf00884 -
23225 -pf03601 -
23230 HMMreg00126_1_mix(pf00126pf03466) EUT2E
23235 HMMreg00486_1_mix(pf00672pf02518pf00512) PDU1F
23240 HMMreg00072_2_pdugrm(pf00072pf00486) EUT2E
23245 -# -
23250 Tsp_orangePink EUT2E
23255 H_russet EUT2E
23260 HMMalddh_2_egra(pf00171) EUT2E
23265 H_fuchsia EUT2E
23270 HMMptac_2_mix(pf06130) EUT2E
23275 HMM02441_05(#) EUT2E EP_NTERM
23280 P_waterBlue EUT2E
23285 Hp_euts_wineRed EUT2E
23290 HMM10662PduV_1_mix(pf10662) EUT2E
23295 HMMreg07568_1_eut2(pf12282pf13581pf07568) EUT2E
23300 HMMreg00072_1_eut2(pf03861pf00072) EUT2E
23305 HMM06277EutA_1_eut(pf06277) EUT2E
23310 HMMetlyEutB_1_EUT1(pf06751) EUT2E
23315 HMMetlyEutC_3_eut2(pf05985) EUT2E
23320 HMM02866_1_pvmeut(pf02866pf00056) EUT2E
23325 -pf01381 -
23330 -pf01551 -
23340 -pf09388 -
BMC-P
BMC-H
BMC-T
Aldehyde dehydrogenase
PTAC
Signature enzyme
Alcohol dehydrogenase
Regulator

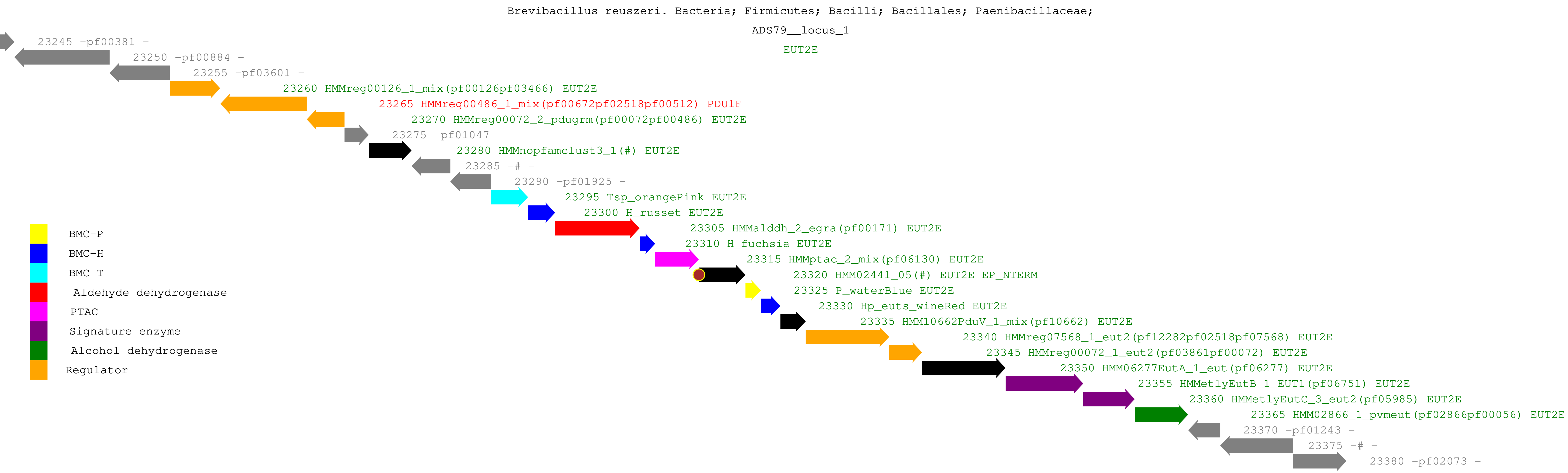
Brevibacillus reuszeri. Bacteria; Firmicutes; Bacilli; Bacillales; Paenibacillaceae;
ADS79__locus_1
EUT2E
23245 -pf00381 -
23250 -pf00884 -
23255 -pf03601 -
23260 HMMreg00126_1_mix(pf00126pf03466) EUT2E
23265 HMMreg00486_1_mix(pf00672pf02518pf00512) PDU1F
23270 HMMreg00072_2_pdugrm(pf00072pf00486) EUT2E
23275 -pf01047 -
23280 HMMnopfamclust3_1(#) EUT2E
23285 -# -
23290 -pf01925 -
23295 Tsp_orangePink EUT2E
23300 H_russet EUT2E
23305 HMMaldh_2_egra(pf00171) EUT2E
23310 H_fuchsia EUT2E
23315 HMMptac_2_mix(pf06130) EUT2E
23320 HMM02441_05(#) EUT2E EP_NTERM
23325 P_waterBlue EUT2E
23330 Hp_euts_wineRed EUT2E
23335 HMM10662PduV_1_mix(pf10662) EUT2E
23340 HMMreg07568_1_eut2(pf12282pf02518pf07568) EUT2E
23345 HMMreg00072_1_eut2(pf03861pf00072) EUT2E
23350 HMM06277EutA_1_eut(pf06277) EUT2E
23355 HMMetlyEutB_1_EUT1(pf06751) EUT2E
23360 HMMetlyEutC_3_eut2(pf05985) EUT2E
23365 HMM02866_1_pvmeut(pf02866pf00056) EUT2E
23370 -pf01243 -
23375 -# -
23380 -pf02073 -
BMC-P
BMC-H
BMC-T
Aldehyde dehydrogenase
PTAC
Signature enzyme
Alcohol dehydrogenase
Regulator

Brevibacillus choshinensis. Bacteria; Firmicutes; Bacilli; Bacillales; Paenibacillaceae;

AN963__locus_2

EUT2E

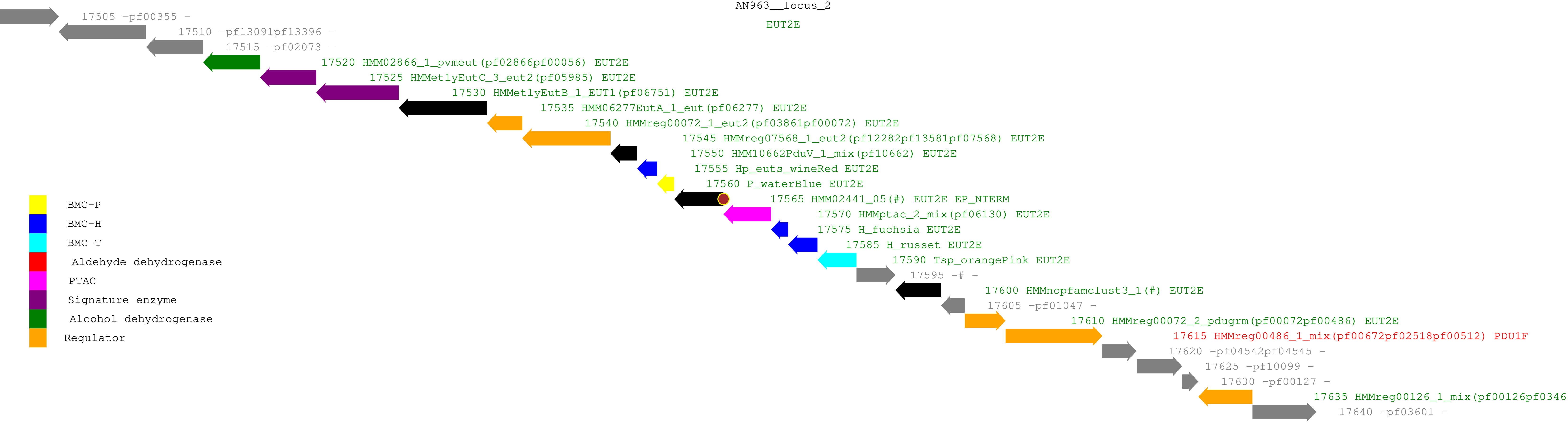

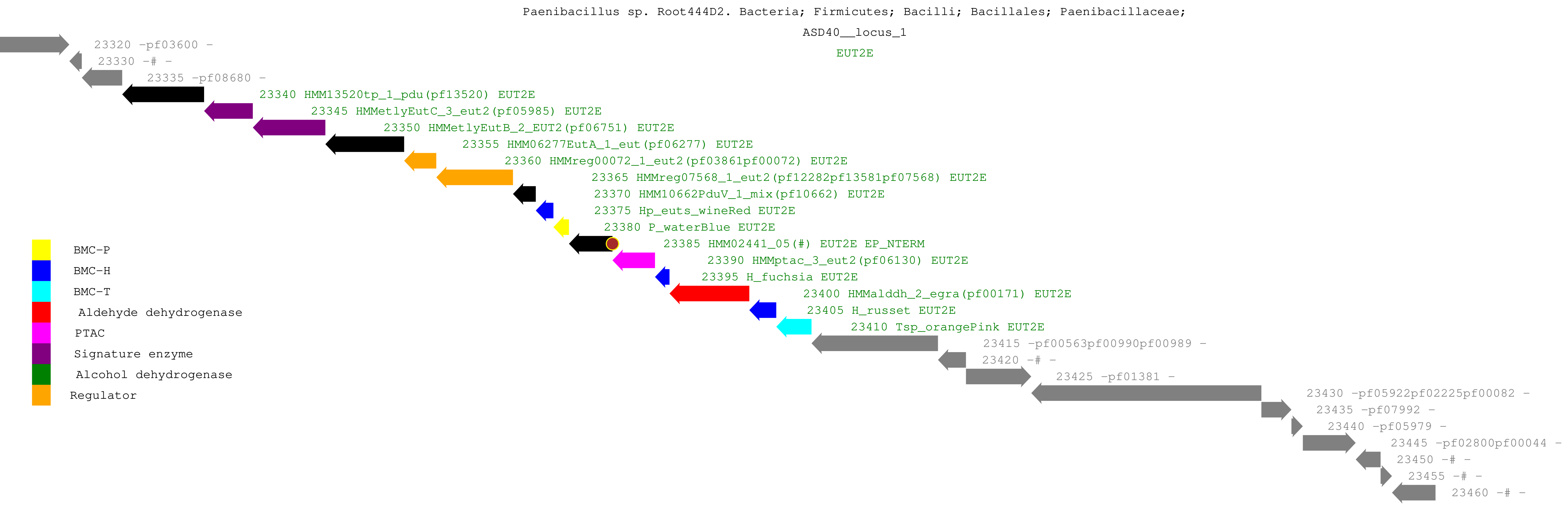

Paenibacillus sp. Root444D2. Bacteria; Firmicutes; Bacilli; Bacillales; Paenibacillaceae;
ASD40__locus_1
EUT2E
23320 -pf03600 -
23330 -# -
23335 -pf08680 -
23340 HMM13520tp_1_pdu(pf13520) EUT2E
23345 HMMetlyEutC_3_eut2(pf05985) EUT2E
23350 HMMetlyEutB_2_EUT2(pf06751) EUT2E
23355 HMM06277EutA_1_eut(pf06277) EUT2E
23360 HMMreg00072_1_eut2(pf03861pf00072) EUT2E
23365 HMMreg07568_1_eut2(pf12282pf13581pf07568) EUT2E
23370 HMM10662PduV_1_mix(pf10662) EUT2E
23375 Hp_euts_wineRed EUT2E
23380 P_waterBlue EUT2E
23385 HMM02441_05(#) EUT2E EP_NTERM
23390 HMMptac_3_eut2(pf06130) EUT2E
23395 H_fuchsia EUT2E
23400 HMMaldh_2_egra(pf00171) EUT2E
23405 H_russet EUT2E
23410 Tsp_orangePink EUT2E
23415 -pf00563pf00990pf00989 -
23420 -# -
23425 -pf01381 -
23430 -pf05922pf02225pf00082 -
23435 -pf07992 -
23440 -pf05979 -
23445 -pf02800pf00044 -
23450 -# -
23455 -# -
23460 -# -
BMC-P
BMC-H
BMC-T
Aldehyde dehydrogenase
PTAC
Signature enzyme
Alcohol dehydrogenase
Regulator

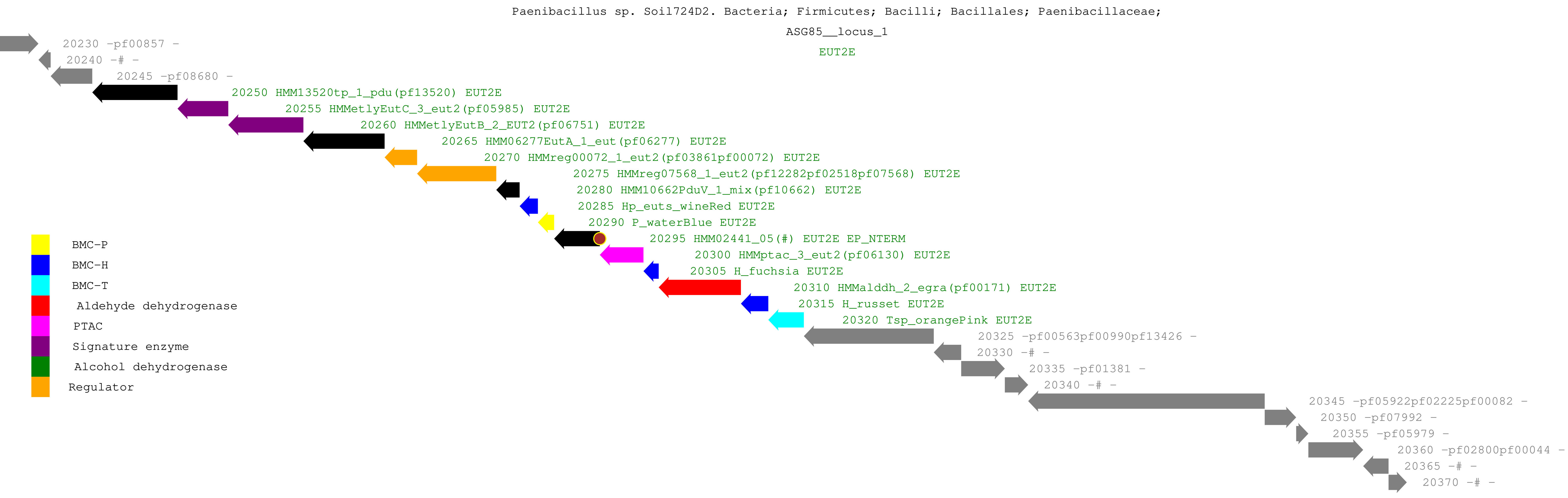

Paenibacillus sp. Soil724D2. Bacteria; Firmicutes; Bacilli; Bacillales; Paenibacillaceae;
ASG85__locus_1
EUT2E
20230 -pf00857 -
20240 -# -
20245 -pf08680 -
20250 HMM13520tp_1_pdu(pf13520) EUT2E
20255 HMMetlyEutC_3_eut2(pf05985) EUT2E
20260 HMMetlyEutB_2_EUT2(pf06751) EUT2E
20265 HMM06277EutA_1_eut(pf06277) EUT2E
20270 HMMreg00072_1_eut2(pf03861pf00072) EUT2E
20275 HMMreg07568_1_eut2(pf12282pf02518pf07568) EUT2E
20280 HMM10662PduV_1_mix(pf10662) EUT2E
20285 Hp_euts_wineRed EUT2E
20290 P_waterBlue EUT2E
20295 HMM02441_05(#) EUT2E EP_NTERM
20300 HMMptac_3_eut2(pf06130) EUT2E
20305 H_fuchsia EUT2E
20310 HMMalddh_2_egra(pf00171) EUT2E
20315 H_russet EUT2E
20320 Tsp_orangePink EUT2E
20325 -pf00563pf00990pf13426 -
20330 -# -
20335 -pf01381 -
20340 -# -
20345 -pf05922pf02225pf00082 -
20350 -pf07992 -
20355 -pf05979 -
20360 -pf02800pf00044 -
20365 -# -
20370 -# -
BMC-P
BMC-H
BMC-T
Aldehyde dehydrogenase
PTAC
Signature enzyme
Alcohol dehydrogenase
Regulator

Brevibacillus parabrevis. Bacteria; Firmicutes; Bacilli; Bacillales; Paenibacillaceae;

AV540__locus_1

EUT2E

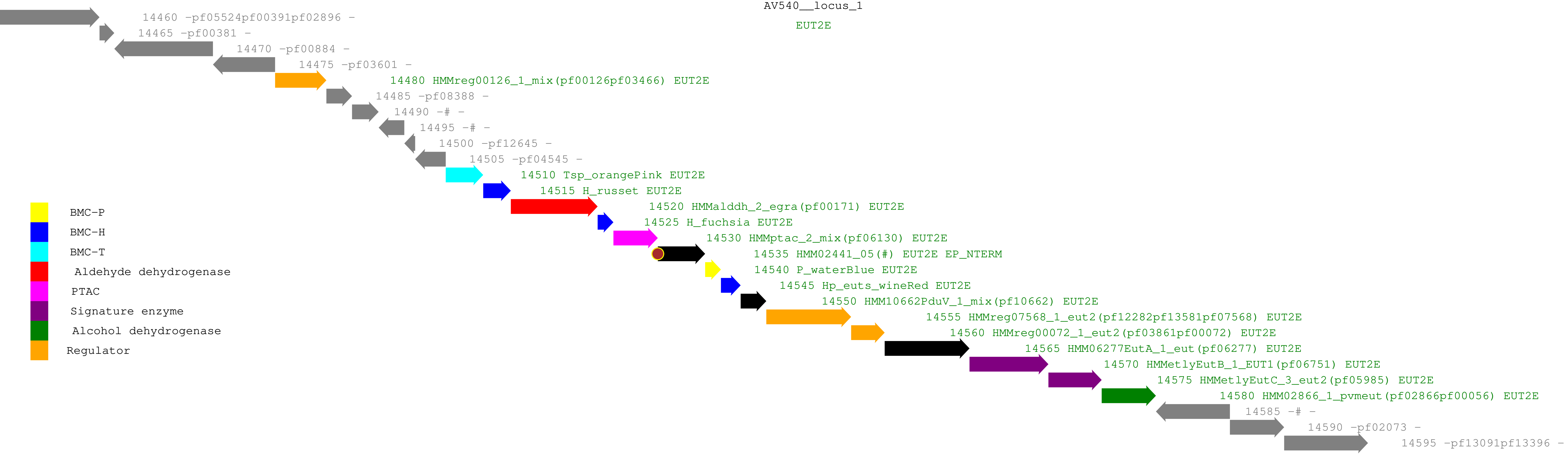

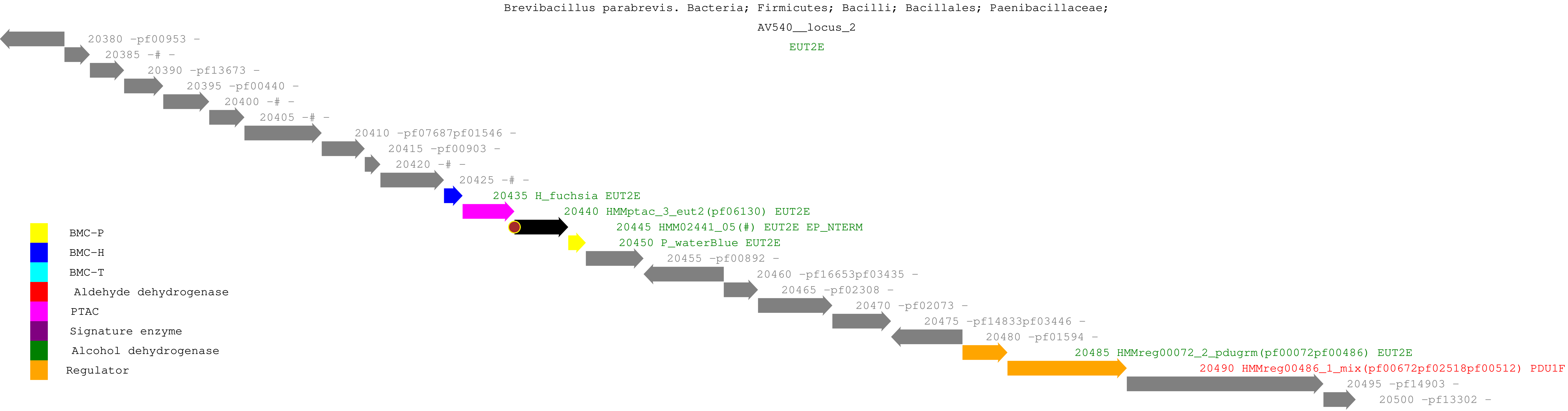
Brevibacillus parabrevis. Bacteria; Firmicutes; Bacilli; Bacillales; Paenibacillaceae;
AV540__locus_2
EUT2E
20380 -pf00953 -
20385 -# -
20390 -pf13673 -
20395 -pf00440 -
20400 -# -
20405 -# -
20410 -pf07687pf01546 -
20415 -pf00903 -
20420 -# -
20425 -# -
20435 H_fuchsia EUT2E
20440 HMMptac_3_eut2(pf06130) EUT2E
20445 HMM02441_05(#) EUT2E EP_NTERM
20450 P_waterBlue EUT2E
20455 -pf00892 -
20460 -pf16653pf03435 -
20465 -pf02308 -
20470 -pf02073 -
20475 -pf14833pf03446 -
20480 -pf01594 -
20485 HMMreg00072_2_pdugrm(pf00072pf00486) EUT2E
20490 HMMreg00486_1_mix(pf00672pf02518pf00512) PDU1F
20495 -pf14903 -
20500 -pf13302 -
BMC-P
BMC-H
BMC-T
Aldehyde dehydrogenase
PTAC
Signature enzyme
Alcohol dehydrogenase
Regulator

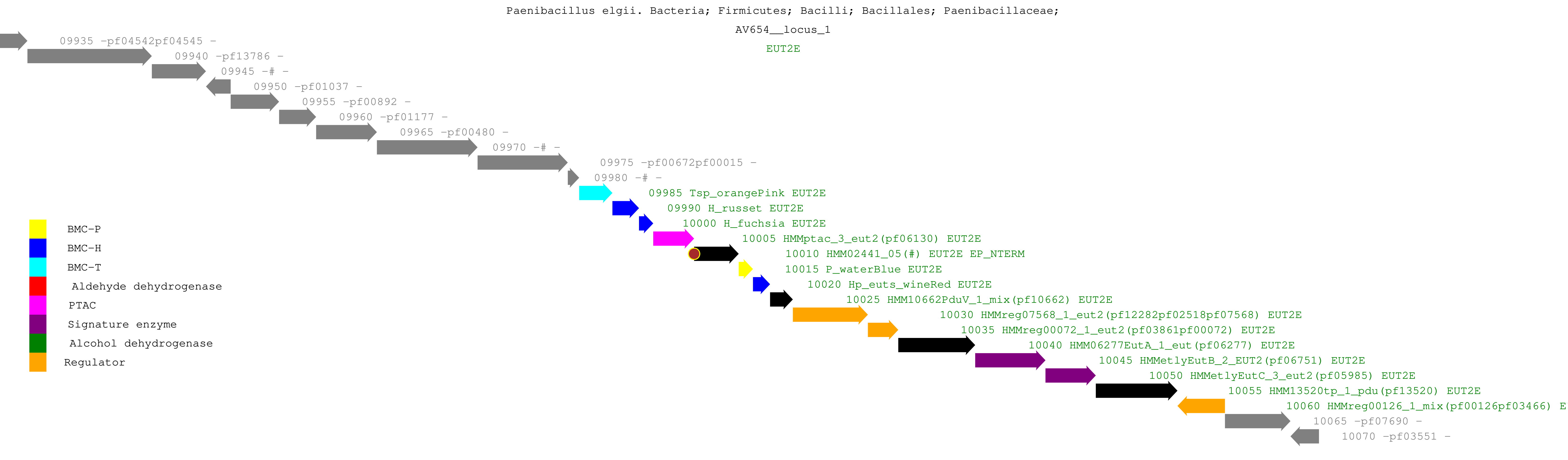
Paenibacillus elgii. Bacteria; Firmicutes; Bacilli; Bacillales; Paenibacillaceae;
AV654__locus_1
EUT2E
09935 -pf04542pf04545 -
09940 -pf13786 -
09945 -# -
09950 -pf01037 -
09955 -pf00892 -
09960 -pf01177 -
09965 -pf00480 -
09970 -# -
09975 -pf00672pf00015 -
09980 -# -
09985 Tsp_orangePink EUT2E
09990 H_russet EUT2E
10000 H_fuchsia EUT2E
10005 HMMptac_3_eut2(pf06130) EUT2E
10010 HMM02441_05(#) EUT2E EP_NTERM
10015 P_waterBlue EUT2E
10020 Hp_euts_wineRed EUT2E
10025 HMM10662PduV_1_mix(pf10662) EUT2E
10030 HMMreg07568_1_eut2(pf12282pf02518pf07568) EUT2E
10035 HMMreg00072_1_eut2(pf03861pf00072) EUT2E
10040 HMM06277EutA_1_eut(pf06277) EUT2E
10045 HMMetlyEutB_2_EUT2(pf06751) EUT2E
10050 HMMetlyEutC_3_eut2(pf05985) EUT2E
10055 HMM13520tp_1_pdu(pf13520) EUT2E
10060 HMMreg00126_1_mix(pf00126pf03466) E
10065 -pf07690 -
10070 -pf03551 -
BMC-P
BMC-H
BMC-T
Aldehyde dehydrogenase
PTAC
Signature enzyme
Alcohol dehydrogenase
Regulator

Parageobacillus caldoxylosilyticus. Bacteria; Firmicutes; Bacilli; Bacillales; Bacillaceae;

B4119__locus_1

EUT2E

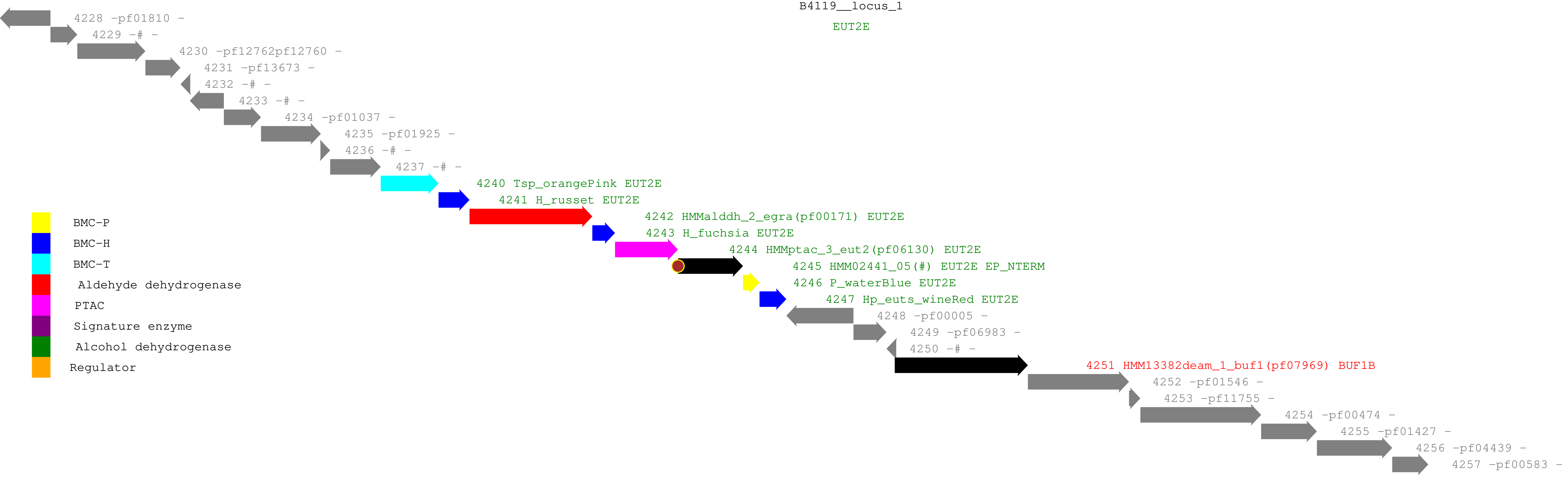

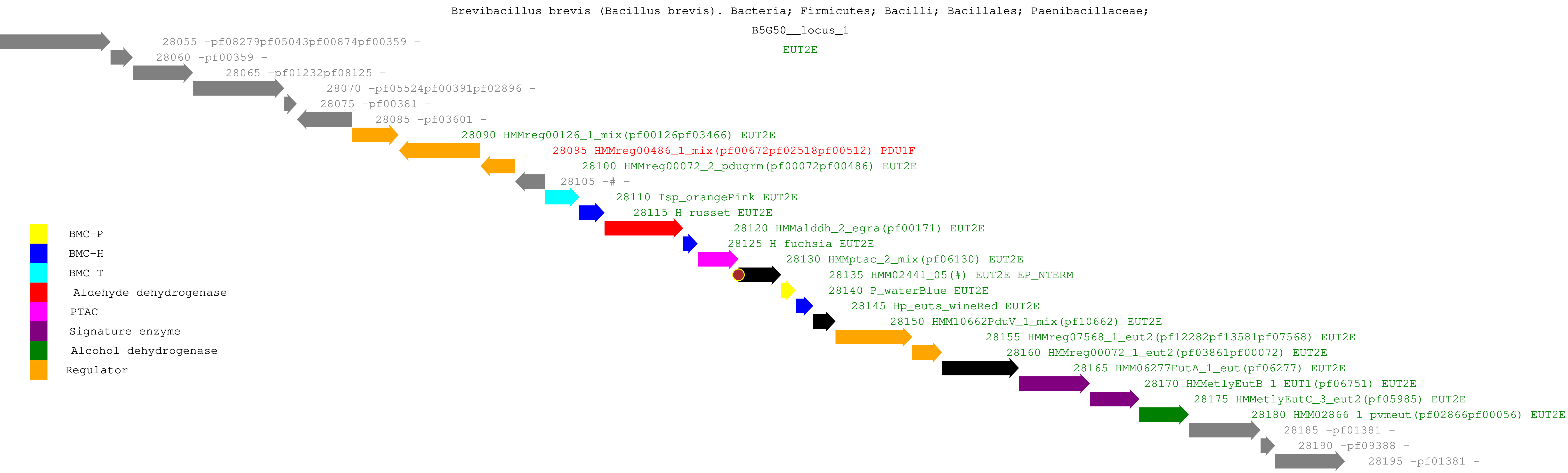
Brevibacillus brevis (Bacillus brevis). Bacteria; Firmicutes; Bacilli; Bacillales; Paenibacillaceae;
B5G50__locus_1
EUT2E
28055 -pf08279pf05043pf00874pf00359 -
28060 -pf00359 -
28065 -pf01232pf08125 -
28070 -pf05524pf00391pf02896 -
28075 -pf00381 -
28085 -pf03601 -
28090 HMMreg00126_1_mix(pf00126pf03466) EUT2E
28095 HMMreg00486_1_mix(pf00672pf02518pf00512) PDU1F
28100 HMMreg00072_2_pdugrm(pf00072pf00486) EUT2E
28105 -# -
28110 Tsp_orangePink EUT2E
28115 H_russet EUT2E
28120 HMMalddh_2_egra(pf00171) EUT2E
28125 H_fuchsia EUT2E
28130 HMMptac_2_mix(pf06130) EUT2E
28135 HMM02441_05(#) EUT2E EP_NTERM
28140 P_waterBlue EUT2E
28145 Hp_euts_wineRed EUT2E
28150 HMM10662PduV_1_mix(pf10662) EUT2E
28155 HMMreg07568_1_eut2(pf12282pf13581pf07568) EUT2E
28160 HMMreg00072_1_eut2(pf03861pf00072) EUT2E
28165 HMM06277EutA_1_eut(pf06277) EUT2E
28170 HMMetlyEutB_1_EUT1(pf06751) EUT2E
28175 HMMetlyEutC_3_eut2(pf05985) EUT2E
28180 HMM02866_1_pvmeut(pf02866pf00056) EUT2E
28185 -pf01381 -
28190 -pf09388 -
28195 -pf01381 -
BMC-P
BMC-H
BMC-T
Aldehyde dehydrogenase
PTAC
Signature enzyme
Alcohol dehydrogenase
Regulator

Geobacillus thermocatenulatus. Bacteria; Firmicutes; Bacilli; Bacillales; Bacillaceae; Geobacillus;

B9L19__locus_1

EUT2E

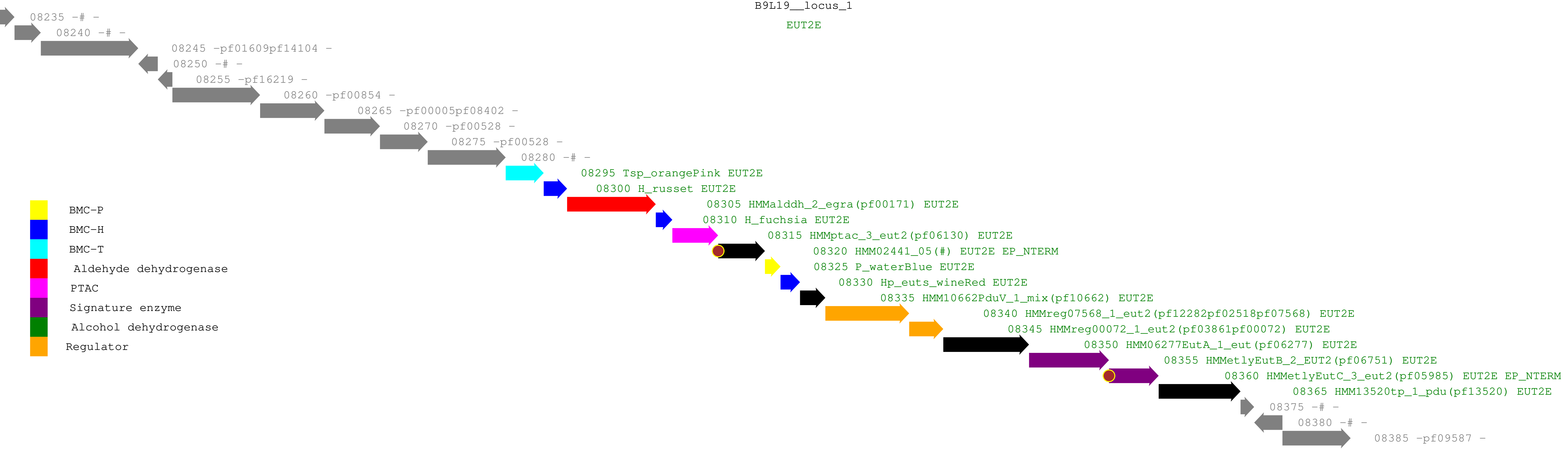

Paenibacillus baekrokdamisoli. Bacteria; Firmicutes; Bacilli; Bacillales; Paenibacillaceae; Paenibacillus.

Back11__locus_1

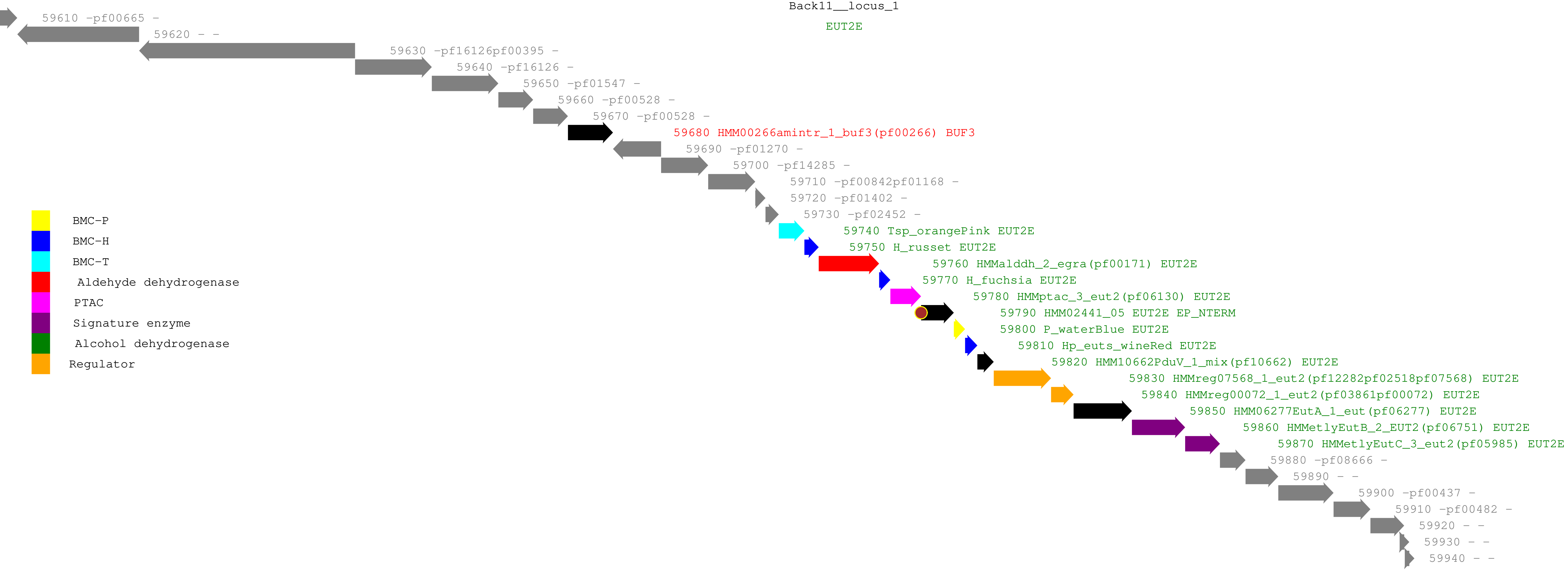

Brevibacillus brevis (strain 47 / JCM 6285 / NBRC 100599). Bacteria; Firmicutes; Bacilli; Bacillales; Paenibacillaceae;

BBR47__locus_1

EUT2E

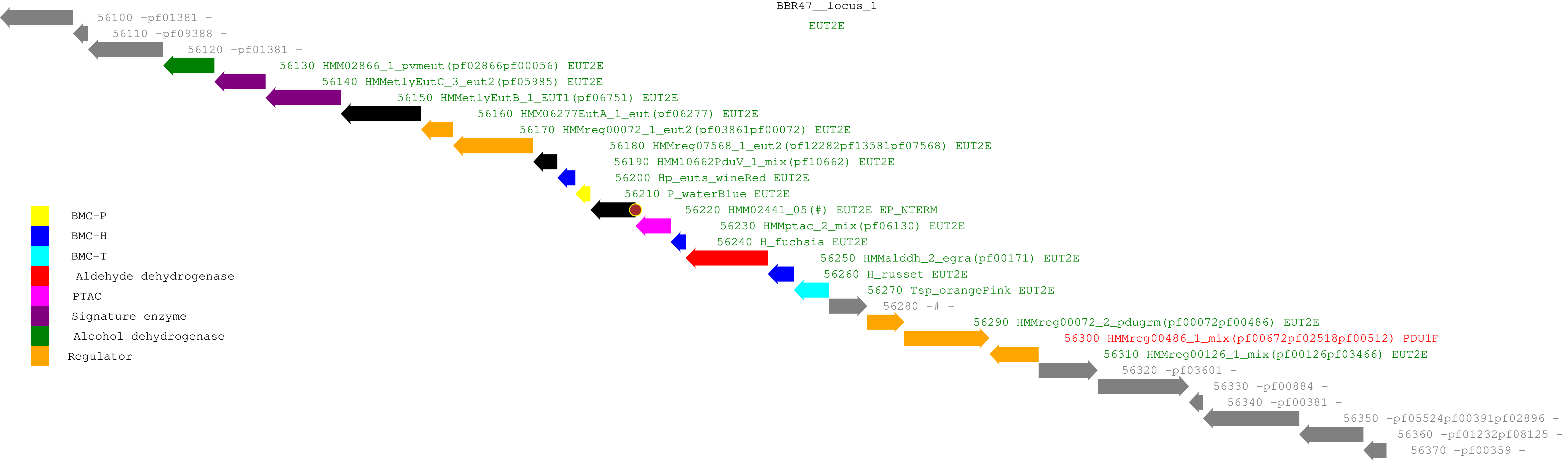

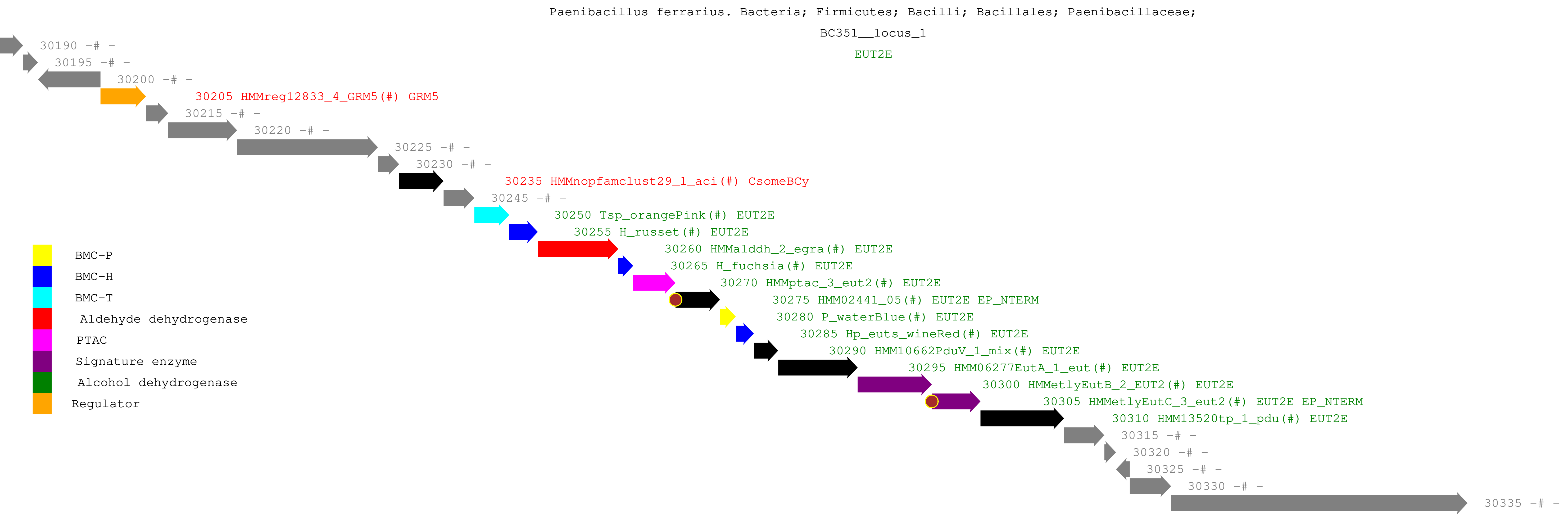

Paenibacillus ferrarius. Bacteria; Firmicutes; Bacilli; Bacillales; Paenibacillaceae;
BC351__locus_1
EUT2E
30190 -# -
30195 -# -
30200 -# -
30205 HMMreg12833_4_GRM5(#) GRM5
30215 -# -
30220 -# -
30225 -# -
30230 -# -
30235 HMMnopfamclust29_1_aci(#) CsomeBCy
30245 -# -
30250 Tsp_orangePink(#) EUT2E
30255 H_russet(#) EUT2E
30260 HMMalddh_2_egra(#) EUT2E
30265 H_fuchsia(#) EUT2E
30270 HMMptac_3_eut2(#) EUT2E
30275 HMM02441_05(#) EUT2E EP_NTERM
30280 P_waterBlue(#) EUT2E
30285 Hp_euts_wineRed(#) EUT2E
30290 HMM10662PduV_1_mix(#) EUT2E
30295 HMM06277EutA_1_eut(#) EUT2E
30300 HMMetlyEutB_2_EUT2(#) EUT2E
30305 HMMetlyEutC_3_eut2(#) EUT2E EP_NTERM
30310 HMM13520tp_1_pdu(#) EUT2E
30315 -# -
30320 -# -
30325 -# -
30330 -# -
30335 -# -
BMC-P
BMC-H
BMC-T
Aldehyde dehydrogenase
PTAC
Signature enzyme
Alcohol dehydrogenase
Regulator

Anoxybacillus amylolyticus. Bacteria; Firmicutes; Bacilli; Bacillales; Bacillaceae; Anoxybacillus.

BDK47__locus_1

EUT2E

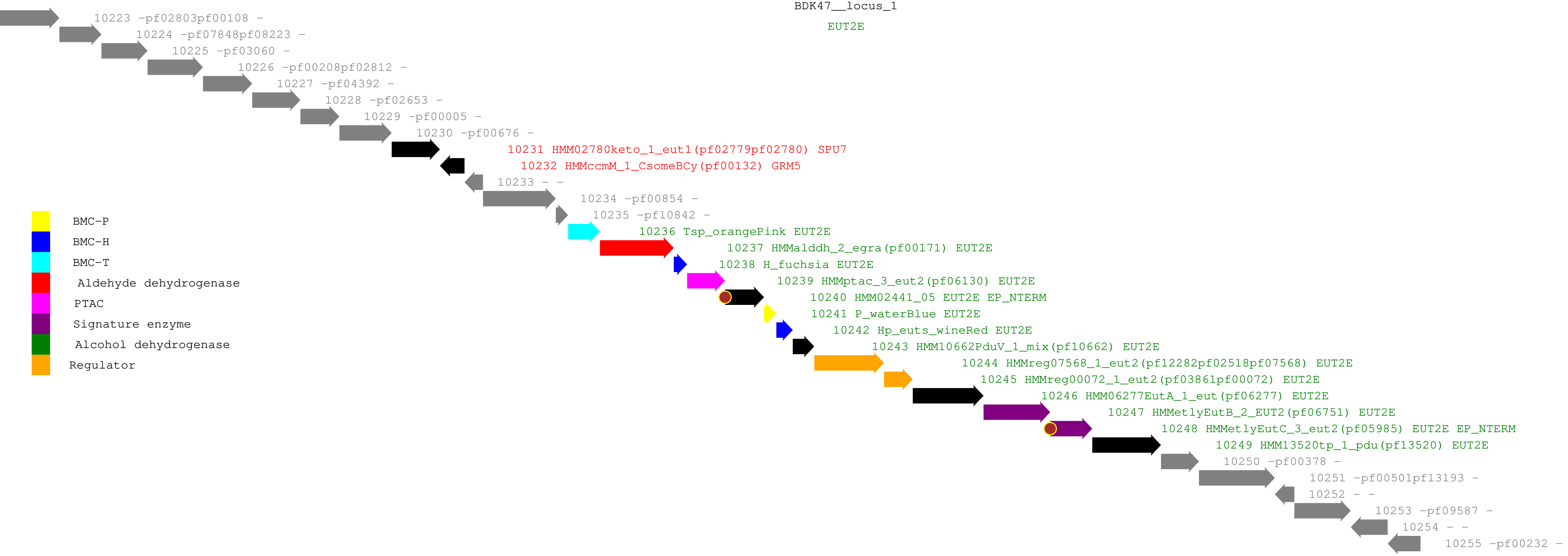

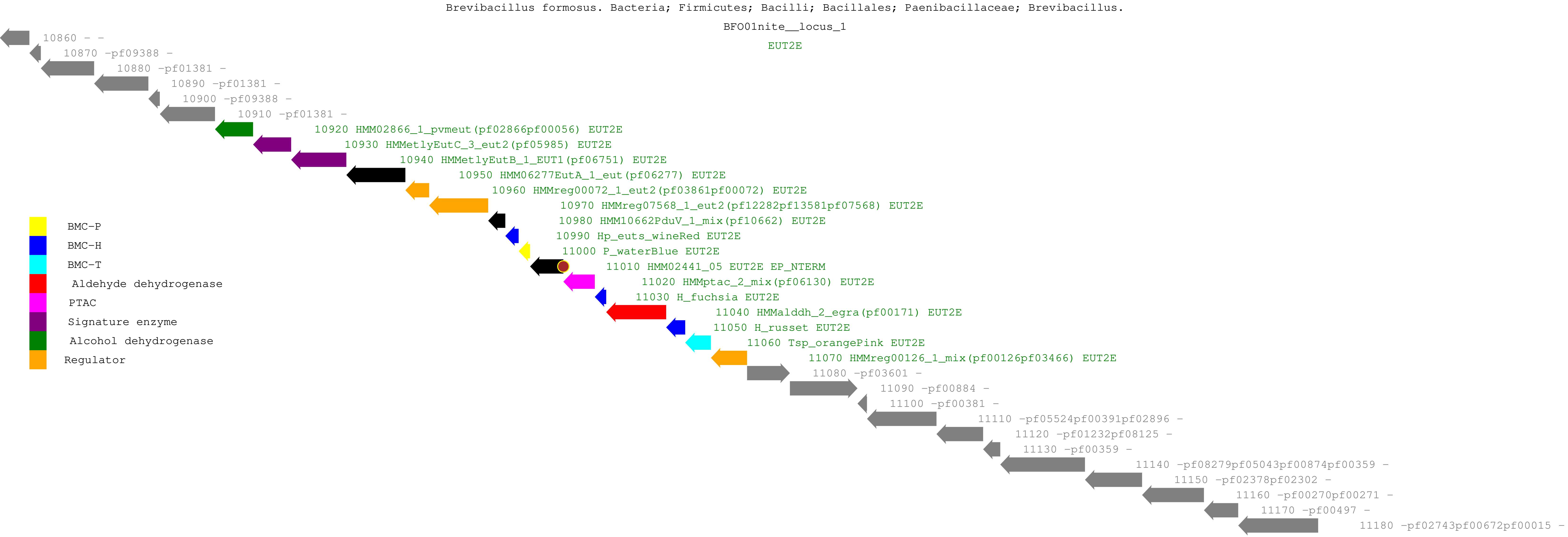
Brevibacillus formosus. Bacteria; Firmicutes; Bacilli; Bacillales; Paenibacillaceae; Brevibacillus.
BFO01nite__locus_1
EUT2E
10860 - -
10870 -pf09388 -
10880 -pf01381 -
10890 -pf01381 -
10900 -pf09388 -
10910 -pf01381 -
10920 HMM02866_1_pvmeut(pf02866pf00056) EUT2E
10930 HMMetlyEutC_3_eut2(pf05985) EUT2E
10940 HMMetlyEutB_1_EUT1(pf06751) EUT2E
10950 HMM06277EutA_1_eut(pf06277) EUT2E
10960 HMMreg00072_1_eut2(pf03861pf00072) EUT2E
10970 HMMreg07568_1_eut2(pf12282pf13581pf07568) EUT2E
10980 HMM10662PduV_1_mix(pf10662) EUT2E
10990 Hp_euts_wineRed EUT2E
11000 P_waterBlue EUT2E
11010 HMM02441_05 EUT2E EP_NTERM
11020 HMMptac_2_mix(pf06130) EUT2E
11030 H_fuchsia EUT2E
11040 HMMalddh_2_egra(pf00171) EUT2E
11050 H_russet EUT2E
11060 Tsp_orangePink EUT2E
11070 HMMreg00126_1_mix(pf00126pf03466) EUT2E
11080 -pf03601 -
11090 -pf00884 -
11100 -pf00381 -
11110 -pf05524pf00391pf02896 -
11120 -pf01232pf08125 -
11130 -pf00359 -
11140 -pf08279pf05043pf00874pf00359 -
11150 -pf02378pf02302 -
11160 -pf00270pf00271 -
11170 -pf00497 -
11180 -pf02743pf00672pf00015 -
BMC-P
BMC-H
BMC-T
Aldehyde dehydrogenase
PTAC
Signature enzyme
Alcohol dehydrogenase
Regulator

Paenibacillus darwinianus. Bacteria; Firmicutes; Bacilli; Bacillales; Paenibacillaceae;

BG52__locus_1_main_sat

EUT2E

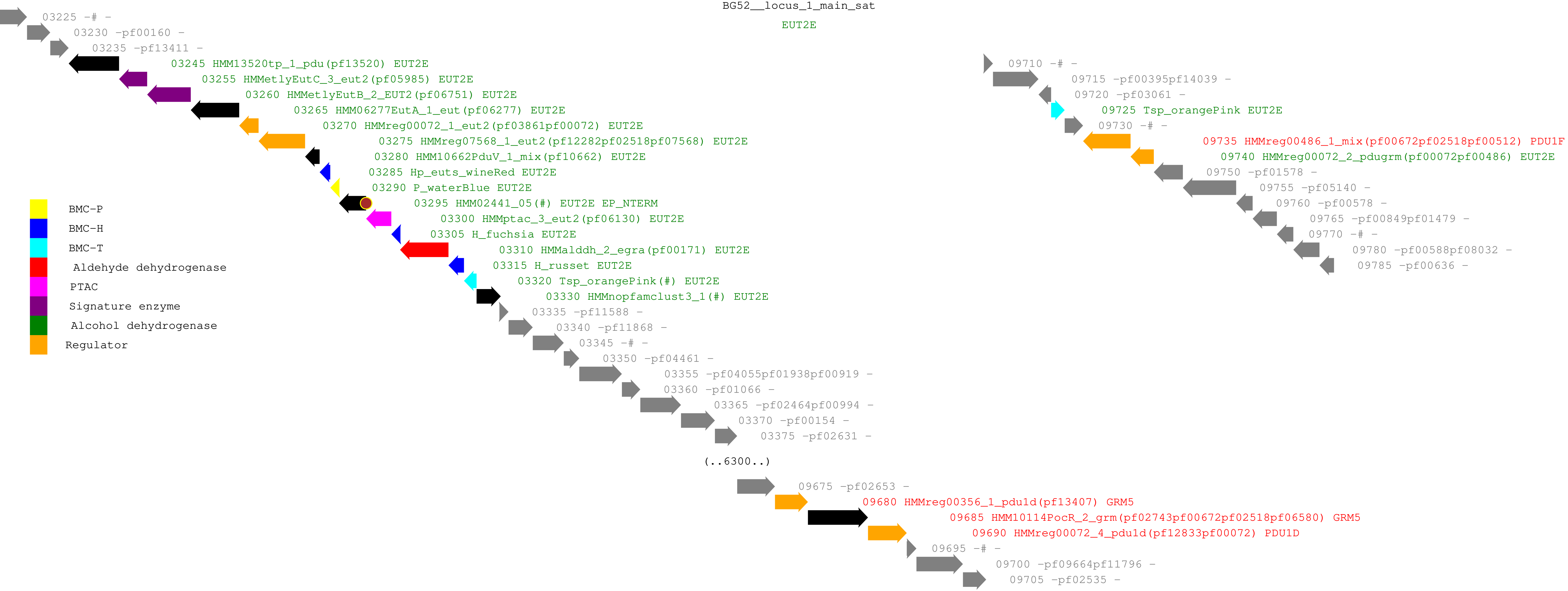

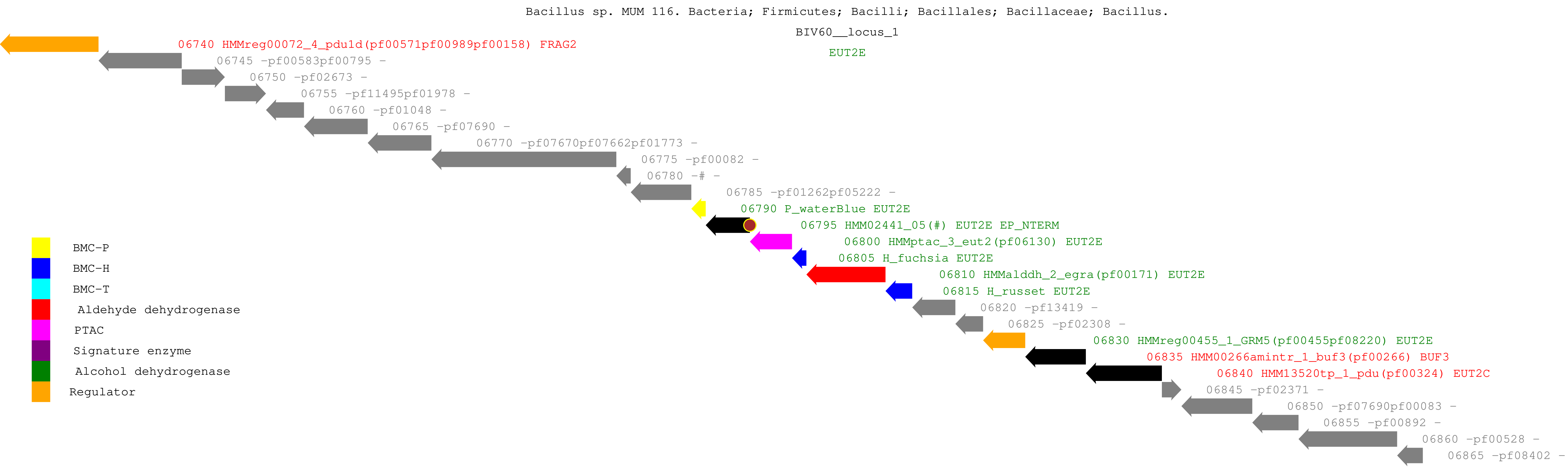

Bacillus sp. MUM 116. Bacteria; Firmicutes; Bacilli; Bacillales; Bacillaceae; Bacillus.
BIV60__locus_1
EUT2E
06740 HMMreg00072_4_pdu1d(pf00571pf00989pf00158) FRAG2
06745 -pf00583pf00795 -
06750 -pf02673 -
06755 -pf11495pf01978 -
06760 -pf01048 -
06765 -pf07690 -
06770 -pf07670pf07662pf01773 -
06775 -pf00082 -
06780 -# -
06785 -pf01262pf05222 -
06790 P_waterBlue EUT2E
06795 HMM02441_05(#) EUT2E EP_NTERM
06800 HMMptac_3_eut2(pf06130) EUT2E
06805 H_fuchsia EUT2E
06810 HMMalddh_2_egra(pf00171) EUT2E
06815 H_russet EUT2E
06820 -pf13419 -
06825 -pf02308 -
06830 HMMreg00455_1_GRM5(pf00455pf08220) EUT2E
06835 HMM00266amintr_1_buf3(pf00266) BUF3
06840 HMM13520tp_1_pdu(pf00324) EUT2C
06845 -pf02371 -
06850 -pf07690pf00083 -
06855 -pf00892 -
06860 -pf00528 -
06865 -pf08402 -
BMC-P
BMC-H
BMC-T
Aldehyde dehydrogenase
PTAC
Signature enzyme
Alcohol dehydrogenase
Regulator

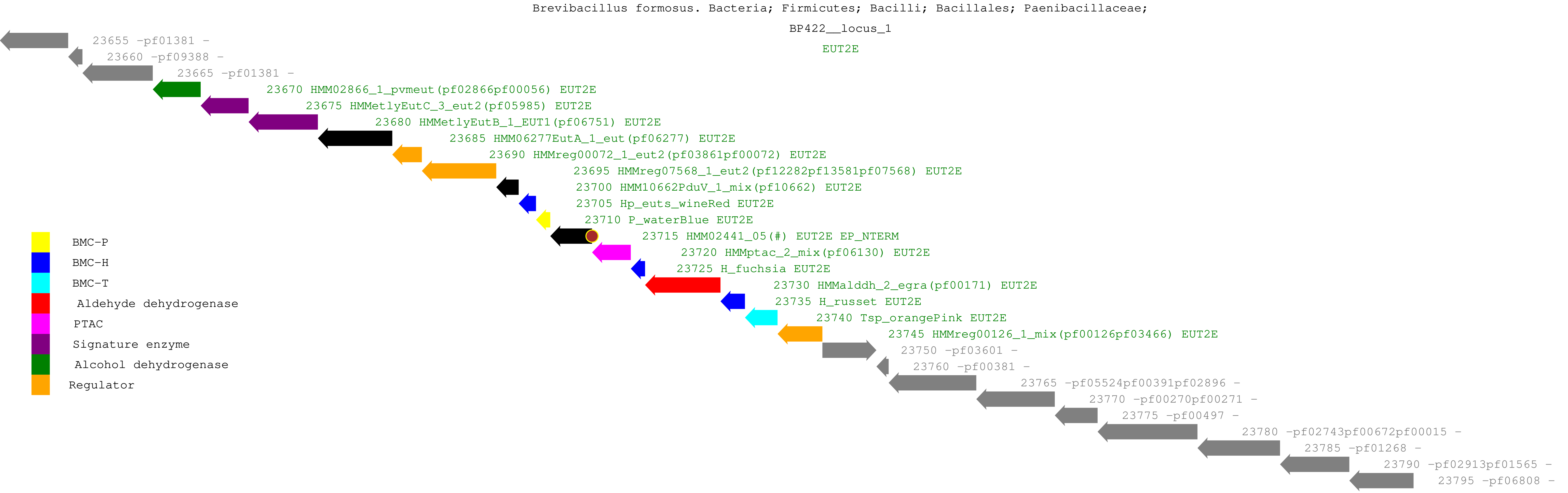

Brevibacillus formosus. Bacteria; Firmicutes; Bacilli; Bacillales; Paenibacillaceae;
BP422__locus_1
EUT2E
23655 -pf01381 -
23660 -pf09388 -
23665 -pf01381 -
23670 HMM02866_1_pvmeut(pf02866pf00056) EUT2E
23675 HMMetlyEutC_3_eut2(pf05985) EUT2E
23680 HMMetlyEutB_1_EUT1(pf06751) EUT2E
23685 HMM06277EutA_1_eut(pf06277) EUT2E
23690 HMMreg00072_1_eut2(pf03861pf00072) EUT2E
23695 HMMreg07568_1_eut2(pf12282pf13581pf07568) EUT2E
23700 HMM10662PduV_1_mix(pf10662) EUT2E
23705 Hp_euts_wineRed EUT2E
23710 P_waterBlue EUT2E
23715 HMM02441_05(#) EUT2E EP_NTERM
23720 HMMptac_2_mix(pf06130) EUT2E
23725 H_fuchsia EUT2E
23730 HMMalddh_2_egra(pf00171) EUT2E
23735 H_russet EUT2E
23740 Tsp_orangePink EUT2E
23745 HMMreg00126_1_mix(pf00126pf03466) EUT2E
23750 -pf03601 -
23760 -pf00381 -
23765 -pf05524pf00391pf02896 -
23770 -pf00270pf00271 -
23775 -pf00497 -
23780 -pf02743pf00672pf00015 -
23785 -pf01268 -
23790 -pf02913pf01565 -
23795 -pf06808 -
BMC-P
BMC-H
BMC-T
Aldehyde dehydrogenase
PTAC
Signature enzyme
Alcohol dehydrogenase
Regulator

Brevibacillus parabrevis. Bacteria; Firmicutes; Bacilli; Bacillales; Paenibacillaceae; Brevibacillus.

BPA01__locus_1

EUT2E

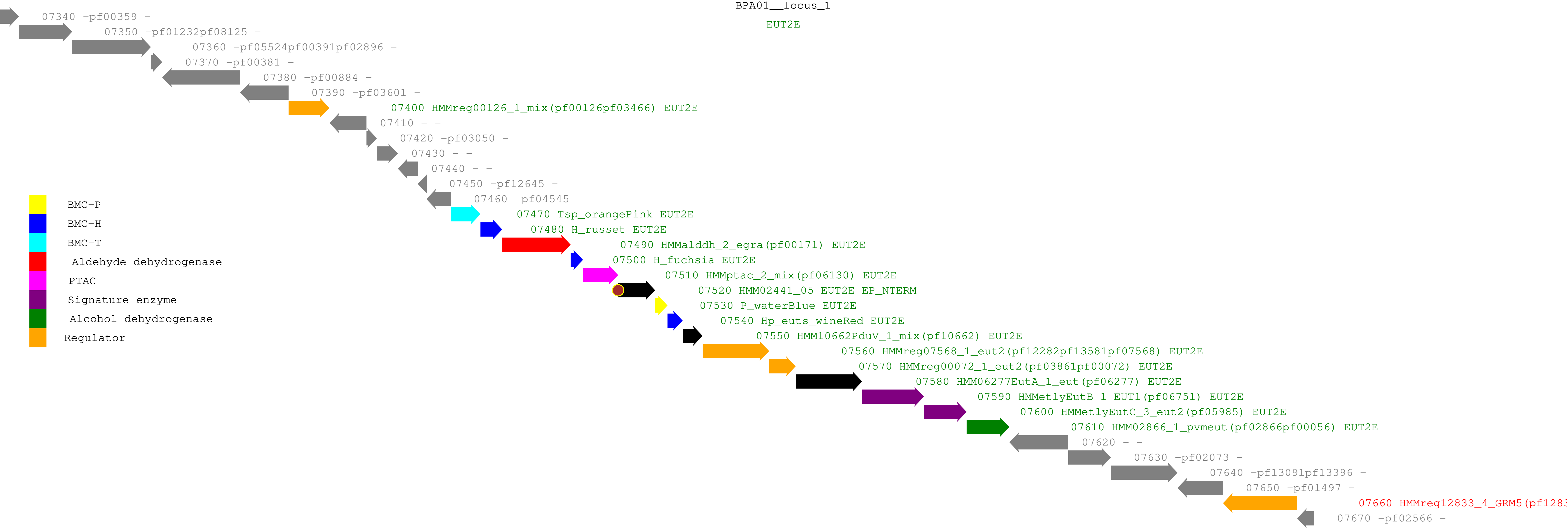

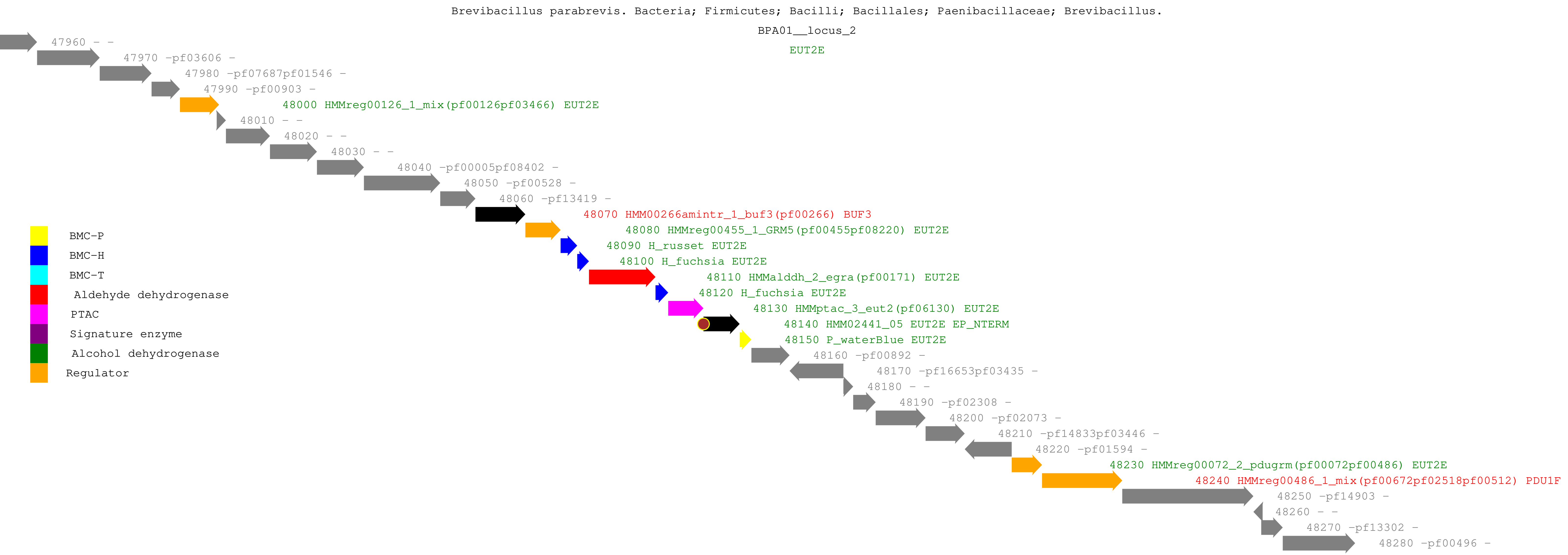
Brevibacillus parabrevis. Bacteria; Firmicutes; Bacilli; Bacillales; Paenibacillaceae; Brevibacillus.
BPA01__locus_2
EUT2E
47960 - -
47970 -pf03606 -
47980 -pf07687pf01546 -
47990 -pf00903 -
48000 HMMreg00126_1_mix(pf00126pf03466) EUT2E
48010 - -
48020 - -
48030 - -
48040 -pf00005pf08402 -
48050 -pf00528 -
48060 -pf13419 -
48070 HMM00266amintr_1_buf3(pf00266) BUF3
48080 HMMreg00455_1_GRM5(pf00455pf08220) EUT2E
48090 H_russet EUT2E
48100 H_fuchsia EUT2E
48110 HMMalddh_2_egra(pf00171) EUT2E
48120 H_fuchsia EUT2E
48130 HMMptac_3_eut2(pf06130) EUT2E
48140 HMM02441_05 EUT2E EP_NTERM
48150 P_waterBlue EUT2E
48160 -pf00892 -
48170 -pf16653pf03435 -
48180 - -
48190 -pf02308 -
48200 -pf02073 -
48210 -pf14833pf03446 -
48220 -pf01594 -
48230 HMMreg00072_2_pdugrm(pf00072pf00486) EUT2E
48240 HMMreg00486_1_mix(pf00672pf02518pf00512) PDU1F
48250 -pf14903 -
48260 - -
48270 -pf13302 -
48280 -pf00496 -
BMC-P
BMC-H
BMC-T
Aldehyde dehydrogenase
PTAC
Signature enzyme
Alcohol dehydrogenase
Regulator

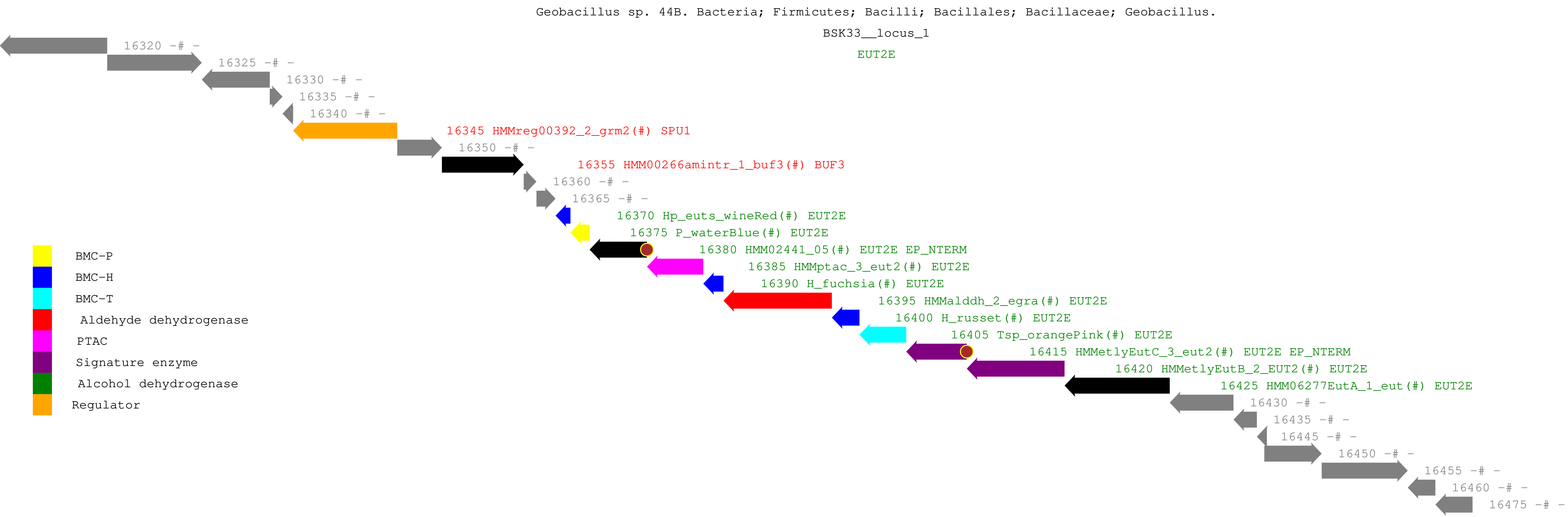
Geobacillus sp. 44B. Bacteria; Firmicutes; Bacilli; Bacillales; Bacillaceae; Geobacillus.
BSK33__locus_1
EUT2E
16320 -# -
16325 -# -
16330 -# -
16335 -# -
16340 -# -
16345 HMMreg00392_2_grm2(#) SPU1
16350 -# -
16355 HMM00266amintr_1_buf3(#) BUF3
16360 -# -
16365 -# -
16370 Hp_euts_wineRed(#) EUT2E
16375 P_waterBlue(#) EUT2E
16380 HMM02441_05(#) EUT2E EP_NTERM
16385 HMMptac_3_eut2(#) EUT2E
16390 H_fuchsia(#) EUT2E
16395 HMMalddh_2_egra(#) EUT2E
16400 H_russet(#) EUT2E
16405 Tsp_orangePink(#) EUT2E
16415 HMMetlyEutC_3_eut2(#) EUT2E EP_NTERM
16420 HMMetlyEutB_2_EUT2(#) EUT2E
16425 HMM06277EutA_1_eut(#) EUT2E
16430 -# -
16435 -# -
16445 -# -
16450 -# -
16455 -# -
16460 -# -
16475 -# -
BMC-P
BMC-H
BMC-T
Aldehyde dehydrogenase
PTAC
Signature enzyme
Alcohol dehydrogenase
Regulator

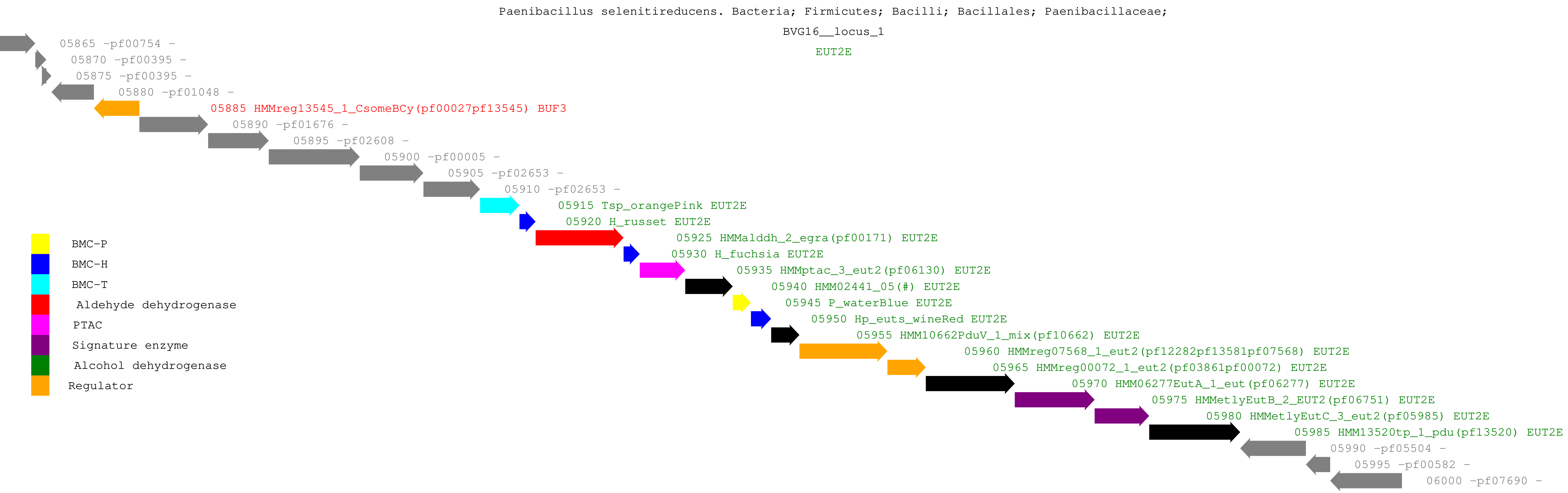
Paenibacillus selenitireducens. Bacteria; Firmicutes; Bacilli; Bacillales; Paenibacillaceae;
BVG16__locus_1
EUT2E
05865 -pf00754 -
05870 -pf00395 -
05875 -pf00395 -
05880 -pf01048 -
05885 HMMreg13545_1_CsomeBCy(pf00027pf13545) BUF3
05890 -pf01676 -
05895 -pf02608 -
05900 -pf00005 -
05905 -pf02653 -
05910 -pf02653 -
05915 Tsp_orangePink EUT2E
05920 H_russet EUT2E
05925 HMMalddh_2_egra(pf00171) EUT2E
05930 H_fuchsia EUT2E
05935 HMMptac_3_eut2(pf06130) EUT2E
05940 HMM02441_05(#) EUT2E
05945 P_waterBlue EUT2E
05950 Hp_euts_wineRed EUT2E
05955 HMM10662PduV_1_mix(pf10662) EUT2E
05960 HMMreg07568_1_eut2(pf12282pf13581pf07568) EUT2E
05965 HMMreg00072_1_eut2(pf03861pf00072) EUT2E
05970 HMM06277EutA_1_eut(pf06277) EUT2E
05975 HMMetlyEutB_2_EUT2(pf06751) EUT2E
05980 HMMetlyEutC_3_eut2(pf05985) EUT2E
05985 HMM13520tp_1_pdu(pf13520) EUT2E
05990 -pf05504 -
05995 -pf00582 -
06000 -pf07690 -
BMC-P
BMC-H
BMC-T
Aldehyde dehydrogenase
PTAC
Signature enzyme
Alcohol dehydrogenase
Regulator

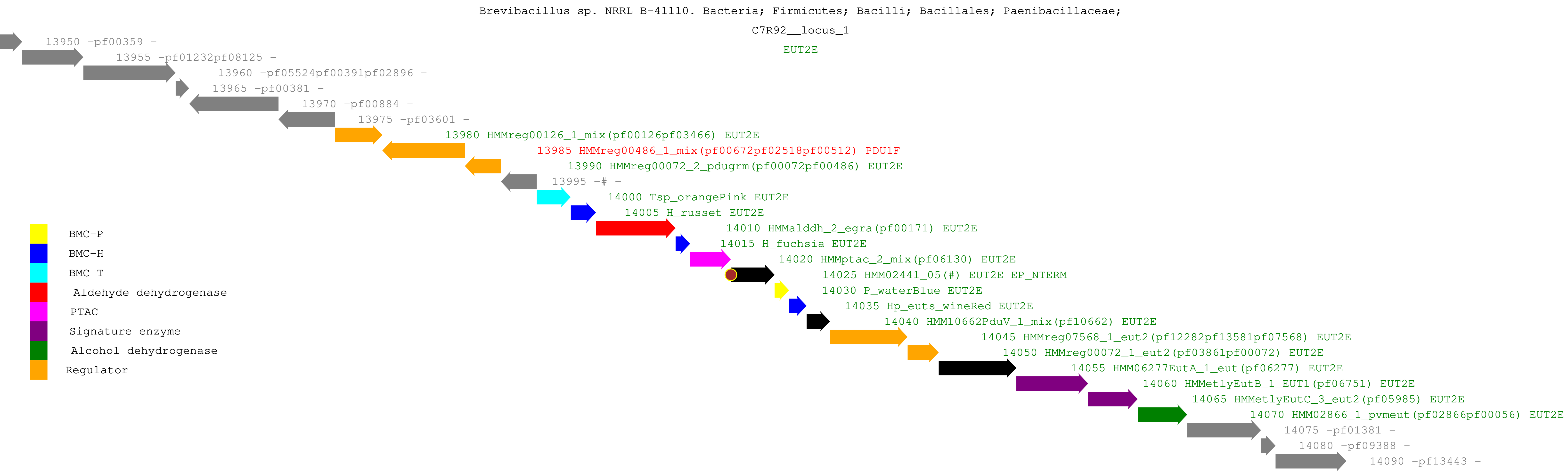
Brevibacillus sp. NRRL B-41110. Bacteria; Firmicutes; Bacilli; Bacillales; Paenibacillaceae;
C7R92__locus_1
EUT2E
13950 -pf00359 -
13955 -pf01232pf08125 -
13960 -pf05524pf00391pf02896 -
13965 -pf00381 -
13970 -pf00884 -
13975 -pf03601 -
13980 HMMreg00126_1_mix(pf00126pf03466) EUT2E
13985 HMMreg00486_1_mix(pf00672pf02518pf00512) PDU1F
13990 HMMreg00072_2_pdugrm(pf00072pf00486) EUT2E
13995 -# -
14000 Tsp_orangePink EUT2E
14005 H_russet EUT2E
14010 HMMalddh_2_egra(pf00171) EUT2E
14015 H_fuchsia EUT2E
14020 HMMptac_2_mix(pf06130) EUT2E
14025 HMM02441_05(#) EUT2E EP_NTERM
14030 P_waterBlue EUT2E
14035 Hp_euts_wineRed EUT2E
14040 HMM10662PduV_1_mix(pf10662) EUT2E
14045 HMMreg07568_1_eut2(pf12282pf13581pf07568) EUT2E
14050 HMMreg00072_1_eut2(pf03861pf00072) EUT2E
14055 HMM06277EutA_1_eut(pf06277) EUT2E
14060 HMMetlyEutB_1_EUT1(pf06751) EUT2E
14065 HMMetlyEutC_3_eut2(pf05985) EUT2E
14070 HMM02866_1_pvmeut(pf02866pf00056) EUT2E
14075 -pf01381 -
14080 -pf09388 -
14090 -pf13443 -
BMC-P
BMC-H
BMC-T
Aldehyde dehydrogenase
PTAC
Signature enzyme
Alcohol dehydrogenase
Regulator

Brevibacillus sp. NRRL NRS-1210. Bacteria; Firmicutes; Bacilli; Bacillales; Paenibacillaceae;

C7R93__locus_1

EUT2E

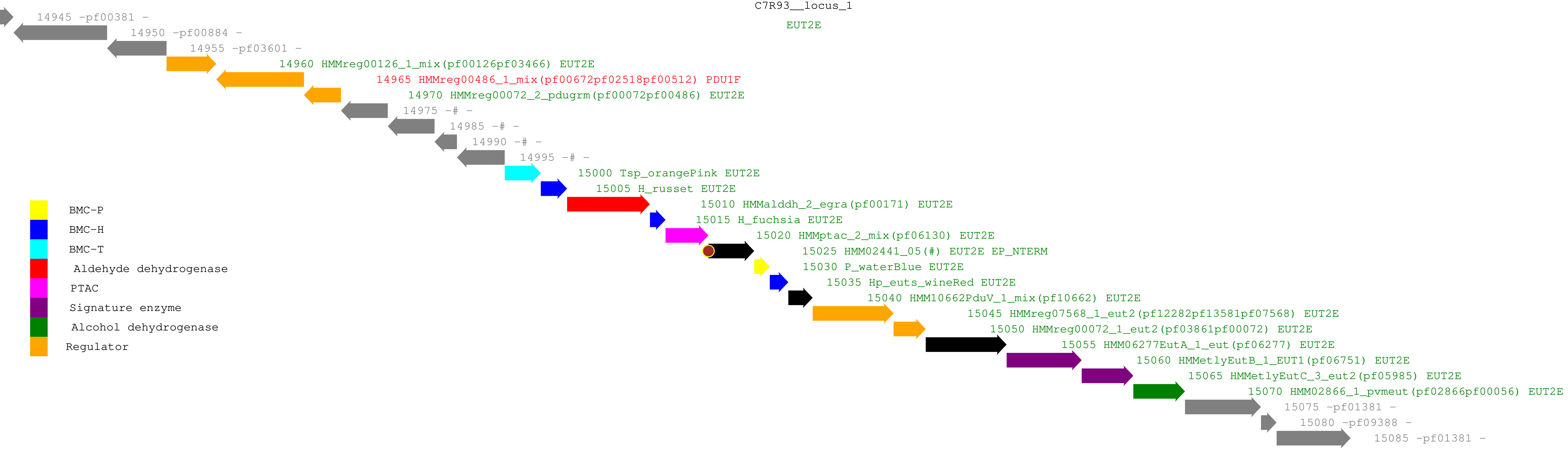

Brevibacillus sp. NRRL NRS-603. Bacteria; Firmicutes; Bacilli; Bacillales; Paenibacillaceae;

C7R94__locus_1

EUT2E

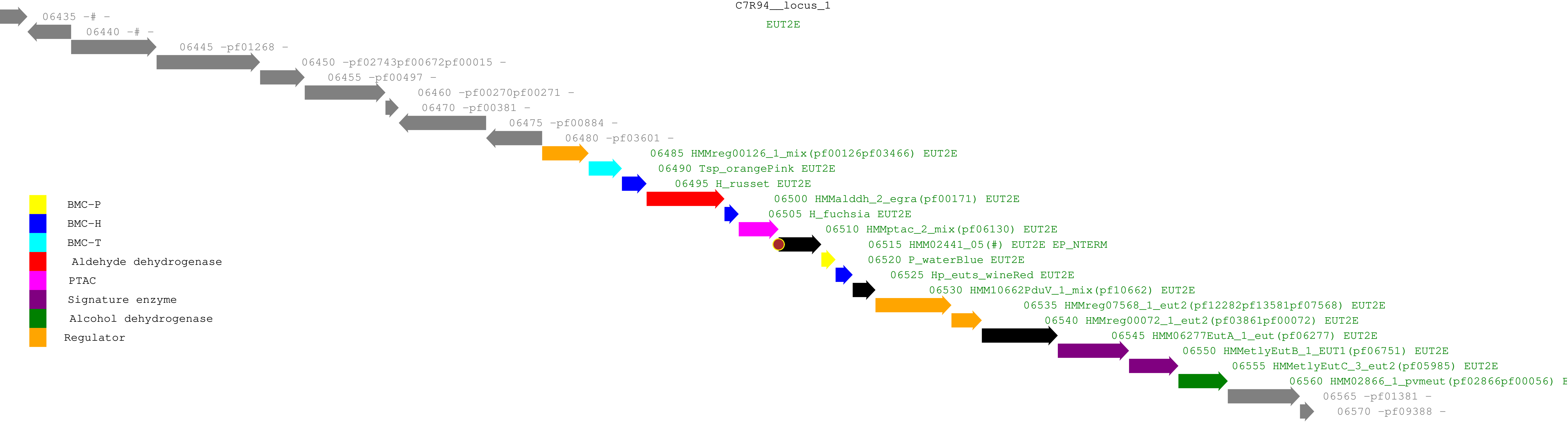

Lysinibacillus sp. SDF0063. Bacteria; Firmicutes; Bacilli; Bacillales; Bacillaceae; Lysinibacillus;

C7Y45__locus_1

EUT2E

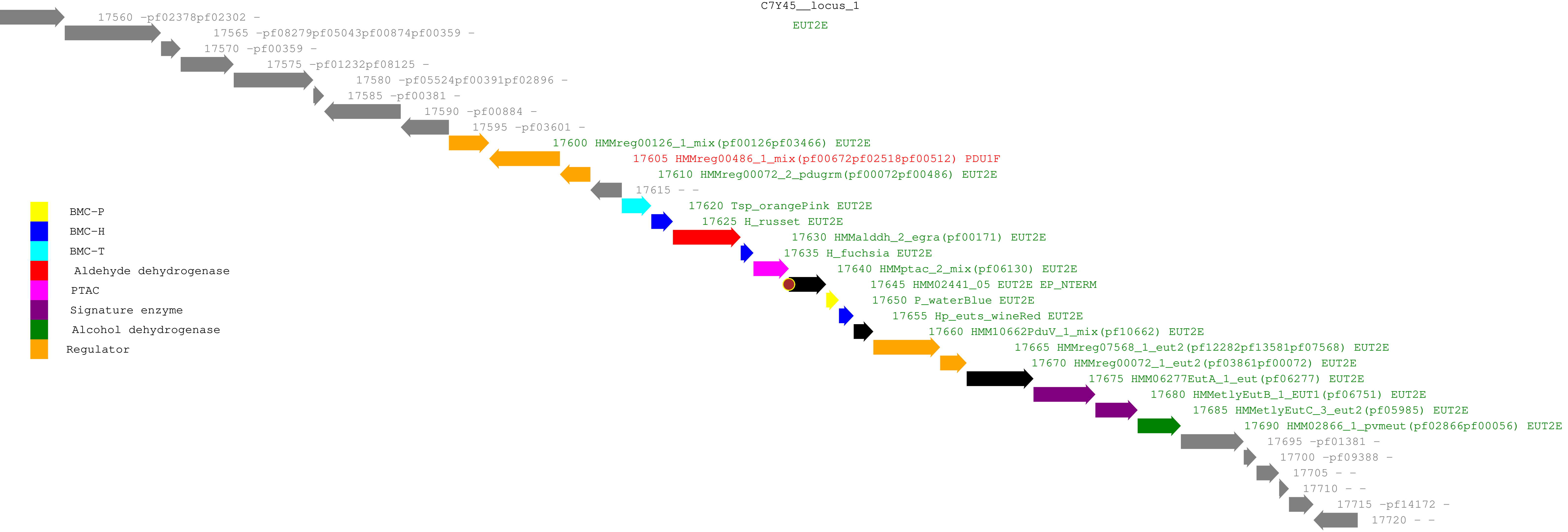

Paenibacillus elgii. Bacteria; Firmicutes; Bacilli; Bacillales; Paenibacillaceae;

C8Z91__locus_1

EUT2E

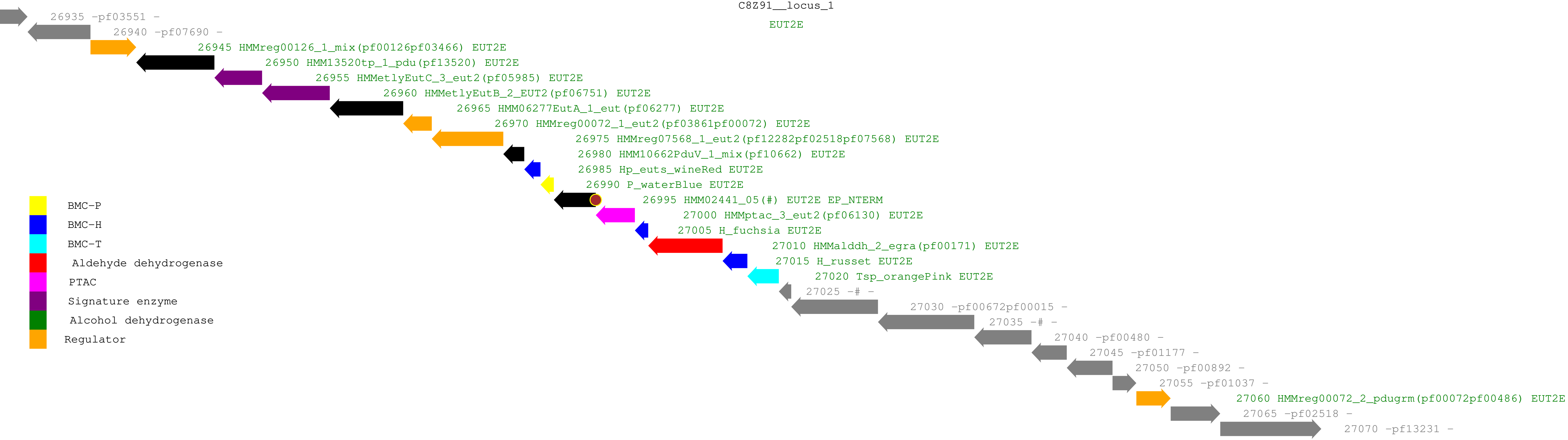

Paenibacillus herberti. Bacteria; Firmicutes; Bacilli; Bacillales; Paenibacillaceae;

CGZ75__locus_1

EUT2E

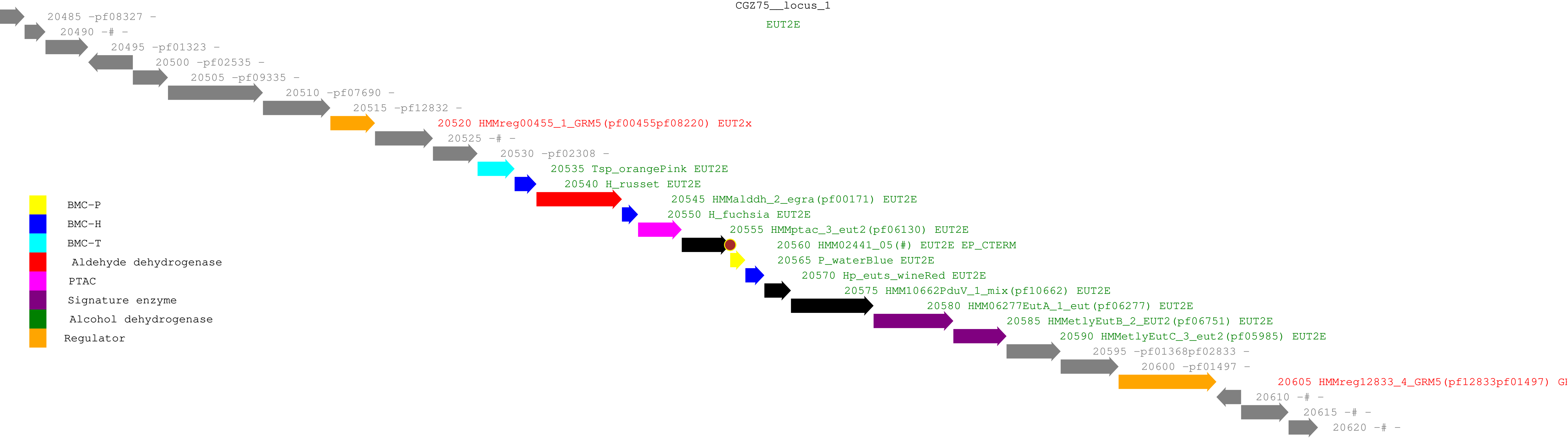

Brevibacillus agri BAB-2500. Bacteria; Firmicutes; Bacilli; Bacillales; Paenibacillaceae;

D478__locus_1

EUT2E

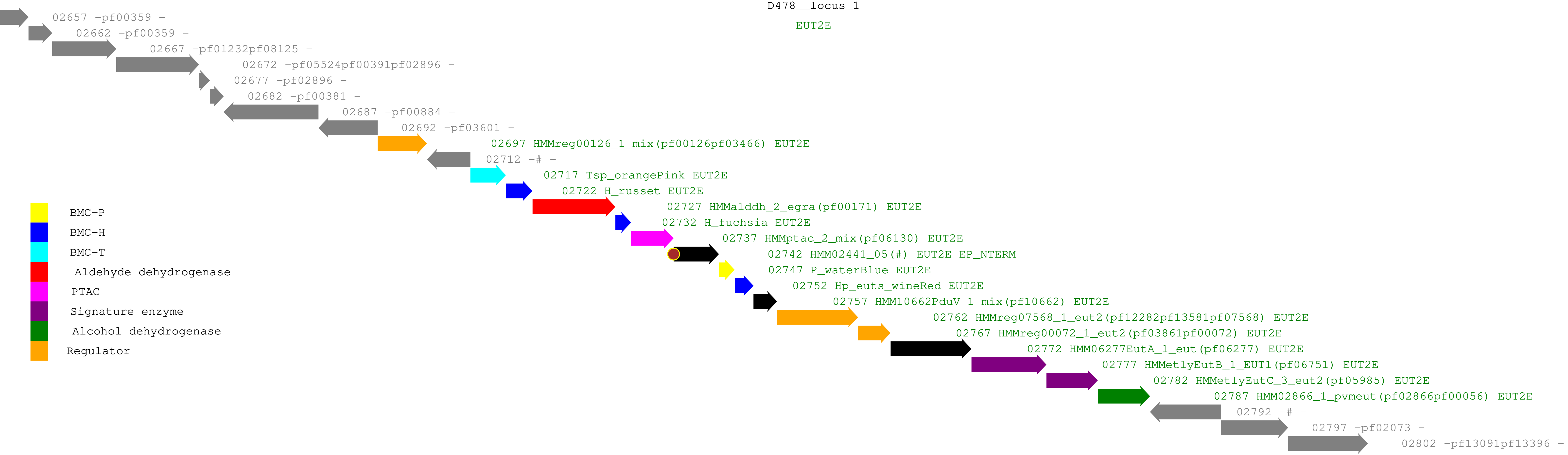

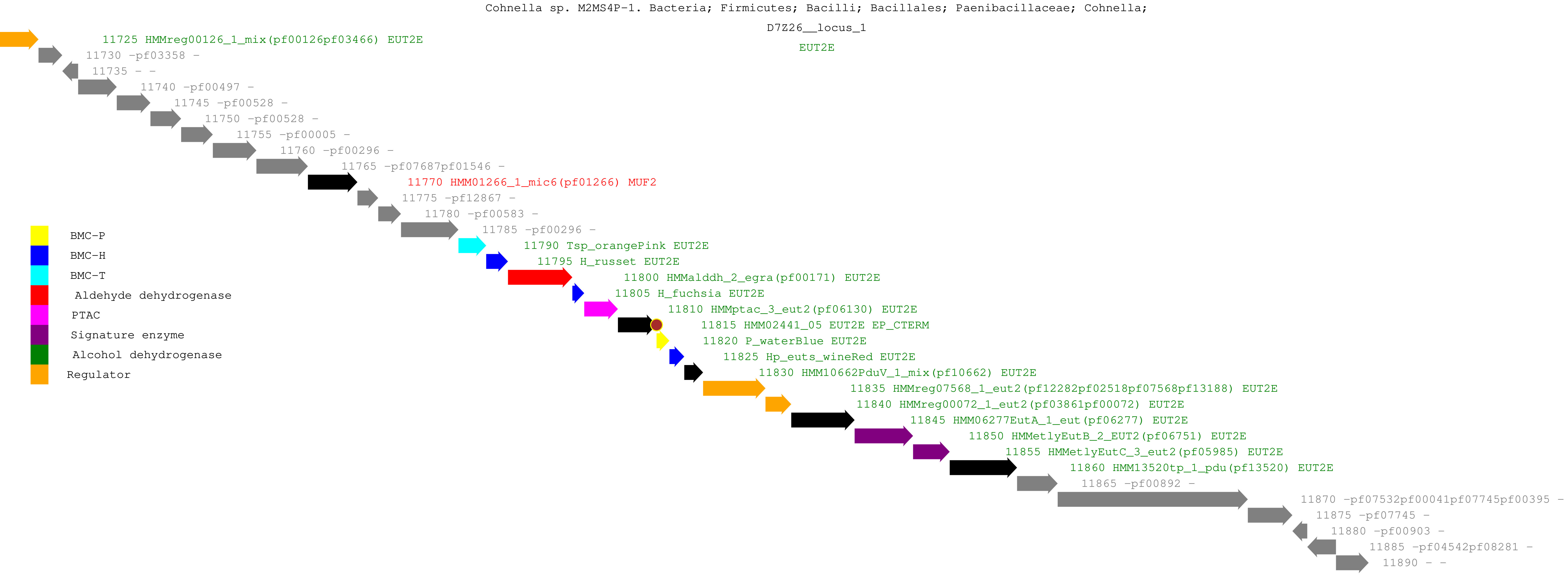
Cohnella sp. M2MS4P-1. Bacteria; Firmicutes; Bacilli; Bacillales; Paenibacillaceae; Cohnella;
D7Z26__locus_1
EUT2E
11725 HMMreg00126_1_mix(pf00126pf03466) EUT2E
11730 -pf03358 -
11735 - -
11740 -pf00497 -
11745 -pf00528 -
11750 -pf00528 -
11755 -pf00005 -
11760 -pf00296 -
11765 -pf07687pf01546 -
11770 HMM01266_1_mic6(pf01266) MUF2
11775 -pf12867 -
11780 -pf00583 -
11785 -pf00296 -
11790 Tsp_orangePink EUT2E
11795 H_russet EUT2E
11800 HMMalddh_2_egra(pf00171) EUT2E
11805 H_fuchsia EUT2E
11810 HMMptac_3_eut2(pf06130) EUT2E
11815 HMM02441_05 EUT2E EP_CTERM
11820 P_waterBlue EUT2E
11825 Hp_euts_wineRed EUT2E
11830 HMM10662PduV_1_mix(pf10662) EUT2E
11835 HMMreg07568_1_eut2(pf12282pf02518pf07568pf13188) EUT2E
11840 HMMreg00072_1_eut2(pf03861pf00072) EUT2E
11845 HMM06277EutA_1_eut(pf06277) EUT2E
11850 HMMetlyEutB_2_EUT2(pf06751) EUT2E
11855 HMMetlyEutC_3_eut2(pf05985) EUT2E
11860 HMM13520tp_1_pdu(pf13520) EUT2E
11865 -pf00892 -
11870 -pf07532pf00041pf07745pf00395 -
11875 -pf07745 -
11880 -pf00903 -
11885 -pf04542pf08281 -
11890 - -
BMC-P
BMC-H
BMC-T
Aldehyde dehydrogenase
PTAC
Signature enzyme
Alcohol dehydrogenase
Regulator

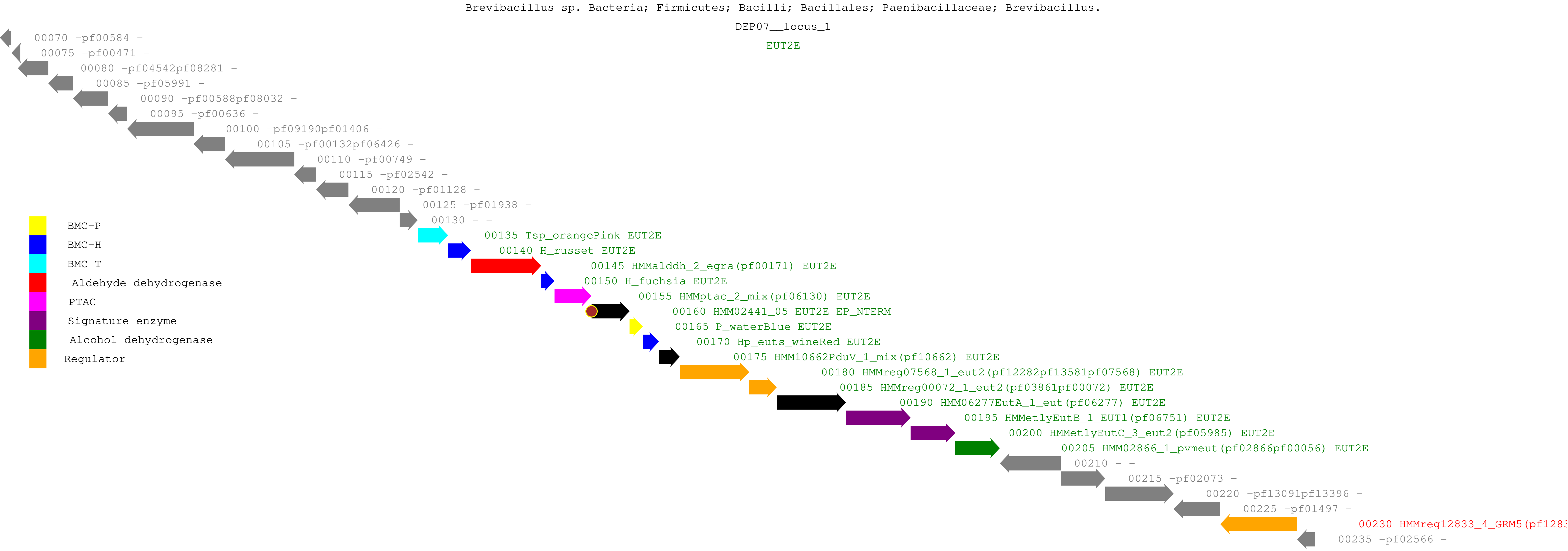
Brevibacillus sp. Bacteria; Firmicutes; Bacilli; Bacillales; Paenibacillaceae; Brevibacillus.
DEP07__locus_1
EUT2E
00070 -pf00584 -
00075 -pf00471 -
00080 -pf04542pf08281 -
00085 -pf05991 -
00090 -pf00588pf08032 -
00095 -pf00636 -
00100 -pf09190pf01406 -
00105 -pf00132pf06426 -
00110 -pf00749 -
00115 -pf02542 -
00120 -pf01128 -
00125 -pf01938 -
00130 - -
00135 Tsp_orangePink EUT2E
00140 H_russet EUT2E
00145 HMMalddh_2_egra(pf00171) EUT2E
00150 H_fuchsia EUT2E
00155 HMMptac_2_mix(pf06130) EUT2E
00160 HMM02441_05 EUT2E EP_NTERM
00165 P_waterBlue EUT2E
00170 Hp_euts_wineRed EUT2E
00175 HMM10662PduV_1_mix(pf10662) EUT2E
00180 HMMreg07568_1_eut2(pf12282pf13581pf07568) EUT2E
00185 HMMreg00072_1_eut2(pf03861pf00072) EUT2E
00190 HMM06277EutA_1_eut(pf06277) EUT2E
00195 HMMetlyEutB_1_EUT1(pf06751) EUT2E
00200 HMMetlyEutC_3_eut2(pf05985) EUT2E
00205 HMM02866_1_pvmeut(pf02866pf00056) EUT2E
00210 - -
00215 -pf02073 -
00220 -pf13091pf13396 -
00225 -pf01497 -
00230 HMMreg12833_4_GRM5(pf1283
00235 -pf02566 -
BMC-P
BMC-H
BMC-T
Aldehyde dehydrogenase
PTAC
Signature enzyme
Alcohol dehydrogenase
Regulator

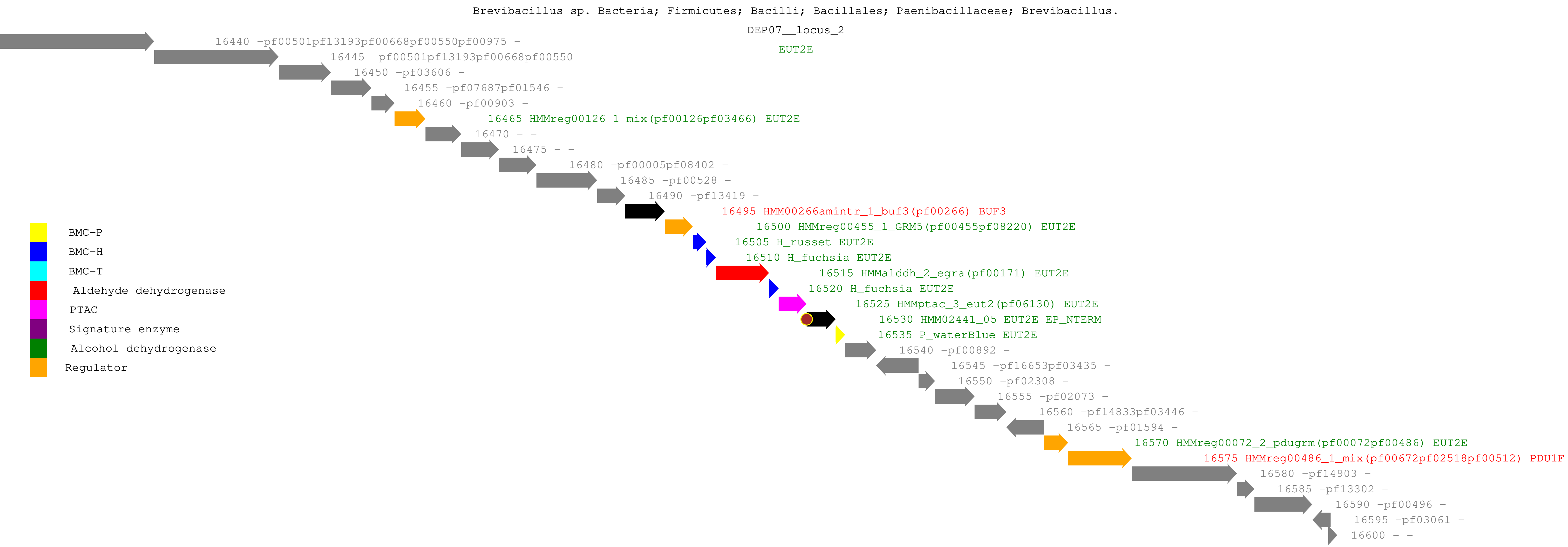
Brevibacillus sp. Bacteria; Firmicutes; Bacilli; Bacillales; Paenibacillaceae; Brevibacillus.
DEP07__locus_2
EUT2E
16440 -pf00501pf13193pf00668pf00550pf00975 -
16445 -pf00501pf13193pf00668pf00550 -
16450 -pf03606 -
16455 -pf07687pf01546 -
16460 -pf00903 -
16465 HMMreg00126_1_mix(pf00126pf03466) EUT2E
16470 - -
16475 - -
16480 -pf00005pf08402 -
16485 -pf00528 -
16490 -pf13419 -
16495 HMM00266amintr_1_buf3(pf00266) BUF3
16500 HMMreg00455_1_GRM5(pf00455pf08220) EUT2E
16505 H_russet EUT2E
16510 H_fuchsia EUT2E
16515 HMMalddh_2_egra(pf00171) EUT2E
16520 H_fuchsia EUT2E
16525 HMMptac_3_eut2(pf06130) EUT2E
16530 HMM02441_05 EUT2E EP_NTERM
16535 P_waterBlue EUT2E
16540 -pf00892 -
16545 -pf16653pf03435 -
16550 -pf02308 -
16555 -pf02073 -
16560 -pf14833pf03446 -
16565 -pf01594 -
16570 HMMreg00072_2_pdugrm(pf00072pf00486) EUT2E
16575 HMMreg00486_1_mix(pf00672pf02518pf00512) PDU1F
16580 -pf14903 -
16585 -pf13302 -
16590 -pf00496 -
16595 -pf03061 -
16600 - -
BMC-P
BMC-H
BMC-T
Aldehyde dehydrogenase
PTAC
Signature enzyme
Alcohol dehydrogenase
Regulator

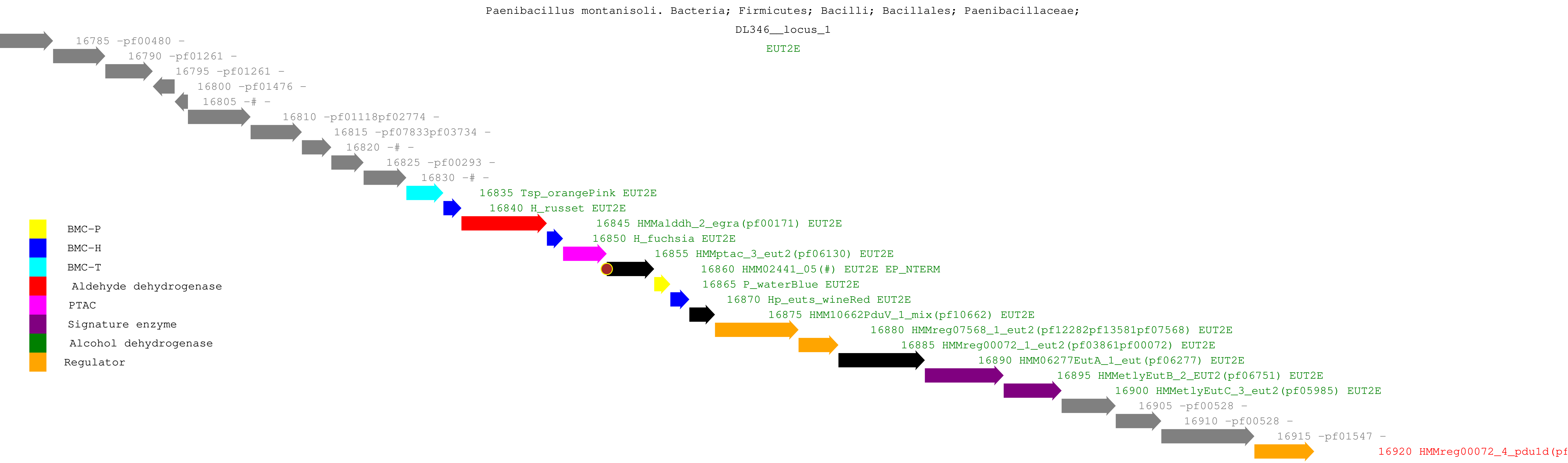

Paenibacillus montanisoli. Bacteria; Firmicutes; Bacilli; Bacillales; Paenibacillaceae;
DL346__locus_1
EUT2E
16785 -pf00480 -
16790 -pf01261 -
16795 -pf01261 -
16800 -pf01476 -
16805 -# -
16810 -pf01118pf02774 -
16815 -pf07833pf03734 -
16820 -# -
16825 -pf00293 -
16830 -# -
16835 Tsp_orangePink EUT2E
16840 H_russet EUT2E
16845 HMMaldd h_2_egra(pf00171) EUT2E
16850 H_fuchsia EUT2E
16855 HMMptac_3_eut2(pf06130) EUT2E
16860 HMM02441_05(#) EUT2E EP_NTERM
16865 P_waterBlue EUT2E
16870 Hp_euts_wineRed EUT2E
16875 HMM10662PduV_1_mix(pf10662) EUT2E
16880 HMMreg07568_1_eut2(pf12282pf13581pf07568) EUT2E
16885 HMMreg00072_1_eut2(pf03861pf00072) EUT2E
16890 HMM06277EutA_1_eut(pf06277) EUT2E
16895 HMMetlyEutB_2_EUT2(pf06751) EUT2E
16900 HMMetlyEutC_3_eut2(pf05985) EUT2E
16905 -pf00528 -
16910 -pf00528 -
16915 -pf01547 -
16920 HMMreg00072_4_pdu1d(pf
BMC-P
BMC-H
BMC-T
Aldehyde dehydrogenase
PTAC
Signature enzyme
Alcohol dehydrogenase
Regulator

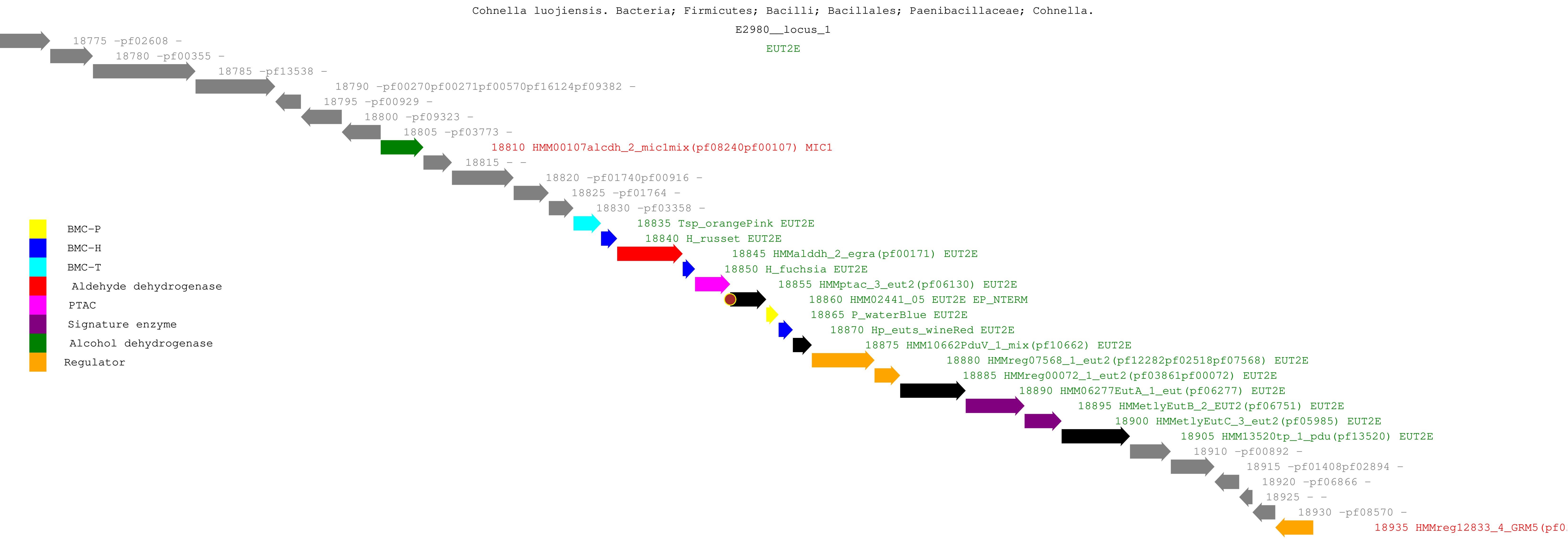

Cohnella luojiensis. Bacteria; Firmicutes; Bacilli; Bacillales; Paenibacillaceae; Cohnella.
E2980__locus_1
EUT2E
18775 -pf02608 -
18780 -pf00355 -
18785 -pf13538 -
18790 -pf00270pf00271pf00570pf16124pf09382 -
18795 -pf00929 -
18800 -pf09323 -
18805 -pf03773 -
18810 HMM00107alcdh_2_mic1mix(pf08240pf00107) MIC1
18815 - -
18820 -pf01740pf00916 -
18825 -pf01764 -
18830 -pf03358 -
18835 Tsp_orangePink EUT2E
18840 H_russet EUT2E
18845 HMMaldh_2_egra(pf00171) EUT2E
18850 H_fuchsia EUT2E
18855 HMMptac_3_eut2(pf06130) EUT2E
18860 HMM02441_05 EUT2E EP_NTERM
18865 P_waterBlue EUT2E
18870 Hp_euts_wineRed EUT2E
18875 HMM10662PduV_1_mix(pf10662) EUT2E
18880 HMMreg07568_1_eut2(pf12282pf02518pf07568) EUT2E
18885 HMMreg00072_1_eut2(pf03861pf00072) EUT2E
18890 HMM06277EutA_1_eut(pf06277) EUT2E
18895 HMMetlyEutB_2_EUT2(pf06751) EUT2E
18900 HMMetlyEutC_3_eut2(pf05985) EUT2E
18905 HMM13520tp_1_pdu(pf13520) EUT2E
18910 -pf00892 -
18915 -pf01408pf02894 -
18920 -pf06866 -
18925 - -
18930 -pf08570 -
18935 HMMreg12833_4_GRM5(pf0
BMC-P
BMC-H
BMC-T
Aldehyde dehydrogenase
PTAC
Signature enzyme
Alcohol dehydrogenase
Regulator

Cohnella sp. CAU 1483. Bacteria; Firmicutes; Bacilli; Bacillales; Paenibacillaceae; Cohnella;

E5161__locus_1

EUT2E

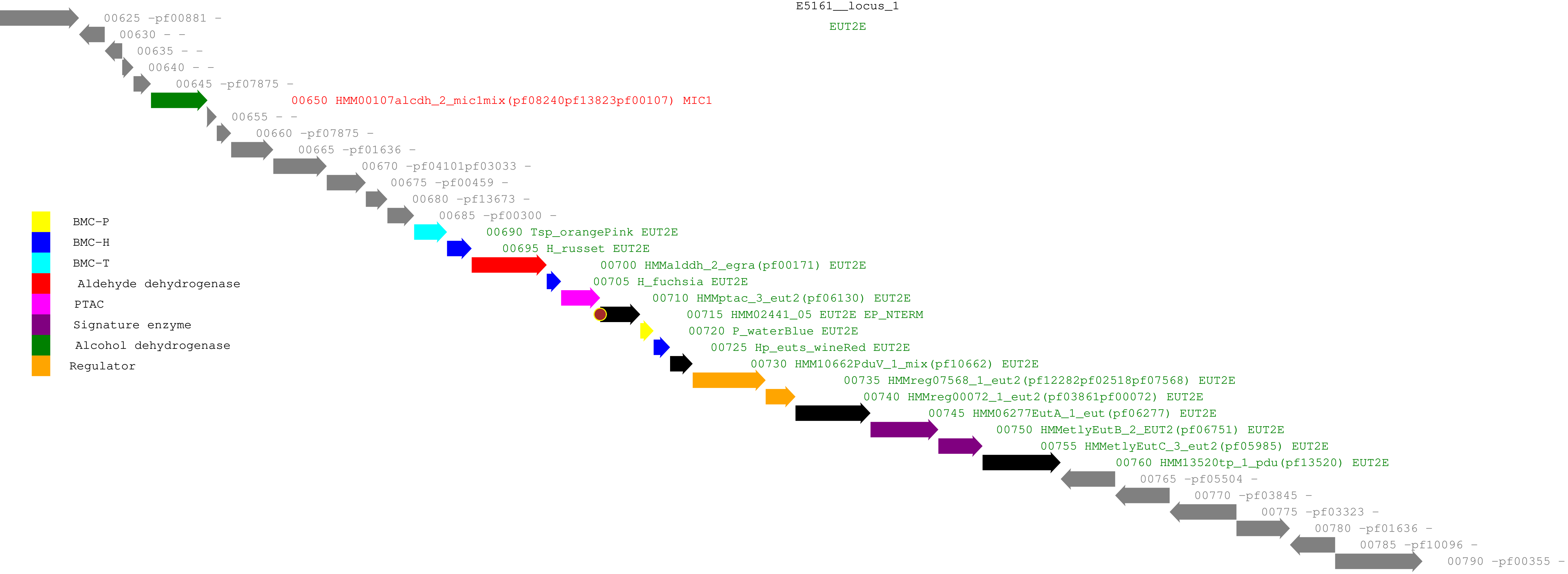

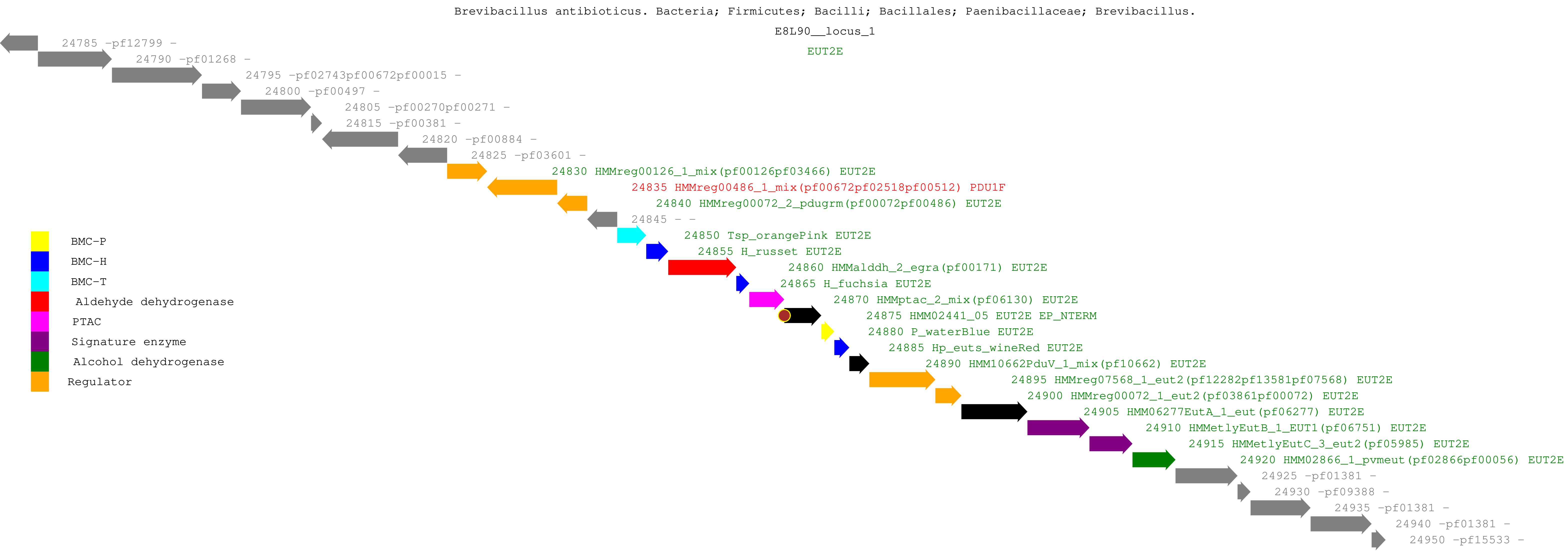

Brevibacillus antibioticus. Bacteria; Firmicutes; Bacilli; Bacillales; Paenibacillaceae; Brevibacillus.
E8L90__locus_1
EUT2E
24785 -pf12799 -
24790 -pf01268 -
24795 -pf02743pf00672pf00015 -
24800 -pf00497 -
24805 -pf00270pf00271 -
24815 -pf00381 -
24820 -pf00884 -
24825 -pf03601 -
24830 HMMreg00126_1_mix(pf00126pf03466) EUT2E
24835 HMMreg00486_1_mix(pf00672pf02518pf00512) PDU1F
24840 HMMreg00072_2_pdugrm(pf00072pf00486) EUT2E
24845 - -
24850 Tsp_orangePink EUT2E
24855 H_russet EUT2E
24860 HMMaldh_2_egra(pf00171) EUT2E
24865 H_fuchsia EUT2E
24870 HMMptac_2_mix(pf06130) EUT2E
24875 HMM02441_05 EUT2E EP_NTERM
24880 P_waterBlue EUT2E
24885 Hp_euts_wineRed EUT2E
24890 HMM10662PduV_1_mix(pf10662) EUT2E
24895 HMMreg07568_1_eut2(pf12282pf13581pf07568) EUT2E
24900 HMMreg00072_1_eut2(pf03861pf00072) EUT2E
24905 HMM06277EutA_1_eut(pf06277) EUT2E
24910 HMMetlyEutB_1_EUT1(pf06751) EUT2E
24915 HMMetlyEutC_3_eut2(pf05985) EUT2E
24920 HMM02866_1_pvmeut(pf02866pf00056) EUT2E
24925 -pf01381 -
24930 -pf09388 -
24935 -pf01381 -
24940 -pf01381 -
24950 -pf15533 -
BMC-P
BMC-H
BMC-T
Aldehyde dehydrogenase
PTAC
Signature enzyme
Alcohol dehydrogenase
Regulator

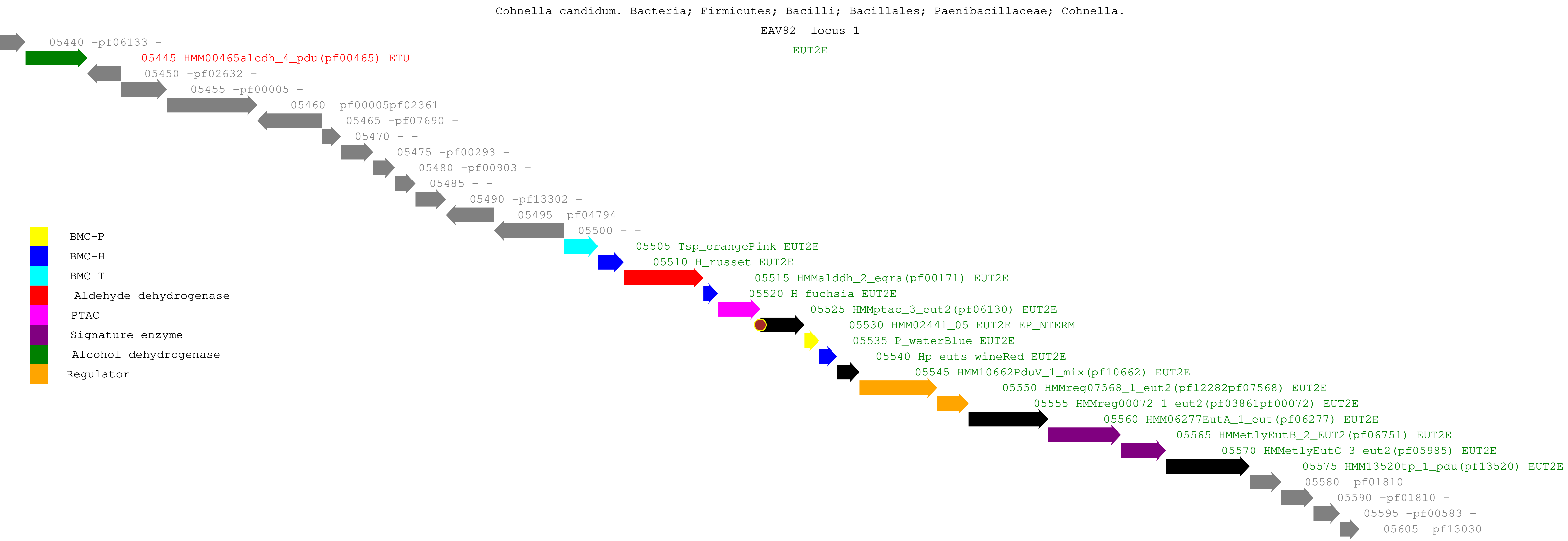
Cohnella candidum. Bacteria; Firmicutes; Bacilli; Bacillales; Paenibacillaceae; Cohnella.
EAV92__locus_1
EUT2E
05440 -pf06133 -
05445 HMM00465alcdh_4_pdu(pf00465) ETU
05450 -pf02632 -
05455 -pf00005 -
05460 -pf00005pf02361 -
05465 -pf07690 -
05470 - -
05475 -pf00293 -
05480 -pf00903 -
05485 - -
05490 -pf13302 -
05495 -pf04794 -
05500 - -
05505 Tsp_orangePink EUT2E
05510 H_russet EUT2E
05515 HMMaldh_2_egra(pf00171) EUT2E
05520 H_fuchsia EUT2E
05525 HMMptac_3_eut2(pf06130) EUT2E
05530 HMM02441_05 EUT2E EP_NTERM
05535 P_waterBlue EUT2E
05540 Hp_euts_wineRed EUT2E
05545 HMM10662PduV_1_mix(pf10662) EUT2E
05550 HMMreg07568_1_eut2(pf12282pf07568) EUT2E
05555 HMMreg00072_1_eut2(pf03861pf00072) EUT2E
05560 HMM06277EutA_1_eut(pf06277) EUT2E
05565 HMMetlyEutB_2_EUT2(pf06751) EUT2E
05570 HMMetlyEutC_3_eut2(pf05985) EUT2E
05575 HMM13520tp_1_pdu(pf13520) EUT2E
05580 -pf01810 -
05590 -pf01810 -
05595 -pf00583 -
05605 -pf13030 -
BMC-P
BMC-H
BMC-T
Aldehyde dehydrogenase
PTAC
Signature enzyme
Alcohol dehydrogenase
Regulator

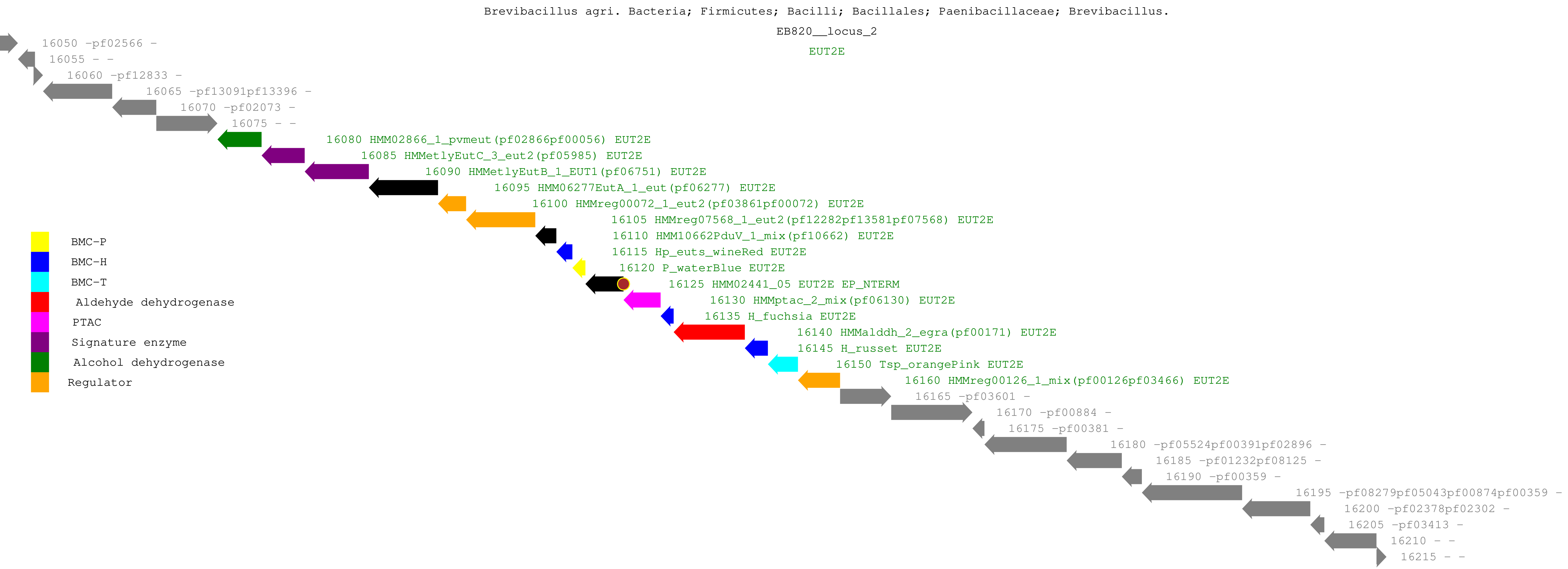
Brevibacillus agri. Bacteria; Firmicutes; Bacilli; Bacillales; Paenibacillaceae; Brevibacillus.
EB820__locus_2
EUT2E
16050 -pf02566 -
16055 - -
16060 -pf12833 -
16065 -pf13091pf13396 -
16070 -pf02073 -
16075 - -
16080 HMM02866_1_pvmeut(pf02866pf00056) EUT2E
16085 HMMetlyEutC_3_eut2(pf05985) EUT2E
16090 HMMetlyEutB_1_EUT1(pf06751) EUT2E
16095 HMM06277EutA_1_eut(pf06277) EUT2E
16100 HMMreg00072_1_eut2(pf03861pf00072) EUT2E
16105 HMMreg07568_1_eut2(pf12282pf13581pf07568) EUT2E
16110 HMM10662PduV_1_mix(pf10662) EUT2E
16115 Hp_euts_wineRed EUT2E
16120 P_waterBlue EUT2E
16125 HMM02441_05 EUT2E EP_NTERM
16130 HMMptac_2_mix(pf06130) EUT2E
16135 H_fuchsia EUT2E
16140 HMMaldh_2_egra(pf00171) EUT2E
16145 H_russet EUT2E
16150 Tsp_orangePink EUT2E
16160 HMMreg00126_1_mix(pf00126pf03466) EUT2E
16165 -pf03601 -
16170 -pf00884 -
16175 -pf00381 -
16180 -pf05524pf00391pf02896 -
16185 -pf01232pf08125 -
16190 -pf00359 -
16195 -pf08279pf05043pf00874pf00359 -
16200 -pf02378pf02302 -
16205 -pf03413 -
16210 - -
16215 - -
BMC-P
BMC-H
BMC-T
Aldehyde dehydrogenase
PTAC
Signature enzyme
Alcohol dehydrogenase
Regulator

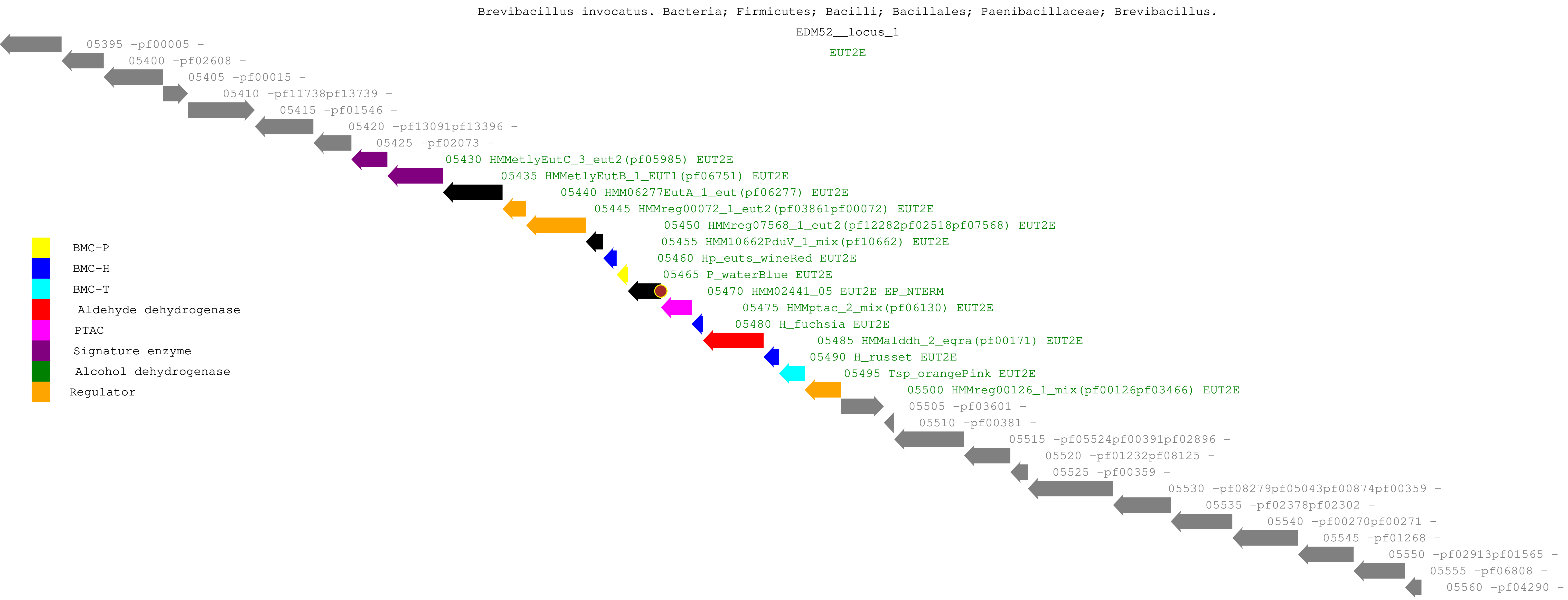
Brevibacillus invocatus. Bacteria; Firmicutes; Bacilli; Bacillales; Paenibacillaceae; Brevibacillus.
EDM52__locus_1
EUT2E
05395 -pf00005 -
05400 -pf02608 -
05405 -pf00015 -
05410 -pf11738pf13739 -
05415 -pf01546 -
05420 -pf13091pf13396 -
05425 -pf02073 -
05430 HMMetlyEutC_3_eut2(pf05985) EUT2E
05435 HMMetlyEutB_1_EUT1(pf06751) EUT2E
05440 HMM06277EutA_1_eut(pf06277) EUT2E
05445 HMMreg00072_1_eut2(pf03861pf00072) EUT2E
05450 HMMreg07568_1_eut2(pf12282pf02518pf07568) EUT2E
05455 HMM10662PduV_1_mix(pf10662) EUT2E
05460 Hp_euts_wineRed EUT2E
05465 P_waterBlue EUT2E
05470 HMM02441_05 EUT2E EP_NTERM
05475 HMMptac_2_mix(pf06130) EUT2E
05480 H_fuchsia EUT2E
05485 HMMalddh_2_egra(pf00171) EUT2E
05490 H_russet EUT2E
05495 Tsp_orangePink EUT2E
05500 HMMreg00126_1_mix(pf00126pf03466) EUT2E
05505 -pf03601 -
05510 -pf00381 -
05515 -pf05524pf00391pf02896 -
05520 -pf01232pf08125 -
05525 -pf00359 -
05530 -pf08279pf05043pf00874pf00359 -
05535 -pf02378pf02302 -
05540 -pf00270pf00271 -
05545 -pf01268 -
05550 -pf02913pf01565 -
05555 -pf06808 -
05560 -pf04290 -
BMC-P
BMC-H
BMC-T
Aldehyde dehydrogenase
PTAC
Signature enzyme
Alcohol dehydrogenase
Regulator

Brevibacillus fluminis. Bacteria; Firmicutes; Bacilli; Bacillales; Paenibacillaceae; Brevibacillus.

EDM56__locus_1

EUT2E

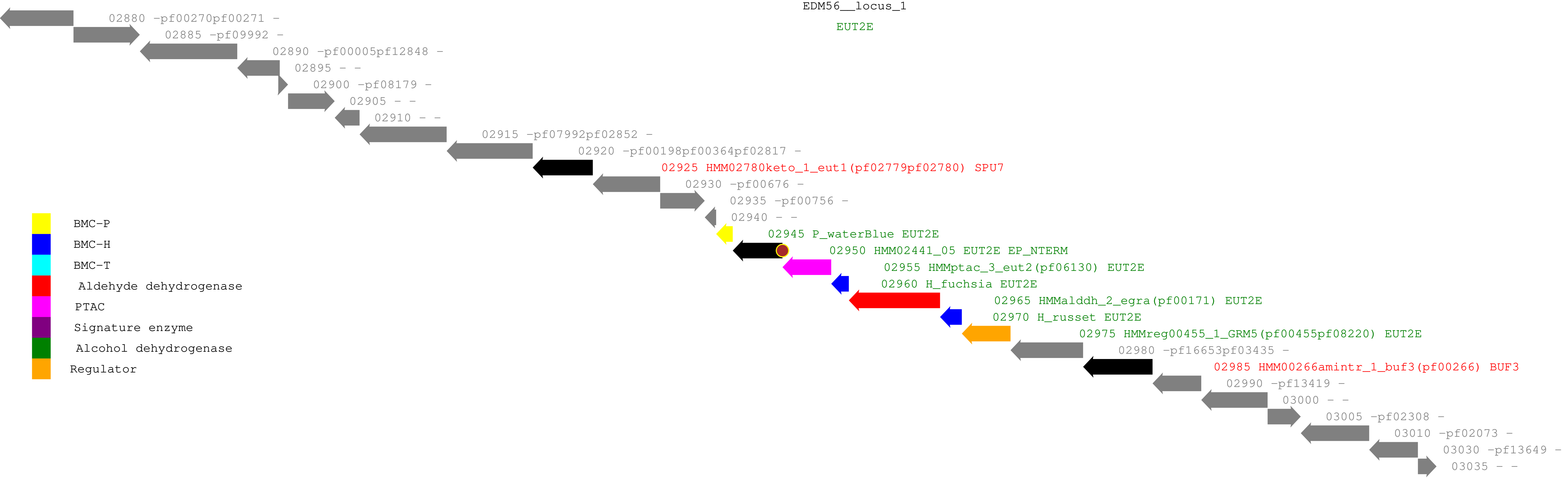

Brevibacillus fluminis. Bacteria; Firmicutes; Bacilli; Bacillales; Paenibacillaceae; Brevibacillus.

EDM56__locus_2

EUT2E

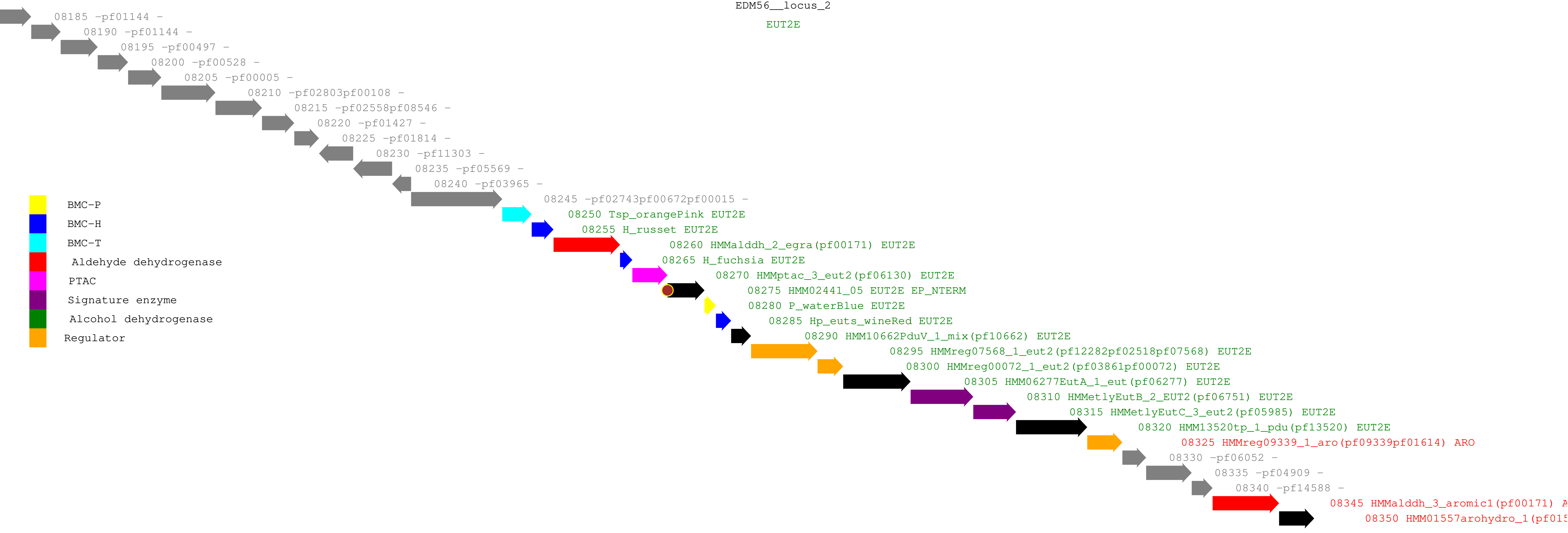

Brevibacillus gelatini. Bacteria; Firmicutes; Bacilli; Bacillales; Paenibacillaceae; Brevibacillus.

EDM57__locus_1

EUT2E

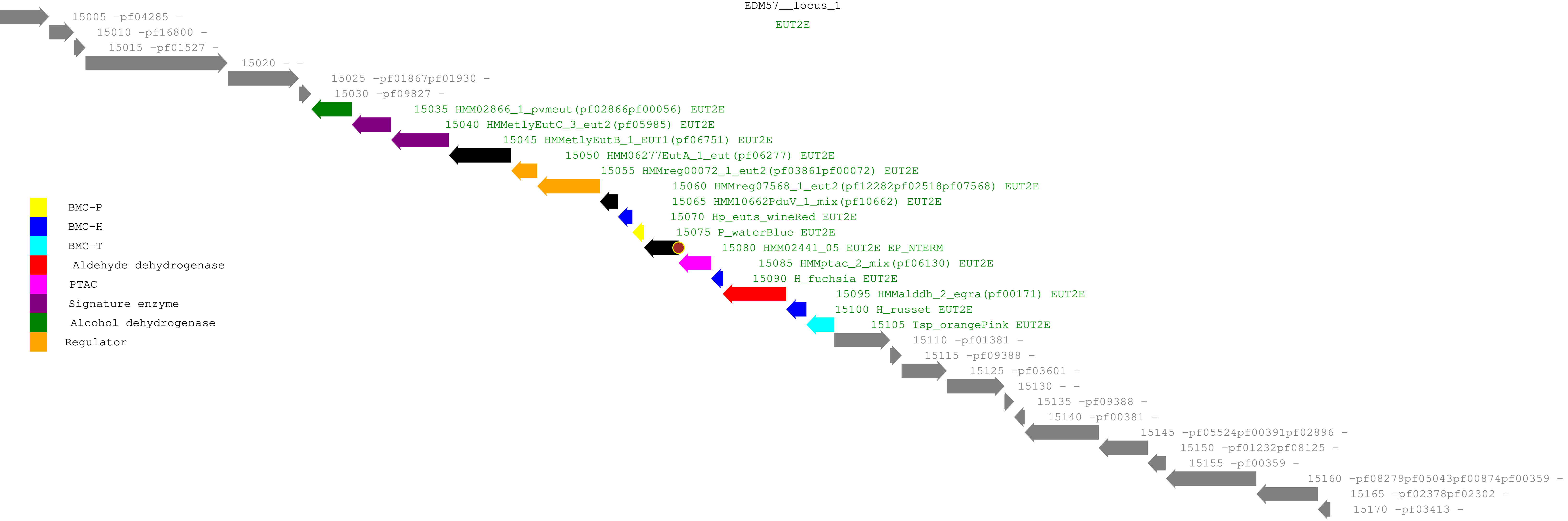

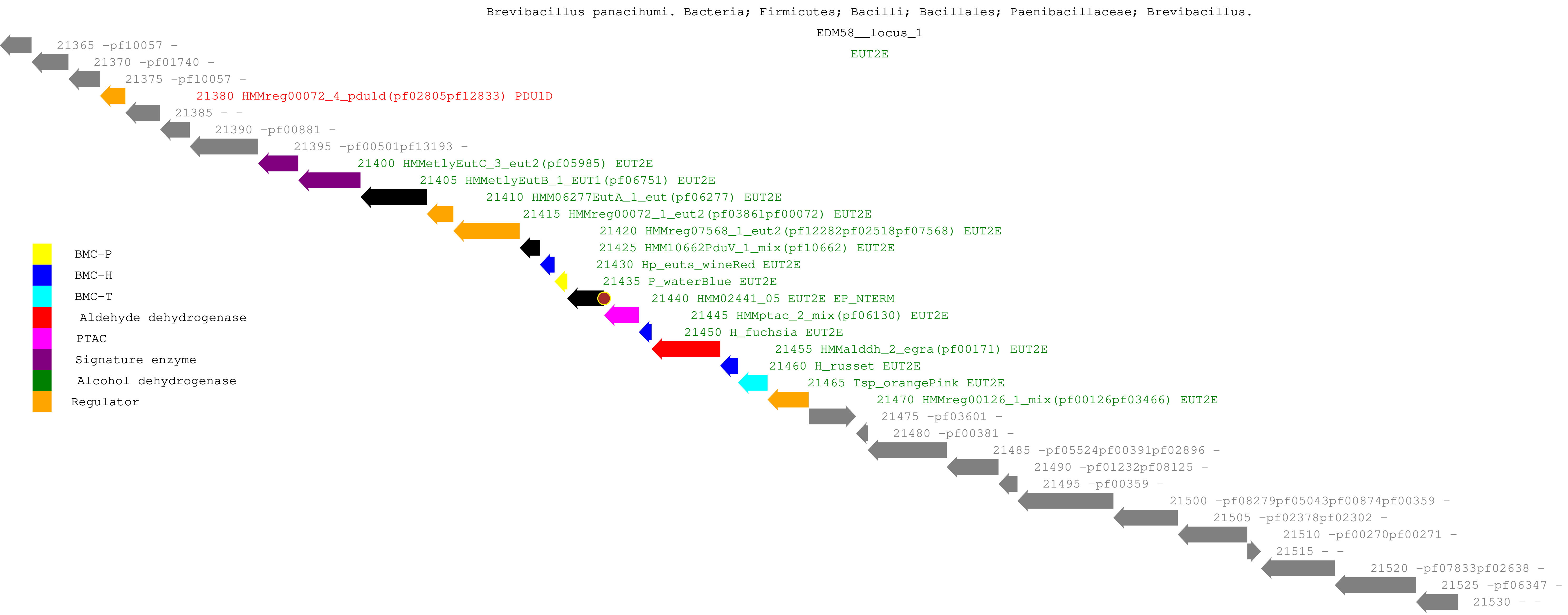
Brevibacillus panacihumi. Bacteria; Firmicutes; Bacilli; Bacillales; Paenibacillaceae; Brevibacillus.
EDM58__locus_1
EUT2E
21365 -pf10057 -
21370 -pf01740 -
21375 -pf10057 -
21380 HMMreg00072_4_pdu1d(pf02805pf12833) PDU1D
21385 - -
21390 -pf00881 -
21395 -pf00501pf13193 -
21400 HMMetlyEutC_3_eut2(pf05985) EUT2E
21405 HMMetlyEutB_1_EUT1(pf06751) EUT2E
21410 HMM06277EutA_1_eut(pf06277) EUT2E
21415 HMMreg00072_1_eut2(pf03861pf00072) EUT2E
21420 HMMreg07568_1_eut2(pf12282pf02518pf07568) EUT2E
21425 HMM10662PduV_1_mix(pf10662) EUT2E
21430 Hp_euts_wineRed EUT2E
21435 P_waterBlue EUT2E
21440 HMM02441_05 EUT2E EP_NTERM
21445 HMMptac_2_mix(pf06130) EUT2E
21450 H_fuchsia EUT2E
21455 HMMalddh_2_egra(pf00171) EUT2E
21460 H_russet EUT2E
21465 Tsp_orangePink EUT2E
21470 HMMreg00126_1_mix(pf00126pf03466) EUT2E
21475 -pf03601 -
21480 -pf00381 -
21485 -pf05524pf00391pf02896 -
21490 -pf01232pf08125 -
21495 -pf00359 -
21500 -pf08279pf05043pf00874pf00359 -
21505 -pf02378pf02302 -
21510 -pf00270pf00271 -
21515 - -
21520 -pf07833pf02638 -
21525 -pf06347 -
21530 - -
BMC-P
BMC-H
BMC-T
Aldehyde dehydrogenase
PTAC
Signature enzyme
Alcohol dehydrogenase
Regulator

Brevibacillus nitrificans. Bacteria; Firmicutes; Bacilli; Bacillales; Paenibacillaceae; Brevibacillus.

EDM59__locus_1

EUT2E

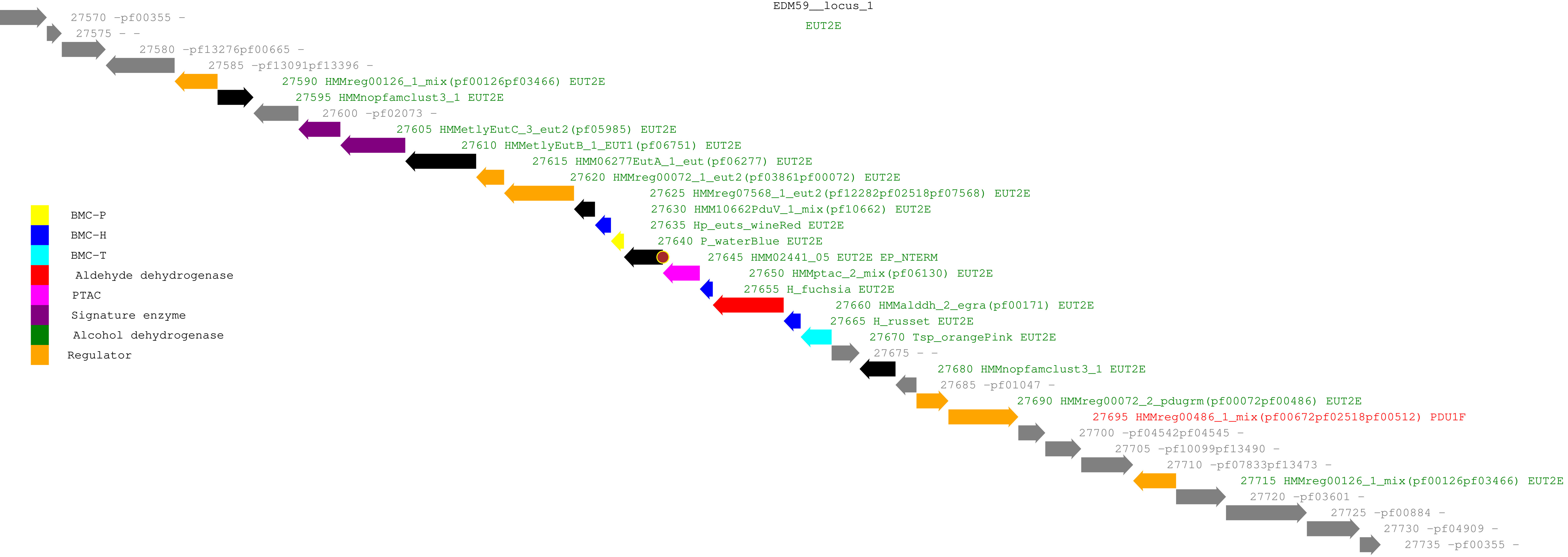

Cohnella sp. AR92. Bacteria; Firmicutes; Bacilli; Bacillales; Paenibacillaceae; Cohnella;

ELR57__locus_2

EUT2E

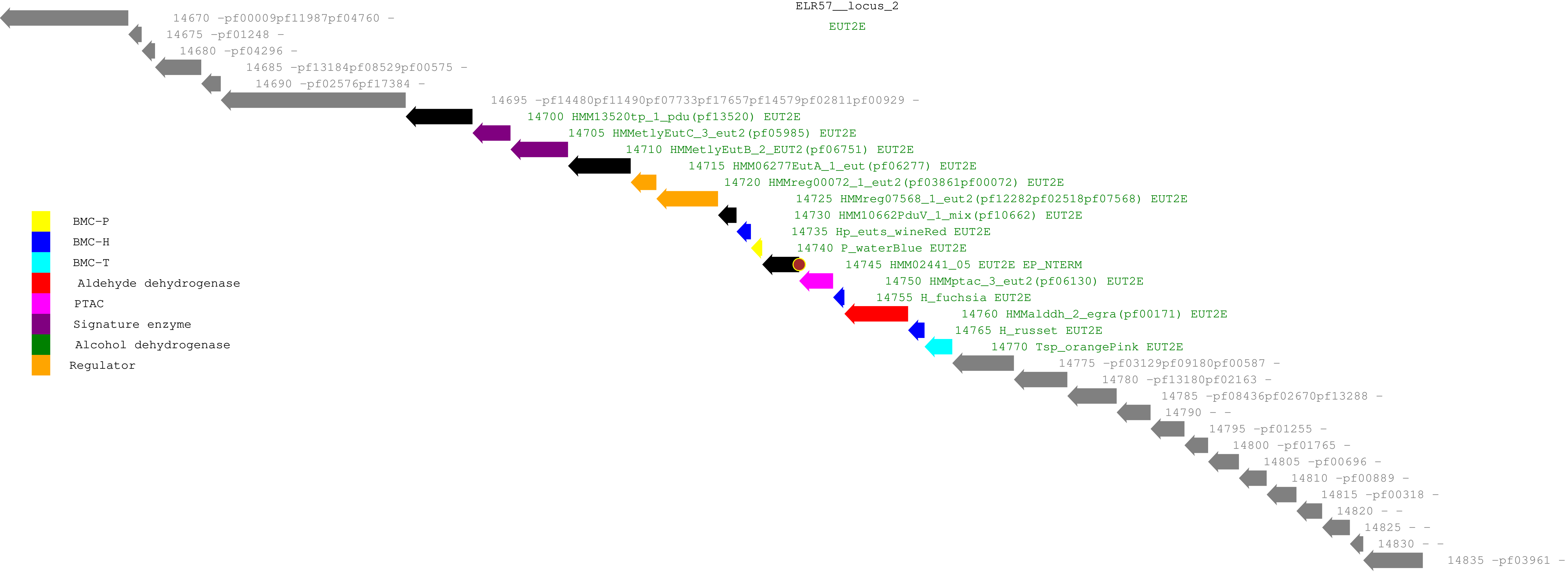

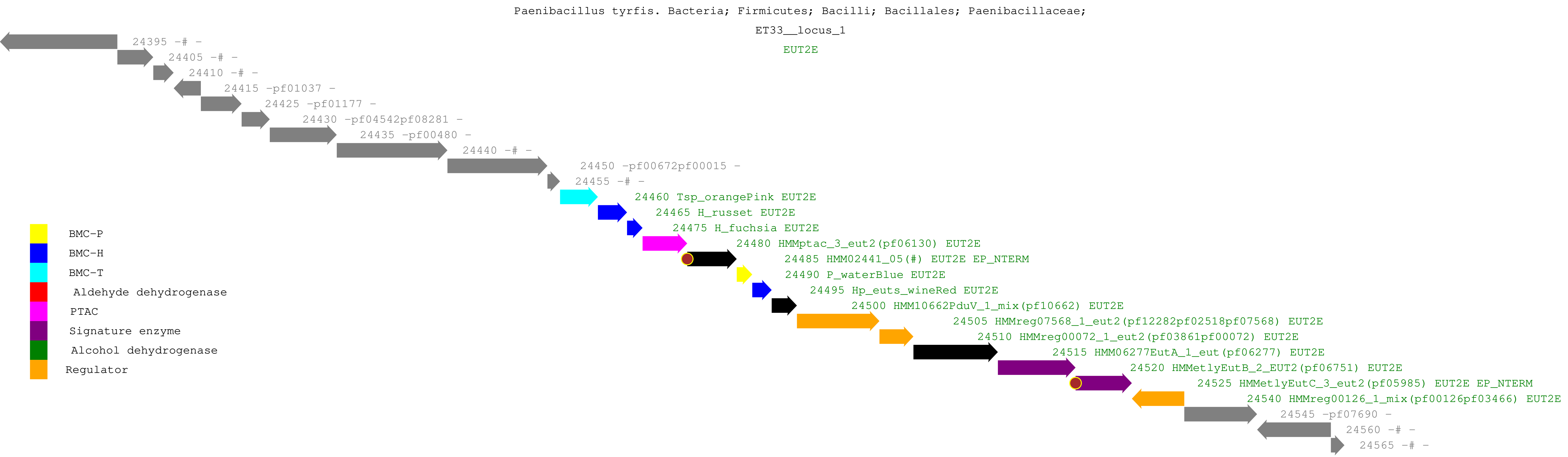
Paenibacillus tyrfis. Bacteria; Firmicutes; Bacilli; Bacillales; Paenibacillaceae;
ET33__locus_1
EUT2E
24395 -# -
24405 -# -
24410 -# -
24415 -pf01037 -
24425 -pf01177 -
24430 -pf04542pf08281 -
24435 -pf00480 -
24440 -# -
24450 -pf00672pf00015 -
24455 -# -
24460 Tsp_orangePink EUT2E
24465 H_russet EUT2E
24475 H_fuchsia EUT2E
24480 HMMptac_3_eut2(pf06130) EUT2E
24485 HMM02441_05(#) EUT2E EP_NTERM
24490 P_waterBlue EUT2E
24495 Hp_euts_wineRed EUT2E
24500 HMM10662PduV_1_mix(pf10662) EUT2E
24505 HMMreg07568_1_eut2(pf12282pf02518pf07568) EUT2E
24510 HMMreg00072_1_eut2(pf03861pf00072) EUT2E
24515 HMM06277EutA_1_eut(pf06277) EUT2E
24520 HMMetlyEutB_2_EUT2(pf06751) EUT2E
24525 HMMetlyEutC_3_eut2(pf05985) EUT2E EP_NTERM
24540 HMMreg00126_1_mix(pf00126pf03466) EUT2E
24545 -pf07690 -
24560 -# -
24565 -# -
BMC-P
BMC-H
BMC-T
Aldehyde dehydrogenase
PTAC
Signature enzyme
Alcohol dehydrogenase
Regulator

Brevibacillus sp. AG162. Bacteria; Firmicutes; Bacilli; Bacillales; Paenibacillaceae; Brevibacillus.

FB479__locus_1

EUT2E

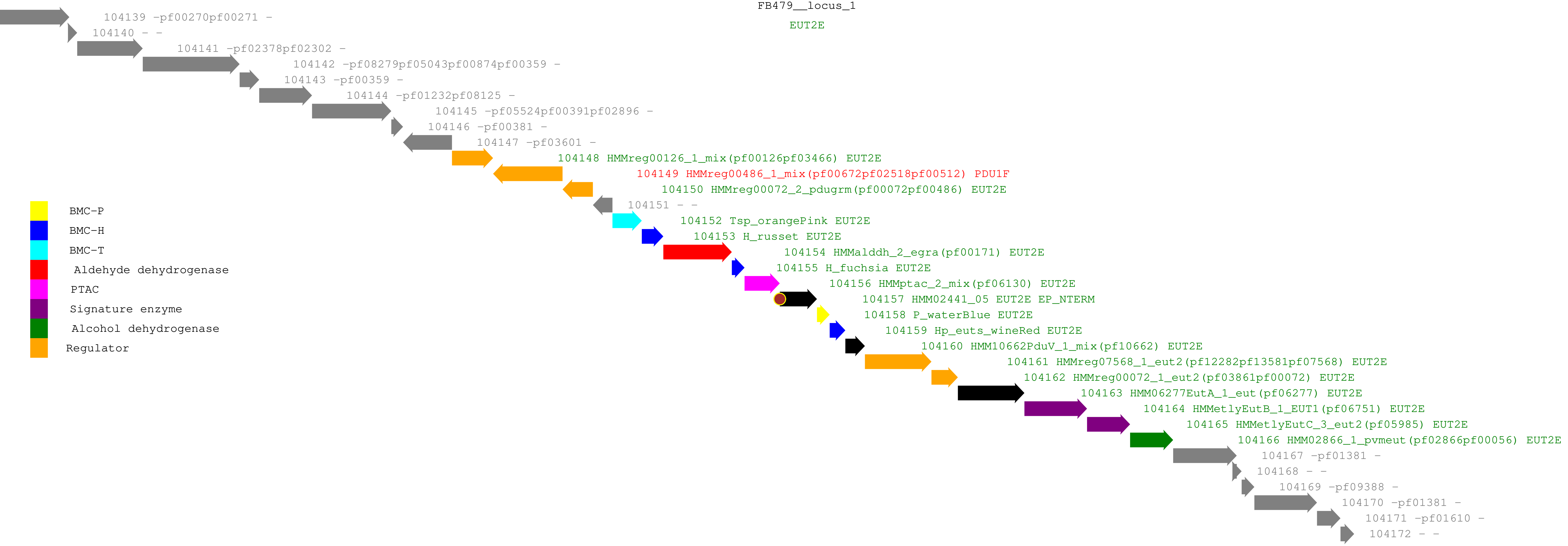

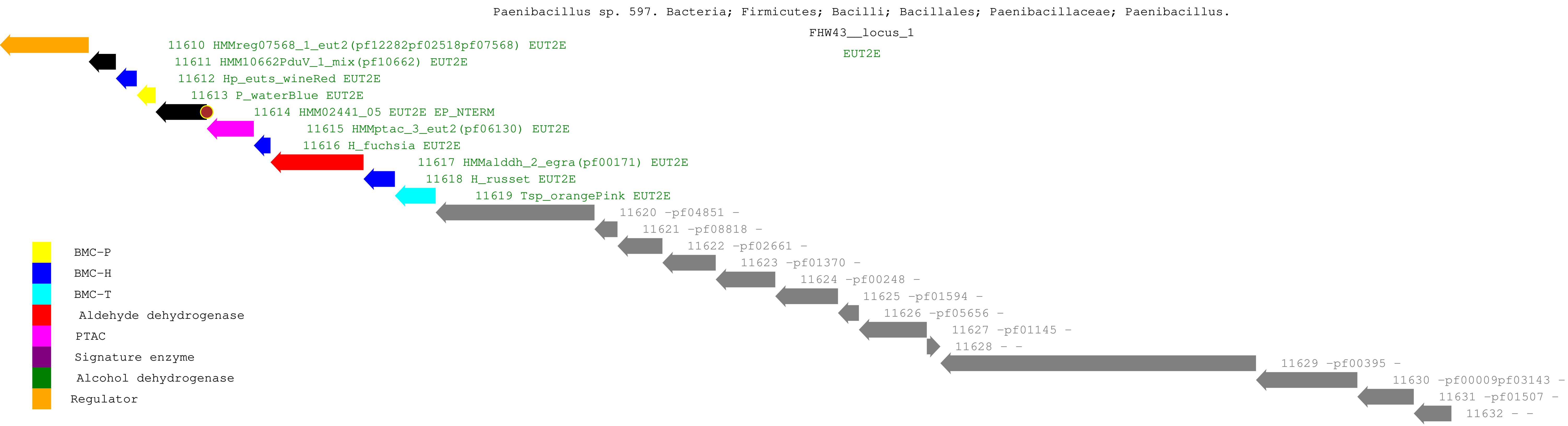

Paenibacillus sp. 597. Bacteria; Firmicutes; Bacilli; Bacillales; Paenibacillaceae; Paenibacillus.
FHW43__locus_1
EUT2E
11610 HMMreg07568_1_eut2(pf12282pf02518pf07568) EUT2E
11611 HMM10662PduV_1_mix(pf10662) EUT2E
11612 Hp_euts_wineRed EUT2E
11613 P_waterBlue EUT2E
11614 HMM02441_05 EUT2E EP_NTERM
11615 HMMptac_3_eut2(pf06130) EUT2E
11616 H_fuchsia EUT2E
11617 HMMalddh_2_egra(pf00171) EUT2E
11618 H_russet EUT2E
11619 Tsp_orangePink EUT2E
11620 -pf04851 -
11621 -pf08818 -
11622 -pf02661 -
11623 -pf01370 -
11624 -pf00248 -
11625 -pf01594 -
11626 -pf05656 -
11627 -pf01145 -
11628 - -
11629 -pf00395 -
11630 -pf00009pf03143 -
11631 -pf01507 -
11632 - -
BMC-P
BMC-H
BMC-T
Aldehyde dehydrogenase
PTAC
Signature enzyme
Alcohol dehydrogenase
Regulator

Geobacillus sp. LEMMJ02. Bacteria; Firmicutes; Bacilli; Bacillales; Bacillaceae; Geobacillus;

FOI67__locus_1

EUT2E

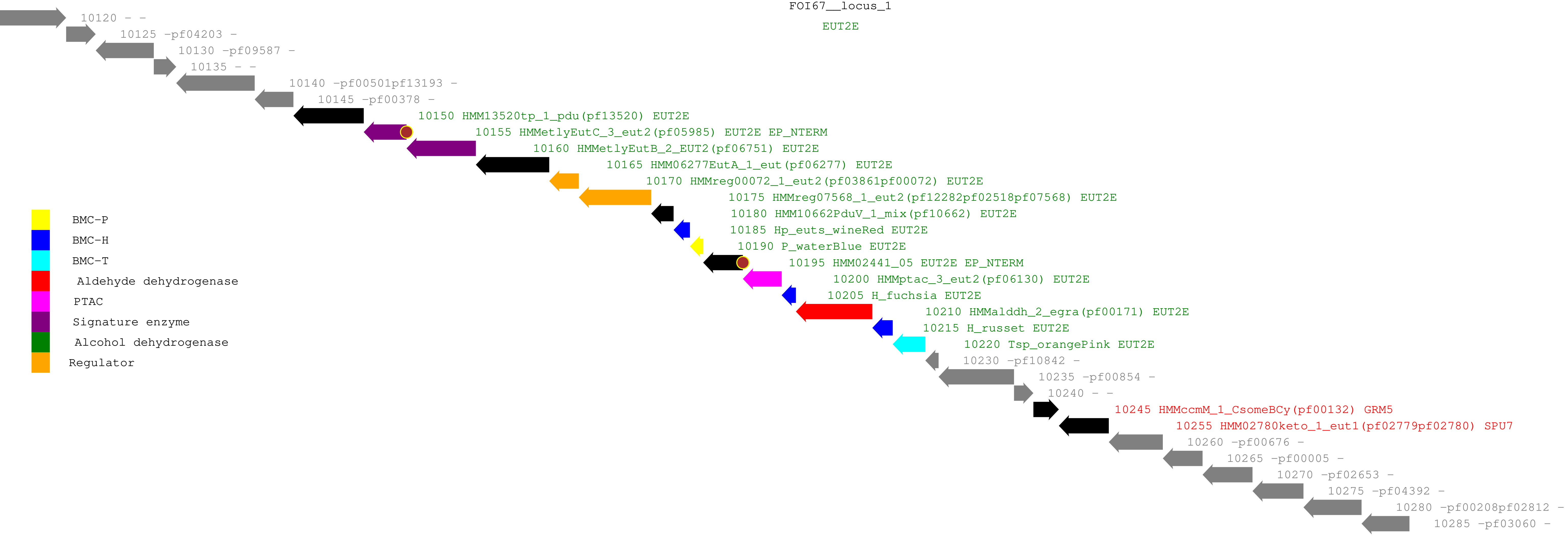

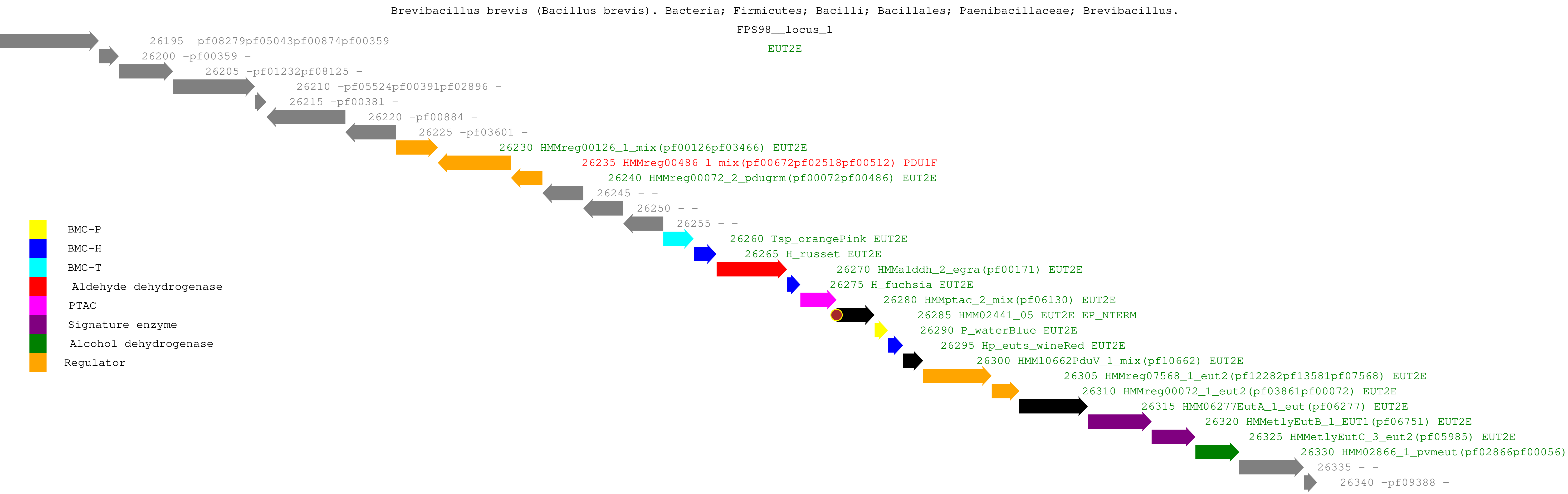
Brevibacillus brevis (Bacillus brevis). Bacteria; Firmicutes; Bacilli; Bacillales; Paenibacillaceae; Brevibacillus.
FPS98__locus_1
EUT2E
26195 -pf08279pf05043pf00874pf00359 -
26200 -pf00359 -
26205 -pf01232pf08125 -
26210 -pf05524pf00391pf02896 -
26215 -pf00381 -
26220 -pf00884 -
26225 -pf03601 -
26230 HMMreg00126_1_mix(pf00126pf03466) EUT2E
26235 HMMreg00486_1_mix(pf00672pf02518pf00512) PDU1F
26240 HMMreg00072_2_pdugrm(pf00072pf00486) EUT2E
26245 - -
26250 - -
26255 - -
26260 Tsp_orangePink EUT2E
26265 H_russet EUT2E
26270 HMMalddh_2_egra(pf00171) EUT2E
26275 H_fuchsia EUT2E
26280 HMMptac_2_mix(pf06130) EUT2E
26285 HMM02441_05 EUT2E EP_NTERM
26290 P_waterBlue EUT2E
26295 Hp_euts_wineRed EUT2E
26300 HMM10662PduV_1_mix(pf10662) EUT2E
26305 HMMreg07568_1_eut2(pf12282pf13581pf07568) EUT2E
26310 HMMreg00072_1_eut2(pf03861pf00072) EUT2E
26315 HMM06277EutA_1_eut(pf06277) EUT2E
26320 HMMetlyEutB_1_EUT1(pf06751) EUT2E
26325 HMMetlyEutC_3_eut2(pf05985) EUT2E
26330 HMM02866_1_pvmeut(pf02866pf00056)
26335 - -
26340 -pf09388 -
BMC-P
BMC-H
BMC-T
Aldehyde dehydrogenase
PTAC
Signature enzyme
Alcohol dehydrogenase
Regulator

Paenibacillus sp. N3.4. Bacteria; Firmicutes; Bacilli; Bacillales; Paenibacillaceae; Paenibacillus.

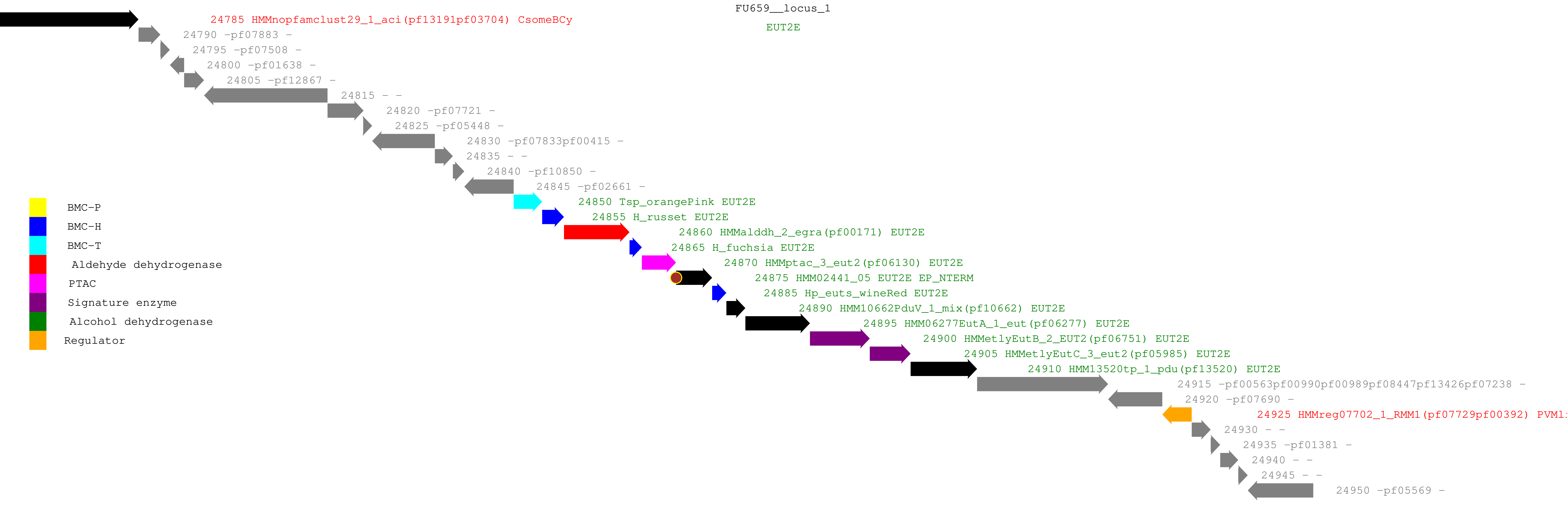

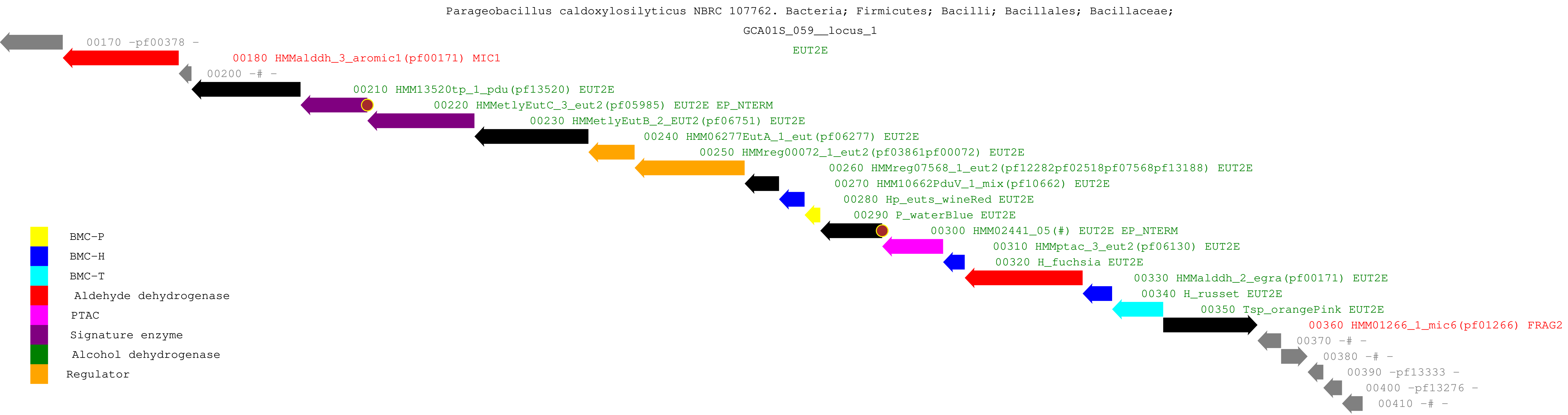

Parageobacillus caldoxylosilyticus NBRC 107762. Bacteria; Firmicutes; Bacilli; Bacillales; Bacillaceae;
GCA01S_059__locus_1
EUT2E
00170 -pf00378 -
00180 HMMalddh_3_aromic1(pf00171) MIC1
00200 -# -
00210 HMM13520tp_1_pdu(pf13520) EUT2E
00220 HMMetlyEutC_3_eut2(pf05985) EUT2E EP_NTERM
00230 HMMetlyEutB_2_EUT2(pf06751) EUT2E
00240 HMM06277EutA_1_eut(pf06277) EUT2E
00250 HMMreg00072_1_eut2(pf03861pf00072) EUT2E
00260 HMMreg07568_1_eut2(pf12282pf02518pf07568pf13188) EUT2E
00270 HMM10662PduV_1_mix(pf10662) EUT2E
00280 Hp_euts_wineRed EUT2E
00290 P_waterBlue EUT2E
00300 HMM02441_05(#) EUT2E EP_NTERM
00310 HMMptac_3_eut2(pf06130) EUT2E
00320 H_fuchsia EUT2E
00330 HMMalddh_2_egra(pf00171) EUT2E
00340 H_russet EUT2E
00350 Tsp_orangePink EUT2E
00360 HMM01266_1_mic6(pf01266) FRAG2
00370 -# -
00380 -# -
00390 -pf13333 -
00400 -pf13276 -
00410 -# -
BMC-P
BMC-H
BMC-T
Aldehyde dehydrogenase
PTAC
Signature enzyme
Alcohol dehydrogenase
Regulator

Brevibacillus borstelensis AK1. Bacteria; Firmicutes; Bacilli; Bacillales; Paenibacillaceae;

I532__locus_1

EUT2E

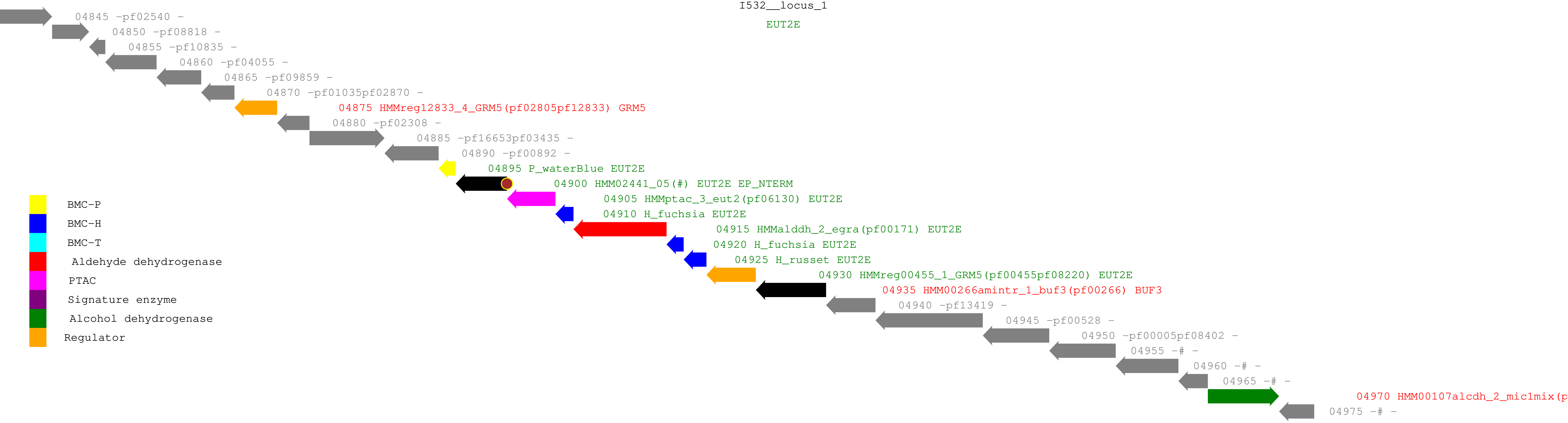

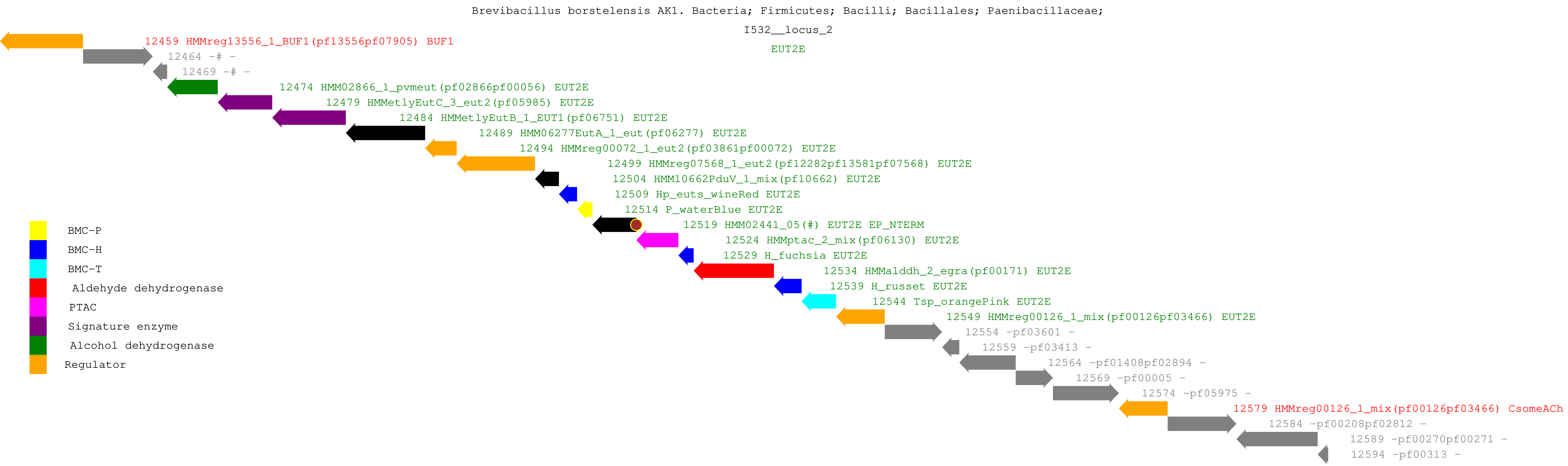
Brevibacillus borstelensis AK1. Bacteria; Firmicutes; Bacilli; Bacillales; Paenibacillaceae;
I532__locus_2
EUT2E
12459 HMMreg13556_1_BUF1(pf13556pf07905) BUF1
12464 -# -
12469 -# -
12474 HMM02866_1_pvmeut(pf02866pf00056) EUT2E
12479 HMMetlyEutC_3_eut2(pf05985) EUT2E
12484 HMMetlyEutB_1_EUT1(pf06751) EUT2E
12489 HMM06277EutA_1_eut(pf06277) EUT2E
12494 HMMreg00072_1_eut2(pf03861pf00072) EUT2E
12499 HMMreg07568_1_eut2(pf12282pf13581pf07568) EUT2E
12504 HMM10662PduV_1_mix(pf10662) EUT2E
12509 Hp_euts_wineRed EUT2E
12514 P_waterBlue EUT2E
12519 HMM02441_05(#) EUT2E EP_NTERM
12524 HMMptac_2_mix(pf06130) EUT2E
12529 H_fuchsia EUT2E
12534 HMMaldh_2_egra(pf00171) EUT2E
12539 H_russet EUT2E
12544 Tsp_orangePink EUT2E
12549 HMMreg00126_1_mix(pf00126pf03466) EUT2E
12554 -pf03601 -
12559 -pf03413 -
12564 -pf01408pf02894 -
12569 -pf00005 -
12574 -pf05975 -
12579 HMMreg00126_1_mix(pf00126pf03466) CsomeACh
12584 -pf00208pf02812 -
12589 -pf00270pf00271 -
12594 -pf00313 -
BMC-P
BMC-H
BMC-T
Aldehyde dehydrogenase
PTAC
Signature enzyme
Alcohol dehydrogenase
Regulator

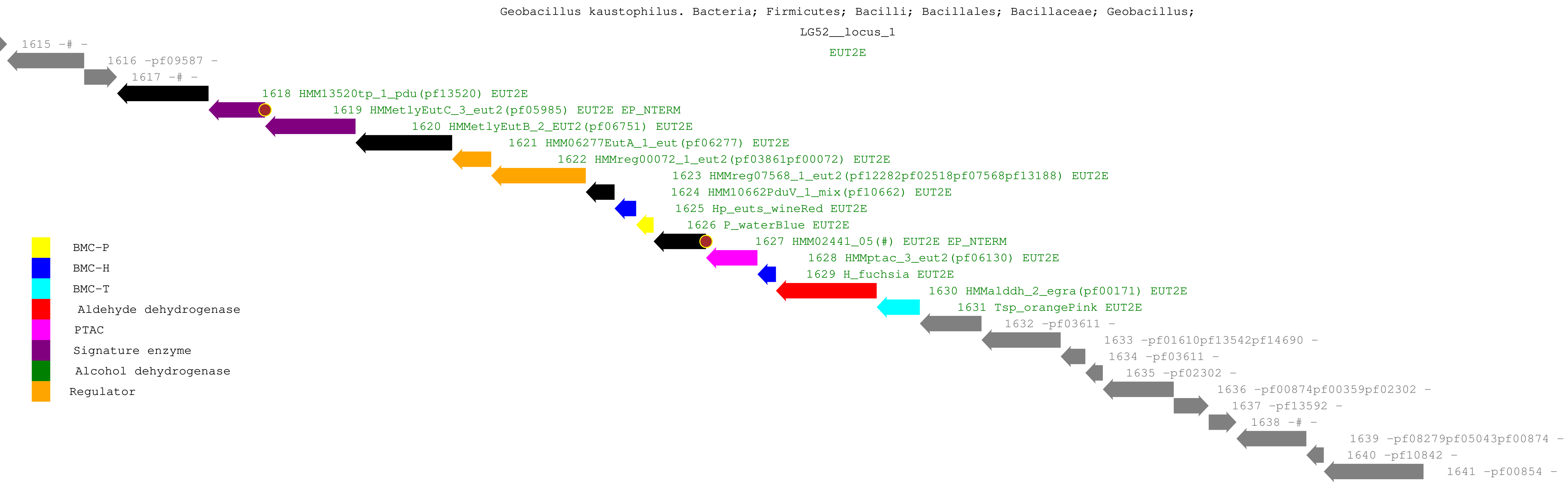
Geobacillus kaustophilus. Bacteria; Firmicutes; Bacilli; Bacillales; Bacillaceae; Geobacillus;
LG52__locus_1
EUT2E
1615 -# -
1616 -pf09587 -
1617 -# -
1618 HMM13520tp_1_pdu(pf13520) EUT2E
1619 HMMetlyEutC_3_eut2(pf05985) EUT2E EP_NTERM
1620 HMMetlyEutB_2_EUT2(pf06751) EUT2E
1621 HMM06277EutA_1_eut(pf06277) EUT2E
1622 HMMreg00072_1_eut2(pf03861pf00072) EUT2E
1623 HMMreg07568_1_eut2(pf12282pf02518pf07568pf13188) EUT2E
1624 HMM10662PduV_1_mix(pf10662) EUT2E
1625 Hp_euts_wineRed EUT2E
1626 P_waterBlue EUT2E
1627 HMM02441_05(#) EUT2E EP_NTERM
1628 HMMptac_3_eut2(pf06130) EUT2E
1629 H_fuchsia EUT2E
1630 HMMalddh_2_egra(pf00171) EUT2E
1631 Tsp_orangePink EUT2E
1632 -pf03611 -
1633 -pf01610pf13542pf14690 -
1634 -pf03611 -
1635 -pf02302 -
1636 -pf00874pf00359pf02302 -
1637 -pf13592 -
1638 -# -
1639 -pf08279pf05043pf00874 -
1640 -pf10842 -
1641 -pf00854 -
BMC-P
BMC-H
BMC-T
Aldehyde dehydrogenase
PTAC
Signature enzyme
Alcohol dehydrogenase
Regulator

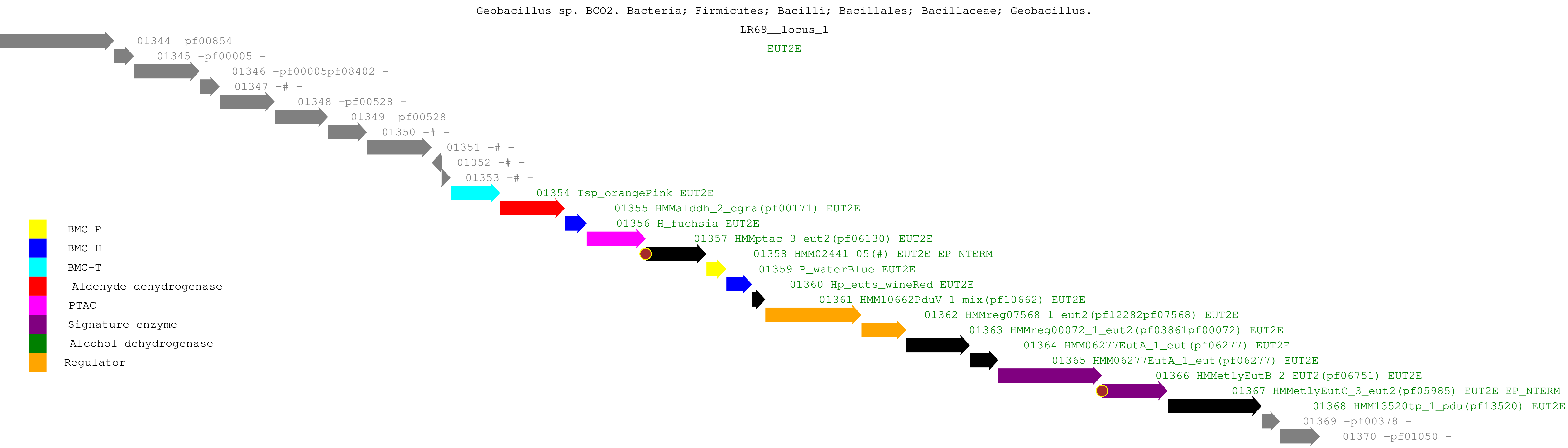

Geobacillus sp. BCO2. Bacteria; Firmicutes; Bacilli; Bacillales; Bacillaceae; Geobacillus.
LR69__locus_1
EUT2E
01344 -pf00854 -
01345 -pf00005 -
01346 -pf00005pf08402 -
01347 -# -
01348 -pf00528 -
01349 -pf00528 -
01350 -# -
01351 -# -
01352 -# -
01353 -# -
01354 Tsp_orangePink EUT2E
01355 HMMalddh_2_egra(pf00171) EUT2E
01356 H_fuchsia EUT2E
01357 HMMptac_3_eut2(pf06130) EUT2E
01358 HMM02441_05(#) EUT2E EP_NTERM
01359 P_waterBlue EUT2E
01360 Hp_euts_wineRed EUT2E
01361 HMM10662PduV_1_mix(pf10662) EUT2E
01362 HMMreg07568_1_eut2(pf12282pf07568) EUT2E
01363 HMMreg00072_1_eut2(pf03861pf00072) EUT2E
01364 HMM06277EutA_1_eut(pf06277) EUT2E
01365 HMM06277EutA_1_eut(pf06277) EUT2E
01366 HMMetlyEutB_2_EUT2(pf06751) EUT2E
01367 HMMetlyEutC_3_eut2(pf05985) EUT2E EP_NTERM
01368 HMM13520tp_1_pdu(pf13520) EUT2E
01369 -pf00378 -
01370 -pf01050 -
BMC-P
BMC-H
BMC-T
Aldehyde dehydrogenase
PTAC
Signature enzyme
Alcohol dehydrogenase
Regulator

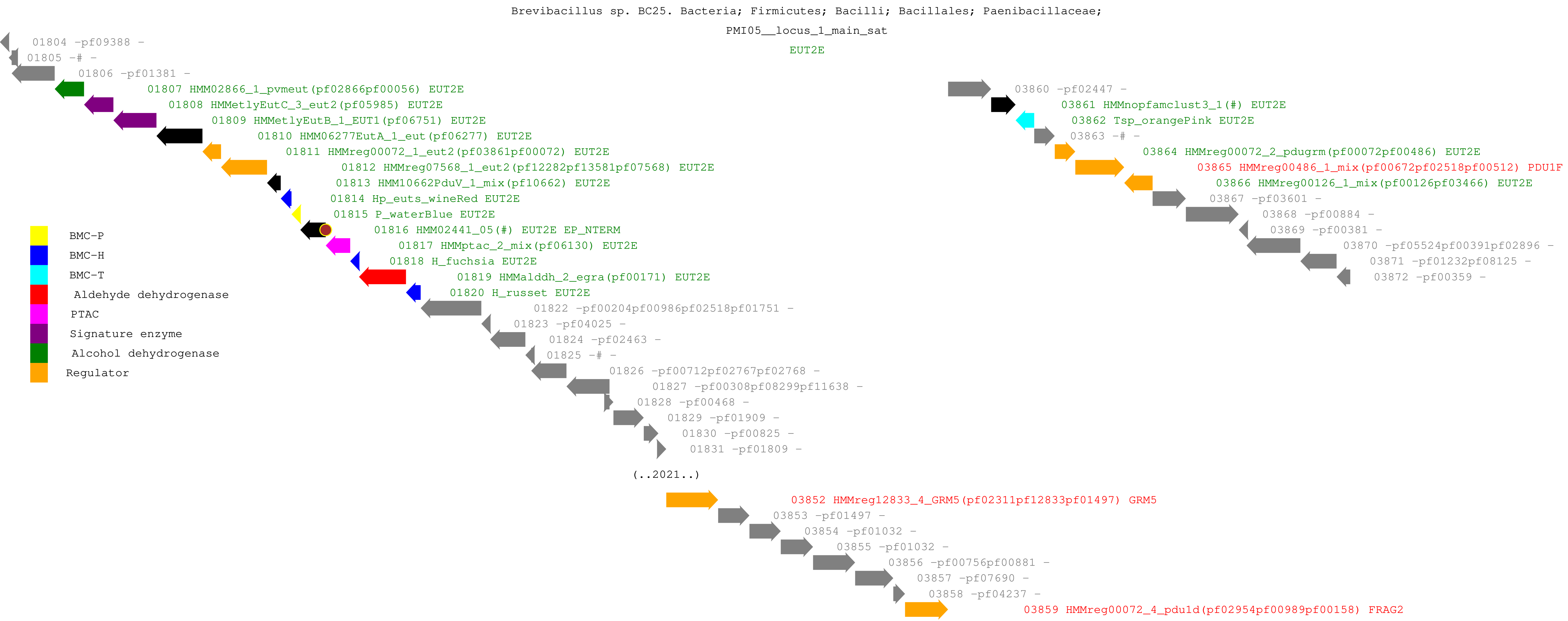
Brevibacillus sp. BC25. Bacteria; Firmicutes; Bacilli; Bacillales; Paenibacillaceae;
PMI05__locus_1_main_sat
EUT2E
01804 -pf09388 -
01805 -# -
01806 -pf01381 -
01807 HMM02866_1_pvmeut(pf02866pf00056) EUT2E
01808 HMMetlyEutC_3_eut2(pf05985) EUT2E
01809 HMMetlyEutB_1_EUT1(pf06751) EUT2E
01810 HMM06277EutA_1_eut(pf06277) EUT2E
01811 HMMreg00072_1_eut2(pf03861pf00072) EUT2E
01812 HMMreg07568_1_eut2(pf12282pf13581pf07568) EUT2E
01813 HMM10662PduV_1_mix(pf10662) EUT2E
01814 Hp_euts_wineRed EUT2E
01815 P_waterBlue EUT2E
01816 HMM02441_05(#) EUT2E EP_NTERM
01817 HMMptac_2_mix(pf06130) EUT2E
01818 H_fuchsia EUT2E
01819 HMMaldh_2_egra(pf00171) EUT2E
01820 H_russet EUT2E
01822 -pf00204pf00986pf02518pf01751 -
01823 -pf04025 -
01824 -pf02463 -
01825 -# -
01826 -pf00712pf02767pf02768 -
01827 -pf00308pf08299pf11638 -
01828 -pf00468 -
01829 -pf01909 -
01830 -pf00825 -
01831 -pf01809 -
(..2021..)
03852 HMMreg12833_4_GRM5(pf02311pf12833pf01497) GRM5
03853 -pf01497 -
03854 -pf01032 -
03855 -pf01032 -
03856 -pf00756pf00881 -
03857 -pf07690 -
03858 -pf04237 -
03859 HMMreg00072_4_pdu1d(pf02954pf00989pf00158) FRAG2
03860 -pf02447 -
03861 HMMnopfamclust3_1(#) EUT2E
03862 Tsp_orangePink EUT2E
03863 -# -
03864 HMMreg00072_2_pdugrm(pf00072pf00486) EUT2E
03865 HMMreg00486_1_mix(pf00672pf02518pf00512) PDU1F
03866 HMMreg00126_1_mix(pf00126pf03466) EUT2E
03867 -pf03601 -
03868 -pf00884 -
03869 -pf00381 -
03870 -pf05524pf00391pf02896 -
03871 -pf01232pf08125 -
03872 -pf00359 -
BMC-P
BMC-H
BMC-T
Aldehyde dehydrogenase
PTAC
Signature enzyme
Alcohol dehydrogenase
Regulator

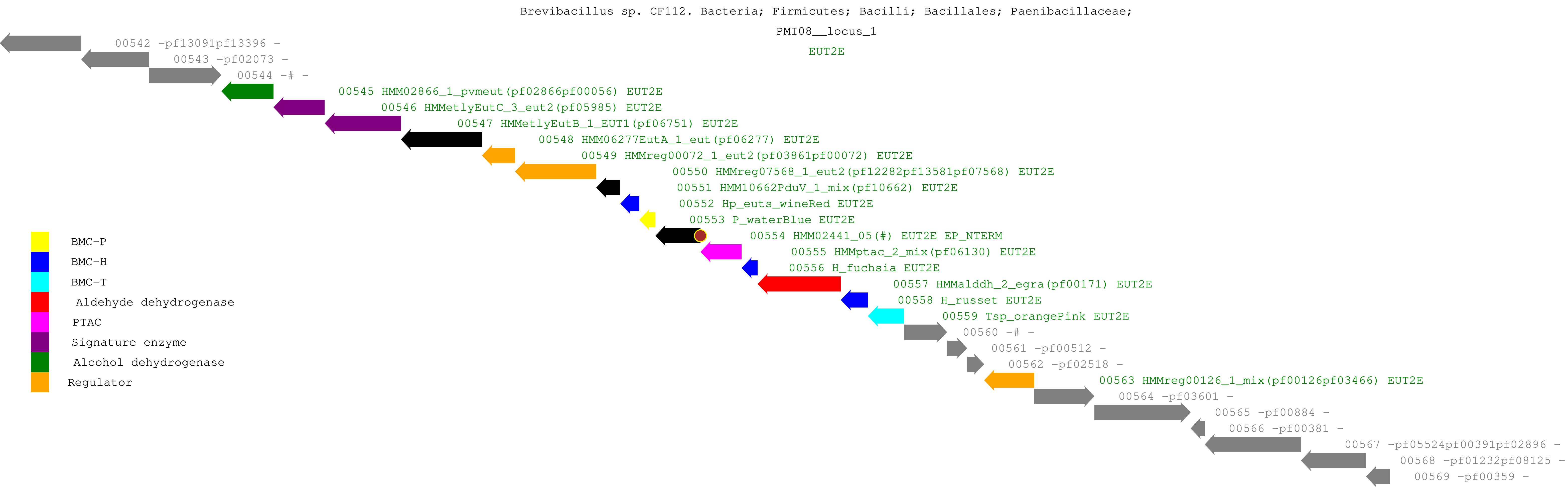
Brevibacillus sp. CF112. Bacteria; Firmicutes; Bacilli; Bacillales; Paenibacillaceae;
PMI08__locus_1
EUT2E
00542 -pf13091pf13396 -
00543 -pf02073 -
00544 -# -
00545 HMM02866_1_pvmeut(pf02866pf00056) EUT2E
00546 HMMetlyEutC_3_eut2(pf05985) EUT2E
00547 HMMetlyEutB_1_EUT1(pf06751) EUT2E
00548 HMM06277EutA_1_eut(pf06277) EUT2E
00549 HMMreg00072_1_eut2(pf03861pf00072) EUT2E
00550 HMMreg07568_1_eut2(pf12282pf13581pf07568) EUT2E
00551 HMM10662PduV_1_mix(pf10662) EUT2E
00552 Hp_euts_wineRed EUT2E
00553 P_waterBlue EUT2E
00554 HMM02441_05(#) EUT2E EP_NTERM
00555 HMMptac_2_mix(pf06130) EUT2E
00556 H_fuchsia EUT2E
00557 HMMalddh_2_egra(pf00171) EUT2E
00558 H_russet EUT2E
00559 Tsp_orangePink EUT2E
00560 -# -
00561 -pf00512 -
00562 -pf02518 -
00563 HMMreg00126_1_mix(pf00126pf03466) EUT2E
00564 -pf03601 -
00565 -pf00884 -
00566 -pf00381 -
00567 -pf05524pf00391pf02896 -
00568 -pf01232pf08125 -
00569 -pf00359 -
BMC-P
BMC-H
BMC-T
Aldehyde dehydrogenase
PTAC
Signature enzyme
Alcohol dehydrogenase
Regulator

Paenibacillus sp. A3. Bacteria; Firmicutes; Bacilli; Bacillales; Paenibacillaceae;

QJ48__locus_1

EUT2E

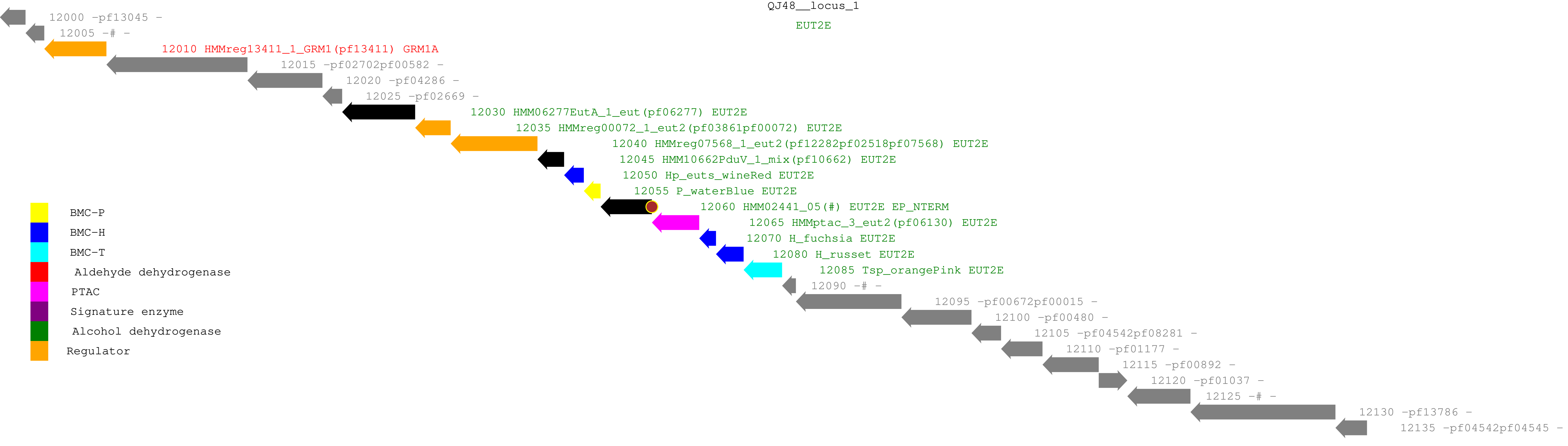

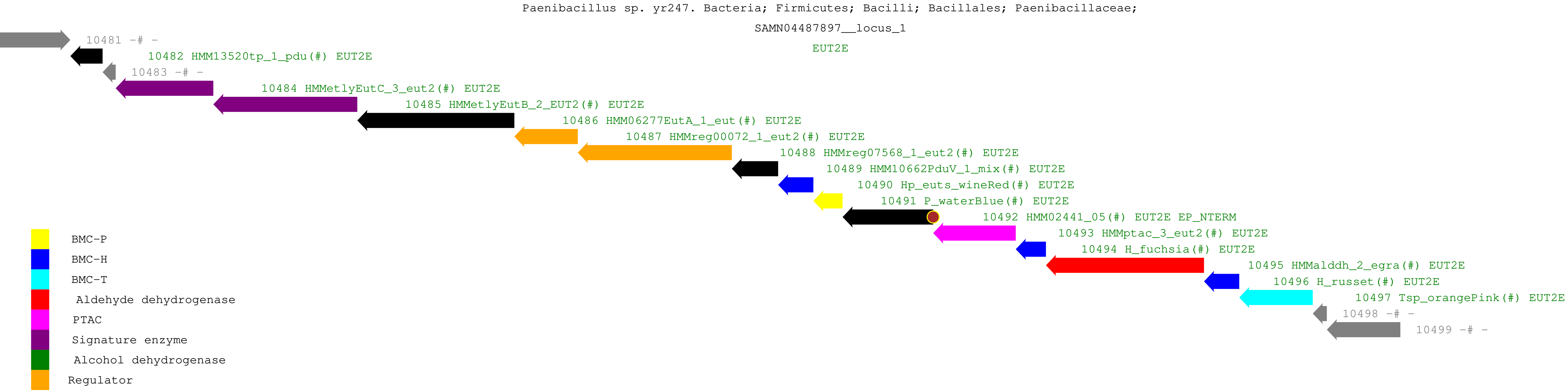

Paenibacillus sp. yr247. Bacteria; Firmicutes; Bacilli; Bacillales; Paenibacillaceae;
SAMN04487897__locus_1
EUT2E
10481 -# -
10482 HMM13520tp_1_pdu(#) EUT2E
10483 -# -
10484 HMMetlyEutC_3_eut2(#) EUT2E
10485 HMMetlyEutB_2_EUT2(#) EUT2E
10486 HMM06277EutA_1_eut(#) EUT2E
10487 HMMreg00072_1_eut2(#) EUT2E
10488 HMMreg07568_1_eut2(#) EUT2E
10489 HMM10662PduV_1_mix(#) EUT2E
10490 Hp_euts_wineRed(#) EUT2E
10491 P_waterBlue(#) EUT2E
10492 HMM02441_05(#) EUT2E EP_NTERM
10493 HMMptac_3_eut2(#) EUT2E
10494 H_fuchsia(#) EUT2E
10495 HMMalddh_2_egra(#) EUT2E
10496 H_russet(#) EUT2E
10497 Tsp_orangePink(#) EUT2E
10498 -# -
10499 -# -
BMC-P
BMC-H
BMC-T
Aldehyde dehydrogenase
PTAC
Signature enzyme
Alcohol dehydrogenase
Regulator

Paenibacillus tianmuensis. Bacteria; Firmicutes; Bacilli; Bacillales; Paenibacillaceae;

SAMN04487970__locus_1

EUT2E

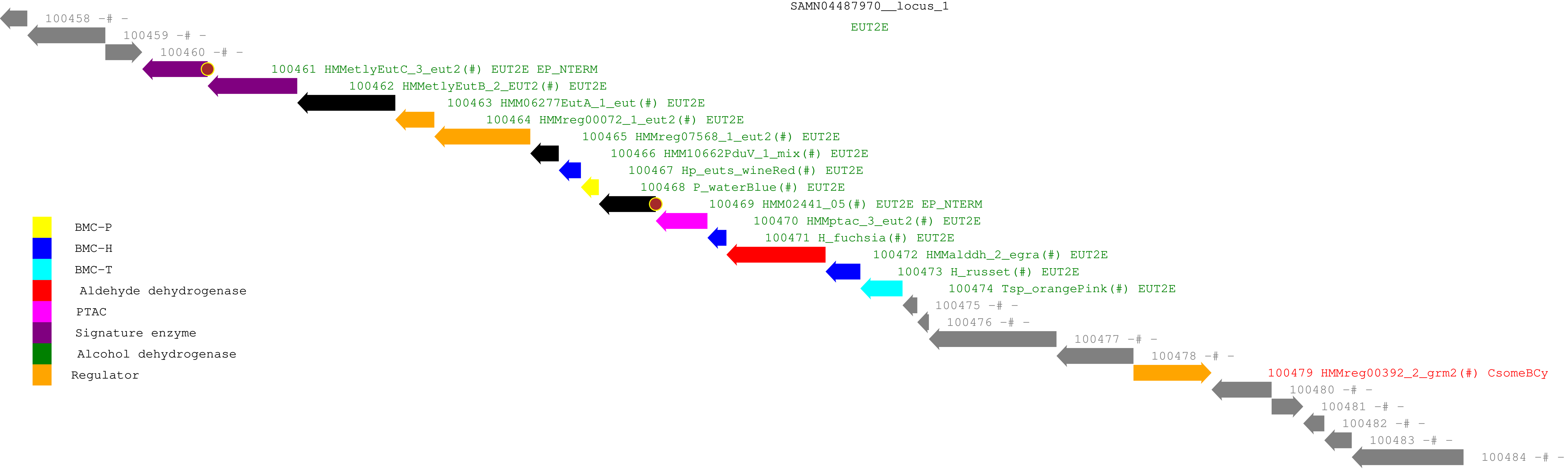

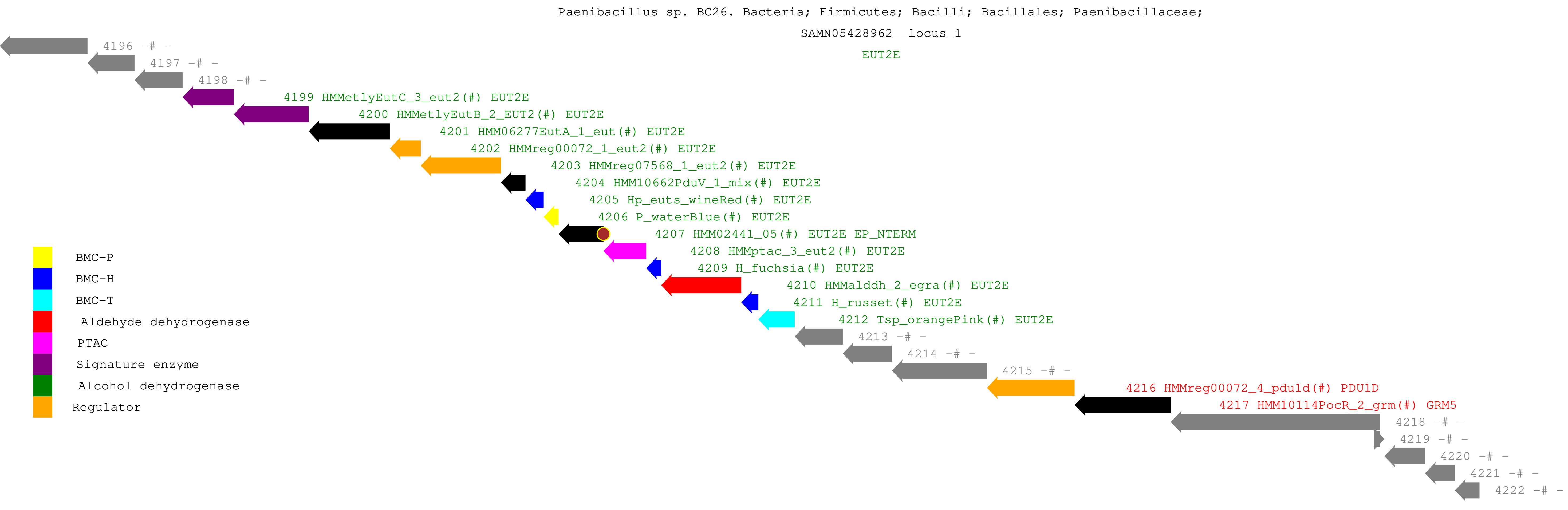
Paenibacillus sp. BC26. Bacteria; Firmicutes; Bacilli; Bacillales; Paenibacillaceae;
SAMN05428962__locus_1
EUT2E
4196 -# -
4197 -# -
4198 -# -
4199 HMMetlyEutC_3_eut2(#) EUT2E
4200 HMMetlyEutB_2_EUT2(#) EUT2E
4201 HMM06277EutA_1_eut(#) EUT2E
4202 HMMreg00072_1_eut2(#) EUT2E
4203 HMMreg07568_1_eut2(#) EUT2E
4204 HMM10662PduV_1_mix(#) EUT2E
4205 Hp_euts_wineRed(#) EUT2E
4206 P_waterBlue(#) EUT2E
4207 HMM02441_05(#) EUT2E EP_NTERM
4208 HMMptac_3_eut2(#) EUT2E
4209 H_fuchsia(#) EUT2E
4210 HMMalddh_2_egra(#) EUT2E
4211 H_russet(#) EUT2E
4212 Tsp_orangePink(#) EUT2E
4213 -# -
4214 -# -
4215 -# -
4216 HMMreg00072_4_pdu1d(#) PDU1D
4217 HMM10114PocR_2_grm(#) GRM5
4218 -# -
4219 -# -
4220 -# -
4221 -# -
4222 -# -
BMC-P
BMC-H
BMC-T
Aldehyde dehydrogenase
PTAC
Signature enzyme
Alcohol dehydrogenase
Regulator

Paenibacillaceae bacterium GAS479. Bacteria; Firmicutes; Bacilli; Bacillales; Paenibacillaceae.

SAMN05444162__locus_1

EUT2E

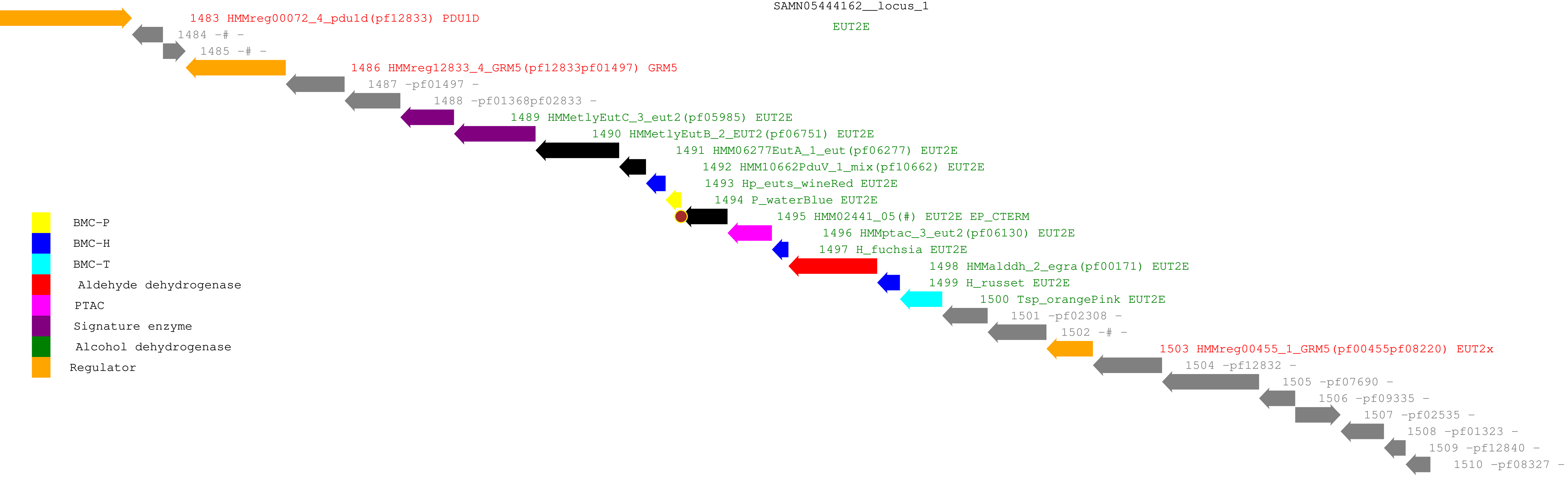

Brevibacillus sp. OK042. Bacteria; Firmicutes; Bacilli; Bacillales; Paenibacillaceae;

SAMN05518846__locus_1

EUT2E

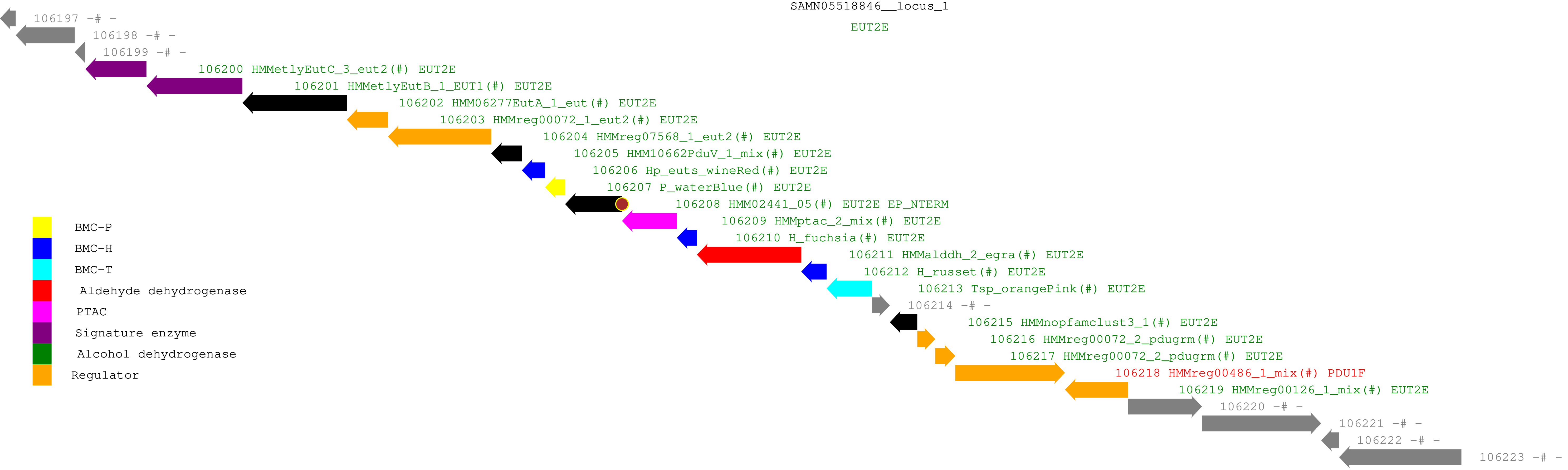

Paenibacillus sp. CF384. Bacteria; Firmicutes; Bacilli; Bacillales; Paenibacillaceae;

SAMN05518855__locus_1

EUT2E

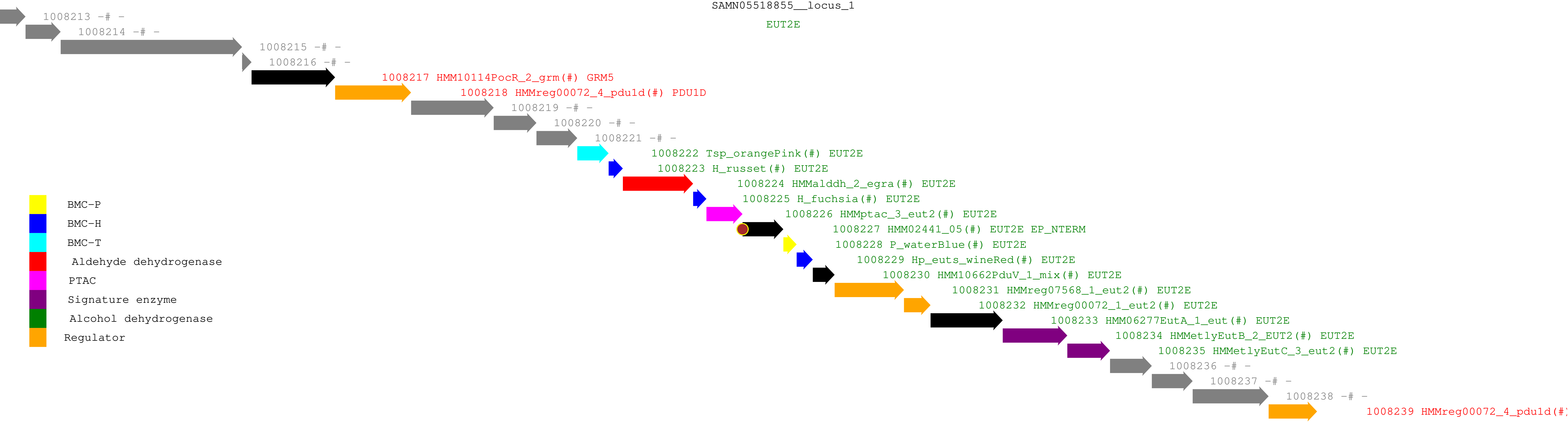

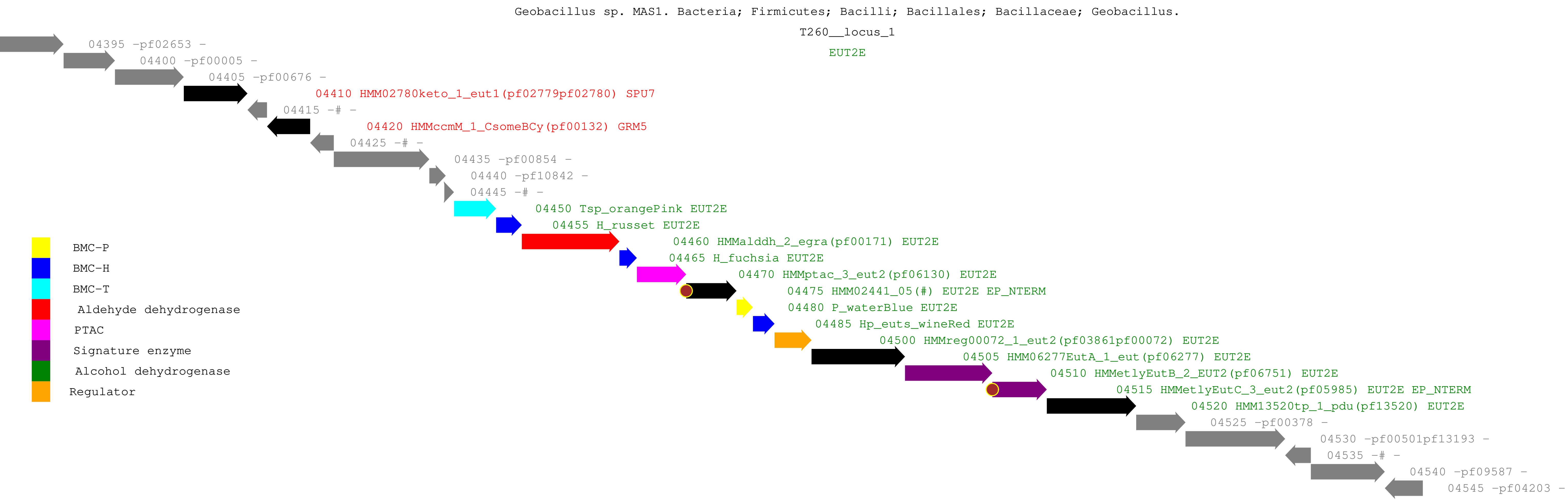

Geobacillus sp. MAS1. Bacteria; Firmicutes; Bacilli; Bacillales; Bacillaceae; Geobacillus.
T260__locus_1
EUT2E
04395 -pf02653 -
04400 -pf00005 -
04405 -pf00676 -
04410 HMM02780keto_1_eut1(pf02779pf02780) SPU7
04415 -# -
04420 HMMccmM_1_CsomeBCy(pf00132) GRM5
04425 -# -
04435 -pf00854 -
04440 -pf10842 -
04445 -# -
04450 Tsp_orangePink EUT2E
04455 H_russet EUT2E
04460 HMMalddh_2_egra(pf00171) EUT2E
04465 H_fuchsia EUT2E
04470 HMMptac_3_eut2(pf06130) EUT2E
04475 HMM02441_05(#) EUT2E EP_NTERM
04480 P_waterBlue EUT2E
04485 Hp_euts_wineRed EUT2E
04500 HMMreg00072_1_eut2(pf03861pf00072) EUT2E
04505 HMM06277EutA_1_eut(pf06277) EUT2E
04510 HMMetlyEutB_2_EUT2(pf06751) EUT2E
04515 HMMetlyEutC_3_eut2(pf05985) EUT2E EP_NTERM
04520 HMM13520tp_1_pdu(pf13520) EUT2E
04525 -pf00378 -
04530 -pf00501pf13193 -
04535 -# -
04540 -pf09587 -
04545 -pf04203 -
BMC-P
BMC-H
BMC-T
Aldehyde dehydrogenase
PTAC
Signature enzyme
Alcohol dehydrogenase
Regulator

Brevibacillus panacihumi W25. Bacteria; Firmicutes; Bacilli; Bacillales; Paenibacillaceae;

T458__locus_1

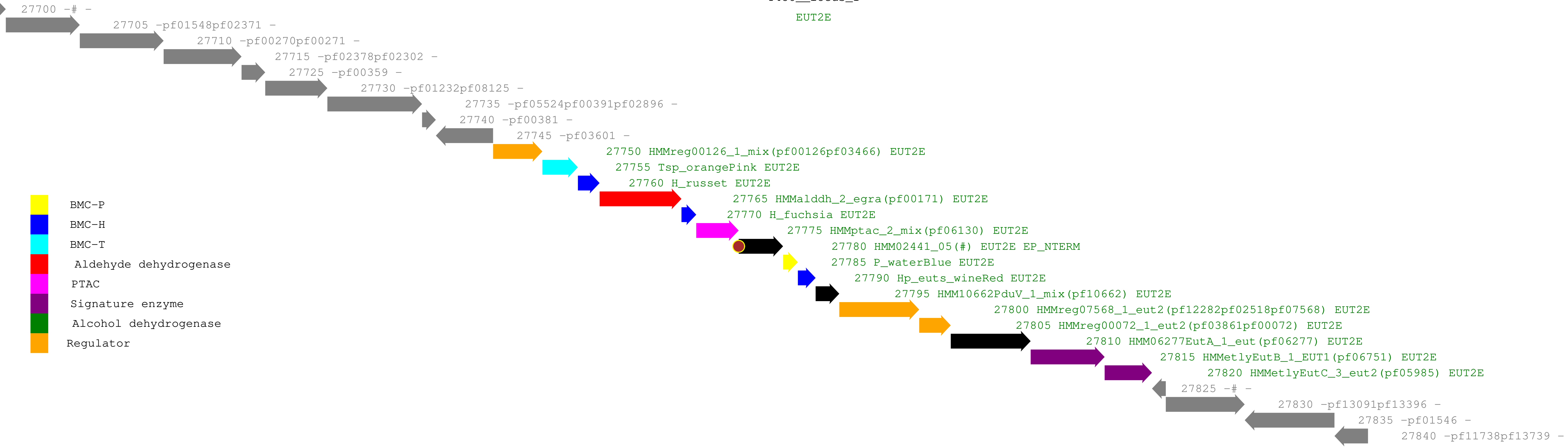

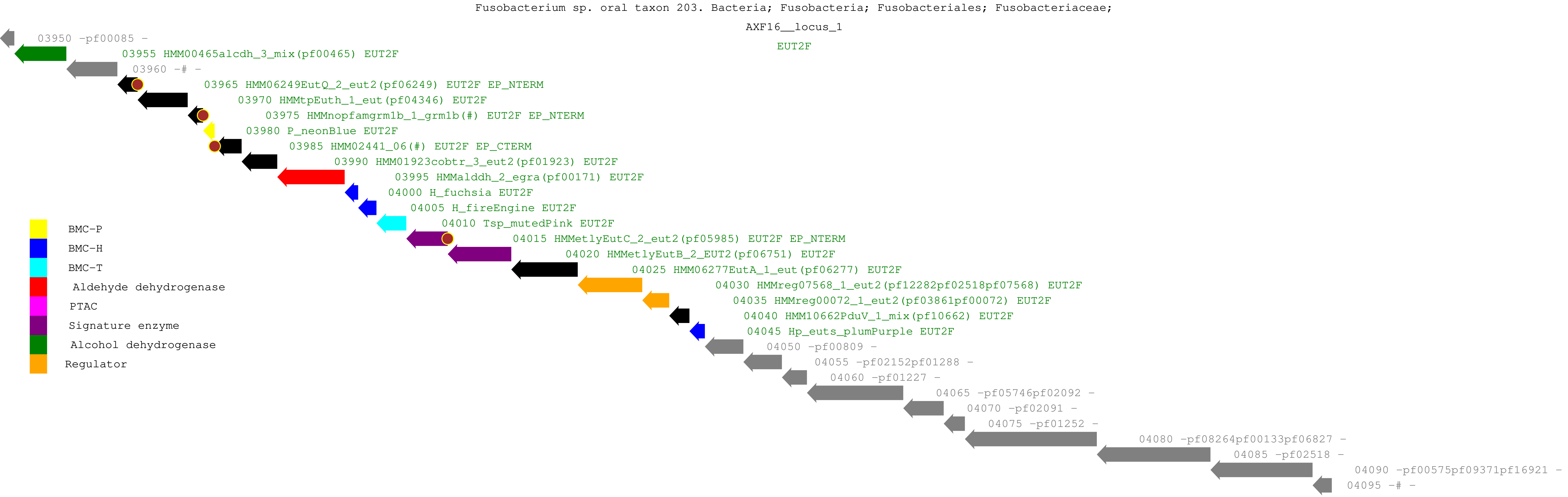
Fusobacterium sp. oral taxon 203. Bacteria; Fusobacteria; Fusobacteriales; Fusobacteriaceae;
AXF16__locus_1
EUT2F
03950 -pf00085 -
03955 HMM00465alcdh_3_mix(pf00465) EUT2F
03960 -# -
03965 HMM06249EutQ_2_eut2(pf06249) EUT2F EP_NTERM
03970 HMMtpEuth_1_eut(pf04346) EUT2F
03975 HMMnopfamgrm1b_1_grm1b(#) EUT2F EP_NTERM
03980 P_neonBlue EUT2F
03985 HMM02441_06(#) EUT2F EP_CTERM
03990 HMM01923cobtr_3_eut2(pf01923) EUT2F
03995 HMMalddh_2_egra(pf00171) EUT2F
04000 H_fuchsia EUT2F
04005 H_fireEngine EUT2F
04010 Tsp_mutedPink EUT2F
04015 HMMetlyEutC_2_eut2(pf05985) EUT2F EP_NTERM
04020 HMMetlyEutB_2_EUT2(pf06751) EUT2F
04025 HMM06277EutA_1_eut(pf06277) EUT2F
04030 HMMreg07568_1_eut2(pf12282pf02518pf07568) EUT2F
04035 HMMreg00072_1_eut2(pf03861pf00072) EUT2F
04040 HMM10662PduV_1_mix(pf10662) EUT2F
04045 Hp_euts_plumPurple EUT2F
04050 -pf00809 -
04055 -pf02152pf01288 -
04060 -pf01227 -
04065 -pf05746pf02092 -
04070 -pf02091 -
04075 -pf01252 -
04080 -pf08264pf00133pf06827 -
04085 -pf02518 -
04090 -pf00575pf09371pf16921 -
04095 -# -
BMC-P
BMC-H
BMC-T
Aldehyde dehydrogenase
PTAC
Signature enzyme
Alcohol dehydrogenase
Regulator

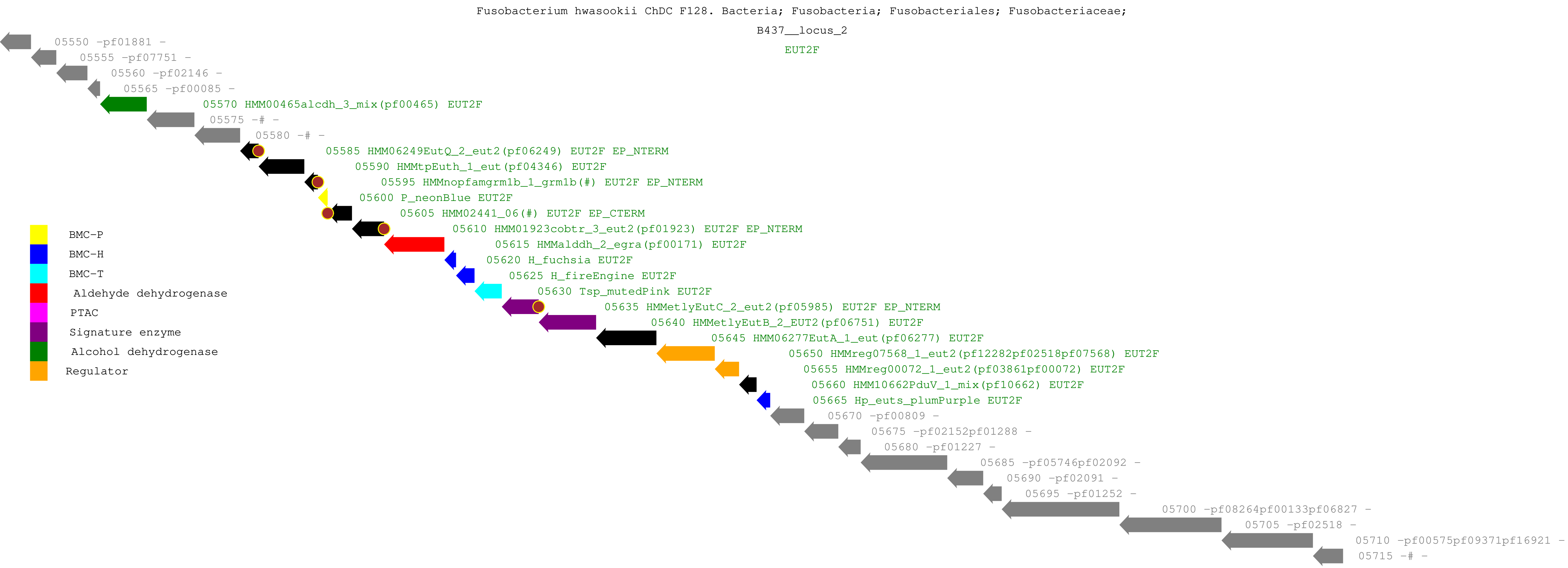
Fusobacterium hwasookii ChDC F128. Bacteria; Fusobacteria; Fusobacteriales; Fusobacteriaceae;
B437__locus_2
EUT2F
05550 -pf01881 -
05555 -pf07751 -
05560 -pf02146 -
05565 -pf00085 -
05570 HMM00465alcdh_3_mix(pf00465) EUT2F
05575 -# -
05580 -# -
05585 HMM06249EutQ_2_eut2(pf06249) EUT2F EP_NTERM
05590 HMMtpEuth_1_eut(pf04346) EUT2F
05595 HMMnopfamgrm1b_1_grm1b(#) EUT2F EP_NTERM
05600 P_neonBlue EUT2F
05605 HMM02441_06(#) EUT2F EP_CTERM
05610 HMM01923cobtr_3_eut2(pf01923) EUT2F EP_NTERM
05615 HMMalddh_2_egra(pf00171) EUT2F
05620 H_fuchsia EUT2F
05625 H_fireEngine EUT2F
05630 Tsp_mutedPink EUT2F
05635 HMMetlyEutC_2_eut2(pf05985) EUT2F EP_NTERM
05640 HMMetlyEutB_2_EUT2(pf06751) EUT2F
05645 HMM06277EutA_1_eut(pf06277) EUT2F
05650 HMMreg07568_1_eut2(pf12282pf02518pf07568) EUT2F
05655 HMMreg00072_1_eut2(pf03861pf00072) EUT2F
05660 HMM10662PduV_1_mix(pf10662) EUT2F
05665 Hp_euts_plumPurple EUT2F
05670 -pf00809 -
05675 -pf02152pf01288 -
05680 -pf01227 -
05685 -pf05746pf02092 -
05690 -pf02091 -
05695 -pf01252 -
05700 -pf08264pf00133pf06827 -
05705 -pf02518 -
05710 -pf00575pf09371pf16921 -
05715 -# -
BMC-P
BMC-H
BMC-T
Aldehyde dehydrogenase
PTAC
Signature enzyme
Alcohol dehydrogenase
Regulator

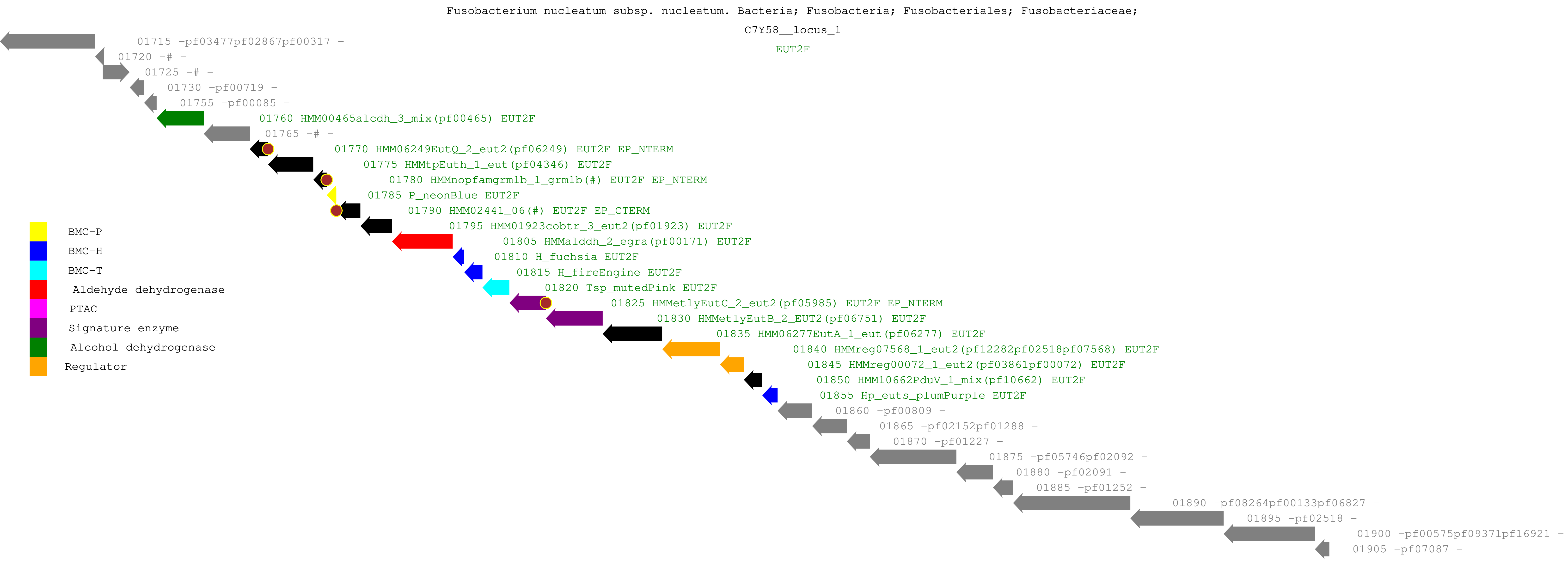

Fusobacterium nucleatum subsp. nucleatum. Bacteria; Fusobacteria; Fusobacteriales; Fusobacteriaceae;
C7Y58__locus_1
EUT2F
01715 -pf03477pf02867pf00317 -
01720 -# -
01725 -# -
01730 -pf00719 -
01755 -pf00085 -
01760 HMM00465alcdh_3_mix(pf00465) EUT2F
01765 -# -
01770 HMM06249EutQ_2_eut2(pf06249) EUT2F EP_NTERM
01775 HMMtpEuth_1_eut(pf04346) EUT2F
01780 HMMnopfamgrm1b_1_grm1b(#) EUT2F EP_NTERM
01785 P_neonBlue EUT2F
01790 HMM02441_06(#) EUT2F EP_CTERM
01795 HMM01923cobtr_3_eut2(pf01923) EUT2F
01805 HMMalddh_2_egra(pf00171) EUT2F
01810 H_fuchsia EUT2F
01815 H_fireEngine EUT2F
01820 Tsp_mutedPink EUT2F
01825 HMMetlyEutC_2_eut2(pf05985) EUT2F EP_NTERM
01830 HMMetlyEutB_2_EUT2(pf06751) EUT2F
01835 HMM06277EutA_1_eut(pf06277) EUT2F
01840 HMMreg07568_1_eut2(pf12282pf02518pf07568) EUT2F
01845 HMMreg00072_1_eut2(pf03861pf00072) EUT2F
01850 HMM10662PduV_1_mix(pf10662) EUT2F
01855 Hp_euts_plumPurple EUT2F
01860 -pf00809 -
01865 -pf02152pf01288 -
01870 -pf01227 -
01875 -pf05746pf02092 -
01880 -pf02091 -
01885 -pf01252 -
01890 -pf08264pf00133pf06827 -
01895 -pf02518 -
01900 -pf00575pf09371pf16921 -
01905 -pf07087 -
BMC-P
BMC-H
BMC-T
Aldehyde dehydrogenase
PTAC
Signature enzyme
Alcohol dehydrogenase
Regulator

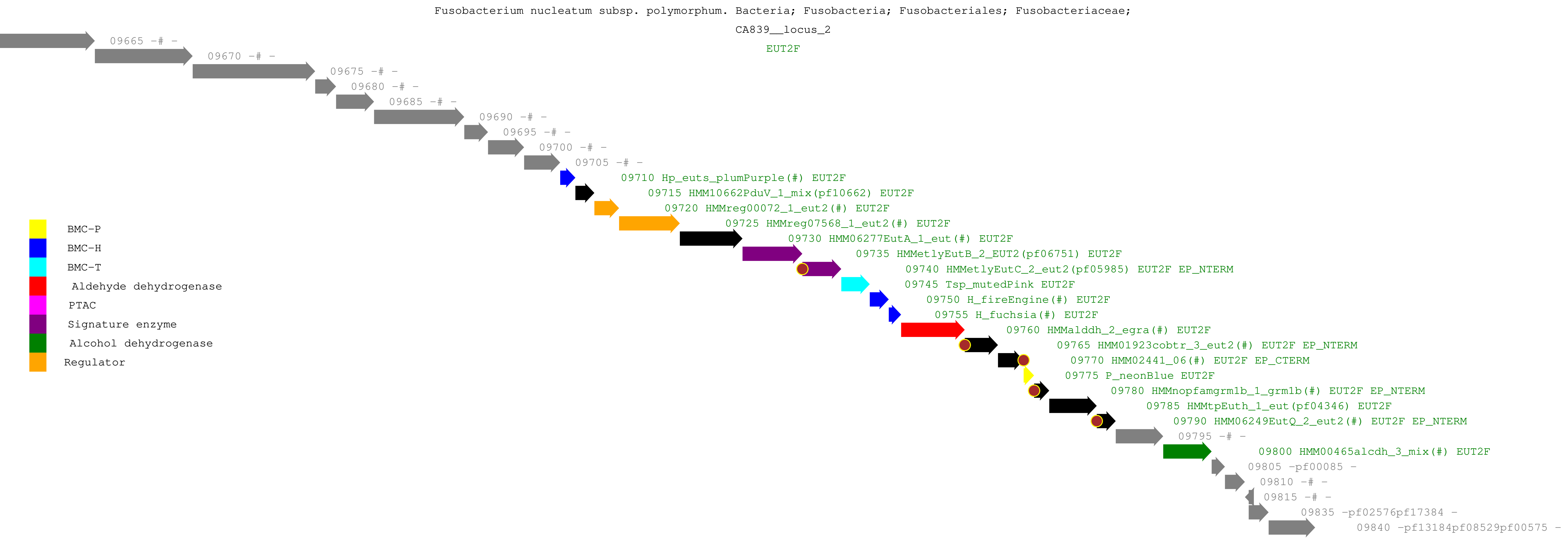
Fusobacterium nucleatum subsp. polymorphum. Bacteria; Fusobacteria; Fusobacteriales; Fusobacteriaceae;
CA839__locus_2
EUT2F
09665 -# -
09670 -# -
09675 -# -
09680 -# -
09685 -# -
09690 -# -
09695 -# -
09700 -# -
09705 -# -
09710 Hp_euts_plumPurple(#) EUT2F
09715 HMM10662PduV_1_mix(pf10662) EUT2F
09720 HMMreg00072_1_eut2(#) EUT2F
09725 HMMreg07568_1_eut2(#) EUT2F
09730 HMM06277EutA_1_eut(#) EUT2F
09735 HMMetlyEutB_2_EUT2(pf06751) EUT2F
09740 HMMetlyEutC_2_eut2(pf05985) EUT2F EP_NTERM
09745 Tsp_mutedPink EUT2F
09750 H_fireEngine(#) EUT2F
09755 H_fuchsia(#) EUT2F
09760 HMMalddh_2_egra(#) EUT2F
09765 HMM01923cobtr_3_eut2(#) EUT2F EP_NTERM
09770 HMM02441_06(#) EUT2F EP_CTERM
09775 P_neonBlue EUT2F
09780 HMMnopfamgrm1b_1_grm1b(#) EUT2F EP_NTERM
09785 HMMtpEuth_1_eut(pf04346) EUT2F
09790 HMM06249EutQ_2_eut2(#) EUT2F EP_NTERM
09795 -# -
09800 HMM00465alcdh_3_mix(#) EUT2F
09805 -pf00085 -
09810 -# -
09815 -# -
09835 -pf02576pf17384 -
09840 -pf13184pf08529pf00575 -
BMC-P
BMC-H
BMC-T
Aldehyde dehydrogenase
PTAC
Signature enzyme
Alcohol dehydrogenase
Regulator

Fusobacterium nucleatum subsp. polymorphum. Bacteria; Fusobacteria; Fusobacteriales; Fusobacteriaceae;

CA840__locus_1

EUT2F

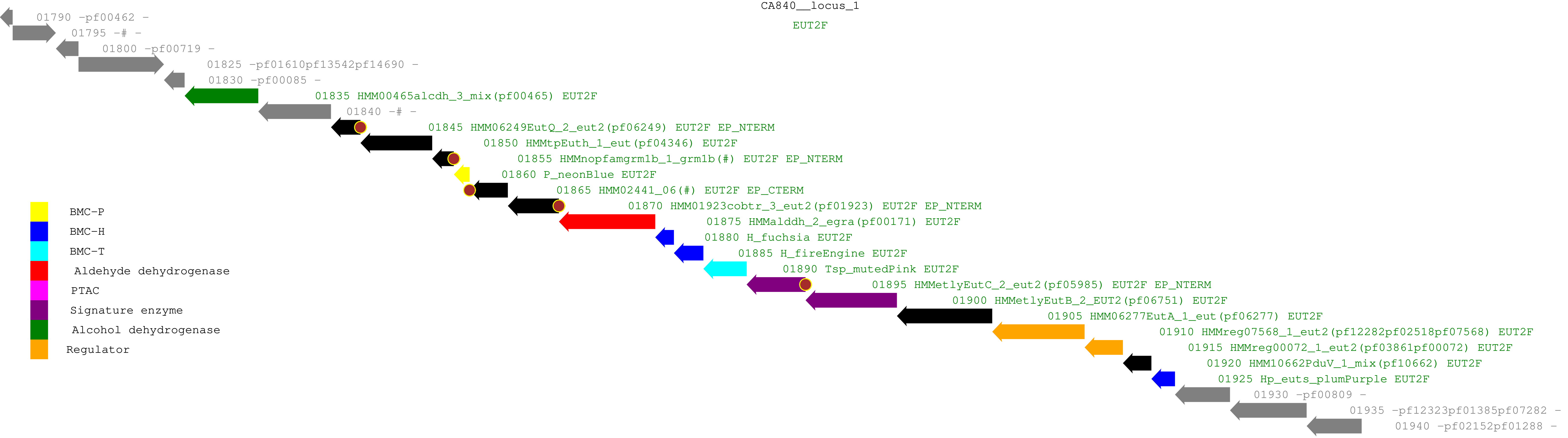

Fusobacterium nucleatum subsp. polymorphum. Bacteria; Fusobacteria; Fusobacteriales; Fusobacteriaceae;

CBG59__locus_1

EUT2F

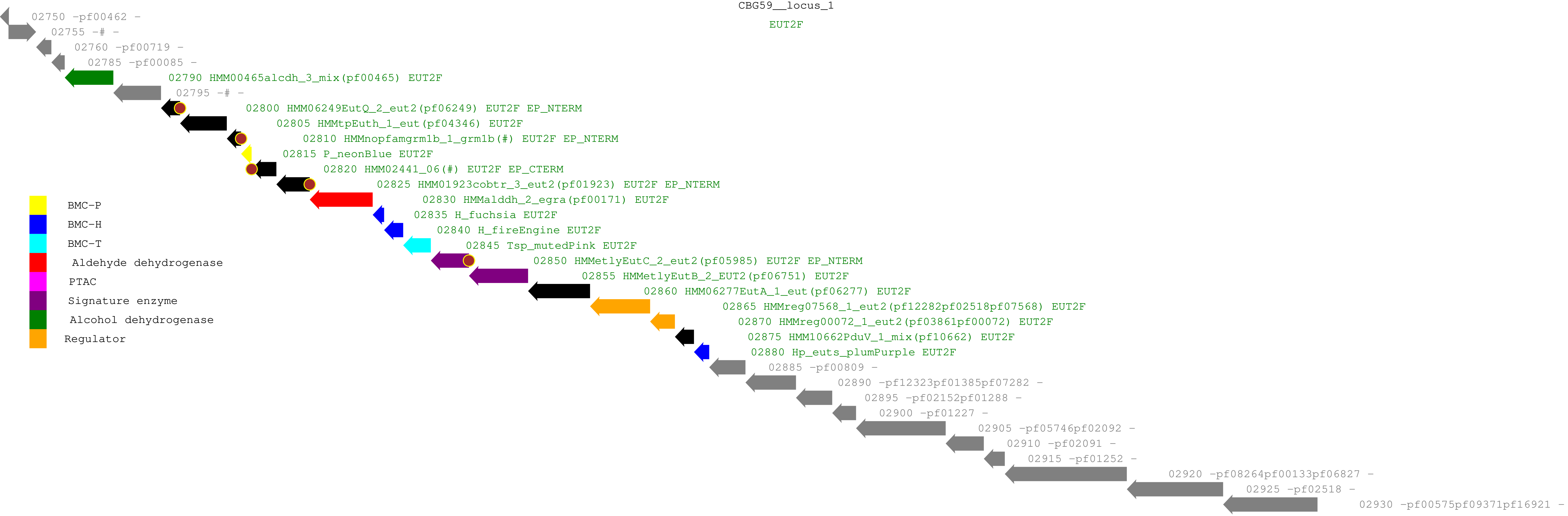

Fusobacterium nucleatum subsp. polymorphum. Bacteria; Fusobacteria; Fusobacteriales; Fusobacteriaceae;

CBG61__locus_1

EUT2F

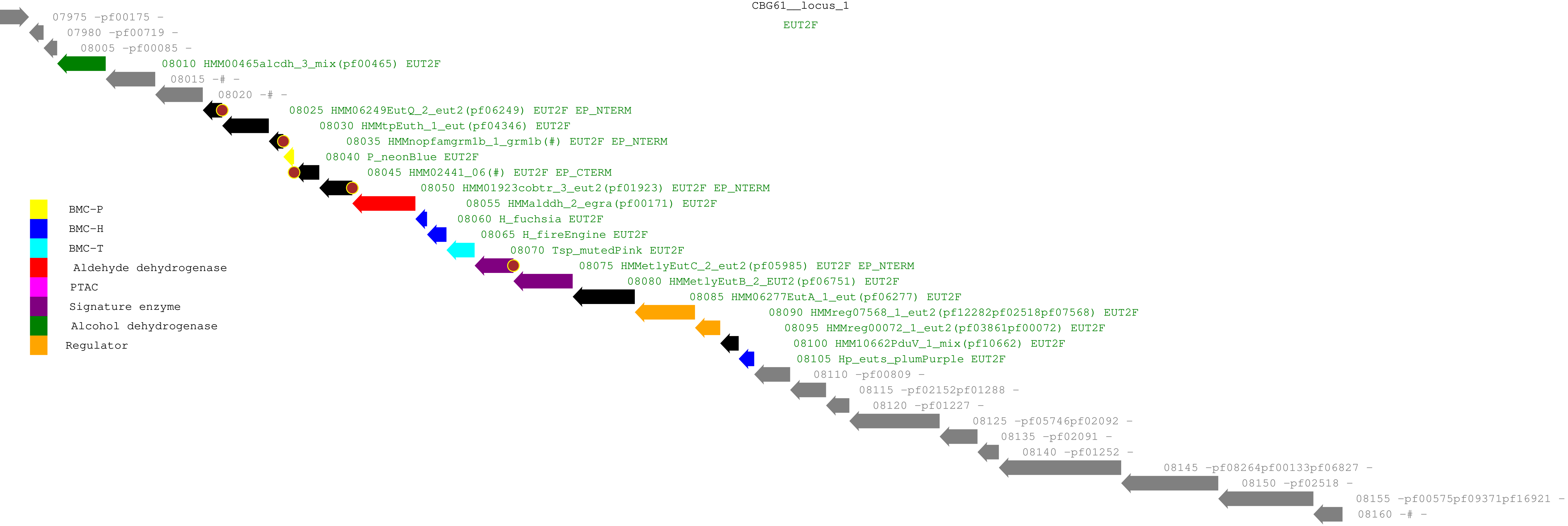

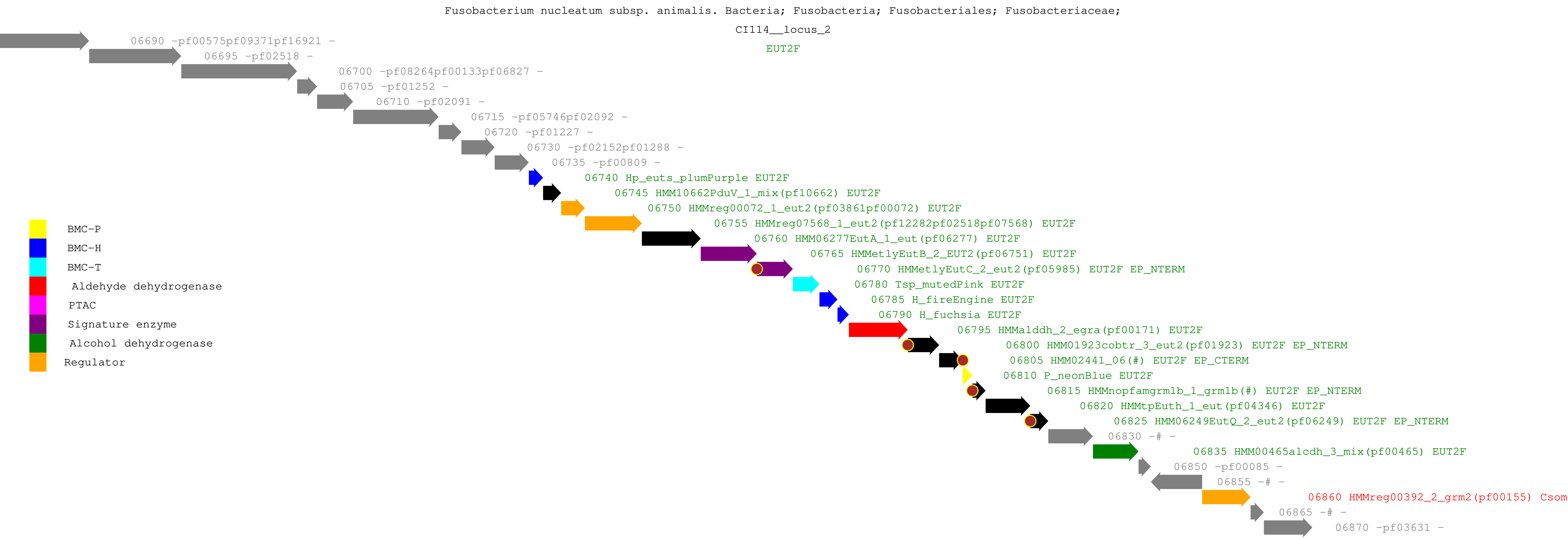
Fusobacterium nucleatum subsp. animalis. Bacteria; Fusobacteria; Fusobacteriales; Fusobacteriaceae;
CI114__locus_2
EUT2F
06690 -pf00575pf09371pf16921 -
06695 -pf02518 -
06700 -pf08264pf00133pf06827 -
06705 -pf01252 -
06710 -pf02091 -
06715 -pf05746pf02092 -
06720 -pf01227 -
06730 -pf02152pf01288 -
06735 -pf00809 -
06740 Hp_euts_plumPurple EUT2F
06745 HMM10662PduV_1_mix(pf10662) EUT2F
06750 HMMreg00072_1_eut2(pf03861pf00072) EUT2F
06755 HMMreg07568_1_eut2(pf12282pf02518pf07568) EUT2F
06760 HMM06277EutA_1_eut(pf06277) EUT2F
06765 HMMetlyEutB_2_EUT2(pf06751) EUT2F
06770 HMMetlyEutC_2_eut2(pf05985) EUT2F EP_NTERM
06780 Tsp_mutedPink EUT2F
06785 H_fireEngine EUT2F
06790 H_fuchsia EUT2F
06795 HMMalddh_2_egra(pf00171) EUT2F
06800 HMM01923cobtr_3_eut2(pf01923) EUT2F EP_NTERM
06805 HMM02441_06(#) EUT2F EP_CTERM
06810 P_neonBlue EUT2F
06815 HMMnopfamgrm1b_1_grm1b(#) EUT2F EP_NTERM
06820 HMMtpEuth_1_eut(pf04346) EUT2F
06825 HMM06249EutQ_2_eut2(pf06249) EUT2F EP_NTERM
06830 -# -
06835 HMM00465alcdh_3_mix(pf00465) EUT2F
06850 -pf00085 -
06855 -# -
06860 HMMreg00392_2_grm2(pf00155) Csom
06865 -# -
06870 -pf03631 -
BMC-P
BMC-H
BMC-T
Aldehyde dehydrogenase
PTAC
Signature enzyme
Alcohol dehydrogenase
Regulator

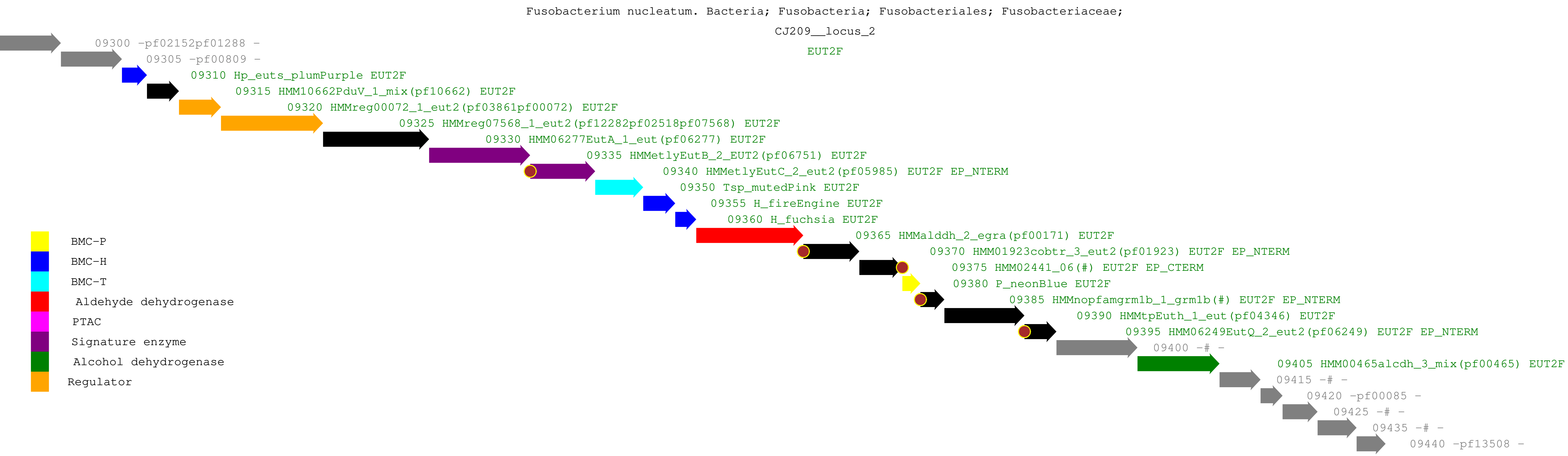
Fusobacterium nucleatum. Bacteria; Fusobacteria; Fusobacteriales; Fusobacteriaceae;
CJ209__locus_2
EUT2F
09300 -pf02152pf01288 -
09305 -pf00809 -
09310 Hp_euts_plumPurple EUT2F
09315 HMM10662PduV_1_mix(pf10662) EUT2F
09320 HMMreg00072_1_eut2(pf03861pf00072) EUT2F
09325 HMMreg07568_1_eut2(pf12282pf02518pf07568) EUT2F
09330 HMM06277EutA_1_eut(pf06277) EUT2F
09335 HMMetlyEutB_2_EUT2(pf06751) EUT2F
09340 HMMetlyEutC_2_eut2(pf05985) EUT2F EP_NTERM
09350 Tsp_mutedPink EUT2F
09355 H_fireEngine EUT2F
09360 H_fuchsia EUT2F
09365 HMMalddh_2_egra(pf00171) EUT2F
09370 HMM01923cobtr_3_eut2(pf01923) EUT2F EP_NTERM
09375 HMM02441_06(#) EUT2F EP_CTERM
09380 P_neonBlue EUT2F
09385 HMMnopfamgrm1b_1_grm1b(#) EUT2F EP_NTERM
09390 HMMtpEuth_1_eut(pf04346) EUT2F
09395 HMM06249EutQ_2_eut2(pf06249) EUT2F EP_NTERM
09400 -# -
09405 HMM00465alcdh_3_mix(pf00465) EUT2F
09415 -# -
09420 -pf00085 -
09425 -# -
09435 -# -
09440 -pf13508 -
BMC-P
BMC-H
BMC-T
Aldehyde dehydrogenase
PTAC
Signature enzyme
Alcohol dehydrogenase
Regulator

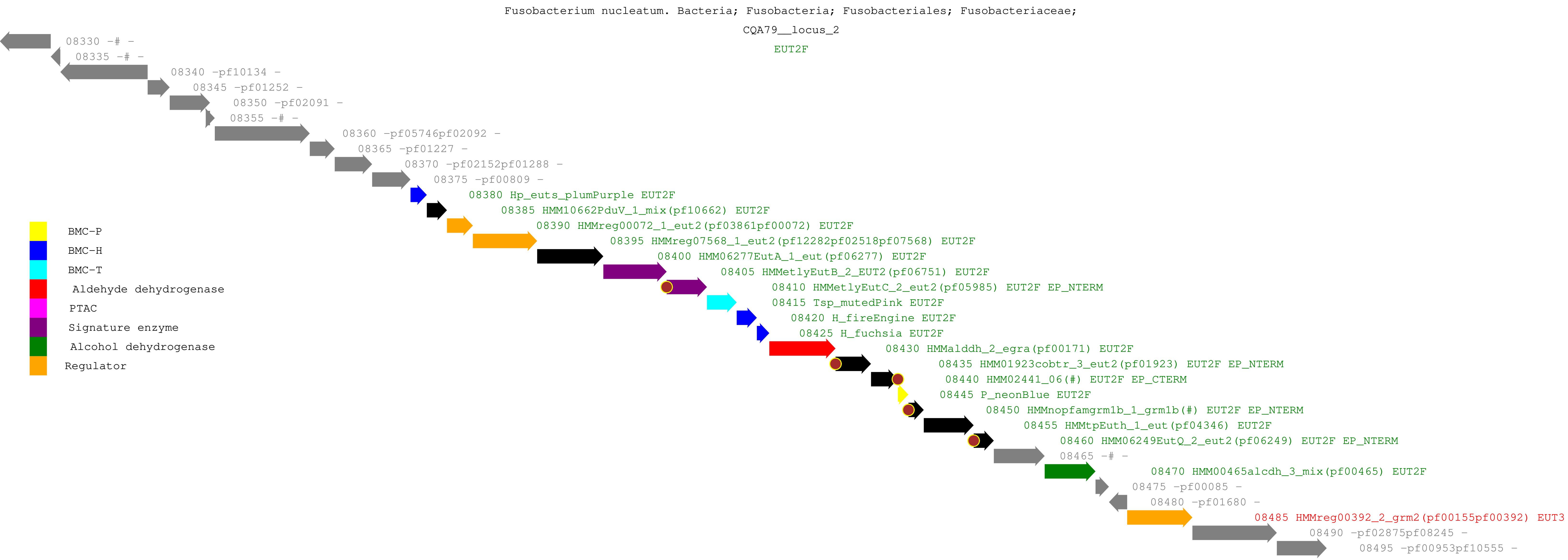
Fusobacterium nucleatum. Bacteria; Fusobacteria; Fusobacteriales; Fusobacteriaceae;
CQA79__locus_2
EUT2F
08330 -# -
08335 -# -
08340 -pf10134 -
08345 -pf01252 -
08350 -pf02091 -
08355 -# -
08360 -pf05746pf02092 -
08365 -pf01227 -
08370 -pf02152pf01288 -
08375 -pf00809 -
08380 Hp_euts_plumPurple EUT2F
08385 HMM10662PduV_1_mix(pf10662) EUT2F
08390 HMMreg00072_1_eut2(pf03861pf00072) EUT2F
08395 HMMreg07568_1_eut2(pf12282pf02518pf07568) EUT2F
08400 HMM06277EutA_1_eut(pf06277) EUT2F
08405 HMMetlyEutB_2_EUT2(pf06751) EUT2F
08410 HMMetlyEutC_2_eut2(pf05985) EUT2F EP_NTERM
08415 Tsp_mutedPink EUT2F
08420 H_fireEngine EUT2F
08425 H_fuchsia EUT2F
08430 HMMalddh_2_egra(pf00171) EUT2F
08435 HMM01923cobtr_3_eut2(pf01923) EUT2F EP_NTERM
08440 HMM02441_06(#) EUT2F EP_CTERM
08445 P_neonBlue EUT2F
08450 HMMnopfamgrm1b_1_grm1b(#) EUT2F EP_NTERM
08455 HMMtpEuth_1_eut(pf04346) EUT2F
08460 HMM06249EutQ_2_eut2(pf06249) EUT2F EP_NTERM
08465 -# -
08470 HMM00465alcdh_3_mix(pf00465) EUT2F
08475 -pf00085 -
08480 -pf01680 -
08485 HMMreg00392_2_grm2(pf00155pf00392) EUT3
08490 -pf02875pf08245 -
08495 -pf00953pf10555 -
BMC-P
BMC-H
BMC-T
Aldehyde dehydrogenase
PTAC
Signature enzyme
Alcohol dehydrogenase
Regulator

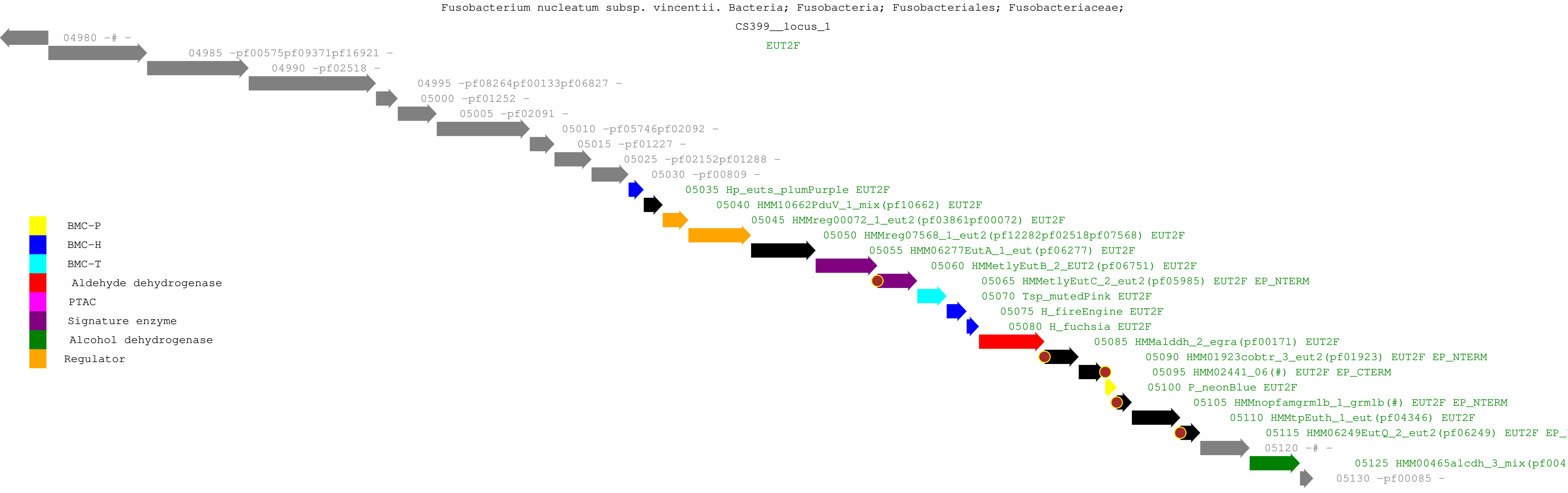
Fusobacterium nucleatum subsp. vincentii. Bacteria; Fusobacteria; Fusobacteriales; Fusobacteriaceae;
CS399__locus_1
EUT2F
04980 -# -
04985 -pf00575pf09371pf16921 -
04990 -pf02518 -
04995 -pf08264pf00133pf06827 -
05000 -pf01252 -
05005 -pf02091 -
05010 -pf05746pf02092 -
05015 -pf01227 -
05025 -pf02152pf01288 -
05030 -pf00809 -
05035 Hp_euts_plumPurple EUT2F
05040 HMM10662PduV_1_mix(pf10662) EUT2F
05045 HMMreg00072_1_eut2(pf03861pf00072) EUT2F
05050 HMMreg07568_1_eut2(pf12282pf02518pf07568) EUT2F
05055 HMM06277EutA_1_eut(pf06277) EUT2F
05060 HMMetlyEutB_2_EUT2(pf06751) EUT2F
05065 HMMetlyEutC_2_eut2(pf05985) EUT2F EP_NTERM
05070 Tsp_mutedPink EUT2F
05075 H_fireEngine EUT2F
05080 H_fuchsia EUT2F
05085 HMMaldh_2_egra(pf00171) EUT2F
05090 HMM01923cobtr_3_eut2(pf01923) EUT2F EP_NTERM
05095 HMM02441_06(#) EUT2F EP_CTERM
05100 P_neonBlue EUT2F
05105 HMMnopfamgrm1b_1_grm1b(#) EUT2F EP_NTERM
05110 HMMtpEuth_1_eut(pf04346) EUT2F
05115 HMM06249EutQ_2_eut2(pf06249) EUT2F EP_
05120 -# -
05125 HMM00465alcdh_3_mix(pf004
05130 -pf00085 -
BMC-P
BMC-H
BMC-T
Aldehyde dehydrogenase
PTAC
Signature enzyme
Alcohol dehydrogenase
Regulator

Fusobacterium periodonticum. Bacteria; Fusobacteria; Fusobacteriales; Fusobacteriaceae;

CTM64__locus_1

EUT2F

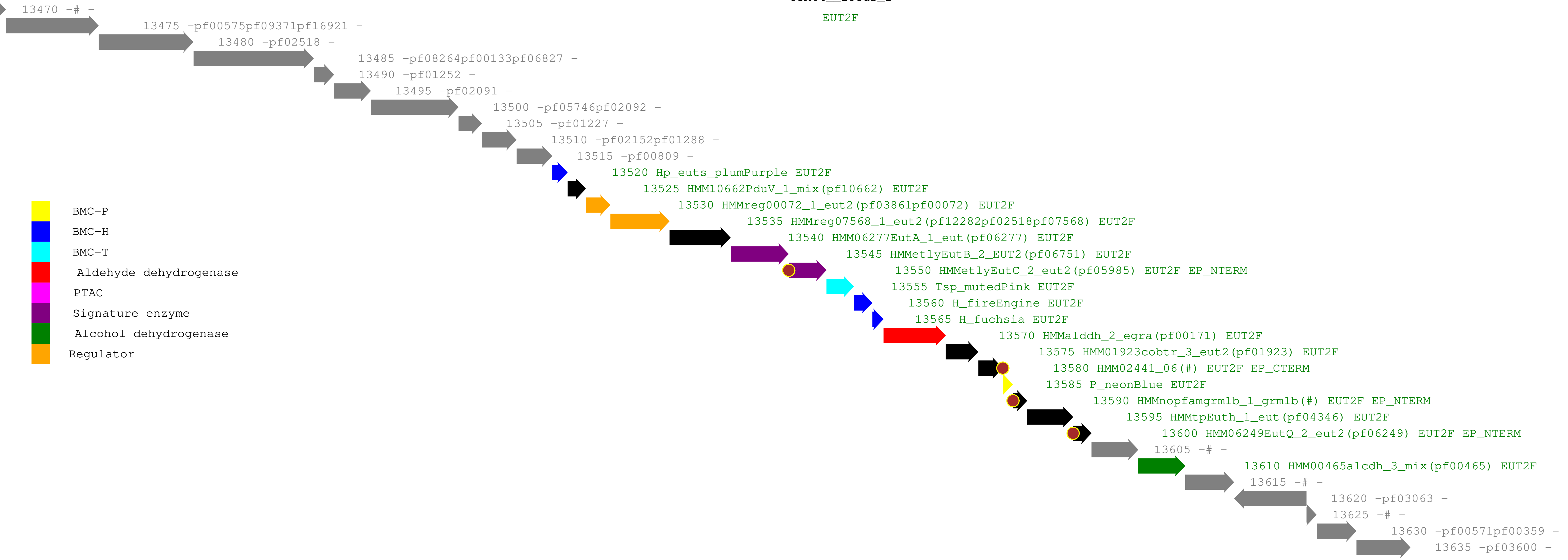

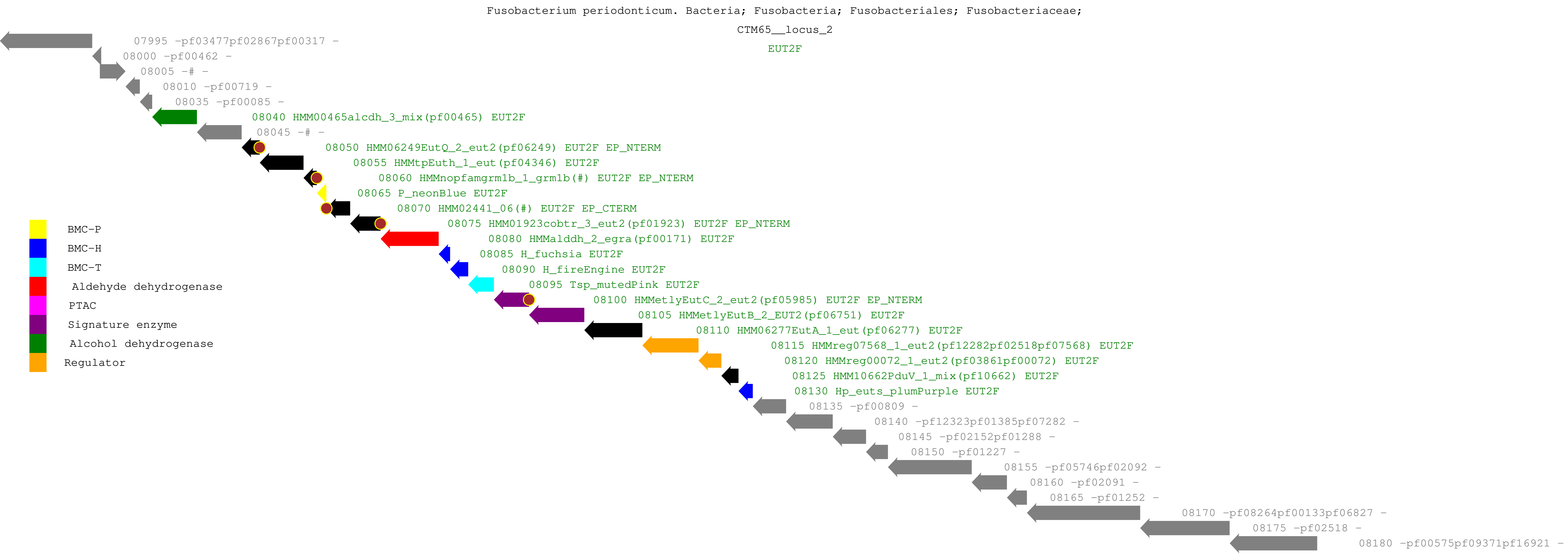
Fusobacterium periodonticum. Bacteria; Fusobacteria; Fusobacteriales; Fusobacteriaceae;
CTM65__locus_2
EUT2F
07995 -pf03477pf02867pf00317 -
08000 -pf00462 -
08005 -# -
08010 -pf00719 -
08035 -pf00085 -
08040 HMM00465alcdh_3_mix(pf00465) EUT2F
08045 -# -
08050 HMM06249EutQ_2_eut2(pf06249) EUT2F EP_NTERM
08055 HMMtpEuth_1_eut(pf04346) EUT2F
08060 HMMnopfamgrm1b_1_grm1b(#) EUT2F EP_NTERM
08065 P_neonBlue EUT2F
08070 HMM02441_06(#) EUT2F EP_CTERM
08075 HMM01923cobtr_3_eut2(pf01923) EUT2F EP_NTERM
08080 HMMalddh_2_egra(pf00171) EUT2F
08085 H_fuchsia EUT2F
08090 H_fireEngine EUT2F
08095 Tsp_mutedPink EUT2F
08100 HMMetlyEutC_2_eut2(pf05985) EUT2F EP_NTERM
08105 HMMetlyEutB_2_EUT2(pf06751) EUT2F
08110 HMM06277EutA_1_eut(pf06277) EUT2F
08115 HMMreg07568_1_eut2(pf12282pf02518pf07568) EUT2F
08120 HMMreg00072_1_eut2(pf03861pf00072) EUT2F
08125 HMM10662PduV_1_mix(pf10662) EUT2F
08130 Hp_euts_plumPurple EUT2F
08135 -pf00809 -
08140 -pf12323pf01385pf07282 -
08145 -pf02152pf01288 -
08150 -pf01227 -
08155 -pf05746pf02092 -
08160 -pf02091 -
08165 -pf01252 -
08170 -pf08264pf00133pf06827 -
08175 -pf02518 -
08180 -pf00575pf09371pf16921 -
BMC-P
BMC-H
BMC-T
Aldehyde dehydrogenase
PTAC
Signature enzyme
Alcohol dehydrogenase
Regulator

Fusobacterium periodonticum. Bacteria; Fusobacteria; Fusobacteriales; Fusobacteriaceae;

CTM69__locus_1

EUT2F

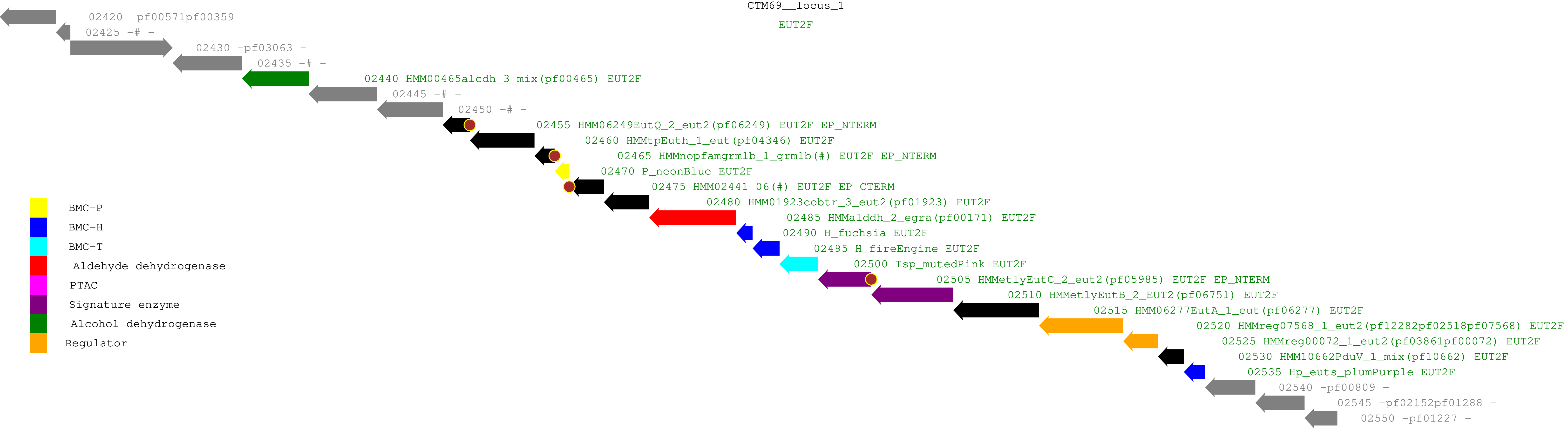

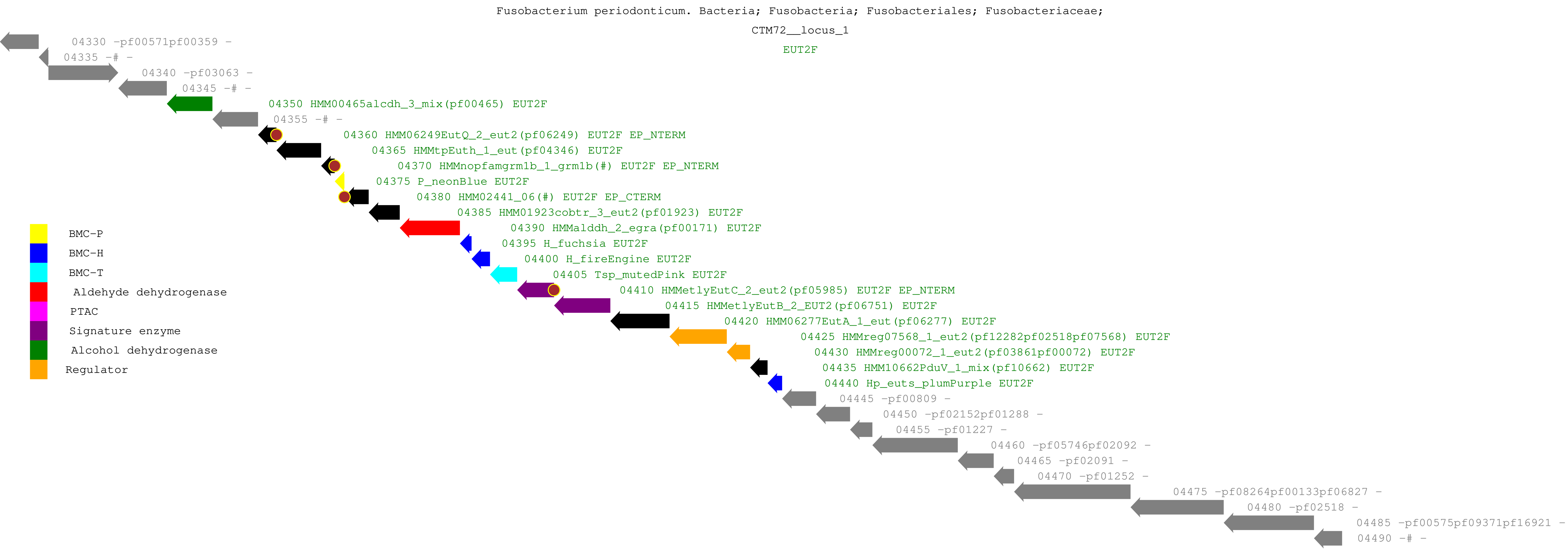
Fusobacterium periodonticum. Bacteria; Fusobacteria; Fusobacteriales; Fusobacteriaceae;
CTM72__locus_1
EUT2F
04330 -pf00571pf00359 -
04335 -# -
04340 -pf03063 -
04345 -# -
04350 HMM00465alcdh_3_mix(pf00465) EUT2F
04355 -# -
04360 HMM06249EutQ_2_eut2(pf06249) EUT2F EP_NTERM
04365 HMMtpEuth_1_eut(pf04346) EUT2F
04370 HMMnopfamgrm1b_1_grm1b(#) EUT2F EP_NTERM
04375 P_neonBlue EUT2F
04380 HMM02441_06(#) EUT2F EP_CTERM
04385 HMM01923cobtr_3_eut2(pf01923) EUT2F
04390 HMMaldh_2_egra(pf00171) EUT2F
04395 H_fuchsia EUT2F
04400 H_fireEngine EUT2F
04405 Tsp_mutedPink EUT2F
04410 HMMetlyEutC_2_eut2(pf05985) EUT2F EP_NTERM
04415 HMMetlyEutB_2_EUT2(pf06751) EUT2F
04420 HMM06277EutA_1_eut(pf06277) EUT2F
04425 HMMreg07568_1_eut2(pf12282pf02518pf07568) EUT2F
04430 HMMreg00072_1_eut2(pf03861pf00072) EUT2F
04435 HMM10662PduV_1_mix(pf10662) EUT2F
04440 Hp_euts_plumPurple EUT2F
04445 -pf00809 -
04450 -pf02152pf01288 -
04455 -pf01227 -
04460 -pf05746pf02092 -
04465 -pf02091 -
04470 -pf01252 -
04475 -pf08264pf00133pf06827 -
04480 -pf02518 -
04485 -pf00575pf09371pf16921 -
04490 -# -
BMC-P
BMC-H
BMC-T
Aldehyde dehydrogenase
PTAC
Signature enzyme
Alcohol dehydrogenase
Regulator

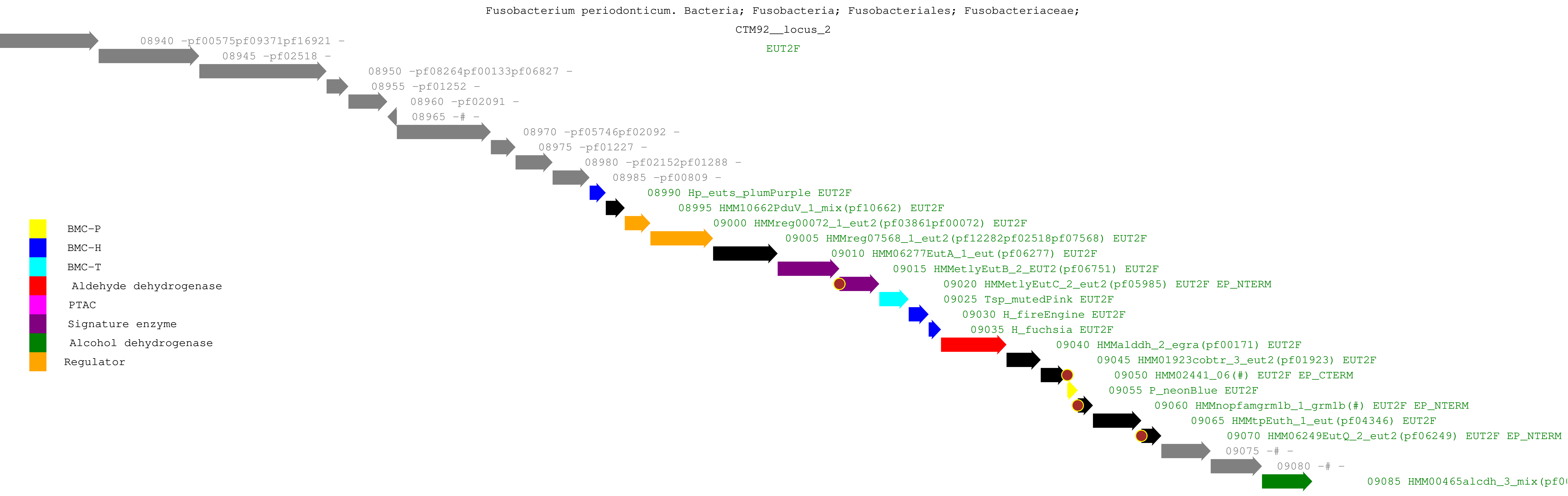
Fusobacterium periodonticum. Bacteria; Fusobacteria; Fusobacteriales; Fusobacteriaceae;
CTM92__locus_2
EUT2F
08940 -pf00575pf09371pf16921 -
08945 -pf02518 -
08950 -pf08264pf00133pf06827 -
08955 -pf01252 -
08960 -pf02091 -
08965 -# -
08970 -pf05746pf02092 -
08975 -pf01227 -
08980 -pf02152pf01288 -
08985 -pf00809 -
08990 Hp_euts_plumPurple EUT2F
08995 HMM10662PduV_1_mix(pf10662) EUT2F
09000 HMMreg00072_1_eut2(pf03861pf00072) EUT2F
09005 HMMreg07568_1_eut2(pf12282pf02518pf07568) EUT2F
09010 HMM06277EutA_1_eut(pf06277) EUT2F
09015 HMMetlyEutB_2_EUT2(pf06751) EUT2F
09020 HMMetlyEutC_2_eut2(pf05985) EUT2F EP_NTERM
09025 Tsp_mutedPink EUT2F
09030 H_fireEngine EUT2F
09035 H_fuchsia EUT2F
09040 HMMalddh_2_egra(pf00171) EUT2F
09045 HMM01923cobtr_3_eut2(pf01923) EUT2F
09050 HMM02441_06(#) EUT2F EP_CTERM
09055 P_neonBlue EUT2F
09060 HMMnopfamgrm1b_1_grm1b(#) EUT2F EP_NTERM
09065 HMMtpEuth_1_eut(pf04346) EUT2F
09070 HMM06249EutQ_2_eut2(pf06249) EUT2F EP_NTERM
09075 -# -
09080 -# -
09085 HMM00465alcdh_3_mix(pf0
BMC-P
BMC-H
BMC-T
Aldehyde dehydrogenase
PTAC
Signature enzyme
Alcohol dehydrogenase
Regulator

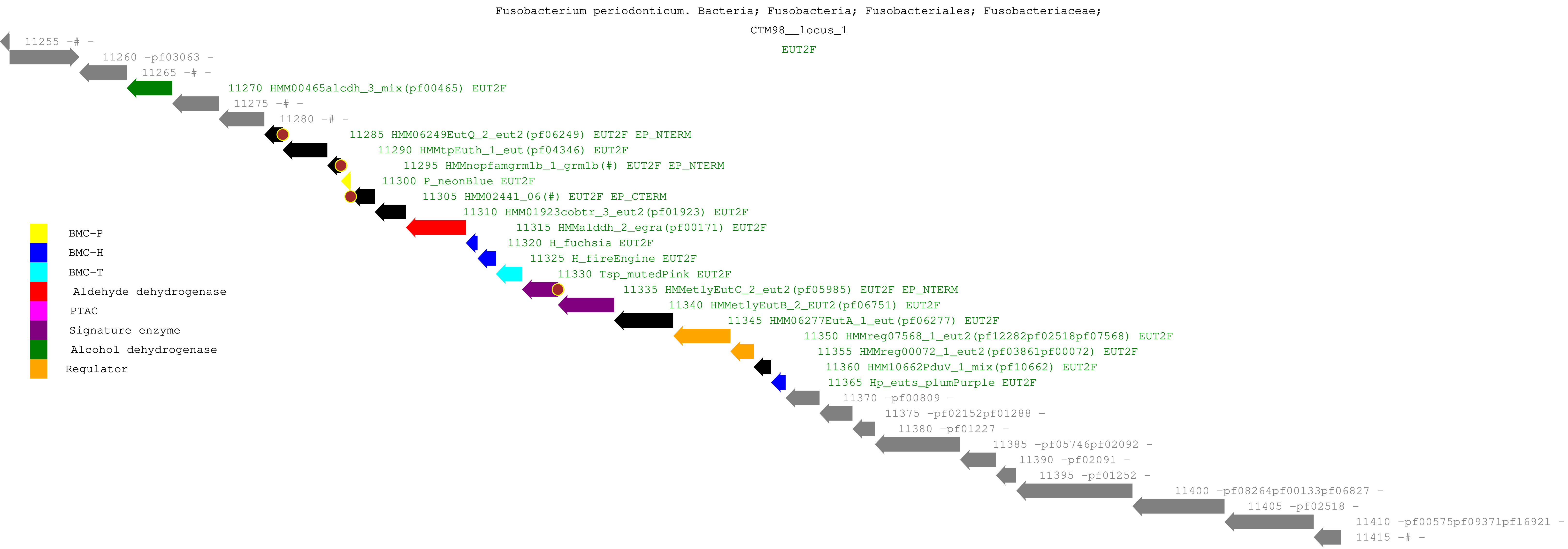

Fusobacterium periodonticum. Bacteria; Fusobacteria; Fusobacteriales; Fusobacteriaceae;
CTM98__locus_1
EUT2F
11255 -# -
11260 -pf03063 -
11265 -# -
11270 HMM00465alcdh_3_mix(pf00465) EUT2F
11275 -# -
11280 -# -
11285 HMM06249EutQ_2_eut2(pf06249) EUT2F EP_NTERM
11290 HMMtpEuth_1_eut(pf04346) EUT2F
11295 HMMnopfamgrm1b_1_grm1b(#) EUT2F EP_NTERM
11300 P_neonBlue EUT2F
11305 HMM02441_06(#) EUT2F EP_CTERM
11310 HMM01923cobtr_3_eut2(pf01923) EUT2F
11315 HMMalddh_2_egra(pf00171) EUT2F
11320 H_fuchsia EUT2F
11325 H_fireEngine EUT2F
11330 Tsp_mutedPink EUT2F
11335 HMMetlyEutC_2_eut2(pf05985) EUT2F EP_NTERM
11340 HMMetlyEutB_2_EUT2(pf06751) EUT2F
11345 HMM06277EutA_1_eut(pf06277) EUT2F
11350 HMMreg07568_1_eut2(pf12282pf02518pf07568) EUT2F
11355 HMMreg00072_1_eut2(pf03861pf00072) EUT2F
11360 HMM10662PduV_1_mix(pf10662) EUT2F
11365 Hp_euts_plumPurple EUT2F
11370 -pf00809 -
11375 -pf02152pf01288 -
11380 -pf01227 -
11385 -pf05746pf02092 -
11390 -pf02091 -
11395 -pf01252 -
11400 -pf08264pf00133pf06827 -
11405 -pf02518 -
11410 -pf00575pf09371pf16921 -
11415 -# -
BMC-P
BMC-H
BMC-T
Aldehyde dehydrogenase
PTAC
Signature enzyme
Alcohol dehydrogenase
Regulator

Fusobacterium periodonticum. Bacteria; Fusobacteria; Fusobacteriales; Fusobacteriaceae;

CTN00__locus_1

EUT2F

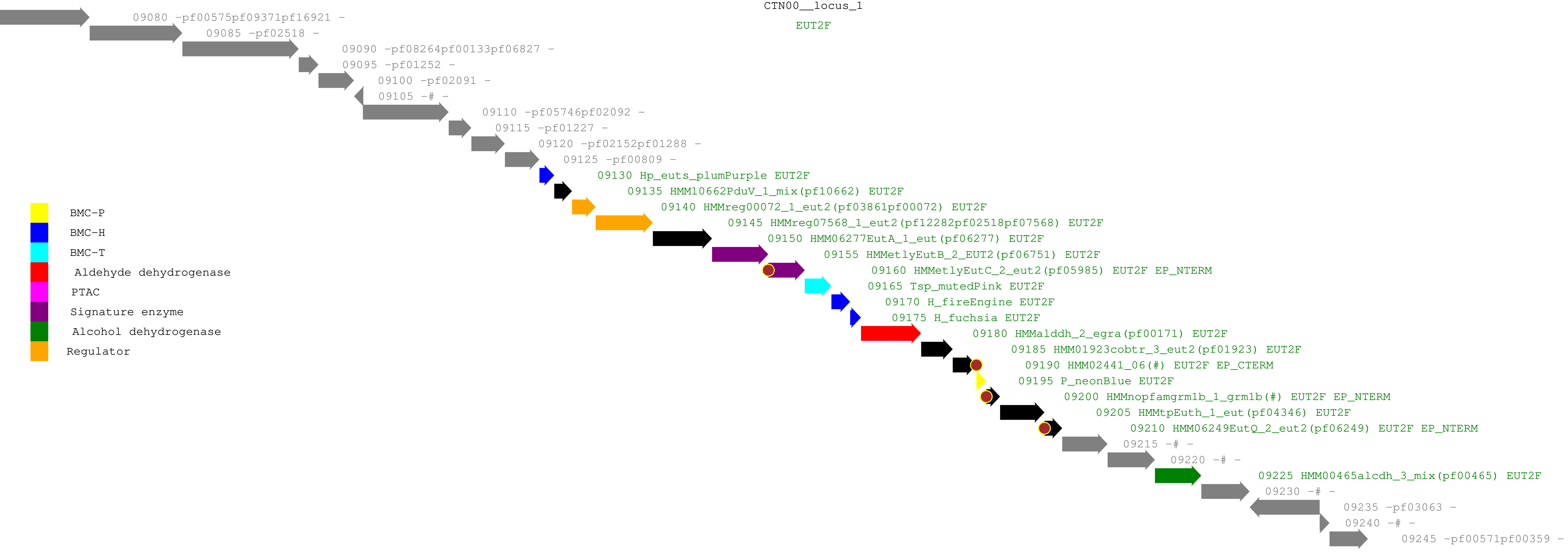

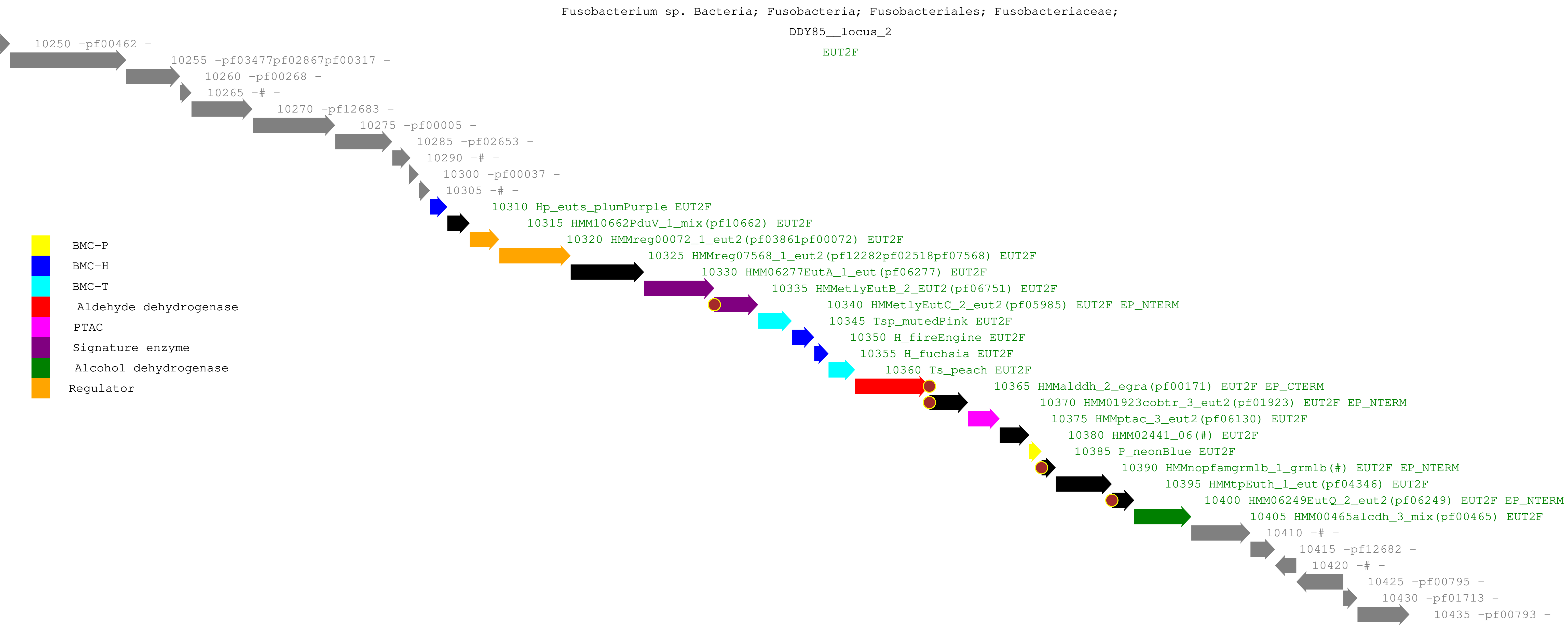

Fusobacterium sp. Bacteria; Fusobacteria; Fusobacteriales; Fusobacteriaceae;
DDY85__locus_2
EUT2F
10250 -pf00462 -
10255 -pf03477pf02867pf00317 -
10260 -pf00268 -
10265 -# -
10270 -pf12683 -
10275 -pf00005 -
10285 -pf02653 -
10290 -# -
10300 -pf00037 -
10305 -# -
10310 Hp_euts_plumPurple EUT2F
10315 HMM10662PduV_1_mix(pf10662) EUT2F
10320 HMMreg00072_1_eut2(pf03861pf00072) EUT2F
10325 HMMreg07568_1_eut2(pf12282pf02518pf07568) EUT2F
10330 HMM06277EutA_1_eut(pf06277) EUT2F
10335 HMMetlyEutB_2_EUT2(pf06751) EUT2F
10340 HMMetlyEutC_2_eut2(pf05985) EUT2F EP_NTERM
10345 Tsp_mutedPink EUT2F
10350 H_fireEngine EUT2F
10355 H_fuchsia EUT2F
10360 Ts_peach EUT2F
10365 HMMaldh_2_egra(pf00171) EUT2F EP_CTERM
10370 HMM01923cobtr_3_eut2(pf01923) EUT2F EP_NTERM
10375 HMMptac_3_eut2(pf06130) EUT2F
10380 HMM02441_06(#) EUT2F
10385 P_neonBlue EUT2F
10390 HMMnopfamgrm1b_1_grm1b(#) EUT2F EP_NTERM
10395 HMMtpEuth_1_eut(pf04346) EUT2F
10400 HMM06249EutQ_2_eut2(pf06249) EUT2F EP_NTERM
10405 HMM00465alcdh_3_mix(pf00465) EUT2F
10410 -# -
10415 -pf12682 -
10420 -# -
10425 -pf00795 -
10430 -pf01713 -
10435 -pf00793 -
BMC-P
BMC-H
BMC-T
Aldehyde dehydrogenase
PTAC
Signature enzyme
Alcohol dehydrogenase
Regulator

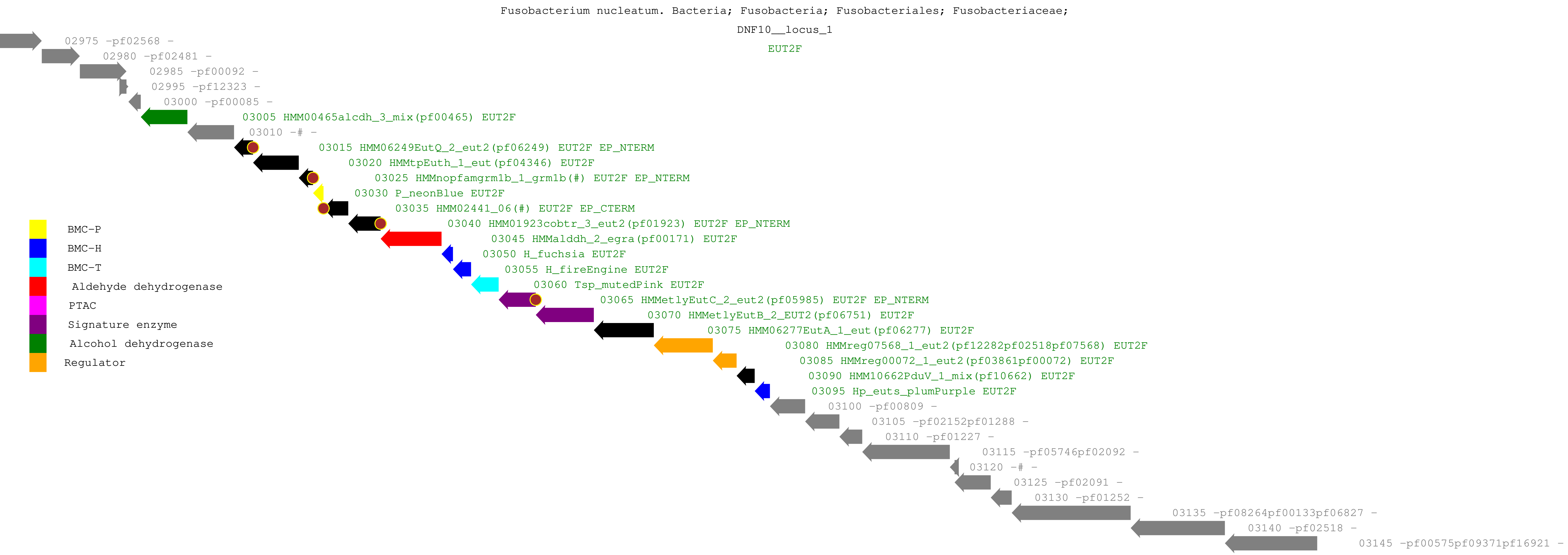
Fusobacterium nucleatum. Bacteria; Fusobacteria; Fusobacteriales; Fusobacteriaceae;
DNF10__locus_1
EUT2F
02975 -pf02568 -
02980 -pf02481 -
02985 -pf00092 -
02995 -pf12323 -
03000 -pf00085 -
03005 HMM00465alcdh_3_mix(pf00465) EUT2F
03010 -# -
03015 HMM06249EutQ_2_eut2(pf06249) EUT2F EP_NTERM
03020 HMMtpEuth_1_eut(pf04346) EUT2F
03025 HMMnopfamgrm1b_1_grm1b(#) EUT2F EP_NTERM
03030 P_neonBlue EUT2F
03035 HMM02441_06(#) EUT2F EP_CTERM
03040 HMM01923cobtr_3_eut2(pf01923) EUT2F EP_NTERM
03045 HMMaldh_2_egra(pf00171) EUT2F
03050 H_fuchsia EUT2F
03055 H_fireEngine EUT2F
03060 Tsp_mutedPink EUT2F
03065 HMMetlyEutC_2_eut2(pf05985) EUT2F EP_NTERM
03070 HMMetlyEutB_2_EUT2(pf06751) EUT2F
03075 HMM06277EutA_1_eut(pf06277) EUT2F
03080 HMMreg07568_1_eut2(pf12282pf02518pf07568) EUT2F
03085 HMMreg00072_1_eut2(pf03861pf00072) EUT2F
03090 HMM10662PduV_1_mix(pf10662) EUT2F
03095 Hp_euts_plumPurple EUT2F
03100 -pf00809 -
03105 -pf02152pf01288 -
03110 -pf01227 -
03115 -pf05746pf02092 -
03120 -# -
03125 -pf02091 -
03130 -pf01252 -
03135 -pf08264pf00133pf06827 -
03140 -pf02518 -
03145 -pf00575pf09371pf16921 -
BMC-P
BMC-H
BMC-T
Aldehyde dehydrogenase
PTAC
Signature enzyme
Alcohol dehydrogenase
Regulator

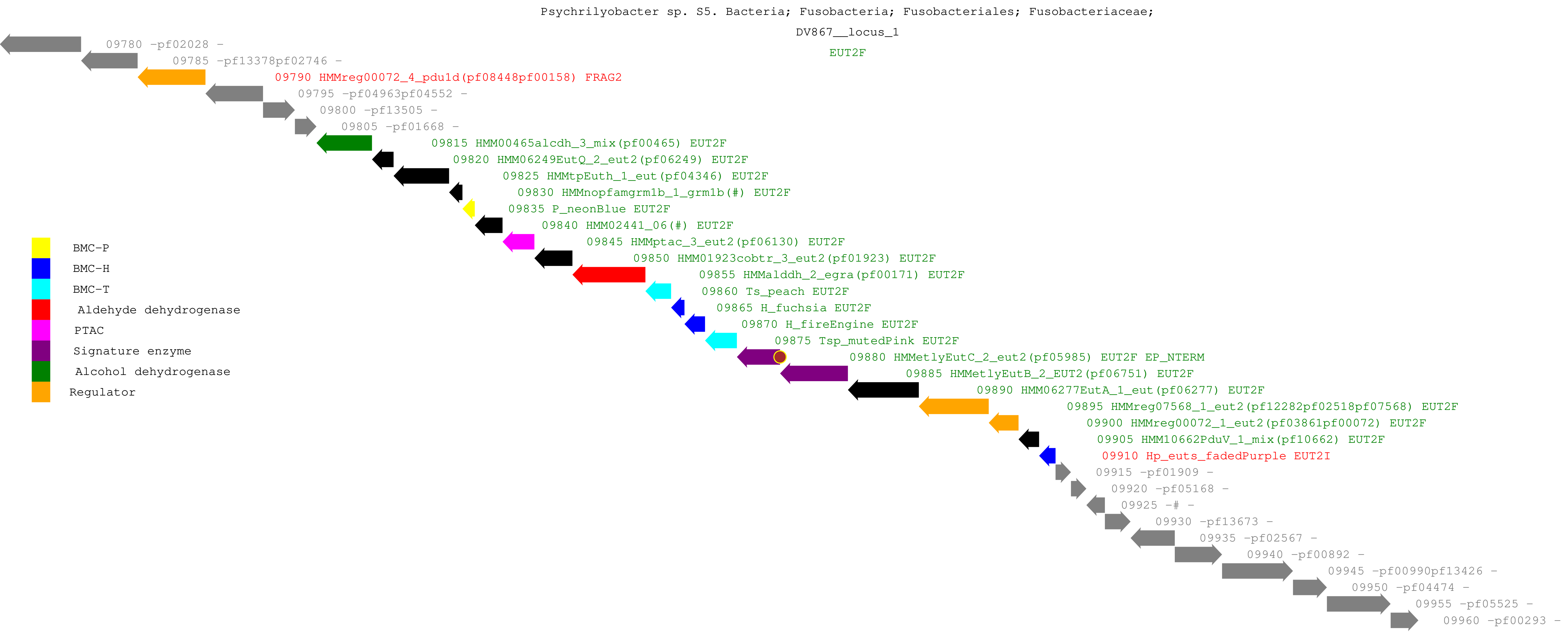
Psychrilyobacter sp. S5. Bacteria; Fusobacteria; Fusobacteriales; Fusobacteriaceae;
DV867__locus_1
EUT2F
09780 -pf02028 -
09785 -pf13378pf02746 -
09790 HMMreg00072_4_pdu1d(pf08448pf00158) FRAG2
09795 -pf04963pf04552 -
09800 -pf13505 -
09805 -pf01668 -
09815 HMM00465alcdh_3_mix(pf00465) EUT2F
09820 HMM06249EutQ_2_eut2(pf06249) EUT2F
09825 HMMtpEuth_1_eut(pf04346) EUT2F
09830 HMMnopfamgrm1b_1_grm1b(#) EUT2F
09835 P_neonBlue EUT2F
09840 HMM02441_06(#) EUT2F
09845 HMMptac_3_eut2(pf06130) EUT2F
09850 HMM01923cobtr_3_eut2(pf01923) EUT2F
09855 HMMalddh_2_egra(pf00171) EUT2F
09860 Ts_peach EUT2F
09865 H_fuchsia EUT2F
09870 H_fireEngine EUT2F
09875 Tsp_mutedPink EUT2F
09880 HMMetlyEutC_2_eut2(pf05985) EUT2F EP_NTERM
09885 HMMetlyEutB_2_EUT2(pf06751) EUT2F
09890 HMM06277EutA_1_eut(pf06277) EUT2F
09895 HMMreg07568_1_eut2(pf12282pf02518pf07568) EUT2F
09900 HMMreg00072_1_eut2(pf03861pf00072) EUT2F
09905 HMM10662PduV_1_mix(pf10662) EUT2F
09910 Hp_euts_fadedPurple EUT2I
09915 -pf01909 -
09920 -pf05168 -
09925 -# -
09930 -pf13673 -
09935 -pf02567 -
09940 -pf00892 -
09945 -pf00990pf13426 -
09950 -pf04474 -
09955 -pf05525 -
09960 -pf00293 -
BMC-P
BMC-H
BMC-T
Aldehyde dehydrogenase
PTAC
Signature enzyme
Alcohol dehydrogenase
Regulator

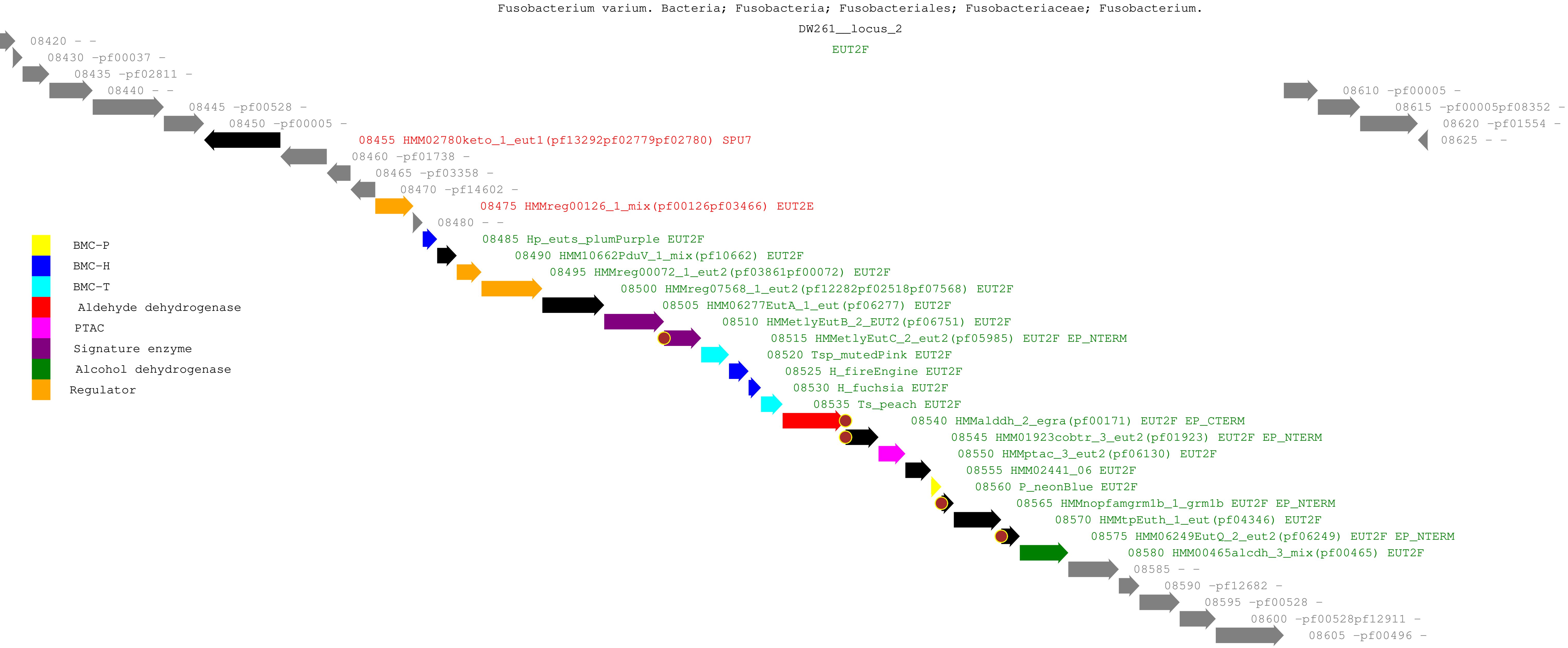
Fusobacterium varium. Bacteria; Fusobacteria; Fusobacteriales; Fusobacteriaceae; Fusobacterium.
DW261__locus_2
EUT2F
08420 - -
08430 -pf00037 -
08435 -pf02811 -
08440 - -
08445 -pf00528 -
08450 -pf00005 -
08455 HMM02780keto_1_eut1(pf13292pf02779pf02780) SPU7
08460 -pf01738 -
08465 -pf03358 -
08470 -pf14602 -
08475 HMMreg00126_1_mix(pf00126pf03466) EUT2E
08480 - -
08485 Hp_euts_plumPurple EUT2F
08490 HMM10662PduV_1_mix(pf10662) EUT2F
08495 HMMreg00072_1_eut2(pf03861pf00072) EUT2F
08500 HMMreg07568_1_eut2(pf12282pf02518pf07568) EUT2F
08505 HMM06277EutA_1_eut(pf06277) EUT2F
08510 HMMetlyEutB_2_EUT2(pf06751) EUT2F
08515 HMMetlyEutC_2_eut2(pf05985) EUT2F EP_NTERM
08520 Tsp_mutedPink EUT2F
08525 H_fireEngine EUT2F
08530 H_fuchsia EUT2F
08535 Ts_peach EUT2F
08540 HMMalddh_2_egra(pf00171) EUT2F EP_CTERM
08545 HMM01923cobtr_3_eut2(pf01923) EUT2F EP_NTERM
08550 HMMptac_3_eut2(pf06130) EUT2F
08555 HMM02441_06 EUT2F
08560 P_neonBlue EUT2F
08565 HMMnopfamgrm1b_1_grm1b EUT2F EP_NTERM
08570 HMMtpEuth_1_eut(pf04346) EUT2F
08575 HMM06249EutQ_2_eut2(pf06249) EUT2F EP_NTERM
08580 HMM00465alcdh_3_mix(pf00465) EUT2F
08585 - -
08590 -pf12682 -
08595 -pf00528 -
08600 -pf00528pf12911 -
08605 -pf00496 -
08610 -pf00005 -
08615 -pf00005pf08352 -
08620 -pf01554 -
08625 - -
BMC-P
BMC-H
BMC-T
Aldehyde dehydrogenase
PTAC
Signature enzyme
Alcohol dehydrogenase
Regulator

Psychrilyobacter sp. SD5. Bacteria; Fusobacteria; Fusobacteriales; Fusobacteriaceae;

DYH56__locus_1

EUT2F

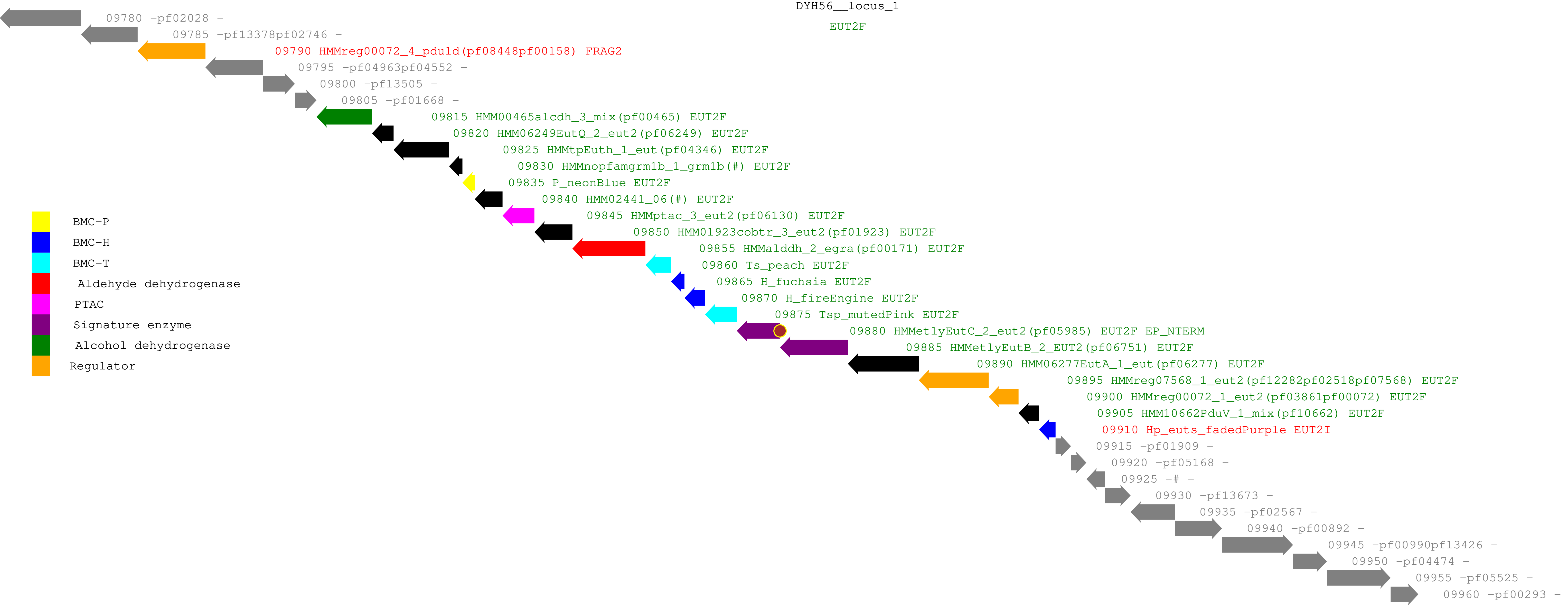

Fusobacterium canifelinum. Bacteria; Fusobacteria; Fusobacteriales; Fusobacteriaceae; Fusobacterium.

EII27__locus_1

EUT2F

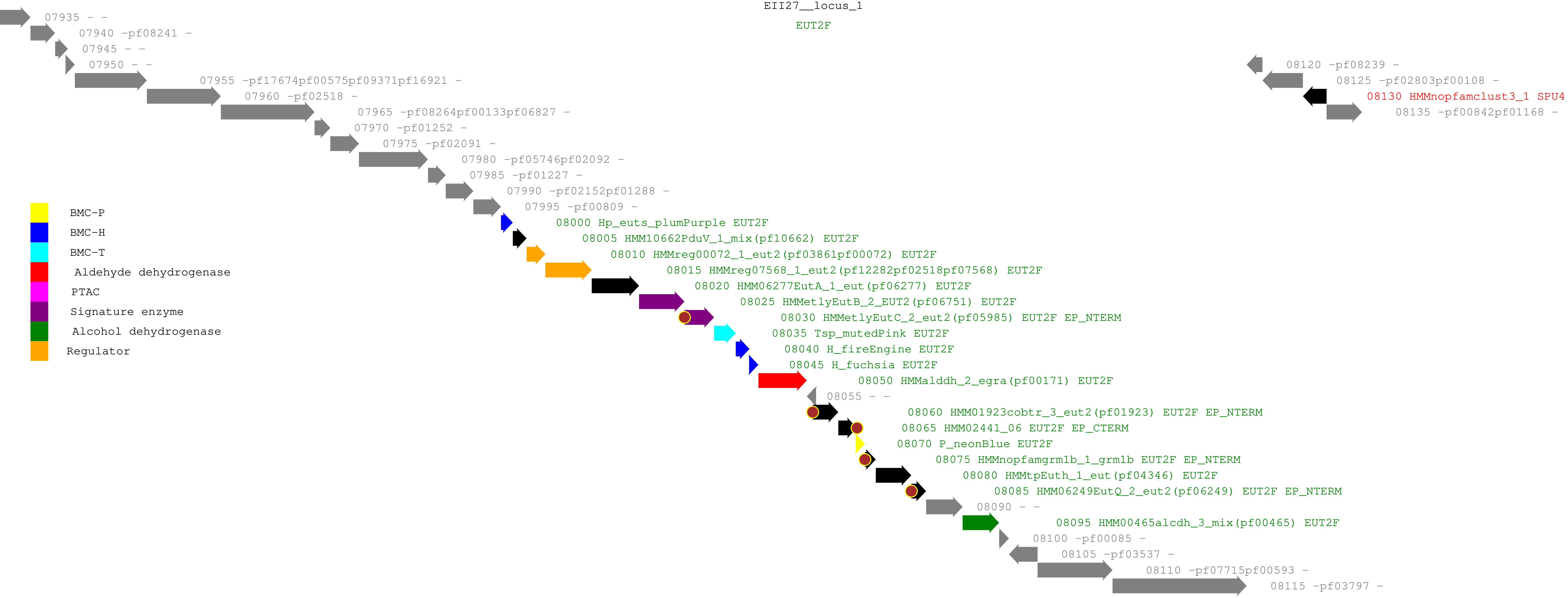

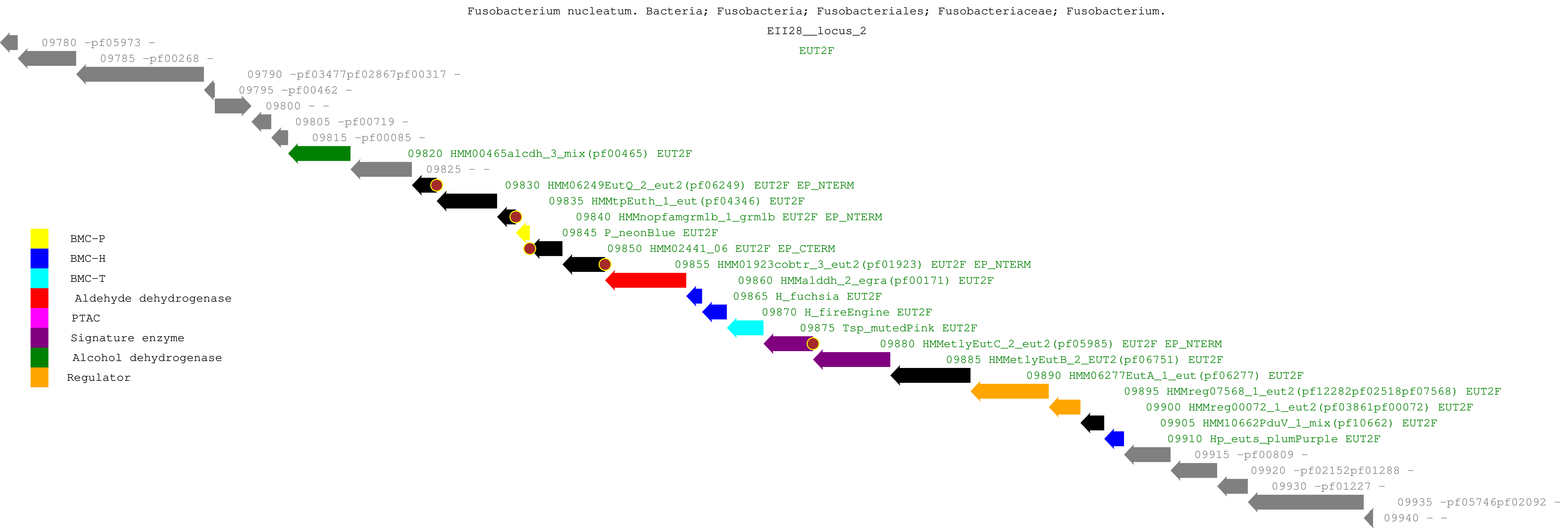
Fusobacterium nucleatum. Bacteria; Fusobacteria; Fusobacteriales; Fusobacteriaceae; Fusobacterium.
EII28__locus_2
EUT2F
09780 -pf05973 -
09785 -pf00268 -
09790 -pf03477pf02867pf00317 -
09795 -pf00462 -
09800 - -
09805 -pf00719 -
09815 -pf00085 -
09820 HMM00465alcdh_3_mix(pf00465) EUT2F
09825 - -
09830 HMM06249EutQ_2_eut2(pf06249) EUT2F EP_NTERM
09835 HMMtpEuth_1_eut(pf04346) EUT2F
09840 HMMnopfamgrm1b_1_grm1b EUT2F EP_NTERM
09845 P_neonBlue EUT2F
09850 HMM02441_06 EUT2F EP_CTERM
09855 HMM01923cobtr_3_eut2(pf01923) EUT2F EP_NTERM
09860 HMMalddh_2_egra(pf00171) EUT2F
09865 H_fuchsia EUT2F
09870 H_fireEngine EUT2F
09875 Tsp_mutedPink EUT2F
09880 HMMetlyEutC_2_eut2(pf05985) EUT2F EP_NTERM
09885 HMMetlyEutB_2_EUT2(pf06751) EUT2F
09890 HMM06277EutA_1_eut(pf06277) EUT2F
09895 HMMreg07568_1_eut2(pf12282pf02518pf07568) EUT2F
09900 HMMreg00072_1_eut2(pf03861pf00072) EUT2F
09905 HMM10662PduV_1_mix(pf10662) EUT2F
09910 Hp_euts_plumPurple EUT2F
09915 -pf00809 -
09920 -pf02152pf01288 -
09930 -pf01227 -
09935 -pf05746pf02092 -
09940 - -
BMC-P
BMC-H
BMC-T
Aldehyde dehydrogenase
PTAC
Signature enzyme
Alcohol dehydrogenase
Regulator

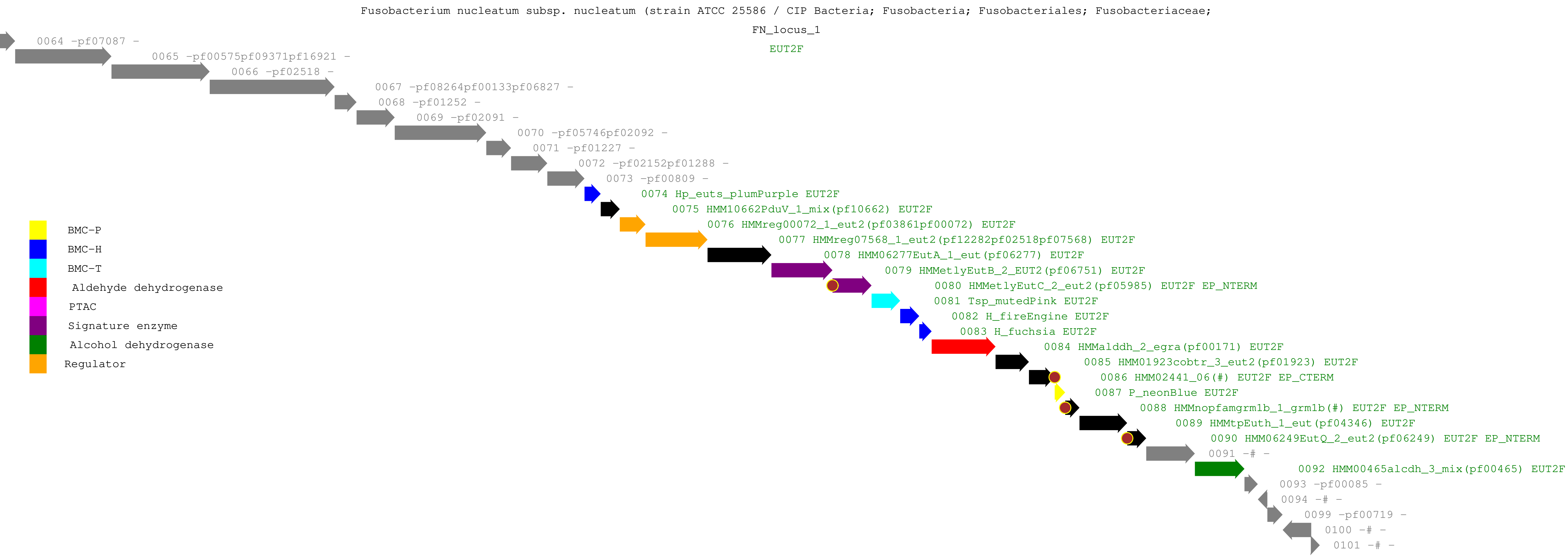
Fusobacterium nucleatum subsp. nucleatum (strain ATCC 25586 / CIP Bacteria; Fusobacteria; Fusobacteriales; Fusobacteriaceae;
FN_locus_1
EUT2F
0064 -pf07087 -
0065 -pf00575pf09371pf16921 -
0066 -pf02518 -
0067 -pf08264pf00133pf06827 -
0068 -pf01252 -
0069 -pf02091 -
0070 -pf05746pf02092 -
0071 -pf01227 -
0072 -pf02152pf01288 -
0073 -pf00809 -
0074 Hp_euts_plumPurple EUT2F
0075 HMM10662PduV_1_mix(pf10662) EUT2F
0076 HMMreg00072_1_eut2(pf03861pf00072) EUT2F
0077 HMMreg07568_1_eut2(pf12282pf02518pf07568) EUT2F
0078 HMM06277EutA_1_eut(pf06277) EUT2F
0079 HMMetlyEutB_2_EUT2(pf06751) EUT2F
0080 HMMetlyEutC_2_eut2(pf05985) EUT2F EP_NTERM
0081 Tsp_mutedPink EUT2F
0082 H_fireEngine EUT2F
0083 H_fuchsia EUT2F
0084 HMMalddh_2_egra(pf00171) EUT2F
0085 HMM01923cobtr_3_eut2(pf01923) EUT2F
0086 HMM02441_06(#) EUT2F EP_CTERM
0087 P_neonBlue EUT2F
0088 HMMnopfamgrm1b_1_grm1b(#) EUT2F EP_NTERM
0089 HMMtpEuth_1_eut(pf04346) EUT2F
0090 HMM06249EutQ_2_eut2(pf06249) EUT2F EP_NTERM
0091 -# -
0092 HMM00465alcdh_3_mix(pf00465) EUT2F
0093 -pf00085 -
0094 -# -
0099 -pf00719 -
0100 -# -
0101 -# -
BMC-P
BMC-H
BMC-T
Aldehyde dehydrogenase
PTAC
Signature enzyme
Alcohol dehydrogenase
Regulator

Fusobacterium nucleatum subsp. polymorphum ATCC 10953. Bacteria; Fusobacteria; Fusobacteriales; Fusobacteriaceae;

FNP__locus_1

EUT2F

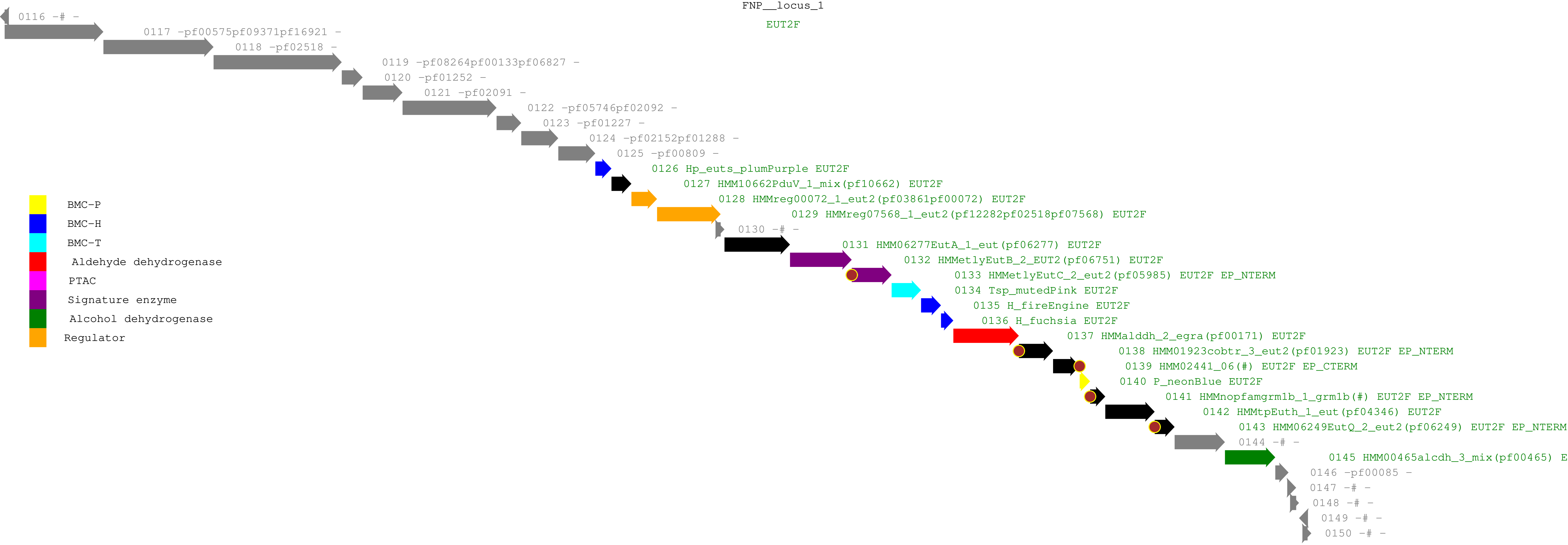

Fusobacterium periodonticum D10. Bacteria; Fusobacteria; Fusobacteriales; Fusobacteriaceae;

FPOG__locus_1

EUT2F

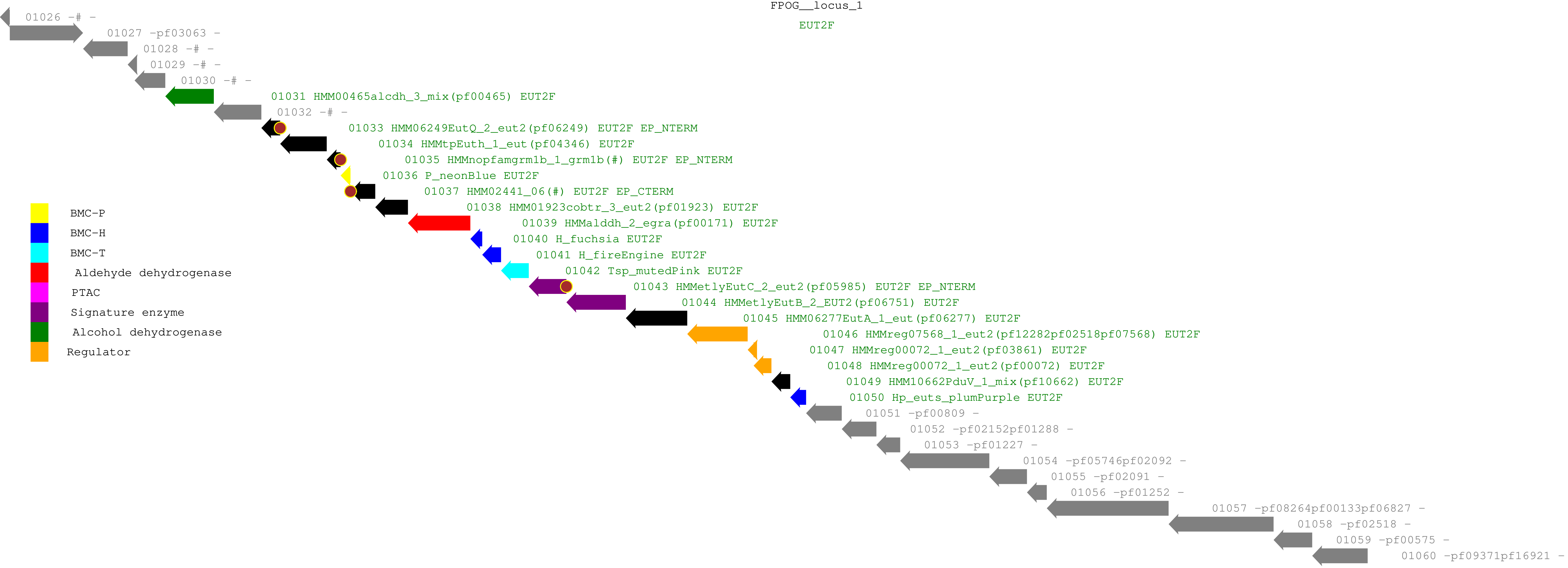

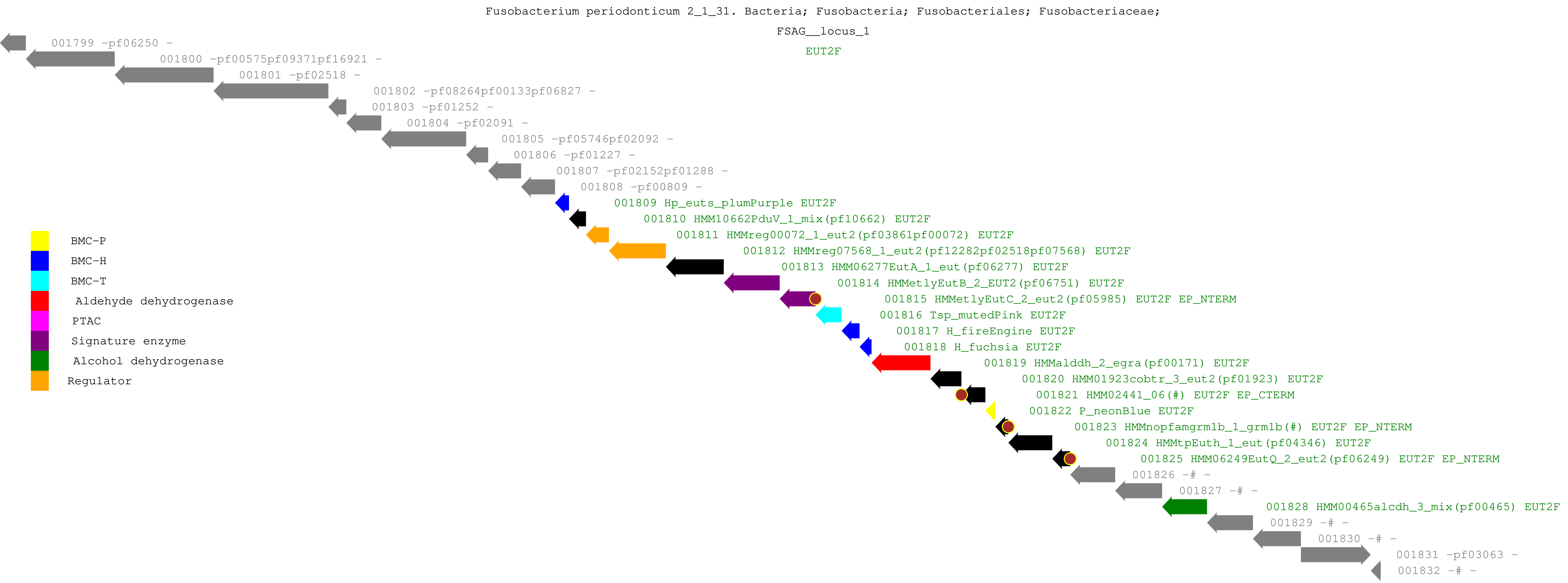

Fusobacterium periodonticum 2_1_31. Bacteria; Fusobacteria; Fusobacteriales; Fusobacteriaceae;
FSAG__locus_1
EUT2F
001799 -pf06250 -
001800 -pf00575pf09371pf16921 -
001801 -pf02518 -
001802 -pf08264pf00133pf06827 -
001803 -pf01252 -
001804 -pf02091 -
001805 -pf05746pf02092 -
001806 -pf01227 -
001807 -pf02152pf01288 -
001808 -pf00809 -
001809 Hp_euts_plumPurple EUT2F
001810 HMM10662PduV_1_mix(pf10662) EUT2F
001811 HMMreg00072_1_eut2(pf03861pf00072) EUT2F
001812 HMMreg07568_1_eut2(pf12282pf02518pf07568) EUT2F
001813 HMM06277EutA_1_eut(pf06277) EUT2F
001814 HMMetlyEutB_2_EUT2(pf06751) EUT2F
001815 HMMetlyEutC_2_eut2(pf05985) EUT2F EP_NTERM
001816 Tsp_mutedPink EUT2F
001817 H_fireEngine EUT2F
001818 H_fuchsia EUT2F
001819 HMMalddh_2_egra(pf00171) EUT2F
001820 HMM01923cobtr_3_eut2(pf01923) EUT2F
001821 HMM02441_06(#) EUT2F EP_CTERM
001822 P_neonBlue EUT2F
001823 HMMnopfamgrm1b_1_grm1b(#) EUT2F EP_NTERM
001824 HMMtpEuth_1_eut(pf04346) EUT2F
001825 HMM06249EutQ_2_eut2(pf06249) EUT2F EP_NTERM
001826 -# -
001827 -# -
001828 HMM00465alcdh_3_mix(pf00465) EUT2F
001829 -# -
001830 -# -
001831 -pf03063 -
001832 -# -
BMC-P
BMC-H
BMC-T
Aldehyde dehydrogenase
PTAC
Signature enzyme
Alcohol dehydrogenase
Regulator

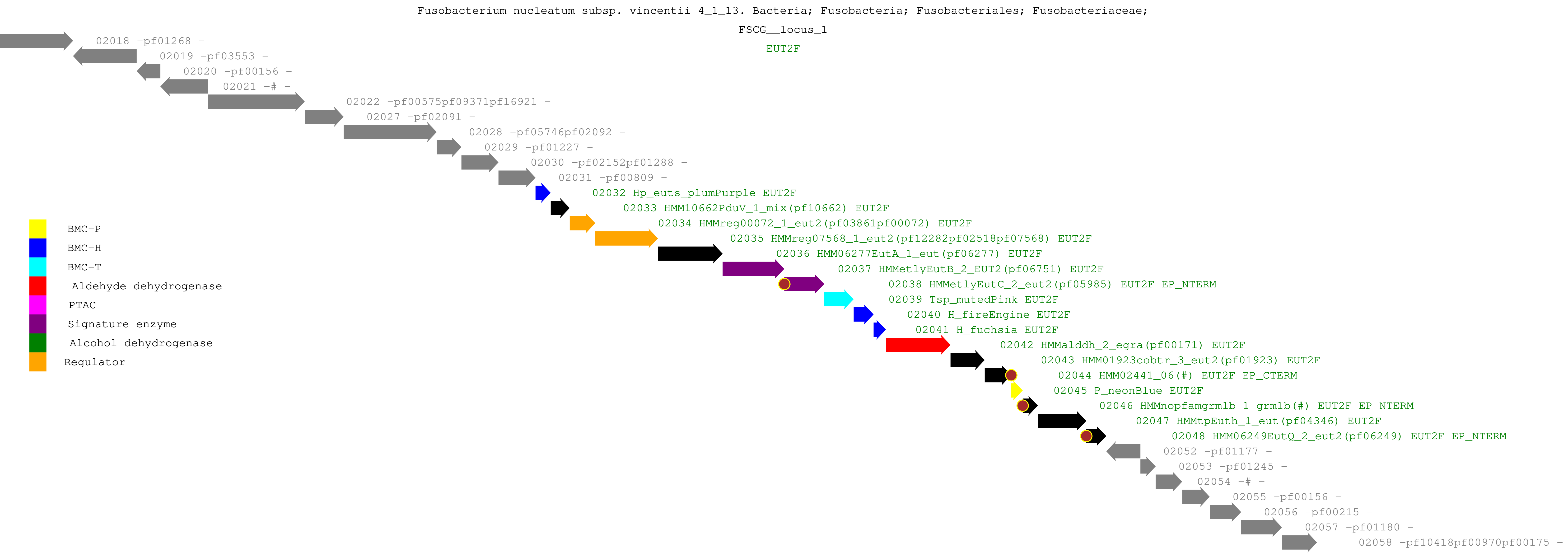

Fusobacterium nucleatum subsp. vincentii 4_1_13. Bacteria; Fusobacteria; Fusobacteriales; Fusobacteriaceae;
FSCG__locus_1
EUT2F
02018 -pf01268 -
02019 -pf03553 -
02020 -pf00156 -
02021 -# -
02022 -pf00575pf09371pf16921 -
02027 -pf02091 -
02028 -pf05746pf02092 -
02029 -pf01227 -
02030 -pf02152pf01288 -
02031 -pf00809 -
02032 Hp_euts_plumPurple EUT2F
02033 HMM10662PduV_1_mix(pf10662) EUT2F
02034 HMMreg00072_1_eut2(pf03861pf00072) EUT2F
02035 HMMreg07568_1_eut2(pf12282pf02518pf07568) EUT2F
02036 HMM06277EutA_1_eut(pf06277) EUT2F
02037 HMMetlyEutB_2_EUT2(pf06751) EUT2F
02038 HMMetlyEutC_2_eut2(pf05985) EUT2F EP_NTERM
02039 Tsp_mutedPink EUT2F
02040 H_fireEngine EUT2F
02041 H_fuchsia EUT2F
02042 HMMaldh_2_egra(pf00171) EUT2F
02043 HMM01923cobtr_3_eut2(pf01923) EUT2F
02044 HMM02441_06(#) EUT2F EP_CTERM
02045 P_neonBlue EUT2F
02046 HMMnopfamgrm1b_1_grm1b(#) EUT2F EP_NTERM
02047 HMMtpEuth_1_eut(pf04346) EUT2F
02048 HMM06249EutQ_2_eut2(pf06249) EUT2F EP_NTERM
02052 -pf01177 -
02053 -pf01245 -
02054 -# -
02055 -pf00156 -
02056 -pf00215 -
02057 -pf01180 -
02058 -pf10418pf00970pf00175 -
BMC-P
BMC-H
BMC-T
Aldehyde dehydrogenase
PTAC
Signature enzyme
Alcohol dehydrogenase
Regulator

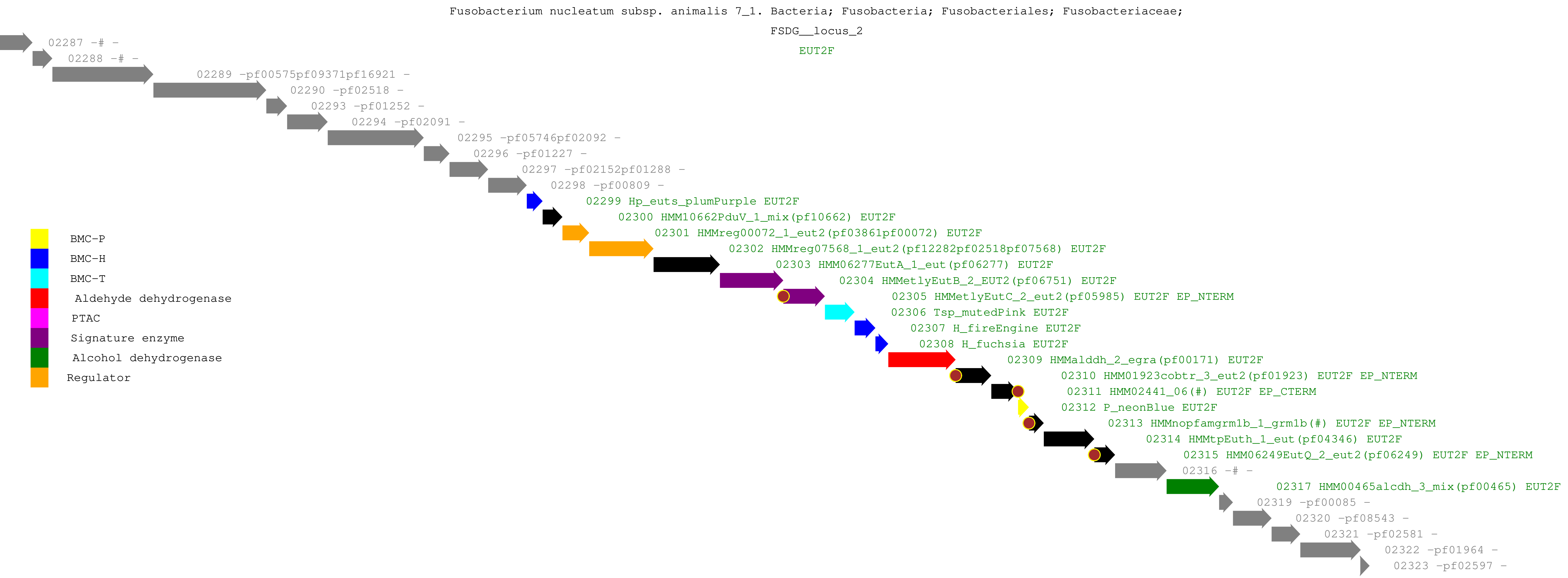
Fusobacterium nucleatum subsp. animalis 7_1. Bacteria; Fusobacteria; Fusobacteriales; Fusobacteriaceae;
FSDG__locus_2
EUT2F
02287 -# -
02288 -# -
02289 -pf00575pf09371pf16921 -
02290 -pf02518 -
02293 -pf01252 -
02294 -pf02091 -
02295 -pf05746pf02092 -
02296 -pf01227 -
02297 -pf02152pf01288 -
02298 -pf00809 -
02299 Hp_euts_plumPurple EUT2F
02300 HMM10662PduV_1_mix(pf10662) EUT2F
02301 HMMreg00072_1_eut2(pf03861pf00072) EUT2F
02302 HMMreg07568_1_eut2(pf12282pf02518pf07568) EUT2F
02303 HMM06277EutA_1_eut(pf06277) EUT2F
02304 HMMetlyEutB_2_EUT2(pf06751) EUT2F
02305 HMMetlyEutC_2_eut2(pf05985) EUT2F EP_NTERM
02306 Tsp_mutedPink EUT2F
02307 H_fireEngine EUT2F
02308 H_fuchsia EUT2F
02309 HMMaldh_2_egra(pf00171) EUT2F
02310 HMM01923cobtr_3_eut2(pf01923) EUT2F EP_NTERM
02311 HMM02441_06(#) EUT2F EP_CTERM
02312 P_neonBlue EUT2F
02313 HMMnopfamgrm1b_1_grm1b(#) EUT2F EP_NTERM
02314 HMMtpEuth_1_eut(pf04346) EUT2F
02315 HMM06249EutQ_2_eut2(pf06249) EUT2F EP_NTERM
02316 -# -
02317 HMM00465alcdh_3_mix(pf00465) EUT2F
02319 -pf00085 -
02320 -pf08543 -
02321 -pf02581 -
02322 -pf01964 -
02323 -pf02597 -
BMC-P
BMC-H
BMC-T
Aldehyde dehydrogenase
PTAC
Signature enzyme
Alcohol dehydrogenase
Regulator

Fusobacterium varium. Bacteria; Fusobacteria; Fusobacteriales; Fusobacteriaceae;

FV113G1__locus_1

EUT2F

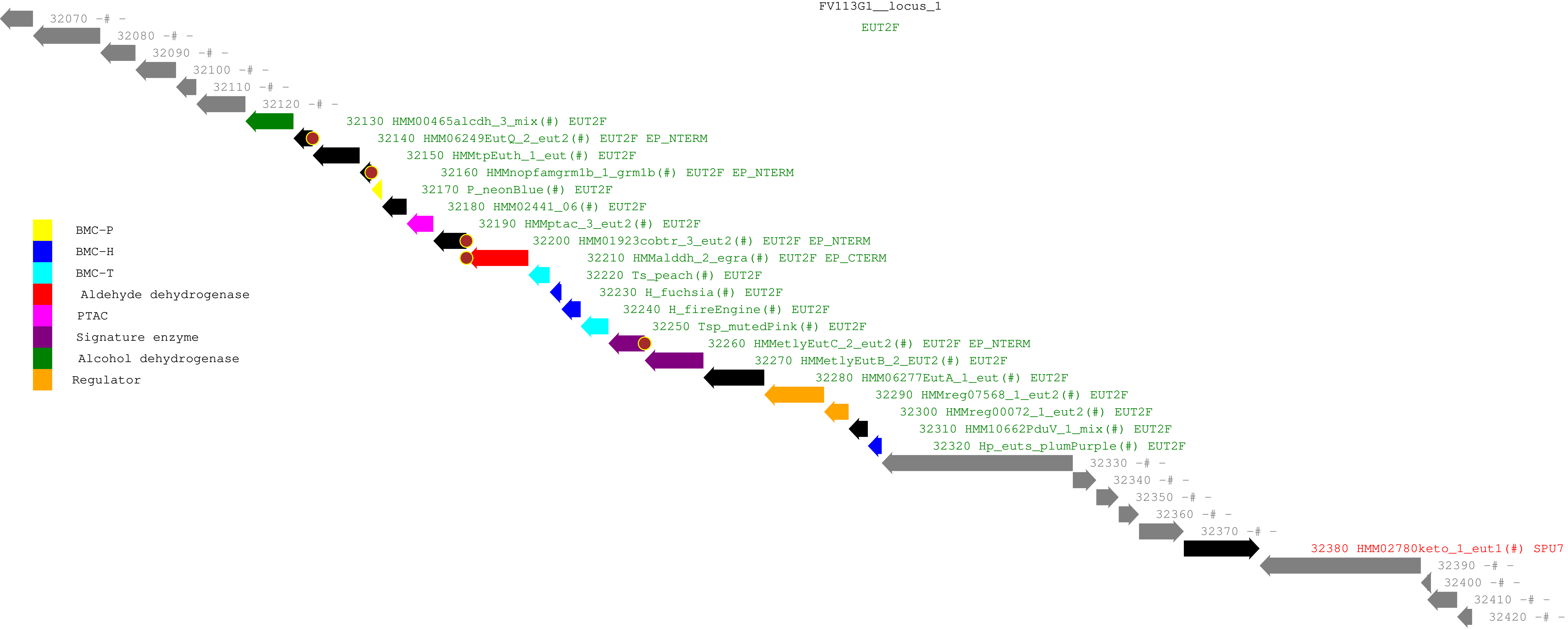

Fusobacterium varium ATCC 27725. Bacteria; Fusobacteria; Fusobacteriales; Fusobacteriaceae;

FVAG__locus_2

EUT2F

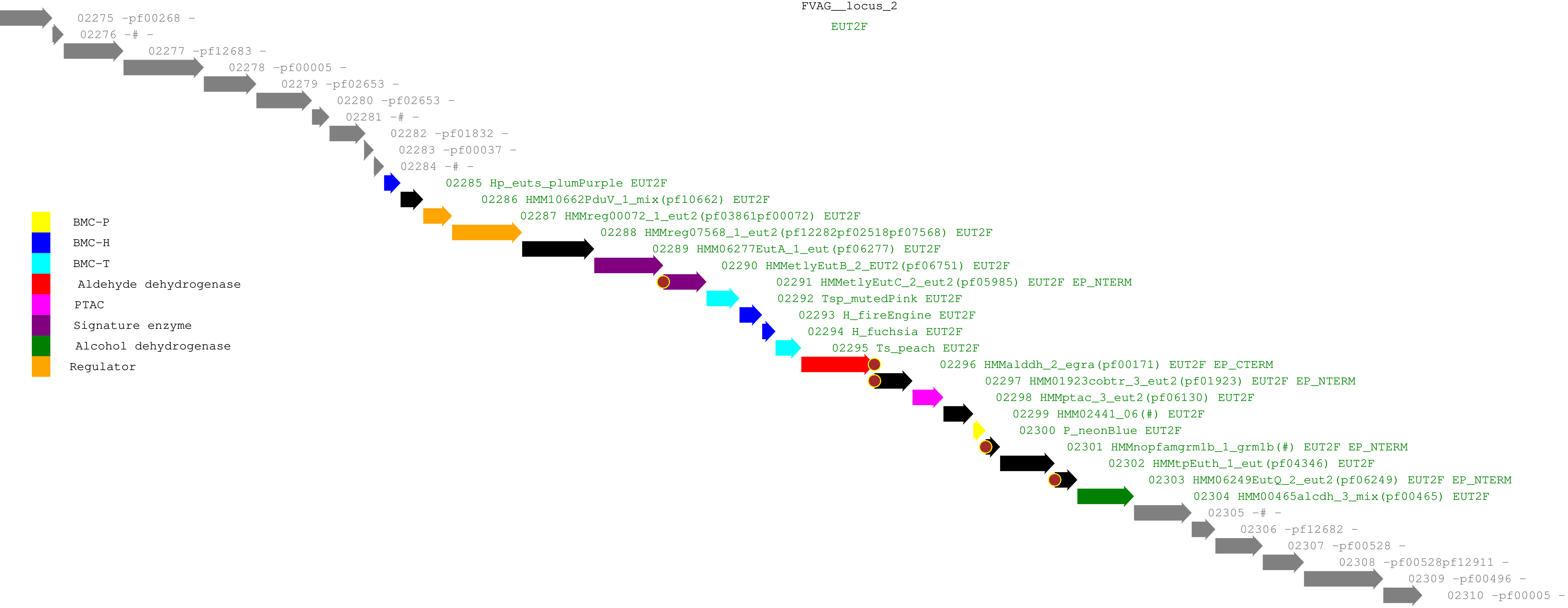

Cetobacterium somerae ATCC BAA-474. Bacteria; Fusobacteria; Fusobacteriales; Fusobacteriaceae;

HMPREF0202__locus_1

EUT2F

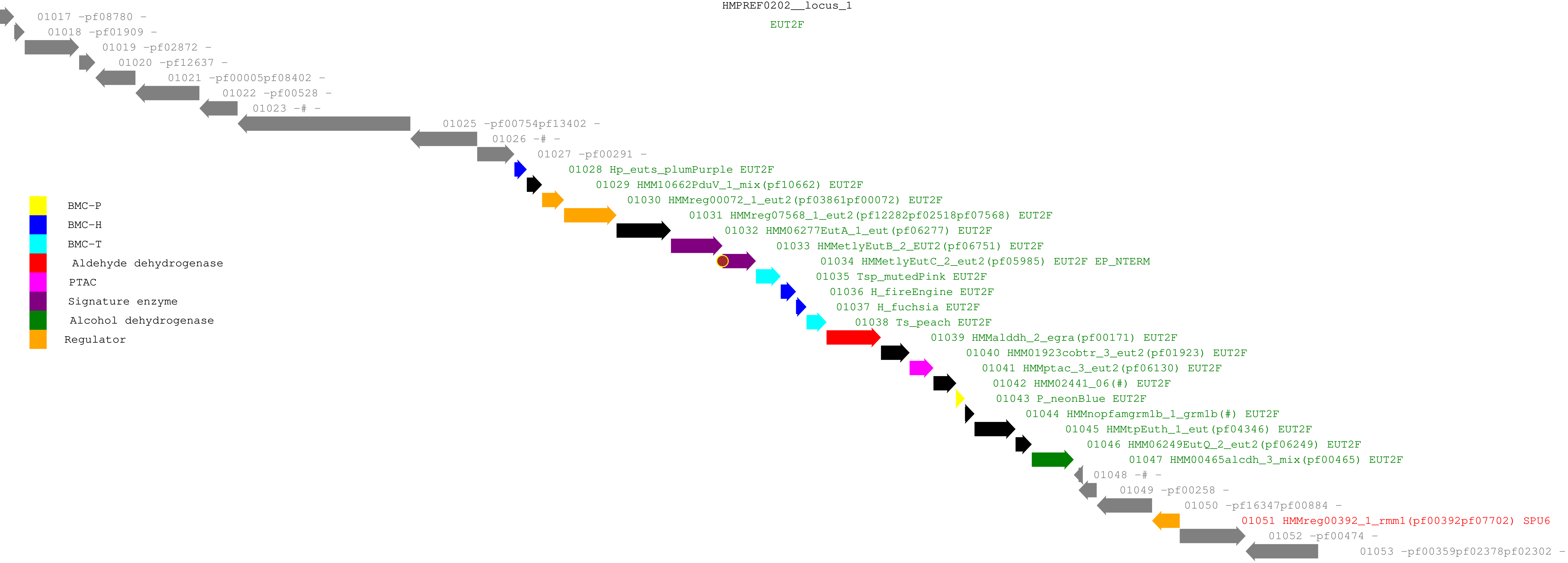

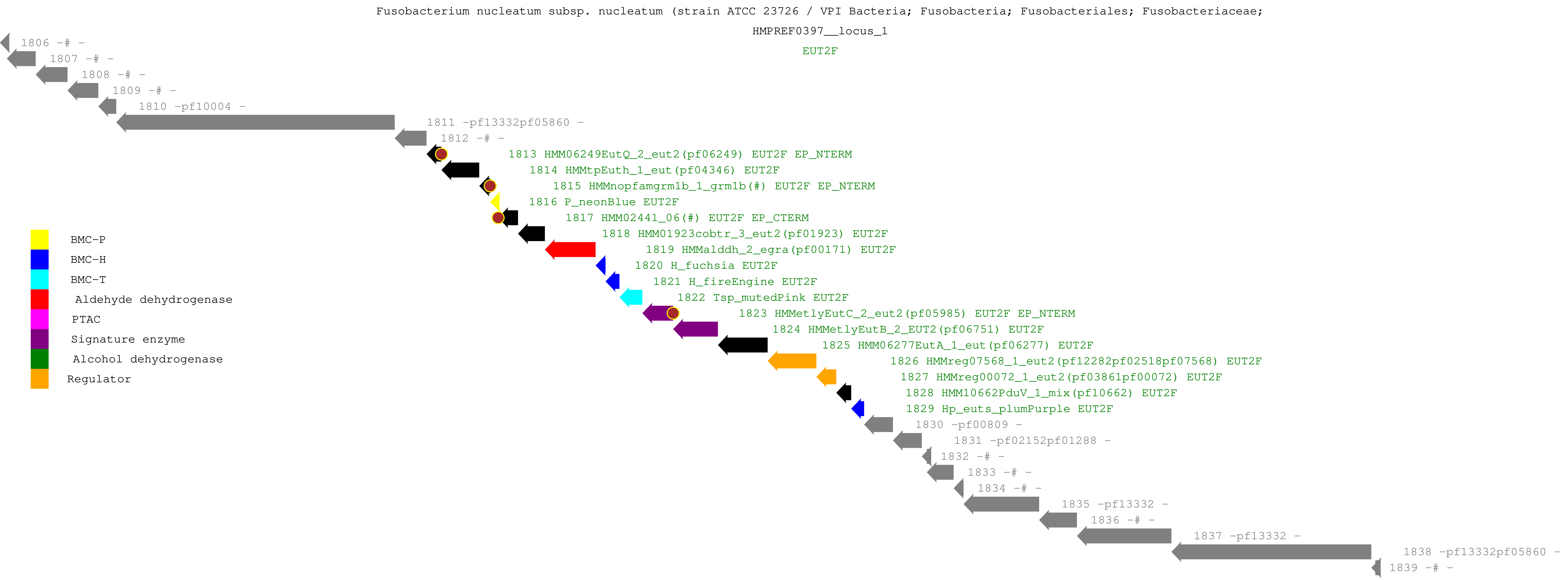
Fusobacterium nucleatum subsp. nucleatum (strain ATCC 23726 / VPI Bacteria; Fusobacteria; Fusobacteriales; Fusobacteriaceae;
HMPREF0397__locus_1
EUT2F
1806 -# -
1807 -# -
1808 -# -
1809 -# -
1810 -pf10004 -
1811 -pf13332pf05860 -
1812 -# -
1813 HMM06249EutQ_2_eut2(pf06249) EUT2F EP_NTERM
1814 HMMtpEuth_1_eut(pf04346) EUT2F
1815 HMMnopfamgrm1b_1_grm1b(#) EUT2F EP_NTERM
1816 P_neonBlue EUT2F
1817 HMM02441_06(#) EUT2F EP_CTERM
1818 HMM01923cobtr_3_eut2(pf01923) EUT2F
1819 HMMalddh_2_egra(pf00171) EUT2F
1820 H_fuchsia EUT2F
1821 H_fireEngine EUT2F
1822 Tsp_mutedPink EUT2F
1823 HMMetlyEutC_2_eut2(pf05985) EUT2F EP_NTERM
1824 HMMetlyEutB_2_EUT2(pf06751) EUT2F
1825 HMM06277EutA_1_eut(pf06277) EUT2F
1826 HMMreg07568_1_eut2(pf12282pf02518pf07568) EUT2F
1827 HMMreg00072_1_eut2(pf03861pf00072) EUT2F
1828 HMM10662PduV_1_mix(pf10662) EUT2F
1829 Hp_euts_plumPurple EUT2F
1830 -pf00809 -
1831 -pf02152pf01288 -
1832 -# -
1833 -# -
1834 -# -
1835 -pf13332 -
1836 -# -
1837 -pf13332 -
1838 -pf13332pf05860 -
1839 -# -
BMC-P
BMC-H
BMC-T
Aldehyde dehydrogenase
PTAC
Signature enzyme
Alcohol dehydrogenase
Regulator

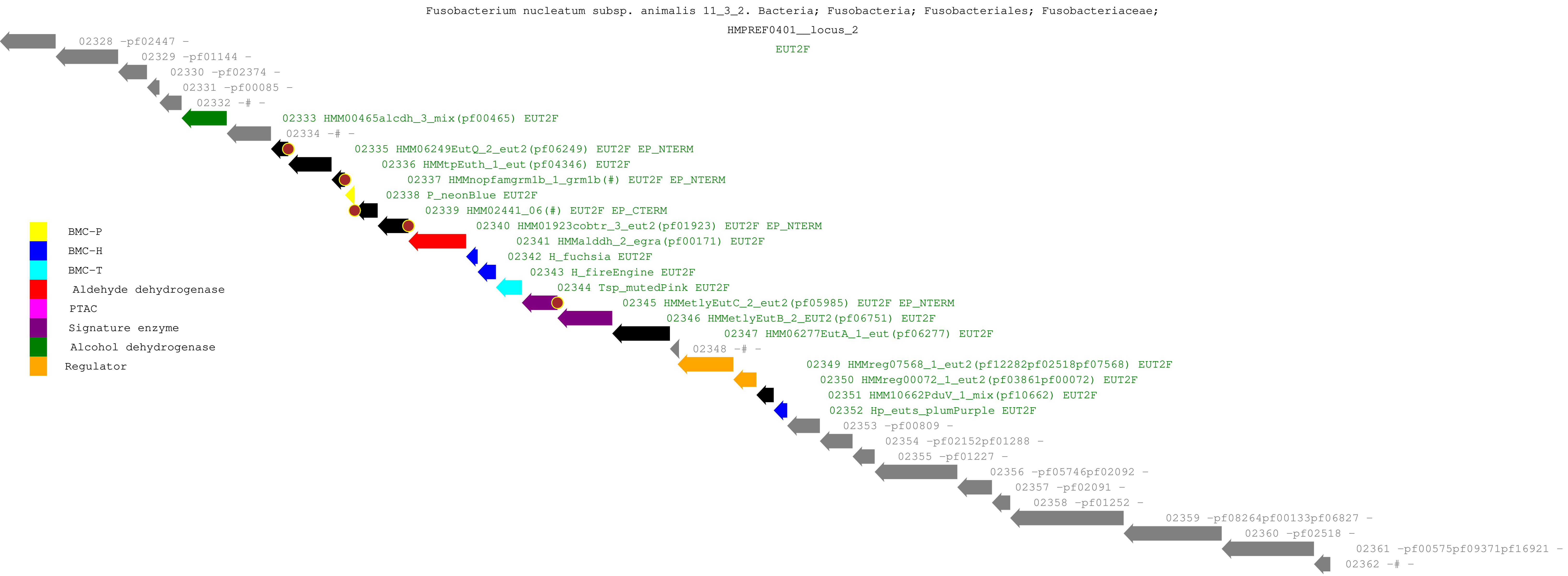
Fusobacterium nucleatum subsp. animalis 11_3_2. Bacteria; Fusobacteria; Fusobacteriales; Fusobacteriaceae;
HMPREF0401__locus_2
EUT2F
02328 -pf02447 -
02329 -pf01144 -
02330 -pf02374 -
02331 -pf00085 -
02332 -# -
02333 HMM00465alcdh_3_mix(pf00465) EUT2F
02334 -# -
02335 HMM06249EutQ_2_eut2(pf06249) EUT2F EP_NTERM
02336 HMMtpEuth_1_eut(pf04346) EUT2F
02337 HMMnopfamgrm1b_1_grm1b(#) EUT2F EP_NTERM
02338 P_neonBlue EUT2F
02339 HMM02441_06(#) EUT2F EP_CTERM
02340 HMM01923cobtr_3_eut2(pf01923) EUT2F EP_NTERM
02341 HMMalddh_2_egra(pf00171) EUT2F
02342 H_fuchsia EUT2F
02343 H_fireEngine EUT2F
02344 Tsp_mutedPink EUT2F
02345 HMMetlyEutC_2_eut2(pf05985) EUT2F EP_NTERM
02346 HMMetlyEutB_2_EUT2(pf06751) EUT2F
02347 HMM06277EutA_1_eut(pf06277) EUT2F
02348 -# -
02349 HMMreg07568_1_eut2(pf12282pf02518pf07568) EUT2F
02350 HMMreg00072_1_eut2(pf03861pf00072) EUT2F
02351 HMM10662PduV_1_mix(pf10662) EUT2F
02352 Hp_euts_plumPurple EUT2F
02353 -pf00809 -
02354 -pf02152pf01288 -
02355 -pf01227 -
02356 -pf05746pf02092 -
02357 -pf02091 -
02358 -pf01252 -
02359 -pf08264pf00133pf06827 -
02360 -pf02518 -
02361 -pf00575pf09371pf16921 -
02362 -# -
BMC-P
BMC-H
BMC-T
Aldehyde dehydrogenase
PTAC
Signature enzyme
Alcohol dehydrogenase
Regulator

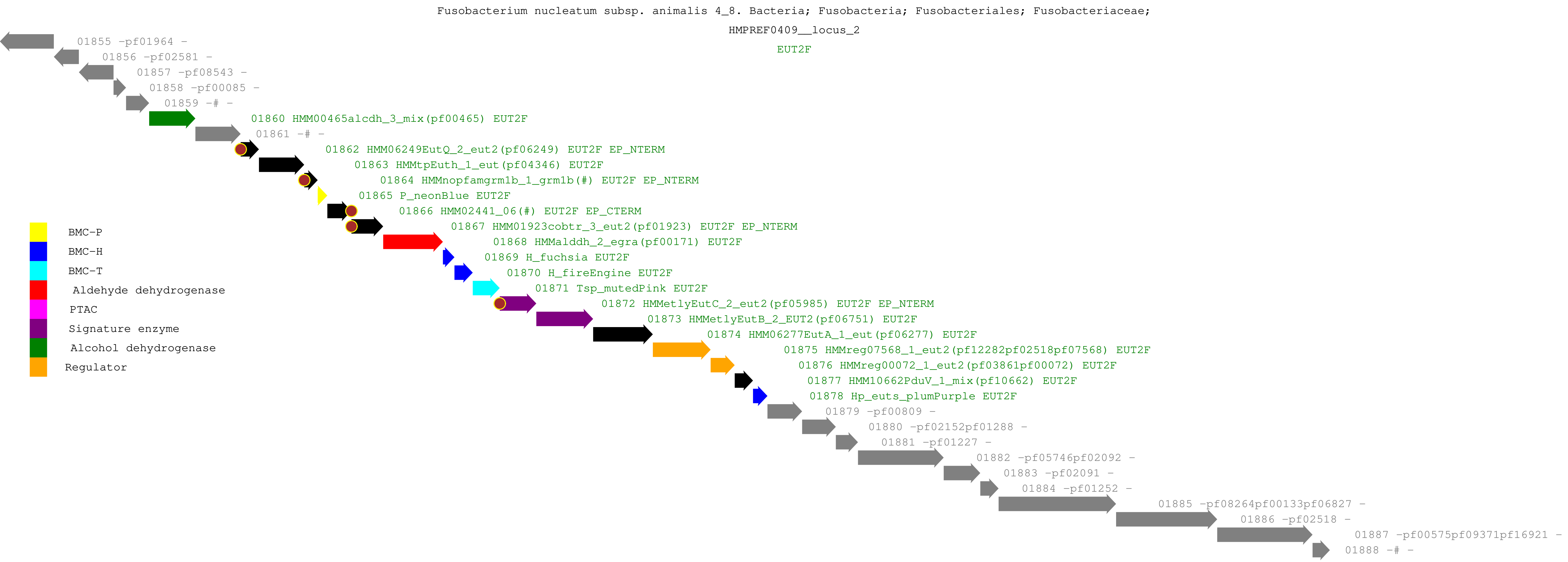
Fusobacterium nucleatum subsp. animalis 4_8. Bacteria; Fusobacteria; Fusobacteriales; Fusobacteriaceae;
HMPREF0409__locus_2
EUT2F
01855 -pf01964 -
01856 -pf02581 -
01857 -pf08543 -
01858 -pf00085 -
01859 -# -
01860 HMM00465alcdh_3_mix(pf00465) EUT2F
01861 -# -
01862 HMM06249EutQ_2_eut2(pf06249) EUT2F EP_NTERM
01863 HMMtpEuth_1_eut(pf04346) EUT2F
01864 HMMnopfamgrm1b_1_grm1b(#) EUT2F EP_NTERM
01865 P_neonBlue EUT2F
01866 HMM02441_06(#) EUT2F EP_CTERM
01867 HMM01923cobtr_3_eut2(pf01923) EUT2F EP_NTERM
01868 HMMalddh_2_egra(pf00171) EUT2F
01869 H_fuchsia EUT2F
01870 H_fireEngine EUT2F
01871 Tsp_mutedPink EUT2F
01872 HMMetlyEutC_2_eut2(pf05985) EUT2F EP_NTERM
01873 HMMetlyEutB_2_EUT2(pf06751) EUT2F
01874 HMM06277EutA_1_eut(pf06277) EUT2F
01875 HMMreg07568_1_eut2(pf12282pf02518pf07568) EUT2F
01876 HMMreg00072_1_eut2(pf03861pf00072) EUT2F
01877 HMM10662PduV_1_mix(pf10662) EUT2F
01878 Hp_euts_plumPurple EUT2F
01879 -pf00809 -
01880 -pf02152pf01288 -
01881 -pf01227 -
01882 -pf05746pf02092 -
01883 -pf02091 -
01884 -pf01252 -
01885 -pf08264pf00133pf06827 -
01886 -pf02518 -
01887 -pf00575pf09371pf16921 -
01888 -# -
BMC-P
BMC-H
BMC-T
Aldehyde dehydrogenase
PTAC
Signature enzyme
Alcohol dehydrogenase
Regulator

Fusobacterium nucleatum subsp. vincentii 3_1_36A2. Bacteria; Fusobacteria; Fusobacteriales; Fusobacteriaceae;

HMPREF0946__locus_1

EUT2F

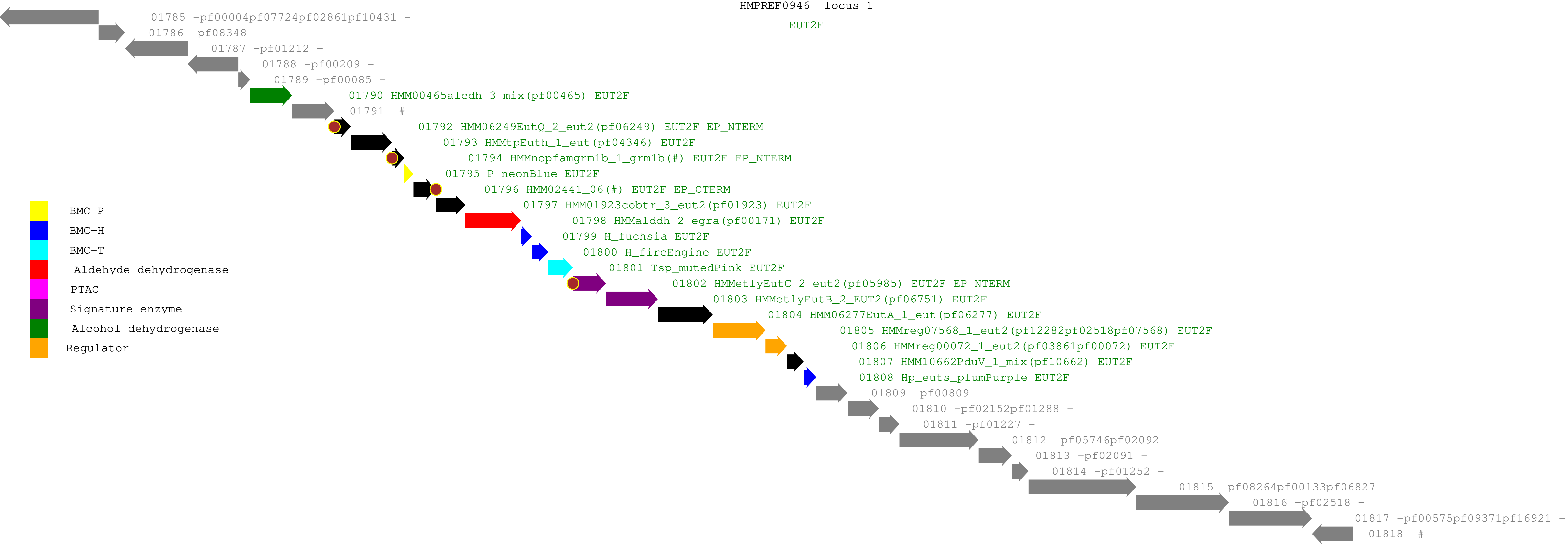

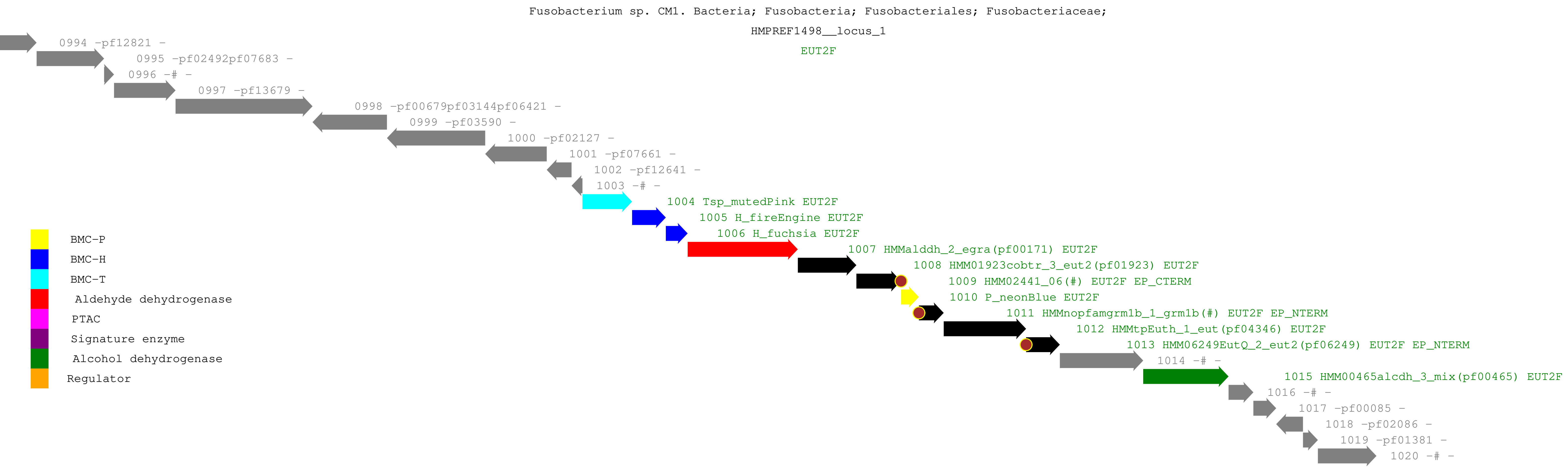
Fusobacterium sp. CM1. Bacteria; Fusobacteria; Fusobacteriales; Fusobacteriaceae;
HMPREF1498__locus_1
EUT2F
0994 -pf12821 -
0995 -pf02492pf07683 -
0996 -# -
0997 -pf13679 -
0998 -pf00679pf03144pf06421 -
0999 -pf03590 -
1000 -pf02127 -
1001 -pf07661 -
1002 -pf12641 -
1003 -# -
1004 Tsp_mutedPink EUT2F
1005 H_fireEngine EUT2F
1006 H_fuchsia EUT2F
1007 HMMalddh_2_egra(pf00171) EUT2F
1008 HMM01923cobtr_3_eut2(pf01923) EUT2F
1009 HMM02441_06(#) EUT2F EP_CTERM
1010 P_neonBlue EUT2F
1011 HMMnopfamgrm1b_1_grm1b(#) EUT2F EP_NTERM
1012 HMMtpEuth_1_eut(pf04346) EUT2F
1013 HMM06249EutQ_2_eut2(pf06249) EUT2F EP_NTERM
1014 -# -
1015 HMM00465alcdh_3_mix(pf00465) EUT2F
1016 -# -
1017 -pf00085 -
1018 -pf02086 -
1019 -pf01381 -
1020 -# -
BMC-P
BMC-H
BMC-T
Aldehyde dehydrogenase
PTAC
Signature enzyme
Alcohol dehydrogenase
Regulator

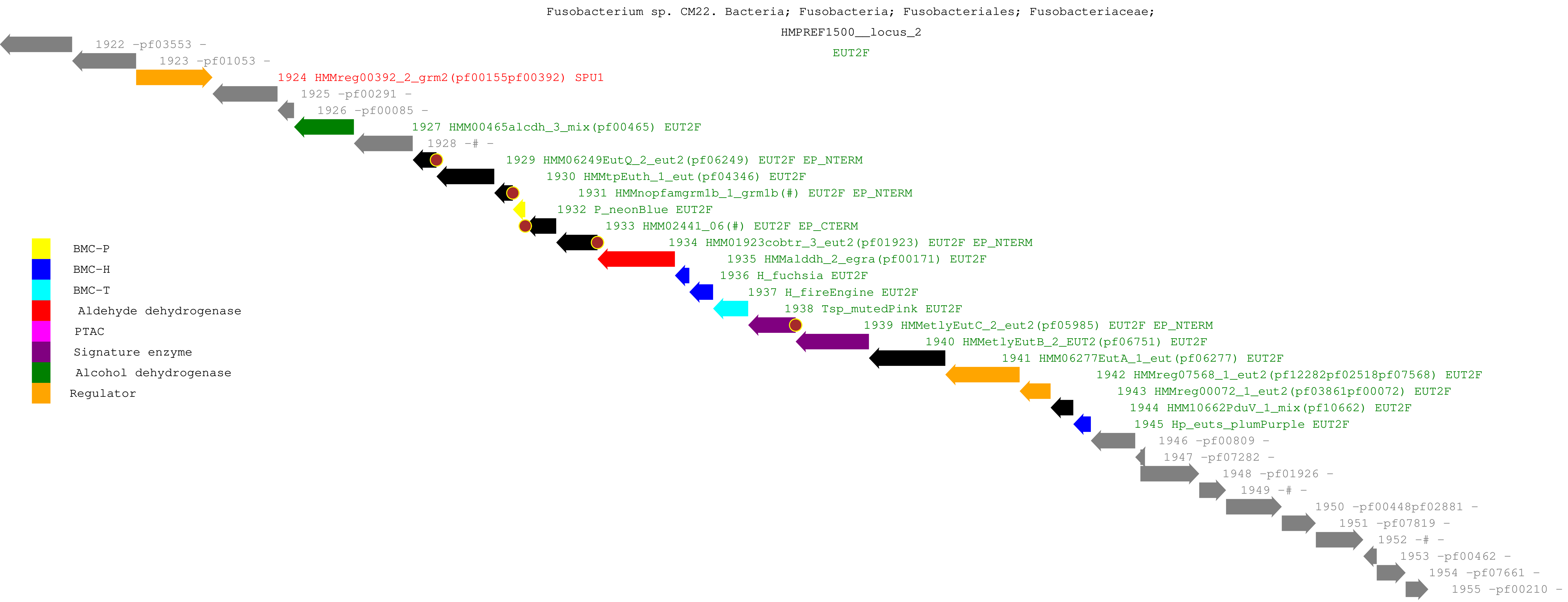
Fusobacterium sp. CM22. Bacteria; Fusobacteria; Fusobacteriales; Fusobacteriaceae;
HMPREF1500__locus_2
EUT2F
1922 -pf03553 -
1923 -pf01053 -
1924 HMMreg00392_2_grm2(pf00155pf00392) SPU1
1925 -pf00291 -
1926 -pf00085 -
1927 HMM00465alcdh_3_mix(pf00465) EUT2F
1928 -# -
1929 HMM06249EutQ_2_eut2(pf06249) EUT2F EP_NTERM
1930 HMMtpEuth_1_eut(pf04346) EUT2F
1931 HMMnopfamgrm1b_1_grm1b(#) EUT2F EP_NTERM
1932 P_neonBlue EUT2F
1933 HMM02441_06(#) EUT2F EP_CTERM
1934 HMM01923cobtr_3_eut2(pf01923) EUT2F EP_NTERM
1935 HMMalddh_2_egra(pf00171) EUT2F
1936 H_fuchsia EUT2F
1937 H_fireEngine EUT2F
1938 Tsp_mutedPink EUT2F
1939 HMMetlyEutC_2_eut2(pf05985) EUT2F EP_NTERM
1940 HMMetlyEutB_2_EUT2(pf06751) EUT2F
1941 HMM06277EutA_1_eut(pf06277) EUT2F
1942 HMMreg07568_1_eut2(pf12282pf02518pf07568) EUT2F
1943 HMMreg00072_1_eut2(pf03861pf00072) EUT2F
1944 HMM10662PduV_1_mix(pf10662) EUT2F
1945 Hp_euts_plumPurple EUT2F
1946 -pf00809 -
1947 -pf07282 -
1948 -pf01926 -
1949 -# -
1950 -pf00448pf02881 -
1951 -pf07819 -
1952 -# -
1953 -pf00462 -
1954 -pf07661 -
1955 -pf00210 -
BMC-P
BMC-H
BMC-T
Aldehyde dehydrogenase
PTAC
Signature enzyme
Alcohol dehydrogenase
Regulator

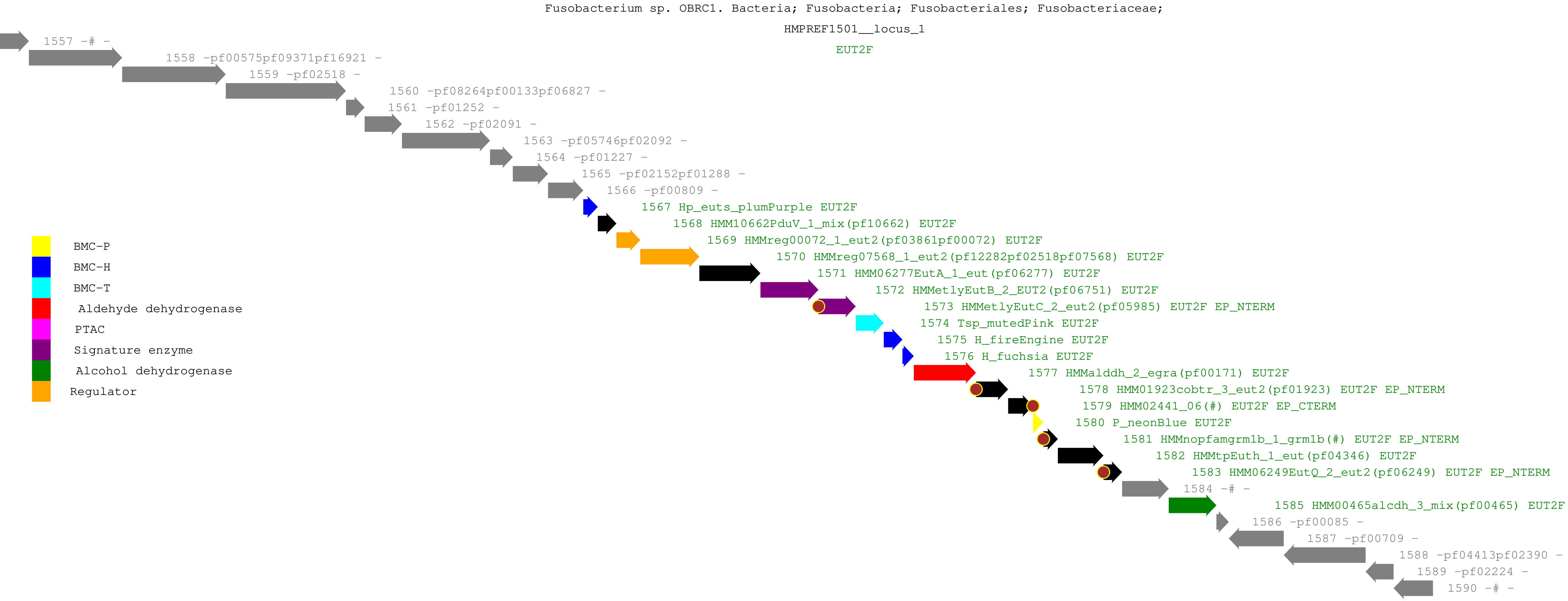
Fusobacterium sp. OBRC1. Bacteria; Fusobacteria; Fusobacteriales; Fusobacteriaceae;
HMPREF1501__locus_1
EUT2F
1557 -# -
1558 -pf00575pf09371pf16921 -
1559 -pf02518 -
1560 -pf08264pf00133pf06827 -
1561 -pf01252 -
1562 -pf02091 -
1563 -pf05746pf02092 -
1564 -pf01227 -
1565 -pf02152pf01288 -
1566 -pf00809 -
1567 Hp_euts_plumPurple EUT2F
1568 HMM10662PduV_1_mix(pf10662) EUT2F
1569 HMMreg00072_1_eut2(pf03861pf00072) EUT2F
1570 HMMreg07568_1_eut2(pf12282pf02518pf07568) EUT2F
1571 HMM06277EutA_1_eut(pf06277) EUT2F
1572 HMMetlyEutB_2_EUT2(pf06751) EUT2F
1573 HMMetlyEutC_2_eut2(pf05985) EUT2F EP_NTERM
1574 Tsp_mutedPink EUT2F
1575 H_fireEngine EUT2F
1576 H_fuchsia EUT2F
1577 HMMalddh_2_egra(pf00171) EUT2F
1578 HMM01923cobtr_3_eut2(pf01923) EUT2F EP_NTERM
1579 HMM02441_06(#) EUT2F EP_CTERM
1580 P_neonBlue EUT2F
1581 HMMnopfamgrm1b_1_grm1b(#) EUT2F EP_NTERM
1582 HMMtpEuth_1_eut(pf04346) EUT2F
1583 HMM06249EutQ_2_eut2(pf06249) EUT2F EP_NTERM
1584 -# -
1585 HMM00465alcdh_3_mix(pf00465) EUT2F
1586 -pf00085 -
1587 -pf00709 -
1588 -pf04413pf02390 -
1589 -pf02224 -
1590 -# -
BMC-P
BMC-H
BMC-T
Aldehyde dehydrogenase
PTAC
Signature enzyme
Alcohol dehydrogenase
Regulator

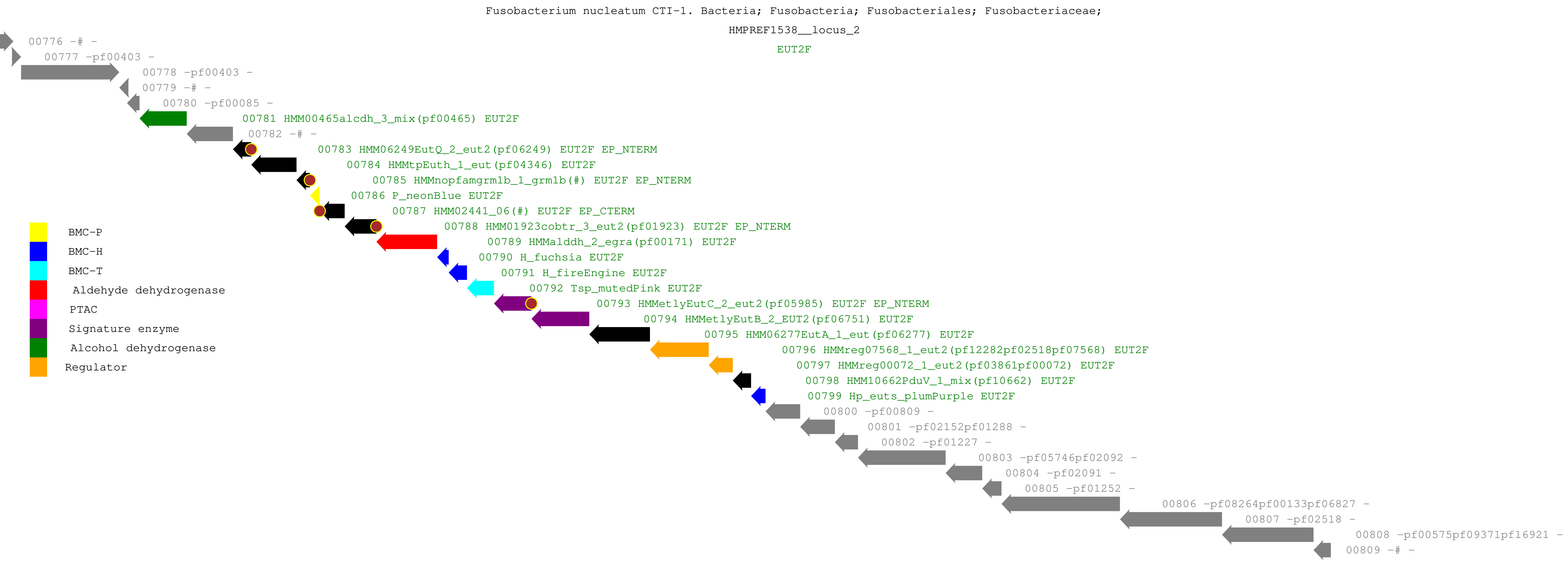

Fusobacterium nucleatum CTI-1. Bacteria; Fusobacteria; Fusobacteriales; Fusobacteriaceae;
HMPREF1538__locus_2
EUT2F
00776 -# -
00777 -pf00403 -
00778 -pf00403 -
00779 -# -
00780 -pf00085 -
00781 HMM00465alcdh_3_mix(pf00465) EUT2F
00782 -# -
00783 HMM06249EutQ_2_eut2(pf06249) EUT2F EP_NTERM
00784 HMMtpEuth_1_eut(pf04346) EUT2F
00785 HMMnopfamgrm1b_1_grm1b(#) EUT2F EP_NTERM
00786 P_neonBlue EUT2F
00787 HMM02441_06(#) EUT2F EP_CTERM
00788 HMM01923cobtr_3_eut2(pf01923) EUT2F EP_NTERM
00789 HMMalddh_2_egra(pf00171) EUT2F
00790 H_fuchsia EUT2F
00791 H_fireEngine EUT2F
00792 Tsp_mutedPink EUT2F
00793 HMMetlyEutC_2_eut2(pf05985) EUT2F EP_NTERM
00794 HMMetlyEutB_2_EUT2(pf06751) EUT2F
00795 HMM06277EutA_1_eut(pf06277) EUT2F
00796 HMMreg07568_1_eut2(pf12282pf02518pf07568) EUT2F
00797 HMMreg00072_1_eut2(pf03861pf00072) EUT2F
00798 HMM10662PduV_1_mix(pf10662) EUT2F
00799 Hp_euts_plumPurple EUT2F
00800 -pf00809 -
00801 -pf02152pf01288 -
00802 -pf01227 -
00803 -pf05746pf02092 -
00804 -pf02091 -
00805 -pf01252 -
00806 -pf08264pf00133pf06827 -
00807 -pf02518 -
00808 -pf00575pf09371pf16921 -
00809 -# -
BMC-P
BMC-H
BMC-T
Aldehyde dehydrogenase
PTAC
Signature enzyme
Alcohol dehydrogenase
Regulator

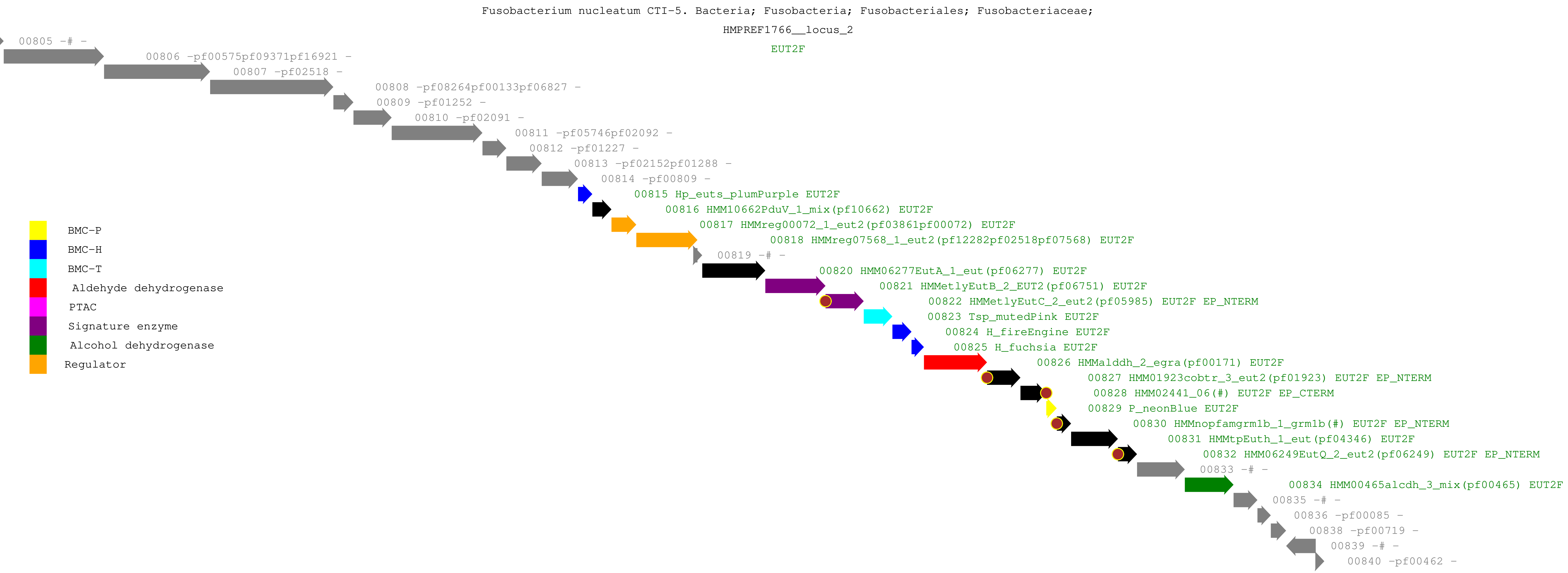
Fusobacterium nucleatum CTI-5. Bacteria; Fusobacteria; Fusobacteriales; Fusobacteriaceae;
HMPREF1766__locus_2
EUT2F
00805 -# -
00806 -pf00575pf09371pf16921 -
00807 -pf02518 -
00808 -pf08264pf00133pf06827 -
00809 -pf01252 -
00810 -pf02091 -
00811 -pf05746pf02092 -
00812 -pf01227 -
00813 -pf02152pf01288 -
00814 -pf00809 -
00815 Hp_euts_plumPurple EUT2F
00816 HMM10662PduV_1_mix(pf10662) EUT2F
00817 HMMreg00072_1_eut2(pf03861pf00072) EUT2F
00818 HMMreg07568_1_eut2(pf12282pf02518pf07568) EUT2F
00819 -# -
00820 HMM06277EutA_1_eut(pf06277) EUT2F
00821 HMMetlyEutB_2_EUT2(pf06751) EUT2F
00822 HMMetlyEutC_2_eut2(pf05985) EUT2F EP_NTERM
00823 Tsp_mutedPink EUT2F
00824 H_fireEngine EUT2F
00825 H_fuchsia EUT2F
00826 HMMalddh_2_egra(pf00171) EUT2F
00827 HMM01923cobtr_3_eut2(pf01923) EUT2F EP_NTERM
00828 HMM02441_06(#) EUT2F EP_CTERM
00829 P_neonBlue EUT2F
00830 HMMnopfamgrm1b_1_grm1b(#) EUT2F EP_NTERM
00831 HMMtpEuth_1_eut(pf04346) EUT2F
00832 HMM06249EutQ_2_eut2(pf06249) EUT2F EP_NTERM
00833 -# -
00834 HMM00465alcdh_3_mix(pf00465) EUT2F
00835 -# -
00836 -pf00085 -
00838 -pf00719 -
00839 -# -
00840 -pf00462 -
BMC-P
BMC-H
BMC-T
Aldehyde dehydrogenase
PTAC
Signature enzyme
Alcohol dehydrogenase
Regulator

Fusobacterium nucleatum CTI-6. Bacteria; Fusobacteria; Fusobacteriales; Fusobacteriaceae;

HMPREF1767__locus_2

EUT2F

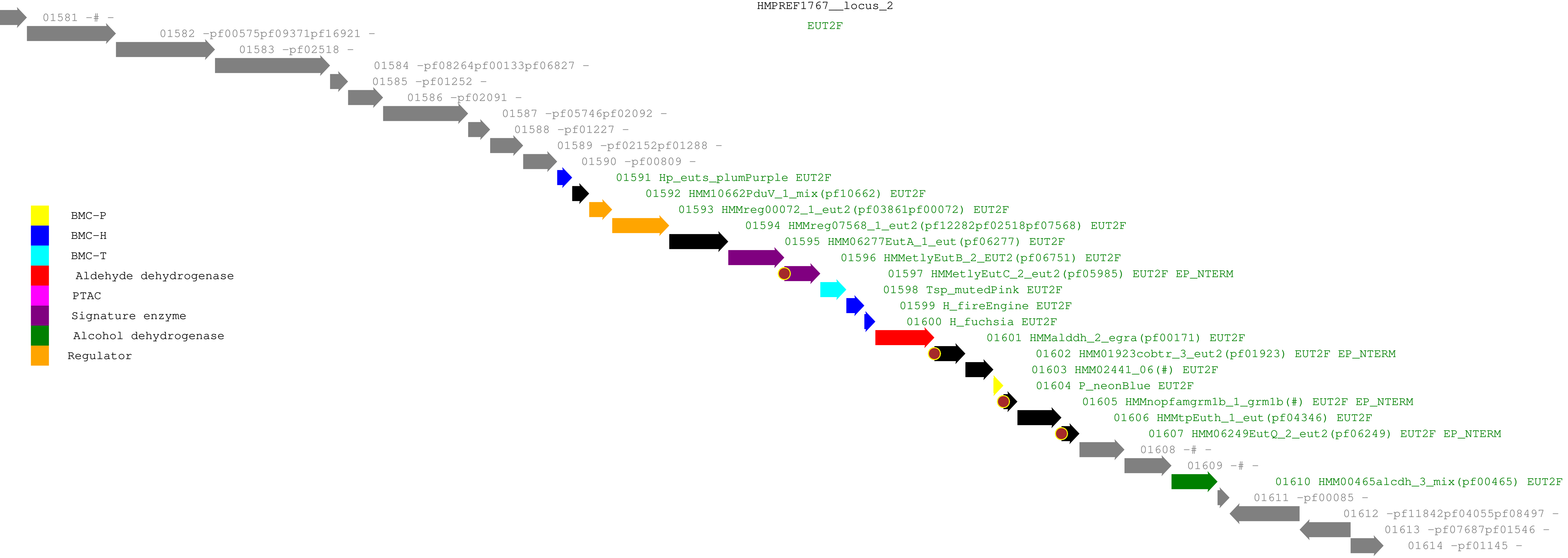

Fusobacterium nucleatum 13_3C. Bacteria; Fusobacteria; Fusobacteriales; Fusobacteriaceae;

HMPREF2085__locus_2

EUT2F

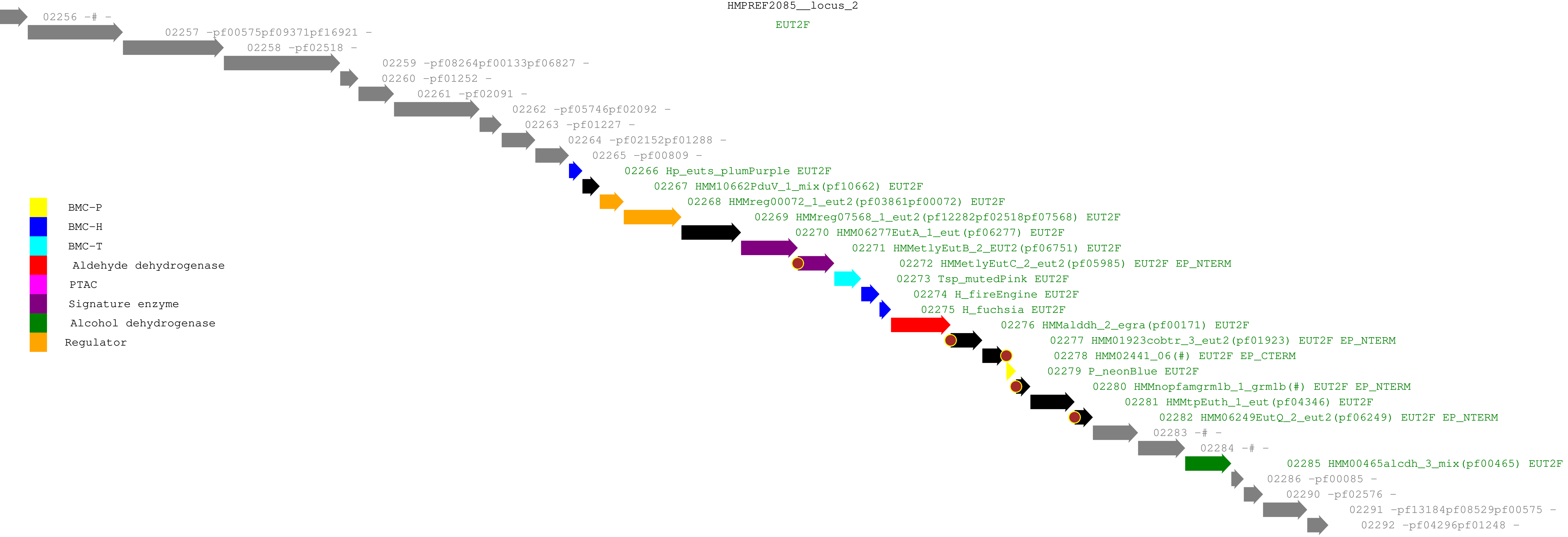

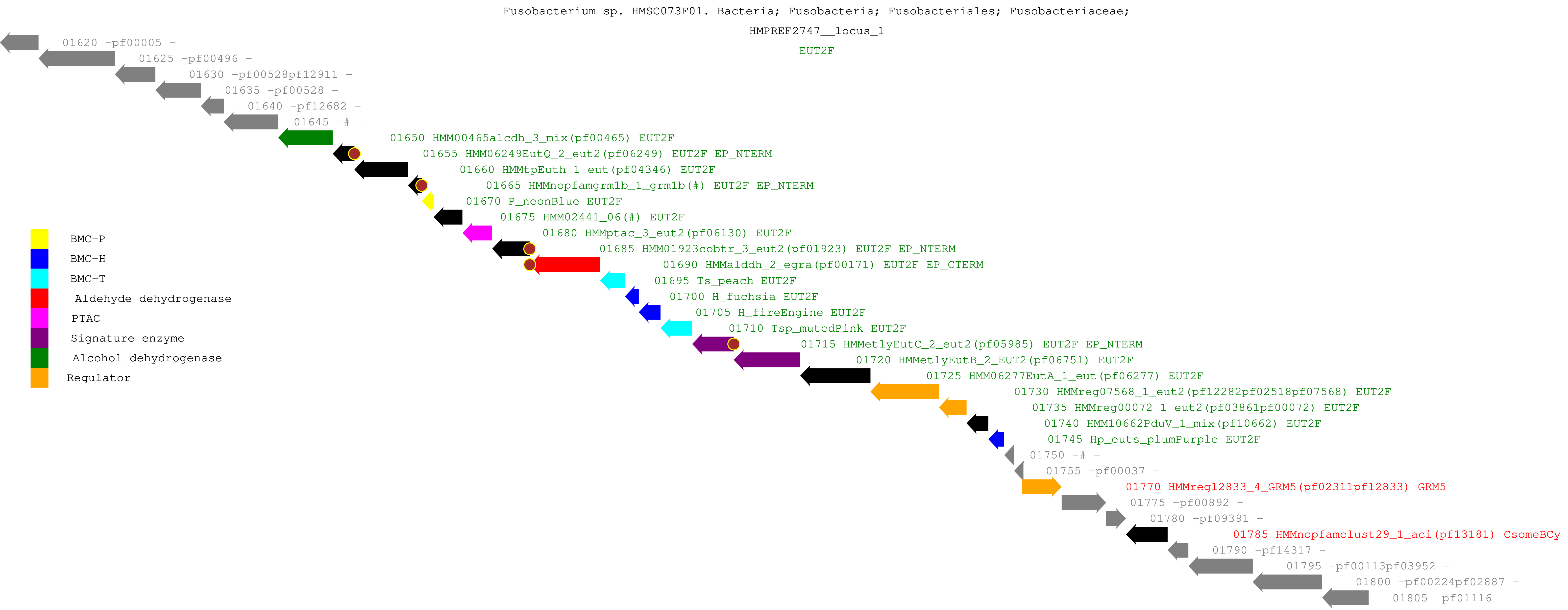
Fusobacterium sp. HMSC073F01. Bacteria; Fusobacteria; Fusobacteriales; Fusobacteriaceae;
HMPREF2747__locus_1
EUT2F
01620 -pf00005 -
01625 -pf00496 -
01630 -pf00528pf12911 -
01635 -pf00528 -
01640 -pf12682 -
01645 -# -
01650 HMM00465alcdh_3_mix(pf00465) EUT2F
01655 HMM06249EutQ_2_eut2(pf06249) EUT2F EP_NTERM
01660 HMMtpEuth_1_eut(pf04346) EUT2F
01665 HMMnopfamgrm1b_1_grm1b(#) EUT2F EP_NTERM
01670 P_neonBlue EUT2F
01675 HMM02441_06(#) EUT2F
01680 HMMptac_3_eut2(pf06130) EUT2F
01685 HMM01923cobtr_3_eut2(pf01923) EUT2F EP_NTERM
01690 HMMalddh_2_egra(pf00171) EUT2F EP_CTERM
01695 Ts_peach EUT2F
01700 H_fuchsia EUT2F
01705 H_fireEngine EUT2F
01710 Tsp_mutedPink EUT2F
01715 HMMetlyEutC_2_eut2(pf05985) EUT2F EP_NTERM
01720 HMMetlyEutB_2_EUT2(pf06751) EUT2F
01725 HMM06277EutA_1_eut(pf06277) EUT2F
01730 HMMreg07568_1_eut2(pf12282pf02518pf07568) EUT2F
01735 HMMreg00072_1_eut2(pf03861pf00072) EUT2F
01740 HMM10662PduV_1_mix(pf10662) EUT2F
01745 Hp_euts_plumPurple EUT2F
01750 -# -
01755 -pf00037 -
01770 HMMreg12833_4_GRM5(pf02311pf12833) GRM5
01775 -pf00892 -
01780 -pf09391 -
01785 HMMnopfamclust29_1_aci(pf13181) CsomeBCy
01790 -pf14317 -
01795 -pf00113pf03952 -
01800 -pf00224pf02887 -
01805 -pf01116 -
BMC-P
BMC-H
BMC-T
Aldehyde dehydrogenase
PTAC
Signature enzyme
Alcohol dehydrogenase
Regulator

Fusobacterium sp. HMSC064B11. Bacteria; Fusobacteria; Fusobacteriales; Fusobacteriaceae;

HMPREF3051__locus_1

EUT2F

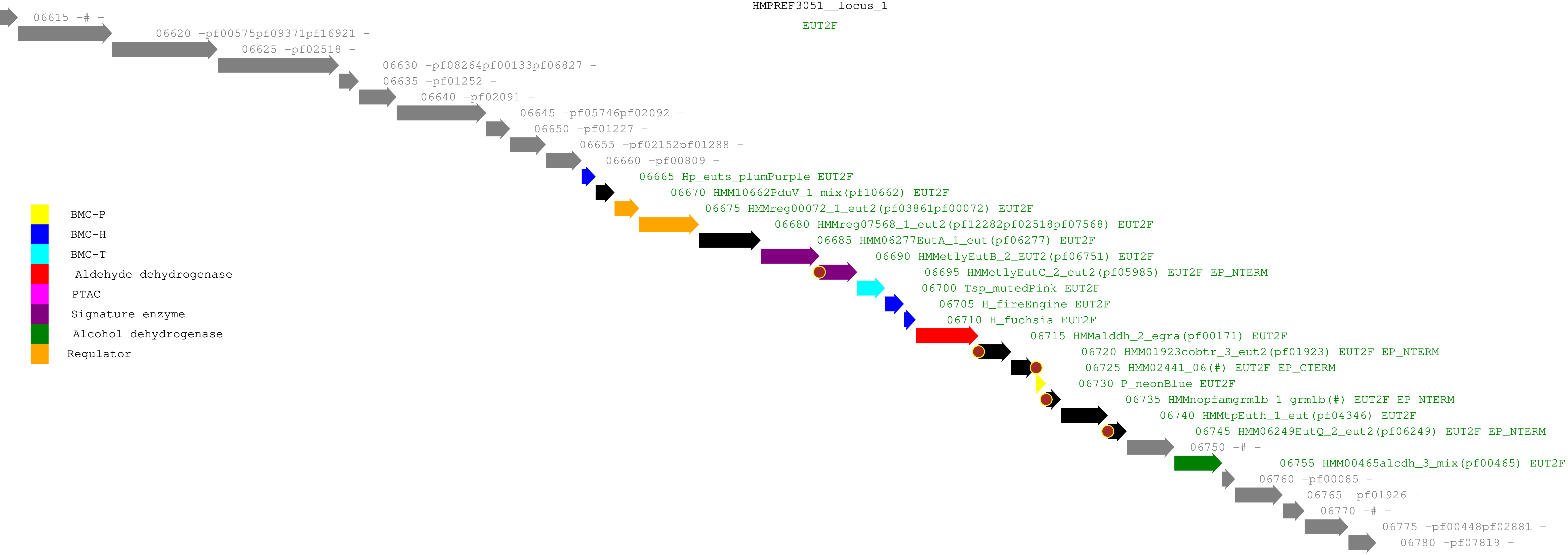

Fusobacterium nucleatum. Bacteria; Fusobacteria; Fusobacteriales; Fusobacteriaceae;

HMPREF3221__locus_1_main_sat

EUT2F

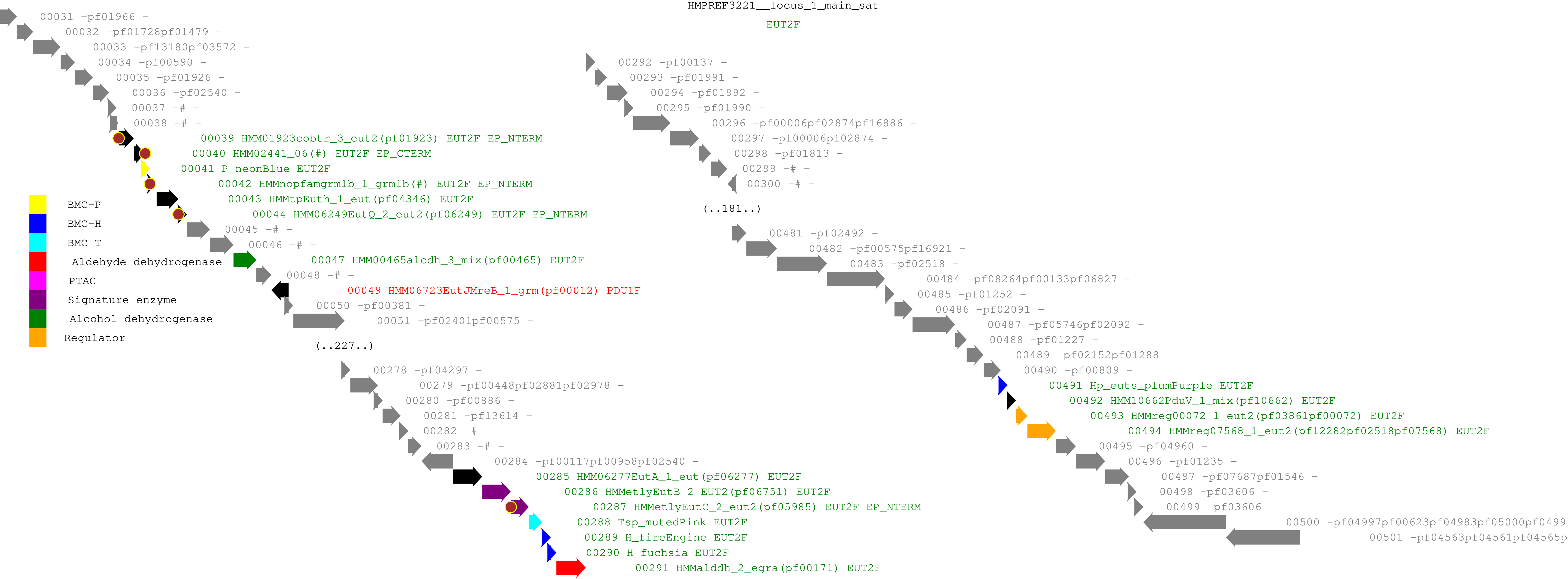

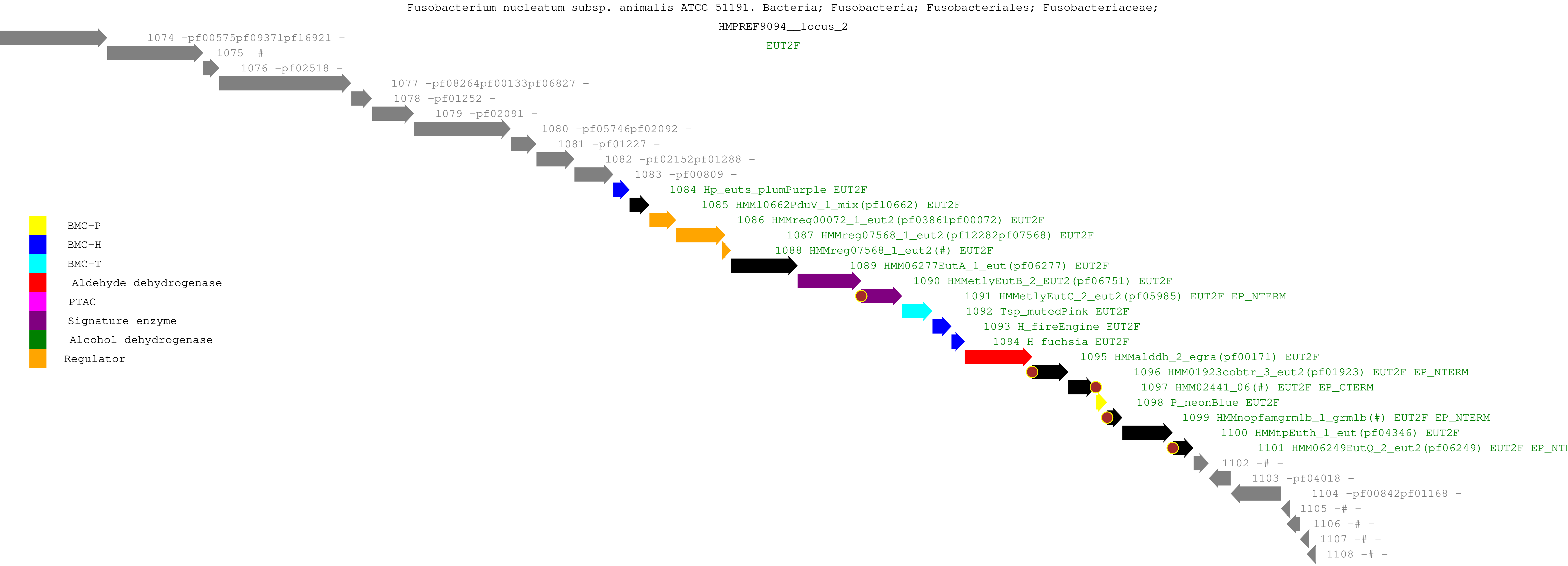

Fusobacterium nucleatum subsp. animalis ATCC 51191. Bacteria; Fusobacteria; Fusobacteriales; Fusobacteriaceae;
HMPREF9094__locus_2
EUT2F
1074 -pf00575pf09371pf16921 -
1075 -# -
1076 -pf02518 -
1077 -pf08264pf00133pf06827 -
1078 -pf01252 -
1079 -pf02091 -
1080 -pf05746pf02092 -
1081 -pf01227 -
1082 -pf02152pf01288 -
1083 -pf00809 -
1084 Hp_euts_plumPurple EUT2F
1085 HMM10662PduV_1_mix(pf10662) EUT2F
1086 HMMreg00072_1_eut2(pf03861pf00072) EUT2F
1087 HMMreg07568_1_eut2(pf12282pf07568) EUT2F
1088 HMMreg07568_1_eut2(#) EUT2F
1089 HMM06277EutA_1_eut(pf06277) EUT2F
1090 HMMetlyEutB_2_EUT2(pf06751) EUT2F
1091 HMMetlyEutC_2_eut2(pf05985) EUT2F EP_NTERM
1092 Tsp_mutedPink EUT2F
1093 H_fireEngine EUT2F
1094 H_fuchsia EUT2F
1095 HMMalddh_2_egra(pf00171) EUT2F
1096 HMM01923cobtr_3_eut2(pf01923) EUT2F EP_NTERM
1097 HMM02441_06(#) EUT2F EP_CTERM
1098 P_neonBlue EUT2F
1099 HMMnopfamgrm1b_1_grm1b(#) EUT2F EP_NTERM
1100 HMMtpEuth_1_eut(pf04346) EUT2F
1101 HMM06249EutQ_2_eut2(pf06249) EUT2F EP_NT
1102 -# -
1103 -pf04018 -
1104 -pf00842pf01168 -
1105 -# -
1106 -# -
1107 -# -
1108 -# -
BMC-P
BMC-H
BMC-T
Aldehyde dehydrogenase
PTAC
Signature enzyme
Alcohol dehydrogenase
Regulator

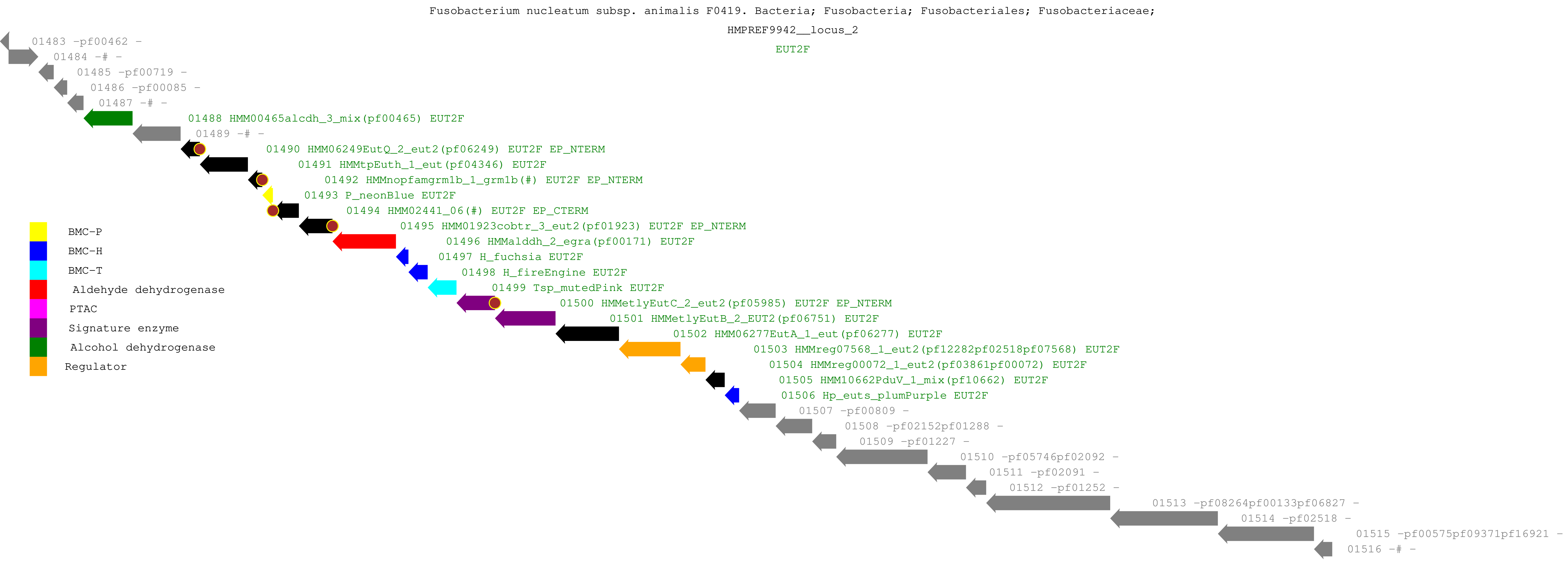
Fusobacterium nucleatum subsp. animalis F0419. Bacteria; Fusobacteria; Fusobacteriales; Fusobacteriaceae;
HMPREF9942__locus_2
EUT2F
01483 -pf00462 -
01484 -# -
01485 -pf00719 -
01486 -pf00085 -
01487 -# -
01488 HMM00465alcdh_3_mix(pf00465) EUT2F
01489 -# -
01490 HMM06249EutQ_2_eut2(pf06249) EUT2F EP_NTERM
01491 HMMtpEuth_1_eut(pf04346) EUT2F
01492 HMMnopfamgrm1b_1_grm1b(#) EUT2F EP_NTERM
01493 P_neonBlue EUT2F
01494 HMM02441_06(#) EUT2F EP_CTERM
01495 HMM01923cobtr_3_eut2(pf01923) EUT2F EP_NTERM
01496 HMMalddh_2_egra(pf00171) EUT2F
01497 H_fuchsia EUT2F
01498 H_fireEngine EUT2F
01499 Tsp_mutedPink EUT2F
01500 HMMetlyEutC_2_eut2(pf05985) EUT2F EP_NTERM
01501 HMMetlyEutB_2_EUT2(pf06751) EUT2F
01502 HMM06277EutA_1_eut(pf06277) EUT2F
01503 HMMreg07568_1_eut2(pf12282pf02518pf07568) EUT2F
01504 HMMreg00072_1_eut2(pf03861pf00072) EUT2F
01505 HMM10662PduV_1_mix(pf10662) EUT2F
01506 Hp_euts_plumPurple EUT2F
01507 -pf00809 -
01508 -pf02152pf01288 -
01509 -pf01227 -
01510 -pf05746pf02092 -
01511 -pf02091 -
01512 -pf01252 -
01513 -pf08264pf00133pf06827 -
01514 -pf02518 -
01515 -pf00575pf09371pf16921 -
01516 -# -
BMC-P
BMC-H
BMC-T
Aldehyde dehydrogenase
PTAC
Signature enzyme
Alcohol dehydrogenase
Regulator

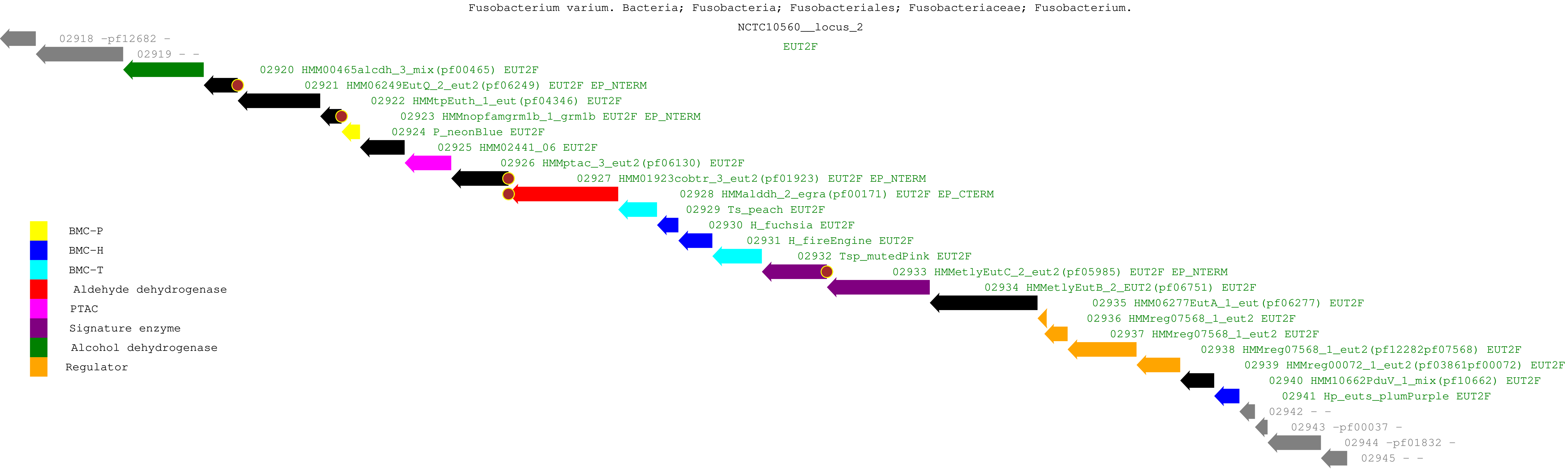
Fusobacterium varium. Bacteria; Fusobacteria; Fusobacteriales; Fusobacteriaceae; Fusobacterium.
NCTC10560__locus_2
EUT2F
02918 -pf12682 -
02919 - -
02920 HMM00465alcdh_3_mix(pf00465) EUT2F
02921 HMM06249EutQ_2_eut2(pf06249) EUT2F EP_NTERM
02922 HMMtpEuth_1_eut(pf04346) EUT2F
02923 HMMnopfamgrm1b_1_grm1b EUT2F EP_NTERM
02924 P_neonBlue EUT2F
02925 HMM02441_06 EUT2F
02926 HMMptac_3_eut2(pf06130) EUT2F
02927 HMM01923cobtr_3_eut2(pf01923) EUT2F EP_NTERM
02928 HMMalddh_2_egra(pf00171) EUT2F EP_CTERM
02929 Ts_peach EUT2F
02930 H_fuchsia EUT2F
02931 H_fireEngine EUT2F
02932 Tsp_mutedPink EUT2F
02933 HMMetlyEutC_2_eut2(pf05985) EUT2F EP_NTERM
02934 HMMetlyEutB_2_EUT2(pf06751) EUT2F
02935 HMM06277EutA_1_eut(pf06277) EUT2F
02936 HMMreg07568_1_eut2 EUT2F
02937 HMMreg07568_1_eut2 EUT2F
02938 HMMreg07568_1_eut2(pf12282pf07568) EUT2F
02939 HMMreg00072_1_eut2(pf03861pf00072) EUT2F
02940 HMM10662PduV_1_mix(pf10662) EUT2F
02941 Hp_euts_plumPurple EUT2F
02942 - -
02943 -pf00037 -
02944 -pf01832 -
02945 - -
BMC-P
BMC-H
BMC-T
Aldehyde dehydrogenase
PTAC
Signature enzyme
Alcohol dehydrogenase
Regulator

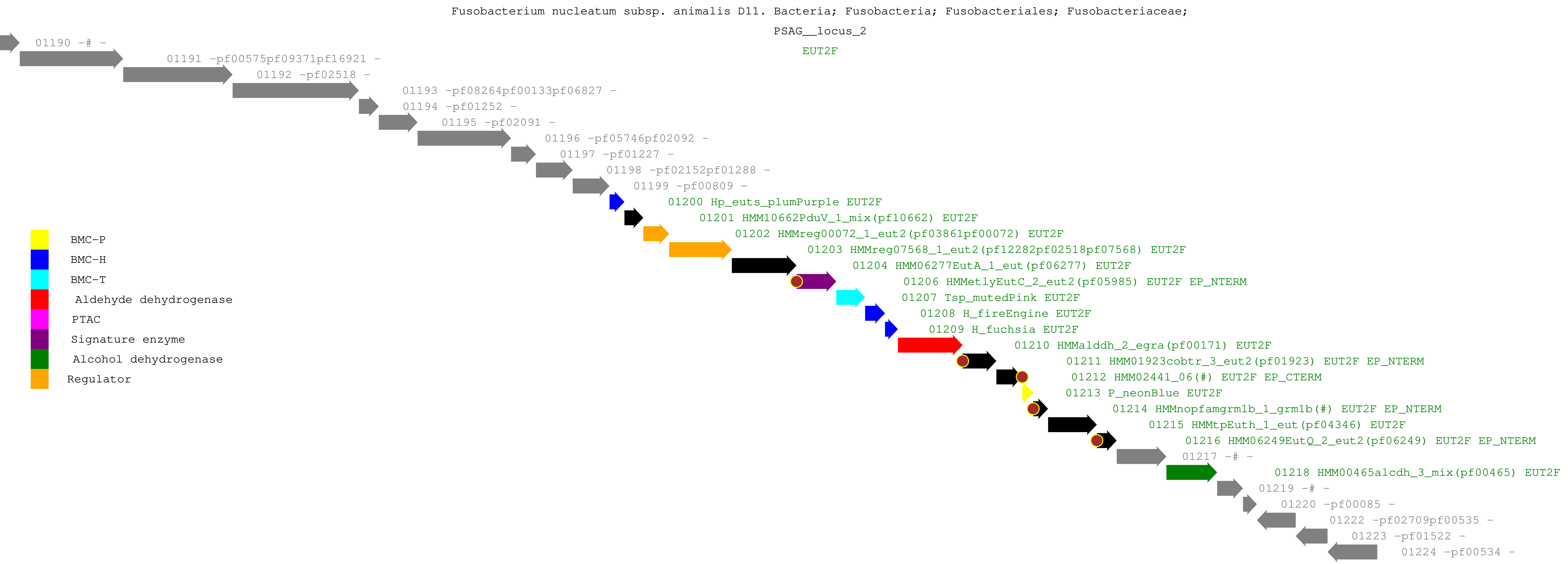
Fusobacterium nucleatum subsp. animalis D11. Bacteria; Fusobacteria; Fusobacteriales; Fusobacteriaceae;
PSAG__locus_2
EUT2F
01190 -# -
01191 -pf00575pf09371pf16921 -
01192 -pf02518 -
01193 -pf08264pf00133pf06827 -
01194 -pf01252 -
01195 -pf02091 -
01196 -pf05746pf02092 -
01197 -pf01227 -
01198 -pf02152pf01288 -
01199 -pf00809 -
01200 Hp_euts_plumPurple EUT2F
01201 HMM10662PduV_1_mix(pf10662) EUT2F
01202 HMMreg00072_1_eut2(pf03861pf00072) EUT2F
01203 HMMreg07568_1_eut2(pf12282pf02518pf07568) EUT2F
01204 HMM06277EutA_1_eut(pf06277) EUT2F
01206 HMMetlyEutC_2_eut2(pf05985) EUT2F EP_NTERM
01207 Tsp_mutedPink EUT2F
01208 H_fireEngine EUT2F
01209 H_fuchsia EUT2F
01210 HMMalddh_2_egra(pf00171) EUT2F
01211 HMM01923cobtr_3_eut2(pf01923) EUT2F EP_NTERM
01212 HMM02441_06(#) EUT2F EP_CTERM
01213 P_neonBlue EUT2F
01214 HMMnopfamgrm1b_1_grm1b(#) EUT2F EP_NTERM
01215 HMMtpEuth_1_eut(pf04346) EUT2F
01216 HMM06249EutQ_2_eut2(pf06249) EUT2F EP_NTERM
01217 -# -
01218 HMM00465alcdh_3_mix(pf00465) EUT2F
01219 -# -
01220 -pf00085 -
01222 -pf02709pf00535 -
01223 -pf01522 -
01224 -pf00534 -
BMC-P
BMC-H
BMC-T
Aldehyde dehydrogenase
PTAC
Signature enzyme
Alcohol dehydrogenase
Regulator

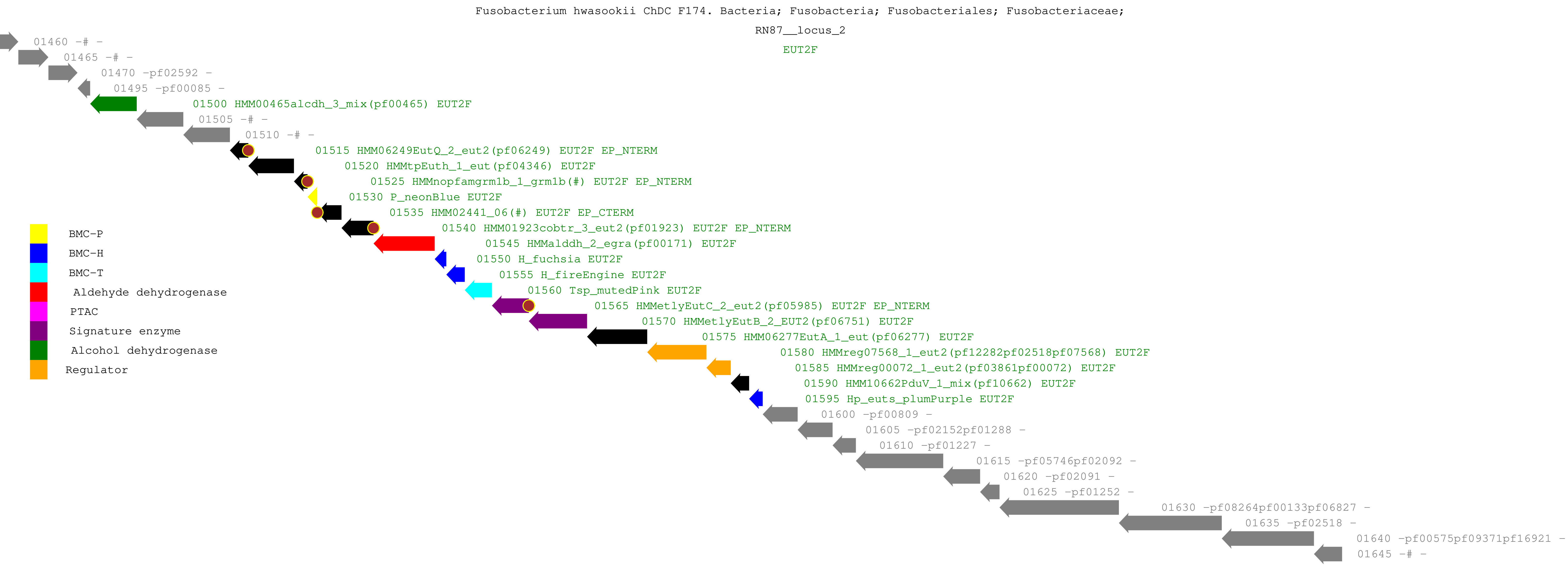
Fusobacterium hwasookii ChDC F174. Bacteria; Fusobacteria; Fusobacteriales; Fusobacteriaceae;
RN87__locus_2
EUT2F
01460 -# -
01465 -# -
01470 -pf02592 -
01495 -pf00085 -
01500 HMM00465alcdh_3_mix(pf00465) EUT2F
01505 -# -
01510 -# -
01515 HMM06249EutQ_2_eut2(pf06249) EUT2F EP_NTERM
01520 HMMtpEuth_1_eut(pf04346) EUT2F
01525 HMMnopfamgrm1b_1_grm1b(#) EUT2F EP_NTERM
01530 P_neonBlue EUT2F
01535 HMM02441_06(#) EUT2F EP_CTERM
01540 HMM01923cobtr_3_eut2(pf01923) EUT2F EP_NTERM
01545 HMMalddh_2_egra(pf00171) EUT2F
01550 H_fuchsia EUT2F
01555 H_fireEngine EUT2F
01560 Tsp_mutedPink EUT2F
01565 HMMetlyEutC_2_eut2(pf05985) EUT2F EP_NTERM
01570 HMMetlyEutB_2_EUT2(pf06751) EUT2F
01575 HMM06277EutA_1_eut(pf06277) EUT2F
01580 HMMreg07568_1_eut2(pf12282pf02518pf07568) EUT2F
01585 HMMreg00072_1_eut2(pf03861pf00072) EUT2F
01590 HMM10662PduV_1_mix(pf10662) EUT2F
01595 Hp_euts_plumPurple EUT2F
01600 -pf00809 -
01605 -pf02152pf01288 -
01610 -pf01227 -
01615 -pf05746pf02092 -
01620 -pf02091 -
01625 -pf01252 -
01630 -pf08264pf00133pf06827 -
01635 -pf02518 -
01640 -pf00575pf09371pf16921 -
01645 -# -
BMC-P
BMC-H
BMC-T
Aldehyde dehydrogenase
PTAC
Signature enzyme
Alcohol dehydrogenase
Regulator

Fusobacterium nucleatum subsp. animalis. Bacteria; Fusobacteria; Fusobacteriales; Fusobacteriaceae;

RN90__locus_2

EUT2F

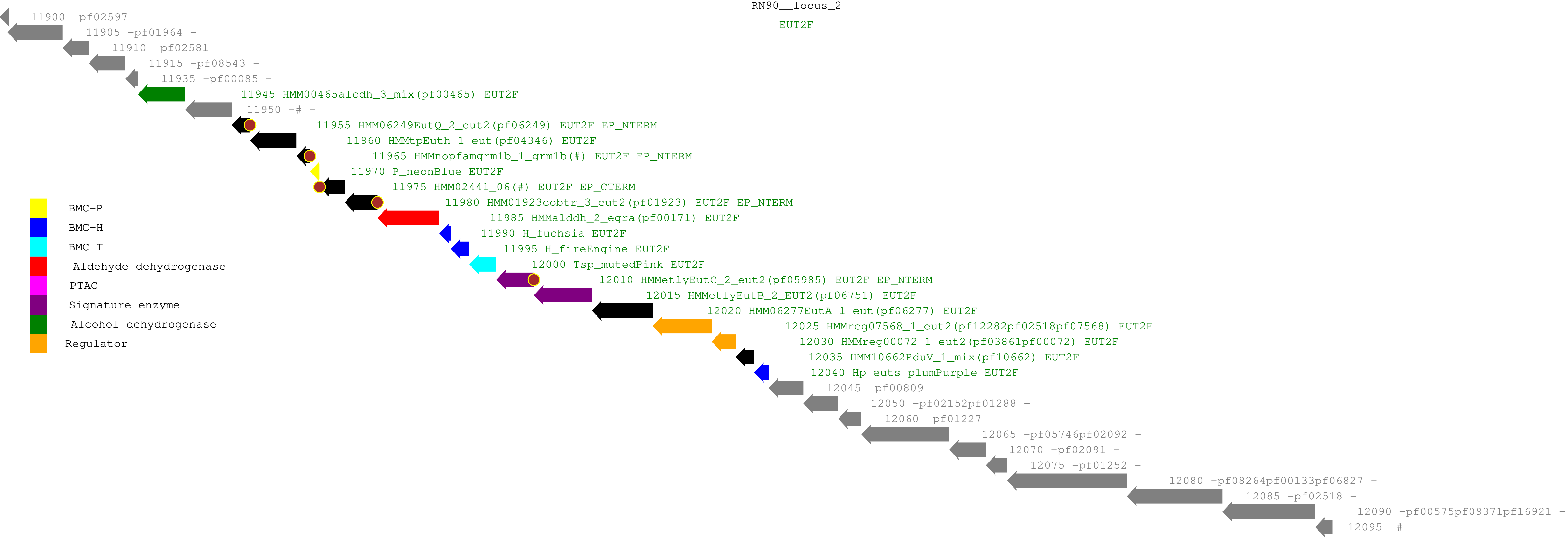

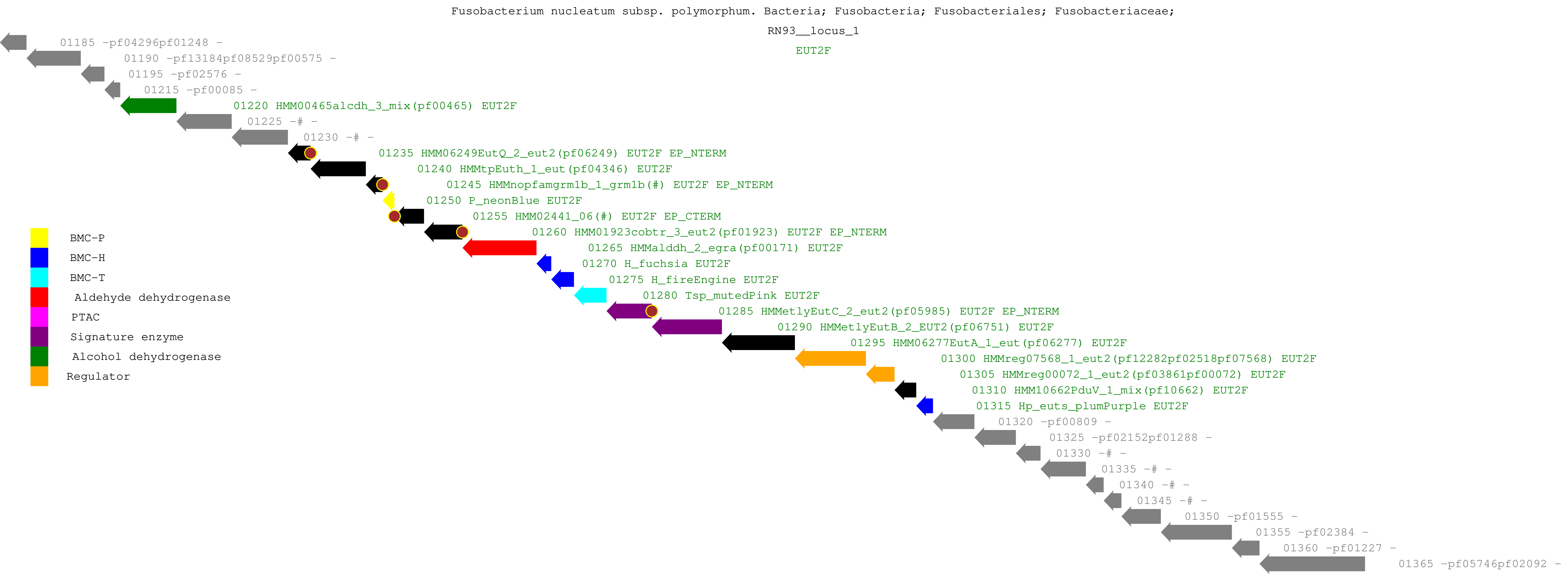

Fusobacterium nucleatum subsp. polymorphum. Bacteria; Fusobacteria; Fusobacteriales; Fusobacteriaceae;
RN93__locus_1
EUT2F
01185 -pf04296pf01248 -
01190 -pf13184pf08529pf00575 -
01195 -pf02576 -
01215 -pf00085 -
01220 HMM00465alcdh_3_mix(pf00465) EUT2F
01225 -# -
01230 -# -
01235 HMM06249EutQ_2_eut2(pf06249) EUT2F EP_NTERM
01240 HMMtpEuth_1_eut(pf04346) EUT2F
01245 HMMnopfamgrm1b_1_grm1b(#) EUT2F EP_NTERM
01250 P_neonBlue EUT2F
01255 HMM02441_06(#) EUT2F EP_CTERM
01260 HMM01923cobtr_3_eut2(pf01923) EUT2F EP_NTERM
01265 HMMalddh_2_egra(pf00171) EUT2F
01270 H_fuchsia EUT2F
01275 H_fireEngine EUT2F
01280 Tsp_mutedPink EUT2F
01285 HMMetlyEutC_2_eut2(pf05985) EUT2F EP_NTERM
01290 HMMetlyEutB_2_EUT2(pf06751) EUT2F
01295 HMM06277EutA_1_eut(pf06277) EUT2F
01300 HMMreg07568_1_eut2(pf12282pf02518pf07568) EUT2F
01305 HMMreg00072_1_eut2(pf03861pf00072) EUT2F
01310 HMM10662PduV_1_mix(pf10662) EUT2F
01315 Hp_euts_plumPurple EUT2F
01320 -pf00809 -
01325 -pf02152pf01288 -
01330 -# -
01335 -# -
01340 -# -
01345 -# -
01350 -pf01555 -
01355 -pf02384 -
01360 -pf01227 -
01365 -pf05746pf02092 -
BMC-P
BMC-H
BMC-T
Aldehyde dehydrogenase
PTAC
Signature enzyme
Alcohol dehydrogenase
Regulator

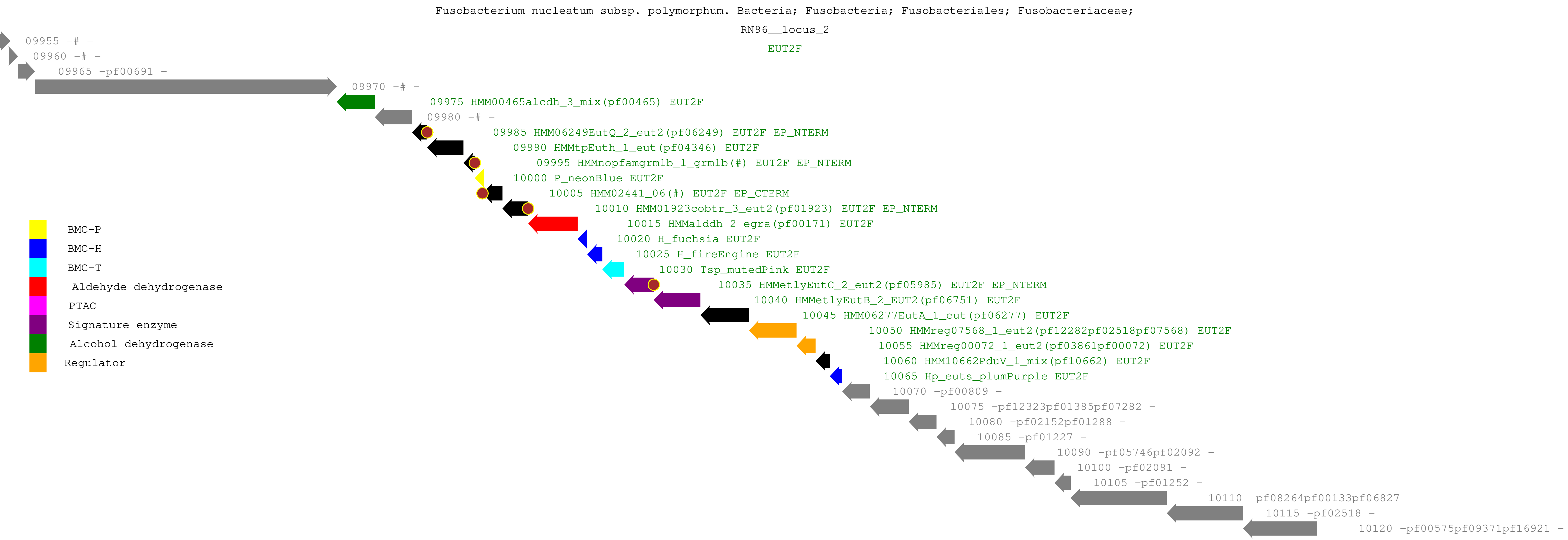

Fusobacterium nucleatum subsp. polymorphum. Bacteria; Fusobacteria; Fusobacteriales; Fusobacteriaceae;
RN96__locus_2
EUT2F
09955 -# -
09960 -# -
09965 -pf00691 -
09970 -# -
09975 HMM00465alcdh_3_mix(pf00465) EUT2F
09980 -# -
09985 HMM06249EutQ_2_eut2(pf06249) EUT2F EP_NTERM
09990 HMMtpEuth_1_eut(pf04346) EUT2F
09995 HMMnopfamgrm1b_1_grm1b(#) EUT2F EP_NTERM
10000 P_neonBlue EUT2F
10005 HMM02441_06(#) EUT2F EP_CTERM
10010 HMM01923cobtr_3_eut2(pf01923) EUT2F EP_NTERM
10015 HMMalddh_2_egra(pf00171) EUT2F
10020 H_fuchsia EUT2F
10025 H_fireEngine EUT2F
10030 Tsp_mutedPink EUT2F
10035 HMMetlyEutC_2_eut2(pf05985) EUT2F EP_NTERM
10040 HMMetlyEutB_2_EUT2(pf06751) EUT2F
10045 HMM06277EutA_1_eut(pf06277) EUT2F
10050 HMMreg07568_1_eut2(pf12282pf02518pf07568) EUT2F
10055 HMMreg00072_1_eut2(pf03861pf00072) EUT2F
10060 HMM10662PduV_1_mix(pf10662) EUT2F
10065 Hp_euts_plumPurple EUT2F
10070 -pf00809 -
10075 -pf12323pf01385pf07282 -
10080 -pf02152pf01288 -
10085 -pf01227 -
10090 -pf05746pf02092 -
10100 -pf02091 -
10105 -pf01252 -
10110 -pf08264pf00133pf06827 -
10115 -pf02518 -
10120 -pf00575pf09371pf16921 -
BMC-P
BMC-H
BMC-T
Aldehyde dehydrogenase
PTAC
Signature enzyme
Alcohol dehydrogenase
Regulator

Fusobacterium hwasookii ChDC F300. Bacteria; Fusobacteria; Fusobacteriales; Fusobacteriaceae;

RN97__locus_1

EUT2F

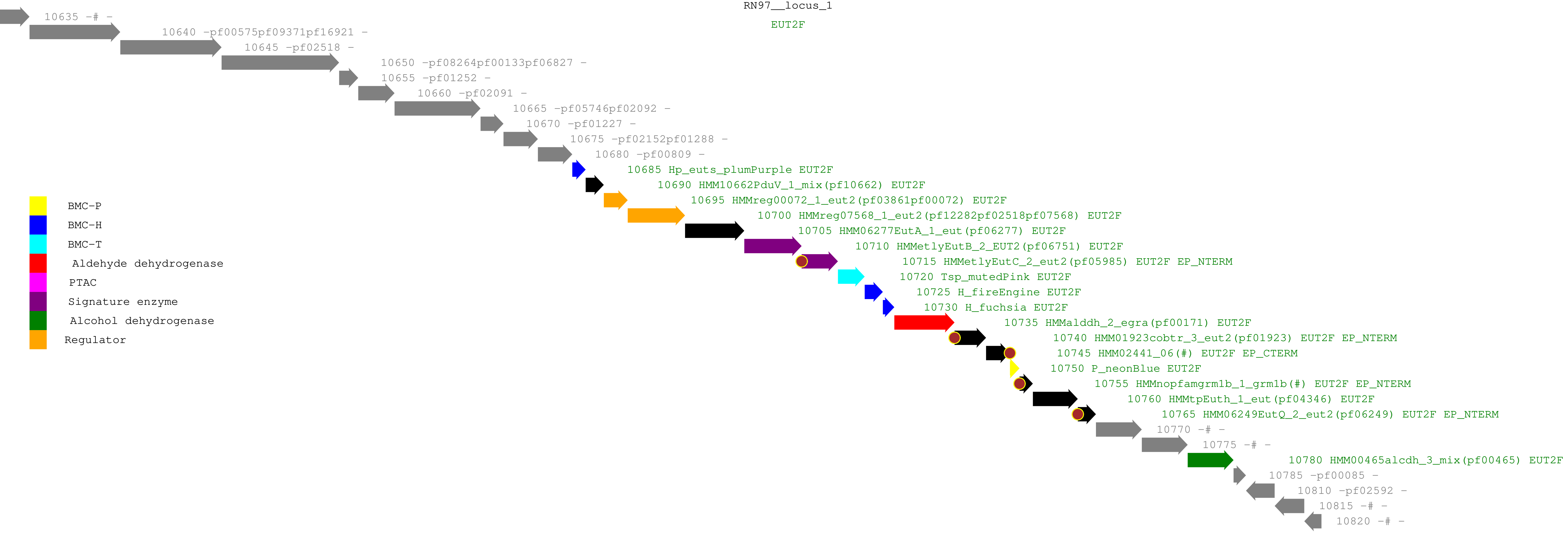

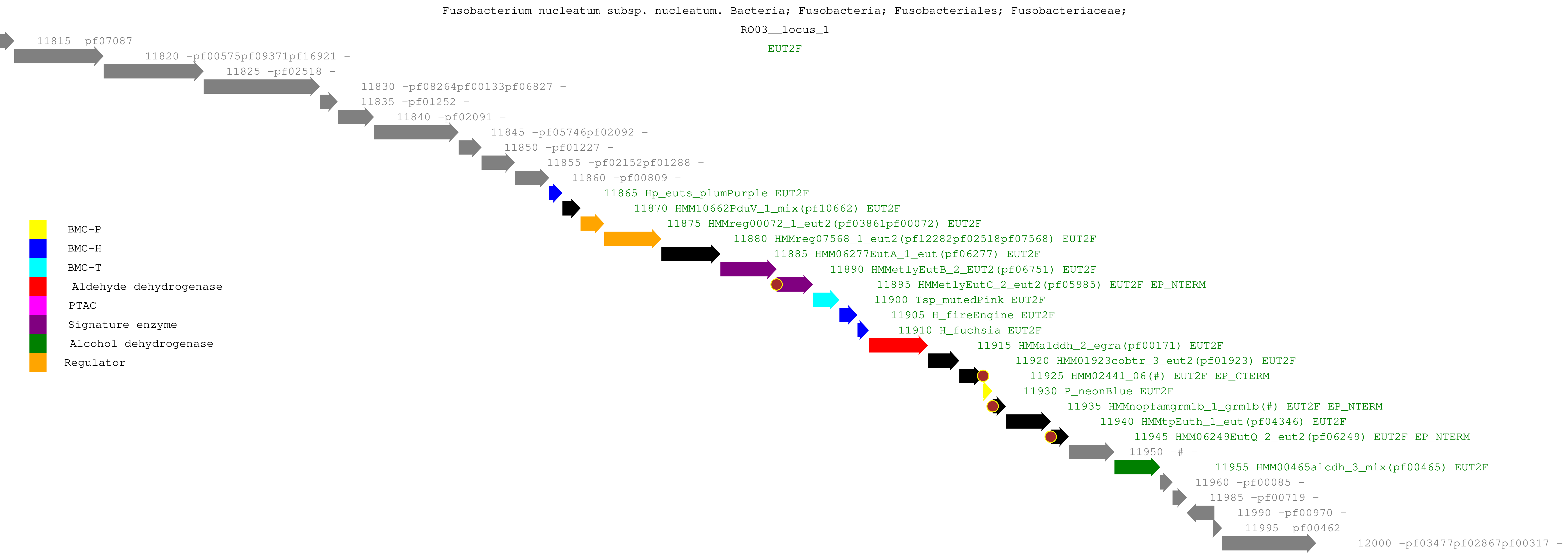

Fusobacterium nucleatum subsp. nucleatum. Bacteria; Fusobacteria; Fusobacteriales; Fusobacteriaceae;
RO03__locus_1
EUT2F
11815 -pf07087 -
11820 -pf00575pf09371pf16921 -
11825 -pf02518 -
11830 -pf08264pf00133pf06827 -
11835 -pf01252 -
11840 -pf02091 -
11845 -pf05746pf02092 -
11850 -pf01227 -
11855 -pf02152pf01288 -
11860 -pf00809 -
11865 Hp_euts_plumPurple EUT2F
11870 HMM10662PduV_1_mix(pf10662) EUT2F
11875 HMMreg00072_1_eut2(pf03861pf00072) EUT2F
11880 HMMreg07568_1_eut2(pf12282pf02518pf07568) EUT2F
11885 HMM06277EutA_1_eut(pf06277) EUT2F
11890 HMMetlyEutB_2_EUT2(pf06751) EUT2F
11895 HMMetlyEutC_2_eut2(pf05985) EUT2F EP_NTERM
11900 Tsp_mutedPink EUT2F
11905 H_fireEngine EUT2F
11910 H_fuchsia EUT2F
11915 HMMaldh_2_egra(pf00171) EUT2F
11920 HMM01923cobtr_3_eut2(pf01923) EUT2F
11925 HMM02441_06(#) EUT2F EP_CTERM
11930 P_neonBlue EUT2F
11935 HMMnopfamgrm1b_1_grm1b(#) EUT2F EP_NTERM
11940 HMMtpEuth_1_eut(pf04346) EUT2F
11945 HMM06249EutQ_2_eut2(pf06249) EUT2F EP_NTERM
11950 -# -
11955 HMM00465alcdh_3_mix(pf00465) EUT2F
11960 -pf00085 -
11985 -pf00719 -
11990 -pf00970 -
11995 -pf00462 -
12000 -pf03477pf02867pf00317 -
BMC-P
BMC-H
BMC-T
Aldehyde dehydrogenase
PTAC
Signature enzyme
Alcohol dehydrogenase
Regulator

Cetobacterium ceti. Bacteria; Fusobacteria; Fusobacteriales; Fusobacteriaceae;

SAMN02745174__locus_1

EUT2F

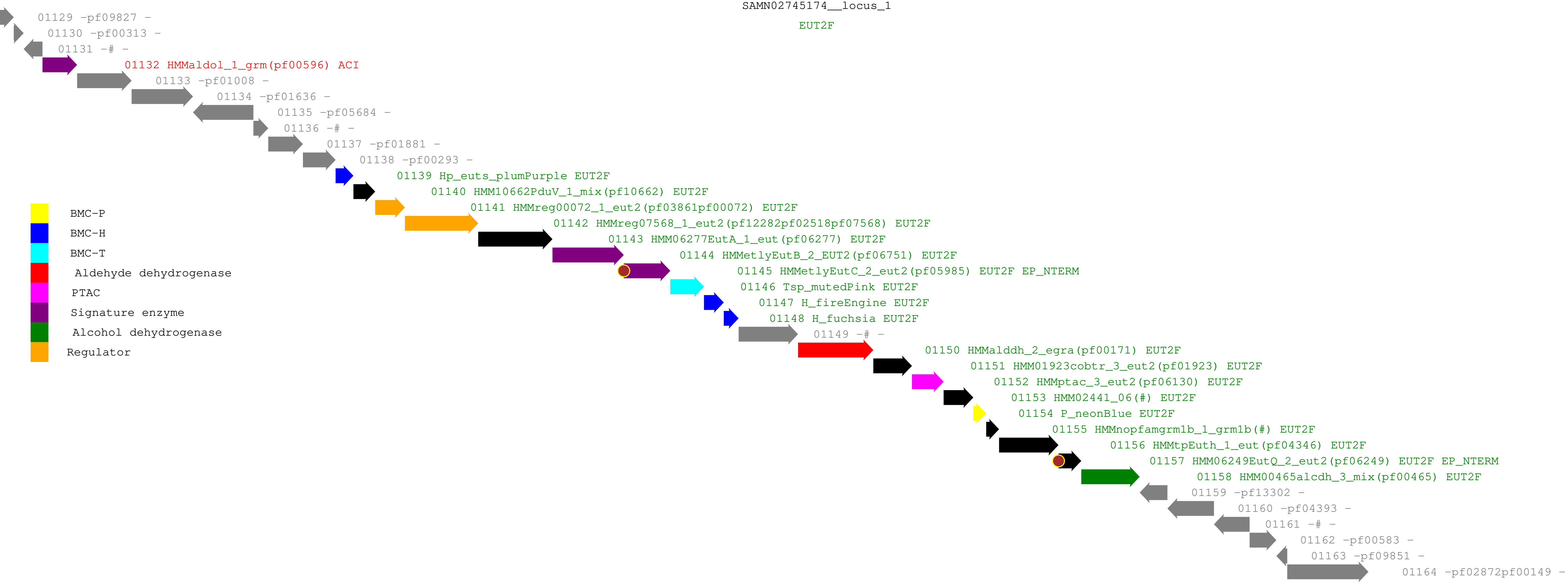

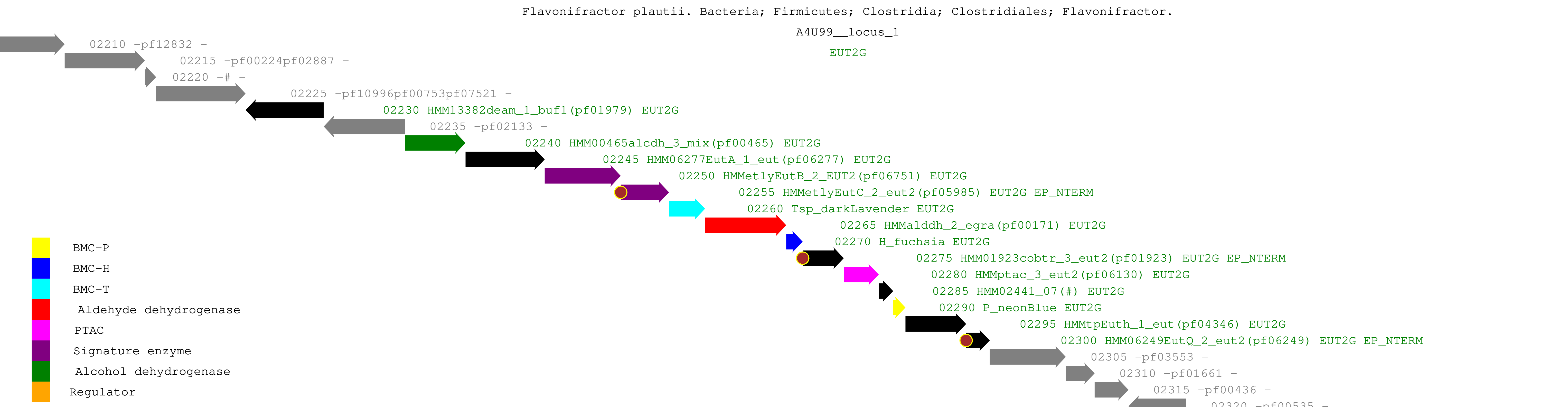

Flavonifractor plautii. Bacteria; Firmicutes; Clostridia; Clostridiales; Flavonifractor.
A4U99__locus_1
EUT2G
02210 -pf12832 -
02215 -pf00224pf02887 -
02220 -# -
02225 -pf10996pf00753pf07521 -
02230 HMM13382deam_1_buf1(pf01979) EUT2G
02235 -pf02133 -
02240 HMM00465alcdh_3_mix(pf00465) EUT2G
02245 HMM06277EutA_1_eut(pf06277) EUT2G
02250 HMMetlyEutB_2_EUT2(pf06751) EUT2G
02255 HMMetlyEutC_2_eut2(pf05985) EUT2G EP_NTERM
02260 Tsp_darkLavender EUT2G
02265 HMMalddh_2_egra(pf00171) EUT2G
02270 H_fuchsia EUT2G
02275 HMM01923cobtr_3_eut2(pf01923) EUT2G EP_NTERM
02280 HMMptac_3_eut2(pf06130) EUT2G
02285 HMM02441_07(#) EUT2G
02290 P_neonBlue EUT2G
02295 HMMtpEuth_1_eut(pf04346) EUT2G
02300 HMM06249EutQ_2_eut2(pf06249) EUT2G EP_NTERM
02305 -pf03553 -
02310 -pf01661 -
02315 -pf00436 -
02320 -pf00535 -
02325 -pf03275 -
02330 -pf00719 -
02335 -pf03965 -
02340 -pf05569 -
BMC-P
BMC-H
BMC-T
Aldehyde dehydrogenase
PTAC
Signature enzyme
Alcohol dehydrogenase
Regulator

Flavonifractor sp. An10. Bacteria; Firmicutes; Clostridia; Clostridiales; Flavonifractor.

B5E42__locus_1

EUT2G

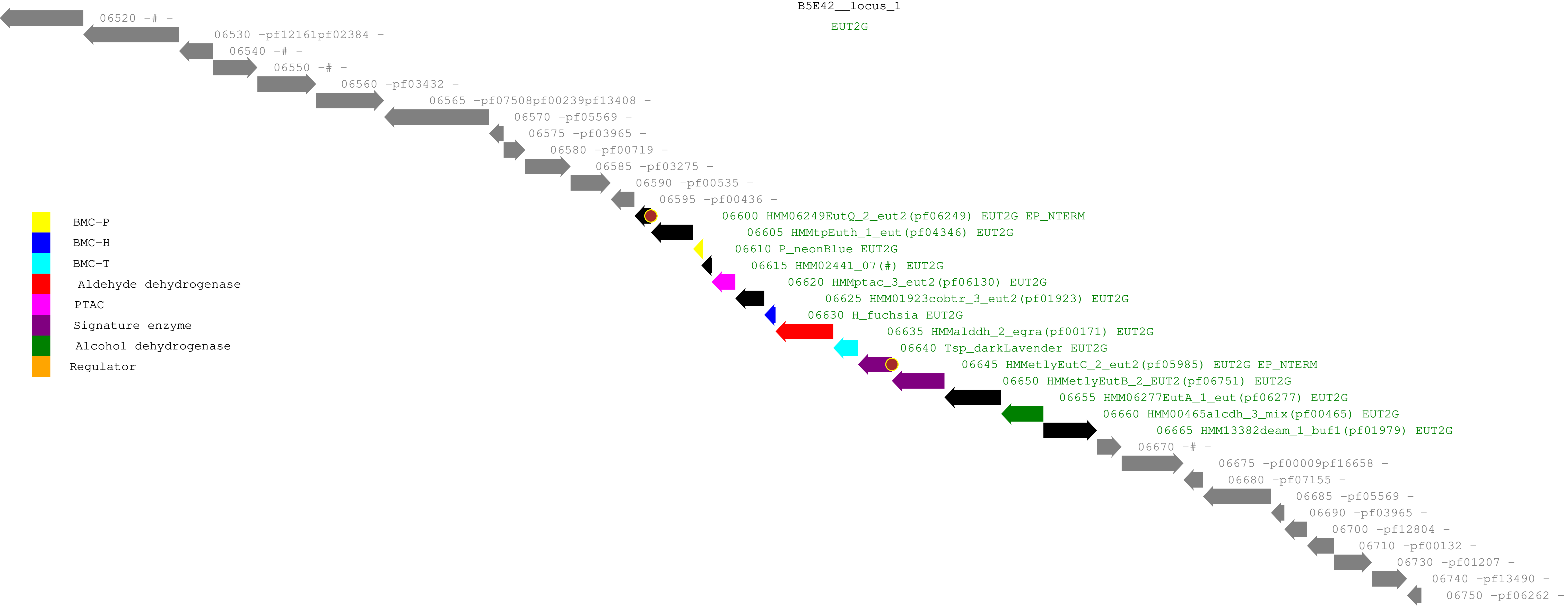

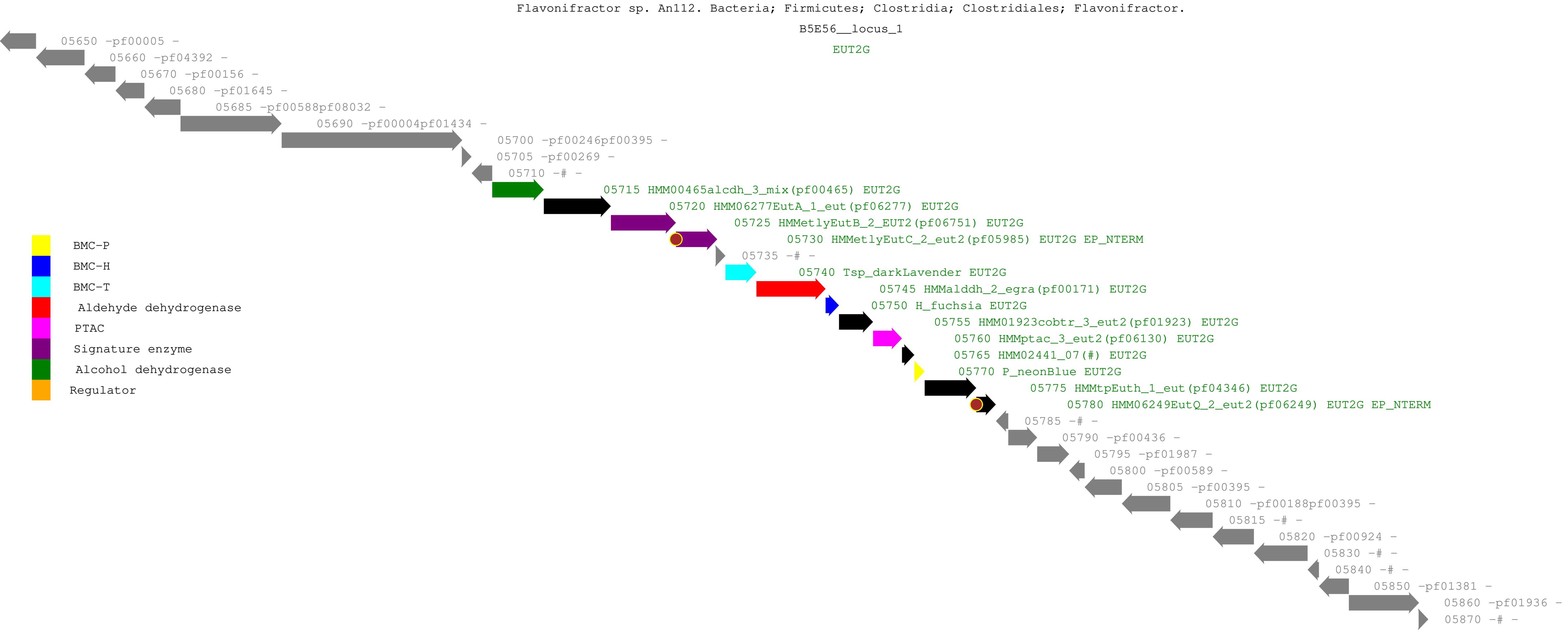
Flavonifractor sp. An112. Bacteria; Firmicutes; Clostridia; Clostridiales; Flavonifractor.
B5E56__locus_1
EUT2G
05650 -pf00005 -
05660 -pf04392 -
05670 -pf00156 -
05680 -pf01645 -
05685 -pf00588pf08032 -
05690 -pf00004pf01434 -
05700 -pf00246pf00395 -
05705 -pf00269 -
05710 -# -
05715 HMM00465alcdh_3_mix(pf00465) EUT2G
05720 HMM06277EutA_1_eut(pf06277) EUT2G
05725 HMMetlyEutB_2_EUT2(pf06751) EUT2G
05730 HMMetlyEutC_2_eut2(pf05985) EUT2G EP_NTERM
05735 -# -
05740 Tsp_darkLavender EUT2G
05745 HMMalddh_2_egra(pf00171) EUT2G
05750 H_fuchsia EUT2G
05755 HMM01923cobtr_3_eut2(pf01923) EUT2G
05760 HMMptac_3_eut2(pf06130) EUT2G
05765 HMM02441_07(#) EUT2G
05770 P_neonBlue EUT2G
05775 HMMtpEuth_1_eut(pf04346) EUT2G
05780 HMM06249EutQ_2_eut2(pf06249) EUT2G EP_NTERM
05785 -# -
05790 -pf00436 -
05795 -pf01987 -
05800 -pf00589 -
05805 -pf00395 -
05810 -pf00188pf00395 -
05815 -# -
05820 -pf00924 -
05830 -# -
05840 -# -
05850 -pf01381 -
05860 -pf01936 -
05870 -# -
BMC-P
BMC-H
BMC-T
Aldehyde dehydrogenase
PTAC
Signature enzyme
Alcohol dehydrogenase
Regulator

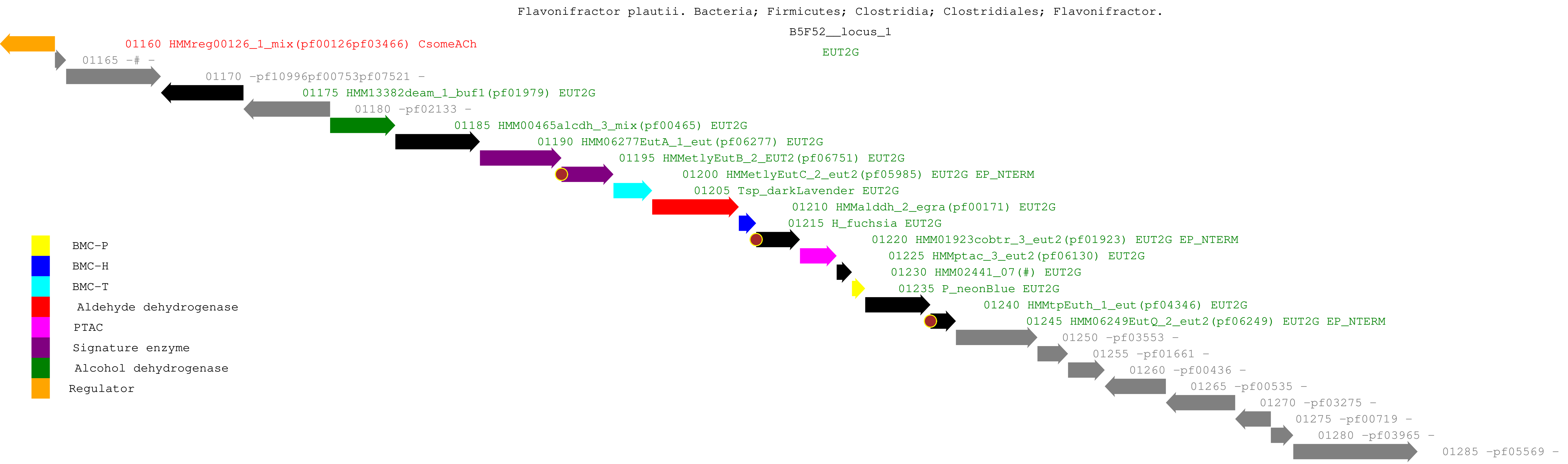

Flavonifractor plautii. Bacteria; Firmicutes; Clostridia; Clostridiales; Flavonifractor.
B5F52__locus_1
EUT2G
01160 HMMreg00126_1_mix(pf00126pf03466) CsomeACh
01165 -# -
01170 -pf10996pf00753pf07521 -
01175 HMM13382deam_1_buf1(pf01979) EUT2G
01180 -pf02133 -
01185 HMM00465alcdh_3_mix(pf00465) EUT2G
01190 HMM06277EutA_1_eut(pf06277) EUT2G
01195 HMMetlyEutB_2_EUT2(pf06751) EUT2G
01200 HMMetlyEutC_2_eut2(pf05985) EUT2G EP_NTERM
01205 Tsp_darkLavender EUT2G
01210 HMMaldh_2_egra(pf00171) EUT2G
01215 H_fuchsia EUT2G
01220 HMM01923cobtr_3_eut2(pf01923) EUT2G EP_NTERM
01225 HMMptac_3_eut2(pf06130) EUT2G
01230 HMM02441_07(#) EUT2G
01235 P_neonBlue EUT2G
01240 HMMtpEuth_1_eut(pf04346) EUT2G
01245 HMM06249EutQ_2_eut2(pf06249) EUT2G EP_NTERM
01250 -pf03553 -
01255 -pf01661 -
01260 -pf00436 -
01265 -pf00535 -
01270 -pf03275 -
01275 -pf00719 -
01280 -pf03965 -
01285 -pf05569 -
BMC-P
BMC-H
BMC-T
Aldehyde dehydrogenase
PTAC
Signature enzyme
Alcohol dehydrogenase
Regulator

Flavonifractor sp. An306. Bacteria; Firmicutes; Clostridia; Clostridiales; Flavonifractor.

B5F88__locus_1

EUT2G

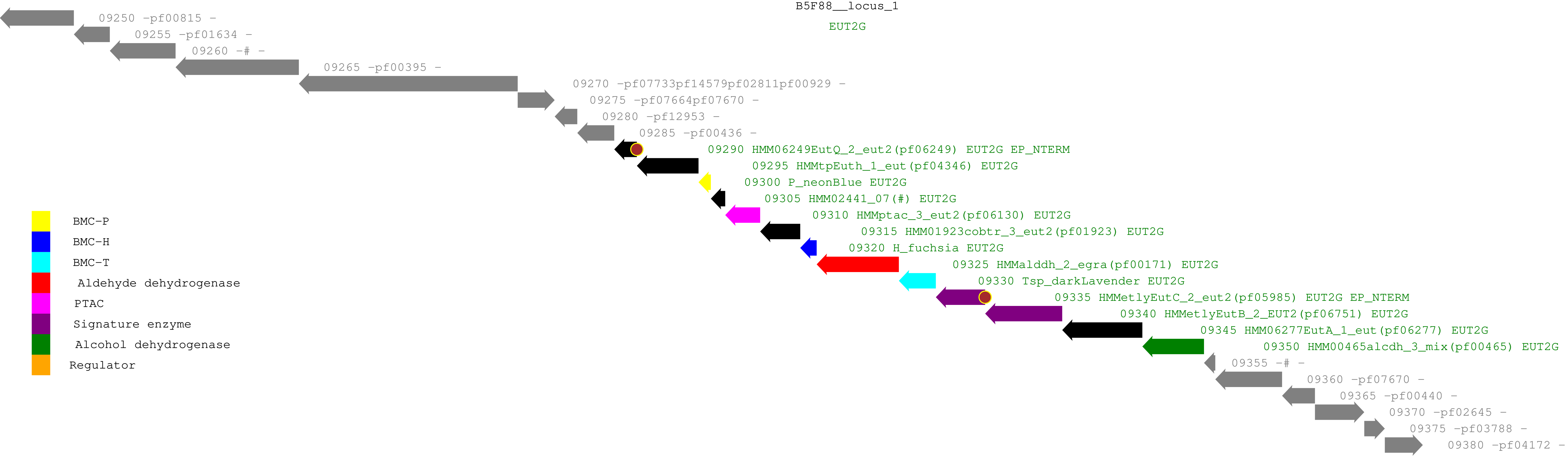

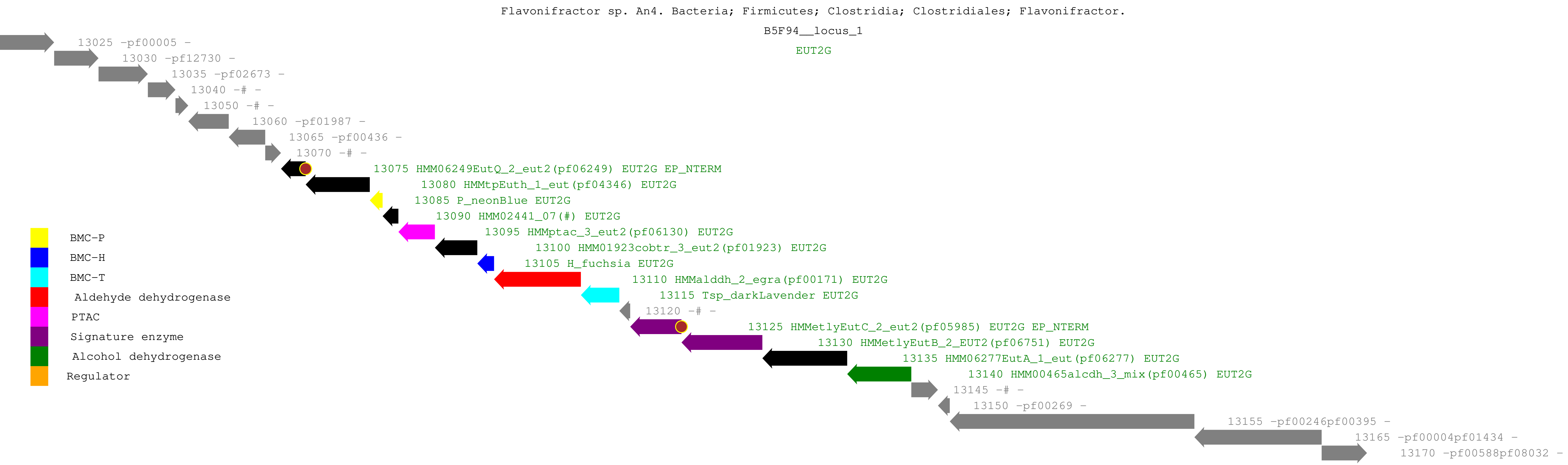

Flavonifractor sp. An4. Bacteria; Firmicutes; Clostridia; Clostridiales; Flavonifractor.
B5F94__locus_1
EUT2G
13025 -pf00005 -
13030 -pf12730 -
13035 -pf02673 -
13040 -# -
13050 -# -
13060 -pf01987 -
13065 -pf00436 -
13070 -# -
13075 HMM06249EutQ_2_eut2(pf06249) EUT2G EP_NTERM
13080 HMMtpEuth_1_eut(pf04346) EUT2G
13085 P_neonBlue EUT2G
13090 HMM02441_07(#) EUT2G
13095 HMMptac_3_eut2(pf06130) EUT2G
13100 HMM01923cobtr_3_eut2(pf01923) EUT2G
13105 H_fuchsia EUT2G
13110 HMMalddh_2_egra(pf00171) EUT2G
13115 Tsp_darkLavender EUT2G
13120 -# -
13125 HMMetlyEutC_2_eut2(pf05985) EUT2G EP_NTERM
13130 HMMetlyEutB_2_EUT2(pf06751) EUT2G
13135 HMM06277EutA_1_eut(pf06277) EUT2G
13140 HMM00465alcdh_3_mix(pf00465) EUT2G
13145 -# -
13150 -pf00269 -
13155 -pf00246pf00395 -
13165 -pf00004pf01434 -
13170 -pf00588pf08032 -
BMC-P
BMC-H
BMC-T
Aldehyde dehydrogenase
PTAC
Signature enzyme
Alcohol dehydrogenase
Regulator

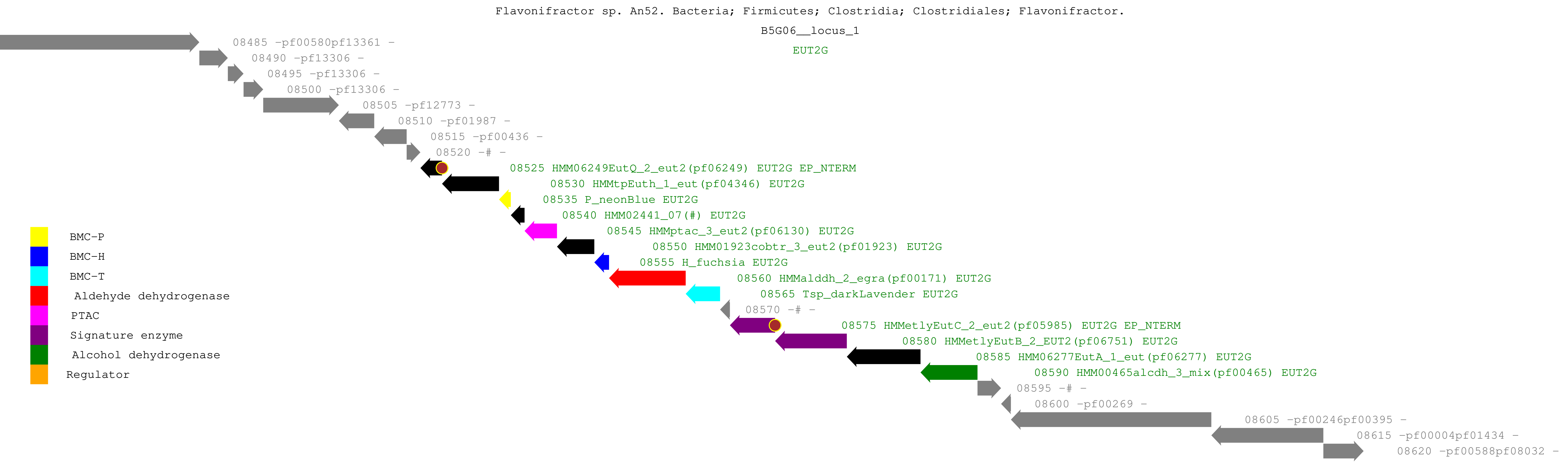
Flavonifractor sp. An52. Bacteria; Firmicutes; Clostridia; Clostridiales; Flavonifractor.
B5G06__locus_1
EUT2G
08485 -pf00580pf13361 -
08490 -pf13306 -
08495 -pf13306 -
08500 -pf13306 -
08505 -pf12773 -
08510 -pf01987 -
08515 -pf00436 -
08520 -# -
08525 HMM06249EutQ_2_eut2(pf06249) EUT2G EP_NTERM
08530 HMMtpEuth_1_eut(pf04346) EUT2G
08535 P_neonBlue EUT2G
08540 HMM02441_07(#) EUT2G
08545 HMMptac_3_eut2(pf06130) EUT2G
08550 HMM01923cobtr_3_eut2(pf01923) EUT2G
08555 H_fuchsia EUT2G
08560 HMMalddh_2_egra(pf00171) EUT2G
08565 Tsp_darkLavender EUT2G
08570 -# -
08575 HMMetlyEutC_2_eut2(pf05985) EUT2G EP_NTERM
08580 HMMetlyEutB_2_EUT2(pf06751) EUT2G
08585 HMM06277EutA_1_eut(pf06277) EUT2G
08590 HMM00465alcdh_3_mix(pf00465) EUT2G
08595 -# -
08600 -pf00269 -
08605 -pf00246pf00395 -
08615 -pf00004pf01434 -
08620 -pf00588pf08032 -
BMC-P
BMC-H
BMC-T
Aldehyde dehydrogenase
PTAC
Signature enzyme
Alcohol dehydrogenase
Regulator

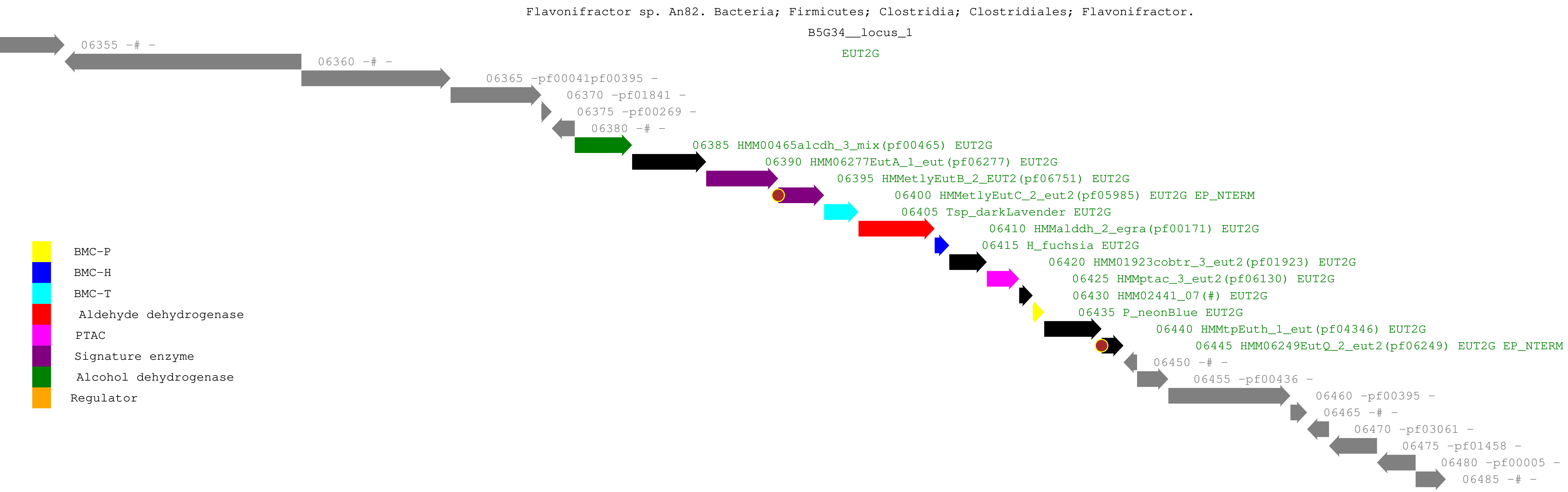
Flavonifractor sp. An82. Bacteria; Firmicutes; Clostridia; Clostridiales; Flavonifractor.
B5G34__locus_1
EUT2G
06355 -# -
06360 -# -
06365 -pf00041pf00395 -
06370 -pf01841 -
06375 -pf00269 -
06380 -# -
06385 HMM00465alcdh_3_mix(pf00465) EUT2G
06390 HMM06277EutA_1_eut(pf06277) EUT2G
06395 HMMetlyEutB_2_EUT2(pf06751) EUT2G
06400 HMMetlyEutC_2_eut2(pf05985) EUT2G EP_NTERM
06405 Tsp_darkLavender EUT2G
06410 HMMaldh_2_egra(pf00171) EUT2G
06415 H_fuchsia EUT2G
06420 HMM01923cobtr_3_eut2(pf01923) EUT2G
06425 HMMptac_3_eut2(pf06130) EUT2G
06430 HMM02441_07(#) EUT2G
06435 P_neonBlue EUT2G
06440 HMMtpEuth_1_eut(pf04346) EUT2G
06445 HMM06249EutQ_2_eut2(pf06249) EUT2G EP_NTERM
06450 -# -
06455 -pf00436 -
06460 -pf00395 -
06465 -# -
06470 -pf03061 -
06475 -pf01458 -
06480 -pf00005 -
06485 -# -
BMC-P
BMC-H
BMC-T
Aldehyde dehydrogenase
PTAC
Signature enzyme
Alcohol dehydrogenase
Regulator

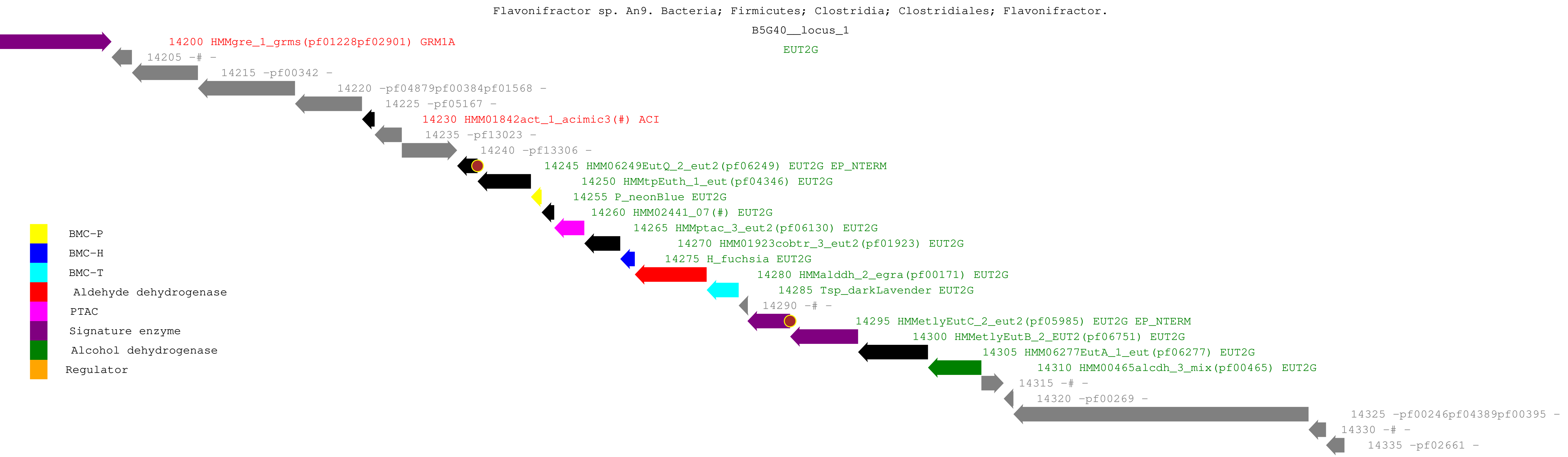
Flavonifractor sp. An9. Bacteria; Firmicutes; Clostridia; Clostridiales; Flavonifractor.
B5G40__locus_1
EUT2G
14200 HMMgre_1_grms(pf01228pf02901) GRM1A
14205 -# -
14215 -pf00342 -
14220 -pf04879pf00384pf01568 -
14225 -pf05167 -
14230 HMM01842act_1_acimic3(#) ACI
14235 -pf13023 -
14240 -pf13306 -
14245 HMM06249EutQ_2_eut2(pf06249) EUT2G EP_NTERM
14250 HMMtpEuth_1_eut(pf04346) EUT2G
14255 P_neonBlue EUT2G
14260 HMM02441_07(#) EUT2G
14265 HMMptac_3_eut2(pf06130) EUT2G
14270 HMM01923cobtr_3_eut2(pf01923) EUT2G
14275 H_fuchsia EUT2G
14280 HMMalddh_2_egra(pf00171) EUT2G
14285 Tsp_darkLavender EUT2G
14290 -# -
14295 HMMetlyEutC_2_eut2(pf05985) EUT2G EP_NTERM
14300 HMMetlyEutB_2_EUT2(pf06751) EUT2G
14305 HMM06277EutA_1_eut(pf06277) EUT2G
14310 HMM00465alcdh_3_mix(pf00465) EUT2G
14315 -# -
14320 -pf00269 -
14325 -pf00246pf04389pf00395 -
14330 -# -
14335 -pf02661 -
BMC-P
BMC-H
BMC-T
Aldehyde dehydrogenase
PTAC
Signature enzyme
Alcohol dehydrogenase
Regulator

Flavonifractor sp. An91. Bacteria; Firmicutes; Clostridia; Clostridiales; Flavonifractor.

B5G42__locus_1

EUT2G

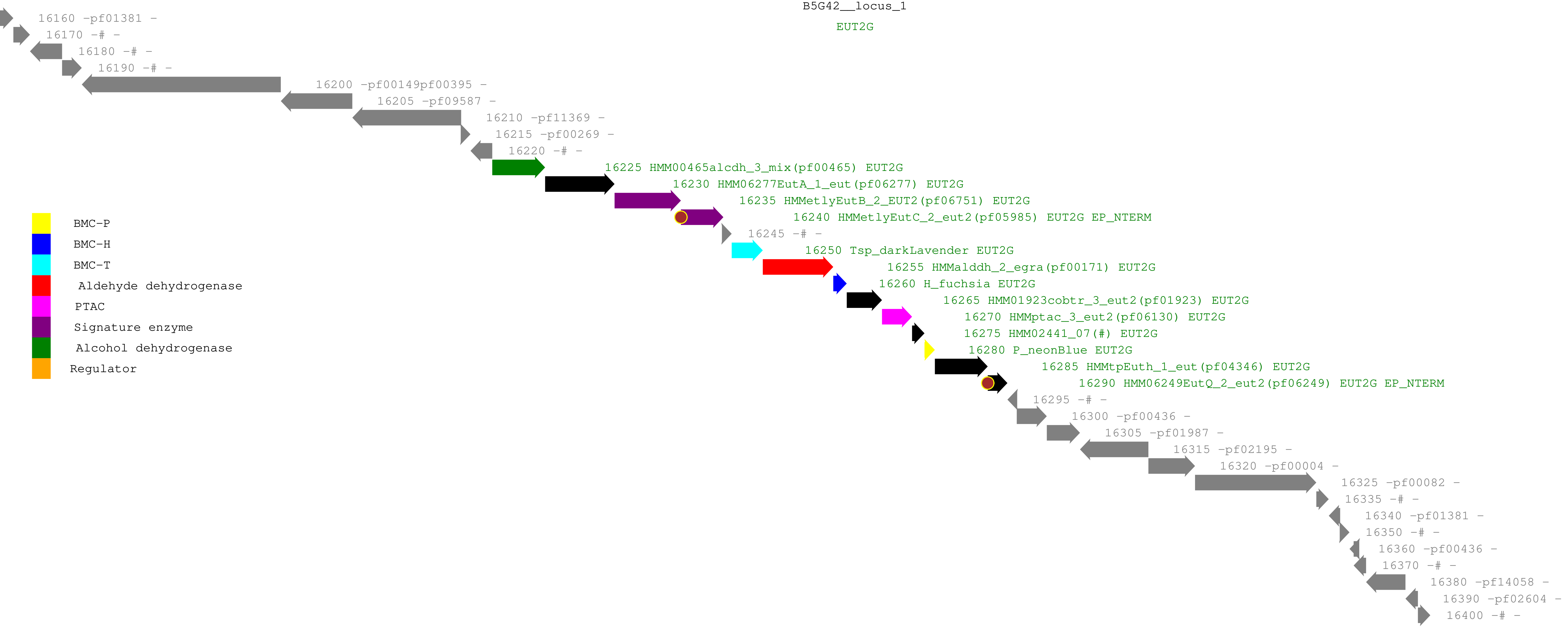

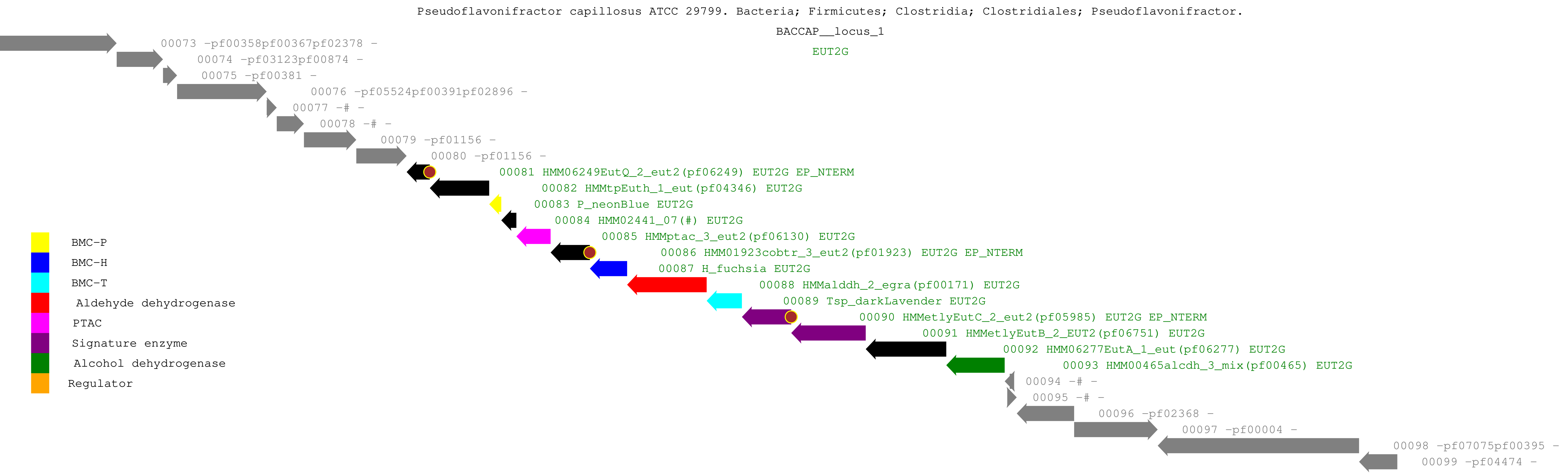
Pseudoflavonifractor capillosus ATCC 29799. Bacteria; Firmicutes; Clostridia; Clostridiales; Pseudoflavonifractor.
BACCAP__locus_1
EUT2G
00073 -pf00358pf00367pf02378 -
00074 -pf03123pf00874 -
00075 -pf00381 -
00076 -pf05524pf00391pf02896 -
00077 -# -
00078 -# -
00079 -pf01156 -
00080 -pf01156 -
00081 HMM06249EutQ_2_eut2(pf06249) EUT2G EP_NTERM
00082 HMMtpEuth_1_eut(pf04346) EUT2G
00083 P_neonBlue EUT2G
00084 HMM02441_07(#) EUT2G
00085 HMMptac_3_eut2(pf06130) EUT2G
00086 HMM01923cobtr_3_eut2(pf01923) EUT2G EP_NTERM
00087 H_fuchsia EUT2G
00088 HMMalddh_2_egra(pf00171) EUT2G
00089 Tsp_darkLavender EUT2G
00090 HMMetlyEutC_2_eut2(pf05985) EUT2G EP_NTERM
00091 HMMetlyEutB_2_EUT2(pf06751) EUT2G
00092 HMM06277EutA_1_eut(pf06277) EUT2G
00093 HMM00465alcdh_3_mix(pf00465) EUT2G
00094 -# -
00095 -# -
00096 -pf02368 -
00097 -pf00004 -
00098 -pf07075pf00395 -
00099 -pf04474 -
BMC-P
BMC-H
BMC-T
Aldehyde dehydrogenase
PTAC
Signature enzyme
Alcohol dehydrogenase
Regulator

Firmicutes bacterium CAG:137_57_8. Bacteria; Firmicutes.

BHW33__locus_1

EUT2G

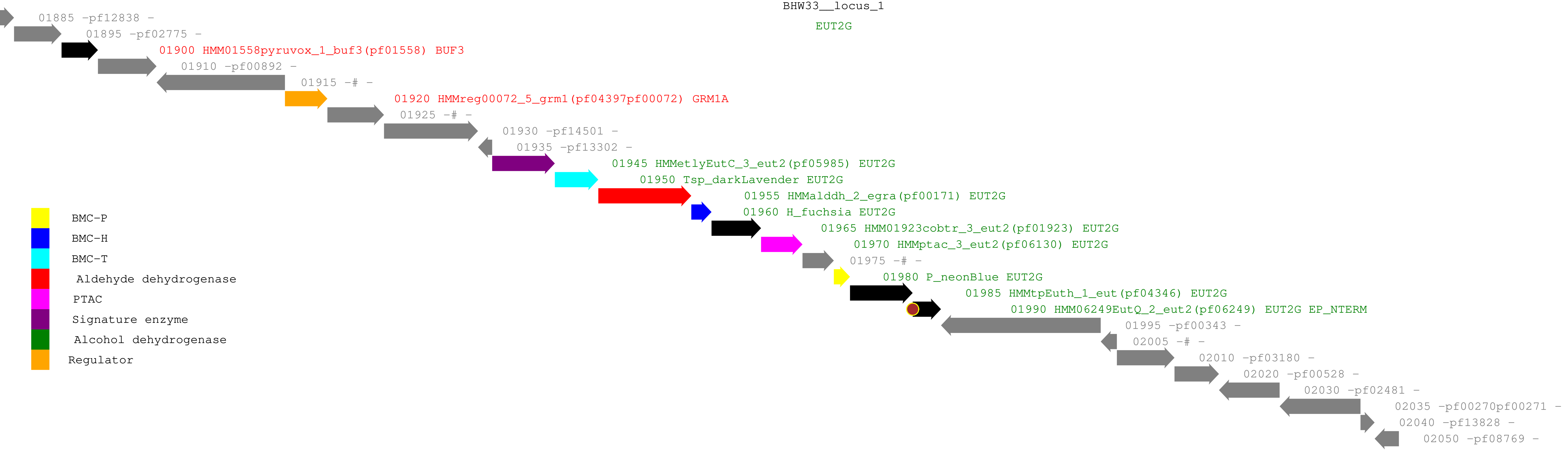

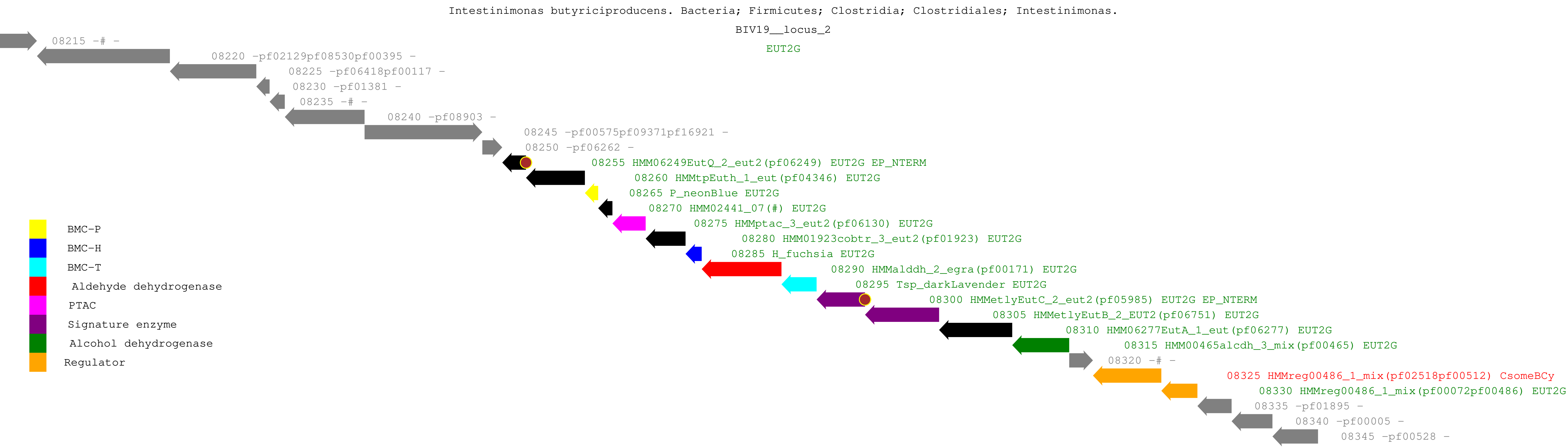
Intestinimonas butyriciproducens. Bacteria; Firmicutes; Clostridia; Clostridiales; Intestinimonas.
BIV19__locus_2
EUT2G
08215 -# -
08220 -pf02129pf08530pf00395 -
08225 -pf06418pf00117 -
08230 -pf01381 -
08235 -# -
08240 -pf08903 -
08245 -pf00575pf09371pf16921 -
08250 -pf06262 -
08255 HMM06249EutQ_2_eut2(pf06249) EUT2G EP_NTERM
08260 HMMtpEuth_1_eut(pf04346) EUT2G
08265 P_neonBlue EUT2G
08270 HMM02441_07(#) EUT2G
08275 HMMptac_3_eut2(pf06130) EUT2G
08280 HMM01923cobtr_3_eut2(pf01923) EUT2G
08285 H_fuchsia EUT2G
08290 HMMalddh_2_egra(pf00171) EUT2G
08295 Tsp_darkLavender EUT2G
08300 HMMetlyEutC_2_eut2(pf05985) EUT2G EP_NTERM
08305 HMMetlyEutB_2_EUT2(pf06751) EUT2G
08310 HMM06277EutA_1_eut(pf06277) EUT2G
08315 HMM00465alcdh_3_mix(pf00465) EUT2G
08320 -# -
08325 HMMreg00486_1_mix(pf02518pf00512) CsomeBCy
08330 HMMreg00486_1_mix(pf00072pf00486) EUT2G
08335 -pf01895 -
08340 -pf00005 -
08345 -pf00528 -
BMC-P
BMC-H
BMC-T
Aldehyde dehydrogenase
PTAC
Signature enzyme
Alcohol dehydrogenase
Regulator

Firmicutes bacterium CAG:110. Bacteria; Firmicutes; environmental samples.

BN466__locus_1

EUT2G

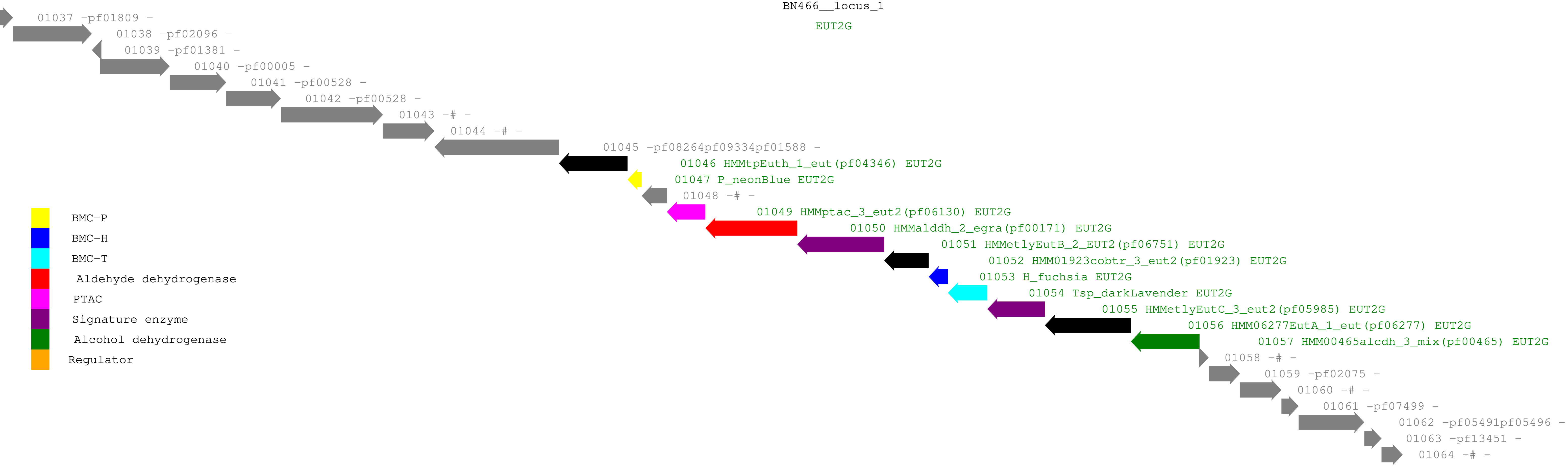

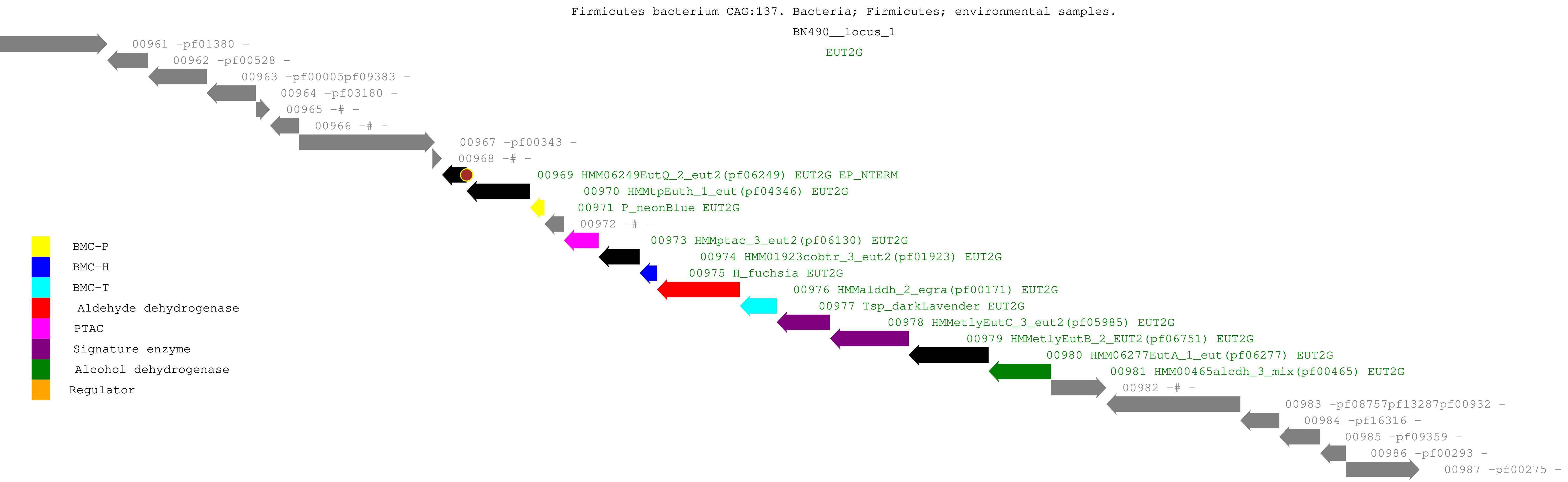
Firmicutes bacterium CAG:137. Bacteria; Firmicutes; environmental samples.
BN490__locus_1
EUT2G
00961 -pf01380 -
00962 -pf00528 -
00963 -pf00005pf09383 -
00964 -pf03180 -
00965 -# -
00966 -# -
00967 -pf00343 -
00968 -# -
00969 HMM06249EutQ_2_eut2(pf06249) EUT2G EP_NTERM
00970 HMMtpEuth_1_eut(pf04346) EUT2G
00971 P_neonBlue EUT2G
00972 -# -
00973 HMMptac_3_eut2(pf06130) EUT2G
00974 HMM01923cobtr_3_eut2(pf01923) EUT2G
00975 H_fuchsia EUT2G
00976 HMMaldh_2_egra(pf00171) EUT2G
00977 Tsp_darkLavender EUT2G
00978 HMMetlyEutC_3_eut2(pf05985) EUT2G
00979 HMMetlyEutB_2_EUT2(pf06751) EUT2G
00980 HMM06277EutA_1_eut(pf06277) EUT2G
00981 HMM00465alcdh_3_mix(pf00465) EUT2G
00982 -# -
00983 -pf08757pf13287pf00932 -
00984 -pf16316 -
00985 -pf09359 -
00986 -pf00293 -
00987 -pf00275 -
BMC-P
BMC-H
BMC-T
Aldehyde dehydrogenase
PTAC
Signature enzyme
Alcohol dehydrogenase
Regulator

Intestinimonas butyriciproducens. Bacteria; Firmicutes; Clostridia; Clostridiales; Intestinimonas.

C7373__locus_1

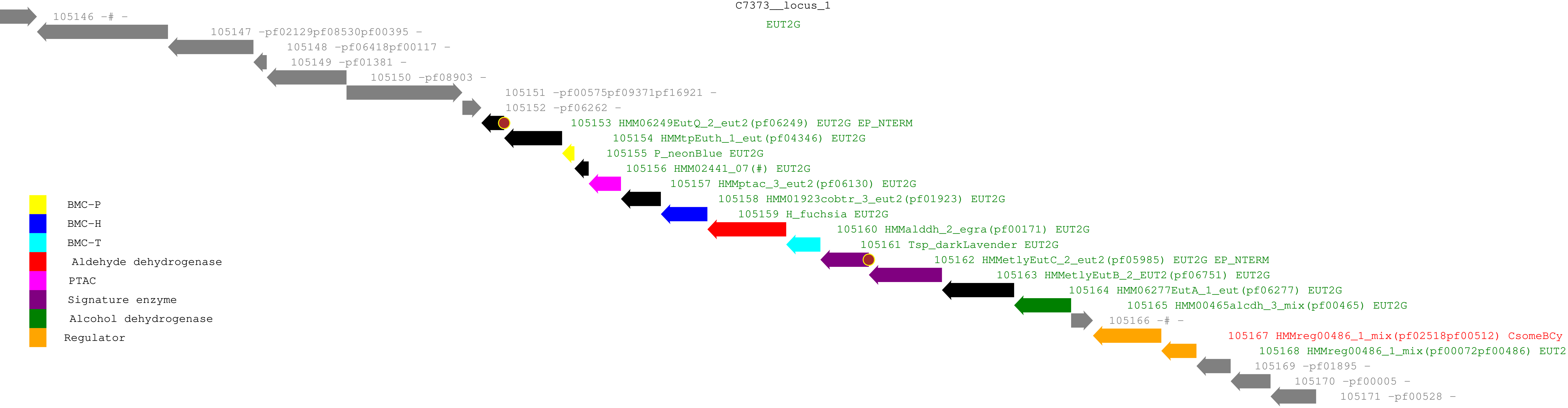

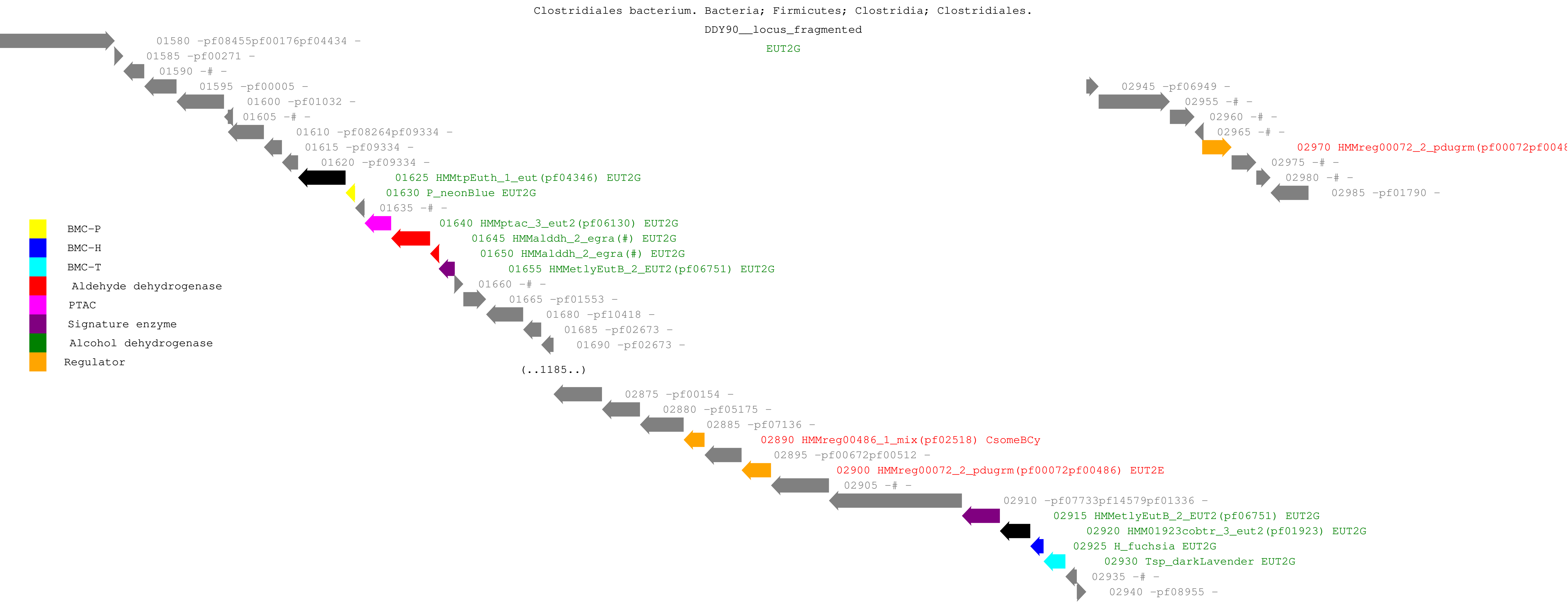
Clostridiales bacterium. Bacteria; Firmicutes; Clostridia; Clostridiales.
DDY90__locus_fragmented
EUT2G
01580 -pf08455pf00176pf04434 -
01585 -pf00271 -
01590 -# -
01595 -pf00005 -
01600 -pf01032 -
01605 -# -
01610 -pf08264pf09334 -
01615 -pf09334 -
01620 -pf09334 -
01625 HMMtpEuth_1_eut(pf04346) EUT2G
01630 P_neonBlue EUT2G
01635 -# -
01640 HMMptac_3_eut2(pf06130) EUT2G
01645 HMMalddh_2_egra(#) EUT2G
01650 HMMalddh_2_egra(#) EUT2G
01655 HMMetlyEutB_2_EUT2(pf06751) EUT2G
01660 -# -
01665 -pf01553 -
01680 -pf10418 -
01685 -pf02673 -
01690 -pf02673 -
(..1185..)
02875 -pf00154 -
02880 -pf05175 -
02885 -pf07136 -
02890 HMMreg00486_1_mix(pf02518) CsomeBCy
02895 -pf00672pf00512 -
02900 HMMreg00072_2_pdugrm(pf00072pf00486) EUT2E
02905 -# -
02910 -pf07733pf14579pf01336 -
02915 HMMetlyEutB_2_EUT2(pf06751) EUT2G
02920 HMM01923cobtr_3_eut2(pf01923) EUT2G
02925 H_fuchsia EUT2G
02930 Tsp_darkLavender EUT2G
02935 -# -
02940 -pf08955 -
02945 -pf06949 -
02955 -# -
02960 -# -
02965 -# -
02970 HMMreg00072_2_pdugrm(pf00072pf0048
02975 -# -
02980 -# -
02985 -pf01790 -
BMC-P
BMC-H
BMC-T
Aldehyde dehydrogenase
PTAC
Signature enzyme
Alcohol dehydrogenase
Regulator

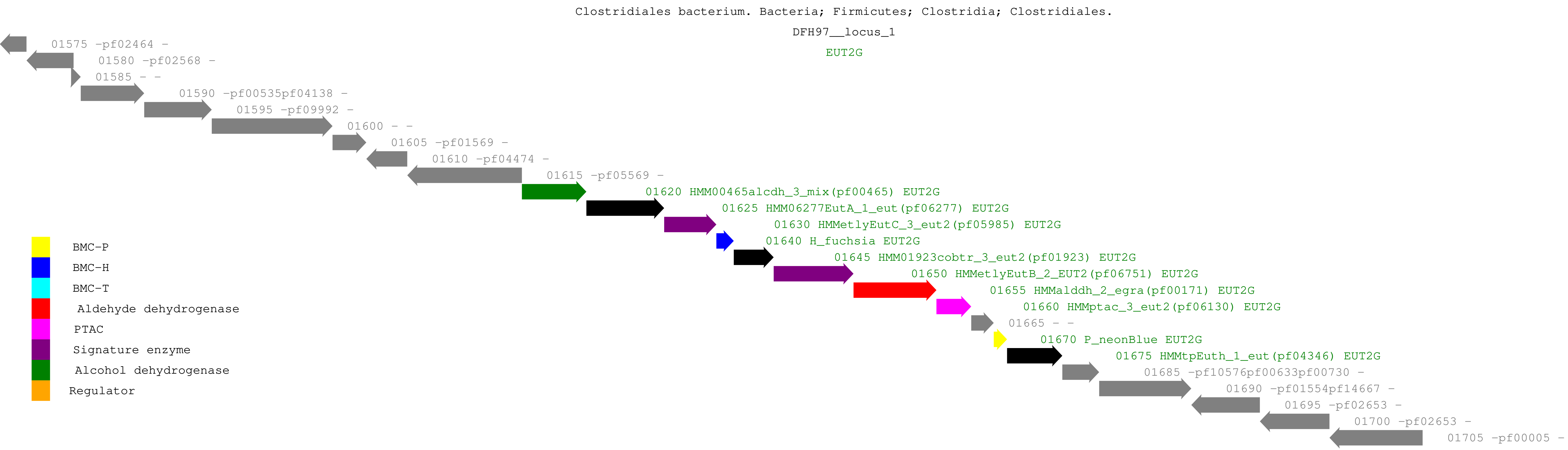
Clostridiales bacterium. Bacteria; Firmicutes; Clostridia; Clostridiales.
DFH97__locus_1
EUT2G
01575 -pf02464 -
01580 -pf02568 -
01585 - -
01590 -pf00535pf04138 -
01595 -pf09992 -
01600 - -
01605 -pf01569 -
01610 -pf04474 -
01615 -pf05569 -
01620 HMM00465alcdh_3_mix(pf00465) EUT2G
01625 HMM06277EutA_1_eut(pf06277) EUT2G
01630 HMMetlyEutC_3_eut2(pf05985) EUT2G
01640 H_fuchsia EUT2G
01645 HMM01923cobtr_3_eut2(pf01923) EUT2G
01650 HMMetlyEutB_2_EUT2(pf06751) EUT2G
01655 HMMalddh_2_egra(pf00171) EUT2G
01660 HMMptac_3_eut2(pf06130) EUT2G
01665 - -
01670 P_neonBlue EUT2G
01675 HMMtpEuth_1_eut(pf04346) EUT2G
01685 -pf10576pf00633pf00730 -
01690 -pf01554pf14667 -
01695 -pf02653 -
01700 -pf02653 -
01705 -pf00005 -
BMC-P
BMC-H
BMC-T
Aldehyde dehydrogenase
PTAC
Signature enzyme
Alcohol dehydrogenase
Regulator

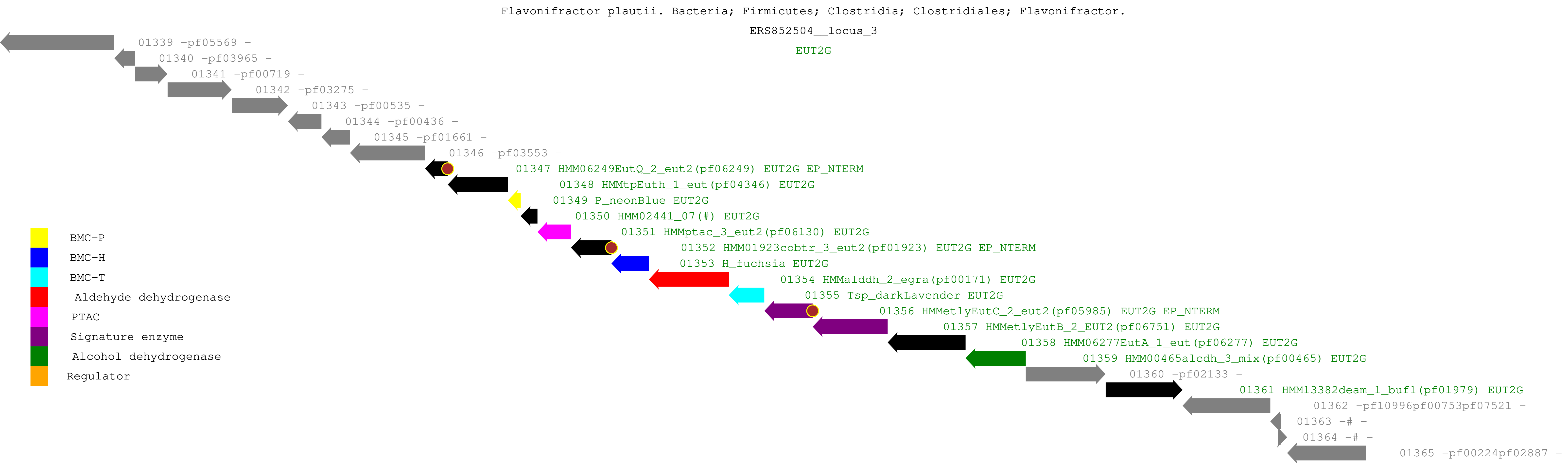
Flavonifractor plautii. Bacteria; Firmicutes; Clostridia; Clostridiales; Flavonifractor.
ERS852504__locus_3
EUT2G
01339 -pf05569 -
01340 -pf03965 -
01341 -pf00719 -
01342 -pf03275 -
01343 -pf00535 -
01344 -pf00436 -
01345 -pf01661 -
01346 -pf03553 -
01347 HMM06249EutQ_2_eut2(pf06249) EUT2G EP_NTERM
01348 HMMtpEuth_1_eut(pf04346) EUT2G
01349 P_neonBlue EUT2G
01350 HMM02441_07(#) EUT2G
01351 HMMptac_3_eut2(pf06130) EUT2G
01352 HMM01923cobtr_3_eut2(pf01923) EUT2G EP_NTERM
01353 H_fuchsia EUT2G
01354 HMMalddh_2_egra(pf00171) EUT2G
01355 Tsp_darkLavender EUT2G
01356 HMMetlyEutC_2_eut2(pf05985) EUT2G EP_NTERM
01357 HMMetlyEutB_2_EUT2(pf06751) EUT2G
01358 HMM06277EutA_1_eut(pf06277) EUT2G
01359 HMM00465alcdh_3_mix(pf00465) EUT2G
01360 -pf02133 -
01361 HMM13382deam_1_buf1(pf01979) EUT2G
01362 -pf10996pf00753pf07521 -
01363 -# -
01364 -# -
01365 -pf00224pf02887 -
BMC-P
BMC-H
BMC-T
Aldehyde dehydrogenase
PTAC
Signature enzyme
Alcohol dehydrogenase
Regulator

Flavonifractor plautii. Bacteria; Firmicutes; Clostridia; Clostridiales; Flavonifractor.

ERS852518__locus_3

EUT2G

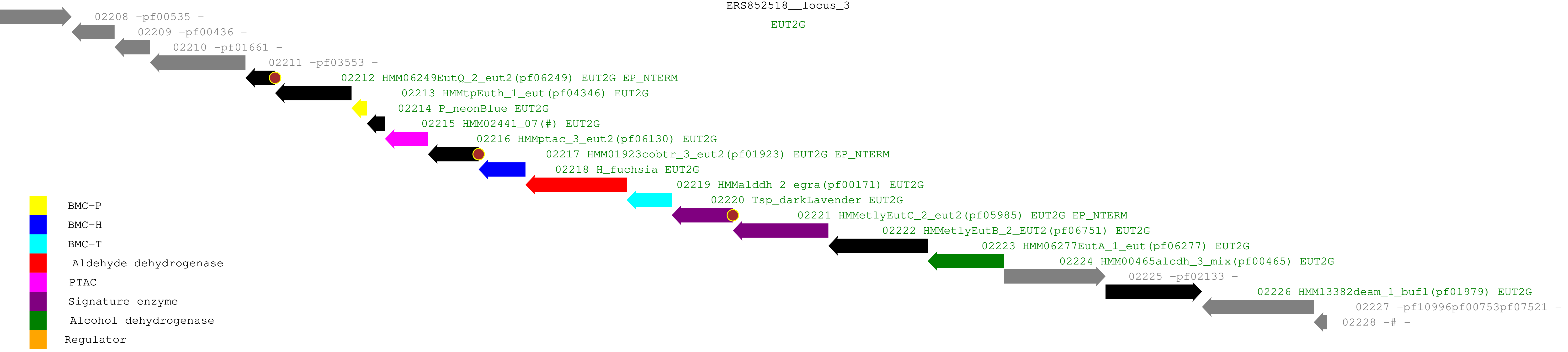

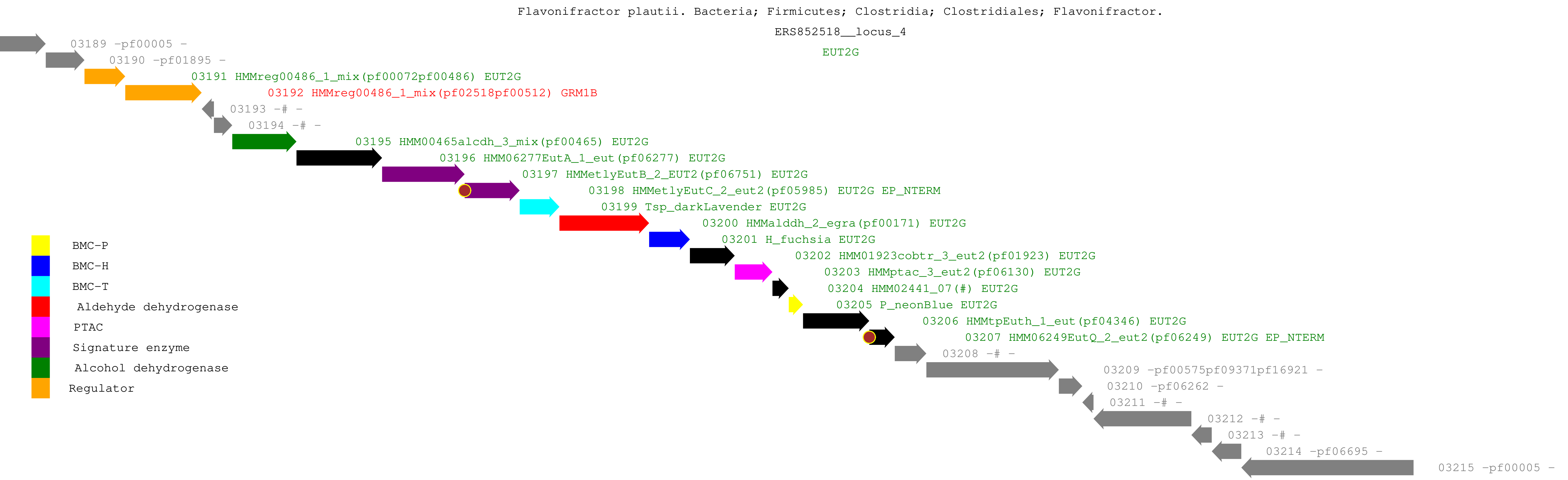
Flavonifractor plautii. Bacteria; Firmicutes; Clostridia; Clostridiales; Flavonifractor.
ERS852518__locus_4
EUT2G
03189 -pf00005 -
03190 -pf01895 -
03191 HMMreg00486_1_mix(pf00072pf00486) EUT2G
03192 HMMreg00486_1_mix(pf02518pf00512) GRM1B
03193 -# -
03194 -# -
03195 HMM00465alcdh_3_mix(pf00465) EUT2G
03196 HMM06277EutA_1_eut(pf06277) EUT2G
03197 HMMetlyEutB_2_EUT2(pf06751) EUT2G
03198 HMMetlyEutC_2_eut2(pf05985) EUT2G EP_NTERM
03199 Tsp_darkLavender EUT2G
03200 HMMalddh_2_egra(pf00171) EUT2G
03201 H_fuchsia EUT2G
03202 HMM01923cobtr_3_eut2(pf01923) EUT2G
03203 HMMptac_3_eut2(pf06130) EUT2G
03204 HMM02441_07(#) EUT2G
03205 P_neonBlue EUT2G
03206 HMMtpEuth_1_eut(pf04346) EUT2G
03207 HMM06249EutQ_2_eut2(pf06249) EUT2G EP_NTERM
03208 -# -
03209 -pf00575pf09371pf16921 -
03210 -pf06262 -
03211 -# -
03212 -# -
03213 -# -
03214 -pf06695 -
03215 -pf00005 -
BMC-P
BMC-H
BMC-T
Aldehyde dehydrogenase
PTAC
Signature enzyme
Alcohol dehydrogenase
Regulator

Flavonifractor plautii DSM 6740. Bacteria; Firmicutes; Clostridia; Clostridiales; Ruminococcaceae;

EV206__locus_2

EUT2G

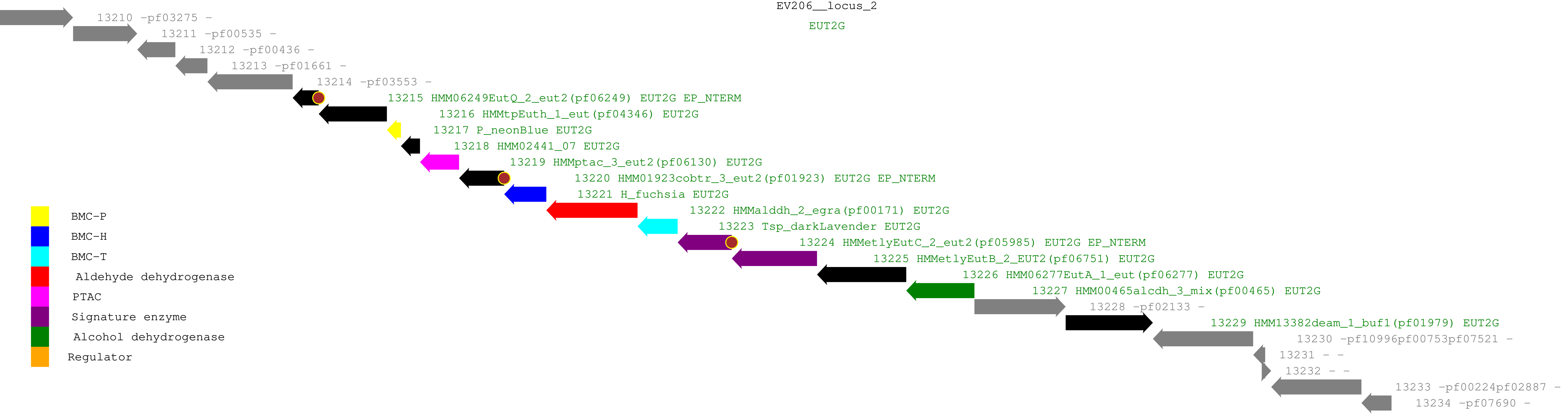

Flavonifractor plautii ATCC 29863. Bacteria; Firmicutes; Clostridia; Clostridiales; Flavonifractor.

HMPREF0372__locus_2

EUT2G

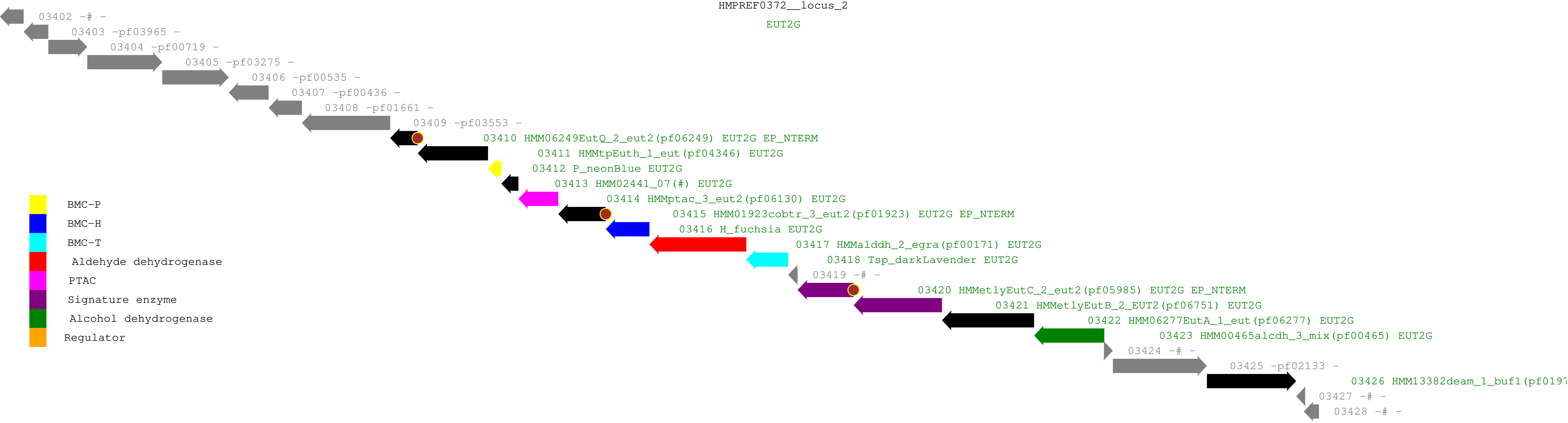

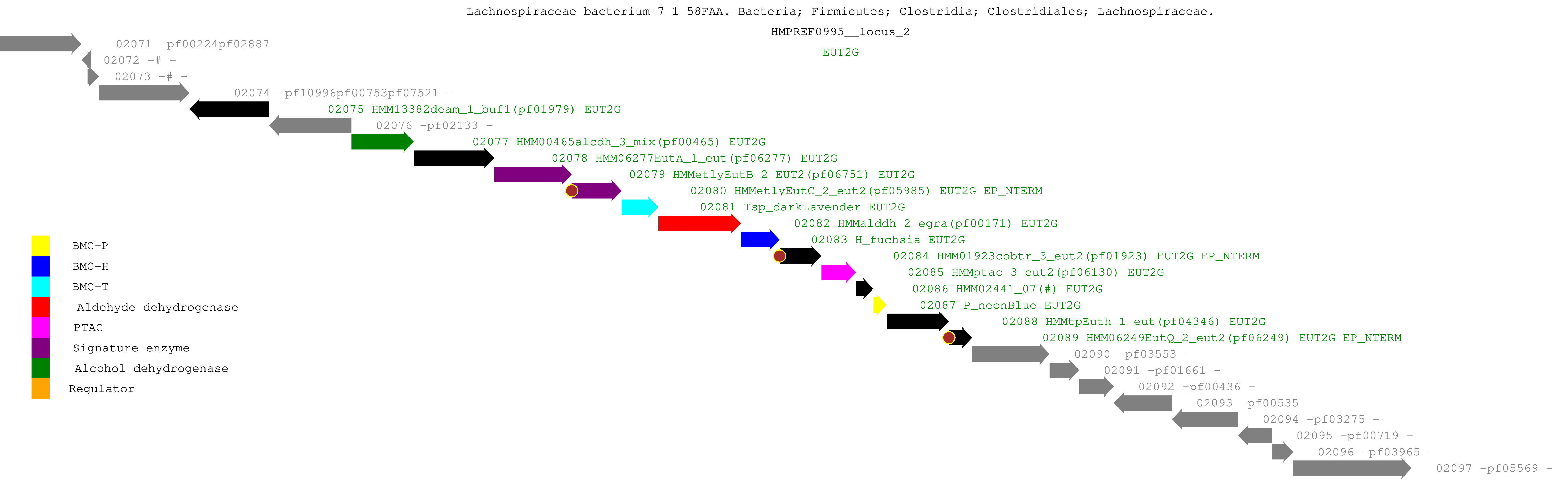

Lachnospiraceae bacterium 7_1_58FAA. Bacteria; Firmicutes; Clostridia; Clostridiales; Lachnospiraceae.
HMPREF0995__locus_2
EUT2G
02071 -pf00224pf02887 -
02072 -# -
02073 -# -
02074 -pf10996pf00753pf07521 -
02075 HMM13382deam_1_buf1(pf01979) EUT2G
02076 -pf02133 -
02077 HMM00465alcdh_3_mix(pf00465) EUT2G
02078 HMM06277EutA_1_eut(pf06277) EUT2G
02079 HMMetlyEutB_2_EUT2(pf06751) EUT2G
02080 HMMetlyEutC_2_eut2(pf05985) EUT2G EP_NTERM
02081 Tsp_darkLavender EUT2G
02082 HMMalddh_2_egra(pf00171) EUT2G
02083 H_fuchsia EUT2G
02084 HMM01923cobtr_3_eut2(pf01923) EUT2G EP_NTERM
02085 HMMptac_3_eut2(pf06130) EUT2G
02086 HMM02441_07(#) EUT2G
02087 P_neonBlue EUT2G
02088 HMMtpEuth_1_eut(pf04346) EUT2G
02089 HMM06249EutQ_2_eut2(pf06249) EUT2G EP_NTERM
02090 -pf03553 -
02091 -pf01661 -
02092 -pf00436 -
02093 -pf00535 -
02094 -pf03275 -
02095 -pf00719 -
02096 -pf03965 -
02097 -pf05569 -
BMC-P
BMC-H
BMC-T
Aldehyde dehydrogenase
PTAC
Signature enzyme
Alcohol dehydrogenase
Regulator

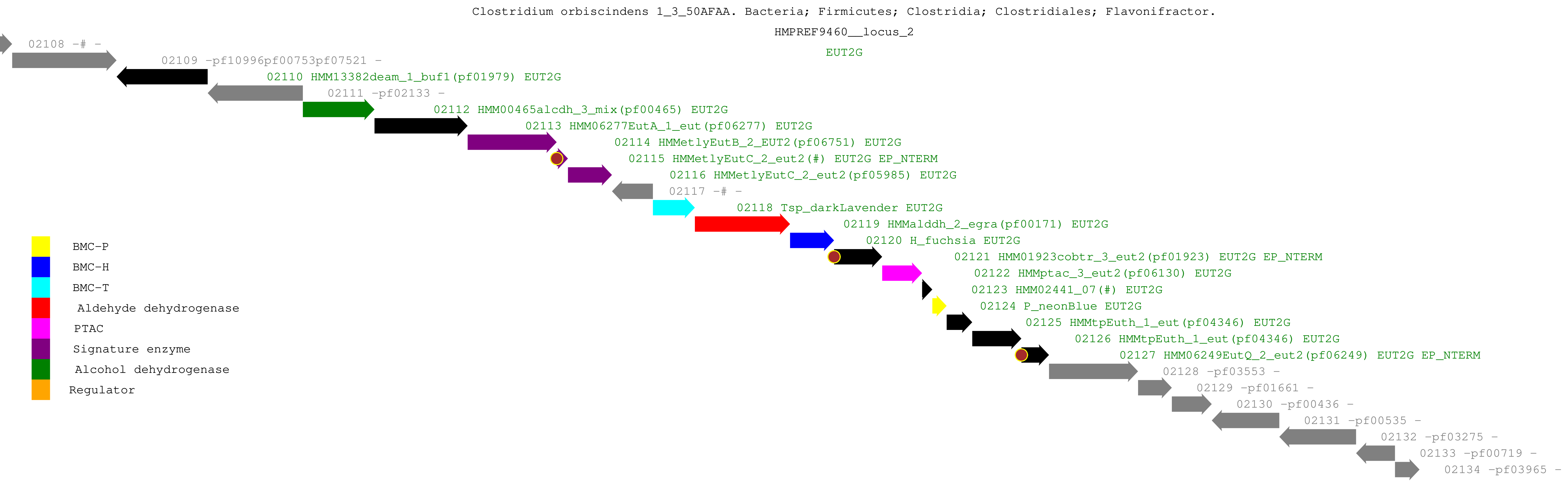

Clostridium orbiscindens 1_3_50AFAA. Bacteria; Firmicutes; Clostridia; Clostridiales; Flavonifractor.
HMPREF9460__locus_2
EUT2G
02108 -# -
02109 -pf10996pf00753pf07521 -
02110 HMM13382deam_1_buf1(pf01979) EUT2G
02111 -pf02133 -
02112 HMM00465alcdh_3_mix(pf00465) EUT2G
02113 HMM06277EutA_1_eut(pf06277) EUT2G
02114 HMMetlyEutB_2_EUT2(pf06751) EUT2G
02115 HMMetlyEutC_2_eut2(#) EUT2G EP_NTERM
02116 HMMetlyEutC_2_eut2(pf05985) EUT2G
02117 -# -
02118 Tsp_darkLavender EUT2G
02119 HMMaldh_2_egra(pf00171) EUT2G
02120 H_fuchsia EUT2G
02121 HMM01923cobtr_3_eut2(pf01923) EUT2G EP_NTERM
02122 HMMptac_3_eut2(pf06130) EUT2G
02123 HMM02441_07(#) EUT2G
02124 P_neonBlue EUT2G
02125 HMMtpEuth_1_eut(pf04346) EUT2G
02126 HMMtpEuth_1_eut(pf04346) EUT2G
02127 HMM06249EutQ_2_eut2(pf06249) EUT2G EP_NTERM
02128 -pf03553 -
02129 -pf01661 -
02130 -pf00436 -
02131 -pf00535 -
02132 -pf03275 -
02133 -pf00719 -
02134 -pf03965 -
BMC-P
BMC-H
BMC-T
Aldehyde dehydrogenase
PTAC
Signature enzyme
Alcohol dehydrogenase
Regulator

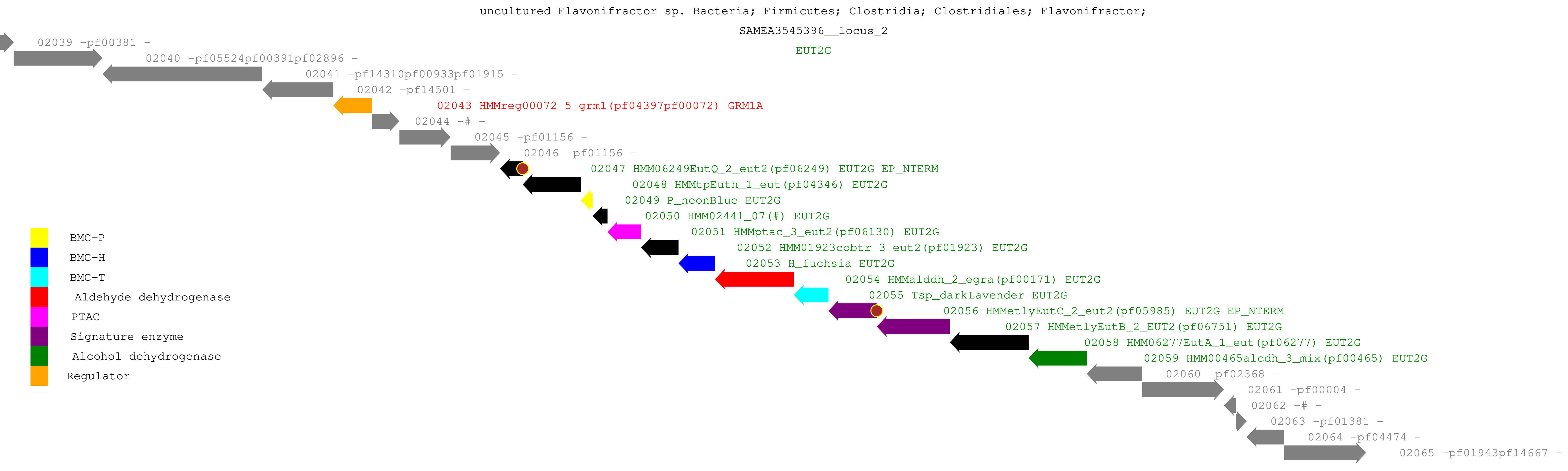
uncultured Flavonifractor sp. Bacteria; Firmicutes; Clostridia; Clostridiales; Flavonifractor;
SAMEA3545396__locus_2
EUT2G
02039 -pf00381 -
02040 -pf05524pf00391pf02896 -
02041 -pf14310pf00933pf01915 -
02042 -pf14501 -
02043 HMMreg00072_5_grm1(pf04397pf00072) GRM1A
02044 -# -
02045 -pf01156 -
02046 -pf01156 -
02047 HMM06249EutQ_2_eut2(pf06249) EUT2G EP_NTERM
02048 HMMtpEuth_1_eut(pf04346) EUT2G
02049 P_neonBlue EUT2G
02050 HMM02441_07(#) EUT2G
02051 HMMptac_3_eut2(pf06130) EUT2G
02052 HMM01923cobtr_3_eut2(pf01923) EUT2G
02053 H_fuchsia EUT2G
02054 HMMalddh_2_egra(pf00171) EUT2G
02055 Tsp_darkLavender EUT2G
02056 HMMetlyEutC_2_eut2(pf05985) EUT2G EP_NTERM
02057 HMMetlyEutB_2_EUT2(pf06751) EUT2G
02058 HMM06277EutA_1_eut(pf06277) EUT2G
02059 HMM00465alcdh_3_mix(pf00465) EUT2G
02060 -pf02368 -
02061 -pf00004 -
02062 -# -
02063 -pf01381 -
02064 -pf04474 -
02065 -pf01943pf14667 -
BMC-P
BMC-H
BMC-T
Aldehyde dehydrogenase
PTAC
Signature enzyme
Alcohol dehydrogenase
Regulator

uncultured Clostridium sp. Bacteria; Firmicutes; Clostridia; Clostridiales; Clostridiaceae;

SAMEA3545399__locus_1

EUT2G

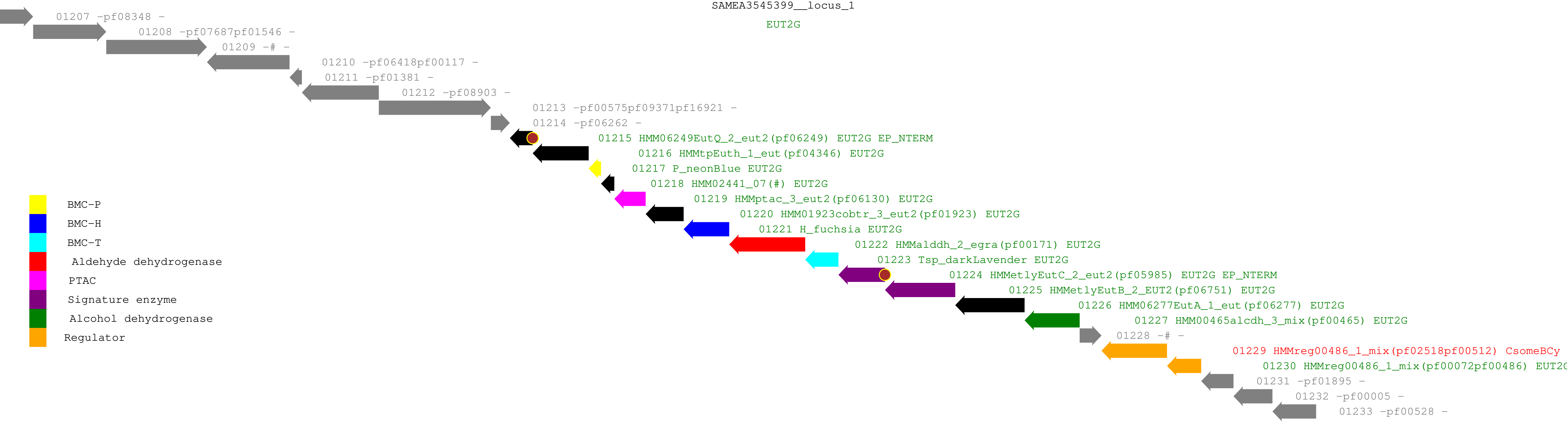

uncultured Flavonifractor sp. Bacteria; Firmicutes; Clostridia; Clostridiales; Flavonifractor;

SAMEA3545403__locus_1_main_sat

EUT2G

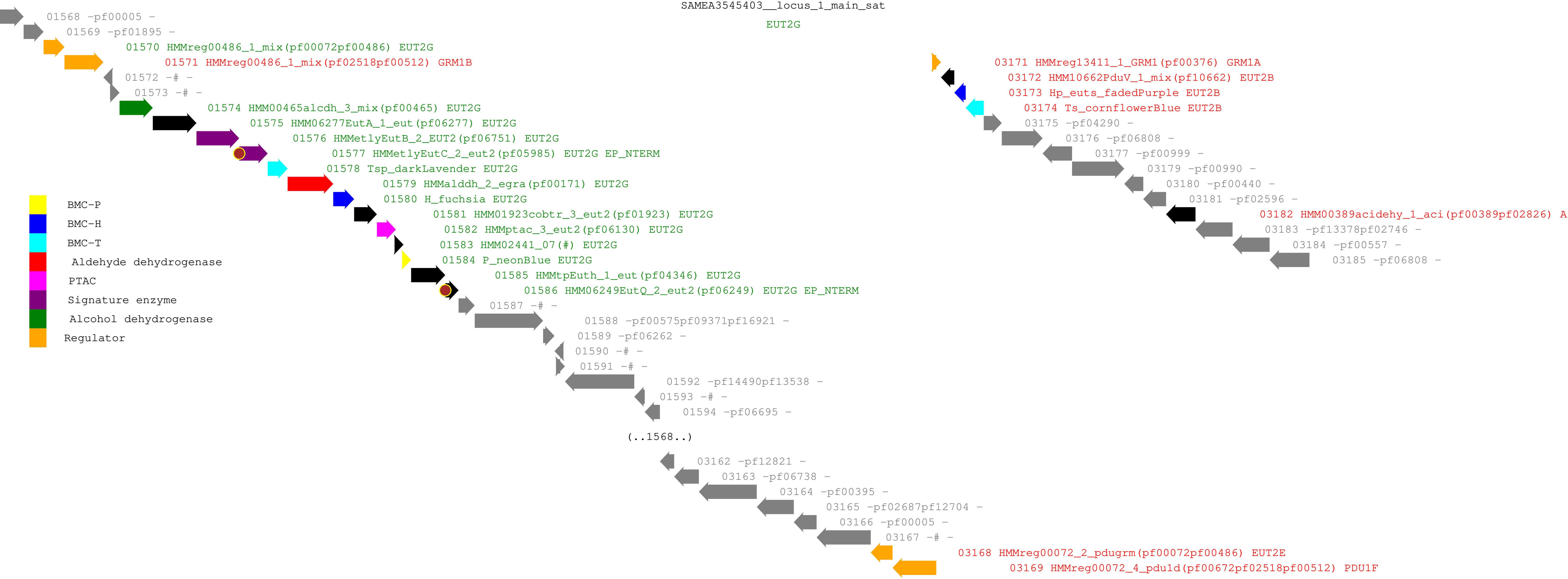

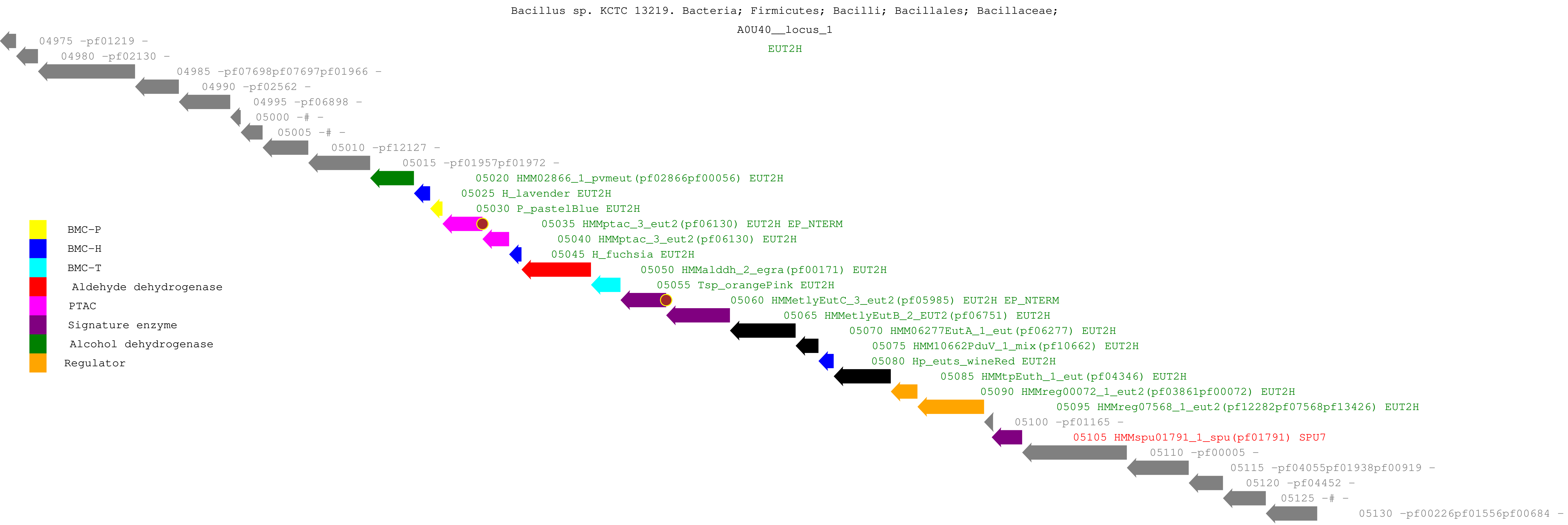
Bacillus sp. KCTC 13219. Bacteria; Firmicutes; Bacilli; Bacillales; Bacillaceae;
A0U40__locus_1
EUT2H
04975 -pf01219 -
04980 -pf02130 -
04985 -pf07698pf07697pf01966 -
04990 -pf02562 -
04995 -pf06898 -
05000 -# -
05005 -# -
05010 -pf12127 -
05015 -pf01957pf01972 -
05020 HMM02866_1_pvmeut(pf02866pf00056) EUT2H
05025 H_lavender EUT2H
05030 P_pastelBlue EUT2H
05035 HMMptac_3_eut2(pf06130) EUT2H EP_NTERM
05040 HMMptac_3_eut2(pf06130) EUT2H
05045 H_fuchsia EUT2H
05050 HMMaldh_2_egra(pf00171) EUT2H
05055 Tsp_orangePink EUT2H
05060 HMMetlyEutC_3_eut2(pf05985) EUT2H EP_NTERM
05065 HMMetlyEutB_2_EUT2(pf06751) EUT2H
05070 HMM06277EutA_1_eut(pf06277) EUT2H
05075 HMM10662PduV_1_mix(pf10662) EUT2H
05080 Hp_euts_wineRed EUT2H
05085 HMMtpEuth_1_eut(pf04346) EUT2H
05090 HMMreg00072_1_eut2(pf03861pf00072) EUT2H
05095 HMMreg07568_1_eut2(pf12282pf07568pf13426) EUT2H
05100 -pf01165 -
05105 HMMspu01791_1_spu(pf01791) SPU7
05110 -pf00005 -
05115 -pf04055pf01938pf00919 -
05120 -pf04452 -
05125 -# -
05130 -pf00226pf01556pf00684 -
BMC-P
BMC-H
BMC-T
Aldehyde dehydrogenase
PTAC
Signature enzyme
Alcohol dehydrogenase
Regulator

Lysinibacillus sp. YR326. Bacteria; Firmicutes; Bacilli; Bacillales; Bacillaceae; Lysinibacillus;

A9A36__locus_1

EUT2H

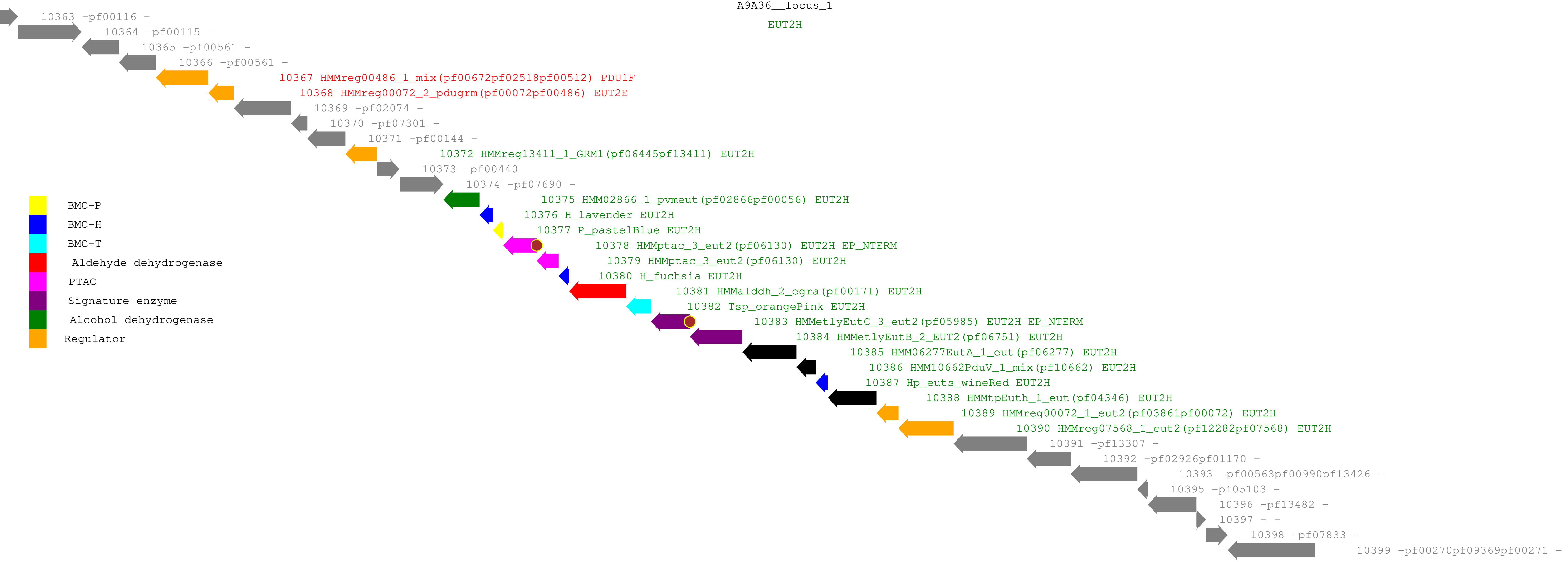

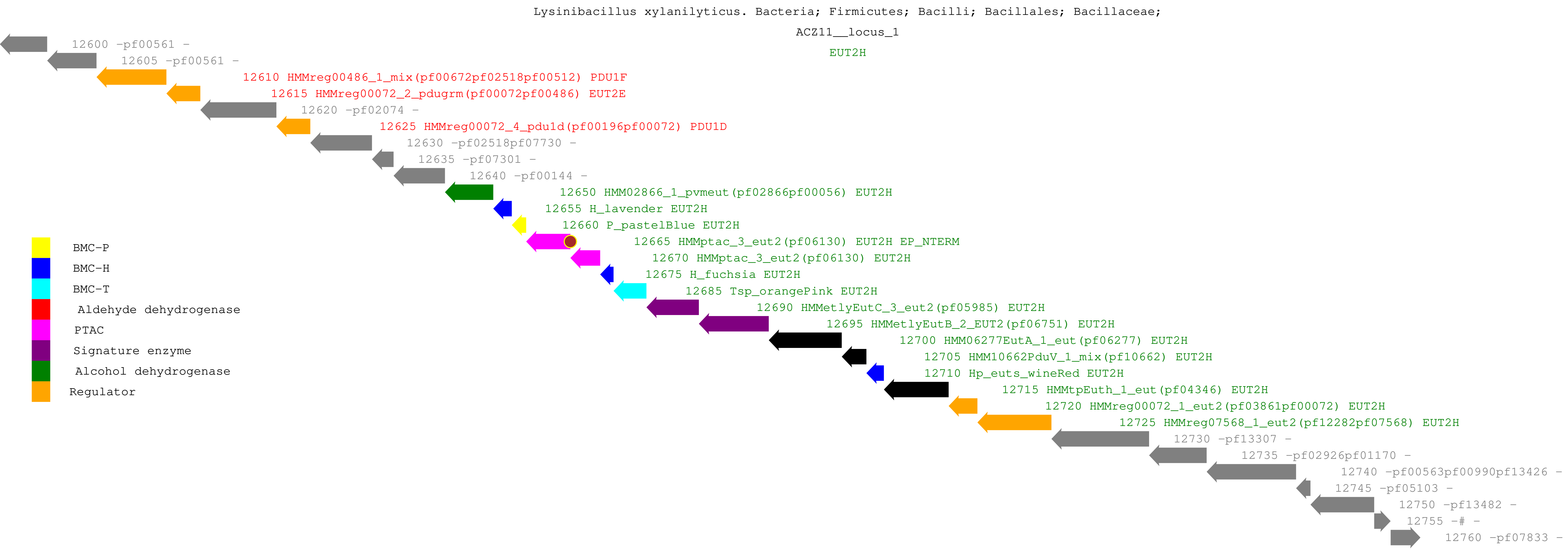
Lysinibacillus xylanilyticus. Bacteria; Firmicutes; Bacilli; Bacillales; Bacillaceae;
ACZ11__locus_1
EUT2H
12600 -pf00561 -
12605 -pf00561 -
12610 HMMreg00486_1_mix(pf00672pf02518pf00512) PDU1F
12615 HMMreg00072_2_pdugrm(pf00072pf00486) EUT2E
12620 -pf02074 -
12625 HMMreg00072_4_pdu1d(pf00196pf00072) PDU1D
12630 -pf02518pf07730 -
12635 -pf07301 -
12640 -pf00144 -
12650 HMM02866_1_pvmeut(pf02866pf00056) EUT2H
12655 H_lavender EUT2H
12660 P_pastelBlue EUT2H
12665 HMMptac_3_eut2(pf06130) EUT2H EP_NTERM
12670 HMMptac_3_eut2(pf06130) EUT2H
12675 H_fuchsia EUT2H
12685 Tsp_orangePink EUT2H
12690 HMMetlyEutC_3_eut2(pf05985) EUT2H
12695 HMMetlyEutB_2_EUT2(pf06751) EUT2H
12700 HMM06277EutA_1_eut(pf06277) EUT2H
12705 HMM10662PduV_1_mix(pf10662) EUT2H
12710 Hp_euts_wineRed EUT2H
12715 HMMtpEuth_1_eut(pf04346) EUT2H
12720 HMMreg00072_1_eut2(pf03861pf00072) EUT2H
12725 HMMreg07568_1_eut2(pf12282pf07568) EUT2H
12730 -pf13307 -
12735 -pf02926pf01170 -
12740 -pf00563pf00990pf13426 -
12745 -pf05103 -
12750 -pf13482 -
12755 -# -
12760 -pf07833 -
BMC-P
BMC-H
BMC-T
Aldehyde dehydrogenase
PTAC
Signature enzyme
Alcohol dehydrogenase
Regulator

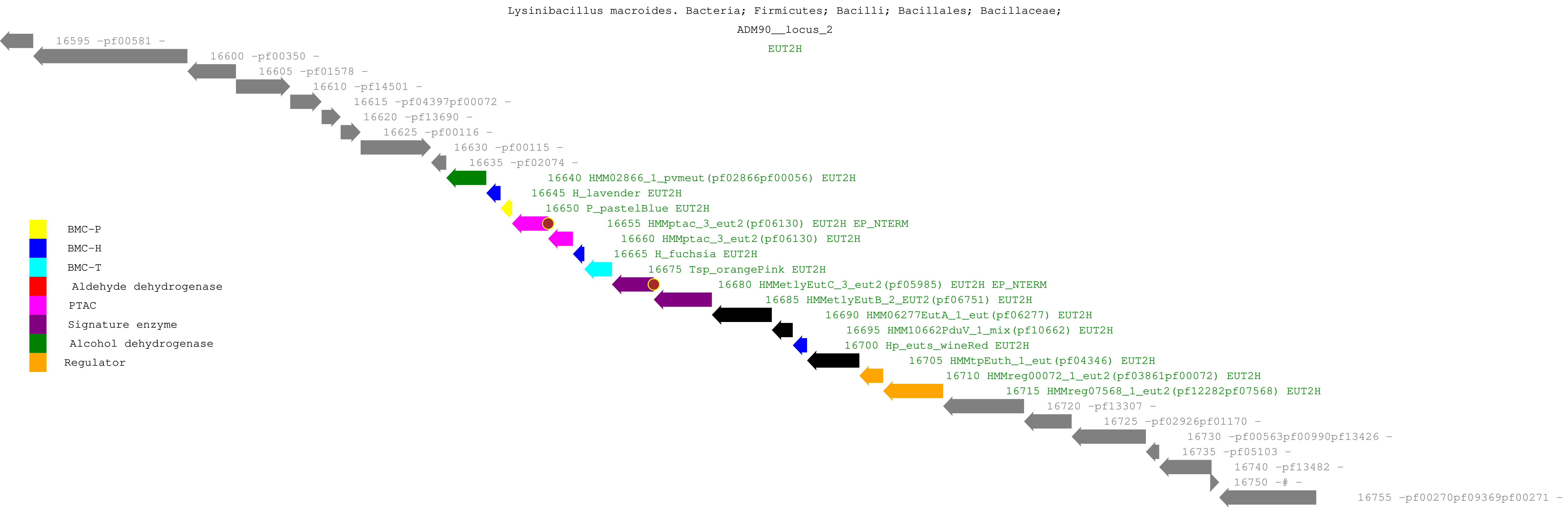

Lysinibacillus macroides. Bacteria; Firmicutes; Bacilli; Bacillales; Bacillaceae;
ADM90__locus_2
EUT2H
16595 -pf00581 -
16600 -pf00350 -
16605 -pf01578 -
16610 -pf14501 -
16615 -pf04397pf00072 -
16620 -pf13690 -
16625 -pf00116 -
16630 -pf00115 -
16635 -pf02074 -
16640 HMM02866_1_pvmeut(pf02866pf00056) EUT2H
16645 H_lavender EUT2H
16650 P_pastelBlue EUT2H
16655 HMMptac_3_eut2(pf06130) EUT2H EP_NTERM
16660 HMMptac_3_eut2(pf06130) EUT2H
16665 H_fuchsia EUT2H
16675 Tsp_orangePink EUT2H
16680 HMMetlyEutC_3_eut2(pf05985) EUT2H EP_NTERM
16685 HMMetlyEutB_2_EUT2(pf06751) EUT2H
16690 HMM06277EutA_1_eut(pf06277) EUT2H
16695 HMM10662PduV_1_mix(pf10662) EUT2H
16700 Hp_euts_wineRed EUT2H
16705 HMMtpEuth_1_eut(pf04346) EUT2H
16710 HMMreg00072_1_eut2(pf03861pf00072) EUT2H
16715 HMMreg07568_1_eut2(pf12282pf07568) EUT2H
16720 -pf13307 -
16725 -pf02926pf01170 -
16730 -pf00563pf00990pf13426 -
16735 -pf05103 -
16740 -pf13482 -
16750 -# -
16755 -pf00270pf09369pf00271 -
BMC-P
BMC-H
BMC-T
Aldehyde dehydrogenase
PTAC
Signature enzyme
Alcohol dehydrogenase
Regulator

Lysinibacillus sp. F5. Bacteria; Firmicutes; Bacilli; Bacillales; Bacillaceae;

AK833__locus_1

EUT2H

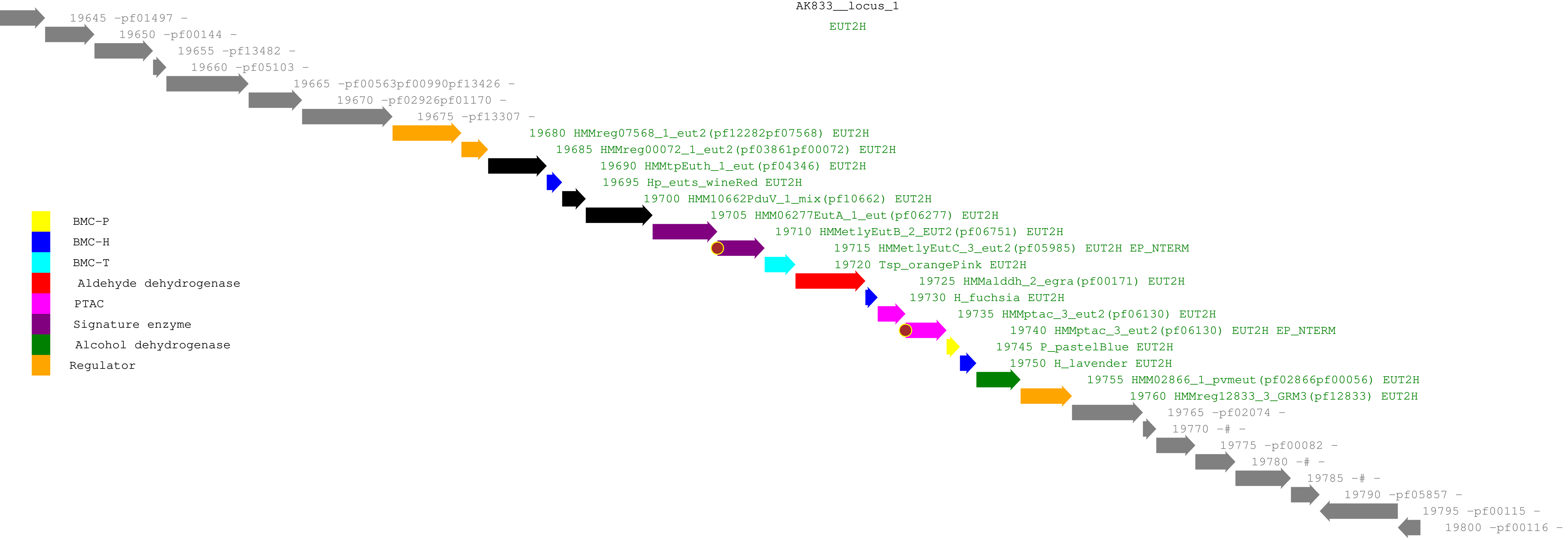

Lysinibacillus sp. FJAT-14745. Bacteria; Firmicutes; Bacilli; Bacillales; Bacillaceae;

AMS59__locus_1

EUT2H

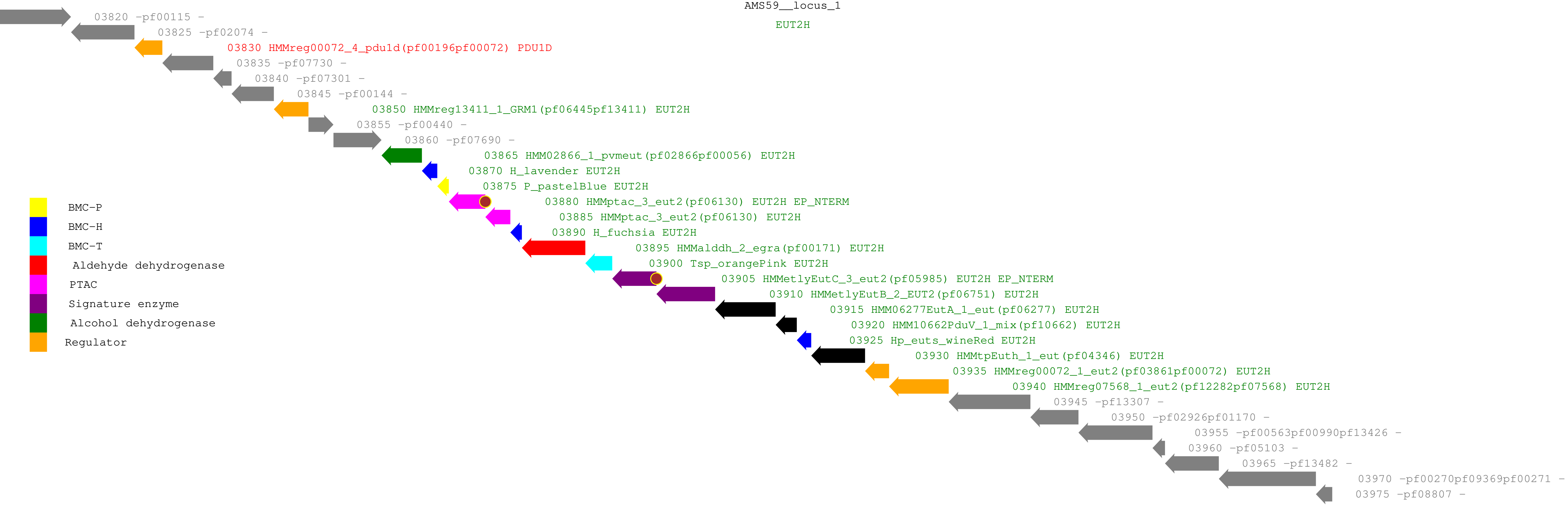

Lysinibacillus sp. FJAT-14222. Bacteria; Firmicutes; Bacilli; Bacillales; Bacillaceae;

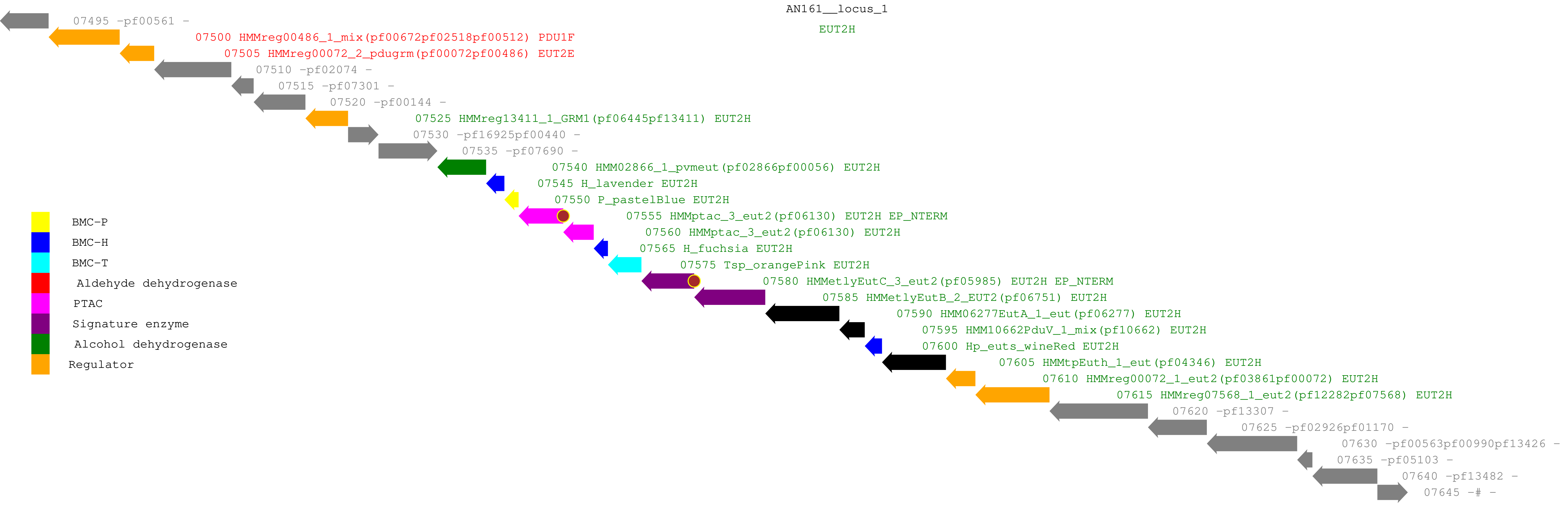

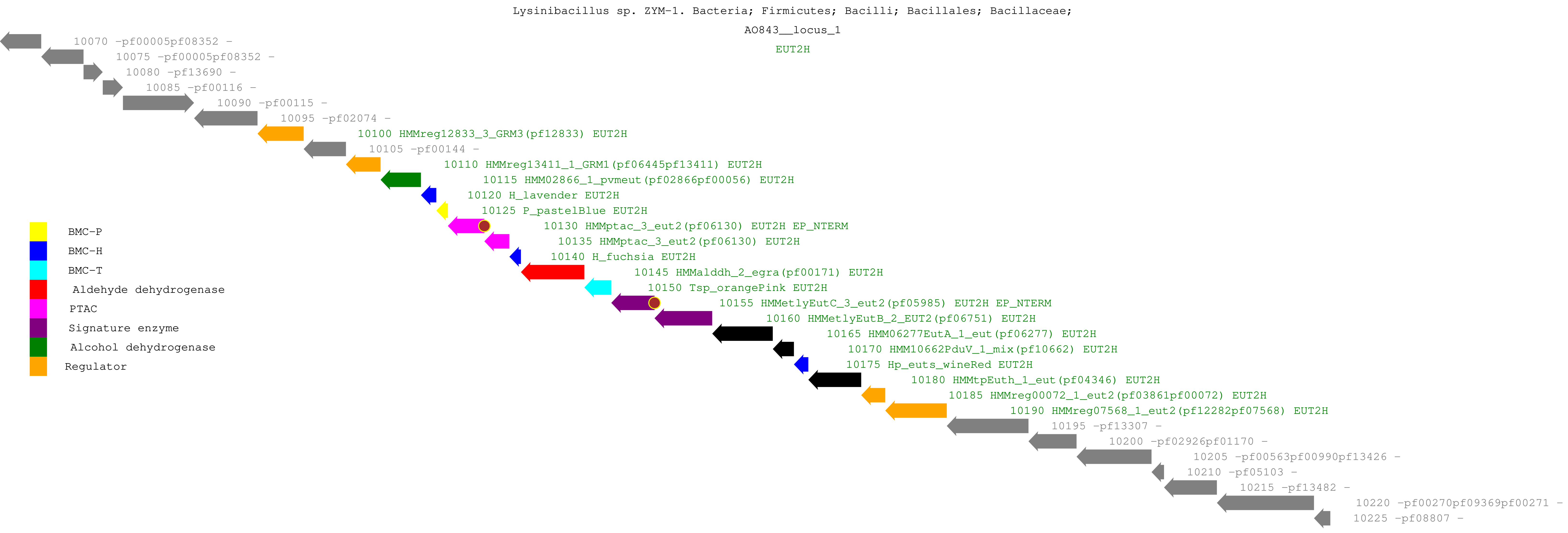

Lysinibacillus sp. ZYM-1. Bacteria; Firmicutes; Bacilli; Bacillales; Bacillaceae;
AO843__locus_1
EUT2H
10070 -pf00005pf08352 -
10075 -pf00005pf08352 -
10080 -pf13690 -
10085 -pf00116 -
10090 -pf00115 -
10095 -pf02074 -
10100 HMMreg12833_3_GRM3(pf12833) EUT2H
10105 -pf00144 -
10110 HMMreg13411_1_GRM1(pf06445pf13411) EUT2H
10115 HMM02866_1_pvmeut(pf02866pf00056) EUT2H
10120 H_lavender EUT2H
10125 P_pastelBlue EUT2H
10130 HMMptac_3_eut2(pf06130) EUT2H EP_NTERM
10135 HMMptac_3_eut2(pf06130) EUT2H
10140 H_fuchsia EUT2H
10145 HMMalddh_2_egra(pf00171) EUT2H
10150 Tsp_orangePink EUT2H
10155 HMMetlyEutC_3_eut2(pf05985) EUT2H EP_NTERM
10160 HMMetlyEutB_2_EUT2(pf06751) EUT2H
10165 HMM06277EutA_1_eut(pf06277) EUT2H
10170 HMM10662PduV_1_mix(pf10662) EUT2H
10175 Hp_euts_wineRed EUT2H
10180 HMMtpEuth_1_eut(pf04346) EUT2H
10185 HMMreg00072_1_eut2(pf03861pf00072) EUT2H
10190 HMMreg07568_1_eut2(pf12282pf07568) EUT2H
10195 -pf13307 -
10200 -pf02926pf01170 -
10205 -pf00563pf00990pf13426 -
10210 -pf05103 -
10215 -pf13482 -
10220 -pf00270pf09369pf00271 -
10225 -pf08807 -
BMC-P
BMC-H
BMC-T
Aldehyde dehydrogenase
PTAC
Signature enzyme
Alcohol dehydrogenase
Regulator

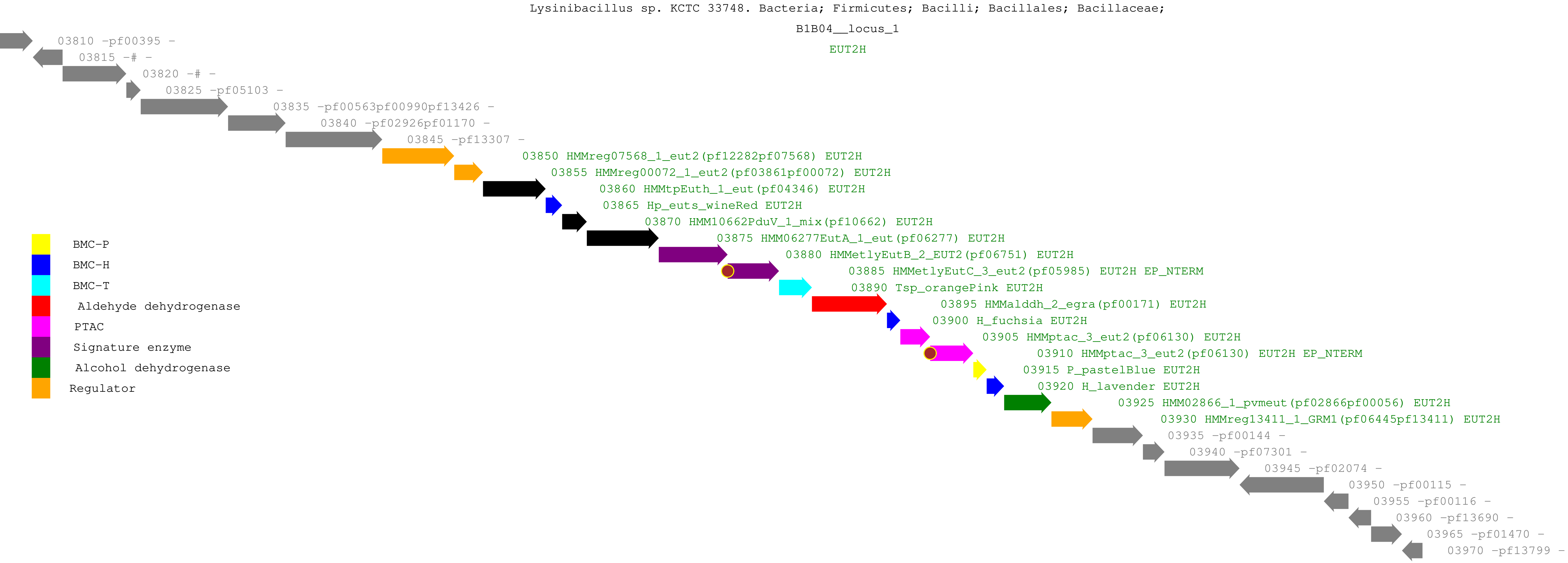
Lysinibacillus sp. KCTC 33748. Bacteria; Firmicutes; Bacilli; Bacillales; Bacillaceae;
B1B04__locus_1
EUT2H
03810 -pf00395 -
03815 -# -
03820 -# -
03825 -pf05103 -
03835 -pf00563pf00990pf13426 -
03840 -pf02926pf01170 -
03845 -pf13307 -
03850 HMMreg07568_1_eut2(pf12282pf07568) EUT2H
03855 HMMreg00072_1_eut2(pf03861pf00072) EUT2H
03860 HMMtpEuth_1_eut(pf04346) EUT2H
03865 Hp_euts_wineRed EUT2H
03870 HMM10662PduV_1_mix(pf10662) EUT2H
03875 HMM06277EutA_1_eut(pf06277) EUT2H
03880 HMMetlyEutB_2_EUT2(pf06751) EUT2H
03885 HMMetlyEutC_3_eut2(pf05985) EUT2H EP_NTERM
03890 Tsp_orangePink EUT2H
03895 HMMalddh_2_egra(pf00171) EUT2H
03900 H_fuchsia EUT2H
03905 HMMptac_3_eut2(pf06130) EUT2H
03910 HMMptac_3_eut2(pf06130) EUT2H EP_NTERM
03915 P_pastelBlue EUT2H
03920 H_lavender EUT2H
03925 HMM02866_1_pvmeut(pf02866pf00056) EUT2H
03930 HMMreg13411_1_GRM1(pf06445pf13411) EUT2H
03935 -pf00144 -
03940 -pf07301 -
03945 -pf02074 -
03950 -pf00115 -
03955 -pf00116 -
03960 -pf13690 -
03965 -pf01470 -
03970 -pf13799 -
BMC-P
BMC-H
BMC-T
Aldehyde dehydrogenase
PTAC
Signature enzyme
Alcohol dehydrogenase
Regulator

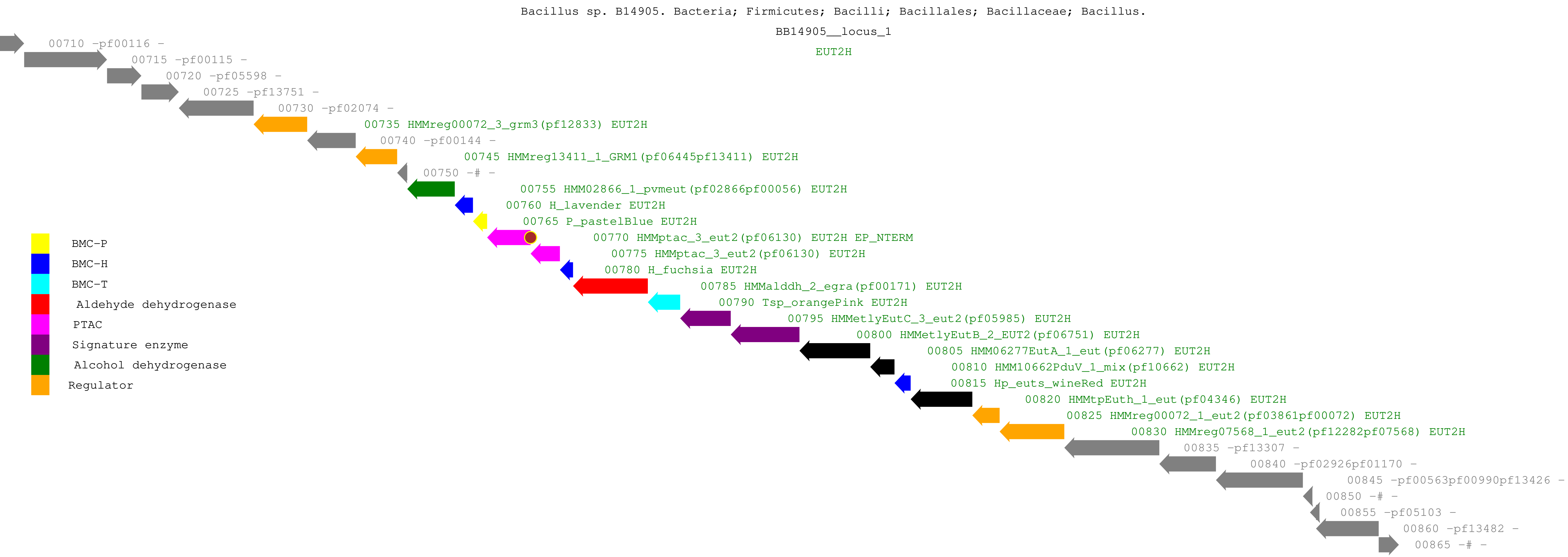

Bacillus sp. B14905. Bacteria; Firmicutes; Bacilli; Bacillales; Bacillaceae; Bacillus.
BB14905__locus_1
EUT2H
00710 -pf00116 -
00715 -pf00115 -
00720 -pf05598 -
00725 -pf13751 -
00730 -pf02074 -
00735 HMMreg00072_3_grm3(pf12833) EUT2H
00740 -pf00144 -
00745 HMMreg13411_1_GRM1(pf06445pf13411) EUT2H
00750 -# -
00755 HMM02866_1_pvmeut(pf02866pf00056) EUT2H
00760 H_lavender EUT2H
00765 P_pastelBlue EUT2H
00770 HMMptac_3_eut2(pf06130) EUT2H EP_NTERM
00775 HMMptac_3_eut2(pf06130) EUT2H
00780 H_fuchsia EUT2H
00785 HMMalddh_2_egra(pf00171) EUT2H
00790 Tsp_orangePink EUT2H
00795 HMMetlyEutC_3_eut2(pf05985) EUT2H
00800 HMMetlyEutB_2_EUT2(pf06751) EUT2H
00805 HMM06277EutA_1_eut(pf06277) EUT2H
00810 HMM10662PduV_1_mix(pf10662) EUT2H
00815 Hp_euts_wineRed EUT2H
00820 HMMtpEuth_1_eut(pf04346) EUT2H
00825 HMMreg00072_1_eut2(pf03861pf00072) EUT2H
00830 HMMreg07568_1_eut2(pf12282pf07568) EUT2H
00835 -pf13307 -
00840 -pf02926pf01170 -
00845 -pf00563pf00990pf13426 -
00850 -# -
00855 -pf05103 -
00860 -pf13482 -
00865 -# -
BMC-P
BMC-H
BMC-T
Aldehyde dehydrogenase
PTAC
Signature enzyme
Alcohol dehydrogenase
Regulator

Lysinibacillus fusiformis ZC1. Bacteria; Firmicutes; Bacilli; Bacillales; Bacillaceae;

BFZC1__locus_1

EUT2H

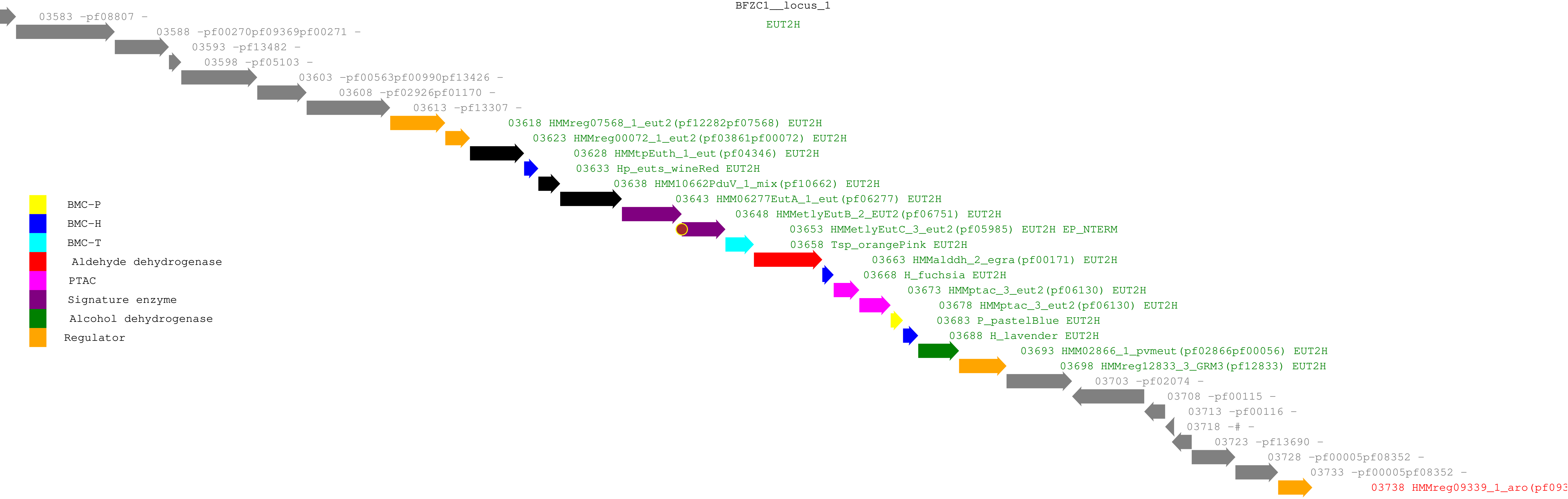

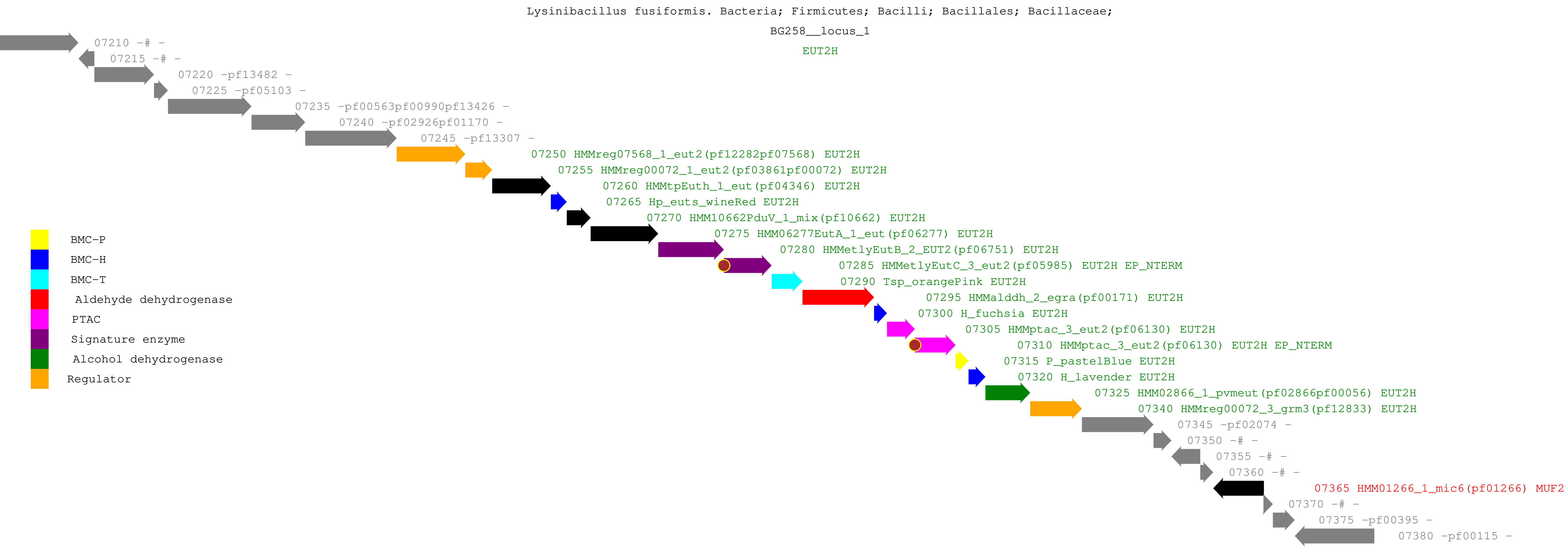
Lysinibacillus fusiformis. Bacteria; Firmicutes; Bacilli; Bacillales; Bacillaceae;
BG258__locus_1
EUT2H
07210 -# -
07215 -# -
07220 -pf13482 -
07225 -pf05103 -
07235 -pf00563pf00990pf13426 -
07240 -pf02926pf01170 -
07245 -pf13307 -
07250 HMMreg07568_1_eut2(pf12282pf07568) EUT2H
07255 HMMreg00072_1_eut2(pf03861pf00072) EUT2H
07260 HMMtpEuth_1_eut(pf04346) EUT2H
07265 Hp_euts_wineRed EUT2H
07270 HMM10662PduV_1_mix(pf10662) EUT2H
07275 HMM06277EutA_1_eut(pf06277) EUT2H
07280 HMMetlyEutB_2_EUT2(pf06751) EUT2H
07285 HMMetlyEutC_3_eut2(pf05985) EUT2H EP_NTERM
07290 Tsp_orangePink EUT2H
07295 HMMalddh_2_egra(pf00171) EUT2H
07300 H_fuchsia EUT2H
07305 HMMptac_3_eut2(pf06130) EUT2H
07310 HMMptac_3_eut2(pf06130) EUT2H EP_NTERM
07315 P_pastelBlue EUT2H
07320 H_lavender EUT2H
07325 HMM02866_1_pvmeut(pf02866pf00056) EUT2H
07340 HMMreg00072_3_grm3(pf12833) EUT2H
07345 -pf02074 -
07350 -# -
07355 -# -
07360 -# -
07365 HMM01266_1_mic6(pf01266) MUF2
07370 -# -
07375 -pf00395 -
07380 -pf00115 -
BMC-P
BMC-H
BMC-T
Aldehyde dehydrogenase
PTAC
Signature enzyme
Alcohol dehydrogenase
Regulator

Lysinibacillus sphaericus (strain C3-41). Bacteria; Firmicutes; Bacilli; Bacillales; Bacillaceae;

Bsph__locus_1

EUT2H

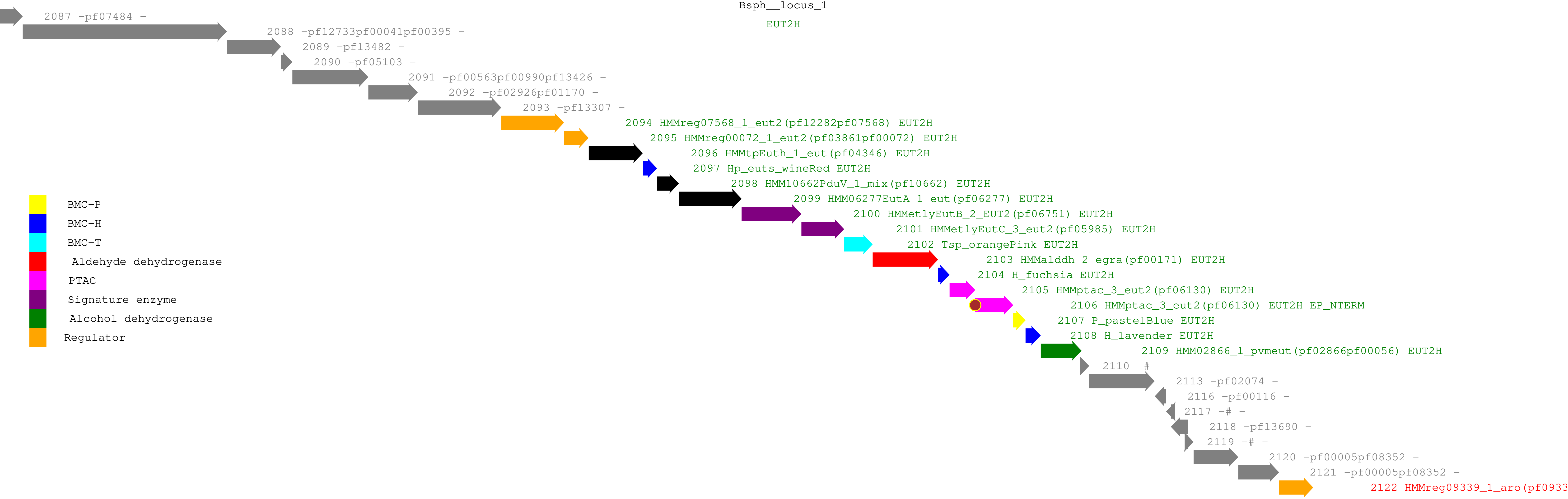

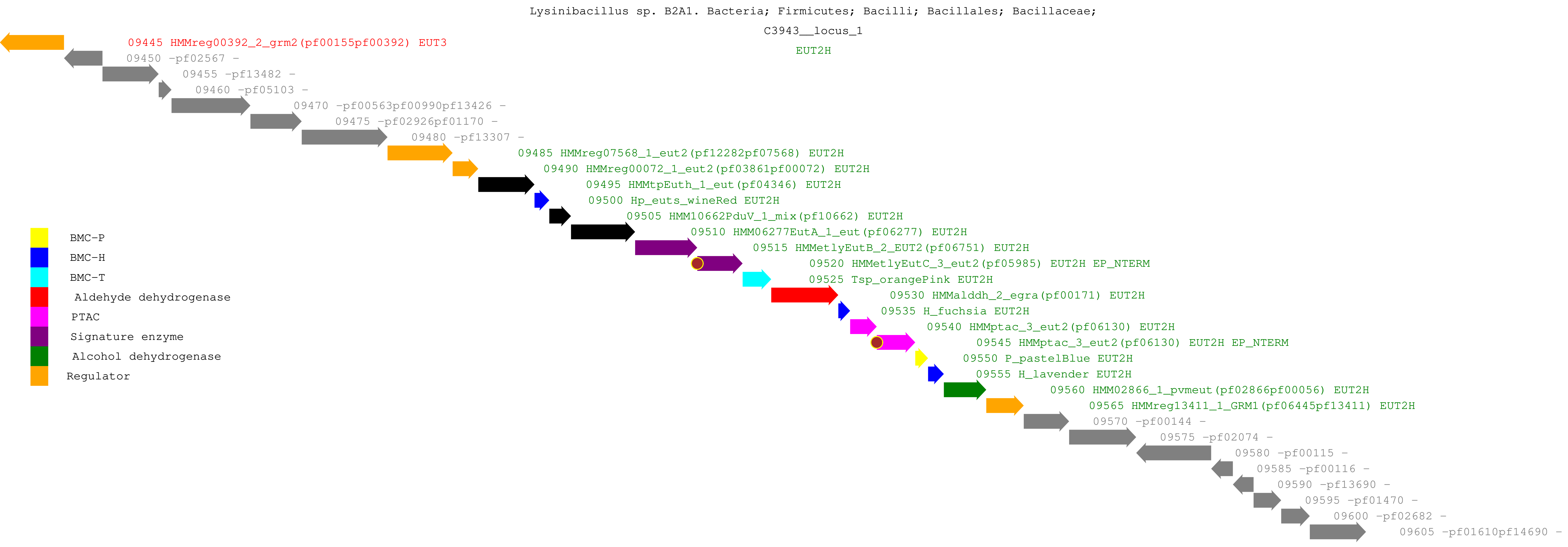
Lysinibacillus sp. B2A1. Bacteria; Firmicutes; Bacilli; Bacillales; Bacillaceae;
C3943__locus_1
EUT2H
09445 HMMreg00392_2_grm2(pf00155pf00392) EUT3
09450 -pf02567 -
09455 -pf13482 -
09460 -pf05103 -
09470 -pf00563pf00990pf13426 -
09475 -pf02926pf01170 -
09480 -pf13307 -
09485 HMMreg07568_1_eut2(pf12282pf07568) EUT2H
09490 HMMreg00072_1_eut2(pf03861pf00072) EUT2H
09495 HMMtpEuth_1_eut(pf04346) EUT2H
09500 Hp_euts_wineRed EUT2H
09505 HMM10662PduV_1_mix(pf10662) EUT2H
09510 HMM06277EutA_1_eut(pf06277) EUT2H
09515 HMMetlyEutB_2_EUT2(pf06751) EUT2H
09520 HMMetlyEutC_3_eut2(pf05985) EUT2H EP_NTERM
09525 Tsp_orangePink EUT2H
09530 HMMalddh_2_egra(pf00171) EUT2H
09535 H_fuchsia EUT2H
09540 HMMptac_3_eut2(pf06130) EUT2H
09545 HMMptac_3_eut2(pf06130) EUT2H EP_NTERM
09550 P_pastelBlue EUT2H
09555 H_lavender EUT2H
09560 HMM02866_1_pvmeut(pf02866pf00056) EUT2H
09565 HMMreg13411_1_GRM1(pf06445pf13411) EUT2H
09570 -pf00144 -
09575 -pf02074 -
09580 -pf00115 -
09585 -pf00116 -
09590 -pf13690 -
09595 -pf01470 -
09600 -pf02682 -
09605 -pf01610pf14690 -
BMC-P
BMC-H
BMC-T
Aldehyde dehydrogenase
PTAC
Signature enzyme
Alcohol dehydrogenase
Regulator

Lysinibacillus sp. SDF0037. Bacteria; Firmicutes; Bacilli; Bacillales; Bacillaceae; Lysinibacillus;

C7Y47__locus_1

EUT2H

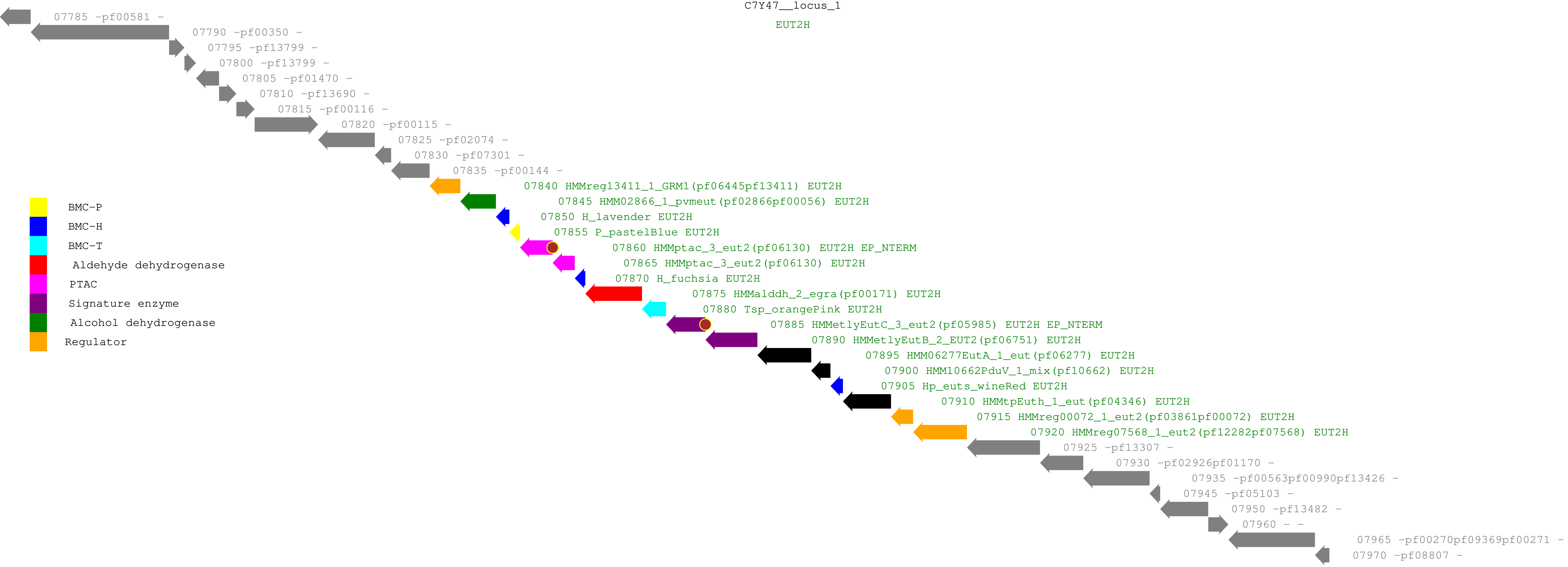

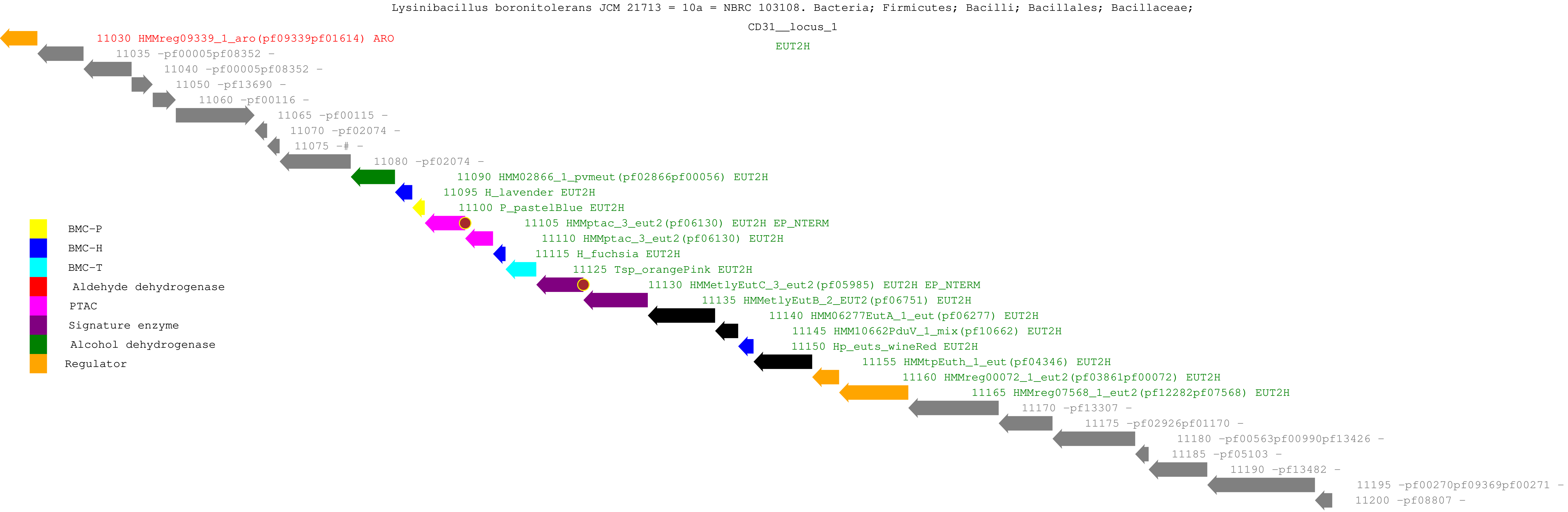
Lysinibacillus boronitolerans JCM 21713 = 10a = NBRC 103108. Bacteria; Firmicutes; Bacilli; Bacillales; Bacillaceae;
CD31__locus_1
EUT2H
11030 HMMreg09339_1_aro(pf09339pf01614) ARO
11035 -pf00005pf08352 -
11040 -pf00005pf08352 -
11050 -pf13690 -
11060 -pf00116 -
11065 -pf00115 -
11070 -pf02074 -
11075 -# -
11080 -pf02074 -
11090 HMM02866_1_pvmeut(pf02866pf00056) EUT2H
11095 H_lavender EUT2H
11100 P_pastelBlue EUT2H
11105 HMMptac_3_eut2(pf06130) EUT2H EP_NTERM
11110 HMMptac_3_eut2(pf06130) EUT2H
11115 H_fuchsia EUT2H
11125 Tsp_orangePink EUT2H
11130 HMMetlyEutC_3_eut2(pf05985) EUT2H EP_NTERM
11135 HMMetlyEutB_2_EUT2(pf06751) EUT2H
11140 HMM06277EutA_1_eut(pf06277) EUT2H
11145 HMM10662PduV_1_mix(pf10662) EUT2H
11150 Hp_euts_wineRed EUT2H
11155 HMMtpEuth_1_eut(pf04346) EUT2H
11160 HMMreg00072_1_eut2(pf03861pf00072) EUT2H
11165 HMMreg07568_1_eut2(pf12282pf07568) EUT2H
11170 -pf13307 -
11175 -pf02926pf01170 -
11180 -pf00563pf00990pf13426 -
11185 -pf05103 -
11190 -pf13482 -
11195 -pf00270pf09369pf00271 -
11200 -pf08807 -
BMC-P
BMC-H
BMC-T
Aldehyde dehydrogenase
PTAC
Signature enzyme
Alcohol dehydrogenase
Regulator

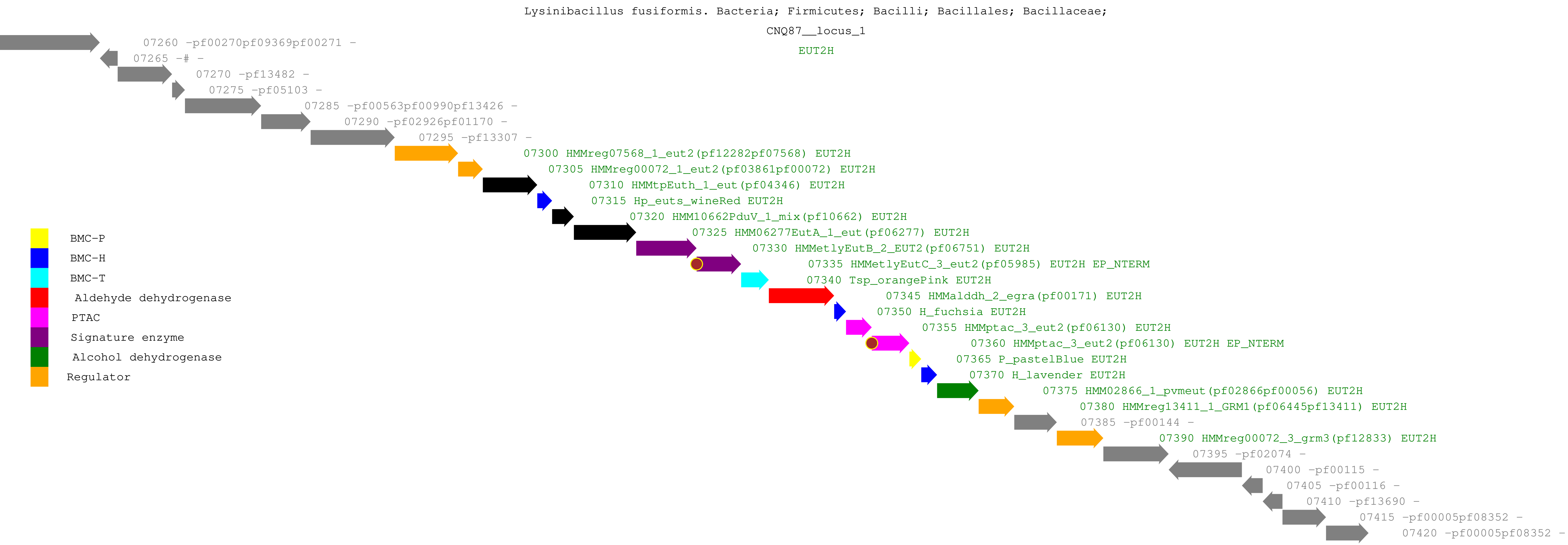
Lysinibacillus fusiformis. Bacteria; Firmicutes; Bacilli; Bacillales; Bacillaceae;
CNQ87__locus_1
EUT2H
07260 -pf00270pf09369pf00271 -
07265 -# -
07270 -pf13482 -
07275 -pf05103 -
07285 -pf00563pf00990pf13426 -
07290 -pf02926pf01170 -
07295 -pf13307 -
07300 HMMreg07568_1_eut2(pf12282pf07568) EUT2H
07305 HMMreg00072_1_eut2(pf03861pf00072) EUT2H
07310 HMMtpEuth_1_eut(pf04346) EUT2H
07315 Hp_euts_wineRed EUT2H
07320 HMM10662PduV_1_mix(pf10662) EUT2H
07325 HMM06277EutA_1_eut(pf06277) EUT2H
07330 HMMetlyEutB_2_EUT2(pf06751) EUT2H
07335 HMMetlyEutC_3_eut2(pf05985) EUT2H EP_NTERM
07340 Tsp_orangePink EUT2H
07345 HMMalddh_2_egra(pf00171) EUT2H
07350 H_fuchsia EUT2H
07355 HMMptac_3_eut2(pf06130) EUT2H
07360 HMMptac_3_eut2(pf06130) EUT2H EP_NTERM
07365 P_pastelBlue EUT2H
07370 H_lavender EUT2H
07375 HMM02866_1_pvmeut(pf02866pf00056) EUT2H
07380 HMMreg13411_1_GRM1(pf06445pf13411) EUT2H
07385 -pf00144 -
07390 HMMreg00072_3_grm3(pf12833) EUT2H
07395 -pf02074 -
07400 -pf00115 -
07405 -pf00116 -
07410 -pf13690 -
07415 -pf00005pf08352 -
07420 -pf00005pf08352 -
BMC-P
BMC-H
BMC-T
Aldehyde dehydrogenase
PTAC
Signature enzyme
Alcohol dehydrogenase
Regulator

Lysinibacillus fusiformis. Bacteria; Firmicutes; Bacilli; Bacillales; Bacillaceae;

CRI88__locus_1

EUT2H

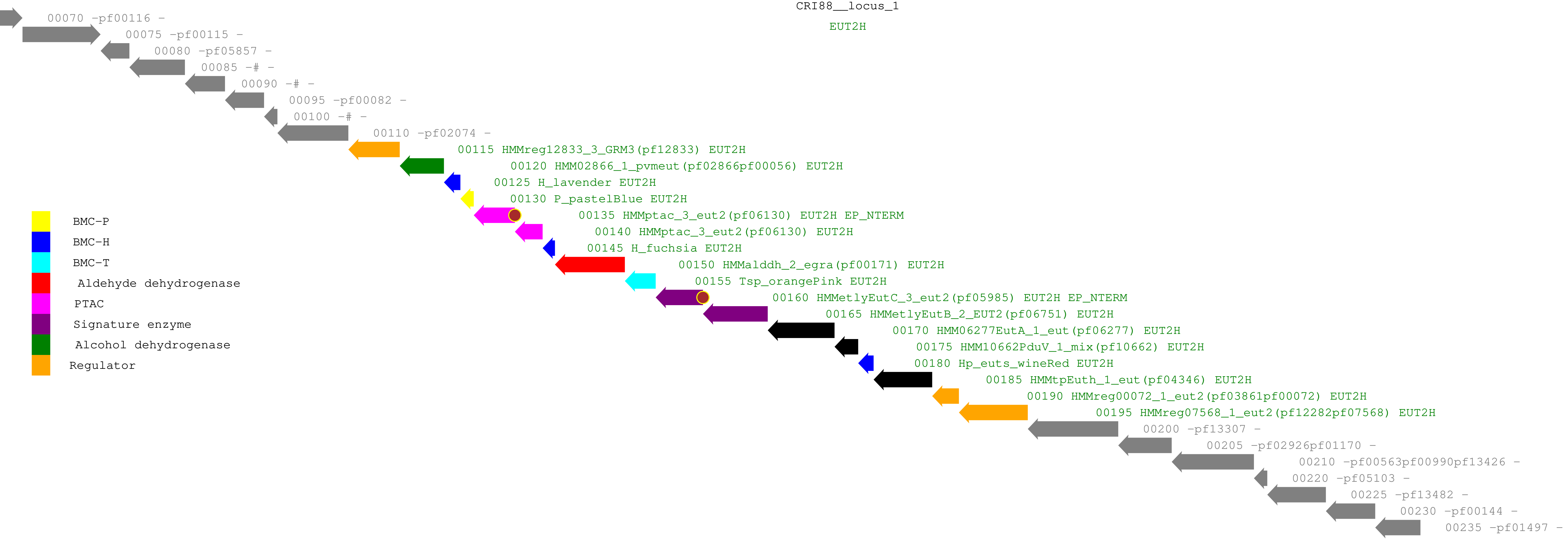

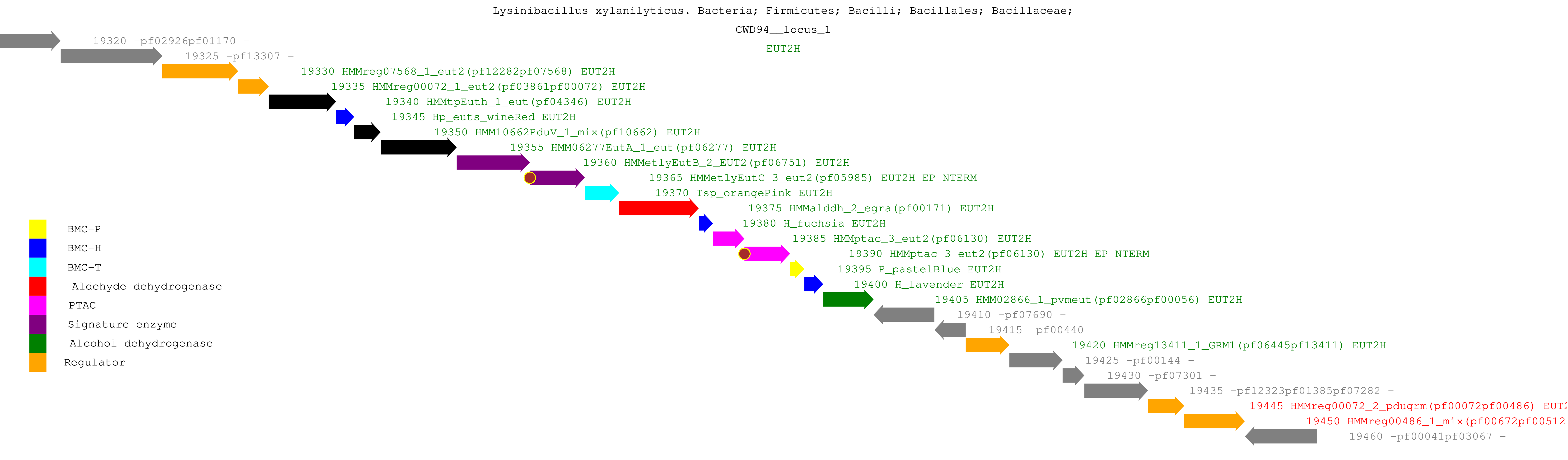

Lysinibacillus xylanilyticus. Bacteria; Firmicutes; Bacilli; Bacillales; Bacillaceae;
CWD94__locus_1
EUT2H
19320 -pf02926pf01170 -
19325 -pf13307 -
19330 HMMreg07568_1_eut2(pf12282pf07568) EUT2H
19335 HMMreg00072_1_eut2(pf03861pf00072) EUT2H
19340 HMMtpEuth_1_eut(pf04346) EUT2H
19345 Hp_euts_wineRed EUT2H
19350 HMM10662PduV_1_mix(pf10662) EUT2H
19355 HMM06277EutA_1_eut(pf06277) EUT2H
19360 HMMetlyEutB_2_EUT2(pf06751) EUT2H
19365 HMMetlyEutC_3_eut2(pf05985) EUT2H EP_NTERM
19370 Tsp_orangePink EUT2H
19375 HMMalddh_2_egra(pf00171) EUT2H
19380 H_fuchsia EUT2H
19385 HMMptac_3_eut2(pf06130) EUT2H
19390 HMMptac_3_eut2(pf06130) EUT2H EP_NTERM
19395 P_pastelBlue EUT2H
19400 H_lavender EUT2H
19405 HMM02866_1_pvmeut(pf02866pf00056) EUT2H
19410 -pf07690 -
19415 -pf00440 -
19420 HMMreg13411_1_GRM1(pf06445pf13411) EUT2H
19425 -pf00144 -
19430 -pf07301 -
19435 -pf12323pf01385pf07282 -
19445 HMMreg00072_2_pdugrm(pf00072pf00486) EUT2
19450 HMMreg00486_1_mix(pf00672pf00512
19460 -pf00041pf03067 -
BMC-P
BMC-H
BMC-T
Aldehyde dehydrogenase
PTAC
Signature enzyme
Alcohol dehydrogenase
Regulator

Lysinibacillus sp. Bacteria; Firmicutes; Bacilli; Bacillales; Bacillaceae;

DCW76__locus_1_main_sat

EUT2H

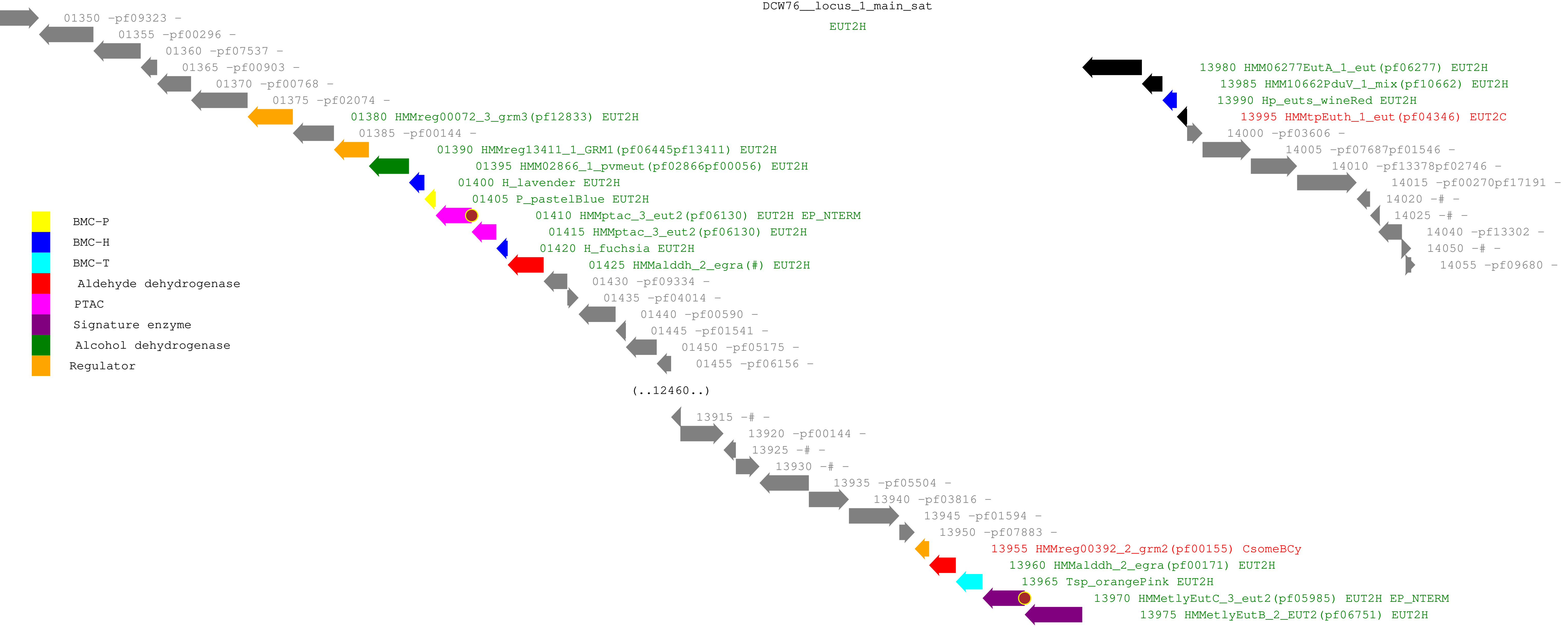

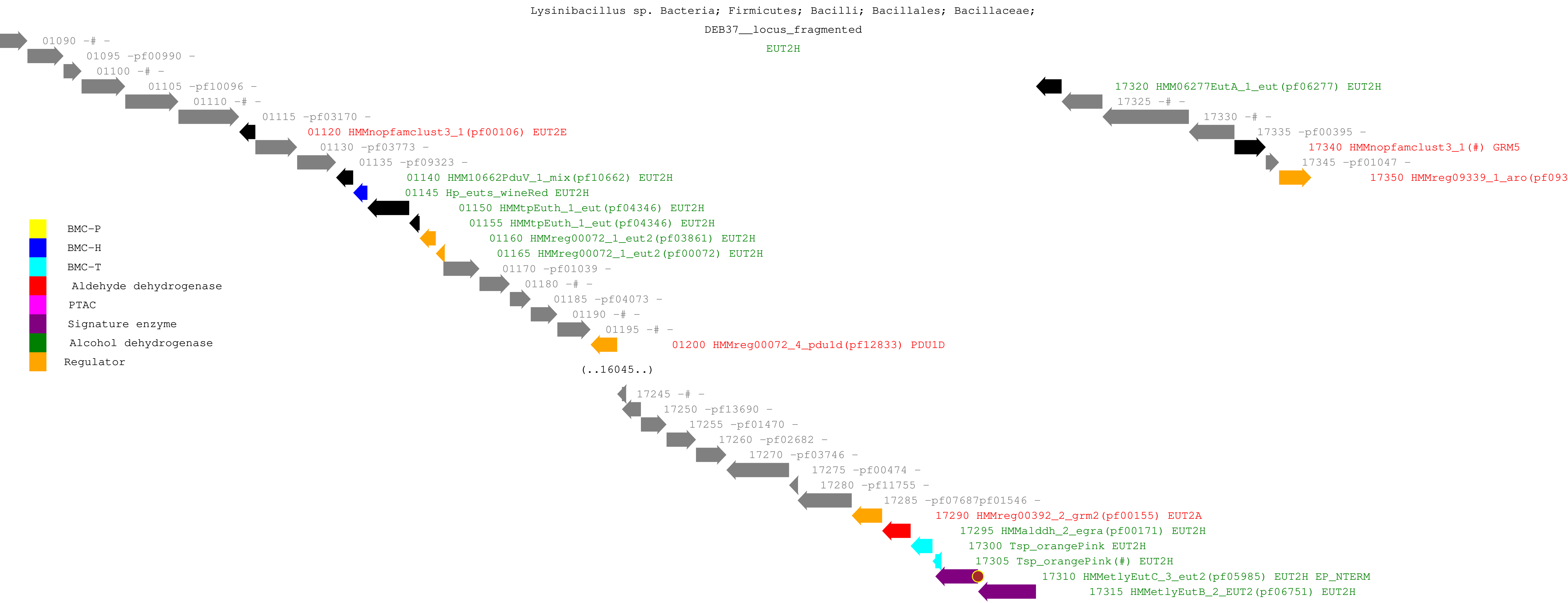
Lysinibacillus sp. Bacteria; Firmicutes; Bacilli; Bacillales; Bacillaceae;
DEB37__locus_fragmented
EUT2H
01090 -# -
01095 -pf00990 -
01100 -# -
01105 -pf10096 -
01110 -# -
01115 -pf03170 -
01120 HMMnopfamclust3_1(pf00106) EUT2E
01130 -pf03773 -
01135 -pf09323 -
01140 HMM10662PduV_1_mix(pf10662) EUT2H
01145 Hp_euts_wineRed EUT2H
01150 HMMtpEuth_1_eut(pf04346) EUT2H
01155 HMMtpEuth_1_eut(pf04346) EUT2H
01160 HMMreg00072_1_eut2(pf03861) EUT2H
01165 HMMreg00072_1_eut2(pf00072) EUT2H
01170 -pf01039 -
01180 -# -
01185 -pf04073 -
01190 -# -
01195 -# -
01200 HMMreg00072_4_pdu1d(pf12833) PDU1D
(..16045..)
17245 -# -
17250 -pf13690 -
17255 -pf01470 -
17260 -pf02682 -
17270 -pf03746 -
17275 -pf00474 -
17280 -pf11755 -
17285 -pf07687pf01546 -
17290 HMMreg00392_2_grm2(pf00155) EUT2A
17295 HMMalddh_2_egra(pf00171) EUT2H
17300 Tsp_orangePink EUT2H
17305 Tsp_orangePink(#) EUT2H
17310 HMMetlyEutC_3_eut2(pf05985) EUT2H EP_NTERM
17315 HMMetlyEutB_2_EUT2(pf06751) EUT2H
17320 HMM06277EutA_1_eut(pf06277) EUT2H
17325 -# -
17330 -# -
17335 -pf00395 -
17340 HMMnopfamclust3_1(#) GRM5
17345 -pf01047 -
17350 HMMreg09339_1_aro(pf093
BMC-P
BMC-H
BMC-T
Aldehyde dehydrogenase
PTAC
Signature enzyme
Alcohol dehydrogenase
Regulator

Lysinibacillus sp. OL1. Bacteria; Firmicutes; Bacilli; Bacillales; Bacillaceae; Lysinibacillus;

EW028__locus_1

EUT2H

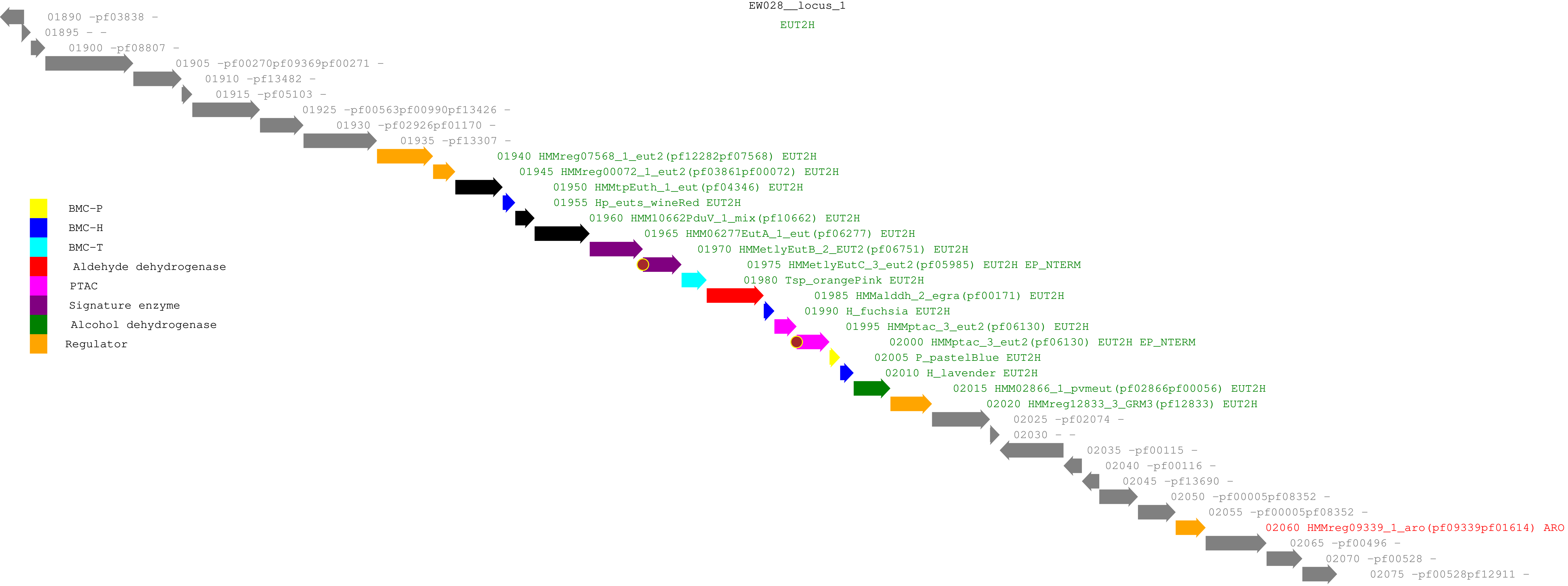

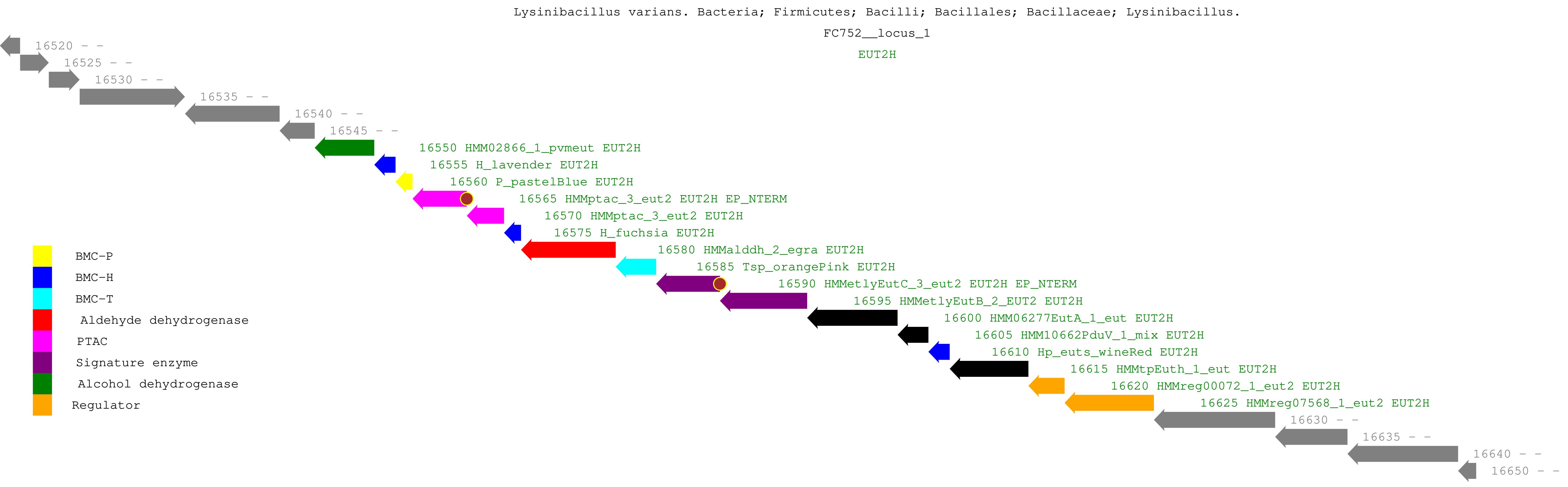
Lysinibacillus varians. Bacteria; Firmicutes; Bacilli; Bacillales; Bacillaceae; Lysinibacillus.
FC752__locus_1
EUT2H
16520 - -
16525 - -
16530 - -
16535 - -
16540 - -
16545 - -
16550 HMM02866_1_pvmeut EUT2H
16555 H_lavender EUT2H
16560 P_pastelBlue EUT2H
16565 HMMptac_3_eut2 EUT2H EP_NTERM
16570 HMMptac_3_eut2 EUT2H
16575 H_fuchsia EUT2H
16580 HMMalddh_2_egra EUT2H
16585 Tsp_orangePink EUT2H
16590 HMMetlyEutC_3_eut2 EUT2H EP_NTERM
16595 HMMetlyEutB_2_EUT2 EUT2H
16600 HMM06277EutA_1_eut EUT2H
16605 HMM10662PduV_1_mix EUT2H
16610 Hp_euts_wineRed EUT2H
16615 HMMtpEuth_1_eut EUT2H
16620 HMMreg00072_1_eut2 EUT2H
16625 HMMreg07568_1_eut2 EUT2H
16630 - -
16635 - -
16640 - -
16650 - -
BMC-P
BMC-H
BMC-T
Aldehyde dehydrogenase
PTAC
Signature enzyme
Alcohol dehydrogenase
Regulator

Lysinibacillus mangiferihumi. Bacteria; Firmicutes; Bacilli; Bacillales; Bacillaceae; Lysinibacillus.

FC756__locus_1

EUT2H

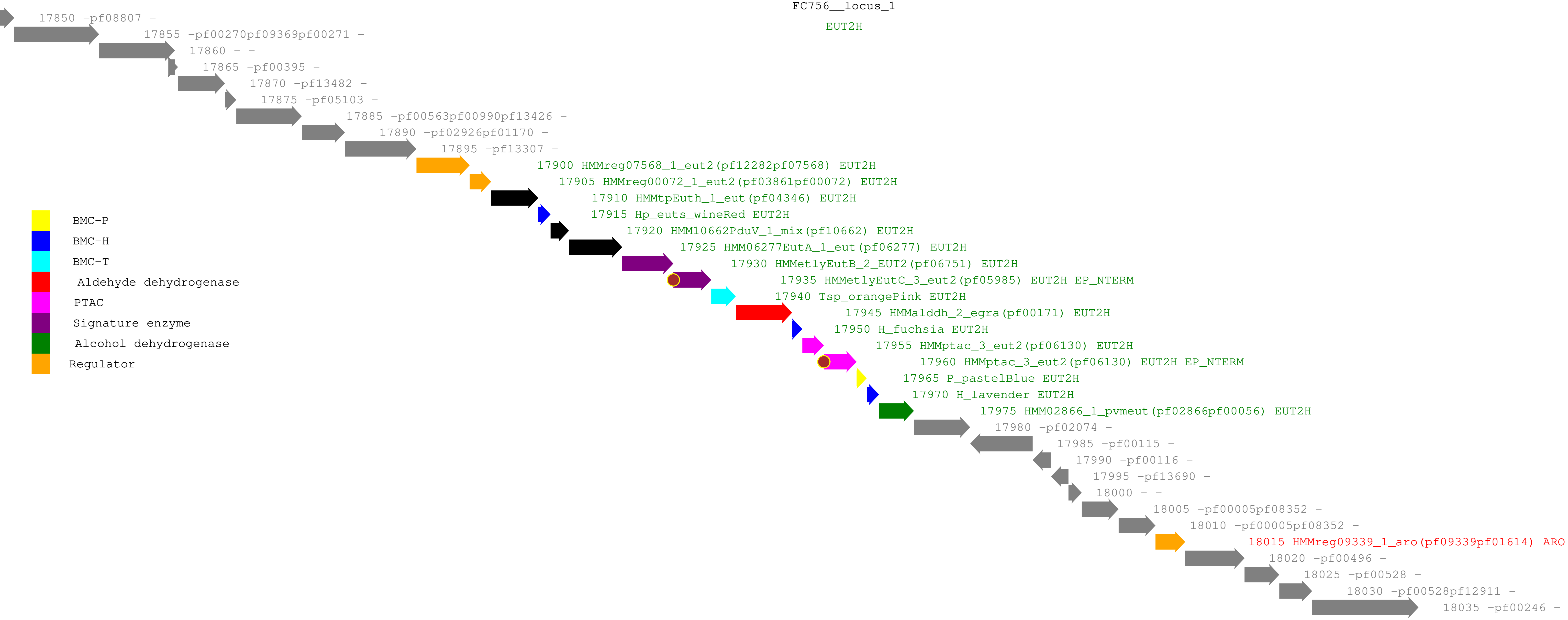

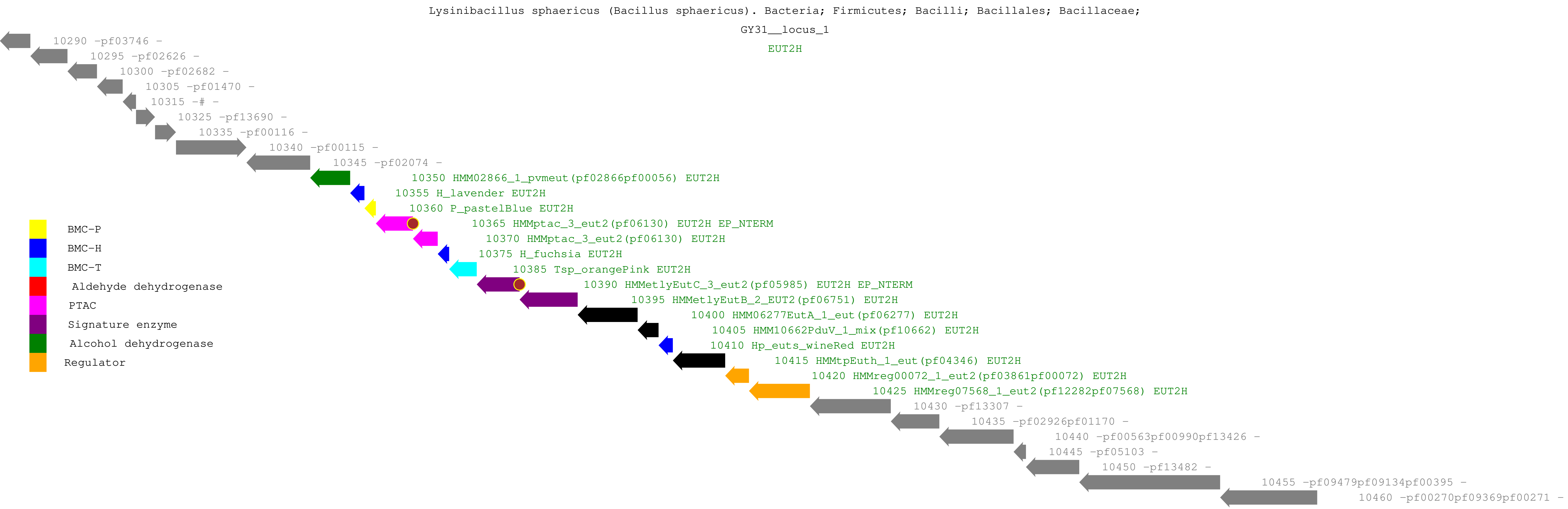
Lysinibacillus sphaericus (Bacillus sphaericus). Bacteria; Firmicutes; Bacilli; Bacillales; Bacillaceae;
GY31__locus_1
EUT2H
10290 -pf03746 -
10295 -pf02626 -
10300 -pf02682 -
10305 -pf01470 -
10315 -# -
10325 -pf13690 -
10335 -pf00116 -
10340 -pf00115 -
10345 -pf02074 -
10350 HMM02866_1_pvmeut(pf02866pf00056) EUT2H
10355 H_lavender EUT2H
10360 P_pastelBlue EUT2H
10365 HMMptac_3_eut2(pf06130) EUT2H EP_NTERM
10370 HMMptac_3_eut2(pf06130) EUT2H
10375 H_fuchsia EUT2H
10385 Tsp_orangePink EUT2H
10390 HMMetlyEutC_3_eut2(pf05985) EUT2H EP_NTERM
10395 HMMetlyEutB_2_EUT2(pf06751) EUT2H
10400 HMM06277EutA_1_eut(pf06277) EUT2H
10405 HMM10662PduV_1_mix(pf10662) EUT2H
10410 Hp_euts_wineRed EUT2H
10415 HMMtpEuth_1_eut(pf04346) EUT2H
10420 HMMreg00072_1_eut2(pf03861pf00072) EUT2H
10425 HMMreg07568_1_eut2(pf12282pf07568) EUT2H
10430 -pf13307 -
10435 -pf02926pf01170 -
10440 -pf00563pf00990pf13426 -
10445 -pf05103 -
10450 -pf13482 -
10455 -pf09479pf09134pf00395 -
10460 -pf00270pf09369pf00271 -
BMC-P
BMC-H
BMC-T
Aldehyde dehydrogenase
PTAC
Signature enzyme
Alcohol dehydrogenase
Regulator

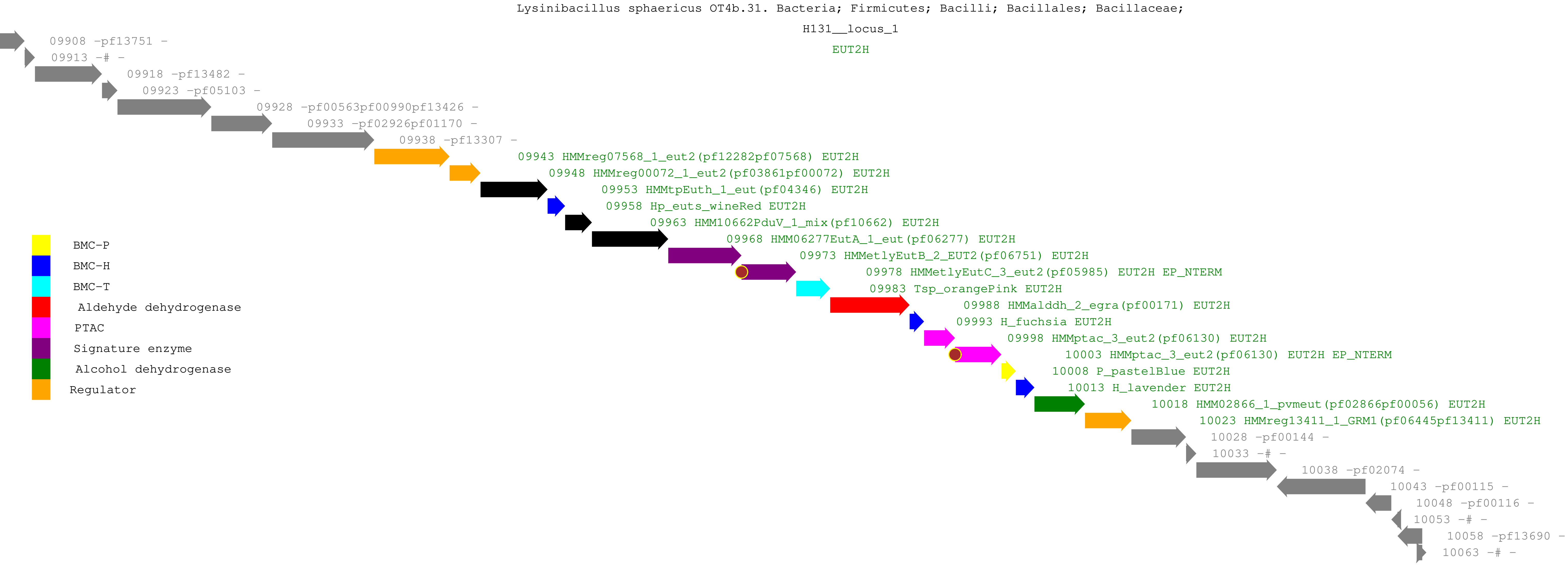

Lysinibacillus sphaericus OT4b.31. Bacteria; Firmicutes; Bacilli; Bacillales; Bacillaceae;
H131__locus_1
EUT2H
09908 -pf13751 -
09913 -# -
09918 -pf13482 -
09923 -pf05103 -
09928 -pf00563pf00990pf13426 -
09933 -pf02926pf01170 -
09938 -pf13307 -
09943 HMMreg07568_1_eut2(pf12282pf07568) EUT2H
09948 HMMreg00072_1_eut2(pf03861pf00072) EUT2H
09953 HMMtpEuth_1_eut(pf04346) EUT2H
09958 Hp_euts_wineRed EUT2H
09963 HMM10662PduV_1_mix(pf10662) EUT2H
09968 HMM06277EutA_1_eut(pf06277) EUT2H
09973 HMMetlyEutB_2_EUT2(pf06751) EUT2H
09978 HMMetlyEutC_3_eut2(pf05985) EUT2H EP_NTERM
09983 Tsp_orangePink EUT2H
09988 HMMaldh_2_egra(pf00171) EUT2H
09993 H_fuchsia EUT2H
09998 HMMptac_3_eut2(pf06130) EUT2H
10003 HMMptac_3_eut2(pf06130) EUT2H EP_NTERM
10008 P_pastelBlue EUT2H
10013 H_lavender EUT2H
10018 HMM02866_1_pvmeut(pf02866pf00056) EUT2H
10023 HMMreg13411_1_GRM1(pf06445pf13411) EUT2H
10028 -pf00144 -
10033 -# -
10038 -pf02074 -
10043 -pf00115 -
10048 -pf00116 -
10053 -# -
10058 -pf13690 -
10063 -# -
BMC-P
BMC-H
BMC-T
Aldehyde dehydrogenase
PTAC
Signature enzyme
Alcohol dehydrogenase
Regulator

Lysinibacillus sp. YS11. Bacteria; Firmicutes; Bacilli; Bacillales; Bacillaceae;

LBYS11__locus_1

EUT2H

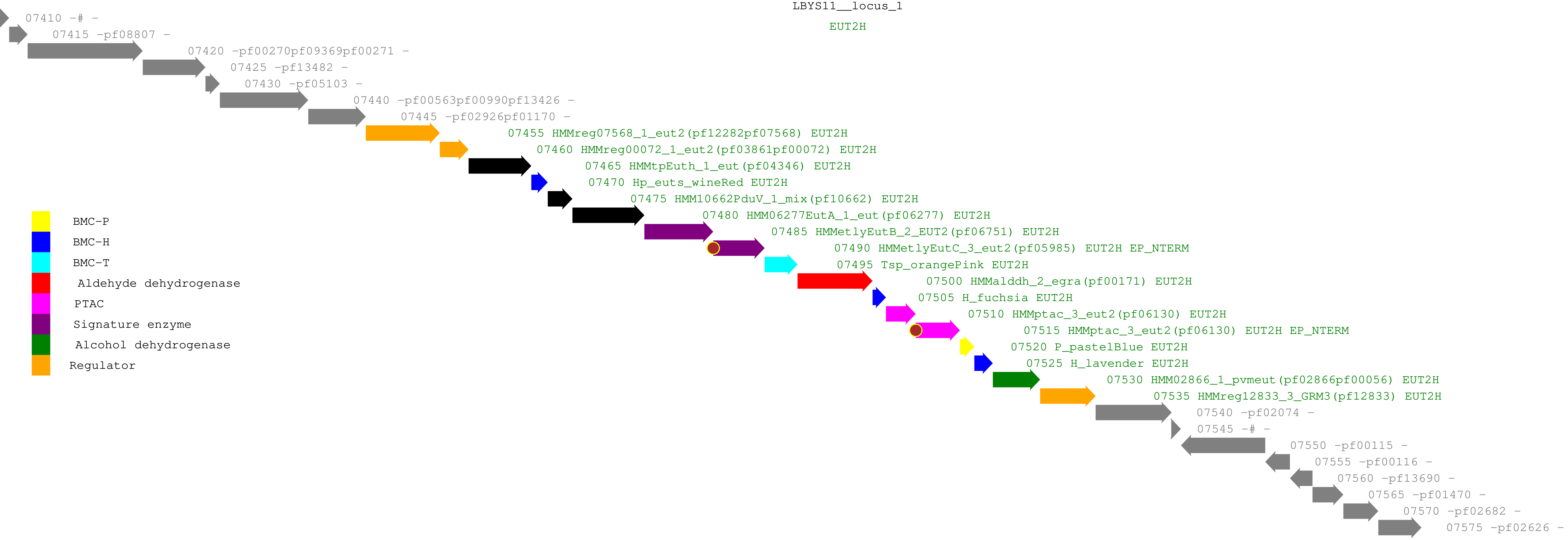

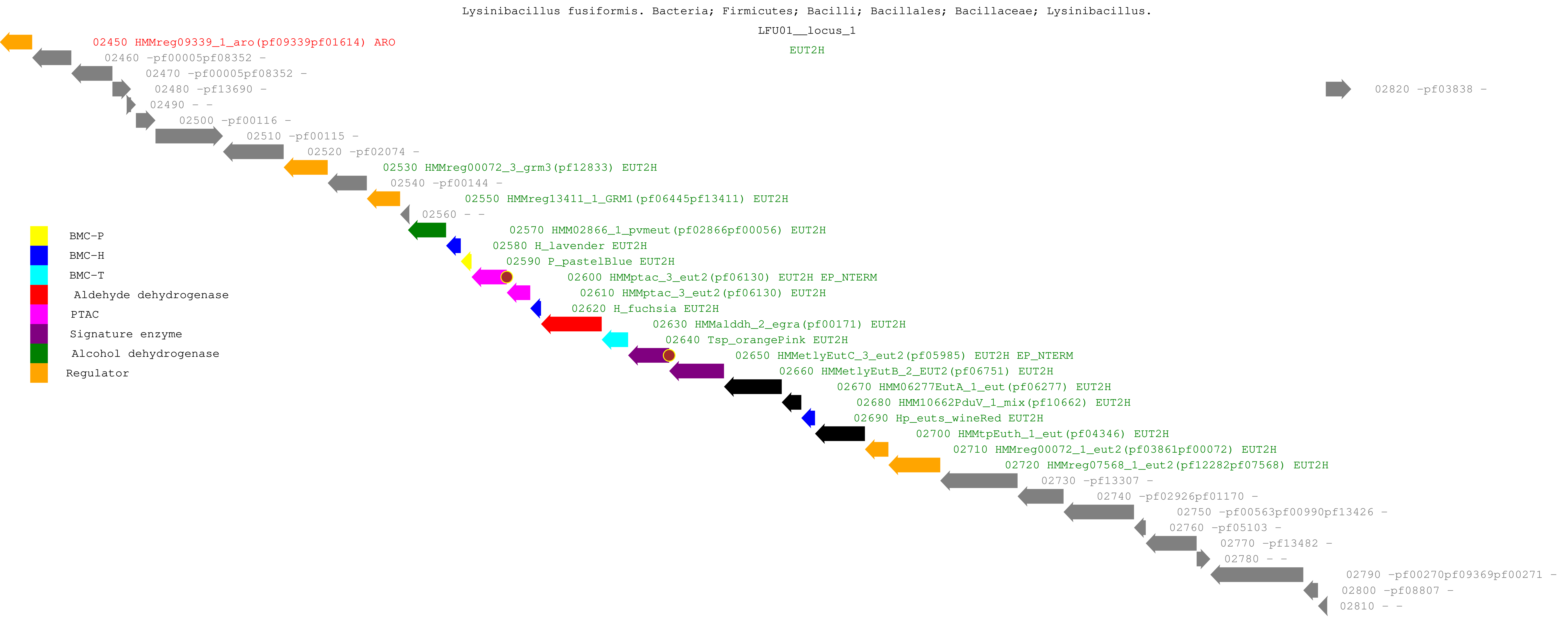

Lysinibacillus fusiformis. Bacteria; Firmicutes; Bacilli; Bacillales; Bacillaceae; Lysinibacillus.
LFU01__locus_1
EUT2H
02450 HMMreg09339_1_aro(pf09339pf01614) ARO
02460 -pf00005pf08352 -
02470 -pf00005pf08352 -
02480 -pf13690 -
02490 - -
02500 -pf00116 -
02510 -pf00115 -
02520 -pf02074 -
02530 HMMreg00072_3_grm3(pf12833) EUT2H
02540 -pf00144 -
02550 HMMreg13411_1_GRM1(pf06445pf13411) EUT2H
02560 - -
02570 HMM02866_1_pvmeut(pf02866pf00056) EUT2H
02580 H_lavender EUT2H
02590 P_pastelBlue EUT2H
02600 HMMptac_3_eut2(pf06130) EUT2H EP_NTERM
02610 HMMptac_3_eut2(pf06130) EUT2H
02620 H_fuchsia EUT2H
02630 HMMalddh_2_egra(pf00171) EUT2H
02640 Tsp_orangePink EUT2H
02650 HMMetlyEutC_3_eut2(pf05985) EUT2H EP_NTERM
02660 HMMetlyEutB_2_EUT2(pf06751) EUT2H
02670 HMM06277EutA_1_eut(pf06277) EUT2H
02680 HMM10662PduV_1_mix(pf10662) EUT2H
02690 Hp_euts_wineRed EUT2H
02700 HMMtpEuth_1_eut(pf04346) EUT2H
02710 HMMreg00072_1_eut2(pf03861pf00072) EUT2H
02720 HMMreg07568_1_eut2(pf12282pf07568) EUT2H
02730 -pf13307 -
02740 -pf02926pf01170 -
02750 -pf00563pf00990pf13426 -
02760 -pf05103 -
02770 -pf13482 -
02780 - -
02790 -pf00270pf09369pf00271 -
02800 -pf08807 -
02810 - -
02820 -pf03838 -
BMC-P
BMC-H
BMC-T
Aldehyde dehydrogenase
PTAC
Signature enzyme
Alcohol dehydrogenase
Regulator

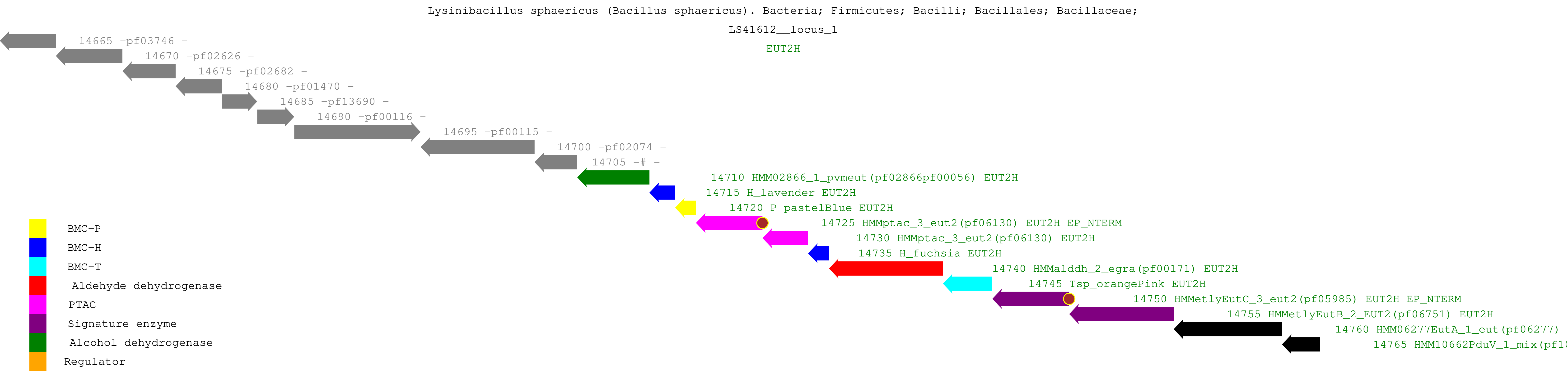
Lysinibacillus sphaericus (Bacillus sphaericus). Bacteria; Firmicutes; Bacilli; Bacillales; Bacillaceae;
LS41612__locus_1
EUT2H
14665 -pf03746 -
14670 -pf02626 -
14675 -pf02682 -
14680 -pf01470 -
14685 -pf13690 -
14690 -pf00116 -
14695 -pf00115 -
14700 -pf02074 -
14705 -# -
14710 HMM02866_1_pvmeut(pf02866pf00056) EUT2H
14715 H_lavender EUT2H
14720 P_pastelBlue EUT2H
14725 HMMptac_3_eut2(pf06130) EUT2H EP_NTERM
14730 HMMptac_3_eut2(pf06130) EUT2H
14735 H_fuchsia EUT2H
14740 HMMalddh_2_egra(pf00171) EUT2H
14745 Tsp_orangePink EUT2H
14750 HMMetlyEutC_3_eut2(pf05985) EUT2H EP_NTERM
14755 HMMetlyEutB_2_EUT2(pf06751) EUT2H
14760 HMM06277EutA_1_eut(pf06277)
14765 HMM10662PduV_1_mix(pf1
BMC-P
BMC-H
BMC-T
Aldehyde dehydrogenase
PTAC
Signature enzyme
Alcohol dehydrogenase
Regulator

Lysinibacillus sphaericus (Bacillus sphaericus). Bacteria; Firmicutes; Bacilli; Bacillales; Bacillaceae; Lysinibacillus.

LSP__locus_1

EUT2H

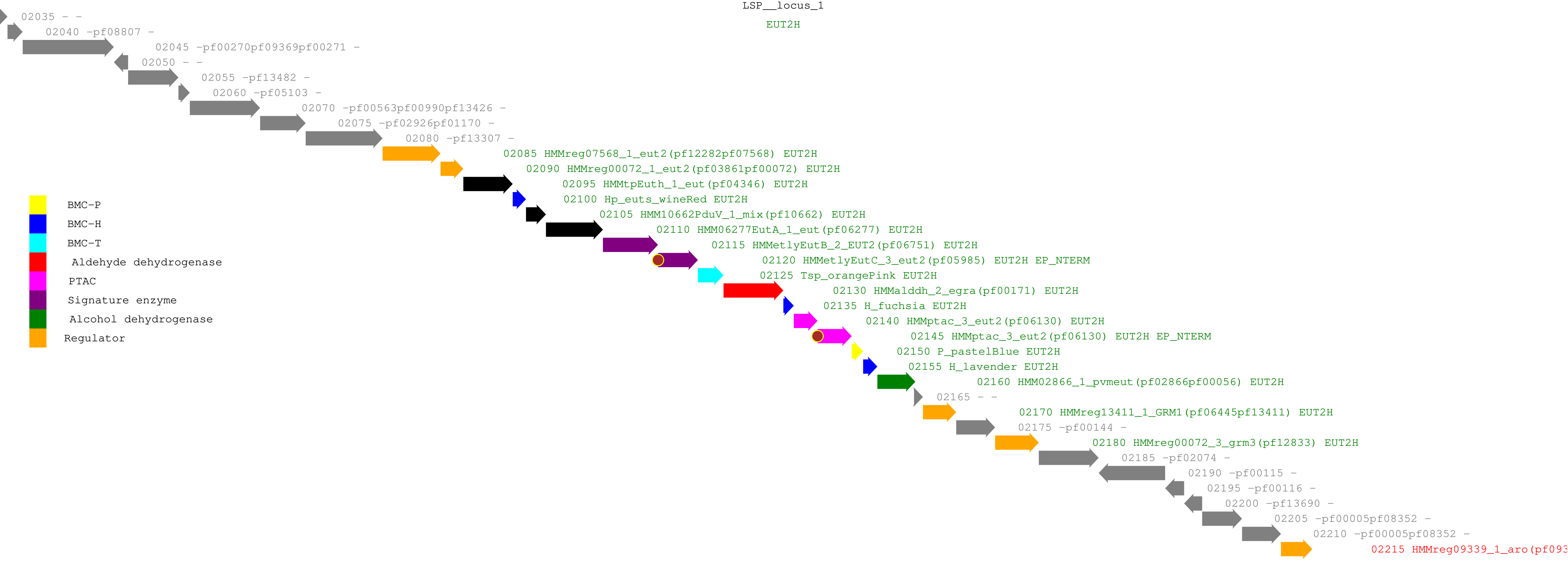

Lysinibacillus sphaericus (Bacillus sphaericus). Bacteria; Firmicutes; Bacilli; Bacillales; Bacillaceae;

LYSIN__locus_1

EUT2H

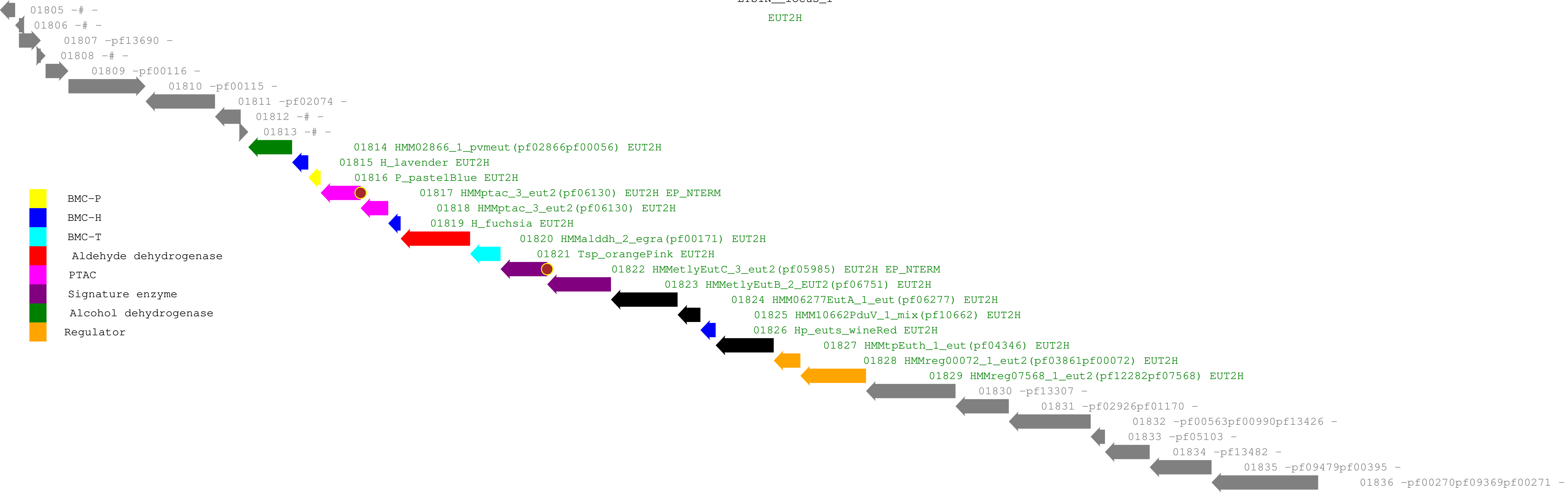

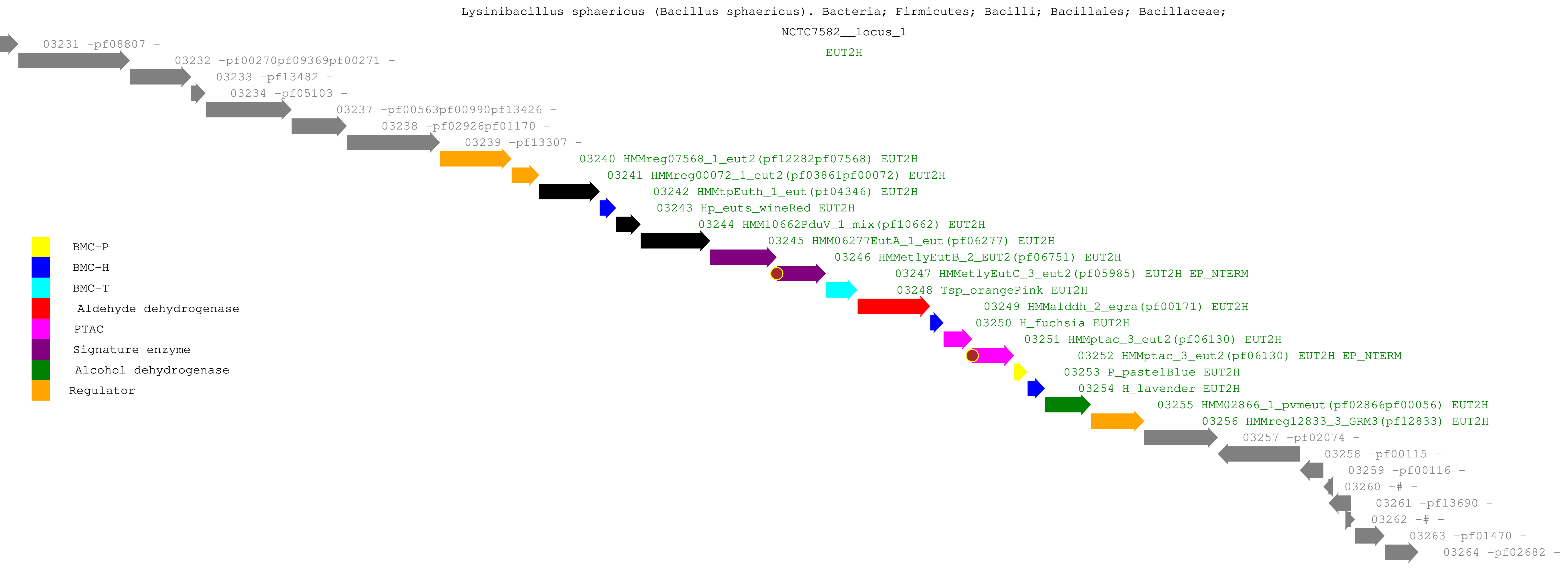
Lysinibacillus sphaericus (Bacillus sphaericus). Bacteria; Firmicutes; Bacilli; Bacillales; Bacillaceae;
NCTC7582__locus_1
EUT2H
03231 -pf08807 -
03232 -pf00270pf09369pf00271 -
03233 -pf13482 -
03234 -pf05103 -
03237 -pf00563pf00990pf13426 -
03238 -pf02926pf01170 -
03239 -pf13307 -
03240 HMMreg07568_1_eut2(pf12282pf07568) EUT2H
03241 HMMreg00072_1_eut2(pf03861pf00072) EUT2H
03242 HMMtpEuth_1_eut(pf04346) EUT2H
03243 Hp_euts_wineRed EUT2H
03244 HMM10662PduV_1_mix(pf10662) EUT2H
03245 HMM06277EutA_1_eut(pf06277) EUT2H
03246 HMMetlyEutB_2_EUT2(pf06751) EUT2H
03247 HMMetlyEutC_3_eut2(pf05985) EUT2H EP_NTERM
03248 Tsp_orangePink EUT2H
03249 HMMalddh_2_egra(pf00171) EUT2H
03250 H_fuchsia EUT2H
03251 HMMptac_3_eut2(pf06130) EUT2H
03252 HMMptac_3_eut2(pf06130) EUT2H EP_NTERM
03253 P_pastelBlue EUT2H
03254 H_lavender EUT2H
03255 HMM02866_1_pvmeut(pf02866pf00056) EUT2H
03256 HMMreg12833_3_GRM3(pf12833) EUT2H
03257 -pf02074 -
03258 -pf00115 -
03259 -pf00116 -
03260 -# -
03261 -pf13690 -
03262 -# -
03263 -pf01470 -
03264 -pf02682 -
BMC-P
BMC-H
BMC-T
Aldehyde dehydrogenase
PTAC
Signature enzyme
Alcohol dehydrogenase
Regulator

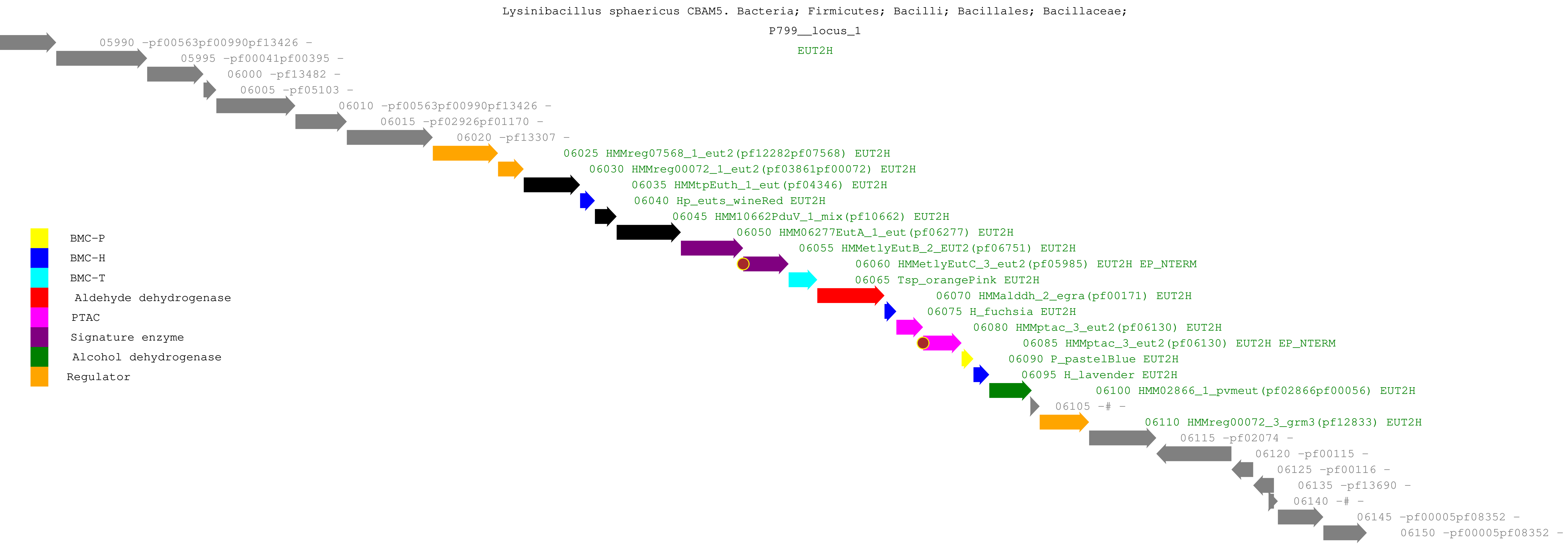
Lysinibacillus sphaericus CBAM5. Bacteria; Firmicutes; Bacilli; Bacillales; Bacillaceae;
P799__locus_1
EUT2H
05990 -pf00563pf00990pf13426 -
05995 -pf00041pf00395 -
06000 -pf13482 -
06005 -pf05103 -
06010 -pf00563pf00990pf13426 -
06015 -pf02926pf01170 -
06020 -pf13307 -
06025 HMMreg07568_1_eut2(pf12282pf07568) EUT2H
06030 HMMreg00072_1_eut2(pf03861pf00072) EUT2H
06035 HMMtpEuth_1_eut(pf04346) EUT2H
06040 Hp_euts_wineRed EUT2H
06045 HMM10662PduV_1_mix(pf10662) EUT2H
06050 HMM06277EutA_1_eut(pf06277) EUT2H
06055 HMMetlyEutB_2_EUT2(pf06751) EUT2H
06060 HMMetlyEutC_3_eut2(pf05985) EUT2H EP_NTERM
06065 Tsp_orangePink EUT2H
06070 HMMalddh_2_egra(pf00171) EUT2H
06075 H_fuchsia EUT2H
06080 HMMptac_3_eut2(pf06130) EUT2H
06085 HMMptac_3_eut2(pf06130) EUT2H EP_NTERM
06090 P_pastelBlue EUT2H
06095 H_lavender EUT2H
06100 HMM02866_1_pvmeut(pf02866pf00056) EUT2H
06105 -# -
06110 HMMreg00072_3_grm3(pf12833) EUT2H
06115 -pf02074 -
06120 -pf00115 -
06125 -pf00116 -
06135 -pf13690 -
06140 -# -
06145 -pf00005pf08352 -
06150 -pf00005pf08352 -
BMC-P
BMC-H
BMC-T
Aldehyde dehydrogenase
PTAC
Signature enzyme
Alcohol dehydrogenase
Regulator

Lysinibacillus sp. A1. Bacteria; Firmicutes; Bacilli; Bacillales; Bacillaceae;

PI85__locus_1

EUT2H

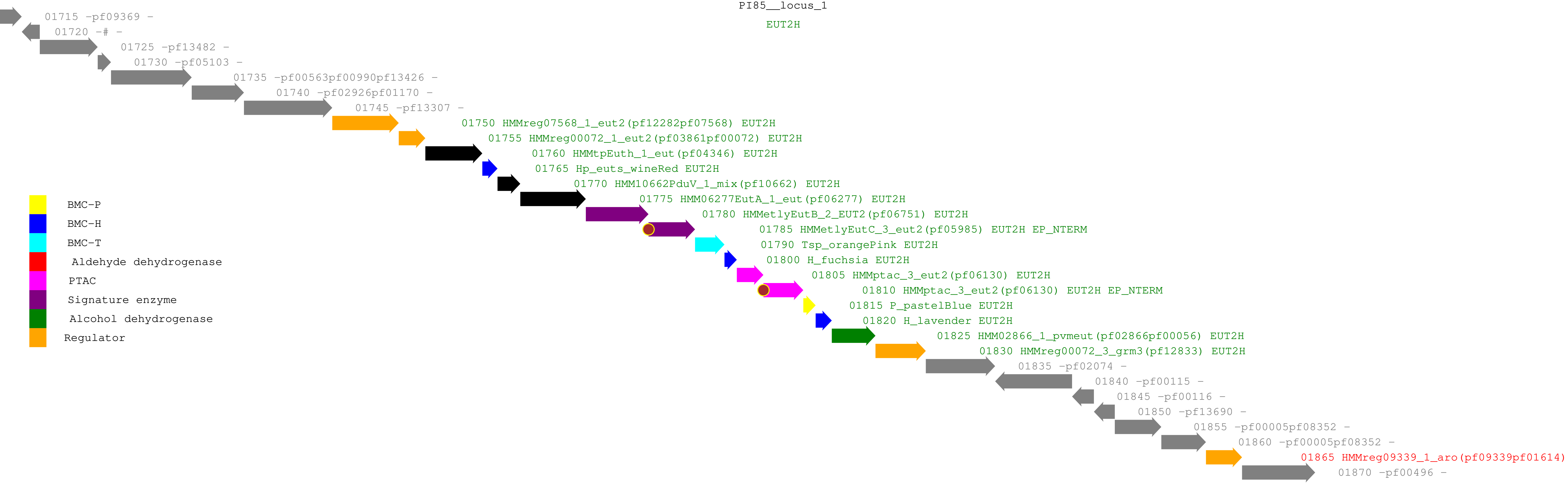

Lysinibacillus sp. SG45. Bacteria; Firmicutes; Bacilli; Bacillales; Bacillaceae;

SAMN02787103__locus_1

EUT2H

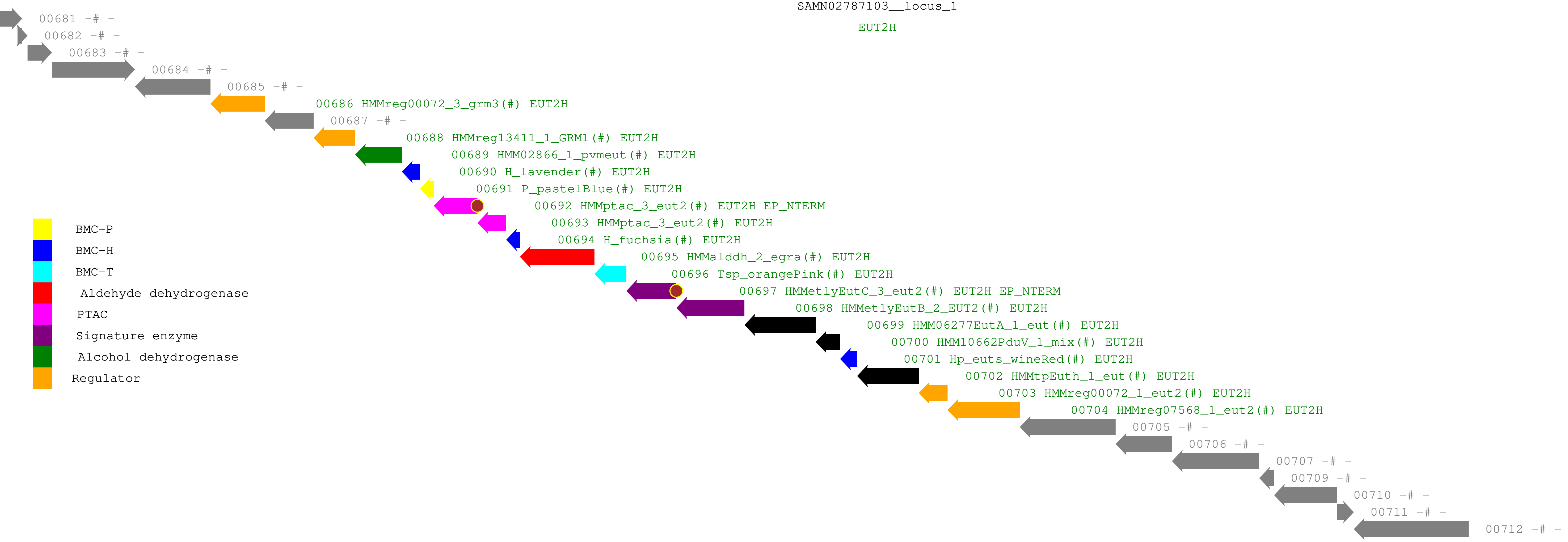

Lysinibacillus sp. AC-3. Bacteria; Firmicutes; Bacilli; Bacillales; Bacillaceae;

SAMN06295926__locus_1

EUT2H

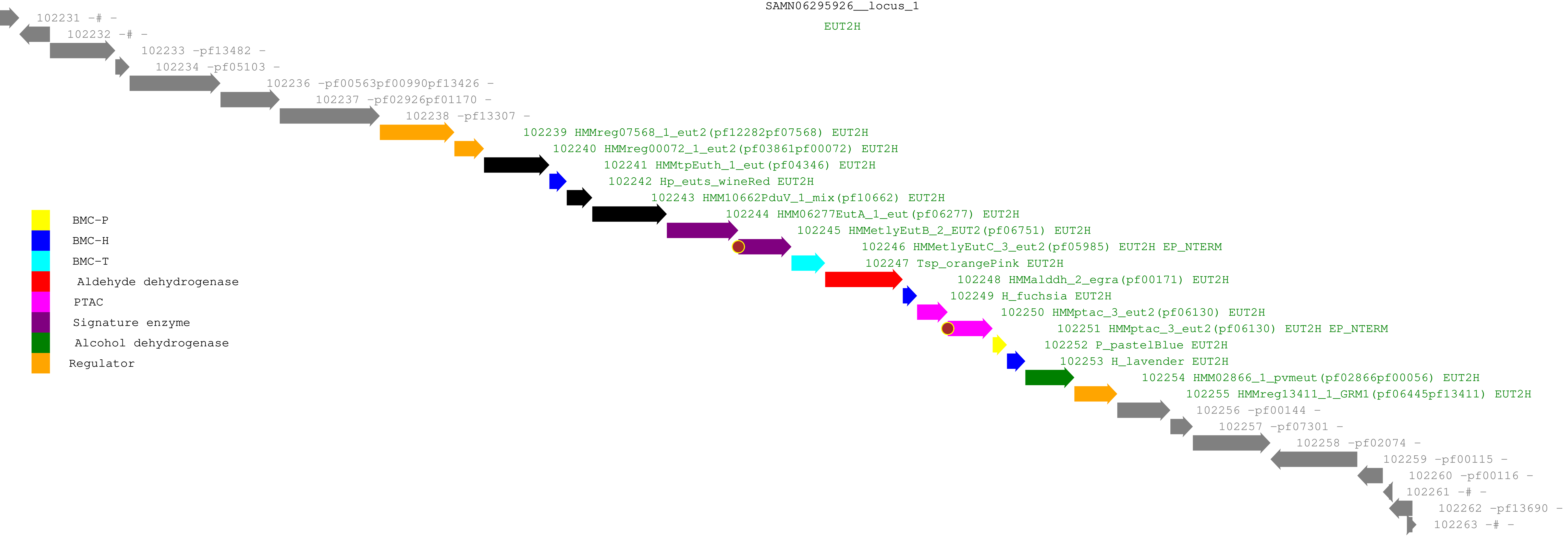

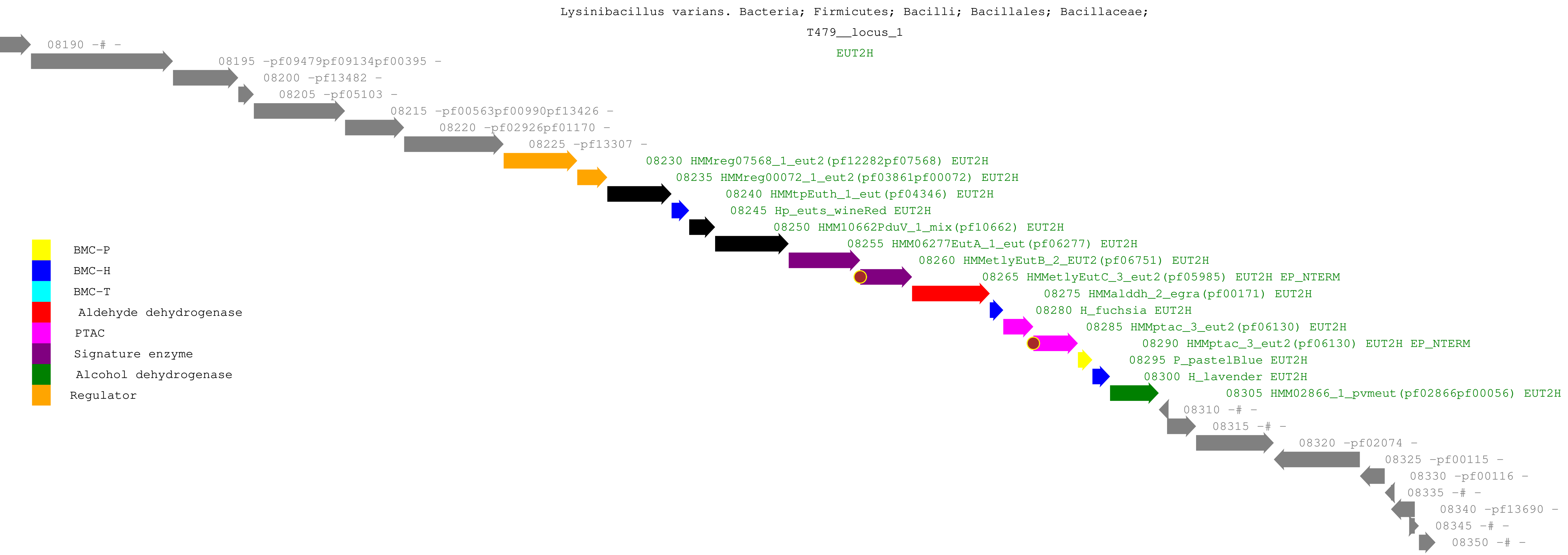

Lysinibacillus varians. Bacteria; Firmicutes; Bacilli; Bacillales; Bacillaceae;
T479__locus_1
EUT2H
08190 -# -
08195 -pf09479pf09134pf00395 -
08200 -pf13482 -
08205 -pf05103 -
08215 -pf00563pf00990pf13426 -
08220 -pf02926pf01170 -
08225 -pf13307 -
08230 HMMreg07568_1_eut2(pf12282pf07568) EUT2H
08235 HMMreg00072_1_eut2(pf03861pf00072) EUT2H
08240 HMMtpEuth_1_eut(pf04346) EUT2H
08245 Hp_euts_wineRed EUT2H
08250 HMM10662PduV_1_mix(pf10662) EUT2H
08255 HMM06277EutA_1_eut(pf06277) EUT2H
08260 HMMetlyEutB_2_EUT2(pf06751) EUT2H
08265 HMMetlyEutC_3_eut2(pf05985) EUT2H EP_NTERM
08275 HMMalddh_2_egra(pf00171) EUT2H
08280 H_fuchsia EUT2H
08285 HMMptac_3_eut2(pf06130) EUT2H
08290 HMMptac_3_eut2(pf06130) EUT2H EP_NTERM
08295 P_pastelBlue EUT2H
08300 H_lavender EUT2H
08305 HMM02866_1_pvmeut(pf02866pf00056) EUT2H
08310 -# -
08315 -# -
08320 -pf02074 -
08325 -pf00115 -
08330 -pf00116 -
08335 -# -
08340 -pf13690 -
08345 -# -
08350 -# -
BMC-P
BMC-H
BMC-T
Aldehyde dehydrogenase
PTAC
Signature enzyme
Alcohol dehydrogenase
Regulator

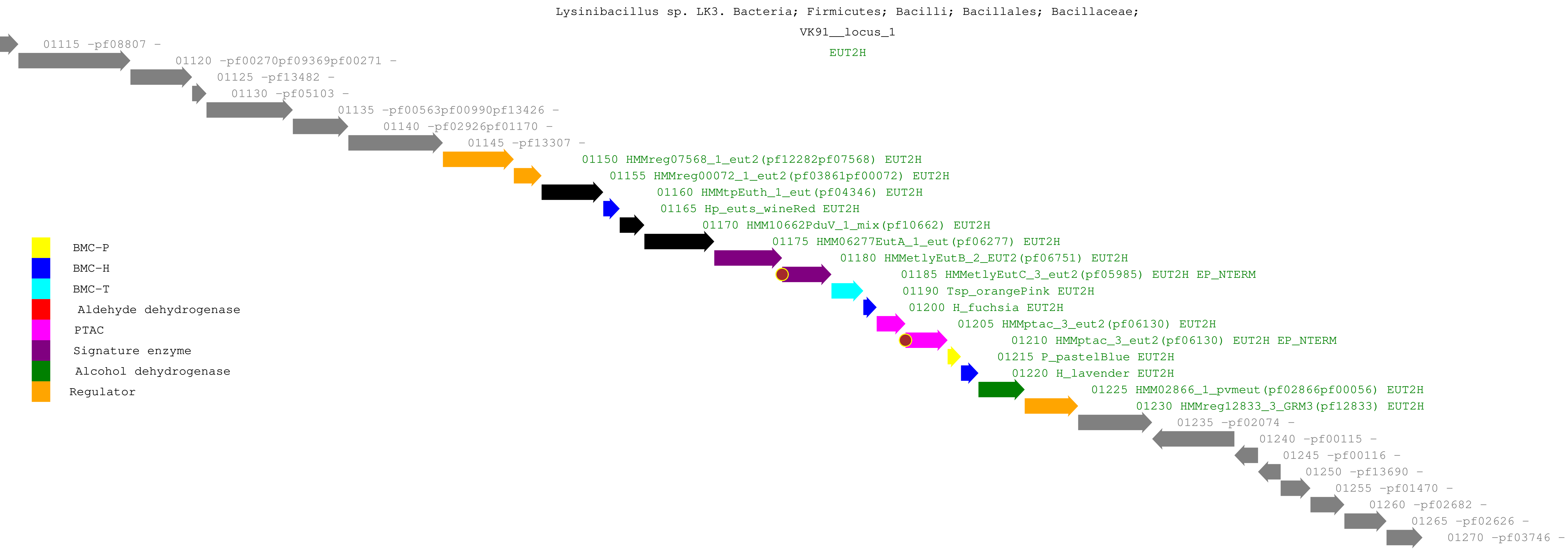
Lysinibacillus sp. LK3. Bacteria; Firmicutes; Bacilli; Bacillales; Bacillaceae;
VK91__locus_1
EUT2H
01115 -pf08807 -
01120 -pf00270pf09369pf00271 -
01125 -pf13482 -
01130 -pf05103 -
01135 -pf00563pf00990pf13426 -
01140 -pf02926pf01170 -
01145 -pf13307 -
01150 HMMreg07568_1_eut2(pf12282pf07568) EUT2H
01155 HMMreg00072_1_eut2(pf03861pf00072) EUT2H
01160 HMMtpEuth_1_eut(pf04346) EUT2H
01165 Hp_euts_wineRed EUT2H
01170 HMM10662PduV_1_mix(pf10662) EUT2H
01175 HMM06277EutA_1_eut(pf06277) EUT2H
01180 HMMetlyEutB_2_EUT2(pf06751) EUT2H
01185 HMMetlyEutC_3_eut2(pf05985) EUT2H EP_NTERM
01190 Tsp_orangePink EUT2H
01200 H_fuchsia EUT2H
01205 HMMptac_3_eut2(pf06130) EUT2H
01210 HMMptac_3_eut2(pf06130) EUT2H EP_NTERM
01215 P_pastelBlue EUT2H
01220 H_lavender EUT2H
01225 HMM02866_1_pvmeut(pf02866pf00056) EUT2H
01230 HMMreg12833_3_GRM3(pf12833) EUT2H
01235 -pf02074 -
01240 -pf00115 -
01245 -pf00116 -
01250 -pf13690 -
01255 -pf01470 -
01260 -pf02682 -
01265 -pf02626 -
01270 -pf03746 -
BMC-P
BMC-H
BMC-T
Aldehyde dehydrogenase
PTAC
Signature enzyme
Alcohol dehydrogenase
Regulator

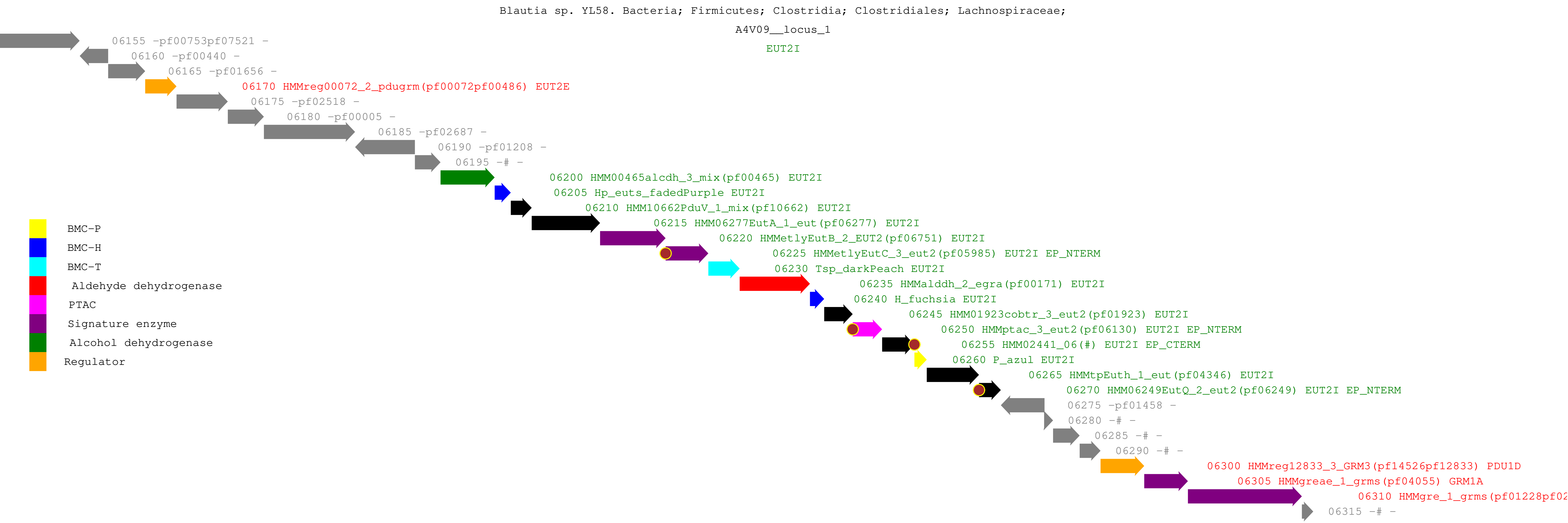

Blautia sp. YL58. Bacteria; Firmicutes; Clostridia; Clostridiales; Lachnospiraceae;
A4V09__locus_1
EUT2I
06155 -pf00753pf07521 -
06160 -pf00440 -
06165 -pf01656 -
06170 HMMreg00072_2_pdugrm(pf00072pf00486) EUT2E
06175 -pf02518 -
06180 -pf00005 -
06185 -pf02687 -
06190 -pf01208 -
06195 -# -
06200 HMM00465alcdh_3_mix(pf00465) EUT2I
06205 Hp_euts_fadedPurple EUT2I
06210 HMM10662PduV_1_mix(pf10662) EUT2I
06215 HMM06277EutA_1_eut(pf06277) EUT2I
06220 HMMetlyEutB_2_EUT2(pf06751) EUT2I
06225 HMMetlyEutC_3_eut2(pf05985) EUT2I EP_NTERM
06230 Tsp_darkPeach EUT2I
06235 HMMalddh_2_egra(pf00171) EUT2I
06240 H_fuchsia EUT2I
06245 HMM01923cobtr_3_eut2(pf01923) EUT2I
06250 HMMptac_3_eut2(pf06130) EUT2I EP_NTERM
06255 HMM02441_06(#) EUT2I EP_CTERM
06260 P_azul EUT2I
06265 HMMtpEuth_1_eut(pf04346) EUT2I
06270 HMM06249EutQ_2_eut2(pf06249) EUT2I EP_NTERM
06275 -pf01458 -
06280 -# -
06285 -# -
06290 -# -
06300 HMMreg12833_3_GRM3(pf14526pf12833) PDU1D
06305 HMMgreae_1_grms(pf04055) GRM1A
06310 HMMgre_1_grms(pf01228pf02
06315 -# -
BMC-P
BMC-H
BMC-T
Aldehyde dehydrogenase
PTAC
Signature enzyme
Alcohol dehydrogenase
Regulator

Clostridium perfringens B str. ATCC 3626. Bacteria; Firmicutes; Clostridia; Clostridiales; Clostridiaceae;

AC1__locus_1

EUT2I

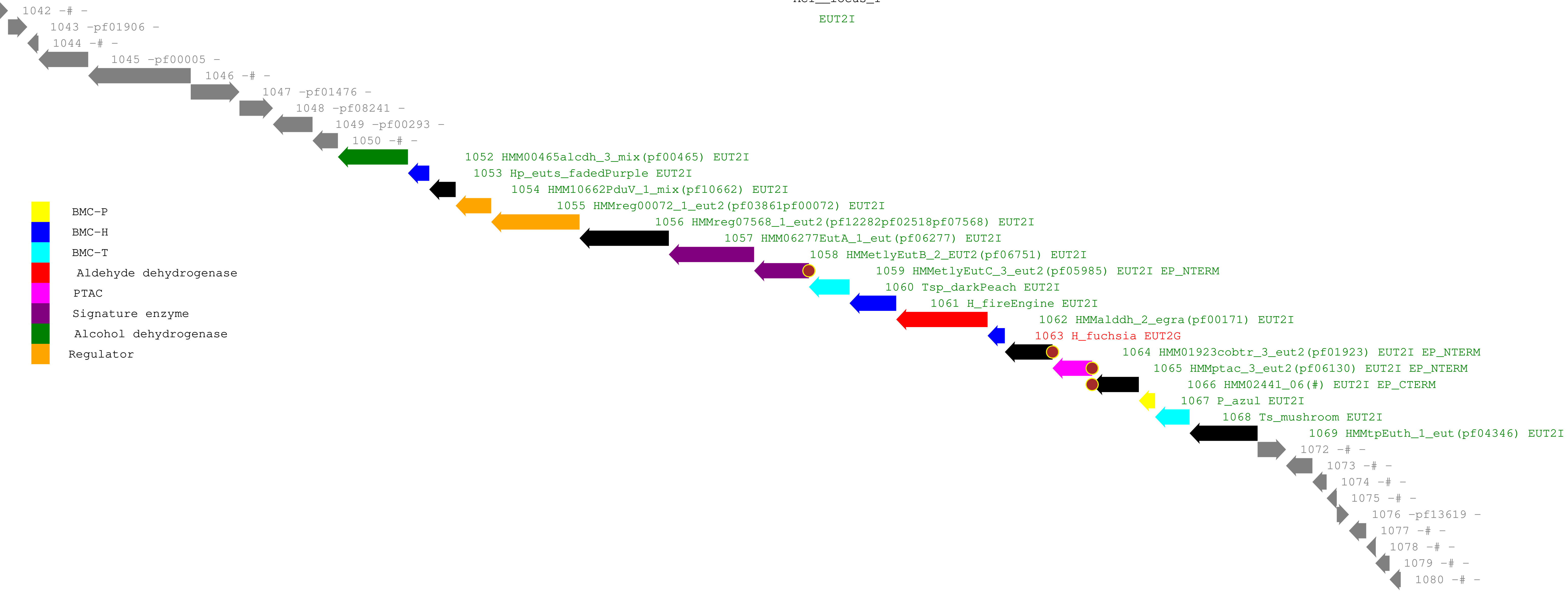

Clostridium perfringens E str. JGS1987. Bacteria; Firmicutes; Clostridia; Clostridiales; Clostridiaceae;

AC3__locus_1

EUT2I

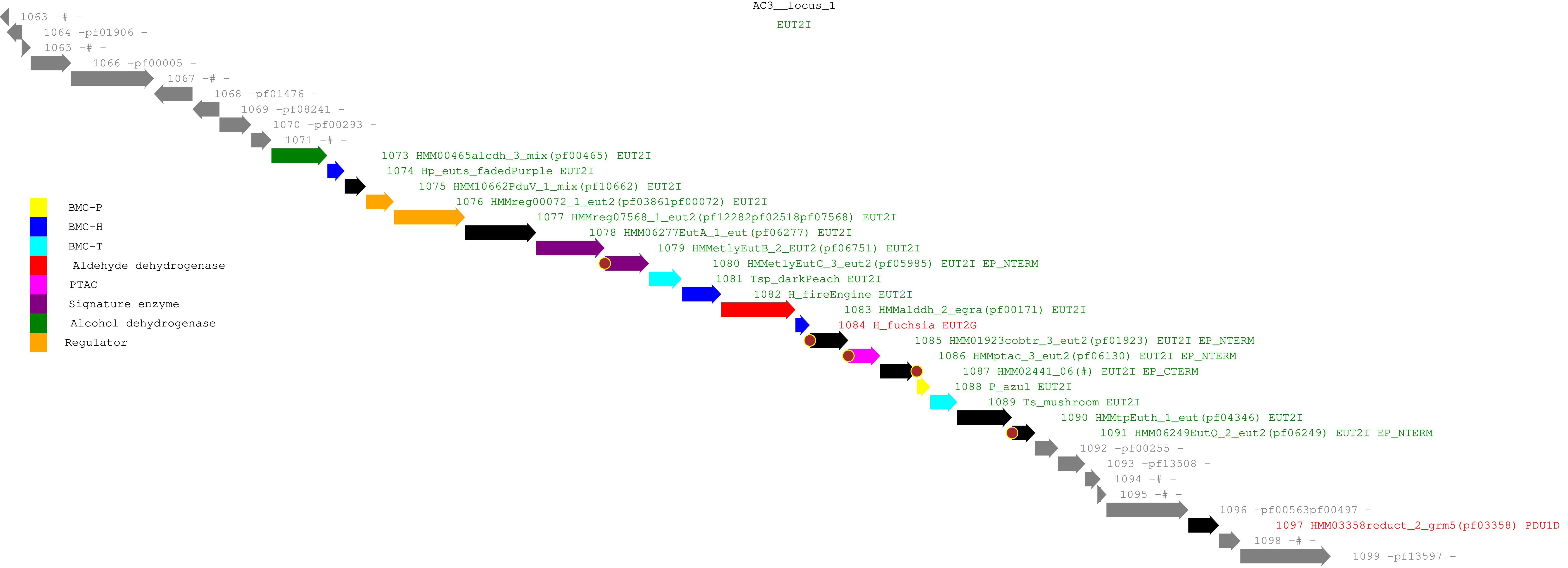

Blautia coccoides. Bacteria; Firmicutes; Clostridia; Clostridiales; Lachnospiraceae;

ADH70__locus_1

EUT2I

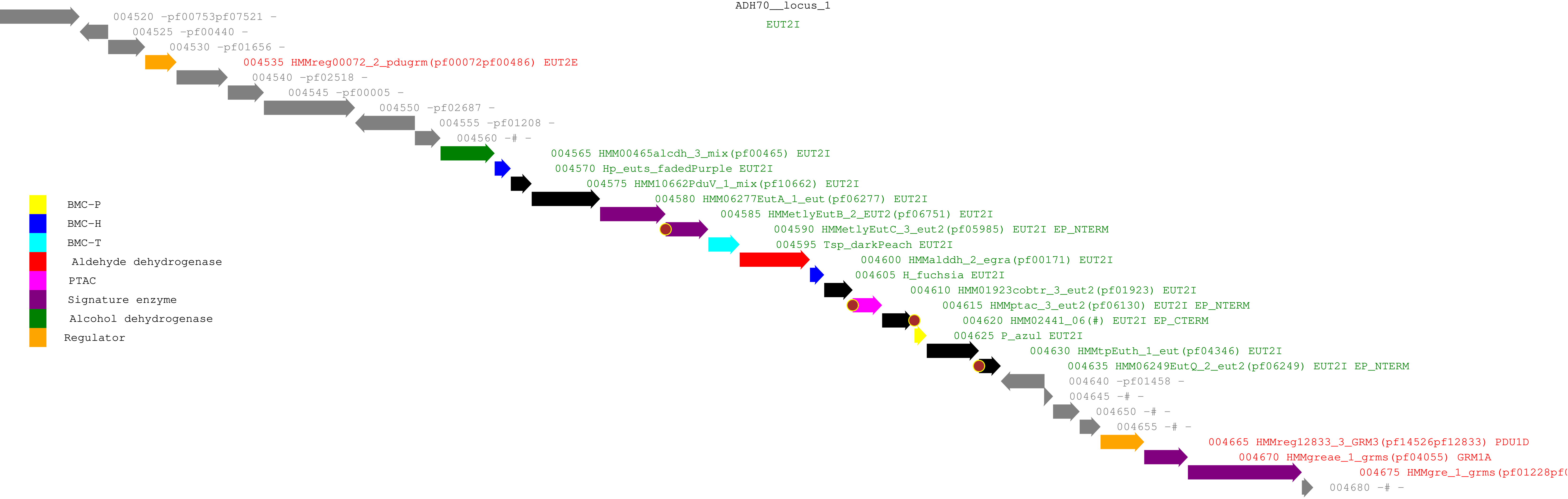

Brevibacillus sp. SKDU10. Bacteria; Firmicutes; Bacilli; Bacillales; Paenibacillaceae;

AYJ08__locus_1_main_sat

EUT2I

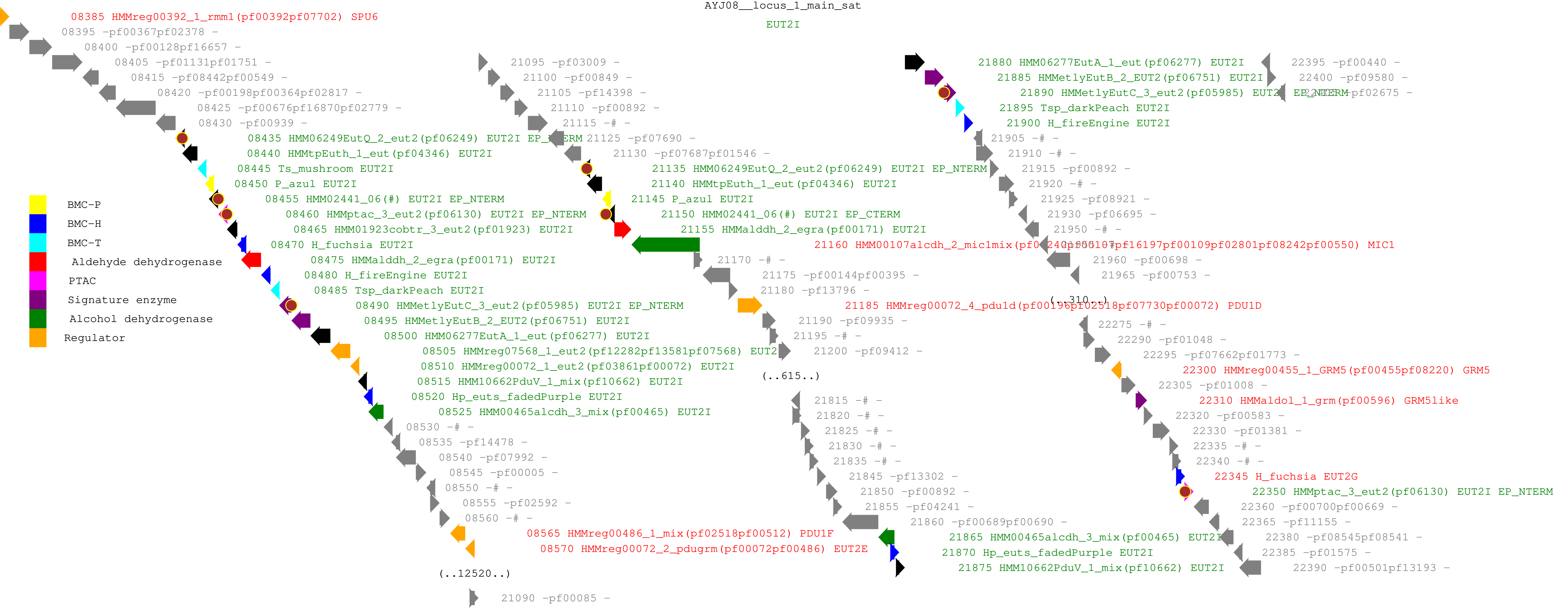

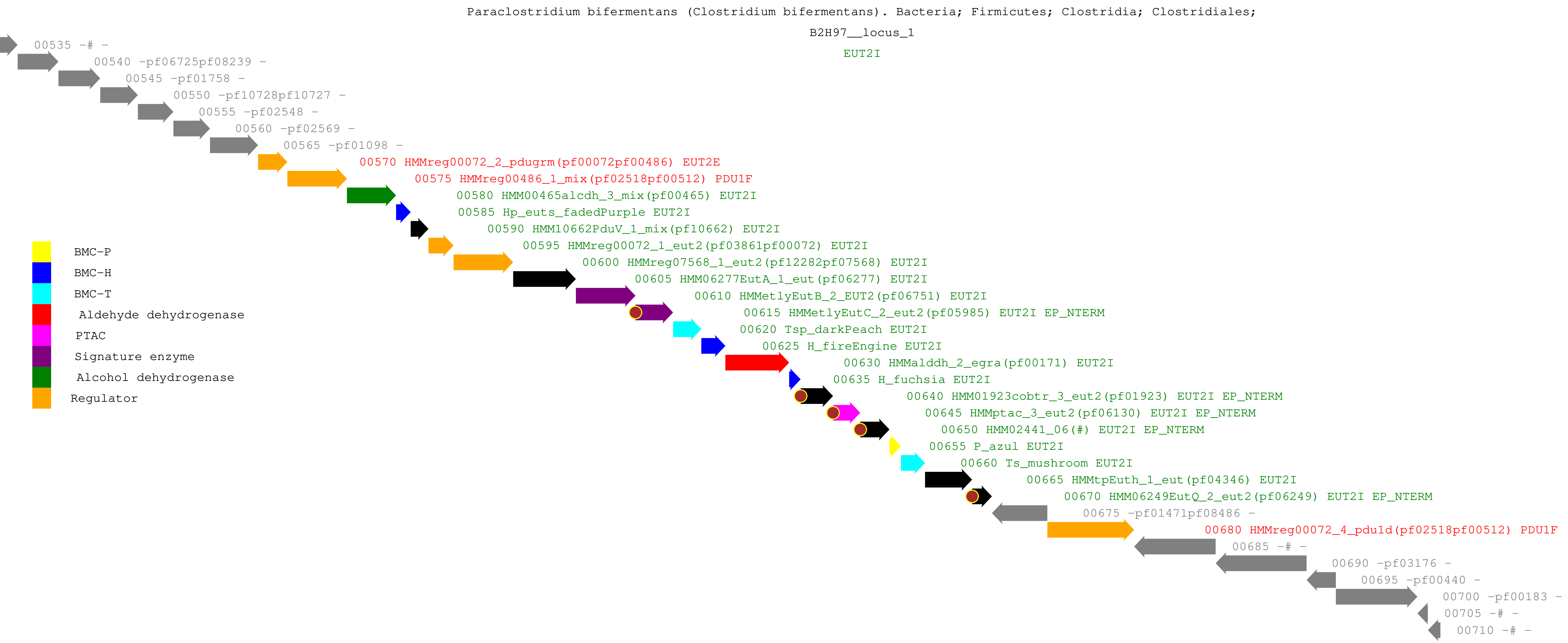
Paraclostridium bifermentans (Clostridium bifermentans). Bacteria; Firmicutes; Clostridia; Clostridiales;
B2H97__locus_1
EUT2I
00535 -# -
00540 -pf06725pf08239 -
00545 -pf01758 -
00550 -pf10728pf10727 -
00555 -pf02548 -
00560 -pf02569 -
00565 -pf01098 -
00570 HMMreg00072_2_pdugrm(pf00072pf00486) EUT2E
00575 HMMreg00486_1_mix(pf02518pf00512) PDU1F
00580 HMM00465alcdh_3_mix(pf00465) EUT2I
00585 Hp_euts_fadedPurple EUT2I
00590 HMM10662PduV_1_mix(pf10662) EUT2I
00595 HMMreg00072_1_eut2(pf03861pf00072) EUT2I
00600 HMMreg07568_1_eut2(pf12282pf07568) EUT2I
00605 HMM06277EutA_1_eut(pf06277) EUT2I
00610 HMMetlyEutB_2_EUT2(pf06751) EUT2I
00615 HMMetlyEutC_2_eut2(pf05985) EUT2I EP_NTERM
00620 Tsp_darkPeach EUT2I
00625 H_fireEngine EUT2I
00630 HMMalddh_2_egra(pf00171) EUT2I
00635 H_fuchsia EUT2I
00640 HMM01923cobtr_3_eut2(pf01923) EUT2I EP_NTERM
00645 HMMptac_3_eut2(pf06130) EUT2I EP_NTERM
00650 HMM02441_06(#) EUT2I EP_NTERM
00655 P_azul EUT2I
00660 Ts_mushroom EUT2I
00665 HMMtpEuth_1_eut(pf04346) EUT2I
00670 HMM06249EutQ_2_eut2(pf06249) EUT2I EP_NTERM
00675 -pf01471pf08486 -
00680 HMMreg00072_4_pdu1d(pf02518pf00512) PDU1F
00685 -# -
00690 -pf03176 -
00695 -pf00440 -
00700 -pf00183 -
00705 -# -
00710 -# -
BMC-P
BMC-H
BMC-T
Aldehyde dehydrogenase
PTAC
Signature enzyme
Alcohol dehydrogenase
Regulator

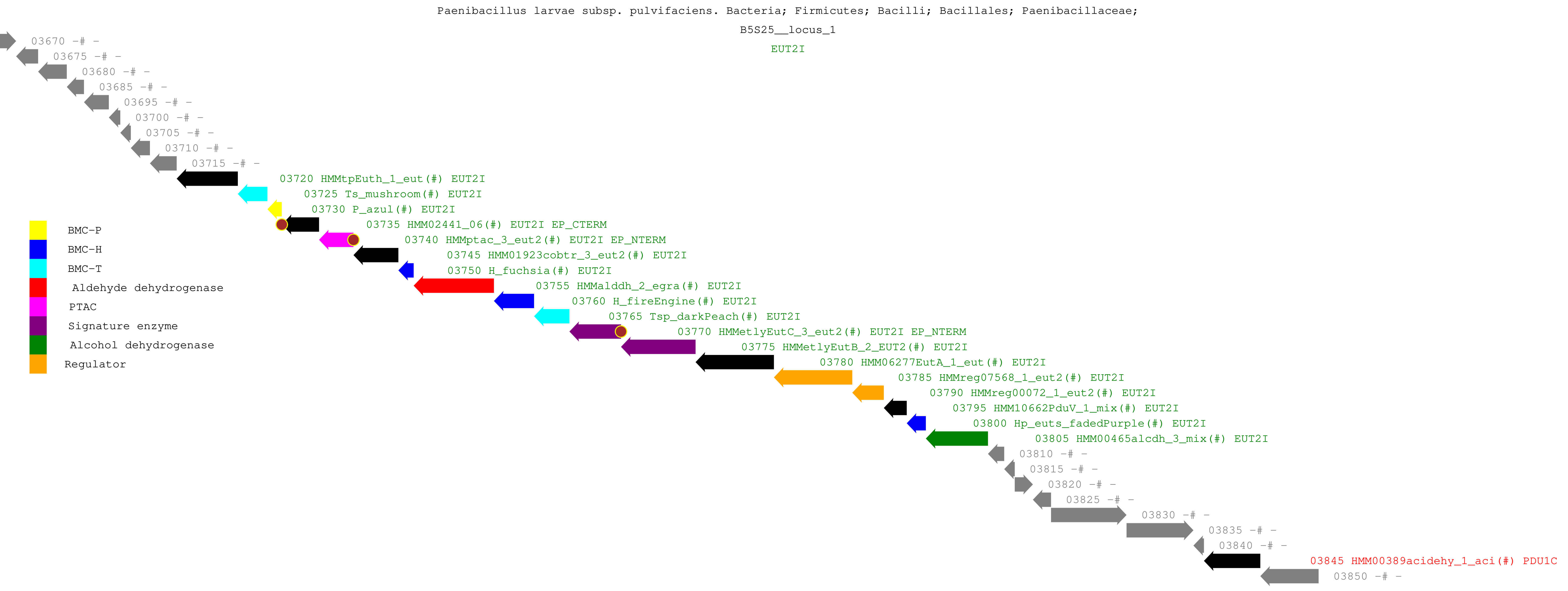
Paenibacillus larvae subsp. pulvifaciens. Bacteria; Firmicutes; Bacilli; Bacillales; Paenibacillaceae;
B5S25__locus_1
EUT2I
03670 -# -
03675 -# -
03680 -# -
03685 -# -
03695 -# -
03700 -# -
03705 -# -
03710 -# -
03715 -# -
03720 HMMtpEuth_1_eut(#) EUT2I
03725 Ts_mushroom(#) EUT2I
03730 P_azul(#) EUT2I
03735 HMM02441_06(#) EUT2I EP_CTERM
03740 HMMptac_3_eut2(#) EUT2I EP_NTERM
03745 HMM01923cobtr_3_eut2(#) EUT2I
03750 H_fuchsia(#) EUT2I
03755 HMMalddh_2_egra(#) EUT2I
03760 H_fireEngine(#) EUT2I
03765 Tsp_darkPeach(#) EUT2I
03770 HMMetlyEutC_3_eut2(#) EUT2I EP_NTERM
03775 HMMetlyEutB_2_EUT2(#) EUT2I
03780 HMM06277EutA_1_eut(#) EUT2I
03785 HMMreg07568_1_eut2(#) EUT2I
03790 HMMreg00072_1_eut2(#) EUT2I
03795 HMM10662PduV_1_mix(#) EUT2I
03800 Hp_euts_fadedPurple(#) EUT2I
03805 HMM00465alcdh_3_mix(#) EUT2I
03810 -# -
03815 -# -
03820 -# -
03825 -# -
03830 -# -
03835 -# -
03840 -# -
03845 HMM00389acidehy_1_aci(#) PDU1C
03850 -# -
BMC-P
BMC-H
BMC-T
Aldehyde dehydrogenase
PTAC
Signature enzyme
Alcohol dehydrogenase
Regulator

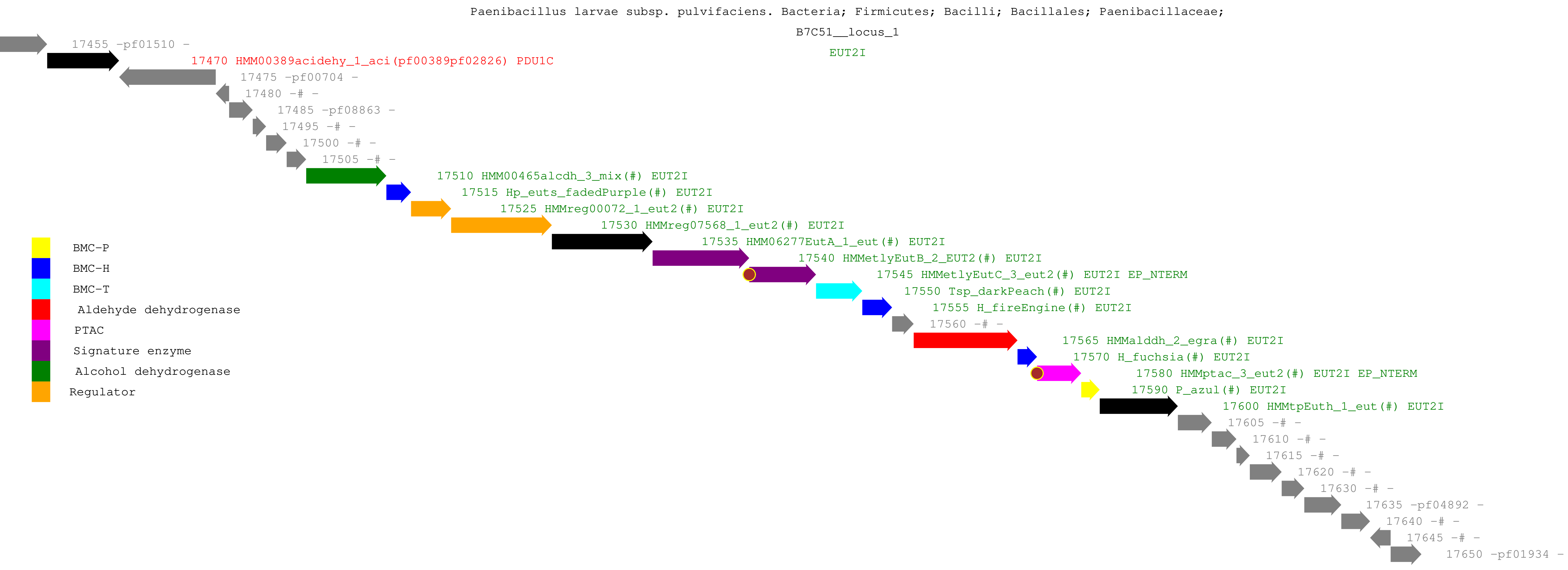

Paenibacillus larvae subsp. pulvifaciens. Bacteria; Firmicutes; Bacilli; Bacillales; Paenibacillaceae;
B7C51__locus_1
EUT2I
17455 -pf01510 -
17470 HMM00389acidehy_1_aci(pf00389pf02826) PDU1C
17475 -pf00704 -
17480 -# -
17485 -pf08863 -
17495 -# -
17500 -# -
17505 -# -
17510 HMM00465alcdh_3_mix(#) EUT2I
17515 Hp_euts_fadedPurple(#) EUT2I
17525 HMMreg00072_1_eut2(#) EUT2I
17530 HMMreg07568_1_eut2(#) EUT2I
17535 HMM06277EutA_1_eut(#) EUT2I
17540 HMMetlyEutB_2_EUT2(#) EUT2I
17545 HMMetlyEutC_3_eut2(#) EUT2I EP_NTERM
17550 Tsp_darkPeach(#) EUT2I
17555 H_fireEngine(#) EUT2I
17560 -# -
17565 HMMalddh_2_egra(#) EUT2I
17570 H_fuchsia(#) EUT2I
17580 HMMptac_3_eut2(#) EUT2I EP_NTERM
17590 P_azul(#) EUT2I
17600 HMMtpEuth_1_eut(#) EUT2I
17605 -# -
17610 -# -
17615 -# -
17620 -# -
17630 -# -
17635 -pf04892 -
17640 -# -
17645 -# -
17650 -pf01934 -
BMC-P
BMC-H
BMC-T
Aldehyde dehydrogenase
PTAC
Signature enzyme
Alcohol dehydrogenase
Regulator

Clostridium drakei. Bacteria; Firmicutes; Clostridia; Clostridiales; Clostridiaceae;

B9W14__locus_1

EUT2I

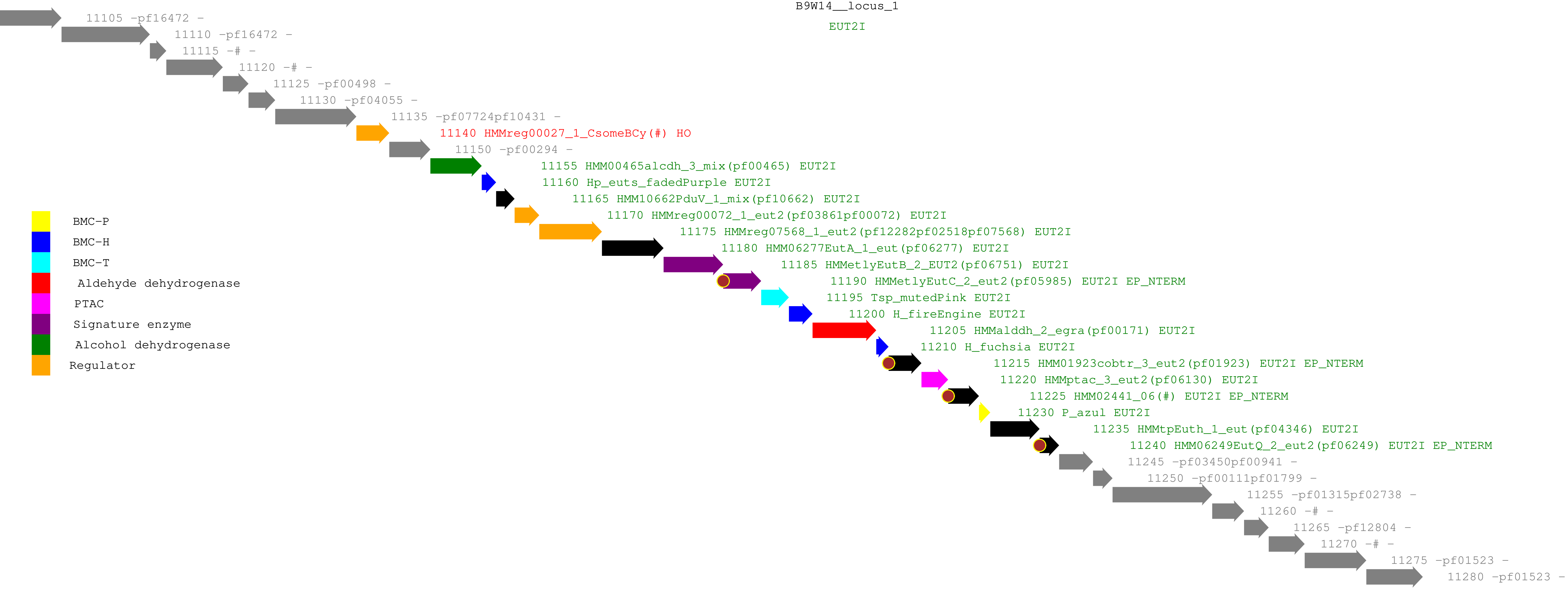

Paenibacillus sp. KS1. Bacteria; Firmicutes; Bacilli; Bacillales; Paenibacillaceae;

BBG47__locus_1

EUT2I

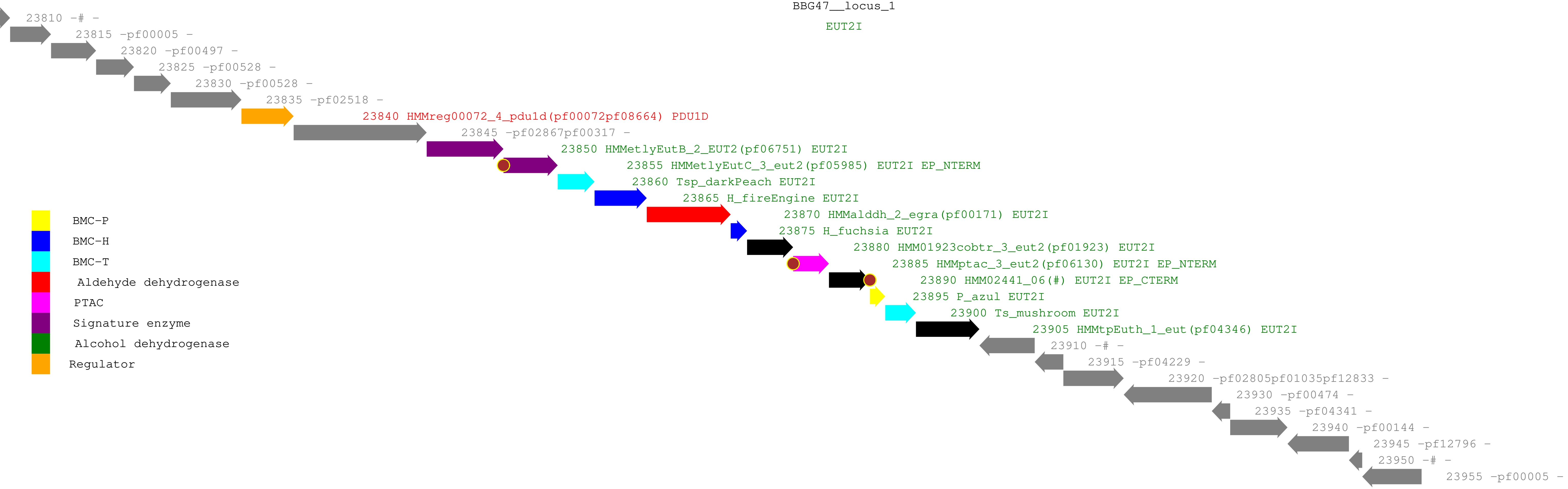

Clostridium algidicarnis DSM 15099. Bacteria; Firmicutes; Clostridia; Clostridiales; Clostridiaceae;

BD821__locus_1

EUT2I

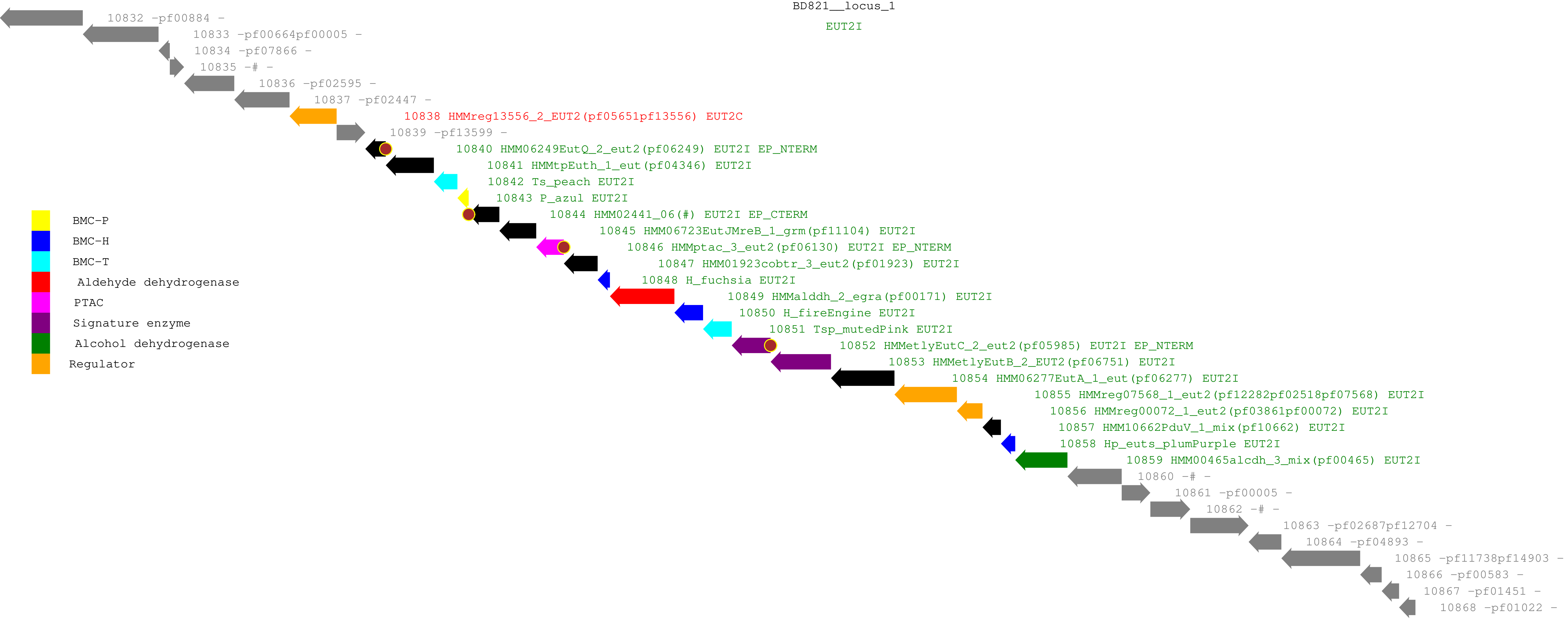

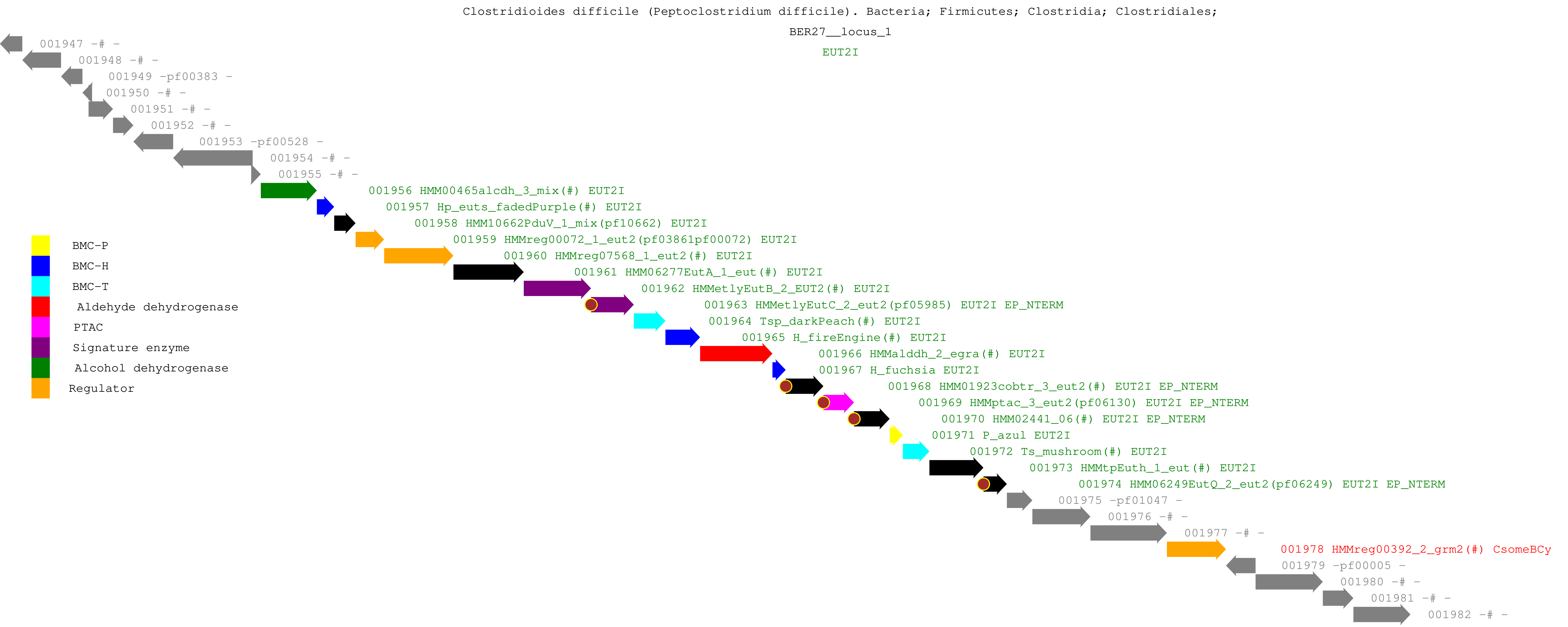
Clostridioides difficile (Peptoclostridium difficile). Bacteria; Firmicutes; Clostridia; Clostridiales;
BER27__locus_1
EUT2I
001947 -# -
001948 -# -
001949 -pf00383 -
001950 -# -
001951 -# -
001952 -# -
001953 -pf00528 -
001954 -# -
001955 -# -
001956 HMM00465alcdh_3_mix(#) EUT2I
001957 Hp_euts_fadedPurple(#) EUT2I
001958 HMM10662PduV_1_mix(pf10662) EUT2I
001959 HMMreg00072_1_eut2(pf03861pf00072) EUT2I
001960 HMMreg07568_1_eut2(#) EUT2I
001961 HMM06277EutA_1_eut(#) EUT2I
001962 HMMetlyEutB_2_EUT2(#) EUT2I
001963 HMMetlyEutC_2_eut2(pf05985) EUT2I EP_NTERM
001964 Tsp_darkPeach(#) EUT2I
001965 H_fireEngine(#) EUT2I
001966 HMMalddh_2_egra(#) EUT2I
001967 H_fuchsia EUT2I
001968 HMM01923cobtr_3_eut2(#) EUT2I EP_NTERM
001969 HMMptac_3_eut2(pf06130) EUT2I EP_NTERM
001970 HMM02441_06(#) EUT2I EP_NTERM
001971 P_azul EUT2I
001972 Ts_mushroom(#) EUT2I
001973 HMMtpEuth_1_eut(#) EUT2I
001974 HMM06249EutQ_2_eut2(pf06249) EUT2I EP_NTERM
001975 -pf01047 -
001976 -# -
001977 -# -
001978 HMMreg00392_2_grm2(#) CsomeBCy
001979 -pf00005 -
001980 -# -
001981 -# -
001982 -# -
BMC-P
BMC-H
BMC-T
Aldehyde dehydrogenase
PTAC
Signature enzyme
Alcohol dehydrogenase
Regulator

Clostridioides difficile (Peptoclostridium difficile). Bacteria; Firmicutes; Clostridia; Clostridiales;

BER46__locus_1

EUT2I

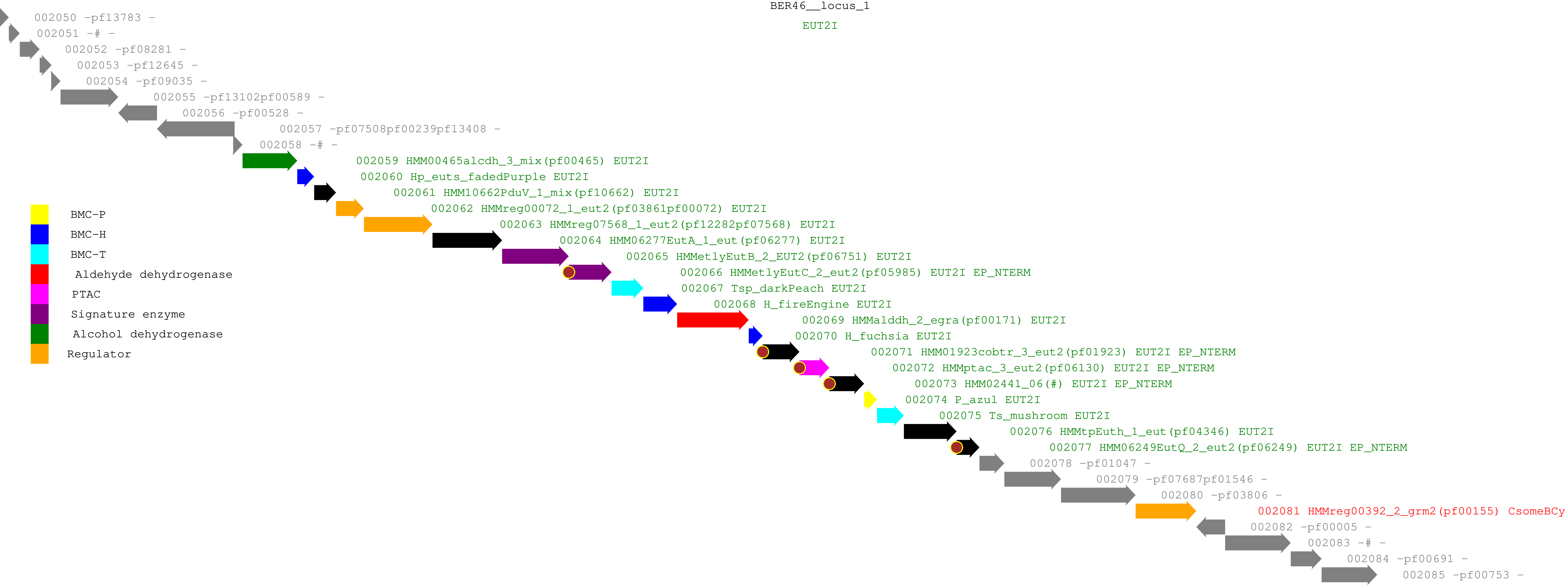

Clostridioides difficile (Peptoclostridium difficile). Bacteria; Firmicutes; Clostridia; Clostridiales;

BGU81__locus_1

EUT2I

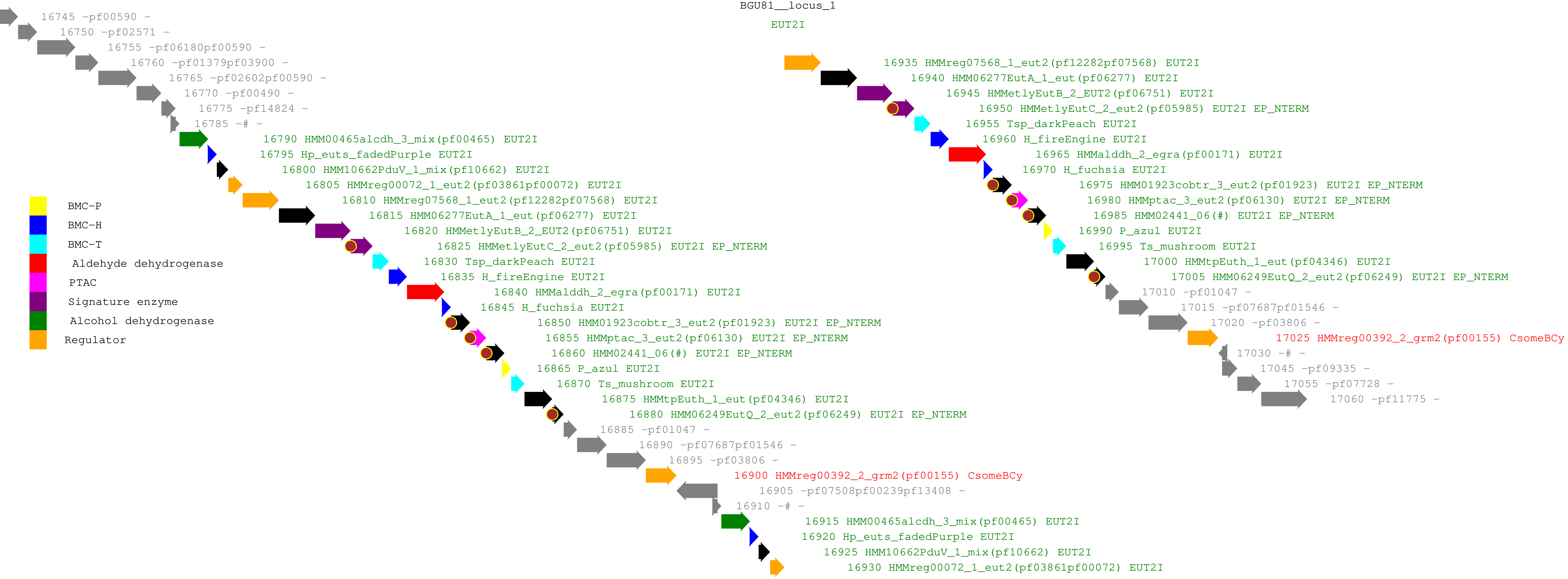

Paenibacillus lautus (Bacillus lautus). Bacteria; Firmicutes; Bacilli; Bacillales; Paenibacillaceae;

BK123__locus_1

EUT2I

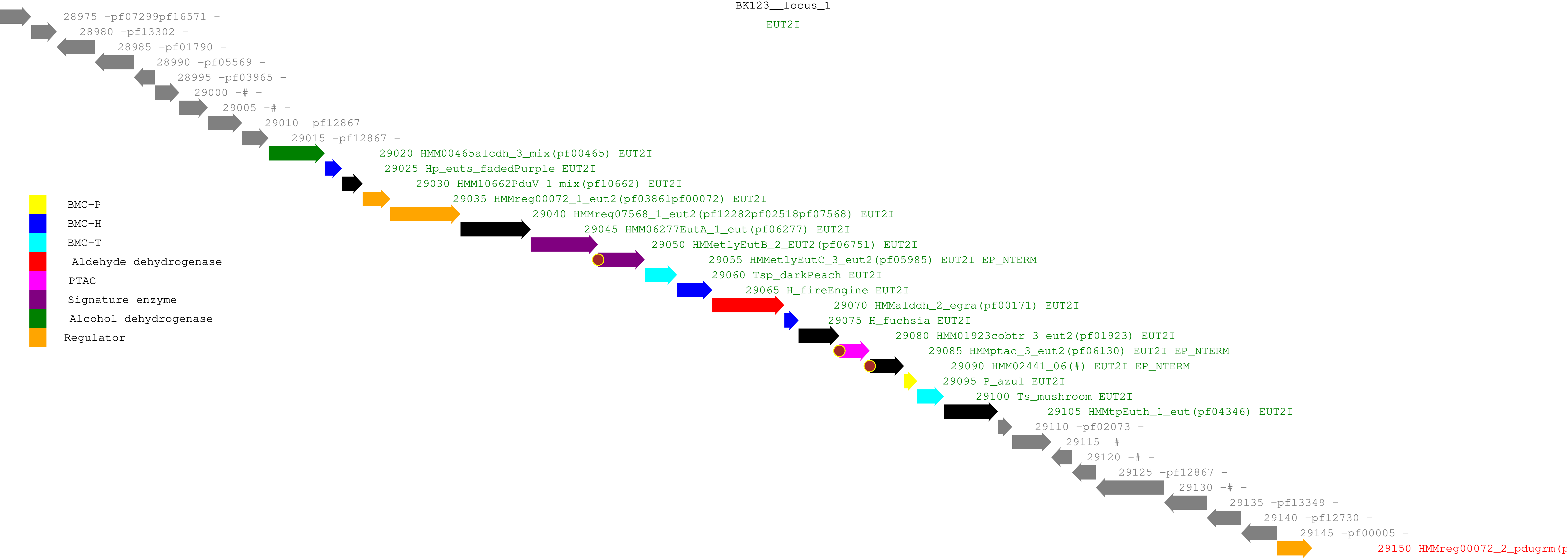

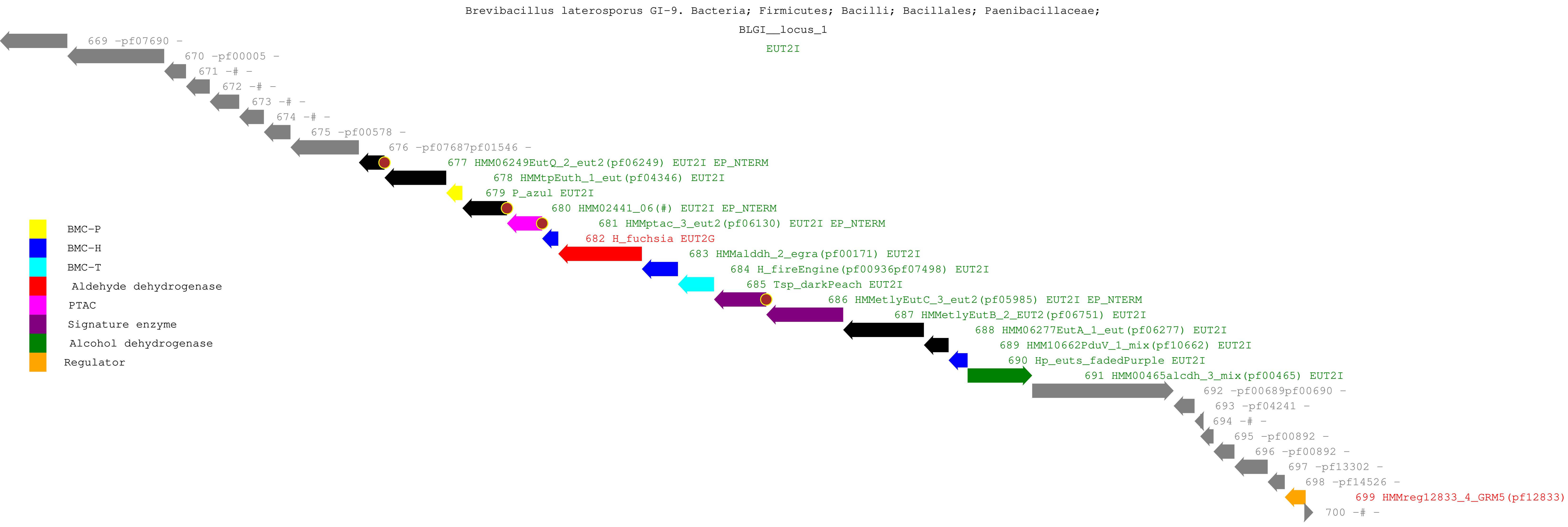
Brevibacillus laterosporus GI-9. Bacteria; Firmicutes; Bacilli; Bacillales; Paenibacillaceae;
BLGI__locus_1
EUT2I
669 -pf07690 -
670 -pf00005 -
671 -# -
672 -# -
673 -# -
674 -# -
675 -pf00578 -
676 -pf07687pf01546 -
677 HMM06249EutQ_2_eut2(pf06249) EUT2I EP_NTERM
678 HMMtpEuth_1_eut(pf04346) EUT2I
679 P_azul EUT2I
680 HMM02441_06(#) EUT2I EP_NTERM
681 HMMptac_3_eut2(pf06130) EUT2I EP_NTERM
682 H_fuchsia EUT2G
683 HMMaldh_2_egra(pf00171) EUT2I
684 H_fireEngine(pf00936pf07498) EUT2I
685 Tsp_darkPeach EUT2I
686 HMMetlyEutC_3_eut2(pf05985) EUT2I EP_NTERM
687 HMMetlyEutB_2_EUT2(pf06751) EUT2I
688 HMM06277EutA_1_eut(pf06277) EUT2I
689 HMM10662PduV_1_mix(pf10662) EUT2I
690 Hp_euts_fadedPurple EUT2I
691 HMM00465alcdh_3_mix(pf00465) EUT2I
692 -pf00689pf00690 -
693 -pf04241 -
694 -# -
695 -pf00892 -
696 -pf00892 -
697 -pf13302 -
698 -pf14526 -
699 HMMreg12833_4_GRM5(pf12833)
700 -# -
BMC-P
BMC-H
BMC-T
Aldehyde dehydrogenase
PTAC
Signature enzyme
Alcohol dehydrogenase
Regulator

Brevibacillus laterosporus GI-9. Bacteria; Firmicutes; Bacilli; Bacillales; Paenibacillaceae;

BLGI__locus_2

EUT2I

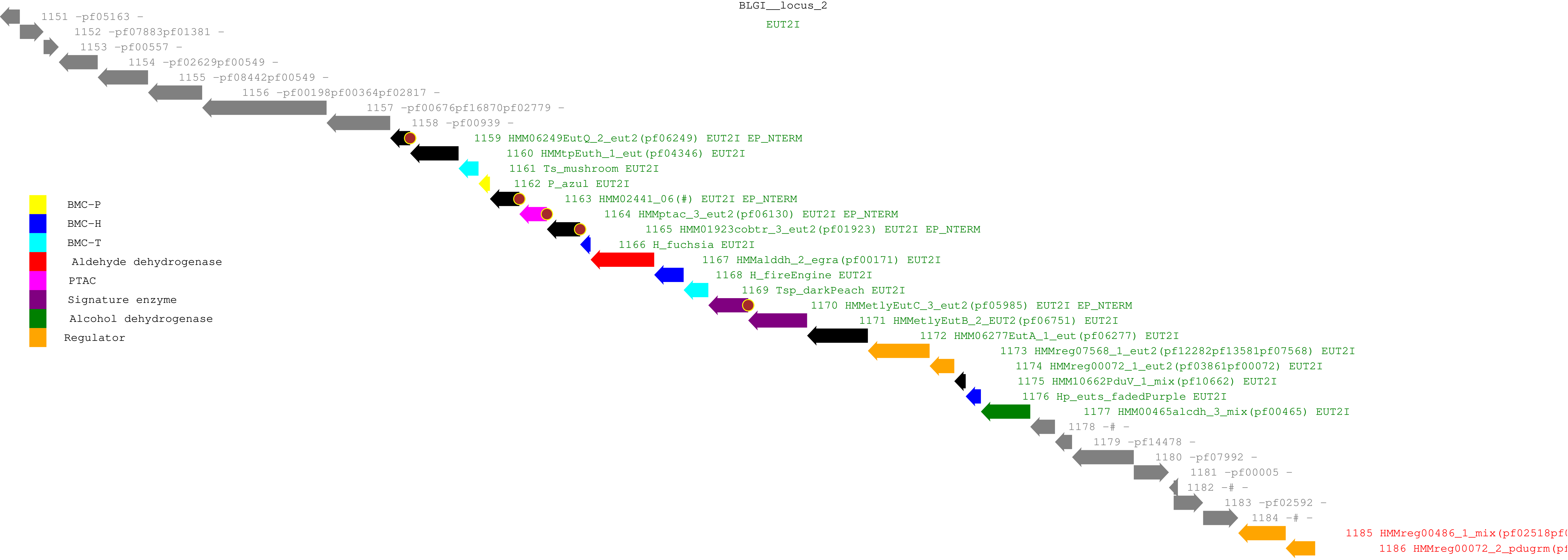

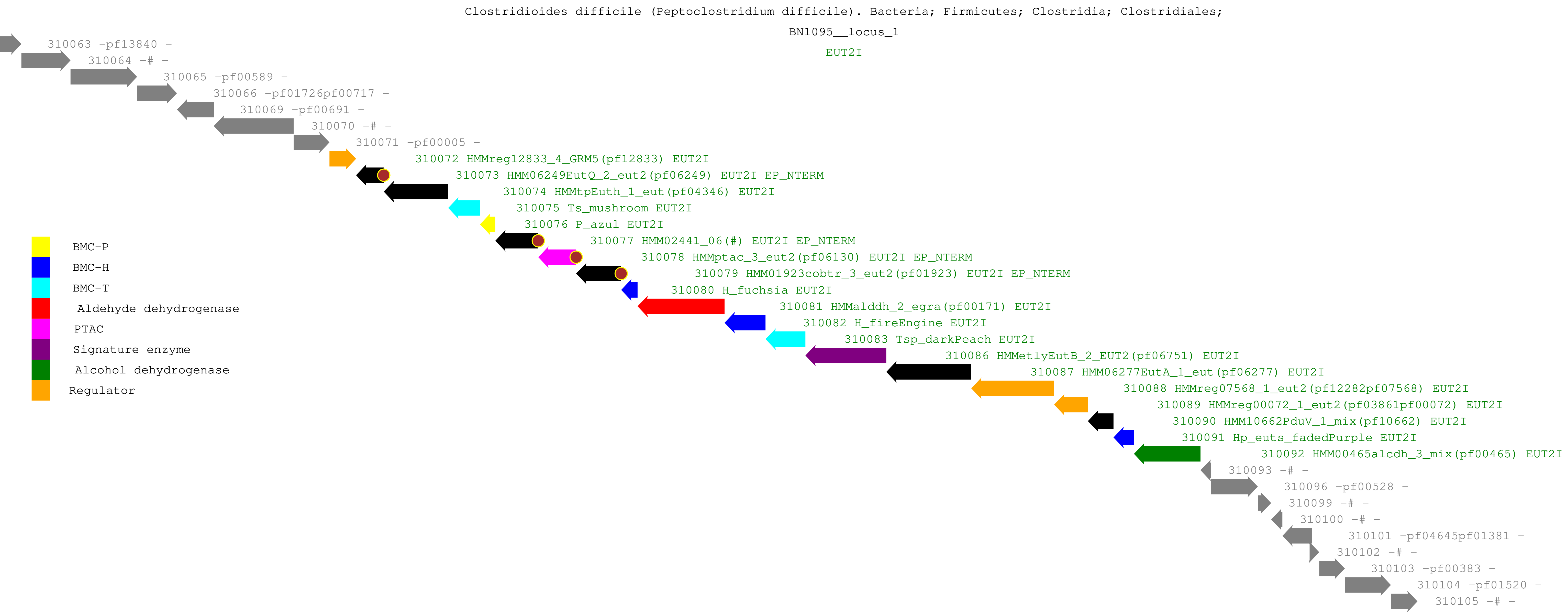
Clostridioides difficile (Peptoclostridium difficile). Bacteria; Firmicutes; Clostridia; Clostridiales;
BN1095__locus_1
EUT2I
310063 -pf13840 -
310064 -# -
310065 -pf00589 -
310066 -pf01726pf00717 -
310069 -pf00691 -
310070 -# -
310071 -pf00005 -
310072 HMMreg12833_4_GRM5(pf12833) EUT2I
310073 HMM06249EutQ_2_eut2(pf06249) EUT2I EP_NTERM
310074 HMMtpEuth_1_eut(pf04346) EUT2I
310075 Ts_mushroom EUT2I
310076 P_azul EUT2I
310077 HMM02441_06(#) EUT2I EP_NTERM
310078 HMMptac_3_eut2(pf06130) EUT2I EP_NTERM
310079 HMM01923cobtr_3_eut2(pf01923) EUT2I EP_NTERM
310080 H_fuchsia EUT2I
310081 HMMalddh_2_egra(pf00171) EUT2I
310082 H_fireEngine EUT2I
310083 Tsp_darkPeach EUT2I
310086 HMMetlyEutB_2_EUT2(pf06751) EUT2I
310087 HMM06277EutA_1_eut(pf06277) EUT2I
310088 HMMreg07568_1_eut2(pf12282pf07568) EUT2I
310089 HMMreg00072_1_eut2(pf03861pf00072) EUT2I
310090 HMM10662PduV_1_mix(pf10662) EUT2I
310091 Hp_euts_fadedPurple EUT2I
310092 HMM00465alcdh_3_mix(pf00465) EUT2I
310093 -# -
310096 -pf00528 -
310099 -# -
310100 -# -
310101 -pf04645pf01381 -
310102 -# -
310103 -pf00383 -
310104 -pf01520 -
310105 -# -
BMC-P
BMC-H
BMC-T
Aldehyde dehydrogenase
PTAC
Signature enzyme
Alcohol dehydrogenase
Regulator

Clostridioides difficile (Peptoclostridium difficile). Bacteria; Firmicutes; Clostridia; Clostridiales;

BN1096__locus_1

EUT2I

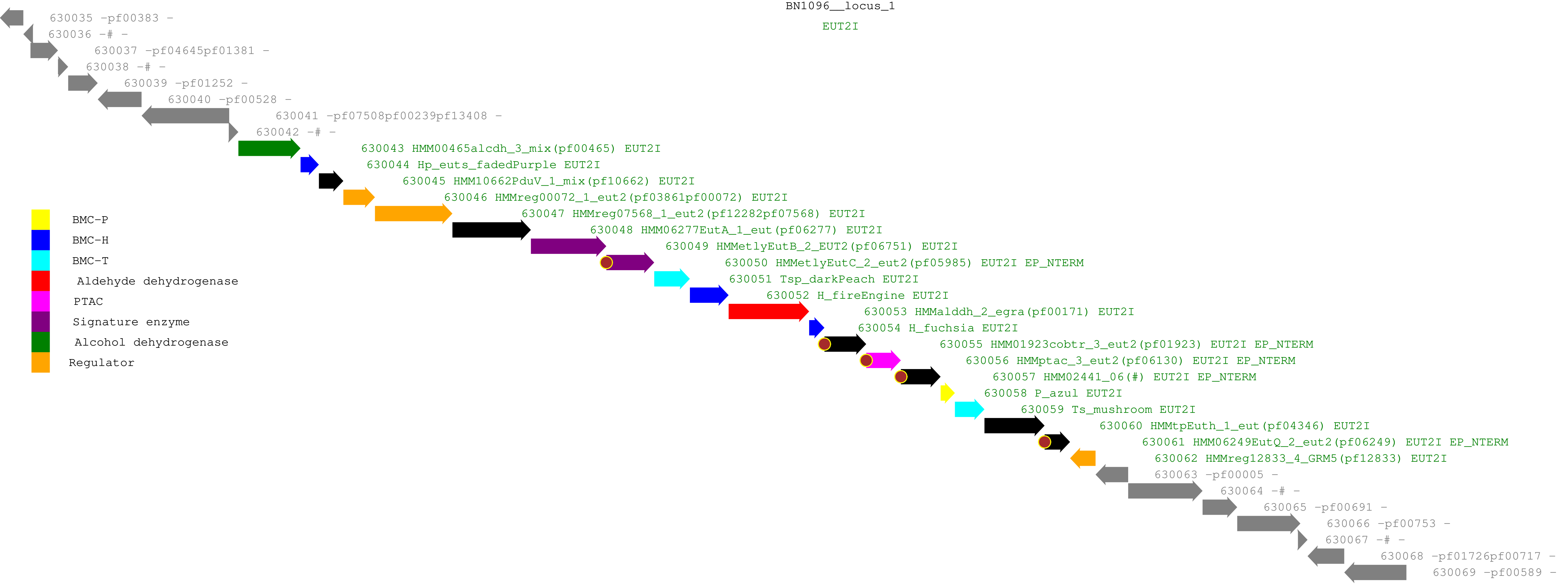

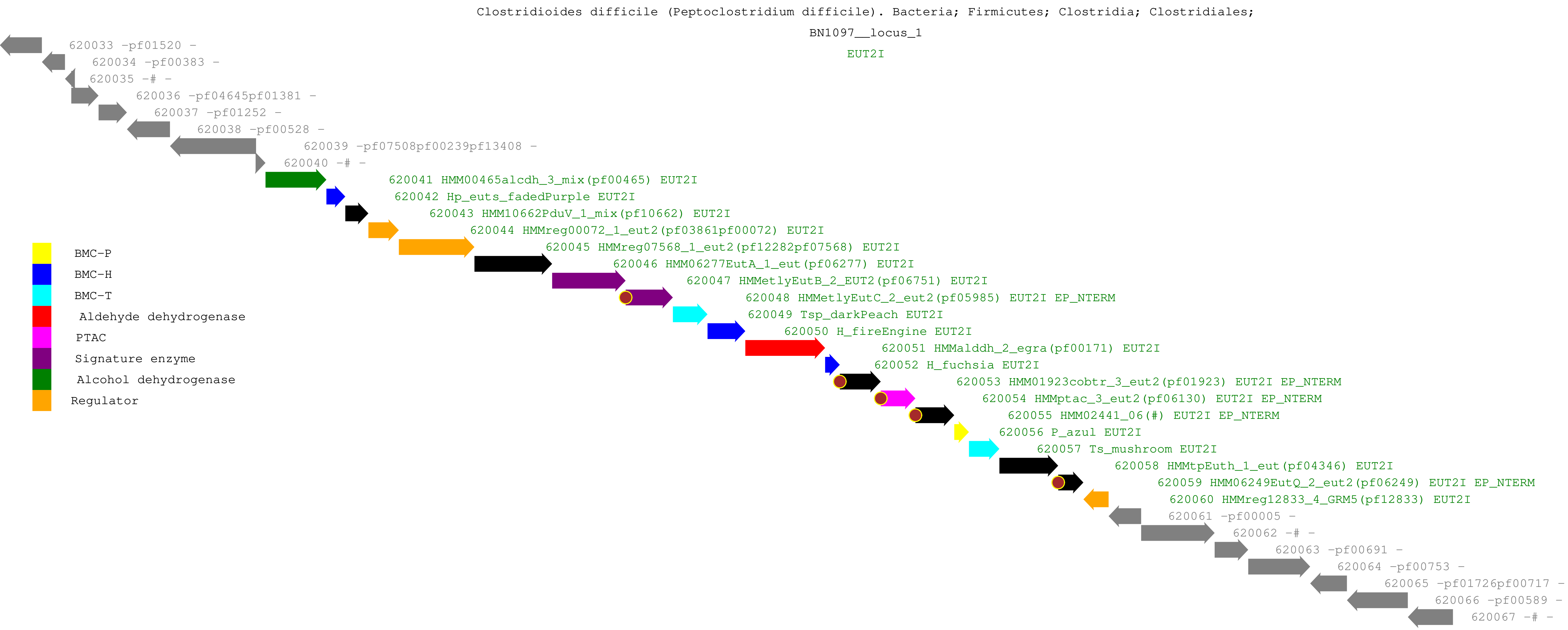
Clostridioides difficile (Peptoclostridium difficile). Bacteria; Firmicutes; Clostridia; Clostridiales;
BN1097__locus_1
EUT2I
620033 -pf01520 -
620034 -pf00383 -
620035 -# -
620036 -pf04645pf01381 -
620037 -pf01252 -
620038 -pf00528 -
620039 -pf07508pf00239pf13408 -
620040 -# -
620041 HMM00465alcdh_3_mix(pf00465) EUT2I
620042 Hp_euts_fadedPurple EUT2I
620043 HMM10662PduV_1_mix(pf10662) EUT2I
620044 HMMreg00072_1_eut2(pf03861pf00072) EUT2I
620045 HMMreg07568_1_eut2(pf12282pf07568) EUT2I
620046 HMM06277EutA_1_eut(pf06277) EUT2I
620047 HMMetlyEutB_2_EUT2(pf06751) EUT2I
620048 HMMetlyEutC_2_eut2(pf05985) EUT2I EP_NTERM
620049 Tsp_darkPeach EUT2I
620050 H_fireEngine EUT2I
620051 HMMalddh_2_egra(pf00171) EUT2I
620052 H_fuchsia EUT2I
620053 HMM01923cobtr_3_eut2(pf01923) EUT2I EP_NTERM
620054 HMMptac_3_eut2(pf06130) EUT2I EP_NTERM
620055 HMM02441_06(#) EUT2I EP_NTERM
620056 P_azul EUT2I
620057 Ts_mushroom EUT2I
620058 HMMtpEuth_1_eut(pf04346) EUT2I
620059 HMM06249EutQ_2_eut2(pf06249) EUT2I EP_NTERM
620060 HMMreg12833_4_GRM5(pf12833) EUT2I
620061 -pf00005 -
620062 -# -
620063 -pf00691 -
620064 -pf00753 -
620065 -pf01726pf00717 -
620066 -pf00589 -
620067 -# -
BMC-P
BMC-H
BMC-T
Aldehyde dehydrogenase
PTAC
Signature enzyme
Alcohol dehydrogenase
Regulator

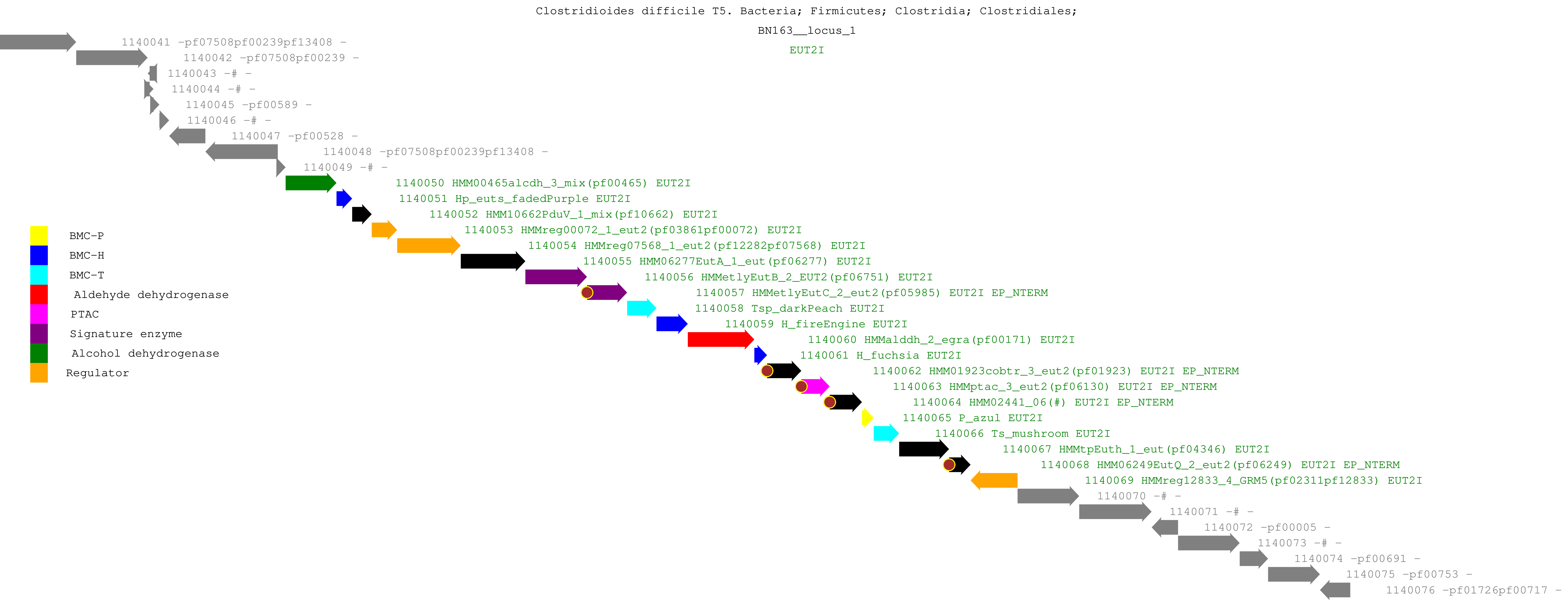

Clostridioides difficile T5. Bacteria; Firmicutes; Clostridia; Clostridiales;
BN163__locus_1
EUT2I
1140041 -pf07508pf00239pf13408 -
1140042 -pf07508pf00239 -
1140043 -# -
1140044 -# -
1140045 -pf00589 -
1140046 -# -
1140047 -pf00528 -
1140048 -pf07508pf00239pf13408 -
1140049 -# -
1140050 HMM00465alcdh_3_mix(pf00465) EUT2I
1140051 Hp_euts_fadedPurple EUT2I
1140052 HMM10662PduV_1_mix(pf10662) EUT2I
1140053 HMMreg00072_1_eut2(pf03861pf00072) EUT2I
1140054 HMMreg07568_1_eut2(pf12282pf07568) EUT2I
1140055 HMM06277EutA_1_eut(pf06277) EUT2I
1140056 HMMetlyEutB_2_EUT2(pf06751) EUT2I
1140057 HMMetlyEutC_2_eut2(pf05985) EUT2I EP_NTERM
1140058 Tsp_darkPeach EUT2I
1140059 H_fireEngine EUT2I
1140060 HMMaldh_2_egra(pf00171) EUT2I
1140061 H_fuchsia EUT2I
1140062 HMM01923cobtr_3_eut2(pf01923) EUT2I EP_NTERM
1140063 HMMptac_3_eut2(pf06130) EUT2I EP_NTERM
1140064 HMM02441_06(#) EUT2I EP_NTERM
1140065 P_azul EUT2I
1140066 Ts_mushroom EUT2I
1140067 HMMtpEuth_1_eut(pf04346) EUT2I
1140068 HMM06249EutQ_2_eut2(pf06249) EUT2I EP_NTERM
1140069 HMMreg12833_4_GRM5(pf02311pf12833) EUT2I
1140070 -# -
1140071 -# -
1140072 -pf00005 -
1140073 -# -
1140074 -pf00691 -
1140075 -pf00753 -
1140076 -pf01726pf00717 -
BMC-P
BMC-H
BMC-T
Aldehyde dehydrogenase
PTAC
Signature enzyme
Alcohol dehydrogenase
Regulator

Clostridioides difficile E1. Bacteria; Firmicutes; Clostridia; Clostridiales;

BN165__locus_1

EUT2I

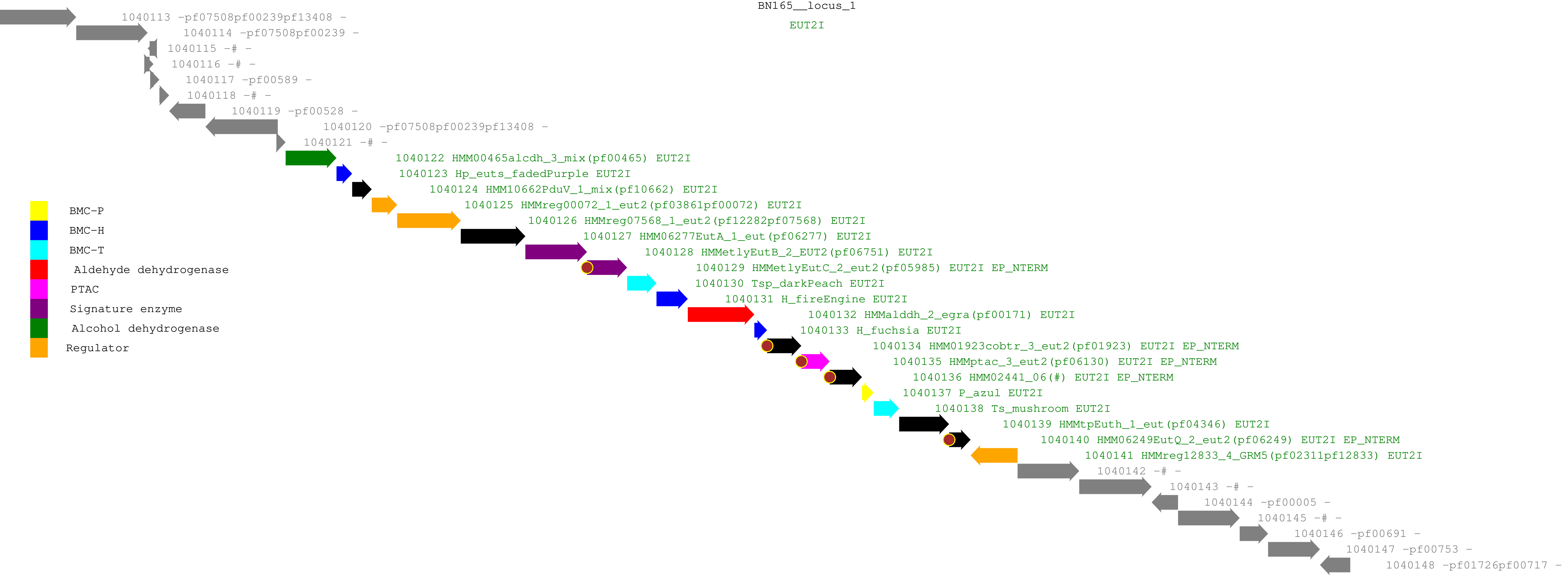

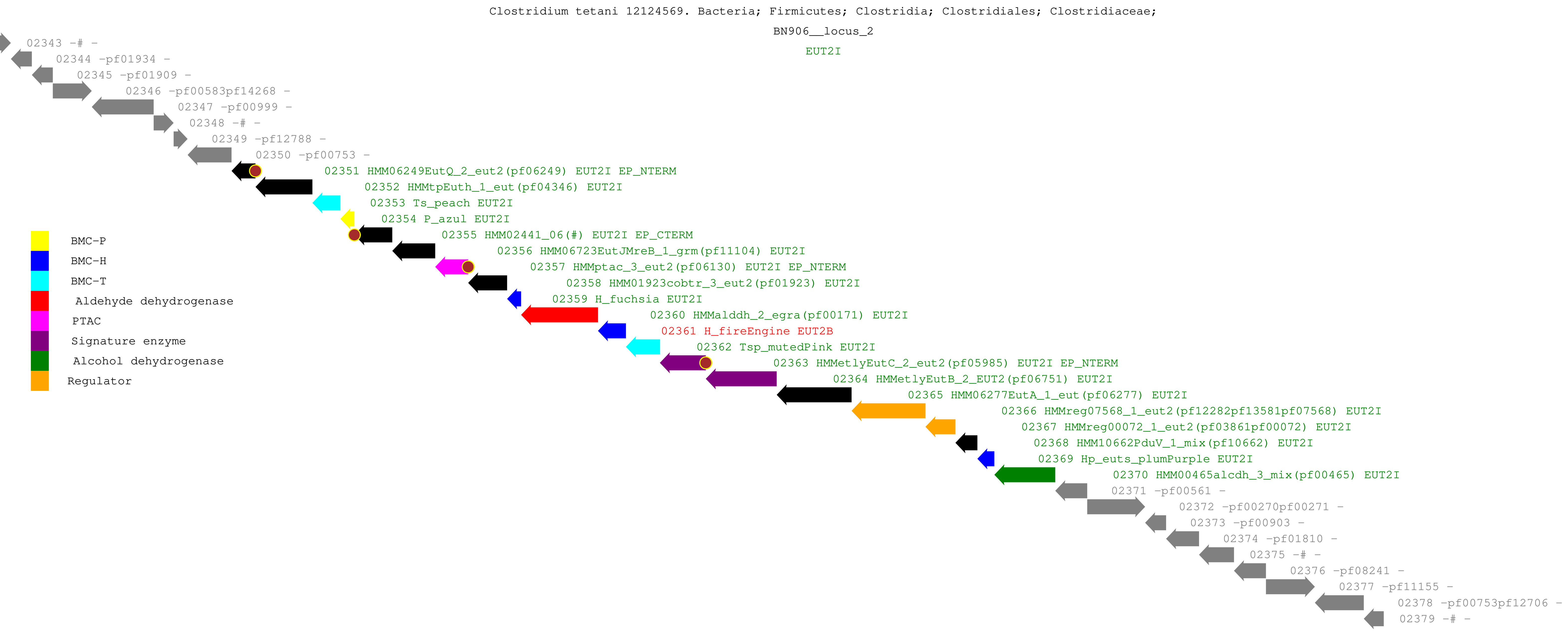

Clostridium tetani 12124569. Bacteria; Firmicutes; Clostridia; Clostridiales; Clostridiaceae;
BN906__locus_2
EUT2I
02343 -# -
02344 -pf01934 -
02345 -pf01909 -
02346 -pf00583pf14268 -
02347 -pf00999 -
02348 -# -
02349 -pf12788 -
02350 -pf00753 -
02351 HMM06249EutQ_2_eut2(pf06249) EUT2I EP_NTERM
02352 HMMtpEuth_1_eut(pf04346) EUT2I
02353 Ts_peach EUT2I
02354 P_azul EUT2I
02355 HMM02441_06(#) EUT2I EP_CTERM
02356 HMM06723EutJMreB_1_grm(pf11104) EUT2I
02357 HMMptac_3_eut2(pf06130) EUT2I EP_NTERM
02358 HMM01923cobtr_3_eut2(pf01923) EUT2I
02359 H_fuchsia EUT2I
02360 HMMalddh_2_egra(pf00171) EUT2I
02361 H_fireEngine EUT2B
02362 Tsp_mutedPink EUT2I
02363 HMMetlyEutC_2_eut2(pf05985) EUT2I EP_NTERM
02364 HMMetlyEutB_2_EUT2(pf06751) EUT2I
02365 HMM06277EutA_1_eut(pf06277) EUT2I
02366 HMMreg07568_1_eut2(pf12282pf13581pf07568) EUT2I
02367 HMMreg00072_1_eut2(pf03861pf00072) EUT2I
02368 HMM10662PduV_1_mix(pf10662) EUT2I
02369 Hp_euts_plumPurple EUT2I
02370 HMM00465alcdh_3_mix(pf00465) EUT2I
02371 -pf00561 -
02372 -pf00270pf00271 -
02373 -pf00903 -
02374 -pf01810 -
02375 -# -
02376 -pf08241 -
02377 -pf11155 -
02378 -pf00753pf12706 -
02379 -# -
BMC-P
BMC-H
BMC-T
Aldehyde dehydrogenase
PTAC
Signature enzyme
Alcohol dehydrogenase
Regulator

Brevibacillus laterosporus DSM 25. Bacteria; Firmicutes; Bacilli; Bacillales; Paenibacillaceae;

BrL25__locus_1

EUT2I

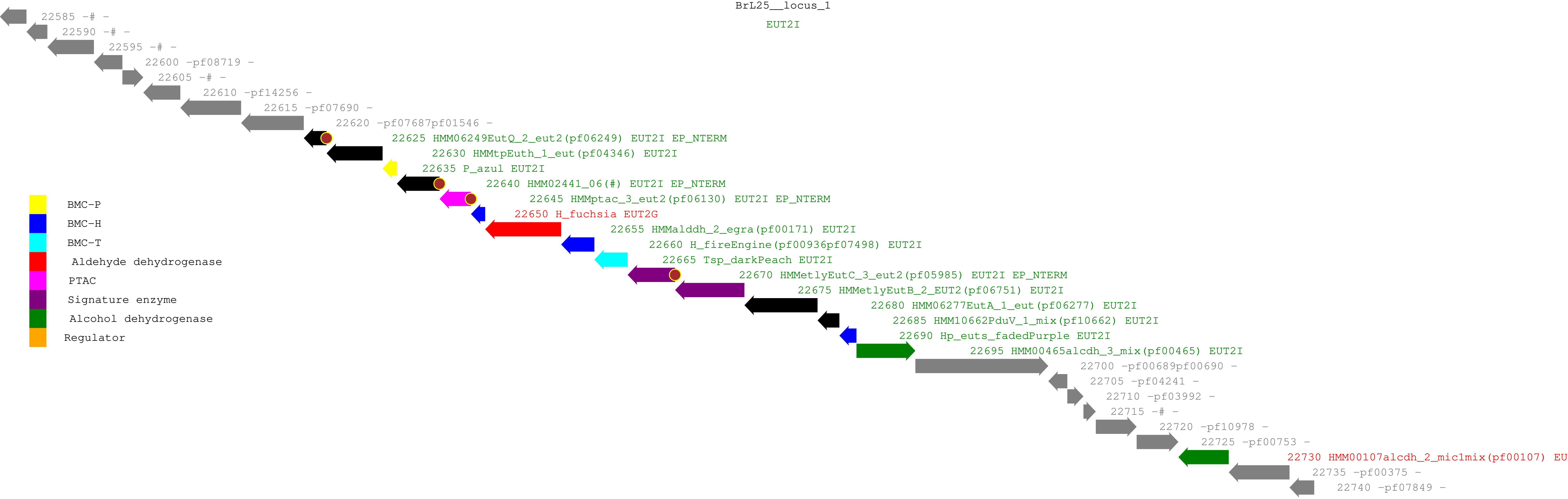

Brevibacillus laterosporus DSM 25. Bacteria; Firmicutes; Bacilli; Bacillales; Paenibacillaceae;

BrL25__locus_2

EUT2I

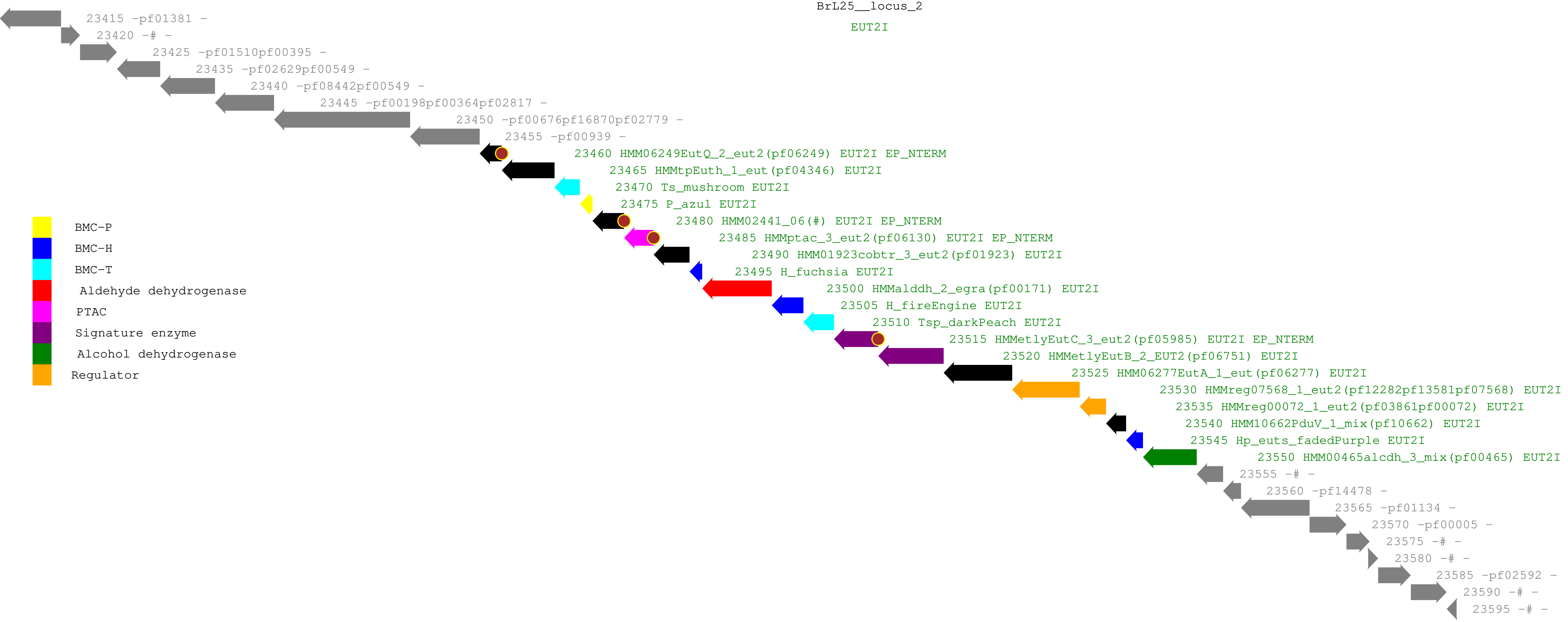

Brevibacillus laterosporus LMG 15441. Bacteria; Firmicutes; Bacilli; Bacillales; Paenibacillaceae;

BRLA_c_locus_1

EUT2I

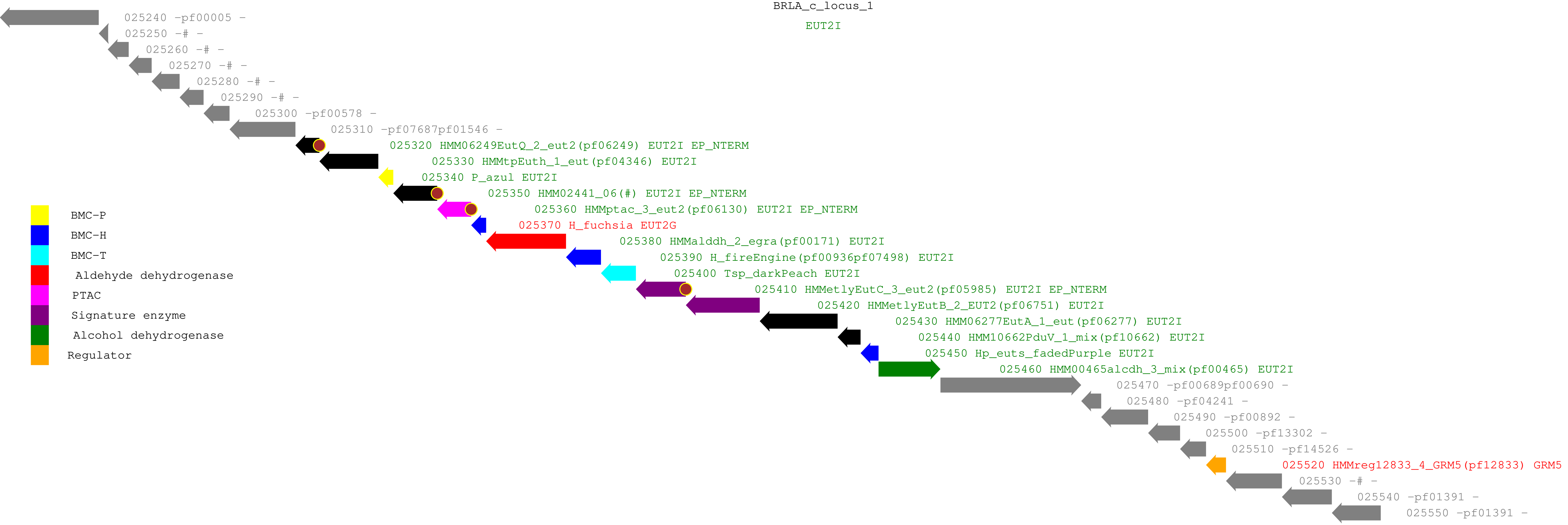

Brevibacillus laterosporus LMG 15441. Bacteria; Firmicutes; Bacilli; Bacillales; Paenibacillaceae;

BRLA_c_locus_2

EUT2I

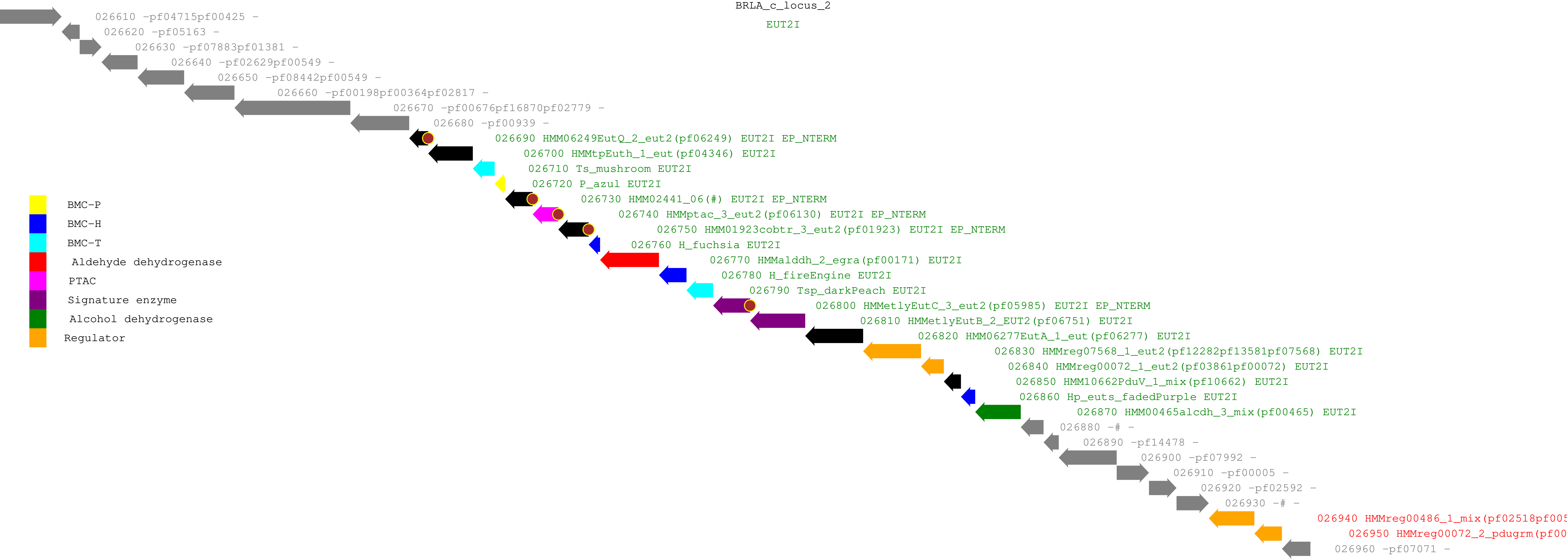

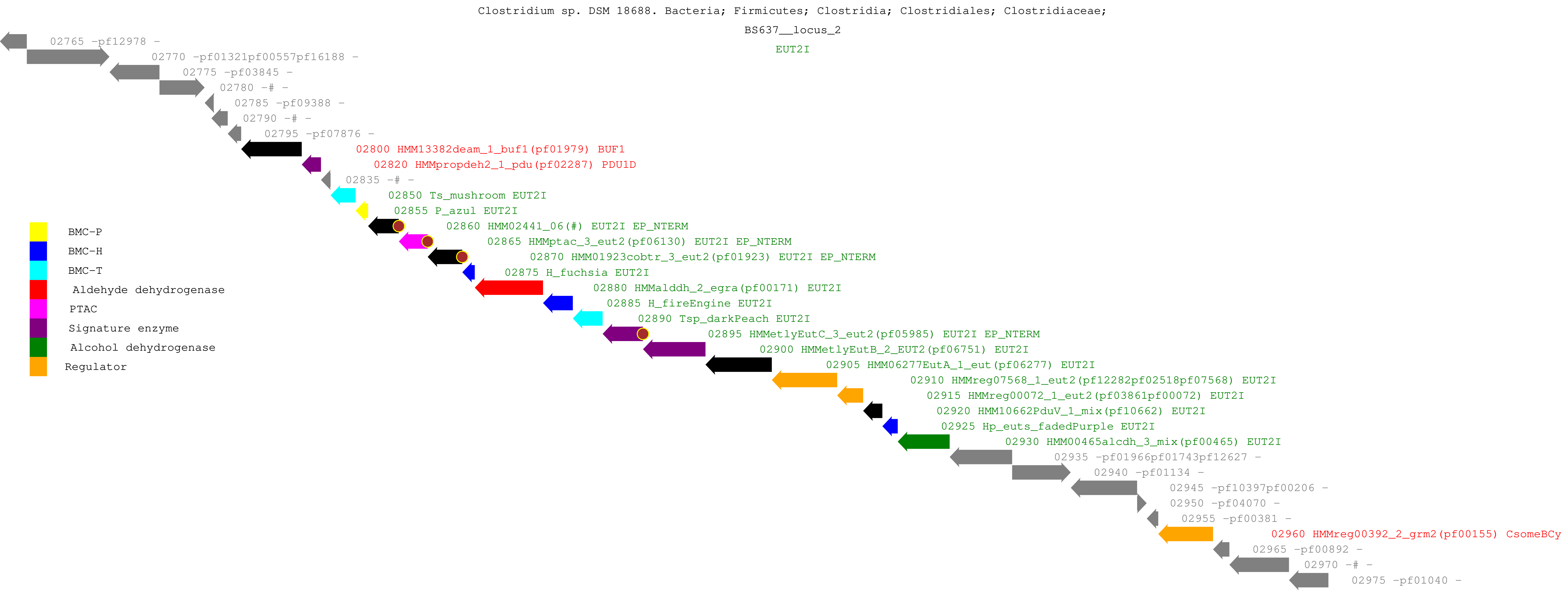
Clostridium sp. DSM 18688. Bacteria; Firmicutes; Clostridia; Clostridiales; Clostridiaceae;
BS637__locus_2
EUT2I
02765 -pf12978 -
02770 -pf01321pf00557pf16188 -
02775 -pf03845 -
02780 -# -
02785 -pf09388 -
02790 -# -
02795 -pf07876 -
02800 HMM13382deam_1_buf1(pf01979) BUF1
02820 HMMpropdeh2_1_pdu(pf02287) PDU1D
02835 -# -
02850 Ts_mushroom EUT2I
02855 P_azul EUT2I
02860 HMM02441_06(#) EUT2I EP_NTERM
02865 HMMptac_3_eut2(pf06130) EUT2I EP_NTERM
02870 HMM01923cobtr_3_eut2(pf01923) EUT2I EP_NTERM
02875 H_fuchsia EUT2I
02880 HMMalddh_2_egra(pf00171) EUT2I
02885 H_fireEngine EUT2I
02890 Tsp_darkPeach EUT2I
02895 HMMetlyEutC_3_eut2(pf05985) EUT2I EP_NTERM
02900 HMMetlyEutB_2_EUT2(pf06751) EUT2I
02905 HMM06277EutA_1_eut(pf06277) EUT2I
02910 HMMreg07568_1_eut2(pf12282pf02518pf07568) EUT2I
02915 HMMreg00072_1_eut2(pf03861pf00072) EUT2I
02920 HMM10662PduV_1_mix(pf10662) EUT2I
02925 Hp_euts_fadedPurple EUT2I
02930 HMM00465alcdh_3_mix(pf00465) EUT2I
02935 -pf01966pf01743pf12627 -
02940 -pf01134 -
02945 -pf10397pf00206 -
02950 -pf04070 -
02955 -pf00381 -
02960 HMMreg00392_2_grm2(pf00155) CsomeBCy
02965 -pf00892 -
02970 -# -
02975 -pf01040 -
BMC-P
BMC-H
BMC-T
Aldehyde dehydrogenase
PTAC
Signature enzyme
Alcohol dehydrogenase
Regulator

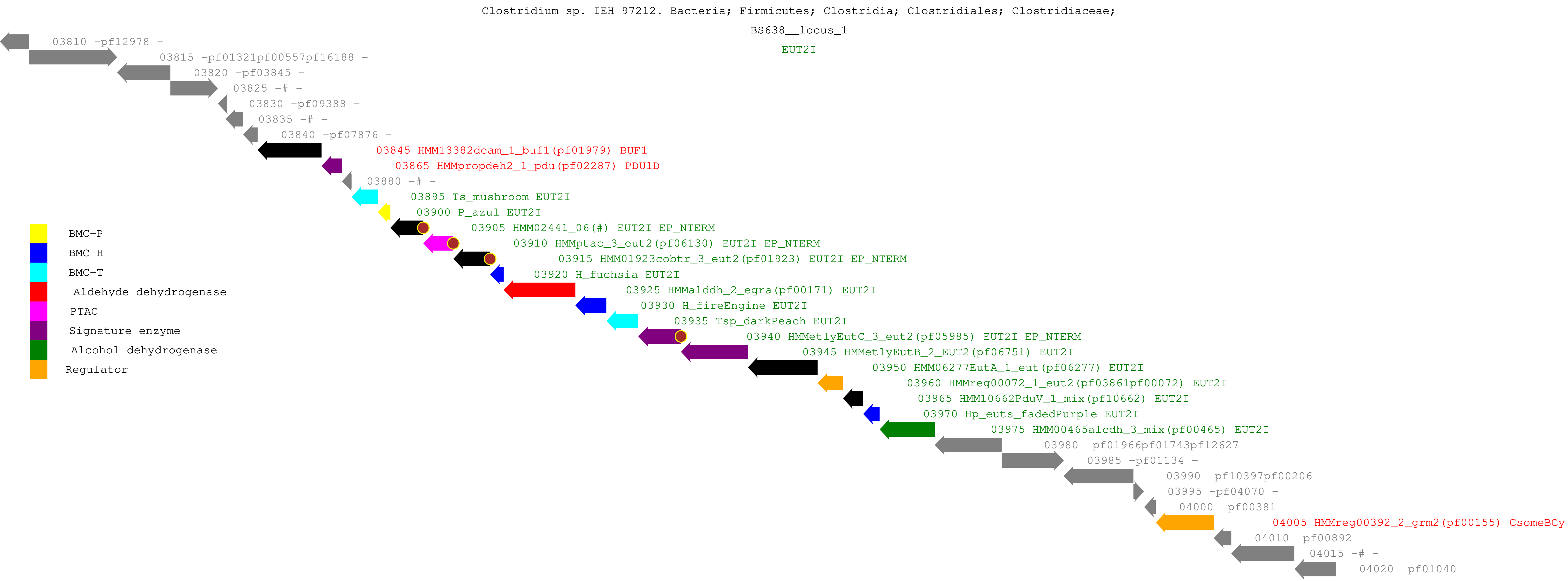
Clostridium sp. IEH 97212. Bacteria; Firmicutes; Clostridia; Clostridiales; Clostridiaceae;
BS638__locus_1
EUT2I
03810 -pf12978 -
03815 -pf01321pf00557pf16188 -
03820 -pf03845 -
03825 -# -
03830 -pf09388 -
03835 -# -
03840 -pf07876 -
03845 HMM13382deam_1_buf1(pf01979) BUF1
03865 HMMpropdeh2_1_pdu(pf02287) PDU1D
03880 -# -
03895 Ts_mushroom EUT2I
03900 P_azul EUT2I
03905 HMM02441_06(#) EUT2I EP_NTERM
03910 HMMptac_3_eut2(pf06130) EUT2I EP_NTERM
03915 HMM01923cobtr_3_eut2(pf01923) EUT2I EP_NTERM
03920 H_fuchsia EUT2I
03925 HMMalddh_2_egra(pf00171) EUT2I
03930 H_fireEngine EUT2I
03935 Tsp_darkPeach EUT2I
03940 HMMetlyEutC_3_eut2(pf05985) EUT2I EP_NTERM
03945 HMMetlyEutB_2_EUT2(pf06751) EUT2I
03950 HMM06277EutA_1_eut(pf06277) EUT2I
03960 HMMreg00072_1_eut2(pf03861pf00072) EUT2I
03965 HMM10662PduV_1_mix(pf10662) EUT2I
03970 Hp_euts_fadedPurple EUT2I
03975 HMM00465alcdh_3_mix(pf00465) EUT2I
03980 -pf01966pf01743pf12627 -
03985 -pf01134 -
03990 -pf10397pf00206 -
03995 -pf04070 -
04000 -pf00381 -
04005 HMMreg00392_2_grm2(pf00155) CsomeBCy
04010 -pf00892 -
04015 -# -
04020 -pf01040 -
BMC-P
BMC-H
BMC-T
Aldehyde dehydrogenase
PTAC
Signature enzyme
Alcohol dehydrogenase
Regulator

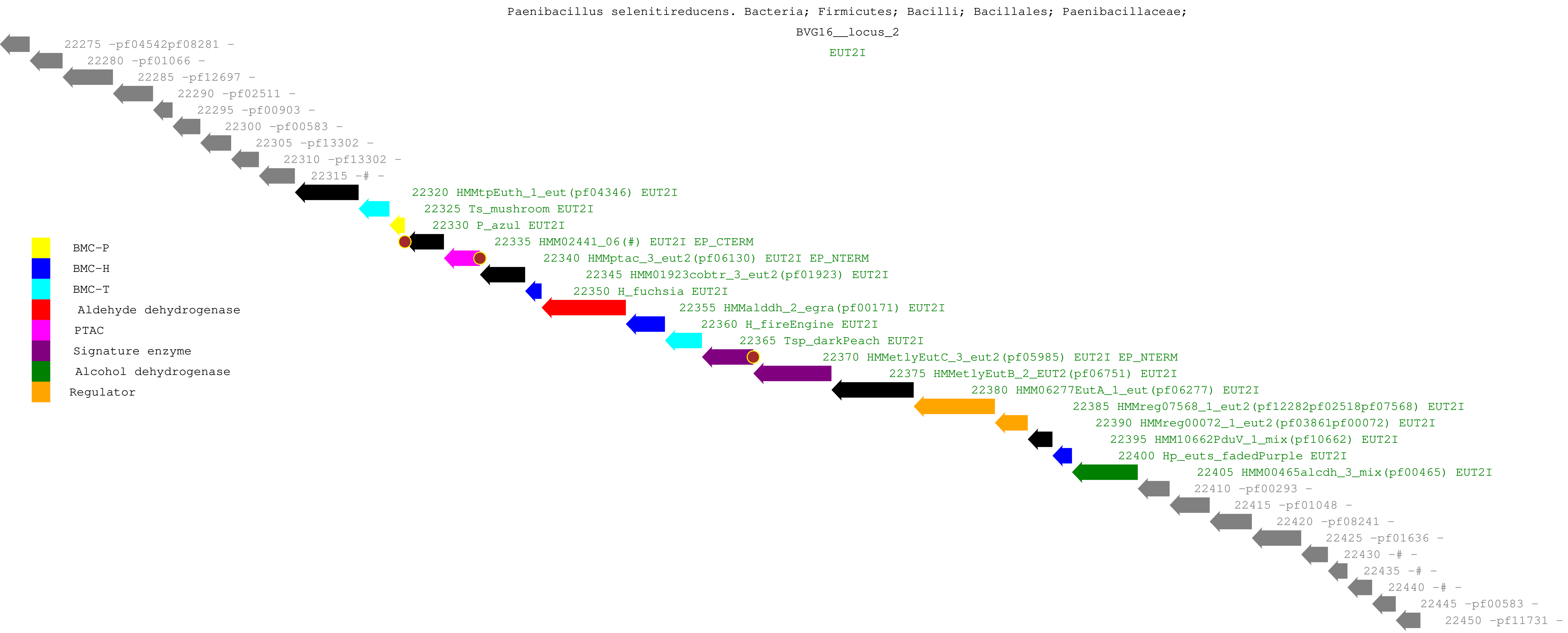
Paenibacillus selenitireducens. Bacteria; Firmicutes; Bacilli; Bacillales; Paenibacillaceae;
BVG16__locus_2
EUT2I
22275 -pf04542pf08281 -
22280 -pf01066 -
22285 -pf12697 -
22290 -pf02511 -
22295 -pf00903 -
22300 -pf00583 -
22305 -pf13302 -
22310 -pf13302 -
22315 -# -
22320 HMMtpEuth_1_eut(pf04346) EUT2I
22325 Ts_mushroom EUT2I
22330 P_azul EUT2I
22335 HMM02441_06(#) EUT2I EP_CTERM
22340 HMMptac_3_eut2(pf06130) EUT2I EP_NTERM
22345 HMM01923cobtr_3_eut2(pf01923) EUT2I
22350 H_fuchsia EUT2I
22355 HMMalddh_2_egra(pf00171) EUT2I
22360 H_fireEngine EUT2I
22365 Tsp_darkPeach EUT2I
22370 HMMetlyEutC_3_eut2(pf05985) EUT2I EP_NTERM
22375 HMMetlyEutB_2_EUT2(pf06751) EUT2I
22380 HMM06277EutA_1_eut(pf06277) EUT2I
22385 HMMreg07568_1_eut2(pf12282pf02518pf07568) EUT2I
22390 HMMreg00072_1_eut2(pf03861pf00072) EUT2I
22395 HMM10662PduV_1_mix(pf10662) EUT2I
22400 Hp_euts_fadedPurple EUT2I
22405 HMM00465alcdh_3_mix(pf00465) EUT2I
22410 -pf00293 -
22415 -pf01048 -
22420 -pf08241 -
22425 -pf01636 -
22430 -# -
22435 -# -
22440 -# -
22445 -pf00583 -
22450 -pf11731 -
BMC-P
BMC-H
BMC-T
Aldehyde dehydrogenase
PTAC
Signature enzyme
Alcohol dehydrogenase
Regulator

Clostridium perfringens. Bacteria; Firmicutes; Clostridia; Clostridiales; Clostridiaceae;

BXT94__locus_1

EUT2I

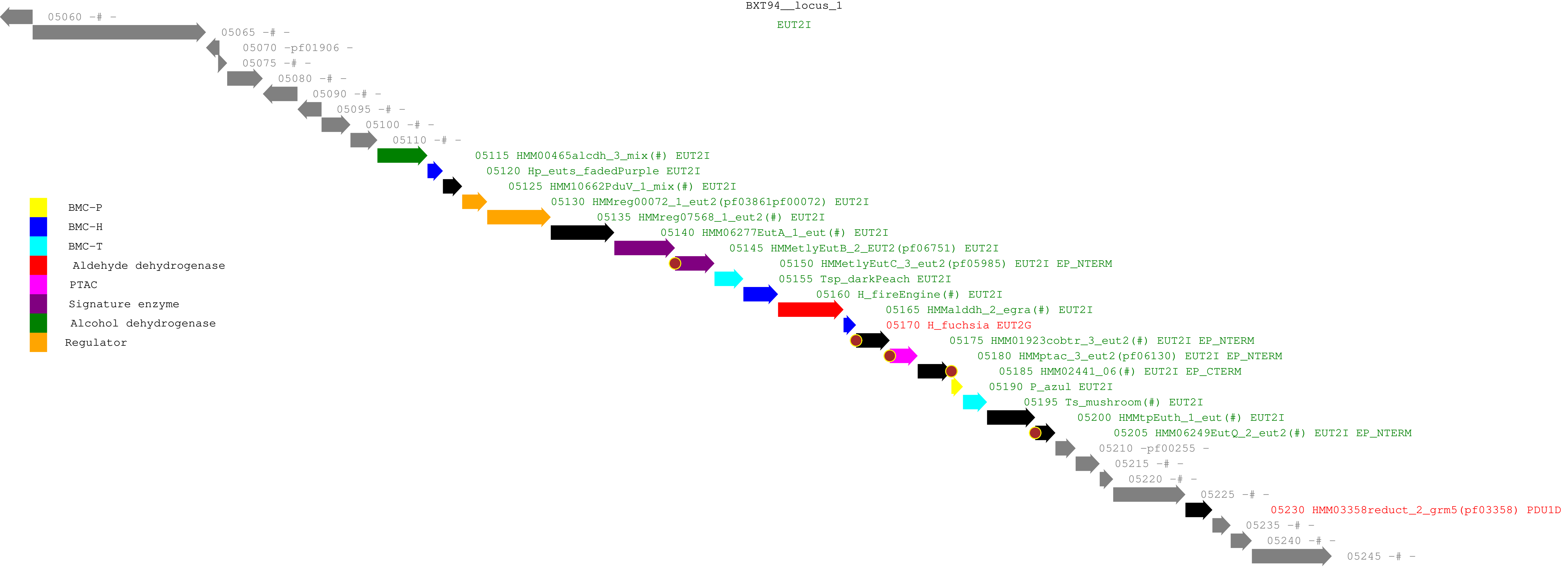

Brevibacillus laterosporus (Bacillus laterosporus). Bacteria; Firmicutes; Bacilli; Bacillales; Paenibacillaceae;

C2W64__locus_1

EUT2I

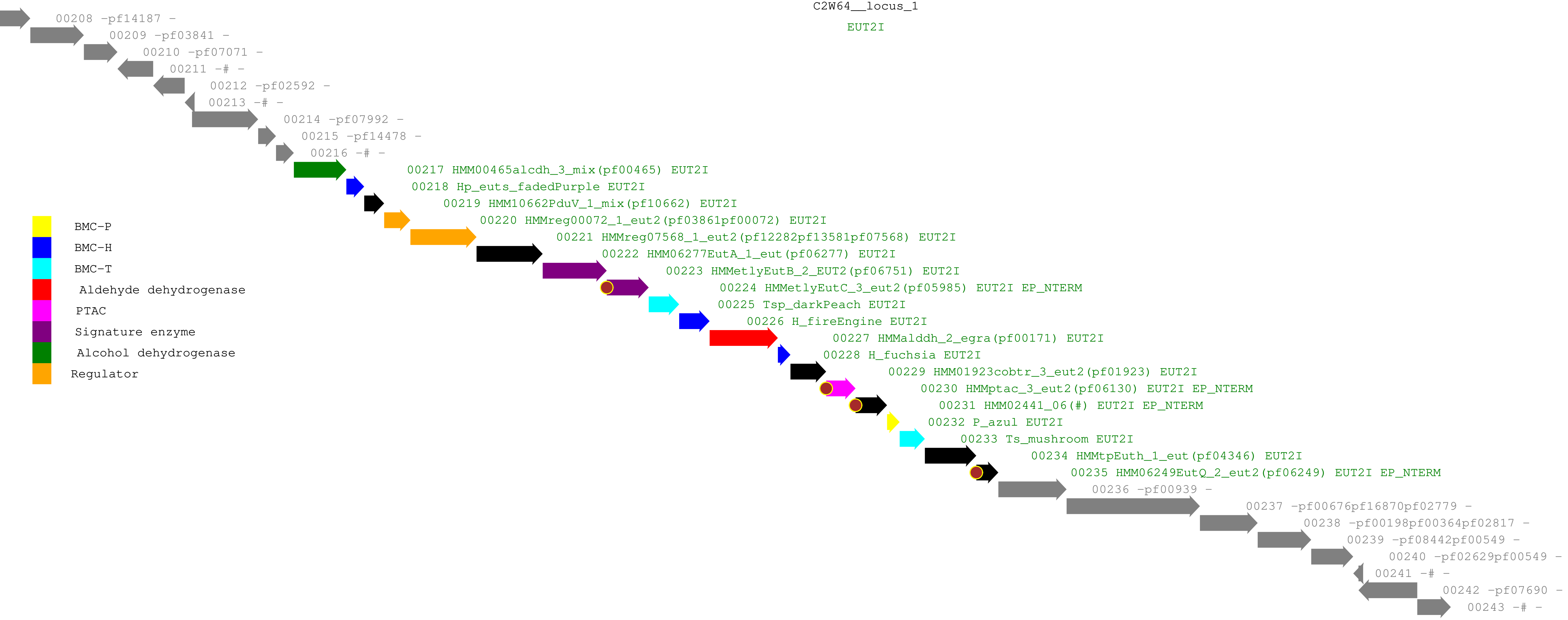

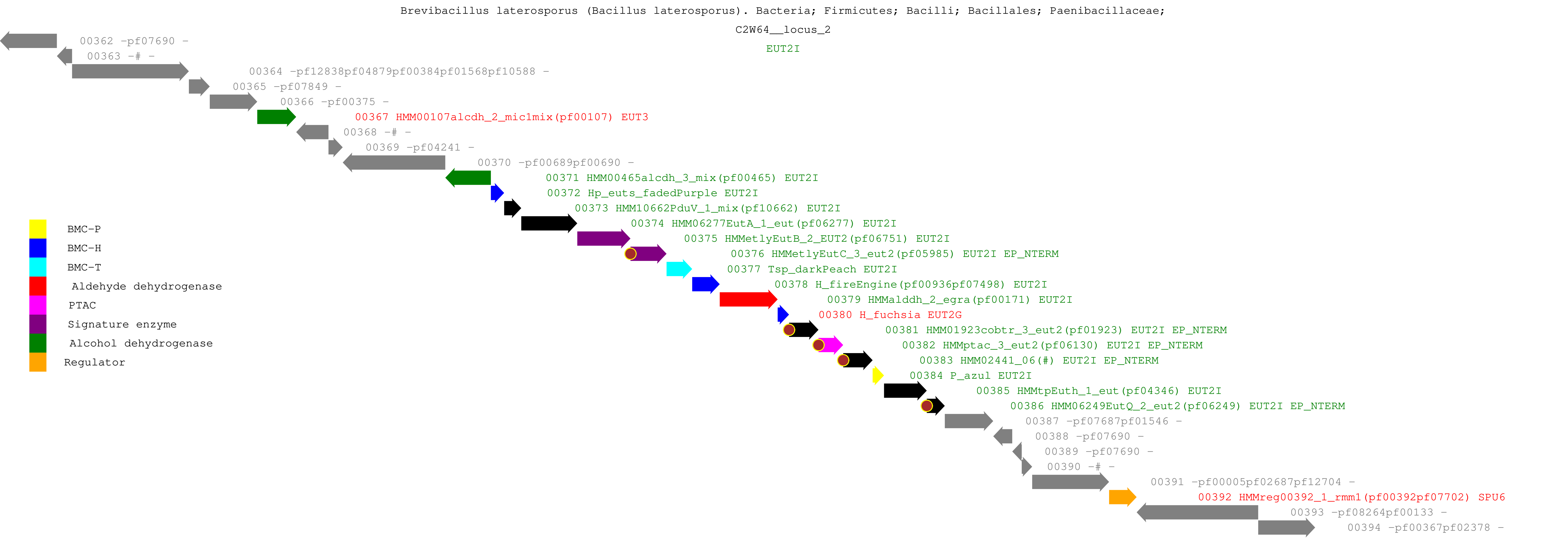
Brevibacillus laterosporus (Bacillus laterosporus). Bacteria; Firmicutes; Bacilli; Bacillales; Paenibacillaceae;
C2W64__locus_2
EUT2I
00362 -pf07690 -
00363 -# -
00364 -pf12838pf04879pf00384pf01568pf10588 -
00365 -pf07849 -
00366 -pf00375 -
00367 HMM00107alcdh_2_mic1mix(pf00107) EUT3
00368 -# -
00369 -pf04241 -
00370 -pf00689pf00690 -
00371 HMM00465alcdh_3_mix(pf00465) EUT2I
00372 Hp_euts_fadedPurple EUT2I
00373 HMM10662PduV_1_mix(pf10662) EUT2I
00374 HMM06277EutA_1_eut(pf06277) EUT2I
00375 HMMetlyEutB_2_EUT2(pf06751) EUT2I
00376 HMMetlyEutC_3_eut2(pf05985) EUT2I EP_NTERM
00377 Tsp_darkPeach EUT2I
00378 H_fireEngine(pf00936pf07498) EUT2I
00379 HMMalddh_2_egra(pf00171) EUT2I
00380 H_fuchsia EUT2G
00381 HMM01923cobtr_3_eut2(pf01923) EUT2I EP_NTERM
00382 HMMptac_3_eut2(pf06130) EUT2I EP_NTERM
00383 HMM02441_06(#) EUT2I EP_NTERM
00384 P_azul EUT2I
00385 HMMtpEuth_1_eut(pf04346) EUT2I
00386 HMM06249EutQ_2_eut2(pf06249) EUT2I EP_NTERM
00387 -pf07687pf01546 -
00388 -pf07690 -
00389 -pf07690 -
00390 -# -
00391 -pf00005pf02687pf12704 -
00392 HMMreg00392_1_rmm1(pf00392pf07702) SPU6
00393 -pf08264pf00133 -
00394 -pf00367pf02378 -
BMC-P
BMC-H
BMC-T
Aldehyde dehydrogenase
PTAC
Signature enzyme
Alcohol dehydrogenase
Regulator

Clostridiaceae bacterium 14S0207. Bacteria; Firmicutes; Clostridia; Clostridiales; Clostridiaceae.

C3495__locus_1

EUT2I

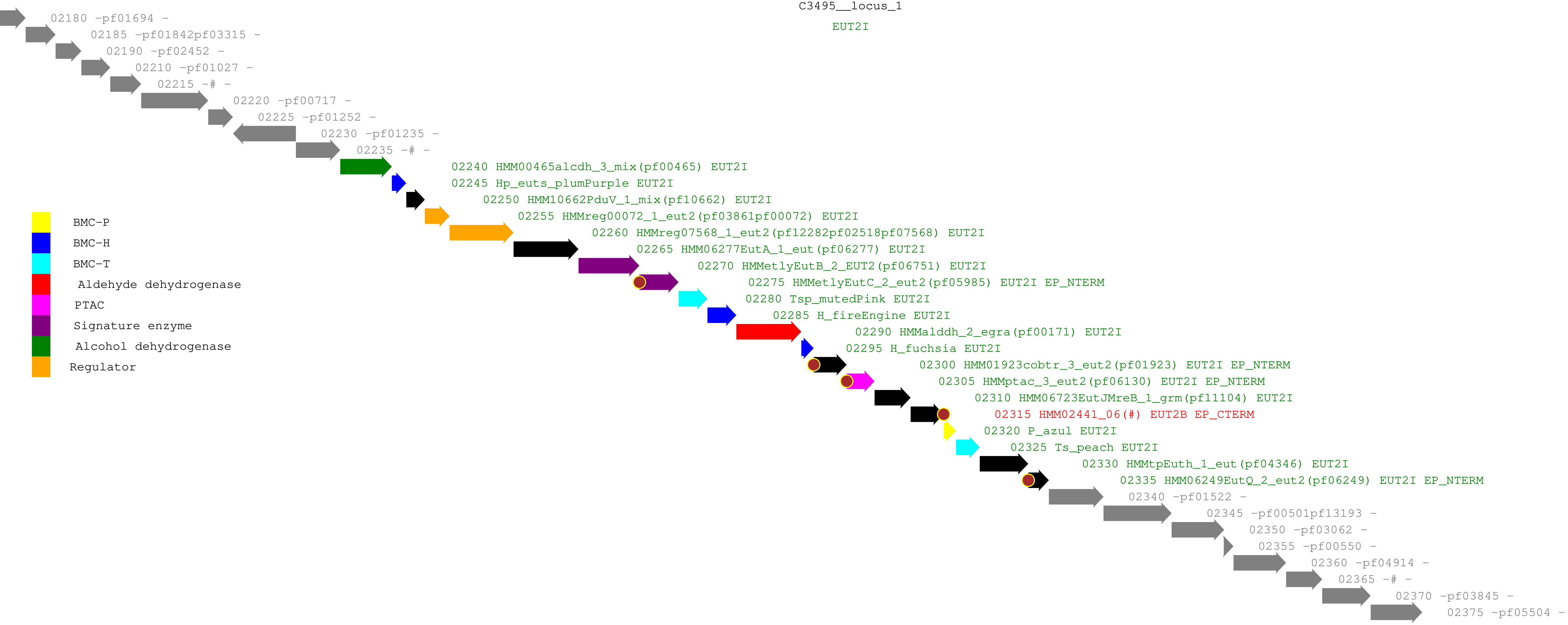

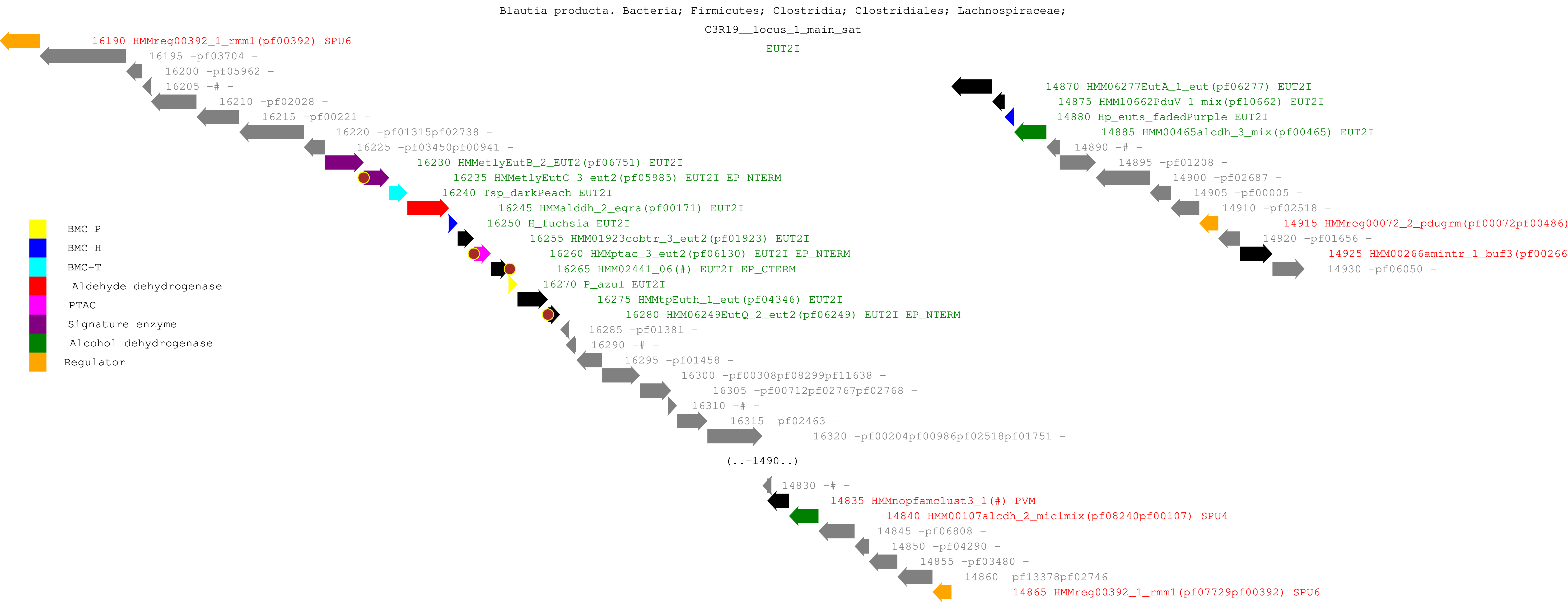

Blautia producta. Bacteria; Firmicutes; Clostridia; Clostridiales; Lachnospiraceae;
C3R19__locus_1_main_sat
EUT2I
16190 HMMreg00392_1_rmm1(pf00392) SPU6
16195 -pf03704 -
16200 -pf05962 -
16205 -# -
16210 -pf02028 -
16215 -pf00221 -
16220 -pf01315pf02738 -
16225 -pf03450pf00941 -
16230 HMMetlyEutB_2_EUT2(pf06751) EUT2I
16235 HMMetlyEutC_3_eut2(pf05985) EUT2I EP_NTERM
16240 Tsp_darkPeach EUT2I
16245 HMMalddh_2_egra(pf00171) EUT2I
16250 H_fuchsia EUT2I
16255 HMM01923cobtr_3_eut2(pf01923) EUT2I
16260 HMMptac_3_eut2(pf06130) EUT2I EP_NTERM
16265 HMM02441_06(#) EUT2I EP_CTERM
16270 P_azul EUT2I
16275 HMMtpEuth_1_eut(pf04346) EUT2I
16280 HMM06249EutQ_2_eut2(pf06249) EUT2I EP_NTERM
16285 -pf01381 -
16290 -# -
16295 -pf01458 -
16300 -pf00308pf08299pf11638 -
16305 -pf00712pf02767pf02768 -
16310 -# -
16315 -pf02463 -
16320 -pf00204pf00986pf02518pf01751 -
(..-1490..)
14830 -# -
14835 HMMnopfamclust3_1(#) PVM
14840 HMM00107alcdh_2_mic1mix(pf08240pf00107) SPU4
14845 -pf06808 -
14850 -pf04290 -
14855 -pf03480 -
14860 -pf13378pf02746 -
14865 HMMreg00392_1_rmm1(pf07729pf00392) SPU6
14870 HMM06277EutA_1_eut(pf06277) EUT2I
14875 HMM10662PduV_1_mix(pf10662) EUT2I
14880 Hp_euts_fadedPurple EUT2I
14885 HMM00465alcdh_3_mix(pf00465) EUT2I
14890 -# -
14895 -pf01208 -
14900 -pf02687 -
14905 -pf00005 -
14910 -pf02518 -
14915 HMMreg00072_2_pdugrm(pf00072pf00486)
14920 -pf01656 -
14925 HMM00266amintr_1_buf3(pf00266
14930 -pf06050 -
BMC-P
BMC-H
BMC-T
Aldehyde dehydrogenase
PTAC
Signature enzyme
Alcohol dehydrogenase
Regulator

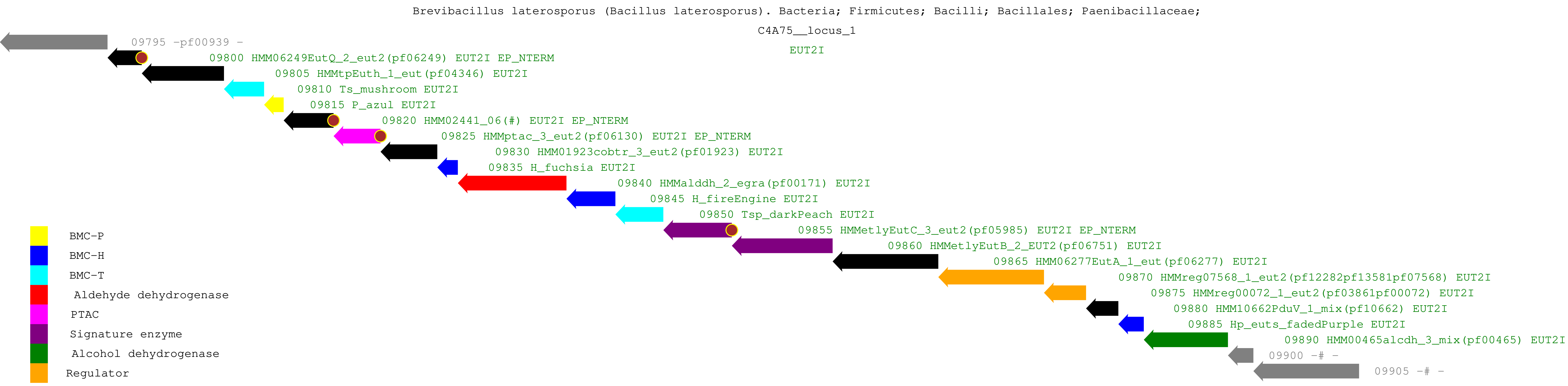
Brevibacillus laterosporus (Bacillus laterosporus). Bacteria; Firmicutes; Bacilli; Bacillales; Paenibacillaceae;
C4A75__locus_1
EUT2I
09795 -pf00939 -
09800 HMM06249EutQ_2_eut2(pf06249) EUT2I EP_NTERM
09805 HMMtpEuth_1_eut(pf04346) EUT2I
09810 Ts_mushroom EUT2I
09815 P_azul EUT2I
09820 HMM02441_06(#) EUT2I EP_NTERM
09825 HMMptac_3_eut2(pf06130) EUT2I EP_NTERM
09830 HMM01923cobtr_3_eut2(pf01923) EUT2I
09835 H_fuchsia EUT2I
09840 HMMalddh_2_egra(pf00171) EUT2I
09845 H_fireEngine EUT2I
09850 Tsp_darkPeach EUT2I
09855 HMMetlyEutC_3_eut2(pf05985) EUT2I EP_NTERM
09860 HMMetlyEutB_2_EUT2(pf06751) EUT2I
09865 HMM06277EutA_1_eut(pf06277) EUT2I
09870 HMMreg07568_1_eut2(pf12282pf13581pf07568) EUT2I
09875 HMMreg00072_1_eut2(pf03861pf00072) EUT2I
09880 HMM10662PduV_1_mix(pf10662) EUT2I
09885 Hp_euts_fadedPurple EUT2I
09890 HMM00465alcdh_3_mix(pf00465) EUT2I
09900 -# -
09905 -# -
BMC-P
BMC-H
BMC-T
Aldehyde dehydrogenase
PTAC
Signature enzyme
Alcohol dehydrogenase
Regulator

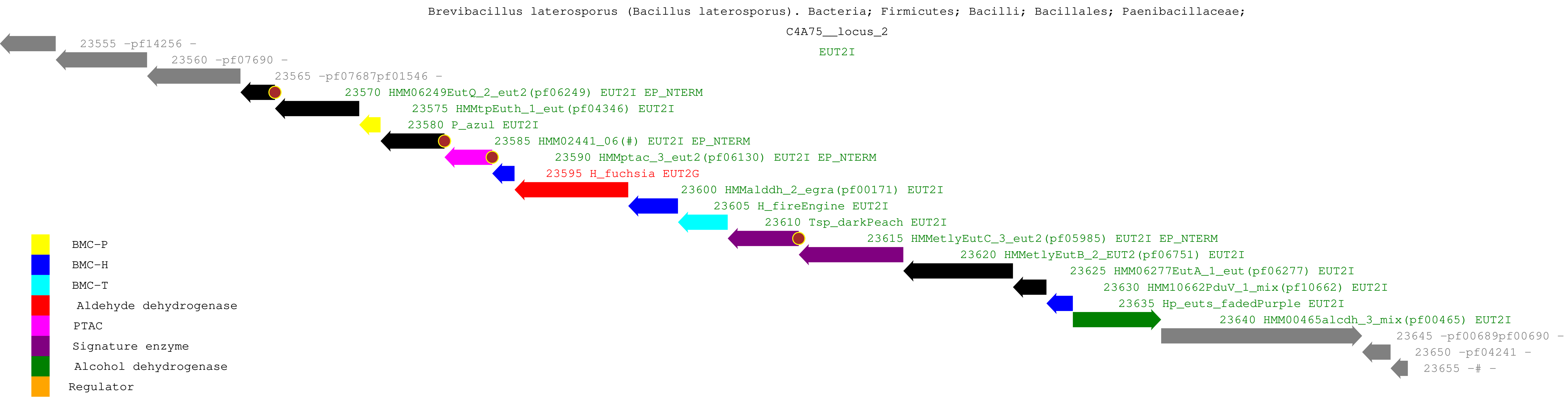
Brevibacillus laterosporus (Bacillus laterosporus). Bacteria; Firmicutes; Bacilli; Bacillales; Paenibacillaceae;
C4A75__locus_2
EUT2I
23555 -pf14256 -
23560 -pf07690 -
23565 -pf07687pf01546 -
23570 HMM06249EutQ_2_eut2(pf06249) EUT2I EP_NTERM
23575 HMMtpEuth_1_eut(pf04346) EUT2I
23580 P_azul EUT2I
23585 HMM02441_06(#) EUT2I EP_NTERM
23590 HMMptac_3_eut2(pf06130) EUT2I EP_NTERM
23595 H_fuchsia EUT2G
23600 HMMalddh_2_egra(pf00171) EUT2I
23605 H_fireEngine EUT2I
23610 Tsp_darkPeach EUT2I
23615 HMMetlyEutC_3_eut2(pf05985) EUT2I EP_NTERM
23620 HMMetlyEutB_2_EUT2(pf06751) EUT2I
23625 HMM06277EutA_1_eut(pf06277) EUT2I
23630 HMM10662PduV_1_mix(pf10662) EUT2I
23635 Hp_euts_fadedPurple EUT2I
23640 HMM00465alcdh_3_mix(pf00465) EUT2I
23645 -pf00689pf00690 -
23650 -pf04241 -
23655 -# -
BMC-P
BMC-H
BMC-T
Aldehyde dehydrogenase
PTAC
Signature enzyme
Alcohol dehydrogenase
Regulator

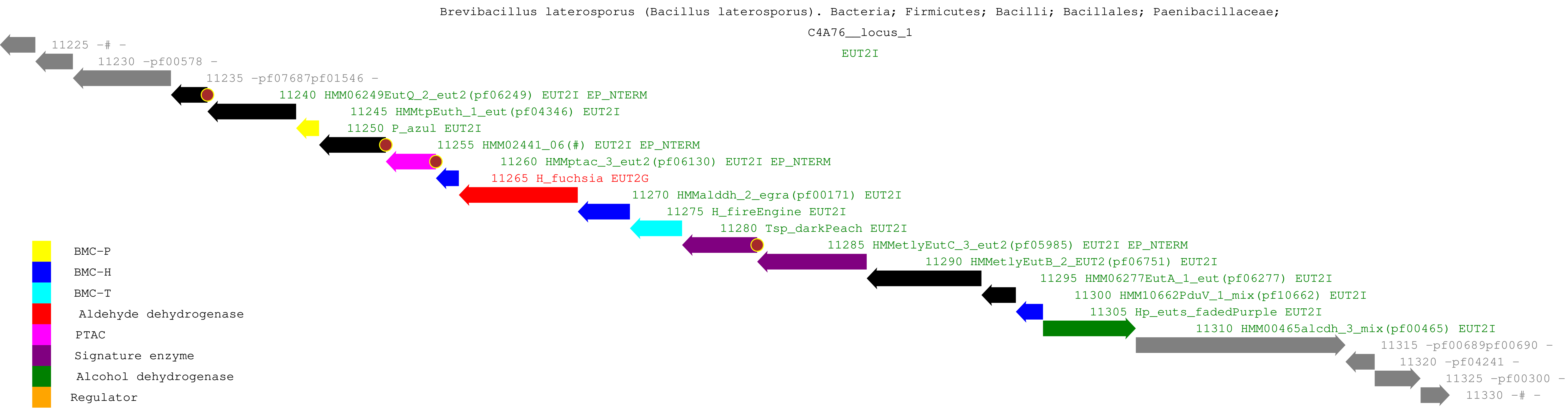

Brevibacillus laterosporus (Bacillus laterosporus). Bacteria; Firmicutes; Bacilli; Bacillales; Paenibacillaceae;
C4A76__locus_1
EUT2I
11225 -# -
11230 -pf00578 -
11235 -pf07687pf01546 -
11240 HMM06249EutQ_2_eut2(pf06249) EUT2I EP_NTERM
11245 HMMtpEuth_1_eut(pf04346) EUT2I
11250 P_azul EUT2I
11255 HMM02441_06(#) EUT2I EP_NTERM
11260 HMMptac_3_eut2(pf06130) EUT2I EP_NTERM
11265 H_fuchsia EUT2G
11270 HMMalddh_2_egra(pf00171) EUT2I
11275 H_fireEngine EUT2I
11280 Tsp_darkPeach EUT2I
11285 HMMetlyEutC_3_eut2(pf05985) EUT2I EP_NTERM
11290 HMMetlyEutB_2_EUT2(pf06751) EUT2I
11295 HMM06277EutA_1_eut(pf06277) EUT2I
11300 HMM10662PduV_1_mix(pf10662) EUT2I
11305 Hp_euts_fadedPurple EUT2I
11310 HMM00465alcdh_3_mix(pf00465) EUT2I
11315 -pf00689pf00690 -
11320 -pf04241 -
11325 -pf00300 -
11330 -# -
BMC-P
BMC-H
BMC-T
Aldehyde dehydrogenase
PTAC
Signature enzyme
Alcohol dehydrogenase
Regulator

Brevibacillus laterosporus (Bacillus laterosporus). Bacteria; Firmicutes; Bacilli; Bacillales; Paenibacillaceae;

C4A77__locus_1

EUT2I

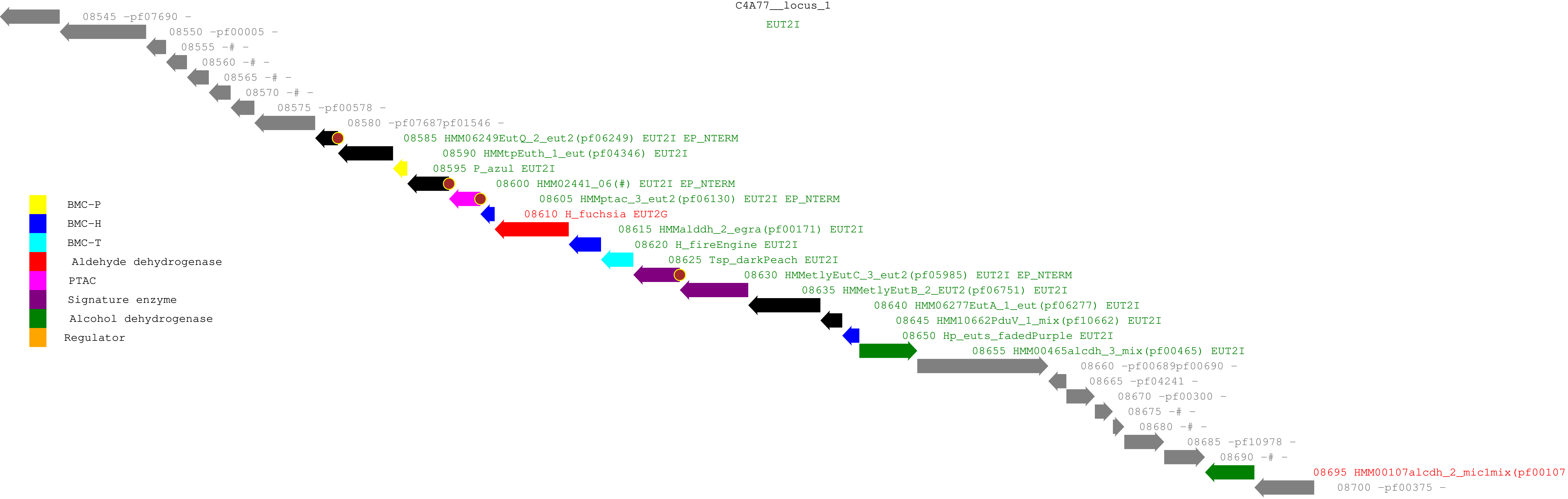

Brevibacillus laterosporus (Bacillus laterosporus). Bacteria; Firmicutes; Bacilli; Bacillales; Paenibacillaceae;

C4A77__locus_2

EUT2I

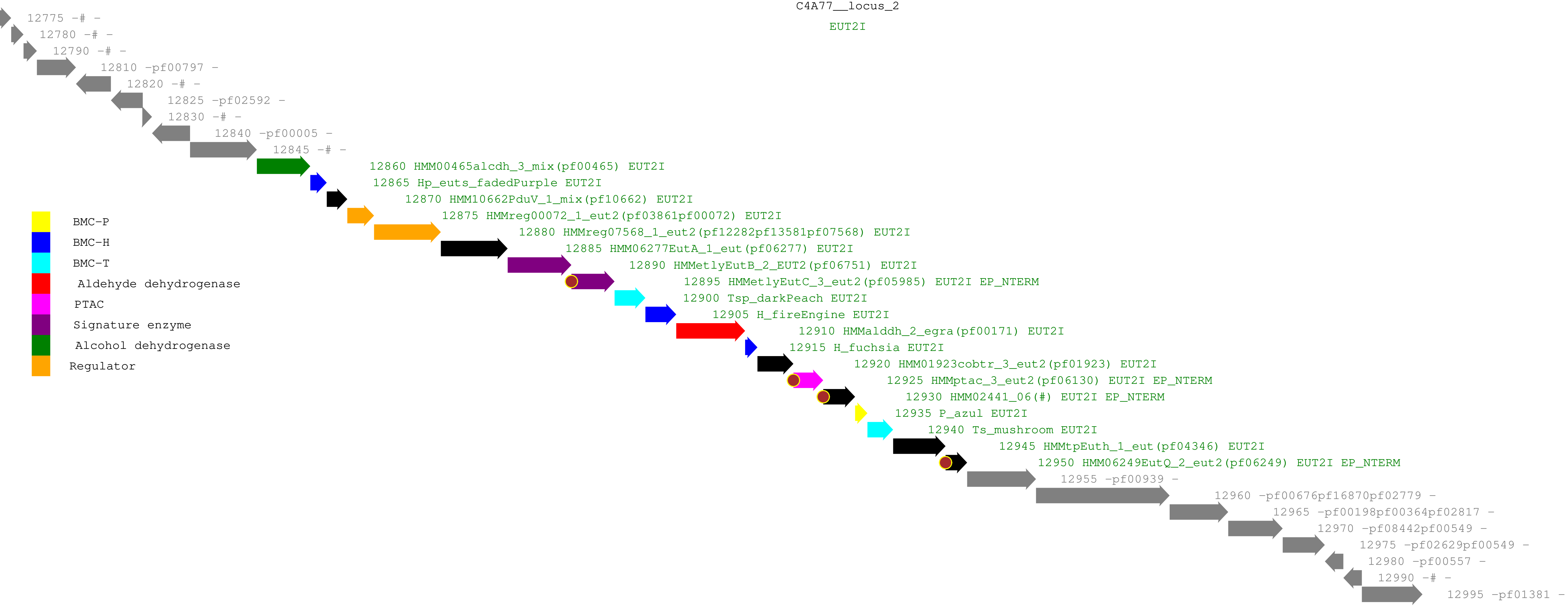

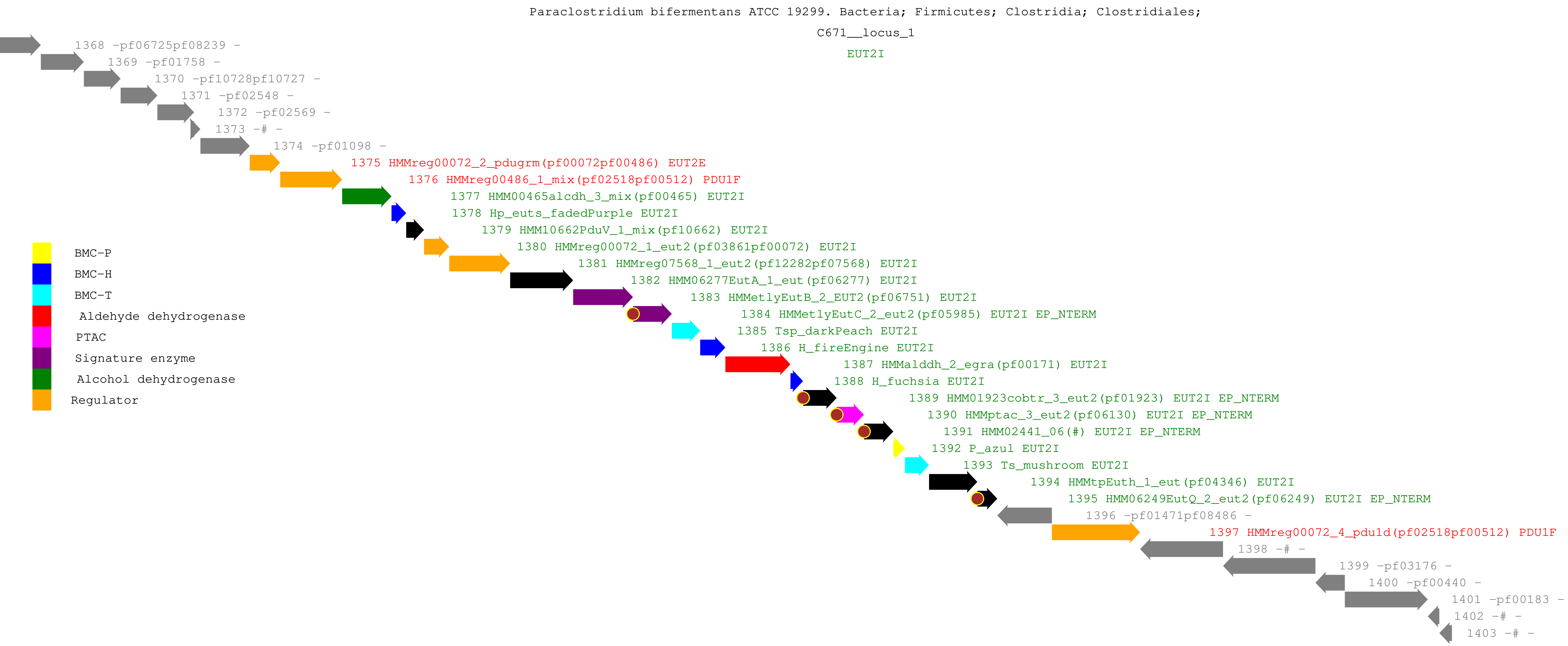
Paraclostridium bifermentans ATCC 19299. Bacteria; Firmicutes; Clostridia; Clostridiales;
C671__locus_1
EUT2I
1368 -pf06725pf08239 -
1369 -pf01758 -
1370 -pf10728pf10727 -
1371 -pf02548 -
1372 -pf02569 -
1373 -# -
1374 -pf01098 -
1375 HMMreg00072_2_pdugrm(pf00072pf00486) EUT2E
1376 HMMreg00486_1_mix(pf02518pf00512) PDU1F
1377 HMM00465alcdh_3_mix(pf00465) EUT2I
1378 Hp_euts_fadedPurple EUT2I
1379 HMM10662PduV_1_mix(pf10662) EUT2I
1380 HMMreg00072_1_eut2(pf03861pf00072) EUT2I
1381 HMMreg07568_1_eut2(pf12282pf07568) EUT2I
1382 HMM06277EutA_1_eut(pf06277) EUT2I
1383 HMMetlyEutB_2_EUT2(pf06751) EUT2I
1384 HMMetlyEutC_2_eut2(pf05985) EUT2I EP_NTERM
1385 Tsp_darkPeach EUT2I
1386 H_fireEngine EUT2I
1387 HMMalddh_2_egra(pf00171) EUT2I
1388 H_fuchsia EUT2I
1389 HMM01923cobtr_3_eut2(pf01923) EUT2I EP_NTERM
1390 HMMptac_3_eut2(pf06130) EUT2I EP_NTERM
1391 HMM02441_06(#) EUT2I EP_NTERM
1392 P_azul EUT2I
1393 Ts_mushroom EUT2I
1394 HMMtpEuth_1_eut(pf04346) EUT2I
1395 HMM06249EutQ_2_eut2(pf06249) EUT2I EP_NTERM
1396 -pf01471pf08486 -
1397 HMMreg00072_4_pdu1d(pf02518pf00512) PDU1F
1398 -# -
1399 -pf03176 -
1400 -pf00440 -
1401 -pf00183 -
1402 -# -
1403 -# -
BMC-P
BMC-H
BMC-T
Aldehyde dehydrogenase
PTAC
Signature enzyme
Alcohol dehydrogenase
Regulator

Paraclostridium bifermentans ATCC 638. Bacteria; Firmicutes; Clostridia; Clostridiales;

C672__locus_1

EUT2I

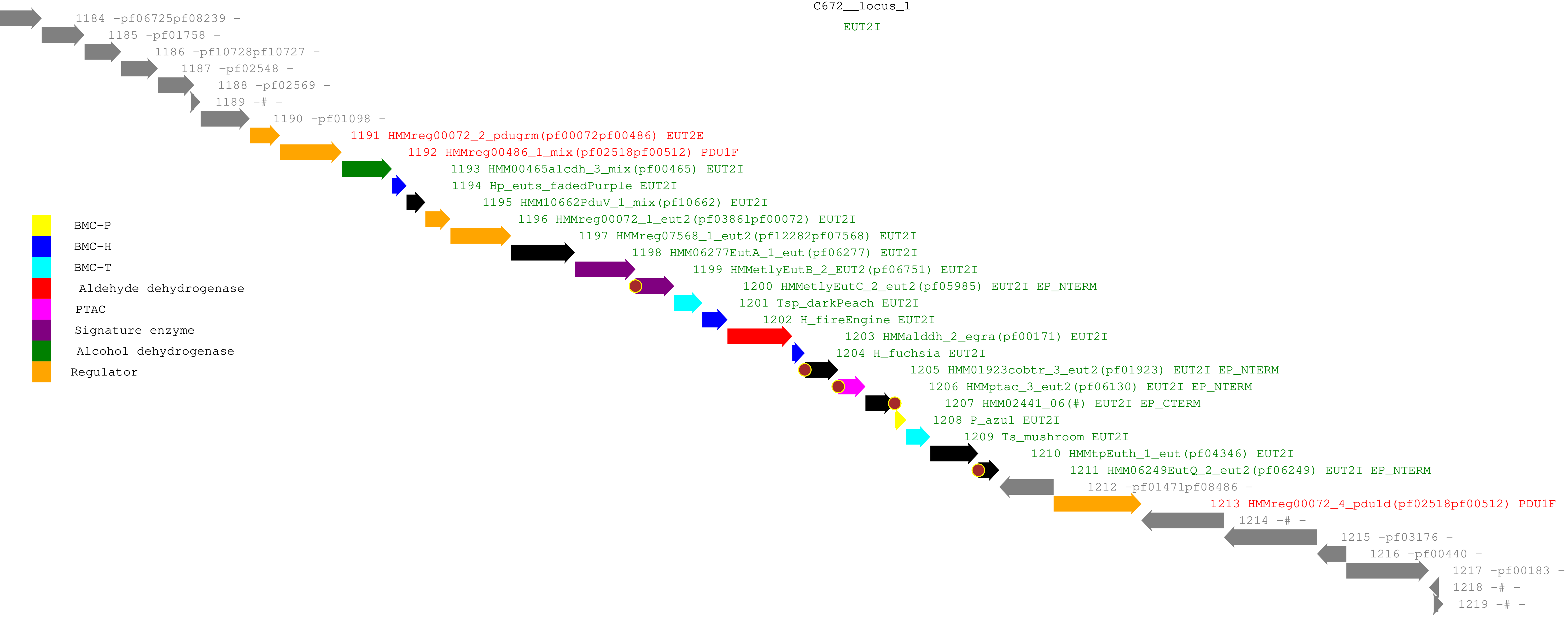

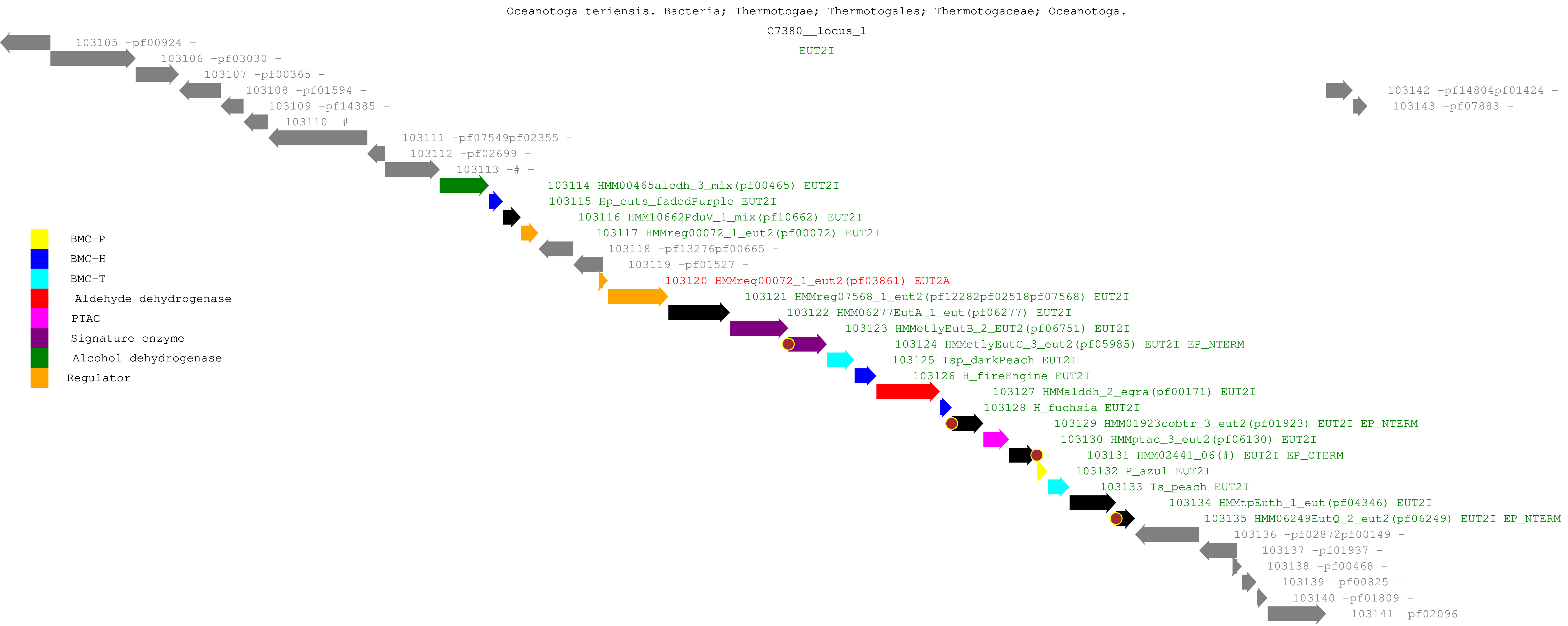
Oceanotoga teriensis. Bacteria; Thermotogae; Thermotogales; Thermotogaceae; Oceanotoga.
C7380__locus_1
EUT2I
103105 -pf00924 -
103106 -pf03030 -
103107 -pf00365 -
103108 -pf01594 -
103109 -pf14385 -
103110 -# -
103111 -pf07549pf02355 -
103112 -pf02699 -
103113 -# -
103114 HMM00465alcdh_3_mix(pf00465) EUT2I
103115 Hp_euts_fadedPurple EUT2I
103116 HMM10662PduV_1_mix(pf10662) EUT2I
103117 HMMreg00072_1_eut2(pf00072) EUT2I
103118 -pf13276pf00665 -
103119 -pf01527 -
103120 HMMreg00072_1_eut2(pf03861) EUT2A
103121 HMMreg07568_1_eut2(pf12282pf02518pf07568) EUT2I
103122 HMM06277EutA_1_eut(pf06277) EUT2I
103123 HMMetlyEutB_2_EUT2(pf06751) EUT2I
103124 HMMetlyEutC_3_eut2(pf05985) EUT2I EP_NTERM
103125 Tsp_darkPeach EUT2I
103126 H_fireEngine EUT2I
103127 HMMalddh_2_egra(pf00171) EUT2I
103128 H_fuchsia EUT2I
103129 HMM01923cobtr_3_eut2(pf01923) EUT2I EP_NTERM
103130 HMMptac_3_eut2(pf06130) EUT2I
103131 HMM02441_06(#) EUT2I EP_CTERM
103132 P_azul EUT2I
103133 Ts_peach EUT2I
103134 HMMtpEuth_1_eut(pf04346) EUT2I
103135 HMM06249EutQ_2_eut2(pf06249) EUT2I EP_NTERM
103136 -pf02872pf00149 -
103137 -pf01937 -
103138 -pf00468 -
103139 -pf00825 -
103140 -pf01809 -
103141 -pf02096 -
103142 -pf14804pf01424 -
103143 -pf07883 -
BMC-P
BMC-H
BMC-T
Aldehyde dehydrogenase
PTAC
Signature enzyme
Alcohol dehydrogenase
Regulator

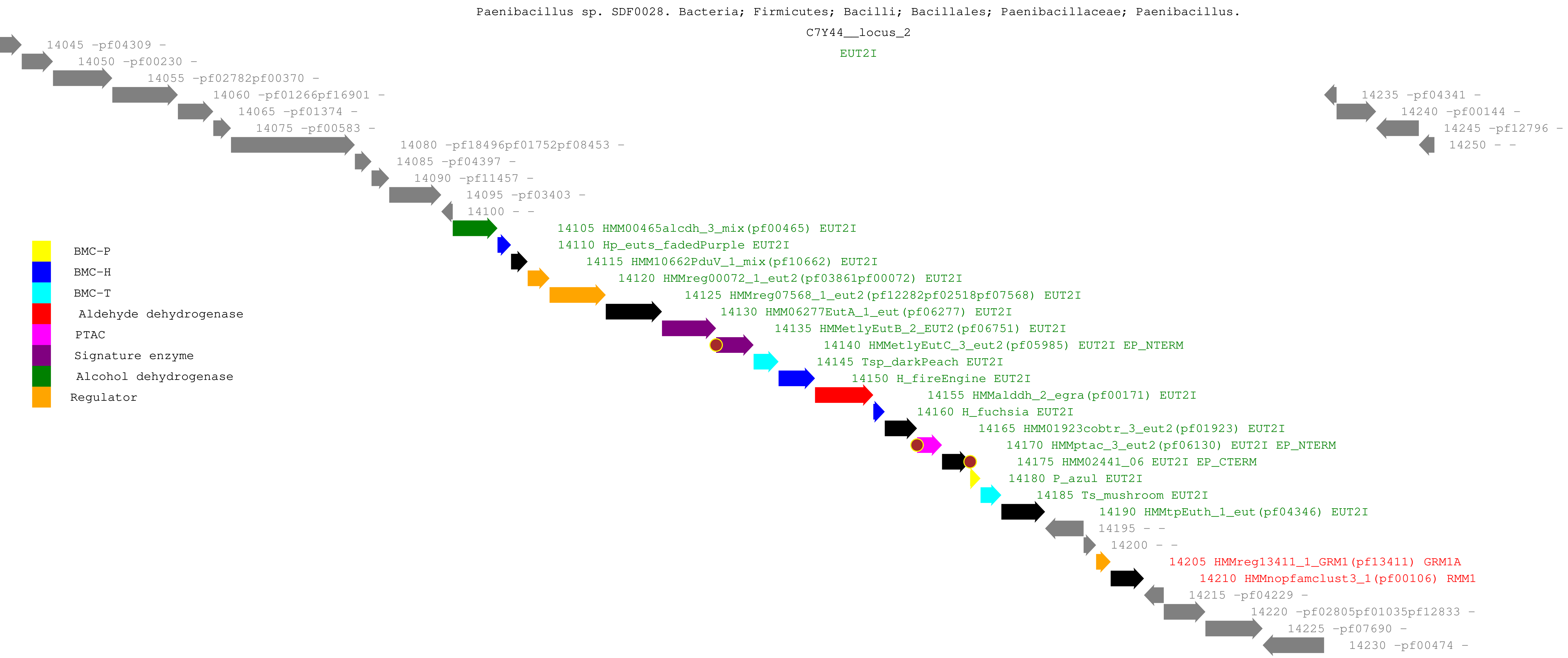

Paenibacillus sp. SDF0028. Bacteria; Firmicutes; Bacilli; Bacillales; Paenibacillaceae; Paenibacillus.
C7Y44__locus_2
EUT2I
14045 -pf04309 -
14050 -pf00230 -
14055 -pf02782pf00370 -
14060 -pf01266pf16901 -
14065 -pf01374 -
14075 -pf00583 -
14080 -pf18496pf01752pf08453 -
14085 -pf04397 -
14090 -pf11457 -
14095 -pf03403 -
14100 - -
14105 HMM00465alcdh_3_mix(pf00465) EUT2I
14110 Hp_euts_fadedPurple EUT2I
14115 HMM10662PduV_1_mix(pf10662) EUT2I
14120 HMMreg00072_1_eut2(pf03861pf00072) EUT2I
14125 HMMreg07568_1_eut2(pf12282pf02518pf07568) EUT2I
14130 HMM06277EutA_1_eut(pf06277) EUT2I
14135 HMMetlyEutB_2_EUT2(pf06751) EUT2I
14140 HMMetlyEutC_3_eut2(pf05985) EUT2I EP_NTERM
14145 Tsp_darkPeach EUT2I
14150 H_fireEngine EUT2I
14155 HMMalddh_2_egra(pf00171) EUT2I
14160 H_fuchsia EUT2I
14165 HMM01923cobtr_3_eut2(pf01923) EUT2I
14170 HMMptac_3_eut2(pf06130) EUT2I EP_NTERM
14175 HMM02441_06 EUT2I EP_CTERM
14180 P_azul EUT2I
14185 Ts_mushroom EUT2I
14190 HMMtpEuth_1_eut(pf04346) EUT2I
14195 - -
14200 - -
14205 HMMreg13411_1_GRM1(pf13411) GRM1A
14210 HMMnopfamclust3_1(pf00106) RMM1
14215 -pf04229 -
14220 -pf02805pf01035pf12833 -
14225 -pf07690 -
14230 -pf00474 -
14235 -pf04341 -
14240 -pf00144 -
14245 -pf12796 -
14250 - -
BMC-P
BMC-H
BMC-T
Aldehyde dehydrogenase
PTAC
Signature enzyme
Alcohol dehydrogenase
Regulator

Peptoclostridium difficile (strain CD196) (Clostridium difficile). Bacteria; Firmicutes; Clostridia; Clostridiales;

CD196__locus_1

EUT2I

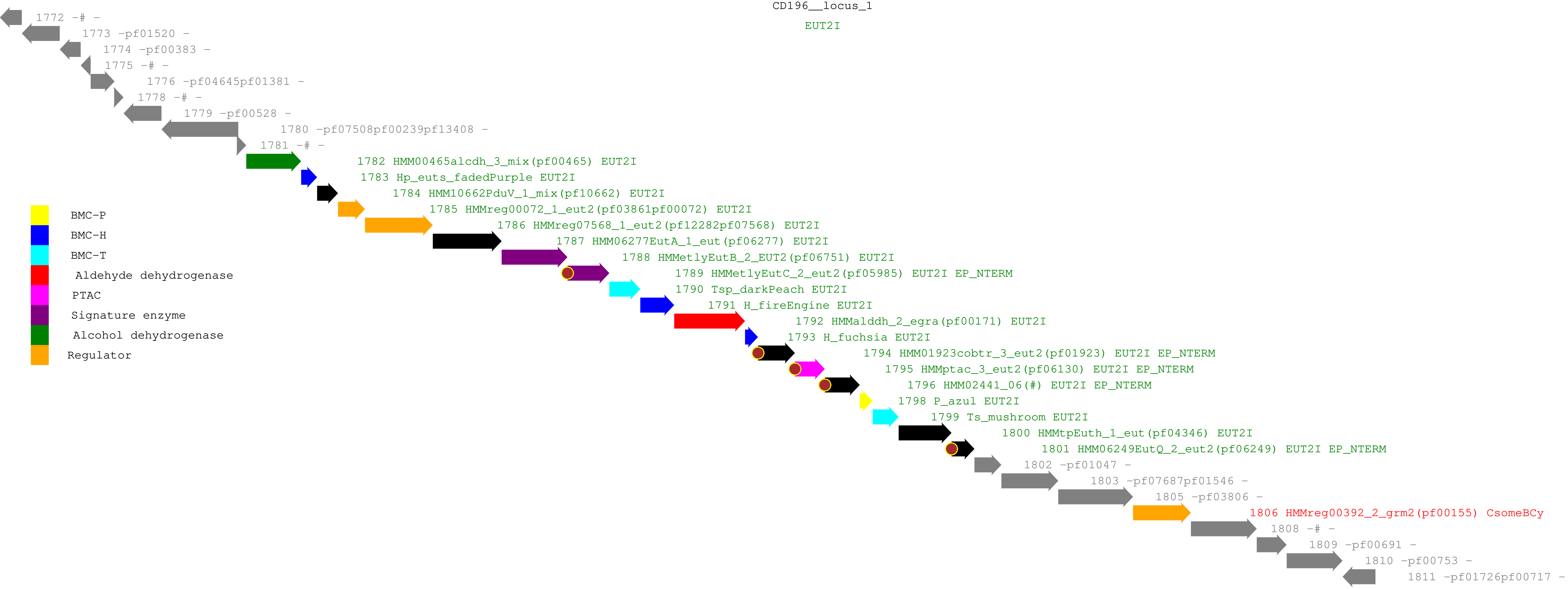

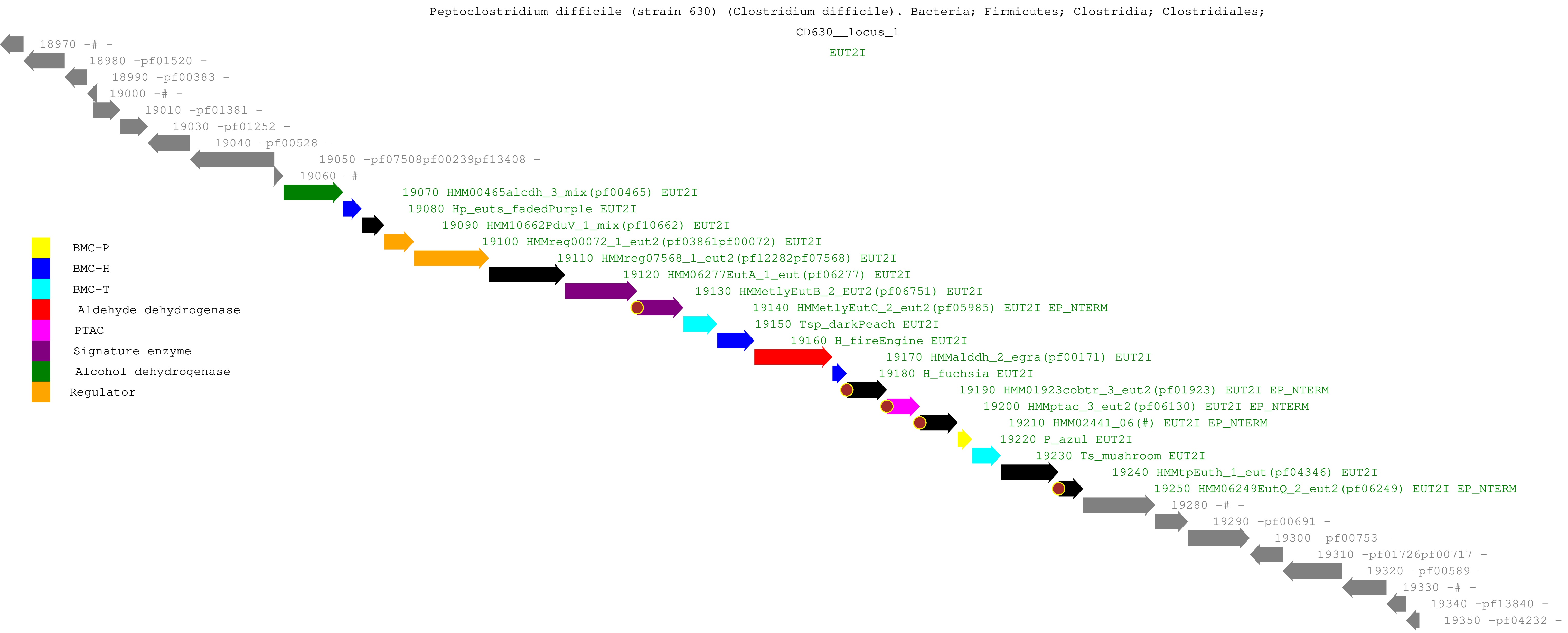
Peptoclostridium difficile (strain 630) (Clostridium difficile). Bacteria; Firmicutes; Clostridia; Clostridiales;
CD630__locus_1
EUT2I
18970 -# -
18980 -pf01520 -
18990 -pf00383 -
19000 -# -
19010 -pf01381 -
19030 -pf01252 -
19040 -pf00528 -
19050 -pf07508pf00239pf13408 -
19060 -# -
19070 HMM00465alcdh_3_mix(pf00465) EUT2I
19080 Hp_euts_fadedPurple EUT2I
19090 HMM10662PduV_1_mix(pf10662) EUT2I
19100 HMMreg00072_1_eut2(pf03861pf00072) EUT2I
19110 HMMreg07568_1_eut2(pf12282pf07568) EUT2I
19120 HMM06277EutA_1_eut(pf06277) EUT2I
19130 HMMetlyEutB_2_EUT2(pf06751) EUT2I
19140 HMMetlyEutC_2_eut2(pf05985) EUT2I EP_NTERM
19150 Tsp_darkPeach EUT2I
19160 H_fireEngine EUT2I
19170 HMMalddh_2_egra(pf00171) EUT2I
19180 H_fuchsia EUT2I
19190 HMM01923cobtr_3_eut2(pf01923) EUT2I EP_NTERM
19200 HMMptac_3_eut2(pf06130) EUT2I EP_NTERM
19210 HMM02441_06(#) EUT2I EP_NTERM
19220 P_azul EUT2I
19230 Ts_mushroom EUT2I
19240 HMMtpEuth_1_eut(pf04346) EUT2I
19250 HMM06249EutQ_2_eut2(pf06249) EUT2I EP_NTERM
19280 -# -
19290 -pf00691 -
19300 -pf00753 -
19310 -pf01726pf00717 -
19320 -pf00589 -
19330 -# -
19340 -pf13840 -
19350 -pf04232 -
BMC-P
BMC-H
BMC-T
Aldehyde dehydrogenase
PTAC
Signature enzyme
Alcohol dehydrogenase
Regulator

Clostridioides difficile (Peptoclostridium difficile). Bacteria; Firmicutes; Clostridia; Clostridiales;

CDFC105__locus_1

EUT2I

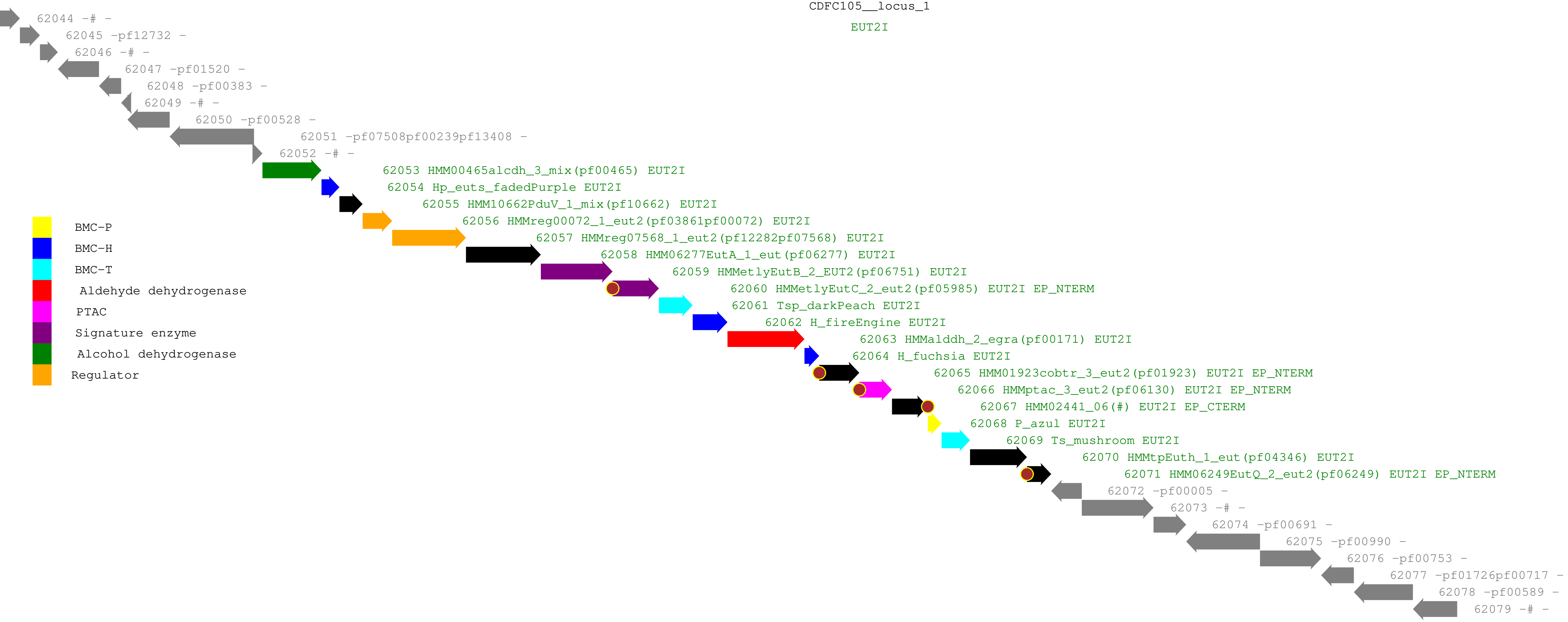

Clostridioides difficile ATCC 9689 = DSM 1296. Bacteria; Firmicutes; Clostridia; Clostridiales;

CDIF1296T__locus_1

EUT2I

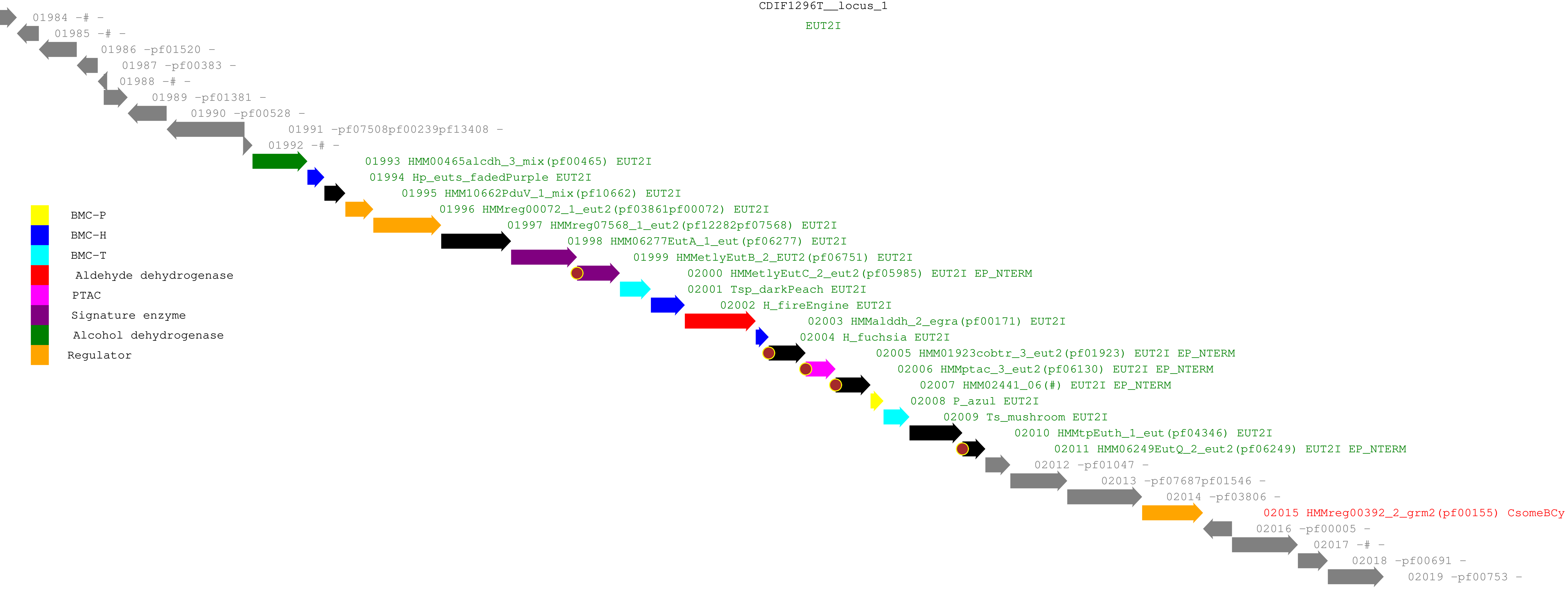

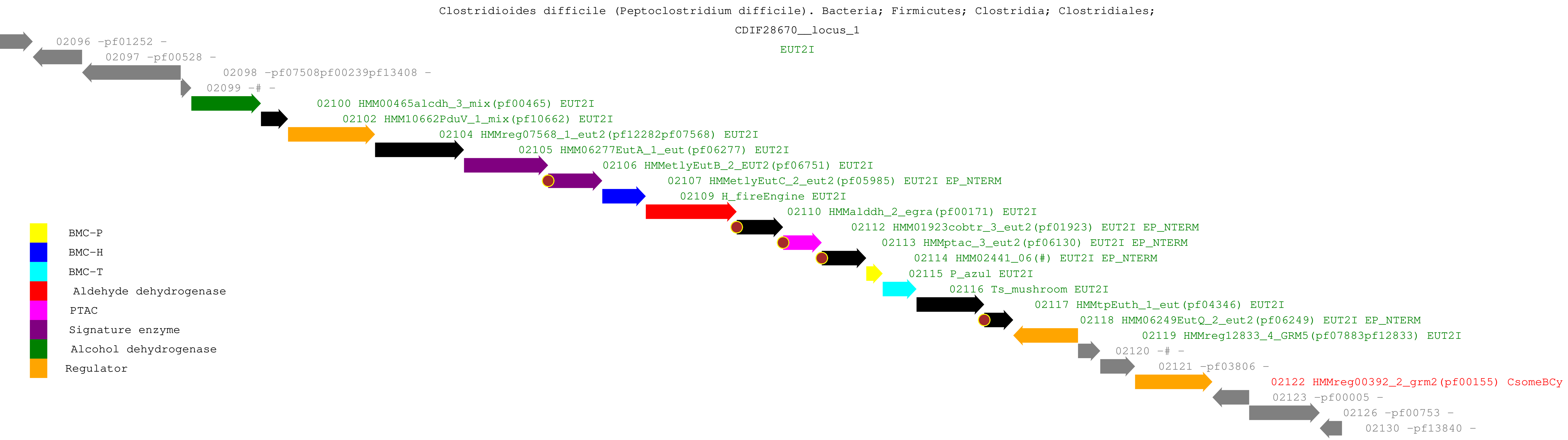
Clostridioides difficile (Peptoclostridium difficile). Bacteria; Firmicutes; Clostridia; Clostridiales;
CDIF28670__locus_1
EUT2I
02096 -pf01252 -
02097 -pf00528 -
02098 -pf07508pf00239pf13408 -
02099 -# -
02100 HMM00465alcdh_3_mix(pf00465) EUT2I
02102 HMM10662PduV_1_mix(pf10662) EUT2I
02104 HMMreg07568_1_eut2(pf12282pf07568) EUT2I
02105 HMM06277EutA_1_eut(pf06277) EUT2I
02106 HMMetlyEutB_2_EUT2(pf06751) EUT2I
02107 HMMetlyEutC_2_eut2(pf05985) EUT2I EP_NTERM
02109 H_fireEngine EUT2I
02110 HMMalddh_2_egra(pf00171) EUT2I
02112 HMM01923cobtr_3_eut2(pf01923) EUT2I EP_NTERM
02113 HMMptac_3_eut2(pf06130) EUT2I EP_NTERM
02114 HMM02441_06(#) EUT2I EP_NTERM
02115 P_azul EUT2I
02116 Ts_mushroom EUT2I
02117 HMMtpEuth_1_eut(pf04346) EUT2I
02118 HMM06249EutQ_2_eut2(pf06249) EUT2I EP_NTERM
02119 HMMreg12833_4_GRM5(pf07883pf12833) EUT2I
02120 -# -
02121 -pf03806 -
02122 HMMreg00392_2_grm2(pf00155) CsomeBCy
02123 -pf00005 -
02126 -pf00753 -
02130 -pf13840 -
BMC-P
BMC-H
BMC-T
Aldehyde dehydrogenase
PTAC
Signature enzyme
Alcohol dehydrogenase
Regulator

Clostridioides difficile (Peptoclostridium difficile). Bacteria; Firmicutes; Clostridia; Clostridiales;

CDIF29627__locus_1

EUT2I

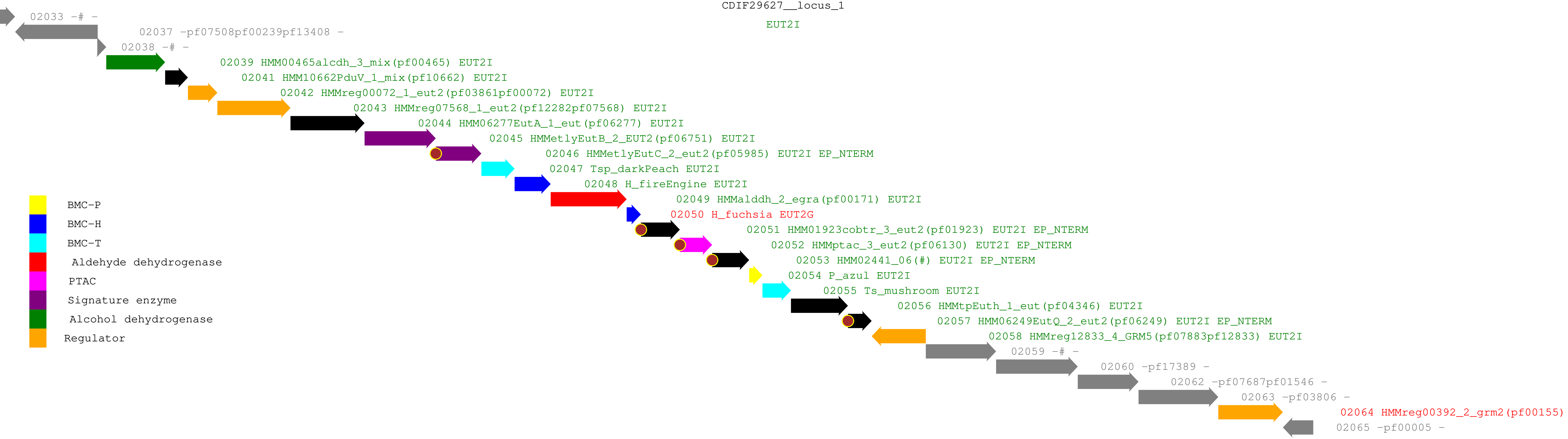

Clostridioides difficile (Peptoclostridium difficile). Bacteria; Firmicutes; Clostridia; Clostridiales;

CDIF29747__locus_1

EUT2I

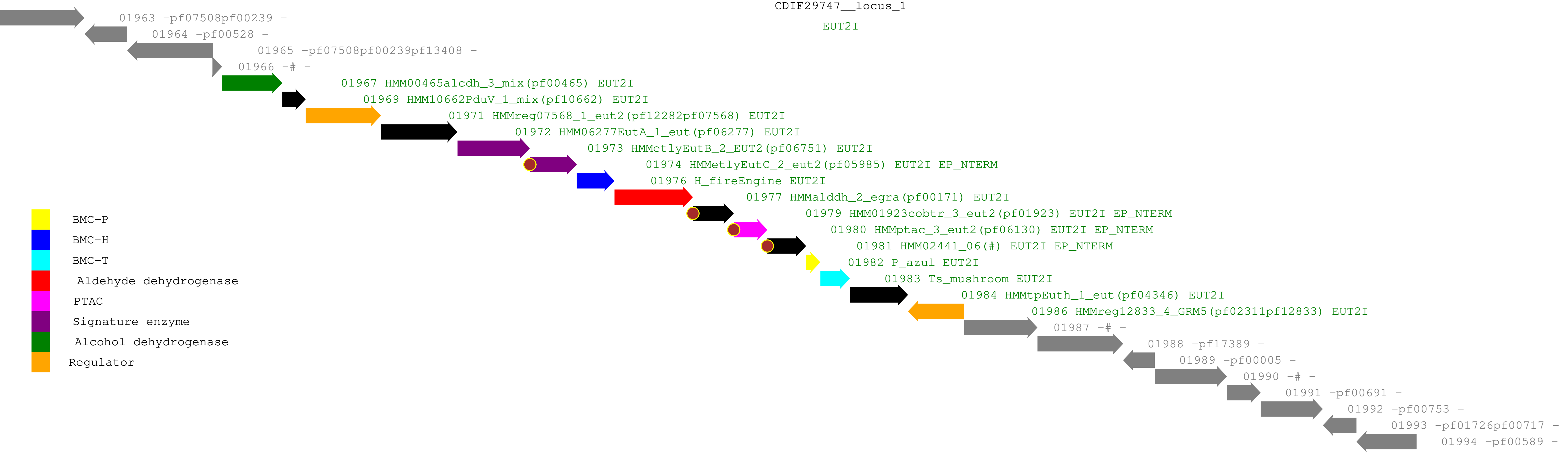

Clostridioides difficile M120. Bacteria; Firmicutes; Clostridia; Clostridiales;

CDIFFM120__locus_1

EUT2I

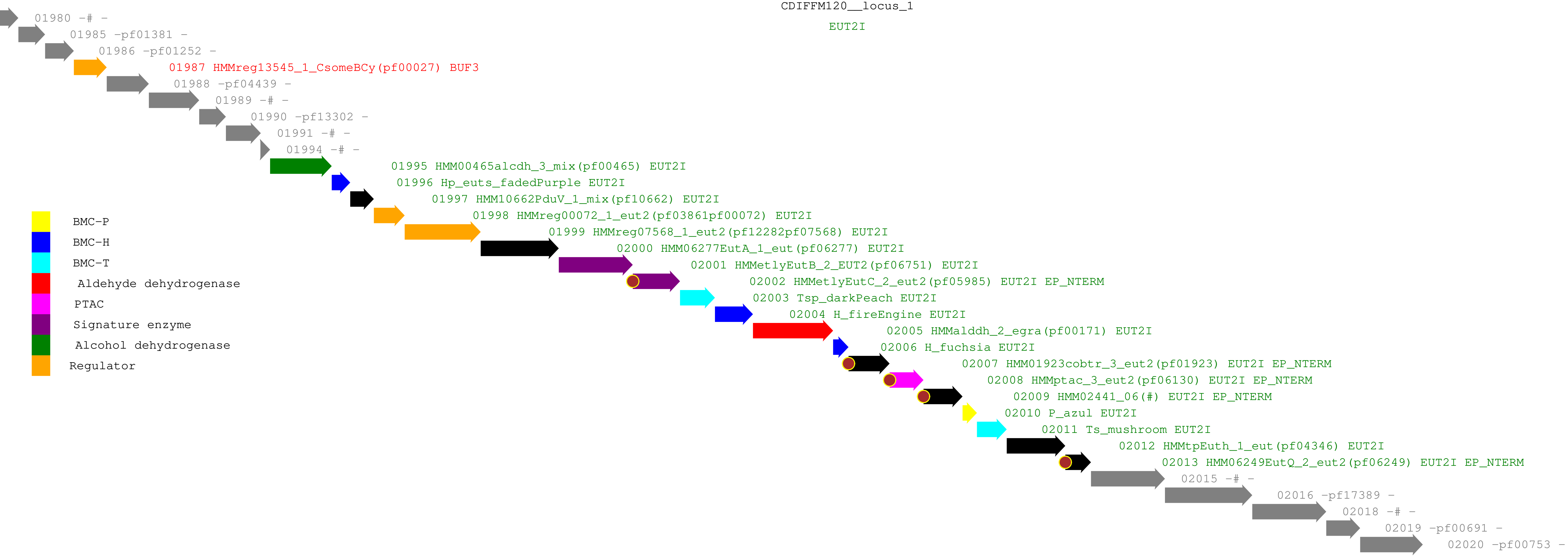

Peptoclostridium difficile (strain R20291) (Clostridium difficile). Bacteria; Firmicutes; Clostridia; Clostridiales;

CDR20291__locus_1

EUT2I

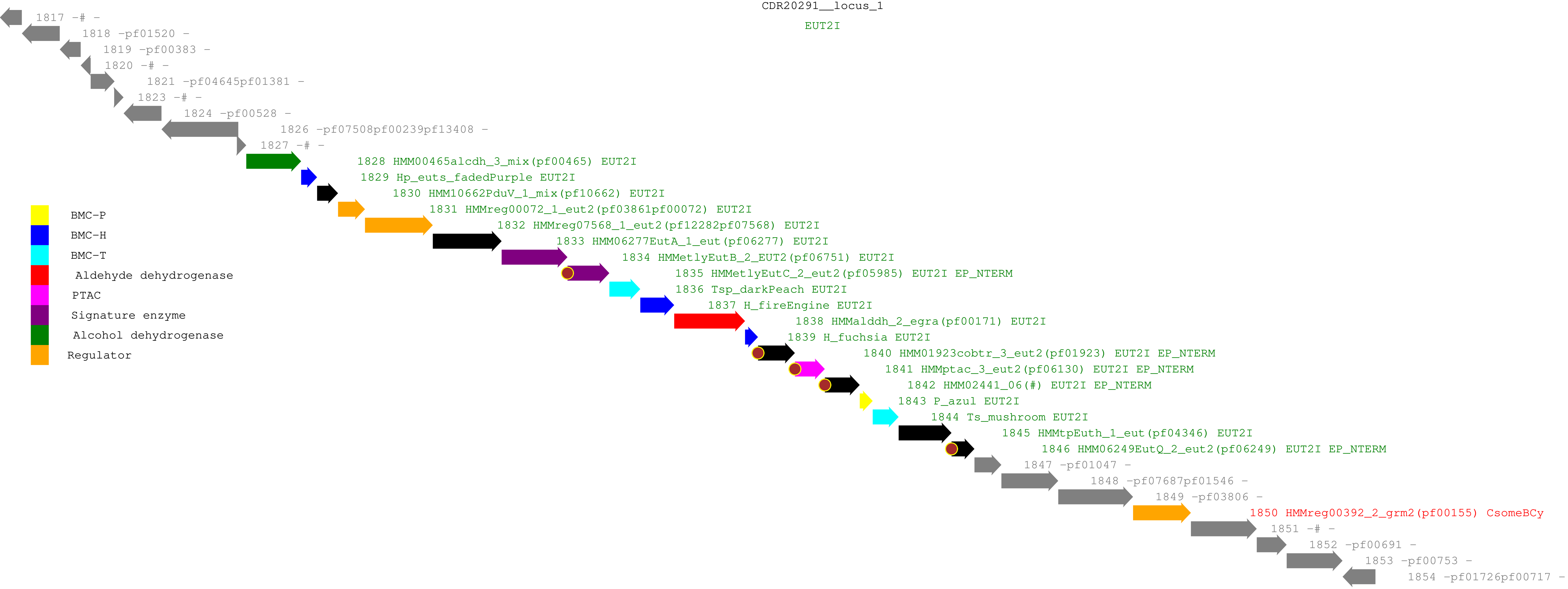

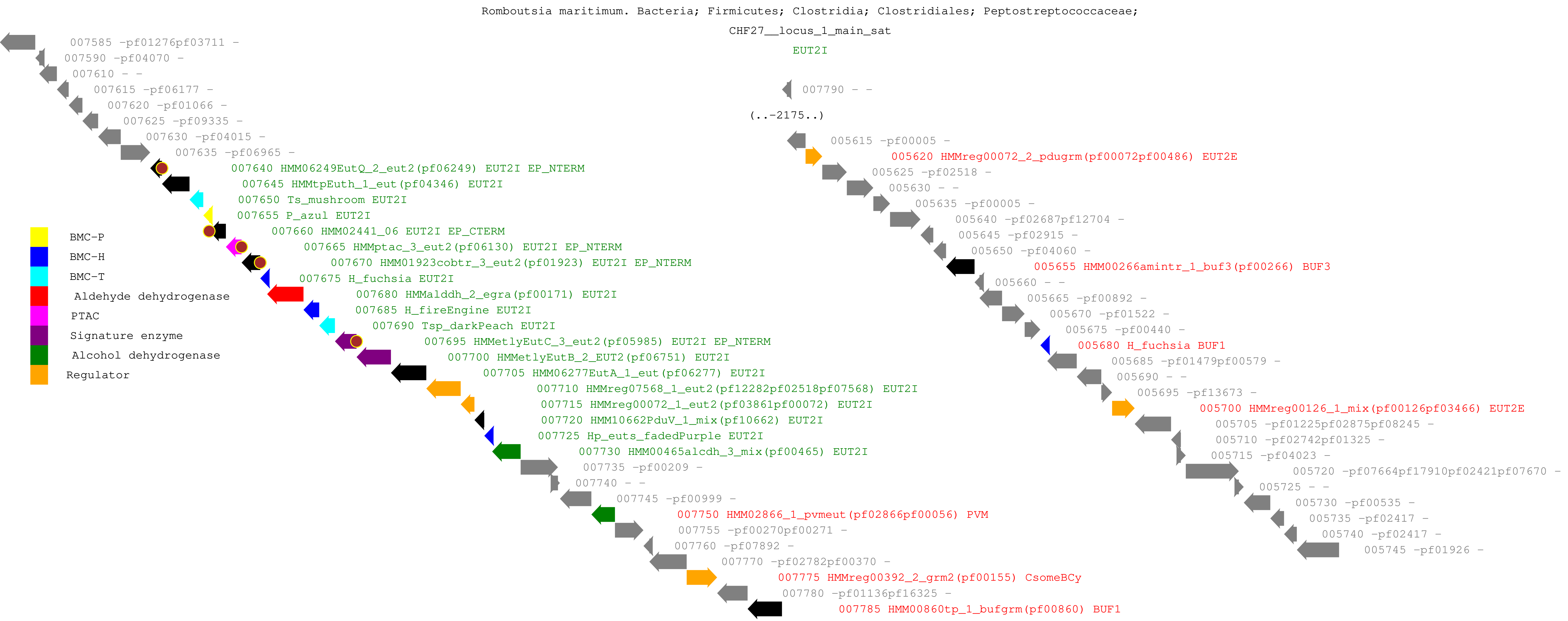
Romboutsia maritimum. Bacteria; Firmicutes; Clostridia; Clostridiales; Peptostreptococcaceae;
CHF27__locus_1_main_sat
EUT2I
007585 -pf01276pf03711 -
007590 -pf04070 -
007610 - -
007615 -pf06177 -
007620 -pf01066 -
007625 -pf09335 -
007630 -pf04015 -
007635 -pf06965 -
007640 HMM06249EutQ_2_eut2(pf06249) EUT2I EP_NTERM
007645 HMMtpEuth_1_eut(pf04346) EUT2I
007650 Ts_mushroom EUT2I
007655 P_azul EUT2I
007660 HMM02441_06 EUT2I EP_CTERM
007665 HMMptac_3_eut2(pf06130) EUT2I EP_NTERM
007670 HMM01923cobtr_3_eut2(pf01923) EUT2I EP_NTERM
007675 H_fuchsia EUT2I
007680 HMMalddh_2_egra(pf00171) EUT2I
007685 H_fireEngine EUT2I
007690 Tsp_darkPeach EUT2I
007695 HMMetlyEutC_3_eut2(pf05985) EUT2I EP_NTERM
007700 HMMetlyEutB_2_EUT2(pf06751) EUT2I
007705 HMM06277EutA_1_eut(pf06277) EUT2I
007710 HMMreg07568_1_eut2(pf12282pf02518pf07568) EUT2I
007715 HMMreg00072_1_eut2(pf03861pf00072) EUT2I
007720 HMM10662PduV_1_mix(pf10662) EUT2I
007725 Hp_euts_fadedPurple EUT2I
007730 HMM00465alcdh_3_mix(pf00465) EUT2I
007735 -pf00209 -
007740 - -
007745 -pf00999 -
007750 HMM02866_1_pvmeut(pf02866pf00056) PVM
007755 -pf00270pf00271 -
007760 -pf07892 -
007770 -pf02782pf00370 -
007775 HMMreg00392_2_grm2(pf00155) CsomeBCy
007780 -pf01136pf16325 -
007785 HMM00860tp_1_bufgrm(pf00860) BUF1
007790 - -
(..-2175..)
005615 -pf00005 -
005620 HMMreg00072_2_pdugrm(pf00072pf00486) EUT2E
005625 -pf02518 -
005630 - -
005635 -pf00005 -
005640 -pf02687pf12704 -
005645 -pf02915 -
005650 -pf04060 -
005655 HMM00266amintr_1_buf3(pf00266) BUF3
005660 - -
005665 -pf00892 -
005670 -pf01522 -
005675 -pf00440 -
005680 H_fuchsia BUF1
005685 -pf01479pf00579 -
005690 - -
005695 -pf13673 -
005700 HMMreg00126_1_mix(pf00126pf03466) EUT2E
005705 -pf01225pf02875pf08245 -
005710 -pf02742pf01325 -
005715 -pf04023 -
005720 -pf07664pf17910pf02421pf07670 -
005725 - -
005730 -pf00535 -
005735 -pf02417 -
005740 -pf02417 -
005745 -pf01926 -
BMC-P
BMC-H
BMC-T
Aldehyde dehydrogenase
PTAC
Signature enzyme
Alcohol dehydrogenase
Regulator

Bacillus sp. 7504-2. Bacteria; Firmicutes; Bacilli; Bacillales; Bacillaceae; Bacillus.

CHH80__locus_2

EUT2I

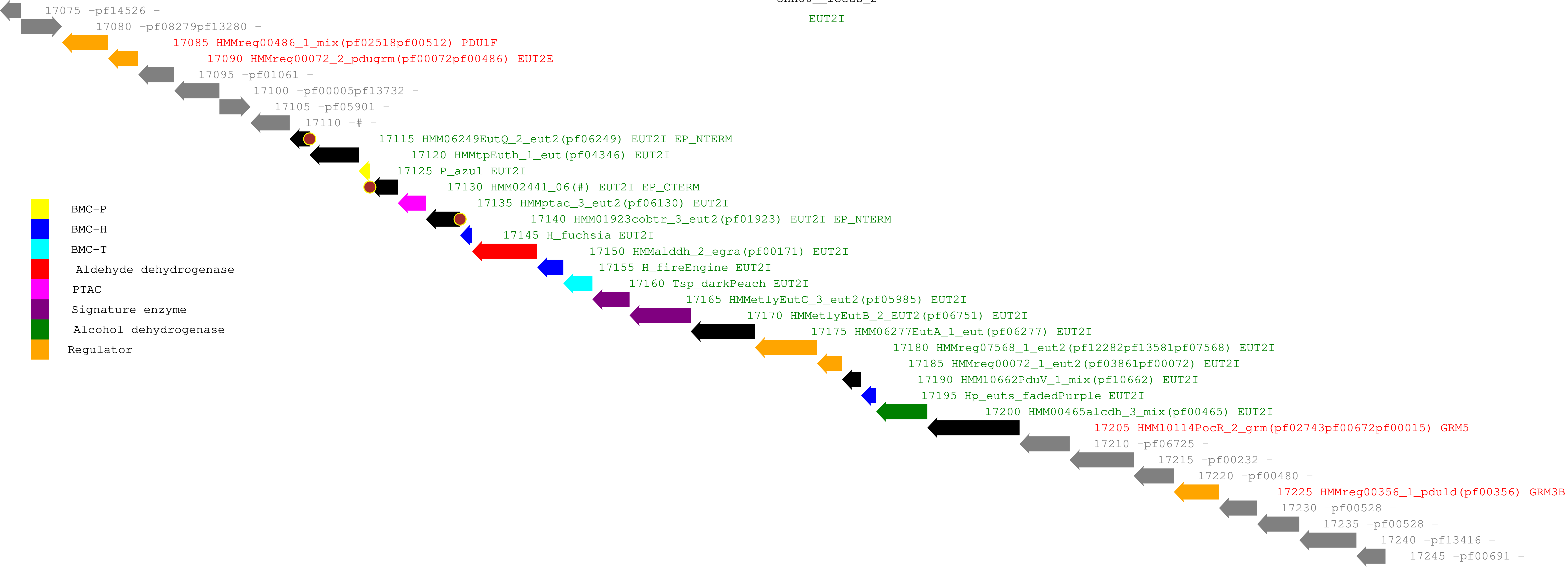

Clostridium perfringens D str. JGS1721. Bacteria; Firmicutes; Clostridia; Clostridiales; Clostridiaceae;

CJD__locus_1

EUT2I

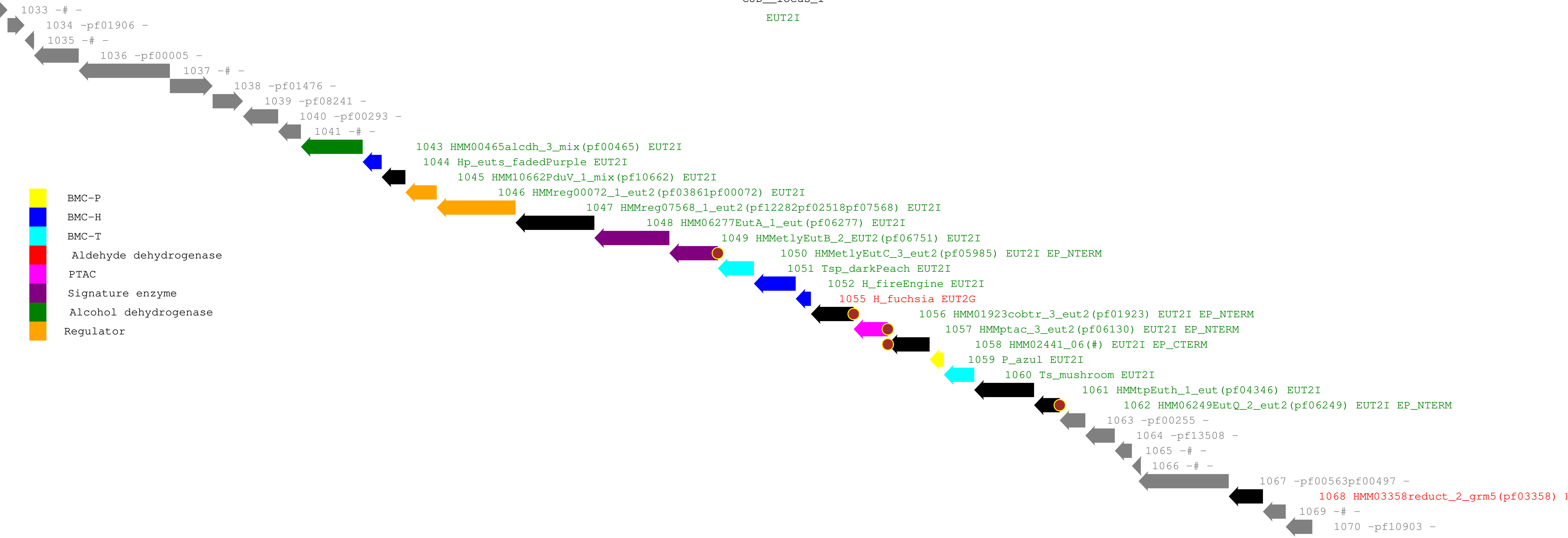

Cellulosilyticum lentocellum (strain ATCC 49066 / DSM 5427 / NCIMB Bacteria; Firmicutes; Clostridia; Clostridiales; Lachnospiraceae;

Clole__locus_1

EUT2I

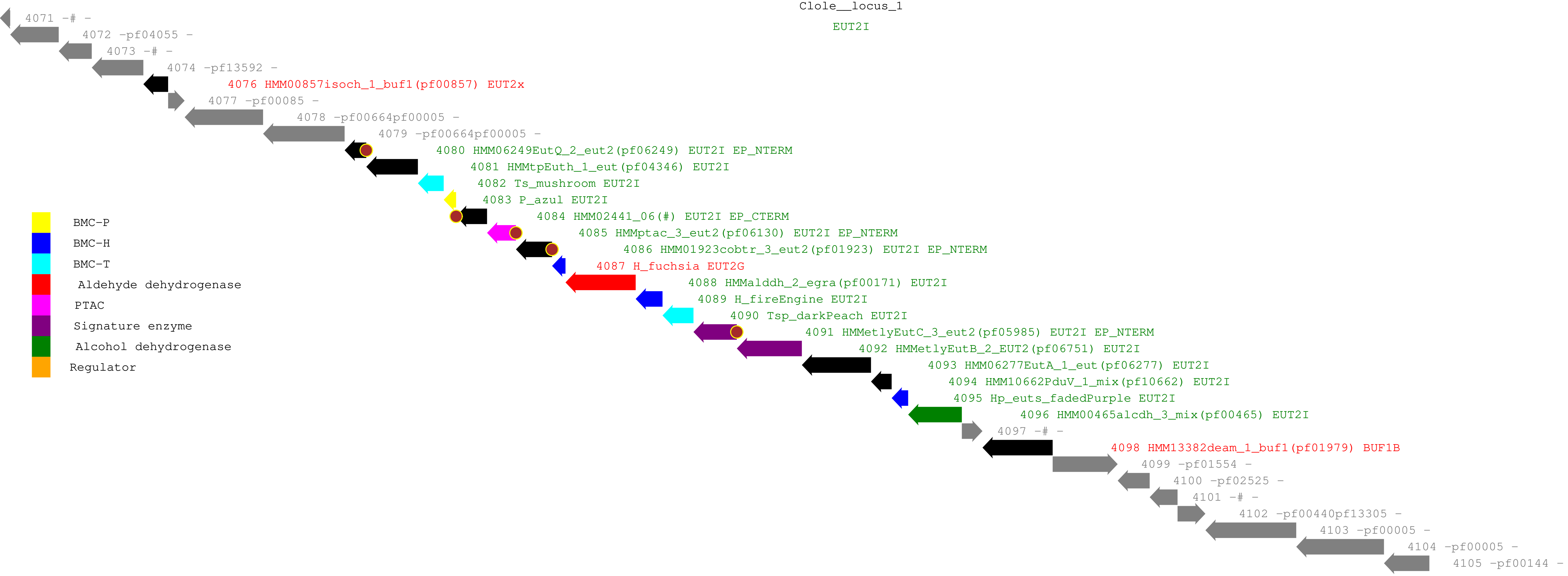

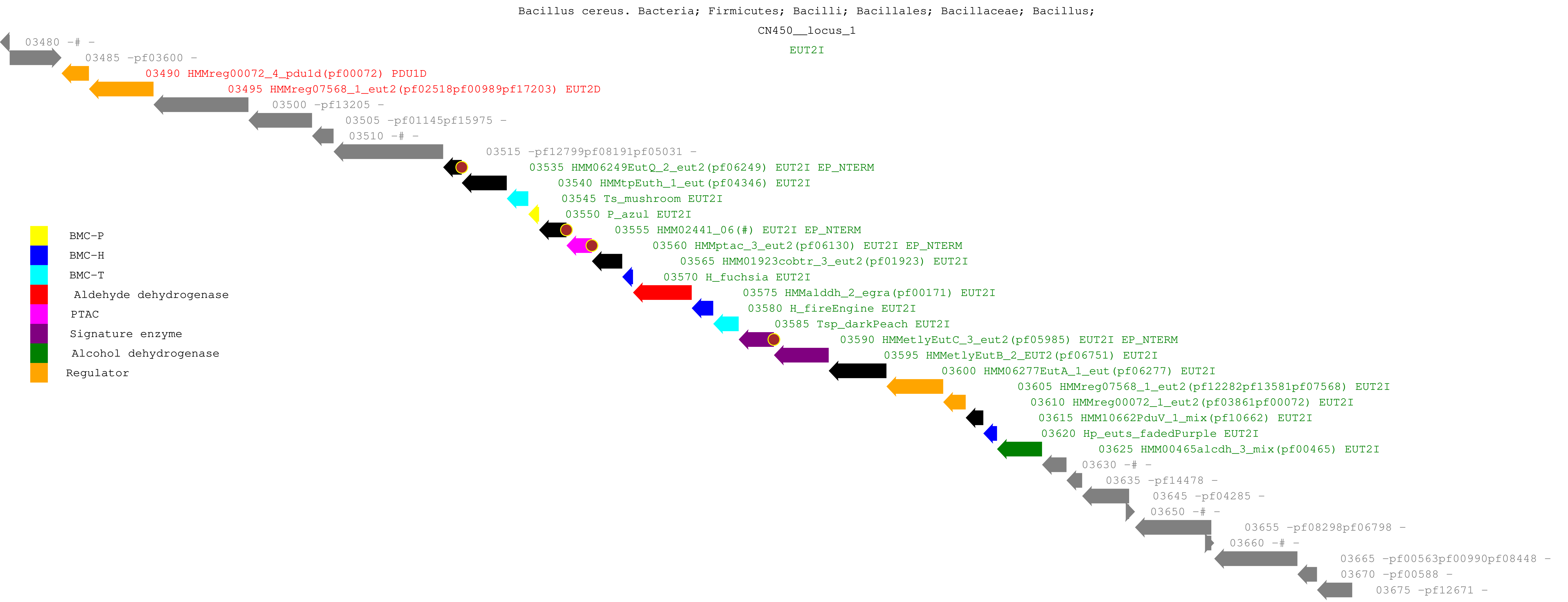

Bacillus cereus. Bacteria; Firmicutes; Bacilli; Bacillales; Bacillaceae; Bacillus;
CN450__locus_1
EUT2I
03480 -# -
03485 -pf03600 -
03490 HMMreg00072_4_pdu1d(pf00072) PDU1D
03495 HMMreg07568_1_eut2(pf02518pf00989pf17203) EUT2D
03500 -pf13205 -
03505 -pf01145pf15975 -
03510 -# -
03515 -pf12799pf08191pf05031 -
03535 HMM06249EutQ_2_eut2(pf06249) EUT2I EP_NTERM
03540 HMMtpEuth_1_eut(pf04346) EUT2I
03545 Ts_mushroom EUT2I
03550 P_azul EUT2I
03555 HMM02441_06(#) EUT2I EP_NTERM
03560 HMMptac_3_eut2(pf06130) EUT2I EP_NTERM
03565 HMM01923cobtr_3_eut2(pf01923) EUT2I
03570 H_fuchsia EUT2I
03575 HMMalddh_2_egra(pf00171) EUT2I
03580 H_fireEngine EUT2I
03585 Tsp_darkPeach EUT2I
03590 HMMetlyEutC_3_eut2(pf05985) EUT2I EP_NTERM
03595 HMMetlyEutB_2_EUT2(pf06751) EUT2I
03600 HMM06277EutA_1_eut(pf06277) EUT2I
03605 HMMreg07568_1_eut2(pf12282pf13581pf07568) EUT2I
03610 HMMreg00072_1_eut2(pf03861pf00072) EUT2I
03615 HMM10662PduV_1_mix(pf10662) EUT2I
03620 Hp_euts_fadedPurple EUT2I
03625 HMM00465alcdh_3_mix(pf00465) EUT2I
03630 -# -
03635 -pf14478 -
03645 -pf04285 -
03650 -# -
03655 -pf08298pf06798 -
03660 -# -
03665 -pf00563pf00990pf08448 -
03670 -pf00588 -
03675 -pf12671 -
BMC-P
BMC-H
BMC-T
Aldehyde dehydrogenase
PTAC
Signature enzyme
Alcohol dehydrogenase
Regulator

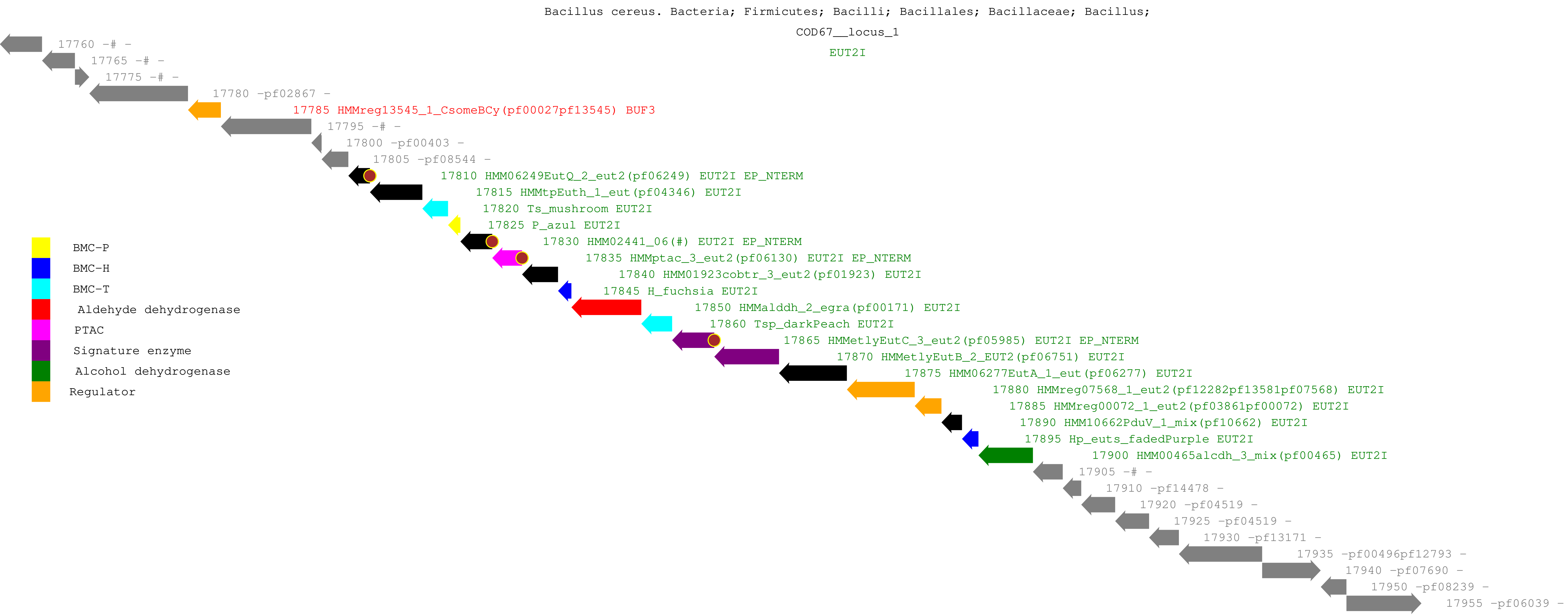
Bacillus cereus. Bacteria; Firmicutes; Bacilli; Bacillales; Bacillaceae; Bacillus;
COD67__locus_1
EUT2I
17760 -# -
17765 -# -
17775 -# -
17780 -pf02867 -
17785 HMMreg13545_1_CsomeBCy(pf00027pf13545) BUF3
17795 -# -
17800 -pf00403 -
17805 -pf08544 -
17810 HMM06249EutQ_2_eut2(pf06249) EUT2I EP_NTERM
17815 HMMtpEuth_1_eut(pf04346) EUT2I
17820 Ts_mushroom EUT2I
17825 P_azul EUT2I
17830 HMM02441_06(#) EUT2I EP_NTERM
17835 HMMptac_3_eut2(pf06130) EUT2I EP_NTERM
17840 HMM01923cobtr_3_eut2(pf01923) EUT2I
17845 H_fuchsia EUT2I
17850 HMMalddh_2_egra(pf00171) EUT2I
17860 Tsp_darkPeach EUT2I
17865 HMMetlyEutC_3_eut2(pf05985) EUT2I EP_NTERM
17870 HMMetlyEutB_2_EUT2(pf06751) EUT2I
17875 HMM06277EutA_1_eut(pf06277) EUT2I
17880 HMMreg07568_1_eut2(pf12282pf13581pf07568) EUT2I
17885 HMMreg00072_1_eut2(pf03861pf00072) EUT2I
17890 HMM10662PduV_1_mix(pf10662) EUT2I
17895 Hp_euts_fadedPurple EUT2I
17900 HMM00465alcdh_3_mix(pf00465) EUT2I
17905 -# -
17910 -pf14478 -
17920 -pf04519 -
17925 -pf04519 -
17930 -pf13171 -
17935 -pf00496pf12793 -
17940 -pf07690 -
17950 -pf08239 -
17955 -pf06039 -
BMC-P
BMC-H
BMC-T
Aldehyde dehydrogenase
PTAC
Signature enzyme
Alcohol dehydrogenase
Regulator

Bacillus cereus. Bacteria; Firmicutes; Bacilli; Bacillales; Bacillaceae; Bacillus;

COI89__locus_1

EUT2I

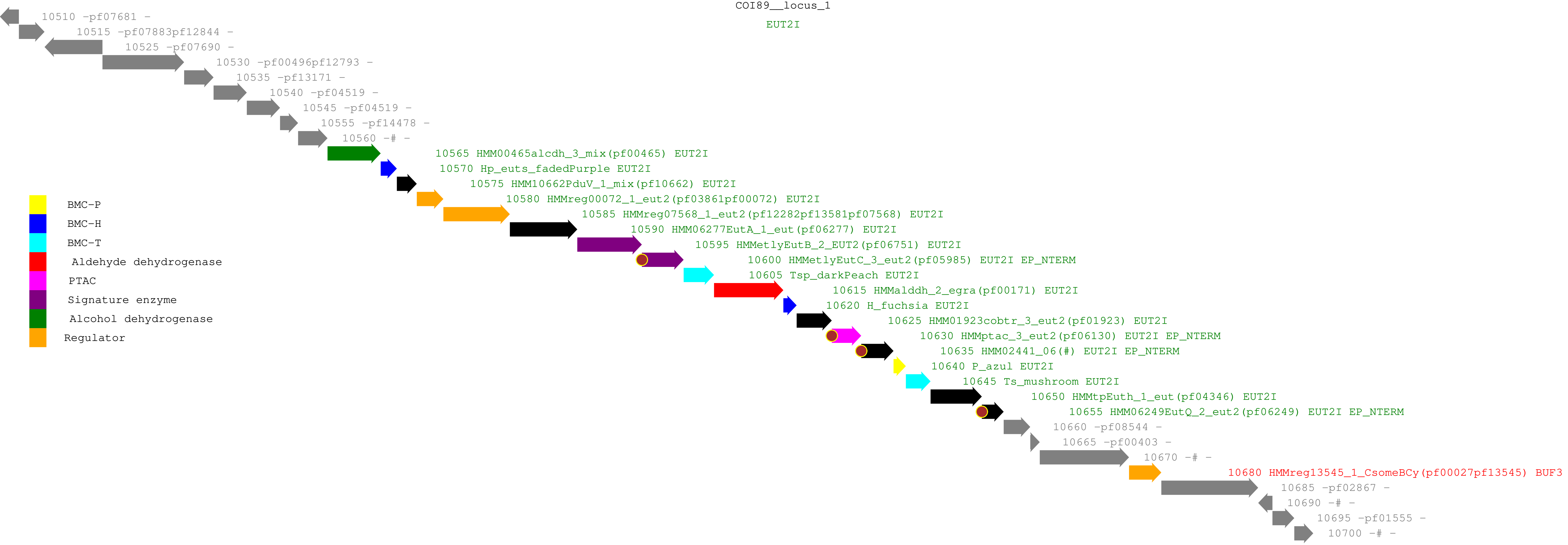

Clostridium perfringens C str. JGS1495. Bacteria; Firmicutes; Clostridia; Clostridiales; Clostridiaceae;

CPC__locus_1

EUT2I

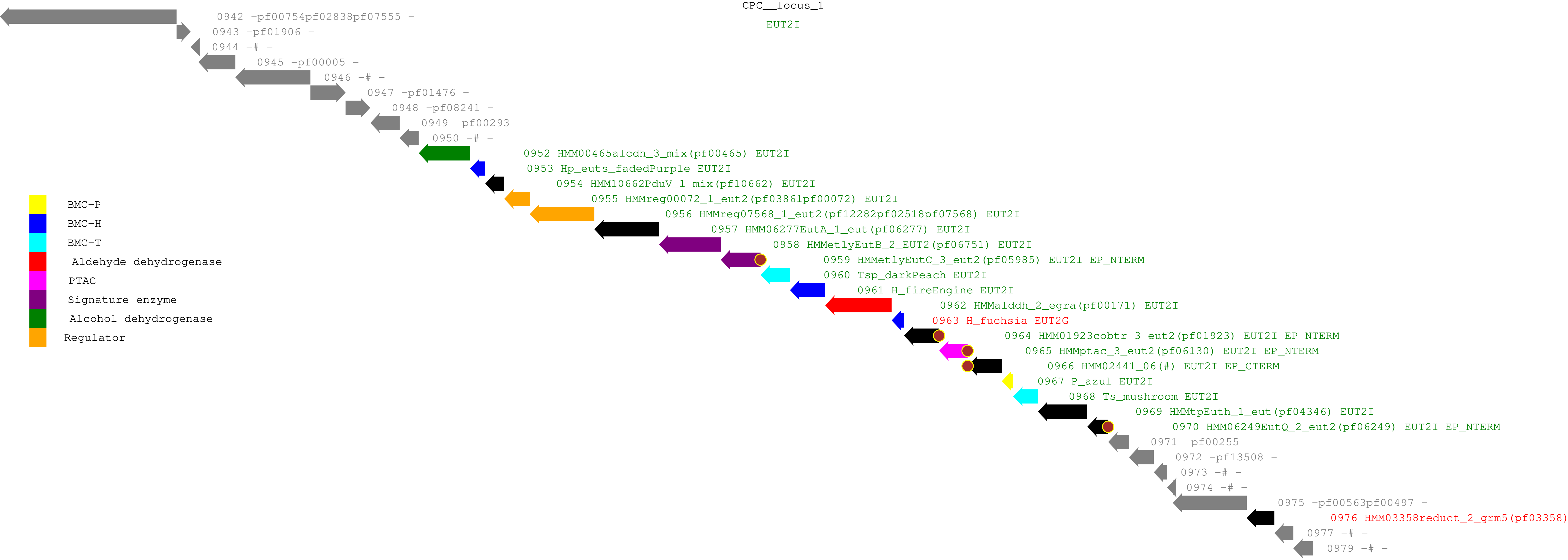

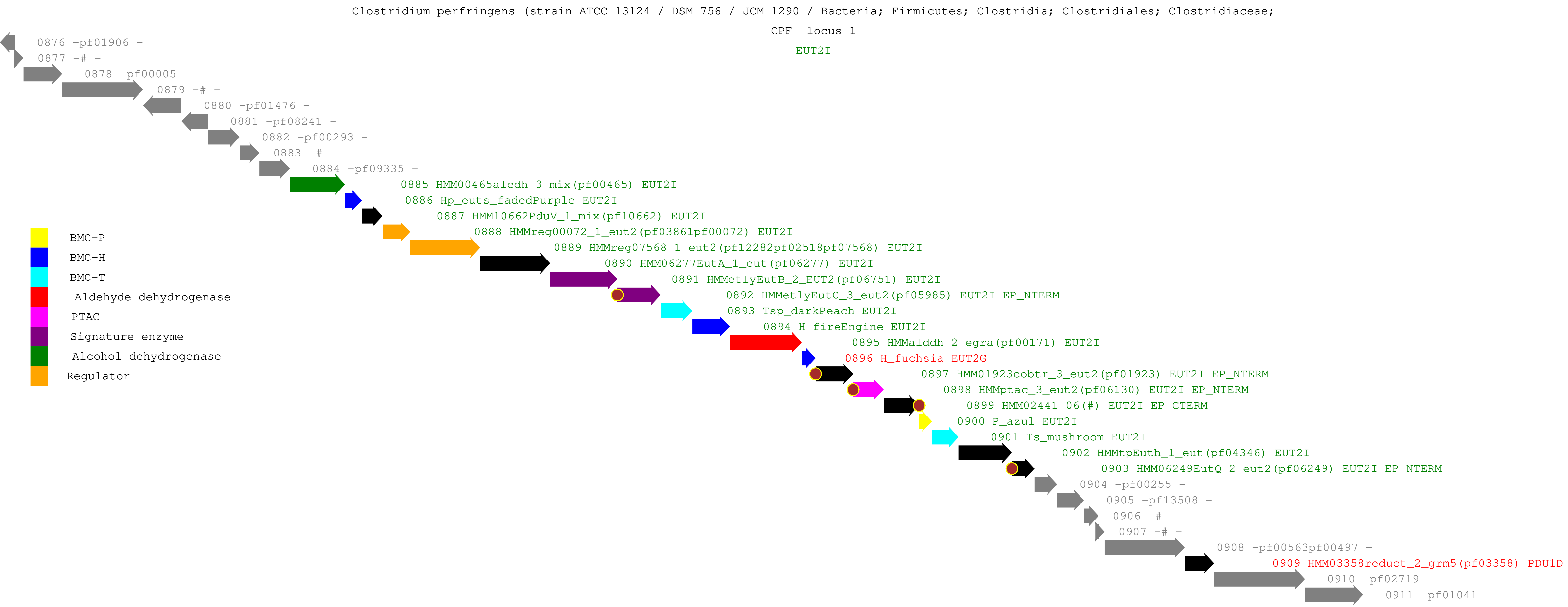
Clostridium perfringens (strain ATCC 13124 / DSM 756 / JCM 1290 / Bacteria; Firmicutes; Clostridia; Clostridiales; Clostridiaceae;
CPF__locus_1
EUT2I
0876 -pf01906 -
0877 -# -
0878 -pf00005 -
0879 -# -
0880 -pf01476 -
0881 -pf08241 -
0882 -pf00293 -
0883 -# -
0884 -pf09335 -
0885 HMM00465alcdh_3_mix(pf00465) EUT2I
0886 Hp_euts_fadedPurple EUT2I
0887 HMM10662PduV_1_mix(pf10662) EUT2I
0888 HMMreg00072_1_eut2(pf03861pf00072) EUT2I
0889 HMMreg07568_1_eut2(pf12282pf02518pf07568) EUT2I
0890 HMM06277EutA_1_eut(pf06277) EUT2I
0891 HMMetlyEutB_2_EUT2(pf06751) EUT2I
0892 HMMetlyEutC_3_eut2(pf05985) EUT2I EP_NTERM
0893 Tsp_darkPeach EUT2I
0894 H_fireEngine EUT2I
0895 HMMaldh_2_egra(pf00171) EUT2I
0896 H_fuchsia EUT2G
0897 HMM01923cobtr_3_eut2(pf01923) EUT2I EP_NTERM
0898 HMMptac_3_eut2(pf06130) EUT2I EP_NTERM
0899 HMM02441_06(#) EUT2I EP_CTERM
0900 P_azul EUT2I
0901 Ts_mushroom EUT2I
0902 HMMtpEuth_1_eut(pf04346) EUT2I
0903 HMM06249EutQ_2_eut2(pf06249) EUT2I EP_NTERM
0904 -pf00255 -
0905 -pf13508 -
0906 -# -
0907 -# -
0908 -pf00563pf00497 -
0909 HMM03358reduct_2_grm5(pf03358) PDU1D
0910 -pf02719 -
0911 -pf01041 -
BMC-P
BMC-H
BMC-T
Aldehyde dehydrogenase
PTAC
Signature enzyme
Alcohol dehydrogenase
Regulator

Lachnoclostridium phytofermentans (strain ATCC 700394 / DSM 18823 / Bacteria; Firmicutes; Clostridia; Clostridiales; Lachnospiraceae.

Cphy__locus_3

EUT2I

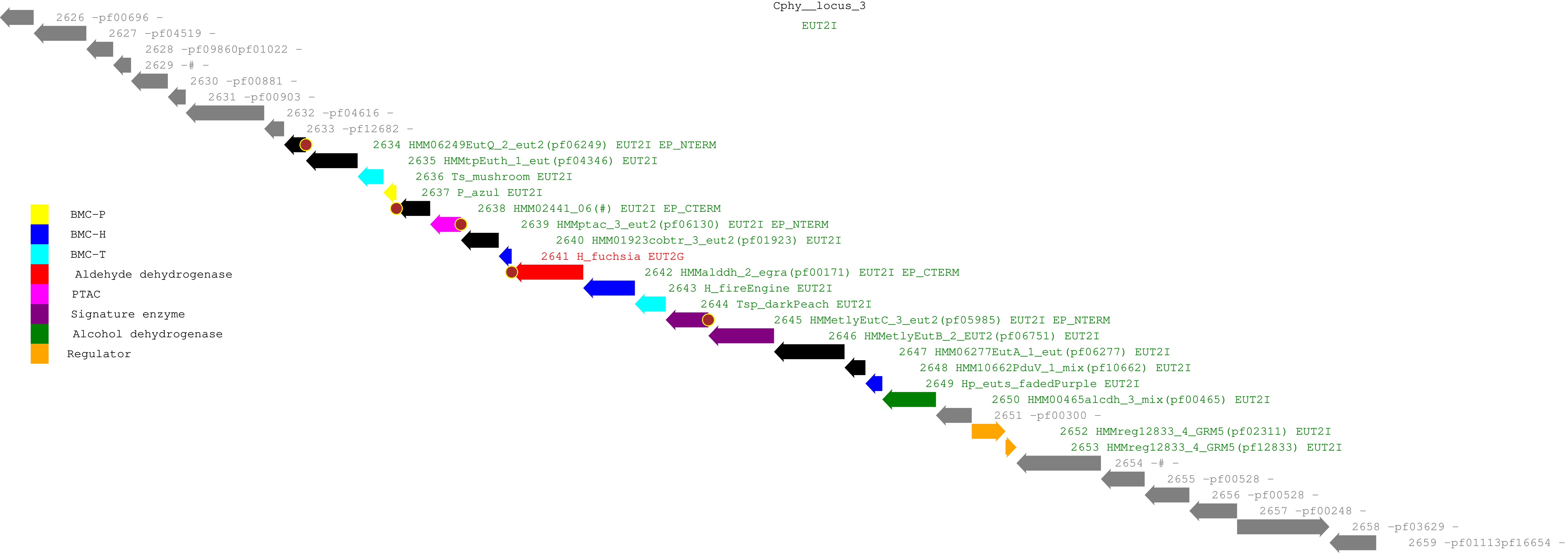

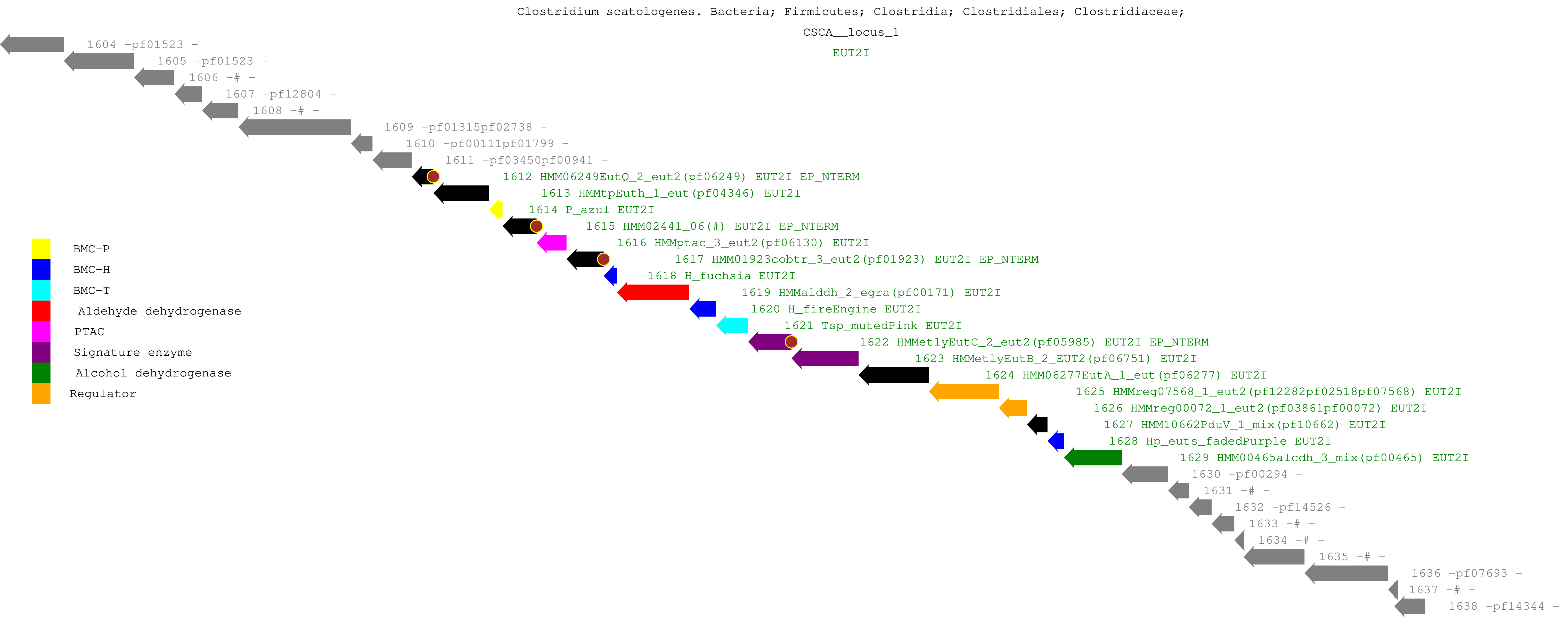
Clostridium scatologenes. Bacteria; Firmicutes; Clostridia; Clostridiales; Clostridiaceae;
CSCA__locus_1
EUT2I
1604 -pf01523 -
1605 -pf01523 -
1606 -# -
1607 -pf12804 -
1608 -# -
1609 -pf01315pf02738 -
1610 -pf00111pf01799 -
1611 -pf03450pf00941 -
1612 HMM06249EutQ_2_eut2(pf06249) EUT2I EP_NTERM
1613 HMMtpEuth_1_eut(pf04346) EUT2I
1614 P_azul EUT2I
1615 HMM02441_06(#) EUT2I EP_NTERM
1616 HMMptac_3_eut2(pf06130) EUT2I
1617 HMM01923cobtr_3_eut2(pf01923) EUT2I EP_NTERM
1618 H_fuchsia EUT2I
1619 HMMaldh_2_egra(pf00171) EUT2I
1620 H_fireEngine EUT2I
1621 Tsp_mutedPink EUT2I
1622 HMMetlyEutC_2_eut2(pf05985) EUT2I EP_NTERM
1623 HMMetlyEutB_2_EUT2(pf06751) EUT2I
1624 HMM06277EutA_1_eut(pf06277) EUT2I
1625 HMMreg07568_1_eut2(pf12282pf02518pf07568) EUT2I
1626 HMMreg00072_1_eut2(pf03861pf00072) EUT2I
1627 HMM10662PduV_1_mix(pf10662) EUT2I
1628 Hp_euts_fadedPurple EUT2I
1629 HMM00465alcdh_3_mix(pf00465) EUT2I
1630 -pf00294 -
1631 -# -
1632 -pf14526 -
1633 -# -
1634 -# -
1635 -# -
1636 -pf07693 -
1637 -# -
1638 -pf14344 -
BMC-P
BMC-H
BMC-T
Aldehyde dehydrogenase
PTAC
Signature enzyme
Alcohol dehydrogenase
Regulator

Clostridium tagluense. Bacteria; Firmicutes; Clostridia; Clostridiales; Clostridiaceae;

Ctaglu__locus_1

EUT2I

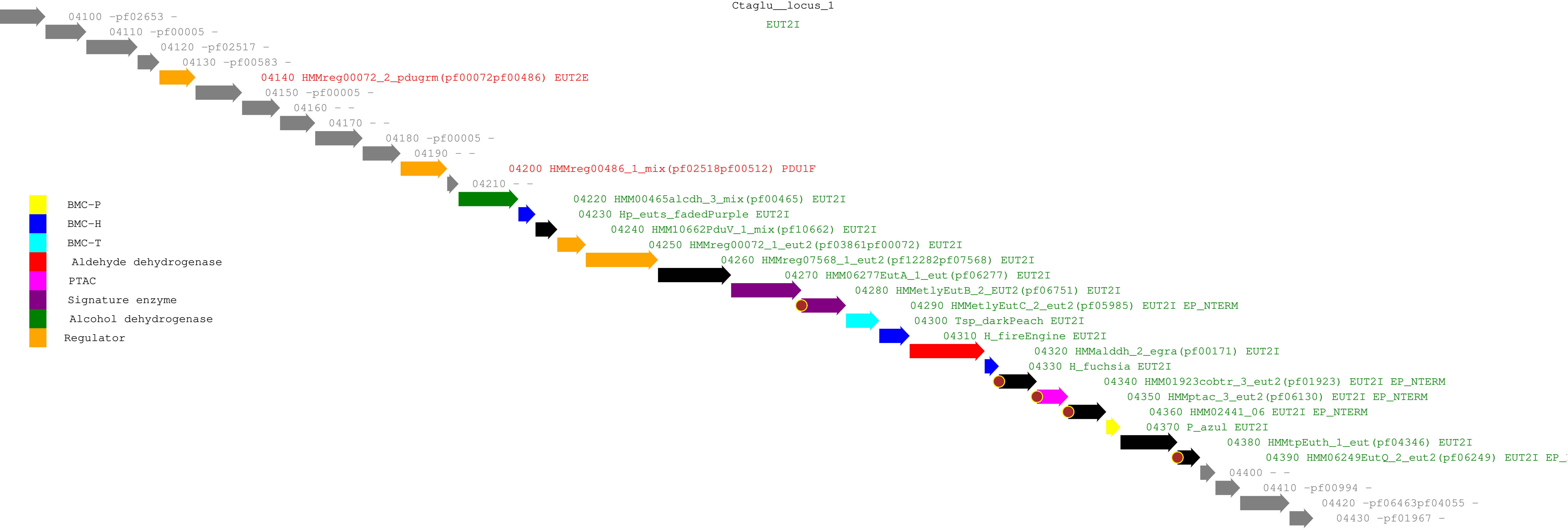

Clostridium tetanomorphum DSM 665. Bacteria; Firmicutes; Clostridia; Clostridiales; Clostridiaceae;

CTM__locus_1

EUT2I

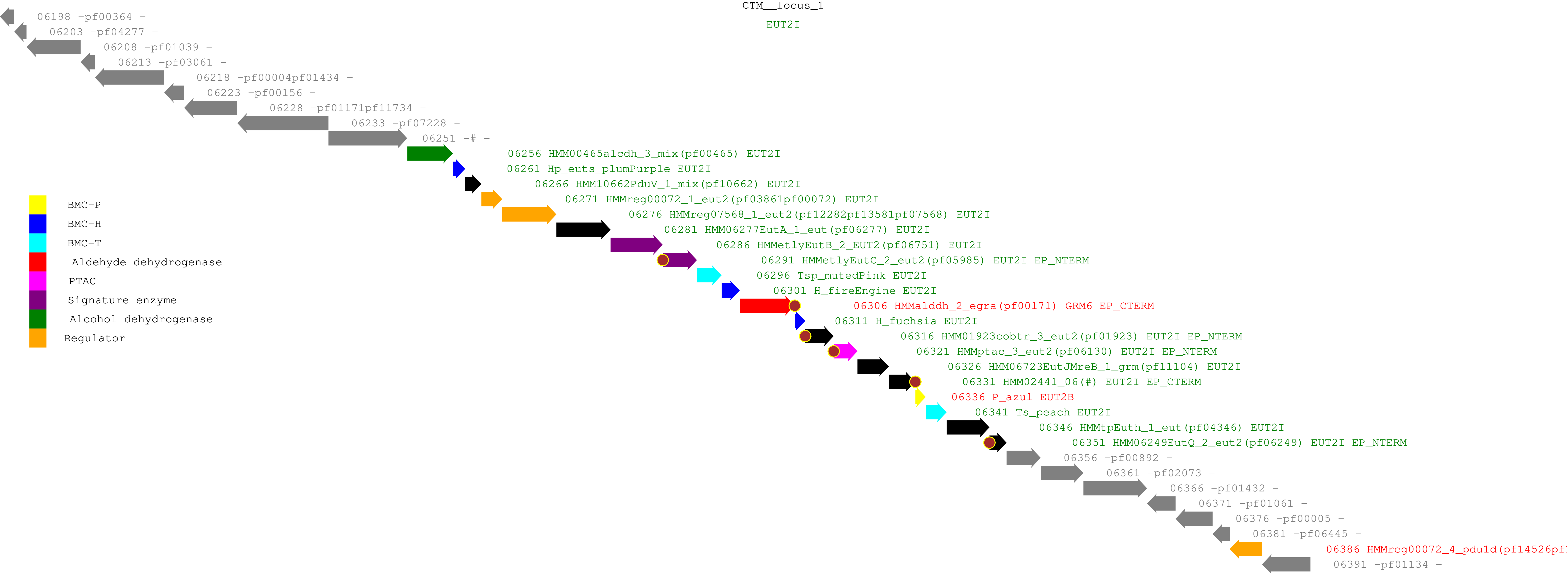

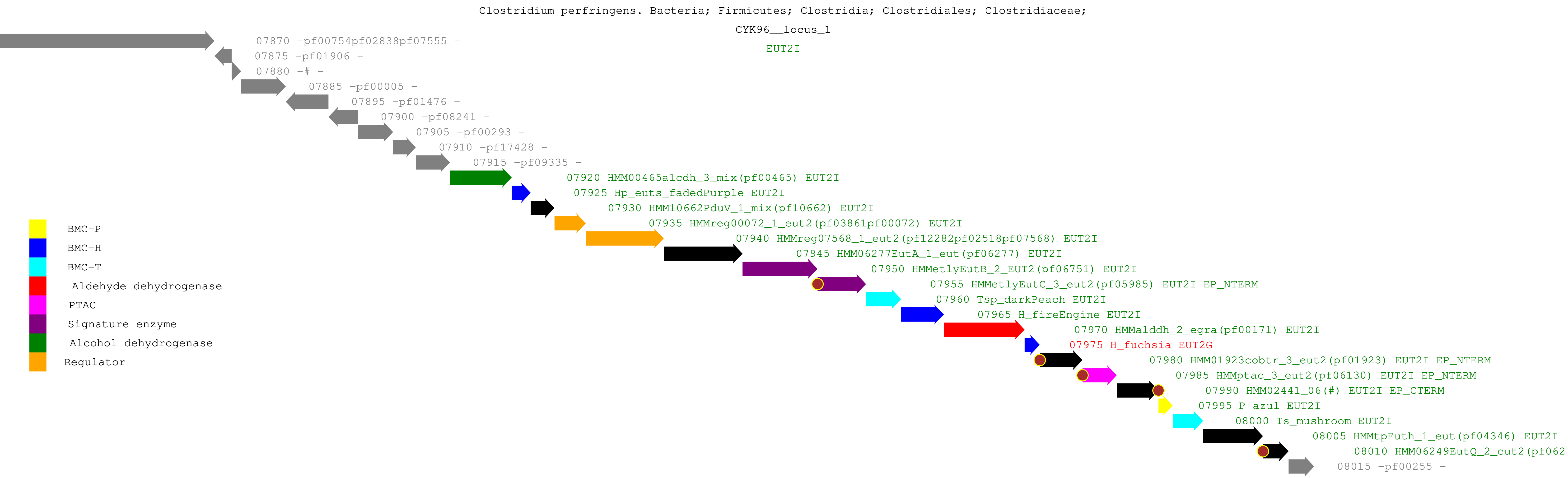
Clostridium perfringens. Bacteria; Firmicutes; Clostridia; Clostridiales; Clostridiaceae;
CYK96__locus_1
EUT2I
07870 -pf00754pf02838pf07555 -
07875 -pf01906 -
07880 -# -
07885 -pf00005 -
07895 -pf01476 -
07900 -pf08241 -
07905 -pf00293 -
07910 -pf17428 -
07915 -pf09335 -
07920 HMM00465alcdh_3_mix(pf00465) EUT2I
07925 Hp_euts_fadedPurple EUT2I
07930 HMM10662PduV_1_mix(pf10662) EUT2I
07935 HMMreg00072_1_eut2(pf03861pf00072) EUT2I
07940 HMMreg07568_1_eut2(pf12282pf02518pf07568) EUT2I
07945 HMM06277EutA_1_eut(pf06277) EUT2I
07950 HMMetlyEutB_2_EUT2(pf06751) EUT2I
07955 HMMetlyEutC_3_eut2(pf05985) EUT2I EP_NTERM
07960 Tsp_darkPeach EUT2I
07965 H_fireEngine EUT2I
07970 HMMalddh_2_egra(pf00171) EUT2I
07975 H_fuchsia EUT2G
07980 HMM01923cobtr_3_eut2(pf01923) EUT2I EP_NTERM
07985 HMMptac_3_eut2(pf06130) EUT2I EP_NTERM
07990 HMM02441_06(#) EUT2I EP_CTERM
07995 P_azul EUT2I
08000 Ts_mushroom EUT2I
08005 HMMtpEuth_1_eut(pf04346) EUT2I
08010 HMM06249EutQ_2_eut2(pf062
08015 -pf00255 -
BMC-P
BMC-H
BMC-T
Aldehyde dehydrogenase
PTAC
Signature enzyme
Alcohol dehydrogenase
Regulator

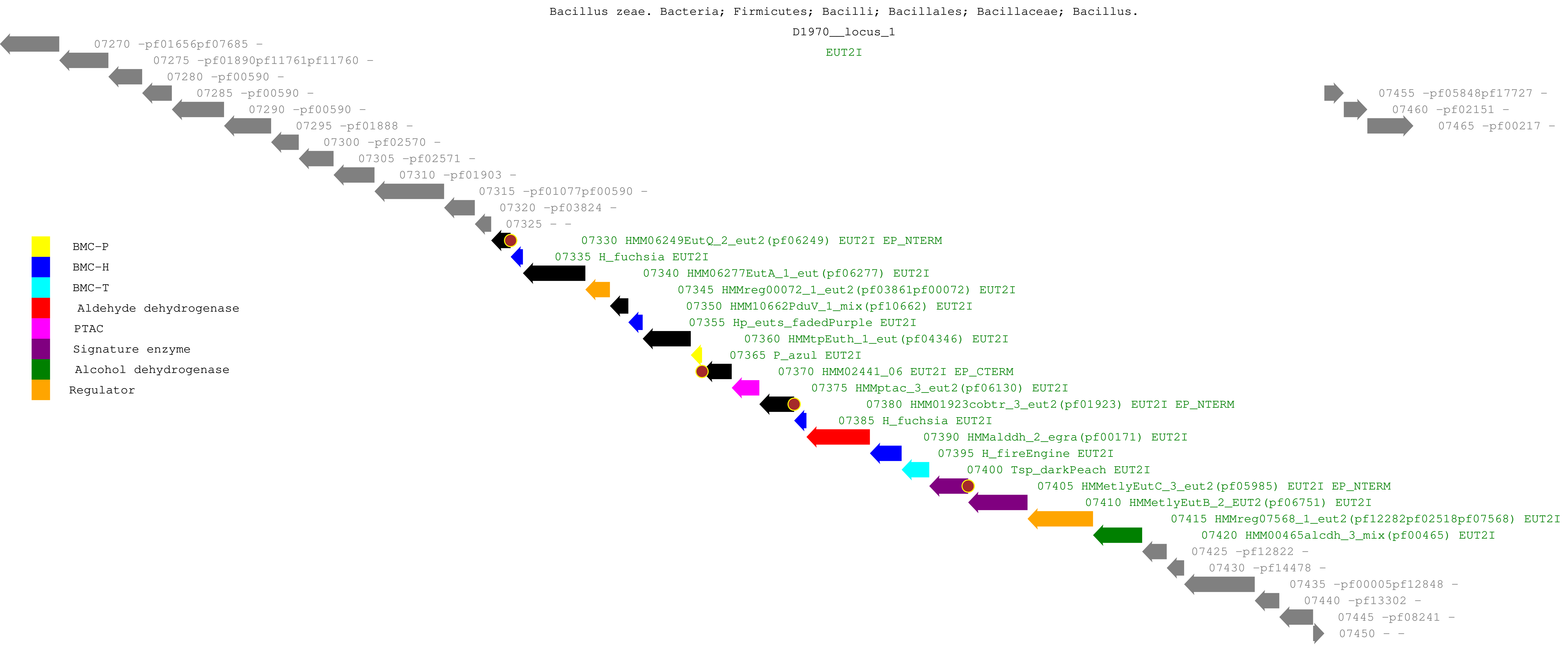
Bacillus zeae. Bacteria; Firmicutes; Bacilli; Bacillales; Bacillaceae; Bacillus.
D1970__locus_1
EUT2I
07270 -pf01656pf07685 -
07275 -pf01890pf11761pf11760 -
07280 -pf00590 -
07285 -pf00590 -
07290 -pf00590 -
07295 -pf01888 -
07300 -pf02570 -
07305 -pf02571 -
07310 -pf01903 -
07315 -pf01077pf00590 -
07320 -pf03824 -
07325 - -
07330 HMM06249EutQ_2_eut2(pf06249) EUT2I EP_NTERM
07335 H_fuchsia EUT2I
07340 HMM06277EutA_1_eut(pf06277) EUT2I
07345 HMMreg00072_1_eut2(pf03861pf00072) EUT2I
07350 HMM10662PduV_1_mix(pf10662) EUT2I
07355 Hp_euts_fadedPurple EUT2I
07360 HMMtpEuth_1_eut(pf04346) EUT2I
07365 P_azul EUT2I
07370 HMM02441_06 EUT2I EP_CTERM
07375 HMMptac_3_eut2(pf06130) EUT2I
07380 HMM01923cobtr_3_eut2(pf01923) EUT2I EP_NTERM
07385 H_fuchsia EUT2I
07390 HMMalddh_2_egra(pf00171) EUT2I
07395 H_fireEngine EUT2I
07400 Tsp_darkPeach EUT2I
07405 HMMetlyEutC_3_eut2(pf05985) EUT2I EP_NTERM
07410 HMMetlyEutB_2_EUT2(pf06751) EUT2I
07415 HMMreg07568_1_eut2(pf12282pf02518pf07568) EUT2I
07420 HMM00465alcdh_3_mix(pf00465) EUT2I
07425 -pf12822 -
07430 -pf14478 -
07435 -pf00005pf12848 -
07440 -pf13302 -
07445 -pf08241 -
07450 - -
07455 -pf05848pf17727 -
07460 -pf02151 -
07465 -pf00217 -
BMC-P
BMC-H
BMC-T
Aldehyde dehydrogenase
PTAC
Signature enzyme
Alcohol dehydrogenase
Regulator

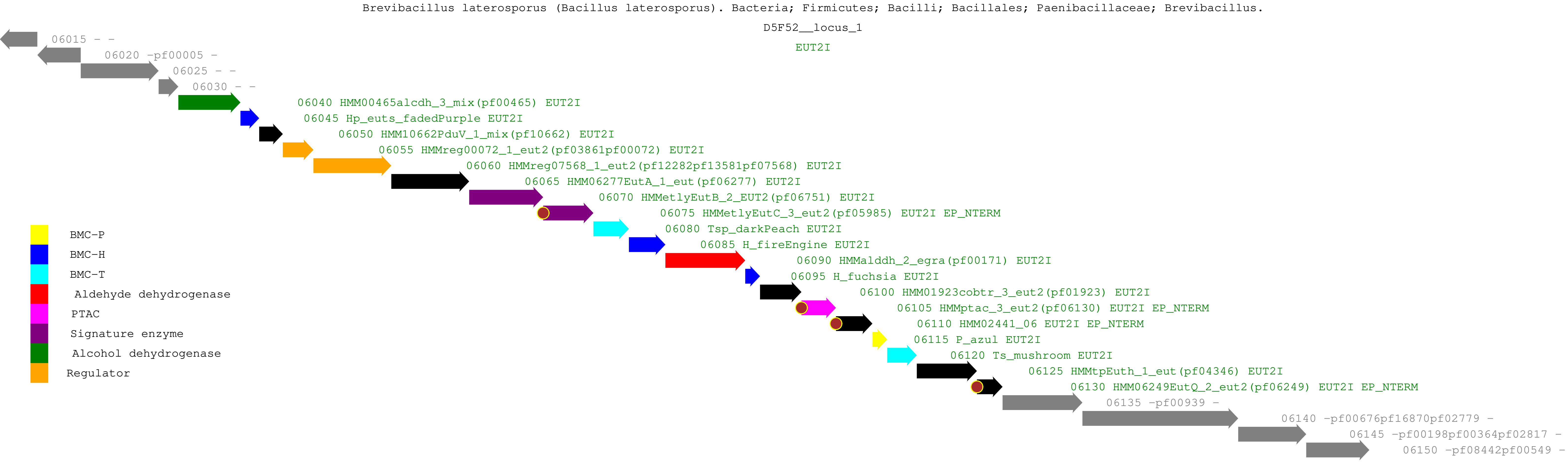

Brevibacillus laterosporus (Bacillus laterosporus). Bacteria; Firmicutes; Bacilli; Bacillales; Paenibacillaceae; Brevibacillus.
D5F52__locus_1
EUT2I
06015 - -
06020 -pf00005 -
06025 - -
06030 - -
06040 HMM00465alcdh_3_mix(pf00465) EUT2I
06045 Hp_euts_fadedPurple EUT2I
06050 HMM10662PduV_1_mix(pf10662) EUT2I
06055 HMMreg00072_1_eut2(pf03861pf00072) EUT2I
06060 HMMreg07568_1_eut2(pf12282pf13581pf07568) EUT2I
06065 HMM06277EutA_1_eut(pf06277) EUT2I
06070 HMMetlyEutB_2_EUT2(pf06751) EUT2I
06075 HMMetlyEutC_3_eut2(pf05985) EUT2I EP_NTERM
06080 Tsp_darkPeach EUT2I
06085 H_fireEngine EUT2I
06090 HMMaldh_2_egra(pf00171) EUT2I
06095 H_fuchsia EUT2I
06100 HMM01923cobtr_3_eut2(pf01923) EUT2I
06105 HMMptac_3_eut2(pf06130) EUT2I EP_NTERM
06110 HMM02441_06 EUT2I EP_NTERM
06115 P_azul EUT2I
06120 Ts_mushroom EUT2I
06125 HMMtpEuth_1_eut(pf04346) EUT2I
06130 HMM06249EutQ_2_eut2(pf06249) EUT2I EP_NTERM
06135 -pf00939 -
06140 -pf00676pf16870pf02779 -
06145 -pf00198pf00364pf02817 -
06150 -pf08442pf00549 -
BMC-P
BMC-H
BMC-T
Aldehyde dehydrogenase
PTAC
Signature enzyme
Alcohol dehydrogenase
Regulator

Brevibacillus laterosporus (Bacillus laterosporus). Bacteria; Firmicutes; Bacilli; Bacillales; Paenibacillaceae; Brevibacillus.

D5F52__locus_2

EUT2I

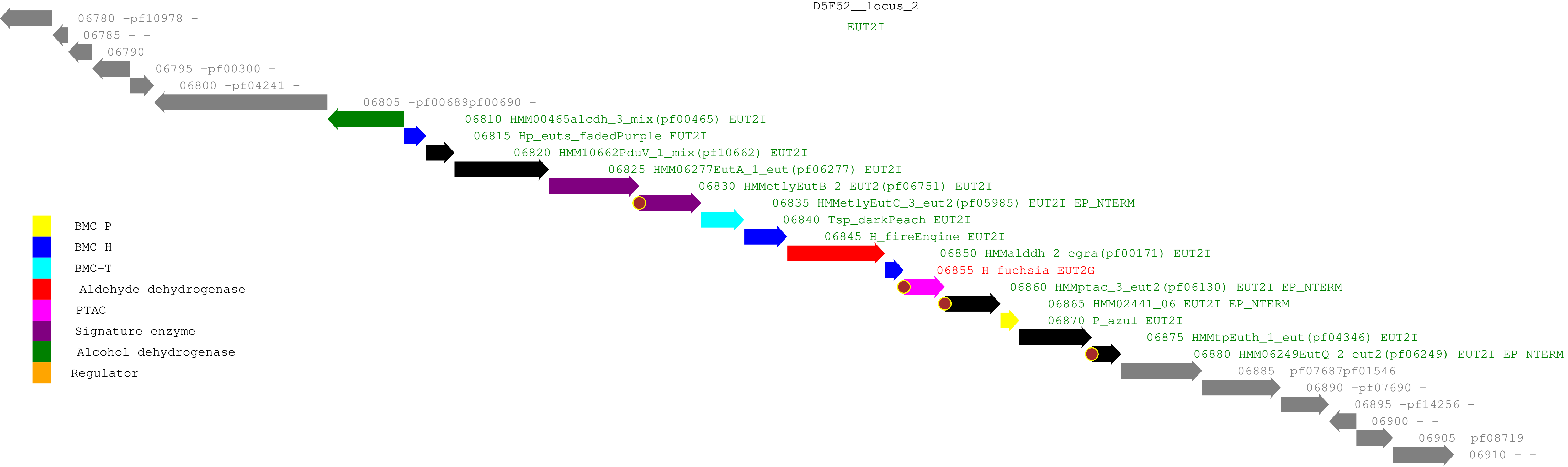

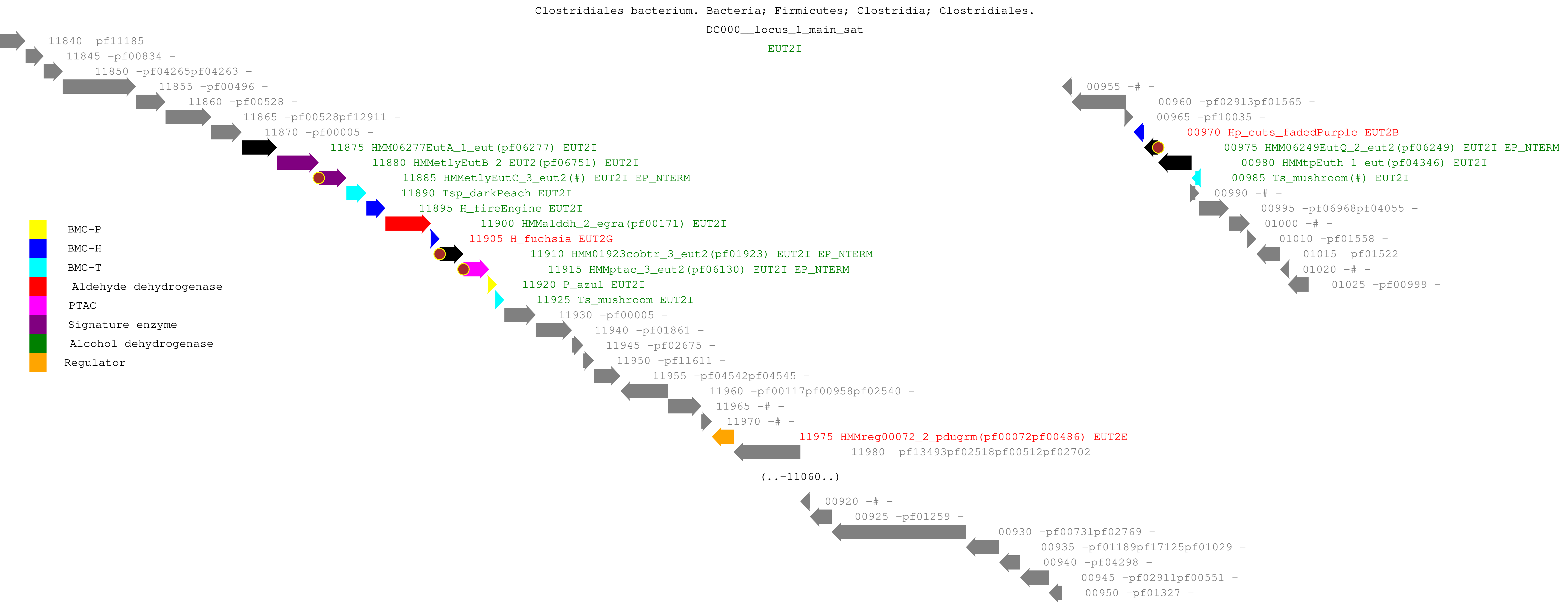
Clostridiales bacterium. Bacteria; Firmicutes; Clostridia; Clostridiales.
DC000__locus_1_main_sat
EUT2I
11840 -pf11185 -
11845 -pf00834 -
11850 -pf04265pf04263 -
11855 -pf00496 -
11860 -pf00528 -
11865 -pf00528pf12911 -
11870 -pf00005 -
11875 HMM06277EutA_1_eut(pf06277) EUT2I
11880 HMMetlyEutB_2_EUT2(pf06751) EUT2I
11885 HMMetlyEutC_3_eut2(#) EUT2I EP_NTERM
11890 Tsp_darkPeach EUT2I
11895 H_fireEngine EUT2I
11900 HMMalddh_2_egra(pf00171) EUT2I
11905 H_fuchsia EUT2G
11910 HMM01923cobtr_3_eut2(pf01923) EUT2I EP_NTERM
11915 HMMptac_3_eut2(pf06130) EUT2I EP_NTERM
11920 P_azul EUT2I
11925 Ts_mushroom EUT2I
11930 -pf00005 -
11940 -pf01861 -
11945 -pf02675 -
11950 -pf11611 -
11955 -pf04542pf04545 -
11960 -pf00117pf00958pf02540 -
11965 -# -
11970 -# -
11975 HMMreg00072_2_pdugrm(pf00072pf00486) EUT2E
11980 -pf13493pf02518pf00512pf02702 -
(..-11060..)
00920 -# -
00925 -pf01259 -
00930 -pf00731pf02769 -
00935 -pf01189pf17125pf01029 -
00940 -pf04298 -
00945 -pf02911pf00551 -
00950 -pf01327 -
00955 -# -
00960 -pf02913pf01565 -
00965 -pf10035 -
00970 Hp_euts_fadedPurple EUT2B
00975 HMM06249EutQ_2_eut2(pf06249) EUT2I EP_NTERM
00980 HMMtpEuth_1_eut(pf04346) EUT2I
00985 Ts_mushroom(#) EUT2I
00990 -# -
00995 -pf06968pf04055 -
01000 -# -
01010 -pf01558 -
01015 -pf01522 -
01020 -# -
01025 -pf00999 -
BMC-P
BMC-H
BMC-T
Aldehyde dehydrogenase
PTAC
Signature enzyme
Alcohol dehydrogenase
Regulator

Clostridium sp. Bacteria; Firmicutes; Clostridia; Clostridiales; Clostridiaceae;

DCL31__locus_1_main_sat

EUT2I

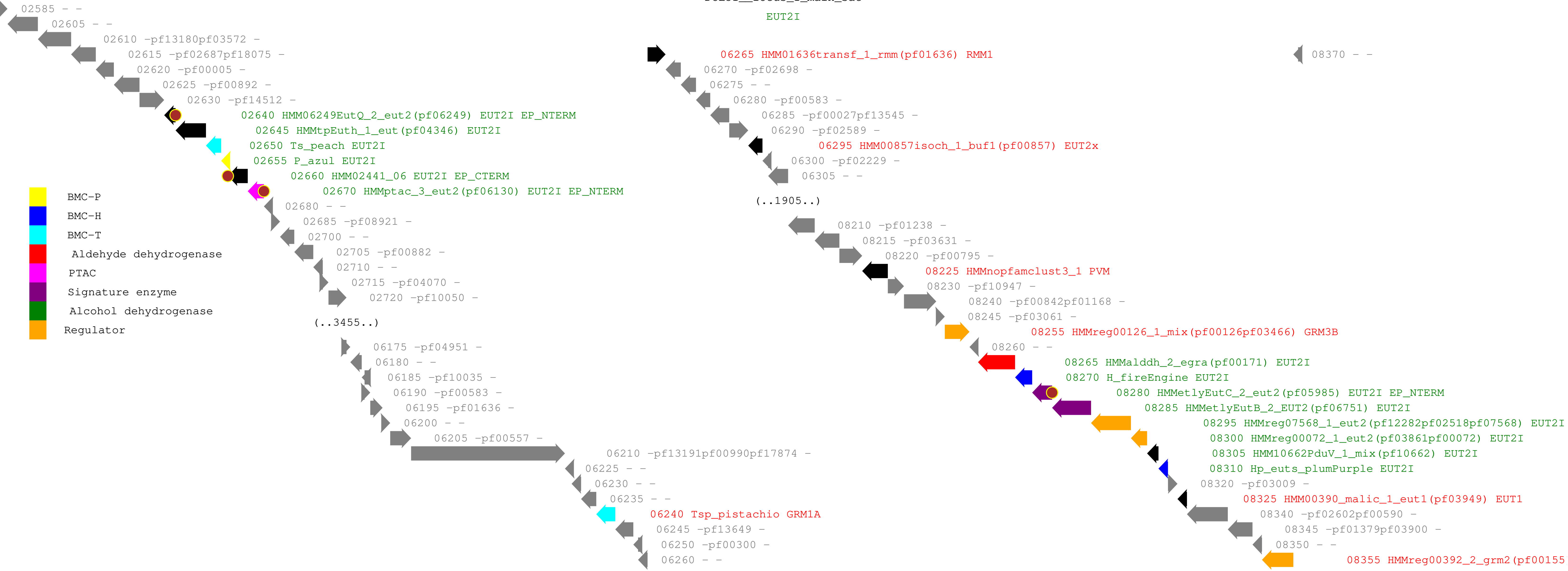

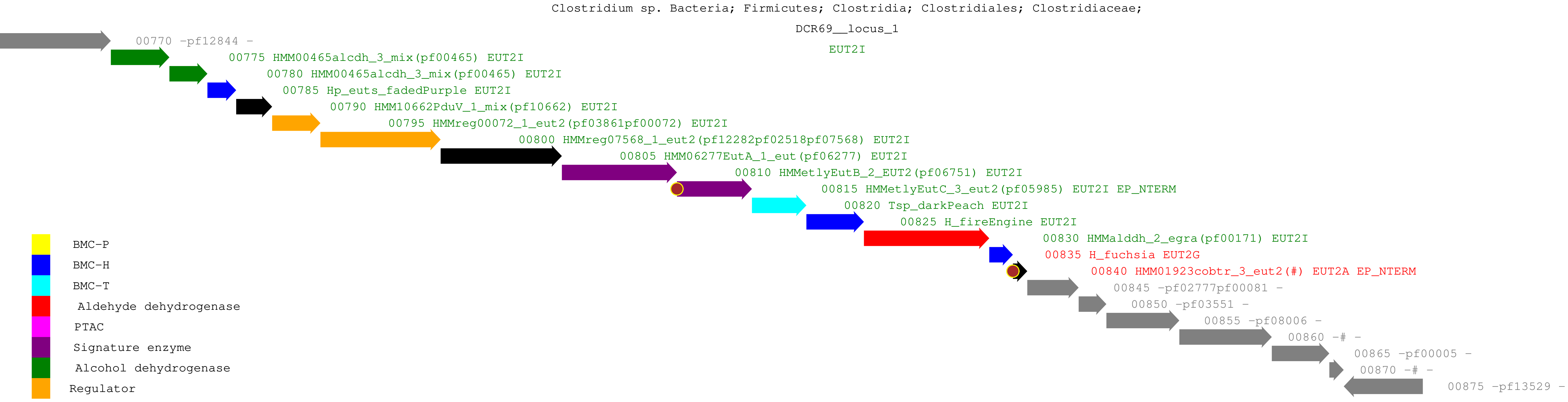
Clostridium sp. Bacteria; Firmicutes; Clostridia; Clostridiales; Clostridiaceae;
DCR69__locus_1
EUT2I
00770 -pf12844 -
00775 HMM00465alcdh_3_mix(pf00465) EUT2I
00780 HMM00465alcdh_3_mix(pf00465) EUT2I
00785 Hp_euts_fadedPurple EUT2I
00790 HMM10662PduV_1_mix(pf10662) EUT2I
00795 HMMreg00072_1_eut2(pf03861pf00072) EUT2I
00800 HMMreg07568_1_eut2(pf12282pf02518pf07568) EUT2I
00805 HMM06277EutA_1_eut(pf06277) EUT2I
00810 HMMetlyEutB_2_EUT2(pf06751) EUT2I
00815 HMMetlyEutC_3_eut2(pf05985) EUT2I EP_NTERM
00820 Tsp_darkPeach EUT2I
00825 H_fireEngine EUT2I
00830 HMMalddh_2_egra(pf00171) EUT2I
00835 H_fuchsia EUT2G
00840 HMM01923cobtr_3_eut2(#) EUT2A EP_NTERM
00845 -pf02777pf00081 -
00850 -pf03551 -
00855 -pf08006 -
00860 -# -
00865 -pf00005 -
00870 -# -
00875 -pf13529 -
BMC-P
BMC-H
BMC-T
Aldehyde dehydrogenase
PTAC
Signature enzyme
Alcohol dehydrogenase
Regulator

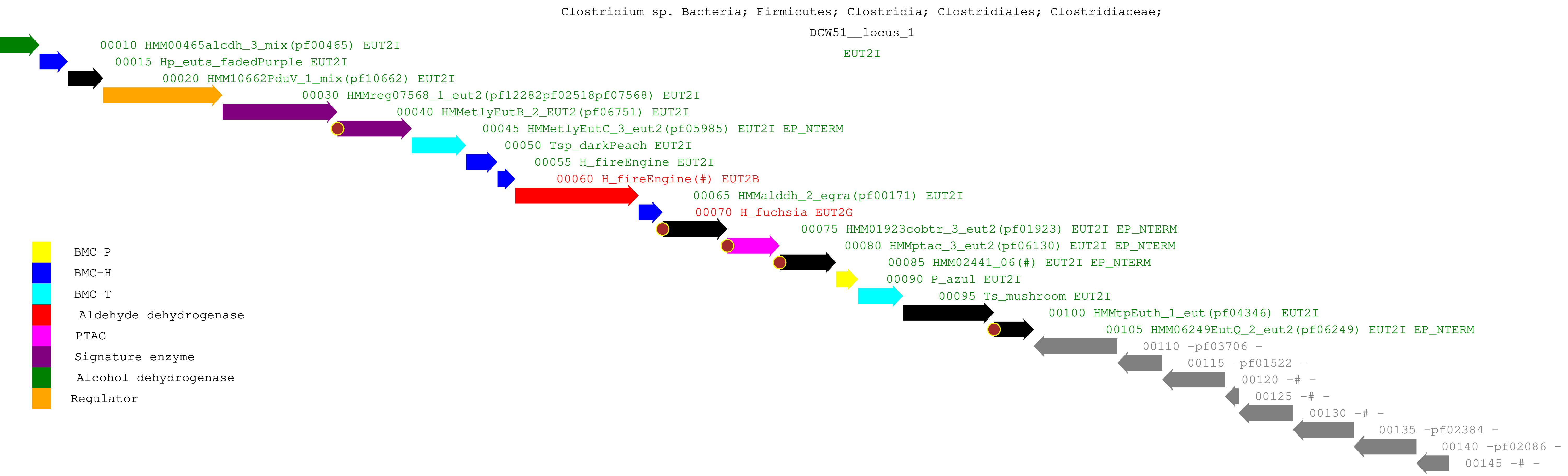
Clostridium sp. Bacteria; Firmicutes; Clostridia; Clostridiales; Clostridiaceae;
DCW51__locus_1
EUT2I
00010 HMM00465alcdh_3_mix(pf00465) EUT2I
00015 Hp_euts_fadedPurple EUT2I
00020 HMM10662PduV_1_mix(pf10662) EUT2I
00030 HMMreg07568_1_eut2(pf12282pf02518pf07568) EUT2I
00040 HMMetlyEutB_2_EUT2(pf06751) EUT2I
00045 HMMetlyEutC_3_eut2(pf05985) EUT2I EP_NTERM
00050 Tsp_darkPeach EUT2I
00055 H_fireEngine EUT2I
00060 H_fireEngine(#) EUT2B
00065 HMMaldh_2_egra(pf00171) EUT2I
00070 H_fuchsia EUT2G
00075 HMM01923cobtr_3_eut2(pf01923) EUT2I EP_NTERM
00080 HMMptac_3_eut2(pf06130) EUT2I EP_NTERM
00085 HMM02441_06(#) EUT2I EP_NTERM
00090 P_azul EUT2I
00095 Ts_mushroom EUT2I
00100 HMMtpEuth_1_eut(pf04346) EUT2I
00105 HMM06249EutQ_2_eut2(pf06249) EUT2I EP_NTERM
00110 -pf03706 -
00115 -pf01522 -
00120 -# -
00125 -# -
00130 -# -
00135 -pf02384 -
00140 -pf02086 -
00145 -# -
BMC-P
BMC-H
BMC-T
Aldehyde dehydrogenase
PTAC
Signature enzyme
Alcohol dehydrogenase
Regulator

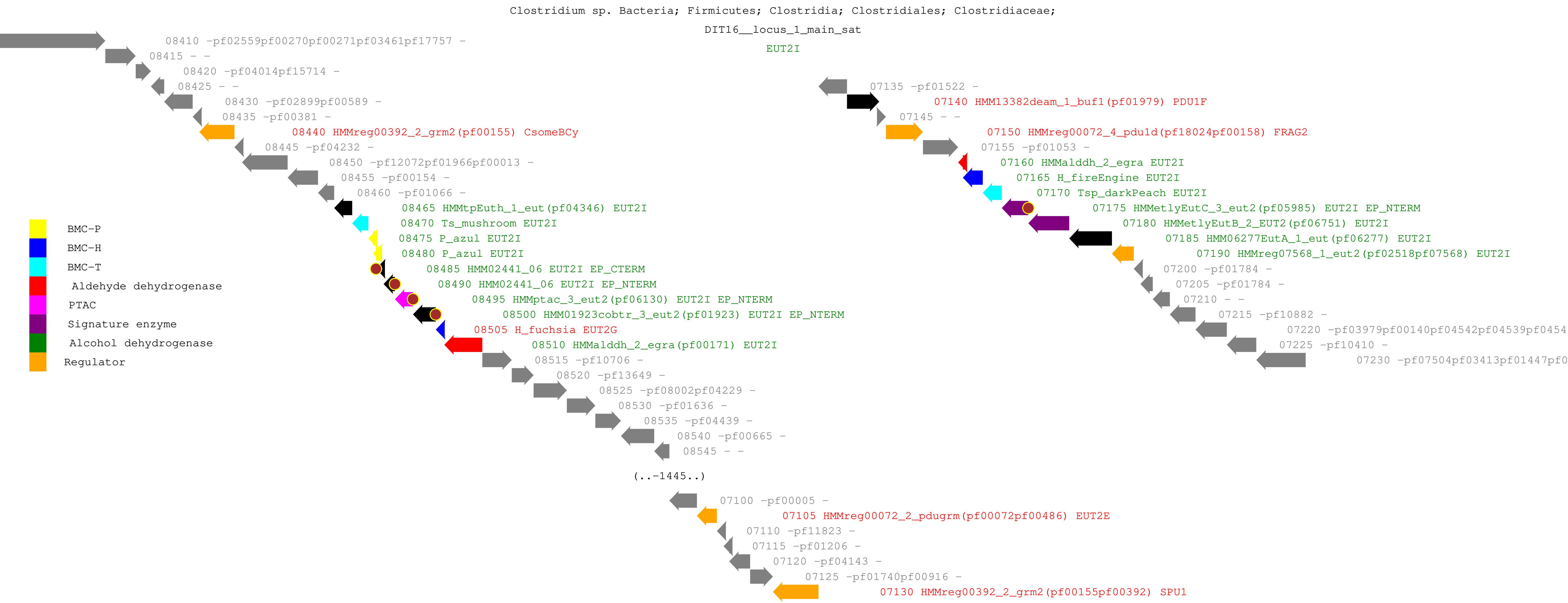
Clostridium sp. Bacteria; Firmicutes; Clostridia; Clostridiales; Clostridiaceae;
DIT16__locus_1_main_sat
EUT2I
08410 -pf02559pf00270pf00271pf03461pf17757 -
08415 - -
08420 -pf04014pf15714 -
08425 - -
08430 -pf02899pf00589 -
08435 -pf00381 -
08440 HMMreg00392_2_grm2(pf00155) CsomeBCy
08445 -pf04232 -
08450 -pf12072pf01966pf00013 -
08455 -pf00154 -
08460 -pf01066 -
08465 HMMtpEuth_1_eut(pf04346) EUT2I
08470 Ts_mushroom EUT2I
08475 P_azul EUT2I
08480 P_azul EUT2I
08485 HMM02441_06 EUT2I EP_CTERM
08490 HMM02441_06 EUT2I EP_NTERM
08495 HMMptac_3_eut2(pf06130) EUT2I EP_NTERM
08500 HMM01923cobtr_3_eut2(pf01923) EUT2I EP_NTERM
08505 H_fuchsia EUT2G
08510 HMMaldh_2_egra(pf00171) EUT2I
08515 -pf10706 -
08520 -pf13649 -
08525 -pf08002pf04229 -
08530 -pf01636 -
08535 -pf04439 -
08540 -pf00665 -
08545 - -
(..-1445..)
07100 -pf00005 -
07105 HMMreg00072_2_pdugrm(pf00072pf00486) EUT2E
07110 -pf11823 -
07115 -pf01206 -
07120 -pf04143 -
07125 -pf01740pf00916 -
07130 HMMreg00392_2_grm2(pf00155pf00392) SPU1
07135 -pf01522 -
07140 HMM13382deam_1_buf1(pf01979) PDU1F
07145 - -
07150 HMMreg00072_4_pdu1d(pf18024pf00158) FRAG2
07155 -pf01053 -
07160 HMMaldh_2_egra EUT2I
07165 H_fireEngine EUT2I
07170 Tsp_darkPeach EUT2I
07175 HMMetlyEutC_3_eut2(pf05985) EUT2I EP_NTERM
07180 HMMetlyEutB_2_EUT2(pf06751) EUT2I
07185 HMM06277EutA_1_eut(pf06277) EUT2I
07190 HMMreg07568_1_eut2(pf02518pf07568) EUT2I
07200 -pf01784 -
07205 -pf01784 -
07210 - -
07215 -pf10882 -
07220 -pf03979pf00140pf04542pf04539pf0454
07225 -pf10410 -
07230 -pf07504pf03413pf01447pf0
BMC-P
BMC-H
BMC-T
Aldehyde dehydrogenase
PTAC
Signature enzyme
Alcohol dehydrogenase
Regulator

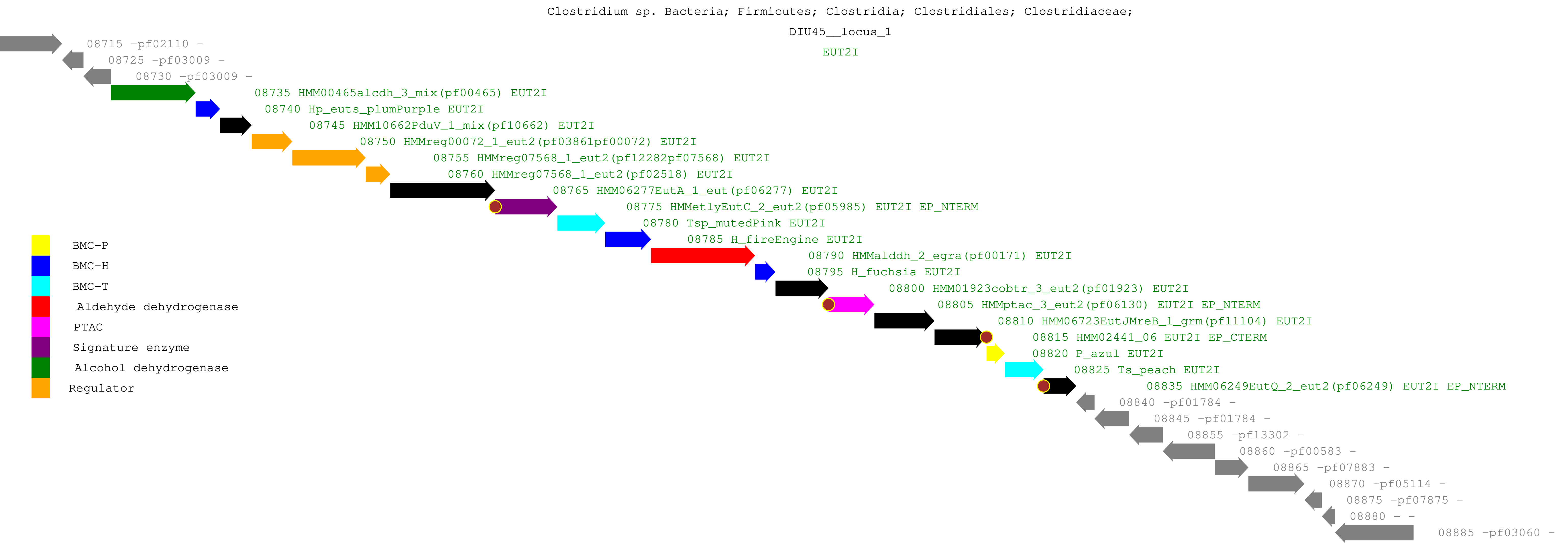
Clostridium sp. Bacteria; Firmicutes; Clostridia; Clostridiales; Clostridiaceae;
DIU45__locus_1
EUT2I
08715 -pf02110 -
08725 -pf03009 -
08730 -pf03009 -
08735 HMM00465alcdh_3_mix(pf00465) EUT2I
08740 Hp_euts_plumPurple EUT2I
08745 HMM10662PduV_1_mix(pf10662) EUT2I
08750 HMMreg00072_1_eut2(pf03861pf00072) EUT2I
08755 HMMreg07568_1_eut2(pf12282pf07568) EUT2I
08760 HMMreg07568_1_eut2(pf02518) EUT2I
08765 HMM06277EutA_1_eut(pf06277) EUT2I
08775 HMMetlyEutC_2_eut2(pf05985) EUT2I EP_NTERM
08780 Tsp_mutedPink EUT2I
08785 H_fireEngine EUT2I
08790 HMMalddh_2_egra(pf00171) EUT2I
08795 H_fuchsia EUT2I
08800 HMM01923cobtr_3_eut2(pf01923) EUT2I
08805 HMMptac_3_eut2(pf06130) EUT2I EP_NTERM
08810 HMM06723EutJMreB_1_grm(pf11104) EUT2I
08815 HMM02441_06 EUT2I EP_CTERM
08820 P_azul EUT2I
08825 Ts_peach EUT2I
08835 HMM06249EutQ_2_eut2(pf06249) EUT2I EP_NTERM
08840 -pf01784 -
08845 -pf01784 -
08855 -pf13302 -
08860 -pf00583 -
08865 -pf07883 -
08870 -pf05114 -
08875 -pf07875 -
08880 - -
08885 -pf03060 -
BMC-P
BMC-H
BMC-T
Aldehyde dehydrogenase
PTAC
Signature enzyme
Alcohol dehydrogenase
Regulator

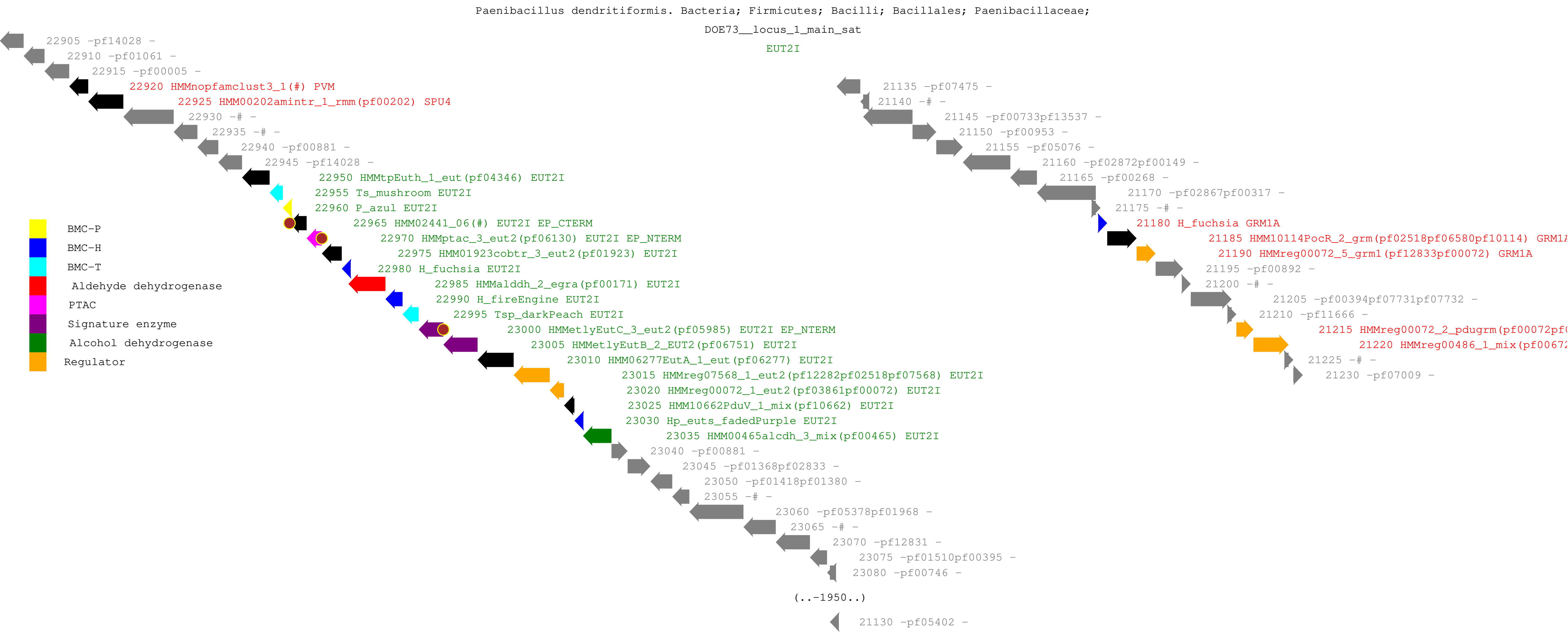

Paenibacillus dendritiformis. Bacteria; Firmicutes; Bacilli; Bacillales; Paenibacillaceae;
DOE73__locus_1_main_sat
EUT2I
22905 -pf14028 -
22910 -pf01061 -
22915 -pf00005 -
22920 HMMnopfamclust3_1(#) PVM
22925 HMM00202amintr_1_rmm(pf00202) SPU4
22930 -# -
22935 -# -
22940 -pf00881 -
22945 -pf14028 -
22950 HMMtpEuth_1_eut(pf04346) EUT2I
22955 Ts_mushroom EUT2I
22960 P_azul EUT2I
22965 HMM02441_06(#) EUT2I EP_CTERM
22970 HMMptac_3_eut2(pf06130) EUT2I EP_NTERM
22975 HMM01923cobtr_3_eut2(pf01923) EUT2I
22980 H_fuchsia EUT2I
22985 HMMalddh_2_egra(pf00171) EUT2I
22990 H_fireEngine EUT2I
22995 Tsp_darkPeach EUT2I
23000 HMMetlyEutC_3_eut2(pf05985) EUT2I EP_NTERM
23005 HMMetlyEutB_2_EUT2(pf06751) EUT2I
23010 HMM06277EutA_1_eut(pf06277) EUT2I
23015 HMMreg07568_1_eut2(pf12282pf02518pf07568) EUT2I
23020 HMMreg00072_1_eut2(pf03861pf00072) EUT2I
23025 HMM10662PduV_1_mix(pf10662) EUT2I
23030 Hp_euts_fadedPurple EUT2I
23035 HMM00465alcdh_3_mix(pf00465) EUT2I
23040 -pf00881 -
23045 -pf01368pf02833 -
23050 -pf01418pf01380 -
23055 -# -
23060 -pf05378pf01968 -
23065 -# -
23070 -pf12831 -
23075 -pf01510pf00395 -
23080 -pf00746 -
(..-1950..)
21130 -pf05402 -
21135 -pf07475 -
21140 -# -
21145 -pf00733pf13537 -
21150 -pf00953 -
21155 -pf05076 -
21160 -pf02872pf00149 -
21165 -pf00268 -
21170 -pf02867pf00317 -
21175 -# -
21180 H_fuchsia GRM1A
21185 HMM10114PocR_2_grm(pf02518pf06580pf10114) GRM1
21190 HMMreg00072_5_grm1(pf12833pf00072) GRM1A
21195 -pf00892 -
21200 -# -
21205 -pf00394pf07731pf07732 -
21210 -pf11666 -
21215 HMMreg00072_2_pdugrm(pf00072pf
21220 HMMreg00486_1_mix(pf0067
21225 -# -
21230 -pf07009 -
BMC-P
BMC-H
BMC-T
Aldehyde dehydrogenase
PTAC
Signature enzyme
Alcohol dehydrogenase
Regulator

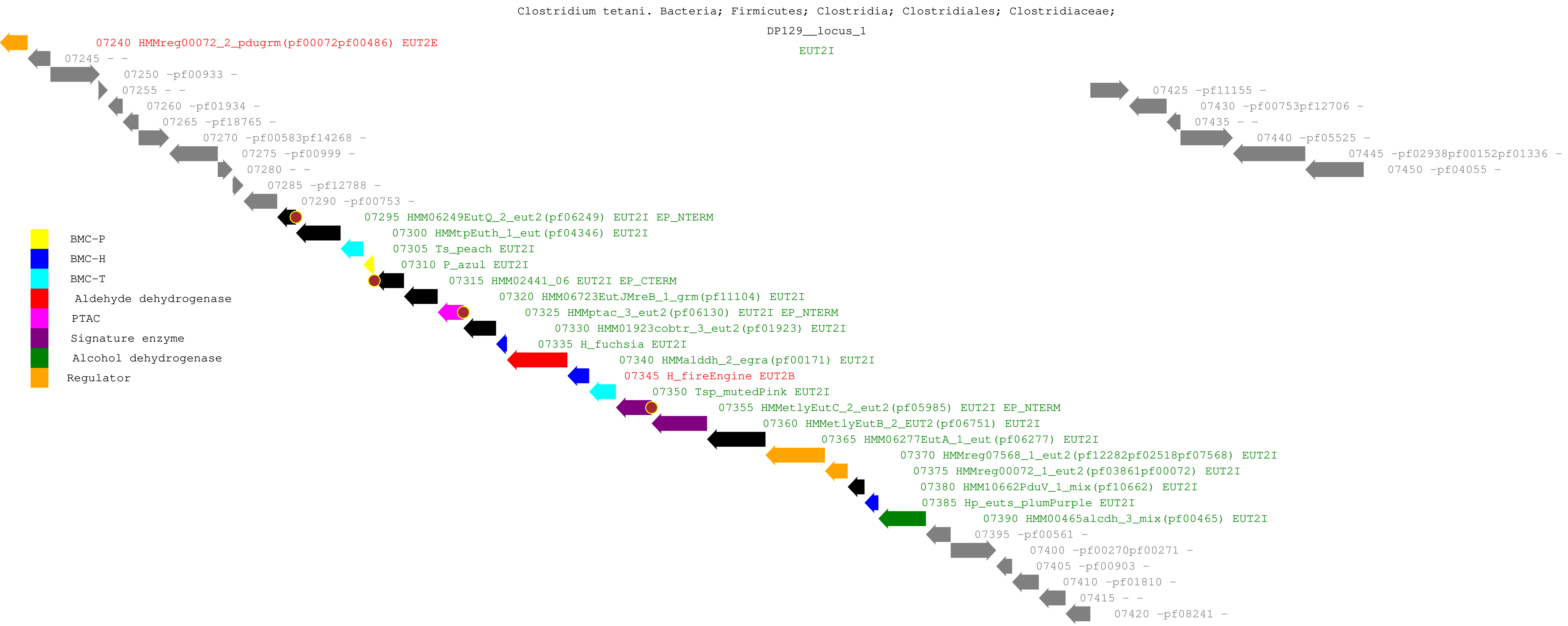
Clostridium tetani. Bacteria; Firmicutes; Clostridia; Clostridiales; Clostridiaceae;
DP129__locus_1
EUT2I
07240 HMMreg00072_2_pdugrm(pf00072pf00486) EUT2E
07245 - -
07250 -pf00933 -
07255 - -
07260 -pf01934 -
07265 -pf18765 -
07270 -pf00583pf14268 -
07275 -pf00999 -
07280 - -
07285 -pf12788 -
07290 -pf00753 -
07295 HMM06249EutQ_2_eut2(pf06249) EUT2I EP_NTERM
07300 HMMtpEuth_1_eut(pf04346) EUT2I
07305 Ts_peach EUT2I
07310 P_azul EUT2I
07315 HMM02441_06 EUT2I EP_CTERM
07320 HMM06723EutJMreB_1_grm(pf11104) EUT2I
07325 HMMptac_3_eut2(pf06130) EUT2I EP_NTERM
07330 HMM01923cobtr_3_eut2(pf01923) EUT2I
07335 H_fuchsia EUT2I
07340 HMMalddh_2_egra(pf00171) EUT2I
07345 H_fireEngine EUT2B
07350 Tsp_mutedPink EUT2I
07355 HMMetlyEutC_2_eut2(pf05985) EUT2I EP_NTERM
07360 HMMetlyEutB_2_EUT2(pf06751) EUT2I
07365 HMM06277EutA_1_eut(pf06277) EUT2I
07370 HMMreg07568_1_eut2(pf12282pf02518pf07568) EUT2I
07375 HMMreg00072_1_eut2(pf03861pf00072) EUT2I
07380 HMM10662PduV_1_mix(pf10662) EUT2I
07385 Hp_euts_plumPurple EUT2I
07390 HMM00465alcdh_3_mix(pf00465) EUT2I
07395 -pf00561 -
07400 -pf00270pf00271 -
07405 -pf00903 -
07410 -pf01810 -
07415 - -
07420 -pf08241 -
07425 -pf11155 -
07430 -pf00753pf12706 -
07435 - -
07440 -pf05525 -
07445 -pf02938pf00152pf01336 -
07450 -pf04055 -
BMC-P
BMC-H
BMC-T
Aldehyde dehydrogenase
PTAC
Signature enzyme
Alcohol dehydrogenase
Regulator

Clostridium tetani. Bacteria; Firmicutes; Clostridia; Clostridiales; Clostridiaceae;

DP130__locus_2

EUT2I

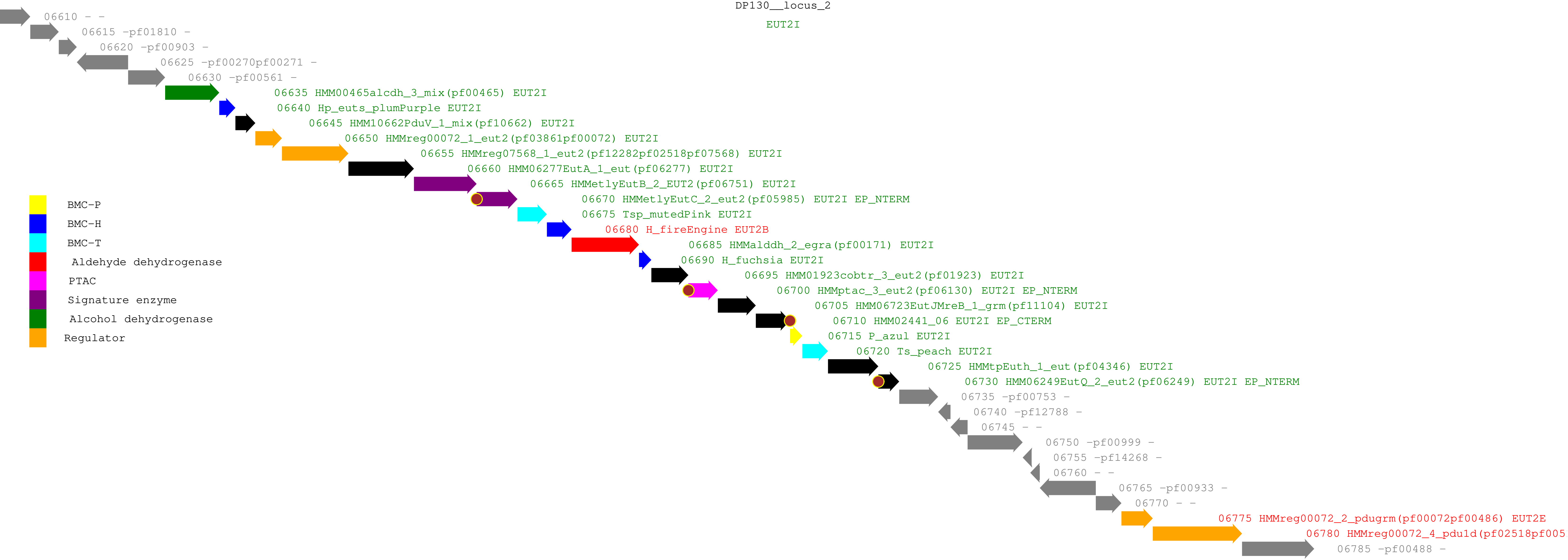

Paenibacillus thiaminolyticus (Bacillus thiaminolyticus). Bacteria; Firmicutes; Bacilli; Bacillales; Paenibacillaceae; Paenibacillus.

DQX05__locus_3

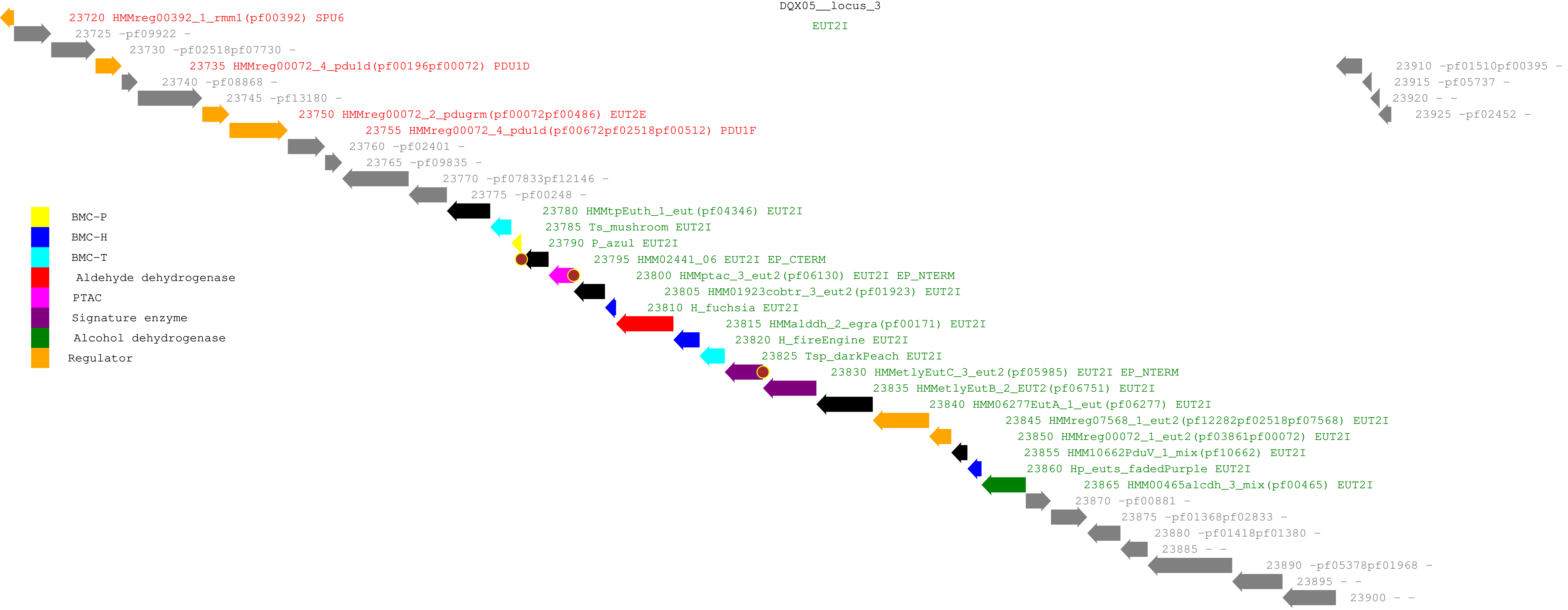

Acinetobacter sp. RIT592. Bacteria; Proteobacteria; Gammaproteobacteria; Pseudomonadales;

DVA85__locus_1

EUT2I

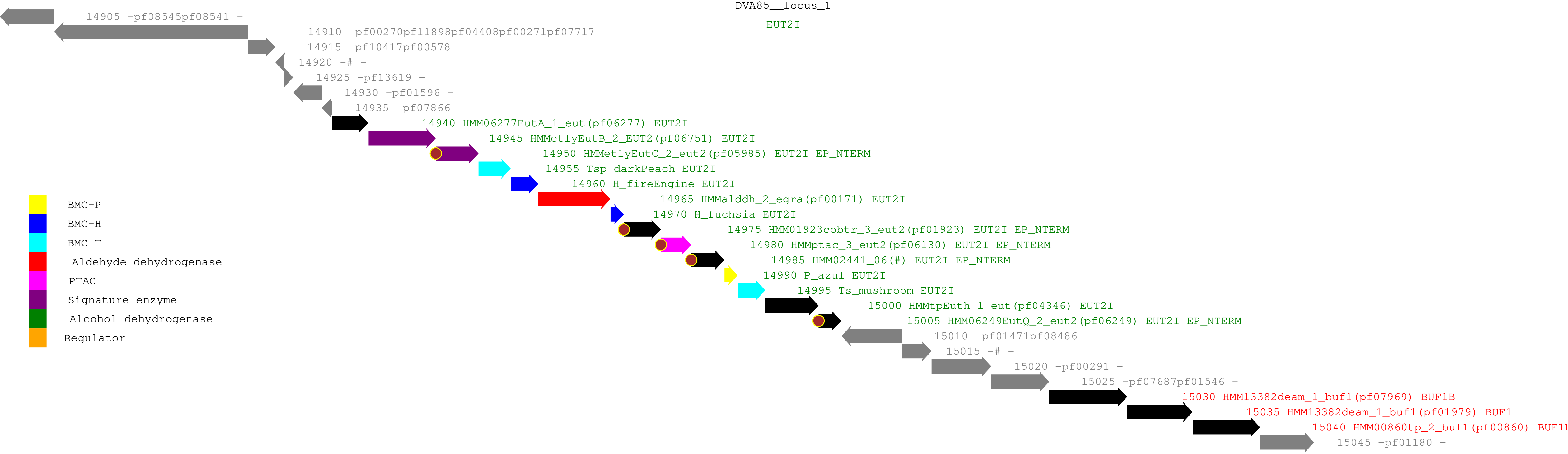

Brevibacillus sp. VP. Bacteria; Firmicutes; Bacilli; Bacillales; Paenibacillaceae;

DZB91__locus_1

EUT2I

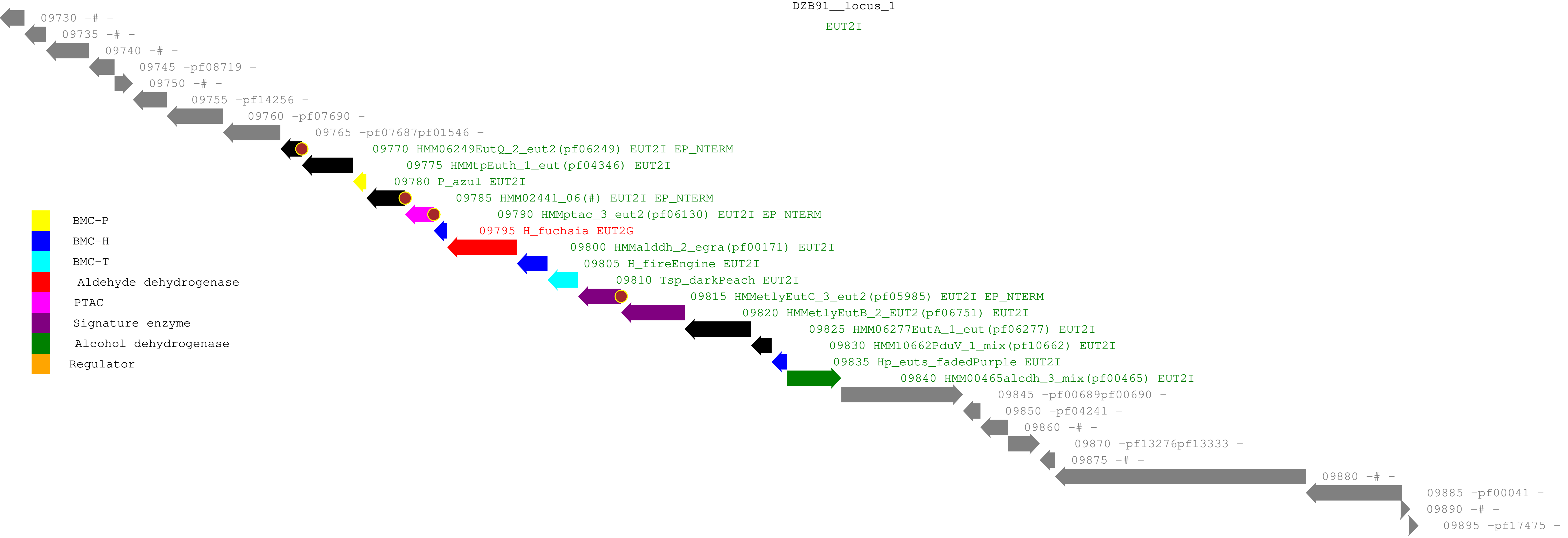

Brevibacillus sp. VP. Bacteria; Firmicutes; Bacilli; Bacillales; Paenibacillaceae;

DZB91__locus_2

EUT2I

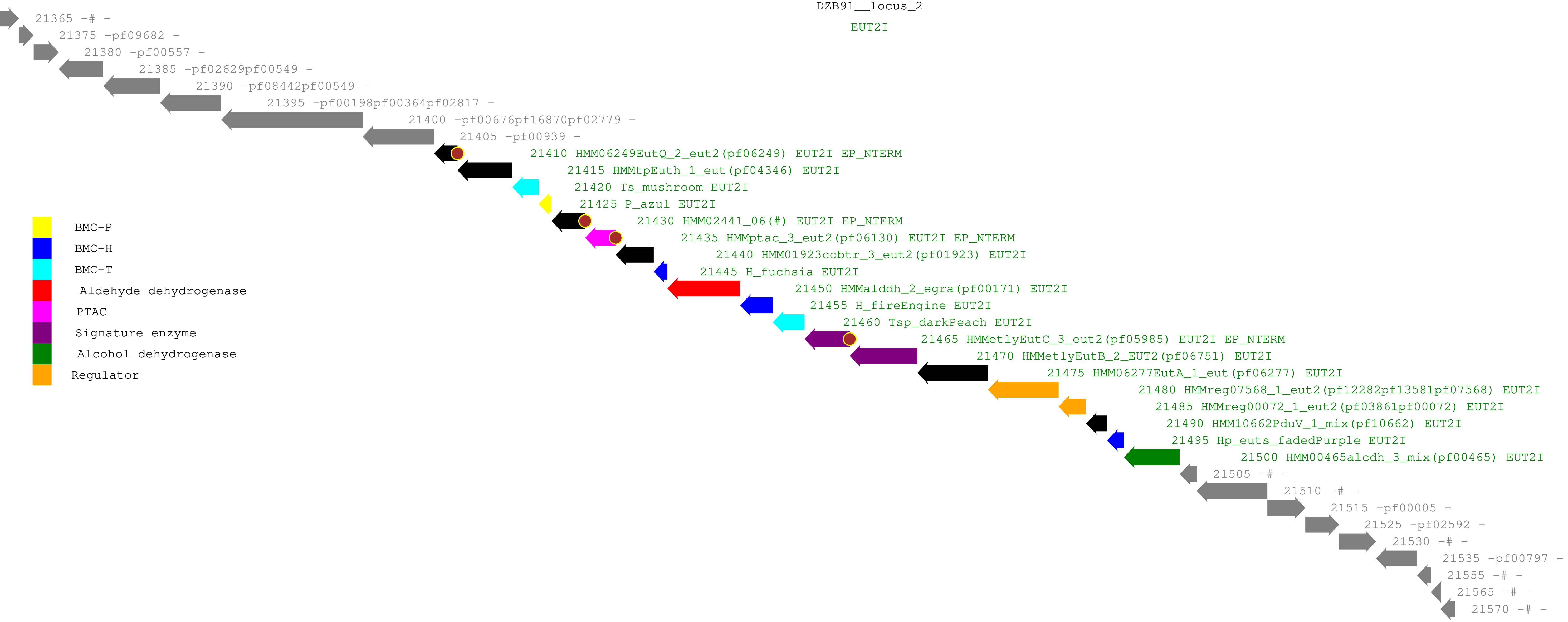

Paenibacillus dendritiformis. Bacteria; Firmicutes; Bacilli; Bacillales; Paenibacillaceae; Paenibacillus.

E2R60__locus_1

EUT2I

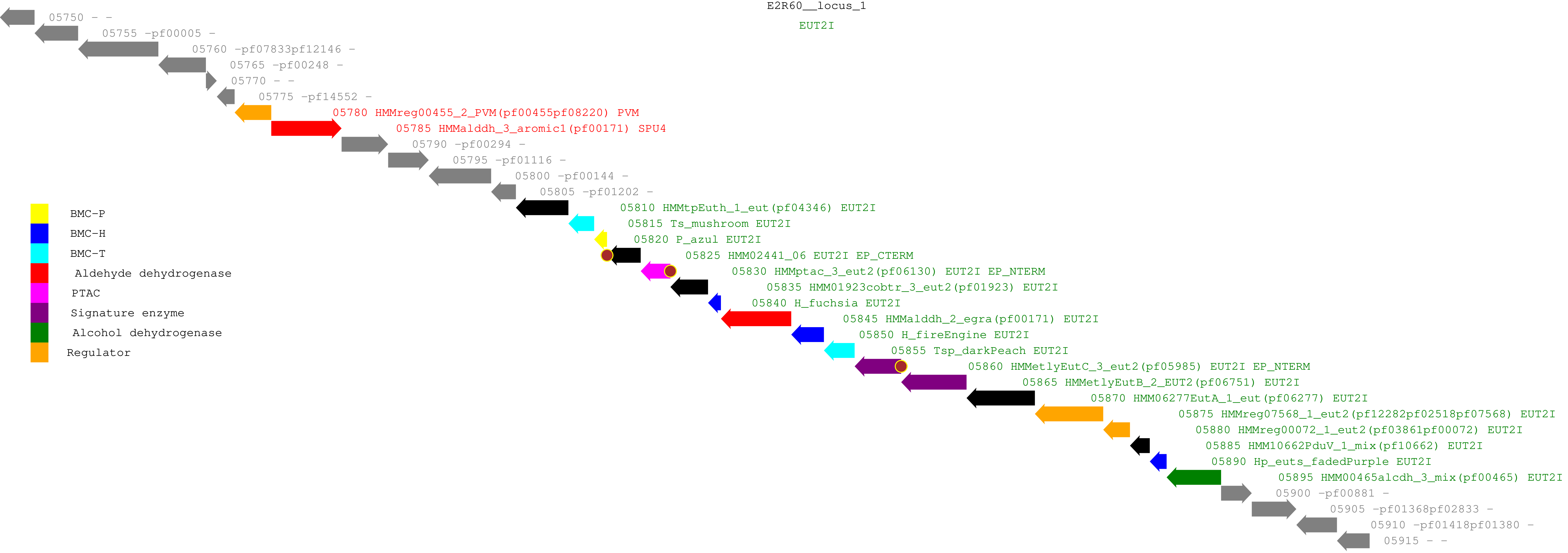

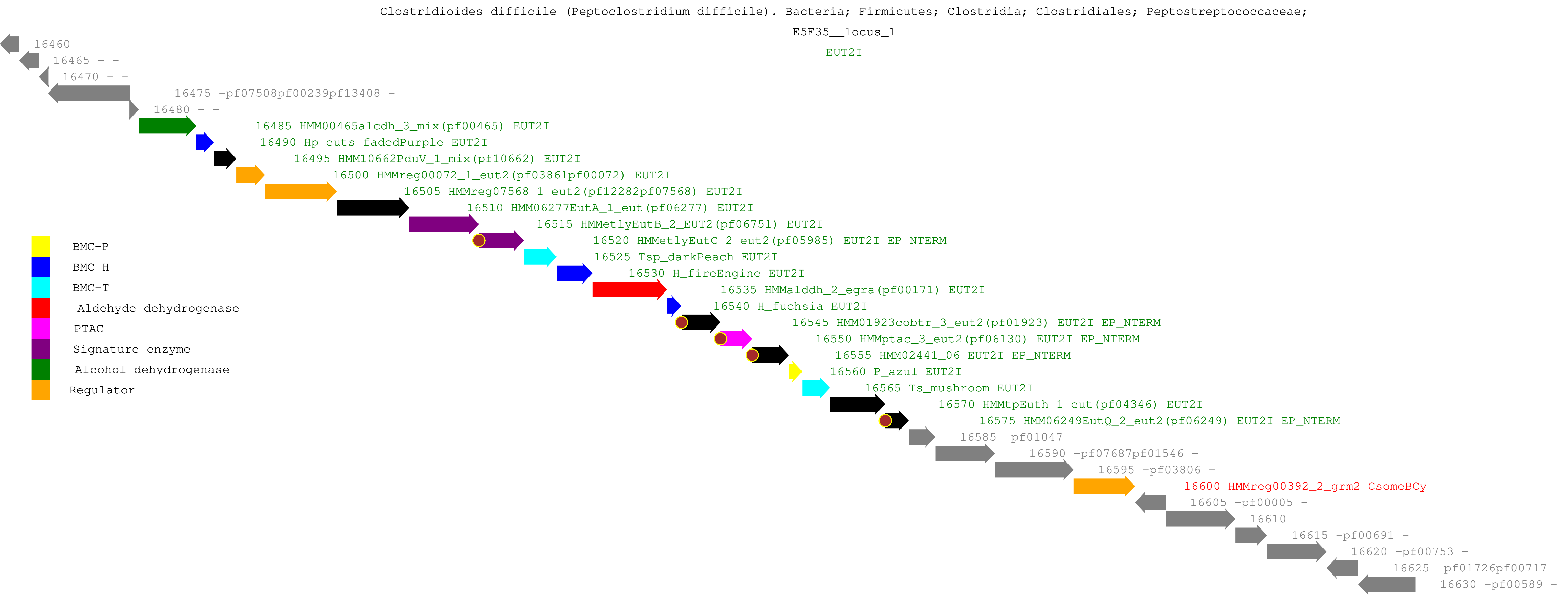

Clostridioides difficile (Peptoclostridium difficile). Bacteria; Firmicutes; Clostridia; Clostridiales; Peptostreptococcaceae;
E5F35__locus_1
EUT2I
16460 - -
16465 - -
16470 - -
16475 -pf07508pf00239pf13408 -
16480 - -
16485 HMM00465alcdh_3_mix(pf00465) EUT2I
16490 Hp_euts_fadedPurple EUT2I
16495 HMM10662PduV_1_mix(pf10662) EUT2I
16500 HMMreg00072_1_eut2(pf03861pf00072) EUT2I
16505 HMMreg07568_1_eut2(pf12282pf07568) EUT2I
16510 HMM06277EutA_1_eut(pf06277) EUT2I
16515 HMMetlyEutB_2_EUT2(pf06751) EUT2I
16520 HMMetlyEutC_2_eut2(pf05985) EUT2I EP_NTERM
16525 Tsp_darkPeach EUT2I
16530 H_fireEngine EUT2I
16535 HMMalddh_2_egra(pf00171) EUT2I
16540 H_fuchsia EUT2I
16545 HMM01923cobtr_3_eut2(pf01923) EUT2I EP_NTERM
16550 HMMptac_3_eut2(pf06130) EUT2I EP_NTERM
16555 HMM02441_06 EUT2I EP_NTERM
16560 P_azul EUT2I
16565 Ts_mushroom EUT2I
16570 HMMtpEuth_1_eut(pf04346) EUT2I
16575 HMM06249EutQ_2_eut2(pf06249) EUT2I EP_NTERM
16585 -pf01047 -
16590 -pf07687pf01546 -
16595 -pf03806 -
16600 HMMreg00392_2_grm2 CsomeBCy
16605 -pf00005 -
16610 - -
16615 -pf00691 -
16620 -pf00753 -
16625 -pf01726pf00717 -
16630 -pf00589 -
BMC-P
BMC-H
BMC-T
Aldehyde dehydrogenase
PTAC
Signature enzyme
Alcohol dehydrogenase
Regulator

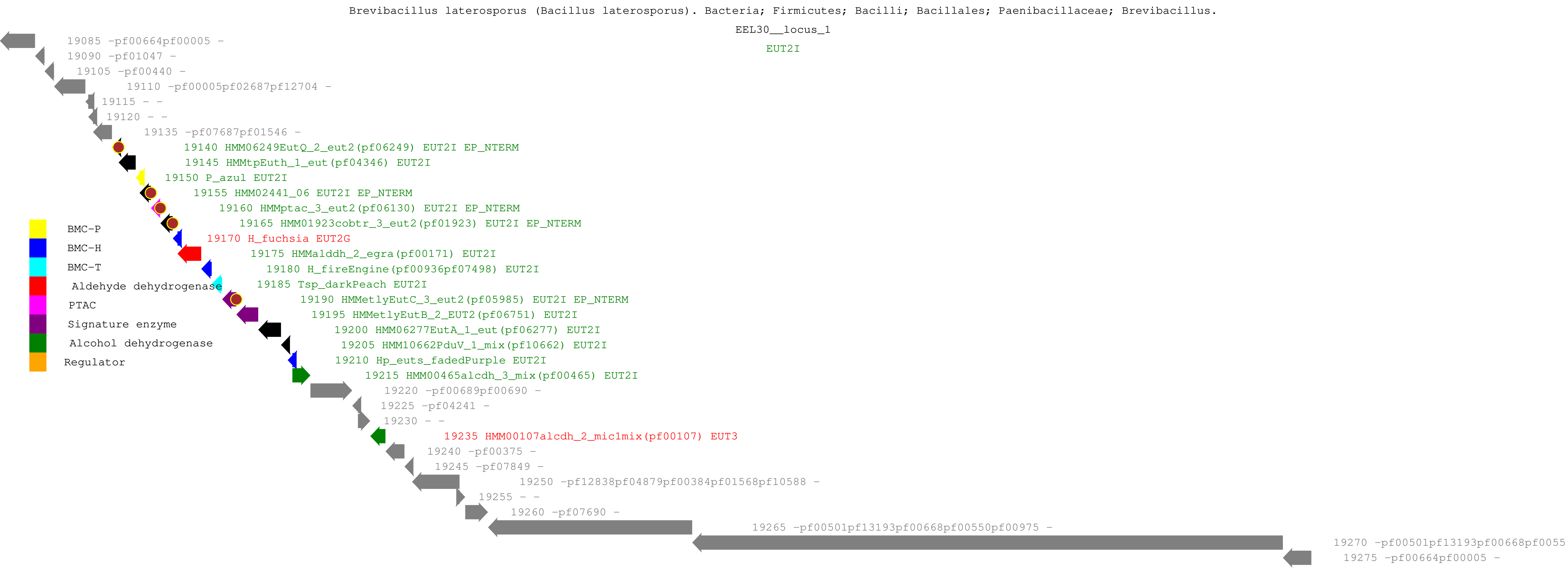
Brevibacillus laterosporus (Bacillus laterosporus). Bacteria; Firmicutes; Bacilli; Bacillales; Paenibacillaceae; Brevibacillus.
EEL30__locus_1
EUT2I
19085 -pf00664pf00005 -
19090 -pf01047 -
19105 -pf00440 -
19110 -pf00005pf02687pf12704 -
19115 - -
19120 - -
19135 -pf07687pf01546 -
19140 HMM06249EutQ_2_eut2(pf06249) EUT2I EP_NTERM
19145 HMMtpEuth_1_eut(pf04346) EUT2I
19150 P_azul EUT2I
19155 HMM02441_06 EUT2I EP_NTERM
19160 HMMptac_3_eut2(pf06130) EUT2I EP_NTERM
19165 HMM01923cobtr_3_eut2(pf01923) EUT2I EP_NTERM
19170 H_fuchsia EUT2G
19175 HMMalddh_2_egra(pf00171) EUT2I
19180 H_fireEngine(pf00936pf07498) EUT2I
19185 Tsp_darkPeach EUT2I
19190 HMMetlyEutC_3_eut2(pf05985) EUT2I EP_NTERM
19195 HMMetlyEutB_2_EUT2(pf06751) EUT2I
19200 HMM06277EutA_1_eut(pf06277) EUT2I
19205 HMM10662PduV_1_mix(pf10662) EUT2I
19210 Hp_euts_fadedPurple EUT2I
19215 HMM00465alcdh_3_mix(pf00465) EUT2I
19220 -pf00689pf00690 -
19225 -pf04241 -
19230 - -
19235 HMM00107alcdh_2_mic1mix(pf00107) EUT3
19240 -pf00375 -
19245 -pf07849 -
19250 -pf12838pf04879pf00384pf01568pf10588 -
19255 - -
19260 -pf07690 -
19265 -pf00501pf13193pf00668pf00550pf00975 -
19270 -pf00501pf13193pf00668pf0055
19275 -pf00664pf00005 -
BMC-P
BMC-H
BMC-T
Aldehyde dehydrogenase
PTAC
Signature enzyme
Alcohol dehydrogenase
Regulator

Brevibacillus laterosporus (Bacillus laterosporus). Bacteria; Firmicutes; Bacilli; Bacillales; Paenibacillaceae; Brevibacillus.

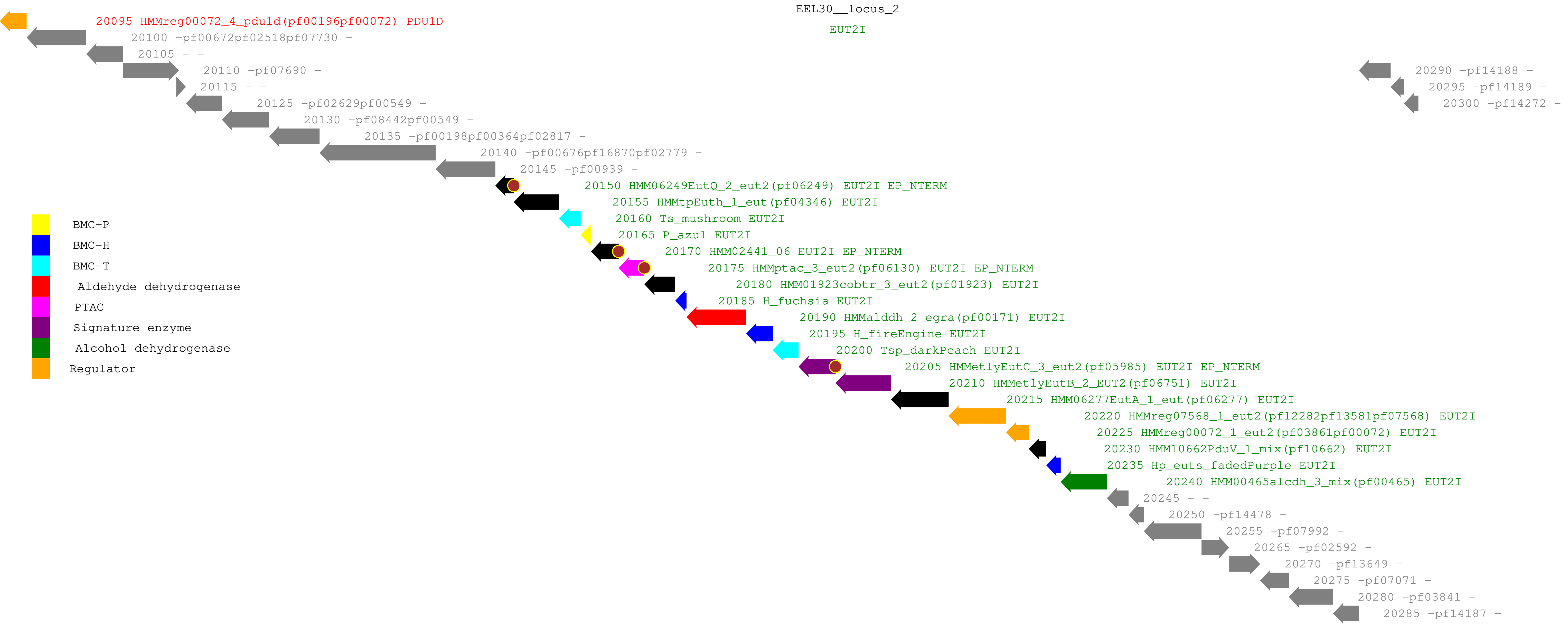

Brevibacillus laterosporus (Bacillus laterosporus). Bacteria; Firmicutes; Bacilli; Bacillales; Paenibacillaceae; Brevibacillus.

EEL32__locus_1

EUT2I

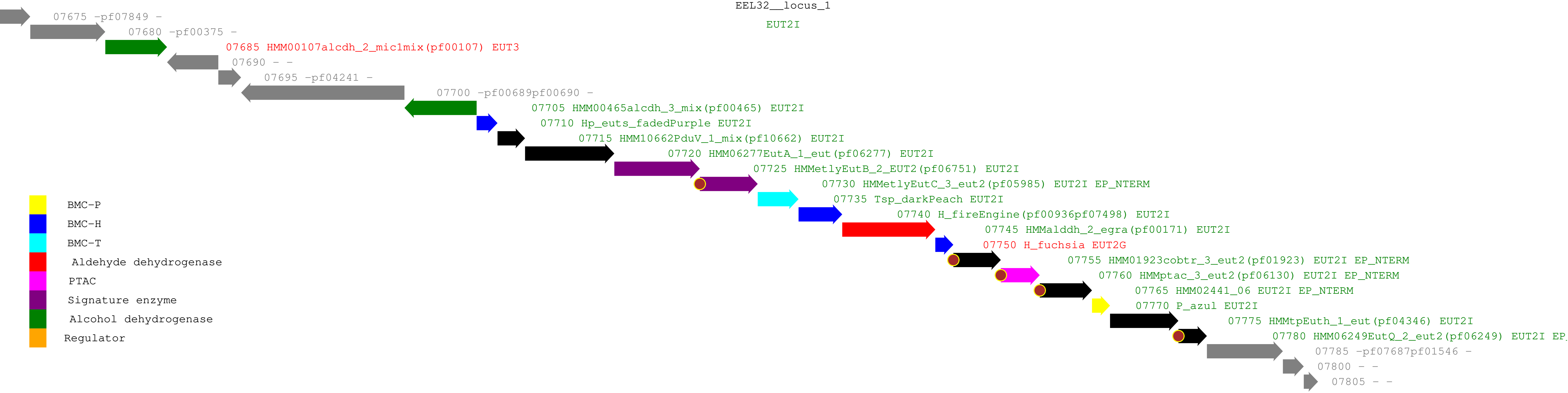

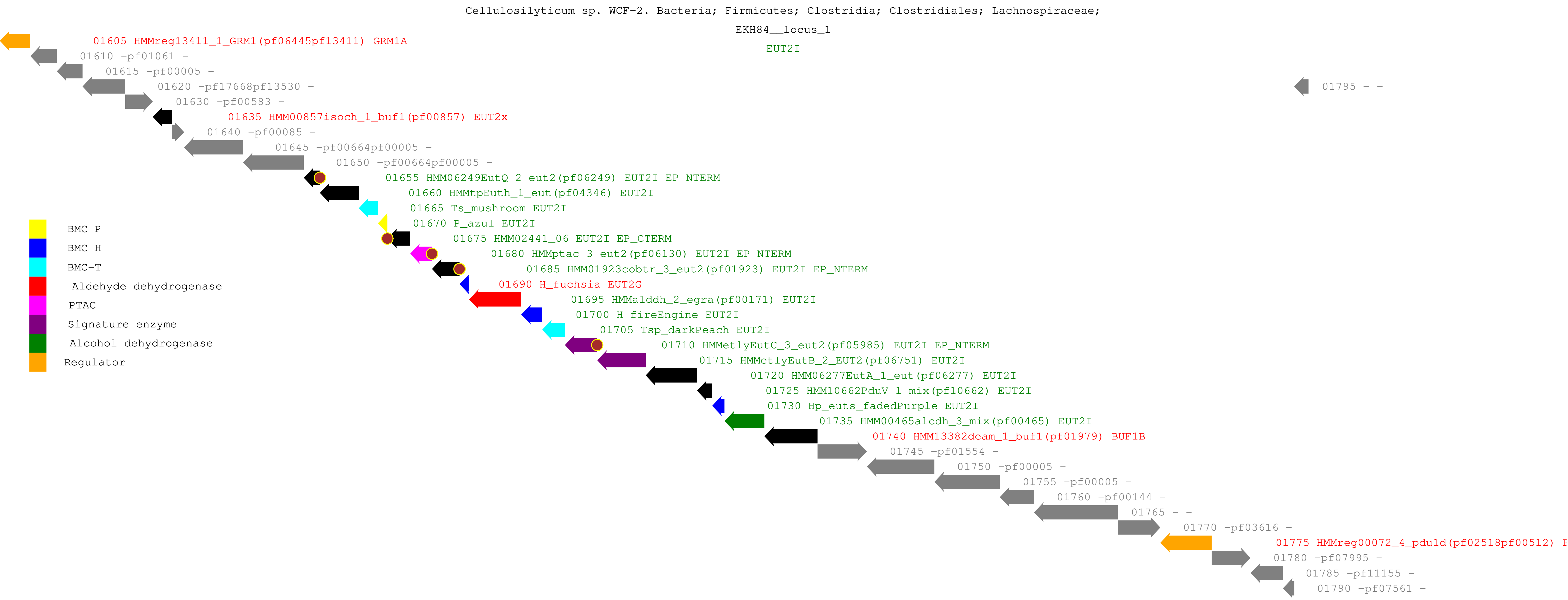
Cellulosilyticum sp. WCF-2. Bacteria; Firmicutes; Clostridia; Clostridiales; Lachnospiraceae;
EKH84__locus_1
EUT2I
01605 HMMreg13411_1_GRM1(pf06445pf13411) GRM1A
01610 -pf01061 -
01615 -pf00005 -
01620 -pf17668pf13530 -
01630 -pf00583 -
01635 HMM00857isoch_1_buf1(pf00857) EUT2x
01640 -pf00085 -
01645 -pf00664pf00005 -
01650 -pf00664pf00005 -
01655 HMM06249EutQ_2_eut2(pf06249) EUT2I EP_NTERM
01660 HMMtpEuth_1_eut(pf04346) EUT2I
01665 Ts_mushroom EUT2I
01670 P_azul EUT2I
01675 HMM02441_06 EUT2I EP_CTERM
01680 HMMptac_3_eut2(pf06130) EUT2I EP_NTERM
01685 HMM01923cobtr_3_eut2(pf01923) EUT2I EP_NTERM
01690 H_fuchsia EUT2G
01695 HMMalddh_2_egra(pf00171) EUT2I
01700 H_fireEngine EUT2I
01705 Tsp_darkPeach EUT2I
01710 HMMetlyEutC_3_eut2(pf05985) EUT2I EP_NTERM
01715 HMMetlyEutB_2_EUT2(pf06751) EUT2I
01720 HMM06277EutA_1_eut(pf06277) EUT2I
01725 HMM10662PduV_1_mix(pf10662) EUT2I
01730 Hp_euts_fadedPurple EUT2I
01735 HMM00465alcdh_3_mix(pf00465) EUT2I
01740 HMM13382deam_1_buf1(pf01979) BUF1B
01745 -pf01554 -
01750 -pf00005 -
01755 -pf00005 -
01760 -pf00144 -
01765 - -
01770 -pf03616 -
01775 HMMreg00072_4_pdu1d(pf02518pf00512)
01780 -pf07995 -
01785 -pf11155 -
01790 -pf07561 -
01795 - -
BMC-P
BMC-H
BMC-T
Aldehyde dehydrogenase
PTAC
Signature enzyme
Alcohol dehydrogenase
Regulator

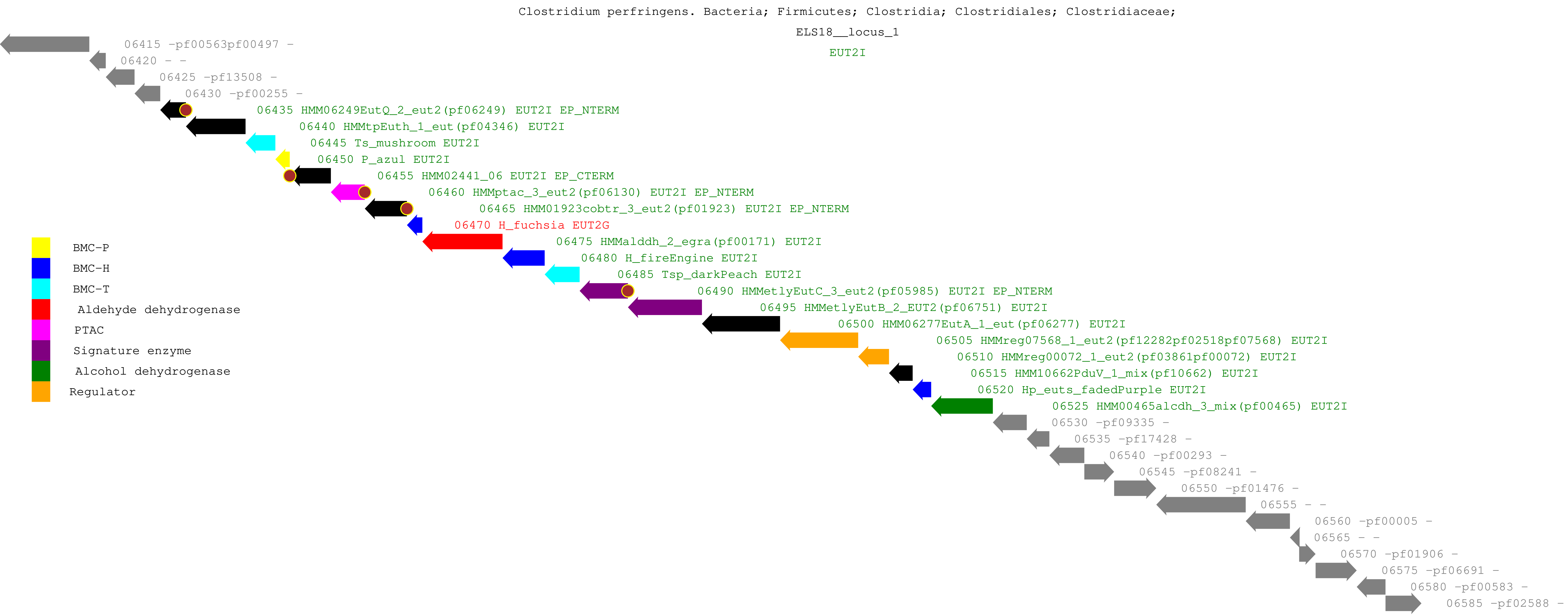
Clostridium perfringens. Bacteria; Firmicutes; Clostridia; Clostridiales; Clostridiaceae;
ELS18__locus_1
EUT2I
06415 -pf00563pf00497 -
06420 - -
06425 -pf13508 -
06430 -pf00255 -
06435 HMM06249EutQ_2_eut2(pf06249) EUT2I EP_NTERM
06440 HMMtpEuth_1_eut(pf04346) EUT2I
06445 Ts_mushroom EUT2I
06450 P_azul EUT2I
06455 HMM02441_06 EUT2I EP_CTERM
06460 HMMptac_3_eut2(pf06130) EUT2I EP_NTERM
06465 HMM01923cobtr_3_eut2(pf01923) EUT2I EP_NTERM
06470 H_fuchsia EUT2G
06475 HMMalddh_2_egra(pf00171) EUT2I
06480 H_fireEngine EUT2I
06485 Tsp_darkPeach EUT2I
06490 HMMetlyEutC_3_eut2(pf05985) EUT2I EP_NTERM
06495 HMMetlyEutB_2_EUT2(pf06751) EUT2I
06500 HMM06277EutA_1_eut(pf06277) EUT2I
06505 HMMreg07568_1_eut2(pf12282pf02518pf07568) EUT2I
06510 HMMreg00072_1_eut2(pf03861pf00072) EUT2I
06515 HMM10662PduV_1_mix(pf10662) EUT2I
06520 Hp_euts_fadedPurple EUT2I
06525 HMM00465alcdh_3_mix(pf00465) EUT2I
06530 -pf09335 -
06535 -pf17428 -
06540 -pf00293 -
06545 -pf08241 -
06550 -pf01476 -
06555 - -
06560 -pf00005 -
06565 - -
06570 -pf01906 -
06575 -pf06691 -
06580 -pf00583 -
06585 -pf02588 -
BMC-P
BMC-H
BMC-T
Aldehyde dehydrogenase
PTAC
Signature enzyme
Alcohol dehydrogenase
Regulator

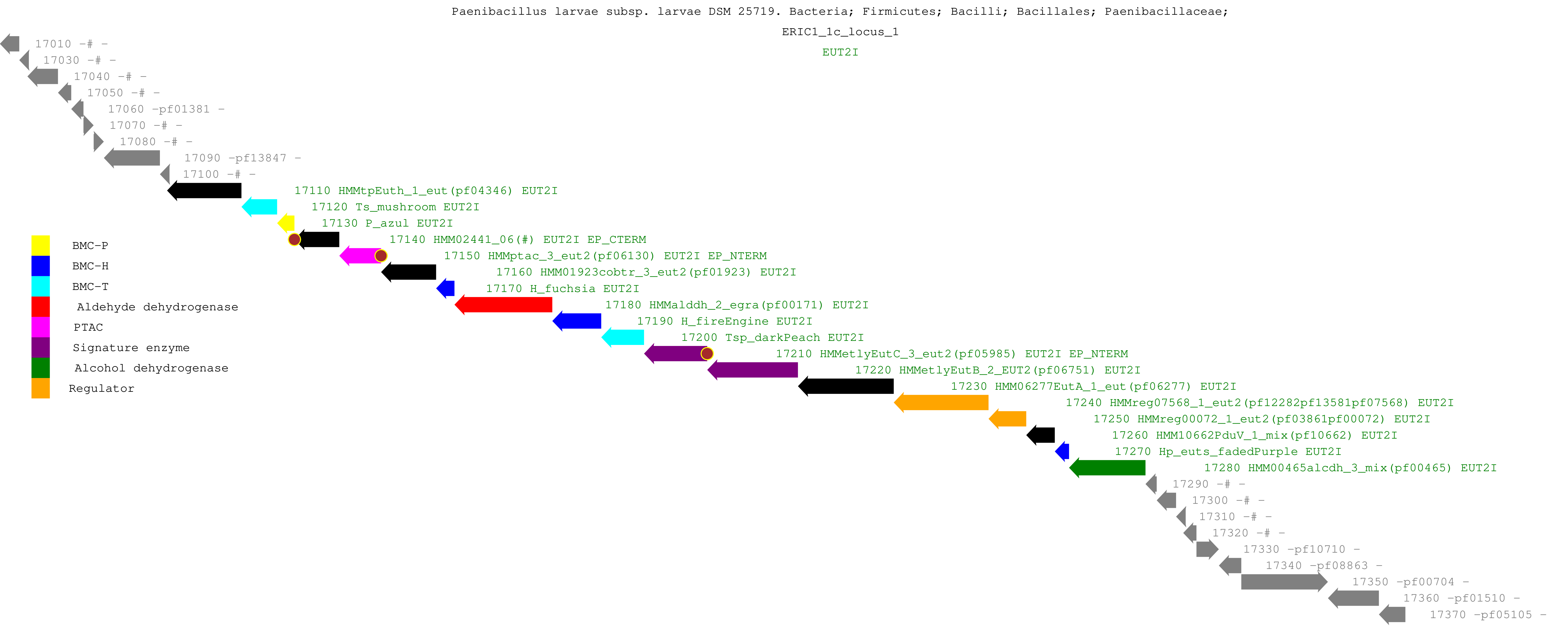
Paenibacillus larvae subsp. larvae DSM 25719. Bacteria; Firmicutes; Bacilli; Bacillales; Paenibacillaceae;
ERIC1_1c_locus_1
EUT2I
17010 -# -
17030 -# -
17040 -# -
17050 -# -
17060 -pf01381 -
17070 -# -
17080 -# -
17090 -pf13847 -
17100 -# -
17110 HMMtpEuth_1_eut(pf04346) EUT2I
17120 Ts_mushroom EUT2I
17130 P_azul EUT2I
17140 HMM02441_06(#) EUT2I EP_CTERM
17150 HMMptac_3_eut2(pf06130) EUT2I EP_NTERM
17160 HMM01923cobtr_3_eut2(pf01923) EUT2I
17170 H_fuchsia EUT2I
17180 HMMalddh_2_egra(pf00171) EUT2I
17190 H_fireEngine EUT2I
17200 Tsp_darkPeach EUT2I
17210 HMMetlyEutC_3_eut2(pf05985) EUT2I EP_NTERM
17220 HMMetlyEutB_2_EUT2(pf06751) EUT2I
17230 HMM06277EutA_1_eut(pf06277) EUT2I
17240 HMMreg07568_1_eut2(pf12282pf13581pf07568) EUT2I
17250 HMMreg00072_1_eut2(pf03861pf00072) EUT2I
17260 HMM10662PduV_1_mix(pf10662) EUT2I
17270 Hp_euts_fadedPurple EUT2I
17280 HMM00465alcdh_3_mix(pf00465) EUT2I
17290 -# -
17300 -# -
17310 -# -
17320 -# -
17330 -pf10710 -
17340 -pf08863 -
17350 -pf00704 -
17360 -pf01510 -
17370 -pf05105 -
BMC-P
BMC-H
BMC-T
Aldehyde dehydrogenase
PTAC
Signature enzyme
Alcohol dehydrogenase
Regulator

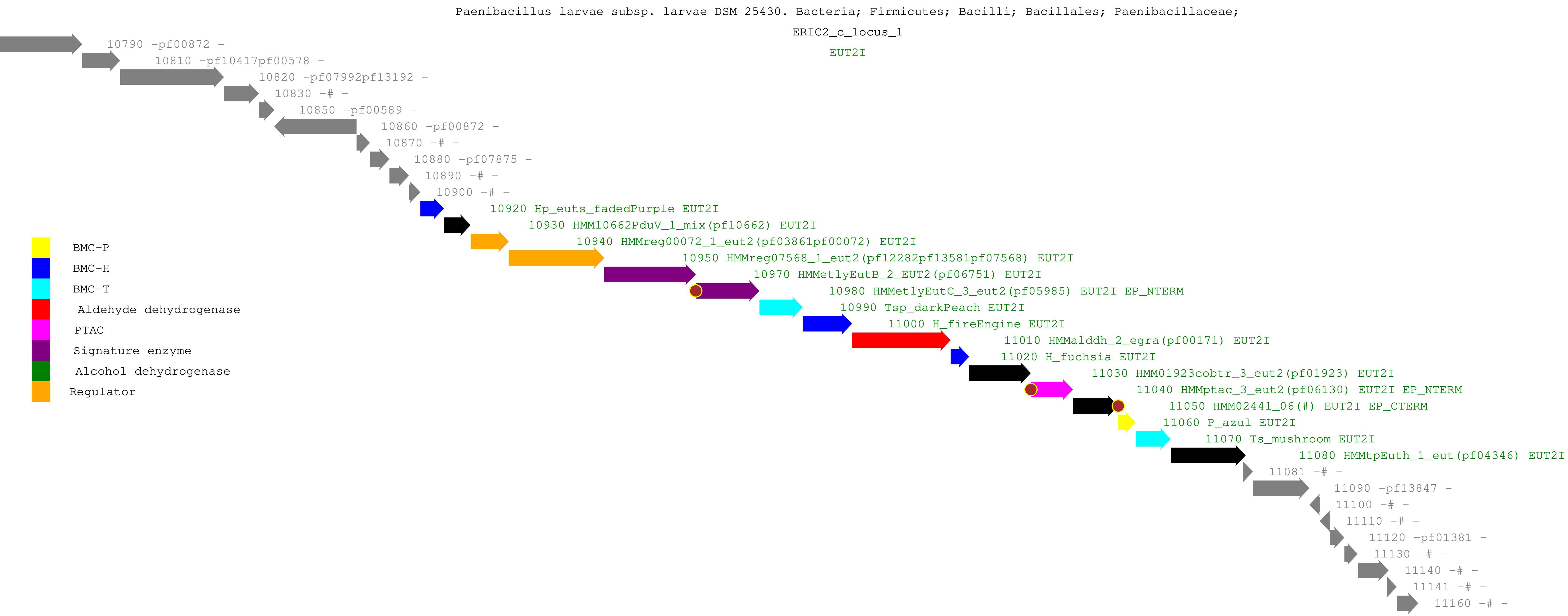
Paenibacillus larvae subsp. larvae DSM 25430. Bacteria; Firmicutes; Bacilli; Bacillales; Paenibacillaceae;
ERIC2_c_locus_1
EUT2I
10790 -pf00872 -
10810 -pf10417pf00578 -
10820 -pf07992pf13192 -
10830 -# -
10850 -pf00589 -
10860 -pf00872 -
10870 -# -
10880 -pf07875 -
10890 -# -
10900 -# -
10920 Hp_euts_fadedPurple EUT2I
10930 HMM10662PduV_1_mix(pf10662) EUT2I
10940 HMMreg00072_1_eut2(pf03861pf00072) EUT2I
10950 HMMreg07568_1_eut2(pf12282pf13581pf07568) EUT2I
10970 HMMetlyEutB_2_EUT2(pf06751) EUT2I
10980 HMMetlyEutC_3_eut2(pf05985) EUT2I EP_NTERM
10990 Tsp_darkPeach EUT2I
11000 H_fireEngine EUT2I
11010 HMMalddh_2_egra(pf00171) EUT2I
11020 H_fuchsia EUT2I
11030 HMM01923cobtr_3_eut2(pf01923) EUT2I
11040 HMMptac_3_eut2(pf06130) EUT2I EP_NTERM
11050 HMM02441_06(#) EUT2I EP_CTERM
11060 P_azul EUT2I
11070 Ts_mushroom EUT2I
11080 HMMtpEuth_1_eut(pf04346) EUT2I
11081 -# -
11090 -pf13847 -
11100 -# -
11110 -# -
11120 -pf01381 -
11130 -# -
11140 -# -
11141 -# -
11160 -# -
BMC-P
BMC-H
BMC-T
Aldehyde dehydrogenase
PTAC
Signature enzyme
Alcohol dehydrogenase
Regulator

Paenibacillus larvae subsp. larvae. Bacteria; Firmicutes; Bacilli; Bacillales; Paenibacillaceae;

ERICIII__locus_1

EUT2I

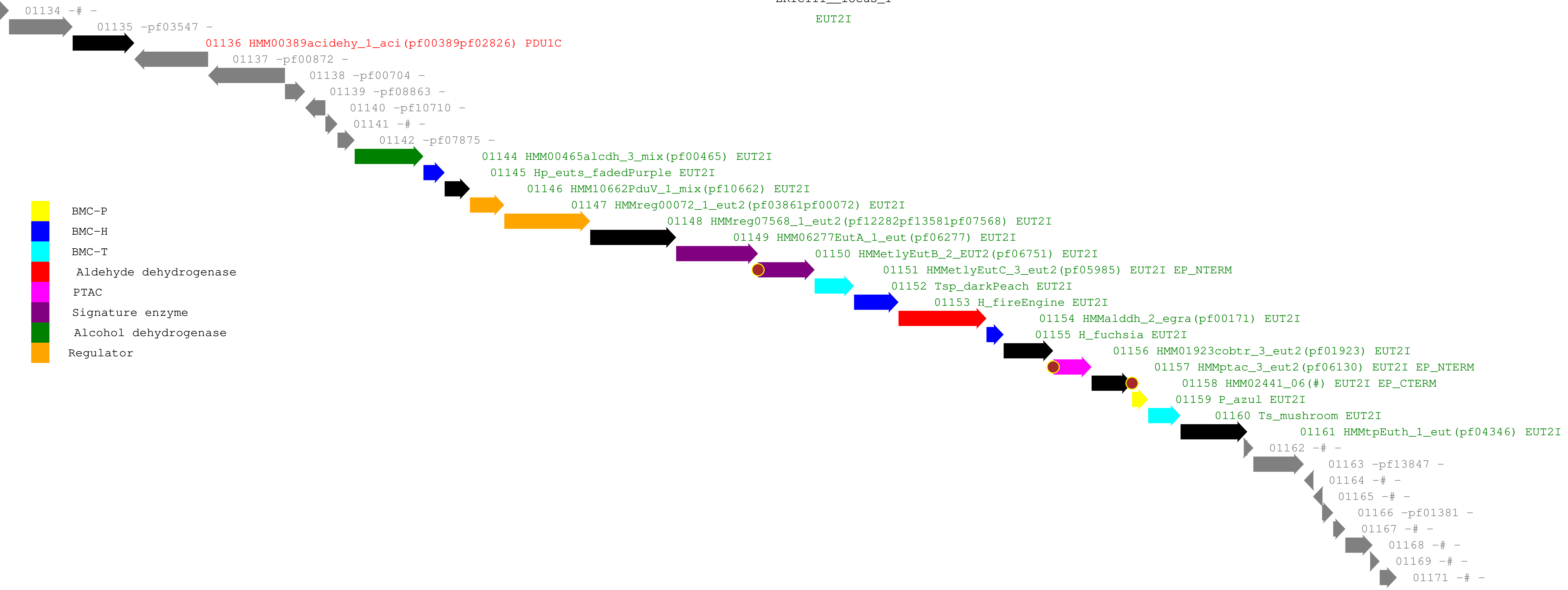

Clostridium perfringens. Bacteria; Firmicutes; Clostridia; Clostridiales; Clostridiaceae;

ERS852446__locus_1

EUT2I

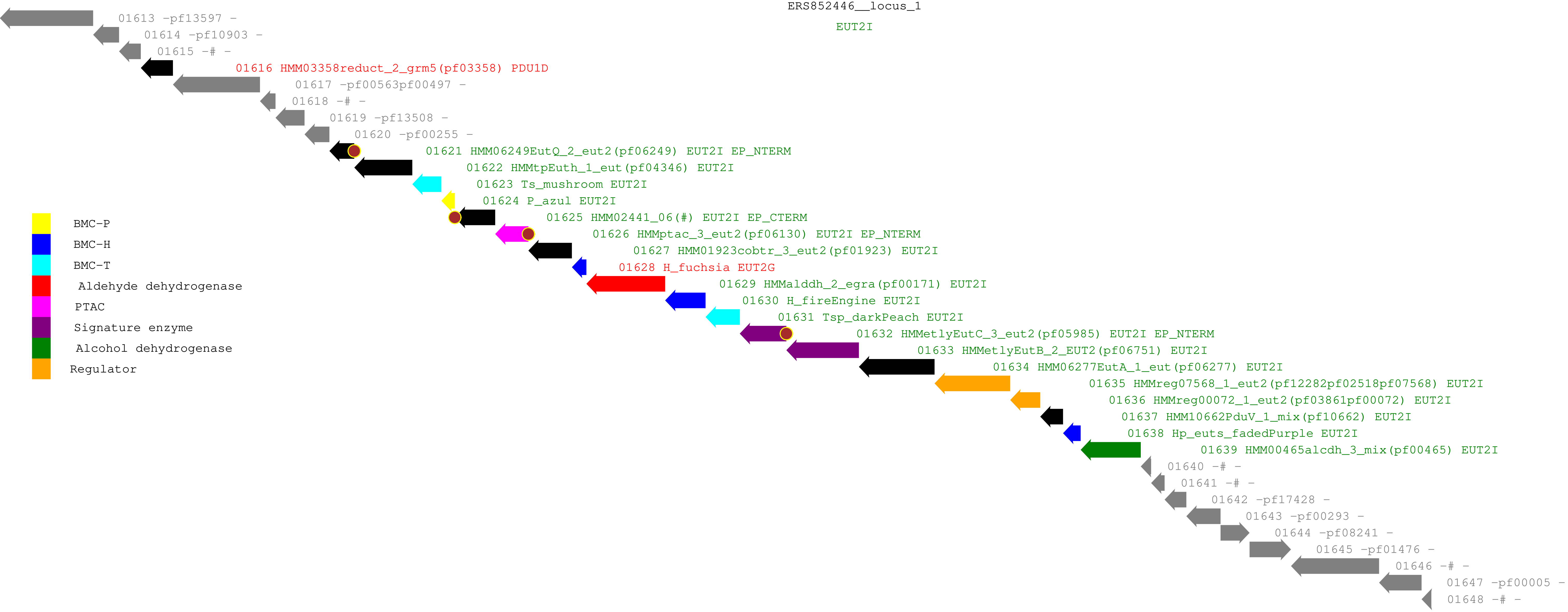

Blautia coccoides. Bacteria; Firmicutes; Clostridia; Clostridiales; Lachnospiraceae; Blautia.

EV205__locus_1_main_sat

EUT2I

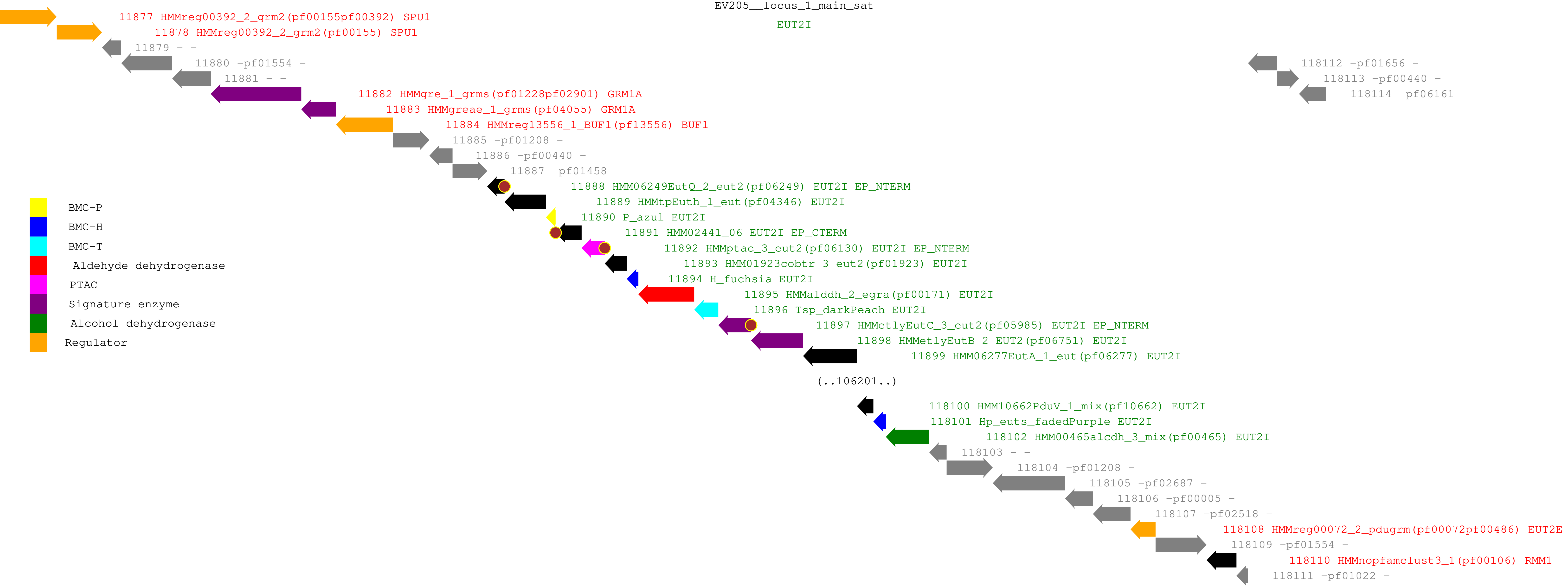

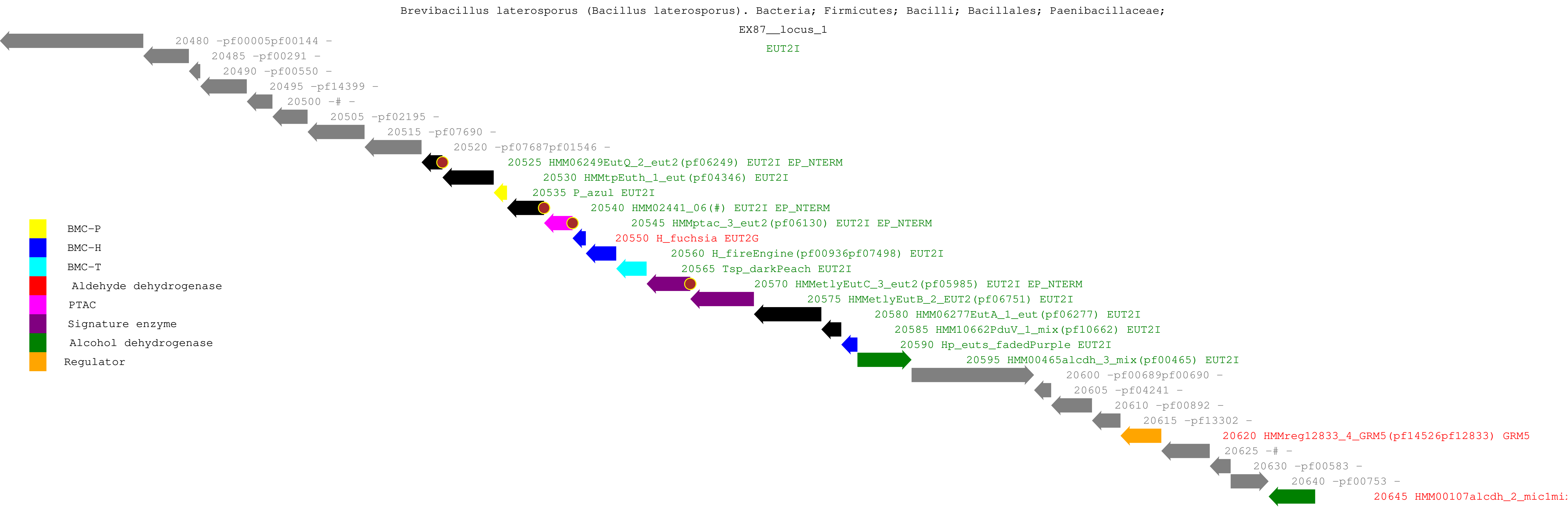
Brevibacillus laterosporus (Bacillus laterosporus). Bacteria; Firmicutes; Bacilli; Bacillales; Paenibacillaceae;
EX87__locus_1
EUT2I
20480 -pf00005pf00144 -
20485 -pf00291 -
20490 -pf00550 -
20495 -pf14399 -
20500 -# -
20505 -pf02195 -
20515 -pf07690 -
20520 -pf07687pf01546 -
20525 HMM06249EutQ_2_eut2(pf06249) EUT2I EP_NTERM
20530 HMMtpEuth_1_eut(pf04346) EUT2I
20535 P_azul EUT2I
20540 HMM02441_06(#) EUT2I EP_NTERM
20545 HMMptac_3_eut2(pf06130) EUT2I EP_NTERM
20550 H_fuchsia EUT2G
20560 H_fireEngine(pf00936pf07498) EUT2I
20565 Tsp_darkPeach EUT2I
20570 HMMetlyEutC_3_eut2(pf05985) EUT2I EP_NTERM
20575 HMMetlyEutB_2_EUT2(pf06751) EUT2I
20580 HMM06277EutA_1_eut(pf06277) EUT2I
20585 HMM10662PduV_1_mix(pf10662) EUT2I
20590 Hp_euts_fadedPurple EUT2I
20595 HMM00465alcdh_3_mix(pf00465) EUT2I
20600 -pf00689pf00690 -
20605 -pf04241 -
20610 -pf00892 -
20615 -pf13302 -
20620 HMMreg12833_4_GRM5(pf14526pf12833) GRM5
20625 -# -
20630 -pf00583 -
20640 -pf00753 -
20645 HMM00107alcdh_2_mic1mi
BMC-P
BMC-H
BMC-T
Aldehyde dehydrogenase
PTAC
Signature enzyme
Alcohol dehydrogenase
Regulator

Brevibacillus laterosporus (Bacillus laterosporus). Bacteria; Firmicutes; Bacilli; Bacillales; Paenibacillaceae;

EX87__locus_2

EUT2I

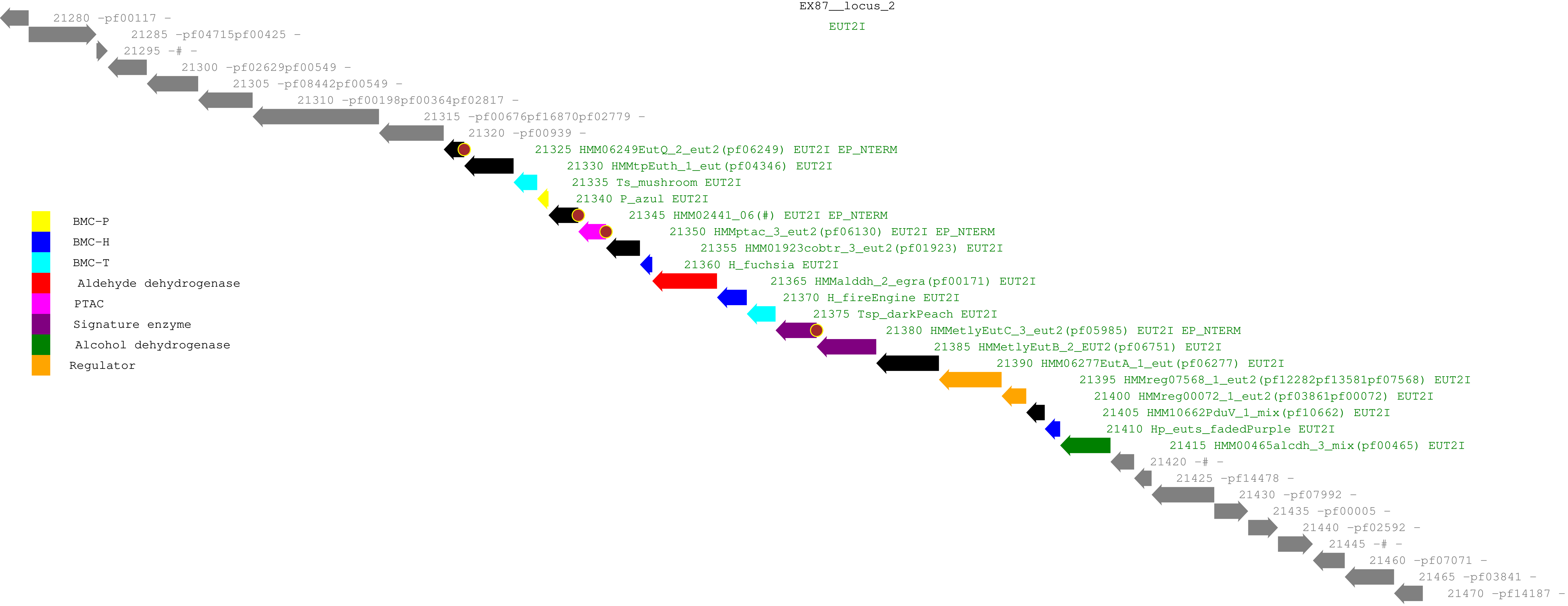

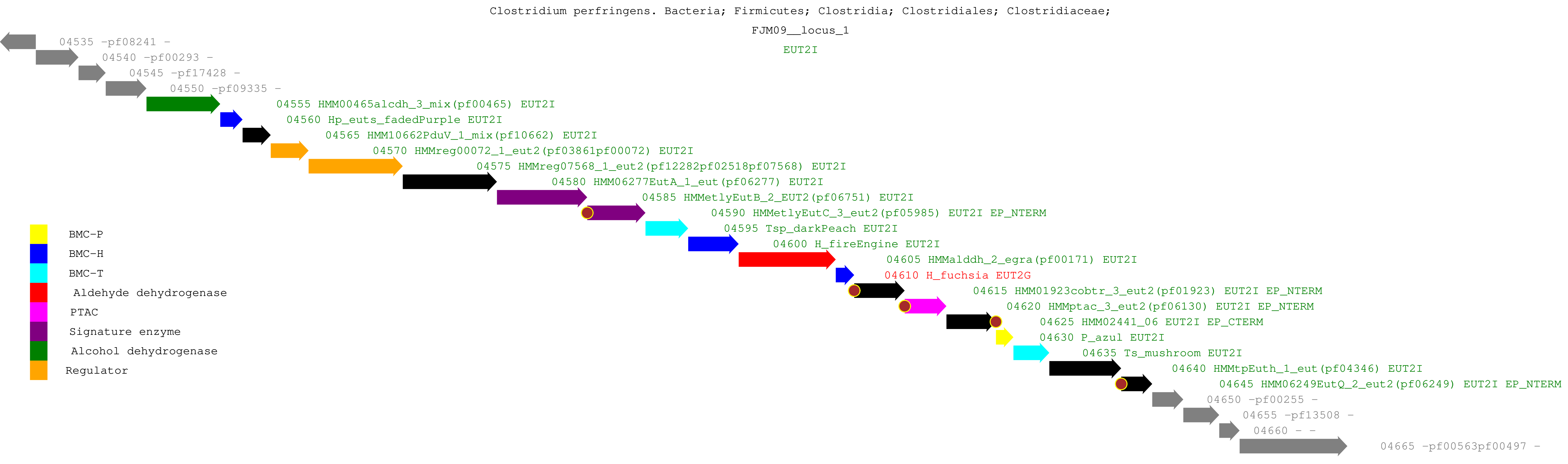
Clostridium perfringens. Bacteria; Firmicutes; Clostridia; Clostridiales; Clostridiaceae;
FJM09__locus_1
EUT2I
04535 -pf08241 -
04540 -pf00293 -
04545 -pf17428 -
04550 -pf09335 -
04555 HMM00465alcdh_3_mix(pf00465) EUT2I
04560 Hp_euts_fadedPurple EUT2I
04565 HMM10662PduV_1_mix(pf10662) EUT2I
04570 HMMreg00072_1_eut2(pf03861pf00072) EUT2I
04575 HMMreg07568_1_eut2(pf12282pf02518pf07568) EUT2I
04580 HMM06277EutA_1_eut(pf06277) EUT2I
04585 HMMetlyEutB_2_EUT2(pf06751) EUT2I
04590 HMMetlyEutC_3_eut2(pf05985) EUT2I EP_NTERM
04595 Tsp_darkPeach EUT2I
04600 H_fireEngine EUT2I
04605 HMMalddh_2_egra(pf00171) EUT2I
04610 H_fuchsia EUT2G
04615 HMM01923cobtr_3_eut2(pf01923) EUT2I EP_NTERM
04620 HMMptac_3_eut2(pf06130) EUT2I EP_NTERM
04625 HMM02441_06 EUT2I EP_CTERM
04630 P_azul EUT2I
04635 Ts_mushroom EUT2I
04640 HMMtpEuth_1_eut(pf04346) EUT2I
04645 HMM06249EutQ_2_eut2(pf06249) EUT2I EP_NTERM
04650 -pf00255 -
04655 -pf13508 -
04660 - -
04665 -pf00563pf00497 -
BMC-P
BMC-H
BMC-T
Aldehyde dehydrogenase
PTAC
Signature enzyme
Alcohol dehydrogenase
Regulator

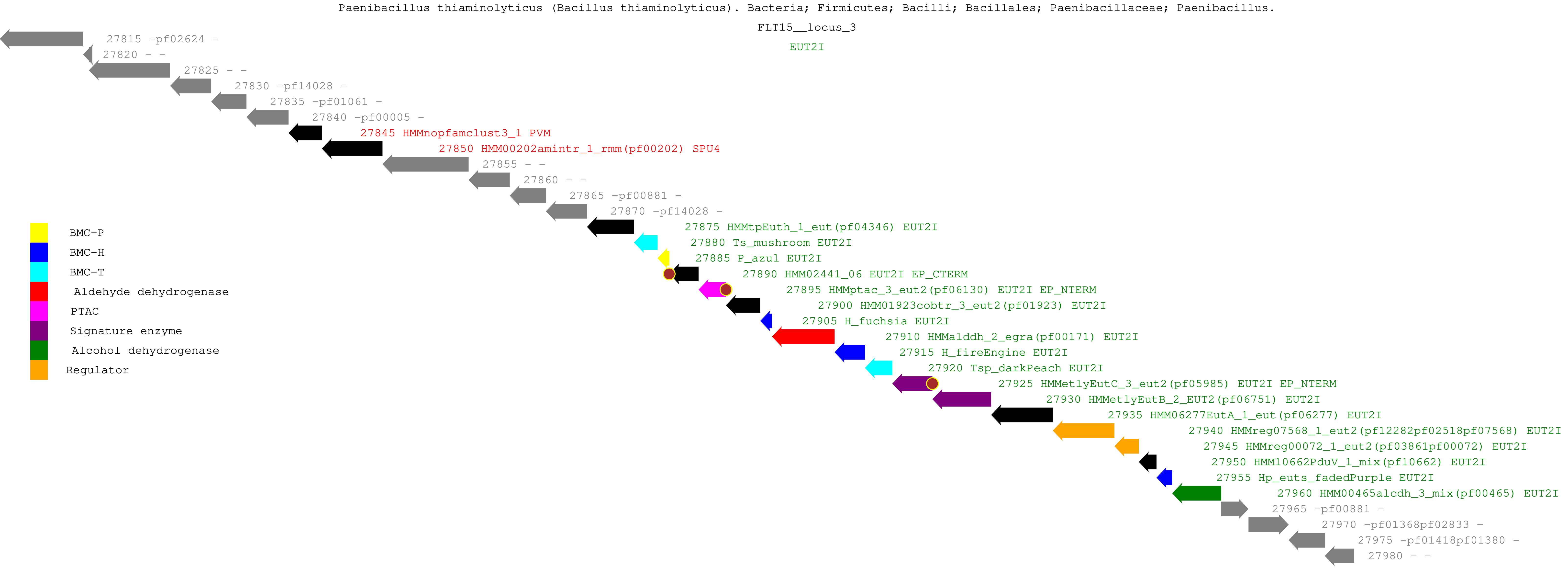
Paenibacillus thiaminolyticus (Bacillus thiaminolyticus). Bacteria; Firmicutes; Bacilli; Bacillales; Paenibacillaceae; Paenibacillus.
FLT15__locus_3
EUT2I
27815 -pf02624 -
27820 - -
27825 - -
27830 -pf14028 -
27835 -pf01061 -
27840 -pf00005 -
27845 HMMnopfamclust3_1 PVM
27850 HMM00202amintr_1_rmm(pf00202) SPU4
27855 - -
27860 - -
27865 -pf00881 -
27870 -pf14028 -
27875 HMMtpEuth_1_eut(pf04346) EUT2I
27880 Ts_mushroom EUT2I
27885 P_azul EUT2I
27890 HMM02441_06 EUT2I EP_CTERM
27895 HMMptac_3_eut2(pf06130) EUT2I EP_NTERM
27900 HMM01923cobtr_3_eut2(pf01923) EUT2I
27905 H_fuchsia EUT2I
27910 HMMalddh_2_egra(pf00171) EUT2I
27915 H_fireEngine EUT2I
27920 Tsp_darkPeach EUT2I
27925 HMMetlyEutC_3_eut2(pf05985) EUT2I EP_NTERM
27930 HMMetlyEutB_2_EUT2(pf06751) EUT2I
27935 HMM06277EutA_1_eut(pf06277) EUT2I
27940 HMMreg07568_1_eut2(pf12282pf02518pf07568) EUT2I
27945 HMMreg00072_1_eut2(pf03861pf00072) EUT2I
27950 HMM10662PduV_1_mix(pf10662) EUT2I
27955 Hp_euts_fadedPurple EUT2I
27960 HMM00465alcdh_3_mix(pf00465) EUT2I
27965 -pf00881 -
27970 -pf01368pf02833 -
27975 -pf01418pf01380 -
27980 - -
BMC-P
BMC-H
BMC-T
Aldehyde dehydrogenase
PTAC
Signature enzyme
Alcohol dehydrogenase
Regulator

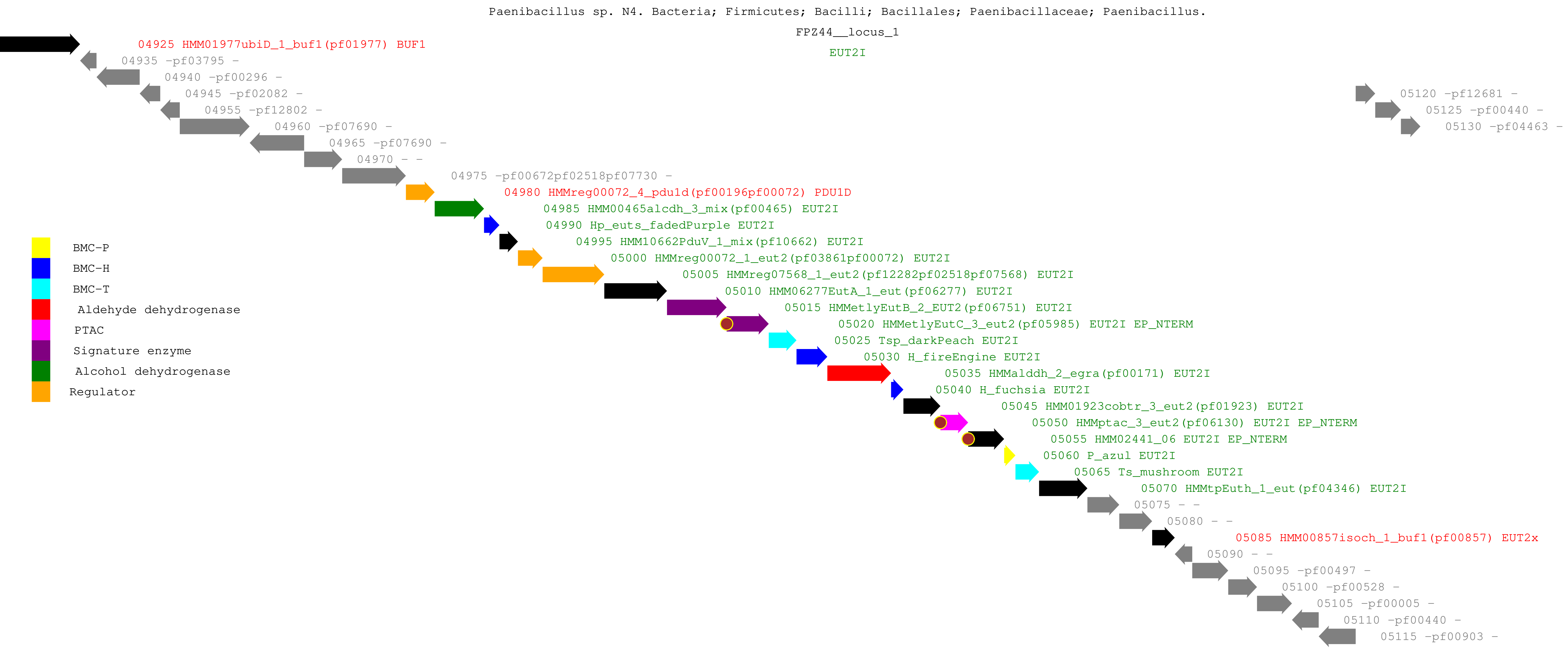
Paenibacillus sp. N4. Bacteria; Firmicutes; Bacilli; Bacillales; Paenibacillaceae; Paenibacillus.
FPZ44__locus_1
EUT2I
04925 HMM01977ubiD_1_buf1(pf01977) BUF1
04935 -pf03795 -
04940 -pf00296 -
04945 -pf02082 -
04955 -pf12802 -
04960 -pf07690 -
04965 -pf07690 -
04970 - -
04975 -pf00672pf02518pf07730 -
04980 HMMreg00072_4_pdu1d(pf00196pf00072) PDU1D
04985 HMM00465alcdh_3_mix(pf00465) EUT2I
04990 Hp_euts_fadedPurple EUT2I
04995 HMM10662PduV_1_mix(pf10662) EUT2I
05000 HMMreg00072_1_eut2(pf03861pf00072) EUT2I
05005 HMMreg07568_1_eut2(pf12282pf02518pf07568) EUT2I
05010 HMM06277EutA_1_eut(pf06277) EUT2I
05015 HMMetlyEutB_2_EUT2(pf06751) EUT2I
05020 HMMetlyEutC_3_eut2(pf05985) EUT2I EP_NTERM
05025 Tsp_darkPeach EUT2I
05030 H_fireEngine EUT2I
05035 HMMalddh_2_egra(pf00171) EUT2I
05040 H_fuchsia EUT2I
05045 HMM01923cobtr_3_eut2(pf01923) EUT2I
05050 HMMptac_3_eut2(pf06130) EUT2I EP_NTERM
05055 HMM02441_06 EUT2I EP_NTERM
05060 P_azul EUT2I
05065 Ts_mushroom EUT2I
05070 HMMtpEuth_1_eut(pf04346) EUT2I
05075 - -
05080 - -
05085 HMM00857isoch_1_buf1(pf00857) EUT2x
05090 - -
05095 -pf00497 -
05100 -pf00528 -
05105 -pf00005 -
05110 -pf00440 -
05115 -pf00903 -
05120 -pf12681 -
05125 -pf00440 -
05130 -pf04463 -
BMC-P
BMC-H
BMC-T
Aldehyde dehydrogenase
PTAC
Signature enzyme
Alcohol dehydrogenase
Regulator

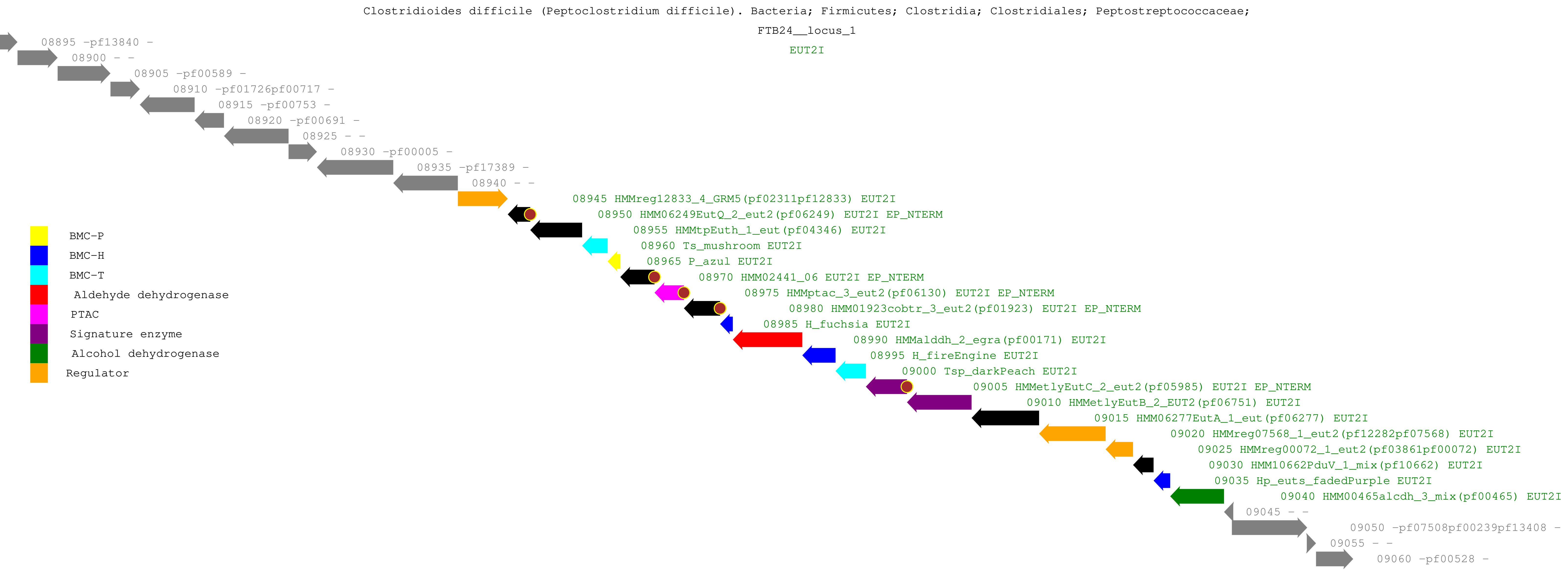
Clostridioides difficile (Peptoclostridium difficile). Bacteria; Firmicutes; Clostridia; Clostridiales; Peptostreptococcaceae;
FTB24__locus_1
EUT2I
08895 -pf13840 -
08900 - -
08905 -pf00589 -
08910 -pf01726pf00717 -
08915 -pf00753 -
08920 -pf00691 -
08925 - -
08930 -pf00005 -
08935 -pf17389 -
08940 - -
08945 HMMreg12833_4_GRM5(pf02311pf12833) EUT2I
08950 HMM06249EutQ_2_eut2(pf06249) EUT2I EP_NTERM
08955 HMMtpEuth_1_eut(pf04346) EUT2I
08960 Ts_mushroom EUT2I
08965 P_azul EUT2I
08970 HMM02441_06 EUT2I EP_NTERM
08975 HMMptac_3_eut2(pf06130) EUT2I EP_NTERM
08980 HMM01923cobtr_3_eut2(pf01923) EUT2I EP_NTERM
08985 H_fuchsia EUT2I
08990 HMMalddh_2_egra(pf00171) EUT2I
08995 H_fireEngine EUT2I
09000 Tsp_darkPeach EUT2I
09005 HMMetlyEutC_2_eut2(pf05985) EUT2I EP_NTERM
09010 HMMetlyEutB_2_EUT2(pf06751) EUT2I
09015 HMM06277EutA_1_eut(pf06277) EUT2I
09020 HMMreg07568_1_eut2(pf12282pf07568) EUT2I
09025 HMMreg00072_1_eut2(pf03861pf00072) EUT2I
09030 HMM10662PduV_1_mix(pf10662) EUT2I
09035 Hp_euts_fadedPurple EUT2I
09040 HMM00465alcdh_3_mix(pf00465) EUT2I
09045 - -
09050 -pf07508pf00239pf13408 -
09055 - -
09060 -pf00528 -
BMC-P
BMC-H
BMC-T
Aldehyde dehydrogenase
PTAC
Signature enzyme
Alcohol dehydrogenase
Regulator

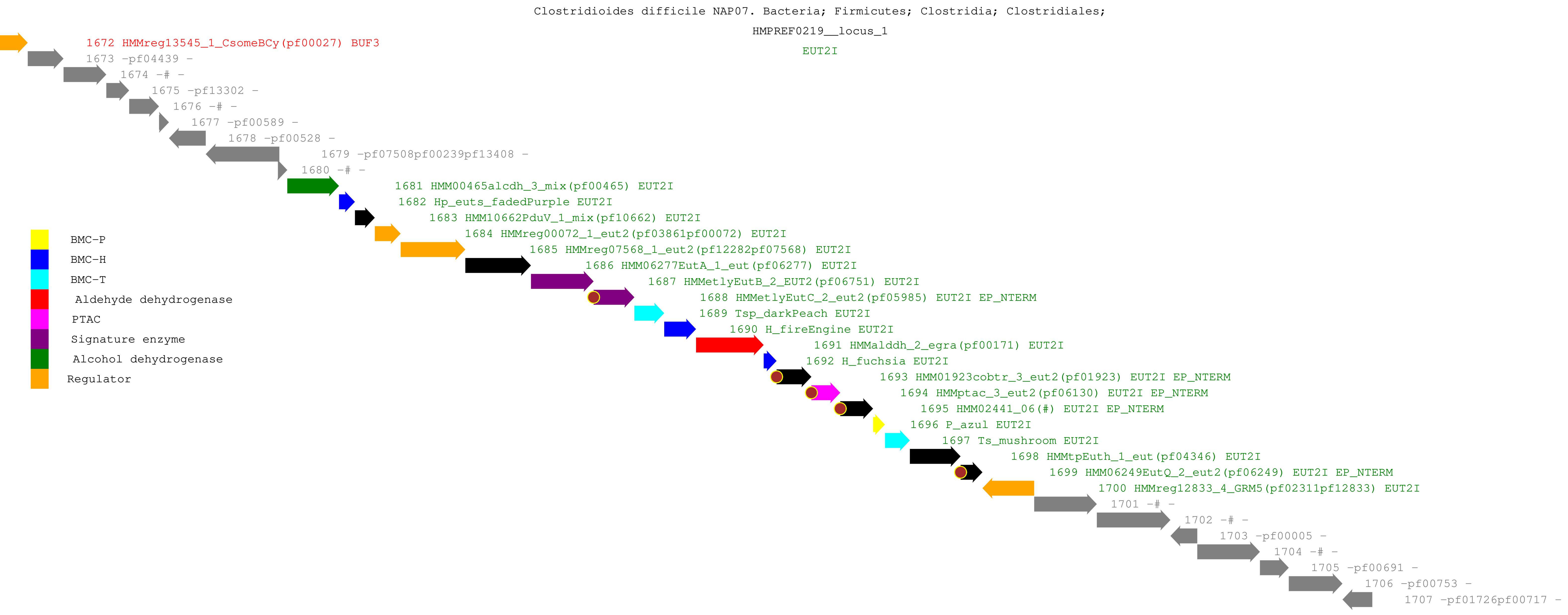
Clostridioides difficile NAP07. Bacteria; Firmicutes; Clostridia; Clostridiales;
HMPREF0219__locus_1
EUT2I
1672 HMMreg13545_1_CsomeBCy(pf00027) BUF3
1673 -pf04439 -
1674 -# -
1675 -pf13302 -
1676 -# -
1677 -pf00589 -
1678 -pf00528 -
1679 -pf07508pf00239pf13408 -
1680 -# -
1681 HMM00465alcdh_3_mix(pf00465) EUT2I
1682 Hp_euts_fadedPurple EUT2I
1683 HMM10662PduV_1_mix(pf10662) EUT2I
1684 HMMreg00072_1_eut2(pf03861pf00072) EUT2I
1685 HMMreg07568_1_eut2(pf12282pf07568) EUT2I
1686 HMM06277EutA_1_eut(pf06277) EUT2I
1687 HMMetlyEutB_2_EUT2(pf06751) EUT2I
1688 HMMetlyEutC_2_eut2(pf05985) EUT2I EP_NTERM
1689 Tsp_darkPeach EUT2I
1690 H_fireEngine EUT2I
1691 HMMalddh_2_egra(pf00171) EUT2I
1692 H_fuchsia EUT2I
1693 HMM01923cobtr_3_eut2(pf01923) EUT2I EP_NTERM
1694 HMMptac_3_eut2(pf06130) EUT2I EP_NTERM
1695 HMM02441_06(#) EUT2I EP_NTERM
1696 P_azul EUT2I
1697 Ts_mushroom EUT2I
1698 HMMtpEuth_1_eut(pf04346) EUT2I
1699 HMM06249EutQ_2_eut2(pf06249) EUT2I EP_NTERM
1700 HMMreg12833_4_GRM5(pf02311pf12833) EUT2I
1701 -# -
1702 -# -
1703 -pf00005 -
1704 -# -
1705 -pf00691 -
1706 -pf00753 -
1707 -pf01726pf00717 -
BMC-P
BMC-H
BMC-T
Aldehyde dehydrogenase
PTAC
Signature enzyme
Alcohol dehydrogenase
Regulator

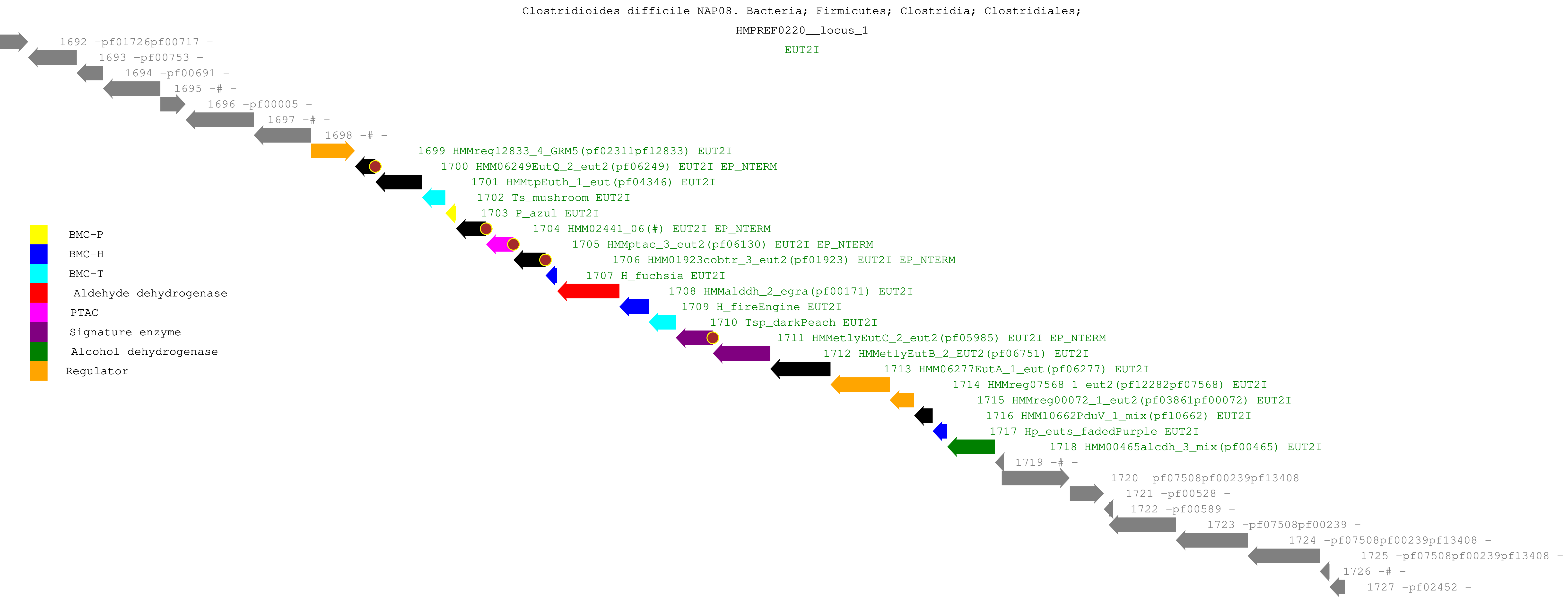
Clostridioides difficile NAP08. Bacteria; Firmicutes; Clostridia; Clostridiales;
HMPREF0220__locus_1
EUT2I
1692 -pf01726pf00717 -
1693 -pf00753 -
1694 -pf00691 -
1695 -# -
1696 -pf00005 -
1697 -# -
1698 -# -
1699 HMMreg12833_4_GRM5(pf02311pf12833) EUT2I
1700 HMM06249EutQ_2_eut2(pf06249) EUT2I EP_NTERM
1701 HMMtpEuth_1_eut(pf04346) EUT2I
1702 Ts_mushroom EUT2I
1703 P_azul EUT2I
1704 HMM02441_06(#) EUT2I EP_NTERM
1705 HMMptac_3_eut2(pf06130) EUT2I EP_NTERM
1706 HMM01923cobtr_3_eut2(pf01923) EUT2I EP_NTERM
1707 H_fuchsia EUT2I
1708 HMMalddh_2_egra(pf00171) EUT2I
1709 H_fireEngine EUT2I
1710 Tsp_darkPeach EUT2I
1711 HMMetlyEutC_2_eut2(pf05985) EUT2I EP_NTERM
1712 HMMetlyEutB_2_EUT2(pf06751) EUT2I
1713 HMM06277EutA_1_eut(pf06277) EUT2I
1714 HMMreg07568_1_eut2(pf12282pf07568) EUT2I
1715 HMMreg00072_1_eut2(pf03861pf00072) EUT2I
1716 HMM10662PduV_1_mix(pf10662) EUT2I
1717 Hp_euts_fadedPurple EUT2I
1718 HMM00465alcdh_3_mix(pf00465) EUT2I
1719 -# -
1720 -pf07508pf00239pf13408 -
1721 -pf00528 -
1722 -pf00589 -
1723 -pf07508pf00239 -
1724 -pf07508pf00239pf13408 -
1725 -pf07508pf00239pf13408 -
1726 -# -
1727 -pf02452 -
BMC-P
BMC-H
BMC-T
Aldehyde dehydrogenase
PTAC
Signature enzyme
Alcohol dehydrogenase
Regulator

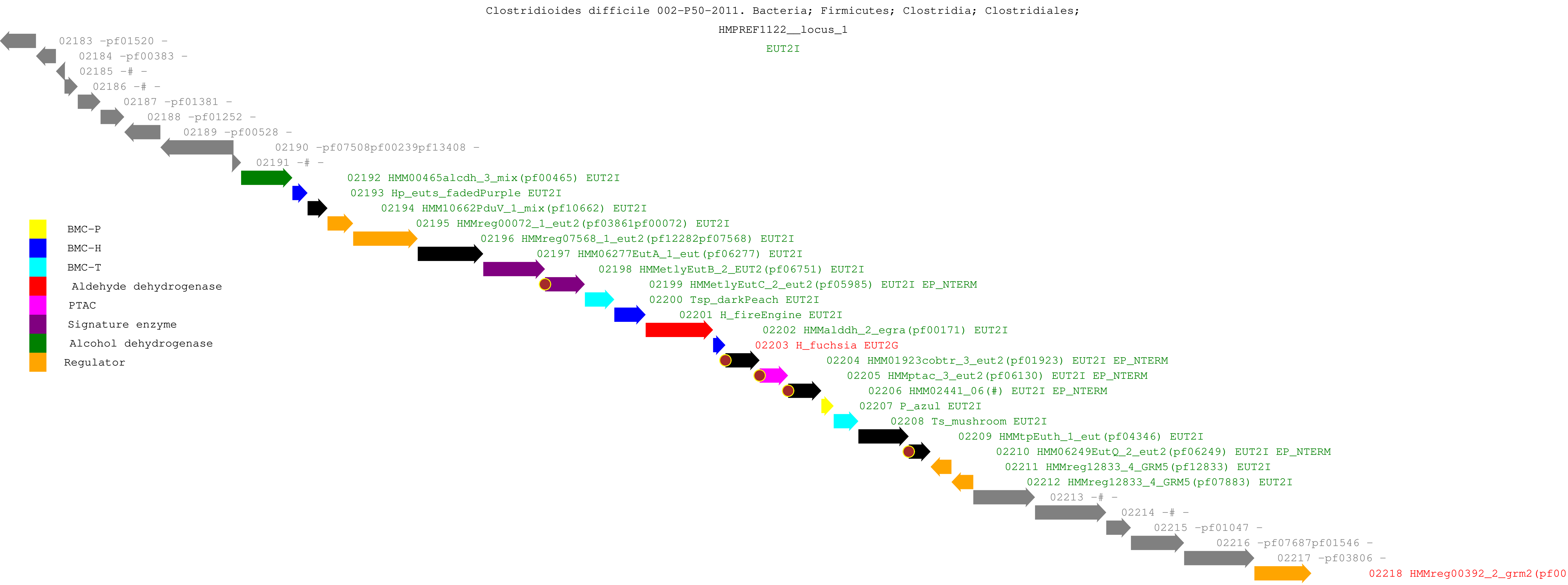
Clostridioides difficile 002-P50-2011. Bacteria; Firmicutes; Clostridia; Clostridiales;
HMPREF1122__locus_1
EUT2I
02183 -pf01520 -
02184 -pf00383 -
02185 -# -
02186 -# -
02187 -pf01381 -
02188 -pf01252 -
02189 -pf00528 -
02190 -pf07508pf00239pf13408 -
02191 -# -
02192 HMM00465alcdh_3_mix(pf00465) EUT2I
02193 Hp_euts_fadedPurple EUT2I
02194 HMM10662PduV_1_mix(pf10662) EUT2I
02195 HMMreg00072_1_eut2(pf03861pf00072) EUT2I
02196 HMMreg07568_1_eut2(pf12282pf07568) EUT2I
02197 HMM06277EutA_1_eut(pf06277) EUT2I
02198 HMMetlyEutB_2_EUT2(pf06751) EUT2I
02199 HMMetlyEutC_2_eut2(pf05985) EUT2I EP_NTERM
02200 Tsp_darkPeach EUT2I
02201 H_fireEngine EUT2I
02202 HMMalddh_2_egra(pf00171) EUT2I
02203 H_fuchsia EUT2G
02204 HMM01923cobtr_3_eut2(pf01923) EUT2I EP_NTERM
02205 HMMptac_3_eut2(pf06130) EUT2I EP_NTERM
02206 HMM02441_06(#) EUT2I EP_NTERM
02207 P_azul EUT2I
02208 Ts_mushroom EUT2I
02209 HMMtpEuth_1_eut(pf04346) EUT2I
02210 HMM06249EutQ_2_eut2(pf06249) EUT2I EP_NTERM
02211 HMMreg12833_4_GRM5(pf12833) EUT2I
02212 HMMreg12833_4_GRM5(pf07883) EUT2I
02213 -# -
02214 -# -
02215 -pf01047 -
02216 -pf07687pf01546 -
02217 -pf03806 -
02218 HMMreg00392_2_grm2(pf00
BMC-P
BMC-H
BMC-T
Aldehyde dehydrogenase
PTAC
Signature enzyme
Alcohol dehydrogenase
Regulator

Clostridioides difficile 050-P50-2011. Bacteria; Firmicutes; Clostridia; Clostridiales;

HMPREF1123__locus_1

EUT2I

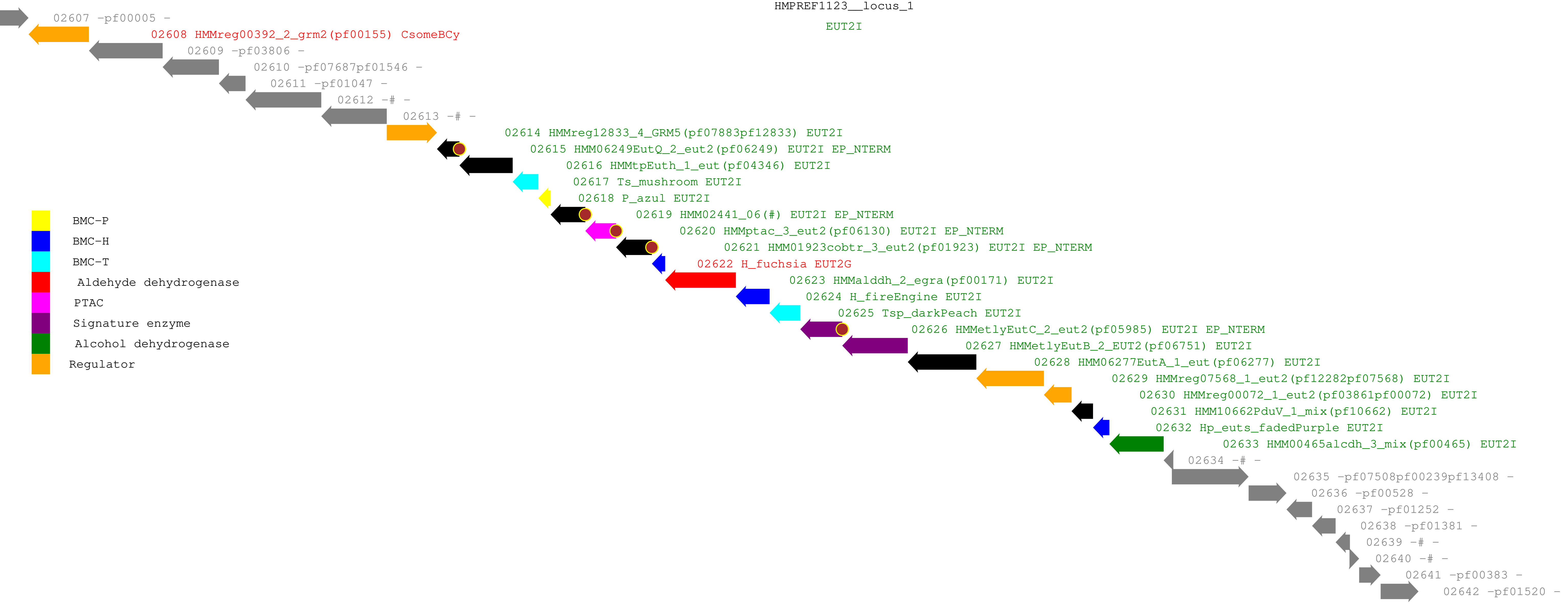

Clostridium sp. HMSC19B10. Bacteria; Firmicutes; Clostridia; Clostridiales; Clostridiaceae;

HMPREF3074__locus_1

EUT2I

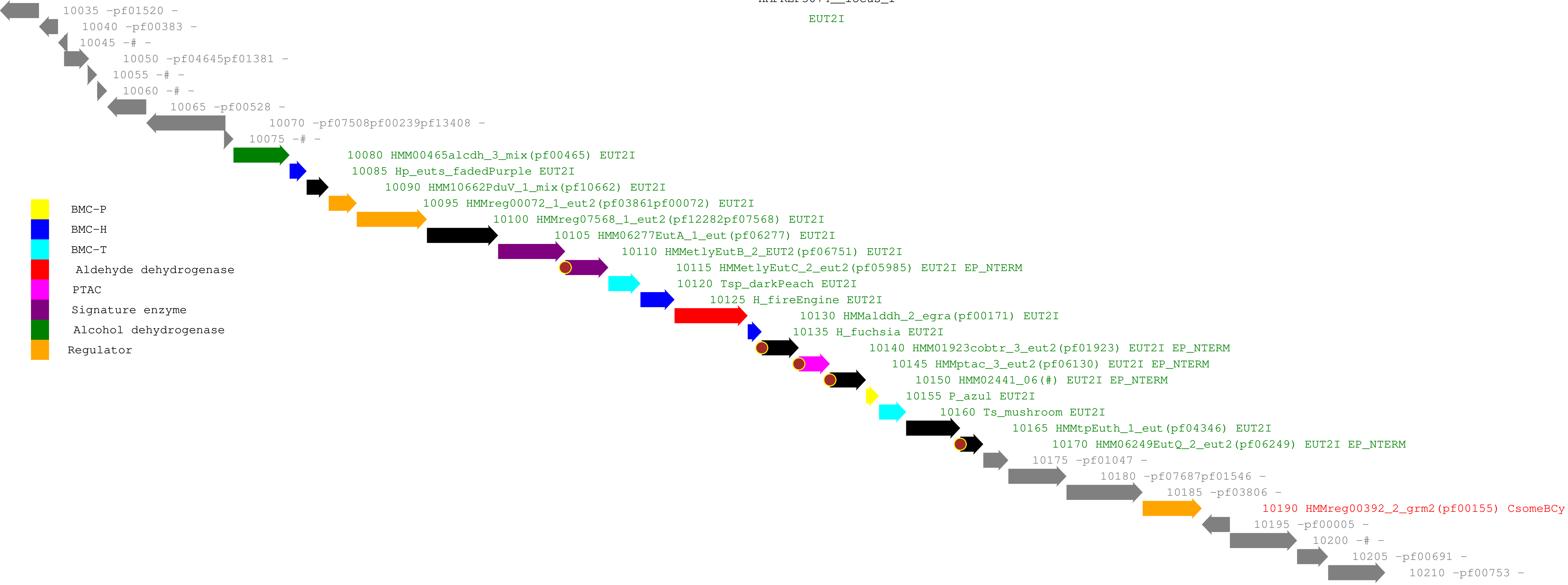

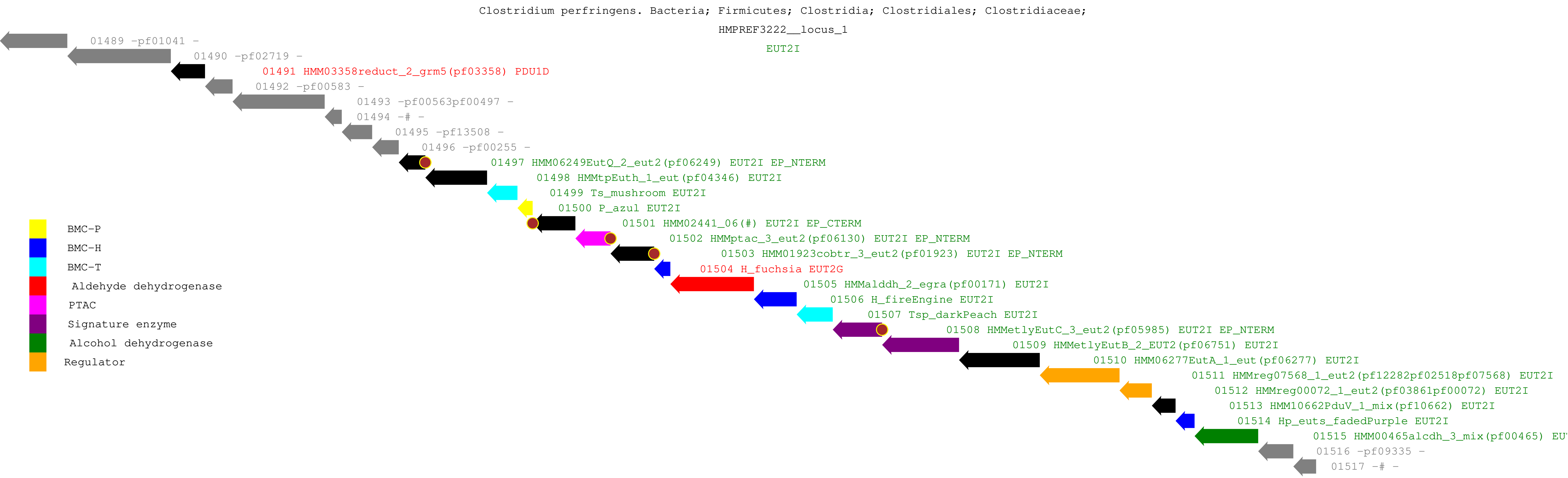
Clostridium perfringens. Bacteria; Firmicutes; Clostridia; Clostridiales; Clostridiaceae;
HMPREF3222__locus_1
EUT2I
01489 -pf01041 -
01490 -pf02719 -
01491 HMM03358reduct_2_grm5(pf03358) PDU1D
01492 -pf00583 -
01493 -pf00563pf00497 -
01494 -# -
01495 -pf13508 -
01496 -pf00255 -
01497 HMM06249EutQ_2_eut2(pf06249) EUT2I EP_NTERM
01498 HMMtpEuth_1_eut(pf04346) EUT2I
01499 Ts_mushroom EUT2I
01500 P_azul EUT2I
01501 HMM02441_06(#) EUT2I EP_CTERM
01502 HMMptac_3_eut2(pf06130) EUT2I EP_NTERM
01503 HMM01923cobtr_3_eut2(pf01923) EUT2I EP_NTERM
01504 H_fuchsia EUT2G
01505 HMMalddh_2_egra(pf00171) EUT2I
01506 H_fireEngine EUT2I
01507 Tsp_darkPeach EUT2I
01508 HMMetlyEutC_3_eut2(pf05985) EUT2I EP_NTERM
01509 HMMetlyEutB_2_EUT2(pf06751) EUT2I
01510 HMM06277EutA_1_eut(pf06277) EUT2I
01511 HMMreg07568_1_eut2(pf12282pf02518pf07568) EUT2I
01512 HMMreg00072_1_eut2(pf03861pf00072) EUT2I
01513 HMM10662PduV_1_mix(pf10662) EUT2I
01514 Hp_euts_fadedPurple EUT2I
01515 HMM00465alcdh_3_mix(pf00465) EU
01516 -pf09335 -
01517 -# -
BMC-P
BMC-H
BMC-T
Aldehyde dehydrogenase
PTAC
Signature enzyme
Alcohol dehydrogenase
Regulator

Clostridioides difficile (Peptoclostridium difficile). Bacteria; Firmicutes; Clostridia; Clostridiales;

IM33__locus_1

EUT2I

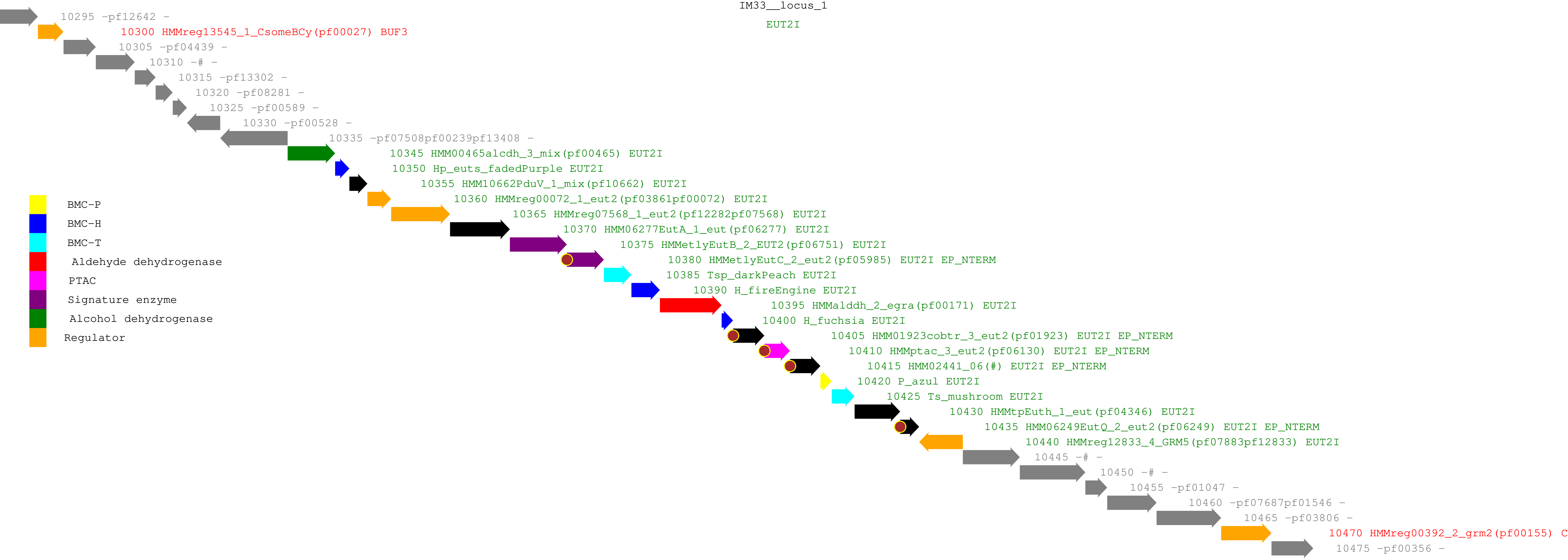

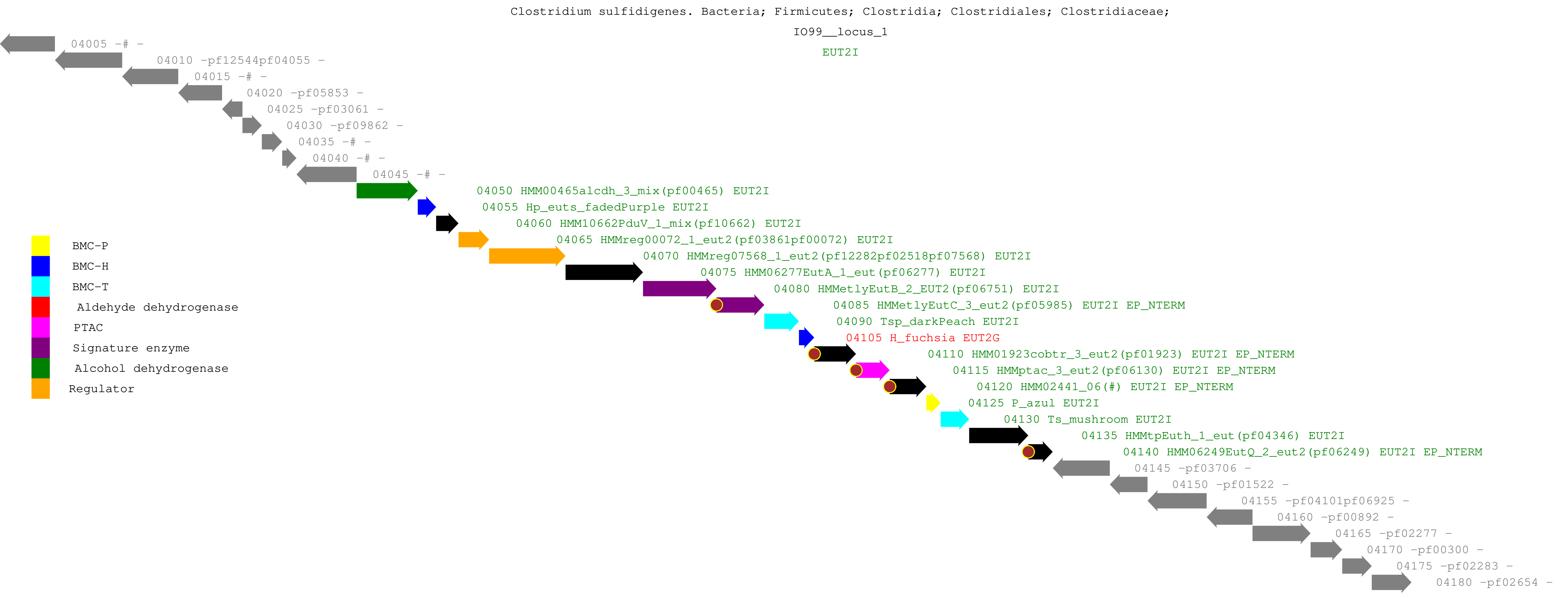
Clostridium sulfidigenes. Bacteria; Firmicutes; Clostridia; Clostridiales; Clostridiaceae;
IO99__locus_1
EUT2I
04005 -# -
04010 -pf12544pf04055 -
04015 -# -
04020 -pf05853 -
04025 -pf03061 -
04030 -pf09862 -
04035 -# -
04040 -# -
04045 -# -
04050 HMM00465alcdh_3_mix(pf00465) EUT2I
04055 Hp_euts_fadedPurple EUT2I
04060 HMM10662PduV_1_mix(pf10662) EUT2I
04065 HMMreg00072_1_eut2(pf03861pf00072) EUT2I
04070 HMMreg07568_1_eut2(pf12282pf02518pf07568) EUT2I
04075 HMM06277EutA_1_eut(pf06277) EUT2I
04080 HMMetlyEutB_2_EUT2(pf06751) EUT2I
04085 HMMetlyEutC_3_eut2(pf05985) EUT2I EP_NTERM
04090 Tsp_darkPeach EUT2I
04105 H_fuchsia EUT2G
04110 HMM01923cobtr_3_eut2(pf01923) EUT2I EP_NTERM
04115 HMMptac_3_eut2(pf06130) EUT2I EP_NTERM
04120 HMM02441_06(#) EUT2I EP_NTERM
04125 P_azul EUT2I
04130 Ts_mushroom EUT2I
04135 HMMtpEuth_1_eut(pf04346) EUT2I
04140 HMM06249EutQ_2_eut2(pf06249) EUT2I EP_NTERM
04145 -pf03706 -
04150 -pf01522 -
04155 -pf04101pf06925 -
04160 -pf00892 -
04165 -pf02277 -
04170 -pf00300 -
04175 -pf02283 -
04180 -pf02654 -
BMC-P
BMC-H
BMC-T
Aldehyde dehydrogenase
PTAC
Signature enzyme
Alcohol dehydrogenase
Regulator

Clostridium perfringens. Bacteria; Firmicutes; Clostridia; Clostridiales; Clostridiaceae;

JFP838__locus_1

EUT2I

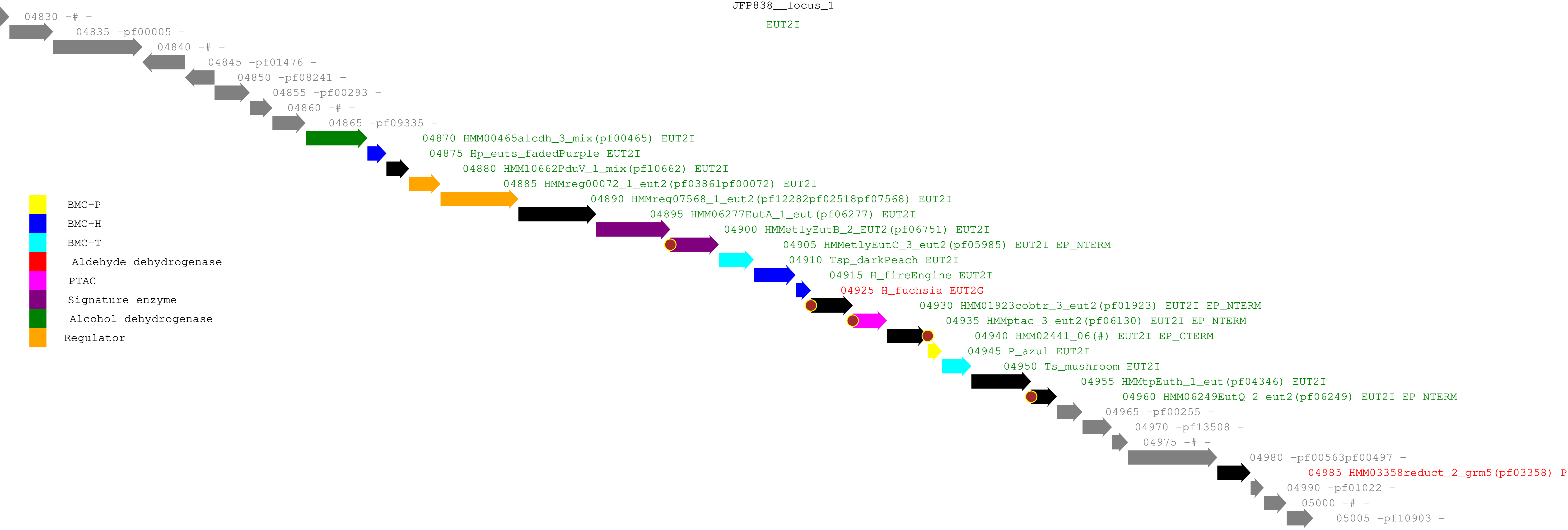

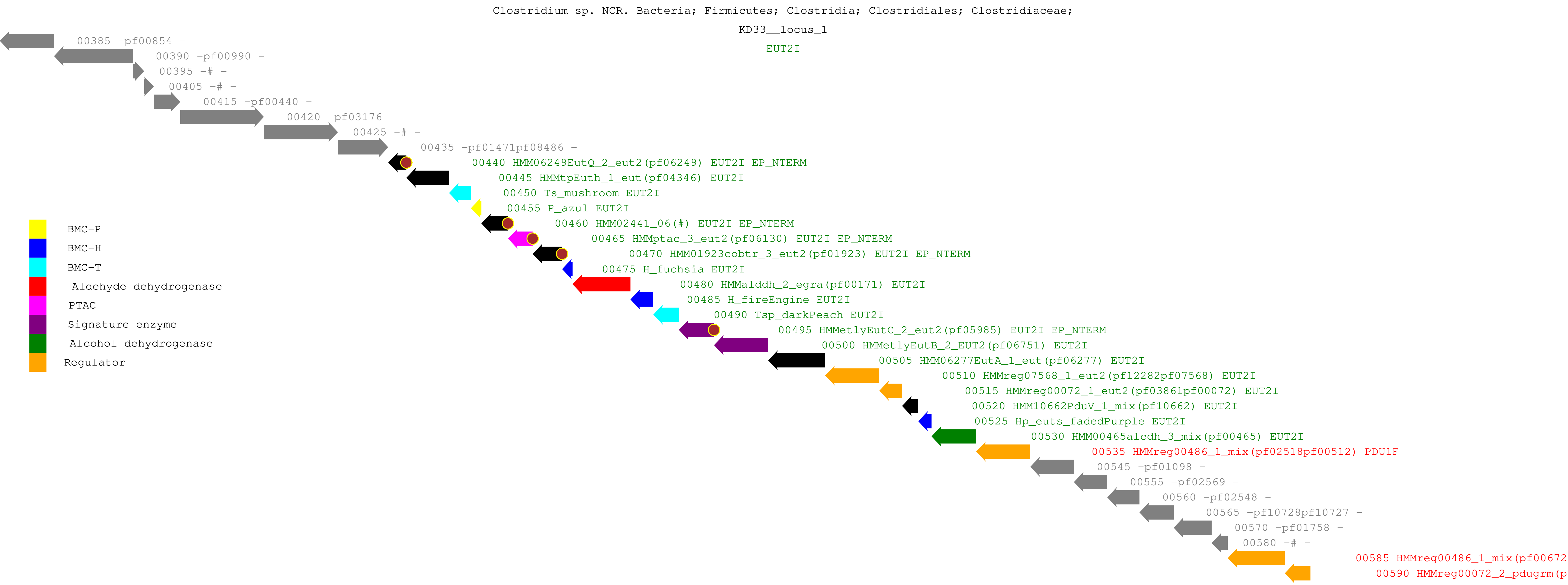
Clostridium sp. NCR. Bacteria; Firmicutes; Clostridia; Clostridiales; Clostridiaceae;
KD33__locus_1
EUT2I
00385 -pf00854 -
00390 -pf00990 -
00395 -# -
00405 -# -
00415 -pf00440 -
00420 -pf03176 -
00425 -# -
00435 -pf01471pf08486 -
00440 HMM06249EutQ_2_eut2(pf06249) EUT2I EP_NTERM
00445 HMMtpEuth_1_eut(pf04346) EUT2I
00450 Ts_mushroom EUT2I
00455 P_azul EUT2I
00460 HMM02441_06(#) EUT2I EP_NTERM
00465 HMMptac_3_eut2(pf06130) EUT2I EP_NTERM
00470 HMM01923cobtr_3_eut2(pf01923) EUT2I EP_NTERM
00475 H_fuchsia EUT2I
00480 HMMalddh_2_egra(pf00171) EUT2I
00485 H_fireEngine EUT2I
00490 Tsp_darkPeach EUT2I
00495 HMMetlyEutC_2_eut2(pf05985) EUT2I EP_NTERM
00500 HMMetlyEutB_2_EUT2(pf06751) EUT2I
00505 HMM06277EutA_1_eut(pf06277) EUT2I
00510 HMMreg07568_1_eut2(pf12282pf07568) EUT2I
00515 HMMreg00072_1_eut2(pf03861pf00072) EUT2I
00520 HMM10662PduV_1_mix(pf10662) EUT2I
00525 Hp_euts_fadedPurple EUT2I
00530 HMM00465alcdh_3_mix(pf00465) EUT2I
00535 HMMreg00486_1_mix(pf02518pf00512) PDU1F
00545 -pf01098 -
00555 -pf02569 -
00560 -pf02548 -
00565 -pf10728pf10727 -
00570 -pf01758 -
00580 -# -
00585 HMMreg00486_1_mix(pf00672
00590 HMMreg00072_2_pdugrm(p
BMC-P
BMC-H
BMC-T
Aldehyde dehydrogenase
PTAC
Signature enzyme
Alcohol dehydrogenase
Regulator

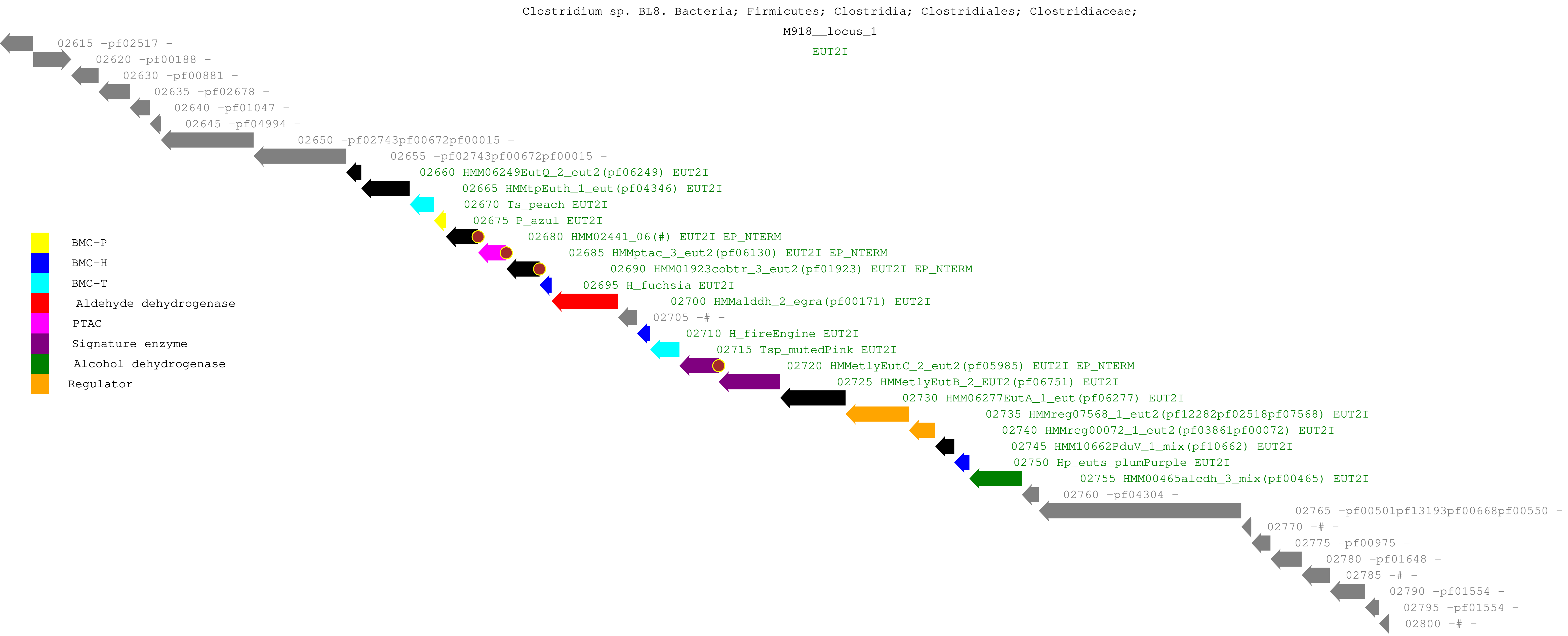

Clostridium sp. BL8. Bacteria; Firmicutes; Clostridia; Clostridiales; Clostridiaceae;
M918__locus_1
EUT2I
02615 -pf02517 -
02620 -pf00188 -
02630 -pf00881 -
02635 -pf02678 -
02640 -pf01047 -
02645 -pf04994 -
02650 -pf02743pf00672pf00015 -
02655 -pf02743pf00672pf00015 -
02660 HMM06249EutQ_2_eut2(pf06249) EUT2I
02665 HMMtpEuth_1_eut(pf04346) EUT2I
02670 Ts_peach EUT2I
02675 P_azul EUT2I
02680 HMM02441_06(#) EUT2I EP_NTERM
02685 HMMptac_3_eut2(pf06130) EUT2I EP_NTERM
02690 HMM01923cobtr_3_eut2(pf01923) EUT2I EP_NTERM
02695 H_fuchsia EUT2I
02700 HMMalddh_2_egra(pf00171) EUT2I
02705 -# -
02710 H_fireEngine EUT2I
02715 Tsp_mutedPink EUT2I
02720 HMMetlyEutC_2_eut2(pf05985) EUT2I EP_NTERM
02725 HMMetlyEutB_2_EUT2(pf06751) EUT2I
02730 HMM06277EutA_1_eut(pf06277) EUT2I
02735 HMMreg07568_1_eut2(pf12282pf02518pf07568) EUT2I
02740 HMMreg00072_1_eut2(pf03861pf00072) EUT2I
02745 HMM10662PduV_1_mix(pf10662) EUT2I
02750 Hp_euts_plumPurple EUT2I
02755 HMM00465alcdh_3_mix(pf00465) EUT2I
02760 -pf04304 -
02765 -pf00501pf13193pf00668pf00550 -
02770 -# -
02775 -pf00975 -
02780 -pf01648 -
02785 -# -
02790 -pf01554 -
02795 -pf01554 -
02800 -# -
BMC-P
BMC-H
BMC-T
Aldehyde dehydrogenase
PTAC
Signature enzyme
Alcohol dehydrogenase
Regulator

Pontibacillus litoralis JSM 072002. Bacteria; Firmicutes; Bacilli; Bacillales; Bacillaceae; Pontibacillus.

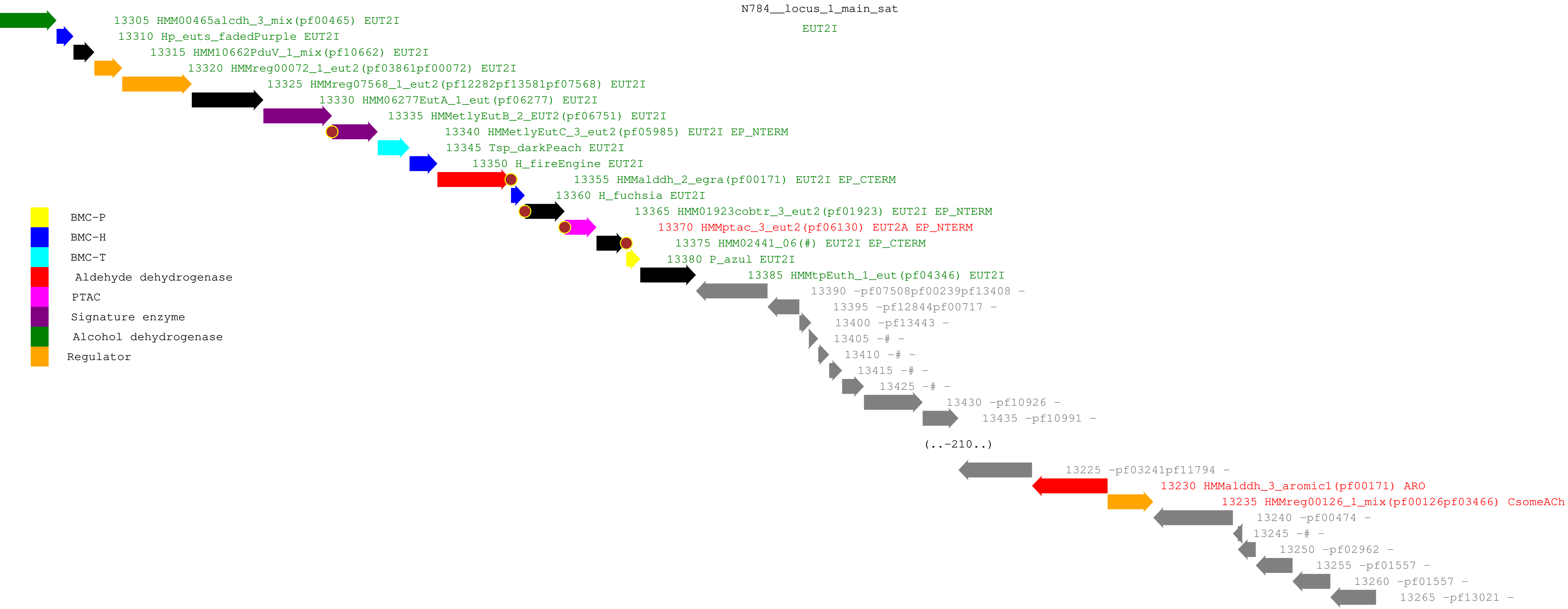

Clostridium perfringens. Bacteria; Firmicutes; Clostridia; Clostridiales; Clostridiaceae;

NCTC10578__locus_1

EUT2I

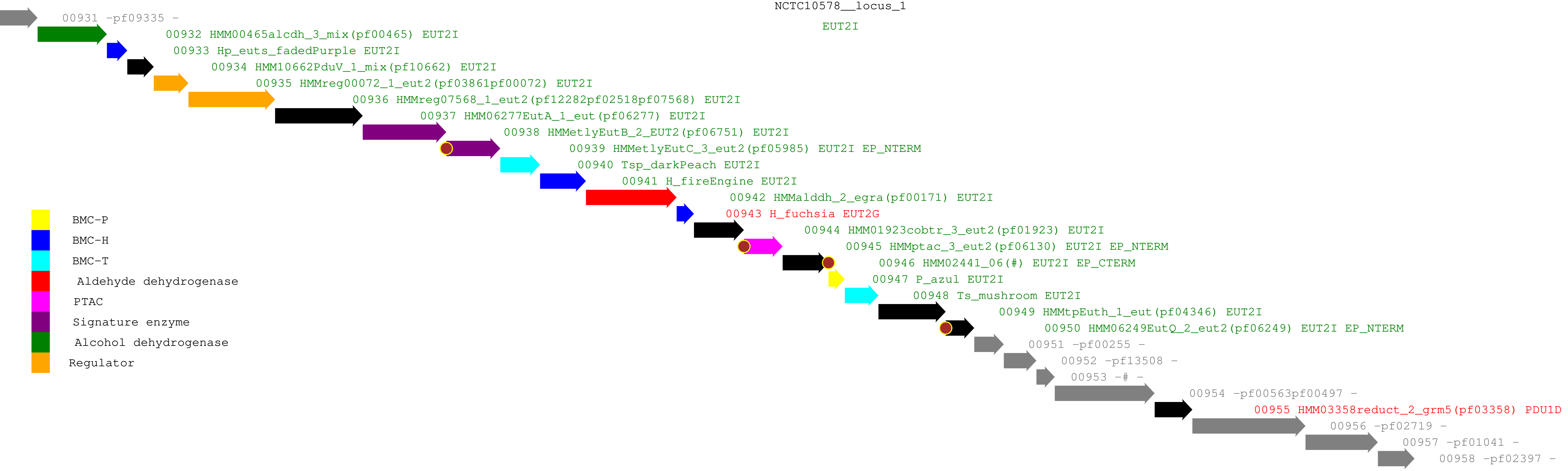

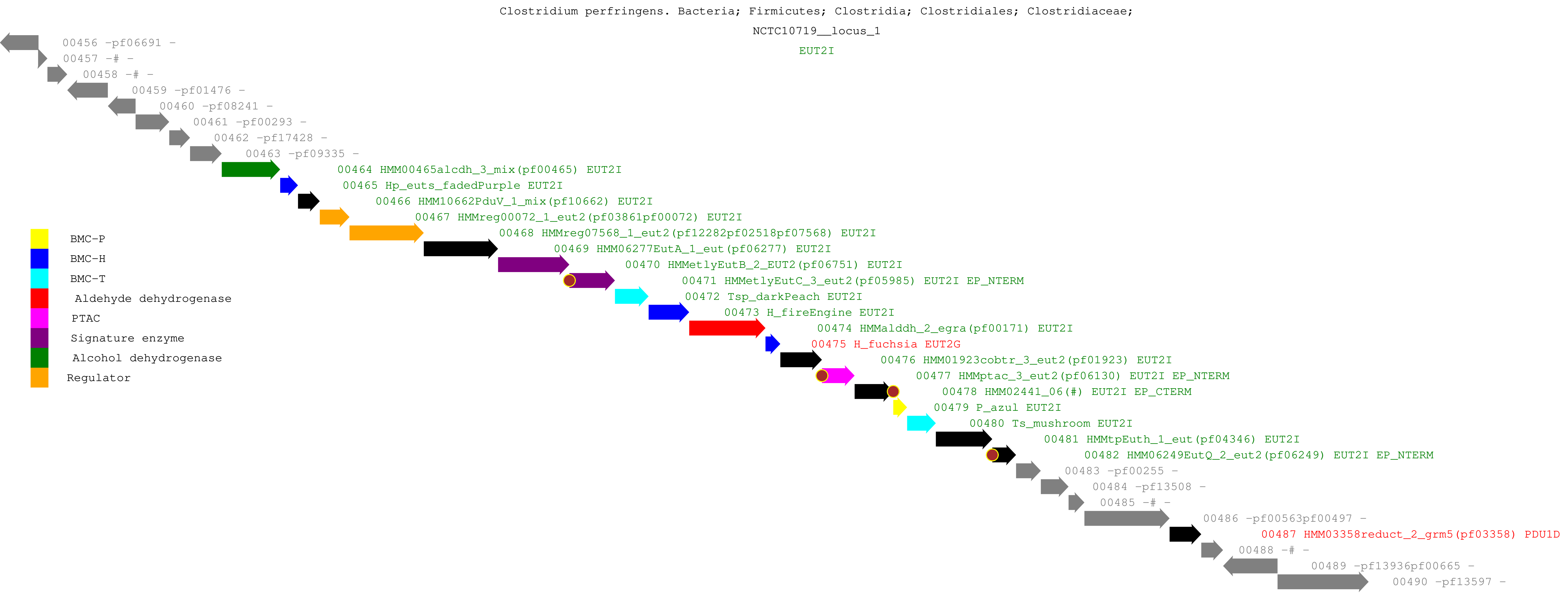

Clostridium perfringens. Bacteria; Firmicutes; Clostridia; Clostridiales; Clostridiaceae;
NCTC10719__locus_1
EUT2I
00456 -pf06691 -
00457 -# -
00458 -# -
00459 -pf01476 -
00460 -pf08241 -
00461 -pf00293 -
00462 -pf17428 -
00463 -pf09335 -
00464 HMM00465alcdh_3_mix(pf00465) EUT2I
00465 Hp_euts_fadedPurple EUT2I
00466 HMM10662PduV_1_mix(pf10662) EUT2I
00467 HMMreg00072_1_eut2(pf03861pf00072) EUT2I
00468 HMMreg07568_1_eut2(pf12282pf02518pf07568) EUT2I
00469 HMM06277EutA_1_eut(pf06277) EUT2I
00470 HMMetlyEutB_2_EUT2(pf06751) EUT2I
00471 HMMetlyEutC_3_eut2(pf05985) EUT2I EP_NTERM
00472 Tsp_darkPeach EUT2I
00473 H_fireEngine EUT2I
00474 HMMaldh_2_egra(pf00171) EUT2I
00475 H_fuchsia EUT2G
00476 HMM01923cobtr_3_eut2(pf01923) EUT2I
00477 HMMptac_3_eut2(pf06130) EUT2I EP_NTERM
00478 HMM02441_06(#) EUT2I EP_CTERM
00479 P_azul EUT2I
00480 Ts_mushroom EUT2I
00481 HMMtpEuth_1_eut(pf04346) EUT2I
00482 HMM06249EutQ_2_eut2(pf06249) EUT2I EP_NTERM
00483 -pf00255 -
00484 -pf13508 -
00485 -# -
00486 -pf00563pf00497 -
00487 HMM03358reduct_2_grm5(pf03358) PDU1D
00488 -# -
00489 -pf13936pf00665 -
00490 -pf13597 -
BMC-P
BMC-H
BMC-T
Aldehyde dehydrogenase
PTAC
Signature enzyme
Alcohol dehydrogenase
Regulator

Paenibacillus thiaminolyticus (Bacillus thiaminolyticus). Bacteria; Firmicutes; Bacilli; Bacillales; Paenibacillaceae;

NCTC11027__locus_2

EUT2I

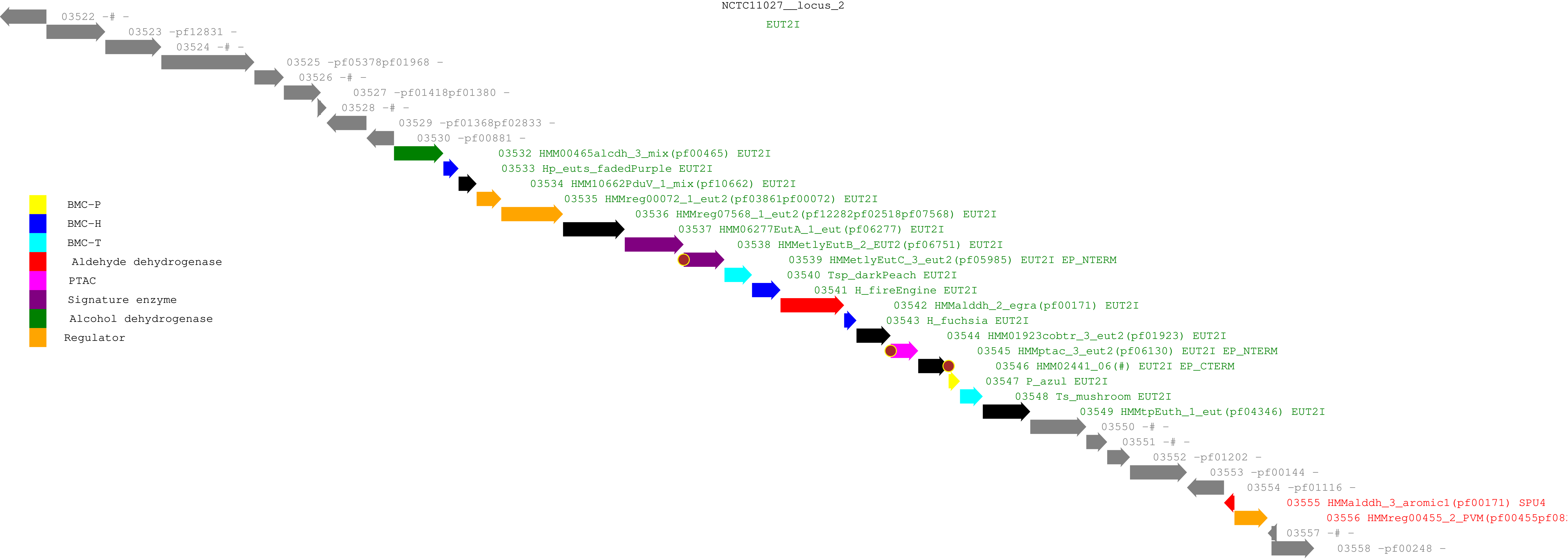

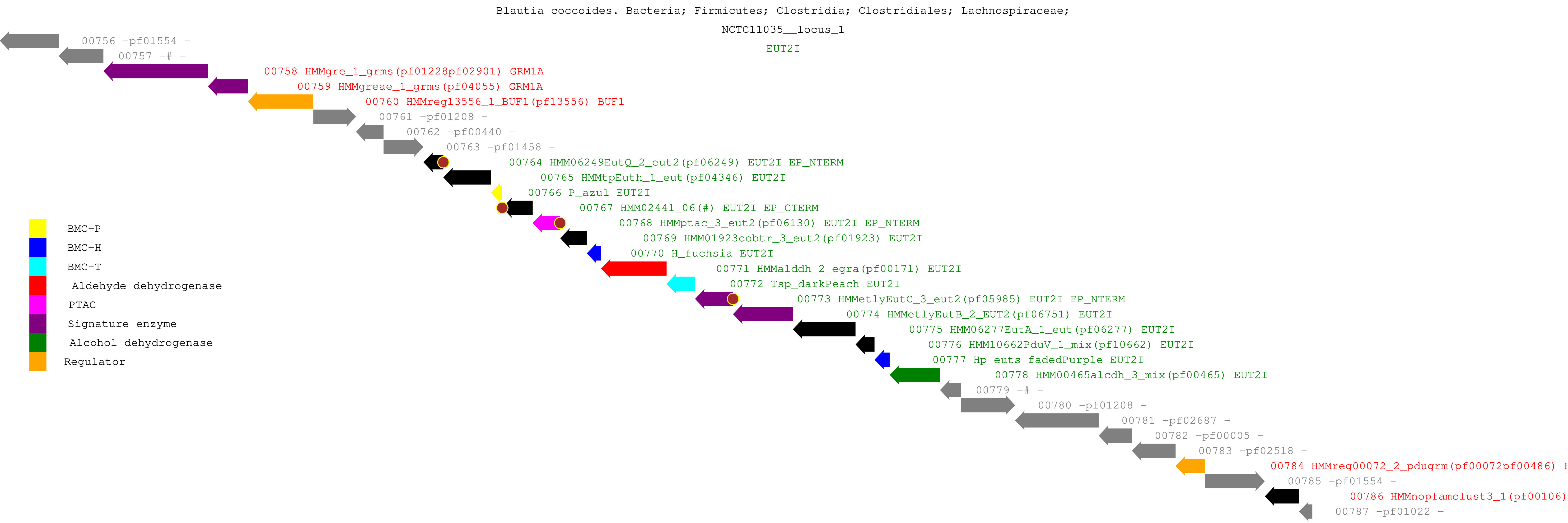
Blautia coccoides. Bacteria; Firmicutes; Clostridia; Clostridiales; Lachnospiraceae;
NCTC11035__locus_1
EUT2I
00756 -pf01554 -
00757 -# -
00758 HMMgre_1_grms(pf01228pf02901) GRM1A
00759 HMMgreae_1_grms(pf04055) GRM1A
00760 HMMreg13556_1_BUF1(pf13556) BUF1
00761 -pf01208 -
00762 -pf00440 -
00763 -pf01458 -
00764 HMM06249EutQ_2_eut2(pf06249) EUT2I EP_NTERM
00765 HMMtpEuth_1_eut(pf04346) EUT2I
00766 P_azul EUT2I
00767 HMM02441_06(#) EUT2I EP_CTERM
00768 HMMptac_3_eut2(pf06130) EUT2I EP_NTERM
00769 HMM01923cobtr_3_eut2(pf01923) EUT2I
00770 H_fuchsia EUT2I
00771 HMMalddh_2_egra(pf00171) EUT2I
00772 Tsp_darkPeach EUT2I
00773 HMMetlyEutC_3_eut2(pf05985) EUT2I EP_NTERM
00774 HMMetlyEutB_2_EUT2(pf06751) EUT2I
00775 HMM06277EutA_1_eut(pf06277) EUT2I
00776 HMM10662PduV_1_mix(pf10662) EUT2I
00777 Hp_euts_fadedPurple EUT2I
00778 HMM00465alcdh_3_mix(pf00465) EUT2I
00779 -# -
00780 -pf01208 -
00781 -pf02687 -
00782 -pf00005 -
00783 -pf02518 -
00784 HMMreg00072_2_pdugrm(pf00072pf00486) I
00785 -pf01554 -
00786 HMMnopfamclust3_1(pf00106)
00787 -pf01022 -
BMC-P
BMC-H
BMC-T
Aldehyde dehydrogenase
PTAC
Signature enzyme
Alcohol dehydrogenase
Regulator

Clostridium perfringens. Bacteria; Firmicutes; Clostridia; Clostridiales; Clostridiaceae;

NCTC3182__locus_1

EUT2I

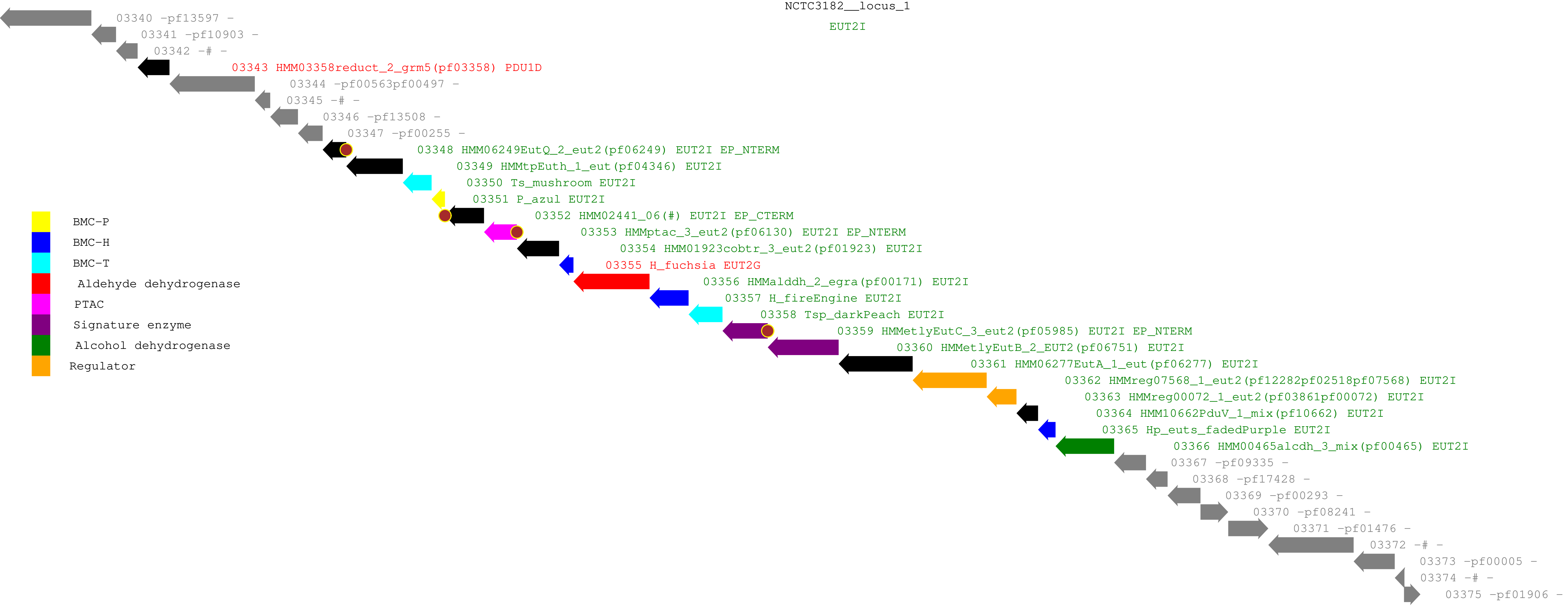

Hathewaya histolytica (Clostridium histolyticum). Bacteria; Firmicutes; Clostridia; Clostridiales; Clostridiaceae; Hathewaya.

NCTC503__locus_1

EUT2I

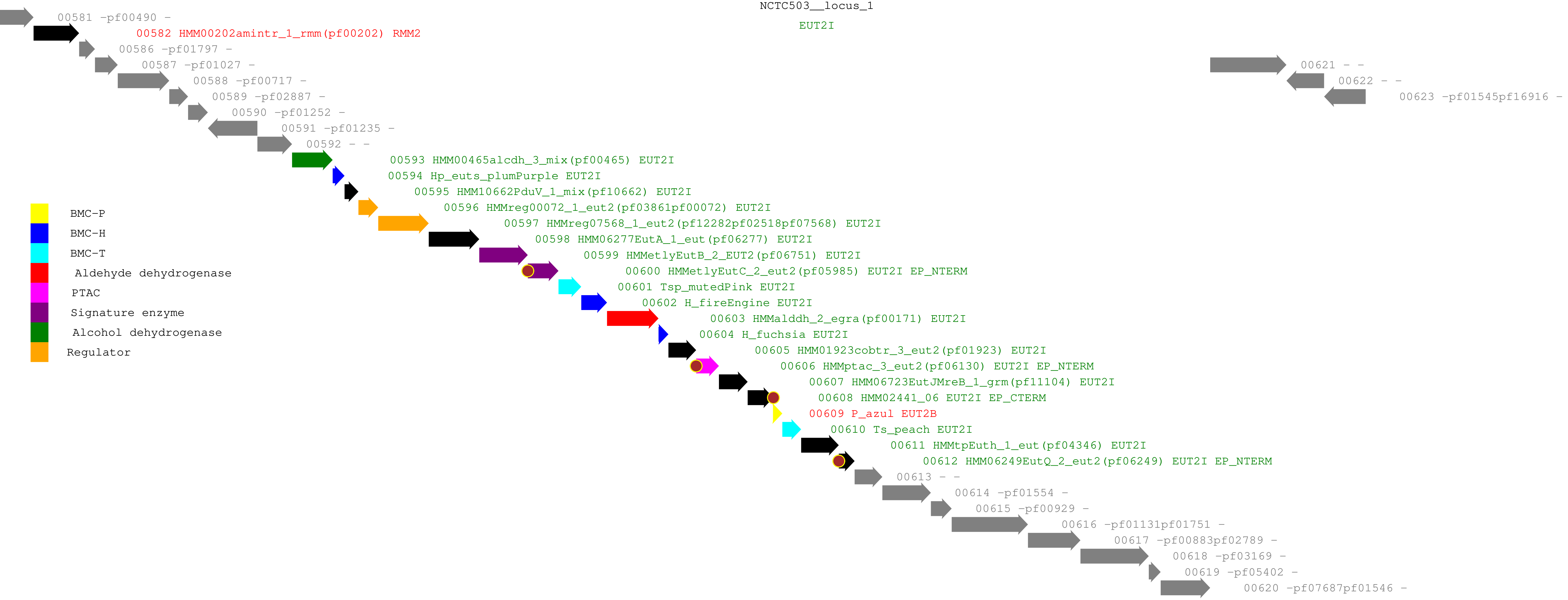

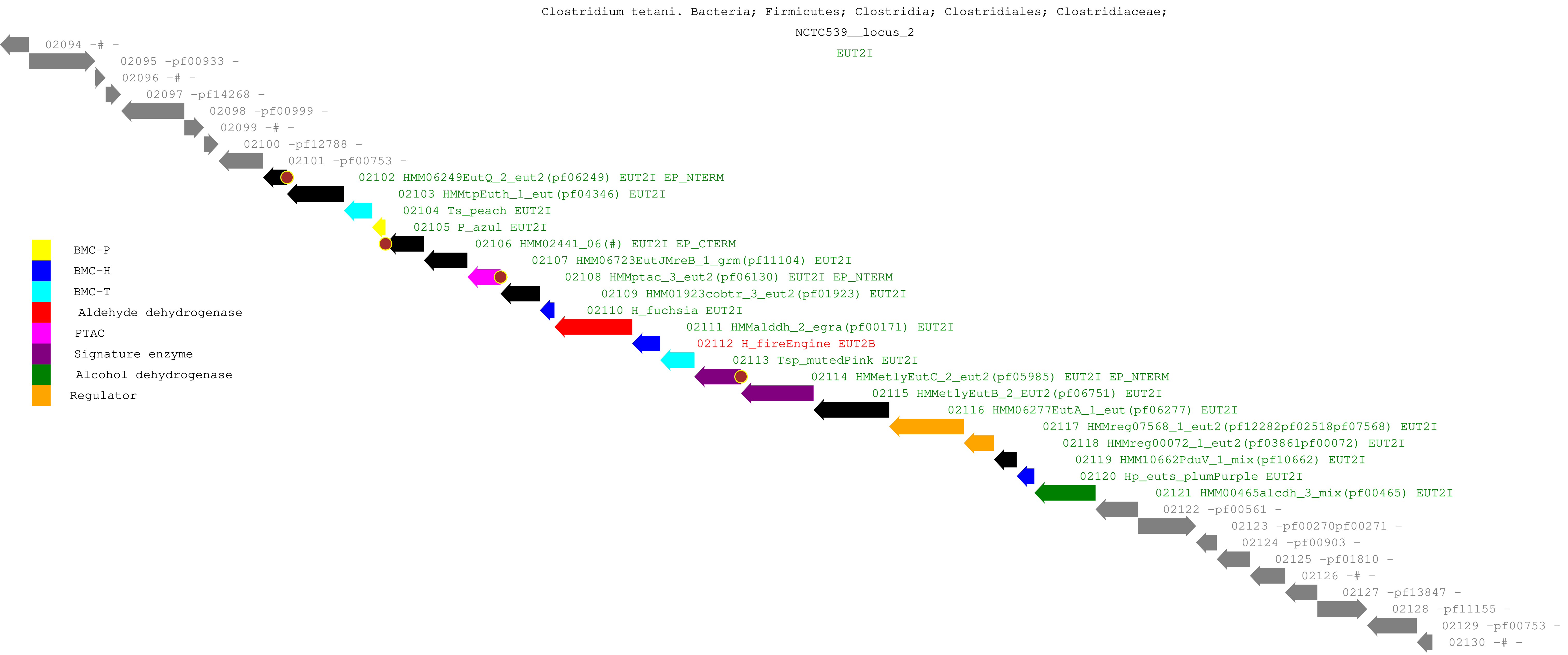
Clostridium tetani. Bacteria; Firmicutes; Clostridia; Clostridiales; Clostridiaceae;
NCTC539__locus_2
EUT2I
02094 -# -
02095 -pf00933 -
02096 -# -
02097 -pf14268 -
02098 -pf00999 -
02099 -# -
02100 -pf12788 -
02101 -pf00753 -
02102 HMM06249EutQ_2_eut2(pf06249) EUT2I EP_NTERM
02103 HMMtpEuth_1_eut(pf04346) EUT2I
02104 Ts_peach EUT2I
02105 P_azul EUT2I
02106 HMM02441_06(#) EUT2I EP_CTERM
02107 HMM06723EutJMreB_1_grm(pf11104) EUT2I
02108 HMMptac_3_eut2(pf06130) EUT2I EP_NTERM
02109 HMM01923cobtr_3_eut2(pf01923) EUT2I
02110 H_fuchsia EUT2I
02111 HMMalddh_2_egra(pf00171) EUT2I
02112 H_fireEngine EUT2B
02113 Tsp_mutedPink EUT2I
02114 HMMetlyEutC_2_eut2(pf05985) EUT2I EP_NTERM
02115 HMMetlyEutB_2_EUT2(pf06751) EUT2I
02116 HMM06277EutA_1_eut(pf06277) EUT2I
02117 HMMreg07568_1_eut2(pf12282pf02518pf07568) EUT2I
02118 HMMreg00072_1_eut2(pf03861pf00072) EUT2I
02119 HMM10662PduV_1_mix(pf10662) EUT2I
02120 Hp_euts_plumPurple EUT2I
02121 HMM00465alcdh_3_mix(pf00465) EUT2I
02122 -pf00561 -
02123 -pf00270pf00271 -
02124 -pf00903 -
02125 -pf01810 -
02126 -# -
02127 -pf13847 -
02128 -pf11155 -
02129 -pf00753 -
02130 -# -
BMC-P
BMC-H
BMC-T
Aldehyde dehydrogenase
PTAC
Signature enzyme
Alcohol dehydrogenase
Regulator

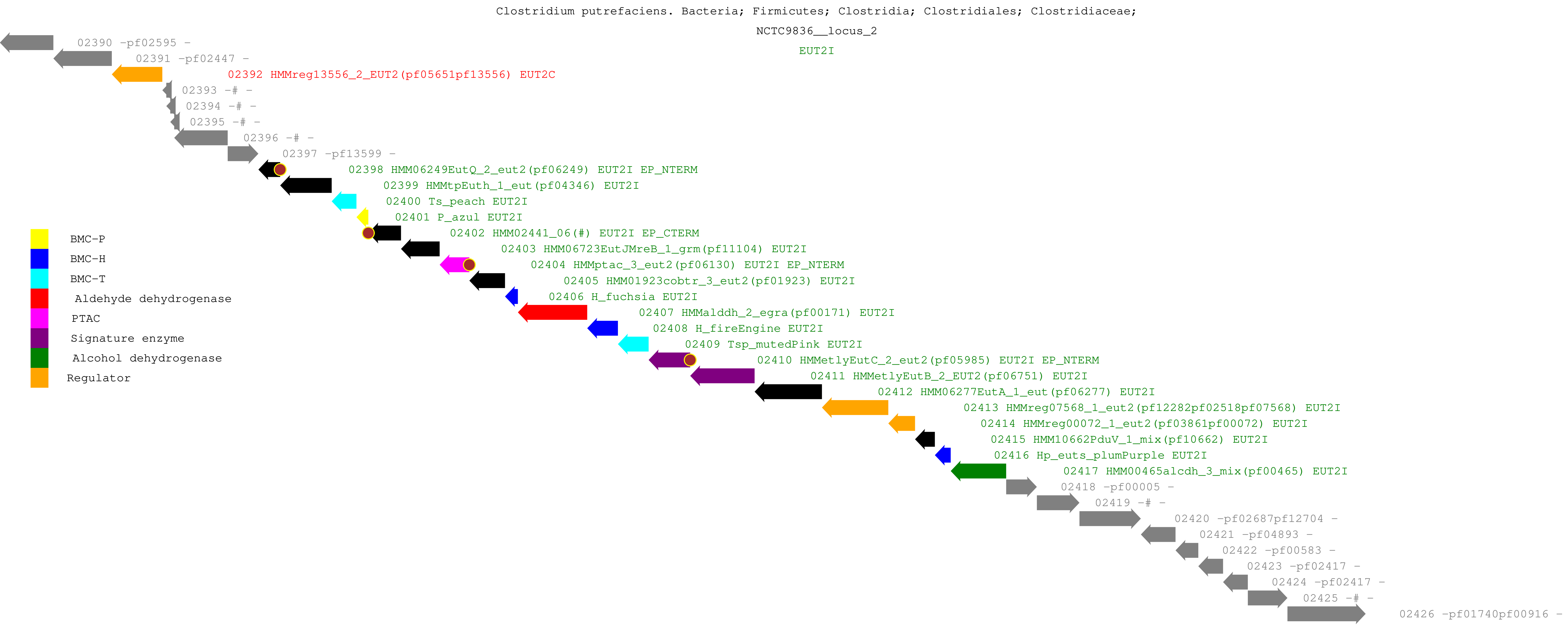
Clostridium putrefaciens. Bacteria; Firmicutes; Clostridia; Clostridiales; Clostridiaceae;
NCTC9836__locus_2
EUT2I
02390 -pf02595 -
02391 -pf02447 -
02392 HMMreg13556_2_EUT2(pf05651pf13556) EUT2C
02393 -# -
02394 -# -
02395 -# -
02396 -# -
02397 -pf13599 -
02398 HMM06249EutQ_2_eut2(pf06249) EUT2I EP_NTERM
02399 HMMtpEuth_1_eut(pf04346) EUT2I
02400 Ts_peach EUT2I
02401 P_azul EUT2I
02402 HMM02441_06(#) EUT2I EP_CTERM
02403 HMM06723EutJMreB_1_grm(pf11104) EUT2I
02404 HMMptac_3_eut2(pf06130) EUT2I EP_NTERM
02405 HMM01923cobtr_3_eut2(pf01923) EUT2I
02406 H_fuchsia EUT2I
02407 HMMalddh_2_egra(pf00171) EUT2I
02408 H_fireEngine EUT2I
02409 Tsp_mutedPink EUT2I
02410 HMMetlyEutC_2_eut2(pf05985) EUT2I EP_NTERM
02411 HMMetlyEutB_2_EUT2(pf06751) EUT2I
02412 HMM06277EutA_1_eut(pf06277) EUT2I
02413 HMMreg07568_1_eut2(pf12282pf02518pf07568) EUT2I
02414 HMMreg00072_1_eut2(pf03861pf00072) EUT2I
02415 HMM10662PduV_1_mix(pf10662) EUT2I
02416 Hp_euts_plumPurple EUT2I
02417 HMM00465alcdh_3_mix(pf00465) EUT2I
02418 -pf00005 -
02419 -# -
02420 -pf02687pf12704 -
02421 -pf04893 -
02422 -pf00583 -
02423 -pf02417 -
02424 -pf02417 -
02425 -# -
02426 -pf01740pf00916 -
BMC-P
BMC-H
BMC-T
Aldehyde dehydrogenase
PTAC
Signature enzyme
Alcohol dehydrogenase
Regulator

Paenibacillus alvei A6-6i-x. Bacteria; Firmicutes; Bacilli; Bacillales; Paenibacillaceae;

PAAL66ix__locus_1

EUT2I

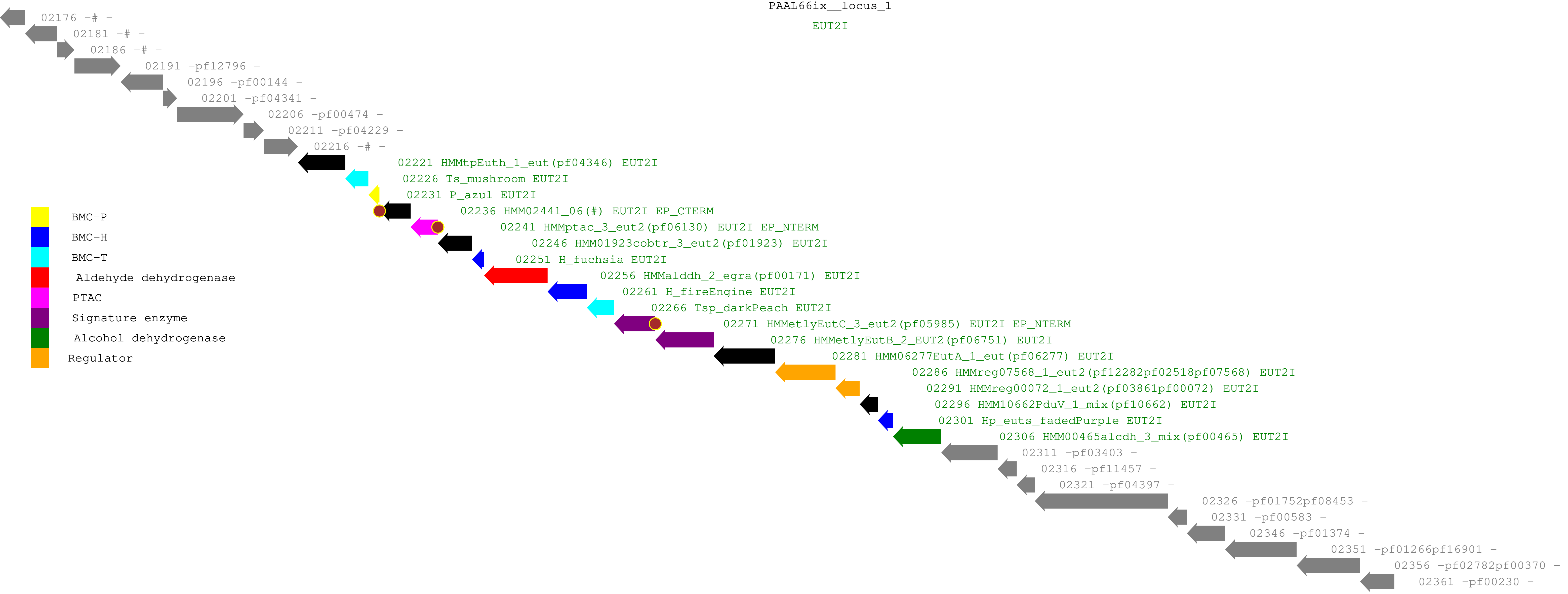

Paenibacillus alvei TS-15. Bacteria; Firmicutes; Bacilli; Bacillales; Paenibacillaceae;

PAALTS15__locus_1

EUT2I

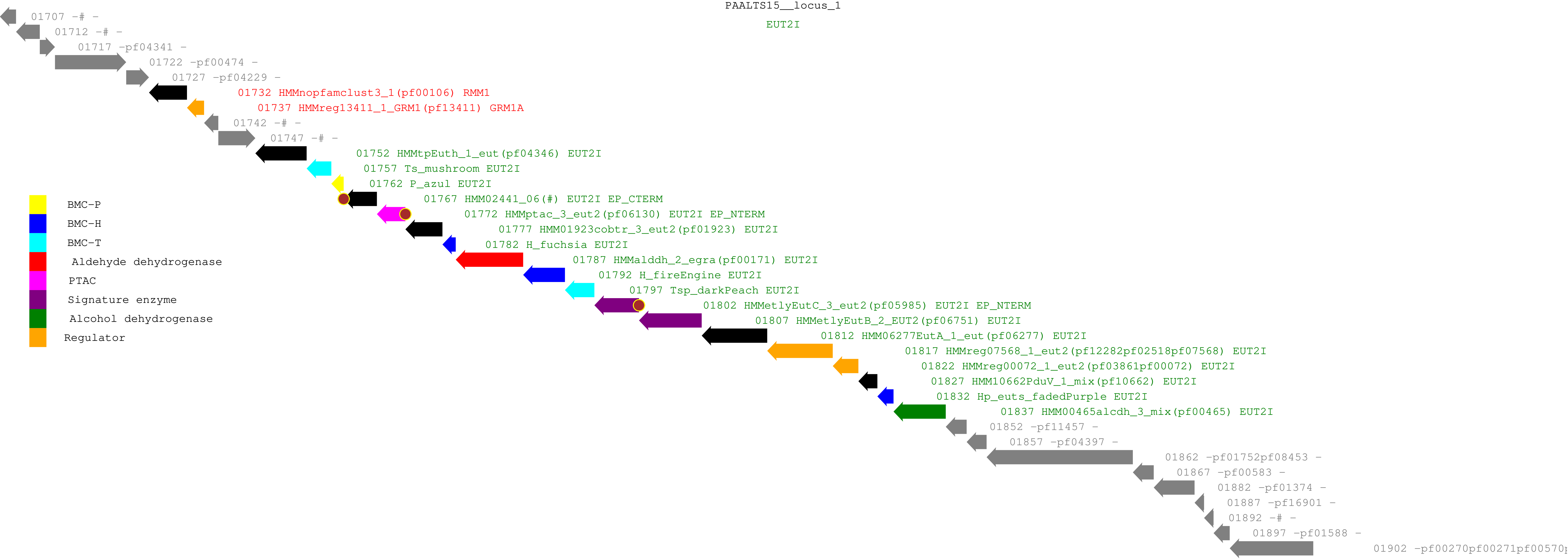

Paenibacillus alvei DSM 29. Bacteria; Firmicutes; Bacilli; Bacillales; Paenibacillaceae;

PAV_6c_locus_1

EUT2I

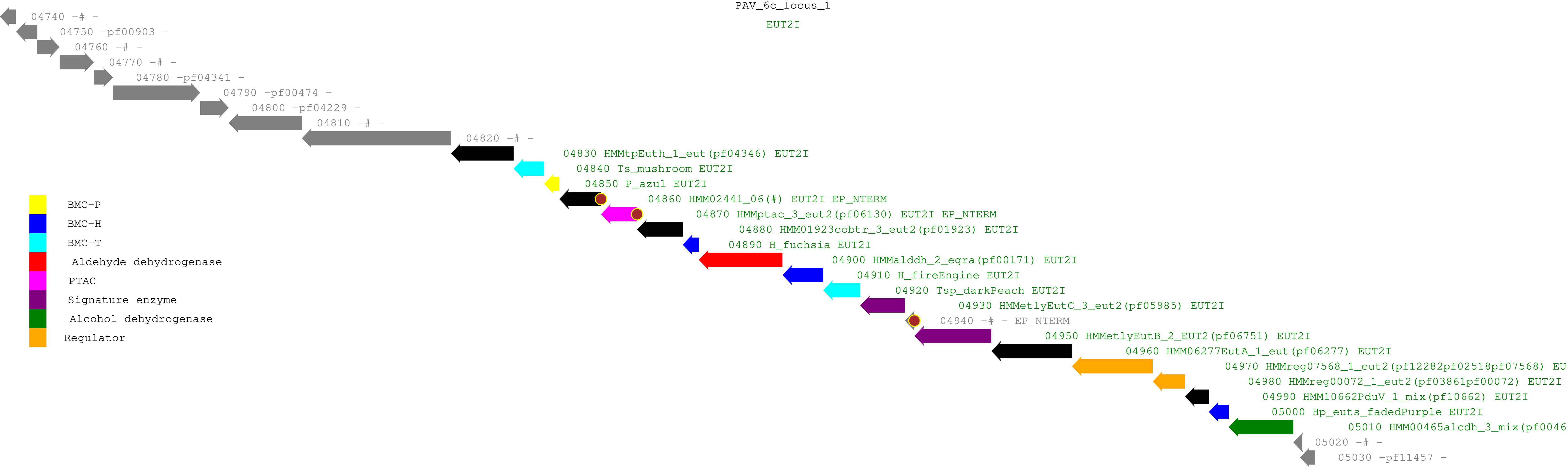

Paenibacillus alvei (Bacillus alvei). Bacteria; Firmicutes; Bacilli; Bacillales; Paenibacillaceae;

PBLR__locus_1

EUT2I

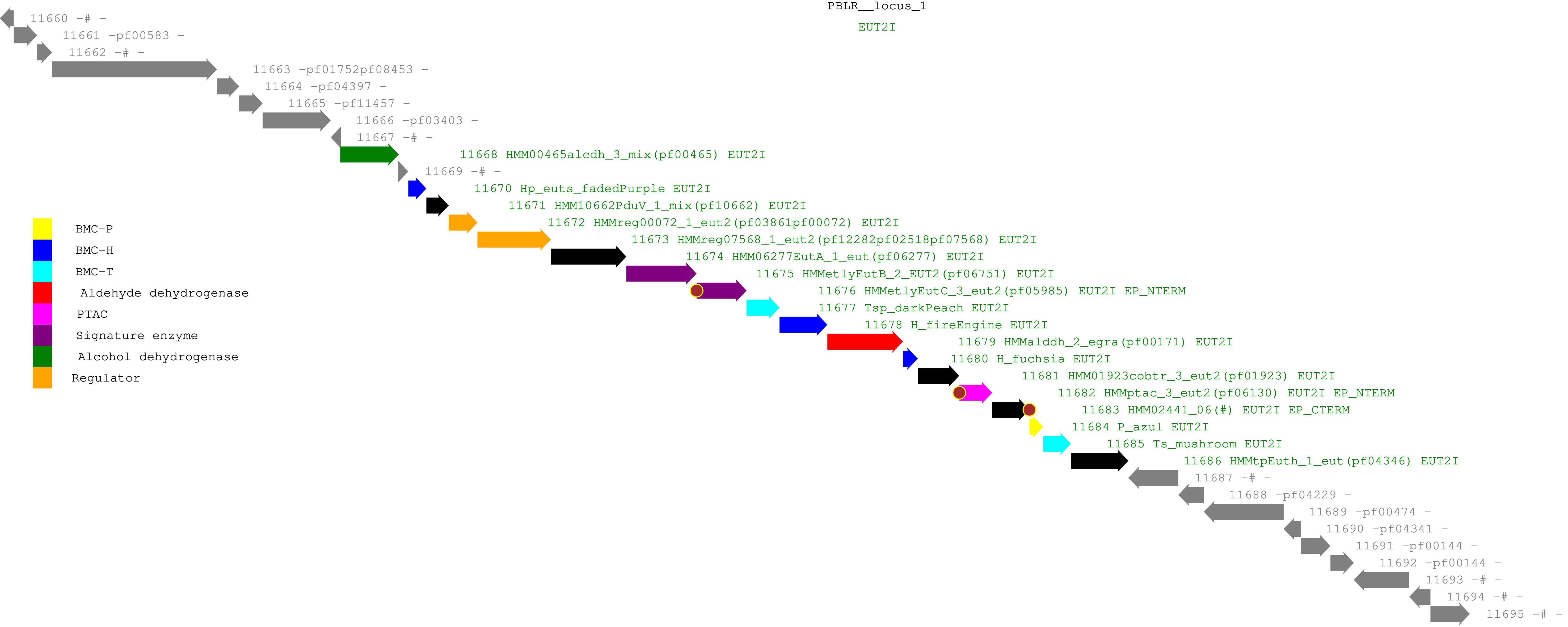

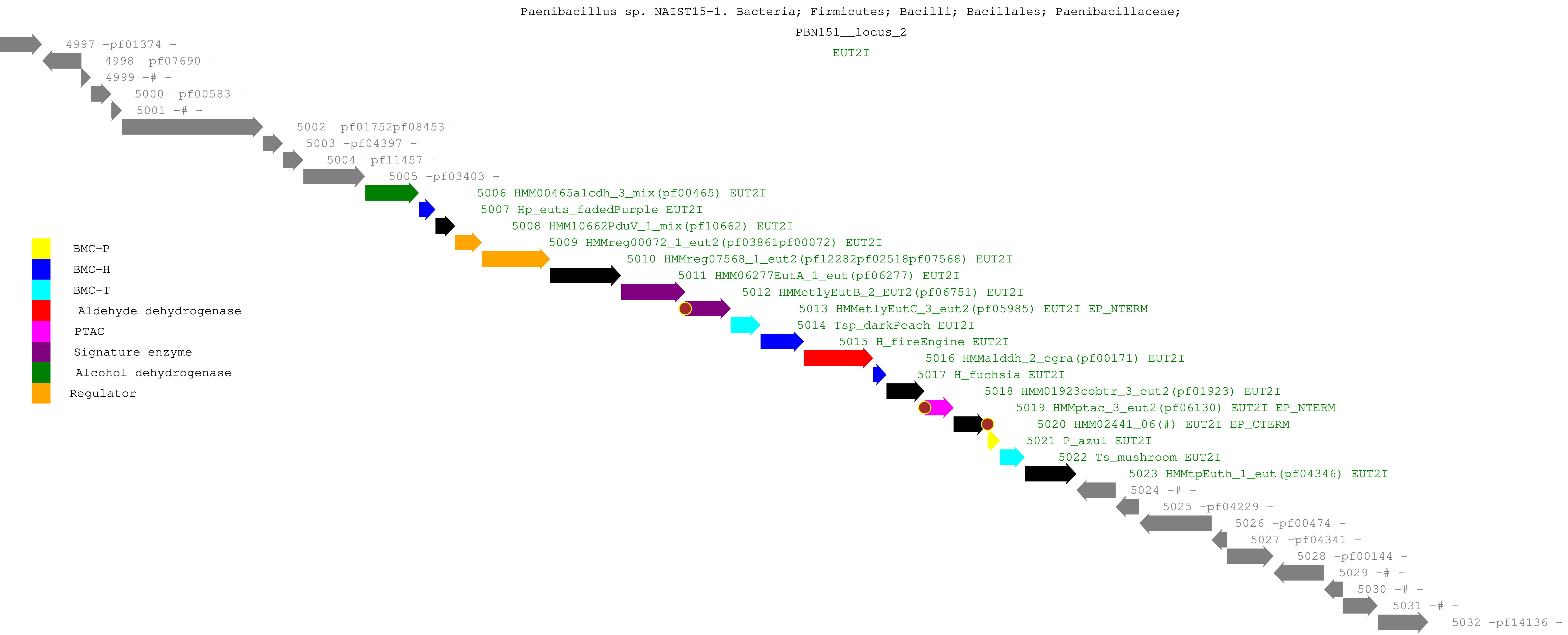

Paenibacillus sp. NAIST15-1. Bacteria; Firmicutes; Bacilli; Bacillales; Paenibacillaceae;
PBN151__locus_2
EUT2I
4997 -pf01374 -
4998 -pf07690 -
4999 -# -
5000 -pf00583 -
5001 -# -
5002 -pf01752pf08453 -
5003 -pf04397 -
5004 -pf11457 -
5005 -pf03403 -
5006 HMM00465alcdh_3_mix(pf00465) EUT2I
5007 Hp_euts_fadedPurple EUT2I
5008 HMM10662PduV_1_mix(pf10662) EUT2I
5009 HMMreg00072_1_eut2(pf03861pf00072) EUT2I
5010 HMMreg07568_1_eut2(pf12282pf02518pf07568) EUT2I
5011 HMM06277EutA_1_eut(pf06277) EUT2I
5012 HMMetlyEutB_2_EUT2(pf06751) EUT2I
5013 HMMetlyEutC_3_eut2(pf05985) EUT2I EP_NTERM
5014 Tsp_darkPeach EUT2I
5015 H_fireEngine EUT2I
5016 HMMalddh_2_egra(pf00171) EUT2I
5017 H_fuchsia EUT2I
5018 HMM01923cobtr_3_eut2(pf01923) EUT2I
5019 HMMptac_3_eut2(pf06130) EUT2I EP_NTERM
5020 HMM02441_06(#) EUT2I EP_CTERM
5021 P_azul EUT2I
5022 Ts_mushroom EUT2I
5023 HMMtpEuth_1_eut(pf04346) EUT2I
5024 -# -
5025 -pf04229 -
5026 -pf00474 -
5027 -pf04341 -
5028 -pf00144 -
5029 -# -
5030 -# -
5031 -# -
5032 -pf14136 -
BMC-P
BMC-H
BMC-T
Aldehyde dehydrogenase
PTAC
Signature enzyme
Alcohol dehydrogenase
Regulator

Paenibacillus dendritiformis C454. Bacteria; Firmicutes; Bacilli; Bacillales; Paenibacillaceae;

PDENDC454__locus_1

EUT2I

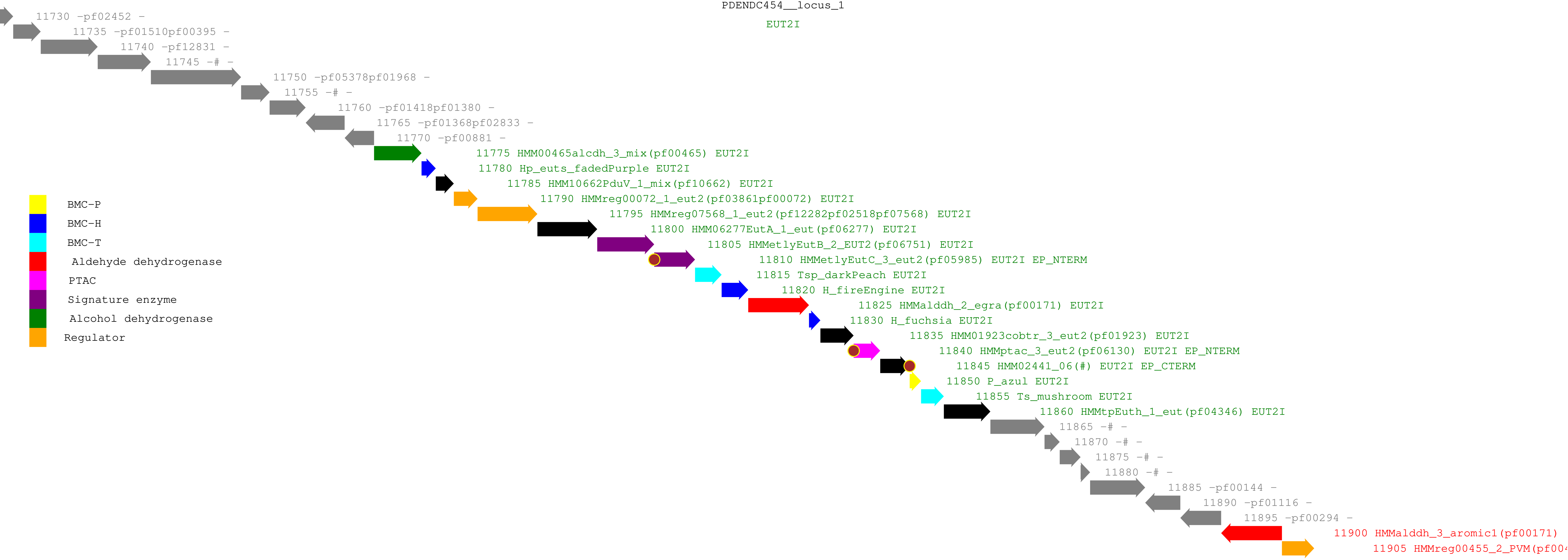

Paenibacillus glacialis. Bacteria; Firmicutes; Bacilli; Bacillales; Paenibacillaceae;

PGLA__locus_1

EUT2I

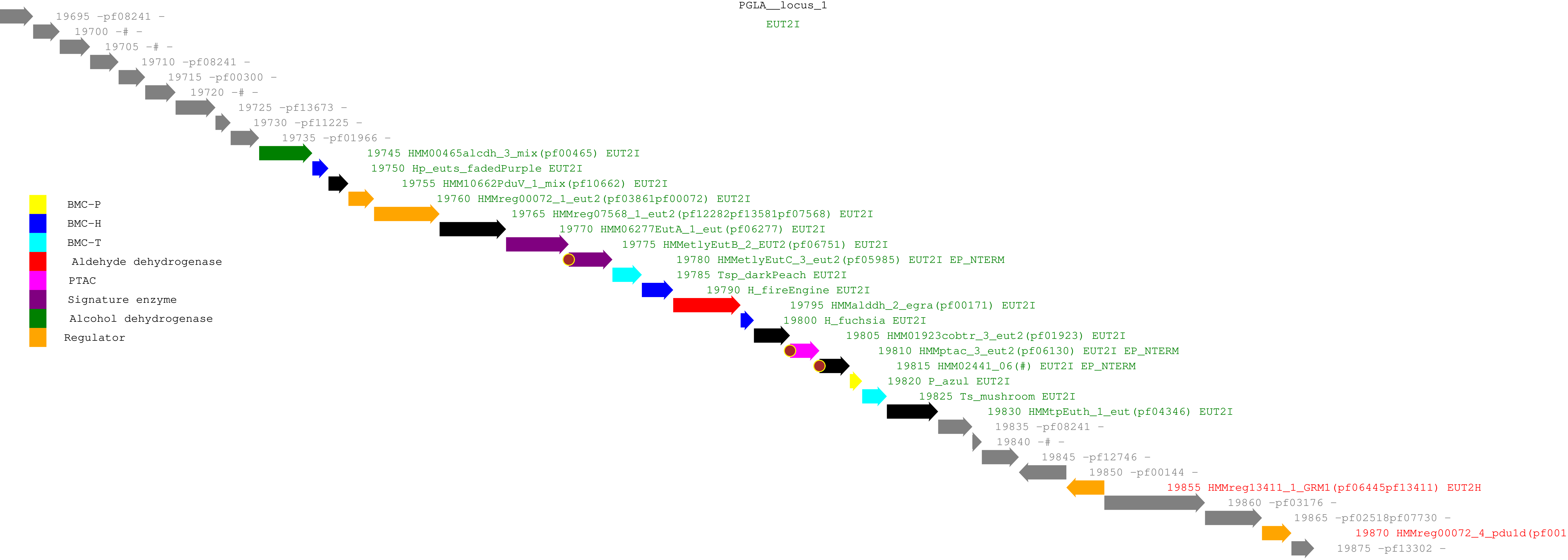

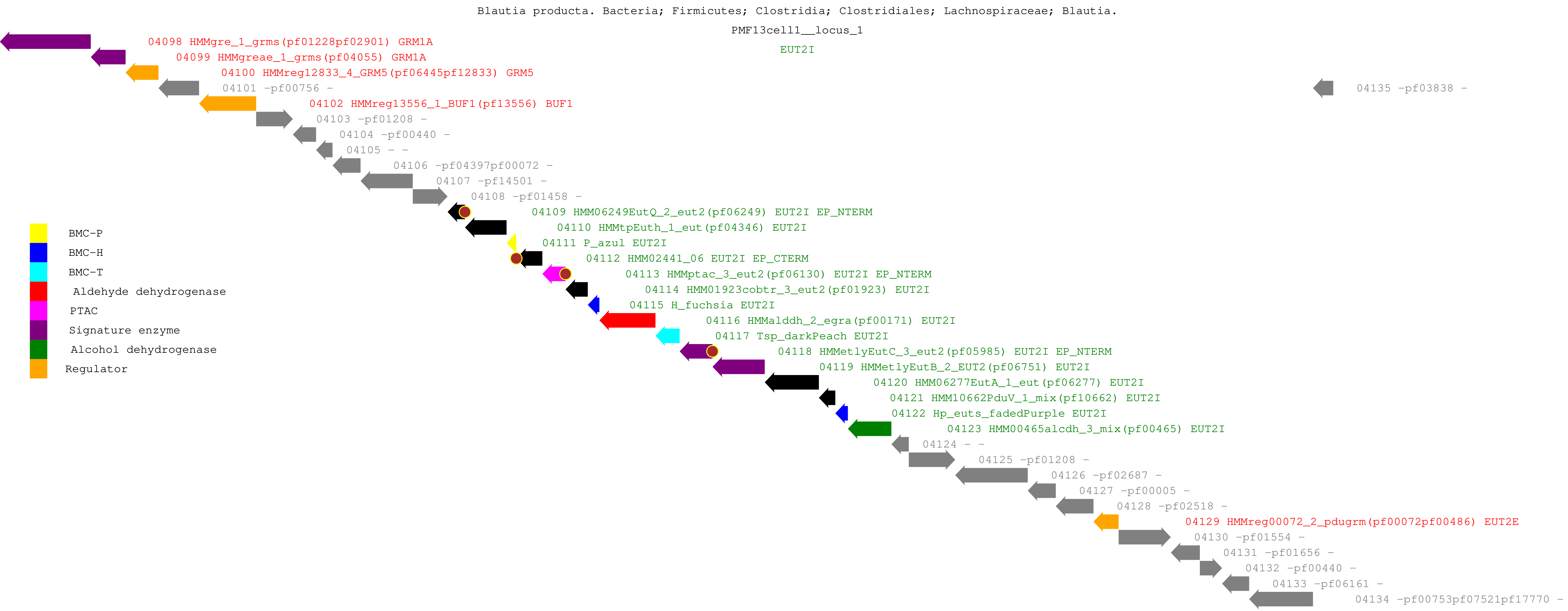
Blautia producta. Bacteria; Firmicutes; Clostridia; Clostridiales; Lachnospiraceae; Blautia.
PMF13cell1__locus_1
EUT2I
04098 HMMgre_1_grms(pf01228pf02901) GRM1A
04099 HMMgreae_1_grms(pf04055) GRM1A
04100 HMMreg12833_4_GRM5(pf06445pf12833) GRM5
04101 -pf00756 -
04102 HMMreg13556_1_BUF1(pf13556) BUF1
04103 -pf01208 -
04104 -pf00440 -
04105 - -
04106 -pf04397pf00072 -
04107 -pf14501 -
04108 -pf01458 -
04109 HMM06249EutQ_2_eut2(pf06249) EUT2I EP_NTERM
04110 HMMtpEuth_1_eut(pf04346) EUT2I
04111 P_azul EUT2I
04112 HMM02441_06 EUT2I EP_CTERM
04113 HMMptac_3_eut2(pf06130) EUT2I EP_NTERM
04114 HMM01923cobtr_3_eut2(pf01923) EUT2I
04115 H_fuchsia EUT2I
04116 HMMalddh_2_egra(pf00171) EUT2I
04117 Tsp_darkPeach EUT2I
04118 HMMetlyEutC_3_eut2(pf05985) EUT2I EP_NTERM
04119 HMMetlyEutB_2_EUT2(pf06751) EUT2I
04120 HMM06277EutA_1_eut(pf06277) EUT2I
04121 HMM10662PduV_1_mix(pf10662) EUT2I
04122 Hp_euts_fadedPurple EUT2I
04123 HMM00465alcdh_3_mix(pf00465) EUT2I
04124 - -
04125 -pf01208 -
04126 -pf02687 -
04127 -pf00005 -
04128 -pf02518 -
04129 HMMreg00072_2_pdugrm(pf00072pf00486) EUT2E
04130 -pf01554 -
04131 -pf01656 -
04132 -pf00440 -
04133 -pf06161 -
04134 -pf00753pf07521pf17770 -
04135 -pf03838 -
BMC-P
BMC-H
BMC-T
Aldehyde dehydrogenase
PTAC
Signature enzyme
Alcohol dehydrogenase
Regulator

Clostridioides difficile CD160. Bacteria; Firmicutes; Clostridia; Clostridiales;

QEW__locus_1

EUT2I

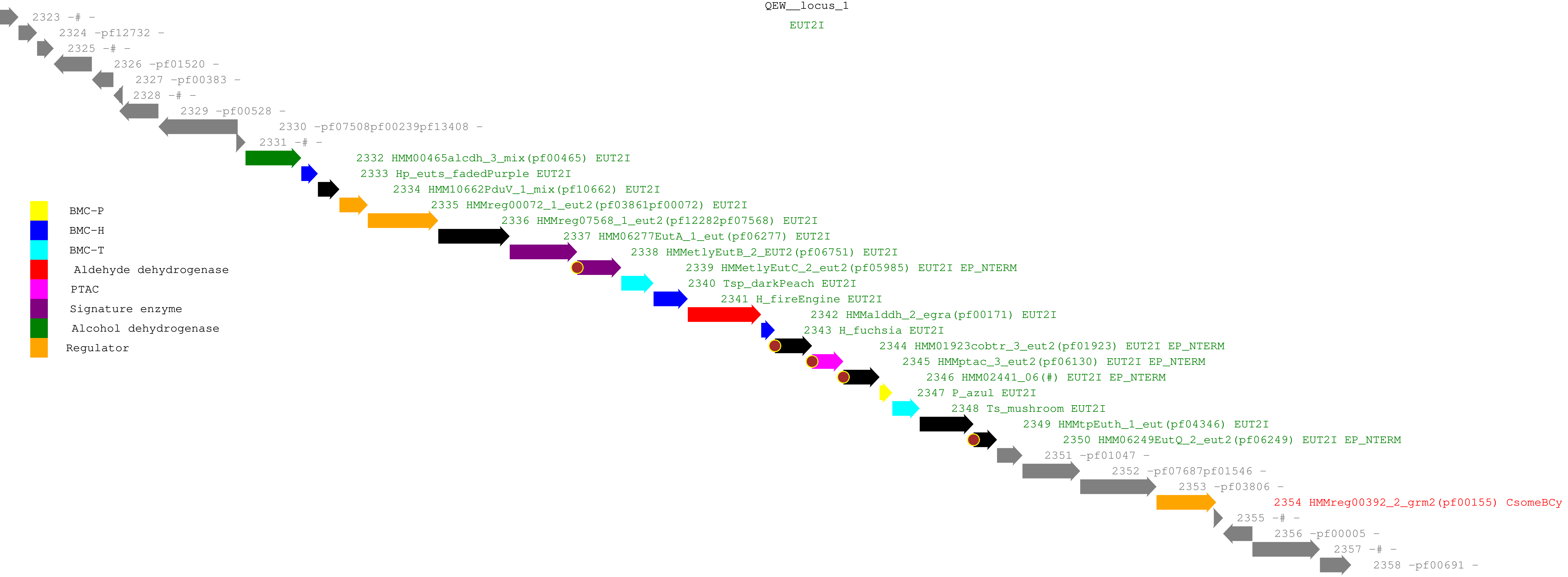

Clostridioides difficile F501. Bacteria; Firmicutes; Clostridia; Clostridiales;

QOE__locus_1

EUT2I

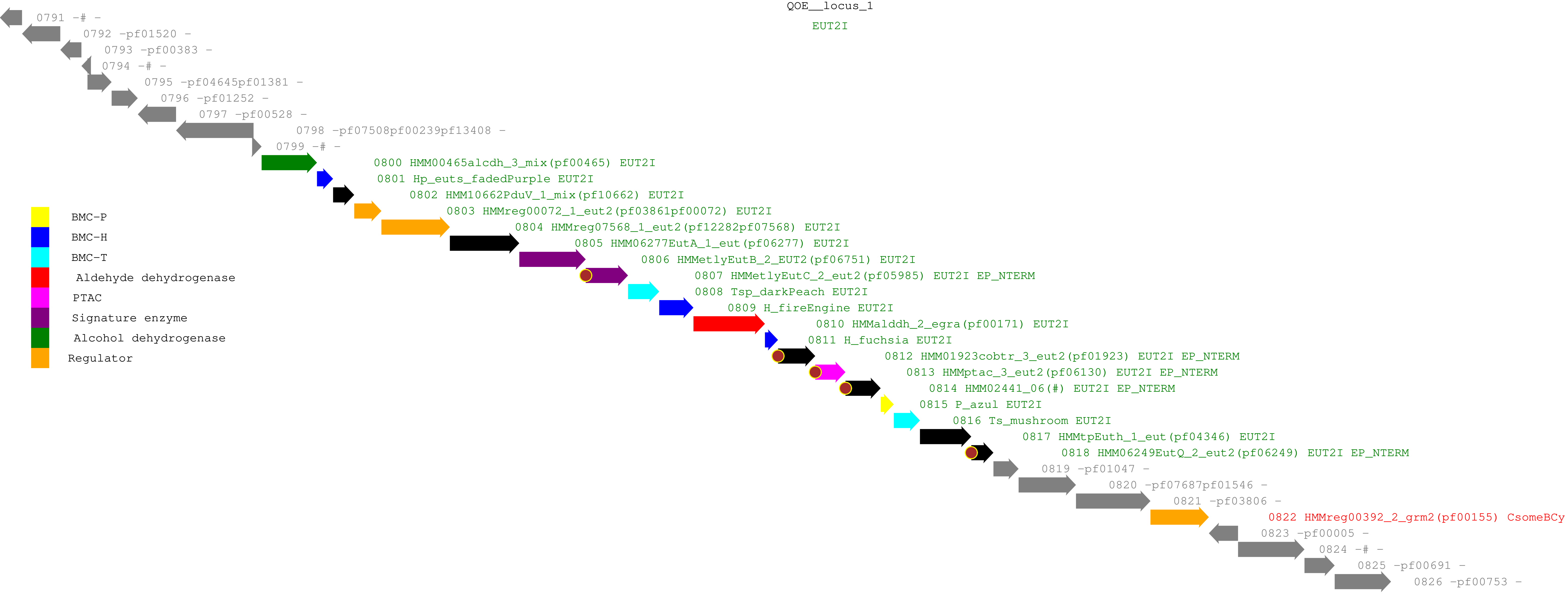

Clostridioides difficile Y384. Bacteria; Firmicutes; Clostridia; Clostridiales;

QQG__locus_1

EUT2I

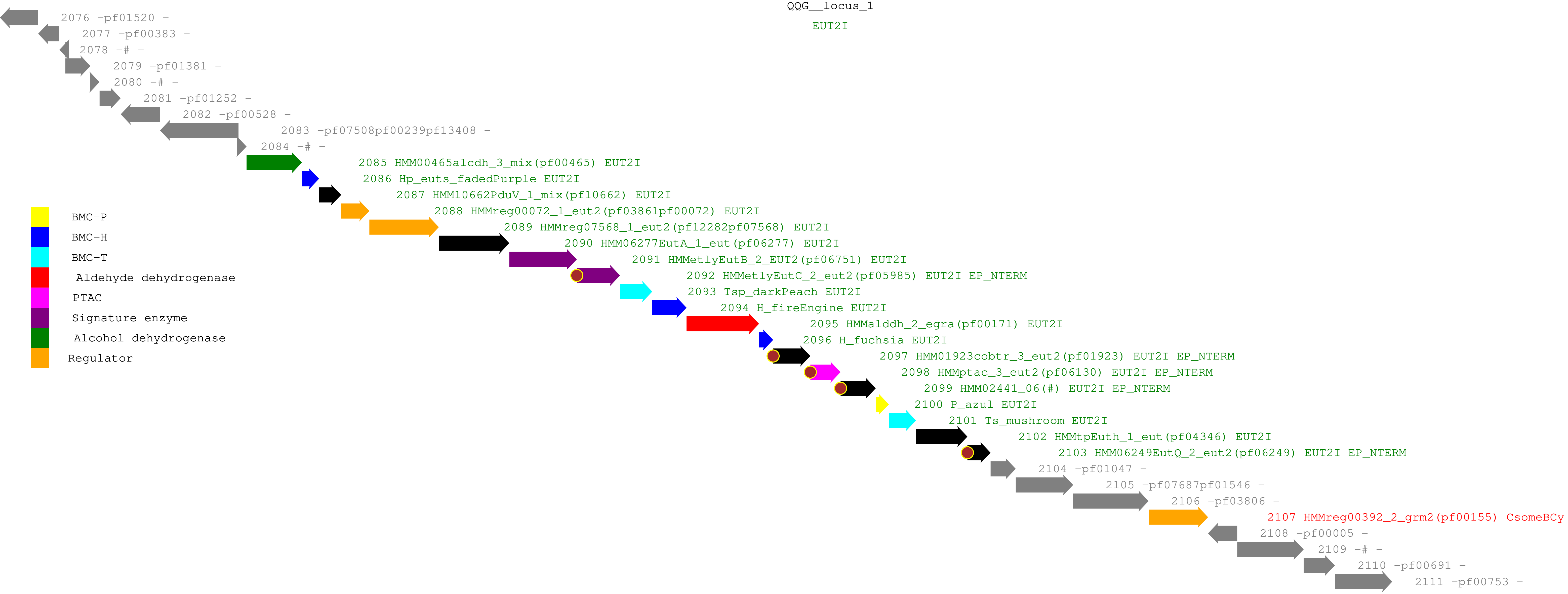

Robinsoniella sp. RHS. Bacteria; Firmicutes; Clostridia; Clostridiales; Lachnospiraceae;

RHS__locus_2

EUT2I

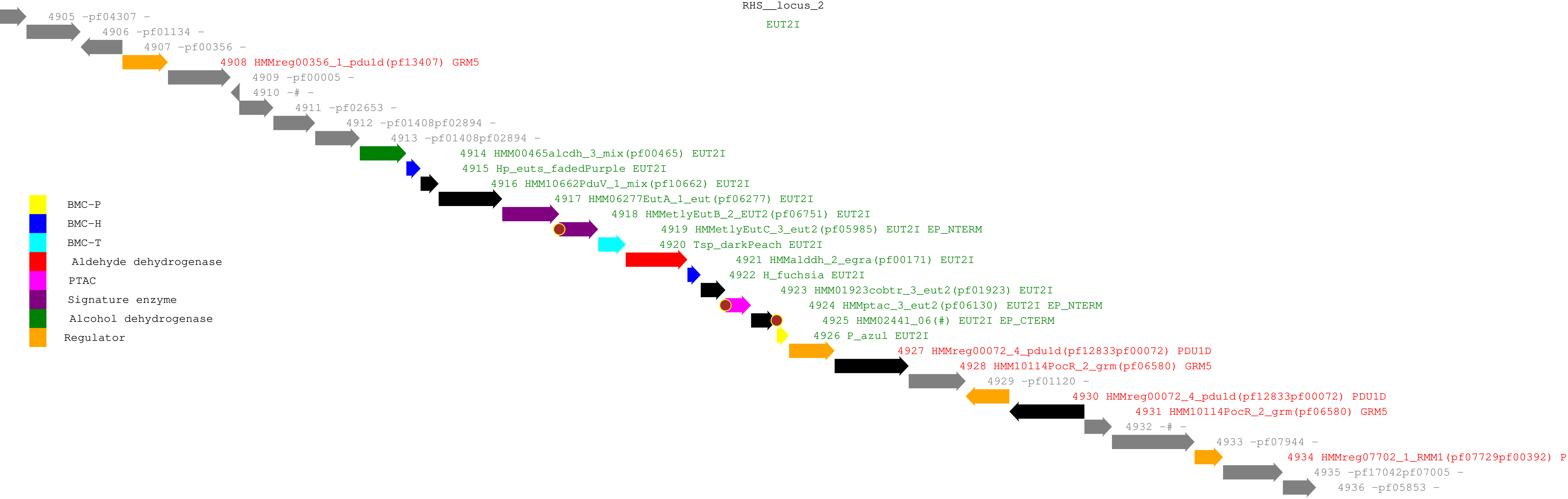

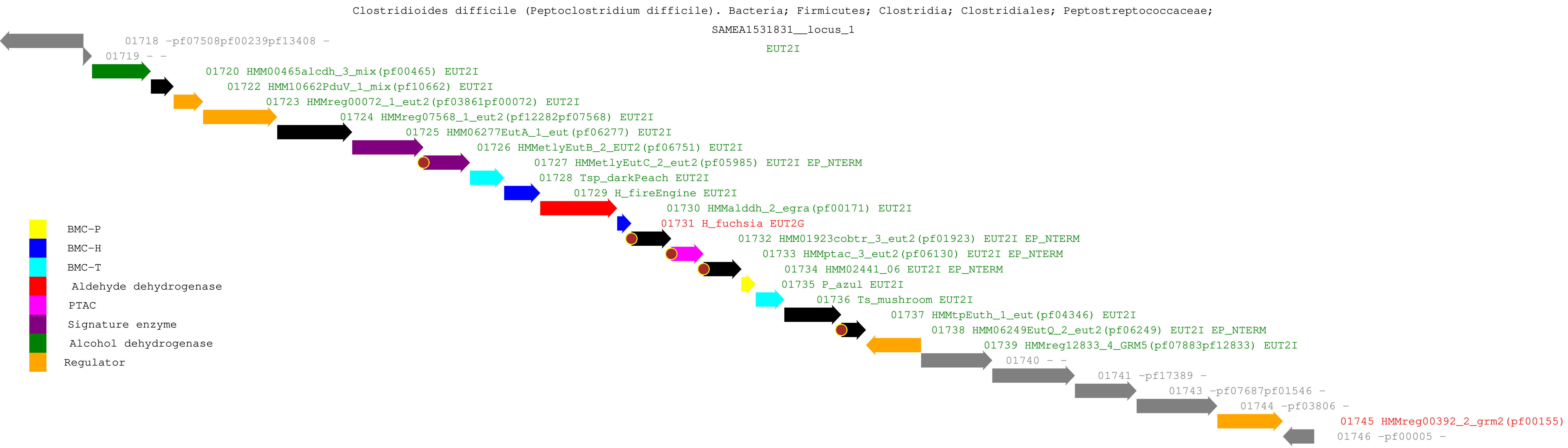
Clostridioides difficile (Peptoclostridium difficile). Bacteria; Firmicutes; Clostridia; Clostridiales; Peptostreptococcaceae;
SAMEA1531831__locus_1
EUT2I
01718 -pf07508pf00239pf13408 -
01719 - -
01720 HMM00465alcdh_3_mix(pf00465) EUT2I
01722 HMM10662PduV_1_mix(pf10662) EUT2I
01723 HMMreg00072_1_eut2(pf03861pf00072) EUT2I
01724 HMMreg07568_1_eut2(pf12282pf07568) EUT2I
01725 HMM06277EutA_1_eut(pf06277) EUT2I
01726 HMMetlyEutB_2_EUT2(pf06751) EUT2I
01727 HMMetlyEutC_2_eut2(pf05985) EUT2I EP_NTERM
01728 Tsp_darkPeach EUT2I
01729 H_fireEngine EUT2I
01730 HMMaldh_2_egra(pf00171) EUT2I
01731 H_fuchsia EUT2G
01732 HMM01923cobtr_3_eut2(pf01923) EUT2I EP_NTERM
01733 HMMptac_3_eut2(pf06130) EUT2I EP_NTERM
01734 HMM02441_06 EUT2I EP_NTERM
01735 P_azul EUT2I
01736 Ts_mushroom EUT2I
01737 HMMtpEuth_1_eut(pf04346) EUT2I
01738 HMM06249EutQ_2_eut2(pf06249) EUT2I EP_NTERM
01739 HMMreg12833_4_GRM5(pf07883pf12833) EUT2I
01740 - -
01741 -pf17389 -
01743 -pf07687pf01546 -
01744 -pf03806 -
01745 HMMreg00392_2_grm2(pf00155)
01746 -pf00005 -
BMC-P
BMC-H
BMC-T
Aldehyde dehydrogenase
PTAC
Signature enzyme
Alcohol dehydrogenase
Regulator

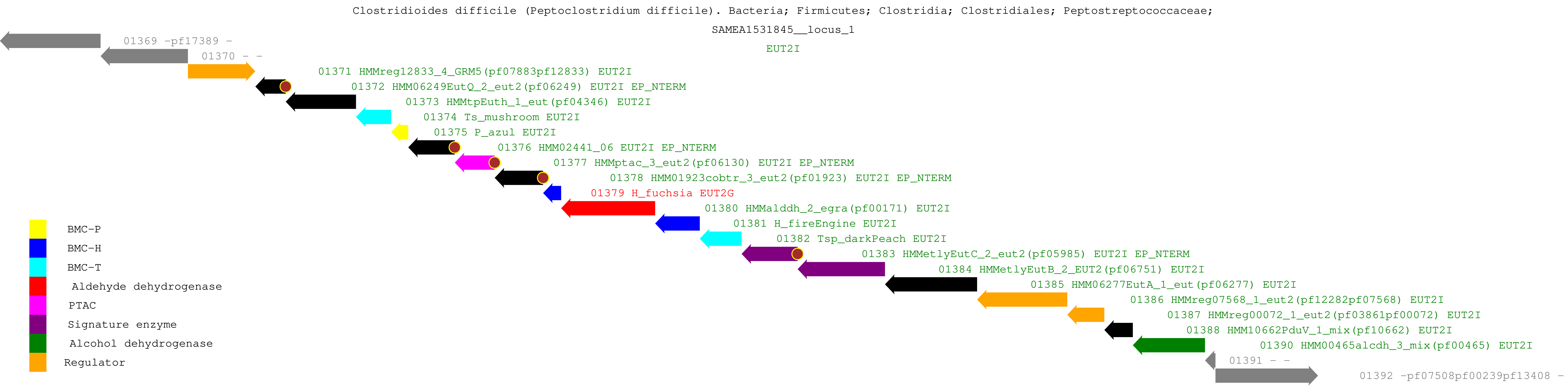

Clostridioides difficile (Peptoclostridium difficile). Bacteria; Firmicutes; Clostridia; Clostridiales; Peptostreptococcaceae;
SAMEA1531845__locus_1
EUT2I
01369 -pf17389 -
01370 - -
01371 HMMreg12833_4_GRM5(pf07883pf12833) EUT2I
01372 HMM06249EutQ_2_eut2(pf06249) EUT2I EP_NTERM
01373 HMMtpEuth_1_eut(pf04346) EUT2I
01374 Ts_mushroom EUT2I
01375 P_azul EUT2I
01376 HMM02441_06 EUT2I EP_NTERM
01377 HMMptac_3_eut2(pf06130) EUT2I EP_NTERM
01378 HMM01923cobtr_3_eut2(pf01923) EUT2I EP_NTERM
01379 H_fuchsia EUT2G
01380 HMMalddh_2_egra(pf00171) EUT2I
01381 H_fireEngine EUT2I
01382 Tsp_darkPeach EUT2I
01383 HMMetlyEutC_2_eut2(pf05985) EUT2I EP_NTERM
01384 HMMetlyEutB_2_EUT2(pf06751) EUT2I
01385 HMM06277EutA_1_eut(pf06277) EUT2I
01386 HMMreg07568_1_eut2(pf12282pf07568) EUT2I
01387 HMMreg00072_1_eut2(pf03861pf00072) EUT2I
01388 HMM10662PduV_1_mix(pf10662) EUT2I
01390 HMM00465alcdh_3_mix(pf00465) EUT2I
01391 - -
01392 -pf07508pf00239pf13408 -
BMC-P
BMC-H
BMC-T
Aldehyde dehydrogenase
PTAC
Signature enzyme
Alcohol dehydrogenase
Regulator

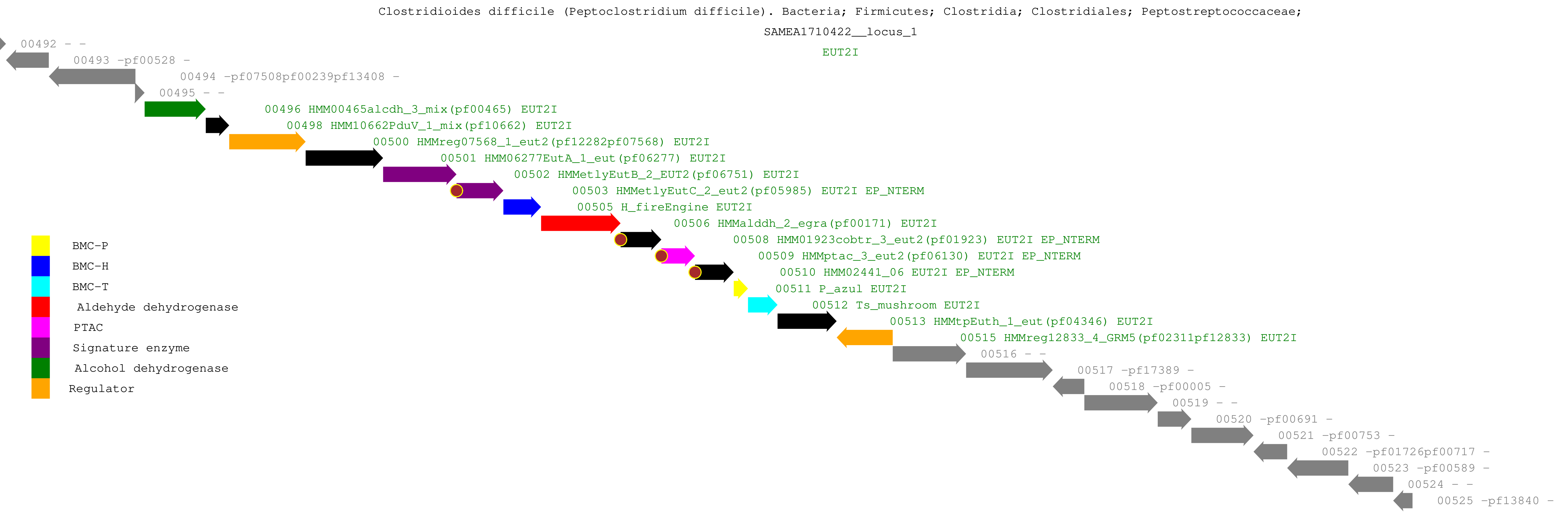

Clostridioides difficile (Peptoclostridium difficile). Bacteria; Firmicutes; Clostridia; Clostridiales; Peptostreptococcaceae;
SAMEA1710422__locus_1
EUT2I
00492 - -
00493 -pf00528 -
00494 -pf07508pf00239pf13408 -
00495 - -
00496 HMM00465alcdh_3_mix(pf00465) EUT2I
00498 HMM10662PduV_1_mix(pf10662) EUT2I
00500 HMMreg07568_1_eut2(pf12282pf07568) EUT2I
00501 HMM06277EutA_1_eut(pf06277) EUT2I
00502 HMMetlyEutB_2_EUT2(pf06751) EUT2I
00503 HMMetlyEutC_2_eut2(pf05985) EUT2I EP_NTERM
00505 H_fireEngine EUT2I
00506 HMMalddh_2_egra(pf00171) EUT2I
00508 HMM01923cobtr_3_eut2(pf01923) EUT2I EP_NTERM
00509 HMMptac_3_eut2(pf06130) EUT2I EP_NTERM
00510 HMM02441_06 EUT2I EP_NTERM
00511 P_azul EUT2I
00512 Ts_mushroom EUT2I
00513 HMMtpEuth_1_eut(pf04346) EUT2I
00515 HMMreg12833_4_GRM5(pf02311pf12833) EUT2I
00516 - -
00517 -pf17389 -
00518 -pf00005 -
00519 - -
00520 -pf00691 -
00521 -pf00753 -
00522 -pf01726pf00717 -
00523 -pf00589 -
00524 - -
00525 -pf13840 -
BMC-P
BMC-H
BMC-T
Aldehyde dehydrogenase
PTAC
Signature enzyme
Alcohol dehydrogenase
Regulator

Clostridioides difficile (Peptoclostridium difficile). Bacteria; Firmicutes; Clostridia; Clostridiales;

SAMEA3374973__locus_1

EUT2I

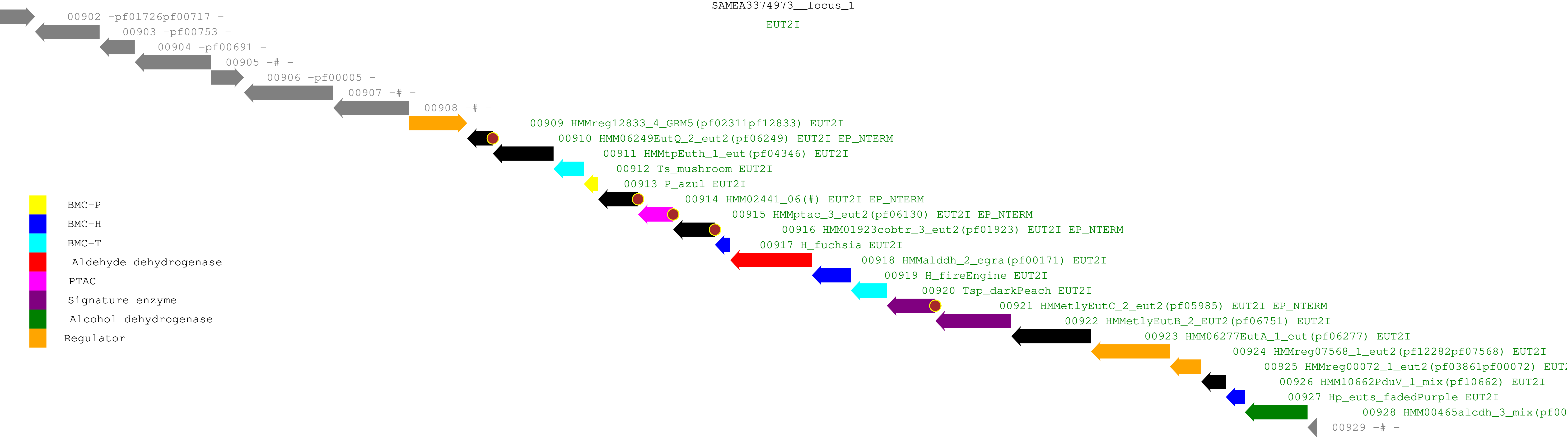

Clostridioides difficile (Peptoclostridium difficile). Bacteria; Firmicutes; Clostridia; Clostridiales;

SAMEA3374989__locus_1

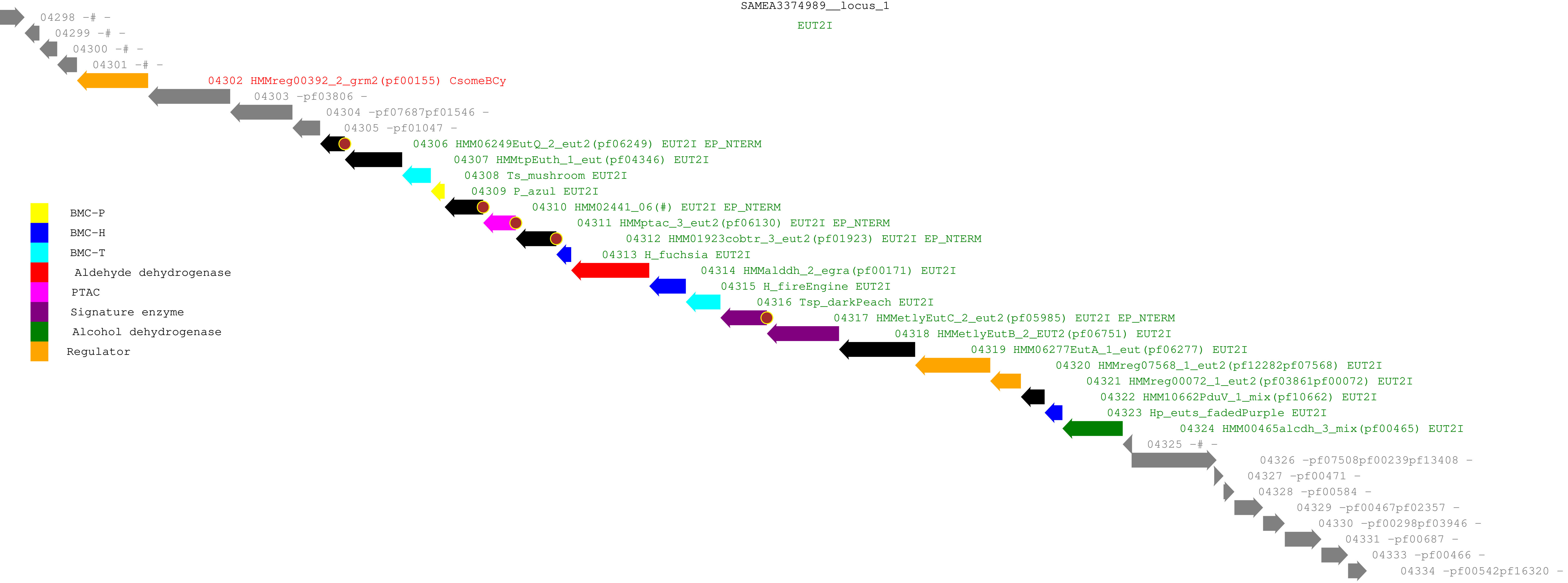

Clostridioides difficile (Peptoclostridium difficile). Bacteria; Firmicutes; Clostridia; Clostridiales;

SAMEA3375004__locus_1

EUT2I

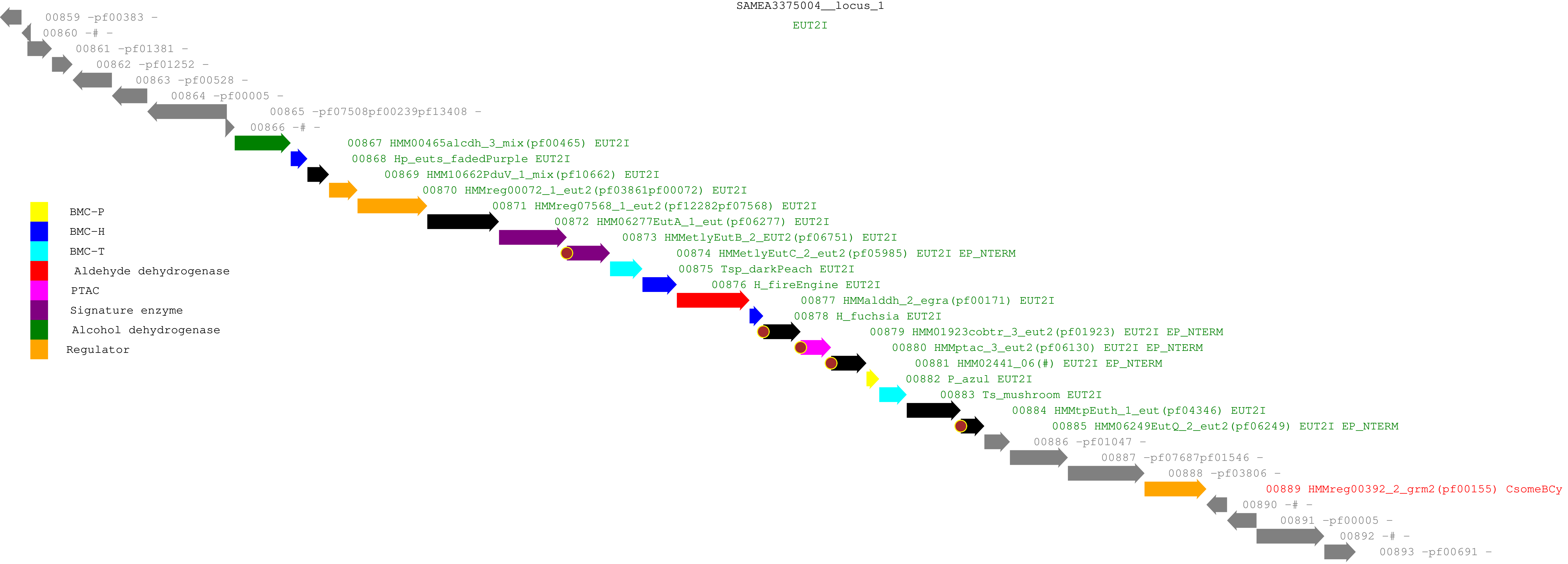

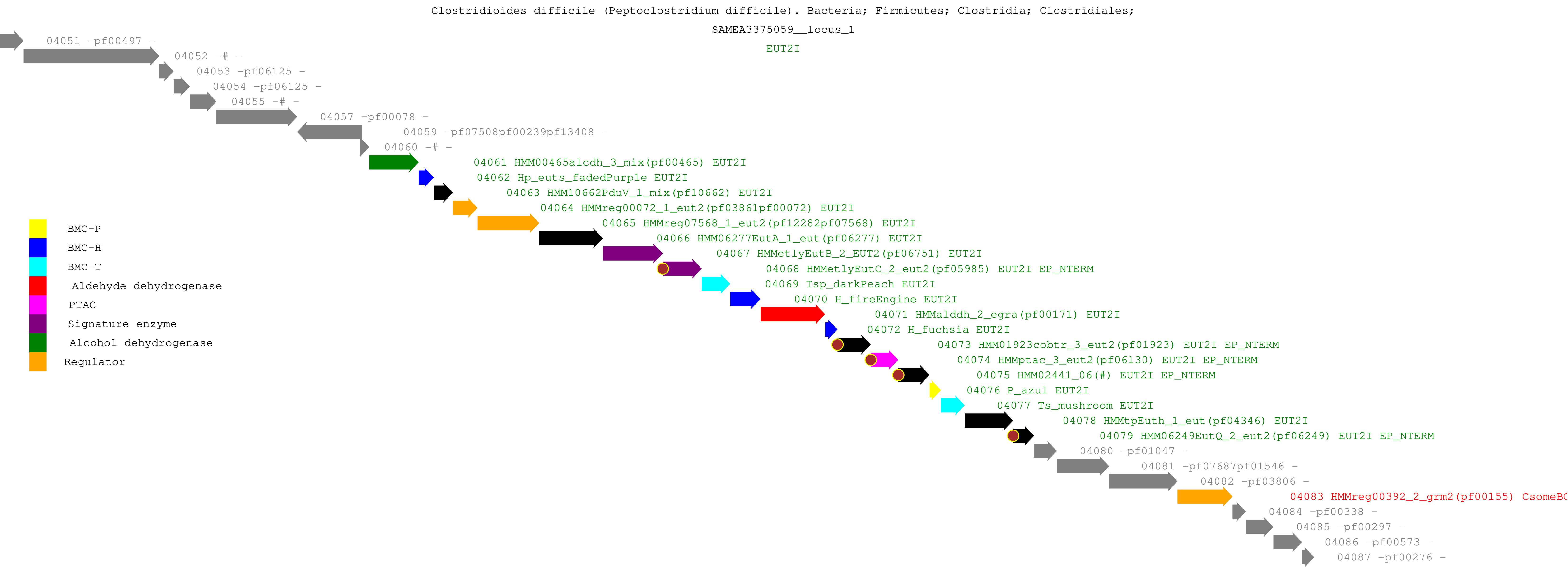
Clostridioides difficile (Peptoclostridium difficile). Bacteria; Firmicutes; Clostridia; Clostridiales;
SAMEA3375059__locus_1
EUT2I
04051 -pf00497 -
04052 -# -
04053 -pf06125 -
04054 -pf06125 -
04055 -# -
04057 -pf00078 -
04059 -pf07508pf00239pf13408 -
04060 -# -
04061 HMM00465alcdh_3_mix(pf00465) EUT2I
04062 Hp_euts_fadedPurple EUT2I
04063 HMM10662PduV_1_mix(pf10662) EUT2I
04064 HMMreg00072_1_eut2(pf03861pf00072) EUT2I
04065 HMMreg07568_1_eut2(pf12282pf07568) EUT2I
04066 HMM06277EutA_1_eut(pf06277) EUT2I
04067 HMMetlyEutB_2_EUT2(pf06751) EUT2I
04068 HMMetlyEutC_2_eut2(pf05985) EUT2I EP_NTERM
04069 Tsp_darkPeach EUT2I
04070 H_fireEngine EUT2I
04071 HMMalddh_2_egra(pf00171) EUT2I
04072 H_fuchsia EUT2I
04073 HMM01923cobtr_3_eut2(pf01923) EUT2I EP_NTERM
04074 HMMptac_3_eut2(pf06130) EUT2I EP_NTERM
04075 HMM02441_06(#) EUT2I EP_NTERM
04076 P_azul EUT2I
04077 Ts_mushroom EUT2I
04078 HMMtpEuth_1_eut(pf04346) EUT2I
04079 HMM06249EutQ_2_eut2(pf06249) EUT2I EP_NTERM
04080 -pf01047 -
04081 -pf07687pf01546 -
04082 -pf03806 -
04083 HMMreg00392_2_grm2(pf00155) CsomeBC
04084 -pf00338 -
04085 -pf00297 -
04086 -pf00573 -
04087 -pf00276 -
BMC-P
BMC-H
BMC-T
Aldehyde dehydrogenase
PTAC
Signature enzyme
Alcohol dehydrogenase
Regulator

Clostridium tetani. Bacteria; Firmicutes; Clostridia; Clostridiales; Clostridiaceae;

SAMN02745112__locus_1

EUT2I

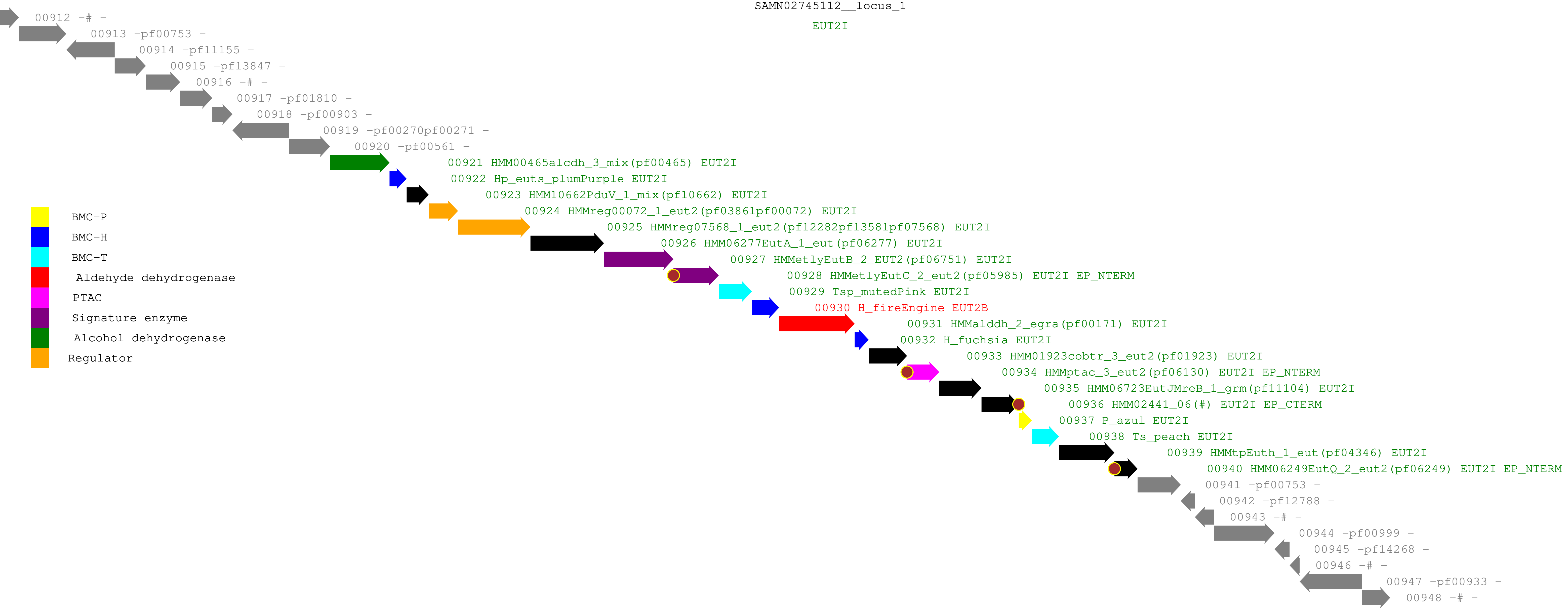

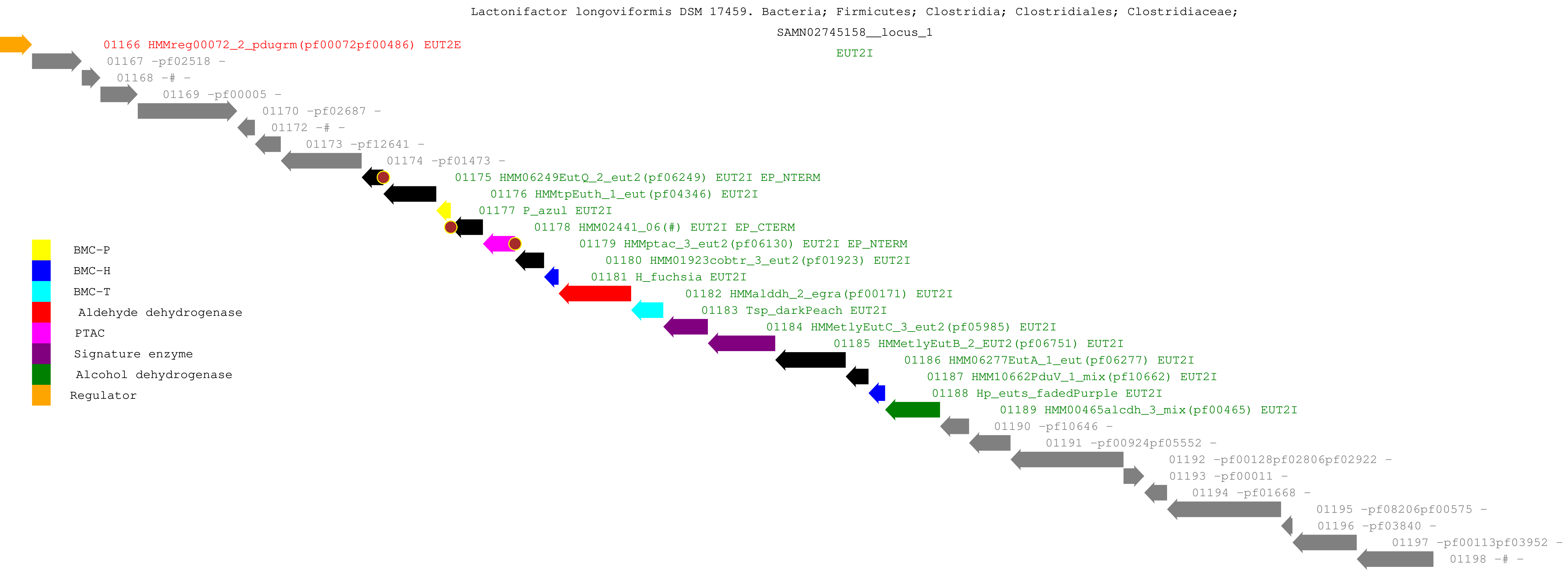
Lactonifactor longoviformis DSM 17459. Bacteria; Firmicutes; Clostridia; Clostridiales; Clostridiaceae;
SAMN02745158__locus_1
EUT2I
01166 HMMreg00072_2_pdugrm(pf00072pf00486) EUT2E
01167 -pf02518 -
01168 -# -
01169 -pf00005 -
01170 -pf02687 -
01172 -# -
01173 -pf12641 -
01174 -pf01473 -
01175 HMM06249EutQ_2_eut2(pf06249) EUT2I EP_NTERM
01176 HMMtpEuth_1_eut(pf04346) EUT2I
01177 P_azul EUT2I
01178 HMM02441_06(#) EUT2I EP_CTERM
01179 HMMptac_3_eut2(pf06130) EUT2I EP_NTERM
01180 HMM01923cobtr_3_eut2(pf01923) EUT2I
01181 H_fuchsia EUT2I
01182 HMMaldh_2_egra(pf00171) EUT2I
01183 Tsp_darkPeach EUT2I
01184 HMMetlyEutC_3_eut2(pf05985) EUT2I
01185 HMMetlyEutB_2_EUT2(pf06751) EUT2I
01186 HMM06277EutA_1_eut(pf06277) EUT2I
01187 HMM10662PduV_1_mix(pf10662) EUT2I
01188 Hp_euts_fadedPurple EUT2I
01189 HMM00465alcdh_3_mix(pf00465) EUT2I
01190 -pf10646 -
01191 -pf00924pf05552 -
01192 -pf00128pf02806pf02922 -
01193 -pf00011 -
01194 -pf01668 -
01195 -pf08206pf00575 -
01196 -pf03840 -
01197 -pf00113pf03952 -
01198 -# -
BMC-P
BMC-H
BMC-T
Aldehyde dehydrogenase
PTAC
Signature enzyme
Alcohol dehydrogenase
Regulator

Clostridium collagenovorans DSM 3089. Bacteria; Firmicutes; Clostridia; Clostridiales; Clostridiaceae;

SAMN02745196__locus_1_main_sat

EUT2I

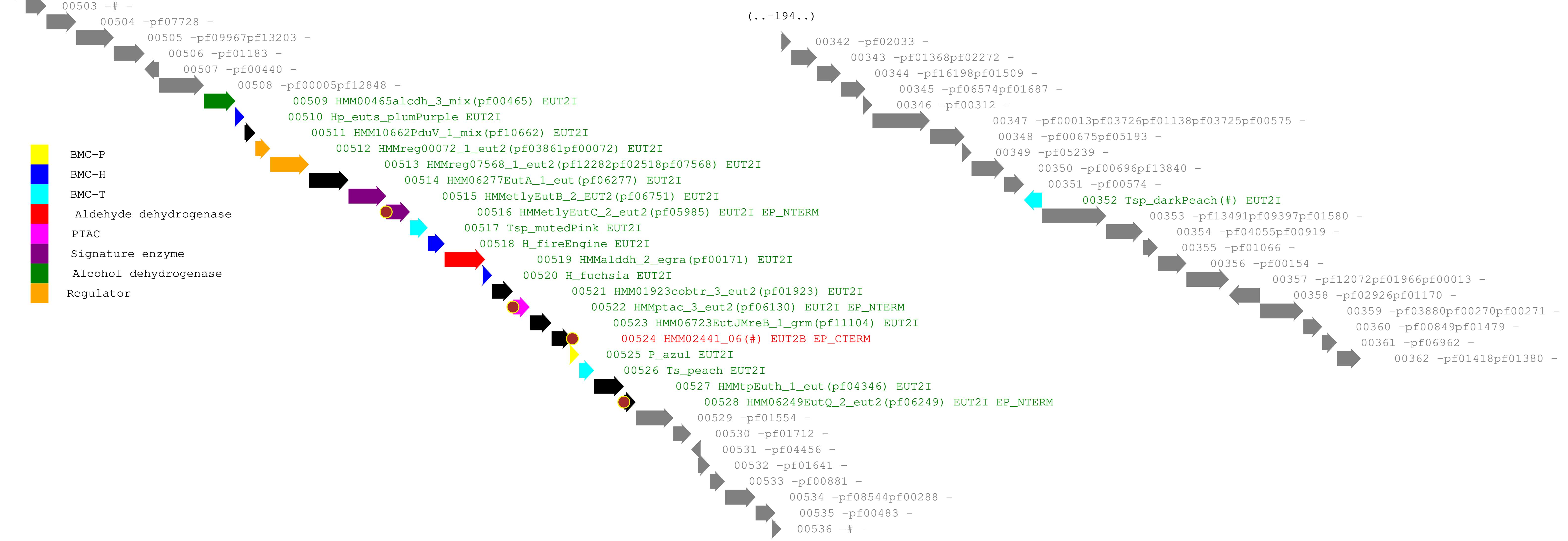

Clostridium proteolyticum DSM 3090. Bacteria; Firmicutes; Clostridia; Clostridiales; Clostridiaceae;

SAMN02745248__locus_1

EUT2I

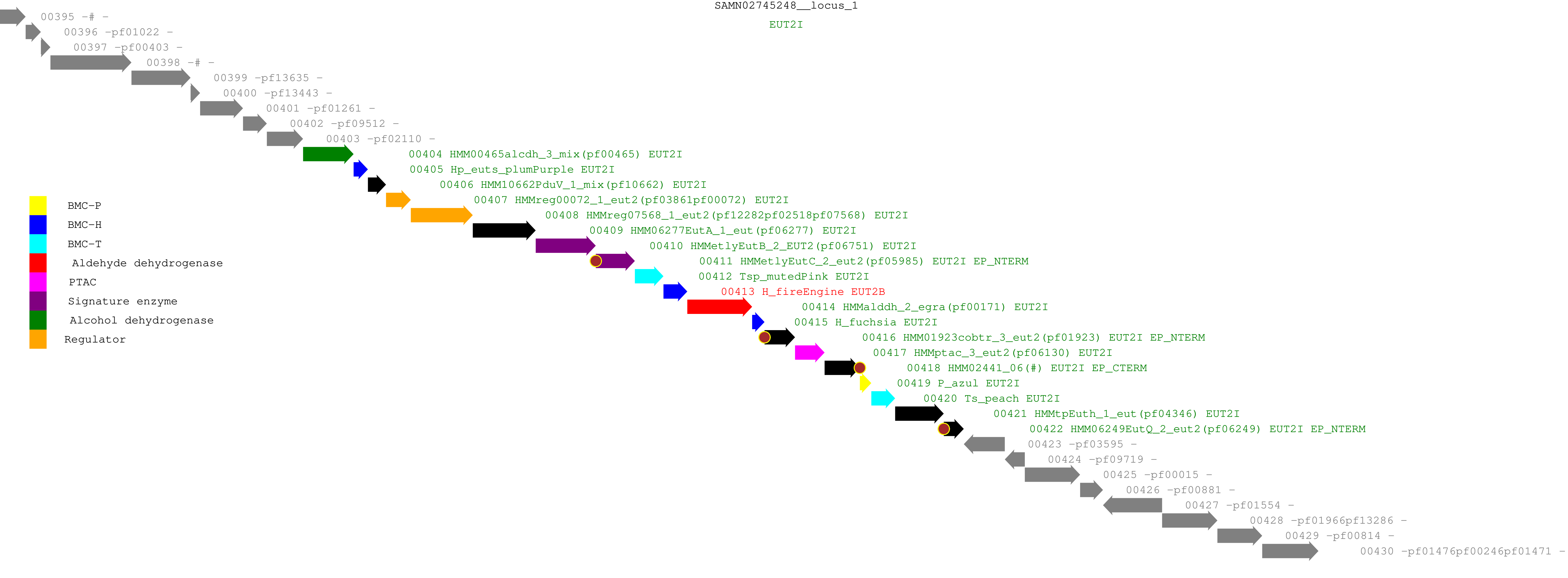

Anaerosporobacter mobilis DSM 15930. Bacteria; Firmicutes; Clostridia; Clostridiales; Lachnospiraceae;

SAMN02746066__locus_1

EUT2I

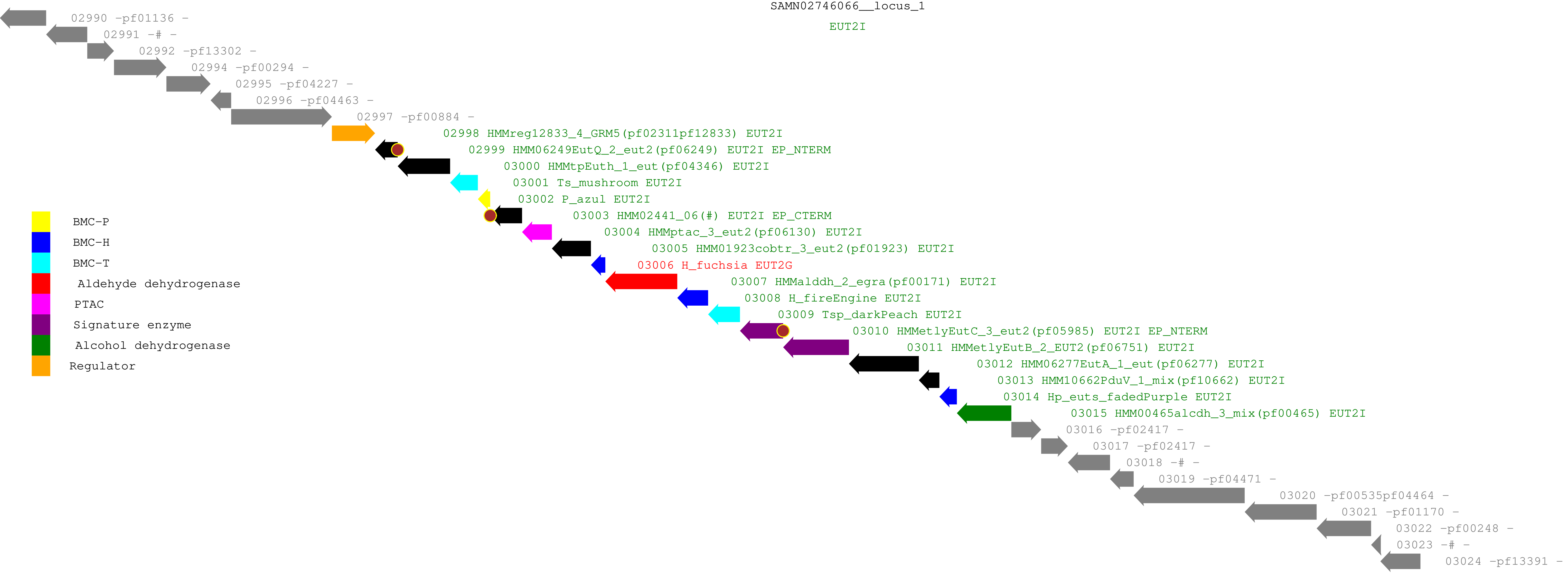

Clostridium cadaveris. Bacteria; Firmicutes; Clostridia; Clostridiales; Clostridiaceae;

SAMN04487885__locus_1

EUT2I

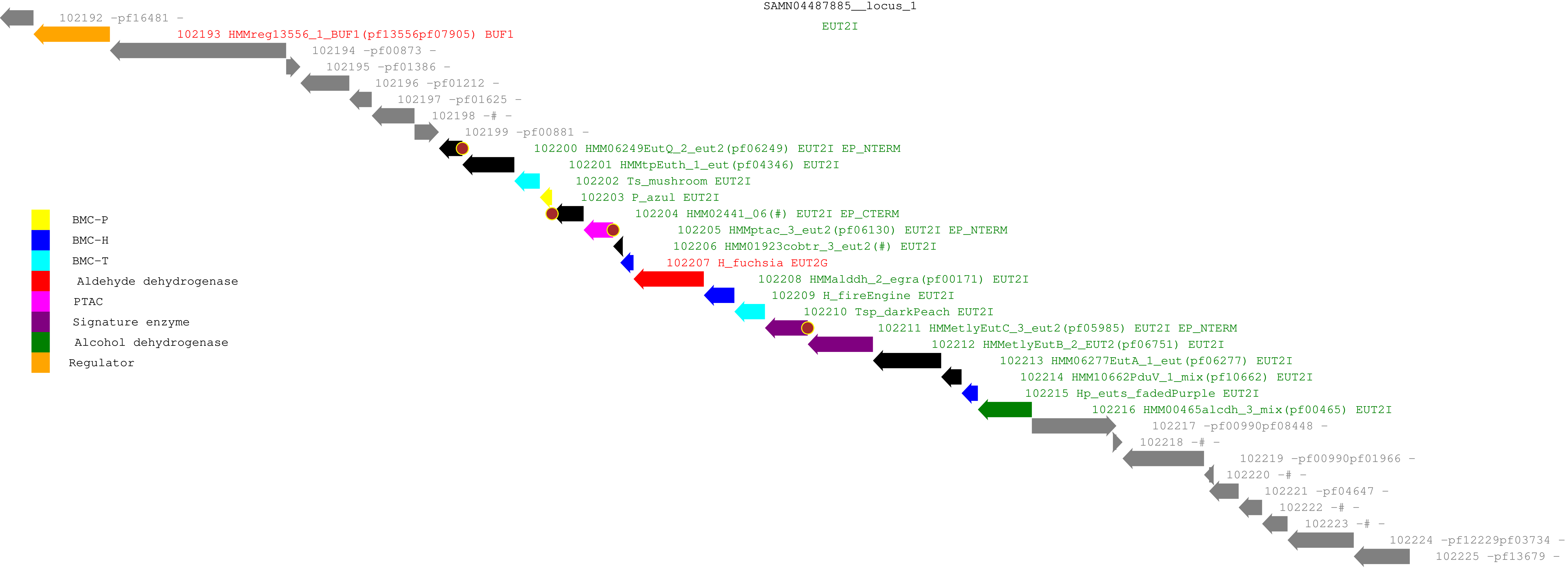

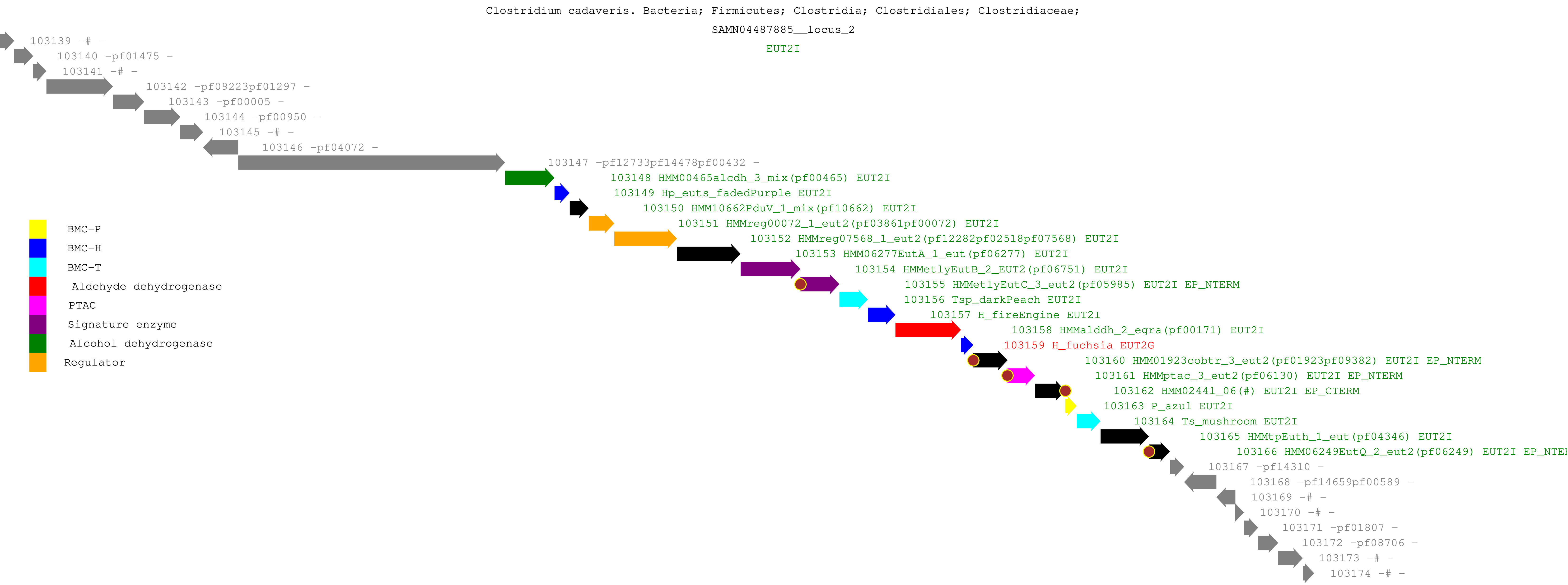

Clostridium cadaveris. Bacteria; Firmicutes; Clostridia; Clostridiales; Clostridiaceae;
SAMN04487885__locus_2
EUT2I
103139 -# -
103140 -pf01475 -
103141 -# -
103142 -pf09223pf01297 -
103143 -pf00005 -
103144 -pf00950 -
103145 -# -
103146 -pf04072 -
103147 -pf12733pf14478pf00432 -
103148 HMM00465alcdh_3_mix(pf00465) EUT2I
103149 Hp_euts_fadedPurple EUT2I
103150 HMM10662PduV_1_mix(pf10662) EUT2I
103151 HMMreg00072_1_eut2(pf03861pf00072) EUT2I
103152 HMMreg07568_1_eut2(pf12282pf02518pf07568) EUT2I
103153 HMM06277EutA_1_eut(pf06277) EUT2I
103154 HMMetlyEutB_2_EUT2(pf06751) EUT2I
103155 HMMetlyEutC_3_eut2(pf05985) EUT2I EP_NTERM
103156 Tsp_darkPeach EUT2I
103157 H_fireEngine EUT2I
103158 HMMalddh_2_egra(pf00171) EUT2I
103159 H_fuchsia EUT2G
103160 HMM01923cobtr_3_eut2(pf01923pf09382) EUT2I EP_NTERM
103161 HMMptac_3_eut2(pf06130) EUT2I EP_NTERM
103162 HMM02441_06(#) EUT2I EP_CTERM
103163 P_azul EUT2I
103164 Ts_mushroom EUT2I
103165 HMMtpEuth_1_eut(pf04346) EUT2I
103166 HMM06249EutQ_2_eut2(pf06249) EUT2I EP_NTE
103167 -pf14310 -
103168 -pf14659pf00589 -
103169 -# -
103170 -# -
103171 -pf01807 -
103172 -pf08706 -
103173 -# -
103174 -# -
BMC-P
BMC-H
BMC-T
Aldehyde dehydrogenase
PTAC
Signature enzyme
Alcohol dehydrogenase
Regulator

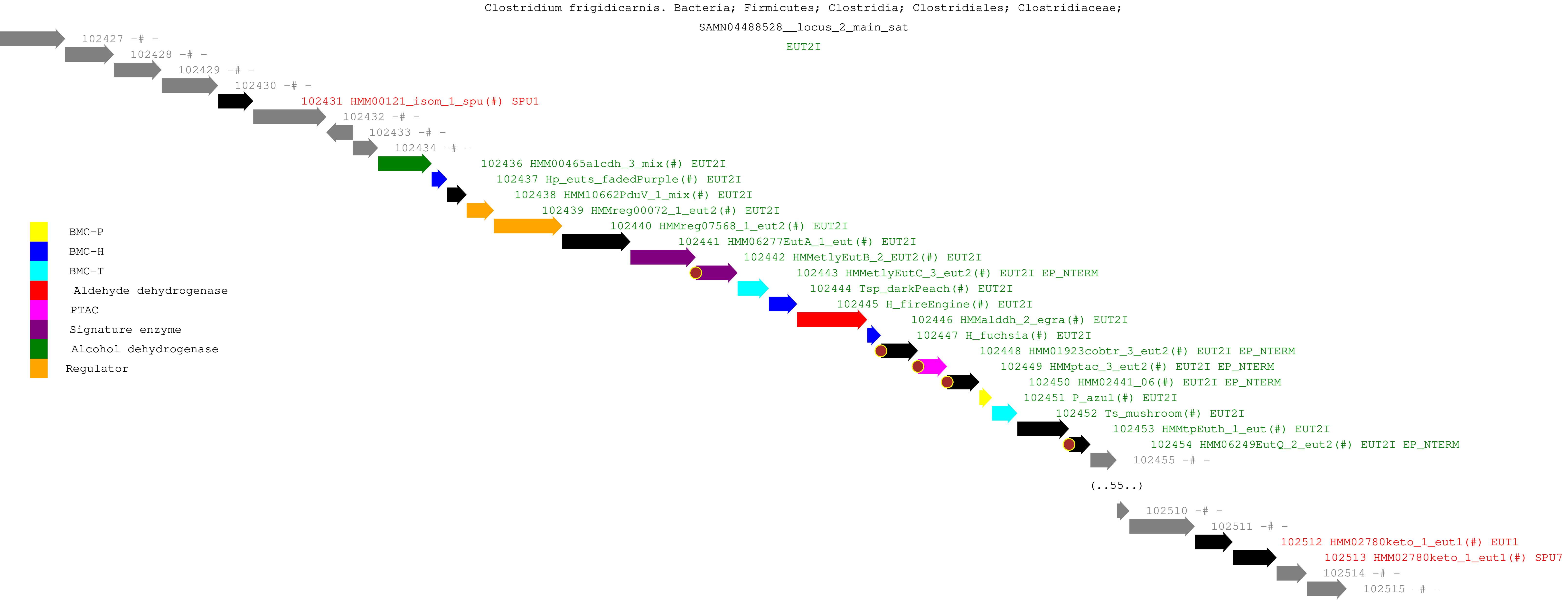

Clostridium frigidicarnis. Bacteria; Firmicutes; Clostridia; Clostridiales; Clostridiaceae;
SAMN04488528__locus_2_main_sat
EUT2I
102427 -# -
102428 -# -
102429 -# -
102430 -# -
102431 HMM00121_isom_1_spu(#) SPU1
102432 -# -
102433 -# -
102434 -# -
102436 HMM00465alcdh_3_mix(#) EUT2I
102437 Hp_euts_fadedPurple(#) EUT2I
102438 HMM10662PduV_1_mix(#) EUT2I
102439 HMMreg00072_1_eut2(#) EUT2I
102440 HMMreg07568_1_eut2(#) EUT2I
102441 HMM06277EutA_1_eut(#) EUT2I
102442 HMMetlyEutB_2_EUT2(#) EUT2I
102443 HMMetlyEutC_3_eut2(#) EUT2I EP_NTERM
102444 Tsp_darkPeach(#) EUT2I
102445 H_fireEngine(#) EUT2I
102446 HMMalddh_2_egra(#) EUT2I
102447 H_fuchsia(#) EUT2I
102448 HMM01923cobtr_3_eut2(#) EUT2I EP_NTERM
102449 HMMptac_3_eut2(#) EUT2I EP_NTERM
102450 HMM02441_06(#) EUT2I EP_NTERM
102451 P_azul(#) EUT2I
102452 Ts_mushroom(#) EUT2I
102453 HMMtpEuth_1_eut(#) EUT2I
102454 HMM06249EutQ_2_eut2(#) EUT2I EP_NTERM
102455 -# -
(..55..)
102510 -# -
102511 -# -
102512 HMM02780keto_1_eut1(#) EUT1
102513 HMM02780keto_1_eut1(#) SPU7
102514 -# -
102515 -# -
BMC-P
BMC-H
BMC-T
Aldehyde dehydrogenase
PTAC
Signature enzyme
Alcohol dehydrogenase
Regulator

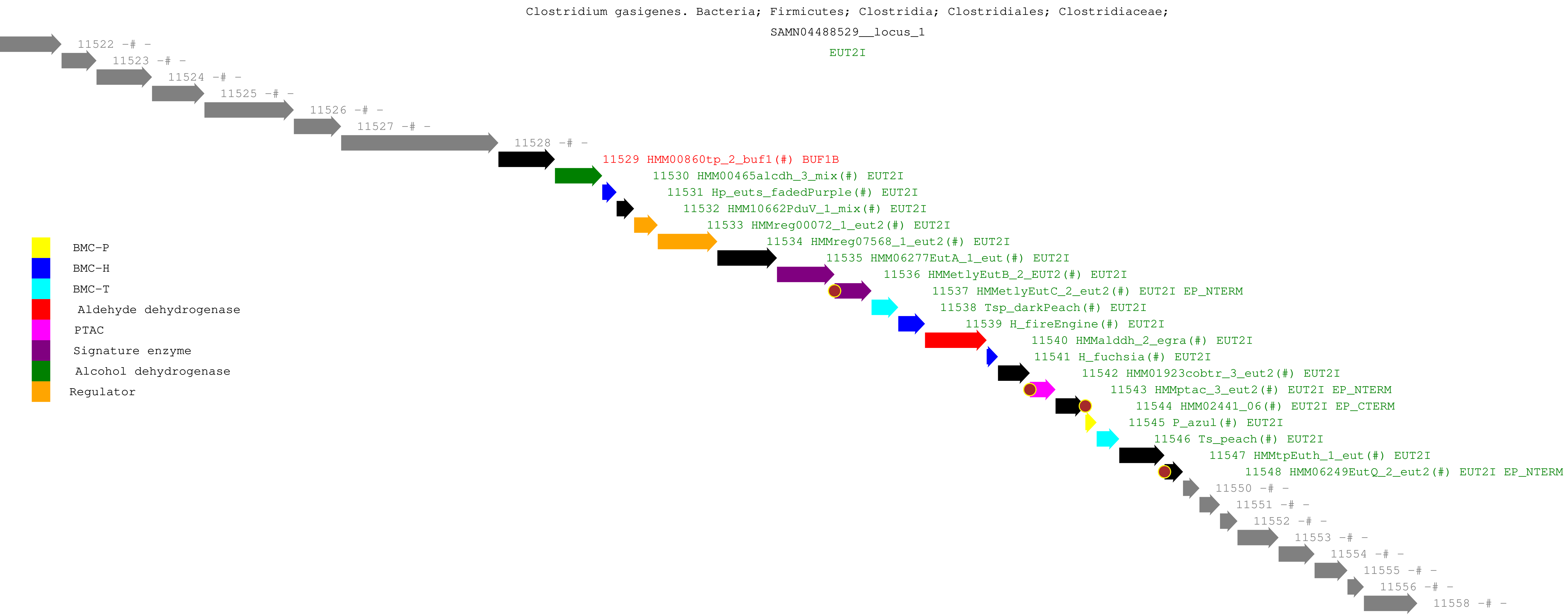

Clostridium gasigenes. Bacteria; Firmicutes; Clostridia; Clostridiales; Clostridiaceae;
SAMN04488529__locus_1
EUT2I
11522 -# -
11523 -# -
11524 -# -
11525 -# -
11526 -# -
11527 -# -
11528 -# -
11529 HMM00860tp_2_buf1(#) BUF1B
11530 HMM00465alcdh_3_mix(#) EUT2I
11531 Hp_euts_fadedPurple(#) EUT2I
11532 HMM10662PduV_1_mix(#) EUT2I
11533 HMMreg00072_1_eut2(#) EUT2I
11534 HMMreg07568_1_eut2(#) EUT2I
11535 HMM06277EutA_1_eut(#) EUT2I
11536 HMMetlyEutB_2_EUT2(#) EUT2I
11537 HMMetlyEutC_2_eut2(#) EUT2I EP_NTERM
11538 Tsp_darkPeach(#) EUT2I
11539 H_fireEngine(#) EUT2I
11540 HMMalddh_2_egra(#) EUT2I
11541 H_fuchsia(#) EUT2I
11542 HMM01923cobtr_3_eut2(#) EUT2I
11543 HMMptac_3_eut2(#) EUT2I EP_NTERM
11544 HMM02441_06(#) EUT2I EP_CTERM
11545 P_azul(#) EUT2I
11546 Ts_peach(#) EUT2I
11547 HMMtpEuth_1_eut(#) EUT2I
11548 HMM06249EutQ_2_eut2(#) EUT2I EP_NTERM
11550 -# -
11551 -# -
11552 -# -
11553 -# -
11554 -# -
11555 -# -
11556 -# -
11558 -# -
BMC-P
BMC-H
BMC-T
Aldehyde dehydrogenase
PTAC
Signature enzyme
Alcohol dehydrogenase
Regulator

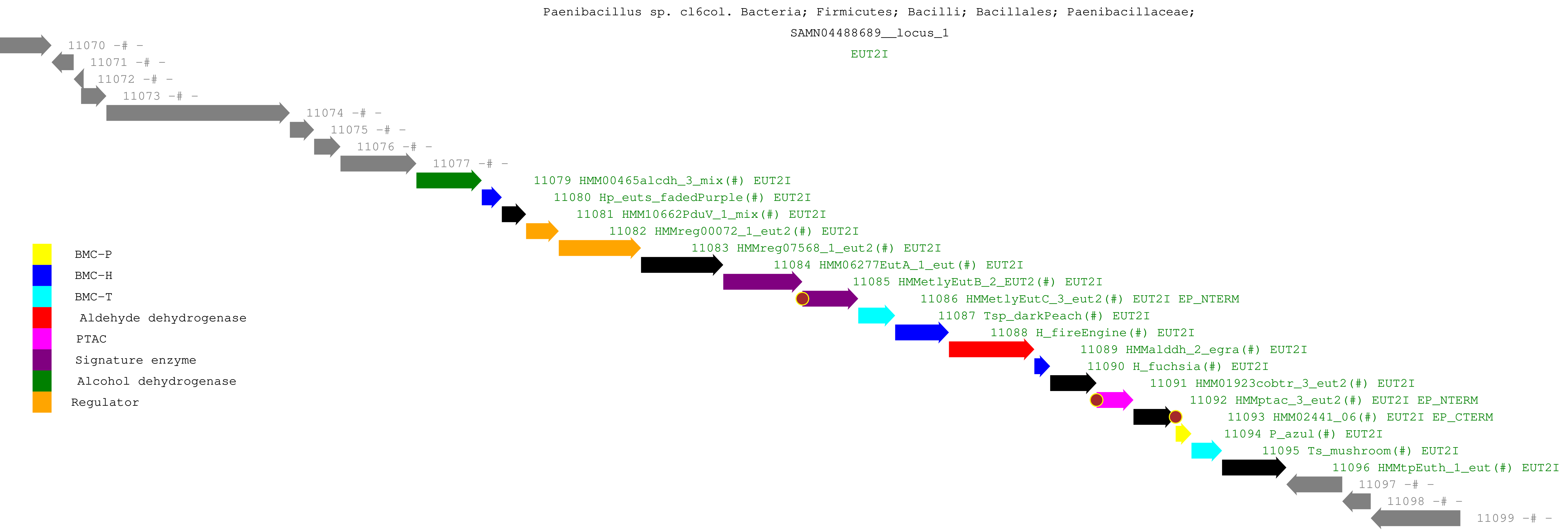
Paenibacillus sp. cl6col. Bacteria; Firmicutes; Bacilli; Bacillales; Paenibacillaceae;
SAMN04488689__locus_1
EUT2I
11070 -# -
11071 -# -
11072 -# -
11073 -# -
11074 -# -
11075 -# -
11076 -# -
11077 -# -
11079 HMM00465alcdh_3_mix(#) EUT2I
11080 Hp_euts_fadedPurple(#) EUT2I
11081 HMM10662PduV_1_mix(#) EUT2I
11082 HMMreg00072_1_eut2(#) EUT2I
11083 HMMreg07568_1_eut2(#) EUT2I
11084 HMM06277EutA_1_eut(#) EUT2I
11085 HMMetlyEutB_2_EUT2(#) EUT2I
11086 HMMetlyEutC_3_eut2(#) EUT2I EP_NTERM
11087 Tsp_darkPeach(#) EUT2I
11088 H_fireEngine(#) EUT2I
11089 HMMalddh_2_egra(#) EUT2I
11090 H_fuchsia(#) EUT2I
11091 HMM01923cobtr_3_eut2(#) EUT2I
11092 HMMptac_3_eut2(#) EUT2I EP_NTERM
11093 HMM02441_06(#) EUT2I EP_CTERM
11094 P_azul(#) EUT2I
11095 Ts_mushroom(#) EUT2I
11096 HMMtpEuth_1_eut(#) EUT2I
11097 -# -
11098 -# -
11099 -# -
BMC-P
BMC-H
BMC-T
Aldehyde dehydrogenase
PTAC
Signature enzyme
Alcohol dehydrogenase
Regulator

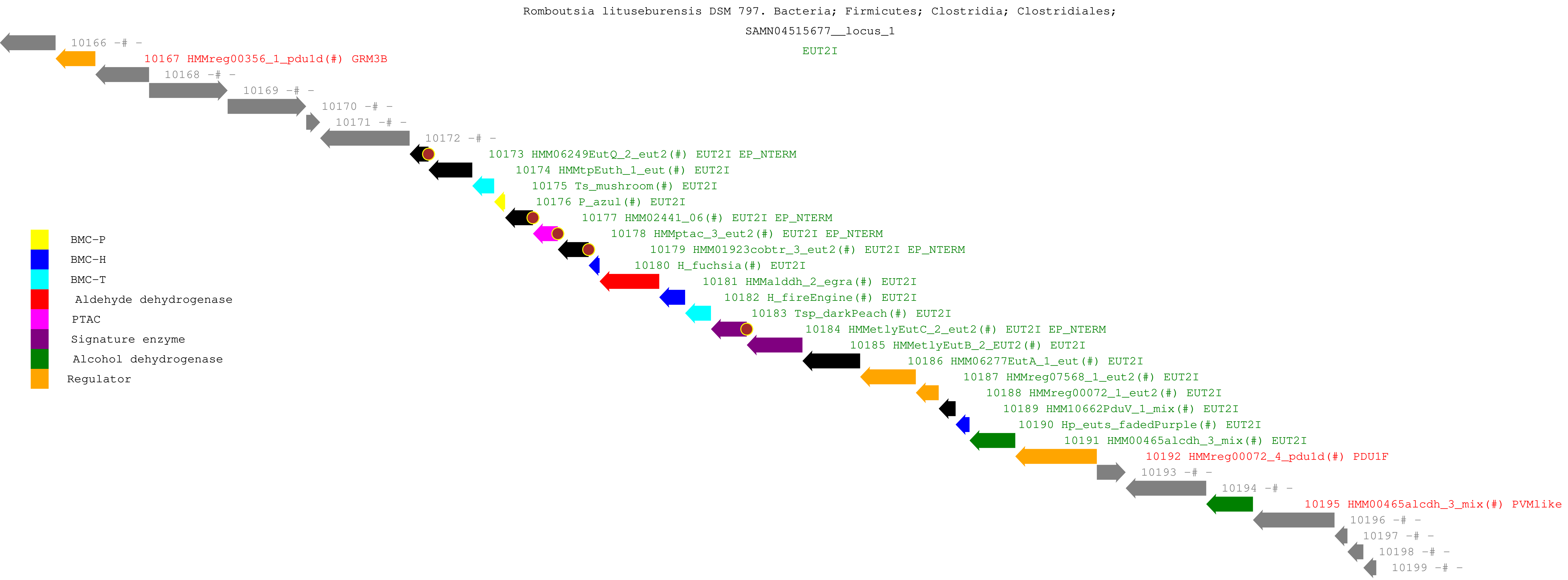
Romboutsia lituseburensis DSM 797. Bacteria; Firmicutes; Clostridia; Clostridiales;
SAMN04515677__locus_1
EUT2I
10166 -# -
10167 HMMreg00356_1_pdu1d(#) GRM3B
10168 -# -
10169 -# -
10170 -# -
10171 -# -
10172 -# -
10173 HMM06249EutQ_2_eut2(#) EUT2I EP_NTERM
10174 HMMtpEuth_1_eut(#) EUT2I
10175 Ts_mushroom(#) EUT2I
10176 P_azul(#) EUT2I
10177 HMM02441_06(#) EUT2I EP_NTERM
10178 HMMptac_3_eut2(#) EUT2I EP_NTERM
10179 HMM01923cobtr_3_eut2(#) EUT2I EP_NTERM
10180 H_fuchsia(#) EUT2I
10181 HMMalddh_2_egra(#) EUT2I
10182 H_fireEngine(#) EUT2I
10183 Tsp_darkPeach(#) EUT2I
10184 HMMetlyEutC_2_eut2(#) EUT2I EP_NTERM
10185 HMMetlyEutB_2_EUT2(#) EUT2I
10186 HMM06277EutA_1_eut(#) EUT2I
10187 HMMreg07568_1_eut2(#) EUT2I
10188 HMMreg00072_1_eut2(#) EUT2I
10189 HMM10662PduV_1_mix(#) EUT2I
10190 Hp_euts_fadedPurple(#) EUT2I
10191 HMM00465alcdh_3_mix(#) EUT2I
10192 HMMreg00072_4_pdu1d(#) PDU1F
10193 -# -
10194 -# -
10195 HMM00465alcdh_3_mix(#) PVMlike
10196 -# -
10197 -# -
10198 -# -
10199 -# -
BMC-P
BMC-H
BMC-T
Aldehyde dehydrogenase
PTAC
Signature enzyme
Alcohol dehydrogenase
Regulator

Paenibacillus uliginis N3/975. Bacteria; Firmicutes; Bacilli; Bacillales; Paenibacillaceae;

SAMN05661091__locus_1

EUT2I

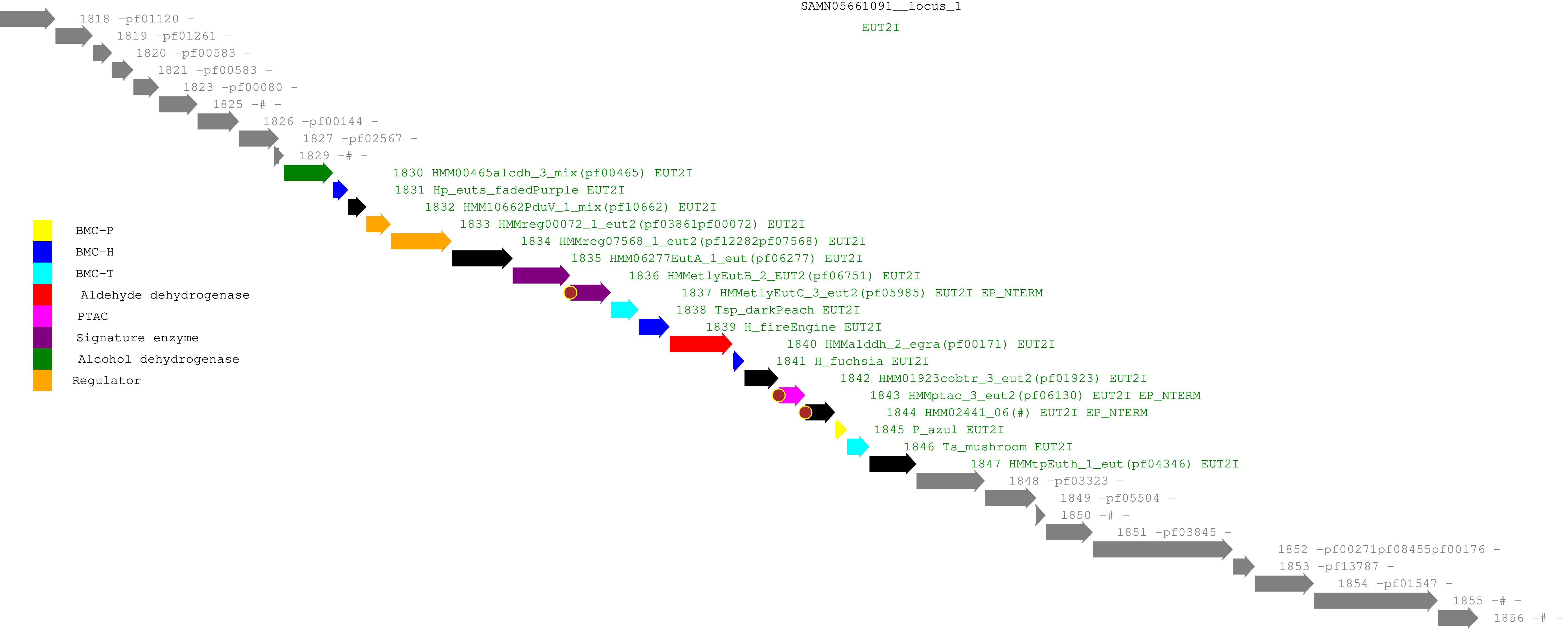

Paenibacillus uliginis N3/975. Bacteria; Firmicutes; Bacilli; Bacillales; Paenibacillaceae;

SAMN05661091__locus_3

EUT2I

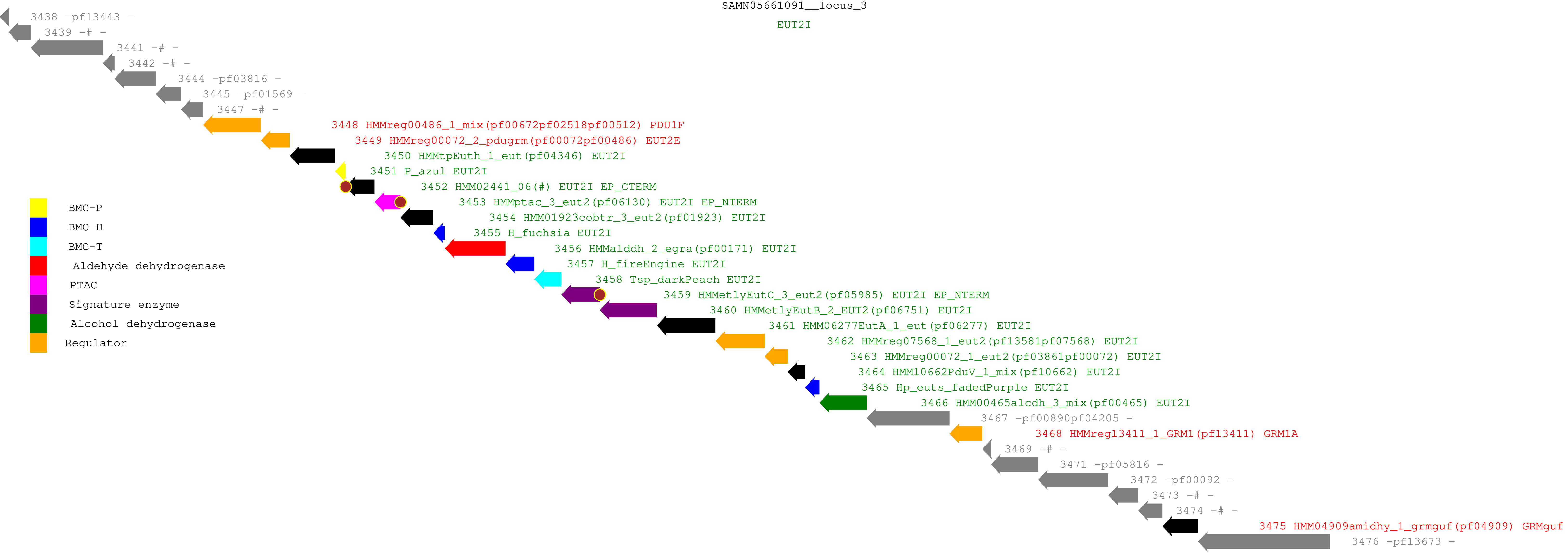

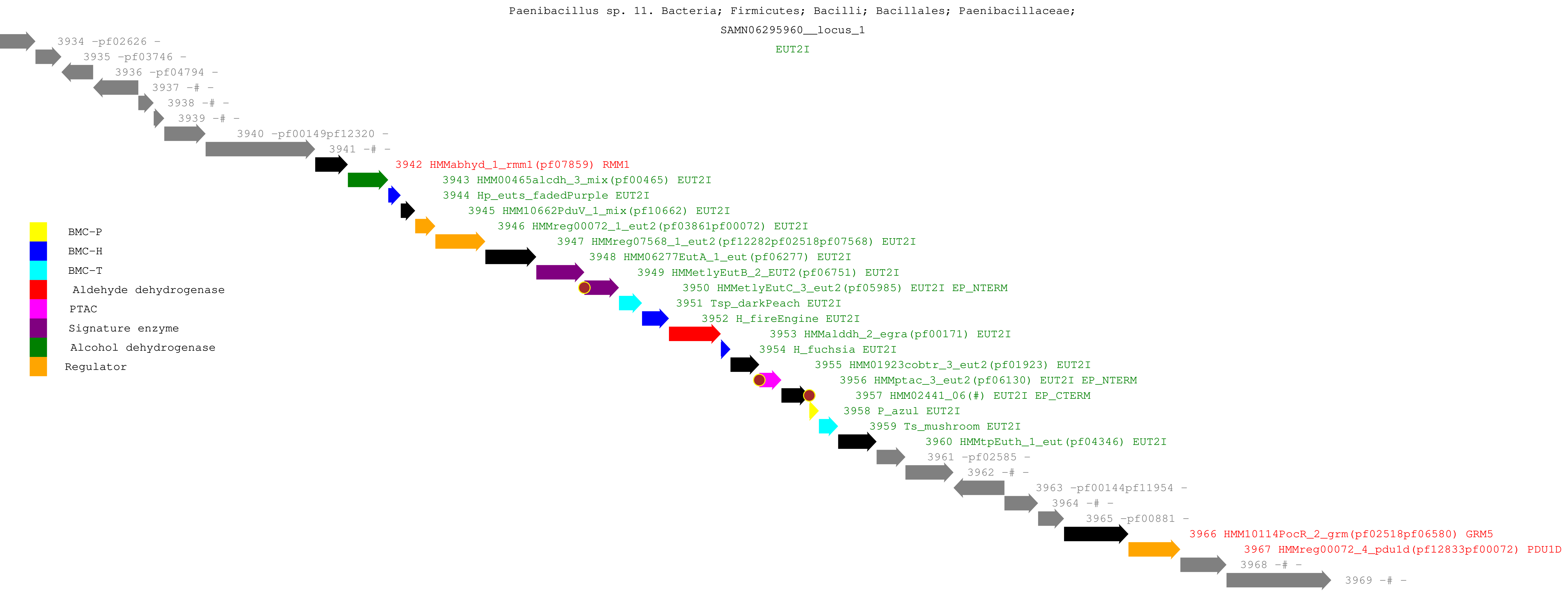
Paenibacillus sp. 11. Bacteria; Firmicutes; Bacilli; Bacillales; Paenibacillaceae;
SAMN06295960__locus_1
EUT2I
3934 -pf02626 -
3935 -pf03746 -
3936 -pf04794 -
3937 -# -
3938 -# -
3939 -# -
3940 -pf00149pf12320 -
3941 -# -
3942 HMMabhyd_1_rmm1(pf07859) RMM1
3943 HMM00465alcdh_3_mix(pf00465) EUT2I
3944 Hp_euts_fadedPurple EUT2I
3945 HMM10662PduV_1_mix(pf10662) EUT2I
3946 HMMreg00072_1_eut2(pf03861pf00072) EUT2I
3947 HMMreg07568_1_eut2(pf12282pf02518pf07568) EUT2I
3948 HMM06277EutA_1_eut(pf06277) EUT2I
3949 HMMetlyEutB_2_EUT2(pf06751) EUT2I
3950 HMMetlyEutC_3_eut2(pf05985) EUT2I EP_NTERM
3951 Tsp_darkPeach EUT2I
3952 H_fireEngine EUT2I
3953 HMMalddh_2_egra(pf00171) EUT2I
3954 H_fuchsia EUT2I
3955 HMM01923cobtr_3_eut2(pf01923) EUT2I
3956 HMMptac_3_eut2(pf06130) EUT2I EP_NTERM
3957 HMM02441_06(#) EUT2I EP_CTERM
3958 P_azul EUT2I
3959 Ts_mushroom EUT2I
3960 HMMtpEuth_1_eut(pf04346) EUT2I
3961 -pf02585 -
3962 -# -
3963 -pf00144pf11954 -
3964 -# -
3965 -pf00881 -
3966 HMM10114PocR_2_grm(pf02518pf06580) GRM5
3967 HMMreg00072_4_pdu1d(pf12833pf00072) PDU1D
3968 -# -
3969 -# -
BMC-P
BMC-H
BMC-T
Aldehyde dehydrogenase
PTAC
Signature enzyme
Alcohol dehydrogenase
Regulator

Clostridium argentinense CDC 2741. Bacteria; Firmicutes; Clostridia; Clostridiales; Clostridiaceae;

U732__locus_1

EUT2I

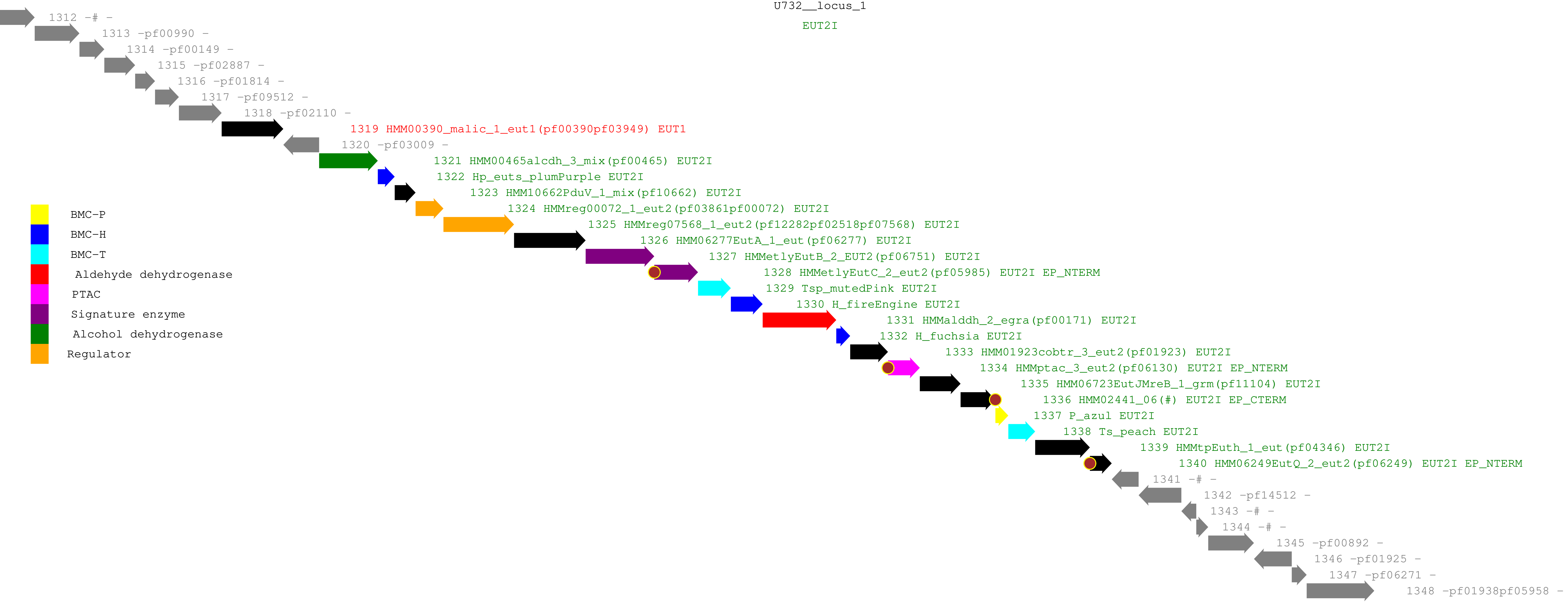

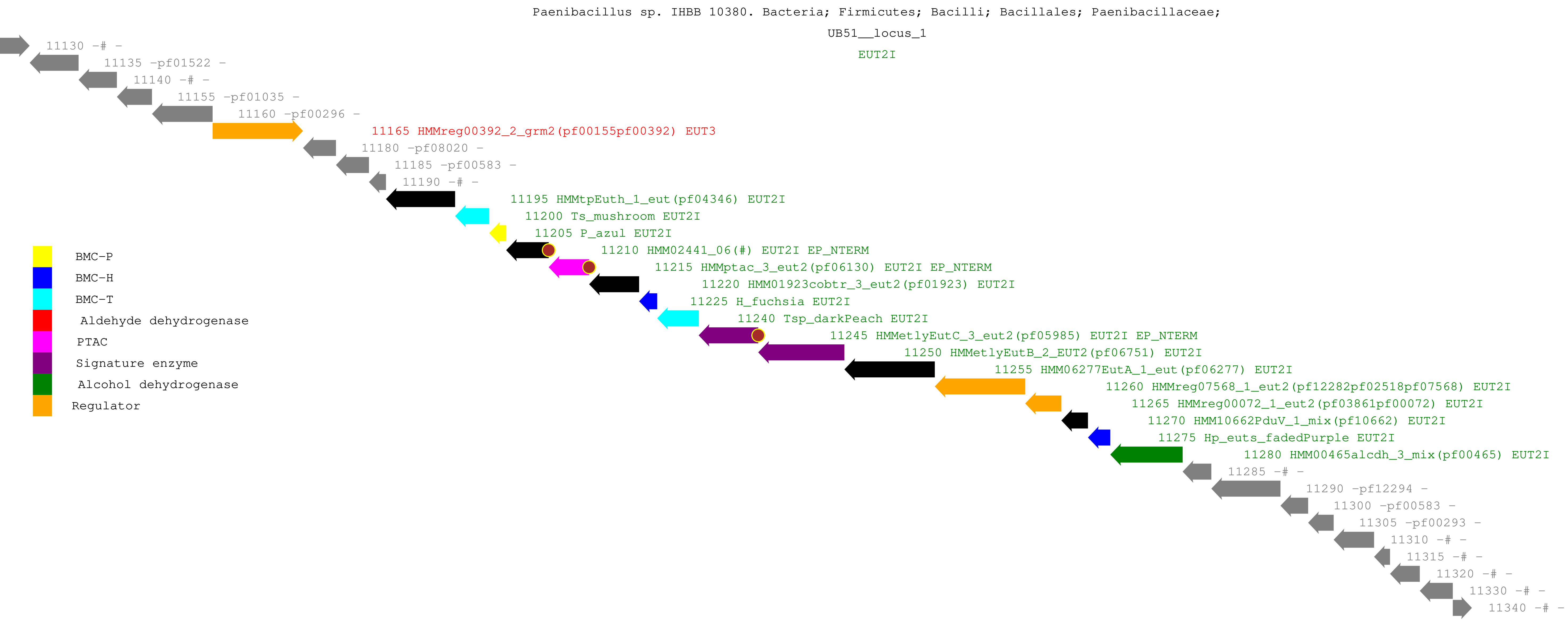
Paenibacillus sp. IHBB 10380. Bacteria; Firmicutes; Bacilli; Bacillales; Paenibacillaceae;
UB51__locus_1
EUT2I
11130 -# -
11135 -pf01522 -
11140 -# -
11155 -pf01035 -
11160 -pf00296 -
11165 HMMreg00392_2_grm2(pf00155pf00392) EUT3
11180 -pf08020 -
11185 -pf00583 -
11190 -# -
11195 HMMtpEuth_1_eut(pf04346) EUT2I
11200 Ts_mushroom EUT2I
11205 P_azul EUT2I
11210 HMM02441_06(#) EUT2I EP_NTERM
11215 HMMptac_3_eut2(pf06130) EUT2I EP_NTERM
11220 HMM01923cobtr_3_eut2(pf01923) EUT2I
11225 H_fuchsia EUT2I
11240 Tsp_darkPeach EUT2I
11245 HMMetlyEutC_3_eut2(pf05985) EUT2I EP_NTERM
11250 HMMetlyEutB_2_EUT2(pf06751) EUT2I
11255 HMM06277EutA_1_eut(pf06277) EUT2I
11260 HMMreg07568_1_eut2(pf12282pf02518pf07568) EUT2I
11265 HMMreg00072_1_eut2(pf03861pf00072) EUT2I
11270 HMM10662PduV_1_mix(pf10662) EUT2I
11275 Hp_euts_fadedPurple EUT2I
11280 HMM00465alcdh_3_mix(pf00465) EUT2I
11285 -# -
11290 -pf12294 -
11300 -pf00583 -
11305 -pf00293 -
11310 -# -
11315 -# -
11320 -# -
11330 -# -
11340 -# -
BMC-P
BMC-H
BMC-T
Aldehyde dehydrogenase
PTAC
Signature enzyme
Alcohol dehydrogenase
Regulator

Paraclostridium benzoelyticum. Bacteria; Firmicutes; Clostridia; Clostridiales;

VN21__locus_2

EUT2I

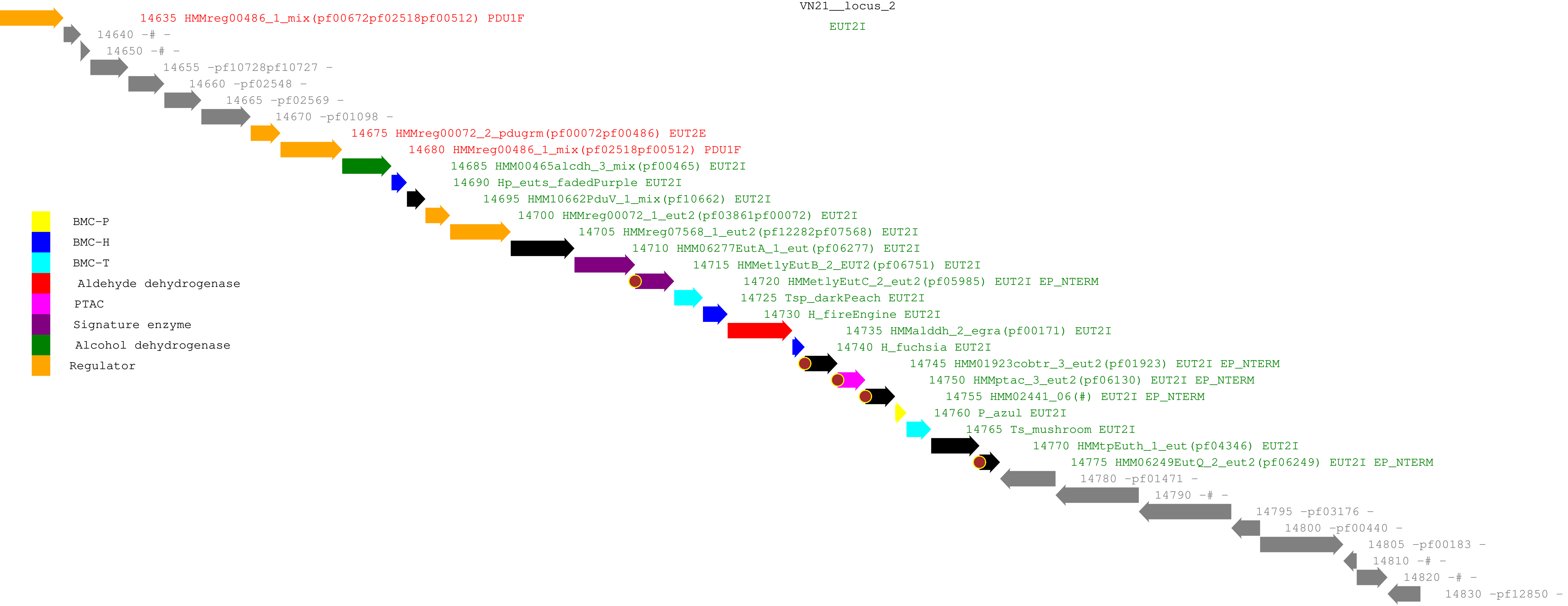

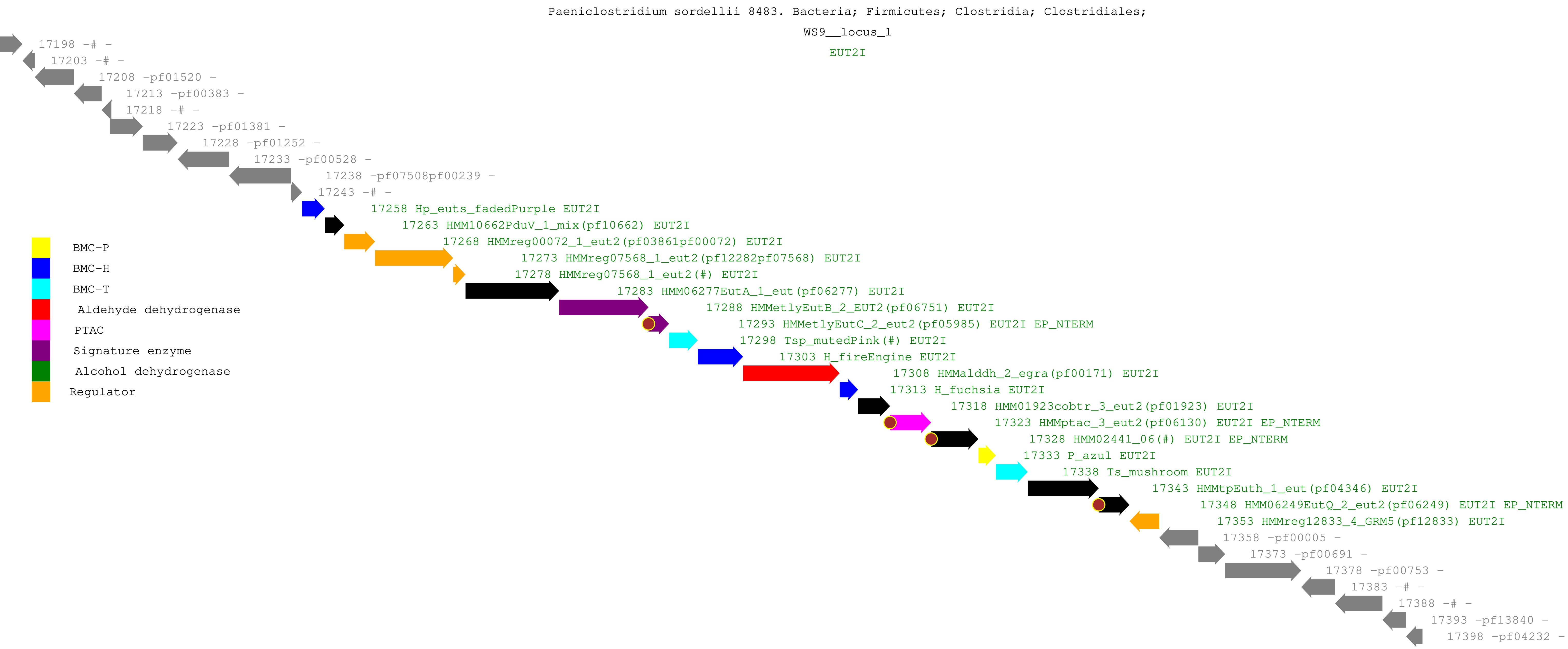
Paeniclostridium sordellii 8483. Bacteria; Firmicutes; Clostridia; Clostridiales;
WS9__locus_1
EUT2I
17198 -# -
17203 -# -
17208 -pf01520 -
17213 -pf00383 -
17218 -# -
17223 -pf01381 -
17228 -pf01252 -
17233 -pf00528 -
17238 -pf07508pf00239 -
17243 -# -
17258 Hp_euts_fadedPurple EUT2I
17263 HMM10662PduV_1_mix(pf10662) EUT2I
17268 HMMreg00072_1_eut2(pf03861pf00072) EUT2I
17273 HMMreg07568_1_eut2(pf12282pf07568) EUT2I
17278 HMMreg07568_1_eut2(#) EUT2I
17283 HMM06277EutA_1_eut(pf06277) EUT2I
17288 HMMetlyEutB_2_EUT2(pf06751) EUT2I
17293 HMMetlyEutC_2_eut2(pf05985) EUT2I EP_NTERM
17298 Tsp_mutedPink(#) EUT2I
17303 H_fireEngine EUT2I
17308 HMMalddh_2_egra(pf00171) EUT2I
17313 H_fuchsia EUT2I
17318 HMM01923cobtr_3_eut2(pf01923) EUT2I
17323 HMMptac_3_eut2(pf06130) EUT2I EP_NTERM
17328 HMM02441_06(#) EUT2I EP_NTERM
17333 P_azul EUT2I
17338 Ts_mushroom EUT2I
17343 HMMtpEuth_1_eut(pf04346) EUT2I
17348 HMM06249EutQ_2_eut2(pf06249) EUT2I EP_NTERM
17353 HMMreg12833_4_GRM5(pf12833) EUT2I
17358 -pf00005 -
17373 -pf00691 -
17378 -pf00753 -
17383 -# -
17388 -# -
17393 -pf13840 -
17398 -pf04232 -
BMC-P
BMC-H
BMC-T
Aldehyde dehydrogenase
PTAC
Signature enzyme
Alcohol dehydrogenase
Regulator

Paenibacillus sp. DMB20. Bacteria; Firmicutes; Bacilli; Bacillales; Paenibacillaceae;

XI25__locus_1

EUT2I

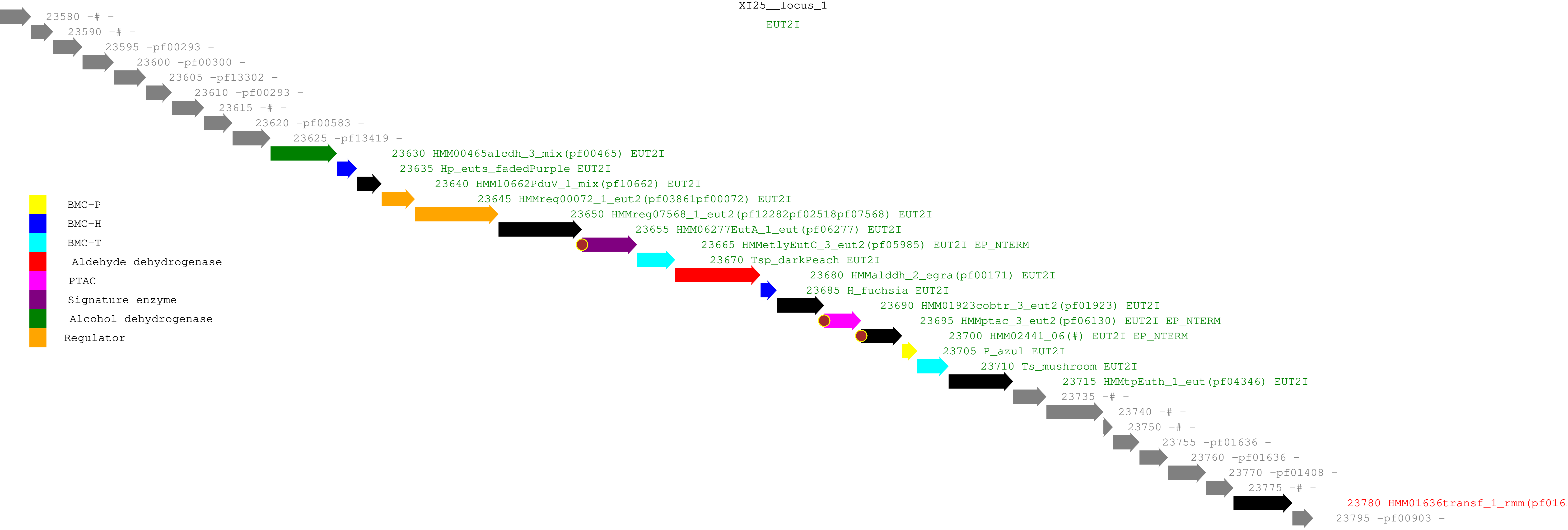

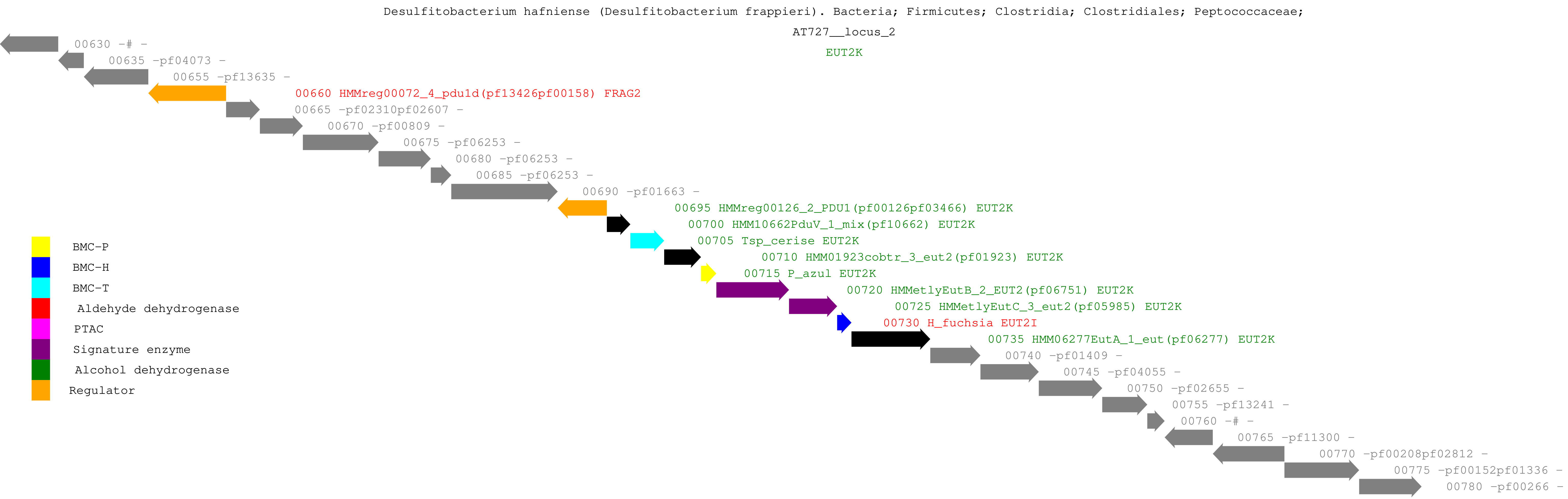
Desulfitobacterium hafniense (Desulfitobacterium frappieri). Bacteria; Firmicutes; Clostridia; Clostridiales; Peptococcaceae;
AT727__locus_2
EUT2K
00630 -# -
00635 -pf04073 -
00655 -pf13635 -
00660 HMMreg00072_4_pdu1d(pf13426pf00158) FRAG2
00665 -pf02310pf02607 -
00670 -pf00809 -
00675 -pf06253 -
00680 -pf06253 -
00685 -pf06253 -
00690 -pf01663 -
00695 HMMreg00126_2_PDU1(pf00126pf03466) EUT2K
00700 HMM10662PduV_1_mix(pf10662) EUT2K
00705 Tsp_cerise EUT2K
00710 HMM01923cobtr_3_eut2(pf01923) EUT2K
00715 P_azul EUT2K
00720 HMMetlyEutB_2_EUT2(pf06751) EUT2K
00725 HMMetlyEutC_3_eut2(pf05985) EUT2K
00730 H_fuchsia EUT2I
00735 HMM06277EutA_1_eut(pf06277) EUT2K
00740 -pf01409 -
00745 -pf04055 -
00750 -pf02655 -
00755 -pf13241 -
00760 -# -
00765 -pf11300 -
00770 -pf00208pf02812 -
00775 -pf00152pf01336 -
00780 -pf00266 -
BMC-P
BMC-H
BMC-T
Aldehyde dehydrogenase
PTAC
Signature enzyme
Alcohol dehydrogenase
Regulator

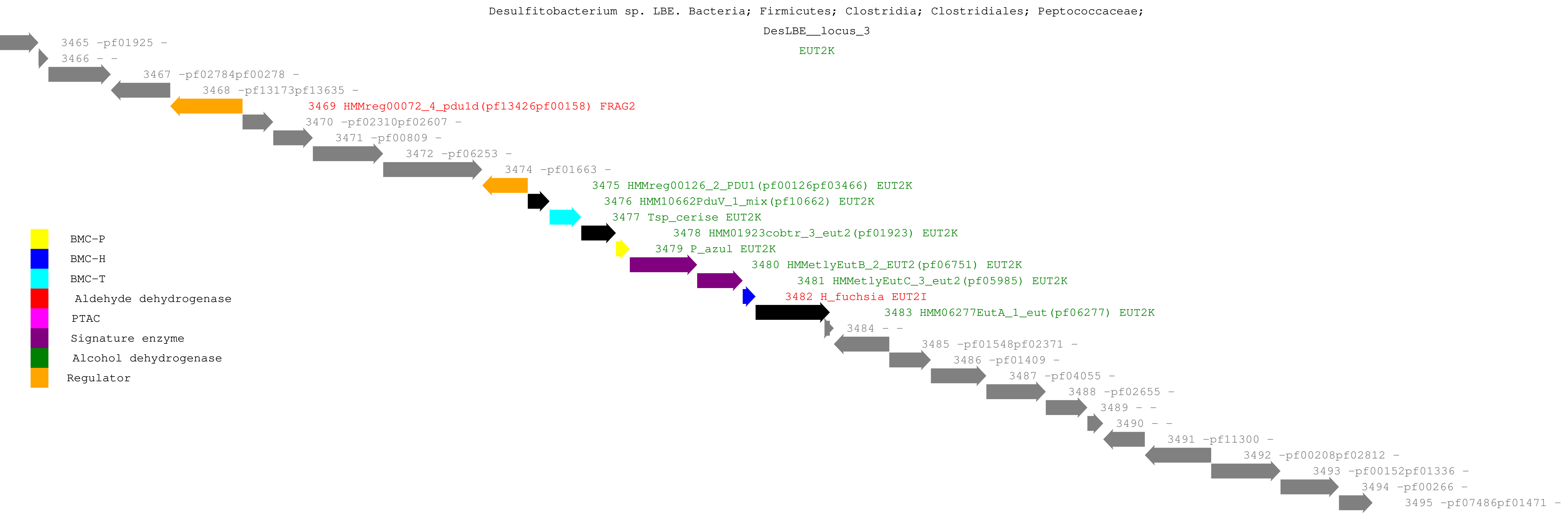
Desulfitobacterium sp. LBE. Bacteria; Firmicutes; Clostridia; Clostridiales; Peptococcaceae;
DesLBE__locus_3
EUT2K
3465 -pf01925 -
3466 - -
3467 -pf02784pf00278 -
3468 -pf13173pf13635 -
3469 HMMreg00072_4_pdu1d(pf13426pf00158) FRAG2
3470 -pf02310pf02607 -
3471 -pf00809 -
3472 -pf06253 -
3474 -pf01663 -
3475 HMMreg00126_2_PDU1(pf00126pf03466) EUT2K
3476 HMM10662PduV_1_mix(pf10662) EUT2K
3477 Tsp_cerise EUT2K
3478 HMM01923cobtr_3_eut2(pf01923) EUT2K
3479 P_azul EUT2K
3480 HMMetlyEutB_2_EUT2(pf06751) EUT2K
3481 HMMetlyEutC_3_eut2(pf05985) EUT2K
3482 H_fuchsia EUT2I
3483 HMM06277EutA_1_eut(pf06277) EUT2K
3484 - -
3485 -pf01548pf02371 -
3486 -pf01409 -
3487 -pf04055 -
3488 -pf02655 -
3489 - -
3490 - -
3491 -pf11300 -
3492 -pf00208pf02812 -
3493 -pf00152pf01336 -
3494 -pf00266 -
3495 -pf07486pf01471 -
BMC-P
BMC-H
BMC-T
Aldehyde dehydrogenase
PTAC
Signature enzyme
Alcohol dehydrogenase
Regulator

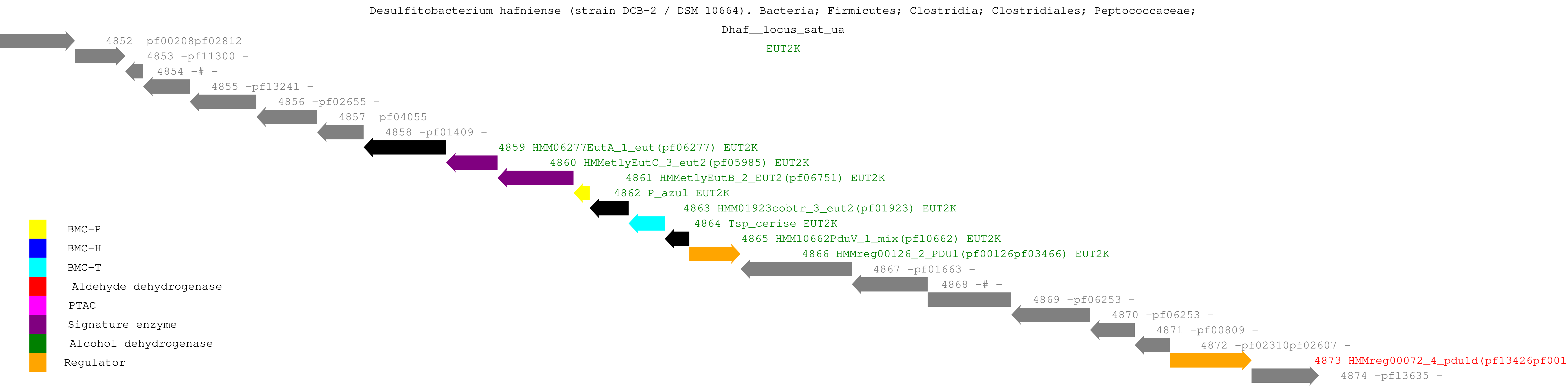
Desulfitobacterium hafniense (strain DCB-2 / DSM 10664). Bacteria; Firmicutes; Clostridia; Clostridiales; Peptococcaceae;
Dhaf__locus_sat_ua
EUT2K
4852 -pf00208pf02812 -
4853 -pf11300 -
4854 -# -
4855 -pf13241 -
4856 -pf02655 -
4857 -pf04055 -
4858 -pf01409 -
4859 HMM06277EutA_1_eut(pf06277) EUT2K
4860 HMMetlyEutC_3_eut2(pf05985) EUT2K
4861 HMMetlyEutB_2_EUT2(pf06751) EUT2K
4862 P_azul EUT2K
4863 HMM01923cobtr_3_eut2(pf01923) EUT2K
4864 Tsp_cerise EUT2K
4865 HMM10662PduV_1_mix(pf10662) EUT2K
4866 HMMreg00126_2_PDU1(pf00126pf03466) EUT2K
4867 -pf01663 -
4868 -# -
4869 -pf06253 -
4870 -pf06253 -
4871 -pf00809 -
4872 -pf02310pf02607 -
4873 HMMreg00072_4_pdu1d(pf13426pf001
4874 -pf13635 -
BMC-P
BMC-H
BMC-T
Aldehyde dehydrogenase
PTAC
Signature enzyme
Alcohol dehydrogenase
Regulator

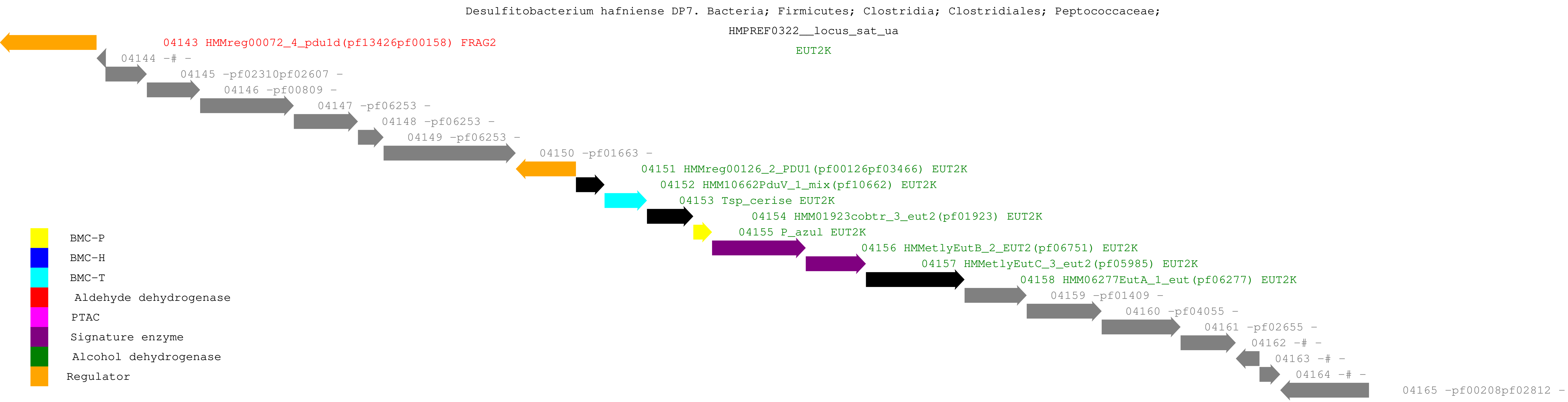
Desulfitobacterium hafniense DP7. Bacteria; Firmicutes; Clostridia; Clostridiales; Peptococcaceae;
HMPREF0322__locus_sat_ua
EUT2K
04143 HMMreg00072_4_pdu1d(pf13426pf00158) FRAG2
04144 -# -
04145 -pf02310pf02607 -
04146 -pf00809 -
04147 -pf06253 -
04148 -pf06253 -
04149 -pf06253 -
04150 -pf01663 -
04151 HMMreg00126_2_PDU1(pf00126pf03466) EUT2K
04152 HMM10662PduV_1_mix(pf10662) EUT2K
04153 Tsp_cerise EUT2K
04154 HMM01923cobtr_3_eut2(pf01923) EUT2K
04155 P_azul EUT2K
04156 HMMetlyEutB_2_EUT2(pf06751) EUT2K
04157 HMMetlyEutC_3_eut2(pf05985) EUT2K
04158 HMM06277EutA_1_eut(pf06277) EUT2K
04159 -pf01409 -
04160 -pf04055 -
04161 -pf02655 -
04162 -# -
04163 -# -
04164 -# -
04165 -pf00208pf02812 -
BMC-P
BMC-H
BMC-T
Aldehyde dehydrogenase
PTAC
Signature enzyme
Alcohol dehydrogenase
Regulator

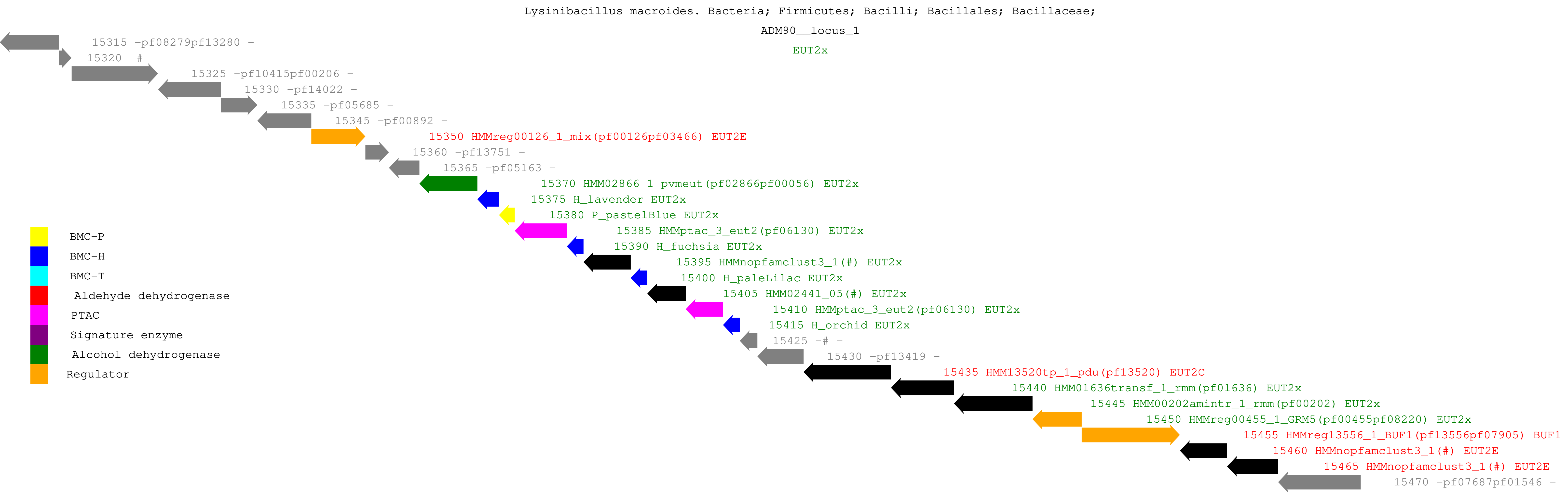
Lysinibacillus macroides. Bacteria; Firmicutes; Bacilli; Bacillales; Bacillaceae;
ADM90__locus_1
EUT2x
15315 -pf08279pf13280 -
15320 -# -
15325 -pf10415pf00206 -
15330 -pf14022 -
15335 -pf05685 -
15345 -pf00892 -
15350 HMMreg00126_1_mix(pf00126pf03466) EUT2E
15360 -pf13751 -
15365 -pf05163 -
15370 HMM02866_1_pvmeut(pf02866pf00056) EUT2x
15375 H_lavender EUT2x
15380 P_pastelBlue EUT2x
15385 HMMptac_3_eut2(pf06130) EUT2x
15390 H_fuchsia EUT2x
15395 HMMnopfamclust3_1(#) EUT2x
15400 H_paleLilac EUT2x
15405 HMM02441_05(#) EUT2x
15410 HMMptac_3_eut2(pf06130) EUT2x
15415 H_orchid EUT2x
15425 -# -
15430 -pf13419 -
15435 HMM13520tp_1_pdu(pf13520) EUT2C
15440 HMM01636transf_1_rmm(pf01636) EUT2x
15445 HMM00202amintr_1_rmm(pf00202) EUT2x
15450 HMMreg00455_1_GRM5(pf00455pf08220) EUT2x
15455 HMMreg13556_1_BUF1(pf13556pf07905) BUF1
15460 HMMnopfamclust3_1(#) EUT2E
15465 HMMnopfamclust3_1(#) EUT2E
15470 -pf07687pf01546 -
BMC-P
BMC-H
BMC-T
Aldehyde dehydrogenase
PTAC
Signature enzyme
Alcohol dehydrogenase
Regulator

Lysinibacillus fusiformis ZC1. Bacteria; Firmicutes; Bacilli; Bacillales; Bacillaceae;

BFZC1__locus_2

EUT2x

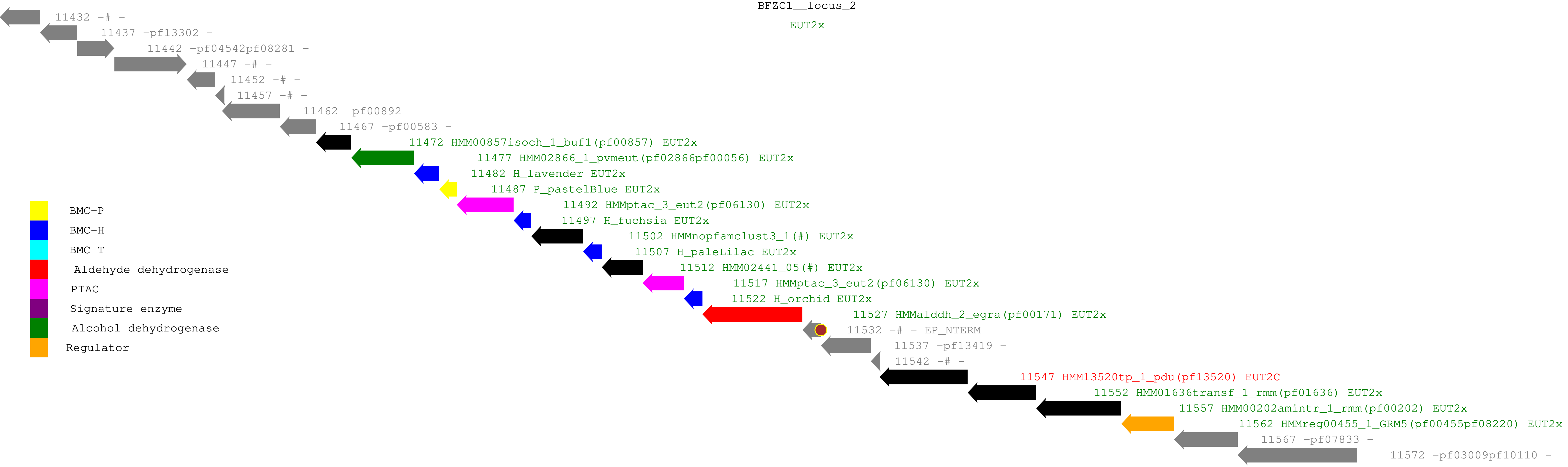

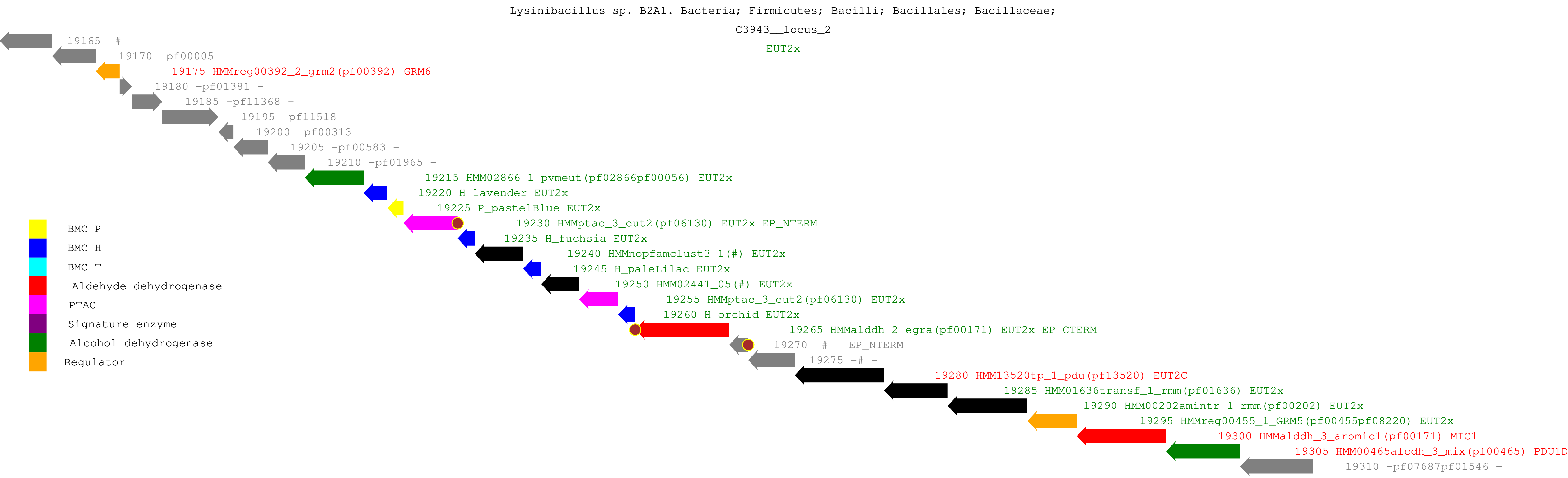
Lysinibacillus sp. B2A1. Bacteria; Firmicutes; Bacilli; Bacillales; Bacillaceae;
C3943__locus_2
EUT2x
19165 -# -
19170 -pf00005 -
19175 HMMreg00392_2_grm2(pf00392) GRM6
19180 -pf01381 -
19185 -pf11368 -
19195 -pf11518 -
19200 -pf00313 -
19205 -pf00583 -
19210 -pf01965 -
19215 HMM02866_1_pvmeut(pf02866pf00056) EUT2x
19220 H_lavender EUT2x
19225 P_pastelBlue EUT2x
19230 HMMptac_3_eut2(pf06130) EUT2x EP_NTERM
19235 H_fuchsia EUT2x
19240 HMMnopfamclust3_1(#) EUT2x
19245 H_paleLilac EUT2x
19250 HMM02441_05(#) EUT2x
19255 HMMptac_3_eut2(pf06130) EUT2x
19260 H_orchid EUT2x
19265 HMMalddh_2_egra(pf00171) EUT2x EP_CTERM
19270 -# - EP_NTERM
19275 -# -
19280 HMM13520tp_1_pdu(pf13520) EUT2C
19285 HMM01636transf_1_rmm(pf01636) EUT2x
19290 HMM00202amintr_1_rmm(pf00202) EUT2x
19295 HMMreg00455_1_GRM5(pf00455pf08220) EUT2x
19300 HMMalddh_3_aromic1(pf00171) MIC1
19305 HMM00465alcdh_3_mix(pf00465) PDU1D
19310 -pf07687pf01546 -
BMC-P
BMC-H
BMC-T
Aldehyde dehydrogenase
PTAC
Signature enzyme
Alcohol dehydrogenase
Regulator

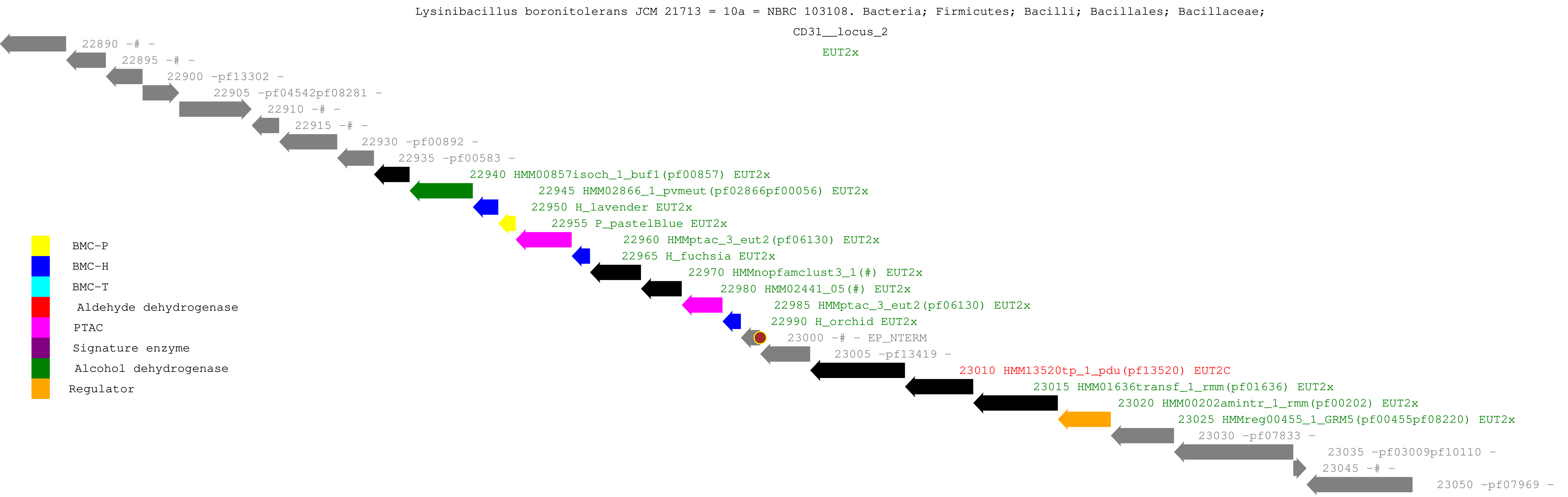
Lysinibacillus boronitolerans JCM 21713 = 10a = NBRC 103108. Bacteria; Firmicutes; Bacilli; Bacillales; Bacillaceae;
CD31__locus_2
EUT2x
22890 -# -
22895 -# -
22900 -pf13302 -
22905 -pf04542pf08281 -
22910 -# -
22915 -# -
22930 -pf00892 -
22935 -pf00583 -
22940 HMM00857isoch_1_buf1(pf00857) EUT2x
22945 HMM02866_1_pvmeut(pf02866pf00056) EUT2x
22950 H_lavender EUT2x
22955 P_pastelBlue EUT2x
22960 HMMptac_3_eut2(pf06130) EUT2x
22965 H_fuchsia EUT2x
22970 HMMnopfamclust3_1(#) EUT2x
22980 HMM02441_05(#) EUT2x
22985 HMMptac_3_eut2(pf06130) EUT2x
22990 H_orchid EUT2x
23000 -# - EP_NTERM
23005 -pf13419 -
23010 HMM13520tp_1_pdu(pf13520) EUT2C
23015 HMM01636transf_1_rmm(pf01636) EUT2x
23020 HMM00202amintr_1_rmm(pf00202) EUT2x
23025 HMMreg00455_1_GRM5(pf00455pf08220) EUT2x
23030 -pf07833 -
23035 -pf03009pf10110 -
23045 -# -
23050 -pf07969 -
BMC-P
BMC-H
BMC-T
Aldehyde dehydrogenase
PTAC
Signature enzyme
Alcohol dehydrogenase
Regulator

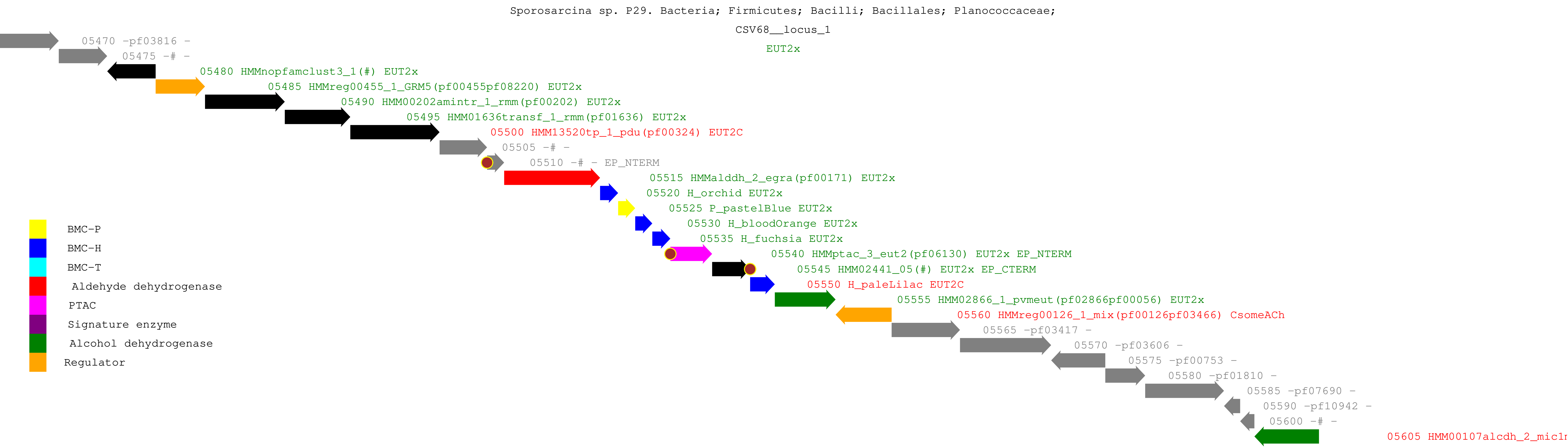
Sporosarcina sp. P29. Bacteria; Firmicutes; Bacilli; Bacillales; Planococcaceae;
CSV68__locus_1
EUT2x
05470 -pf03816 -
05475 -# -
05480 HMMnopfamclust3_1(#) EUT2x
05485 HMMreg00455_1_GRM5(pf00455pf08220) EUT2x
05490 HMM00202amintr_1_rmm(pf00202) EUT2x
05495 HMM01636transf_1_rmm(pf01636) EUT2x
05500 HMM13520tp_1_pdu(pf00324) EUT2C
05505 -# -
05510 -# - EP_NTERM
05515 HMMalddh_2_egra(pf00171) EUT2x
05520 H_orchid EUT2x
05525 P_pastelBlue EUT2x
05530 H_bloodOrange EUT2x
05535 H_fuchsia EUT2x
05540 HMMptac_3_eut2(pf06130) EUT2x EP_NTERM
05545 HMM02441_05(#) EUT2x EP_CTERM
05550 H_paleLilac EUT2C
05555 HMM02866_1_pvmeut(pf02866pf00056) EUT2x
05560 HMMreg00126_1_mix(pf00126pf03466) CsomeACh
05565 -pf03417 -
05570 -pf03606 -
05575 -pf00753 -
05580 -pf01810 -
05585 -pf07690 -
05590 -pf10942 -
05600 -# -
05605 HMM00107alcdh_2_mic1r
BMC-P
BMC-H
BMC-T
Aldehyde dehydrogenase
PTAC
Signature enzyme
Alcohol dehydrogenase
Regulator

Sporosarcina sp. P16b. Bacteria; Firmicutes; Bacilli; Bacillales; Planococcaceae;

CSV77__locus_1

EUT2x

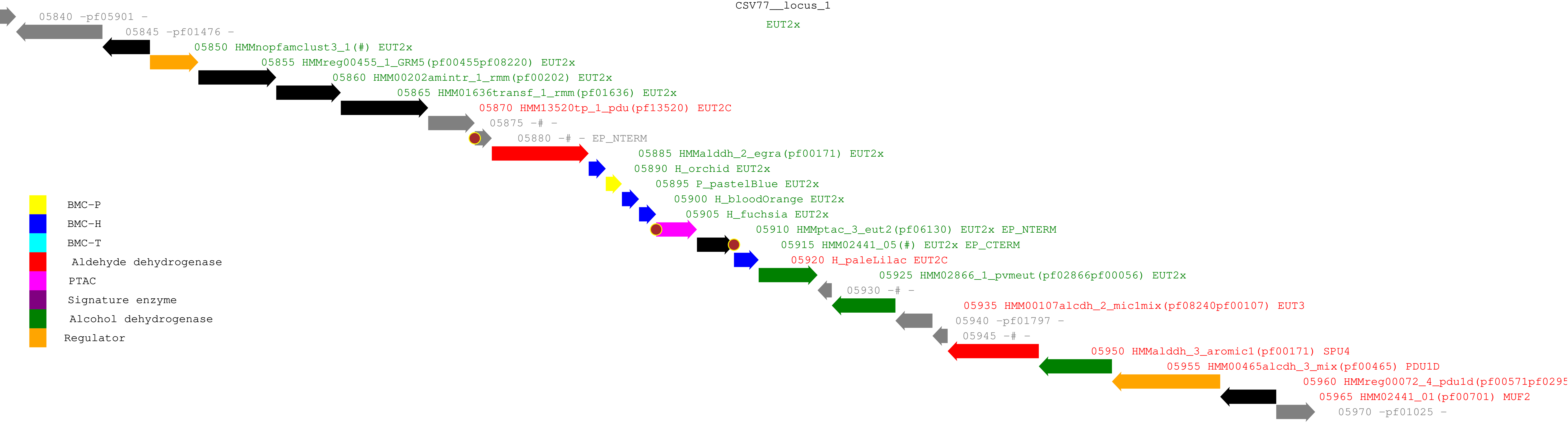

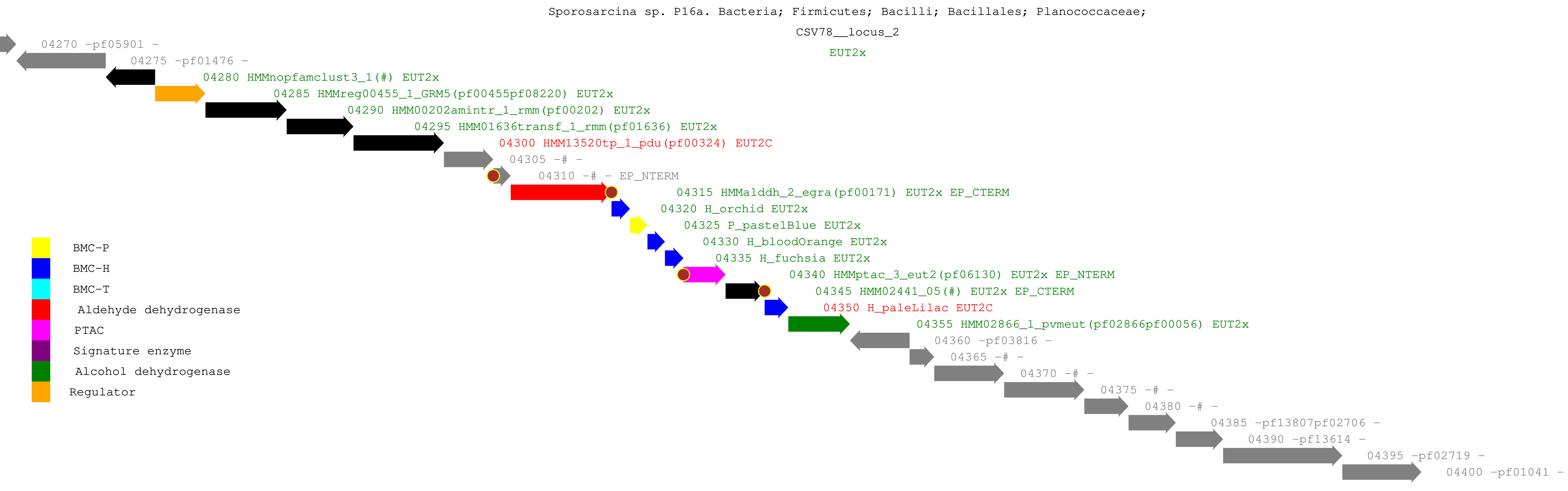

Sporosarcina sp. P16a. Bacteria; Firmicutes; Bacilli; Bacillales; Planococcaceae;
CSV78__locus_2
EUT2x
04270 -pf05901 -
04275 -pf01476 -
04280 HMMnopfamclust3_1(#) EUT2x
04285 HMMreg00455_1_GRM5(pf00455pf08220) EUT2x
04290 HMM00202amintr_1_rmm(pf00202) EUT2x
04295 HMM01636transf_1_rmm(pf01636) EUT2x
04300 HMM13520tp_1_pdu(pf00324) EUT2C
04305 -# -
04310 -# - EP_NTERM
04315 HMMaldd h_2_egra(pf00171) EUT2x EP_CTERM
04320 H_orchid EUT2x
04325 P_pastelBlue EUT2x
04330 H_bloodOrange EUT2x
04335 H_fuchsia EUT2x
04340 HMMptac_3_eut2(pf06130) EUT2x EP_NTERM
04345 HMM02441_05(#) EUT2x EP_CTERM
04350 H_paleLilac EUT2C
04355 HMM02866_1_pvmeut(pf02866pf00056) EUT2x
04360 -pf03816 -
04365 -# -
04370 -# -
04375 -# -
04380 -# -
04385 -pf13807pf02706 -
04390 -pf13614 -
04395 -pf02719 -
04400 -pf01041 -
BMC-P
BMC-H
BMC-T
Aldehyde dehydrogenase
PTAC
Signature enzyme
Alcohol dehydrogenase
Regulator

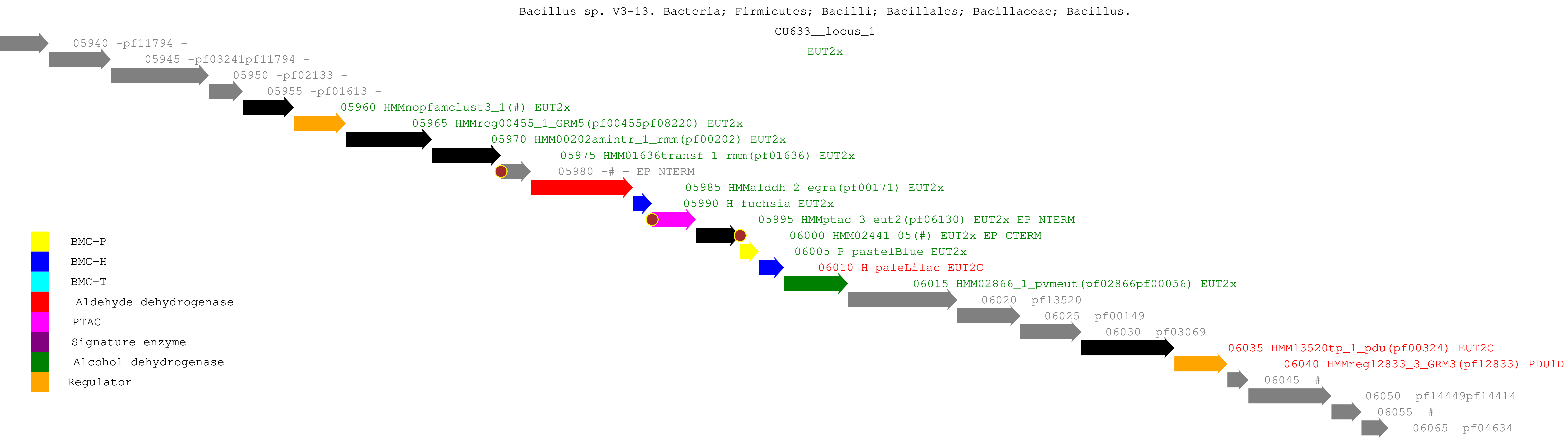

Bacillus sp. V3-13. Bacteria; Firmicutes; Bacilli; Bacillales; Bacillaceae; Bacillus.
CU633__locus_1
EUT2x
05940 -pf11794 -
05945 -pf03241pf11794 -
05950 -pf02133 -
05955 -pf01613 -
05960 HMMnopfamclust3_1(#) EUT2x
05965 HMMreg00455_1_GRM5(pf00455pf08220) EUT2x
05970 HMM00202amintr_1_rmm(pf00202) EUT2x
05975 HMM01636transf_1_rmm(pf01636) EUT2x
05980 -# - EP_NTERM
05985 HMMaldh_2_egra(pf00171) EUT2x
05990 H_fuchsia EUT2x
05995 HMMptac_3_eut2(pf06130) EUT2x EP_NTERM
06000 HMM02441_05(#) EUT2x EP_CTERM
06005 P_pastelBlue EUT2x
06010 H_paleLilac EUT2C
06015 HMM02866_1_pvmeut(pf02866pf00056) EUT2x
06020 -pf13520 -
06025 -pf00149 -
06030 -pf03069 -
06035 HMM13520tp_1_pdu(pf00324) EUT2C
06040 HMMreg12833_3_GRM3(pf12833) PDU1D
06045 -# -
06050 -pf14449pf14414 -
06055 -# -
06065 -pf04634 -
BMC-P
BMC-H
BMC-T
Aldehyde dehydrogenase
PTAC
Signature enzyme
Alcohol dehydrogenase
Regulator

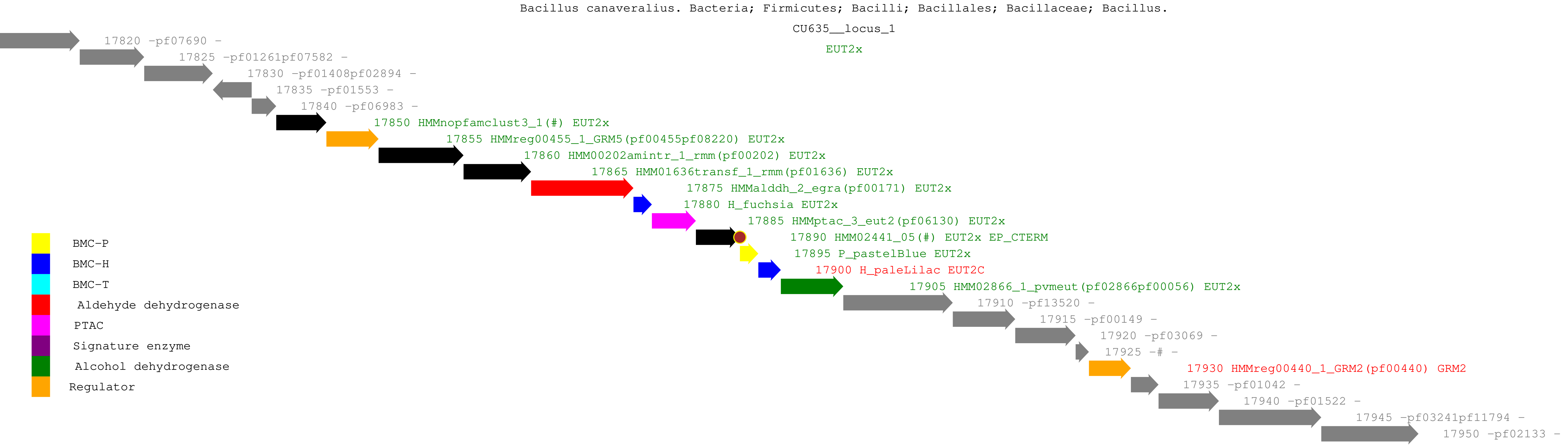

Bacillus canaveralius. Bacteria; Firmicutes; Bacilli; Bacillales; Bacillaceae; Bacillus.
CU635__locus_1
EUT2x
17820 -pf07690 -
17825 -pf01261pf07582 -
17830 -pf01408pf02894 -
17835 -pf01553 -
17840 -pf06983 -
17850 HMMnopfamclust3_1(#) EUT2x
17855 HMMreg00455_1_GRM5(pf00455pf08220) EUT2x
17860 HMM00202amintr_1_rmm(pf00202) EUT2x
17865 HMM01636transf_1_rmm(pf01636) EUT2x
17875 HMMalddh_2_egra(pf00171) EUT2x
17880 H_fuchsia EUT2x
17885 HMMptac_3_eut2(pf06130) EUT2x
17890 HMM02441_05(#) EUT2x EP_CTERM
17895 P_pastelBlue EUT2x
17900 H_paleLilac EUT2C
17905 HMM02866_1_pvmeut(pf02866pf00056) EUT2x
17910 -pf13520 -
17915 -pf00149 -
17920 -pf03069 -
17925 -# -
17930 HMMreg00440_1_GRM2(pf00440) GRM2
17935 -pf01042 -
17940 -pf01522 -
17945 -pf03241pf11794 -
17950 -pf02133 -
BMC-P
BMC-H
BMC-T
Aldehyde dehydrogenase
PTAC
Signature enzyme
Alcohol dehydrogenase
Regulator

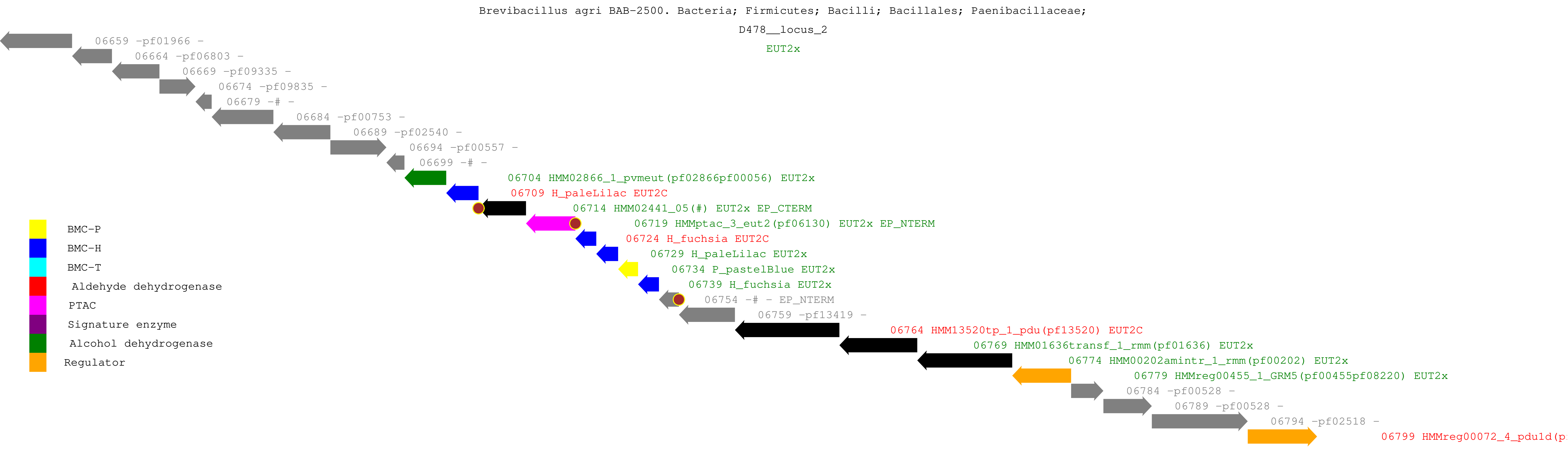
Brevibacillus agri BAB-2500. Bacteria; Firmicutes; Bacilli; Bacillales; Paenibacillaceae;
D478__locus_2
EUT2x
06659 -pf01966 -
06664 -pf06803 -
06669 -pf09335 -
06674 -pf09835 -
06679 -# -
06684 -pf00753 -
06689 -pf02540 -
06694 -pf00557 -
06699 -# -
06704 HMM02866_1_pvmeut(pf02866pf00056) EUT2x
06709 H_paleLilac EUT2C
06714 HMM02441_05(#) EUT2x EP_CTERM
06719 HMMptac_3_eut2(pf06130) EUT2x EP_NTERM
06724 H_fuchsia EUT2C
06729 H_paleLilac EUT2x
06734 P_pastelBlue EUT2x
06739 H_fuchsia EUT2x
06754 -# - EP_NTERM
06759 -pf13419 -
06764 HMM13520tp_1_pdu(pf13520) EUT2C
06769 HMM01636transf_1_rmm(pf01636) EUT2x
06774 HMM00202amintr_1_rmm(pf00202) EUT2x
06779 HMMreg00455_1_GRM5(pf00455pf08220) EUT2x
06784 -pf00528 -
06789 -pf00528 -
06794 -pf02518 -
06799 HMMreg00072_4_pdu1d(p
BMC-P
BMC-H
BMC-T
Aldehyde dehydrogenase
PTAC
Signature enzyme
Alcohol dehydrogenase
Regulator

Pueribacillus theae. Bacteria; Firmicutes; Bacilli; Bacillales; Bacillaceae; Pueribacillus.

DCC39__locus_1

EUT2x

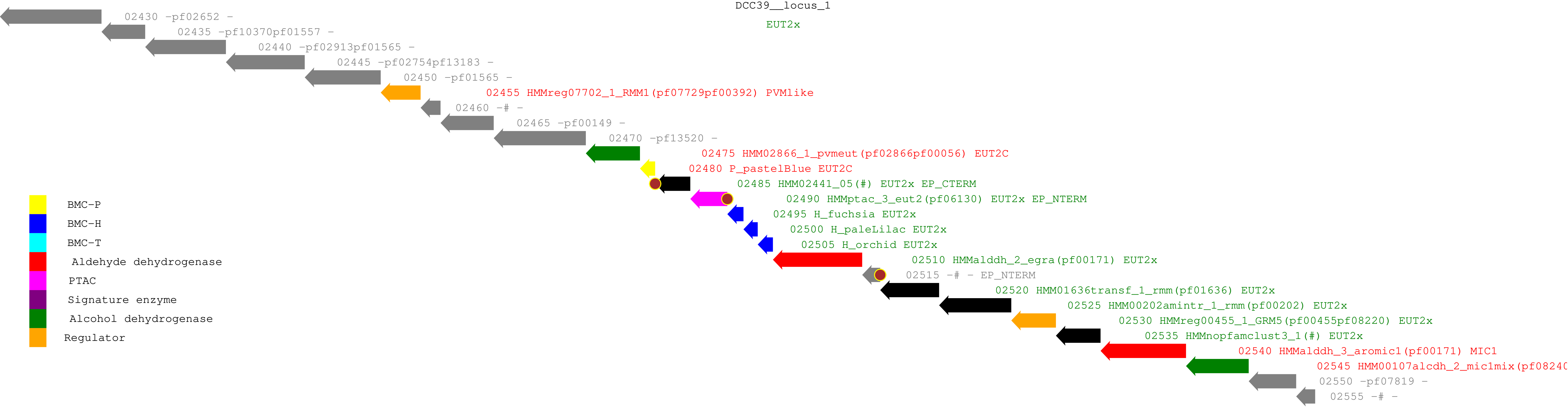

Pseudogracilibacillus auburnensis. Bacteria; Firmicutes; Bacilli; Bacillales; Bacillaceae;

DFR56__locus_3

EUT2x

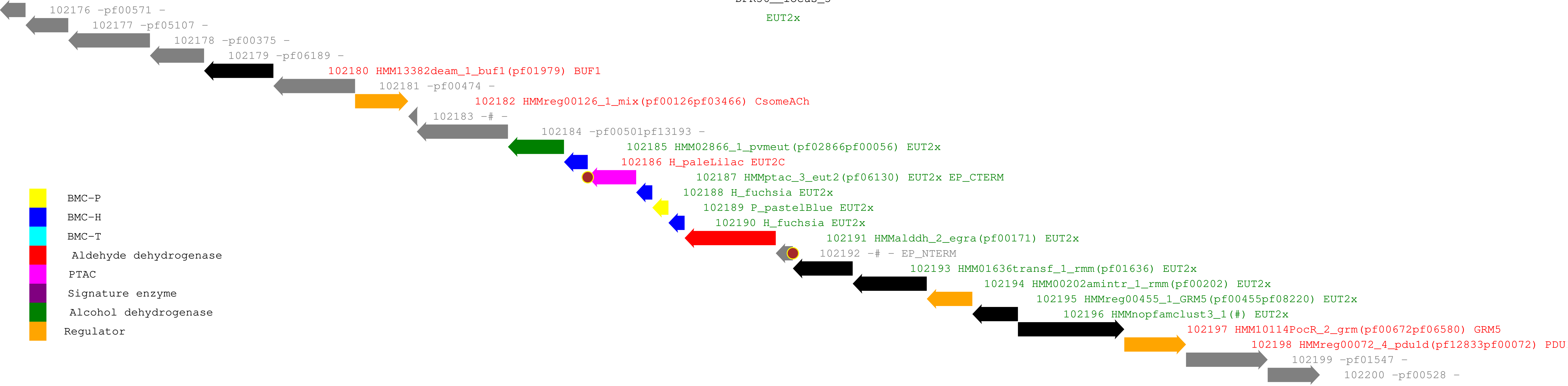

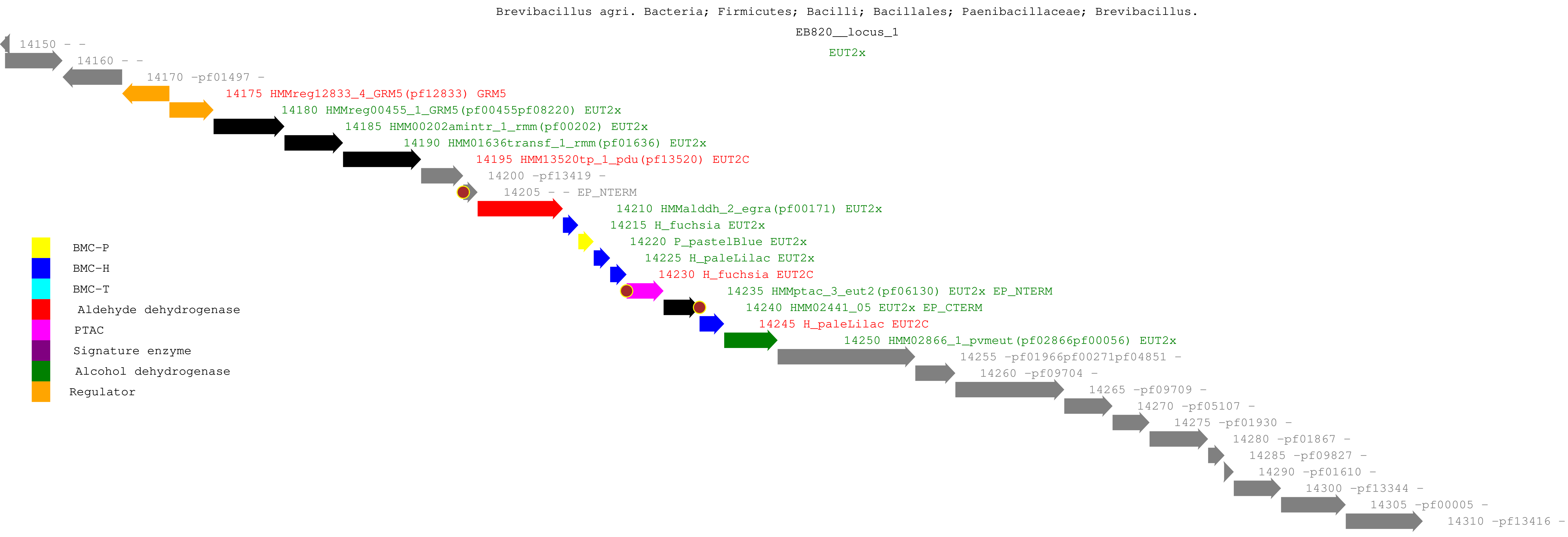
Brevibacillus agri. Bacteria; Firmicutes; Bacilli; Bacillales; Paenibacillaceae; Brevibacillus.
EB820__locus_1
EUT2x
14150 - -
14160 - -
14170 -pf01497 -
14175 HMMreg12833_4_GRM5(pf12833) GRM5
14180 HMMreg00455_1_GRM5(pf00455pf08220) EUT2x
14185 HMM00202amintr_1_rmm(pf00202) EUT2x
14190 HMM01636transf_1_rmm(pf01636) EUT2x
14195 HMM13520tp_1_pdu(pf13520) EUT2C
14200 -pf13419 -
14205 - - EP_NTERM
14210 HMMalddh_2_egra(pf00171) EUT2x
14215 H_fuchsia EUT2x
14220 P_pastelBlue EUT2x
14225 H_paleLilac EUT2x
14230 H_fuchsia EUT2C
14235 HMMptac_3_eut2(pf06130) EUT2x EP_NTERM
14240 HMM02441_05 EUT2x EP_CTERM
14245 H_paleLilac EUT2C
14250 HMM02866_1_pvmeut(pf02866pf00056) EUT2x
14255 -pf01966pf00271pf04851 -
14260 -pf09704 -
14265 -pf09709 -
14270 -pf05107 -
14275 -pf01930 -
14280 -pf01867 -
14285 -pf09827 -
14290 -pf01610 -
14300 -pf13344 -
14305 -pf00005 -
14310 -pf13416 -
BMC-P
BMC-H
BMC-T
Aldehyde dehydrogenase
PTAC
Signature enzyme
Alcohol dehydrogenase
Regulator

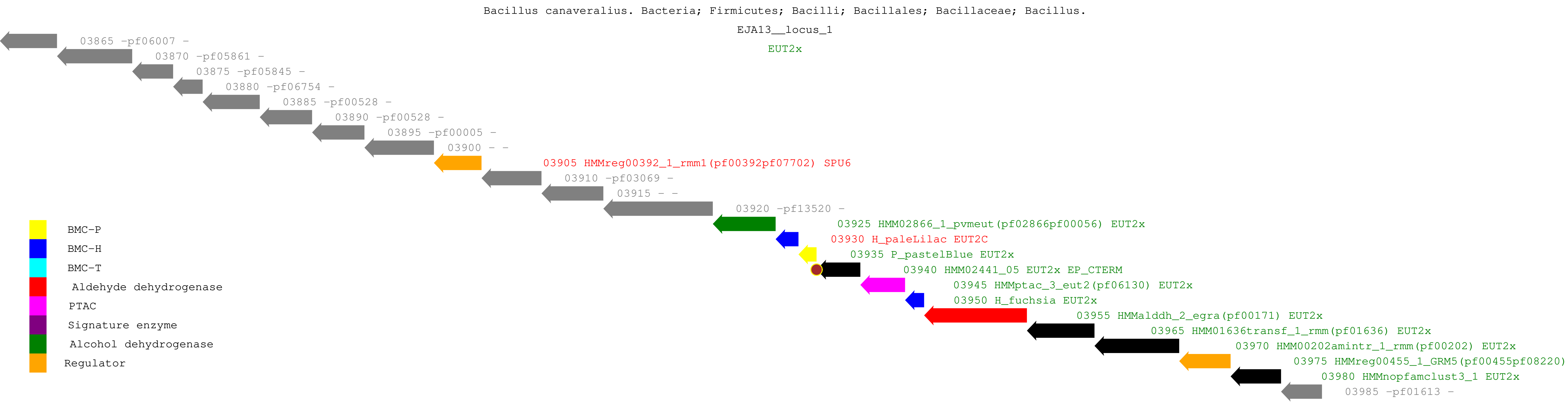

Bacillus canaveralius. Bacteria; Firmicutes; Bacilli; Bacillales; Bacillaceae; Bacillus.
EJA13__locus_1
EUT2x
03865 -pf06007 -
03870 -pf05861 -
03875 -pf05845 -
03880 -pf06754 -
03885 -pf00528 -
03890 -pf00528 -
03895 -pf00005 -
03900 - -
03905 HMMreg00392_1_rmm1(pf00392pf07702) SPU6
03910 -pf03069 -
03915 - -
03920 -pf13520 -
03925 HMM02866_1_pvmeut(pf02866pf00056) EUT2x
03930 H_paleLilac EUT2C
03935 P_pastelBlue EUT2x
03940 HMM02441_05 EUT2x EP_CTERM
03945 HMMptac_3_eut2(pf06130) EUT2x
03950 H_fuchsia EUT2x
03955 HMMaldh_2_egra(pf00171) EUT2x
03965 HMM01636transf_1_rmm(pf01636) EUT2x
03970 HMM00202amintr_1_rmm(pf00202) EUT2x
03975 HMMreg00455_1_GRM5(pf00455pf08220)
03980 HMMnopfamclust3_1 EUT2x
03985 -pf01613 -
BMC-P
BMC-H
BMC-T
Aldehyde dehydrogenase
PTAC
Signature enzyme
Alcohol dehydrogenase
Regulator

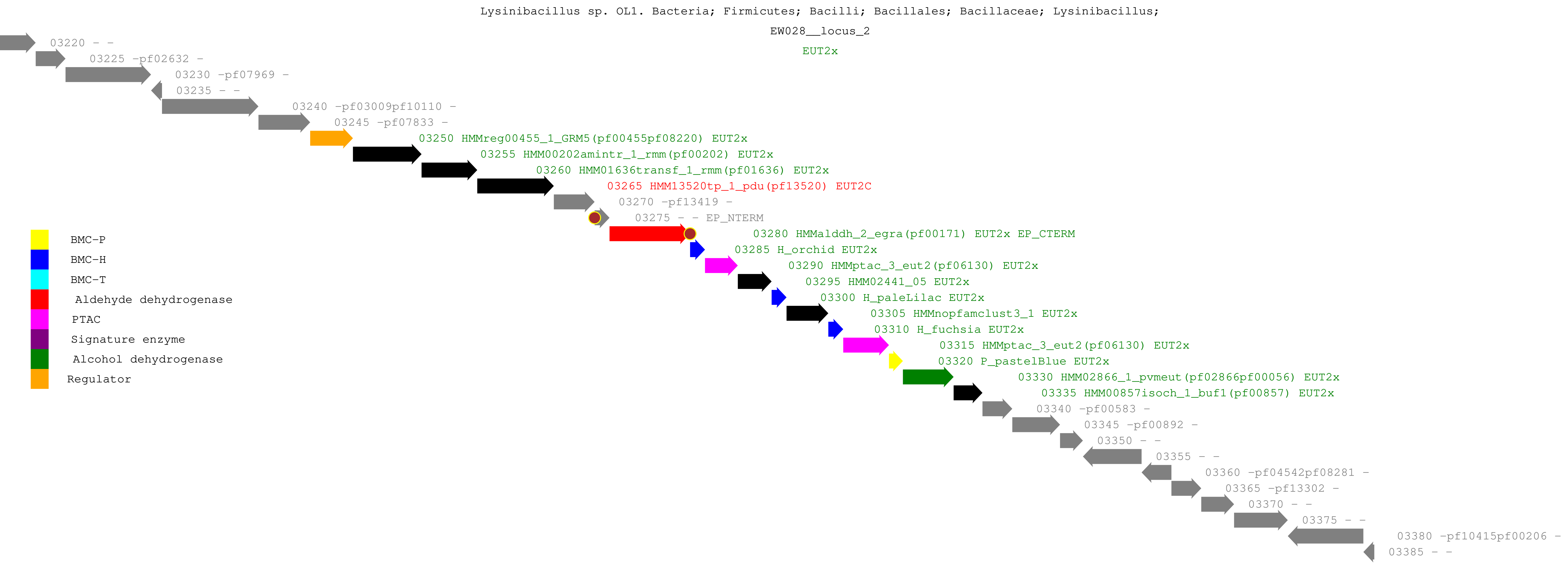

Lysinibacillus sp. OL1. Bacteria; Firmicutes; Bacilli; Bacillales; Bacillaceae; Lysinibacillus;
EW028__locus_2
EUT2x
03220 - -
03225 -pf02632 -
03230 -pf07969 -
03235 - -
03240 -pf03009pf10110 -
03245 -pf07833 -
03250 HMMreg00455_1_GRM5(pf00455pf08220) EUT2x
03255 HMM00202amintr_1_rmm(pf00202) EUT2x
03260 HMM01636transf_1_rmm(pf01636) EUT2x
03265 HMM13520tp_1_pdu(pf13520) EUT2C
03270 -pf13419 -
03275 - - EP_NTERM
03280 HMMalddh_2_egra(pf00171) EUT2x EP_CTERM
03285 H_orchid EUT2x
03290 HMMptac_3_eut2(pf06130) EUT2x
03295 HMM02441_05 EUT2x
03300 H_paleLilac EUT2x
03305 HMMnopfamclust3_1 EUT2x
03310 H_fuchsia EUT2x
03315 HMMptac_3_eut2(pf06130) EUT2x
03320 P_pastelBlue EUT2x
03330 HMM02866_1_pvmeut(pf02866pf00056) EUT2x
03335 HMM00857isoch_1_buf1(pf00857) EUT2x
03340 -pf00583 -
03345 -pf00892 -
03350 - -
03355 - -
03360 -pf04542pf08281 -
03365 -pf13302 -
03370 - -
03375 - -
03380 -pf10415pf00206 -
03385 - -
BMC-P
BMC-H
BMC-T
Aldehyde dehydrogenase
PTAC
Signature enzyme
Alcohol dehydrogenase
Regulator

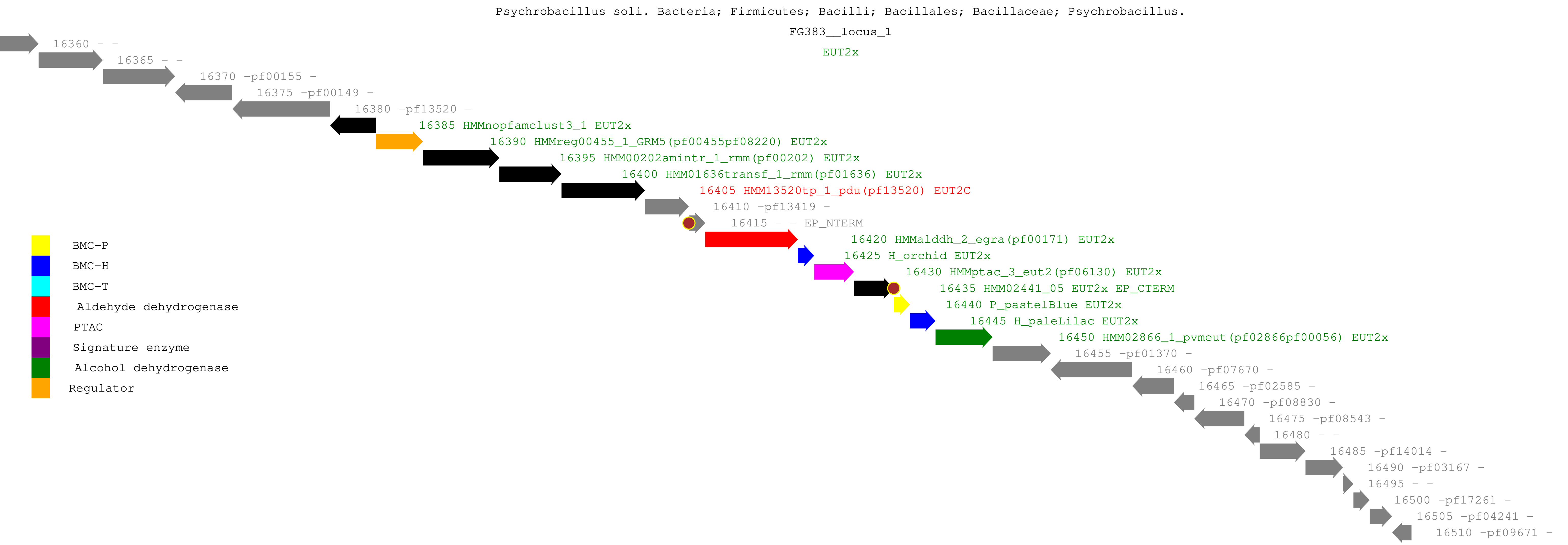

Psychrobacillus soli. Bacteria; Firmicutes; Bacilli; Bacillales; Bacillaceae; Psychrobacillus.
FG383__locus_1
EUT2x
16360 - -
16365 - -
16370 -pf00155 -
16375 -pf00149 -
16380 -pf13520 -
16385 HMMnopfamclust3_1 EUT2x
16390 HMMreg00455_1_GRM5(pf00455pf08220) EUT2x
16395 HMM00202amintr_1_rmm(pf00202) EUT2x
16400 HMM01636transf_1_rmm(pf01636) EUT2x
16405 HMM13520tp_1_pdu(pf13520) EUT2C
16410 -pf13419 -
16415 - - EP_NTERM
16420 HMMaldh_2_egra(pf00171) EUT2x
16425 H_orchid EUT2x
16430 HMMptac_3_eut2(pf06130) EUT2x
16435 HMM02441_05 EUT2x EP_CTERM
16440 P_pastelBlue EUT2x
16445 H_paleLilac EUT2x
16450 HMM02866_1_pvmeut(pf02866pf00056) EUT2x
16455 -pf01370 -
16460 -pf07670 -
16465 -pf02585 -
16470 -pf08830 -
16475 -pf08543 -
16480 - -
16485 -pf14014 -
16490 -pf03167 -
16495 - -
16500 -pf17261 -
16505 -pf04241 -
16510 -pf09671 -
BMC-P
BMC-H
BMC-T
Aldehyde dehydrogenase
PTAC
Signature enzyme
Alcohol dehydrogenase
Regulator

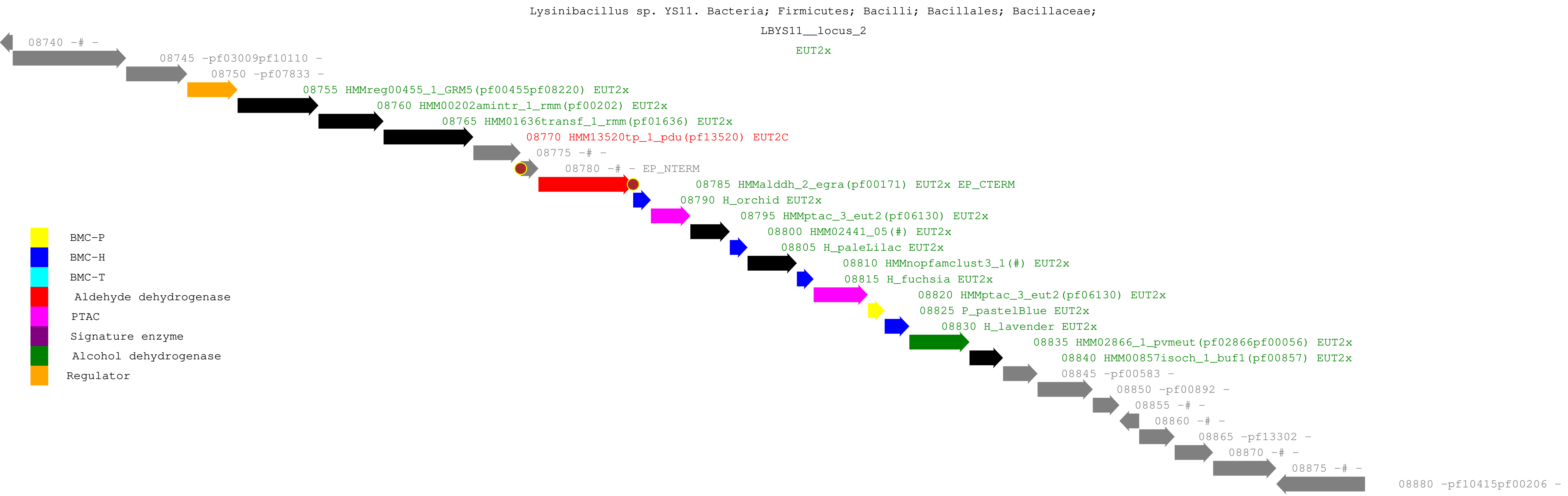

Lysinibacillus sp. YS11. Bacteria; Firmicutes; Bacilli; Bacillales; Bacillaceae;
LBYS11__locus_2
EUT2x
08740 -# -
08745 -pf03009pf10110 -
08750 -pf07833 -
08755 HMMreg00455_1_GRM5(pf00455pf08220) EUT2x
08760 HMM00202amintr_1_rmm(pf00202) EUT2x
08765 HMM01636transf_1_rmm(pf01636) EUT2x
08770 HMM13520tp_1_pdu(pf13520) EUT2C
08775 -# -
08780 -# - EP_NTERM
08785 HMMalddh_2_egra(pf00171) EUT2x EP_CTERM
08790 H_orchid EUT2x
08795 HMMptac_3_eut2(pf06130) EUT2x
08800 HMM02441_05(#) EUT2x
08805 H_paleLilac EUT2x
08810 HMMnopfamclust3_1(#) EUT2x
08815 H_fuchsia EUT2x
08820 HMMptac_3_eut2(pf06130) EUT2x
08825 P_pastelBlue EUT2x
08830 H_lavender EUT2x
08835 HMM02866_1_pvmeut(pf02866pf00056) EUT2x
08840 HMM00857isoch_1_buf1(pf00857) EUT2x
08845 -pf00583 -
08850 -pf00892 -
08855 -# -
08860 -# -
08865 -pf13302 -
08870 -# -
08875 -# -
08880 -pf10415pf00206 -
BMC-P
BMC-H
BMC-T
Aldehyde dehydrogenase
PTAC
Signature enzyme
Alcohol dehydrogenase
Regulator

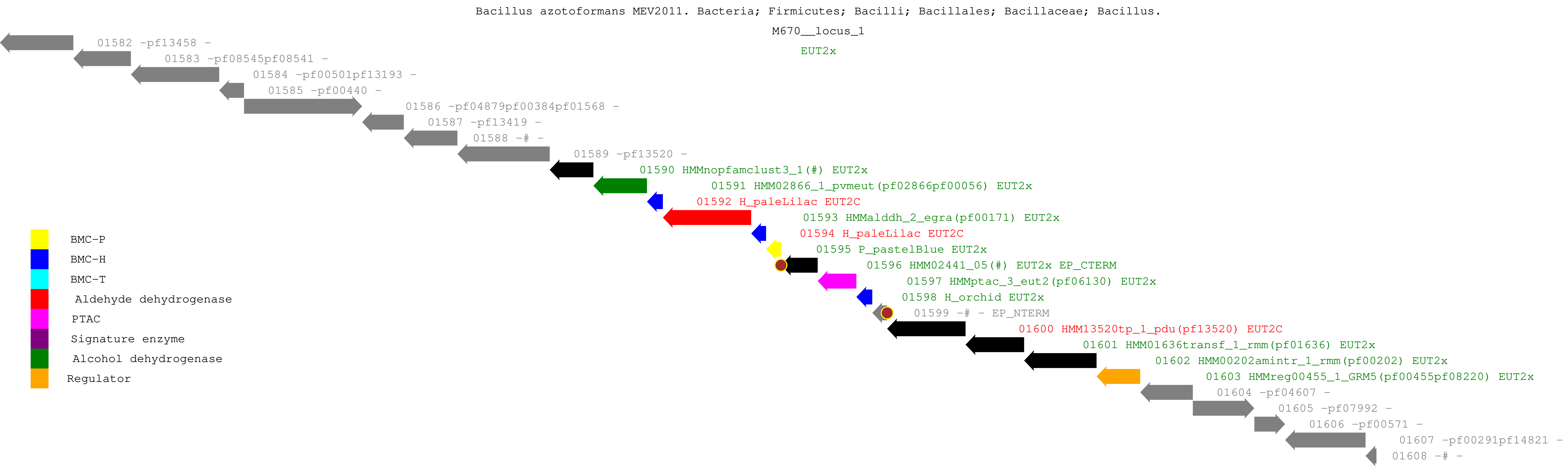
Bacillus azotoformans MEV2011. Bacteria; Firmicutes; Bacilli; Bacillales; Bacillaceae; Bacillus.
M670__locus_1
EUT2x
01582 -pf13458 -
01583 -pf08545pf08541 -
01584 -pf00501pf13193 -
01585 -pf00440 -
01586 -pf04879pf00384pf01568 -
01587 -pf13419 -
01588 -# -
01589 -pf13520 -
01590 HMMnopfamclust3_1(#) EUT2x
01591 HMM02866_1_pvmeut(pf02866pf00056) EUT2x
01592 H_paleLilac EUT2C
01593 HMMaldh_2_egra(pf00171) EUT2x
01594 H_paleLilac EUT2C
01595 P_pastelBlue EUT2x
01596 HMM02441_05(#) EUT2x EP_CTERM
01597 HMMptac_3_eut2(pf06130) EUT2x
01598 H_orchid EUT2x
01599 -# - EP_NTERM
01600 HMM13520tp_1_pdu(pf13520) EUT2C
01601 HMM01636transf_1_rmm(pf01636) EUT2x
01602 HMM00202amintr_1_rmm(pf00202) EUT2x
01603 HMMreg00455_1_GRM5(pf00455pf08220) EUT2x
01604 -pf04607 -
01605 -pf07992 -
01606 -pf00571 -
01607 -pf00291pf14821 -
01608 -# -
BMC-P
BMC-H
BMC-T
Aldehyde dehydrogenase
PTAC
Signature enzyme
Alcohol dehydrogenase
Regulator

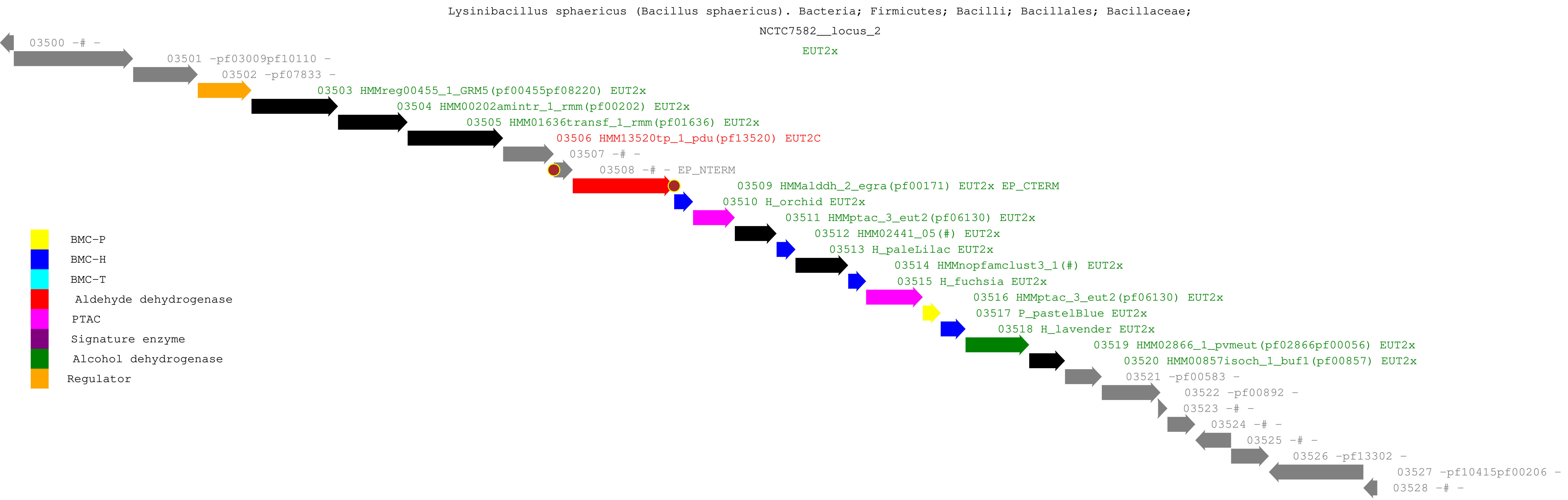
Lysinibacillus sphaericus (Bacillus sphaericus). Bacteria; Firmicutes; Bacilli; Bacillales; Bacillaceae;
NCTC7582__locus_2
EUT2x
03500 -# -
03501 -pf03009pf10110 -
03502 -pf07833 -
03503 HMMreg00455_1_GRM5(pf00455pf08220) EUT2x
03504 HMM00202amintr_1_rmm(pf00202) EUT2x
03505 HMM01636transf_1_rmm(pf01636) EUT2x
03506 HMM13520tp_1_pdu(pf13520) EUT2C
03507 -# -
03508 -# - EP_NTERM
03509 HMMaldd_2_egra(pf00171) EUT2x EP_CTERM
03510 H_orchid EUT2x
03511 HMMptac_3_eut2(pf06130) EUT2x
03512 HMM02441_05(#) EUT2x
03513 H_paleLilac EUT2x
03514 HMMnopfamclust3_1(#) EUT2x
03515 H_fuchsia EUT2x
03516 HMMptac_3_eut2(pf06130) EUT2x
03517 P_pastelBlue EUT2x
03518 H_lavender EUT2x
03519 HMM02866_1_pvmeut(pf02866pf00056) EUT2x
03520 HMM00857isoch_1_buf1(pf00857) EUT2x
03521 -pf00583 -
03522 -pf00892 -
03523 -# -
03524 -# -
03525 -# -
03526 -pf13302 -
03527 -pf10415pf00206 -
03528 -# -
BMC-P
BMC-H
BMC-T
Aldehyde dehydrogenase
PTAC
Signature enzyme
Alcohol dehydrogenase
Regulator

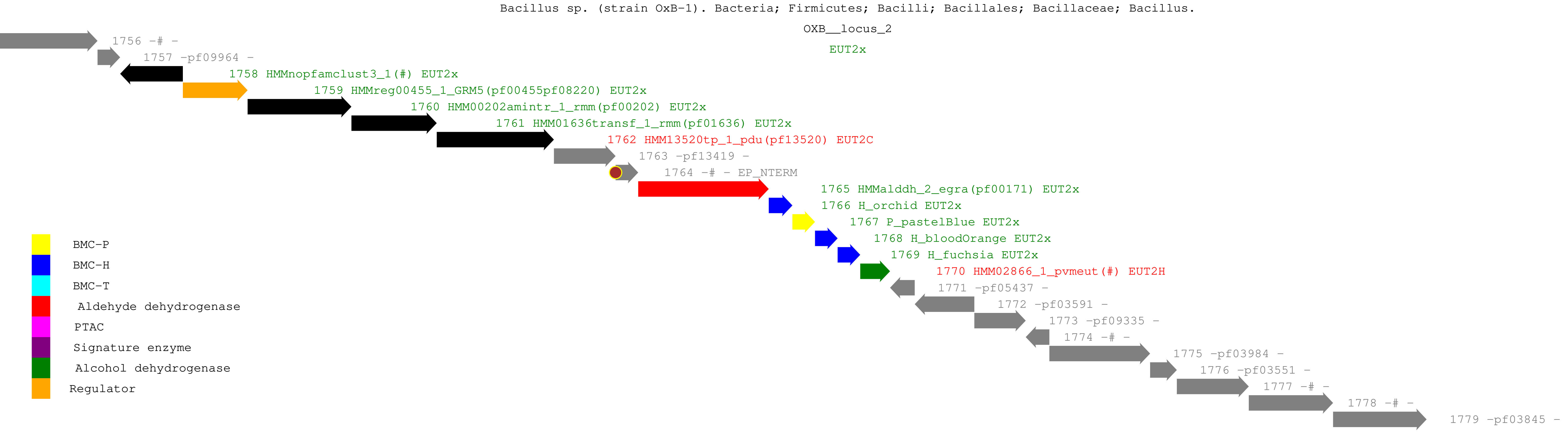
Bacillus sp. (strain OxB-1). Bacteria; Firmicutes; Bacilli; Bacillales; Bacillaceae; Bacillus.
OXB__locus_2
EUT2x
1756 -# -
1757 -pf09964 -
1758 HMMnopfamclust3_1(#) EUT2x
1759 HMMreg00455_1_GRM5(pf00455pf08220) EUT2x
1760 HMM00202amintr_1_rmm(pf00202) EUT2x
1761 HMM01636transf_1_rmm(pf01636) EUT2x
1762 HMM13520tp_1_pdu(pf13520) EUT2C
1763 -pf13419 -
1764 -# - EP_NTERM
1765 HMMaldd_2_egra(pf00171) EUT2x
1766 H_orchid EUT2x
1767 P_pastelBlue EUT2x
1768 H_bloodOrange EUT2x
1769 H_fuchsia EUT2x
1770 HMM02866_1_pvmeut(#) EUT2H
1771 -pf05437 -
1772 -pf03591 -
1773 -pf09335 -
1774 -# -
1775 -pf03984 -
1776 -pf03551 -
1777 -# -
1778 -# -
1779 -pf03845 -
BMC-P
BMC-H
BMC-T
Aldehyde dehydrogenase
PTAC
Signature enzyme
Alcohol dehydrogenase
Regulator

Brevibacillus sp. CF112. Bacteria; Firmicutes; Bacilli; Bacillales; Paenibacillaceae;

PMI08__locus_2

EUT2x

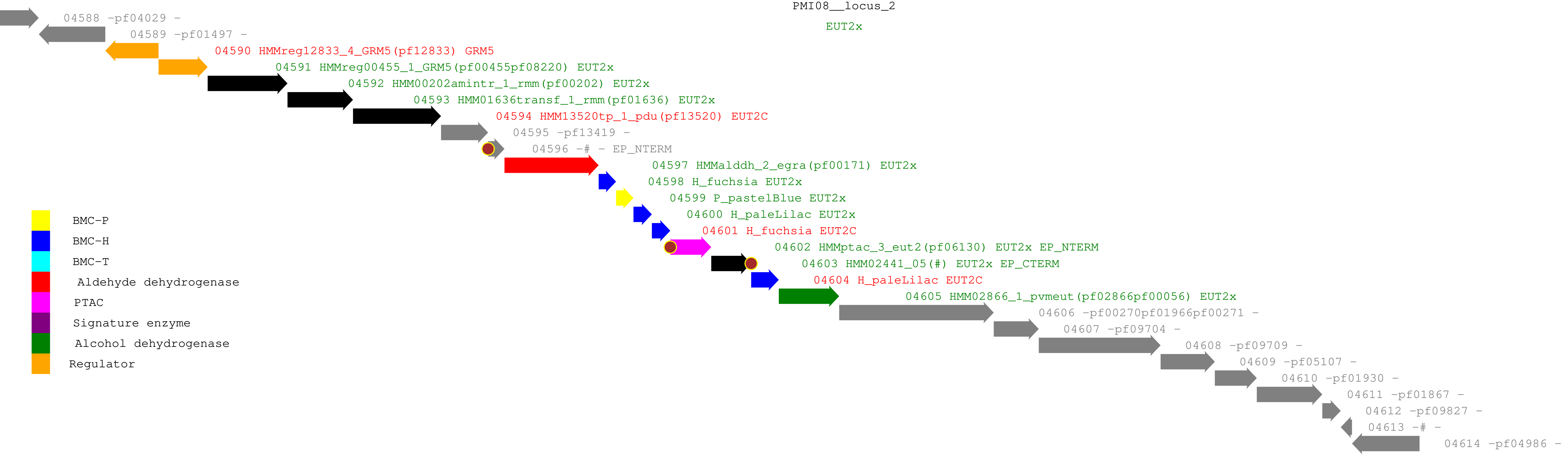

Sporosarcina ureae. Bacteria; Firmicutes; Bacilli; Bacillales; Planococcaceae;

SporoP8__locus_2

EUT2x

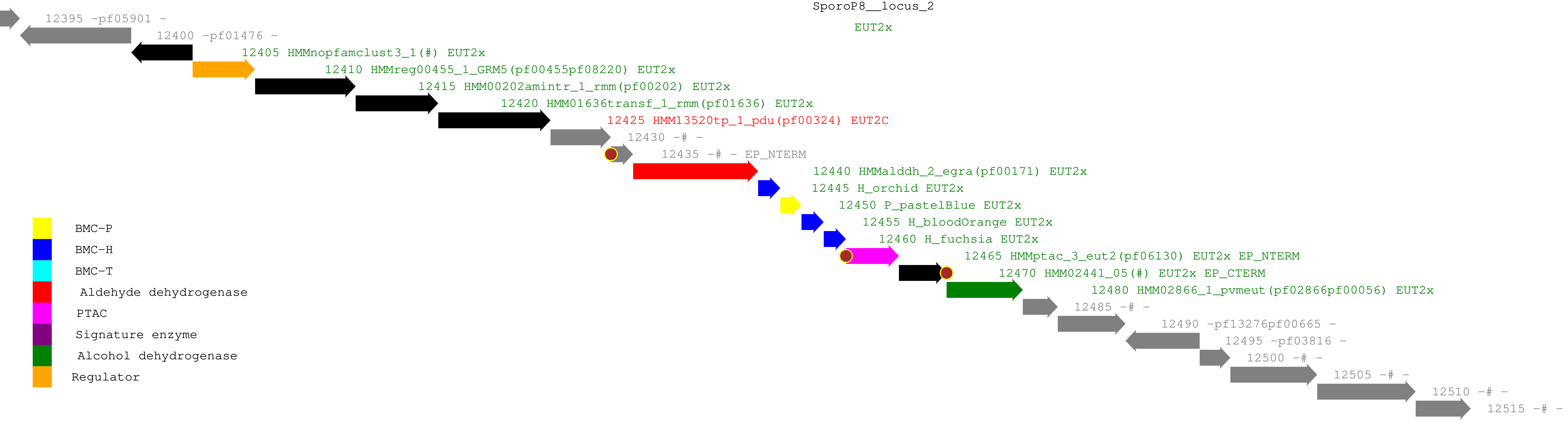

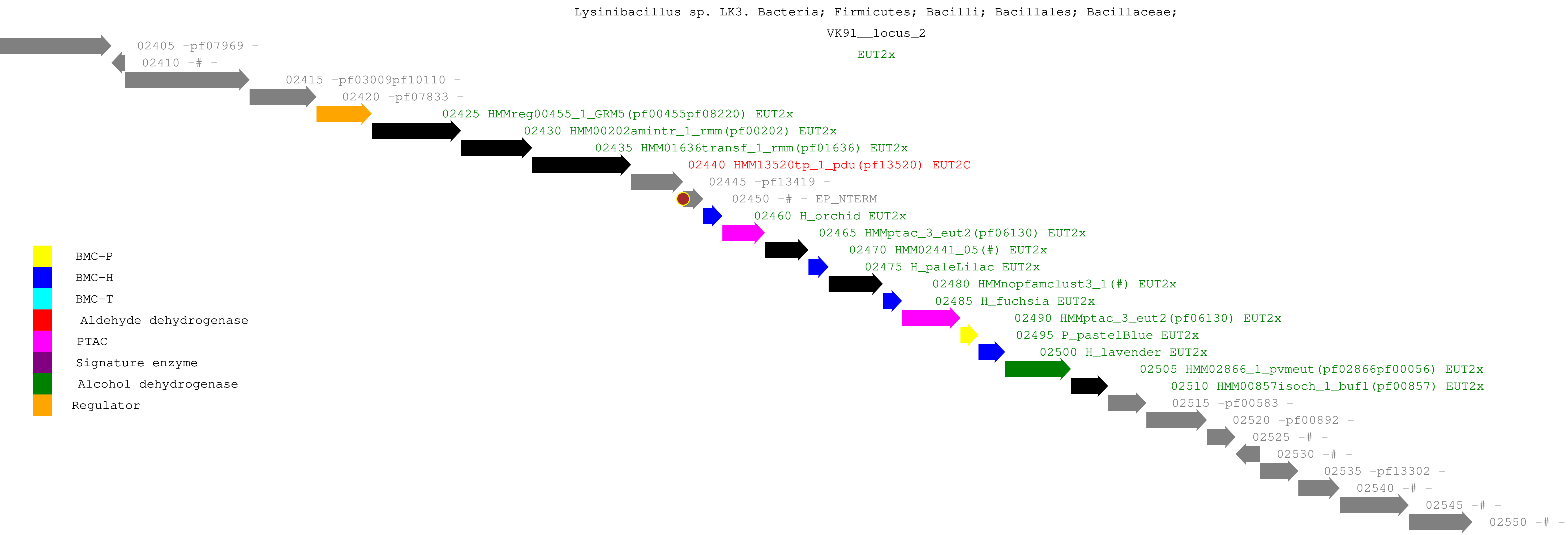
Lysinibacillus sp. LK3. Bacteria; Firmicutes; Bacilli; Bacillales; Bacillaceae;
VK91__locus_2
EUT2x
02405 -pf07969 -
02410 -# -
02415 -pf03009pf10110 -
02420 -pf07833 -
02425 HMMreg00455_1_GRM5(pf00455pf08220) EUT2x
02430 HMM00202amintr_1_rmm(pf00202) EUT2x
02435 HMM01636transf_1_rmm(pf01636) EUT2x
02440 HMM13520tp_1_pdu(pf13520) EUT2C
02445 -pf13419 -
02450 -# - EP_NTERM
02460 H_orchid EUT2x
02465 HMMptac_3_eut2(pf06130) EUT2x
02470 HMM02441_05(#) EUT2x
02475 H_paleLilac EUT2x
02480 HMMnopfamclust3_1(#) EUT2x
02485 H_fuchsia EUT2x
02490 HMMptac_3_eut2(pf06130) EUT2x
02495 P_pastelBlue EUT2x
02500 H_lavender EUT2x
02505 HMM02866_1_pvmeut(pf02866pf00056) EUT2x
02510 HMM00857isoch_1_buf1(pf00857) EUT2x
02515 -pf00583 -
02520 -pf00892 -
02525 -# -
02530 -# -
02535 -pf13302 -
02540 -# -
02545 -# -
02550 -# -
BMC-P
BMC-H
BMC-T
Aldehyde dehydrogenase
PTAC
Signature enzyme
Alcohol dehydrogenase
Regulator

Anaerolinea thermophila (strain DSM 14523 / JCM 11388 / NBRC 100420 / Bacteria; Chloroflexi; Anaerolineae; Anaerolineales; Anaerolineaceae;

ANT__locus_1

EUT3

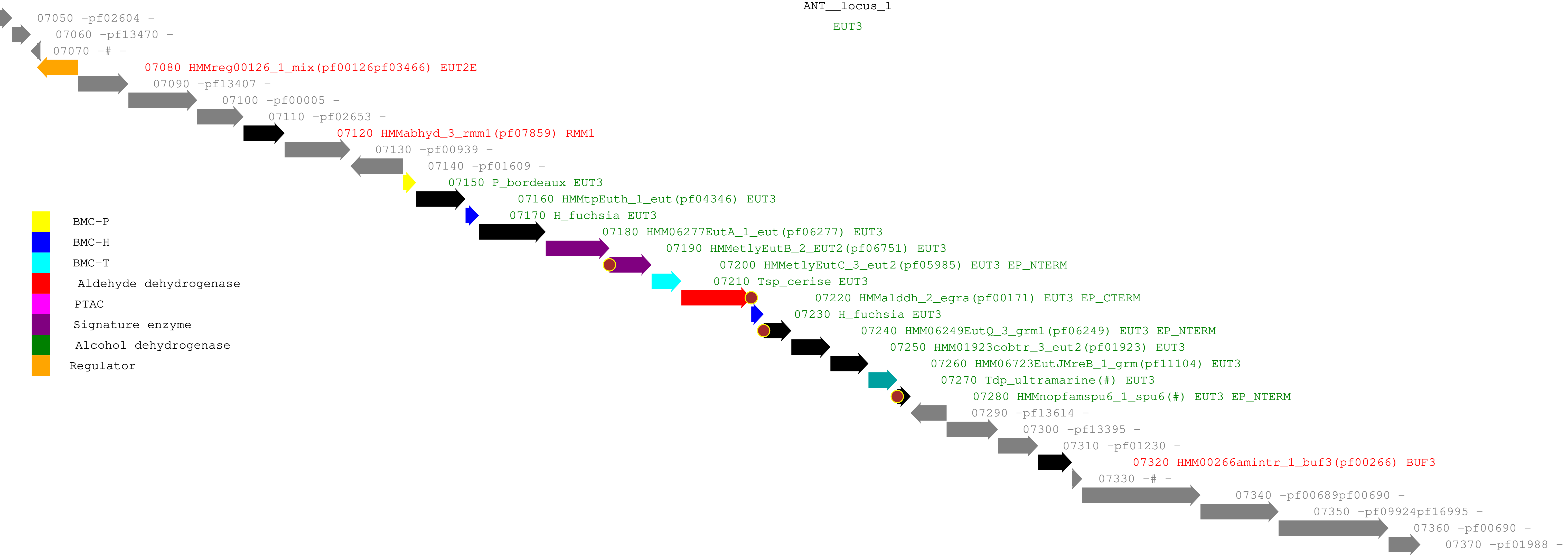

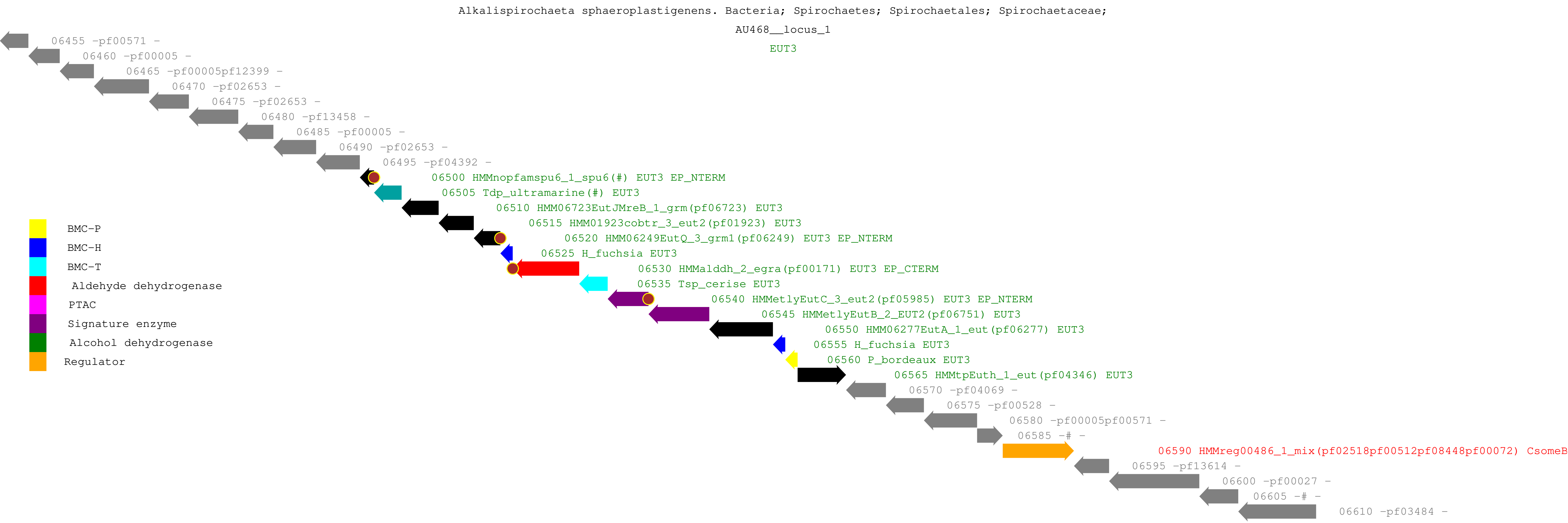

Alkalispirochaeta sphaeroplastigenens. Bacteria; Spirochaetes; Spirochaetales; Spirochaetaceae;
AU468__locus_1
EUT3
06455 -pf00571 -
06460 -pf00005 -
06465 -pf00005pf12399 -
06470 -pf02653 -
06475 -pf02653 -
06480 -pf13458 -
06485 -pf00005 -
06490 -pf02653 -
06495 -pf04392 -
06500 HMMnopfamspu6_1_spu6(#) EUT3 EP_NTERM
06505 Tdp_ultramarine(#) EUT3
06510 HMM06723EutJMreB_1_grm(pf06723) EUT3
06515 HMM01923cobtr_3_eut2(pf01923) EUT3
06520 HMM06249EutQ_3_grm1(pf06249) EUT3 EP_NTERM
06525 H_fuchsia EUT3
06530 HMMalddh_2_egra(pf00171) EUT3 EP_CTERM
06535 Tsp_cerise EUT3
06540 HMMetlyEutC_3_eut2(pf05985) EUT3 EP_NTERM
06545 HMMetlyEutB_2_EUT2(pf06751) EUT3
06550 HMM06277EutA_1_eut(pf06277) EUT3
06555 H_fuchsia EUT3
06560 P_bordeaux EUT3
06565 HMMtpEuth_1_eut(pf04346) EUT3
06570 -pf04069 -
06575 -pf00528 -
06580 -pf00005pf00571 -
06585 -# -
06590 HMMreg00486_1_mix(pf02518pf00512pf08448pf00072) CsomeB
06595 -pf13614 -
06600 -pf00027 -
06605 -# -
06610 -pf03484 -
BMC-P
BMC-H
BMC-T
Aldehyde dehydrogenase
PTAC
Signature enzyme
Alcohol dehydrogenase
Regulator

Tessaracoccus lapidicaptus. Bacteria; Actinobacteria; Propionibacteriales; Propionibacteriaceae;

BCR15__locus_1

EUT3

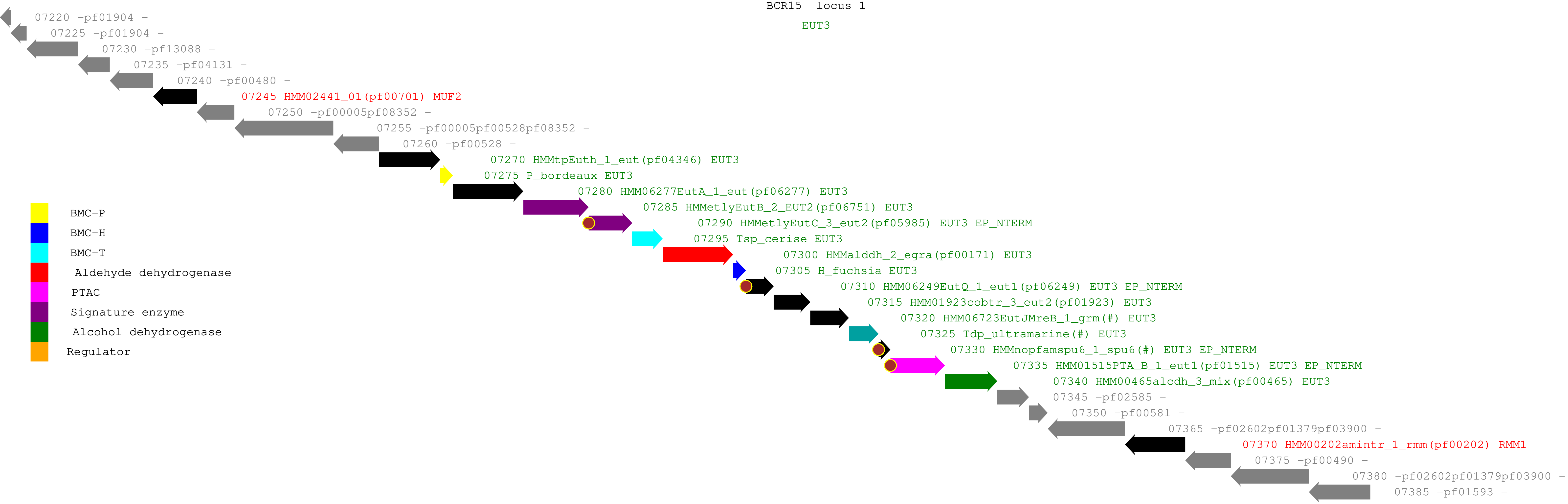

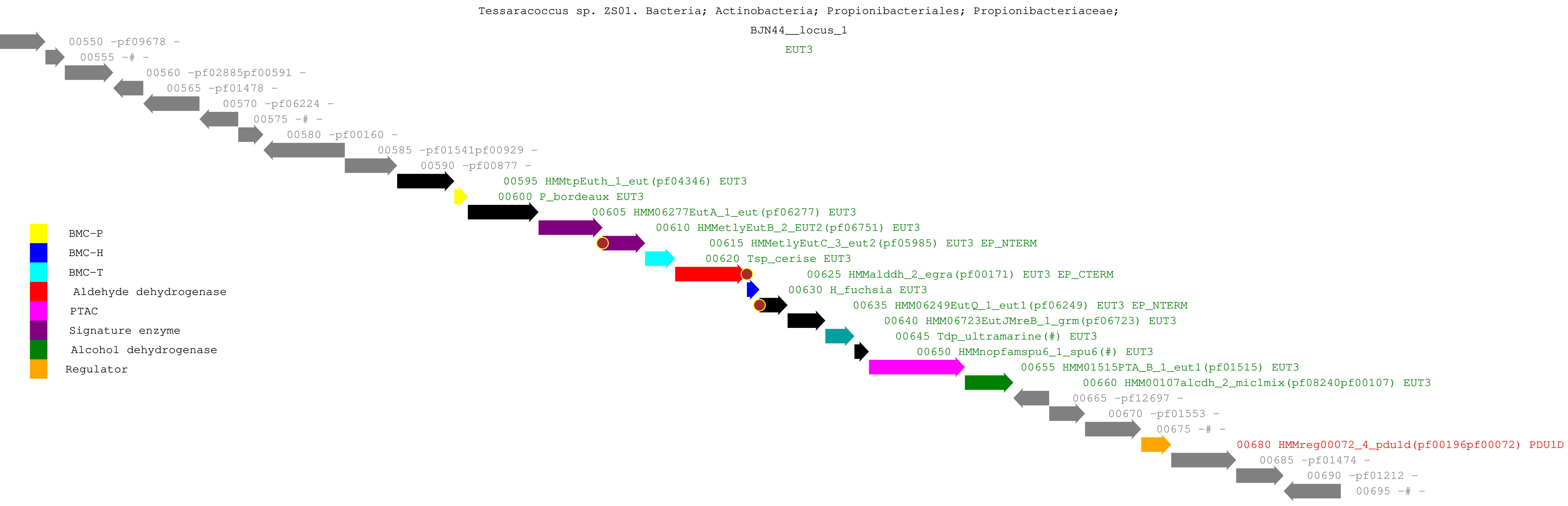

Tessaracoccus sp. ZS01. Bacteria; Actinobacteria; Propionibacteriales; Propionibacteriaceae;
BJN44__locus_1
EUT3
00550 -pf09678 -
00555 -# -
00560 -pf02885pf00591 -
00565 -pf01478 -
00570 -pf06224 -
00575 -# -
00580 -pf00160 -
00585 -pf01541pf00929 -
00590 -pf00877 -
00595 HMMtpEuth_1_eut(pf04346) EUT3
00600 P_bordeaux EUT3
00605 HMM06277EutA_1_eut(pf06277) EUT3
00610 HMMetlyEutB_2_EUT2(pf06751) EUT3
00615 HMMetlyEutC_3_eut2(pf05985) EUT3 EP_NTERM
00620 Tsp_cerise EUT3
00625 HMMalddh_2_egra(pf00171) EUT3 EP_CTERM
00630 H_fuchsia EUT3
00635 HMM06249EutQ_1_eut1(pf06249) EUT3 EP_NTERM
00640 HMM06723EutJMreB_1_grm(pf06723) EUT3
00645 Tdp_ultramarine(#) EUT3
00650 HMMnopfamspu6_1_spu6(#) EUT3
00655 HMM01515PTA_B_1_eut1(pf01515) EUT3
00660 HMM00107alcdh_2_mic1mix(pf08240pf00107) EUT3
00665 -pf12697 -
00670 -pf01553 -
00675 -# -
00680 HMMreg00072_4_pdu1d(pf00196pf00072) PDU1D
00685 -pf01474 -
00690 -pf01212 -
00695 -# -
BMC-P
BMC-H
BMC-T
Aldehyde dehydrogenase
PTAC
Signature enzyme
Alcohol dehydrogenase
Regulator

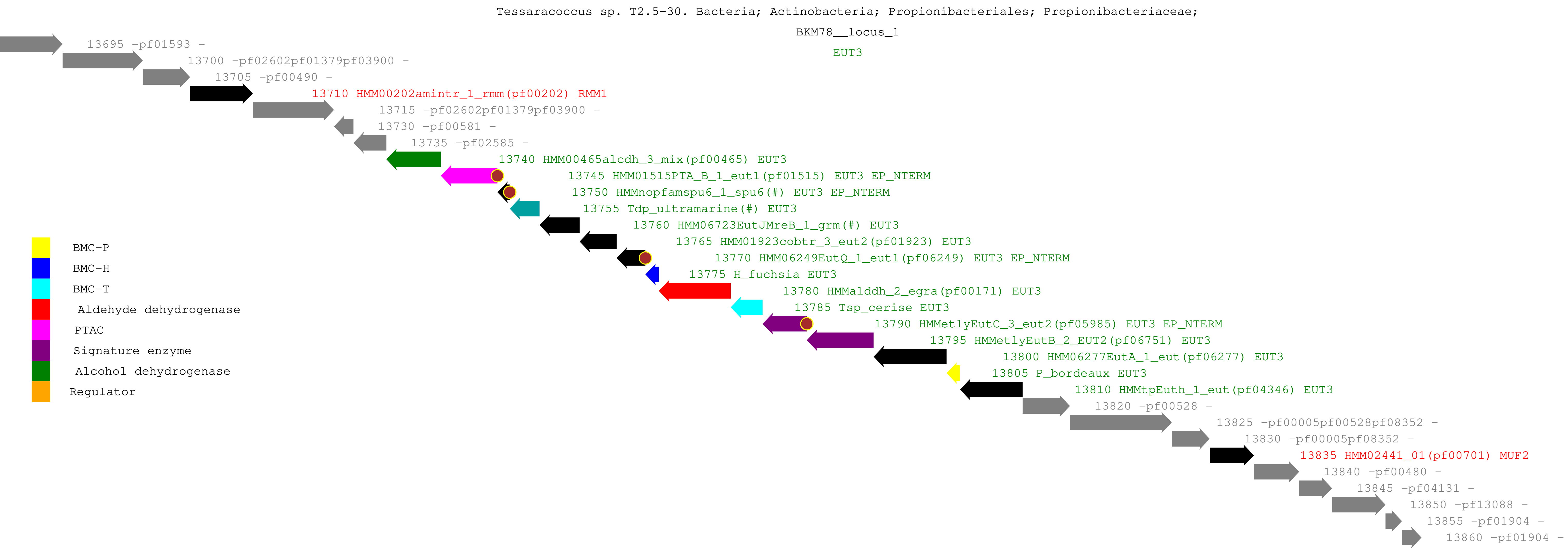

Tessaracoccus sp. T2.5-30. Bacteria; Actinobacteria; Propionibacteriales; Propionibacteriaceae;
BKM78__locus_1
EUT3
13695 -pf01593 -
13700 -pf02602pf01379pf03900 -
13705 -pf00490 -
13710 HMM00202amintr_1_rmm(pf00202) RMM1
13715 -pf02602pf01379pf03900 -
13730 -pf00581 -
13735 -pf02585 -
13740 HMM00465alcdh_3_mix(pf00465) EUT3
13745 HMM01515PTA_B_1_eut1(pf01515) EUT3 EP_NTERM
13750 HMMnopfamspu6_1_spu6(#) EUT3 EP_NTERM
13755 Tdp_ultramarine(#) EUT3
13760 HMM06723EutJMreB_1_grm(#) EUT3
13765 HMM01923cobtr_3_eut2(pf01923) EUT3
13770 HMM06249EutQ_1_eut1(pf06249) EUT3 EP_NTERM
13775 H_fuchsia EUT3
13780 HMMaldh_2_egra(pf00171) EUT3
13785 Tsp_cerise EUT3
13790 HMMetlyEutC_3_eut2(pf05985) EUT3 EP_NTERM
13795 HMMetlyEutB_2_EUT2(pf06751) EUT3
13800 HMM06277EutA_1_eut(pf06277) EUT3
13805 P_bordeaux EUT3
13810 HMMtpEuth_1_eut(pf04346) EUT3
13820 -pf00528 -
13825 -pf00005pf00528pf08352 -
13830 -pf00005pf08352 -
13835 HMM02441_01(pf00701) MUF2
13840 -pf00480 -
13845 -pf04131 -
13850 -pf13088 -
13855 -pf01904 -
13860 -pf01904 -
BMC-P
BMC-H
BMC-T
Aldehyde dehydrogenase
PTAC
Signature enzyme
Alcohol dehydrogenase
Regulator

Tessaracoccus sp. NSG39. Bacteria; Actinobacteria; Propionibacteriales; Propionibacteriaceae;

BW730__locus_1

EUT3

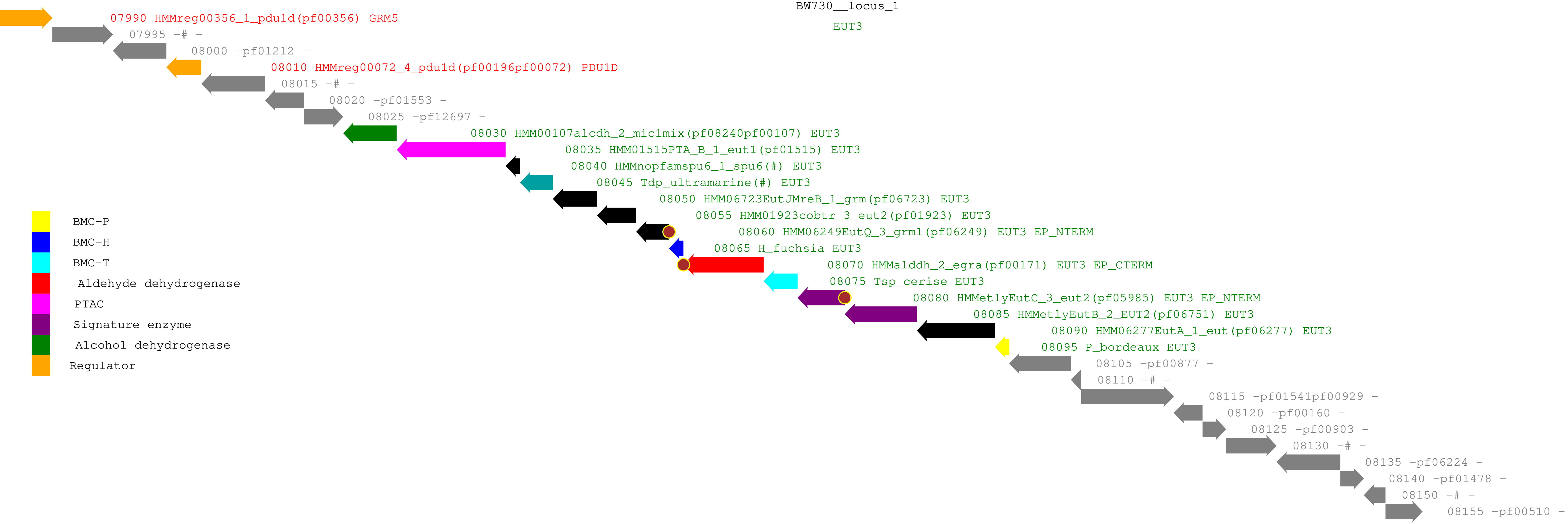

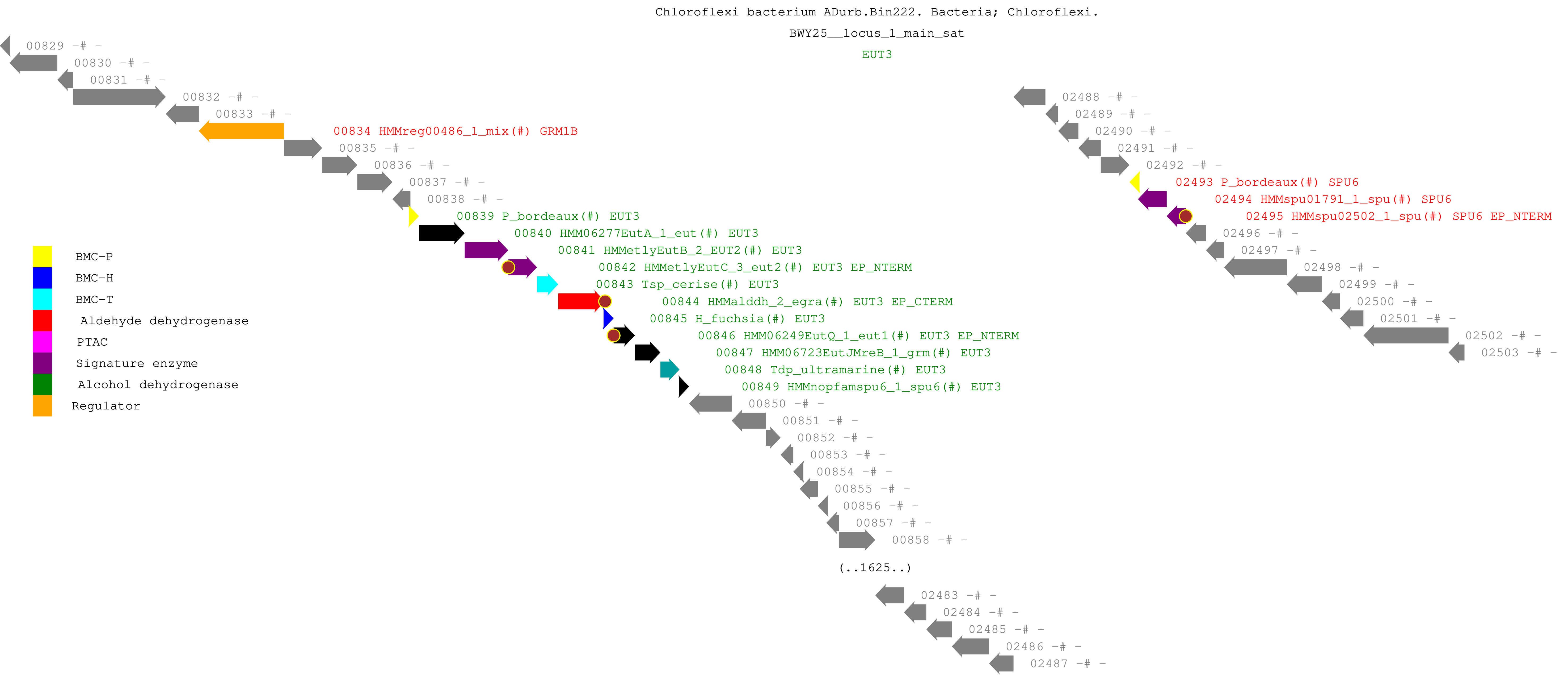
Chloroflexi bacterium ADurb.Bin222. Bacteria; Chloroflexi.
BWY25__locus_1_main_sat
EUT3
00829 -# -
00830 -# -
00831 -# -
00832 -# -
00833 -# -
00834 HMMreg00486_1_mix(#) GRM1B
00835 -# -
00836 -# -
00837 -# -
00838 -# -
00839 P_bordeaux(#) EUT3
00840 HMM06277EutA_1_eut(#) EUT3
00841 HMMetlyEutB_2_EUT2(#) EUT3
00842 HMMetlyEutC_3_eut2(#) EUT3 EP_NTERM
00843 Tsp_cerise(#) EUT3
00844 HMMalddh_2_egra(#) EUT3 EP_CTERM
00845 H_fuchsia(#) EUT3
00846 HMM06249EutQ_1_eut1(#) EUT3 EP_NTERM
00847 HMM06723EutJMreB_1_grm(#) EUT3
00848 Tdp_ultramarine(#) EUT3
00849 HMMnopfamspu6_1_spu6(#) EUT3
00850 -# -
00851 -# -
00852 -# -
00853 -# -
00854 -# -
00855 -# -
00856 -# -
00857 -# -
00858 -# -
(..1625..)
02483 -# -
02484 -# -
02485 -# -
02486 -# -
02487 -# -
02488 -# -
02489 -# -
02490 -# -
02491 -# -
02492 -# -
02493 P_bordeaux(#) SPU6
02494 HMMspu01791_1_spu(#) SPU6
02495 HMMspu02502_1_spu(#) SPU6 EP_NTERM
02496 -# -
02497 -# -
02498 -# -
02499 -# -
02500 -# -
02501 -# -
02502 -# -
02503 -# -
BMC-P
BMC-H
BMC-T
Aldehyde dehydrogenase
PTAC
Signature enzyme
Alcohol dehydrogenase
Regulator

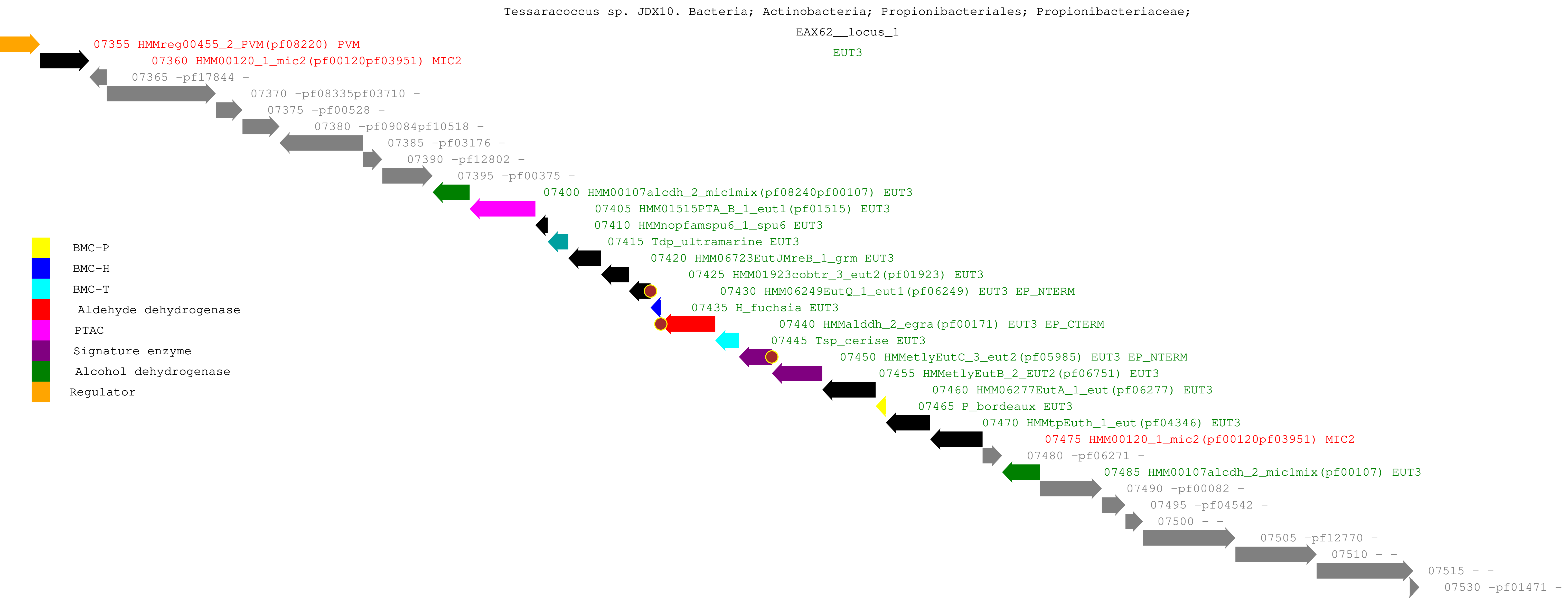
Tessaracoccus sp. JDX10. Bacteria; Actinobacteria; Propionibacteriales; Propionibacteriaceae;
EAX62__locus_1
EUT3
07355 HMMreg00455_2_PVM(pf08220) PVM
07360 HMM00120_1_mic2(pf00120pf03951) MIC2
07365 -pf17844 -
07370 -pf08335pf03710 -
07375 -pf00528 -
07380 -pf09084pf10518 -
07385 -pf03176 -
07390 -pf12802 -
07395 -pf00375 -
07400 HMM00107alcdh_2_mic1mix(pf08240pf00107) EUT3
07405 HMM01515PTA_B_1_eut1(pf01515) EUT3
07410 HMMnopfamspu6_1_spu6 EUT3
07415 Tdp_ultramarine EUT3
07420 HMM06723EutJMreB_1_grm EUT3
07425 HMM01923cobtr_3_eut2(pf01923) EUT3
07430 HMM06249EutQ_1_eut1(pf06249) EUT3 EP_NTERM
07435 H_fuchsia EUT3
07440 HMMalddh_2_egra(pf00171) EUT3 EP_CTERM
07445 Tsp_cerise EUT3
07450 HMMetlyEutC_3_eut2(pf05985) EUT3 EP_NTERM
07455 HMMetlyEutB_2_EUT2(pf06751) EUT3
07460 HMM06277EutA_1_eut(pf06277) EUT3
07465 P_bordeaux EUT3
07470 HMMtpEuth_1_eut(pf04346) EUT3
07475 HMM00120_1_mic2(pf00120pf03951) MIC2
07480 -pf06271 -
07485 HMM00107alcdh_2_mic1mix(pf00107) EUT3
07490 -pf00082 -
07495 -pf04542 -
07500 - -
07505 -pf12770 -
07510 - -
07515 - -
07530 -pf01471 -
BMC-P
BMC-H
BMC-T
Aldehyde dehydrogenase
PTAC
Signature enzyme
Alcohol dehydrogenase
Regulator

Pseudopropionibacterium propionicum. Bacteria; Actinobacteria; Propionibacteriales; Propionibacteriaceae;

EII34__locus_1_main_sat

EUT3

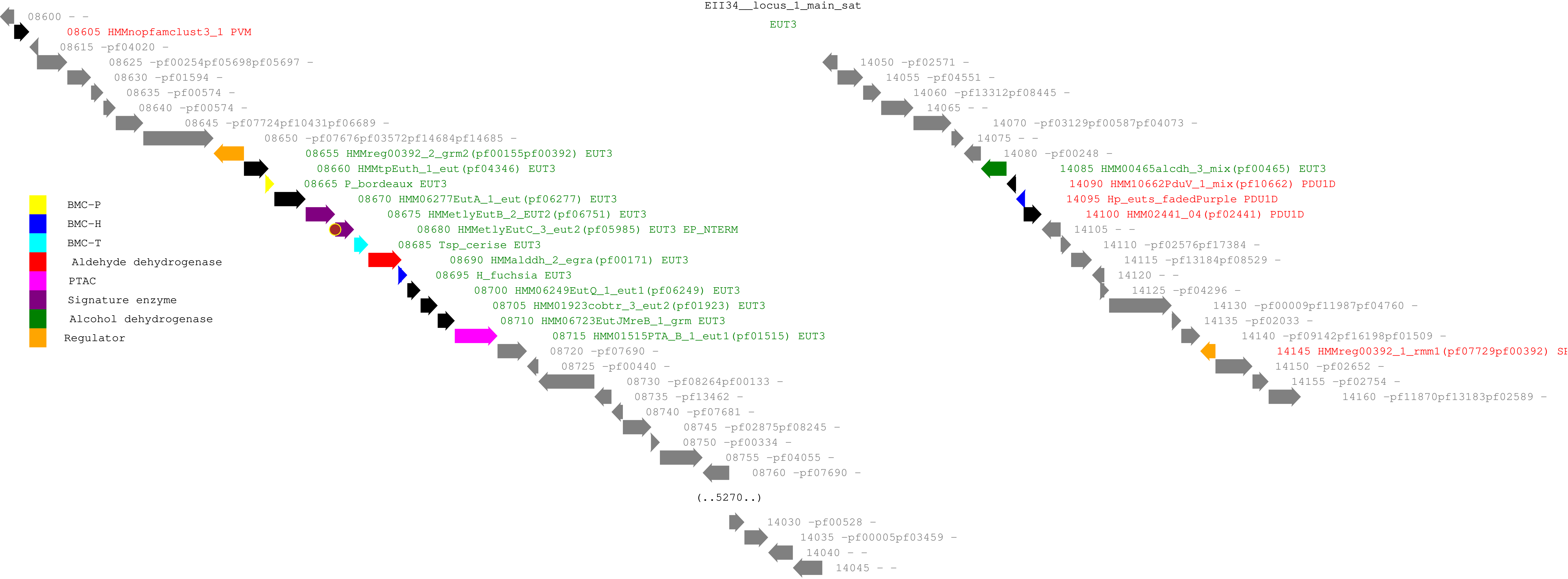

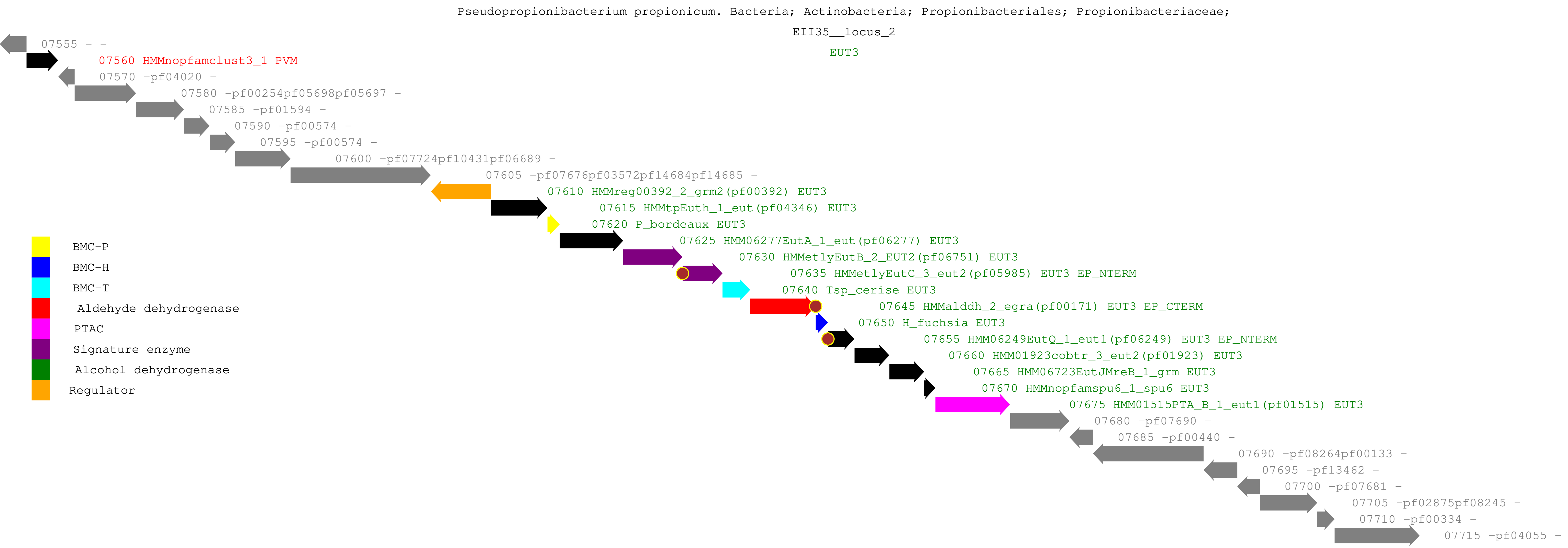
Pseudopropionibacterium propionicum. Bacteria; Actinobacteria; Propionibacteriales; Propionibacteriaceae;
EII35__locus_2
EUT3
07555 - -
07560 HMMnopfamclust3_1 PVM
07570 -pf04020 -
07580 -pf00254pf05698pf05697 -
07585 -pf01594 -
07590 -pf00574 -
07595 -pf00574 -
07600 -pf07724pf10431pf06689 -
07605 -pf07676pf03572pf14684pf14685 -
07610 HMMreg00392_2_grm2(pf00392) EUT3
07615 HMMtpEuth_1_eut(pf04346) EUT3
07620 P_bordeaux EUT3
07625 HMM06277EutA_1_eut(pf06277) EUT3
07630 HMMetlyEutB_2_EUT2(pf06751) EUT3
07635 HMMetlyEutC_3_eut2(pf05985) EUT3 EP_NTERM
07640 Tsp_cerise EUT3
07645 HMMalddh_2_egra(pf00171) EUT3 EP_CTERM
07650 H_fuchsia EUT3
07655 HMM06249EutQ_1_eut1(pf06249) EUT3 EP_NTERM
07660 HMM01923cobtr_3_eut2(pf01923) EUT3
07665 HMM06723EutJMreB_1_grm EUT3
07670 HMMnopfamspu6_1_spu6 EUT3
07675 HMM01515PTA_B_1_eut1(pf01515) EUT3
07680 -pf07690 -
07685 -pf00440 -
07690 -pf08264pf00133 -
07695 -pf13462 -
07700 -pf07681 -
07705 -pf02875pf08245 -
07710 -pf00334 -
07715 -pf04055 -
BMC-P
BMC-H
BMC-T
Aldehyde dehydrogenase
PTAC
Signature enzyme
Alcohol dehydrogenase
Regulator

Xylanibacterium sp. 2JSPR-7. Bacteria; Actinobacteria; Micrococcales; Promicromonosporaceae;

ET495__locus_1

EUT3

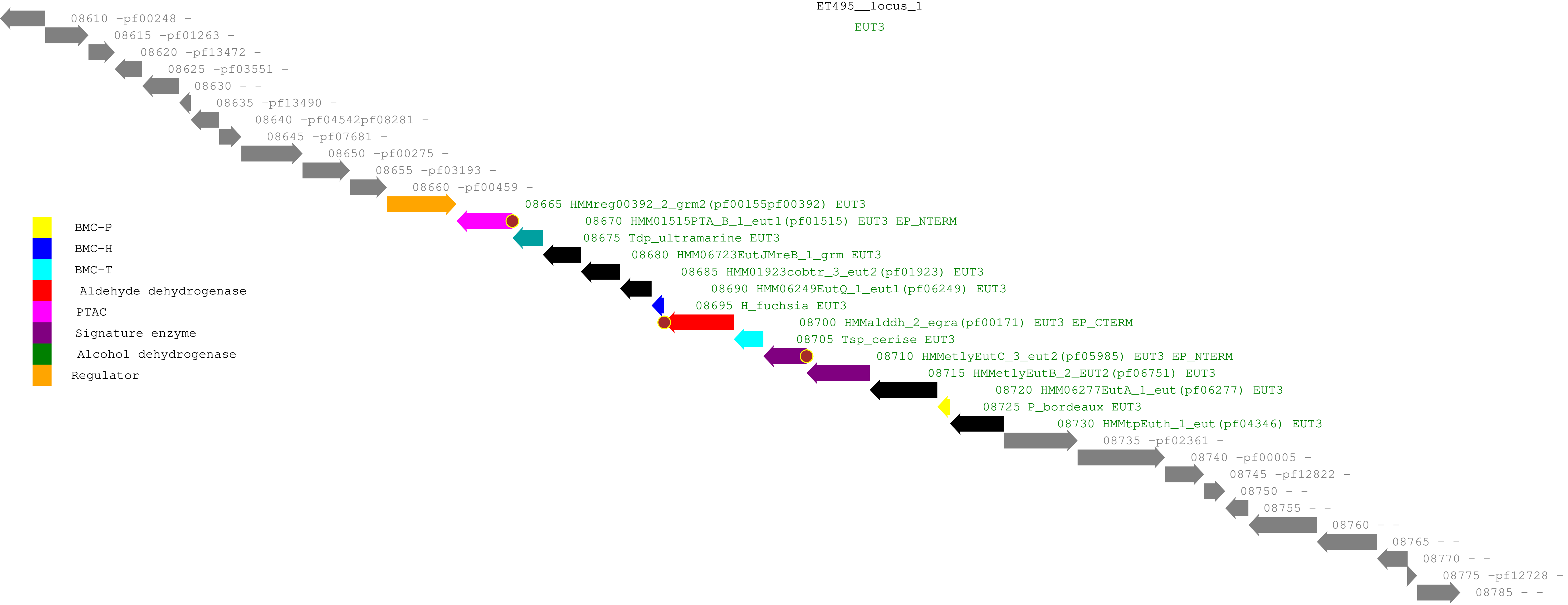

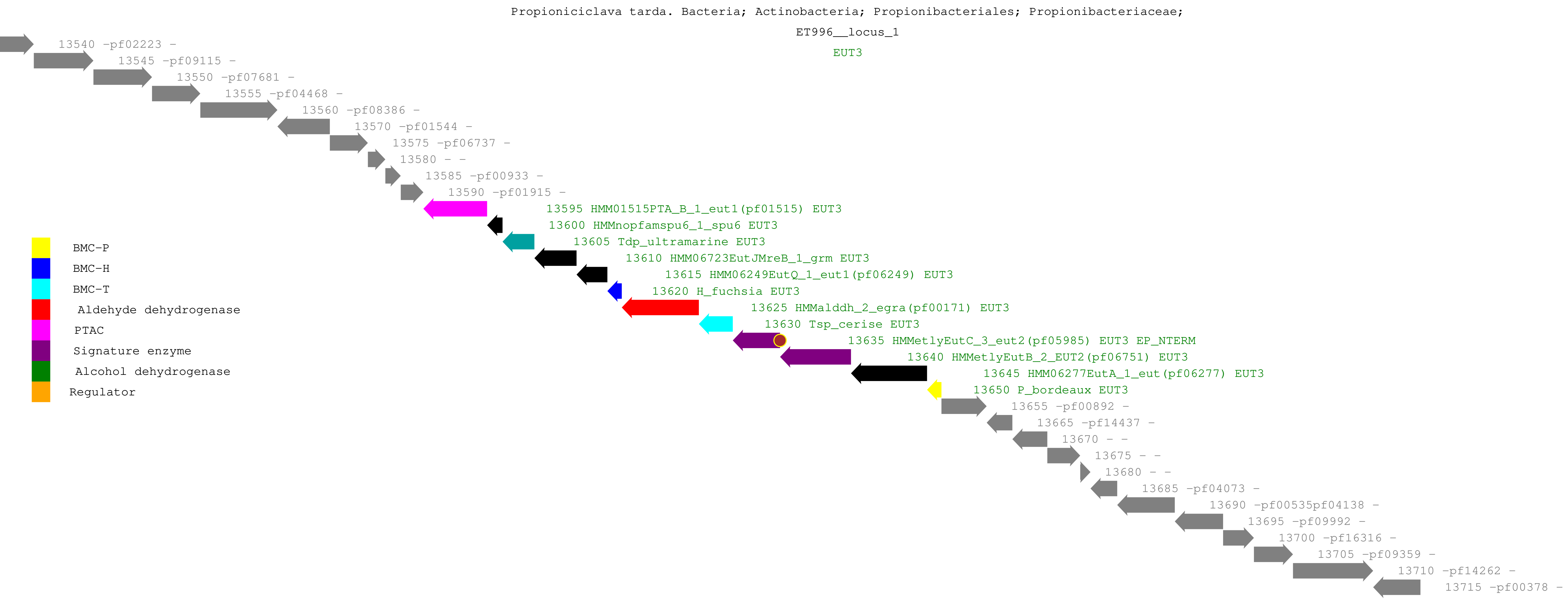

Propioniciclava tarda. Bacteria; Actinobacteria; Propionibacteriales; Propionibacteriaceae;
ET996__locus_1
EUT3
13540 -pf02223 -
13545 -pf09115 -
13550 -pf07681 -
13555 -pf04468 -
13560 -pf08386 -
13570 -pf01544 -
13575 -pf06737 -
13580 - -
13585 -pf00933 -
13590 -pf01915 -
13595 HMM01515PTA_B_1_eut1(pf01515) EUT3
13600 HMMnopfamspu6_1_spu6 EUT3
13605 Tdp_ultramarine EUT3
13610 HMM06723EutJMreB_1_grm EUT3
13615 HMM06249EutQ_1_eut1(pf06249) EUT3
13620 H_fuchsia EUT3
13625 HMMaldh_2_egra(pf00171) EUT3
13630 Tsp_cerise EUT3
13635 HMMetlyEutC_3_eut2(pf05985) EUT3 EP_NTERM
13640 HMMetlyEutB_2_EUT2(pf06751) EUT3
13645 HMM06277EutA_1_eut(pf06277) EUT3
13650 P_bordeaux EUT3
13655 -pf00892 -
13665 -pf14437 -
13670 - -
13675 - -
13680 - -
13685 -pf04073 -
13690 -pf00535pf04138 -
13695 -pf09992 -
13700 -pf16316 -
13705 -pf09359 -
13710 -pf14262 -
13715 -pf00378 -
BMC-P
BMC-H
BMC-T
Aldehyde dehydrogenase
PTAC
Signature enzyme
Alcohol dehydrogenase
Regulator

Xylanibacterium ulmi. Bacteria; Actinobacteria; Micrococcales; Promicromonosporaceae;

EV386__locus_1

EUT3

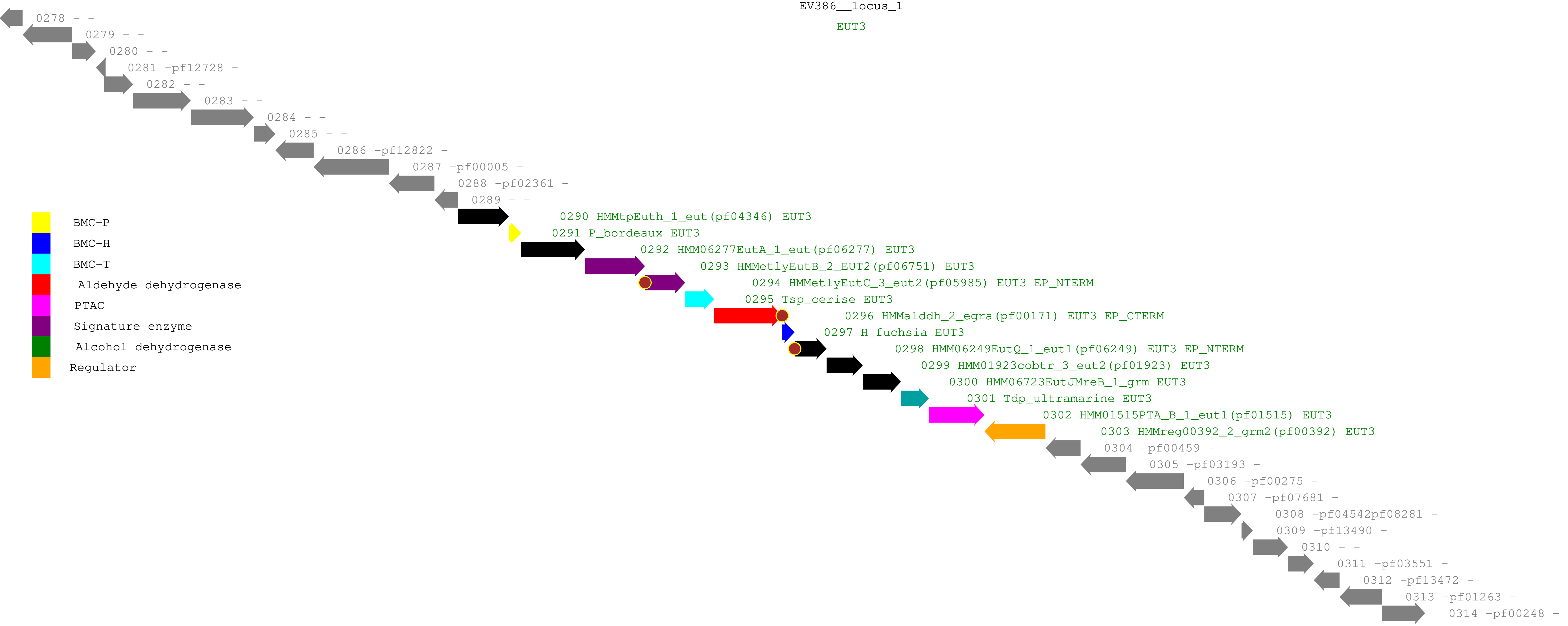

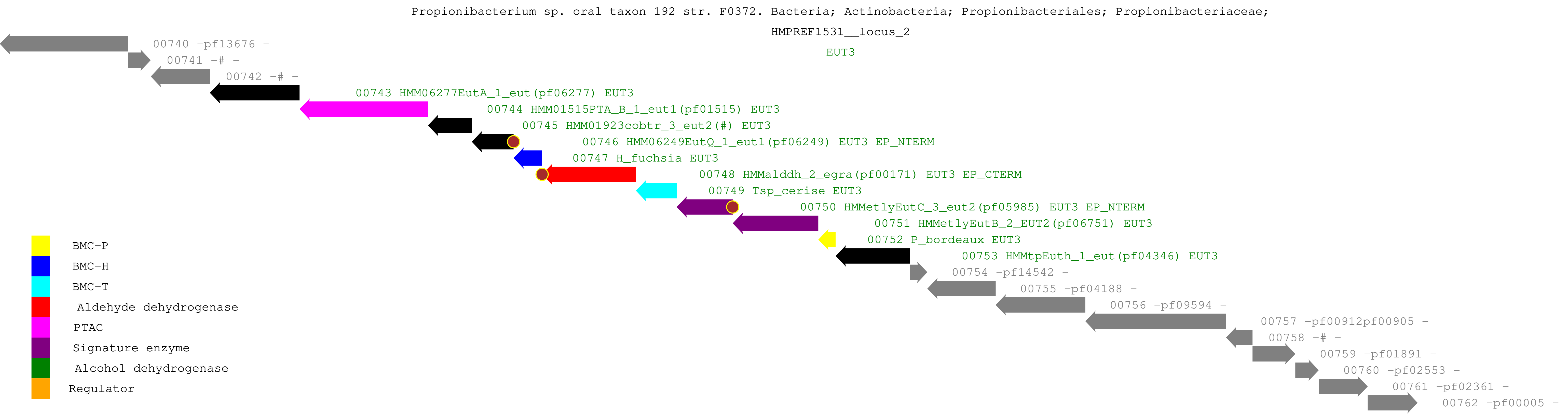
Propionibacterium sp. oral taxon 192 str. F0372. Bacteria; Actinobacteria; Propionibacteriales; Propionibacteriaceae;
HMPREF1531__locus_2
EUT3
00740 -pf13676 -
00741 -# -
00742 -# -
00743 HMM06277EutA_1_eut(pf06277) EUT3
00744 HMM01515PTA_B_1_eut1(pf01515) EUT3
00745 HMM01923cobtr_3_eut2(#) EUT3
00746 HMM06249EutQ_1_eut1(pf06249) EUT3 EP_NTERM
00747 H_fuchsia EUT3
00748 HMMalddh_2_egra(pf00171) EUT3 EP_CTERM
00749 Tsp_cerise EUT3
00750 HMMetlyEutC_3_eut2(pf05985) EUT3 EP_NTERM
00751 HMMetlyEutB_2_EUT2(pf06751) EUT3
00752 P_bordeaux EUT3
00753 HMMtpEuth_1_eut(pf04346) EUT3
00754 -pf14542 -
00755 -pf04188 -
00756 -pf09594 -
00757 -pf00912pf00905 -
00758 -# -
00759 -pf01891 -
00760 -pf02553 -
00761 -pf02361 -
00762 -pf00005 -
BMC-P
BMC-H
BMC-T
Aldehyde dehydrogenase
PTAC
Signature enzyme
Alcohol dehydrogenase
Regulator

Kineosphaera limosa NBRC 100340. Bacteria; Actinobacteria; Micrococcales; Dermatophilaceae;

KILIM_086__locus_1

EUT3

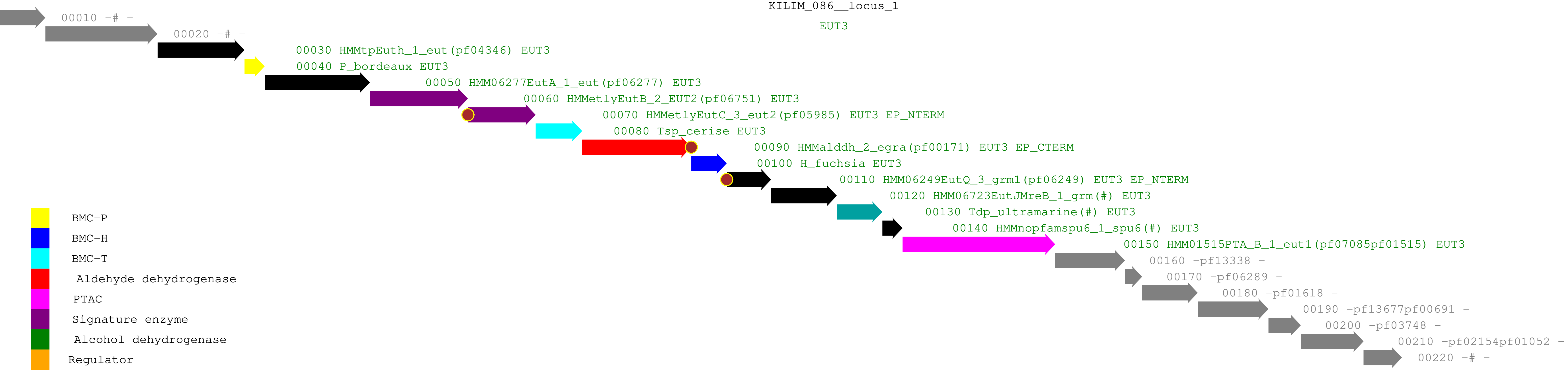

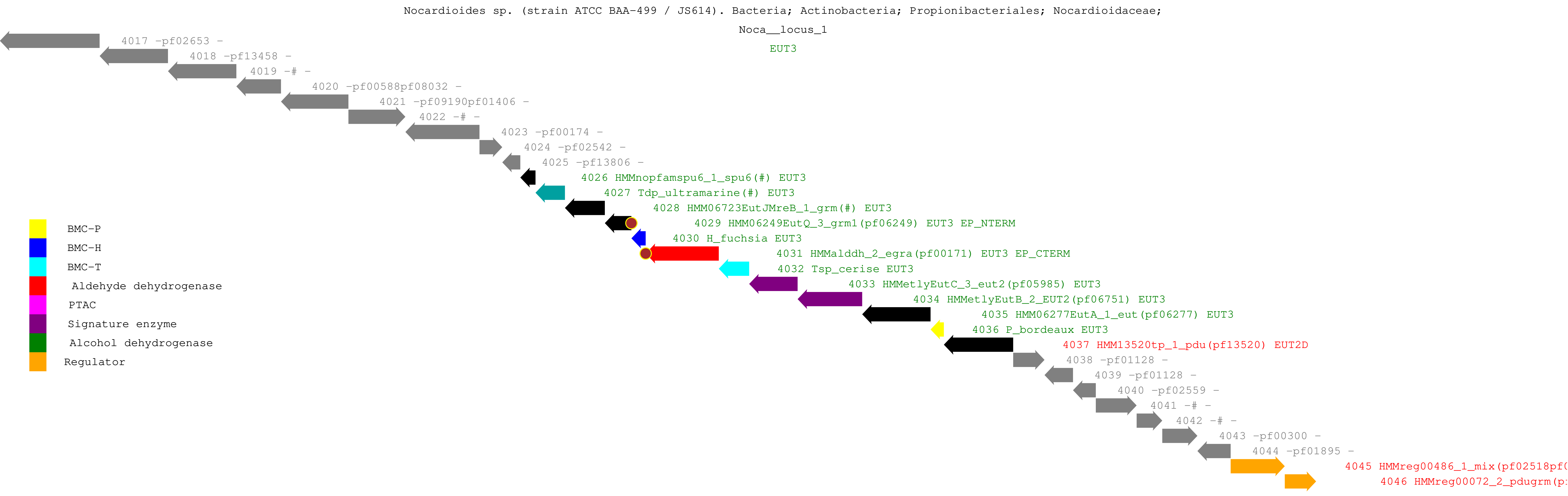

Nocardioides sp. (strain ATCC BAA-499 / JS614). Bacteria; Actinobacteria; Propionibacteriales; Nocardioidaceae;
Noca__locus_1
EUT3
4017 -pf02653 -
4018 -pf13458 -
4019 -# -
4020 -pf00588pf08032 -
4021 -pf09190pf01406 -
4022 -# -
4023 -pf00174 -
4024 -pf02542 -
4025 -pf13806 -
4026 HMMnopfamspu6_1_spu6(#) EUT3
4027 Tdp_ultramarine(#) EUT3
4028 HMM06723EutJMreB_1_grm(#) EUT3
4029 HMM06249EutQ_3_grm1(pf06249) EUT3 EP_NTERM
4030 H_fuchsia EUT3
4031 HMMalddh_2_egra(pf00171) EUT3 EP_CTERM
4032 Tsp_cerise EUT3
4033 HMMetlyEutC_3_eut2(pf05985) EUT3
4034 HMMetlyEutB_2_EUT2(pf06751) EUT3
4035 HMM06277EutA_1_eut(pf06277) EUT3
4036 P_bordeaux EUT3
4037 HMM13520tp_1_pdu(pf13520) EUT2D
4038 -pf01128 -
4039 -pf01128 -
4040 -pf02559 -
4041 -# -
4042 -# -
4043 -pf00300 -
4044 -pf01895 -
4045 HMMreg00486_1_mix(pf02518pf0
4046 HMMreg00072_2_pdugrm(p
BMC-P
BMC-H
BMC-T
Aldehyde dehydrogenase
PTAC
Signature enzyme
Alcohol dehydrogenase
Regulator

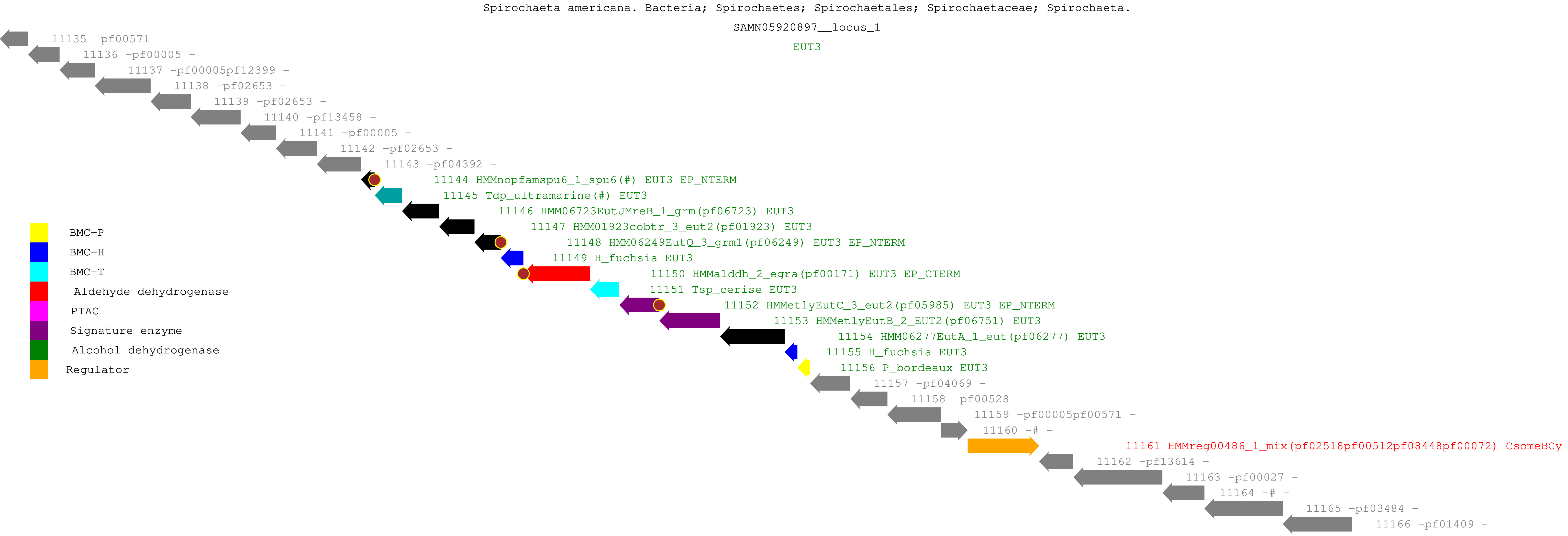

Spirochaeta americana. Bacteria; Spirochaetes; Spirochaetales; Spirochaetaceae; Spirochaeta.
SAMN05920897__locus_1
EUT3
11135 -pf00571 -
11136 -pf00005 -
11137 -pf00005pf12399 -
11138 -pf02653 -
11139 -pf02653 -
11140 -pf13458 -
11141 -pf00005 -
11142 -pf02653 -
11143 -pf04392 -
11144 HMMnopfamspu6_1_spu6(#) EUT3 EP_NTERM
11145 Tdp_ultramarine(#) EUT3
11146 HMM06723EutJMreB_1_grm(pf06723) EUT3
11147 HMM01923cobtr_3_eut2(pf01923) EUT3
11148 HMM06249EutQ_3_grm1(pf06249) EUT3 EP_NTERM
11149 H_fuchsia EUT3
11150 HMMalddh_2_egra(pf00171) EUT3 EP_CTERM
11151 Tsp_cerise EUT3
11152 HMMetlyEutC_3_eut2(pf05985) EUT3 EP_NTERM
11153 HMMetlyEutB_2_EUT2(pf06751) EUT3
11154 HMM06277EutA_1_eut(pf06277) EUT3
11155 H_fuchsia EUT3
11156 P_bordeaux EUT3
11157 -pf04069 -
11158 -pf00528 -
11159 -pf00005pf00571 -
11160 -# -
11161 HMMreg00486_1_mix(pf02518pf00512pf08448pf00072) CsomeBCy
11162 -pf13614 -
11163 -pf00027 -
11164 -# -
11165 -pf03484 -
11166 -pf01409 -
BMC-P
BMC-H
BMC-T
Aldehyde dehydrogenase
PTAC
Signature enzyme
Alcohol dehydrogenase
Regulator

Ignavibacteria bacterium GWA2_35_9. Bacteria; Ignavibacteriae; Ignavibacteria.

A2057__locus_fragmented

FRAG1

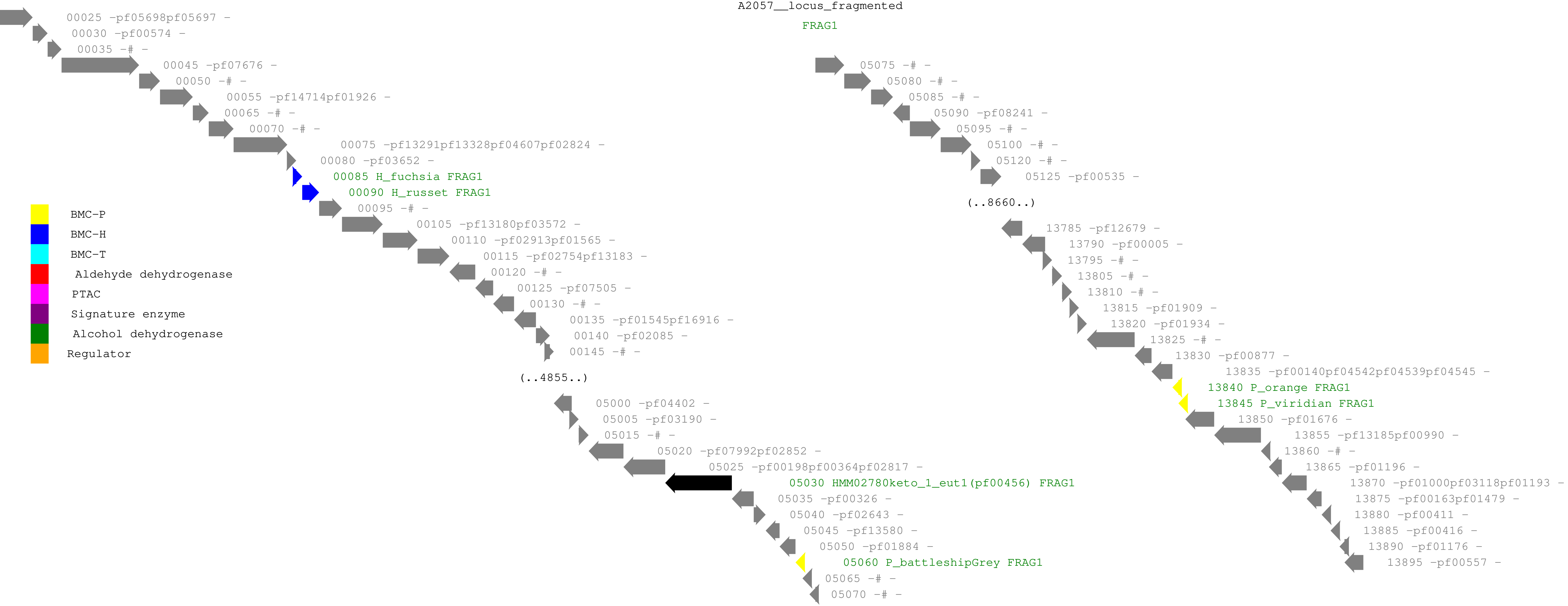

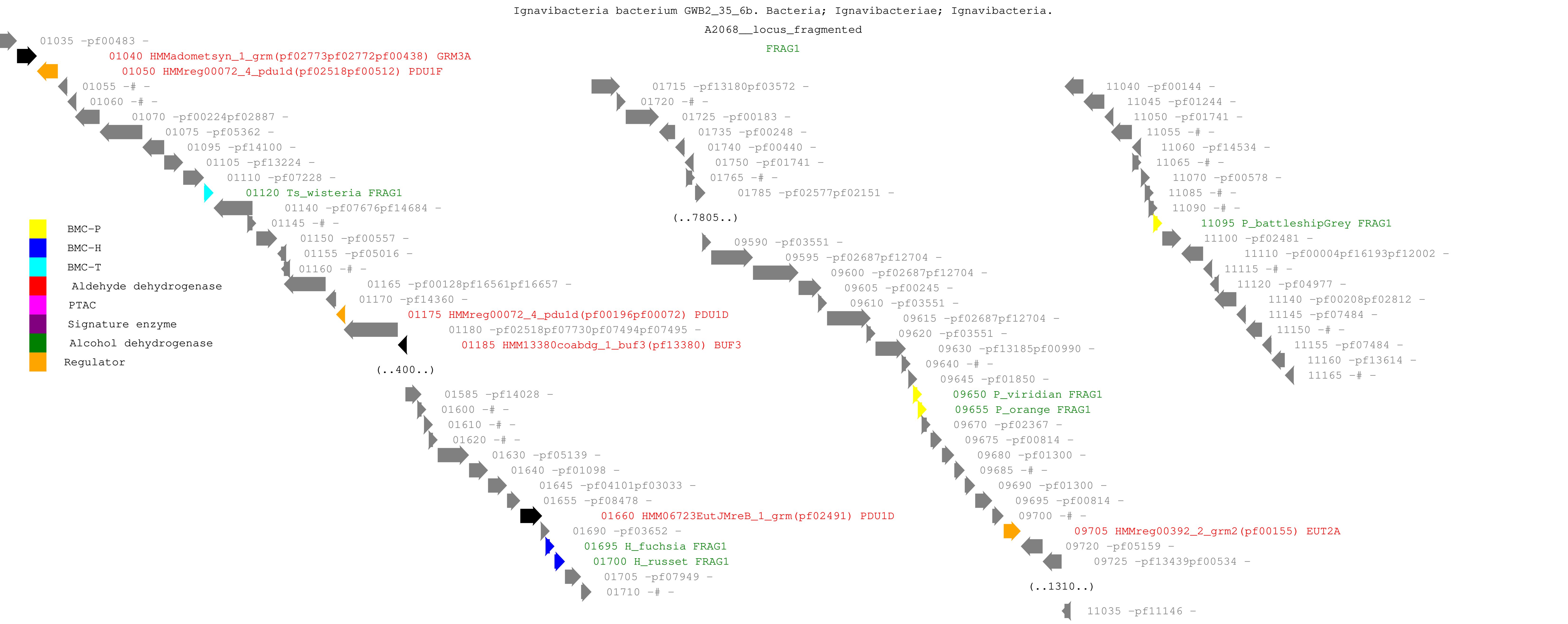

Ignavibacteria bacterium GWB2_35_6b. Bacteria; Ignavibacteriae; Ignavibacteria.
A2068__locus_fragmented
FRAG1
01035 -pf00483 -
01040 HMMadometsyn_1_grm(pf02773pf02772pf00438) GRM3A
01050 HMMreg00072_4_pdu1d(pf02518pf00512) PDU1F
01055 -# -
01060 -# -
01070 -pf00224pf02887 -
01075 -pf05362 -
01095 -pf14100 -
01105 -pf13224 -
01110 -pf07228 -
01120 Ts_wisteria FRAG1
01140 -pf07676pf14684 -
01145 -# -
01150 -pf00557 -
01155 -pf05016 -
01160 -# -
01165 -pf00128pf16561pf16657 -
01170 -pf14360 -
01175 HMMreg00072_4_pdu1d(pf00196pf00072) PDU1D
01180 -pf02518pf07730pf07494pf07495 -
01185 HMM13380coabdg_1_buf3(pf13380) BUF3
(..400..)
01585 -pf14028 -
01600 -# -
01610 -# -
01620 -# -
01630 -pf05139 -
01640 -pf01098 -
01645 -pf04101pf03033 -
01655 -pf08478 -
01660 HMM06723EutJMreB_1_grm(pf02491) PDU1D
01690 -pf03652 -
01695 H_fuchsia FRAG1
01700 H_russet FRAG1
01705 -pf07949 -
01710 -# -
01715 -pf13180pf03572 -
01720 -# -
01725 -pf00183 -
01735 -pf00248 -
01740 -pf00440 -
01750 -pf01741 -
01765 -# -
01785 -pf02577pf02151 -
(..7805..)
09590 -pf03551 -
09595 -pf02687pf12704 -
09600 -pf02687pf12704 -
09605 -pf00245 -
09610 -pf03551 -
09615 -pf02687pf12704 -
09620 -pf03551 -
09630 -pf13185pf00990 -
09640 -# -
09645 -pf01850 -
09650 P_viridian FRAG1
09655 P_orange FRAG1
09670 -pf02367 -
09675 -pf00814 -
09680 -pf01300 -
09685 -# -
09690 -pf01300 -
09695 -pf00814 -
09700 -# -
09705 HMMreg00392_2_grm2(pf00155) EUT2A
09720 -pf05159 -
09725 -pf13439pf00534 -
(..1310..)
11035 -pf11146 -
11040 -pf00144 -
11045 -pf01244 -
11050 -pf01741 -
11055 -# -
11060 -pf14534 -
11065 -# -
11070 -pf00578 -
11085 -# -
11090 -# -
11095 P_battleshipGrey FRAG1
11100 -pf02481 -
11110 -pf00004pf16193pf12002 -
11115 -# -
11120 -pf04977 -
11140 -pf00208pf02812 -
11145 -pf07484 -
11150 -# -
11155 -pf07484 -
11160 -pf13614 -
11165 -# -
BMC-P
BMC-H
BMC-T
Aldehyde dehydrogenase
PTAC
Signature enzyme
Alcohol dehydrogenase
Regulator

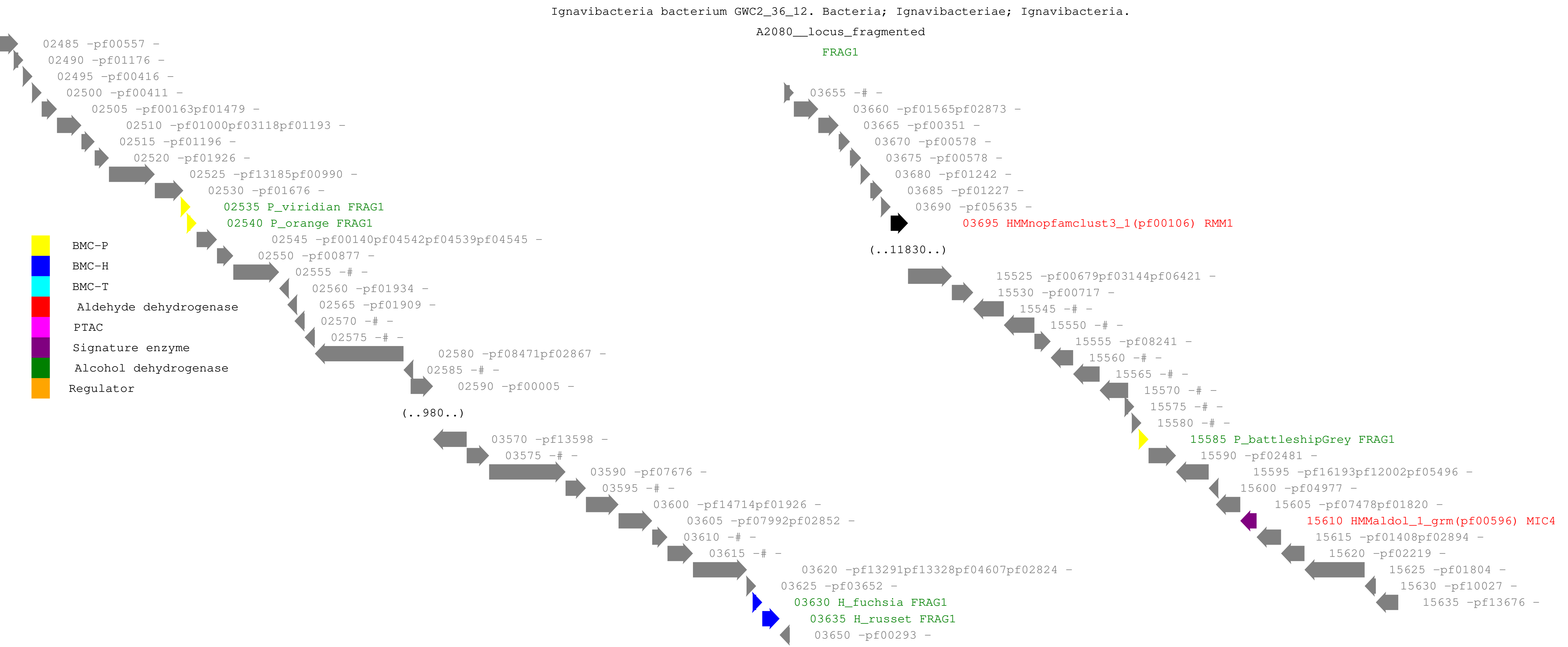

Ignavibacteria bacterium GWC2_36_12. Bacteria; Ignavibacteriae; Ignavibacteria.
A2080__locus_fragmented
FRAG1
02485 -pf00557 -
02490 -pf01176 -
02495 -pf00416 -
02500 -pf00411 -
02505 -pf00163pf01479 -
02510 -pf01000pf03118pf01193 -
02515 -pf01196 -
02520 -pf01926 -
02525 -pf13185pf00990 -
02530 -pf01676 -
02535 P_viridian FRAG1
02540 P_orange FRAG1
02545 -pf00140pf04542pf04539pf04545 -
02550 -pf00877 -
02555 -# -
02560 -pf01934 -
02565 -pf01909 -
02570 -# -
02575 -# -
02580 -pf08471pf02867 -
02585 -# -
02590 -pf00005 -
(..980..)
03570 -pf13598 -
03575 -# -
03590 -pf07676 -
03595 -# -
03600 -pf14714pf01926 -
03605 -pf07992pf02852 -
03610 -# -
03615 -# -
03620 -pf13291pf13328pf04607pf02824 -
03625 -pf03652 -
03630 H_fuchsia FRAG1
03635 H_russet FRAG1
03650 -pf00293 -
03655 -# -
03660 -pf01565pf02873 -
03665 -pf00351 -
03670 -pf00578 -
03675 -pf00578 -
03680 -pf01242 -
03685 -pf01227 -
03690 -pf05635 -
03695 HMMnopfamclust3_1(pf00106) RMM1
(..11830..)
15525 -pf00679pf03144pf06421 -
15530 -pf00717 -
15545 -# -
15550 -# -
15555 -pf08241 -
15560 -# -
15565 -# -
15570 -# -
15575 -# -
15580 -# -
15585 P_battleshipGrey FRAG1
15590 -pf02481 -
15595 -pf16193pf12002pf05496 -
15600 -pf04977 -
15605 -pf07478pf01820 -
15610 HMMaldol_1_grm(pf00596) MIC4
15615 -pf01408pf02894 -
15620 -pf02219 -
15625 -pf01804 -
15630 -pf10027 -
15635 -pf13676 -
BMC-P
BMC-H
BMC-T
Aldehyde dehydrogenase
PTAC
Signature enzyme
Alcohol dehydrogenase
Regulator

Ignavibacteria bacterium RIFOXYB2_FULL_37_11. Bacteria; Ignavibacteriae; Ignavibacteria.

A2299__locus_fragmented

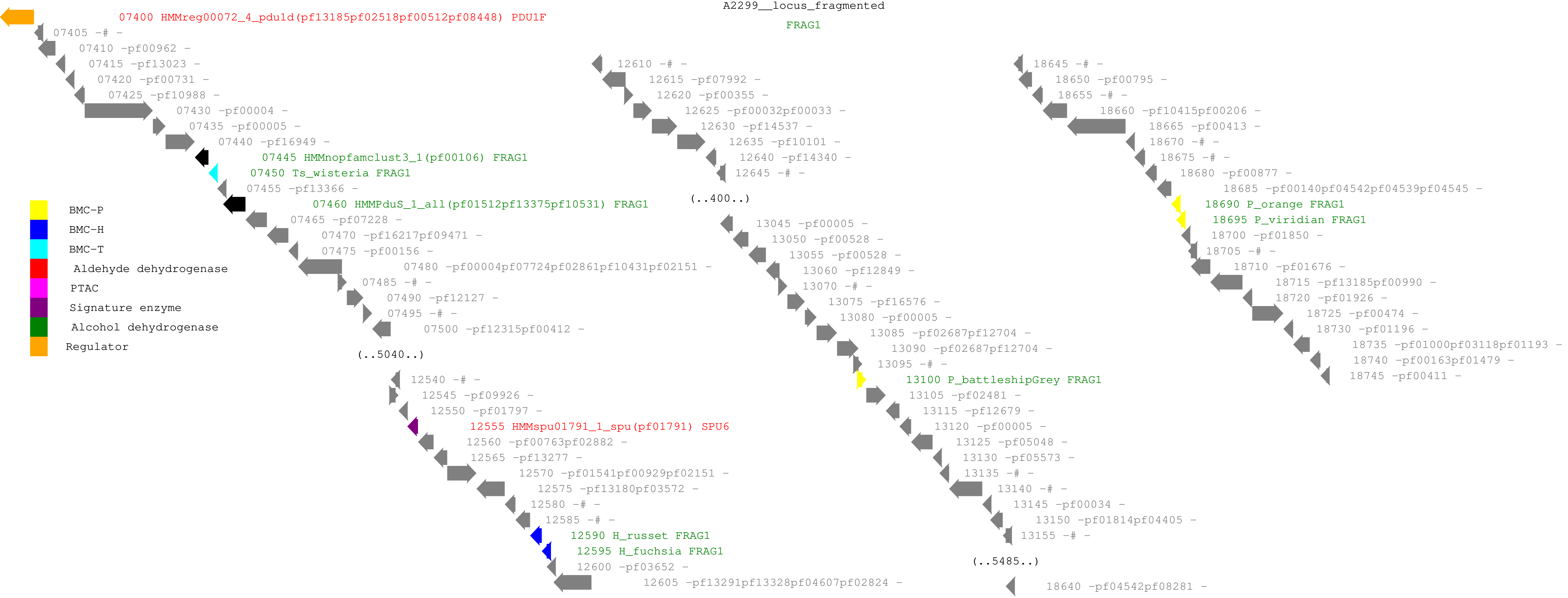

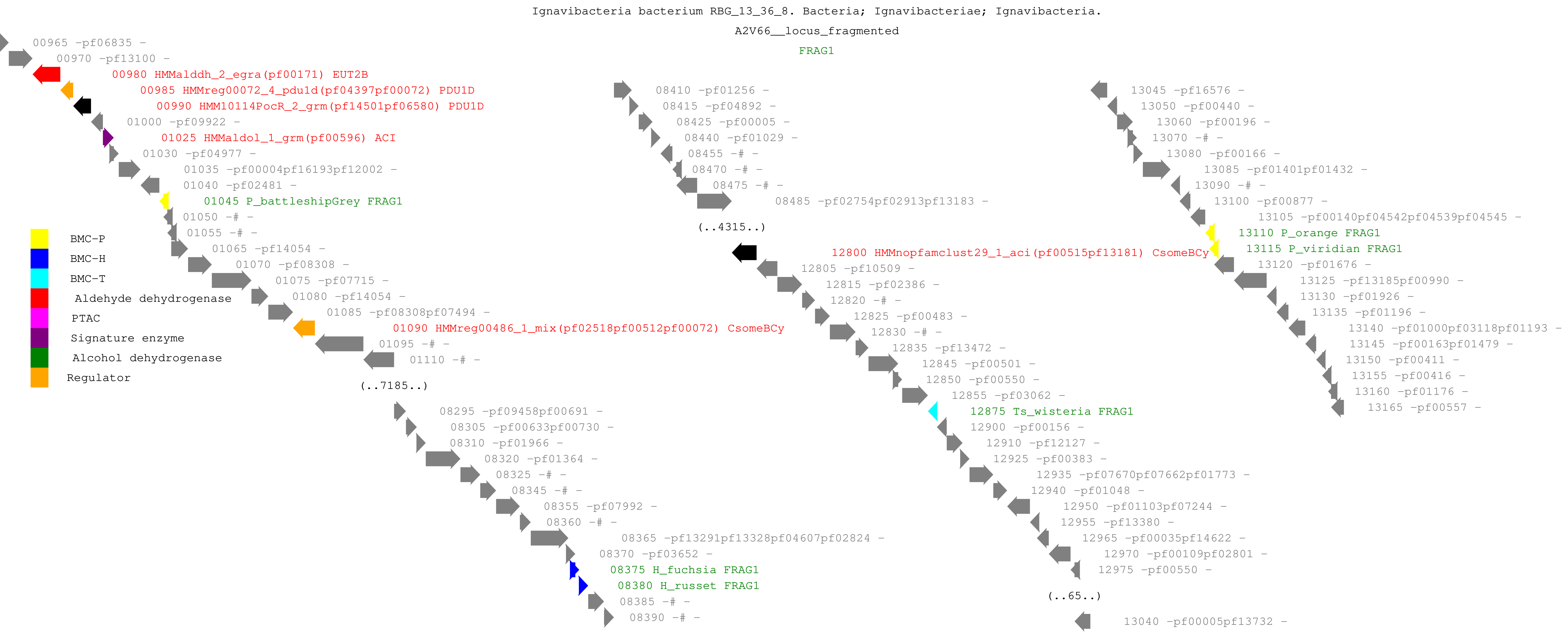
Ignavibacteria bacterium RBG_13_36_8. Bacteria; Ignavibacteriae; Ignavibacteria.
A2V66__locus_fragmented
FRAG1
00965 -pf06835 -
00970 -pf13100 -
00980 HMMalddh_2_egra(pf00171) EUT2B
00985 HMMreg00072_4_pdu1d(pf04397pf00072) PDU1D
00990 HMM10114PocR_2_grm(pf14501pf06580) PDU1D
01000 -pf09922 -
01025 HMMaldol_1_grm(pf00596) ACI
01030 -pf04977 -
01035 -pf00004pf16193pf12002 -
01040 -pf02481 -
01045 P_battleshipGrey FRAG1
01050 -# -
01055 -# -
01065 -pf14054 -
01070 -pf08308 -
01075 -pf07715 -
01080 -pf14054 -
01085 -pf08308pf07494 -
01090 HMMreg00486_1_mix(pf02518pf00512pf00072) CsomeBCy
01095 -# -
01110 -# -
(..7185..)
08295 -pf09458pf00691 -
08305 -pf00633pf00730 -
08310 -pf01966 -
08320 -pf01364 -
08325 -# -
08345 -# -
08355 -pf07992 -
08360 -# -
08365 -pf13291pf13328pf04607pf02824 -
08370 -pf03652 -
08375 H_fuchsia FRAG1
08380 H_russet FRAG1
08385 -# -
08390 -# -
BMC-P
BMC-H
BMC-T
Aldehyde dehydrogenase
PTAC
Signature enzyme
Alcohol dehydrogenase
Regulator
08410 -pf01256 -
08415 -pf04892 -
08425 -pf00005 -
08440 -pf01029 -
08455 -# -
08470 -# -
08475 -# -
08485 -pf02754pf02913pf13183 -
(..4315..)
12800 HMMnopfamclust29_1_aci(pf00515pf13181) CsomeBCy
12805 -pf10509 -
12815 -pf02386 -
12820 -# -
12825 -pf00483 -
12830 -# -
12835 -pf13472 -
12845 -pf00501 -
12850 -pf00550 -
12855 -pf03062 -
12875 Ts_wisteria FRAG1
12900 -pf00156 -
12910 -pf12127 -
12925 -pf00383 -
12935 -pf07670pf07662pf01773 -
12940 -pf01048 -
12950 -pf01103pf07244 -
12955 -pf13380 -
12965 -pf00035pf14622 -
12970 -pf00109pf02801 -
12975 -pf00550 -
(..65..)
13040 -pf00005pf13732 -
13045 -pf16576 -
13050 -pf00440 -
13060 -pf00196 -
13070 -# -
13080 -pf00166 -
13085 -pf01401pf01432 -
13090 -# -
13100 -pf00877 -
13105 -pf00140pf04542pf04539pf04545 -
13110 P_orange FRAG1
13115 P_viridian FRAG1
13120 -pf01676 -
13125 -pf13185pf00990 -
13130 -pf01926 -
13135 -pf01196 -
13140 -pf01000pf03118pf01193 -
13145 -pf00163pf01479 -
13150 -pf00411 -
13155 -pf00416 -
13160 -pf01176 -
13165 -pf00557 -

Ignavibacteria bacterium RBG_16_36_9. Bacteria; Ignavibacteriae; Ignavibacteria.

A2W30__locus_fragmented

FRAG1

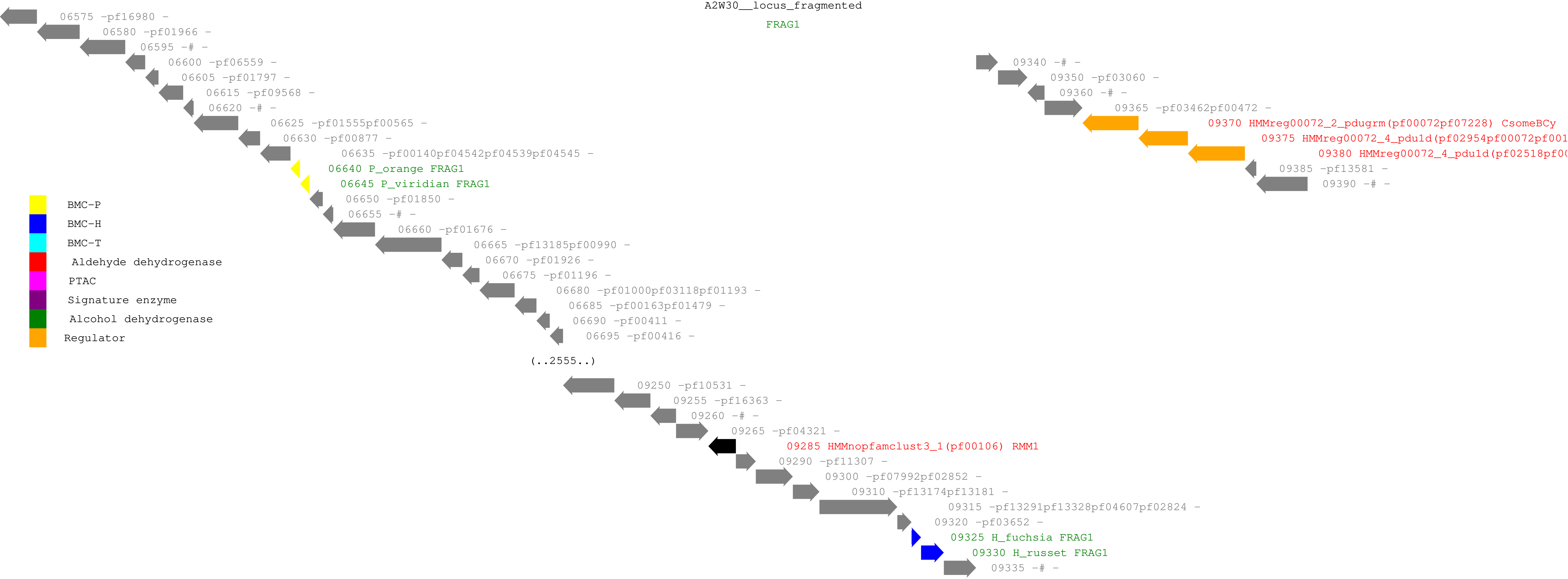

Melioribacter sp. GWF2_38_21. Bacteria; Ignavibacteriae; Ignavibacteria; Ignavibacteriales;

A2X65__locus_fragmented

FRAG1

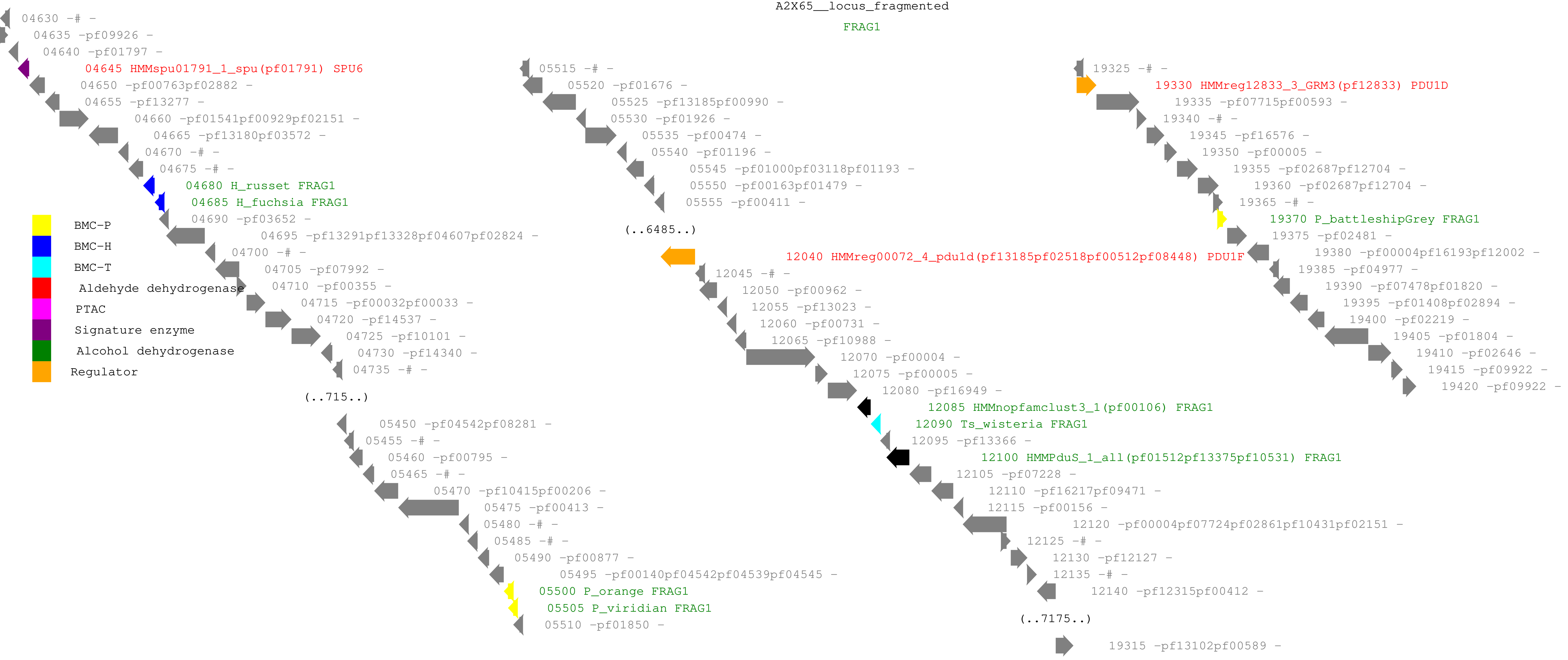

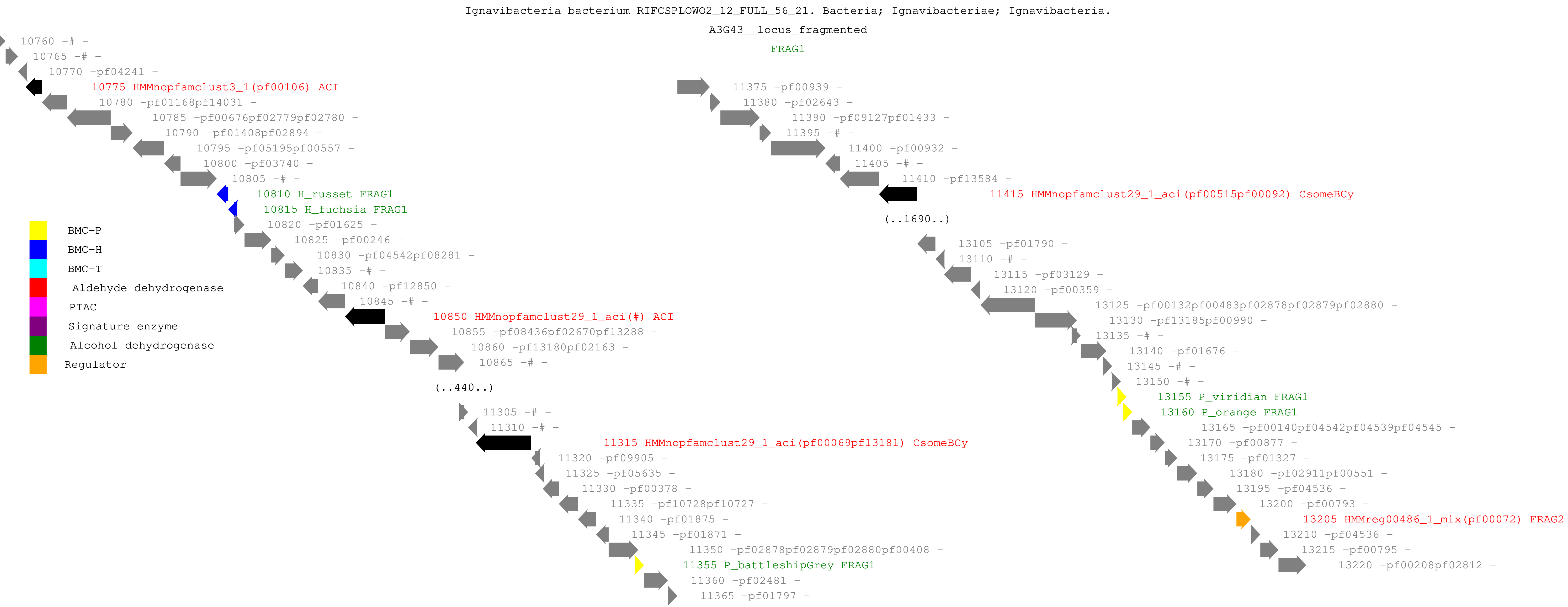

Ignavibacteria bacterium RIFCSPLOWO2_12_FULL_56_21. Bacteria; Ignavibacteriae; Ignavibacteria.
A3G43__locus_fragmented
FRAG1
10760 -# -
10765 -# -
10770 -pf04241 -
10775 HMMnopfamclust3_1(pf00106) ACI
10780 -pf01168pf14031 -
10785 -pf00676pf02779pf02780 -
10790 -pf01408pf02894 -
10795 -pf05195pf00557 -
10800 -pf03740 -
10805 -# -
10810 H_russet FRAG1
10815 H_fuchsia FRAG1
10820 -pf01625 -
10825 -pf00246 -
10830 -pf04542pf08281 -
10835 -# -
10840 -pf12850 -
10845 -# -
10850 HMMnopfamclust29_1_aci(#) ACI
10855 -pf08436pf02670pf13288 -
10860 -pf13180pf02163 -
10865 -# -
(..440..)
11305 -# -
11310 -# -
11315 HMMnopfamclust29_1_aci(pf00069pf13181) CsomeBCy
11320 -pf09905 -
11325 -pf05635 -
11330 -pf00378 -
11335 -pf10728pf10727 -
11340 -pf01875 -
11345 -pf01871 -
11350 -pf02878pf02879pf02880pf00408 -
11355 P_battleshipGrey FRAG1
11360 -pf02481 -
11365 -pf01797 -
11375 -pf00939 -
11380 -pf02643 -
11390 -pf09127pf01433 -
11395 -# -
11400 -pf00932 -
11405 -# -
11410 -pf13584 -
11415 HMMnopfamclust29_1_aci(pf00515pf00092) CsomeBCy
(..1690..)
13105 -pf01790 -
13110 -# -
13115 -pf03129 -
13120 -pf00359 -
13125 -pf00132pf00483pf02878pf02879pf02880 -
13130 -pf13185pf00990 -
13135 -# -
13140 -pf01676 -
13145 -# -
13150 -# -
13155 P_viridian FRAG1
13160 P_orange FRAG1
13165 -pf00140pf04542pf04539pf04545 -
13170 -pf00877 -
13175 -pf01327 -
13180 -pf02911pf00551 -
13195 -pf04536 -
13200 -pf00793 -
13205 HMMreg00486_1_mix(pf00072) FRAG2
13210 -pf04536 -
13215 -pf00795 -
13220 -pf00208pf02812 -
BMC-P
BMC-H
BMC-T
Aldehyde dehydrogenase
PTAC
Signature enzyme
Alcohol dehydrogenase
Regulator

Ignavibacteria bacterium RIFOXYA2_FULL_37_17. Bacteria; Ignavibacteriae; Ignavibacteria.

A3J84__locus_fragmented

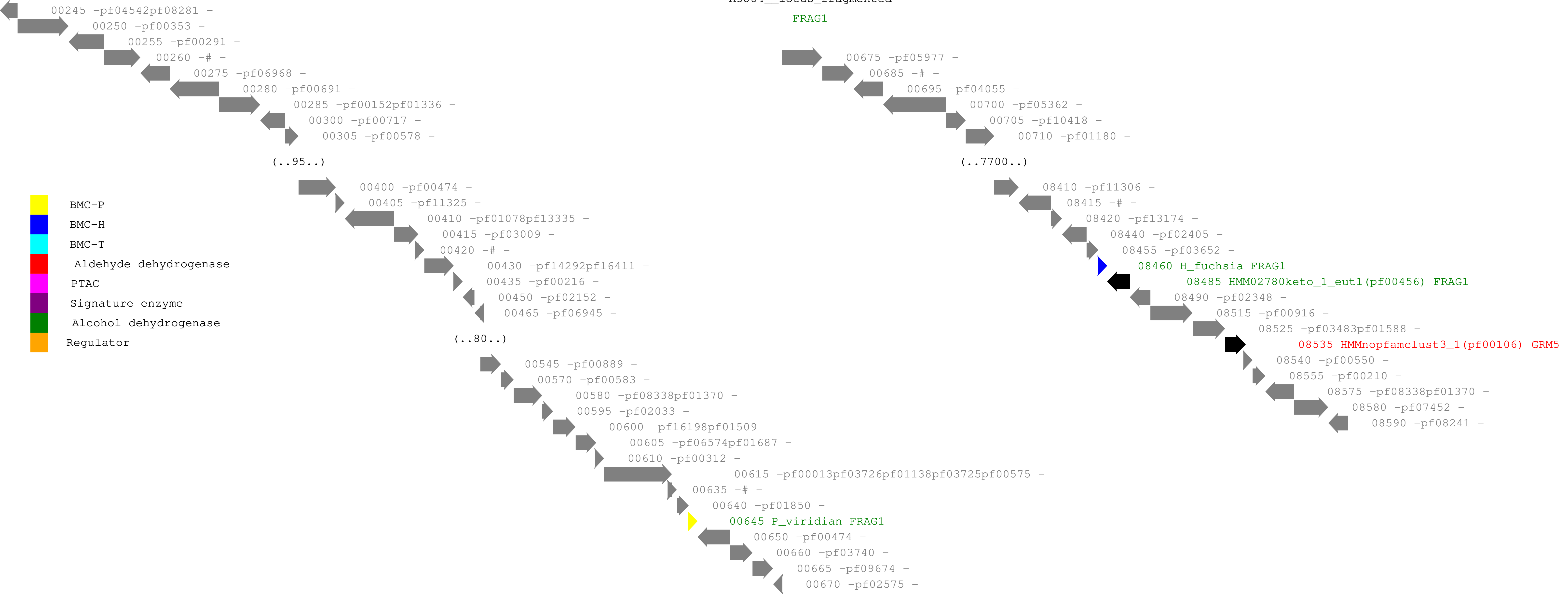

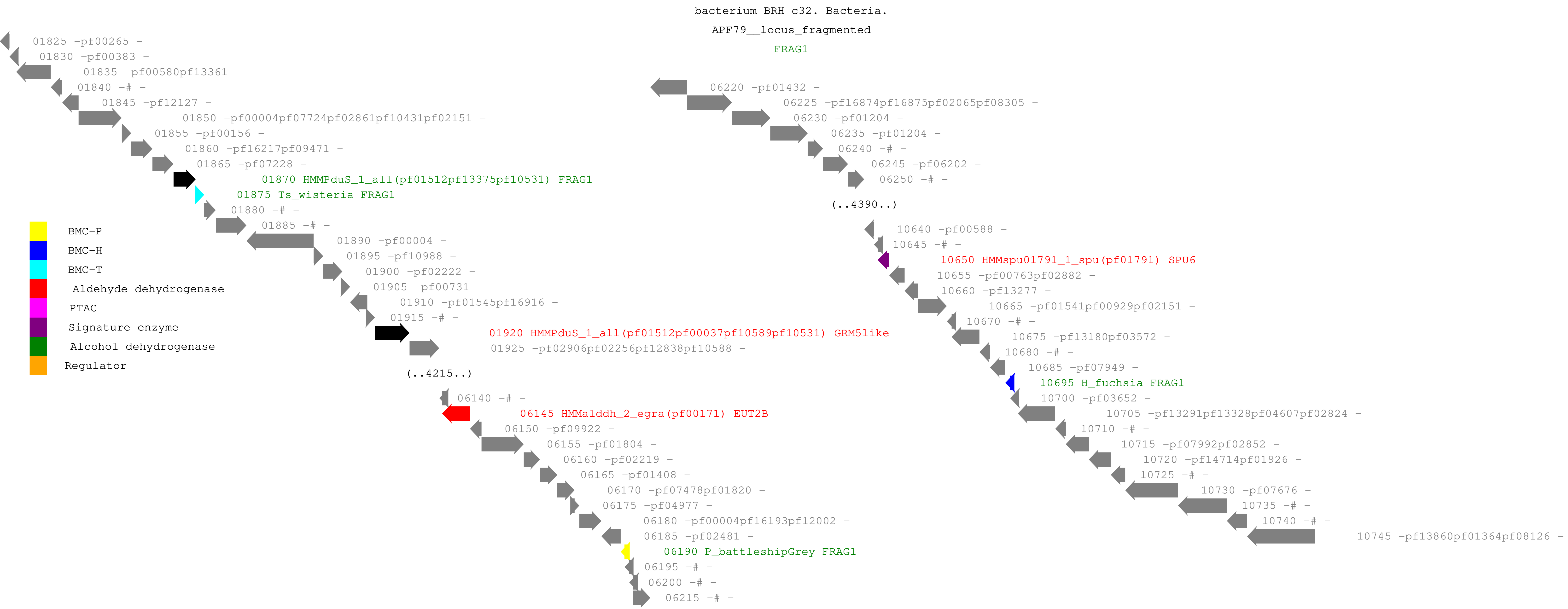
bacterium BRH_c32. Bacteria.
APF79__locus_fragmented
FRAG1
01825 -pf00265 -
01830 -pf00383 -
01835 -pf00580pf13361 -
01840 -# -
01845 -pf12127 -
01850 -pf00004pf07724pf02861pf10431pf02151 -
01855 -pf00156 -
01860 -pf16217pf09471 -
01865 -pf07228 -
01870 HMMPduS_1_all(pf01512pf13375pf10531) FRAG1
01875 Ts_wisteria FRAG1
01880 -# -
01885 -# -
01890 -pf00004 -
01895 -pf10988 -
01900 -pf02222 -
01905 -pf00731 -
01910 -pf01545pf16916 -
01915 -# -
01920 HMMPduS_1_all(pf01512pf00037pf10589pf10531) GRM5like
01925 -pf02906pf02256pf12838pf10588 -
(..4215..)
06140 -# -
06145 HMMalddh_2_egra(pf00171) EUT2B
06150 -pf09922 -
06155 -pf01804 -
06160 -pf02219 -
06165 -pf01408 -
06170 -pf07478pf01820 -
06175 -pf04977 -
06180 -pf00004pf16193pf12002 -
06185 -pf02481 -
06190 P_battleshipGrey FRAG1
06195 -# -
06200 -# -
06215 -# -
06220 -pf01432 -
06225 -pf16874pf16875pf02065pf08305 -
06230 -pf01204 -
06235 -pf01204 -
06240 -# -
06245 -pf06202 -
06250 -# -
(..4390..)
10640 -pf00588 -
10645 -# -
10650 HMMspu01791_1_spu(pf01791) SPU6
10655 -pf00763pf02882 -
10660 -pf13277 -
10665 -pf01541pf00929pf02151 -
10670 -# -
10675 -pf13180pf03572 -
10680 -# -
10685 -pf07949 -
10695 H_fuchsia FRAG1
10700 -pf03652 -
10705 -pf13291pf13328pf04607pf02824 -
10710 -# -
10715 -pf07992pf02852 -
10720 -pf14714pf01926 -
10725 -# -
10730 -pf07676 -
10735 -# -
10740 -# -
10745 -pf13860pf01364pf08126 -
BMC-P
BMC-H
BMC-T
Aldehyde dehydrogenase
PTAC
Signature enzyme
Alcohol dehydrogenase
Regulator

hydrocarbon metagenome. unclassified sequences; metagenomes; ecological metagenomes.

ASZ90__locus_fragmented

FRAG1

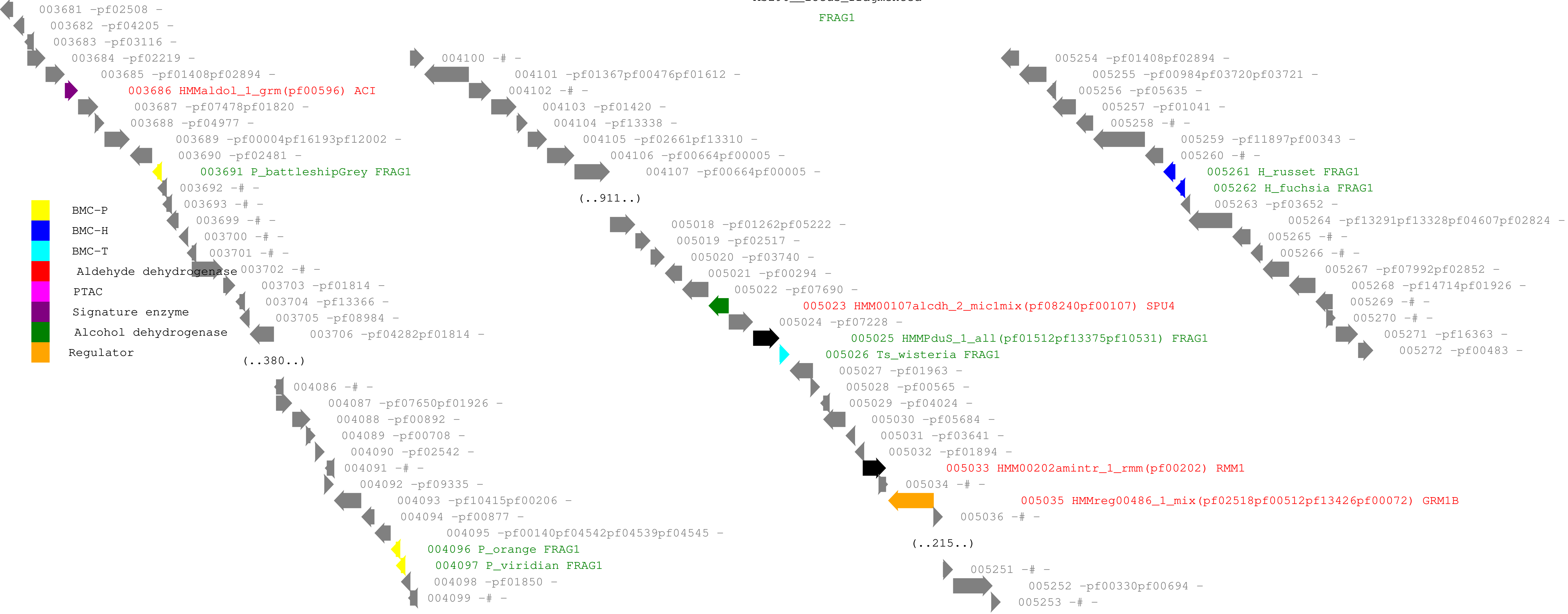

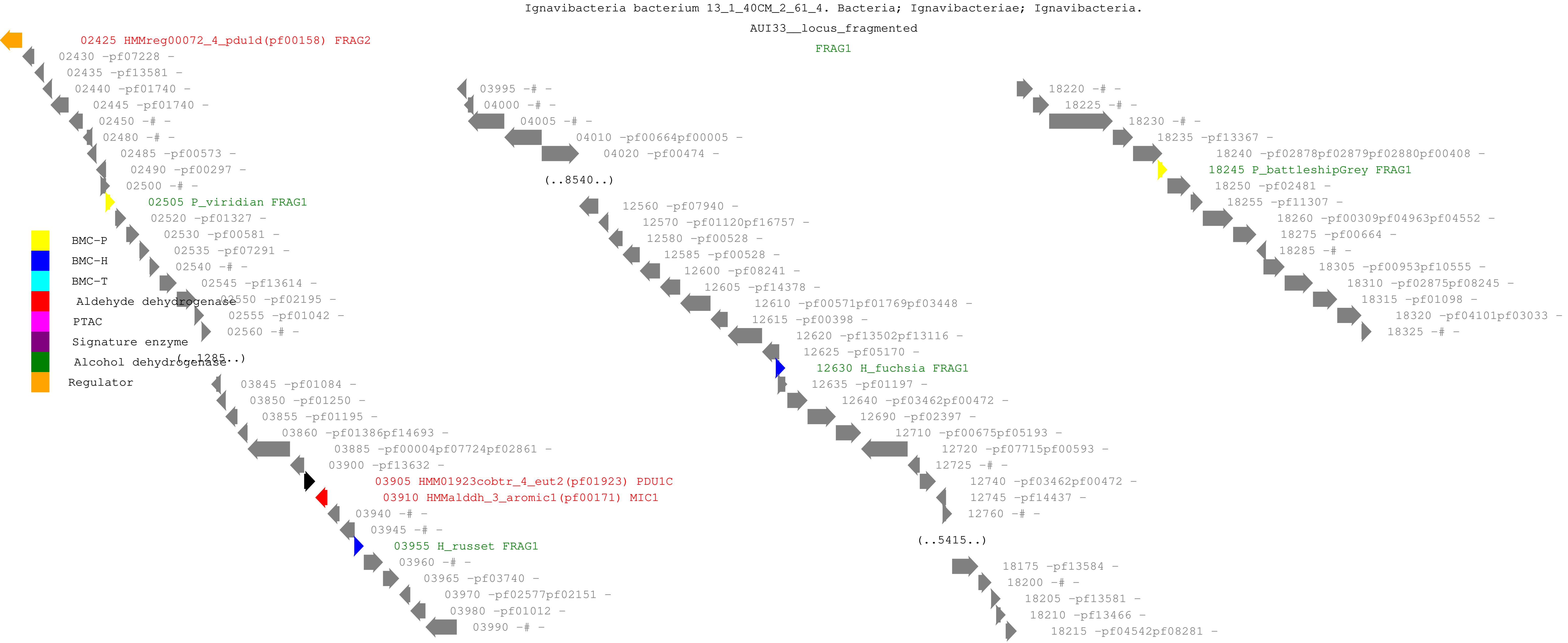

Ignavibacteria bacterium 13_1_40CM_2_61_4. Bacteria; Ignavibacteriae; Ignavibacteria.
AUI33__locus_fragmented
FRAG1
02425 HMMreg00072_4_pdu1d(pf00158) FRAG2
02430 -pf07228 -
02435 -pf13581 -
02440 -pf01740 -
02445 -pf01740 -
02450 -# -
02480 -# -
02485 -pf00573 -
02490 -pf00297 -
02500 -# -
02505 P_viridian FRAG1
02520 -pf01327 -
02530 -pf00581 -
02535 -pf07291 -
02540 -# -
02545 -pf13614 -
02550 -pf02195 -
02555 -pf01042 -
02560 -# -
(..1285..)
03845 -pf01084 -
03850 -pf01250 -
03855 -pf01195 -
03860 -pf01386pf14693 -
03885 -pf00004pf07724pf02861 -
03900 -pf13632 -
03905 HMM01923cobtr_4_eut2(pf01923) PDU1C
03910 HMMalddh_3_aromic1(pf00171) MIC1
03940 -# -
03945 -# -
03955 H_russet FRAG1
03960 -# -
03965 -pf03740 -
03970 -pf02577pf02151 -
03980 -pf01012 -
03990 -# -
BMC-P
BMC-H
BMC-T
Aldehyde dehydrogenase
PTAC
Signature enzyme
Alcohol dehydrogenase
Regulator
03995 -# -
04000 -# -
04005 -# -
04010 -pf00664pf00005 -
04020 -pf00474 -
(..8540..)
12560 -pf07940 -
12570 -pf01120pf16757 -
12580 -pf00528 -
12585 -pf00528 -
12600 -pf08241 -
12605 -pf14378 -
12610 -pf00571pf01769pf03448 -
12615 -pf00398 -
12620 -pf13502pf13116 -
12625 -pf05170 -
12630 H_fuchsia FRAG1
12635 -pf01197 -
12640 -pf03462pf00472 -
12690 -pf02397 -
12710 -pf00675pf05193 -
12720 -pf07715pf00593 -
12725 -# -
12740 -pf03462pf00472 -
12745 -pf14437 -
12760 -# -
(..5415..)
18175 -pf13584 -
18200 -# -
18205 -pf13581 -
18210 -pf13466 -
18215 -pf04542pf08281 -
18220 -# -
18225 -# -
18230 -# -
18235 -pf13367 -
18240 -pf02878pf02879pf02880pf00408 -
18245 P_battleshipGrey FRAG1
18250 -pf02481 -
18255 -pf11307 -
18260 -pf00309pf04963pf04552 -
18275 -pf00664 -
18285 -# -
18305 -pf00953pf10555 -
18310 -pf02875pf08245 -
18315 -pf01098 -
18320 -pf04101pf03033 -
18325 -# -

Ignavibacteria bacterium CG1_02_37_35. Bacteria; Ignavibacteriae; Ignavibacteria.

AUJ54__locus_fragmented

FRAG1

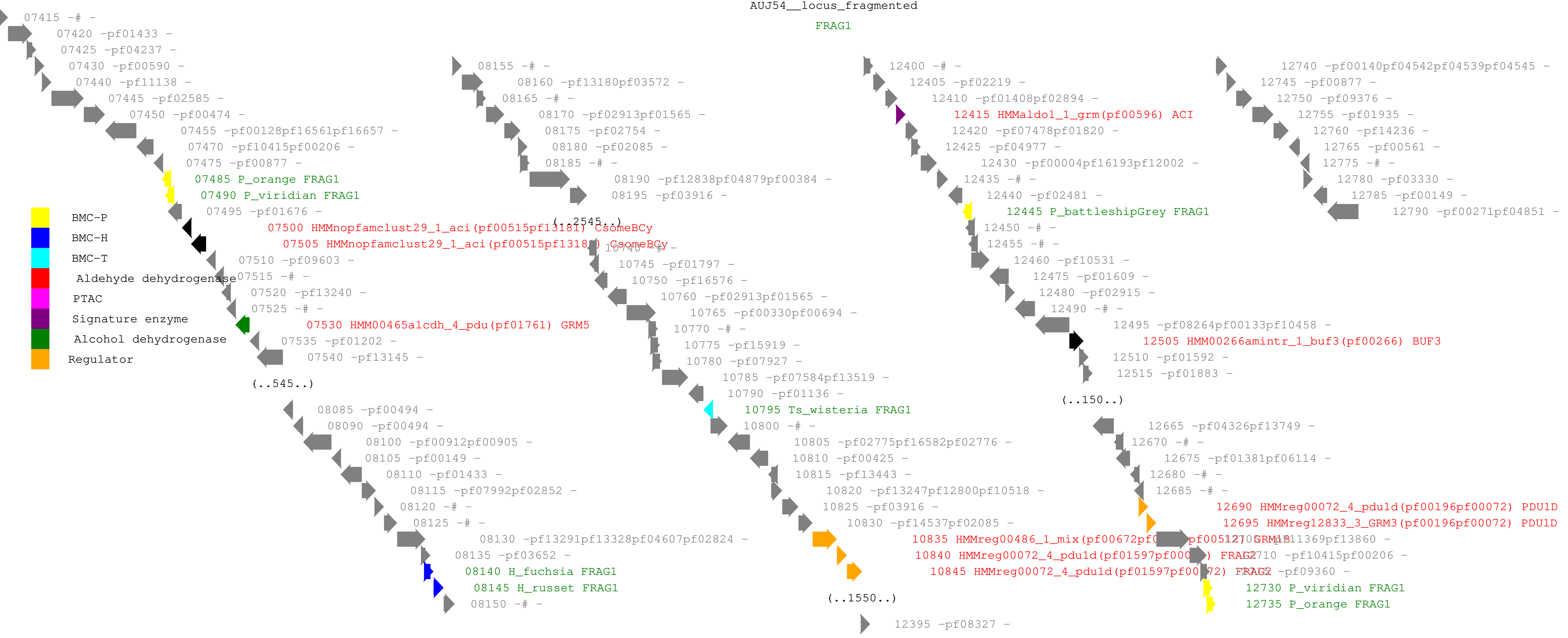

Rubrivirga sp. SAORIC476. Bacteria; Rhodothermaeota; Rhodothermia; Rhodothermales;

B1759__locus_fragmented

FRAG1

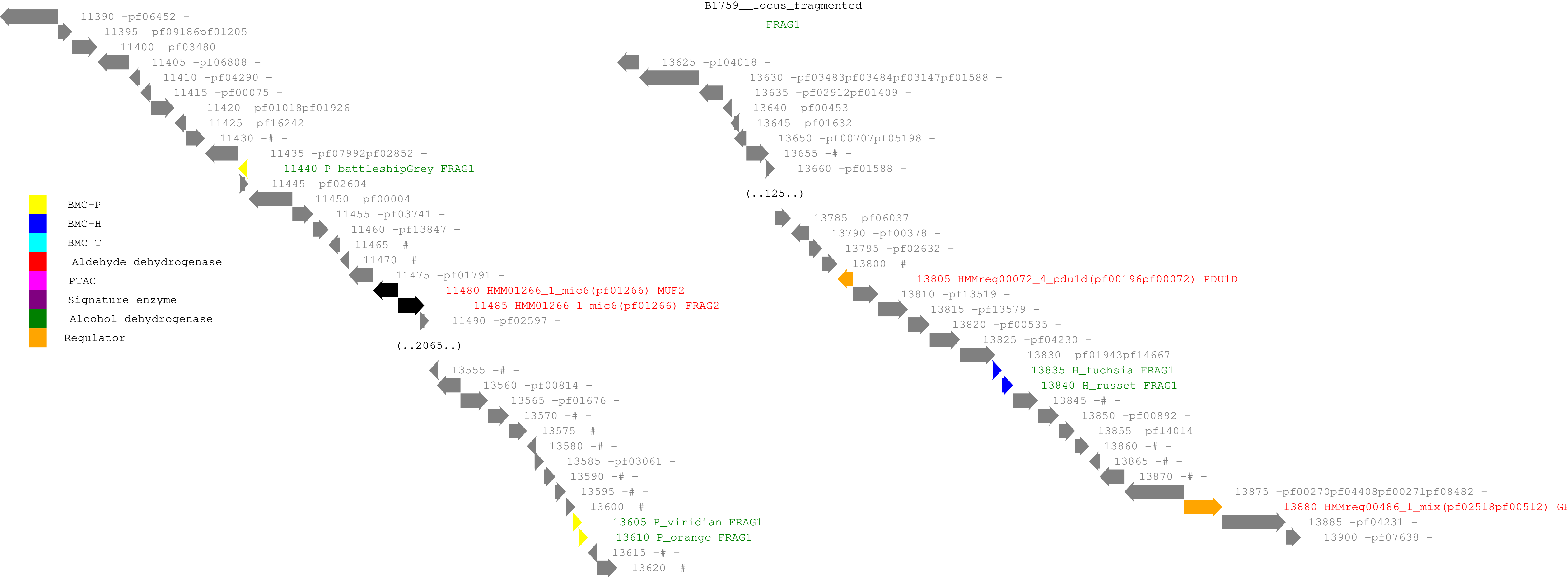

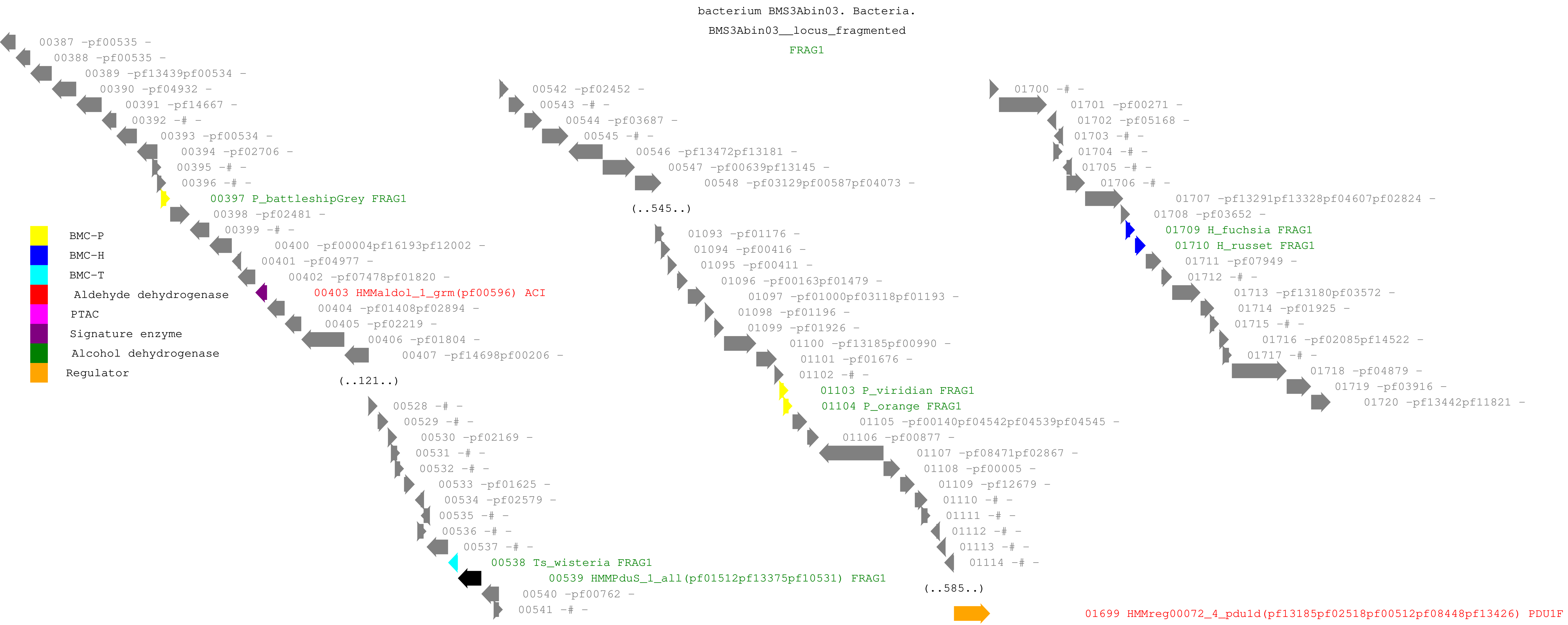
bacterium BMS3Abin03. Bacteria.
BMS3Abin03__locus_fragmented
FRAG1
00387 -pf00535 -
00388 -pf00535 -
00389 -pf13439pf00534 -
00390 -pf04932 -
00391 -pf14667 -
00392 -# -
00393 -pf00534 -
00394 -pf02706 -
00395 -# -
00396 -# -
00397 P_battleshipGrey FRAG1
00398 -pf02481 -
00399 -# -
00400 -pf00004pf16193pf12002 -
00401 -pf04977 -
00402 -pf07478pf01820 -
00403 HMMaldol_1_grm(pf00596) ACI
00404 -pf01408pf02894 -
00405 -pf02219 -
00406 -pf01804 -
00407 -pf14698pf00206 -
(..121..)
00528 -# -
00529 -# -
00530 -pf02169 -
00531 -# -
00532 -# -
00533 -pf01625 -
00534 -pf02579 -
00535 -# -
00536 -# -
00537 -# -
00538 Ts_wisteria FRAG1
00539 HMMPduS_1_all(pf01512pf13375pf10531) FRAG1
00540 -pf00762 -
00541 -# -
00542 -pf02452 -
00543 -# -
00544 -pf03687 -
00545 -# -
00546 -pf13472pf13181 -
00547 -pf00639pf13145 -
00548 -pf03129pf00587pf04073 -
(..545..)
01093 -pf01176 -
01094 -pf00416 -
01095 -pf00411 -
01096 -pf00163pf01479 -
01097 -pf01000pf03118pf01193 -
01098 -pf01196 -
01099 -pf01926 -
01100 -pf13185pf00990 -
01101 -pf01676 -
01102 -# -
01103 P_viridian FRAG1
01104 P_orange FRAG1
01105 -pf00140pf04542pf04539pf04545 -
01106 -pf00877 -
01107 -pf08471pf02867 -
01108 -pf00005 -
01109 -pf12679 -
01110 -# -
01111 -# -
01112 -# -
01113 -# -
01114 -# -
(..585..)
01699 HMMreg00072_4_pdu1d(pf13185pf02518pf00512pf08448pf13426) PDU1F
01700 -# -
01701 -pf00271 -
01702 -pf05168 -
01703 -# -
01704 -# -
01705 -# -
01706 -# -
01707 -pf13291pf13328pf04607pf02824 -
01708 -pf03652 -
01709 H_fuchsia FRAG1
01710 H_russet FRAG1
01711 -pf07949 -
01712 -# -
01713 -pf13180pf03572 -
01714 -pf01925 -
01715 -# -
01716 -pf02085pf14522 -
01717 -# -
01718 -pf04879 -
01719 -pf03916 -
01720 -pf13442pf11821 -
BMC-P
BMC-H
BMC-T
Aldehyde dehydrogenase
PTAC
Signature enzyme
Alcohol dehydrogenase
Regulator

bacterium BMS3Abin04. Bacteria.
BMS3Abin04__locus_fragmented
FRAG1

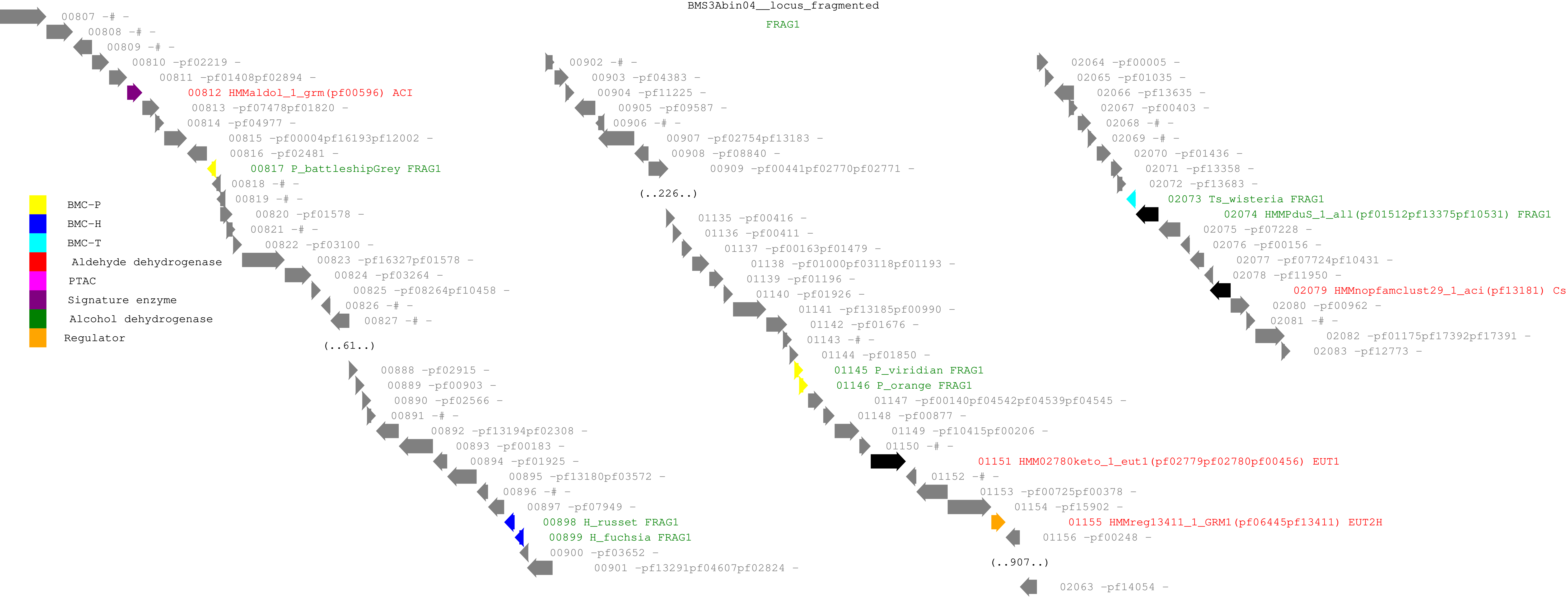

Rubricoccus marinus. Bacteria; Rhodothermaeota; Rhodothermia; Rhodothermales;

BSZ36__locus_fragmented

FRAG1

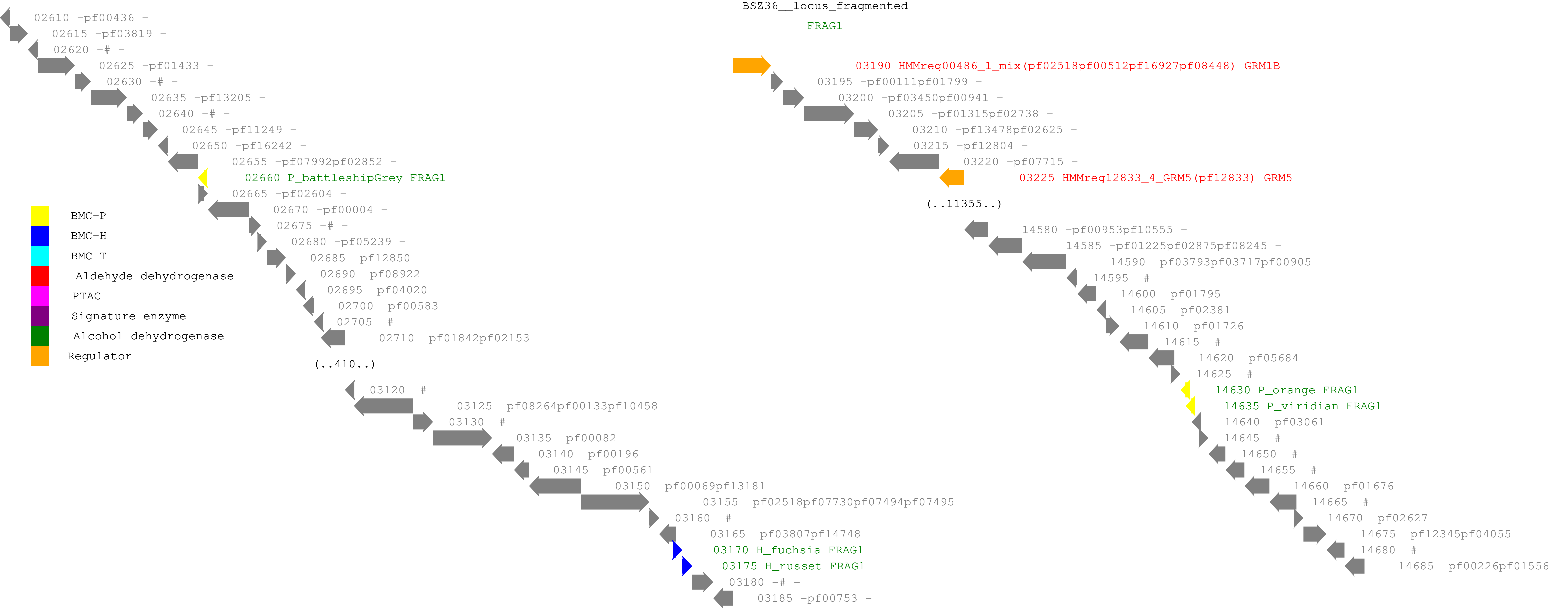

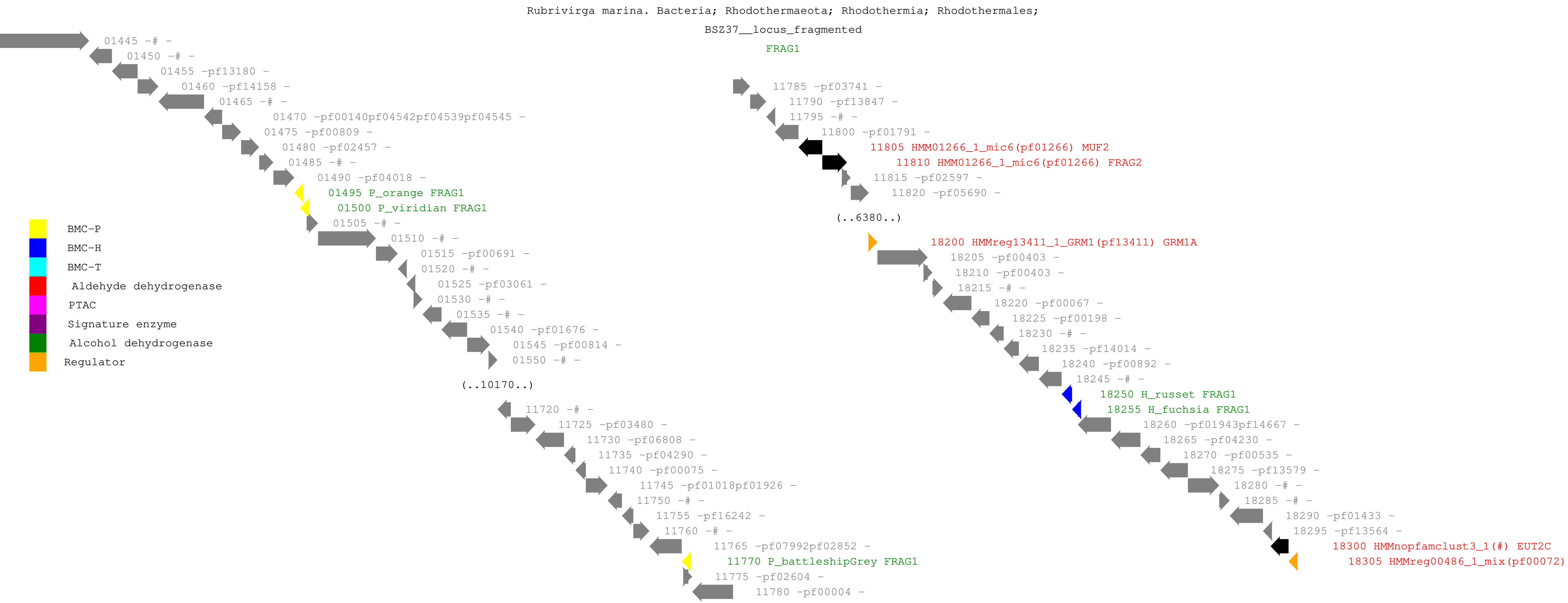
Rubrivirga marina. Bacteria; Rhodothermaeota; Rhodothermia; Rhodothermales;
BSZ37__locus_fragmented
FRAG1
01445 -# -
01450 -# -
01455 -pf13180 -
01460 -pf14158 -
01465 -# -
01470 -pf00140pf04542pf04539pf04545 -
01475 -pf00809 -
01480 -pf02457 -
01485 -# -
01490 -pf04018 -
01495 P_orange FRAG1
01500 P_viridian FRAG1
01505 -# -
01510 -# -
01515 -pf00691 -
01520 -# -
01525 -pf03061 -
01530 -# -
01535 -# -
01540 -pf01676 -
01545 -pf00814 -
01550 -# -
(..10170..)
11720 -# -
11725 -pf03480 -
11730 -pf06808 -
11735 -pf04290 -
11740 -pf00075 -
11745 -pf01018pf01926 -
11750 -# -
11755 -pf16242 -
11760 -# -
11765 -pf07992pf02852 -
11770 P_battleshipGrey FRAG1
11775 -pf02604 -
11780 -pf00004 -
11785 -pf03741 -
11790 -pf13847 -
11795 -# -
11800 -pf01791 -
11805 HMM01266_1_mic6(pf01266) MUF2
11810 HMM01266_1_mic6(pf01266) FRAG2
11815 -pf02597 -
11820 -pf05690 -
(..6380..)
18200 HMMreg13411_1_GRM1(pf13411) GRM1A
18205 -pf00403 -
18210 -pf00403 -
18215 -# -
18220 -pf00067 -
18225 -pf00198 -
18230 -# -
18235 -pf14014 -
18240 -pf00892 -
18245 -# -
18250 H_russet FRAG1
18255 H_fuchsia FRAG1
18260 -pf01943pf14667 -
18265 -pf04230 -
18270 -pf00535 -
18275 -pf13579 -
18280 -# -
18285 -# -
18290 -pf01433 -
18295 -pf13564 -
18300 HMMnopfamclust3_1(#) EUT2C
18305 HMMreg00486_1_mix(pf00072)
BMC-P
BMC-H
BMC-T
Aldehyde dehydrogenase
PTAC
Signature enzyme
Alcohol dehydrogenase
Regulator

Ignavibacteriales bacterium. Bacteria; Ignavibacteriae; Ignavibacteria; Ignavibacteriales.

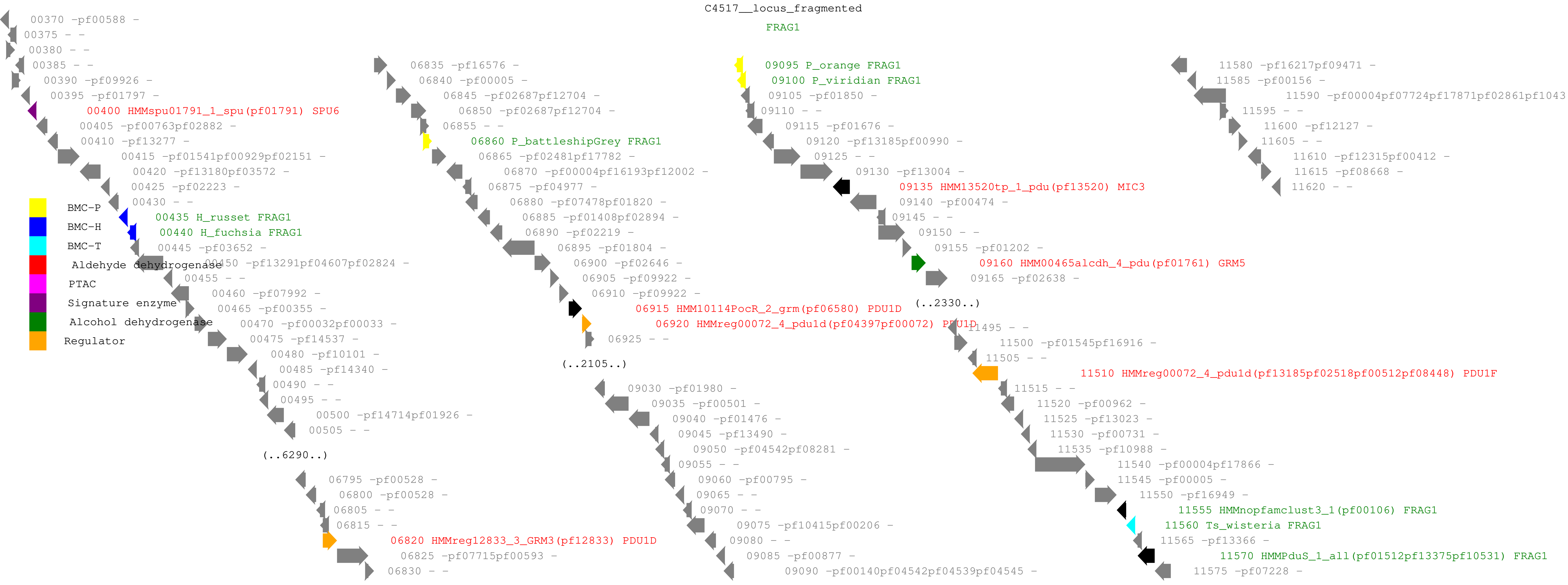

Ignavibacteriales bacterium. Bacteria; Ignavibacteriae; Ignavibacteria; Ignavibacteriales.

C4543__locus_fragmented

FRAG1

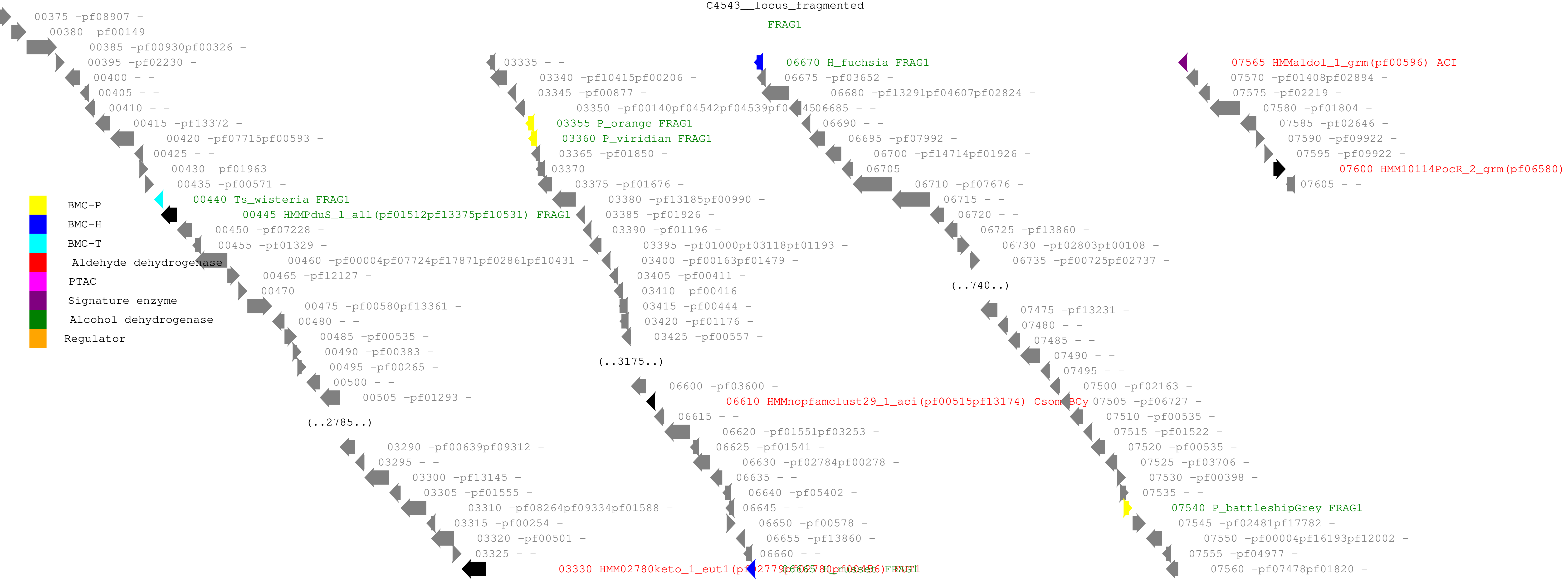

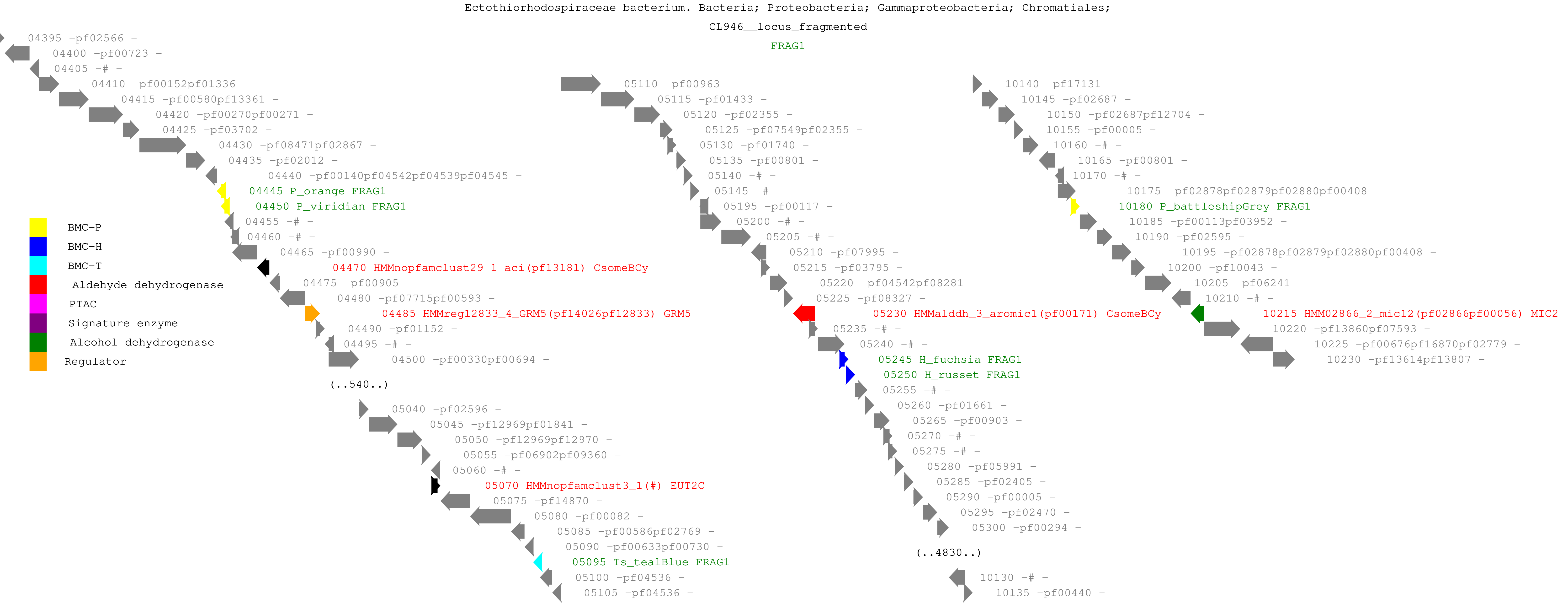

Ectothiorhodospiraceae bacterium. Bacteria; Proteobacteria; Gammaproteobacteria; Chromatiales;
CL946__locus_fragmented
FRAG1
04395 -pf02566 -
04400 -pf00723 -
04405 -# -
04410 -pf00152pf01336 -
04415 -pf00580pf13361 -
04420 -pf00270pf00271 -
04425 -pf03702 -
04430 -pf08471pf02867 -
04435 -pf02012 -
04440 -pf00140pf04542pf04539pf04545 -
04445 P_orange FRAG1
04450 P_viridian FRAG1
04455 -# -
04460 -# -
04465 -pf00990 -
04470 HMMnopfamclust29_1_aci(pf13181) CsomeBCy
04475 -pf00905 -
04480 -pf07715pf00593 -
04485 HMMreg12833_4_GRM5(pf14026pf12833) GRM5
04490 -pf01152 -
04495 -# -
04500 -pf00330pf00694 -
(..540..)
05040 -pf02596 -
05045 -pf12969pf01841 -
05050 -pf12969pf12970 -
05055 -pf06902pf09360 -
05060 -# -
05070 HMMnopfamclust3_1(#) EUT2C
05075 -pf14870 -
05080 -pf00082 -
05085 -pf00586pf02769 -
05090 -pf00633pf00730 -
05095 Ts_tealBlue FRAG1
05100 -pf04536 -
05105 -pf04536 -
05110 -pf00963 -
05115 -pf01433 -
05120 -pf02355 -
05125 -pf07549pf02355 -
05130 -pf01740 -
05135 -pf00801 -
05140 -# -
05145 -# -
05195 -pf00117 -
05200 -# -
05205 -# -
05210 -pf07995 -
05215 -pf03795 -
05220 -pf04542pf08281 -
05225 -pf08327 -
05230 HMMalddh_3_aromic1(pf00171) CsomeBCy
05235 -# -
05240 -# -
05245 H_fuchsia FRAG1
05250 H_russet FRAG1
05255 -# -
05260 -pf01661 -
05265 -pf00903 -
05270 -# -
05275 -# -
05280 -pf05991 -
05285 -pf02405 -
05290 -pf00005 -
05295 -pf02470 -
05300 -pf00294 -
(..4830..)
10130 -# -
10135 -pf00440 -
10140 -pf17131 -
10145 -pf02687 -
10150 -pf02687pf12704 -
10155 -pf00005 -
10160 -# -
10165 -pf00801 -
10170 -# -
10175 -pf02878pf02879pf02880pf00408 -
10180 P_battleshipGrey FRAG1
10185 -pf00113pf03952 -
10190 -pf02595 -
10195 -pf02878pf02879pf02880pf00408 -
10200 -pf10043 -
10205 -pf06241 -
10210 -# -
10215 HMM02866_2_mic12(pf02866pf00056) MIC2
10220 -pf13860pf07593 -
10225 -pf00676pf16870pf02779 -
10230 -pf13614pf13807 -
BMC-P
BMC-H
BMC-T
Aldehyde dehydrogenase
PTAC
Signature enzyme
Alcohol dehydrogenase
Regulator

Ignavibacteriae bacterium. Bacteria; Ignavibacteriae.

CMF23__locus_fragmented

FRAG1

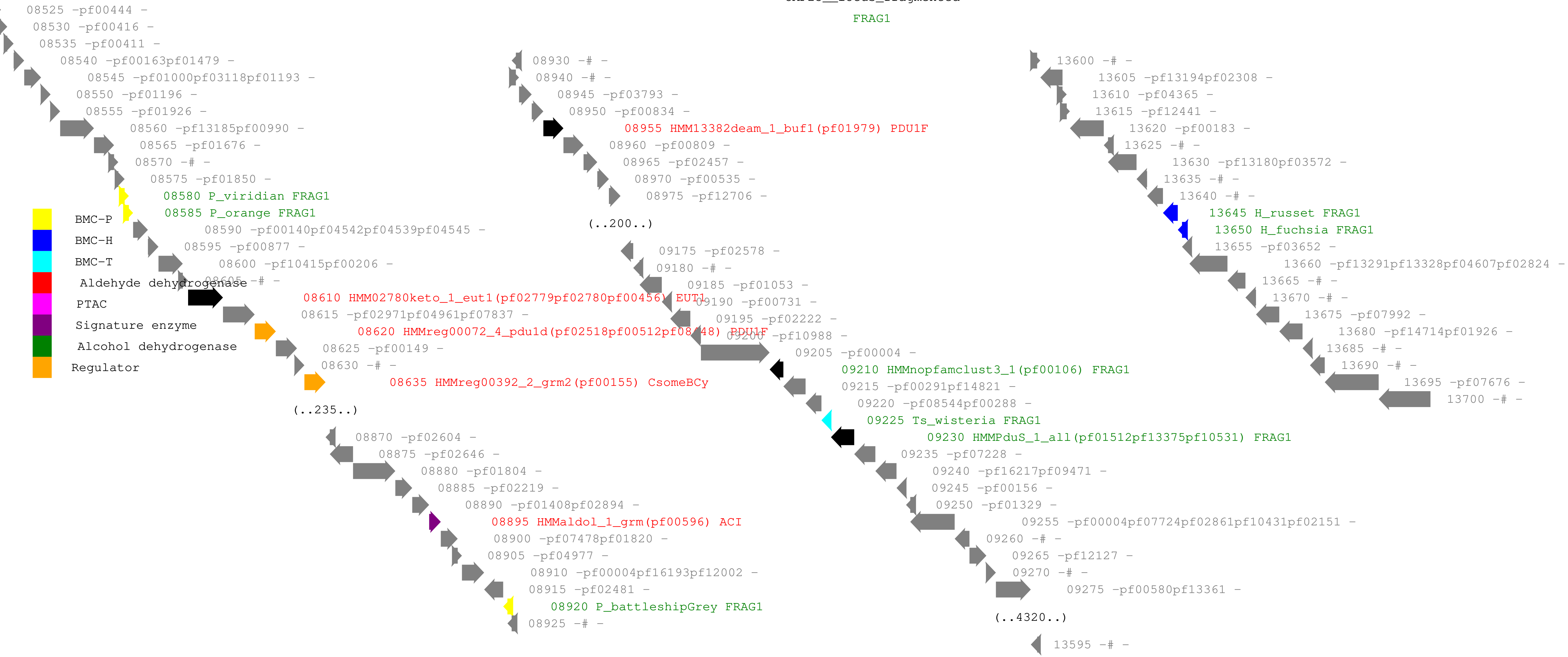

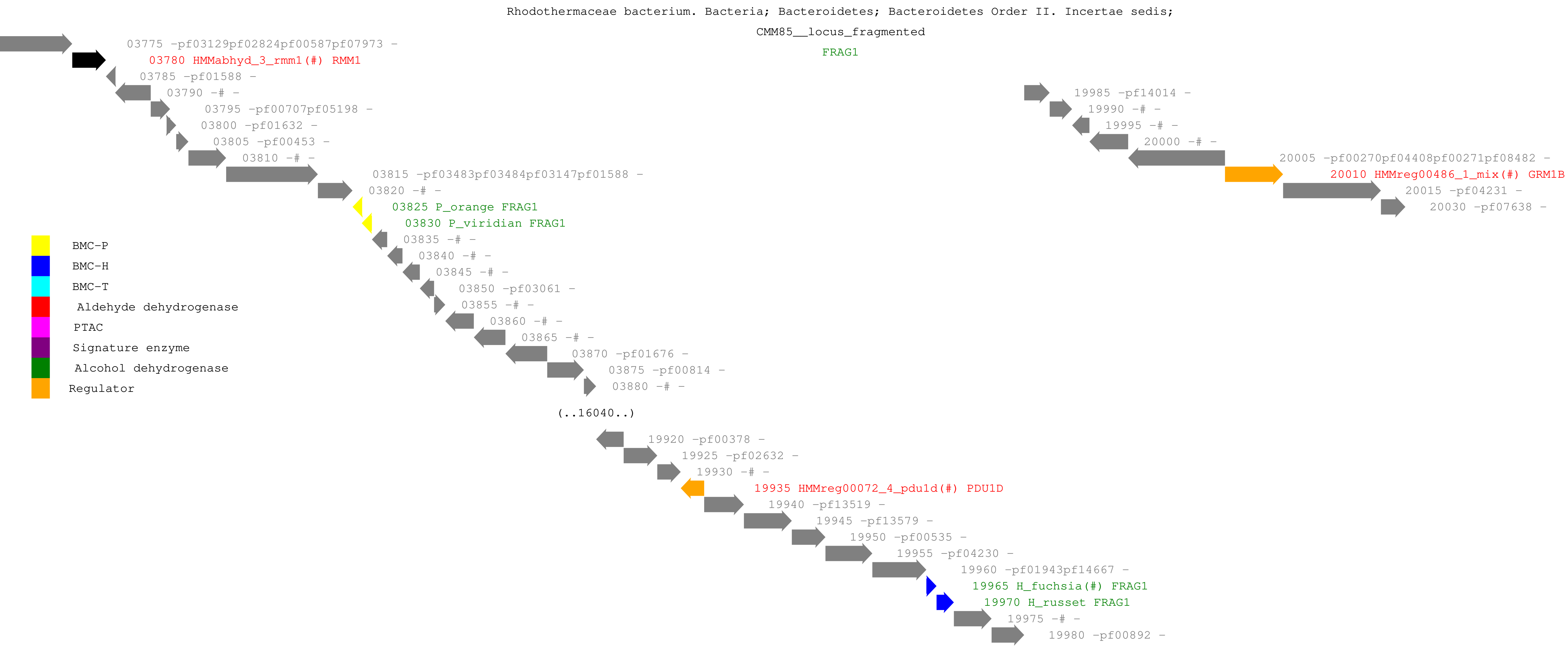

Rhodothermaceae bacterium. Bacteria; Bacteroidetes; Bacteroidetes Order II. Incertae sedis;
CMM85__locus_fragmented
FRAG1
03775 -pf03129pf02824pf00587pf07973 -
03780 HMMabhyd_3_rmm1(#) RMM1
03785 -pf01588 -
03790 -# -
03795 -pf00707pf05198 -
03800 -pf01632 -
03805 -pf00453 -
03810 -# -
03815 -pf03483pf03484pf03147pf01588 -
03820 -# -
03825 P_orange FRAG1
03830 P_viridian FRAG1
03835 -# -
03840 -# -
03845 -# -
03850 -pf03061 -
03855 -# -
03860 -# -
03865 -# -
03870 -pf01676 -
03875 -pf00814 -
03880 -# -
(..16040..)
19920 -pf00378 -
19925 -pf02632 -
19930 -# -
19935 HMMreg00072_4_pdu1d(#) PDU1D
19940 -pf13519 -
19945 -pf13579 -
19950 -pf00535 -
19955 -pf04230 -
19960 -pf01943pf14667 -
19965 H_fuchsia(#) FRAG1
19970 H_russet FRAG1
19975 -# -
19980 -pf00892 -
19985 -pf14014 -
19990 -# -
19995 -# -
20000 -# -
20005 -pf00270pf04408pf00271pf08482 -
20010 HMMreg00486_1_mix(#) GRM1B
20015 -pf04231 -
20030 -pf07638 -
BMC-P
BMC-H
BMC-T
Aldehyde dehydrogenase
PTAC
Signature enzyme
Alcohol dehydrogenase
Regulator

Ignavibacteria bacterium CG_4_9_14_0_2_um_filter_37_13. Bacteria; Ignavibacteriae; Ignavibacteria.

CO025__locus_fragmented

FRAG1

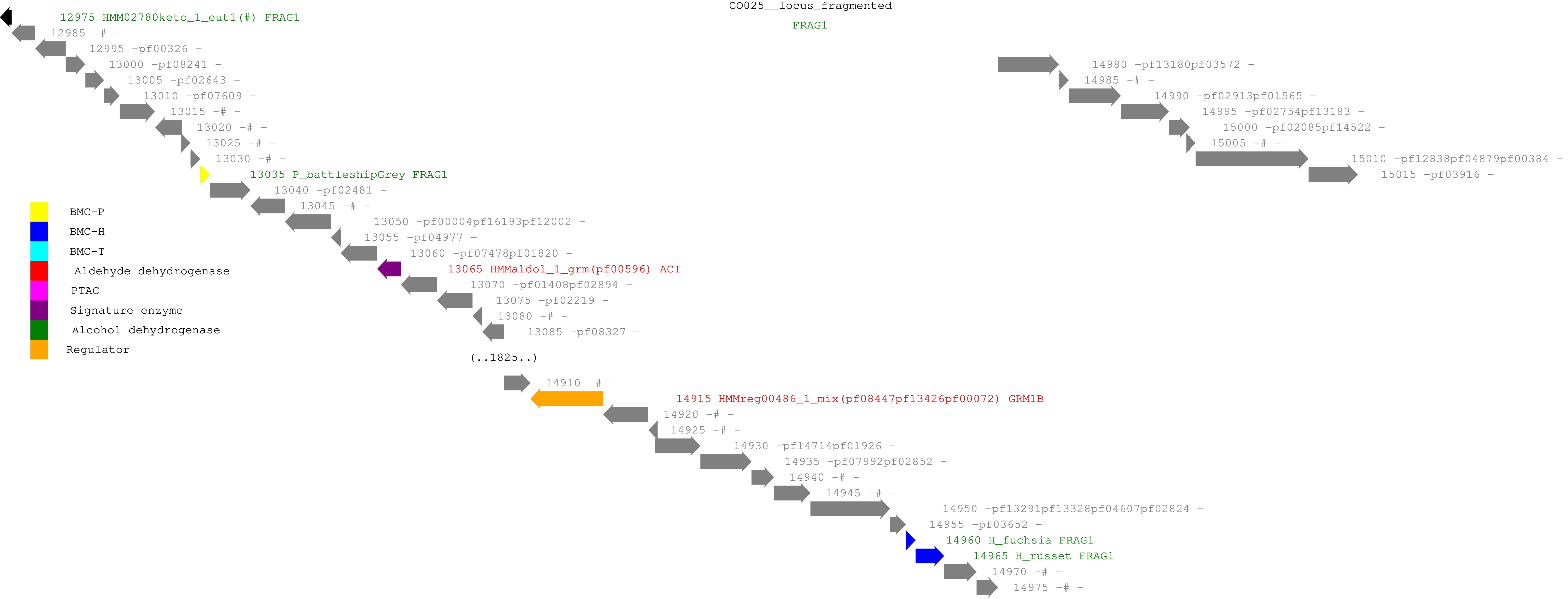

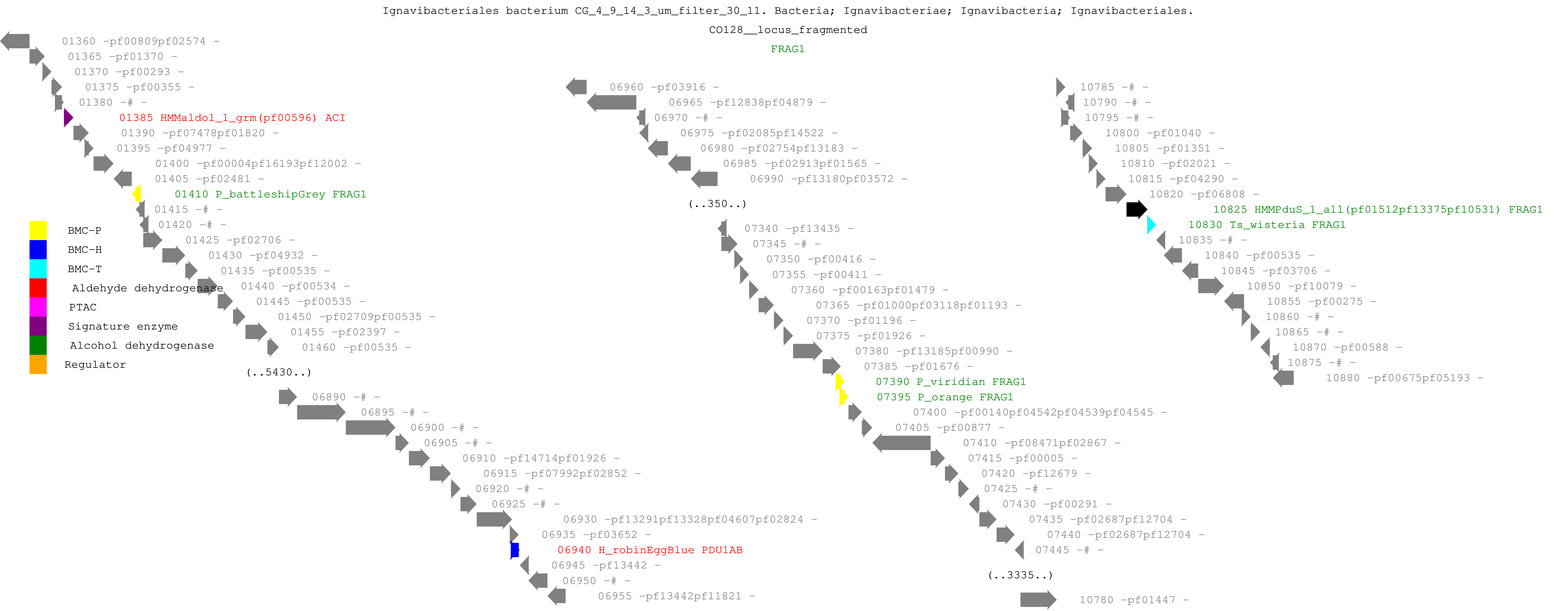
Ignavibacteriales bacterium CG_4_9_14_3_um_filter_30_11. Bacteria; Ignavibacteriae; Ignavibacteria; Ignavibacteriales.
CO128__locus_fragmented
FRAG1
01360 -pf00809pf02574 -
01365 -pf01370 -
01370 -pf00293 -
01375 -pf00355 -
01380 -# -
01385 HMMaldol_1_grm(pf00596) ACI
01390 -pf07478pf01820 -
01395 -pf04977 -
01400 -pf00004pf16193pf12002 -
01405 -pf02481 -
01410 P_battleshipGrey FRAG1
01415 -# -
01420 -# -
01425 -pf02706 -
01430 -pf04932 -
01435 -pf00535 -
01440 -pf00534 -
01445 -pf00535 -
01450 -pf02709pf00535 -
01455 -pf02397 -
01460 -pf00535 -
(..5430..)
06890 -# -
06895 -# -
06900 -# -
06905 -# -
06910 -pf14714pf01926 -
06915 -pf07992pf02852 -
06920 -# -
06925 -# -
06930 -pf13291pf13328pf04607pf02824 -
06935 -pf03652 -
06940 H_robinEggBlue PDU1AB
06945 -pf13442 -
06950 -# -
06955 -pf13442pf11821 -
BMC-P
BMC-H
BMC-T
Aldehyde dehydrogenase
PTAC
Signature enzyme
Alcohol dehydrogenase
Regulator
06960 -pf03916 -
06965 -pf12838pf04879 -
06970 -# -
06975 -pf02085pf14522 -
06980 -pf02754pf13183 -
06985 -pf02913pf01565 -
06990 -pf13180pf03572 -
(..350..)
07340 -pf13435 -
07345 -# -
07350 -pf00416 -
07355 -pf00411 -
07360 -pf00163pf01479 -
07365 -pf01000pf03118pf01193 -
07370 -pf01196 -
07375 -pf01926 -
07380 -pf13185pf00990 -
07385 -pf01676 -
07390 P_viridian FRAG1
07395 P_orange FRAG1
07400 -pf00140pf04542pf04539pf04545 -
07405 -pf00877 -
07410 -pf08471pf02867 -
07415 -pf00005 -
07420 -pf12679 -
07425 -# -
07430 -pf00291 -
07435 -pf02687pf12704 -
07440 -pf02687pf12704 -
07445 -# -
(..3335..)
10780 -pf01447 -
10785 -# -
10790 -# -
10795 -# -
10800 -pf01040 -
10805 -pf01351 -
10810 -pf02021 -
10815 -pf04290 -
10820 -pf06808 -
10825 HMMPduS_1_all(pf01512pf13375pf10531) FRAG1
10830 Ts_wisteria FRAG1
10835 -# -
10840 -pf00535 -
10845 -pf03706 -
10850 -pf10079 -
10855 -pf00275 -
10860 -# -
10865 -# -
10870 -pf00588 -
10875 -# -
10880 -pf00675pf05193 -

Ignavibacteriales bacterium CG_4_9_14_3_um_filter_34_10. Bacteria; Ignavibacteriae; Ignavibacteria; Ignavibacteriales.

CO129__locus_fragmented

FRAG1

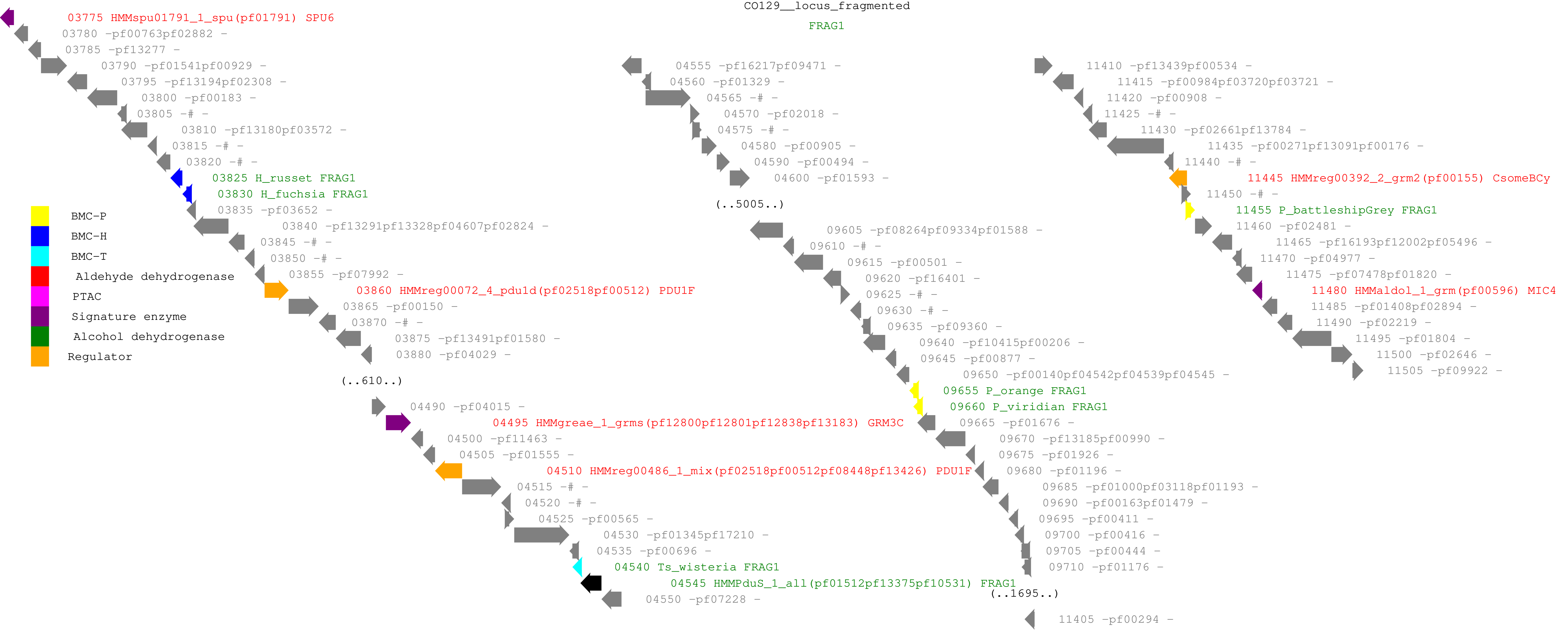

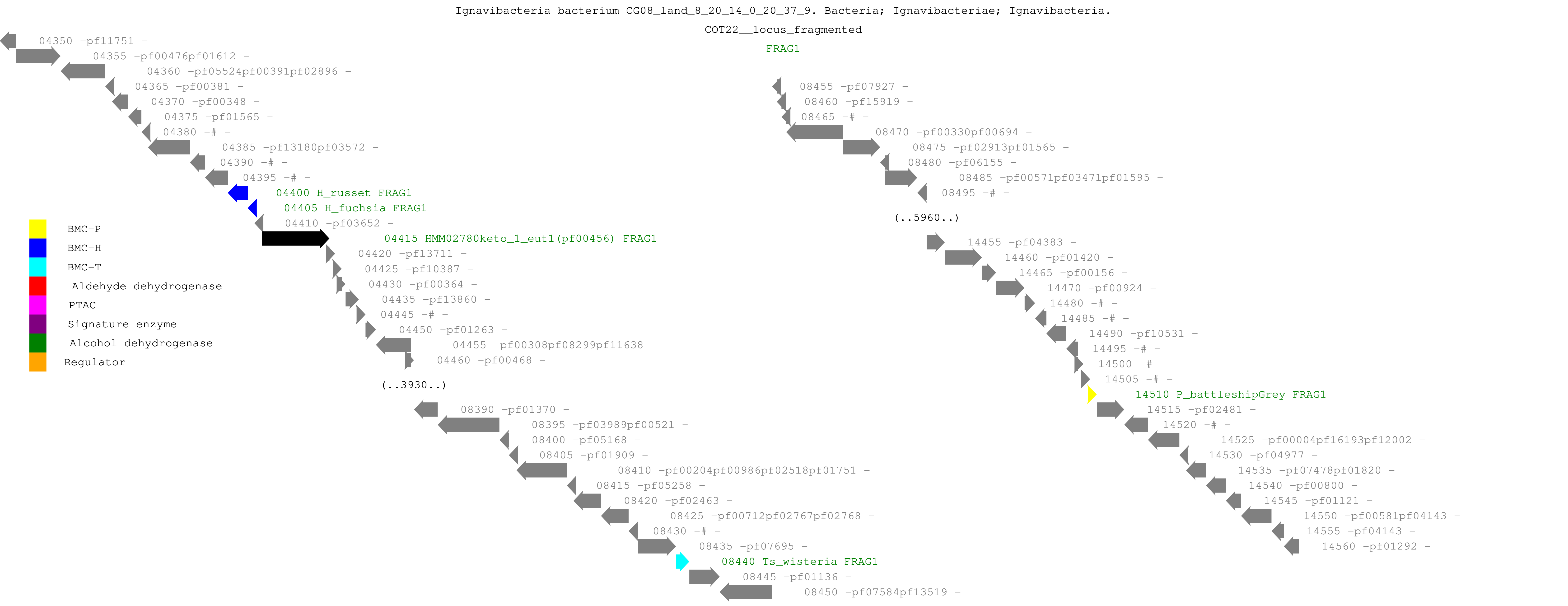
Ignavibacteria bacterium CG08_land_8_20_14_0_20_37_9. Bacteria; Ignavibacteriae; Ignavibacteria.
COT22__locus_fragmented
FRAG1
04350 -pf11751 -
04355 -pf00476pf01612 -
04360 -pf05524pf00391pf02896 -
04365 -pf00381 -
04370 -pf00348 -
04375 -pf01565 -
04380 -# -
04385 -pf13180pf03572 -
04390 -# -
04395 -# -
04400 H_russet FRAG1
04405 H_fuchsia FRAG1
04410 -pf03652 -
04415 HMM02780keto_1_eut1(pf00456) FRAG1
04420 -pf13711 -
04425 -pf10387 -
04430 -pf00364 -
04435 -pf13860 -
04445 -# -
04450 -pf01263 -
04455 -pf00308pf08299pf11638 -
04460 -pf00468 -
(..3930..)
08390 -pf01370 -
08395 -pf03989pf00521 -
08400 -pf05168 -
08405 -pf01909 -
08410 -pf00204pf00986pf02518pf01751 -
08415 -pf05258 -
08420 -pf02463 -
08425 -pf00712pf02767pf02768 -
08430 -# -
08435 -pf07695 -
08440 Ts_wisteria FRAG1
08445 -pf01136 -
08450 -pf07584pf13519 -
08455 -pf07927 -
08460 -pf15919 -
08465 -# -
08470 -pf00330pf00694 -
08475 -pf02913pf01565 -
08480 -pf06155 -
08485 -pf00571pf03471pf01595 -
08495 -# -
(..5960..)
14455 -pf04383 -
14460 -pf01420 -
14465 -pf00156 -
14470 -pf00924 -
14480 -# -
14485 -# -
14490 -pf10531 -
14495 -# -
14500 -# -
14505 -# -
14510 P_battleshipGrey FRAG1
14515 -pf02481 -
14520 -# -
14525 -pf00004pf16193pf12002 -
14530 -pf04977 -
14535 -pf07478pf01820 -
14540 -pf00800 -
14545 -pf01121 -
14550 -pf00581pf04143 -
14555 -pf04143 -
14560 -pf01292 -
BMC-P
BMC-H
BMC-T
Aldehyde dehydrogenase
PTAC
Signature enzyme
Alcohol dehydrogenase
Regulator

Bacteroidetes bacterium CG12_big_fil_rev_8_21_14_0_65_60_17. Bacteria; Bacteroidetes.

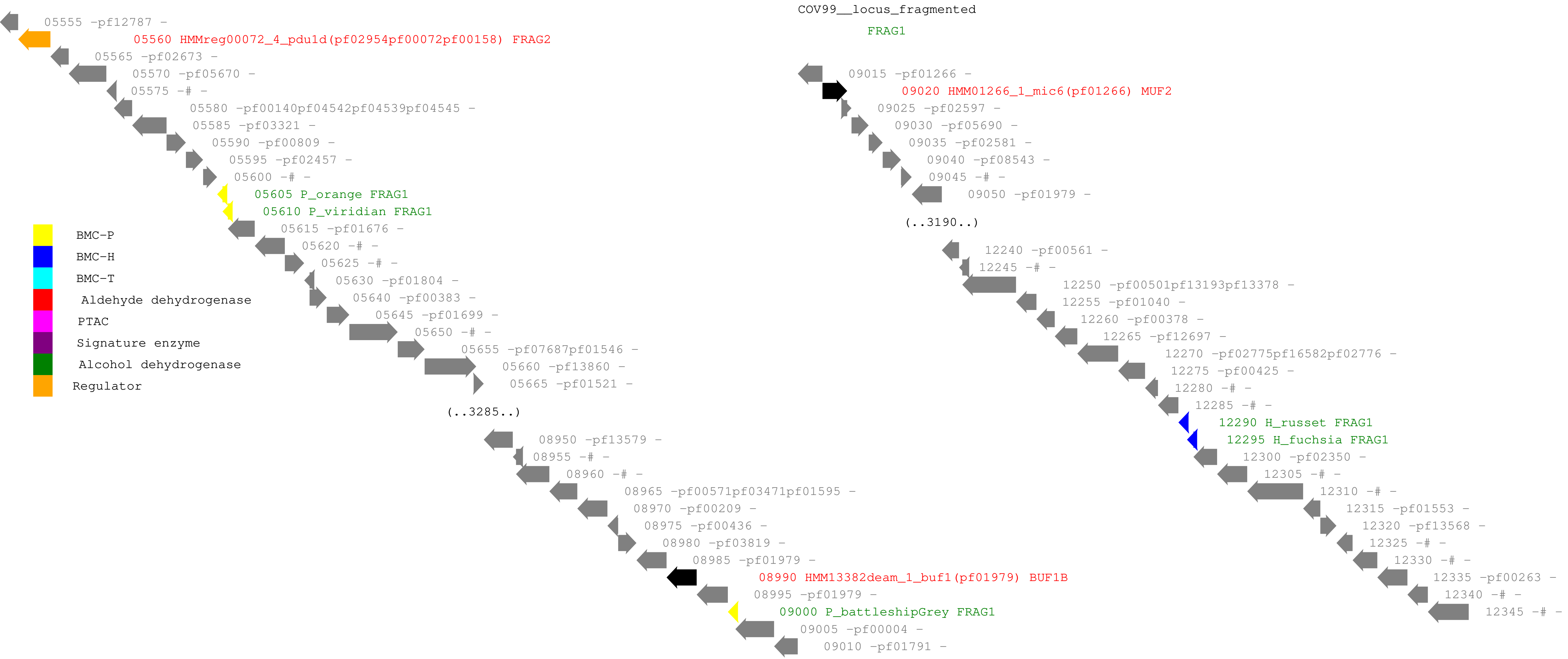

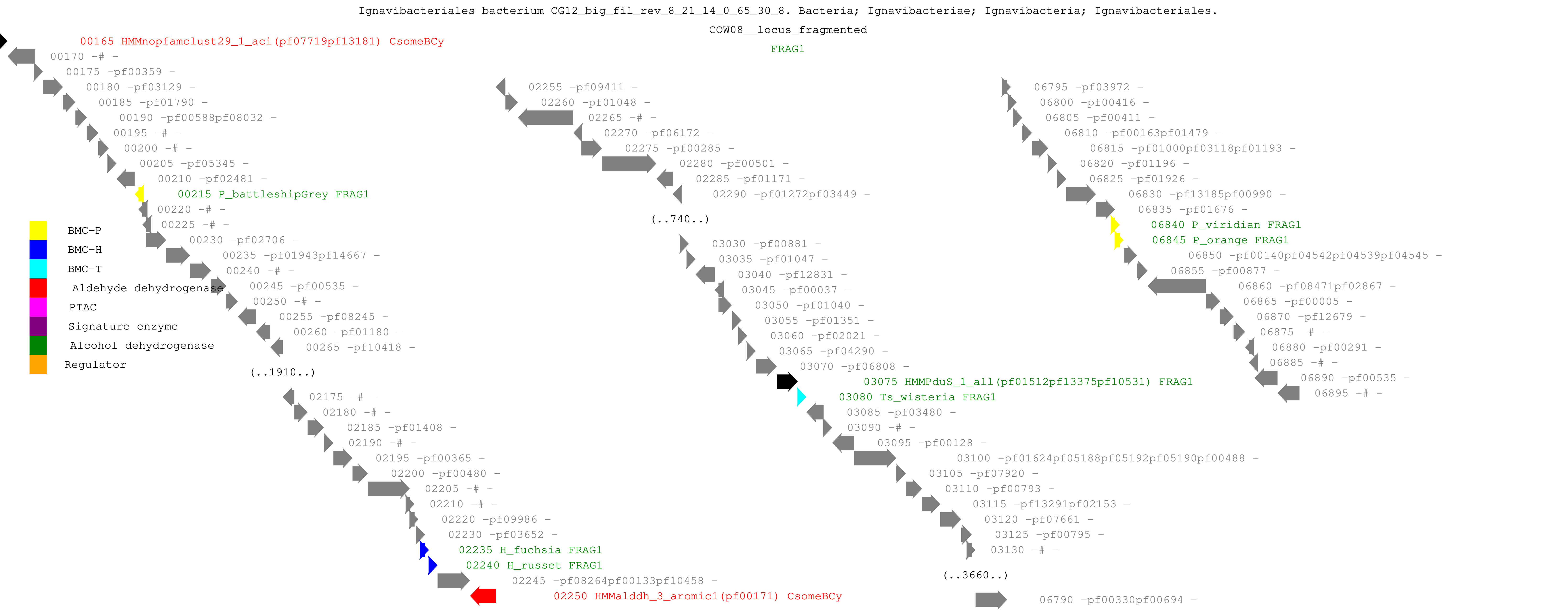
Ignavibacteriales bacterium CG12_big_fil_rev_8_21_14_0_65_30_8. Bacteria; Ignavibacteriae; Ignavibacteria; Ignavibacteriales.
COW08__locus_fragmented
FRAG1
00165 HMMnopfamclust29_1_aci(pf07719pf13181) CsomeBCy
00170 -# -
00175 -pf00359 -
00180 -pf03129 -
00185 -pf01790 -
00190 -pf00588pf08032 -
00195 -# -
00200 -# -
00205 -pf05345 -
00210 -pf02481 -
00215 P_battleshipGrey FRAG1
00220 -# -
00225 -# -
00230 -pf02706 -
00235 -pf01943pf14667 -
00240 -# -
00245 -pf00535 -
00250 -# -
00255 -pf08245 -
00260 -pf01180 -
00265 -pf10418 -
(..1910..)
02175 -# -
02180 -# -
02185 -pf01408 -
02190 -# -
02195 -pf00365 -
02200 -pf00480 -
02205 -# -
02210 -# -
02220 -pf09986 -
02230 -pf03652 -
02235 H_fuchsia FRAG1
02240 H_russet FRAG1
02245 -pf08264pf00133pf10458 -
02250 HMMaldh_3_aromic1(pf00171) CsomeBCy
BMC-P
BMC-H
BMC-T
Aldehyde dehydrogenase
PTAC
Signature enzyme
Alcohol dehydrogenase
Regulator
02255 -pf09411 -
02260 -pf01048 -
02265 -# -
02270 -pf06172 -
02275 -pf00285 -
02280 -pf00501 -
02285 -pf01171 -
02290 -pf01272pf03449 -
(..740..)
03030 -pf00881 -
03035 -pf01047 -
03040 -pf12831 -
03045 -pf00037 -
03050 -pf01040 -
03055 -pf01351 -
03060 -pf02021 -
03065 -pf04290 -
03070 -pf06808 -
03075 HMMPduS_1_all(pf01512pf13375pf10531) FRAG1
03080 Ts_wisteria FRAG1
03085 -pf03480 -
03090 -# -
03095 -pf00128 -
03100 -pf01624pf05188pf05192pf05190pf00488 -
03105 -pf07920 -
03110 -pf00793 -
03115 -pf13291pf02153 -
03120 -pf07661 -
03125 -pf00795 -
03130 -# -
(..3660..)
06790 -pf00330pf00694 -
06795 -pf03972 -
06800 -pf00416 -
06805 -pf00411 -
06810 -pf00163pf01479 -
06815 -pf01000pf03118pf01193 -
06820 -pf01196 -
06825 -pf01926 -
06830 -pf13185pf00990 -
06835 -pf01676 -
06840 P_viridian FRAG1
06845 P_orange FRAG1
06850 -pf00140pf04542pf04539pf04545 -
06855 -pf00877 -
06860 -pf08471pf02867 -
06865 -pf00005 -
06870 -pf12679 -
06875 -# -
06880 -pf00291 -
06885 -# -
06890 -pf00535 -
06895 -# -

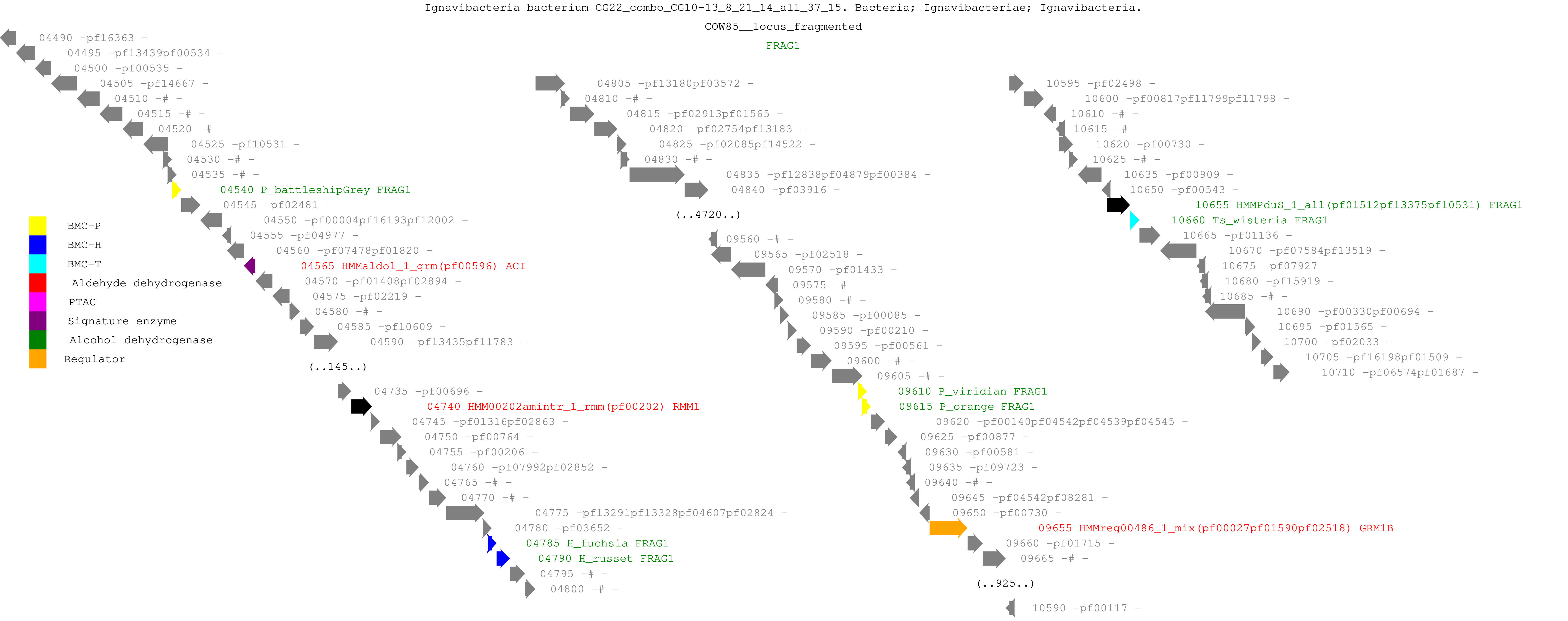

Ignavibacteria bacterium CG22_combo_CG10-13_8_21_14_all_37_15. Bacteria; Ignavibacteriae; Ignavibacteria.
COW85__locus_fragmented
FRAG1
04490 -pf16363 -
04495 -pf13439pf00534 -
04500 -pf00535 -
04505 -pf14667 -
04510 -# -
04515 -# -
04520 -# -
04525 -pf10531 -
04530 -# -
04535 -# -
04540 P_battleshipGrey FRAG1
04545 -pf02481 -
04550 -pf00004pf16193pf12002 -
04555 -pf04977 -
04560 -pf07478pf01820 -
04565 HMMaldol_1_grm(pf00596) ACI
04570 -pf01408pf02894 -
04575 -pf02219 -
04580 -# -
04585 -pf10609 -
04590 -pf13435pf11783 -
(..145..)
04735 -pf00696 -
04740 HMM00202amintr_1_rmm(pf00202) RMM1
04745 -pf01316pf02863 -
04750 -pf00764 -
04755 -pf00206 -
04760 -pf07992pf02852 -
04765 -# -
04770 -# -
04775 -pf13291pf13328pf04607pf02824 -
04780 -pf03652 -
04785 H_fuchsia FRAG1
04790 H_russet FRAG1
04795 -# -
04800 -# -
04805 -pf13180pf03572 -
04810 -# -
04815 -pf02913pf01565 -
04820 -pf02754pf13183 -
04825 -pf02085pf14522 -
04830 -# -
04835 -pf12838pf04879pf00384 -
04840 -pf03916 -
(..4720..)
09560 -# -
09565 -pf02518 -
09570 -pf01433 -
09575 -# -
09580 -# -
09585 -pf00085 -
09590 -pf00210 -
09595 -pf00561 -
09600 -# -
09605 -# -
09610 P_viridian FRAG1
09615 P_orange FRAG1
09620 -pf00140pf04542pf04539pf04545 -
09625 -pf00877 -
09630 -pf00581 -
09635 -pf09723 -
09640 -# -
09645 -pf04542pf08281 -
09650 -pf00730 -
09655 HMMreg00486_1_mix(pf00027pf01590pf02518) GRM1B
09660 -pf01715 -
09665 -# -
(..925..)
10590 -pf00117 -
10595 -pf02498 -
10600 -pf00817pf11799pf11798 -
10610 -# -
10615 -# -
10620 -pf00730 -
10625 -# -
10635 -pf00909 -
10650 -pf00543 -
10655 HMMPduS_1_all(pf01512pf13375pf10531) FRAG1
10660 Ts_wisteria FRAG1
10665 -pf01136 -
10670 -pf07584pf13519 -
10675 -pf07927 -
10680 -pf15919 -
10685 -# -
10690 -pf00330pf00694 -
10695 -pf01565 -
10700 -pf02033 -
10705 -pf16198pf01509 -
10710 -pf06574pf01687 -
BMC-P
BMC-H
BMC-T
Aldehyde dehydrogenase
PTAC
Signature enzyme
Alcohol dehydrogenase
Regulator

Ignavibacteria bacterium CG_4_10_14_3_um_filter_37_18. Bacteria; Ignavibacteriae; Ignavibacteria.

COZ25__locus_fragmented

FRAG1

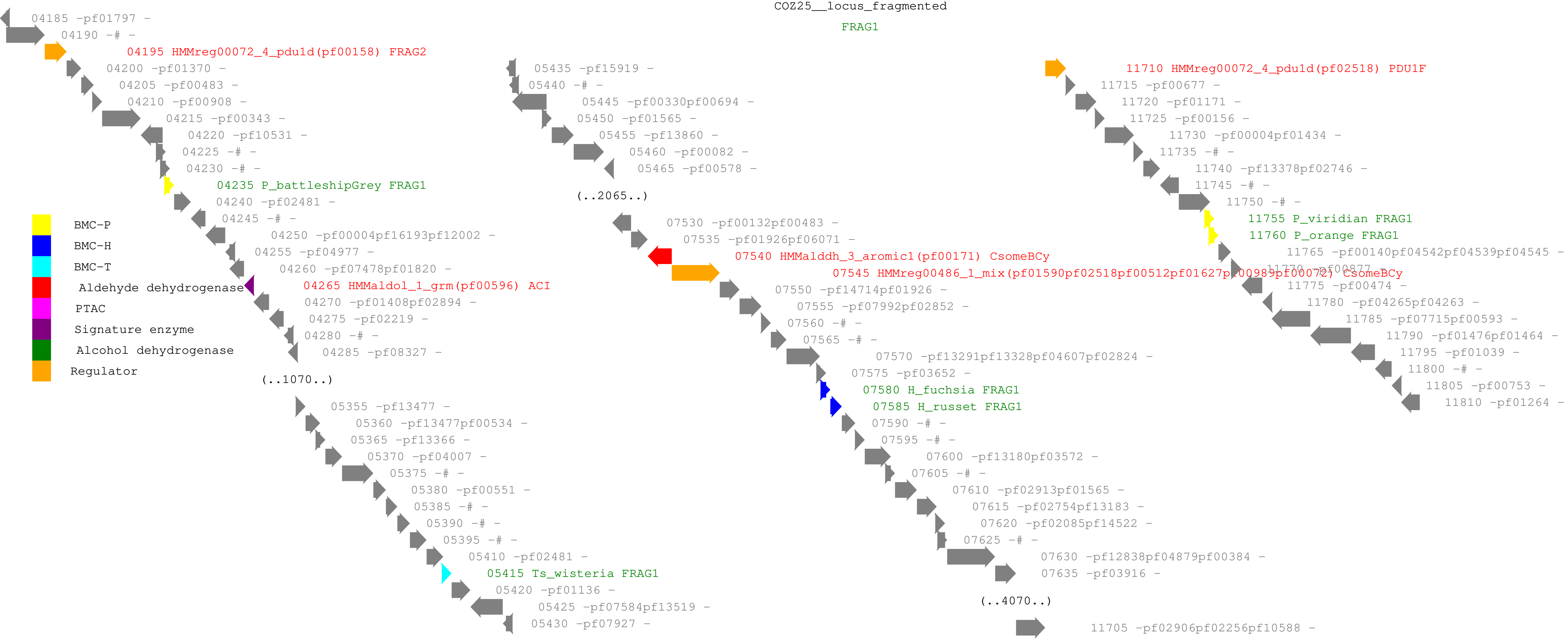

Ignavibacteria bacterium CG_4_8_14_3_um_filter_37_9. Bacteria; Ignavibacteriae; Ignavibacteria.

COZ80__locus_fragmented

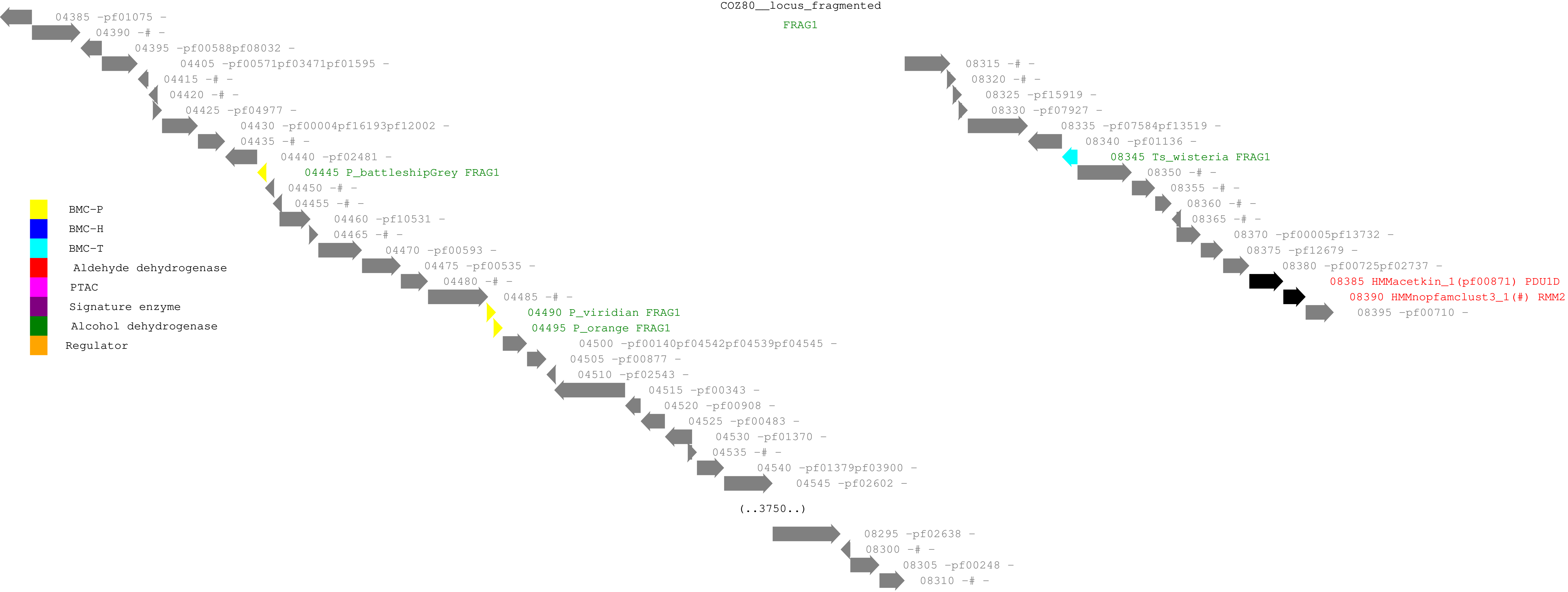

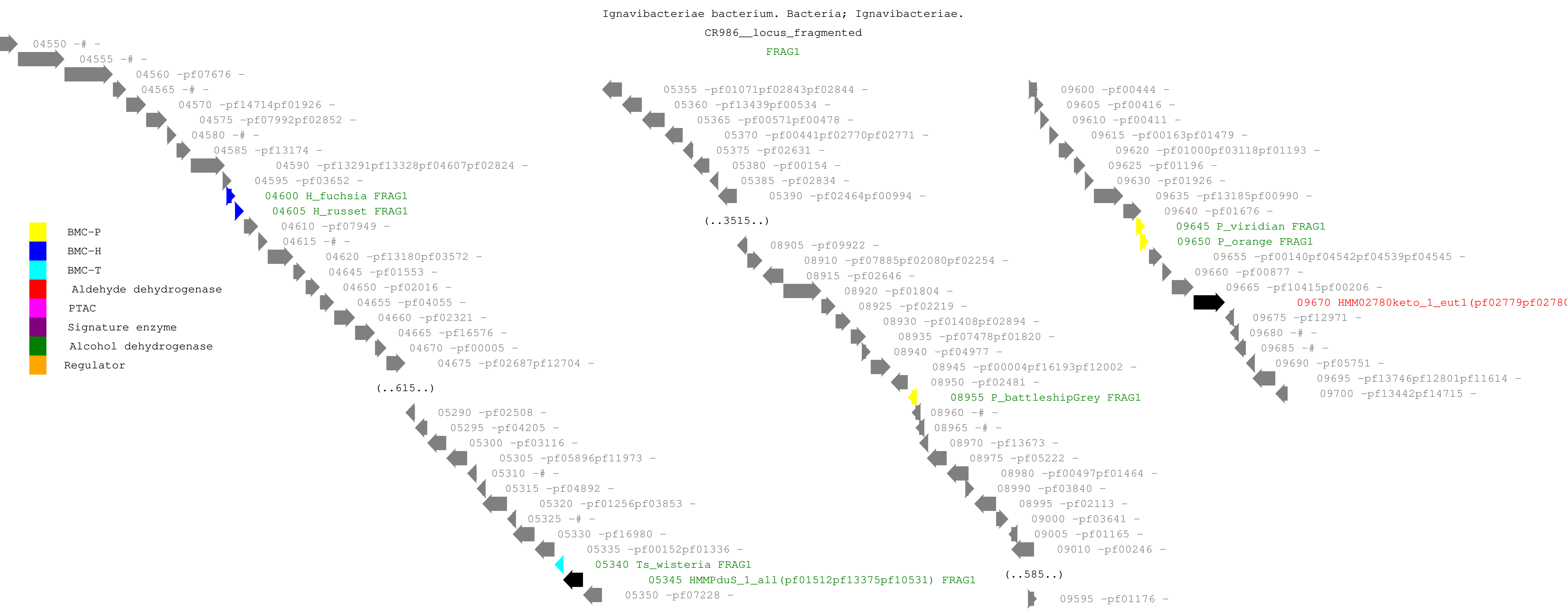

Ignavibacteriae bacterium. Bacteria; Ignavibacteriae.
CR986__locus_fragmented
FRAG1
04550 -# -
04555 -# -
04560 -pf07676 -
04565 -# -
04570 -pf14714pf01926 -
04575 -pf07992pf02852 -
04580 -# -
04585 -pf13174 -
04590 -pf13291pf13328pf04607pf02824 -
04595 -pf03652 -
04600 H_fuchsia FRAG1
04605 H_russet FRAG1
04610 -pf07949 -
04615 -# -
04620 -pf13180pf03572 -
04645 -pf01553 -
04650 -pf02016 -
04655 -pf04055 -
04660 -pf02321 -
04665 -pf16576 -
04670 -pf00005 -
04675 -pf02687pf12704 -
(..615..)
05290 -pf02508 -
05295 -pf04205 -
05300 -pf03116 -
05305 -pf05896pf11973 -
05310 -# -
05315 -pf04892 -
05320 -pf01256pf03853 -
05325 -# -
05330 -pf16980 -
05335 -pf00152pf01336 -
05340 Ts_wisteria FRAG1
05345 HMMPduS_1_all(pf01512pf13375pf10531) FRAG1
05350 -pf07228 -
BMC-P
BMC-H
BMC-T
Aldehyde dehydrogenase
PTAC
Signature enzyme
Alcohol dehydrogenase
Regulator
05355 -pf01071pf02843pf02844 -
05360 -pf13439pf00534 -
05365 -pf00571pf00478 -
05370 -pf00441pf02770pf02771 -
05375 -pf02631 -
05380 -pf00154 -
05385 -pf02834 -
05390 -pf02464pf00994 -
(..3515..)
08905 -pf09922 -
08910 -pf07885pf02080pf02254 -
08915 -pf02646 -
08920 -pf01804 -
08925 -pf02219 -
08930 -pf01408pf02894 -
08935 -pf07478pf01820 -
08940 -pf04977 -
08945 -pf00004pf16193pf12002 -
08950 -pf02481 -
08955 P_battleshipGrey FRAG1
08960 -# -
08965 -# -
08970 -pf13673 -
08975 -pf05222 -
08980 -pf00497pf01464 -
08990 -pf03840 -
08995 -pf02113 -
09000 -pf03641 -
09005 -pf01165 -
09010 -pf00246 -
(..585..)
09595 -pf01176 -
09600 -pf00444 -
09605 -pf00416 -
09610 -pf00411 -
09615 -pf00163pf01479 -
09620 -pf01000pf03118pf01193 -
09625 -pf01196 -
09630 -pf01926 -
09635 -pf13185pf00990 -
09640 -pf01676 -
09645 P_viridian FRAG1
09650 P_orange FRAG1
09655 -pf00140pf04542pf04539pf04545 -
09660 -pf00877 -
09665 -pf10415pf00206 -
09670 HMM02780keto_1_eut1(pf02779pf02780
09675 -pf12971 -
09680 -# -
09685 -# -
09690 -pf05751 -
09695 -pf13746pf12801pf11614 -
09700 -pf13442pf14715 -

Chloroherpeton thalassium (strain ATCC 35110 / GB-78). Bacteria; Chlorobi; Chlorobia; Chlorobiales; Chlorobiaceae;

Ctha__locus_fragmented

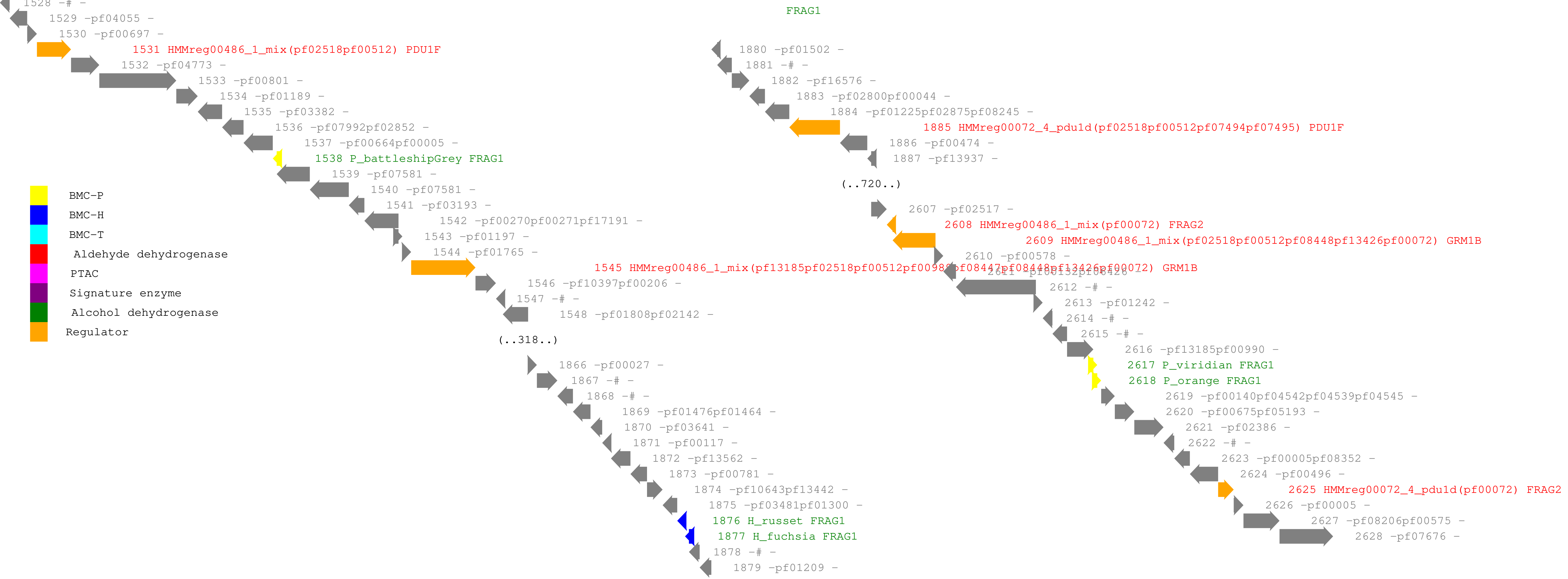

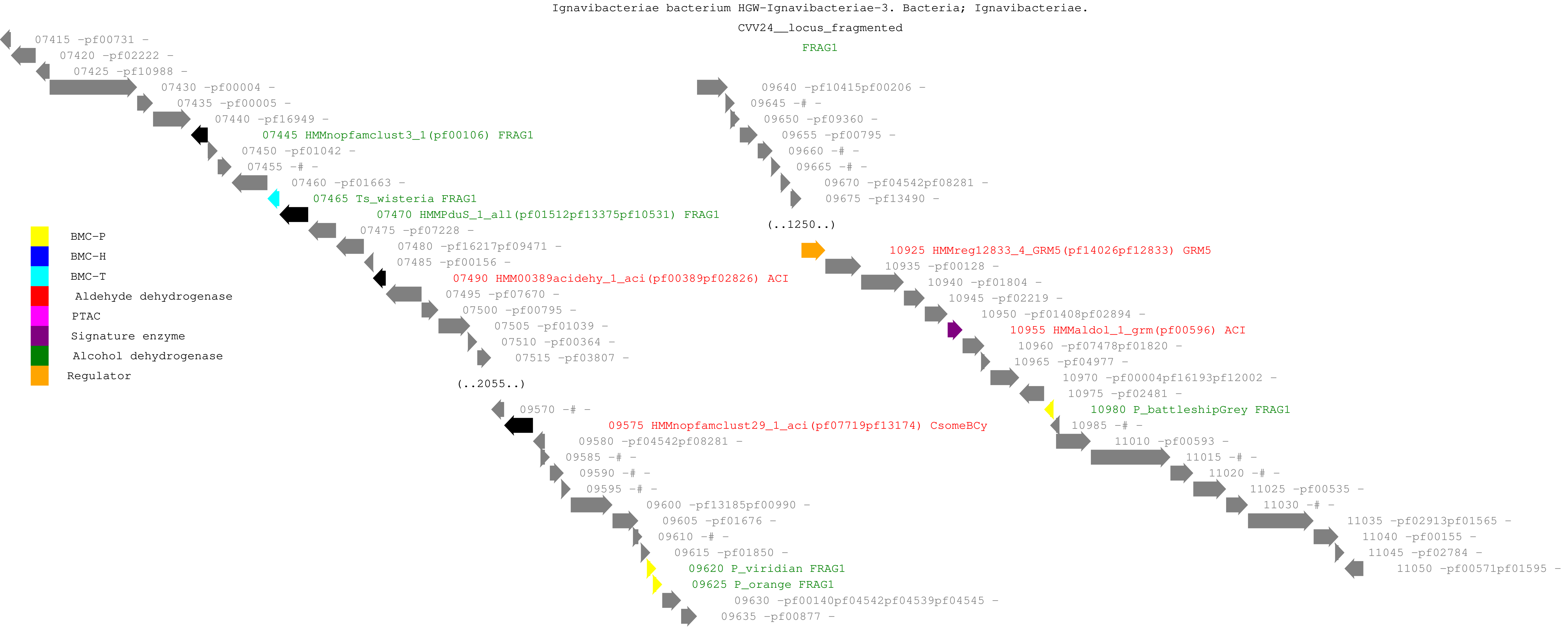

Ignavibacteriae bacterium HGW-Ignavibacteriae-3. Bacteria; Ignavibacteriae.
CVV24__locus_fragmented
FRAG1
07415 -pf00731 -
07420 -pf02222 -
07425 -pf10988 -
07430 -pf00004 -
07435 -pf00005 -
07440 -pf16949 -
07445 HMMnopfamclust3_1(pf00106) FRAG1
07450 -pf01042 -
07455 -# -
07460 -pf01663 -
07465 Ts_wisteria FRAG1
07470 HMMPduS_1_all(pf01512pf13375pf10531) FRAG1
07475 -pf07228 -
07480 -pf16217pf09471 -
07485 -pf00156 -
07490 HMM00389acidehy_1_aci(pf00389pf02826) ACI
07495 -pf07670 -
07500 -pf00795 -
07505 -pf01039 -
07510 -pf00364 -
07515 -pf03807 -
(..2055..)
09570 -# -
09575 HMMnopfamclust29_1_aci(pf07719pf13174) CsomeBCy
09580 -pf04542pf08281 -
09585 -# -
09590 -# -
09595 -# -
09600 -pf13185pf00990 -
09605 -pf01676 -
09610 -# -
09615 -pf01850 -
09620 P_viridian FRAG1
09625 P_orange FRAG1
09630 -pf00140pf04542pf04539pf04545 -
09635 -pf00877 -
09640 -pf10415pf00206 -
09645 -# -
09650 -pf09360 -
09655 -pf00795 -
09660 -# -
09665 -# -
09670 -pf04542pf08281 -
09675 -pf13490 -
(..1250..)
10925 HMMreg12833_4_GRM5(pf14026pf12833) GRM5
10935 -pf00128 -
10940 -pf01804 -
10945 -pf02219 -
10950 -pf01408pf02894 -
10955 HMMaldol_1_grm(pf00596) ACI
10960 -pf07478pf01820 -
10965 -pf04977 -
10970 -pf00004pf16193pf12002 -
10975 -pf02481 -
10980 P_battleshipGrey FRAG1
10985 -# -
11010 -pf00593 -
11015 -# -
11020 -# -
11025 -pf00535 -
11030 -# -
11035 -pf02913pf01565 -
11040 -pf00155 -
11045 -pf02784 -
11050 -pf00571pf01595 -
BMC-P
BMC-H
BMC-T
Aldehyde dehydrogenase
PTAC
Signature enzyme
Alcohol dehydrogenase
Regulator

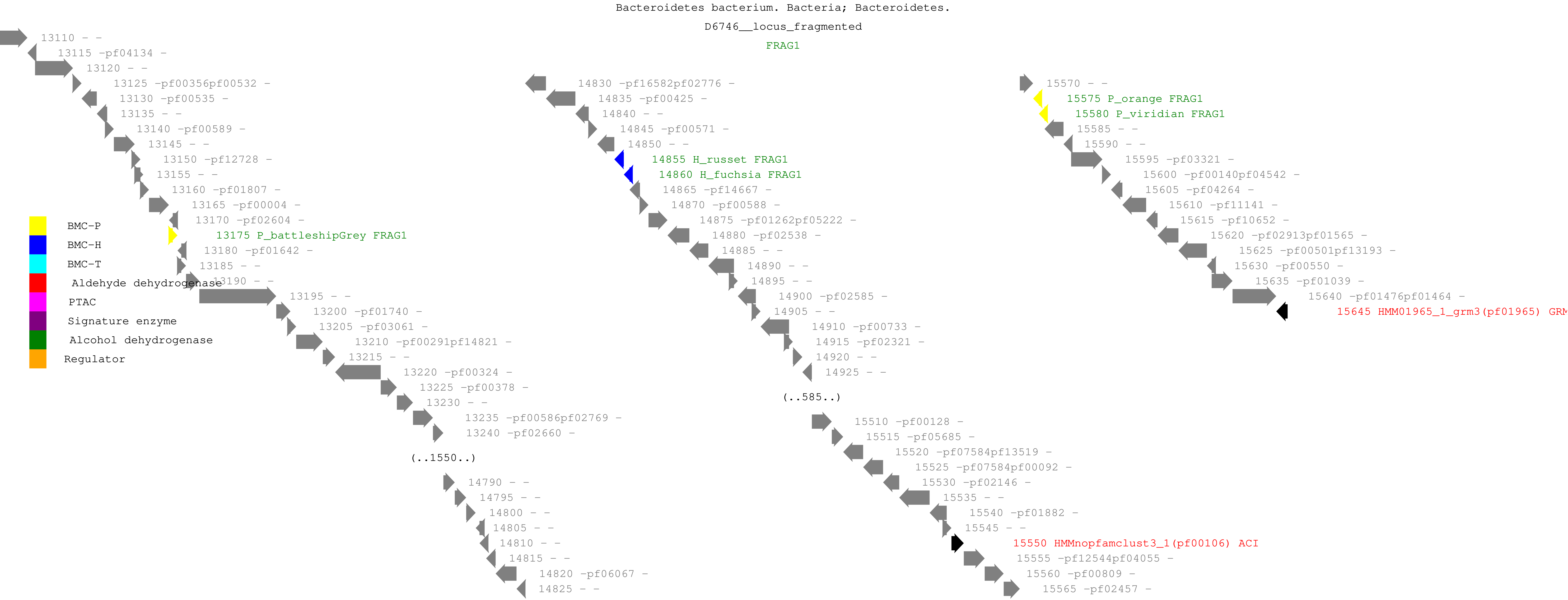

Bacteroidetes bacterium. Bacteria; Bacteroidetes.
D6746__locus_fragmented
FRAG1
13110 - -
13115 -pf04134 -
13120 - -
13125 -pf00356pf00532 -
13130 -pf00535 -
13135 - -
13140 -pf00589 -
13145 - -
13150 -pf12728 -
13155 - -
13160 -pf01807 -
13165 -pf00004 -
13170 -pf02604 -
13175 P_battleshipGrey FRAG1
13180 -pf01642 -
13185 - -
13190 - -
13195 - -
13200 -pf01740 -
13205 -pf03061 -
13210 -pf00291pf14821 -
13215 - -
13220 -pf00324 -
13225 -pf00378 -
13230 - -
13235 -pf00586pf02769 -
13240 -pf02660 -
(..1550..)
14790 - -
14795 - -
14800 - -
14805 - -
14810 - -
14815 - -
14820 -pf06067 -
14825 - -
14830 -pf16582pf02776 -
14835 -pf00425 -
14840 - -
14845 -pf00571 -
14850 - -
14855 H_russet FRAG1
14860 H_fuchsia FRAG1
14865 -pf14667 -
14870 -pf00588 -
14875 -pf01262pf05222 -
14880 -pf02538 -
14885 - -
14890 - -
14895 - -
14900 -pf02585 -
14905 - -
14910 -pf00733 -
14915 -pf02321 -
14920 - -
14925 - -
(..585..)
15510 -pf00128 -
15515 -pf05685 -
15520 -pf07584pf13519 -
15525 -pf07584pf00092 -
15530 -pf02146 -
15535 - -
15540 -pf01882 -
15545 - -
15550 HMMnopfamclust3_1(pf00106) ACI
15555 -pf12544pf04055 -
15560 -pf00809 -
15565 -pf02457 -
15570 - -
15575 P_orange FRAG1
15580 P_viridian FRAG1
15585 - -
15590 - -
15595 -pf03321 -
15600 -pf00140pf04542 -
15605 -pf04264 -
15610 -pf11141 -
15615 -pf10652 -
15620 -pf02913pf01565 -
15625 -pf00501pf13193 -
15630 -pf00550 -
15635 -pf01039 -
15640 -pf01476pf01464 -
15645 HMM01965_1_grm3(pf01965) GRM
BMC-P
BMC-H
BMC-T
Aldehyde dehydrogenase
PTAC
Signature enzyme
Alcohol dehydrogenase
Regulator

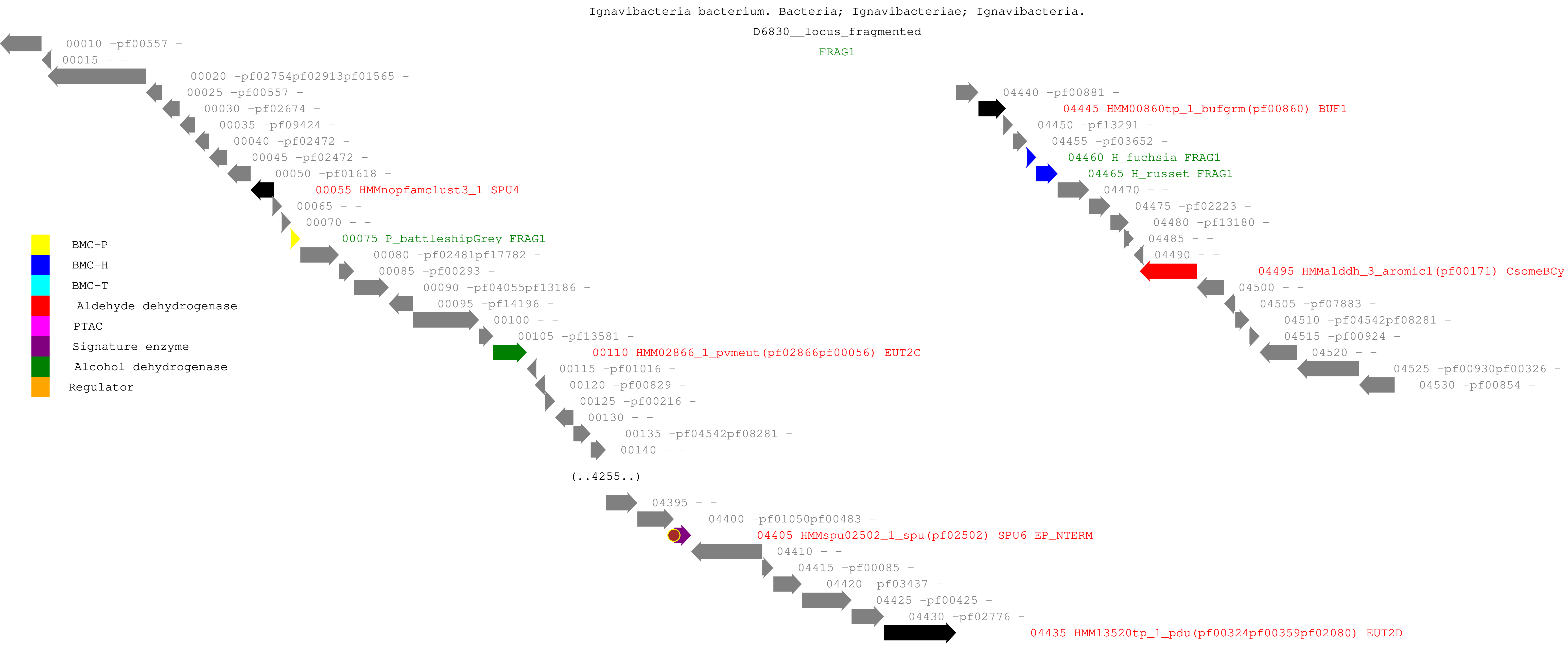
Ignavibacteria bacterium. Bacteria; Ignavibacteriae; Ignavibacteria.
D6830__locus_fragmented
FRAG1
00010 -pf00557 -
00015 - -
00020 -pf02754pf02913pf01565 -
00025 -pf00557 -
00030 -pf02674 -
00035 -pf09424 -
00040 -pf02472 -
00045 -pf02472 -
00050 -pf01618 -
00055 HMMnopfamclust3_1 SPU4
00065 - -
00070 - -
00075 P_battleshipGrey FRAG1
00080 -pf02481pf17782 -
00085 -pf00293 -
00090 -pf04055pf13186 -
00095 -pf14196 -
00100 - -
00105 -pf13581 -
00110 HMM02866_1_pvmeut(pf02866pf00056) EUT2C
00115 -pf01016 -
00120 -pf00829 -
00125 -pf00216 -
00130 - -
00135 -pf04542pf08281 -
00140 - -
(..4255..)
04395 - -
04400 -pf01050pf00483 -
04405 HMMspu02502_1_spu(pf02502) SPU6 EP_NTERM
04410 - -
04415 -pf00085 -
04420 -pf03437 -
04425 -pf00425 -
04430 -pf02776 -
04435 HMM13520tp_1_pdu(pf00324pf00359pf02080) EUT2D
04440 -pf00881 -
04445 HMM00860tp_1_bufgrm(pf00860) BUF1
04450 -pf13291 -
04455 -pf03652 -
04460 H_fuchsia FRAG1
04465 H_russet FRAG1
04470 - -
04475 -pf02223 -
04480 -pf13180 -
04485 - -
04490 - -
04495 HMMalddh_3_aromic1(pf00171) CsomeBCy
04500 - -
04505 -pf07883 -
04510 -pf04542pf08281 -
04515 -pf00924 -
04520 - -
04525 -pf00930pf00326 -
04530 -pf00854 -
BMC-P
BMC-H
BMC-T
Aldehyde dehydrogenase
PTAC
Signature enzyme
Alcohol dehydrogenase
Regulator

Bacteroidetes bacterium. Bacteria; Bacteroidetes.

DCX46__locus_fragmented

FRAG1

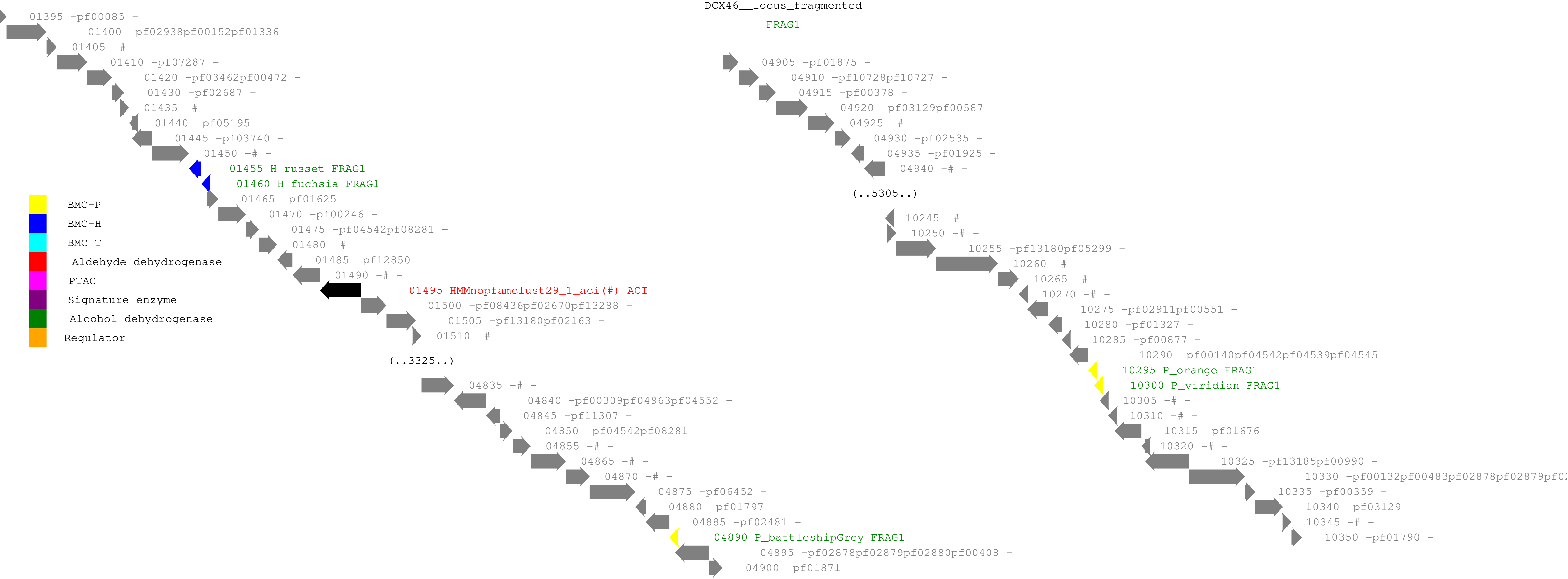

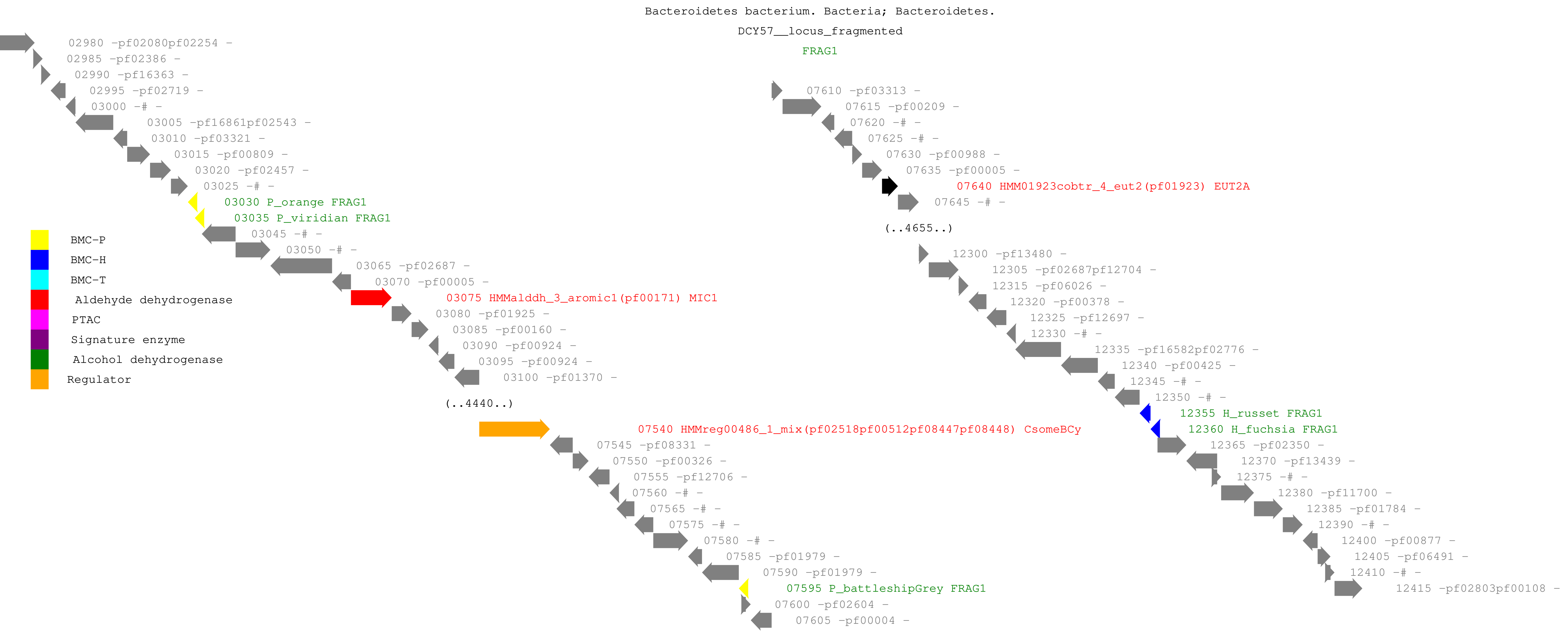

Bacteroidetes bacterium. Bacteria; Bacteroidetes.
DCY57__locus_fragmented
FRAG1
02980 -pf02080pf02254 -
02985 -pf02386 -
02990 -pf16363 -
02995 -pf02719 -
03000 -# -
03005 -pf16861pf02543 -
03010 -pf03321 -
03015 -pf00809 -
03020 -pf02457 -
03025 -# -
03030 P_orange FRAG1
03035 P_viridian FRAG1
03045 -# -
03050 -# -
03065 -pf02687 -
03070 -pf00005 -
03075 HMMaldDH_3_aromic1(pf00171) MIC1
03080 -pf01925 -
03085 -pf00160 -
03090 -pf00924 -
03095 -pf00924 -
03100 -pf01370 -
(..4440..)
07540 HMMreg00486_1_mix(pf02518pf00512pf08447pf08448) CsomeBCy
07545 -pf08331 -
07550 -pf00326 -
07555 -pf12706 -
07560 -# -
07565 -# -
07575 -# -
07580 -# -
07585 -pf01979 -
07590 -pf01979 -
07595 P_battleshipGrey FRAG1
07600 -pf02604 -
07605 -pf00004 -
07610 -pf03313 -
07615 -pf00209 -
07620 -# -
07625 -# -
07630 -pf00988 -
07635 -pf00005 -
07640 HMM01923cobtr_4_eut2(pf01923) EUT2A
07645 -# -
(..4655..)
12300 -pf13480 -
12305 -pf02687pf12704 -
12315 -pf06026 -
12320 -pf00378 -
12325 -pf12697 -
12330 -# -
12335 -pf16582pf02776 -
12340 -pf00425 -
12345 -# -
12350 -# -
12355 H_russet FRAG1
12360 H_fuchsia FRAG1
12365 -pf02350 -
12370 -pf13439 -
12375 -# -
12380 -pf11700 -
12385 -pf01784 -
12390 -# -
12400 -pf00877 -
12405 -pf06491 -
12410 -# -
12415 -pf02803pf00108 -
BMC-P
BMC-H
BMC-T
Aldehyde dehydrogenase
PTAC
Signature enzyme
Alcohol dehydrogenase
Regulator

Bacteroidetes bacterium. Bacteria; Bacteroidetes.

DGH68__locus_fragmented

FRAG1

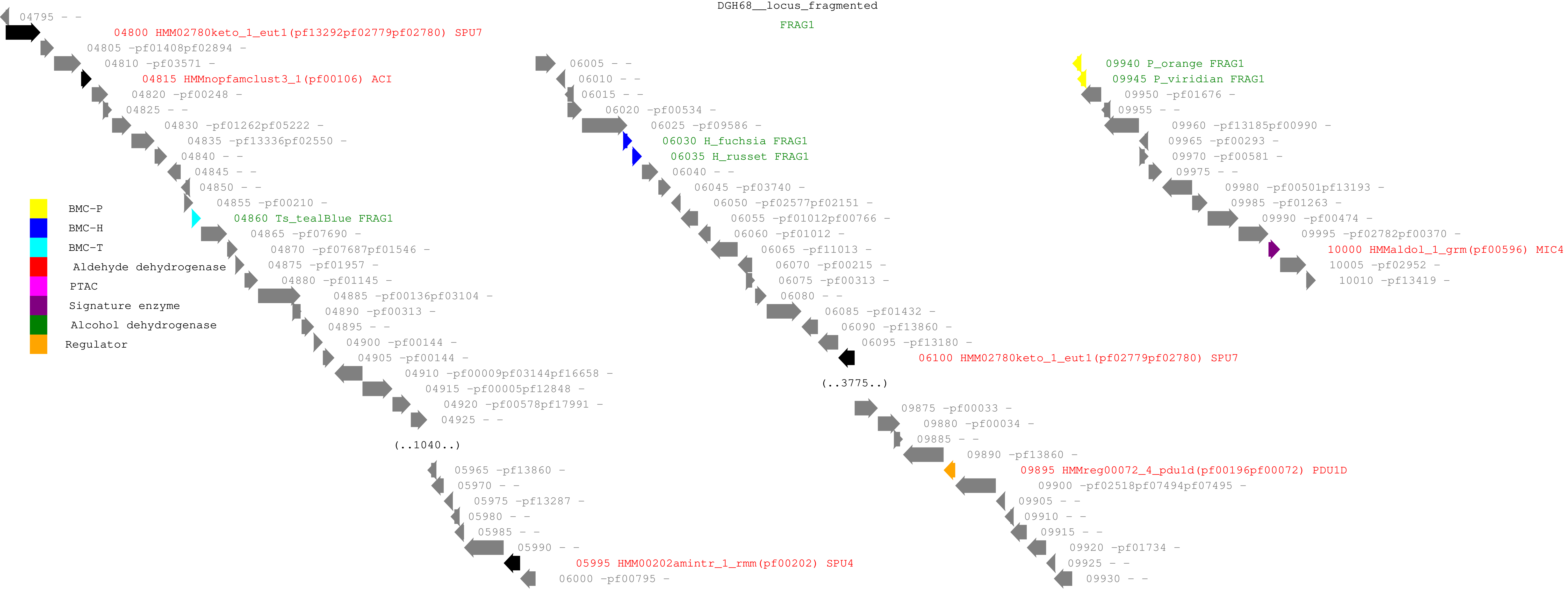

Ignavibacteria bacterium. Bacteria; Ignavibacteriae; Ignavibacteria.

E2046__locus_fragmented

FRAG1

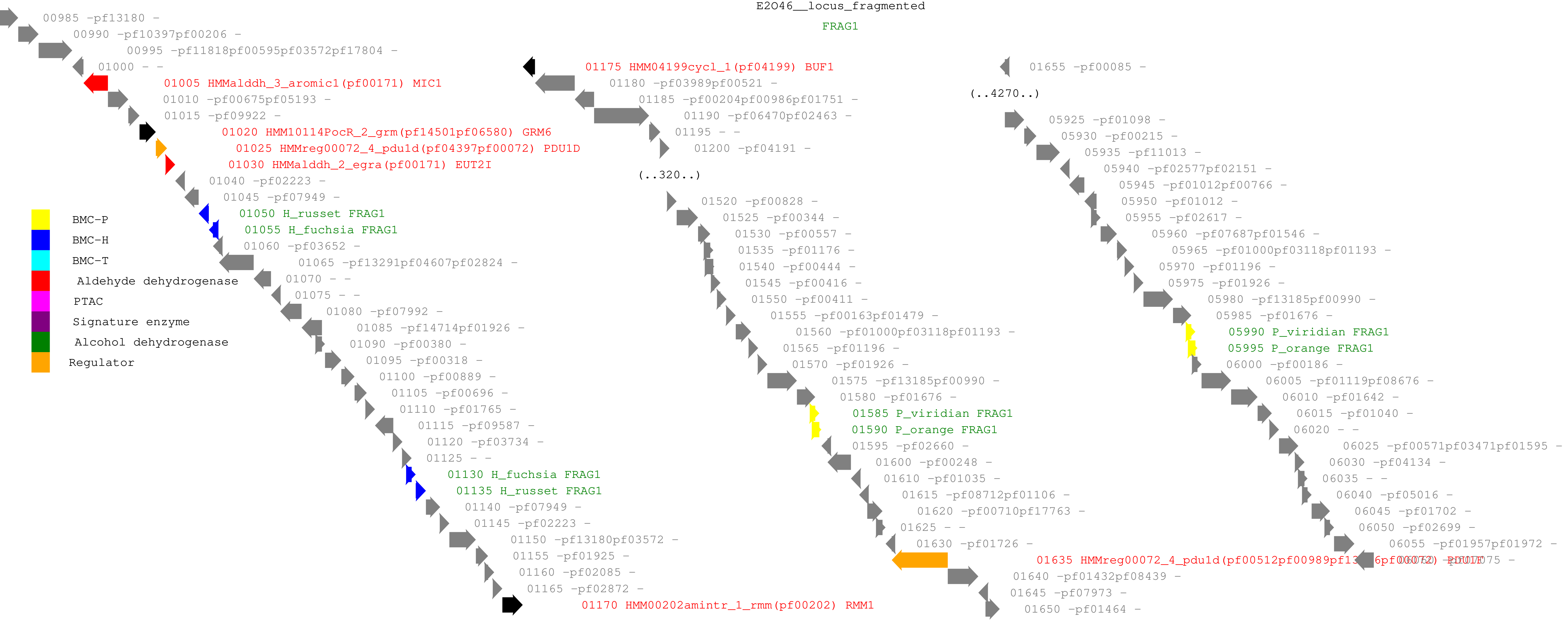

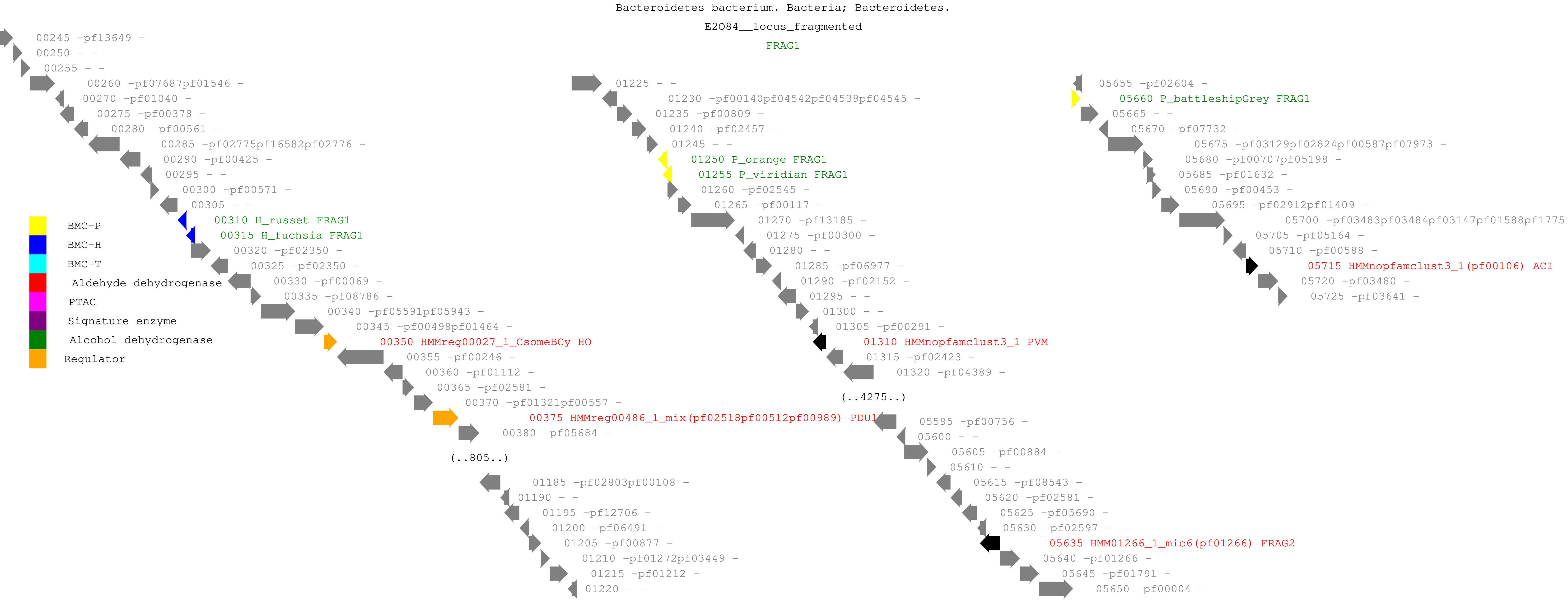

Bacteroidetes bacterium. Bacteria; Bacteroidetes.
E2084__locus_fragmented
FRAG1
00245 -pf13649 -
00250 - -
00255 - -
00260 -pf07687pf01546 -
00270 -pf01040 -
00275 -pf00378 -
00280 -pf00561 -
00285 -pf02775pf16582pf02776 -
00290 -pf00425 -
00295 - -
00300 -pf00571 -
00305 - -
00310 H_russet FRAG1
00315 H_fuchsia FRAG1
00320 -pf02350 -
00325 -pf02350 -
00330 -pf00069 -
00335 -pf08786 -
00340 -pf05591pf05943 -
00345 -pf00498pf01464 -
00350 HMMreg00027_1_CsomeBCy HO
00355 -pf00246 -
00360 -pf01112 -
00365 -pf02581 -
00370 -pf01321pf00557 -
00375 HMMreg00486_1_mix(pf02518pf00512pf00989) PDU1
00380 -pf05684 -
(..805..)
01185 -pf02803pf00108 -
01190 - -
01195 -pf12706 -
01200 -pf06491 -
01205 -pf00877 -
01210 -pf01272pf03449 -
01215 -pf01212 -
01220 - -
01225 - -
01230 -pf00140pf04542pf04539pf04545 -
01235 -pf00809 -
01240 -pf02457 -
01245 - -
01250 P_orange FRAG1
01255 P_viridian FRAG1
01260 -pf02545 -
01265 -pf00117 -
01270 -pf13185 -
01275 -pf00300 -
01280 - -
01285 -pf06977 -
01290 -pf02152 -
01295 - -
01300 - -
01305 -pf00291 -
01310 HMMnopfamclust3_1 PVM
01315 -pf02423 -
01320 -pf04389 -
(..4275..)
05595 -pf00756 -
05600 - -
05605 -pf00884 -
05610 - -
05615 -pf08543 -
05620 -pf02581 -
05625 -pf05690 -
05630 -pf02597 -
05635 HMM01266_1_mic6(pf01266) FRAG2
05640 -pf01266 -
05645 -pf01791 -
05650 -pf00004 -
05655 -pf02604 -
05660 P_battleshipGrey FRAG1
05665 - -
05670 -pf07732 -
05675 -pf03129pf02824pf00587pf07973 -
05680 -pf00707pf05198 -
05685 -pf01632 -
05690 -pf00453 -
05695 -pf02912pf01409 -
05700 -pf03483pf03484pf03147pf01588pf1775
05705 -pf05164 -
05710 -pf00588 -
05715 HMMnopfamclust3_1(pf00106) ACI
05720 -pf03480 -
05725 -pf03641 -
BMC-P
BMC-H
BMC-T
Aldehyde dehydrogenase
PTAC
Signature enzyme
Alcohol dehydrogenase
Regulator

Ignavibacteria bacterium. Bacteria; Ignavibacteriae; Ignavibacteria.

E6K56__locus_fragmented

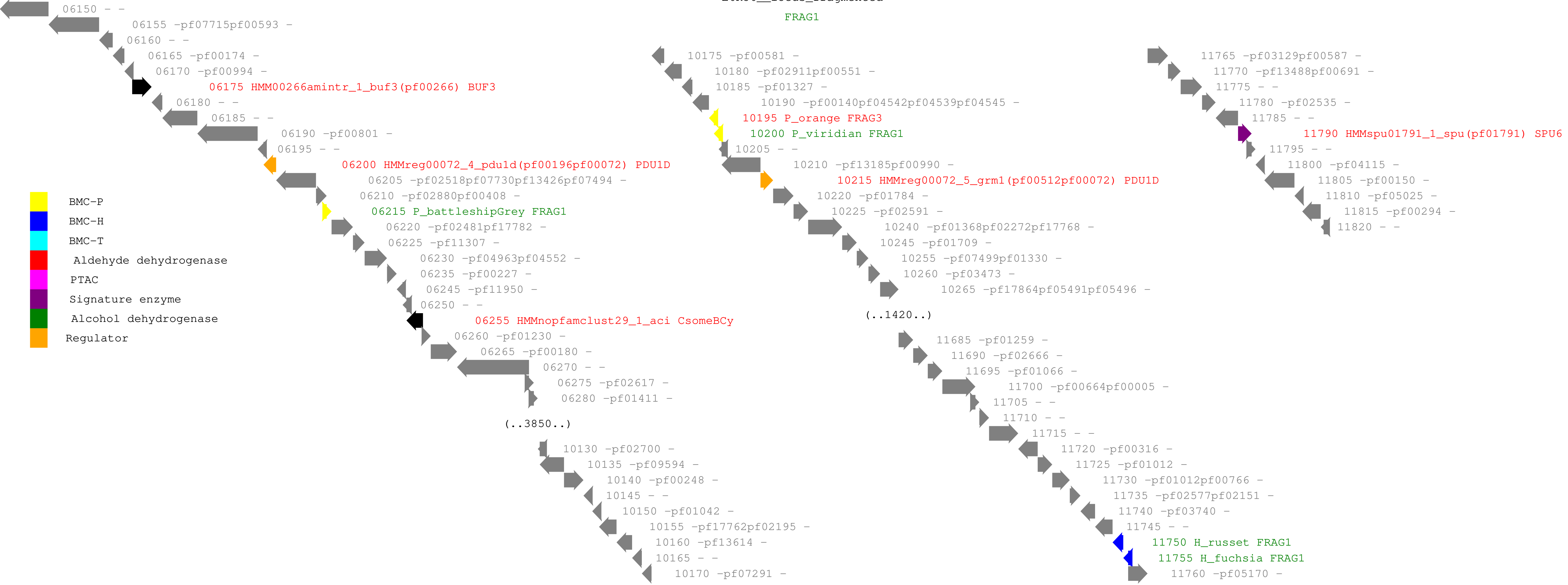

Ignavibacteriales bacterium. Bacteria; Ignavibacteriae; Ignavibacteria; Ignavibacteriales.

EHM47__locus_fragmented

FRAG1

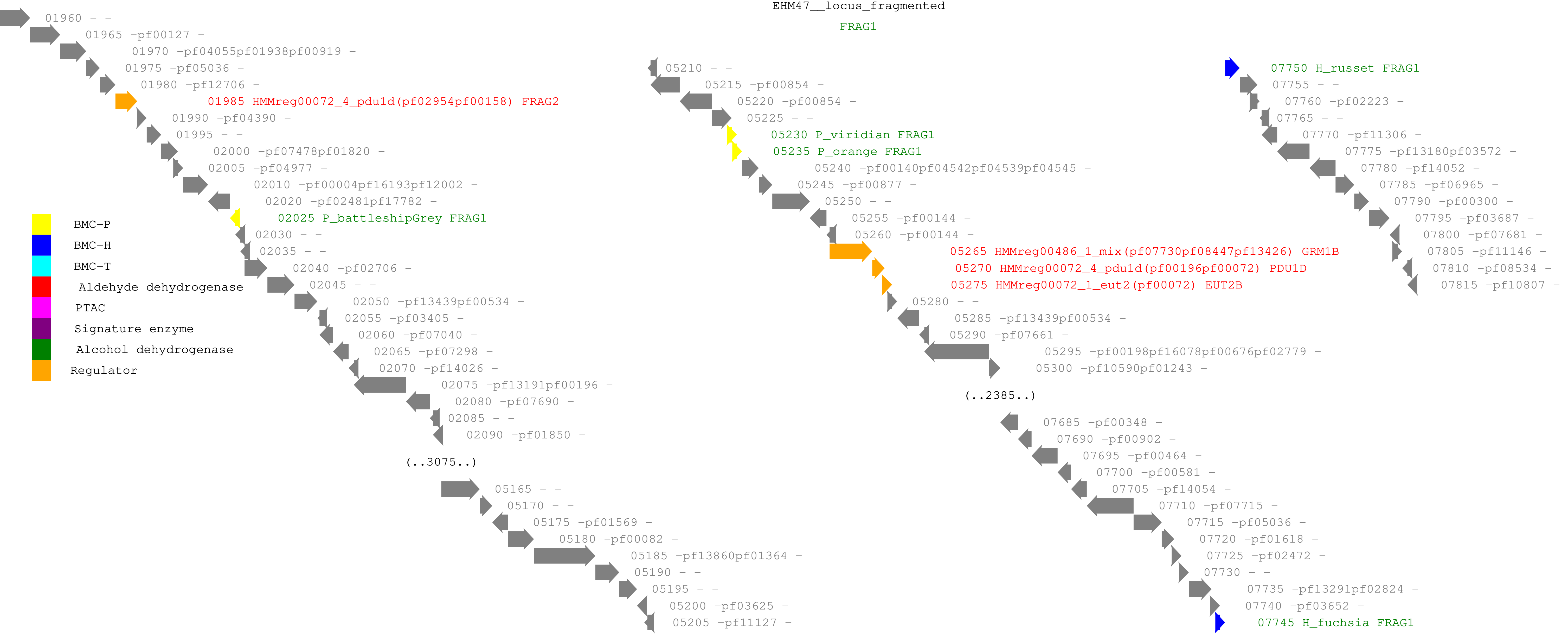

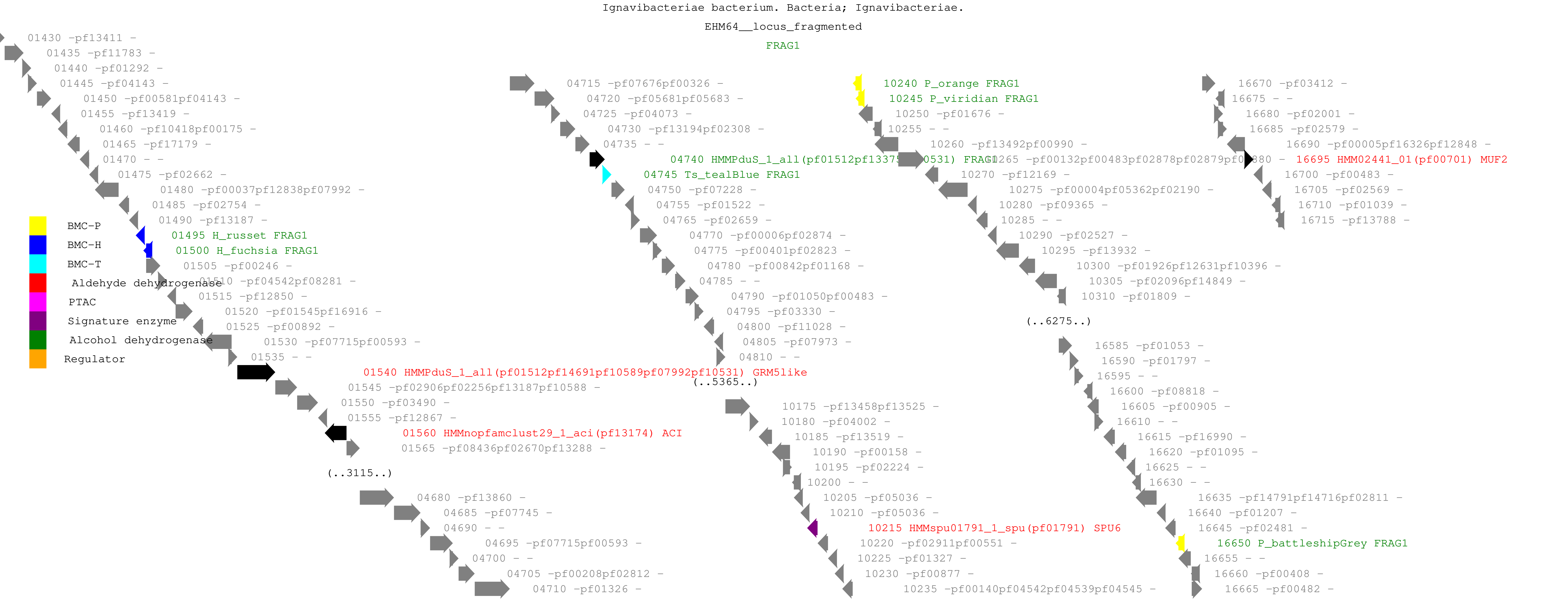

Ignavibacteriae bacterium. Bacteria; Ignavibacteriae.
EHM64__locus_fragmented
FRAG1
01430 -pf13411 -
01435 -pf11783 -
01440 -pf01292 -
01445 -pf04143 -
01450 -pf00581pf04143 -
01455 -pf13419 -
01460 -pf10418pf00175 -
01465 -pf17179 -
01470 - -
01475 -pf02662 -
01480 -pf00037pf12838pf07992 -
01485 -pf02754 -
01490 -pf13187 -
01495 H_russet FRAG1
01500 H_fuchsia FRAG1
01505 -pf00246 -
01510 -pf04542pf08281 -
01515 -pf12850 -
01520 -pf01545pf16916 -
01525 -pf00892 -
01530 -pf07715pf00593 -
01535 - -
01540 HMMPduS_1_all(pf01512pf14691pf10589pf07992pf10531) GRM5like
01545 -pf02906pf02256pf13187pf10588 -
01550 -pf03490 -
01555 -pf12867 -
01560 HMMnopfamclust29_1_aci(pf13174) ACI
01565 -pf08436pf02670pf13288 -
(..3115..)
04680 -pf13860 -
04685 -pf07745 -
04690 - -
04695 -pf07715pf00593 -
04700 - -
04705 -pf00208pf02812 -
04710 -pf01326 -
04715 -pf07676pf00326 -
04720 -pf05681pf05683 -
04725 -pf04073 -
04730 -pf13194pf02308 -
04735 - -
04740 HMMPduS_1_all(pf01512pf13375pf10531) FRAG1
04745 Ts_tealBlue FRAG1
04750 -pf07228 -
04755 -pf01522 -
04765 -pf02659 -
04770 -pf00006pf02874 -
04775 -pf00401pf02823 -
04780 -pf00842pf01168 -
04785 - -
04790 -pf01050pf00483 -
04795 -pf03330 -
04800 -pf11028 -
04805 -pf07973 -
04810 - -
(..5365..)
10175 -pf13458pf13525 -
10180 -pf04002 -
10185 -pf13519 -
10190 -pf00158 -
10195 -pf02224 -
10200 - -
10205 -pf05036 -
10210 -pf05036 -
10215 HMMspu01791_1_spu(pf01791) SPU6
10220 -pf02911pf00551 -
10225 -pf01327 -
10230 -pf00877 -
10235 -pf00140pf04542pf04539pf04545 -
10240 P_orange FRAG1
10245 P_viridian FRAG1
10250 -pf01676 -
10255 - -
10260 -pf13492pf00990 -
10265 -pf00132pf00483pf02878pf02879pf0[illegible]880 -
10270 -pf12169 -
10275 -pf00004pf05362pf02190 -
10280 -pf09365 -
10285 - -
10290 -pf02527 -
10295 -pf13932 -
10300 -pf01926pf12631pf10396 -
10305 -pf02096pf14849 -
10310 -pf01809 -
(..6275..)
16585 -pf01053 -
16590 -pf01797 -
16595 - -
16600 -pf08818 -
16605 -pf00905 -
16610 - -
16615 -pf16990 -
16620 -pf01095 -
16625 - -
16630 - -
16635 -pf14791pf14716pf02811 -
16640 -pf01207 -
16645 -pf02481 -
16650 P_battleshipGrey FRAG1
16655 - -
16660 -pf00408 -
16665 -pf00482 -
16670 -pf03412 -
16675 - -
16680 -pf02001 -
16685 -pf02579 -
16690 -pf00005pf16326pf12848 -
16695 HMM02441_01(pf00701) MUF2
16700 -pf00483 -
16705 -pf02569 -
16710 -pf01039 -
16715 -pf13788 -
BMC-P
BMC-H
BMC-T
Aldehyde dehydrogenase
PTAC
Signature enzyme
Alcohol dehydrogenase
Regulator

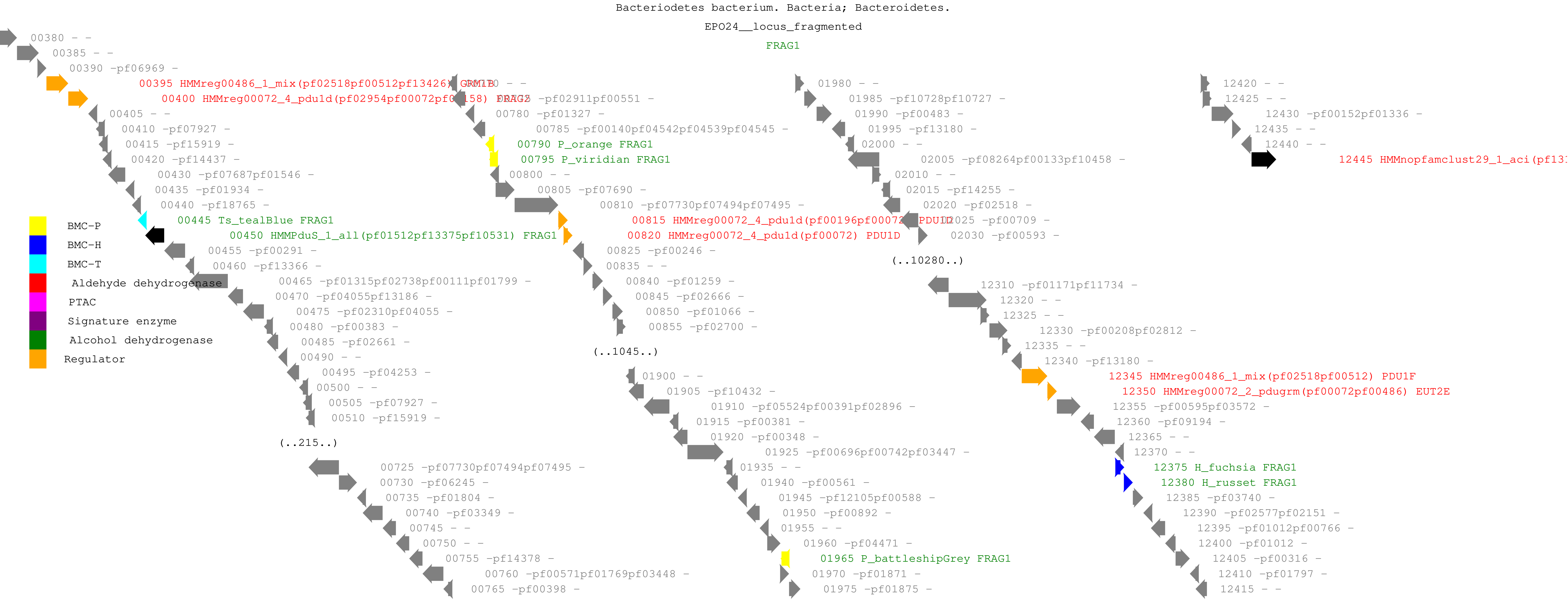

Bacteriodetes bacterium. Bacteria; Bacteroidetes.
EPO24__locus_fragmented
FRAG1
00380 - -
00385 - -
00390 -pf06969 -
00395 HMMreg00486_1_mix(pf02518pf00512pf13426) GRM1B
00400 HMMreg00072_4_pdu1d(pf02954pf00072pf0
158) FRAG2
00405 - -
00410 -pf07927 -
00415 -pf15919 -
00420 -pf14437 -
00430 -pf07687pf01546 -
00435 -pf01934 -
00440 -pf18765 -
00445 Ts_tealBlue FRAG1
00450 HMMPduS_1_all(pf01512pf13375pf10531) FRAG1
00455 -pf00291 -
00460 -pf13366 -
00465 -pf01315pf02738pf00111pf01799 -
00470 -pf04055pf13186 -
00475 -pf02310pf04055 -
00480 -pf00383 -
00485 -pf02661 -
00490 - -
00495 -pf04253 -
00500 - -
00505 -pf07927 -
00510 -pf15919 -
(..215..)
00725 -pf07730pf07494pf07495 -
00730 -pf06245 -
00735 -pf01804 -
00740 -pf03349 -
00745 - -
00750 - -
00755 -pf14378 -
00760 -pf00571pf01769pf03448 -
00765 -pf00398 -
- -
-pf02911pf00551 -
00780 -pf01327 -
00785 -pf00140pf04542pf04539pf04545 -
00790 P_orange FRAG1
00795 P_viridian FRAG1
00800 - -
00805 -pf07690 -
00810 -pf07730pf07494pf07495 -
00815 HMMreg00072_4_pdu1d(pf00196pf00072
PDU1D
00820 HMMreg00072_4_pdu1d(pf00072) PDU1D
00825 -pf00246 -
00835 - -
00840 -pf01259 -
00845 -pf02666 -
00850 -pf01066 -
00855 -pf02700 -
(..1045..)
01900 - -
01905 -pf10432 -
01910 -pf05524pf00391pf02896 -
01915 -pf00381 -
01920 -pf00348 -
01925 -pf00696pf00742pf03447 -
01935 - -
01940 -pf00561 -
01945 -pf12105pf00588 -
01950 -pf00892 -
01955 - -
01960 -pf04471 -
01965 P_battleshipGrey FRAG1
01970 -pf01871 -
01975 -pf01875 -
01980 - -
01985 -pf10728pf10727 -
01990 -pf00483 -
01995 -pf13180 -
02000 - -
02005 -pf08264pf00133pf10458 -
02010 - -
02015 -pf14255 -
02020 -pf02518 -
02025 -pf00709 -
02030 -pf00593 -
(..10280..)
12310 -pf01171pf11734 -
12320 - -
12325 - -
12330 -pf00208pf02812 -
12335 - -
12340 -pf13180 -
12345 HMMreg00486_1_mix(pf02518pf00512) PDU1F
12350 HMMreg00072_2_pdugrm(pf00072pf00486) EUT2E
12355 -pf00595pf03572 -
12360 -pf09194 -
12365 - -
12370 - -
12375 H_fuchsia FRAG1
12380 H_russet FRAG1
12385 -pf03740 -
12390 -pf02577pf02151 -
12395 -pf01012pf00766 -
12400 -pf01012 -
12405 -pf00316 -
12410 -pf01797 -
12415 - -
12420 - -
12425 - -
12430 -pf00152pf01336 -
12435 - -
12440 - -
12445 HMMnopfamclust29_1_aci(pf13
BMC-P
BMC-H
BMC-T
Aldehyde dehydrogenase
PTAC
Signature enzyme
Alcohol dehydrogenase
Regulator

Chlorobium sp. GBChlB. Bacteria; Chlorobi; Chlorobia; Chlorobiales; Chlorobiaceae;

HY22__locus_fragmented

FRAG1

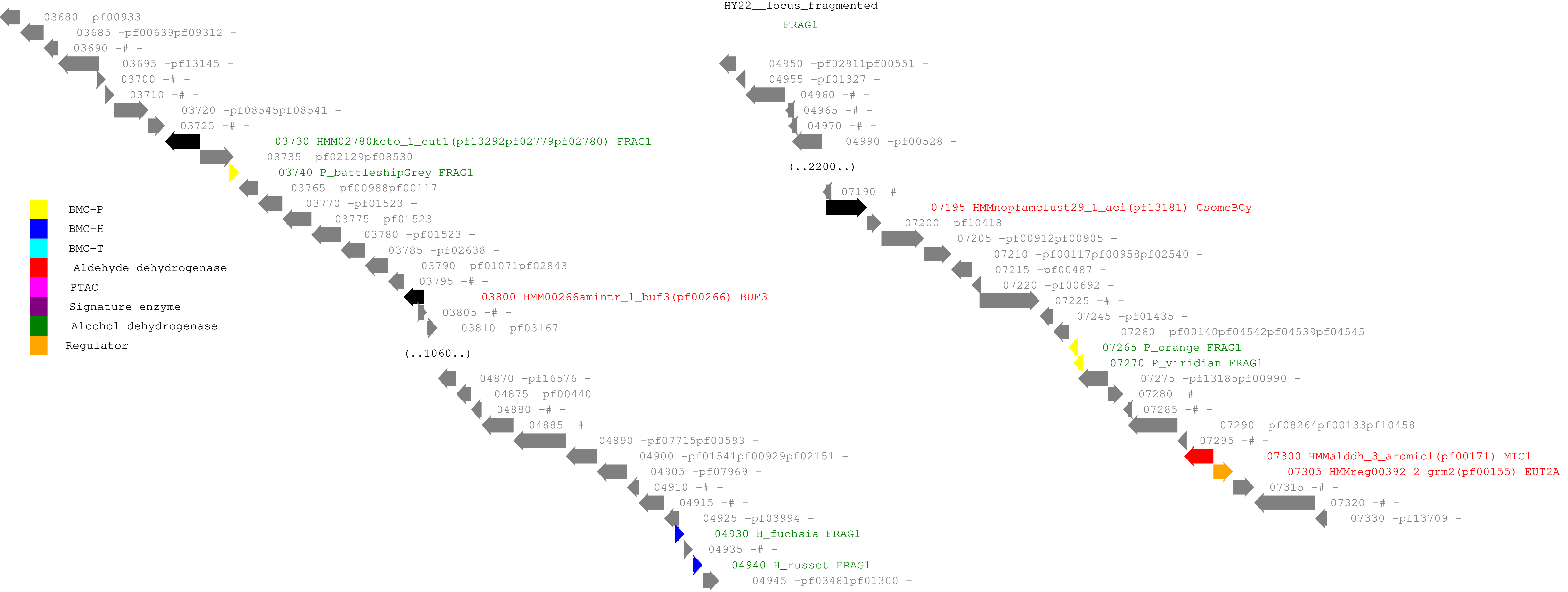

Ignavibacteriae bacterium. Bacteria; Ignavibacteriae.

IGBAC__locus_fragmented

FRAG1

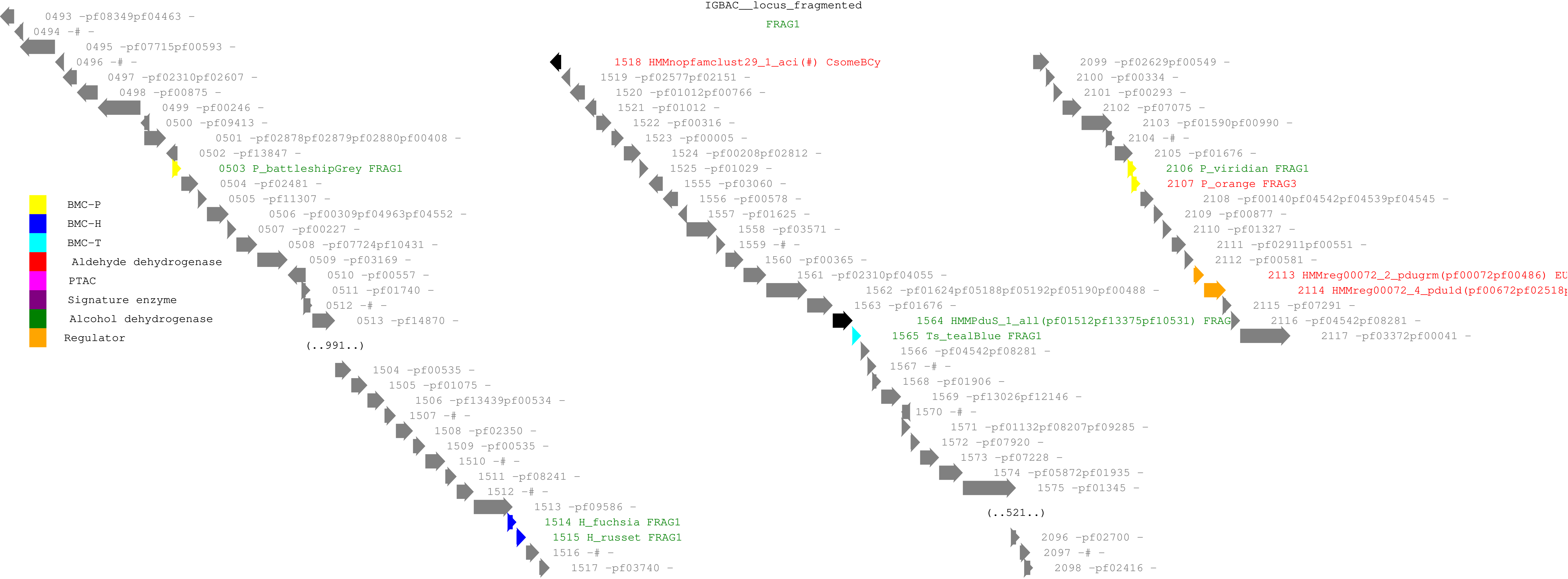

Candidatus Kryptonium thompsoni. Bacteria; Candidatus Kryptonia; Candidatus Kryptonium.

JGI10__locus_fragmented

FRAG1

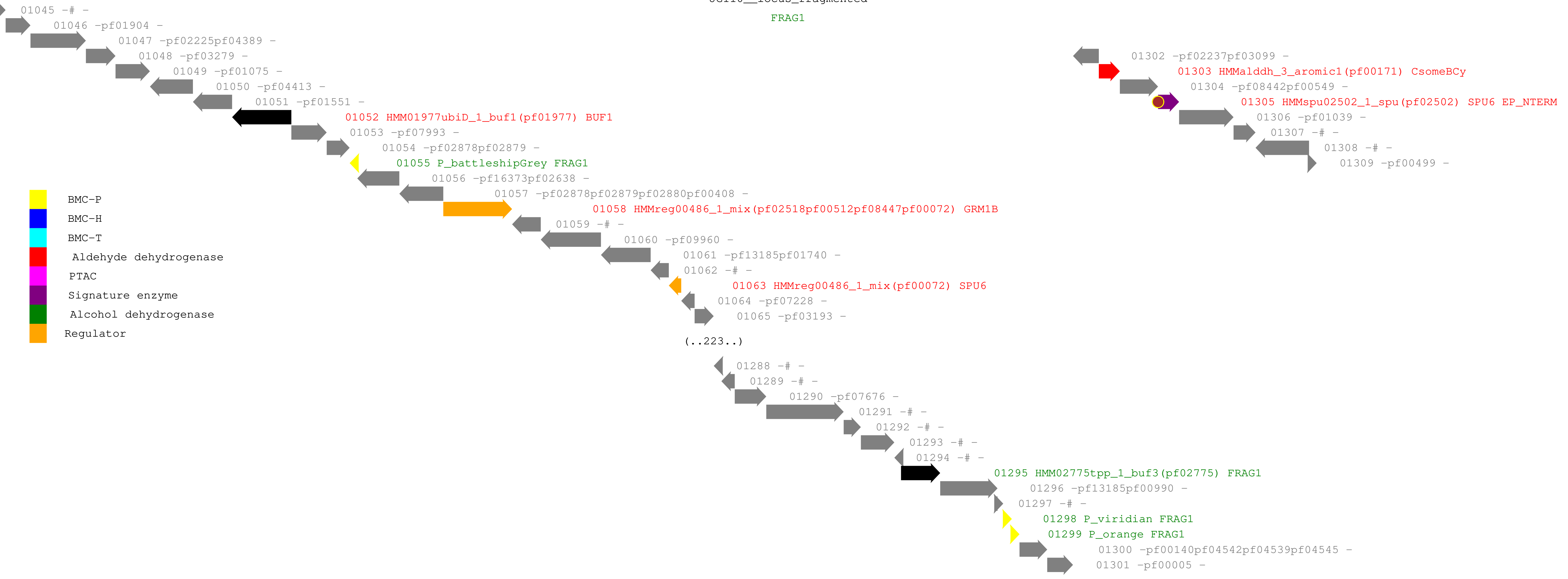

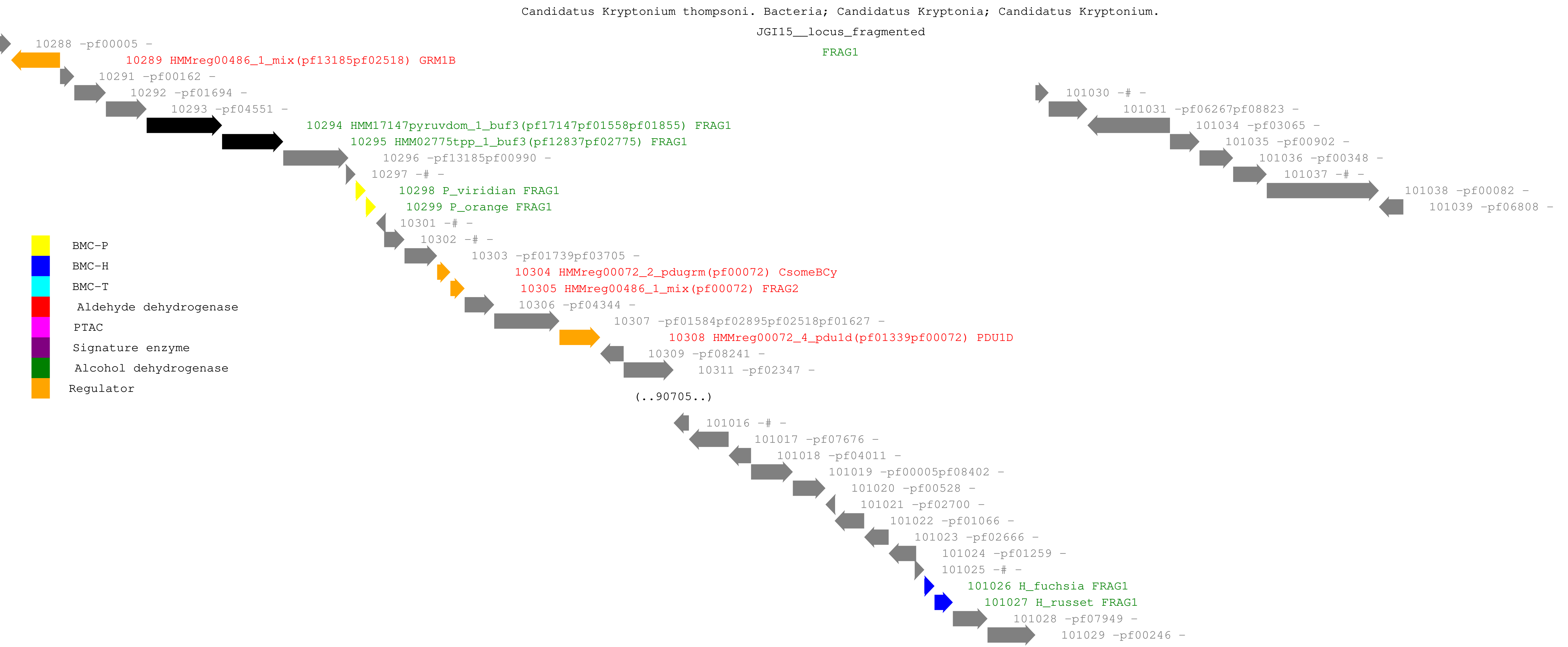
Candidatus Kryptonium thompsoni. Bacteria; Candidatus Kryptonia; Candidatus Kryptonium.
JGI15__locus_fragmented
FRAG1
10288 -pf00005 -
10289 HMMreg00486_1_mix(pf13185pf02518) GRM1B
10291 -pf00162 -
10292 -pf01694 -
10293 -pf04551 -
10294 HMM17147pyruvdom_1_buf3(pf17147pf01558pf01855) FRAG1
10295 HMM02775tpp_1_buf3(pf12837pf02775) FRAG1
10296 -pf13185pf00990 -
10297 -# -
10298 P_viridian FRAG1
10299 P_orange FRAG1
10301 -# -
10302 -# -
10303 -pf01739pf03705 -
10304 HMMreg00072_2_pdugrm(pf00072) CsomeBCy
10305 HMMreg00486_1_mix(pf00072) FRAG2
10306 -pf04344 -
10307 -pf01584pf02895pf02518pf01627 -
10308 HMMreg00072_4_pdu1d(pf01339pf00072) PDU1D
10309 -pf08241 -
10311 -pf02347 -
(..90705..)
101016 -# -
101017 -pf07676 -
101018 -pf04011 -
101019 -pf00005pf08402 -
101020 -pf00528 -
101021 -pf02700 -
101022 -pf01066 -
101023 -pf02666 -
101024 -pf01259 -
101025 -# -
101026 H_fuchsia FRAG1
101027 H_russet FRAG1
101028 -pf07949 -
101029 -pf00246 -
101030 -# -
101031 -pf06267pf08823 -
101034 -pf03065 -
101035 -pf00902 -
101036 -pf00348 -
101037 -# -
101038 -pf00082 -
101039 -pf06808 -
BMC-P
BMC-H
BMC-T
Aldehyde dehydrogenase
PTAC
Signature enzyme
Alcohol dehydrogenase
Regulator

Candidatus Thermokryptus mobilis. Bacteria; Candidatus Kryptonia; Candidatus Thermokryptus.

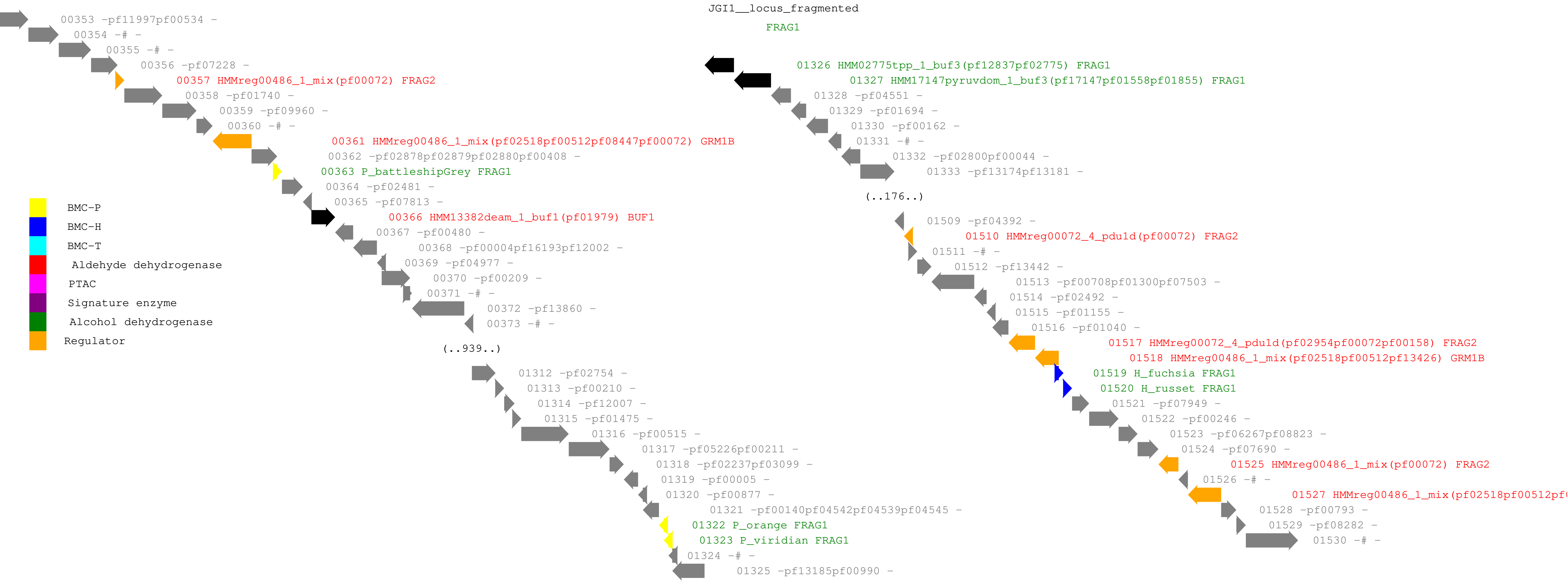

Candidatus Kryptobacter tengchongensis. Bacteria; Candidatus Kryptonia; Candidatus Kryptobacter.

JGI20__locus_fragmented

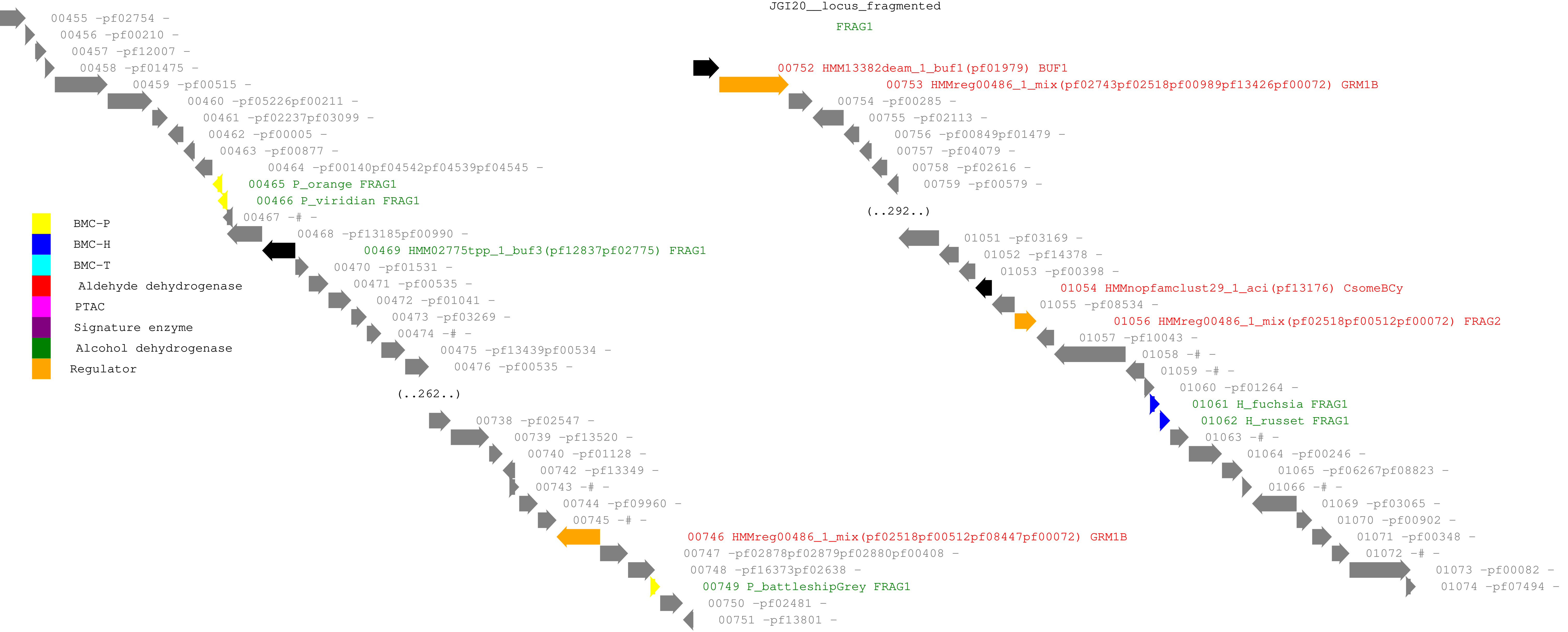

Candidatus Kryptobacter tengchongensis. Bacteria; Candidatus Kryptonia; Candidatus Kryptobacter.

JGI3__locus_fragmented

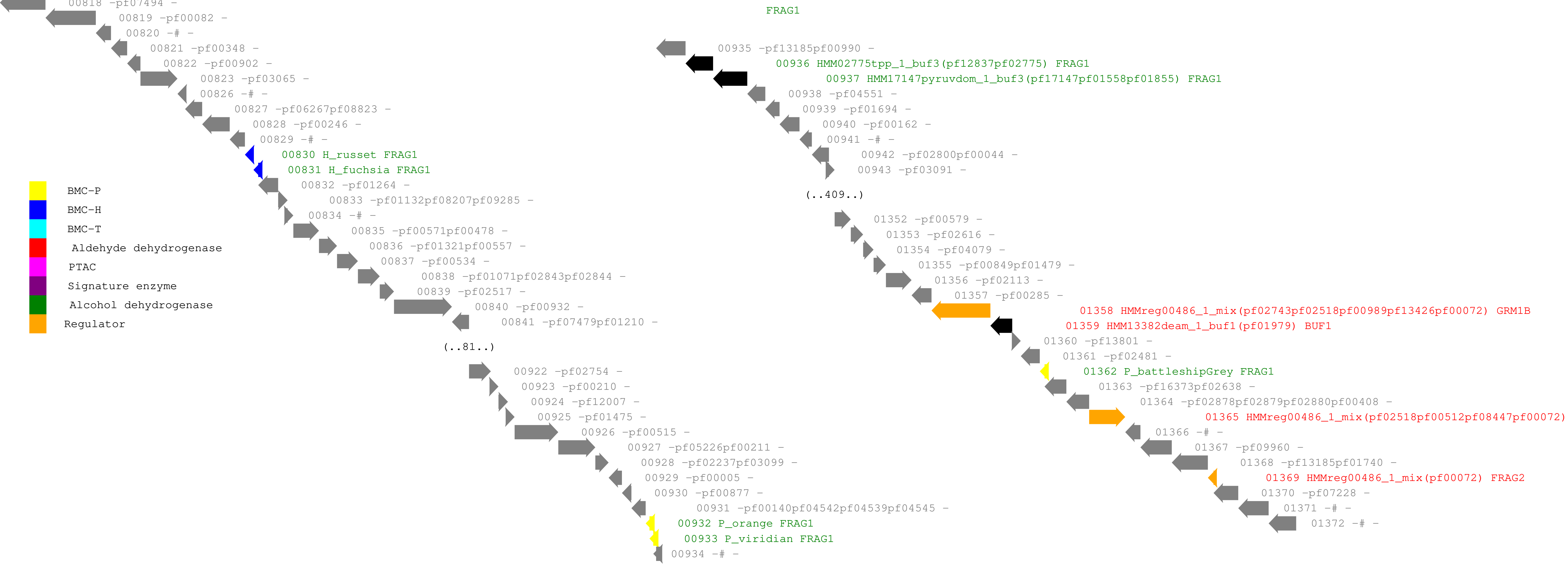

Candidatus Kryptonium thompsoni. Bacteria; Candidatus Kryptonia; Candidatus Kryptonium.

JGI4__locus_fragmented

FRAG1

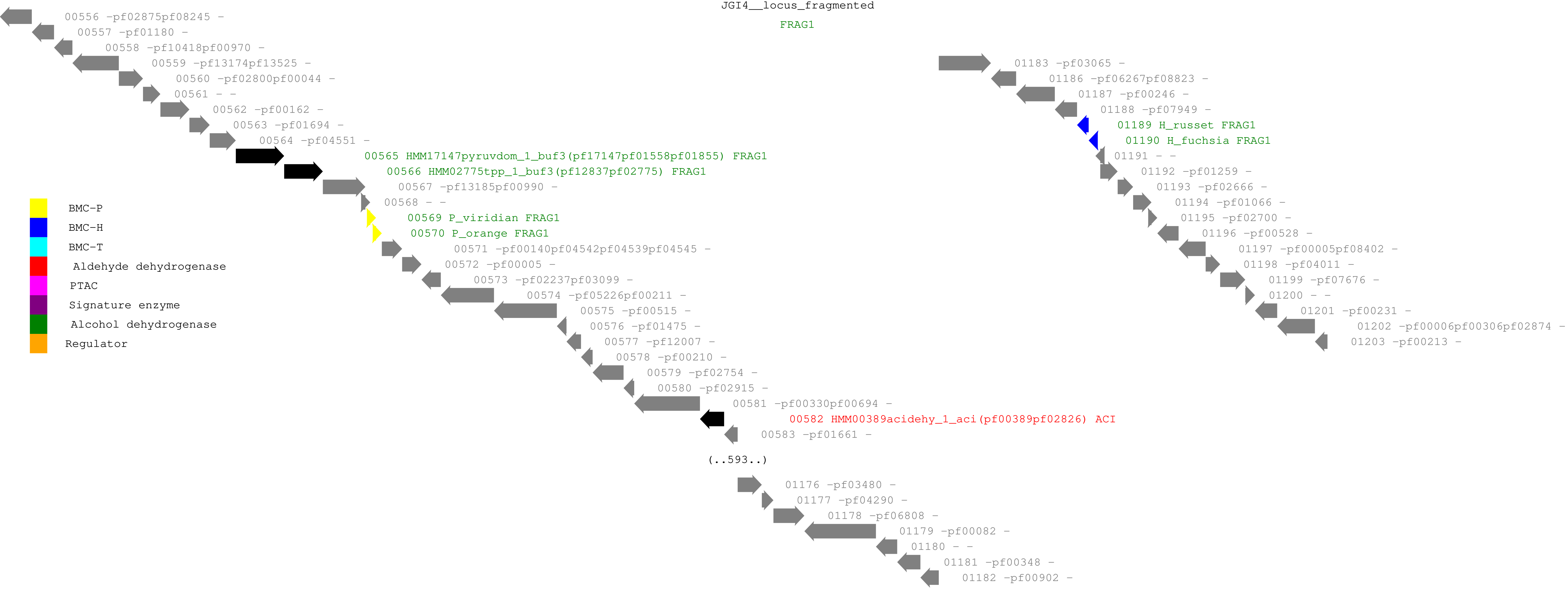

hydrothermal vent metagenome. unclassified sequences; metagenomes; ecological metagenomes.

MNBD_IGNAVI01-_locus_fragmented

FRAG1

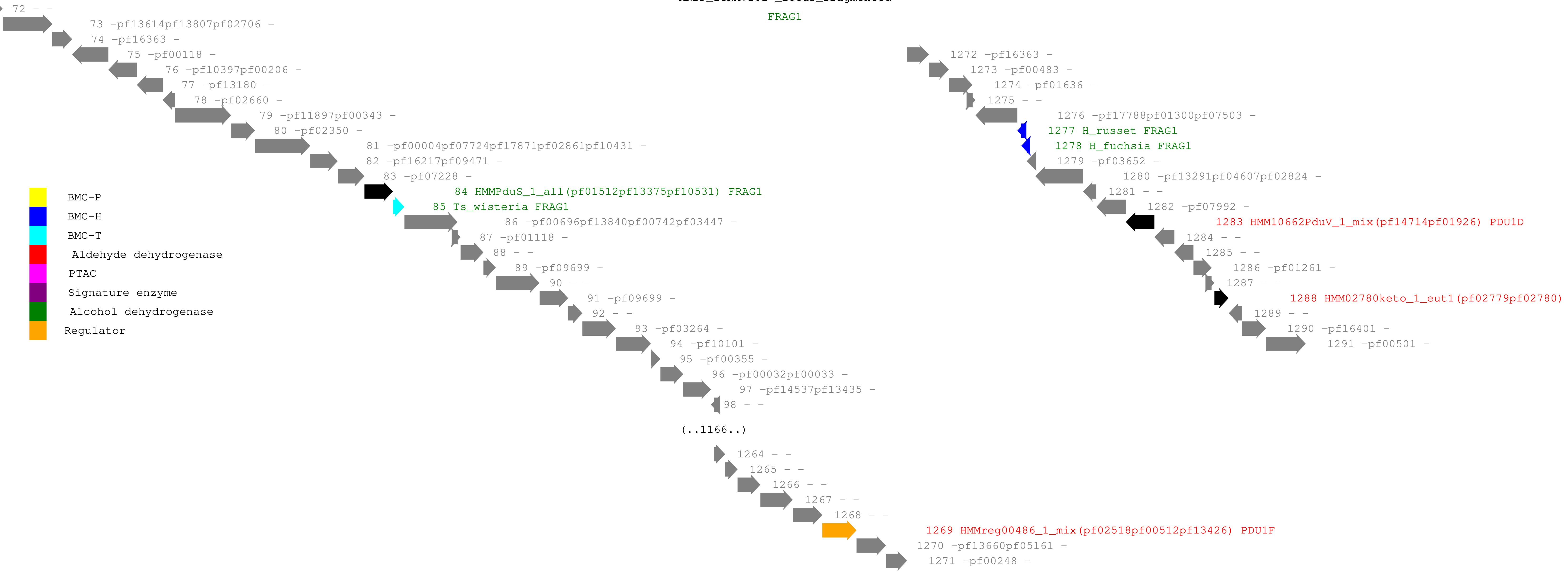

Melioribacter roseus (strain JCM 17771 / P3M-2). Bacteria; Ignavibacteriae; Ignavibacteria; Ignavibacteriales;

MROS__locus_fragmented

FRAG1

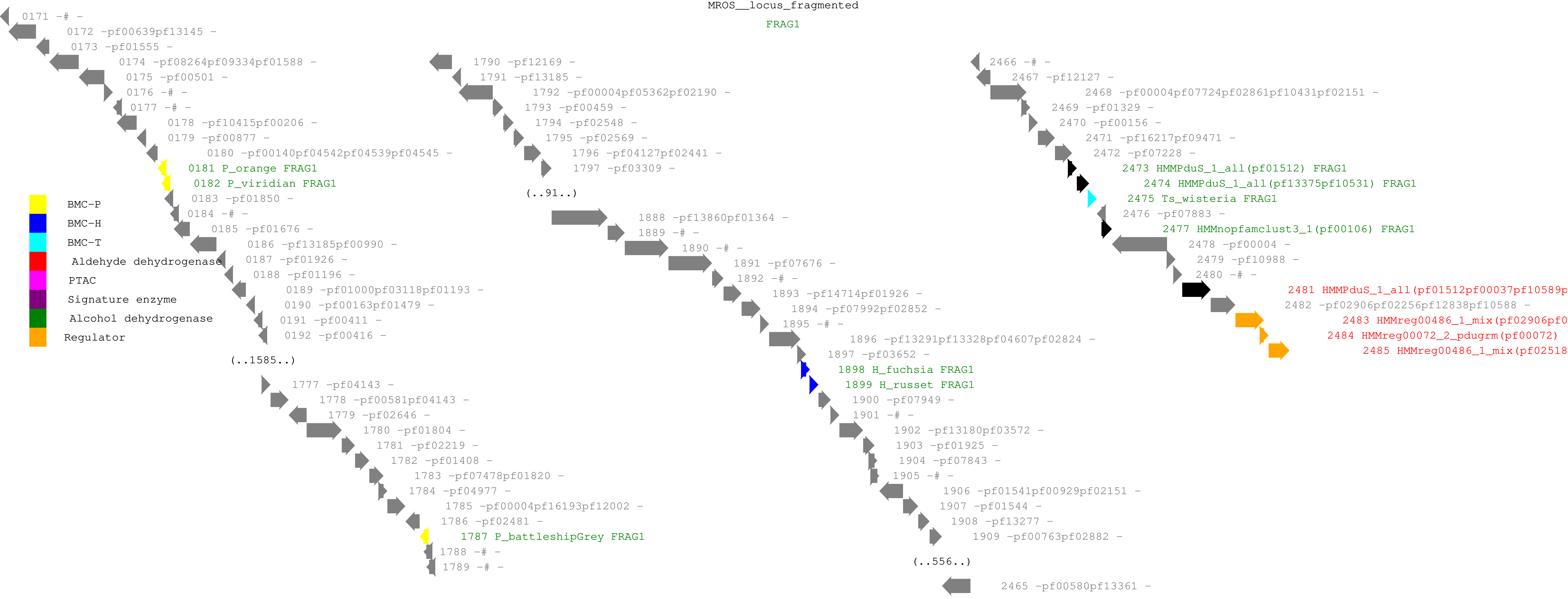

Candidatus Latescibacteria bacterium 4484_107. Bacteria; Candidatus Latescibacteria.

B1H02__locus_1_main_sat

FRAG2

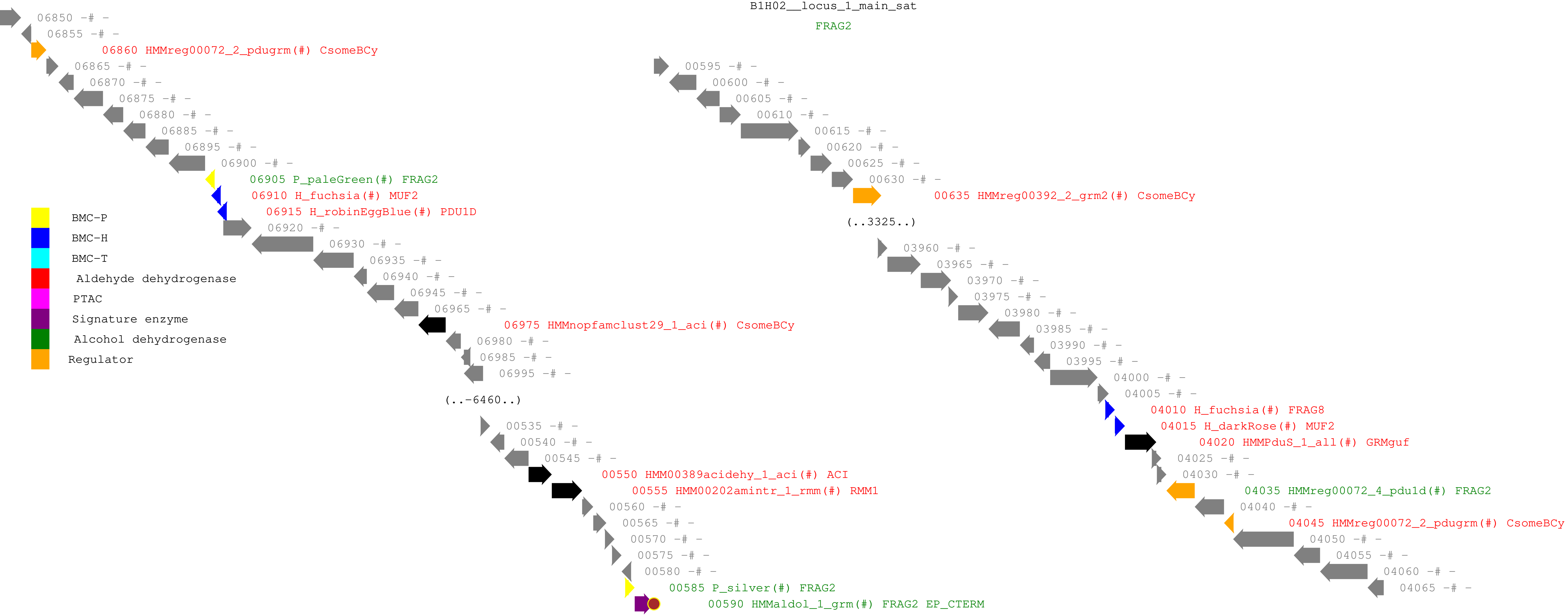

Candidatus Omnitrophica bacterium. Bacteria; Candidatus Omnitrophica.

C4527__locus_fragmented

FRAG2

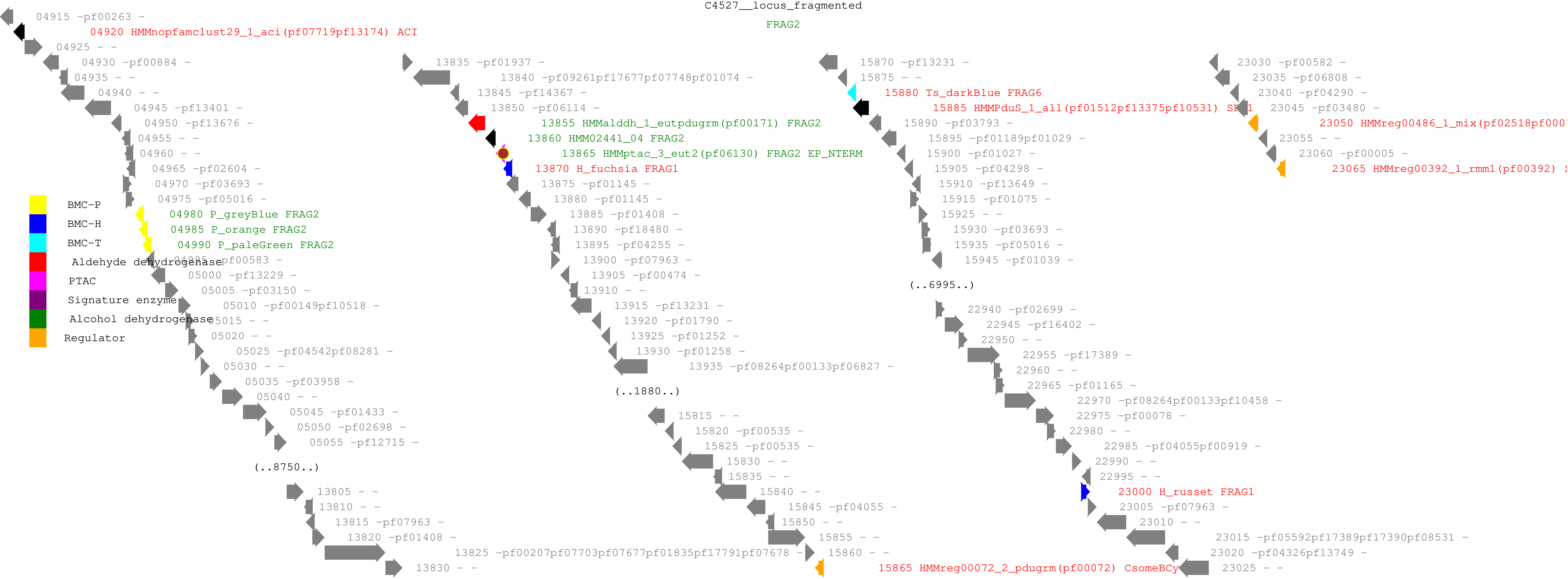

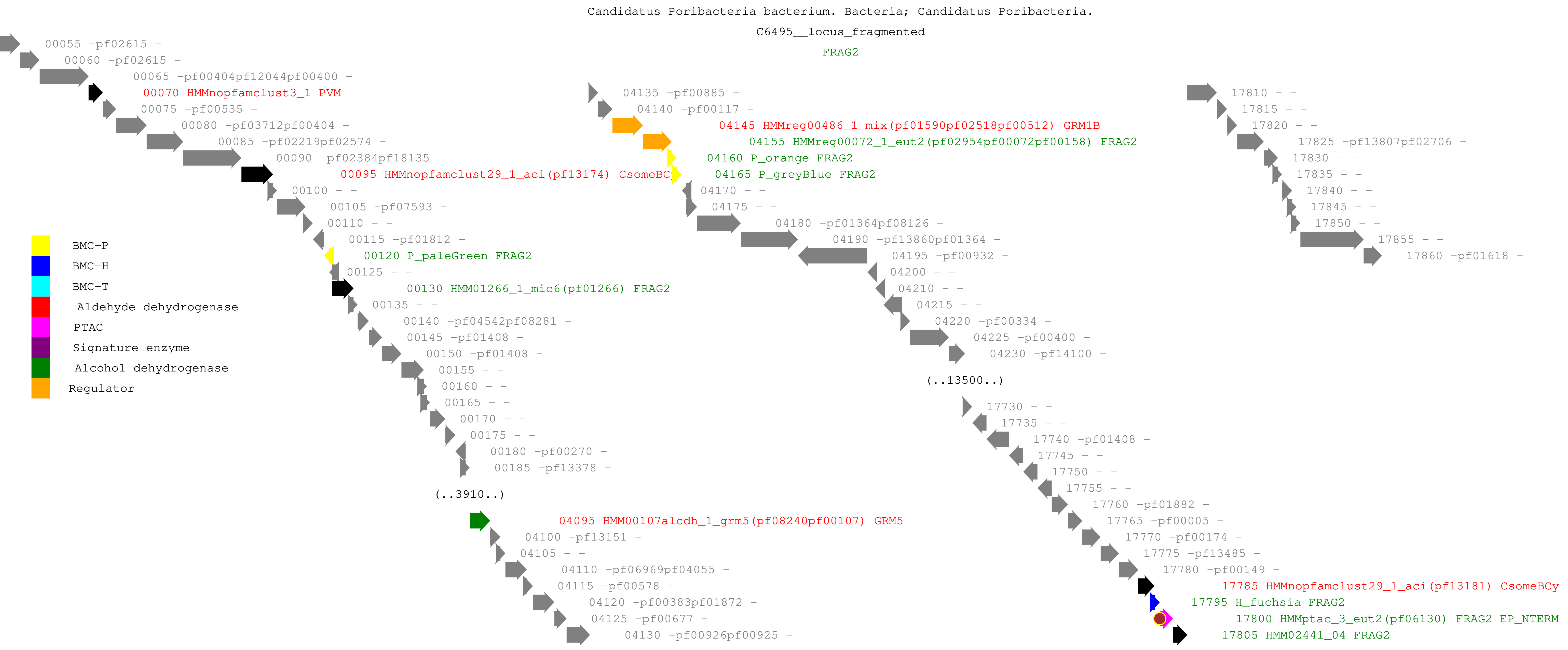

Candidatus Poribacteria bacterium. Bacteria; Candidatus Poribacteria.
C6495__locus_fragmented
FRAG2
00055 -pf02615 -
00060 -pf02615 -
00065 -pf00404pf12044pf00400 -
00070 HMMnopfamclust3_1 PVM
00075 -pf00535 -
00080 -pf03712pf00404 -
00085 -pf02219pf02574 -
00090 -pf02384pf18135 -
00095 HMMnopfamclust29_1_aci(pf13174) CsomeBCy
00100 - -
00105 -pf07593 -
00110 - -
00115 -pf01812 -
00120 P_paleGreen FRAG2
00125 - -
00130 HMM01266_1_mic6(pf01266) FRAG2
00135 - -
00140 -pf04542pf08281 -
00145 -pf01408 -
00150 -pf01408 -
00155 - -
00160 - -
00165 - -
00170 - -
00175 - -
00180 -pf00270 -
00185 -pf13378 -
(..3910..)
04095 HMM00107alcdh_1_grm5(pf08240pf00107) GRM5
04100 -pf13151 -
04105 - -
04110 -pf06969pf04055 -
04115 -pf00578 -
04120 -pf00383pf01872 -
04125 -pf00677 -
04130 -pf00926pf00925 -
04135 -pf00885 -
04140 -pf00117 -
04145 HMMreg00486_1_mix(pf01590pf02518pf00512) GRM1B
04155 HMMreg00072_1_eut2(pf02954pf00072pf00158) FRAG2
04160 P_orange FRAG2
04165 P_greyBlue FRAG2
04170 - -
04175 - -
04180 -pf01364pf08126 -
04190 -pf13860pf01364 -
04195 -pf00932 -
04200 - -
04210 - -
04215 - -
04220 -pf00334 -
04225 -pf00400 -
04230 -pf14100 -
(..13500..)
17730 - -
17735 - -
17740 -pf01408 -
17745 - -
17750 - -
17755 - -
17760 -pf01882 -
17765 -pf00005 -
17770 -pf00174 -
17775 -pf13485 -
17780 -pf00149 -
17785 HMMnopfamclust29_1_aci(pf13181) CsomeBCy
17795 H_fuchsia FRAG2
17800 HMMptac_3_eut2(pf06130) FRAG2 EP_NTERM
17805 HMM02441_04 FRAG2
17810 - -
17815 - -
17820 - -
17825 -pf13807pf02706 -
17830 - -
17835 - -
17840 - -
17845 - -
17850 - -
17855 - -
17860 -pf01618 -
BMC-P
BMC-H
BMC-T
Aldehyde dehydrogenase
PTAC
Signature enzyme
Alcohol dehydrogenase
Regulator

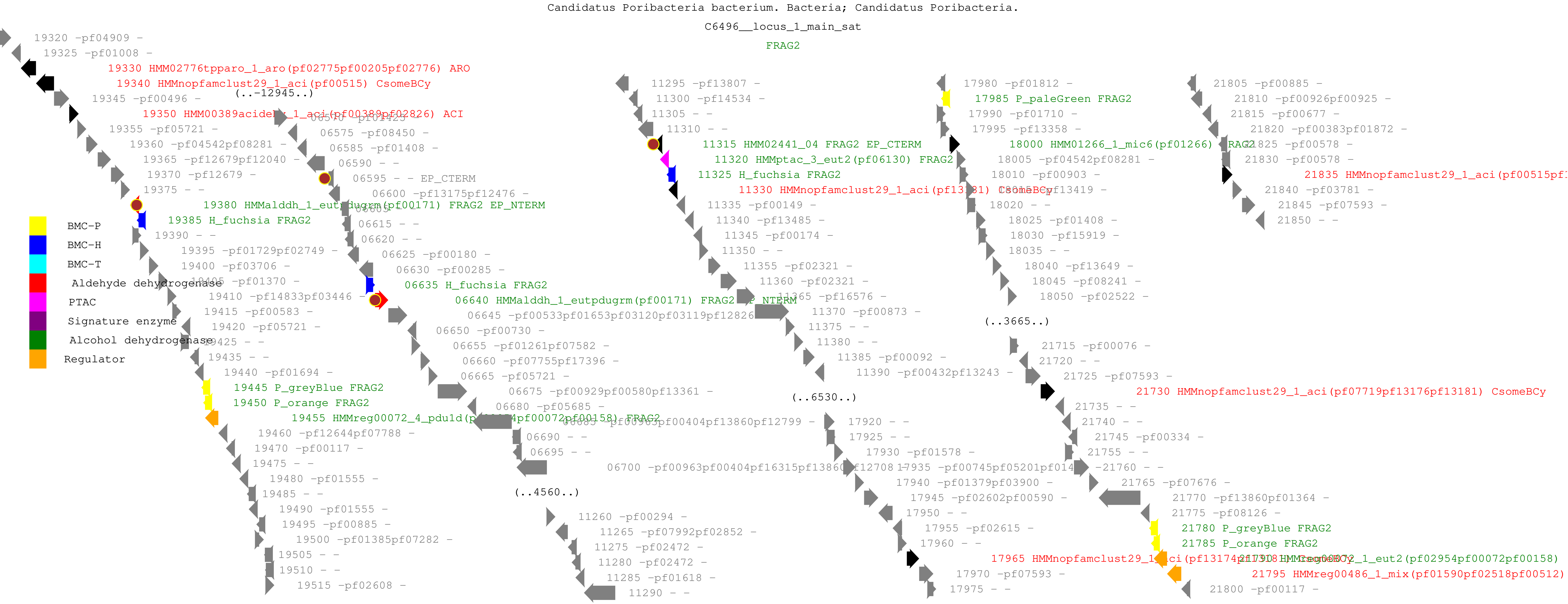
Candidatus Poribacteria bacterium. Bacteria; Candidatus Poribacteria.
C6496__locus_1_main_sat
FRAG2
19320 -pf04909 -
19325 -pf01008 -
19330 HMM02776tpparo_1_aro(pf02775pf00205pf02776) ARO
19340 HMMnopfamclust29_1_aci(pf00515) CsomeBCy
19345 -pf00496 -
(..-12945..)
19350 HMM00389acide _1_aci(pf00389pf02826) ACI
19355 -pf05721 -
19360 -pf04542pf08281 -
19365 -pf12679pf12040 -
19370 -pf12679 -
19375 - -
19380 HMMalddh_1_eutpdugrm(pf00171) FRAG2 EP_NTERM
19385 H_fuchsia FRAG2
19390 - -
19395 -pf01729pf02749 -
19400 -pf03706 -
19405 -pf01370 -
19410 -pf14833pf03446 -
19415 -pf00583 -
19420 -pf05721 -
19425 - -
19435 - -
19440 -pf01694 -
19445 P_greyBlue FRAG2
19450 P_orange FRAG2
19455 HMMreg00072_4_pdu1d(p 4pf00072pf00158) FRAG2
19460 -pf12644pf07788 -
19470 -pf00117 -
19475 - -
19480 -pf01555 -
19485 - -
19490 -pf01555 -
19495 -pf00885 -
19500 -pf01385pf07282 -
19505 - -
19510 - -
19515 -pf02608 -
BMC-P
BMC-H
BMC-T
Aldehyde dehydrogenase
PTAC
Signature enzyme
Alcohol dehydrogenase
Regulator
06570 -pf01425 -
06575 -pf08450 -
06585 -pf01408 -
06590 - -
06595 - - EP_CTERM
06600 -pf13175pf12476 -
06605
06615 - -
06620 - -
06625 -pf00180 -
06630 -pf00285 -
06635 H_fuchsia FRAG2
06640 HMMalddh_1_eutpdugrm(pf00171) FRAG2 EP_NTERM
06645 -pf00533pf01653pf03120pf03119pf12826
06650 -pf00730 -
06655 -pf01261pf07582 -
06660 -pf07755pf17396 -
06665 -pf05721 -
06675 -pf00929pf00580pf13361 -
06680 -pf05685 -
06685 -pf00563pf00404pf13860pf12799 -
06690 - -
06695 - -
06700 -pf00963pf00404pf16315pf1386 pf12708
(..4560..)
11260 -pf00294 -
11265 -pf07992pf02852 -
11275 -pf02472 -
11280 -pf02472 -
11285 -pf01618 -
11290 - -
11295 -pf13807 -
11300 -pf14534 -
11305 - -
11310 - -
11315 HMM02441_04 FRAG2 EP_CTERM
11320 HMMptac_3_eut2(pf06130) FRAG2
11325 H_fuchsia FRAG2
11330 HMMnopfamclust29_1_aci(pf13181) CsomeBCy
11335 -pf00149 -
11340 -pf13485 -
11345 -pf00174 -
11350 - -
11355 -pf02321 -
11360 -pf02321 -
11365 -pf16576 -
11370 -pf00873 -
11375 - -
11380 - -
11385 -pf00092 -
11390 -pf00432pf13243 -
(..6530..)
17920 - -
17925 - -
17930 -pf01578 -
17935 -pf00745pf05201pf014
17940 -pf01379pf03900 -
17945 -pf02602pf00590 -
17950 - -
17955 -pf02615 -
17960 - -
17965 HMMnopfamclust29_1_aci(pf13174pf13181) CsomeBCy
17970 -pf07593 -
17975 - -
17980 -pf01812 -
17985 P_paleGreen FRAG2
17990 -pf01710 -
17995 -pf13358 -
18000 HMM01266_1_mic6(pf01266) FRAG2
18005 -pf04542pf08281 -
18010 -pf00903 -
18015 -pf13419 -
18020 - -
18025 -pf01408 -
18030 -pf15919 -
18035 - -
18040 -pf13649 -
18045 -pf08241 -
18050 -pf02522 -
(..3665..)
21715 -pf00076 -
21720 - -
21725 -pf07593 -
21730 HMMnopfamclust29_1_aci(pf07719pf13176pf13181) CsomeBCy
21735 - -
21740 - -
21745 -pf00334 -
21755 - -
21760 - -
21765 -pf07676 -
21770 -pf13860pf01364 -
21775 -pf08126 -
21780 P_greyBlue FRAG2
21785 P_orange FRAG2
21790 HMMreg00072_1_eut2(pf02954pf00072pf00158)
21795 HMMreg00486_1_mix(pf01590pf02518pf00512)
21800 -pf00117 -
21805 -pf00885 -
21810 -pf00926pf00925 -
21815 -pf00677 -
21820 -pf00383pf01872 -
21825 -pf00578 -
21830 -pf00578 -
21835 HMMnopfamclust29_1_aci(pf00515pf
21840 -pf03781 -
21845 -pf07593 -
21850 - -

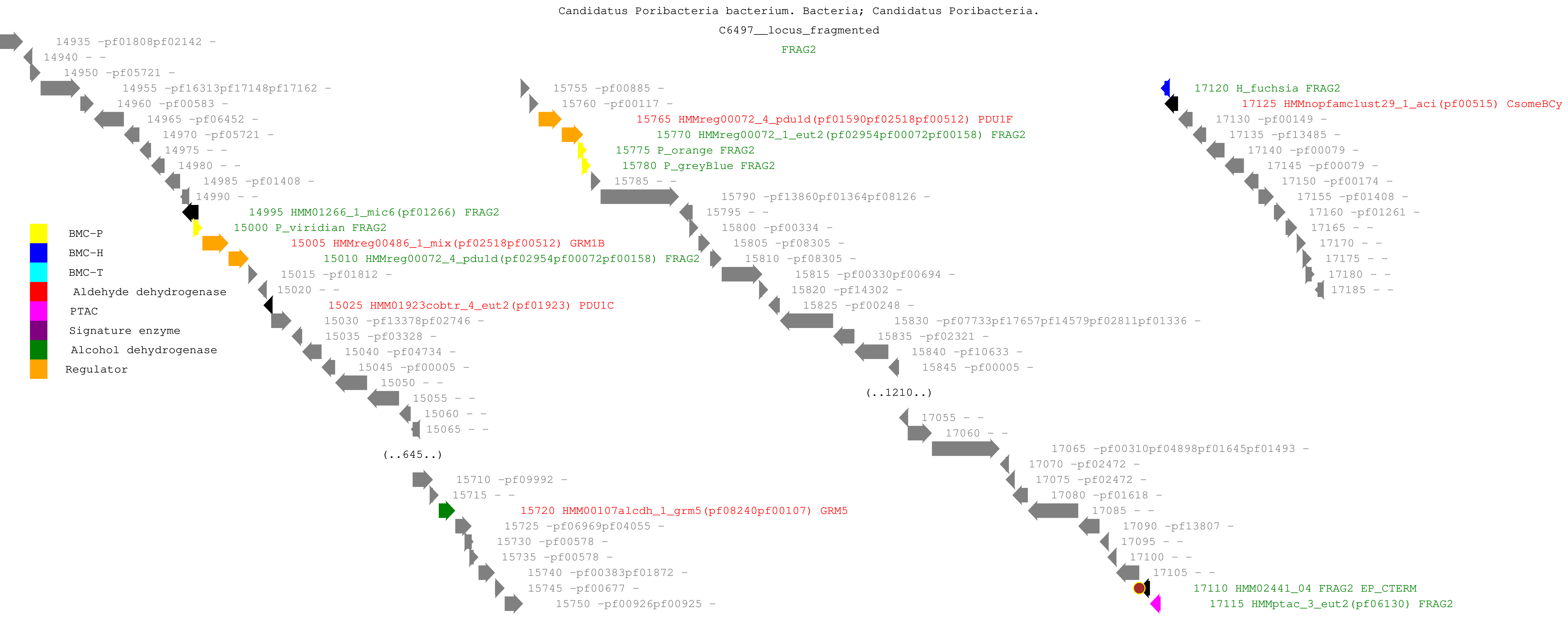

Candidatus Poribacteria bacterium. Bacteria; Candidatus Poribacteria.
C6497__locus_fragmented
FRAG2
14935 -pf01808pf02142 -
14940 - -
14950 -pf05721 -
14955 -pf16313pf17148pf17162 -
14960 -pf00583 -
14965 -pf06452 -
14970 -pf05721 -
14975 - -
14980 - -
14985 -pf01408 -
14990 - -
14995 HMM01266_1_mic6(pf01266) FRAG2
15000 P_viridian FRAG2
15005 HMMreg00486_1_mix(pf02518pf00512) GRM1B
15010 HMMreg00072_4_pdu1d(pf02954pf00072pf00158) FRAG2
15015 -pf01812 -
15020 - -
15025 HMM01923cobtr_4_eut2(pf01923) PDU1C
15030 -pf13378pf02746 -
15035 -pf03328 -
15040 -pf04734 -
15045 -pf00005 -
15050 - -
15055 - -
15060 - -
15065 - -
(..645..)
15710 -pf09992 -
15715 - -
15720 HMM00107alcdh_1_grm5(pf08240pf00107) GRM5
15725 -pf06969pf04055 -
15730 -pf00578 -
15735 -pf00578 -
15740 -pf00383pf01872 -
15745 -pf00677 -
15750 -pf00926pf00925 -
15755 -pf00885 -
15760 -pf00117 -
15765 HMMreg00072_4_pdu1d(pf01590pf02518pf00512) PDU1F
15770 HMMreg00072_1_eut2(pf02954pf00072pf00158) FRAG2
15775 P_orange FRAG2
15780 P_greyBlue FRAG2
15785 - -
15790 -pf13860pf01364pf08126 -
15795 - -
15800 -pf00334 -
15805 -pf08305 -
15810 -pf08305 -
15815 -pf00330pf00694 -
15820 -pf14302 -
15825 -pf00248 -
15830 -pf07733pf17657pf14579pf02811pf01336 -
15835 -pf02321 -
15840 -pf10633 -
15845 -pf00005 -
(..1210..)
17055 - -
17060 - -
17065 -pf00310pf04898pf01645pf01493 -
17070 -pf02472 -
17075 -pf02472 -
17080 -pf01618 -
17085 - -
17090 -pf13807 -
17095 - -
17100 - -
17105 - -
17110 HMM02441_04 FRAG2 EP_CTERM
17115 HMMptac_3_eut2(pf06130) FRAG2
17120 H_fuchsia FRAG2
17125 HMMnopfamclust29_1_aci(pf00515) CsomeBCy
17130 -pf00149 -
17135 -pf13485 -
17140 -pf00079 -
17145 -pf00079 -
17150 -pf00174 -
17155 -pf01408 -
17160 -pf01261 -
17165 - -
17170 - -
17175 - -
17180 - -
17185 - -
BMC-P
BMC-H
BMC-T
Aldehyde dehydrogenase
PTAC
Signature enzyme
Alcohol dehydrogenase
Regulator

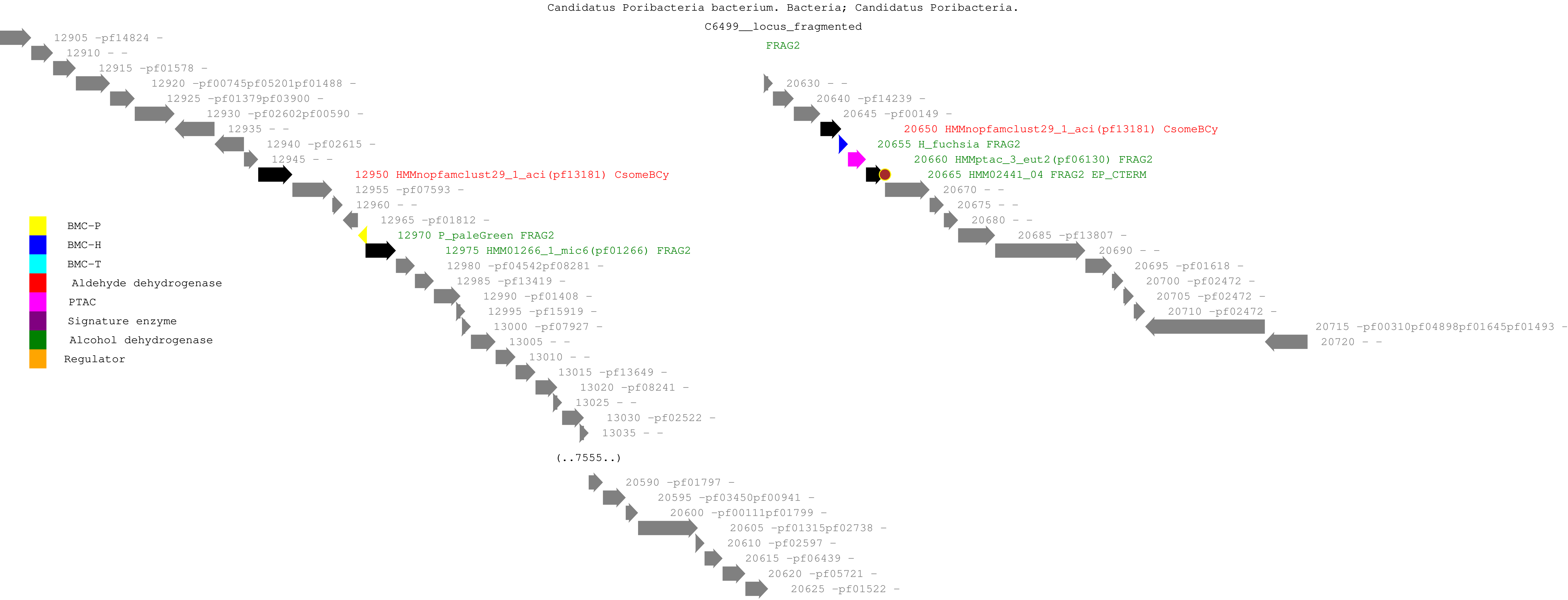

Candidatus Poribacteria bacterium. Bacteria; Candidatus Poribacteria.
C6499__locus_fragmented
FRAG2
12905 -pf14824 -
12910 - -
12915 -pf01578 -
12920 -pf00745pf05201pf01488 -
12925 -pf01379pf03900 -
12930 -pf02602pf00590 -
12935 - -
12940 -pf02615 -
12945 - -
12950 HMMnopfamclust29_1_aci(pf13181) CsomeBCy
12955 -pf07593 -
12960 - -
12965 -pf01812 -
12970 P_paleGreen FRAG2
12975 HMM01266_1_mic6(pf01266) FRAG2
12980 -pf04542pf08281 -
12985 -pf13419 -
12990 -pf01408 -
12995 -pf15919 -
13000 -pf07927 -
13005 - -
13010 - -
13015 -pf13649 -
13020 -pf08241 -
13025 - -
13030 -pf02522 -
13035 - -
(..7555..)
20590 -pf01797 -
20595 -pf03450pf00941 -
20600 -pf00111pf01799 -
20605 -pf01315pf02738 -
20610 -pf02597 -
20615 -pf06439 -
20620 -pf05721 -
20625 -pf01522 -
20630 - -
20640 -pf14239 -
20645 -pf00149 -
20650 HMMnopfamclust29_1_aci(pf13181) CsomeBCy
20655 H_fuchsia FRAG2
20660 HMMptac_3_eut2(pf06130) FRAG2
20665 HMM02441_04 FRAG2 EP_CTERM
20670 - -
20675 - -
20680 - -
20685 -pf13807 -
20690 - -
20695 -pf01618 -
20700 -pf02472 -
20705 -pf02472 -
20710 -pf02472 -
20715 -pf00310pf04898pf01645pf01493 -
20720 - -
BMC-P
BMC-H
BMC-T
Aldehyde dehydrogenase
PTAC
Signature enzyme
Alcohol dehydrogenase
Regulator

Candidatus Poribacteria bacterium. Bacteria; Candidatus Poribacteria.

C6500__locus_fragmented

FRAG2

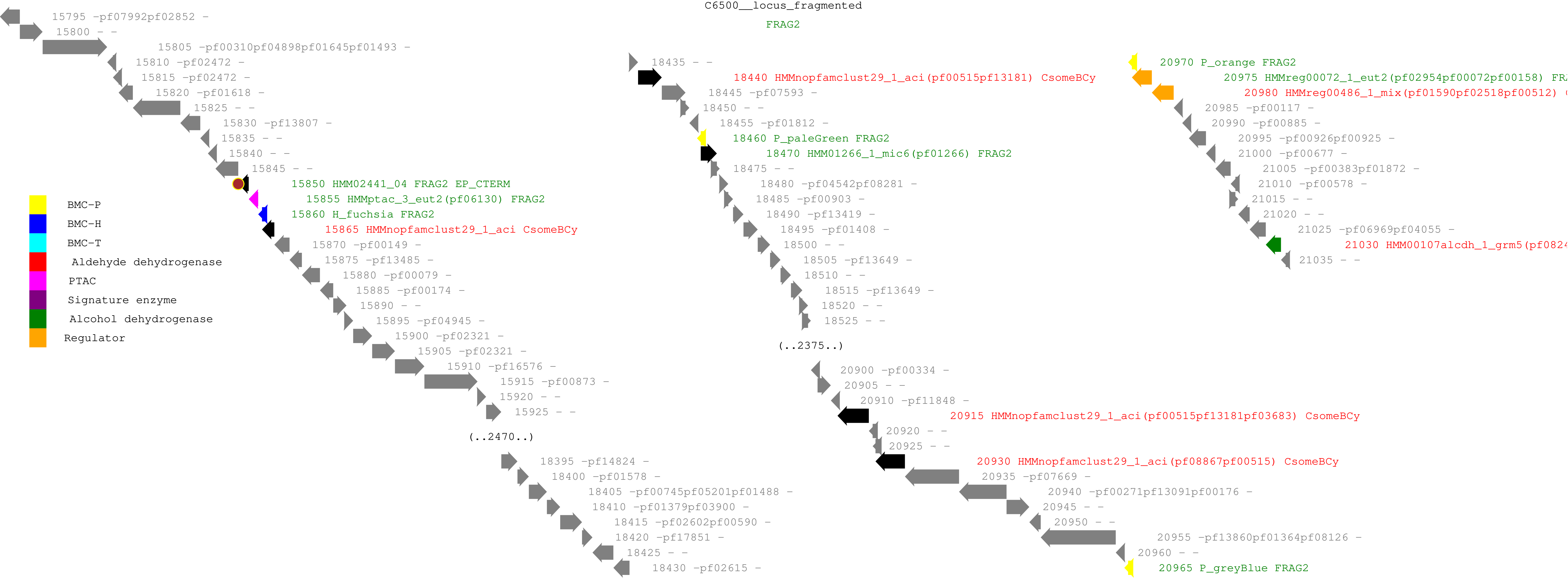

Candidatus Poribacteria bacterium. Bacteria; Candidatus Poribacteria.

C6501__locus_fragmented

FRAG2

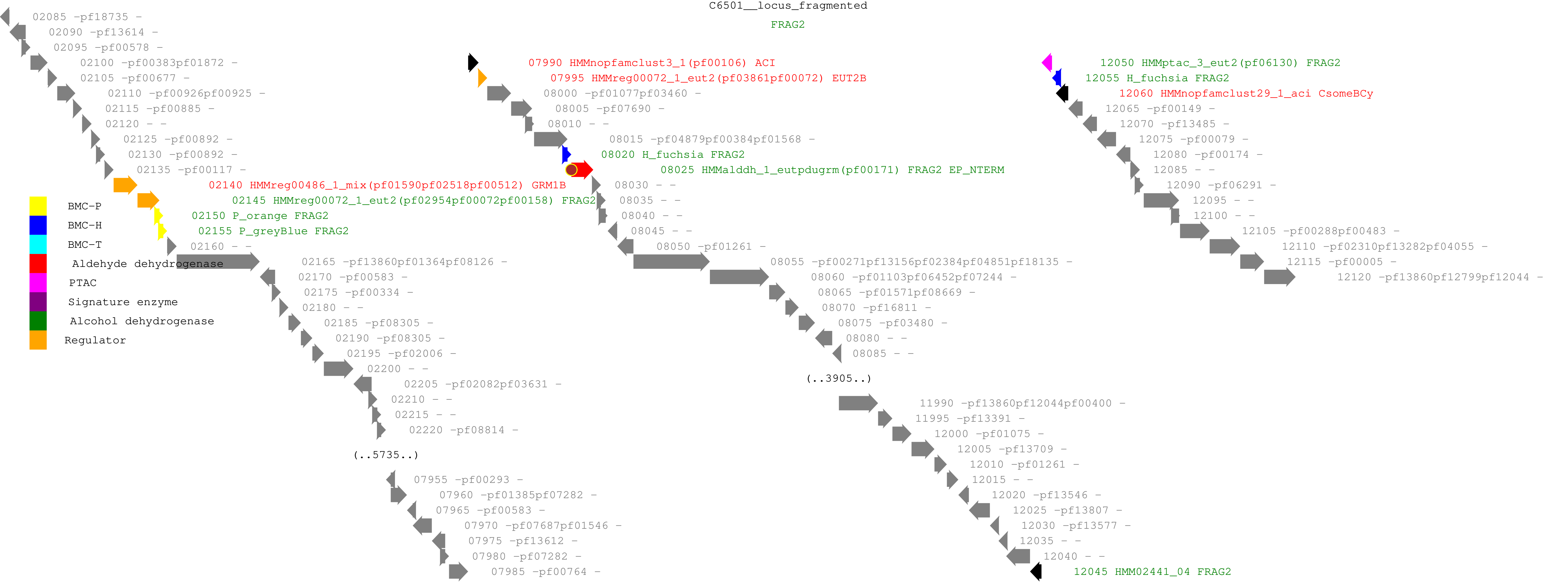

Candidatus Poribacteria bacterium. Bacteria; Candidatus Poribacteria.

C6502__locus_fragmented

FRAG2

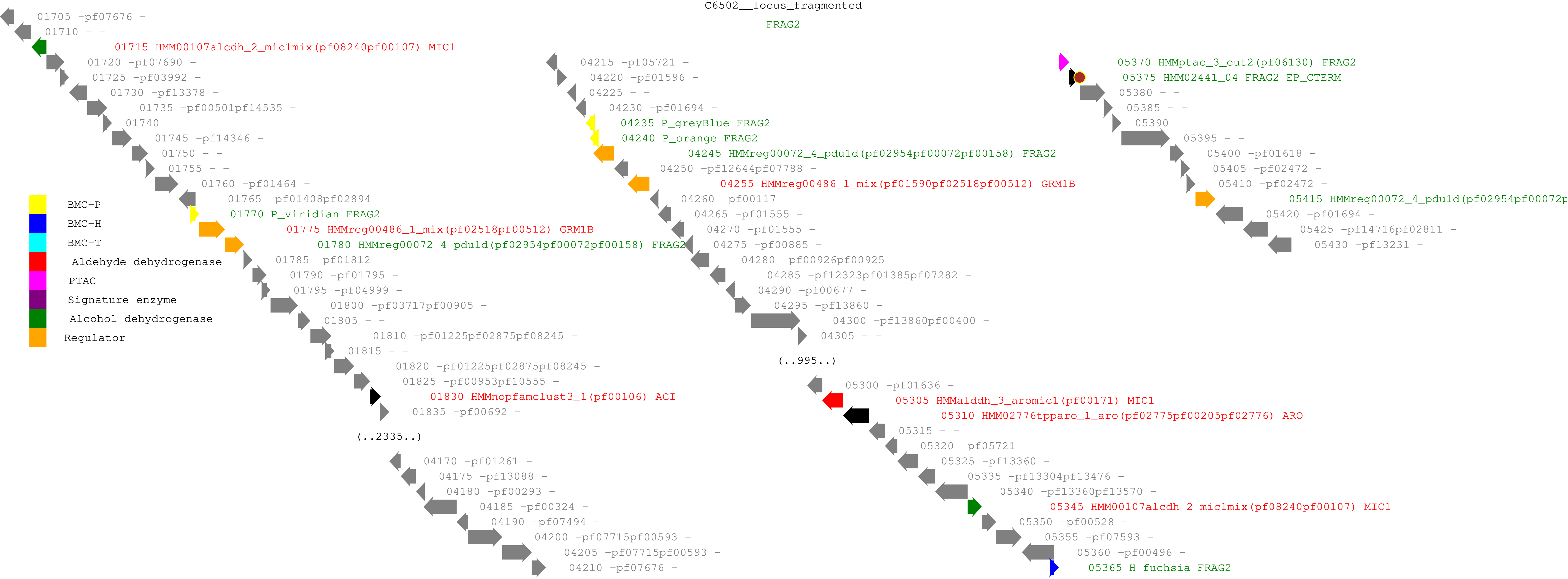

Candidatus Poribacteria bacterium. Bacteria; Candidatus Poribacteria.

C6503__locus_fragmented

FRAG2

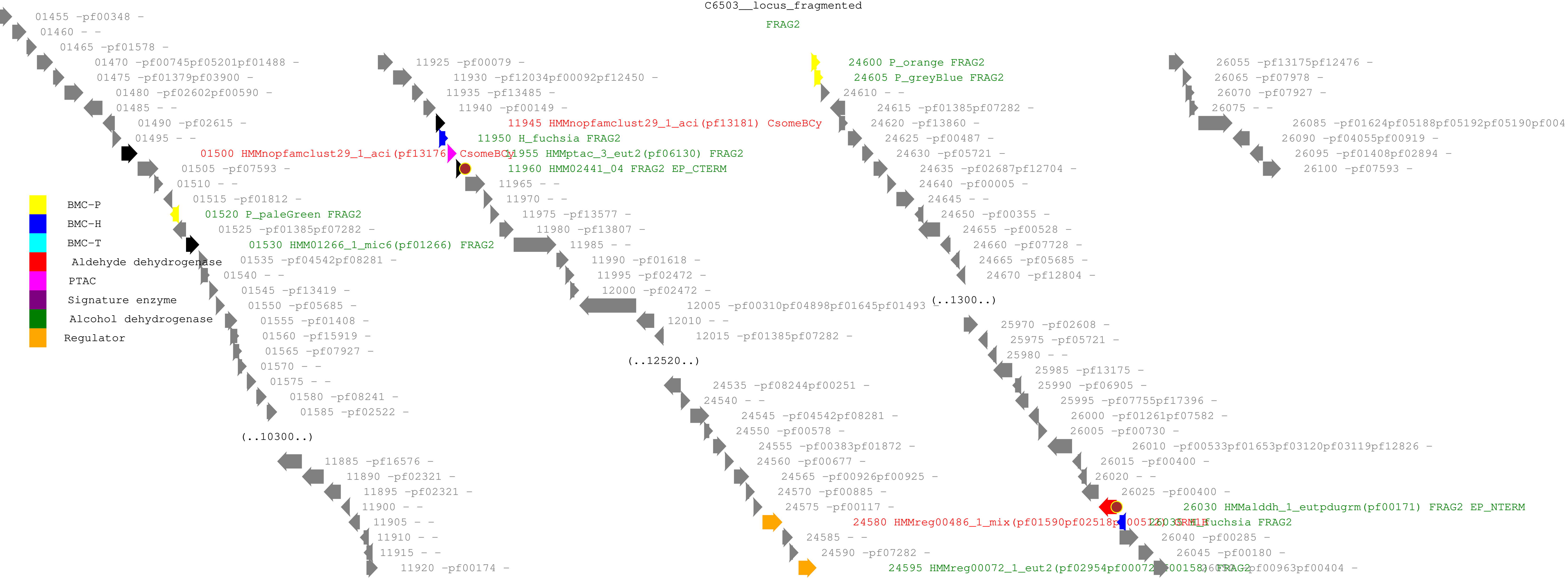

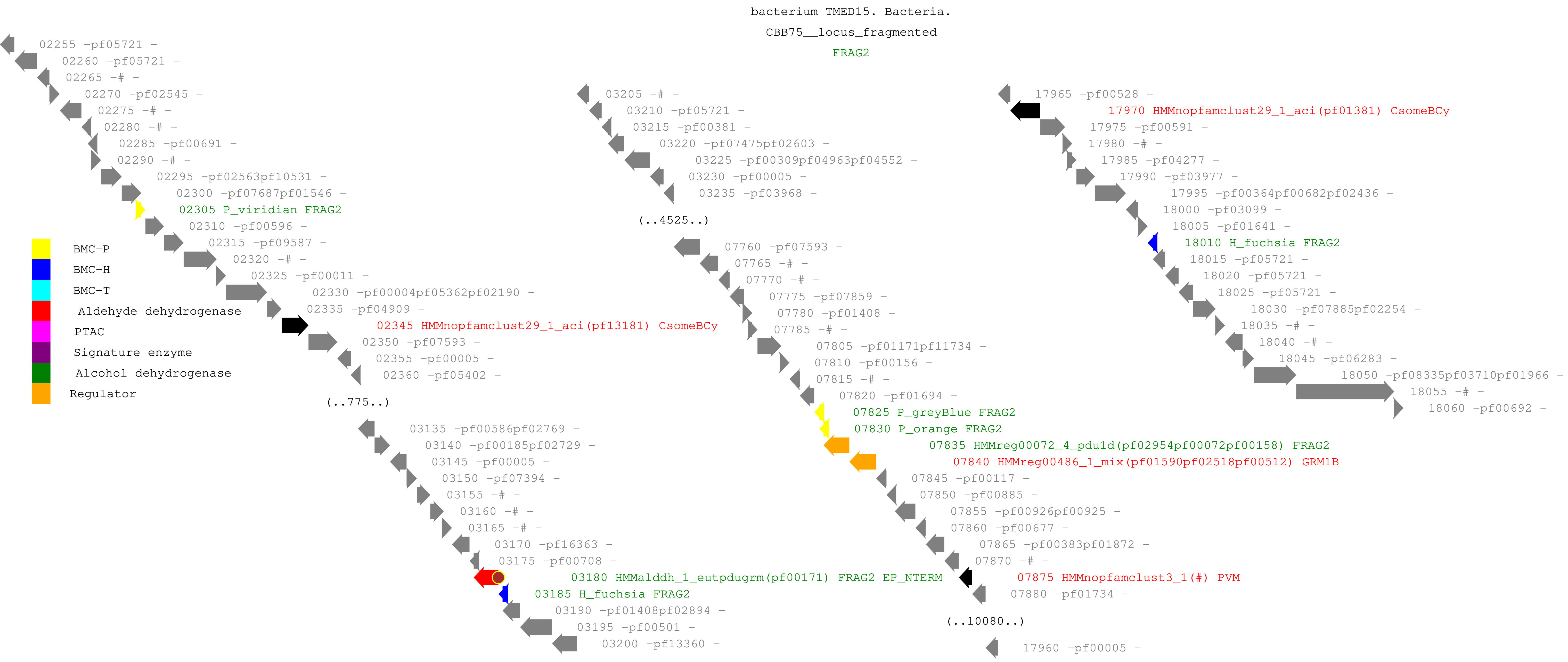

bacterium TMED15. Bacteria.
CBB75__locus_fragmented
FRAG2
02255 -pf05721 -
02260 -pf05721 -
02265 -# -
02270 -pf02545 -
02275 -# -
02280 -# -
02285 -pf00691 -
02290 -# -
02295 -pf02563pf10531 -
02300 -pf07687pf01546 -
02305 P_viridian FRAG2
02310 -pf00596 -
02315 -pf09587 -
02320 -# -
02325 -pf00011 -
02330 -pf00004pf05362pf02190 -
02335 -pf04909 -
02345 HMMnopfamclust29_1_aci(pf13181) CsomeBCy
02350 -pf07593 -
02355 -pf00005 -
02360 -pf05402 -
(..775..)
03135 -pf00586pf02769 -
03140 -pf00185pf02729 -
03145 -pf00005 -
03150 -pf07394 -
03155 -# -
03160 -# -
03165 -# -
03170 -pf16363 -
03175 -pf00708 -
03180 HMMalddh_1_eutpdugrm(pf00171) FRAG2 EP_NTERM
03185 H_fuchsia FRAG2
03190 -pf01408pf02894 -
03195 -pf00501 -
03200 -pf13360 -
03205 -# -
03210 -pf05721 -
03215 -pf00381 -
03220 -pf07475pf02603 -
03225 -pf00309pf04963pf04552 -
03230 -pf00005 -
03235 -pf03968 -
(..4525..)
07760 -pf07593 -
07765 -# -
07770 -# -
07775 -pf07859 -
07780 -pf01408 -
07785 -# -
07805 -pf01171pf11734 -
07810 -pf00156 -
07815 -# -
07820 -pf01694 -
07825 P_greyBlue FRAG2
07830 P_orange FRAG2
07835 HMMreg00072_4_pdu1d(pf02954pf00072pf00158) FRAG2
07840 HMMreg00486_1_mix(pf01590pf02518pf00512) GRM1B
07845 -pf00117 -
07850 -pf00885 -
07855 -pf00926pf00925 -
07860 -pf00677 -
07865 -pf00383pf01872 -
07870 -# -
07875 HMMnopfamclust3_1(#) PVM
07880 -pf01734 -
(..10080..)
17960 -pf00005 -
17965 -pf00528 -
17970 HMMnopfamclust29_1_aci(pf01381) CsomeBCy
17975 -pf00591 -
17980 -# -
17985 -pf04277 -
17990 -pf03977 -
17995 -pf00364pf00682pf02436 -
18000 -pf03099 -
18005 -pf01641 -
18010 H_fuchsia FRAG2
18015 -pf05721 -
18020 -pf05721 -
18025 -pf05721 -
18030 -pf07885pf02254 -
18035 -# -
18040 -# -
18045 -pf06283 -
18050 -pf08335pf03710pf01966 -
18055 -# -
18060 -pf00692 -
BMC-P
BMC-H
BMC-T
Aldehyde dehydrogenase
PTAC
Signature enzyme
Alcohol dehydrogenase
Regulator

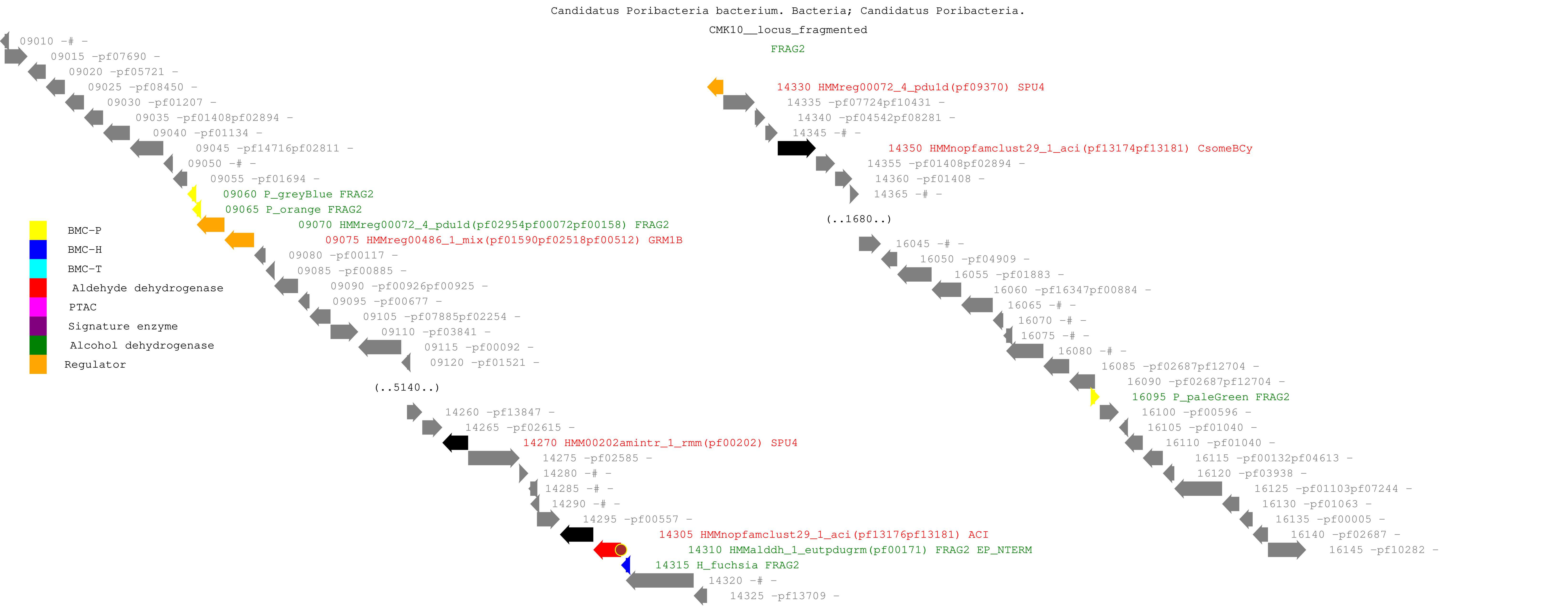
Candidatus Poribacteria bacterium. Bacteria; Candidatus Poribacteria.
CMK10__locus_fragmented
FRAG2
09010 -# -
09015 -pf07690 -
09020 -pf05721 -
09025 -pf08450 -
09030 -pf01207 -
09035 -pf01408pf02894 -
09040 -pf01134 -
09045 -pf14716pf02811 -
09050 -# -
09055 -pf01694 -
09060 P_greyBlue FRAG2
09065 P_orange FRAG2
09070 HMMreg00072_4_pdu1d(pf02954pf00072pf00158) FRAG2
09075 HMMreg00486_1_mix(pf01590pf02518pf00512) GRM1B
09080 -pf00117 -
09085 -pf00885 -
09090 -pf00926pf00925 -
09095 -pf00677 -
09105 -pf07885pf02254 -
09110 -pf03841 -
09115 -pf00092 -
09120 -pf01521 -
(..5140..)
14260 -pf13847 -
14265 -pf02615 -
14270 HMM00202amintr_1_rmm(pf00202) SPU4
14275 -pf02585 -
14280 -# -
14285 -# -
14290 -# -
14295 -pf00557 -
14305 HMMnopfamclust29_1_aci(pf13176pf13181) ACI
14310 HMMalddh_1_eutpdugrm(pf00171) FRAG2 EP_NTERM
14315 H_fuchsia FRAG2
14320 -# -
14325 -pf13709 -
14330 HMMreg00072_4_pdu1d(pf09370) SPU4
14335 -pf07724pf10431 -
14340 -pf04542pf08281 -
14345 -# -
14350 HMMnopfamclust29_1_aci(pf13174pf13181) CsomeBCy
14355 -pf01408pf02894 -
14360 -pf01408 -
14365 -# -
(..1680..)
16045 -# -
16050 -pf04909 -
16055 -pf01883 -
16060 -pf16347pf00884 -
16065 -# -
16070 -# -
16075 -# -
16080 -# -
16085 -pf02687pf12704 -
16090 -pf02687pf12704 -
16095 P_paleGreen FRAG2
16100 -pf00596 -
16105 -pf01040 -
16110 -pf01040 -
16115 -pf00132pf04613 -
16120 -pf03938 -
16125 -pf01103pf07244 -
16130 -pf01063 -
16135 -pf00005 -
16140 -pf02687 -
16145 -pf10282 -
BMC-P
BMC-H
BMC-T
Aldehyde dehydrogenase
PTAC
Signature enzyme
Alcohol dehydrogenase
Regulator

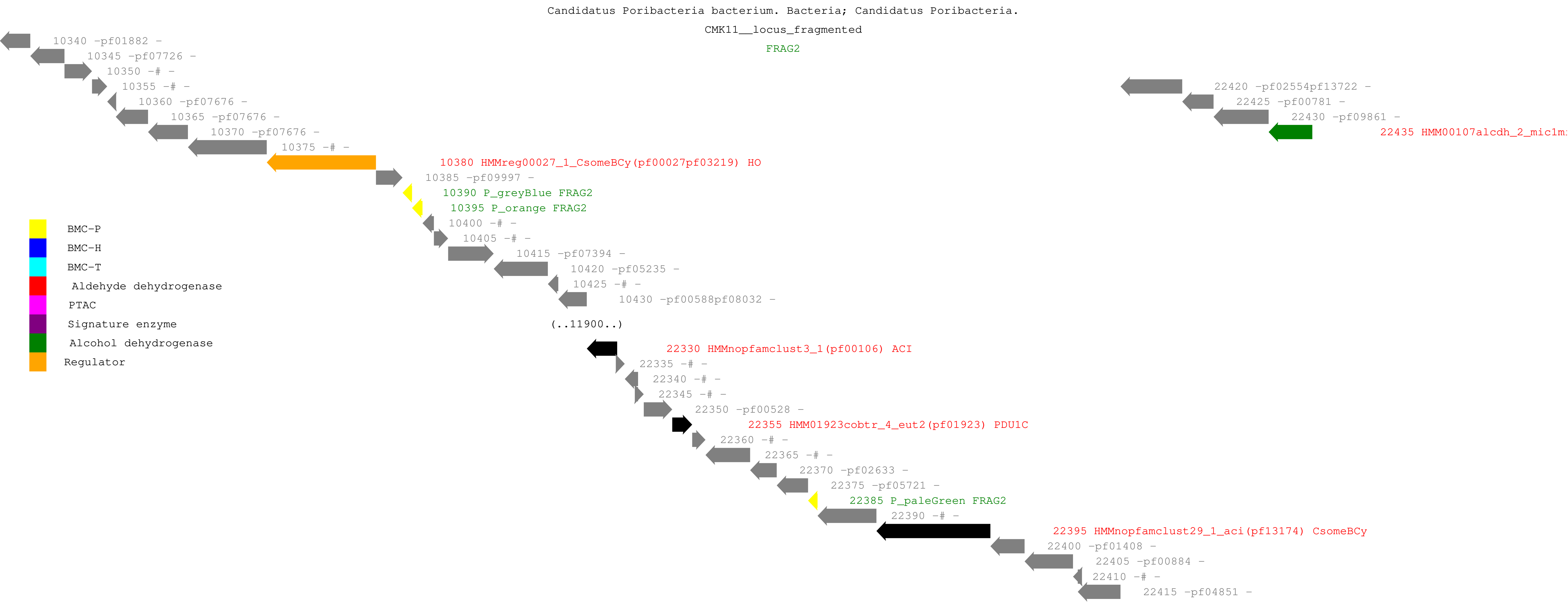
Candidatus Poribacteria bacterium. Bacteria; Candidatus Poribacteria.
CMK11__locus_fragmented
FRAG2
10340 -pf01882 -
10345 -pf07726 -
10350 -# -
10355 -# -
10360 -pf07676 -
10365 -pf07676 -
10370 -pf07676 -
10375 -# -
10380 HMMreg00027_1_CsomeBCy(pf00027pf03219) HO
10385 -pf09997 -
10390 P_greyBlue FRAG2
10395 P_orange FRAG2
10400 -# -
10405 -# -
10415 -pf07394 -
10420 -pf05235 -
10425 -# -
10430 -pf00588pf08032 -
(..11900..)
22330 HMMnopfamclust3_1(pf00106) ACI
22335 -# -
22340 -# -
22345 -# -
22350 -pf00528 -
22355 HMM01923cobtr_4_eut2(pf01923) PDU1C
22360 -# -
22365 -# -
22370 -pf02633 -
22375 -pf05721 -
22385 P_paleGreen FRAG2
22390 -# -
22395 HMMnopfamclust29_1_aci(pf13174) CsomeBCy
22400 -pf01408 -
22405 -pf00884 -
22410 -# -
22415 -pf04851 -
22420 -pf02554pf13722 -
22425 -pf00781 -
22430 -pf09861 -
22435 HMM00107alcdh_2_mic1m
BMC-P
BMC-H
BMC-T
Aldehyde dehydrogenase
PTAC
Signature enzyme
Alcohol dehydrogenase
Regulator

Candidatus Poribacteria bacterium. Bacteria; Candidatus Poribacteria.

CMK13__locus_fragmented

FRAG2

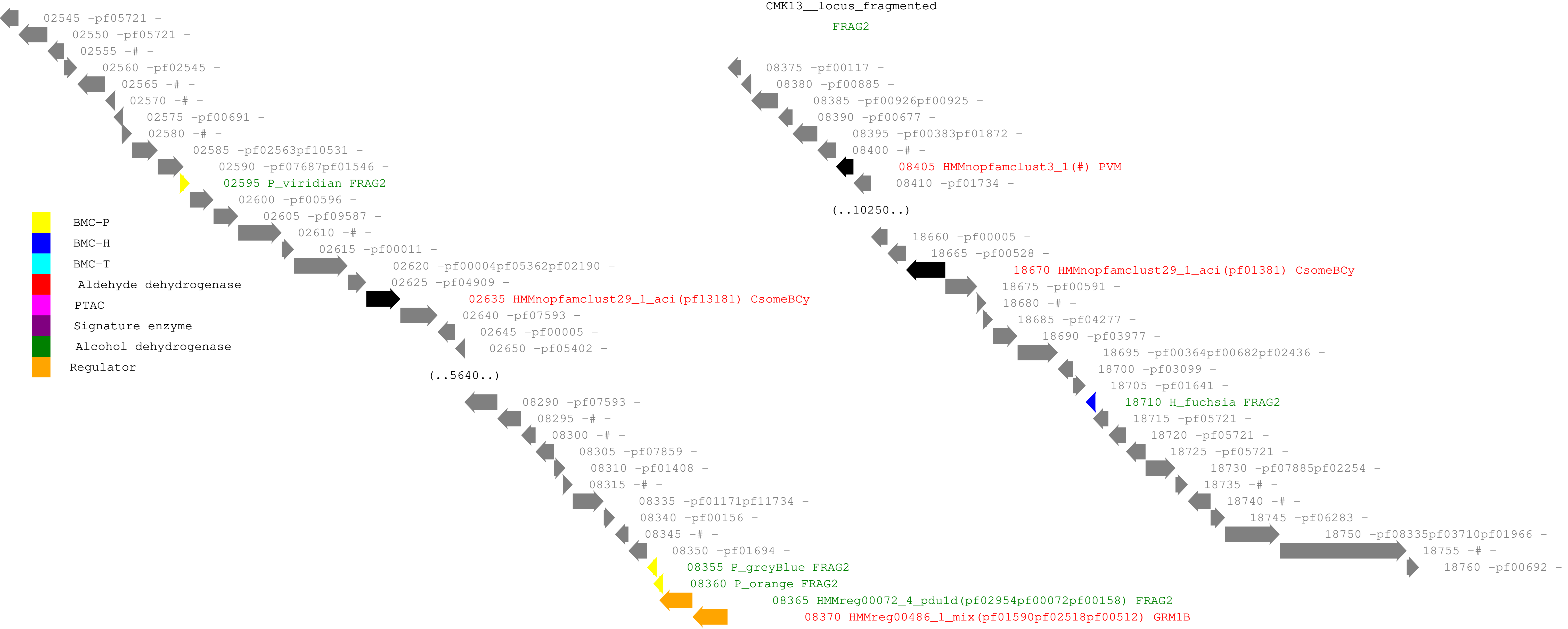

Candidatus Poribacteria bacterium. Bacteria; Candidatus Poribacteria.

CMK14__locus_fragmented

FRAG2

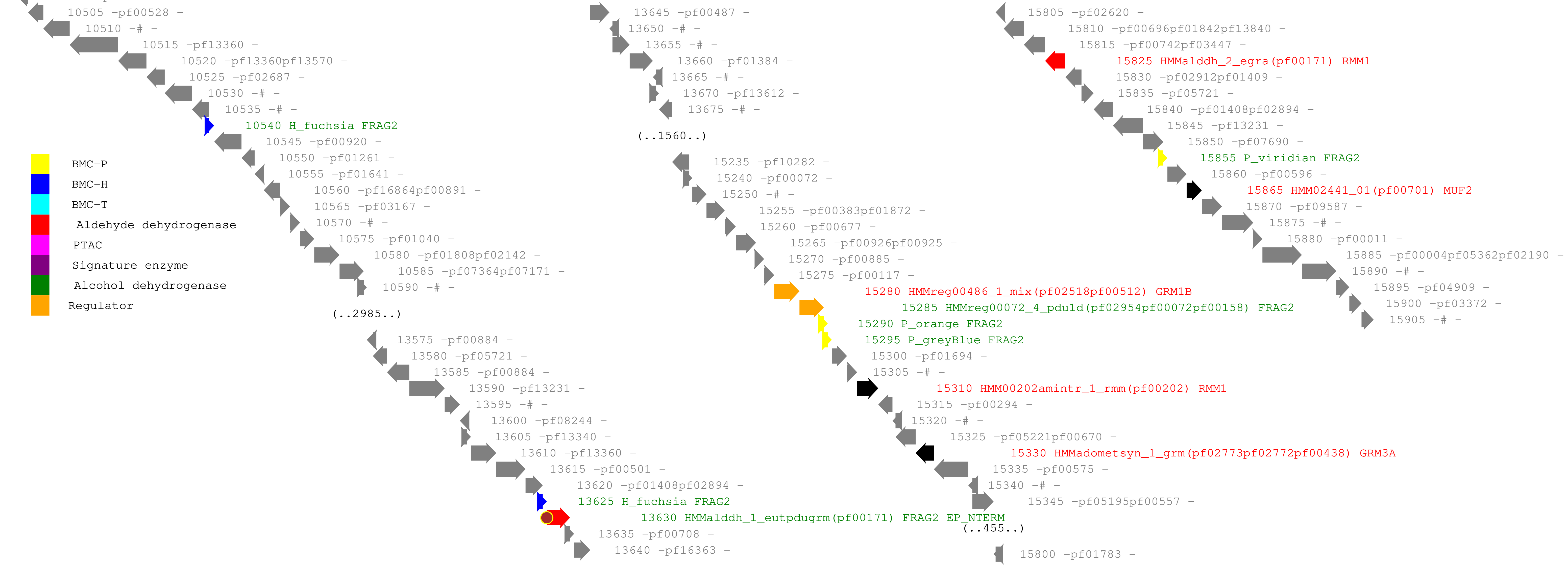

Candidatus Poribacteria bacterium. Bacteria; Candidatus Poribacteria.

CMK16__locus_fragmented

FRAG2

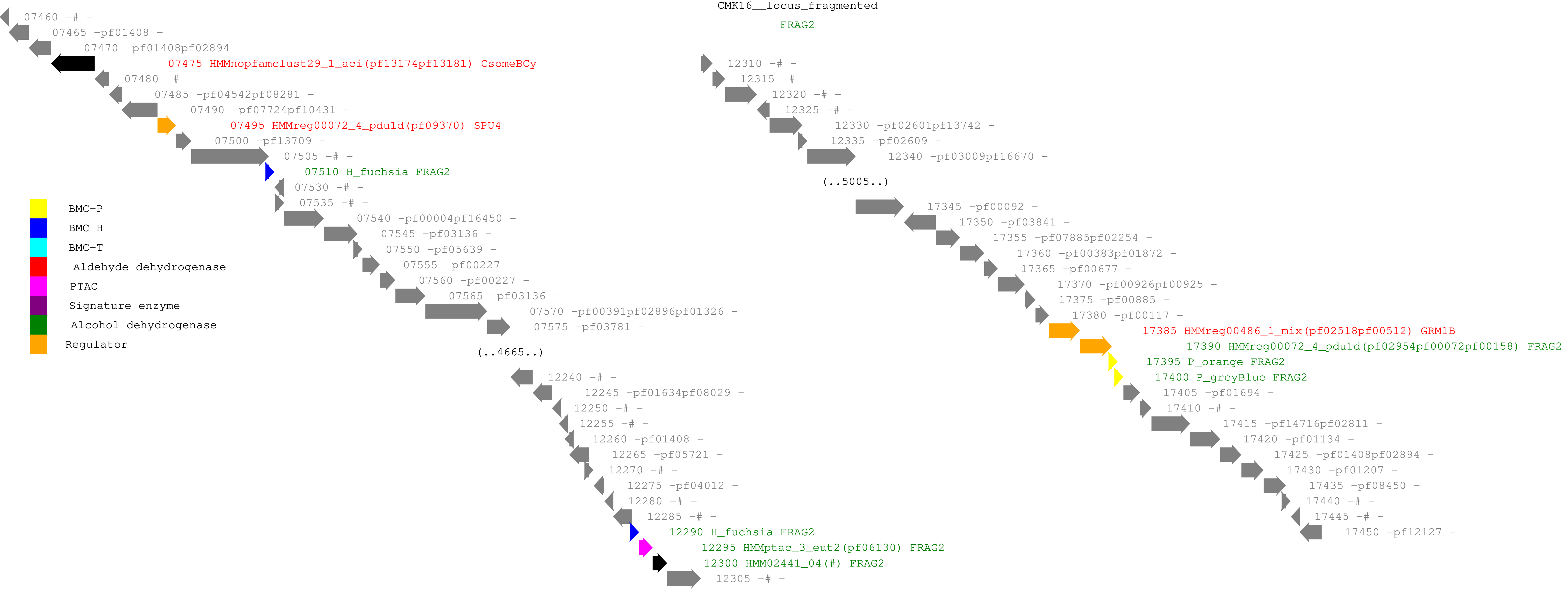

Candidatus Poribacteria bacterium. Bacteria; Candidatus Poribacteria.

CMK17__locus_fragmented

FRAG2

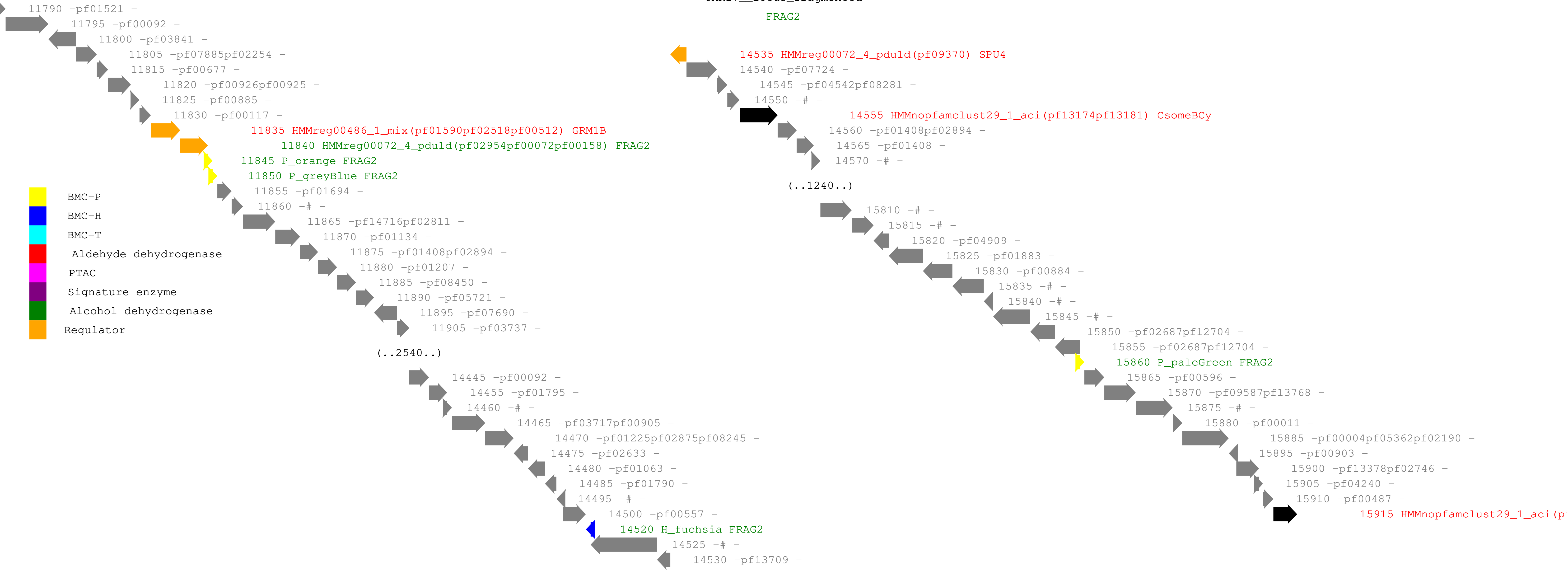

Candidatus Poribacteria bacterium. Bacteria; Candidatus Poribacteria.

CMK18__locus_fragmented

FRAG2

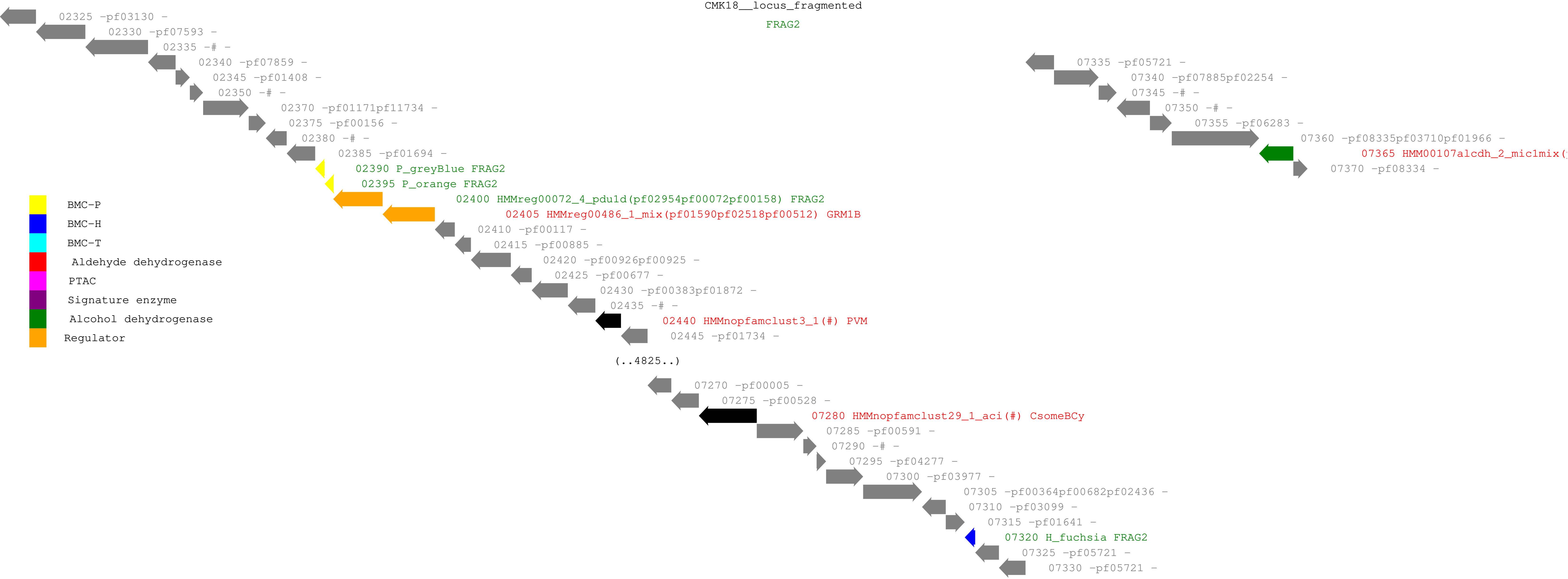

Candidatus Poribacteria bacterium. Bacteria; Candidatus Poribacteria.

CMK19__locus_fragmented

FRAG2

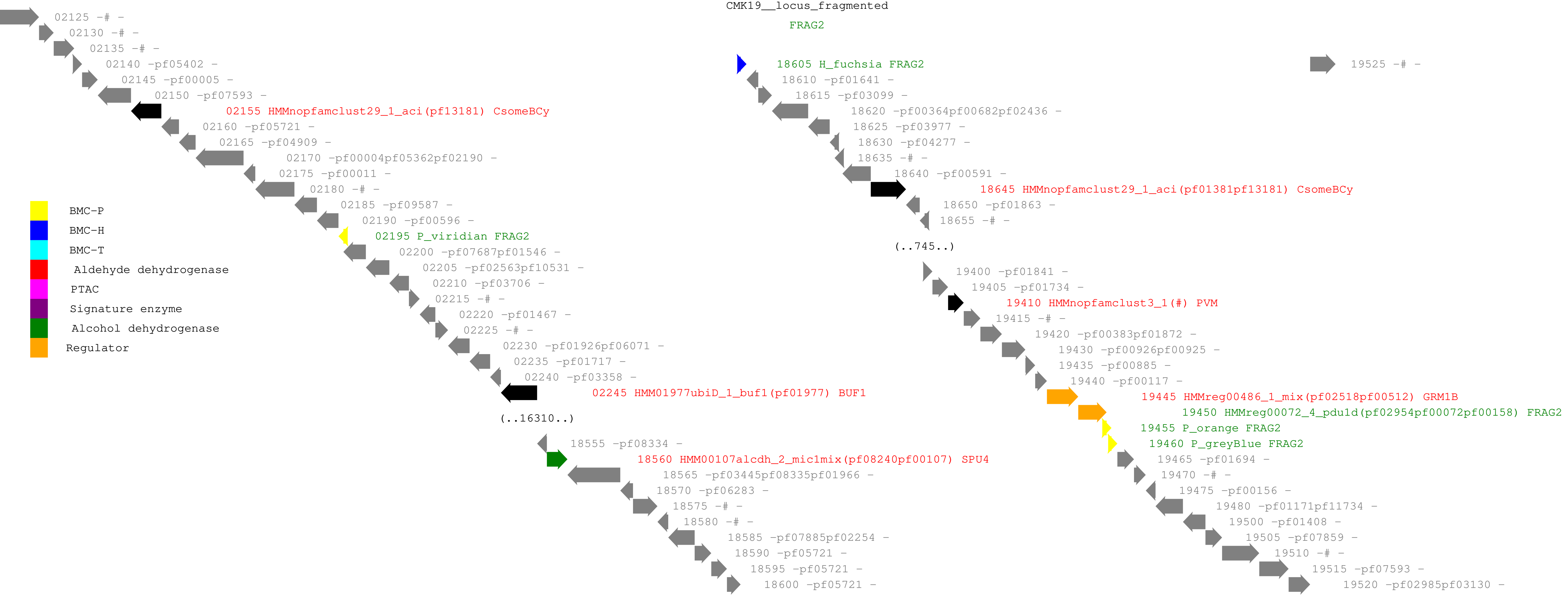

Candidatus Poribacteria bacterium. Bacteria; Candidatus Poribacteria.

CMK20__locus_fragmented

FRAG2

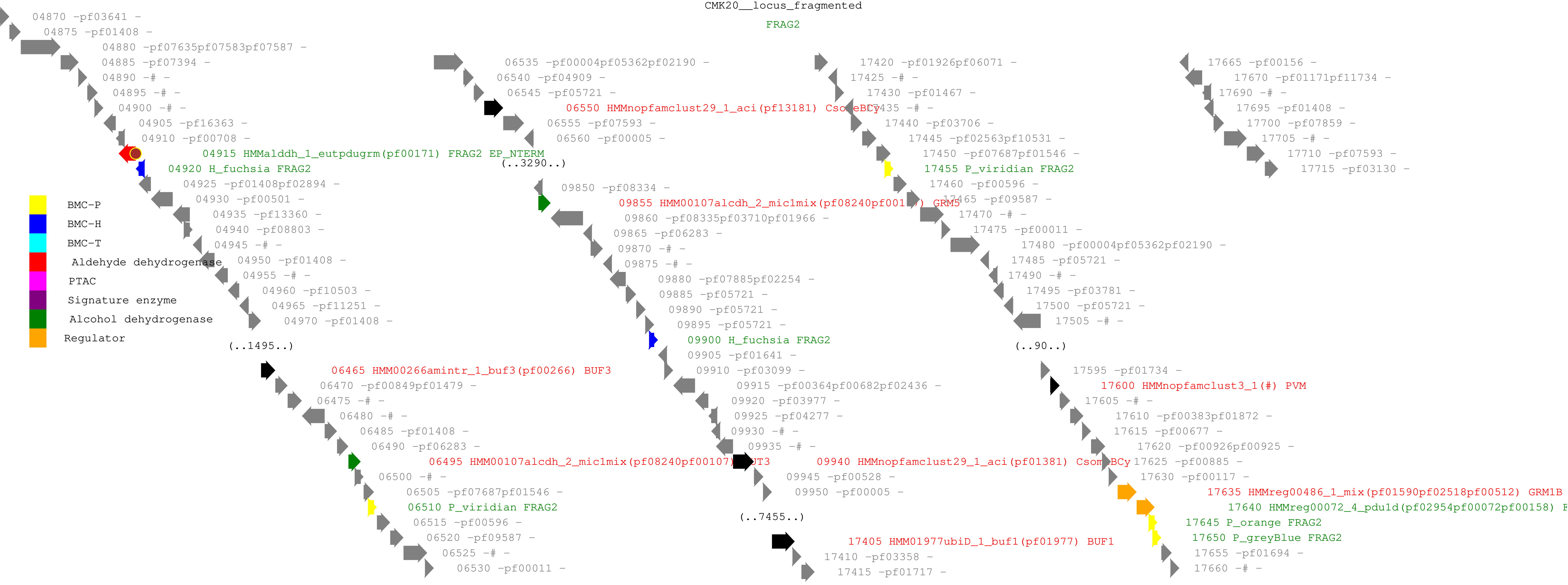

Candidatus Poribacteria bacterium. Bacteria; Candidatus Poribacteria.

CMK21__locus_fragmented

FRAG2

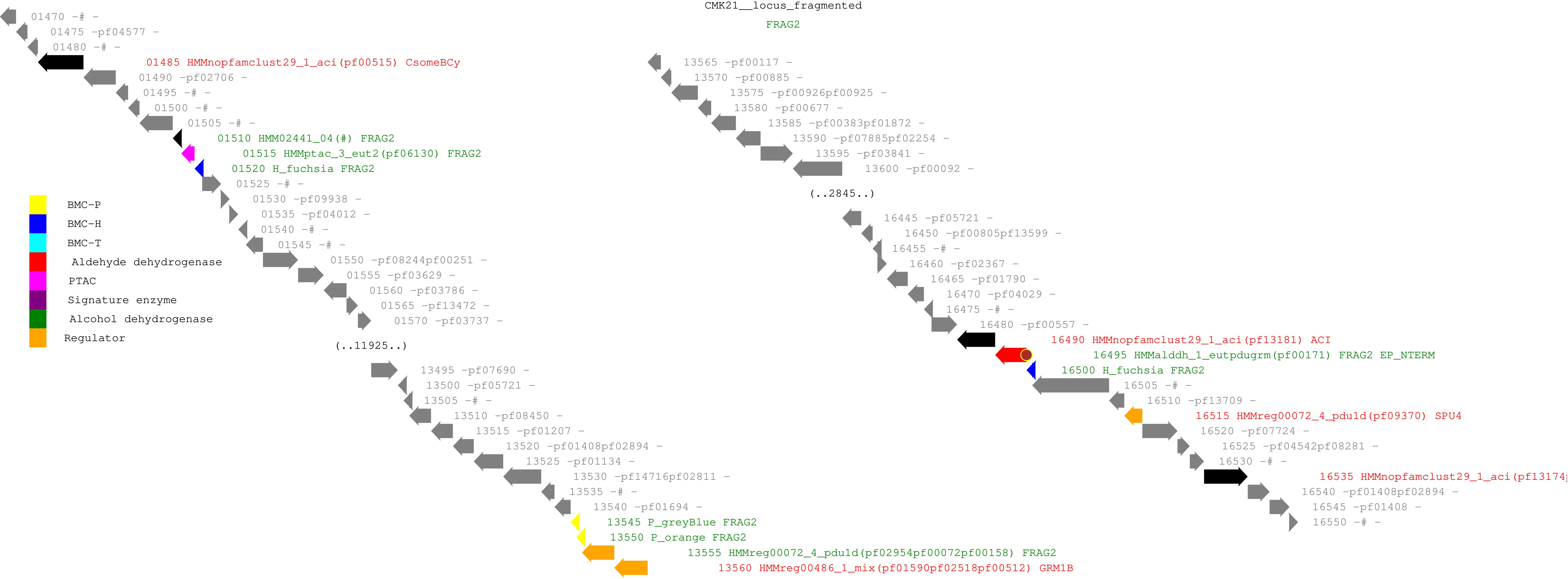

Candidatus Poribacteria bacterium. Bacteria; Candidatus Poribacteria.

CMK22__locus_fragmented

FRAG2

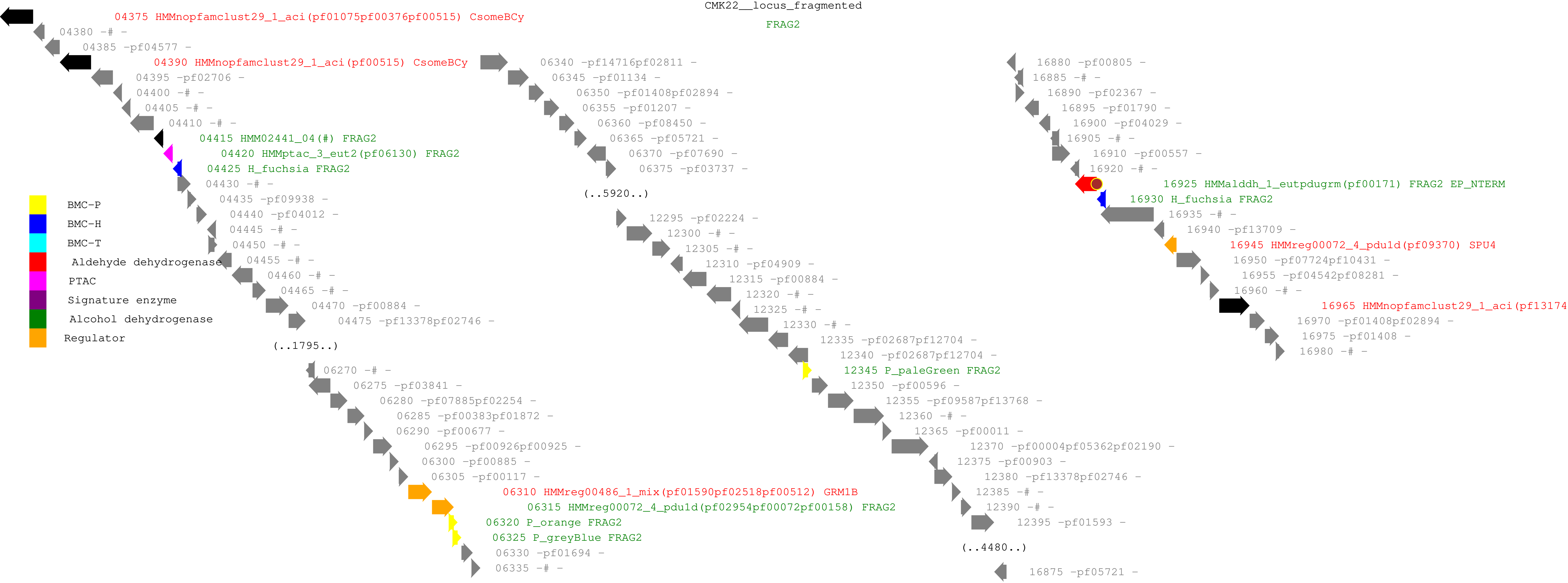

bacterium (Candidatus Blackallbacteria) Bacteria.

COW36__locus_1_main_sat

FRAG2

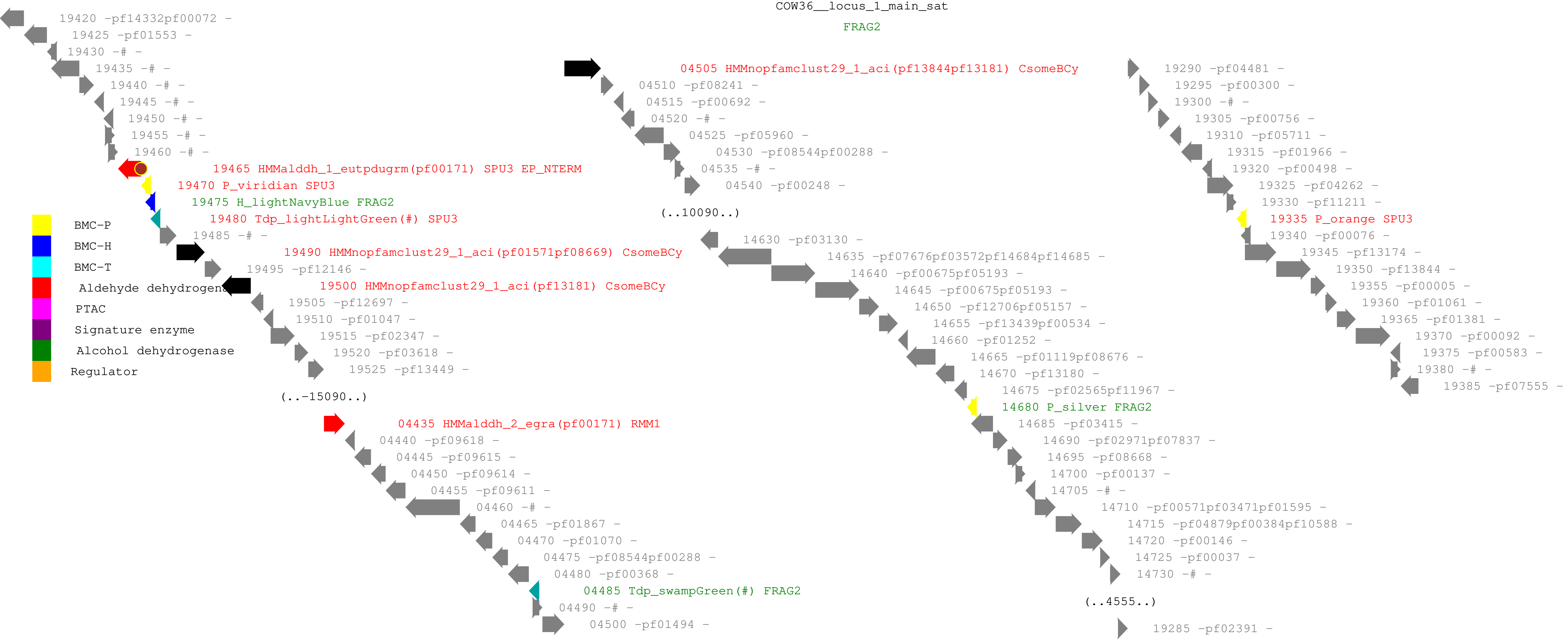

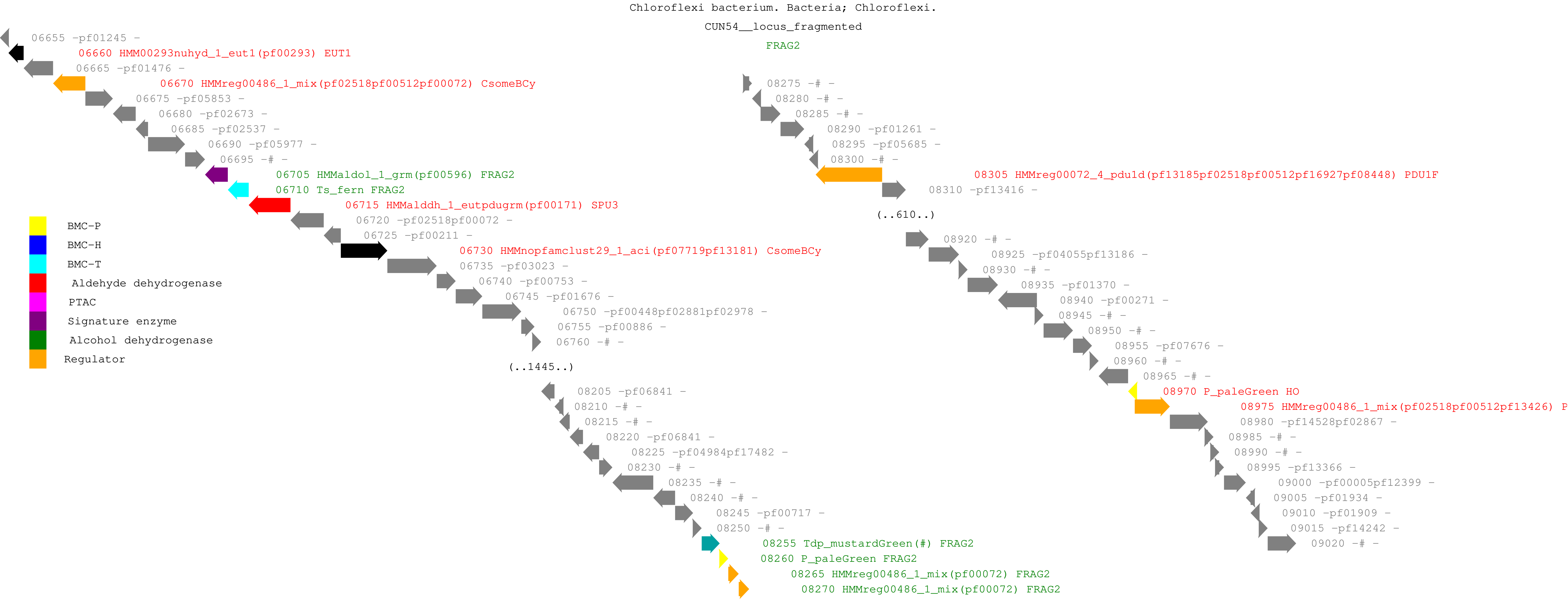

Chloroflexi bacterium. Bacteria; Chloroflexi.
CUN54__locus_fragmented
FRAG2
06655 -pf01245 -
06660 HMM00293nuhyd_1_eut1(pf00293) EUT1
06665 -pf01476 -
06670 HMMreg00486_1_mix(pf02518pf00512pf00072) CsomeBCy
06675 -pf05853 -
06680 -pf02673 -
06685 -pf02537 -
06690 -pf05977 -
06695 -# -
06705 HMMaldol_1_grm(pf00596) FRAG2
06710 Ts_fern FRAG2
06715 HMMalddh_1_eutpdugrm(pf00171) SPU3
06720 -pf02518pf00072 -
06725 -pf00211 -
06730 HMMnopfamclust29_1_aci(pf07719pf13181) CsomeBCy
06735 -pf03023 -
06740 -pf00753 -
06745 -pf01676 -
06750 -pf00448pf02881pf02978 -
06755 -pf00886 -
06760 -# -
(..1445..)
08205 -pf06841 -
08210 -# -
08215 -# -
08220 -pf06841 -
08225 -pf04984pf17482 -
08230 -# -
08235 -# -
08240 -# -
08245 -pf00717 -
08250 -# -
08255 Tdp_mustardGreen(#) FRAG2
08260 P_paleGreen FRAG2
08265 HMMreg00486_1_mix(pf00072) FRAG2
08270 HMMreg00486_1_mix(pf00072) FRAG2
08275 -# -
08280 -# -
08285 -# -
08290 -pf01261 -
08295 -pf05685 -
08300 -# -
08305 HMMreg00072_4_pdu1d(pf13185pf02518pf00512pf16927pf08448) PDU1F
08310 -pf13416 -
(..610..)
08920 -# -
08925 -pf04055pf13186 -
08930 -# -
08935 -pf01370 -
08940 -pf00271 -
08945 -# -
08950 -# -
08955 -pf07676 -
08960 -# -
08965 -# -
08970 P_paleGreen HO
08975 HMMreg00486_1_mix(pf02518pf00512pf13426) P
08980 -pf14528pf02867 -
08985 -# -
08990 -# -
08995 -pf13366 -
09000 -pf00005pf12399 -
09005 -pf01934 -
09010 -pf01909 -
09015 -pf14242 -
09020 -# -
BMC-P
BMC-H
BMC-T
Aldehyde dehydrogenase
PTAC
Signature enzyme
Alcohol dehydrogenase
Regulator

Calditrichaeota bacterium. Bacteria; Calditrichaeota.

D6677__locus_fragmented

FRAG2

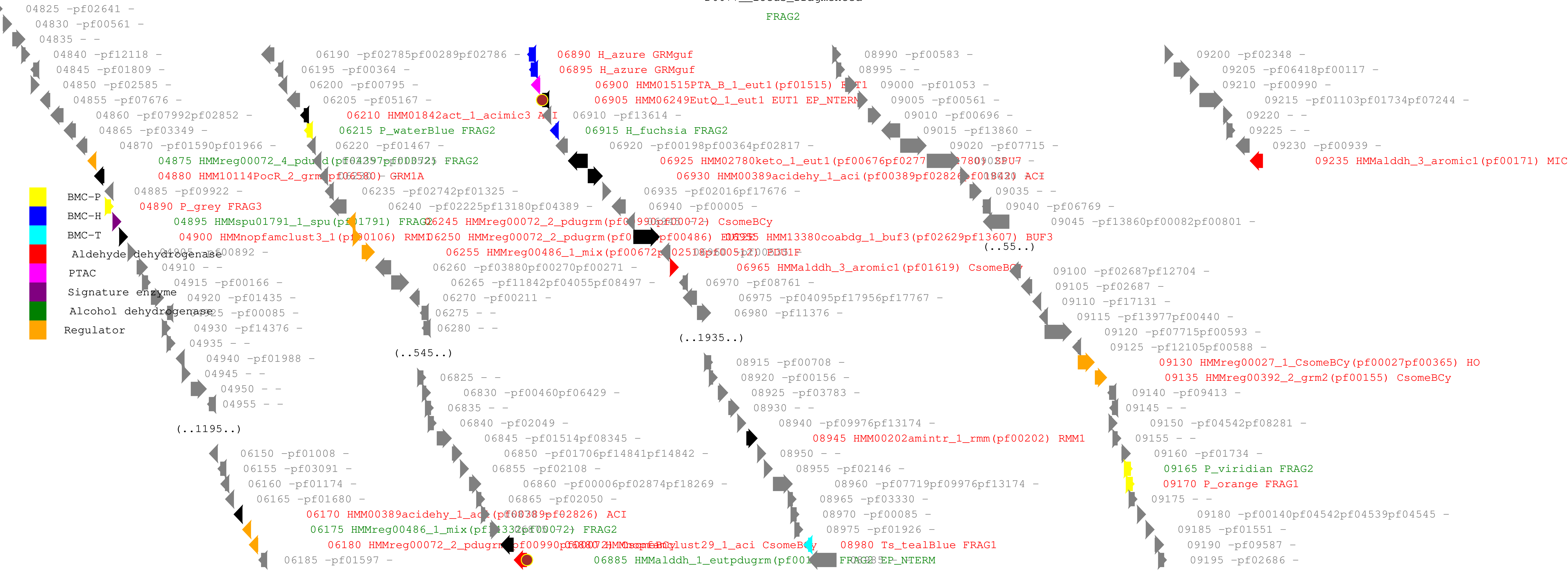

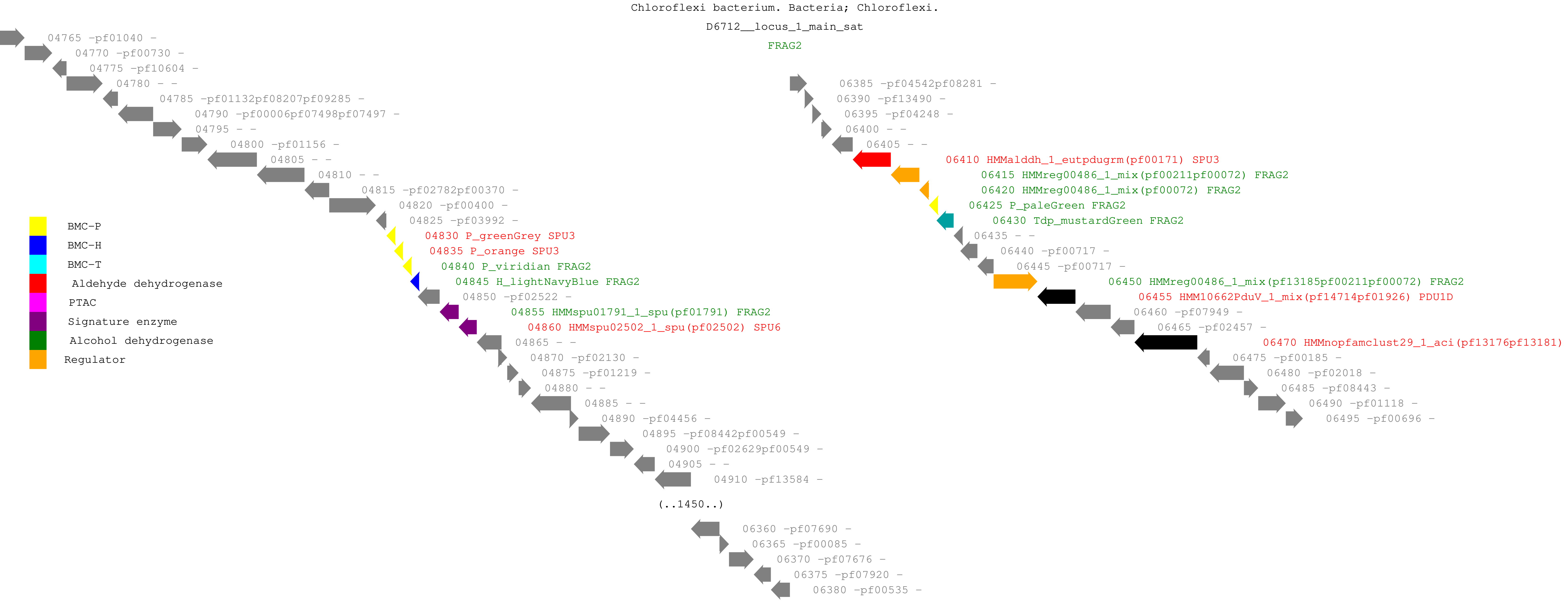

Chloroflexi bacterium. Bacteria; Chloroflexi.
D6712__locus_1_main_sat
FRAG2
04765 -pf01040 -
04770 -pf00730 -
04775 -pf10604 -
04780 - -
04785 -pf01132pf08207pf09285 -
04790 -pf00006pf07498pf07497 -
04795 - -
04800 -pf01156 -
04805 - -
04810 - -
04815 -pf02782pf00370 -
04820 -pf00400 -
04825 -pf03992 -
04830 P_greenGrey SPU3
04835 P_orange SPU3
04840 P_viridian FRAG2
04845 H_lightNavyBlue FRAG2
04850 -pf02522 -
04855 HMMspu01791_1_spu(pf01791) FRAG2
04860 HMMspu02502_1_spu(pf02502) SPU6
04865 - -
04870 -pf02130 -
04875 -pf01219 -
04880 - -
04885 - -
04890 -pf04456 -
04895 -pf08442pf00549 -
04900 -pf02629pf00549 -
04905 - -
04910 -pf13584 -
(..1450..)
06360 -pf07690 -
06365 -pf00085 -
06370 -pf07676 -
06375 -pf07920 -
06380 -pf00535 -
06385 -pf04542pf08281 -
06390 -pf13490 -
06395 -pf04248 -
06400 - -
06405 - -
06410 HMMalddh_1_eutpdugrm(pf00171) SPU3
06415 HMMreg00486_1_mix(pf00211pf00072) FRAG2
06420 HMMreg00486_1_mix(pf00072) FRAG2
06425 P_paleGreen FRAG2
06430 Tdp_mustardGreen FRAG2
06435 - -
06440 -pf00717 -
06445 -pf00717 -
06450 HMMreg00486_1_mix(pf13185pf00211pf00072) FRAG2
06455 HMM10662PduV_1_mix(pf14714pf01926) PDU1D
06460 -pf07949 -
06465 -pf02457 -
06470 HMMnopfamclust29_1_aci(pf13176pf13181)
06475 -pf00185 -
06480 -pf02018 -
06485 -pf08443 -
06490 -pf01118 -
06495 -pf00696 -
BMC-P
BMC-H
BMC-T
Aldehyde dehydrogenase
PTAC
Signature enzyme
Alcohol dehydrogenase
Regulator

Chloroflexi bacterium. Bacteria; Chloroflexi.

D6737__locus_1_main_sat

FRAG2

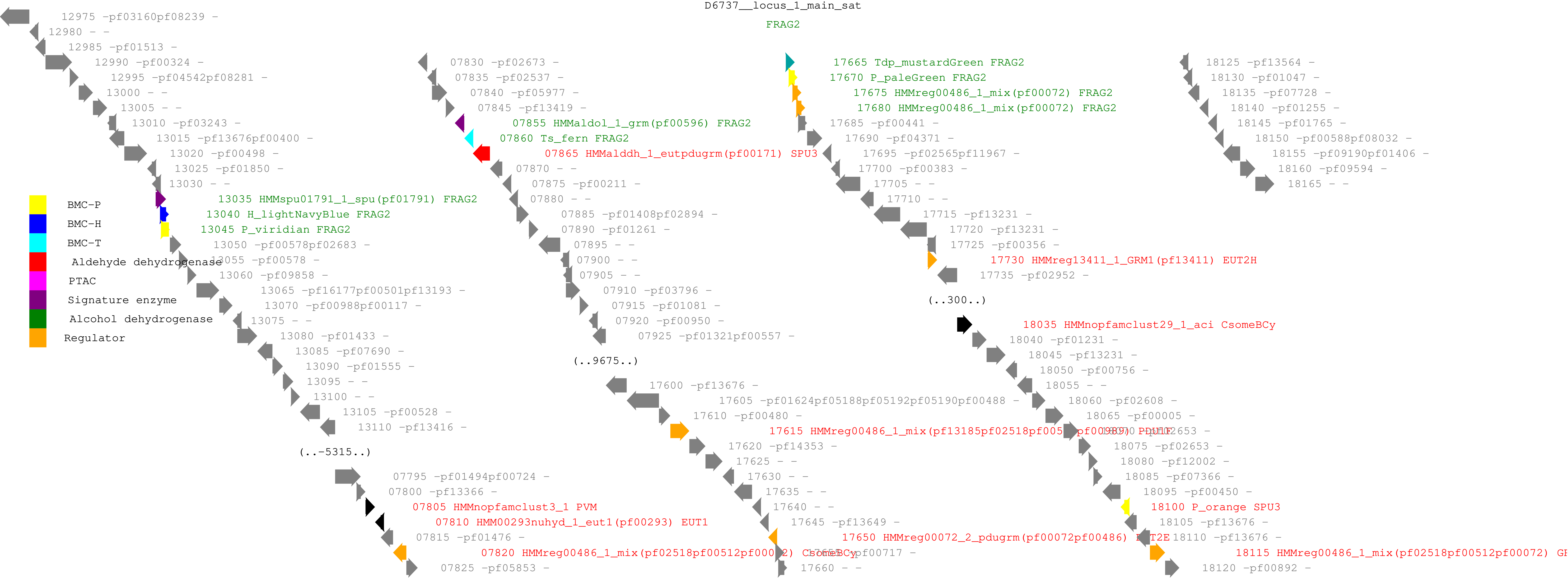

Planctomycetes bacterium. Bacteria; Planctomycetes.

D6785__locus_1_main_sat

FRAG2

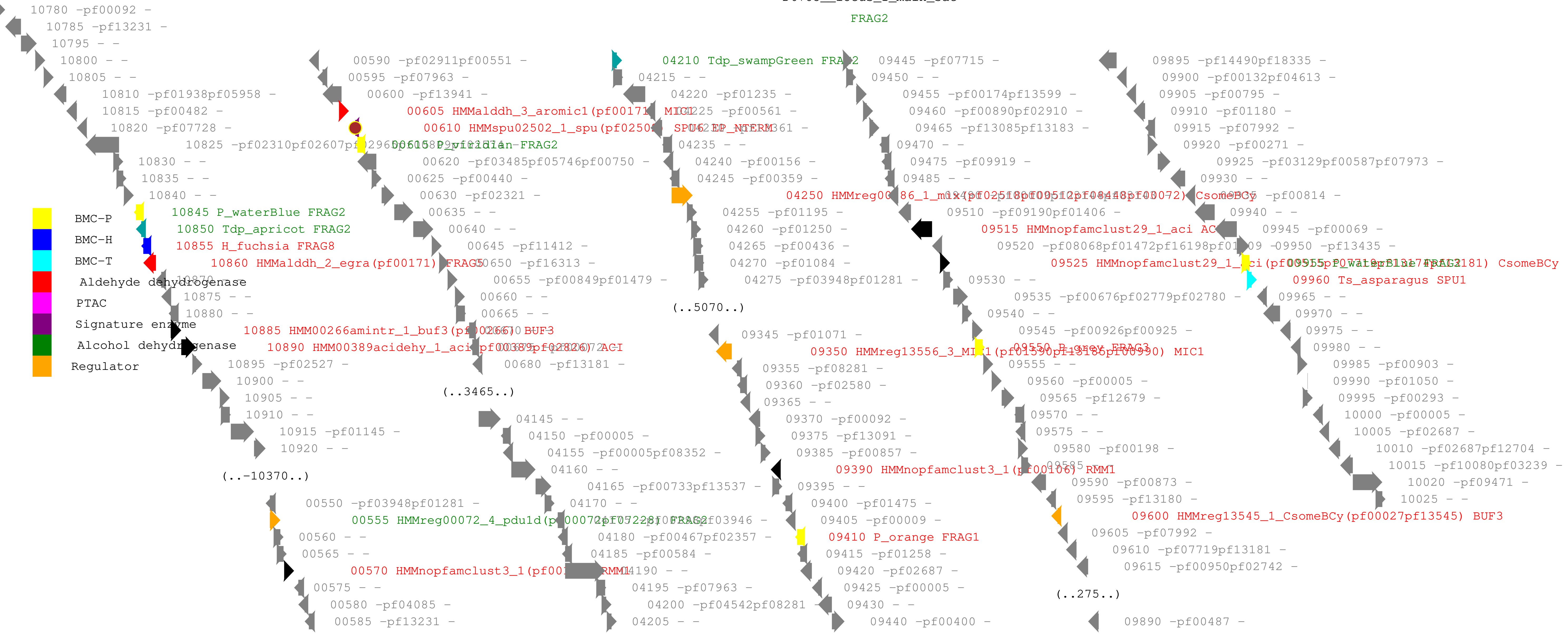

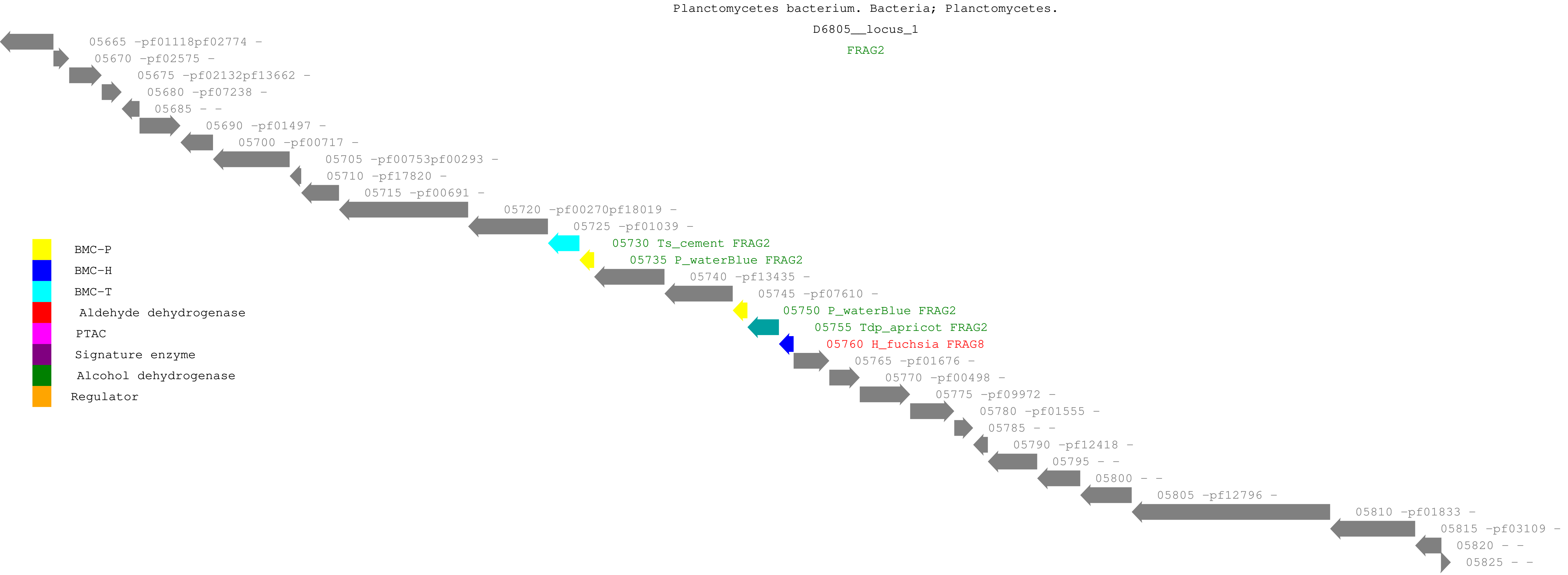

Planctomycetes bacterium. Bacteria; Planctomycetes.
D6805__locus_1
FRAG2
05665 -pf01118pf02774 -
05670 -pf02575 -
05675 -pf02132pf13662 -
05680 -pf07238 -
05685 - -
05690 -pf01497 -
05700 -pf00717 -
05705 -pf00753pf00293 -
05710 -pf17820 -
05715 -pf00691 -
05720 -pf00270pf18019 -
05725 -pf01039 -
05730 Ts_cement FRAG2
05735 P_waterBlue FRAG2
05740 -pf13435 -
05745 -pf07610 -
05750 P_waterBlue FRAG2
05755 Tdp_apricot FRAG2
05760 H_fuchsia FRAG8
05765 -pf01676 -
05770 -pf00498 -
05775 -pf09972 -
05780 -pf01555 -
05785 - -
05790 -pf12418 -
05795 - -
05800 - -
05805 -pf12796 -
05810 -pf01833 -
05815 -pf03109 -
05820 - -
05825 - -
BMC-P
BMC-H
BMC-T
Aldehyde dehydrogenase
PTAC
Signature enzyme
Alcohol dehydrogenase
Regulator

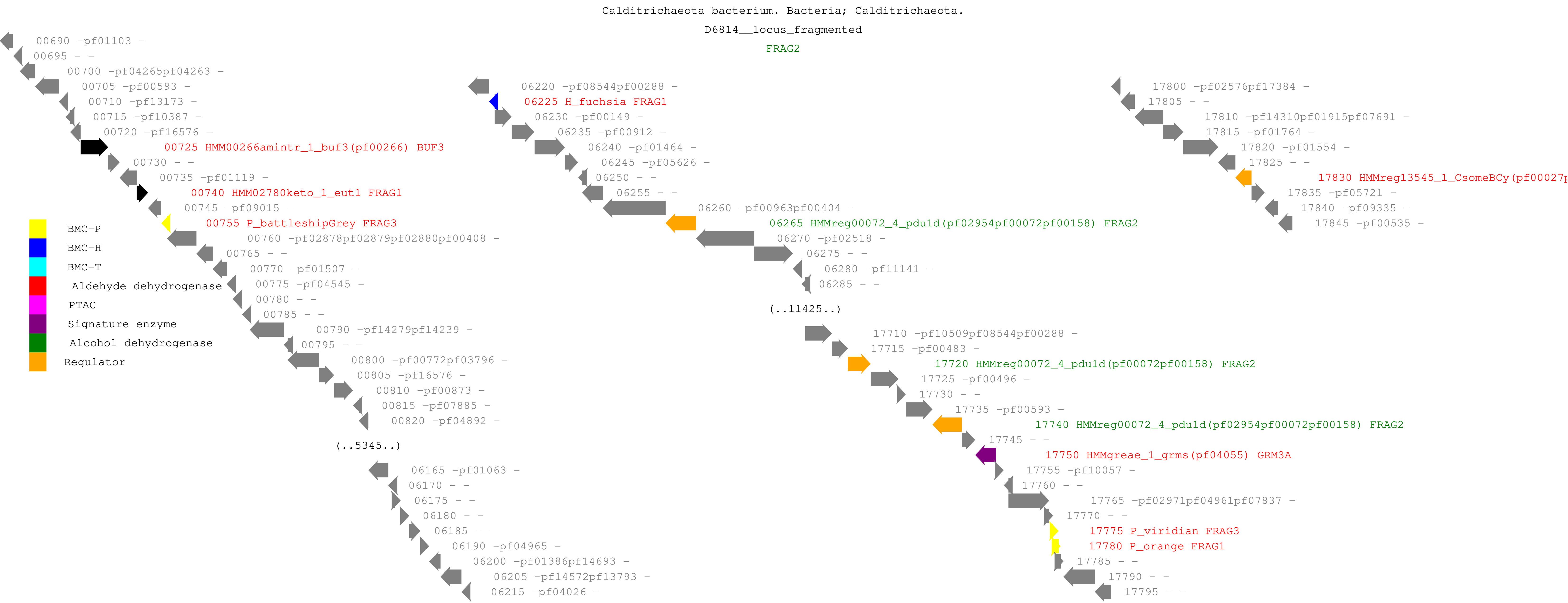
Calditrichaeota bacterium. Bacteria; Calditrichaeota.
D6814__locus_fragmented
FRAG2
00690 -pf01103 -
00695 - -
00700 -pf04265pf04263 -
00705 -pf00593 -
00710 -pf13173 -
00715 -pf10387 -
00720 -pf16576 -
00725 HMM00266amintr_1_buf3(pf00266) BUF3
00730 - -
00735 -pf01119 -
00740 HMM02780keto_1_eut1 FRAG1
00745 -pf09015 -
00755 P_battleshipGrey FRAG3
00760 -pf02878pf02879pf02880pf00408 -
00765 - -
00770 -pf01507 -
00775 -pf04545 -
00780 - -
00785 - -
00790 -pf14279pf14239 -
00795 - -
00800 -pf00772pf03796 -
00805 -pf16576 -
00810 -pf00873 -
00815 -pf07885 -
00820 -pf04892 -
(..5345..)
06165 -pf01063 -
06170 - -
06175 - -
06180 - -
06185 - -
06190 -pf04965 -
06200 -pf01386pf14693 -
06205 -pf14572pf13793 -
06215 -pf04026 -
06220 -pf08544pf00288 -
06225 H_fuchsia FRAG1
06230 -pf00149 -
06235 -pf00912 -
06240 -pf01464 -
06245 -pf05626 -
06250 - -
06255 - -
06260 -pf00963pf00404 -
06265 HMMreg00072_4_pdu1d(pf02954pf00072pf00158) FRAG2
06270 -pf02518 -
06275 - -
06280 -pf11141 -
06285 - -
(..11425..)
17710 -pf10509pf08544pf00288 -
17715 -pf00483 -
17720 HMMreg00072_4_pdu1d(pf00072pf00158) FRAG2
17725 -pf00496 -
17730 - -
17735 -pf00593 -
17740 HMMreg00072_4_pdu1d(pf02954pf00072pf00158) FRAG2
17745 - -
17750 HMMgreae_1_grms(pf04055) GRM3A
17755 -pf10057 -
17760 - -
17765 -pf02971pf04961pf07837 -
17770 - -
17775 P_viridian FRAG3
17780 P_orange FRAG1
17785 - -
17790 - -
17795 - -
17800 -pf02576pf17384 -
17805 - -
17810 -pf14310pf01915pf07691 -
17815 -pf01764 -
17820 -pf01554 -
17825 - -
17830 HMMreg13545_1_CsomeBCy(pf00027
17835 -pf05721 -
17840 -pf09335 -
17845 -pf00535 -
BMC-P
BMC-H
BMC-T
Aldehyde dehydrogenase
PTAC
Signature enzyme
Alcohol dehydrogenase
Regulator

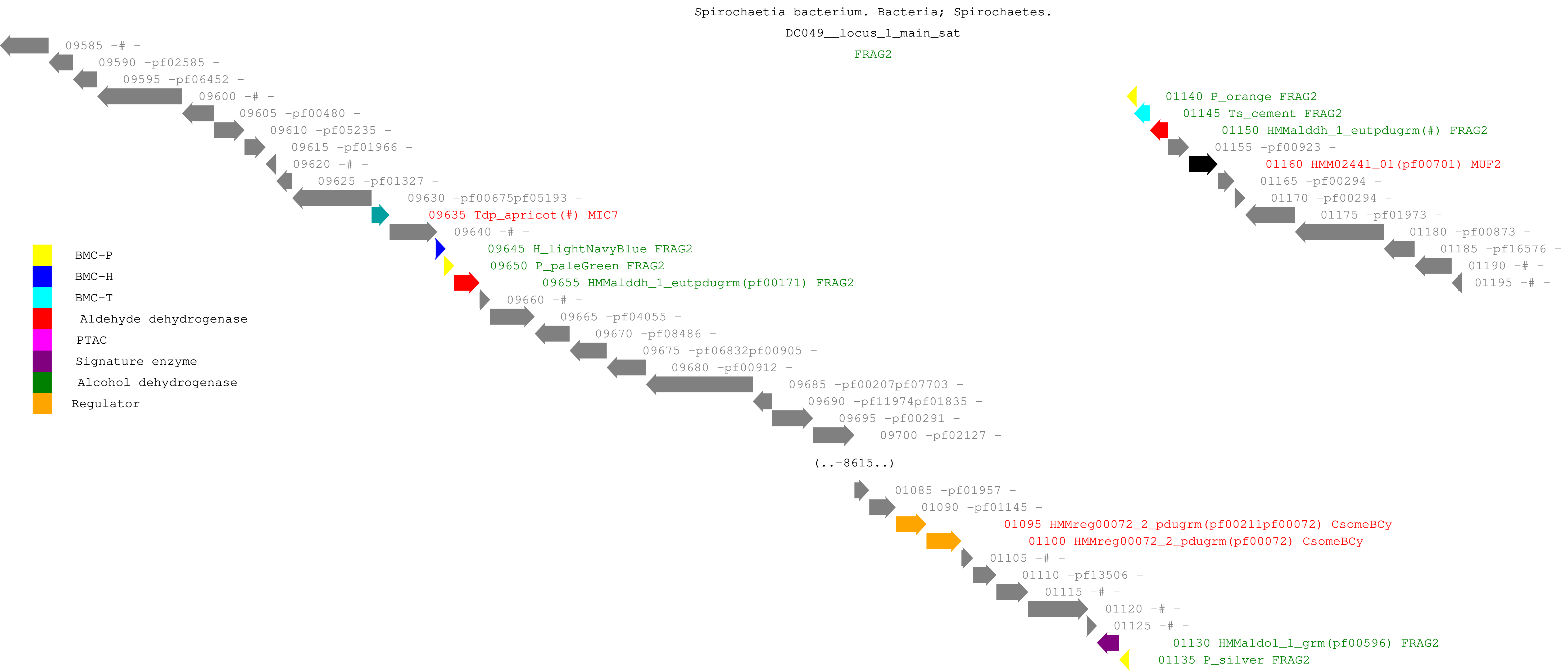
Spirochaetia bacterium. Bacteria; Spirochaetes.
DC049__locus_1_main_sat
FRAG2
09585 -# -
09590 -pf02585 -
09595 -pf06452 -
09600 -# -
09605 -pf00480 -
09610 -pf05235 -
09615 -pf01966 -
09620 -# -
09625 -pf01327 -
09630 -pf00675pf05193 -
09635 Tdp_apricot(#) MIC7
09640 -# -
09645 H_lightNavyBlue FRAG2
09650 P_paleGreen FRAG2
09655 HMMaldh_1_eutpdugrm(pf00171) FRAG2
09660 -# -
09665 -pf04055 -
09670 -pf08486 -
09675 -pf06832pf00905 -
09680 -pf00912 -
09685 -pf00207pf07703 -
09690 -pf11974pf01835 -
09695 -pf00291 -
09700 -pf02127 -
(..-8615..)
01085 -pf01957 -
01090 -pf01145 -
01095 HMMreg00072_2_pdugrm(pf00211pf00072) CsomeBCy
01100 HMMreg00072_2_pdugrm(pf00072) CsomeBCy
01105 -# -
01110 -pf13506 -
01115 -# -
01120 -# -
01125 -# -
01130 HMMaldol_1_grm(pf00596) FRAG2
01135 P_silver FRAG2
01140 P_orange FRAG2
01145 Ts_cement FRAG2
01150 HMMalddh_1_eutpdugrm(#) FRAG2
01155 -pf00923 -
01160 HMM02441_01(pf00701) MUF2
01165 -pf00294 -
01170 -pf00294 -
01175 -pf01973 -
01180 -pf00873 -
01185 -pf16576 -
01190 -# -
01195 -# -
BMC-P
BMC-H
BMC-T
Aldehyde dehydrogenase
PTAC
Signature enzyme
Alcohol dehydrogenase
Regulator

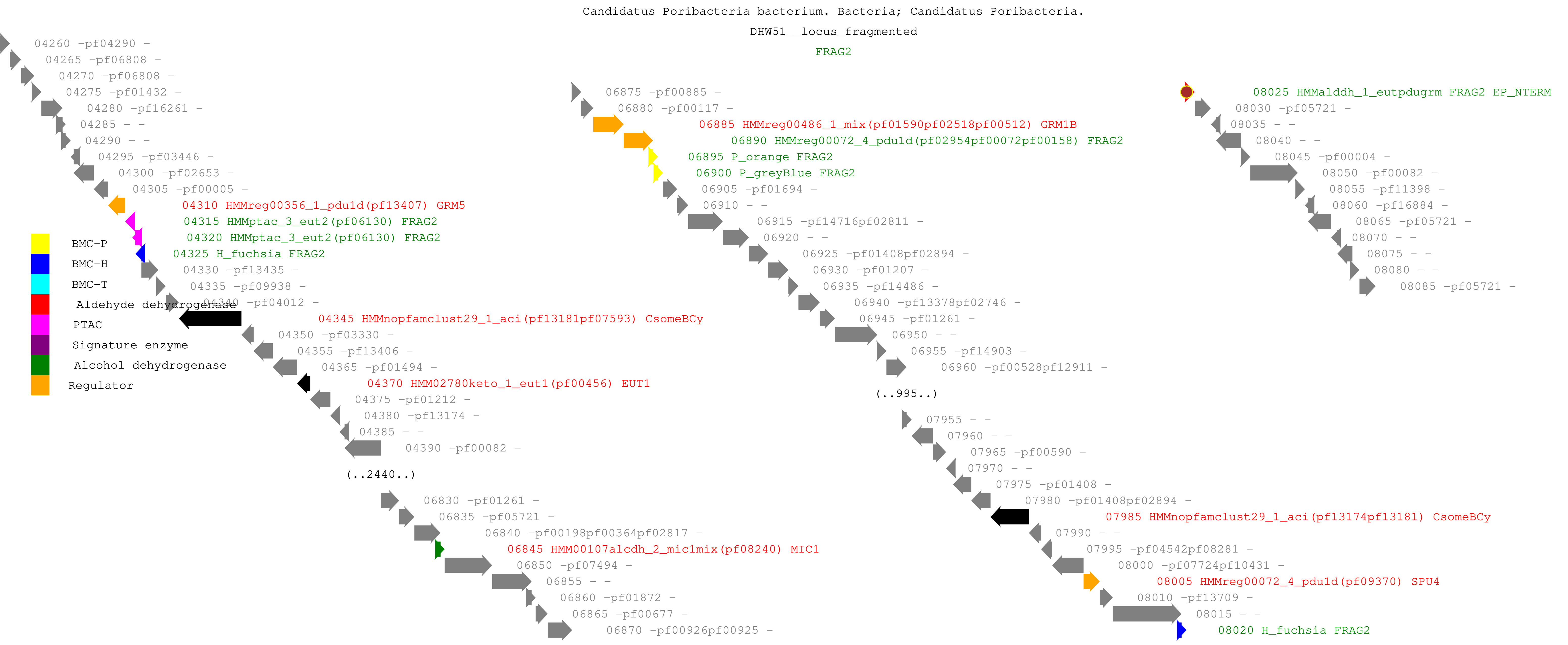
Candidatus Poribacteria bacterium. Bacteria; Candidatus Poribacteria.
DHW51__locus_fragmented
FRAG2
04260 -pf04290 -
04265 -pf06808 -
04270 -pf06808 -
04275 -pf01432 -
04280 -pf16261 -
04285 - -
04290 - -
04295 -pf03446 -
04300 -pf02653 -
04305 -pf00005 -
04310 HMMreg00356_1_pdu1d(pf13407) GRM5
04315 HMMptac_3_eut2(pf06130) FRAG2
04320 HMMptac_3_eut2(pf06130) FRAG2
04325 H_fuchsia FRAG2
04330 -pf13435 -
04335 -pf09938 -
04340 -pf04012 -
04345 HMMnopfamclust29_1_aci(pf13181pf07593) CsomeBCy
04350 -pf03330 -
04355 -pf13406 -
04365 -pf01494 -
04370 HMM02780keto_1_eut1(pf00456) EUT1
04375 -pf01212 -
04380 -pf13174 -
04385 - -
04390 -pf00082 -
(..2440..)
06830 -pf01261 -
06835 -pf05721 -
06840 -pf00198pf00364pf02817 -
06845 HMM00107alcdh_2_mic1mix(pf08240) MIC1
06850 -pf07494 -
06855 - -
06860 -pf01872 -
06865 -pf00677 -
06870 -pf00926pf00925 -
06875 -pf00885 -
06880 -pf00117 -
06885 HMMreg00486_1_mix(pf01590pf02518pf00512) GRM1B
06890 HMMreg00072_4_pdu1d(pf02954pf00072pf00158) FRAG2
06895 P_orange FRAG2
06900 P_greyBlue FRAG2
06905 -pf01694 -
06910 - -
06915 -pf14716pf02811 -
06920 - -
06925 -pf01408pf02894 -
06930 -pf01207 -
06935 -pf14486 -
06940 -pf13378pf02746 -
06945 -pf01261 -
06950 - -
06955 -pf14903 -
06960 -pf00528pf12911 -
(..995..)
07955 - -
07960 - -
07965 -pf00590 -
07970 - -
07975 -pf01408 -
07980 -pf01408pf02894 -
07985 HMMnopfamclust29_1_aci(pf13174pf13181) CsomeBCy
07990 - -
07995 -pf04542pf08281 -
08000 -pf07724pf10431 -
08005 HMMreg00072_4_pdu1d(pf09370) SPU4
08010 -pf13709 -
08015 - -
08020 H_fuchsia FRAG2
08025 HMMaldh_1_eutpdugrm FRAG2 EP_NTERM
08030 -pf05721 -
08035 - -
08040 - -
08045 -pf00004 -
08050 -pf00082 -
08055 -pf11398 -
08060 -pf16884 -
08065 -pf05721 -
08070 - -
08075 - -
08080 - -
08085 -pf05721 -
BMC-P
BMC-H
BMC-T
Aldehyde dehydrogenase
PTAC
Signature enzyme
Alcohol dehydrogenase
Regulator

candidate division Zixibacteria bacterium. Bacteria.

C4524__locus_fragmented

FRAG3

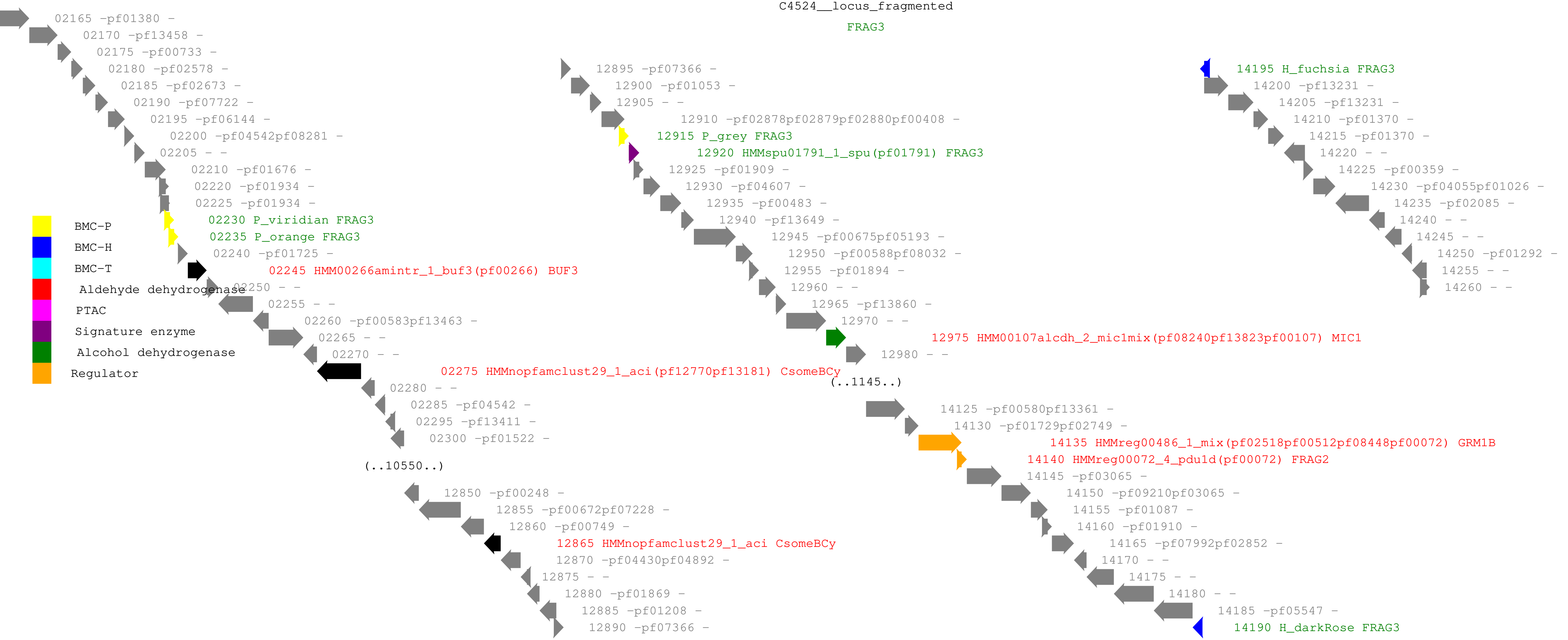

candidate division LCP-89 bacterium B3_LCP. Bacteria; candidate division LCP-89.

CEE37__locus_fragmented

FRAG3

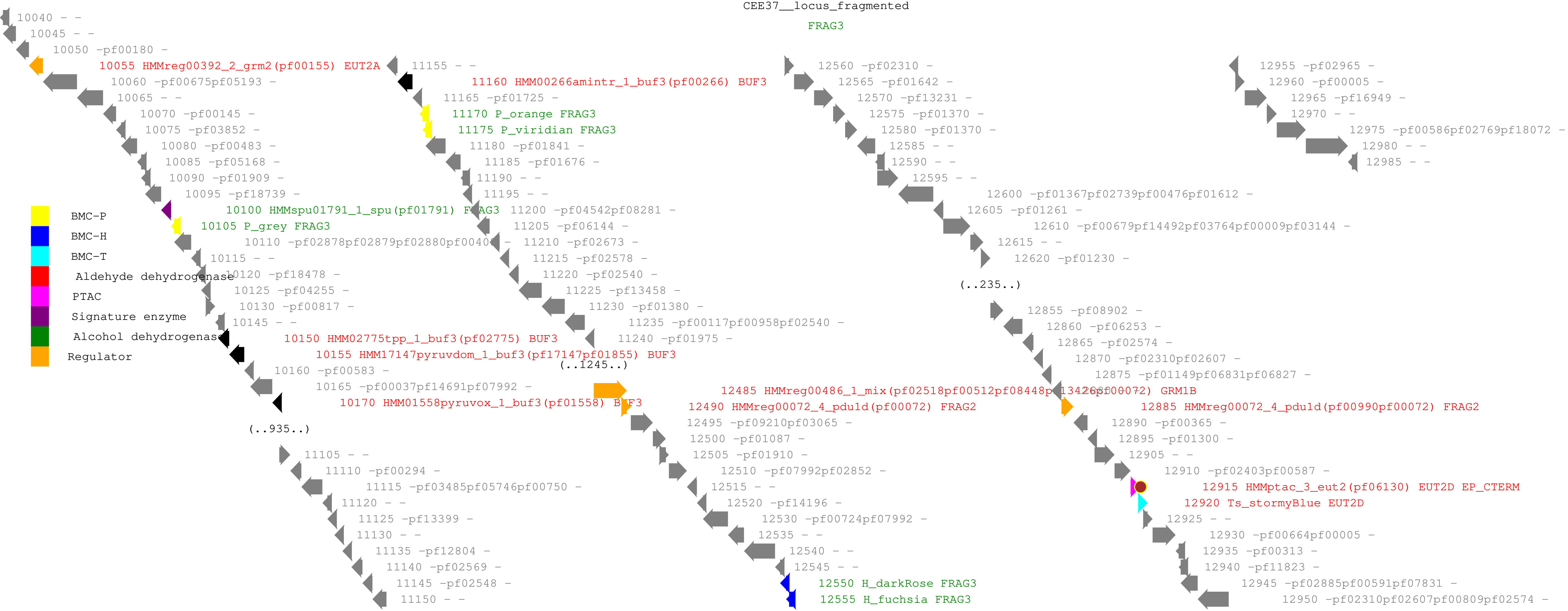

Calditrichaeota bacterium. Bacteria; Calditrichaeota.

D6743__locus_fragmented

FRAG3

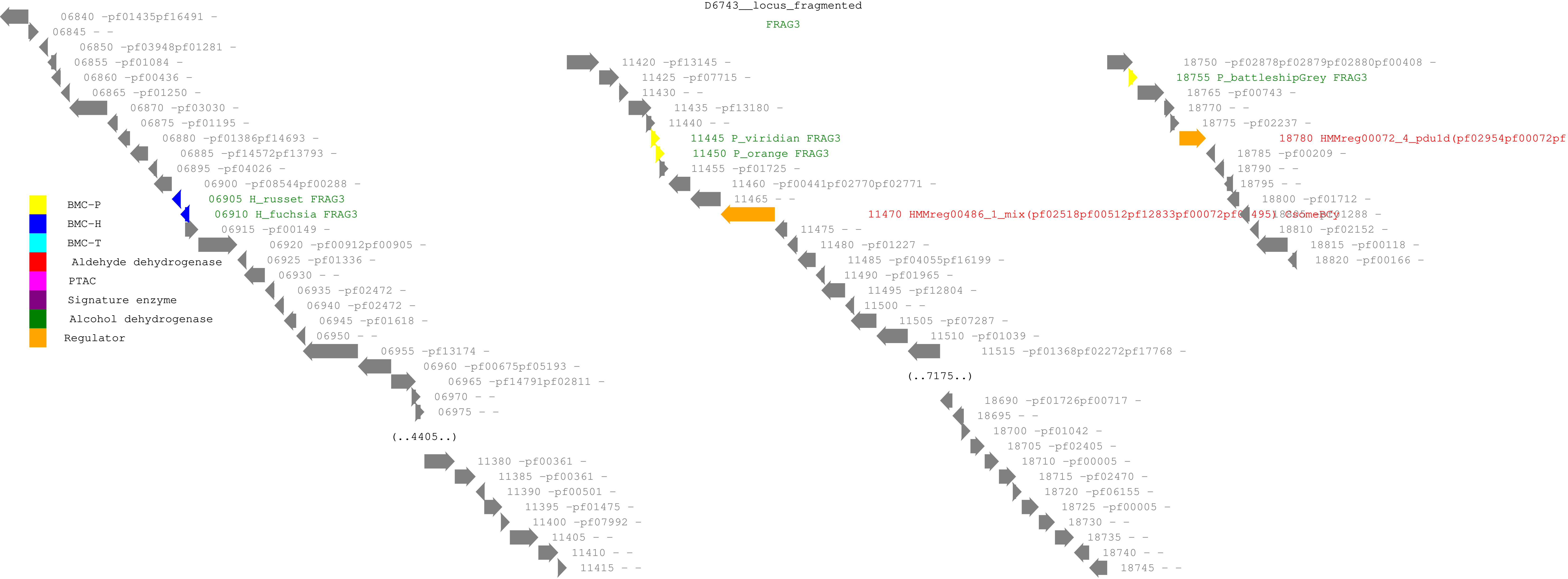

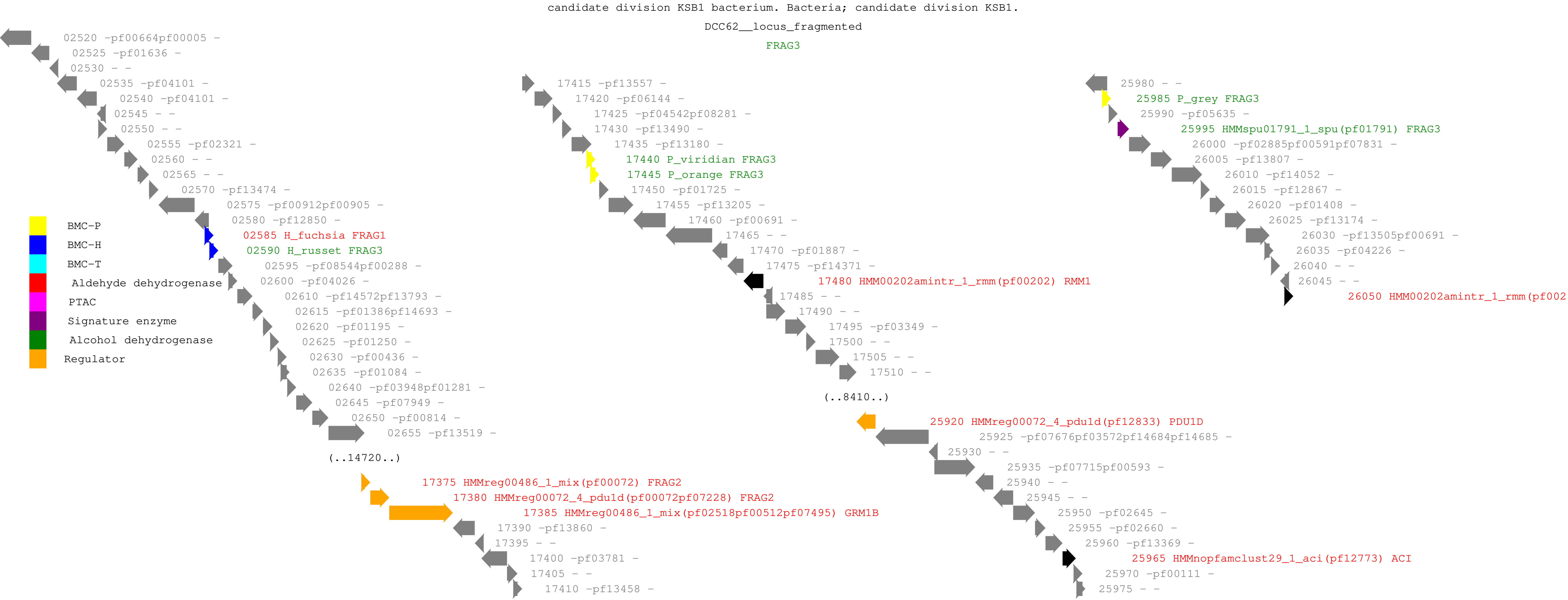

candidate division KSB1 bacterium. Bacteria; candidate division KSB1.
DCC62__locus_fragmented
FRAG3
02520 -pf00664pf00005 -
02525 -pf01636 -
02530 - -
02535 -pf04101 -
02540 -pf04101 -
02545 - -
02550 - -
02555 -pf02321 -
02560 - -
02565 - -
02570 -pf13474 -
02575 -pf00912pf00905 -
02580 -pf12850 -
02585 H_fuchsia FRAG1
02590 H_russet FRAG3
02595 -pf08544pf00288 -
02600 -pf04026 -
02610 -pf14572pf13793 -
02615 -pf01386pf14693 -
02620 -pf01195 -
02625 -pf01250 -
02630 -pf00436 -
02635 -pf01084 -
02640 -pf03948pf01281 -
02645 -pf07949 -
02650 -pf00814 -
02655 -pf13519 -
(..14720..)
17375 HMMreg00486_1_mix(pf00072) FRAG2
17380 HMMreg00072_4_pdu1d(pf00072pf07228) FRAG2
17385 HMMreg00486_1_mix(pf02518pf00512pf07495) GRM1B
17390 -pf13860 -
17395 - -
17400 -pf03781 -
17405 - -
17410 -pf13458 -
17415 -pf13557 -
17420 -pf06144 -
17425 -pf04542pf08281 -
17430 -pf13490 -
17435 -pf13180 -
17440 P_viridian FRAG3
17445 P_orange FRAG3
17450 -pf01725 -
17455 -pf13205 -
17460 -pf00691 -
17465 - -
17470 -pf01887 -
17475 -pf14371 -
17480 HMM00202amintr_1_rmm(pf00202) RMM1
17485 - -
17490 - -
17495 -pf03349 -
17500 - -
17505 - -
17510 - -
(..8410..)
25920 HMMreg00072_4_pdu1d(pf12833) PDU1D
25925 -pf07676pf03572pf14684pf14685 -
25930 - -
25935 -pf07715pf00593 -
25940 - -
25945 - -
25950 -pf02645 -
25955 -pf02660 -
25960 -pf13369 -
25965 HMMnopfamclust29_1_aci(pf12773) ACI
25970 -pf00111 -
25975 - -
25980 - -
25985 P_grey FRAG3
25990 -pf05635 -
25995 HMMspu01791_1_spu(pf01791) FRAG3
26000 -pf02885pf00591pf07831 -
26005 -pf13807 -
26010 -pf14052 -
26015 -pf12867 -
26020 -pf01408 -
26025 -pf13174 -
26030 -pf13505pf00691 -
26035 -pf04226 -
26040 - -
26045 - -
26050 HMM00202amintr_1_rmm(pf002
BMC-P
BMC-H
BMC-T
Aldehyde dehydrogenase
PTAC
Signature enzyme
Alcohol dehydrogenase
Regulator

Caldithrix sp. Bacteria; Calditrichaeota; Calditrichae; Calditrichales; Calditrichaceae;

E2076__locus_fragmented

FRAG3

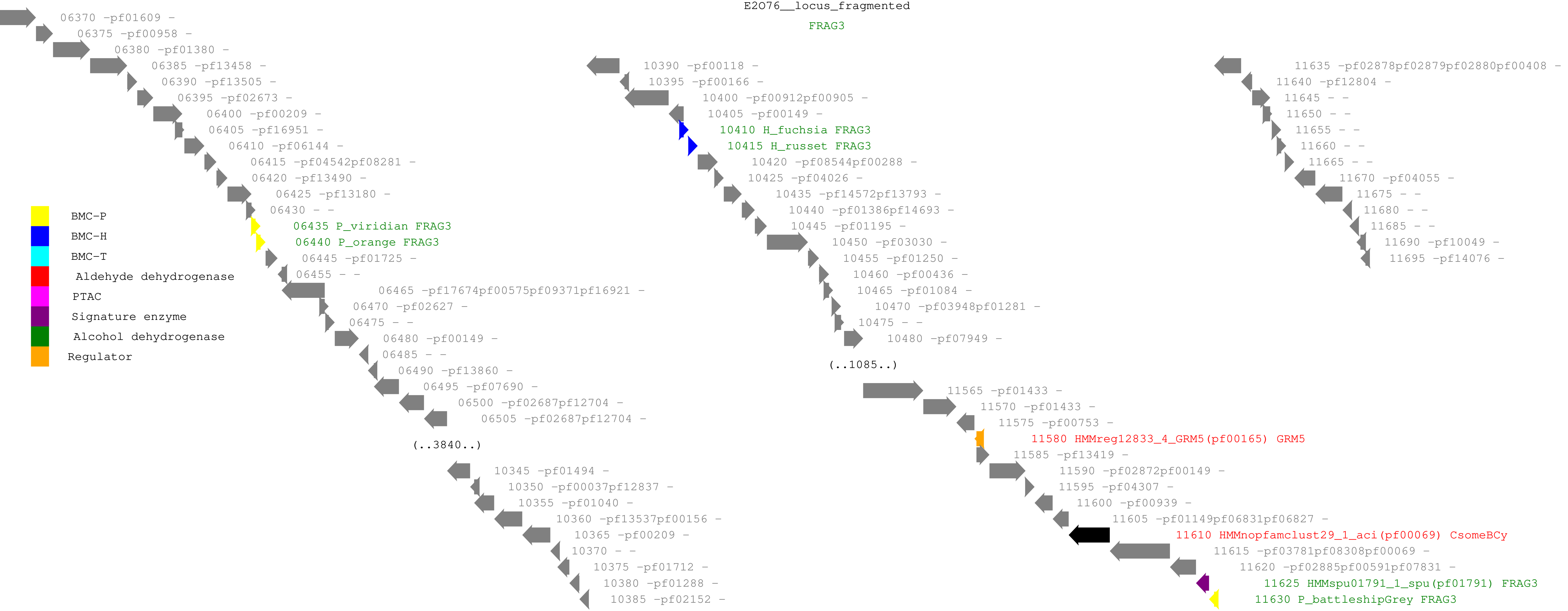

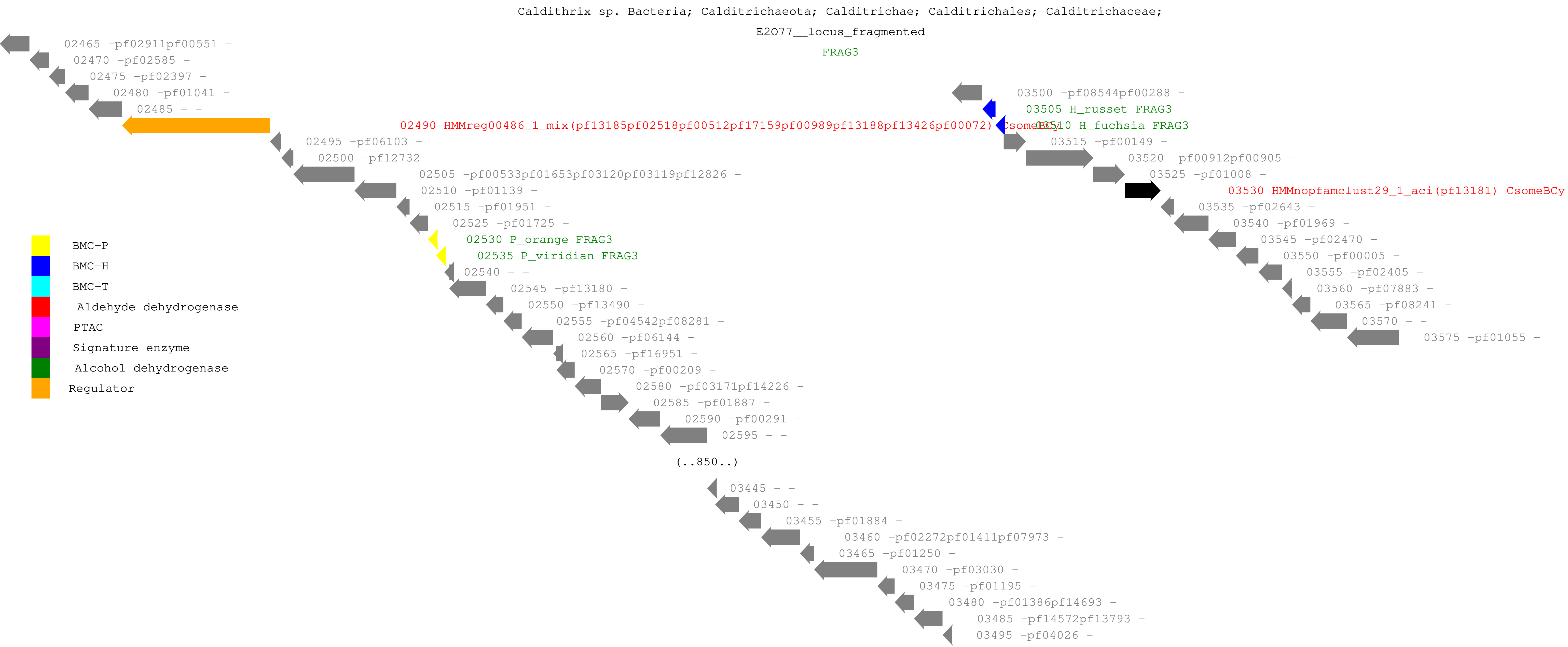
Caldithrix sp. Bacteria; Calditrichaeota; Calditrichae; Calditrichales; Calditrichaceae;
E2077__locus_fragmented
FRAG3
02465 -pf02911pf00551 -
02470 -pf02585 -
02475 -pf02397 -
02480 -pf01041 -
02485 - -
02490 HMMreg00486_1_mix(pf13185pf02518pf00512pf17159pf00989pf13188pf13426pf00072) CsomeBCy
02495 -pf06103 -
02500 -pf12732 -
02505 -pf00533pf01653pf03120pf03119pf12826 -
02510 -pf01139 -
02515 -pf01951 -
02525 -pf01725 -
02530 P_orange FRAG3
02535 P_viridian FRAG3
02540 - -
02545 -pf13180 -
02550 -pf13490 -
02555 -pf04542pf08281 -
02560 -pf06144 -
02565 -pf16951 -
02570 -pf00209 -
02580 -pf03171pf14226 -
02585 -pf01887 -
02590 -pf00291 -
02595 - -
(..850..)
03445 - -
03450 - -
03455 -pf01884 -
03460 -pf02272pf01411pf07973 -
03465 -pf01250 -
03470 -pf03030 -
03475 -pf01195 -
03480 -pf01386pf14693 -
03485 -pf14572pf13793 -
03495 -pf04026 -
03500 -pf08544pf00288 -
03505 H_russet FRAG3
03510 H_fuchsia FRAG3
03515 -pf00149 -
03520 -pf00912pf00905 -
03525 -pf01008 -
03530 HMMnopfamclust29_1_aci(pf13181) CsomeBCy
03535 -pf02643 -
03540 -pf01969 -
03545 -pf02470 -
03550 -pf00005 -
03555 -pf02405 -
03560 -pf07883 -
03565 -pf08241 -
03570 - -
03575 -pf01055 -
BMC-P
BMC-H
BMC-T
Aldehyde dehydrogenase
PTAC
Signature enzyme
Alcohol dehydrogenase
Regulator

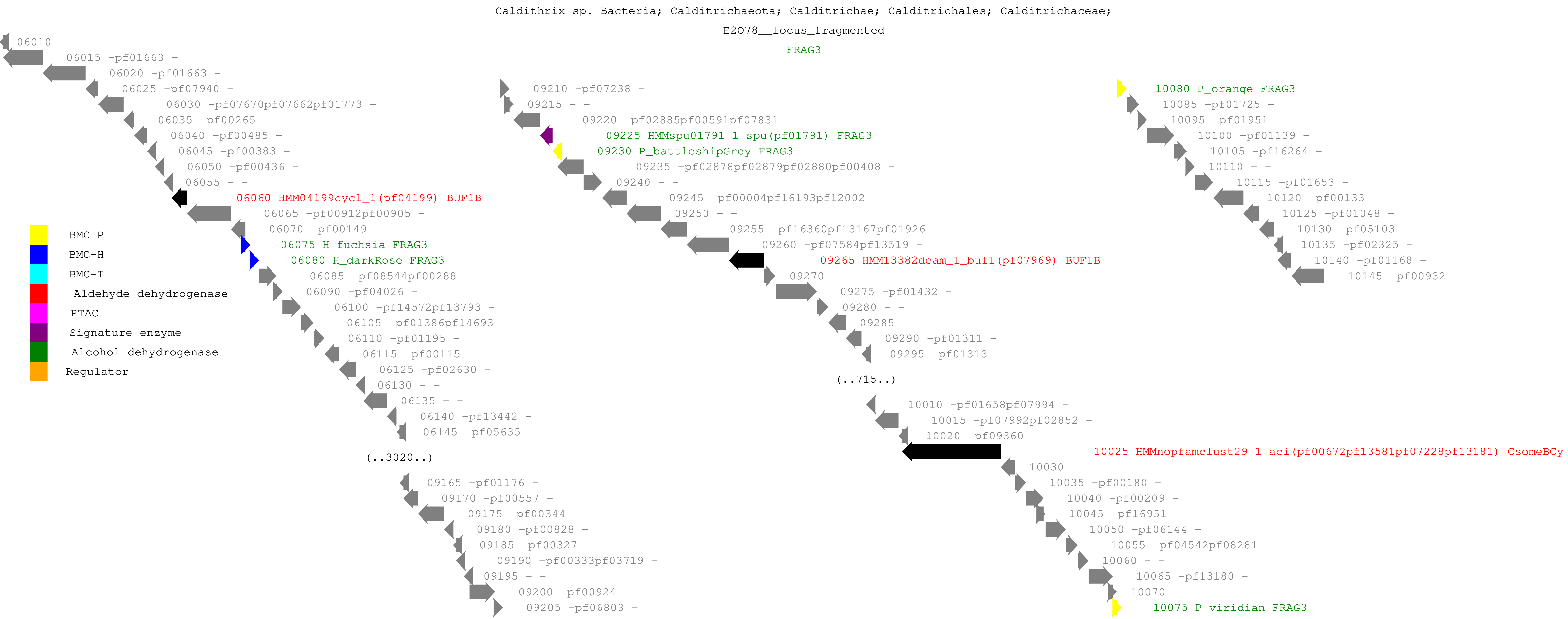

Caldithrix sp. Bacteria; Calditrichaeota; Calditrichae; Calditrichales; Calditrichaceae;
E2078__locus_fragmented
FRAG3
06010 - -
06015 -pf01663 -
06020 -pf01663 -
06025 -pf07940 -
06030 -pf07670pf07662pf01773 -
06035 -pf00265 -
06040 -pf00485 -
06045 -pf00383 -
06050 -pf00436 -
06055 - -
06060 HMM04199cycl_1(pf04199) BUF1B
06065 -pf00912pf00905 -
06070 -pf00149 -
06075 H_fuchsia FRAG3
06080 H_darkRose FRAG3
06085 -pf08544pf00288 -
06090 -pf04026 -
06100 -pf14572pf13793 -
06105 -pf01386pf14693 -
06110 -pf01195 -
06115 -pf00115 -
06125 -pf02630 -
06130 - -
06135 - -
06140 -pf13442 -
06145 -pf05635 -
(..3020..)
09165 -pf01176 -
09170 -pf00557 -
09175 -pf00344 -
09180 -pf00828 -
09185 -pf00327 -
09190 -pf00333pf03719 -
09195 - -
09200 -pf00924 -
09205 -pf06803 -
09210 -pf07238 -
09215 - -
09220 -pf02885pf00591pf07831 -
09225 HMMspu01791_1_spu(pf01791) FRAG3
09230 P_battleshipGrey FRAG3
09235 -pf02878pf02879pf02880pf00408 -
09240 - -
09245 -pf00004pf16193pf12002 -
09250 - -
09255 -pf16360pf13167pf01926 -
09260 -pf07584pf13519 -
09265 HMM13382deam_1_buf1(pf07969) BUF1B
09270 - -
09275 -pf01432 -
09280 - -
09285 - -
09290 -pf01311 -
09295 -pf01313 -
(..715..)
10010 -pf01658pf07994 -
10015 -pf07992pf02852 -
10020 -pf09360 -
10025 HMMnopfamclust29_1_aci(pf00672pf13581pf07228pf13181) CsomeBCy
10030 - -
10035 -pf00180 -
10040 -pf00209 -
10045 -pf16951 -
10050 -pf06144 -
10055 -pf04542pf08281 -
10060 - -
10065 -pf13180 -
10070 - -
10075 P_viridian FRAG3
10080 P_orange FRAG3
10085 -pf01725 -
10095 -pf01951 -
10100 -pf01139 -
10105 -pf16264 -
10110 - -
10115 -pf01653 -
10120 -pf00133 -
10125 -pf01048 -
10130 -pf05103 -
10135 -pf02325 -
10140 -pf01168 -
10145 -pf00932 -
BMC-P
BMC-H
BMC-T
Aldehyde dehydrogenase
PTAC
Signature enzyme
Alcohol dehydrogenase
Regulator

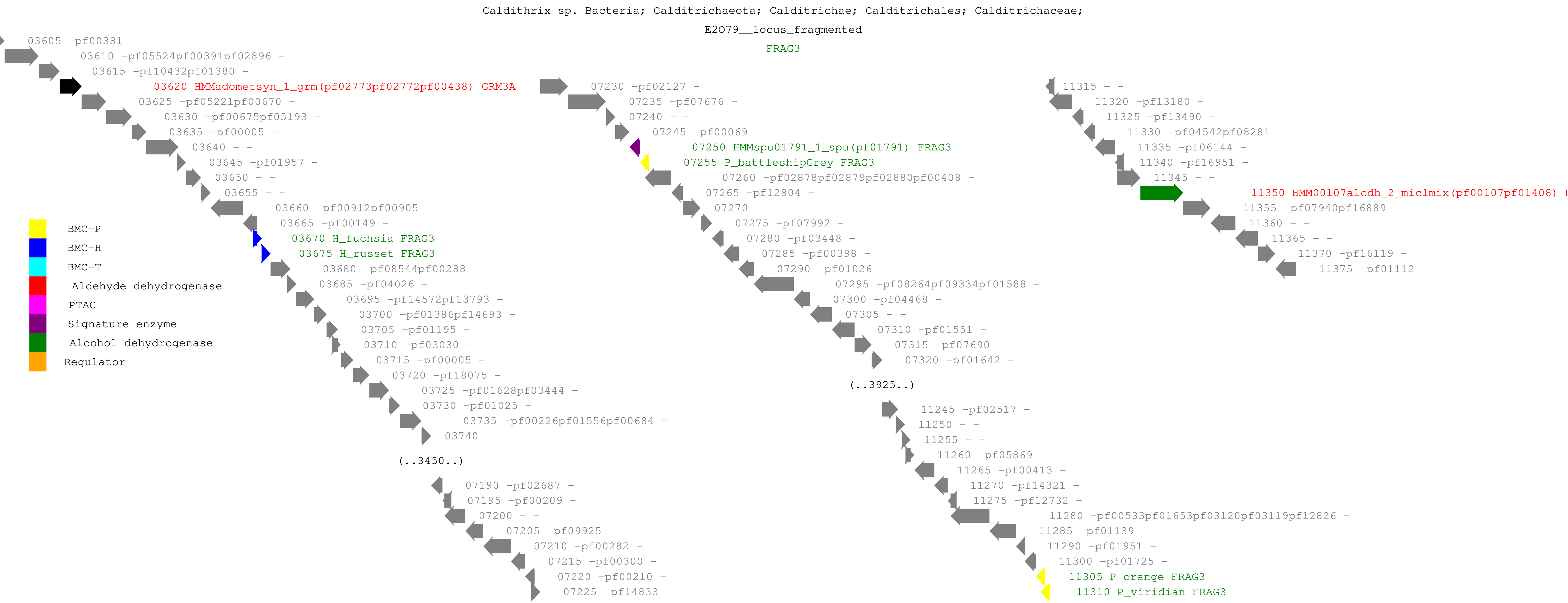
Caldithrix sp. Bacteria; Calditrichaeota; Calditrichae; Calditrichales; Calditrichaceae;
E2079__locus_fragmented
FRAG3
03605 -pf00381 -
03610 -pf05524pf00391pf02896 -
03615 -pf10432pf01380 -
03620 HMMadometsyn_1_grm(pf02773pf02772pf00438) GRM3A
03625 -pf05221pf00670 -
03630 -pf00675pf05193 -
03635 -pf00005 -
03640 - -
03645 -pf01957 -
03650 - -
03655 - -
03660 -pf00912pf00905 -
03665 -pf00149 -
03670 H_fuchsia FRAG3
03675 H_russet FRAG3
03680 -pf08544pf00288 -
03685 -pf04026 -
03695 -pf14572pf13793 -
03700 -pf01386pf14693 -
03705 -pf01195 -
03710 -pf03030 -
03715 -pf00005 -
03720 -pf18075 -
03725 -pf01628pf03444 -
03730 -pf01025 -
03735 -pf00226pf01556pf00684 -
03740 - -
(..3450..)
07190 -pf02687 -
07195 -pf00209 -
07200 - -
07205 -pf09925 -
07210 -pf00282 -
07215 -pf00300 -
07220 -pf00210 -
07225 -pf14833 -
07230 -pf02127 -
07235 -pf07676 -
07240 - -
07245 -pf00069 -
07250 HMMspu01791_1_spu(pf01791) FRAG3
07255 P_battleshipGrey FRAG3
07260 -pf02878pf02879pf02880pf00408 -
07265 -pf12804 -
07270 - -
07275 -pf07992 -
07280 -pf03448 -
07285 -pf00398 -
07290 -pf01026 -
07295 -pf08264pf09334pf01588 -
07300 -pf04468 -
07305 - -
07310 -pf01551 -
07315 -pf07690 -
07320 -pf01642 -
(..3925..)
11245 -pf02517 -
11250 - -
11255 - -
11260 -pf05869 -
11265 -pf00413 -
11270 -pf14321 -
11275 -pf12732 -
11280 -pf00533pf01653pf03120pf03119pf12826 -
11285 -pf01139 -
11290 -pf01951 -
11300 -pf01725 -
11305 P_orange FRAG3
11310 P_viridian FRAG3
11315 - -
11320 -pf13180 -
11325 -pf13490 -
11330 -pf04542pf08281 -
11335 -pf06144 -
11340 -pf16951 -
11345 - -
11350 HMM00107alcdh_2_mic1mix(pf00107pf01408) I
11355 -pf07940pf16889 -
11360 - -
11365 - -
11370 -pf16119 -
11375 -pf01112 -
BMC-P
BMC-H
BMC-T
Aldehyde dehydrogenase
PTAC
Signature enzyme
Alcohol dehydrogenase
Regulator

Candidatus Marinimicrobia bacterium CG1_02_48_14. Bacteria; Candidatus Marinimicrobia.

AUJ47__locus_fragmented

FRAG4

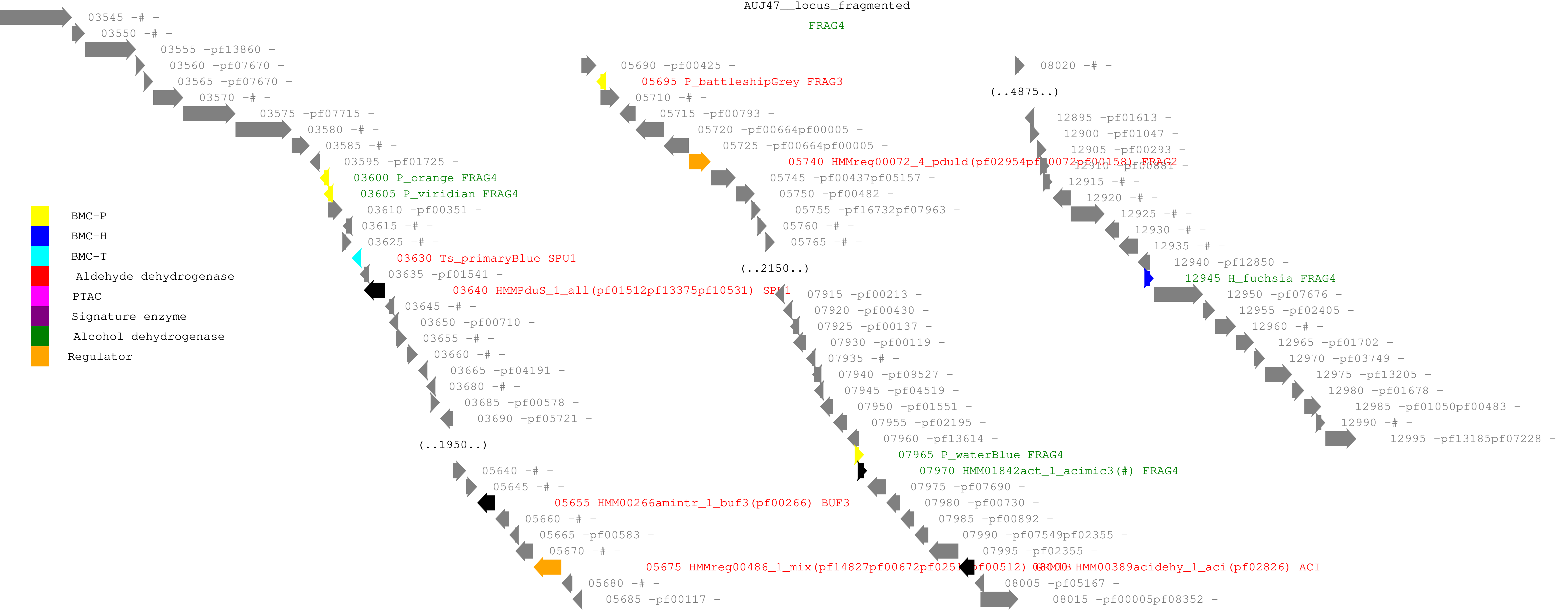

Candidatus Marinimicrobia bacterium CG_4_9_14_3_um_filter_48_9. Bacteria; Candidatus Marinimicrobia.

CO167__locus_fragmented

FRAG4

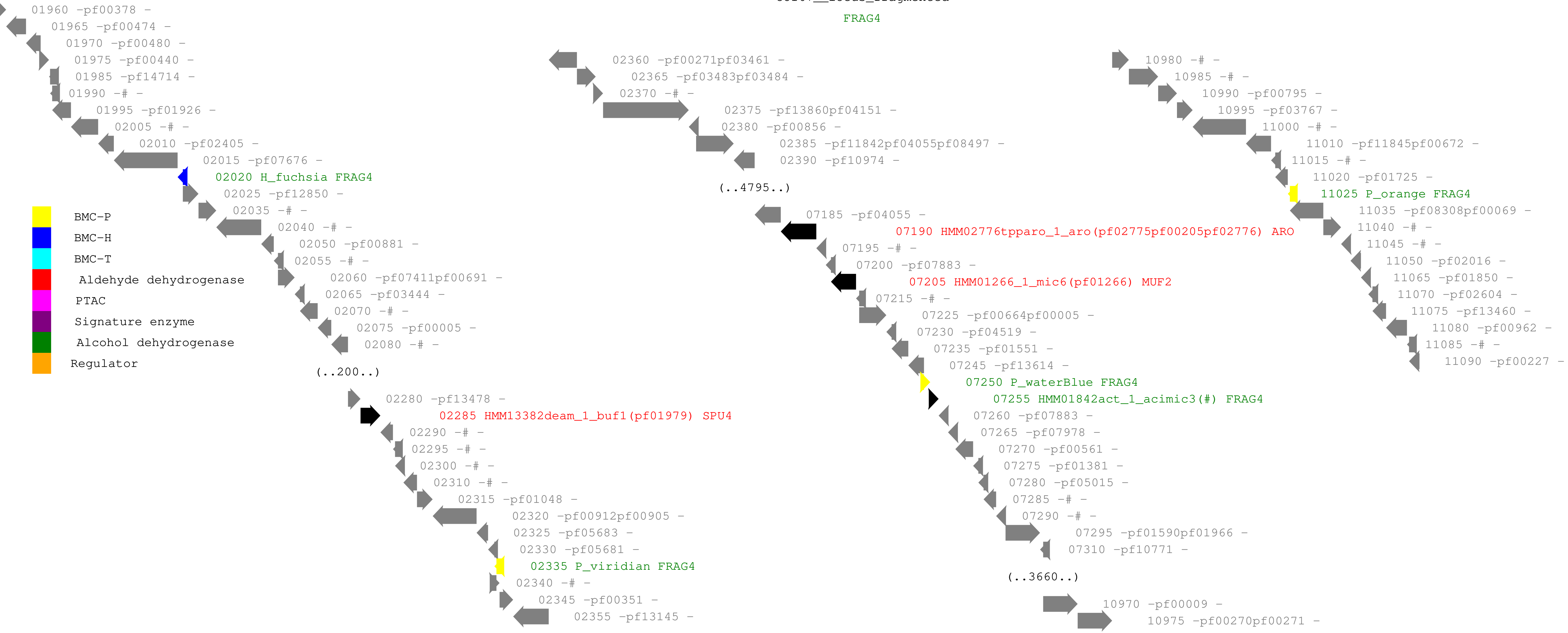

Candidatus Marinimicrobia bacterium CG_4_10_14_0_2_um_filter_48_9. Bacteria; Candidatus Marinimicrobia.

COY19__locus_fragmented

FRAG4

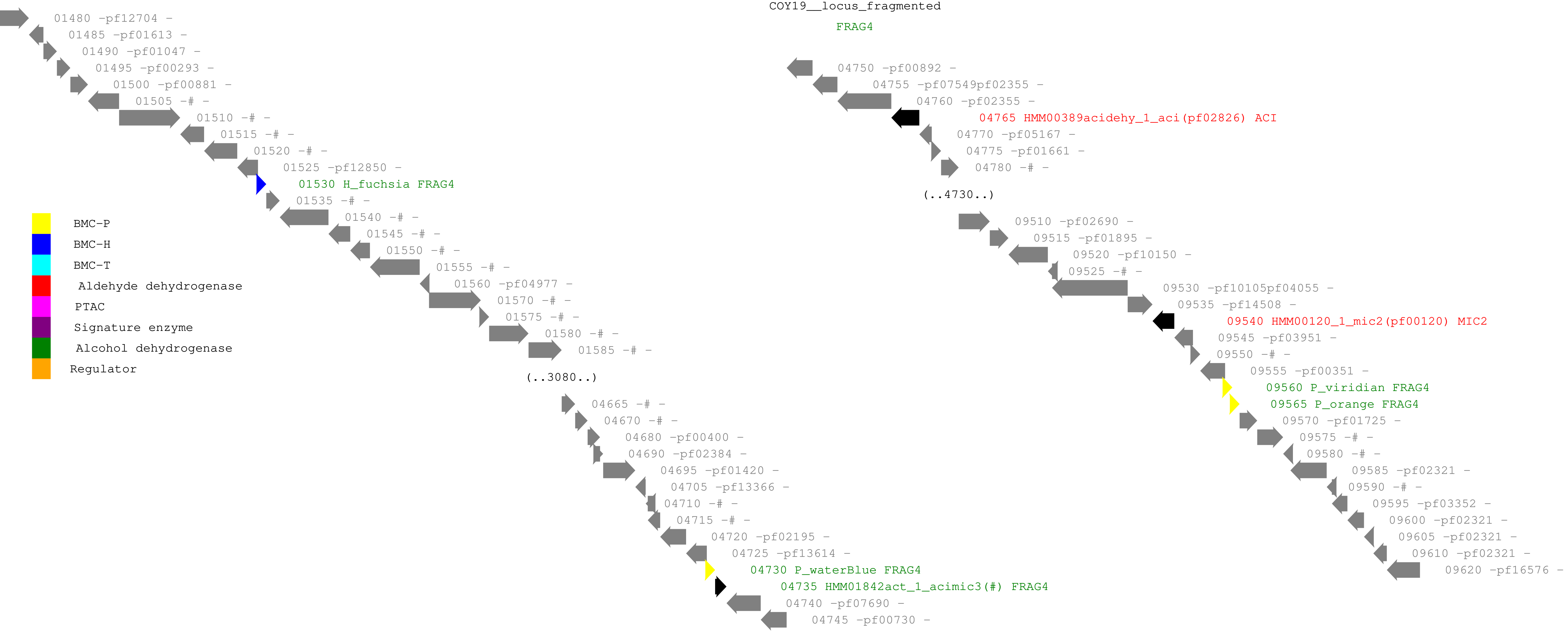

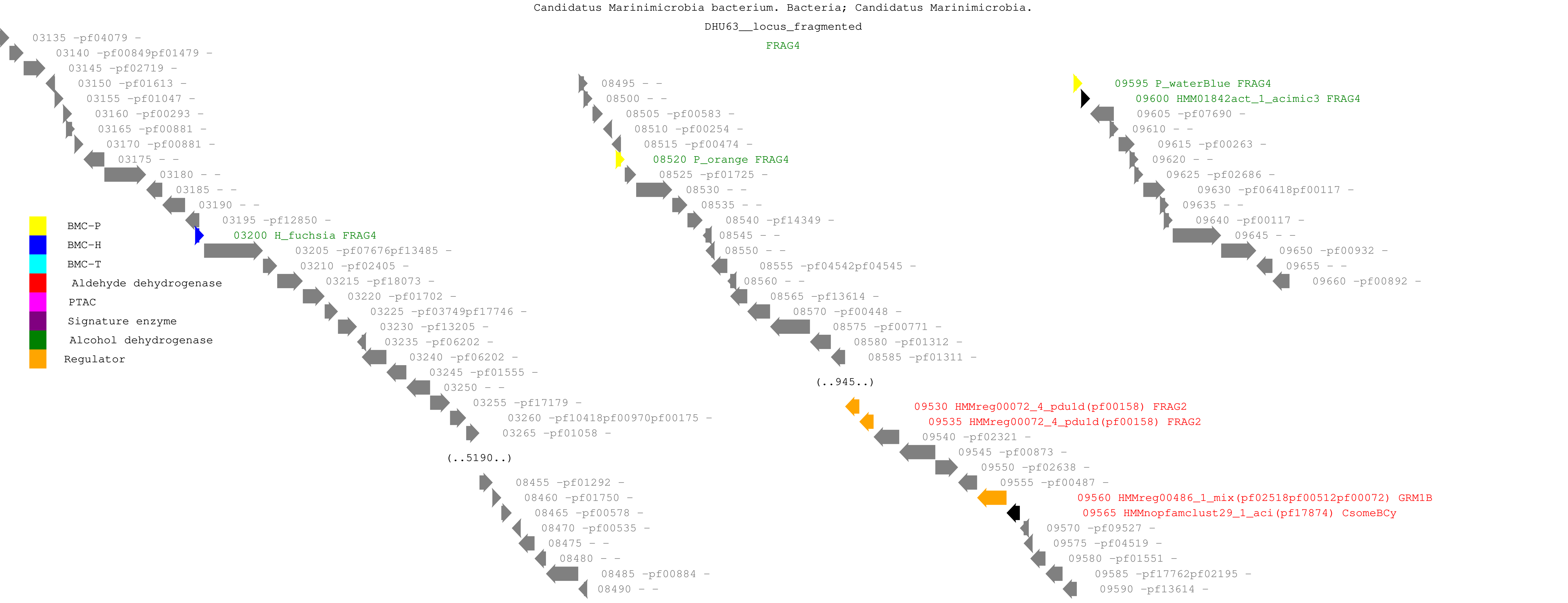

Candidatus Marinimicrobia bacterium. Bacteria; Candidatus Marinimicrobia.
DHU63__locus_fragmented
FRAG4
03135 -pf04079 -
03140 -pf00849pf01479 -
03145 -pf02719 -
03150 -pf01613 -
03155 -pf01047 -
03160 -pf00293 -
03165 -pf00881 -
03170 -pf00881 -
03175 - -
03180 - -
03185 - -
03190 - -
03195 -pf12850 -
03200 H_fuchsia FRAG4
03205 -pf07676pf13485 -
03210 -pf02405 -
03215 -pf18073 -
03220 -pf01702 -
03225 -pf03749pf17746 -
03230 -pf13205 -
03235 -pf06202 -
03240 -pf06202 -
03245 -pf01555 -
03250 - -
03255 -pf17179 -
03260 -pf10418pf00970pf00175 -
03265 -pf01058 -
(..5190..)
08455 -pf01292 -
08460 -pf01750 -
08465 -pf00578 -
08470 -pf00535 -
08475 - -
08480 - -
08485 -pf00884 -
08490 - -
08495 - -
08500 - -
08505 -pf00583 -
08510 -pf00254 -
08515 -pf00474 -
08520 P_orange FRAG4
08525 -pf01725 -
08530 - -
08535 - -
08540 -pf14349 -
08545 - -
08550 - -
08555 -pf04542pf04545 -
08560 - -
08565 -pf13614 -
08570 -pf00448 -
08575 -pf00771 -
08580 -pf01312 -
08585 -pf01311 -
(..945..)
09530 HMMreg00072_4_pdu1d(pf00158) FRAG2
09535 HMMreg00072_4_pdu1d(pf00158) FRAG2
09540 -pf02321 -
09545 -pf00873 -
09550 -pf02638 -
09555 -pf00487 -
09560 HMMreg00486_1_mix(pf02518pf00512pf00072) GRM1B
09565 HMMnopfamclust29_1_aci(pf17874) CsomeBCy
09570 -pf09527 -
09575 -pf04519 -
09580 -pf01551 -
09585 -pf17762pf02195 -
09590 -pf13614 -
09595 P_waterBlue FRAG4
09600 HMM01842act_1_acimic3 FRAG4
09605 -pf07690 -
09610 - -
09615 -pf00263 -
09620 - -
09625 -pf02686 -
09630 -pf06418pf00117 -
09635 - -
09640 -pf00117 -
09645 - -
09650 -pf00932 -
09655 - -
09660 -pf00892 -
BMC-P
BMC-H
BMC-T
Aldehyde dehydrogenase
PTAC
Signature enzyme
Alcohol dehydrogenase
Regulator

Planctomycetes bacterium. Bacteria; Planctomycetes.

D6731__locus_fragmented

FRAG5

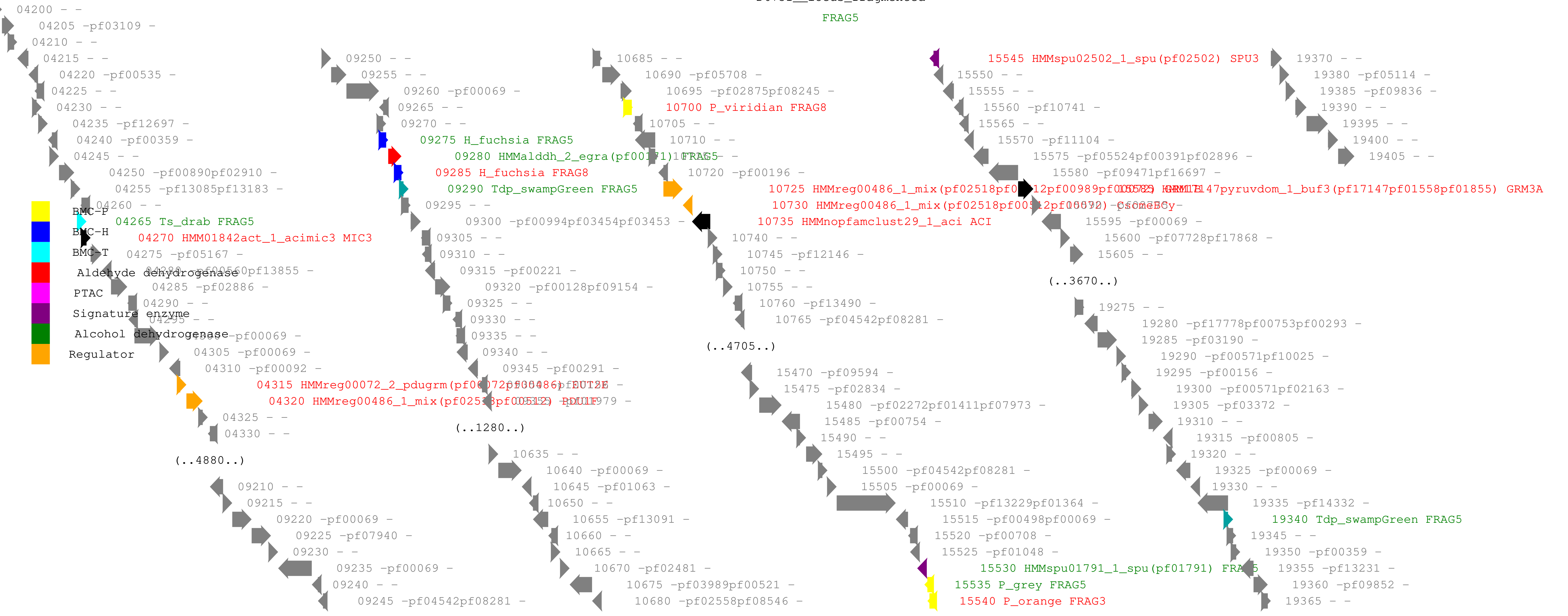

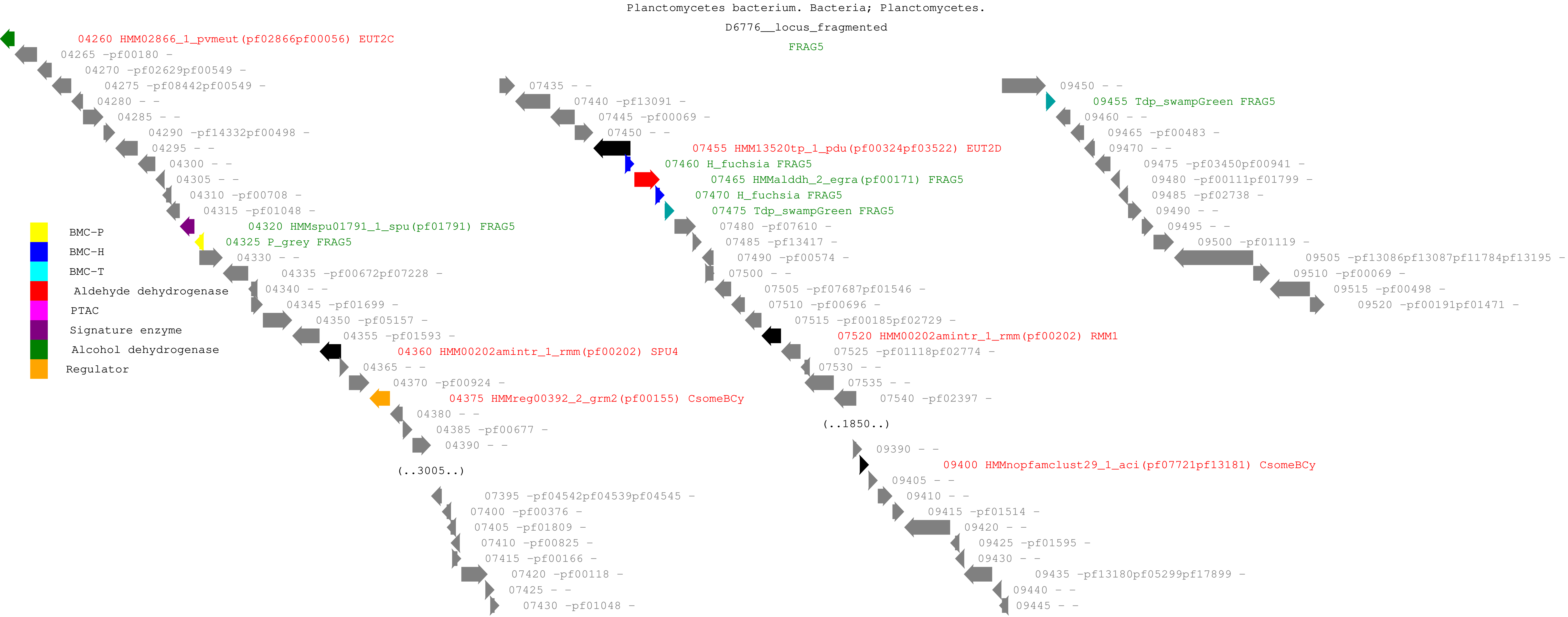

Planctomycetes bacterium. Bacteria; Planctomycetes.
D6776__locus_fragmented
FRAG5
04260 HMM02866_1_pvmeut(pf02866pf00056) EUT2C
04265 -pf00180 -
04270 -pf02629pf00549 -
04275 -pf08442pf00549 -
04280 - -
04285 - -
04290 -pf14332pf00498 -
04295 - -
04300 - -
04305 - -
04310 -pf00708 -
04315 -pf01048 -
04320 HMMspu01791_1_spu(pf01791) FRAG5
04325 P_grey FRAG5
04330 - -
04335 -pf00672pf07228 -
04340 - -
04345 -pf01699 -
04350 -pf05157 -
04355 -pf01593 -
04360 HMM00202amintr_1_rmm(pf00202) SPU4
04365 - -
04370 -pf00924 -
04375 HMMreg00392_2_grm2(pf00155) CsomeBCy
04380 - -
04385 -pf00677 -
04390 - -
(..3005..)
07395 -pf04542pf04539pf04545 -
07400 -pf00376 -
07405 -pf01809 -
07410 -pf00825 -
07415 -pf00166 -
07420 -pf00118 -
07425 - -
07430 -pf01048 -
07435 - -
07440 -pf13091 -
07445 -pf00069 -
07450 - -
07455 HMM13520tp_1_pdu(pf00324pf03522) EUT2D
07460 H_fuchsia FRAG5
07465 HMMalddh_2_egra(pf00171) FRAG5
07470 H_fuchsia FRAG5
07475 Tdp_swampGreen FRAG5
07480 -pf07610 -
07485 -pf13417 -
07490 -pf00574 -
07500 - -
07505 -pf07687pf01546 -
07510 -pf00696 -
07515 -pf00185pf02729 -
07520 HMM00202amintr_1_rmm(pf00202) RMM1
07525 -pf01118pf02774 -
07530 - -
07535 - -
07540 -pf02397 -
(..1850..)
09390 - -
09400 HMMnopfamclust29_1_aci(pf07721pf13181) CsomeBCy
09405 - -
09410 - -
09415 -pf01514 -
09420 - -
09425 -pf01595 -
09430 - -
09435 -pf13180pf05299pf17899 -
09440 - -
09445 - -
09450 - -
09455 Tdp_swampGreen FRAG5
09460 - -
09465 -pf00483 -
09470 - -
09475 -pf03450pf00941 -
09480 -pf00111pf01799 -
09485 -pf02738 -
09490 - -
09495 - -
09500 -pf01119 -
09505 -pf13086pf13087pf11784pf13195 -
09510 -pf00069 -
09515 -pf00498 -
09520 -pf00191pf01471 -
BMC-P
BMC-H
BMC-T
Aldehyde dehydrogenase
PTAC
Signature enzyme
Alcohol dehydrogenase
Regulator

Planctomycetes bacterium. Bacteria; Planctomycetes.

DEA08__locus_fragmented

FRAG5

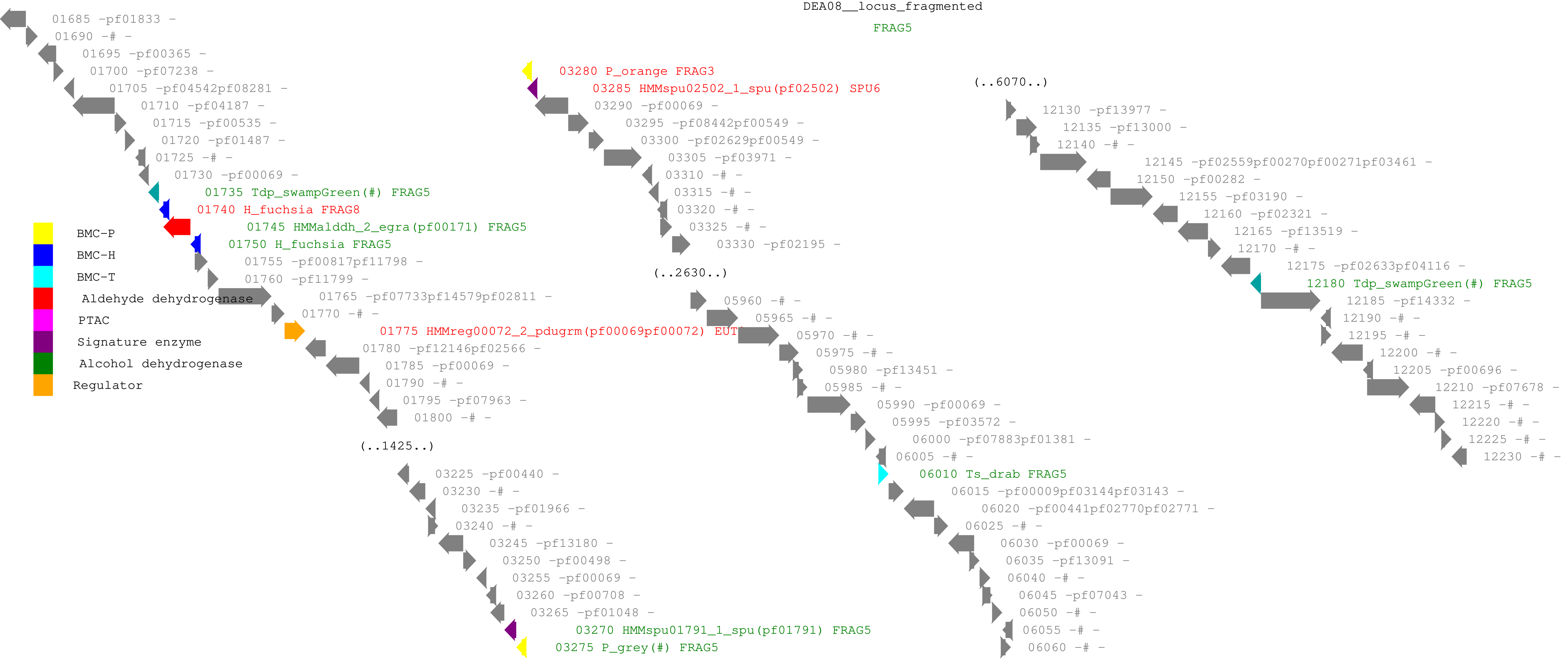

Candidatus Sumerlaea chitinovorans. Bacteria; Candidatus Sumerlaeota; Candidatus Sumerlaeia;

BRCON__locus_fragmented

FRAG6

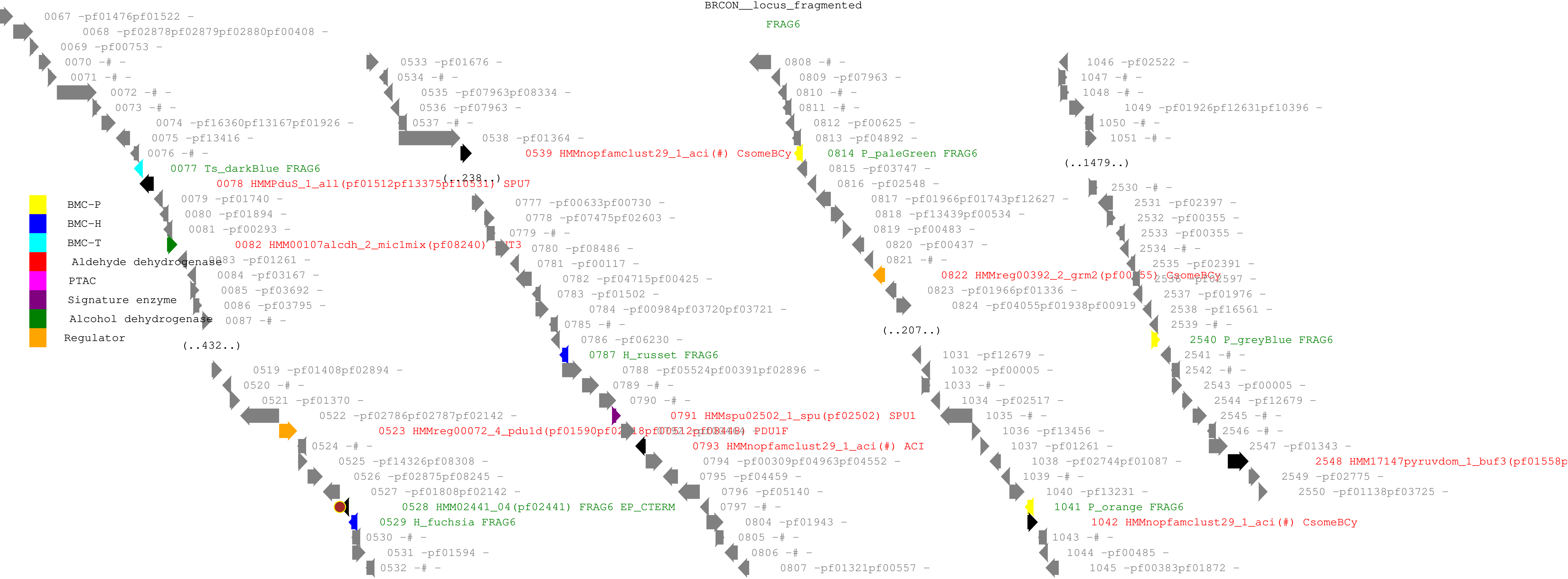

candidate division BRC1 bacterium ADurb.Bin183. Bacteria; candidate division BRC1.

BWY12__locus_fragmented

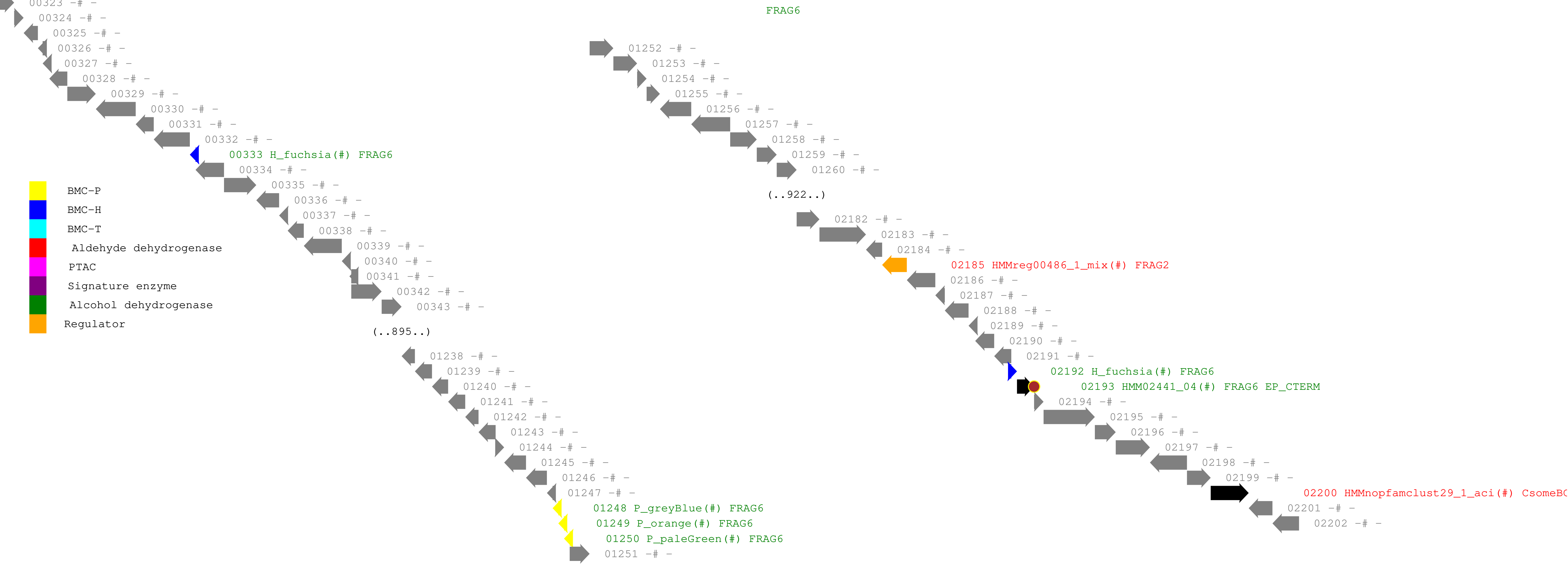

candidate division BRC1 bacterium ADurb.BinA364. Bacteria; candidate division BRC1.

BWZ10__locus_fragmented

FRAG6

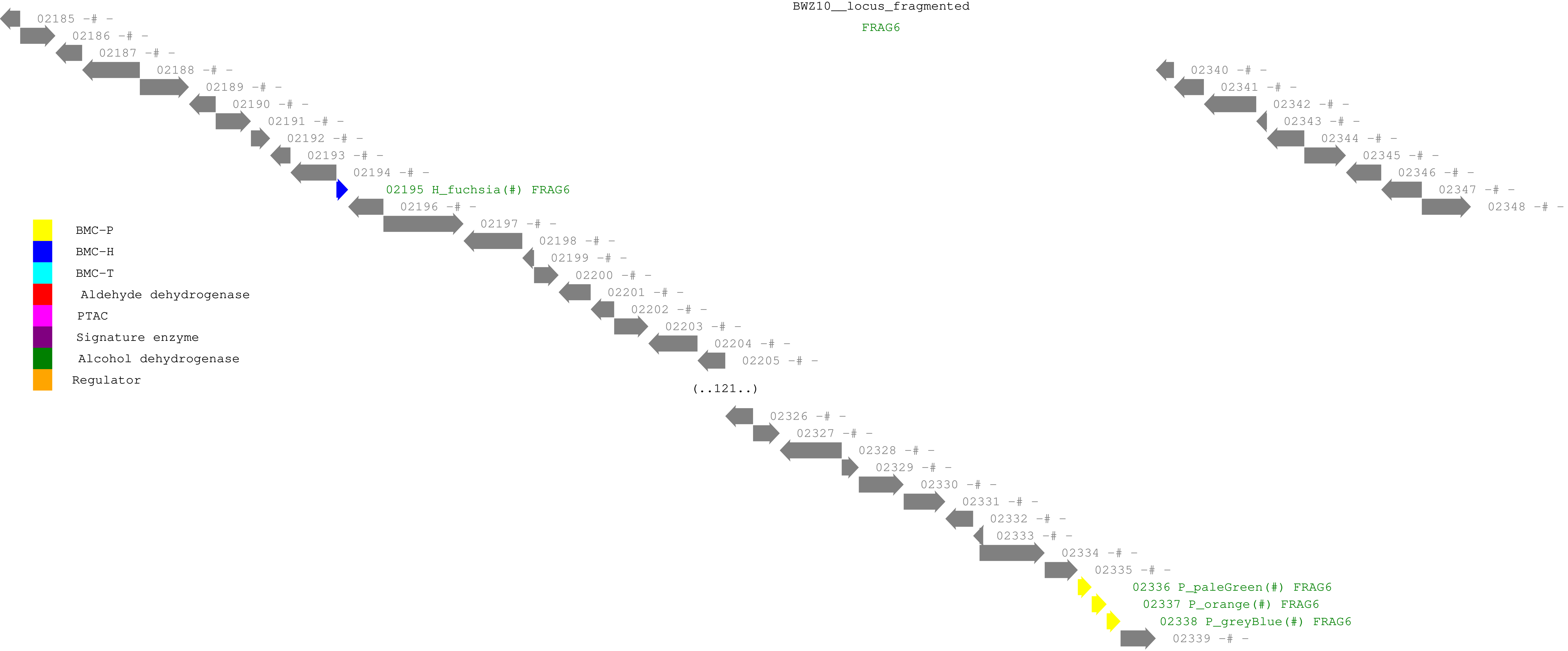

candidate division BRC1 bacterium HGW-BRC1-1. Bacteria; candidate division BRC1.

CVU37__locus_1_main_sat

FRAG6

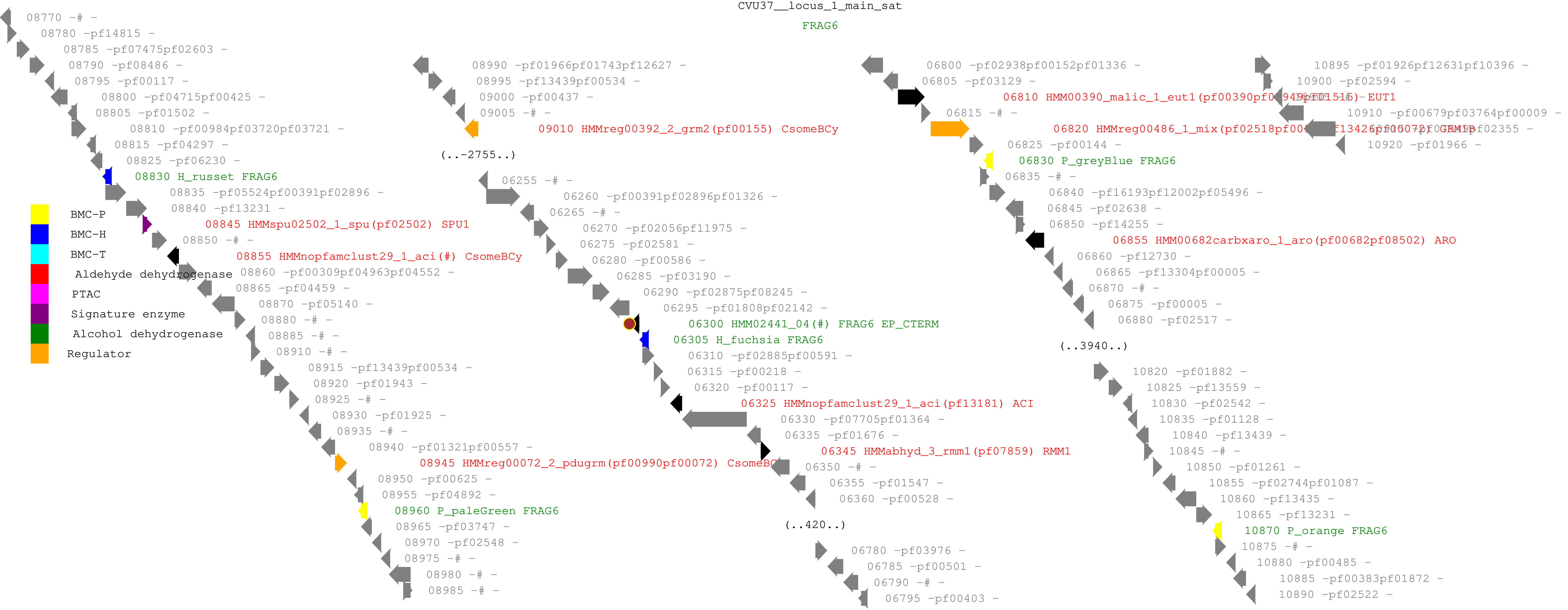

Candidatus Hydrogenedentes bacterium. Bacteria; Candidatus Hydrogenedentes.

D6691__locus_fragmented

FRAG6

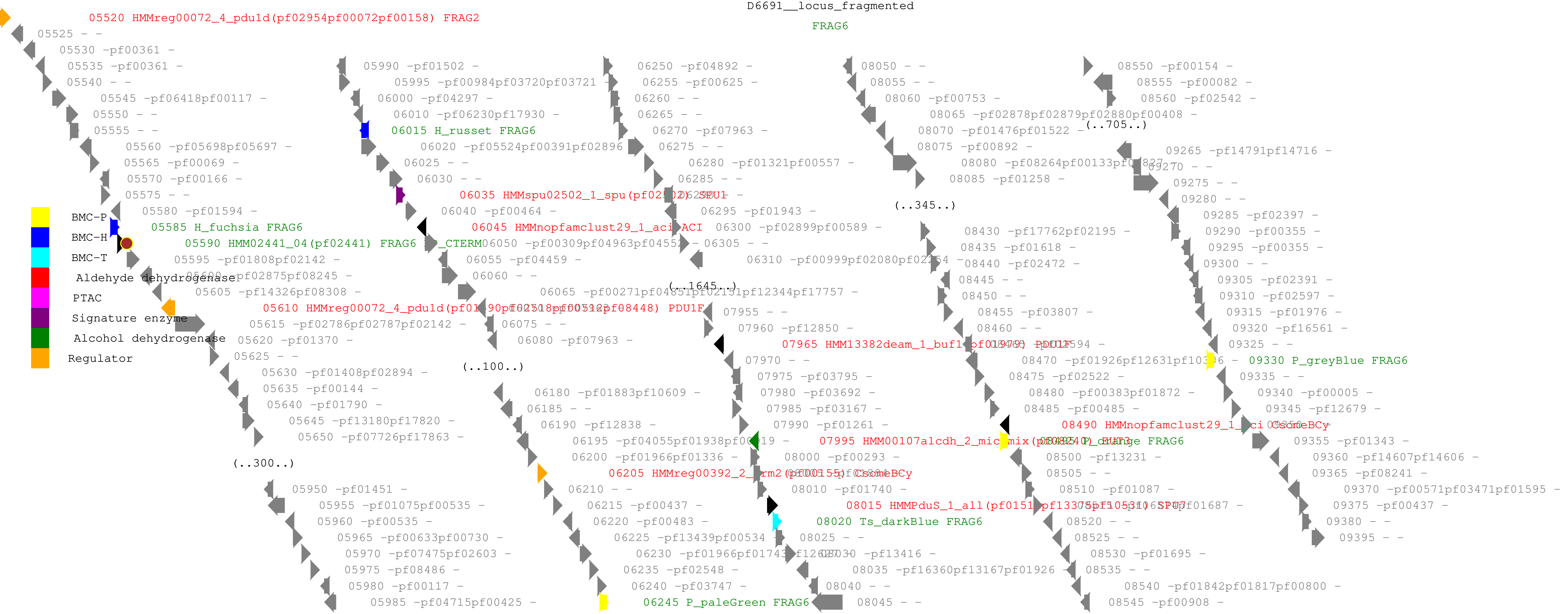

Candidatus Hydrogenedentes bacterium CG1_02_42_14. Bacteria; Candidatus Hydrogenedentes.

AUJ18__locus_fragmented

FRAG7

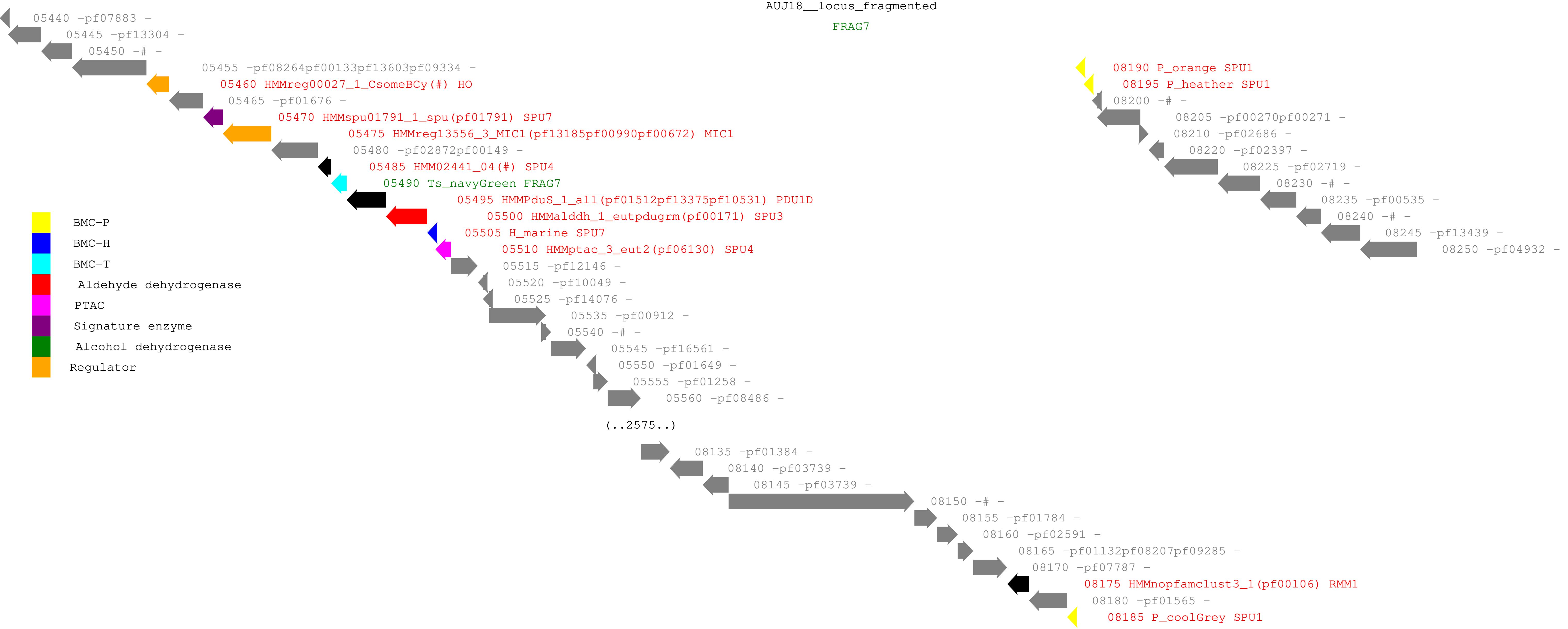

Candidatus Hydrogenedentes bacterium CG07_land_8_20_14_0_80_42_17. Bacteria; Candidatus Hydrogenedentes.

COS94__locus_fragmented

FRAG7

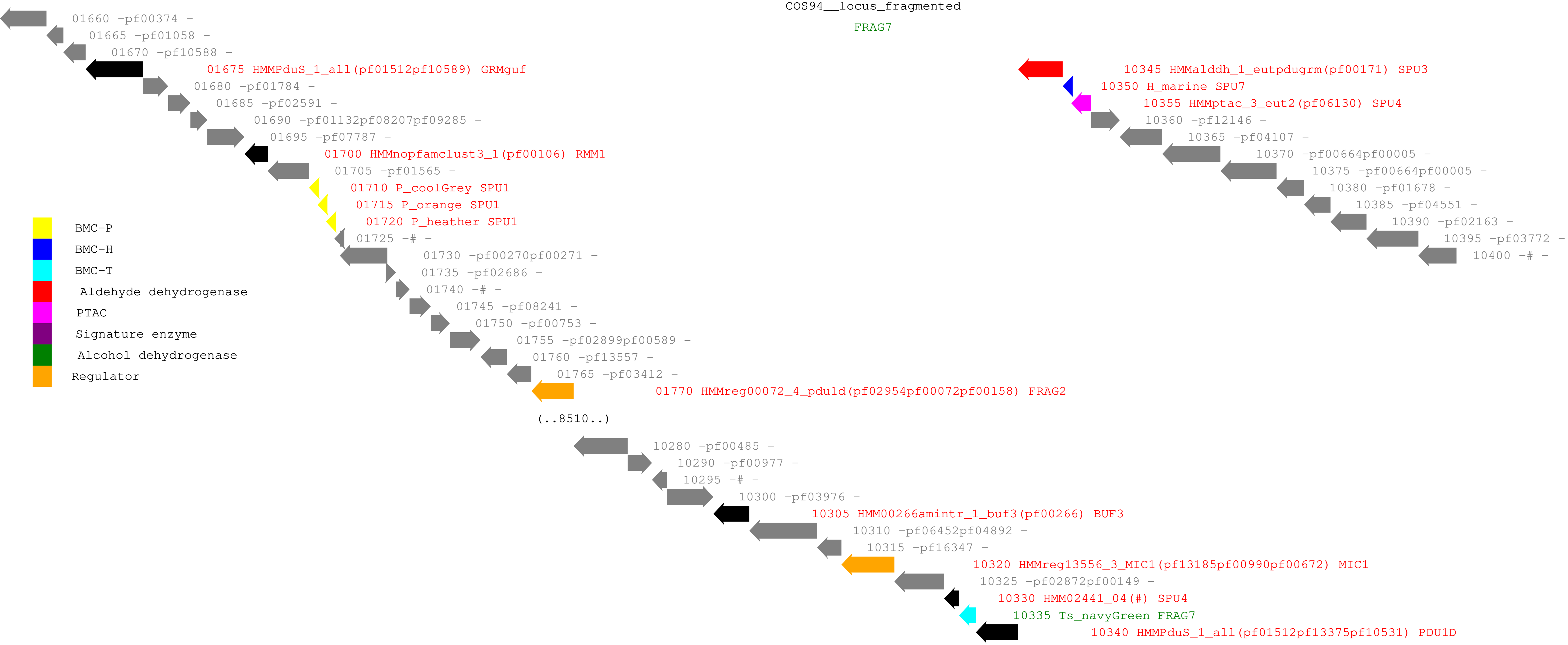

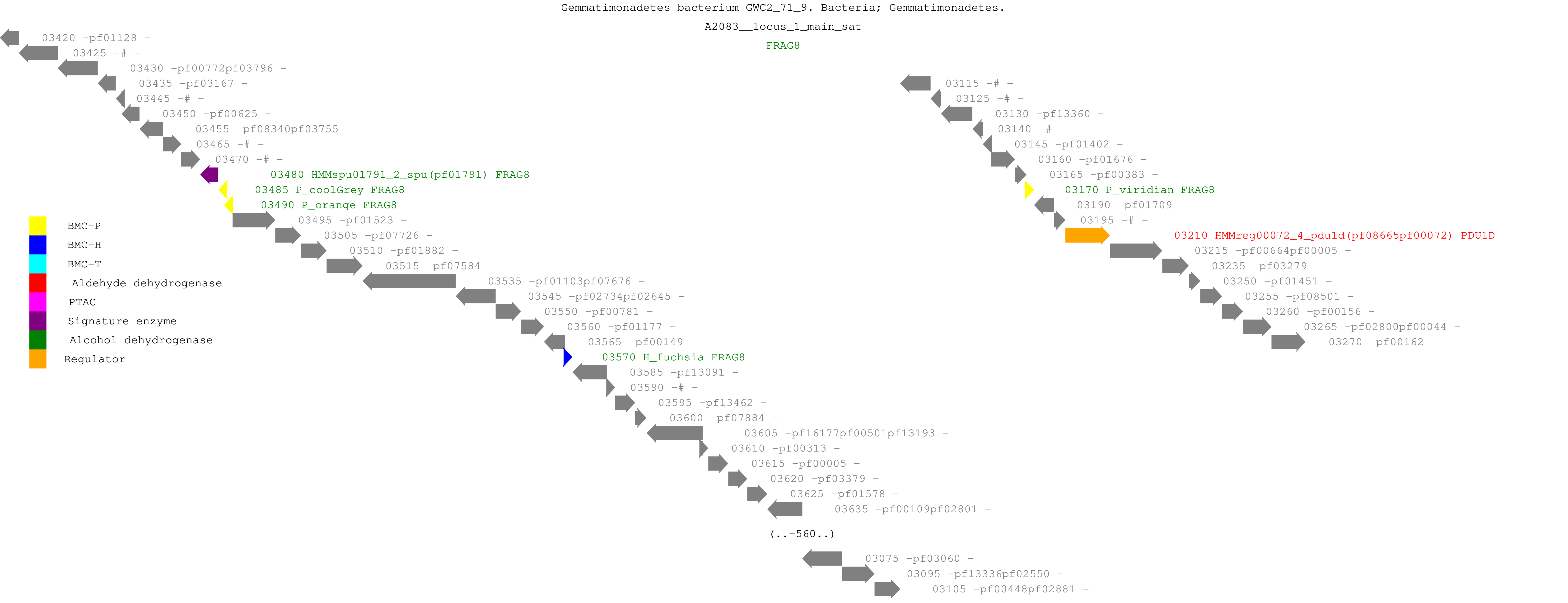
Gemmatimonadetes bacterium GWC2_71_9. Bacteria; Gemmatimonadetes.
A2083__locus_1_main_sat
FRAG8
03420 -pf01128 -
03425 -# -
03430 -pf00772pf03796 -
03435 -pf03167 -
03445 -# -
03450 -pf00625 -
03455 -pf08340pf03755 -
03465 -# -
03470 -# -
03480 HMMspu01791_2_spu(pf01791) FRAG8
03485 P_coolGrey FRAG8
03490 P_orange FRAG8
03495 -pf01523 -
03505 -pf07726 -
03510 -pf01882 -
03515 -pf07584 -
03535 -pf01103pf07676 -
03545 -pf02734pf02645 -
03550 -pf00781 -
03560 -pf01177 -
03565 -pf00149 -
03570 H_fuchsia FRAG8
03585 -pf13091 -
03590 -# -
03595 -pf13462 -
03600 -pf07884 -
03605 -pf16177pf00501pf13193 -
03610 -pf00313 -
03615 -pf00005 -
03620 -pf03379 -
03625 -pf01578 -
03635 -pf00109pf02801 -
(..-560..)
03075 -pf03060 -
03095 -pf13336pf02550 -
03105 -pf00448pf02881 -
03115 -# -
03125 -# -
03130 -pf13360 -
03140 -# -
03145 -pf01402 -
03160 -pf01676 -
03165 -pf00383 -
03170 P_viridian FRAG8
03190 -pf01709 -
03195 -# -
03210 HMMreg00072_4_pdu1d(pf08665pf00072) PDU1D
03215 -pf00664pf00005 -
03235 -pf03279 -
03250 -pf01451 -
03255 -pf08501 -
03260 -pf00156 -
03265 -pf02800pf00044 -
03270 -pf00162 -
BMC-P
BMC-H
BMC-T
Aldehyde dehydrogenase
PTAC
Signature enzyme
Alcohol dehydrogenase
Regulator

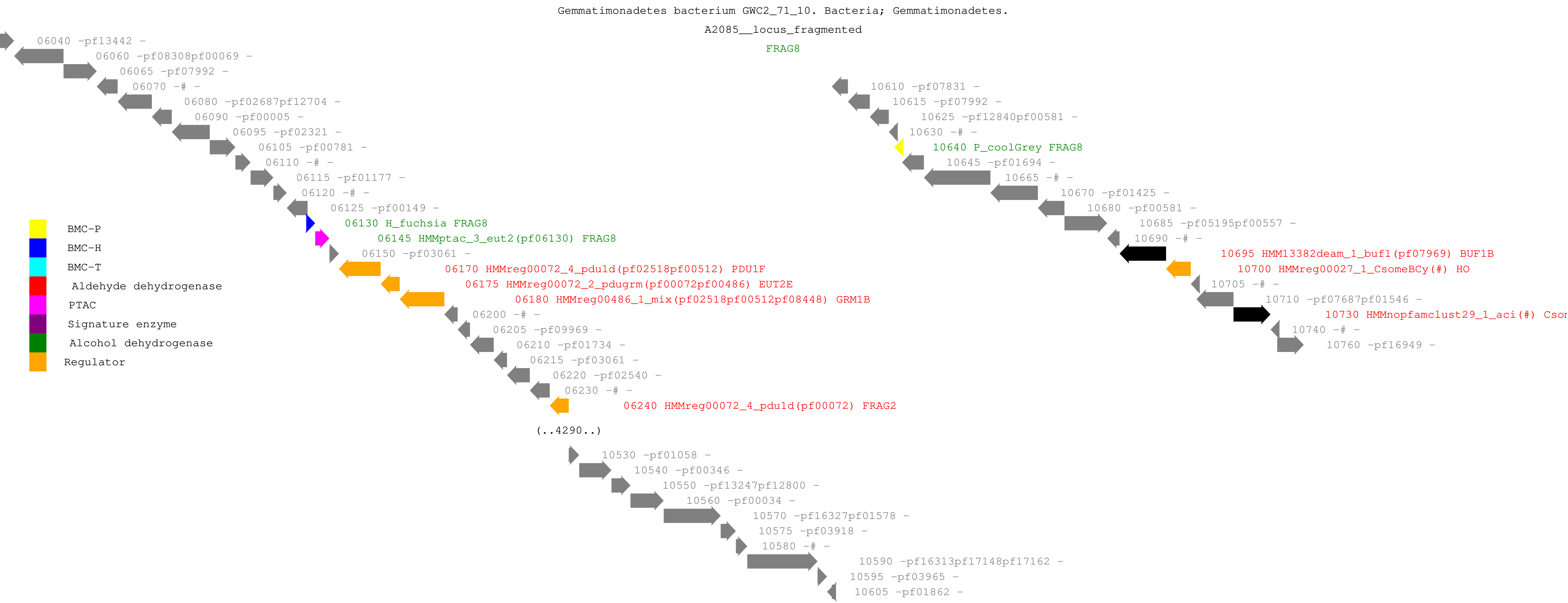

Gemmatimonadetes bacterium GWC2_71_10. Bacteria; Gemmatimonadetes.
A2085__locus_fragmented
FRAG8
06040 -pf13442 -
06060 -pf08308pf00069 -
06065 -pf07992 -
06070 -# -
06080 -pf02687pf12704 -
06090 -pf00005 -
06095 -pf02321 -
06105 -pf00781 -
06110 -# -
06115 -pf01177 -
06120 -# -
06125 -pf00149 -
06130 H_fuchsia FRAG8
06145 HMMptac_3_eut2(pf06130) FRAG8
06150 -pf03061 -
06170 HMMreg00072_4_pdu1d(pf02518pf00512) PDU1F
06175 HMMreg00072_2_pdugrm(pf00072pf00486) EUT2E
06180 HMMreg00486_1_mix(pf02518pf00512pf08448) GRM1B
06200 -# -
06205 -pf09969 -
06210 -pf01734 -
06215 -pf03061 -
06220 -pf02540 -
06230 -# -
06240 HMMreg00072_4_pdu1d(pf00072) FRAG2
(..4290..)
10530 -pf01058 -
10540 -pf00346 -
10550 -pf13247pf12800 -
10560 -pf00034 -
10570 -pf16327pf01578 -
10575 -pf03918 -
10580 -# -
10590 -pf16313pf17148pf17162 -
10595 -pf03965 -
10605 -pf01862 -
10610 -pf07831 -
10615 -pf07992 -
10625 -pf12840pf00581 -
10630 -# -
10640 P_coolGrey FRAG8
10645 -pf01694 -
10665 -# -
10670 -pf01425 -
10680 -pf00581 -
10685 -pf05195pf00557 -
10690 -# -
10695 HMM13382deam_1_buf1(pf07969) BUF1B
10700 HMMreg00027_1_CsomeBCy(#) HO
10705 -# -
10710 -pf07687pf01546 -
10730 HMMnopfamclust29_1_aci(#) Csor
10740 -# -
10760 -pf16949 -
BMC-P
BMC-H
BMC-T
Aldehyde dehydrogenase
PTAC
Signature enzyme
Alcohol dehydrogenase
Regulator

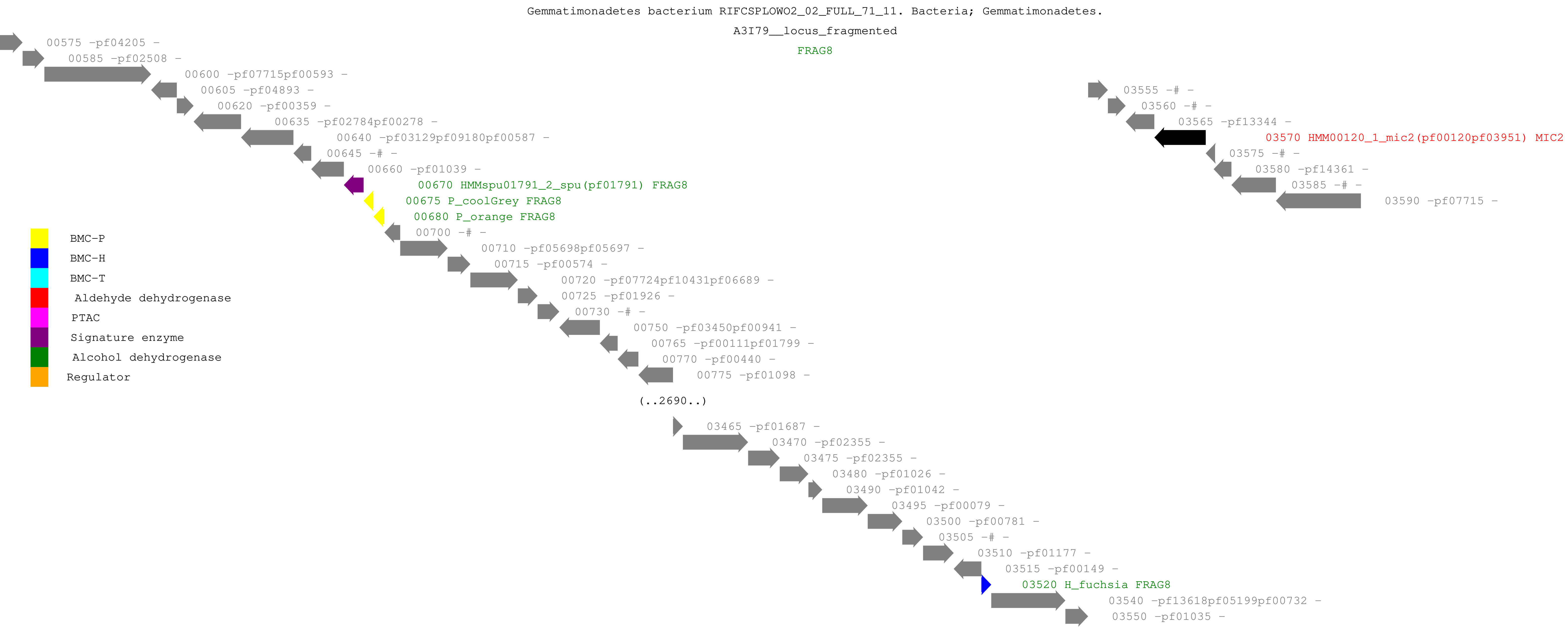
Gemmatimonadetes bacterium RIFCSPLOWO2_02_FULL_71_11. Bacteria; Gemmatimonadetes.
A3I79__locus_fragmented
FRAG8
00575 -pf04205 -
00585 -pf02508 -
00600 -pf07715pf00593 -
00605 -pf04893 -
00620 -pf00359 -
00635 -pf02784pf00278 -
00640 -pf03129pf09180pf00587 -
00645 -# -
00660 -pf01039 -
00670 HMMspu01791_2_spu(pf01791) FRAG8
00675 P_coolGrey FRAG8
00680 P_orange FRAG8
00700 -# -
00710 -pf05698pf05697 -
00715 -pf00574 -
00720 -pf07724pf10431pf06689 -
00725 -pf01926 -
00730 -# -
00750 -pf03450pf00941 -
00765 -pf00111pf01799 -
00770 -pf00440 -
00775 -pf01098 -
(..2690..)
03465 -pf01687 -
03470 -pf02355 -
03475 -pf02355 -
03480 -pf01026 -
03490 -pf01042 -
03495 -pf00079 -
03500 -pf00781 -
03505 -# -
03510 -pf01177 -
03515 -pf00149 -
03520 H_fuchsia FRAG8
03540 -pf13618pf05199pf00732 -
03550 -pf01035 -
03555 -# -
03560 -# -
03565 -pf13344 -
03570 HMM00120_1_mic2(pf00120pf03951) MIC2
03575 -# -
03580 -pf14361 -
03585 -# -
03590 -pf07715 -
BMC-P
BMC-H
BMC-T
Aldehyde dehydrogenase
PTAC
Signature enzyme
Alcohol dehydrogenase
Regulator

Gemmatimonadetes bacterium SCN 70-22. Bacteria; Gemmatimonadetes.

ABS52__locus_fragmented

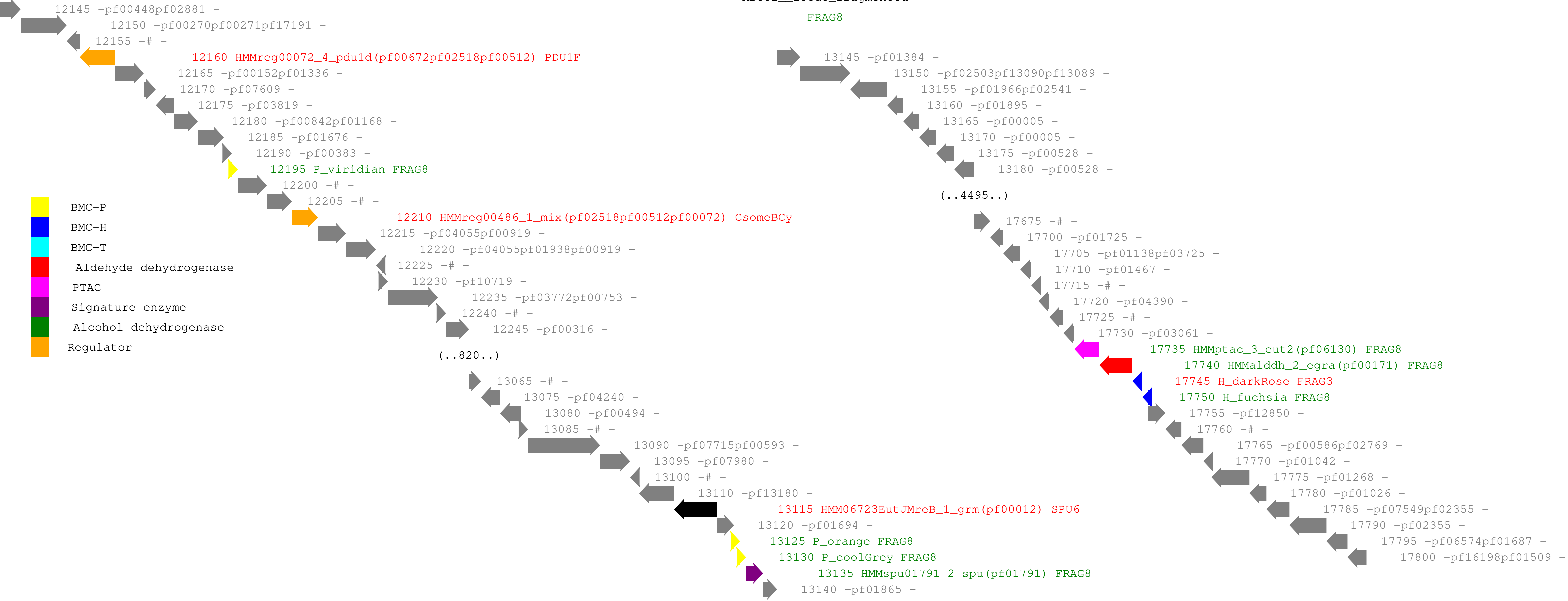

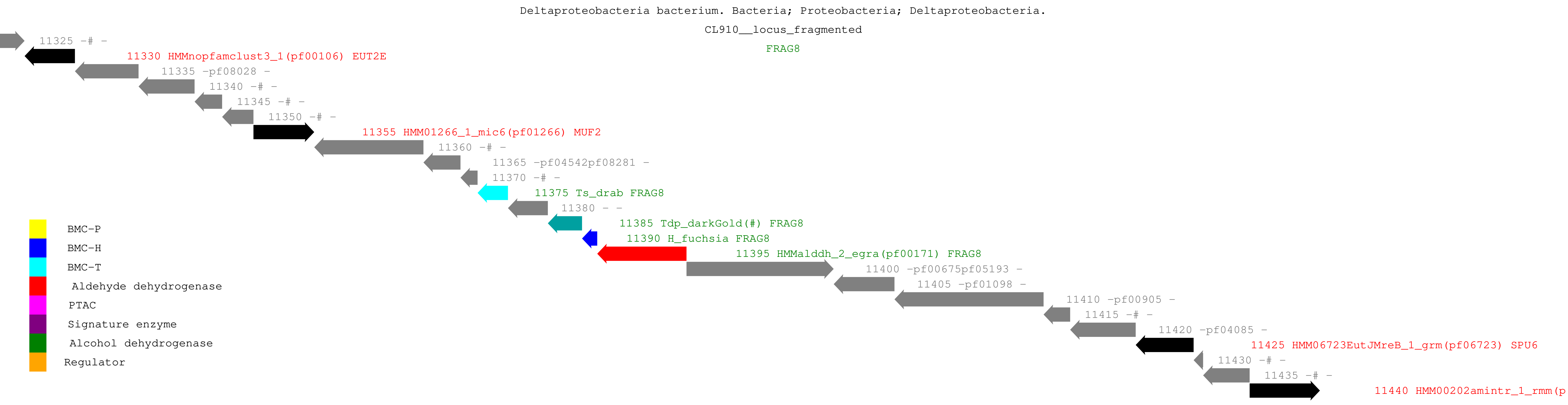
Deltaproteobacteria bacterium. Bacteria; Proteobacteria; Deltaproteobacteria.
CL910__locus_fragmented
FRAG8
11325 -# -
11330 HMMnopfamclust3_1(pf00106) EUT2E
11335 -pf08028 -
11340 -# -
11345 -# -
11350 -# -
11355 HMM01266_1_mic6(pf01266) MUF2
11360 -# -
11365 -pf04542pf08281 -
11370 -# -
11375 Ts_drab FRAG8
11380 - -
11385 Tdp_darkGold(#) FRAG8
11390 H_fuchsia FRAG8
11395 HMMalddh_2_egra(pf00171) FRAG8
11400 -pf00675pf05193 -
11405 -pf01098 -
11410 -pf00905 -
11415 -# -
11420 -pf04085 -
11425 HMM06723EutJMreB_1_grm(pf06723) SPU6
11430 -# -
11435 -# -
11440 HMM00202amintr_1_rmm(p
BMC-P
BMC-H
BMC-T
Aldehyde dehydrogenase
PTAC
Signature enzyme
Alcohol dehydrogenase
Regulator

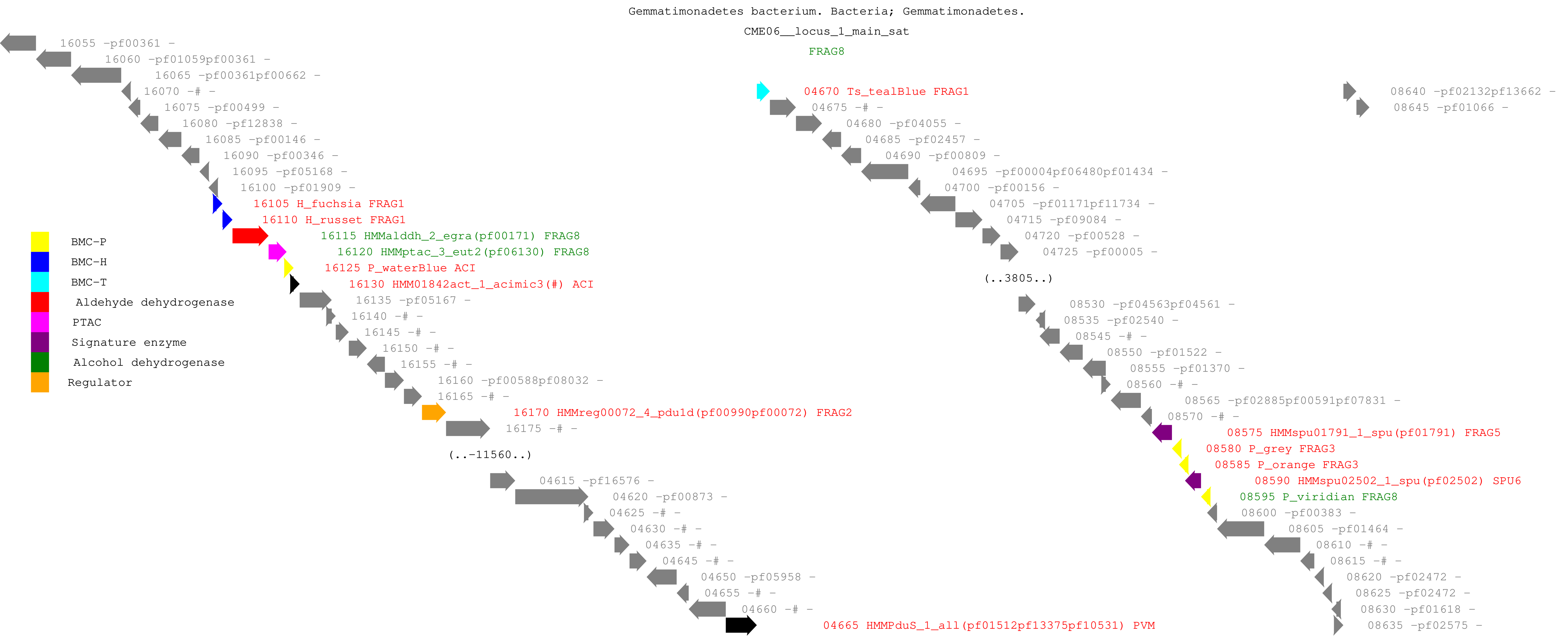
Gemmatimonadetes bacterium. Bacteria; Gemmatimonadetes.
CME06__locus_1_main_sat
FRAG8
16055 -pf00361 -
16060 -pf01059pf00361 -
16065 -pf00361pf00662 -
16070 -# -
16075 -pf00499 -
16080 -pf12838 -
16085 -pf00146 -
16090 -pf00346 -
16095 -pf05168 -
16100 -pf01909 -
16105 H_fuchsia FRAG1
16110 H_russet FRAG1
16115 HMMalddh_2_egra(pf00171) FRAG8
16120 HMMptac_3_eut2(pf06130) FRAG8
16125 P_waterBlue ACI
16130 HMM01842act_1_acimic3(#) ACI
16135 -pf05167 -
16140 -# -
16145 -# -
16150 -# -
16155 -# -
16160 -pf00588pf08032 -
16165 -# -
16170 HMMreg00072_4_pduld(pf00990pf00072) FRAG2
16175 -# -
(..-11560..)
04615 -pf16576 -
04620 -pf00873 -
04625 -# -
04630 -# -
04635 -# -
04645 -# -
04650 -pf05958 -
04655 -# -
04660 -# -
04665 HMMPduS_1_all(pf01512pf13375pf10531) PVM
04670 Ts_tealBlue FRAG1
04675 -# -
04680 -pf04055 -
04685 -pf02457 -
04690 -pf00809 -
04695 -pf00004pf06480pf01434 -
04700 -pf00156 -
04705 -pf01171pf11734 -
04715 -pf09084 -
04720 -pf00528 -
04725 -pf00005 -
(..3805..)
08530 -pf04563pf04561 -
08535 -pf02540 -
08545 -# -
08550 -pf01522 -
08555 -pf01370 -
08560 -# -
08565 -pf02885pf00591pf07831 -
08570 -# -
08575 HMMspu01791_1_spu(pf01791) FRAG5
08580 P_grey FRAG3
08585 P_orange FRAG3
08590 HMMspu02502_1_spu(pf02502) SPU6
08595 P_viridian FRAG8
08600 -pf00383 -
08605 -pf01464 -
08610 -# -
08615 -# -
08620 -pf02472 -
08625 -pf02472 -
08630 -pf01618 -
08635 -pf02575 -
08640 -pf02132pf13662 -
08645 -pf01066 -
BMC-P
BMC-H
BMC-T
Aldehyde dehydrogenase
PTAC
Signature enzyme
Alcohol dehydrogenase
Regulator

Planctomycetes bacterium. Bacteria; Planctomycetes.

CMJ90__locus_fragmented

FRAG8

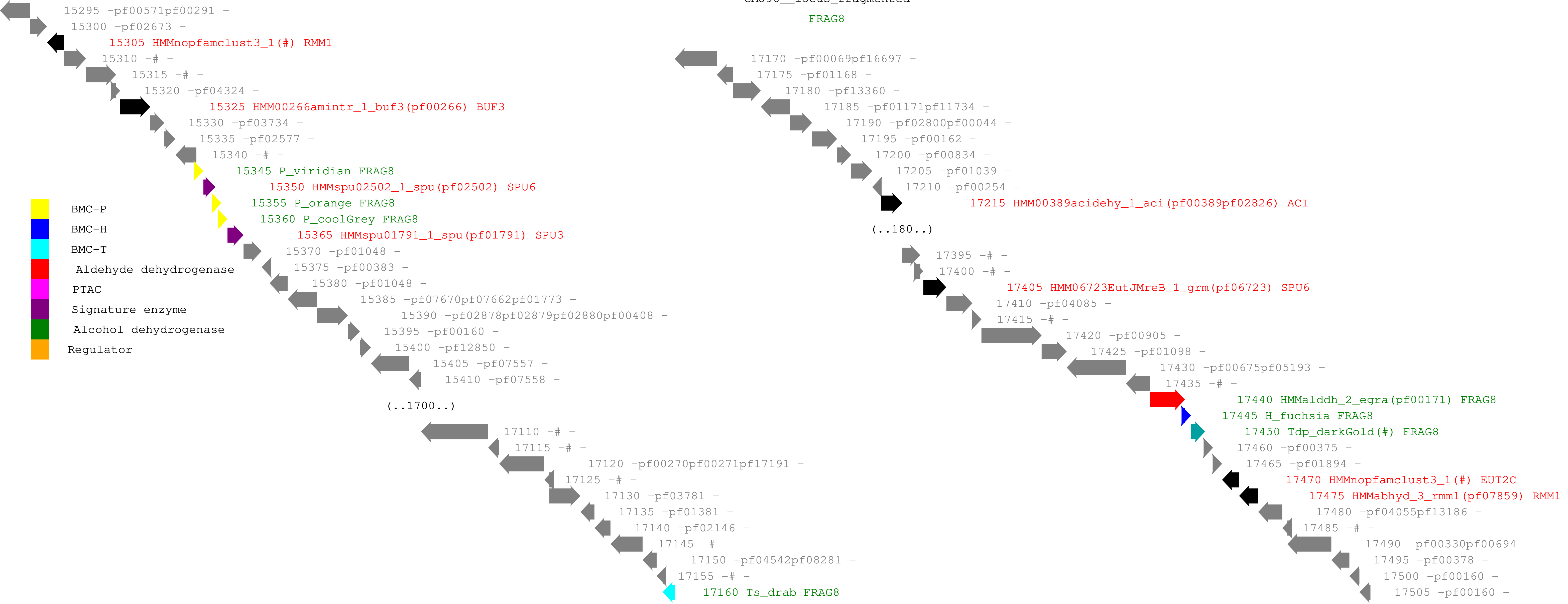

Lachnoclostridium sp. YL32. Bacteria; Firmicutes; Clostridia; Clostridiales; Lachnospiraceae.

A4V08__locus_2

GRM1A

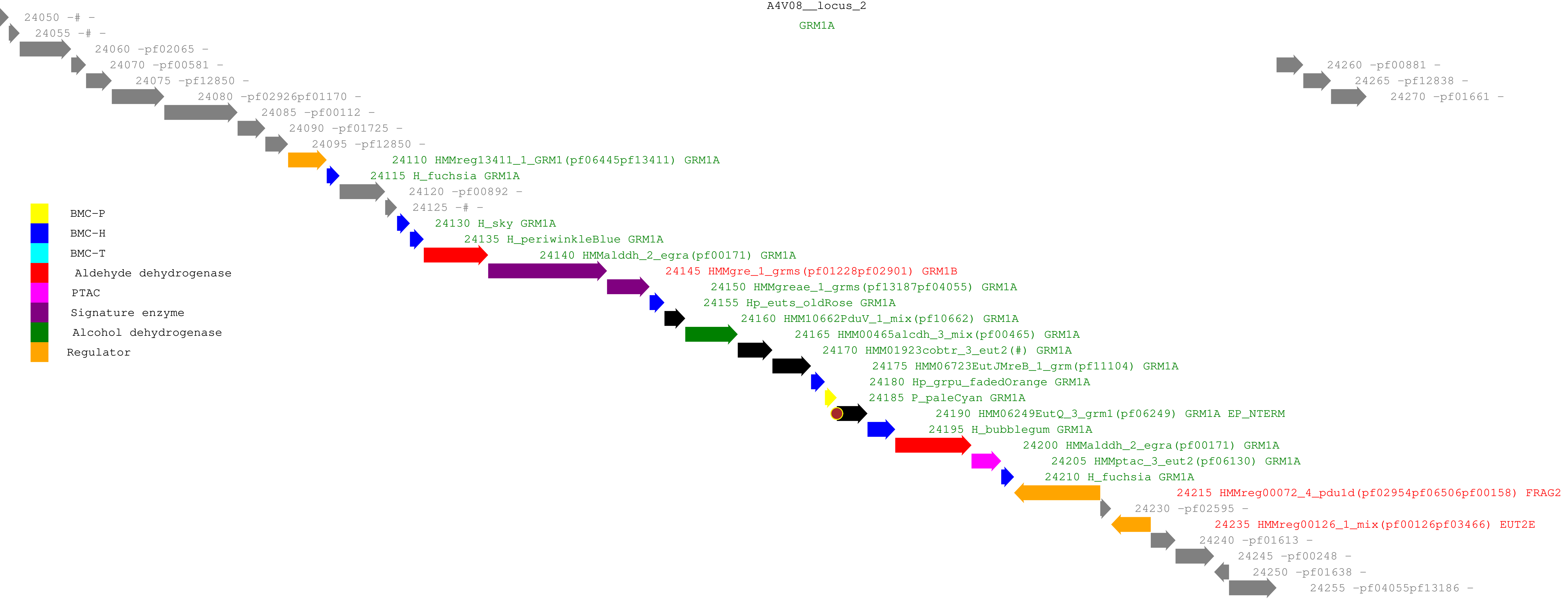

Clostridium sartagoforme AAU1. Bacteria; Firmicutes; Clostridia; Clostridiales; Clostridiaceae;

A500__locus_1

GRM1A

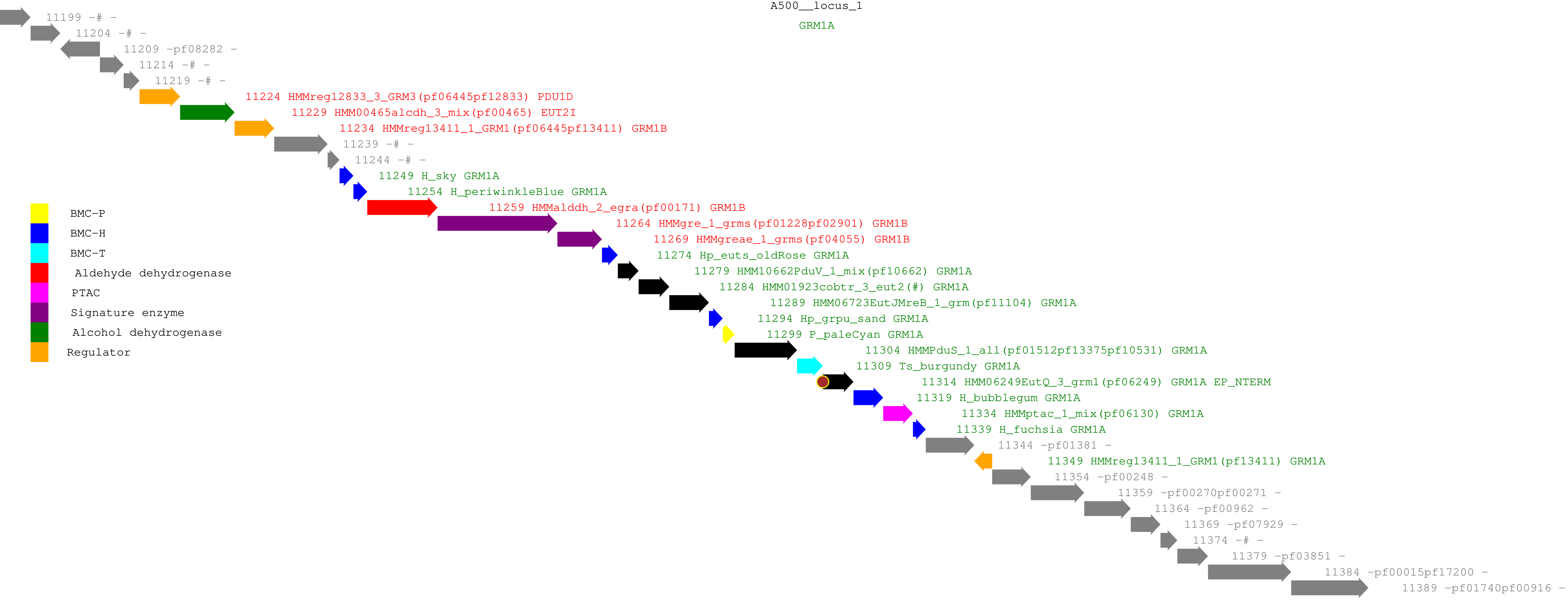

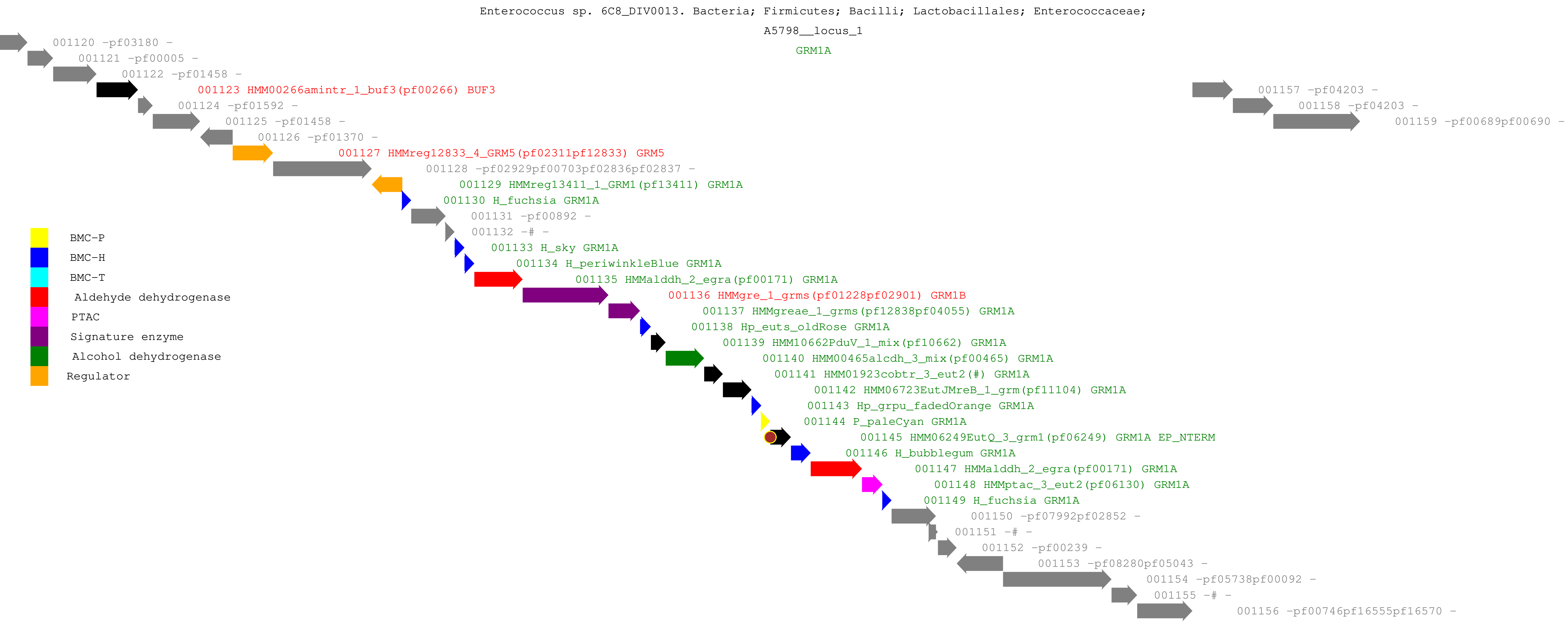
Enterococcus sp. 6C8_DIV0013. Bacteria; Firmicutes; Bacilli; Lactobacillales; Enterococcaceae;
A5798__locus_1
GRM1A
001120 -pf03180 -
001121 -pf00005 -
001122 -pf01458 -
001123 HMM00266amintr_1_buf3(pf00266) BUF3
001124 -pf01592 -
001125 -pf01458 -
001126 -pf01370 -
001127 HMMreg12833_4_GRM5(pf02311pf12833) GRM5
001128 -pf02929pf00703pf02836pf02837 -
001129 HMMreg13411_1_GRM1(pf13411) GRM1A
001130 H_fuchsia GRM1A
001131 -pf00892 -
001132 -# -
001133 H_sky GRM1A
001134 H_periwinkleBlue GRM1A
001135 HMMalddh_2_egra(pf00171) GRM1A
001136 HMMgre_1_grms(pf01228pf02901) GRM1B
001137 HMMgreae_1_grms(pf12838pf04055) GRM1A
001138 Hp_euts_oldRose GRM1A
001139 HMM10662PduV_1_mix(pf10662) GRM1A
001140 HMM00465alcdh_3_mix(pf00465) GRM1A
001141 HMM01923cobtr_3_eut2(#) GRM1A
001142 HMM06723EutJMreB_1_grm(pf11104) GRM1A
001143 Hp_grpu_fadedOrange GRM1A
001144 P_paleCyan GRM1A
001145 HMM06249EutQ_3_grm1(pf06249) GRM1A EP_NTERM
001146 H_bubblegum GRM1A
001147 HMMalddh_2_egra(pf00171) GRM1A
001148 HMMptac_3_eut2(pf06130) GRM1A
001149 H_fuchsia GRM1A
001150 -pf07992pf02852 -
001151 -# -
001152 -pf00239 -
001153 -pf08280pf05043 -
001154 -pf05738pf00092 -
001155 -# -
001156 -pf00746pf16555pf16570 -
001157 -pf04203 -
001158 -pf04203 -
001159 -pf00689pf00690 -
BMC-P
BMC-H
BMC-T
Aldehyde dehydrogenase
PTAC
Signature enzyme
Alcohol dehydrogenase
Regulator

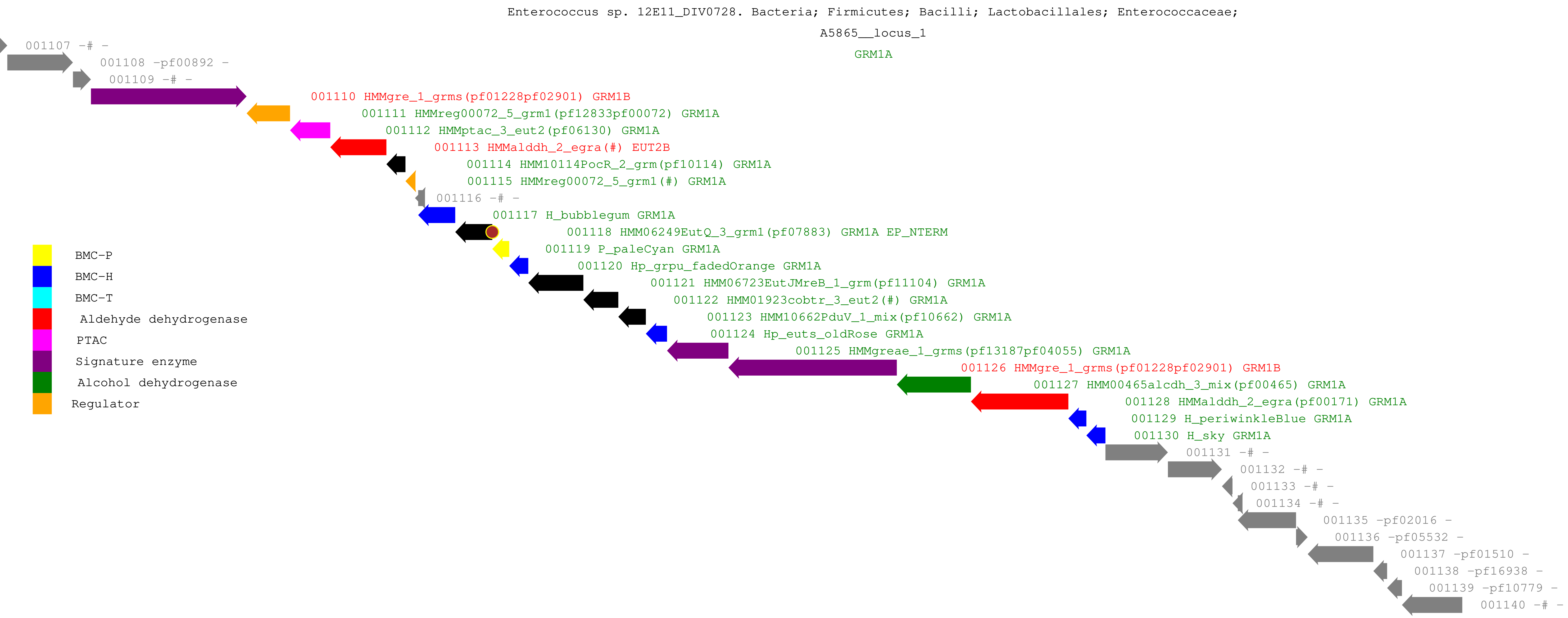
Enterococcus sp. 12E11_DIV0728. Bacteria; Firmicutes; Bacilli; Lactobacillales; Enterococcaceae;
A5865__locus_1
GRM1A
001107 -# -
001108 -pf00892 -
001109 -# -
001110 HMMgre_1_grms(pf01228pf02901) GRM1B
001111 HMMreg00072_5_grm1(pf12833pf00072) GRM1A
001112 HMMptac_3_eut2(pf06130) GRM1A
001113 HMMalddh_2_egra(#) EUT2B
001114 HMM10114PocR_2_grm(pf10114) GRM1A
001115 HMMreg00072_5_grm1(#) GRM1A
001116 -# -
001117 H_bubblegum GRM1A
001118 HMM06249EutQ_3_grm1(pf07883) GRM1A EP_NTERM
001119 P_paleCyan GRM1A
001120 Hp_grpu_fadedOrange GRM1A
001121 HMM06723EutJMreB_1_grm(pf11104) GRM1A
001122 HMM01923cobtr_3_eut2(#) GRM1A
001123 HMM10662PduV_1_mix(pf10662) GRM1A
001124 Hp_euts_oldRose GRM1A
001125 HMMgreae_1_grms(pf13187pf04055) GRM1A
001126 HMMgre_1_grms(pf01228pf02901) GRM1B
001127 HMM00465alcdh_3_mix(pf00465) GRM1A
001128 HMMaldd h_2_egra(pf00171) GRM1A
001129 H_periwinkleBlue GRM1A
001130 H_sky GRM1A
001131 -# -
001132 -# -
001133 -# -
001134 -# -
001135 -pf02016 -
001136 -pf05532 -
001137 -pf01510 -
001138 -pf16938 -
001139 -pf10779 -
001140 -# -
BMC-P
BMC-H
BMC-T
Aldehyde dehydrogenase
PTAC
Signature enzyme
Alcohol dehydrogenase
Regulator

Enterococcus sp. 12F9_DIV0723. Bacteria; Firmicutes; Bacilli; Lactobacillales; Enterococcaceae;

A5868__locus_1

GRM1A

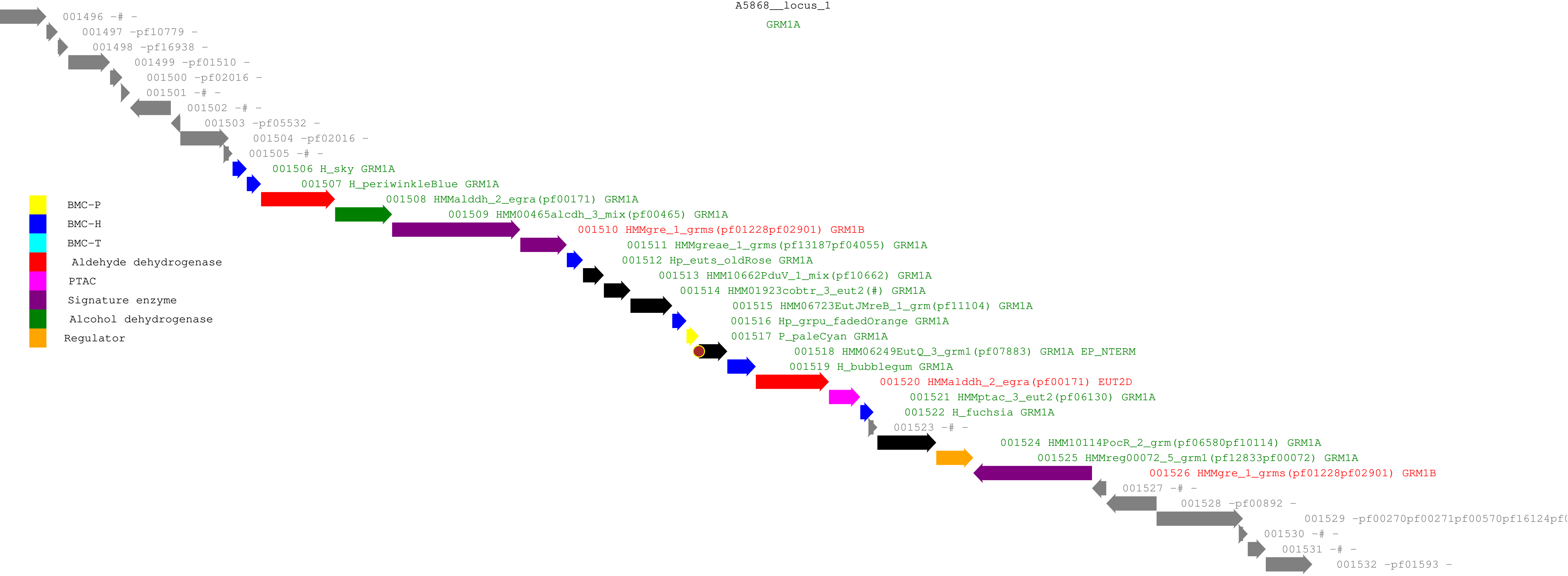

Enterococcus sp. 3H8_DIV0648. Bacteria; Firmicutes; Bacilli; Lactobacillales; Enterococcaceae;

A5875__locus_2

GRM1A

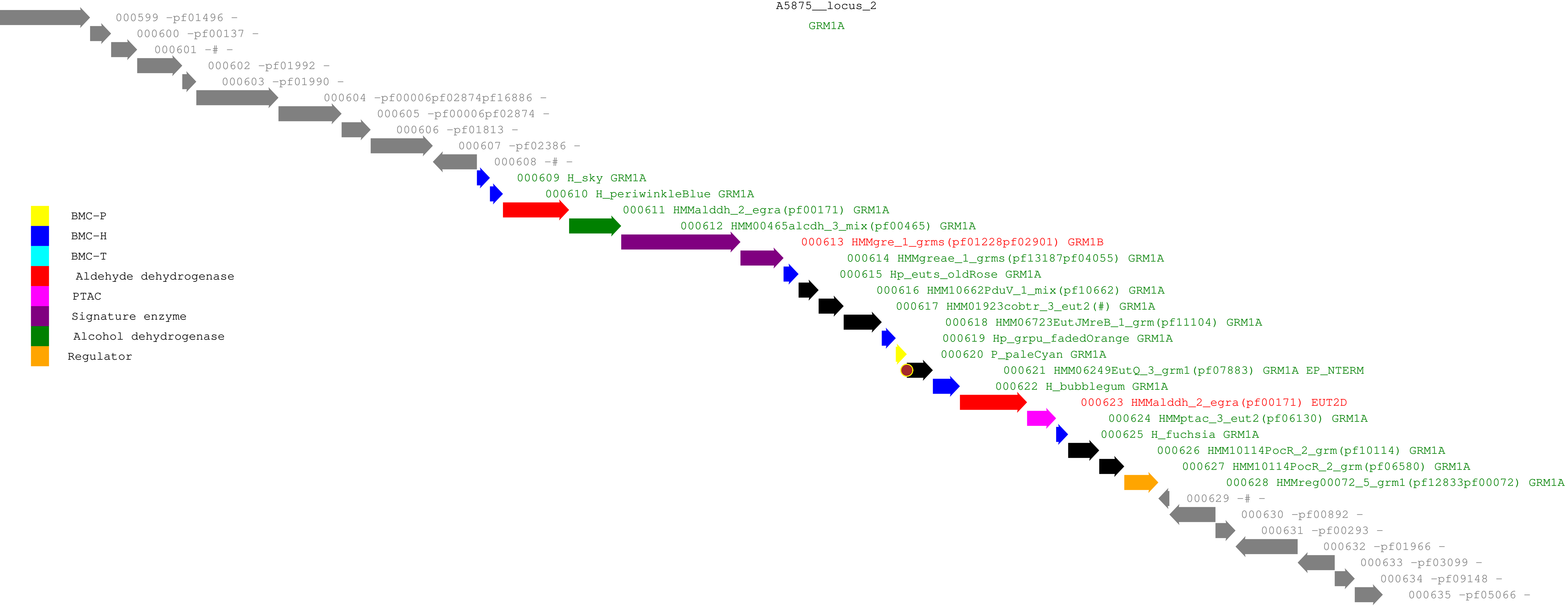

Enterococcus termitis. Bacteria; Firmicutes; Bacilli; Lactobacillales; Enterococcaceae;

A5881__locus_1

GRM1A

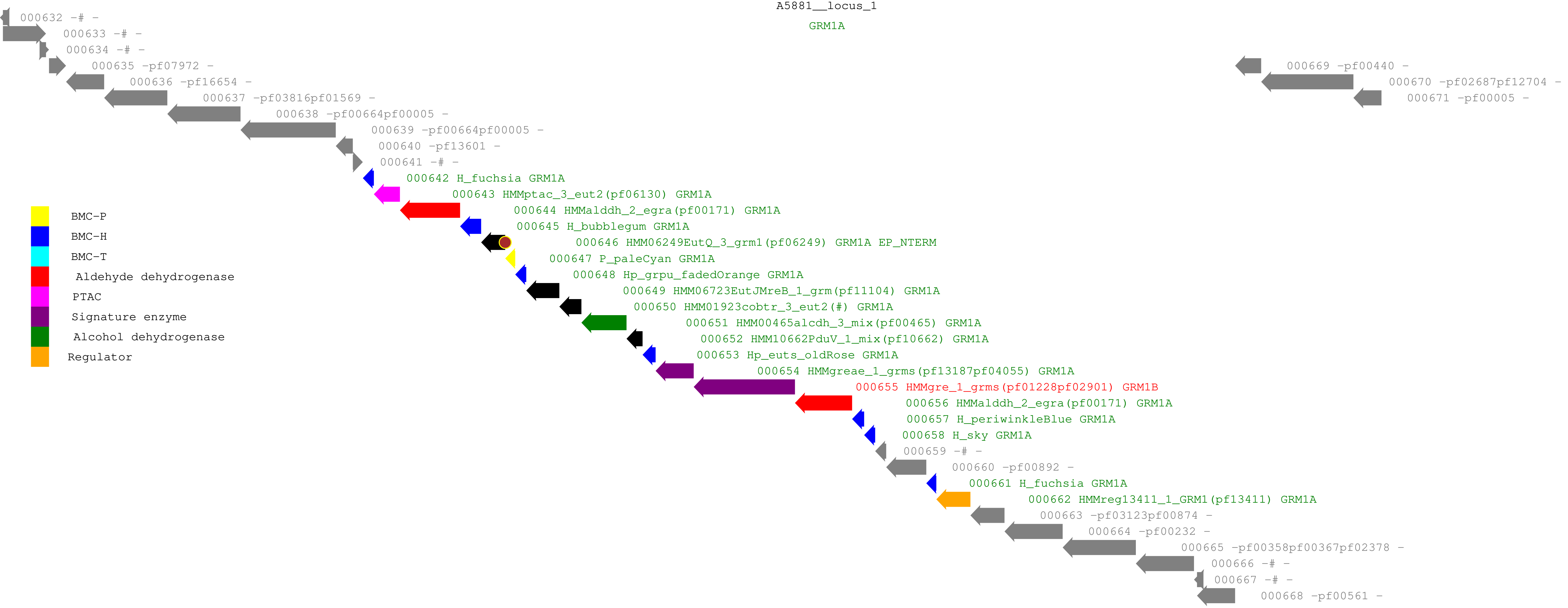

Enterococcus sp. 9E7_DIV0242. Bacteria; Firmicutes; Bacilli; Lactobacillales; Enterococcaceae;

A5888__locus_1_main_sat

GRM1A

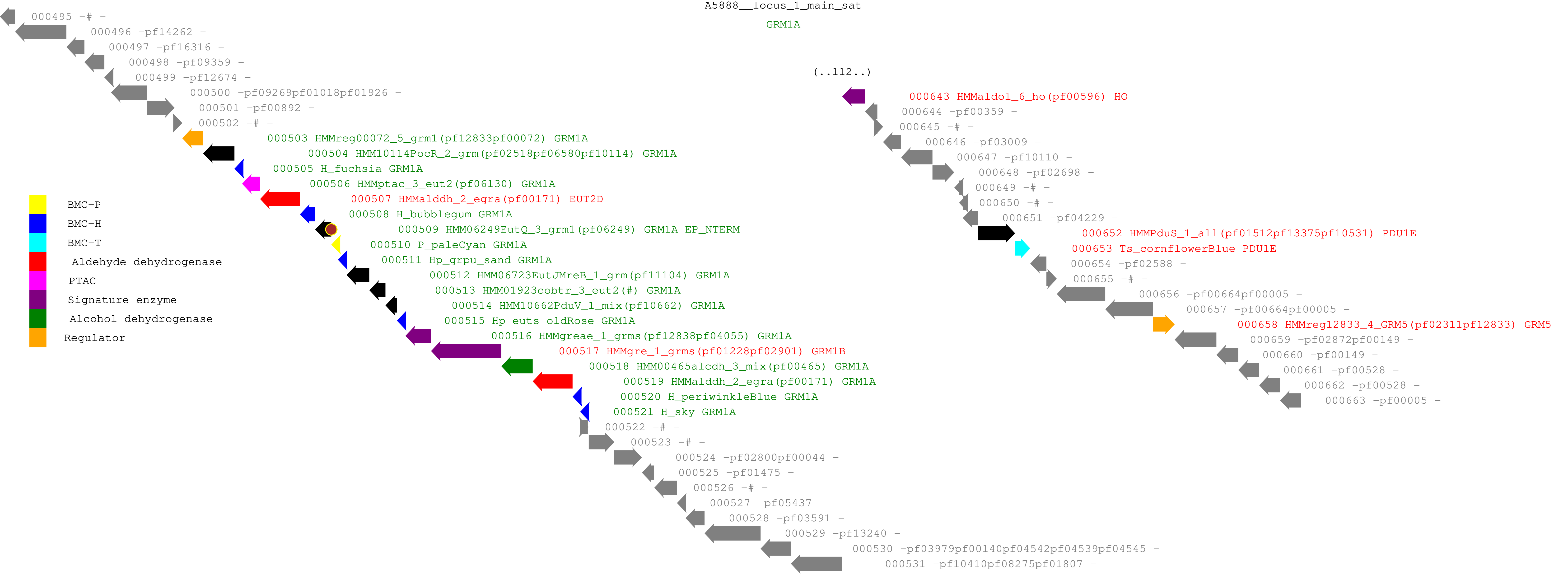

Enterococcus sp. 9D6_DIV0238. Bacteria; Firmicutes; Bacilli; Lactobacillales; Enterococcaceae;

A5889__locus_1

GRM1A

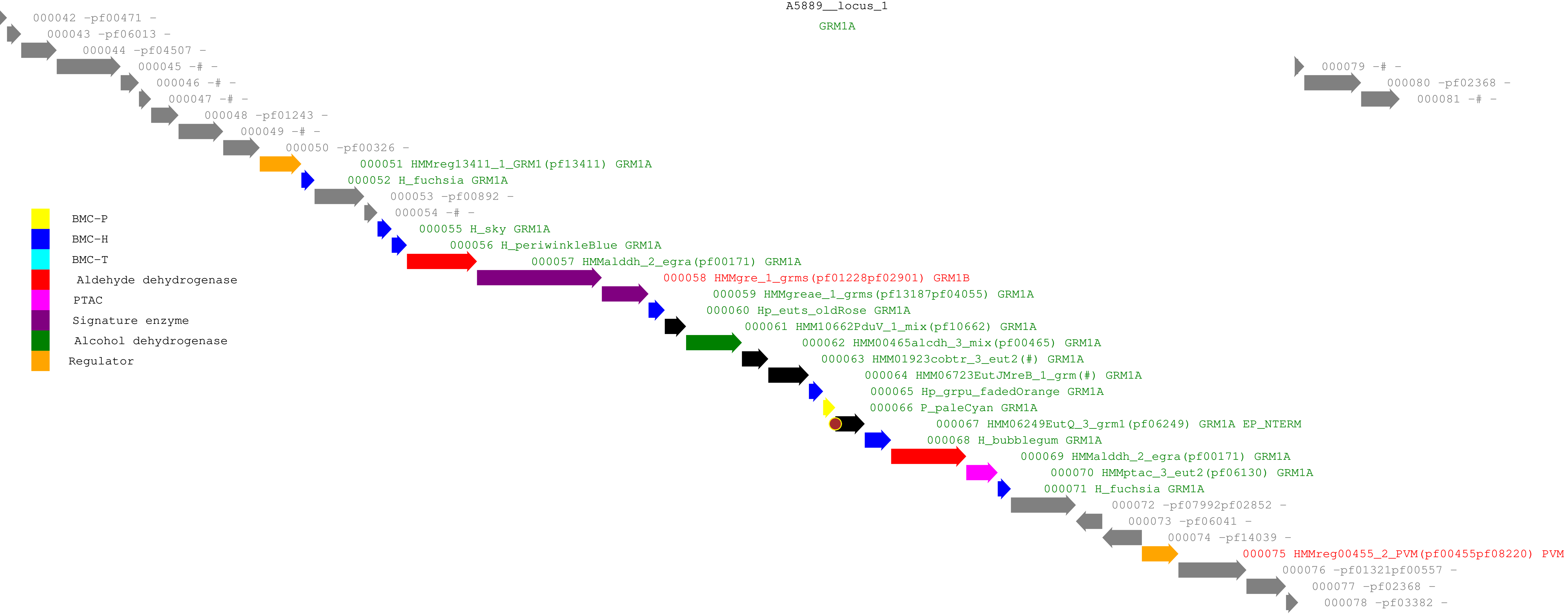

Clostridium sp. C8. Bacteria; Firmicutes; Clostridia; Clostridiales; Clostridiaceae;

AAT22__locus_1

GRM1A

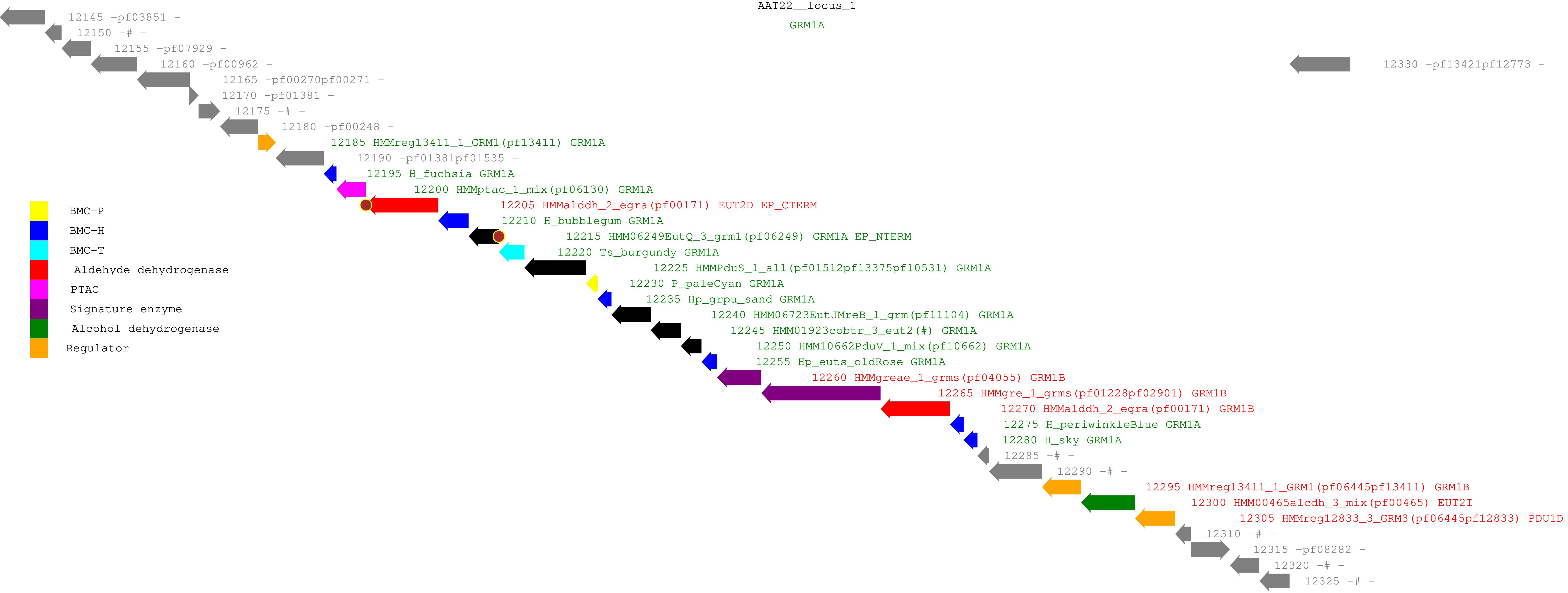

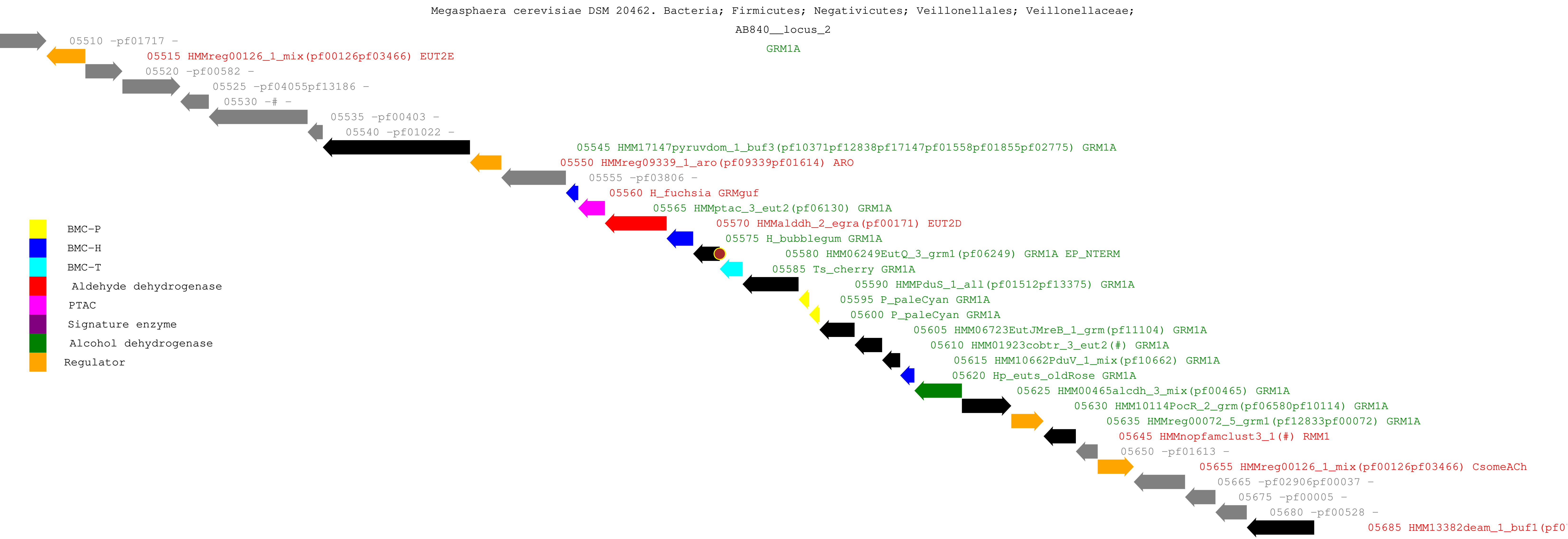
Megasphaera cerevisiae DSM 20462. Bacteria; Firmicutes; Negativicutes; Veillonellales; Veillonellaceae;
AB840__locus_2
GRM1A
05510 -pf01717 -
05515 HMMreg00126_1_mix(pf00126pf03466) EUT2E
05520 -pf00582 -
05525 -pf04055pf13186 -
05530 -# -
05535 -pf00403 -
05540 -pf01022 -
05545 HMM17147pyruvdom_1_buf3(pf10371pf12838pf17147pf01558pf01855pf02775) GRM1A
05550 HMMreg09339_1_aro(pf09339pf01614) ARO
05555 -pf03806 -
05560 H_fuchsia GRMguf
05565 HMMptac_3_eut2(pf06130) GRM1A
05570 HMMalddh_2_egra(pf00171) EUT2D
05575 H_bubblegum GRM1A
05580 HMM06249EutQ_3_grm1(pf06249) GRM1A EP_NTERM
05585 Ts_cherry GRM1A
05590 HMMPduS_1_all(pf01512pf13375) GRM1A
05595 P_paleCyan GRM1A
05600 P_paleCyan GRM1A
05605 HMM06723EutJMreB_1_grm(pf11104) GRM1A
05610 HMM01923cobtr_3_eut2(#) GRM1A
05615 HMM10662PduV_1_mix(pf10662) GRM1A
05620 Hp_euts_oldRose GRM1A
05625 HMM00465alcdh_3_mix(pf00465) GRM1A
05630 HMM10114PocR_2_grm(pf06580pf10114) GRM1A
05635 HMMreg00072_5_grm1(pf12833pf00072) GRM1A
05645 HMMnopfamclust3_1(#) RMM1
05650 -pf01613 -
05655 HMMreg00126_1_mix(pf00126pf03466) CsomeACh
05665 -pf02906pf00037 -
05675 -pf00005 -
05680 -pf00528 -
05685 HMM13382deam_1_buf1(pf0
BMC-P
BMC-H
BMC-T
Aldehyde dehydrogenase
PTAC
Signature enzyme
Alcohol dehydrogenase
Regulator

Clostridium botulinum. Bacteria; Firmicutes; Clostridia; Clostridiales; Clostridiaceae;

ACP50__locus_2

GRM1A

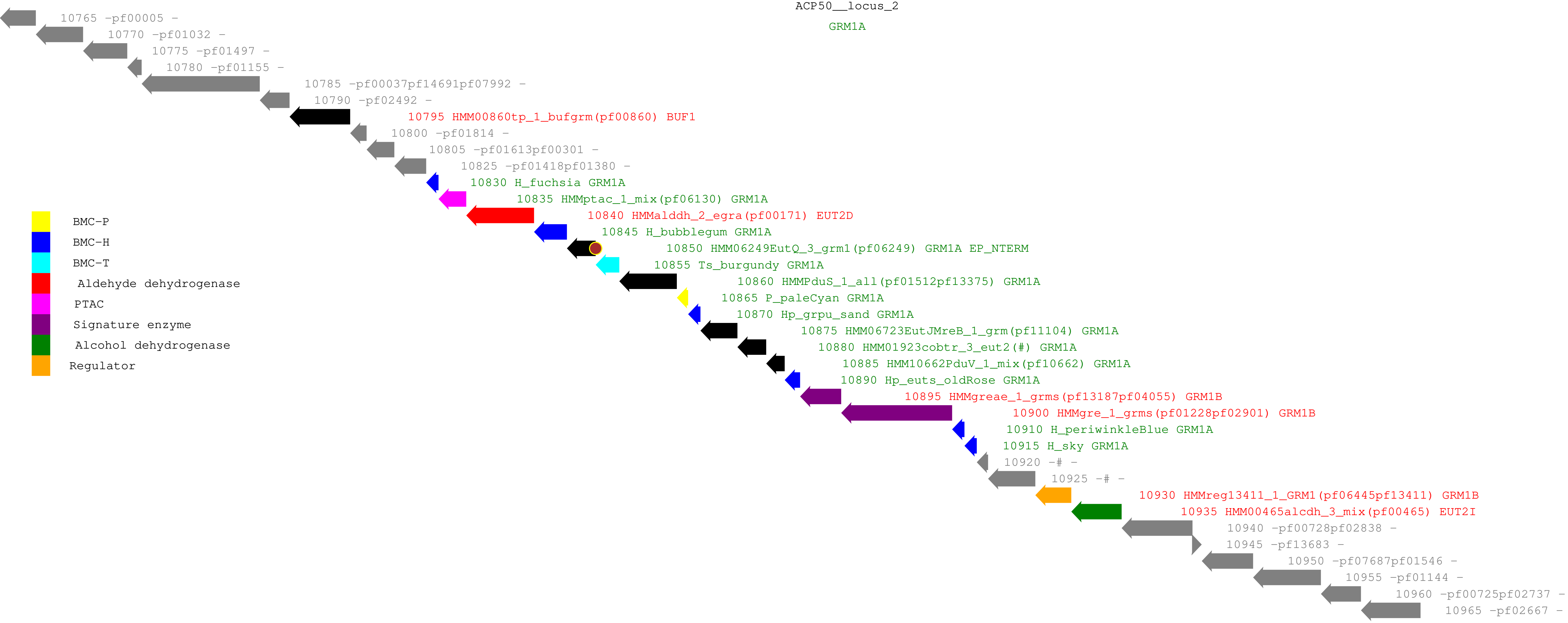

Clostridium botulinum. Bacteria; Firmicutes; Clostridia; Clostridiales; Clostridiaceae;

ACP52__locus_1

GRM1A

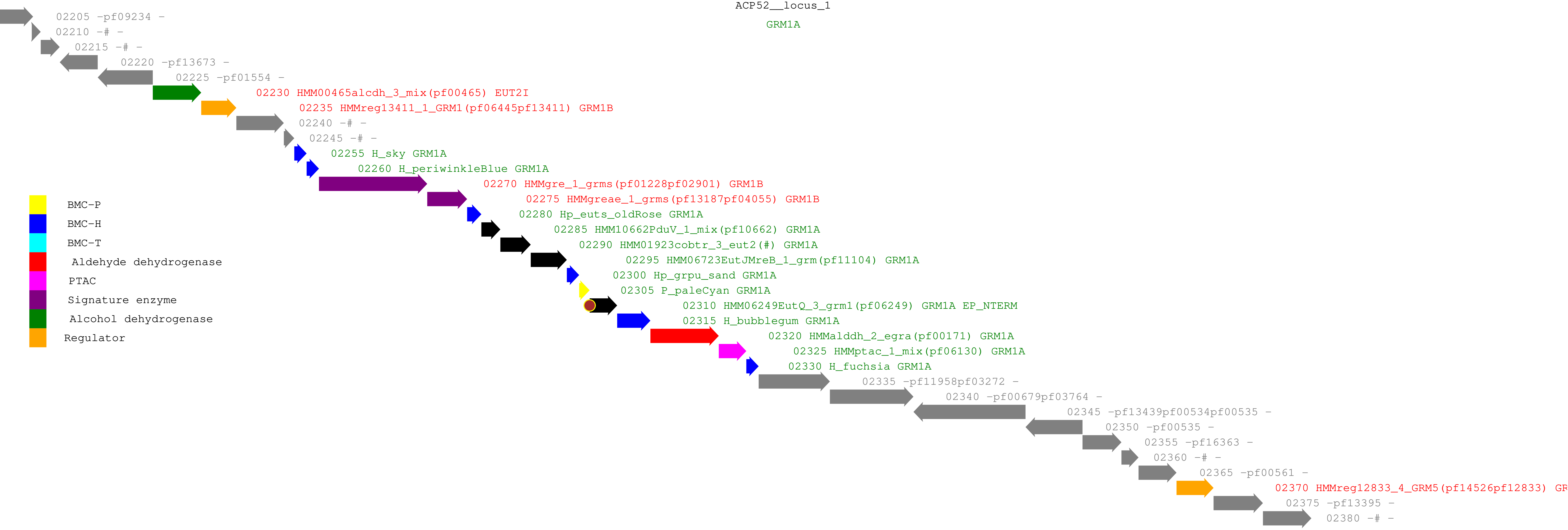

Megasphaera hexanoica. Bacteria; Firmicutes; Negativicutes; Veillonellales; Veillonellaceae;

ACT01__locus_1

GRM1A

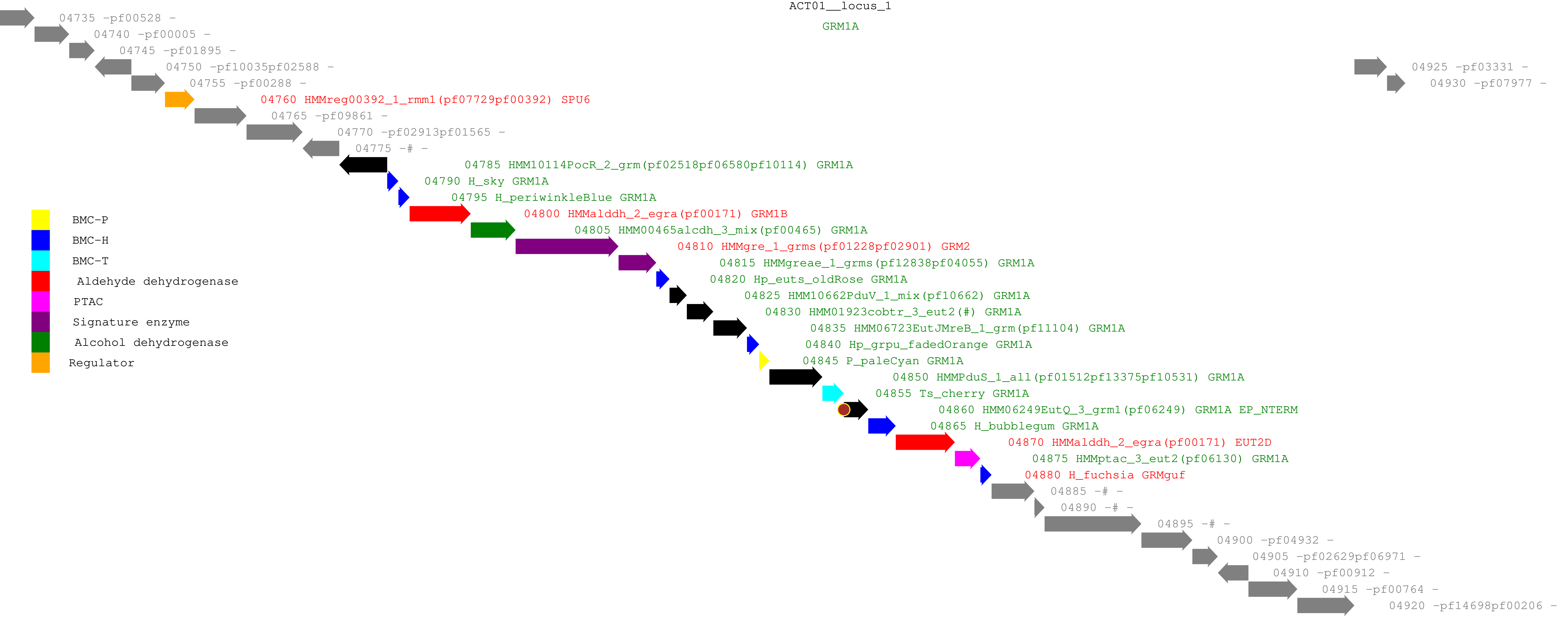

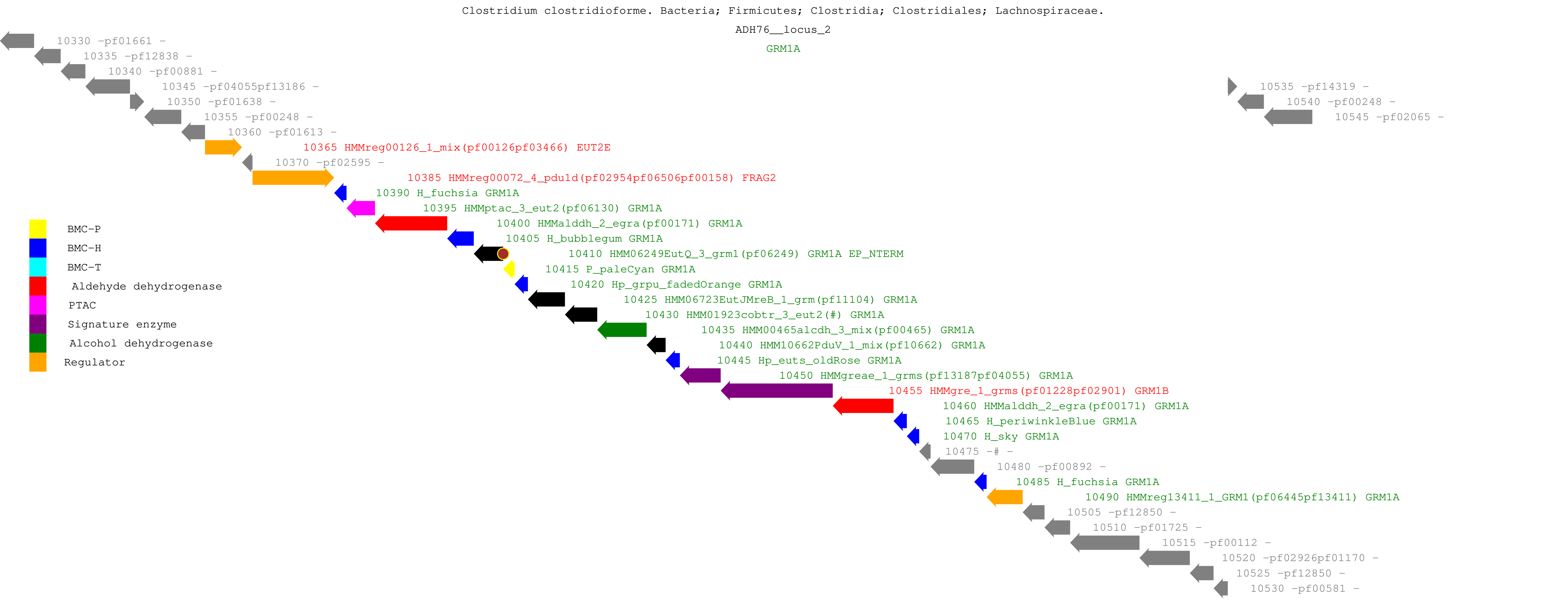

Clostridium clostridioforme. Bacteria; Firmicutes; Clostridia; Clostridiales; Lachnospiraceae.
ADH76__locus_2
GRM1A
10330 -pf01661 -
10335 -pf12838 -
10340 -pf00881 -
10345 -pf04055pf13186 -
10350 -pf01638 -
10355 -pf00248 -
10360 -pf01613 -
10365 HMMreg00126_1_mix(pf00126pf03466) EUT2E
10370 -pf02595 -
10385 HMMreg00072_4_pdu1d(pf02954pf06506pf00158) FRAG2
10390 H_fuchsia GRM1A
10395 HMMptac_3_eut2(pf06130) GRM1A
10400 HMMaldh_2_egra(pf00171) GRM1A
10405 H_bubblegum GRM1A
10410 HMM06249EutQ_3_grm1(pf06249) GRM1A EP_NTERM
10415 P_paleCyan GRM1A
10420 Hp_grpu_fadedOrange GRM1A
10425 HMM06723EutJMreB_1_grm(pf11104) GRM1A
10430 HMM01923cobtr_3_eut2(#) GRM1A
10435 HMM00465alcdh_3_mix(pf00465) GRM1A
10440 HMM10662PduV_1_mix(pf10662) GRM1A
10445 Hp_euts_oldRose GRM1A
10450 HMMgreae_1_grms(pf13187pf04055) GRM1A
10455 HMMgre_1_grms(pf01228pf02901) GRM1B
10460 HMMaldh_2_egra(pf00171) GRM1A
10465 H_periwinkleBlue GRM1A
10470 H_sky GRM1A
10475 -# -
10480 -pf00892 -
10485 H_fuchsia GRM1A
10490 HMMreg13411_1_GRM1(pf06445pf13411) GRM1A
10505 -pf12850 -
10510 -pf01725 -
10515 -pf00112 -
10520 -pf02926pf01170 -
10525 -pf12850 -
10530 -pf00581 -
10535 -pf14319 -
10540 -pf00248 -
10545 -pf02065 -
BMC-P
BMC-H
BMC-T
Aldehyde dehydrogenase
PTAC
Signature enzyme
Alcohol dehydrogenase
Regulator

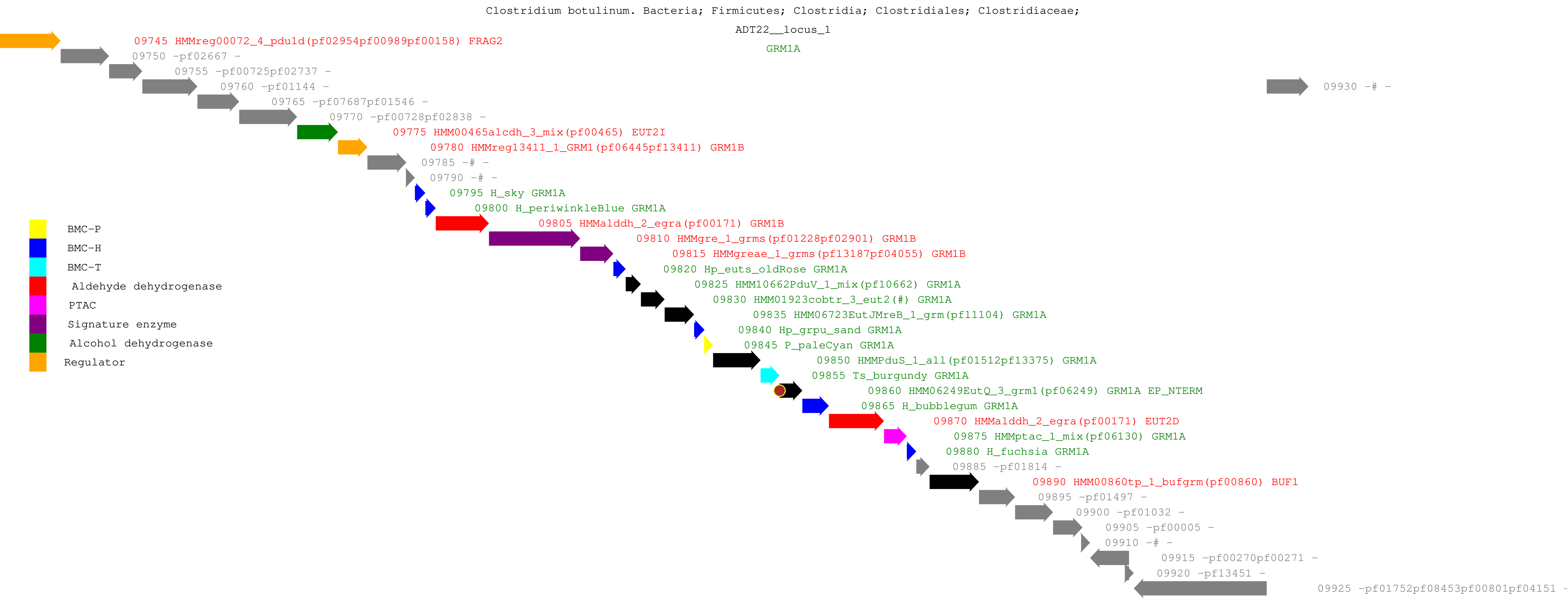
Clostridium botulinum. Bacteria; Firmicutes; Clostridia; Clostridiales; Clostridiaceae;
ADT22__locus_1
GRM1A
09745 HMMreg00072_4_pdu1d(pf02954pf00989pf00158) FRAG2
09750 -pf02667 -
09755 -pf00725pf02737 -
09760 -pf01144 -
09765 -pf07687pf01546 -
09770 -pf00728pf02838 -
09775 HMM00465alcdh_3_mix(pf00465) EUT2I
09780 HMMreg13411_1_GRM1(pf06445pf13411) GRM1B
09785 -# -
09790 -# -
09795 H_sky GRM1A
09800 H_periwinkleBlue GRM1A
09805 HMMalddh_2_egra(pf00171) GRM1B
09810 HMMgre_1_grms(pf01228pf02901) GRM1B
09815 HMMgreae_1_grms(pf13187pf04055) GRM1B
09820 Hp_euts_oldRose GRM1A
09825 HMM10662PduV_1_mix(pf10662) GRM1A
09830 HMM01923cobtr_3_eut2(#) GRM1A
09835 HMM06723EutJMreB_1_grm(pf11104) GRM1A
09840 Hp_grpu_sand GRM1A
09845 P_paleCyan GRM1A
09850 HMMPduS_1_all(pf01512pf13375) GRM1A
09855 Ts_burgundy GRM1A
09860 HMM06249EutQ_3_grm1(pf06249) GRM1A EP_NTERM
09865 H_bubblegum GRM1A
09870 HMMalddh_2_egra(pf00171) EUT2D
09875 HMMptac_1_mix(pf06130) GRM1A
09880 H_fuchsia GRM1A
09885 -pf01814 -
09890 HMM00860tp_1_bufgrm(pf00860) BUF1
09895 -pf01497 -
09900 -pf01032 -
09905 -pf00005 -
09910 -# -
09915 -pf00270pf00271 -
09920 -pf13451 -
09925 -pf01752pf08453pf00801pf04151 -
09930 -# -
BMC-P
BMC-H
BMC-T
Aldehyde dehydrogenase
PTAC
Signature enzyme
Alcohol dehydrogenase
Regulator

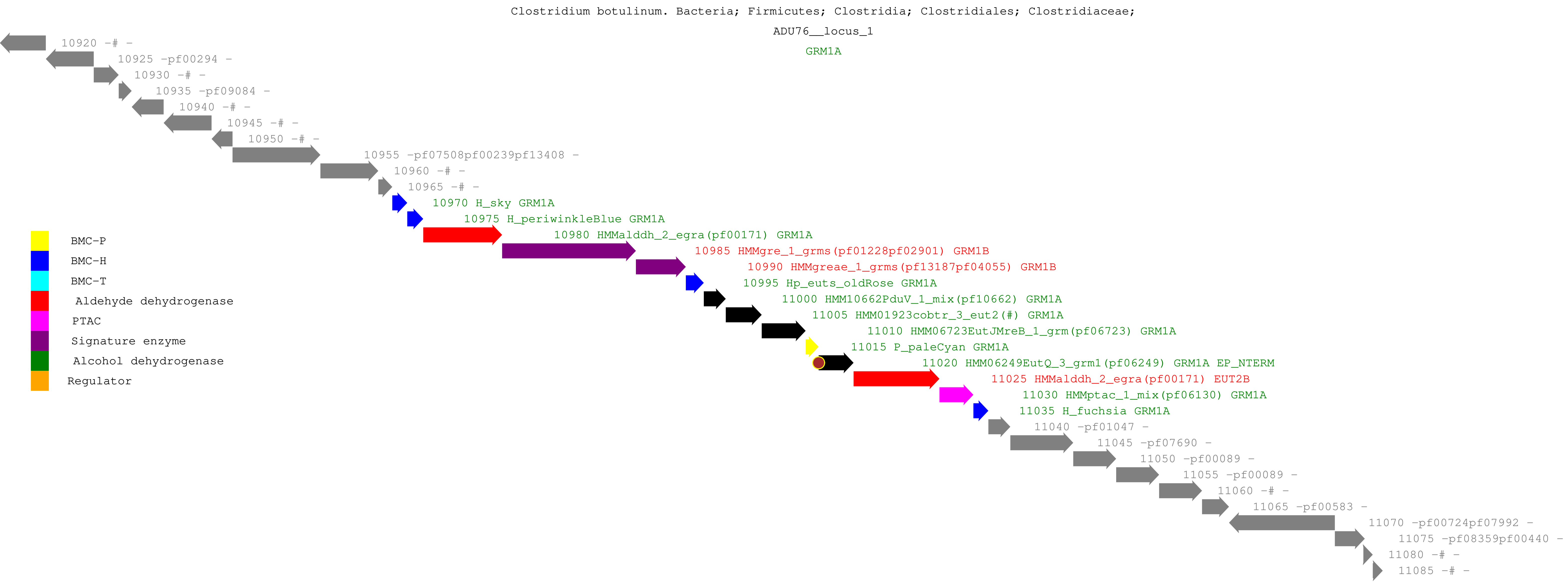
Clostridium botulinum. Bacteria; Firmicutes; Clostridia; Clostridiales; Clostridiaceae;
ADU76__locus_1
GRM1A
10920 -# -
10925 -pf00294 -
10930 -# -
10935 -pf09084 -
10940 -# -
10945 -# -
10950 -# -
10955 -pf07508pf00239pf13408 -
10960 -# -
10965 -# -
10970 H_sky GRM1A
10975 H_periwinkleBlue GRM1A
10980 HMMalddh_2_egra(pf00171) GRM1A
10985 HMMgre_1_grms(pf01228pf02901) GRM1B
10990 HMMgreae_1_grms(pf13187pf04055) GRM1B
10995 Hp_euts_oldRose GRM1A
11000 HMM10662PduV_1_mix(pf10662) GRM1A
11005 HMM01923cobtr_3_eut2(#) GRM1A
11010 HMM06723EutJMreB_1_grm(pf06723) GRM1A
11015 P_paleCyan GRM1A
11020 HMM06249EutQ_3_grm1(pf06249) GRM1A EP_NTERM
11025 HMMalddh_2_egra(pf00171) EUT2B
11030 HMMptac_1_mix(pf06130) GRM1A
11035 H_fuchsia GRM1A
11040 -pf01047 -
11045 -pf07690 -
11050 -pf00089 -
11055 -pf00089 -
11060 -# -
11065 -pf00583 -
11070 -pf00724pf07992 -
11075 -pf08359pf00440 -
11080 -# -
11085 -# -
BMC-P
BMC-H
BMC-T
Aldehyde dehydrogenase
PTAC
Signature enzyme
Alcohol dehydrogenase
Regulator

Alkaliphilus metalliredigens (strain QYMF). Bacteria; Firmicutes; Clostridia; Clostridiales; Clostridiaceae;

Amet__locus_2

GRM1A

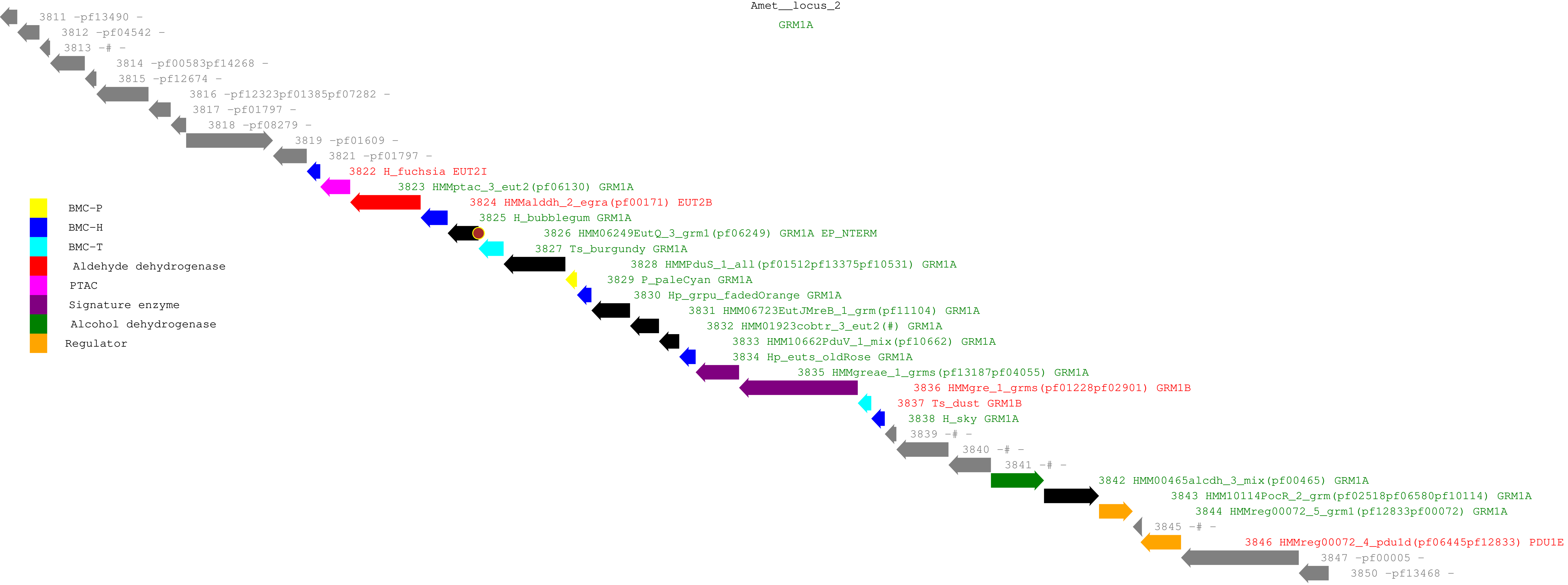

Anaerococcus hydrogenalis DSM 7454. Bacteria; Firmicutes; Tissierellia; Tissierellales; Peptoniphilaceae;

ANHYDRO__locus_1

GRM1A

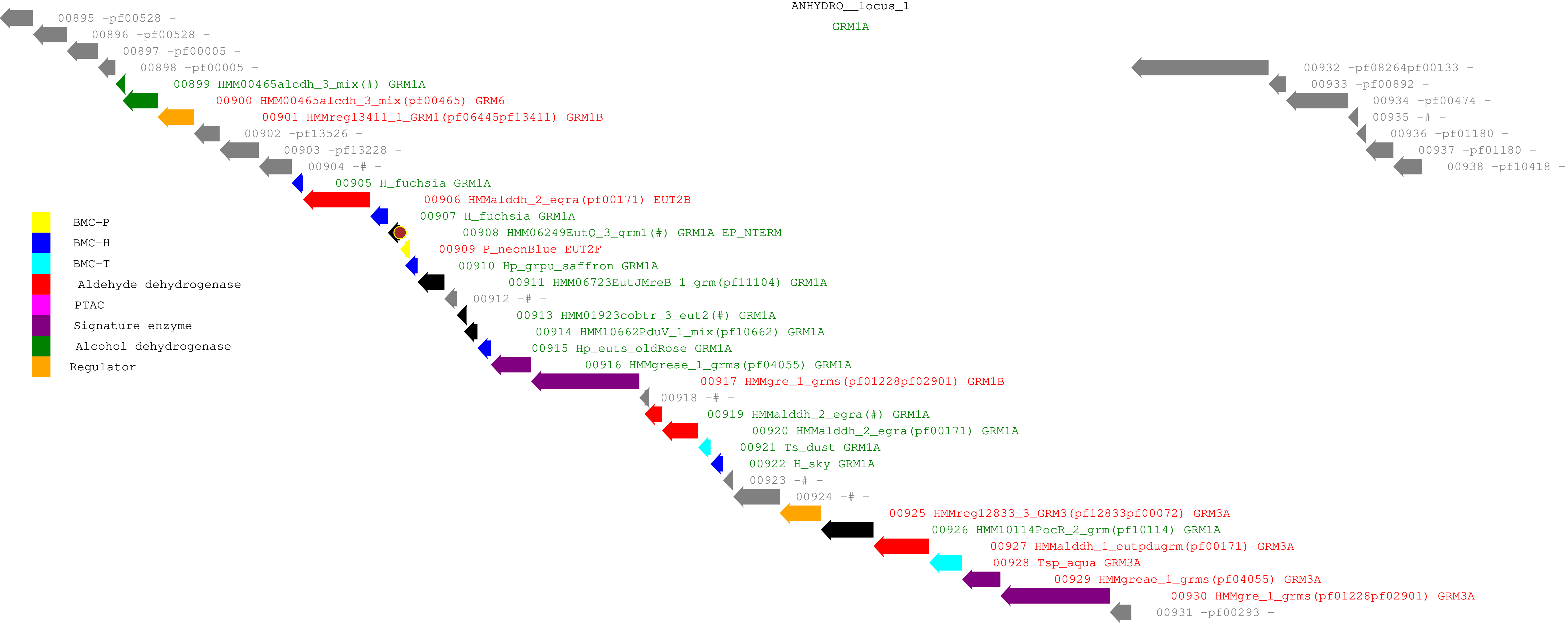

Collinsella aerofaciens. Bacteria; Actinobacteria; Coriobacteriia; Coriobacteriales;

AOBCEHNH__locus_1

GRM1A

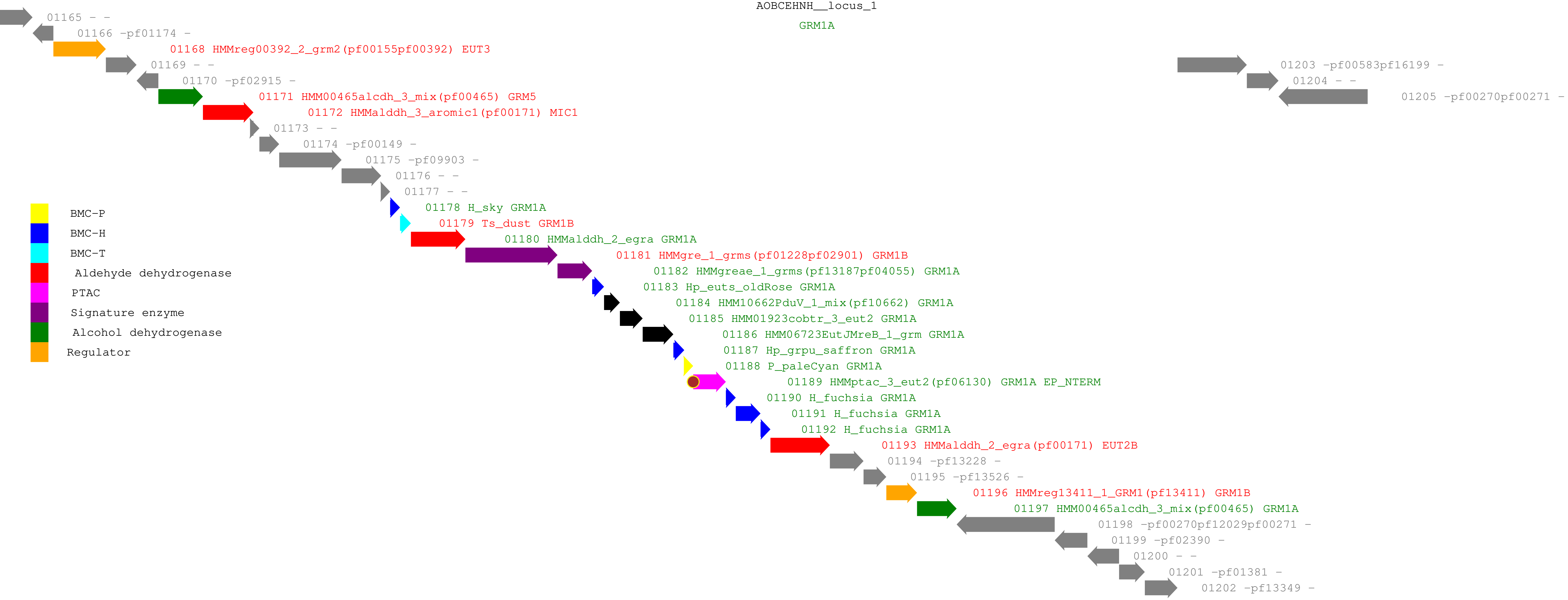

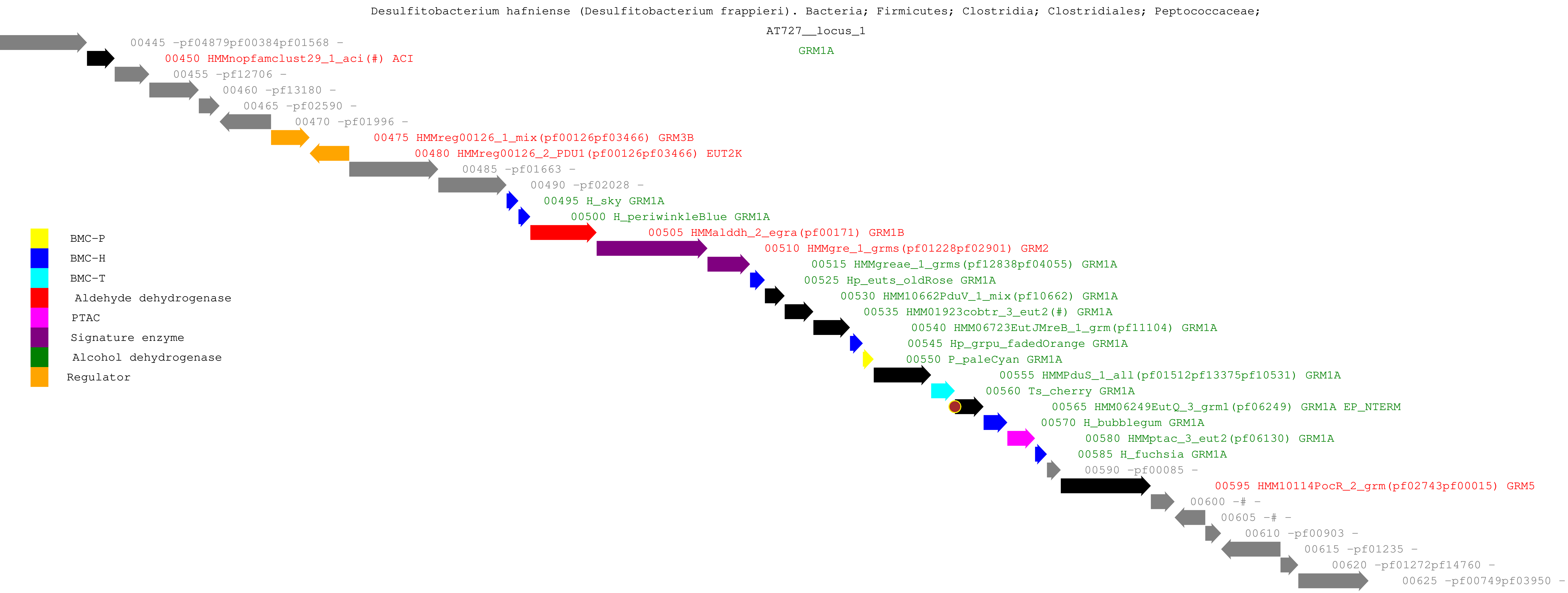
Desulfitobacterium hafniense (Desulfitobacterium frappieri). Bacteria; Firmicutes; Clostridia; Clostridiales; Peptococcaceae;
AT727__locus_1
GRM1A
00445 -pf04879pf00384pf01568 -
00450 HMMnopfamclust29_1_aci(#) ACI
00455 -pf12706 -
00460 -pf13180 -
00465 -pf02590 -
00470 -pf01996 -
00475 HMMreg00126_1_mix(pf00126pf03466) GRM3B
00480 HMMreg00126_2_PDU1(pf00126pf03466) EUT2K
00485 -pf01663 -
00490 -pf02028 -
00495 H_sky GRM1A
00500 H_periwinkleBlue GRM1A
00505 HMMalddh_2_egra(pf00171) GRM1B
00510 HMMgre_1_grms(pf01228pf02901) GRM2
00515 HMMgreae_1_grms(pf12838pf04055) GRM1A
00525 Hp_euts_oldRose GRM1A
00530 HMM10662PduV_1_mix(pf10662) GRM1A
00535 HMM01923cobtr_3_eut2(#) GRM1A
00540 HMM06723EutJMreB_1_grm(pf11104) GRM1A
00545 Hp_grpu_fadedOrange GRM1A
00550 P_paleCyan GRM1A
00555 HMMPduS_1_all(pf01512pf13375pf10531) GRM1A
00560 Ts_cherry GRM1A
00565 HMM06249EutQ_3_grm1(pf06249) GRM1A EP_NTERM
00570 H_bubblegum GRM1A
00580 HMMptac_3_eut2(pf06130) GRM1A
00585 H_fuchsia GRM1A
00590 -pf00085 -
00595 HMM10114PocR_2_grm(pf02743pf00015) GRM5
00600 -# -
00605 -# -
00610 -pf00903 -
00615 -pf01235 -
00620 -pf01272pf14760 -
00625 -pf00749pf03950 -
BMC-P
BMC-H
BMC-T
Aldehyde dehydrogenase
PTAC
Signature enzyme
Alcohol dehydrogenase
Regulator

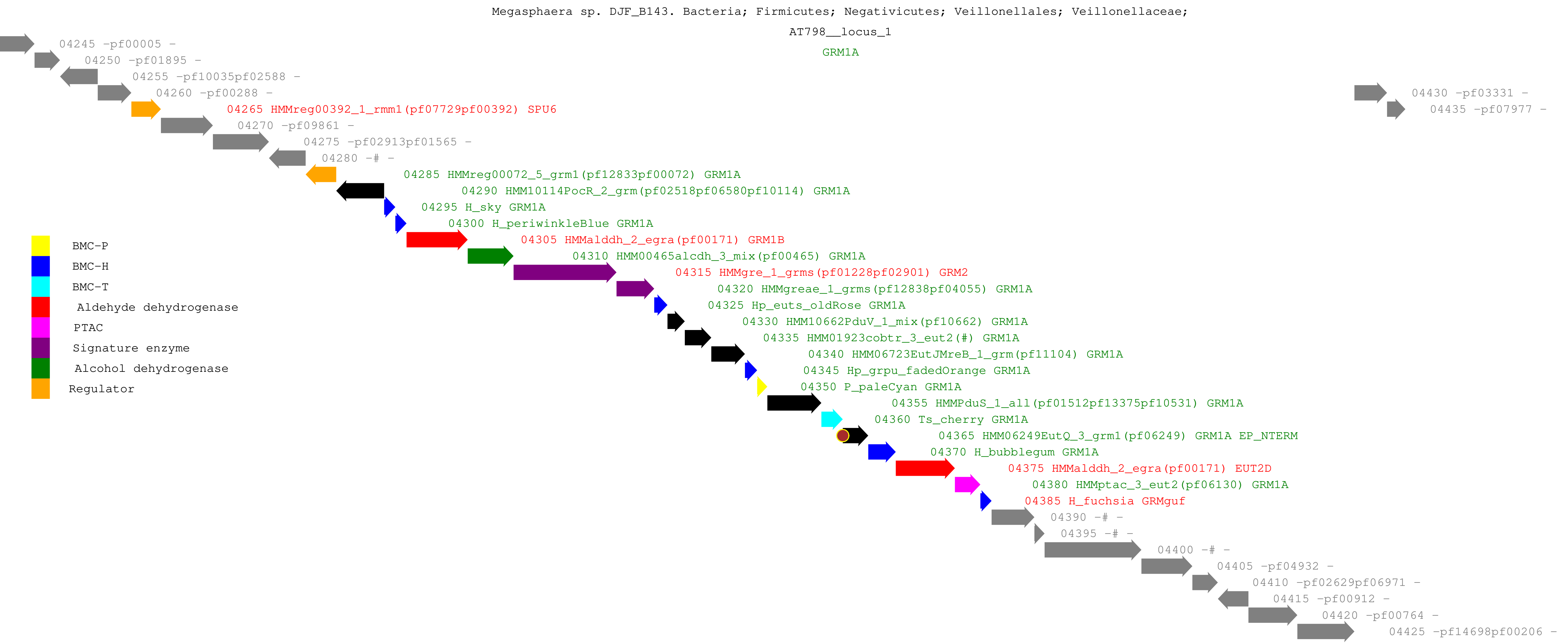
Megasphaera sp. DJF_B143. Bacteria; Firmicutes; Negativicutes; Veillonellales; Veillonellaceae;
AT798__locus_1
GRM1A
04245 -pf00005 -
04250 -pf01895 -
04255 -pf10035pf02588 -
04260 -pf00288 -
04265 HMMreg00392_1_rmm1(pf07729pf00392) SPU6
04270 -pf09861 -
04275 -pf02913pf01565 -
04280 -# -
04285 HMMreg00072_5_grm1(pf12833pf00072) GRM1A
04290 HMM10114PocR_2_grm(pf02518pf06580pf10114) GRM1A
04295 H_sky GRM1A
04300 H_periwinkleBlue GRM1A
04305 HMMalddh_2_egra(pf00171) GRM1B
04310 HMM00465alcdh_3_mix(pf00465) GRM1A
04315 HMMgre_1_grms(pf01228pf02901) GRM2
04320 HMMgreae_1_grms(pf12838pf04055) GRM1A
04325 Hp_euts_oldRose GRM1A
04330 HMM10662PduV_1_mix(pf10662) GRM1A
04335 HMM01923cobtr_3_eut2(#) GRM1A
04340 HMM06723EutJMreB_1_grm(pf11104) GRM1A
04345 Hp_grpu_fadedOrange GRM1A
04350 P_paleCyan GRM1A
04355 HMMPduS_1_all(pf01512pf13375pf10531) GRM1A
04360 Ts_cherry GRM1A
04365 HMM06249EutQ_3_grm1(pf06249) GRM1A EP_NTERM
04370 H_bubblegum GRM1A
04375 HMMalddh_2_egra(pf00171) EUT2D
04380 HMMptac_3_eut2(pf06130) GRM1A
04385 H_fuchsia GRMguf
04390 -# -
04395 -# -
04400 -# -
04405 -pf04932 -
04410 -pf02629pf06971 -
04415 -pf00912 -
04420 -pf00764 -
04425 -pf14698pf00206 -
04430 -pf03331 -
04435 -pf07977 -
BMC-P
BMC-H
BMC-T
Aldehyde dehydrogenase
PTAC
Signature enzyme
Alcohol dehydrogenase
Regulator

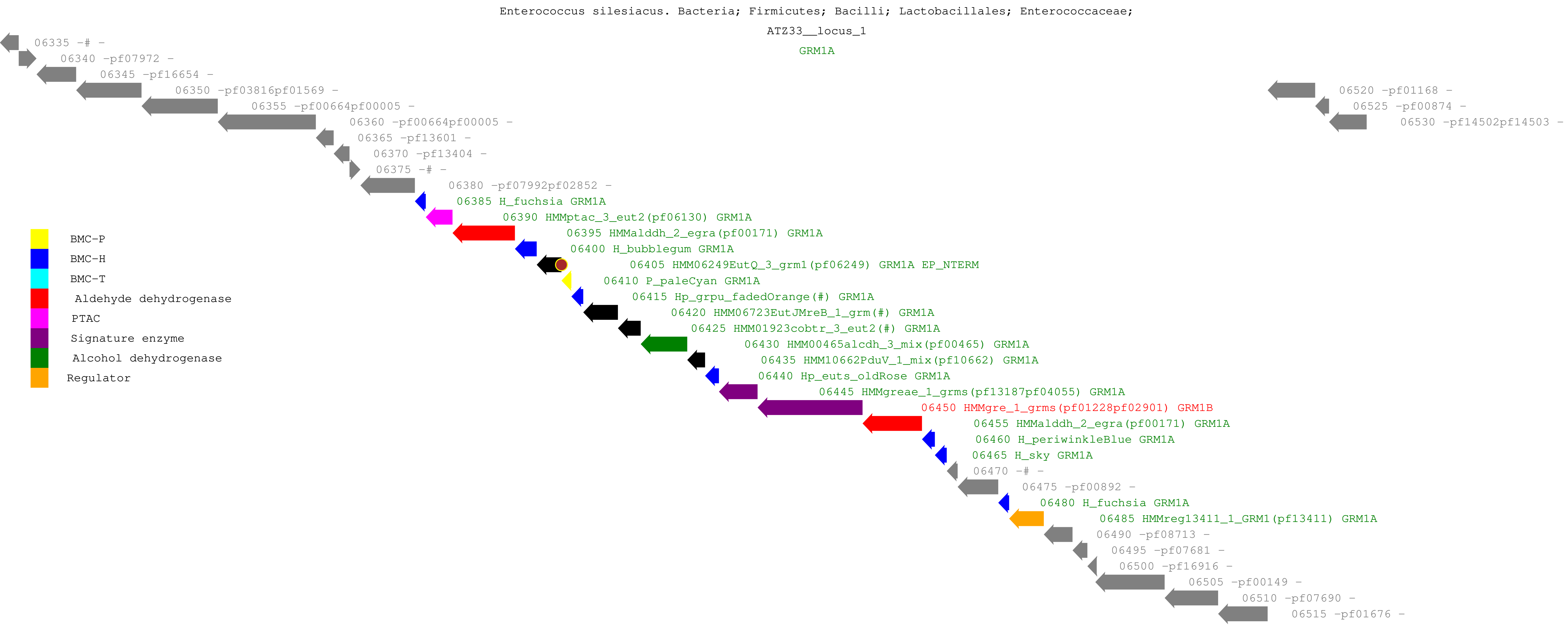

Enterococcus silesiacus. Bacteria; Firmicutes; Bacilli; Lactobacillales; Enterococcaceae;
ATZ33__locus_1
GRM1A
06335 -# -
06340 -pf07972 -
06345 -pf16654 -
06350 -pf03816pf01569 -
06355 -pf00664pf00005 -
06360 -pf00664pf00005 -
06365 -pf13601 -
06370 -pf13404 -
06375 -# -
06380 -pf07992pf02852 -
06385 H_fuchsia GRM1A
06390 HMMptac_3_eut2(pf06130) GRM1A
06395 HMMalddh_2_egra(pf00171) GRM1A
06400 H_bubblegum GRM1A
06405 HMM06249EutQ_3_grm1(pf06249) GRM1A EP_NTERM
06410 P_paleCyan GRM1A
06415 Hp_grpu_fadedOrange(#) GRM1A
06420 HMM06723EutJMreB_1_grm(#) GRM1A
06425 HMM01923cobtr_3_eut2(#) GRM1A
06430 HMM00465alcdh_3_mix(pf00465) GRM1A
06435 HMM10662PduV_1_mix(pf10662) GRM1A
06440 Hp_euts_oldRose GRM1A
06445 HMMgreae_1_grms(pf13187pf04055) GRM1A
06450 HMMgre_1_grms(pf01228pf02901) GRM1B
06455 HMMalddh_2_egra(pf00171) GRM1A
06460 H_periwinkleBlue GRM1A
06465 H_sky GRM1A
06470 -# -
06475 -pf00892 -
06480 H_fuchsia GRM1A
06485 HMMreg13411_1_GRM1(pf13411) GRM1A
06490 -pf08713 -
06495 -pf07681 -
06500 -pf16916 -
06505 -pf00149 -
06510 -pf07690 -
06515 -pf01676 -
06520 -pf01168 -
06525 -pf00874 -
06530 -pf14502pf14503 -
BMC-P
BMC-H
BMC-T
Aldehyde dehydrogenase
PTAC
Signature enzyme
Alcohol dehydrogenase
Regulator

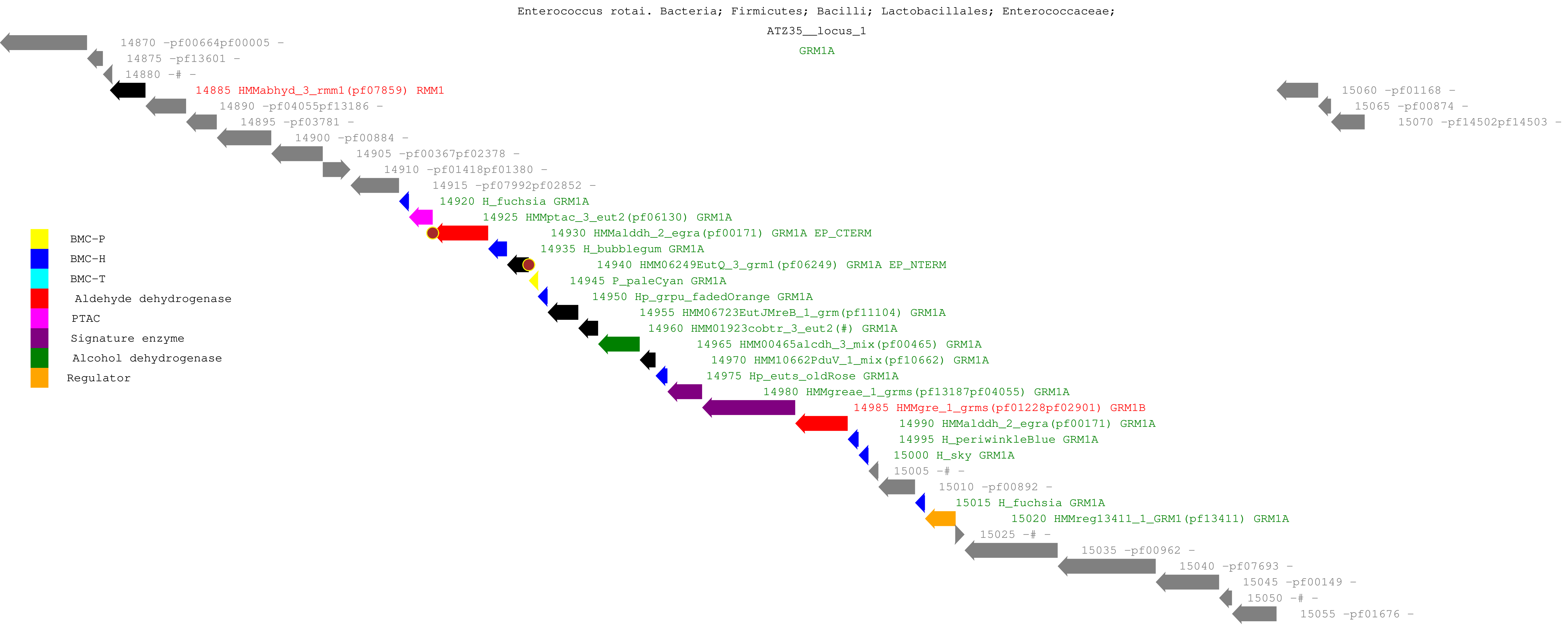
Enterococcus rotai. Bacteria; Firmicutes; Bacilli; Lactobacillales; Enterococcaceae;
ATZ35__locus_1
GRM1A
14870 -pf00664pf00005 -
14875 -pf13601 -
14880 -# -
14885 HMMabhyd_3_rmm1(pf07859) RMM1
14890 -pf04055pf13186 -
14895 -pf03781 -
14900 -pf00884 -
14905 -pf00367pf02378 -
14910 -pf01418pf01380 -
14915 -pf07992pf02852 -
14920 H_fuchsia GRM1A
14925 HMMptac_3_eut2(pf06130) GRM1A
14930 HMMalddh_2_egra(pf00171) GRM1A EP_CTERM
14935 H_bubblegum GRM1A
14940 HMM06249EutQ_3_grm1(pf06249) GRM1A EP_NTERM
14945 P_paleCyan GRM1A
14950 Hp_grpu_fadedOrange GRM1A
14955 HMM06723EutJMreB_1_grm(pf11104) GRM1A
14960 HMM01923cobtr_3_eut2(#) GRM1A
14965 HMM00465alcdh_3_mix(pf00465) GRM1A
14970 HMM10662PduV_1_mix(pf10662) GRM1A
14975 Hp_euts_oldRose GRM1A
14980 HMMgreae_1_grms(pf13187pf04055) GRM1A
14985 HMMgre_1_grms(pf01228pf02901) GRM1B
14990 HMMalddh_2_egra(pf00171) GRM1A
14995 H_periwinkleBlue GRM1A
15000 H_sky GRM1A
15005 -# -
15010 -pf00892 -
15015 H_fuchsia GRM1A
15020 HMMreg13411_1_GRM1(pf13411) GRM1A
15025 -# -
15035 -pf00962 -
15040 -pf07693 -
15045 -pf00149 -
15050 -# -
15055 -pf01676 -
15060 -pf01168 -
15065 -pf00874 -
15070 -pf14502pf14503 -
BMC-P
BMC-H
BMC-T
Aldehyde dehydrogenase
PTAC
Signature enzyme
Alcohol dehydrogenase
Regulator

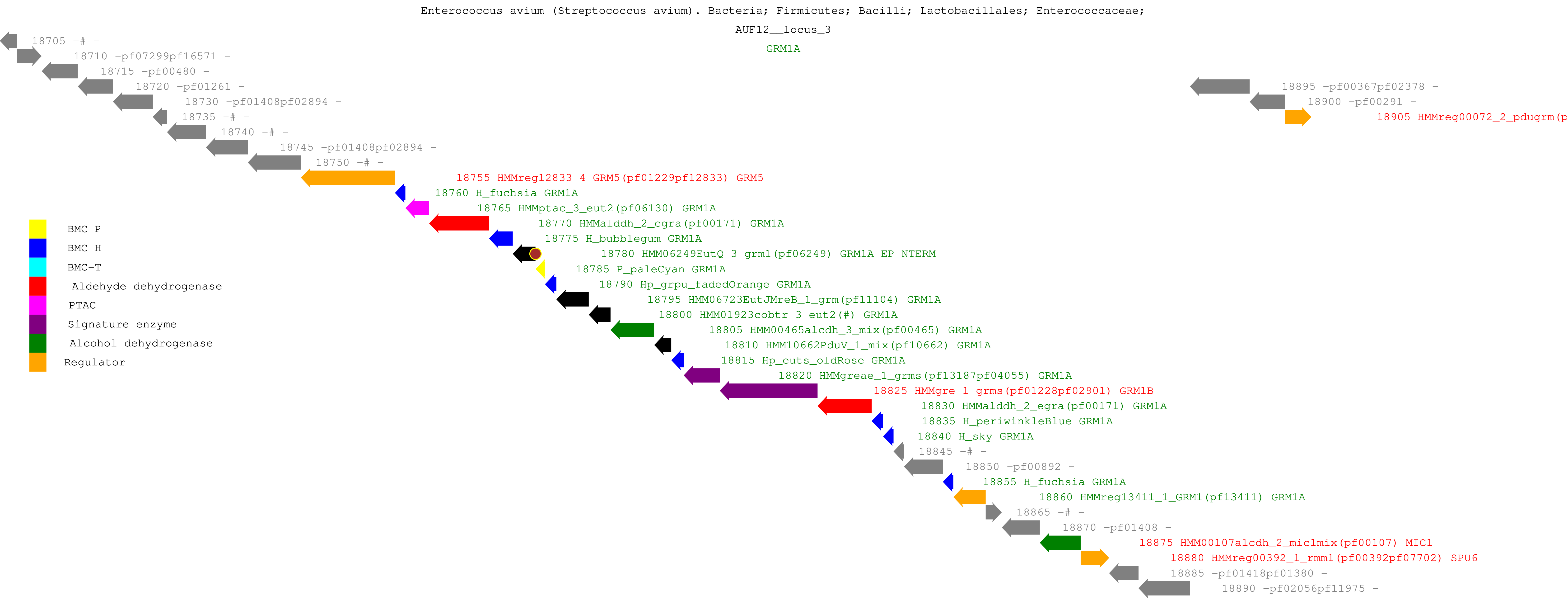
Enterococcus avium (Streptococcus avium). Bacteria; Firmicutes; Bacilli; Lactobacillales; Enterococcaceae;
AUF12__locus_3
GRM1A
18705 -# -
18710 -pf07299pf16571 -
18715 -pf00480 -
18720 -pf01261 -
18730 -pf01408pf02894 -
18735 -# -
18740 -# -
18745 -pf01408pf02894 -
18750 -# -
18755 HMMreg12833_4_GRM5(pf01229pf12833) GRM5
18760 H_fuchsia GRM1A
18765 HMMptac_3_eut2(pf06130) GRM1A
18770 HMMalddh_2_egra(pf00171) GRM1A
18775 H_bubblegum GRM1A
18780 HMM06249EutQ_3_grm1(pf06249) GRM1A EP_NTERM
18785 P_paleCyan GRM1A
18790 Hp_grpu_fadedOrange GRM1A
18795 HMM06723EutJMreB_1_grm(pf11104) GRM1A
18800 HMM01923cobtr_3_eut2(#) GRM1A
18805 HMM00465alcdh_3_mix(pf00465) GRM1A
18810 HMM10662PduV_1_mix(pf10662) GRM1A
18815 Hp_euts_oldRose GRM1A
18820 HMMgreae_1_grms(pf13187pf04055) GRM1A
18825 HMMgre_1_grms(pf01228pf02901) GRM1B
18830 HMMalddh_2_egra(pf00171) GRM1A
18835 H_periwinkleBlue GRM1A
18840 H_sky GRM1A
18845 -# -
18850 -pf00892 -
18855 H_fuchsia GRM1A
18860 HMMreg13411_1_GRM1(pf13411) GRM1A
18865 -# -
18870 -pf01408 -
18875 HMM00107alcdh_2_mic1mix(pf00107) MIC1
18880 HMMreg00392_1_rmm1(pf00392pf07702) SPU6
18885 -pf01418pf01380 -
18890 -pf02056pf11975 -
18895 -pf00367pf02378 -
18900 -pf00291 -
18905 HMMreg00072_2_pdugrm(p
BMC-P
BMC-H
BMC-T
Aldehyde dehydrogenase
PTAC
Signature enzyme
Alcohol dehydrogenase
Regulator

Clostridium haemolyticum. Bacteria; Firmicutes; Clostridia; Clostridiales; Clostridiaceae;

AXF41__locus_1

GRM1A

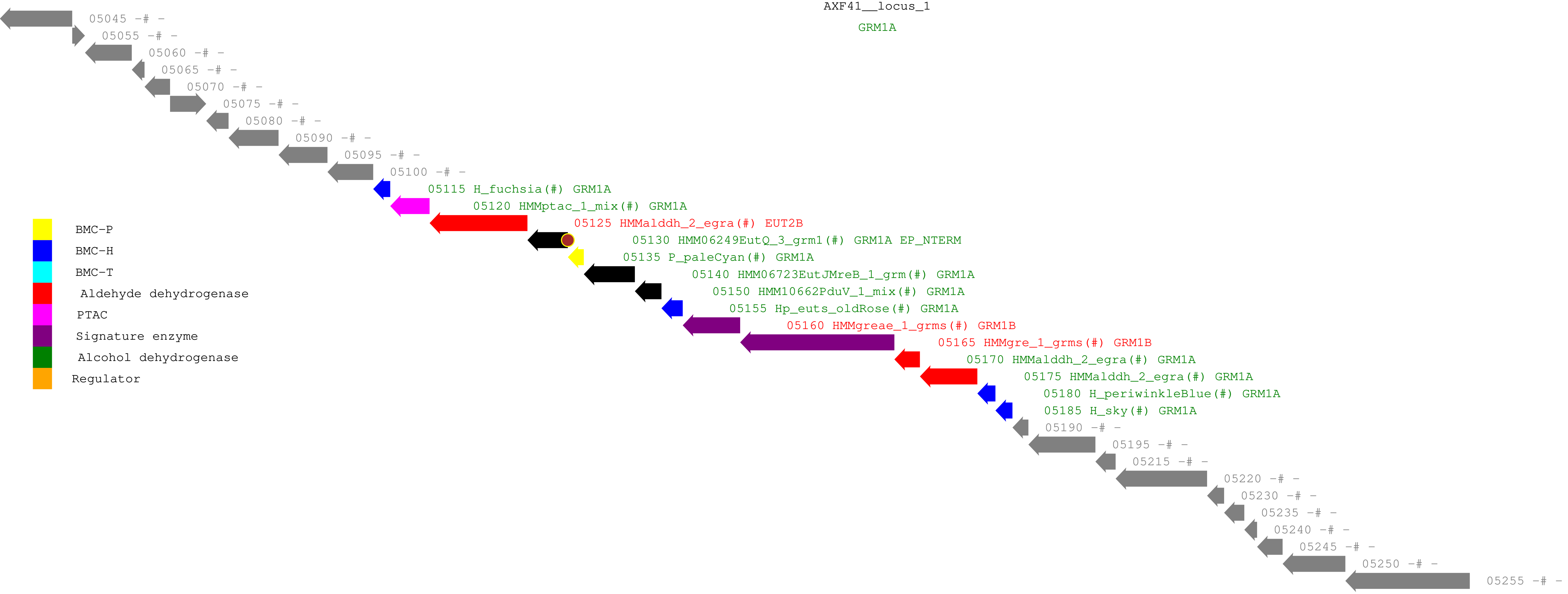

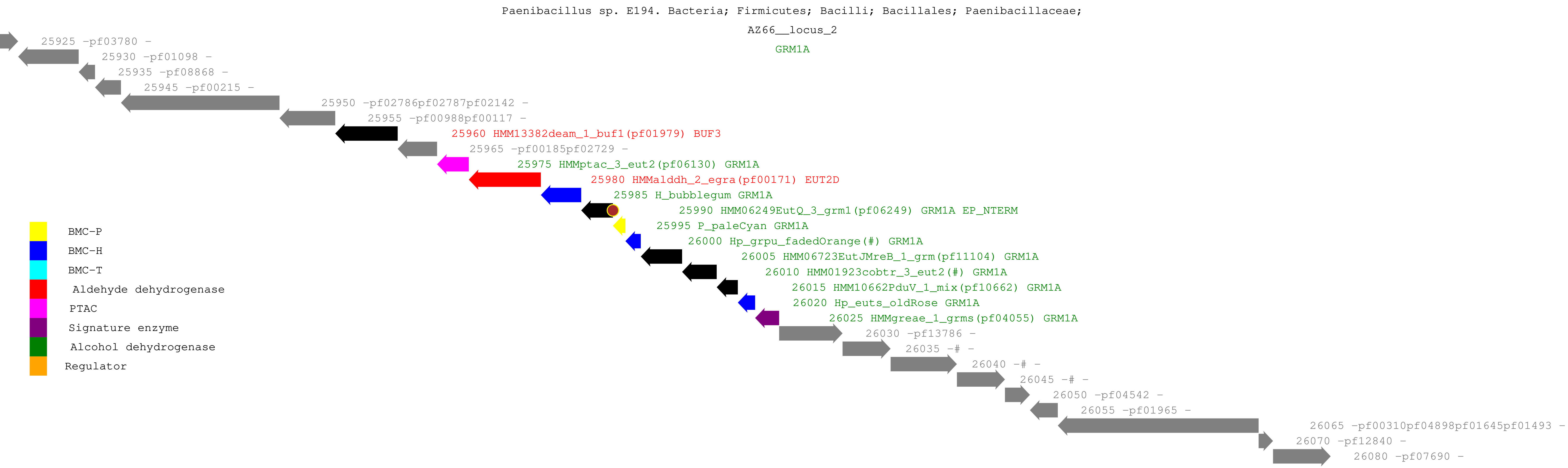

Paenibacillus sp. E194. Bacteria; Firmicutes; Bacilli; Bacillales; Paenibacillaceae;
AZ66__locus_2
GRM1A
25925 -pf03780 -
25930 -pf01098 -
25935 -pf08868 -
25945 -pf00215 -
25950 -pf02786pf02787pf02142 -
25955 -pf00988pf00117 -
25960 HMM13382deam_1_buf1(pf01979) BUF3
25965 -pf00185pf02729 -
25975 HMMptac_3_eut2(pf06130) GRM1A
25980 HMMalddh_2_egra(pf00171) EUT2D
25985 H_bubblegum GRM1A
25990 HMM06249EutQ_3_grm1(pf06249) GRM1A EP_NTERM
25995 P_paleCyan GRM1A
26000 Hp_grpu_fadedOrange(#) GRM1A
26005 HMM06723EutJMreB_1_grm(pf11104) GRM1A
26010 HMM01923cobtr_3_eut2(#) GRM1A
26015 HMM10662PduV_1_mix(pf10662) GRM1A
26020 Hp_euts_oldRose GRM1A
26025 HMMgreae_1_grms(pf04055) GRM1A
26030 -pf13786 -
26035 -# -
26040 -# -
26045 -# -
26050 -pf04542 -
26055 -pf01965 -
26065 -pf00310pf04898pf01645pf01493 -
26070 -pf12840 -
26080 -pf07690 -
BMC-P
BMC-H
BMC-T
Aldehyde dehydrogenase
PTAC
Signature enzyme
Alcohol dehydrogenase
Regulator

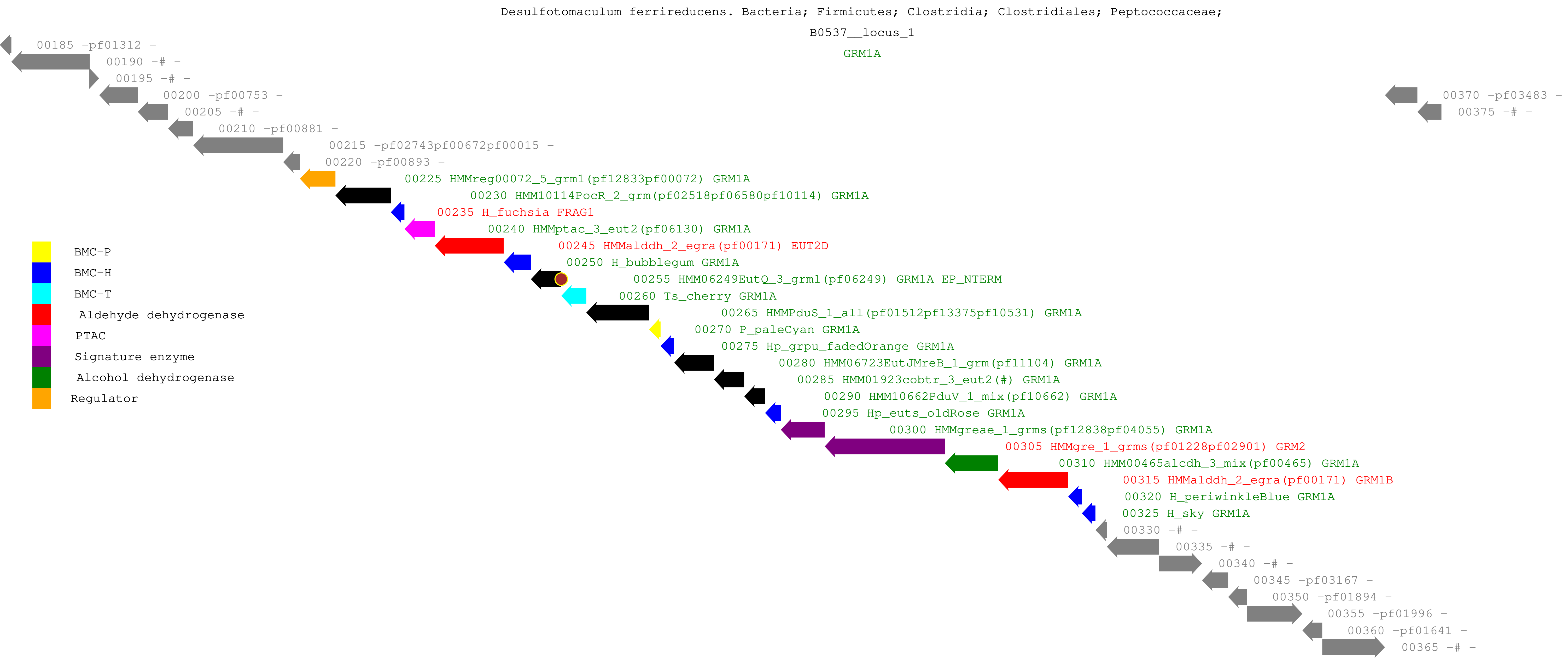
Desulfotomaculum ferrireducens. Bacteria; Firmicutes; Clostridia; Clostridiales; Peptococcaceae;
B0537__locus_1
GRM1A
00185 -pf01312 -
00190 -# -
00195 -# -
00200 -pf00753 -
00205 -# -
00210 -pf00881 -
00215 -pf02743pf00672pf00015 -
00220 -pf00893 -
00225 HMMreg00072_5_grm1(pf12833pf00072) GRM1A
00230 HMM10114PocR_2_grm(pf02518pf06580pf10114) GRM1A
00235 H_fuchsia FRAG1
00240 HMMptac_3_eut2(pf06130) GRM1A
00245 HMMaldh_2_egra(pf00171) EUT2D
00250 H_bubblegum GRM1A
00255 HMM06249EutQ_3_grm1(pf06249) GRM1A EP_NTERM
00260 Ts_cherry GRM1A
00265 HMMPduS_1_all(pf01512pf13375pf10531) GRM1A
00270 P_paleCyan GRM1A
00275 Hp_grpu_fadedOrange GRM1A
00280 HMM06723EutJMreB_1_grm(pf11104) GRM1A
00285 HMM01923cobtr_3_eut2(#) GRM1A
00290 HMM10662PduV_1_mix(pf10662) GRM1A
00295 Hp_euts_oldRose GRM1A
00300 HMMgreae_1_grms(pf12838pf04055) GRM1A
00305 HMMgre_1_grms(pf01228pf02901) GRM2
00310 HMM00465alcdh_3_mix(pf00465) GRM1A
00315 HMMaldh_2_egra(pf00171) GRM1B
00320 H_periwinkleBlue GRM1A
00325 H_sky GRM1A
00330 -# -
00335 -# -
00340 -# -
00345 -pf03167 -
00350 -pf01894 -
00355 -pf01996 -
00360 -pf01641 -
00365 -# -
00370 -pf03483 -
00375 -# -
BMC-P
BMC-H
BMC-T
Aldehyde dehydrogenase
PTAC
Signature enzyme
Alcohol dehydrogenase
Regulator

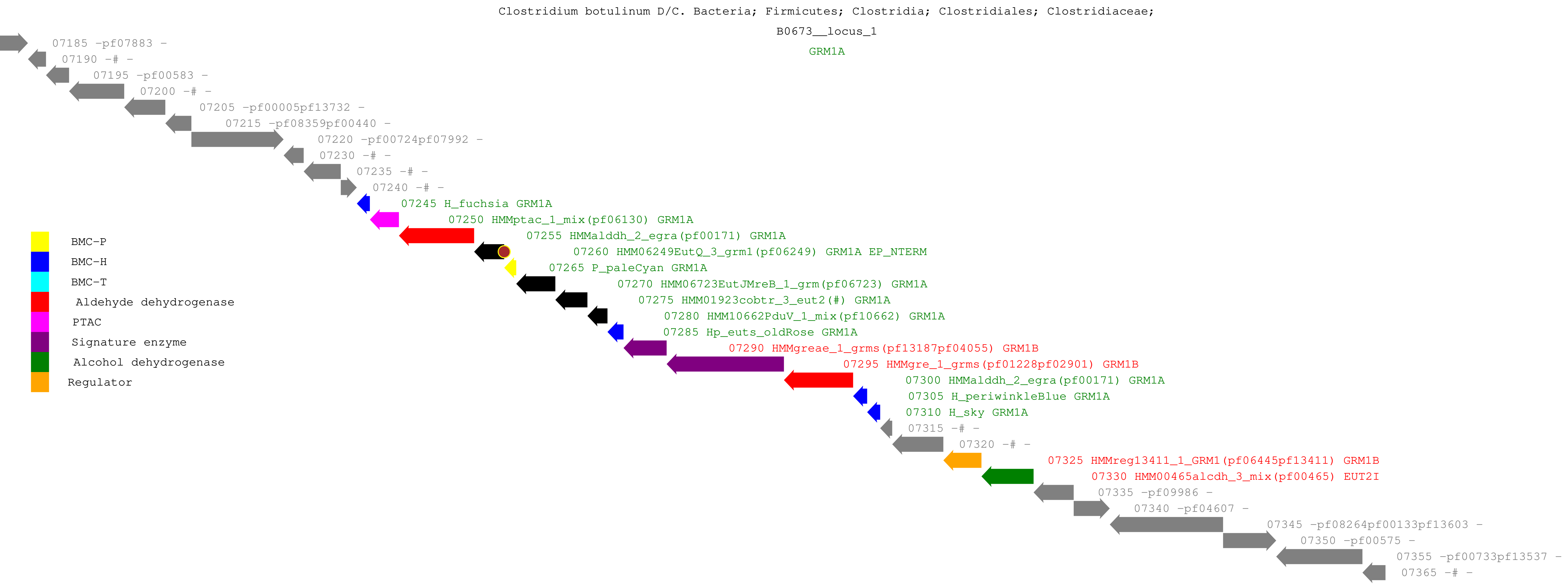
Clostridium botulinum D/C. Bacteria; Firmicutes; Clostridia; Clostridiales; Clostridiaceae;
B0673__locus_1
GRM1A
07185 -pf07883 -
07190 -# -
07195 -pf00583 -
07200 -# -
07205 -pf00005pf13732 -
07215 -pf08359pf00440 -
07220 -pf00724pf07992 -
07230 -# -
07235 -# -
07240 -# -
07245 H_fuchsia GRM1A
07250 HMMptac_1_mix(pf06130) GRM1A
07255 HMMaldh_2_egra(pf00171) GRM1A
07260 HMM06249EutQ_3_grm1(pf06249) GRM1A EP_NTERM
07265 P_paleCyan GRM1A
07270 HMM06723EutJMreB_1_grm(pf06723) GRM1A
07275 HMM01923cobtr_3_eut2(#) GRM1A
07280 HMM10662PduV_1_mix(pf10662) GRM1A
07285 Hp_euts_oldRose GRM1A
07290 HMMgreae_1_grms(pf13187pf04055) GRM1B
07295 HMMgre_1_grms(pf01228pf02901) GRM1B
07300 HMMaldh_2_egra(pf00171) GRM1A
07305 H_periwinkleBlue GRM1A
07310 H_sky GRM1A
07315 -# -
07320 -# -
07325 HMMreg13411_1_GRM1(pf06445pf13411) GRM1B
07330 HMM00465alcdh_3_mix(pf00465) EUT2I
07335 -pf09986 -
07340 -pf04607 -
07345 -pf08264pf00133pf13603 -
07350 -pf00575 -
07355 -pf00733pf13537 -
07365 -# -
BMC-P
BMC-H
BMC-T
Aldehyde dehydrogenase
PTAC
Signature enzyme
Alcohol dehydrogenase
Regulator

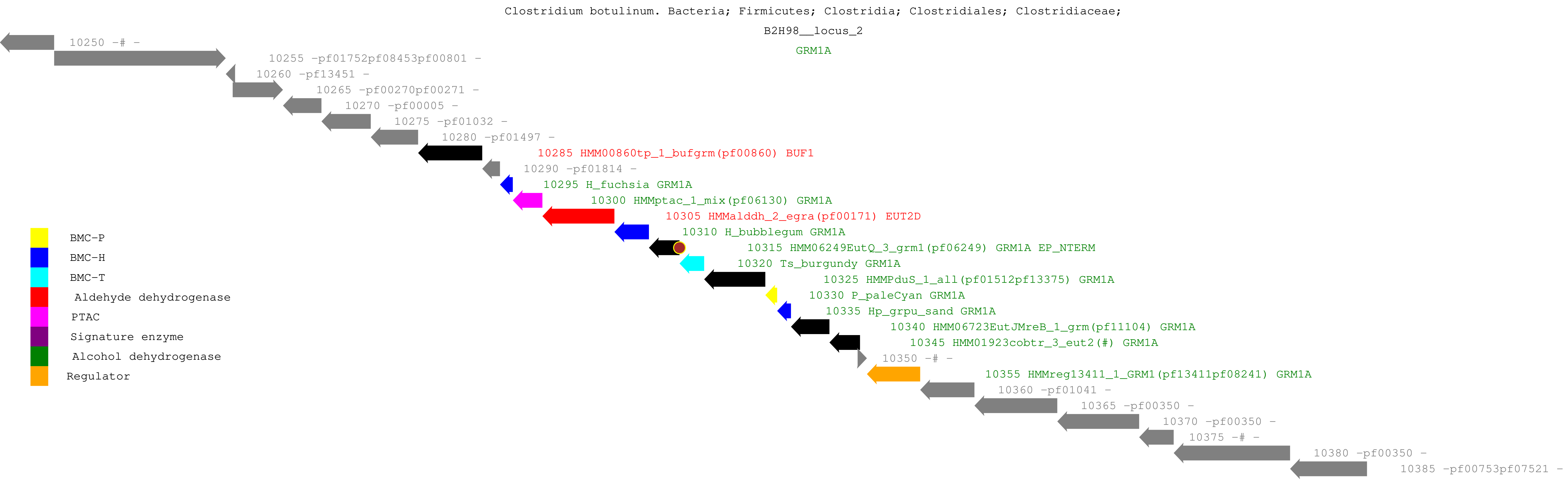
Clostridium botulinum. Bacteria; Firmicutes; Clostridia; Clostridiales; Clostridiaceae;
B2H98__locus_2
GRM1A
10250 -# -
10255 -pf01752pf08453pf00801 -
10260 -pf13451 -
10265 -pf00270pf00271 -
10270 -pf00005 -
10275 -pf01032 -
10280 -pf01497 -
10285 HMM00860tp_1_bufgrm(pf00860) BUF1
10290 -pf01814 -
10295 H_fuchsia GRM1A
10300 HMMptac_1_mix(pf06130) GRM1A
10305 HMMaldh_2_egra(pf00171) EUT2D
10310 H_bubblegum GRM1A
10315 HMM06249EutQ_3_grm1(pf06249) GRM1A EP_NTERM
10320 Ts_burgundy GRM1A
10325 HMMPduS_1_all(pf01512pf13375) GRM1A
10330 P_paleCyan GRM1A
10335 Hp_grpu_sand GRM1A
10340 HMM06723EutJMreB_1_grm(pf11104) GRM1A
10345 HMM01923cobtr_3_eut2(#) GRM1A
10350 -# -
10355 HMMreg13411_1_GRM1(pf13411pf08241) GRM1A
10360 -pf01041 -
10365 -pf00350 -
10370 -pf00350 -
10375 -# -
10380 -pf00350 -
10385 -pf00753pf07521 -
BMC-P
BMC-H
BMC-T
Aldehyde dehydrogenase
PTAC
Signature enzyme
Alcohol dehydrogenase
Regulator

Gemmiger sp. An194. Bacteria; Firmicutes; Clostridia; Clostridiales; Ruminococcaceae;

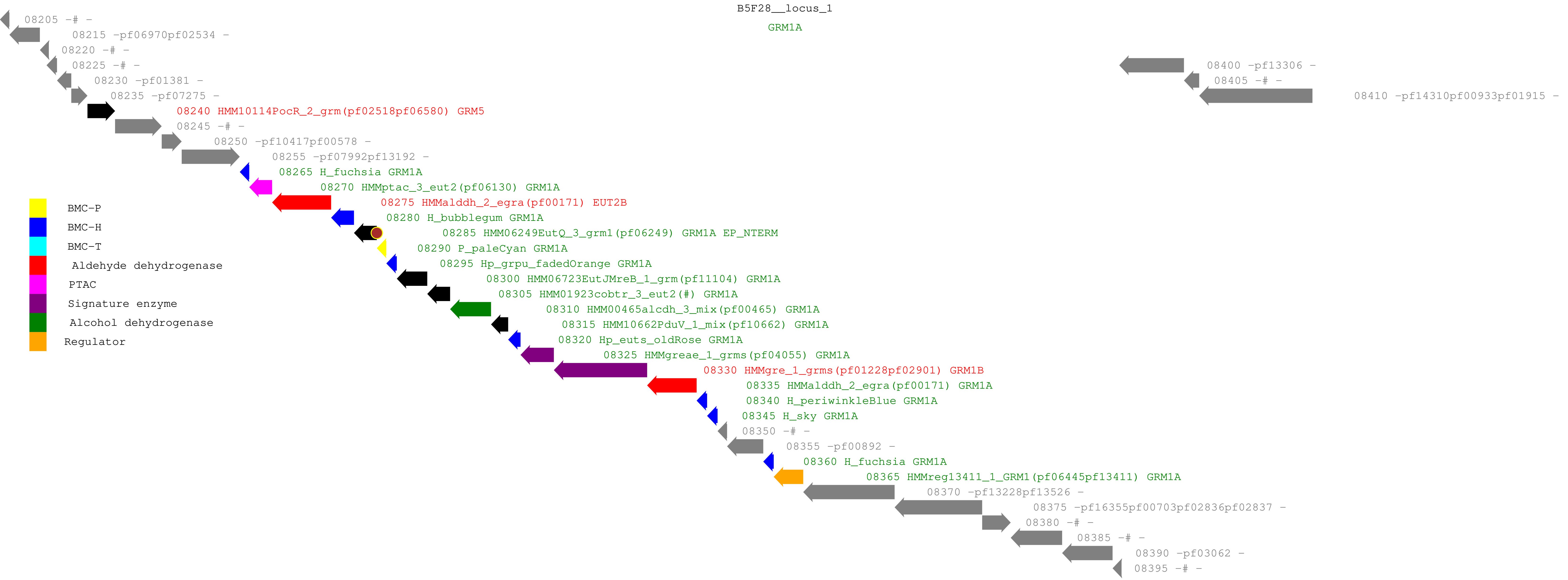

Collinsella sp. An271. Bacteria; Actinobacteria; Coriobacteriia; Coriobacteriales;

B5F74__locus_1_main_sat

GRM1A

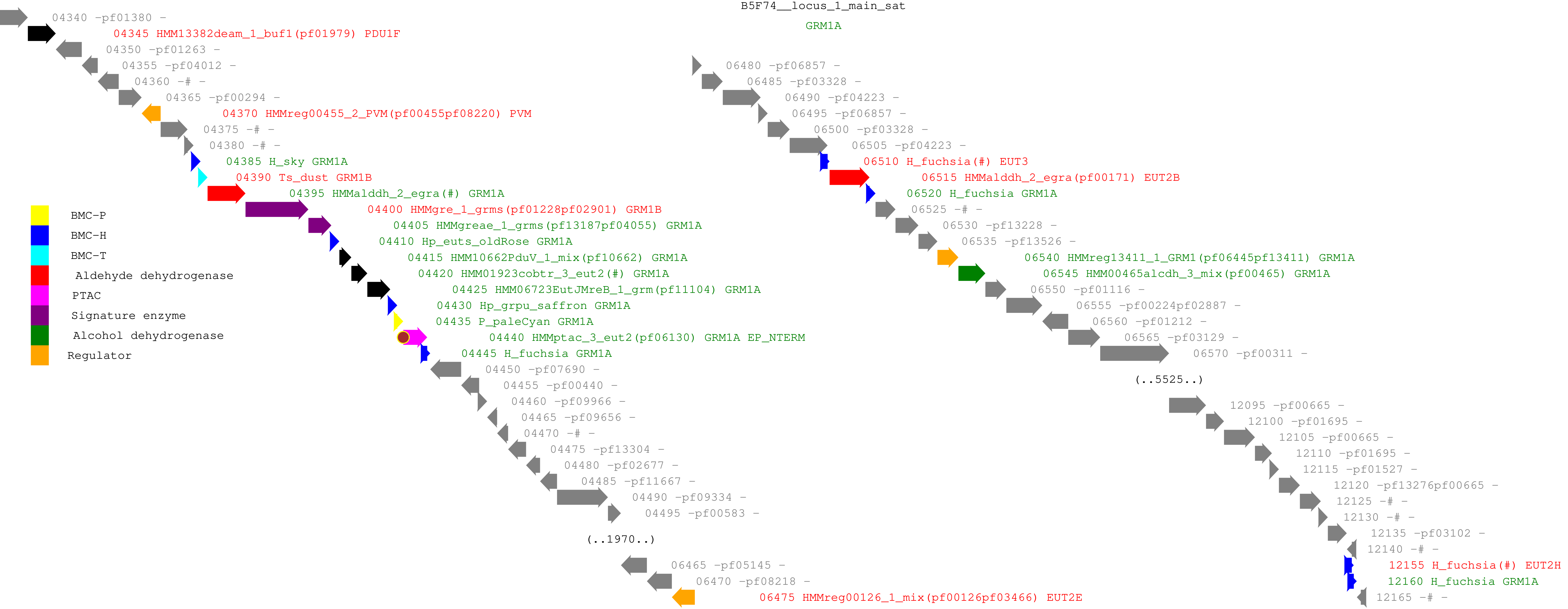

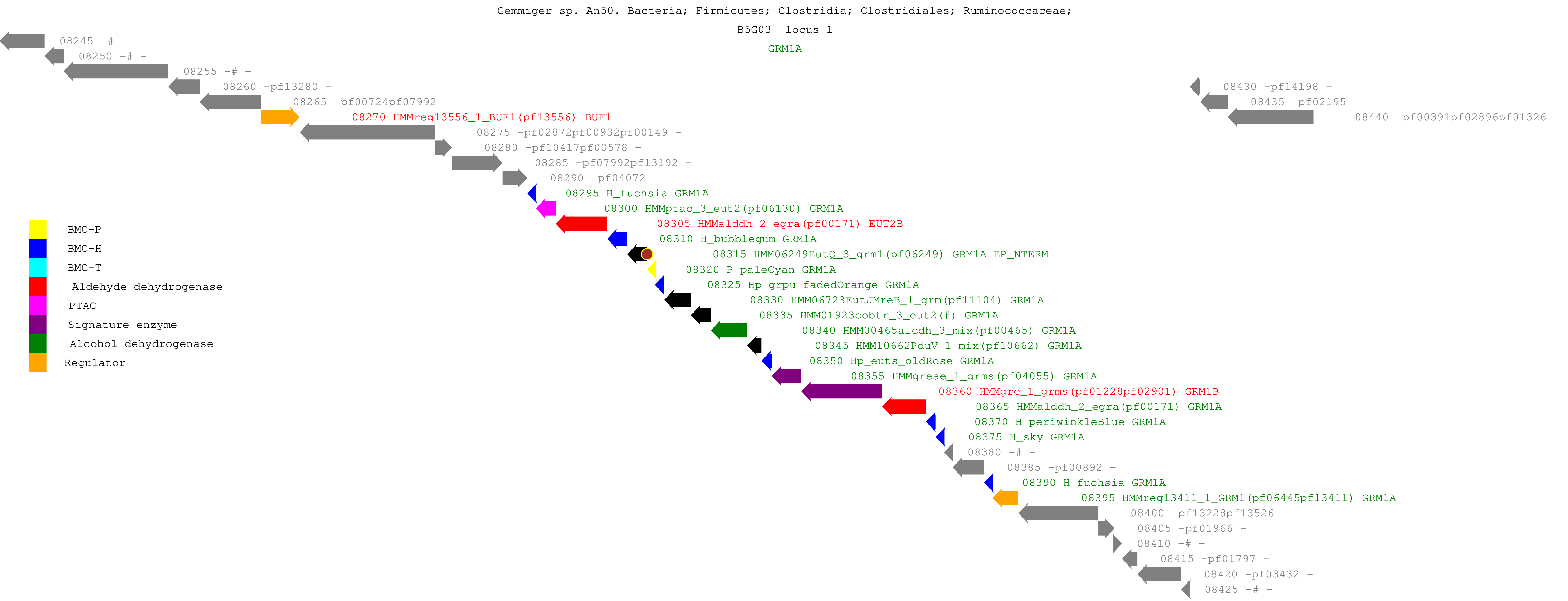

Gemmiger sp. An50. Bacteria; Firmicutes; Clostridia; Clostridiales; Ruminococcaceae;
B5G03__locus_1
GRM1A
08245 -# -
08250 -# -
08255 -# -
08260 -pf13280 -
08265 -pf00724pf07992 -
08270 HMMreg13556_1_BUF1(pf13556) BUF1
08275 -pf02872pf00932pf00149 -
08280 -pf10417pf00578 -
08285 -pf07992pf13192 -
08290 -pf04072 -
08295 H_fuchsia GRM1A
08300 HMMptac_3_eut2(pf06130) GRM1A
08305 HMMaldдh_2_egra(pf00171) EUT2B
08310 H_bubblegum GRM1A
08315 HMM06249EutQ_3_grm1(pf06249) GRM1A EP_NTERM
08320 P_paleCyan GRM1A
08325 Hp_grpu_fadedOrange GRM1A
08330 HMM06723EutJMreB_1_grm(pf11104) GRM1A
08335 HMM01923cobtr_3_eut2(#) GRM1A
08340 HMM00465alcdh_3_mix(pf00465) GRM1A
08345 HMM10662PduV_1_mix(pf10662) GRM1A
08350 Hp_euts_oldRose GRM1A
08355 HMMgreae_1_grms(pf04055) GRM1A
08360 HMMgre_1_grms(pf01228pf02901) GRM1B
08365 HMMaldдh_2_egra(pf00171) GRM1A
08370 H_periwinkleBlue GRM1A
08375 H_sky GRM1A
08380 -# -
08385 -pf00892 -
08390 H_fuchsia GRM1A
08395 HMMreg13411_1_GRM1(pf06445pf13411) GRM1A
08400 -pf13228pf13526 -
08405 -pf01966 -
08410 -# -
08415 -pf01797 -
08420 -pf03432 -
08425 -# -
08430 -pf14198 -
08435 -pf02195 -
08440 -pf00391pf02896pf01326 -
BMC-P
BMC-H
BMC-T
Aldehyde dehydrogenase
PTAC
Signature enzyme
Alcohol dehydrogenase
Regulator

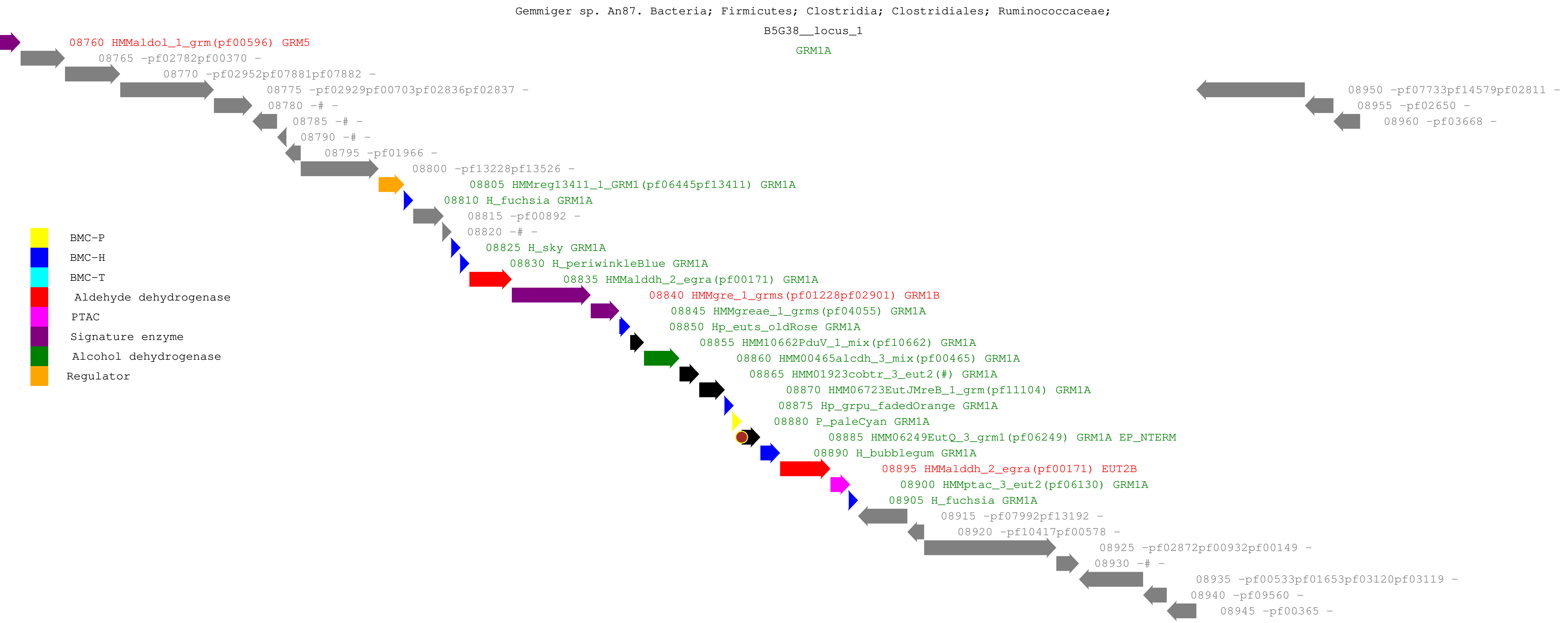
Gemmiger sp. An87. Bacteria; Firmicutes; Clostridia; Clostridiales; Ruminococcaceae;
B5G38__locus_1
GRM1A
08760 HMMaldol_1_grm(pf00596) GRM5
08765 -pf02782pf00370 -
08770 -pf02952pf07881pf07882 -
08775 -pf02929pf00703pf02836pf02837 -
08780 -# -
08785 -# -
08790 -# -
08795 -pf01966 -
08800 -pf13228pf13526 -
08805 HMMreg13411_1_GRM1(pf06445pf13411) GRM1A
08810 H_fuchsia GRM1A
08815 -pf00892 -
08820 -# -
08825 H_sky GRM1A
08830 H_periwinkleBlue GRM1A
08835 HMMaldh_2_egra(pf00171) GRM1A
08840 HMMgre_1_grms(pf01228pf02901) GRM1B
08845 HMMgreae_1_grms(pf04055) GRM1A
08850 Hp_euts_oldRose GRM1A
08855 HMM10662PduV_1_mix(pf10662) GRM1A
08860 HMM00465alcdh_3_mix(pf00465) GRM1A
08865 HMM01923cobtr_3_eut2(#) GRM1A
08870 HMM06723EutJMreB_1_grm(pf11104) GRM1A
08875 Hp_grpu_fadedOrange GRM1A
08880 P_paleCyan GRM1A
08885 HMM06249EutQ_3_grm1(pf06249) GRM1A EP_NTERM
08890 H_bubblegum GRM1A
08895 HMMaldh_2_egra(pf00171) EUT2B
08900 HMMptac_3_eut2(pf06130) GRM1A
08905 H_fuchsia GRM1A
08915 -pf07992pf13192 -
08920 -pf10417pf00578 -
08925 -pf02872pf00932pf00149 -
08930 -# -
08935 -pf00533pf01653pf03120pf03119 -
08940 -pf09560 -
08945 -pf00365 -
08950 -pf07733pf14579pf02811 -
08955 -pf02650 -
08960 -pf03668 -
BMC-P
BMC-H
BMC-T
Aldehyde dehydrogenase
PTAC
Signature enzyme
Alcohol dehydrogenase
Regulator

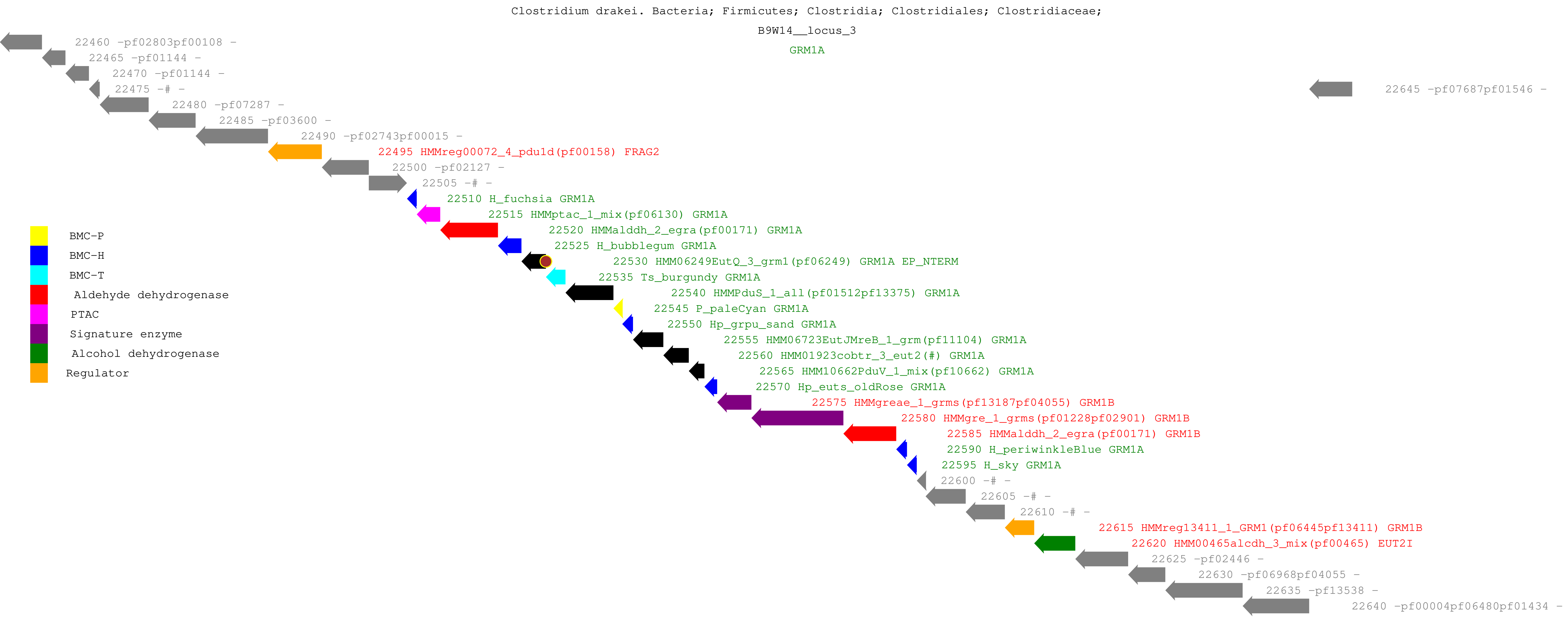
Clostridium drakei. Bacteria; Firmicutes; Clostridia; Clostridiales; Clostridiaceae;
B9W14__locus_3
GRM1A
22460 -pf02803pf00108 -
22465 -pf01144 -
22470 -pf01144 -
22475 -# -
22480 -pf07287 -
22485 -pf03600 -
22490 -pf02743pf00015 -
22495 HMMreg00072_4_pdu1d(pf00158) FRAG2
22500 -pf02127 -
22505 -# -
22510 H_fuchsia GRM1A
22515 HMMptac_1_mix(pf06130) GRM1A
22520 HMMalddh_2_egra(pf00171) GRM1A
22525 H_bubblegum GRM1A
22530 HMM06249EutQ_3_grm1(pf06249) GRM1A EP_NTERM
22535 Ts_burgundy GRM1A
22540 HMMPduS_1_all(pf01512pf13375) GRM1A
22545 P_paleCyan GRM1A
22550 Hp_grpu_sand GRM1A
22555 HMM06723EutJMreB_1_grm(pf11104) GRM1A
22560 HMM01923cobtr_3_eut2(#) GRM1A
22565 HMM10662PduV_1_mix(pf10662) GRM1A
22570 Hp_euts_oldRose GRM1A
22575 HMMgreae_1_grms(pf13187pf04055) GRM1B
22580 HMMgre_1_grms(pf01228pf02901) GRM1B
22585 HMMalddh_2_egra(pf00171) GRM1B
22590 H_periwinkleBlue GRM1A
22595 H_sky GRM1A
22600 -# -
22605 -# -
22610 -# -
22615 HMMreg13411_1_GRM1(pf06445pf13411) GRM1B
22620 HMM00465alcdh_3_mix(pf00465) EUT2I
22625 -pf02446 -
22630 -pf06968pf04055 -
22635 -pf13538 -
22640 -pf00004pf06480pf01434 -
22645 -pf07687pf01546 -
BMC-P
BMC-H
BMC-T
Aldehyde dehydrogenase
PTAC
Signature enzyme
Alcohol dehydrogenase
Regulator

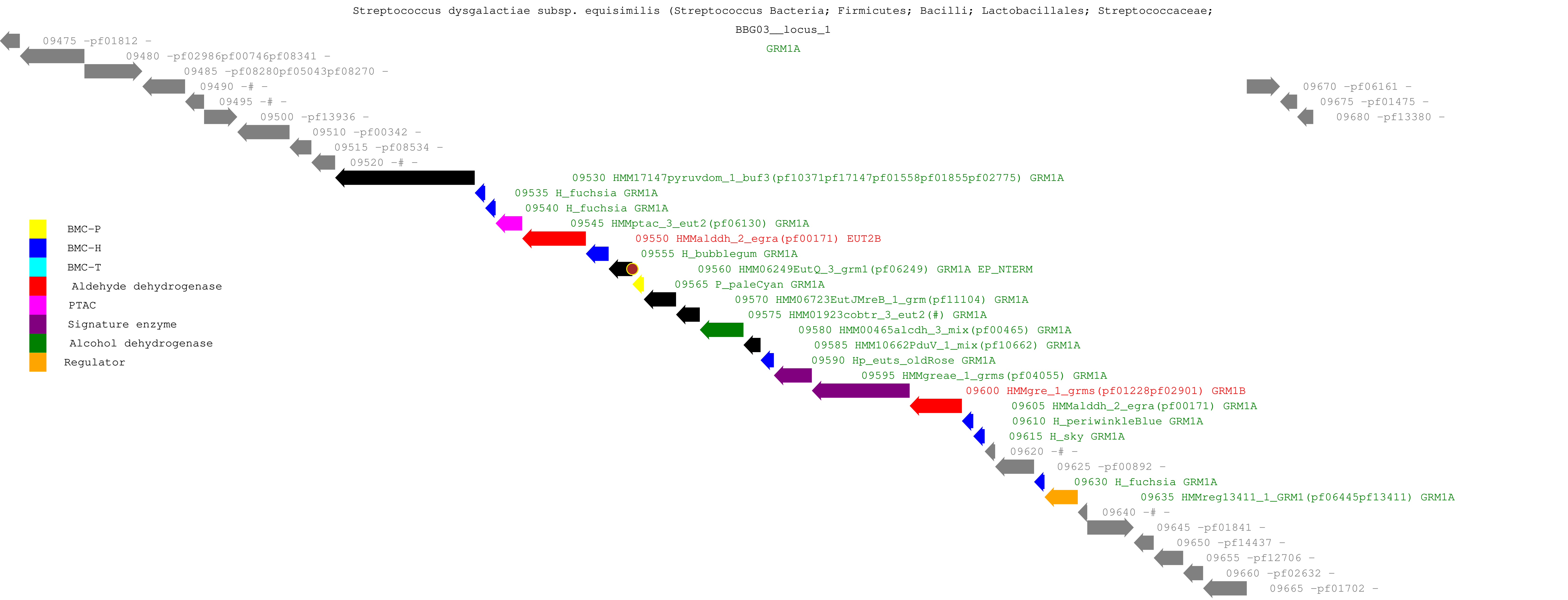
Streptococcus dysgalactiae subsp. equisimilis (Streptococcus Bacteria; Firmicutes; Bacilli; Lactobacillales; Streptococcaceae;
BBG03__locus_1
GRM1A
09475 -pf01812 -
09480 -pf02986pf00746pf08341 -
09485 -pf08280pf05043pf08270 -
09490 -# -
09495 -# -
09500 -pf13936 -
09510 -pf00342 -
09515 -pf08534 -
09520 -# -
09530 HMM17147pyruvdom_1_buf3(pf10371pf17147pf01558pf01855pf02775) GRM1A
09535 H_fuchsia GRM1A
09540 H_fuchsia GRM1A
09545 HMMptac_3_eut2(pf06130) GRM1A
09550 HMMalddh_2_egra(pf00171) EUT2B
09555 H_bubblegum GRM1A
09560 HMM06249EutQ_3_grm1(pf06249) GRM1A EP_NTERM
09565 P_paleCyan GRM1A
09570 HMM06723EutJMreB_1_grm(pf11104) GRM1A
09575 HMM01923cobtr_3_eut2(#) GRM1A
09580 HMM00465alcdh_3_mix(pf00465) GRM1A
09585 HMM10662PduV_1_mix(pf10662) GRM1A
09590 Hp_euts_oldRose GRM1A
09595 HMMgreae_1_grms(pf04055) GRM1A
09600 HMMgre_1_grms(pf01228pf02901) GRM1B
09605 HMMalddh_2_egra(pf00171) GRM1A
09610 H_periwinkleBlue GRM1A
09615 H_sky GRM1A
09620 -# -
09625 -pf00892 -
09630 H_fuchsia GRM1A
09635 HMMreg13411_1_GRM1(pf06445pf13411) GRM1A
09640 -# -
09645 -pf01841 -
09650 -pf14437 -
09655 -pf12706 -
09660 -pf02632 -
09665 -pf01702 -
09670 -pf06161 -
09675 -pf01475 -
09680 -pf13380 -
BMC-P
BMC-H
BMC-T
Aldehyde dehydrogenase
PTAC
Signature enzyme
Alcohol dehydrogenase
Regulator

Paenibacillus sp. KS1. Bacteria; Firmicutes; Bacilli; Bacillales; Paenibacillaceae;

BBG47__locus_2

GRM1A

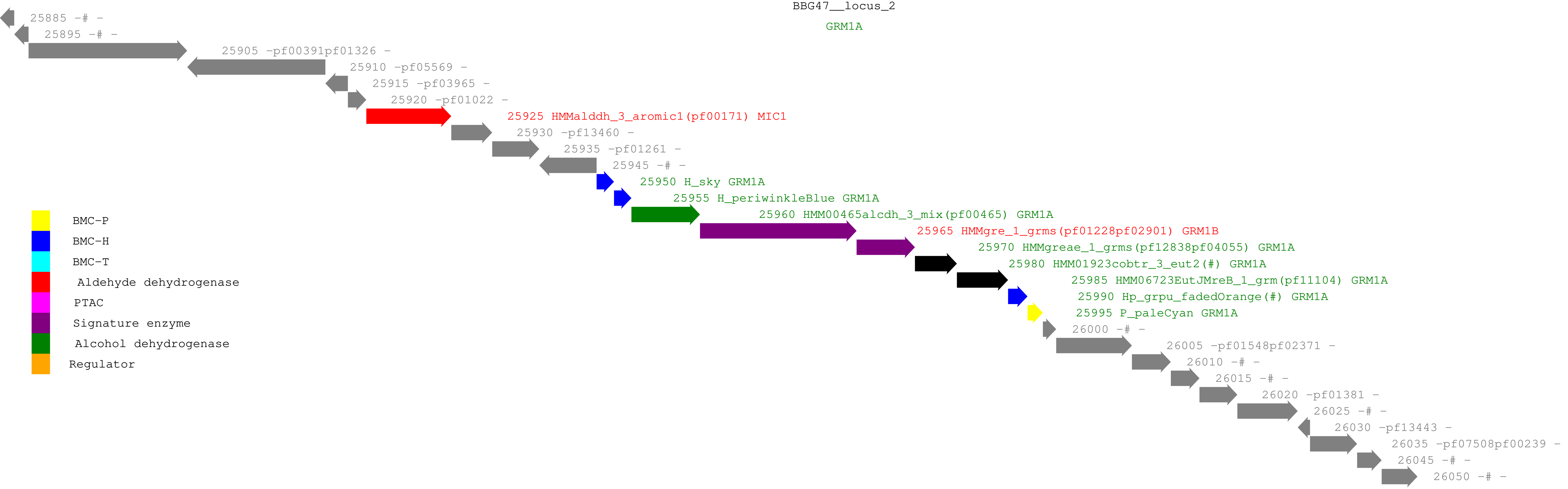

Enterococcus plantarum. Bacteria; Firmicutes; Bacilli; Lactobacillales; Enterococcaceae;

BCR22__locus_1

GRM1A

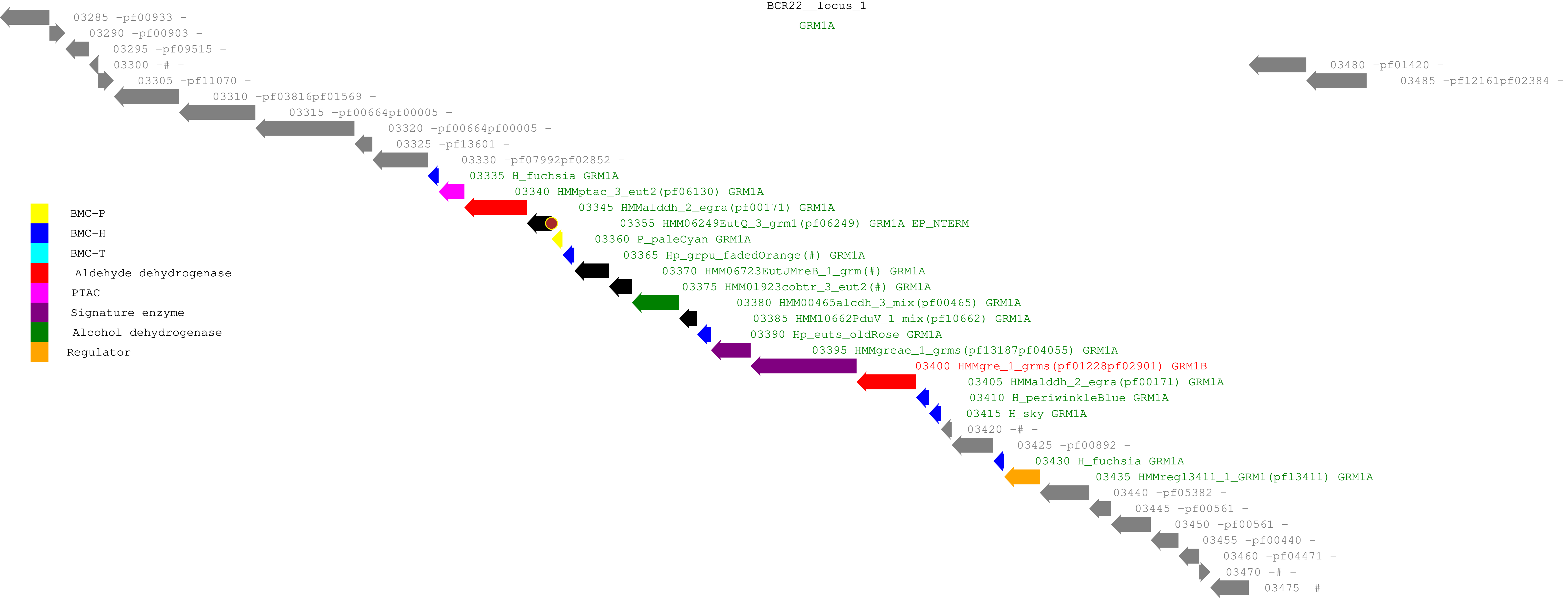

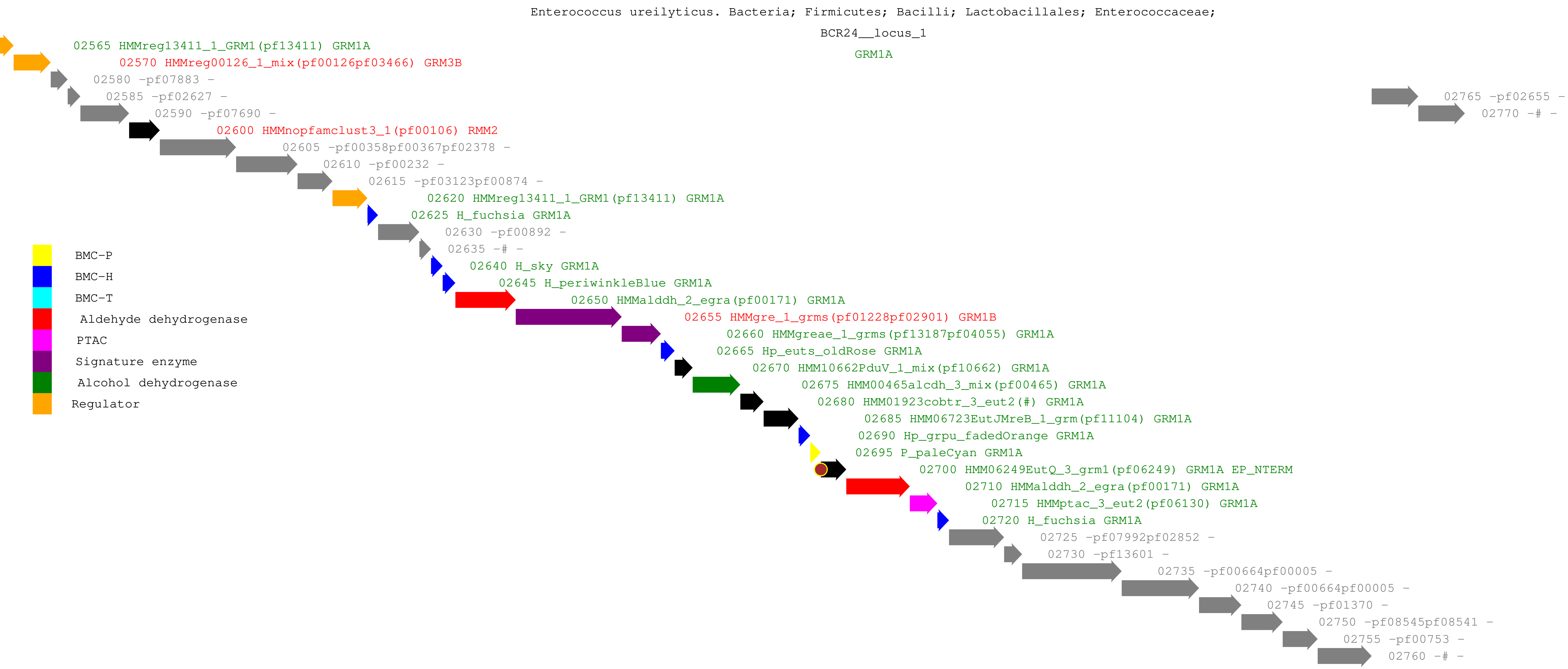
Enterococcus ureilyticus. Bacteria; Firmicutes; Bacilli; Lactobacillales; Enterococcaceae;
BCR24__locus_1
GRM1A
02565 HMMreg13411_1_GRM1(pf13411) GRM1A
02570 HMMreg00126_1_mix(pf00126pf03466) GRM3B
02580 -pf07883 -
02585 -pf02627 -
02590 -pf07690 -
02600 HMMnopfamclust3_1(pf00106) RMM2
02605 -pf00358pf00367pf02378 -
02610 -pf00232 -
02615 -pf03123pf00874 -
02620 HMMreg13411_1_GRM1(pf13411) GRM1A
02625 H_fuchsia GRM1A
02630 -pf00892 -
02635 -# -
02640 H_sky GRM1A
02645 H_periwinkleBlue GRM1A
02650 HMMalddh_2_egra(pf00171) GRM1A
02655 HMMgre_1_grms(pf01228pf02901) GRM1B
02660 HMMgreae_1_grms(pf13187pf04055) GRM1A
02665 Hp_euts_oldRose GRM1A
02670 HMM10662PduV_1_mix(pf10662) GRM1A
02675 HMM00465alcdh_3_mix(pf00465) GRM1A
02680 HMM01923cobtr_3_eut2(#) GRM1A
02685 HMM06723EutJMreB_1_grm(pf11104) GRM1A
02690 Hp_grpu_fadedOrange GRM1A
02695 P_paleCyan GRM1A
02700 HMM06249EutQ_3_grm1(pf06249) GRM1A EP_NTERM
02710 HMMalddh_2_egra(pf00171) GRM1A
02715 HMMptac_3_eut2(pf06130) GRM1A
02720 H_fuchsia GRM1A
02725 -pf07992pf02852 -
02730 -pf13601 -
02735 -pf00664pf00005 -
02740 -pf00664pf00005 -
02745 -pf01370 -
02750 -pf08545pf08541 -
02755 -pf00753 -
02760 -# -
02765 -pf02655 -
02770 -# -
BMC-P
BMC-H
BMC-T
Aldehyde dehydrogenase
PTAC
Signature enzyme
Alcohol dehydrogenase
Regulator

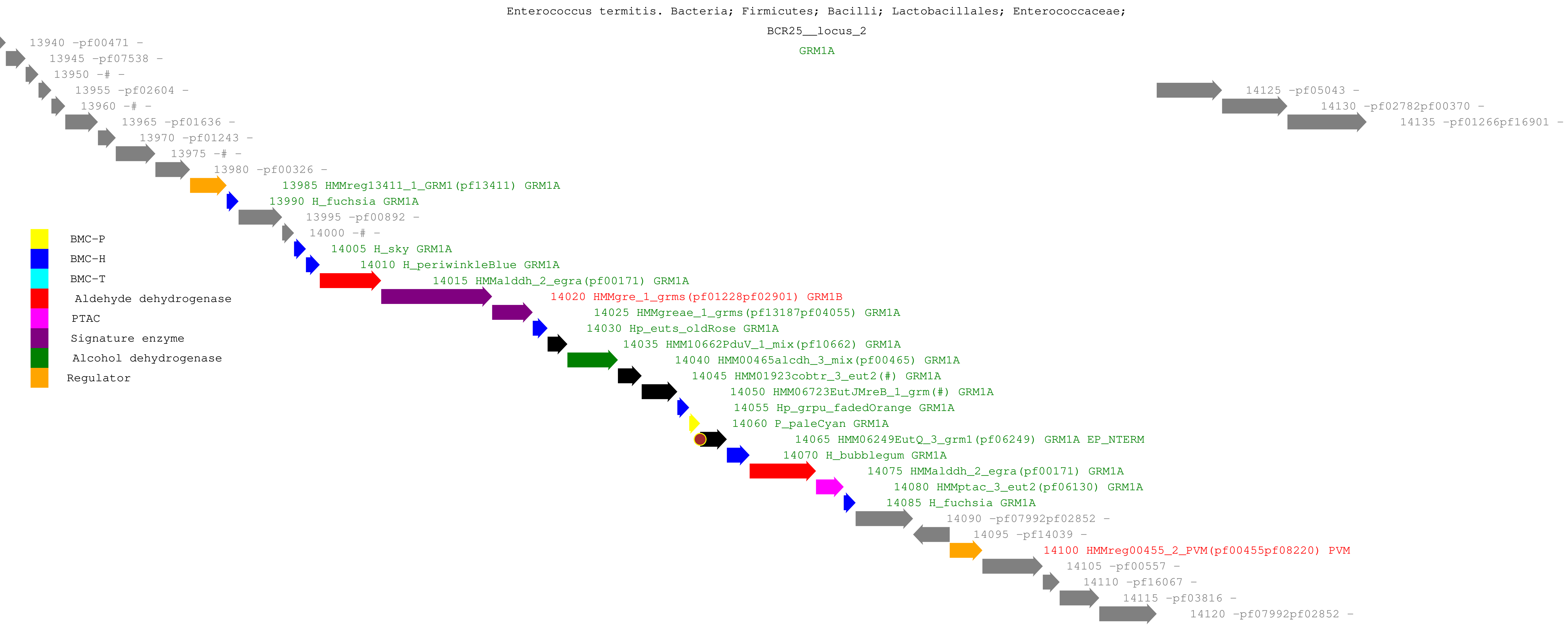
Enterococcus termitis. Bacteria; Firmicutes; Bacilli; Lactobacillales; Enterococcaceae;
BCR25__locus_2
GRM1A
13940 -pf00471 -
13945 -pf07538 -
13950 -# -
13955 -pf02604 -
13960 -# -
13965 -pf01636 -
13970 -pf01243 -
13975 -# -
13980 -pf00326 -
13985 HMMreg13411_1_GRM1(pf13411) GRM1A
13990 H_fuchsia GRM1A
13995 -pf00892 -
14000 -# -
14005 H_sky GRM1A
14010 H_periwinkleBlue GRM1A
14015 HMMalddh_2_egra(pf00171) GRM1A
14020 HMMgre_1_grms(pf01228pf02901) GRM1B
14025 HMMgreae_1_grms(pf13187pf04055) GRM1A
14030 Hp_euts_oldRose GRM1A
14035 HMM10662PduV_1_mix(pf10662) GRM1A
14040 HMM00465alcdh_3_mix(pf00465) GRM1A
14045 HMM01923cobtr_3_eut2(#) GRM1A
14050 HMM06723EutJMreB_1_grm(#) GRM1A
14055 Hp_grpu_fadedOrange GRM1A
14060 P_paleCyan GRM1A
14065 HMM06249EutQ_3_grm1(pf06249) GRM1A EP_NTERM
14070 H_bubblegum GRM1A
14075 HMMalddh_2_egra(pf00171) GRM1A
14080 HMMptac_3_eut2(pf06130) GRM1A
14085 H_fuchsia GRM1A
14090 -pf07992pf02852 -
14095 -pf14039 -
14100 HMMreg00455_2_PVM(pf00455pf08220) PVM
14105 -pf00557 -
14110 -pf16067 -
14115 -pf03816 -
14120 -pf07992pf02852 -
14125 -pf05043 -
14130 -pf02782pf00370 -
14135 -pf01266pf16901 -
BMC-P
BMC-H
BMC-T
Aldehyde dehydrogenase
PTAC
Signature enzyme
Alcohol dehydrogenase
Regulator

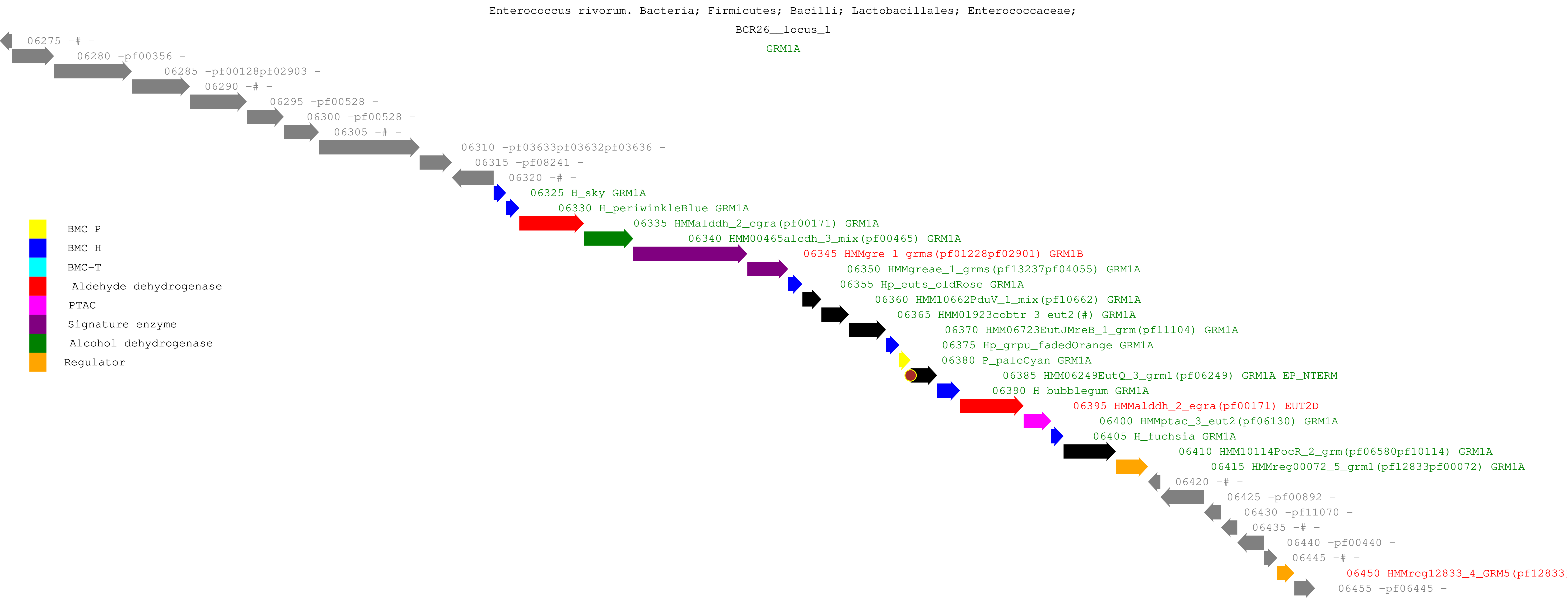
Enterococcus rivorum. Bacteria; Firmicutes; Bacilli; Lactobacillales; Enterococcaceae;
BCR26__locus_1
GRM1A
06275 -# -
06280 -pf00356 -
06285 -pf00128pf02903 -
06290 -# -
06295 -pf00528 -
06300 -pf00528 -
06305 -# -
06310 -pf03633pf03632pf03636 -
06315 -pf08241 -
06320 -# -
06325 H_sky GRM1A
06330 H_periwinkleBlue GRM1A
06335 HMMaldh_2_egra(pf00171) GRM1A
06340 HMM00465alcdh_3_mix(pf00465) GRM1A
06345 HMMgre_1_grms(pf01228pf02901) GRM1B
06350 HMMgreae_1_grms(pf13237pf04055) GRM1A
06355 Hp_euts_oldRose GRM1A
06360 HMM10662PduV_1_mix(pf10662) GRM1A
06365 HMM01923cobtr_3_eut2(#) GRM1A
06370 HMM06723EutJMreB_1_grm(pf11104) GRM1A
06375 Hp_grpu_fadedOrange GRM1A
06380 P_paleCyan GRM1A
06385 HMM06249EutQ_3_grm1(pf06249) GRM1A EP_NTERM
06390 H_bubblegum GRM1A
06395 HMMaldh_2_egra(pf00171) EUT2D
06400 HMMptac_3_eut2(pf06130) GRM1A
06405 H_fuchsia GRM1A
06410 HMM10114PocR_2_grm(pf06580pf10114) GRM1A
06415 HMMreg00072_5_grm1(pf12833pf00072) GRM1A
06420 -# -
06425 -pf00892 -
06430 -pf11070 -
06435 -# -
06440 -pf00440 -
06445 -# -
06450 HMMreg12833_4_GRM5(pf12833
06455 -pf06445 -
BMC-P
BMC-H
BMC-T
Aldehyde dehydrogenase
PTAC
Signature enzyme
Alcohol dehydrogenase
Regulator

Clostridium algidicarnis DSM 15099. Bacteria; Firmicutes; Clostridia; Clostridiales; Clostridiaceae;

BD821__locus_2

GRM1A

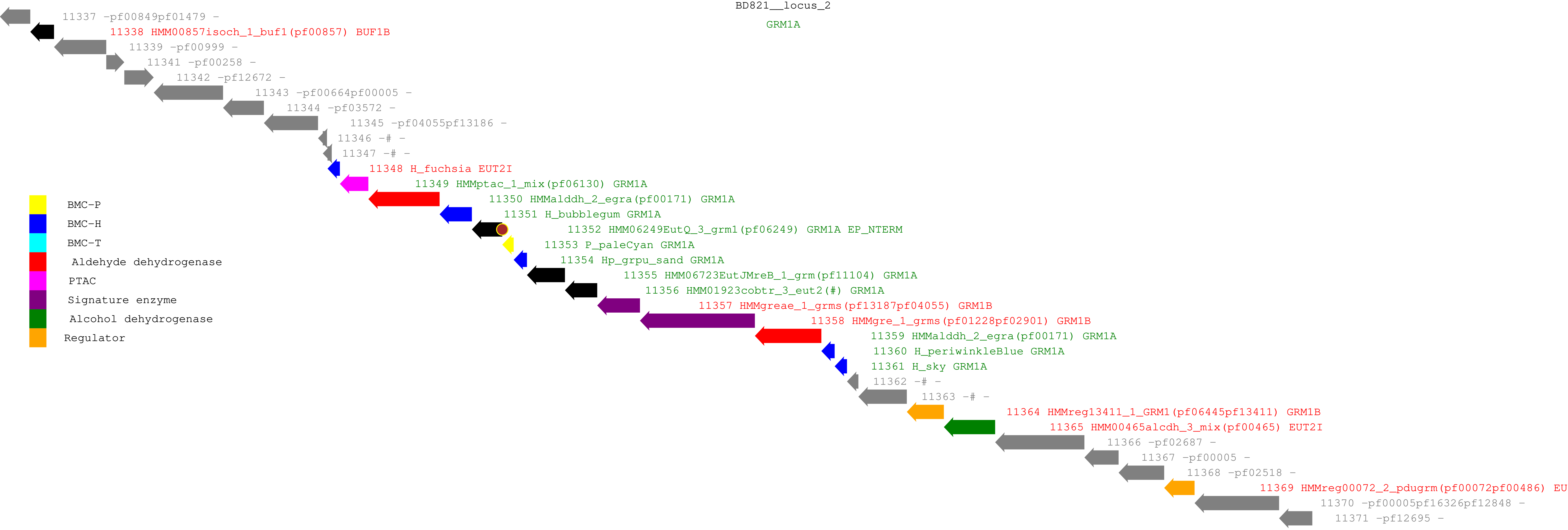

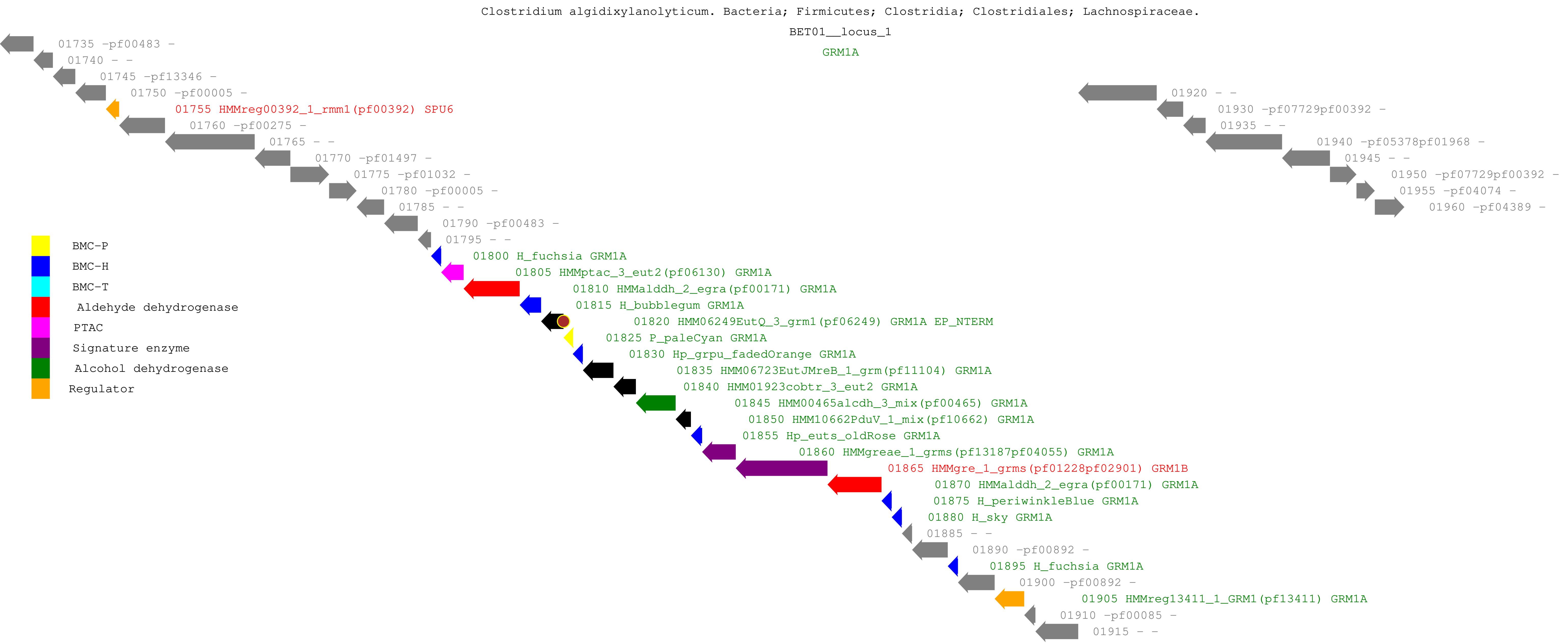
Clostridium algidixylanolyticum. Bacteria; Firmicutes; Clostridia; Clostridiales; Lachnospiraceae.
BET01__locus_1
GRM1A
01735 -pf00483 -
01740 - -
01745 -pf13346 -
01750 -pf00005 -
01755 HMMreg00392_1_rmm1(pf00392) SPU6
01760 -pf00275 -
01765 - -
01770 -pf01497 -
01775 -pf01032 -
01780 -pf00005 -
01785 - -
01790 -pf00483 -
01795 - -
01800 H_fuchsia GRM1A
01805 HMMptac_3_eut2(pf06130) GRM1A
01810 HMMaldh_2_egra(pf00171) GRM1A
01815 H_bubblegum GRM1A
01820 HMM06249EutQ_3_grm1(pf06249) GRM1A EP_NTERM
01825 P_paleCyan GRM1A
01830 Hp_grpu_fadedOrange GRM1A
01835 HMM06723EutJMreB_1_grm(pf11104) GRM1A
01840 HMM01923cobtr_3_eut2 GRM1A
01845 HMM00465alcdh_3_mix(pf00465) GRM1A
01850 HMM10662PduV_1_mix(pf10662) GRM1A
01855 Hp_euts_oldRose GRM1A
01860 HMMgreae_1_grms(pf13187pf04055) GRM1A
01865 HMMgre_1_grms(pf01228pf02901) GRM1B
01870 HMMaldh_2_egra(pf00171) GRM1A
01875 H_periwinkleBlue GRM1A
01880 H_sky GRM1A
01885 - -
01890 -pf00892 -
01895 H_fuchsia GRM1A
01900 -pf00892 -
01905 HMMreg13411_1_GRM1(pf13411) GRM1A
01910 -pf00085 -
01915 - -
01920 - -
01930 -pf07729pf00392 -
01935 - -
01940 -pf05378pf01968 -
01945 - -
01950 -pf07729pf00392 -
01955 -pf04074 -
01960 -pf04389 -
BMC-P
BMC-H
BMC-T
Aldehyde dehydrogenase
PTAC
Signature enzyme
Alcohol dehydrogenase
Regulator

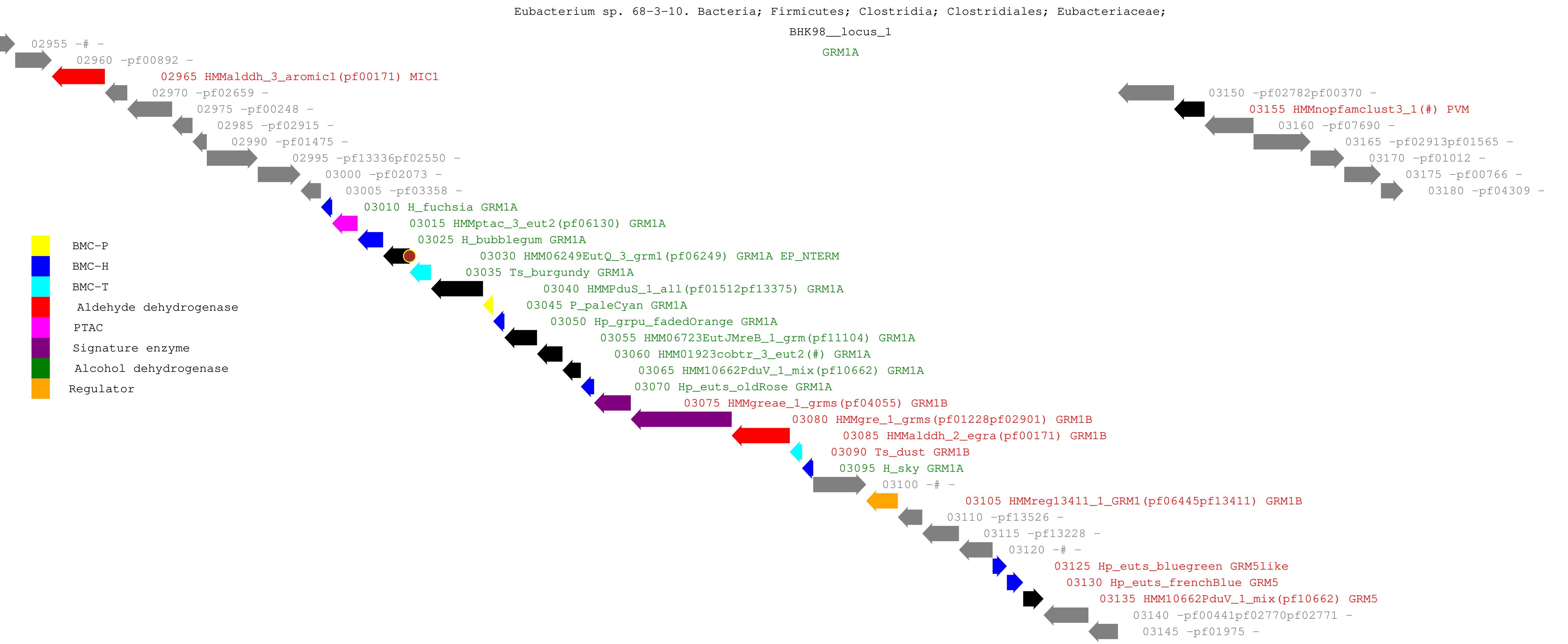
Eubacterium sp. 68-3-10. Bacteria; Firmicutes; Clostridia; Clostridiales; Eubacteriaceae;
BHK98__locus_1
GRM1A
02955 -# -
02960 -pf00892 -
02965 HMMalddh_3_aromic1(pf00171) MIC1
02970 -pf02659 -
02975 -pf00248 -
02985 -pf02915 -
02990 -pf01475 -
02995 -pf13336pf02550 -
03000 -pf02073 -
03005 -pf03358 -
03010 H_fuchsia GRM1A
03015 HMMptac_3_eut2(pf06130) GRM1A
03025 H_bubblegum GRM1A
03030 HMM06249EutQ_3_grm1(pf06249) GRM1A EP_NTERM
03035 Ts_burgundy GRM1A
03040 HMMPduS_1_all(pf01512pf13375) GRM1A
03045 P_paleCyan GRM1A
03050 Hp_grpu_fadedOrange GRM1A
03055 HMM06723EutJMreB_1_grm(pf11104) GRM1A
03060 HMM01923cobtr_3_eut2(#) GRM1A
03065 HMM10662PduV_1_mix(pf10662) GRM1A
03070 Hp_euts_oldRose GRM1A
03075 HMMgreae_1_grms(pf04055) GRM1B
03080 HMMgre_1_grms(pf01228pf02901) GRM1B
03085 HMMalddh_2_egra(pf00171) GRM1B
03090 Ts_dust GRM1B
03095 H_sky GRM1A
03100 -# -
03105 HMMreg13411_1_GRM1(pf06445pf13411) GRM1B
03110 -pf13526 -
03115 -pf13228 -
03120 -# -
03125 Hp_euts_bluegreen GRM5like
03130 Hp_euts_frenchBlue GRM5
03135 HMM10662PduV_1_mix(pf10662) GRM5
03140 -pf00441pf02770pf02771 -
03145 -pf01975 -
03150 -pf02782pf00370 -
03155 HMMnopfamclust3_1(#) PVM
03160 -pf07690 -
03165 -pf02913pf01565 -
03170 -pf01012 -
03175 -pf00766 -
03180 -pf04309 -
BMC-P
BMC-H
BMC-T
Aldehyde dehydrogenase
PTAC
Signature enzyme
Alcohol dehydrogenase
Regulator

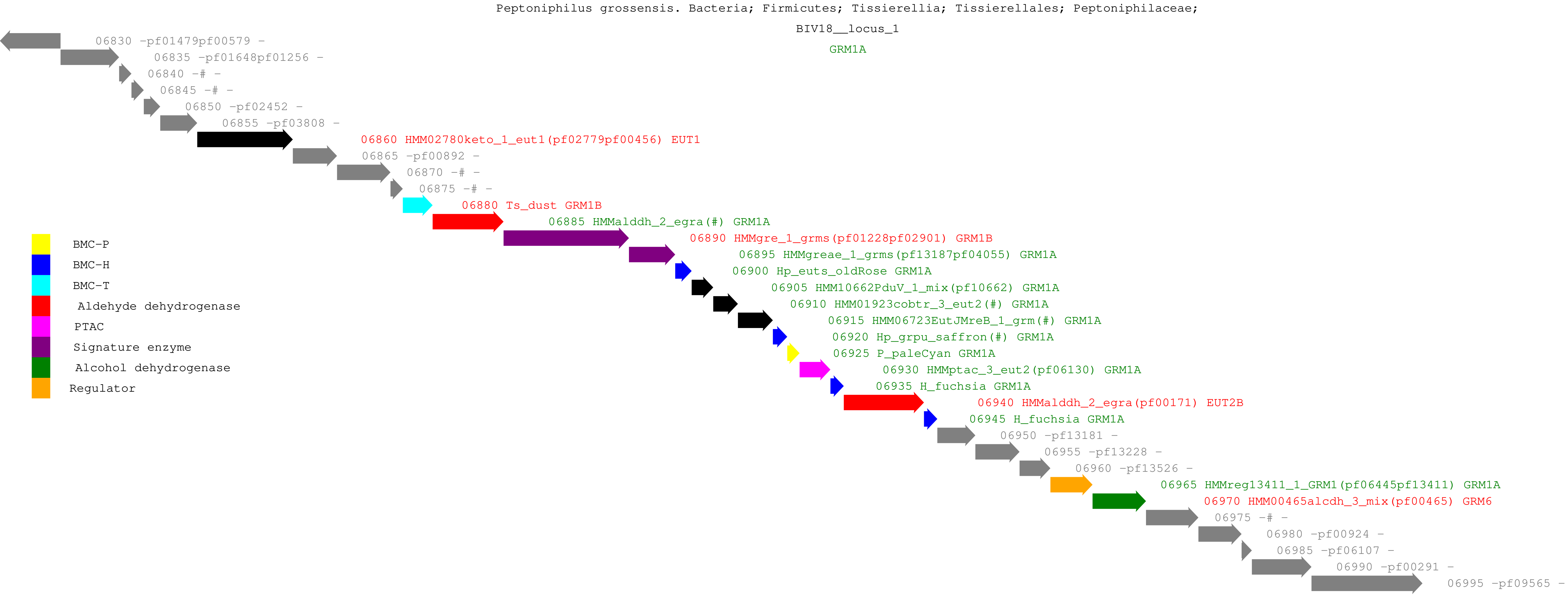
Peptoniphilus grossensis. Bacteria; Firmicutes; Tissierellia; Tissierellales; Peptoniphilaceae;
BIV18__locus_1
GRM1A
06830 -pf01479pf00579 -
06835 -pf01648pf01256 -
06840 -# -
06845 -# -
06850 -pf02452 -
06855 -pf03808 -
06860 HMM02780keto_1_eut1(pf02779pf00456) EUT1
06865 -pf00892 -
06870 -# -
06875 -# -
06880 Ts_dust GRM1B
06885 HMMalddh_2_egra(#) GRM1A
06890 HMMgre_1_grms(pf01228pf02901) GRM1B
06895 HMMgreae_1_grms(pf13187pf04055) GRM1A
06900 Hp_euts_oldRose GRM1A
06905 HMM10662PduV_1_mix(pf10662) GRM1A
06910 HMM01923cobtr_3_eut2(#) GRM1A
06915 HMM06723EutJMreB_1_grm(#) GRM1A
06920 Hp_grpu_saffron(#) GRM1A
06925 P_paleCyan GRM1A
06930 HMMptac_3_eut2(pf06130) GRM1A
06935 H_fuchsia GRM1A
06940 HMMaldddh_2_egra(pf00171) EUT2B
06945 H_fuchsia GRM1A
06950 -pf13181 -
06955 -pf13228 -
06960 -pf13526 -
06965 HMMreg13411_1_GRM1(pf06445pf13411) GRM1A
06970 HMM00465alcdh_3_mix(pf00465) GRM6
06975 -# -
06980 -pf00924 -
06985 -pf06107 -
06990 -pf00291 -
06995 -pf09565 -
BMC-P
BMC-H
BMC-T
Aldehyde dehydrogenase
PTAC
Signature enzyme
Alcohol dehydrogenase
Regulator

Clostridium formicaceticum. Bacteria; Firmicutes; Clostridia; Clostridiales; Clostridiaceae;

BJL90__locus_1

GRM1A

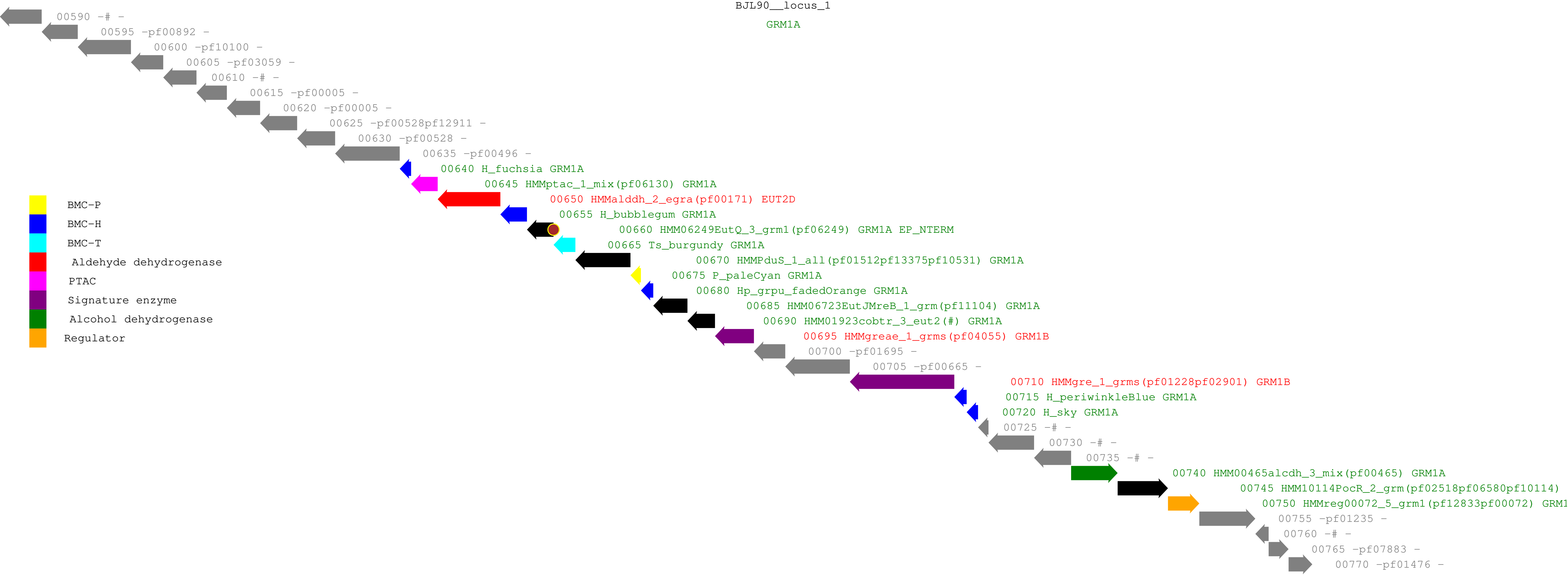

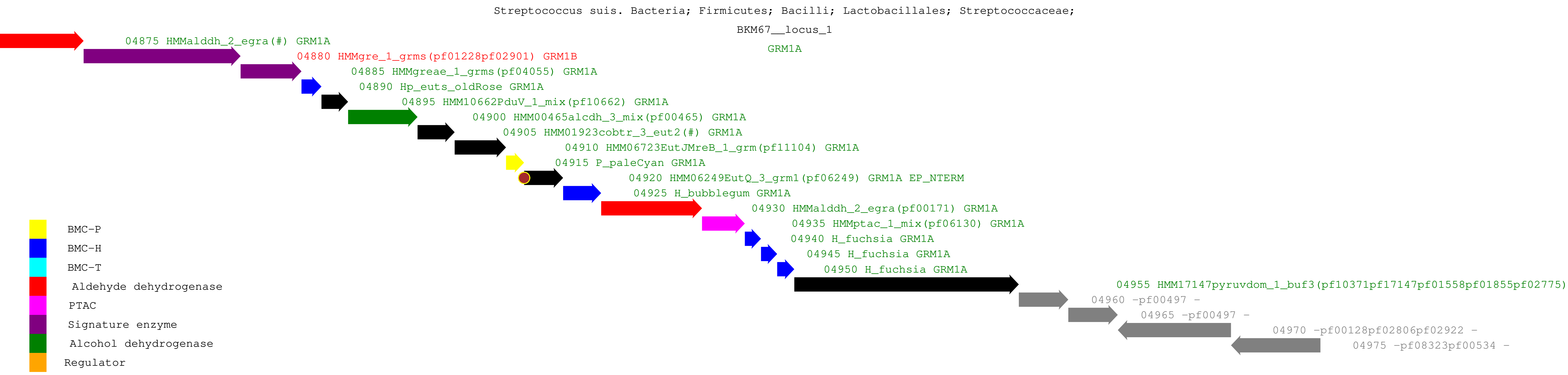
Streptococcus suis. Bacteria; Firmicutes; Bacilli; Lactobacillales; Streptococcaceae;
BKM67__locus_1
GRM1A
04875 HMMalddh_2_egra(#) GRM1A
04880 HMMgre_1_grms(pf01228pf02901) GRM1B
04885 HMMgreae_1_grms(pf04055) GRM1A
04890 Hp_euts_oldRose GRM1A
04895 HMM10662PduV_1_mix(pf10662) GRM1A
04900 HMM00465alcdh_3_mix(pf00465) GRM1A
04905 HMM01923cobtr_3_eut2(#) GRM1A
04910 HMM06723EutJMreB_1_grm(pf11104) GRM1A
04915 P_paleCyan GRM1A
04920 HMM06249EutQ_3_grm1(pf06249) GRM1A EP_NTERM
04925 H_bubblegum GRM1A
04930 HMMalddh_2_egra(pf00171) GRM1A
04935 HMMptac_1_mix(pf06130) GRM1A
04940 H_fuchsia GRM1A
04945 H_fuchsia GRM1A
04950 H_fuchsia GRM1A
04955 HMM17147pyruvdom_1_buf3(pf10371pf17147pf01558pf01855pf02775)
04960 -pf00497 -
04965 -pf00497 -
04970 -pf00128pf02806pf02922 -
04975 -pf08323pf00534 -
BMC-P
BMC-H
BMC-T
Aldehyde dehydrogenase
PTAC
Signature enzyme
Alcohol dehydrogenase
Regulator

Streptococcus iniae. Bacteria; Firmicutes; Bacilli; Lactobacillales; Streptococcaceae;

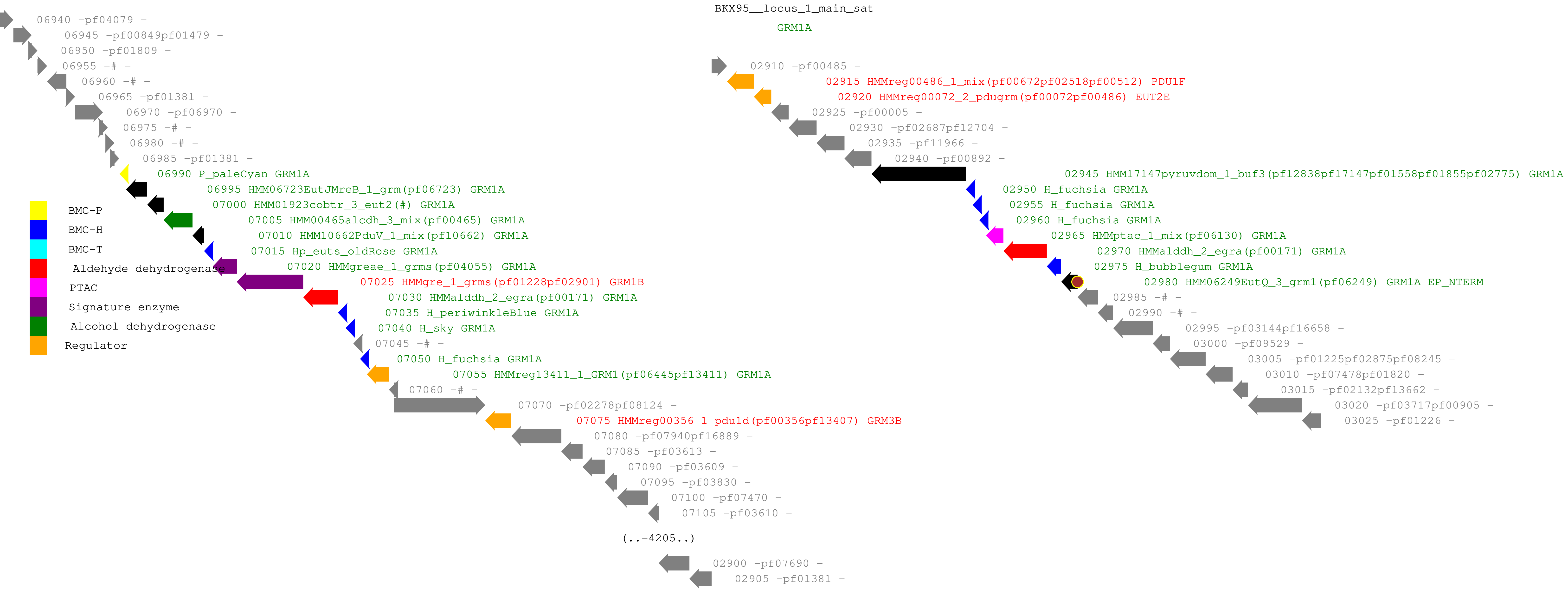

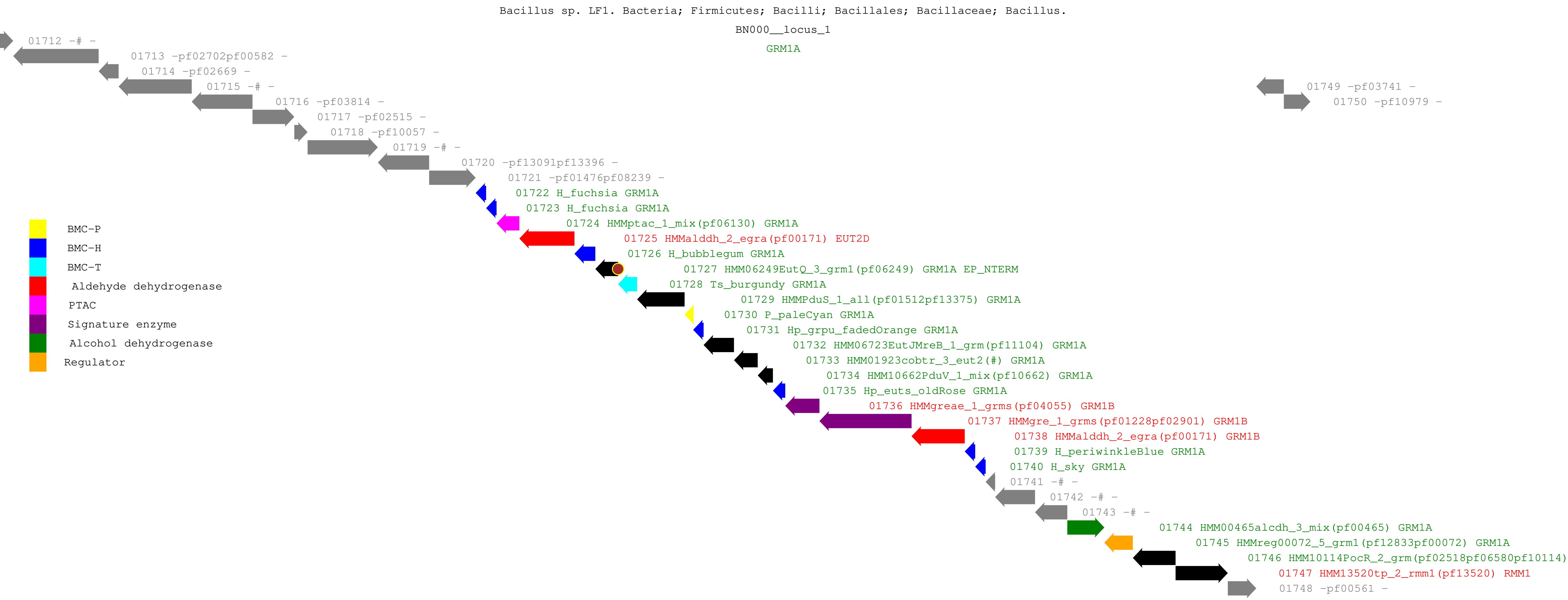

Bacillus sp. LF1. Bacteria; Firmicutes; Bacilli; Bacillales; Bacillaceae; Bacillus.
BN000__locus_1
GRM1A
01712 -# -
01713 -pf02702pf00582 -
01714 -pf02669 -
01715 -# -
01716 -pf03814 -
01717 -pf02515 -
01718 -pf10057 -
01719 -# -
01720 -pf13091pf13396 -
01721 -pf01476pf08239 -
01722 H_fuchsia GRM1A
01723 H_fuchsia GRM1A
01724 HMMptac_1_mix(pf06130) GRM1A
01725 HMMaldh_2_egra(pf00171) EUT2D
01726 H_bubblegum GRM1A
01727 HMM06249EutQ_3_grm1(pf06249) GRM1A EP_NTERM
01728 Ts_burgundy GRM1A
01729 HMMPduS_1_all(pf01512pf13375) GRM1A
01730 P_paleCyan GRM1A
01731 Hp_grpu_fadedOrange GRM1A
01732 HMM06723EutJMreB_1_grm(pf11104) GRM1A
01733 HMM01923cobtr_3_eut2(#) GRM1A
01734 HMM10662PduV_1_mix(pf10662) GRM1A
01735 Hp_euts_oldRose GRM1A
01736 HMMgreae_1_grms(pf04055) GRM1B
01737 HMMgre_1_grms(pf01228pf02901) GRM1B
01738 HMMaldh_2_egra(pf00171) GRM1B
01739 H_periwinkleBlue GRM1A
01740 H_sky GRM1A
01741 -# -
01742 -# -
01743 -# -
01744 HMM00465alcdh_3_mix(pf00465) GRM1A
01745 HMMreg00072_5_grm1(pf12833pf00072) GRM1A
01746 HMM10114PocR_2_grm(pf02518pf06580pf10114)
01747 HMM13520tp_2_rmm1(pf13520) RMM1
01748 -pf00561 -
01749 -pf03741 -
01750 -pf10979 -
BMC-P
BMC-H
BMC-T
Aldehyde dehydrogenase
PTAC
Signature enzyme
Alcohol dehydrogenase
Regulator

Clostridium sp. C105KSO15. Bacteria; Firmicutes; Clostridia; Clostridiales; Clostridiaceae;

BN3590__locus_2

GRM1A

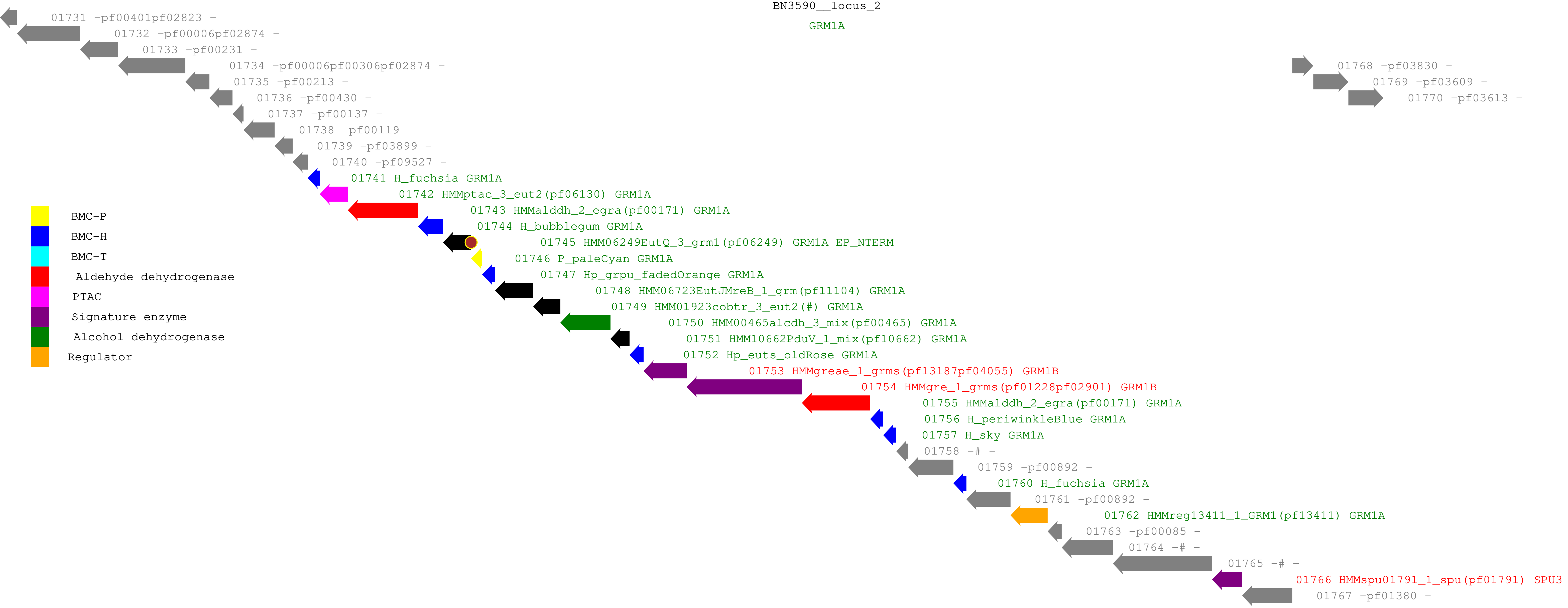

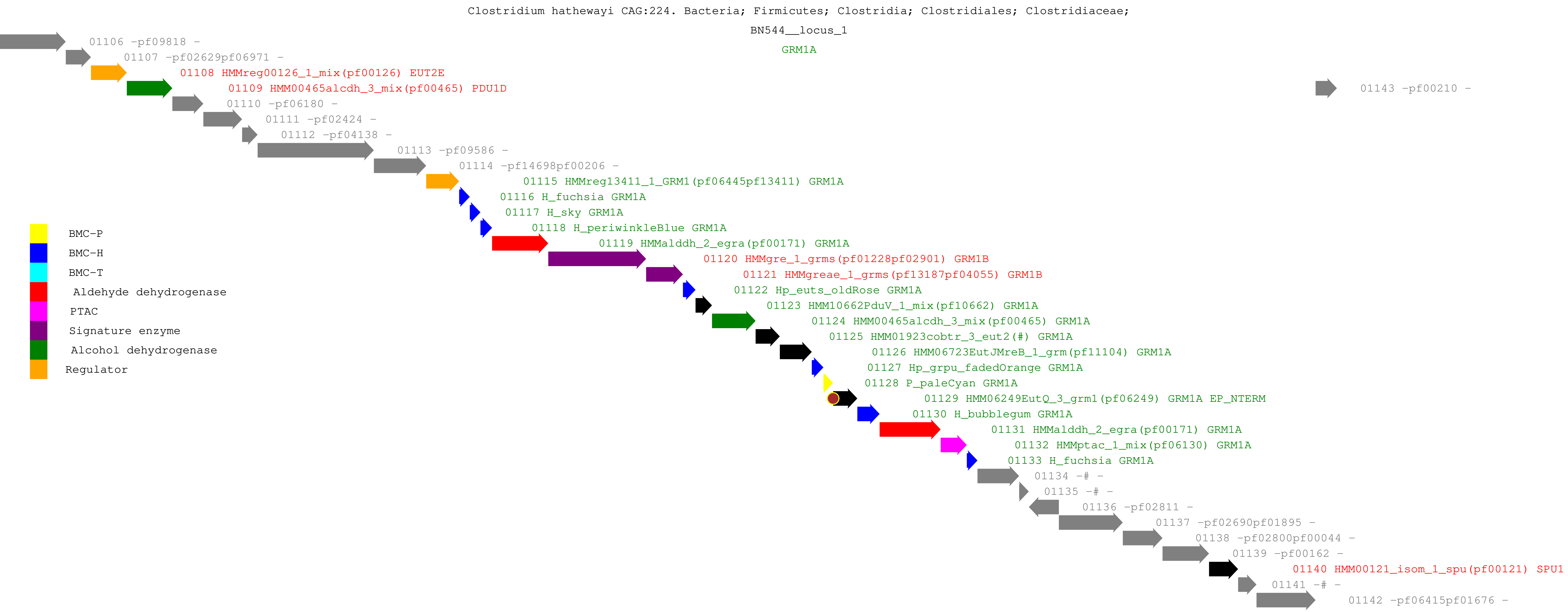
Clostridium hathewayi CAG:224. Bacteria; Firmicutes; Clostridia; Clostridiales; Clostridiaceae;
BN544__locus_1
GRM1A
01106 -pf09818 -
01107 -pf02629pf06971 -
01108 HMMreg00126_1_mix(pf00126) EUT2E
01109 HMM00465alcdh_3_mix(pf00465) PDU1D
01110 -pf06180 -
01111 -pf02424 -
01112 -pf04138 -
01113 -pf09586 -
01114 -pf14698pf00206 -
01115 HMMreg13411_1_GRM1(pf06445pf13411) GRM1A
01116 H_fuchsia GRM1A
01117 H_sky GRM1A
01118 H_periwinkleBlue GRM1A
01119 HMMalddh_2_egra(pf00171) GRM1A
01120 HMMgre_1_grms(pf01228pf02901) GRM1B
01121 HMMgreae_1_grms(pf13187pf04055) GRM1B
01122 Hp_euts_oldRose GRM1A
01123 HMM10662PduV_1_mix(pf10662) GRM1A
01124 HMM00465alcdh_3_mix(pf00465) GRM1A
01125 HMM01923cobtr_3_eut2(#) GRM1A
01126 HMM06723EutJMreB_1_grm(pf11104) GRM1A
01127 Hp_grpu_fadedOrange GRM1A
01128 P_paleCyan GRM1A
01129 HMM06249EutQ_3_grm1(pf06249) GRM1A EP_NTERM
01130 H_bubblegum GRM1A
01131 HMMalddh_2_egra(pf00171) GRM1A
01132 HMMptac_1_mix(pf06130) GRM1A
01133 H_fuchsia GRM1A
01134 -# -
01135 -# -
01136 -pf02811 -
01137 -pf02690pf01895 -
01138 -pf02800pf00044 -
01139 -pf00162 -
01140 HMM00121_isom_1_spu(pf00121) SPU1
01141 -# -
01142 -pf06415pf01676 -
01143 -pf00210 -
BMC-P
BMC-H
BMC-T
Aldehyde dehydrogenase
PTAC
Signature enzyme
Alcohol dehydrogenase
Regulator

Collinsella sp. CAG:289. Bacteria; Actinobacteria; Coriobacteriia; Coriobacteriales;

BN589__locus_1

GRM1A

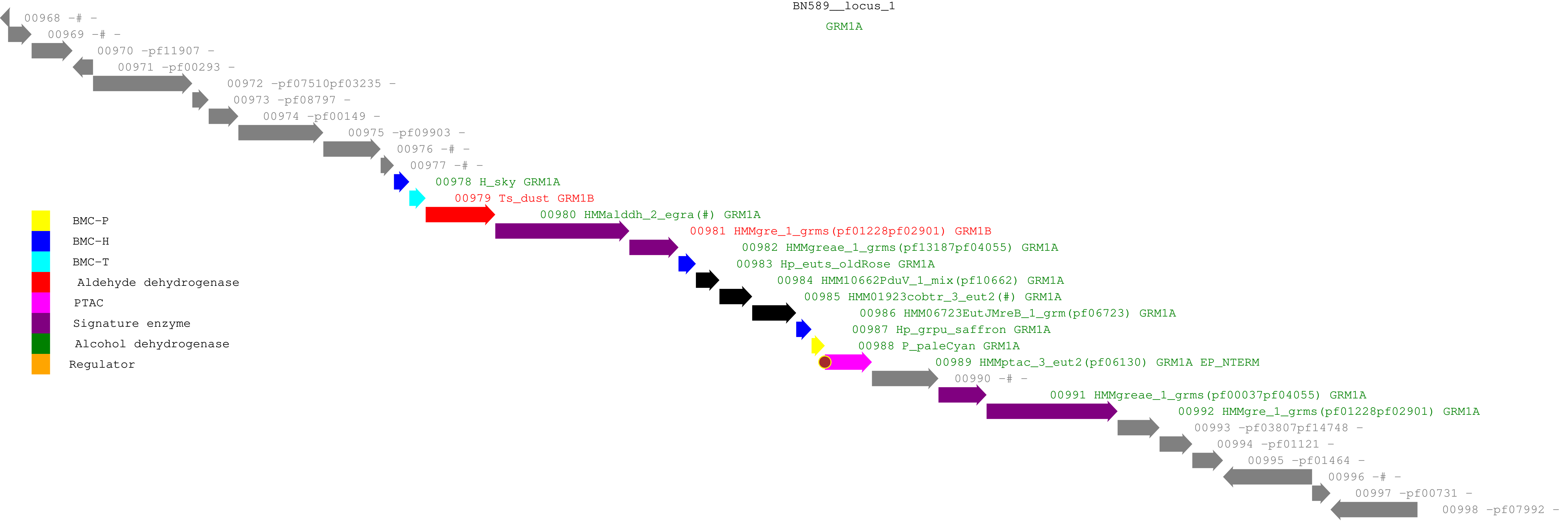

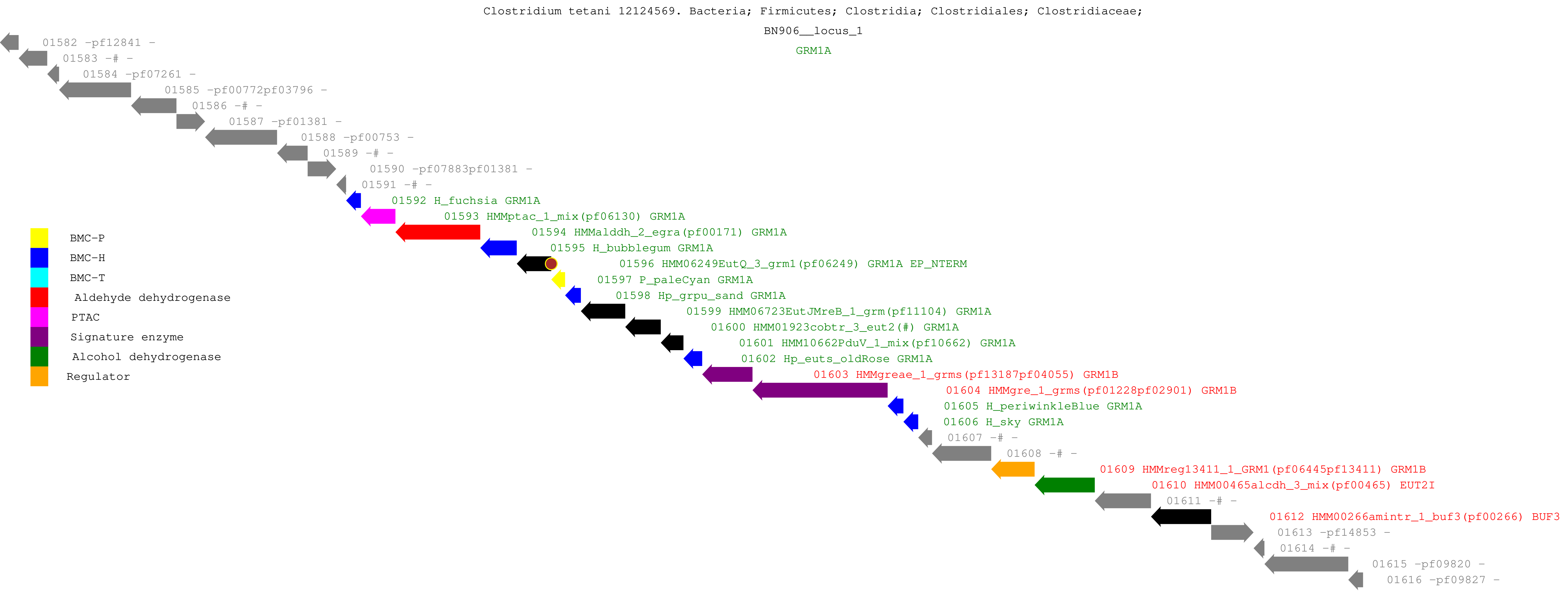
Clostridium tetani 12124569. Bacteria; Firmicutes; Clostridia; Clostridiales; Clostridiaceae;
BN906__locus_1
GRM1A
01582 -pf12841 -
01583 -# -
01584 -pf07261 -
01585 -pf00772pf03796 -
01586 -# -
01587 -pf01381 -
01588 -pf00753 -
01589 -# -
01590 -pf07883pf01381 -
01591 -# -
01592 H_fuchsia GRM1A
01593 HMMptac_1_mix(pf06130) GRM1A
01594 HMMalddh_2_egra(pf00171) GRM1A
01595 H_bubblegum GRM1A
01596 HMM06249EutQ_3_grm1(pf06249) GRM1A EP_NTERM
01597 P_paleCyan GRM1A
01598 Hp_grpu_sand GRM1A
01599 HMM06723EutJMreB_1_grm(pf11104) GRM1A
01600 HMM01923cobtr_3_eut2(#) GRM1A
01601 HMM10662PduV_1_mix(pf10662) GRM1A
01602 Hp_euts_oldRose GRM1A
01603 HMMgreae_1_grms(pf13187pf04055) GRM1B
01604 HMMgre_1_grms(pf01228pf02901) GRM1B
01605 H_periwinkleBlue GRM1A
01606 H_sky GRM1A
01607 -# -
01608 -# -
01609 HMMreg13411_1_GRM1(pf06445pf13411) GRM1B
01610 HMM00465alcdh_3_mix(pf00465) EUT2I
01611 -# -
01612 HMM00266amintr_1_buf3(pf00266) BUF3
01613 -pf14853 -
01614 -# -
01615 -pf09820 -
01616 -pf09827 -
BMC-P
BMC-H
BMC-T
Aldehyde dehydrogenase
PTAC
Signature enzyme
Alcohol dehydrogenase
Regulator

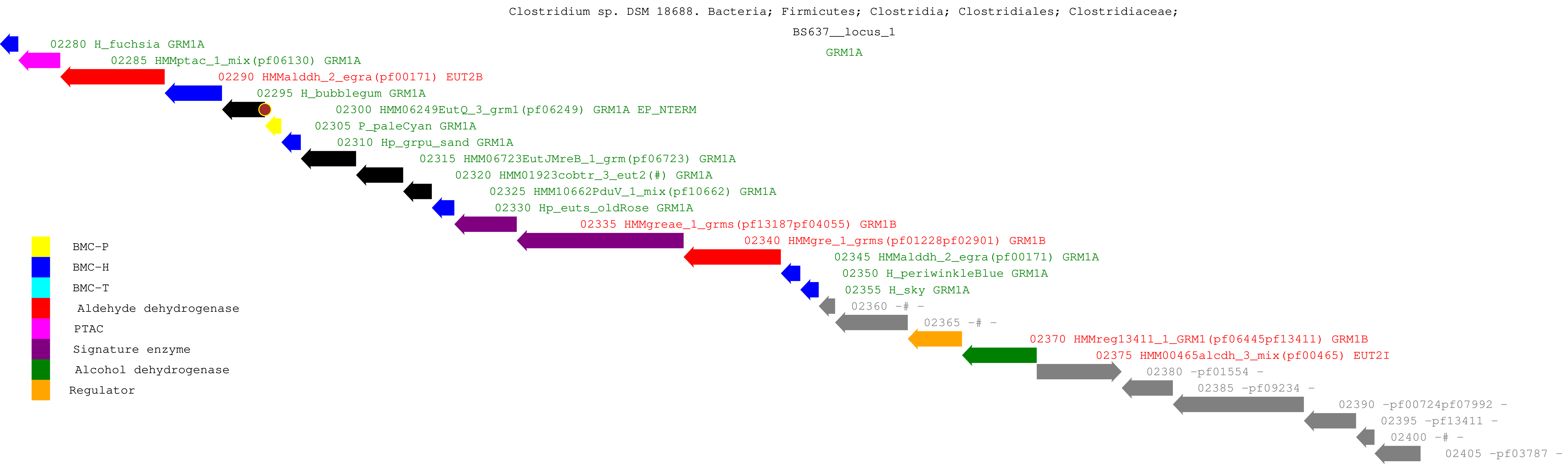
Clostridium sp. DSM 18688. Bacteria; Firmicutes; Clostridia; Clostridiales; Clostridiaceae;
BS637__locus_1
GRM1A
02280 H_fuchsia GRM1A
02285 HMMptac_1_mix(pf06130) GRM1A
02290 HMMaldh_2_egra(pf00171) EUT2B
02295 H_bubblegum GRM1A
02300 HMM06249EutQ_3_grm1(pf06249) GRM1A EP_NTERM
02305 P_paleCyan GRM1A
02310 Hp_grpu_sand GRM1A
02315 HMM06723EutJMreB_1_grm(pf06723) GRM1A
02320 HMM01923cobtr_3_eut2(#) GRM1A
02325 HMM10662PduV_1_mix(pf10662) GRM1A
02330 Hp_euts_oldRose GRM1A
02335 HMMgreae_1_grms(pf13187pf04055) GRM1B
02340 HMMgre_1_grms(pf01228pf02901) GRM1B
02345 HMMaldh_2_egra(pf00171) GRM1A
02350 H_periwinkleBlue GRM1A
02355 H_sky GRM1A
02360 -# -
02365 -# -
02370 HMMreg13411_1_GRM1(pf06445pf13411) GRM1B
02375 HMM00465alcdh_3_mix(pf00465) EUT2I
02380 -pf01554 -
02385 -pf09234 -
02390 -pf00724pf07992 -
02395 -pf13411 -
02400 -# -
02405 -pf03787 -
BMC-P
BMC-H
BMC-T
Aldehyde dehydrogenase
PTAC
Signature enzyme
Alcohol dehydrogenase
Regulator

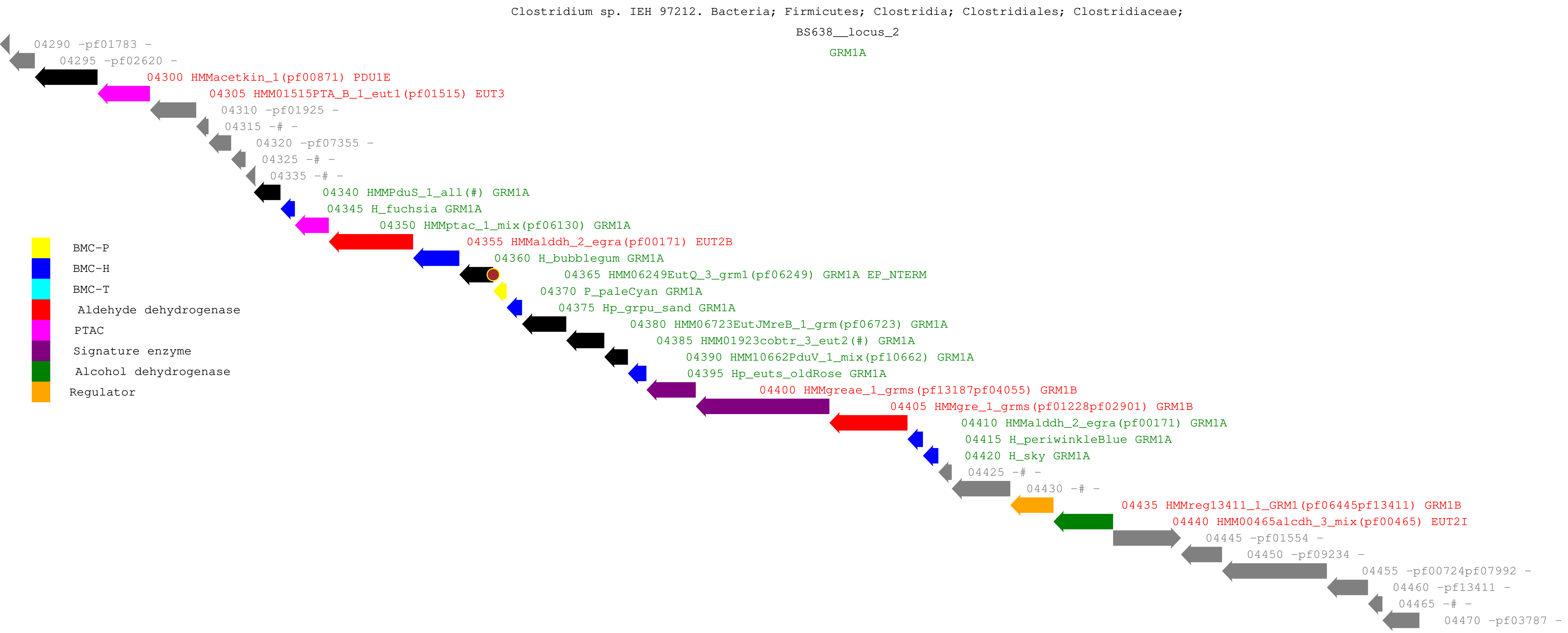
Clostridium sp. IEH 97212. Bacteria; Firmicutes; Clostridia; Clostridiales; Clostridiaceae;
BS638__locus_2
GRM1A
04290 -pf01783 -
04295 -pf02620 -
04300 HMMacetkin_1(pf00871) PDU1E
04305 HMM01515PTA_B_1_eut1(pf01515) EUT3
04310 -pf01925 -
04315 -# -
04320 -pf07355 -
04325 -# -
04335 -# -
04340 HMMPduS_1_all(#) GRM1A
04345 H_fuchsia GRM1A
04350 HMMptac_1_mix(pf06130) GRM1A
04355 HMMalddh_2_egra(pf00171) EUT2B
04360 H_bubblegum GRM1A
04365 HMM06249EutQ_3_grm1(pf06249) GRM1A EP_NTERM
04370 P_paleCyan GRM1A
04375 Hp_grpu_sand GRM1A
04380 HMM06723EutJMreB_1_grm(pf06723) GRM1A
04385 HMM01923cobtr_3_eut2(#) GRM1A
04390 HMM10662PduV_1_mix(pf10662) GRM1A
04395 Hp_euts_oldRose GRM1A
04400 HMMgreae_1_grms(pf13187pf04055) GRM1B
04405 HMMgre_1_grms(pf01228pf02901) GRM1B
04410 HMMalddh_2_egra(pf00171) GRM1A
04415 H_periwinkleBlue GRM1A
04420 H_sky GRM1A
04425 -# -
04430 -# -
04435 HMMreg13411_1_GRM1(pf06445pf13411) GRM1B
04440 HMM00465alcdh_3_mix(pf00465) EUT2I
04445 -pf01554 -
04450 -pf09234 -
04455 -pf00724pf07992 -
04460 -pf13411 -
04465 -# -
04470 -pf03787 -
BMC-P
BMC-H
BMC-T
Aldehyde dehydrogenase
PTAC
Signature enzyme
Alcohol dehydrogenase
Regulator

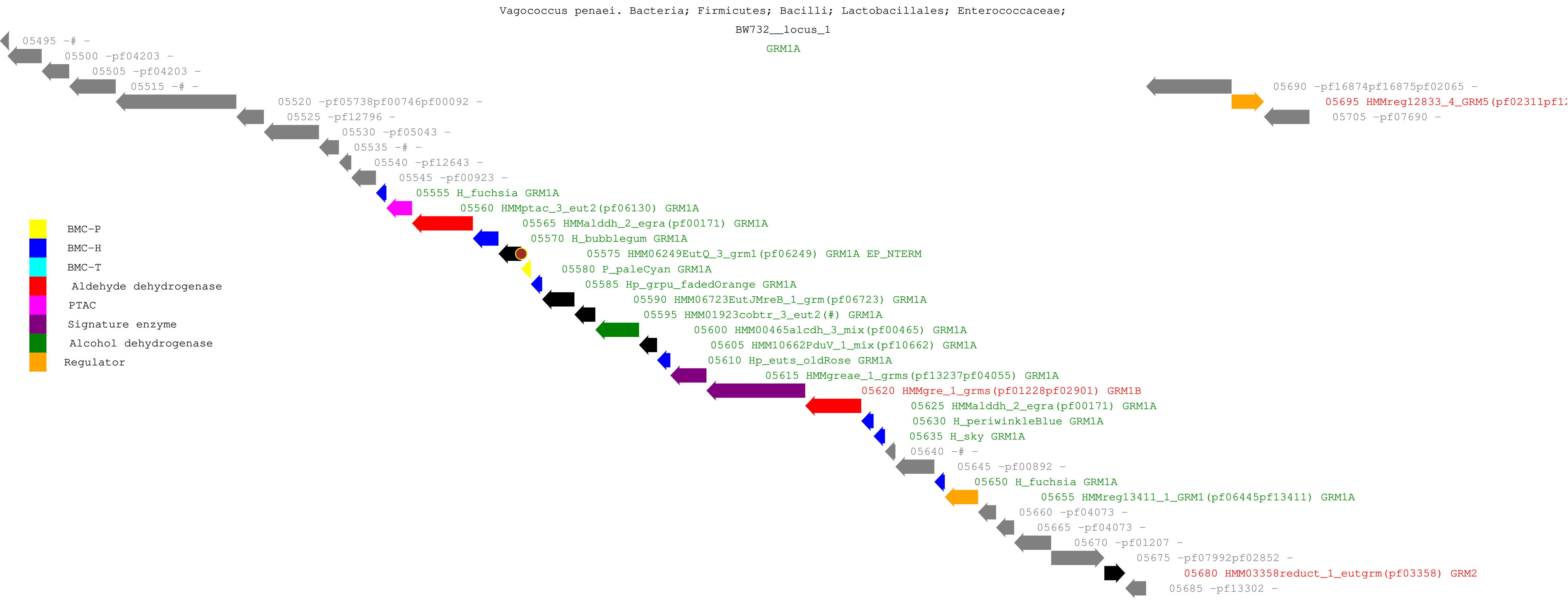
Vagococcus penaei. Bacteria; Firmicutes; Bacilli; Lactobacillales; Enterococcaceae;
BW732__locus_1
GRM1A
05495 -# -
05500 -pf04203 -
05505 -pf04203 -
05515 -# -
05520 -pf05738pf00746pf00092 -
05525 -pf12796 -
05530 -pf05043 -
05535 -# -
05540 -pf12643 -
05545 -pf00923 -
05555 H_fuchsia GRM1A
05560 HMMptac_3_eut2(pf06130) GRM1A
05565 HMMaldh_2_egra(pf00171) GRM1A
05570 H_bubblegum GRM1A
05575 HMM06249EutQ_3_grm1(pf06249) GRM1A EP_NTERM
05580 P_paleCyan GRM1A
05585 Hp_grpu_fadedOrange GRM1A
05590 HMM06723EutJMreB_1_grm(pf06723) GRM1A
05595 HMM01923cobtr_3_eut2(#) GRM1A
05600 HMM00465alcdh_3_mix(pf00465) GRM1A
05605 HMM10662PduV_1_mix(pf10662) GRM1A
05610 Hp_euts_oldRose GRM1A
05615 HMMgreae_1_grms(pf13237pf04055) GRM1A
05620 HMMgre_1_grms(pf01228pf02901) GRM1B
05625 HMMaldh_2_egra(pf00171) GRM1A
05630 H_periwinkleBlue GRM1A
05635 H_sky GRM1A
05640 -# -
05645 -pf00892 -
05650 H_fuchsia GRM1A
05655 HMMreg13411_1_GRM1(pf06445pf13411) GRM1A
05660 -pf04073 -
05665 -pf04073 -
05670 -pf01207 -
05675 -pf07992pf02852 -
05680 HMM03358reduct_1_eutgrm(pf03358) GRM2
05685 -pf13302 -
05690 -pf16874pf16875pf02065 -
05695 HMMreg12833_4_GRM5(pf02311pf1
05705 -pf07690 -
BMC-P
BMC-H
BMC-T
Aldehyde dehydrogenase
PTAC
Signature enzyme
Alcohol dehydrogenase
Regulator

Bacteroides xylanolyticus. Bacteria; Bacteroidetes; Bacteroidia; Bacteroidales; Bacteroidaceae;

BXY41__locus_1

GRM1A

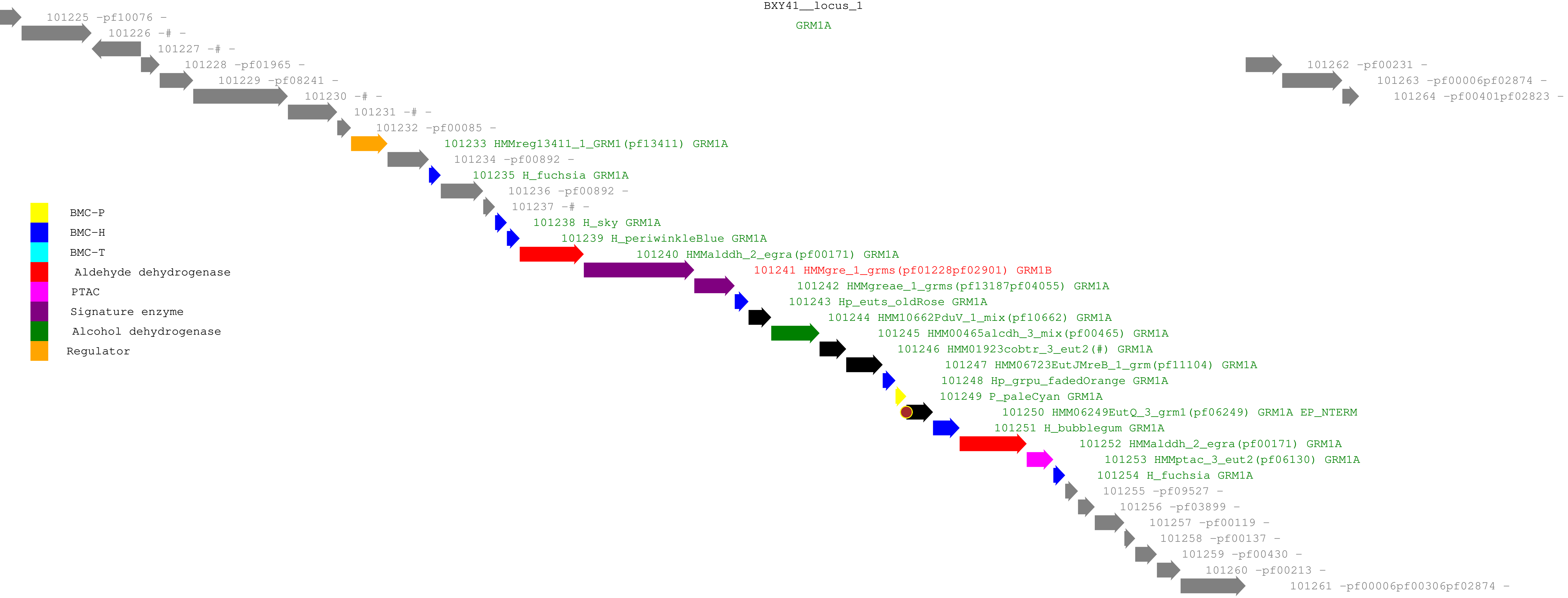

Clostridium botulinum. Bacteria; Firmicutes; Clostridia; Clostridiales; Clostridiaceae;

C1149__locus_1

GRM1A

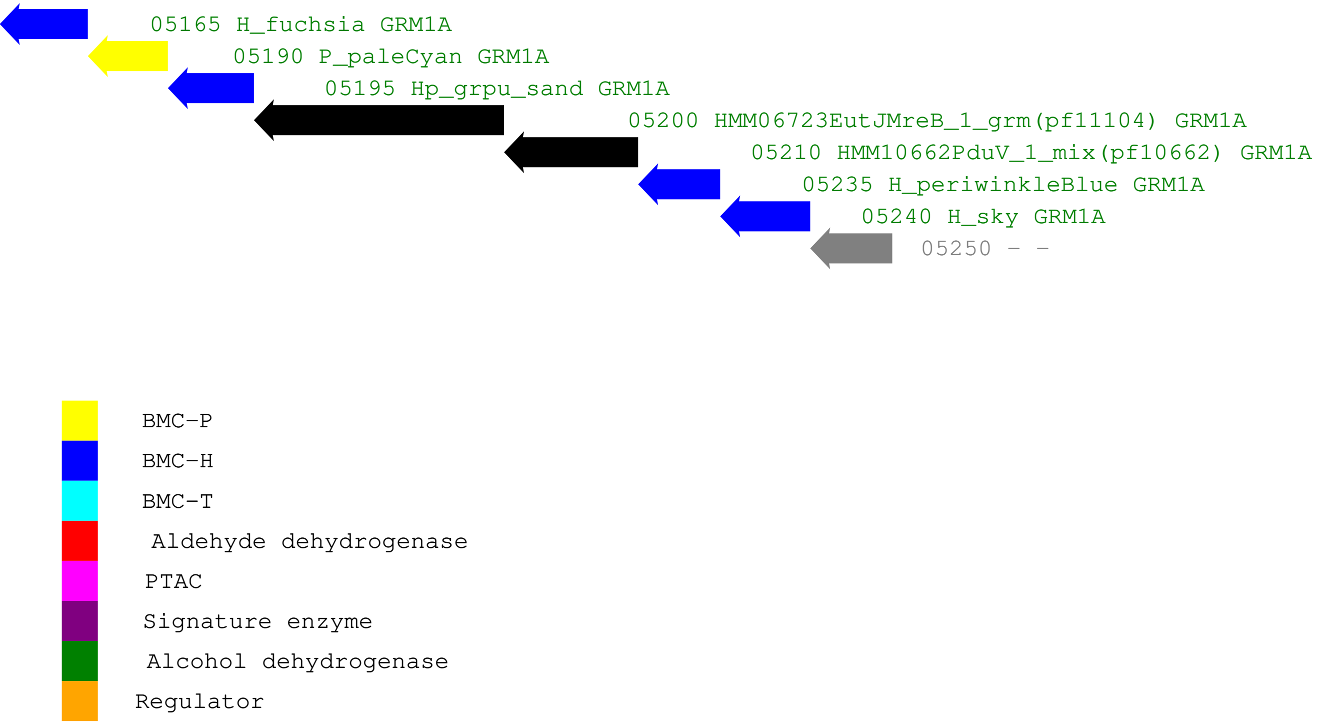

Clostridiaceae bacterium 14S0207. Bacteria; Firmicutes; Clostridia; Clostridiales; Clostridiaceae.

C3495__locus_2

GRM1A

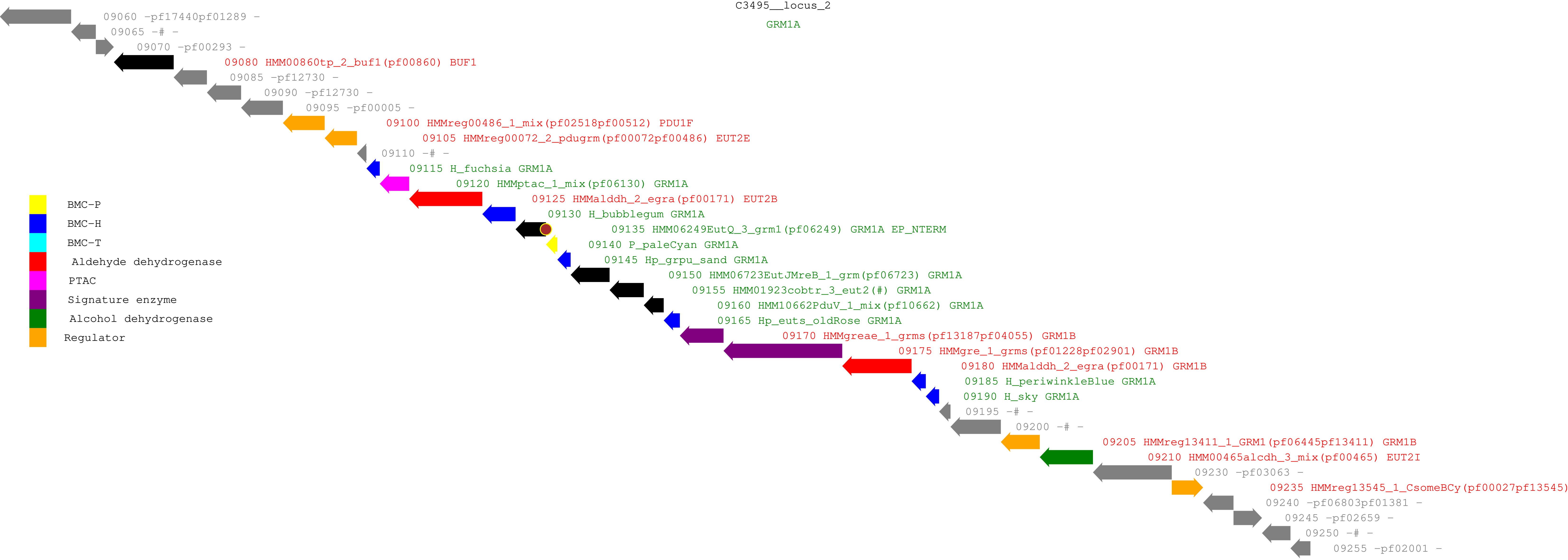

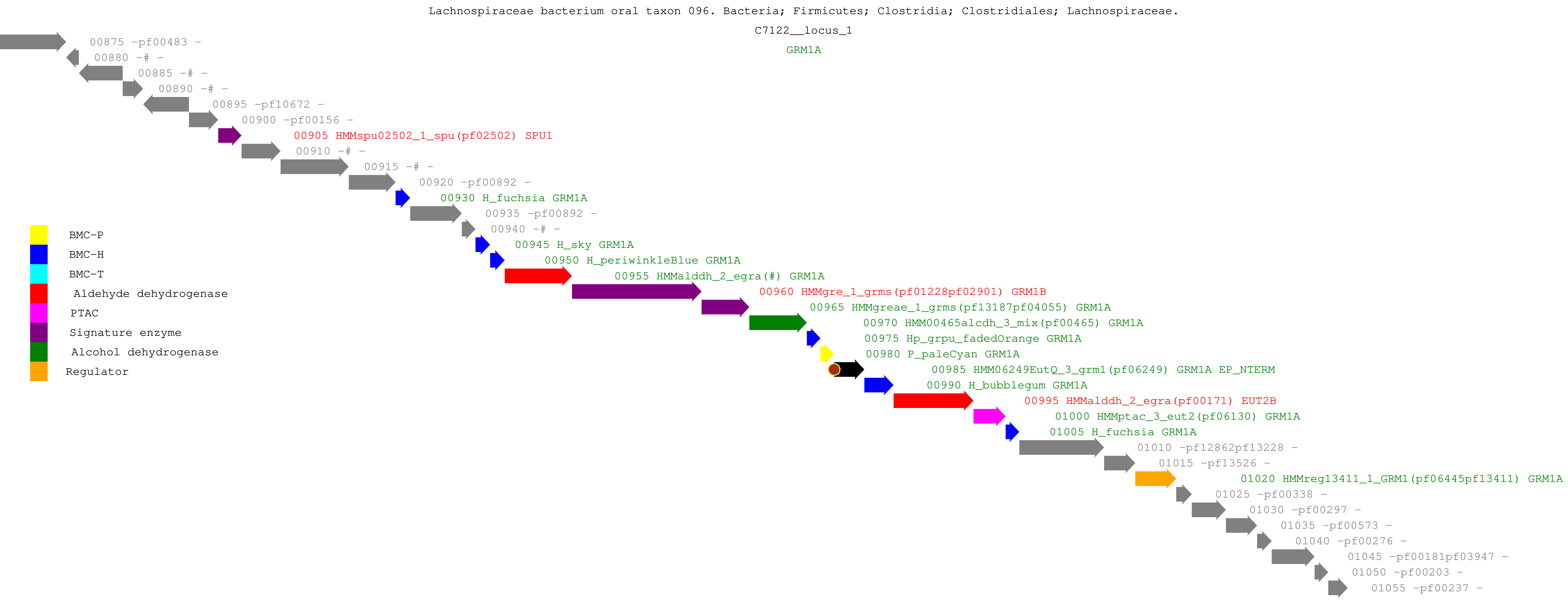
Lachnospiraceae bacterium oral taxon 096. Bacteria; Firmicutes; Clostridia; Clostridiales; Lachnospiraceae.
C7122__locus_1
GRM1A
00875 -pf00483 -
00880 -# -
00885 -# -
00890 -# -
00895 -pf10672 -
00900 -pf00156 -
00905 HMMspu02502_1_spu(pf02502) SPU1
00910 -# -
00915 -# -
00920 -pf00892 -
00930 H_fuchsia GRM1A
00935 -pf00892 -
00940 -# -
00945 H_sky GRM1A
00950 H_periwinkleBlue GRM1A
00955 HMMaldh_2_egra(#) GRM1A
00960 HMMgre_1_grms(pf01228pf02901) GRM1B
00965 HMMgreae_1_grms(pf13187pf04055) GRM1A
00970 HMM00465alcdh_3_mix(pf00465) GRM1A
00975 Hp_grpu_fadedOrange GRM1A
00980 P_paleCyan GRM1A
00985 HMM06249EutQ_3_grm1(pf06249) GRM1A EP_NTERM
00990 H_bubblegum GRM1A
00995 HMMaldh_2_egra(pf00171) EUT2B
01000 HMMptac_3_eut2(pf06130) GRM1A
01005 H_fuchsia GRM1A
01010 -pf12862pf13228 -
01015 -pf13526 -
01020 HMMreg13411_1_GRM1(pf06445pf13411) GRM1A
01025 -pf00338 -
01030 -pf00297 -
01035 -pf00573 -
01040 -pf00276 -
01045 -pf00181pf03947 -
01050 -pf00203 -
01055 -pf00237 -
BMC-P
BMC-H
BMC-T
Aldehyde dehydrogenase
PTAC
Signature enzyme
Alcohol dehydrogenase
Regulator

Clostridium lavalense. Bacteria; Firmicutes; Clostridia; Clostridiales; Lachnospiraceae.

C7256__locus_2

GRM1A

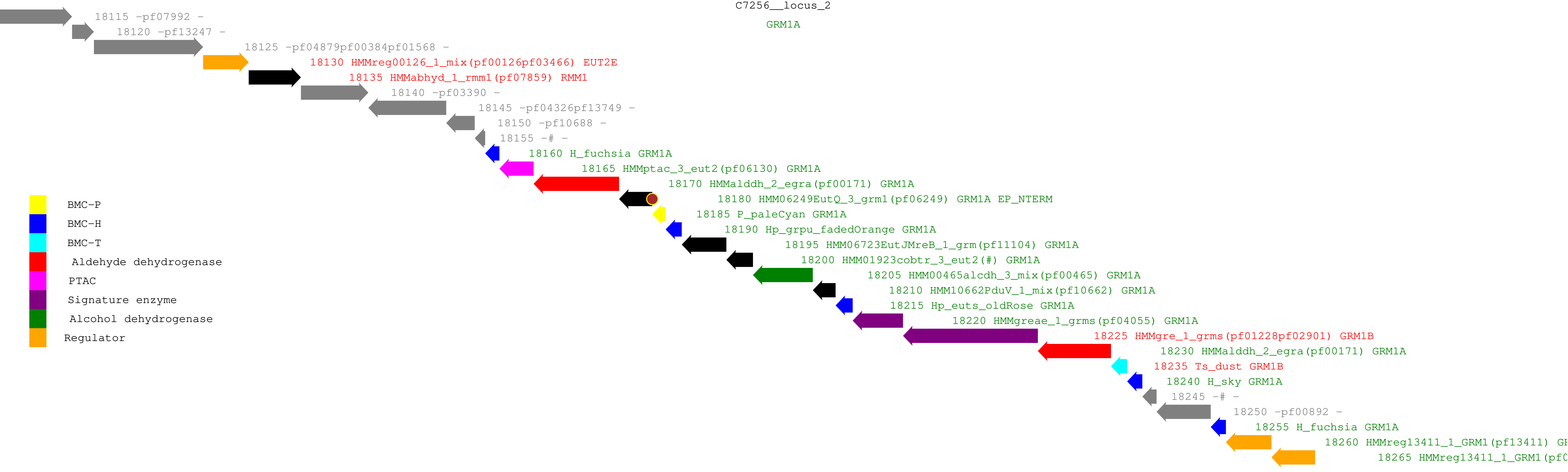

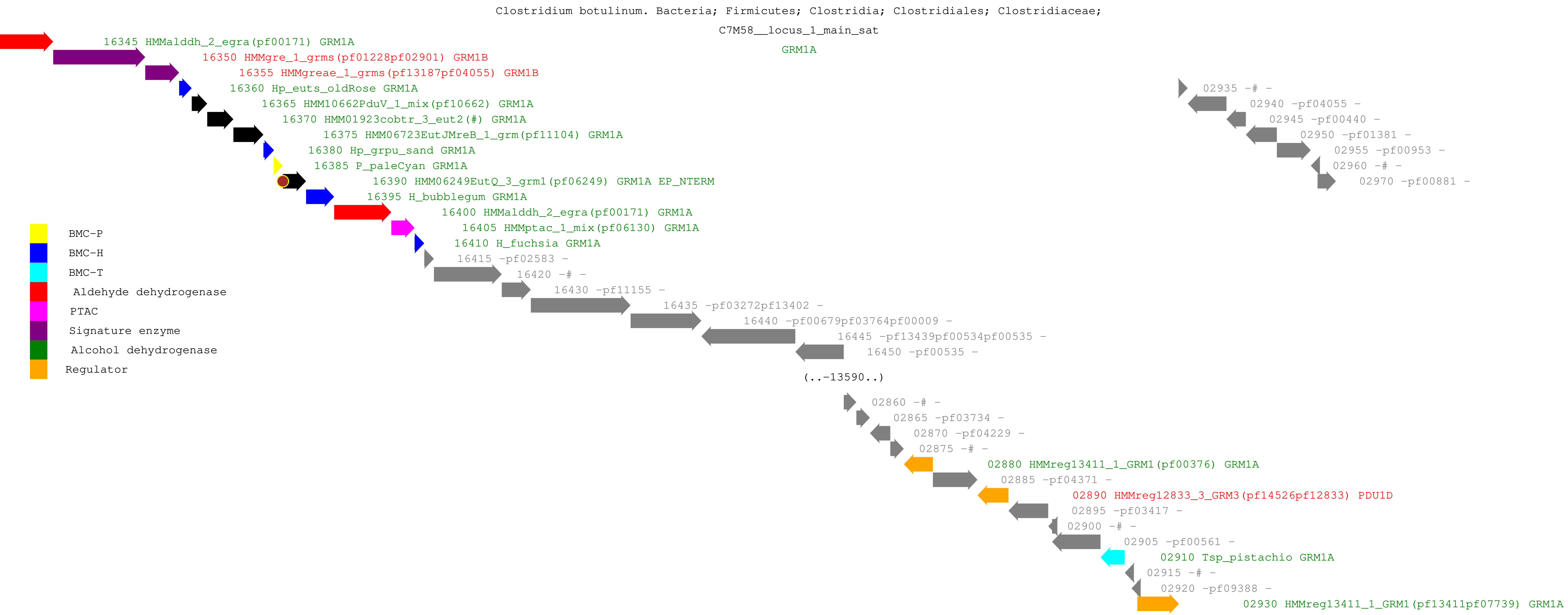
Clostridium botulinum. Bacteria; Firmicutes; Clostridia; Clostridiales; Clostridiaceae;
C7M58__locus_1_main_sat
GRM1A
16345 HMMalddh_2_egra(pf00171) GRM1A
16350 HMMgre_1_grms(pf01228pf02901) GRM1B
16355 HMMgreae_1_grms(pf13187pf04055) GRM1B
16360 Hp_euts_oldRose GRM1A
16365 HMM10662PduV_1_mix(pf10662) GRM1A
16370 HMM01923cobtr_3_eut2(#) GRM1A
16375 HMM06723EutJMreB_1_grm(pf11104) GRM1A
16380 Hp_grpu_sand GRM1A
16385 P_paleCyan GRM1A
16390 HMM06249EutQ_3_grm1(pf06249) GRM1A EP_NTERM
16395 H_bubblegum GRM1A
16400 HMMalddh_2_egra(pf00171) GRM1A
16405 HMMptac_1_mix(pf06130) GRM1A
16410 H_fuchsia GRM1A
16415 -pf02583 -
16420 -# -
16430 -pf11155 -
16435 -pf03272pf13402 -
16440 -pf00679pf03764pf00009 -
16445 -pf13439pf00534pf00535 -
16450 -pf00535 -
(..-13590..)
02860 -# -
02865 -pf03734 -
02870 -pf04229 -
02875 -# -
02880 HMMreg13411_1_GRM1(pf00376) GRM1A
02885 -pf04371 -
02890 HMMreg12833_3_GRM3(pf14526pf12833) PDU1D
02895 -pf03417 -
02900 -# -
02905 -pf00561 -
02910 Tsp_pistachio GRM1A
02915 -# -
02920 -pf09388 -
02930 HMMreg13411_1_GRM1(pf13411pf07739) GRM1A
02935 -# -
02940 -pf04055 -
02945 -pf00440 -
02950 -pf01381 -
02955 -pf00953 -
02960 -# -
02970 -pf00881 -
BMC-P
BMC-H
BMC-T
Aldehyde dehydrogenase
PTAC
Signature enzyme
Alcohol dehydrogenase
Regulator

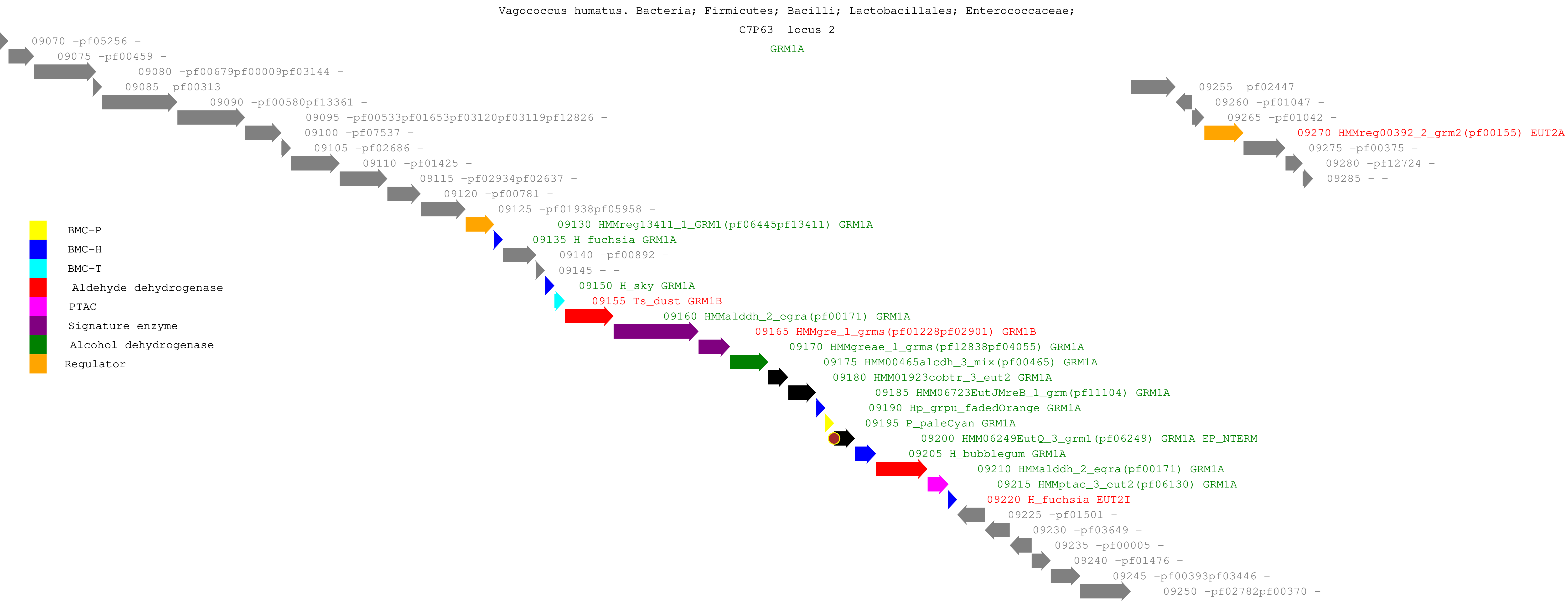

Vagococcus humatus. Bacteria; Firmicutes; Bacilli; Lactobacillales; Enterococcaceae;
C7P63__locus_2
GRM1A
09070 -pf05256 -
09075 -pf00459 -
09080 -pf00679pf00009pf03144 -
09085 -pf00313 -
09090 -pf00580pf13361 -
09095 -pf00533pf01653pf03120pf03119pf12826 -
09100 -pf07537 -
09105 -pf02686 -
09110 -pf01425 -
09115 -pf02934pf02637 -
09120 -pf00781 -
09125 -pf01938pf05958 -
09130 HMMreg13411_1_GRM1(pf06445pf13411) GRM1A
09135 H_fuchsia GRM1A
09140 -pf00892 -
09145 - -
09150 H_sky GRM1A
09155 Ts_dust GRM1B
09160 HMMaldh_2_egra(pf00171) GRM1A
09165 HMMgre_1_grms(pf01228pf02901) GRM1B
09170 HMMgreae_1_grms(pf12838pf04055) GRM1A
09175 HMM00465alcdh_3_mix(pf00465) GRM1A
09180 HMM01923cobtr_3_eut2 GRM1A
09185 HMM06723EutJMreB_1_grm(pf11104) GRM1A
09190 Hp_grpu_fadedOrange GRM1A
09195 P_paleCyan GRM1A
09200 HMM06249EutQ_3_grm1(pf06249) GRM1A EP_NTERM
09205 H_bubblegum GRM1A
09210 HMMaldh_2_egra(pf00171) GRM1A
09215 HMMptac_3_eut2(pf06130) GRM1A
09220 H_fuchsia EUT2I
09225 -pf01501 -
09230 -pf03649 -
09235 -pf00005 -
09240 -pf01476 -
09245 -pf00393pf03446 -
09250 -pf02782pf00370 -
09255 -pf02447 -
09260 -pf01047 -
09265 -pf01042 -
09270 HMMreg00392_2_grm2(pf00155) EUT2A
09275 -pf00375 -
09280 -pf12724 -
09285 - -
BMC-P
BMC-H
BMC-T
Aldehyde dehydrogenase
PTAC
Signature enzyme
Alcohol dehydrogenase
Regulator

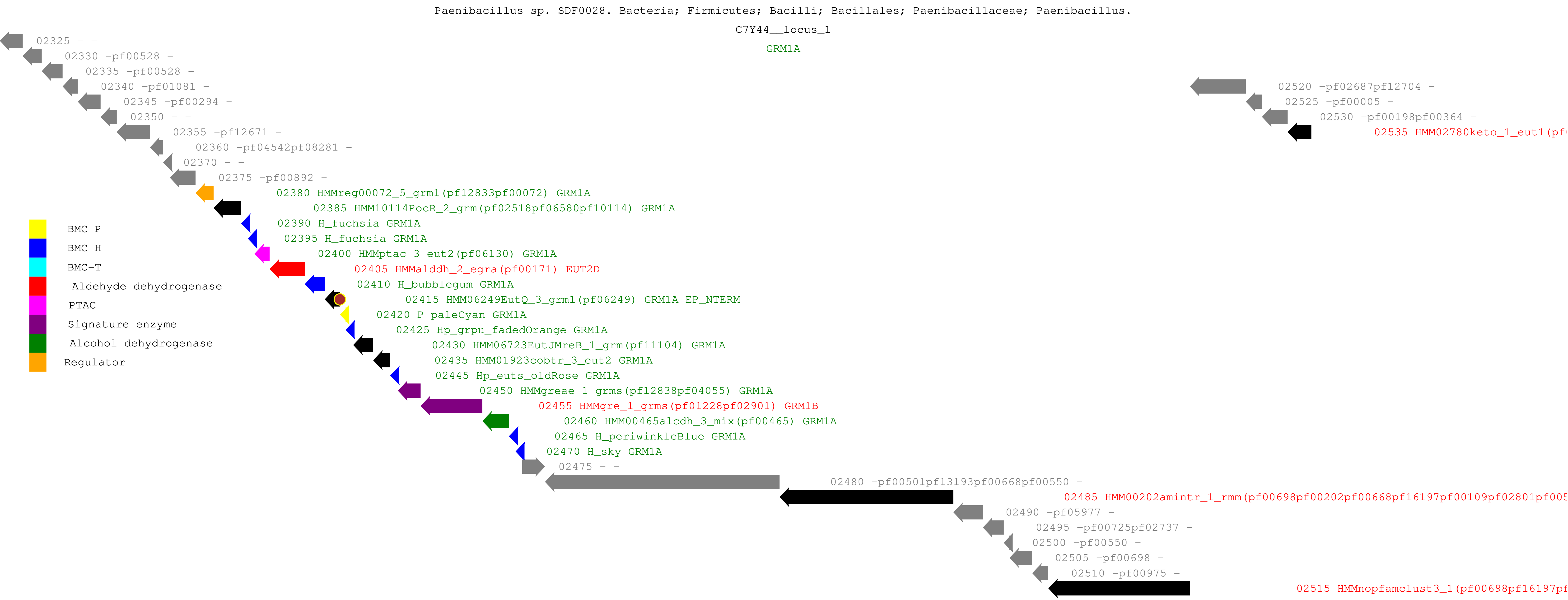
Paenibacillus sp. SDF0028. Bacteria; Firmicutes; Bacilli; Bacillales; Paenibacillaceae; Paenibacillus.
C7Y44__locus_1
GRM1A
02325 - -
02330 -pf00528 -
02335 -pf00528 -
02340 -pf01081 -
02345 -pf00294 -
02350 - -
02355 -pf12671 -
02360 -pf04542pf08281 -
02370 - -
02375 -pf00892 -
02380 HMMreg00072_5_grm1(pf12833pf00072) GRM1A
02385 HMM10114PocR_2_grm(pf02518pf06580pf10114) GRM1A
02390 H_fuchsia GRM1A
02395 H_fuchsia GRM1A
02400 HMMptac_3_eut2(pf06130) GRM1A
02405 HMMalddh_2_egra(pf00171) EUT2D
02410 H_bubblegum GRM1A
02415 HMM06249EutQ_3_grm1(pf06249) GRM1A EP_NTERM
02420 P_paleCyan GRM1A
02425 Hp_grpu_fadedOrange GRM1A
02430 HMM06723EutJMreB_1_grm(pf11104) GRM1A
02435 HMM01923cobtr_3_eut2 GRM1A
02445 Hp_euts_oldRose GRM1A
02450 HMMgreae_1_grms(pf12838pf04055) GRM1A
02455 HMMgre_1_grms(pf01228pf02901) GRM1B
02460 HMM00465alcdh_3_mix(pf00465) GRM1A
02465 H_periwinkleBlue GRM1A
02470 H_sky GRM1A
02475 - -
02480 -pf00501pf13193pf00668pf00550 -
02485 HMM00202amintr_1_rmm(pf00698pf00202pf00668pf16197pf00109pf02801pf005
02490 -pf05977 -
02495 -pf00725pf02737 -
02500 -pf00550 -
02505 -pf00698 -
02510 -pf00975 -
02515 HMMnopfamclust3_1(pf00698pf16197pf
02520 -pf02687pf12704 -
02525 -pf00005 -
02530 -pf00198pf00364 -
02535 HMM02780keto_1_eut1(pf
BMC-P
BMC-H
BMC-T
Aldehyde dehydrogenase
PTAC
Signature enzyme
Alcohol dehydrogenase
Regulator

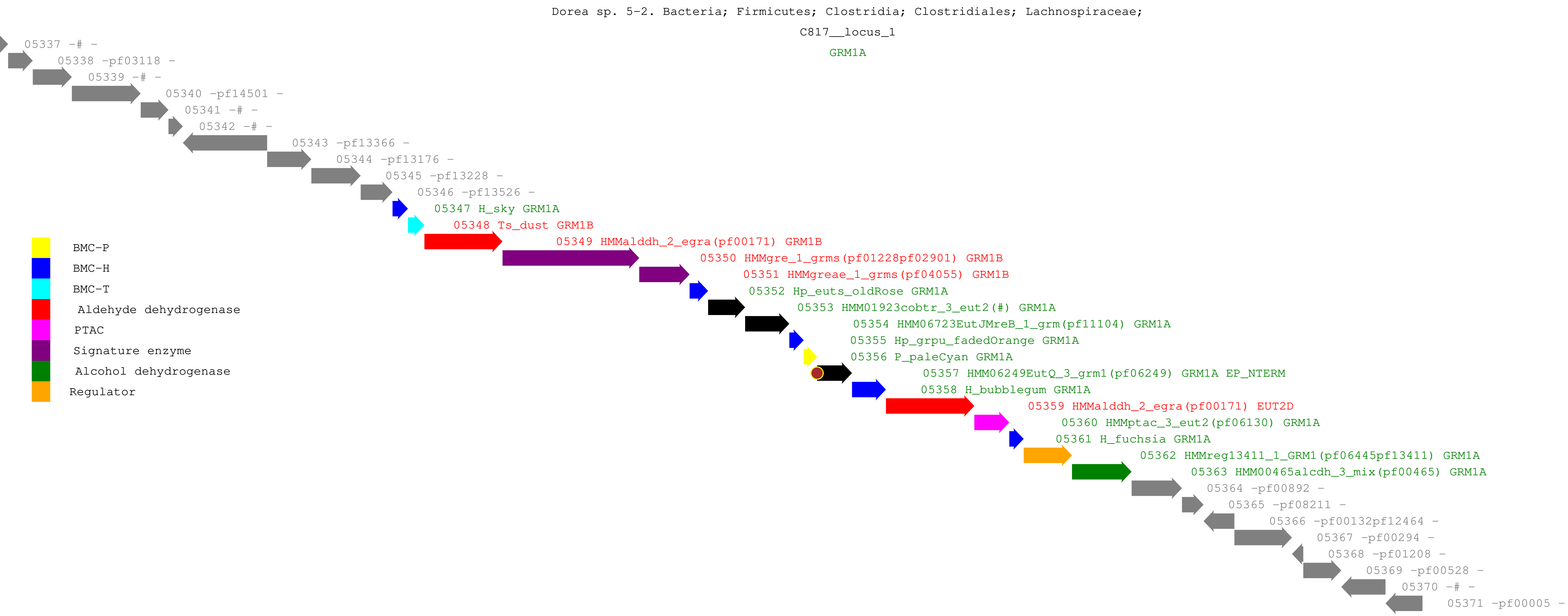
Dorea sp. 5-2. Bacteria; Firmicutes; Clostridia; Clostridiales; Lachnospiraceae;
C817__locus_1
GRM1A
05337 -# -
05338 -pf03118 -
05339 -# -
05340 -pf14501 -
05341 -# -
05342 -# -
05343 -pf13366 -
05344 -pf13176 -
05345 -pf13228 -
05346 -pf13526 -
05347 H_sky GRM1A
05348 Ts_dust GRM1B
05349 HMMalddh_2_egra(pf00171) GRM1B
05350 HMMgre_1_grms(pf01228pf02901) GRM1B
05351 HMMgreae_1_grms(pf04055) GRM1B
05352 Hp_euts_oldRose GRM1A
05353 HMM01923cobtr_3_eut2(#) GRM1A
05354 HMM06723EutJMreB_1_grm(pf11104) GRM1A
05355 Hp_grpu_fadedOrange GRM1A
05356 P_paleCyan GRM1A
05357 HMM06249EutQ_3_grm1(pf06249) GRM1A EP_NTERM
05358 H_bubblegum GRM1A
05359 HMMalddh_2_egra(pf00171) EUT2D
05360 HMMptac_3_eut2(pf06130) GRM1A
05361 H_fuchsia GRM1A
05362 HMMreg13411_1_GRM1(pf06445pf13411) GRM1A
05363 HMM00465alcdh_3_mix(pf00465) GRM1A
05364 -pf00892 -
05365 -pf08211 -
05366 -pf00132pf12464 -
05367 -pf00294 -
05368 -pf01208 -
05369 -pf00528 -
05370 -# -
05371 -pf00005 -
BMC-P
BMC-H
BMC-T
Aldehyde dehydrogenase
PTAC
Signature enzyme
Alcohol dehydrogenase
Regulator

Clostridium aceticum. Bacteria; Firmicutes; Clostridia; Clostridiales; Clostridiaceae;

CACET_c_locus_2

GRM1A

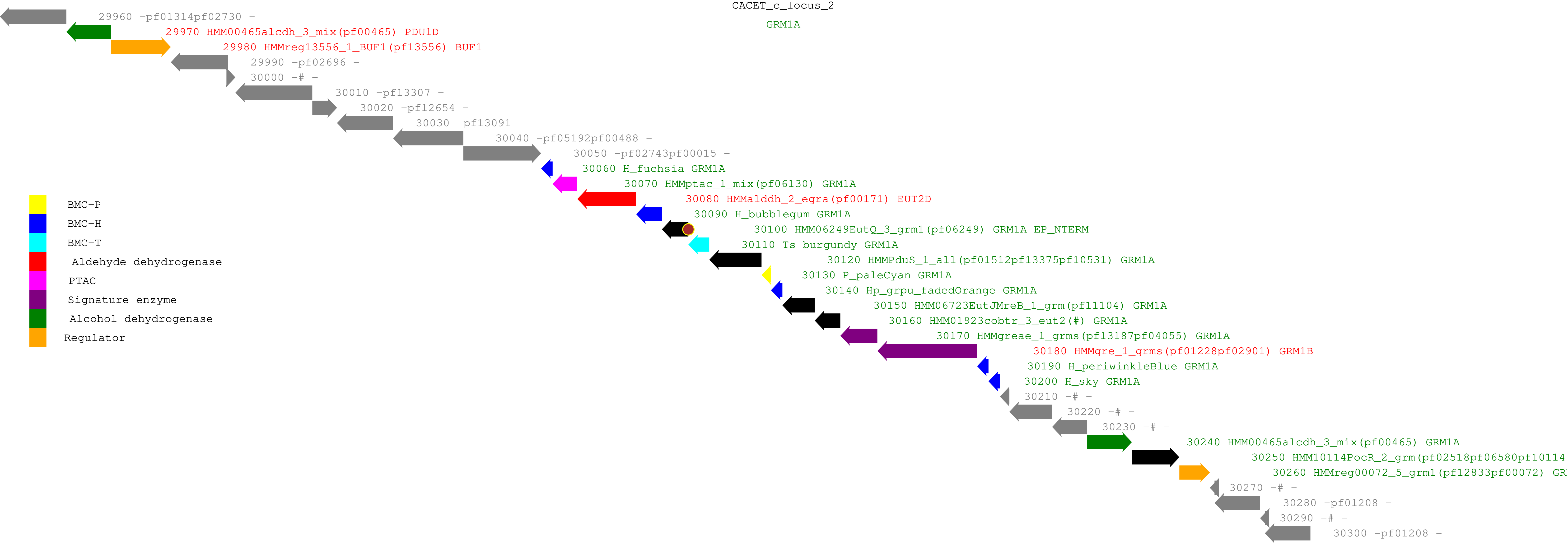

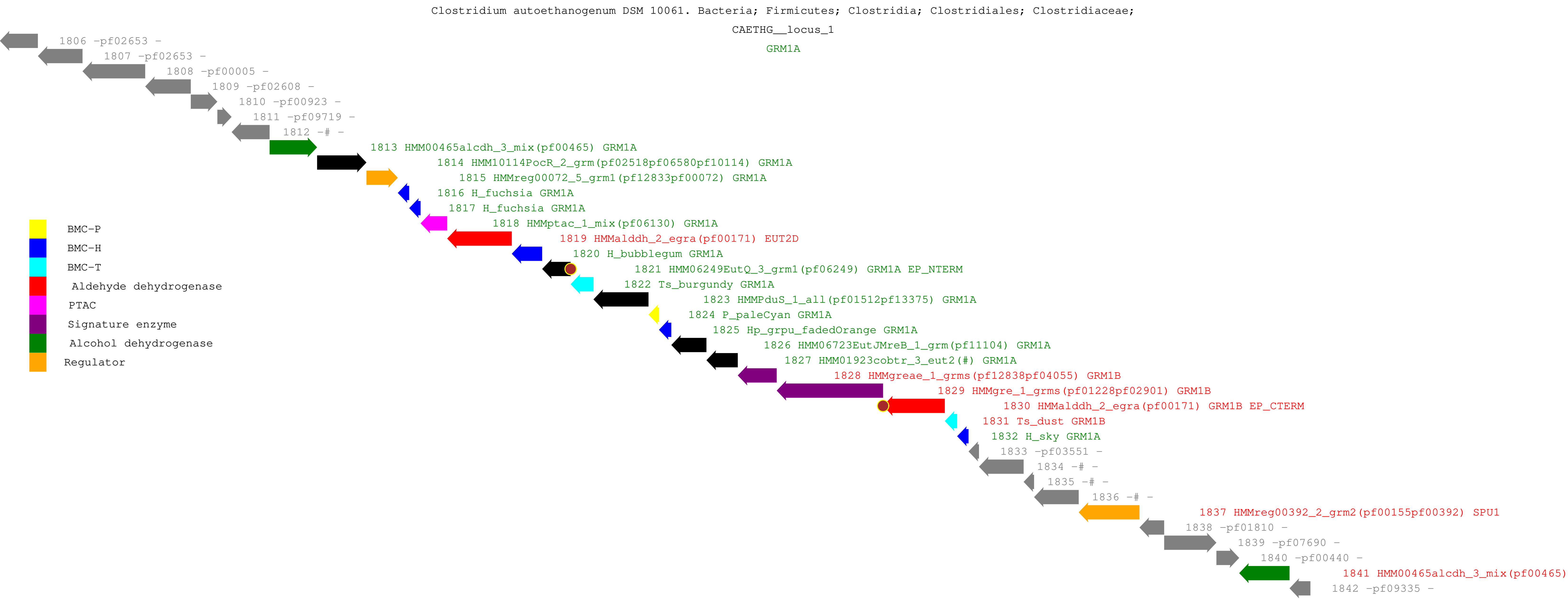
Clostridium autoethanogenum DSM 10061. Bacteria; Firmicutes; Clostridia; Clostridiales; Clostridiaceae;
CAETHG__locus_1
GRM1A
1806 -pf02653 -
1807 -pf02653 -
1808 -pf00005 -
1809 -pf02608 -
1810 -pf00923 -
1811 -pf09719 -
1812 -# -
1813 HMM00465alcdh_3_mix(pf00465) GRM1A
1814 HMM10114PocR_2_grm(pf02518pf06580pf10114) GRM1A
1815 HMMreg00072_5_grm1(pf12833pf00072) GRM1A
1816 H_fuchsia GRM1A
1817 H_fuchsia GRM1A
1818 HMMptac_1_mix(pf06130) GRM1A
1819 HMMaldh_2_egra(pf00171) EUT2D
1820 H_bubblegum GRM1A
1821 HMM06249EutQ_3_grm1(pf06249) GRM1A EP_NTERM
1822 Ts_burgundy GRM1A
1823 HMMPduS_1_all(pf01512pf13375) GRM1A
1824 P_paleCyan GRM1A
1825 Hp_grpu_fadedOrange GRM1A
1826 HMM06723EutJMreB_1_grm(pf11104) GRM1A
1827 HMM01923cobtr_3_eut2(#) GRM1A
1828 HMMgreae_1_grms(pf12838pf04055) GRM1B
1829 HMMgre_1_grms(pf01228pf02901) GRM1B
1830 HMMaldh_2_egra(pf00171) GRM1B EP_CTERM
1831 Ts_dust GRM1B
1832 H_sky GRM1A
1833 -pf03551 -
1834 -# -
1835 -# -
1836 -# -
1837 HMMreg00392_2_grm2(pf00155pf00392) SPU1
1838 -pf01810 -
1839 -pf07690 -
1840 -pf00440 -
1841 HMM00465alcdh_3_mix(pf00465)
1842 -pf09335 -
BMC-P
BMC-H
BMC-T
Aldehyde dehydrogenase
PTAC
Signature enzyme
Alcohol dehydrogenase
Regulator

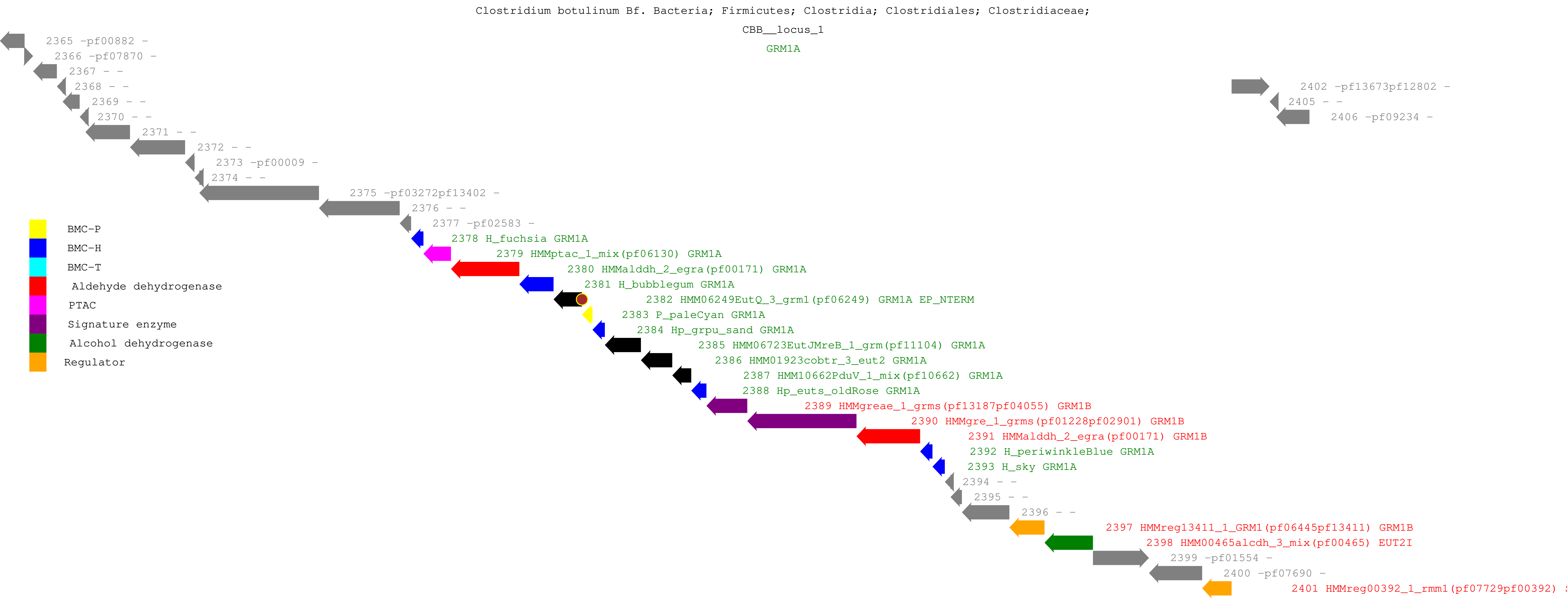
Clostridium botulinum Bf. Bacteria; Firmicutes; Clostridia; Clostridiales; Clostridiaceae;
CBB__locus_1
GRM1A
2365 -pf00882 -
2366 -pf07870 -
2367 - -
2368 - -
2369 - -
2370 - -
2371 - -
2372 - -
2373 -pf00009 -
2374 - -
2375 -pf03272pf13402 -
2376 - -
2377 -pf02583 -
2378 H_fuchsia GRM1A
2379 HMMptac_1_mix(pf06130) GRM1A
2380 HMMaldдh_2_egra(pf00171) GRM1A
2381 H_bubblegum GRM1A
2382 HMM06249EutQ_3_grm1(pf06249) GRM1A EP_NTERM
2383 P_paleCyan GRM1A
2384 Hp_grpu_sand GRM1A
2385 HMM06723EutJMreB_1_grm(pf11104) GRM1A
2386 HMM01923cobtr_3_eut2 GRM1A
2387 HMM10662PduV_1_mix(pf10662) GRM1A
2388 Hp_euts_oldRose GRM1A
2389 HMMgreae_1_grms(pf13187pf04055) GRM1B
2390 HMMgre_1_grms(pf01228pf02901) GRM1B
2391 HMMaldдh_2_egra(pf00171) GRM1B
2392 H_periwinkleBlue GRM1A
2393 H_sky GRM1A
2394 - -
2395 - -
2396 - -
2397 HMMreg13411_1_GRM1(pf06445pf13411) GRM1B
2398 HMM00465alcdh_3_mix(pf00465) EUT2I
2399 -pf01554 -
2400 -pf07690 -
2401 HMMreg00392_1_rmm1(pf07729pf00392)
2402 -pf13673pf12802 -
2405 - -
2406 -pf09234 -
BMC-P
BMC-H
BMC-T
Aldehyde dehydrogenase
PTAC
Signature enzyme
Alcohol dehydrogenase
Regulator

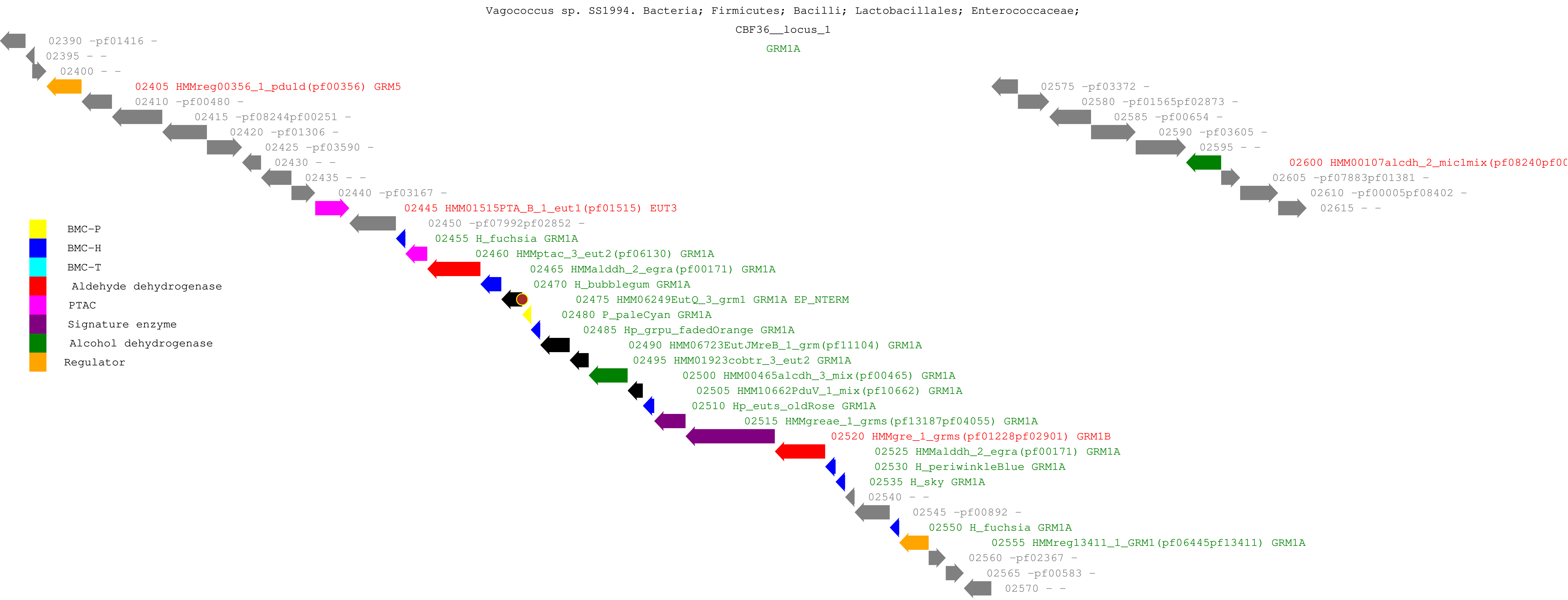

Vagococcus sp. SS1994. Bacteria; Firmicutes; Bacilli; Lactobacillales; Enterococcaceae;
CBF36__locus_1
GRM1A
02390 -pf01416 -
02395 - -
02400 - -
02405 HMMreg00356_1_pdu1d(pf00356) GRM5
02410 -pf00480 -
02415 -pf08244pf00251 -
02420 -pf01306 -
02425 -pf03590 -
02430 - -
02435 - -
02440 -pf03167 -
02445 HMM01515PTA_B_1_eut1(pf01515) EUT3
02450 -pf07992pf02852 -
02455 H_fuchsia GRM1A
02460 HMMptac_3_eut2(pf06130) GRM1A
02465 HMMalddh_2_egra(pf00171) GRM1A
02470 H_bubblegum GRM1A
02475 HMM06249EutQ_3_grm1 GRM1A EP_NTERM
02480 P_paleCyan GRM1A
02485 Hp_grpu_fadedOrange GRM1A
02490 HMM06723EutJMreB_1_grm(pf11104) GRM1A
02495 HMM01923cobtr_3_eut2 GRM1A
02500 HMM00465alcdh_3_mix(pf00465) GRM1A
02505 HMM10662PduV_1_mix(pf10662) GRM1A
02510 Hp_euts_oldRose GRM1A
02515 HMMgreae_1_grms(pf13187pf04055) GRM1A
02520 HMMgre_1_grms(pf01228pf02901) GRM1B
02525 HMMalddh_2_egra(pf00171) GRM1A
02530 H_periwinkleBlue GRM1A
02535 H_sky GRM1A
02540 - -
02545 -pf00892 -
02550 H_fuchsia GRM1A
02555 HMMreg13411_1_GRM1(pf06445pf13411) GRM1A
02560 -pf02367 -
02565 -pf00583 -
02570 - -
02575 -pf03372 -
02580 -pf01565pf02873 -
02585 -pf00654 -
02590 -pf03605 -
02595 - -
02600 HMM00107alcdh_2_mic1mix(pf08240pf00
02605 -pf07883pf01381 -
02610 -pf00005pf08402 -
02615 - -
BMC-P
BMC-H
BMC-T
Aldehyde dehydrogenase
PTAC
Signature enzyme
Alcohol dehydrogenase
Regulator

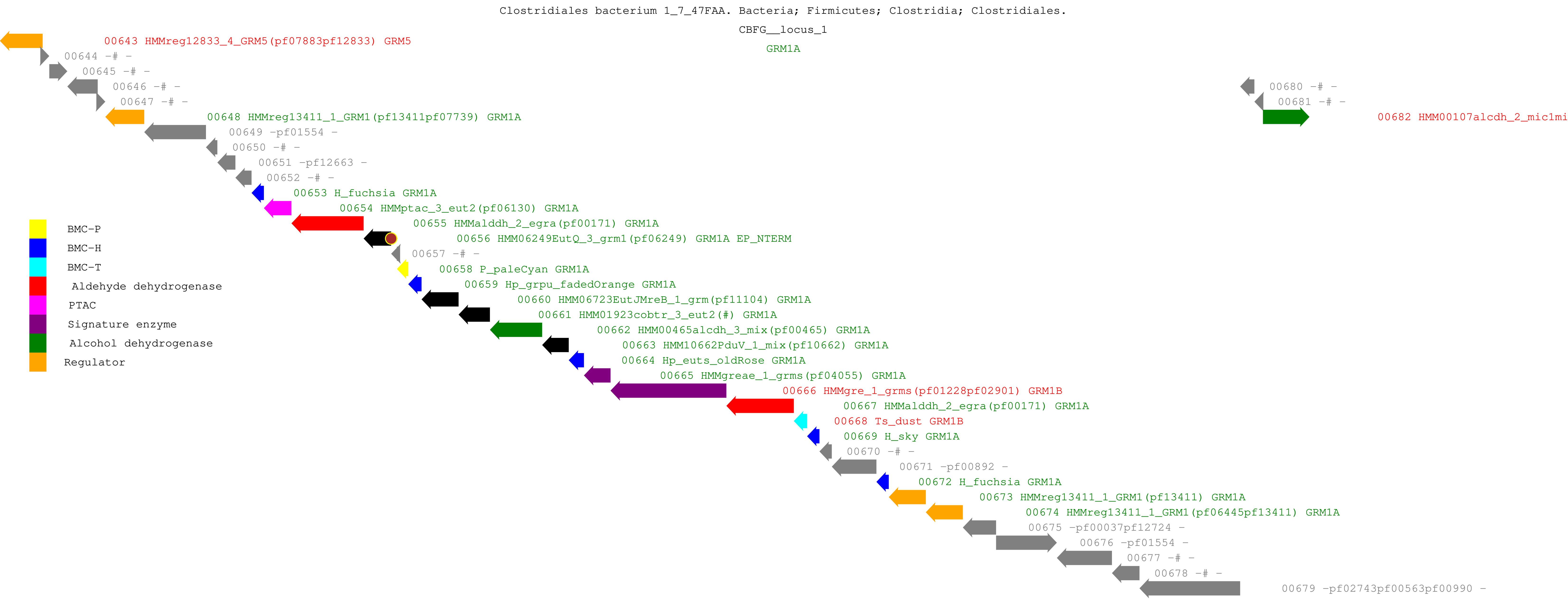

Clostridiales bacterium 1_7_47FAA. Bacteria; Firmicutes; Clostridia; Clostridiales.
CBFG__locus_1
GRM1A
00643 HMMreg12833_4_GRM5(pf07883pf12833) GRM5
00644 -# -
00645 -# -
00646 -# -
00647 -# -
00648 HMMreg13411_1_GRM1(pf13411pf07739) GRM1A
00649 -pf01554 -
00650 -# -
00651 -pf12663 -
00652 -# -
00653 H_fuchsia GRM1A
00654 HMMptac_3_eut2(pf06130) GRM1A
00655 HMMaldh_2_egra(pf00171) GRM1A
00656 HMM06249EutQ_3_grm1(pf06249) GRM1A EP_NTERM
00657 -# -
00658 P_paleCyan GRM1A
00659 Hp_grpu_fadedOrange GRM1A
00660 HMM06723EutJMreB_1_grm(pf11104) GRM1A
00661 HMM01923cobtr_3_eut2(#) GRM1A
00662 HMM00465alcdh_3_mix(pf00465) GRM1A
00663 HMM10662PduV_1_mix(pf10662) GRM1A
00664 Hp_euts_oldRose GRM1A
00665 HMMgreae_1_grms(pf04055) GRM1A
00666 HMMgre_1_grms(pf01228pf02901) GRM1B
00667 HMMaldh_2_egra(pf00171) GRM1A
00668 Ts_dust GRM1B
00669 H_sky GRM1A
00670 -# -
00671 -pf00892 -
00672 H_fuchsia GRM1A
00673 HMMreg13411_1_GRM1(pf13411) GRM1A
00674 HMMreg13411_1_GRM1(pf06445pf13411) GRM1A
00675 -pf00037pf12724 -
00676 -pf01554 -
00677 -# -
00678 -# -
00679 -pf02743pf00563pf00990 -
00680 -# -
00681 -# -
00682 HMM00107alcdh_2_mic1mi
BMC-P
BMC-H
BMC-T
Aldehyde dehydrogenase
PTAC
Signature enzyme
Alcohol dehydrogenase
Regulator

Clostridium botulinum B str. Osaka05. Bacteria; Firmicutes; Clostridia; Clostridiales; Clostridiaceae;

CBO05C__locus_1_main_sat

GRM1A

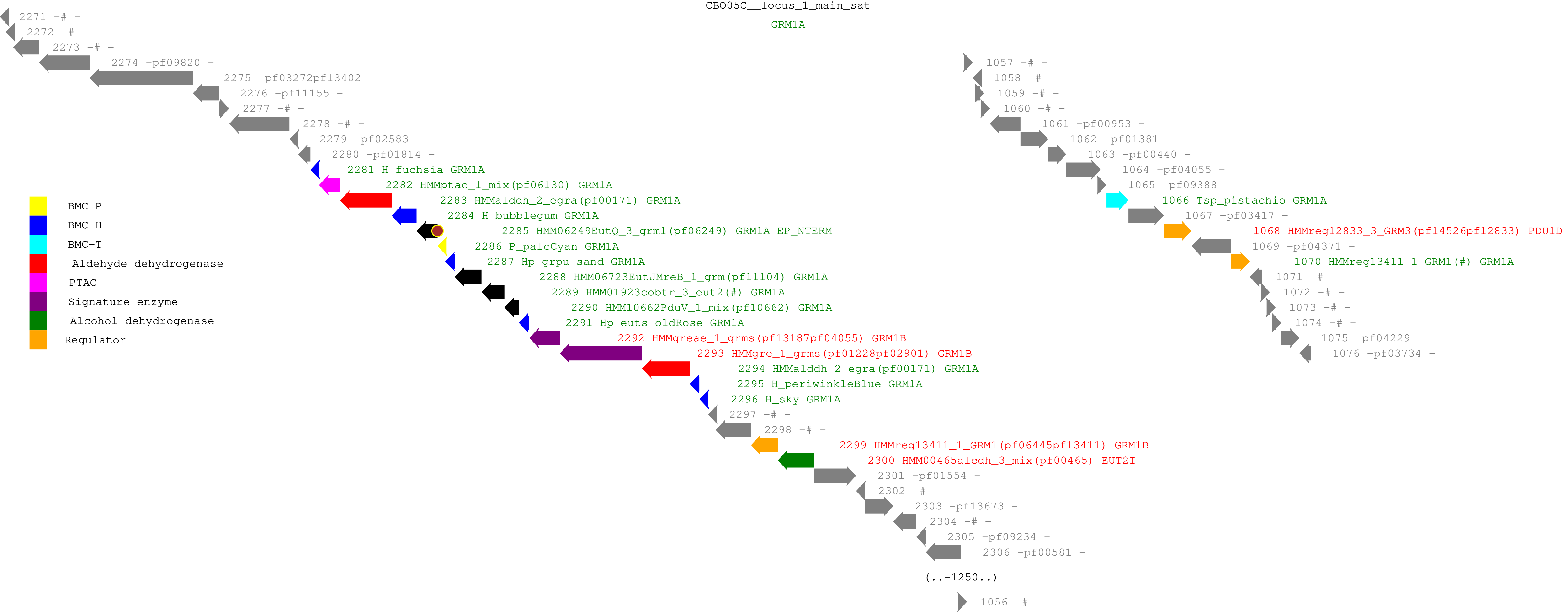

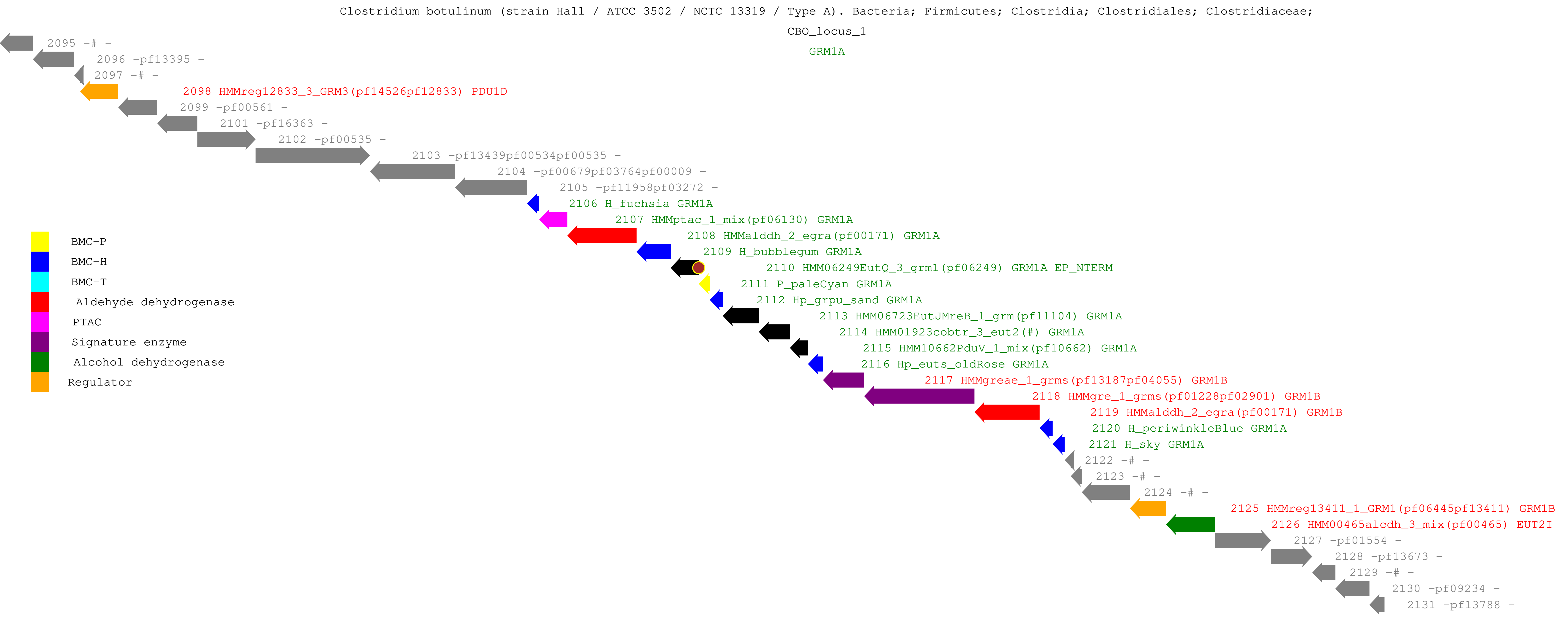

Clostridium botulinum (strain Hall / ATCC 3502 / NCTC 13319 / Type A). Bacteria; Firmicutes; Clostridia; Clostridiales; Clostridiaceae;
CBO_locus_1
GRM1A
2095 -# -
2096 -pf13395 -
2097 -# -
2098 HMMreg12833_3_GRM3(pf14526pf12833) PDU1D
2099 -pf00561 -
2101 -pf16363 -
2102 -pf00535 -
2103 -pf13439pf00534pf00535 -
2104 -pf00679pf03764pf00009 -
2105 -pf11958pf03272 -
2106 H_fuchsia GRM1A
2107 HMMptac_1_mix(pf06130) GRM1A
2108 HMMaldh_2_egra(pf00171) GRM1A
2109 H_bubblegum GRM1A
2110 HMM06249EutQ_3_grm1(pf06249) GRM1A EP_NTERM
2111 P_paleCyan GRM1A
2112 Hp_grpu_sand GRM1A
2113 HMM06723EutJMreB_1_grm(pf11104) GRM1A
2114 HMM01923cobtr_3_eut2(#) GRM1A
2115 HMM10662PduV_1_mix(pf10662) GRM1A
2116 Hp_euts_oldRose GRM1A
2117 HMMgreae_1_grms(pf13187pf04055) GRM1B
2118 HMMgre_1_grms(pf01228pf02901) GRM1B
2119 HMMaldh_2_egra(pf00171) GRM1B
2120 H_periwinkleBlue GRM1A
2121 H_sky GRM1A
2122 -# -
2123 -# -
2124 -# -
2125 HMMreg13411_1_GRM1(pf06445pf13411) GRM1B
2126 HMM00465alcdh_3_mix(pf00465) EUT2I
2127 -pf01554 -
2128 -pf13673 -
2129 -# -
2130 -pf09234 -
2131 -pf13788 -
BMC-P
BMC-H
BMC-T
Aldehyde dehydrogenase
PTAC
Signature enzyme
Alcohol dehydrogenase
Regulator

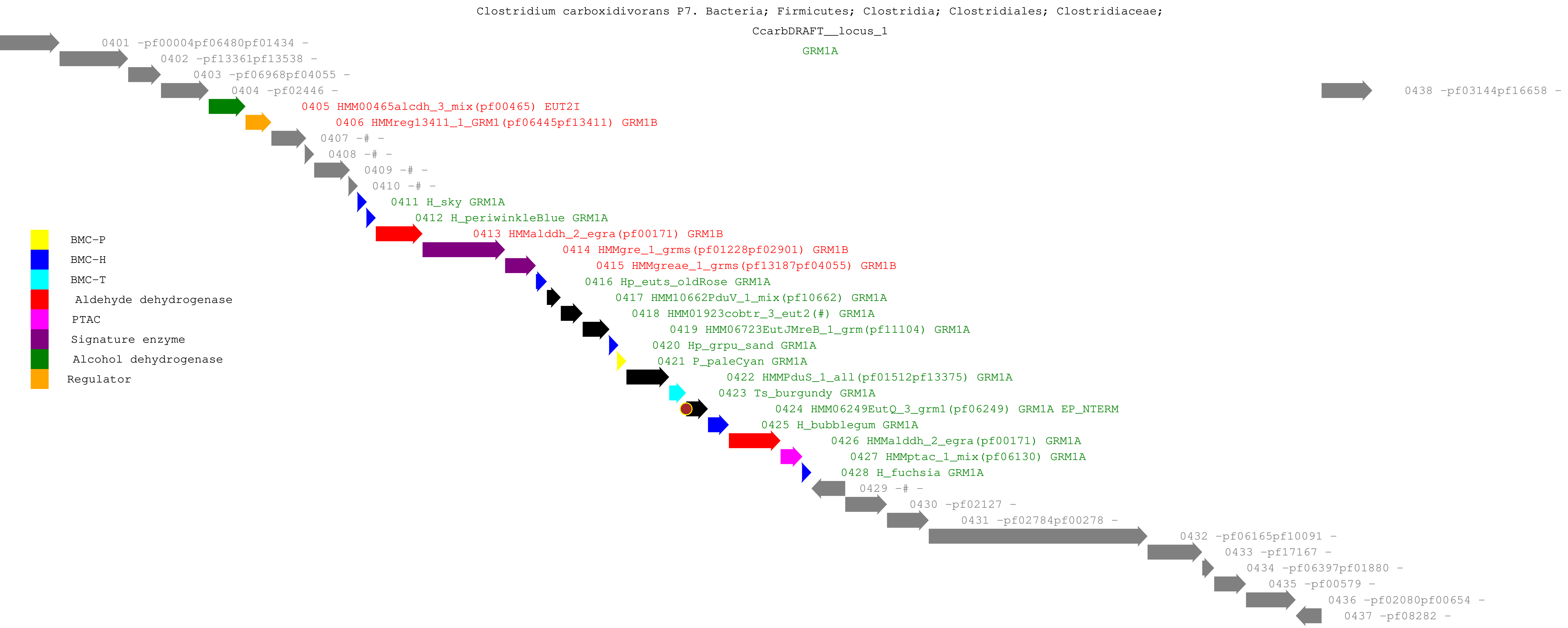

Clostridium carboxidivorans P7. Bacteria; Firmicutes; Clostridia; Clostridiales; Clostridiaceae;
CcarbDRAFT__locus_1
GRM1A
0401 -pf00004pf06480pf01434 -
0402 -pf13361pf13538 -
0403 -pf06968pf04055 -
0404 -pf02446 -
0438 -pf03144pf16658 -
0405 HMM00465alcdh_3_mix(pf00465) EUT2I
0406 HMMreg13411_1_GRM1(pf06445pf13411) GRM1B
0407 -# -
0408 -# -
0409 -# -
0410 -# -
0411 H_sky GRM1A
0412 H_periwinkleBlue GRM1A
0413 HMMaldh_2_egra(pf00171) GRM1B
0414 HMMgre_1_grms(pf01228pf02901) GRM1B
0415 HMMgreae_1_grms(pf13187pf04055) GRM1B
0416 Hp_euts_oldRose GRM1A
0417 HMM10662PduV_1_mix(pf10662) GRM1A
0418 HMM01923cobtr_3_eut2(#) GRM1A
0419 HMM06723EutJMreB_1_grm(pf11104) GRM1A
0420 Hp_grpu_sand GRM1A
0421 P_paleCyan GRM1A
0422 HMMPduS_1_all(pf01512pf13375) GRM1A
0423 Ts_burgundy GRM1A
0424 HMM06249EutQ_3_grm1(pf06249) GRM1A EP_NTERM
0425 H_bubblegum GRM1A
0426 HMMaldh_2_egra(pf00171) GRM1A
0427 HMMptac_1_mix(pf06130) GRM1A
0428 H_fuchsia GRM1A
0429 -# -
0430 -pf02127 -
0431 -pf02784pf00278 -
0432 -pf06165pf10091 -
0433 -pf17167 -
0434 -pf06397pf01880 -
0435 -pf00579 -
0436 -pf02080pf00654 -
0437 -pf08282 -
BMC-P
BMC-H
BMC-T
Aldehyde dehydrogenase
PTAC
Signature enzyme
Alcohol dehydrogenase
Regulator

Clostridium chauvoei JF4335. Bacteria; Firmicutes; Clostridia; Clostridiales; Clostridiaceae;

CCH01__locus_2

GRM1A

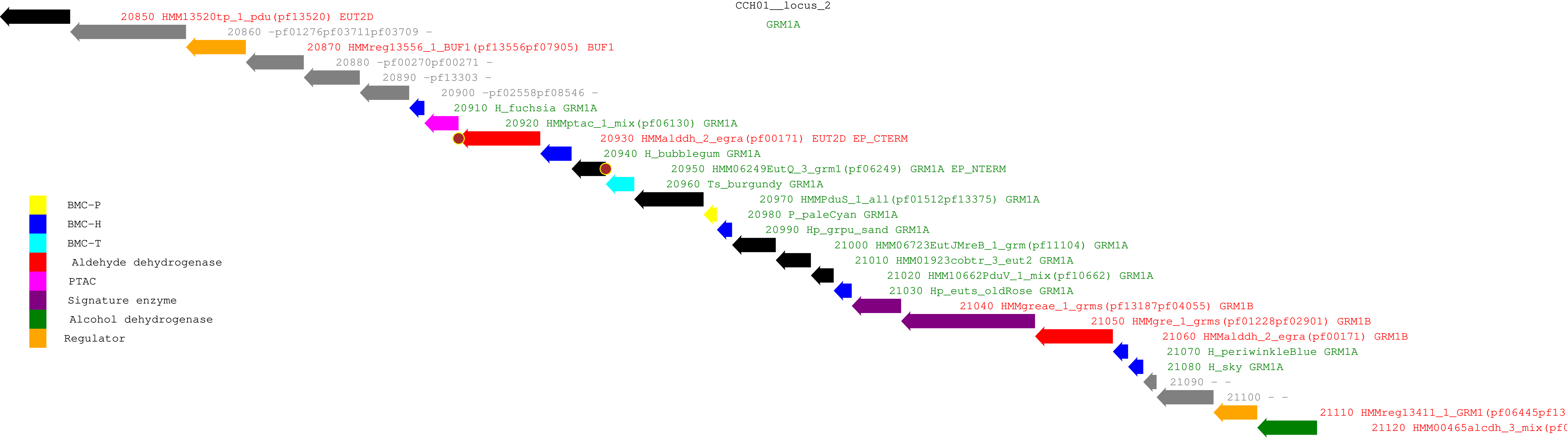

Clostridium botulinum CFSAN001627. Bacteria; Firmicutes; Clostridia; Clostridiales; Clostridiaceae;

CFSAN001627__locus_1_main_sat

GRM1A

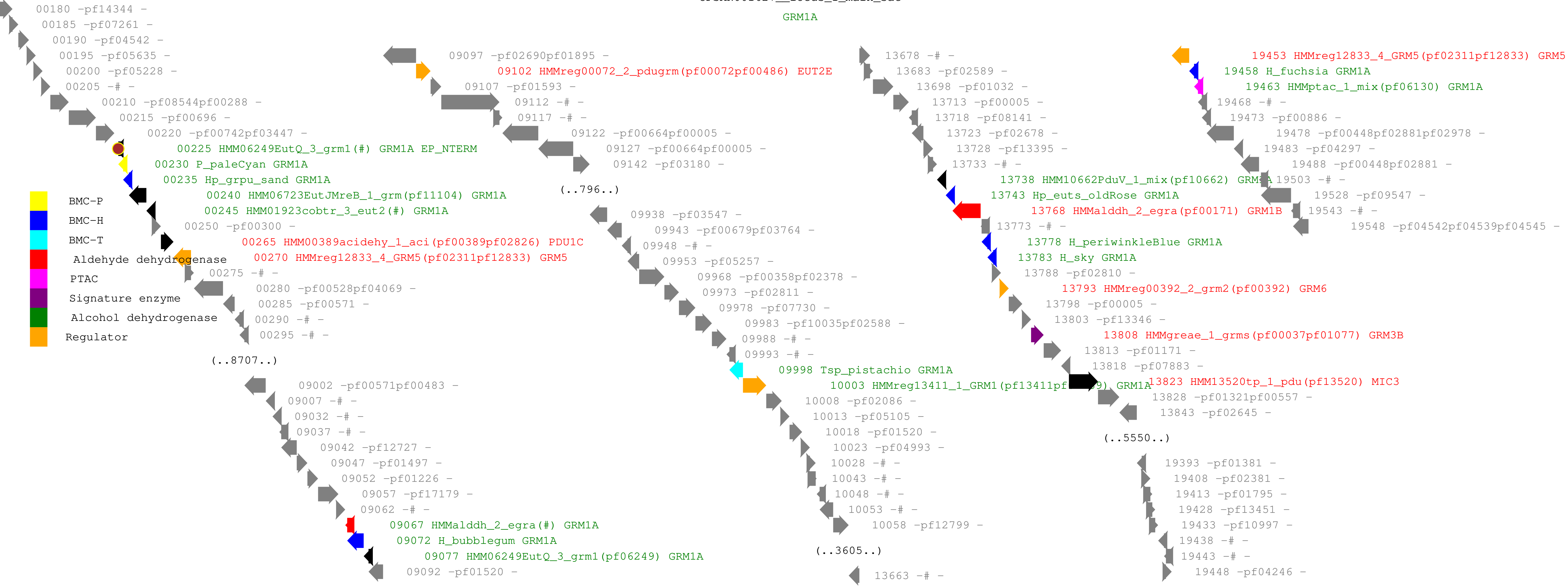

Enterococcus plantarum. Bacteria; Firmicutes; Bacilli; Lactobacillales; Enterococcaceae;

CI088__locus_1

GRM1A

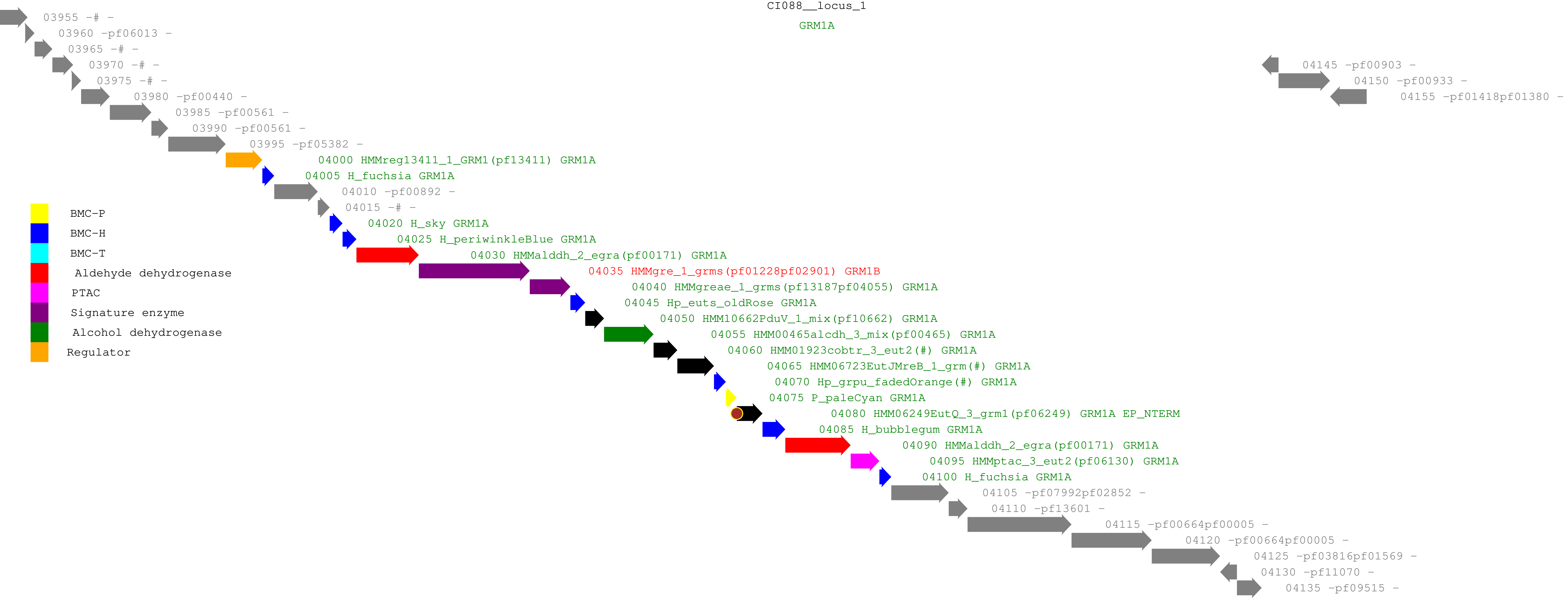

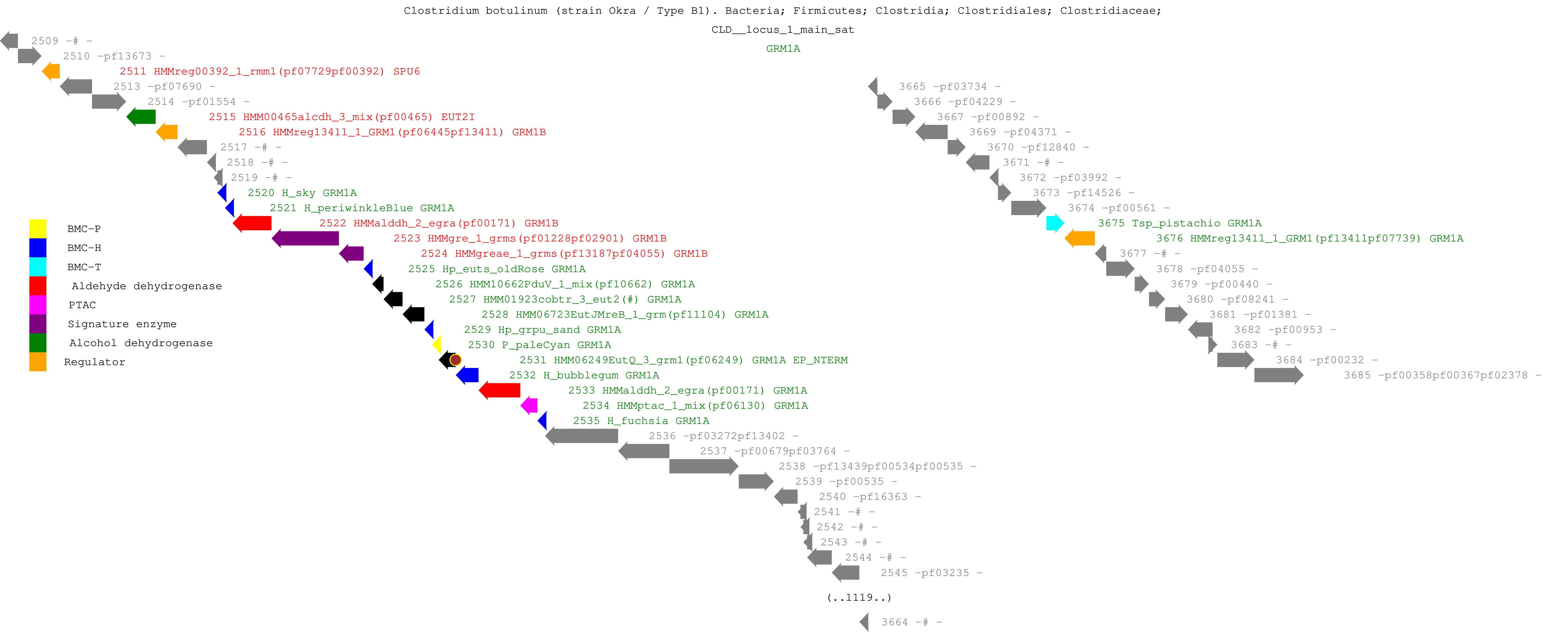

Clostridium botulinum (strain Okra / Type B1). Bacteria; Firmicutes; Clostridia; Clostridiales; Clostridiaceae;
CLD__locus_1_main_sat
GRM1A
2509 -# -
2510 -pf13673 -
2511 HMMreg00392_1_rmm1(pf07729pf00392) SPU6
2513 -pf07690 -
2514 -pf01554 -
2515 HMM00465alcdh_3_mix(pf00465) EUT2I
2516 HMMreg13411_1_GRM1(pf06445pf13411) GRM1B
2517 -# -
2518 -# -
2519 -# -
2520 H_sky GRM1A
2521 H_periwinkleBlue GRM1A
2522 HMMaldh_2_egra(pf00171) GRM1B
2523 HMMgre_1_grms(pf01228pf02901) GRM1B
2524 HMMgreae_1_grms(pf13187pf04055) GRM1B
2525 Hp_euts_oldRose GRM1A
2526 HMM10662PduV_1_mix(pf10662) GRM1A
2527 HMM01923cobtr_3_eut2(#) GRM1A
2528 HMM06723EutJMreB_1_grm(pf11104) GRM1A
2529 Hp_grpu_sand GRM1A
2530 P_paleCyan GRM1A
2531 HMM06249EutQ_3_grm1(pf06249) GRM1A EP_NTERM
2532 H_bubblegum GRM1A
2533 HMMaldh_2_egra(pf00171) GRM1A
2534 HMMptac_1_mix(pf06130) GRM1A
2535 H_fuchsia GRM1A
2536 -pf03272pf13402 -
2537 -pf00679pf03764 -
2538 -pf13439pf00534pf00535 -
2539 -pf00535 -
2540 -pf16363 -
2541 -# -
2542 -# -
2543 -# -
2544 -# -
2545 -pf03235 -
(..1119..)
3664 -# -
3665 -pf03734 -
3666 -pf04229 -
3667 -pf00892 -
3669 -pf04371 -
3670 -pf12840 -
3671 -# -
3672 -pf03992 -
3673 -pf14526 -
3674 -pf00561 -
3675 Tsp_pistachio GRM1A
3676 HMMreg13411_1_GRM1(pf13411pf07739) GRM1A
3677 -# -
3678 -pf04055 -
3679 -pf00440 -
3680 -pf08241 -
3681 -pf01381 -
3682 -pf00953 -
3683 -# -
3684 -pf00232 -
3685 -pf00358pf00367pf02378 -
BMC-P
BMC-H
BMC-T
Aldehyde dehydrogenase
PTAC
Signature enzyme
Alcohol dehydrogenase
Regulator

Clostridium sp. FS41. Bacteria; Firmicutes; Clostridia; Clostridiales; Clostridiaceae;

CLFS41__locus_1

GRM1A

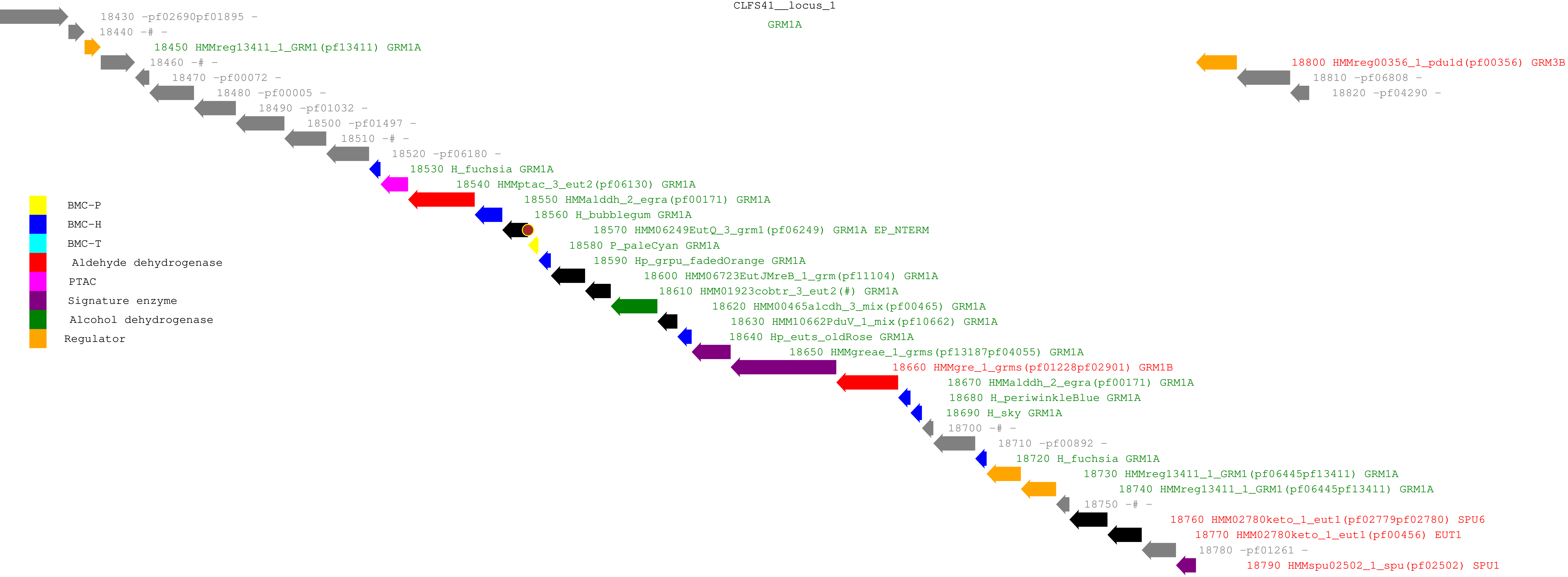

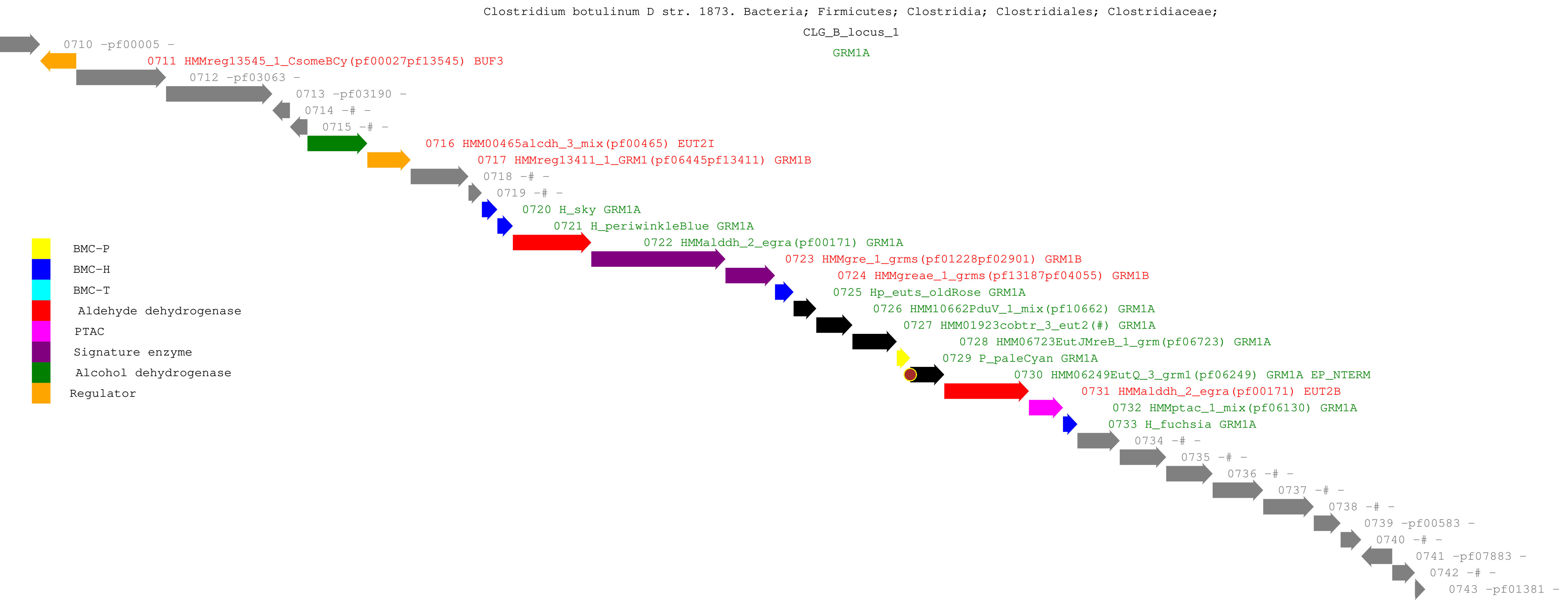
Clostridium botulinum D str. 1873. Bacteria; Firmicutes; Clostridia; Clostridiales; Clostridiaceae;
CLG_B_locus_1
GRM1A
0710 -pf00005 -
0711 HMMreg13545_1_CsomeBCy(pf00027pf13545) BUF3
0712 -pf03063 -
0713 -pf03190 -
0714 -# -
0715 -# -
0716 HMM00465alcdh_3_mix(pf00465) EUT2I
0717 HMMreg13411_1_GRM1(pf06445pf13411) GRM1B
0718 -# -
0719 -# -
0720 H_sky GRM1A
0721 H_periwinkleBlue GRM1A
0722 HMMalddh_2_egra(pf00171) GRM1A
0723 HMMgre_1_grms(pf01228pf02901) GRM1B
0724 HMMgreae_1_grms(pf13187pf04055) GRM1B
0725 Hp_euts_oldRose GRM1A
0726 HMM10662PduV_1_mix(pf10662) GRM1A
0727 HMM01923cobtr_3_eut2(#) GRM1A
0728 HMM06723EutJMreB_1_grm(pf06723) GRM1A
0729 P_paleCyan GRM1A
0730 HMM06249EutQ_3_grm1(pf06249) GRM1A EP_NTERM
0731 HMMalddh_2_egra(pf00171) EUT2B
0732 HMMptac_1_mix(pf06130) GRM1A
0733 H_fuchsia GRM1A
0734 -# -
0735 -# -
0736 -# -
0737 -# -
0738 -# -
0739 -pf00583 -
0740 -# -
0741 -pf07883 -
0742 -# -
0743 -pf01381 -
BMC-P
BMC-H
BMC-T
Aldehyde dehydrogenase
PTAC
Signature enzyme
Alcohol dehydrogenase
Regulator

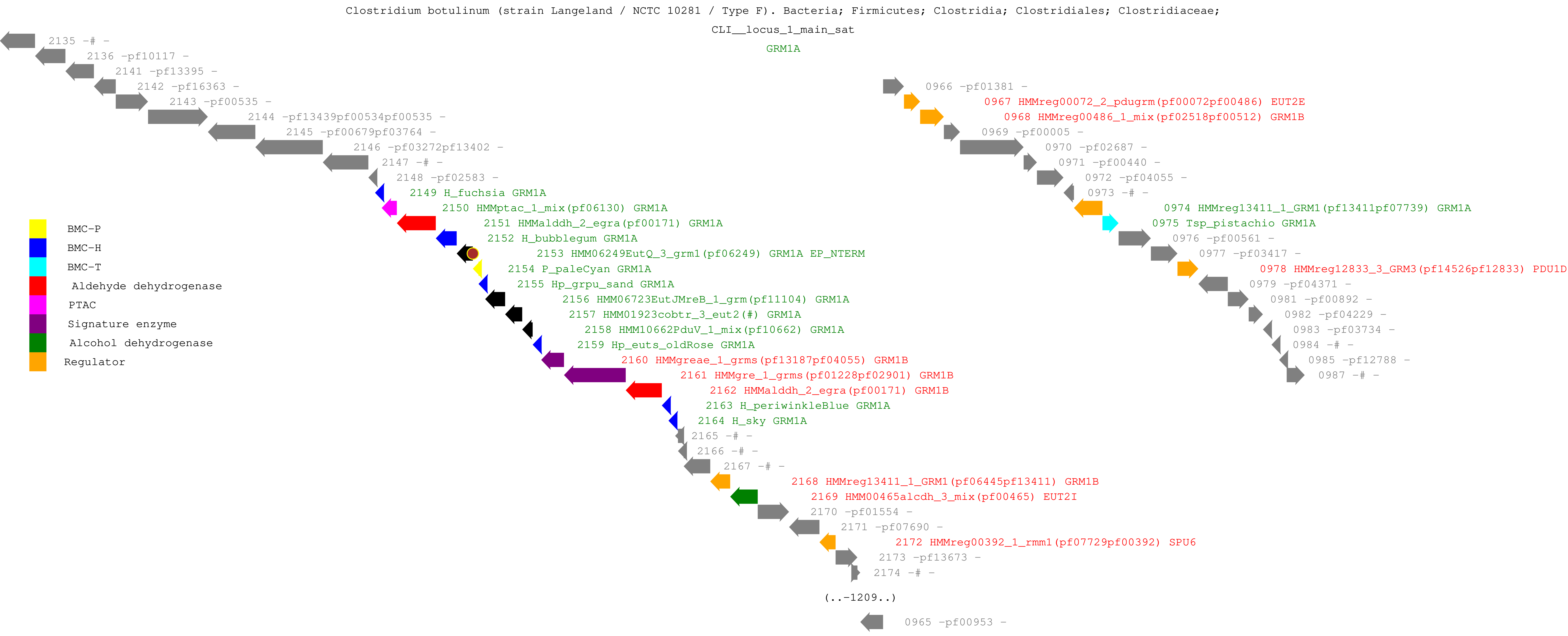

Clostridium botulinum (strain Langeland / NCTC 10281 / Type F). Bacteria; Firmicutes; Clostridia; Clostridiales; Clostridiaceae;
CLI__locus_1_main_sat
GRM1A
2135 -# -
2136 -pf10117 -
2141 -pf13395 -
2142 -pf16363 -
2143 -pf00535 -
2144 -pf13439pf00534pf00535 -
2145 -pf00679pf03764 -
2146 -pf03272pf13402 -
2147 -# -
2148 -pf02583 -
2149 H_fuchsia GRM1A
2150 HMMptac_1_mix(pf06130) GRM1A
2151 HMMalddh_2_egra(pf00171) GRM1A
2152 H_bubblegum GRM1A
2153 HMM06249EutQ_3_grm1(pf06249) GRM1A EP_NTERM
2154 P_paleCyan GRM1A
2155 Hp_grpu_sand GRM1A
2156 HMM06723EutJMreB_1_grm(pf11104) GRM1A
2157 HMM01923cobtr_3_eut2(#) GRM1A
2158 HMM10662PduV_1_mix(pf10662) GRM1A
2159 Hp_euts_oldRose GRM1A
2160 HMMgreae_1_grms(pf13187pf04055) GRM1B
2161 HMMgre_1_grms(pf01228pf02901) GRM1B
2162 HMMalddh_2_egra(pf00171) GRM1B
2163 H_periwinkleBlue GRM1A
2164 H_sky GRM1A
2165 -# -
2166 -# -
2167 -# -
2168 HMMreg13411_1_GRM1(pf06445pf13411) GRM1B
2169 HMM00465alcdh_3_mix(pf00465) EUT2I
2170 -pf01554 -
2171 -pf07690 -
2172 HMMreg00392_1_rmm1(pf07729pf00392) SPU6
2173 -pf13673 -
2174 -# -
(..-1209..)
0965 -pf00953 -
0966 -pf01381 -
0967 HMMreg00072_2_pdugrm(pf00072pf00486) EUT2E
0968 HMMreg00486_1_mix(pf02518pf00512) GRM1B
0969 -pf00005 -
0970 -pf02687 -
0971 -pf00440 -
0972 -pf04055 -
0973 -# -
0974 HMMreg13411_1_GRM1(pf13411pf07739) GRM1A
0975 Tsp_pistachio GRM1A
0976 -pf00561 -
0977 -pf03417 -
0978 HMMreg12833_3_GRM3(pf14526pf12833) PDU1D
0979 -pf04371 -
0981 -pf00892 -
0982 -pf04229 -
0983 -pf03734 -
0984 -# -
0985 -pf12788 -
0987 -# -
BMC-P
BMC-H
BMC-T
Aldehyde dehydrogenase
PTAC
Signature enzyme
Alcohol dehydrogenase
Regulator

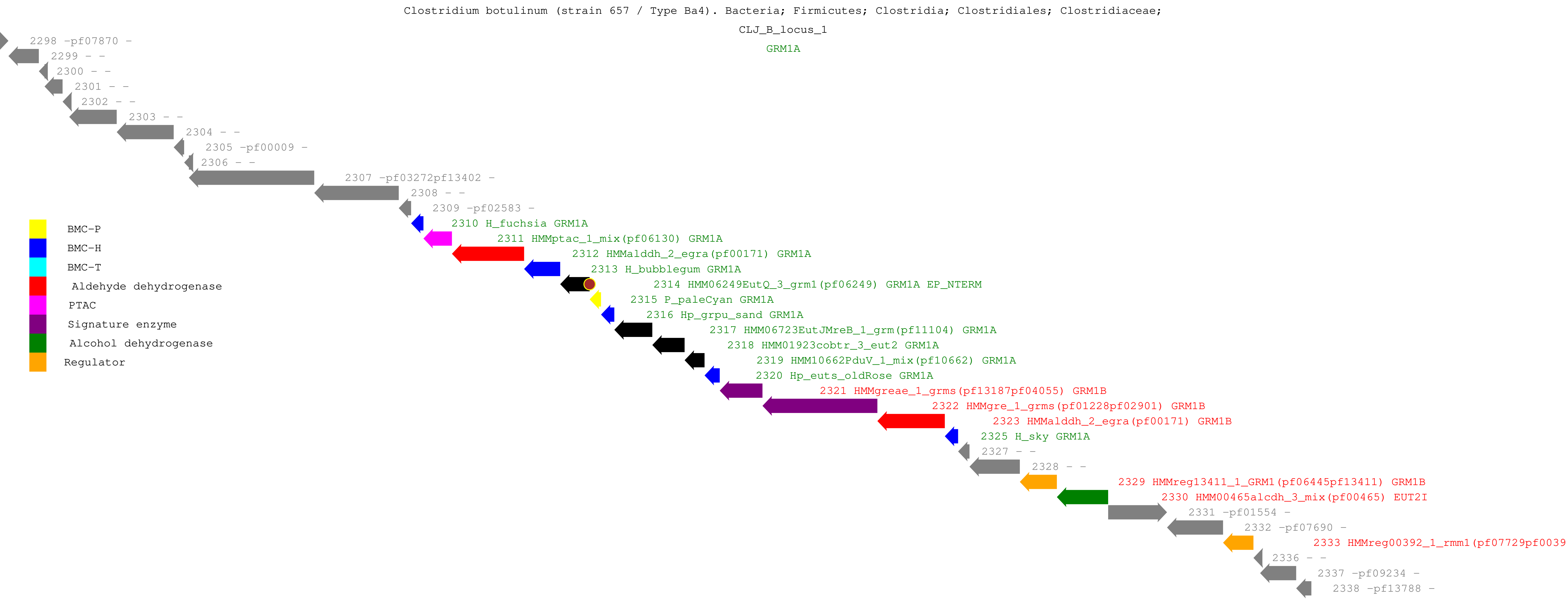

Clostridium botulinum (strain 657 / Type Ba4). Bacteria; Firmicutes; Clostridia; Clostridiales; Clostridiaceae;
CLJ_B_locus_1
GRM1A
2298 -pf07870 -
2299 - -
2300 - -
2301 - -
2302 - -
2303 - -
2304 - -
2305 -pf00009 -
2306 - -
2307 -pf03272pf13402 -
2308 - -
2309 -pf02583 -
2310 H_fuchsia GRM1A
2311 HMMptac_1_mix(pf06130) GRM1A
2312 HMMalddh_2_egra(pf00171) GRM1A
2313 H_bubblegum GRM1A
2314 HMM06249EutQ_3_grm1(pf06249) GRM1A EP_NTERM
2315 P_paleCyan GRM1A
2316 Hp_grpu_sand GRM1A
2317 HMM06723EutJMreB_1_grm(pf11104) GRM1A
2318 HMM01923cobtr_3_eut2 GRM1A
2319 HMM10662PduV_1_mix(pf10662) GRM1A
2320 Hp_euts_oldRose GRM1A
2321 HMMgreae_1_grms(pf13187pf04055) GRM1B
2322 HMMgre_1_grms(pf01228pf02901) GRM1B
2323 HMMalddh_2_egra(pf00171) GRM1B
2325 H_sky GRM1A
2327 - -
2328 - -
2329 HMMreg13411_1_GRM1(pf06445pf13411) GRM1B
2330 HMM00465alcdh_3_mix(pf00465) EUT2I
2331 -pf01554 -
2332 -pf07690 -
2333 HMMreg00392_1_rmm1(pf07729pf0039
2336 - -
2337 -pf09234 -
2338 -pf13788 -
BMC-P
BMC-H
BMC-T
Aldehyde dehydrogenase
PTAC
Signature enzyme
Alcohol dehydrogenase
Regulator

Clostridium ljungdahlii (strain ATCC 55383 / DSM 13528 / PETC). Bacteria; Firmicutes; Clostridia; Clostridiales; Clostridiaceae;

CLJU_c_locus_2

GRM1A

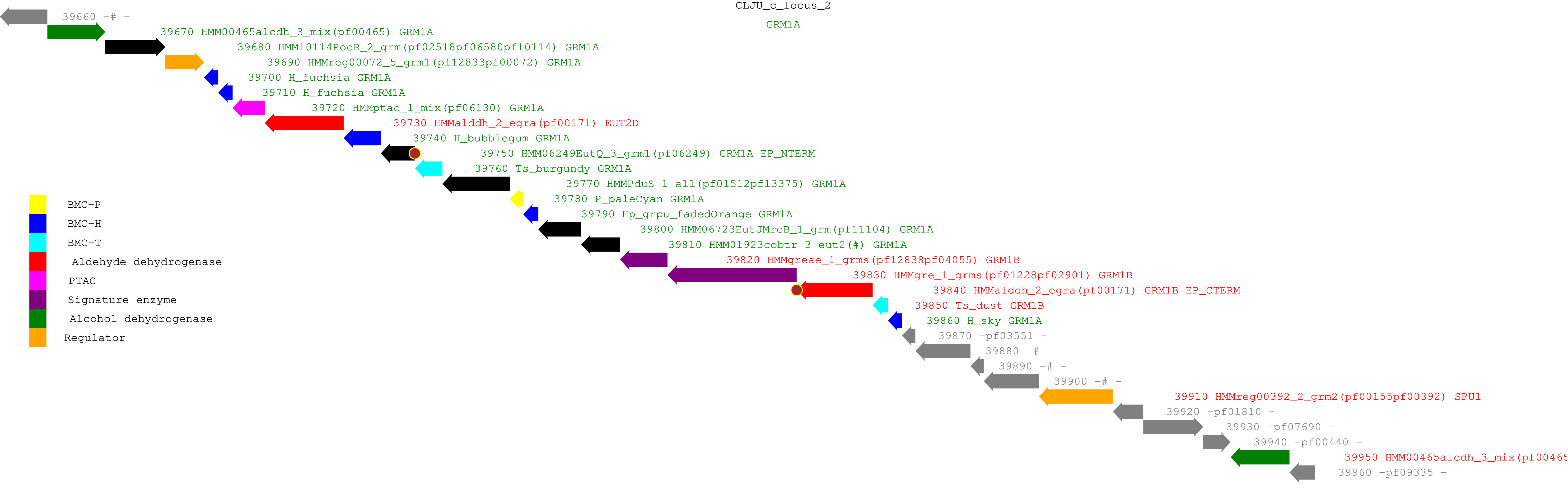

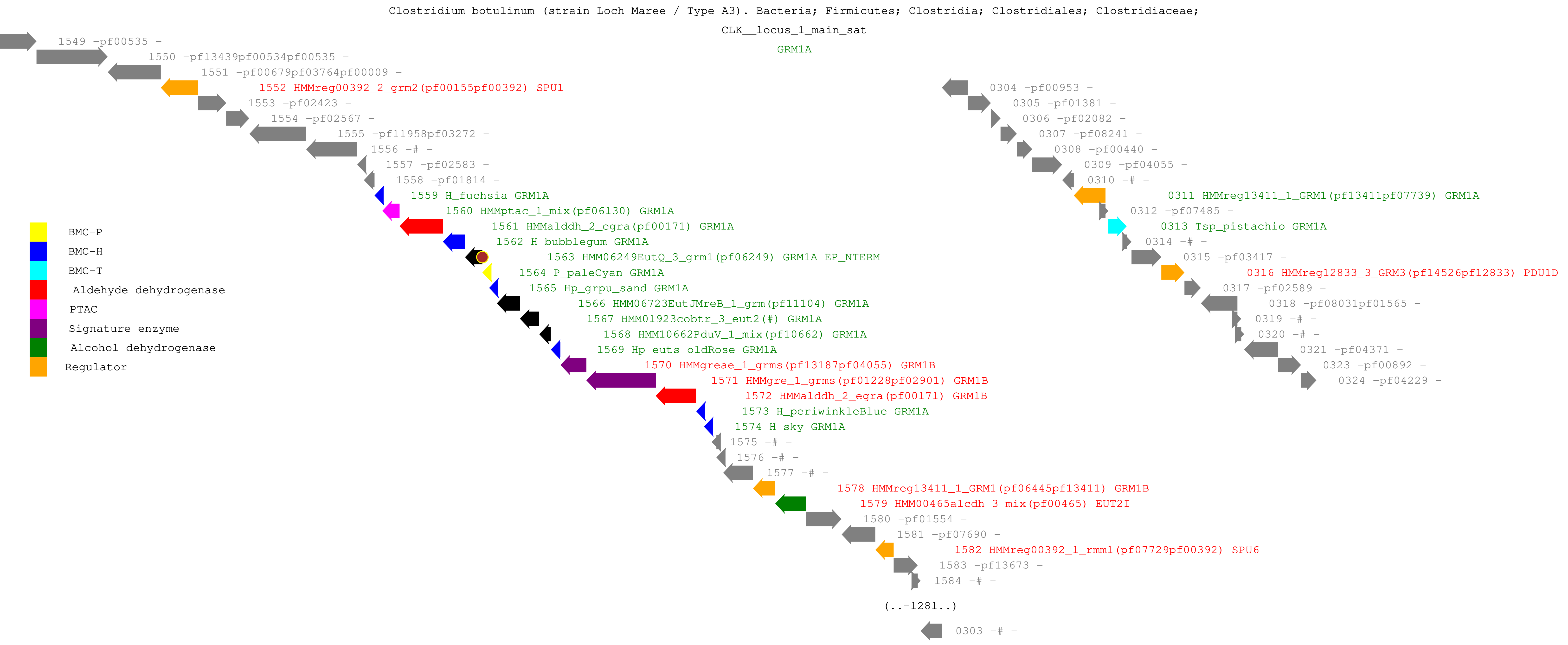
Clostridium botulinum (strain Loch Maree / Type A3). Bacteria; Firmicutes; Clostridia; Clostridiales; Clostridiaceae;
CLK__locus_1_main_sat
GRM1A
1549 -pf00535 -
1550 -pf13439pf00534pf00535 -
1551 -pf00679pf03764pf00009 -
1552 HMMreg00392_2_grm2(pf00155pf00392) SPU1
1553 -pf02423 -
1554 -pf02567 -
1555 -pf11958pf03272 -
1556 -# -
1557 -pf02583 -
1558 -pf01814 -
1559 H_fuchsia GRM1A
1560 HMMptac_1_mix(pf06130) GRM1A
1561 HMMaldh_2_egra(pf00171) GRM1A
1562 H_bubblegum GRM1A
1563 HMM06249EutQ_3_grm1(pf06249) GRM1A EP_NTERM
1564 P_paleCyan GRM1A
1565 Hp_grpu_sand GRM1A
1566 HMM06723EutJMreB_1_grm(pf11104) GRM1A
1567 HMM01923cobtr_3_eut2(#) GRM1A
1568 HMM10662PduV_1_mix(pf10662) GRM1A
1569 Hp_euts_oldRose GRM1A
1570 HMMgreae_1_grms(pf13187pf04055) GRM1B
1571 HMMgre_1_grms(pf01228pf02901) GRM1B
1572 HMMaldh_2_egra(pf00171) GRM1B
1573 H_periwinkleBlue GRM1A
1574 H_sky GRM1A
1575 -# -
1576 -# -
1577 -# -
1578 HMMreg13411_1_GRM1(pf06445pf13411) GRM1B
1579 HMM00465alcdh_3_mix(pf00465) EUT2I
1580 -pf01554 -
1581 -pf07690 -
1582 HMMreg00392_1_rmm1(pf07729pf00392) SPU6
1583 -pf13673 -
1584 -# -
(..-1281..)
0303 -# -
0304 -pf00953 -
0305 -pf01381 -
0306 -pf02082 -
0307 -pf08241 -
0308 -pf00440 -
0309 -pf04055 -
0310 -# -
0311 HMMreg13411_1_GRM1(pf13411pf07739) GRM1A
0312 -pf07485 -
0313 Tsp_pistachio GRM1A
0314 -# -
0315 -pf03417 -
0316 HMMreg12833_3_GRM3(pf14526pf12833) PDU1D
0317 -pf02589 -
0318 -pf08031pf01565 -
0319 -# -
0320 -# -
0321 -pf04371 -
0323 -pf00892 -
0324 -pf04229 -
BMC-P
BMC-H
BMC-T
Aldehyde dehydrogenase
PTAC
Signature enzyme
Alcohol dehydrogenase
Regulator

Clostridium botulinum (strain Eklund 17B / Type B). Bacteria; Firmicutes; Clostridia; Clostridiales; Clostridiaceae;

CLL_A_locus_1

GRM1A

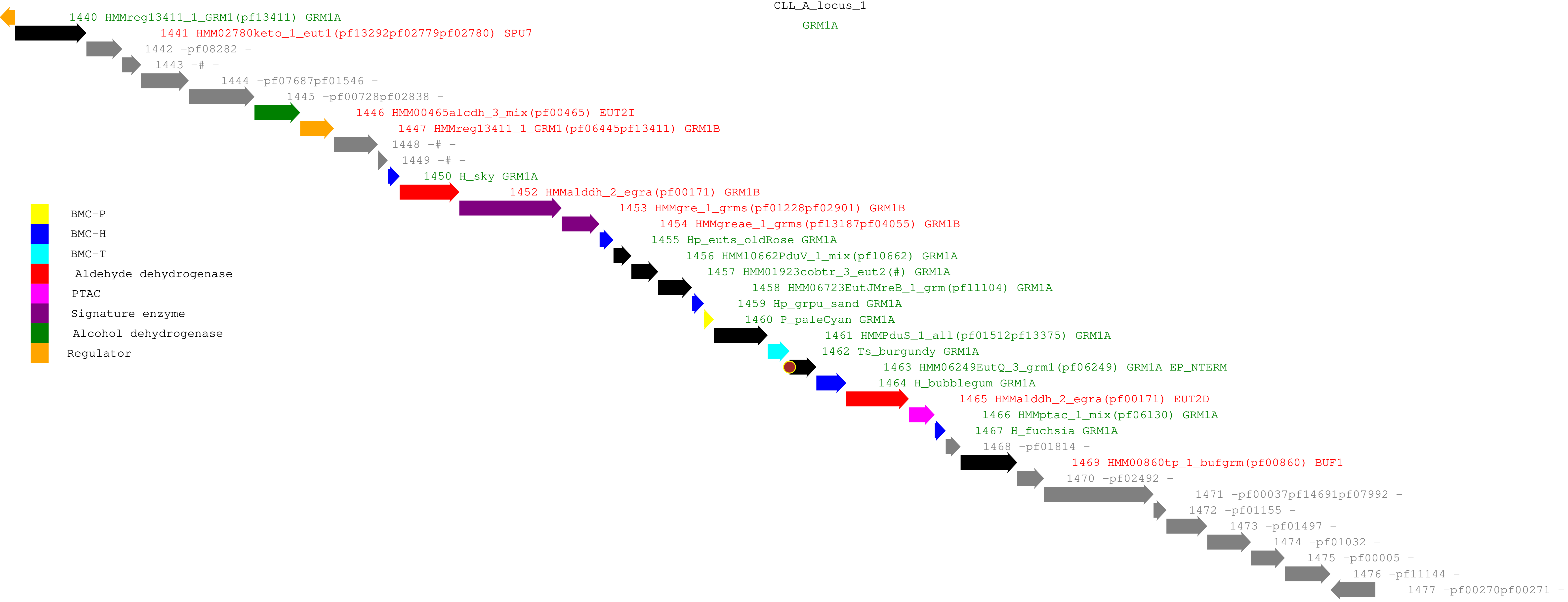

Clostridium magnum DSM 2767. Bacteria; Firmicutes; Clostridia; Clostridiales; Clostridiaceae;

CLMAG__locus_2

GRM1A

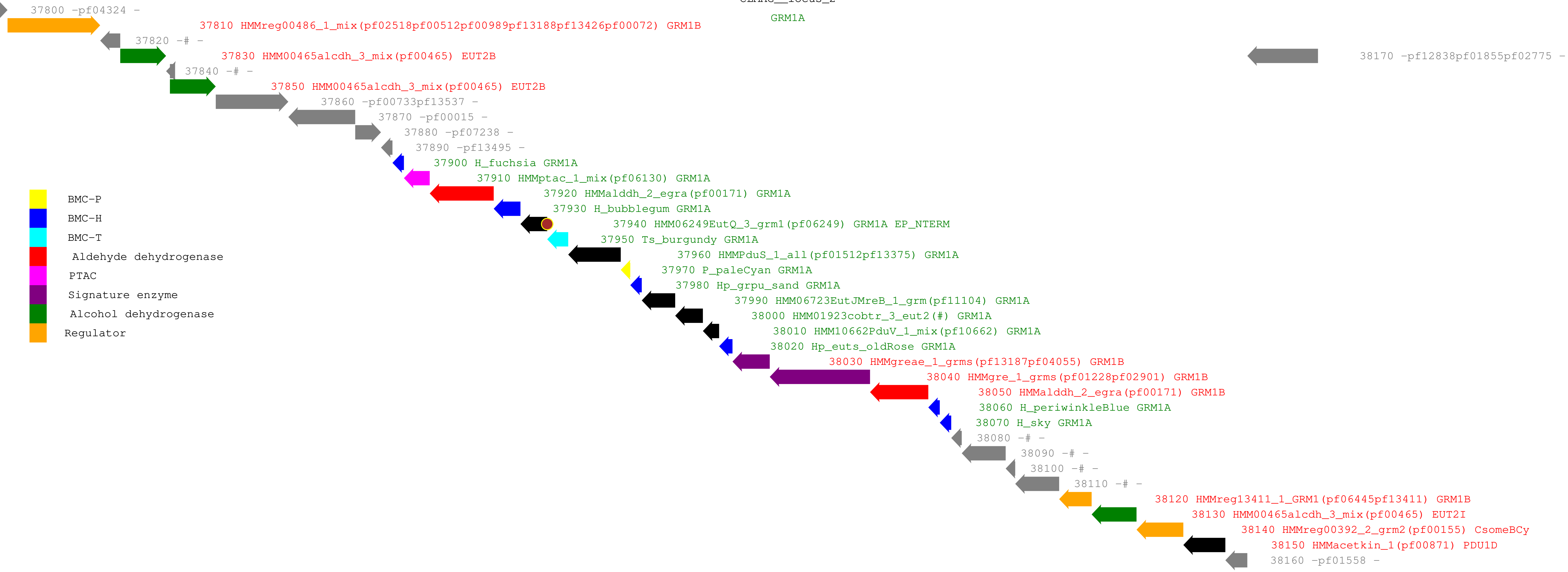

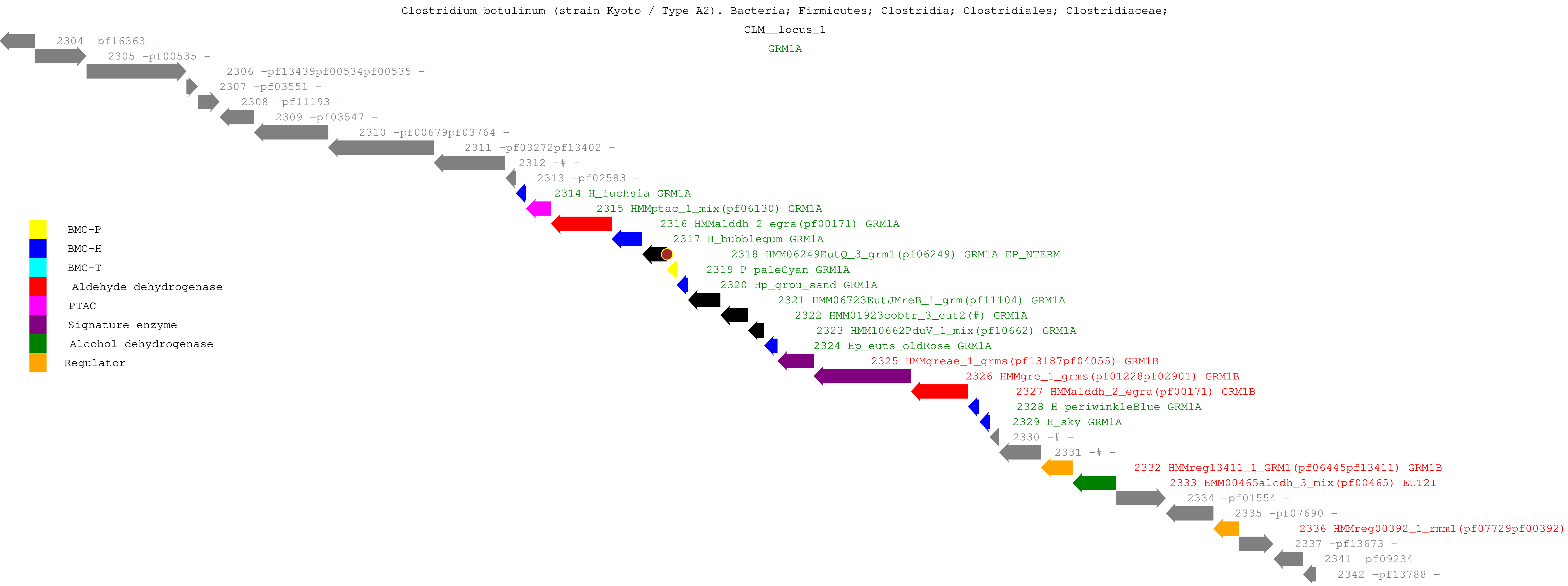
Clostridium botulinum (strain Kyoto / Type A2). Bacteria; Firmicutes; Clostridia; Clostridiales; Clostridiaceae;
CLM__locus_1
GRM1A
2304 -pf16363 -
2305 -pf00535 -
2306 -pf13439pf00534pf00535 -
2307 -pf03551 -
2308 -pf11193 -
2309 -pf03547 -
2310 -pf00679pf03764 -
2311 -pf03272pf13402 -
2312 -# -
2313 -pf02583 -
2314 H_fuchsia GRM1A
2315 HMMptac_1_mix(pf06130) GRM1A
2316 HMMaldh_2_egra(pf00171) GRM1A
2317 H_bubblegum GRM1A
2318 HMM06249EutQ_3_grm1(pf06249) GRM1A EP_NTERM
2319 P_paleCyan GRM1A
2320 Hp_grpu_sand GRM1A
2321 HMM06723EutJMreB_1_grm(pf11104) GRM1A
2322 HMM01923cobtr_3_eut2(#) GRM1A
2323 HMM10662PduV_1_mix(pf10662) GRM1A
2324 Hp_euts_oldRose GRM1A
2325 HMMgreae_1_grms(pf13187pf04055) GRM1B
2326 HMMgre_1_grms(pf01228pf02901) GRM1B
2327 HMMaldh_2_egra(pf00171) GRM1B
2328 H_periwinkleBlue GRM1A
2329 H_sky GRM1A
2330 -# -
2331 -# -
2332 HMMreg13411_1_GRM1(pf06445pf13411) GRM1B
2333 HMM00465alcdh_3_mix(pf00465) EUT2I
2334 -pf01554 -
2335 -pf07690 -
2336 HMMreg00392_1_rmm1(pf07729pf00392)
2337 -pf13673 -
2341 -pf09234 -
2342 -pf13788 -
BMC-P
BMC-H
BMC-T
Aldehyde dehydrogenase
PTAC
Signature enzyme
Alcohol dehydrogenase
Regulator

Clostridium botulinum E1 str. 'BoNT E Beluga'. Bacteria; Firmicutes; Clostridia; Clostridiales; Clostridiaceae;

CLO__locus_2

GRM1A

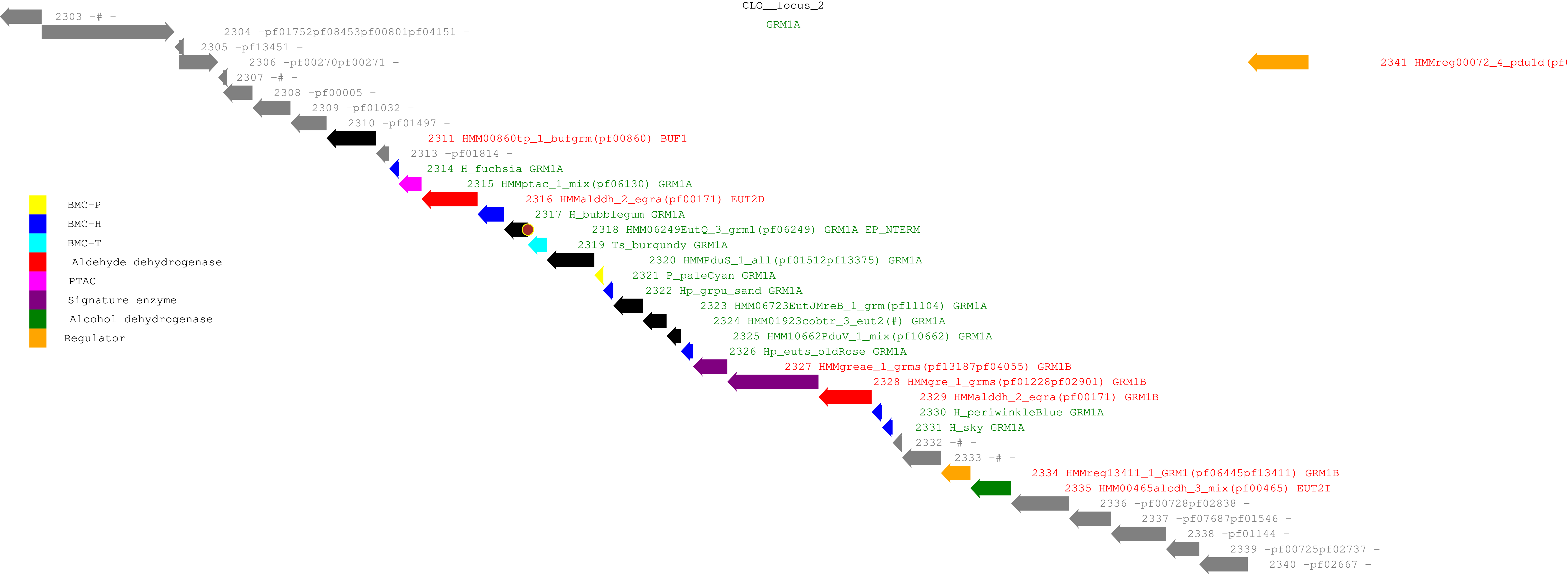

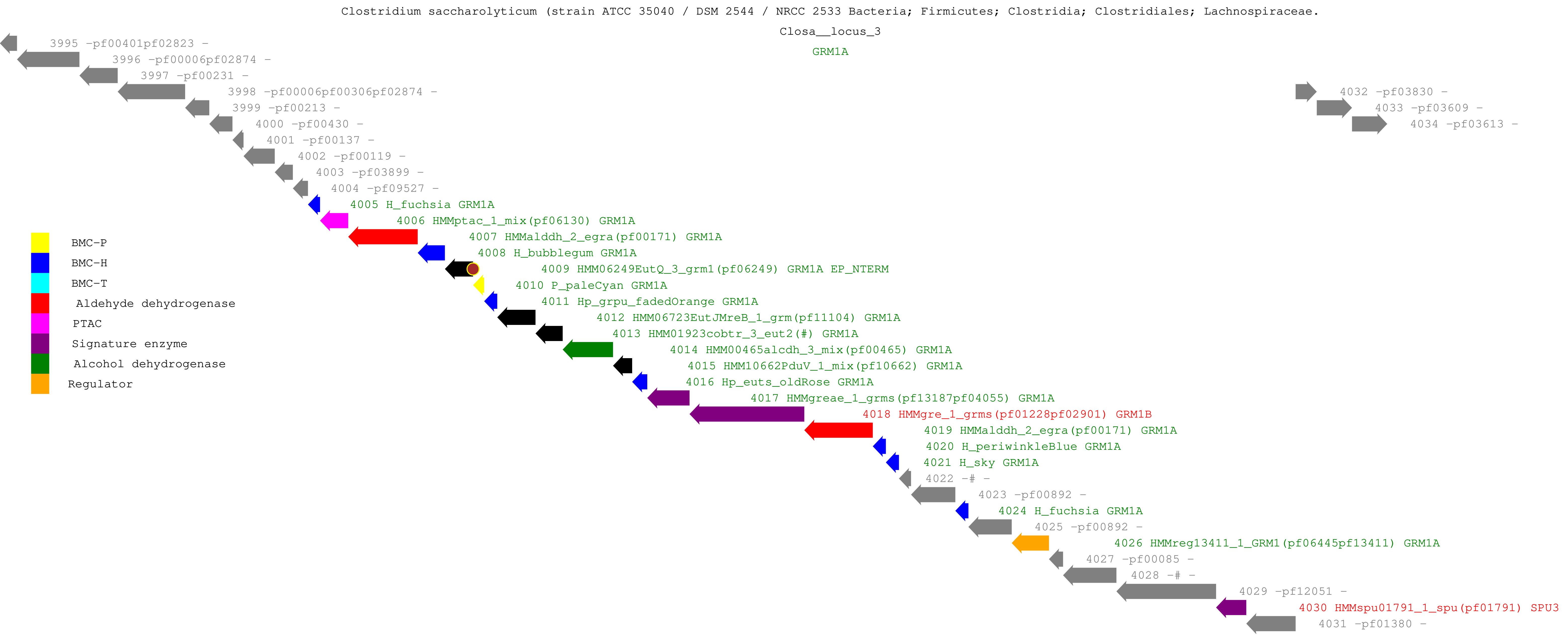
Clostridium saccharolyticum (strain ATCC 35040 / DSM 2544 / NRCC 2533 Bacteria; Firmicutes; Clostridia; Clostridiales; Lachnospiraceae.
Closa__locus_3
GRM1A
3995 -pf00401pf02823 -
3996 -pf00006pf02874 -
3997 -pf00231 -
3998 -pf00006pf00306pf02874 -
3999 -pf00213 -
4000 -pf00430 -
4001 -pf00137 -
4002 -pf00119 -
4003 -pf03899 -
4004 -pf09527 -
4005 H_fuchsia GRM1A
4006 HMMptac_1_mix(pf06130) GRM1A
4007 HMMalddh_2_egra(pf00171) GRM1A
4008 H_bubblegum GRM1A
4009 HMM06249EutQ_3_grm1(pf06249) GRM1A EP_NTERM
4010 P_paleCyan GRM1A
4011 Hp_grpu_fadedOrange GRM1A
4012 HMM06723EutJMreB_1_grm(pf11104) GRM1A
4013 HMM01923cobtr_3_eut2(#) GRM1A
4014 HMM00465alcdh_3_mix(pf00465) GRM1A
4015 HMM10662PduV_1_mix(pf10662) GRM1A
4016 Hp_euts_oldRose GRM1A
4017 HMMgreae_1_grms(pf13187pf04055) GRM1A
4018 HMMgre_1_grms(pf01228pf02901) GRM1B
4019 HMMalddh_2_egra(pf00171) GRM1A
4020 H_periwinkleBlue GRM1A
4021 H_sky GRM1A
4022 -# -
4023 -pf00892 -
4024 H_fuchsia GRM1A
4025 -pf00892 -
4026 HMMreg13411_1_GRM1(pf06445pf13411) GRM1A
4027 -pf00085 -
4028 -# -
4029 -pf12051 -
4030 HMMspu01791_1_spu(pf01791) SPU3
4031 -pf01380 -
4032 -pf03830 -
4033 -pf03609 -
4034 -pf03613 -
BMC-P
BMC-H
BMC-T
Aldehyde dehydrogenase
PTAC
Signature enzyme
Alcohol dehydrogenase
Regulator

Alkaliphilus oremlandii (strain OhILAs) (Clostridium oremlandii Bacteria; Firmicutes; Clostridia; Clostridiales; Clostridiaceae;

Clos__locus_3

GRM1A

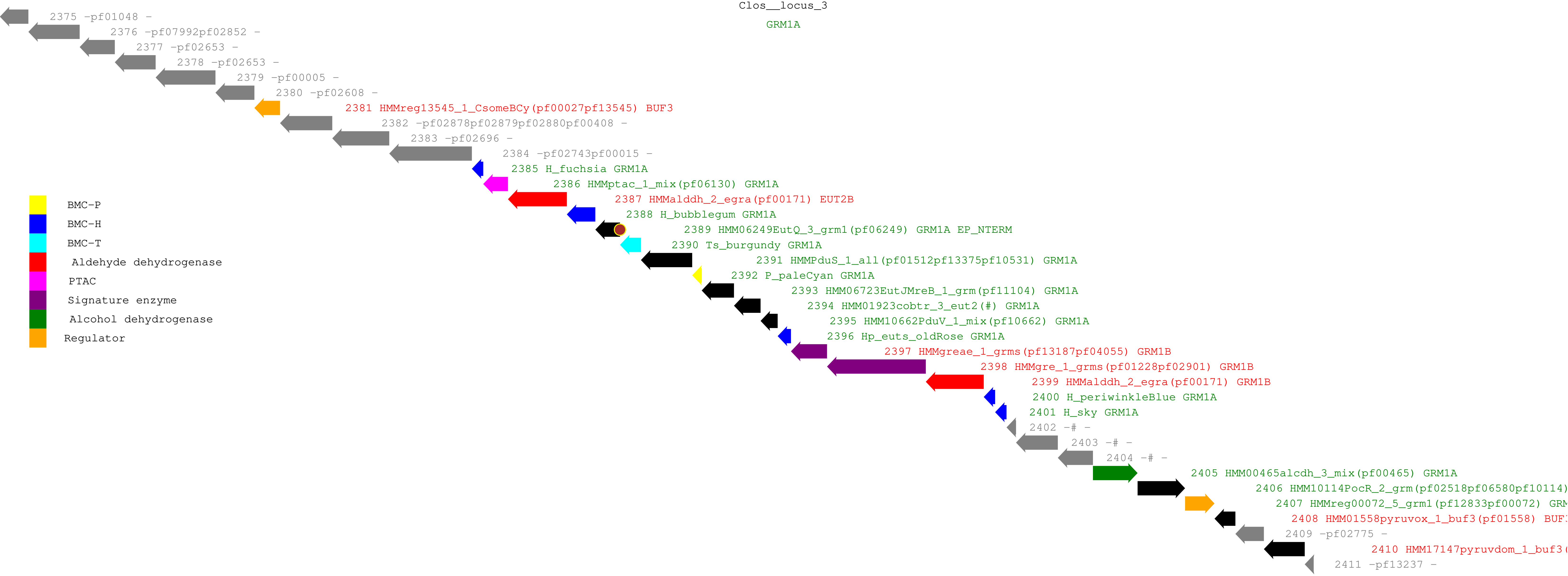

Clostridium sporogenes ATCC 15579. Bacteria; Firmicutes; Clostridia; Clostridiales; Clostridiaceae;

CLOSPO__locus_1_main_sat

GRM1A

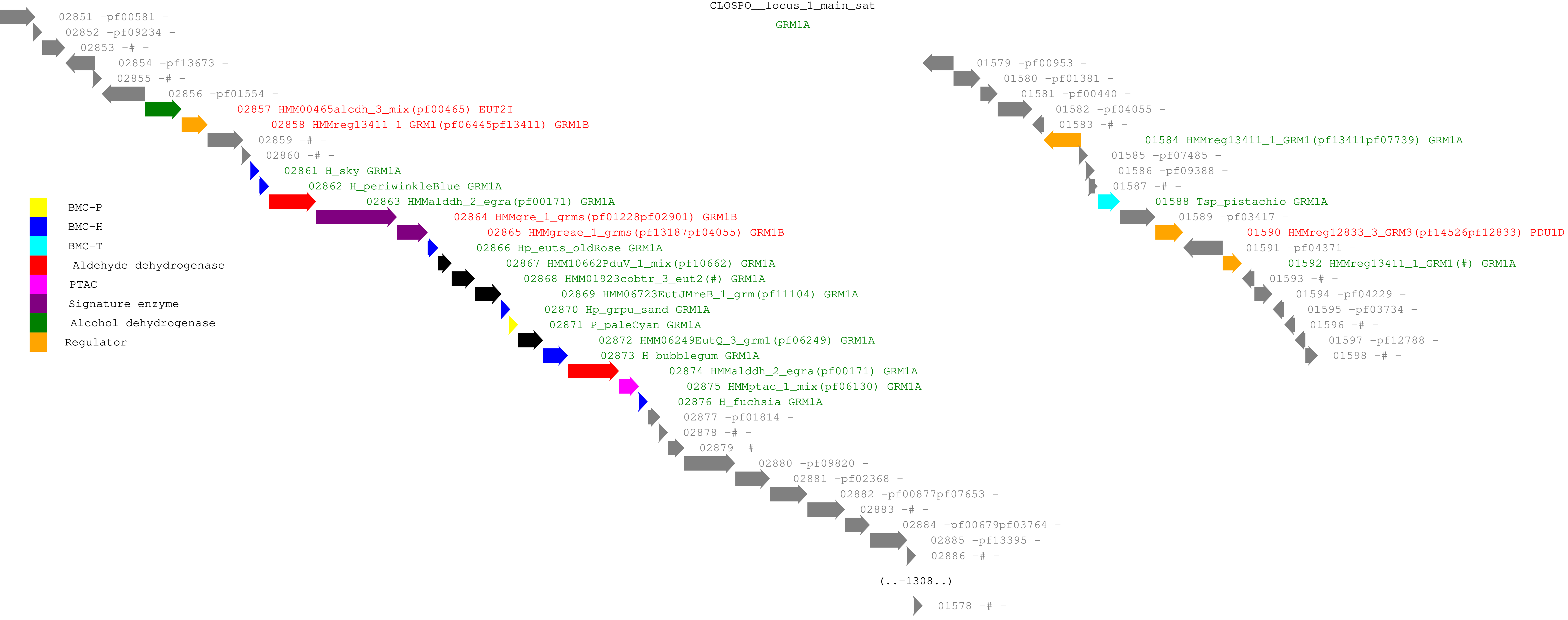

Clostridium asparagiforme DSM 15981. Bacteria; Firmicutes; Clostridia; Clostridiales; Lachnospiraceae.

CLOSTASPAR__locus_3

GRM1A

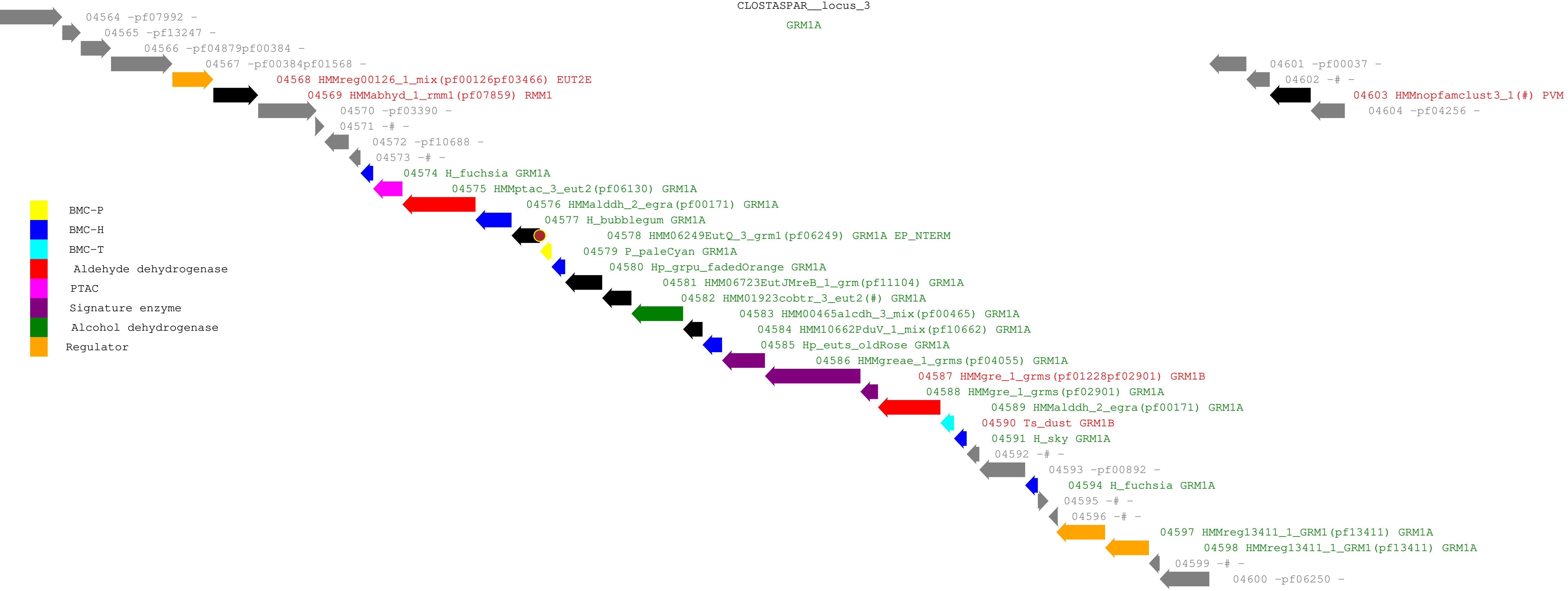

Hungatella hathewayi DSM 13479. Bacteria; Firmicutes; Clostridia; Clostridiales; Clostridiaceae;

CLOSTHATH__locus_1

GRM1A

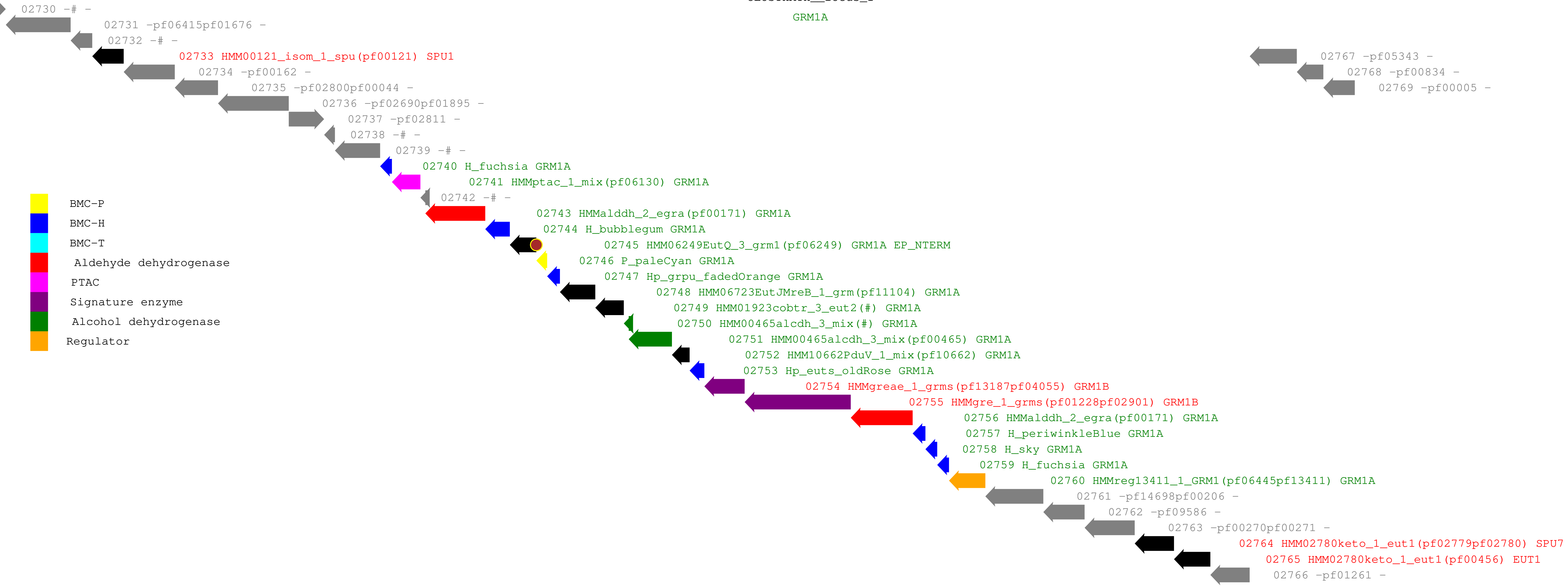

Clostridium ragsdalei P11. Bacteria; Firmicutes; Clostridia; Clostridiales; Clostridiaceae;

CLRAG__locus_2

GRM1A

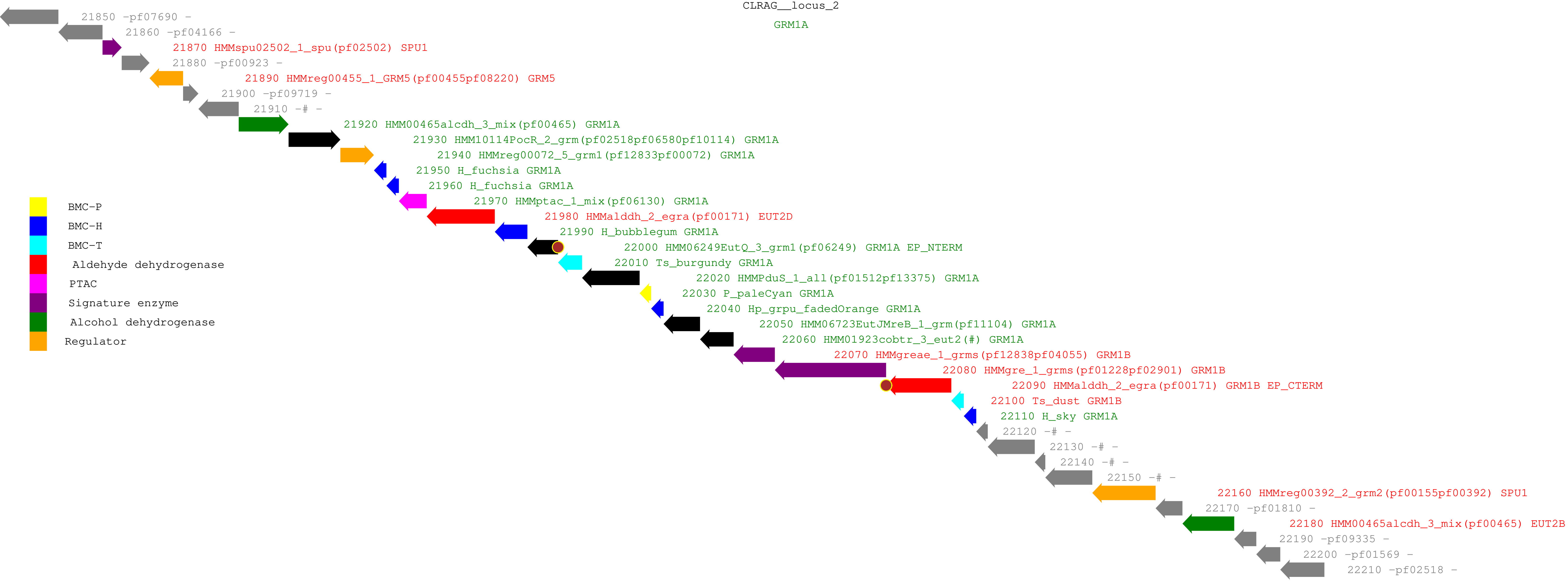

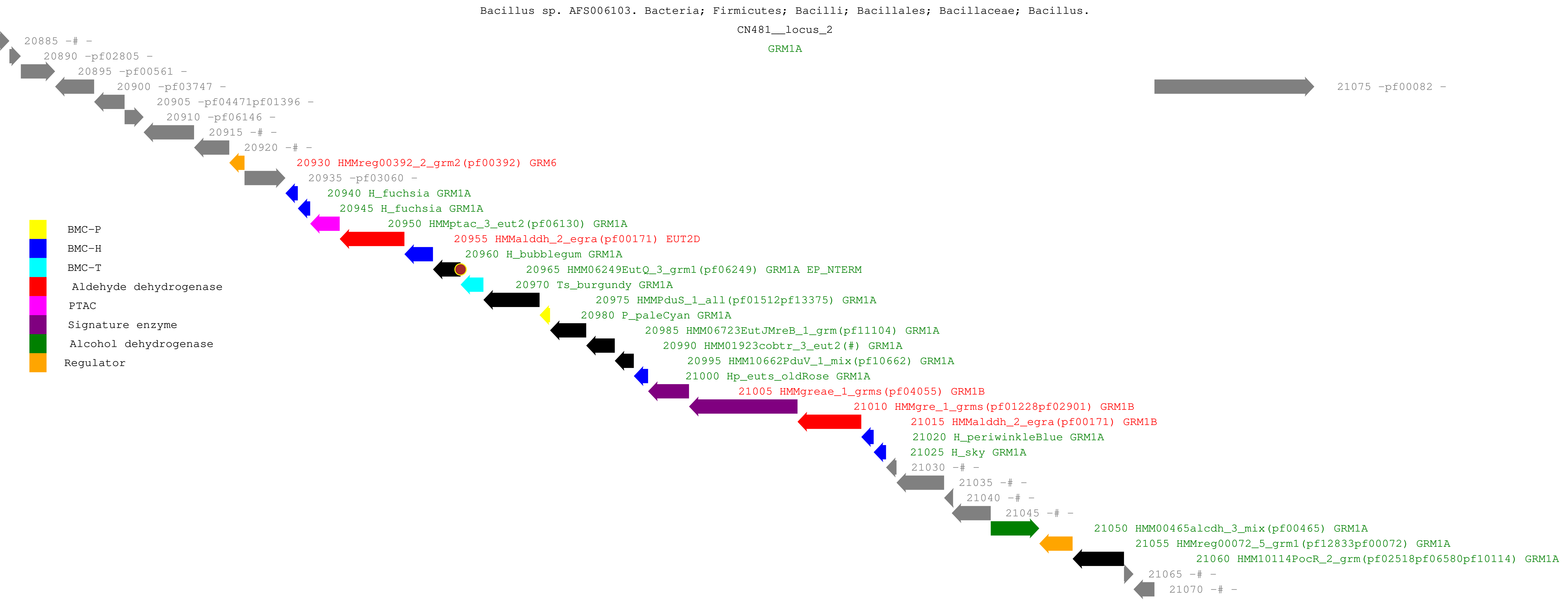
Bacillus sp. AFS006103. Bacteria; Firmicutes; Bacilli; Bacillales; Bacillaceae; Bacillus.
CN481__locus_2
GRM1A
20885 -# -
20890 -pf02805 -
20895 -pf00561 -
20900 -pf03747 -
20905 -pf04471pf01396 -
20910 -pf06146 -
20915 -# -
20920 -# -
20930 HMMreg00392_2_grm2(pf00392) GRM6
20935 -pf03060 -
20940 H_fuchsia GRM1A
20945 H_fuchsia GRM1A
20950 HMMptac_3_eut2(pf06130) GRM1A
20955 HMMaldh_2_egra(pf00171) EUT2D
20960 H_bubblegum GRM1A
20965 HMM06249EutQ_3_grm1(pf06249) GRM1A EP_NTERM
20970 Ts_burgundy GRM1A
20975 HMMPduS_1_all(pf01512pf13375) GRM1A
20980 P_paleCyan GRM1A
20985 HMM06723EutJMreB_1_grm(pf11104) GRM1A
20990 HMM01923cobtr_3_eut2(#) GRM1A
20995 HMM10662PduV_1_mix(pf10662) GRM1A
21000 Hp_euts_oldRose GRM1A
21005 HMMgreae_1_grms(pf04055) GRM1B
21010 HMMgre_1_grms(pf01228pf02901) GRM1B
21015 HMMaldh_2_egra(pf00171) GRM1B
21020 H_periwinkleBlue GRM1A
21025 H_sky GRM1A
21030 -# -
21035 -# -
21040 -# -
21045 -# -
21050 HMM00465alcdh_3_mix(pf00465) GRM1A
21055 HMMreg00072_5_grm1(pf12833pf00072) GRM1A
21060 HMM10114PocR_2_grm(pf02518pf06580pf10114) GRM1A
21065 -# -
21070 -# -
21075 -pf00082 -
BMC-P
BMC-H
BMC-T
Aldehyde dehydrogenase
PTAC
Signature enzyme
Alcohol dehydrogenase
Regulator

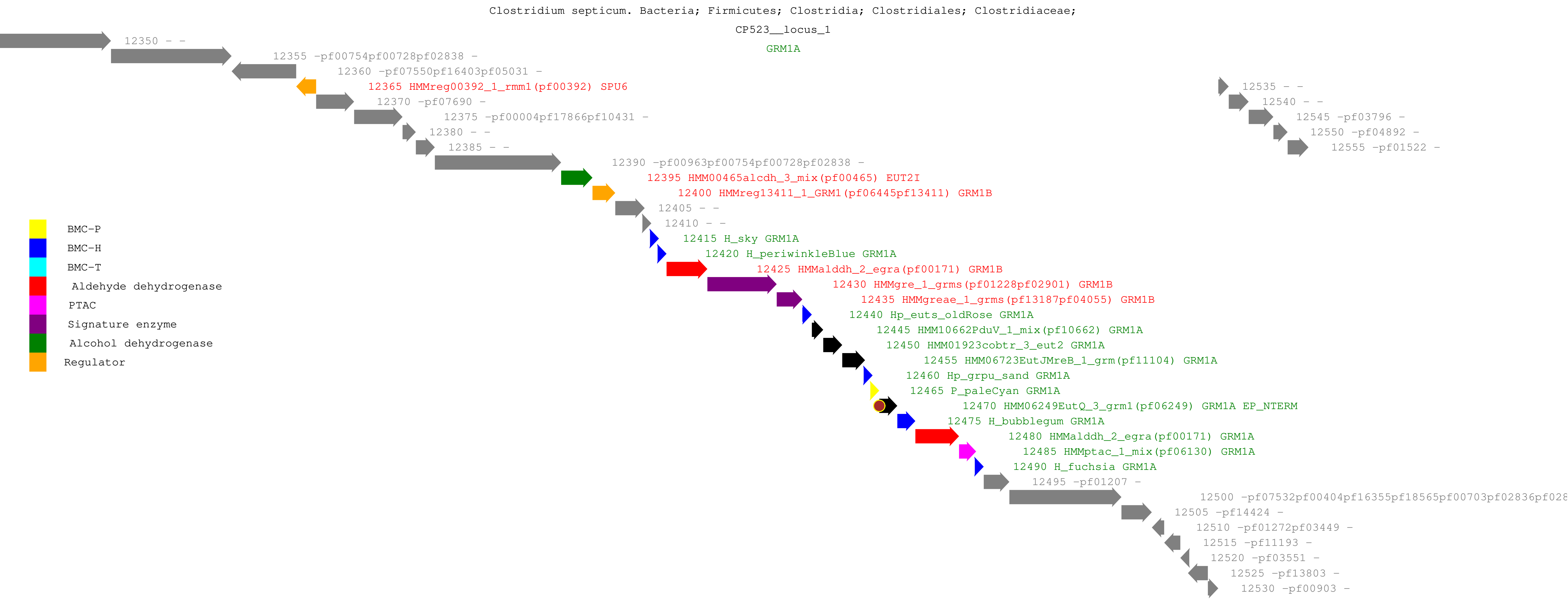

Clostridium septicum. Bacteria; Firmicutes; Clostridia; Clostridiales; Clostridiaceae;
CP523__locus_1
GRM1A
12350 - -
12355 -pf00754pf00728pf02838 -
12360 -pf07550pf16403pf05031 -
12365 HMMreg00392_1_rmm1(pf00392) SPU6
12370 -pf07690 -
12375 -pf00004pf17866pf10431 -
12380 - -
12385 - -
12390 -pf00963pf00754pf00728pf02838 -
12395 HMM00465alcdh_3_mix(pf00465) EUT2I
12400 HMMreg13411_1_GRM1(pf06445pf13411) GRM1B
12405 - -
12410 - -
12415 H_sky GRM1A
12420 H_periwinkleBlue GRM1A
12425 HMMaldh_2_egra(pf00171) GRM1B
12430 HMMgre_1_grms(pf01228pf02901) GRM1B
12435 HMMgreae_1_grms(pf13187pf04055) GRM1B
12440 Hp_euts_oldRose GRM1A
12445 HMM10662PduV_1_mix(pf10662) GRM1A
12450 HMM01923cobtr_3_eut2 GRM1A
12455 HMM06723EutJMreB_1_grm(pf11104) GRM1A
12460 Hp_grpu_sand GRM1A
12465 P_paleCyan GRM1A
12470 HMM06249EutQ_3_grm1(pf06249) GRM1A EP_NTERM
12475 H_bubblegum GRM1A
12480 HMMaldh_2_egra(pf00171) GRM1A
12485 HMMptac_1_mix(pf06130) GRM1A
12490 H_fuchsia GRM1A
12495 -pf01207 -
12500 -pf07532pf00404pf16355pf18565pf00703pf02836pf028
12505 -pf14424 -
12510 -pf01272pf03449 -
12515 -pf11193 -
12520 -pf03551 -
12525 -pf13803 -
12530 -pf00903 -
12535 - -
12540 - -
12545 -pf03796 -
12550 -pf04892 -
12555 -pf01522 -
BMC-P
BMC-H
BMC-T
Aldehyde dehydrogenase
PTAC
Signature enzyme
Alcohol dehydrogenase
Regulator

Lachnoclostridium phytofermentans (strain ATCC 700394 / DSM 18823 / Bacteria; Firmicutes; Clostridia; Clostridiales; Lachnospiraceae.

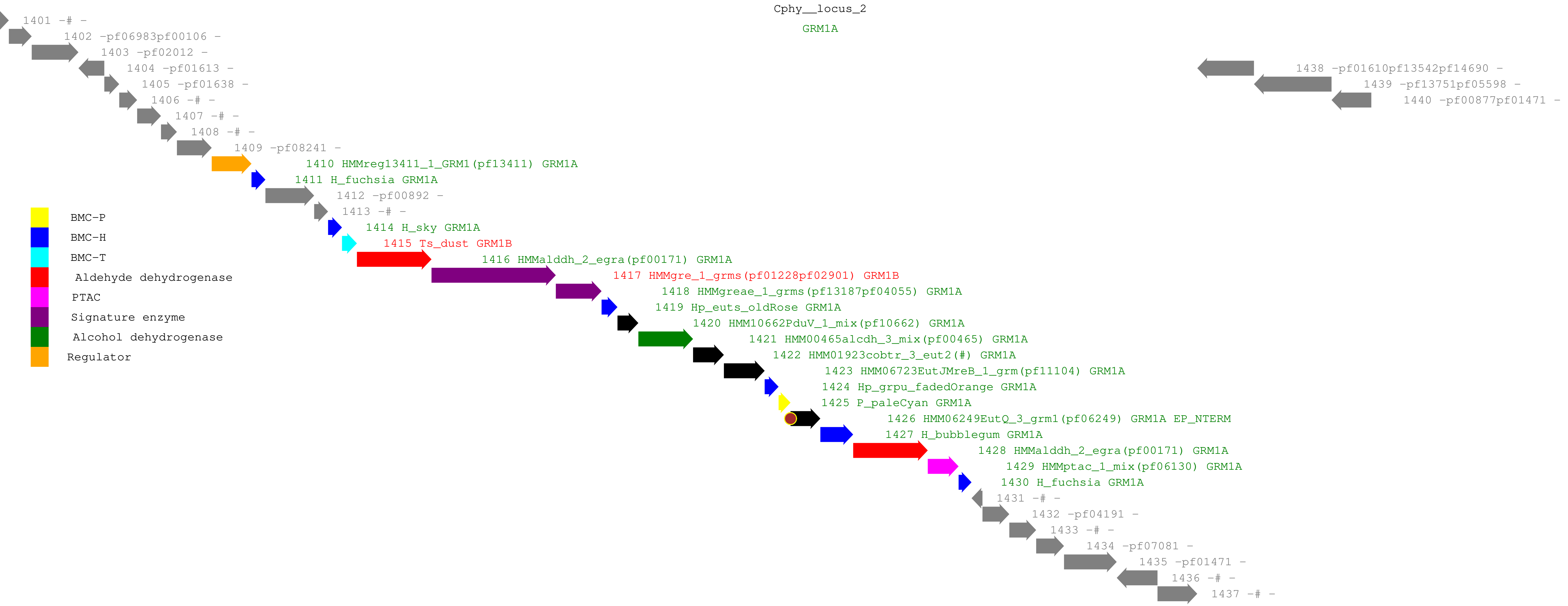

Eubacterium combesii. Bacteria; Firmicutes; Clostridia; Clostridiales; Eubacteriaceae;

CS538__locus_1_main_sat

GRM1A

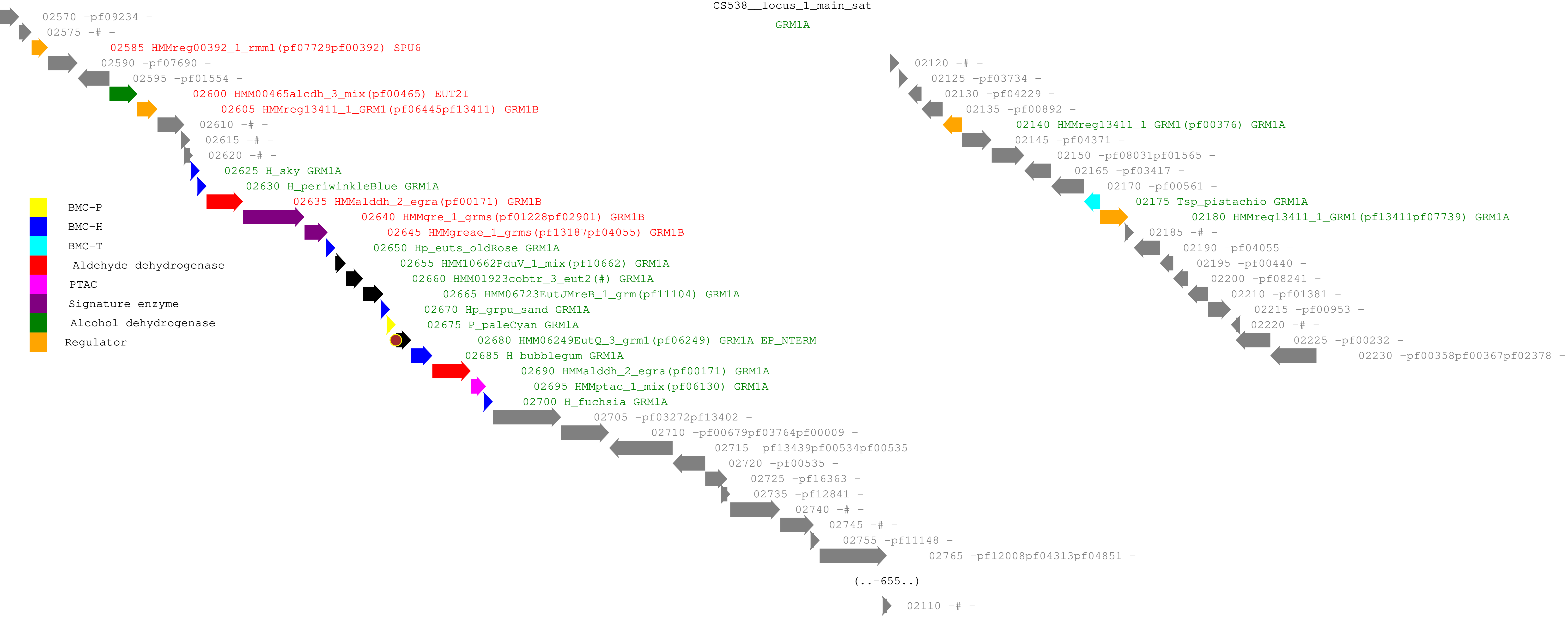

Clostridium scatologenes. Bacteria; Firmicutes; Clostridia; Clostridiales; Clostridiaceae;

CSCA__locus_2

GRM1A

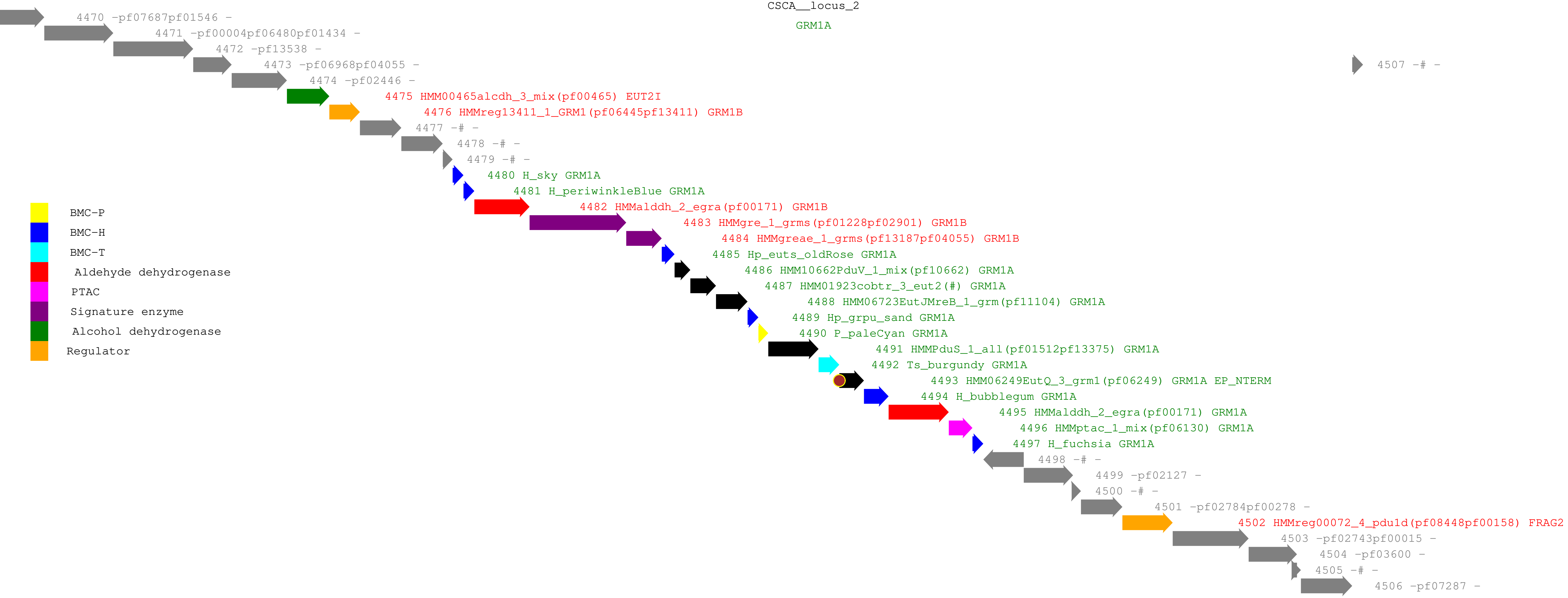

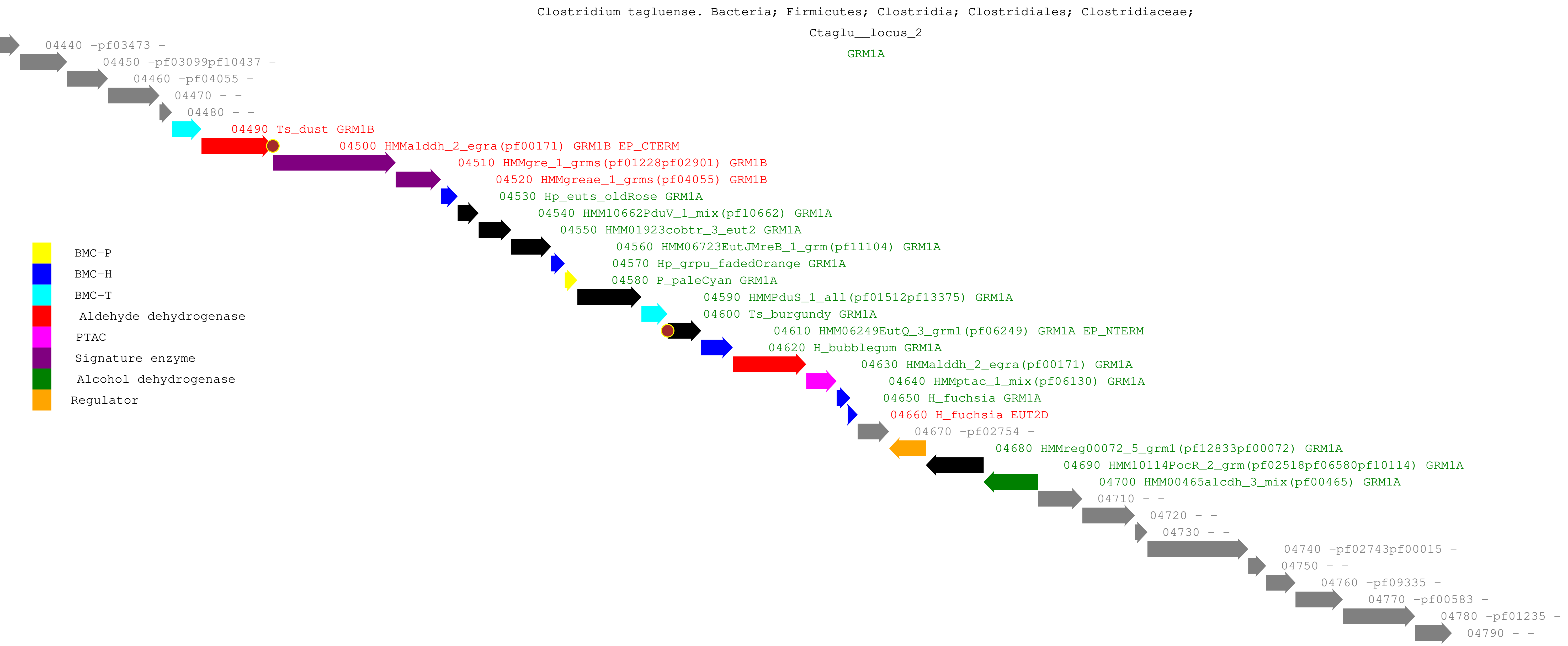
Clostridium tagluense. Bacteria; Firmicutes; Clostridia; Clostridiales; Clostridiaceae;
Ctaglu__locus_2
GRM1A
04440 -pf03473 -
04450 -pf03099pf10437 -
04460 -pf04055 -
04470 - -
04480 - -
04490 Ts_dust GRM1B
04500 HMMalddh_2_egra(pf00171) GRM1B EP_CTERM
04510 HMMgre_1_grms(pf01228pf02901) GRM1B
04520 HMMgreae_1_grms(pf04055) GRM1B
04530 Hp_euts_oldRose GRM1A
04540 HMM10662PduV_1_mix(pf10662) GRM1A
04550 HMM01923cobtr_3_eut2 GRM1A
04560 HMM06723EutJMreB_1_grm(pf11104) GRM1A
04570 Hp_grpu_fadedOrange GRM1A
04580 P_paleCyan GRM1A
04590 HMMPduS_1_all(pf01512pf13375) GRM1A
04600 Ts_burgundy GRM1A
04610 HMM06249EutQ_3_grm1(pf06249) GRM1A EP_NTERM
04620 H_bubblegum GRM1A
04630 HMMalddh_2_egra(pf00171) GRM1A
04640 HMMptac_1_mix(pf06130) GRM1A
04650 H_fuchsia GRM1A
04660 H_fuchsia EUT2D
04670 -pf02754 -
04680 HMMreg00072_5_grm1(pf12833pf00072) GRM1A
04690 HMM10114PocR_2_grm(pf02518pf06580pf10114) GRM1A
04700 HMM00465alcdh_3_mix(pf00465) GRM1A
04710 - -
04720 - -
04730 - -
04740 -pf02743pf00015 -
04750 - -
04760 -pf09335 -
04770 -pf00583 -
04780 -pf01235 -
04790 - -
BMC-P
BMC-H
BMC-T
Aldehyde dehydrogenase
PTAC
Signature enzyme
Alcohol dehydrogenase
Regulator

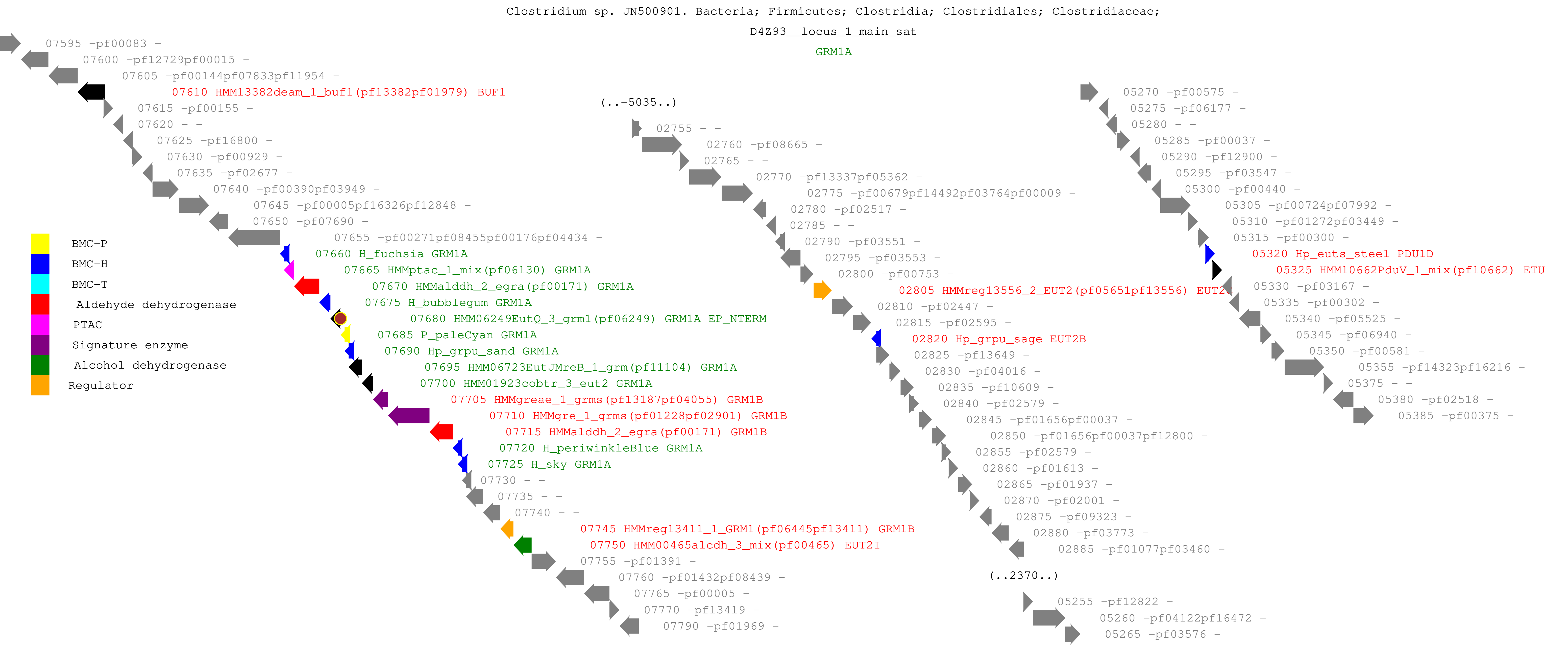

Clostridium sp. JN500901. Bacteria; Firmicutes; Clostridia; Clostridiales; Clostridiaceae;
D4Z93__locus_1_main_sat
GRM1A
BMC-P
BMC-H
BMC-T
Aldehyde dehydrogenase
PTAC
Signature enzyme
Alcohol dehydrogenase
Regulator
07595 -pf00083 -
07600 -pf12729pf00015 -
07605 -pf00144pf07833pf11954 -
07610 HMM13382deam_1_buf1(pf13382pf01979) BUF1
07615 -pf00155 -
07620 - -
07625 -pf16800 -
07630 -pf00929 -
07635 -pf02677 -
07640 -pf00390pf03949 -
07645 -pf00005pf16326pf12848 -
07650 -pf07690 -
07655 -pf00271pf08455pf00176pf04434 -
07660 H_fuchsia GRM1A
07665 HMMptac_1_mix(pf06130) GRM1A
07670 HMMalddh_2_egra(pf00171) GRM1A
07675 H_bubblegum GRM1A
07680 HMM06249EutQ_3_grm1(pf06249) GRM1A EP_NTERM
07685 P_paleCyan GRM1A
07690 Hp_grpu_sand GRM1A
07695 HMM06723EutJMreB_1_grm(pf11104) GRM1A
07700 HMM01923cobtr_3_eut2 GRM1A
07705 HMMgreae_1_grms(pf13187pf04055) GRM1B
07710 HMMgre_1_grms(pf01228pf02901) GRM1B
07715 HMMalddh_2_egra(pf00171) GRM1B
07720 H_periwinkleBlue GRM1A
07725 H_sky GRM1A
07730 - -
07735 - -
07740 - -
07745 HMMreg13411_1_GRM1(pf06445pf13411) GRM1B
07750 HMM00465alcdh_3_mix(pf00465) EUT2I
07755 -pf01391 -
07760 -pf01432pf08439 -
07765 -pf00005 -
07770 -pf13419 -
07790 -pf01969 -
(..-5035..)
02755 - -
02760 -pf08665 -
02765 - -
02770 -pf13337pf05362 -
02775 -pf00679pf14492pf03764pf00009 -
02780 -pf02517 -
02785 - -
02790 -pf03551 -
02795 -pf03553 -
02800 -pf00753 -
02805 HMMreg13556_2_EUT2(pf05651pf13556) EUT2
02810 -pf02447 -
02815 -pf02595 -
02820 Hp_grpu_sage EUT2B
02825 -pf13649 -
02830 -pf04016 -
02835 -pf10609 -
02840 -pf02579 -
02845 -pf01656pf00037 -
02850 -pf01656pf00037pf12800 -
02855 -pf02579 -
02860 -pf01613 -
02865 -pf01937 -
02870 -pf02001 -
02875 -pf09323 -
02880 -pf03773 -
02885 -pf01077pf03460 -
(..2370..)
05255 -pf12822 -
05260 -pf04122pf16472 -
05265 -pf03576 -
05270 -pf00575 -
05275 -pf06177 -
05280 - -
05285 -pf00037 -
05290 -pf12900 -
05295 -pf03547 -
05300 -pf00440 -
05305 -pf00724pf07992 -
05310 -pf01272pf03449 -
05315 -pf00300 -
05320 Hp_euts_steel PDU1D
05325 HMM10662PduV_1_mix(pf10662) ETU
05330 -pf03167 -
05335 -pf00302 -
05340 -pf05525 -
05345 -pf06940 -
05350 -pf00581 -
05355 -pf14323pf16216 -
05375 - -
05380 -pf02518 -
05385 -pf00375 -

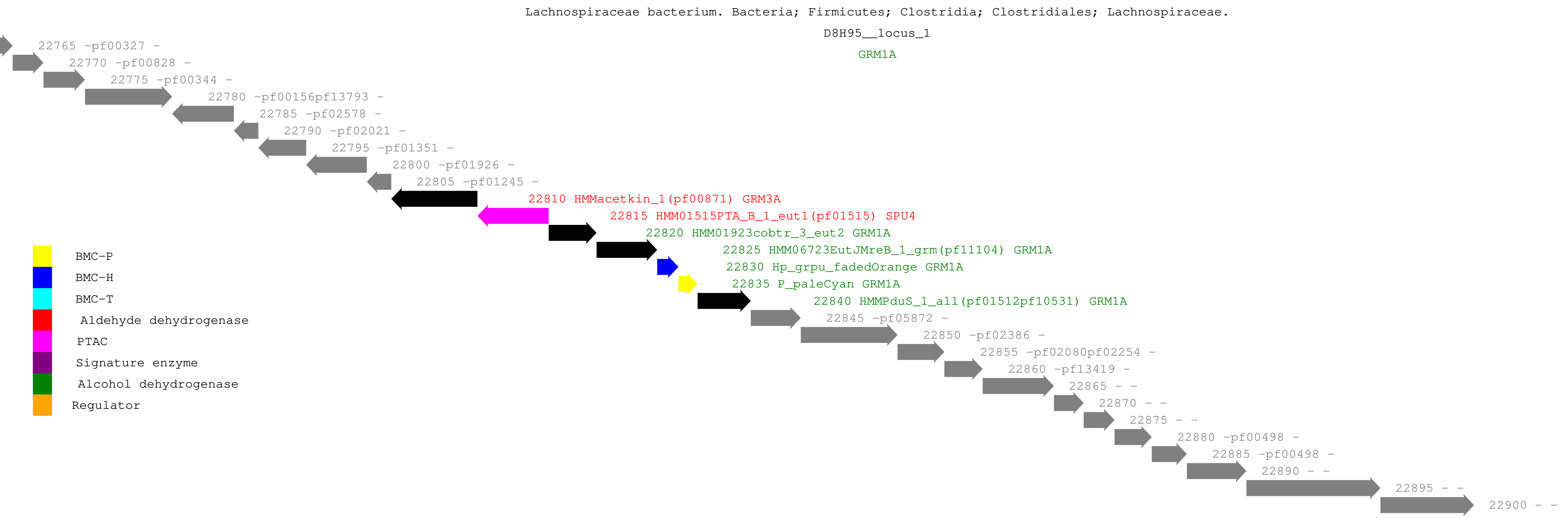
Lachnospiraceae bacterium. Bacteria; Firmicutes; Clostridia; Clostridiales; Lachnospiraceae.
D8H95__locus_1
GRM1A
22765 -pf00327 -
22770 -pf00828 -
22775 -pf00344 -
22780 -pf00156pf13793 -
22785 -pf02578 -
22790 -pf02021 -
22795 -pf01351 -
22800 -pf01926 -
22805 -pf01245 -
22810 HMMacetkin_1(pf00871) GRM3A
22815 HMM01515PTA_B_1_eut1(pf01515) SPU4
22820 HMM01923cobtr_3_eut2 GRM1A
22825 HMM06723EutJMreB_1_grm(pf11104) GRM1A
22830 Hp_grpu_fadedOrange GRM1A
22835 P_paleCyan GRM1A
22840 HMMPduS_1_all(pf01512pf10531) GRM1A
22845 -pf05872 -
22850 -pf02386 -
22855 -pf02080pf02254 -
22860 -pf13419 -
22865 - -
22870 - -
22875 - -
22880 -pf00498 -
22885 -pf00498 -
22890 - -
22895 - -
22900 - -
BMC-P
BMC-H
BMC-T
Aldehyde dehydrogenase
PTAC
Signature enzyme
Alcohol dehydrogenase
Regulator

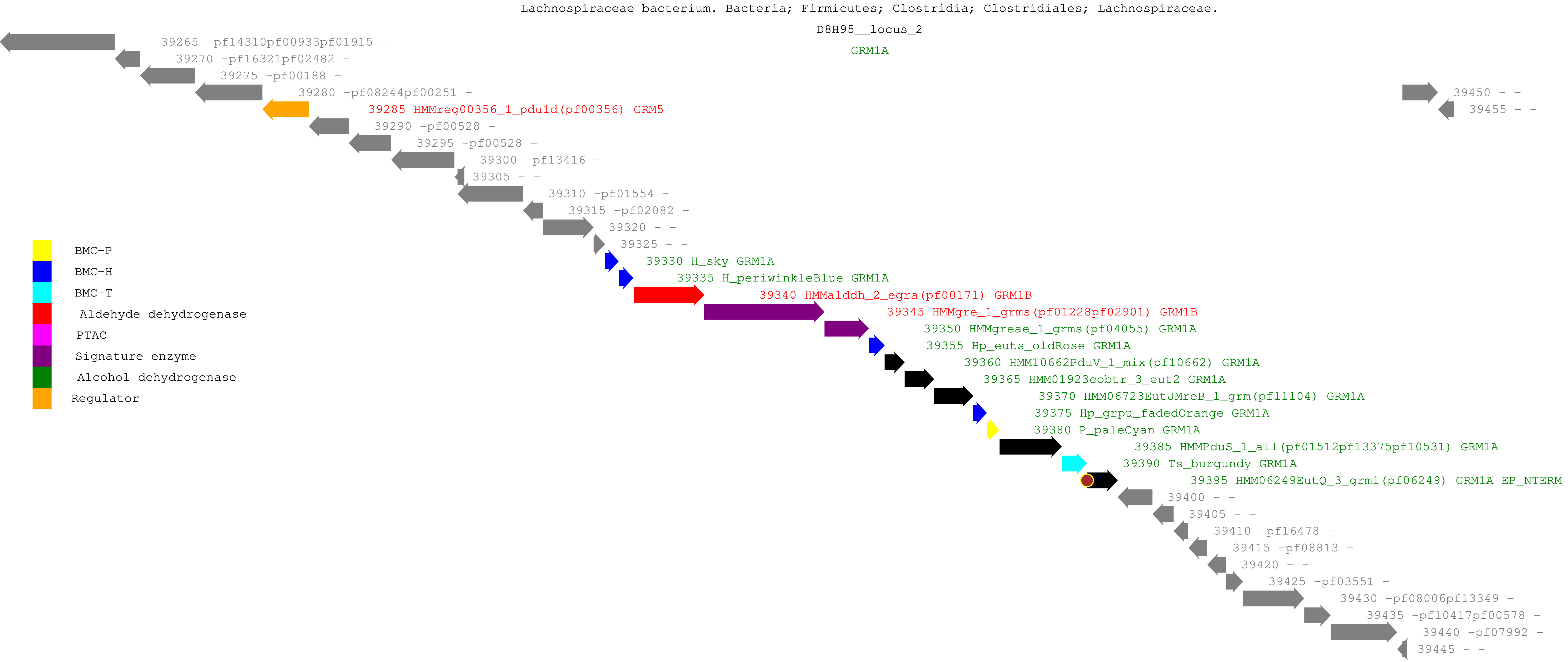

Lachnospiraceae bacterium. Bacteria; Firmicutes; Clostridia; Clostridiales; Lachnospiraceae.
D8H95__locus_2
GRM1A
39265 -pf14310pf00933pf01915 -
39270 -pf16321pf02482 -
39275 -pf00188 -
39280 -pf08244pf00251 -
39285 HMMreg00356_1_pdu1d(pf00356) GRM5
39290 -pf00528 -
39295 -pf00528 -
39300 -pf13416 -
39305 - -
39310 -pf01554 -
39315 -pf02082 -
39320 - -
39325 - -
39330 H_sky GRM1A
39335 H_periwinkleBlue GRM1A
39340 HMMalddh_2_egra(pf00171) GRM1B
39345 HMMgre_1_grms(pf01228pf02901) GRM1B
39350 HMMgreae_1_grms(pf04055) GRM1A
39355 Hp_euts_oldRose GRM1A
39360 HMM10662PduV_1_mix(pf10662) GRM1A
39365 HMM01923cobtr_3_eut2 GRM1A
39370 HMM06723EutJMreB_1_grm(pf11104) GRM1A
39375 Hp_grpu_fadedOrange GRM1A
39380 P_paleCyan GRM1A
39385 HMMPduS_1_all(pf01512pf13375pf10531) GRM1A
39390 Ts_burgundy GRM1A
39395 HMM06249EutQ_3_grm1(pf06249) GRM1A EP_NTERM
39400 - -
39405 - -
39410 -pf16478 -
39415 -pf08813 -
39420 - -
39425 -pf03551 -
39430 -pf08006pf13349 -
39435 -pf10417pf00578 -
39440 -pf07992 -
39445 - -
39450 - -
39455 - -
BMC-P
BMC-H
BMC-T
Aldehyde dehydrogenase
PTAC
Signature enzyme
Alcohol dehydrogenase
Regulator

Streptococcus sp. Bacteria; Firmicutes; Bacilli; Lactobacillales; Streptococcaceae;

D8H99__locus_2

GRM1A

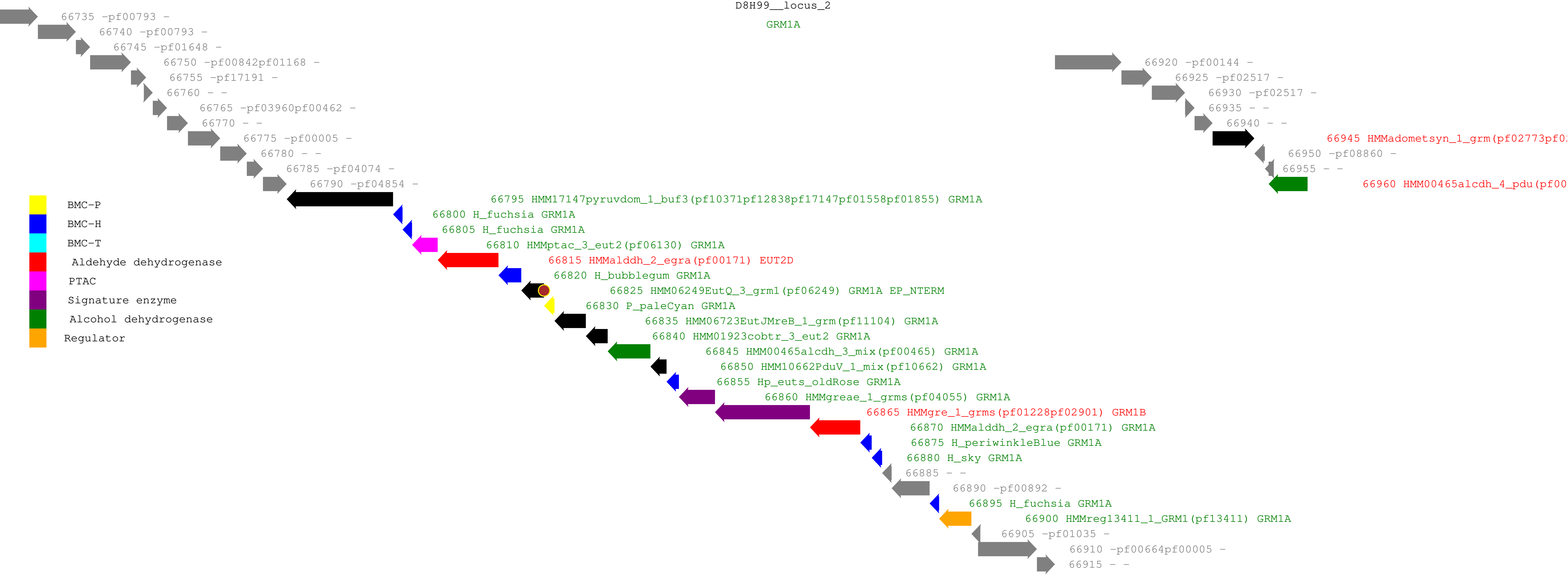

Clostridium autoethanogenum. Bacteria; Firmicutes; Clostridia; Clostridiales; Clostridiaceae;

D9040__locus_2

GRM1A

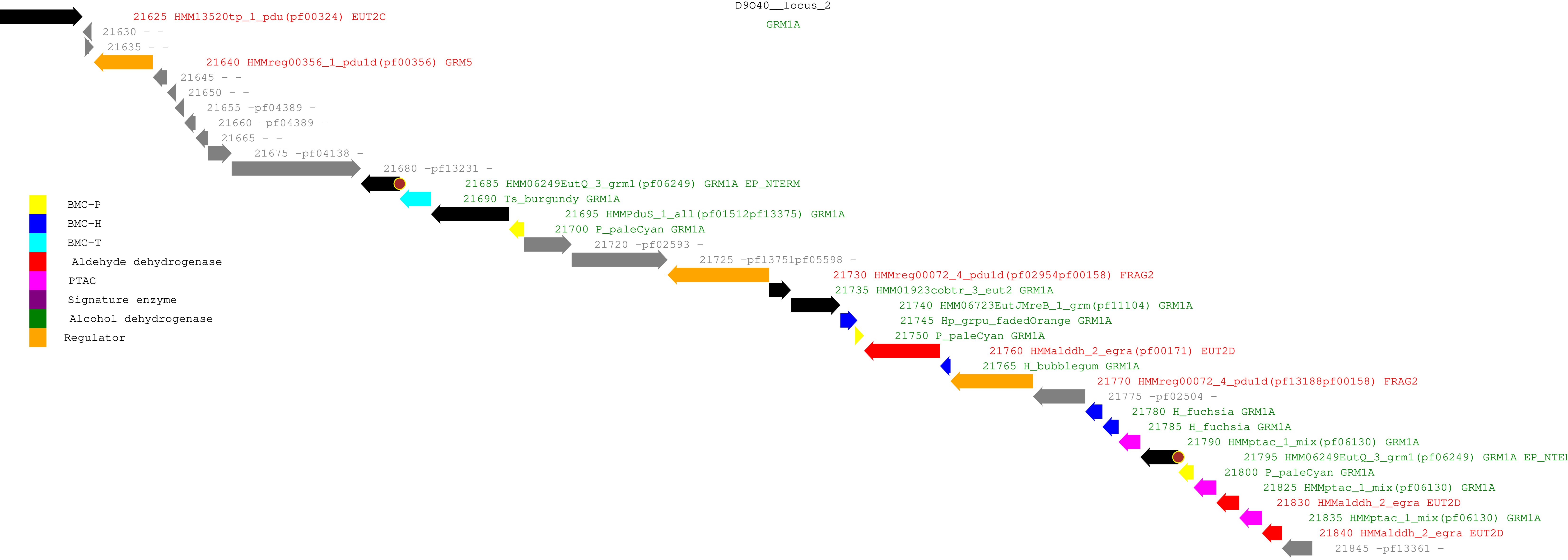

Lachnoclostridium sp. Bacteria; Firmicutes; Clostridia; Clostridiales; Lachnospiraceae.

DC053__locus_3

GRM1A

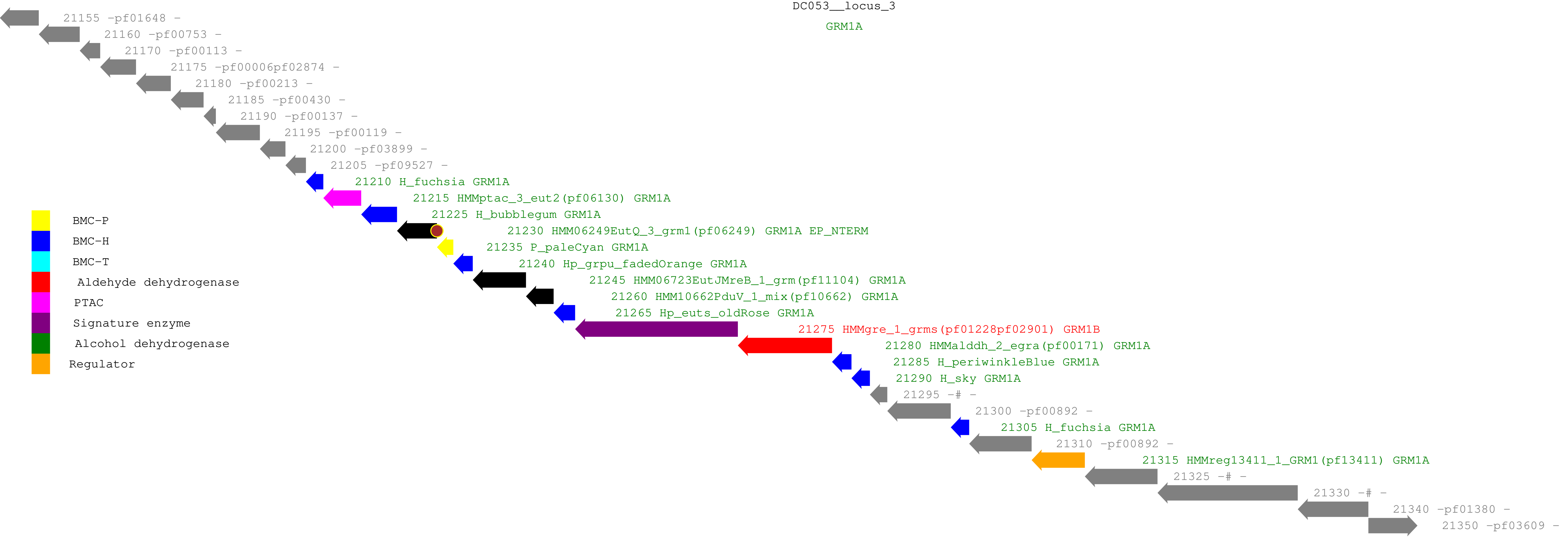

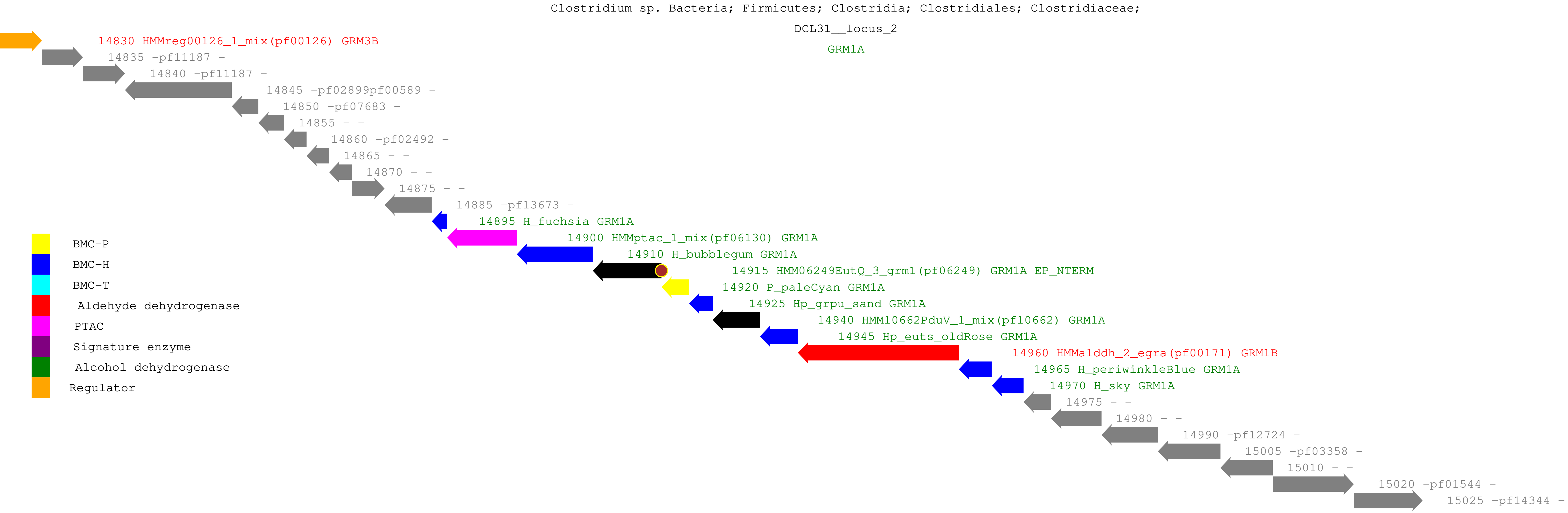
Clostridium sp. Bacteria; Firmicutes; Clostridia; Clostridiales; Clostridiaceae;
DCL31__locus_2
GRM1A
14830 HMMreg00126_1_mix(pf00126) GRM3B
14835 -pf11187 -
14840 -pf11187 -
14845 -pf02899pf00589 -
14850 -pf07683 -
14855 - -
14860 -pf02492 -
14865 - -
14870 - -
14875 - -
14885 -pf13673 -
14895 H_fuchsia GRM1A
14900 HMMptac_1_mix(pf06130) GRM1A
14910 H_bubblegum GRM1A
14915 HMM06249EutQ_3_grm1(pf06249) GRM1A EP_NTERM
14920 P_paleCyan GRM1A
14925 Hp_grpu_sand GRM1A
14940 HMM10662PduV_1_mix(pf10662) GRM1A
14945 Hp_euts_oldRose GRM1A
14960 HMMalddh_2_egra(pf00171) GRM1B
14965 H_periwinkleBlue GRM1A
14970 H_sky GRM1A
14975 - -
14980 - -
14990 -pf12724 -
15005 -pf03358 -
15010 - -
15020 -pf01544 -
15025 -pf14344 -
BMC-P
BMC-H
BMC-T
Aldehyde dehydrogenase
PTAC
Signature enzyme
Alcohol dehydrogenase
Regulator

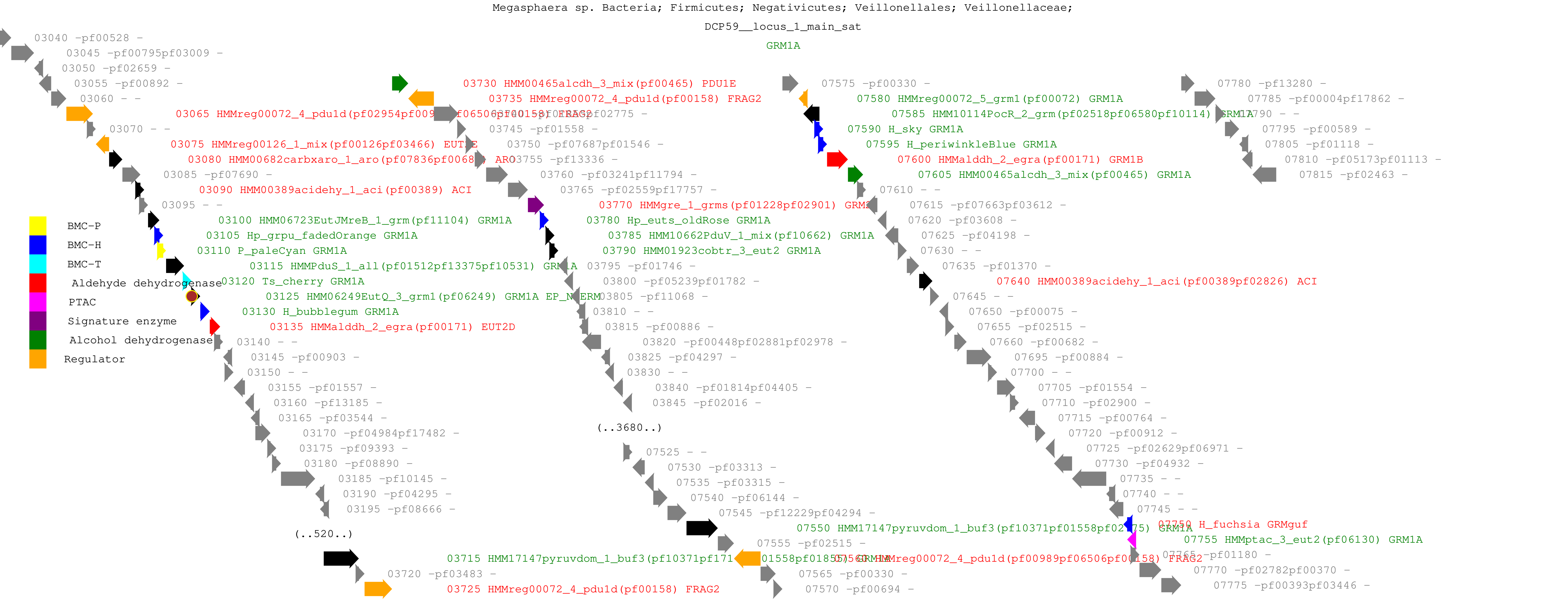

Megasphaera sp. Bacteria; Firmicutes; Negativicutes; Veillonellales; Veillonellaceae;
DCP59__locus_1_main_sat
GRM1A
BMC-P
BMC-H
BMC-T
Aldehyde dehydrogenase
PTAC
Signature enzyme
Alcohol dehydrogenase
Regulator
03040 -pf00528 -
03045 -pf00795pf03009 -
03050 -pf02659 -
03055 -pf00892 -
03060 - -
03065 HMMreg00072_4_pdu1d(pf02954pf00989pf06506pf00158) FRAG2
03070 - -
03075 HMMreg00126_1_mix(pf00126pf03466) EUT2E
03080 HMM00682carbxaro_1_aro(pf07836pf00682) ARO
03085 -pf07690 -
03090 HMM00389acidehy_1_aci(pf00389) ACI
03095 - -
03100 HMM06723EutJMreB_1_grm(pf11104) GRM1A
03105 Hp_grpu_fadedOrange GRM1A
03110 P_paleCyan GRM1A
03115 HMMPduS_1_all(pf01512pf13375pf10531) GRM1A
03120 Ts_cherry GRM1A
03125 HMM06249EutQ_3_grm1(pf06249) GRM1A EP_N_TERM
03130 H_bubblegum GRM1A
03135 HMMaldh_2_egra(pf00171) EUT2D
03140 - -
03145 -pf00903 -
03150 - -
03155 -pf01557 -
03160 -pf13185 -
03165 -pf03544 -
03170 -pf04984pf17482 -
03175 -pf09393 -
03180 -pf08890 -
03185 -pf10145 -
03190 -pf04295 -
03195 -pf08666 -
(..520..)
03715 HMM17147pyruvdom_1_buf3(pf10371pf01558pf01855) GRM1A
03720 -pf03483 -
03725 HMMreg00072_4_pdu1d(pf00158) FRAG2
03730 HMM00465alcdh_3_mix(pf00465) PDU1E
03735 HMMreg00072_4_pdu1d(pf00158) FRAG2
03740 -pf00171pf02775 -
03745 -pf01558 -
03750 -pf07687pf01546 -
03755 -pf13336 -
03760 -pf03241pf11794 -
03765 -pf02559pf17757 -
03770 HMMgre_1_grms(pf01228pf02901) GRM1A
03780 Hp_euts_oldRose GRM1A
03785 HMM10662PduV_1_mix(pf10662) GRM1A
03790 HMM01923cobtr_3_eut2 GRM1A
03795 -pf01746 -
03800 -pf05239pf01782 -
03805 -pf11068 -
03810 - -
03815 -pf00886 -
03820 -pf00448pf02881pf02978 -
03825 -pf04297 -
03830 - -
03840 -pf01814pf04405 -
03845 -pf02016 -
(..3680..)
07525 - -
07530 -pf03313 -
07535 -pf03315 -
07540 -pf06144 -
07545 -pf12229pf04294 -
07550 HMM17147pyruvdom_1_buf3(pf10371pf01558pf01855) GRM1A
07555 -pf02515 -
07560 HMMreg00072_4_pdu1d(pf00989pf06506pf00158) FRAG2
07565 -pf00330 -
07570 -pf00694 -
07575 -pf00330 -
07580 HMMreg00072_5_grm1(pf00072) GRM1A
07585 HMM10114PocR_2_grm(pf02518pf06580pf10114) GRM1A
07590 H_sky GRM1A
07595 H_periwinkleBlue GRM1A
07600 HMMaldh_2_egra(pf00171) GRM1B
07605 HMM00465alcdh_3_mix(pf00465) GRM1A
07610 - -
07615 -pf07663pf03612 -
07620 -pf03608 -
07625 -pf04198 -
07630 - -
07635 -pf01370 -
07640 HMM00389acidehy_1_aci(pf00389pf02826) ACI
07645 - -
07650 -pf00075 -
07655 -pf02515 -
07660 -pf00682 -
07695 -pf00884 -
07700 - -
07705 -pf01554 -
07710 -pf02900 -
07715 -pf00764 -
07720 -pf00912 -
07725 -pf02629pf06971 -
07730 -pf04932 -
07735 - -
07740 - -
07745 - -
07750 H_fuchsia GRMguf
07755 HMMptac_3_eut2(pf06130) GRM1A
07765 -pf01180 -
07770 -pf02782pf00370 -
07775 -pf00393pf03446 -
07780 -pf13280 -
07785 -pf00004pf17862 -
07790 - -
07795 -pf00589 -
07805 -pf01118 -
07810 -pf05173pf01113 -
07815 -pf02463 -

Clostridium sp. Bacteria; Firmicutes; Clostridia; Clostridiales; Clostridiaceae;

DCR69__locus_2

GRM1A

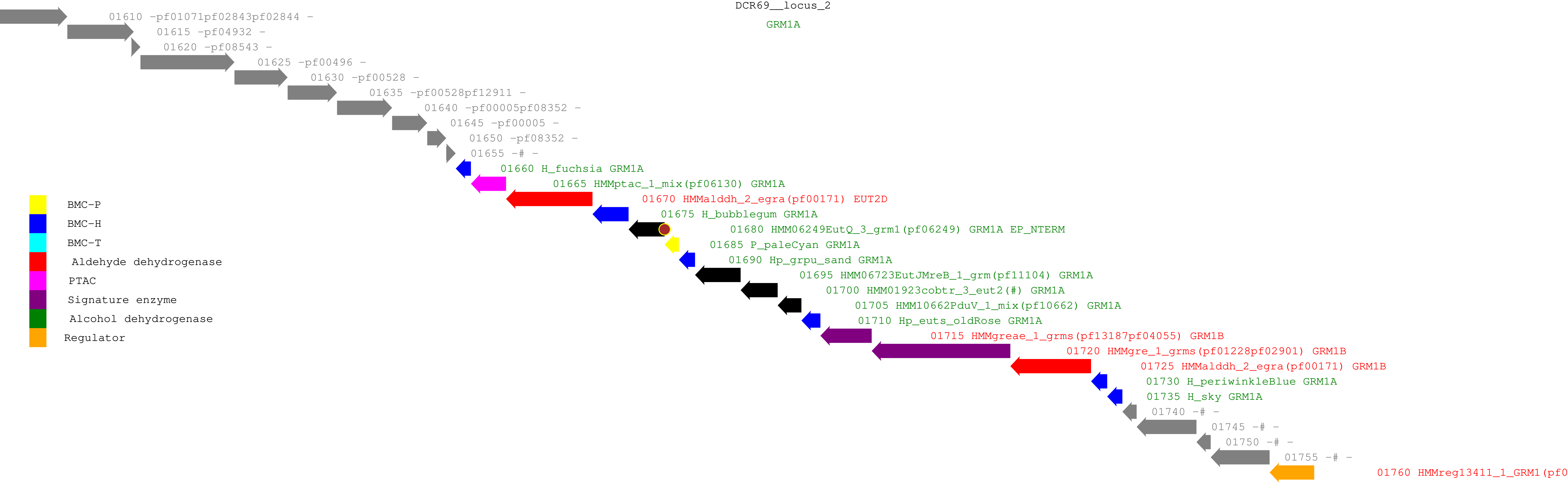

Clostridium sp. Bacteria; Firmicutes; Clostridia; Clostridiales; Clostridiaceae;

DCW51__locus_2

GRM1A

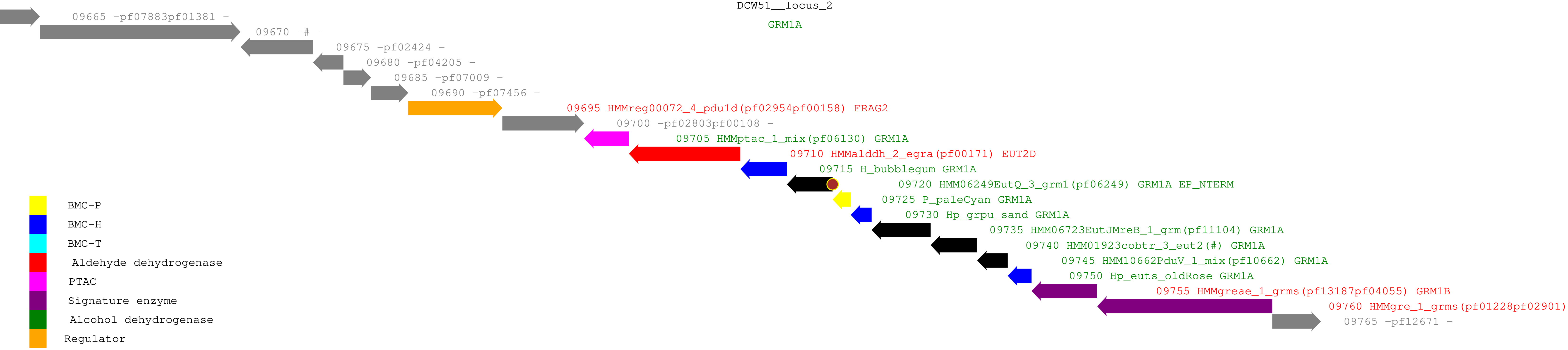

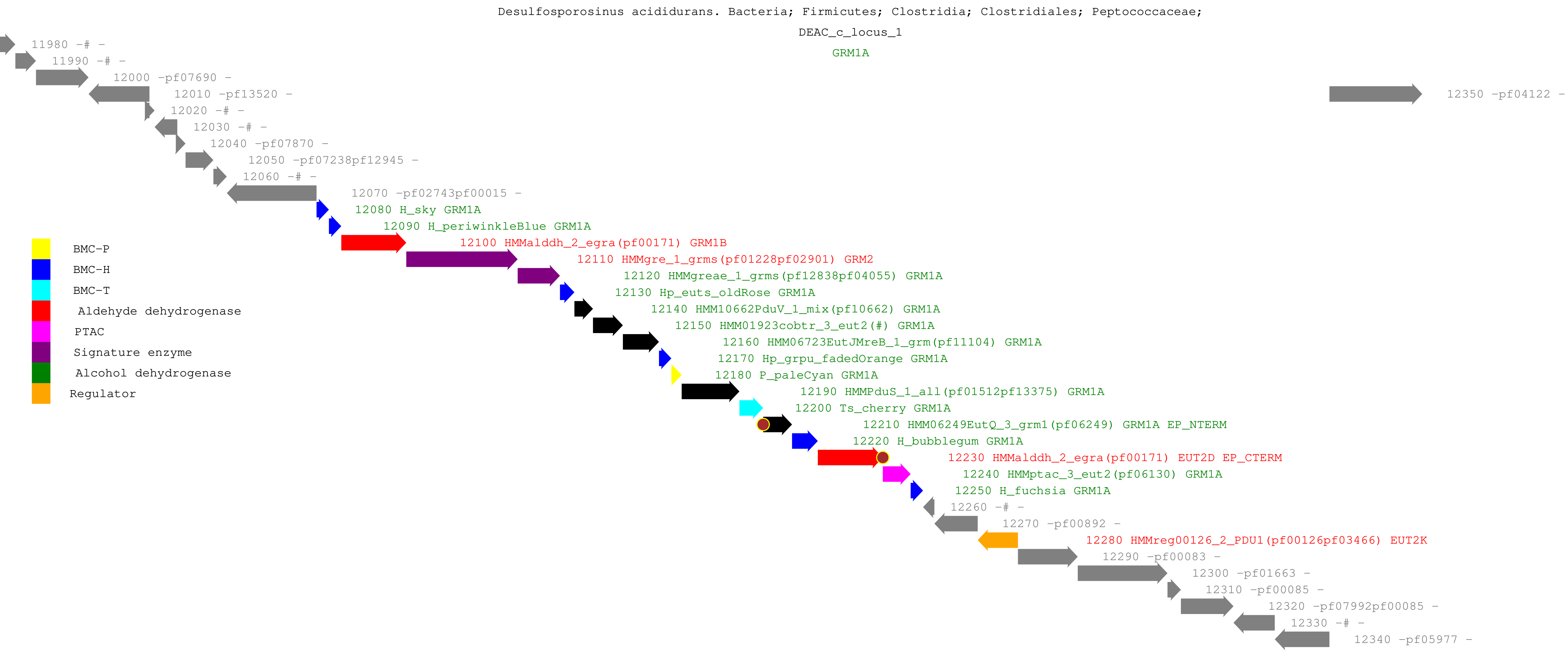
Desulfosporosinus acididurans. Bacteria; Firmicutes; Clostridia; Clostridiales; Peptococcaceae;
DEAC_c_locus_1
GRM1A
11980 -# -
11990 -# -
12000 -pf07690 -
12010 -pf13520 -
12020 -# -
12030 -# -
12040 -pf07870 -
12050 -pf07238pf12945 -
12060 -# -
12070 -pf02743pf00015 -
12080 H_sky GRM1A
12090 H_periwinkleBlue GRM1A
12100 HMMalddh_2_egra(pf00171) GRM1B
12110 HMMgre_1_grms(pf01228pf02901) GRM2
12120 HMMgreae_1_grms(pf12838pf04055) GRM1A
12130 Hp_euts_oldRose GRM1A
12140 HMM10662PduV_1_mix(pf10662) GRM1A
12150 HMM01923cobtr_3_eut2(#) GRM1A
12160 HMM06723EutJMreB_1_grm(pf11104) GRM1A
12170 Hp_grpu_fadedOrange GRM1A
12180 P_paleCyan GRM1A
12190 HMMPduS_1_all(pf01512pf13375) GRM1A
12200 Ts_cherry GRM1A
12210 HMM06249EutQ_3_grm1(pf06249) GRM1A EP_NTERM
12220 H_bubblegum GRM1A
12230 HMMalddh_2_egra(pf00171) EUT2D EP_CTERM
12240 HMMptac_3_eut2(pf06130) GRM1A
12250 H_fuchsia GRM1A
12260 -# -
12270 -pf00892 -
12280 HMMreg00126_2_PDU1(pf00126pf03466) EUT2K
12290 -pf00083 -
12300 -pf01663 -
12310 -pf00085 -
12320 -pf07992pf00085 -
12330 -# -
12340 -pf05977 -
12350 -pf04122 -
BMC-P
BMC-H
BMC-T
Aldehyde dehydrogenase
PTAC
Signature enzyme
Alcohol dehydrogenase
Regulator

Desulfosporosinus sp. Bacteria; Firmicutes; Clostridia; Clostridiales; Peptococcaceae;

DEF42__locus_3

GRM1A

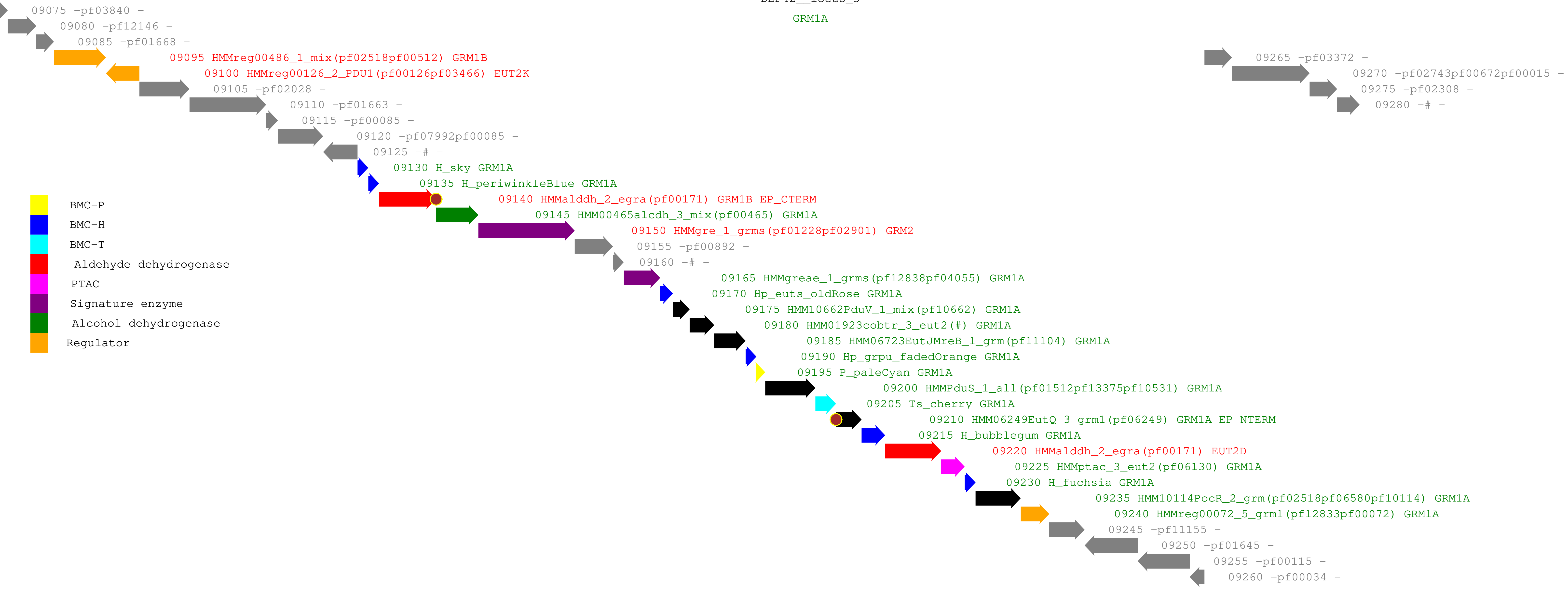

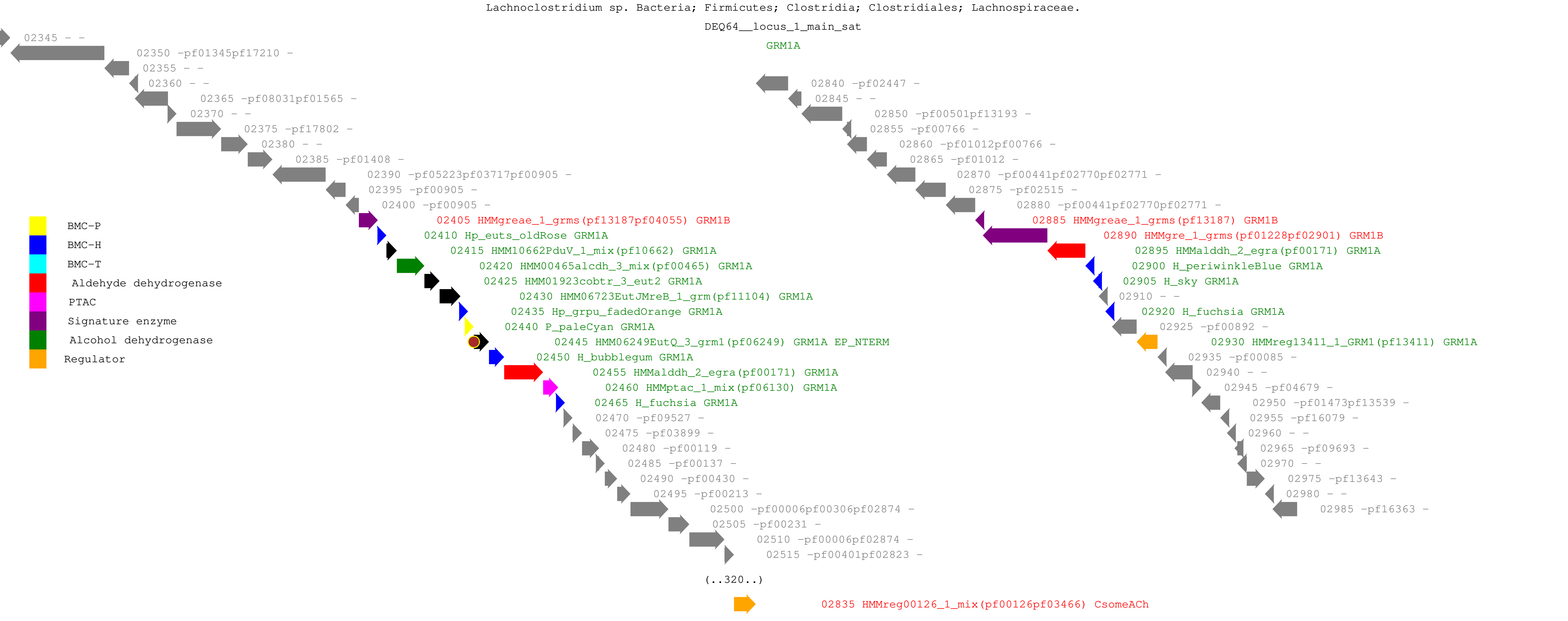
Lachnoclostridium sp. Bacteria; Firmicutes; Clostridia; Clostridiales; Lachnospiraceae.
DEQ64__locus_1_main_sat
GRM1A
02345 - -
02350 -pf01345pf17210 -
02355 - -
02360 - -
02365 -pf08031pf01565 -
02370 - -
02375 -pf17802 -
02380 - -
02385 -pf01408 -
02390 -pf05223pf03717pf00905 -
02395 -pf00905 -
02400 -pf00905 -
02405 HMMgreae_1_grms(pf13187pf04055) GRM1B
02410 Hp_euts_oldRose GRM1A
02415 HMM10662PduV_1_mix(pf10662) GRM1A
02420 HMM00465alcdh_3_mix(pf00465) GRM1A
02425 HMM01923cobtr_3_eut2 GRM1A
02430 HMM06723EutJMreB_1_grm(pf11104) GRM1A
02435 Hp_grpu_fadedOrange GRM1A
02440 P_paleCyan GRM1A
02445 HMM06249EutQ_3_grm1(pf06249) GRM1A EP_NTERM
02450 H_bubblegum GRM1A
02455 HMMalddh_2_egra(pf00171) GRM1A
02460 HMMptac_1_mix(pf06130) GRM1A
02465 H_fuchsia GRM1A
02470 -pf09527 -
02475 -pf03899 -
02480 -pf00119 -
02485 -pf00137 -
02490 -pf00430 -
02495 -pf00213 -
02500 -pf00006pf00306pf02874 -
02505 -pf00231 -
02510 -pf00006pf02874 -
02515 -pf00401pf02823 -
(..320..)
02835 HMMreg00126_1_mix(pf00126pf03466) CsomeACh
02840 -pf02447 -
02845 - -
02850 -pf00501pf13193 -
02855 -pf00766 -
02860 -pf01012pf00766 -
02865 -pf01012 -
02870 -pf00441pf02770pf02771 -
02875 -pf02515 -
02880 -pf00441pf02770pf02771 -
02885 HMMgreae_1_grms(pf13187) GRM1B
02890 HMMgre_1_grms(pf01228pf02901) GRM1B
02895 HMMalddh_2_egra(pf00171) GRM1A
02900 H_periwinkleBlue GRM1A
02905 H_sky GRM1A
02910 - -
02920 H_fuchsia GRM1A
02925 -pf00892 -
02930 HMMreg13411_1_GRM1(pf13411) GRM1A
02935 -pf00085 -
02940 - -
02945 -pf04679 -
02950 -pf01473pf13539 -
02955 -pf16079 -
02960 - -
02965 -pf09693 -
02970 - -
02975 -pf13643 -
02980 - -
02985 -pf16363 -
BMC-P
BMC-H
BMC-T
Aldehyde dehydrogenase
PTAC
Signature enzyme
Alcohol dehydrogenase
Regulator

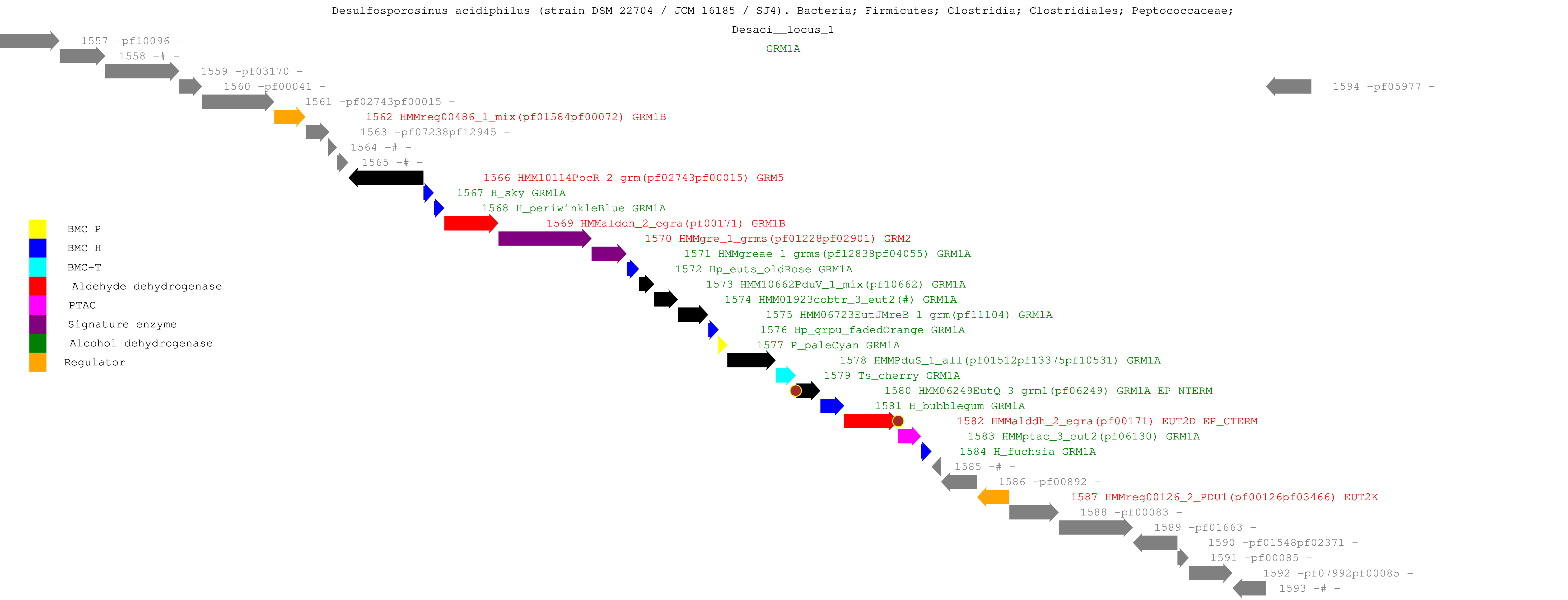
Desulfosporosinus acidiphilus (strain DSM 22704 / JCM 16185 / SJ4). Bacteria; Firmicutes; Clostridia; Clostridiales; Peptococcaceae;
Desaci__locus_1
GRM1A
1557 -pf10096 -
1558 -# -
1559 -pf03170 -
1560 -pf00041 -
1561 -pf02743pf00015 -
1562 HMMreg00486_1_mix(pf01584pf00072) GRM1B
1563 -pf07238pf12945 -
1564 -# -
1565 -# -
1566 HMM10114PocR_2_grm(pf02743pf00015) GRM5
1567 H_sky GRM1A
1568 H_periwinkleBlue GRM1A
1569 HMMaldh_2_egra(pf00171) GRM1B
1570 HMMgre_1_grms(pf01228pf02901) GRM2
1571 HMMgreae_1_grms(pf12838pf04055) GRM1A
1572 Hp_euts_oldRose GRM1A
1573 HMM10662PduV_1_mix(pf10662) GRM1A
1574 HMM01923cobtr_3_eut2(#) GRM1A
1575 HMM06723EutJMreB_1_grm(pf11104) GRM1A
1576 Hp_grpu_fadedOrange GRM1A
1577 P_paleCyan GRM1A
1578 HMMPduS_1_all(pf01512pf13375pf10531) GRM1A
1579 Ts_cherry GRM1A
1580 HMM06249EutQ_3_grm1(pf06249) GRM1A EP_NTERM
1581 H_bubblegum GRM1A
1582 HMMaldh_2_egra(pf00171) EUT2D EP_CTERM
1583 HMMptac_3_eut2(pf06130) GRM1A
1584 H_fuchsia GRM1A
1585 -# -
1586 -pf00892 -
1587 HMMreg00126_2_PDU1(pf00126pf03466) EUT2K
1588 -pf00083 -
1589 -pf01663 -
1590 -pf01548pf02371 -
1591 -pf00085 -
1592 -pf07992pf00085 -
1593 -# -
1594 -pf05977 -
BMC-P
BMC-H
BMC-T
Aldehyde dehydrogenase
PTAC
Signature enzyme
Alcohol dehydrogenase
Regulator

Desulfitobacterium dehalogenans (strain ATCC 51507 / DSM 9161 / Bacteria; Firmicutes; Clostridia; Clostridiales; Peptococcaceae;

Desde__locus_2

GRM1A

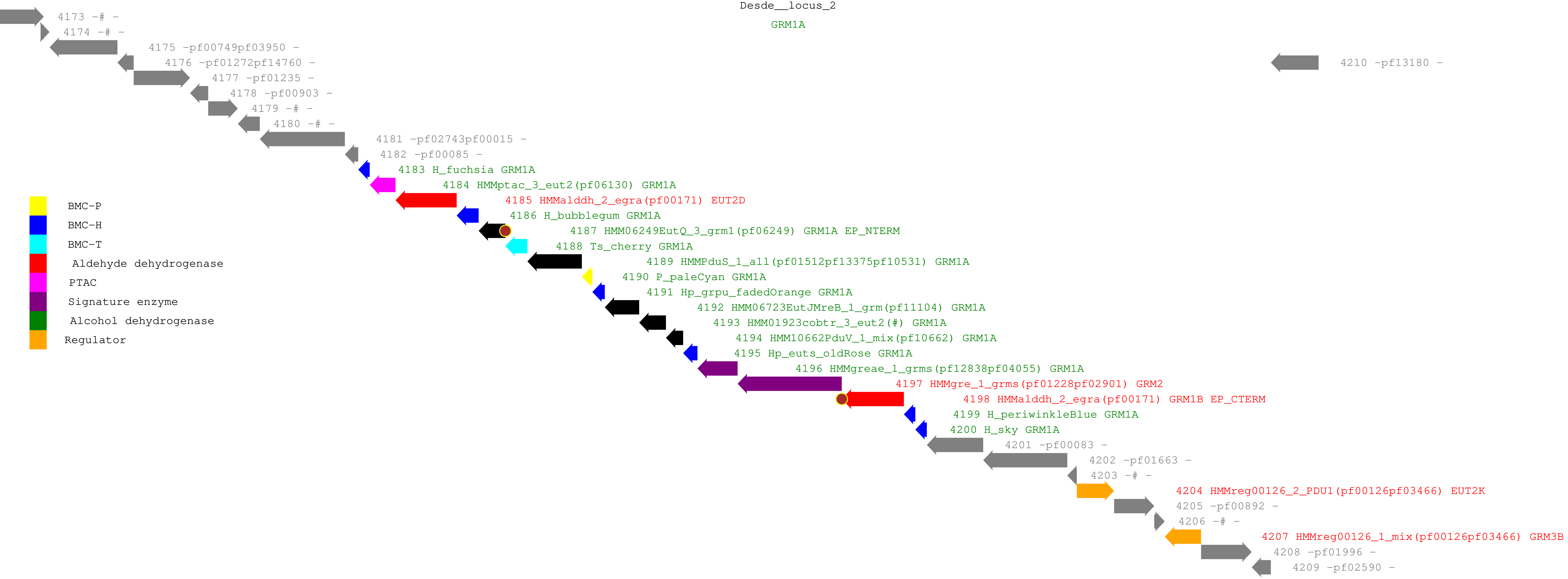

Desulfotomaculum hydrothermale Lam5 = DSM 18033. Bacteria; Firmicutes; Clostridia; Clostridiales; Peptococcaceae;

DESHY__locus_1

GRM1A

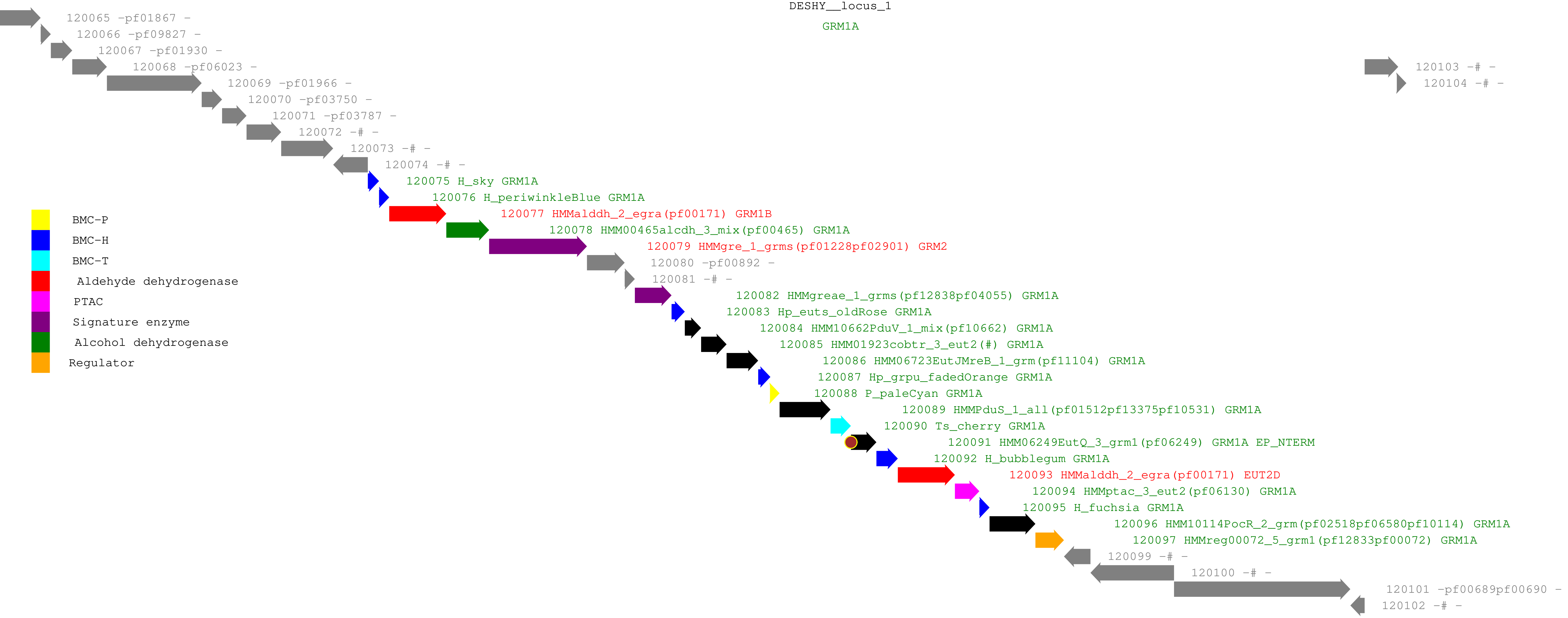

Desulfitobacterium sp. LBE. Bacteria; Firmicutes; Clostridia; Clostridiales; Peptococcaceae;

DesLBE__locus_2

GRM1A

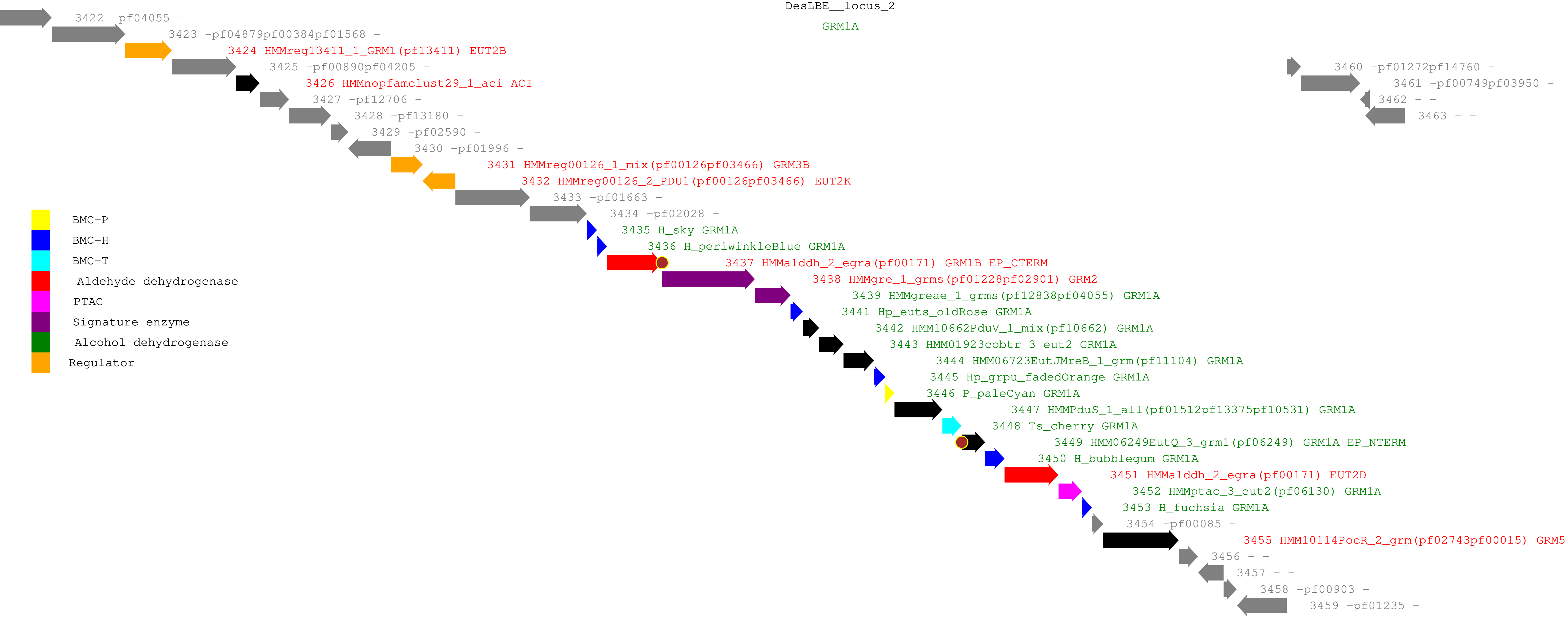

Desulfosporosinus meridiei (strain ATCC BAA-275 / DSM 13257 / NCIMB Bacteria; Firmicutes; Clostridia; Clostridiales; Peptococcaceae;

Desmer__locus_3

GRM1A

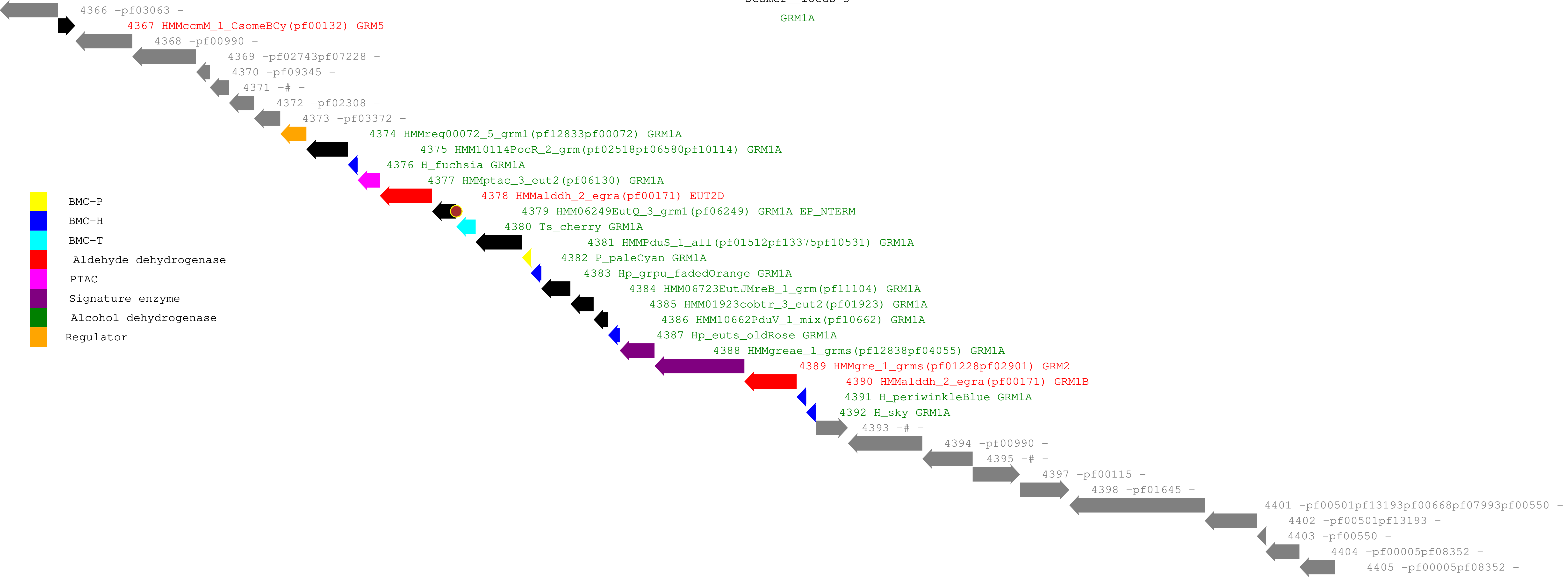

Desulfosporosinus orientis (strain ATCC 19365 / DSM 765 / NCIMB 8382 / Bacteria; Firmicutes; Clostridia; Clostridiales; Peptococcaceae;

Desor__locus_3

GRM1A

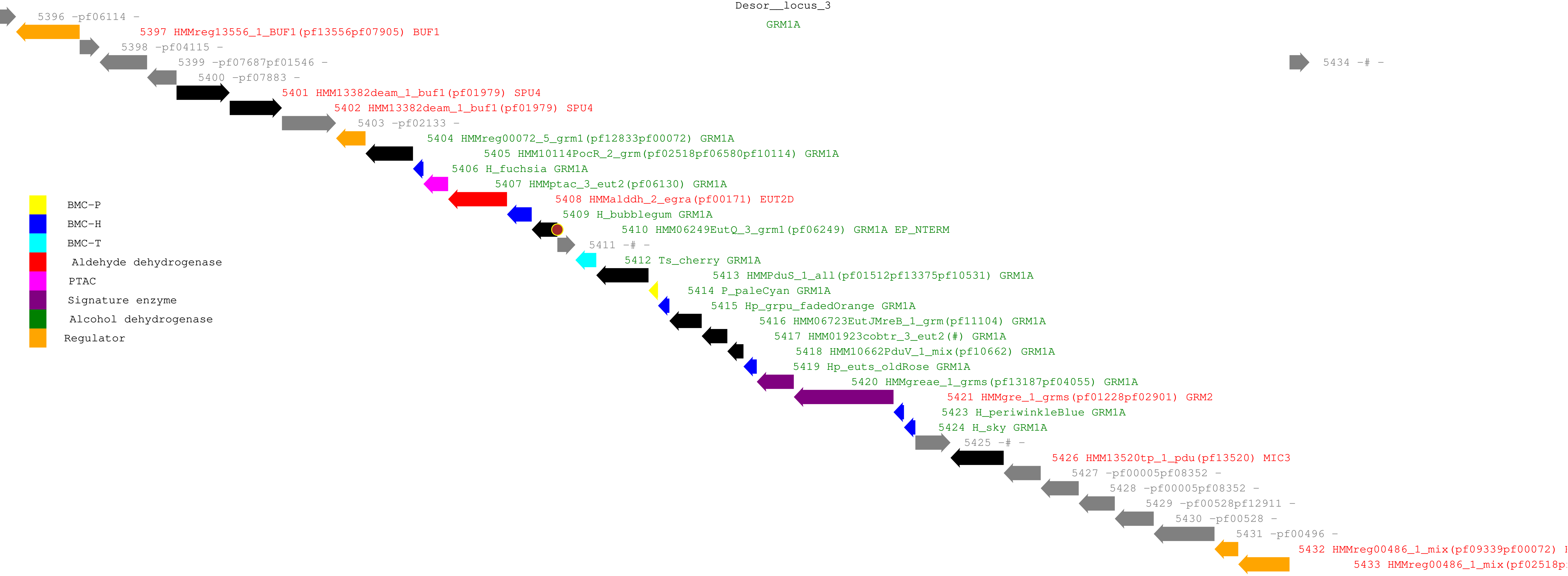

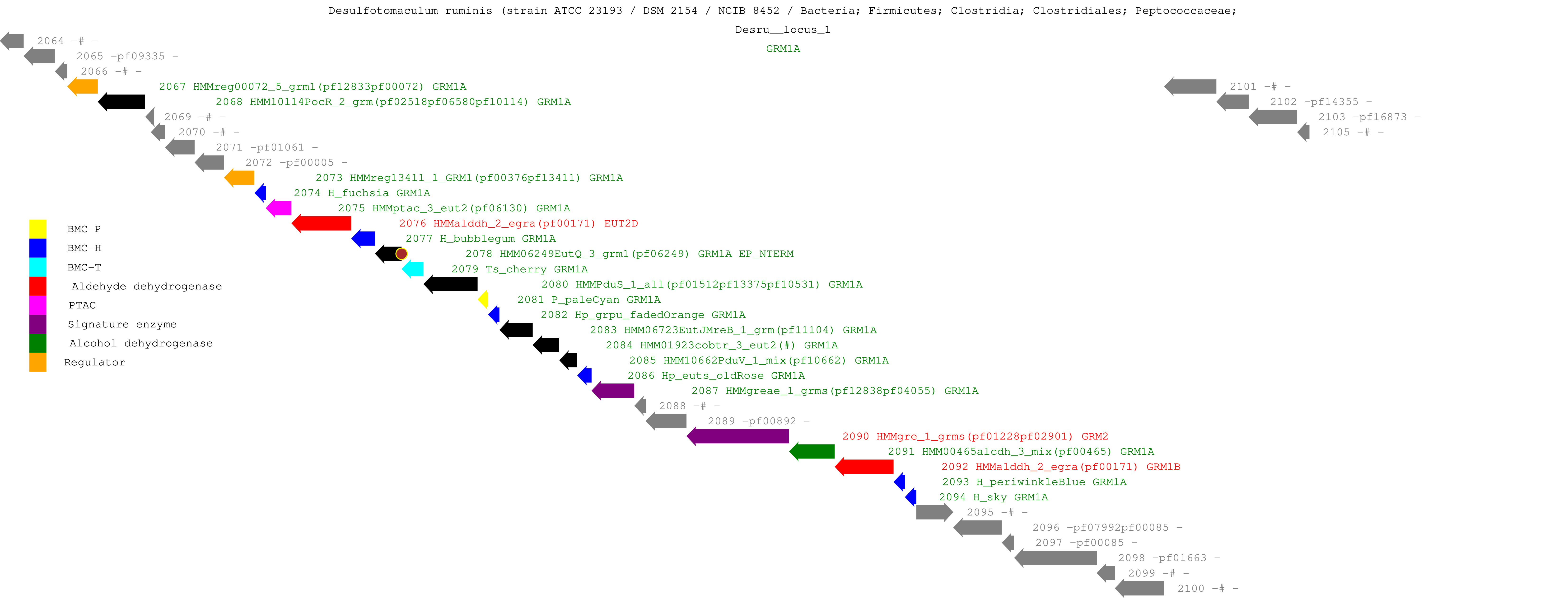

Desulfotomaculum ruminis (strain ATCC 23193 / DSM 2154 / NCIB 8452 / Bacteria; Firmicutes; Clostridia; Clostridiales; Peptococcaceae;
Desru__locus_1
GRM1A
2064 -# -
2065 -pf09335 -
2066 -# -
2067 HMMreg00072_5_grm1(pf12833pf00072) GRM1A
2068 HMM10114PocR_2_grm(pf02518pf06580pf10114) GRM1A
2069 -# -
2070 -# -
2071 -pf01061 -
2072 -pf00005 -
2073 HMMreg13411_1_GRM1(pf00376pf13411) GRM1A
2074 H_fuchsia GRM1A
2075 HMMptac_3_eut2(pf06130) GRM1A
2076 HMMalddh_2_egra(pf00171) EUT2D
2077 H_bubblegum GRM1A
2078 HMM06249EutQ_3_grm1(pf06249) GRM1A EP_NTERM
2079 Ts_cherry GRM1A
2080 HMMPduS_1_all(pf01512pf13375pf10531) GRM1A
2081 P_paleCyan GRM1A
2082 Hp_grpu_fadedOrange GRM1A
2083 HMM06723EutJMreB_1_grm(pf11104) GRM1A
2084 HMM01923cobtr_3_eut2(#) GRM1A
2085 HMM10662PduV_1_mix(pf10662) GRM1A
2086 Hp_euts_oldRose GRM1A
2087 HMMgreae_1_grms(pf12838pf04055) GRM1A
2088 -# -
2089 -pf00892 -
2090 HMMgre_1_grms(pf01228pf02901) GRM2
2091 HMM00465alcdh_3_mix(pf00465) GRM1A
2092 HMMalddh_2_egra(pf00171) GRM1B
2093 H_periwinkleBlue GRM1A
2094 H_sky GRM1A
2095 -# -
2096 -pf07992pf00085 -
2097 -pf00085 -
2098 -pf01663 -
2099 -# -
2100 -# -
2101 -# -
2102 -pf14355 -
2103 -pf16873 -
2105 -# -
BMC-P
BMC-H
BMC-T
Aldehyde dehydrogenase
PTAC
Signature enzyme
Alcohol dehydrogenase
Regulator

Desulfosporosinus youngiae DSM 17734. Bacteria; Firmicutes; Clostridia; Clostridiales; Peptococcaceae;

DesyoDRAFT__locus_3

GRM1A

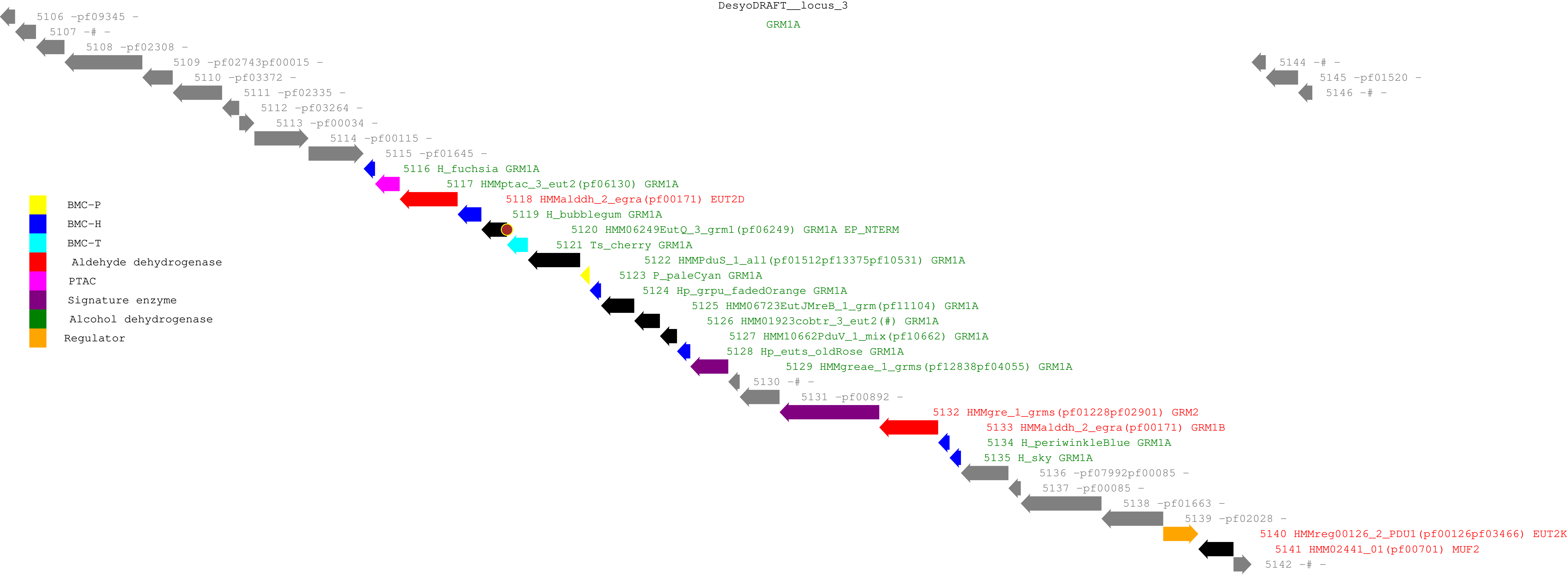

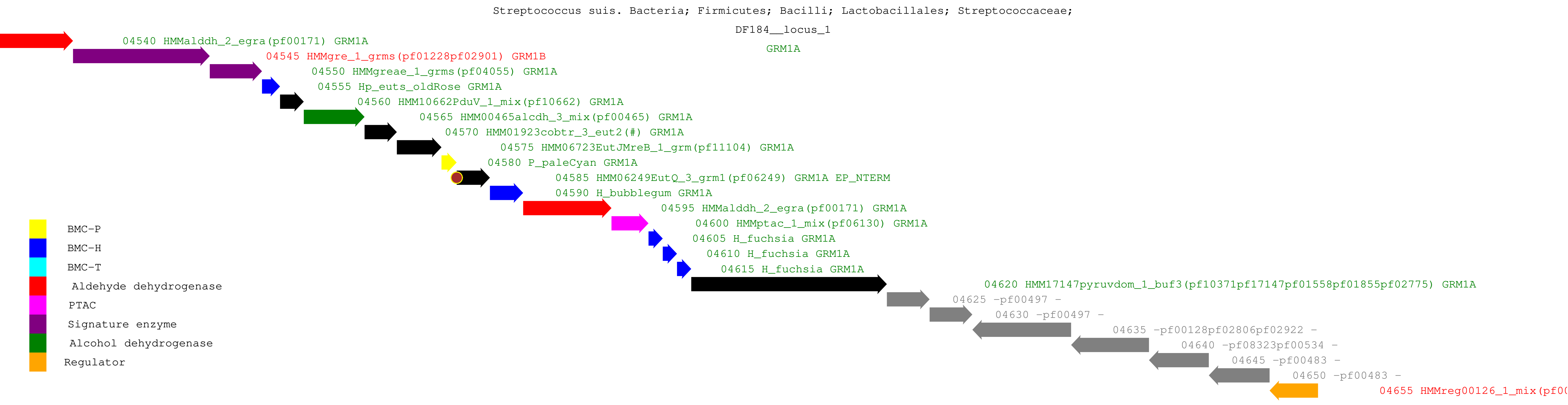
Streptococcus suis. Bacteria; Firmicutes; Bacilli; Lactobacillales; Streptococcaceae;
DF184__locus_1
GRM1A
04540 HMMalddh_2_egra(pf00171) GRM1A
04545 HMMgre_1_grms(pf01228pf02901) GRM1B
04550 HMMgreae_1_grms(pf04055) GRM1A
04555 Hp_euts_oldRose GRM1A
04560 HMM10662PduV_1_mix(pf10662) GRM1A
04565 HMM00465alcdh_3_mix(pf00465) GRM1A
04570 HMM01923cobtr_3_eut2(#) GRM1A
04575 HMM06723EutJMreB_1_grm(pf11104) GRM1A
04580 P_paleCyan GRM1A
04585 HMM06249EutQ_3_grm1(pf06249) GRM1A EP_NTERM
04590 H_bubblegum GRM1A
04595 HMMalddh_2_egra(pf00171) GRM1A
04600 HMMptac_1_mix(pf06130) GRM1A
04605 H_fuchsia GRM1A
04610 H_fuchsia GRM1A
04615 H_fuchsia GRM1A
04620 HMM17147pyruvdom_1_buf3(pf10371pf17147pf01558pf01855pf02775) GRM1A
04625 -pf00497 -
04630 -pf00497 -
04635 -pf00128pf02806pf02922 -
04640 -pf08323pf00534 -
04645 -pf00483 -
04650 -pf00483 -
04655 HMMreg00126_1_mix(pf0
BMC-P
BMC-H
BMC-T
Aldehyde dehydrogenase
PTAC
Signature enzyme
Alcohol dehydrogenase
Regulator

Clostridium novyi. Bacteria; Firmicutes; Clostridia; Clostridiales; Clostridiaceae;

DFH04__locus_1

GRM1A

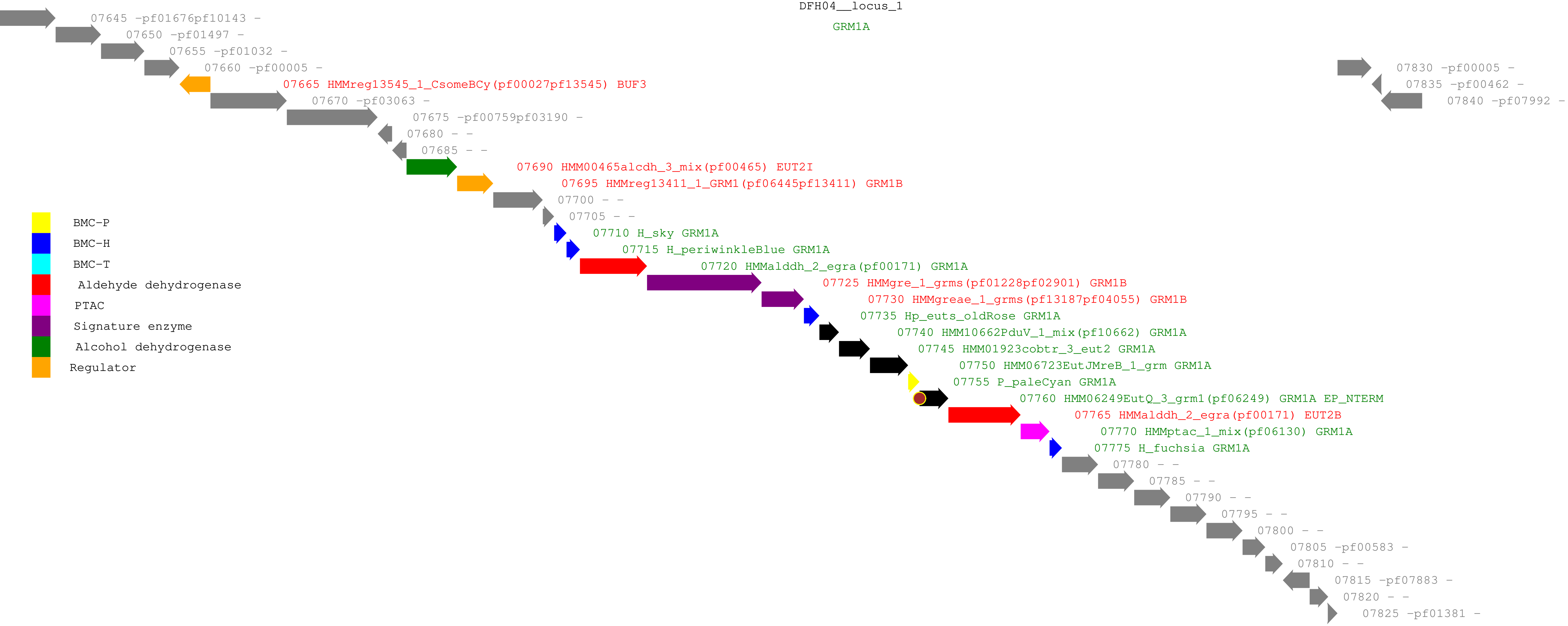

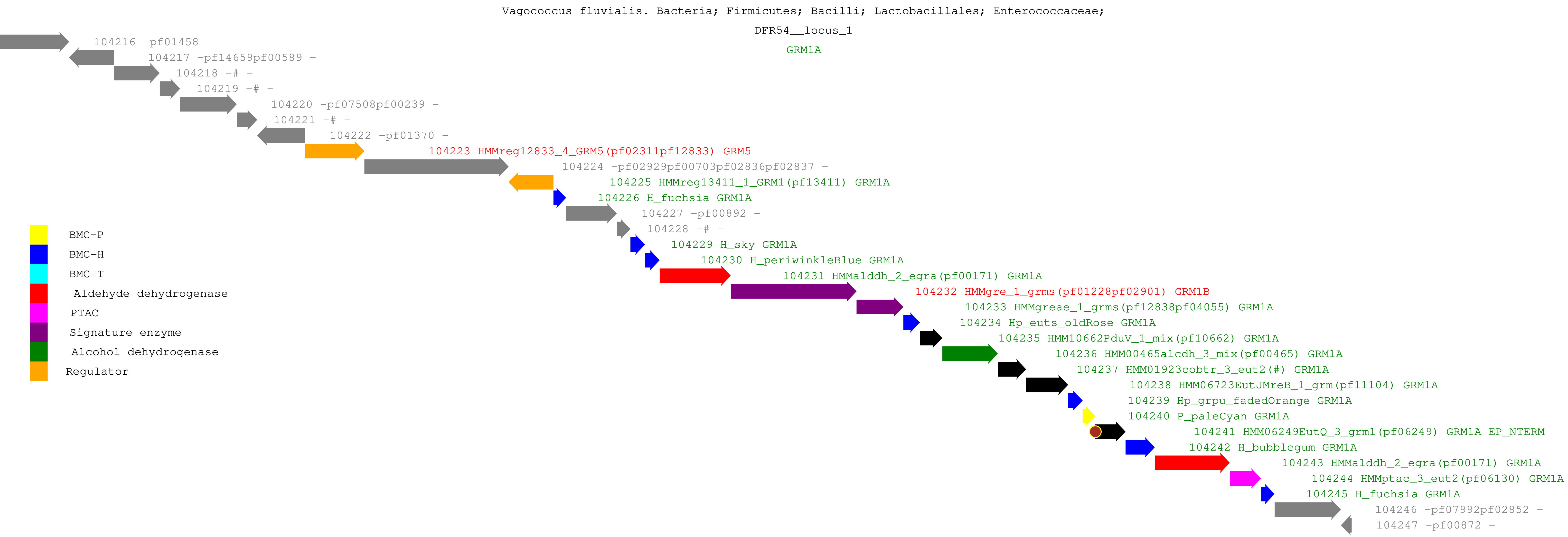
Vagococcus fluvialis. Bacteria; Firmicutes; Bacilli; Lactobacillales; Enterococcaceae;
DFR54__locus_1
GRM1A
104216 -pf01458 -
104217 -pf14659pf00589 -
104218 -# -
104219 -# -
104220 -pf07508pf00239 -
104221 -# -
104222 -pf01370 -
104223 HMMreg12833_4_GRM5(pf02311pf12833) GRM5
104224 -pf02929pf00703pf02836pf02837 -
104225 HMMreg13411_1_GRM1(pf13411) GRM1A
104226 H_fuchsia GRM1A
104227 -pf00892 -
104228 -# -
104229 H_sky GRM1A
104230 H_periwinkleBlue GRM1A
104231 HMMaldd h_2_egra(pf00171) GRM1A
104232 HMMgre_1_grms(pf01228pf02901) GRM1B
104233 HMMgreae_1_grms(pf12838pf04055) GRM1A
104234 Hp_euts_oldRose GRM1A
104235 HMM10662PduV_1_mix(pf10662) GRM1A
104236 HMM00465alcdh_3_mix(pf00465) GRM1A
104237 HMM01923cobtr_3_eut2(#) GRM1A
104238 HMM06723EutJMreB_1_grm(pf11104) GRM1A
104239 Hp_grpu_fadedOrange GRM1A
104240 P_paleCyan GRM1A
104241 HMM06249EutQ_3_grm1(pf06249) GRM1A EP_NTERM
104242 H_bubblegum GRM1A
104243 HMMaldd h_2_egra(pf00171) GRM1A
104244 HMMptac_3_eut2(pf06130) GRM1A
104245 H_fuchsia GRM1A
104246 -pf07992pf02852 -
104247 -pf00872 -
BMC-P
BMC-H
BMC-T
Aldehyde dehydrogenase
PTAC
Signature enzyme
Alcohol dehydrogenase
Regulator

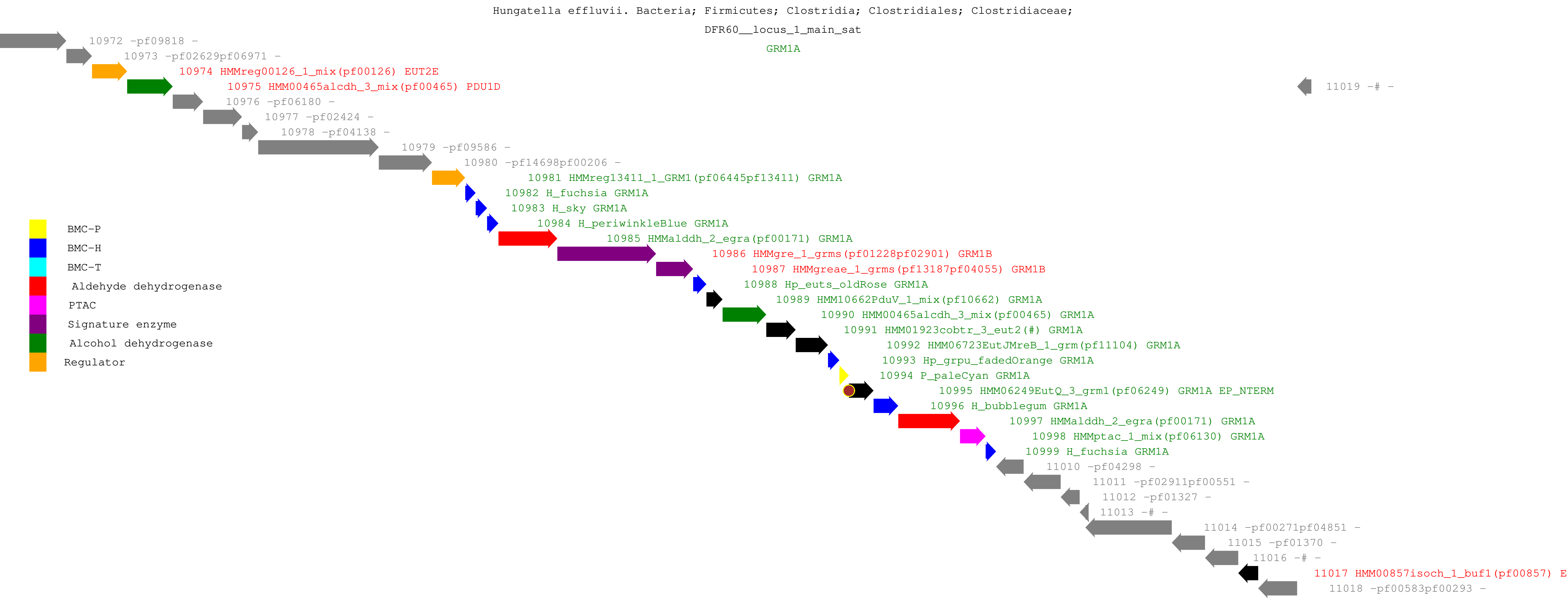

Hungatella effluvii. Bacteria; Firmicutes; Clostridia; Clostridiales; Clostridiaceae;
DFR60__locus_1_main_sat
GRM1A
10972 -pf09818 -
10973 -pf02629pf06971 -
10974 HMMreg00126_1_mix(pf00126) EUT2E
10975 HMM00465alcdh_3_mix(pf00465) PDU1D
10976 -pf06180 -
10977 -pf02424 -
10978 -pf04138 -
10979 -pf09586 -
10980 -pf14698pf00206 -
10981 HMMreg13411_1_GRM1(pf06445pf13411) GRM1A
10982 H_fuchsia GRM1A
10983 H_sky GRM1A
10984 H_periwinkleBlue GRM1A
10985 HMMalddh_2_egra(pf00171) GRM1A
10986 HMMgre_1_grms(pf01228pf02901) GRM1B
10987 HMMgreae_1_grms(pf13187pf04055) GRM1B
10988 Hp_euts_oldRose GRM1A
10989 HMM10662PduV_1_mix(pf10662) GRM1A
10990 HMM00465alcdh_3_mix(pf00465) GRM1A
10991 HMM01923cobtr_3_eut2(#) GRM1A
10992 HMM06723EutJMreB_1_grm(pf11104) GRM1A
10993 Hp_grpu_fadedOrange GRM1A
10994 P_paleCyan GRM1A
10995 HMM06249EutQ_3_grm1(pf06249) GRM1A EP_NTERM
10996 H_bubblegum GRM1A
10997 HMMalddh_2_egra(pf00171) GRM1A
10998 HMMptac_1_mix(pf06130) GRM1A
10999 H_fuchsia GRM1A
11010 -pf04298 -
11011 -pf02911pf00551 -
11012 -pf01327 -
11013 -# -
11014 -pf00271pf04851 -
11015 -pf01370 -
11016 -# -
11017 HMM00857isoch_1_buf1(pf00857) E
11018 -pf00583pf00293 -
11019 -# -
BMC-P
BMC-H
BMC-T
Aldehyde dehydrogenase
PTAC
Signature enzyme
Alcohol dehydrogenase
Regulator

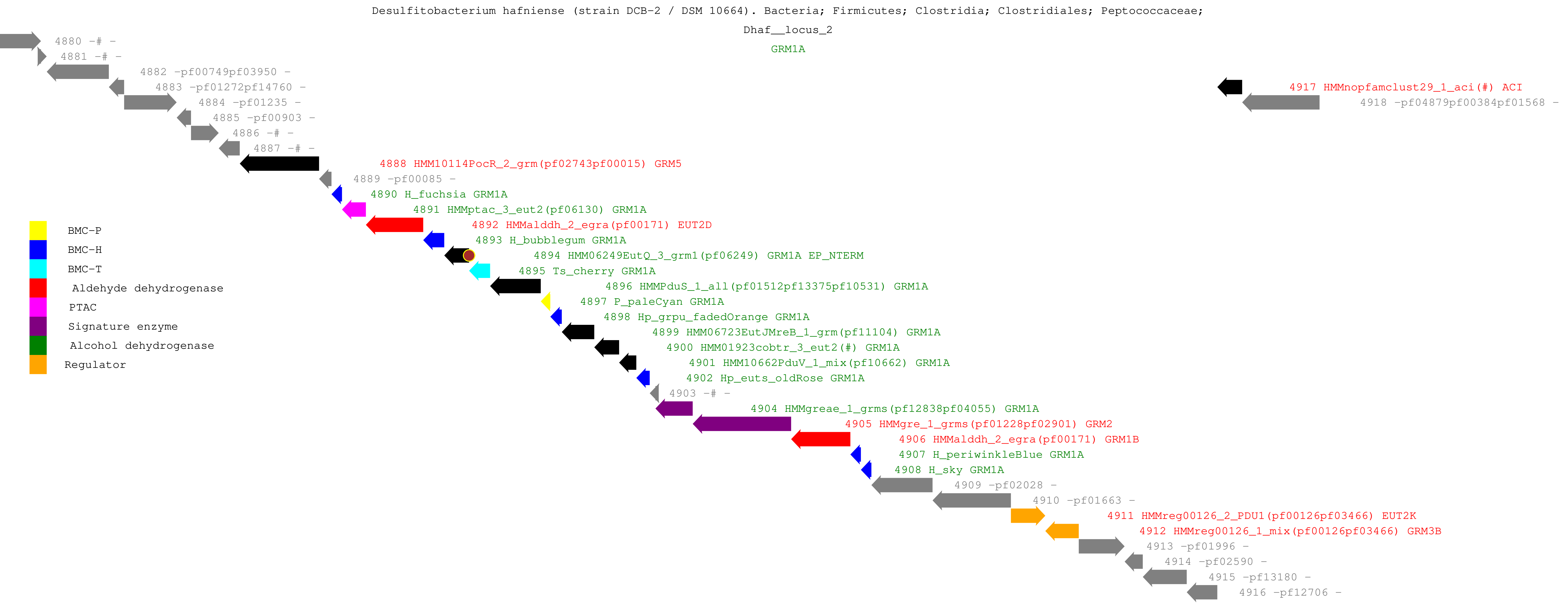
Desulfitobacterium hafniense (strain DCB-2 / DSM 10664). Bacteria; Firmicutes; Clostridia; Clostridiales; Peptococcaceae;
Dhaf__locus_2
GRM1A
4880 -# -
4881 -# -
4882 -pf00749pf03950 -
4883 -pf01272pf14760 -
4884 -pf01235 -
4885 -pf00903 -
4886 -# -
4887 -# -
4888 HMM10114PocR_2_grm(pf02743pf00015) GRM5
4889 -pf00085 -
4890 H_fuchsia GRM1A
4891 HMMptac_3_eut2(pf06130) GRM1A
4892 HMMalddh_2_egra(pf00171) EUT2D
4893 H_bubblegum GRM1A
4894 HMM06249EutQ_3_grm1(pf06249) GRM1A EP_NTERM
4895 Ts_cherry GRM1A
4896 HMMPduS_1_all(pf01512pf13375pf10531) GRM1A
4897 P_paleCyan GRM1A
4898 Hp_grpu_fadedOrange GRM1A
4899 HMM06723EutJMreB_1_grm(pf11104) GRM1A
4900 HMM01923cobtr_3_eut2(#) GRM1A
4901 HMM10662PduV_1_mix(pf10662) GRM1A
4902 Hp_euts_oldRose GRM1A
4903 -# -
4904 HMMgreae_1_grms(pf12838pf04055) GRM1A
4905 HMMgre_1_grms(pf01228pf02901) GRM2
4906 HMMalddh_2_egra(pf00171) GRM1B
4907 H_periwinkleBlue GRM1A
4908 H_sky GRM1A
4909 -pf02028 -
4910 -pf01663 -
4911 HMMreg00126_2_PDU1(pf00126pf03466) EUT2K
4912 HMMreg00126_1_mix(pf00126pf03466) GRM3B
4913 -pf01996 -
4914 -pf02590 -
4915 -pf13180 -
4916 -pf12706 -
4917 HMMnopfamclust29_1_aci(#) ACI
4918 -pf04879pf00384pf01568 -
BMC-P
BMC-H
BMC-T
Aldehyde dehydrogenase
PTAC
Signature enzyme
Alcohol dehydrogenase
Regulator

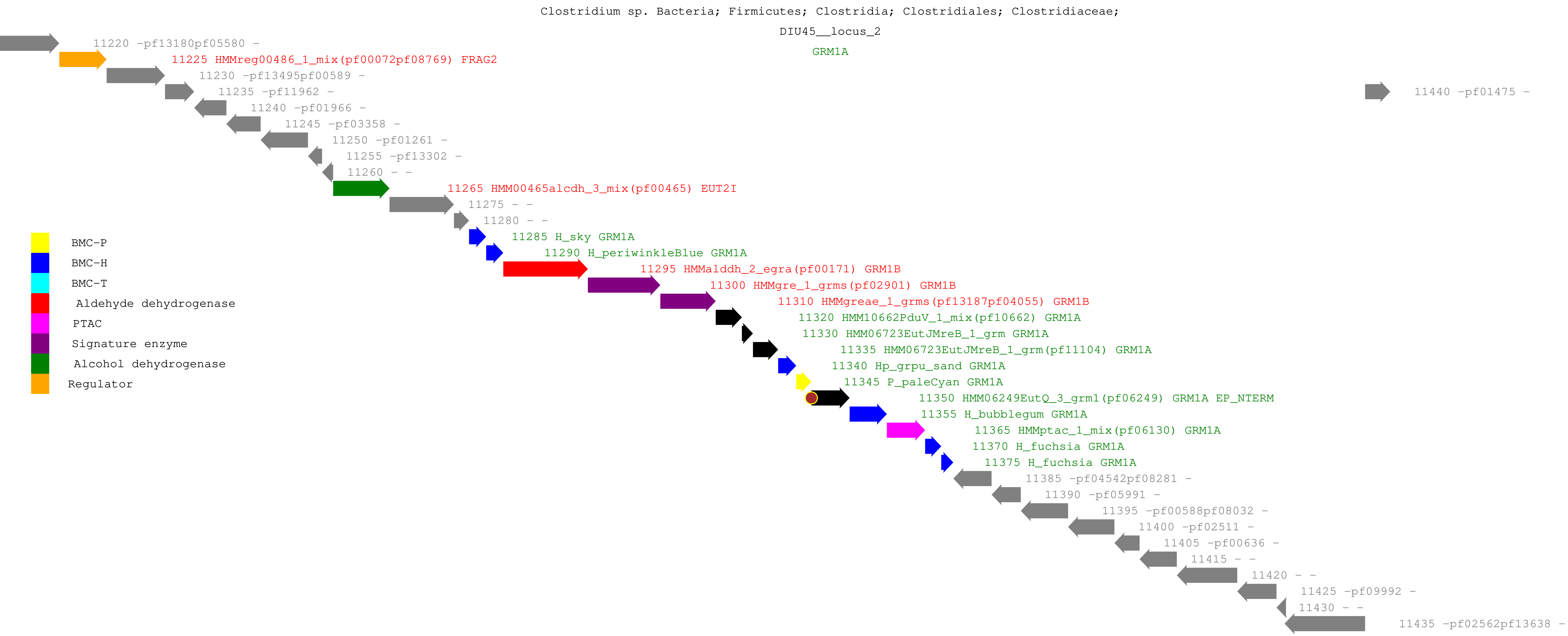
Clostridium sp. Bacteria; Firmicutes; Clostridia; Clostridiales; Clostridiaceae;
DIU45__locus_2
GRM1A
11220 -pf13180pf05580 -
11225 HMMreg00486_1_mix(pf00072pf08769) FRAG2
11230 -pf13495pf00589 -
11235 -pf11962 -
11240 -pf01966 -
11245 -pf03358 -
11250 -pf01261 -
11255 -pf13302 -
11260 - -
11265 HMM00465alcdh_3_mix(pf00465) EUT2I
11275 - -
11280 - -
11285 H_sky GRM1A
11290 H_periwinkleBlue GRM1A
11295 HMMaldh_2_egra(pf00171) GRM1B
11300 HMMgre_1_grms(pf02901) GRM1B
11310 HMMgreae_1_grms(pf13187pf04055) GRM1B
11320 HMM10662PduV_1_mix(pf10662) GRM1A
11330 HMM06723EutJMreB_1_grm GRM1A
11335 HMM06723EutJMreB_1_grm(pf11104) GRM1A
11340 Hp_grpu_sand GRM1A
11345 P_paleCyan GRM1A
11350 HMM06249EutQ_3_grm1(pf06249) GRM1A EP_NTERM
11355 H_bubblegum GRM1A
11365 HMMptac_1_mix(pf06130) GRM1A
11370 H_fuchsia GRM1A
11375 H_fuchsia GRM1A
11385 -pf04542pf08281 -
11390 -pf05991 -
11395 -pf00588pf08032 -
11400 -pf02511 -
11405 -pf00636 -
11415 - -
11420 - -
11425 -pf09992 -
11430 - -
11435 -pf02562pf13638 -
11440 -pf01475 -
BMC-P
BMC-H
BMC-T
Aldehyde dehydrogenase
PTAC
Signature enzyme
Alcohol dehydrogenase
Regulator

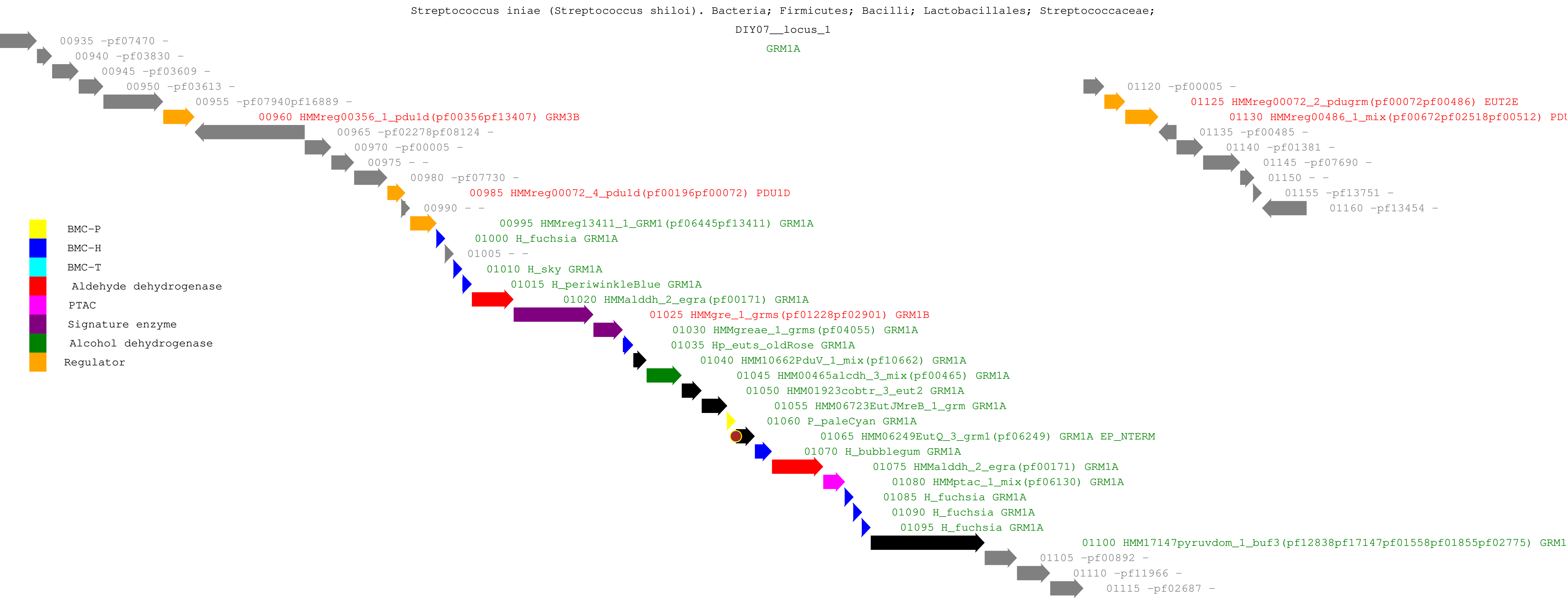

Streptococcus iniae (Streptococcus shiloi). Bacteria; Firmicutes; Bacilli; Lactobacillales; Streptococcaceae;
DIY07__locus_1
GRM1A
00935 -pf07470 -
00940 -pf03830 -
00945 -pf03609 -
00950 -pf03613 -
00955 -pf07940pf16889 -
00960 HMMreg00356_1_pdu1d(pf00356pf13407) GRM3B
00965 -pf02278pf08124 -
00970 -pf00005 -
00975 - -
00980 -pf07730 -
00985 HMMreg00072_4_pdu1d(pf00196pf00072) PDU1D
00990 - -
00995 HMMreg13411_1_GRM1(pf06445pf13411) GRM1A
01000 H_fuchsia GRM1A
01005 - -
01010 H_sky GRM1A
01015 H_periwinkleBlue GRM1A
01020 HMMaldh_2_egra(pf00171) GRM1A
01025 HMMgre_1_grms(pf01228pf02901) GRM1B
01030 HMMgreae_1_grms(pf04055) GRM1A
01035 Hp_euts_oldRose GRM1A
01040 HMM10662PduV_1_mix(pf10662) GRM1A
01045 HMM00465alcdh_3_mix(pf00465) GRM1A
01050 HMM01923cobtr_3_eut2 GRM1A
01055 HMM06723EutJMreB_1_grm GRM1A
01060 P_paleCyan GRM1A
01065 HMM06249EutQ_3_grm1(pf06249) GRM1A EP_NTERM
01070 H_bubblegum GRM1A
01075 HMMaldh_2_egra(pf00171) GRM1A
01080 HMMptac_1_mix(pf06130) GRM1A
01085 H_fuchsia GRM1A
01090 H_fuchsia GRM1A
01095 H_fuchsia GRM1A
01100 HMM17147pyruvdom_1_buf3(pf12838pf17147pf01558pf01855pf02775) GRM1
01105 -pf00892 -
01110 -pf11966 -
01115 -pf02687 -
01120 -pf00005 -
01125 HMMreg00072_2_pdugrm(pf00072pf00486) EUT2E
01130 HMMreg00486_1_mix(pf00672pf02518pf00512) PDU
01135 -pf00485 -
01140 -pf01381 -
01145 -pf07690 -
01150 - -
01155 -pf13751 -
01160 -pf13454 -
BMC-P
BMC-H
BMC-T
Aldehyde dehydrogenase
PTAC
Signature enzyme
Alcohol dehydrogenase
Regulator

Clostridium sp. AWRP. Bacteria; Firmicutes; Clostridia; Clostridiales; Clostridiaceae;

DMR38__locus_1

GRM1A

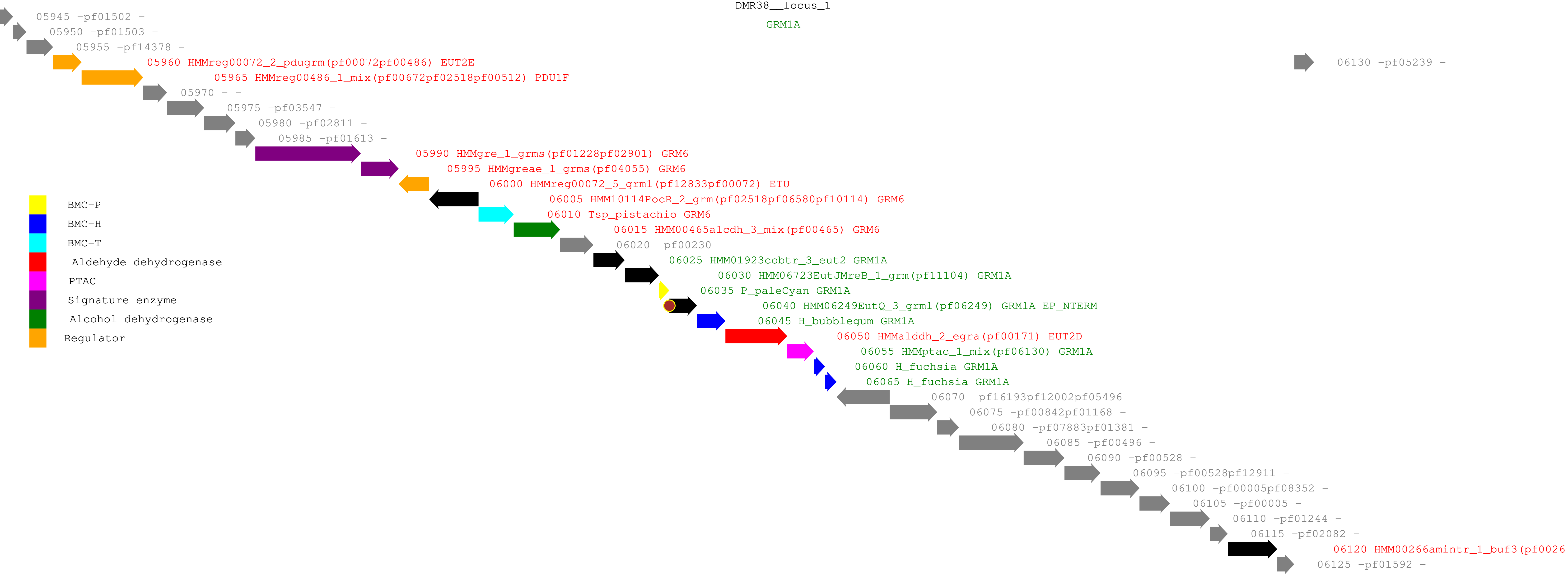

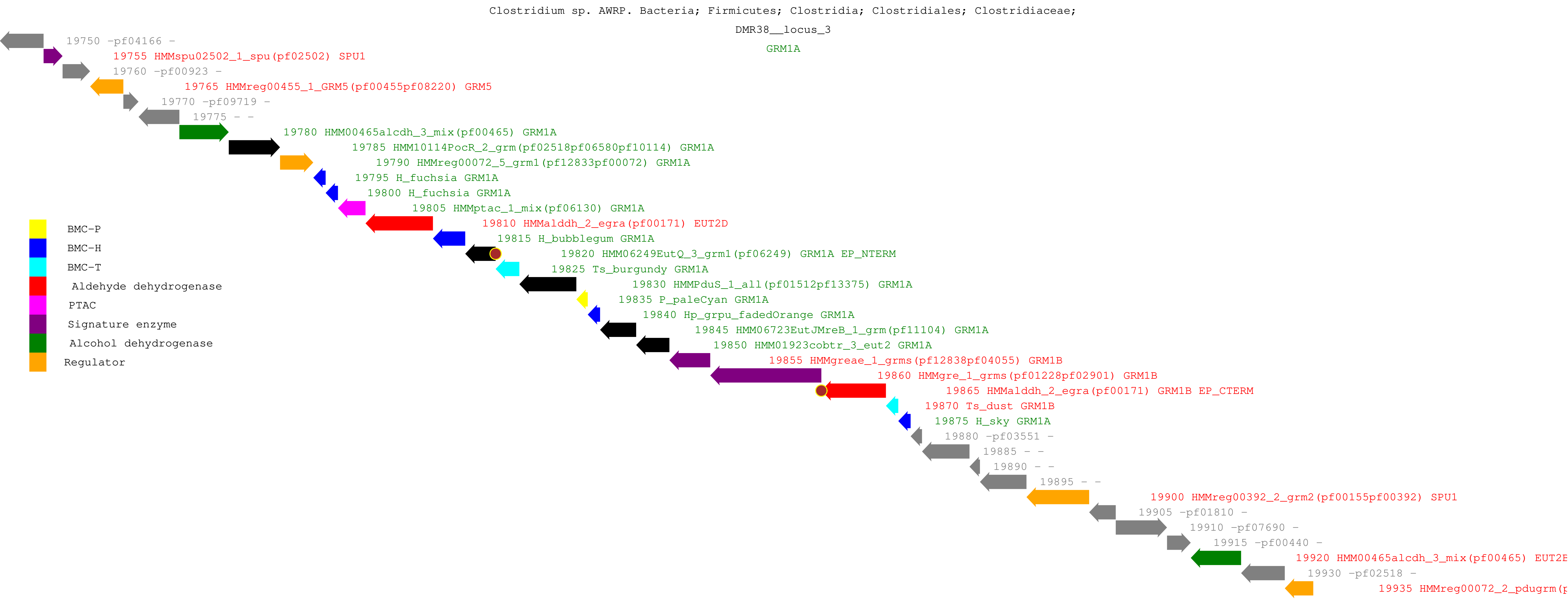
Clostridium sp. AWRP. Bacteria; Firmicutes; Clostridia; Clostridiales; Clostridiaceae;
DMR38__locus_3
GRM1A
19750 -pf04166 -
19755 HMMspu02502_1_spu(pf02502) SPU1
19760 -pf00923 -
19765 HMMreg00455_1_GRM5(pf00455pf08220) GRM5
19770 -pf09719 -
19775 - -
19780 HMM00465alcdh_3_mix(pf00465) GRM1A
19785 HMM10114PocR_2_grm(pf02518pf06580pf10114) GRM1A
19790 HMMreg00072_5_grm1(pf12833pf00072) GRM1A
19795 H_fuchsia GRM1A
19800 H_fuchsia GRM1A
19805 HMMptac_1_mix(pf06130) GRM1A
19810 HMMaldh_2_egra(pf00171) EUT2D
19815 H_bubblegum GRM1A
19820 HMM06249EutQ_3_grm1(pf06249) GRM1A EP_NTERM
19825 Ts_burgundy GRM1A
19830 HMMPduS_1_all(pf01512pf13375) GRM1A
19835 P_paleCyan GRM1A
19840 Hp_grpu_fadedOrange GRM1A
19845 HMM06723EutJMreB_1_grm(pf11104) GRM1A
19850 HMM01923cobtr_3_eut2 GRM1A
19855 HMMgreae_1_grms(pf12838pf04055) GRM1B
19860 HMMgre_1_grms(pf01228pf02901) GRM1B
19865 HMMaldh_2_egra(pf00171) GRM1B EP_CTERM
19870 Ts_dust GRM1B
19875 H_sky GRM1A
19880 -pf03551 -
19885 - -
19890 - -
19895 - -
19900 HMMreg00392_2_grm2(pf00155pf00392) SPU1
19905 -pf01810 -
19910 -pf07690 -
19915 -pf00440 -
19920 HMM00465alcdh_3_mix(pf00465) EUT2E
19930 -pf02518 -
19935 HMMreg00072_2_pdugrm(p
BMC-P
BMC-H
BMC-T
Aldehyde dehydrogenase
PTAC
Signature enzyme
Alcohol dehydrogenase
Regulator

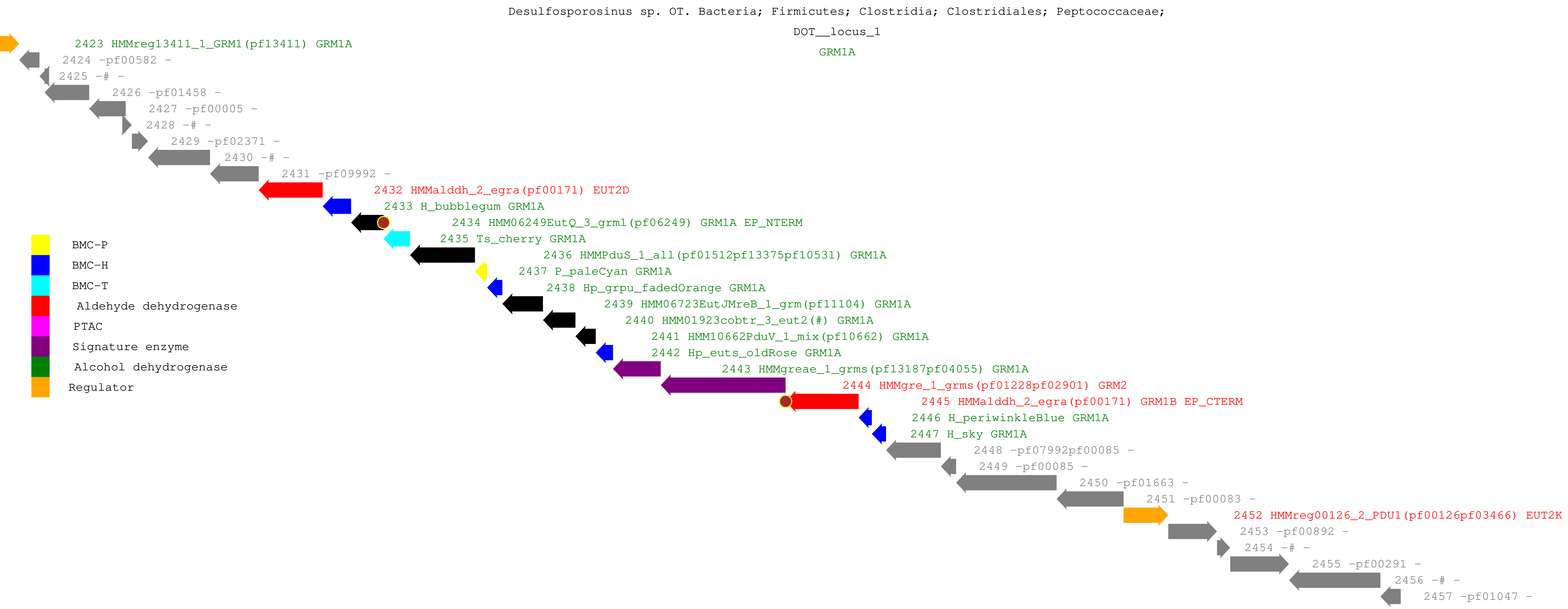

Desulfosporosinus sp. OT. Bacteria; Firmicutes; Clostridia; Clostridiales; Peptococcaceae;
DOT__locus_1
GRM1A
2423 HMMreg13411_1_GRM1(pf13411) GRM1A
2424 -pf00582 -
2425 -# -
2426 -pf01458 -
2427 -pf00005 -
2428 -# -
2429 -pf02371 -
2430 -# -
2431 -pf09992 -
2432 HMMaldh_2_egra(pf00171) EUT2D
2433 H_bubblegum GRM1A
2434 HMM06249EutQ_3_grm1(pf06249) GRM1A EP_NTERM
2435 Ts_cherry GRM1A
2436 HMMPduS_1_all(pf01512pf13375pf10531) GRM1A
2437 P_paleCyan GRM1A
2438 Hp_grpu_fadedOrange GRM1A
2439 HMM06723EutJMreB_1_grm(pf11104) GRM1A
2440 HMM01923cobtr_3_eut2(#) GRM1A
2441 HMM10662PduV_1_mix(pf10662) GRM1A
2442 Hp_euts_oldRose GRM1A
2443 HMMgreae_1_grms(pf13187pf04055) GRM1A
2444 HMMgre_1_grms(pf01228pf02901) GRM2
2445 HMMaldh_2_egra(pf00171) GRM1B EP_CTERM
2446 H_periwinkleBlue GRM1A
2447 H_sky GRM1A
2448 -pf07992pf00085 -
2449 -pf00085 -
2450 -pf01663 -
2451 -pf00083 -
2452 HMMreg00126_2_PDU1(pf00126pf03466) EUT2K
2453 -pf00892 -
2454 -# -
2455 -pf00291 -
2456 -# -
2457 -pf01047 -
BMC-P
BMC-H
BMC-T
Aldehyde dehydrogenase
PTAC
Signature enzyme
Alcohol dehydrogenase
Regulator

Streptococcus suis. Bacteria; Firmicutes; Bacilli; Lactobacillales; Streptococcaceae;

DP112__locus_1

GRM1A

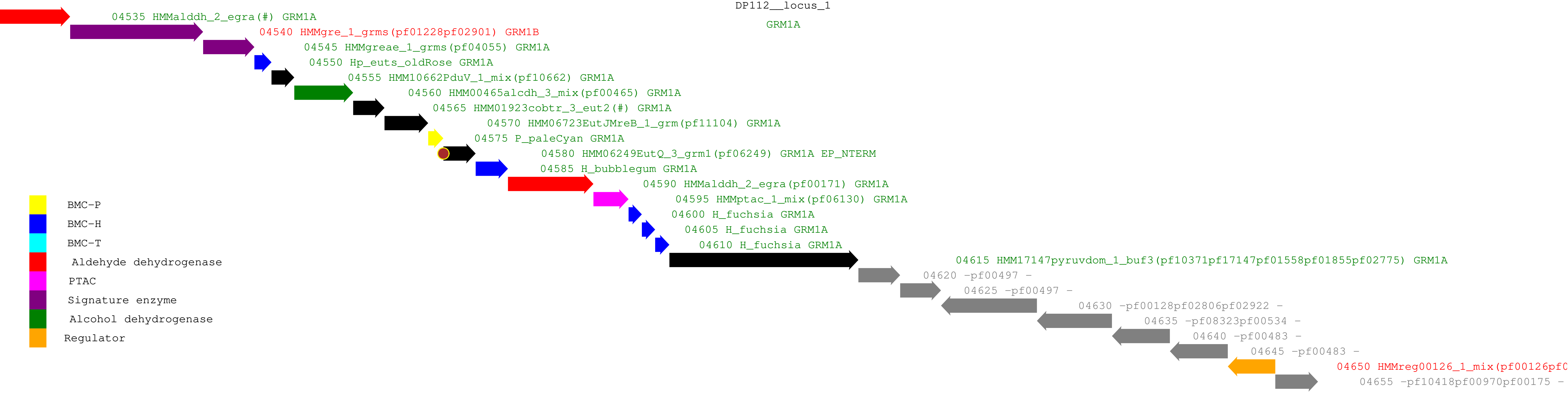

Clostridium tetani. Bacteria; Firmicutes; Clostridia; Clostridiales; Clostridiaceae;

DP129__locus_2

GRM1A

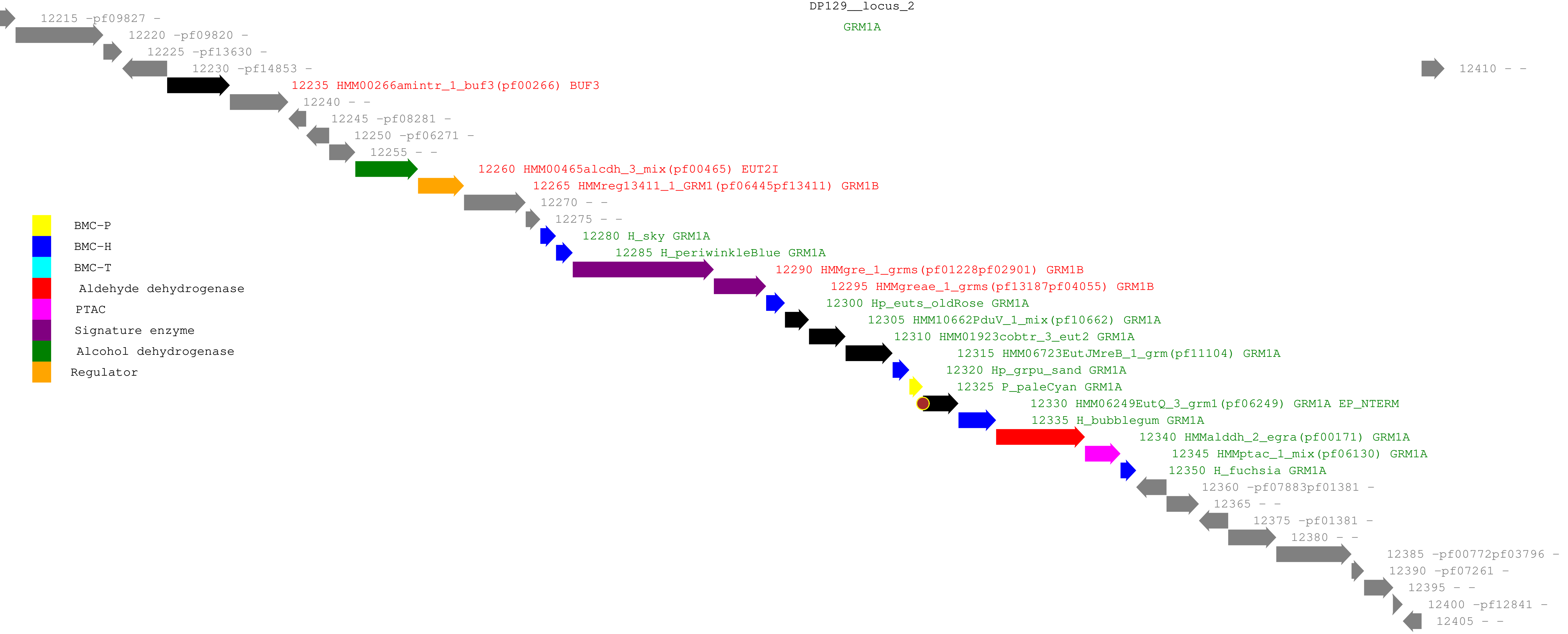

Clostridium tetani. Bacteria; Firmicutes; Clostridia; Clostridiales; Clostridiaceae;

DP130__locus_1

GRM1A

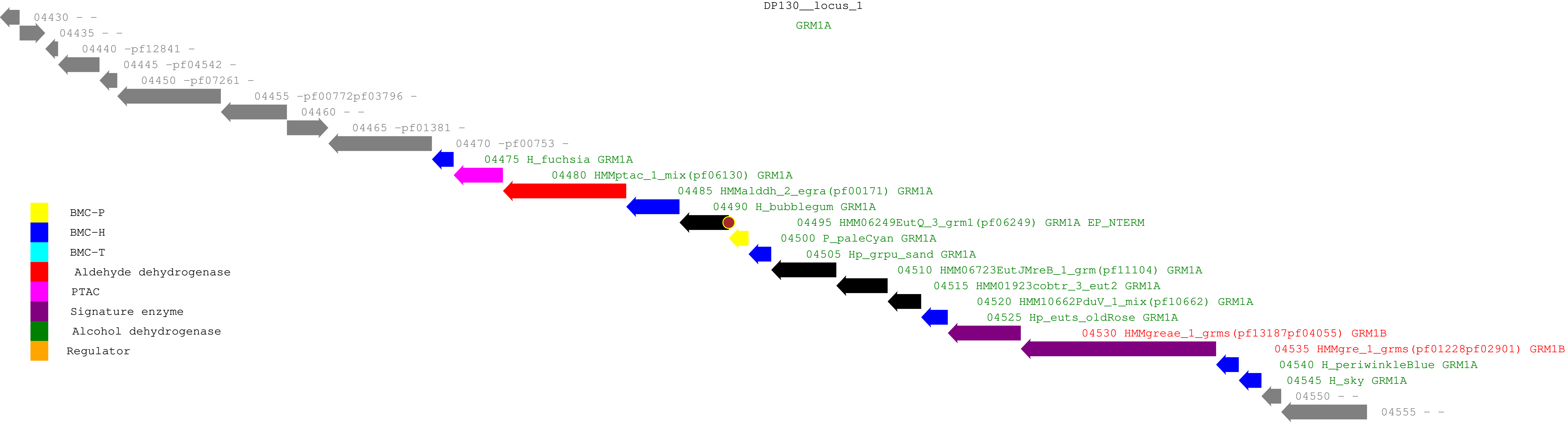

Desulfosporosinus sp. HMP52. Bacteria; Firmicutes; Clostridia; Clostridiales; Peptococcaceae;

DP73__locus_1

GRM1A

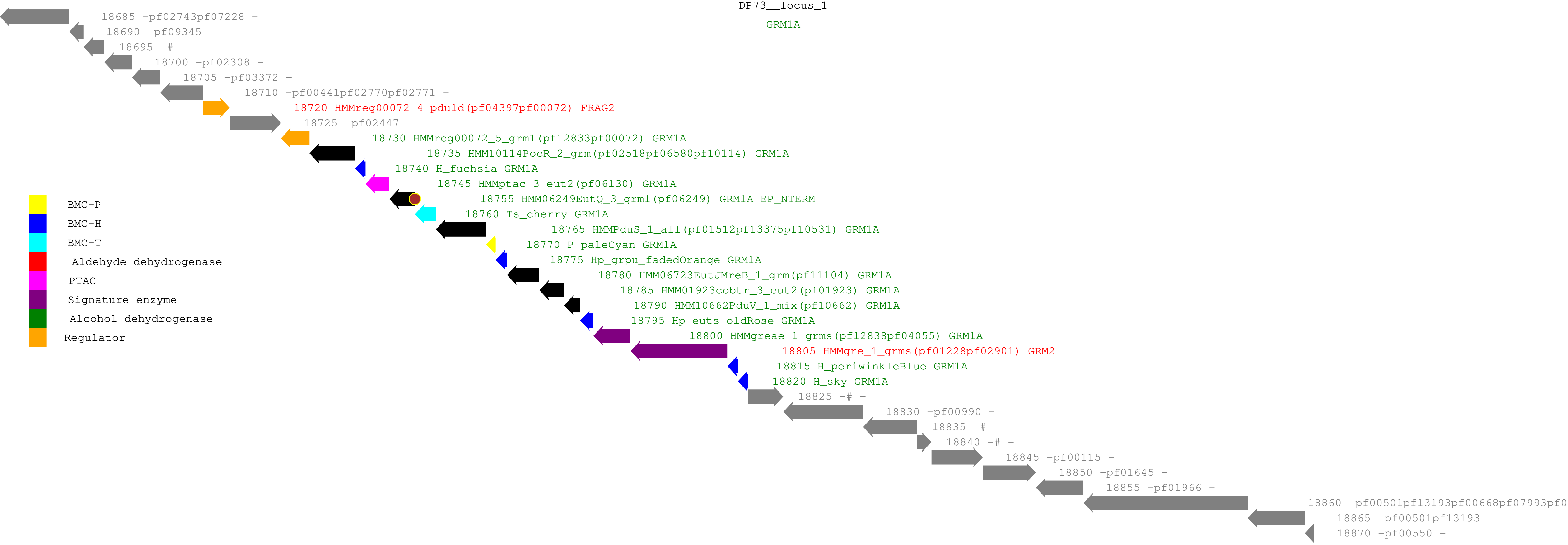

Desulfitobacterium hafniense (Desulfitobacterium frappieri). Bacteria; Firmicutes; Clostridia; Clostridiales; Peptococcaceae;

DPCES__locus_1

GRM1A

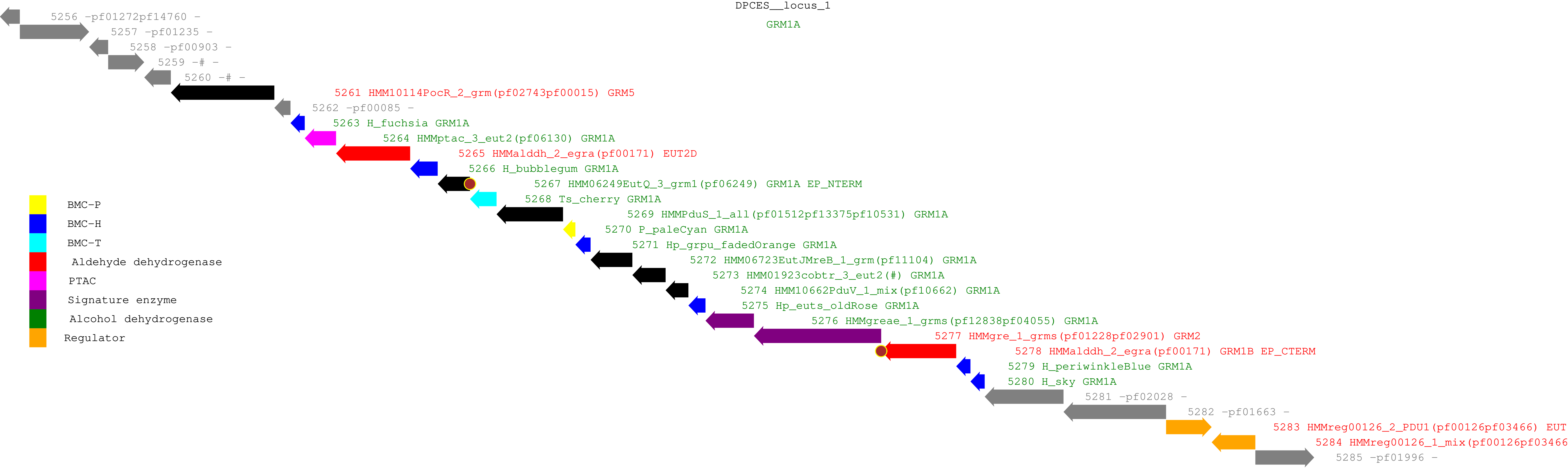

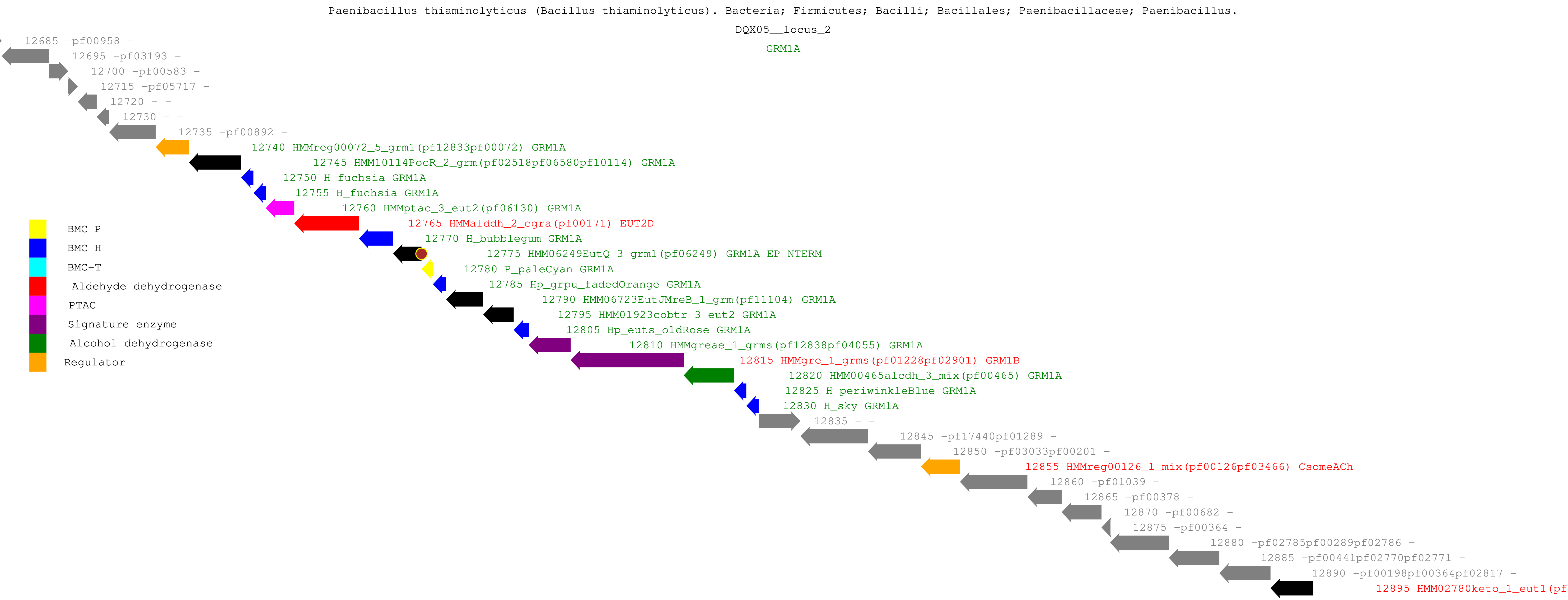
Paenibacillus thiaminolyticus (Bacillus thiaminolyticus). Bacteria; Firmicutes; Bacilli; Bacillales; Paenibacillaceae; Paenibacillus.
DQX05__locus_2
GRM1A
12685 -pf00958 -
12695 -pf03193 -
12700 -pf00583 -
12715 -pf05717 -
12720 - -
12730 - -
12735 -pf00892 -
12740 HMMreg00072_5_grm1(pf12833pf00072) GRM1A
12745 HMM10114PocR_2_grm(pf02518pf06580pf10114) GRM1A
12750 H_fuchsia GRM1A
12755 H_fuchsia GRM1A
12760 HMMptac_3_eut2(pf06130) GRM1A
12765 HMMalddh_2_egra(pf00171) EUT2D
12770 H_bubblegum GRM1A
12775 HMM06249EutQ_3_grm1(pf06249) GRM1A EP_NTERM
12780 P_paleCyan GRM1A
12785 Hp_grpu_fadedOrange GRM1A
12790 HMM06723EutJMreB_1_grm(pf11104) GRM1A
12795 HMM01923cobtr_3_eut2 GRM1A
12805 Hp_euts_oldRose GRM1A
12810 HMMgreae_1_grms(pf12838pf04055) GRM1A
12815 HMMgre_1_grms(pf01228pf02901) GRM1B
12820 HMM00465alcdh_3_mix(pf00465) GRM1A
12825 H_periwinkleBlue GRM1A
12830 H_sky GRM1A
12835 - -
12845 -pf17440pf01289 -
12850 -pf03033pf00201 -
12855 HMMreg00126_1_mix(pf00126pf03466) CsomeACh
12860 -pf01039 -
12865 -pf00378 -
12870 -pf00682 -
12875 -pf00364 -
12880 -pf02785pf00289pf02786 -
12885 -pf00441pf02770pf02771 -
12890 -pf00198pf00364pf02817 -
12895 HMM02780keto_1_eut1(pf
BMC-P
BMC-H
BMC-T
Aldehyde dehydrogenase
PTAC
Signature enzyme
Alcohol dehydrogenase
Regulator

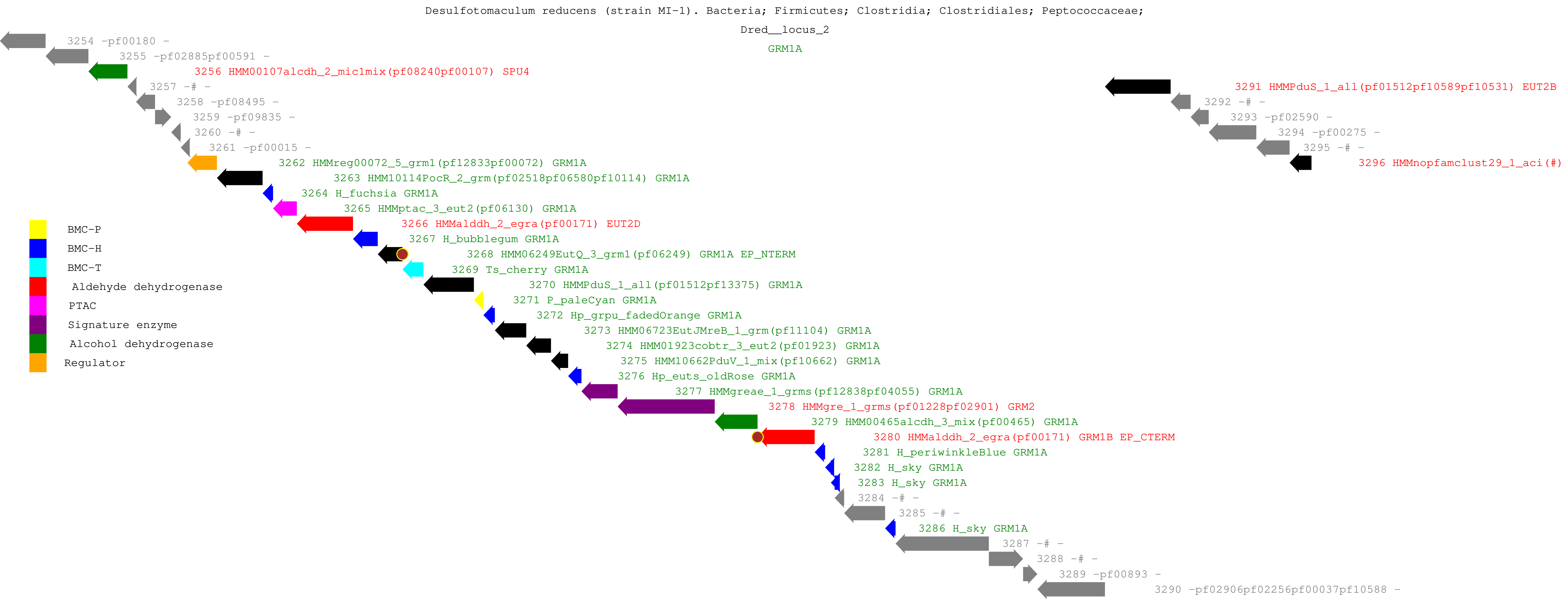

Desulfotomaculum reducens (strain MI-1). Bacteria; Firmicutes; Clostridia; Clostridiales; Peptococcaceae;
Dred__locus_2
GRM1A
3254 -pf00180 -
3255 -pf02885pf00591 -
3256 HMM00107alcdh_2_mic1mix(pf08240pf00107) SPU4
3257 -# -
3258 -pf08495 -
3259 -pf09835 -
3260 -# -
3261 -pf00015 -
3262 HMMreg00072_5_grm1(pf12833pf00072) GRM1A
3263 HMM10114PocR_2_grm(pf02518pf06580pf10114) GRM1A
3264 H_fuchsia GRM1A
3265 HMMptac_3_eut2(pf06130) GRM1A
3266 HMMalddh_2_egra(pf00171) EUT2D
3267 H_bubblegum GRM1A
3268 HMM06249EutQ_3_grm1(pf06249) GRM1A EP_NTERM
3269 Ts_cherry GRM1A
3270 HMMPduS_1_all(pf01512pf13375) GRM1A
3271 P_paleCyan GRM1A
3272 Hp_grpu_fadedOrange GRM1A
3273 HMM06723EutJMreB_1_grm(pf11104) GRM1A
3274 HMM01923cobtr_3_eut2(pf01923) GRM1A
3275 HMM10662PduV_1_mix(pf10662) GRM1A
3276 Hp_euts_oldRose GRM1A
3277 HMMgreae_1_grms(pf12838pf04055) GRM1A
3278 HMMgre_1_grms(pf01228pf02901) GRM2
3279 HMM00465alcdh_3_mix(pf00465) GRM1A
3280 HMMaldh_2_egra(pf00171) GRM1B EP_CTERM
3281 H_periwinkleBlue GRM1A
3282 H_sky GRM1A
3283 H_sky GRM1A
3284 -# -
3285 -# -
3286 H_sky GRM1A
3287 -# -
3288 -# -
3289 -pf00893 -
3290 -pf02906pf02256pf00037pf10588 -
3291 HMMPduS_1_all(pf01512pf10589pf10531) EUT2B
3292 -# -
3293 -pf02590 -
3294 -pf00275 -
3295 -# -
3296 HMMnopfamclust29_1_aci(#)
BMC-P
BMC-H
BMC-T
Aldehyde dehydrogenase
PTAC
Signature enzyme
Alcohol dehydrogenase
Regulator

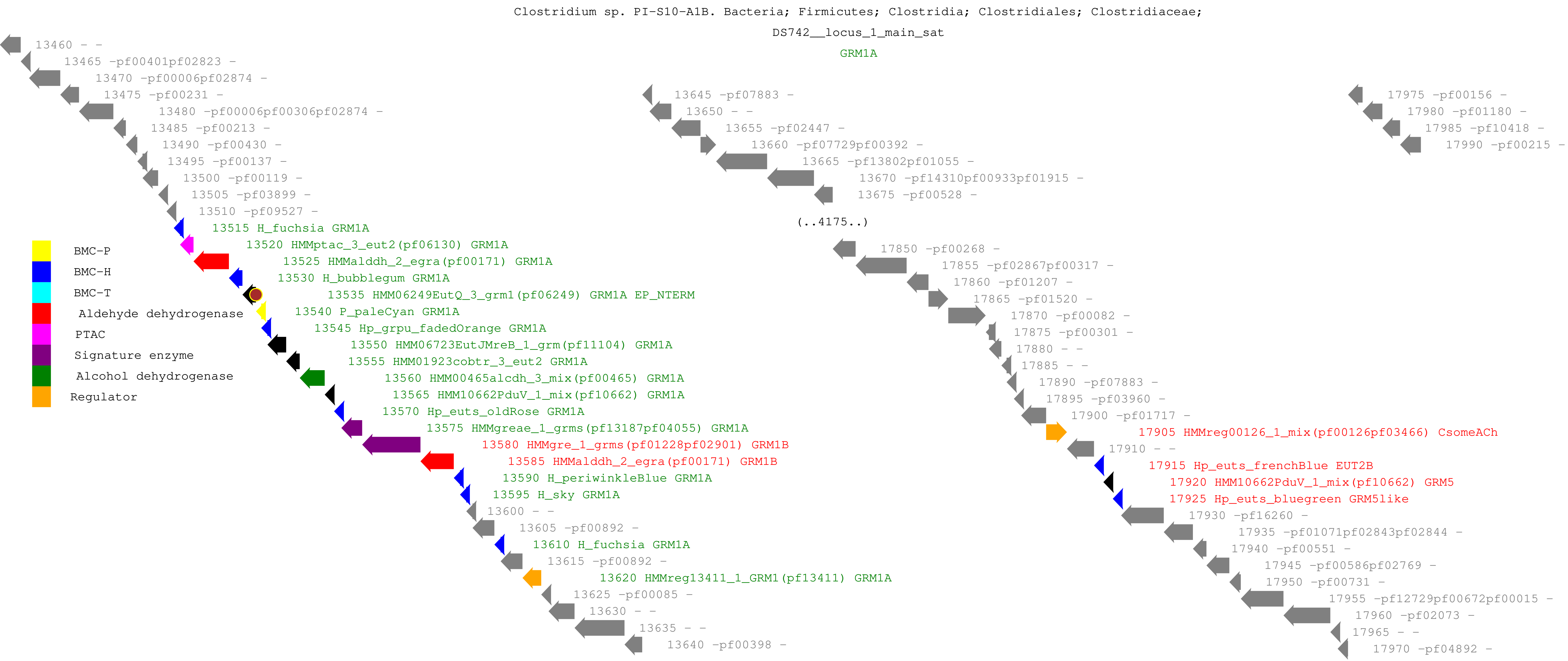
Clostridium sp. PI-S10-A1B. Bacteria; Firmicutes; Clostridia; Clostridiales; Clostridiaceae;
DS742__locus_1_main_sat
GRM1A
13460 - -
13465 -pf00401pf02823 -
13470 -pf00006pf02874 -
13475 -pf00231 -
13480 -pf00006pf00306pf02874 -
13485 -pf00213 -
13490 -pf00430 -
13495 -pf00137 -
13500 -pf00119 -
13505 -pf03899 -
13510 -pf09527 -
13515 H_fuchsia GRM1A
13520 HMMptac_3_eut2(pf06130) GRM1A
13525 HMMalddh_2_egra(pf00171) GRM1A
13530 H_bubblegum GRM1A
13535 HMM06249EutQ_3_grm1(pf06249) GRM1A EP_NTERM
13540 P_paleCyan GRM1A
13545 Hp_grpu_fadedOrange GRM1A
13550 HMM06723EutJMreB_1_grm(pf11104) GRM1A
13555 HMM01923cobtr_3_eut2 GRM1A
13560 HMM00465alcdh_3_mix(pf00465) GRM1A
13565 HMM10662PduV_1_mix(pf10662) GRM1A
13570 Hp_euts_oldRose GRM1A
13575 HMMgreae_1_grms(pf13187pf04055) GRM1A
13580 HMMgre_1_grms(pf01228pf02901) GRM1B
13585 HMMalddh_2_egra(pf00171) GRM1B
13590 H_periwinkleBlue GRM1A
13595 H_sky GRM1A
13600 - -
13605 -pf00892 -
13610 H_fuchsia GRM1A
13615 -pf00892 -
13620 HMMreg13411_1_GRM1(pf13411) GRM1A
13625 -pf00085 -
13630 - -
13635 - -
13640 -pf00398 -
13645 -pf07883 -
13650 - -
13655 -pf02447 -
13660 -pf07729pf00392 -
13665 -pf13802pf01055 -
13670 -pf14310pf00933pf01915 -
13675 -pf00528 -
(..4175..)
17850 -pf00268 -
17855 -pf02867pf00317 -
17860 -pf01207 -
17865 -pf01520 -
17870 -pf00082 -
17875 -pf00301 -
17880 - -
17885 - -
17890 -pf07883 -
17895 -pf03960 -
17900 -pf01717 -
17905 HMMreg00126_1_mix(pf00126pf03466) CsomeACh
17910 - -
17915 Hp_euts_frenchBlue EUT2B
17920 HMM10662PduV_1_mix(pf10662) GRM5
17925 Hp_euts_bluegreen GRM5like
17930 -pf16260 -
17935 -pf01071pf02843pf02844 -
17940 -pf00551 -
17945 -pf00586pf02769 -
17950 -pf00731 -
17955 -pf12729pf00672pf00015 -
17960 -pf02073 -
17965 - -
17970 -pf04892 -
17975 -pf00156 -
17980 -pf01180 -
17985 -pf10418 -
17990 -pf00215 -
BMC-P
BMC-H
BMC-T
Aldehyde dehydrogenase
PTAC
Signature enzyme
Alcohol dehydrogenase
Regulator

Desulfosporosinus sp. BG. Bacteria; Firmicutes; Clostridia; Clostridiales; Peptococcaceae;

DSBG__locus_1

GRM1A

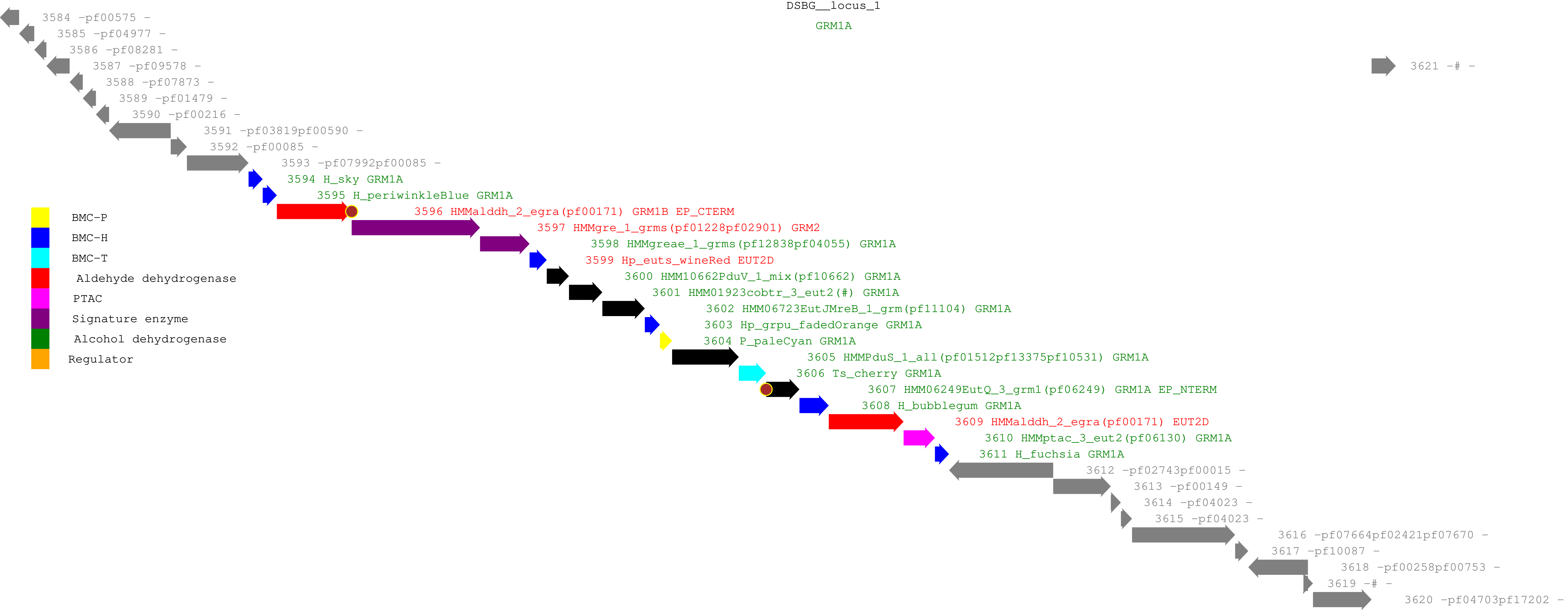

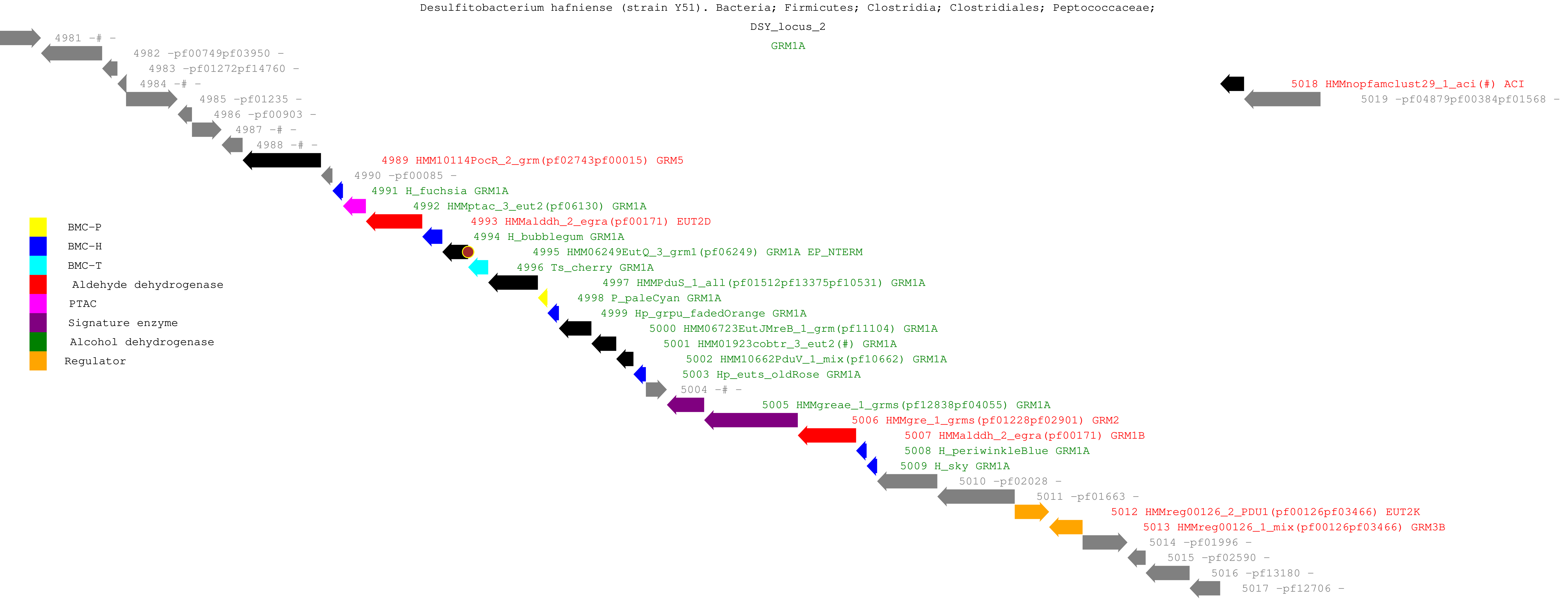
Desulfitobacterium hafniense (strain Y51). Bacteria; Firmicutes; Clostridia; Clostridiales; Peptococcaceae;
DSY_locus_2
GRM1A
4981 -# -
4982 -pf00749pf03950 -
4983 -pf01272pf14760 -
4984 -# -
4985 -pf01235 -
4986 -pf00903 -
4987 -# -
4988 -# -
4989 HMM10114PocR_2_grm(pf02743pf00015) GRM5
4990 -pf00085 -
4991 H_fuchsia GRM1A
4992 HMMptac_3_eut2(pf06130) GRM1A
4993 HMMalddh_2_egra(pf00171) EUT2D
4994 H_bubblegum GRM1A
4995 HMM06249EutQ_3_grm1(pf06249) GRM1A EP_NTERM
4996 Ts_cherry GRM1A
4997 HMMPduS_1_all(pf01512pf13375pf10531) GRM1A
4998 P_paleCyan GRM1A
4999 Hp_grpu_fadedOrange GRM1A
5000 HMM06723EutJMreB_1_grm(pf11104) GRM1A
5001 HMM01923cobtr_3_eut2(#) GRM1A
5002 HMM10662PduV_1_mix(pf10662) GRM1A
5003 Hp_euts_oldRose GRM1A
5004 -# -
5005 HMMgreae_1_grms(pf12838pf04055) GRM1A
5006 HMMgre_1_grms(pf01228pf02901) GRM2
5007 HMMalddh_2_egra(pf00171) GRM1B
5008 H_periwinkleBlue GRM1A
5009 H_sky GRM1A
5010 -pf02028 -
5011 -pf01663 -
5012 HMMreg00126_2_PDU1(pf00126pf03466) EUT2K
5013 HMMreg00126_1_mix(pf00126pf03466) GRM3B
5014 -pf01996 -
5015 -pf02590 -
5016 -pf13180 -
5017 -pf12706 -
5018 HMMnopfamclust29_1_aci(#) ACI
5019 -pf04879pf00384pf01568 -
BMC-P
BMC-H
BMC-T
Aldehyde dehydrogenase
PTAC
Signature enzyme
Alcohol dehydrogenase
Regulator

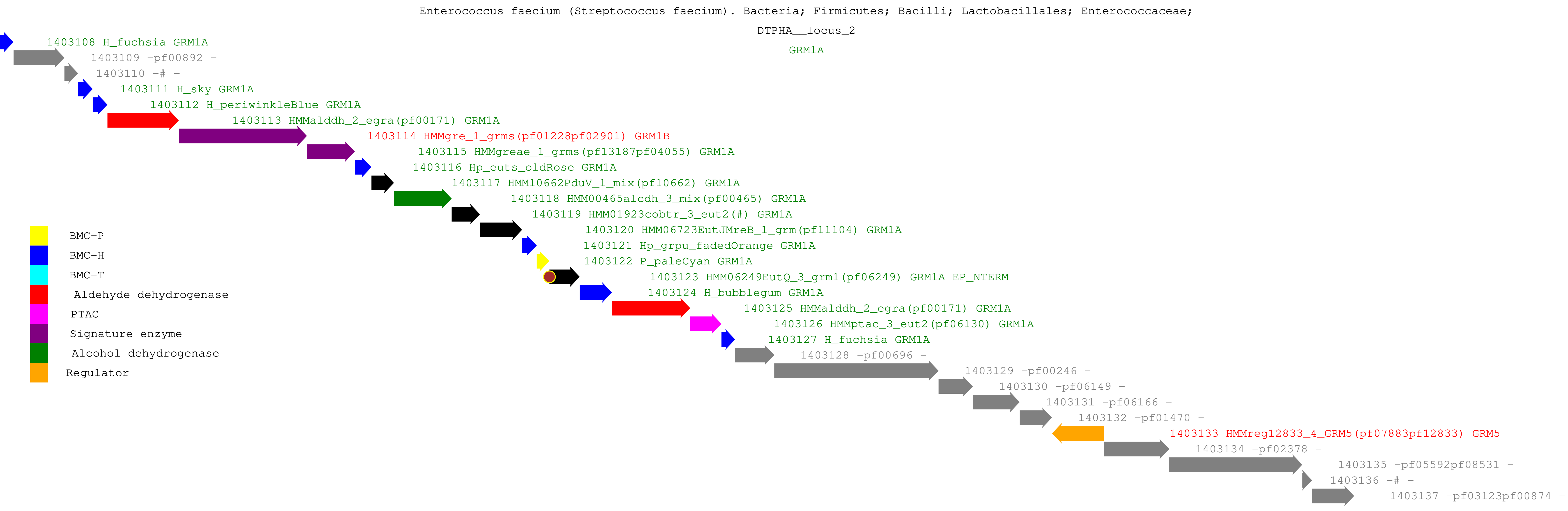

Enterococcus faecium (Streptococcus faecium). Bacteria; Firmicutes; Bacilli; Lactobacillales; Enterococcaceae;
DTPHA__locus_2
GRM1A
1403108 H_fuchsia GRM1A
1403109 -pf00892 -
1403110 -# -
1403111 H_sky GRM1A
1403112 H_periwinkleBlue GRM1A
1403113 HMMalddh_2_egra(pf00171) GRM1A
1403114 HMMgre_1_grms(pf01228pf02901) GRM1B
1403115 HMMgreae_1_grms(pf13187pf04055) GRM1A
1403116 Hp_euts_oldRose GRM1A
1403117 HMM10662PduV_1_mix(pf10662) GRM1A
1403118 HMM00465alcdh_3_mix(pf00465) GRM1A
1403119 HMM01923cobtr_3_eut2(#) GRM1A
1403120 HMM06723EutJMreB_1_grm(pf11104) GRM1A
1403121 Hp_grpu_fadedOrange GRM1A
1403122 P_paleCyan GRM1A
1403123 HMM06249EutQ_3_grm1(pf06249) GRM1A EP_NTERM
1403124 H_bubblegum GRM1A
1403125 HMMalddh_2_egra(pf00171) GRM1A
1403126 HMMptac_3_eut2(pf06130) GRM1A
1403127 H_fuchsia GRM1A
1403128 -pf00696 -
1403129 -pf00246 -
1403130 -pf06149 -
1403131 -pf06166 -
1403132 -pf01470 -
1403133 HMMreg12833_4_GRM5(pf07883pf12833) GRM5
1403134 -pf02378 -
1403135 -pf05592pf08531 -
1403136 -# -
1403137 -pf03123pf00874 -
BMC-P
BMC-H
BMC-T
Aldehyde dehydrogenase
PTAC
Signature enzyme
Alcohol dehydrogenase
Regulator

Coriobacteriaceae bacterium. Bacteria; Actinobacteria; Coriobacteriia; Coriobacteriales;

DUD33__locus_1_main_sat

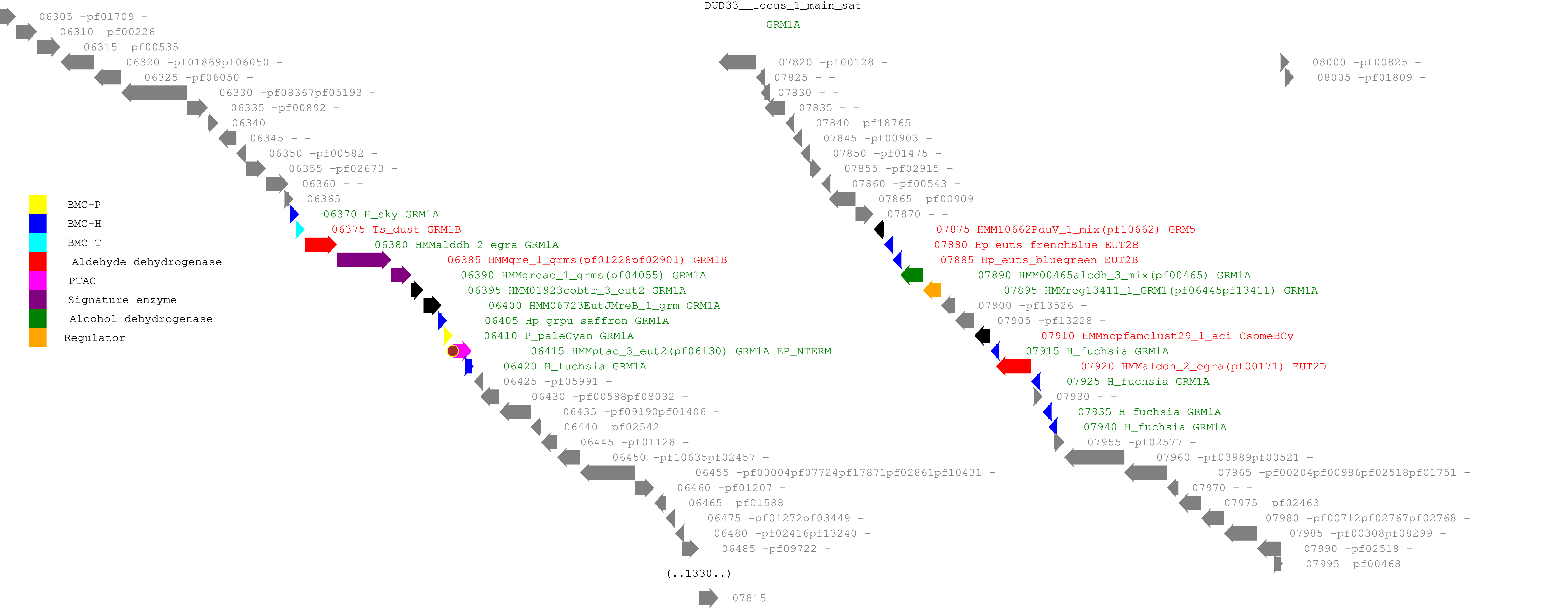

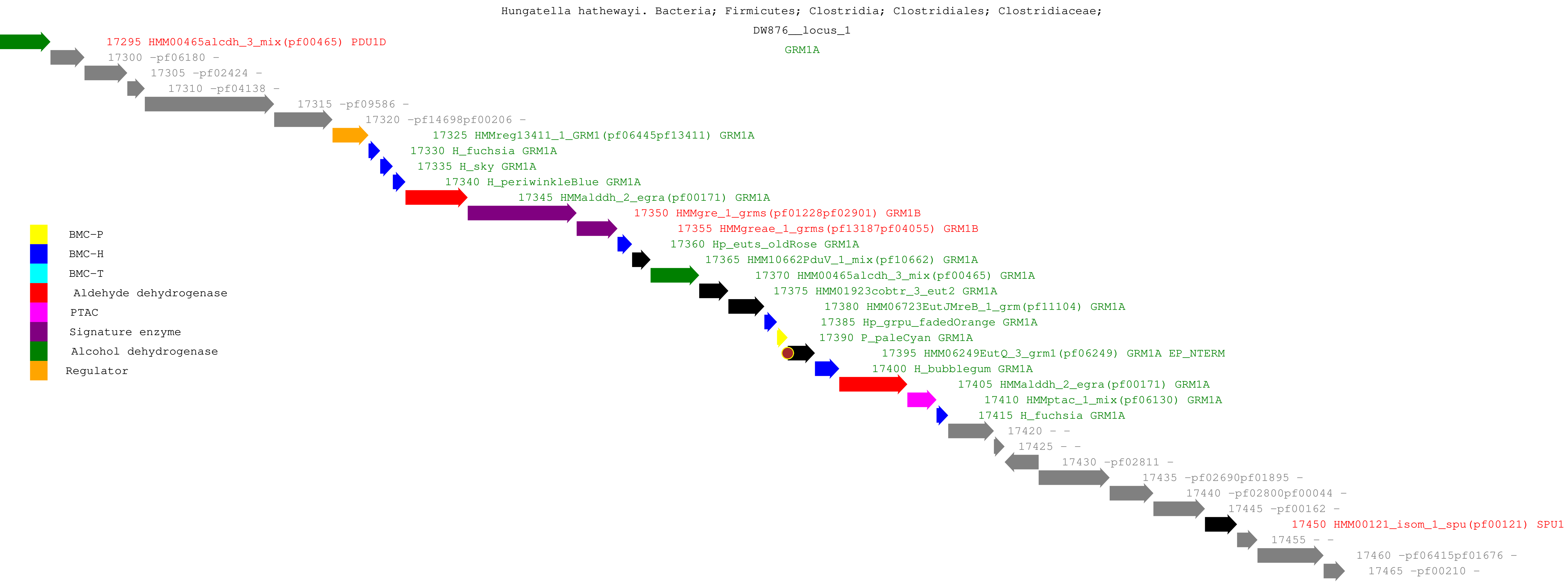
Hungatella hathewayi. Bacteria; Firmicutes; Clostridia; Clostridiales; Clostridiaceae;
DW876__locus_1
GRM1A
17295 HMM00465alcdh_3_mix(pf00465) PDU1D
17300 -pf06180 -
17305 -pf02424 -
17310 -pf04138 -
17315 -pf09586 -
17320 -pf14698pf00206 -
17325 HMMreg13411_1_GRM1(pf06445pf13411) GRM1A
17330 H_fuchsia GRM1A
17335 H_sky GRM1A
17340 H_periwinkleBlue GRM1A
17345 HMMalddh_2_egra(pf00171) GRM1A
17350 HMMgre_1_grms(pf01228pf02901) GRM1B
17355 HMMgreae_1_grms(pf13187pf04055) GRM1B
17360 Hp_euts_oldRose GRM1A
17365 HMM10662PduV_1_mix(pf10662) GRM1A
17370 HMM00465alcdh_3_mix(pf00465) GRM1A
17375 HMM01923cobtr_3_eut2 GRM1A
17380 HMM06723EutJMreB_1_grm(pf11104) GRM1A
17385 Hp_grpu_fadedOrange GRM1A
17390 P_paleCyan GRM1A
17395 HMM06249EutQ_3_grm1(pf06249) GRM1A EP_NTERM
17400 H_bubblegum GRM1A
17405 HMMalddh_2_egra(pf00171) GRM1A
17410 HMMptac_1_mix(pf06130) GRM1A
17415 H_fuchsia GRM1A
17420 - -
17425 - -
17430 -pf02811 -
17435 -pf02690pf01895 -
17440 -pf02800pf00044 -
17445 -pf00162 -
17450 HMM00121_isom_1_spu(pf00121) SPU1
17455 - -
17460 -pf06415pf01676 -
17465 -pf00210 -
BMC-P
BMC-H
BMC-T
Aldehyde dehydrogenase
PTAC
Signature enzyme
Alcohol dehydrogenase
Regulator

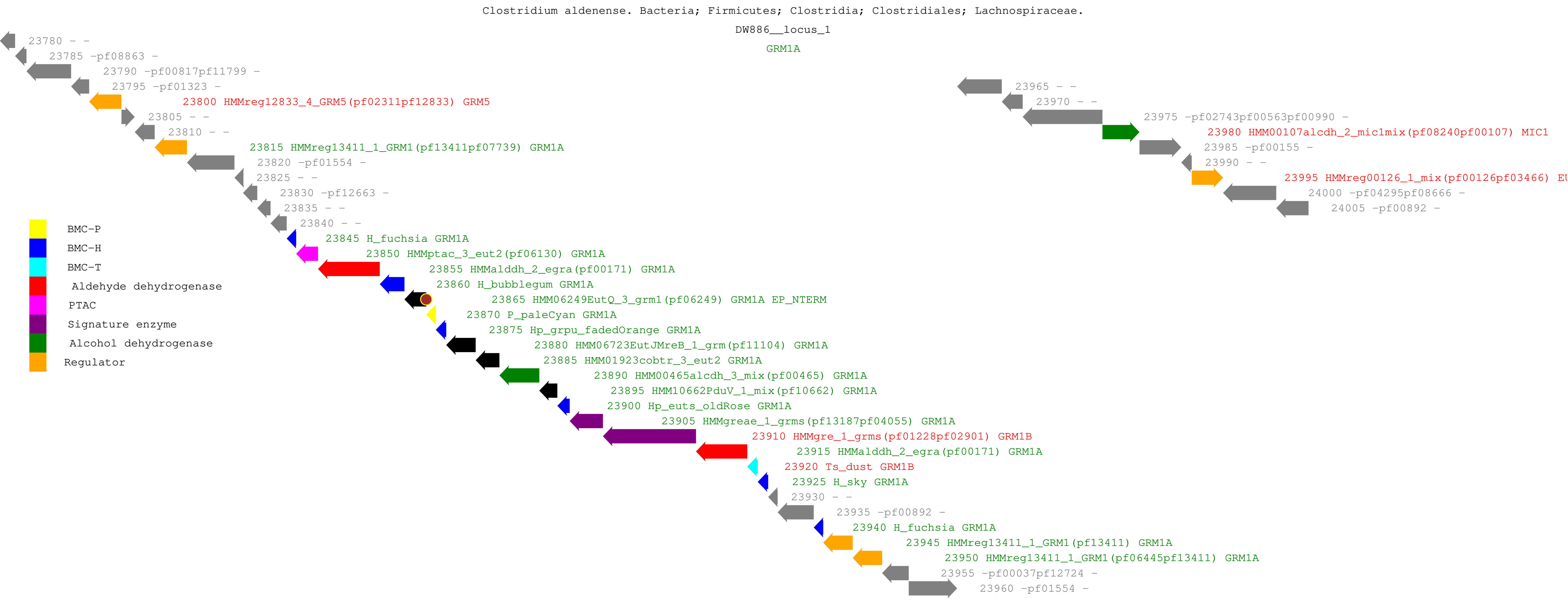
Clostridium aldenense. Bacteria; Firmicutes; Clostridia; Clostridiales; Lachnospiraceae.
DW886__locus_1
GRM1A
23780 - -
23785 -pf08863 -
23790 -pf00817pf11799 -
23795 -pf01323 -
23800 HMMreg12833_4_GRM5(pf02311pf12833) GRM5
23805 - -
23810 - -
23815 HMMreg13411_1_GRM1(pf13411pf07739) GRM1A
23820 -pf01554 -
23825 - -
23830 -pf12663 -
23835 - -
23840 - -
23845 H_fuchsia GRM1A
23850 HMMptac_3_eut2(pf06130) GRM1A
23855 HMMalddh_2_egra(pf00171) GRM1A
23860 H_bubblegum GRM1A
23865 HMM06249EutQ_3_grm1(pf06249) GRM1A EP_NTERM
23870 P_paleCyan GRM1A
23875 Hp_grpu_fadedOrange GRM1A
23880 HMM06723EutJMreB_1_grm(pf11104) GRM1A
23885 HMM01923cobtr_3_eut2 GRM1A
23890 HMM00465alcdh_3_mix(pf00465) GRM1A
23895 HMM10662PduV_1_mix(pf10662) GRM1A
23900 Hp_euts_oldRose GRM1A
23905 HMMgreae_1_grms(pf13187pf04055) GRM1A
23910 HMMgre_1_grms(pf01228pf02901) GRM1B
23915 HMMalddh_2_egra(pf00171) GRM1A
23920 Ts_dust GRM1B
23925 H_sky GRM1A
23930 - -
23935 -pf00892 -
23940 H_fuchsia GRM1A
23945 HMMreg13411_1_GRM1(pf13411) GRM1A
23950 HMMreg13411_1_GRM1(pf06445pf13411) GRM1A
23955 -pf00037pf12724 -
23960 -pf01554 -
23965 - -
23970 - -
23975 -pf02743pf00563pf00990 -
23980 HMM00107alcdh_2_mic1mix(pf08240pf00107) MIC1
23985 -pf00155 -
23990 - -
23995 HMMreg00126_1_mix(pf00126pf03466) EU
24000 -pf04295pf08666 -
24005 -pf00892 -
BMC-P
BMC-H
BMC-T
Aldehyde dehydrogenase
PTAC
Signature enzyme
Alcohol dehydrogenase
Regulator

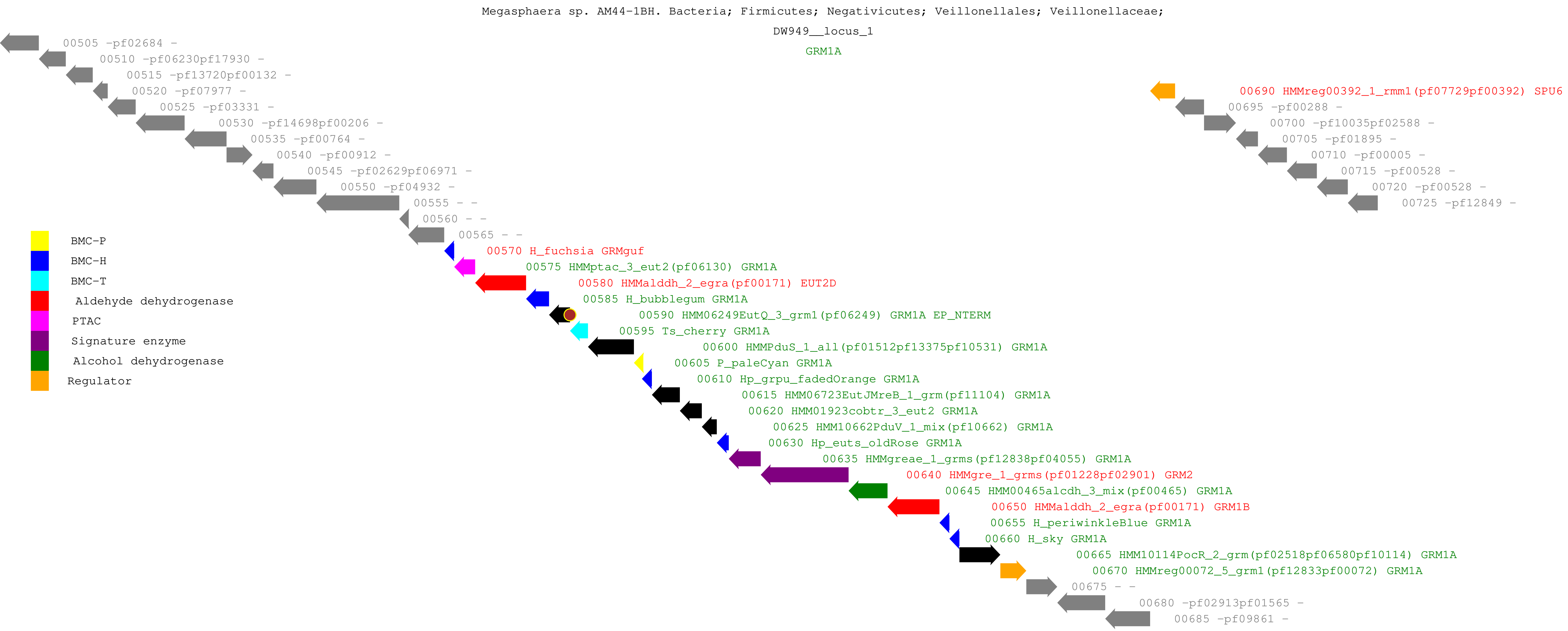
Megasphaera sp. AM44-1BH. Bacteria; Firmicutes; Negativicutes; Veillonellales; Veillonellaceae;
DW949__locus_1
GRM1A
00505 -pf02684 -
00510 -pf06230pf17930 -
00515 -pf13720pf00132 -
00520 -pf07977 -
00525 -pf03331 -
00530 -pf14698pf00206 -
00535 -pf00764 -
00540 -pf00912 -
00545 -pf02629pf06971 -
00550 -pf04932 -
00555 - -
00560 - -
00565 - -
00570 H_fuchsia GRMguf
00575 HMMptac_3_eut2(pf06130) GRM1A
00580 HMMalddh_2_egra(pf00171) EUT2D
00585 H_bubblegum GRM1A
00590 HMM06249EutQ_3_grm1(pf06249) GRM1A EP_NTERM
00595 Ts_cherry GRM1A
00600 HMMPduS_1_all(pf01512pf13375pf10531) GRM1A
00605 P_paleCyan GRM1A
00610 Hp_grpu_fadedOrange GRM1A
00615 HMM06723EutJMreB_1_grm(pf11104) GRM1A
00620 HMM01923cobtr_3_eut2 GRM1A
00625 HMM10662PduV_1_mix(pf10662) GRM1A
00630 Hp_euts_oldRose GRM1A
00635 HMMgreae_1_grms(pf12838pf04055) GRM1A
00640 HMMgre_1_grms(pf01228pf02901) GRM2
00645 HMM00465alcdh_3_mix(pf00465) GRM1A
00650 HMMalddh_2_egra(pf00171) GRM1B
00655 H_periwinkleBlue GRM1A
00660 H_sky GRM1A
00665 HMM10114PocR_2_grm(pf02518pf06580pf10114) GRM1A
00670 HMMreg00072_5_grm1(pf12833pf00072) GRM1A
00675 - -
00680 -pf02913pf01565 -
00685 -pf09861 -
00690 HMMreg00392_1_rmm1(pf07729pf00392) SPU6
00695 -pf00288 -
00700 -pf10035pf02588 -
00705 -pf01895 -
00710 -pf00005 -
00715 -pf00528 -
00720 -pf00528 -
00725 -pf12849 -
BMC-P
BMC-H
BMC-T
Aldehyde dehydrogenase
PTAC
Signature enzyme
Alcohol dehydrogenase
Regulator

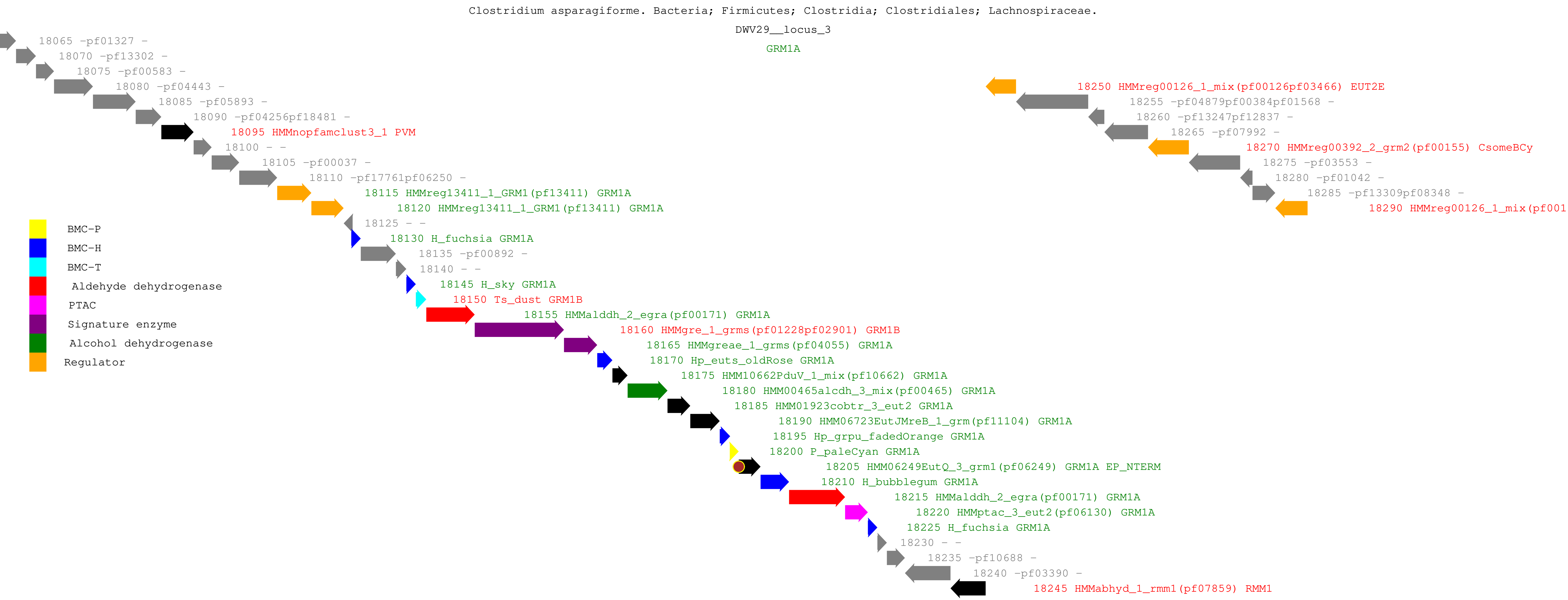
Clostridium asparagiforme. Bacteria; Firmicutes; Clostridia; Clostridiales; Lachnospiraceae.
DWV29__locus_3
GRM1A
18065 -pf01327 -
18070 -pf13302 -
18075 -pf00583 -
18080 -pf04443 -
18085 -pf05893 -
18090 -pf04256pf18481 -
18095 HMMnopfamclust3_1 PVM
18100 - -
18105 -pf00037 -
18110 -pf17761pf06250 -
18115 HMMreg13411_1_GRM1(pf13411) GRM1A
18120 HMMreg13411_1_GRM1(pf13411) GRM1A
18125 - -
18130 H_fuchsia GRM1A
18135 -pf00892 -
18140 - -
18145 H_sky GRM1A
18150 Ts_dust GRM1B
18155 HMMaldh_2_egra(pf00171) GRM1A
18160 HMMgre_1_grms(pf01228pf02901) GRM1B
18165 HMMgreae_1_grms(pf04055) GRM1A
18170 Hp_euts_oldRose GRM1A
18175 HMM10662PduV_1_mix(pf10662) GRM1A
18180 HMM00465alcdh_3_mix(pf00465) GRM1A
18185 HMM01923cobtr_3_eut2 GRM1A
18190 HMM06723EutJMreB_1_grm(pf11104) GRM1A
18195 Hp_grpu_fadedOrange GRM1A
18200 P_paleCyan GRM1A
18205 HMM06249EutQ_3_grm1(pf06249) GRM1A EP_NTERM
18210 H_bubblegum GRM1A
18215 HMMaldh_2_egra(pf00171) GRM1A
18220 HMMptac_3_eut2(pf06130) GRM1A
18225 H_fuchsia GRM1A
18230 - -
18235 -pf10688 -
18240 -pf03390 -
18245 HMMabhyd_1_rmm1(pf07859) RMM1
18250 HMMreg00126_1_mix(pf00126pf03466) EUT2E
18255 -pf04879pf00384pf01568 -
18260 -pf13247pf12837 -
18265 -pf07992 -
18270 HMMreg00392_2_grm2(pf00155) CsomeBCy
18275 -pf03553 -
18280 -pf01042 -
18285 -pf13309pf08348 -
18290 HMMreg00126_1_mix(pf001
BMC-P
BMC-H
BMC-T
Aldehyde dehydrogenase
PTAC
Signature enzyme
Alcohol dehydrogenase
Regulator

Enterococcus asini. Bacteria; Firmicutes; Bacilli; Lactobacillales; Enterococcaceae;
DWV91__locus_1
GRM1A

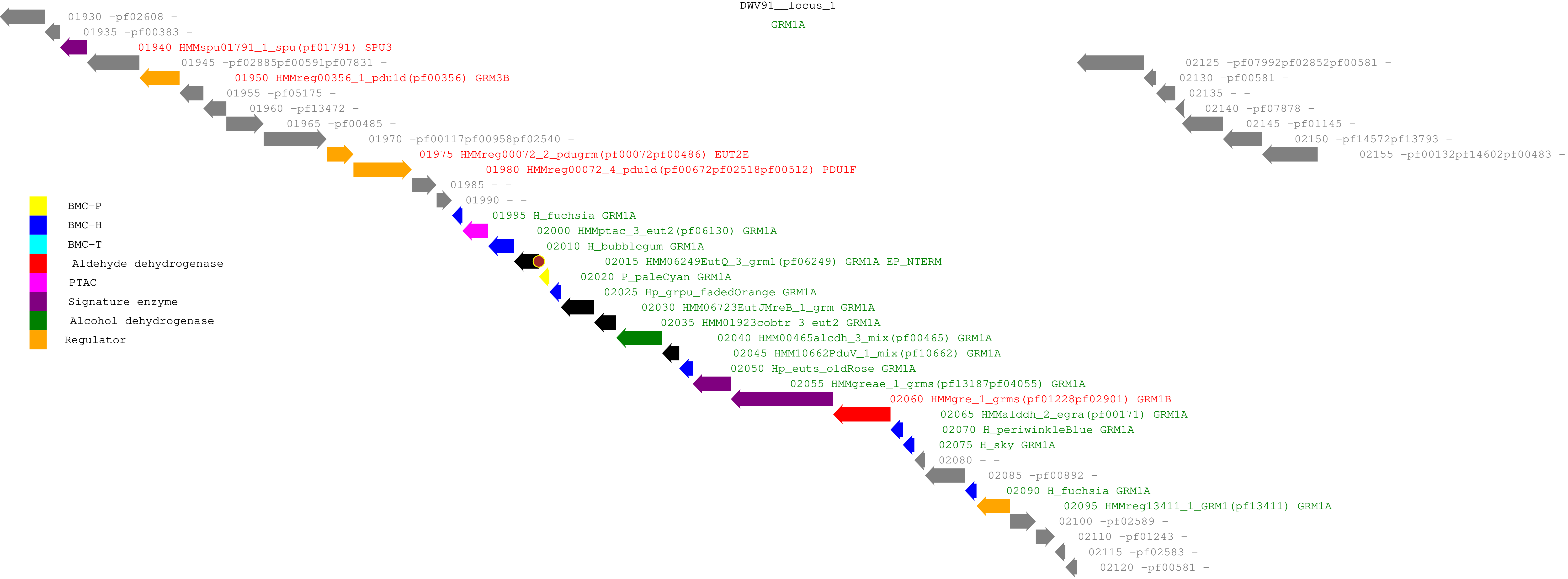

Clostridium sp. AF18-27. Bacteria; Firmicutes; Clostridia; Clostridiales; Clostridiaceae;

DWX10__locus_3

GRM1A

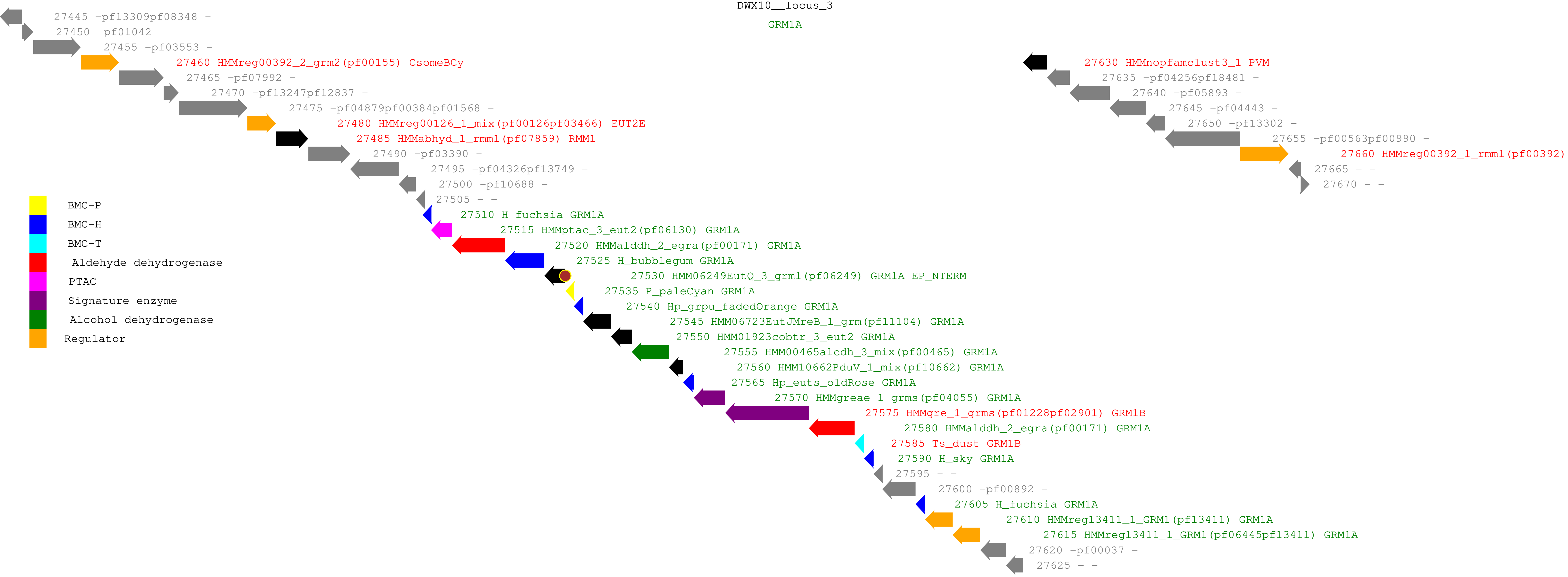

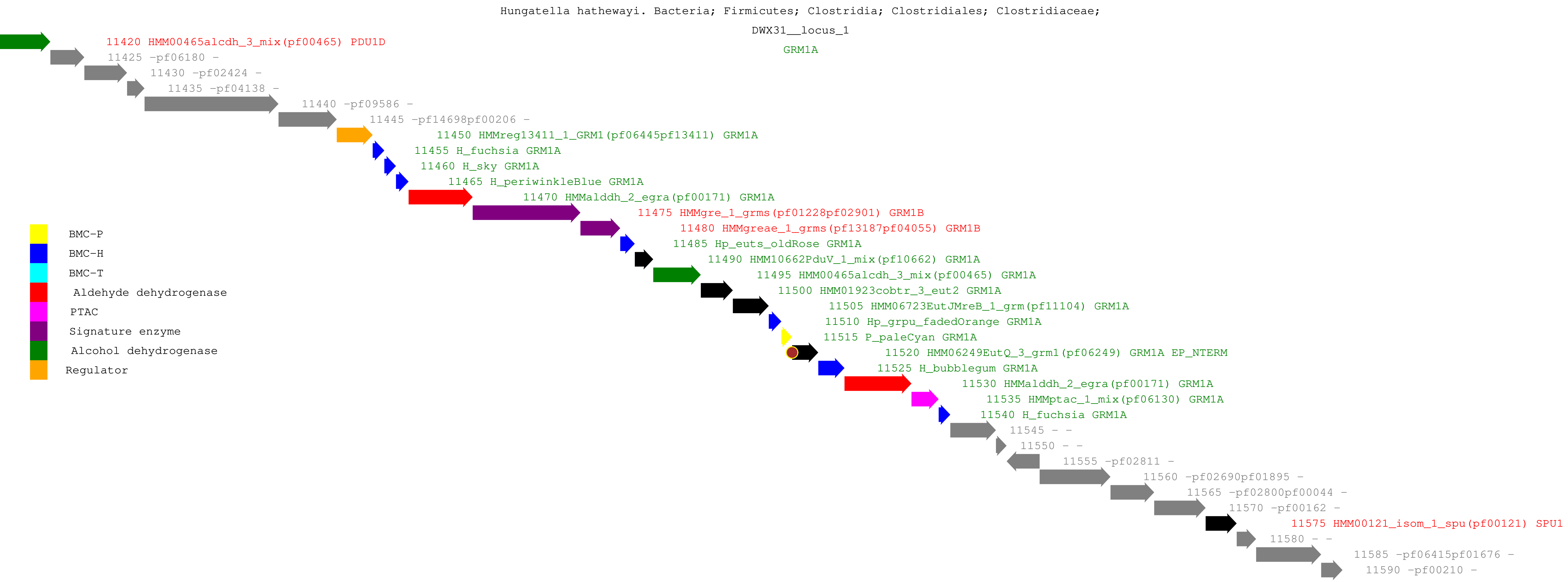
Hungatella hathewayi. Bacteria; Firmicutes; Clostridia; Clostridiales; Clostridiaceae;
DWX31__locus_1
GRM1A
11420 HMM00465alcdh_3_mix(pf00465) PDU1D
11425 -pf06180 -
11430 -pf02424 -
11435 -pf04138 -
11440 -pf09586 -
11445 -pf14698pf00206 -
11450 HMMreg13411_1_GRM1(pf06445pf13411) GRM1A
11455 H_fuchsia GRM1A
11460 H_sky GRM1A
11465 H_periwinkleBlue GRM1A
11470 HMMalddh_2_egra(pf00171) GRM1A
11475 HMMgre_1_grms(pf01228pf02901) GRM1B
11480 HMMgreae_1_grms(pf13187pf04055) GRM1B
11485 Hp_euts_oldRose GRM1A
11490 HMM10662PduV_1_mix(pf10662) GRM1A
11495 HMM00465alcdh_3_mix(pf00465) GRM1A
11500 HMM01923cobtr_3_eut2 GRM1A
11505 HMM06723EutJMreB_1_grm(pf11104) GRM1A
11510 Hp_grpu_fadedOrange GRM1A
11515 P_paleCyan GRM1A
11520 HMM06249EutQ_3_grm1(pf06249) GRM1A EP_NTERM
11525 H_bubblegum GRM1A
11530 HMMalddh_2_egra(pf00171) GRM1A
11535 HMMptac_1_mix(pf06130) GRM1A
11540 H_fuchsia GRM1A
11545 - -
11550 - -
11555 -pf02811 -
11560 -pf02690pf01895 -
11565 -pf02800pf00044 -
11570 -pf00162 -
11575 HMM00121_isom_1_spu(pf00121) SPU1
11580 - -
11585 -pf06415pf01676 -
11590 -pf00210 -
BMC-P
BMC-H
BMC-T
Aldehyde dehydrogenase
PTAC
Signature enzyme
Alcohol dehydrogenase
Regulator

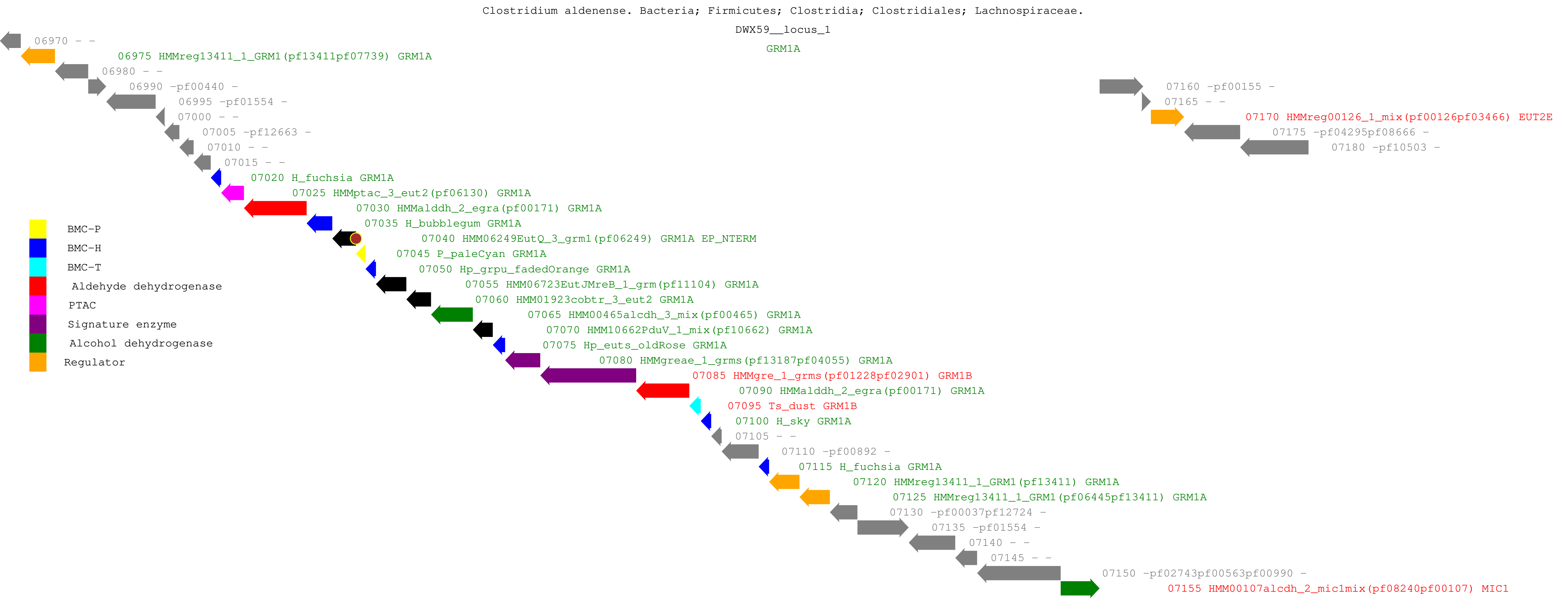
Clostridium aldenense. Bacteria; Firmicutes; Clostridia; Clostridiales; Lachnospiraceae.
DWX59__locus_1
GRM1A
06970 - -
06975 HMMreg13411_1_GRM1(pf13411pf07739) GRM1A
06980 - -
06990 -pf00440 -
06995 -pf01554 -
07000 - -
07005 -pf12663 -
07010 - -
07015 - -
07020 H_fuchsia GRM1A
07025 HMMptac_3_eut2(pf06130) GRM1A
07030 HMMaldh_2_egra(pf00171) GRM1A
07035 H_bubblegum GRM1A
07040 HMM06249EutQ_3_grm1(pf06249) GRM1A EP_NTERM
07045 P_paleCyan GRM1A
07050 Hp_grpu_fadedOrange GRM1A
07055 HMM06723EutJMreB_1_grm(pf11104) GRM1A
07060 HMM01923cobtr_3_eut2 GRM1A
07065 HMM00465alcdh_3_mix(pf00465) GRM1A
07070 HMM10662PduV_1_mix(pf10662) GRM1A
07075 Hp_euts_oldRose GRM1A
07080 HMMgreae_1_grms(pf13187pf04055) GRM1A
07085 HMMgre_1_grms(pf01228pf02901) GRM1B
07090 HMMaldh_2_egra(pf00171) GRM1A
07095 Ts_dust GRM1B
07100 H_sky GRM1A
07105 - -
07110 -pf00892 -
07115 H_fuchsia GRM1A
07120 HMMreg13411_1_GRM1(pf13411) GRM1A
07125 HMMreg13411_1_GRM1(pf06445pf13411) GRM1A
07130 -pf00037pf12724 -
07135 -pf01554 -
07140 - -
07145 - -
07150 -pf02743pf00563pf00990 -
07155 HMM00107alcdh_2_mic1mix(pf08240pf00107) MIC1
07160 -pf00155 -
07165 - -
07170 HMMreg00126_1_mix(pf00126pf03466) EUT2E
07175 -pf04295pf08666 -
07180 -pf10503 -
BMC-P
BMC-H
BMC-T
Aldehyde dehydrogenase
PTAC
Signature enzyme
Alcohol dehydrogenase
Regulator

Collinsella sp. AF20-14LB. Bacteria; Actinobacteria; Coriobacteriia; Coriobacteriales;
DWX63__locus_1
GRM1A

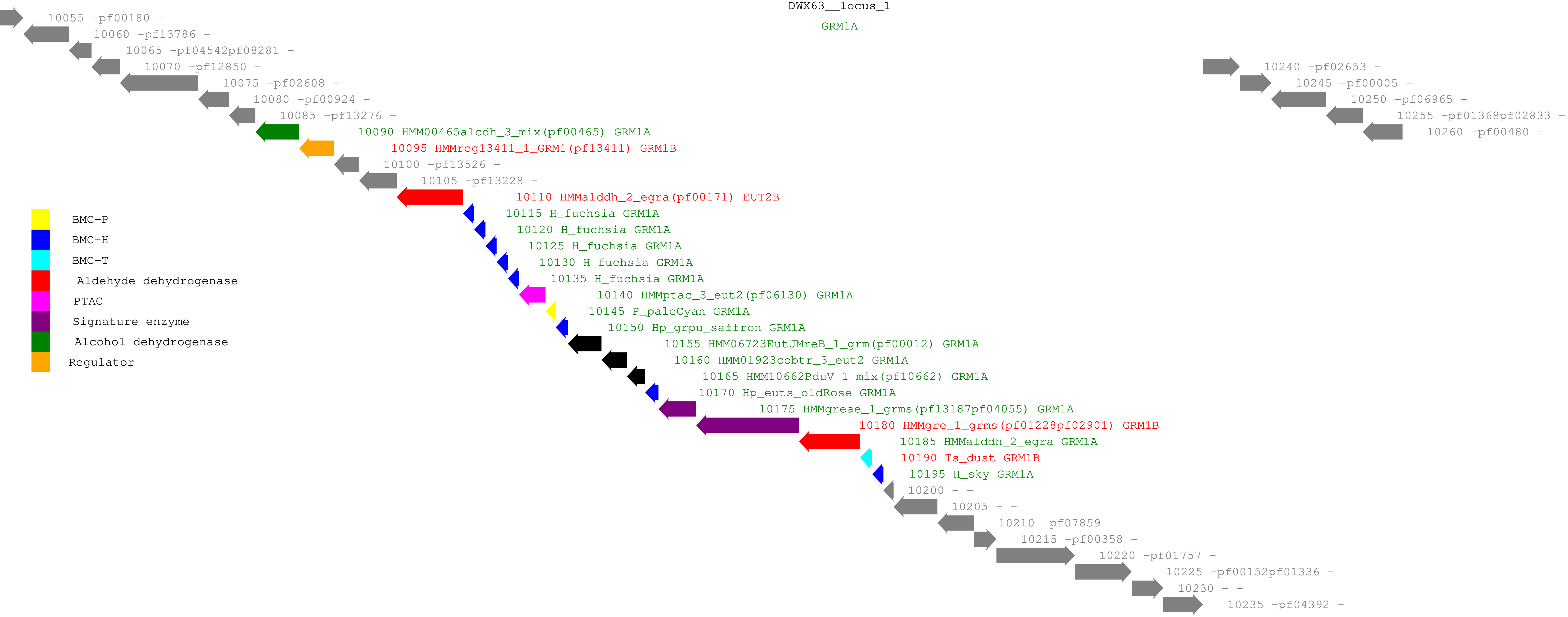

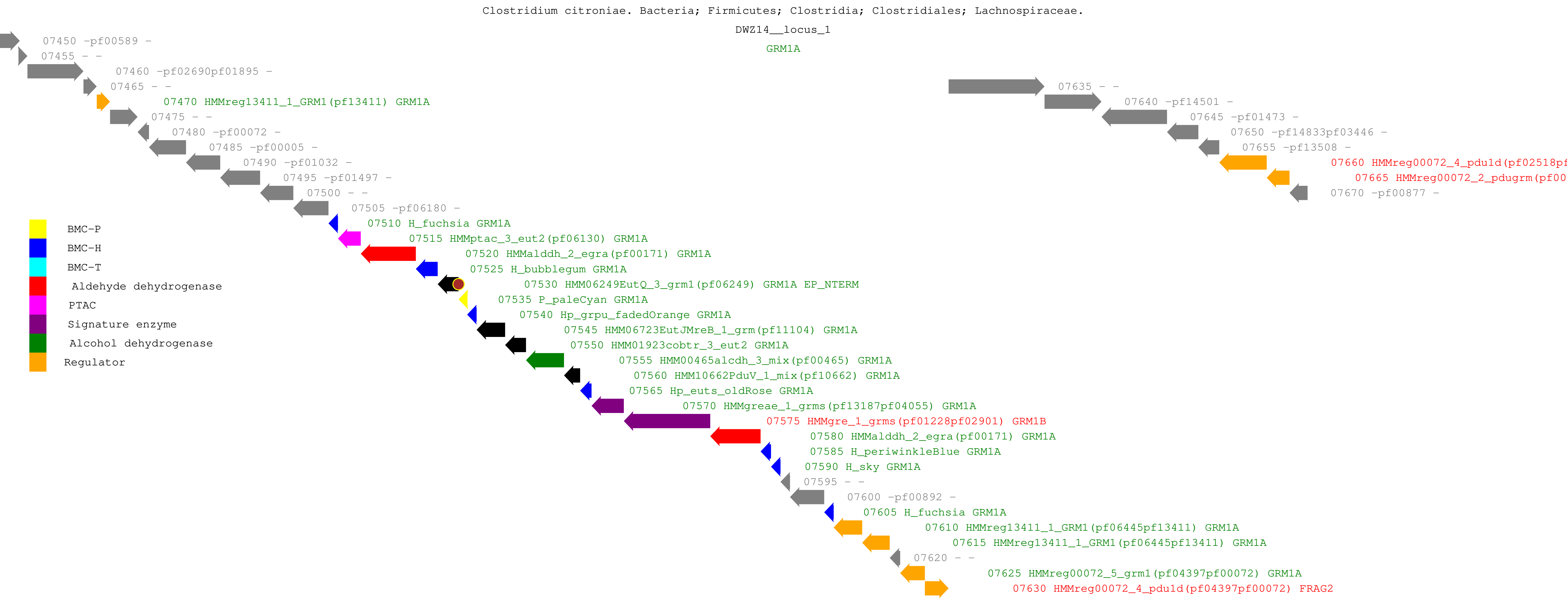
Clostridium citroniae. Bacteria; Firmicutes; Clostridia; Clostridiales; Lachnospiraceae.
DWZ14__locus_1
GRM1A
07450 -pf00589 -
07455 - -
07460 -pf02690pf01895 -
07465 - -
07470 HMMreg13411_1_GRM1(pf13411) GRM1A
07475 - -
07480 -pf00072 -
07485 -pf00005 -
07490 -pf01032 -
07495 -pf01497 -
07500 - -
07505 -pf06180 -
07510 H_fuchsia GRM1A
07515 HMMptac_3_eut2(pf06130) GRM1A
07520 HMMalddh_2_egra(pf00171) GRM1A
07525 H_bubblegum GRM1A
07530 HMM06249EutQ_3_grm1(pf06249) GRM1A EP_NTERM
07535 P_paleCyan GRM1A
07540 Hp_grpu_fadedOrange GRM1A
07545 HMM06723EutJMreB_1_grm(pf11104) GRM1A
07550 HMM01923cobtr_3_eut2 GRM1A
07555 HMM00465alcdh_3_mix(pf00465) GRM1A
07560 HMM10662PduV_1_mix(pf10662) GRM1A
07565 Hp_euts_oldRose GRM1A
07570 HMMgreae_1_grms(pf13187pf04055) GRM1A
07575 HMMgre_1_grms(pf01228pf02901) GRM1B
07580 HMMaldlh_2_egra(pf00171) GRM1A
07585 H_periwinkleBlue GRM1A
07590 H_sky GRM1A
07595 - -
07600 -pf00892 -
07605 H_fuchsia GRM1A
07610 HMMreg13411_1_GRM1(pf06445pf13411) GRM1A
07615 HMMreg13411_1_GRM1(pf06445pf13411) GRM1A
07620 - -
07625 HMMreg00072_5_grm1(pf04397pf00072) GRM1A
07630 HMMreg00072_4_pdu1d(pf04397pf00072) FRAG2
07635 - -
07640 -pf14501 -
07645 -pf01473 -
07650 -pf14833pf03446 -
07655 -pf13508 -
07660 HMMreg00072_4_pdu1d(pf02518pf
07665 HMMreg00072_2_pdugrm(pf00
07670 -pf00877 -
BMC-P
BMC-H
BMC-T
Aldehyde dehydrogenase
PTAC
Signature enzyme
Alcohol dehydrogenase
Regulator

Melissococcus sp. OM08-11BH. Bacteria; Firmicutes; Bacilli; Lactobacillales; Enterococcaceae;

DXC12__locus_1

GRM1A

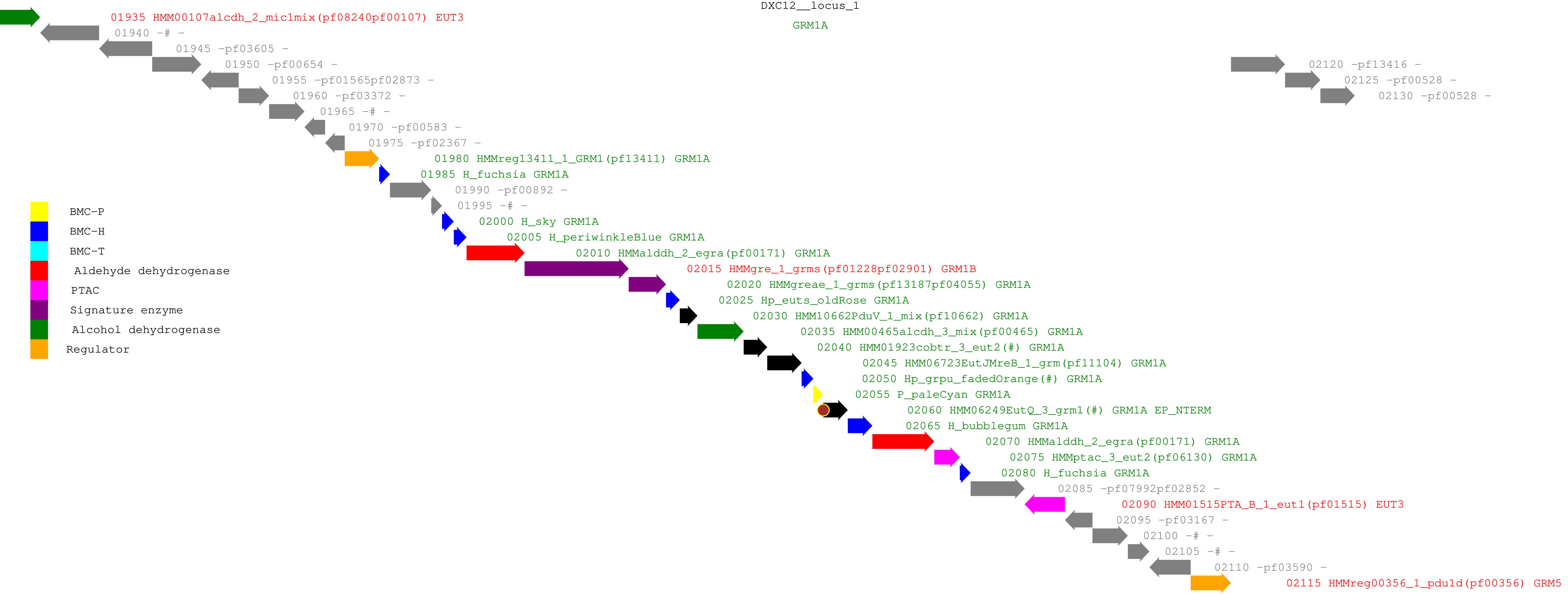

Hungatella hathewayi. Bacteria; Firmicutes; Clostridia; Clostridiales; Clostridiaceae;

DXC39__locus_1

GRM1A

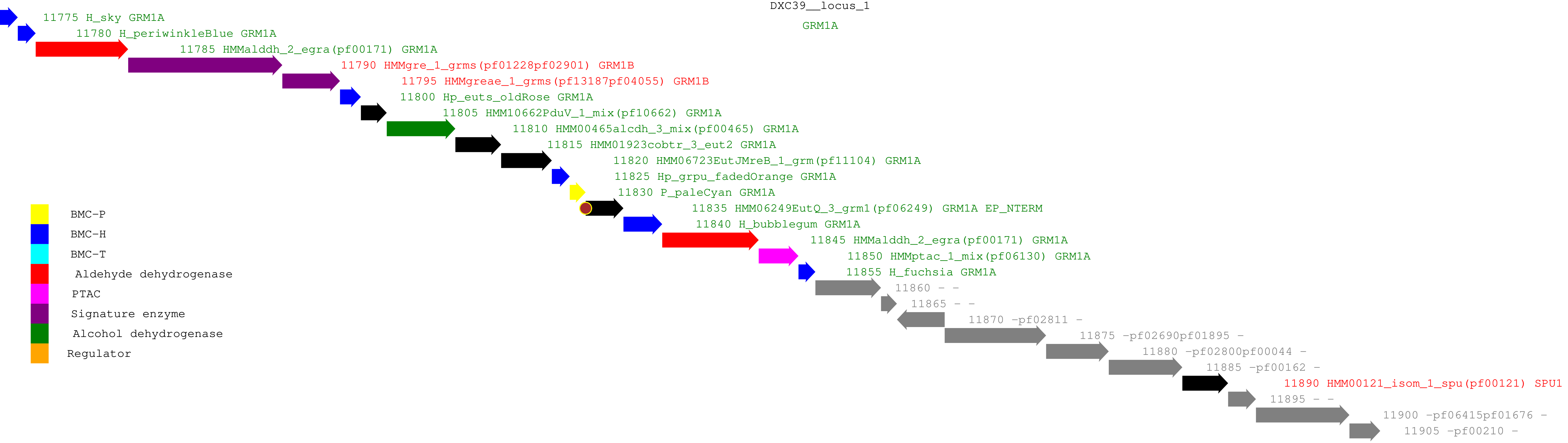

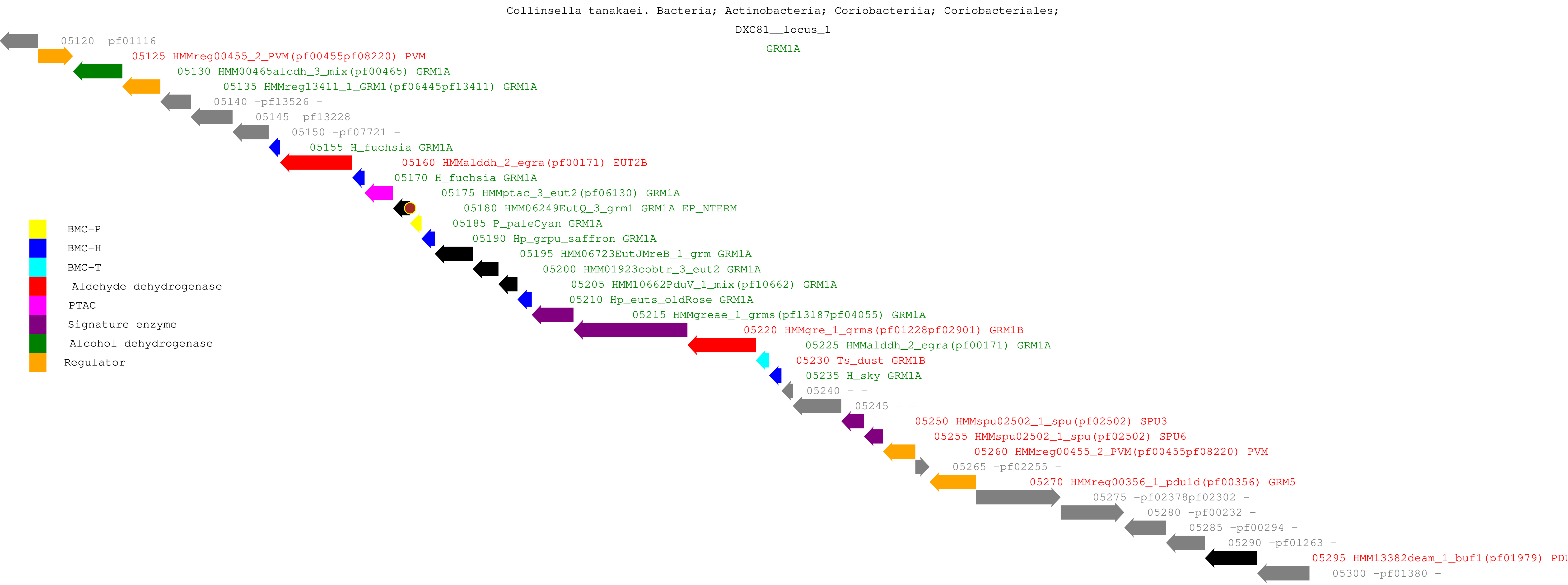
Collinsella tanakaei. Bacteria; Actinobacteria; Coriobacteriia; Coriobacteriales;
DXC81__locus_1
GRM1A
05120 -pf01116 -
05125 HMMreg00455_2_PVM(pf00455pf08220) PVM
05130 HMM00465alcdh_3_mix(pf00465) GRM1A
05135 HMMreg13411_1_GRM1(pf06445pf13411) GRM1A
05140 -pf13526 -
05145 -pf13228 -
05150 -pf07721 -
05155 H_fuchsia GRM1A
05160 HMMaldh_2_egra(pf00171) EUT2B
05170 H_fuchsia GRM1A
05175 HMMptac_3_eut2(pf06130) GRM1A
05180 HMM06249EutQ_3_grm1 GRM1A EP_NTERM
05185 P_paleCyan GRM1A
05190 Hp_grpu_saffron GRM1A
05195 HMM06723EutJMreB_1_grm GRM1A
05200 HMM01923cobtr_3_eut2 GRM1A
05205 HMM10662PduV_1_mix(pf10662) GRM1A
05210 Hp_euts_oldRose GRM1A
05215 HMMgreae_1_grms(pf13187pf04055) GRM1A
05220 HMMgre_1_grms(pf01228pf02901) GRM1B
05225 HMMaldh_2_egra(pf00171) GRM1A
05230 Ts_dust GRM1B
05235 H_sky GRM1A
05240 - -
05245 - -
05250 HMMspu02502_1_spu(pf02502) SPU3
05255 HMMspu02502_1_spu(pf02502) SPU6
05260 HMMreg00455_2_PVM(pf00455pf08220) PVM
05265 -pf02255 -
05270 HMMreg00356_1_pdu1d(pf00356) GRM5
05275 -pf02378pf02302 -
05280 -pf00232 -
05285 -pf00294 -
05290 -pf01263 -
05295 HMM13382deam_1_buf1(pf01979) PDU
05300 -pf01380 -
BMC-P
BMC-H
BMC-T
Aldehyde dehydrogenase
PTAC
Signature enzyme
Alcohol dehydrogenase
Regulator

Clostridiales bacterium TF09-2AC. Bacteria; Firmicutes; Clostridia; Clostridiales.

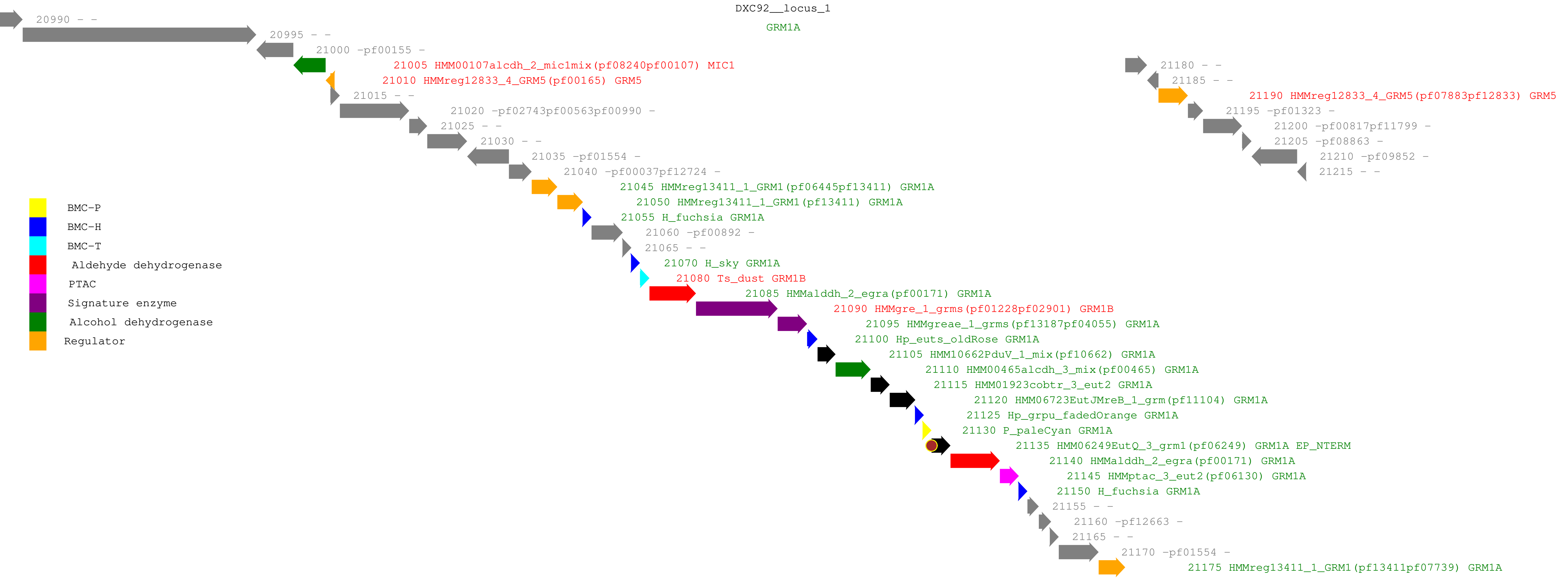

Clostridium botulinum. Bacteria; Firmicutes; Clostridia; Clostridiales; Clostridiaceae;

DZC34__locus_1_main_sat

GRM1A

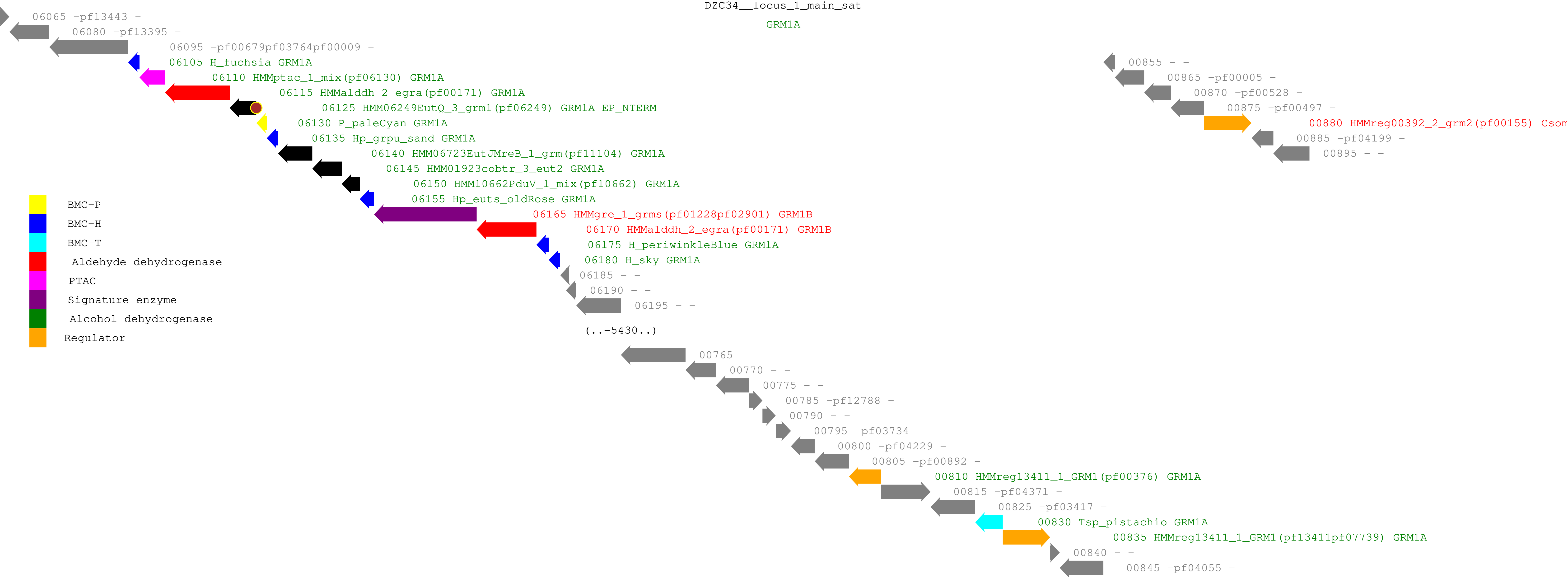

Paenibacillus dendritiformis. Bacteria; Firmicutes; Bacilli; Bacillales; Paenibacillaceae; Paenibacillus.

E2R60__locus_2

GRM1A

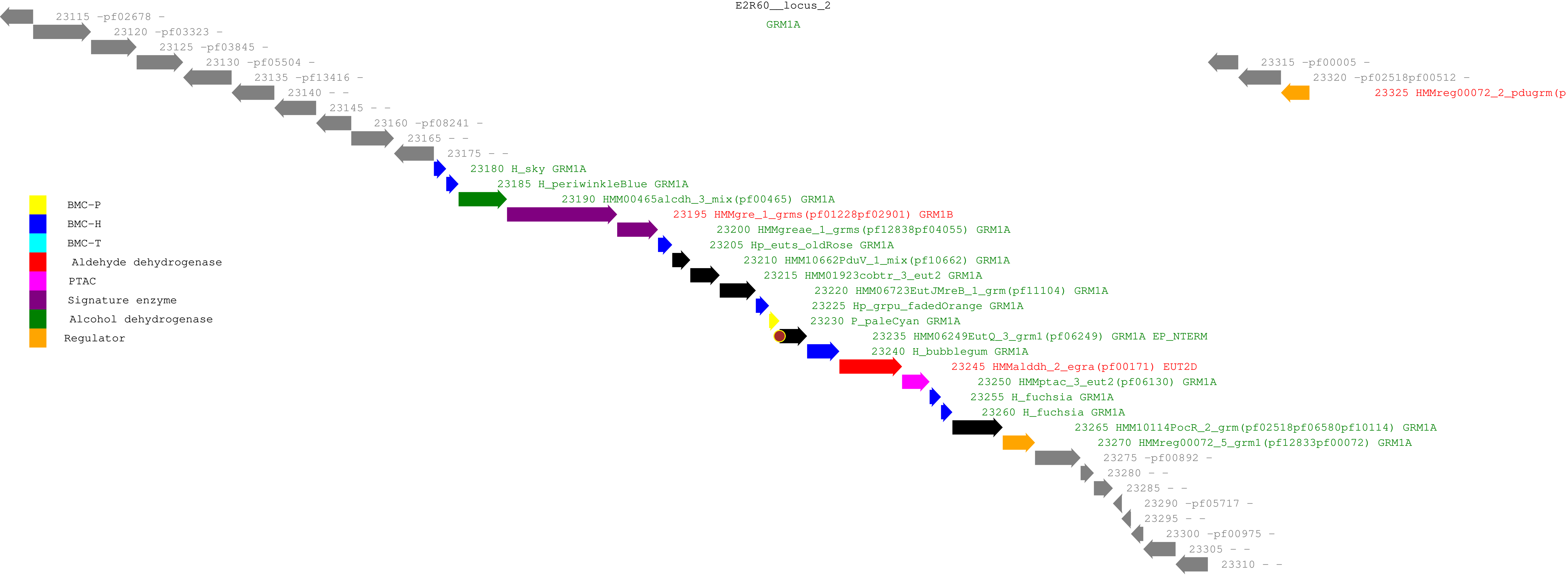

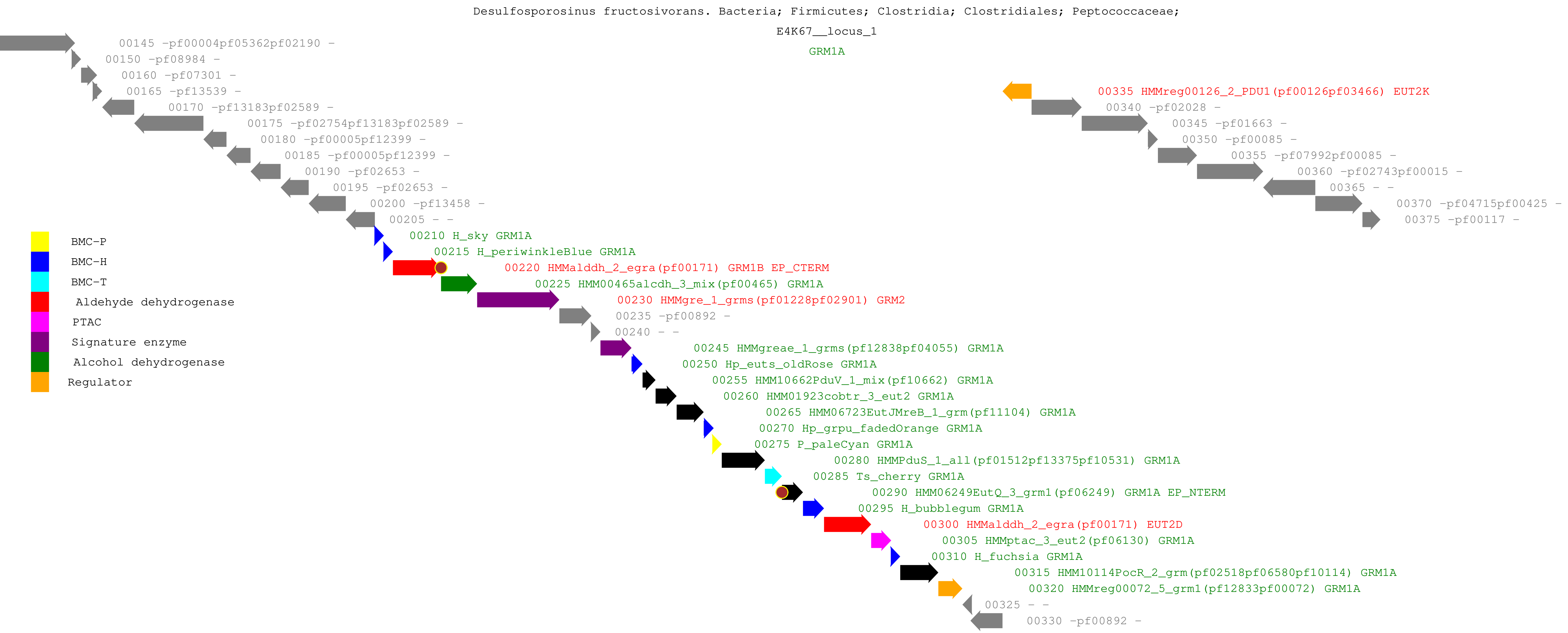
Desulfosporosinus fructosivorans. Bacteria; Firmicutes; Clostridia; Clostridiales; Peptococcaceae;
E4K67__locus_1
GRM1A
00145 -pf00004pf05362pf02190 -
00150 -pf08984 -
00160 -pf07301 -
00165 -pf13539 -
00170 -pf13183pf02589 -
00175 -pf02754pf13183pf02589 -
00180 -pf00005pf12399 -
00185 -pf00005pf12399 -
00190 -pf02653 -
00195 -pf02653 -
00200 -pf13458 -
00205 - -
00210 H_sky GRM1A
00215 H_periwinkleBlue GRM1A
00220 HMMaldddh_2_egra(pf00171) GRM1B EP_CTERM
00225 HMM00465alcdh_3_mix(pf00465) GRM1A
00230 HMMgre_1_grms(pf01228pf02901) GRM2
00235 -pf00892 -
00240 - -
00245 HMMgreae_1_grms(pf12838pf04055) GRM1A
00250 Hp_euts_oldRose GRM1A
00255 HMM10662PduV_1_mix(pf10662) GRM1A
00260 HMM01923cobtr_3_eut2 GRM1A
00265 HMM06723EutJMreB_1_grm(pf11104) GRM1A
00270 Hp_grpu_fadedOrange GRM1A
00275 P_paleCyan GRM1A
00280 HMMPduS_1_all(pf01512pf13375pf10531) GRM1A
00285 Ts_cherry GRM1A
00290 HMM06249EutQ_3_grm1(pf06249) GRM1A EP_NTERM
00295 H_bubblegum GRM1A
00300 HMMaldddh_2_egra(pf00171) EUT2D
00305 HMMptac_3_eut2(pf06130) GRM1A
00310 H_fuchsia GRM1A
00315 HMM10114PocR_2_grm(pf02518pf06580pf10114) GRM1A
00320 HMMreg00072_5_grm1(pf12833pf00072) GRM1A
00325 - -
00330 -pf00892 -
00335 HMMreg00126_2_PDU1(pf00126pf03466) EUT2K
00340 -pf02028 -
00345 -pf01663 -
00350 -pf00085 -
00355 -pf07992pf00085 -
00360 -pf02743pf00015 -
00365 - -
00370 -pf04715pf00425 -
00375 -pf00117 -
BMC-P
BMC-H
BMC-T
Aldehyde dehydrogenase
PTAC
Signature enzyme
Alcohol dehydrogenase
Regulator

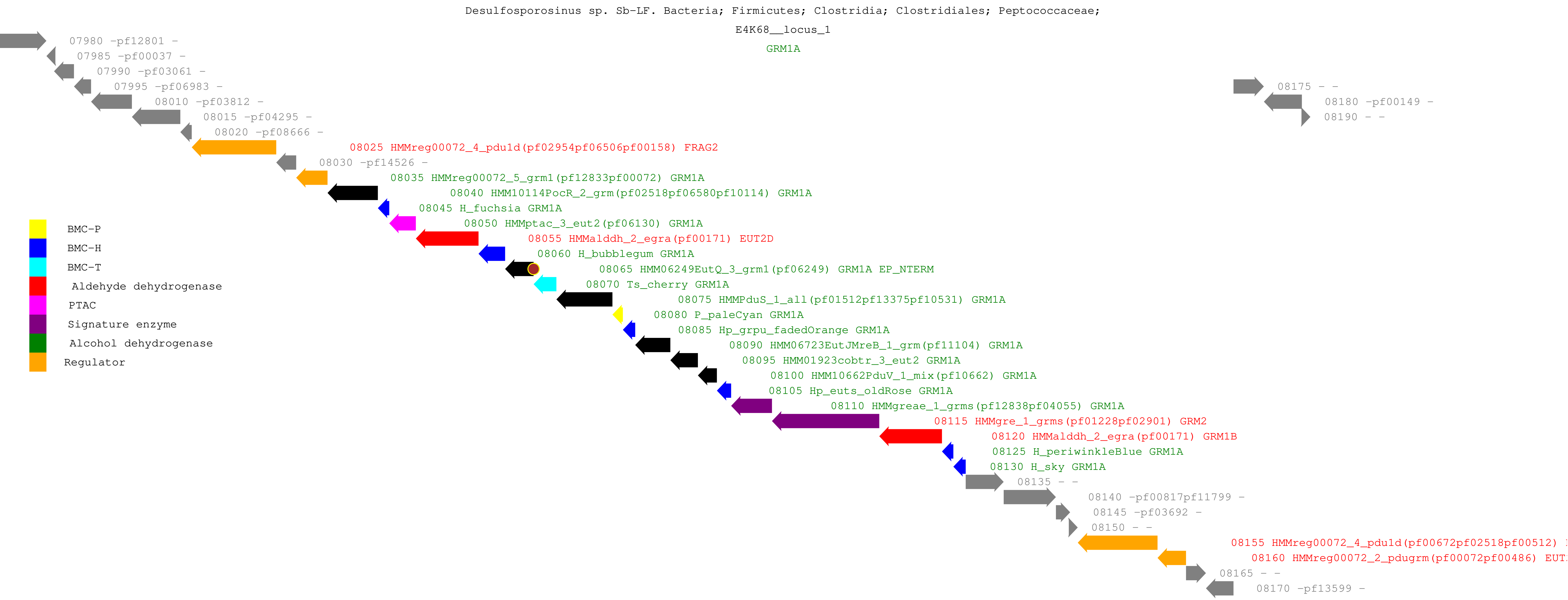

Desulfosporosinus sp. Sb-LF. Bacteria; Firmicutes; Clostridia; Clostridiales; Peptococcaceae;
E4K68__locus_1
GRM1A
07980 -pf12801 -
07985 -pf00037 -
07990 -pf03061 -
07995 -pf06983 -
08010 -pf03812 -
08015 -pf04295 -
08020 -pf08666 -
08025 HMMreg00072_4_pdu1d(pf02954pf06506pf00158) FRAG2
08030 -pf14526 -
08035 HMMreg00072_5_grm1(pf12833pf00072) GRM1A
08040 HMM10114PocR_2_grm(pf02518pf06580pf10114) GRM1A
08045 H_fuchsia GRM1A
08050 HMMptac_3_eut2(pf06130) GRM1A
08055 HMMaldh_2_egra(pf00171) EUT2D
08060 H_bubblegum GRM1A
08065 HMM06249EutQ_3_grm1(pf06249) GRM1A EP_NTERM
08070 Ts_cherry GRM1A
08075 HMMPduS_1_all(pf01512pf13375pf10531) GRM1A
08080 P_paleCyan GRM1A
08085 Hp_grpu_fadedOrange GRM1A
08090 HMM06723EutJMreB_1_grm(pf11104) GRM1A
08095 HMM01923cobtr_3_eut2 GRM1A
08100 HMM10662PduV_1_mix(pf10662) GRM1A
08105 Hp_euts_oldRose GRM1A
08110 HMMgreae_1_grms(pf12838pf04055) GRM1A
08115 HMMgre_1_grms(pf01228pf02901) GRM2
08120 HMMaldh_2_egra(pf00171) GRM1B
08125 H_periwinkleBlue GRM1A
08130 H_sky GRM1A
08135 - -
08140 -pf00817pf11799 -
08145 -pf03692 -
08150 - -
08155 HMMreg00072_4_pdu1d(pf00672pf02518pf00512)
08160 HMMreg00072_2_pdugrm(pf00072pf00486) EUT
08165 - -
08170 -pf13599 -
08175 - -
08180 -pf00149 -
08190 - -
BMC-P
BMC-H
BMC-T
Aldehyde dehydrogenase
PTAC
Signature enzyme
Alcohol dehydrogenase
Regulator

Streptococcus sp. LYSM12. Bacteria; Firmicutes; Bacilli; Lactobacillales; Streptococcaceae;

E4T79__locus_1

GRM1A

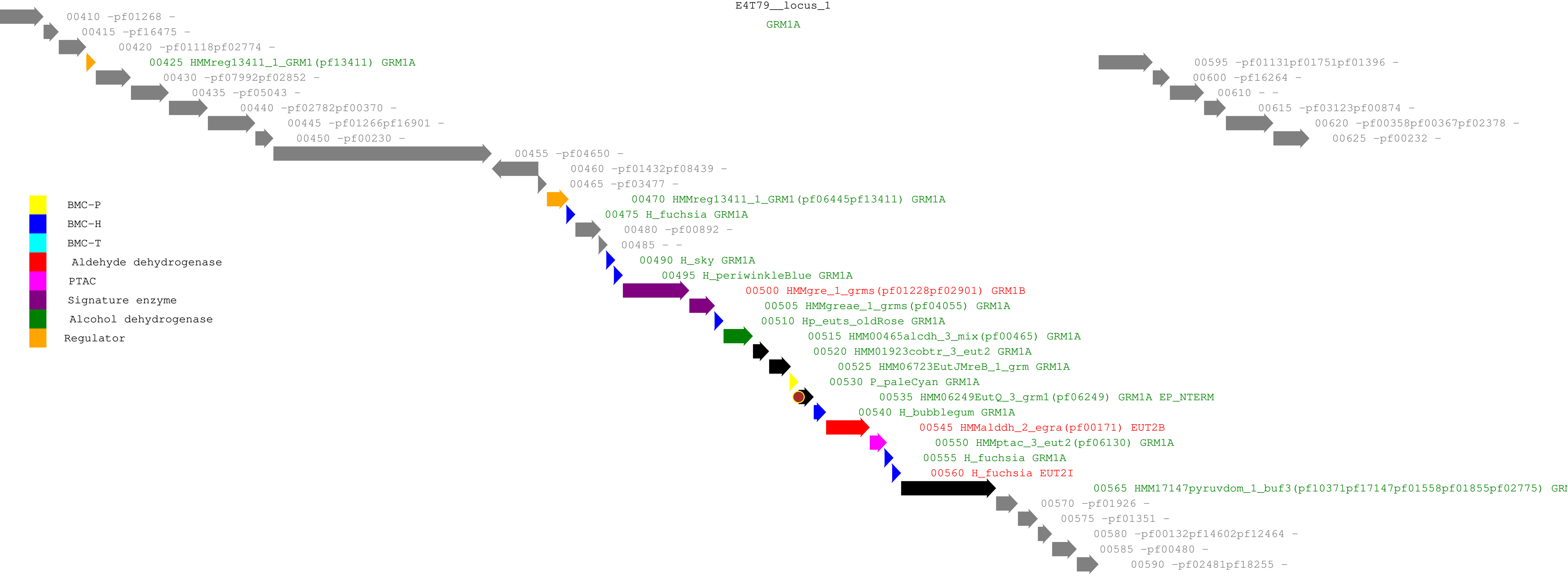

Fournierella massiliensis. Bacteria; Firmicutes; Clostridia; Clostridiales; Ruminococcaceae;

EDD77__locus_1_main_sat

GRM1A

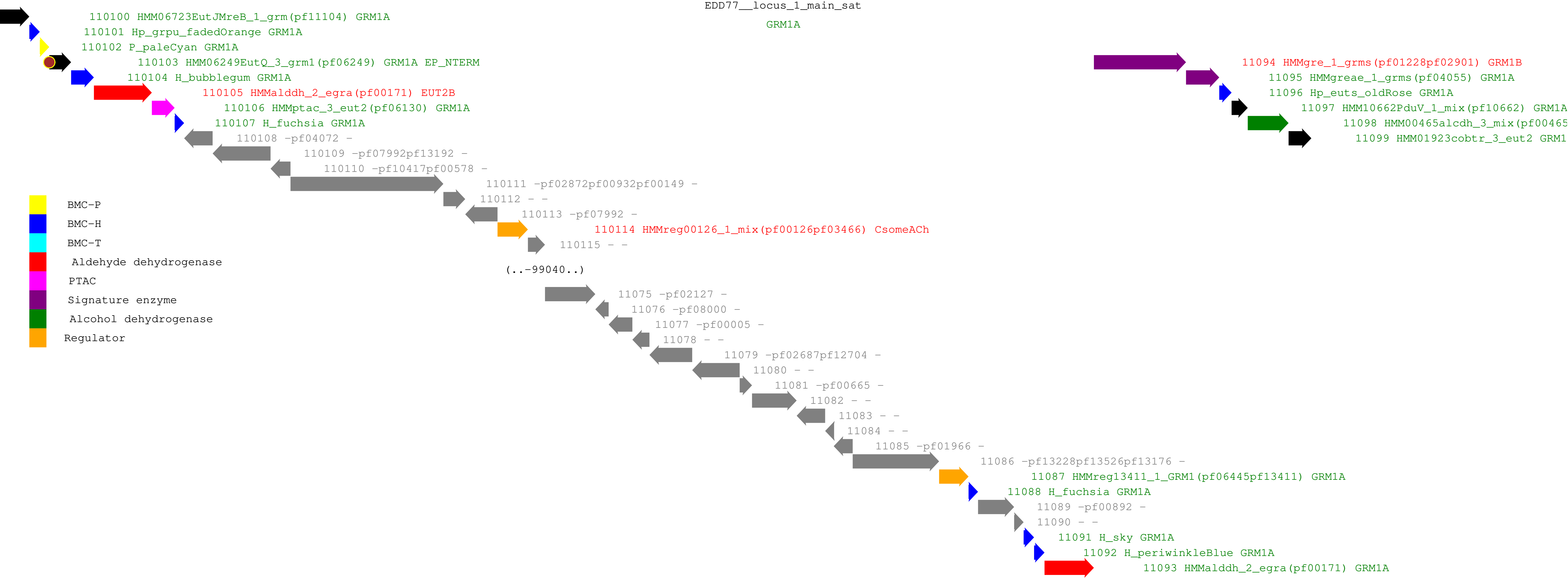

Enterococcus gilvus. Bacteria; Firmicutes; Bacilli; Lactobacillales; Enterococcaceae;

EGCR1__locus_1

GRM1A

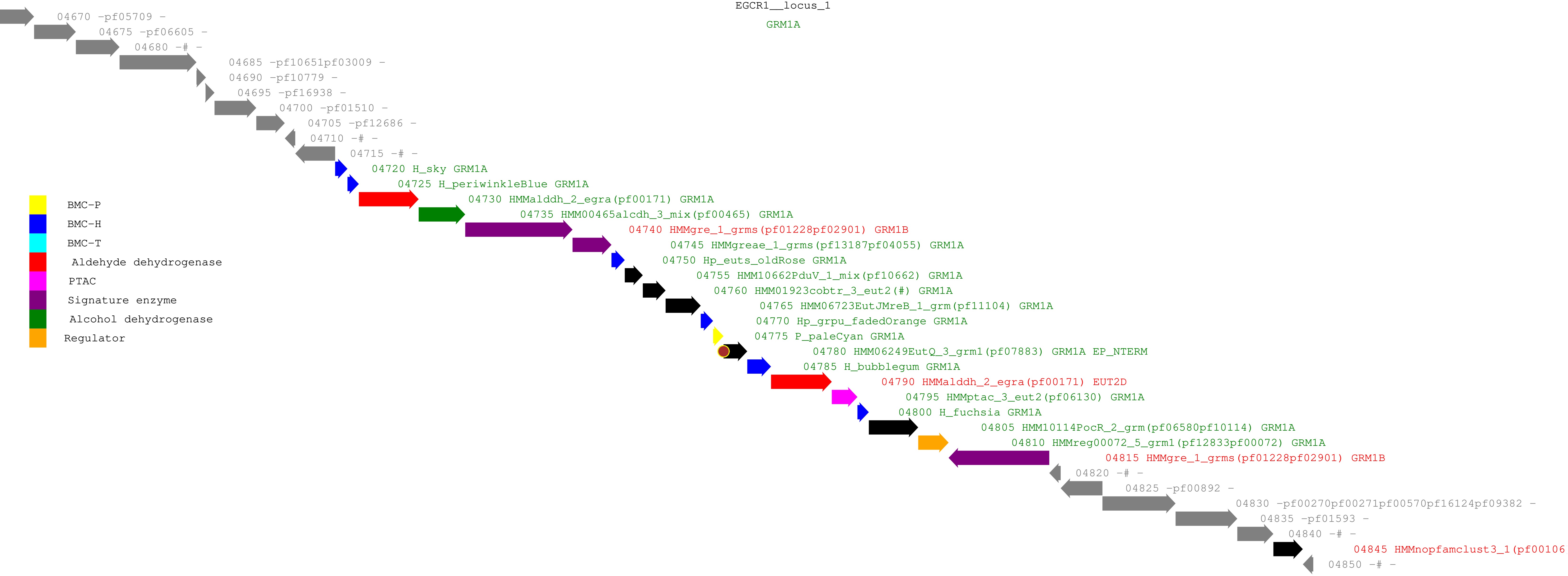

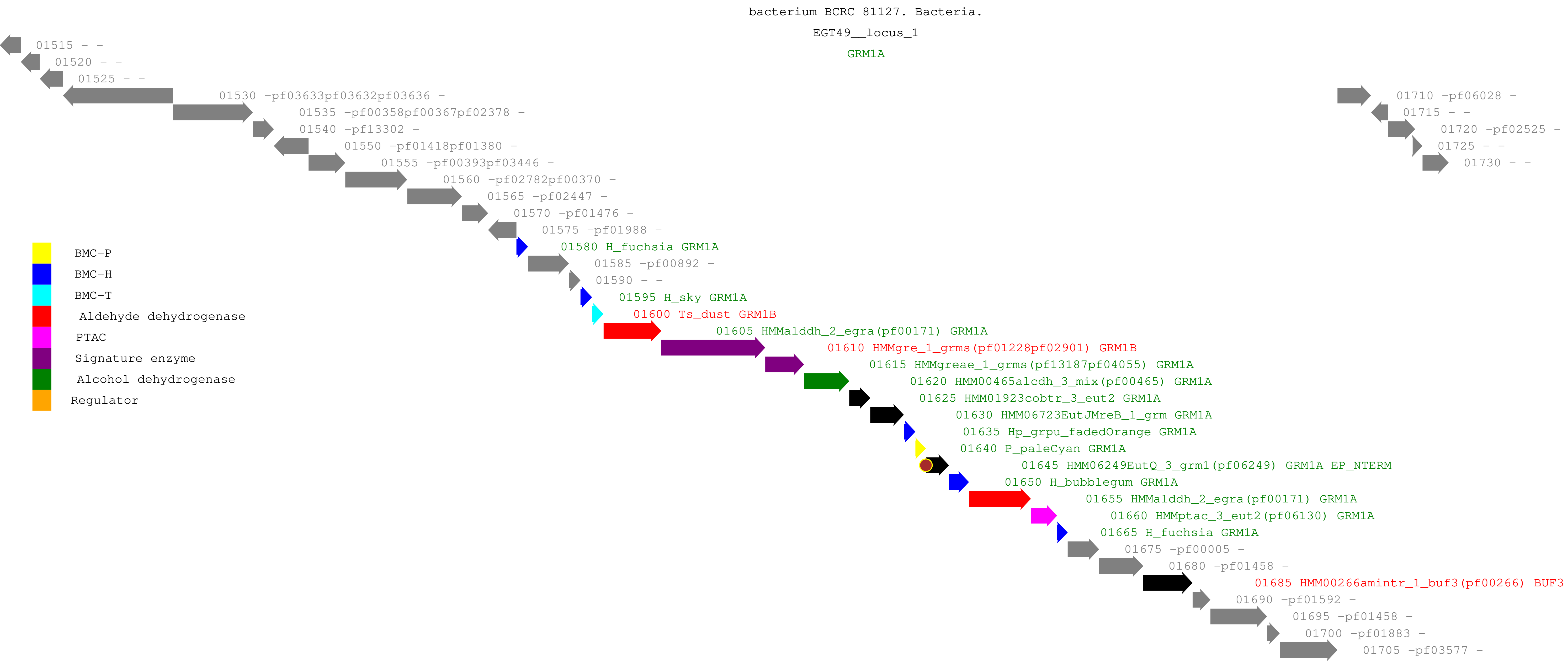
bacterium BCRC 81127. Bacteria.
EGT49__locus_1
GRM1A
01515 - -
01520 - -
01525 - -
01530 -pf03633pf03632pf03636 -
01535 -pf00358pf00367pf02378 -
01540 -pf13302 -
01550 -pf01418pf01380 -
01555 -pf00393pf03446 -
01560 -pf02782pf00370 -
01565 -pf02447 -
01570 -pf01476 -
01575 -pf01988 -
01580 H_fuchsia GRM1A
01585 -pf00892 -
01590 - -
01595 H_sky GRM1A
01600 Ts_dust GRM1B
01605 HMMalddh_2_egra(pf00171) GRM1A
01610 HMMgre_1_grms(pf01228pf02901) GRM1B
01615 HMMgreae_1_grms(pf13187pf04055) GRM1A
01620 HMM00465alcdh_3_mix(pf00465) GRM1A
01625 HMM01923cobtr_3_eut2 GRM1A
01630 HMM06723EutJMreB_1_grm GRM1A
01635 Hp_grpu_fadedOrange GRM1A
01640 P_paleCyan GRM1A
01645 HMM06249EutQ_3_grm1(pf06249) GRM1A EP_NTERM
01650 H_bubblegum GRM1A
01655 HMMalddh_2_egra(pf00171) GRM1A
01660 HMMptac_3_eut2(pf06130) GRM1A
01665 H_fuchsia GRM1A
01675 -pf00005 -
01680 -pf01458 -
01685 HMM00266amintr_1_buf3(pf00266) BUF3
01690 -pf01592 -
01695 -pf01458 -
01700 -pf01883 -
01705 -pf03577 -
01710 -pf06028 -
01715 - -
01720 -pf02525 -
01725 - -
01730 - -
BMC-P
BMC-H
BMC-T
Aldehyde dehydrogenase
PTAC
Signature enzyme
Alcohol dehydrogenase
Regulator

Enterococcus avium (Streptococcus avium). Bacteria; Firmicutes; Bacilli; Lactobacillales; Enterococcaceae;

EH197__locus_1

GRM1A

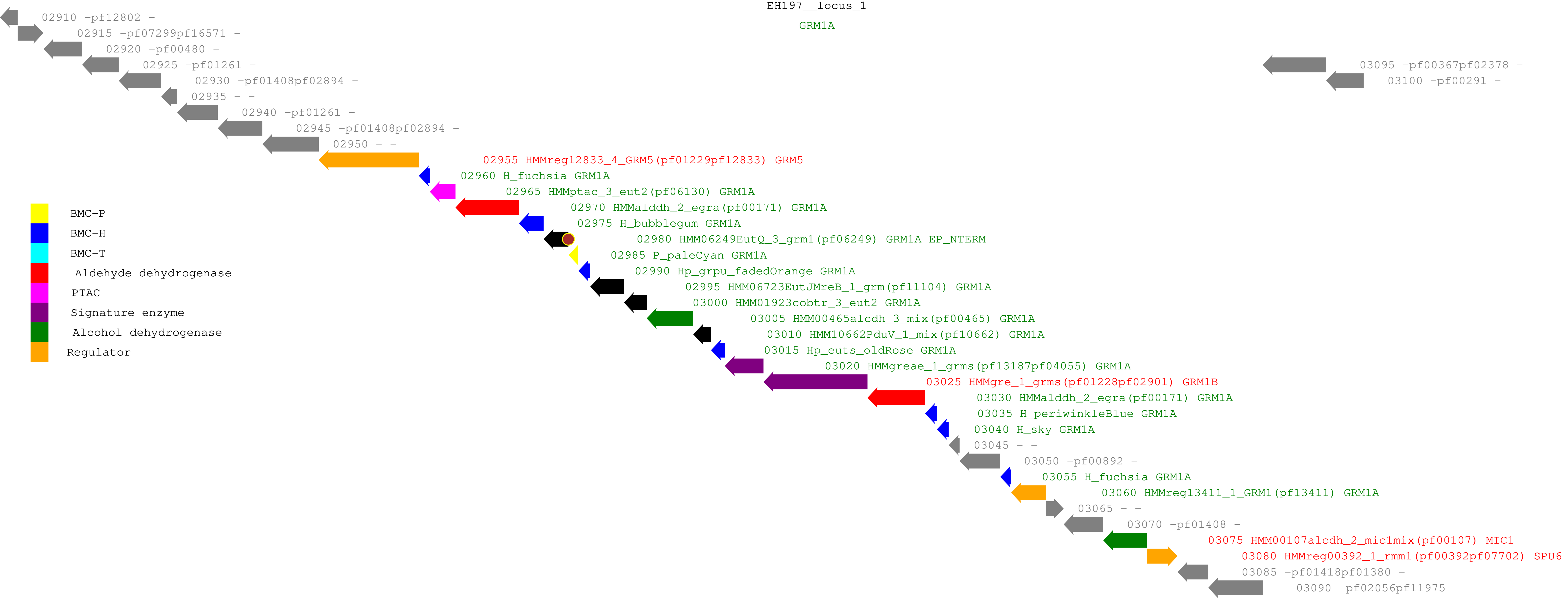

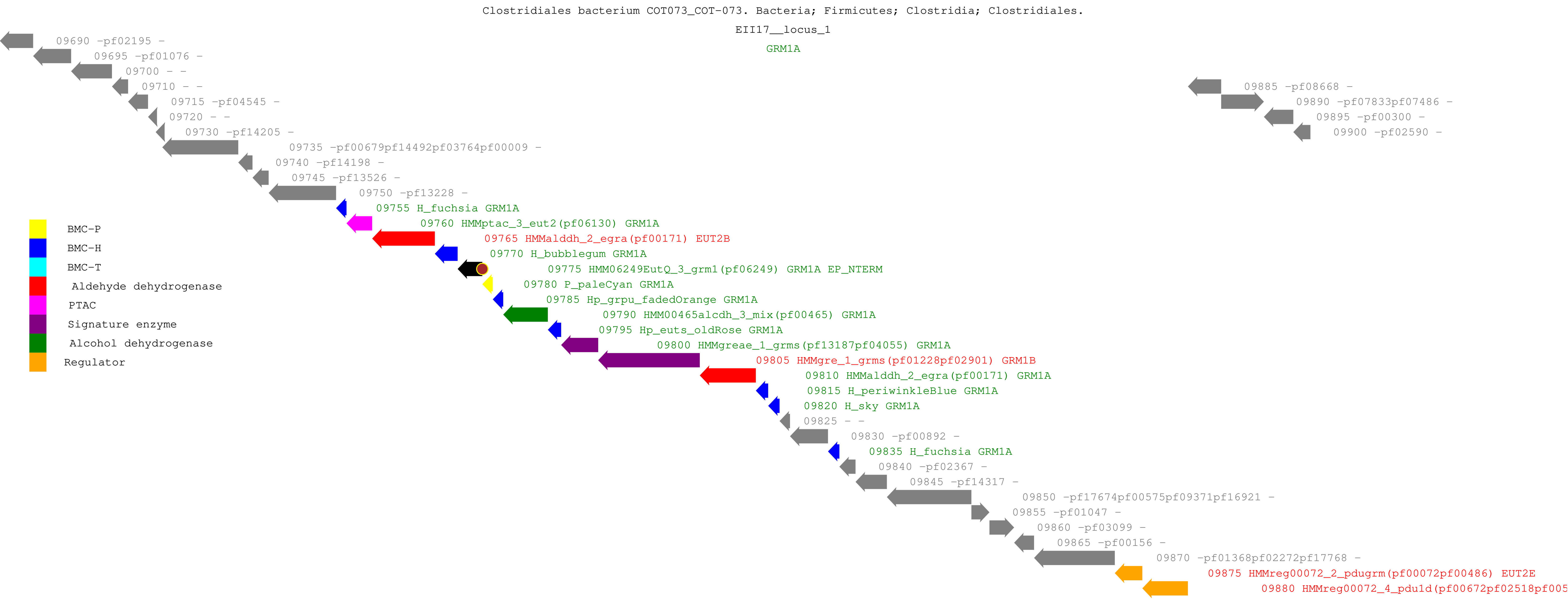
Clostridiales bacterium COT073_COT-073. Bacteria; Firmicutes; Clostridia; Clostridiales.
EII17__locus_1
GRM1A
09690 -pf02195 -
09695 -pf01076 -
09700 - -
09710 - -
09715 -pf04545 -
09720 - -
09730 -pf14205 -
09735 -pf00679pf14492pf03764pf00009 -
09740 -pf14198 -
09745 -pf13526 -
09750 -pf13228 -
09755 H_fuchsia GRM1A
09760 HMMptac_3_eut2(pf06130) GRM1A
09765 HMMalddh_2_egra(pf00171) EUT2B
09770 H_bubblegum GRM1A
09775 HMM06249EutQ_3_grm1(pf06249) GRM1A EP_NTERM
09780 P_paleCyan GRM1A
09785 Hp_grpu_fadedOrange GRM1A
09790 HMM00465alcdh_3_mix(pf00465) GRM1A
09795 Hp_euts_oldRose GRM1A
09800 HMMgreae_1_grms(pf13187pf04055) GRM1A
09805 HMMgre_1_grms(pf01228pf02901) GRM1B
09810 HMMalddh_2_egra(pf00171) GRM1A
09815 H_periwinkleBlue GRM1A
09820 H_sky GRM1A
09825 - -
09830 -pf00892 -
09835 H_fuchsia GRM1A
09840 -pf02367 -
09845 -pf14317 -
09850 -pf17674pf00575pf09371pf16921 -
09855 -pf01047 -
09860 -pf03099 -
09865 -pf00156 -
09870 -pf01368pf02272pf17768 -
09875 HMMreg00072_2_pdugrm(pf00072pf00486) EUT2E
09880 HMMreg00072_4_pdu1d(pf00672pf02518pf005
09885 -pf08668 -
09890 -pf07833pf07486 -
09895 -pf00300 -
09900 -pf02590 -
BMC-P
BMC-H
BMC-T
Aldehyde dehydrogenase
PTAC
Signature enzyme
Alcohol dehydrogenase
Regulator

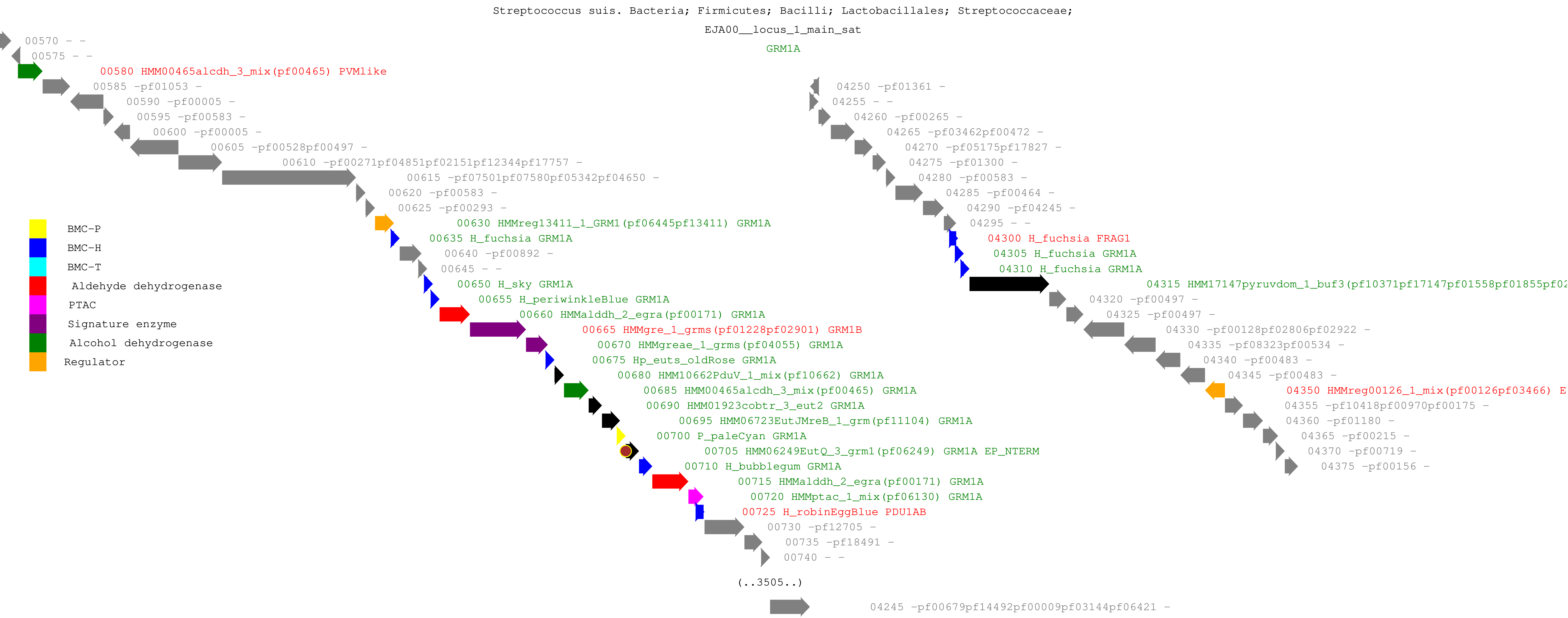
Streptococcus suis. Bacteria; Firmicutes; Bacilli; Lactobacillales; Streptococcaceae;
EJA00__locus_1_main_sat
GRM1A
00570 - -
00575 - -
00580 HMM00465alcdh_3_mix(pf00465) PVMlike
00585 -pf01053 -
00590 -pf00005 -
00595 -pf00583 -
00600 -pf00005 -
00605 -pf00528pf00497 -
00610 -pf00271pf04851pf02151pf12344pf17757 -
00615 -pf07501pf07580pf05342pf04650 -
00620 -pf00583 -
00625 -pf00293 -
00630 HMMreg13411_1_GRM1(pf06445pf13411) GRM1A
00635 H_fuchsia GRM1A
00640 -pf00892 -
00645 - -
00650 H_sky GRM1A
00655 H_periwinkleBlue GRM1A
00660 HMMalddh_2_egra(pf00171) GRM1A
00665 HMMgre_1_grms(pf01228pf02901) GRM1B
00670 HMMgreae_1_grms(pf04055) GRM1A
00675 Hp_euts_oldRose GRM1A
00680 HMM10662PduV_1_mix(pf10662) GRM1A
00685 HMM00465alcdh_3_mix(pf00465) GRM1A
00690 HMM01923cobtr_3_eut2 GRM1A
00695 HMM06723EutJMreB_1_grm(pf11104) GRM1A
00700 P_paleCyan GRM1A
00705 HMM06249EutQ_3_grm1(pf06249) GRM1A EP_NTERM
00710 H_bubblegum GRM1A
00715 HMMalddh_2_egra(pf00171) GRM1A
00720 HMMptac_1_mix(pf06130) GRM1A
00725 H_robinEggBlue PDU1AB
00730 -pf12705 -
00735 -pf18491 -
00740 - -
(..3505..)
04245 -pf00679pf14492pf00009pf03144pf06421 -
04250 -pf01361 -
04255 - -
04260 -pf00265 -
04265 -pf03462pf00472 -
04270 -pf05175pf17827 -
04275 -pf01300 -
04280 -pf00583 -
04285 -pf00464 -
04290 -pf04245 -
04295 - -
04300 H_fuchsia FRAG1
04305 H_fuchsia GRM1A
04310 H_fuchsia GRM1A
04315 HMM17147pyruvdom_1_buf3(pf10371pf17147pf01558pf01855pf02
04320 -pf00497 -
04325 -pf00497 -
04330 -pf00128pf02806pf02922 -
04335 -pf08323pf00534 -
04340 -pf00483 -
04345 -pf00483 -
04350 HMMreg00126_1_mix(pf00126pf03466) E
04355 -pf10418pf00970pf00175 -
04360 -pf01180 -
04365 -pf00215 -
04370 -pf00719 -
04375 -pf00156 -
BMC-P
BMC-H
BMC-T
Aldehyde dehydrogenase
PTAC
Signature enzyme
Alcohol dehydrogenase
Regulator

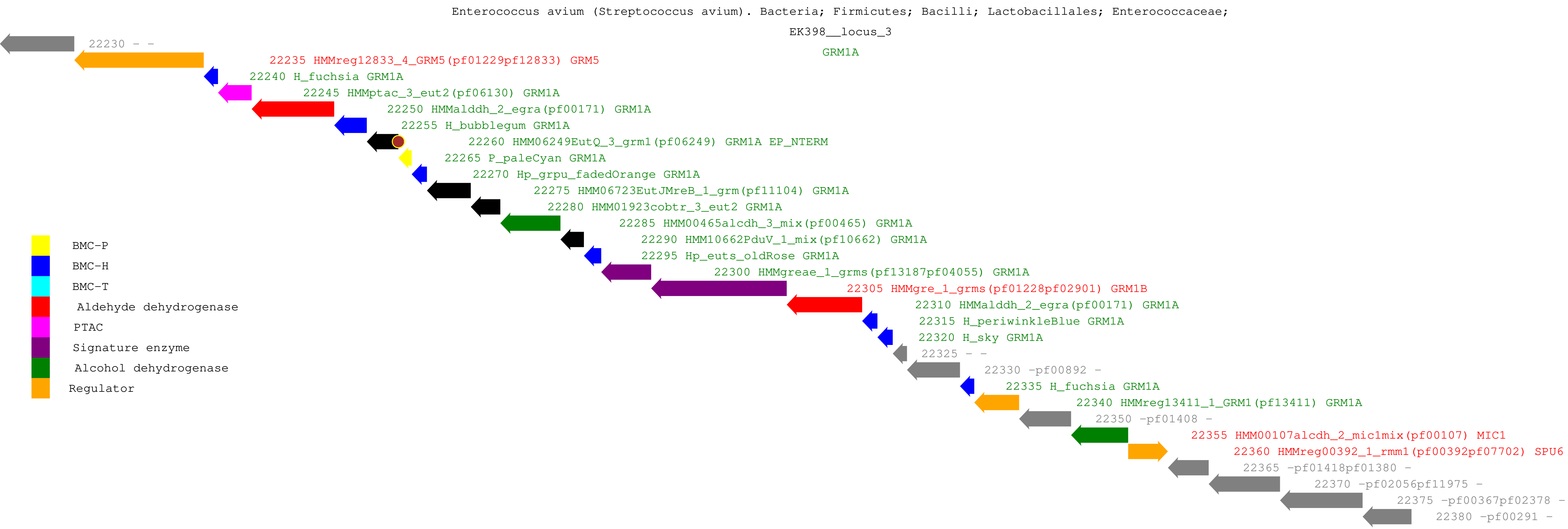

Enterococcus avium (Streptococcus avium). Bacteria; Firmicutes; Bacilli; Lactobacillales; Enterococcaceae;
EK398__locus_3
GRM1A
22230 - -
22235 HMMreg12833_4_GRM5(pf01229pf12833) GRM5
22240 H_fuchsia GRM1A
22245 HMMptac_3_eut2(pf06130) GRM1A
22250 HMMalddh_2_egra(pf00171) GRM1A
22255 H_bubblegum GRM1A
22260 HMM06249EutQ_3_grm1(pf06249) GRM1A EP_NTERM
22265 P_paleCyan GRM1A
22270 Hp_grpu_fadedOrange GRM1A
22275 HMM06723EutJMreB_1_grm(pf11104) GRM1A
22280 HMM01923cobtr_3_eut2 GRM1A
22285 HMM00465alcdh_3_mix(pf00465) GRM1A
22290 HMM10662PduV_1_mix(pf10662) GRM1A
22295 Hp_euts_oldRose GRM1A
22300 HMMgreae_1_grms(pf13187pf04055) GRM1A
22305 HMMgre_1_grms(pf01228pf02901) GRM1B
22310 HMMalddh_2_egra(pf00171) GRM1A
22315 H_periwinkleBlue GRM1A
22320 H_sky GRM1A
22325 - -
22330 -pf00892 -
22335 H_fuchsia GRM1A
22340 HMMreg13411_1_GRM1(pf13411) GRM1A
22350 -pf01408 -
22355 HMM00107alcdh_2_mic1mix(pf00107) MIC1
22360 HMMreg00392_1_rmm1(pf00392pf07702) SPU6
22365 -pf01418pf01380 -
22370 -pf02056pf11975 -
22375 -pf00367pf02378 -
22380 -pf00291 -
BMC-P
BMC-H
BMC-T
Aldehyde dehydrogenase
PTAC
Signature enzyme
Alcohol dehydrogenase
Regulator

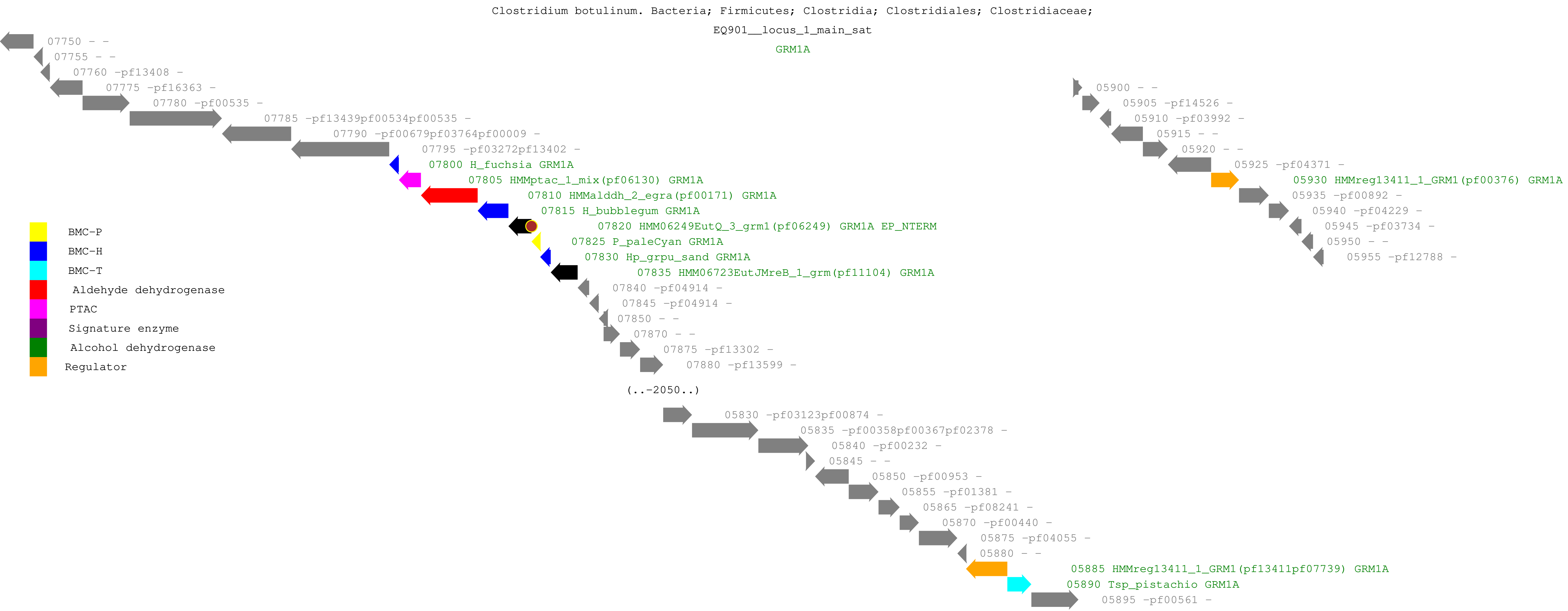
Clostridium botulinum. Bacteria; Firmicutes; Clostridia; Clostridiales; Clostridiaceae;
EQ901__locus_1_main_sat
GRM1A
07750 - -
07755 - -
07760 -pf13408 -
07775 -pf16363 -
07780 -pf00535 -
07785 -pf13439pf00534pf00535 -
07790 -pf00679pf03764pf00009 -
07795 -pf03272pf13402 -
07800 H_fuchsia GRM1A
07805 HMMptac_1_mix(pf06130) GRM1A
07810 HMMalddh_2_egra(pf00171) GRM1A
07815 H_bubblegum GRM1A
07820 HMM06249EutQ_3_grm1(pf06249) GRM1A EP_NTERM
07825 P_paleCyan GRM1A
07830 Hp_grpu_sand GRM1A
07835 HMM06723EutJMreB_1_grm(pf11104) GRM1A
07840 -pf04914 -
07845 -pf04914 -
07850 - -
07870 - -
07875 -pf13302 -
07880 -pf13599 -
(..-2050..)
05830 -pf03123pf00874 -
05835 -pf00358pf00367pf02378 -
05840 -pf00232 -
05845 - -
05850 -pf00953 -
05855 -pf01381 -
05865 -pf08241 -
05870 -pf00440 -
05875 -pf04055 -
05880 - -
05885 HMMreg13411_1_GRM1(pf13411pf07739) GRM1A
05890 Tsp_pistachio GRM1A
05895 -pf00561 -
05900 - -
05905 -pf14526 -
05910 -pf03992 -
05915 - -
05920 - -
05925 -pf04371 -
05930 HMMreg13411_1_GRM1(pf00376) GRM1A
05935 -pf00892 -
05940 -pf04229 -
05945 -pf03734 -
05950 - -
05955 -pf12788 -
BMC-P
BMC-H
BMC-T
Aldehyde dehydrogenase
PTAC
Signature enzyme
Alcohol dehydrogenase
Regulator

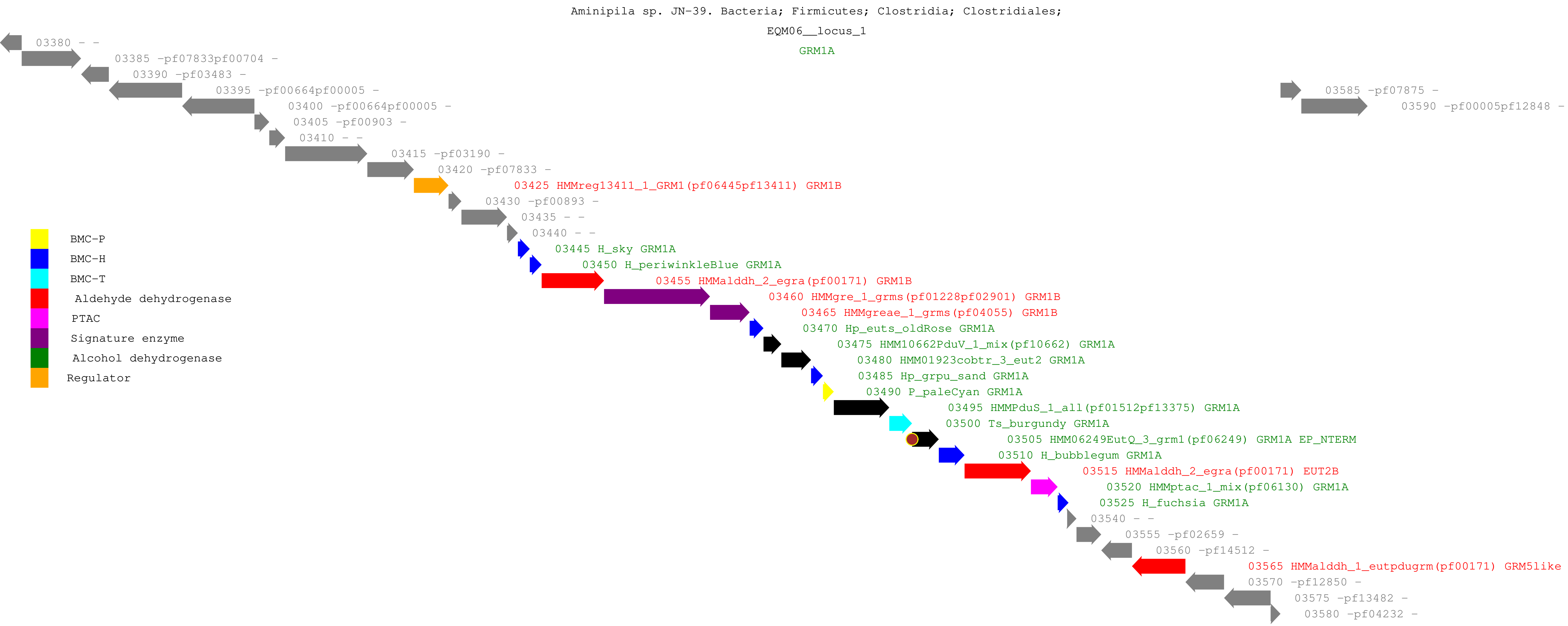
Aminipila sp. JN-39. Bacteria; Firmicutes; Clostridia; Clostridiales;
EQM06__locus_1
GRM1A
03380 - -
03385 -pf07833pf00704 -
03390 -pf03483 -
03395 -pf00664pf00005 -
03400 -pf00664pf00005 -
03405 -pf00903 -
03410 - -
03415 -pf03190 -
03420 -pf07833 -
03425 HMMreg13411_1_GRM1(pf06445pf13411) GRM1B
03430 -pf00893 -
03435 - -
03440 - -
03445 H_sky GRM1A
03450 H_periwinkleBlue GRM1A
03455 HMMaldh_2_egra(pf00171) GRM1B
03460 HMMgre_1_grms(pf01228pf02901) GRM1B
03465 HMMgreae_1_grms(pf04055) GRM1B
03470 Hp_euts_oldRose GRM1A
03475 HMM10662PduV_1_mix(pf10662) GRM1A
03480 HMM01923cobtr_3_eut2 GRM1A
03485 Hp_grpu_sand GRM1A
03490 P_paleCyan GRM1A
03495 HMMPduS_1_all(pf01512pf13375) GRM1A
03500 Ts_burgundy GRM1A
03505 HMM06249EutQ_3_grm1(pf06249) GRM1A EP_NTERM
03510 H_bubblegum GRM1A
03515 HMMaldh_2_egra(pf00171) EUT2B
03520 HMMptac_1_mix(pf06130) GRM1A
03525 H_fuchsia GRM1A
03540 - -
03555 -pf02659 -
03560 -pf14512 -
03565 HMMaldh_1_eutpdugrm(pf00171) GRM5like
03570 -pf12850 -
03575 -pf13482 -
03580 -pf04232 -
03585 -pf07875 -
03590 -pf00005pf12848 -
BMC-P
BMC-H
BMC-T
Aldehyde dehydrogenase
PTAC
Signature enzyme
Alcohol dehydrogenase
Regulator

Streptococcus suis. Bacteria; Firmicutes; Bacilli; Lactobacillales; Streptococcaceae;

ERS132399__locus_1_main_sat

GRM1A

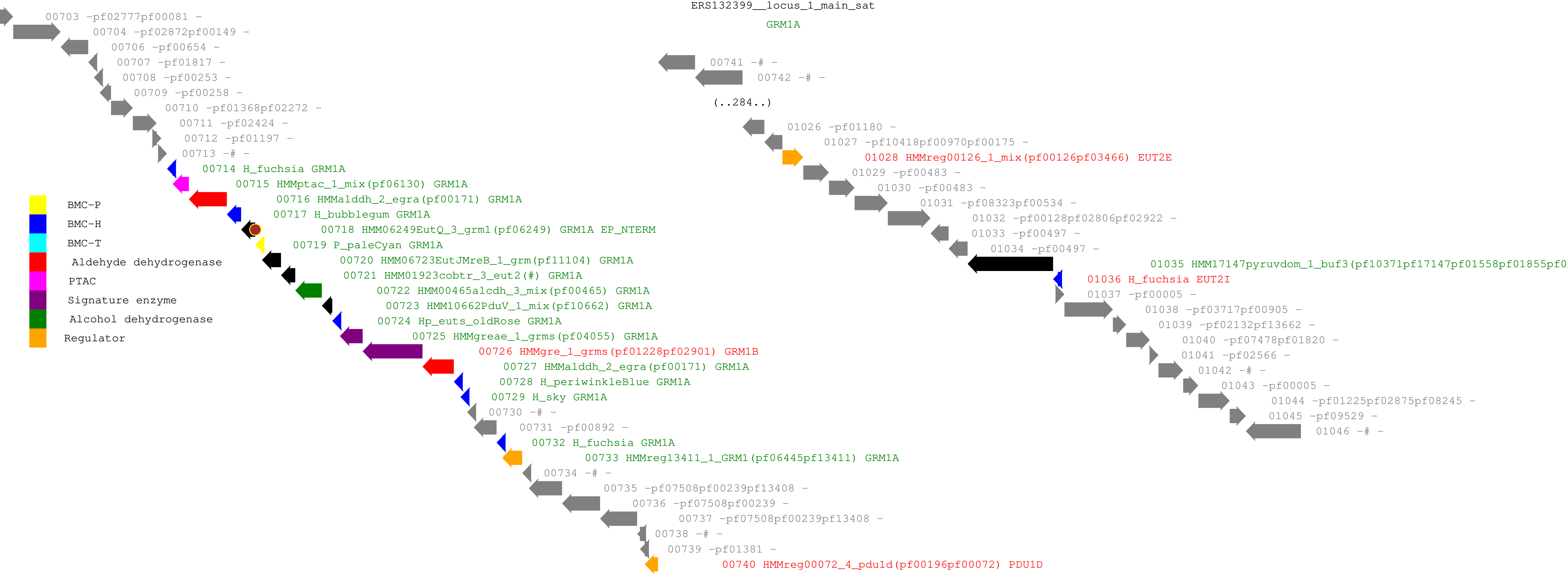

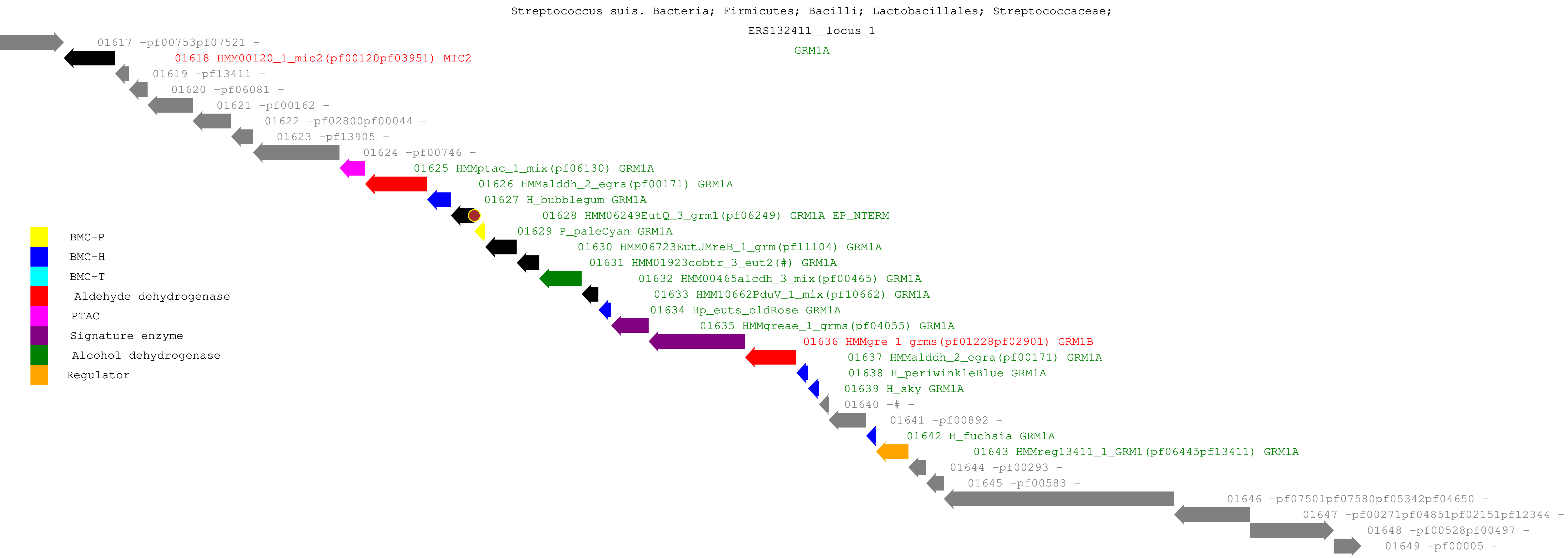
Streptococcus suis. Bacteria; Firmicutes; Bacilli; Lactobacillales; Streptococcaceae;
ERS132411__locus_1
GRM1A
01617 -pf00753pf07521 -
01618 HMM00120_1_mic2(pf00120pf03951) MIC2
01619 -pf13411 -
01620 -pf06081 -
01621 -pf00162 -
01622 -pf02800pf00044 -
01623 -pf13905 -
01624 -pf00746 -
01625 HMMptac_1_mix(pf06130) GRM1A
01626 HMMalddh_2_egra(pf00171) GRM1A
01627 H_bubblegum GRM1A
01628 HMM06249EutQ_3_grm1(pf06249) GRM1A EP_NTERM
01629 P_paleCyan GRM1A
01630 HMM06723EutJMreB_1_grm(pf11104) GRM1A
01631 HMM01923cobtr_3_eut2(#) GRM1A
01632 HMM00465alcdh_3_mix(pf00465) GRM1A
01633 HMM10662PduV_1_mix(pf10662) GRM1A
01634 Hp_euts_oldRose GRM1A
01635 HMMgreae_1_grms(pf04055) GRM1A
01636 HMMgre_1_grms(pf01228pf02901) GRM1B
01637 HMMalddh_2_egra(pf00171) GRM1A
01638 H_periwinkleBlue GRM1A
01639 H_sky GRM1A
01640 -# -
01641 -pf00892 -
01642 H_fuchsia GRM1A
01643 HMMreg13411_1_GRM1(pf06445pf13411) GRM1A
01644 -pf00293 -
01645 -pf00583 -
01646 -pf07501pf07580pf05342pf04650 -
01647 -pf00271pf04851pf02151pf12344 -
01648 -pf00528pf00497 -
01649 -pf00005 -
BMC-P
BMC-H
BMC-T
Aldehyde dehydrogenase
PTAC
Signature enzyme
Alcohol dehydrogenase
Regulator

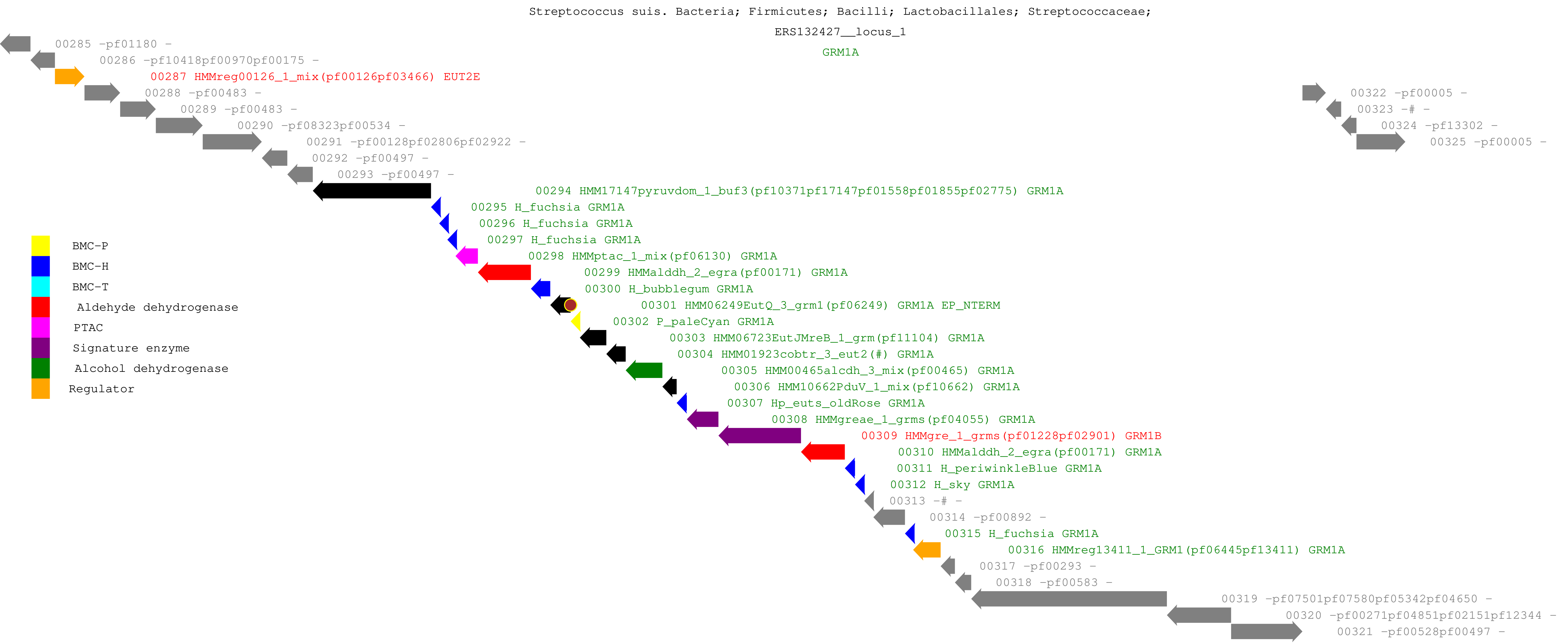
Streptococcus suis. Bacteria; Firmicutes; Bacilli; Lactobacillales; Streptococcaceae;
ERS132427__locus_1
GRM1A
00285 -pf01180 -
00286 -pf10418pf00970pf00175 -
00287 HMMreg00126_1_mix(pf00126pf03466) EUT2E
00288 -pf00483 -
00289 -pf00483 -
00290 -pf08323pf00534 -
00291 -pf00128pf02806pf02922 -
00292 -pf00497 -
00293 -pf00497 -
00294 HMM17147pyruvdom_1_buf3(pf10371pf17147pf01558pf01855pf02775) GRM1A
00295 H_fuchsia GRM1A
00296 H_fuchsia GRM1A
00297 H_fuchsia GRM1A
00298 HMMptac_1_mix(pf06130) GRM1A
00299 HMMalddh_2_egra(pf00171) GRM1A
00300 H_bubblegum GRM1A
00301 HMM06249EutQ_3_grm1(pf06249) GRM1A EP_NTERM
00302 P_paleCyan GRM1A
00303 HMM06723EutJMreB_1_grm(pf11104) GRM1A
00304 HMM01923cobtr_3_eut2(#) GRM1A
00305 HMM00465alcdh_3_mix(pf00465) GRM1A
00306 HMM10662PduV_1_mix(pf10662) GRM1A
00307 Hp_euts_oldRose GRM1A
00308 HMMgreae_1_grms(pf04055) GRM1A
00309 HMMgre_1_grms(pf01228pf02901) GRM1B
00310 HMMalddh_2_egra(pf00171) GRM1A
00311 H_periwinkleBlue GRM1A
00312 H_sky GRM1A
00313 -# -
00314 -pf00892 -
00315 H_fuchsia GRM1A
00316 HMMreg13411_1_GRM1(pf06445pf13411) GRM1A
00317 -pf00293 -
00318 -pf00583 -
00319 -pf07501pf07580pf05342pf04650 -
00320 -pf00271pf04851pf02151pf12344 -
00321 -pf00528pf00497 -
00322 -pf00005 -
00323 -# -
00324 -pf13302 -
00325 -pf00005 -
BMC-P
BMC-H
BMC-T
Aldehyde dehydrogenase
PTAC
Signature enzyme
Alcohol dehydrogenase
Regulator

Streptococcus suis. Bacteria; Firmicutes; Bacilli; Lactobacillales; Streptococcaceae;

ERS132452__locus_1

GRM1A

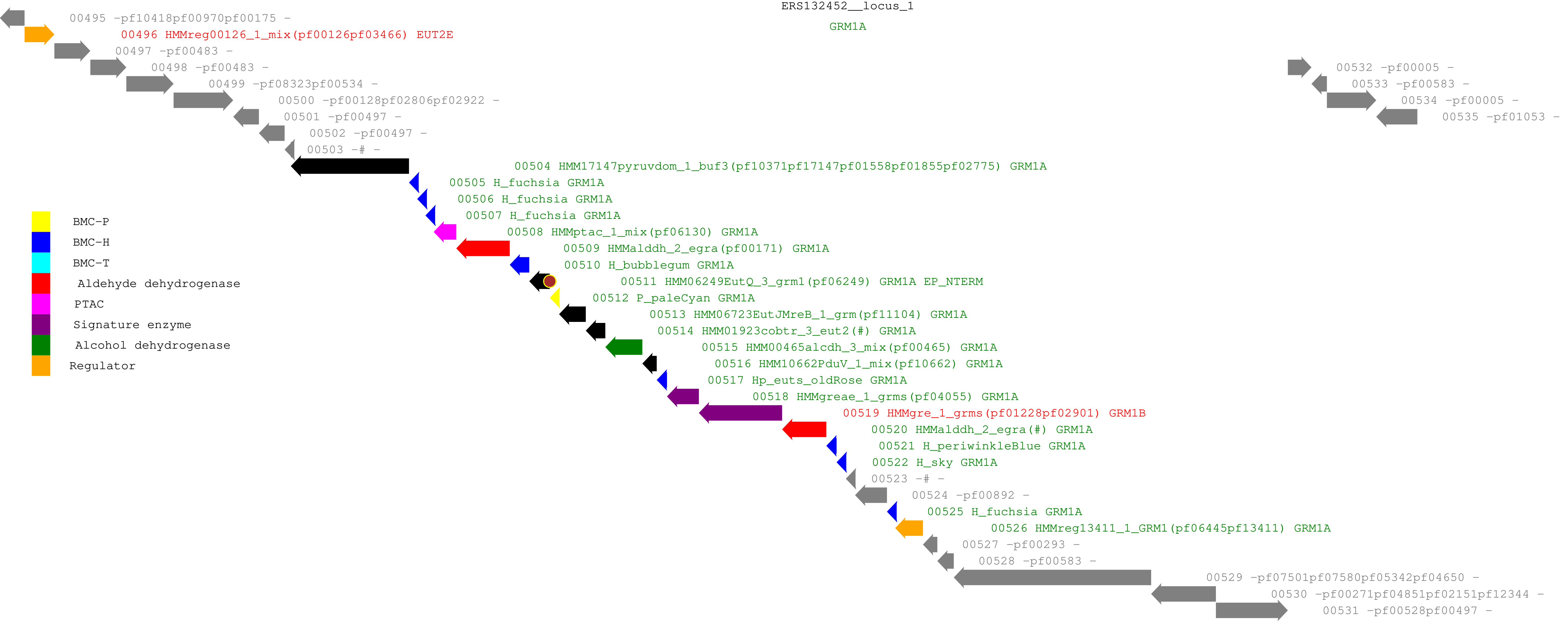

Streptococcus suis. Bacteria; Firmicutes; Bacilli; Lactobacillales; Streptococcaceae;

ERS132457__locus_1

GRM1A

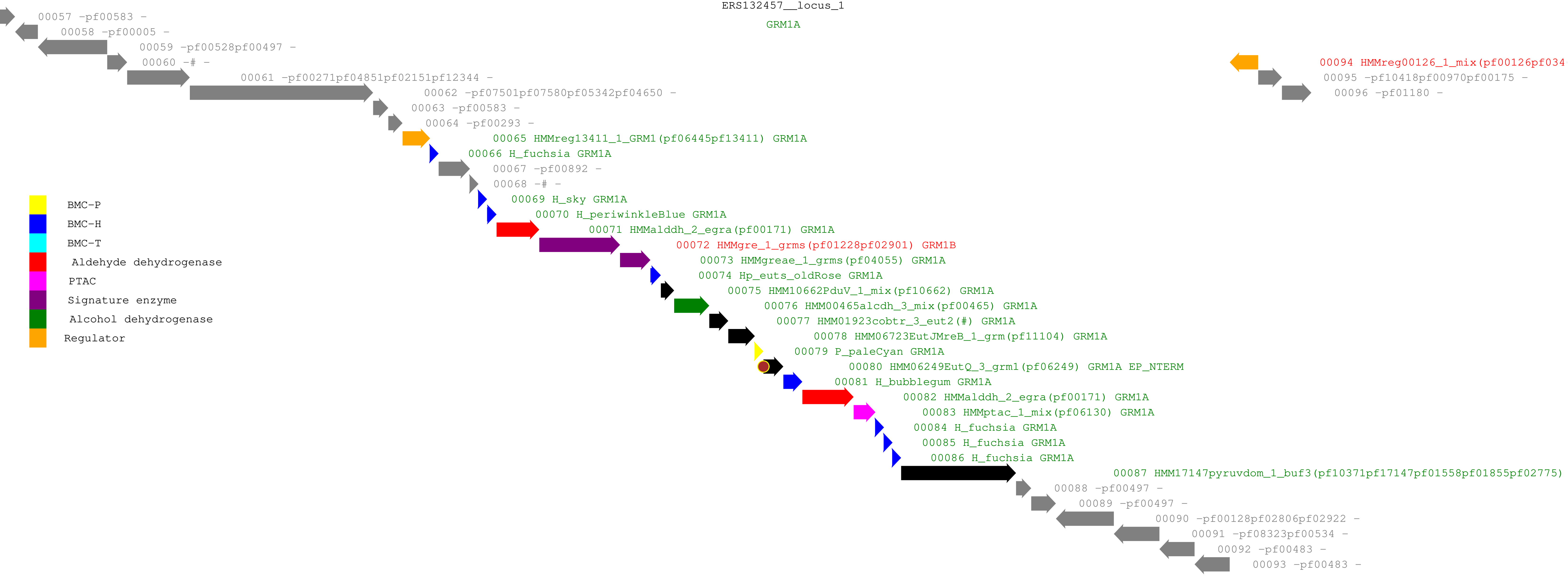

Chlamydia trachomatis. Bacteria; Chlamydiae; Chlamydiales; Chlamydiaceae;

ERS133248__locus_fragmented

GRM1A

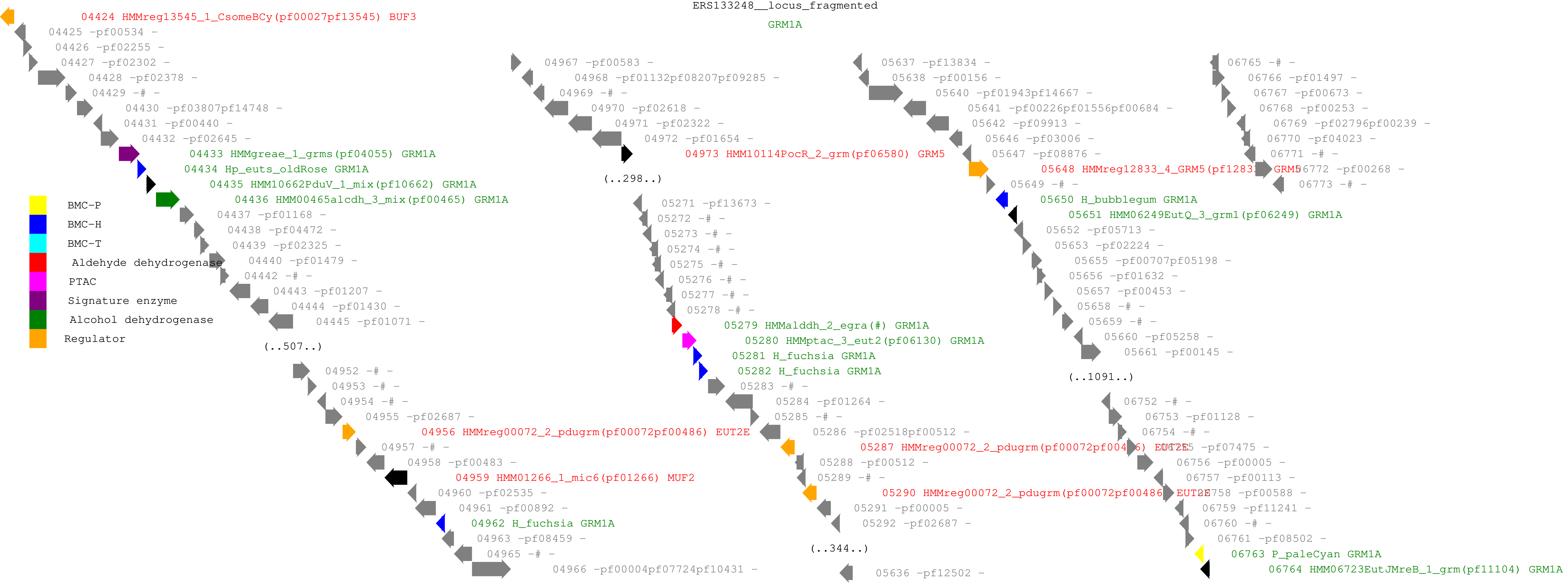

Hungatella hathewayi. Bacteria; Firmicutes; Clostridia; Clostridiales; Clostridiaceae;

ERS852407__locus_1

GRM1A

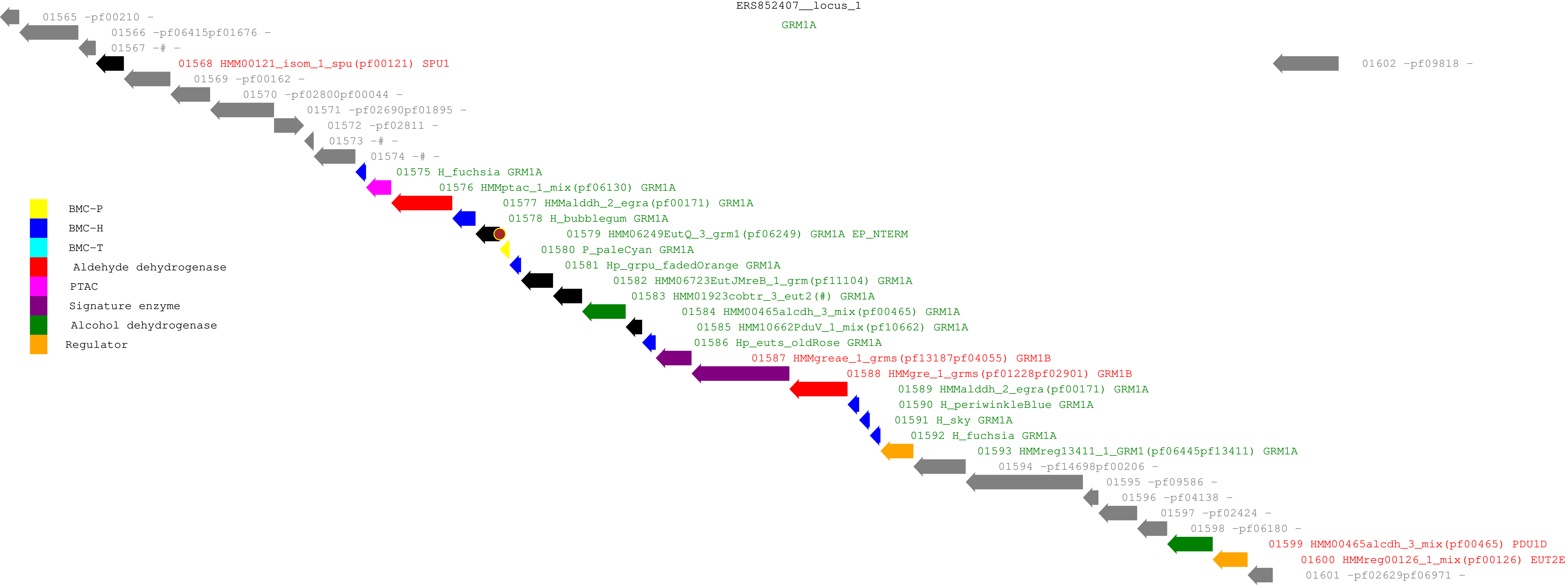

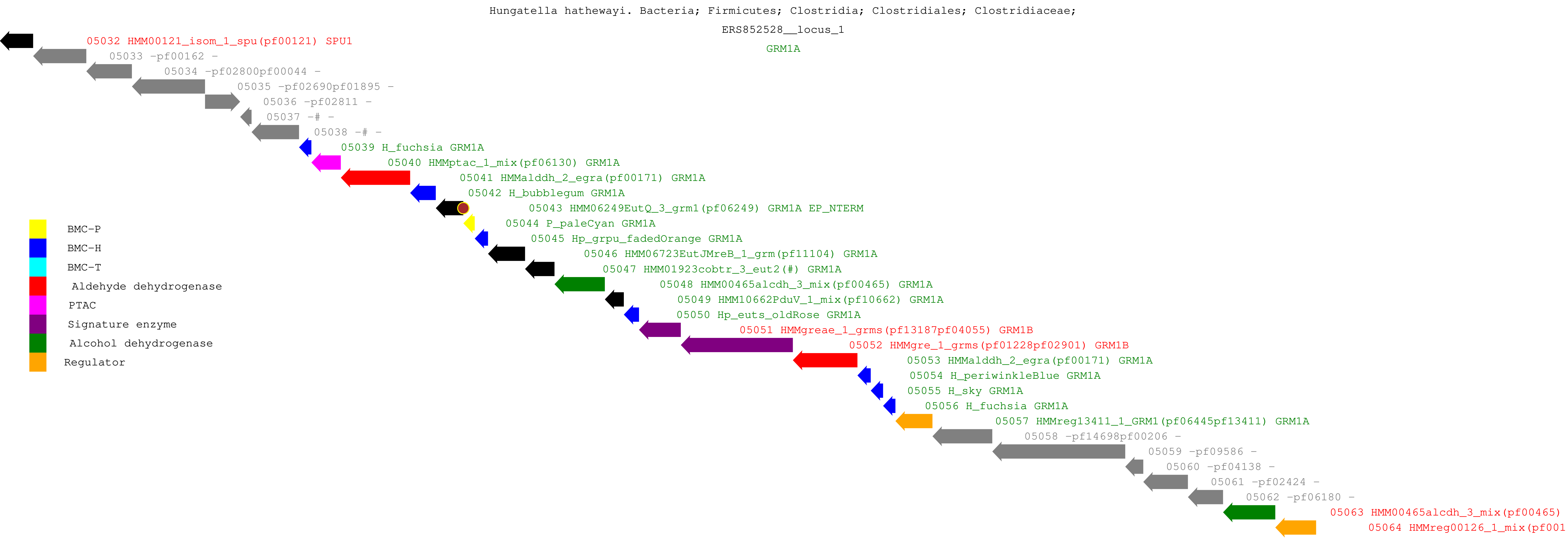

Hungatella hathewayi. Bacteria; Firmicutes; Clostridia; Clostridiales; Clostridiaceae;
ERS852528__locus_1
GRM1A
05032 HMM00121_isom_1_spu(pf00121) SPU1
05033 -pf00162 -
05034 -pf02800pf00044 -
05035 -pf02690pf01895 -
05036 -pf02811 -
05037 -# -
05038 -# -
05039 H_fuchsia GRM1A
05040 HMMptac_1_mix(pf06130) GRM1A
05041 HMMalddh_2_egra(pf00171) GRM1A
05042 H_bubblegum GRM1A
05043 HMM06249EutQ_3_grm1(pf06249) GRM1A EP_NTERM
05044 P_paleCyan GRM1A
05045 Hp_grpu_fadedOrange GRM1A
05046 HMM06723EutJMreB_1_grm(pf11104) GRM1A
05047 HMM01923cobtr_3_eut2(#) GRM1A
05048 HMM00465alcdh_3_mix(pf00465) GRM1A
05049 HMM10662PduV_1_mix(pf10662) GRM1A
05050 Hp_euts_oldRose GRM1A
05051 HMMgreae_1_grms(pf13187pf04055) GRM1B
05052 HMMgre_1_grms(pf01228pf02901) GRM1B
05053 HMMalddh_2_egra(pf00171) GRM1A
05054 H_periwinkleBlue GRM1A
05055 H_sky GRM1A
05056 H_fuchsia GRM1A
05057 HMMreg13411_1_GRM1(pf06445pf13411) GRM1A
05058 -pf14698pf00206 -
05059 -pf09586 -
05060 -pf04138 -
05061 -pf02424 -
05062 -pf06180 -
05063 HMM00465alcdh_3_mix(pf00465)
05064 HMMreg00126_1_mix(pf001
BMC-P
BMC-H
BMC-T
Aldehyde dehydrogenase
PTAC
Signature enzyme
Alcohol dehydrogenase
Regulator

Clostridium baratii. Bacteria; Firmicutes; Clostridia; Clostridiales; Clostridiaceae;

ERS852568__locus_1

GRM1A

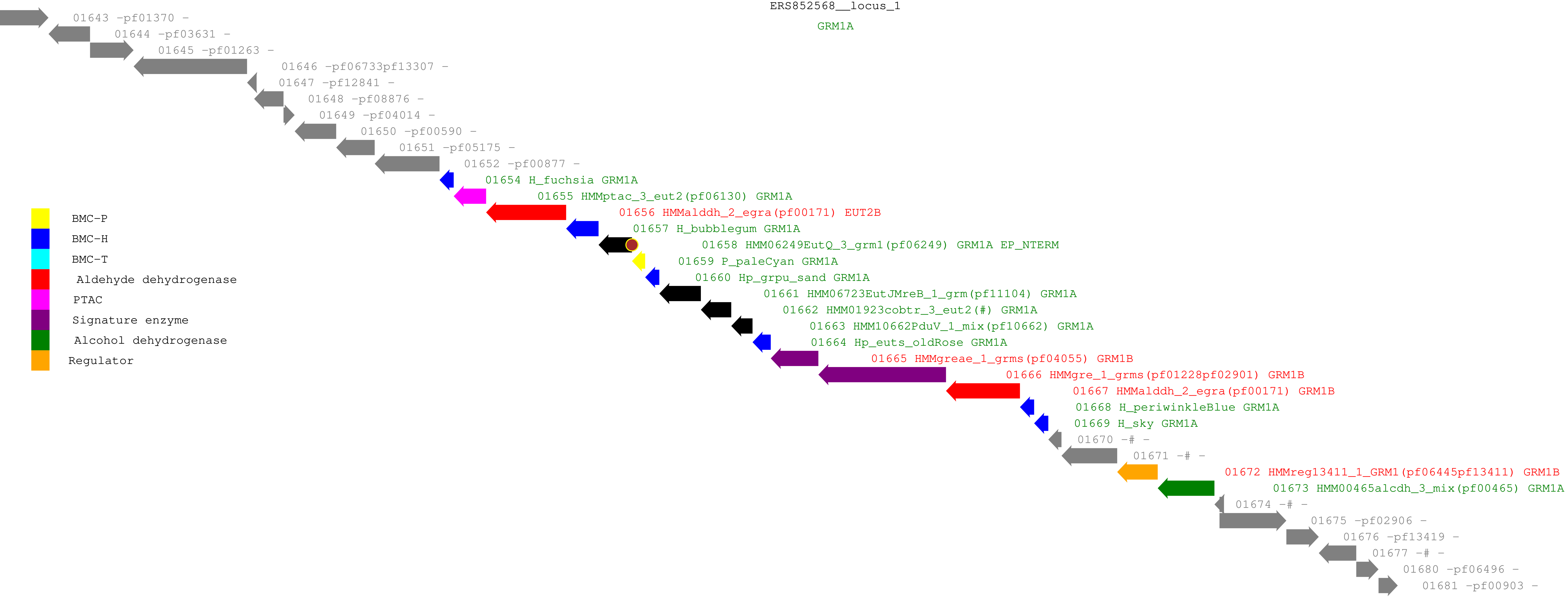

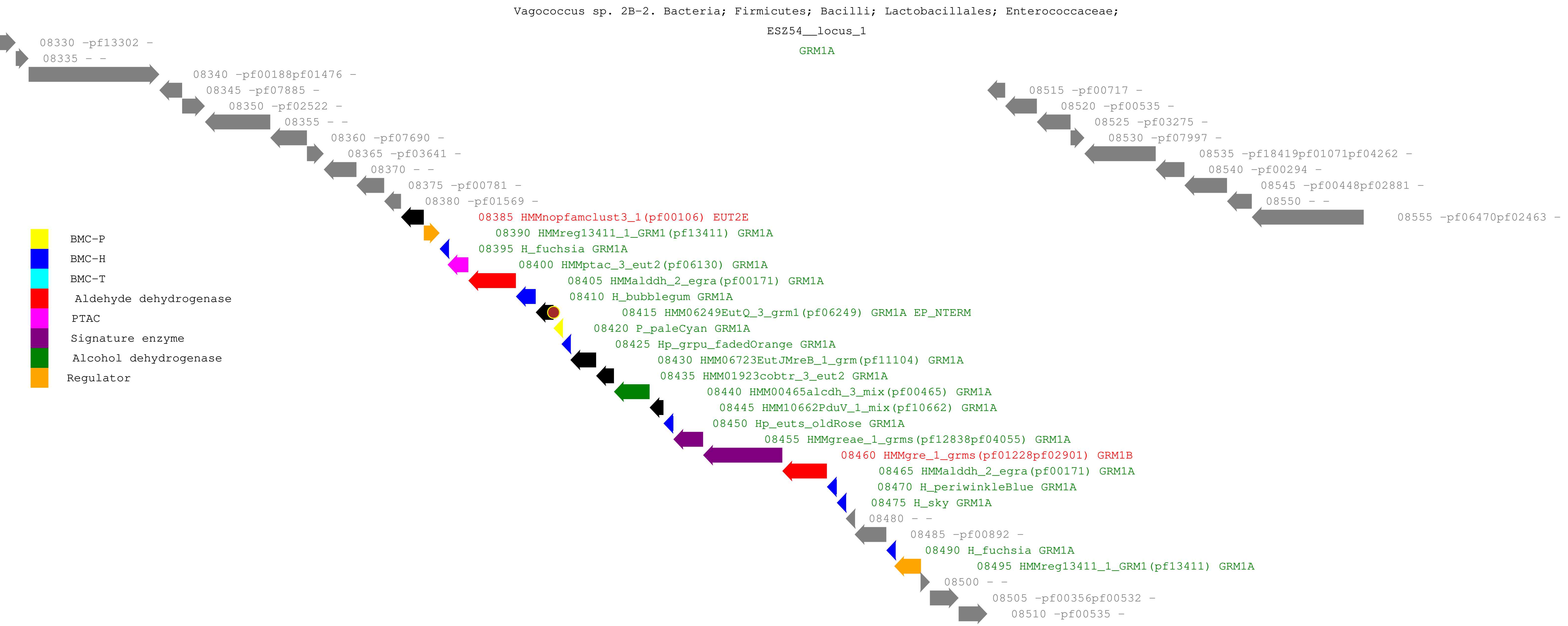

Vagococcus sp. 2B-2. Bacteria; Firmicutes; Bacilli; Lactobacillales; Enterococcaceae;
ESZ54__locus_1
GRM1A
08330 -pf13302 -
08335 - -
08340 -pf00188pf01476 -
08345 -pf07885 -
08350 -pf02522 -
08355 - -
08360 -pf07690 -
08365 -pf03641 -
08370 - -
08375 -pf00781 -
08380 -pf01569 -
08385 HMMnopfamclust3_1(pf00106) EUT2E
08390 HMMreg13411_1_GRM1(pf13411) GRM1A
08395 H_fuchsia GRM1A
08400 HMMptac_3_eut2(pf06130) GRM1A
08405 HMMaldDH_2_egra(pf00171) GRM1A
08410 H_bubblegum GRM1A
08415 HMM06249EutQ_3_grm1(pf06249) GRM1A EP_NTERM
08420 P_paleCyan GRM1A
08425 Hp_grpu_fadedOrange GRM1A
08430 HMM06723EutJMreB_1_grm(pf11104) GRM1A
08435 HMM01923cobtr_3_eut2 GRM1A
08440 HMM00465alcdh_3_mix(pf00465) GRM1A
08445 HMM10662PduV_1_mix(pf10662) GRM1A
08450 Hp_euts_oldRose GRM1A
08455 HMMgreae_1_grms(pf12838pf04055) GRM1A
08460 HMMgre_1_grms(pf01228pf02901) GRM1B
08465 HMMaldDH_2_egra(pf00171) GRM1A
08470 H_periwinkleBlue GRM1A
08475 H_sky GRM1A
08480 - -
08485 -pf00892 -
08490 H_fuchsia GRM1A
08495 HMMreg13411_1_GRM1(pf13411) GRM1A
08500 - -
08505 -pf00356pf00532 -
08510 -pf00535 -
08515 -pf00717 -
08520 -pf00535 -
08525 -pf03275 -
08530 -pf07997 -
08535 -pf18419pf01071pf04262 -
08540 -pf00294 -
08545 -pf00448pf02881 -
08550 - -
08555 -pf06470pf02463 -
BMC-P
BMC-H
BMC-T
Aldehyde dehydrogenase
PTAC
Signature enzyme
Alcohol dehydrogenase
Regulator

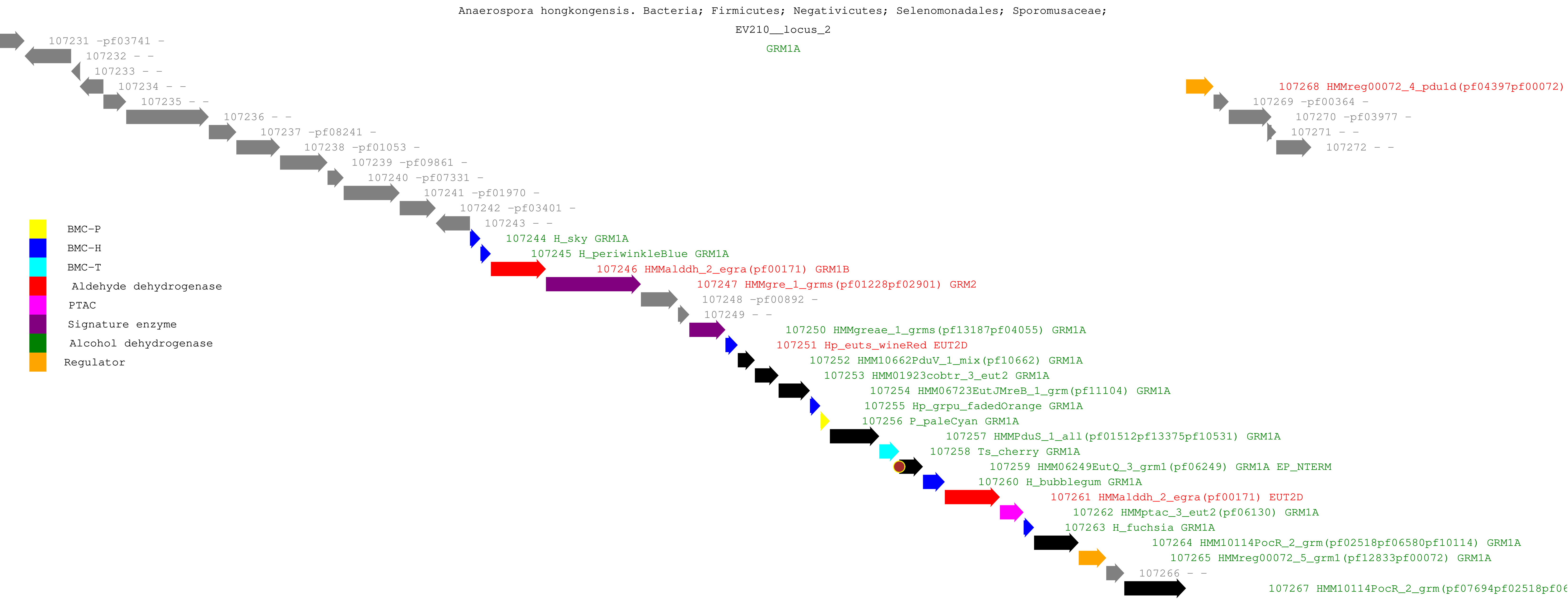

Anaerospora hongkongensis. Bacteria; Firmicutes; Negativicutes; Selenomonadales; Sporomusaceae;
EV210__locus_2
GRM1A
107231 -pf03741 -
107232 - -
107233 - -
107234 - -
107235 - -
107236 - -
107237 -pf08241 -
107238 -pf01053 -
107239 -pf09861 -
107240 -pf07331 -
107241 -pf01970 -
107242 -pf03401 -
107243 - -
107244 H_sky GRM1A
107245 H_periwinkleBlue GRM1A
107246 HMMaldh_2_egra(pf00171) GRM1B
107247 HMMgre_1_grms(pf01228pf02901) GRM2
107248 -pf00892 -
107249 - -
107250 HMMgreae_1_grms(pf13187pf04055) GRM1A
107251 Hp_euts_wineRed EUT2D
107252 HMM10662PduV_1_mix(pf10662) GRM1A
107253 HMM01923cobtr_3_eut2 GRM1A
107254 HMM06723EutJMreB_1_grm(pf11104) GRM1A
107255 Hp_grpu_fadedOrange GRM1A
107256 P_paleCyan GRM1A
107257 HMMPduS_1_all(pf01512pf13375pf10531) GRM1A
107258 Ts_cherry GRM1A
107259 HMM06249EutQ_3_grm1(pf06249) GRM1A EP_NTERM
107260 H_bubblegum GRM1A
107261 HMMaldh_2_egra(pf00171) EUT2D
107262 HMMptac_3_eut2(pf06130) GRM1A
107263 H_fuchsia GRM1A
107264 HMM10114PocR_2_grm(pf02518pf06580pf10114) GRM1A
107265 HMMreg00072_5_grm1(pf12833pf00072) GRM1A
107266 - -
107267 HMM10114PocR_2_grm(pf07694pf02518pf06
107268 HMMreg00072_4_pdu1d(pf04397pf00072)
107269 -pf00364 -
107270 -pf03977 -
107271 - -
107272 - -
BMC-P
BMC-H
BMC-T
Aldehyde dehydrogenase
PTAC
Signature enzyme
Alcohol dehydrogenase
Regulator

Lactobacillus paralimentarius DSM 19674. Bacteria; Firmicutes; Bacilli; Lactobacillales; Lactobacillaceae;

FC78_GL_locus_1

GRM1A

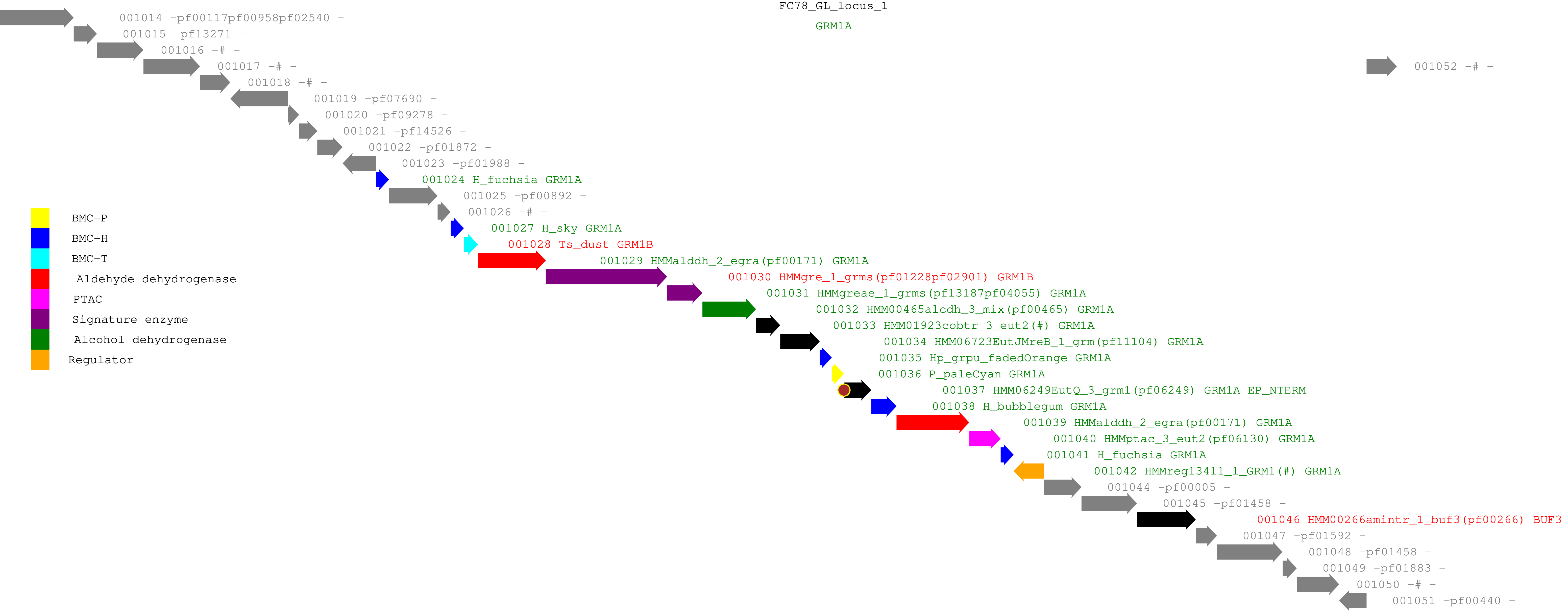

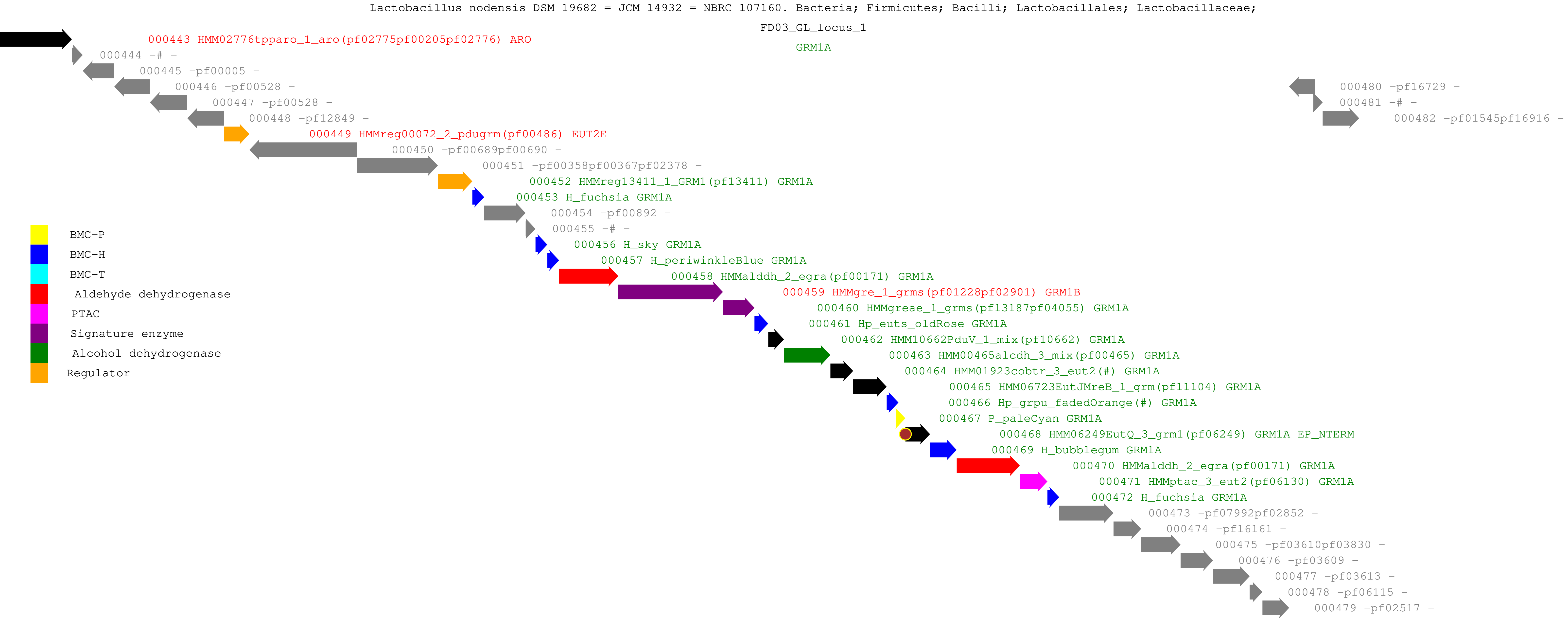

Lactobacillus nodensis DSM 19682 = JCM 14932 = NBRC 107160. Bacteria; Firmicutes; Bacilli; Lactobacillales; Lactobacillaceae;
FD03_GL_locus_1
GRM1A
000443 HMM02776tpparo_1_aro(pf02775pf00205pf02776) ARO
000444 -# -
000445 -pf00005 -
000446 -pf00528 -
000447 -pf00528 -
000448 -pf12849 -
000449 HMMreg00072_2_pdugrm(pf00486) EUT2E
000450 -pf00689pf00690 -
000451 -pf00358pf00367pf02378 -
000452 HMMreg13411_1_GRM1(pf13411) GRM1A
000453 H_fuchsia GRM1A
000454 -pf00892 -
000455 -# -
000456 H_sky GRM1A
000457 H_periwinkleBlue GRM1A
000458 HMMalddh_2_egra(pf00171) GRM1A
000459 HMMgre_1_grms(pf01228pf02901) GRM1B
000460 HMMgreae_1_grms(pf13187pf04055) GRM1A
000461 Hp_euts_oldRose GRM1A
000462 HMM10662PduV_1_mix(pf10662) GRM1A
000463 HMM00465alcdh_3_mix(pf00465) GRM1A
000464 HMM01923cobtr_3_eut2(#) GRM1A
000465 HMM06723EutJMreB_1_grm(pf11104) GRM1A
000466 Hp_grpu_fadedOrange(#) GRM1A
000467 P_paleCyan GRM1A
000468 HMM06249EutQ_3_grm1(pf06249) GRM1A EP_NTERM
000469 H_bubblegum GRM1A
000470 HMMalddh_2_egra(pf00171) GRM1A
000471 HMMptac_3_eut2(pf06130) GRM1A
000472 H_fuchsia GRM1A
000473 -pf07992pf02852 -
000474 -pf16161 -
000475 -pf03610pf03830 -
000476 -pf03609 -
000477 -pf03613 -
000478 -pf06115 -
000479 -pf02517 -
000480 -pf16729 -
000481 -# -
000482 -pf01545pf16916 -
BMC-P
BMC-H
BMC-T
Aldehyde dehydrogenase
PTAC
Signature enzyme
Alcohol dehydrogenase
Regulator

Lactobacillus paralimentarius DSM 13238 = JCM 10415. Bacteria; Firmicutes; Bacilli; Lactobacillales; Lactobacillaceae;

FD33_GL_locus_1

GRM1A

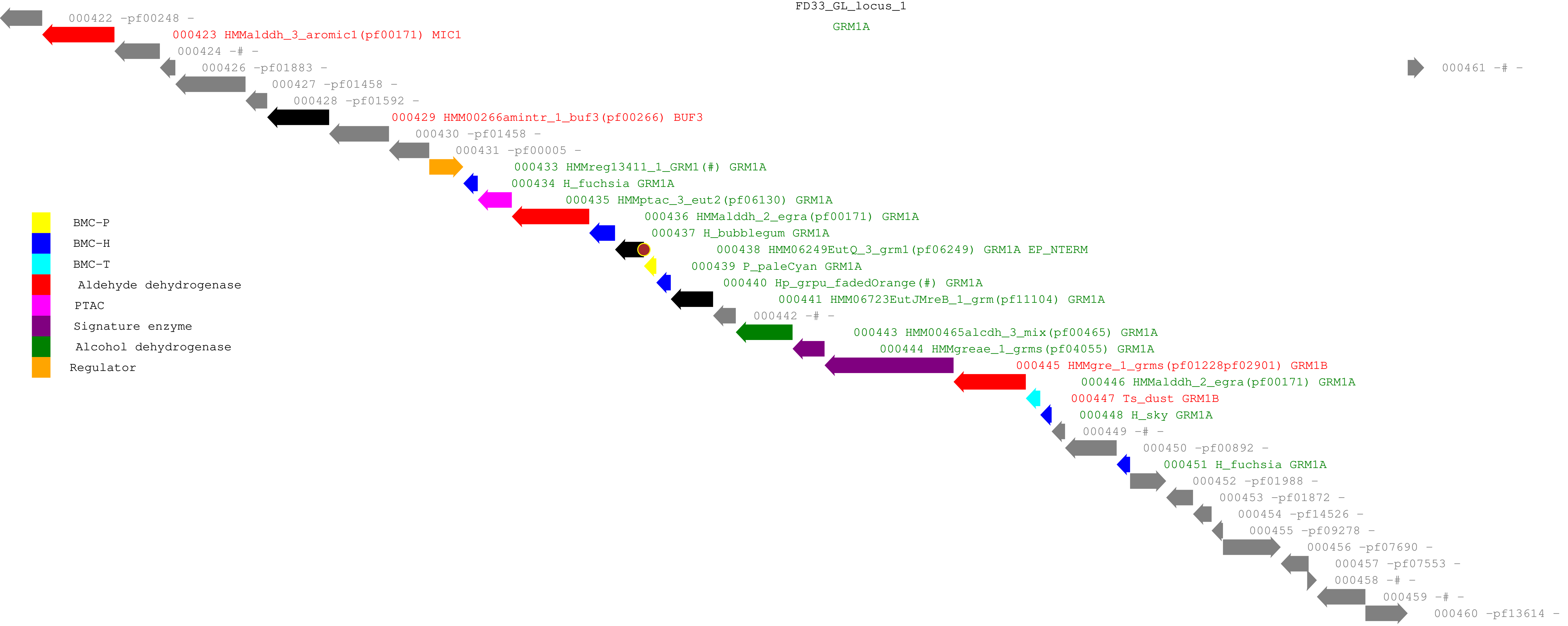

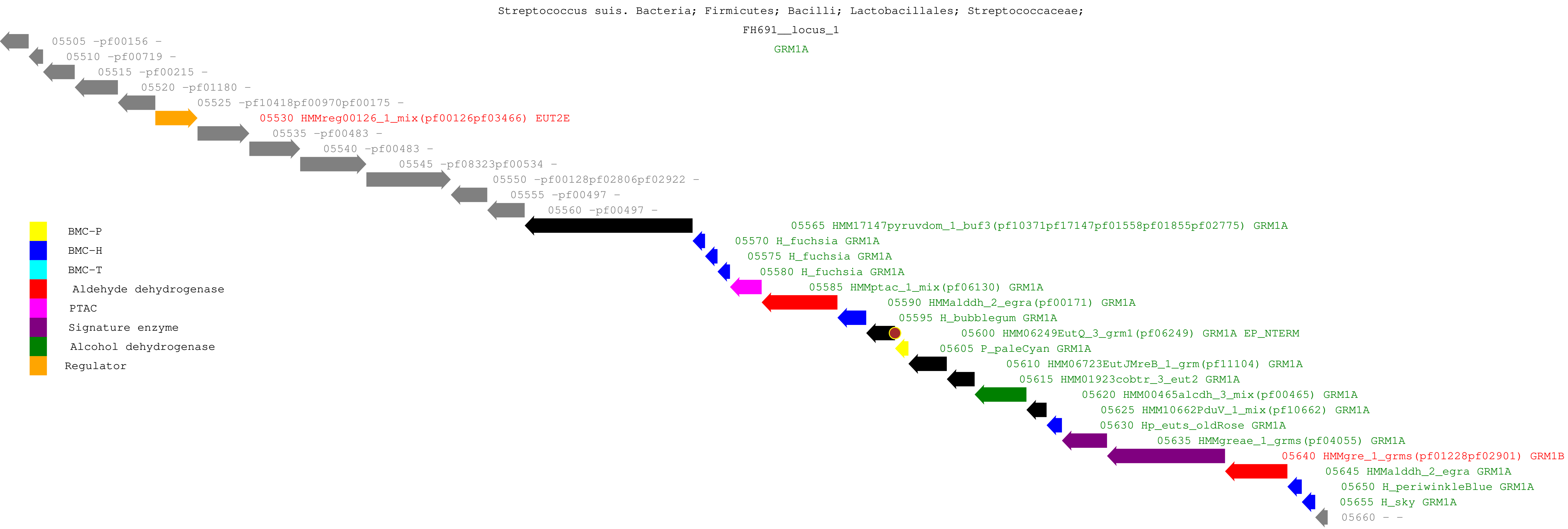
Streptococcus suis. Bacteria; Firmicutes; Bacilli; Lactobacillales; Streptococcaceae;
FH691__locus_1
GRM1A
05505 -pf00156 -
05510 -pf00719 -
05515 -pf00215 -
05520 -pf01180 -
05525 -pf10418pf00970pf00175 -
05530 HMMreg00126_1_mix(pf00126pf03466) EUT2E
05535 -pf00483 -
05540 -pf00483 -
05545 -pf08323pf00534 -
05550 -pf00128pf02806pf02922 -
05555 -pf00497 -
05560 -pf00497 -
05565 HMM17147pyruvdom_1_buf3(pf10371pf17147pf01558pf01855pf02775) GRM1A
05570 H_fuchsia GRM1A
05575 H_fuchsia GRM1A
05580 H_fuchsia GRM1A
05585 HMMptac_1_mix(pf06130) GRM1A
05590 HMMaldddh_2_egra(pf00171) GRM1A
05595 H_bubblegum GRM1A
05600 HMM06249EutQ_3_grm1(pf06249) GRM1A EP_NTERM
05605 P_paleCyan GRM1A
05610 HMM06723EutJMreB_1_grm(pf11104) GRM1A
05615 HMM01923cobtr_3_eut2 GRM1A
05620 HMM00465alcdh_3_mix(pf00465) GRM1A
05625 HMM10662PduV_1_mix(pf10662) GRM1A
05630 Hp_euts_oldRose GRM1A
05635 HMMgreae_1_grms(pf04055) GRM1A
05640 HMMgre_1_grms(pf01228pf02901) GRM1B
05645 HMMaldddh_2_egra GRM1A
05650 H_periwinkleBlue GRM1A
05655 H_sky GRM1A
05660 - -
BMC-P
BMC-H
BMC-T
Aldehyde dehydrogenase
PTAC
Signature enzyme
Alcohol dehydrogenase
Regulator

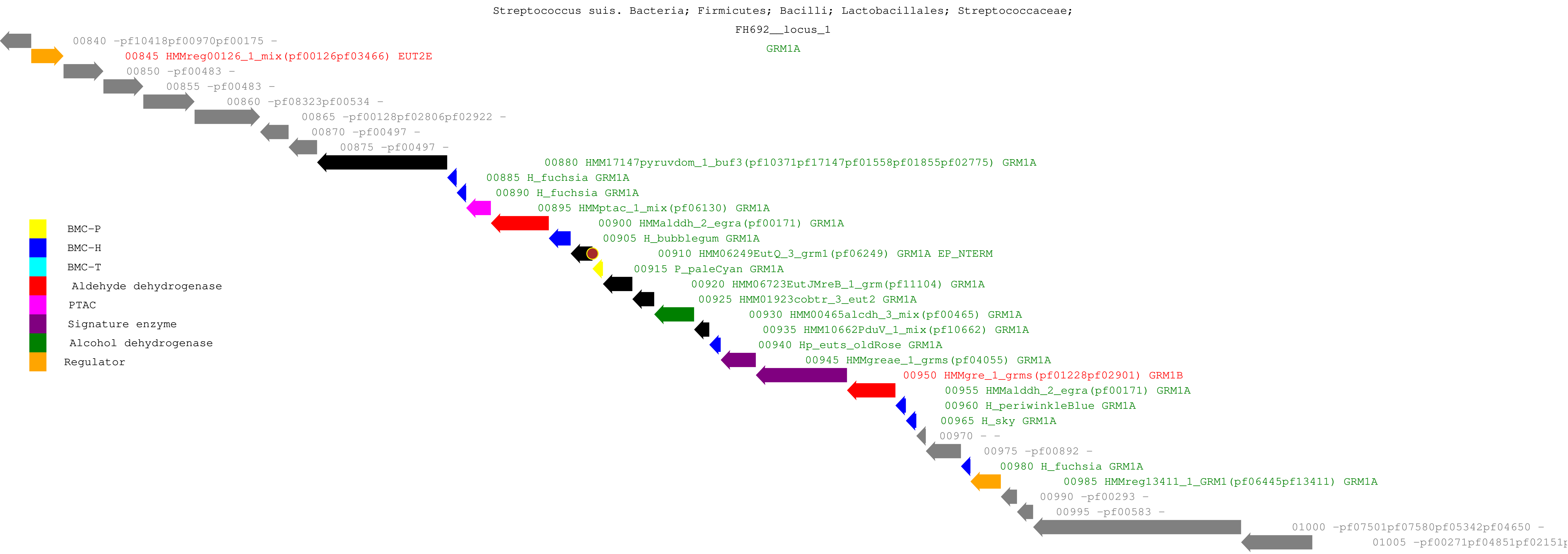
Streptococcus suis. Bacteria; Firmicutes; Bacilli; Lactobacillales; Streptococcaceae;
FH692__locus_1
GRM1A
00840 -pf10418pf00970pf00175 -
00845 HMMreg00126_1_mix(pf00126pf03466) EUT2E
00850 -pf00483 -
00855 -pf00483 -
00860 -pf08323pf00534 -
00865 -pf00128pf02806pf02922 -
00870 -pf00497 -
00875 -pf00497 -
00880 HMM17147pyruvdom_1_buf3(pf10371pf17147pf01558pf01855pf02775) GRM1A
00885 H_fuchsia GRM1A
00890 H_fuchsia GRM1A
00895 HMMptac_1_mix(pf06130) GRM1A
00900 HMMaldh_2_egra(pf00171) GRM1A
00905 H_bubblegum GRM1A
00910 HMM06249EutQ_3_grm1(pf06249) GRM1A EP_NTERM
00915 P_paleCyan GRM1A
00920 HMM06723EutJMreB_1_grm(pf11104) GRM1A
00925 HMM01923cobtr_3_eut2 GRM1A
00930 HMM00465alcdh_3_mix(pf00465) GRM1A
00935 HMM10662PduV_1_mix(pf10662) GRM1A
00940 Hp_euts_oldRose GRM1A
00945 HMMgreae_1_grms(pf04055) GRM1A
00950 HMMgre_1_grms(pf01228pf02901) GRM1B
00955 HMMaldh_2_egra(pf00171) GRM1A
00960 H_periwinkleBlue GRM1A
00965 H_sky GRM1A
00970 - -
00975 -pf00892 -
00980 H_fuchsia GRM1A
00985 HMMreg13411_1_GRM1(pf06445pf13411) GRM1A
00990 -pf00293 -
00995 -pf00583 -
01000 -pf07501pf07580pf05342pf04650 -
01005 -pf00271pf04851pf02151p
BMC-P
BMC-H
BMC-T
Aldehyde dehydrogenase
PTAC
Signature enzyme
Alcohol dehydrogenase
Regulator

Paenibacillus thiaminolyticus (Bacillus thiaminolyticus). Bacteria; Firmicutes; Bacilli; Bacillales; Paenibacillaceae; Paenibacillus.

FLT15__locus_2

GRM1A

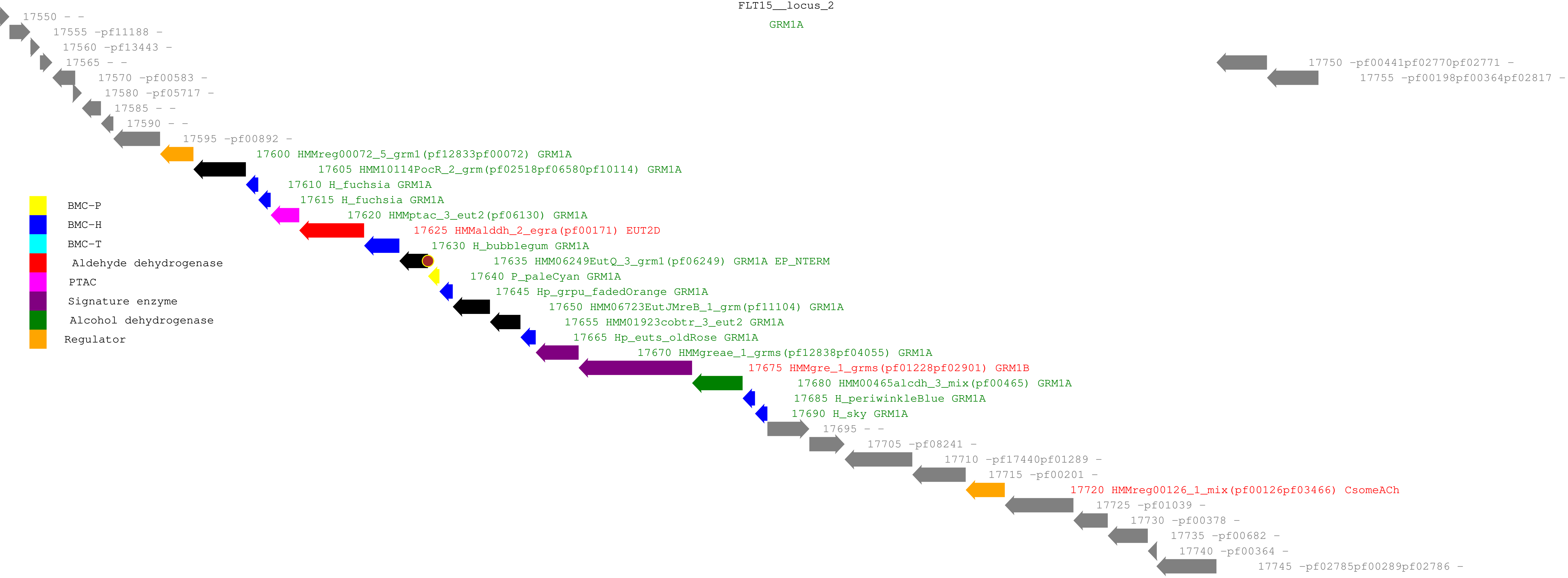

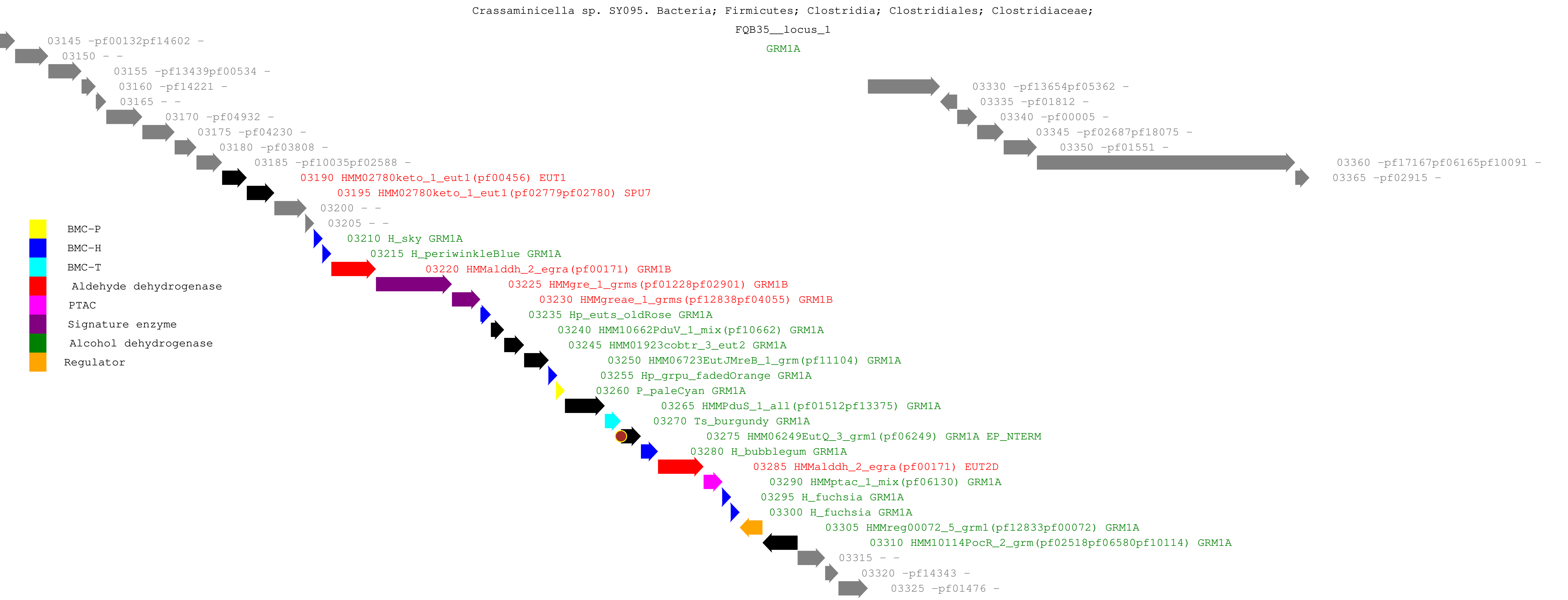
Crassaminicella sp. SY095. Bacteria; Firmicutes; Clostridia; Clostridiales; Clostridiaceae;
FQB35__locus_1
GRM1A
03145 -pf00132pf14602 -
03150 - -
03155 -pf13439pf00534 -
03160 -pf14221 -
03165 - -
03170 -pf04932 -
03175 -pf04230 -
03180 -pf03808 -
03185 -pf10035pf02588 -
03190 HMM02780keto_1_eut1(pf00456) EUT1
03195 HMM02780keto_1_eut1(pf02779pf02780) SPU7
03200 - -
03205 - -
03210 H_sky GRM1A
03215 H_periwinkleBlue GRM1A
03220 HMMalddh_2_egra(pf00171) GRM1B
03225 HMMgre_1_grms(pf01228pf02901) GRM1B
03230 HMMgreae_1_grms(pf12838pf04055) GRM1B
03235 Hp_euts_oldRose GRM1A
03240 HMM10662PduV_1_mix(pf10662) GRM1A
03245 HMM01923cobtr_3_eut2 GRM1A
03250 HMM06723EutJMreB_1_grm(pf11104) GRM1A
03255 Hp_grpu_fadedOrange GRM1A
03260 P_paleCyan GRM1A
03265 HMMPduS_1_all(pf01512pf13375) GRM1A
03270 Ts_burgundy GRM1A
03275 HMM06249EutQ_3_grm1(pf06249) GRM1A EP_NTERM
03280 H_bubblegum GRM1A
03285 HMMaldh_2_egra(pf00171) EUT2D
03290 HMMptac_1_mix(pf06130) GRM1A
03295 H_fuchsia GRM1A
03300 H_fuchsia GRM1A
03305 HMMreg00072_5_grm1(pf12833pf00072) GRM1A
03310 HMM10114PocR_2_grm(pf02518pf06580pf10114) GRM1A
03315 - -
03320 -pf14343 -
03325 -pf01476 -
03330 -pf13654pf05362 -
03335 -pf01812 -
03340 -pf00005 -
03345 -pf02687pf18075 -
03350 -pf01551 -
03360 -pf17167pf06165pf10091 -
03365 -pf02915 -
BMC-P
BMC-H
BMC-T
Aldehyde dehydrogenase
PTAC
Signature enzyme
Alcohol dehydrogenase
Regulator

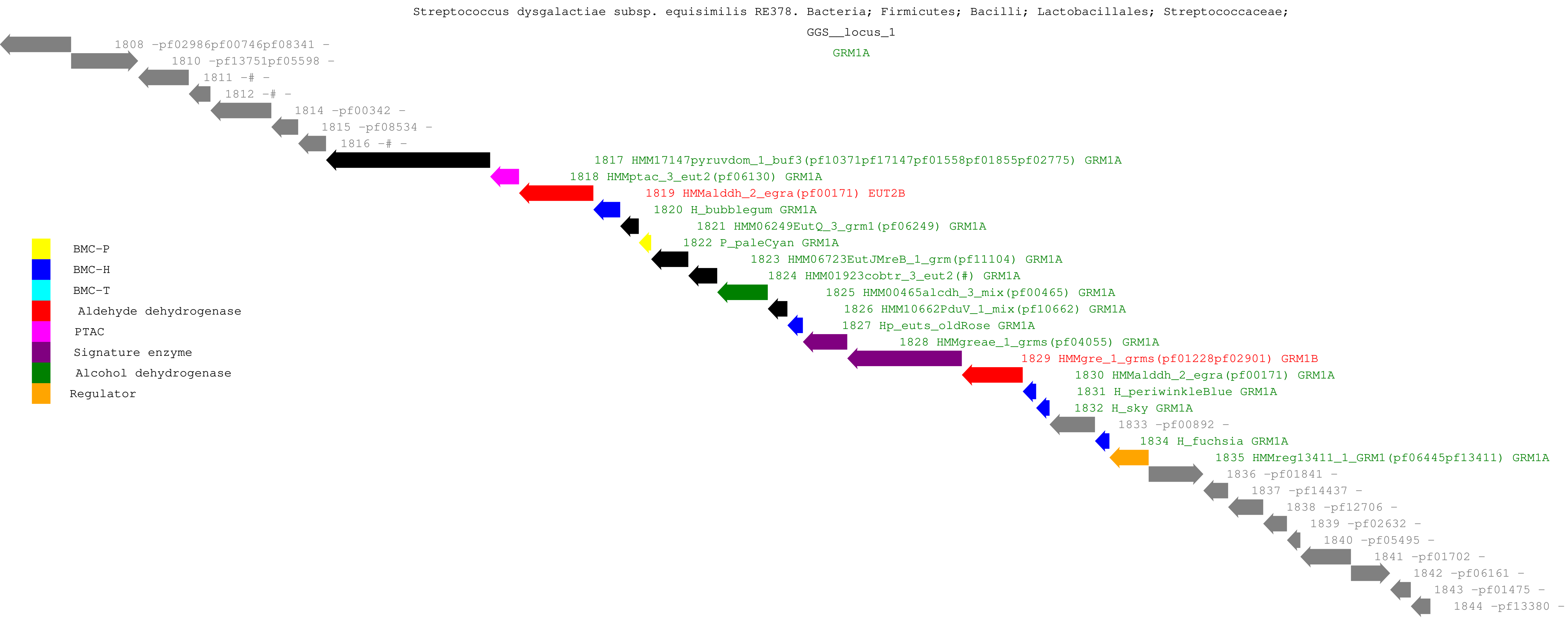
Streptococcus dysgalactiae subsp. equisimilis RE378. Bacteria; Firmicutes; Bacilli; Lactobacillales; Streptococcaceae;
GGS__locus_1
GRM1A
1808 -pf02986pf00746pf08341 -
1810 -pf13751pf05598 -
1811 -# -
1812 -# -
1814 -pf00342 -
1815 -pf08534 -
1816 -# -
1817 HMM17147pyruvdom_1_buf3(pf10371pf17147pf01558pf01855pf02775) GRM1A
1818 HMMptac_3_eut2(pf06130) GRM1A
1819 HMMaldh_2_egra(pf00171) EUT2B
1820 H_bubblegum GRM1A
1821 HMM06249EutQ_3_grm1(pf06249) GRM1A
1822 P_paleCyan GRM1A
1823 HMM06723EutJMreB_1_grm(pf11104) GRM1A
1824 HMM01923cobtr_3_eut2(#) GRM1A
1825 HMM00465alcdh_3_mix(pf00465) GRM1A
1826 HMM10662PduV_1_mix(pf10662) GRM1A
1827 Hp_euts_oldRose GRM1A
1828 HMMgreae_1_grms(pf04055) GRM1A
1829 HMMgre_1_grms(pf01228pf02901) GRM1B
1830 HMMaldh_2_egra(pf00171) GRM1A
1831 H_periwinkleBlue GRM1A
1832 H_sky GRM1A
1833 -pf00892 -
1834 H_fuchsia GRM1A
1835 HMMreg13411_1_GRM1(pf06445pf13411) GRM1A
1836 -pf01841 -
1837 -pf14437 -
1838 -pf12706 -
1839 -pf02632 -
1840 -pf05495 -
1841 -pf01702 -
1842 -pf06161 -
1843 -pf01475 -
1844 -pf13380 -
BMC-P
BMC-H
BMC-T
Aldehyde dehydrogenase
PTAC
Signature enzyme
Alcohol dehydrogenase
Regulator

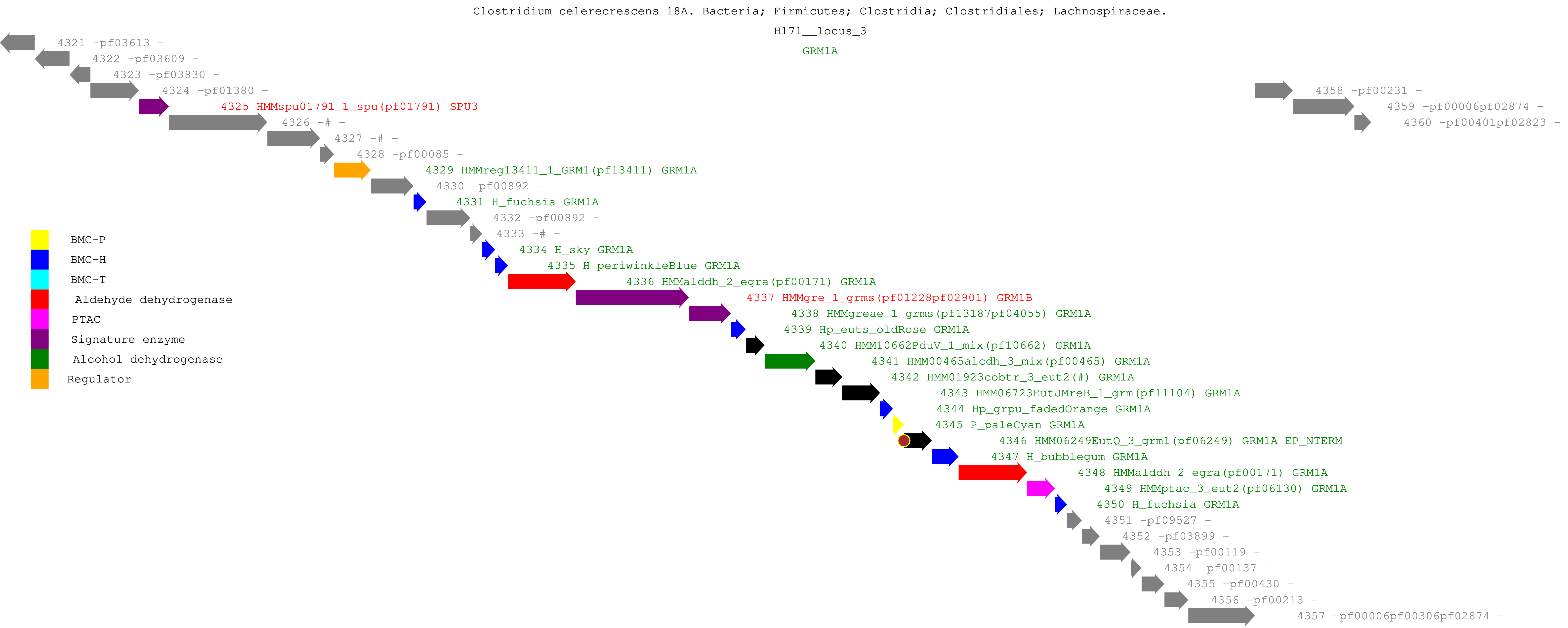
Clostridium celerecrescens 18A. Bacteria; Firmicutes; Clostridia; Clostridiales; Lachnospiraceae.
H171__locus_3
GRM1A
4321 -pf03613 -
4322 -pf03609 -
4323 -pf03830 -
4324 -pf01380 -
4325 HMMspu01791_1_spu(pf01791) SPU3
4326 -# -
4327 -# -
4328 -pf00085 -
4329 HMMreg13411_1_GRM1(pf13411) GRM1A
4330 -pf00892 -
4331 H_fuchsia GRM1A
4332 -pf00892 -
4333 -# -
4334 H_sky GRM1A
4335 H_periwinkleBlue GRM1A
4336 HMMalddh_2_egra(pf00171) GRM1A
4337 HMMgre_1_grms(pf01228pf02901) GRM1B
4338 HMMgreae_1_grms(pf13187pf04055) GRM1A
4339 Hp_euts_oldRose GRM1A
4340 HMM10662PduV_1_mix(pf10662) GRM1A
4341 HMM00465alcdh_3_mix(pf00465) GRM1A
4342 HMM01923cobtr_3_eut2(#) GRM1A
4343 HMM06723EutJMreB_1_grm(pf11104) GRM1A
4344 Hp_grpu_fadedOrange GRM1A
4345 P_paleCyan GRM1A
4346 HMM06249EutQ_3_grm1(pf06249) GRM1A EP_NTERM
4347 H_bubblegum GRM1A
4348 HMMalddh_2_egra(pf00171) GRM1A
4349 HMMptac_3_eut2(pf06130) GRM1A
4350 H_fuchsia GRM1A
4351 -pf09527 -
4352 -pf03899 -
4353 -pf00119 -
4354 -pf00137 -
4355 -pf00430 -
4356 -pf00213 -
4357 -pf00006pf00306pf02874 -
4358 -pf00231 -
4359 -pf00006pf02874 -
4360 -pf00401pf02823 -
BMC-P
BMC-H
BMC-T
Aldehyde dehydrogenase
PTAC
Signature enzyme
Alcohol dehydrogenase
Regulator

Anaerococcus tetradius ATCC 35098. Bacteria; Firmicutes; Tissierellia; Tissierellales; Peptoniphilaceae;

HMPREF0077__locus_1

GRM1A

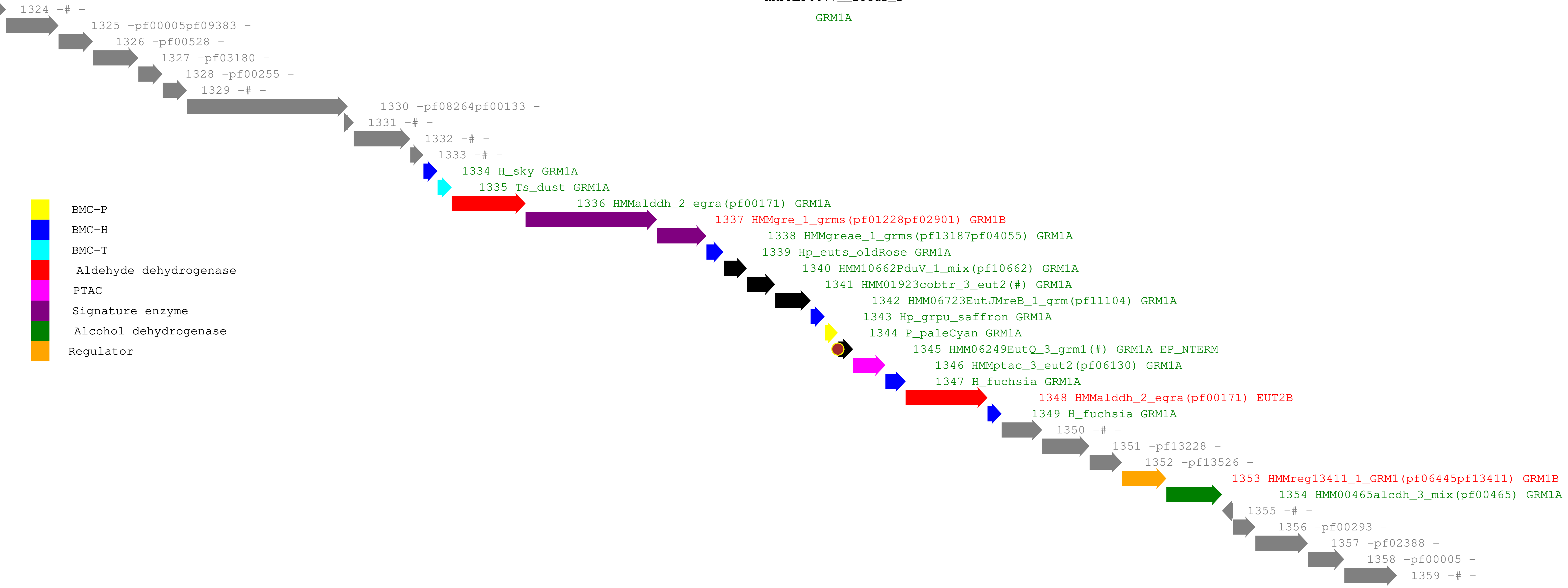

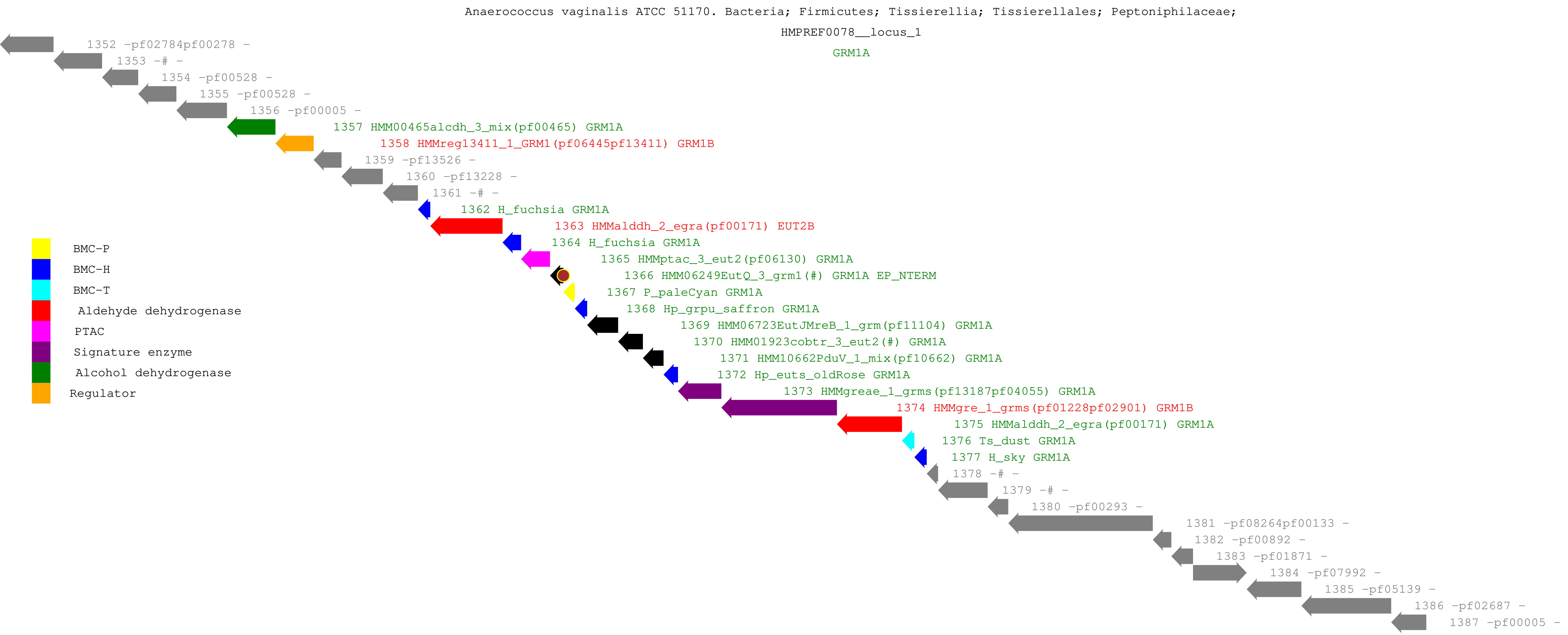
Anaerococcus vaginalis ATCC 51170. Bacteria; Firmicutes; Tissierellia; Tissierellales; Peptoniphilaceae;
HMPREF0078__locus_1
GRM1A
1352 -pf02784pf00278 -
1353 -# -
1354 -pf00528 -
1355 -pf00528 -
1356 -pf00005 -
1357 HMM00465alcdh_3_mix(pf00465) GRM1A
1358 HMMreg13411_1_GRM1(pf06445pf13411) GRM1B
1359 -pf13526 -
1360 -pf13228 -
1361 -# -
1362 H_fuchsia GRM1A
1363 HMMaldh_2_egra(pf00171) EUT2B
1364 H_fuchsia GRM1A
1365 HMMptac_3_eut2(pf06130) GRM1A
1366 HMM06249EutQ_3_grm1(#) GRM1A EP_NTERM
1367 P_paleCyan GRM1A
1368 Hp_grpu_saffron GRM1A
1369 HMM06723EutJMreB_1_grm(pf11104) GRM1A
1370 HMM01923cobtr_3_eut2(#) GRM1A
1371 HMM10662PduV_1_mix(pf10662) GRM1A
1372 Hp_euts_oldRose GRM1A
1373 HMMgreae_1_grms(pf13187pf04055) GRM1A
1374 HMMgre_1_grms(pf01228pf02901) GRM1B
1375 HMMaldh_2_egra(pf00171) GRM1A
1376 Ts_dust GRM1A
1377 H_sky GRM1A
1378 -# -
1379 -# -
1380 -pf00293 -
1381 -pf08264pf00133 -
1382 -pf00892 -
1383 -pf01871 -
1384 -pf07992 -
1385 -pf05139 -
1386 -pf02687 -
1387 -pf00005 -
BMC-P
BMC-H
BMC-T
Aldehyde dehydrogenase
PTAC
Signature enzyme
Alcohol dehydrogenase
Regulator

Desulfitobacterium hafniense DP7. Bacteria; Firmicutes; Clostridia; Clostridiales; Peptococcaceae;

HMPREF0322__locus_2

GRM1A

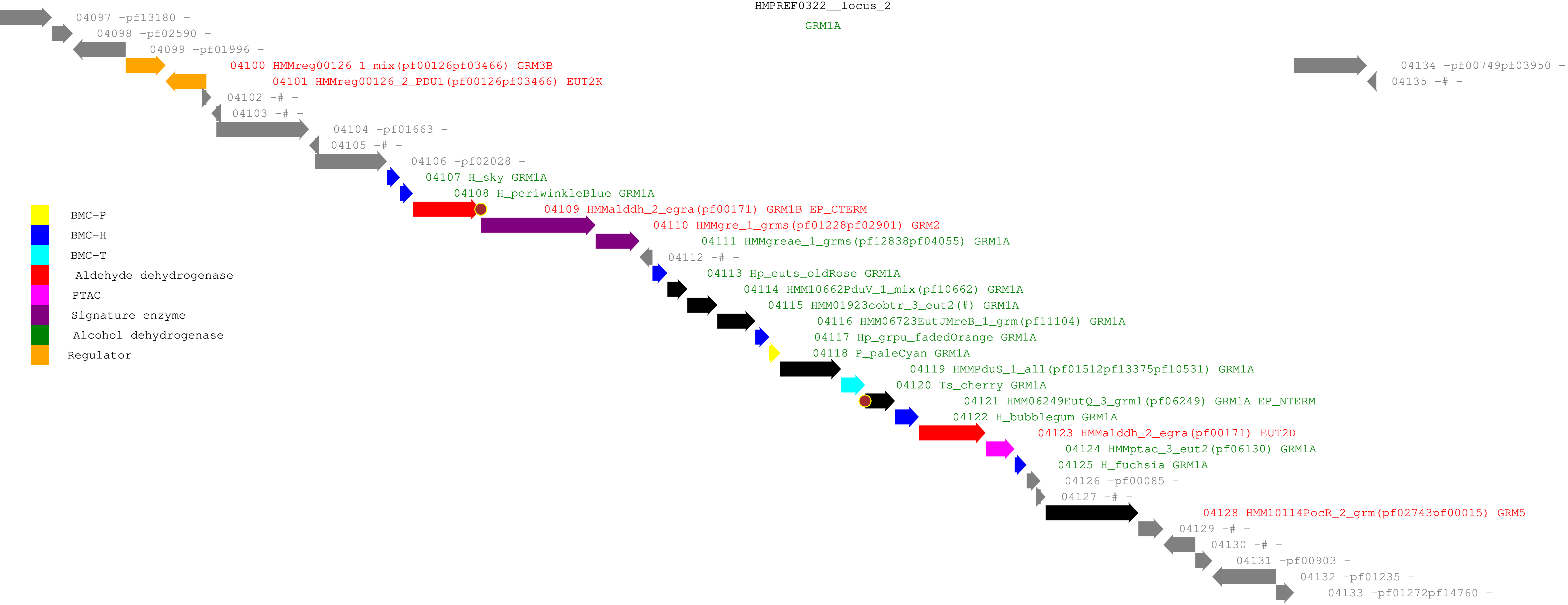

Atopobium minutum 10063974. Bacteria; Actinobacteria; Coriobacteriia; Coriobacteriales;

HMPREF1091__locus_1

GRM1A

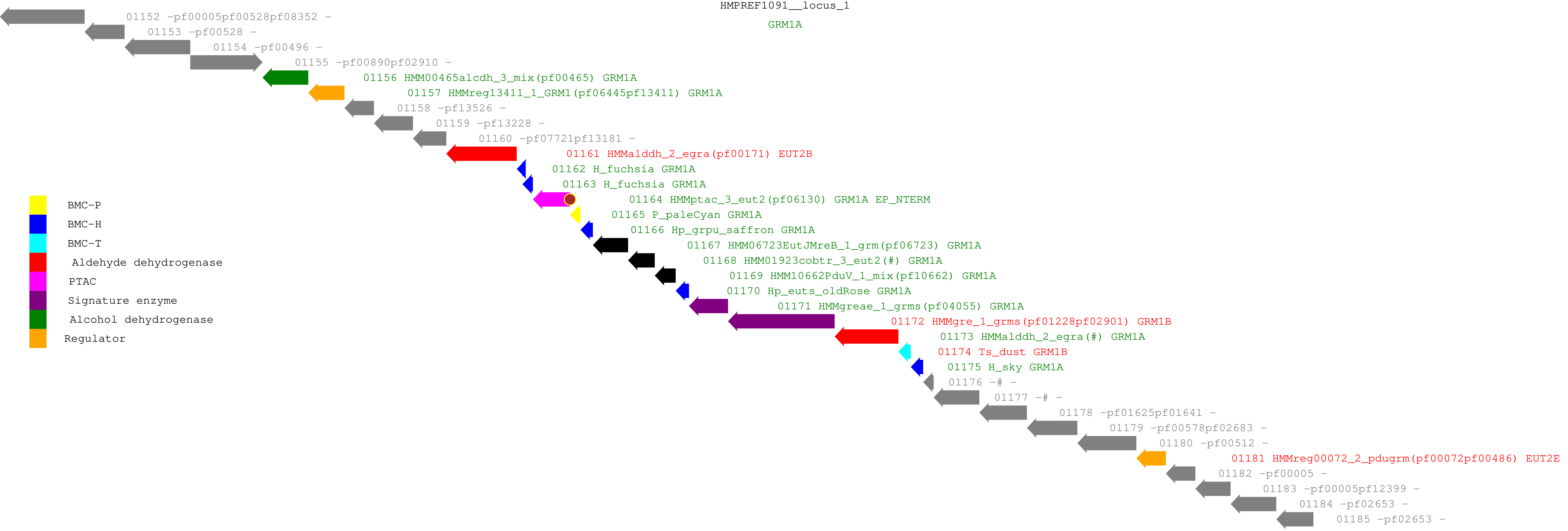

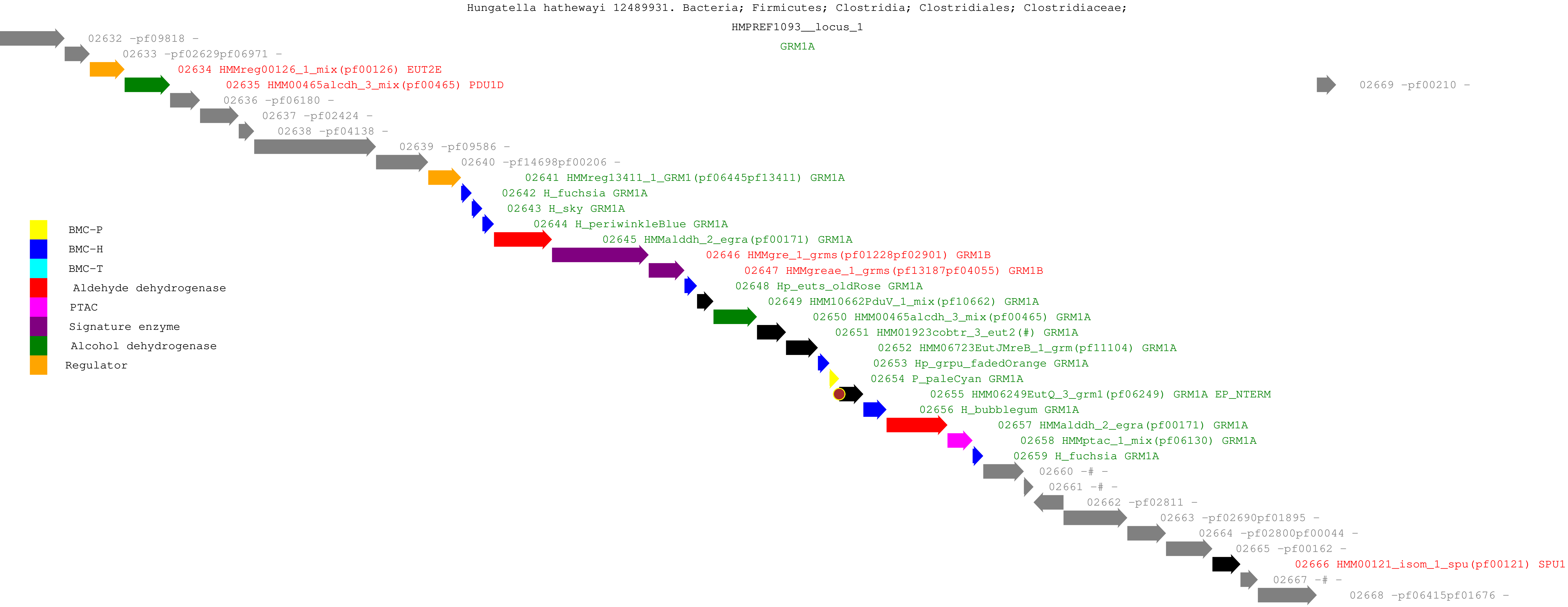
Hungatella hathewayi 12489931. Bacteria; Firmicutes; Clostridia; Clostridiales; Clostridiaceae;
HMPREF1093__locus_1
GRM1A
02632 -pf09818 -
02633 -pf02629pf06971 -
02634 HMMreg00126_1_mix(pf00126) EUT2E
02635 HMM00465alcdh_3_mix(pf00465) PDU1D
02669 -pf00210 -
02636 -pf06180 -
02637 -pf02424 -
02638 -pf04138 -
02639 -pf09586 -
02640 -pf14698pf00206 -
02641 HMMreg13411_1_GRM1(pf06445pf13411) GRM1A
02642 H_fuchsia GRM1A
02643 H_sky GRM1A
02644 H_periwinkleBlue GRM1A
02645 HMMaldh_2_egra(pf00171) GRM1A
02646 HMMgre_1_grms(pf01228pf02901) GRM1B
02647 HMMgreae_1_grms(pf13187pf04055) GRM1B
02648 Hp_euts_oldRose GRM1A
02649 HMM10662PduV_1_mix(pf10662) GRM1A
02650 HMM00465alcdh_3_mix(pf00465) GRM1A
02651 HMM01923cobtr_3_eut2(#) GRM1A
02652 HMM06723EutJMreB_1_grm(pf11104) GRM1A
02653 Hp_grpu_fadedOrange GRM1A
02654 P_paleCyan GRM1A
02655 HMM06249EutQ_3_grm1(pf06249) GRM1A EP_NTERM
02656 H_bubblegum GRM1A
02657 HMMaldh_2_egra(pf00171) GRM1A
02658 HMMptac_1_mix(pf06130) GRM1A
02659 H_fuchsia GRM1A
02660 -# -
02661 -# -
02662 -pf02811 -
02663 -pf02690pf01895 -
02664 -pf02800pf00044 -
02665 -pf00162 -
02666 HMM00121_isom_1_spu(pf00121) SPU1
02667 -# -
02668 -pf06415pf01676 -
BMC-P
BMC-H
BMC-T
Aldehyde dehydrogenase
PTAC
Signature enzyme
Alcohol dehydrogenase
Regulator

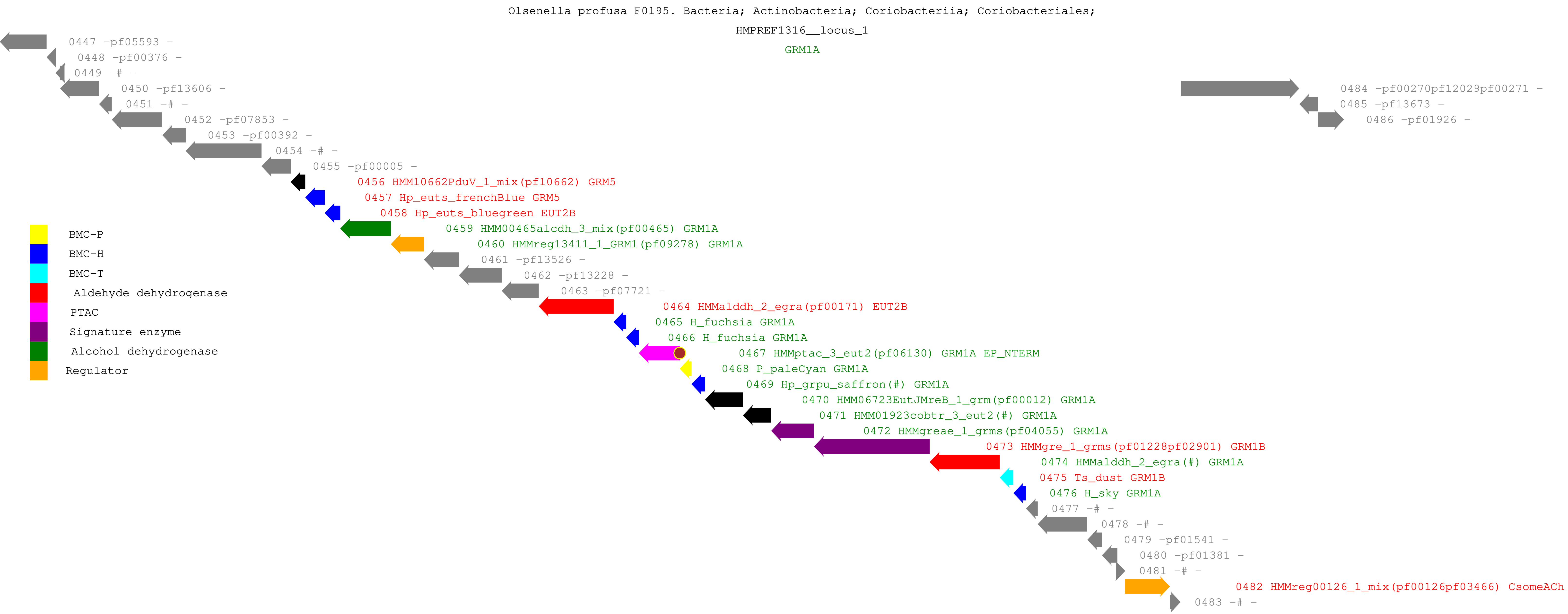

Olsenella profusa F0195. Bacteria; Actinobacteria; Coriobacteriia; Coriobacteriales;
HMPREF1316__locus_1
GRM1A
0447 -pf05593 -
0448 -pf00376 -
0449 -# -
0450 -pf13606 -
0451 -# -
0452 -pf07853 -
0453 -pf00392 -
0454 -# -
0455 -pf00005 -
0456 HMM10662PduV_1_mix(pf10662) GRM5
0457 Hp_euts_frenchBlue GRM5
0458 Hp_euts_bluegreen EUT2B
0459 HMM00465alcdh_3_mix(pf00465) GRM1A
0460 HMMreg13411_1_GRM1(pf09278) GRM1A
0461 -pf13526 -
0462 -pf13228 -
0463 -pf07721 -
0464 HMMalddh_2_egra(pf00171) EUT2B
0465 H_fuchsia GRM1A
0466 H_fuchsia GRM1A
0467 HMMptac_3_eut2(pf06130) GRM1A EP_NTERM
0468 P_paleCyan GRM1A
0469 Hp_grpu_saffron(#) GRM1A
0470 HMM06723EutJMreB_1_grm(pf00012) GRM1A
0471 HMM01923cobtr_3_eut2(#) GRM1A
0472 HMMgreae_1_grms(pf04055) GRM1A
0473 HMMgre_1_grms(pf01228pf02901) GRM1B
0474 HMMalddh_2_egra(#) GRM1A
0475 Ts_dust GRM1B
0476 H_sky GRM1A
0477 -# -
0478 -# -
0479 -pf01541 -
0480 -pf01381 -
0481 -# -
0482 HMMreg00126_1_mix(pf00126pf03466) CsomeACh
0483 -# -
0484 -pf00270pf12029pf00271 -
0485 -pf13673 -
0486 -pf01926 -
BMC-P
BMC-H
BMC-T
Aldehyde dehydrogenase
PTAC
Signature enzyme
Alcohol dehydrogenase
Regulator

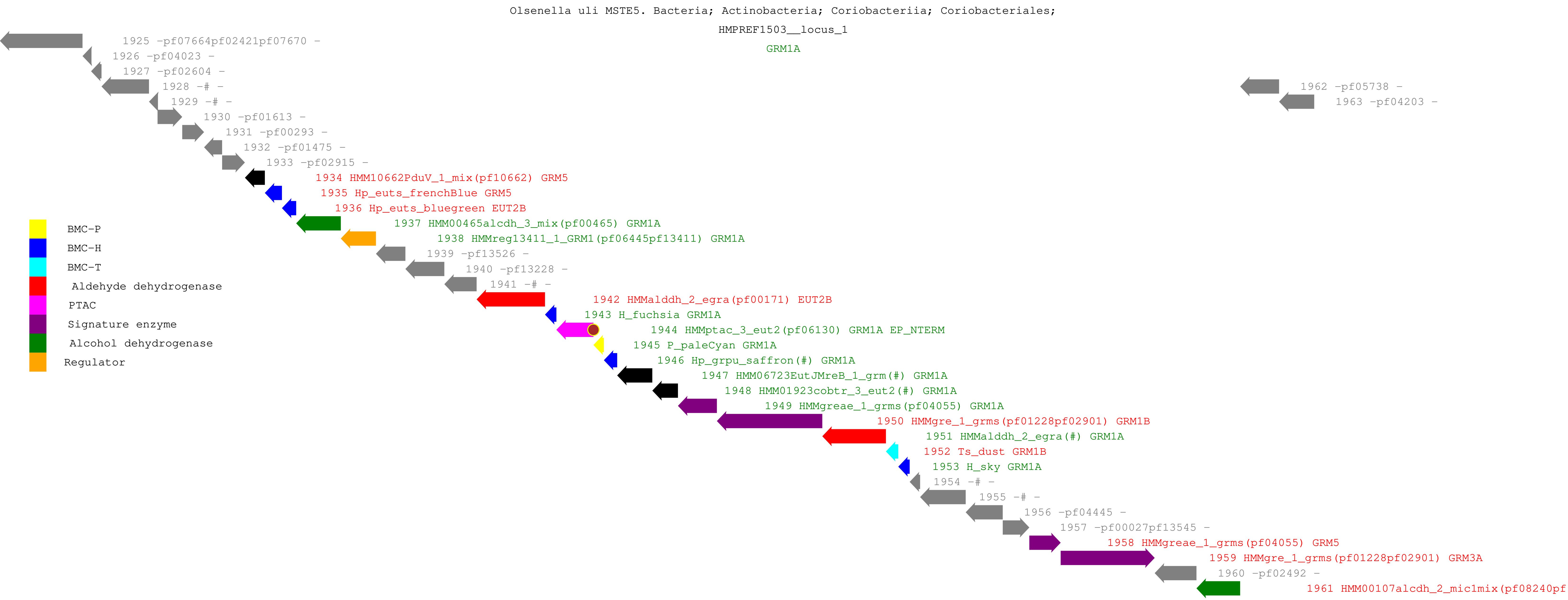
Olsenella uli MSTE5. Bacteria; Actinobacteria; Coriobacteriia; Coriobacteriales;
HMPREF1503__locus_1
GRM1A
1925 -pf07664pf02421pf07670 -
1926 -pf04023 -
1927 -pf02604 -
1928 -# -
1929 -# -
1930 -pf01613 -
1931 -pf00293 -
1932 -pf01475 -
1933 -pf02915 -
1934 HMM10662PduV_1_mix(pf10662) GRM5
1935 Hp_euts_frenchBlue GRM5
1936 Hp_euts_bluegreen EUT2B
1937 HMM00465alcdh_3_mix(pf00465) GRM1A
1938 HMMreg13411_1_GRM1(pf06445pf13411) GRM1A
1939 -pf13526 -
1940 -pf13228 -
1941 -# -
1942 HMMalddh_2_egra(pf00171) EUT2B
1943 H_fuchsia GRM1A
1944 HMMptac_3_eut2(pf06130) GRM1A EP_NTERM
1945 P_paleCyan GRM1A
1946 Hp_grpu_saffron(#) GRM1A
1947 HMM06723EutJMreB_1_grm(#) GRM1A
1948 HMM01923cobtr_3_eut2(#) GRM1A
1949 HMMgreae_1_grms(pf04055) GRM1A
1950 HMMgre_1_grms(pf01228pf02901) GRM1B
1951 HMMalddh_2_egra(#) GRM1A
1952 Ts_dust GRM1B
1953 H_sky GRM1A
1954 -# -
1955 -# -
1956 -pf04445 -
1957 -pf00027pf13545 -
1958 HMMgreae_1_grms(pf04055) GRM5
1959 HMMgre_1_grms(pf01228pf02901) GRM3A
1960 -pf02492 -
1961 HMM00107alcdh_2_mic1mix(pf08240pf
1962 -pf05738 -
1963 -pf04203 -
BMC-P
BMC-H
BMC-T
Aldehyde dehydrogenase
PTAC
Signature enzyme
Alcohol dehydrogenase
Regulator

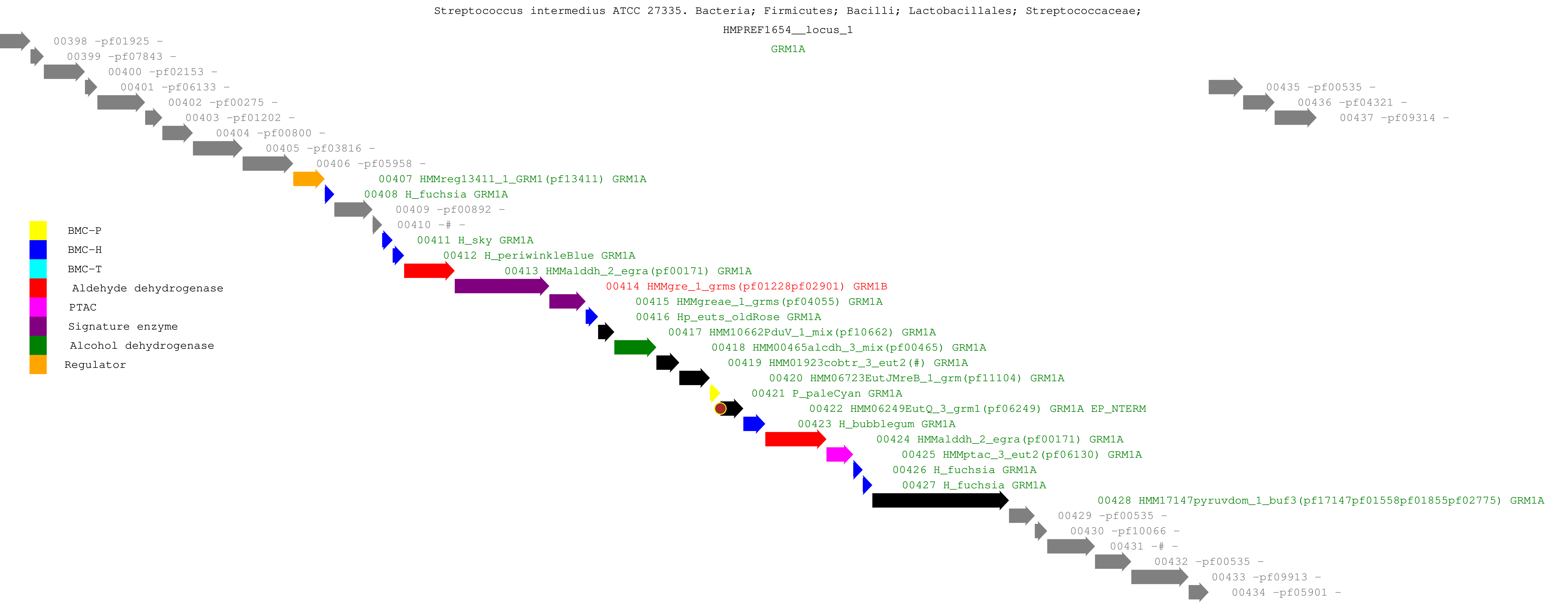
Streptococcus intermedius ATCC 27335. Bacteria; Firmicutes; Bacilli; Lactobacillales; Streptococcaceae;
HMPREF1654__locus_1
GRM1A
00398 -pf01925 -
00399 -pf07843 -
00400 -pf02153 -
00401 -pf06133 -
00402 -pf00275 -
00403 -pf01202 -
00404 -pf00800 -
00405 -pf03816 -
00406 -pf05958 -
00407 HMMreg13411_1_GRM1(pf13411) GRM1A
00408 H_fuchsia GRM1A
00409 -pf00892 -
00410 -# -
00411 H_sky GRM1A
00412 H_periwinkleBlue GRM1A
00413 HMMaldh_2_egra(pf00171) GRM1A
00414 HMMgre_1_grms(pf01228pf02901) GRM1B
00415 HMMgreae_1_grms(pf04055) GRM1A
00416 Hp_euts_oldRose GRM1A
00417 HMM10662PduV_1_mix(pf10662) GRM1A
00418 HMM00465alcdh_3_mix(pf00465) GRM1A
00419 HMM01923cobtr_3_eut2(#) GRM1A
00420 HMM06723EutJMreB_1_grm(pf11104) GRM1A
00421 P_paleCyan GRM1A
00422 HMM06249EutQ_3_grm1(pf06249) GRM1A EP_NTERM
00423 H_bubblegum GRM1A
00424 HMMaldh_2_egra(pf00171) GRM1A
00425 HMMptac_3_eut2(pf06130) GRM1A
00426 H_fuchsia GRM1A
00427 H_fuchsia GRM1A
00428 HMM17147pyruvdom_1_buf3(pf17147pf01558pf01855pf02775) GRM1A
00429 -pf00535 -
00430 -pf10066 -
00431 -# -
00432 -pf00535 -
00433 -pf09913 -
00434 -pf05901 -
00435 -pf00535 -
00436 -pf04321 -
00437 -pf09314 -
BMC-P
BMC-H
BMC-T
Aldehyde dehydrogenase
PTAC
Signature enzyme
Alcohol dehydrogenase
Regulator

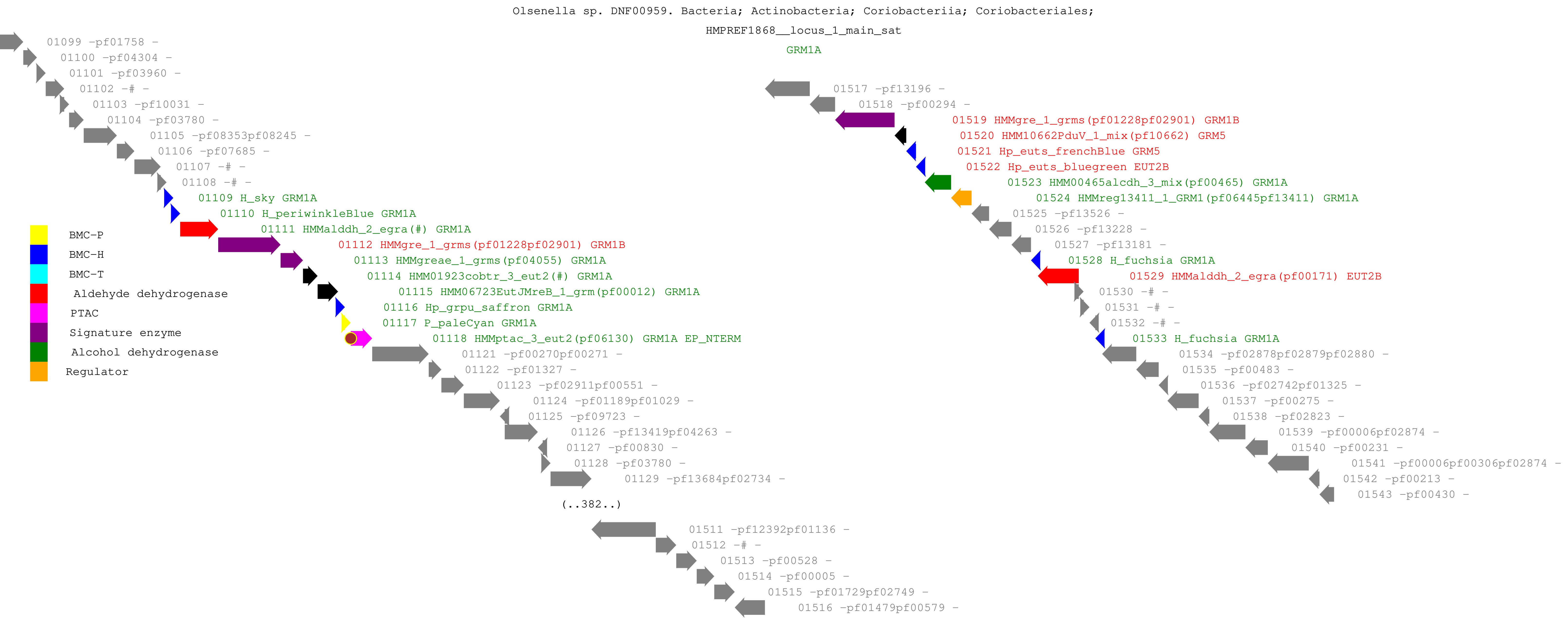

Olsenella sp. DNF00959. Bacteria; Actinobacteria; Coriobacteriia; Coriobacteriales;
HMPREF1868__locus_1_main_sat
GRM1A
01099 -pf01758 -
01100 -pf04304 -
01101 -pf03960 -
01102 -# -
01103 -pf10031 -
01104 -pf03780 -
01105 -pf08353pf08245 -
01106 -pf07685 -
01107 -# -
01108 -# -
01109 H_sky GRM1A
01110 H_periwinkleBlue GRM1A
01111 HMMaldh_2_egra(#) GRM1A
01112 HMMgre_1_grms(pf01228pf02901) GRM1B
01113 HMMgreae_1_grms(pf04055) GRM1A
01114 HMM01923cobtr_3_eut2(#) GRM1A
01115 HMM06723EutJMreB_1_grm(pf00012) GRM1A
01116 Hp_grpu_saffron GRM1A
01117 P_paleCyan GRM1A
01118 HMMptac_3_eut2(pf06130) GRM1A EP_NTERM
01121 -pf00270pf00271 -
01122 -pf01327 -
01123 -pf02911pf00551 -
01124 -pf01189pf01029 -
01125 -pf09723 -
01126 -pf13419pf04263 -
01127 -pf00830 -
01128 -pf03780 -
01129 -pf13684pf02734 -
(..382..)
01511 -pf12392pf01136 -
01512 -# -
01513 -pf00528 -
01514 -pf00005 -
01515 -pf01729pf02749 -
01516 -pf01479pf00579 -
01517 -pf13196 -
01518 -pf00294 -
01519 HMMgre_1_grms(pf01228pf02901) GRM1B
01520 HMM10662PduV_1_mix(pf10662) GRM5
01521 Hp_euts_frenchBlue GRM5
01522 Hp_euts_bluegreen EUT2B
01523 HMM00465alcdh_3_mix(pf00465) GRM1A
01524 HMMreg13411_1_GRM1(pf06445pf13411) GRM1A
01525 -pf13526 -
01526 -pf13228 -
01527 -pf13181 -
01528 H_fuchsia GRM1A
01529 HMMaldh_2_egra(pf00171) EUT2B
01530 -# -
01531 -# -
01532 -# -
01533 H_fuchsia GRM1A
01534 -pf02878pf02879pf02880 -
01535 -pf00483 -
01536 -pf02742pf01325 -
01537 -pf00275 -
01538 -pf02823 -
01539 -pf00006pf02874 -
01540 -pf00231 -
01541 -pf00006pf00306pf02874 -
01542 -pf00213 -
01543 -pf00430 -
BMC-P
BMC-H
BMC-T
Aldehyde dehydrogenase
PTAC
Signature enzyme
Alcohol dehydrogenase
Regulator

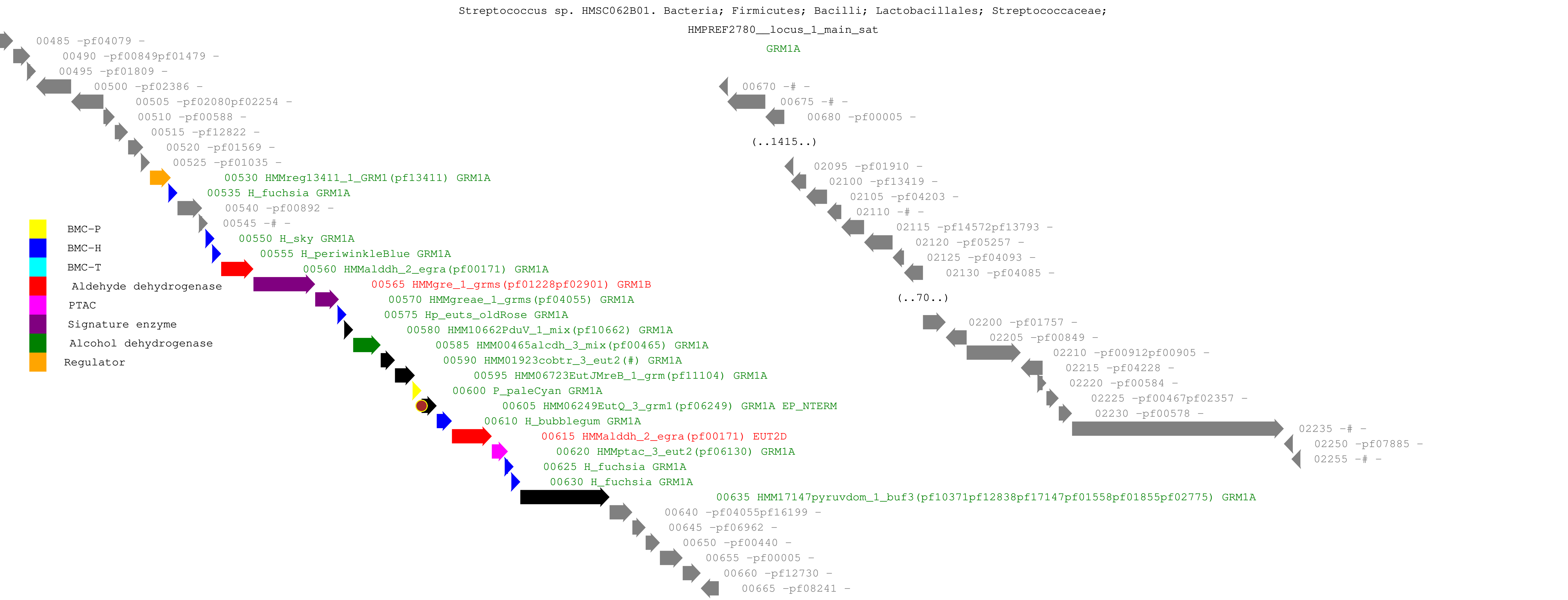
Streptococcus sp. HMSC062B01. Bacteria; Firmicutes; Bacilli; Lactobacillales; Streptococcaceae;
HMPREF2780__locus_1_main_sat
GRM1A
00485 -pf04079 -
00490 -pf00849pf01479 -
00495 -pf01809 -
00500 -pf02386 -
00505 -pf02080pf02254 -
00510 -pf00588 -
00515 -pf12822 -
00520 -pf01569 -
00525 -pf01035 -
00530 HMMreg13411_1_GRM1(pf13411) GRM1A
00535 H_fuchsia GRM1A
00540 -pf00892 -
00545 -# -
00550 H_sky GRM1A
00555 H_periwinkleBlue GRM1A
00560 HMMaldh_2_egra(pf00171) GRM1A
00565 HMMgre_1_grms(pf01228pf02901) GRM1B
00570 HMMgreae_1_grms(pf04055) GRM1A
00575 Hp_euts_oldRose GRM1A
00580 HMM10662PduV_1_mix(pf10662) GRM1A
00585 HMM00465alcdh_3_mix(pf00465) GRM1A
00590 HMM01923cobtr_3_eut2(#) GRM1A
00595 HMM06723EutJMreB_1_grm(pf11104) GRM1A
00600 P_paleCyan GRM1A
00605 HMM06249EutQ_3_grm1(pf06249) GRM1A EP_NTERM
00610 H_bubblegum GRM1A
00615 HMMaldh_2_egra(pf00171) EUT2D
00620 HMMptac_3_eut2(pf06130) GRM1A
00625 H_fuchsia GRM1A
00630 H_fuchsia GRM1A
00635 HMM17147pyruvdom_1_buf3(pf10371pf12838pf17147pf01558pf01855pf02775) GRM1A
00640 -pf04055pf16199 -
00645 -pf06962 -
00650 -pf00440 -
00655 -pf00005 -
00660 -pf12730 -
00665 -pf08241 -
00670 -# -
00675 -# -
00680 -pf00005 -
(..1415..)
02095 -pf01910 -
02100 -pf13419 -
02105 -pf04203 -
02110 -# -
02115 -pf14572pf13793 -
02120 -pf05257 -
02125 -pf04093 -
02130 -pf04085 -
(..70..)
02200 -pf01757 -
02205 -pf00849 -
02210 -pf00912pf00905 -
02215 -pf04228 -
02220 -pf00584 -
02225 -pf00467pf02357 -
02230 -pf00578 -
02235 -# -
02250 -pf07885 -
02255 -# -
BMC-P
BMC-H
BMC-T
Aldehyde dehydrogenase
PTAC
Signature enzyme
Alcohol dehydrogenase
Regulator

Anaerococcus sp. HMSC065G05. Bacteria; Firmicutes; Tissierellia; Tissierellales; Peptoniphilaceae;

HMPREF2852__locus_1

GRM1A

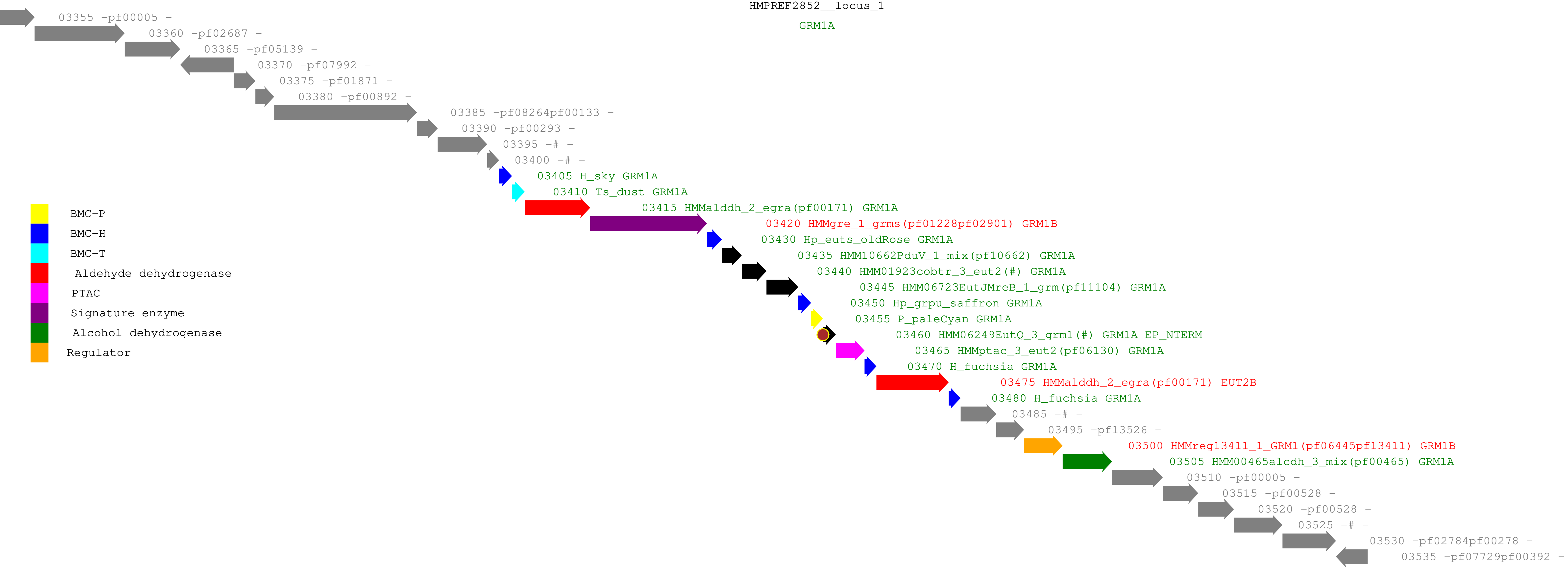

Anaerococcus tetradius. Bacteria; Firmicutes; Tissierellia; Tissierellales; Peptoniphilaceae;

HMPREF3200__locus_1

GRM1A

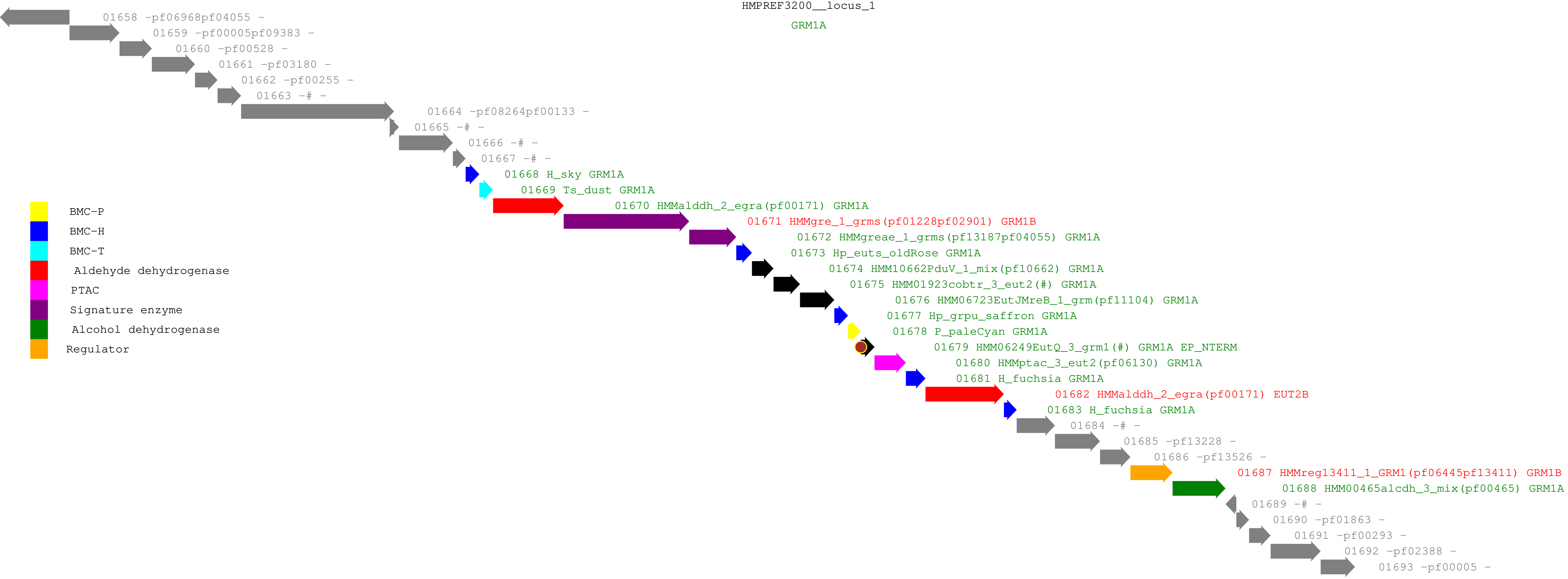

Anaerococcus hydrogenalis ACS-025-V-Sch4. Bacteria; Firmicutes; Tissierellia; Tissierellales; Peptoniphilaceae;

HMPREF9246__locus_1

GRM1A

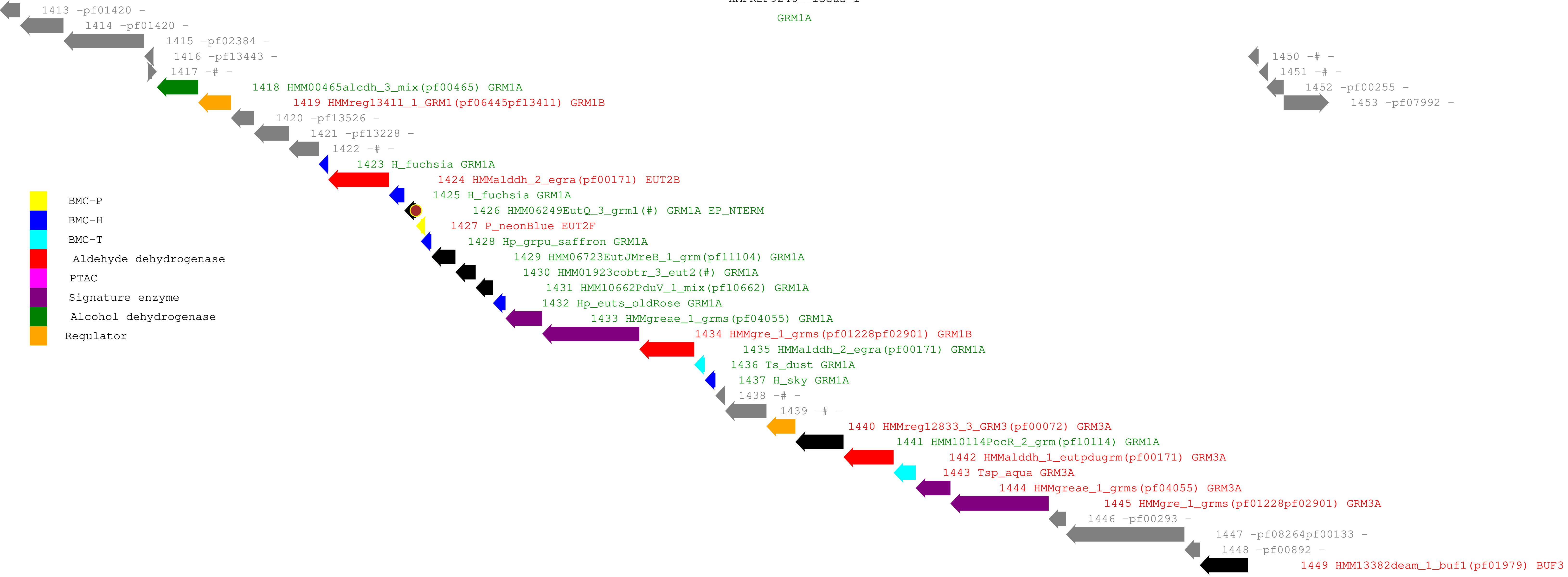

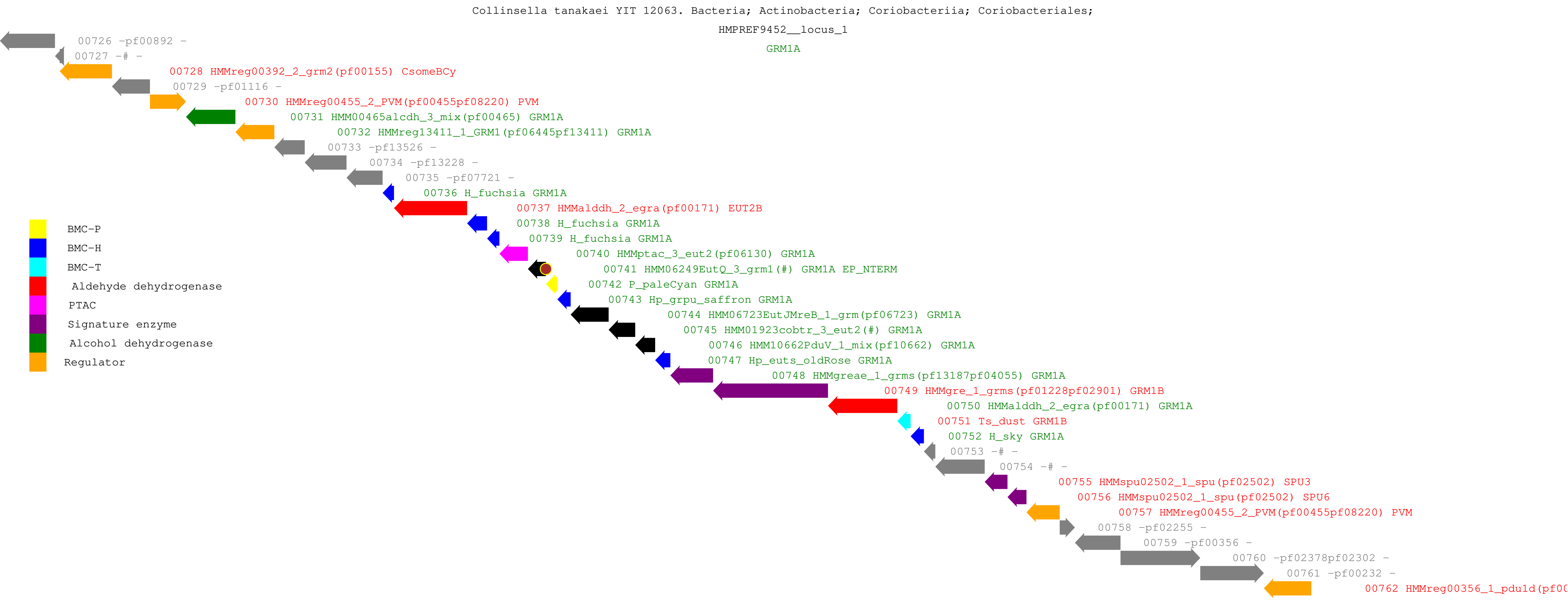
Collinsella tanakaei YIT 12063. Bacteria; Actinobacteria; Coriobacteriia; Coriobacteriales;
HMPREF9452__locus_1
GRM1A
00726 -pf00892 -
00727 -# -
00728 HMMreg00392_2_grm2(pf00155) CsomeBCy
00729 -pf01116 -
00730 HMMreg00455_2_PVM(pf00455pf08220) PVM
00731 HMM00465alcdh_3_mix(pf00465) GRM1A
00732 HMMreg13411_1_GRM1(pf06445pf13411) GRM1A
00733 -pf13526 -
00734 -pf13228 -
00735 -pf07721 -
00736 H_fuchsia GRM1A
00737 HMMalddh_2_egra(pf00171) EUT2B
00738 H_fuchsia GRM1A
00739 H_fuchsia GRM1A
00740 HMMptac_3_eut2(pf06130) GRM1A
00741 HMM06249EutQ_3_grm1(#) GRM1A EP_NTERM
00742 P_paleCyan GRM1A
00743 Hp_grpu_saffron GRM1A
00744 HMM06723EutJMreB_1_grm(pf06723) GRM1A
00745 HMM01923cobtr_3_eut2(#) GRM1A
00746 HMM10662PduV_1_mix(pf10662) GRM1A
00747 Hp_euts_oldRose GRM1A
00748 HMMgreae_1_grms(pf13187pf04055) GRM1A
00749 HMMgre_1_grms(pf01228pf02901) GRM1B
00750 HMMalddh_2_egra(pf00171) GRM1A
00751 Ts_dust GRM1B
00752 H_sky GRM1A
00753 -# -
00754 -# -
00755 HMMspu02502_1_spu(pf02502) SPU3
00756 HMMspu02502_1_spu(pf02502) SPU6
00757 HMMreg00455_2_PVM(pf00455pf08220) PVM
00758 -pf02255 -
00759 -pf00356 -
00760 -pf02378pf02302 -
00761 -pf00232 -
00762 HMMreg00356_1_pdu1d(pf00
BMC-P
BMC-H
BMC-T
Aldehyde dehydrogenase
PTAC
Signature enzyme
Alcohol dehydrogenase
Regulator

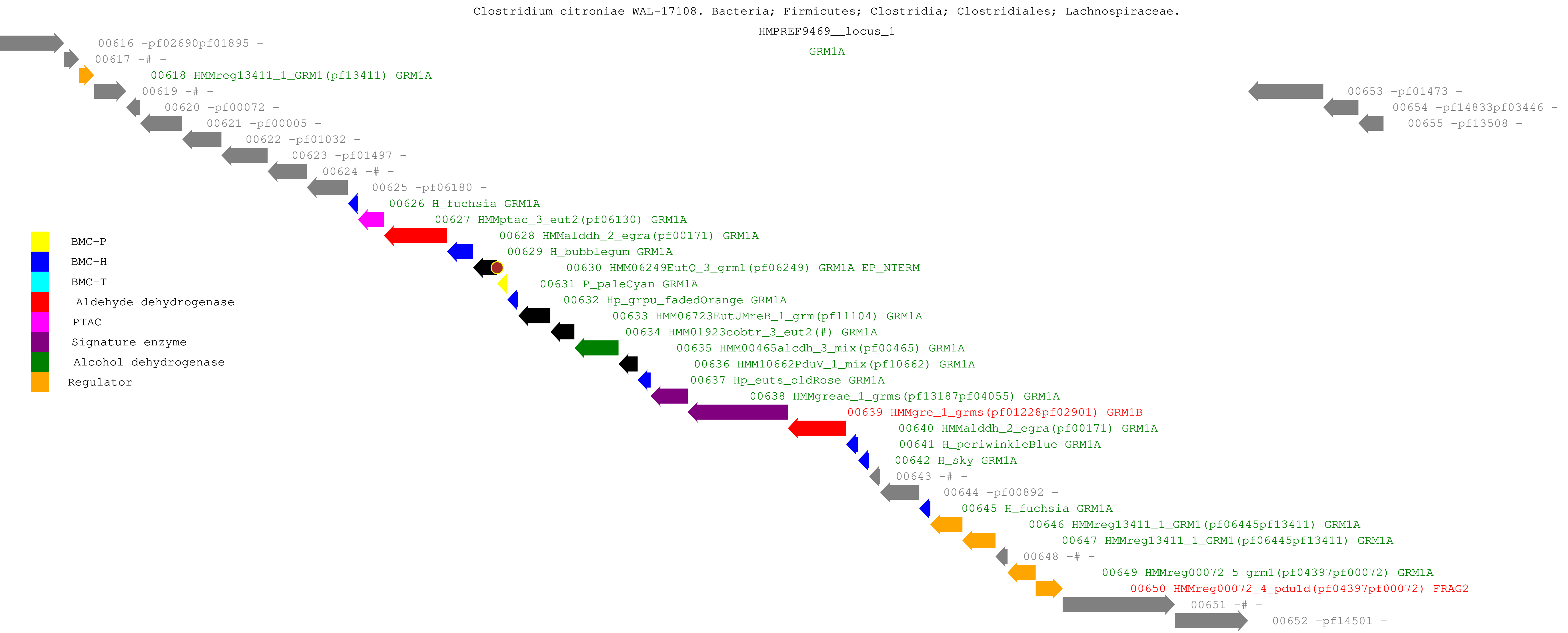

Clostridium citroniae WAL-17108. Bacteria; Firmicutes; Clostridia; Clostridiales; Lachnospiraceae.
HMPREF9469__locus_1
GRM1A
00616 -pf02690pf01895 -
00617 -# -
00618 HMMreg13411_1_GRM1(pf13411) GRM1A
00619 -# -
00620 -pf00072 -
00621 -pf00005 -
00622 -pf01032 -
00623 -pf01497 -
00624 -# -
00625 -pf06180 -
00626 H_fuchsia GRM1A
00627 HMMptac_3_eut2(pf06130) GRM1A
00628 HMMaldd h_2_egra(pf00171) GRM1A
00629 H_bubblegum GRM1A
00630 HMM06249EutQ_3_grm1(pf06249) GRM1A EP_NTERM
00631 P_paleCyan GRM1A
00632 Hp_grpu_fadedOrange GRM1A
00633 HMM06723EutJMreB_1_grm(pf11104) GRM1A
00634 HMM01923cobtr_3_eut2(#) GRM1A
00635 HMM00465alcdh_3_mix(pf00465) GRM1A
00636 HMM10662PduV_1_mix(pf10662) GRM1A
00637 Hp_euts_oldRose GRM1A
00638 HMMgreae_1_grms(pf13187pf04055) GRM1A
00639 HMMgre_1_grms(pf01228pf02901) GRM1B
00640 HMMaldd h_2_egra(pf00171) GRM1A
00641 H_periwinkleBlue GRM1A
00642 H_sky GRM1A
00643 -# -
00644 -pf00892 -
00645 H_fuchsia GRM1A
00646 HMMreg13411_1_GRM1(pf06445pf13411) GRM1A
00647 HMMreg13411_1_GRM1(pf06445pf13411) GRM1A
00648 -# -
00649 HMMreg00072_5_grm1(pf04397pf00072) GRM1A
00650 HMMreg00072_4_pdu1d(pf04397pf00072) FRAG2
00651 -# -
00652 -pf14501 -
00653 -pf01473 -
00654 -pf14833pf03446 -
00655 -pf13508 -
BMC-P
BMC-H
BMC-T
Aldehyde dehydrogenase
PTAC
Signature enzyme
Alcohol dehydrogenase
Regulator

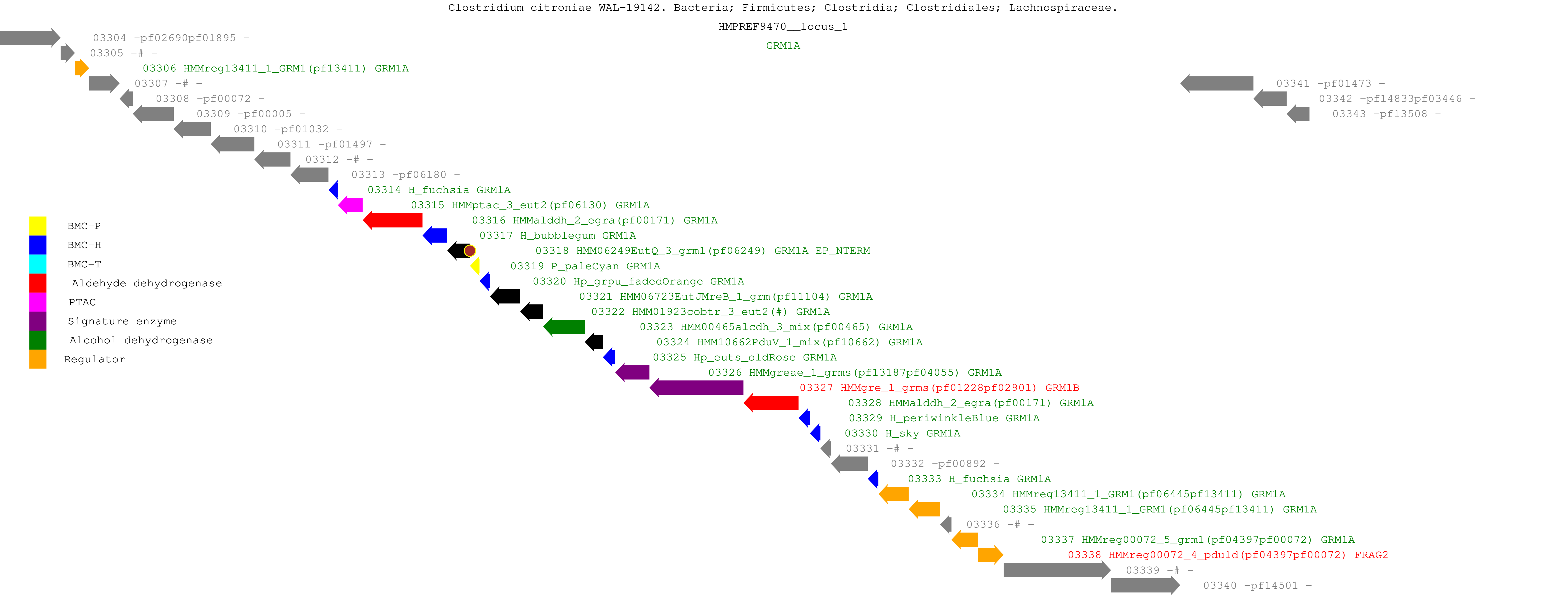

Clostridium citroniae WAL-19142. Bacteria; Firmicutes; Clostridia; Clostridiales; Lachnospiraceae.
HMPREF9470__locus_1
GRM1A
03304 -pf02690pf01895 -
03305 -# -
03306 HMMreg13411_1_GRM1(pf13411) GRM1A
03307 -# -
03308 -pf00072 -
03309 -pf00005 -
03310 -pf01032 -
03311 -pf01497 -
03312 -# -
03313 -pf06180 -
03314 H_fuchsia GRM1A
03315 HMMptac_3_eut2(pf06130) GRM1A
03316 HMMaldh_2_egra(pf00171) GRM1A
03317 H_bubblegum GRM1A
03318 HMM06249EutQ_3_grm1(pf06249) GRM1A EP_NTERM
03319 P_paleCyan GRM1A
03320 Hp_grpu_fadedOrange GRM1A
03321 HMM06723EutJMreB_1_grm(pf11104) GRM1A
03322 HMM01923cobtr_3_eut2(#) GRM1A
03323 HMM00465alcdh_3_mix(pf00465) GRM1A
03324 HMM10662PduV_1_mix(pf10662) GRM1A
03325 Hp_euts_oldRose GRM1A
03326 HMMgreae_1_grms(pf13187pf04055) GRM1A
03327 HMMgre_1_grms(pf01228pf02901) GRM1B
03328 HMMaldh_2_egra(pf00171) GRM1A
03329 H_periwinkleBlue GRM1A
03330 H_sky GRM1A
03331 -# -
03332 -pf00892 -
03333 H_fuchsia GRM1A
03334 HMMreg13411_1_GRM1(pf06445pf13411) GRM1A
03335 HMMreg13411_1_GRM1(pf06445pf13411) GRM1A
03336 -# -
03337 HMMreg00072_5_grm1(pf04397pf00072) GRM1A
03338 HMMreg00072_4_pdu1d(pf04397pf00072) FRAG2
03339 -# -
03340 -pf14501 -
03341 -pf01473 -
03342 -pf14833pf03446 -
03343 -pf13508 -
BMC-P
BMC-H
BMC-T
Aldehyde dehydrogenase
PTAC
Signature enzyme
Alcohol dehydrogenase
Regulator

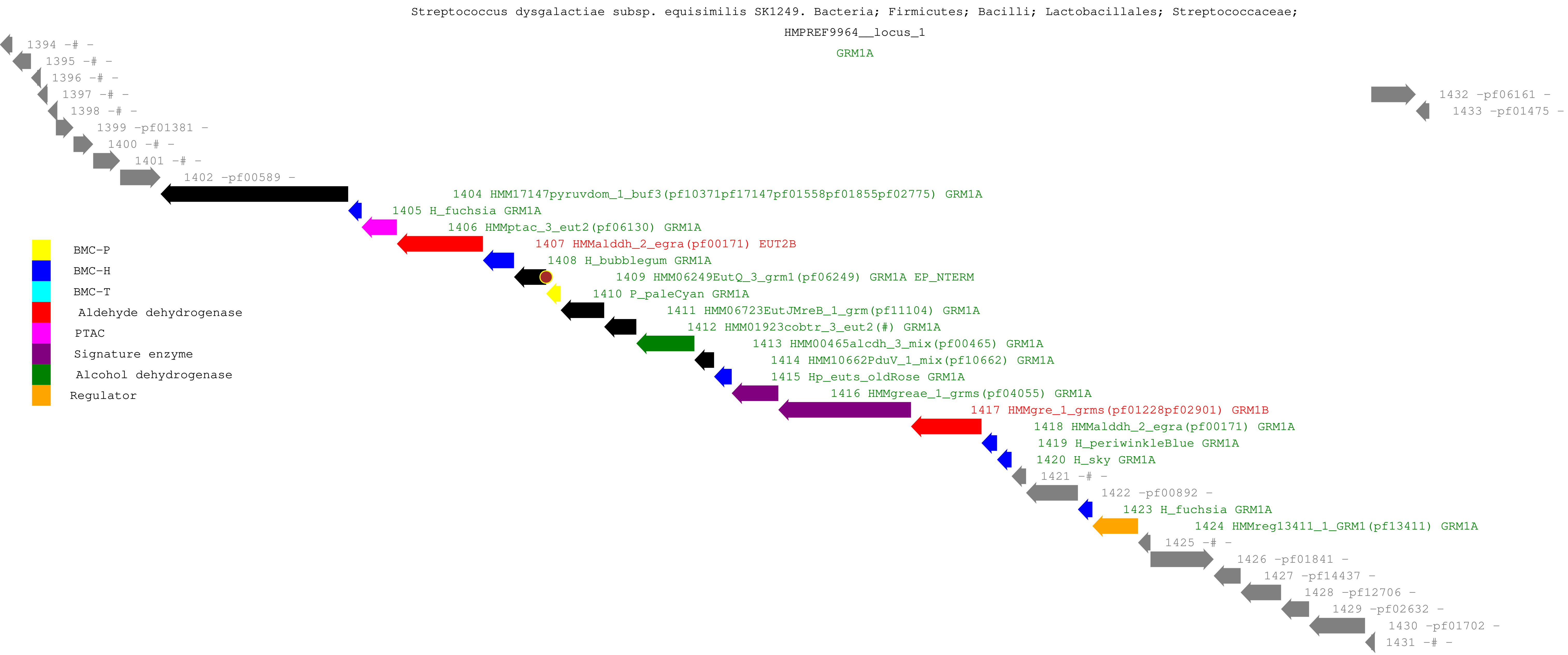

Streptococcus dysgalactiae subsp. equisimilis SK1249. Bacteria; Firmicutes; Bacilli; Lactobacillales; Streptococcaceae;
HMPREF9964__locus_1
GRM1A
1394 -# -
1395 -# -
1396 -# -
1397 -# -
1398 -# -
1399 -pf01381 -
1400 -# -
1401 -# -
1402 -pf00589 -
1404 HMM17147pyruvdom_1_buf3(pf10371pf17147pf01558pf01855pf02775) GRM1A
1405 H_fuchsia GRM1A
1406 HMMptac_3_eut2(pf06130) GRM1A
1407 HMMalddh_2_egra(pf00171) EUT2B
1408 H_bubblegum GRM1A
1409 HMM06249EutQ_3_grm1(pf06249) GRM1A EP_NTERM
1410 P_paleCyan GRM1A
1411 HMM06723EutJMreB_1_grm(pf11104) GRM1A
1412 HMM01923cobtr_3_eut2(#) GRM1A
1413 HMM00465alcdh_3_mix(pf00465) GRM1A
1414 HMM10662PduV_1_mix(pf10662) GRM1A
1415 Hp_euts_oldRose GRM1A
1416 HMMgreae_1_grms(pf04055) GRM1A
1417 HMMgre_1_grms(pf01228pf02901) GRM1B
1418 HMMalddh_2_egra(pf00171) GRM1A
1419 H_periwinkleBlue GRM1A
1420 H_sky GRM1A
1421 -# -
1422 -pf00892 -
1423 H_fuchsia GRM1A
1424 HMMreg13411_1_GRM1(pf13411) GRM1A
1425 -# -
1426 -pf01841 -
1427 -pf14437 -
1428 -pf12706 -
1429 -pf02632 -
1430 -pf01702 -
1431 -# -
1432 -pf06161 -
1433 -pf01475 -
BMC-P
BMC-H
BMC-T
Aldehyde dehydrogenase
PTAC
Signature enzyme
Alcohol dehydrogenase
Regulator

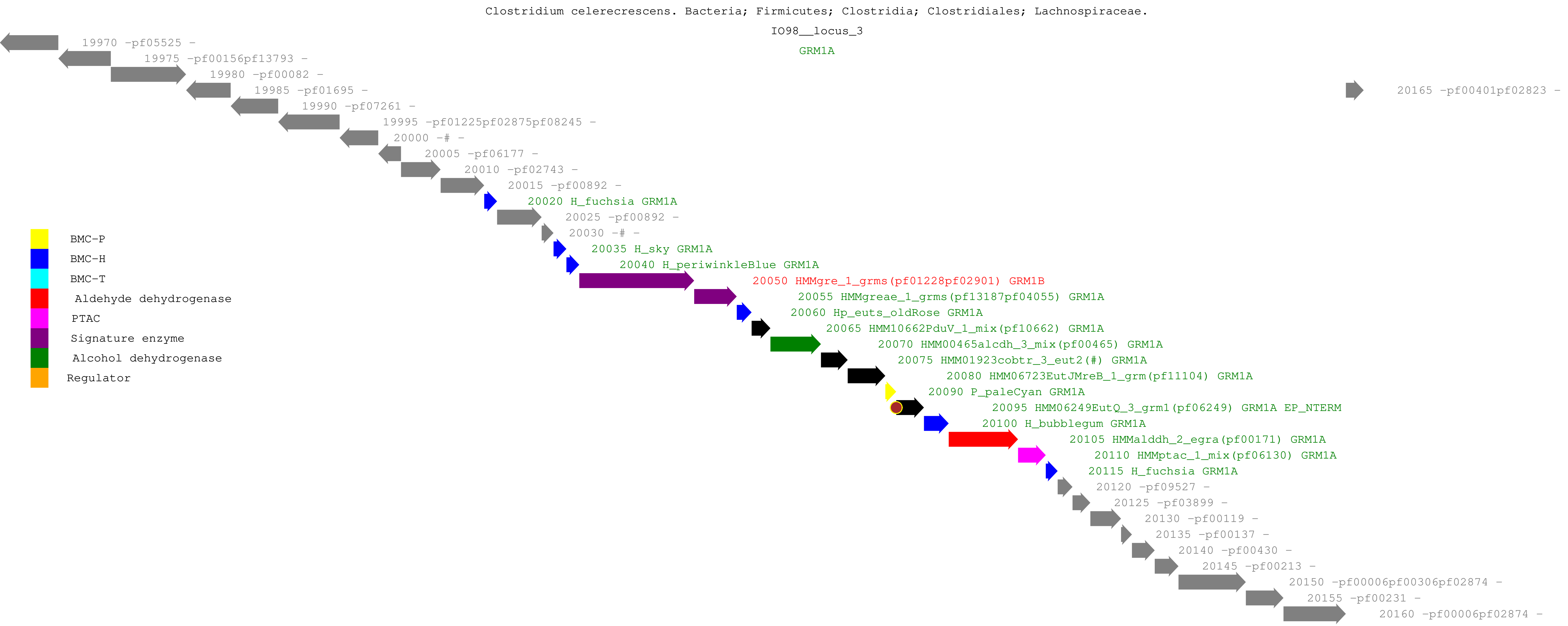
Clostridium celerecrescens. Bacteria; Firmicutes; Clostridia; Clostridiales; Lachnospiraceae.
IO98__locus_3
GRM1A
19970 -pf05525 -
19975 -pf00156pf13793 -
19980 -pf00082 -
19985 -pf01695 -
19990 -pf07261 -
19995 -pf01225pf02875pf08245 -
20000 -# -
20005 -pf06177 -
20010 -pf02743 -
20015 -pf00892 -
20020 H_fuchsia GRM1A
20025 -pf00892 -
20030 -# -
20035 H_sky GRM1A
20040 H_periwinkleBlue GRM1A
20050 HMMgre_1_grms(pf01228pf02901) GRM1B
20055 HMMgreae_1_grms(pf13187pf04055) GRM1A
20060 Hp_euts_oldRose GRM1A
20065 HMM10662PduV_1_mix(pf10662) GRM1A
20070 HMM00465alcdh_3_mix(pf00465) GRM1A
20075 HMM01923cobtr_3_eut2(#) GRM1A
20080 HMM06723EutJMreB_1_grm(pf11104) GRM1A
20090 P_paleCyan GRM1A
20095 HMM06249EutQ_3_grm1(pf06249) GRM1A EP_NTERM
20100 H_bubblegum GRM1A
20105 HMMalddh_2_egra(pf00171) GRM1A
20110 HMMptac_1_mix(pf06130) GRM1A
20115 H_fuchsia GRM1A
20120 -pf09527 -
20125 -pf03899 -
20130 -pf00119 -
20135 -pf00137 -
20140 -pf00430 -
20145 -pf00213 -
20150 -pf00006pf00306pf02874 -
20155 -pf00231 -
20160 -pf00006pf02874 -
20165 -pf00401pf02823 -
BMC-P
BMC-H
BMC-T
Aldehyde dehydrogenase
PTAC
Signature enzyme
Alcohol dehydrogenase
Regulator

Clostridium sulfidigenes. Bacteria; Firmicutes; Clostridia; Clostridiales; Clostridiaceae;

IO99__locus_2

GRM1A

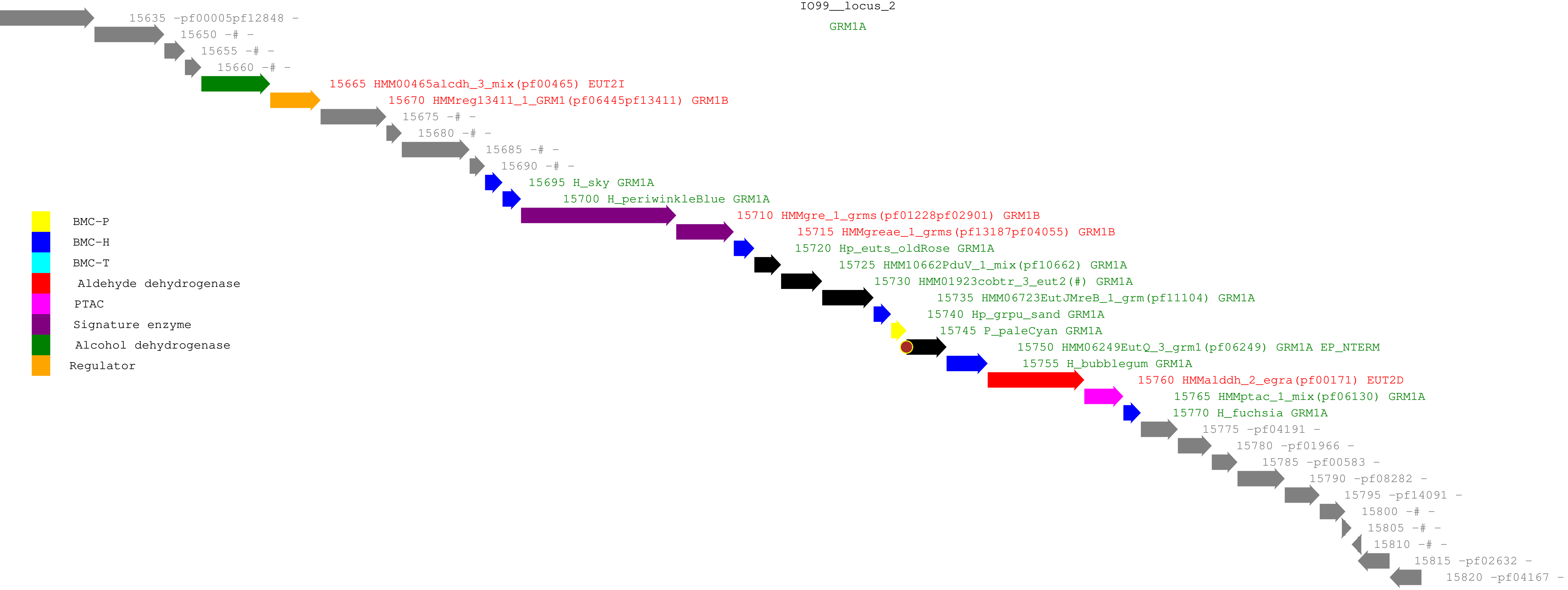

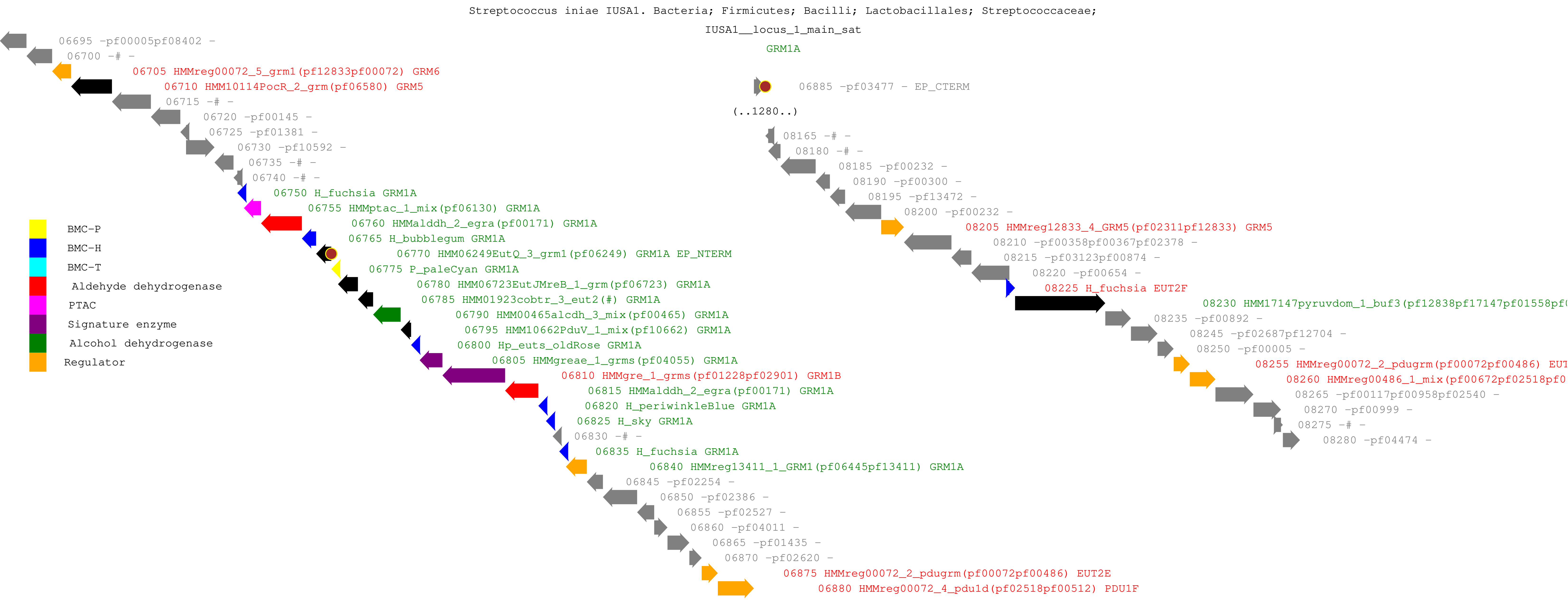
Streptococcus iniae IUSA1. Bacteria; Firmicutes; Bacilli; Lactobacillales; Streptococcaceae;
IUSA1__locus_1_main_sat
GRM1A
06695 -pf00005pf08402 -
06700 -# -
06705 HMMreg00072_5_grm1(pf12833pf00072) GRM6
06710 HMM10114PocR_2_grm(pf06580) GRM5
06715 -# -
06720 -pf00145 -
06725 -pf01381 -
06730 -pf10592 -
06735 -# -
06740 -# -
06750 H_fuchsia GRM1A
06755 HMMptac_1_mix(pf06130) GRM1A
06760 HMMaldh_2_egra(pf00171) GRM1A
06765 H_bubblegum GRM1A
06770 HMM06249EutQ_3_grm1(pf06249) GRM1A EP_NTERM
06775 P_paleCyan GRM1A
06780 HMM06723EutJMreB_1_grm(pf06723) GRM1A
06785 HMM01923cobtr_3_eut2(#) GRM1A
06790 HMM00465alcdh_3_mix(pf00465) GRM1A
06795 HMM10662PduV_1_mix(pf10662) GRM1A
06800 Hp_euts_oldRose GRM1A
06805 HMMgreae_1_grms(pf04055) GRM1A
06810 HMMgre_1_grms(pf01228pf02901) GRM1B
06815 HMMaldh_2_egra(pf00171) GRM1A
06820 H_periwinkleBlue GRM1A
06825 H_sky GRM1A
06830 -# -
06835 H_fuchsia GRM1A
06840 HMMreg13411_1_GRM1(pf06445pf13411) GRM1A
06845 -pf02254 -
06850 -pf02386 -
06855 -pf02527 -
06860 -pf04011 -
06865 -pf01435 -
06870 -pf02620 -
06875 HMMreg00072_2_pdugrm(pf00072pf00486) EUT2E
06880 HMMreg00072_4_pdu1d(pf02518pf00512) PDU1F
BMC-P
BMC-H
BMC-T
Aldehyde dehydrogenase
PTAC
Signature enzyme
Alcohol dehydrogenase
Regulator
06885 -pf03477 - EP_CTERM
(..1280..)
08165 -# -
08180 -# -
08185 -pf00232 -
08190 -pf00300 -
08195 -pf13472 -
08200 -pf00232 -
08205 HMMreg12833_4_GRM5(pf02311pf12833) GRM5
08210 -pf00358pf00367pf02378 -
08215 -pf03123pf00874 -
08220 -pf00654 -
08225 H_fuchsia EUT2F
08230 HMM17147pyruvdom_1_buf3(pf12838pf17147pf01558pf(
08235 -pf00892 -
08245 -pf02687pf12704 -
08250 -pf00005 -
08255 HMMreg00072_2_pdugrm(pf00072pf00486) EUT
08260 HMMreg00486_1_mix(pf00672pf02518pf0
08265 -pf00117pf00958pf02540 -
08270 -pf00999 -
08275 -# -
08280 -pf04474 -

Clostridium sp. ASBs410. Bacteria; Firmicutes; Clostridia; Clostridiales; Clostridiaceae;

K413DRAFT__locus_1

GRM1A

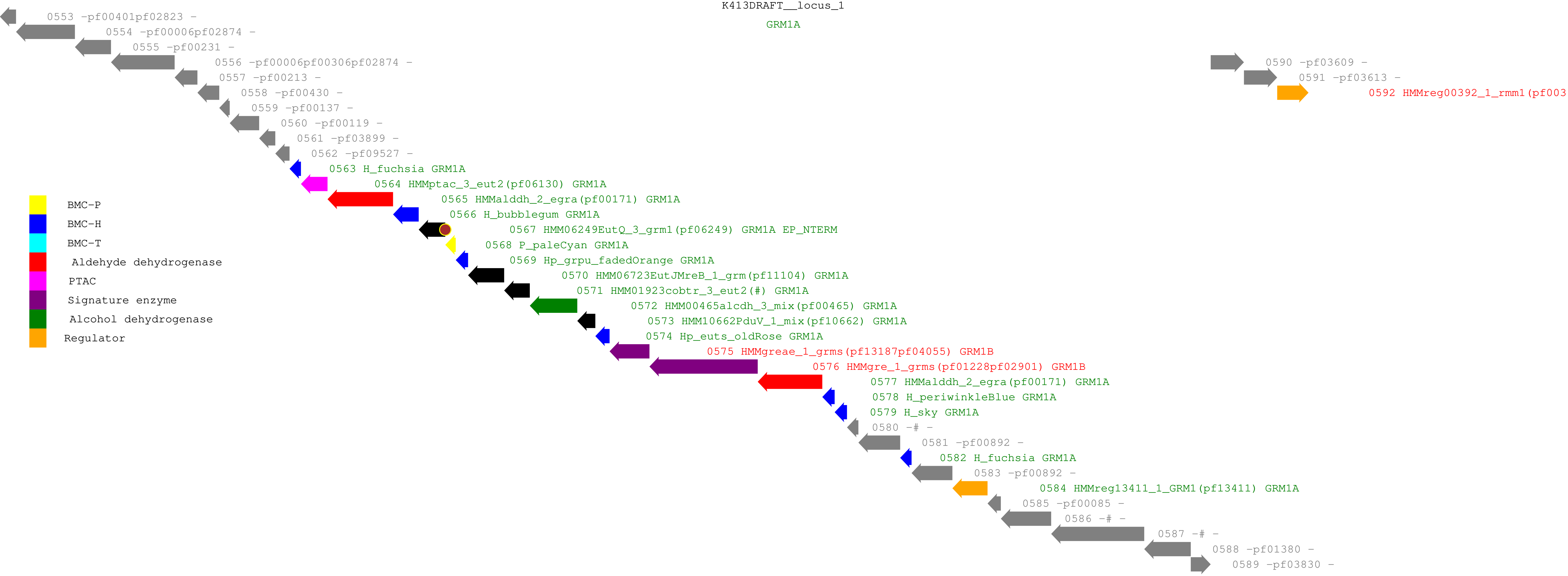

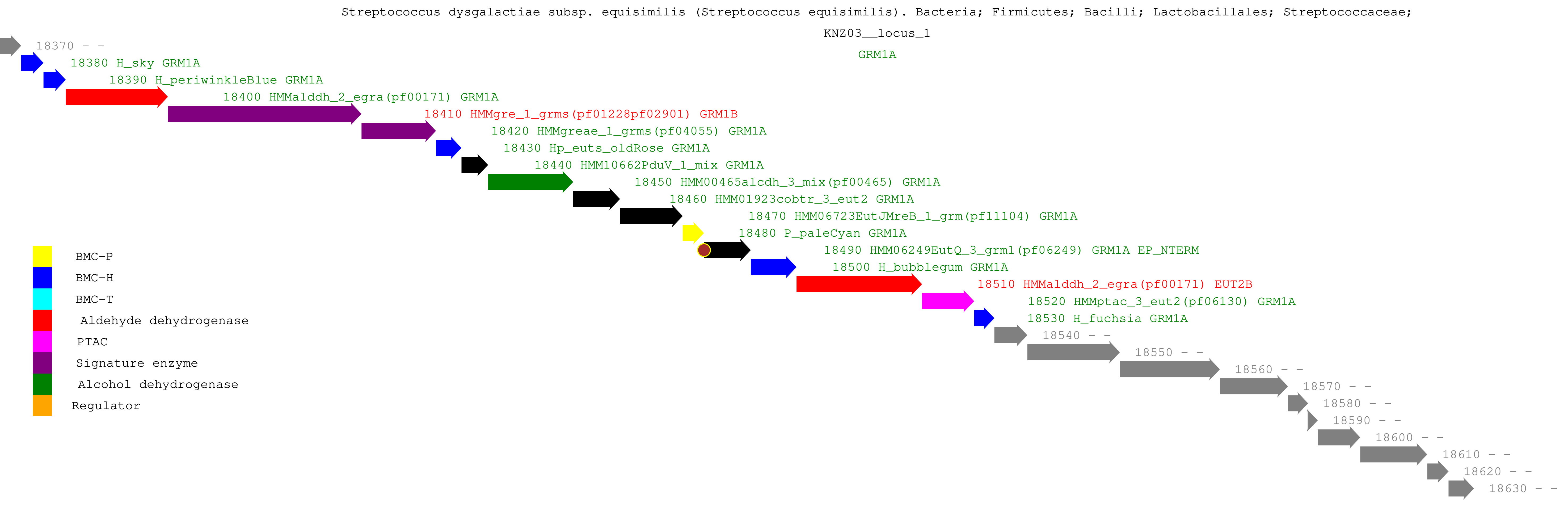
Streptococcus dysgalactiae subsp. equisimilis (Streptococcus equisimilis). Bacteria; Firmicutes; Bacilli; Lactobacillales; Streptococcaceae;
KNZ03__locus_1
GRM1A
18370 - -
18380 H_sky GRM1A
18390 H_periwinkleBlue GRM1A
18400 HMMalddh_2_egra(pf00171) GRM1A
18410 HMMgre_1_grms(pf01228pf02901) GRM1B
18420 HMMgreae_1_grms(pf04055) GRM1A
18430 Hp_euts_oldRose GRM1A
18440 HMM10662PduV_1_mix GRM1A
18450 HMM00465alcdh_3_mix(pf00465) GRM1A
18460 HMM01923cobtr_3_eut2 GRM1A
18470 HMM06723EutJMreB_1_grm(pf11104) GRM1A
18480 P_paleCyan GRM1A
18490 HMM06249EutQ_3_grm1(pf06249) GRM1A EP_NTERM
18500 H_bubblegum GRM1A
18510 HMMalddh_2_egra(pf00171) EUT2B
18520 HMMptac_3_eut2(pf06130) GRM1A
18530 H_fuchsia GRM1A
18540 - -
18550 - -
18560 - -
18570 - -
18580 - -
18590 - -
18600 - -
18610 - -
18620 - -
18630 - -
BMC-P
BMC-H
BMC-T
Aldehyde dehydrogenase
PTAC
Signature enzyme
Alcohol dehydrogenase
Regulator

Streptococcus dysgalactiae subsp. equisimilis (Streptococcus equisimilis). Bacteria; Firmicutes; Bacilli; Lactobacillales; Streptococcaceae;

KNZ07__locus_1

GRM1A

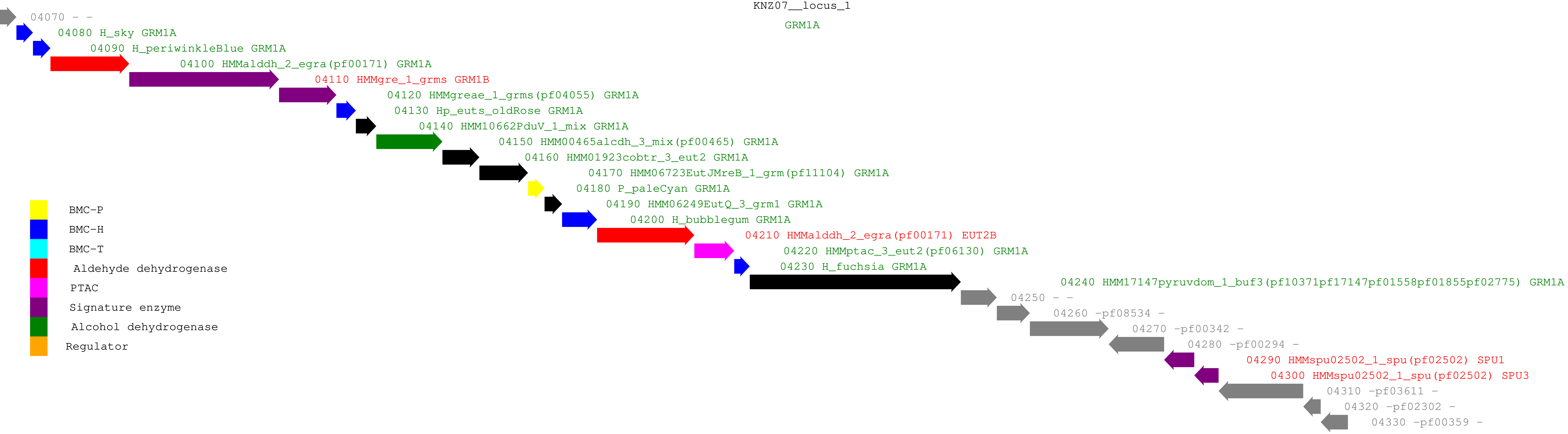

Collinsella aerofaciens. Bacteria; Actinobacteria; Coriobacteriia; Coriobacteriales;

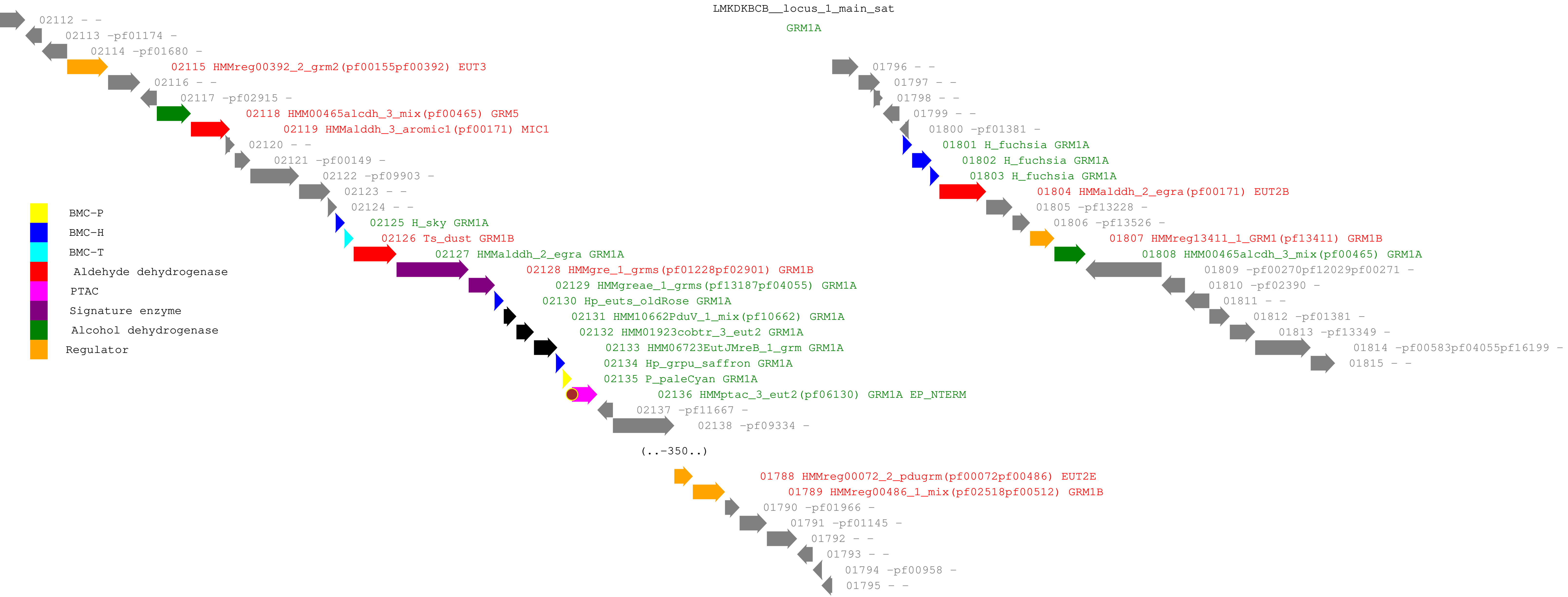

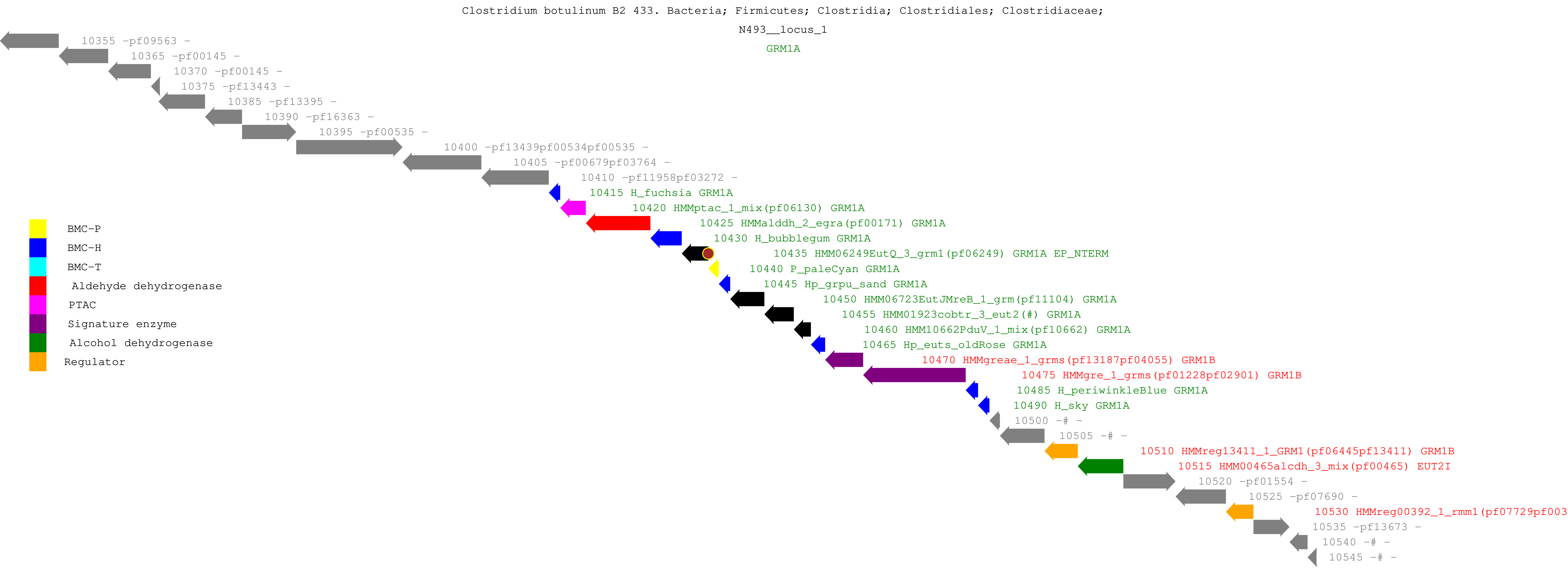

Clostridium botulinum B2 433. Bacteria; Firmicutes; Clostridia; Clostridiales; Clostridiaceae;
N493__locus_1
GRM1A
10355 -pf09563 -
10365 -pf00145 -
10370 -pf00145 -
10375 -pf13443 -
10385 -pf13395 -
10390 -pf16363 -
10395 -pf00535 -
10400 -pf13439pf00534pf00535 -
10405 -pf00679pf03764 -
10410 -pf11958pf03272 -
10415 H_fuchsia GRM1A
10420 HMMptac_1_mix(pf06130) GRM1A
10425 HMMalddh_2_egra(pf00171) GRM1A
10430 H_bubblegum GRM1A
10435 HMM06249EutQ_3_grm1(pf06249) GRM1A EP_NTERM
10440 P_paleCyan GRM1A
10445 Hp_grpu_sand GRM1A
10450 HMM06723EutJMreB_1_grm(pf11104) GRM1A
10455 HMM01923cobtr_3_eut2(#) GRM1A
10460 HMM10662PduV_1_mix(pf10662) GRM1A
10465 Hp_euts_oldRose GRM1A
10470 HMMgreae_1_grms(pf13187pf04055) GRM1B
10475 HMMgre_1_grms(pf01228pf02901) GRM1B
10485 H_periwinkleBlue GRM1A
10490 H_sky GRM1A
10500 -# -
10505 -# -
10510 HMMreg13411_1_GRM1(pf06445pf13411) GRM1B
10515 HMM00465alcdh_3_mix(pf00465) EUT2I
10520 -pf01554 -
10525 -pf07690 -
10530 HMMreg00392_1_rmm1(pf07729pf003
10535 -pf13673 -
10540 -# -
10545 -# -
BMC-P
BMC-H
BMC-T
Aldehyde dehydrogenase
PTAC
Signature enzyme
Alcohol dehydrogenase
Regulator

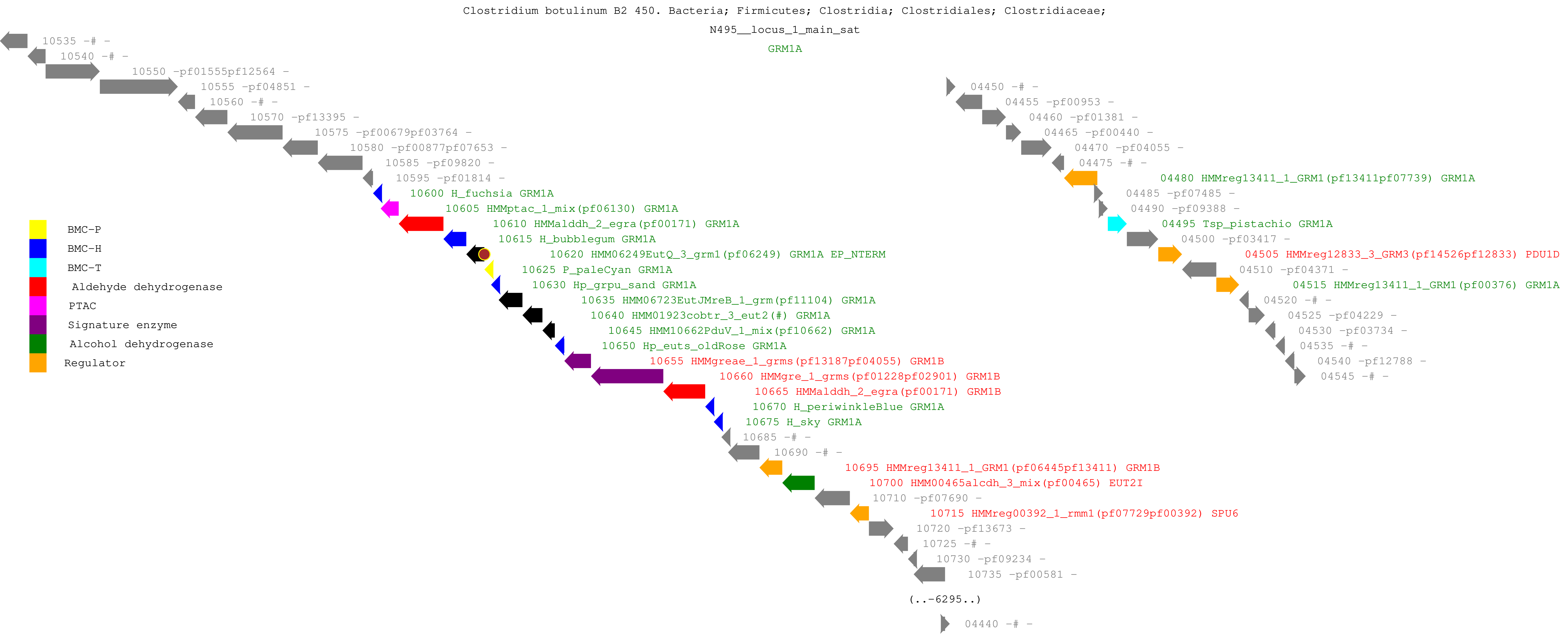
Clostridium botulinum B2 450. Bacteria; Firmicutes; Clostridia; Clostridiales; Clostridiaceae;
N495__locus_1_main_sat
GRM1A
10535 -# -
10540 -# -
10550 -pf01555pf12564 -
10555 -pf04851 -
10560 -# -
10570 -pf13395 -
10575 -pf00679pf03764 -
10580 -pf00877pf07653 -
10585 -pf09820 -
10595 -pf01814 -
10600 H_fuchsia GRM1A
10605 HMMptac_1_mix(pf06130) GRM1A
10610 HMMalddh_2_egra(pf00171) GRM1A
10615 H_bubblegum GRM1A
10620 HMM06249EutQ_3_grm1(pf06249) GRM1A EP_NTERM
10625 P_paleCyan GRM1A
10630 Hp_grpu_sand GRM1A
10635 HMM06723EutJMreB_1_grm(pf11104) GRM1A
10640 HMM01923cobtr_3_eut2(#) GRM1A
10645 HMM10662PduV_1_mix(pf10662) GRM1A
10650 Hp_euts_oldRose GRM1A
10655 HMMgreae_1_grms(pf13187pf04055) GRM1B
10660 HMMgre_1_grms(pf01228pf02901) GRM1B
10665 HMMalddh_2_egra(pf00171) GRM1B
10670 H_periwinkleBlue GRM1A
10675 H_sky GRM1A
10685 -# -
10690 -# -
10695 HMMreg13411_1_GRM1(pf06445pf13411) GRM1B
10700 HMM00465alcdh_3_mix(pf00465) EUT2I
10710 -pf07690 -
10715 HMMreg00392_1_rmm1(pf07729pf00392) SPU6
10720 -pf13673 -
10725 -# -
10730 -pf09234 -
10735 -pf00581 -
(..-6295..)
04440 -# -
04450 -# -
04455 -pf00953 -
04460 -pf01381 -
04465 -pf00440 -
04470 -pf04055 -
04475 -# -
04480 HMMreg13411_1_GRM1(pf13411pf07739) GRM1A
04485 -pf07485 -
04490 -pf09388 -
04495 Tsp_pistachio GRM1A
04500 -pf03417 -
04505 HMMreg12833_3_GRM3(pf14526pf12833) PDU1D
04510 -pf04371 -
04515 HMMreg13411_1_GRM1(pf00376) GRM1A
04520 -# -
04525 -pf04229 -
04530 -pf03734 -
04535 -# -
04540 -pf12788 -
04545 -# -
BMC-P
BMC-H
BMC-T
Aldehyde dehydrogenase
PTAC
Signature enzyme
Alcohol dehydrogenase
Regulator

Desulfotomaculum aquiferis. Bacteria; Firmicutes; Clostridia; Clostridiales; Peptococcaceae;

N752__locus_1

GRM1A

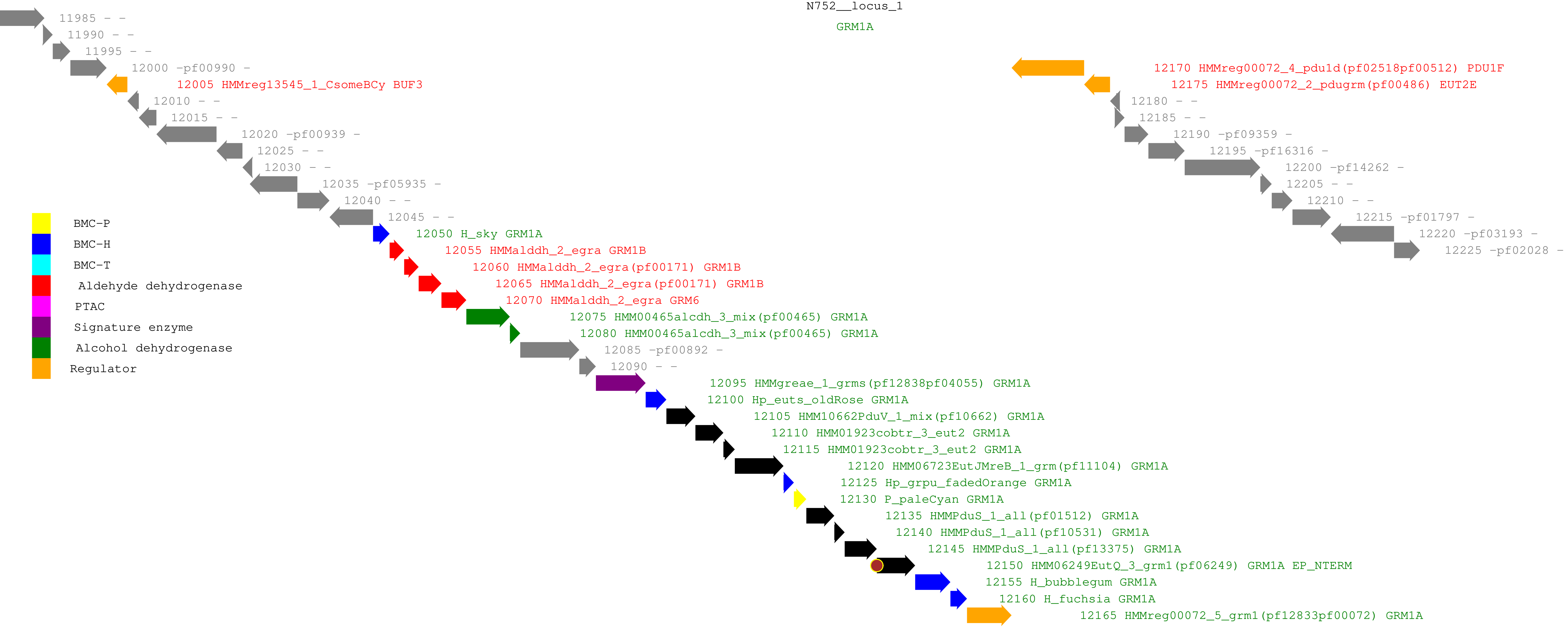

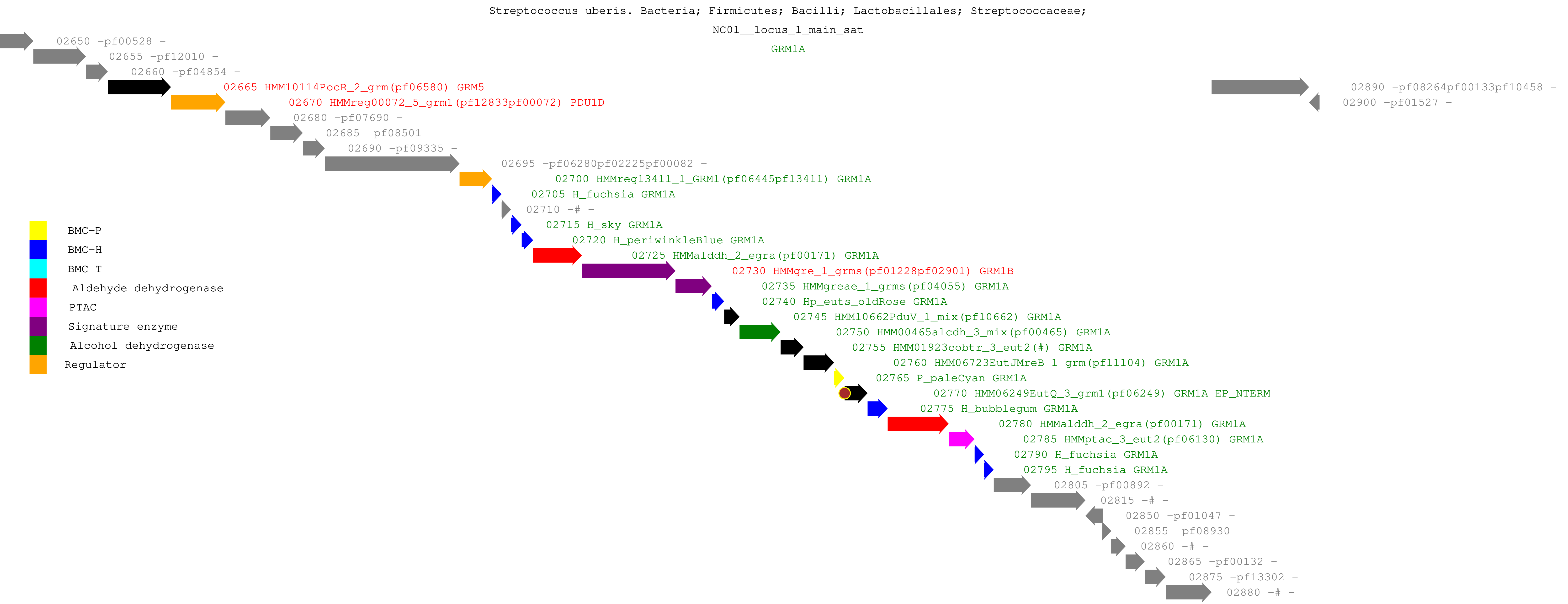

Streptococcus uberis. Bacteria; Firmicutes; Bacilli; Lactobacillales; Streptococcaceae;
NC01__locus_1_main_sat
GRM1A
02650 -pf00528 -
02655 -pf12010 -
02660 -pf04854 -
02665 HMM10114PocR_2_grm(pf06580) GRM5
02670 HMMreg00072_5_grm1(pf12833pf00072) PDU1D
02680 -pf07690 -
02685 -pf08501 -
02690 -pf09335 -
02695 -pf06280pf02225pf00082 -
02700 HMMreg13411_1_GRM1(pf06445pf13411) GRM1A
02705 H_fuchsia GRM1A
02710 -# -
02715 H_sky GRM1A
02720 H_periwinkleBlue GRM1A
02725 HMMalddh_2_egra(pf00171) GRM1A
02730 HMMgre_1_grms(pf01228pf02901) GRM1B
02735 HMMgreae_1_grms(pf04055) GRM1A
02740 Hp_euts_oldRose GRM1A
02745 HMM10662PduV_1_mix(pf10662) GRM1A
02750 HMM00465alcdh_3_mix(pf00465) GRM1A
02755 HMM01923cobtr_3_eut2(#) GRM1A
02760 HMM06723EutJMreB_1_grm(pf11104) GRM1A
02765 P_paleCyan GRM1A
02770 HMM06249EutQ_3_grm1(pf06249) GRM1A EP_NTERM
02775 H_bubblegum GRM1A
02780 HMMalddh_2_egra(pf00171) GRM1A
02785 HMMptac_3_eut2(pf06130) GRM1A
02790 H_fuchsia GRM1A
02795 H_fuchsia GRM1A
02805 -pf00892 -
02815 -# -
02850 -pf01047 -
02855 -pf08930 -
02860 -# -
02865 -pf00132 -
02875 -pf13302 -
02880 -# -
02890 -pf08264pf00133pf10458 -
02900 -pf01527 -
BMC-P
BMC-H
BMC-T
Aldehyde dehydrogenase
PTAC
Signature enzyme
Alcohol dehydrogenase
Regulator

Paenibacillus thiaminolyticus (Bacillus thiaminolyticus). Bacteria; Firmicutes; Bacilli; Bacillales; Paenibacillaceae;

NCTC11027__locus_3

GRM1A

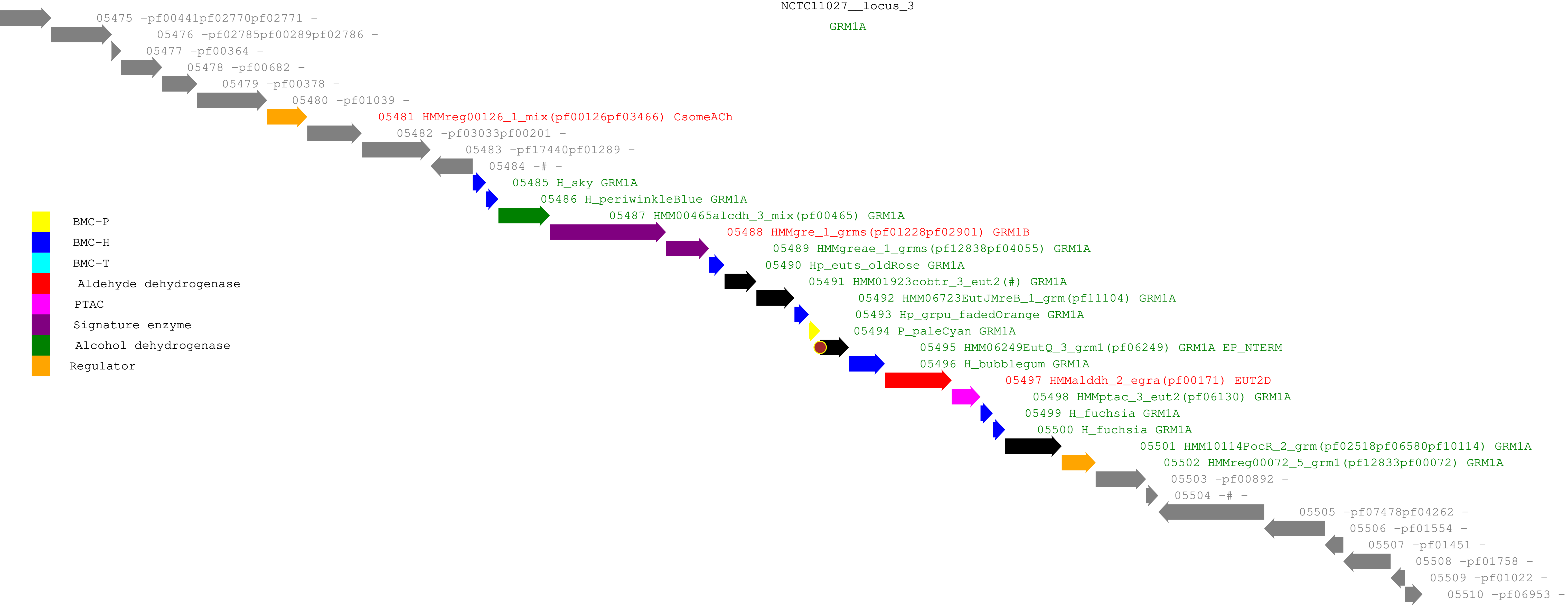

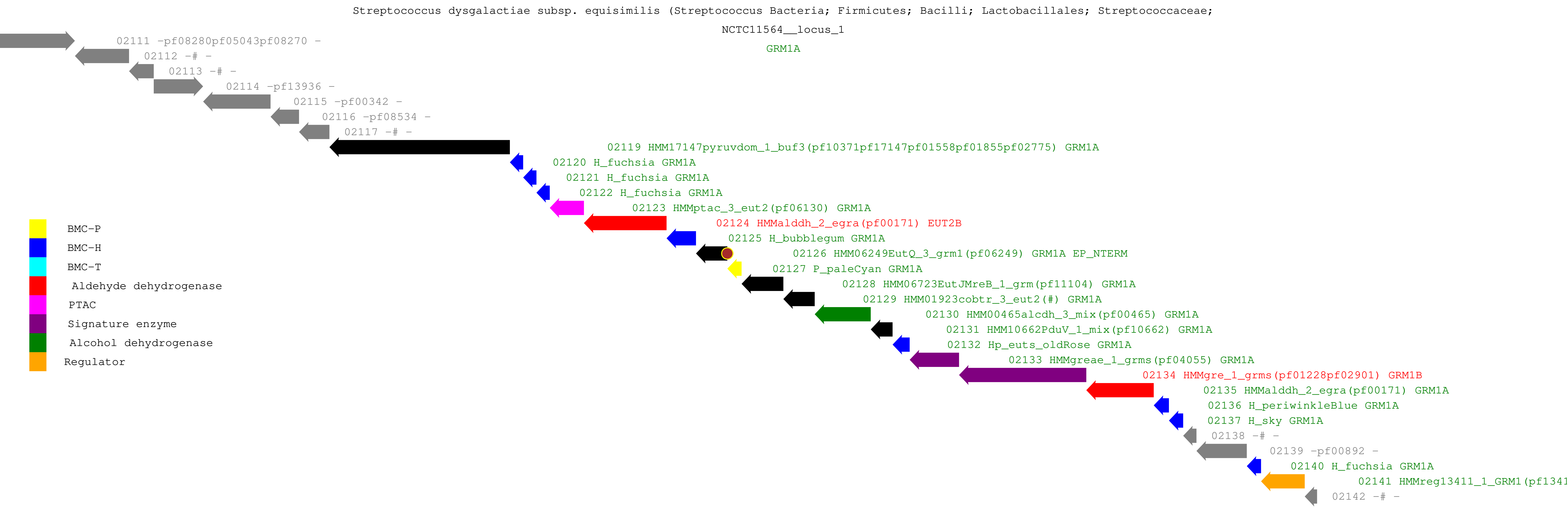

Streptococcus dysgalactiae subsp. equisimilis (Streptococcus Bacteria; Firmicutes; Bacilli; Lactobacillales; Streptococcaceae;
NCTC11564__locus_1
GRM1A
02111 -pf08280pf05043pf08270 -
02112 -# -
02113 -# -
02114 -pf13936 -
02115 -pf00342 -
02116 -pf08534 -
02117 -# -
02119 HMM17147pyruvdom_1_buf3(pf10371pf17147pf01558pf01855pf02775) GRM1A
02120 H_fuchsia GRM1A
02121 H_fuchsia GRM1A
02122 H_fuchsia GRM1A
02123 HMMptac_3_eut2(pf06130) GRM1A
02124 HMMalddh_2_egra(pf00171) EUT2B
02125 H_bubblegum GRM1A
02126 HMM06249EutQ_3_grm1(pf06249) GRM1A EP_NTERM
02127 P_paleCyan GRM1A
02128 HMM06723EutJMreB_1_grm(pf11104) GRM1A
02129 HMM01923cobtr_3_eut2(#) GRM1A
02130 HMM00465alcdh_3_mix(pf00465) GRM1A
02131 HMM10662PduV_1_mix(pf10662) GRM1A
02132 Hp_euts_oldRose GRM1A
02133 HMMgreae_1_grms(pf04055) GRM1A
02134 HMMgre_1_grms(pf01228pf02901) GRM1B
02135 HMMalddh_2_egra(pf00171) GRM1A
02136 H_periwinkleBlue GRM1A
02137 H_sky GRM1A
02138 -# -
02139 -pf00892 -
02140 H_fuchsia GRM1A
02141 HMMreg13411_1_GRM1(pf1341
02142 -# -
BMC-P
BMC-H
BMC-T
Aldehyde dehydrogenase
PTAC
Signature enzyme
Alcohol dehydrogenase
Regulator

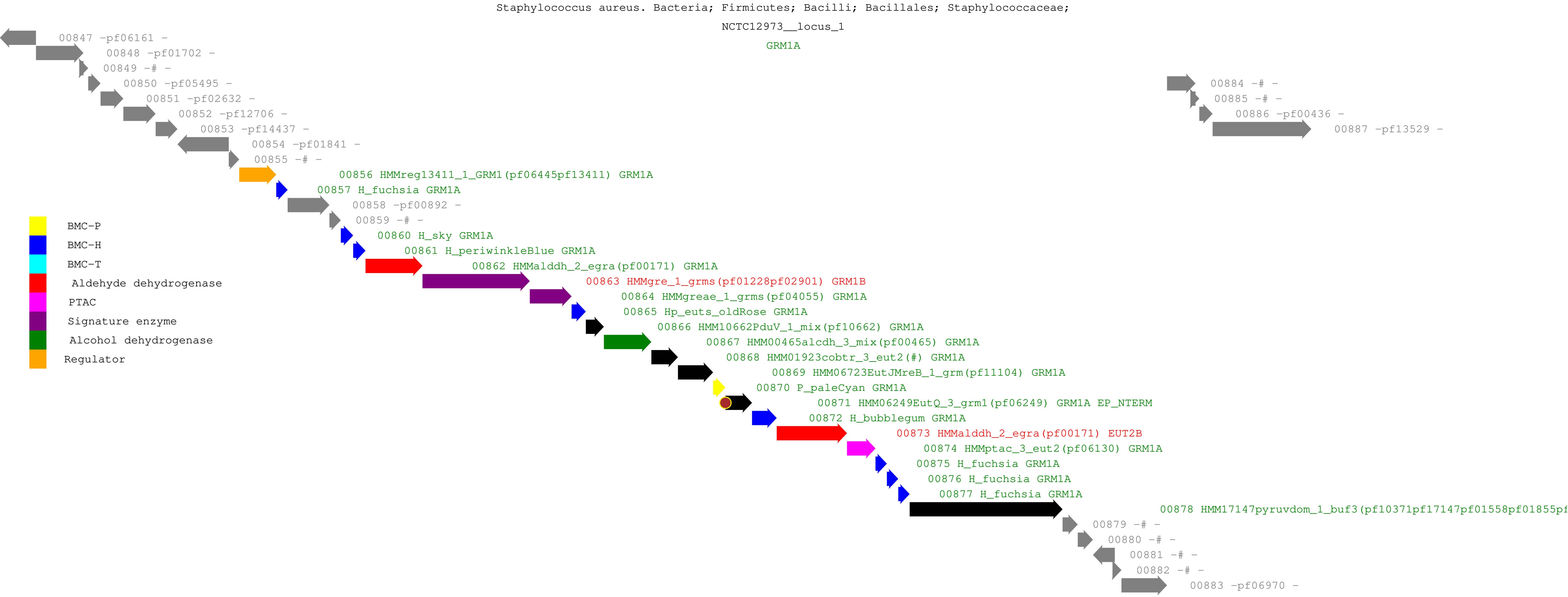
Staphylococcus aureus. Bacteria; Firmicutes; Bacilli; Bacillales; Staphylococcaceae;
NCTC12973__locus_1
GRM1A
00847 -pf06161 -
00848 -pf01702 -
00849 -# -
00850 -pf05495 -
00851 -pf02632 -
00852 -pf12706 -
00853 -pf14437 -
00854 -pf01841 -
00855 -# -
00856 HMMreg13411_1_GRM1(pf06445pf13411) GRM1A
00857 H_fuchsia GRM1A
00858 -pf00892 -
00859 -# -
00860 H_sky GRM1A
00861 H_periwinkleBlue GRM1A
00862 HMMaldh_2_egra(pf00171) GRM1A
00863 HMMgre_1_grms(pf01228pf02901) GRM1B
00864 HMMgreae_1_grms(pf04055) GRM1A
00865 Hp_euts_oldRose GRM1A
00866 HMM10662PduV_1_mix(pf10662) GRM1A
00867 HMM00465alcdh_3_mix(pf00465) GRM1A
00868 HMM01923cobtr_3_eut2(#) GRM1A
00869 HMM06723EutJMreB_1_grm(pf11104) GRM1A
00870 P_paleCyan GRM1A
00871 HMM06249EutQ_3_grm1(pf06249) GRM1A EP_NTERM
00872 H_bubblegum GRM1A
00873 HMMaldh_2_egra(pf00171) EUT2B
00874 HMMptac_3_eut2(pf06130) GRM1A
00875 H_fuchsia GRM1A
00876 H_fuchsia GRM1A
00877 H_fuchsia GRM1A
00878 HMM17147pyruvdom_1_buf3(pf10371pf17147pf01558pf01855pf
00879 -# -
00880 -# -
00881 -# -
00882 -# -
00883 -pf06970 -
00884 -# -
00885 -# -
00886 -pf00436 -
00887 -pf13529 -
BMC-P
BMC-H
BMC-T
Aldehyde dehydrogenase
PTAC
Signature enzyme
Alcohol dehydrogenase
Regulator

Clostridium sporogenes. Bacteria; Firmicutes; Clostridia; Clostridiales; Clostridiaceae;

NCTC13020__locus_1_main_sat

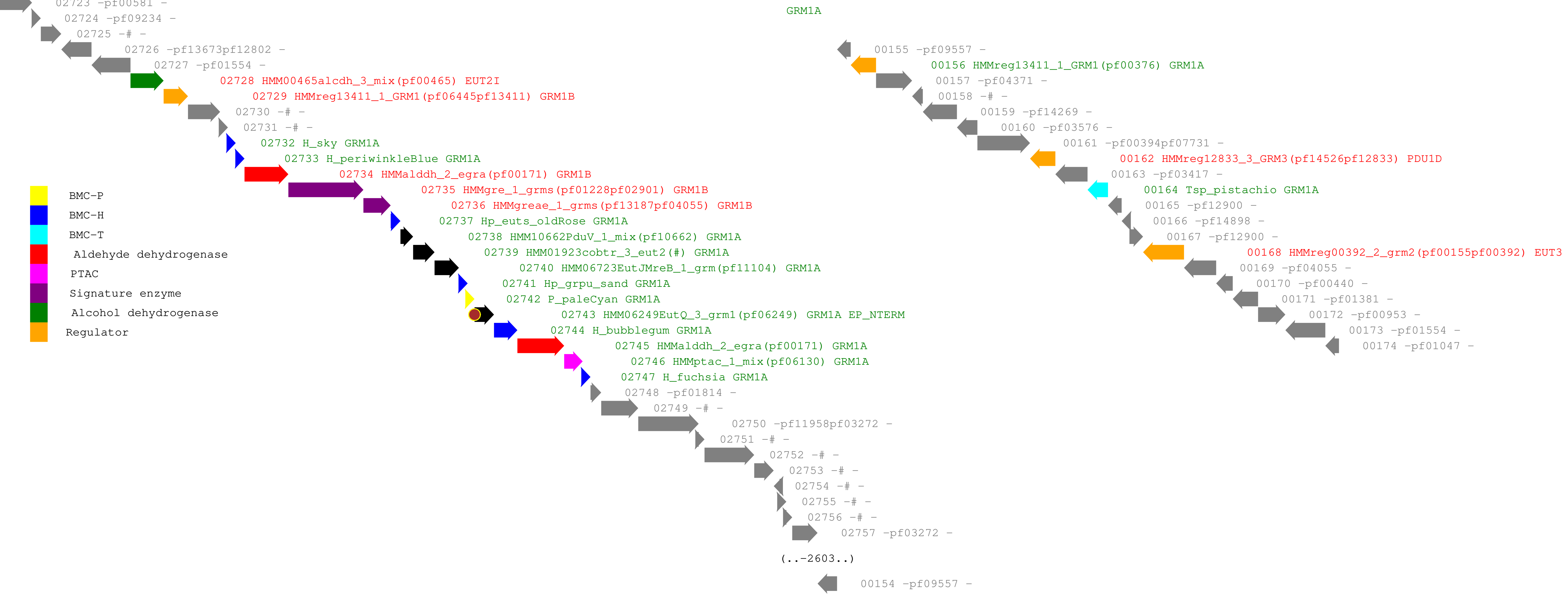

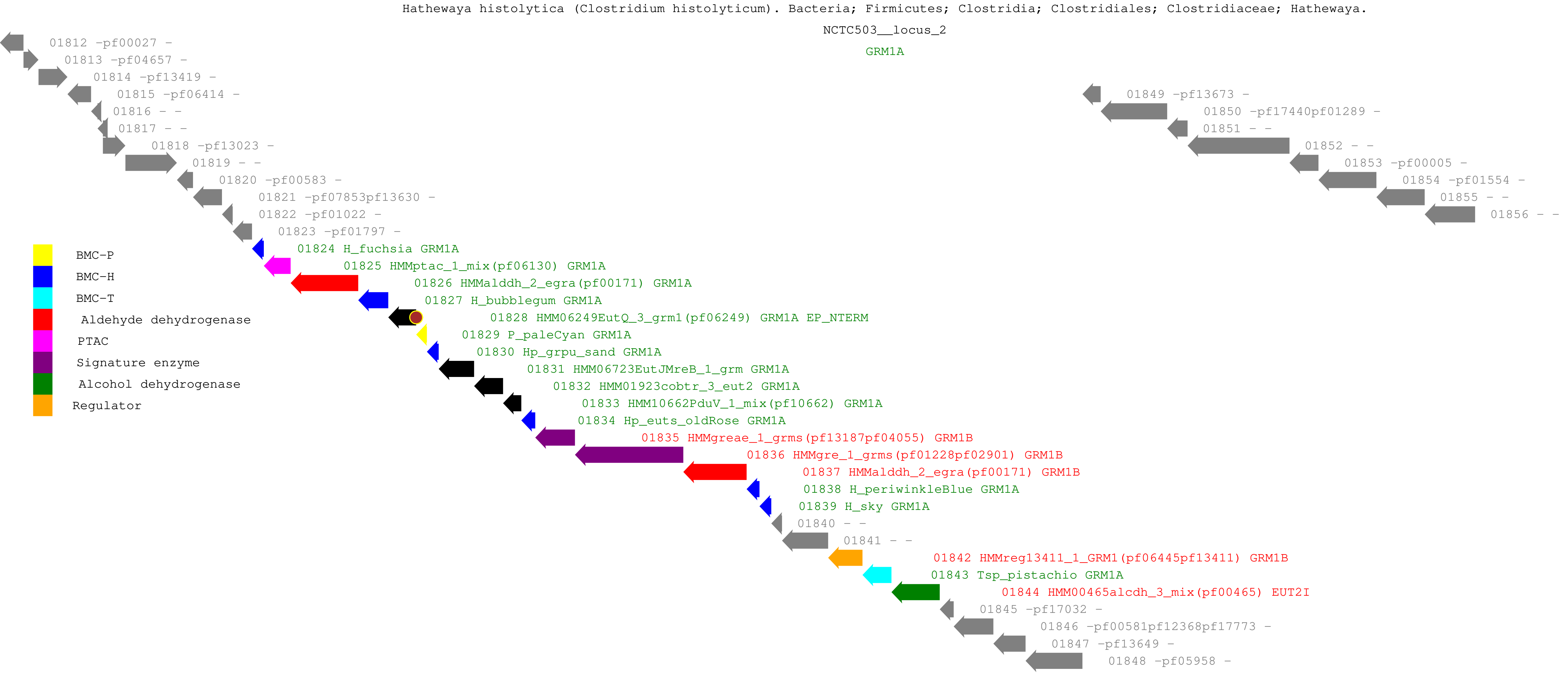
Hathewaya histolytica (Clostridium histolyticum). Bacteria; Firmicutes; Clostridia; Clostridiales; Clostridiaceae; Hathewaya.
NCTC503__locus_2
GRM1A
01812 -pf00027 -
01813 -pf04657 -
01814 -pf13419 -
01815 -pf06414 -
01816 - -
01817 - -
01818 -pf13023 -
01819 - -
01820 -pf00583 -
01821 -pf07853pf13630 -
01822 -pf01022 -
01823 -pf01797 -
01824 H_fuchsia GRM1A
01825 HMMptac_1_mix(pf06130) GRM1A
01826 HMMalddh_2_egra(pf00171) GRM1A
01827 H_bubblegum GRM1A
01828 HMM06249EutQ_3_grm1(pf06249) GRM1A EP_NTERM
01829 P_paleCyan GRM1A
01830 Hp_grpu_sand GRM1A
01831 HMM06723EutJMreB_1_grm GRM1A
01832 HMM01923cobtr_3_eut2 GRM1A
01833 HMM10662PduV_1_mix(pf10662) GRM1A
01834 Hp_euts_oldRose GRM1A
01835 HMMgreae_1_grms(pf13187pf04055) GRM1B
01836 HMMgre_1_grms(pf01228pf02901) GRM1B
01837 HMMalddh_2_egra(pf00171) GRM1B
01838 H_periwinkleBlue GRM1A
01839 H_sky GRM1A
01840 - -
01841 - -
01842 HMMreg13411_1_GRM1(pf06445pf13411) GRM1B
01843 Tsp_pistachio GRM1A
01844 HMM00465alcdh_3_mix(pf00465) EUT2I
01845 -pf17032 -
01846 -pf00581pf12368pf17773 -
01847 -pf13649 -
01848 -pf05958 -
01849 -pf13673 -
01850 -pf17440pf01289 -
01851 - -
01852 - -
01853 -pf00005 -
01854 -pf01554 -
01855 - -
01856 - -
BMC-P
BMC-H
BMC-T
Aldehyde dehydrogenase
PTAC
Signature enzyme
Alcohol dehydrogenase
Regulator

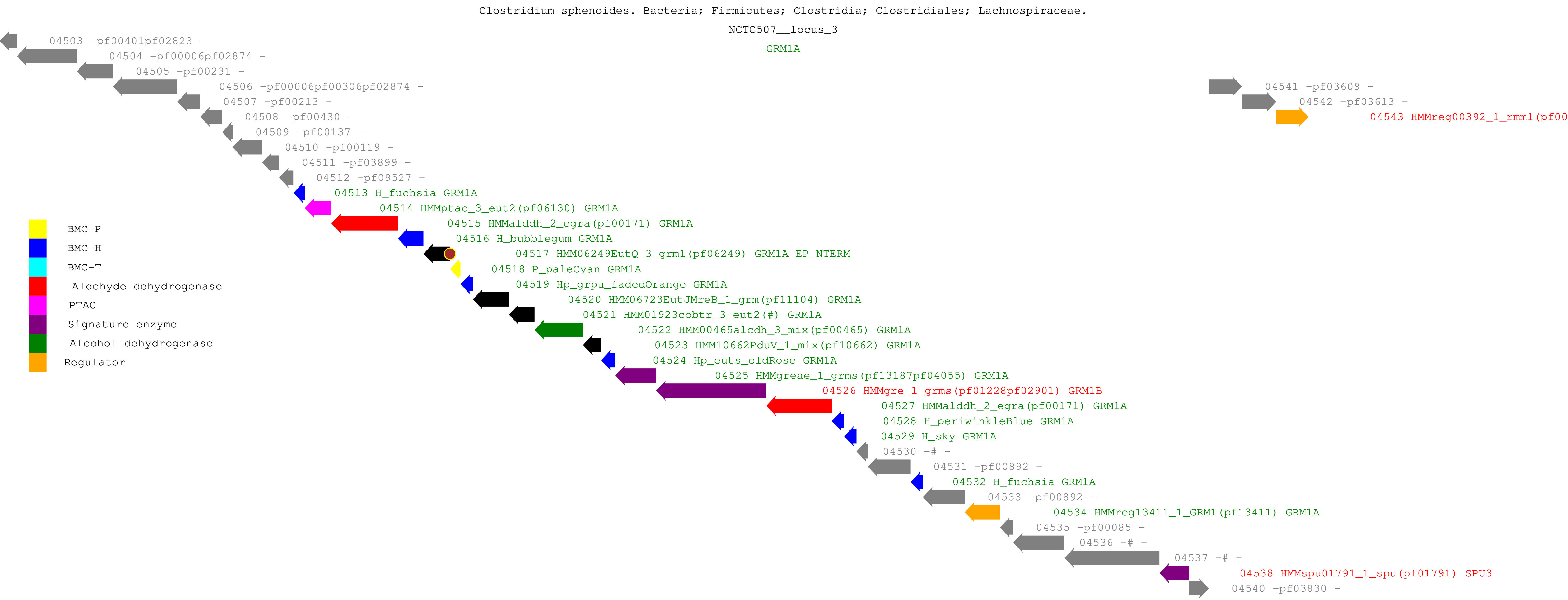

Clostridium sphenoides. Bacteria; Firmicutes; Clostridia; Clostridiales; Lachnospiraceae.
NCTC507__locus_3
GRM1A
04503 -pf00401pf02823 -
04504 -pf00006pf02874 -
04505 -pf00231 -
04506 -pf00006pf00306pf02874 -
04507 -pf00213 -
04508 -pf00430 -
04509 -pf00137 -
04510 -pf00119 -
04511 -pf03899 -
04512 -pf09527 -
04513 H_fuchsia GRM1A
04514 HMMptac_3_eut2(pf06130) GRM1A
04515 HMMaldh_2_egra(pf00171) GRM1A
04516 H_bubblegum GRM1A
04517 HMM06249EutQ_3_grm1(pf06249) GRM1A EP_NTERM
04518 P_paleCyan GRM1A
04519 Hp_grpu_fadedOrange GRM1A
04520 HMM06723EutJMreB_1_grm(pf11104) GRM1A
04521 HMM01923cobtr_3_eut2(#) GRM1A
04522 HMM00465alcdh_3_mix(pf00465) GRM1A
04523 HMM10662PduV_1_mix(pf10662) GRM1A
04524 Hp_euts_oldRose GRM1A
04525 HMMgreae_1_grms(pf13187pf04055) GRM1A
04526 HMMgre_1_grms(pf01228pf02901) GRM1B
04527 HMMaldh_2_egra(pf00171) GRM1A
04528 H_periwinkleBlue GRM1A
04529 H_sky GRM1A
04530 -# -
04531 -pf00892 -
04532 H_fuchsia GRM1A
04533 -pf00892 -
04534 HMMreg13411_1_GRM1(pf13411) GRM1A
04535 -pf00085 -
04536 -# -
04537 -# -
04538 HMMspu01791_1_spu(pf01791) SPU3
04540 -pf03830 -
04541 -pf03609 -
04542 -pf03613 -
04543 HMMreg00392_1_rmm1(pf00
BMC-P
BMC-H
BMC-T
Aldehyde dehydrogenase
PTAC
Signature enzyme
Alcohol dehydrogenase
Regulator

Clostridium tetani. Bacteria; Firmicutes; Clostridia; Clostridiales; Clostridiaceae;

NCTC539__locus_1

GRM1A

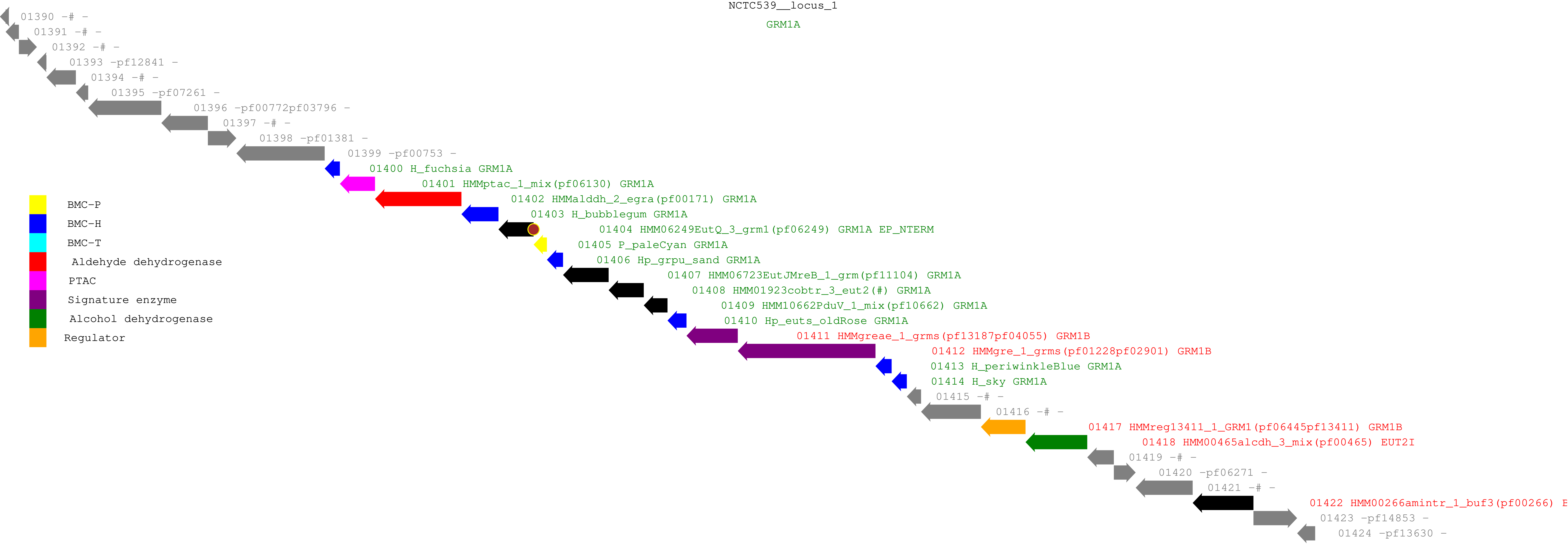

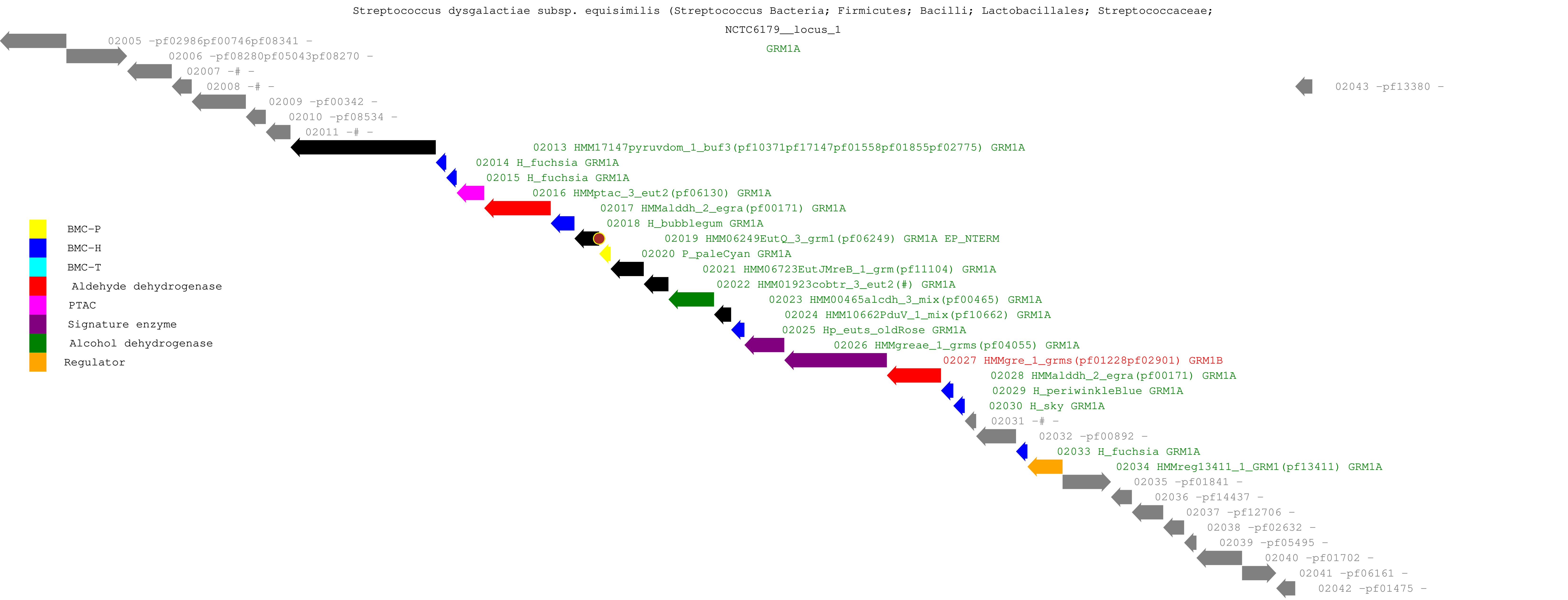
Streptococcus dysgalactiae subsp. equisimilis (Streptococcus Bacteria; Firmicutes; Bacilli; Lactobacillales; Streptococcaceae;
NCTC6179__locus_1
GRM1A
02005 -pf02986pf00746pf08341 -
02006 -pf08280pf05043pf08270 -
02007 -# -
02008 -# -
02009 -pf00342 -
02010 -pf08534 -
02011 -# -
02013 HMM17147pyruvdom_1_buf3(pf10371pf17147pf01558pf01855pf02775) GRM1A
02014 H_fuchsia GRM1A
02015 H_fuchsia GRM1A
02016 HMMptac_3_eut2(pf06130) GRM1A
02017 HMMalddh_2_egra(pf00171) GRM1A
02018 H_bubblegum GRM1A
02019 HMM06249EutQ_3_grm1(pf06249) GRM1A EP_NTERM
02020 P_paleCyan GRM1A
02021 HMM06723EutJMreB_1_grm(pf11104) GRM1A
02022 HMM01923cobtr_3_eut2(#) GRM1A
02023 HMM00465alcdh_3_mix(pf00465) GRM1A
02024 HMM10662PduV_1_mix(pf10662) GRM1A
02025 Hp_euts_oldRose GRM1A
02026 HMMgreae_1_grms(pf04055) GRM1A
02027 HMMgre_1_grms(pf01228pf02901) GRM1B
02028 HMMalddh_2_egra(pf00171) GRM1A
02029 H_periwinkleBlue GRM1A
02030 H_sky GRM1A
02031 -# -
02032 -pf00892 -
02033 H_fuchsia GRM1A
02034 HMMreg13411_1_GRM1(pf13411) GRM1A
02035 -pf01841 -
02036 -pf14437 -
02037 -pf12706 -
02038 -pf02632 -
02039 -pf05495 -
02040 -pf01702 -
02041 -pf06161 -
02042 -pf01475 -
02043 -pf13380 -
BMC-P
BMC-H
BMC-T
Aldehyde dehydrogenase
PTAC
Signature enzyme
Alcohol dehydrogenase
Regulator

Streptococcus dysgalactiae subsp. equisimilis (Streptococcus equisimilis). Bacteria; Firmicutes; Bacilli; Lactobacillales; Streptococcaceae;

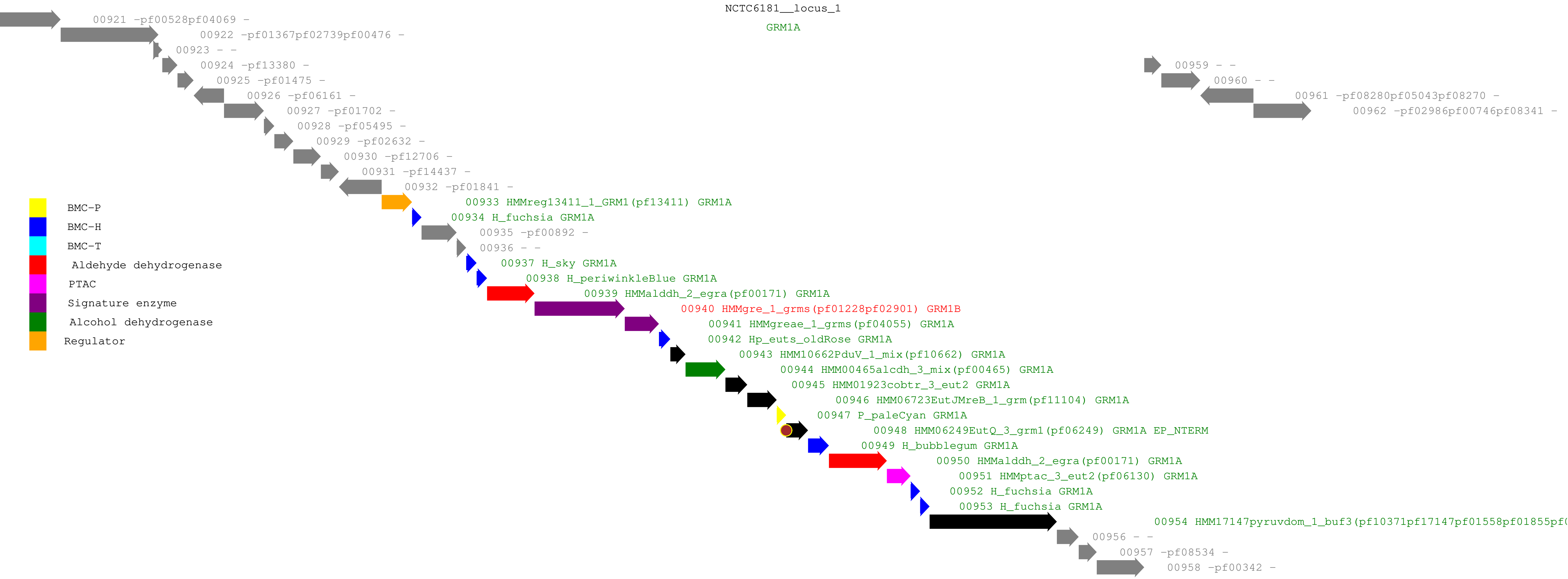

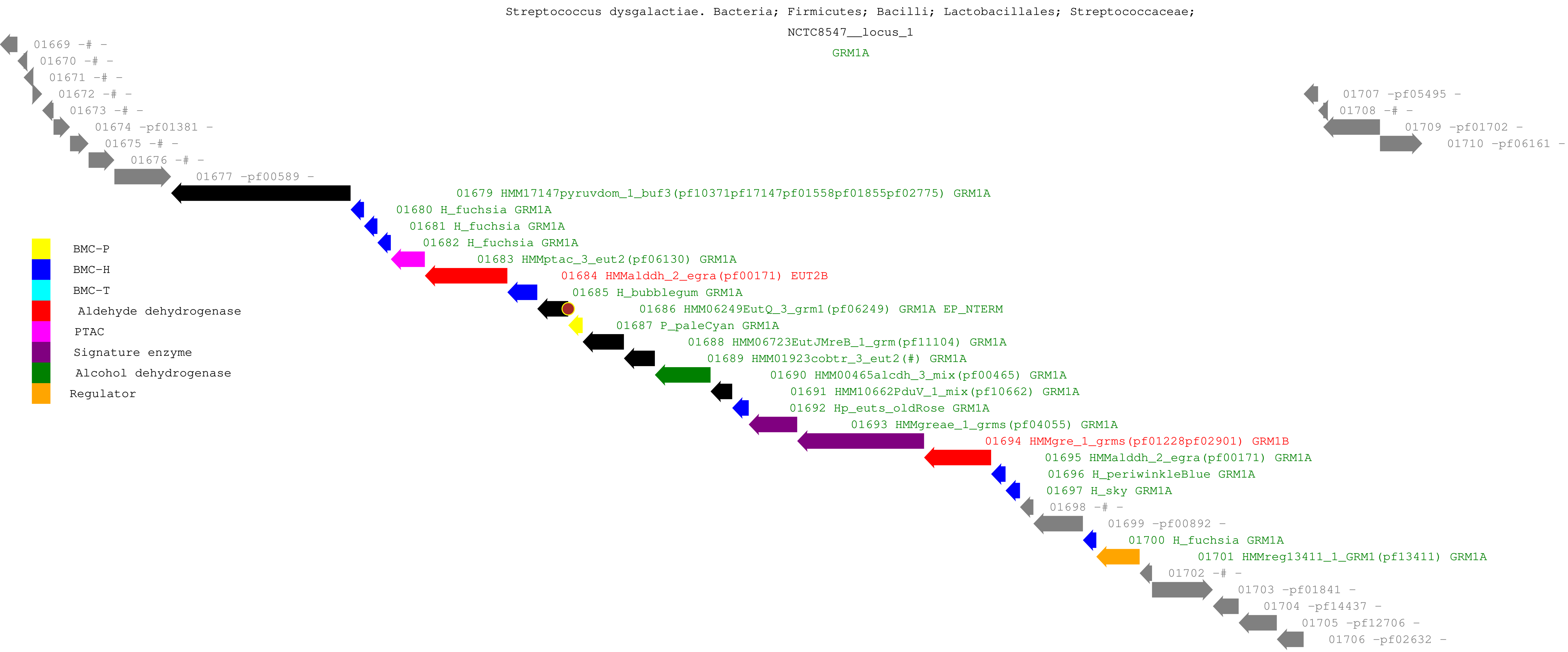
Streptococcus dysgalactiae. Bacteria; Firmicutes; Bacilli; Lactobacillales; Streptococcaceae;
NCTC8547__locus_1
GRM1A
01669 -# -
01670 -# -
01671 -# -
01672 -# -
01673 -# -
01674 -pf01381 -
01675 -# -
01676 -# -
01677 -pf00589 -
01679 HMM17147pyruvdom_1_buf3(pf10371pf17147pf01558pf01855pf02775) GRM1A
01680 H_fuchsia GRM1A
01681 H_fuchsia GRM1A
01682 H_fuchsia GRM1A
01683 HMMptac_3_eut2(pf06130) GRM1A
01684 HMMaldh_2_egra(pf00171) EUT2B
01685 H_bubblegum GRM1A
01686 HMM06249EutQ_3_grm1(pf06249) GRM1A EP_NTERM
01687 P_paleCyan GRM1A
01688 HMM06723EutJMreB_1_grm(pf11104) GRM1A
01689 HMM01923cobtr_3_eut2(#) GRM1A
01690 HMM00465alcdh_3_mix(pf00465) GRM1A
01691 HMM10662PduV_1_mix(pf10662) GRM1A
01692 Hp_euts_oldRose GRM1A
01693 HMMgreae_1_grms(pf04055) GRM1A
01694 HMMgre_1_grms(pf01228pf02901) GRM1B
01695 HMMaldh_2_egra(pf00171) GRM1A
01696 H_periwinkleBlue GRM1A
01697 H_sky GRM1A
01698 -# -
01699 -pf00892 -
01700 H_fuchsia GRM1A
01701 HMMreg13411_1_GRM1(pf13411) GRM1A
01702 -# -
01703 -pf01841 -
01704 -pf14437 -
01705 -pf12706 -
01706 -pf02632 -
01707 -pf05495 -
01708 -# -
01709 -pf01702 -
01710 -pf06161 -
BMC-P
BMC-H
BMC-T
Aldehyde dehydrogenase
PTAC
Signature enzyme
Alcohol dehydrogenase
Regulator

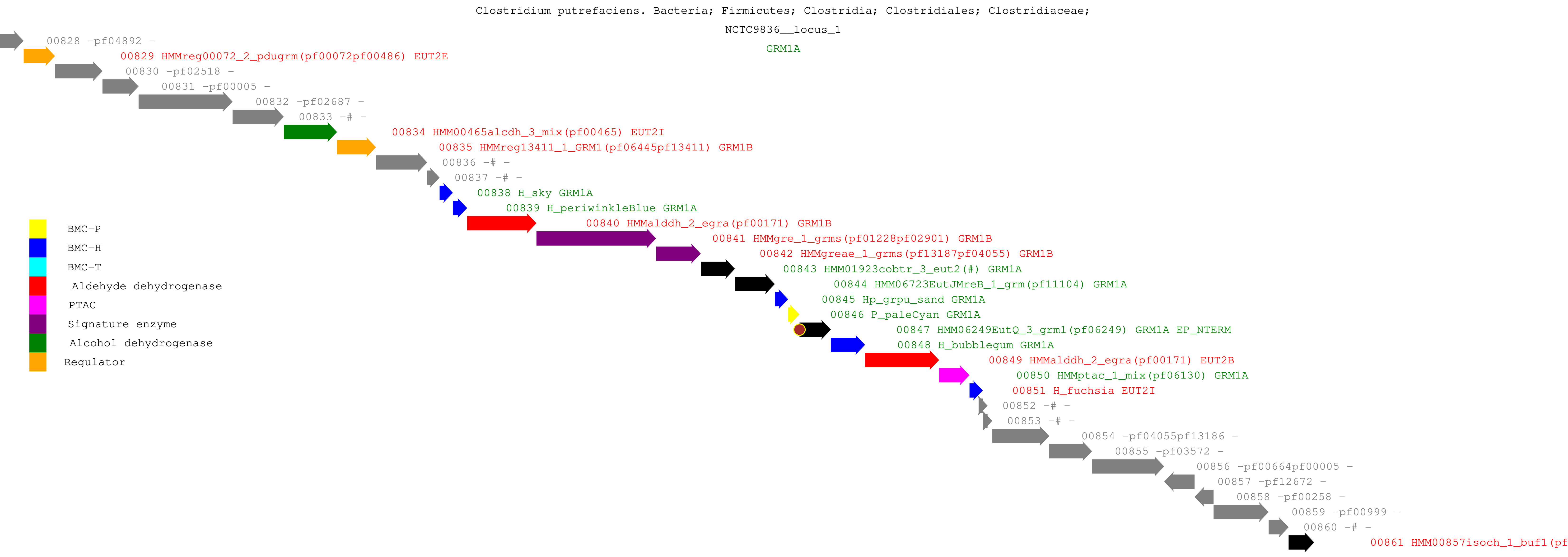
Clostridium putrefaciens. Bacteria; Firmicutes; Clostridia; Clostridiales; Clostridiaceae;
NCTC9836__locus_1
GRM1A
00828 -pf04892 -
00829 HMMreg00072_2_pdugrm(pf00072pf00486) EUT2E
00830 -pf02518 -
00831 -pf00005 -
00832 -pf02687 -
00833 -# -
00834 HMM00465alcdh_3_mix(pf00465) EUT2I
00835 HMMreg13411_1_GRM1(pf06445pf13411) GRM1B
00836 -# -
00837 -# -
00838 H_sky GRM1A
00839 H_periwinkleBlue GRM1A
00840 HMMalddh_2_egra(pf00171) GRM1B
00841 HMMgre_1_grms(pf01228pf02901) GRM1B
00842 HMMgreae_1_grms(pf13187pf04055) GRM1B
00843 HMM01923cobtr_3_eut2(#) GRM1A
00844 HMM06723EutJMreB_1_grm(pf11104) GRM1A
00845 Hp_grpu_sand GRM1A
00846 P_paleCyan GRM1A
00847 HMM06249EutQ_3_grm1(pf06249) GRM1A EP_NTERM
00848 H_bubblegum GRM1A
00849 HMMalddh_2_egra(pf00171) EUT2B
00850 HMMptac_1_mix(pf06130) GRM1A
00851 H_fuchsia EUT2I
00852 -# -
00853 -# -
00854 -pf04055pf13186 -
00855 -pf03572 -
00856 -pf00664pf00005 -
00857 -pf12672 -
00858 -pf00258 -
00859 -pf00999 -
00860 -# -
00861 HMM00857isoch_1_buf1(pf
BMC-P
BMC-H
BMC-T
Aldehyde dehydrogenase
PTAC
Signature enzyme
Alcohol dehydrogenase
Regulator

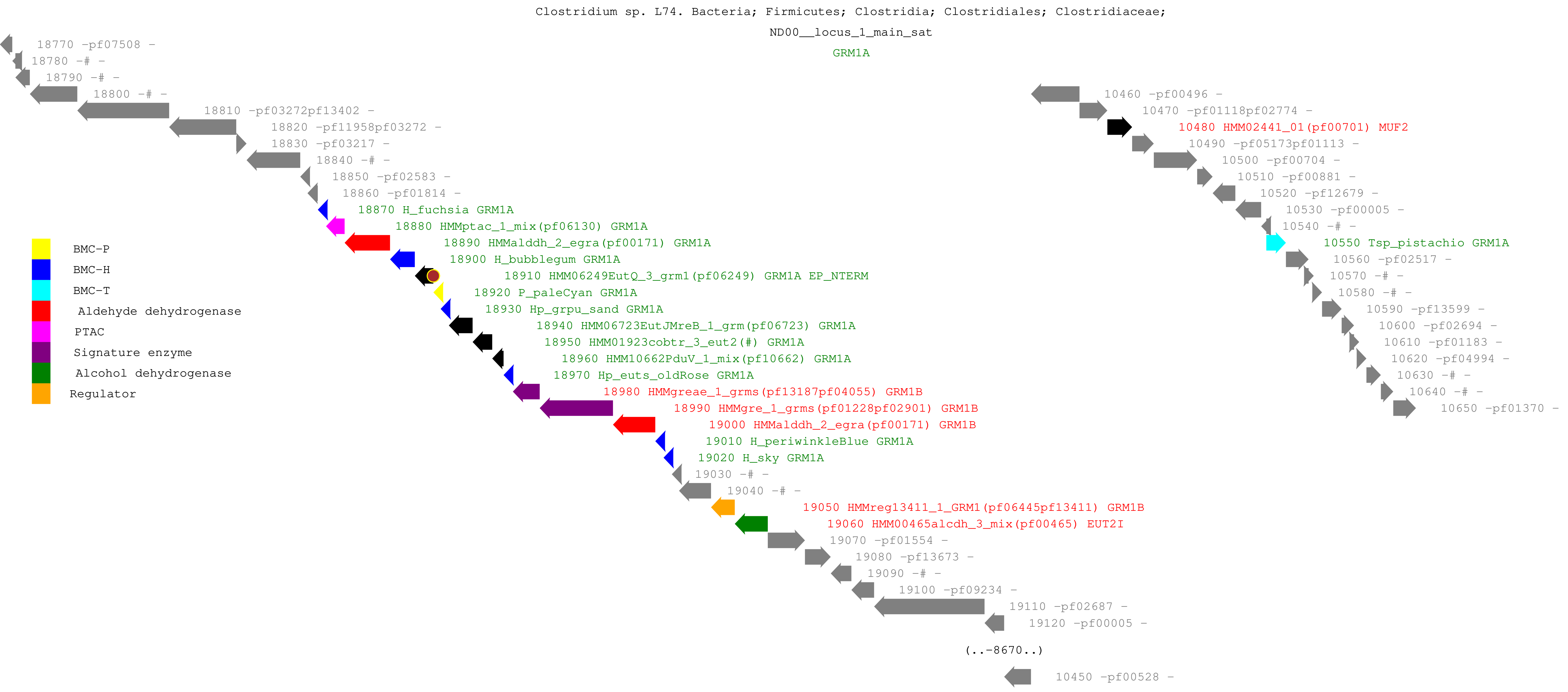
Clostridium sp. L74. Bacteria; Firmicutes; Clostridia; Clostridiales; Clostridiaceae;
ND00__locus_1_main_sat
GRM1A
18770 -pf07508 -
18780 -# -
18790 -# -
18800 -# -
18810 -pf03272pf13402 -
18820 -pf11958pf03272 -
18830 -pf03217 -
18840 -# -
18850 -pf02583 -
18860 -pf01814 -
18870 H_fuchsia GRM1A
18880 HMMptac_1_mix(pf06130) GRM1A
18890 HMMalddh_2_egra(pf00171) GRM1A
18900 H_bubblegum GRM1A
18910 HMM06249EutQ_3_grm1(pf06249) GRM1A EP_NTERM
18920 P_paleCyan GRM1A
18930 Hp_grpu_sand GRM1A
18940 HMM06723EutJMreB_1_grm(pf06723) GRM1A
18950 HMM01923cobtr_3_eut2(#) GRM1A
18960 HMM10662PduV_1_mix(pf10662) GRM1A
18970 Hp_euts_oldRose GRM1A
18980 HMMgreae_1_grms(pf13187pf04055) GRM1B
18990 HMMgre_1_grms(pf01228pf02901) GRM1B
19000 HMMalddh_2_egra(pf00171) GRM1B
19010 H_periwinkleBlue GRM1A
19020 H_sky GRM1A
19030 -# -
19040 -# -
19050 HMMreg13411_1_GRM1(pf06445pf13411) GRM1B
19060 HMM00465alcdh_3_mix(pf00465) EUT2I
19070 -pf01554 -
19080 -pf13673 -
19090 -# -
19100 -pf09234 -
19110 -pf02687 -
19120 -pf00005 -
(..-8670..)
10450 -pf00528 -
10460 -pf00496 -
10470 -pf01118pf02774 -
10480 HMM02441_01(pf00701) MUF2
10490 -pf05173pf01113 -
10500 -pf00704 -
10510 -pf00881 -
10520 -pf12679 -
10530 -pf00005 -
10540 -# -
10550 Tsp_pistachio GRM1A
10560 -pf02517 -
10570 -# -
10580 -# -
10590 -pf13599 -
10600 -pf02694 -
10610 -pf01183 -
10620 -pf04994 -
10630 -# -
10640 -# -
10650 -pf01370 -
BMC-P
BMC-H
BMC-T
Aldehyde dehydrogenase
PTAC
Signature enzyme
Alcohol dehydrogenase
Regulator

Clostridium botulinum. Bacteria; Firmicutes; Clostridia; Clostridiales; Clostridiaceae;

NPD5__locus_1

GRM1A

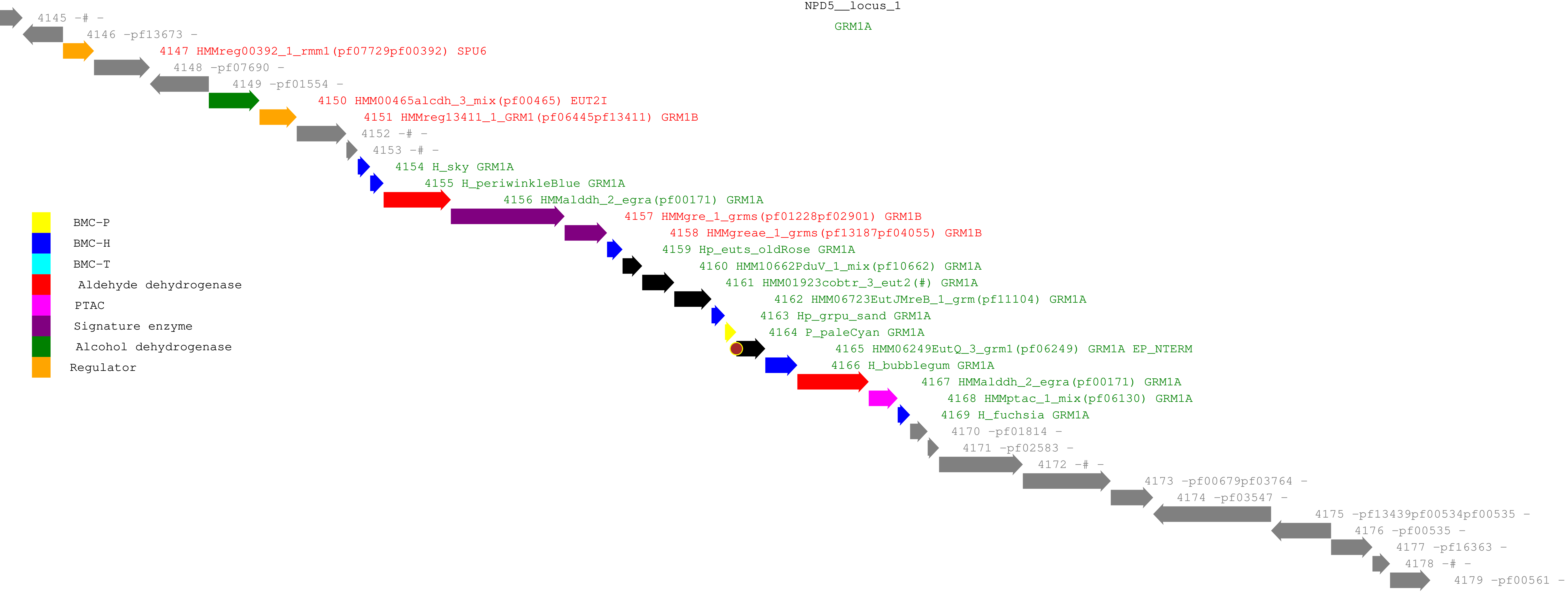

Clostridium botulinum. Bacteria; Firmicutes; Clostridia; Clostridiales; Clostridiaceae;

NPD7__locus_1_main_sat

GRM1A

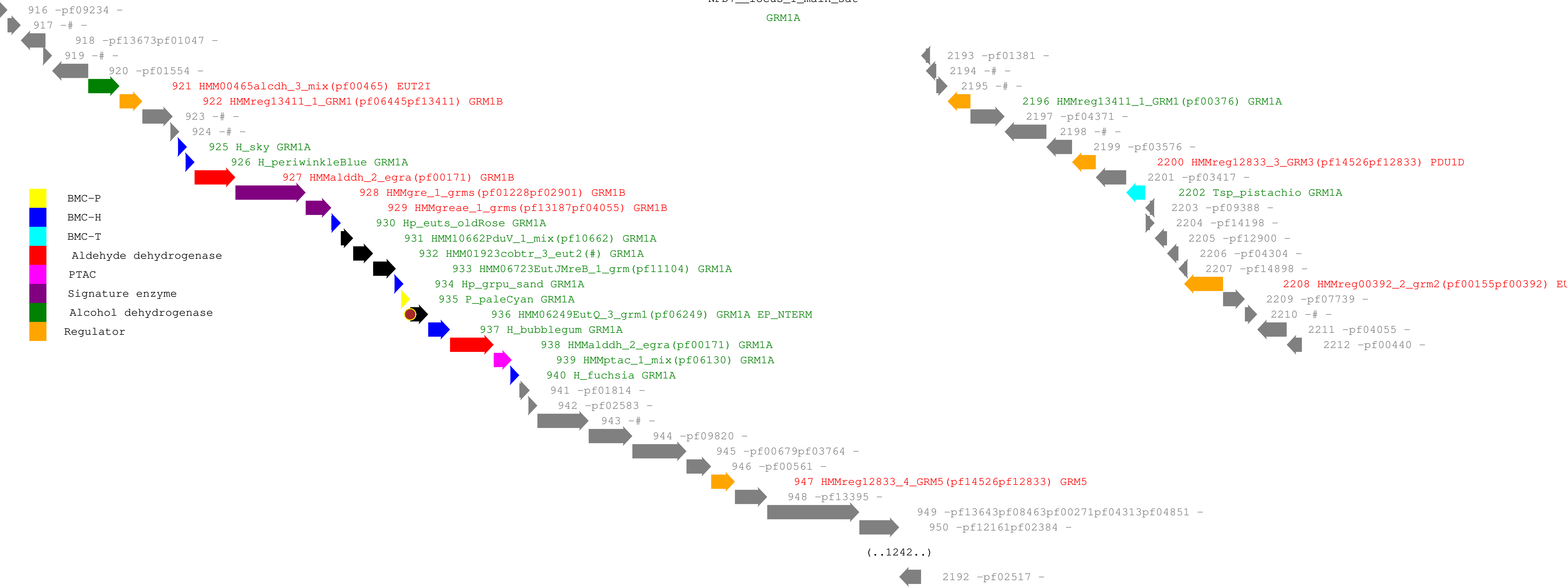

Olsenella uli (strain ATCC 49627 / DSM 7084 / CIP 109912 / JCM 12494 / Bacteria; Actinobacteria; Coriobacteriia; Coriobacteriales;

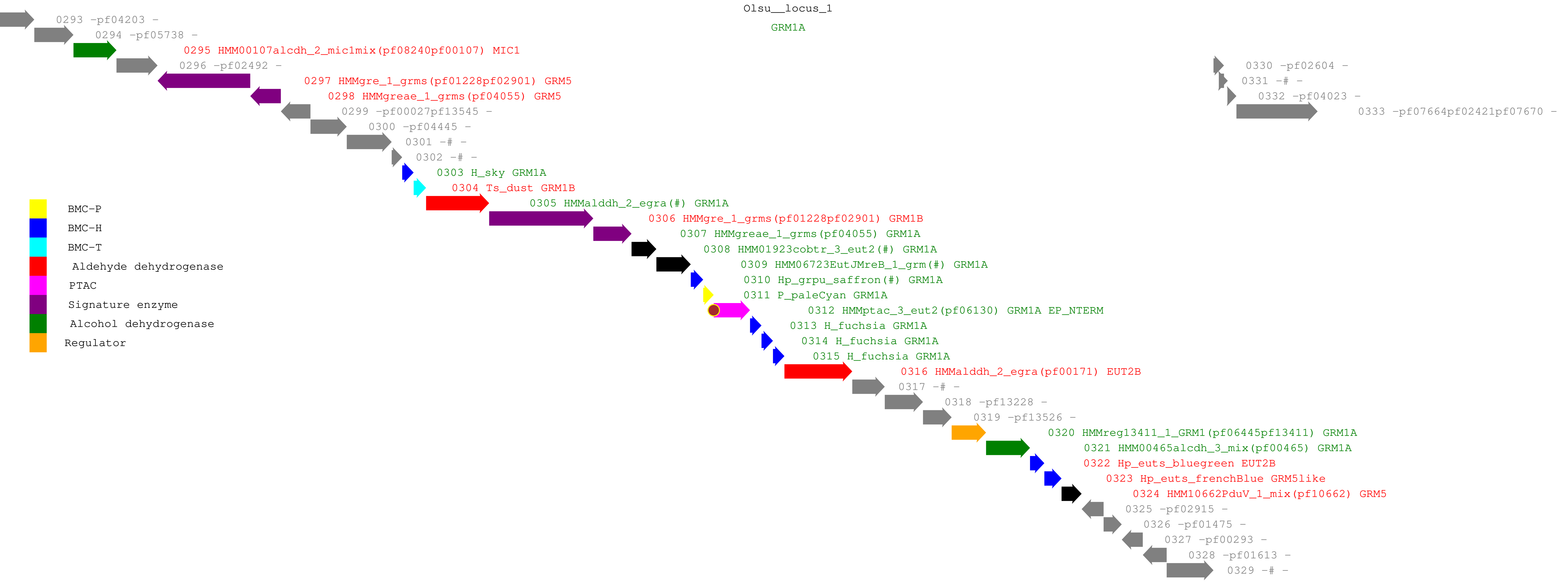

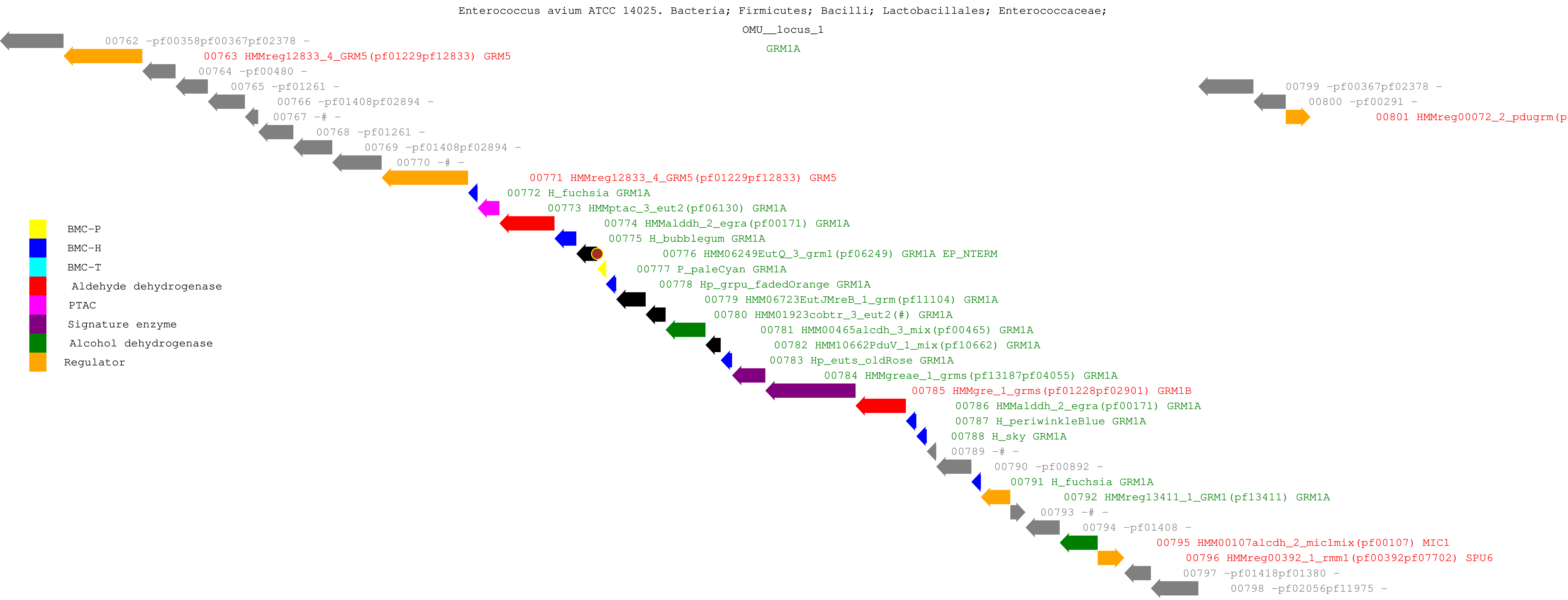
Enterococcus avium ATCC 14025. Bacteria; Firmicutes; Bacilli; Lactobacillales; Enterococcaceae;
OMU__locus_1
GRM1A
00762 -pf00358pf00367pf02378 -
00763 HMMreg12833_4_GRM5(pf01229pf12833) GRM5
00764 -pf00480 -
00765 -pf01261 -
00766 -pf01408pf02894 -
00767 -# -
00768 -pf01261 -
00769 -pf01408pf02894 -
00770 -# -
00771 HMMreg12833_4_GRM5(pf01229pf12833) GRM5
00772 H_fuchsia GRM1A
00773 HMMptac_3_eut2(pf06130) GRM1A
00774 HMMalddh_2_egra(pf00171) GRM1A
00775 H_bubblegum GRM1A
00776 HMM06249EutQ_3_grm1(pf06249) GRM1A EP_NTERM
00777 P_paleCyan GRM1A
00778 Hp_grpu_fadedOrange GRM1A
00779 HMM06723EutJMreB_1_grm(pf11104) GRM1A
00780 HMM01923cobtr_3_eut2(#) GRM1A
00781 HMM00465alcdh_3_mix(pf00465) GRM1A
00782 HMM10662PduV_1_mix(pf10662) GRM1A
00783 Hp_euts_oldRose GRM1A
00784 HMMgreae_1_grms(pf13187pf04055) GRM1A
00785 HMMgre_1_grms(pf01228pf02901) GRM1B
00786 HMMalddh_2_egra(pf00171) GRM1A
00787 H_periwinkleBlue GRM1A
00788 H_sky GRM1A
00789 -# -
00790 -pf00892 -
00791 H_fuchsia GRM1A
00792 HMMreg13411_1_GRM1(pf13411) GRM1A
00793 -# -
00794 -pf01408 -
00795 HMM00107alcdh_2_mic1mix(pf00107) MIC1
00796 HMMreg00392_1_rmm1(pf00392pf07702) SPU6
00797 -pf01418pf01380 -
00798 -pf02056pf11975 -
00799 -pf00367pf02378 -
00800 -pf00291 -
00801 HMMreg00072_2_pdugrm(p
BMC-P
BMC-H
BMC-T
Aldehyde dehydrogenase
PTAC
Signature enzyme
Alcohol dehydrogenase
Regulator

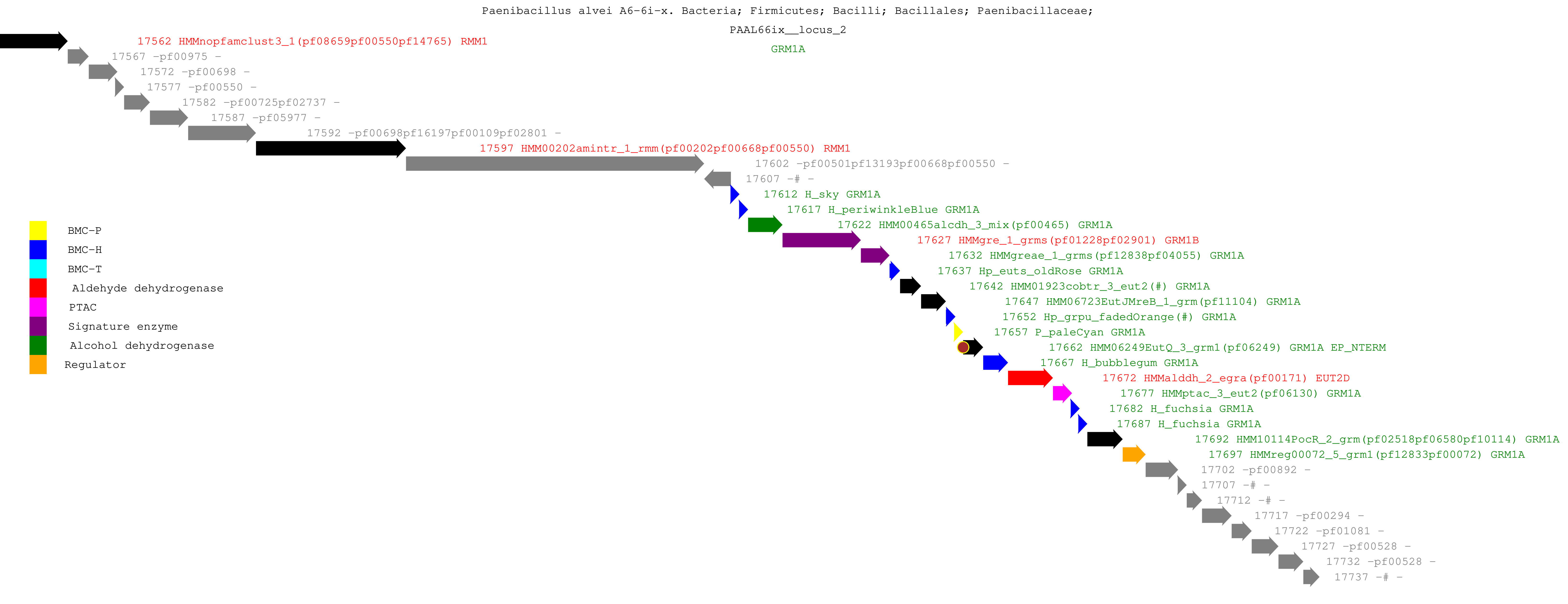
Paenibacillus alvei A6-6i-x. Bacteria; Firmicutes; Bacilli; Bacillales; Paenibacillaceae;
PAAL66ix__locus_2
GRM1A
17562 HMMnopfamclust3_1(pf08659pf00550pf14765) RMM1
17567 -pf00975 -
17572 -pf00698 -
17577 -pf00550 -
17582 -pf00725pf02737 -
17587 -pf05977 -
17592 -pf00698pf16197pf00109pf02801 -
17597 HMM00202amintr_1_rmm(pf00202pf00668pf00550) RMM1
17602 -pf00501pf13193pf00668pf00550 -
17607 -# -
17612 H_sky GRM1A
17617 H_periwinkleBlue GRM1A
17622 HMM00465alcdh_3_mix(pf00465) GRM1A
17627 HMMgre_1_grms(pf01228pf02901) GRM1B
17632 HMMgreae_1_grms(pf12838pf04055) GRM1A
17637 Hp_euts_oldRose GRM1A
17642 HMM01923cobtr_3_eut2(#) GRM1A
17647 HMM06723EutJMreB_1_grm(pf11104) GRM1A
17652 Hp_grpu_fadedOrange(#) GRM1A
17657 P_paleCyan GRM1A
17662 HMM06249EutQ_3_grm1(pf06249) GRM1A EP_NTERM
17667 H_bubblegum GRM1A
17672 HMMalddh_2_egra(pf00171) EUT2D
17677 HMMptac_3_eut2(pf06130) GRM1A
17682 H_fuchsia GRM1A
17687 H_fuchsia GRM1A
17692 HMM10114PocR_2_grm(pf02518pf06580pf10114) GRM1A
17697 HMMreg00072_5_grm1(pf12833pf00072) GRM1A
17702 -pf00892 -
17707 -# -
17712 -# -
17717 -pf00294 -
17722 -pf01081 -
17727 -pf00528 -
17732 -pf00528 -
17737 -# -
BMC-P
BMC-H
BMC-T
Aldehyde dehydrogenase
PTAC
Signature enzyme
Alcohol dehydrogenase
Regulator

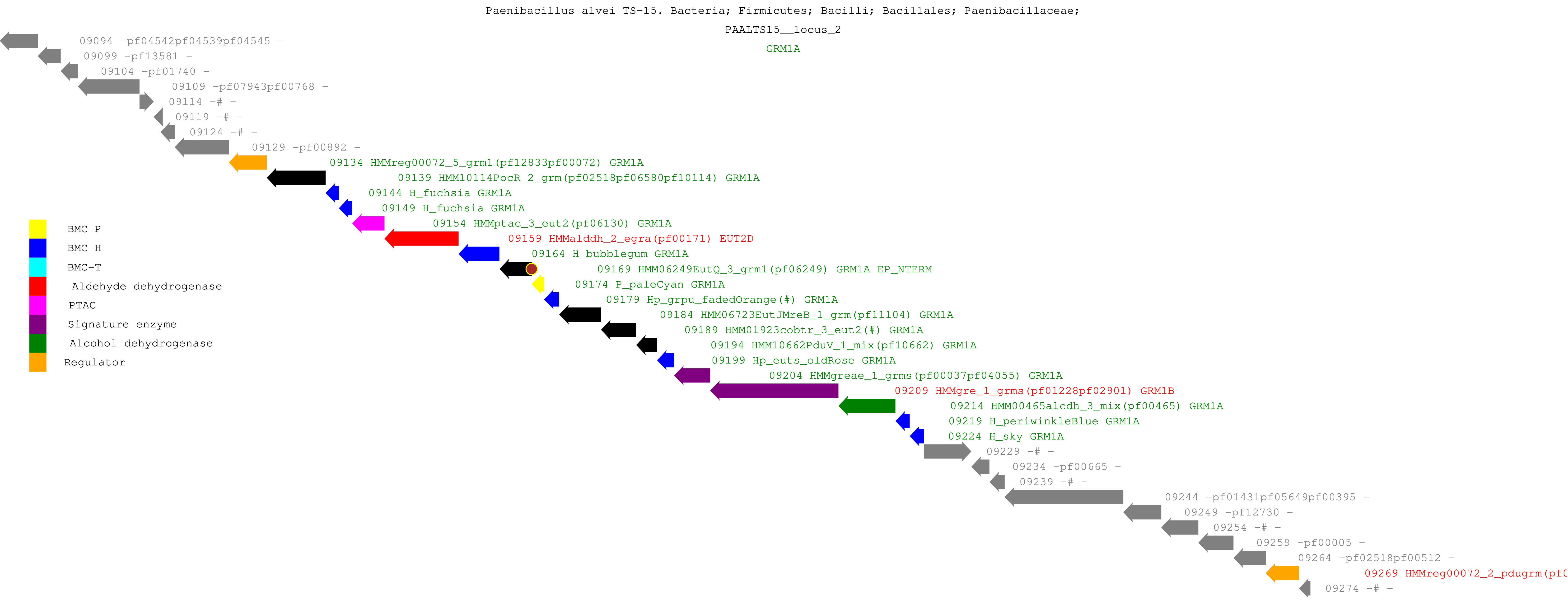
Paenibacillus alvei TS-15. Bacteria; Firmicutes; Bacilli; Bacillales; Paenibacillaceae;
PAALTS15__locus_2
GRM1A
09094 -pf04542pf04539pf04545 -
09099 -pf13581 -
09104 -pf01740 -
09109 -pf07943pf00768 -
09114 -# -
09119 -# -
09124 -# -
09129 -pf00892 -
09134 HMMreg00072_5_grm1(pf12833pf00072) GRM1A
09139 HMM10114PocR_2_grm(pf02518pf06580pf10114) GRM1A
09144 H_fuchsia GRM1A
09149 H_fuchsia GRM1A
09154 HMMptac_3_eut2(pf06130) GRM1A
09159 HMMaldddh_2_egra(pf00171) EUT2D
09164 H_bubblegum GRM1A
09169 HMM06249EutQ_3_grm1(pf06249) GRM1A EP_NTERM
09174 P_paleCyan GRM1A
09179 Hp_grpu_fadedOrange(#) GRM1A
09184 HMM06723EutJMreB_1_grm(pf11104) GRM1A
09189 HMM01923cobtr_3_eut2(#) GRM1A
09194 HMM10662PduV_1_mix(pf10662) GRM1A
09199 Hp_euts_oldRose GRM1A
09204 HMMgreae_1_grms(pf00037pf04055) GRM1A
09209 HMMgre_1_grms(pf01228pf02901) GRM1B
09214 HMM00465alcdh_3_mix(pf00465) GRM1A
09219 H_periwinkleBlue GRM1A
09224 H_sky GRM1A
09229 -# -
09234 -pf00665 -
09239 -# -
09244 -pf01431pf05649pf00395 -
09249 -pf12730 -
09254 -# -
09259 -pf00005 -
09264 -pf02518pf00512 -
09269 HMMreg00072_2_pdugrm(pf0
09274 -# -
BMC-P
BMC-H
BMC-T
Aldehyde dehydrogenase
PTAC
Signature enzyme
Alcohol dehydrogenase
Regulator

Paenibacillus alvei DSM 29. Bacteria; Firmicutes; Bacilli; Bacillales; Paenibacillaceae;

PAV_7c_locus_1

GRM1A

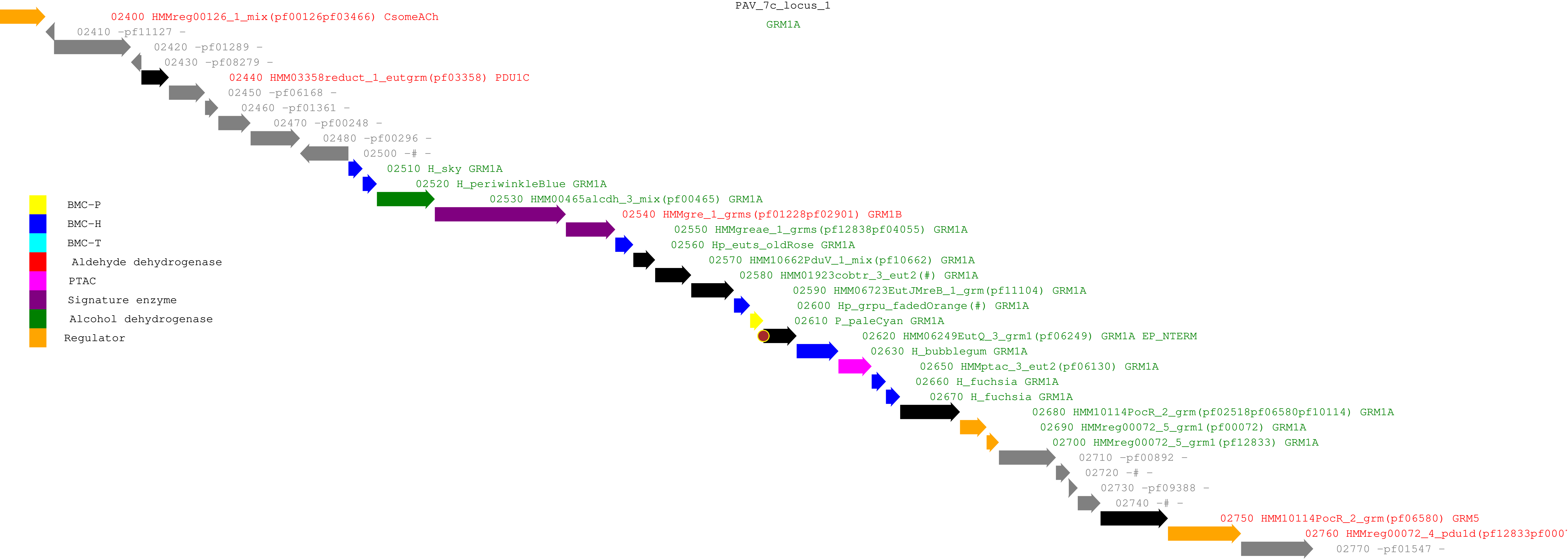

Paenibacillus alvei (Bacillus alvei). Bacteria; Firmicutes; Bacilli; Bacillales; Paenibacillaceae;

PBLR__locus_2

GRM1A

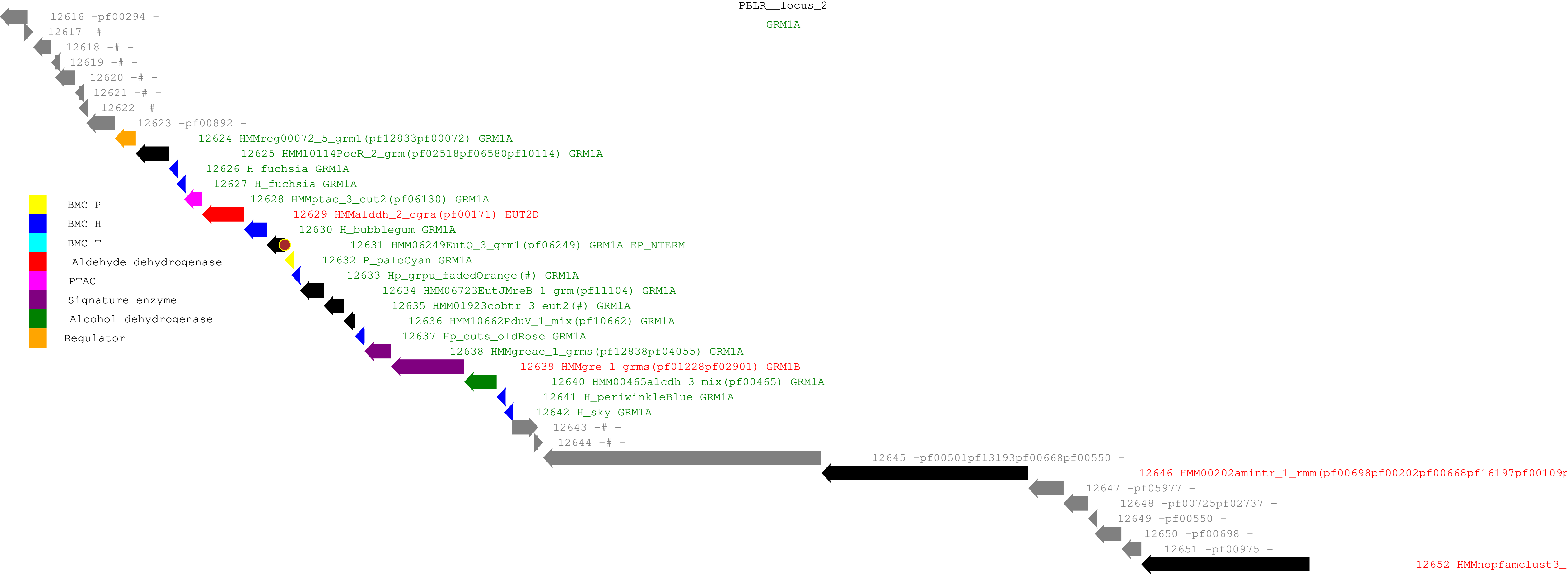

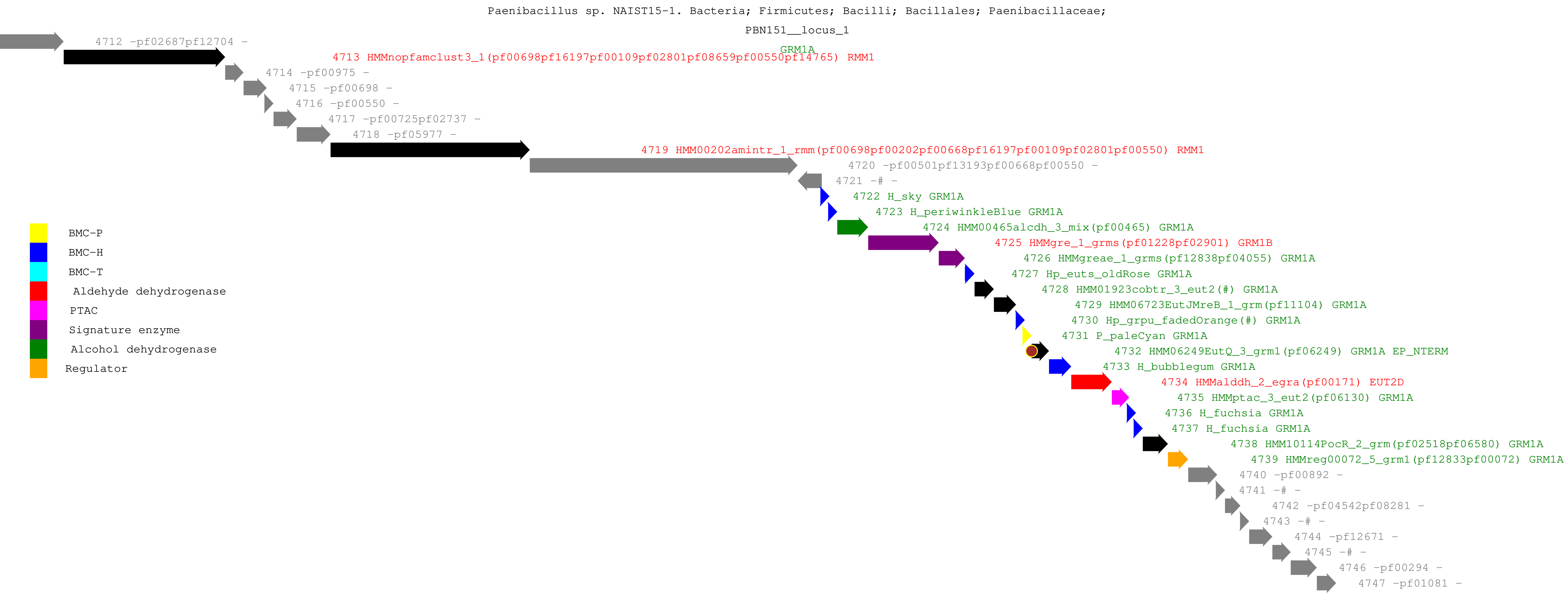
Paenibacillus sp. NAIST15-1. Bacteria; Firmicutes; Bacilli; Bacillales; Paenibacillaceae;
PBN151__locus_1
GRM1A
4712 -pf02687pf12704 -
4713 HMMnopfamclust3_1(pf00698pf16197pf00109pf02801pf08659pf00550pf14765) RMM1
4714 -pf00975 -
4715 -pf00698 -
4716 -pf00550 -
4717 -pf00725pf02737 -
4718 -pf05977 -
4719 HMM00202amintr_1_rmm(pf00698pf00202pf00668pf16197pf00109pf02801pf00550) RMM1
4720 -pf00501pf13193pf00668pf00550 -
4721 -# -
4722 H_sky GRM1A
4723 H_periwinkleBlue GRM1A
4724 HMM00465alcdh_3_mix(pf00465) GRM1A
4725 HMMgre_1_grms(pf01228pf02901) GRM1B
4726 HMMgreae_1_grms(pf12838pf04055) GRM1A
4727 Hp_euts_oldRose GRM1A
4728 HMM01923cobtr_3_eut2(#) GRM1A
4729 HMM06723EutJMreB_1_grm(pf11104) GRM1A
4730 Hp_grpu_fadedOrange(#) GRM1A
4731 P_paleCyan GRM1A
4732 HMM06249EutQ_3_grm1(pf06249) GRM1A EP_NTERM
4733 H_bubblegum GRM1A
4734 HMMalddh_2_egra(pf00171) EUT2D
4735 HMMptac_3_eut2(pf06130) GRM1A
4736 H_fuchsia GRM1A
4737 H_fuchsia GRM1A
4738 HMM10114PocR_2_grm(pf02518pf06580) GRM1A
4739 HMMreg00072_5_grm1(pf12833pf00072) GRM1A
4740 -pf00892 -
4741 -# -
4742 -pf04542pf08281 -
4743 -# -
4744 -pf12671 -
4745 -# -
4746 -pf00294 -
4747 -pf01081 -
BMC-P
BMC-H
BMC-T
Aldehyde dehydrogenase
PTAC
Signature enzyme
Alcohol dehydrogenase
Regulator

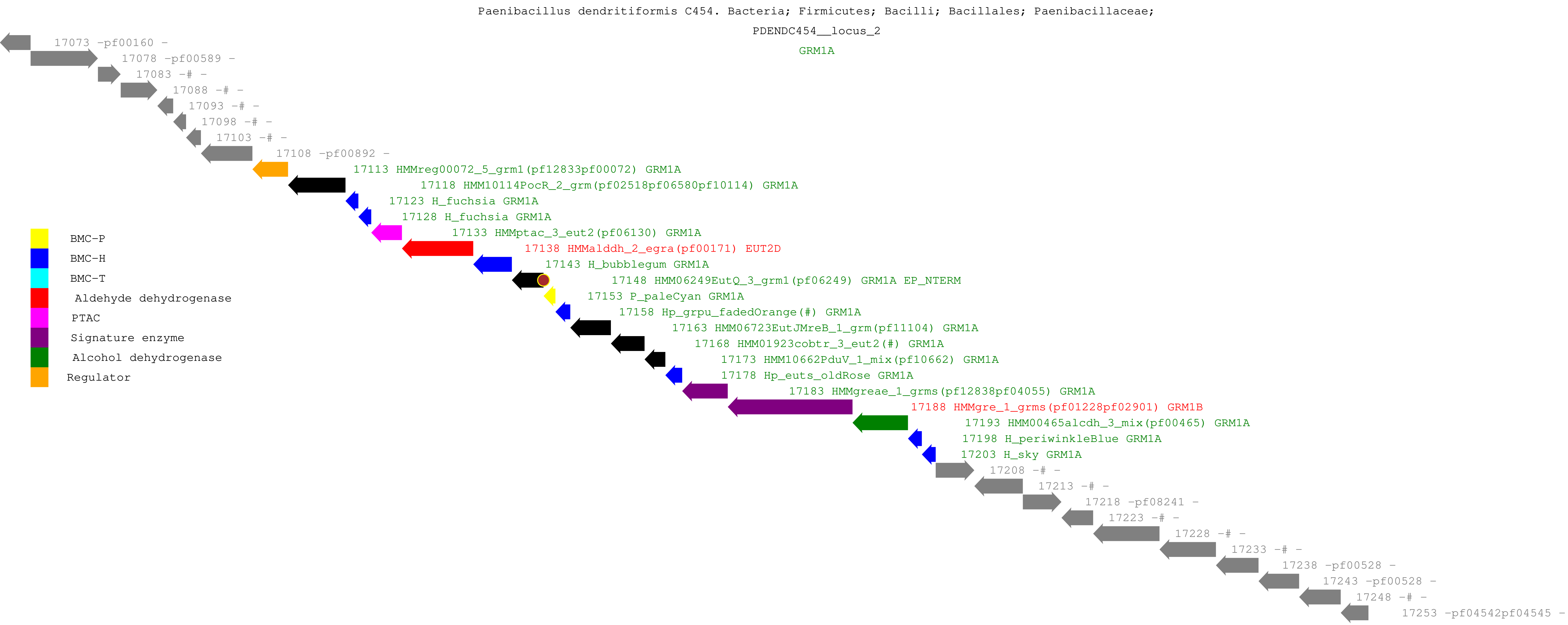
Paenibacillus dendritiformis C454. Bacteria; Firmicutes; Bacilli; Bacillales; Paenibacillaceae;
PDENDC454__locus_2
GRM1A
17073 -pf00160 -
17078 -pf00589 -
17083 -# -
17088 -# -
17093 -# -
17098 -# -
17103 -# -
17108 -pf00892 -
17113 HMMreg00072_5_grm1(pf12833pf00072) GRM1A
17118 HMM10114PocR_2_grm(pf02518pf06580pf10114) GRM1A
17123 H_fuchsia GRM1A
17128 H_fuchsia GRM1A
17133 HMMptac_3_eut2(pf06130) GRM1A
17138 HMMaldh_2_egra(pf00171) EUT2D
17143 H_bubblegum GRM1A
17148 HMM06249EutQ_3_grm1(pf06249) GRM1A EP_NTERM
17153 P_paleCyan GRM1A
17158 Hp_grpu_fadedOrange(#) GRM1A
17163 HMM06723EutJMreB_1_grm(pf11104) GRM1A
17168 HMM01923cobtr_3_eut2(#) GRM1A
17173 HMM10662PduV_1_mix(pf10662) GRM1A
17178 Hp_euts_oldRose GRM1A
17183 HMMgreae_1_grms(pf12838pf04055) GRM1A
17188 HMMgre_1_grms(pf01228pf02901) GRM1B
17193 HMM00465alcdh_3_mix(pf00465) GRM1A
17198 H_periwinkleBlue GRM1A
17203 H_sky GRM1A
17208 -# -
17213 -# -
17218 -pf08241 -
17223 -# -
17228 -# -
17233 -# -
17238 -pf00528 -
17243 -pf00528 -
17248 -# -
17253 -pf04542pf04545 -
BMC-P
BMC-H
BMC-T
Aldehyde dehydrogenase
PTAC
Signature enzyme
Alcohol dehydrogenase
Regulator

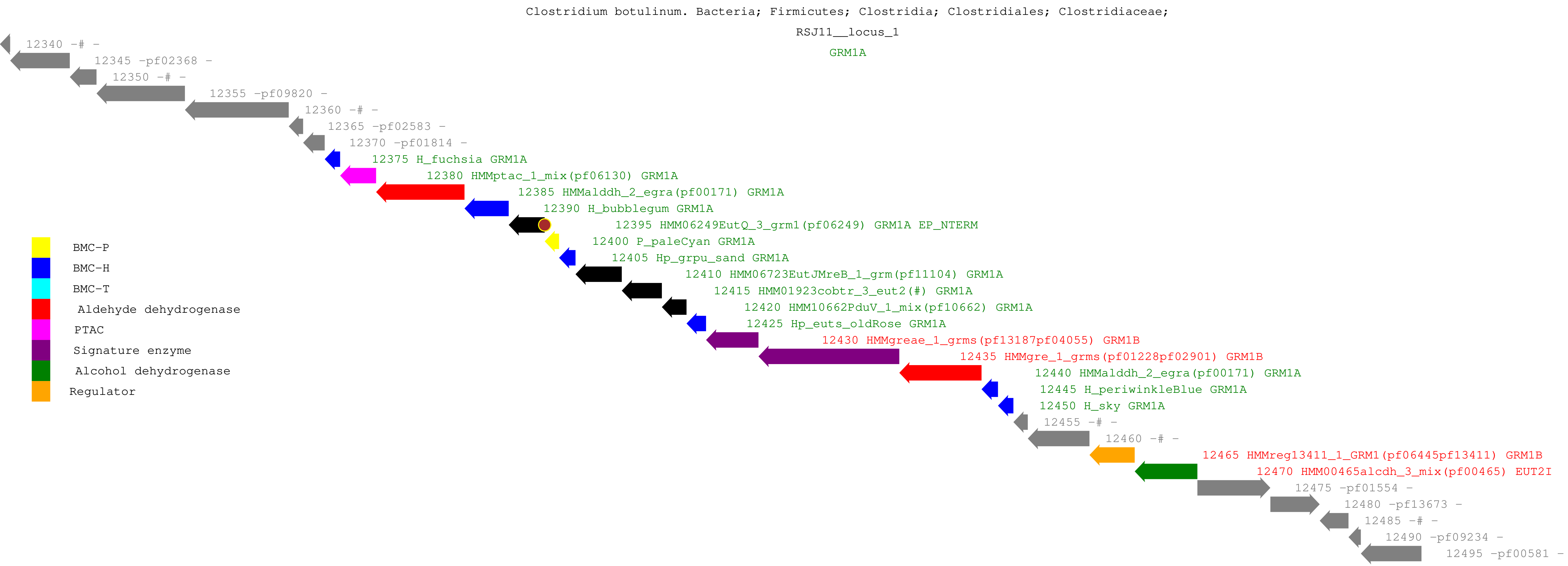
Clostridium botulinum. Bacteria; Firmicutes; Clostridia; Clostridiales; Clostridiaceae;
RSJ11__locus_1
GRM1A
12340 -# -
12345 -pf02368 -
12350 -# -
12355 -pf09820 -
12360 -# -
12365 -pf02583 -
12370 -pf01814 -
12375 H_fuchsia GRM1A
12380 HMMptac_1_mix(pf06130) GRM1A
12385 HMMaldh_2_egra(pf00171) GRM1A
12390 H_bubblegum GRM1A
12395 HMM06249EutQ_3_grm1(pf06249) GRM1A EP_NTERM
12400 P_paleCyan GRM1A
12405 Hp_grpu_sand GRM1A
12410 HMM06723EutJMreB_1_grm(pf11104) GRM1A
12415 HMM01923cobtr_3_eut2(#) GRM1A
12420 HMM10662PduV_1_mix(pf10662) GRM1A
12425 Hp_euts_oldRose GRM1A
12430 HMMgreae_1_grms(pf13187pf04055) GRM1B
12435 HMMgre_1_grms(pf01228pf02901) GRM1B
12440 HMMaldh_2_egra(pf00171) GRM1A
12445 H_periwinkleBlue GRM1A
12450 H_sky GRM1A
12455 -# -
12460 -# -
12465 HMMreg13411_1_GRM1(pf06445pf13411) GRM1B
12470 HMM00465alcdh_3_mix(pf00465) EUT2I
12475 -pf01554 -
12480 -pf13673 -
12485 -# -
12490 -pf09234 -
12495 -pf00581 -
BMC-P
BMC-H
BMC-T
Aldehyde dehydrogenase
PTAC
Signature enzyme
Alcohol dehydrogenase
Regulator

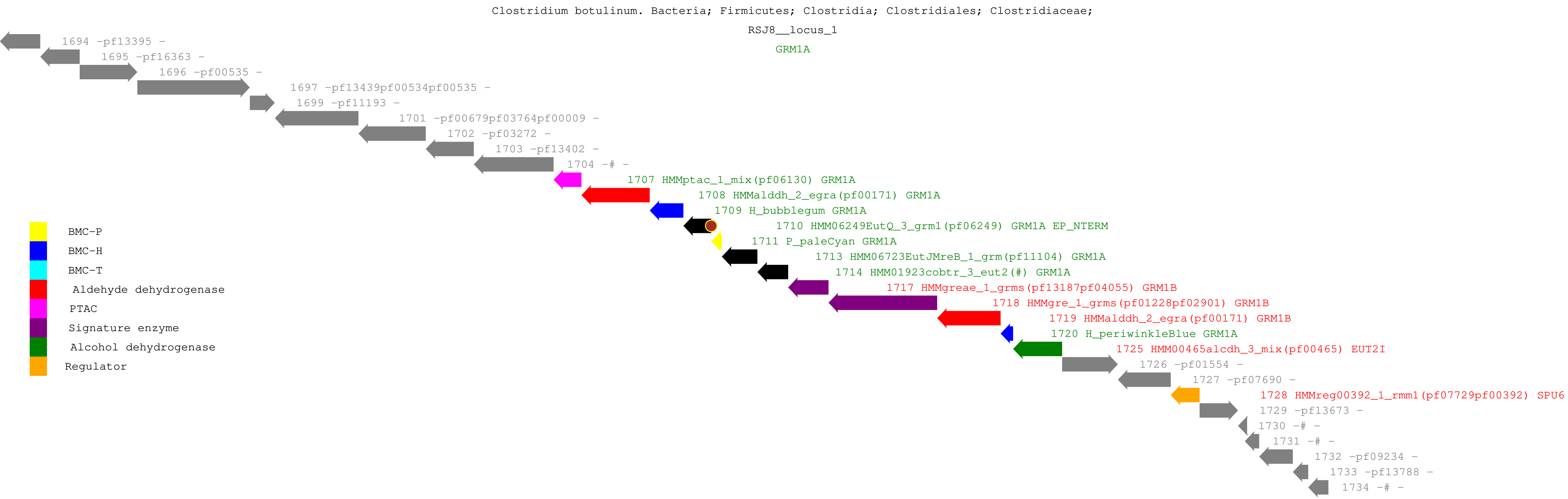

Clostridium botulinum. Bacteria; Firmicutes; Clostridia; Clostridiales; Clostridiaceae;
RSJ8__locus_1
GRM1A
1694 -pf13395 -
1695 -pf16363 -
1696 -pf00535 -
1697 -pf13439pf00534pf00535 -
1699 -pf11193 -
1701 -pf00679pf03764pf00009 -
1702 -pf03272 -
1703 -pf13402 -
1704 -# -
1707 HMMptac_1_mix(pf06130) GRM1A
1708 HMMaldh_2_egra(pf00171) GRM1A
1709 H_bubblegum GRM1A
1710 HMM06249EutQ_3_grm1(pf06249) GRM1A EP_NTERM
1711 P_paleCyan GRM1A
1713 HMM06723EutJMreB_1_grm(pf11104) GRM1A
1714 HMM01923cobtr_3_eut2(#) GRM1A
1717 HMMgreae_1_grms(pf13187pf04055) GRM1B
1718 HMMgre_1_grms(pf01228pf02901) GRM1B
1719 HMMaldh_2_egra(pf00171) GRM1B
1720 H_periwinkleBlue GRM1A
1725 HMM00465alcdh_3_mix(pf00465) EUT2I
1726 -pf01554 -
1727 -pf07690 -
1728 HMMreg00392_1_rmm1(pf07729pf00392) SPU6
1729 -pf13673 -
1730 -# -
1731 -# -
1732 -pf09234 -
1733 -pf13788 -
1734 -# -
BMC-P
BMC-H
BMC-T
Aldehyde dehydrogenase
PTAC
Signature enzyme
Alcohol dehydrogenase
Regulator

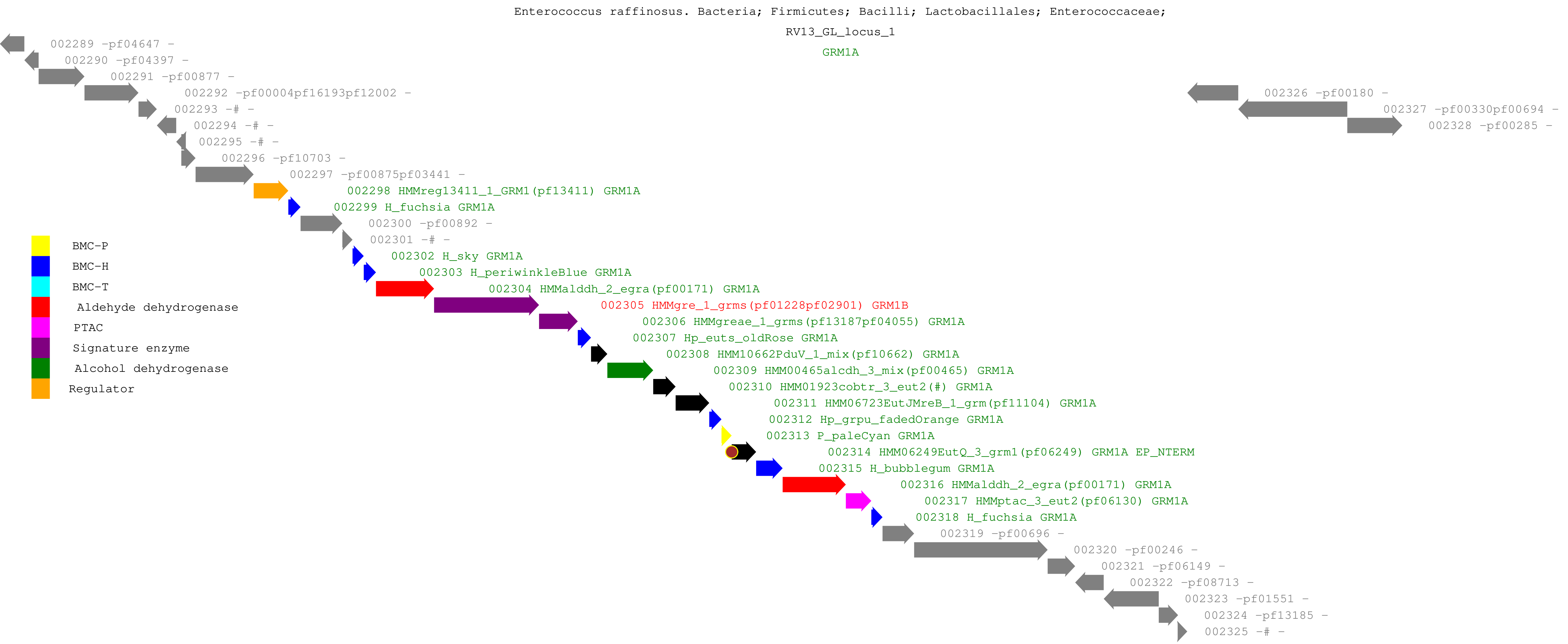
Enterococcus raffinosus. Bacteria; Firmicutes; Bacilli; Lactobacillales; Enterococcaceae;
RV13_GL_locus_1
GRM1A
002289 -pf04647 -
002290 -pf04397 -
002291 -pf00877 -
002292 -pf00004pf16193pf12002 -
002293 -# -
002294 -# -
002295 -# -
002296 -pf10703 -
002297 -pf00875pf03441 -
002298 HMMreg13411_1_GRM1(pf13411) GRM1A
002299 H_fuchsia GRM1A
002300 -pf00892 -
002301 -# -
002302 H_sky GRM1A
002303 H_periwinkleBlue GRM1A
002304 HMMalddh_2_egra(pf00171) GRM1A
002305 HMMgre_1_grms(pf01228pf02901) GRM1B
002306 HMMgreae_1_grms(pf13187pf04055) GRM1A
002307 Hp_euts_oldRose GRM1A
002308 HMM10662PduV_1_mix(pf10662) GRM1A
002309 HMM00465alcdh_3_mix(pf00465) GRM1A
002310 HMM01923cobtr_3_eut2(#) GRM1A
002311 HMM06723EutJMreB_1_grm(pf11104) GRM1A
002312 Hp_grpu_fadedOrange GRM1A
002313 P_paleCyan GRM1A
002314 HMM06249EutQ_3_grm1(pf06249) GRM1A EP_NTERM
002315 H_bubblegum GRM1A
002316 HMMalddh_2_egra(pf00171) GRM1A
002317 HMMptac_3_eut2(pf06130) GRM1A
002318 H_fuchsia GRM1A
002319 -pf00696 -
002320 -pf00246 -
002321 -pf06149 -
002322 -pf08713 -
002323 -pf01551 -
002324 -pf13185 -
002325 -# -
002326 -pf00180 -
002327 -pf00330pf00694 -
002328 -pf00285 -
BMC-P
BMC-H
BMC-T
Aldehyde dehydrogenase
PTAC
Signature enzyme
Alcohol dehydrogenase
Regulator

uncultured Eubacterium sp. Bacteria; Firmicutes; Clostridia; Clostridiales; Eubacteriaceae;

SAMEA3545284__locus_1

GRM1A

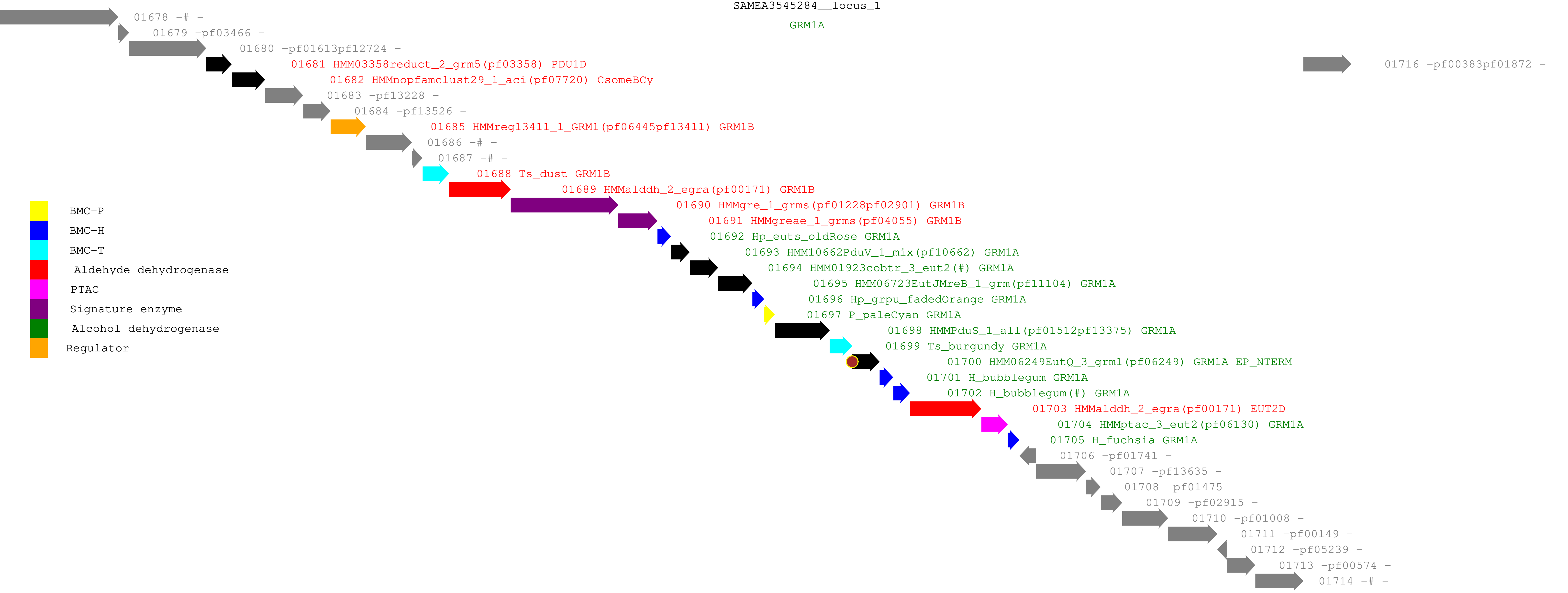

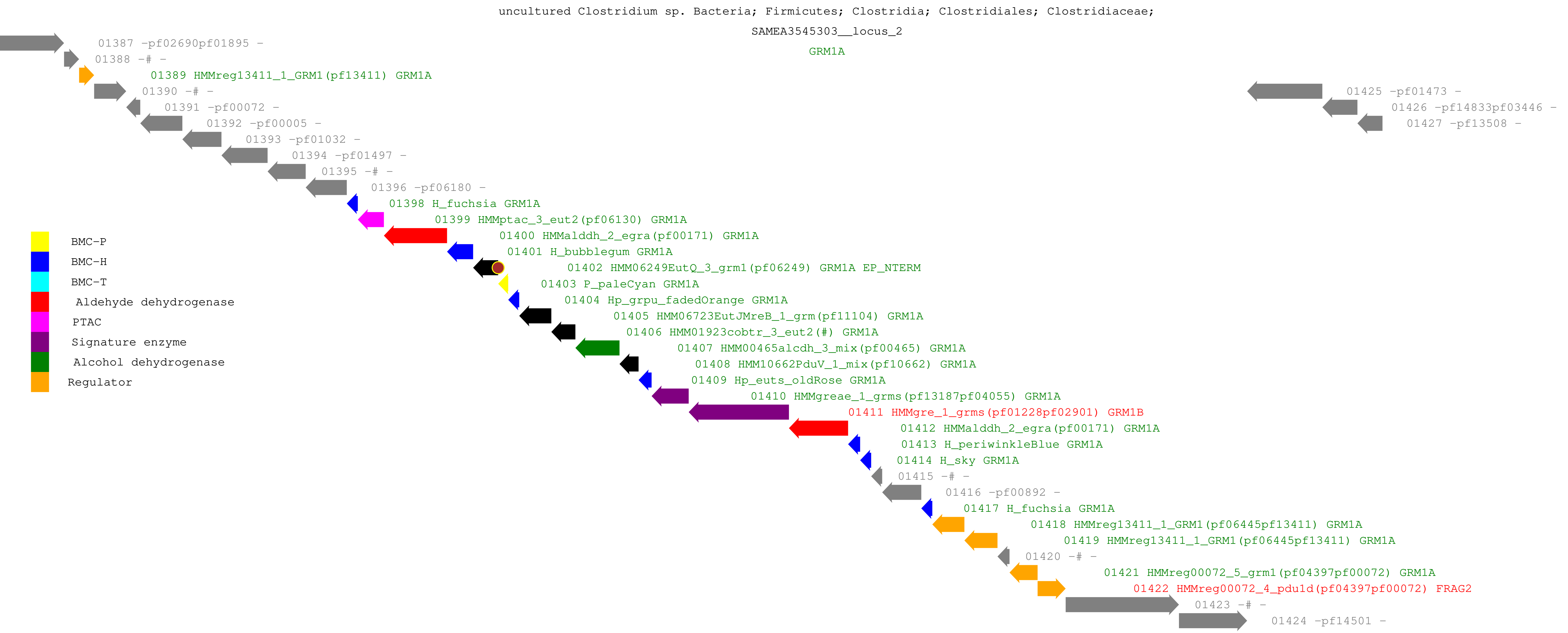
uncultured Clostridium sp. Bacteria; Firmicutes; Clostridia; Clostridiales; Clostridiaceae;
SAMEA3545303__locus_2
GRM1A
01387 -pf02690pf01895 -
01388 -# -
01389 HMMreg13411_1_GRM1(pf13411) GRM1A
01390 -# -
01391 -pf00072 -
01392 -pf00005 -
01393 -pf01032 -
01394 -pf01497 -
01395 -# -
01396 -pf06180 -
01398 H_fuchsia GRM1A
01399 HMMptac_3_eut2(pf06130) GRM1A
01400 HMMaldh_2_egra(pf00171) GRM1A
01401 H_bubblegum GRM1A
01402 HMM06249EutQ_3_grm1(pf06249) GRM1A EP_NTERM
01403 P_paleCyan GRM1A
01404 Hp_grpu_fadedOrange GRM1A
01405 HMM06723EutJMreB_1_grm(pf11104) GRM1A
01406 HMM01923cobtr_3_eut2(#) GRM1A
01407 HMM00465alcdh_3_mix(pf00465) GRM1A
01408 HMM10662PduV_1_mix(pf10662) GRM1A
01409 Hp_euts_oldRose GRM1A
01410 HMMgreae_1_grms(pf13187pf04055) GRM1A
01411 HMMgre_1_grms(pf01228pf02901) GRM1B
01412 HMMaldh_2_egra(pf00171) GRM1A
01413 H_periwinkleBlue GRM1A
01414 H_sky GRM1A
01415 -# -
01416 -pf00892 -
01417 H_fuchsia GRM1A
01418 HMMreg13411_1_GRM1(pf06445pf13411) GRM1A
01419 HMMreg13411_1_GRM1(pf06445pf13411) GRM1A
01420 -# -
01421 HMMreg00072_5_grm1(pf04397pf00072) GRM1A
01422 HMMreg00072_4_pdu1d(pf04397pf00072) FRAG2
01423 -# -
01424 -pf14501 -
01425 -pf01473 -
01426 -pf14833pf03446 -
01427 -pf13508 -
BMC-P
BMC-H
BMC-T
Aldehyde dehydrogenase
PTAC
Signature enzyme
Alcohol dehydrogenase
Regulator

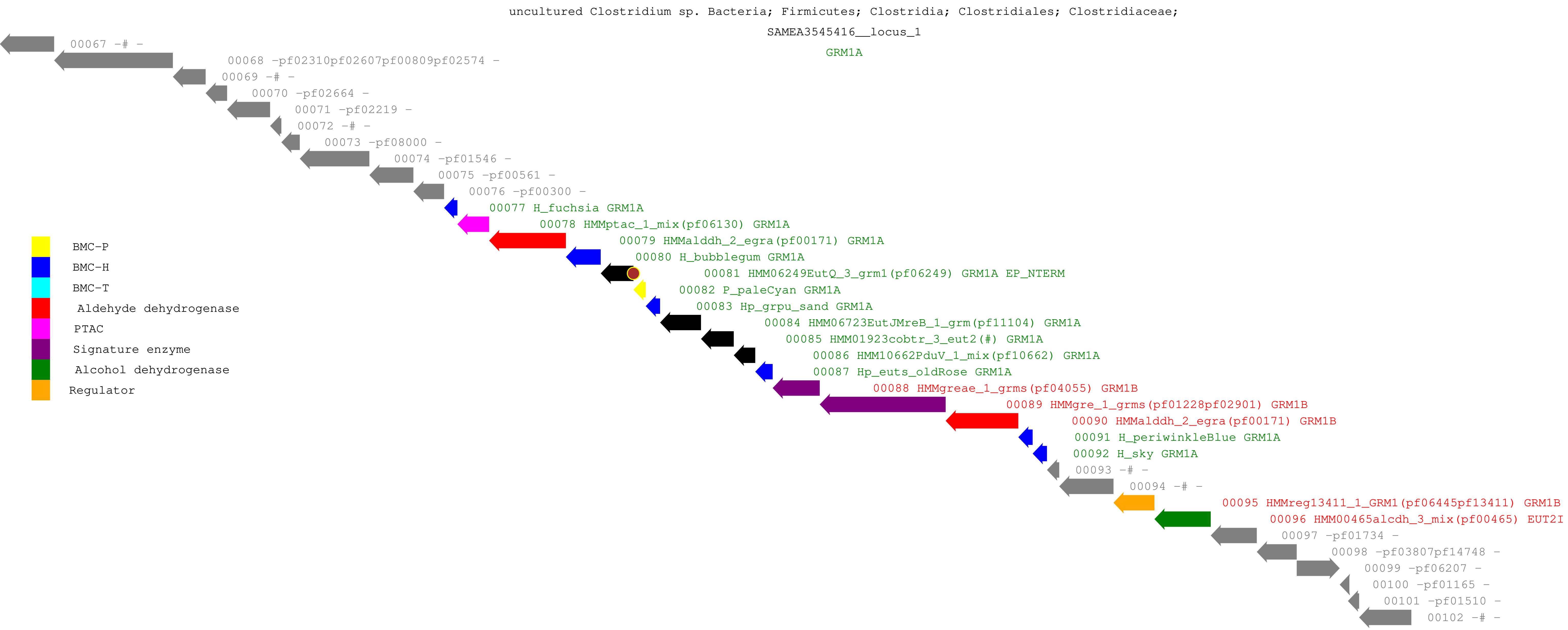
uncultured Clostridium sp. Bacteria; Firmicutes; Clostridia; Clostridiales; Clostridiaceae;
SAMEA3545416__locus_1
GRM1A
00067 -# -
00068 -pf02310pf02607pf00809pf02574 -
00069 -# -
00070 -pf02664 -
00071 -pf02219 -
00072 -# -
00073 -pf08000 -
00074 -pf01546 -
00075 -pf00561 -
00076 -pf00300 -
00077 H_fuchsia GRM1A
00078 HMMptac_1_mix(pf06130) GRM1A
00079 HMMalddh_2_egra(pf00171) GRM1A
00080 H_bubblegum GRM1A
00081 HMM06249EutQ_3_grm1(pf06249) GRM1A EP_NTERM
00082 P_paleCyan GRM1A
00083 Hp_grpu_sand GRM1A
00084 HMM06723EutJMreB_1_grm(pf11104) GRM1A
00085 HMM01923cobtr_3_eut2(#) GRM1A
00086 HMM10662PduV_1_mix(pf10662) GRM1A
00087 Hp_euts_oldRose GRM1A
00088 HMMgreae_1_grms(pf04055) GRM1B
00089 HMMgre_1_grms(pf01228pf02901) GRM1B
00090 HMMalddh_2_egra(pf00171) GRM1B
00091 H_periwinkleBlue GRM1A
00092 H_sky GRM1A
00093 -# -
00094 -# -
00095 HMMreg13411_1_GRM1(pf06445pf13411) GRM1B
00096 HMM00465alcdh_3_mix(pf00465) EUT2I
00097 -pf01734 -
00098 -pf03807pf14748 -
00099 -pf06207 -
00100 -pf01165 -
00101 -pf01510 -
00102 -# -
BMC-P
BMC-H
BMC-T
Aldehyde dehydrogenase
PTAC
Signature enzyme
Alcohol dehydrogenase
Regulator

Streptococcus merionis. Bacteria; Firmicutes; Bacilli; Lactobacillales; Streptococcaceae;

SAMEA4412692__locus_2

GRM1A

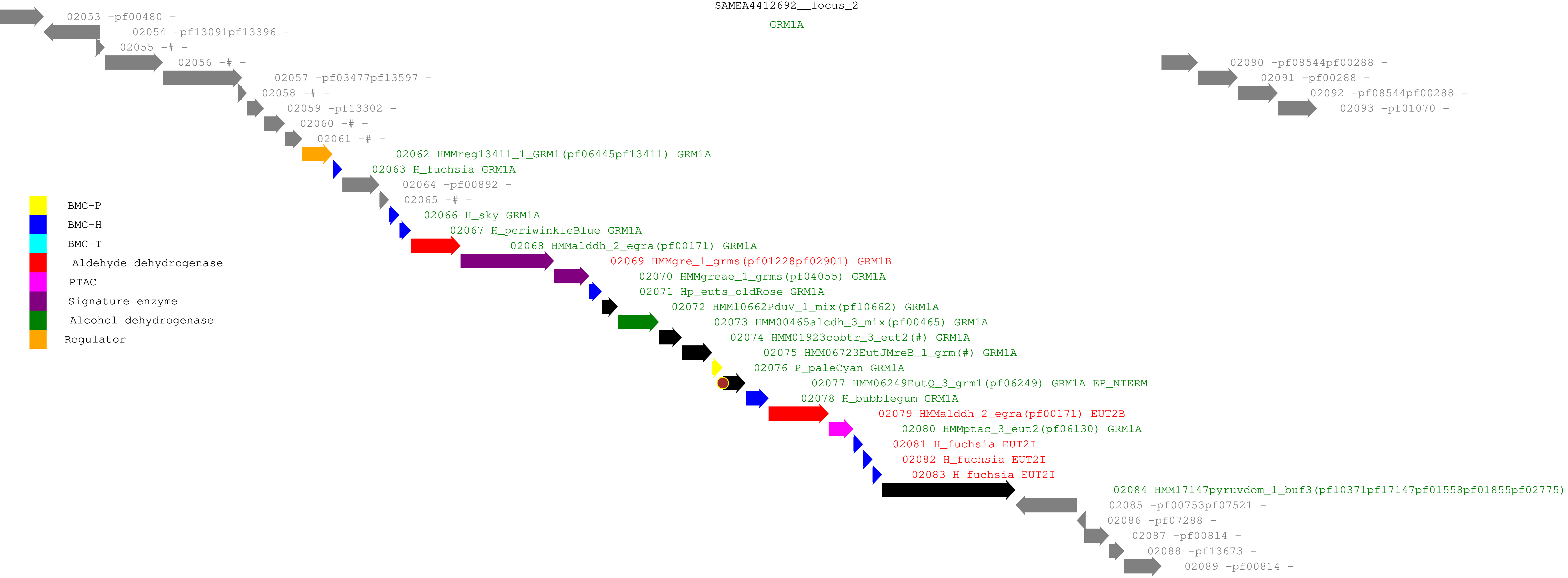

Maledivibacter halophilus. Bacteria; Firmicutes; Clostridia; Clostridiales; Clostridiaceae;

SAMN02194393__locus_5

GRM1A

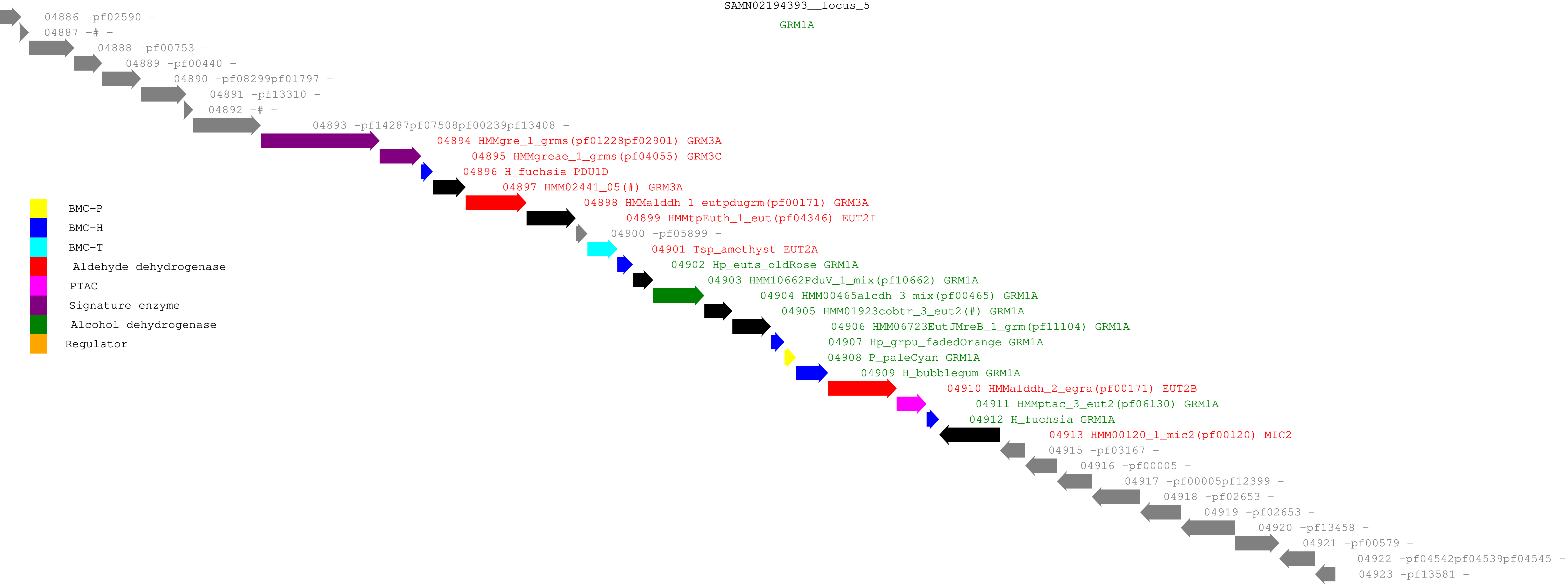

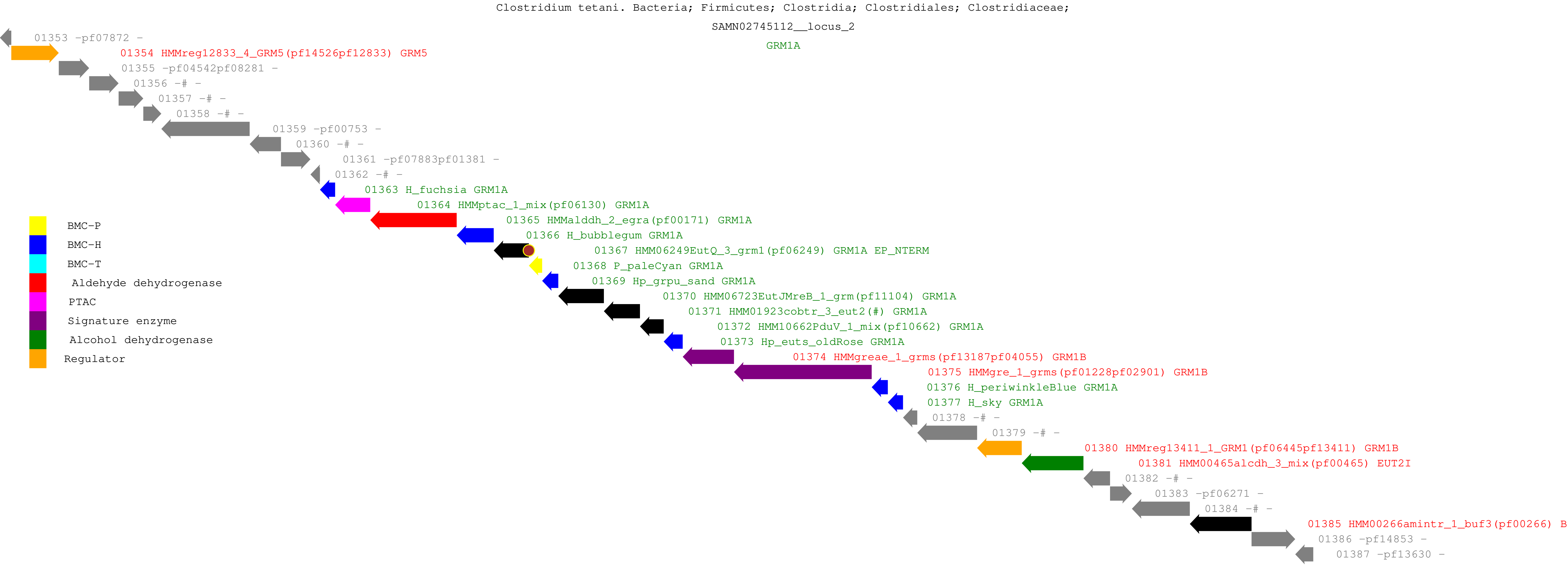
Clostridium tetani. Bacteria; Firmicutes; Clostridia; Clostridiales; Clostridiaceae;
SAMN02745112__locus_2
GRM1A
01353 -pf07872 -
01354 HMMreg12833_4_GRM5(pf14526pf12833) GRM5
01355 -pf04542pf08281 -
01356 -# -
01357 -# -
01358 -# -
01359 -pf00753 -
01360 -# -
01361 -pf07883pf01381 -
01362 -# -
01363 H_fuchsia GRM1A
01364 HMMptac_1_mix(pf06130) GRM1A
01365 HMMalddh_2_egra(pf00171) GRM1A
01366 H_bubblegum GRM1A
01367 HMM06249EutQ_3_grm1(pf06249) GRM1A EP_NTERM
01368 P_paleCyan GRM1A
01369 Hp_grpu_sand GRM1A
01370 HMM06723EutJMreB_1_grm(pf11104) GRM1A
01371 HMM01923cobtr_3_eut2(#) GRM1A
01372 HMM10662PduV_1_mix(pf10662) GRM1A
01373 Hp_euts_oldRose GRM1A
01374 HMMgreae_1_grms(pf13187pf04055) GRM1B
01375 HMMgre_1_grms(pf01228pf02901) GRM1B
01376 H_periwinkleBlue GRM1A
01377 H_sky GRM1A
01378 -# -
01379 -# -
01380 HMMreg13411_1_GRM1(pf06445pf13411) GRM1B
01381 HMM00465alcdh_3_mix(pf00465) EUT2I
01382 -# -
01383 -pf06271 -
01384 -# -
01385 HMM00266amintr_1_buf3(pf00266) B
01386 -pf14853 -
01387 -pf13630 -
BMC-P
BMC-H
BMC-T
Aldehyde dehydrogenase
PTAC
Signature enzyme
Alcohol dehydrogenase
Regulator

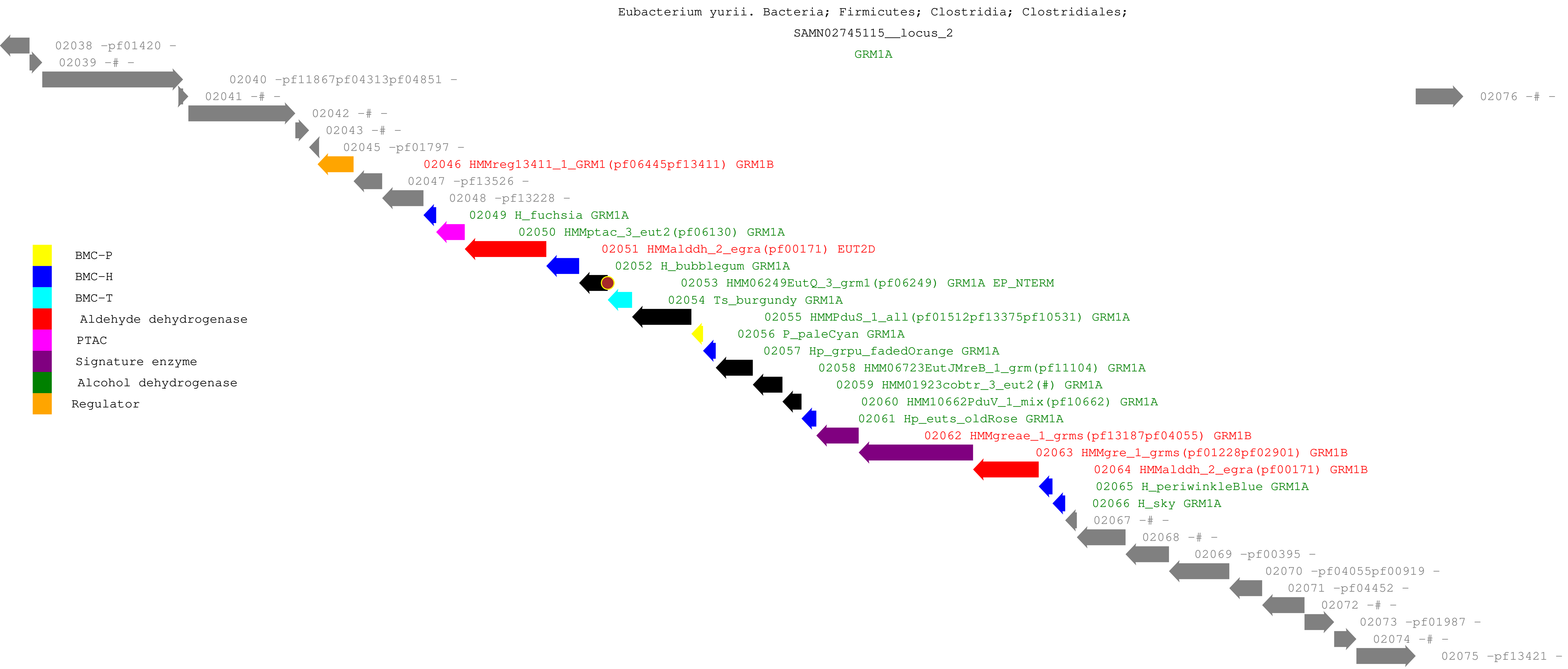
Eubacterium yurii. Bacteria; Firmicutes; Clostridia; Clostridiales;
SAMN02745115__locus_2
GRM1A
02038 -pf01420 -
02039 -# -
02040 -pf11867pf04313pf04851 -
02041 -# -
02042 -# -
02043 -# -
02045 -pf01797 -
02046 HMMreg13411_1_GRM1(pf06445pf13411) GRM1B
02047 -pf13526 -
02048 -pf13228 -
02049 H_fuchsia GRM1A
02050 HMMptac_3_eut2(pf06130) GRM1A
02051 HMMaldh_2_egra(pf00171) EUT2D
02052 H_bubblegum GRM1A
02053 HMM06249EutQ_3_grm1(pf06249) GRM1A EP_NTERM
02054 Ts_burgundy GRM1A
02055 HMMPduS_1_all(pf01512pf13375pf10531) GRM1A
02056 P_paleCyan GRM1A
02057 Hp_grpu_fadedOrange GRM1A
02058 HMM06723EutJMreB_1_grm(pf11104) GRM1A
02059 HMM01923cobtr_3_eut2(#) GRM1A
02060 HMM10662PduV_1_mix(pf10662) GRM1A
02061 Hp_euts_oldRose GRM1A
02062 HMMgreae_1_grms(pf13187pf04055) GRM1B
02063 HMMgre_1_grms(pf01228pf02901) GRM1B
02064 HMMaldh_2_egra(pf00171) GRM1B
02065 H_periwinkleBlue GRM1A
02066 H_sky GRM1A
02067 -# -
02068 -# -
02069 -pf00395 -
02070 -pf04055pf00919 -
02071 -pf04452 -
02072 -# -
02073 -pf01987 -
02074 -# -
02075 -pf13421 -
02076 -# -
BMC-P
BMC-H
BMC-T
Aldehyde dehydrogenase
PTAC
Signature enzyme
Alcohol dehydrogenase
Regulator

Desulfotomaculum aeronauticum DSM 10349. Bacteria; Firmicutes; Clostridia; Clostridiales; Peptococcaceae;

SAMN02745123__locus_1

GRM1A

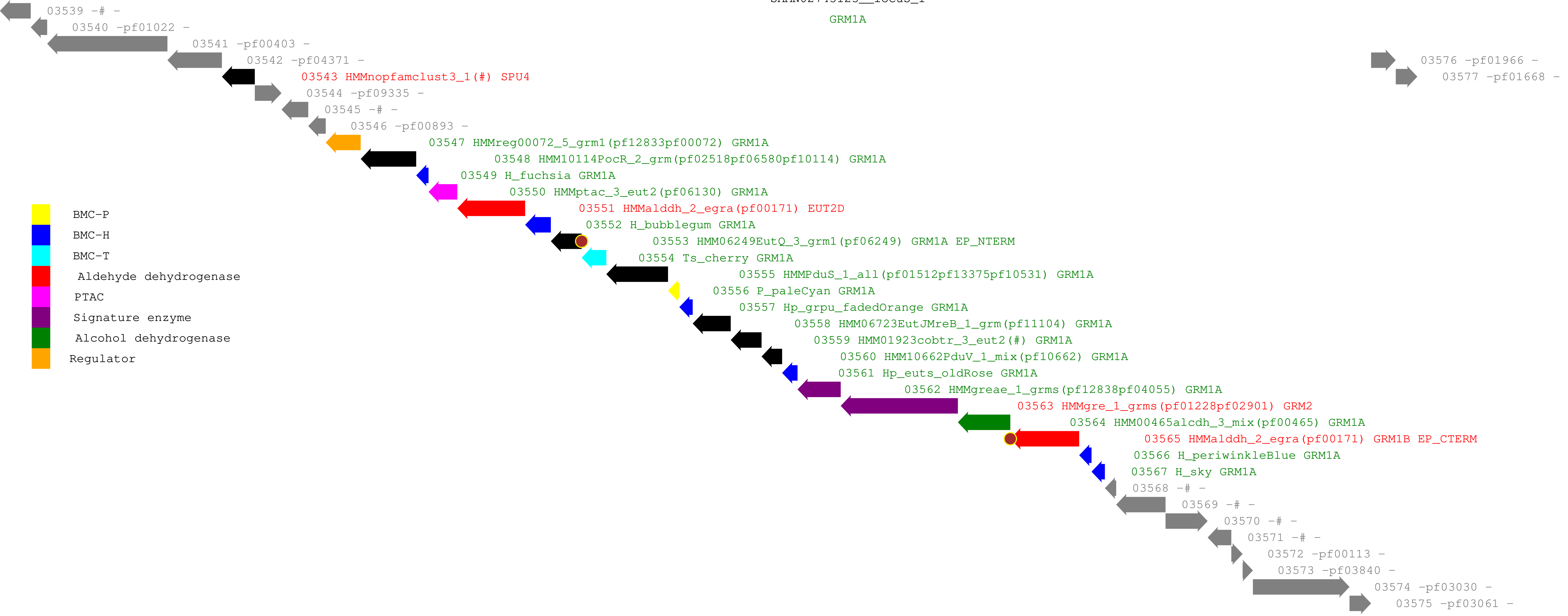

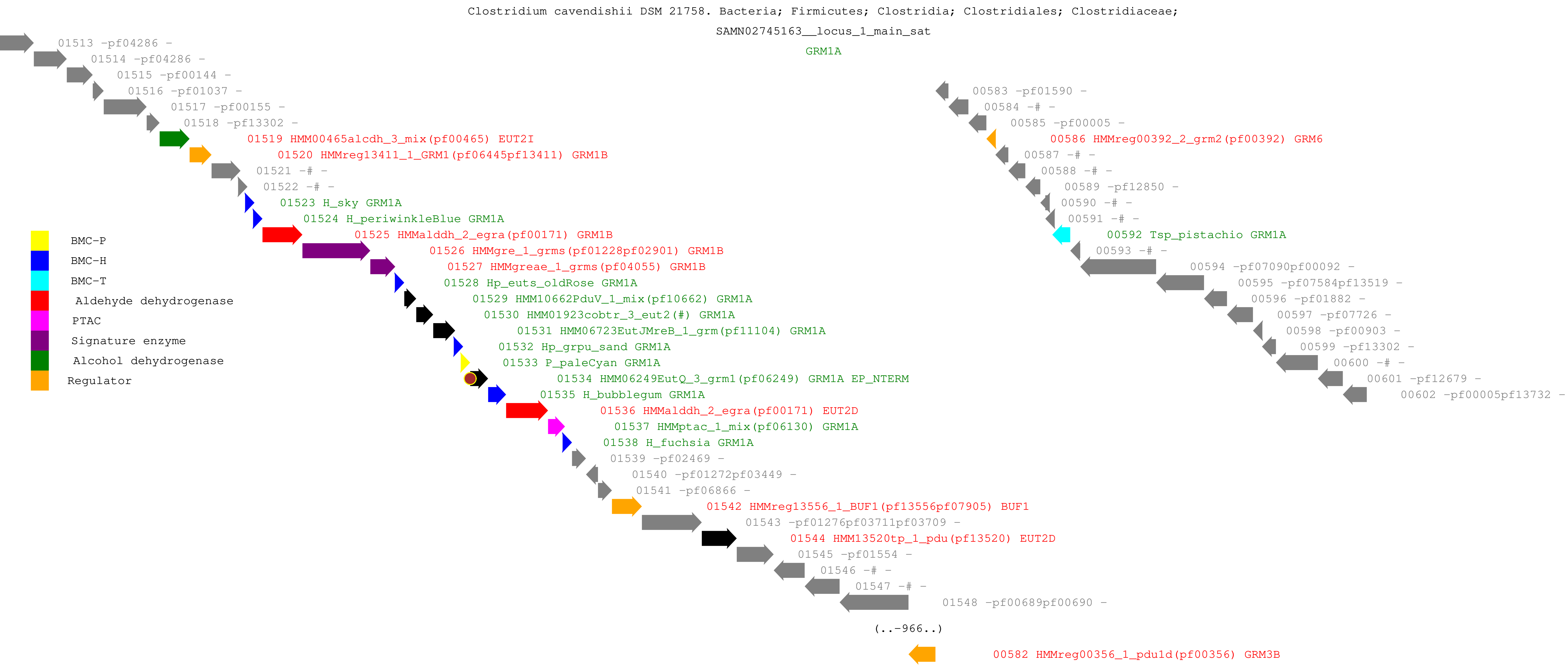

Clostridium cavendishii DSM 21758. Bacteria; Firmicutes; Clostridia; Clostridiales; Clostridiaceae;
SAMN02745163__locus_1_main_sat
GRM1A
01513 -pf04286 -
01514 -pf04286 -
01515 -pf00144 -
01516 -pf01037 -
01517 -pf00155 -
01518 -pf13302 -
01519 HMM00465alcdh_3_mix(pf00465) EUT2I
01520 HMMreg13411_1_GRM1(pf06445pf13411) GRM1B
01521 -# -
01522 -# -
01523 H_sky GRM1A
01524 H_periwinkleBlue GRM1A
01525 HMMalddh_2_egra(pf00171) GRM1B
01526 HMMgre_1_grms(pf01228pf02901) GRM1B
01527 HMMgreae_1_grms(pf04055) GRM1B
01528 Hp_euts_oldRose GRM1A
01529 HMM10662PduV_1_mix(pf10662) GRM1A
01530 HMM01923cobtr_3_eut2(#) GRM1A
01531 HMM06723EutJMreB_1_grm(pf11104) GRM1A
01532 Hp_grpu_sand GRM1A
01533 P_paleCyan GRM1A
01534 HMM06249EutQ_3_grm1(pf06249) GRM1A EP_NTERM
01535 H_bubblegum GRM1A
01536 HMMalddh_2_egra(pf00171) EUT2D
01537 HMMptac_1_mix(pf06130) GRM1A
01538 H_fuchsia GRM1A
01539 -pf02469 -
01540 -pf01272pf03449 -
01541 -pf06866 -
01542 HMMreg13556_1_BUF1(pf13556pf07905) BUF1
01543 -pf01276pf03711pf03709 -
01544 HMM13520tp_1_pdu(pf13520) EUT2D
01545 -pf01554 -
01546 -# -
01547 -# -
01548 -pf00689pf00690 -
(..-966..)
00582 HMMreg00356_1_pdu1d(pf00356) GRM3B
00583 -pf01590 -
00584 -# -
00585 -pf00005 -
00586 HMMreg00392_2_grm2(pf00392) GRM6
00587 -# -
00588 -# -
00589 -pf12850 -
00590 -# -
00591 -# -
00592 Tsp_pistachio GRM1A
00593 -# -
00594 -pf07090pf00092 -
00595 -pf07584pf13519 -
00596 -pf01882 -
00597 -pf07726 -
00598 -pf00903 -
00599 -pf13302 -
00600 -# -
00601 -pf12679 -
00602 -pf00005pf13732 -
BMC-P
BMC-H
BMC-T
Aldehyde dehydrogenase
PTAC
Signature enzyme
Alcohol dehydrogenase
Regulator

Cetobacterium ceti. Bacteria; Fusobacteria; Fusobacteriales; Fusobacteriaceae;

SAMN02745174__locus_2

GRM1A

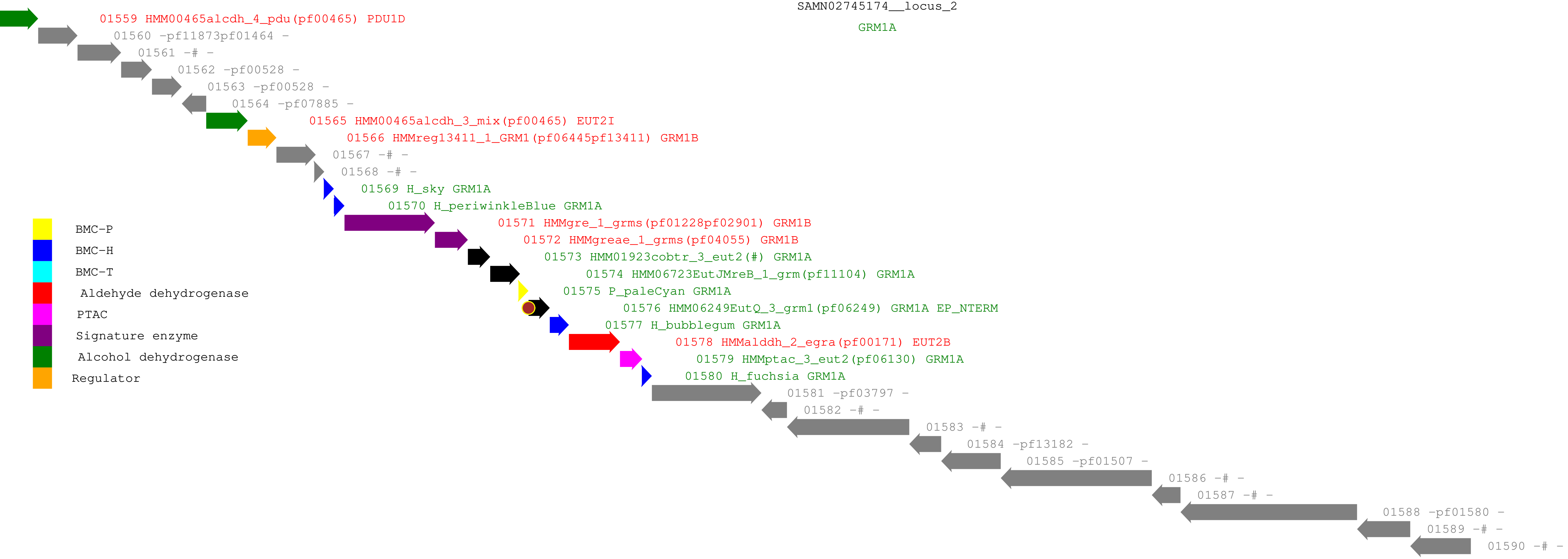

Clostridium grantii DSM 8605. Bacteria; Firmicutes; Clostridia; Clostridiales; Clostridiaceae;

SAMN02745207__locus_2

GRM1A

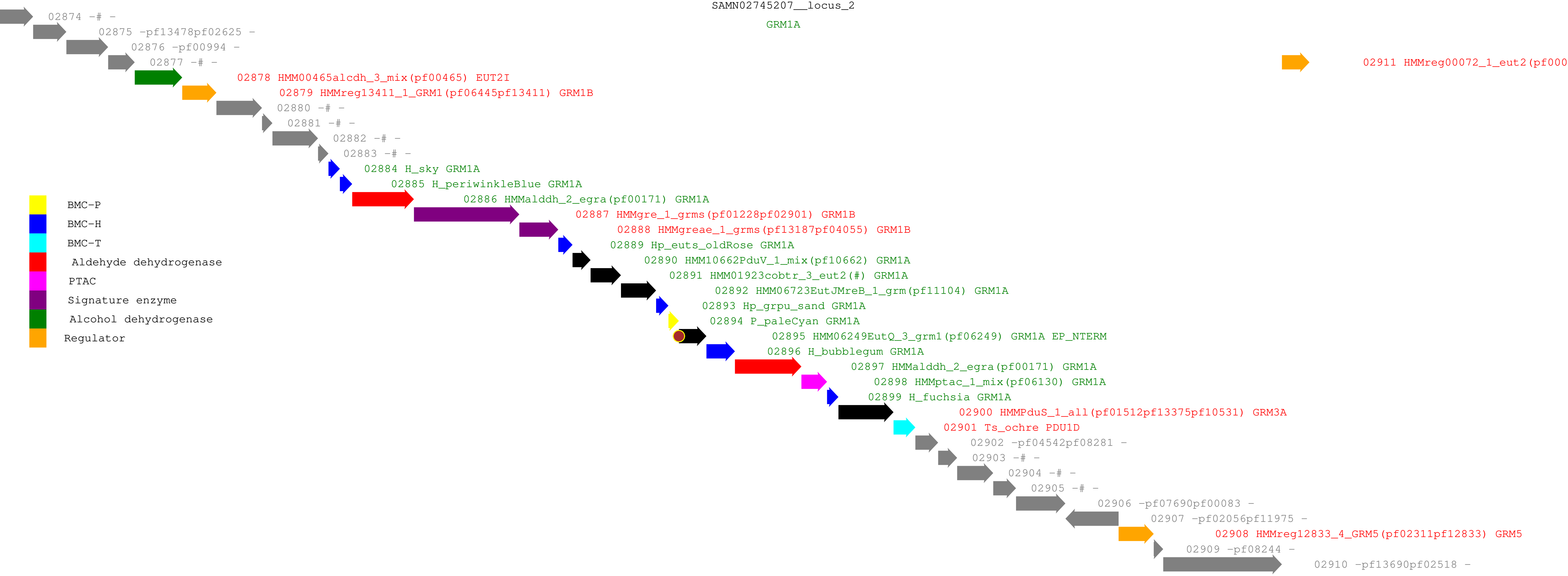

Desulfitobacterium chlororespirans DSM 11544. Bacteria; Firmicutes; Clostridia; Clostridiales; Peptococcaceae;

SAMN02745215__locus_2

GRM1A

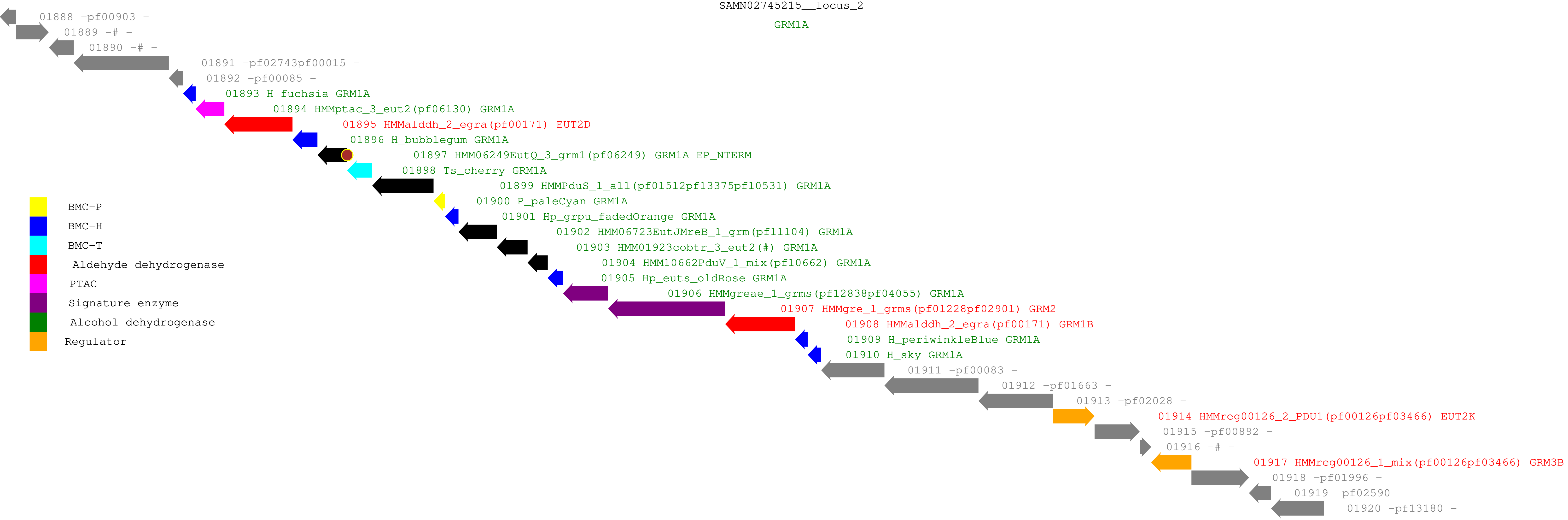

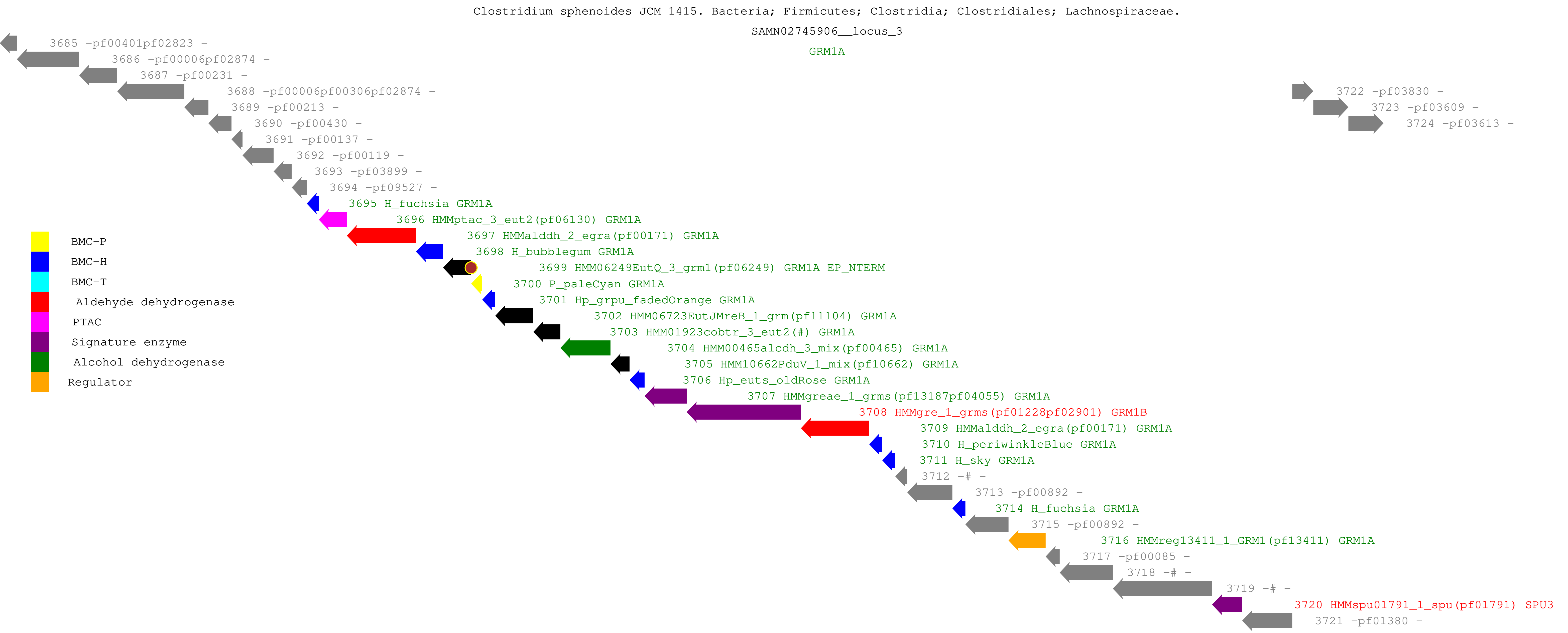

Clostridium sphenoides JCM 1415. Bacteria; Firmicutes; Clostridia; Clostridiales; Lachnospiraceae.
SAMN02745906__locus_3
GRM1A
3685 -pf00401pf02823 -
3686 -pf00006pf02874 -
3687 -pf00231 -
3688 -pf00006pf00306pf02874 -
3689 -pf00213 -
3690 -pf00430 -
3691 -pf00137 -
3692 -pf00119 -
3693 -pf03899 -
3694 -pf09527 -
3695 H_fuchsia GRM1A
3696 HMMptac_3_eut2(pf06130) GRM1A
3697 HMMalddh_2_egra(pf00171) GRM1A
3698 H_bubblegum GRM1A
3699 HMM06249EutQ_3_grm1(pf06249) GRM1A EP_NTERM
3700 P_paleCyan GRM1A
3701 Hp_grpu_fadedOrange GRM1A
3702 HMM06723EutJMreB_1_grm(pf11104) GRM1A
3703 HMM01923cobtr_3_eut2(#) GRM1A
3704 HMM00465alcdh_3_mix(pf00465) GRM1A
3705 HMM10662PduV_1_mix(pf10662) GRM1A
3706 Hp_euts_oldRose GRM1A
3707 HMMgreae_1_grms(pf13187pf04055) GRM1A
3708 HMMgre_1_grms(pf01228pf02901) GRM1B
3709 HMMalddh_2_egra(pf00171) GRM1A
3710 H_periwinkleBlue GRM1A
3711 H_sky GRM1A
3712 -# -
3713 -pf00892 -
3714 H_fuchsia GRM1A
3715 -pf00892 -
3716 HMMreg13411_1_GRM1(pf13411) GRM1A
3717 -pf00085 -
3718 -# -
3719 -# -
3720 HMMspu01791_1_spu(pf01791) SPU3
3721 -pf01380 -
3722 -pf03830 -
3723 -pf03609 -
3724 -pf03613 -
BMC-P
BMC-H
BMC-T
Aldehyde dehydrogenase
PTAC
Signature enzyme
Alcohol dehydrogenase
Regulator

Desulfosporosinus lacus DSM 15449. Bacteria; Firmicutes; Clostridia; Clostridiales; Peptococcaceae;

SAMN02746098__locus_1

GRM1A

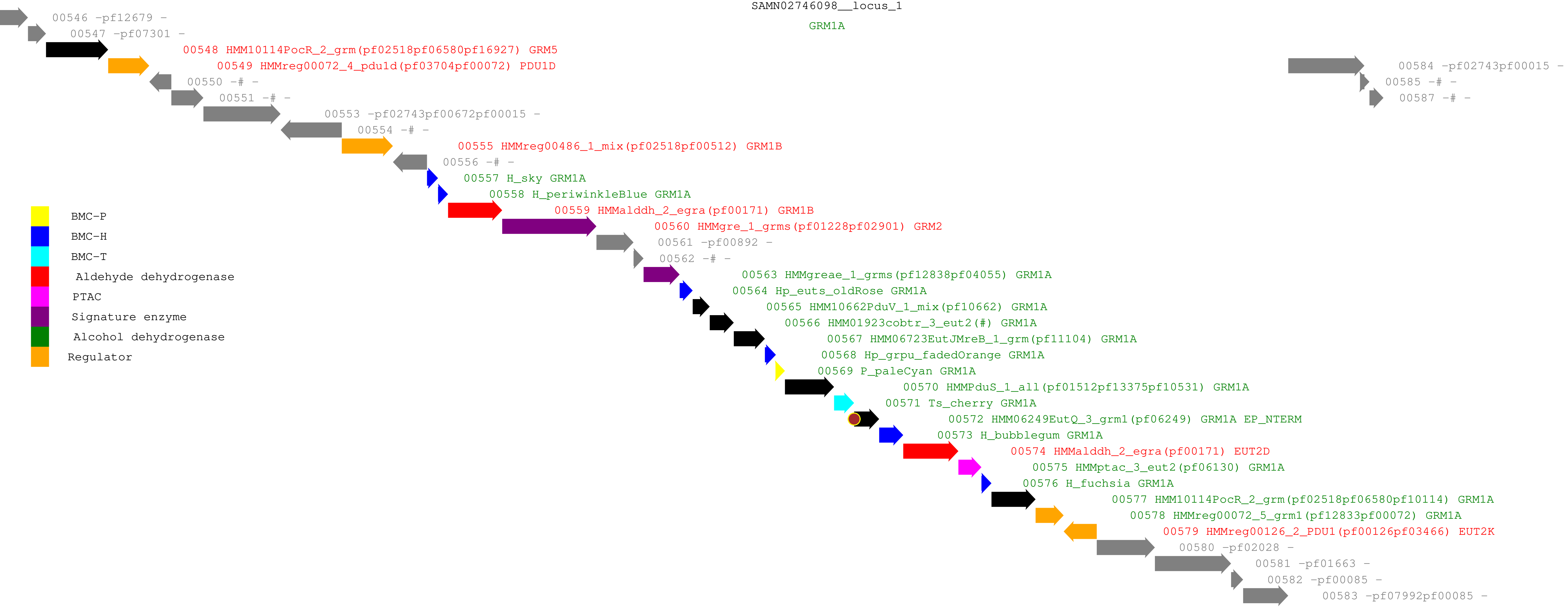

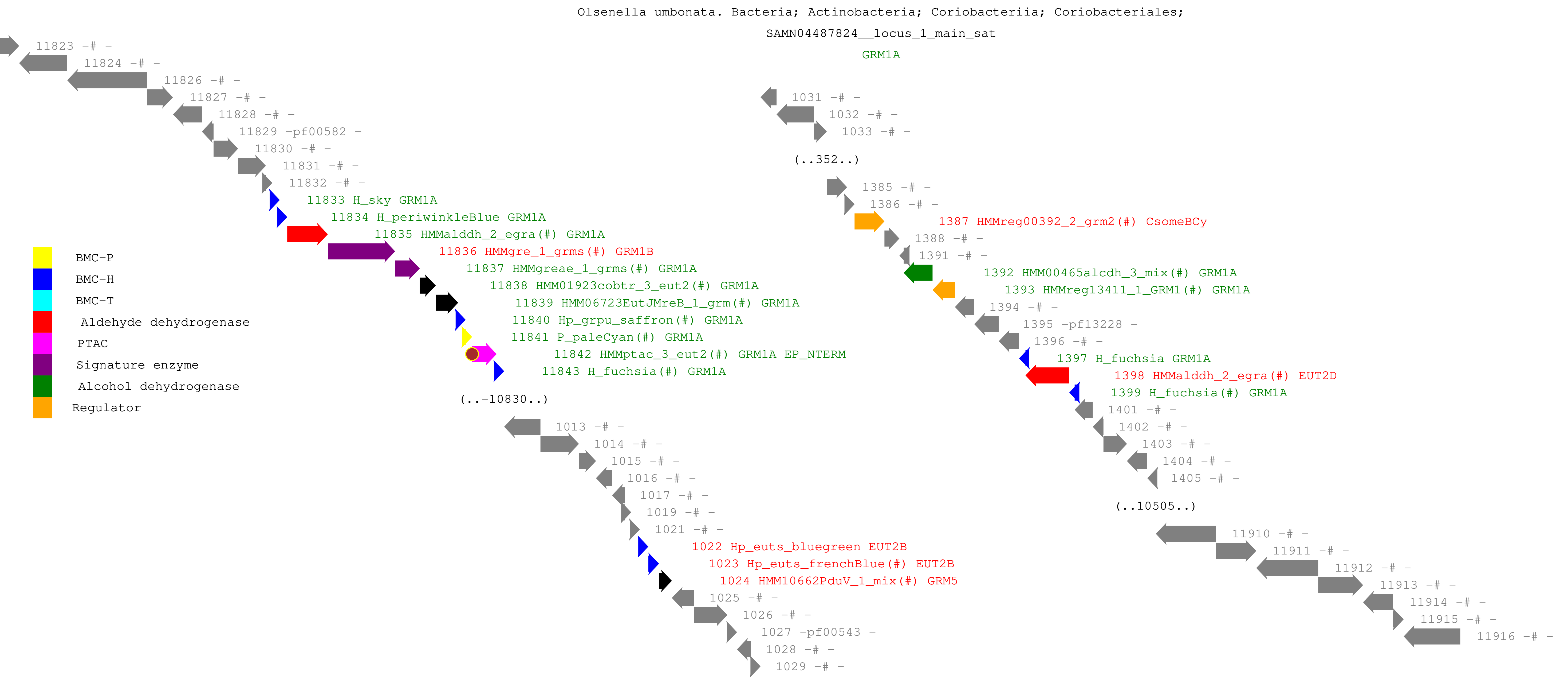
Olsenella umbonata. Bacteria; Actinobacteria; Coriobacteriia; Coriobacteriales;
SAMN04487824__locus_1_main_sat
GRM1A
11823 -# -
11824 -# -
11826 -# -
11827 -# -
11828 -# -
11829 -pf00582 -
11830 -# -
11831 -# -
11832 -# -
11833 H_sky GRM1A
11834 H_periwinkleBlue GRM1A
11835 HMMalddh_2_egra(#) GRM1A
11836 HMMgre_1_grms(#) GRM1B
11837 HMMgreae_1_grms(#) GRM1A
11838 HMM01923cobtr_3_eut2(#) GRM1A
11839 HMM06723EutJMreB_1_grm(#) GRM1A
11840 Hp_grpu_saffron(#) GRM1A
11841 P_paleCyan(#) GRM1A
11842 HMMptac_3_eut2(#) GRM1A EP_NTERM
11843 H_fuchsia(#) GRM1A
(..-10830..)
1013 -# -
1014 -# -
1015 -# -
1016 -# -
1017 -# -
1019 -# -
1021 -# -
1022 Hp_euts_bluegreen EUT2B
1023 Hp_euts_frenchBlue(#) EUT2B
1024 HMM10662PduV_1_mix(#) GRM5
1025 -# -
1026 -# -
1027 -pf00543 -
1028 -# -
1029 -# -
1031 -# -
1032 -# -
1033 -# -
(..352..)
1385 -# -
1386 -# -
1387 HMMreg00392_2_grm2(#) CsomeBCy
1388 -# -
1391 -# -
1392 HMM00465alcdh_3_mix(#) GRM1A
1393 HMMreg13411_1_GRM1(#) GRM1A
1394 -# -
1395 -pf13228 -
1396 -# -
1397 H_fuchsia GRM1A
1398 HMMaldh_2_egra(#) EUT2D
1399 H_fuchsia(#) GRM1A
1401 -# -
1402 -# -
1403 -# -
1404 -# -
1405 -# -
(..10505..)
11910 -# -
11911 -# -
11912 -# -
11913 -# -
11914 -# -
11915 -# -
11916 -# -
BMC-P
BMC-H
BMC-T
Aldehyde dehydrogenase
PTAC
Signature enzyme
Alcohol dehydrogenase
Regulator

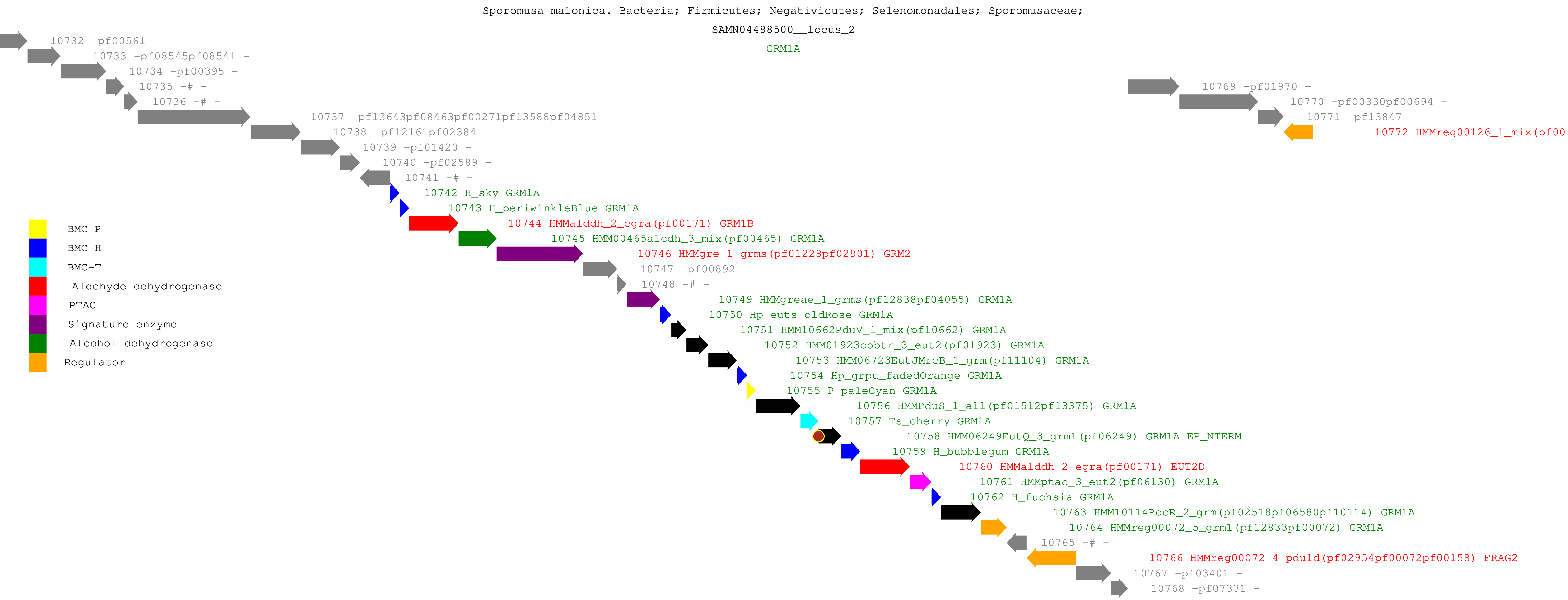
Sporomusa malonica. Bacteria; Firmicutes; Negativicutes; Selenomonadales; Sporomusaceae;
SAMN04488500__locus_2
GRM1A
10732 -pf00561 -
10733 -pf08545pf08541 -
10734 -pf00395 -
10735 -# -
10736 -# -
10737 -pf13643pf08463pf00271pf13588pf04851 -
10738 -pf12161pf02384 -
10739 -pf01420 -
10740 -pf02589 -
10741 -# -
10742 H_sky GRM1A
10743 H_periwinkleBlue GRM1A
10744 HMMalddh_2_egra(pf00171) GRM1B
10745 HMM00465alcdh_3_mix(pf00465) GRM1A
10746 HMMgre_1_grms(pf01228pf02901) GRM2
10747 -pf00892 -
10748 -# -
10749 HMMgreae_1_grms(pf12838pf04055) GRM1A
10750 Hp_euts_oldRose GRM1A
10751 HMM10662PduV_1_mix(pf10662) GRM1A
10752 HMM01923cobtr_3_eut2(pf01923) GRM1A
10753 HMM06723EutJMreB_1_grm(pf11104) GRM1A
10754 Hp_grpu_fadedOrange GRM1A
10755 P_paleCyan GRM1A
10756 HMMPduS_1_all(pf01512pf13375) GRM1A
10757 Ts_cherry GRM1A
10758 HMM06249EutQ_3_grm1(pf06249) GRM1A EP_NTERM
10759 H_bubblegum GRM1A
10760 HMMalddh_2_egra(pf00171) EUT2D
10761 HMMptac_3_eut2(pf06130) GRM1A
10762 H_fuchsia GRM1A
10763 HMM10114PocR_2_grm(pf02518pf06580pf10114) GRM1A
10764 HMMreg00072_5_grm1(pf12833pf00072) GRM1A
10765 -# -
10766 HMMreg00072_4_pdu1d(pf02954pf00072pf00158) FRAG2
10767 -pf03401 -
10768 -pf07331 -
10769 -pf01970 -
10770 -pf00330pf00694 -
10771 -pf13847 -
10772 HMMreg00126_1_mix(pf00
BMC-P
BMC-H
BMC-T
Aldehyde dehydrogenase
PTAC
Signature enzyme
Alcohol dehydrogenase
Regulator

Paenibacillus sp. cl6col. Bacteria; Firmicutes; Bacilli; Bacillales; Paenibacillaceae;

SAMN04488689__locus_2

GRM1A

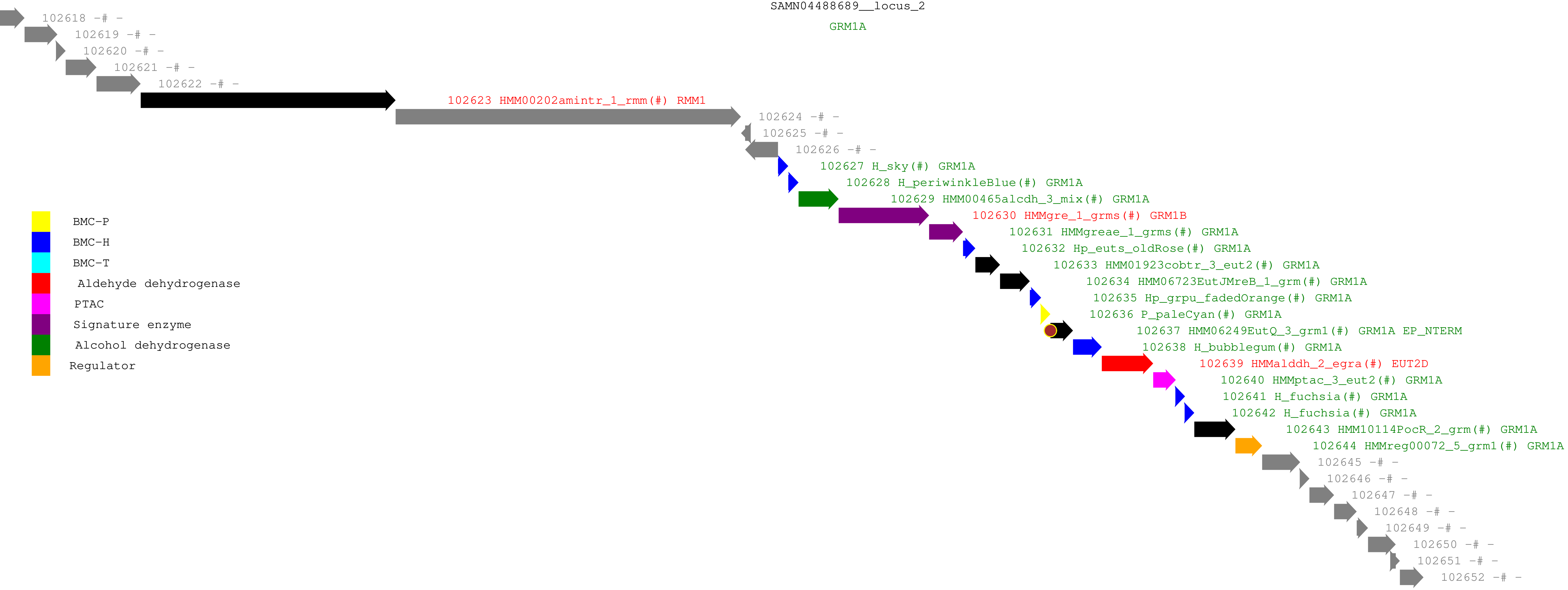

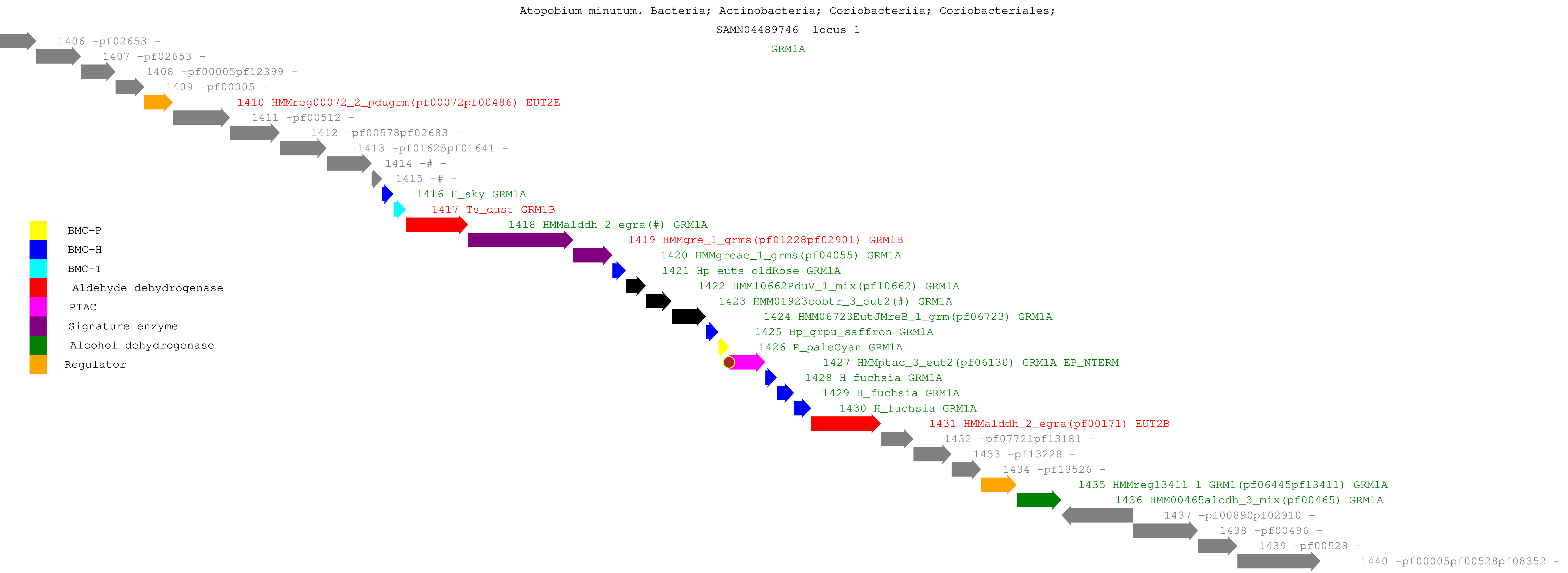
Atopobium minutum. Bacteria; Actinobacteria; Coriobacteriia; Coriobacteriales;
SAMN04489746__locus_1
GRM1A
1406 -pf02653 -
1407 -pf02653 -
1408 -pf00005pf12399 -
1409 -pf00005 -
1410 HMMreg00072_2_pdugrm(pf00072pf00486) EUT2E
1411 -pf00512 -
1412 -pf00578pf02683 -
1413 -pf01625pf01641 -
1414 -# -
1415 -# -
1416 H_sky GRM1A
1417 Ts_dust GRM1B
1418 HMMalddh_2_egra(#) GRM1A
1419 HMMgre_1_grms(pf01228pf02901) GRM1B
1420 HMMgreae_1_grms(pf04055) GRM1A
1421 Hp_euts_oldRose GRM1A
1422 HMM10662PduV_1_mix(pf10662) GRM1A
1423 HMM01923cobtr_3_eut2(#) GRM1A
1424 HMM06723EutJMreB_1_grm(pf06723) GRM1A
1425 Hp_grpu_saffron GRM1A
1426 P_paleCyan GRM1A
1427 HMMptac_3_eut2(pf06130) GRM1A EP_NTERM
1428 H_fuchsia GRM1A
1429 H_fuchsia GRM1A
1430 H_fuchsia GRM1A
1431 HMMalddh_2_egra(pf00171) EUT2B
1432 -pf07721pf13181 -
1433 -pf13228 -
1434 -pf13526 -
1435 HMMreg13411_1_GRM1(pf06445pf13411) GRM1A
1436 HMM00465alcdh_3_mix(pf00465) GRM1A
1437 -pf00890pf02910 -
1438 -pf00496 -
1439 -pf00528 -
1440 -pf00005pf00528pf08352 -
BMC-P
BMC-H
BMC-T
Aldehyde dehydrogenase
PTAC
Signature enzyme
Alcohol dehydrogenase
Regulator

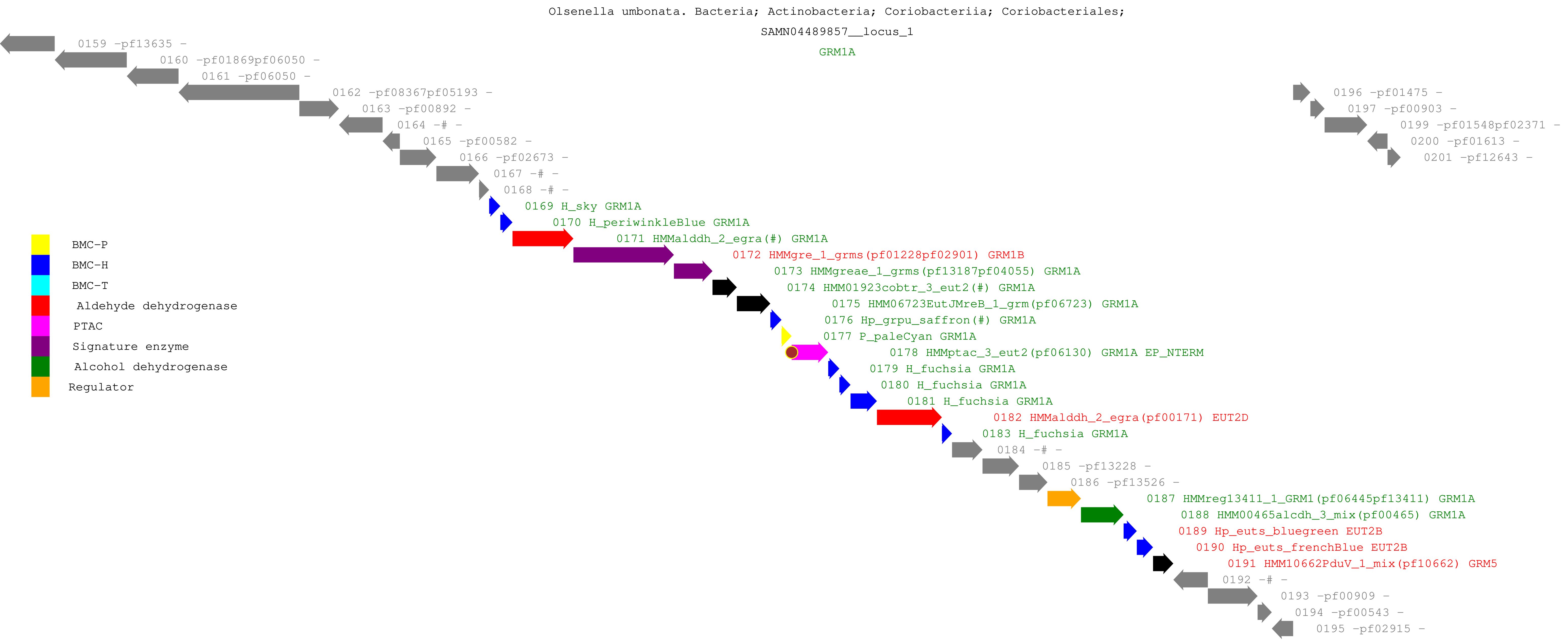
Olsenella umbonata. Bacteria; Actinobacteria; Coriobacteriia; Coriobacteriales;
SAMN04489857__locus_1
GRM1A
0159 -pf13635 -
0160 -pf01869pf06050 -
0161 -pf06050 -
0162 -pf08367pf05193 -
0163 -pf00892 -
0164 -# -
0165 -pf00582 -
0166 -pf02673 -
0167 -# -
0168 -# -
0169 H_sky GRM1A
0170 H_periwinkleBlue GRM1A
0171 HMMalddh_2_egra(#) GRM1A
0172 HMMgre_1_grms(pf01228pf02901) GRM1B
0173 HMMgreae_1_grms(pf13187pf04055) GRM1A
0174 HMM01923cobtr_3_eut2(#) GRM1A
0175 HMM06723EutJMreB_1_grm(pf06723) GRM1A
0176 Hp_grpu_saffron(#) GRM1A
0177 P_paleCyan GRM1A
0178 HMMptac_3_eut2(pf06130) GRM1A EP_NTERM
0179 H_fuchsia GRM1A
0180 H_fuchsia GRM1A
0181 H_fuchsia GRM1A
0182 HMMalddh_2_egra(pf00171) EUT2D
0183 H_fuchsia GRM1A
0184 -# -
0185 -pf13228 -
0186 -pf13526 -
0187 HMMreg13411_1_GRM1(pf06445pf13411) GRM1A
0188 HMM00465alcdh_3_mix(pf00465) GRM1A
0189 Hp_euts_bluegreen EUT2B
0190 Hp_euts_frenchBlue EUT2B
0191 HMM10662PduV_1_mix(pf10662) GRM5
0192 -# -
0193 -pf00909 -
0194 -pf00543 -
0195 -pf02915 -
0196 -pf01475 -
0197 -pf00903 -
0199 -pf01548pf02371 -
0200 -pf01613 -
0201 -pf12643 -
BMC-P
BMC-H
BMC-T
Aldehyde dehydrogenase
PTAC
Signature enzyme
Alcohol dehydrogenase
Regulator

Peptococcus niger. Bacteria; Firmicutes; Clostridia; Clostridiales; Peptococcaceae;

SAMN04489866__locus_1

GRM1A

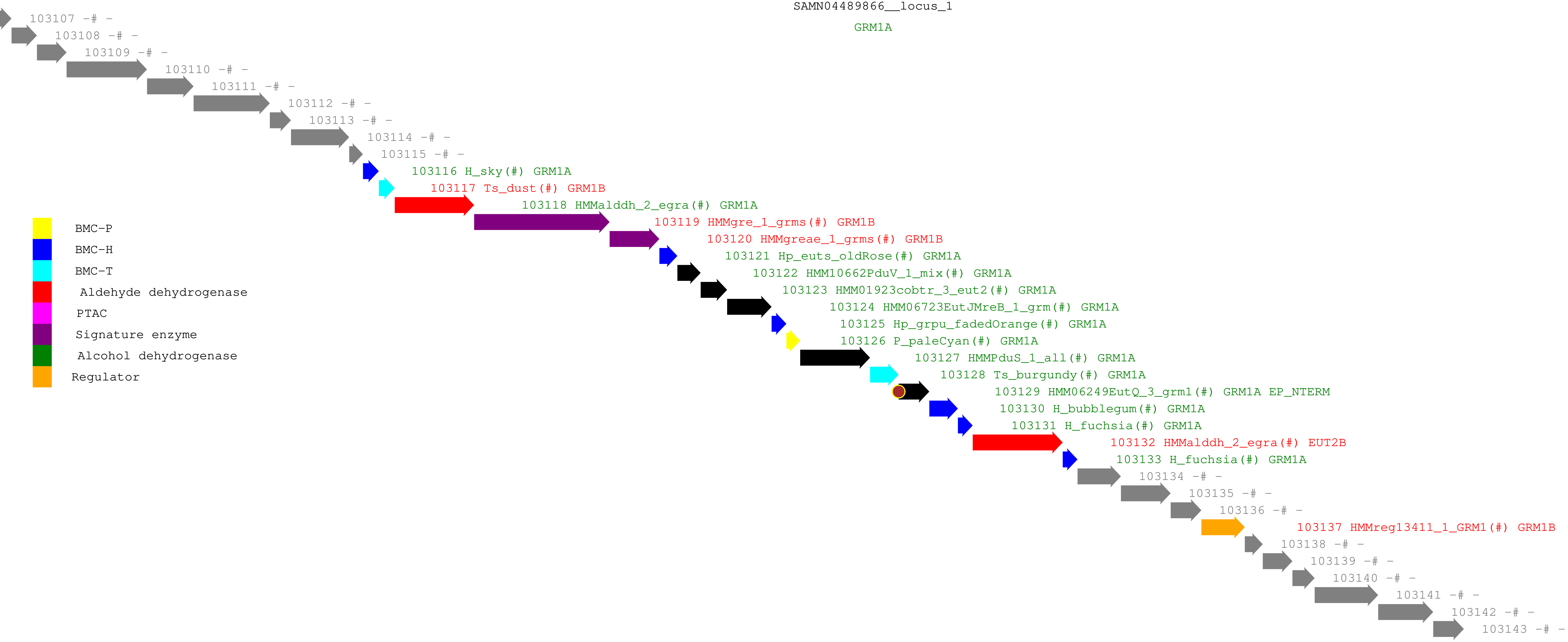

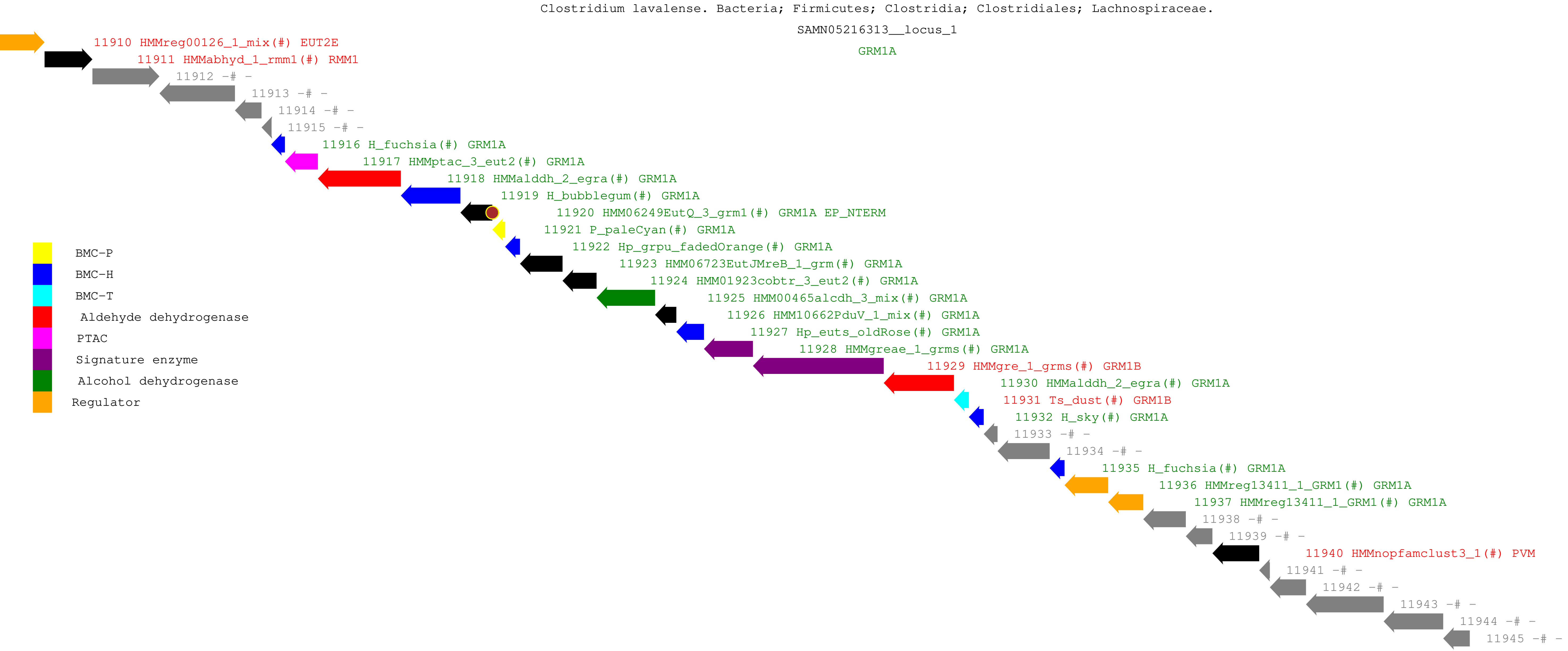

Clostridium lavalense. Bacteria; Firmicutes; Clostridia; Clostridiales; Lachnospiraceae.
SAMN05216313__locus_1
GRM1A
11910 HMMreg00126_1_mix(#) EUT2E
11911 HMMabhyd_1_rmm1(#) RMM1
11912 -# -
11913 -# -
11914 -# -
11915 -# -
11916 H_fuchsia(#) GRM1A
11917 HMMptac_3_eut2(#) GRM1A
11918 HMMalddh_2_egra(#) GRM1A
11919 H_bubblegum(#) GRM1A
11920 HMM06249EutQ_3_grm1(#) GRM1A EP_NTERM
11921 P_paleCyan(#) GRM1A
11922 Hp_grpu_fadedOrange(#) GRM1A
11923 HMM06723EutJMreB_1_grm(#) GRM1A
11924 HMM01923cobtr_3_eut2(#) GRM1A
11925 HMM00465alcdh_3_mix(#) GRM1A
11926 HMM10662PduV_1_mix(#) GRM1A
11927 Hp_euts_oldRose(#) GRM1A
11928 HMMgreae_1_grms(#) GRM1A
11929 HMMgre_1_grms(#) GRM1B
11930 HMMalddh_2_egra(#) GRM1A
11931 Ts_dust(#) GRM1B
11932 H_sky(#) GRM1A
11933 -# -
11934 -# -
11935 H_fuchsia(#) GRM1A
11936 HMMreg13411_1_GRM1(#) GRM1A
11937 HMMreg13411_1_GRM1(#) GRM1A
11938 -# -
11939 -# -
11940 HMMnopfamclust3_1(#) PVM
11941 -# -
11942 -# -
11943 -# -
11944 -# -
11945 -# -
BMC-P
BMC-H
BMC-T
Aldehyde dehydrogenase
PTAC
Signature enzyme
Alcohol dehydrogenase
Regulator

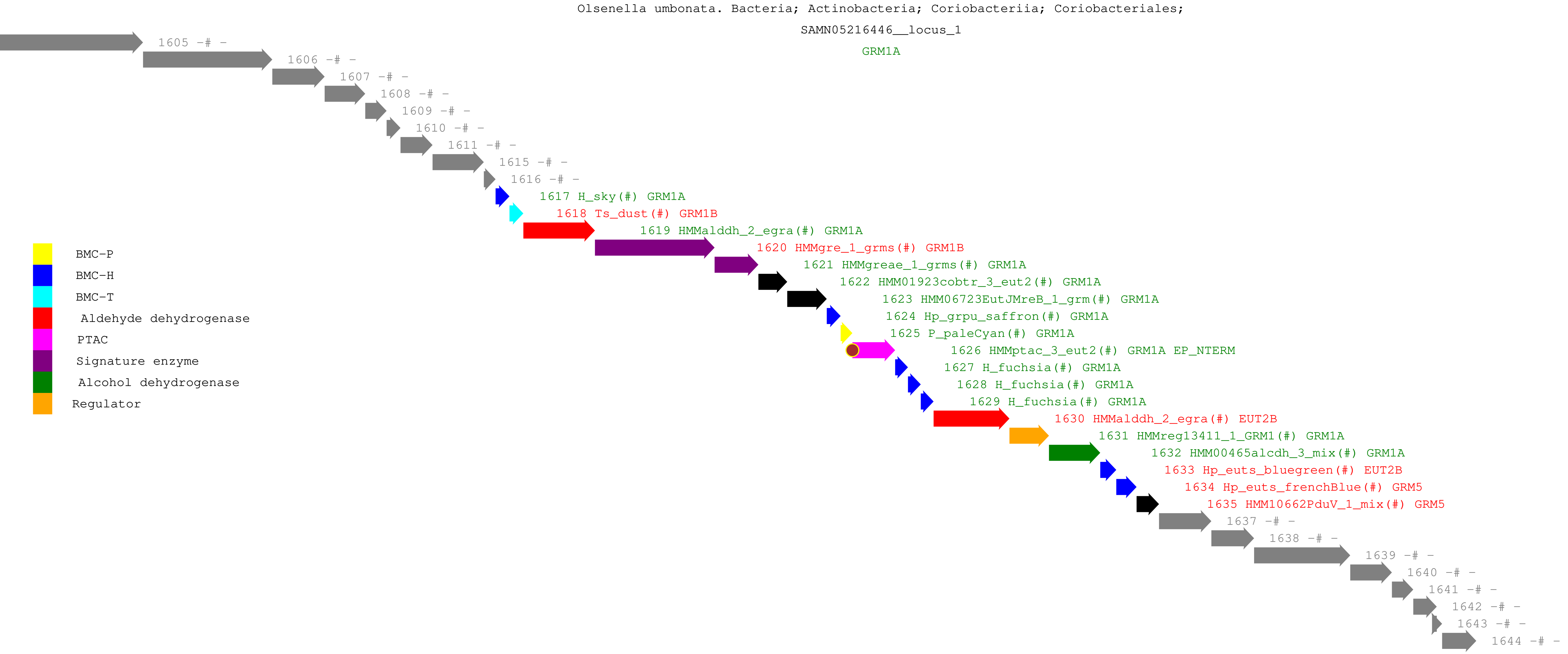

Olsenella umbonata. Bacteria; Actinobacteria; Coriobacteriia; Coriobacteriales;
SAMN05216446__locus_1
GRM1A
1605 -# -
1606 -# -
1607 -# -
1608 -# -
1609 -# -
1610 -# -
1611 -# -
1615 -# -
1616 -# -
1617 H_sky(#) GRM1A
1618 Ts_dust(#) GRM1B
1619 HMMalddh_2_egra(#) GRM1A
1620 HMMgre_1_grms(#) GRM1B
1621 HMMgreae_1_grms(#) GRM1A
1622 HMM01923cobtr_3_eut2(#) GRM1A
1623 HMM06723EutJMreB_1_grm(#) GRM1A
1624 Hp_grpu_saffron(#) GRM1A
1625 P_paleCyan(#) GRM1A
1626 HMMptac_3_eut2(#) GRM1A EP_NTERM
1627 H_fuchsia(#) GRM1A
1628 H_fuchsia(#) GRM1A
1629 H_fuchsia(#) GRM1A
1630 HMMalddh_2_egra(#) EUT2B
1631 HMMreg13411_1_GRM1(#) GRM1A
1632 HMM00465alcdh_3_mix(#) GRM1A
1633 Hp_euts_bluegreen(#) EUT2B
1634 Hp_euts_frenchBlue(#) GRM5
1635 HMM10662PduV_1_mix(#) GRM5
1637 -# -
1638 -# -
1639 -# -
1640 -# -
1641 -# -
1642 -# -
1643 -# -
1644 -# -
BMC-P
BMC-H
BMC-T
Aldehyde dehydrogenase
PTAC
Signature enzyme
Alcohol dehydrogenase
Regulator

Granulicatella balaenopterae. Bacteria; Firmicutes; Bacilli; Lactobacillales; Carnobacteriaceae;

SAMN05421767__locus_1

GRM1A

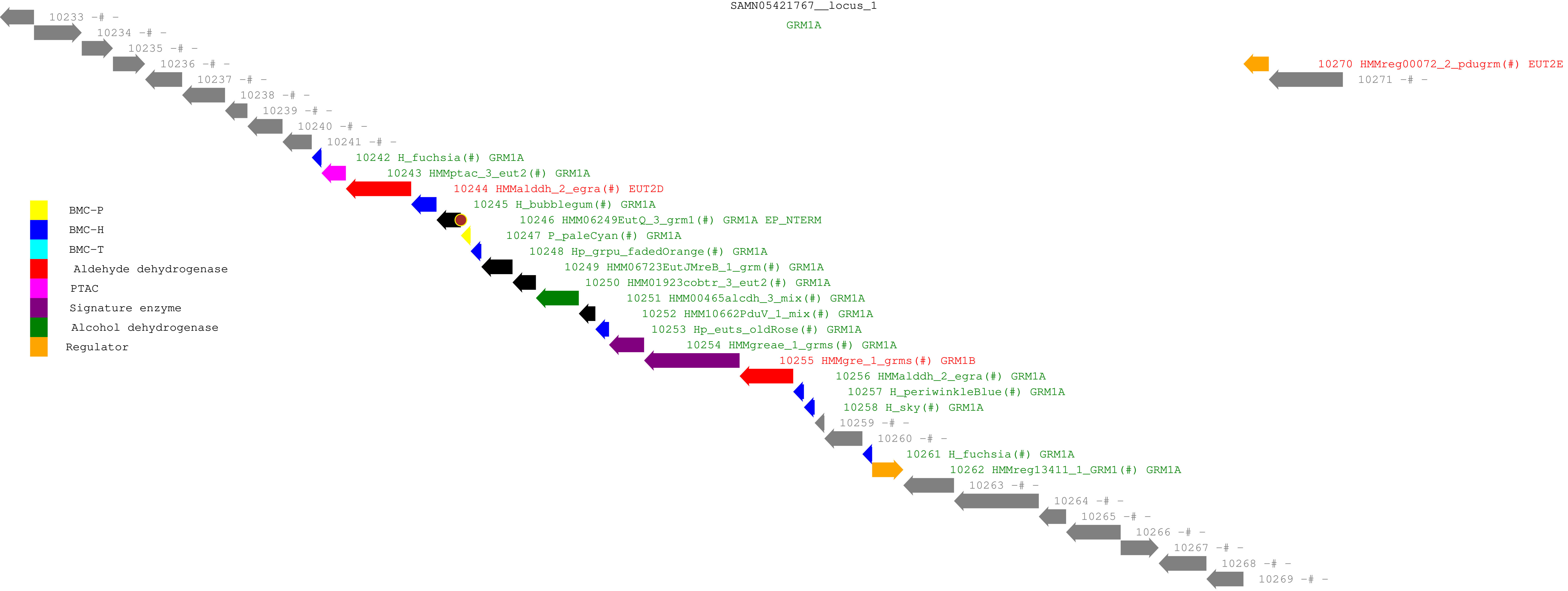

Desulfotomaculum guttoideum. Bacteria; Firmicutes; Clostridia; Clostridiales; Lachnospiraceae.

SAMN05443270__locus_1

GRM1A

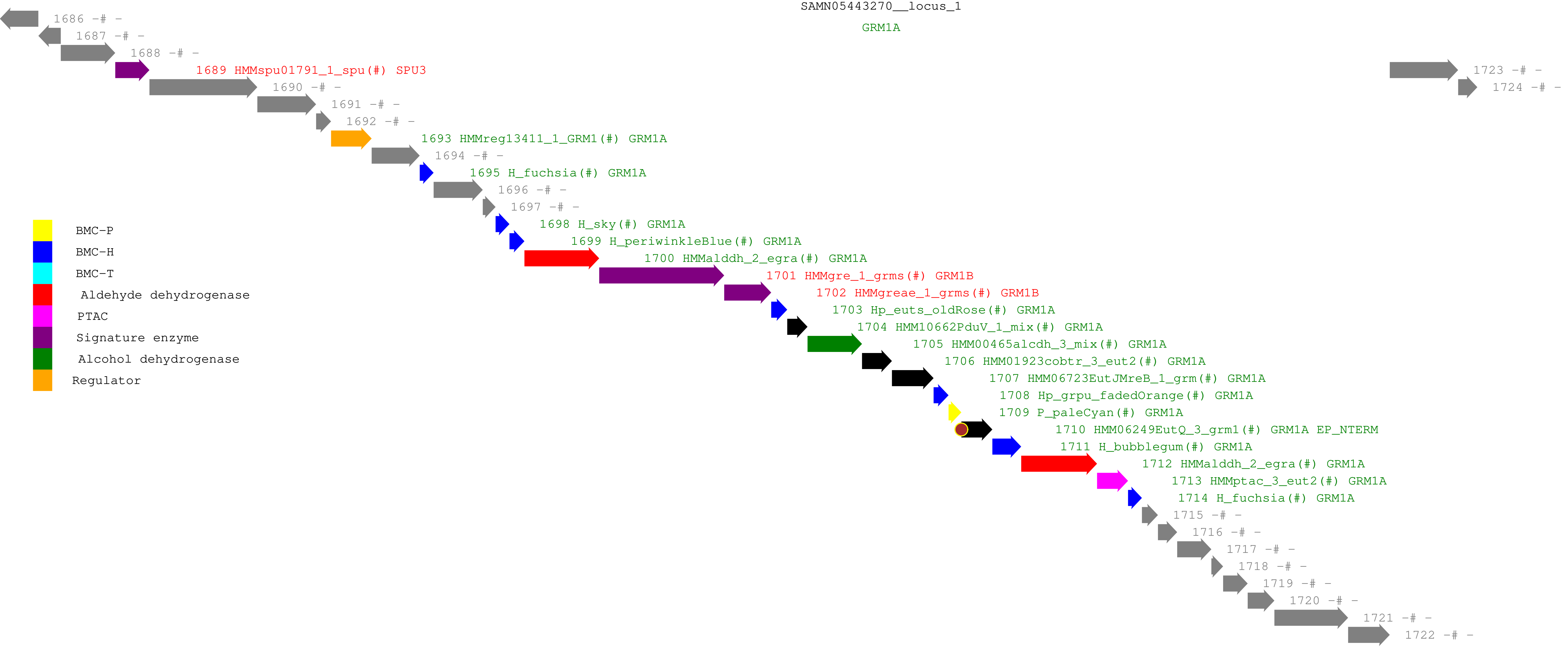

Caloramator quimbayensis. Bacteria; Firmicutes; Clostridia; Clostridiales; Clostridiaceae;

SAMN05443428__locus_1

GRM1A

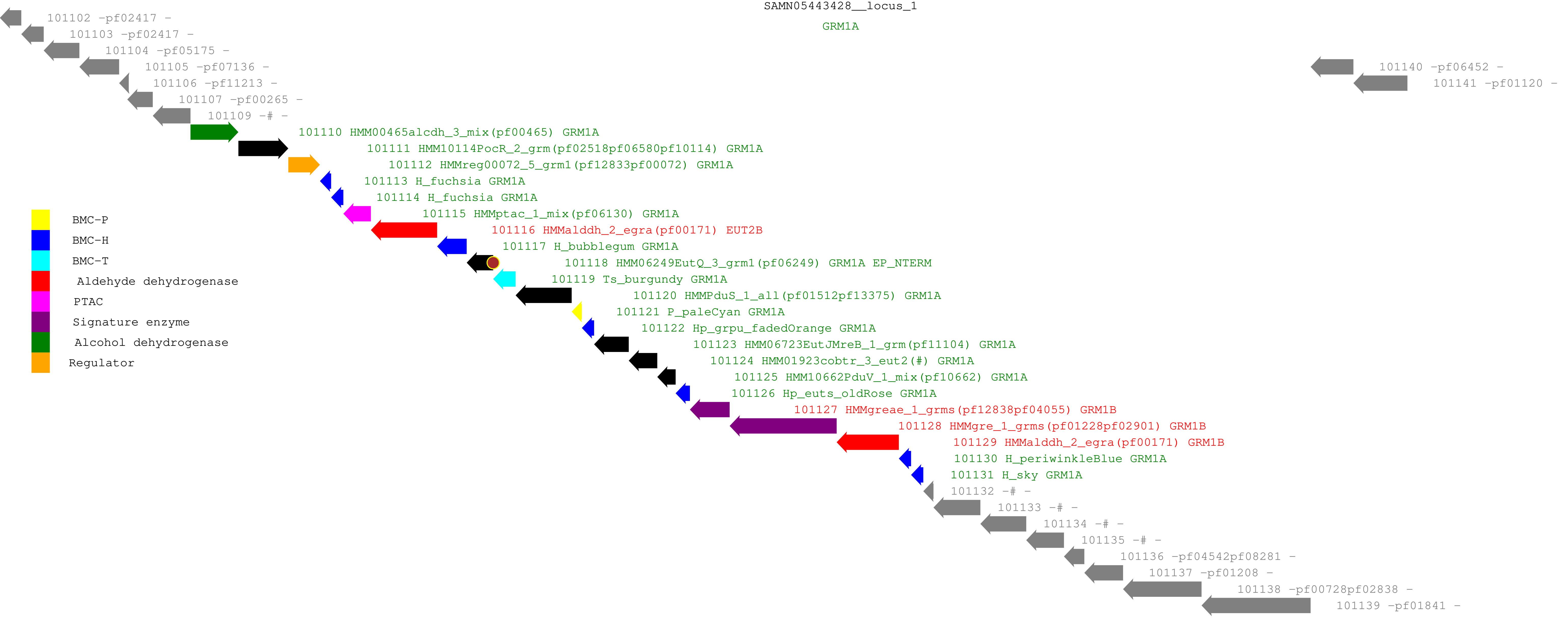

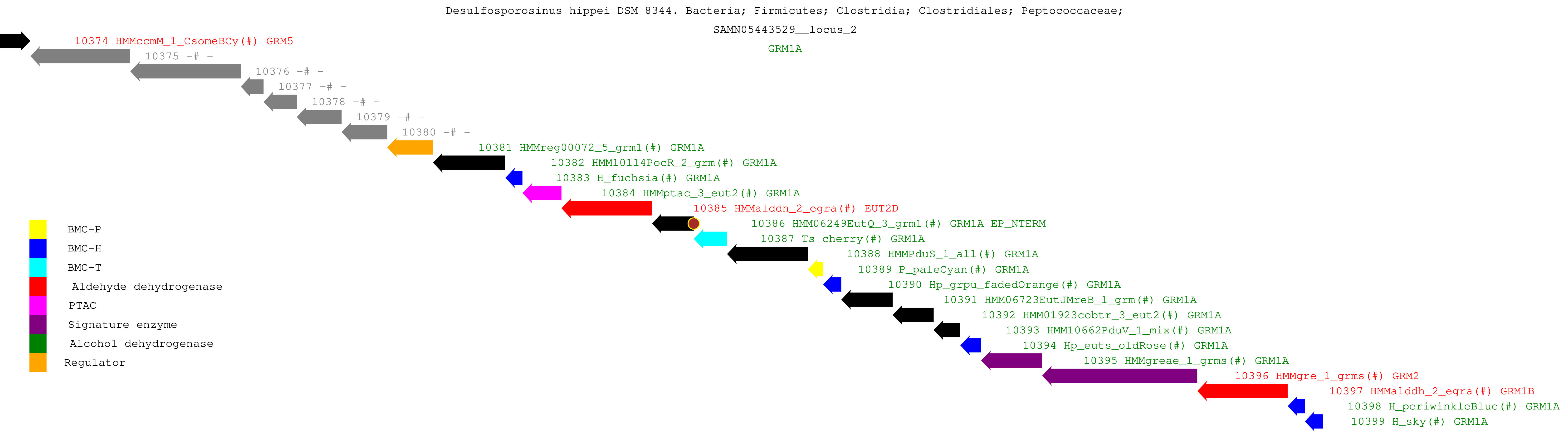
Desulfosporosinus hippei DSM 8344. Bacteria; Firmicutes; Clostridia; Clostridiales; Peptococcaceae;
SAMN05443529__locus_2
GRM1A
10374 HMMccmM_1_CsomeBCy(#) GRM5
10375 -# -
10376 -# -
10377 -# -
10378 -# -
10379 -# -
10380 -# -
10381 HMMreg00072_5_grm1(#) GRM1A
10382 HMM10114PocR_2_grm(#) GRM1A
10383 H_fuchsia(#) GRM1A
10384 HMMptac_3_eut2(#) GRM1A
10385 HMMalddh_2_egra(#) EUT2D
10386 HMM06249EutQ_3_grm1(#) GRM1A EP_NTERM
10387 Ts_cherry(#) GRM1A
10388 HMMPduS_1_all(#) GRM1A
10389 P_paleCyan(#) GRM1A
10390 Hp_grpu_fadedOrange(#) GRM1A
10391 HMM06723EutJMreB_1_grm(#) GRM1A
10392 HMM01923cobtr_3_eut2(#) GRM1A
10393 HMM10662PduV_1_mix(#) GRM1A
10394 Hp_euts_oldRose(#) GRM1A
10395 HMMgreae_1_grms(#) GRM1A
10396 HMMgre_1_grms(#) GRM2
10397 HMMalddh_2_egra(#) GRM1B
10398 H_periwinkleBlue(#) GRM1A
10399 H_sky(#) GRM1A
BMC-P
BMC-H
BMC-T
Aldehyde dehydrogenase
PTAC
Signature enzyme
Alcohol dehydrogenase
Regulator

Desulfovibrio desulfuricans. Bacteria; Proteobacteria; Deltaproteobacteria; Desulfovibrionales;

SAMN05660830__locus_2

GRM1A

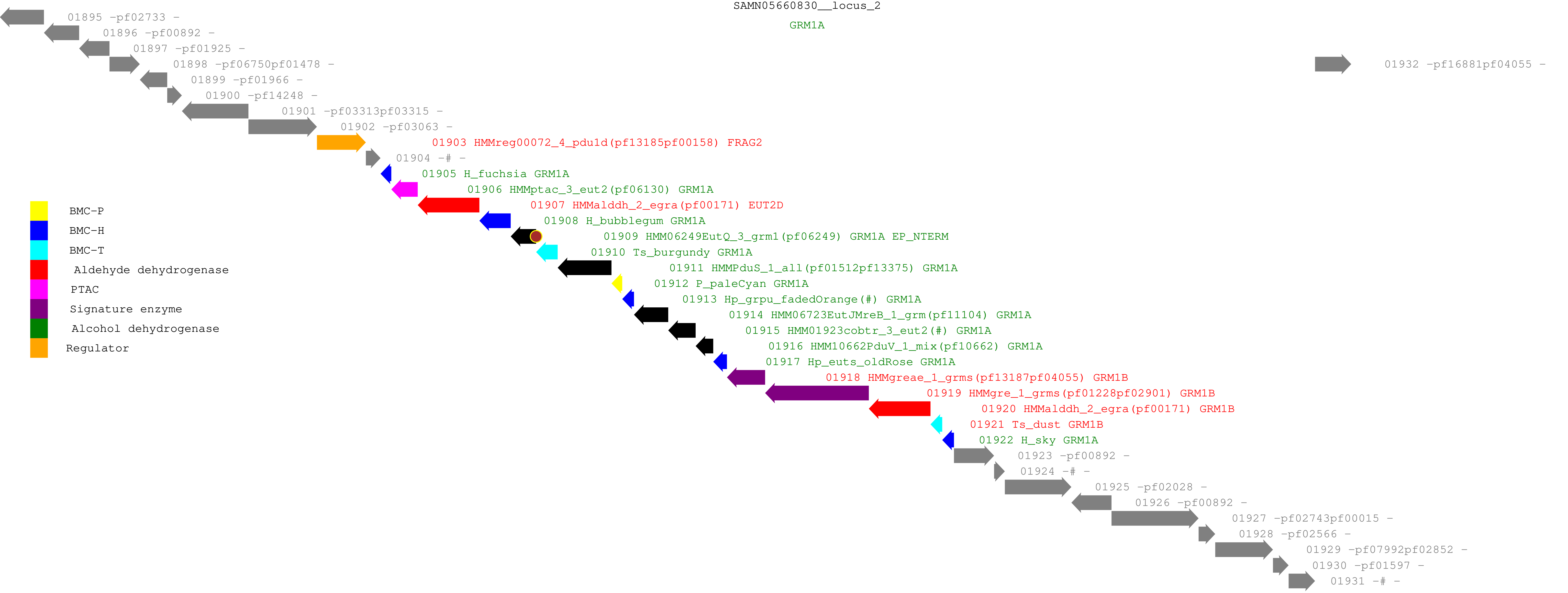

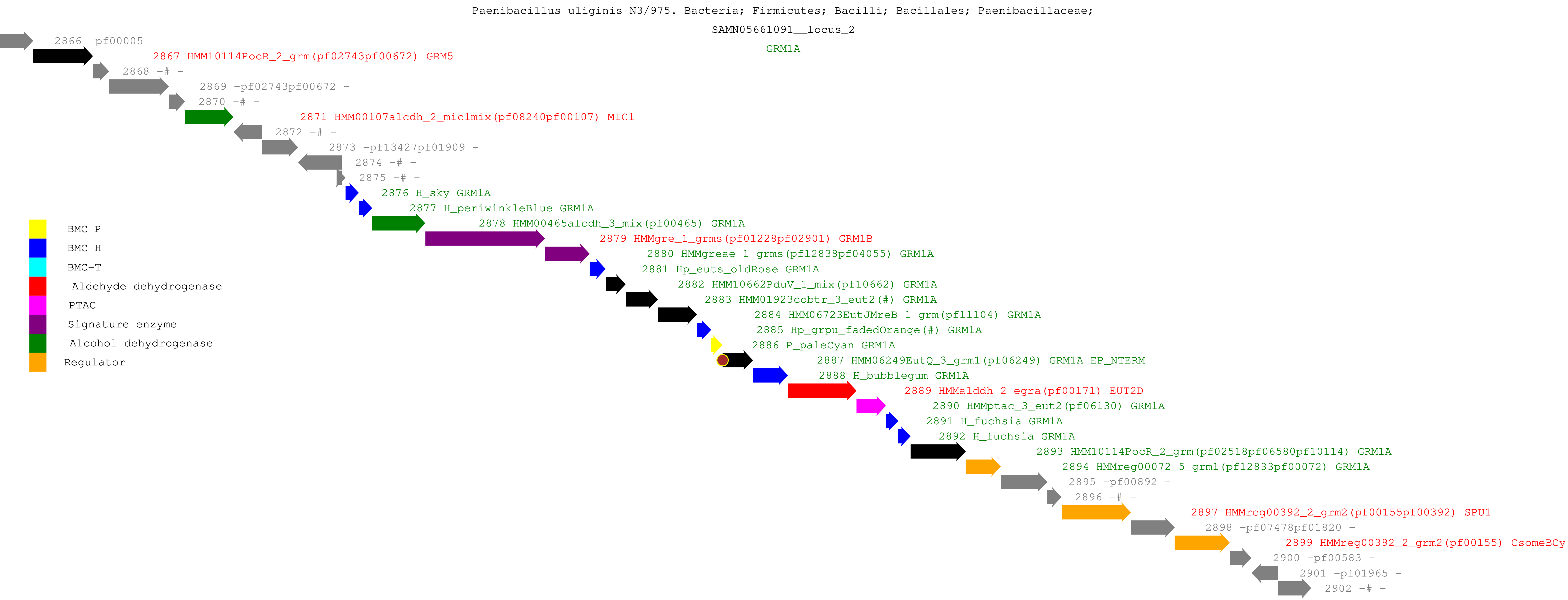

Paenibacillus uliginis N3/975. Bacteria; Firmicutes; Bacilli; Bacillales; Paenibacillaceae;
SAMN05661091__locus_2
GRM1A
2866 -pf00005 -
2867 HMM10114PocR_2_grm(pf02743pf00672) GRM5
2868 -# -
2869 -pf02743pf00672 -
2870 -# -
2871 HMM00107alcdh_2_mic1mix(pf08240pf00107) MIC1
2872 -# -
2873 -pf13427pf01909 -
2874 -# -
2875 -# -
2876 H_sky GRM1A
2877 H_periwinkleBlue GRM1A
2878 HMM00465alcdh_3_mix(pf00465) GRM1A
2879 HMMgre_1_grms(pf01228pf02901) GRM1B
2880 HMMgreae_1_grms(pf12838pf04055) GRM1A
2881 Hp_euts_oldRose GRM1A
2882 HMM10662PduV_1_mix(pf10662) GRM1A
2883 HMM01923cobtr_3_eut2(#) GRM1A
2884 HMM06723EutJMreB_1_grm(pf11104) GRM1A
2885 Hp_grpu_fadedOrange(#) GRM1A
2886 P_paleCyan GRM1A
2887 HMM06249EutQ_3_grm1(pf06249) GRM1A EP_NTERM
2888 H_bubblegum GRM1A
2889 HMMaldh_2_egra(pf00171) EUT2D
2890 HMMptac_3_eut2(pf06130) GRM1A
2891 H_fuchsia GRM1A
2892 H_fuchsia GRM1A
2893 HMM10114PocR_2_grm(pf02518pf06580pf10114) GRM1A
2894 HMMreg00072_5_grm1(pf12833pf00072) GRM1A
2895 -pf00892 -
2896 -# -
2897 HMMreg00392_2_grm2(pf00155pf00392) SPU1
2898 -pf07478pf01820 -
2899 HMMreg00392_2_grm2(pf00155) CsomeBCy
2900 -pf00583 -
2901 -pf01965 -
2902 -# -
BMC-P
BMC-H
BMC-T
Aldehyde dehydrogenase
PTAC
Signature enzyme
Alcohol dehydrogenase
Regulator

Clostridium botulinum. Bacteria; Firmicutes; Clostridia; Clostridiales; Clostridiaceae;

SD74__locus_1

GRM1A

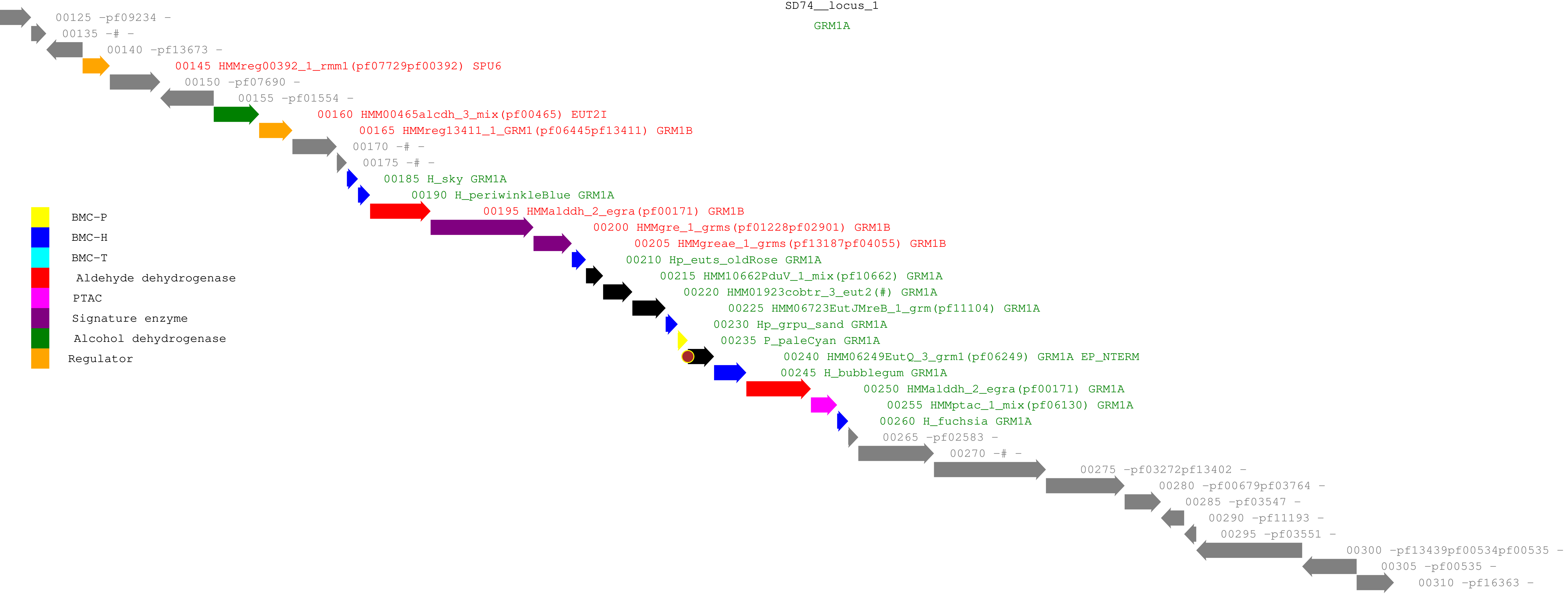

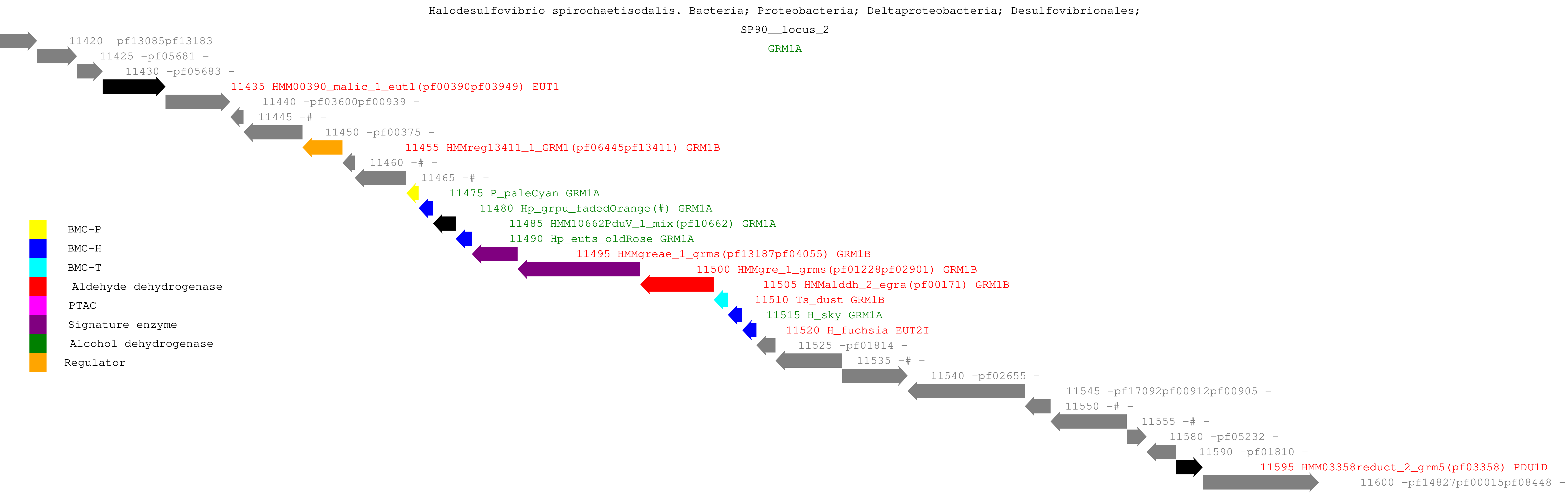

Halodesulfovibrio spirochaetisodalis. Bacteria; Proteobacteria; Deltaproteobacteria; Desulfovibrionales;
SP90__locus_2
GRM1A
11420 -pf13085pf13183 -
11425 -pf05681 -
11430 -pf05683 -
11435 HMM00390_malic_1_eut1(pf00390pf03949) EUT1
11440 -pf03600pf00939 -
11445 -# -
11450 -pf00375 -
11455 HMMreg13411_1_GRM1(pf06445pf13411) GRM1B
11460 -# -
11465 -# -
11475 P_paleCyan GRM1A
11480 Hp_grpu_fadedOrange(#) GRM1A
11485 HMM10662PduV_1_mix(pf10662) GRM1A
11490 Hp_euts_oldRose GRM1A
11495 HMMgreae_1_grms(pf13187pf04055) GRM1B
11500 HMMgre_1_grms(pf01228pf02901) GRM1B
11505 HMMalddh_2_egra(pf00171) GRM1B
11510 Ts_dust GRM1B
11515 H_sky GRM1A
11520 H_fuchsia EUT2I
11525 -pf01814 -
11535 -# -
11540 -pf02655 -
11545 -pf17092pf00912pf00905 -
11550 -# -
11555 -# -
11580 -pf05232 -
11590 -pf01810 -
11595 HMM03358reduct_2_grm5(pf03358) PDU1D
11600 -pf14827pf00015pf08448 -
BMC-P
BMC-H
BMC-T
Aldehyde dehydrogenase
PTAC
Signature enzyme
Alcohol dehydrogenase
Regulator

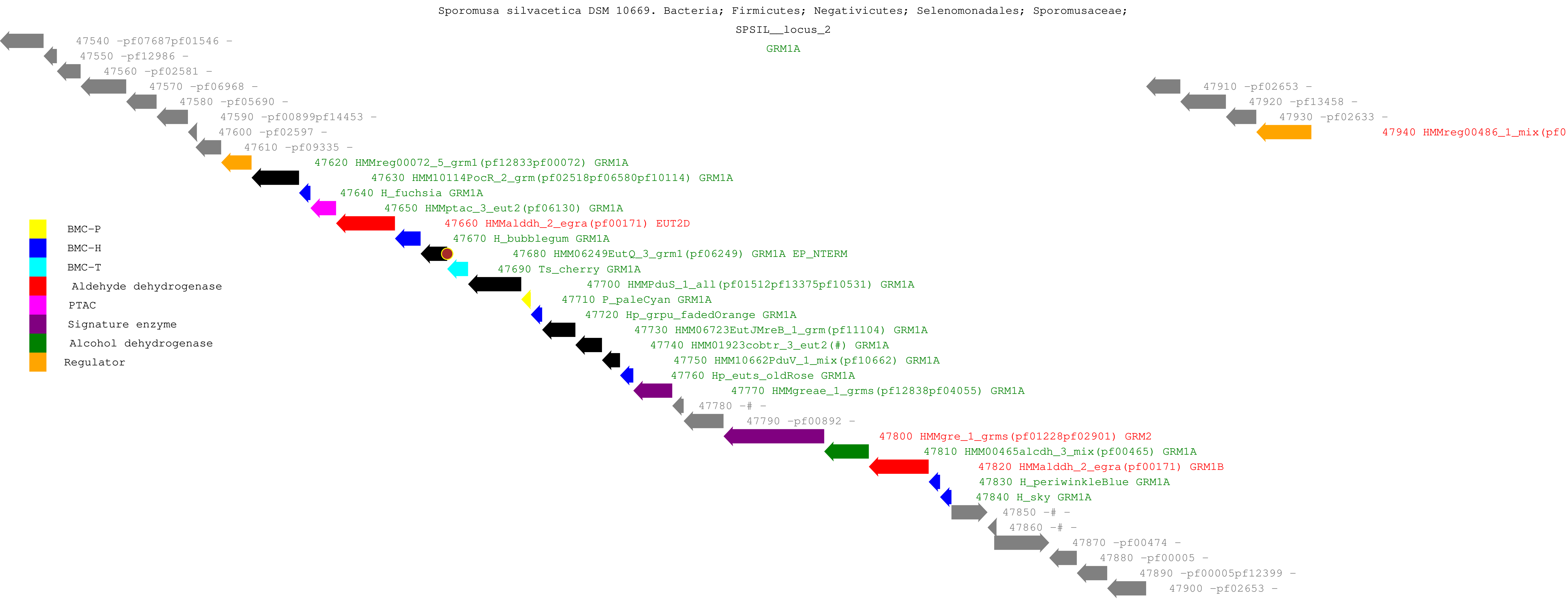
Sporomusa silvacetica DSM 10669. Bacteria; Firmicutes; Negativicutes; Selenomonadales; Sporomusaceae;
SPSIL__locus_2
GRM1A
47540 -pf07687pf01546 -
47550 -pf12986 -
47560 -pf02581 -
47570 -pf06968 -
47580 -pf05690 -
47590 -pf00899pf14453 -
47600 -pf02597 -
47610 -pf09335 -
47620 HMMreg00072_5_grm1(pf12833pf00072) GRM1A
47630 HMM10114PocR_2_grm(pf02518pf06580pf10114) GRM1A
47640 H_fuchsia GRM1A
47650 HMMptac_3_eut2(pf06130) GRM1A
47660 HMMaldDH_2_egra(pf00171) EUT2D
47670 H_bubblegum GRM1A
47680 HMM06249EutQ_3_grm1(pf06249) GRM1A EP_NTERM
47690 Ts_cherry GRM1A
47700 HMMPduS_1_all(pf01512pf13375pf10531) GRM1A
47710 P_paleCyan GRM1A
47720 Hp_grpu_fadedOrange GRM1A
47730 HMM06723EutJMreB_1_grm(pf11104) GRM1A
47740 HMM01923cobtr_3_eut2(#) GRM1A
47750 HMM10662PduV_1_mix(pf10662) GRM1A
47760 Hp_euts_oldRose GRM1A
47770 HMMgreae_1_grms(pf12838pf04055) GRM1A
47780 -# -
47790 -pf00892 -
47800 HMMgre_1_grms(pf01228pf02901) GRM2
47810 HMM00465alcdh_3_mix(pf00465) GRM1A
47820 HMMaldDH_2_egra(pf00171) GRM1B
47830 H_periwinkleBlue GRM1A
47840 H_sky GRM1A
47850 -# -
47860 -# -
47870 -pf00474 -
47880 -pf00005 -
47890 -pf00005pf12399 -
47900 -pf02653 -
47910 -pf02653 -
47920 -pf13458 -
47930 -pf02633 -
47940 HMMreg00486_1_mix(pf0
BMC-P
BMC-H
BMC-T
Aldehyde dehydrogenase
PTAC
Signature enzyme
Alcohol dehydrogenase
Regulator

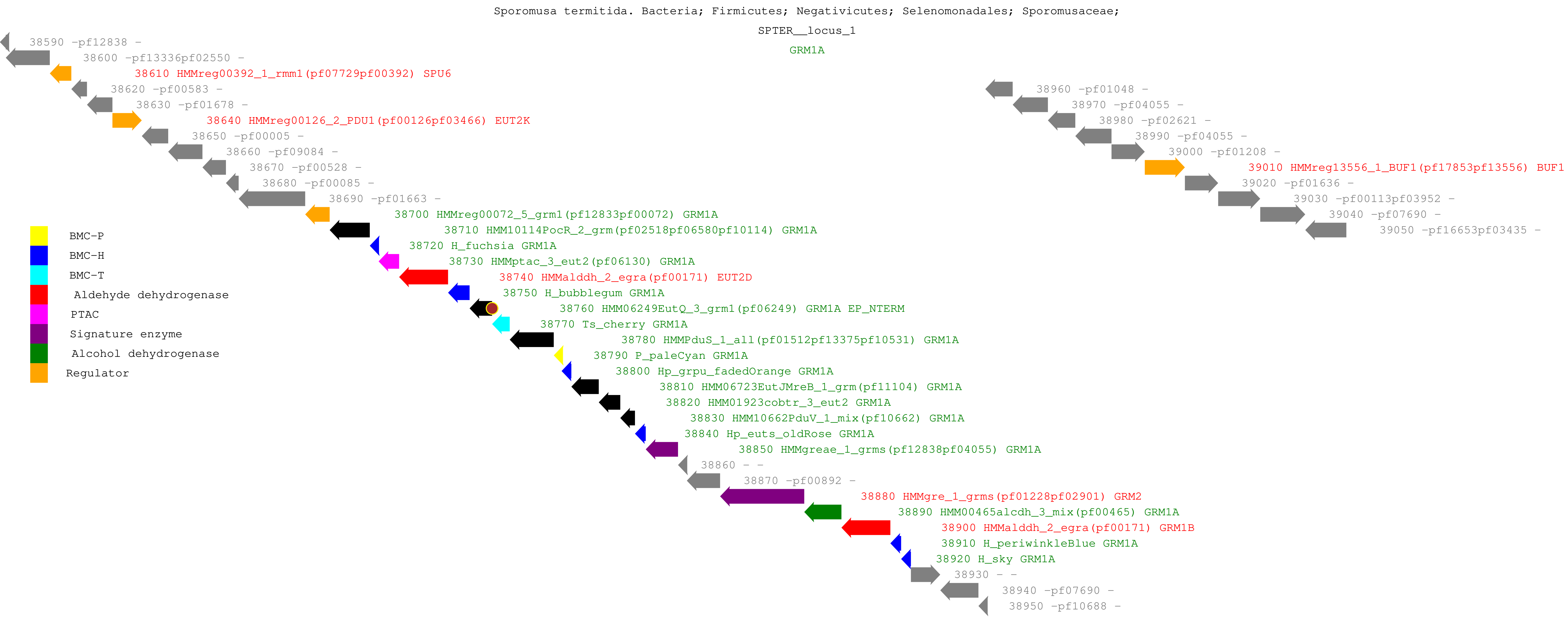
Sporomusa termitida. Bacteria; Firmicutes; Negativicutes; Selenomonadales; Sporomusaceae;
SPTER__locus_1
GRM1A
38590 -pf12838 -
38600 -pf13336pf02550 -
38610 HMMreg00392_1_rmm1(pf07729pf00392) SPU6
38620 -pf00583 -
38630 -pf01678 -
38640 HMMreg00126_2_PDU1(pf00126pf03466) EUT2K
38650 -pf00005 -
38660 -pf09084 -
38670 -pf00528 -
38680 -pf00085 -
38690 -pf01663 -
38700 HMMreg00072_5_grm1(pf12833pf00072) GRM1A
38710 HMM10114PocR_2_grm(pf02518pf06580pf10114) GRM1A
38720 H_fuchsia GRM1A
38730 HMMptac_3_eut2(pf06130) GRM1A
38740 HMMaldh_2_egra(pf00171) EUT2D
38750 H_bubblegum GRM1A
38760 HMM06249EutQ_3_grm1(pf06249) GRM1A EP_NTERM
38770 Ts_cherry GRM1A
38780 HMMPduS_1_all(pf01512pf13375pf10531) GRM1A
38790 P_paleCyan GRM1A
38800 Hp_grpu_fadedOrange GRM1A
38810 HMM06723EutJMreB_1_grm(pf11104) GRM1A
38820 HMM01923cobtr_3_eut2 GRM1A
38830 HMM10662PduV_1_mix(pf10662) GRM1A
38840 Hp_euts_oldRose GRM1A
38850 HMMgreae_1_grms(pf12838pf04055) GRM1A
38860 - -
38870 -pf00892 -
38880 HMMgre_1_grms(pf01228pf02901) GRM2
38890 HMM00465alcdh_3_mix(pf00465) GRM1A
38900 HMMaldh_2_egra(pf00171) GRM1B
38910 H_periwinkleBlue GRM1A
38920 H_sky GRM1A
38930 - -
38940 -pf07690 -
38950 -pf10688 -
38960 -pf01048 -
38970 -pf04055 -
38980 -pf02621 -
38990 -pf04055 -
39000 -pf01208 -
39010 HMMreg13556_1_BUF1(pf17853pf13556) BUF1
39020 -pf01636 -
39030 -pf00113pf03952 -
39040 -pf07690 -
39050 -pf16653pf03435 -
BMC-P
BMC-H
BMC-T
Aldehyde dehydrogenase
PTAC
Signature enzyme
Alcohol dehydrogenase
Regulator

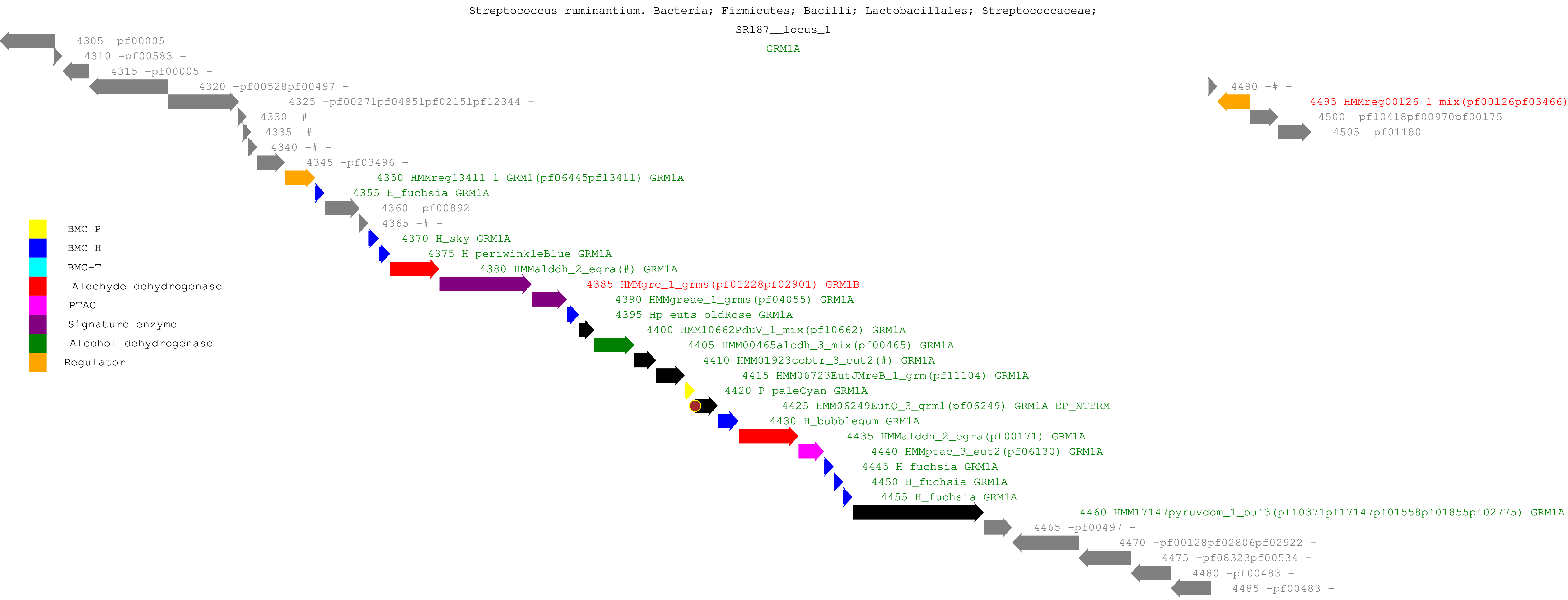

Streptococcus ruminantium. Bacteria; Firmicutes; Bacilli; Lactobacillales; Streptococcaceae;
SR187__locus_1
GRM1A
4305 -pf00005 -
4310 -pf00583 -
4315 -pf00005 -
4320 -pf00528pf00497 -
4325 -pf00271pf04851pf02151pf12344 -
4330 -# -
4335 -# -
4340 -# -
4345 -pf03496 -
4350 HMMreg13411_1_GRM1(pf06445pf13411) GRM1A
4355 H_fuchsia GRM1A
4360 -pf00892 -
4365 -# -
4370 H_sky GRM1A
4375 H_periwinkleBlue GRM1A
4380 HMMalddh_2_egra(#) GRM1A
4385 HMMgre_1_grms(pf01228pf02901) GRM1B
4390 HMMgreae_1_grms(pf04055) GRM1A
4395 Hp_euts_oldRose GRM1A
4400 HMM10662PduV_1_mix(pf10662) GRM1A
4405 HMM00465alcdh_3_mix(pf00465) GRM1A
4410 HMM01923cobtr_3_eut2(#) GRM1A
4415 HMM06723EutJMreB_1_grm(pf11104) GRM1A
4420 P_paleCyan GRM1A
4425 HMM06249EutQ_3_grm1(pf06249) GRM1A EP_NTERM
4430 H_bubblegum GRM1A
4435 HMMalddh_2_egra(pf00171) GRM1A
4440 HMMptac_3_eut2(pf06130) GRM1A
4445 H_fuchsia GRM1A
4450 H_fuchsia GRM1A
4455 H_fuchsia GRM1A
4460 HMM17147pyruvdom_1_buf3(pf10371pf17147pf01558pf01855pf02775) GRM1A
4465 -pf00497 -
4470 -pf00128pf02806pf02922 -
4475 -pf08323pf00534 -
4480 -pf00483 -
4485 -pf00483 -
4490 -# -
4495 HMMreg00126_1_mix(pf00126pf03466)
4500 -pf10418pf00970pf00175 -
4505 -pf01180 -
BMC-P
BMC-H
BMC-T
Aldehyde dehydrogenase
PTAC
Signature enzyme
Alcohol dehydrogenase
Regulator

Clostridium botulinum Prevot_594. Bacteria; Firmicutes; Clostridia; Clostridiales; Clostridiaceae;

T258__locus_1_main_sat

GRM1A

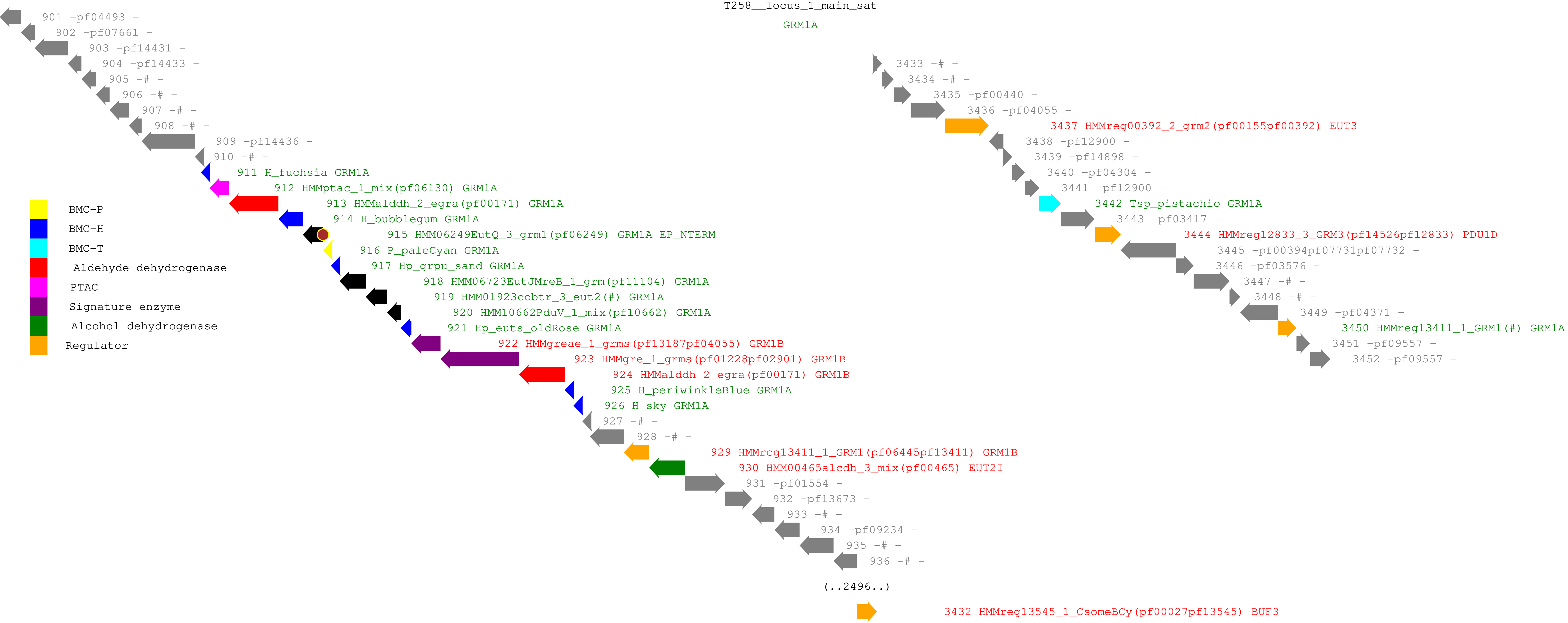

Clostridium botulinum CDC_1436. Bacteria; Firmicutes; Clostridia; Clostridiales; Clostridiaceae;

T259__locus_1

GRM1A

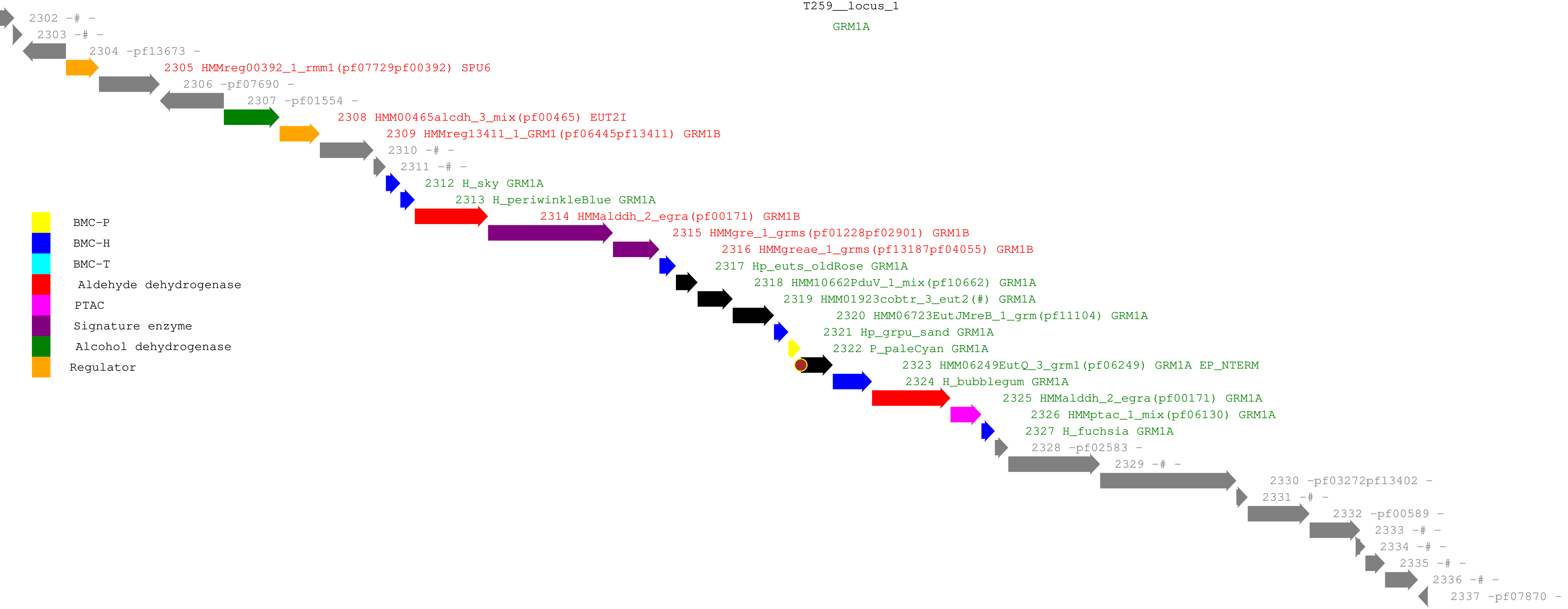

Clostridium baratii str. Sullivan. Bacteria; Firmicutes; Clostridia; Clostridiales; Clostridiaceae;

U729__locus_1

GRM1A

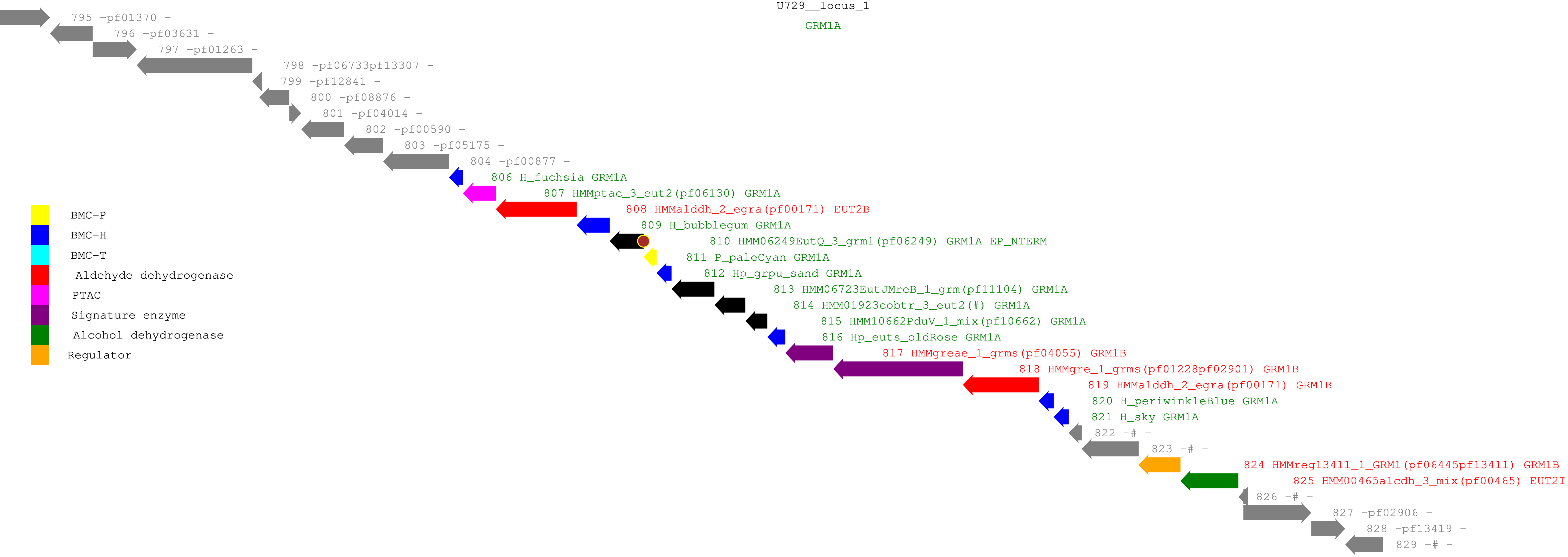

Clostridium argentinense CDC 2741. Bacteria; Firmicutes; Clostridia; Clostridiales; Clostridiaceae;

U732__locus_2

GRM1A

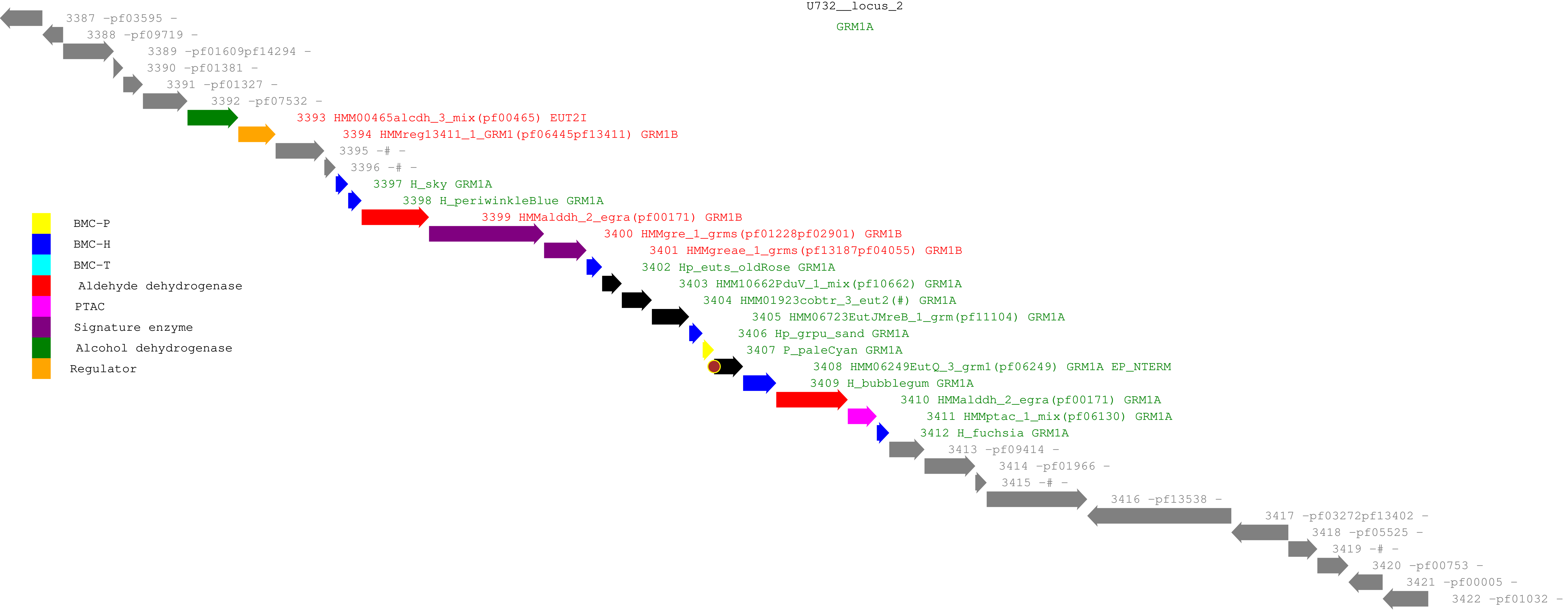

Enterococcus raffinosus ATCC 49464. Bacteria; Firmicutes; Bacilli; Lactobacillales; Enterococcaceae;

UAK__locus_1

GRM1A

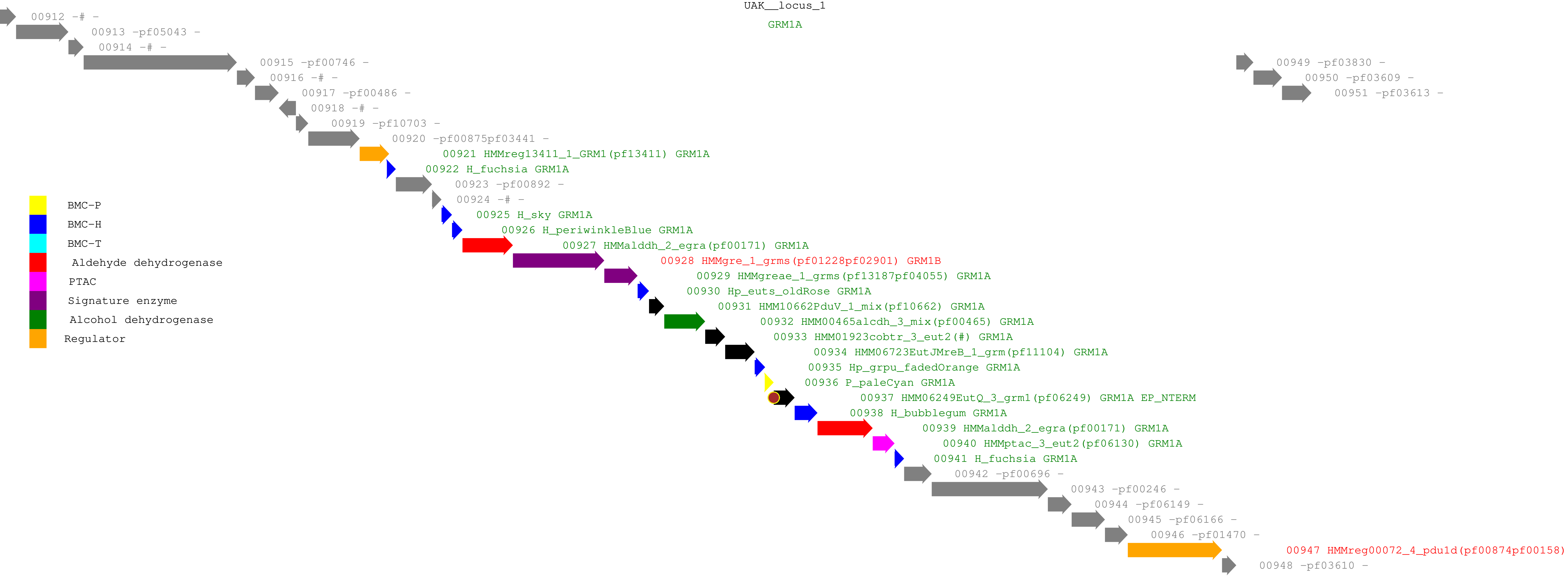

Enterococcus asini ATCC 700915. Bacteria; Firmicutes; Bacilli; Lactobacillales; Enterococcaceae;

UAS__locus_1

GRM1A

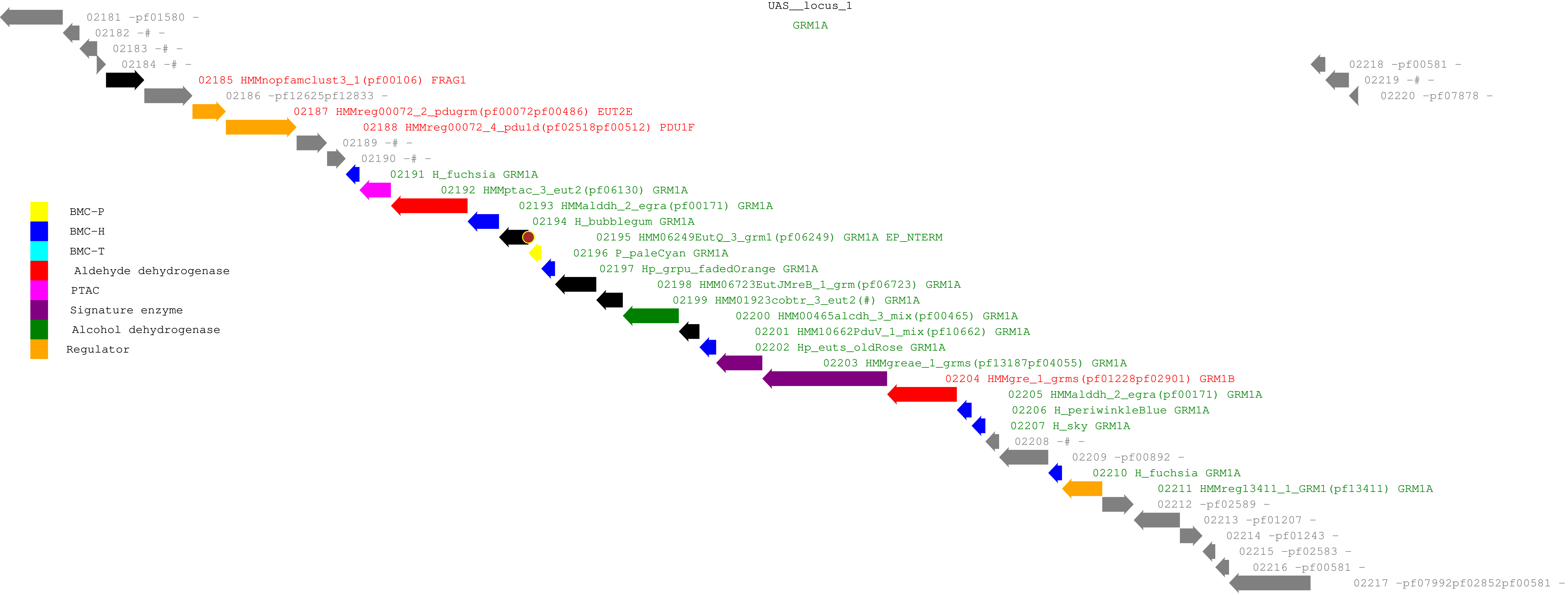

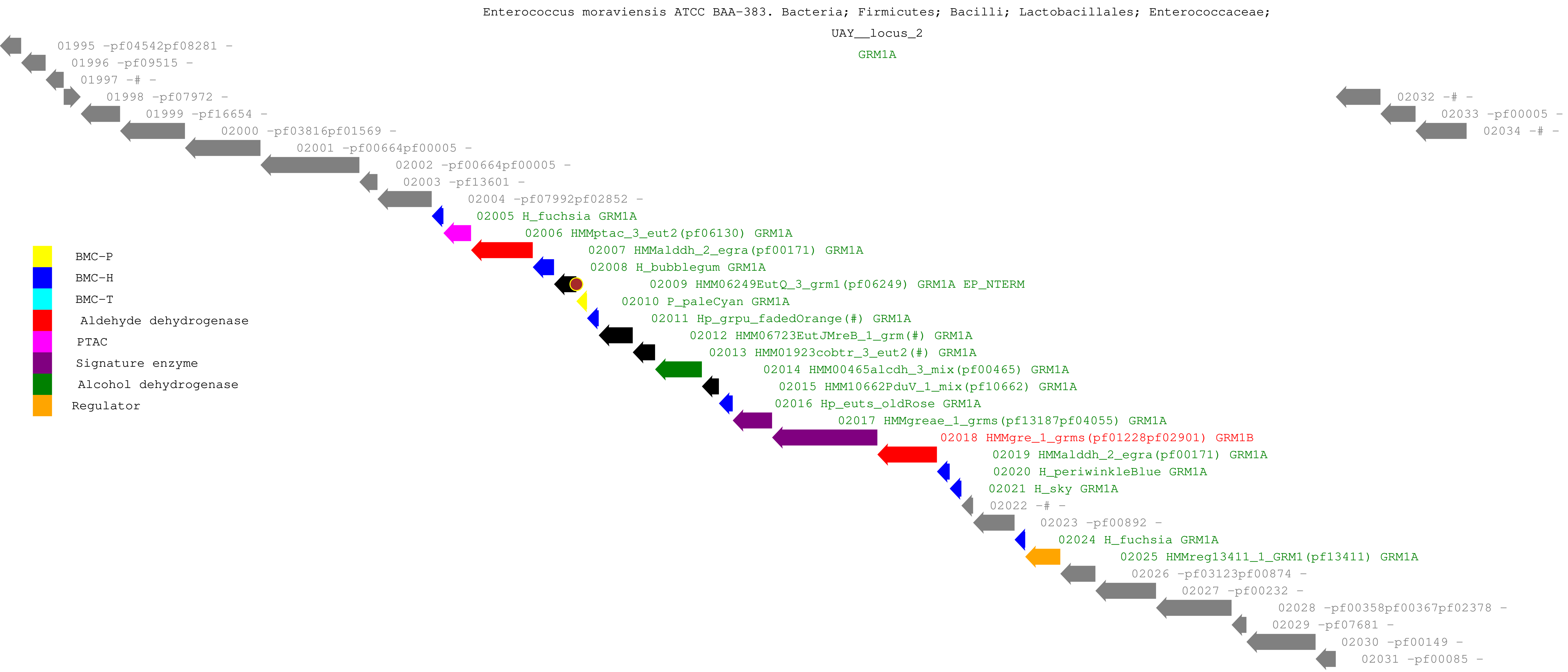
Enterococcus moraviensis ATCC BAA-383. Bacteria; Firmicutes; Bacilli; Lactobacillales; Enterococcaceae;
UAY__locus_2
GRM1A
01995 -pf04542pf08281 -
01996 -pf09515 -
01997 -# -
01998 -pf07972 -
01999 -pf16654 -
02000 -pf03816pf01569 -
02001 -pf00664pf00005 -
02002 -pf00664pf00005 -
02003 -pf13601 -
02004 -pf07992pf02852 -
02005 H_fuchsia GRM1A
02006 HMMptac_3_eut2(pf06130) GRM1A
02007 HMMalddh_2_egra(pf00171) GRM1A
02008 H_bubblegum GRM1A
02009 HMM06249EutQ_3_grm1(pf06249) GRM1A EP_NTERM
02010 P_paleCyan GRM1A
02011 Hp_grpu_fadedOrange(#) GRM1A
02012 HMM06723EutJMreB_1_grm(#) GRM1A
02013 HMM01923cobtr_3_eut2(#) GRM1A
02014 HMM00465alcdh_3_mix(pf00465) GRM1A
02015 HMM10662PduV_1_mix(pf10662) GRM1A
02016 Hp_euts_oldRose GRM1A
02017 HMMgreae_1_grms(pf13187pf04055) GRM1A
02018 HMMgre_1_grms(pf01228pf02901) GRM1B
02019 HMMalddh_2_egra(pf00171) GRM1A
02020 H_periwinkleBlue GRM1A
02021 H_sky GRM1A
02022 -# -
02023 -pf00892 -
02024 H_fuchsia GRM1A
02025 HMMreg13411_1_GRM1(pf13411) GRM1A
02026 -pf03123pf00874 -
02027 -pf00232 -
02028 -pf00358pf00367pf02378 -
02029 -pf07681 -
02030 -pf00149 -
02031 -pf00085 -
02032 -# -
02033 -pf00005 -
02034 -# -
BMC-P
BMC-H
BMC-T
Aldehyde dehydrogenase
PTAC
Signature enzyme
Alcohol dehydrogenase
Regulator

Enterococcus phoeniculicola ATCC BAA-412. Bacteria; Firmicutes; Bacilli; Lactobacillales; Enterococcaceae;

UC3__locus_1

GRM1A

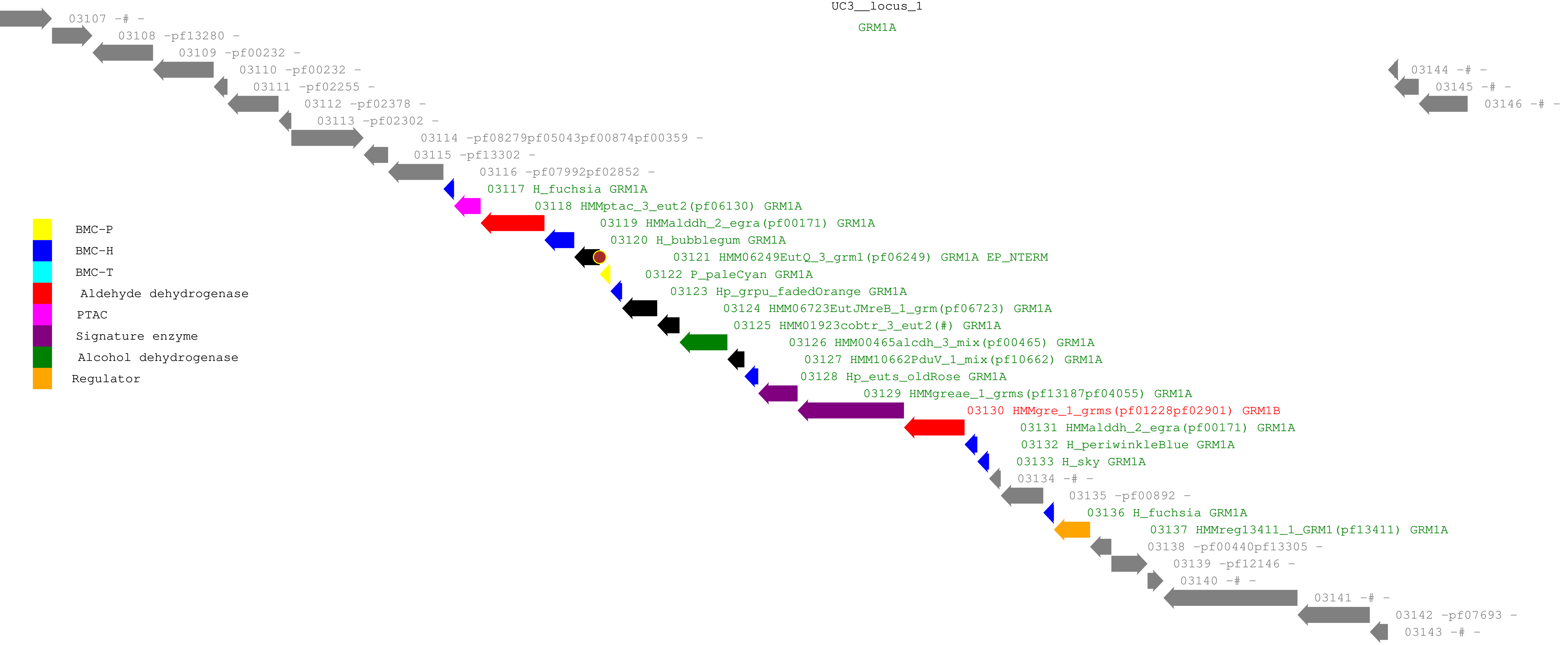

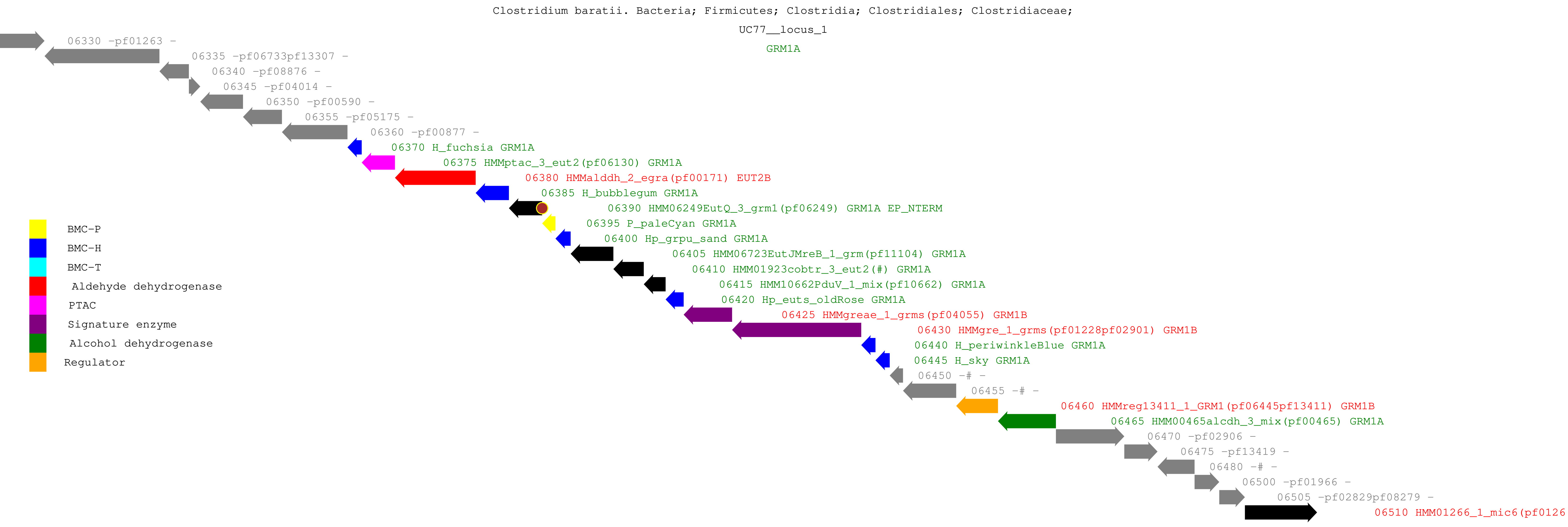
Clostridium baratii. Bacteria; Firmicutes; Clostridia; Clostridiales; Clostridiaceae;
UC77__locus_1
GRM1A
06330 -pf01263 -
06335 -pf06733pf13307 -
06340 -pf08876 -
06345 -pf04014 -
06350 -pf00590 -
06355 -pf05175 -
06360 -pf00877 -
06370 H_fuchsia GRM1A
06375 HMMptac_3_eut2(pf06130) GRM1A
06380 HMMaldh_2_egra(pf00171) EUT2B
06385 H_bubblegum GRM1A
06390 HMM06249EutQ_3_grm1(pf06249) GRM1A EP_NTERM
06395 P_paleCyan GRM1A
06400 Hp_grpu_sand GRM1A
06405 HMM06723EutJMreB_1_grm(pf11104) GRM1A
06410 HMM01923cobtr_3_eut2(#) GRM1A
06415 HMM10662PduV_1_mix(pf10662) GRM1A
06420 Hp_euts_oldRose GRM1A
06425 HMMgreae_1_grms(pf04055) GRM1B
06430 HMMgre_1_grms(pf01228pf02901) GRM1B
06440 H_periwinkleBlue GRM1A
06445 H_sky GRM1A
06450 -# -
06455 -# -
06460 HMMreg13411_1_GRM1(pf06445pf13411) GRM1B
06465 HMM00465alcdh_3_mix(pf00465) GRM1A
06470 -pf02906 -
06475 -pf13419 -
06480 -# -
06500 -pf01966 -
06505 -pf02829pf08279 -
06510 HMM01266_1_mic6(pf0126
BMC-P
BMC-H
BMC-T
Aldehyde dehydrogenase
PTAC
Signature enzyme
Alcohol dehydrogenase
Regulator

Enterococcus caccae ATCC BAA-1240. Bacteria; Firmicutes; Bacilli; Lactobacillales; Enterococcaceae;

UC7__locus_1

GRM1A

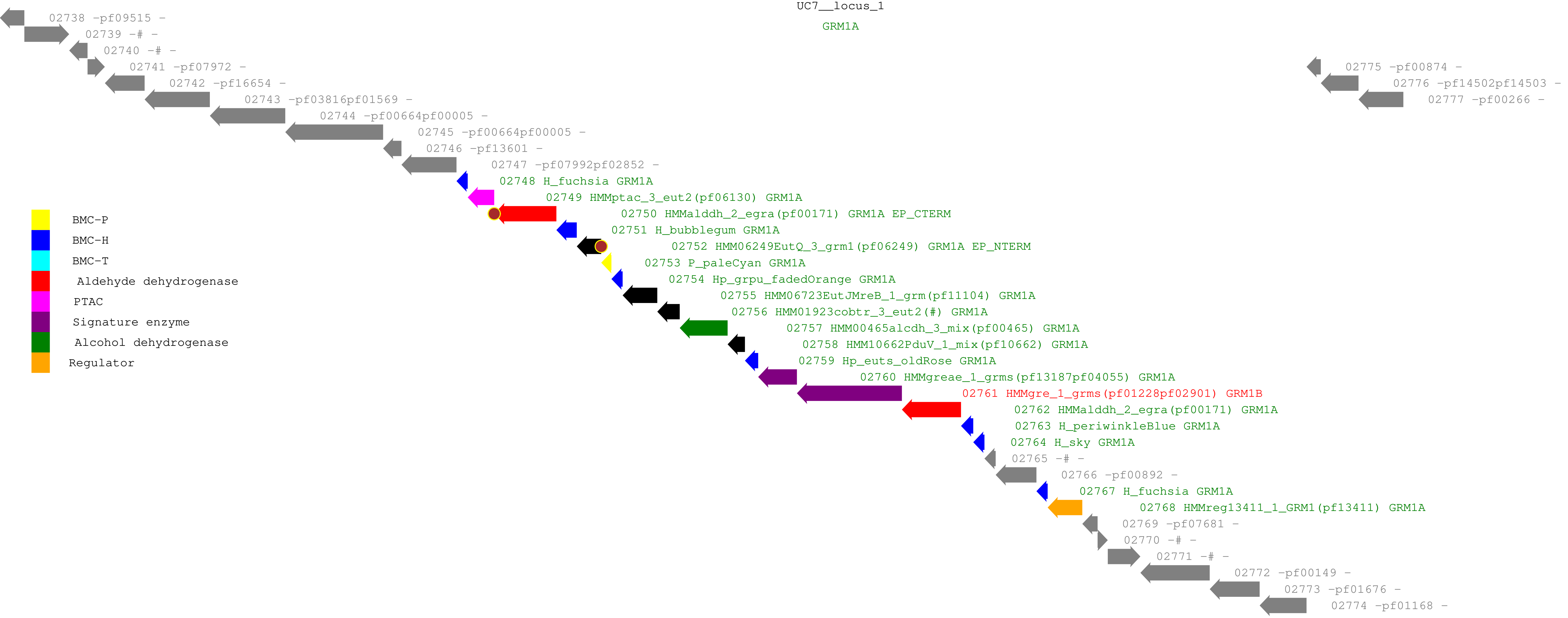

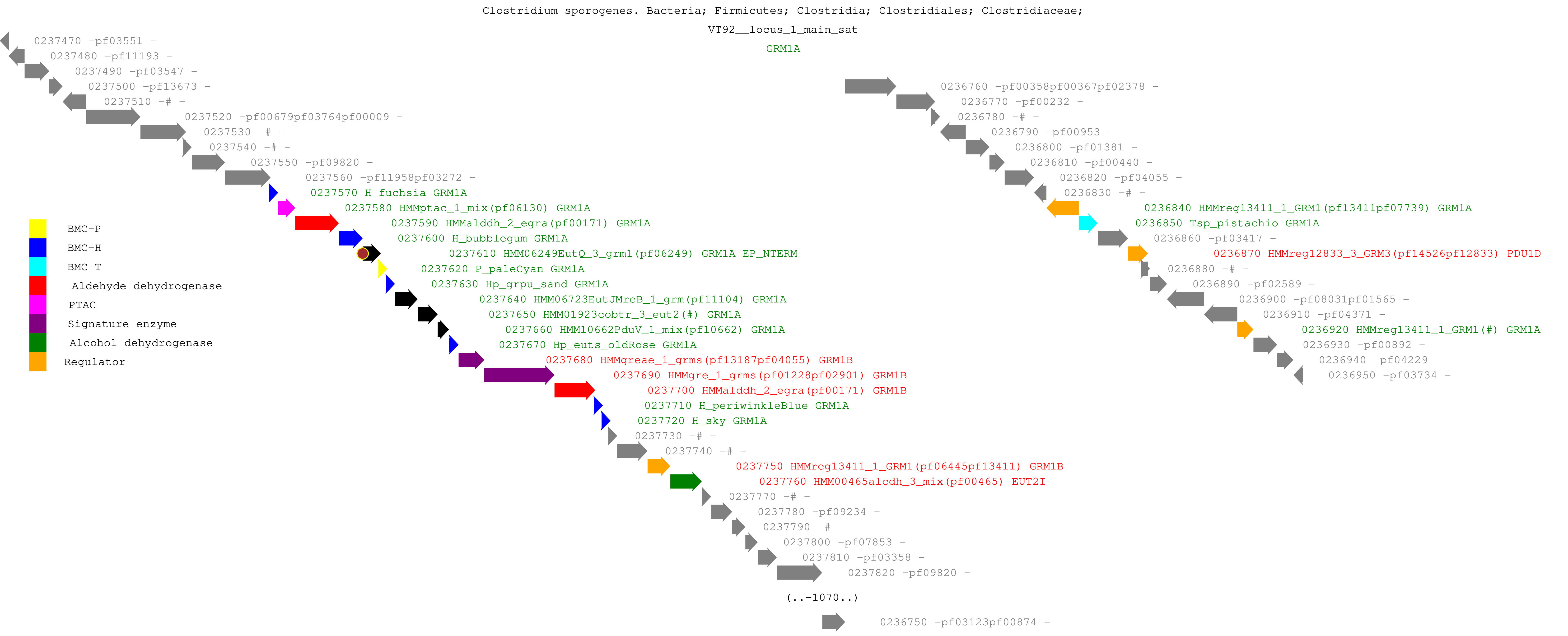
Clostridium sporogenes. Bacteria; Firmicutes; Clostridia; Clostridiales; Clostridiaceae;
VT92__locus_1_main_sat
GRM1A
0237470 -pf03551 -
0237480 -pf11193 -
0237490 -pf03547 -
0237500 -pf13673 -
0237510 -# -
0237520 -pf00679pf03764pf00009 -
0237530 -# -
0237540 -# -
0237550 -pf09820 -
0237560 -pf11958pf03272 -
0237570 H_fuchsia GRM1A
0237580 HMMptac_1_mix(pf06130) GRM1A
0237590 HMMaldd h_2_egra(pf00171) GRM1A
0237600 H_bubblegum GRM1A
0237610 HMM06249EutQ_3_grm1(pf06249) GRM1A EP_NTERM
0237620 P_paleCyan GRM1A
0237630 Hp_grpu_sand GRM1A
0237640 HMM06723EutJMreB_1_grm(pf11104) GRM1A
0237650 HMM01923cobtr_3_eut2(#) GRM1A
0237660 HMM10662PduV_1_mix(pf10662) GRM1A
0237670 Hp_euts_oldRose GRM1A
0237680 HMMgreae_1_grms(pf13187pf04055) GRM1B
0237690 HMMgre_1_grms(pf01228pf02901) GRM1B
0237700 HMMaldd h_2_egra(pf00171) GRM1B
0237710 H_periwinkleBlue GRM1A
0237720 H_sky GRM1A
0237730 -# -
0237740 -# -
0237750 HMMreg13411_1_GRM1(pf06445pf13411) GRM1B
0237760 HMM00465alcdh_3_mix(pf00465) EUT2I
0237770 -# -
0237780 -pf09234 -
0237790 -# -
0237800 -pf07853 -
0237810 -pf03358 -
0237820 -pf09820 -
(..-1070..)
0236750 -pf03123pf00874 -
0236760 -pf00358pf00367pf02378 -
0236770 -pf00232 -
0236780 -# -
0236790 -pf00953 -
0236800 -pf01381 -
0236810 -pf00440 -
0236820 -pf04055 -
0236830 -# -
0236840 HMMreg13411_1_GRM1(pf13411pf07739) GRM1A
0236850 Tsp_pistachio GRM1A
0236860 -pf03417 -
0236870 HMMreg12833_3_GRM3(pf14526pf12833) PDU1D
0236880 -# -
0236890 -pf02589 -
0236900 -pf08031pf01565 -
0236910 -pf04371 -
0236920 HMMreg13411_1_GRM1(#) GRM1A
0236930 -pf00892 -
0236940 -pf04229 -
0236950 -pf03734 -
BMC-P
BMC-H
BMC-T
Aldehyde dehydrogenase
PTAC
Signature enzyme
Alcohol dehydrogenase
Regulator

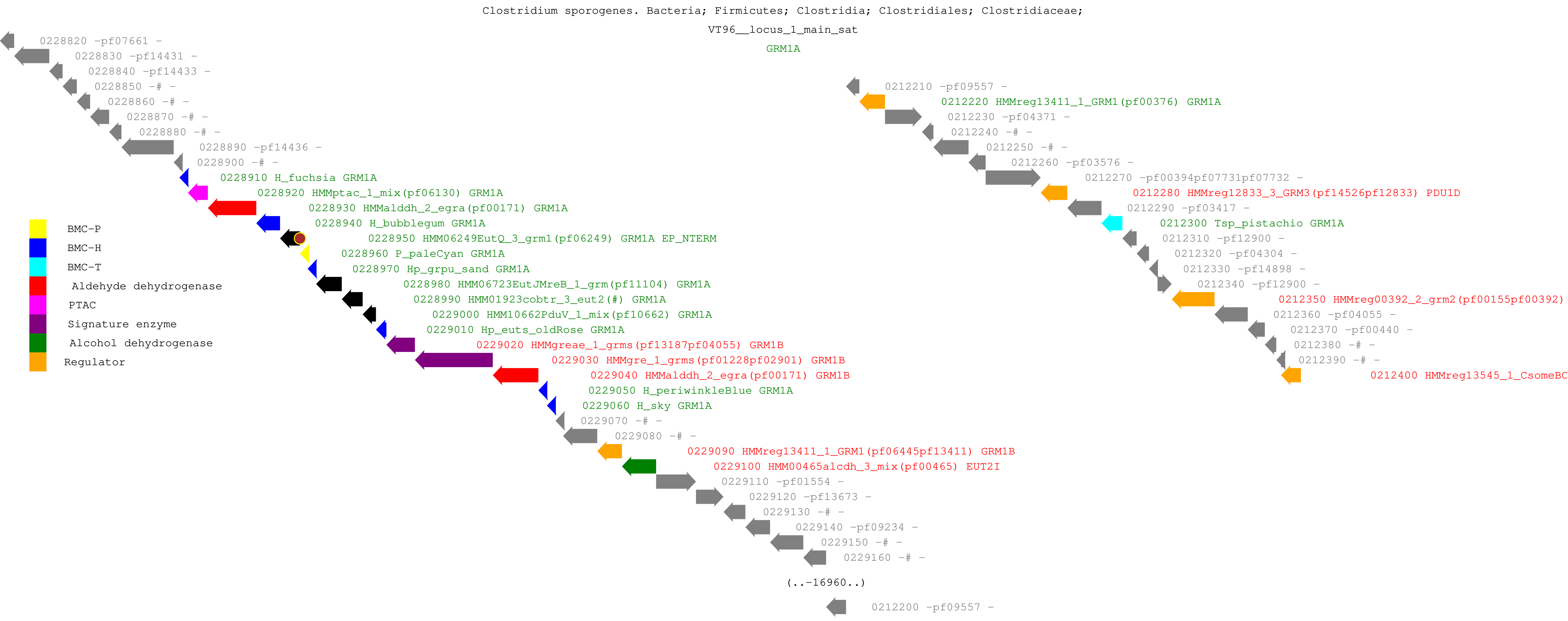
Clostridium sporogenes. Bacteria; Firmicutes; Clostridia; Clostridiales; Clostridiaceae;
VT96__locus_1_main_sat
GRM1A
0228820 -pf07661 -
0228830 -pf14431 -
0228840 -pf14433 -
0228850 -# -
0228860 -# -
0228870 -# -
0228880 -# -
0228890 -pf14436 -
0228900 -# -
0228910 H_fuchsia GRM1A
0228920 HMMptac_1_mix(pf06130) GRM1A
0228930 HMMaldh_2_egra(pf00171) GRM1A
0228940 H_bubblegum GRM1A
0228950 HMM06249EutQ_3_grm1(pf06249) GRM1A EP_NTERM
0228960 P_paleCyan GRM1A
0228970 Hp_grpu_sand GRM1A
0228980 HMM06723EutJMreB_1_grm(pf11104) GRM1A
0228990 HMM01923cobtr_3_eut2(#) GRM1A
0229000 HMM10662PduV_1_mix(pf10662) GRM1A
0229010 Hp_euts_oldRose GRM1A
0229020 HMMgreae_1_grms(pf13187pf04055) GRM1B
0229030 HMMgre_1_grms(pf01228pf02901) GRM1B
0229040 HMMaldh_2_egra(pf00171) GRM1B
0229050 H_periwinkleBlue GRM1A
0229060 H_sky GRM1A
0229070 -# -
0229080 -# -
0229090 HMMreg13411_1_GRM1(pf06445pf13411) GRM1B
0229100 HMM00465alcdh_3_mix(pf00465) EUT2I
0229110 -pf01554 -
0229120 -pf13673 -
0229130 -# -
0229140 -pf09234 -
0229150 -# -
0229160 -# -
(..-16960..)
0212200 -pf09557 -
0212210 -pf09557 -
0212220 HMMreg13411_1_GRM1(pf00376) GRM1A
0212230 -pf04371 -
0212240 -# -
0212250 -# -
0212260 -pf03576 -
0212270 -pf00394pf07731pf07732 -
0212280 HMMreg12833_3_GRM3(pf14526pf12833) PDU1D
0212290 -pf03417 -
0212300 Tsp_pistachio GRM1A
0212310 -pf12900 -
0212320 -pf04304 -
0212330 -pf14898 -
0212340 -pf12900 -
0212350 HMMreg00392_2_grm2(pf00155pf00392)
0212360 -pf04055 -
0212370 -pf00440 -
0212380 -# -
0212390 -# -
0212400 HMMreg13545_1_CsomeBC
BMC-P
BMC-H
BMC-T
Aldehyde dehydrogenase
PTAC
Signature enzyme
Alcohol dehydrogenase
Regulator

Paenibacillus sp. DMB20. Bacteria; Firmicutes; Bacilli; Bacillales; Paenibacillaceae;

XI25__locus_2

GRM1A

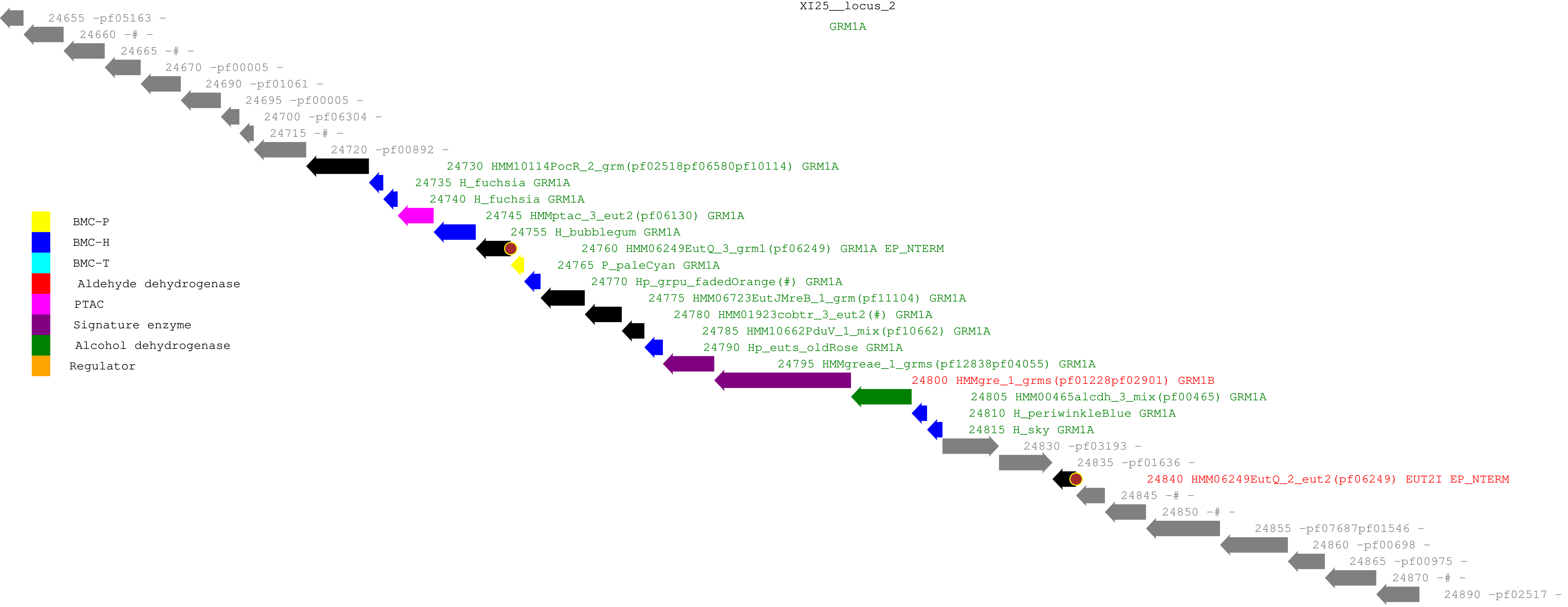

Clostridium botulinum C/D str. DC5. Bacteria; Firmicutes; Clostridia; Clostridiales; Clostridiaceae;

Z955__locus_1

GRM1A

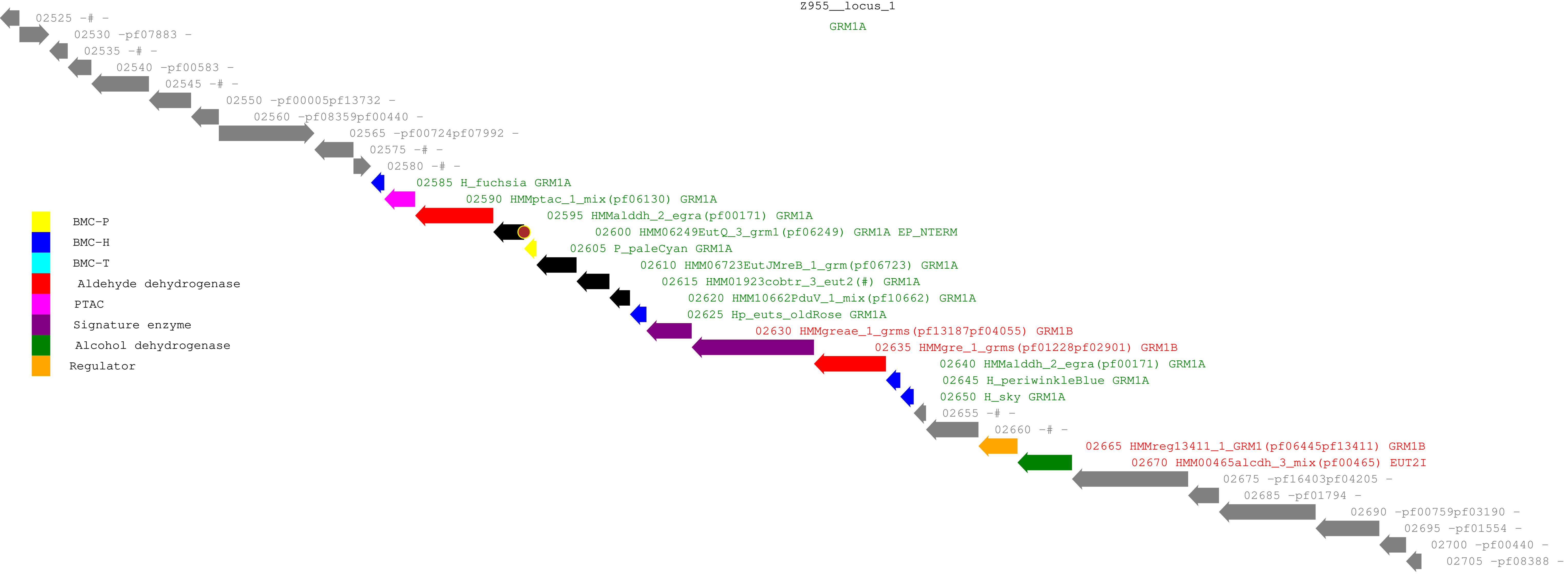

Clostridium sp. K25. Bacteria; Firmicutes; Clostridia; Clostridiales; Clostridiaceae;

Z957__locus_1

GRM1A

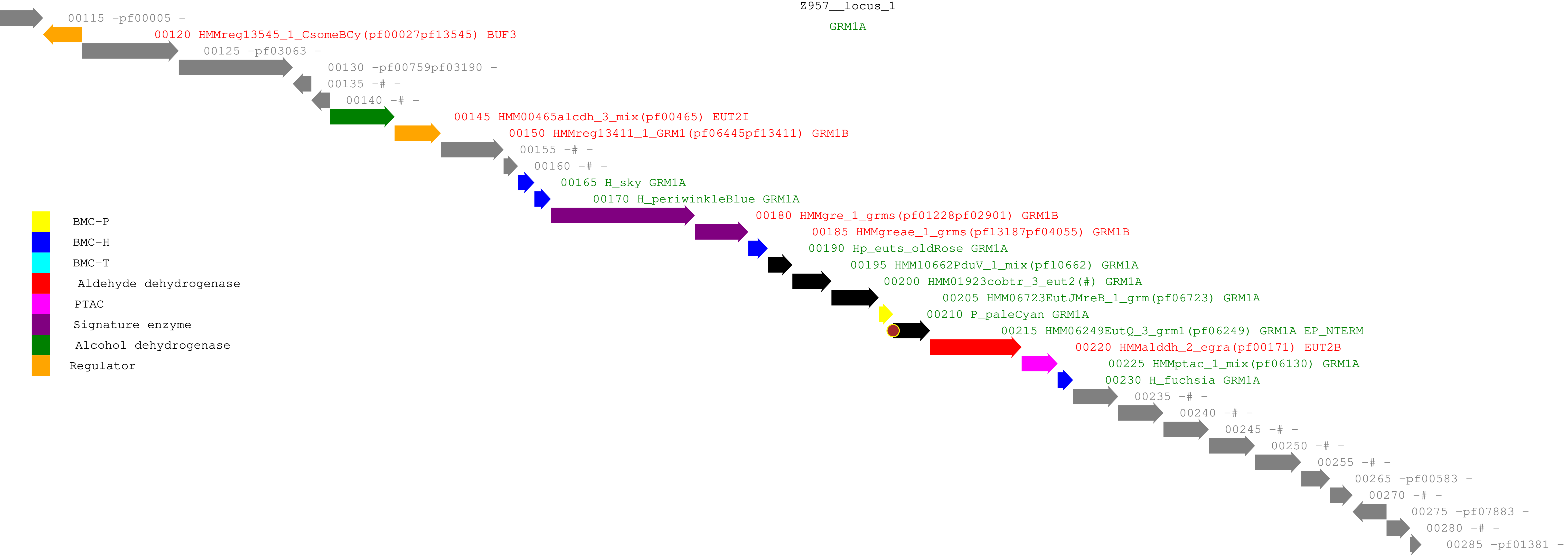

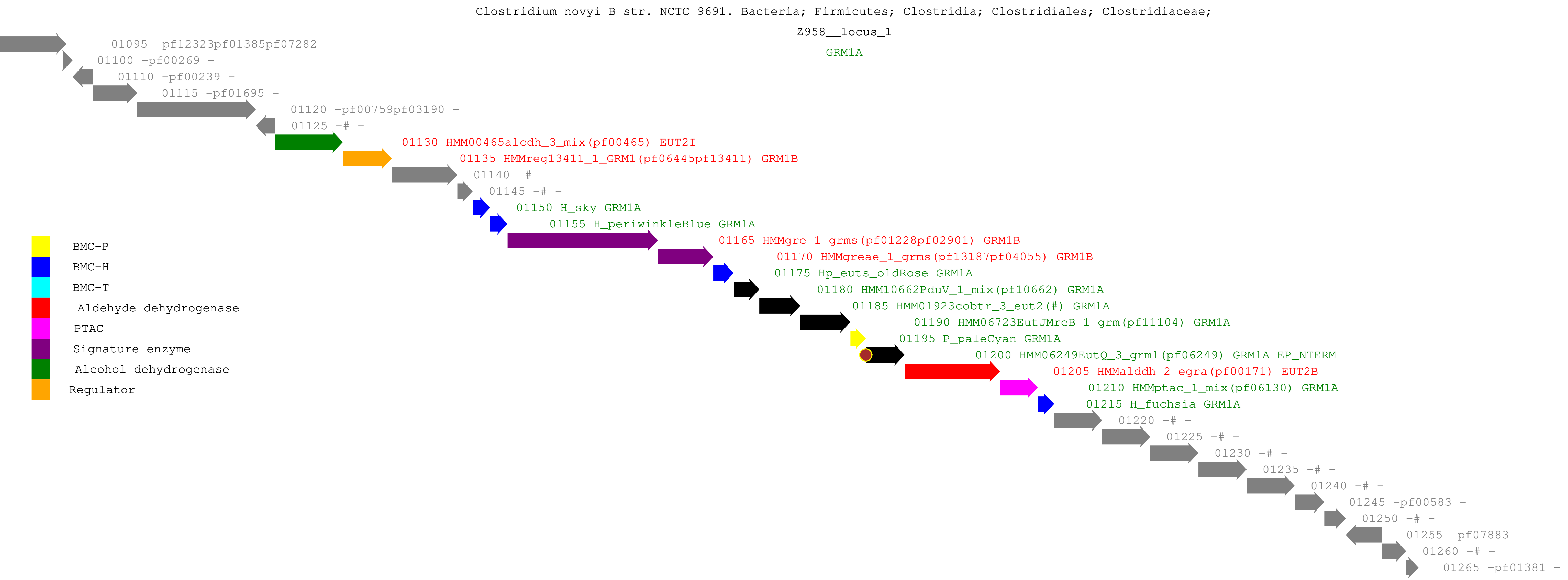
Clostridium novyi B str. NCTC 9691. Bacteria; Firmicutes; Clostridia; Clostridiales; Clostridiaceae;
Z958__locus_1
GRM1A
01095 -pf12323pf01385pf07282 -
01100 -pf00269 -
01110 -pf00239 -
01115 -pf01695 -
01120 -pf00759pf03190 -
01125 -# -
01130 HMM00465alcdh_3_mix(pf00465) EUT2I
01135 HMMreg13411_1_GRM1(pf06445pf13411) GRM1B
01140 -# -
01145 -# -
01150 H_sky GRM1A
01155 H_periwinkleBlue GRM1A
01165 HMMgre_1_grms(pf01228pf02901) GRM1B
01170 HMMgreae_1_grms(pf13187pf04055) GRM1B
01175 Hp_euts_oldRose GRM1A
01180 HMM10662PduV_1_mix(pf10662) GRM1A
01185 HMM01923cobtr_3_eut2(#) GRM1A
01190 HMM06723EutJMreB_1_grm(pf11104) GRM1A
01195 P_paleCyan GRM1A
01200 HMM06249EutQ_3_grm1(pf06249) GRM1A EP_NTERM
01205 HMMalddh_2_egra(pf00171) EUT2B
01210 HMMptac_1_mix(pf06130) GRM1A
01215 H_fuchsia GRM1A
01220 -# -
01225 -# -
01230 -# -
01235 -# -
01240 -# -
01245 -pf00583 -
01250 -# -
01255 -pf07883 -
01260 -# -
01265 -pf01381 -
BMC-P
BMC-H
BMC-T
Aldehyde dehydrogenase
PTAC
Signature enzyme
Alcohol dehydrogenase
Regulator

Clostridium haemolyticum NCTC 8350. Bacteria; Firmicutes; Clostridia; Clostridiales; Clostridiaceae;

Z961__locus_1

GRM1A

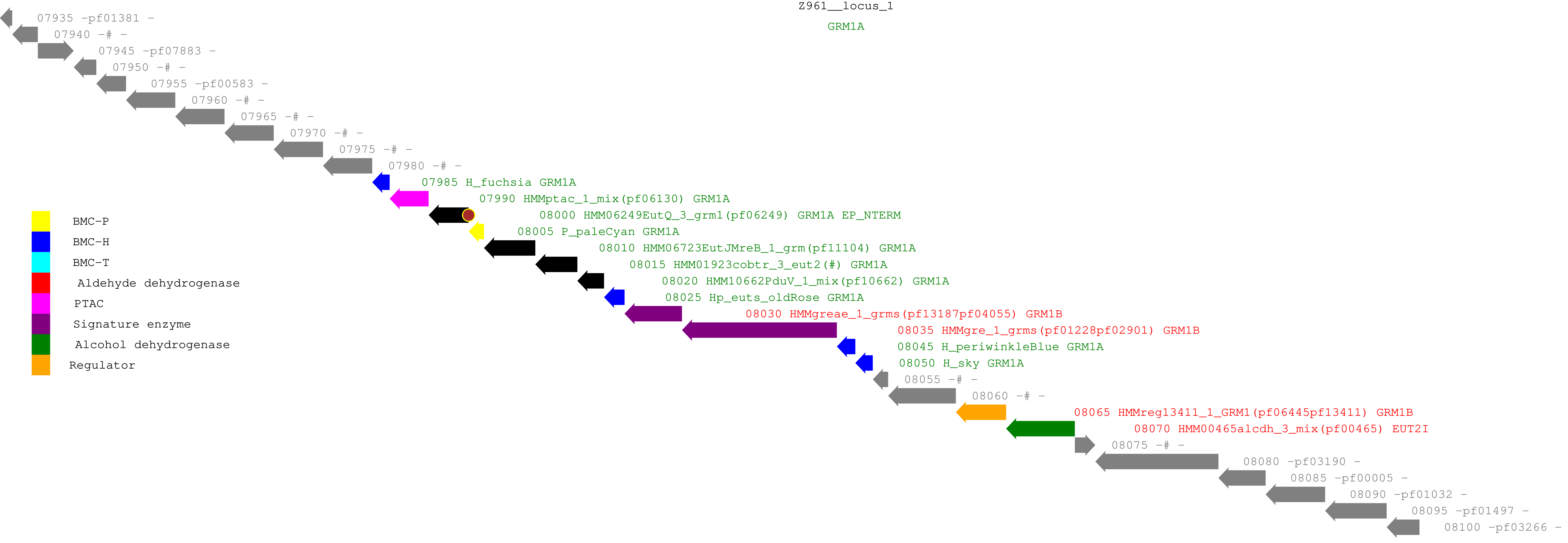

Desulfovibrio sp. MES5. Bacteria; Proteobacteria; Deltaproteobacteria; Desulfovibrionales;

BCS36__locus_1

GRM1B

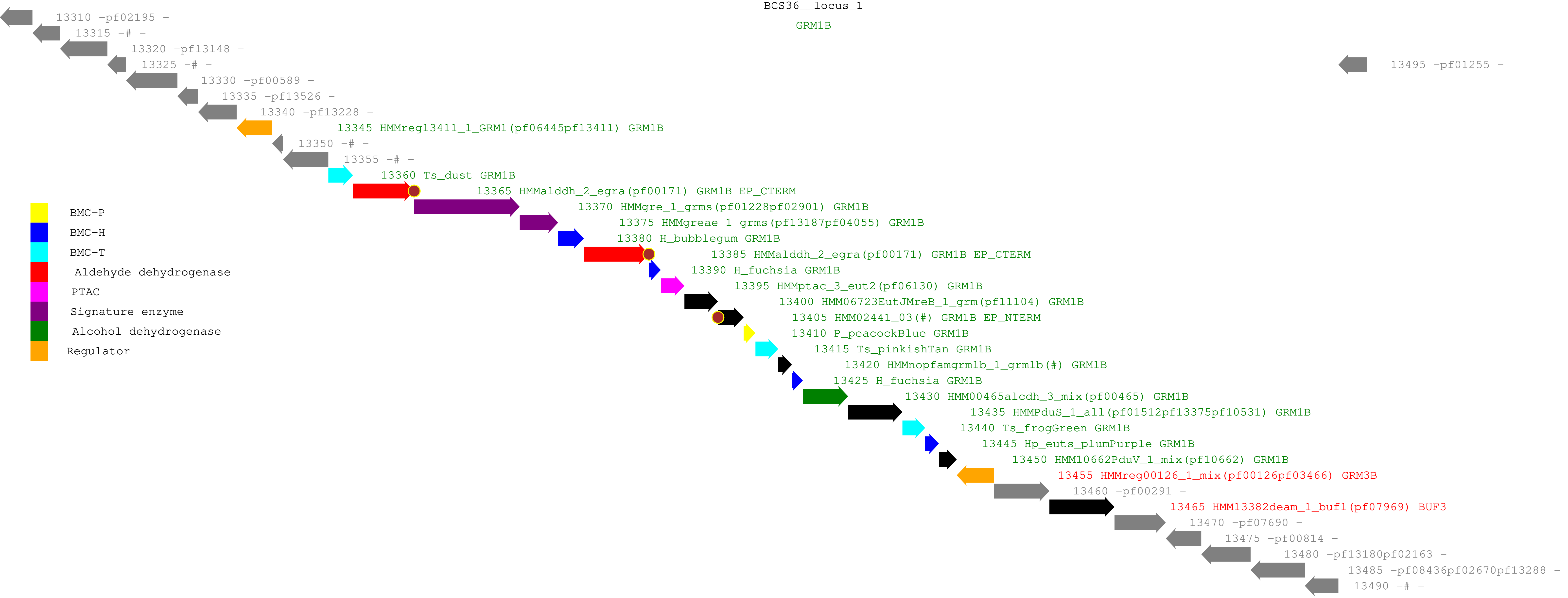

Photobacterium sp. 13-12. Bacteria; Proteobacteria; Gammaproteobacteria; Vibrionales;

BIT28__locus_2

GRM1B

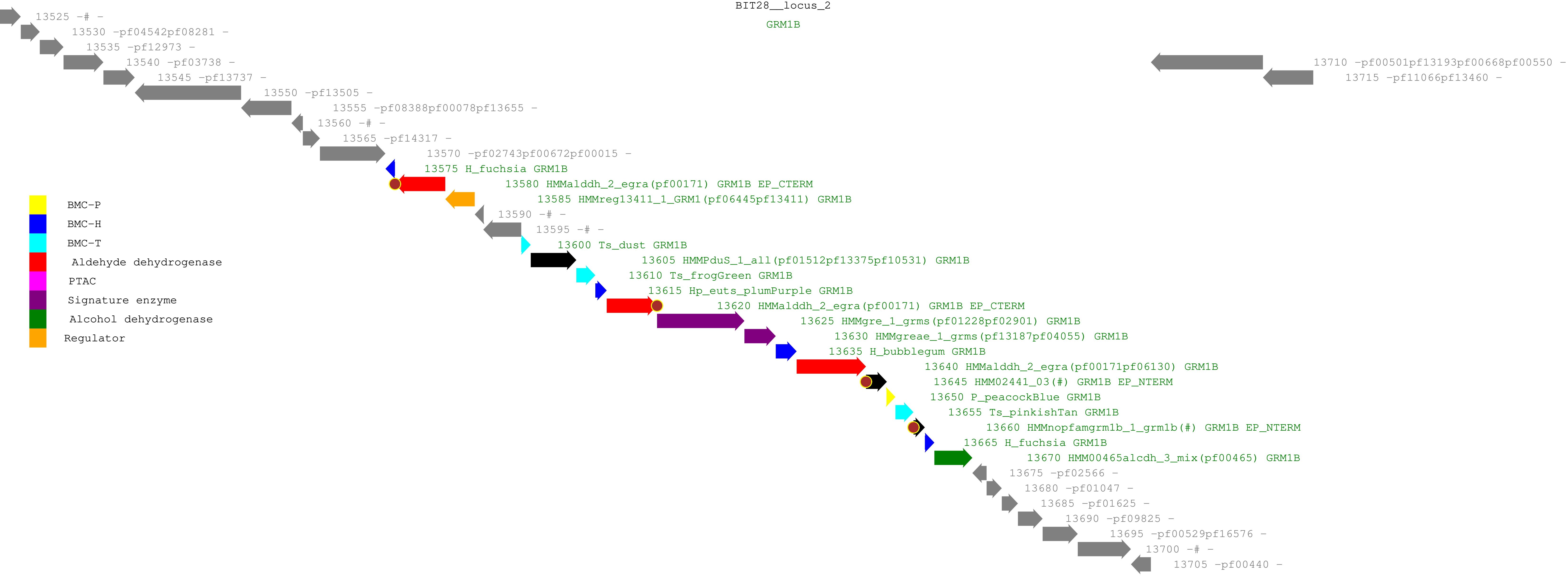

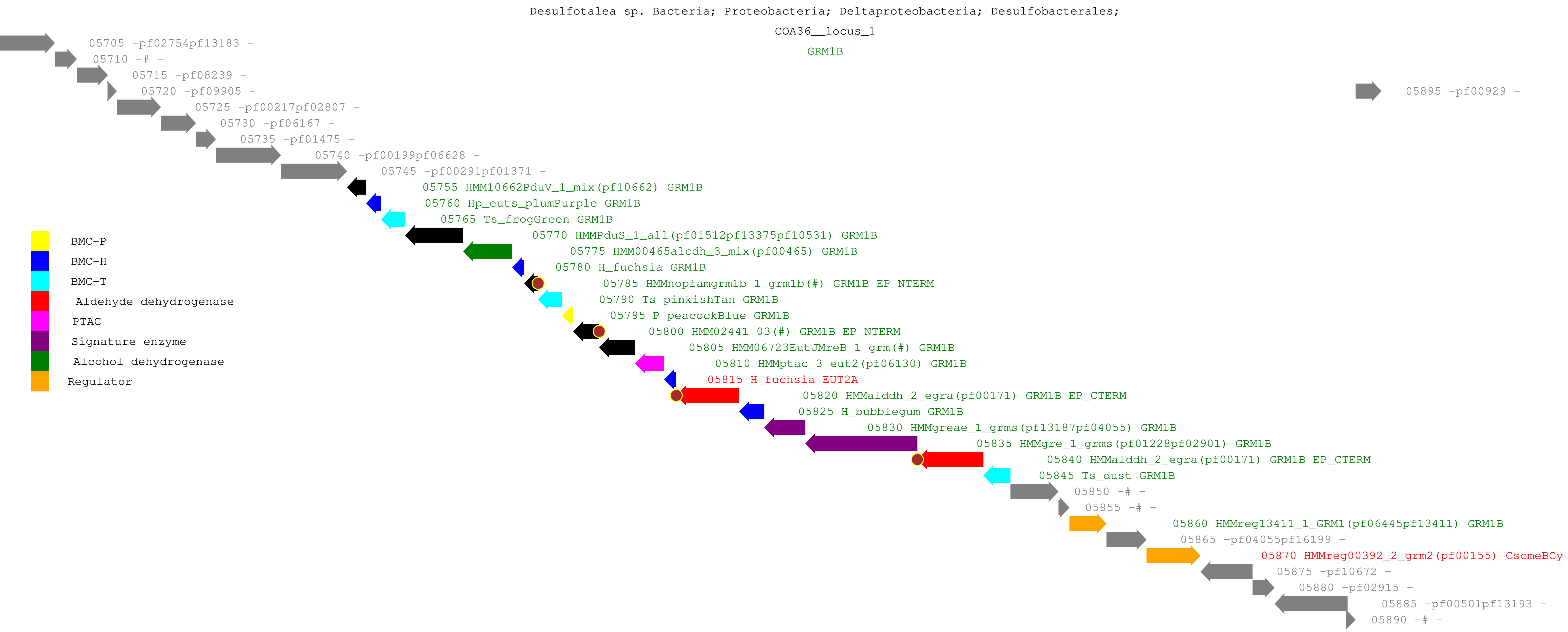

Desulfotalea sp. Bacteria; Proteobacteria; Deltaproteobacteria; Desulfobacterales;
COA36__locus_1
GRM1B
05705 -pf02754pf13183 -
05710 -# -
05715 -pf08239 -
05720 -pf09905 -
05725 -pf00217pf02807 -
05730 -pf06167 -
05735 -pf01475 -
05740 -pf00199pf06628 -
05745 -pf00291pf01371 -
05755 HMM10662PduV_1_mix(pf10662) GRM1B
05760 Hp_euts_plumPurple GRM1B
05765 Ts_frogGreen GRM1B
05770 HMMPduS_1_all(pf01512pf13375pf10531) GRM1B
05775 HMM00465alcdh_3_mix(pf00465) GRM1B
05780 H_fuchsia GRM1B
05785 HMMnopfamgrm1b_1_grm1b(#) GRM1B EP_NTERM
05790 Ts_pinkishTan GRM1B
05795 P_peacockBlue GRM1B
05800 HMM02441_03(#) GRM1B EP_NTERM
05805 HMM06723EutJMreB_1_grm(#) GRM1B
05810 HMMptac_3_eut2(pf06130) GRM1B
05815 H_fuchsia EUT2A
05820 HMMalddh_2_egra(pf00171) GRM1B EP_CTERM
05825 H_bubblegum GRM1B
05830 HMMgreae_1_grms(pf13187pf04055) GRM1B
05835 HMMgre_1_grms(pf01228pf02901) GRM1B
05840 HMMalddh_2_egra(pf00171) GRM1B EP_CTERM
05845 Ts_dust GRM1B
05850 -# -
05855 -# -
05860 HMMreg13411_1_GRM1(pf06445pf13411) GRM1B
05865 -pf04055pf16199 -
05870 HMMreg00392_2_grm2(pf00155) CsomeBCy
05875 -pf10672 -
05880 -pf02915 -
05885 -pf00501pf13193 -
05890 -# -
05895 -pf00929 -
BMC-P
BMC-H
BMC-T
Aldehyde dehydrogenase
PTAC
Signature enzyme
Alcohol dehydrogenase
Regulator

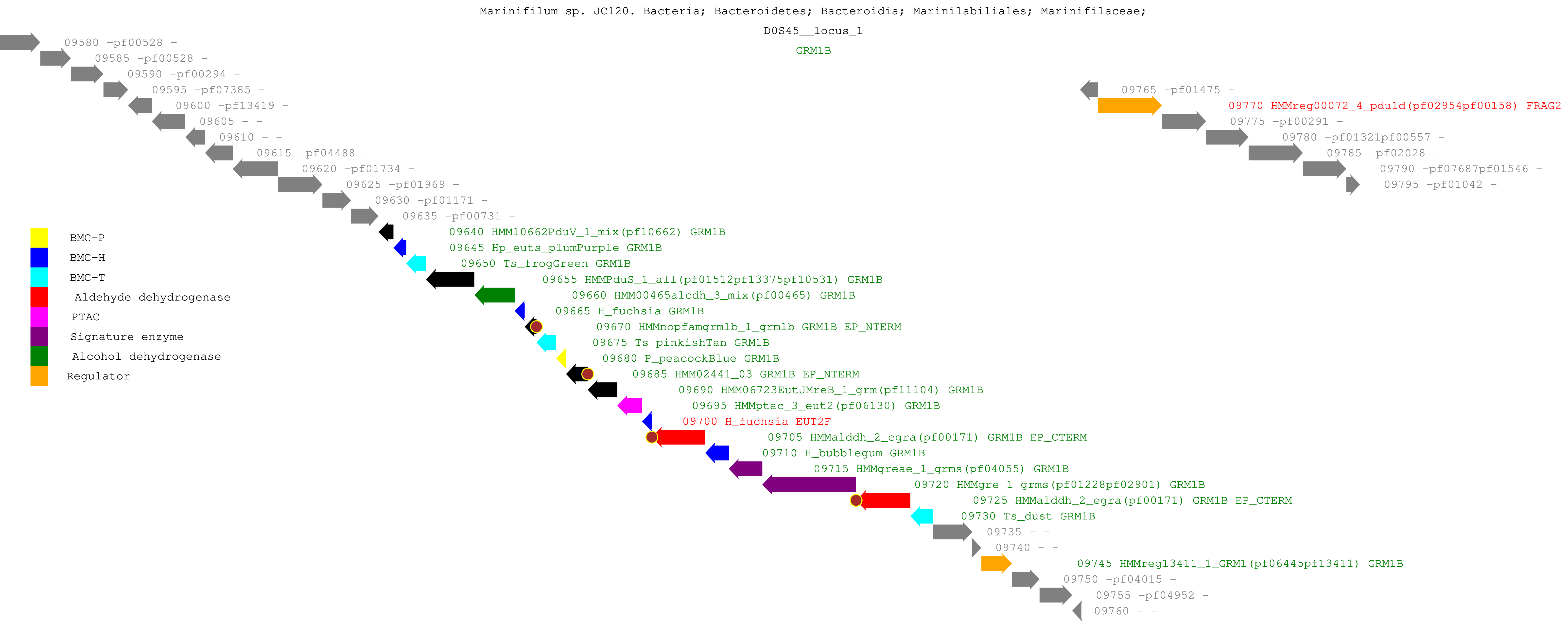
Marinifilum sp. JC120. Bacteria; Bacteroidetes; Bacteroidia; Marinilabiliales; Marinifilaceae;
D0S45__locus_1
GRM1B
09580 -pf00528 -
09585 -pf00528 -
09590 -pf00294 -
09595 -pf07385 -
09600 -pf13419 -
09605 - -
09610 - -
09615 -pf04488 -
09620 -pf01734 -
09625 -pf01969 -
09630 -pf01171 -
09635 -pf00731 -
09640 HMM10662PduV_1_mix(pf10662) GRM1B
09645 Hp_euts_plumPurple GRM1B
09650 Ts_frogGreen GRM1B
09655 HMMPduS_1_all(pf01512pf13375pf10531) GRM1B
09660 HMM00465alcdh_3_mix(pf00465) GRM1B
09665 H_fuchsia GRM1B
09670 HMMnopfamgrm1b_1_grm1b GRM1B EP_NTERM
09675 Ts_pinkishTan GRM1B
09680 P_peacockBlue GRM1B
09685 HMM02441_03 GRM1B EP_NTERM
09690 HMM06723EutJMreB_1_grm(pf11104) GRM1B
09695 HMMptac_3_eut2(pf06130) GRM1B
09700 H_fuchsia EUT2F
09705 HMMalddh_2_egra(pf00171) GRM1B EP_CTERM
09710 H_bubblegum GRM1B
09715 HMMgreae_1_grms(pf04055) GRM1B
09720 HMMgre_1_grms(pf01228pf02901) GRM1B
09725 HMMalddh_2_egra(pf00171) GRM1B EP_CTERM
09730 Ts_dust GRM1B
09735 - -
09740 - -
09745 HMMreg13411_1_GRM1(pf06445pf13411) GRM1B
09750 -pf04015 -
09755 -pf04952 -
09760 - -
09765 -pf01475 -
09770 HMMreg00072_4_pdu1d(pf02954pf00158) FRAG2
09775 -pf00291 -
09780 -pf01321pf00557 -
09785 -pf02028 -
09790 -pf07687pf01546 -
09795 -pf01042 -
BMC-P
BMC-H
BMC-T
Aldehyde dehydrogenase
PTAC
Signature enzyme
Alcohol dehydrogenase
Regulator

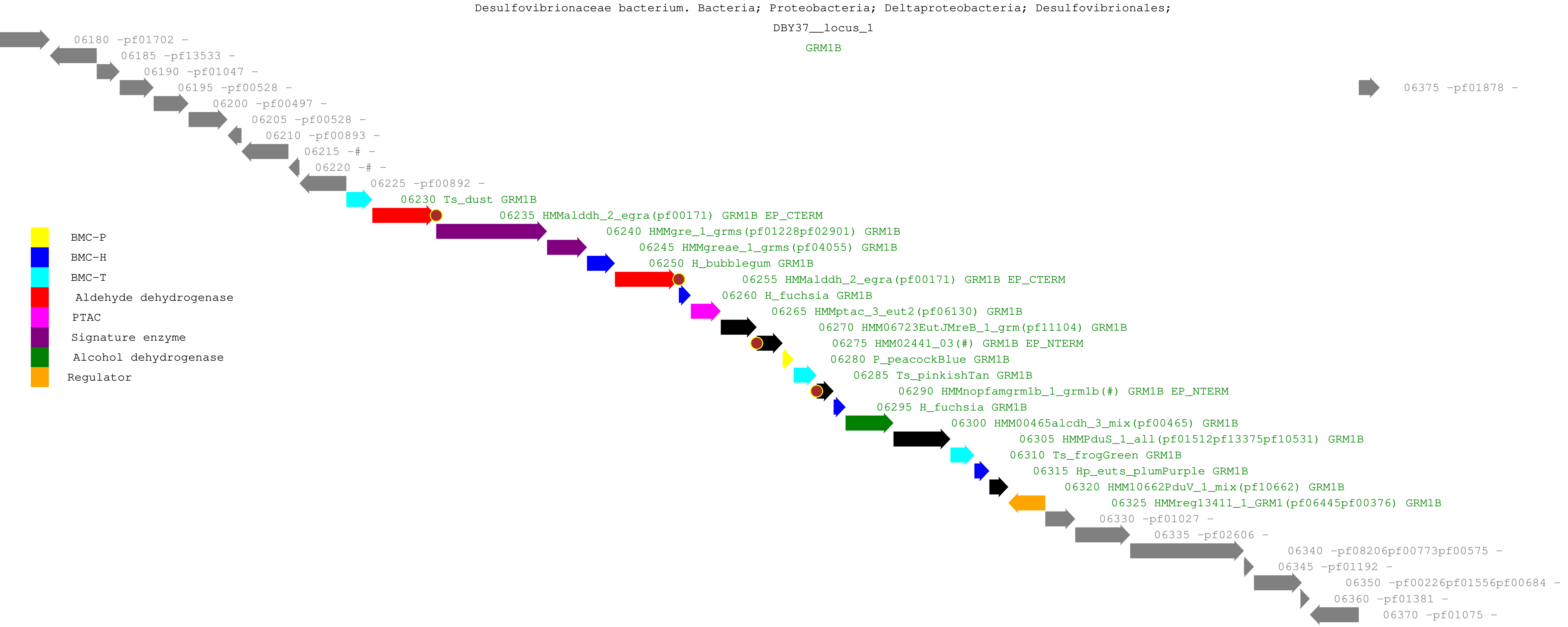
Desulfovibrionaceae bacterium. Bacteria; Proteobacteria; Deltaproteobacteria; Desulfovibrionales;
DBY37__locus_1
GRM1B
06180 -pf01702 -
06185 -pf13533 -
06190 -pf01047 -
06195 -pf00528 -
06200 -pf00497 -
06205 -pf00528 -
06210 -pf00893 -
06215 -# -
06220 -# -
06225 -pf00892 -
06230 Ts_dust GRM1B
06235 HMMaldh_2_egra(pf00171) GRM1B EP_CTERM
06240 HMMgre_1_grms(pf01228pf02901) GRM1B
06245 HMMgreae_1_grms(pf04055) GRM1B
06250 H_bubblegum GRM1B
06255 HMMaldh_2_egra(pf00171) GRM1B EP_CTERM
06260 H_fuchsia GRM1B
06265 HMMptac_3_eut2(pf06130) GRM1B
06270 HMM06723EutJMreB_1_grm(pf11104) GRM1B
06275 HMM02441_03(#) GRM1B EP_NTERM
06280 P_peacockBlue GRM1B
06285 Ts_pinkishTan GRM1B
06290 HMMnopfamgrm1b_1_grm1b(#) GRM1B EP_NTERM
06295 H_fuchsia GRM1B
06300 HMM00465alcdh_3_mix(pf00465) GRM1B
06305 HMMPduS_1_all(pf01512pf13375pf10531) GRM1B
06310 Ts_frogGreen GRM1B
06315 Hp_euts_plumPurple GRM1B
06320 HMM10662PduV_1_mix(pf10662) GRM1B
06325 HMMreg13411_1_GRM1(pf06445pf00376) GRM1B
06330 -pf01027 -
06335 -pf02606 -
06340 -pf08206pf00773pf00575 -
06345 -pf01192 -
06350 -pf00226pf01556pf00684 -
06360 -pf01381 -
06370 -pf01075 -
06375 -pf01878 -
BMC-P
BMC-H
BMC-T
Aldehyde dehydrogenase
PTAC
Signature enzyme
Alcohol dehydrogenase
Regulator

Desulfovibrio sp. Bacteria; Proteobacteria; Deltaproteobacteria; Desulfovibrionales;

DCS48__locus_1_main_sat

GRM1B

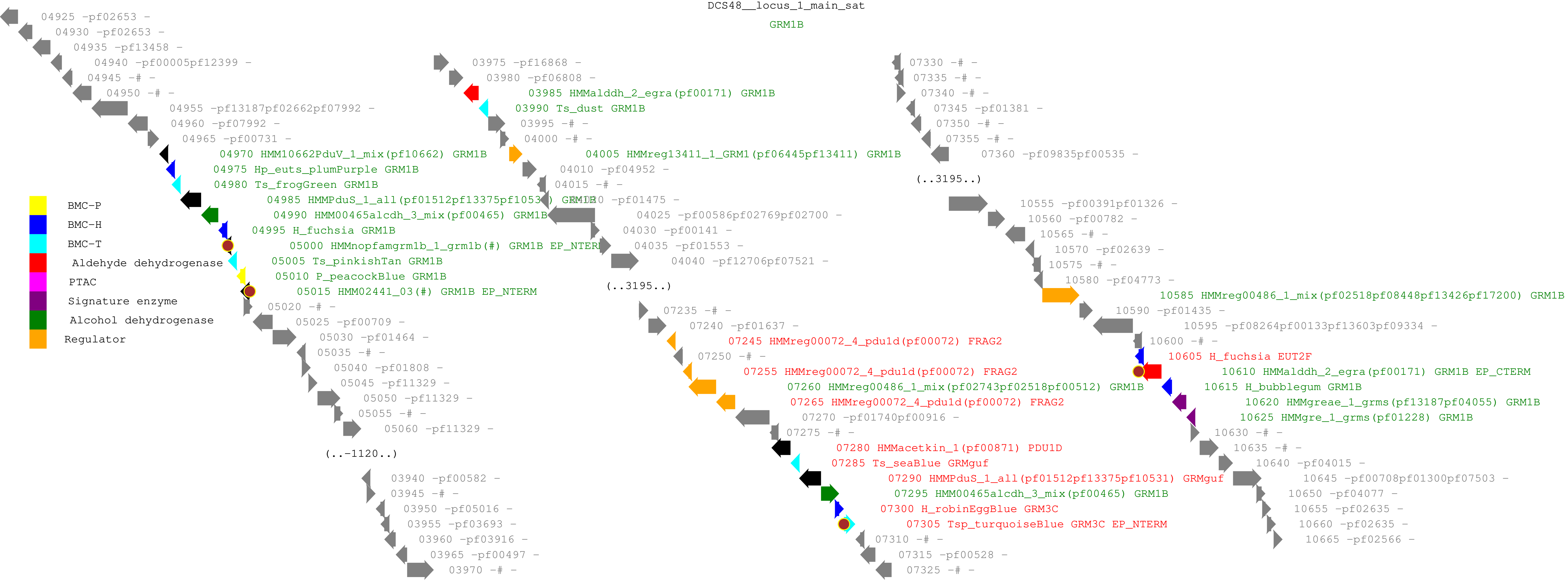

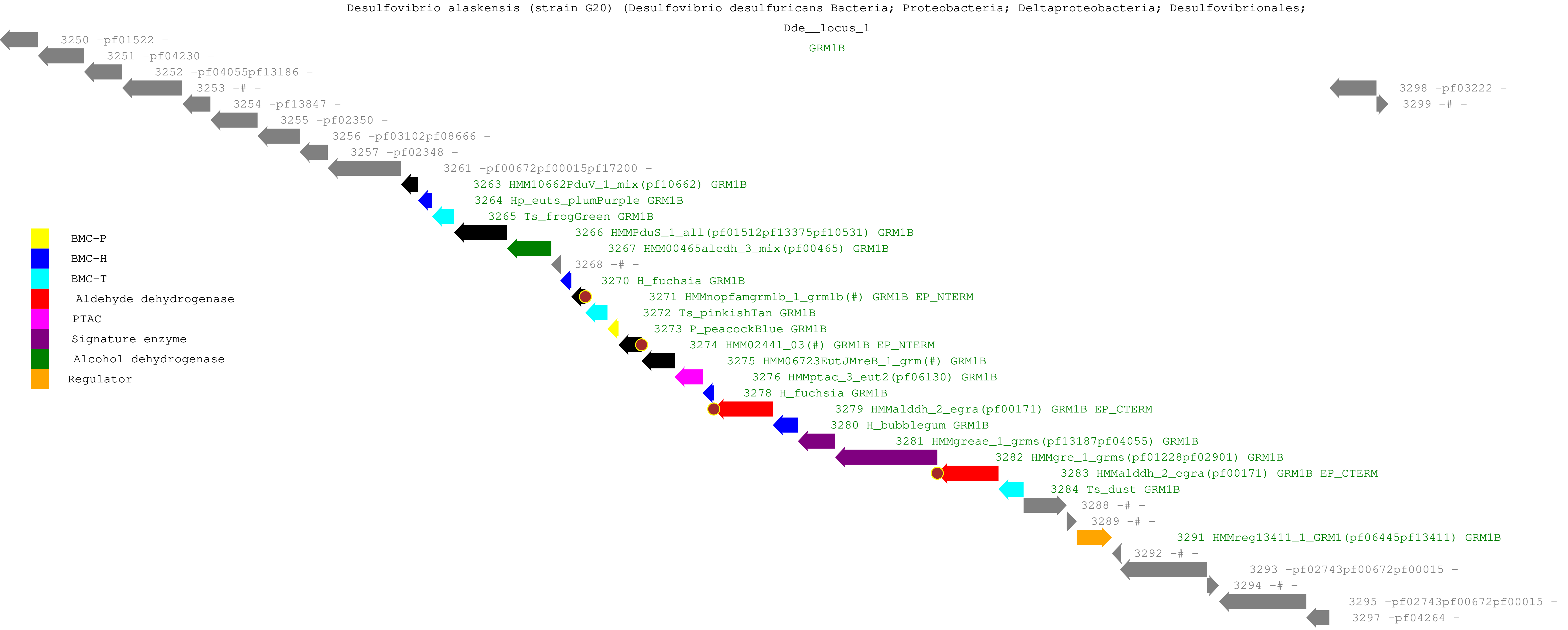
Desulfovibrio alaskensis (strain G20) (Desulfovibrio desulfuricans Bacteria; Proteobacteria; Deltaproteobacteria; Desulfovibrionales;
Dde__locus_1
GRM1B
3250 -pf01522 -
3251 -pf04230 -
3252 -pf04055pf13186 -
3253 -# -
3254 -pf13847 -
3255 -pf02350 -
3256 -pf03102pf08666 -
3257 -pf02348 -
3261 -pf00672pf00015pf17200 -
3263 HMM10662PduV_1_mix(pf10662) GRM1B
3264 Hp_euts_plumPurple GRM1B
3265 Ts_frogGreen GRM1B
3266 HMMPduS_1_all(pf01512pf13375pf10531) GRM1B
3267 HMM00465alcdh_3_mix(pf00465) GRM1B
3268 -# -
3270 H_fuchsia GRM1B
3271 HMMnopfamgrm1b_1_grm1b(#) GRM1B EP_NTERM
3272 Ts_pinkishTan GRM1B
3273 P_peacockBlue GRM1B
3274 HMM02441_03(#) GRM1B EP_NTERM
3275 HMM06723EutJMreB_1_grm(#) GRM1B
3276 HMMptac_3_eut2(pf06130) GRM1B
3278 H_fuchsia GRM1B
3279 HMMalddh_2_egra(pf00171) GRM1B EP_CTERM
3280 H_bubblegum GRM1B
3281 HMMgreae_1_grms(pf13187pf04055) GRM1B
3282 HMMgre_1_grms(pf01228pf02901) GRM1B
3283 HMMalddh_2_egra(pf00171) GRM1B EP_CTERM
3284 Ts_dust GRM1B
3288 -# -
3289 -# -
3291 HMMreg13411_1_GRM1(pf06445pf13411) GRM1B
3292 -# -
3293 -pf02743pf00672pf00015 -
3294 -# -
3295 -pf02743pf00672pf00015 -
3297 -pf04264 -
3298 -pf03222 -
3299 -# -
BMC-P
BMC-H
BMC-T
Aldehyde dehydrogenase
PTAC
Signature enzyme
Alcohol dehydrogenase
Regulator

Desulfovibrio desulfuricans (strain ATCC 27774 / DSM 6949). Bacteria; Proteobacteria; Deltaproteobacteria; Desulfovibrionales;

Ddes__locus_1

GRM1B

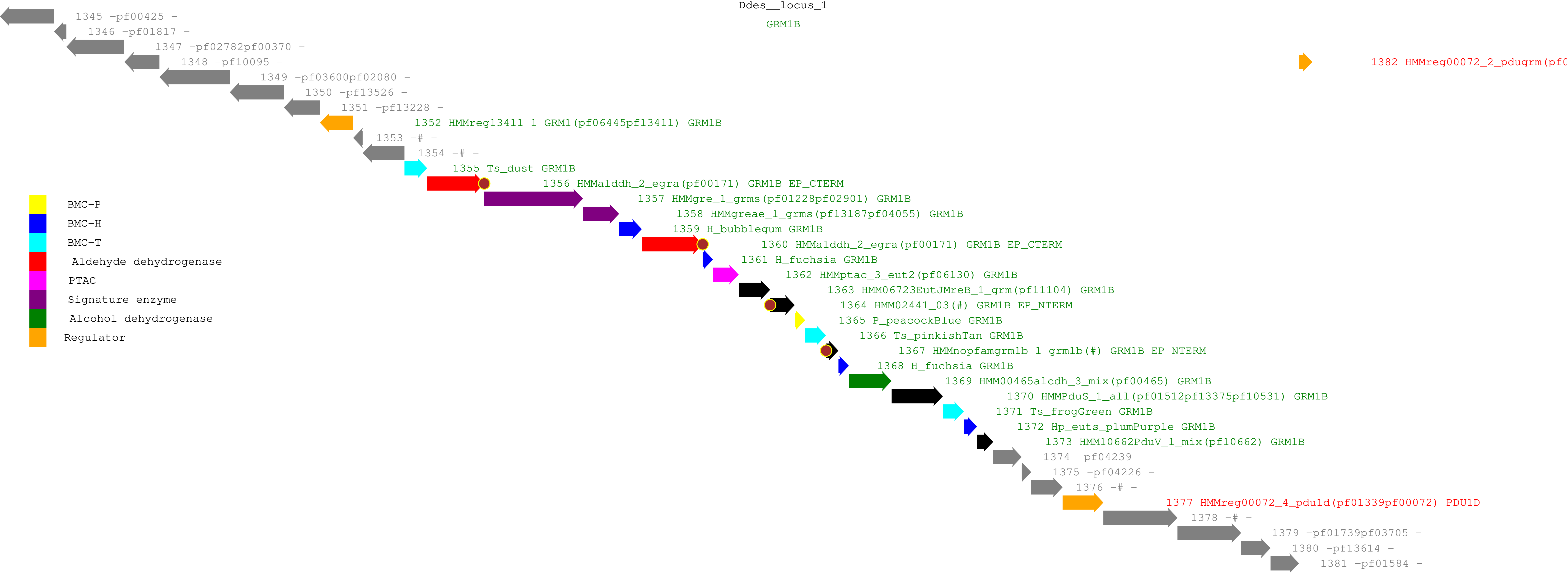

Desulfovibrio salexigens (strain ATCC 14822 / DSM 2638 / NCIB 8403 / Bacteria; Proteobacteria; Deltaproteobacteria; Desulfovibrionales;

Desal__locus_1

GRM1B

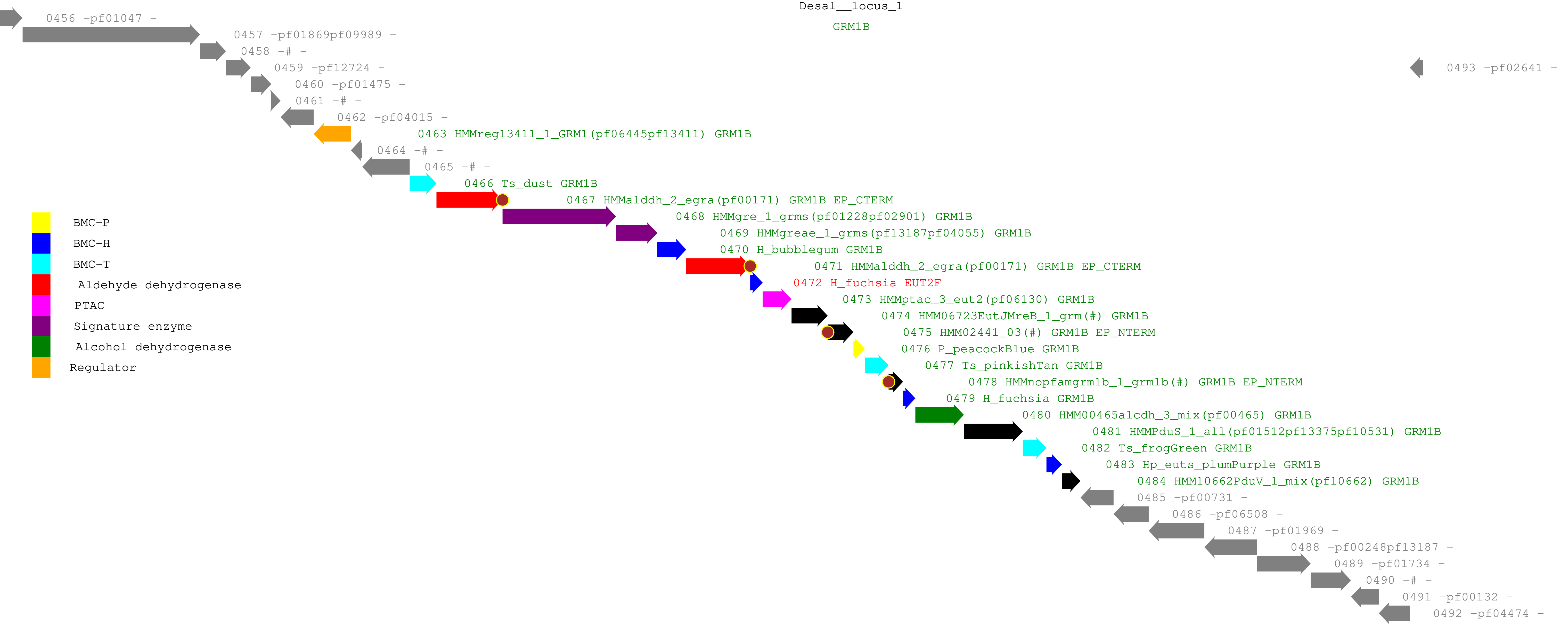

Desulfovibrio hydrothermalis AM13 = DSM 14728. Bacteria; Proteobacteria; Deltaproteobacteria; Desulfovibrionales;

DESAM__locus_1

GRM1B

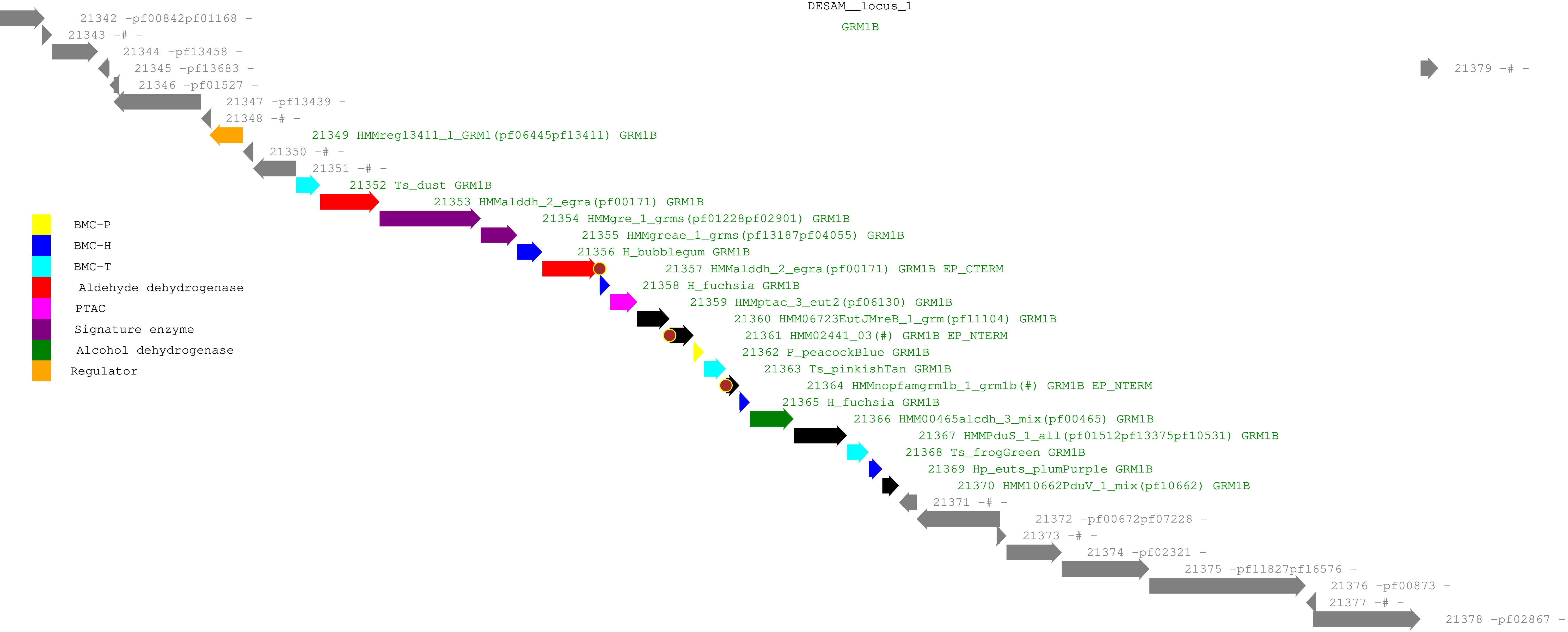

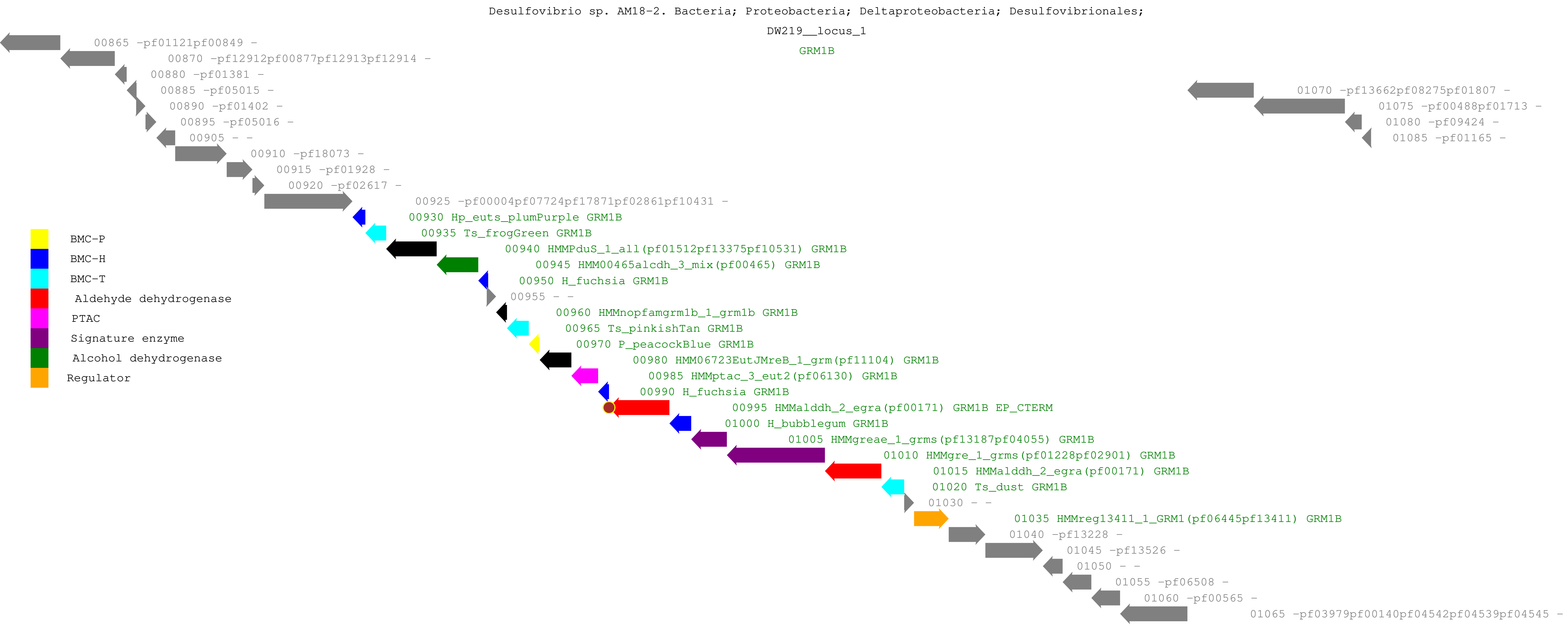
Desulfovibrio sp. AM18-2. Bacteria; Proteobacteria; Deltaproteobacteria; Desulfovibrionales;
DW219__locus_1
GRM1B
00865 -pf01121pf00849 -
00870 -pf12912pf00877pf12913pf12914 -
00880 -pf01381 -
00885 -pf05015 -
00890 -pf01402 -
00895 -pf05016 -
00905 - -
00910 -pf18073 -
00915 -pf01928 -
00920 -pf02617 -
00925 -pf00004pf07724pf17871pf02861pf10431 -
00930 Hp_euts_plumPurple GRM1B
00935 Ts_frogGreen GRM1B
00940 HMMPduS_1_all(pf01512pf13375pf10531) GRM1B
00945 HMM00465alcdh_3_mix(pf00465) GRM1B
00950 H_fuchsia GRM1B
00955 - -
00960 HMMnopfamgrm1b_1_grm1b GRM1B
00965 Ts_pinkishTan GRM1B
00970 P_peacockBlue GRM1B
00980 HMM06723EutJMreB_1_grm(pf11104) GRM1B
00985 HMMptac_3_eut2(pf06130) GRM1B
00990 H_fuchsia GRM1B
00995 HMMaldddh_2_egra(pf00171) GRM1B EP_CTERM
01000 H_bubblegum GRM1B
01005 HMMgreae_1_grms(pf13187pf04055) GRM1B
01010 HMMgre_1_grms(pf01228pf02901) GRM1B
01015 HMMaldddh_2_egra(pf00171) GRM1B
01020 Ts_dust GRM1B
01030 - -
01035 HMMreg13411_1_GRM1(pf06445pf13411) GRM1B
01040 -pf13228 -
01045 -pf13526 -
01050 - -
01055 -pf06508 -
01060 -pf00565 -
01065 -pf03979pf00140pf04542pf04539pf04545 -
01070 -pf13662pf08275pf01807 -
01075 -pf00488pf01713 -
01080 -pf09424 -
01085 -pf01165 -
BMC-P
BMC-H
BMC-T
Aldehyde dehydrogenase
PTAC
Signature enzyme
Alcohol dehydrogenase
Regulator

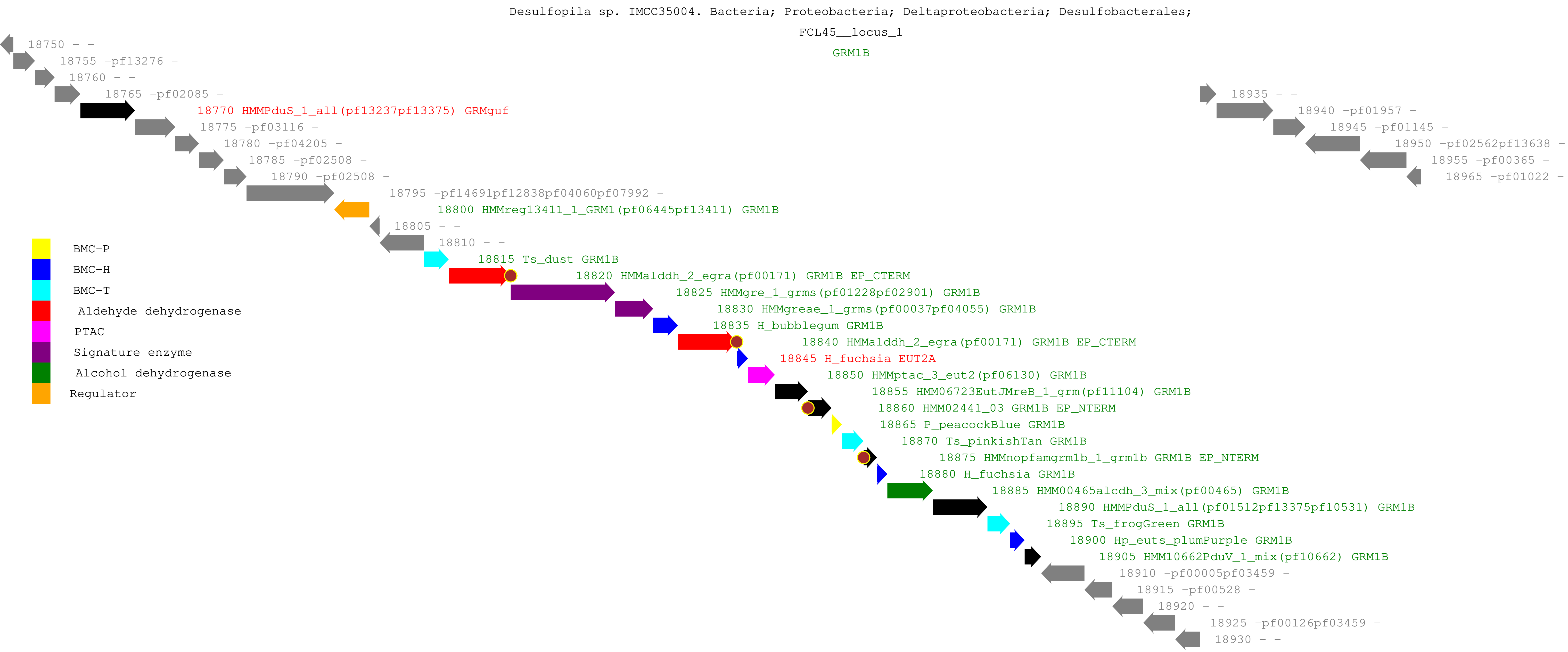
Desulfopila sp. IMCC35004. Bacteria; Proteobacteria; Deltaproteobacteria; Desulfobacterales;
FCL45__locus_1
GRM1B
18750 - -
18755 -pf13276 -
18760 - -
18765 -pf02085 -
18770 HMMPduS_1_all(pf13237pf13375) GRMguf
18775 -pf03116 -
18780 -pf04205 -
18785 -pf02508 -
18790 -pf02508 -
18795 -pf14691pf12838pf04060pf07992 -
18800 HMMreg13411_1_GRM1(pf06445pf13411) GRM1B
18805 - -
18810 - -
18815 Ts_dust GRM1B
18820 HMMalddh_2_egra(pf00171) GRM1B EP_CTERM
18825 HMMgre_1_grms(pf01228pf02901) GRM1B
18830 HMMgreae_1_grms(pf00037pf04055) GRM1B
18835 H_bubblegum GRM1B
18840 HMMalddh_2_egra(pf00171) GRM1B EP_CTERM
18845 H_fuchsia EUT2A
18850 HMMptac_3_eut2(pf06130) GRM1B
18855 HMM06723EutJMreB_1_grm(pf11104) GRM1B
18860 HMM02441_03 GRM1B EP_NTERM
18865 P_peacockBlue GRM1B
18870 Ts_pinkishTan GRM1B
18875 HMMnopfamgrm1b_1_grm1b GRM1B EP_NTERM
18880 H_fuchsia GRM1B
18885 HMM00465alcdh_3_mix(pf00465) GRM1B
18890 HMMPduS_1_all(pf01512pf13375pf10531) GRM1B
18895 Ts_frogGreen GRM1B
18900 Hp_euts_plumPurple GRM1B
18905 HMM10662PduV_1_mix(pf10662) GRM1B
18910 -pf00005pf03459 -
18915 -pf00528 -
18920 - -
18925 -pf00126pf03459 -
18930 - -
18935 - -
18940 -pf01957 -
18945 -pf01145 -
18950 -pf02562pf13638 -
18955 -pf00365 -
18965 -pf01022 -
BMC-P
BMC-H
BMC-T
Aldehyde dehydrogenase
PTAC
Signature enzyme
Alcohol dehydrogenase
Regulator

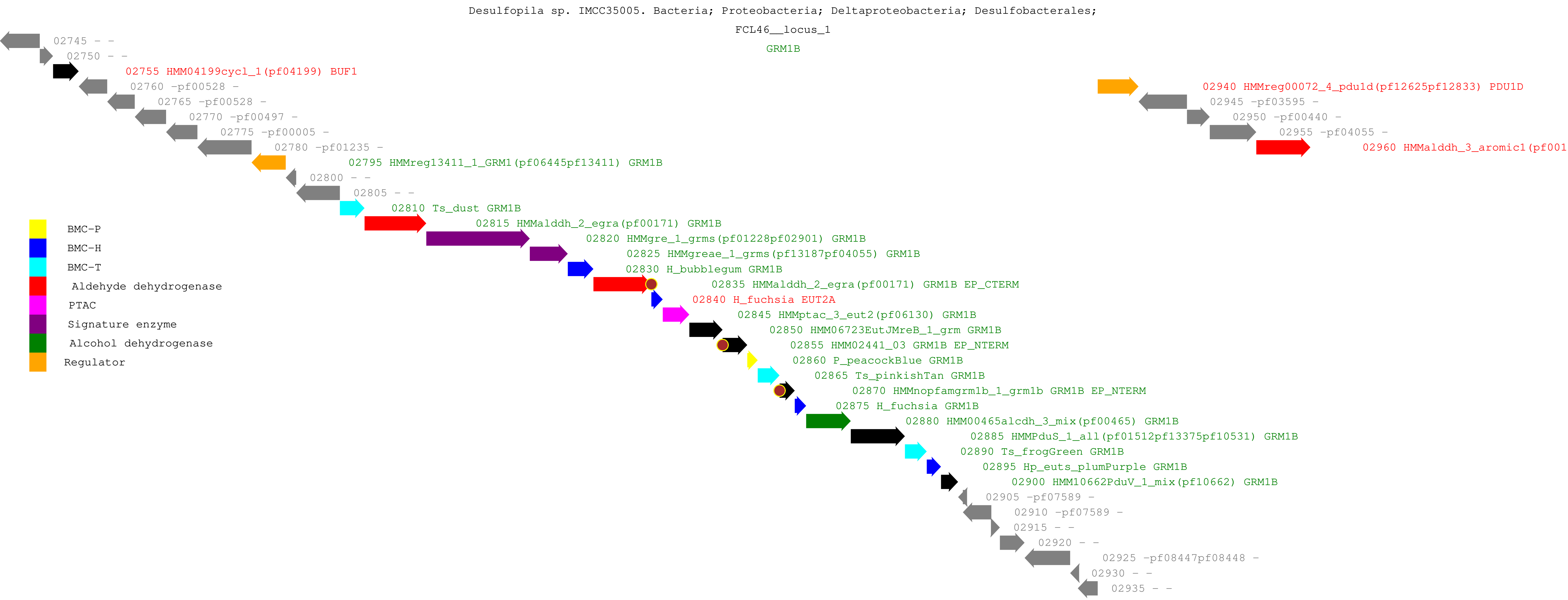
Desulfopila sp. IMCC35005. Bacteria; Proteobacteria; Deltaproteobacteria; Desulfobacterales;
FCL46__locus_1
GRM1B
02745 - -
02750 - -
02755 HMM04199cycl_1(pf04199) BUF1
02760 -pf00528 -
02765 -pf00528 -
02770 -pf00497 -
02775 -pf00005 -
02780 -pf01235 -
02795 HMMreg13411_1_GRM1(pf06445pf13411) GRM1B
02800 - -
02805 - -
02810 Ts_dust GRM1B
02815 HMMalddh_2_egra(pf00171) GRM1B
02820 HMMgre_1_grms(pf01228pf02901) GRM1B
02825 HMMgreae_1_grms(pf13187pf04055) GRM1B
02830 H_bubblegum GRM1B
02835 HMMalddh_2_egra(pf00171) GRM1B EP_CTERM
02840 H_fuchsia EUT2A
02845 HMMptac_3_eut2(pf06130) GRM1B
02850 HMM06723EutJMreB_1_grm GRM1B
02855 HMM02441_03 GRM1B EP_NTERM
02860 P_peacockBlue GRM1B
02865 Ts_pinkishTan GRM1B
02870 HMMnopfamgrm1b_1_grm1b GRM1B EP_NTERM
02875 H_fuchsia GRM1B
02880 HMM00465alcdh_3_mix(pf00465) GRM1B
02885 HMMPduS_1_all(pf01512pf13375pf10531) GRM1B
02890 Ts_frogGreen GRM1B
02895 Hp_euts_plumPurple GRM1B
02900 HMM10662PduV_1_mix(pf10662) GRM1B
02905 -pf07589 -
02910 -pf07589 -
02915 - -
02920 - -
02925 -pf08447pf08448 -
02930 - -
02935 - -
02940 HMMreg00072_4_pdu1d(pf12625pf12833) PDU1D
02945 -pf03595 -
02950 -pf00440 -
02955 -pf04055 -
02960 HMMalddh_3_aromic1(pf001
BMC-P
BMC-H
BMC-T
Aldehyde dehydrogenase
PTAC
Signature enzyme
Alcohol dehydrogenase
Regulator

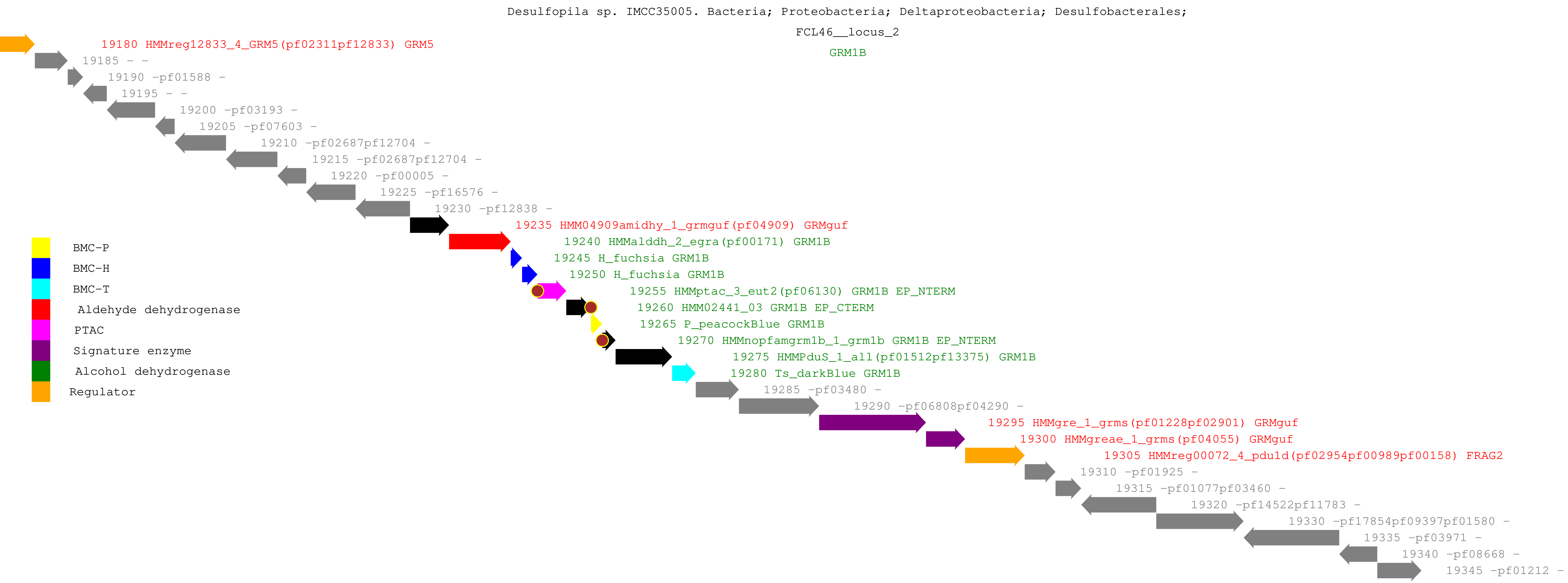
Desulfopila sp. IMCC35005. Bacteria; Proteobacteria; Deltaproteobacteria; Desulfobacterales;
FCL46__locus_2
GRM1B
19180 HMMreg12833_4_GRM5(pf02311pf12833) GRM5
19185 - -
19190 -pf01588 -
19195 - -
19200 -pf03193 -
19205 -pf07603 -
19210 -pf02687pf12704 -
19215 -pf02687pf12704 -
19220 -pf00005 -
19225 -pf16576 -
19230 -pf12838 -
19235 HMM04909amidhy_1_grmguf(pf04909) GRMguf
19240 HMMalddh_2_egra(pf00171) GRM1B
19245 H_fuchsia GRM1B
19250 H_fuchsia GRM1B
19255 HMMptac_3_eut2(pf06130) GRM1B EP_NTERM
19260 HMM02441_03 GRM1B EP_CTERM
19265 P_peacockBlue GRM1B
19270 HMMnopfamgrm1b_1_grm1b GRM1B EP_NTERM
19275 HMMPduS_1_all(pf01512pf13375) GRM1B
19280 Ts_darkBlue GRM1B
19285 -pf03480 -
19290 -pf06808pf04290 -
19295 HMMgre_1_grms(pf01228pf02901) GRMguf
19300 HMMgreae_1_grms(pf04055) GRMguf
19305 HMMreg00072_4_pduld(pf02954pf00989pf00158) FRAG2
19310 -pf01925 -
19315 -pf01077pf03460 -
19320 -pf14522pf11783 -
19330 -pf17854pf09397pf01580 -
19335 -pf03971 -
19340 -pf08668 -
19345 -pf01212 -
BMC-P
BMC-H
BMC-T
Aldehyde dehydrogenase
PTAC
Signature enzyme
Alcohol dehydrogenase
Regulator

Desulforhopalus sp. IMCC35007. Bacteria; Proteobacteria; Deltaproteobacteria; Desulfobacterales;

FCL48__locus_1

GRM1B

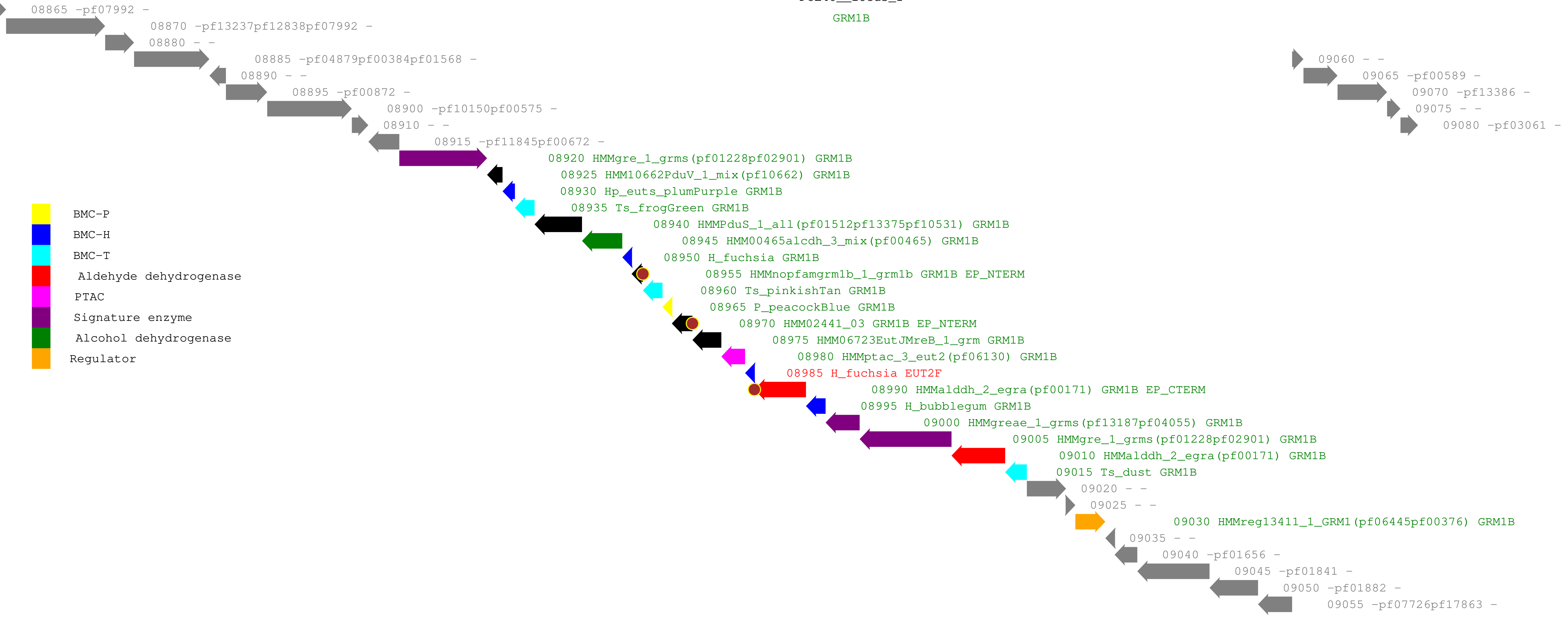

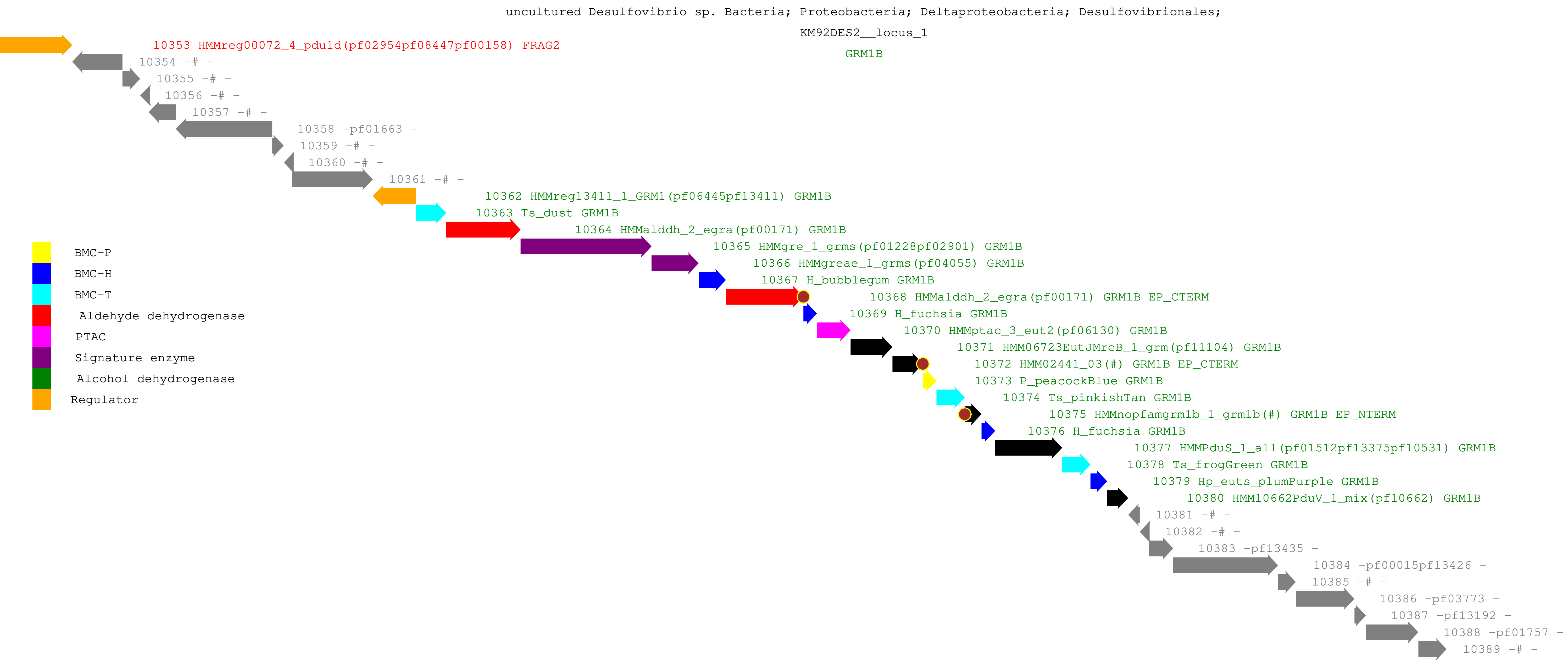
uncultured Desulfovibrio sp. Bacteria; Proteobacteria; Deltaproteobacteria; Desulfovibrionales;
KM92DES2__locus_1
GRM1B
10353 HMMreg00072_4_pdu1d(pf02954pf08447pf00158) FRAG2
10354 -# -
10355 -# -
10356 -# -
10357 -# -
10358 -pf01663 -
10359 -# -
10360 -# -
10361 -# -
10362 HMMreg13411_1_GRM1(pf06445pf13411) GRM1B
10363 Ts_dust GRM1B
10364 HMMalddh_2_egra(pf00171) GRM1B
10365 HMMgre_1_grms(pf01228pf02901) GRM1B
10366 HMMgreae_1_grms(pf04055) GRM1B
10367 H_bubblegum GRM1B
10368 HMMalddh_2_egra(pf00171) GRM1B EP_CTERM
10369 H_fuchsia GRM1B
10370 HMMptac_3_eut2(pf06130) GRM1B
10371 HMM06723EutJMreB_1_grm(pf11104) GRM1B
10372 HMM02441_03(#) GRM1B EP_CTERM
10373 P_peacockBlue GRM1B
10374 Ts_pinkishTan GRM1B
10375 HMMnopfamgrm1b_1_grm1b(#) GRM1B EP_NTERM
10376 H_fuchsia GRM1B
10377 HMMPduS_1_all(pf01512pf13375pf10531) GRM1B
10378 Ts_frogGreen GRM1B
10379 Hp_euts_plumPurple GRM1B
10380 HMM10662PduV_1_mix(pf10662) GRM1B
10381 -# -
10382 -# -
10383 -pf13435 -
10384 -pf00015pf13426 -
10385 -# -
10386 -pf03773 -
10387 -pf13192 -
10388 -pf01757 -
10389 -# -
BMC-P
BMC-H
BMC-T
Aldehyde dehydrogenase
PTAC
Signature enzyme
Alcohol dehydrogenase
Regulator

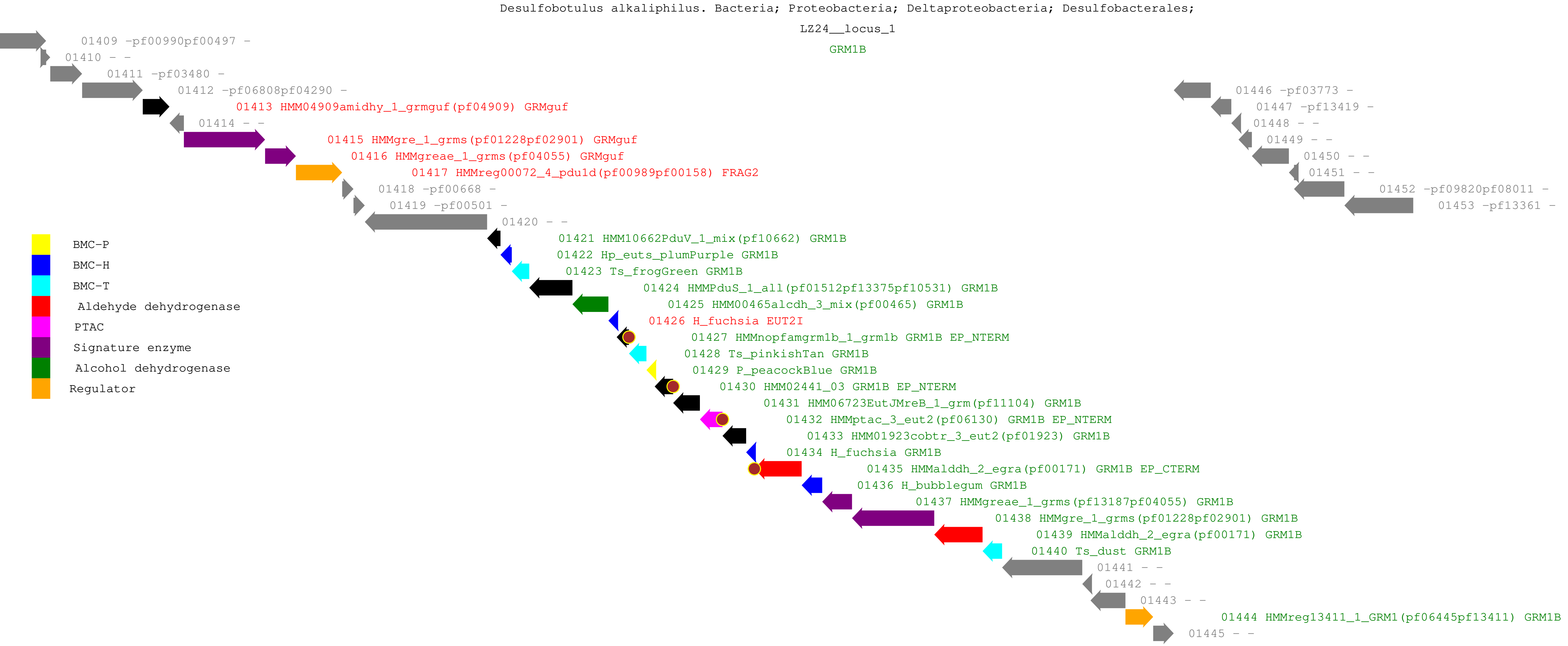
Desulfobotulus alkaliphilus. Bacteria; Proteobacteria; Deltaproteobacteria; Desulfobacterales;
LZ24__locus_1
GRM1B
01409 -pf00990pf00497 -
01410 - -
01411 -pf03480 -
01412 -pf06808pf04290 -
01413 HMM04909amidhy_1_grmguf(pf04909) GRMguf
01414 - -
01415 HMMgre_1_grms(pf01228pf02901) GRMguf
01416 HMMgreae_1_grms(pf04055) GRMguf
01417 HMMreg00072_4_pdu1d(pf00989pf00158) FRAG2
01418 -pf00668 -
01419 -pf00501 -
01420 - -
01421 HMM10662PduV_1_mix(pf10662) GRM1B
01422 Hp_euts_plumPurple GRM1B
01423 Ts_frogGreen GRM1B
01424 HMMPduS_1_all(pf01512pf13375pf10531) GRM1B
01425 HMM00465alcdh_3_mix(pf00465) GRM1B
01426 H_fuchsia EUT2I
01427 HMMnopfamgrm1b_1_grm1b GRM1B EP_NTERM
01428 Ts_pinkishTan GRM1B
01429 P_peacockBlue GRM1B
01430 HMM02441_03 GRM1B EP_NTERM
01431 HMM06723EutJMreB_1_grm(pf11104) GRM1B
01432 HMMptac_3_eut2(pf06130) GRM1B EP_NTERM
01433 HMM01923cobtr_3_eut2(pf01923) GRM1B
01434 H_fuchsia GRM1B
01435 HMMaldh_2_egra(pf00171) GRM1B EP_CTERM
01436 H_bubblegum GRM1B
01437 HMMgreae_1_grms(pf13187pf04055) GRM1B
01438 HMMgre_1_grms(pf01228pf02901) GRM1B
01439 HMMaldh_2_egra(pf00171) GRM1B
01440 Ts_dust GRM1B
01441 - -
01442 - -
01443 - -
01444 HMMreg13411_1_GRM1(pf06445pf13411) GRM1B
01445 - -
01446 -pf03773 -
01447 -pf13419 -
01448 - -
01449 - -
01450 - -
01451 - -
01452 -pf09820pf08011 -
01453 -pf13361 -
BMC-P
BMC-H
BMC-T
Aldehyde dehydrogenase
PTAC
Signature enzyme
Alcohol dehydrogenase
Regulator

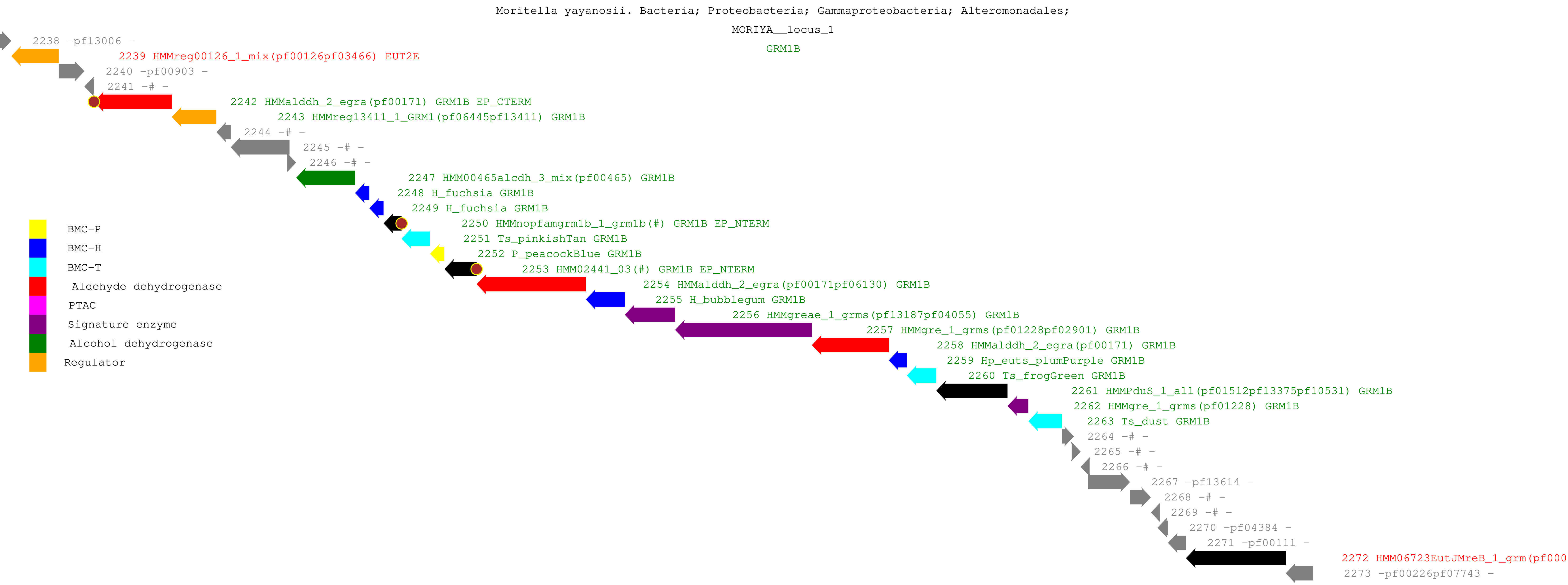

Moritella yayanosii. Bacteria; Proteobacteria; Gammaproteobacteria; Alteromonadales;
MORIYA__locus_1
GRM1B
2238 -pf13006 -
2239 HMMreg00126_1_mix(pf00126pf03466) EUT2E
2240 -pf00903 -
2241 -# -
2242 HMMaldh_2_egra(pf00171) GRM1B EP_CTERM
2243 HMMreg13411_1_GRM1(pf06445pf13411) GRM1B
2244 -# -
2245 -# -
2246 -# -
2247 HMM00465alcdh_3_mix(pf00465) GRM1B
2248 H_fuchsia GRM1B
2249 H_fuchsia GRM1B
2250 HMMnopfamgrm1b_1_grm1b(#) GRM1B EP_NTERM
2251 Ts_pinkishTan GRM1B
2252 P_peacockBlue GRM1B
2253 HMM02441_03(#) GRM1B EP_NTERM
2254 HMMaldh_2_egra(pf00171pf06130) GRM1B
2255 H_bubblegum GRM1B
2256 HMMgreae_1_grms(pf13187pf04055) GRM1B
2257 HMMgre_1_grms(pf01228pf02901) GRM1B
2258 HMMaldh_2_egra(pf00171) GRM1B
2259 Hp_euts_plumPurple GRM1B
2260 Ts_frogGreen GRM1B
2261 HMMPduS_1_all(pf01512pf13375pf10531) GRM1B
2262 HMMgre_1_grms(pf01228) GRM1B
2263 Ts_dust GRM1B
2264 -# -
2265 -# -
2266 -# -
2267 -pf13614 -
2268 -# -
2269 -# -
2270 -pf04384 -
2271 -pf00111 -
2272 HMM06723EutJMreB_1_grm(pf000
2273 -pf00226pf07743 -
BMC-P
BMC-H
BMC-T
Aldehyde dehydrogenase
PTAC
Signature enzyme
Alcohol dehydrogenase
Regulator

Desulfoluna butyratoxydans. Bacteria; Proteobacteria; Deltaproteobacteria; Desulfobacterales;

MSL71__locus_1

GRM1B

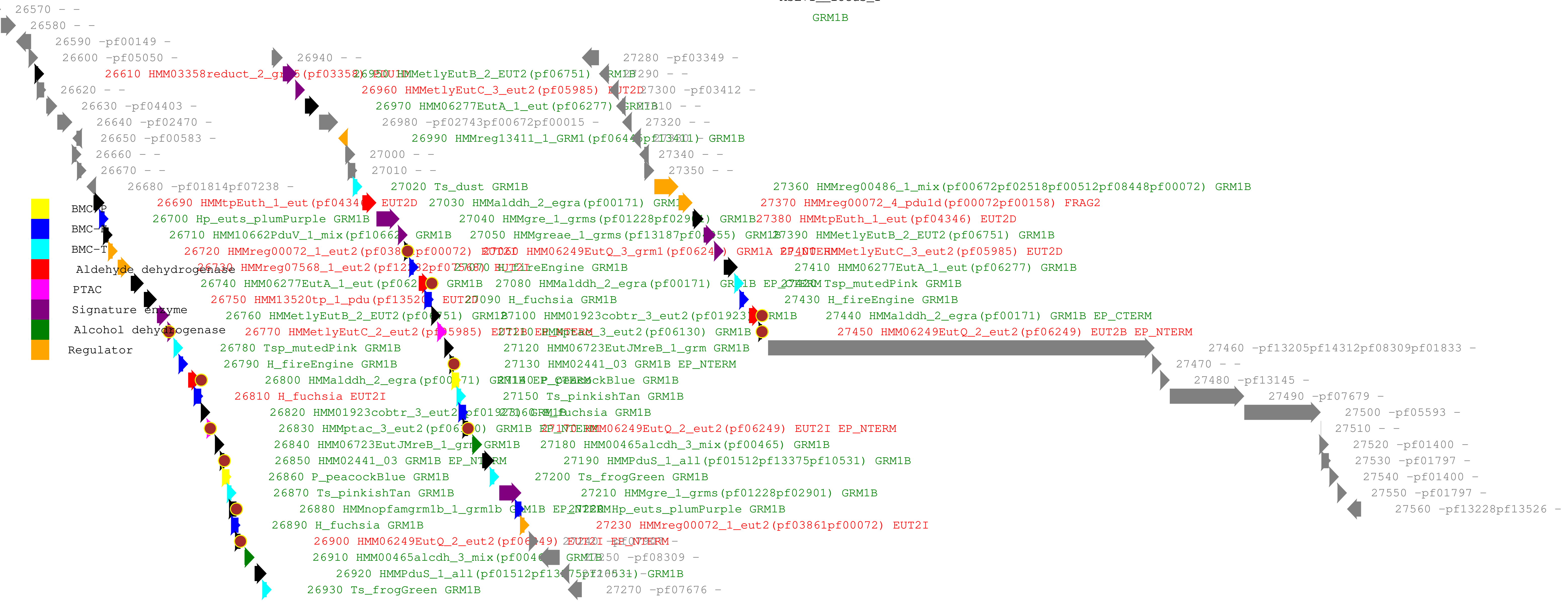

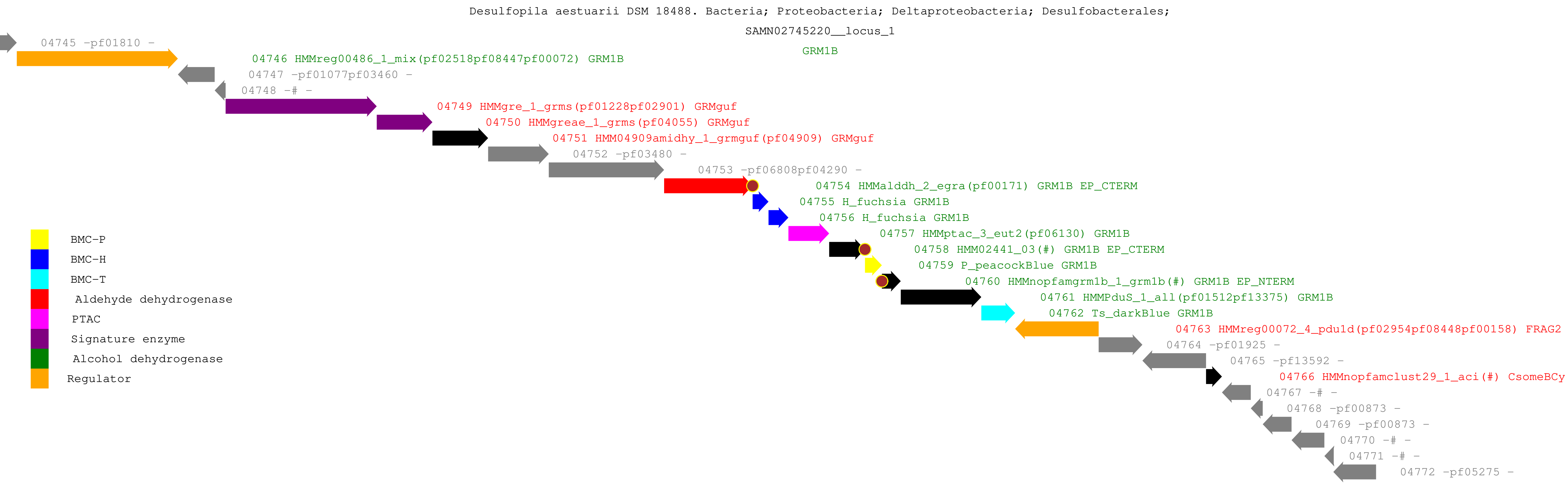
Desulfopila aestuarii DSM 18488. Bacteria; Proteobacteria; Deltaproteobacteria; Desulfobacterales;
SAMN02745220__locus_1
GRM1B
04745 -pf01810 -
04746 HMMreg00486_1_mix(pf02518pf08447pf00072) GRM1B
04747 -pf01077pf03460 -
04748 -# -
04749 HMMgre_1_grms(pf01228pf02901) GRMguf
04750 HMMgreae_1_grms(pf04055) GRMguf
04751 HMM04909amidhy_1_grmguf(pf04909) GRMguf
04752 -pf03480 -
04753 -pf06808pf04290 -
04754 HMMalddh_2_egra(pf00171) GRM1B EP_CTERM
04755 H_fuchsia GRM1B
04756 H_fuchsia GRM1B
04757 HMMptac_3_eut2(pf06130) GRM1B
04758 HMM02441_03(#) GRM1B EP_CTERM
04759 P_peacockBlue GRM1B
04760 HMMnopfamgrm1b_1_grm1b(#) GRM1B EP_NTERM
04761 HMMPduS_1_all(pf01512pf13375) GRM1B
04762 Ts_darkBlue GRM1B
04763 HMMreg00072_4_pduld(pf02954pf08448pf00158) FRAG2
04764 -pf01925 -
04765 -pf13592 -
04766 HMMnopfamclust29_1_aci(#) CsomeBCy
04767 -# -
04768 -pf00873 -
04769 -pf00873 -
04770 -# -
04771 -# -
04772 -pf05275 -
BMC-P
BMC-H
BMC-T
Aldehyde dehydrogenase
PTAC
Signature enzyme
Alcohol dehydrogenase
Regulator

Desulfovibrio bizertensis DSM 18034. Bacteria; Proteobacteria; Deltaproteobacteria; Desulfovibrionales;

SAMN02745702__locus_1

GRM1B

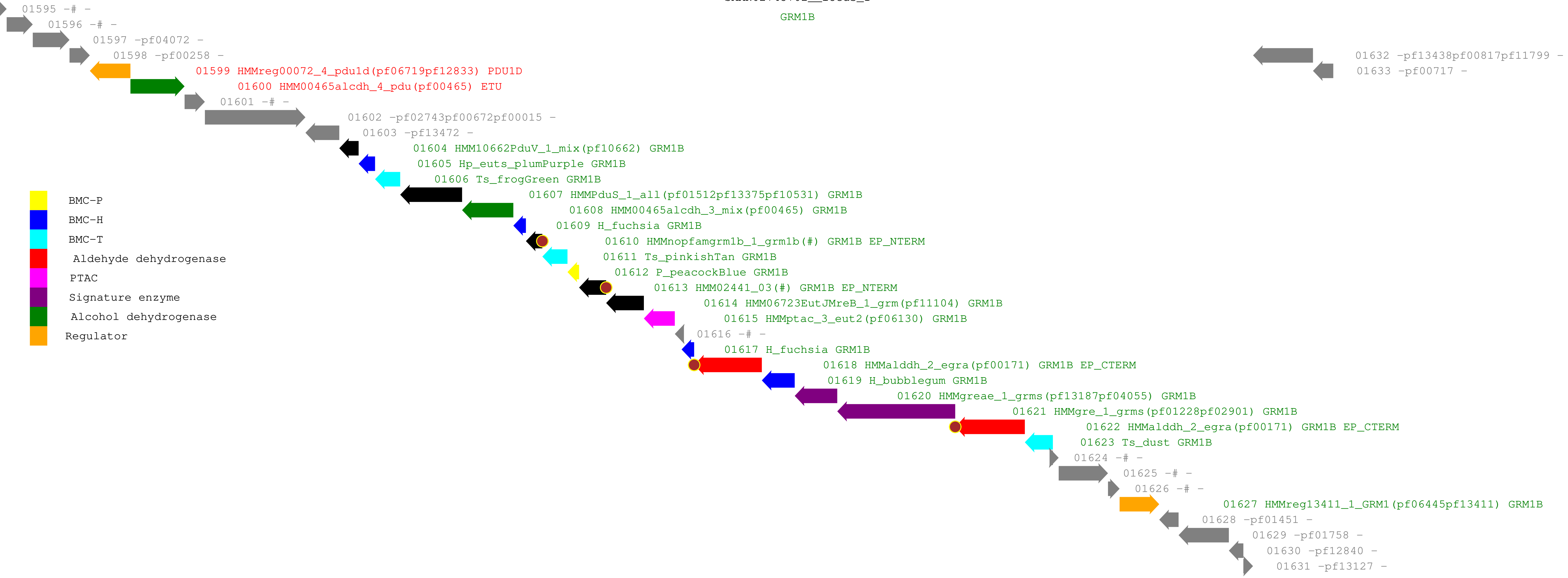

Desulfovibrio desulfuricans. Bacteria; Proteobacteria; Deltaproteobacteria; Desulfovibrionales;

SAMN02910291__locus_1

GRM1B

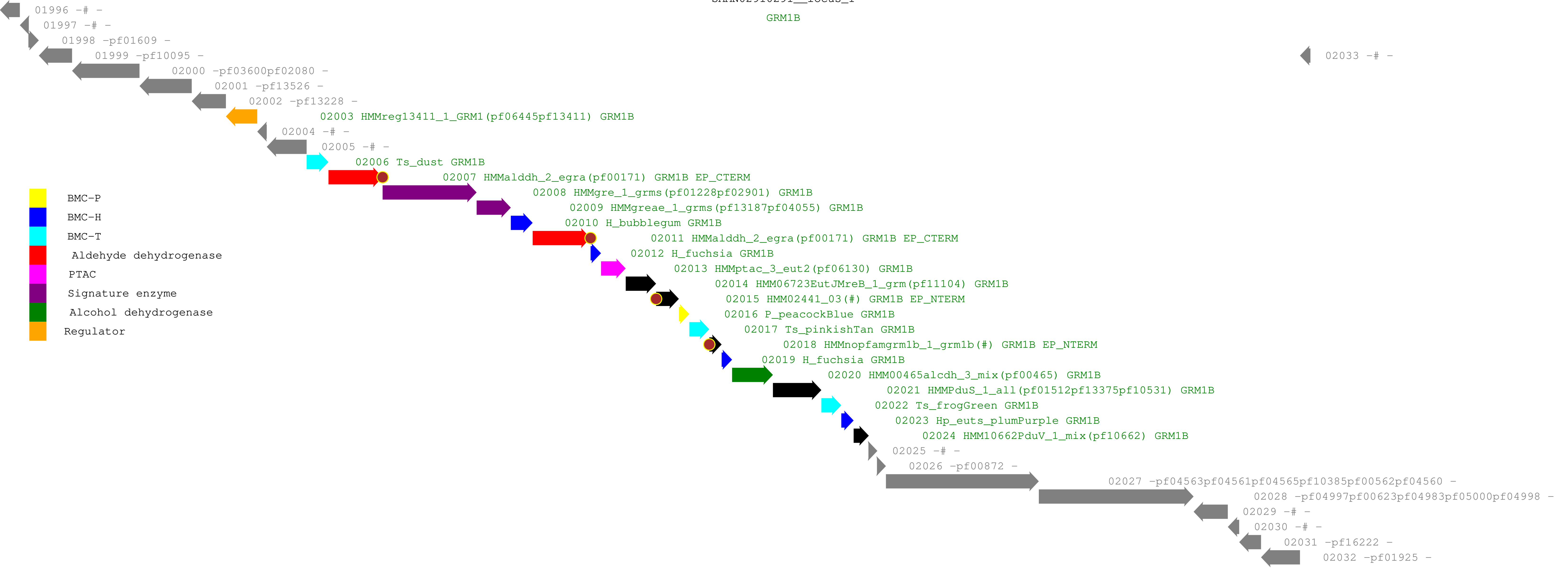

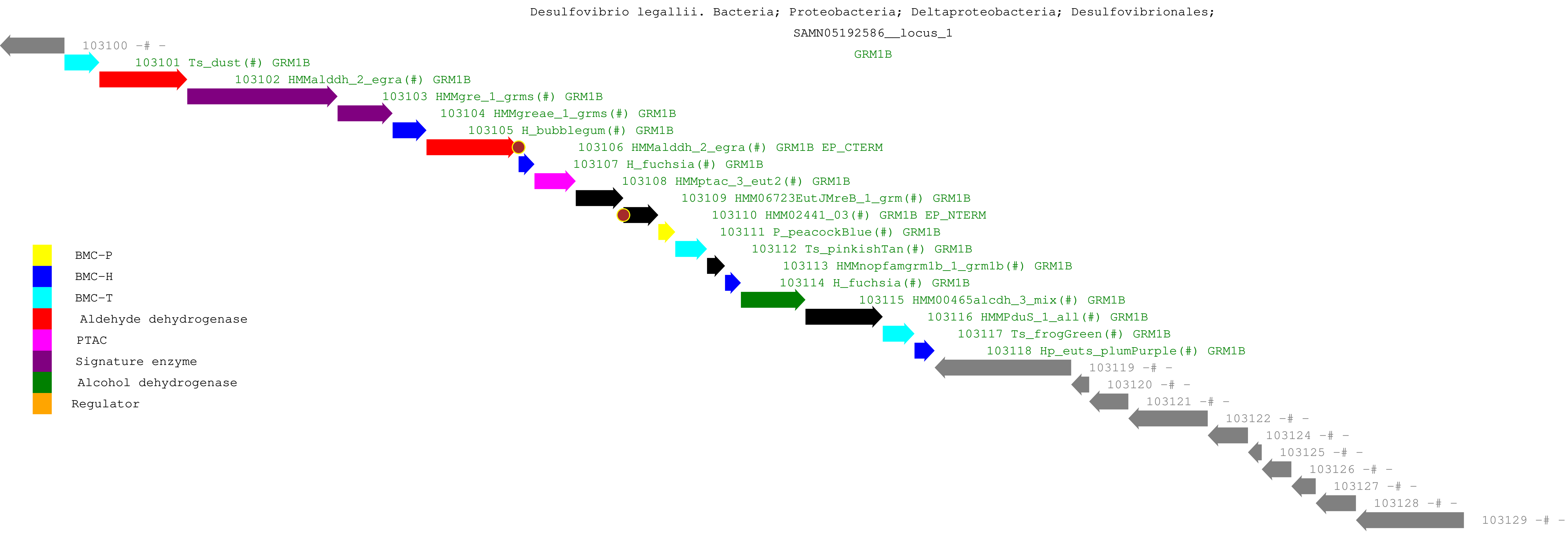
Desulfovibrio legallii. Bacteria; Proteobacteria; Deltaproteobacteria; Desulfovibrionales;
SAMN05192586__locus_1
GRM1B
103100 -# -
103101 Ts_dust(#) GRM1B
103102 HMMalddh_2_egra(#) GRM1B
103103 HMMgre_1_grms(#) GRM1B
103104 HMMgreae_1_grms(#) GRM1B
103105 H_bubblegum(#) GRM1B
103106 HMMalddh_2_egra(#) GRM1B EP_CTERM
103107 H_fuchsia(#) GRM1B
103108 HMMptac_3_eut2(#) GRM1B
103109 HMM06723EutJMreB_1_grm(#) GRM1B
103110 HMM02441_03(#) GRM1B EP_NTERM
103111 P_peacockBlue(#) GRM1B
103112 Ts_pinkishTan(#) GRM1B
103113 HMMnopfamgrm1b_1_grm1b(#) GRM1B
103114 H_fuchsia(#) GRM1B
103115 HMM00465alcdh_3_mix(#) GRM1B
103116 HMMPduS_1_all(#) GRM1B
103117 Ts_frogGreen(#) GRM1B
103118 Hp_euts_plumPurple(#) GRM1B
103119 -# -
103120 -# -
103121 -# -
103122 -# -
103124 -# -
103125 -# -
103126 -# -
103127 -# -
103128 -# -
103129 -# -
BMC-P
BMC-H
BMC-T
Aldehyde dehydrogenase
PTAC
Signature enzyme
Alcohol dehydrogenase
Regulator

Desulfoluna spongiiphila. Bacteria; Proteobacteria; Deltaproteobacteria; Desulfobacterales;

SAMN05216233__locus_1

GRM1B

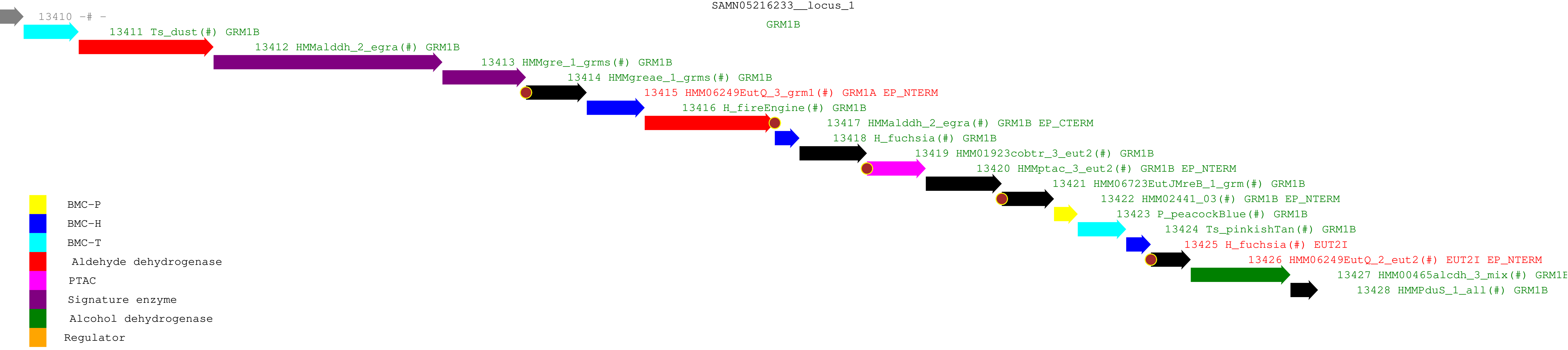

Desulfovibrio sp. K3S. Bacteria; Proteobacteria; Deltaproteobacteria; Desulfovibrionales;

SAMN06295933__locus_1

GRM1B

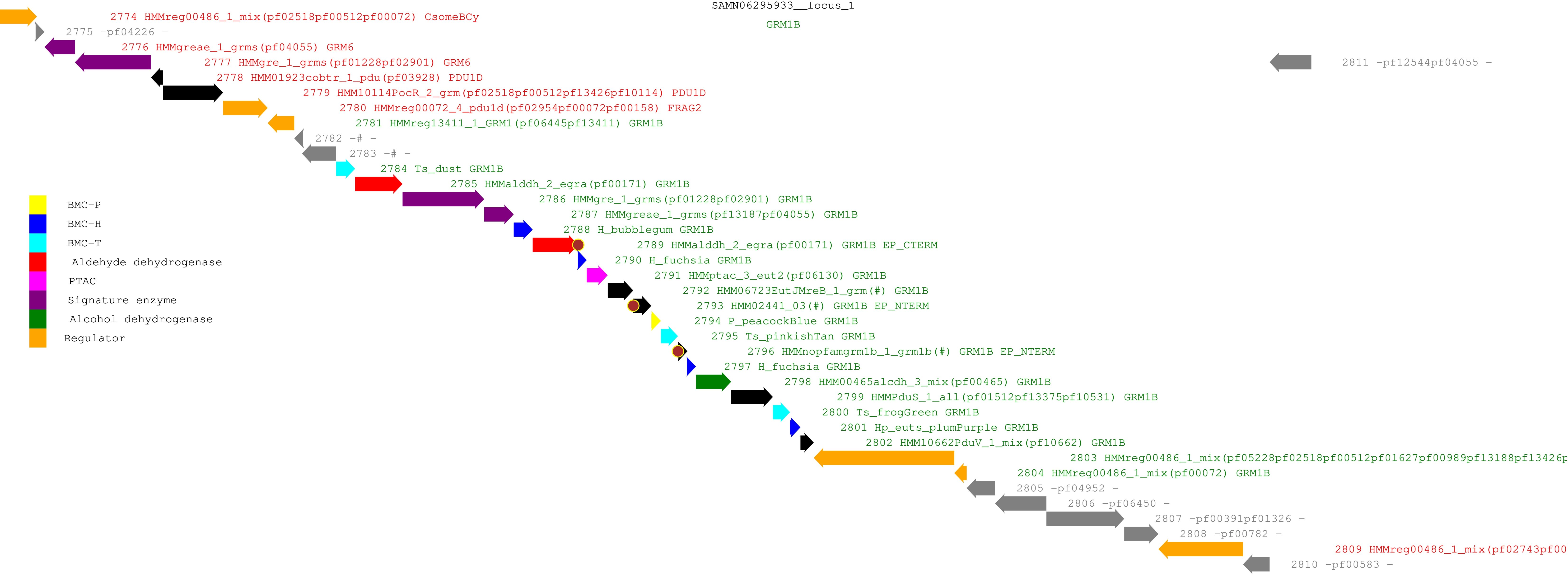

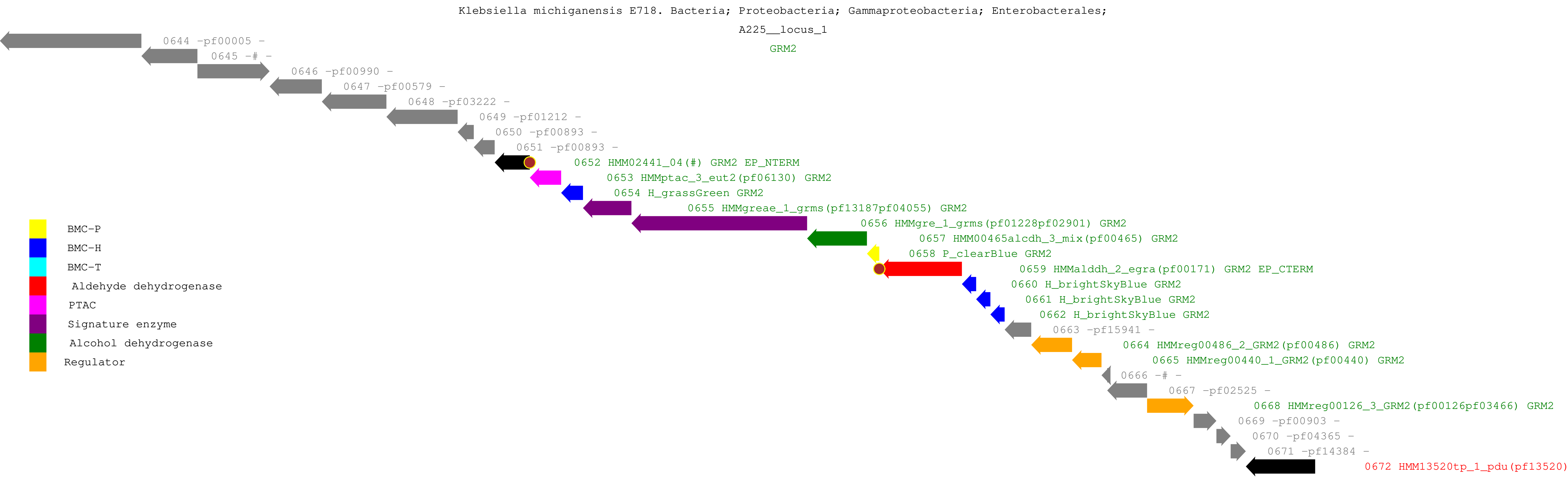

Klebsiella michiganensis E718. Bacteria; Proteobacteria; Gammaproteobacteria; Enterobacterales;
A225__locus_1
GRM2
0644 -pf00005 -
0645 -# -
0646 -pf00990 -
0647 -pf00579 -
0648 -pf03222 -
0649 -pf01212 -
0650 -pf00893 -
0651 -pf00893 -
0652 HMM02441_04(#) GRM2 EP_NTERM
0653 HMMptac_3_eut2(pf06130) GRM2
0654 H_grassGreen GRM2
0655 HMMgreae_1_grms(pf13187pf04055) GRM2
0656 HMMgre_1_grms(pf01228pf02901) GRM2
0657 HMM00465alcdh_3_mix(pf00465) GRM2
0658 P_clearBlue GRM2
0659 HMMalddh_2_egra(pf00171) GRM2 EP_CTERM
0660 H_brightSkyBlue GRM2
0661 H_brightSkyBlue GRM2
0662 H_brightSkyBlue GRM2
0663 -pf15941 -
0664 HMMreg00486_2_GRM2(pf00486) GRM2
0665 HMMreg00440_1_GRM2(pf00440) GRM2
0666 -# -
0667 -pf02525 -
0668 HMMreg00126_3_GRM2(pf00126pf03466) GRM2
0669 -pf00903 -
0670 -pf04365 -
0671 -pf14384 -
0672 HMM13520tp_1_pdu(pf13520)
BMC-P
BMC-H
BMC-T
Aldehyde dehydrogenase
PTAC
Signature enzyme
Alcohol dehydrogenase
Regulator

Photobacterium jeanii. Bacteria; Proteobacteria; Gammaproteobacteria; Vibrionales;

A3K86__locus_1

GRM2

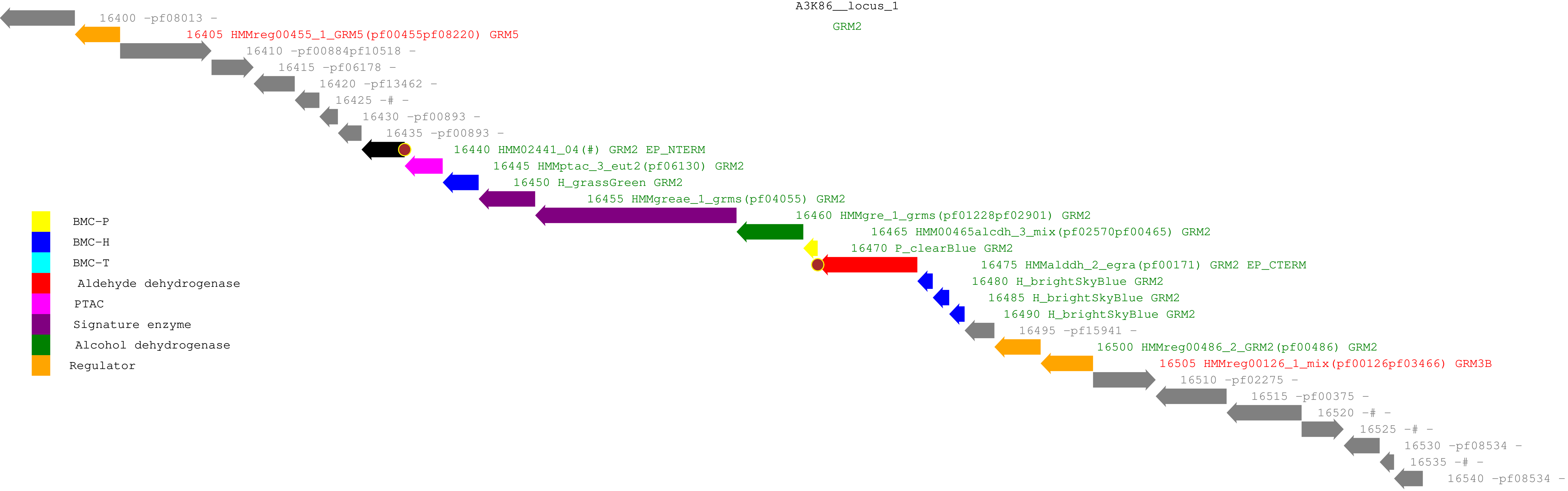

Escherichia coli. Bacteria; Proteobacteria; Gammaproteobacteria; Enterobacterales;

A4T50__locus_2

GRM2

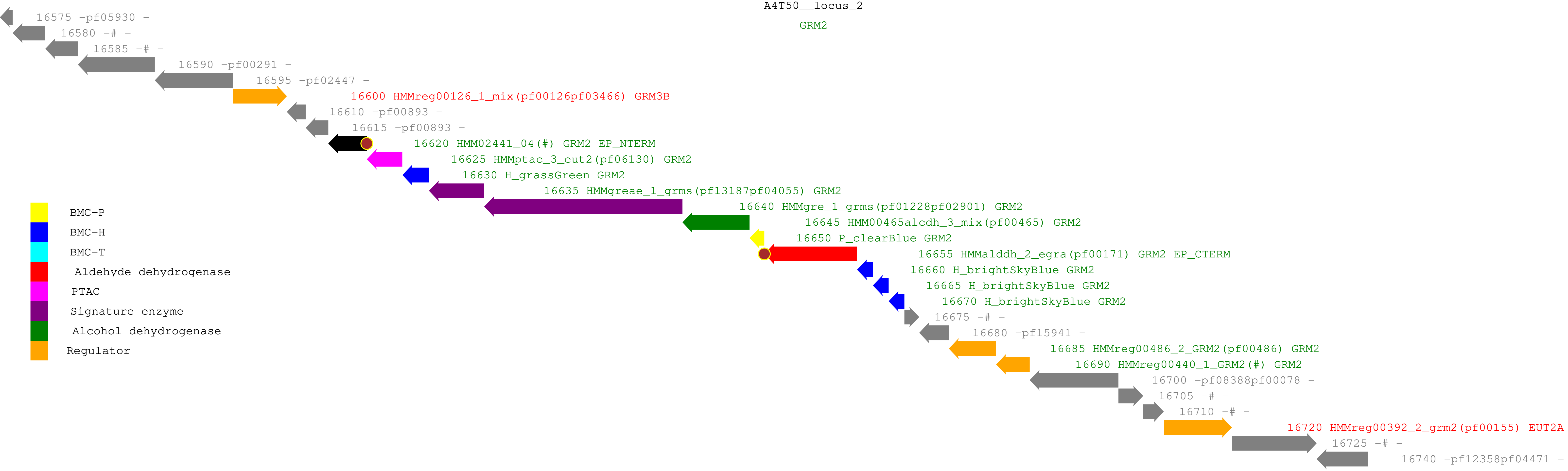

Klebsiella michiganensis. Bacteria; Proteobacteria; Gammaproteobacteria; Enterobacterales;

A6A30__locus_1

GRM2

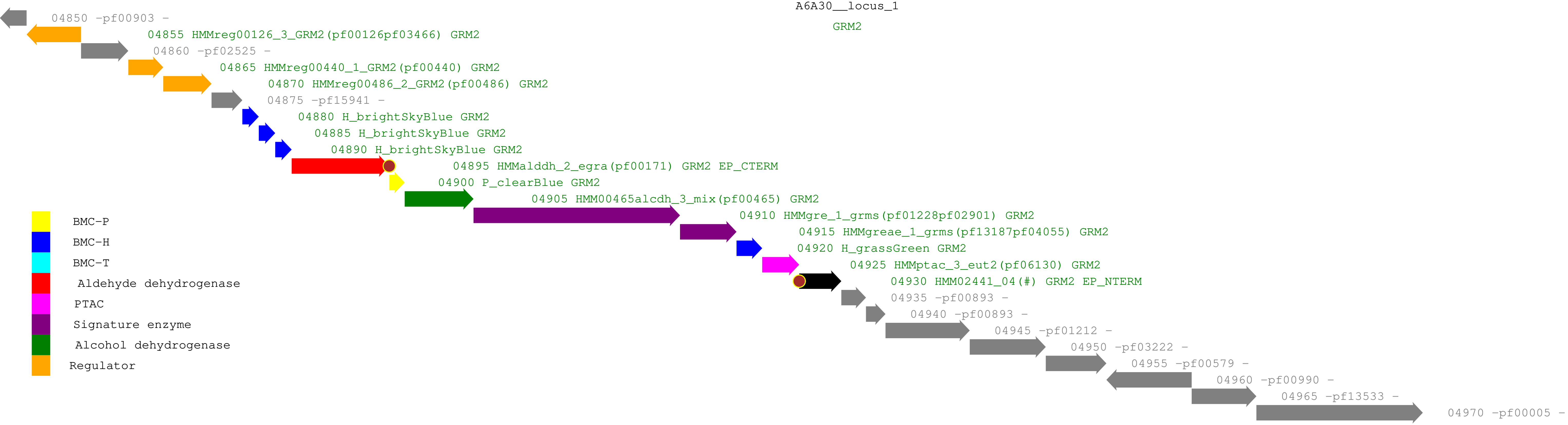

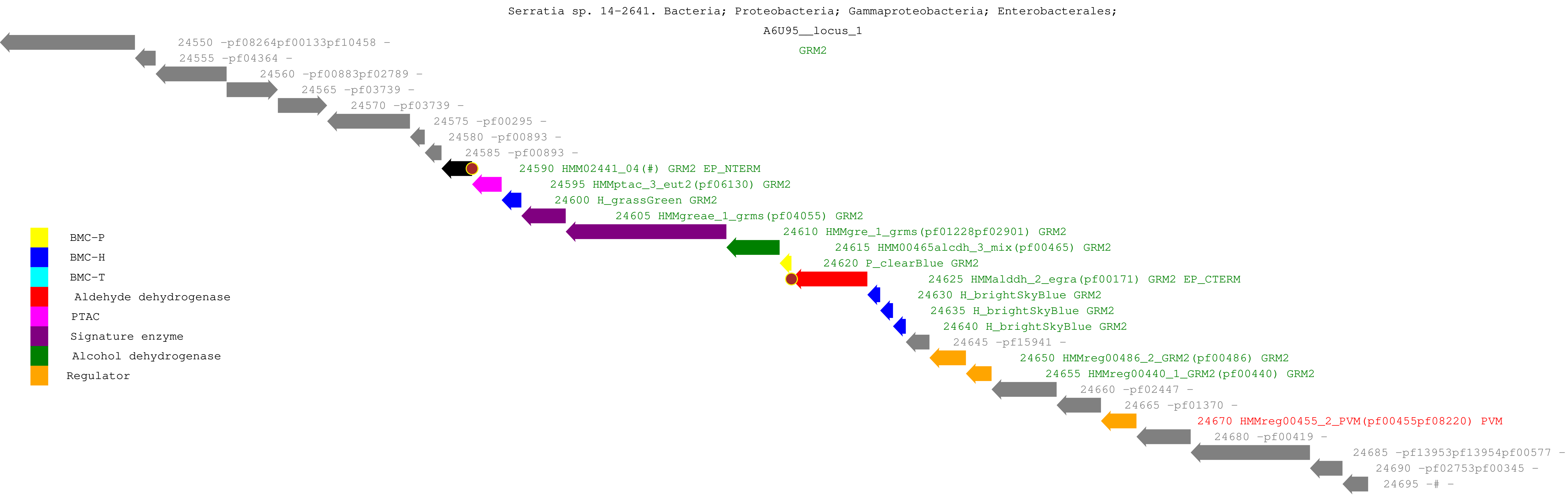
Serratia sp. 14-2641. Bacteria; Proteobacteria; Gammaproteobacteria; Enterobacterales;
A6U95__locus_1
GRM2
24550 -pf08264pf00133pf10458 -
24555 -pf04364 -
24560 -pf00883pf02789 -
24565 -pf03739 -
24570 -pf03739 -
24575 -pf00295 -
24580 -pf00893 -
24585 -pf00893 -
24590 HMM02441_04(#) GRM2 EP_NTERM
24595 HMMptac_3_eut2(pf06130) GRM2
24600 H_grassGreen GRM2
24605 HMMgreae_1_grms(pf04055) GRM2
24610 HMMgre_1_grms(pf01228pf02901) GRM2
24615 HMM00465alcdh_3_mix(pf00465) GRM2
24620 P_clearBlue GRM2
24625 HMMalddh_2_egra(pf00171) GRM2 EP_CTERM
24630 H_brightSkyBlue GRM2
24635 H_brightSkyBlue GRM2
24640 H_brightSkyBlue GRM2
24645 -pf15941 -
24650 HMMreg00486_2_GRM2(pf00486) GRM2
24655 HMMreg00440_1_GRM2(pf00440) GRM2
24660 -pf02447 -
24665 -pf01370 -
24670 HMMreg00455_2_PVM(pf00455pf08220) PVM
24680 -pf00419 -
24685 -pf13953pf13954pf00577 -
24690 -pf02753pf00345 -
24695 -# -
BMC-P
BMC-H
BMC-T
Aldehyde dehydrogenase
PTAC
Signature enzyme
Alcohol dehydrogenase
Regulator

Escherichia coli. Bacteria; Proteobacteria; Gammaproteobacteria; Enterobacterales;

A8M42__locus_2

GRM2

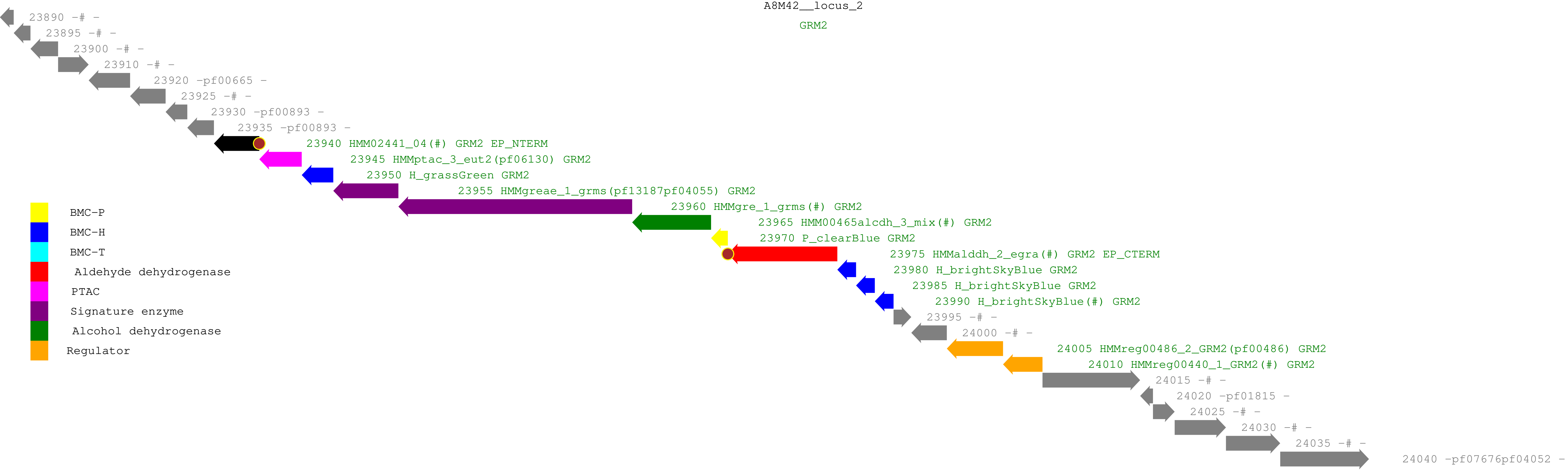

Aeromonas dhakensis. Bacteria; Proteobacteria; Gammaproteobacteria; Aeromonadales;

A9196__locus_1

GRM2

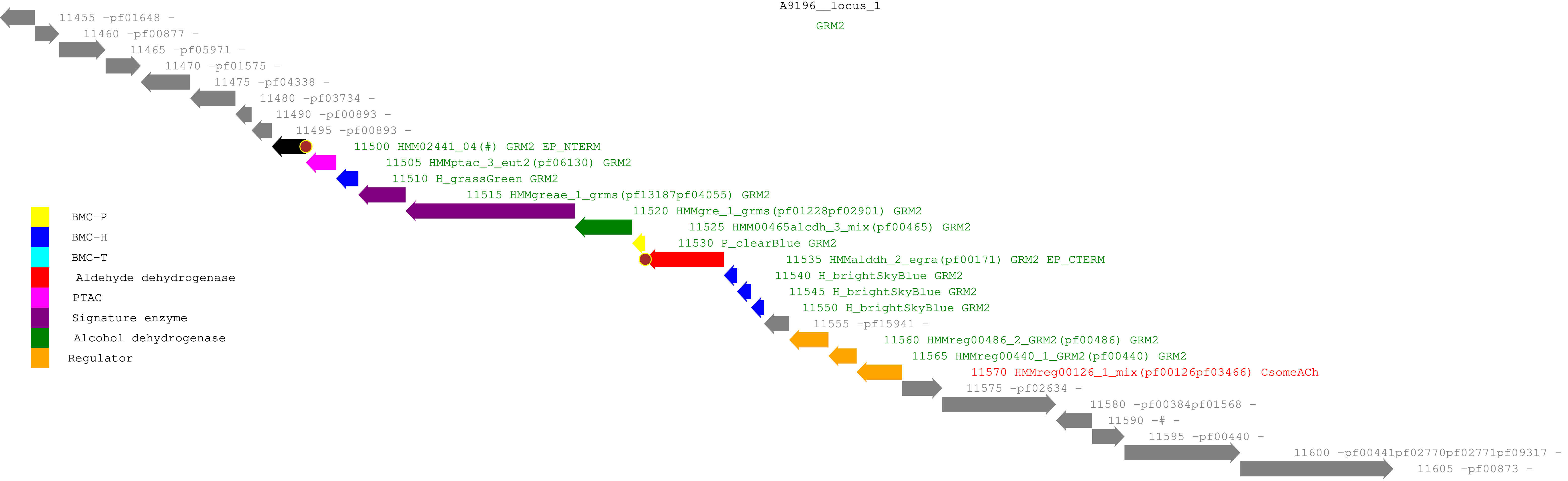

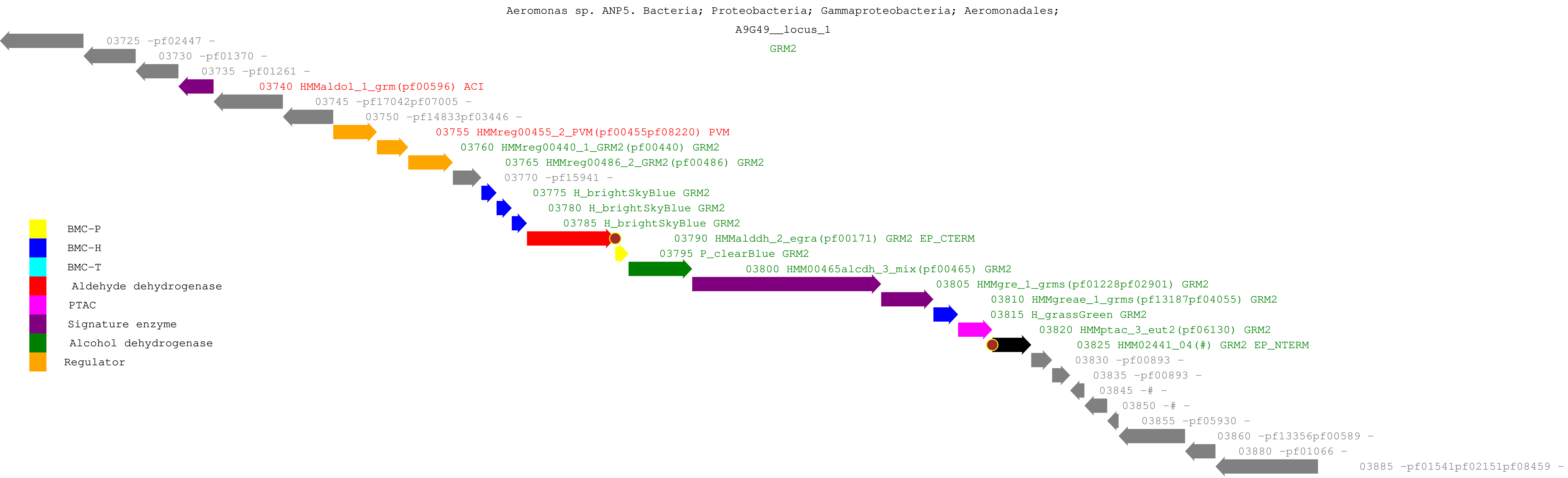
Aeromonas sp. ANP5. Bacteria; Proteobacteria; Gammaproteobacteria; Aeromonadales;
A9G49__locus_1
GRM2
03725 -pf02447 -
03730 -pf01370 -
03735 -pf01261 -
03740 HMMaldol_1_grm(pf00596) ACI
03745 -pf17042pf07005 -
03750 -pf14833pf03446 -
03755 HMMreg00455_2_PVM(pf00455pf08220) PVM
03760 HMMreg00440_1_GRM2(pf00440) GRM2
03765 HMMreg00486_2_GRM2(pf00486) GRM2
03770 -pf15941 -
03775 H_brightSkyBlue GRM2
03780 H_brightSkyBlue GRM2
03785 H_brightSkyBlue GRM2
03790 HMMalddh_2_egra(pf00171) GRM2 EP_CTERM
03795 P_clearBlue GRM2
03800 HMM00465alcdh_3_mix(pf00465) GRM2
03805 HMMgre_1_grms(pf01228pf02901) GRM2
03810 HMMgreae_1_grms(pf13187pf04055) GRM2
03815 H_grassGreen GRM2
03820 HMMptac_3_eut2(pf06130) GRM2
03825 HMM02441_04(#) GRM2 EP_NTERM
03830 -pf00893 -
03835 -pf00893 -
03845 -# -
03850 -# -
03855 -pf05930 -
03860 -pf13356pf00589 -
03880 -pf01066 -
03885 -pf01541pf02151pf08459 -
BMC-P
BMC-H
BMC-T
Aldehyde dehydrogenase
PTAC
Signature enzyme
Alcohol dehydrogenase
Regulator

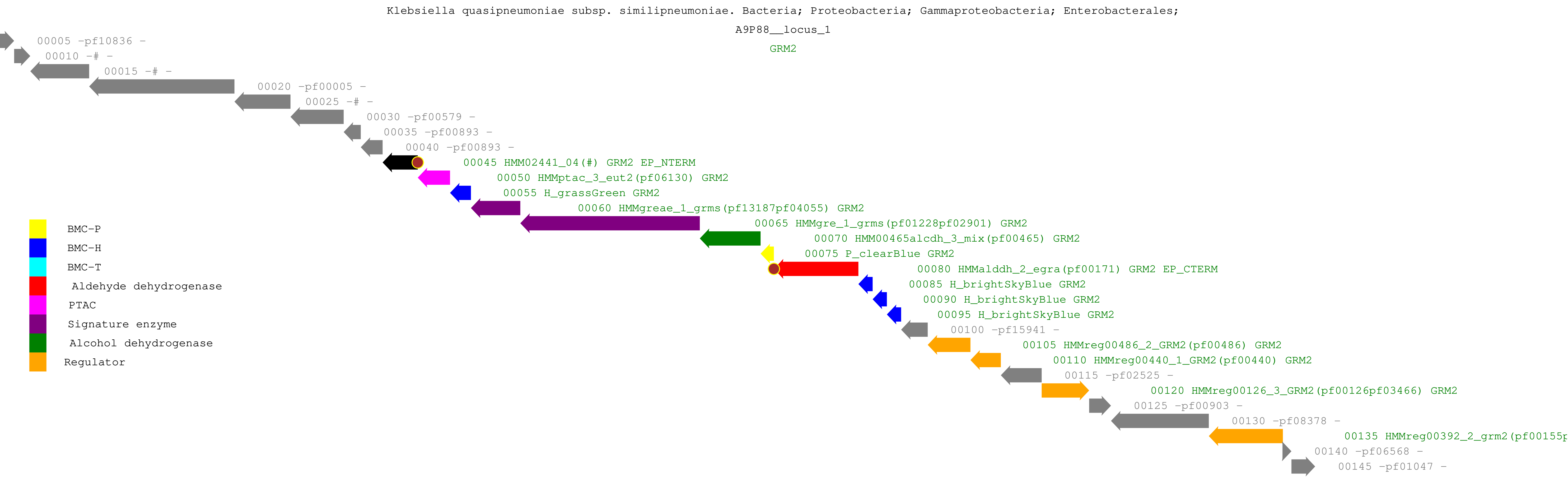

Klebsiella quasipneumoniae subsp. similipneumoniae. Bacteria; Proteobacteria; Gammaproteobacteria; Enterobacterales;
A9P88__locus_1
GRM2
00005 -pf10836 -
00010 -# -
00015 -# -
00020 -pf00005 -
00025 -# -
00030 -pf00579 -
00035 -pf00893 -
00040 -pf00893 -
00045 HMM02441_04(#) GRM2 EP_NTERM
00050 HMMptac_3_eut2(pf06130) GRM2
00055 H_grassGreen GRM2
00060 HMMgreae_1_grms(pf13187pf04055) GRM2
00065 HMMgre_1_grms(pf01228pf02901) GRM2
00070 HMM00465alcdh_3_mix(pf00465) GRM2
00075 P_clearBlue GRM2
00080 HMMaldh_2_egra(pf00171) GRM2 EP_CTERM
00085 H_brightSkyBlue GRM2
00090 H_brightSkyBlue GRM2
00095 H_brightSkyBlue GRM2
00100 -pf15941 -
00105 HMMreg00486_2_GRM2(pf00486) GRM2
00110 HMMreg00440_1_GRM2(pf00440) GRM2
00115 -pf02525 -
00120 HMMreg00126_3_GRM2(pf00126pf03466) GRM2
00125 -pf00903 -
00130 -pf08378 -
00135 HMMreg00392_2_grm2(pf00155p
00140 -pf06568 -
00145 -pf01047 -
BMC-P
BMC-H
BMC-T
Aldehyde dehydrogenase
PTAC
Signature enzyme
Alcohol dehydrogenase
Regulator

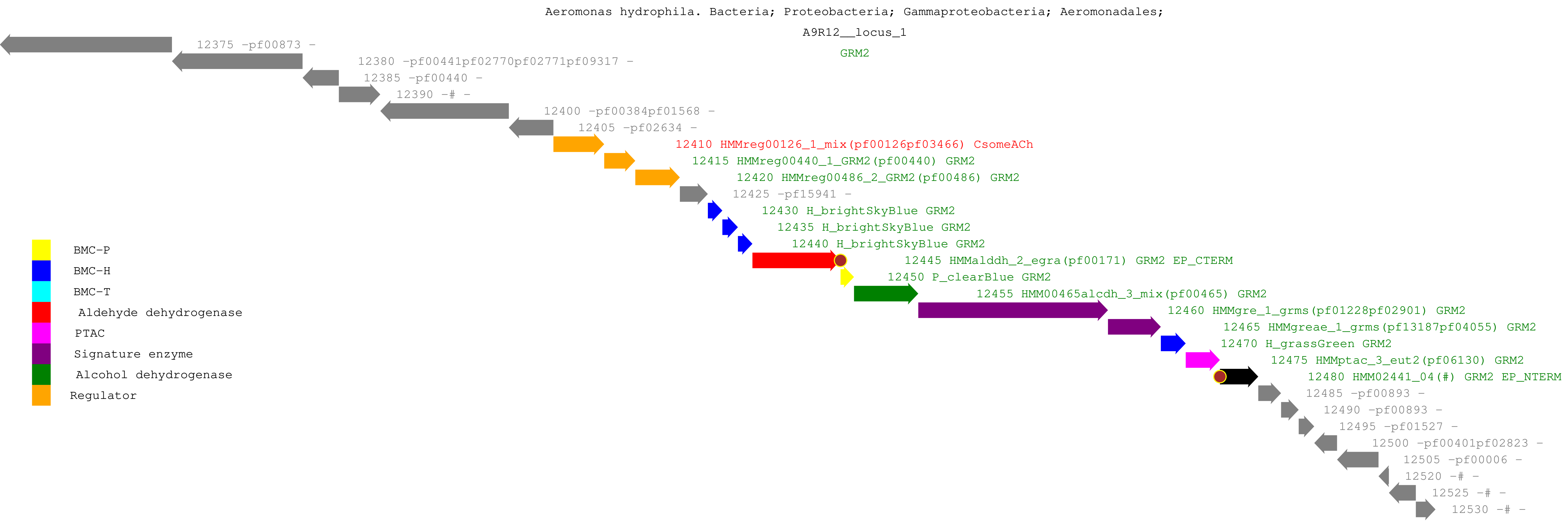
Aeromonas hydrophila. Bacteria; Proteobacteria; Gammaproteobacteria; Aeromonadales;
A9R12__locus_1
GRM2
12375 -pf00873 -
12380 -pf00441pf02770pf02771pf09317 -
12385 -pf00440 -
12390 -# -
12400 -pf00384pf01568 -
12405 -pf02634 -
12410 HMMreg00126_1_mix(pf00126pf03466) CsomeACh
12415 HMMreg00440_1_GRM2(pf00440) GRM2
12420 HMMreg00486_2_GRM2(pf00486) GRM2
12425 -pf15941 -
12430 H_brightSkyBlue GRM2
12435 H_brightSkyBlue GRM2
12440 H_brightSkyBlue GRM2
12445 HMMaldh_2_egra(pf00171) GRM2 EP_CTERM
12450 P_clearBlue GRM2
12455 HMM00465alcdh_3_mix(pf00465) GRM2
12460 HMMgre_1_grms(pf01228pf02901) GRM2
12465 HMMgreae_1_grms(pf13187pf04055) GRM2
12470 H_grassGreen GRM2
12475 HMMptac_3_eut2(pf06130) GRM2
12480 HMM02441_04(#) GRM2 EP_NTERM
12485 -pf00893 -
12490 -pf00893 -
12495 -pf01527 -
12500 -pf00401pf02823 -
12505 -pf00006 -
12520 -# -
12525 -# -
12530 -# -
BMC-P
BMC-H
BMC-T
Aldehyde dehydrogenase
PTAC
Signature enzyme
Alcohol dehydrogenase
Regulator

Providencia rettgeri. Bacteria; Proteobacteria; Gammaproteobacteria; Enterobacterales;

AAY77__locus_1

GRM2

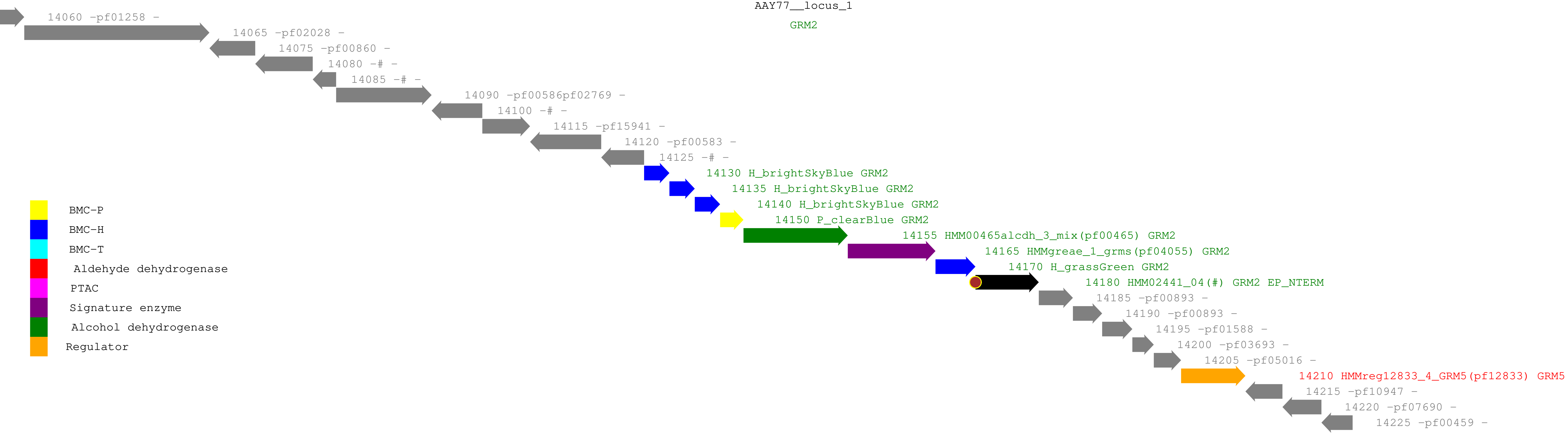

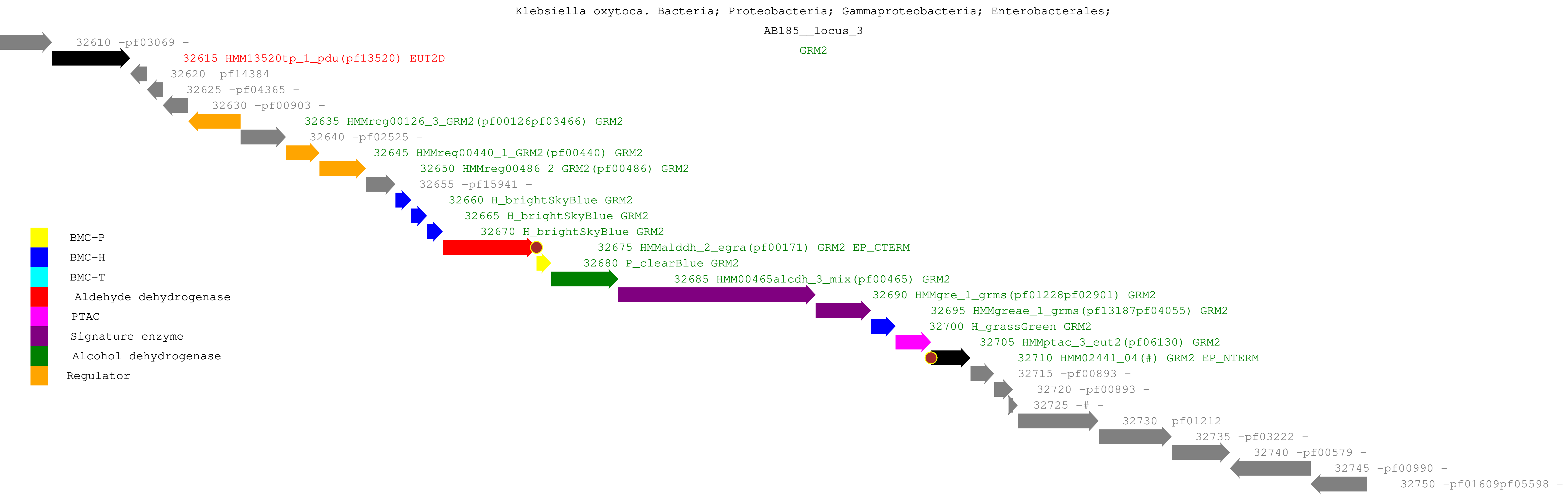
Klebsiella oxytoca. Bacteria; Proteobacteria; Gammaproteobacteria; Enterobacterales;
AB185__locus_3
GRM2
32610 -pf03069 -
32615 HMM13520tp_1_pdu(pf13520) EUT2D
32620 -pf14384 -
32625 -pf04365 -
32630 -pf00903 -
32635 HMMreg00126_3_GRM2(pf00126pf03466) GRM2
32640 -pf02525 -
32645 HMMreg00440_1_GRM2(pf00440) GRM2
32650 HMMreg00486_2_GRM2(pf00486) GRM2
32655 -pf15941 -
32660 H_brightSkyBlue GRM2
32665 H_brightSkyBlue GRM2
32670 H_brightSkyBlue GRM2
32675 HMMaldddh_2_egra(pf00171) GRM2 EP_CTERM
32680 P_clearBlue GRM2
32685 HMM00465alcdh_3_mix(pf00465) GRM2
32690 HMMgre_1_grms(pf01228pf02901) GRM2
32695 HMMgreae_1_grms(pf13187pf04055) GRM2
32700 H_grassGreen GRM2
32705 HMMptac_3_eut2(pf06130) GRM2
32710 HMM02441_04(#) GRM2 EP_NTERM
32715 -pf00893 -
32720 -pf00893 -
32725 -# -
32730 -pf01212 -
32735 -pf03222 -
32740 -pf00579 -
32745 -pf00990 -
32750 -pf01609pf05598 -
BMC-P
BMC-H
BMC-T
Aldehyde dehydrogenase
PTAC
Signature enzyme
Alcohol dehydrogenase
Regulator

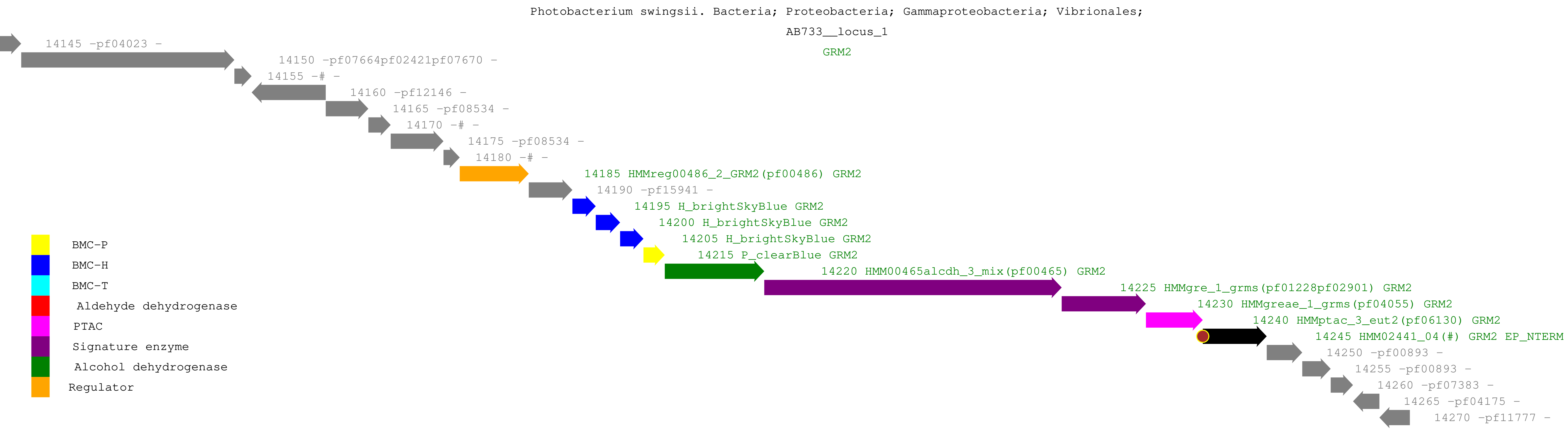
Photobacterium swingsii. Bacteria; Proteobacteria; Gammaproteobacteria; Vibrionales;
AB733__locus_1
GRM2
14145 -pf04023 -
14150 -pf07664pf02421pf07670 -
14155 -# -
14160 -pf12146 -
14165 -pf08534 -
14170 -# -
14175 -pf08534 -
14180 -# -
14185 HMMreg00486_2_GRM2(pf00486) GRM2
14190 -pf15941 -
14195 H_brightSkyBlue GRM2
14200 H_brightSkyBlue GRM2
14205 H_brightSkyBlue GRM2
14215 P_clearBlue GRM2
14220 HMM00465alcdh_3_mix(pf00465) GRM2
14225 HMMgre_1_grms(pf01228pf02901) GRM2
14230 HMMgreae_1_grms(pf04055) GRM2
14240 HMMptac_3_eut2(pf06130) GRM2
14245 HMM02441_04(#) GRM2 EP_NTERM
14250 -pf00893 -
14255 -pf00893 -
14260 -pf07383 -
14265 -pf04175 -
14270 -pf11777 -
BMC-P
BMC-H
BMC-T
Aldehyde dehydrogenase
PTAC
Signature enzyme
Alcohol dehydrogenase
Regulator

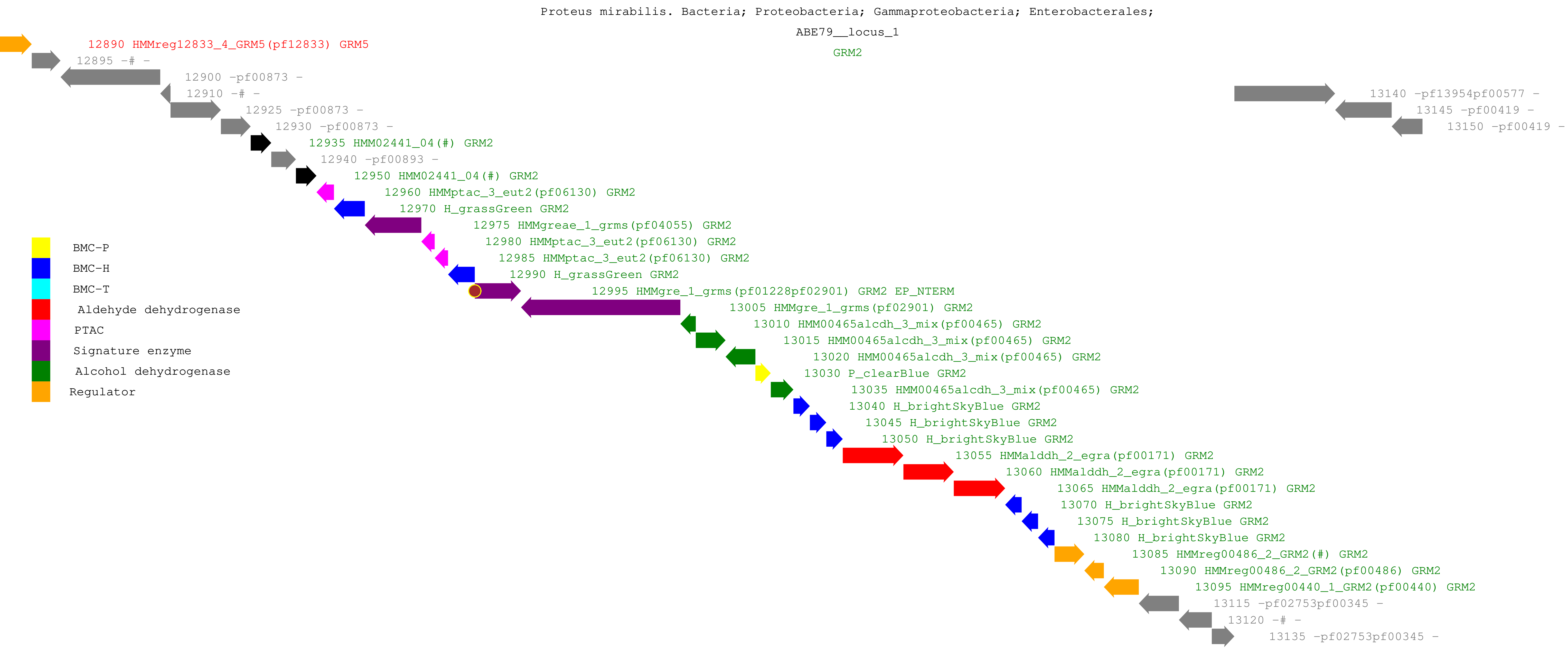
Proteus mirabilis. Bacteria; Proteobacteria; Gammaproteobacteria; Enterobacterales;
ABE79__locus_1
GRM2
12890 HMMreg12833_4_GRM5(pf12833) GRM5
12895 -# -
12900 -pf00873 -
12910 -# -
12925 -pf00873 -
12930 -pf00873 -
12935 HMM02441_04(#) GRM2
12940 -pf00893 -
12950 HMM02441_04(#) GRM2
12960 HMMptac_3_eut2(pf06130) GRM2
12970 H_grassGreen GRM2
12975 HMMgreae_1_grms(pf04055) GRM2
12980 HMMptac_3_eut2(pf06130) GRM2
12985 HMMptac_3_eut2(pf06130) GRM2
12990 H_grassGreen GRM2
12995 HMMgre_1_grms(pf01228pf02901) GRM2 EP_NTERM
13005 HMMgre_1_grms(pf02901) GRM2
13010 HMM00465alcdh_3_mix(pf00465) GRM2
13015 HMM00465alcdh_3_mix(pf00465) GRM2
13020 HMM00465alcdh_3_mix(pf00465) GRM2
13030 P_clearBlue GRM2
13035 HMM00465alcdh_3_mix(pf00465) GRM2
13040 H_brightSkyBlue GRM2
13045 H_brightSkyBlue GRM2
13050 H_brightSkyBlue GRM2
13055 HMMalddh_2_egra(pf00171) GRM2
13060 HMMalddh_2_egra(pf00171) GRM2
13065 HMMalddh_2_egra(pf00171) GRM2
13070 H_brightSkyBlue GRM2
13075 H_brightSkyBlue GRM2
13080 H_brightSkyBlue GRM2
13085 HMMreg00486_2_GRM2(#) GRM2
13090 HMMreg00486_2_GRM2(pf00486) GRM2
13095 HMMreg00440_1_GRM2(pf00440) GRM2
13115 -pf02753pf00345 -
13120 -# -
13135 -pf02753pf00345 -
13140 -pf13954pf00577 -
13145 -pf00419 -
13150 -pf00419 -
BMC-P
BMC-H
BMC-T
Aldehyde dehydrogenase
PTAC
Signature enzyme
Alcohol dehydrogenase
Regulator

Photobacterium ganghwense. Bacteria; Proteobacteria; Gammaproteobacteria; Vibrionales;

ABT57__locus_1

GRM2

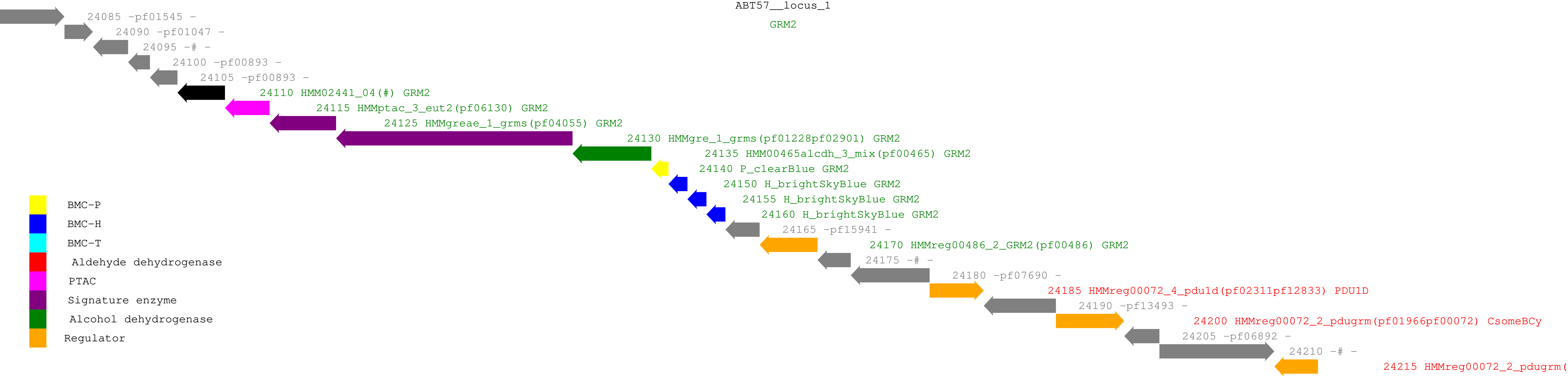

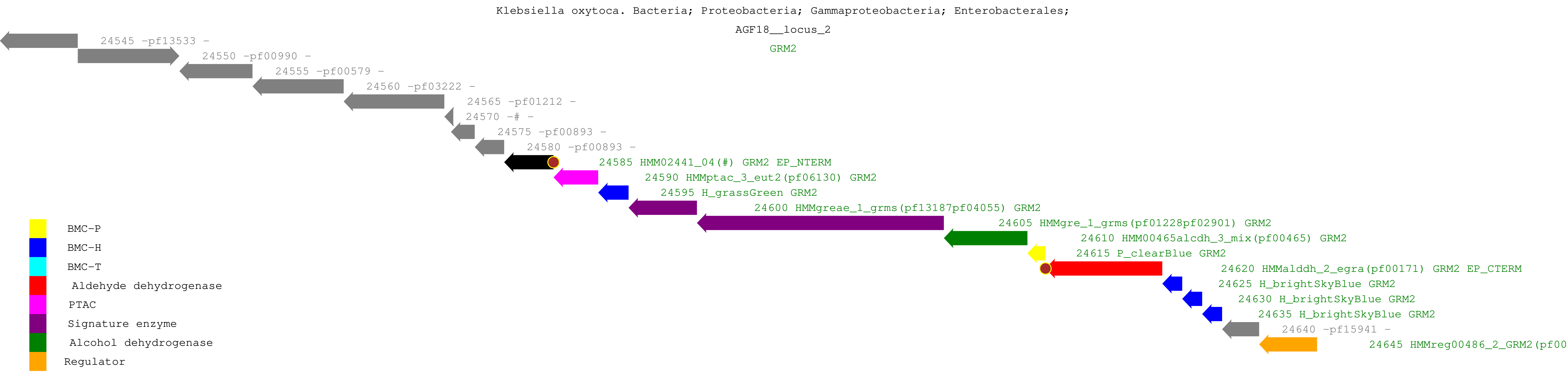
Klebsiella oxytoca. Bacteria; Proteobacteria; Gammaproteobacteria; Enterobacterales;
AGF18__locus_2
GRM2
24545 -pf13533 -
24550 -pf00990 -
24555 -pf00579 -
24560 -pf03222 -
24565 -pf01212 -
24570 -# -
24575 -pf00893 -
24580 -pf00893 -
24585 HMM02441_04(#) GRM2 EP_NTERM
24590 HMMptac_3_eut2(pf06130) GRM2
24595 H_grassGreen GRM2
24600 HMMgreae_1_grms(pf13187pf04055) GRM2
24605 HMMgre_1_grms(pf01228pf02901) GRM2
24610 HMM00465alcdh_3_mix(pf00465) GRM2
24615 P_clearBlue GRM2
24620 HMMalddh_2_egra(pf00171) GRM2 EP_CTERM
24625 H_brightSkyBlue GRM2
24630 H_brightSkyBlue GRM2
24635 H_brightSkyBlue GRM2
24640 -pf15941 -
24645 HMMreg00486_2_GRM2(pf00
BMC-P
BMC-H
BMC-T
Aldehyde dehydrogenase
PTAC
Signature enzyme
Alcohol dehydrogenase
Regulator

Klebsiella michiganensis. Bacteria; Proteobacteria; Gammaproteobacteria; Enterobacterales;

AGG97__locus_1

GRM2

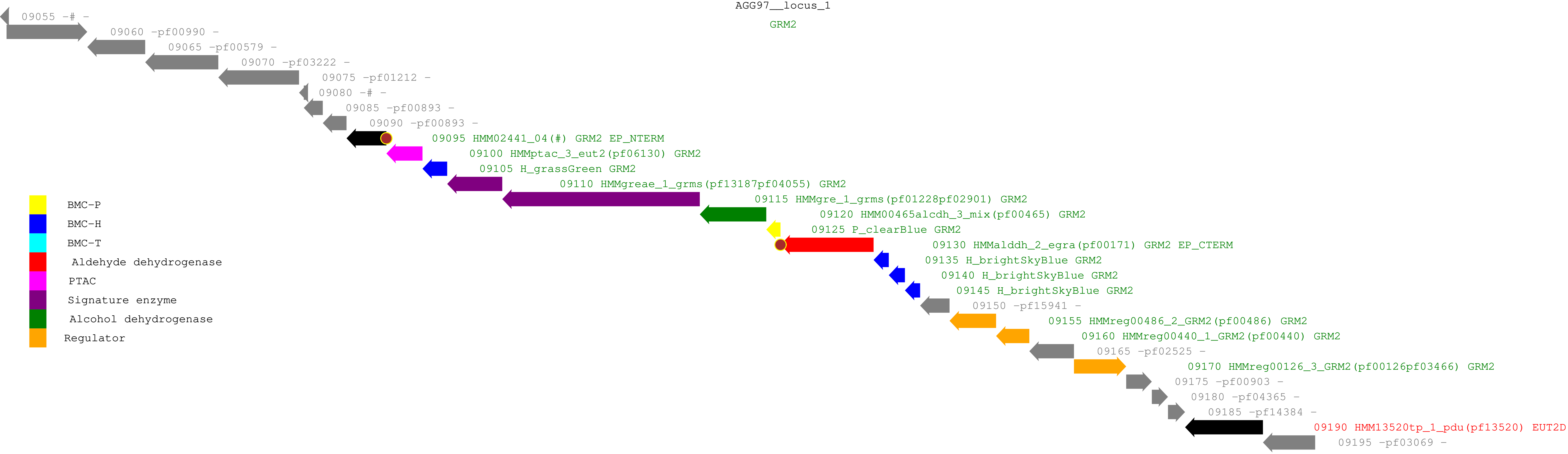

Aeromonas hydrophila subsp. hydrophila (strain ATCC 7966 / DSM 30187 / Bacteria; Proteobacteria; Gammaproteobacteria; Aeromonadales;

AHA__locus_1

GRM2

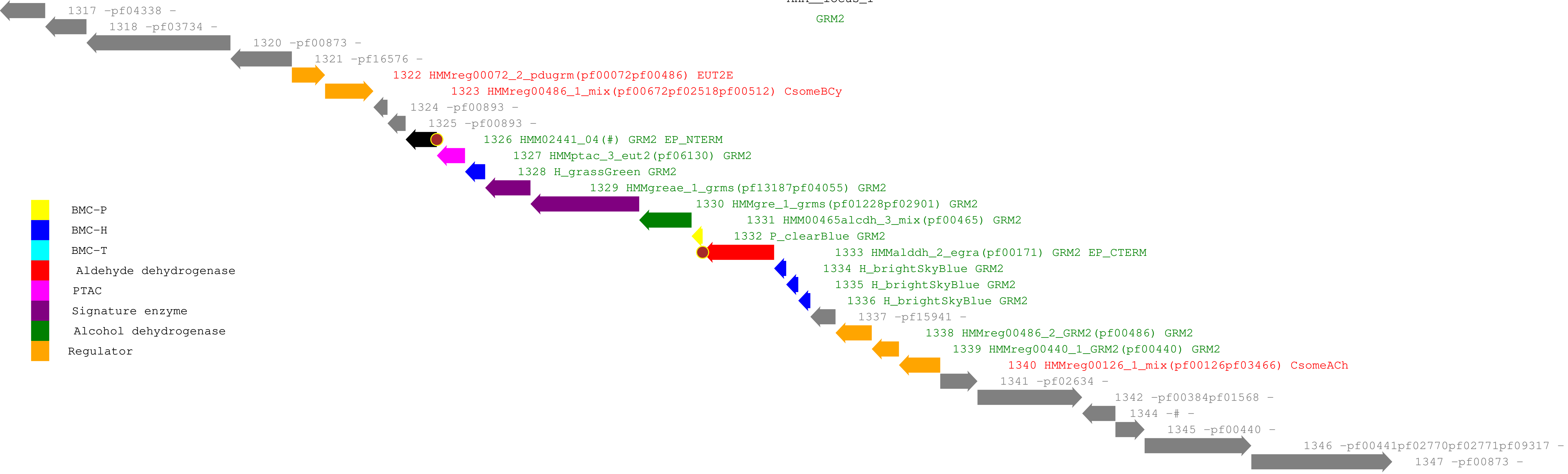

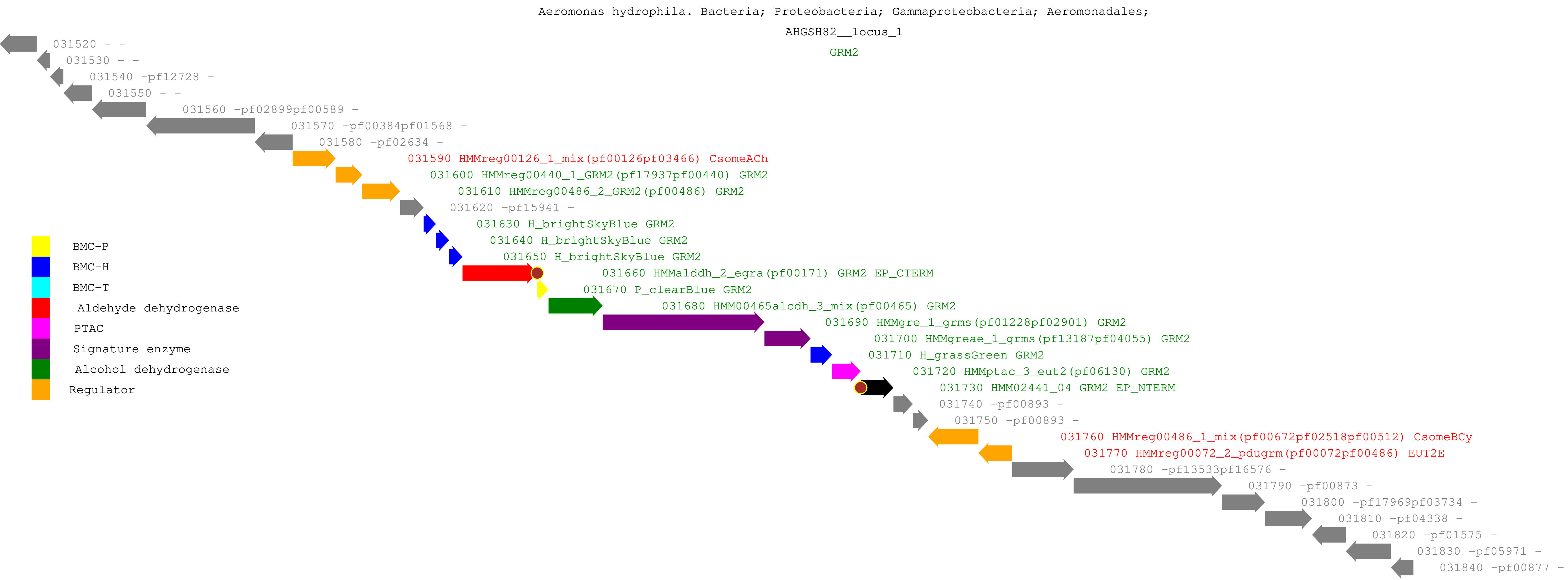
Aeromonas hydrophila. Bacteria; Proteobacteria; Gammaproteobacteria; Aeromonadales;
AHGSH82__locus_1
GRM2
031520 - -
031530 - -
031540 -pf12728 -
031550 - -
031560 -pf02899pf00589 -
031570 -pf00384pf01568 -
031580 -pf02634 -
031590 HMMreg00126_1_mix(pf00126pf03466) CsomeACh
031600 HMMreg00440_1_GRM2(pf17937pf00440) GRM2
031610 HMMreg00486_2_GRM2(pf00486) GRM2
031620 -pf15941 -
031630 H_brightSkyBlue GRM2
031640 H_brightSkyBlue GRM2
031650 H_brightSkyBlue GRM2
031660 HMMaldh_2_egra(pf00171) GRM2 EP_CTERM
031670 P_clearBlue GRM2
031680 HMM00465alcdh_3_mix(pf00465) GRM2
031690 HMMgre_1_grms(pf01228pf02901) GRM2
031700 HMMgreae_1_grms(pf13187pf04055) GRM2
031710 H_grassGreen GRM2
031720 HMMptac_3_eut2(pf06130) GRM2
031730 HMM02441_04 GRM2 EP_NTERM
031740 -pf00893 -
031750 -pf00893 -
031760 HMMreg00486_1_mix(pf00672pf02518pf00512) CsomeBCy
031770 HMMreg00072_2_pdugrm(pf00072pf00486) EUT2E
031780 -pf13533pf16576 -
031790 -pf00873 -
031800 -pf17969pf03734 -
031810 -pf04338 -
031820 -pf01575 -
031830 -pf05971 -
031840 -pf00877 -
BMC-P
BMC-H
BMC-T
Aldehyde dehydrogenase
PTAC
Signature enzyme
Alcohol dehydrogenase
Regulator

Proteus mirabilis. Bacteria; Proteobacteria; Gammaproteobacteria; Enterobacterales;

AM402__locus_1

GRM2

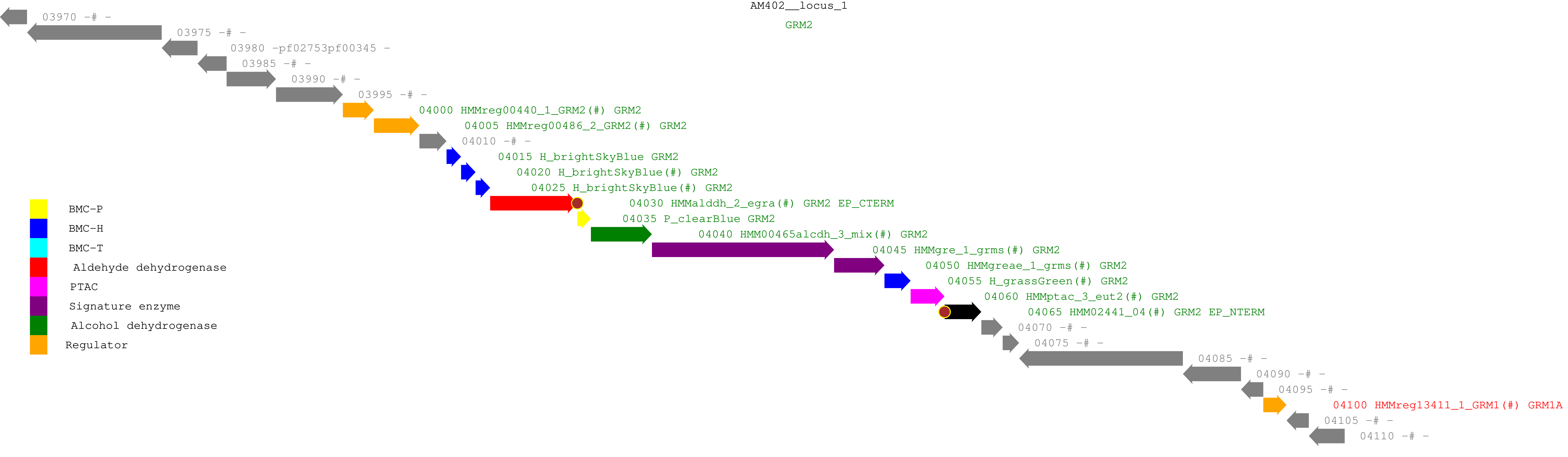

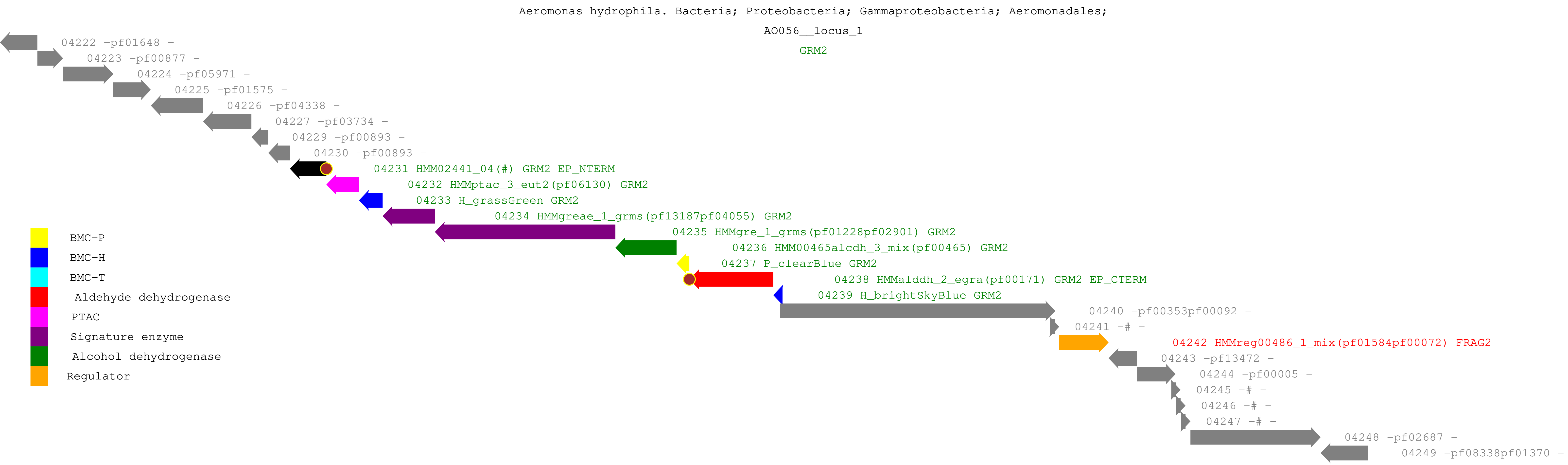
Aeromonas hydrophila. Bacteria; Proteobacteria; Gammaproteobacteria; Aeromonadales;
AO056__locus_1
GRM2
04222 -pf01648 -
04223 -pf00877 -
04224 -pf05971 -
04225 -pf01575 -
04226 -pf04338 -
04227 -pf03734 -
04229 -pf00893 -
04230 -pf00893 -
04231 HMM02441_04(#) GRM2 EP_NTERM
04232 HMMptac_3_eut2(pf06130) GRM2
04233 H_grassGreen GRM2
04234 HMMgreae_1_grms(pf13187pf04055) GRM2
04235 HMMgre_1_grms(pf01228pf02901) GRM2
04236 HMM00465alcdh_3_mix(pf00465) GRM2
04237 P_clearBlue GRM2
04238 HMMaldd_2_egra(pf00171) GRM2 EP_CTERM
04239 H_brightSkyBlue GRM2
04240 -pf00353pf00092 -
04241 -# -
04242 HMMreg00486_1_mix(pf01584pf00072) FRAG2
04243 -pf13472 -
04244 -pf00005 -
04245 -# -
04246 -# -
04247 -# -
04248 -pf02687 -
04249 -pf08338pf01370 -
BMC-P
BMC-H
BMC-T
Aldehyde dehydrogenase
PTAC
Signature enzyme
Alcohol dehydrogenase
Regulator

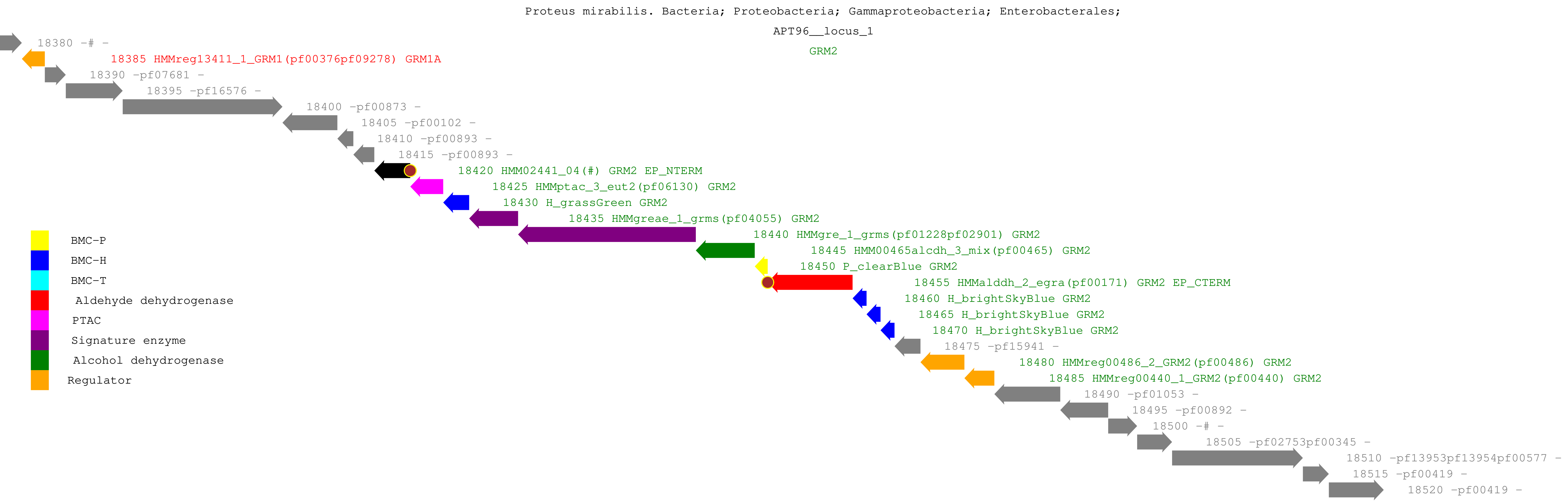
Proteus mirabilis. Bacteria; Proteobacteria; Gammaproteobacteria; Enterobacterales;
APT96__locus_1
GRM2
18380 -# -
18385 HMMreg13411_1_GRM1(pf00376pf09278) GRM1A
18390 -pf07681 -
18395 -pf16576 -
18400 -pf00873 -
18405 -pf00102 -
18410 -pf00893 -
18415 -pf00893 -
18420 HMM02441_04(#) GRM2 EP_NTERM
18425 HMMptac_3_eut2(pf06130) GRM2
18430 H_grassGreen GRM2
18435 HMMgreae_1_grms(pf04055) GRM2
18440 HMMgre_1_grms(pf01228pf02901) GRM2
18445 HMM00465alcdh_3_mix(pf00465) GRM2
18450 P_clearBlue GRM2
18455 HMMalddh_2_egra(pf00171) GRM2 EP_CTERM
18460 H_brightSkyBlue GRM2
18465 H_brightSkyBlue GRM2
18470 H_brightSkyBlue GRM2
18475 -pf15941 -
18480 HMMreg00486_2_GRM2(pf00486) GRM2
18485 HMMreg00440_1_GRM2(pf00440) GRM2
18490 -pf01053 -
18495 -pf00892 -
18500 -# -
18505 -pf02753pf00345 -
18510 -pf13953pf13954pf00577 -
18515 -pf00419 -
18520 -pf00419 -
BMC-P
BMC-H
BMC-T
Aldehyde dehydrogenase
PTAC
Signature enzyme
Alcohol dehydrogenase
Regulator

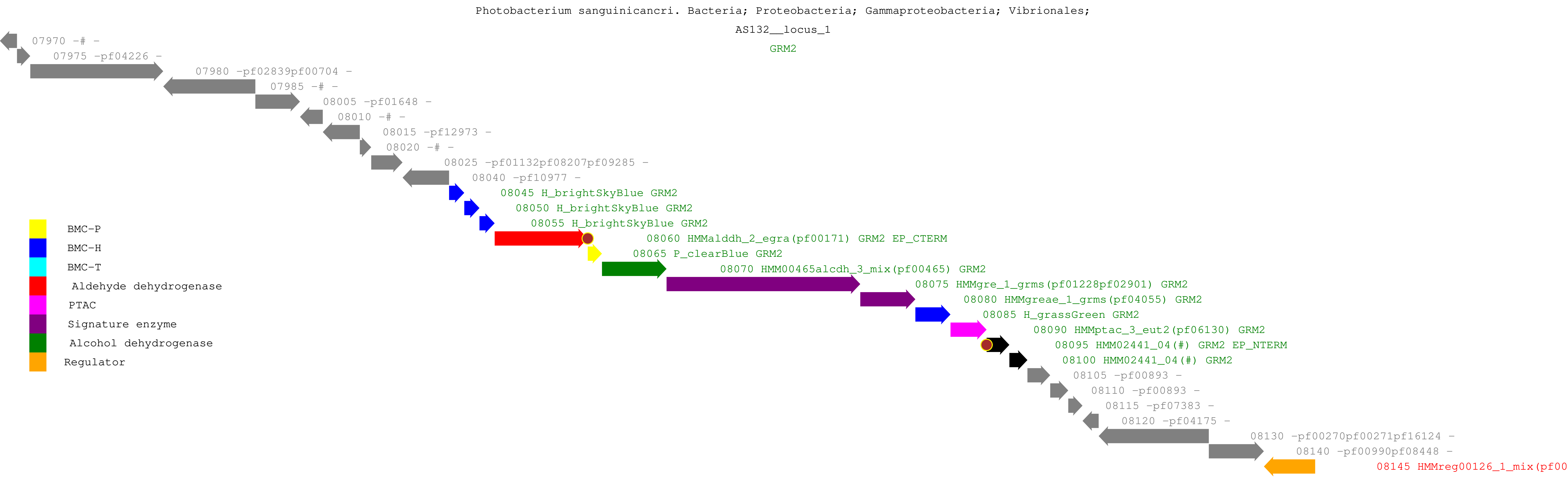
Photobacterium sanguinicancri. Bacteria; Proteobacteria; Gammaproteobacteria; Vibrionales;
AS132__locus_1
GRM2
07970 -# -
07975 -pf04226 -
07980 -pf02839pf00704 -
07985 -# -
08005 -pf01648 -
08010 -# -
08015 -pf12973 -
08020 -# -
08025 -pf01132pf08207pf09285 -
08040 -pf10977 -
08045 H_brightSkyBlue GRM2
08050 H_brightSkyBlue GRM2
08055 H_brightSkyBlue GRM2
08060 HMMalddh_2_egra(pf00171) GRM2 EP_CTERM
08065 P_clearBlue GRM2
08070 HMM00465alcdh_3_mix(pf00465) GRM2
08075 HMMgre_1_grms(pf01228pf02901) GRM2
08080 HMMgreae_1_grms(pf04055) GRM2
08085 H_grassGreen GRM2
08090 HMMptac_3_eut2(pf06130) GRM2
08095 HMM02441_04(#) GRM2 EP_NTERM
08100 HMM02441_04(#) GRM2
08105 -pf00893 -
08110 -pf00893 -
08115 -pf07383 -
08120 -pf04175 -
08130 -pf00270pf00271pf16124 -
08140 -pf00990pf08448 -
08145 HMMreg00126_1_mix(pf00
BMC-P
BMC-H
BMC-T
Aldehyde dehydrogenase
PTAC
Signature enzyme
Alcohol dehydrogenase
Regulator

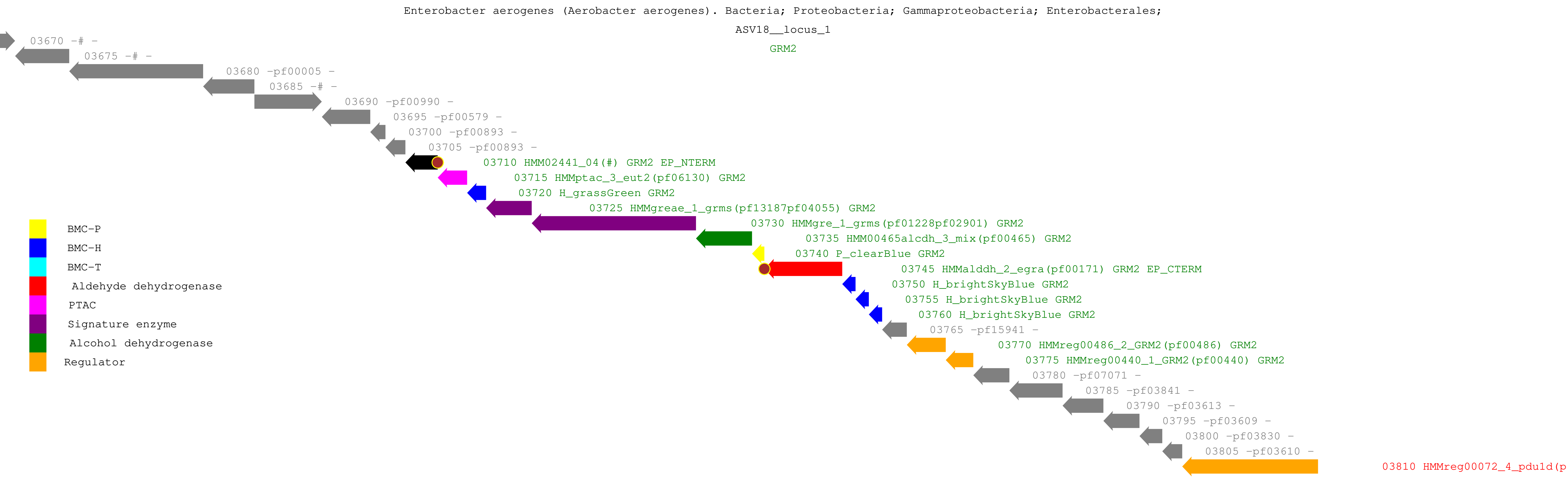
Enterobacter aerogenes (Aerobacter aerogenes). Bacteria; Proteobacteria; Gammaproteobacteria; Enterobacterales;
ASV18__locus_1
GRM2
03670 -# -
03675 -# -
03680 -pf00005 -
03685 -# -
03690 -pf00990 -
03695 -pf00579 -
03700 -pf00893 -
03705 -pf00893 -
03710 HMM02441_04(#) GRM2 EP_NTERM
03715 HMMptac_3_eut2(pf06130) GRM2
03720 H_grassGreen GRM2
03725 HMMgreae_1_grms(pf13187pf04055) GRM2
03730 HMMgre_1_grms(pf01228pf02901) GRM2
03735 HMM00465alcdh_3_mix(pf00465) GRM2
03740 P_clearBlue GRM2
03745 HMMalddh_2_egra(pf00171) GRM2 EP_CTERM
03750 H_brightSkyBlue GRM2
03755 H_brightSkyBlue GRM2
03760 H_brightSkyBlue GRM2
03765 -pf15941 -
03770 HMMreg00486_2_GRM2(pf00486) GRM2
03775 HMMreg00440_1_GRM2(pf00440) GRM2
03780 -pf07071 -
03785 -pf03841 -
03790 -pf03613 -
03795 -pf03609 -
03800 -pf03830 -
03805 -pf03610 -
03810 HMMreg00072_4_pdu1d(p
BMC-P
BMC-H
BMC-T
Aldehyde dehydrogenase
PTAC
Signature enzyme
Alcohol dehydrogenase
Regulator

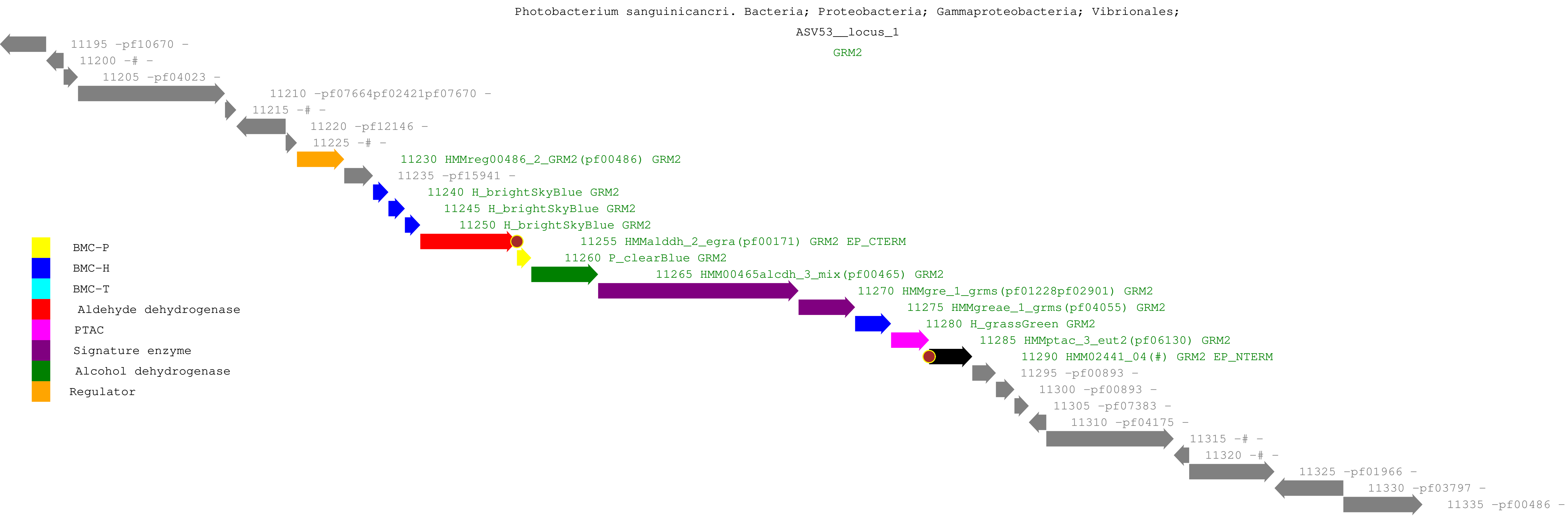

Photobacterium sanguinicancri. Bacteria; Proteobacteria; Gammaproteobacteria; Vibrionales;
ASV53__locus_1
GRM2
11195 -pf10670 -
11200 -# -
11205 -pf04023 -
11210 -pf07664pf02421pf07670 -
11215 -# -
11220 -pf12146 -
11225 -# -
11230 HMMreg00486_2_GRM2(pf00486) GRM2
11235 -pf15941 -
11240 H_brightSkyBlue GRM2
11245 H_brightSkyBlue GRM2
11250 H_brightSkyBlue GRM2
11255 HMMaldh_2_egra(pf00171) GRM2 EP_CTERM
11260 P_clearBlue GRM2
11265 HMM00465alcdh_3_mix(pf00465) GRM2
11270 HMMgre_1_grms(pf01228pf02901) GRM2
11275 HMMgreae_1_grms(pf04055) GRM2
11280 H_grassGreen GRM2
11285 HMMptac_3_eut2(pf06130) GRM2
11290 HMM02441_04(#) GRM2 EP_NTERM
11295 -pf00893 -
11300 -pf00893 -
11305 -pf07383 -
11310 -pf04175 -
11315 -# -
11320 -# -
11325 -pf01966 -
11330 -pf03797 -
11335 -pf00486 -
BMC-P
BMC-H
BMC-T
Aldehyde dehydrogenase
PTAC
Signature enzyme
Alcohol dehydrogenase
Regulator

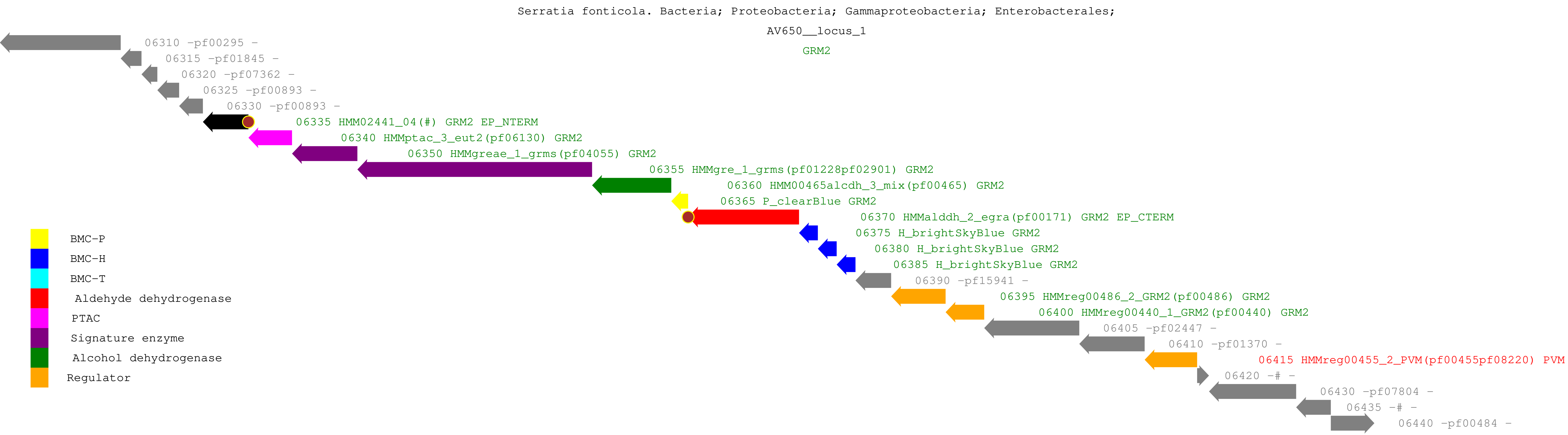
Serratia fonticola. Bacteria; Proteobacteria; Gammaproteobacteria; Enterobacterales;
AV650__locus_1
GRM2
06310 -pf00295 -
06315 -pf01845 -
06320 -pf07362 -
06325 -pf00893 -
06330 -pf00893 -
06335 HMM02441_04(#) GRM2 EP_NTERM
06340 HMMptac_3_eut2(pf06130) GRM2
06350 HMMgreae_1_grms(pf04055) GRM2
06355 HMMgre_1_grms(pf01228pf02901) GRM2
06360 HMM00465alcdh_3_mix(pf00465) GRM2
06365 P_clearBlue GRM2
06370 HMMalddh_2_egra(pf00171) GRM2 EP_CTERM
06375 H_brightSkyBlue GRM2
06380 H_brightSkyBlue GRM2
06385 H_brightSkyBlue GRM2
06390 -pf15941 -
06395 HMMreg00486_2_GRM2(pf00486) GRM2
06400 HMMreg00440_1_GRM2(pf00440) GRM2
06405 -pf02447 -
06410 -pf01370 -
06415 HMMreg00455_2_PVM(pf00455pf08220) PVM
06420 -# -
06430 -pf07804 -
06435 -# -
06440 -pf00484 -
BMC-P
BMC-H
BMC-T
Aldehyde dehydrogenase
PTAC
Signature enzyme
Alcohol dehydrogenase
Regulator

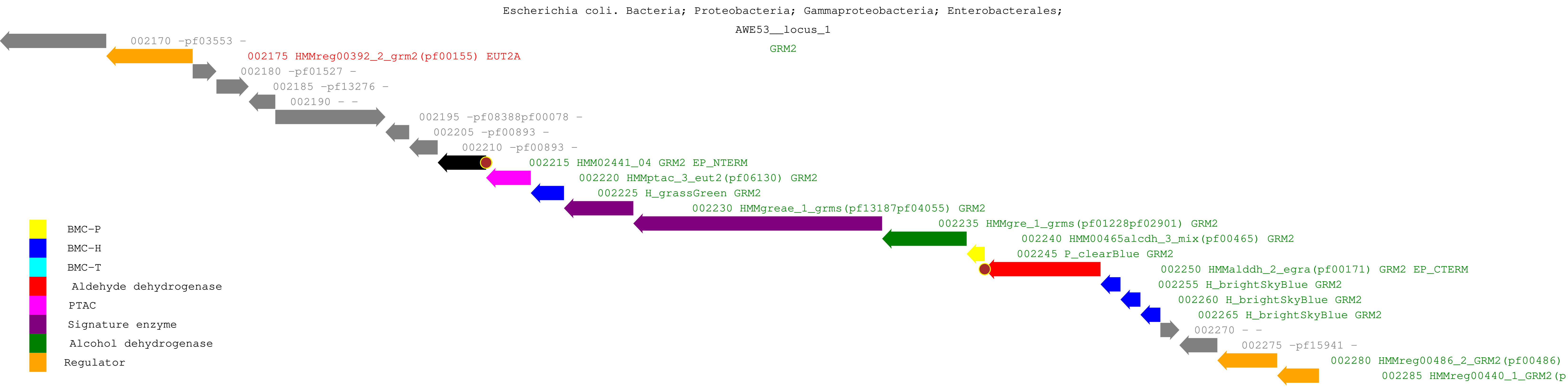

Escherichia coli. Bacteria; Proteobacteria; Gammaproteobacteria; Enterobacterales;
AWE53__locus_1
GRM2
002170 -pf03553 -
002175 HMMreg00392_2_grm2(pf00155) EUT2A
002180 -pf01527 -
002185 -pf13276 -
002190 - -
002195 -pf08388pf00078 -
002205 -pf00893 -
002210 -pf00893 -
002215 HMM02441_04 GRM2 EP_NTERM
002220 HMMptac_3_eut2(pf06130) GRM2
002225 H_grassGreen GRM2
002230 HMMgreae_1_grms(pf13187pf04055) GRM2
002235 HMMgre_1_grms(pf01228pf02901) GRM2
002240 HMM00465alcdh_3_mix(pf00465) GRM2
002245 P_clearBlue GRM2
002250 HMMalddh_2_egra(pf00171) GRM2 EP_CTERM
002255 H_brightSkyBlue GRM2
002260 H_brightSkyBlue GRM2
002265 H_brightSkyBlue GRM2
002270 - -
002275 -pf15941 -
002280 HMMreg00486_2_GRM2(pf00486)
002285 HMMreg00440_1_GRM2(p
BMC-P
BMC-H
BMC-T
Aldehyde dehydrogenase
PTAC
Signature enzyme
Alcohol dehydrogenase
Regulator

Klebsiella variicola. Bacteria; Proteobacteria; Gammaproteobacteria; Enterobacterales;

AWN63__locus_2

GRM2

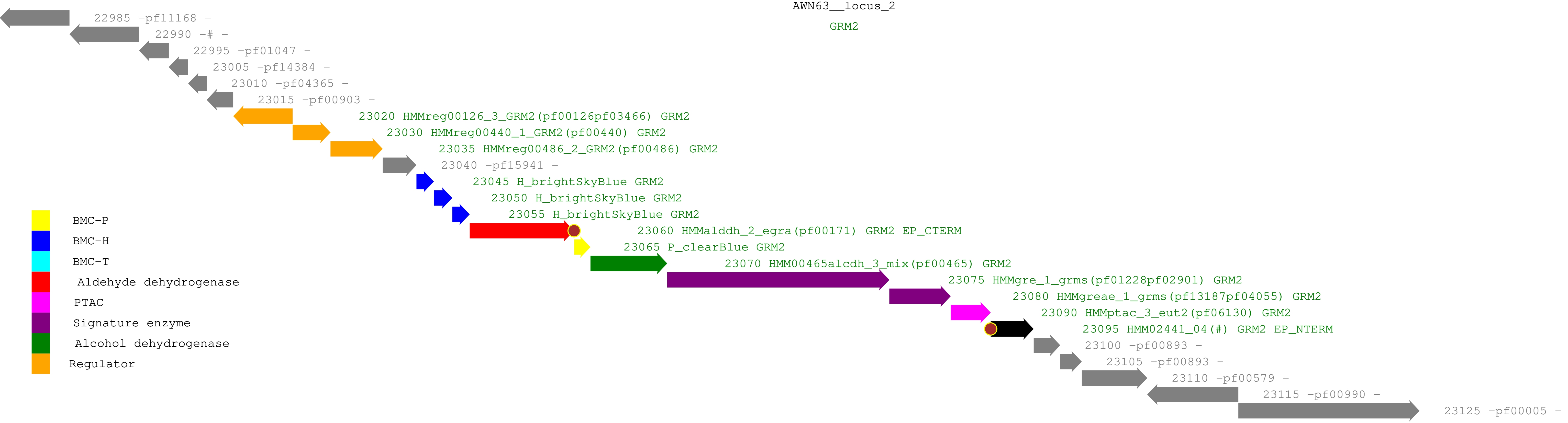

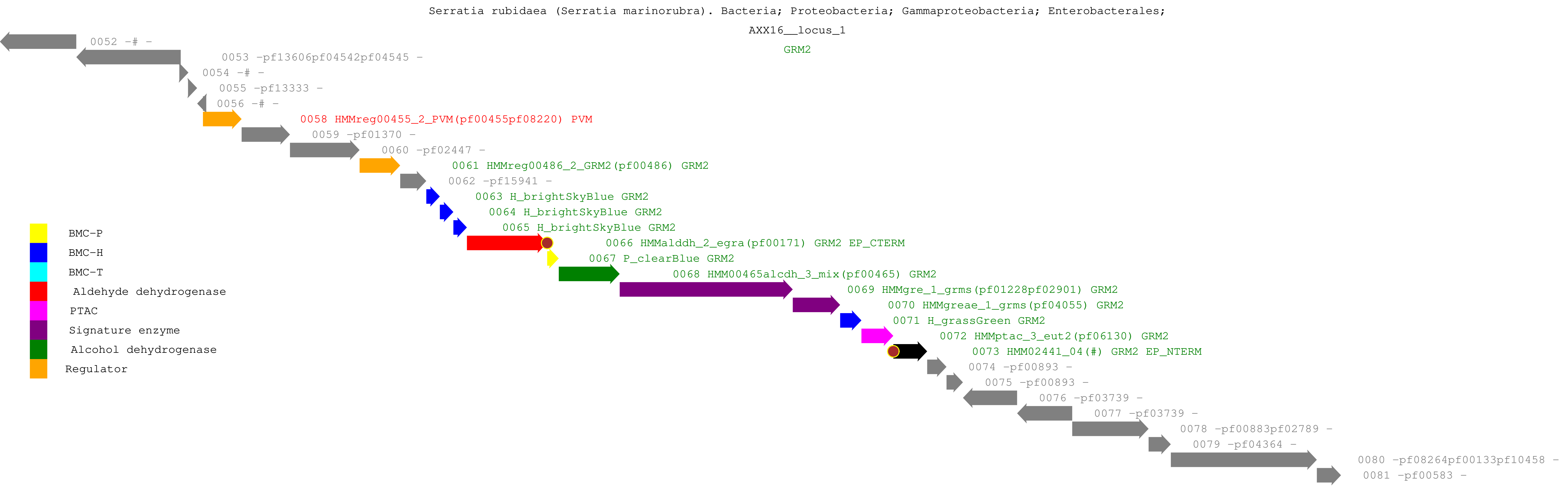

Serratia rubidaea (Serratia marinorubra). Bacteria; Proteobacteria; Gammaproteobacteria; Enterobacterales;
AXX16__locus_1
GRM2
0052 -# -
0053 -pf13606pf04542pf04545 -
0054 -# -
0055 -pf13333 -
0056 -# -
0058 HMMreg00455_2_PVM(pf00455pf08220) PVM
0059 -pf01370 -
0060 -pf02447 -
0061 HMMreg00486_2_GRM2(pf00486) GRM2
0062 -pf15941 -
0063 H_brightSkyBlue GRM2
0064 H_brightSkyBlue GRM2
0065 H_brightSkyBlue GRM2
0066 HMMalddh_2_egra(pf00171) GRM2 EP_CTERM
0067 P_clearBlue GRM2
0068 HMM00465alcdh_3_mix(pf00465) GRM2
0069 HMMgre_1_grms(pf01228pf02901) GRM2
0070 HMMgreae_1_grms(pf04055) GRM2
0071 H_grassGreen GRM2
0072 HMMptac_3_eut2(pf06130) GRM2
0073 HMM02441_04(#) GRM2 EP_NTERM
0074 -pf00893 -
0075 -pf00893 -
0076 -pf03739 -
0077 -pf03739 -
0078 -pf00883pf02789 -
0079 -pf04364 -
0080 -pf08264pf00133pf10458 -
0081 -pf00583 -
BMC-P
BMC-H
BMC-T
Aldehyde dehydrogenase
PTAC
Signature enzyme
Alcohol dehydrogenase
Regulator

Raoultella planticola (Klebsiella planticola). Bacteria; Proteobacteria; Gammaproteobacteria; Enterobacterales;

AYO04__locus_1

GRM2

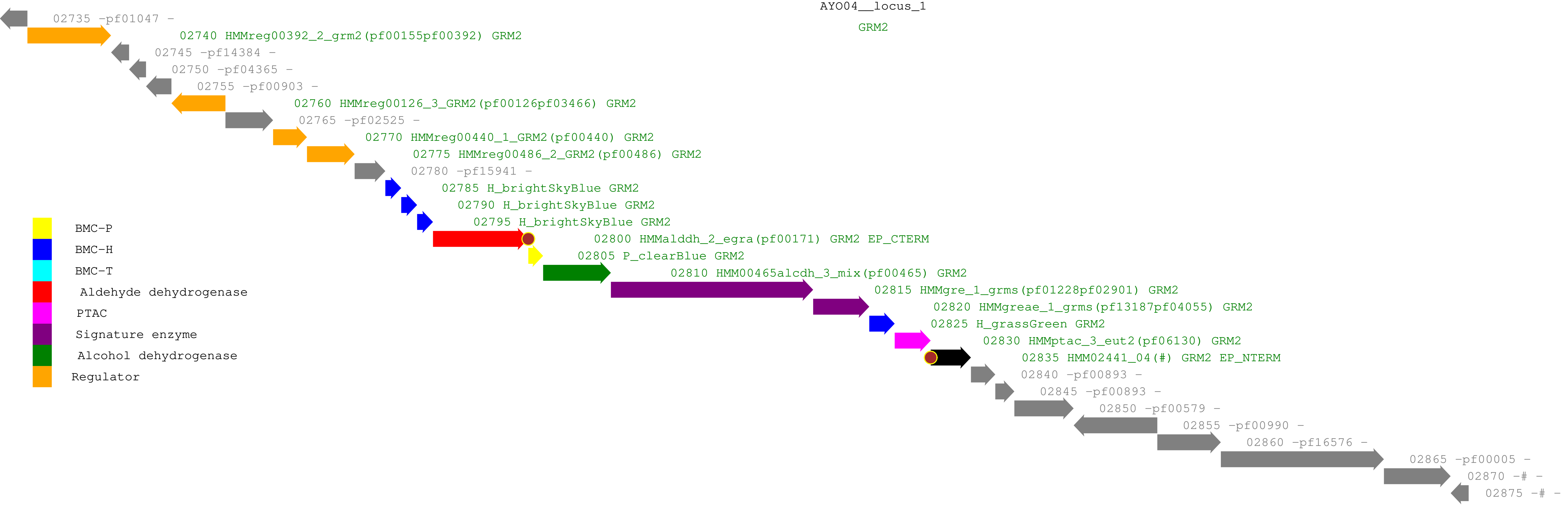

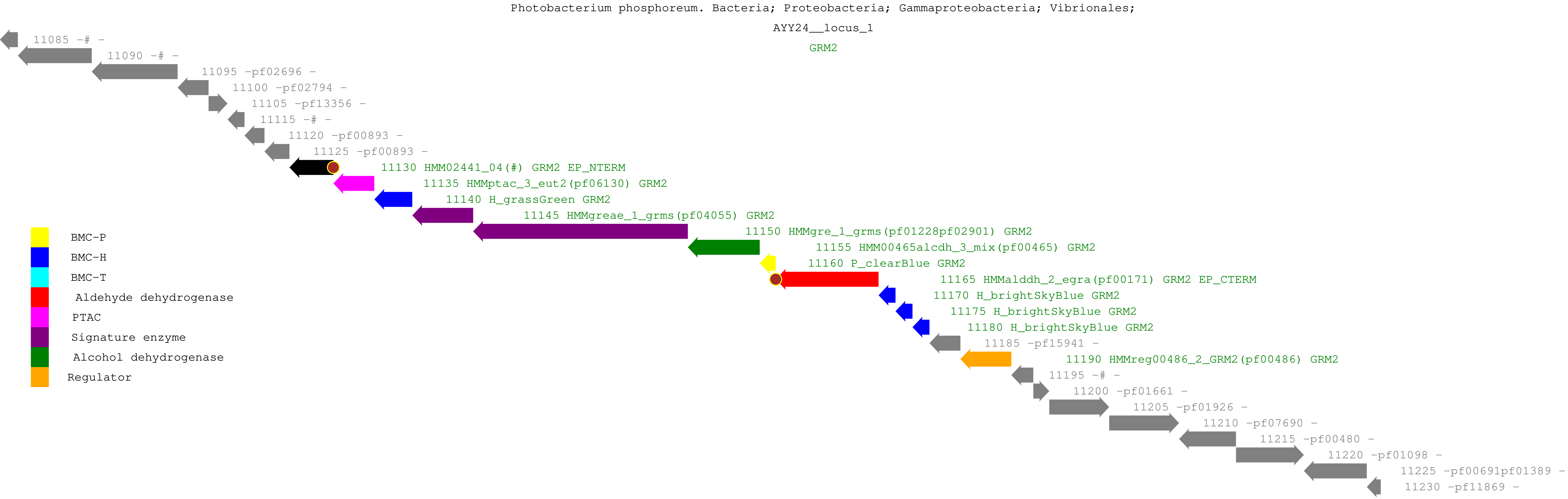
Photobacterium phosphoreum. Bacteria; Proteobacteria; Gammaproteobacteria; Vibrionales;
AYY24__locus_1
GRM2
11085 -# -
11090 -# -
11095 -pf02696 -
11100 -pf02794 -
11105 -pf13356 -
11115 -# -
11120 -pf00893 -
11125 -pf00893 -
11130 HMM02441_04(#) GRM2 EP_NTERM
11135 HMMptac_3_eut2(pf06130) GRM2
11140 H_grassGreen GRM2
11145 HMMgreae_1_grms(pf04055) GRM2
11150 HMMgre_1_grms(pf01228pf02901) GRM2
11155 HMM00465alcdh_3_mix(pf00465) GRM2
11160 P_clearBlue GRM2
11165 HMMalddh_2_egra(pf00171) GRM2 EP_CTERM
11170 H_brightSkyBlue GRM2
11175 H_brightSkyBlue GRM2
11180 H_brightSkyBlue GRM2
11185 -pf15941 -
11190 HMMreg00486_2_GRM2(pf00486) GRM2
11195 -# -
11200 -pf01661 -
11205 -pf01926 -
11210 -pf07690 -
11215 -pf00480 -
11220 -pf01098 -
11225 -pf00691pf01389 -
11230 -pf11869 -
BMC-P
BMC-H
BMC-T
Aldehyde dehydrogenase
PTAC
Signature enzyme
Alcohol dehydrogenase
Regulator

Klebsiella aerogenes (Enterobacter aerogenes). Bacteria; Proteobacteria; Gammaproteobacteria; Enterobacterales;

AZZ82__locus_1

GRM2

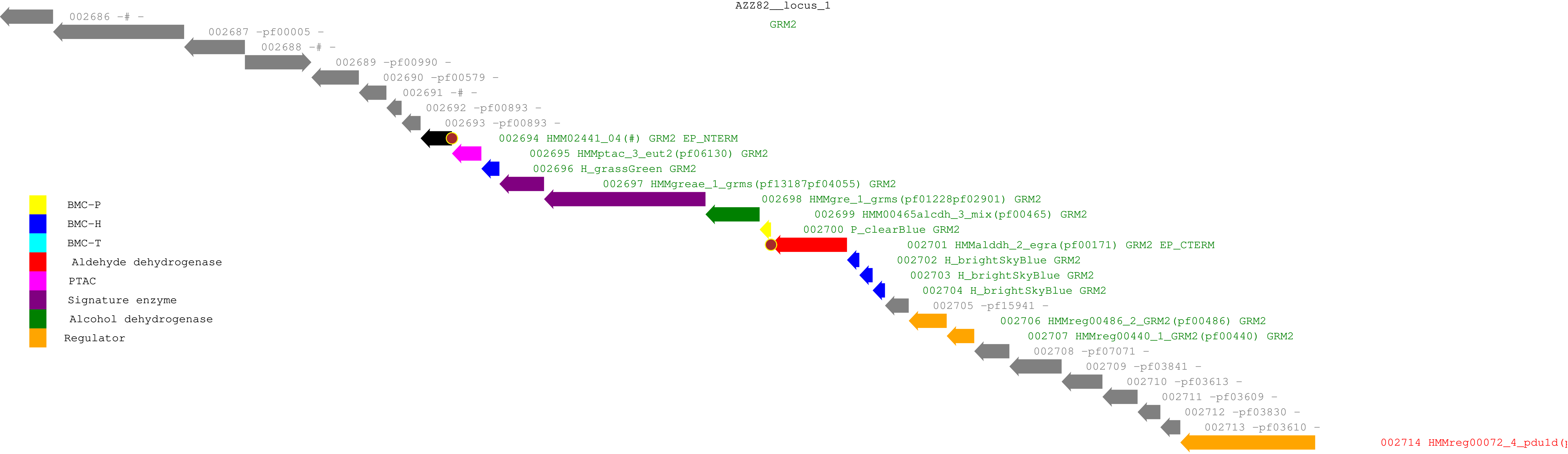

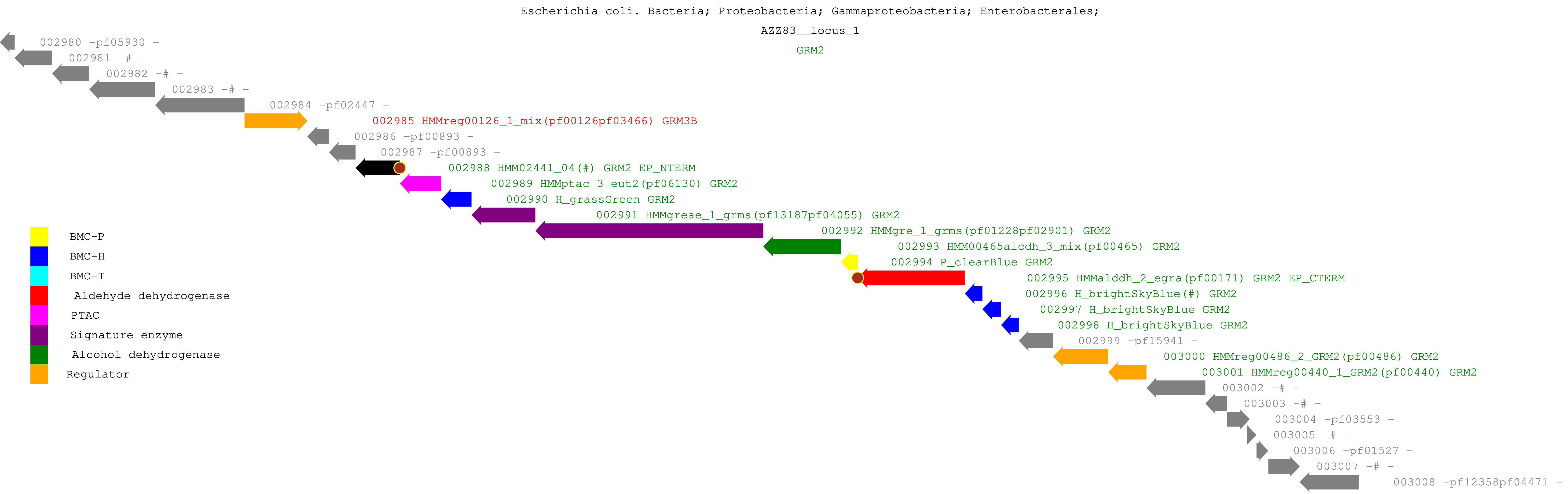

Escherichia coli. Bacteria; Proteobacteria; Gammaproteobacteria; Enterobacterales;
AZZ83__locus_1
GRM2
002980 -pf05930 -
002981 -# -
002982 -# -
002983 -# -
002984 -pf02447 -
002985 HMMreg00126_1_mix(pf00126pf03466) GRM3B
002986 -pf00893 -
002987 -pf00893 -
002988 HMM02441_04(#) GRM2 EP_NTERM
002989 HMMptac_3_eut2(pf06130) GRM2
002990 H_grassGreen GRM2
002991 HMMgreae_1_grms(pf13187pf04055) GRM2
002992 HMMgre_1_grms(pf01228pf02901) GRM2
002993 HMM00465alcdh_3_mix(pf00465) GRM2
002994 P_clearBlue GRM2
002995 HMMalddh_2_egra(pf00171) GRM2 EP_CTERM
002996 H_brightSkyBlue(#) GRM2
002997 H_brightSkyBlue GRM2
002998 H_brightSkyBlue GRM2
002999 -pf15941 -
003000 HMMreg00486_2_GRM2(pf00486) GRM2
003001 HMMreg00440_1_GRM2(pf00440) GRM2
003002 -# -
003003 -# -
003004 -pf03553 -
003005 -# -
003006 -pf01527 -
003007 -# -
003008 -pf12358pf04471 -
BMC-P
BMC-H
BMC-T
Aldehyde dehydrogenase
PTAC
Signature enzyme
Alcohol dehydrogenase
Regulator

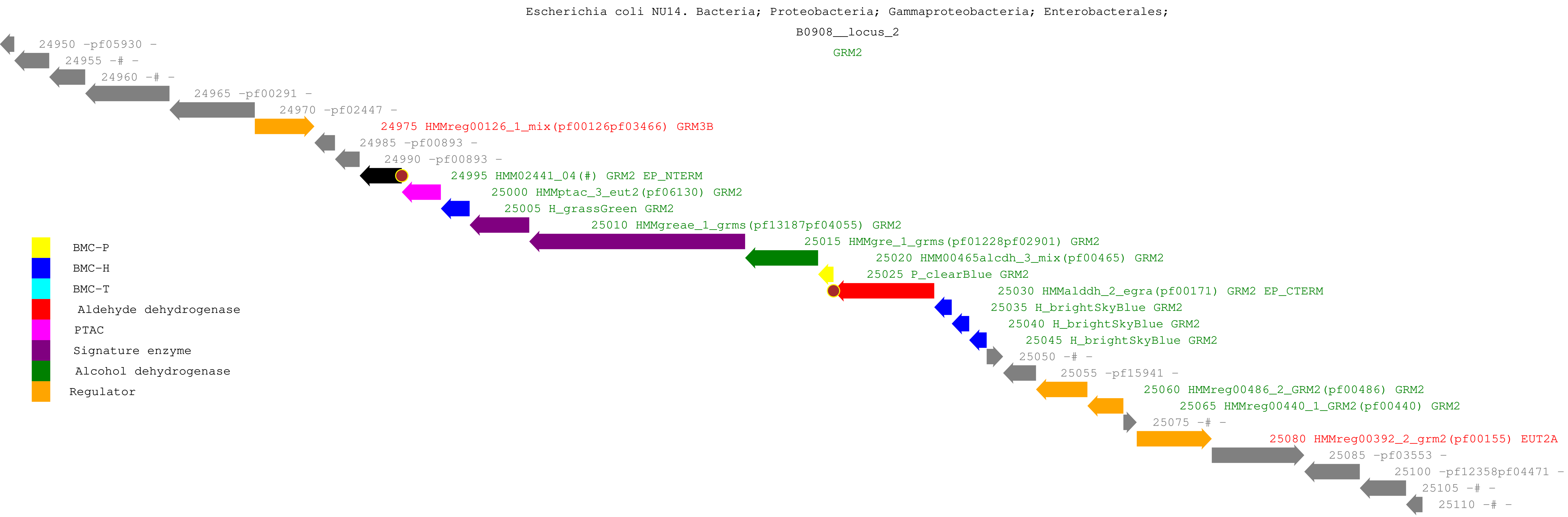
Escherichia coli NU14. Bacteria; Proteobacteria; Gammaproteobacteria; Enterobacterales;
B0908__locus_2
GRM2
24950 -pf05930 -
24955 -# -
24960 -# -
24965 -pf00291 -
24970 -pf02447 -
24975 HMMreg00126_1_mix(pf00126pf03466) GRM3B
24985 -pf00893 -
24990 -pf00893 -
24995 HMM02441_04(#) GRM2 EP_NTERM
25000 HMMptac_3_eut2(pf06130) GRM2
25005 H_grassGreen GRM2
25010 HMMgreae_1_grms(pf13187pf04055) GRM2
25015 HMMgre_1_grms(pf01228pf02901) GRM2
25020 HMM00465alcdh_3_mix(pf00465) GRM2
25025 P_clearBlue GRM2
25030 HMMalddh_2_egra(pf00171) GRM2 EP_CTERM
25035 H_brightSkyBlue GRM2
25040 H_brightSkyBlue GRM2
25045 H_brightSkyBlue GRM2
25050 -# -
25055 -pf15941 -
25060 HMMreg00486_2_GRM2(pf00486) GRM2
25065 HMMreg00440_1_GRM2(pf00440) GRM2
25075 -# -
25080 HMMreg00392_2_grm2(pf00155) EUT2A
25085 -pf03553 -
25100 -pf12358pf04471 -
25105 -# -
25110 -# -
BMC-P
BMC-H
BMC-T
Aldehyde dehydrogenase
PTAC
Signature enzyme
Alcohol dehydrogenase
Regulator

Aeromonas hydrophila. Bacteria; Proteobacteria; Gammaproteobacteria; Aeromonadales;

B7E00__locus_1

GRM2

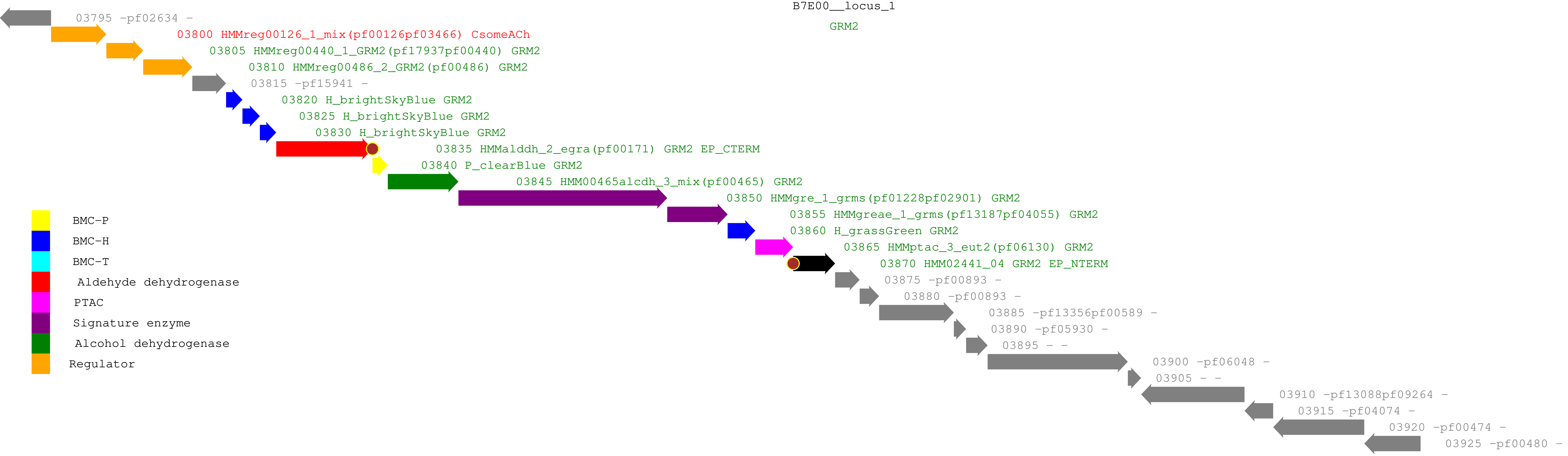

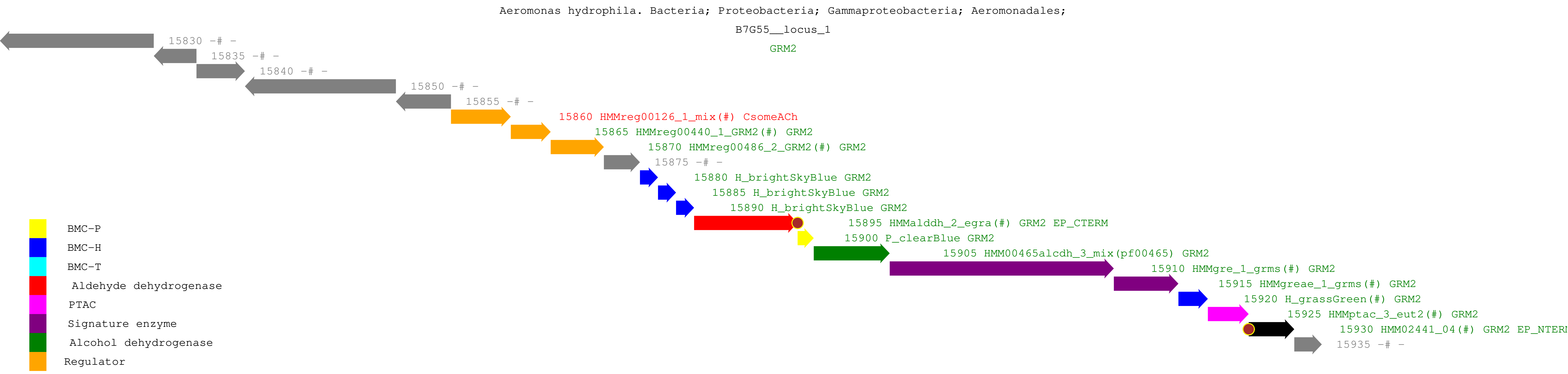

Aeromonas hydrophila. Bacteria; Proteobacteria; Gammaproteobacteria; Aeromonadales;
B7G55__locus_1
GRM2
15830 -# -
15835 -# -
15840 -# -
15850 -# -
15855 -# -
15860 HMMreg00126_1_mix(#) CsomeACh
15865 HMMreg00440_1_GRM2(#) GRM2
15870 HMMreg00486_2_GRM2(#) GRM2
15875 -# -
15880 H_brightSkyBlue GRM2
15885 H_brightSkyBlue GRM2
15890 H_brightSkyBlue GRM2
15895 HMMaldh_2_egra(#) GRM2 EP_CTERM
15900 P_clearBlue GRM2
15905 HMM00465alcdh_3_mix(pf00465) GRM2
15910 HMMgre_1_grms(#) GRM2
15915 HMMgreae_1_grms(#) GRM2
15920 H_grassGreen(#) GRM2
15925 HMMptac_3_eut2(#) GRM2
15930 HMM02441_04(#) GRM2 EP_NTERI
15935 -# -
BMC-P
BMC-H
BMC-T
Aldehyde dehydrogenase
PTAC
Signature enzyme
Alcohol dehydrogenase
Regulator

Proteus mirabilis. Bacteria; Proteobacteria; Gammaproteobacteria; Enterobacterales;

B9475__locus_1

GRM2

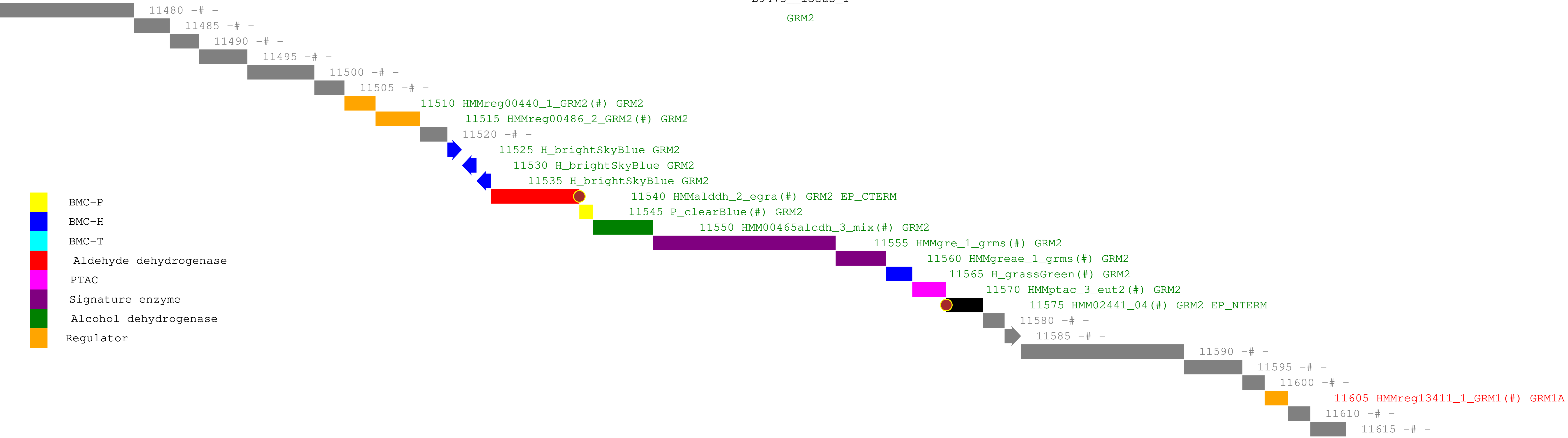

Klebsiella variicola. Bacteria; Proteobacteria; Gammaproteobacteria; Enterobacterales;

BBD63__locus_1

GRM2

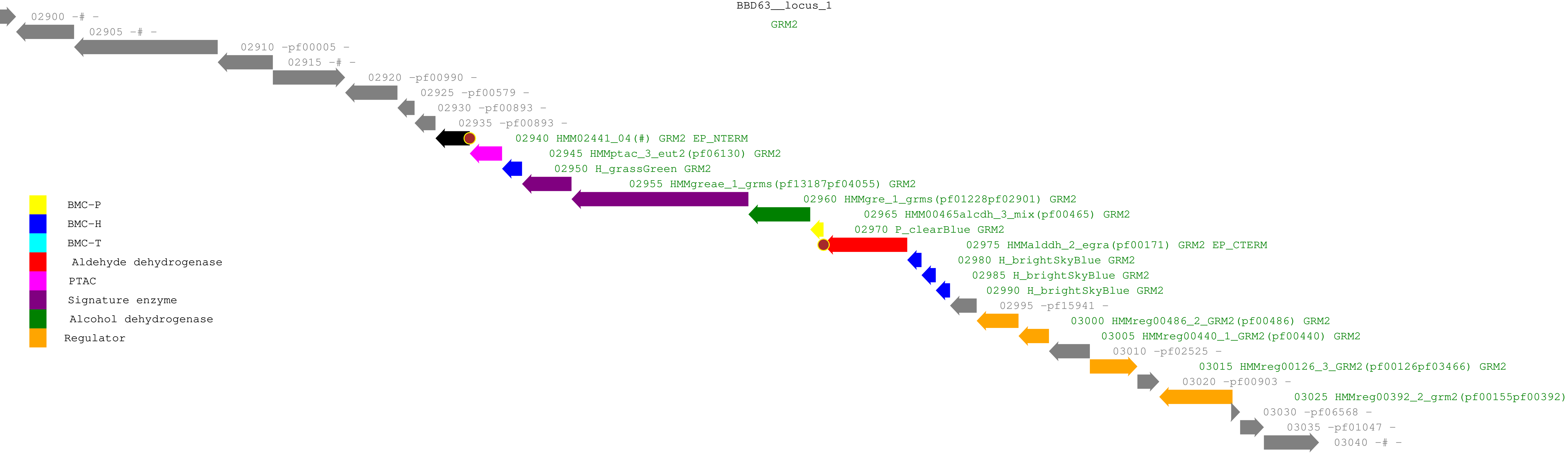

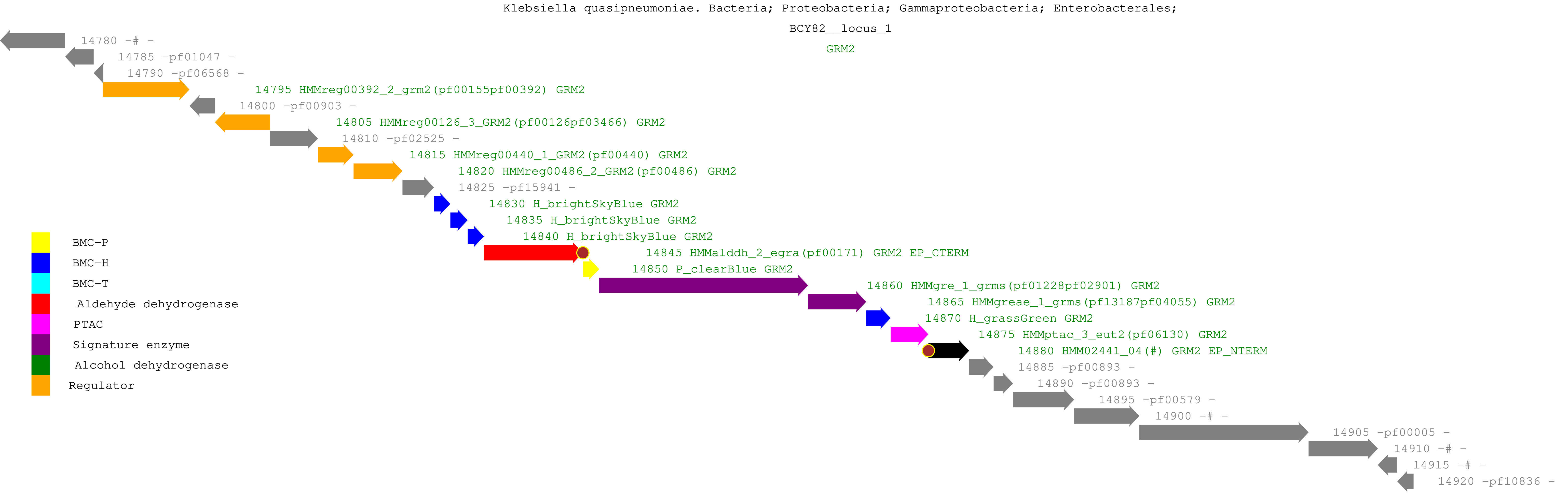
Klebsiella quasipneumoniae. Bacteria; Proteobacteria; Gammaproteobacteria; Enterobacterales;
BCY82__locus_1
GRM2
14780 -# -
14785 -pf01047 -
14790 -pf06568 -
14795 HMMreg00392_2_grm2(pf00155pf00392) GRM2
14800 -pf00903 -
14805 HMMreg00126_3_GRM2(pf00126pf03466) GRM2
14810 -pf02525 -
14815 HMMreg00440_1_GRM2(pf00440) GRM2
14820 HMMreg00486_2_GRM2(pf00486) GRM2
14825 -pf15941 -
14830 H_brightSkyBlue GRM2
14835 H_brightSkyBlue GRM2
14840 H_brightSkyBlue GRM2
14845 HMMalddh_2_egra(pf00171) GRM2 EP_CTERM
14850 P_clearBlue GRM2
14860 HMMgre_1_grms(pf01228pf02901) GRM2
14865 HMMgreae_1_grms(pf13187pf04055) GRM2
14870 H_grassGreen GRM2
14875 HMMptac_3_eut2(pf06130) GRM2
14880 HMM02441_04(#) GRM2 EP_NTERM
14885 -pf00893 -
14890 -pf00893 -
14895 -pf00579 -
14900 -# -
14905 -pf00005 -
14910 -# -
14915 -# -
14920 -pf10836 -
BMC-P
BMC-H
BMC-T
Aldehyde dehydrogenase
PTAC
Signature enzyme
Alcohol dehydrogenase
Regulator

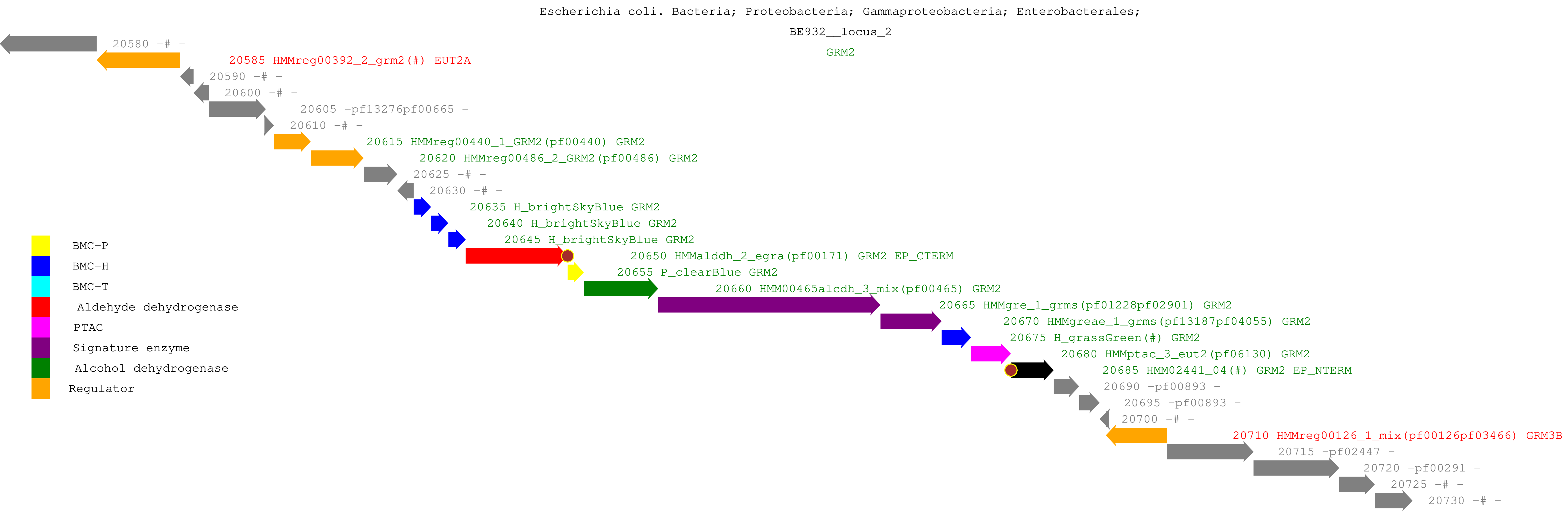
Escherichia coli. Bacteria; Proteobacteria; Gammaproteobacteria; Enterobacterales;
BE932__locus_2
GRM2
20580 -# -
20585 HMMreg00392_2_grm2(#) EUT2A
20590 -# -
20600 -# -
20605 -pf13276pf00665 -
20610 -# -
20615 HMMreg00440_1_GRM2(pf00440) GRM2
20620 HMMreg00486_2_GRM2(pf00486) GRM2
20625 -# -
20630 -# -
20635 H_brightSkyBlue GRM2
20640 H_brightSkyBlue GRM2
20645 H_brightSkyBlue GRM2
20650 HMMalddh_2_egra(pf00171) GRM2 EP_CTERM
20655 P_clearBlue GRM2
20660 HMM00465alcdh_3_mix(pf00465) GRM2
20665 HMMgre_1_grms(pf01228pf02901) GRM2
20670 HMMgreae_1_grms(pf13187pf04055) GRM2
20675 H_grassGreen(#) GRM2
20680 HMMptac_3_eut2(pf06130) GRM2
20685 HMM02441_04(#) GRM2 EP_NTERM
20690 -pf00893 -
20695 -pf00893 -
20700 -# -
20710 HMMreg00126_1_mix(pf00126pf03466) GRM3B
20715 -pf02447 -
20720 -pf00291 -
20725 -# -
20730 -# -
BMC-P
BMC-H
BMC-T
Aldehyde dehydrogenase
PTAC
Signature enzyme
Alcohol dehydrogenase
Regulator

Shigella sp. FC2928. Bacteria; Proteobacteria; Gammaproteobacteria; Enterobacterales;

BFF50__locus_2

GRM2

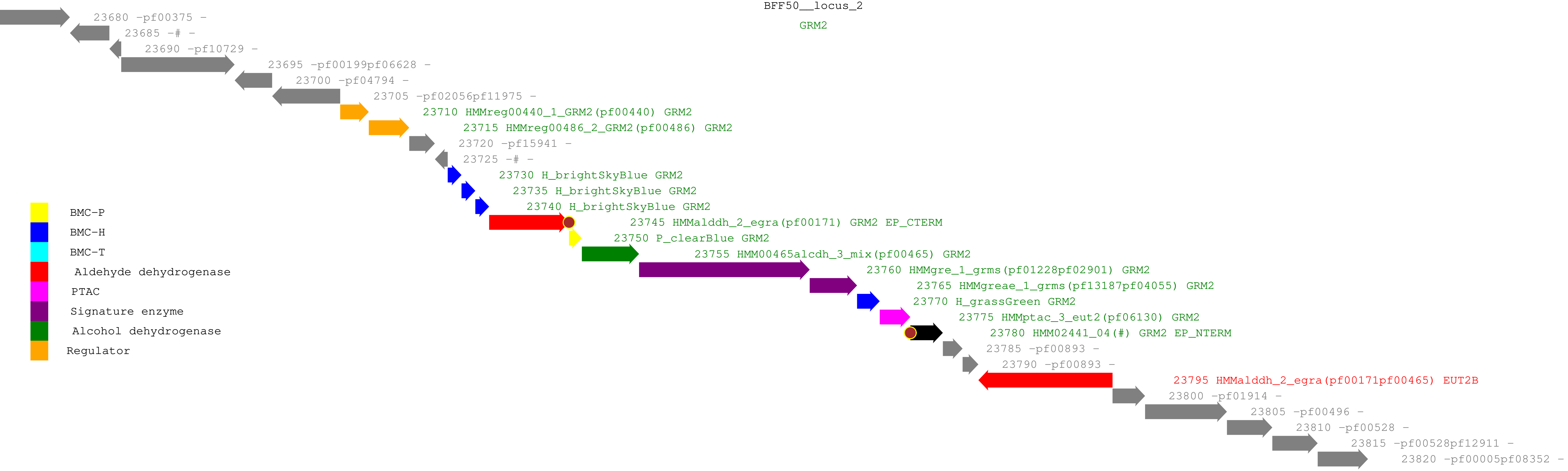

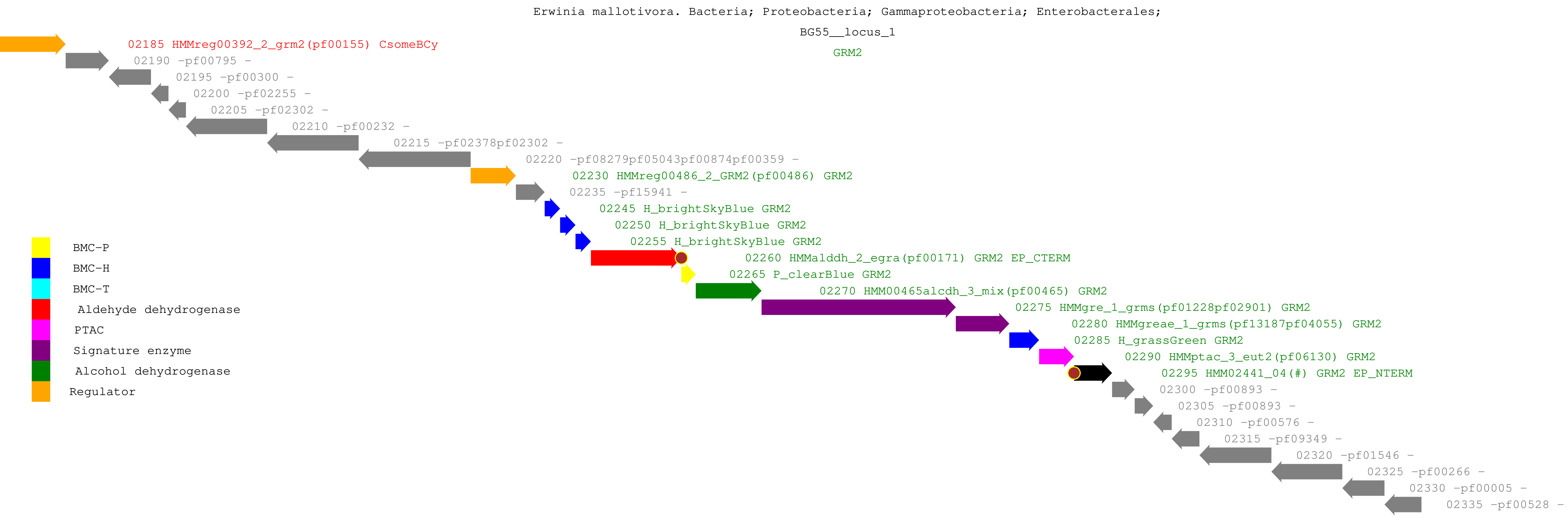
Erwinia mallotivora. Bacteria; Proteobacteria; Gammaproteobacteria; Enterobacterales;
BG55__locus_1
GRM2
02185 HMMreg00392_2_grm2(pf00155) CsomeBCy
02190 -pf00795 -
02195 -pf00300 -
02200 -pf02255 -
02205 -pf02302 -
02210 -pf00232 -
02215 -pf02378pf02302 -
02220 -pf08279pf05043pf00874pf00359 -
02230 HMMreg00486_2_GRM2(pf00486) GRM2
02235 -pf15941 -
02245 H_brightSkyBlue GRM2
02250 H_brightSkyBlue GRM2
02255 H_brightSkyBlue GRM2
02260 HMMalddh_2_egra(pf00171) GRM2 EP_CTERM
02265 P_clearBlue GRM2
02270 HMM00465alcdh_3_mix(pf00465) GRM2
02275 HMMgre_1_grms(pf01228pf02901) GRM2
02280 HMMgreae_1_grms(pf13187pf04055) GRM2
02285 H_grassGreen GRM2
02290 HMMptac_3_eut2(pf06130) GRM2
02295 HMM02441_04(#) GRM2 EP_NTERM
02300 -pf00893 -
02305 -pf00893 -
02310 -pf00576 -
02315 -pf09349 -
02320 -pf01546 -
02325 -pf00266 -
02330 -pf00005 -
02335 -pf00528 -
BMC-P
BMC-H
BMC-T
Aldehyde dehydrogenase
PTAC
Signature enzyme
Alcohol dehydrogenase
Regulator

Shigella sp. FC130. Bacteria; Proteobacteria; Gammaproteobacteria; Enterobacterales;

BGK50__locus_1

GRM2

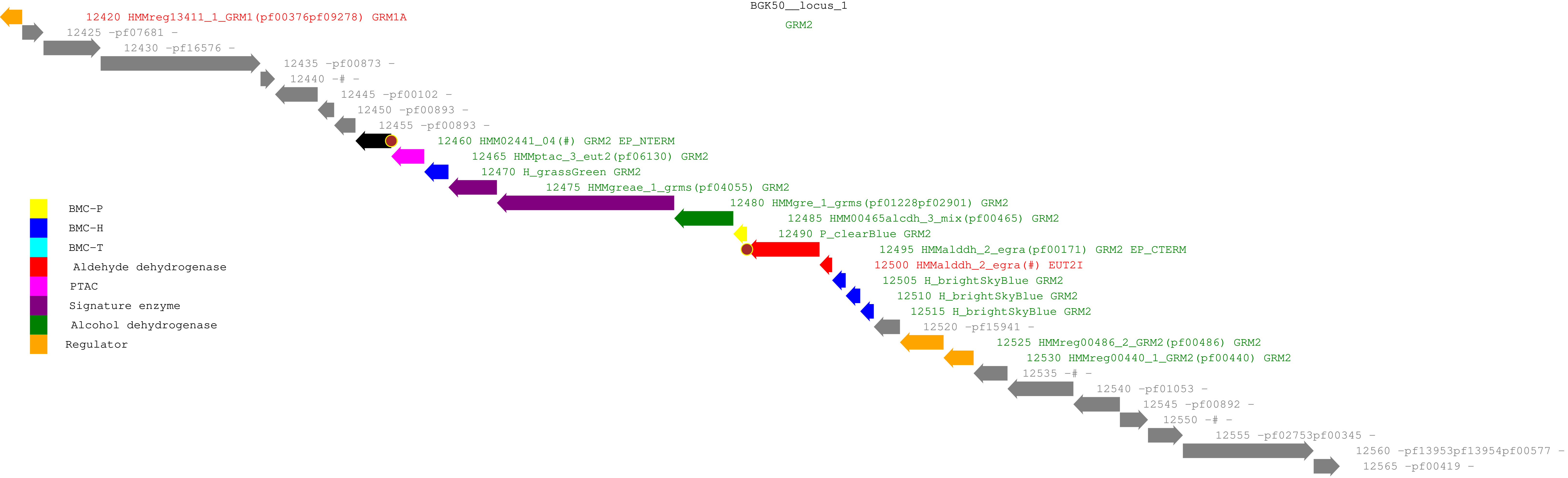

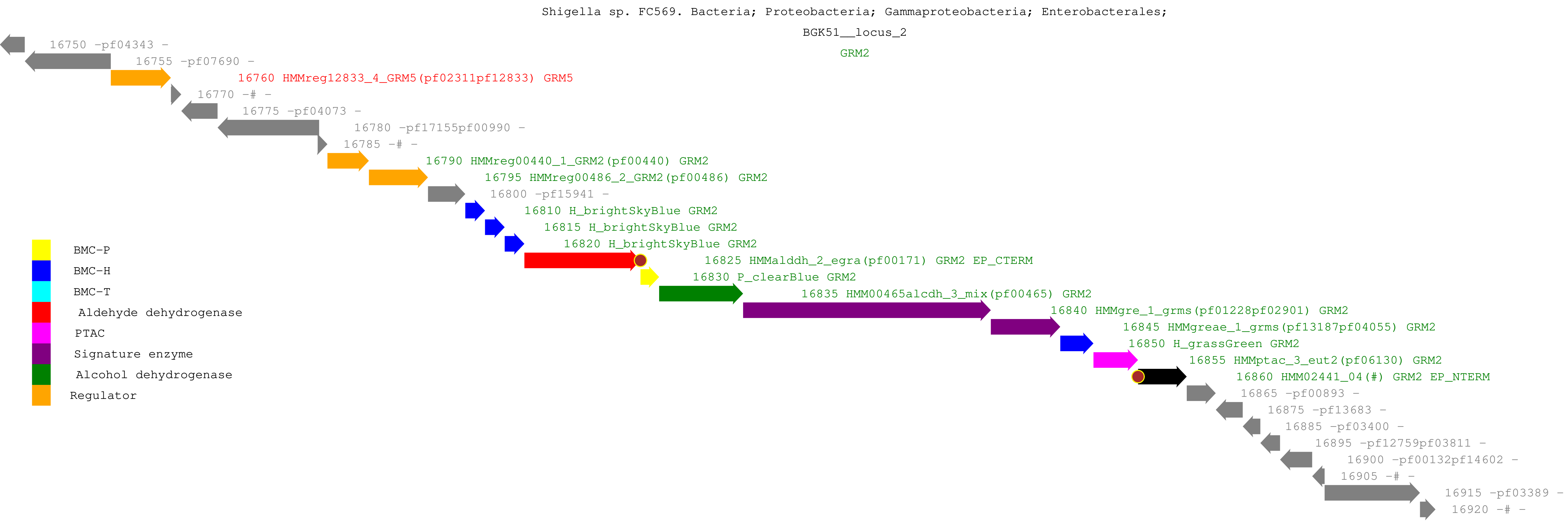

Shigella sp. FC569. Bacteria; Proteobacteria; Gammaproteobacteria; Enterobacterales;
BGK51__locus_2
GRM2
16750 -pf04343 -
16755 -pf07690 -
16760 HMMreg12833_4_GRM5(pf02311pf12833) GRM5
16770 -# -
16775 -pf04073 -
16780 -pf17155pf00990 -
16785 -# -
16790 HMMreg00440_1_GRM2(pf00440) GRM2
16795 HMMreg00486_2_GRM2(pf00486) GRM2
16800 -pf15941 -
16810 H_brightSkyBlue GRM2
16815 H_brightSkyBlue GRM2
16820 H_brightSkyBlue GRM2
16825 HMMaldh_2_egra(pf00171) GRM2 EP_CTERM
16830 P_clearBlue GRM2
16835 HMM00465alcdh_3_mix(pf00465) GRM2
16840 HMMgre_1_grms(pf01228pf02901) GRM2
16845 HMMgreae_1_grms(pf13187pf04055) GRM2
16850 H_grassGreen GRM2
16855 HMMptac_3_eut2(pf06130) GRM2
16860 HMM02441_04(#) GRM2 EP_NTERM
16865 -pf00893 -
16875 -pf13683 -
16885 -pf03400 -
16895 -pf12759pf03811 -
16900 -pf00132pf14602 -
16905 -# -
16915 -pf03389 -
16920 -# -
BMC-P
BMC-H
BMC-T
Aldehyde dehydrogenase
PTAC
Signature enzyme
Alcohol dehydrogenase
Regulator

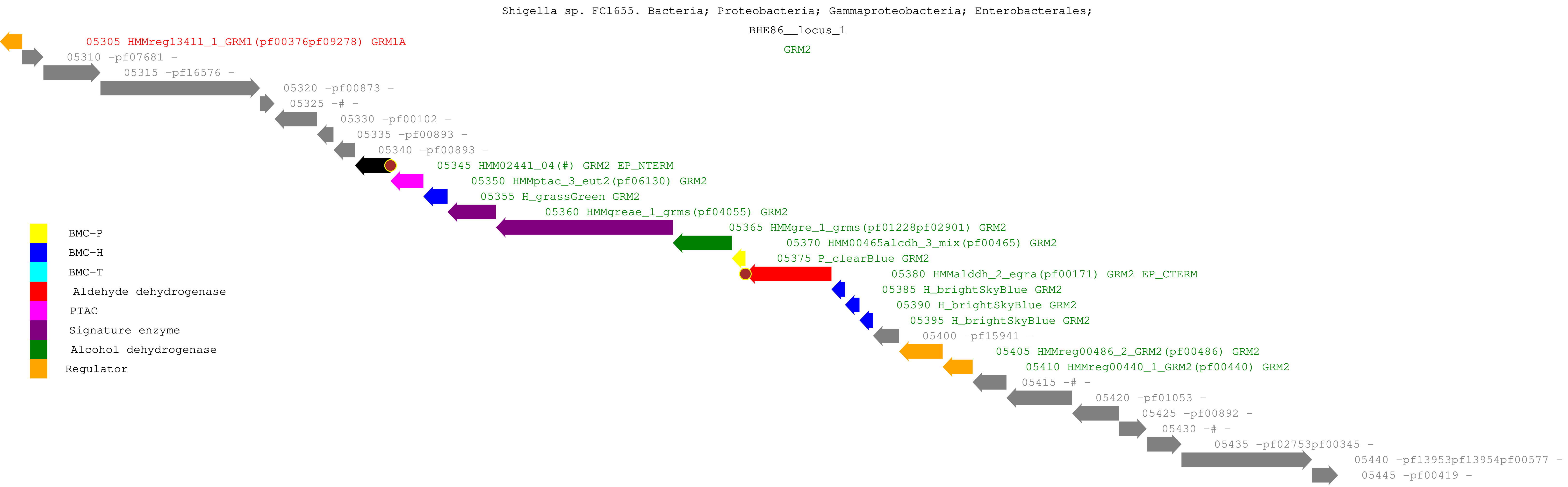
Shigella sp. FC1655. Bacteria; Proteobacteria; Gammaproteobacteria; Enterobacterales;
BHE86__locus_1
GRM2
05305 HMMreg13411_1_GRM1(pf00376pf09278) GRM1A
05310 -pf07681 -
05315 -pf16576 -
05320 -pf00873 -
05325 -# -
05330 -pf00102 -
05335 -pf00893 -
05340 -pf00893 -
05345 HMM02441_04(#) GRM2 EP_NTERM
05350 HMMptac_3_eut2(pf06130) GRM2
05355 H_grassGreen GRM2
05360 HMMgreae_1_grms(pf04055) GRM2
05365 HMMgre_1_grms(pf01228pf02901) GRM2
05370 HMM00465alcdh_3_mix(pf00465) GRM2
05375 P_clearBlue GRM2
05380 HMMalddh_2_egra(pf00171) GRM2 EP_CTERM
05385 H_brightSkyBlue GRM2
05390 H_brightSkyBlue GRM2
05395 H_brightSkyBlue GRM2
05400 -pf15941 -
05405 HMMreg00486_2_GRM2(pf00486) GRM2
05410 HMMreg00440_1_GRM2(pf00440) GRM2
05415 -# -
05420 -pf01053 -
05425 -pf00892 -
05430 -# -
05435 -pf02753pf00345 -
05440 -pf13953pf13954pf00577 -
05445 -pf00419 -
BMC-P
BMC-H
BMC-T
Aldehyde dehydrogenase
PTAC
Signature enzyme
Alcohol dehydrogenase
Regulator

Shigella sp. FC1967. Bacteria; Proteobacteria; Gammaproteobacteria; Enterobacterales;

BHE89__locus_1_main_sat

GRM2

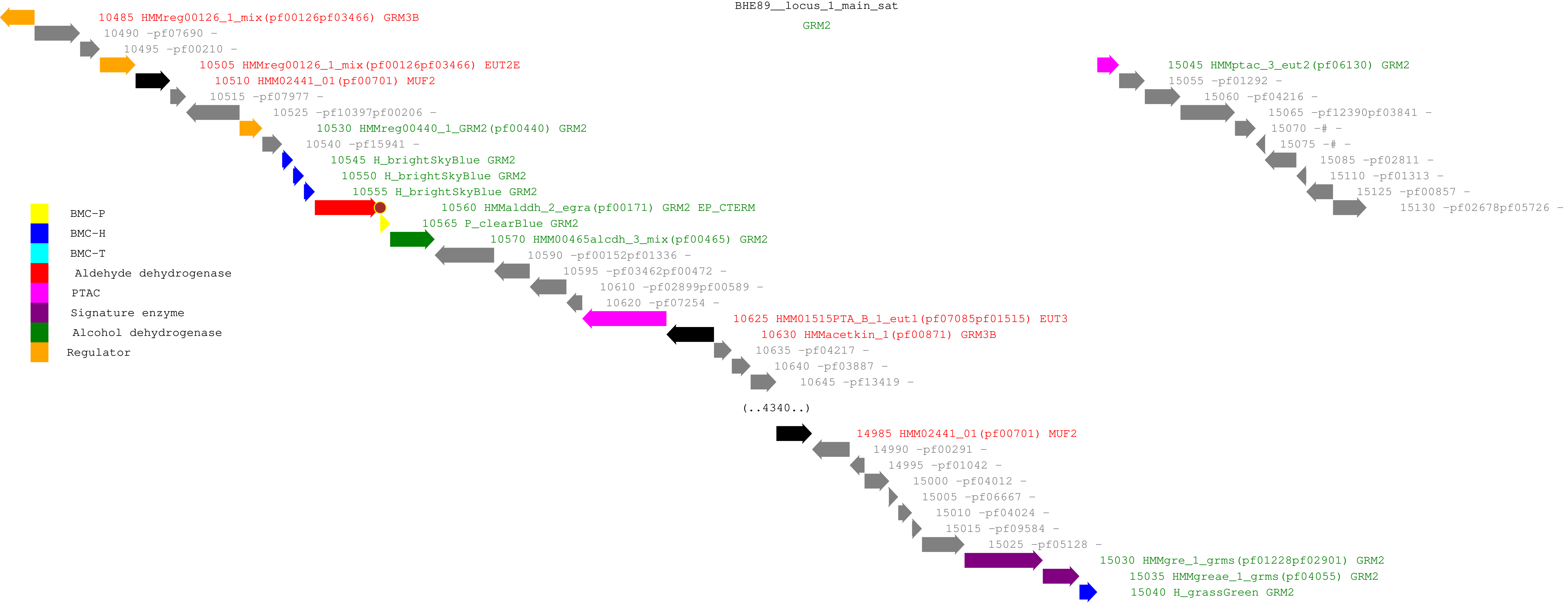

Photobacterium sp. 13-12. Bacteria; Proteobacteria; Gammaproteobacteria; Vibrionales;

BIT28__locus_1

GRM2

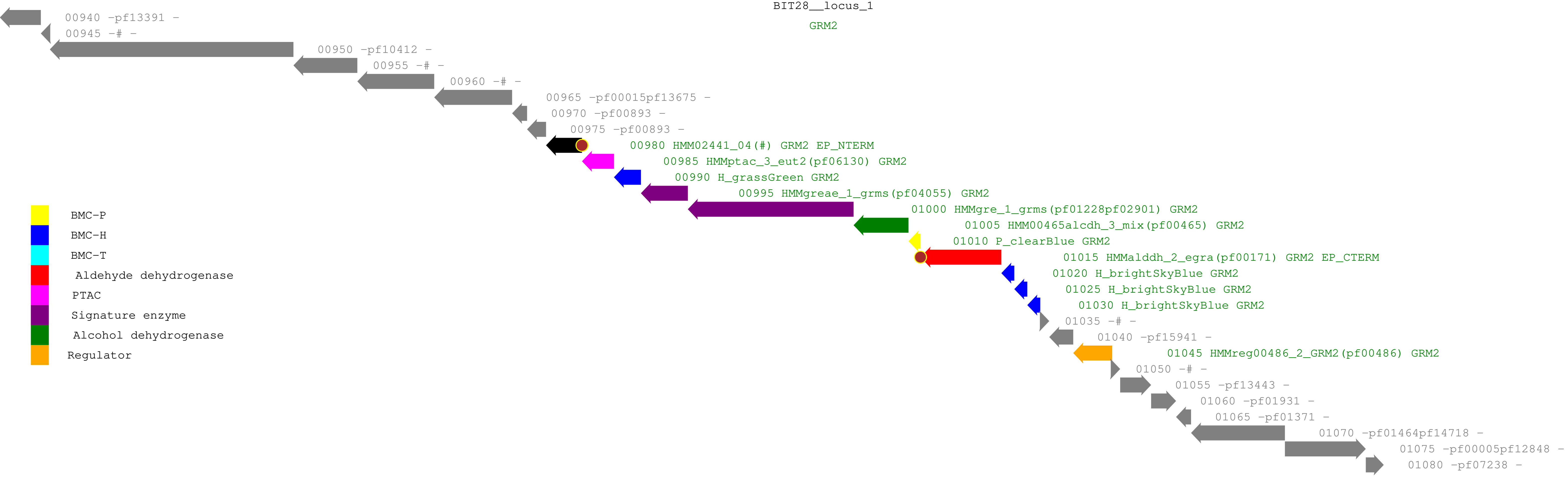

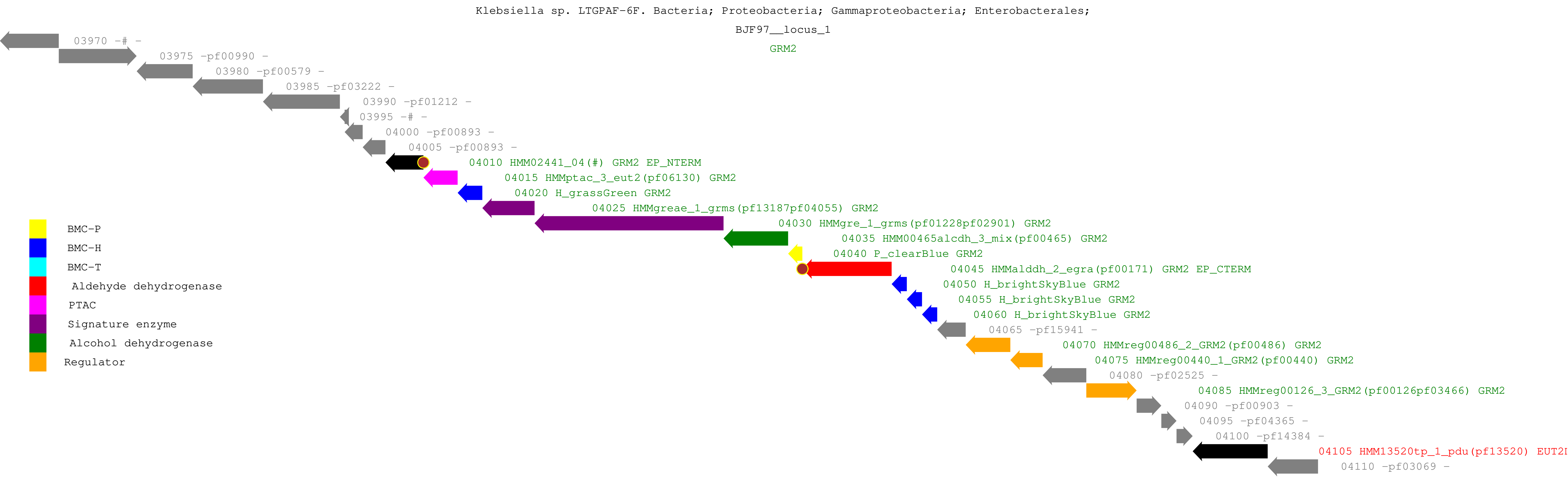

Klebsiella sp. LTGPAF-6F. Bacteria; Proteobacteria; Gammaproteobacteria; Enterobacterales;
BJF97__locus_1
GRM2
03970 -# -
03975 -pf00990 -
03980 -pf00579 -
03985 -pf03222 -
03990 -pf01212 -
03995 -# -
04000 -pf00893 -
04005 -pf00893 -
04010 HMM02441_04(#) GRM2 EP_NTERM
04015 HMMptac_3_eut2(pf06130) GRM2
04020 H_grassGreen GRM2
04025 HMMgreae_1_grms(pf13187pf04055) GRM2
04030 HMMgre_1_grms(pf01228pf02901) GRM2
04035 HMM00465alcdh_3_mix(pf00465) GRM2
04040 P_clearBlue GRM2
04045 HMMaldh_2_egra(pf00171) GRM2 EP_CTERM
04050 H_brightSkyBlue GRM2
04055 H_brightSkyBlue GRM2
04060 H_brightSkyBlue GRM2
04065 -pf15941 -
04070 HMMreg00486_2_GRM2(pf00486) GRM2
04075 HMMreg00440_1_GRM2(pf00440) GRM2
04080 -pf02525 -
04085 HMMreg00126_3_GRM2(pf00126pf03466) GRM2
04090 -pf00903 -
04095 -pf04365 -
04100 -pf14384 -
04105 HMM13520tp_1_pdu(pf13520) EUT2I
04110 -pf03069 -
BMC-P
BMC-H
BMC-T
Aldehyde dehydrogenase
PTAC
Signature enzyme
Alcohol dehydrogenase
Regulator

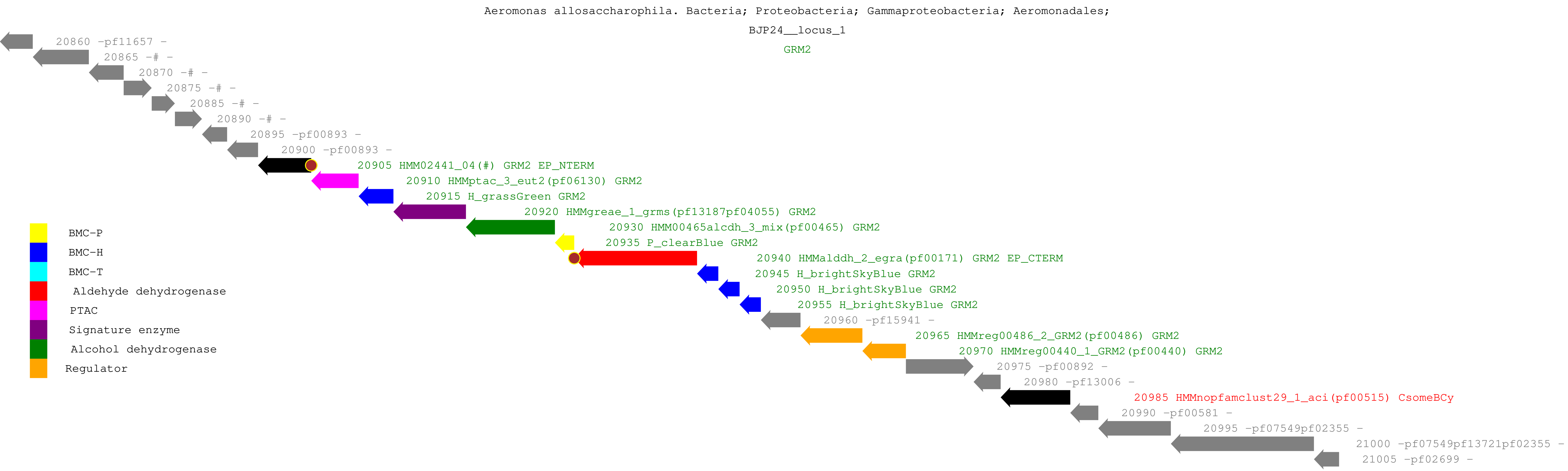
Aeromonas allosaccharophila. Bacteria; Proteobacteria; Gammaproteobacteria; Aeromonadales;
BJP24__locus_1
GRM2
20860 -pf11657 -
20865 -# -
20870 -# -
20875 -# -
20885 -# -
20890 -# -
20895 -pf00893 -
20900 -pf00893 -
20905 HMM02441_04(#) GRM2 EP_NTERM
20910 HMMptac_3_eut2(pf06130) GRM2
20915 H_grassGreen GRM2
20920 HMMgreae_1_grms(pf13187pf04055) GRM2
20930 HMM00465alcdh_3_mix(pf00465) GRM2
20935 P_clearBlue GRM2
20940 HMMalddh_2_egra(pf00171) GRM2 EP_CTERM
20945 H_brightSkyBlue GRM2
20950 H_brightSkyBlue GRM2
20955 H_brightSkyBlue GRM2
20960 -pf15941 -
20965 HMMreg00486_2_GRM2(pf00486) GRM2
20970 HMMreg00440_1_GRM2(pf00440) GRM2
20975 -pf00892 -
20980 -pf13006 -
20985 HMMnopfamclust29_1_aci(pf00515) CsomeBCy
20990 -pf00581 -
20995 -pf07549pf02355 -
21000 -pf07549pf13721pf02355 -
21005 -pf02699 -
BMC-P
BMC-H
BMC-T
Aldehyde dehydrogenase
PTAC
Signature enzyme
Alcohol dehydrogenase
Regulator

Raoultella ornithinolytica (Klebsiella ornithinolytica). Bacteria; Proteobacteria; Gammaproteobacteria; Enterobacterales;

BK817__locus_1

GRM2

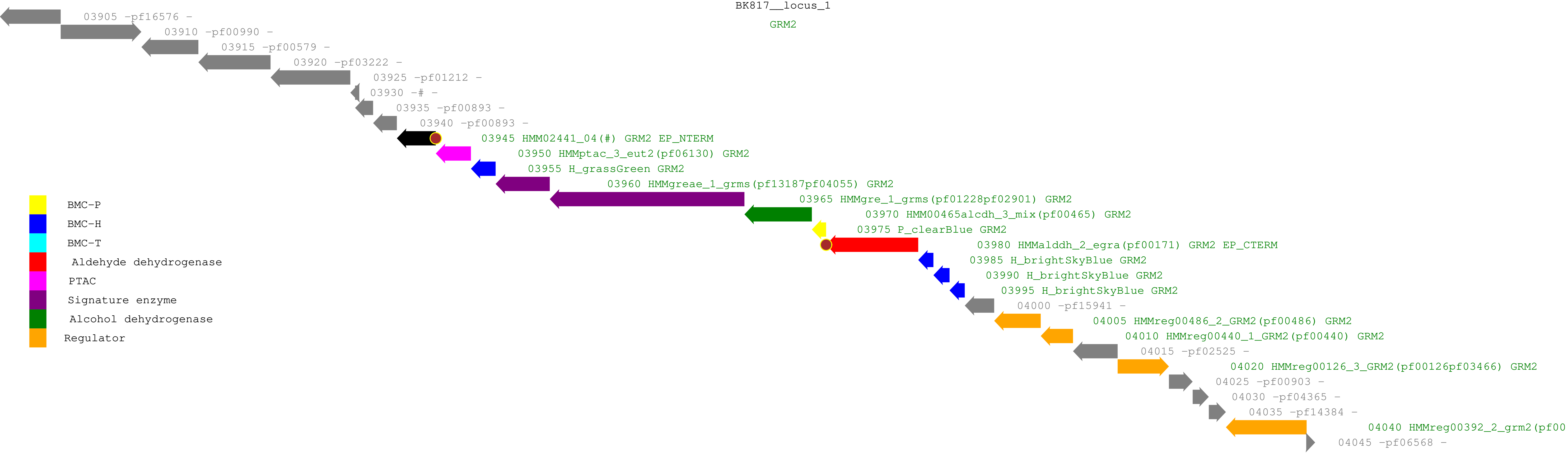

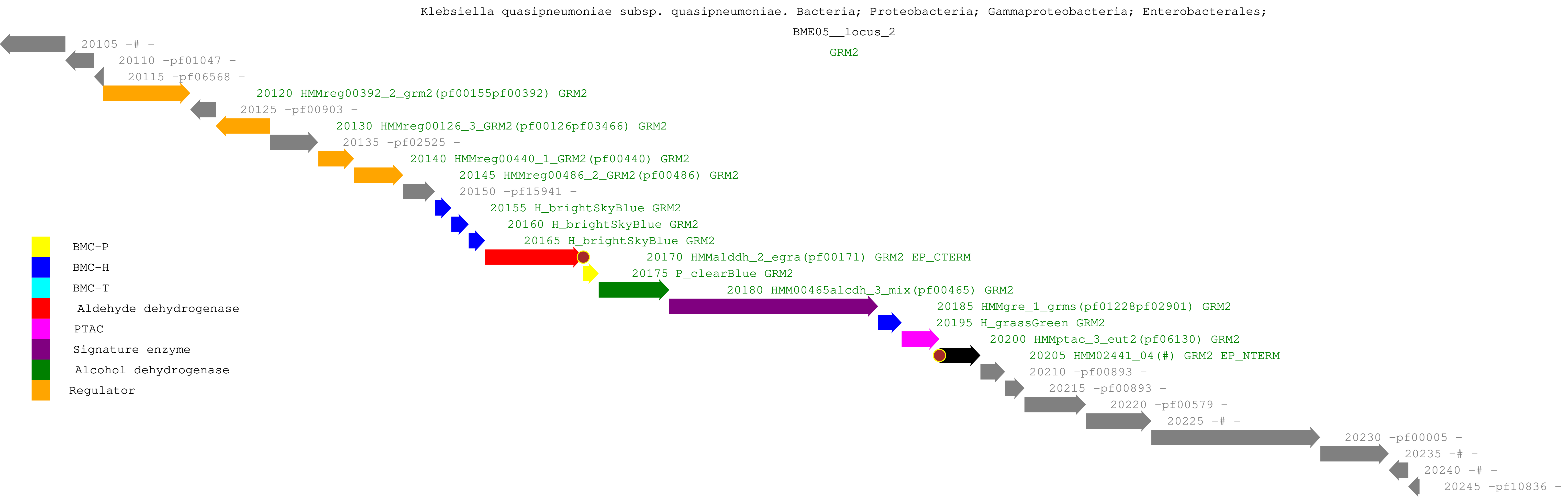
Klebsiella quasipneumoniae subsp. quasipneumoniae. Bacteria; Proteobacteria; Gammaproteobacteria; Enterobacterales;
BME05__locus_2
GRM2
20105 -# -
20110 -pf01047 -
20115 -pf06568 -
20120 HMMreg00392_2_grm2(pf00155pf00392) GRM2
20125 -pf00903 -
20130 HMMreg00126_3_GRM2(pf00126pf03466) GRM2
20135 -pf02525 -
20140 HMMreg00440_1_GRM2(pf00440) GRM2
20145 HMMreg00486_2_GRM2(pf00486) GRM2
20150 -pf15941 -
20155 H_brightSkyBlue GRM2
20160 H_brightSkyBlue GRM2
20165 H_brightSkyBlue GRM2
20170 HMMalddh_2_egra(pf00171) GRM2 EP_CTERM
20175 P_clearBlue GRM2
20180 HMM00465alcdh_3_mix(pf00465) GRM2
20185 HMMgre_1_grms(pf01228pf02901) GRM2
20195 H_grassGreen GRM2
20200 HMMptac_3_eut2(pf06130) GRM2
20205 HMM02441_04(#) GRM2 EP_NTERM
20210 -pf00893 -
20215 -pf00893 -
20220 -pf00579 -
20225 -# -
20230 -pf00005 -
20235 -# -
20240 -# -
20245 -pf10836 -
BMC-P
BMC-H
BMC-T
Aldehyde dehydrogenase
PTAC
Signature enzyme
Alcohol dehydrogenase
Regulator

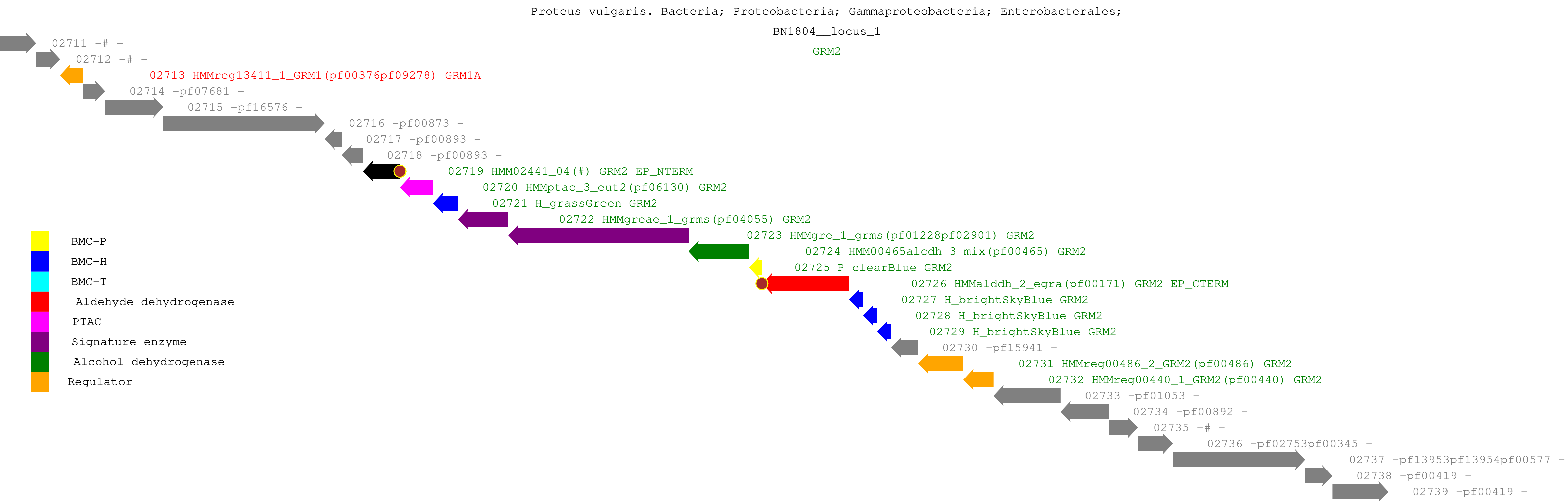
Proteus vulgaris. Bacteria; Proteobacteria; Gammaproteobacteria; Enterobacterales;
BN1804__locus_1
GRM2
02711 -# -
02712 -# -
02713 HMMreg13411_1_GRM1(pf00376pf09278) GRM1A
02714 -pf07681 -
02715 -pf16576 -
02716 -pf00873 -
02717 -pf00893 -
02718 -pf00893 -
02719 HMM02441_04(#) GRM2 EP_NTERM
02720 HMMptac_3_eut2(pf06130) GRM2
02721 H_grassGreen GRM2
02722 HMMgreae_1_grms(pf04055) GRM2
02723 HMMgre_1_grms(pf01228pf02901) GRM2
02724 HMM00465alcdh_3_mix(pf00465) GRM2
02725 P_clearBlue GRM2
02726 HMMalddh_2_egra(pf00171) GRM2 EP_CTERM
02727 H_brightSkyBlue GRM2
02728 H_brightSkyBlue GRM2
02729 H_brightSkyBlue GRM2
02730 -pf15941 -
02731 HMMreg00486_2_GRM2(pf00486) GRM2
02732 HMMreg00440_1_GRM2(pf00440) GRM2
02733 -pf01053 -
02734 -pf00892 -
02735 -# -
02736 -pf02753pf00345 -
02737 -pf13953pf13954pf00577 -
02738 -pf00419 -
02739 -pf00419 -
BMC-P
BMC-H
BMC-T
Aldehyde dehydrogenase
PTAC
Signature enzyme
Alcohol dehydrogenase
Regulator

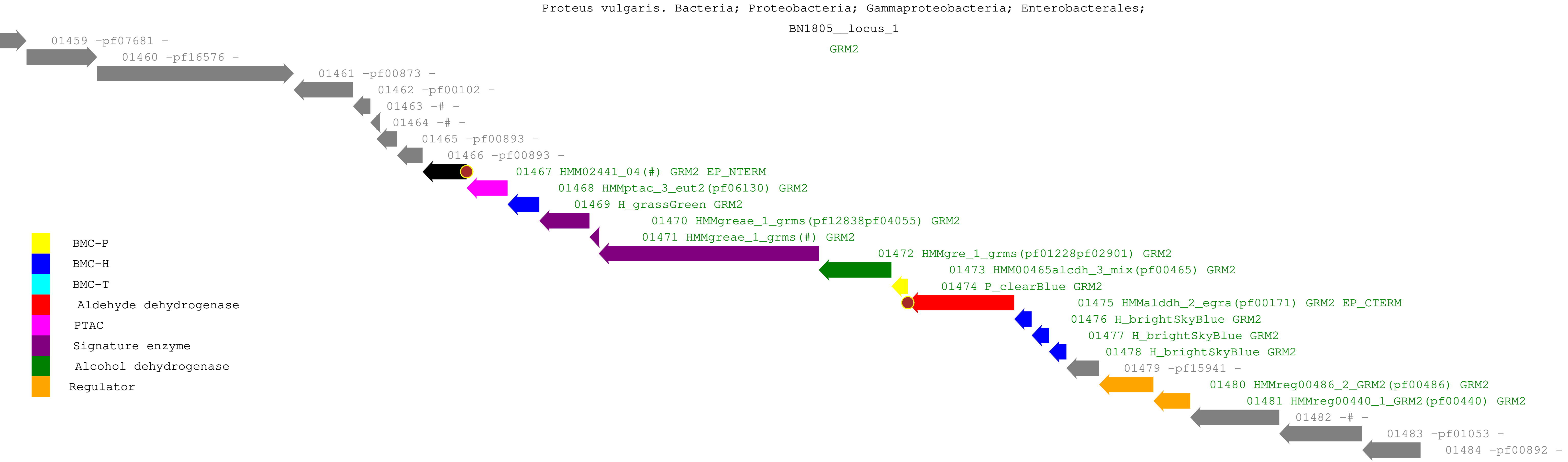
Proteus vulgaris. Bacteria; Proteobacteria; Gammaproteobacteria; Enterobacterales;
BN1805__locus_1
GRM2
01459 -pf07681 -
01460 -pf16576 -
01461 -pf00873 -
01462 -pf00102 -
01463 -# -
01464 -# -
01465 -pf00893 -
01466 -pf00893 -
01467 HMM02441_04(#) GRM2 EP_NTERM
01468 HMMptac_3_eut2(pf06130) GRM2
01469 H_grassGreen GRM2
01470 HMMgreae_1_grms(pf12838pf04055) GRM2
01471 HMMgreae_1_grms(#) GRM2
01472 HMMgre_1_grms(pf01228pf02901) GRM2
01473 HMM00465alcdh_3_mix(pf00465) GRM2
01474 P_clearBlue GRM2
01475 HMMaldh_2_egra(pf00171) GRM2 EP_CTERM
01476 H_brightSkyBlue GRM2
01477 H_brightSkyBlue GRM2
01478 H_brightSkyBlue GRM2
01479 -pf15941 -
01480 HMMreg00486_2_GRM2(pf00486) GRM2
01481 HMMreg00440_1_GRM2(pf00440) GRM2
01482 -# -
01483 -pf01053 -
01484 -pf00892 -
BMC-P
BMC-H
BMC-T
Aldehyde dehydrogenase
PTAC
Signature enzyme
Alcohol dehydrogenase
Regulator

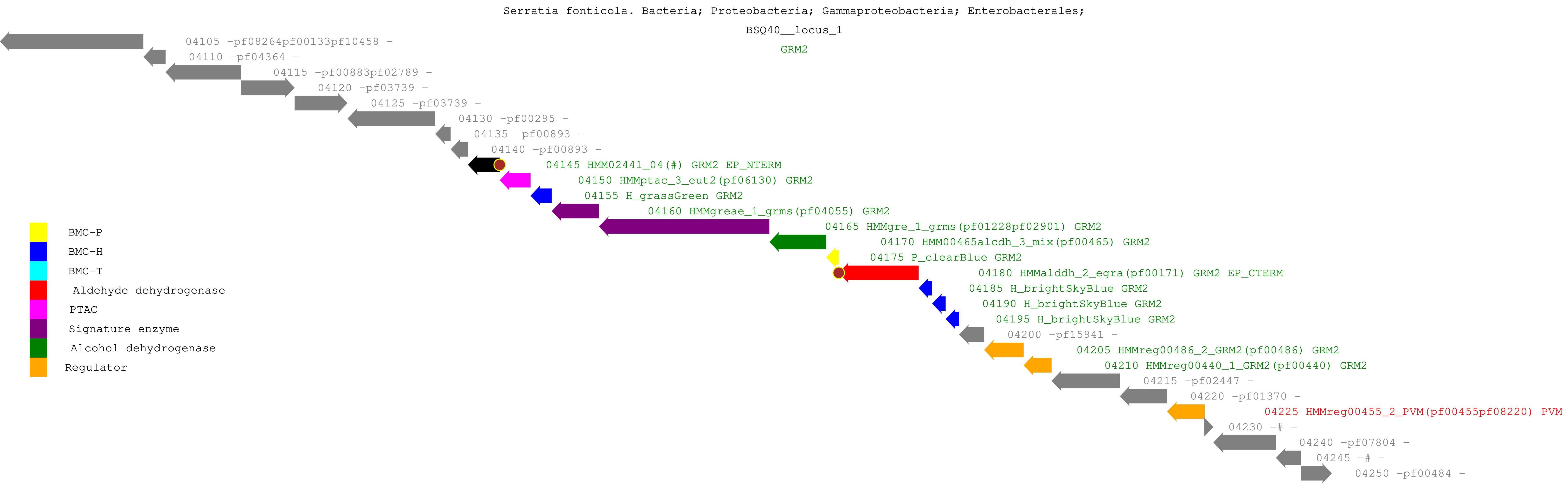

Serratia fonticola. Bacteria; Proteobacteria; Gammaproteobacteria; Enterobacterales;
BSQ40__locus_1
GRM2
04105 -pf08264pf00133pf10458 -
04110 -pf04364 -
04115 -pf00883pf02789 -
04120 -pf03739 -
04125 -pf03739 -
04130 -pf00295 -
04135 -pf00893 -
04140 -pf00893 -
04145 HMM02441_04(#) GRM2 EP_NTERM
04150 HMMptac_3_eut2(pf06130) GRM2
04155 H_grassGreen GRM2
04160 HMMgreae_1_grms(pf04055) GRM2
04165 HMMgre_1_grms(pf01228pf02901) GRM2
04170 HMM00465alcdh_3_mix(pf00465) GRM2
04175 P_clearBlue GRM2
04180 HMMaldh_2_egra(pf00171) GRM2 EP_CTERM
04185 H_brightSkyBlue GRM2
04190 H_brightSkyBlue GRM2
04195 H_brightSkyBlue GRM2
04200 -pf15941 -
04205 HMMreg00486_2_GRM2(pf00486) GRM2
04210 HMMreg00440_1_GRM2(pf00440) GRM2
04215 -pf02447 -
04220 -pf01370 -
04225 HMMreg00455_2_PVM(pf00455pf08220) PVM
04230 -# -
04240 -pf07804 -
04245 -# -
04250 -pf00484 -
BMC-P
BMC-H
BMC-T
Aldehyde dehydrogenase
PTAC
Signature enzyme
Alcohol dehydrogenase
Regulator

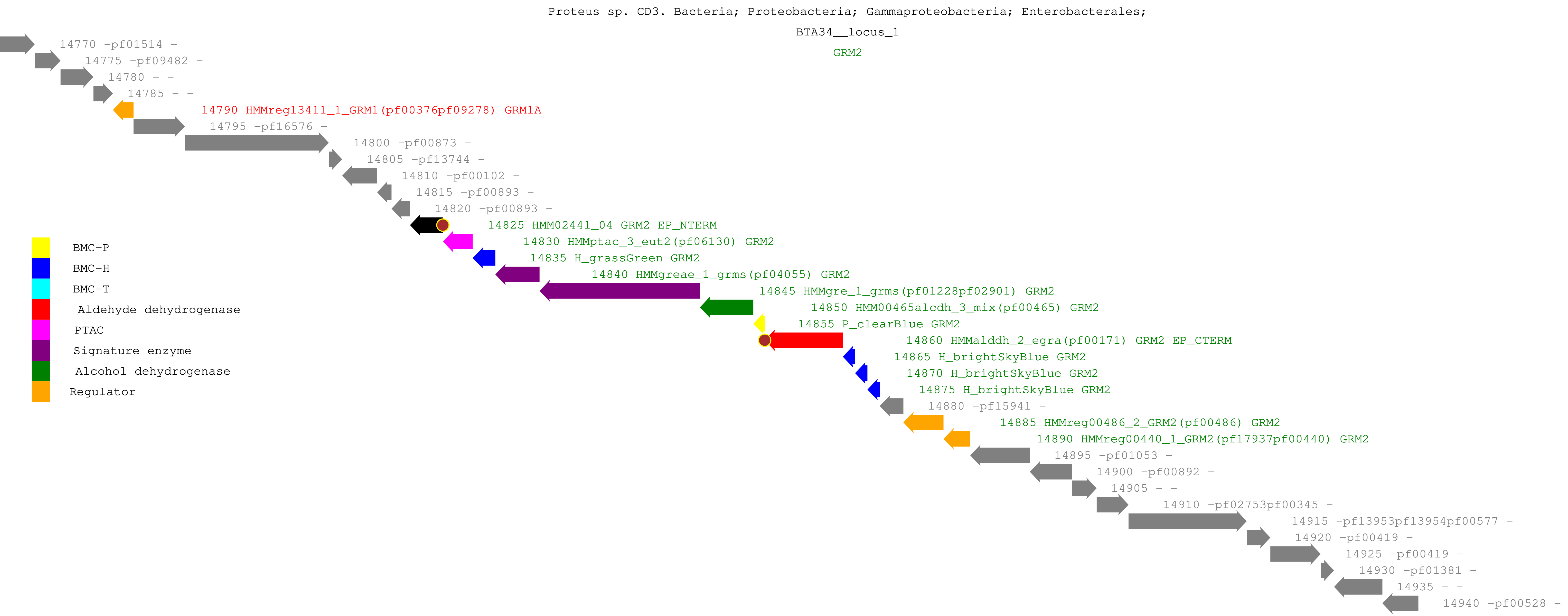
Proteus sp. CD3. Bacteria; Proteobacteria; Gammaproteobacteria; Enterobacterales;
BTA34__locus_1
GRM2
14770 -pf01514 -
14775 -pf09482 -
14780 - -
14785 - -
14790 HMMreg13411_1_GRM1(pf00376pf09278) GRM1A
14795 -pf16576 -
14800 -pf00873 -
14805 -pf13744 -
14810 -pf00102 -
14815 -pf00893 -
14820 -pf00893 -
14825 HMM02441_04 GRM2 EP_NTERM
14830 HMMptac_3_eut2(pf06130) GRM2
14835 H_grassGreen GRM2
14840 HMMgreae_1_grms(pf04055) GRM2
14845 HMMgre_1_grms(pf01228pf02901) GRM2
14850 HMM00465alcdh_3_mix(pf00465) GRM2
14855 P_clearBlue GRM2
14860 HMMaldh_2_egra(pf00171) GRM2 EP_CTERM
14865 H_brightSkyBlue GRM2
14870 H_brightSkyBlue GRM2
14875 H_brightSkyBlue GRM2
14880 -pf15941 -
14885 HMMreg00486_2_GRM2(pf00486) GRM2
14890 HMMreg00440_1_GRM2(pf17937pf00440) GRM2
14895 -pf01053 -
14900 -pf00892 -
14905 - -
14910 -pf02753pf00345 -
14915 -pf13953pf13954pf00577 -
14920 -pf00419 -
14925 -pf00419 -
14930 -pf01381 -
14935 - -
14940 -pf00528 -
BMC-P
BMC-H
BMC-T
Aldehyde dehydrogenase
PTAC
Signature enzyme
Alcohol dehydrogenase
Regulator

Klebsiella pneumoniae. Bacteria; Proteobacteria; Gammaproteobacteria; Enterobacterales;

BU230__locus_1

GRM2

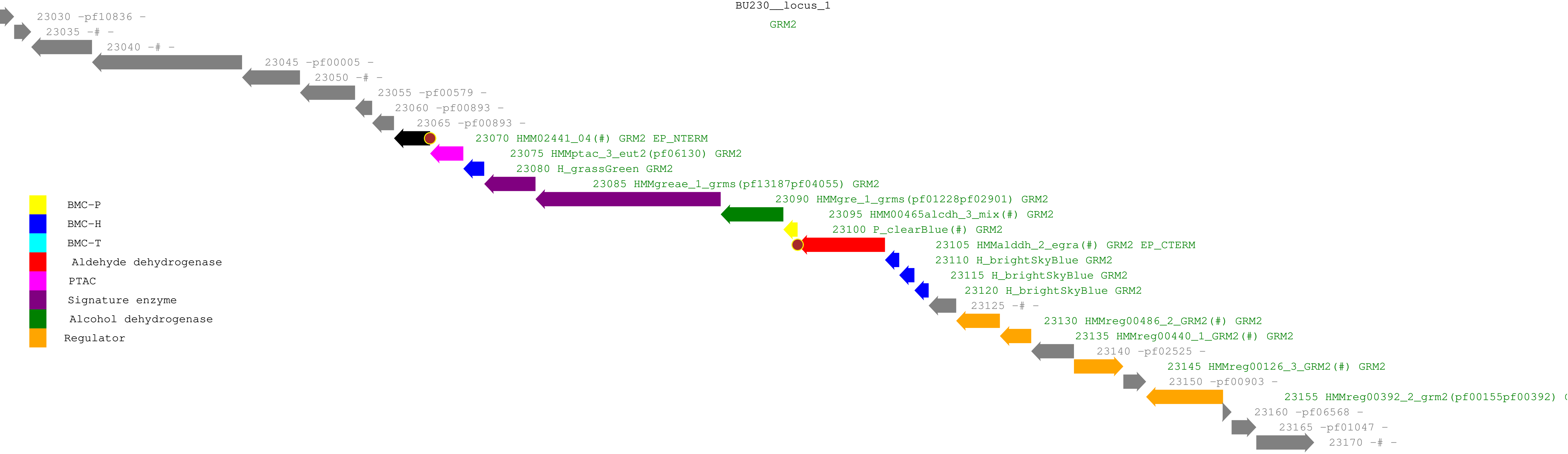

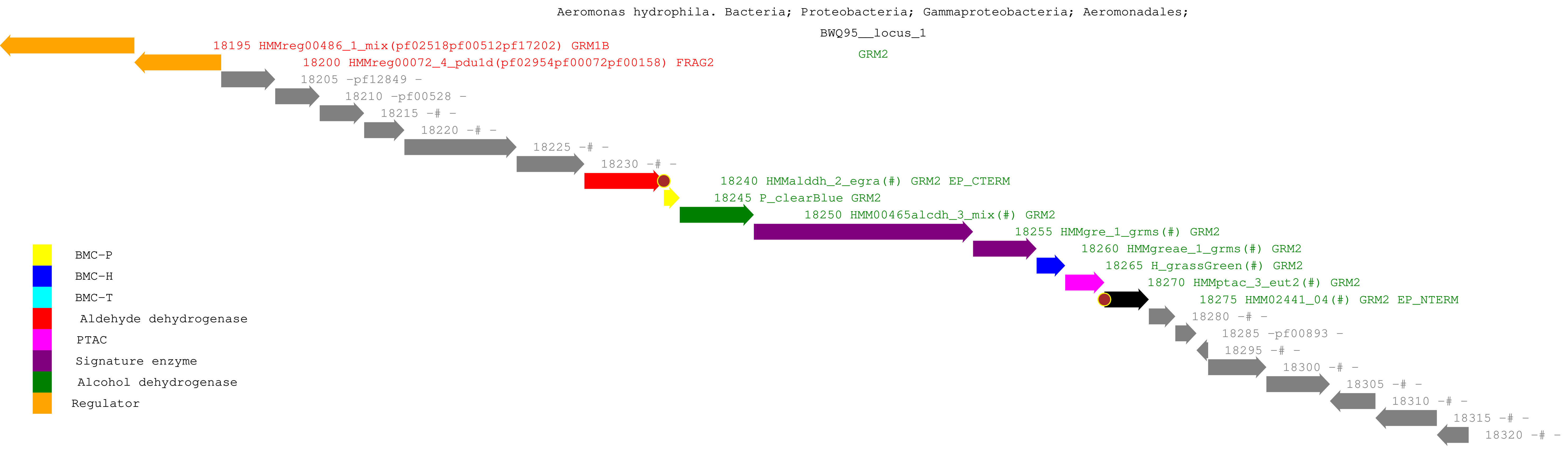

Aeromonas hydrophila. Bacteria; Proteobacteria; Gammaproteobacteria; Aeromonadales;
BWQ95__locus_1
GRM2
18195 HMMreg00486_1_mix(pf02518pf00512pf17202) GRM1B
18200 HMMreg00072_4_pdu1d(pf02954pf00072pf00158) FRAG2
18205 -pf12849 -
18210 -pf00528 -
18215 -# -
18220 -# -
18225 -# -
18230 -# -
18240 HMMalddh_2_egra(#) GRM2 EP_CTERM
18245 P_clearBlue GRM2
18250 HMM00465alcdh_3_mix(#) GRM2
18255 HMMgre_1_grms(#) GRM2
18260 HMMgreae_1_grms(#) GRM2
18265 H_grassGreen(#) GRM2
18270 HMMptac_3_eut2(#) GRM2
18275 HMM02441_04(#) GRM2 EP_NTERM
18280 -# -
18285 -pf00893 -
18295 -# -
18300 -# -
18305 -# -
18310 -# -
18315 -# -
18320 -# -
BMC-P
BMC-H
BMC-T
Aldehyde dehydrogenase
PTAC
Signature enzyme
Alcohol dehydrogenase
Regulator

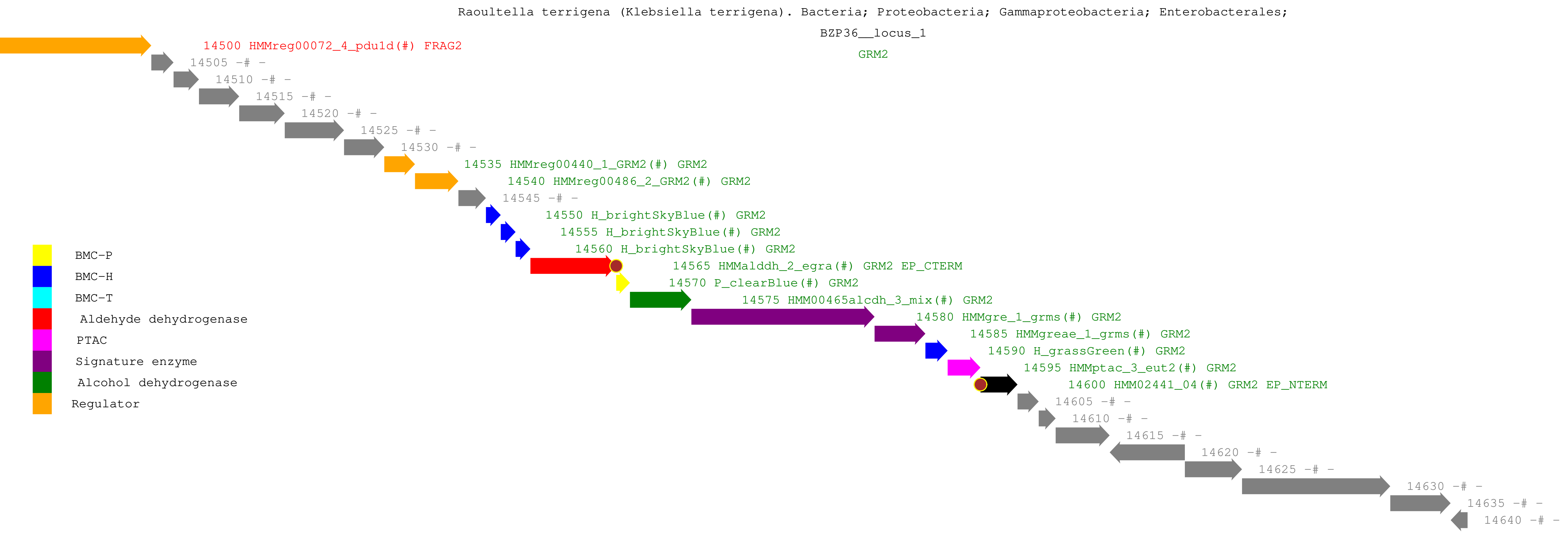
Raoultella terrigena (Klebsiella terrigena). Bacteria; Proteobacteria; Gammaproteobacteria; Enterobacterales;
BZP36__locus_1
GRM2
14500 HMMreg00072_4_pdu1d(#) FRAG2
14505 -# -
14510 -# -
14515 -# -
14520 -# -
14525 -# -
14530 -# -
14535 HMMreg00440_1_GRM2(#) GRM2
14540 HMMreg00486_2_GRM2(#) GRM2
14545 -# -
14550 H_brightSkyBlue(#) GRM2
14555 H_brightSkyBlue(#) GRM2
14560 H_brightSkyBlue(#) GRM2
14565 HMMalddh_2_egra(#) GRM2 EP_CTERM
14570 P_clearBlue(#) GRM2
14575 HMM00465alcdh_3_mix(#) GRM2
14580 HMMgre_1_grms(#) GRM2
14585 HMMgreae_1_grms(#) GRM2
14590 H_grassGreen(#) GRM2
14595 HMMptac_3_eut2(#) GRM2
14600 HMM02441_04(#) GRM2 EP_NTERM
14605 -# -
14610 -# -
14615 -# -
14620 -# -
14625 -# -
14630 -# -
14635 -# -
14640 -# -
BMC-P
BMC-H
BMC-T
Aldehyde dehydrogenase
PTAC
Signature enzyme
Alcohol dehydrogenase
Regulator

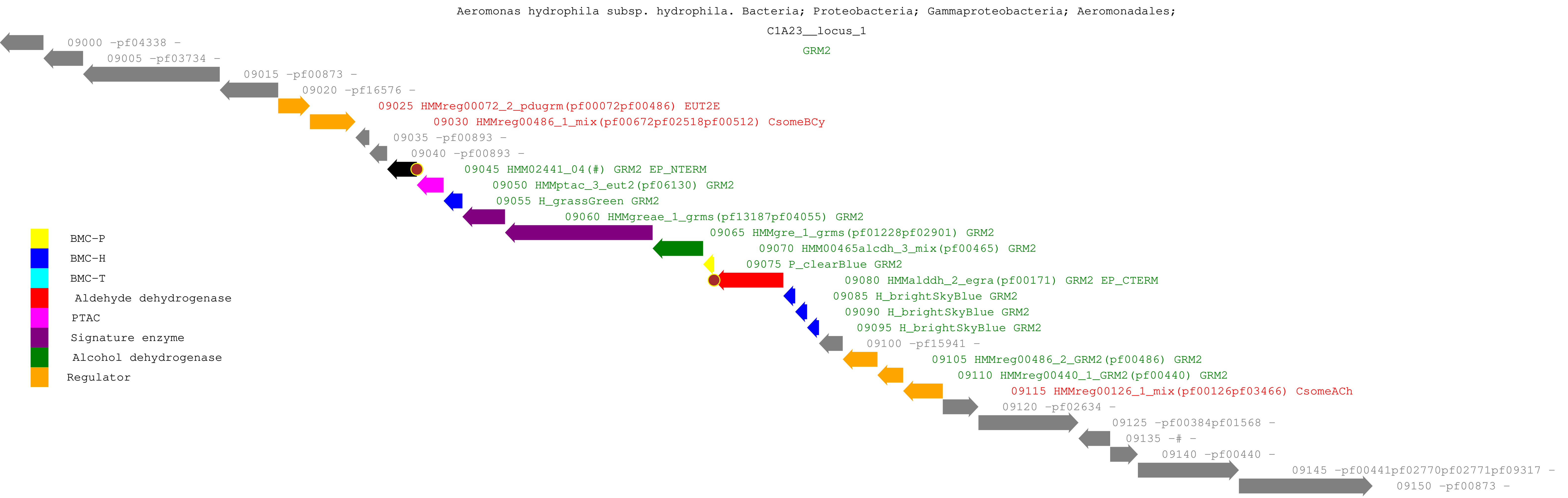
Aeromonas hydrophila subsp. hydrophila. Bacteria; Proteobacteria; Gammaproteobacteria; Aeromonadales;
C1A23__locus_1
GRM2
09000 -pf04338 -
09005 -pf03734 -
09015 -pf00873 -
09020 -pf16576 -
09025 HMMreg00072_2_pdugrm(pf00072pf00486) EUT2E
09030 HMMreg00486_1_mix(pf00672pf02518pf00512) CsomeBCy
09035 -pf00893 -
09040 -pf00893 -
09045 HMM02441_04(#) GRM2 EP_NTERM
09050 HMMptac_3_eut2(pf06130) GRM2
09055 H_grassGreen GRM2
09060 HMMgreae_1_grms(pf13187pf04055) GRM2
09065 HMMgre_1_grms(pf01228pf02901) GRM2
09070 HMM00465alcdh_3_mix(pf00465) GRM2
09075 P_clearBlue GRM2
09080 HMMalddh_2_egra(pf00171) GRM2 EP_CTERM
09085 H_brightSkyBlue GRM2
09090 H_brightSkyBlue GRM2
09095 H_brightSkyBlue GRM2
09100 -pf15941 -
09105 HMMreg00486_2_GRM2(pf00486) GRM2
09110 HMMreg00440_1_GRM2(pf00440) GRM2
09115 HMMreg00126_1_mix(pf00126pf03466) CsomeACh
09120 -pf02634 -
09125 -pf00384pf01568 -
09135 -# -
09140 -pf00440 -
09145 -pf00441pf02770pf02771pf09317 -
09150 -pf00873 -
BMC-P
BMC-H
BMC-T
Aldehyde dehydrogenase
PTAC
Signature enzyme
Alcohol dehydrogenase
Regulator

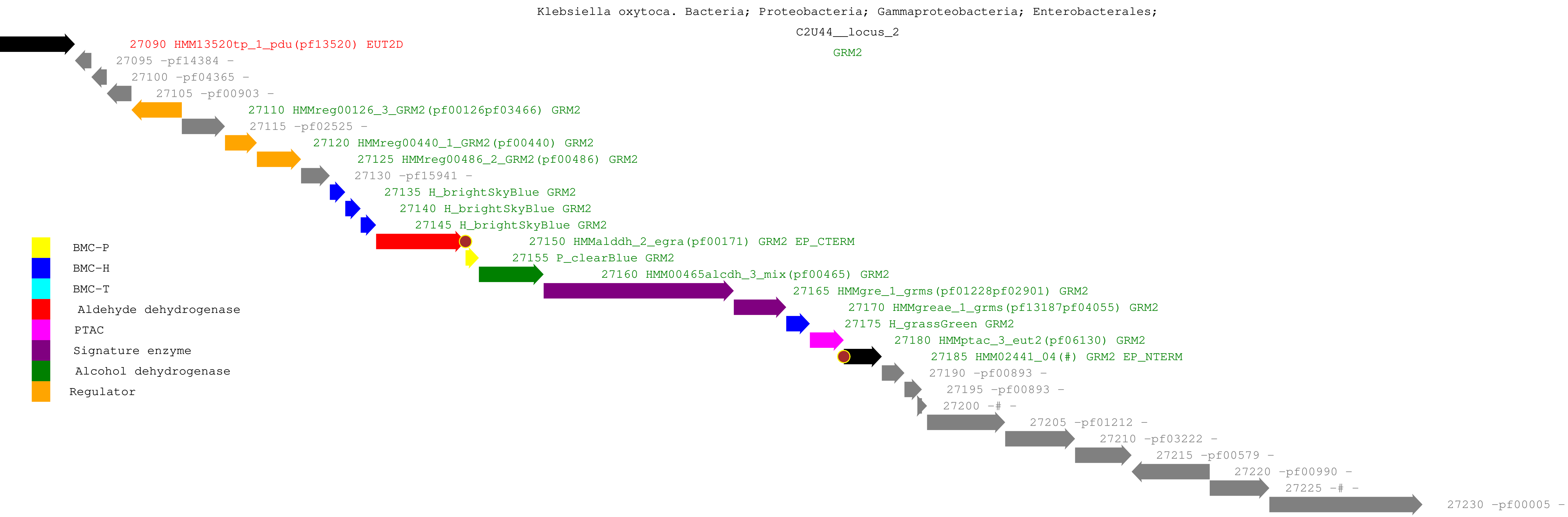
Klebsiella oxytoca. Bacteria; Proteobacteria; Gammaproteobacteria; Enterobacterales;
C2U44__locus_2
GRM2
27090 HMM13520tp_1_pdu(pf13520) EUT2D
27095 -pf14384 -
27100 -pf04365 -
27105 -pf00903 -
27110 HMMreg00126_3_GRM2(pf00126pf03466) GRM2
27115 -pf02525 -
27120 HMMreg00440_1_GRM2(pf00440) GRM2
27125 HMMreg00486_2_GRM2(pf00486) GRM2
27130 -pf15941 -
27135 H_brightSkyBlue GRM2
27140 H_brightSkyBlue GRM2
27145 H_brightSkyBlue GRM2
27150 HMMalddh_2_egra(pf00171) GRM2 EP_CTERM
27155 P_clearBlue GRM2
27160 HMM00465alcdh_3_mix(pf00465) GRM2
27165 HMMgre_1_grms(pf01228pf02901) GRM2
27170 HMMgreae_1_grms(pf13187pf04055) GRM2
27175 H_grassGreen GRM2
27180 HMMptac_3_eut2(pf06130) GRM2
27185 HMM02441_04(#) GRM2 EP_NTERM
27190 -pf00893 -
27195 -pf00893 -
27200 -# -
27205 -pf01212 -
27210 -pf03222 -
27215 -pf00579 -
27220 -pf00990 -
27225 -# -
27230 -pf00005 -
BMC-P
BMC-H
BMC-T
Aldehyde dehydrogenase
PTAC
Signature enzyme
Alcohol dehydrogenase
Regulator

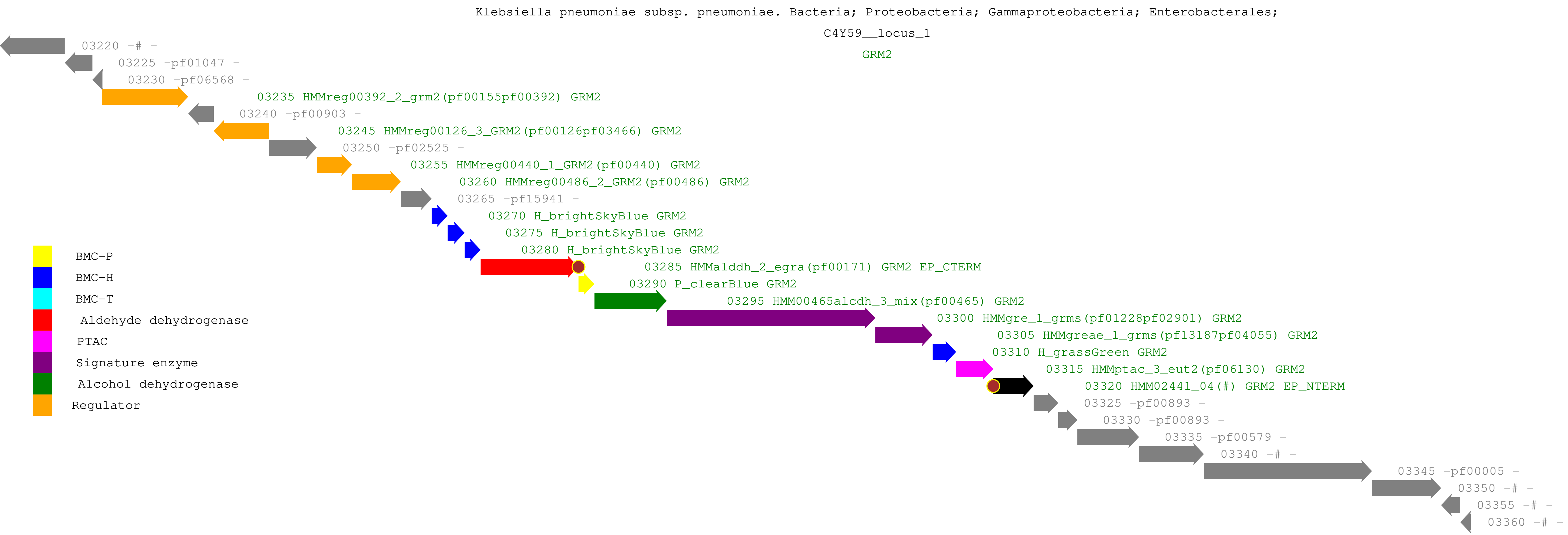
Klebsiella pneumoniae subsp. pneumoniae. Bacteria; Proteobacteria; Gammaproteobacteria; Enterobacterales;
C4Y59__locus_1
GRM2
03220 -# -
03225 -pf01047 -
03230 -pf06568 -
03235 HMMreg00392_2_grm2(pf00155pf00392) GRM2
03240 -pf00903 -
03245 HMMreg00126_3_GRM2(pf00126pf03466) GRM2
03250 -pf02525 -
03255 HMMreg00440_1_GRM2(pf00440) GRM2
03260 HMMreg00486_2_GRM2(pf00486) GRM2
03265 -pf15941 -
03270 H_brightSkyBlue GRM2
03275 H_brightSkyBlue GRM2
03280 H_brightSkyBlue GRM2
03285 HMMaldd h_2_egra(pf00171) GRM2 EP_CTERM
03290 P_clearBlue GRM2
03295 HMM00465alcdh_3_mix(pf00465) GRM2
03300 HMMgre_1_grms(pf01228pf02901) GRM2
03305 HMMgreae_1_grms(pf13187pf04055) GRM2
03310 H_grassGreen GRM2
03315 HMMptac_3_eut2(pf06130) GRM2
03320 HMM02441_04(#) GRM2 EP_NTERM
03325 -pf00893 -
03330 -pf00893 -
03335 -pf00579 -
03340 -# -
03345 -pf00005 -
03350 -# -
03355 -# -
03360 -# -
BMC-P
BMC-H
BMC-T
Aldehyde dehydrogenase
PTAC
Signature enzyme
Alcohol dehydrogenase
Regulator

Klebsiella quasipneumoniae. Bacteria; Proteobacteria; Gammaproteobacteria; Enterobacterales;

C5672__locus_2

GRM2

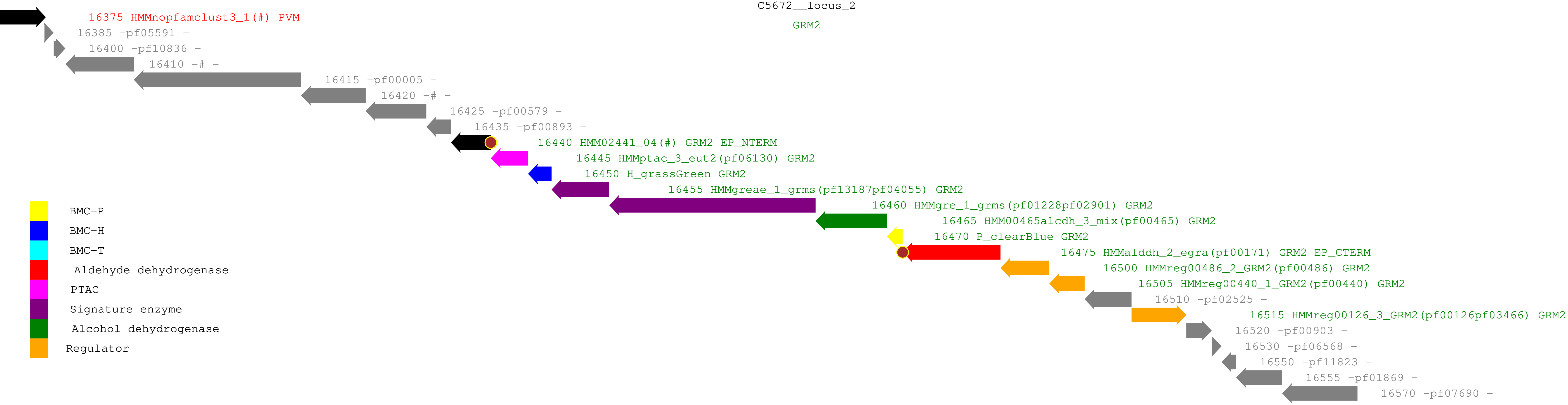

Proteus mirabilis. Bacteria; Proteobacteria; Gammaproteobacteria; Enterobacterales;

C6A90__locus_1

GRM2

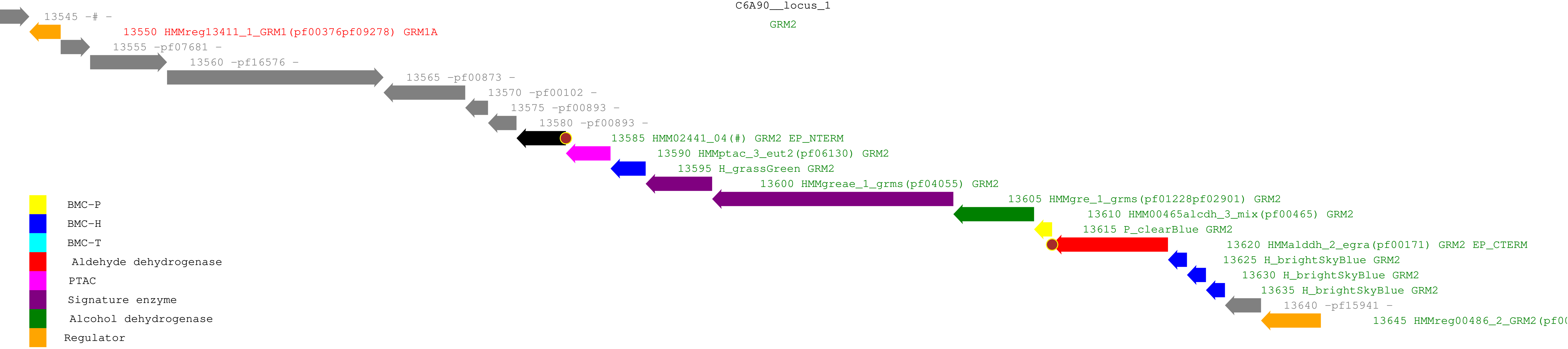

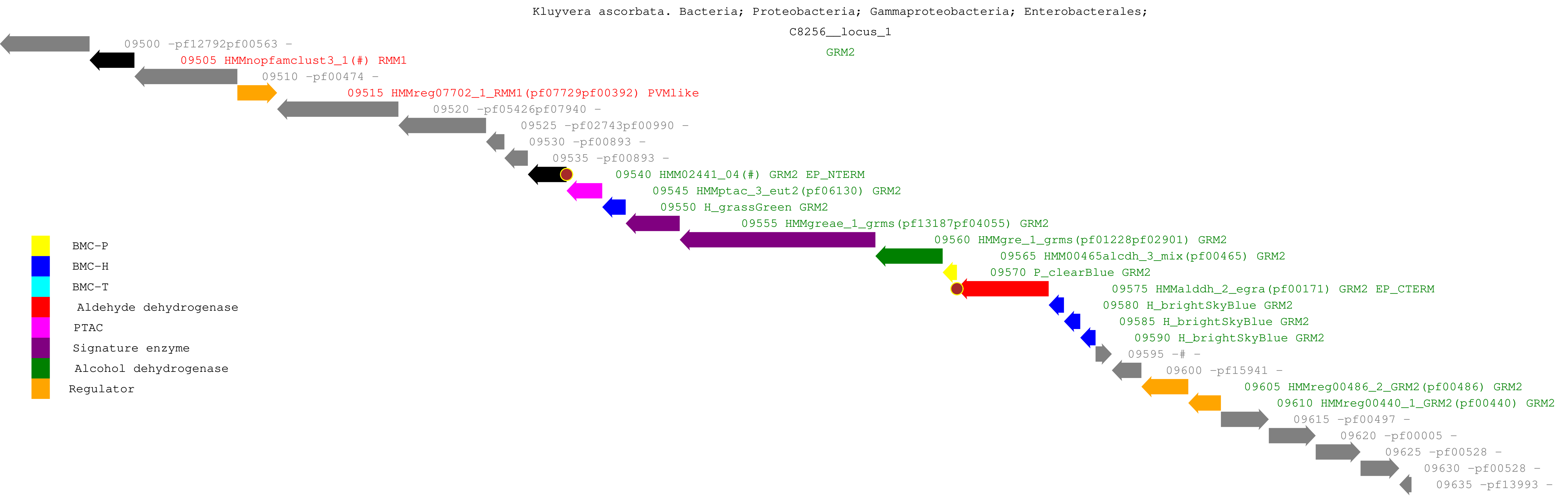
Kluyvera ascorbata. Bacteria; Proteobacteria; Gammaproteobacteria; Enterobacterales;
C8256__locus_1
GRM2
09500 -pf12792pf00563 -
09505 HMMnopfamclust3_1(#) RMM1
09510 -pf00474 -
09515 HMMreg07702_1_RMM1(pf07729pf00392) PVMlike
09520 -pf05426pf07940 -
09525 -pf02743pf00990 -
09530 -pf00893 -
09535 -pf00893 -
09540 HMM02441_04(#) GRM2 EP_NTERM
09545 HMMptac_3_eut2(pf06130) GRM2
09550 H_grassGreen GRM2
09555 HMMgreae_1_grms(pf13187pf04055) GRM2
09560 HMMgre_1_grms(pf01228pf02901) GRM2
09565 HMM00465alcdh_3_mix(pf00465) GRM2
09570 P_clearBlue GRM2
09575 HMMaldh_2_egra(pf00171) GRM2 EP_CTERM
09580 H_brightSkyBlue GRM2
09585 H_brightSkyBlue GRM2
09590 H_brightSkyBlue GRM2
09595 -# -
09600 -pf15941 -
09605 HMMreg00486_2_GRM2(pf00486) GRM2
09610 HMMreg00440_1_GRM2(pf00440) GRM2
09615 -pf00497 -
09620 -pf00005 -
09625 -pf00528 -
09630 -pf00528 -
09635 -pf13993 -
BMC-P
BMC-H
BMC-T
Aldehyde dehydrogenase
PTAC
Signature enzyme
Alcohol dehydrogenase
Regulator

Enterobacillus tribolii. Bacteria; Proteobacteria; Gammaproteobacteria; Enterobacterales;

C8D90__locus_1

GRM2

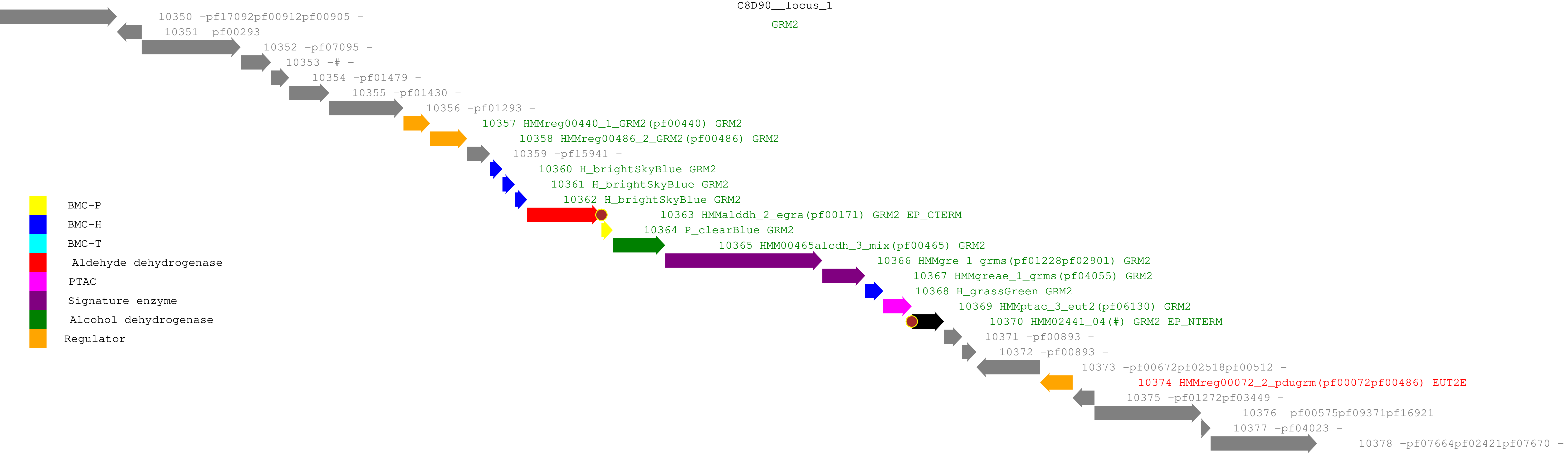

Providencia heimbachae. Bacteria; Proteobacteria; Gammaproteobacteria; Enterobacterales;

C9446__locus_1

GRM2

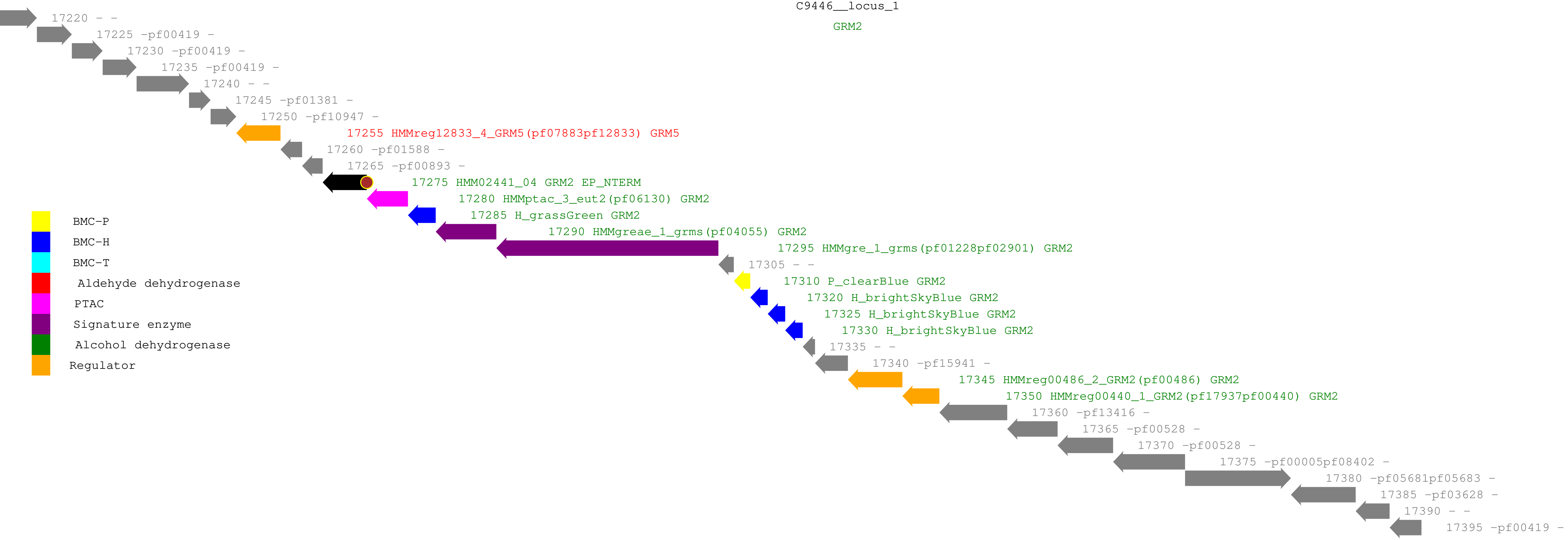

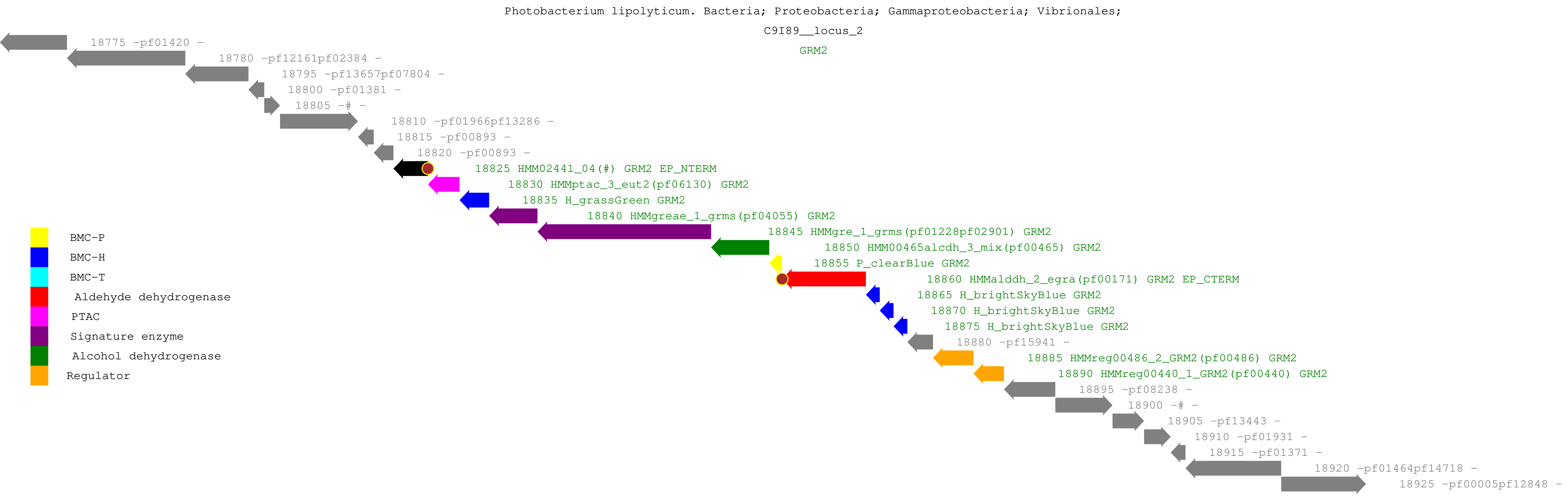
Photobacterium lipolyticum. Bacteria; Proteobacteria; Gammaproteobacteria; Vibrionales;
C9I89__locus_2
GRM2
18775 -pf01420 -
18780 -pf12161pf02384 -
18795 -pf13657pf07804 -
18800 -pf01381 -
18805 -# -
18810 -pf01966pf13286 -
18815 -pf00893 -
18820 -pf00893 -
18825 HMM02441_04(#) GRM2 EP_NTERM
18830 HMMptac_3_eut2(pf06130) GRM2
18835 H_grassGreen GRM2
18840 HMMgreae_1_grms(pf04055) GRM2
18845 HMMgre_1_grms(pf01228pf02901) GRM2
18850 HMM00465alcdh_3_mix(pf00465) GRM2
18855 P_clearBlue GRM2
18860 HMMalddh_2_egra(pf00171) GRM2 EP_CTERM
18865 H_brightSkyBlue GRM2
18870 H_brightSkyBlue GRM2
18875 H_brightSkyBlue GRM2
18880 -pf15941 -
18885 HMMreg00486_2_GRM2(pf00486) GRM2
18890 HMMreg00440_1_GRM2(pf00440) GRM2
18895 -pf08238 -
18900 -# -
18905 -pf13443 -
18910 -pf01931 -
18915 -pf01371 -
18920 -pf01464pf14718 -
18925 -pf00005pf12848 -
BMC-P
BMC-H
BMC-T
Aldehyde dehydrogenase
PTAC
Signature enzyme
Alcohol dehydrogenase
Regulator

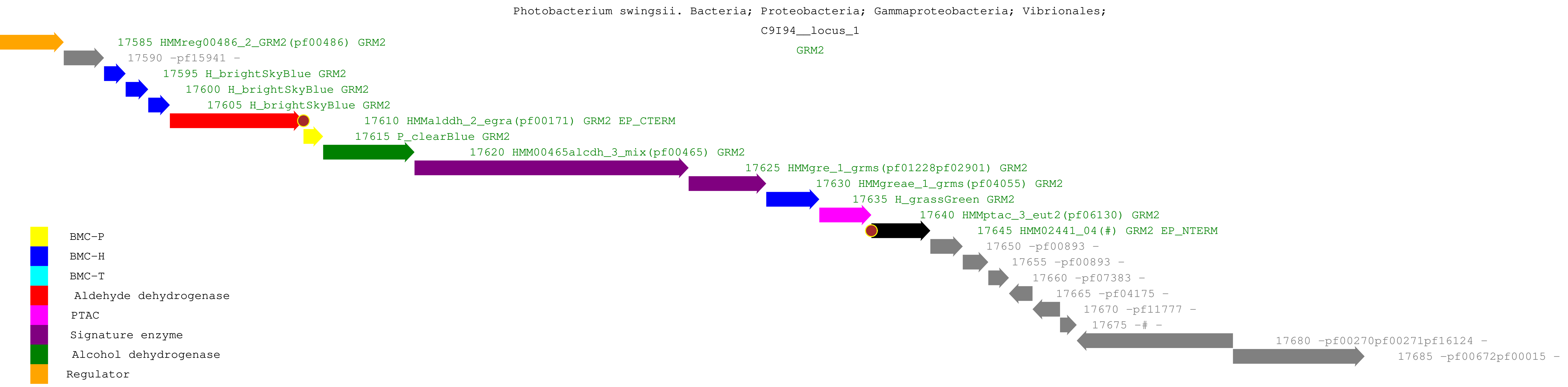

Photobacterium swingsii. Bacteria; Proteobacteria; Gammaproteobacteria; Vibrionales;
C9I94__locus_1
GRM2
17585 HMMreg00486_2_GRM2(pf00486) GRM2
17590 -pf15941 -
17595 H_brightSkyBlue GRM2
17600 H_brightSkyBlue GRM2
17605 H_brightSkyBlue GRM2
17610 HMMalddh_2_egra(pf00171) GRM2 EP_CTERM
17615 P_clearBlue GRM2
17620 HMM00465alcdh_3_mix(pf00465) GRM2
17625 HMMgre_1_grms(pf01228pf02901) GRM2
17630 HMMgreae_1_grms(pf04055) GRM2
17635 H_grassGreen GRM2
17640 HMMptac_3_eut2(pf06130) GRM2
17645 HMM02441_04(#) GRM2 EP_NTERM
17650 -pf00893 -
17655 -pf00893 -
17660 -pf07383 -
17665 -pf04175 -
17670 -pf11777 -
17675 -# -
17680 -pf00270pf00271pf16124 -
17685 -pf00672pf00015 -
BMC-P
BMC-H
BMC-T
Aldehyde dehydrogenase
PTAC
Signature enzyme
Alcohol dehydrogenase
Regulator

Raoultella ornithinolytica (Klebsiella ornithinolytica). Bacteria; Proteobacteria; Gammaproteobacteria; Enterobacterales;

CAC00__locus_2

GRM2

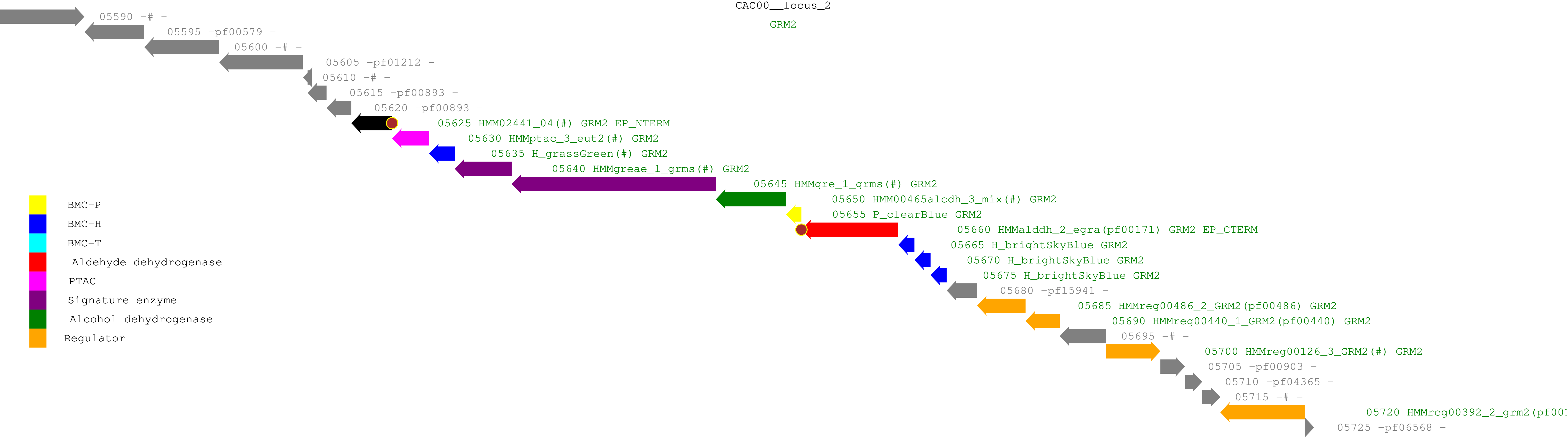

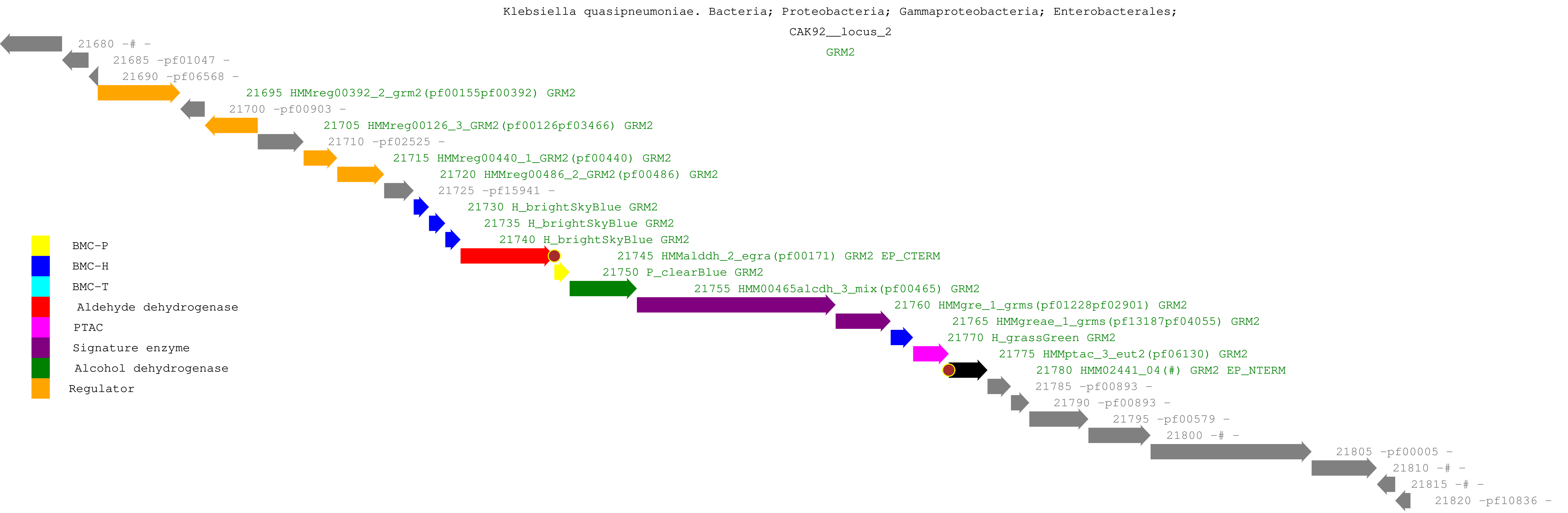
Klebsiella quasipneumoniae. Bacteria; Proteobacteria; Gammaproteobacteria; Enterobacterales;
CAK92__locus_2
GRM2
21680 -# -
21685 -pf01047 -
21690 -pf06568 -
21695 HMMreg00392_2_grm2(pf00155pf00392) GRM2
21700 -pf00903 -
21705 HMMreg00126_3_GRM2(pf00126pf03466) GRM2
21710 -pf02525 -
21715 HMMreg00440_1_GRM2(pf00440) GRM2
21720 HMMreg00486_2_GRM2(pf00486) GRM2
21725 -pf15941 -
21730 H_brightSkyBlue GRM2
21735 H_brightSkyBlue GRM2
21740 H_brightSkyBlue GRM2
21745 HMMaldh_2_egra(pf00171) GRM2 EP_CTERM
21750 P_clearBlue GRM2
21755 HMM00465alcdh_3_mix(pf00465) GRM2
21760 HMMgre_1_grms(pf01228pf02901) GRM2
21765 HMMgreae_1_grms(pf13187pf04055) GRM2
21770 H_grassGreen GRM2
21775 HMMptac_3_eut2(pf06130) GRM2
21780 HMM02441_04(#) GRM2 EP_NTERM
21785 -pf00893 -
21790 -pf00893 -
21795 -pf00579 -
21800 -# -
21805 -pf00005 -
21810 -# -
21815 -# -
21820 -pf10836 -
BMC-P
BMC-H
BMC-T
Aldehyde dehydrogenase
PTAC
Signature enzyme
Alcohol dehydrogenase
Regulator

Klebsiella pneumoniae. Bacteria; Proteobacteria; Gammaproteobacteria; Enterobacterales;

CBL45__locus_1

GRM2

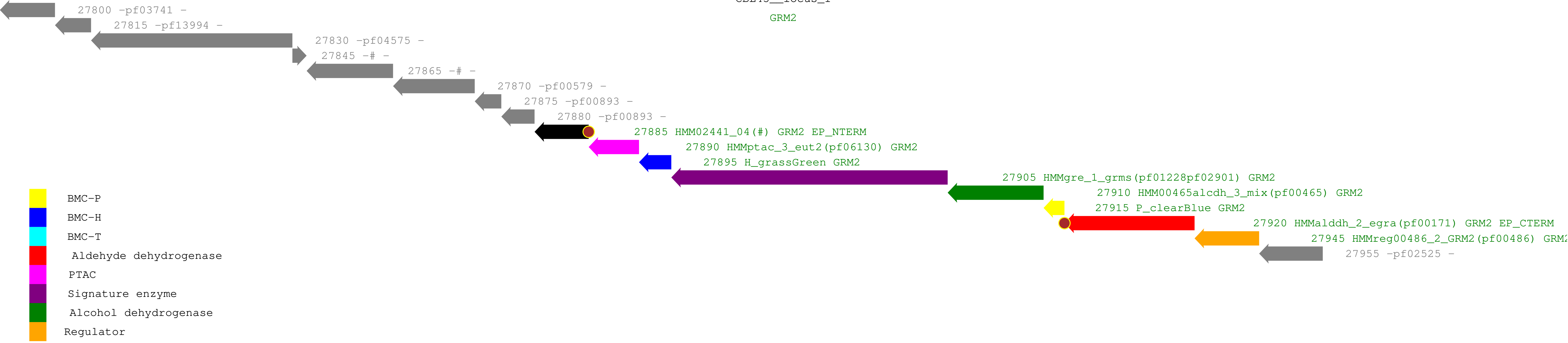

Enterobacter sp. 10-1. Bacteria; Proteobacteria; Gammaproteobacteria; Enterobacterales;

CD006__locus_2

GRM2

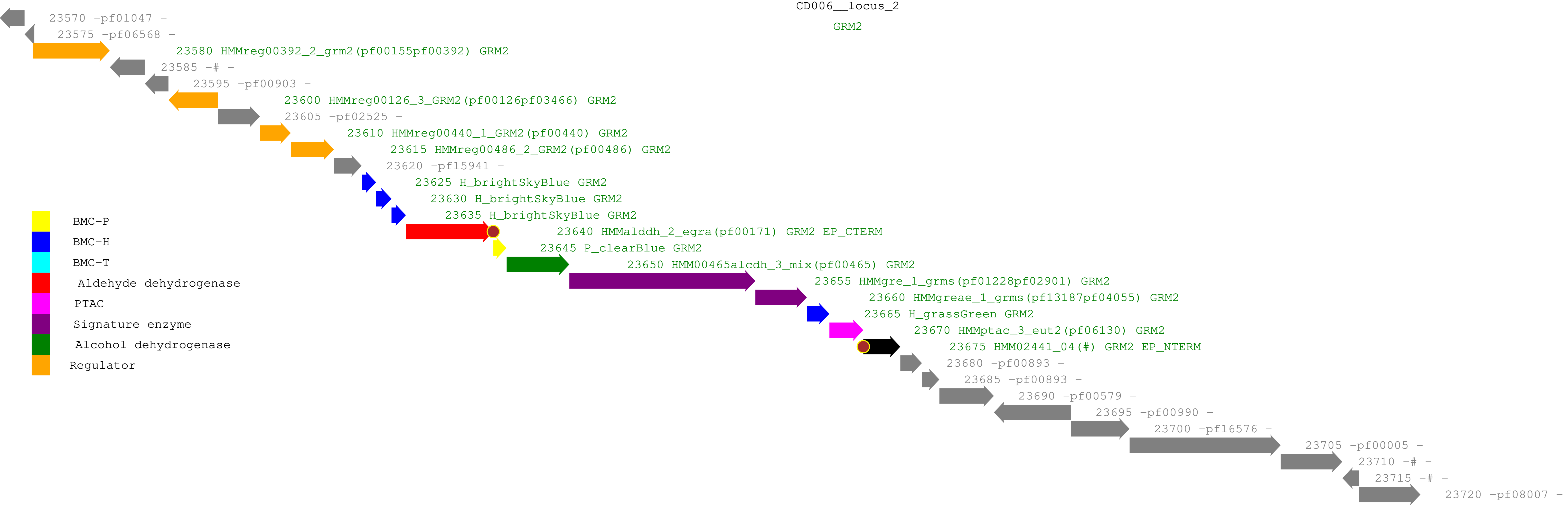

Escherichia coli O7:K1 str. CE10. Bacteria; Proteobacteria; Gammaproteobacteria; Enterobacterales;

CE10__locus_2

GRM2

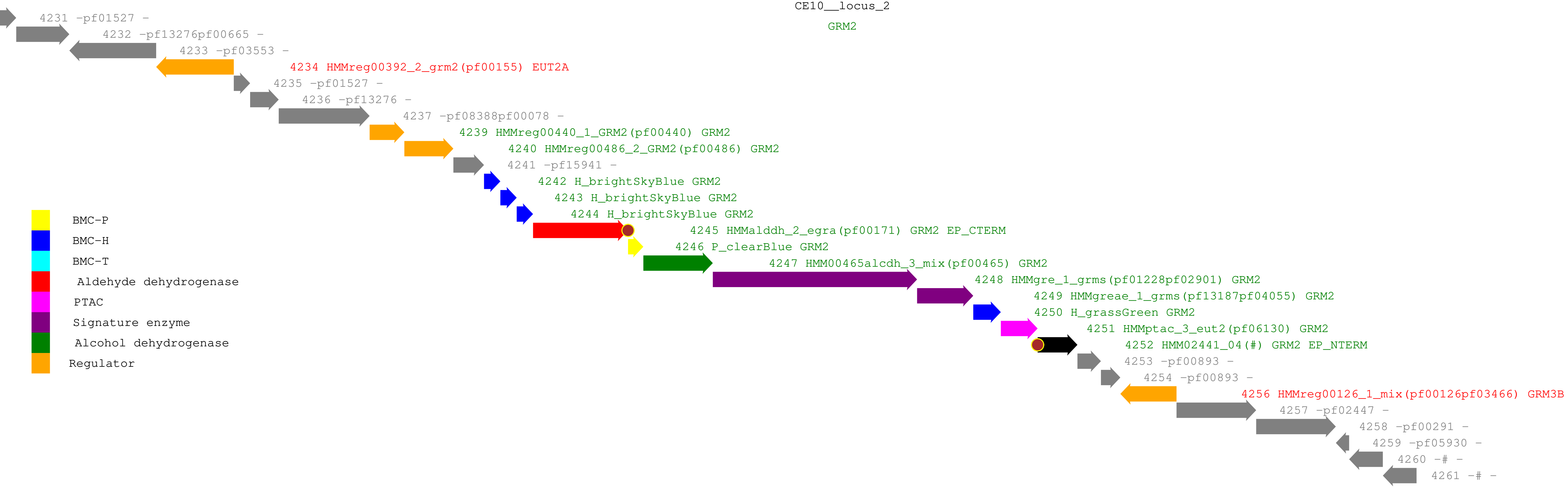

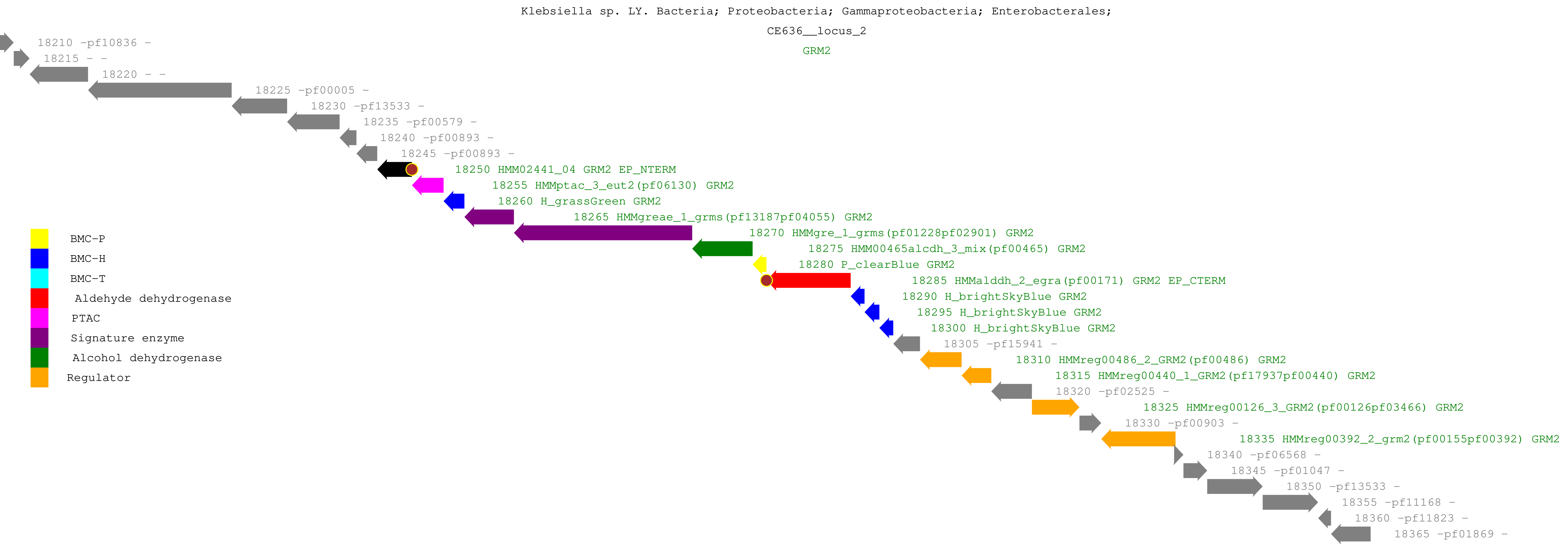
Klebsiella sp. LY. Bacteria; Proteobacteria; Gammaproteobacteria; Enterobacterales;
CE636__locus_2
GRM2
18210 -pf10836 -
18215 - -
18220 - -
18225 -pf00005 -
18230 -pf13533 -
18235 -pf00579 -
18240 -pf00893 -
18245 -pf00893 -
18250 HMM02441_04 GRM2 EP_NTERM
18255 HMMptac_3_eut2(pf06130) GRM2
18260 H_grassGreen GRM2
18265 HMMgreae_1_grms(pf13187pf04055) GRM2
18270 HMMgre_1_grms(pf01228pf02901) GRM2
18275 HMM00465alcdh_3_mix(pf00465) GRM2
18280 P_clearBlue GRM2
18285 HMMalddh_2_egra(pf00171) GRM2 EP_CTERM
18290 H_brightSkyBlue GRM2
18295 H_brightSkyBlue GRM2
18300 H_brightSkyBlue GRM2
18305 -pf15941 -
18310 HMMreg00486_2_GRM2(pf00486) GRM2
18315 HMMreg00440_1_GRM2(pf17937pf00440) GRM2
18320 -pf02525 -
18325 HMMreg00126_3_GRM2(pf00126pf03466) GRM2
18330 -pf00903 -
18335 HMMreg00392_2_grm2(pf00155pf00392) GRM2
18340 -pf06568 -
18345 -pf01047 -
18350 -pf13533 -
18355 -pf11168 -
18360 -pf11823 -
18365 -pf01869 -
BMC-P
BMC-H
BMC-T
Aldehyde dehydrogenase
PTAC
Signature enzyme
Alcohol dehydrogenase
Regulator

Proteus mirabilis. Bacteria; Proteobacteria; Gammaproteobacteria; Enterobacterales;

CEP63__locus_1

GRM2

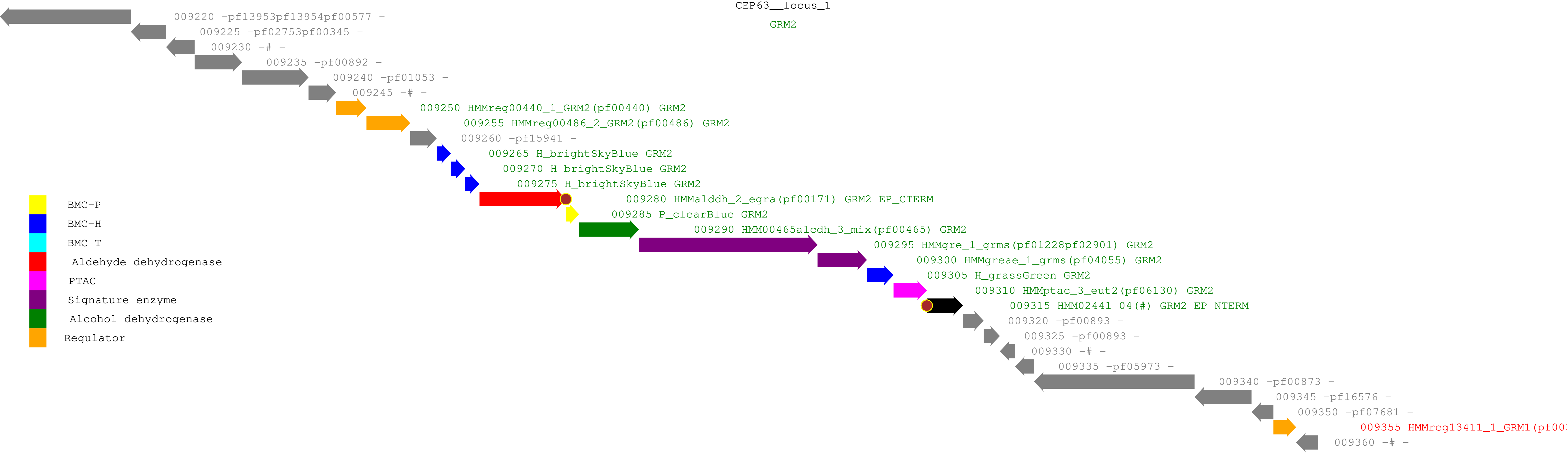

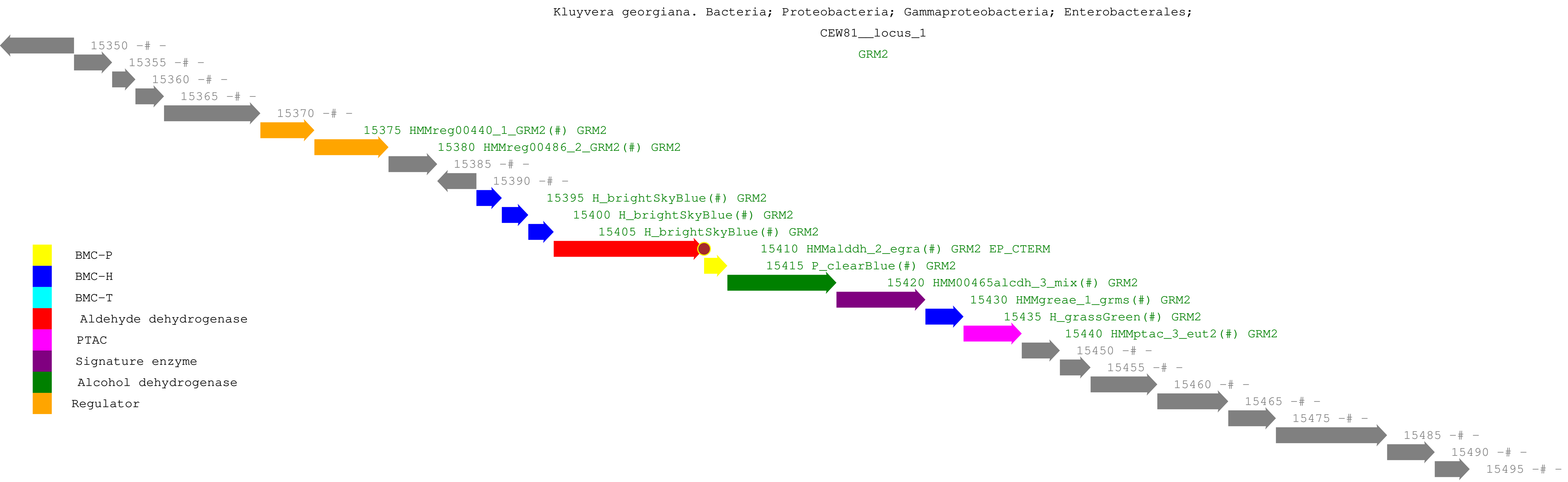

Kluyvera georgiana. Bacteria; Proteobacteria; Gammaproteobacteria; Enterobacterales;
CEW81__locus_1
GRM2
15350 -# -
15355 -# -
15360 -# -
15365 -# -
15370 -# -
15375 HMMreg00440_1_GRM2(#) GRM2
15380 HMMreg00486_2_GRM2(#) GRM2
15385 -# -
15390 -# -
15395 H_brightSkyBlue(#) GRM2
15400 H_brightSkyBlue(#) GRM2
15405 H_brightSkyBlue(#) GRM2
15410 HMMalddh_2_egra(#) GRM2 EP_CTERM
15415 P_clearBlue(#) GRM2
15420 HMM00465alcdh_3_mix(#) GRM2
15430 HMMgreae_1_grms(#) GRM2
15435 H_grassGreen(#) GRM2
15440 HMMptac_3_eut2(#) GRM2
15450 -# -
15455 -# -
15460 -# -
15465 -# -
15475 -# -
15485 -# -
15490 -# -
15495 -# -
BMC-P
BMC-H
BMC-T
Aldehyde dehydrogenase
PTAC
Signature enzyme
Alcohol dehydrogenase
Regulator

Klebsiella quasipneumoniae subsp. similipneumoniae. Bacteria; Proteobacteria; Gammaproteobacteria; Enterobacterales;

CI612__locus_2

GRM2

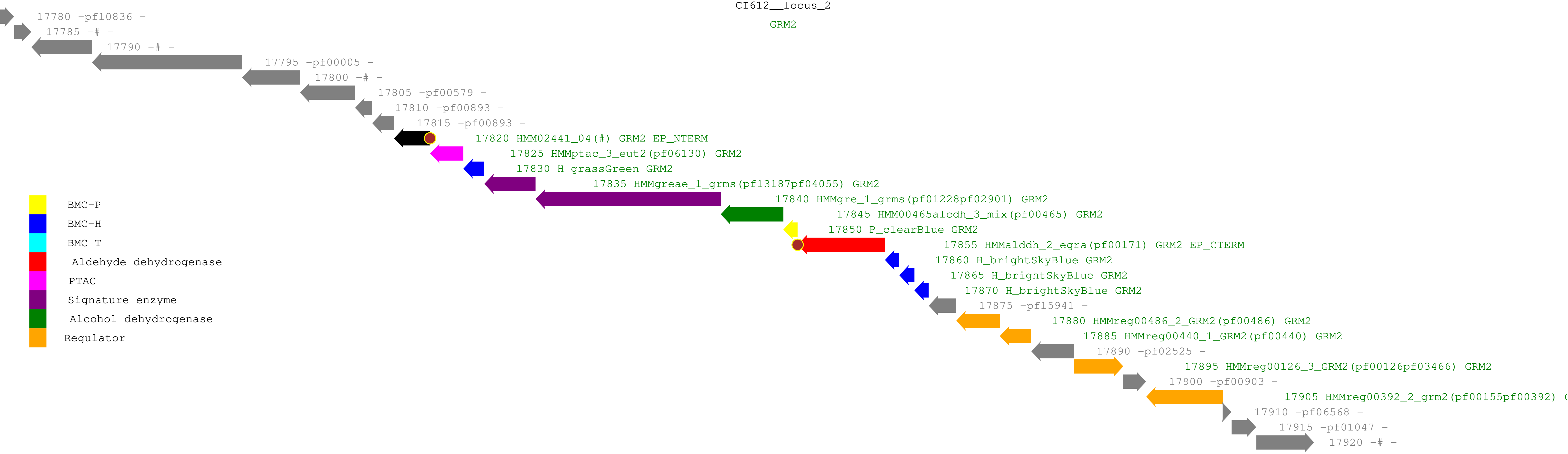

Klebsiella quasipneumoniae subsp. similipneumoniae. Bacteria; Proteobacteria; Gammaproteobacteria; Enterobacterales;

CI664__locus_1

GRM2

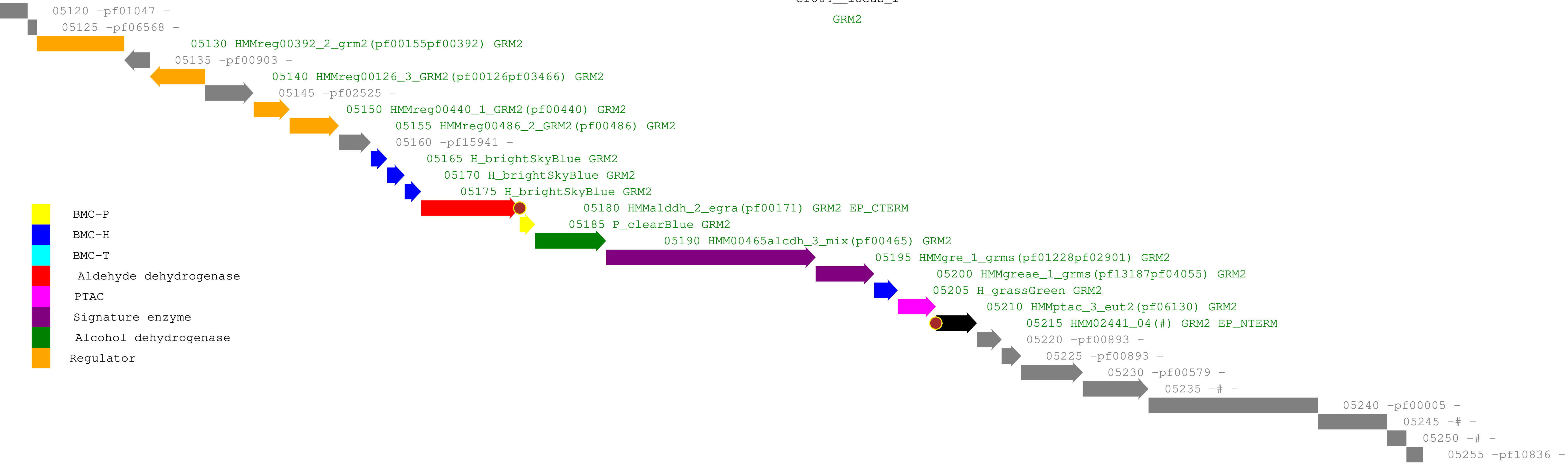

Klebsiella pneumoniae subsp. pneumoniae. Bacteria; Proteobacteria; Gammaproteobacteria; Enterobacterales;

CI740__locus_1

GRM2

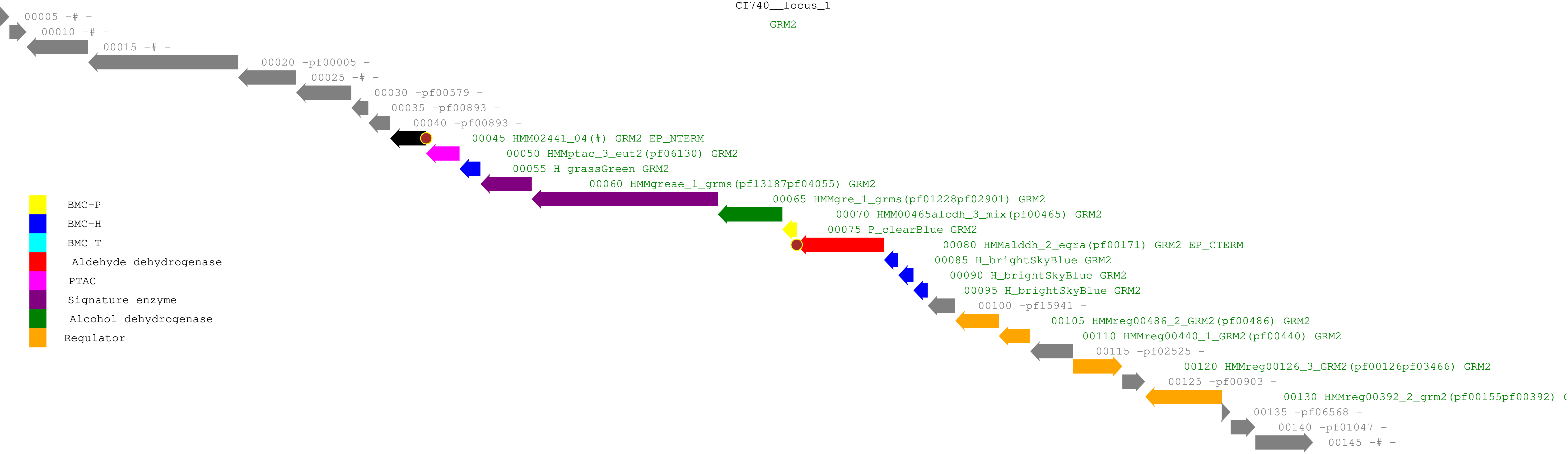

Raoultella planticola (Klebsiella planticola). Bacteria; Proteobacteria; Gammaproteobacteria; Enterobacterales;

CIG23__locus_2

GRM2

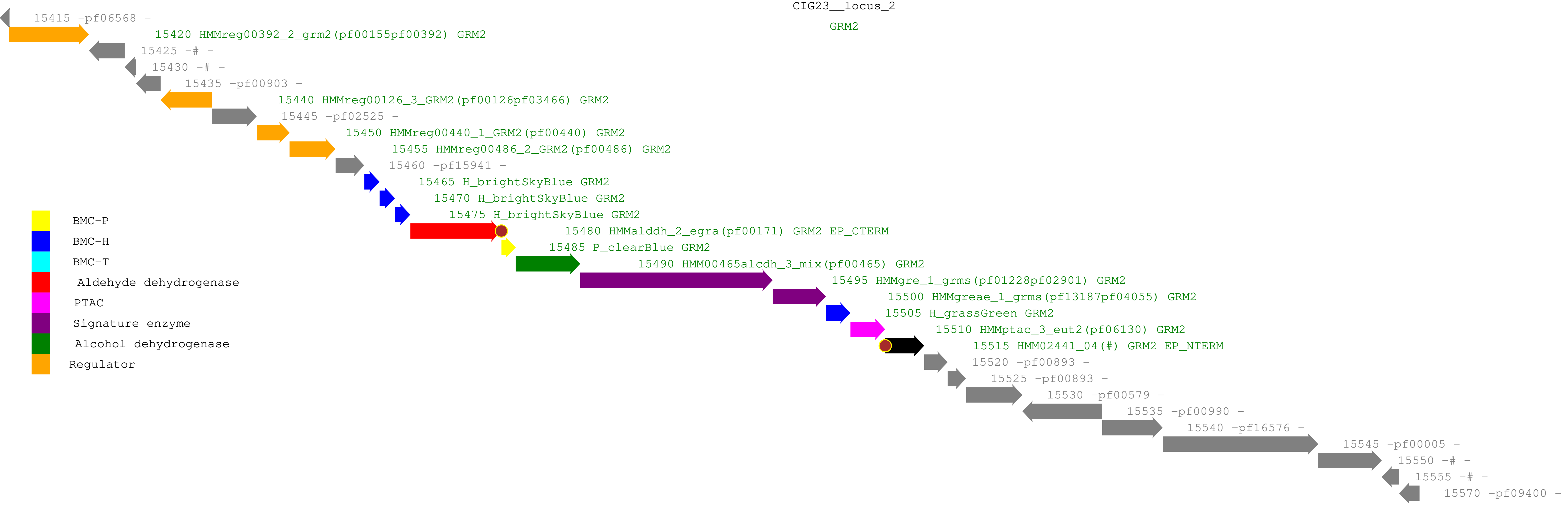

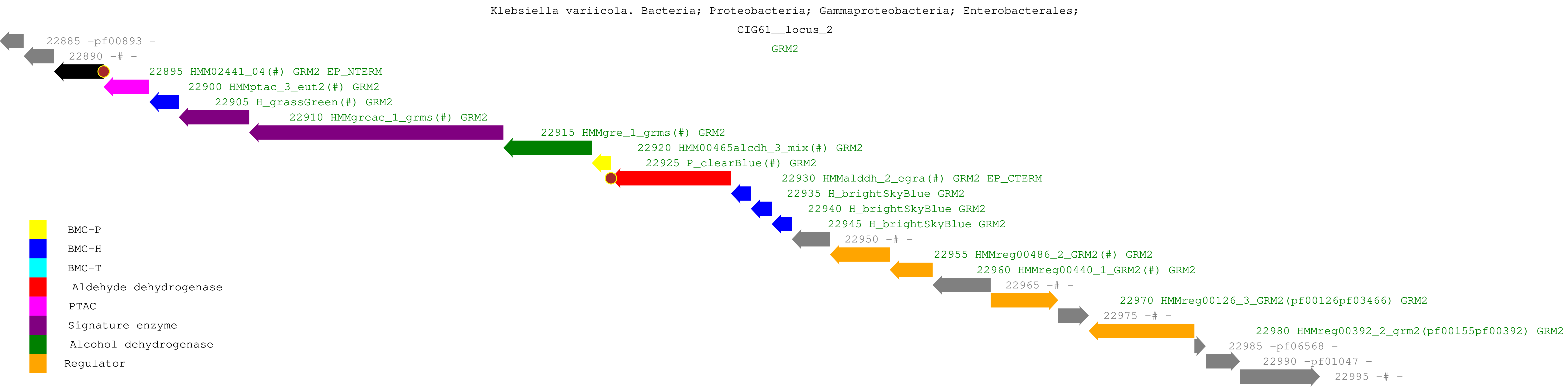

Klebsiella variicola. Bacteria; Proteobacteria; Gammaproteobacteria; Enterobacterales;
CIG61__locus_2
GRM2
22885 -pf00893 -
22890 -# -
22895 HMM02441_04(#) GRM2 EP_NTERM
22900 HMMptac_3_eut2(#) GRM2
22905 H_grassGreen(#) GRM2
22910 HMMgreae_1_grms(#) GRM2
22915 HMMgre_1_grms(#) GRM2
22920 HMM00465alcdh_3_mix(#) GRM2
22925 P_clearBlue(#) GRM2
22930 HMMalddh_2_egra(#) GRM2 EP_CTERM
22935 H_brightSkyBlue GRM2
22940 H_brightSkyBlue GRM2
22945 H_brightSkyBlue GRM2
22950 -# -
22955 HMMreg00486_2_GRM2(#) GRM2
22960 HMMreg00440_1_GRM2(#) GRM2
22965 -# -
22970 HMMreg00126_3_GRM2(pf00126pf03466) GRM2
22975 -# -
22980 HMMreg00392_2_grm2(pf00155pf00392) GRM2
22985 -pf06568 -
22990 -pf01047 -
22995 -# -
BMC-P
BMC-H
BMC-T
Aldehyde dehydrogenase
PTAC
Signature enzyme
Alcohol dehydrogenase
Regulator

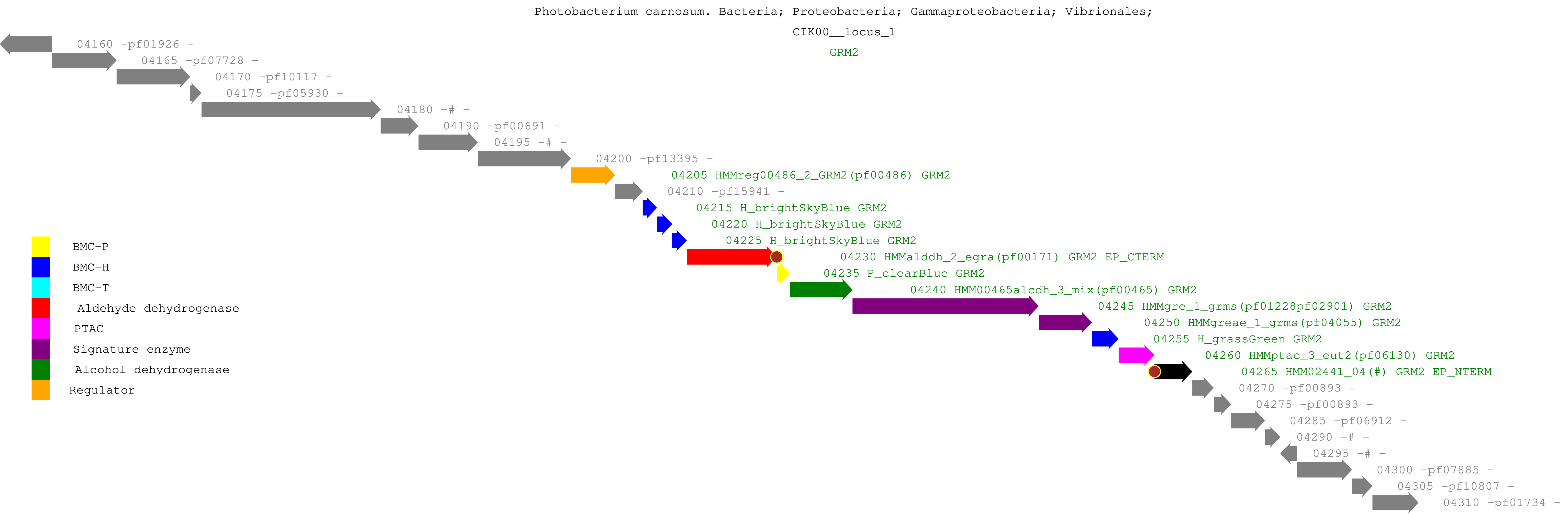

Photobacterium carnosum. Bacteria; Proteobacteria; Gammaproteobacteria; Vibrionales;
CIK00__locus_1
GRM2
04160 -pf01926 -
04165 -pf07728 -
04170 -pf10117 -
04175 -pf05930 -
04180 -# -
04190 -pf00691 -
04195 -# -
04200 -pf13395 -
04205 HMMreg00486_2_GRM2(pf00486) GRM2
04210 -pf15941 -
04215 H_brightSkyBlue GRM2
04220 H_brightSkyBlue GRM2
04225 H_brightSkyBlue GRM2
04230 HMMalddh_2_egra(pf00171) GRM2 EP_CTERM
04235 P_clearBlue GRM2
04240 HMM00465alcdh_3_mix(pf00465) GRM2
04245 HMMgre_1_grms(pf01228pf02901) GRM2
04250 HMMgreae_1_grms(pf04055) GRM2
04255 H_grassGreen GRM2
04260 HMMptac_3_eut2(pf06130) GRM2
04265 HMM02441_04(#) GRM2 EP_NTERM
04270 -pf00893 -
04275 -pf00893 -
04285 -pf06912 -
04290 -# -
04295 -# -
04300 -pf07885 -
04305 -pf10807 -
04310 -pf01734 -
BMC-P
BMC-H
BMC-T
Aldehyde dehydrogenase
PTAC
Signature enzyme
Alcohol dehydrogenase
Regulator

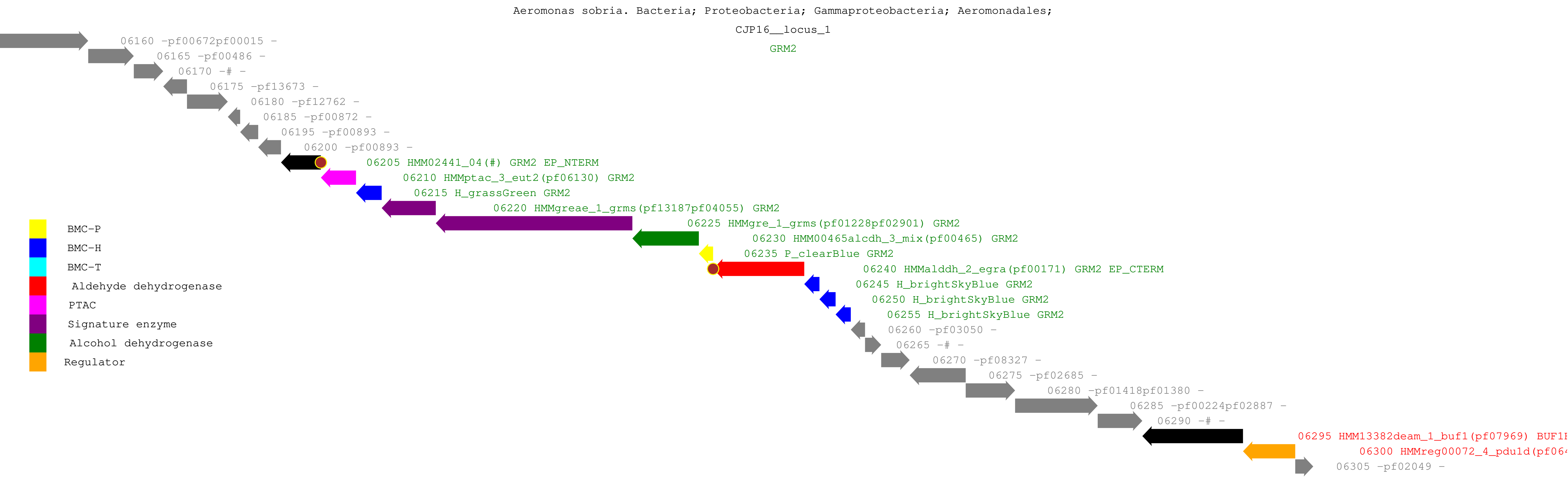

Aeromonas sobria. Bacteria; Proteobacteria; Gammaproteobacteria; Aeromonadales;
CJP16__locus_1
GRM2
06160 -pf00672pf00015 -
06165 -pf00486 -
06170 -# -
06175 -pf13673 -
06180 -pf12762 -
06185 -pf00872 -
06195 -pf00893 -
06200 -pf00893 -
06205 HMM02441_04(#) GRM2 EP_NTERM
06210 HMMptac_3_eut2(pf06130) GRM2
06215 H_grassGreen GRM2
06220 HMMgreae_1_grms(pf13187pf04055) GRM2
06225 HMMgre_1_grms(pf01228pf02901) GRM2
06230 HMM00465alcdh_3_mix(pf00465) GRM2
06235 P_clearBlue GRM2
06240 HMMalddh_2_egra(pf00171) GRM2 EP_CTERM
06245 H_brightSkyBlue GRM2
06250 H_brightSkyBlue GRM2
06255 H_brightSkyBlue GRM2
06260 -pf03050 -
06265 -# -
06270 -pf08327 -
06275 -pf02685 -
06280 -pf01418pf01380 -
06285 -pf00224pf02887 -
06290 -# -
06295 HMM13382deam_1_buf1(pf07969) BUF1
06300 HMMreg00072_4_pdu1d(pf06
06305 -pf02049 -
BMC-P
BMC-H
BMC-T
Aldehyde dehydrogenase
PTAC
Signature enzyme
Alcohol dehydrogenase
Regulator

Aeromonas sp. CA23. Bacteria; Proteobacteria; Gammaproteobacteria; Aeromonadales;

CK910__locus_1

GRM2

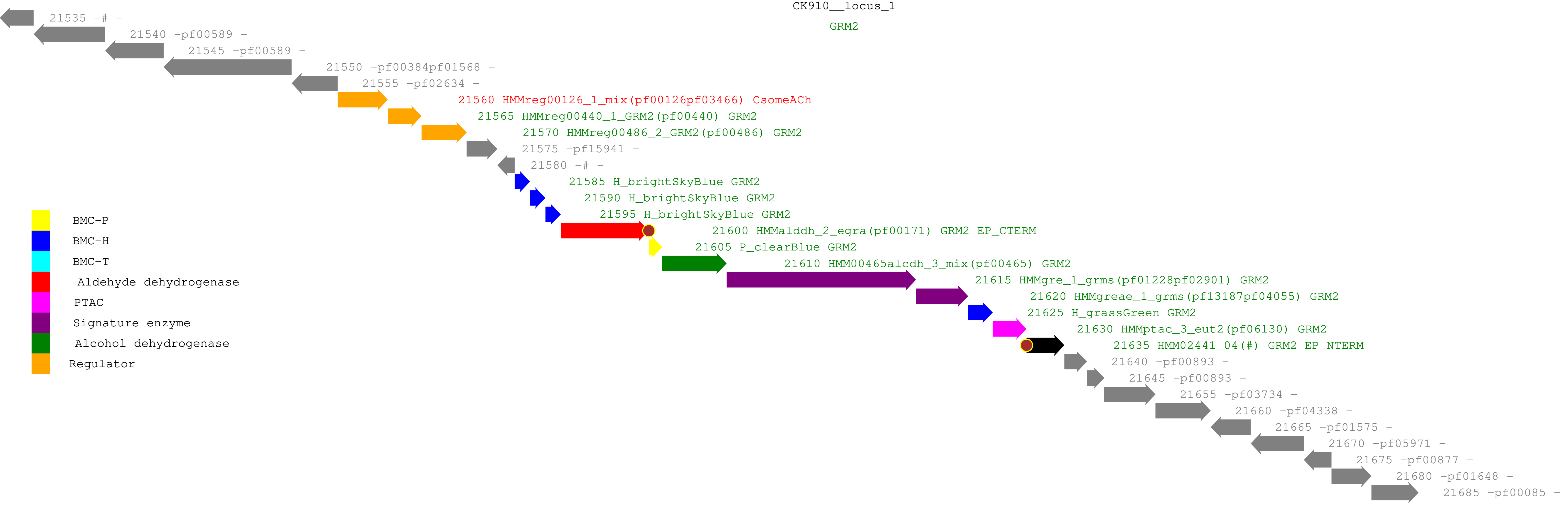

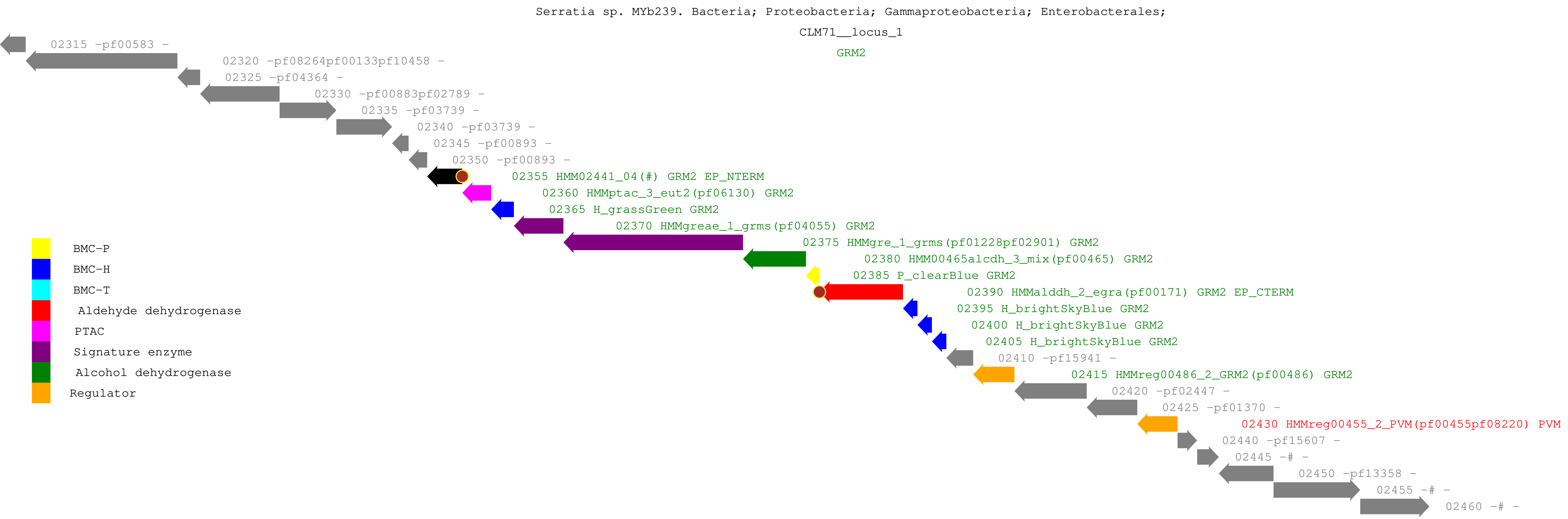
Serratia sp. MYb239. Bacteria; Proteobacteria; Gammaproteobacteria; Enterobacterales;
CLM71__locus_1
GRM2
02315 -pf00583 -
02320 -pf08264pf00133pf10458 -
02325 -pf04364 -
02330 -pf00883pf02789 -
02335 -pf03739 -
02340 -pf03739 -
02345 -pf00893 -
02350 -pf00893 -
02355 HMM02441_04(#) GRM2 EP_NTERM
02360 HMMptac_3_eut2(pf06130) GRM2
02365 H_grassGreen GRM2
02370 HMMgreae_1_grms(pf04055) GRM2
02375 HMMgre_1_grms(pf01228pf02901) GRM2
02380 HMM00465alcdh_3_mix(pf00465) GRM2
02385 P_clearBlue GRM2
02390 HMMaldh_2_egra(pf00171) GRM2 EP_CTERM
02395 H_brightSkyBlue GRM2
02400 H_brightSkyBlue GRM2
02405 H_brightSkyBlue GRM2
02410 -pf15941 -
02415 HMMreg00486_2_GRM2(pf00486) GRM2
02420 -pf02447 -
02425 -pf01370 -
02430 HMMreg00455_2_PVM(pf00455pf08220) PVM
02440 -pf15607 -
02445 -# -
02450 -pf13358 -
02455 -# -
02460 -# -
BMC-P
BMC-H
BMC-T
Aldehyde dehydrogenase
PTAC
Signature enzyme
Alcohol dehydrogenase
Regulator

Providencia alcalifaciens. Bacteria; Proteobacteria; Gammaproteobacteria; Enterobacterales;

CO695__locus_1

GRM2

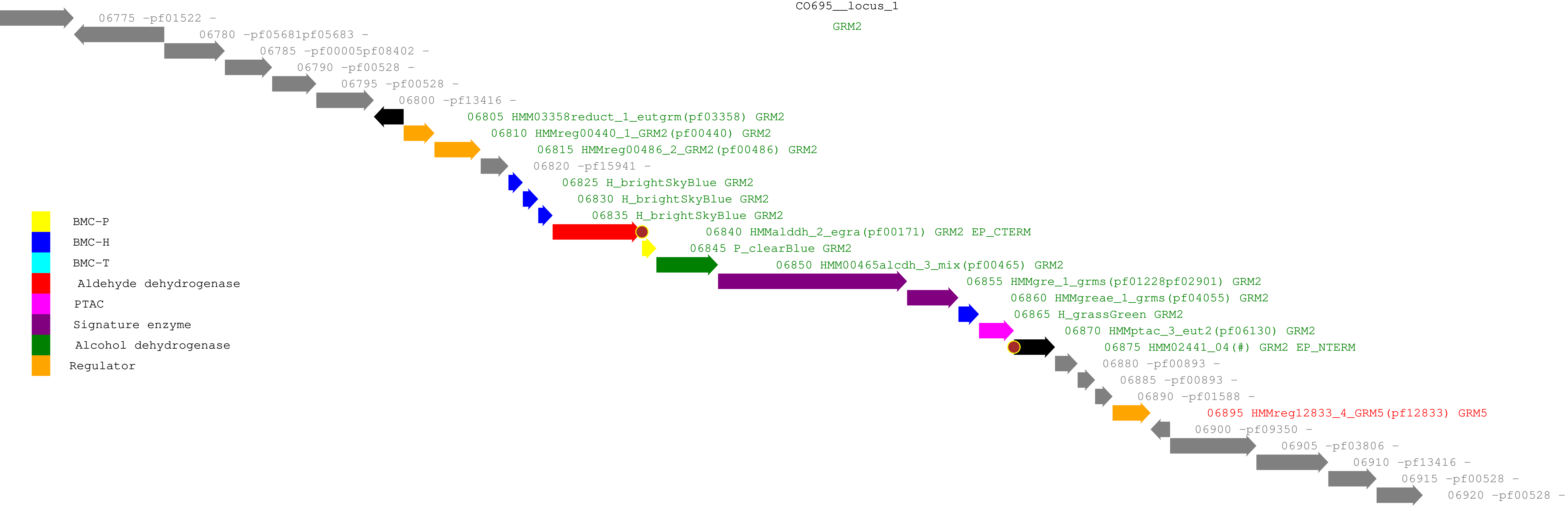

Klebsiella quasipneumoniae. Bacteria; Proteobacteria; Gammaproteobacteria; Enterobacterales;

CP911__locus_2

GRM2

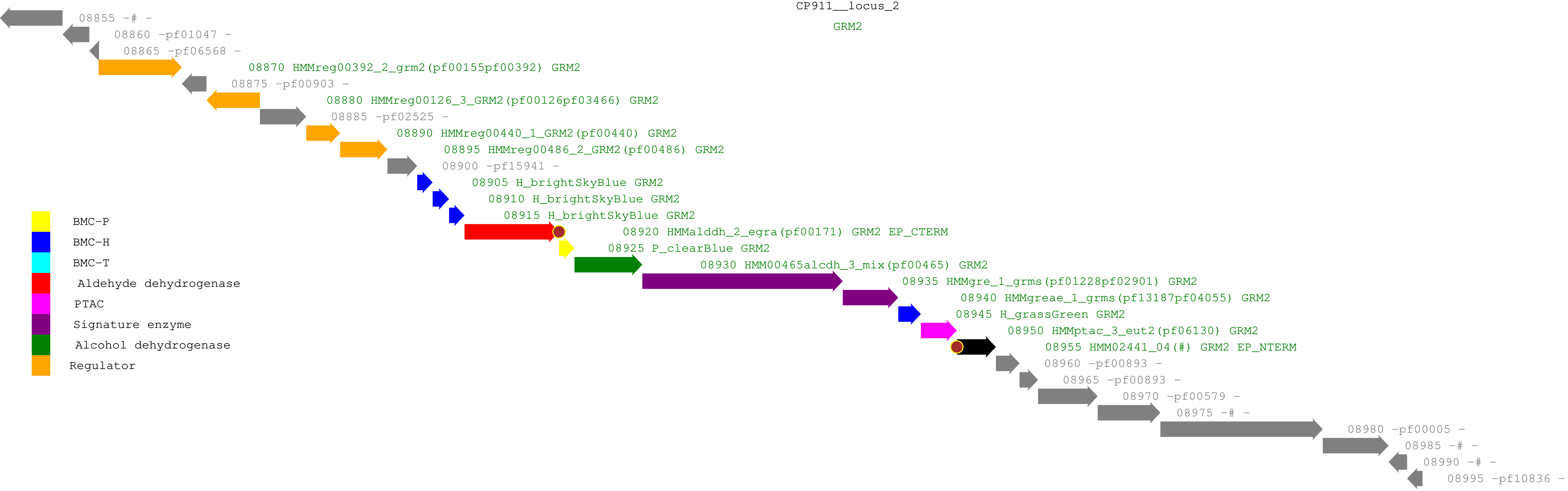

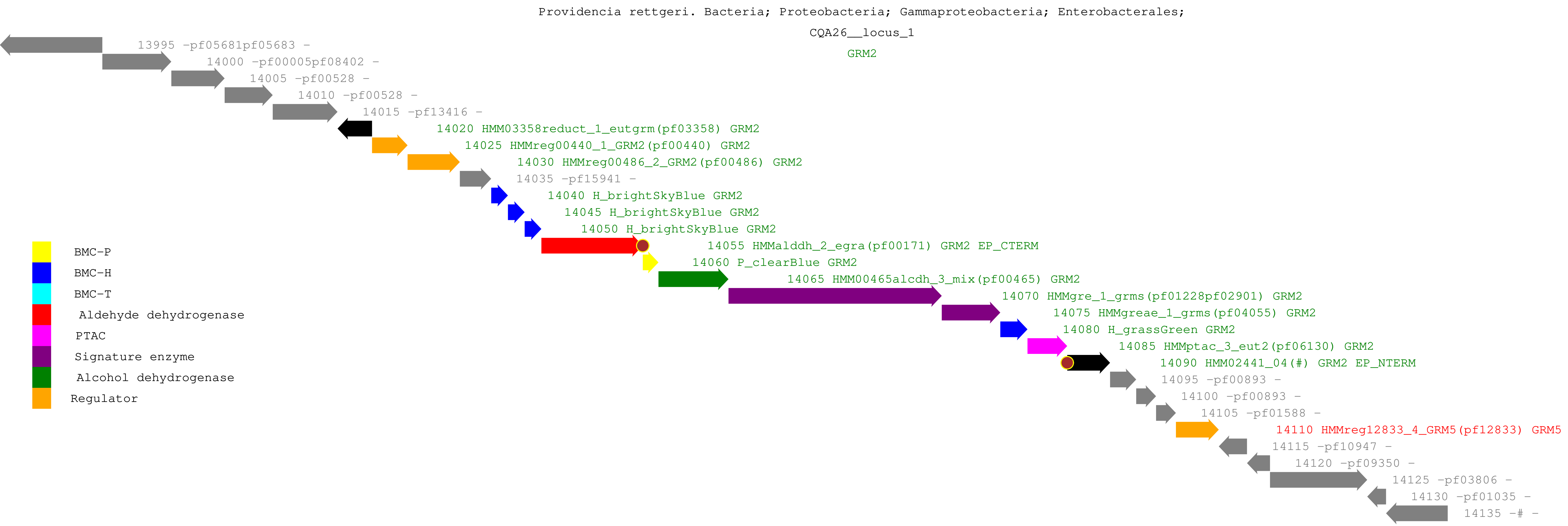
Providencia rettgeri. Bacteria; Proteobacteria; Gammaproteobacteria; Enterobacterales;
CQA26__locus_1
GRM2
13995 -pf05681pf05683 -
14000 -pf00005pf08402 -
14005 -pf00528 -
14010 -pf00528 -
14015 -pf13416 -
14020 HMM03358reduct_1_eutgrm(pf03358) GRM2
14025 HMMreg00440_1_GRM2(pf00440) GRM2
14030 HMMreg00486_2_GRM2(pf00486) GRM2
14035 -pf15941 -
14040 H_brightSkyBlue GRM2
14045 H_brightSkyBlue GRM2
14050 H_brightSkyBlue GRM2
14055 HMMalddh_2_egra(pf00171) GRM2 EP_CTERM
14060 P_clearBlue GRM2
14065 HMM00465alcdh_3_mix(pf00465) GRM2
14070 HMMgre_1_grms(pf01228pf02901) GRM2
14075 HMMgreae_1_grms(pf04055) GRM2
14080 H_grassGreen GRM2
14085 HMMptac_3_eut2(pf06130) GRM2
14090 HMM02441_04(#) GRM2 EP_NTERM
14095 -pf00893 -
14100 -pf00893 -
14105 -pf01588 -
14110 HMMreg12833_4_GRM5(pf12833) GRM5
14115 -pf10947 -
14120 -pf09350 -
14125 -pf03806 -
14130 -pf01035 -
14135 -# -
BMC-P
BMC-H
BMC-T
Aldehyde dehydrogenase
PTAC
Signature enzyme
Alcohol dehydrogenase
Regulator

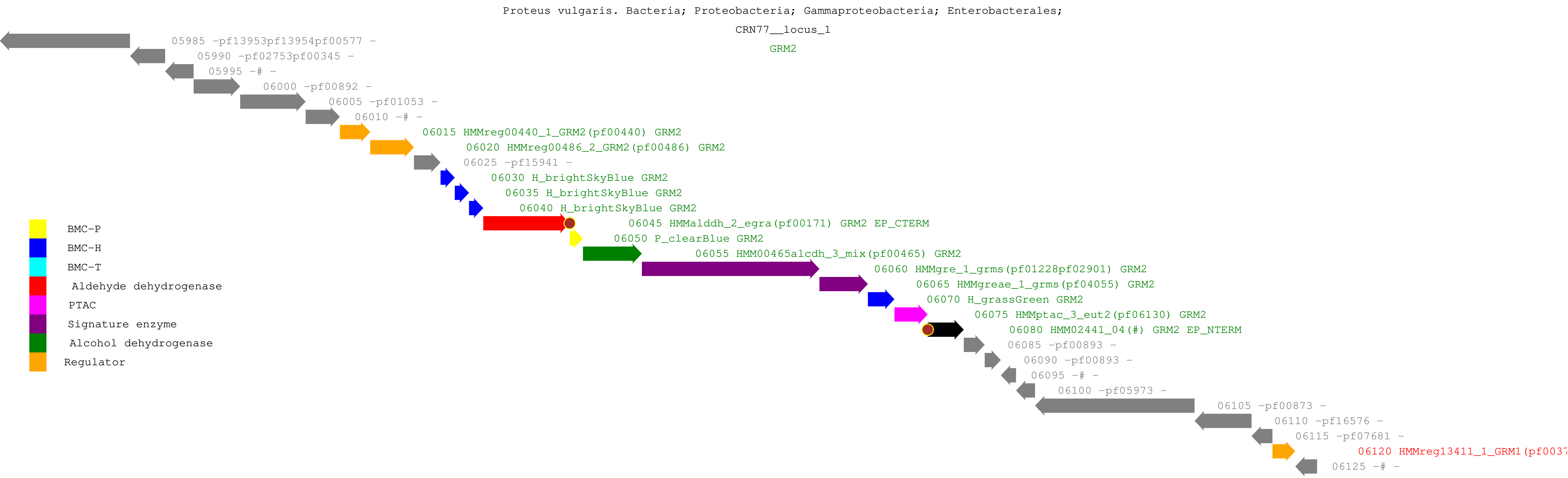

Proteus vulgaris. Bacteria; Proteobacteria; Gammaproteobacteria; Enterobacterales;
CRN77__locus_1
GRM2
05985 -pf13953pf13954pf00577 -
05990 -pf02753pf00345 -
05995 -# -
06000 -pf00892 -
06005 -pf01053 -
06010 -# -
06015 HMMreg00440_1_GRM2(pf00440) GRM2
06020 HMMreg00486_2_GRM2(pf00486) GRM2
06025 -pf15941 -
06030 H_brightSkyBlue GRM2
06035 H_brightSkyBlue GRM2
06040 H_brightSkyBlue GRM2
06045 HMMaldh_2_egra(pf00171) GRM2 EP_CTERM
06050 P_clearBlue GRM2
06055 HMM00465alcdh_3_mix(pf00465) GRM2
06060 HMMgre_1_grms(pf01228pf02901) GRM2
06065 HMMgreae_1_grms(pf04055) GRM2
06070 H_grassGreen GRM2
06075 HMMptac_3_eut2(pf06130) GRM2
06080 HMM02441_04(#) GRM2 EP_NTERM
06085 -pf00893 -
06090 -pf00893 -
06095 -# -
06100 -pf05973 -
06105 -pf00873 -
06110 -pf16576 -
06115 -pf07681 -
06120 HMMreg13411_1_GRM1(pf0037
06125 -# -
BMC-P
BMC-H
BMC-T
Aldehyde dehydrogenase
PTAC
Signature enzyme
Alcohol dehydrogenase
Regulator

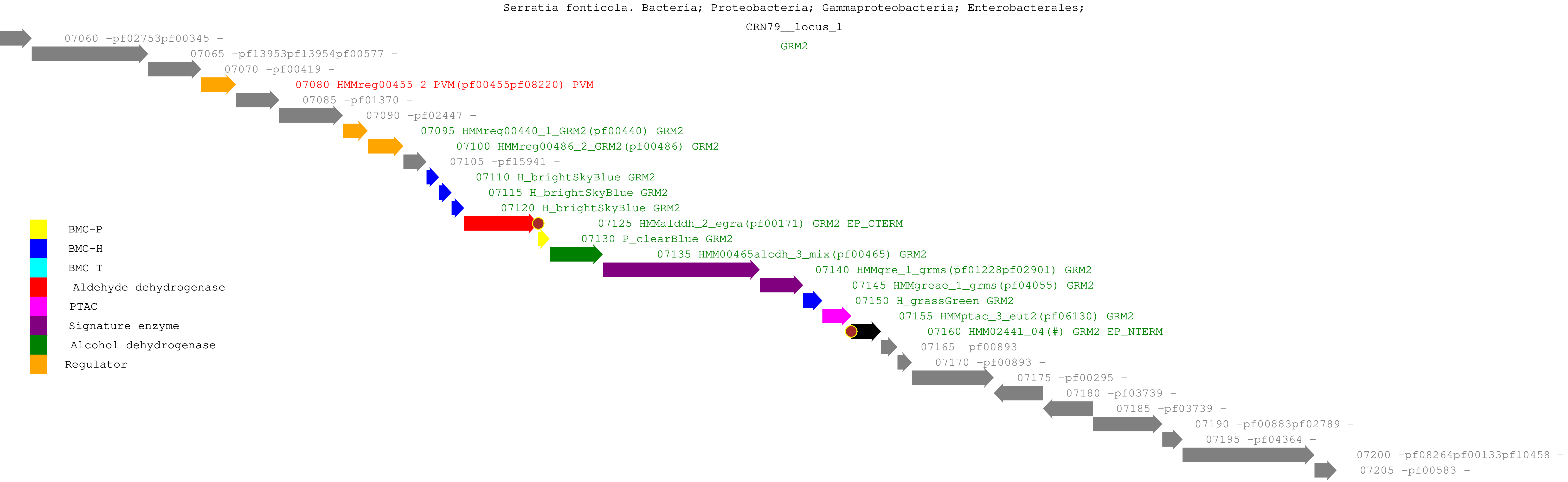

Serratia fonticola. Bacteria; Proteobacteria; Gammaproteobacteria; Enterobacterales;
CRN79__locus_1
GRM2
07060 -pf02753pf00345 -
07065 -pf13953pf13954pf00577 -
07070 -pf00419 -
07080 HMMreg00455_2_PVM(pf00455pf08220) PVM
07085 -pf01370 -
07090 -pf02447 -
07095 HMMreg00440_1_GRM2(pf00440) GRM2
07100 HMMreg00486_2_GRM2(pf00486) GRM2
07105 -pf15941 -
07110 H_brightSkyBlue GRM2
07115 H_brightSkyBlue GRM2
07120 H_brightSkyBlue GRM2
07125 HMMalddh_2_egra(pf00171) GRM2 EP_CTERM
07130 P_clearBlue GRM2
07135 HMM00465alcdh_3_mix(pf00465) GRM2
07140 HMMgre_1_grms(pf01228pf02901) GRM2
07145 HMMgreae_1_grms(pf04055) GRM2
07150 H_grassGreen GRM2
07155 HMMptac_3_eut2(pf06130) GRM2
07160 HMM02441_04(#) GRM2 EP_NTERM
07165 -pf00893 -
07170 -pf00893 -
07175 -pf00295 -
07180 -pf03739 -
07185 -pf03739 -
07190 -pf00883pf02789 -
07195 -pf04364 -
07200 -pf08264pf00133pf10458 -
07205 -pf00583 -
BMC-P
BMC-H
BMC-T
Aldehyde dehydrogenase
PTAC
Signature enzyme
Alcohol dehydrogenase
Regulator

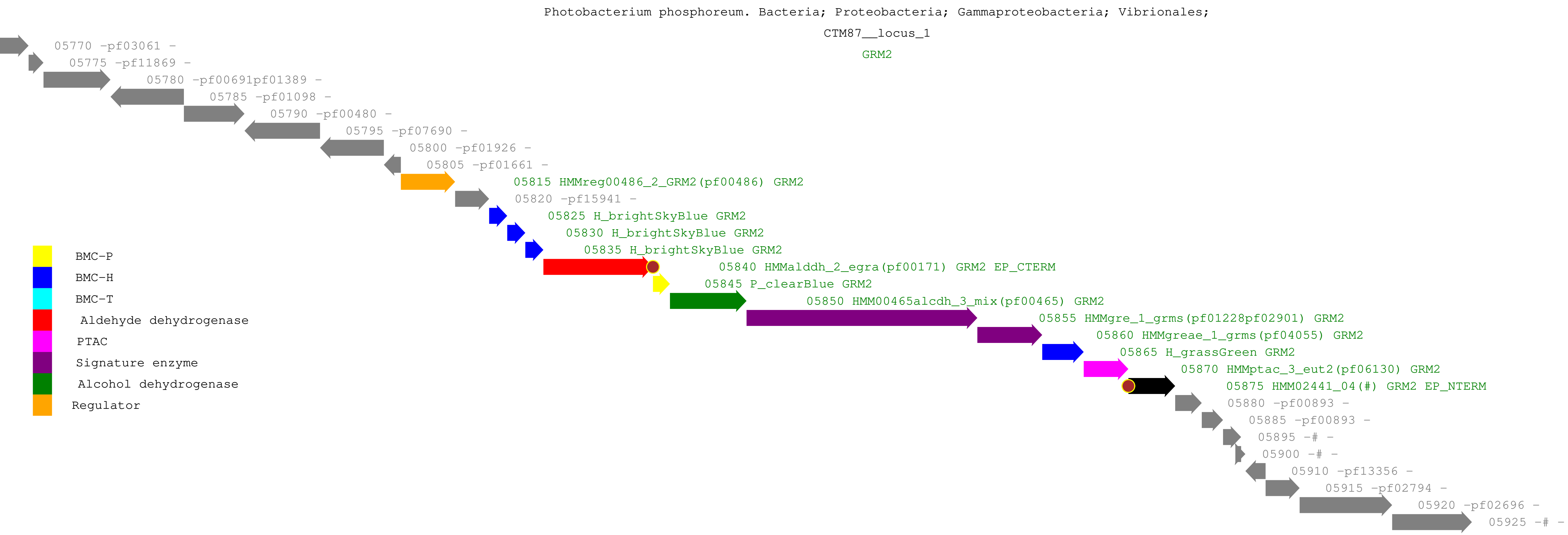
Photobacterium phosphoreum. Bacteria; Proteobacteria; Gammaproteobacteria; Vibrionales;
CTM87__locus_1
GRM2
05770 -pf03061 -
05775 -pf11869 -
05780 -pf00691pf01389 -
05785 -pf01098 -
05790 -pf00480 -
05795 -pf07690 -
05800 -pf01926 -
05805 -pf01661 -
05815 HMMreg00486_2_GRM2(pf00486) GRM2
05820 -pf15941 -
05825 H_brightSkyBlue GRM2
05830 H_brightSkyBlue GRM2
05835 H_brightSkyBlue GRM2
05840 HMMaldh_2_egra(pf00171) GRM2 EP_CTERM
05845 P_clearBlue GRM2
05850 HMM00465alcdh_3_mix(pf00465) GRM2
05855 HMMgre_1_grms(pf01228pf02901) GRM2
05860 HMMgreae_1_grms(pf04055) GRM2
05865 H_grassGreen GRM2
05870 HMMptac_3_eut2(pf06130) GRM2
05875 HMM02441_04(#) GRM2 EP_NTERM
05880 -pf00893 -
05885 -pf00893 -
05895 -# -
05900 -# -
05910 -pf13356 -
05915 -pf02794 -
05920 -pf02696 -
05925 -# -
BMC-P
BMC-H
BMC-T
Aldehyde dehydrogenase
PTAC
Signature enzyme
Alcohol dehydrogenase
Regulator

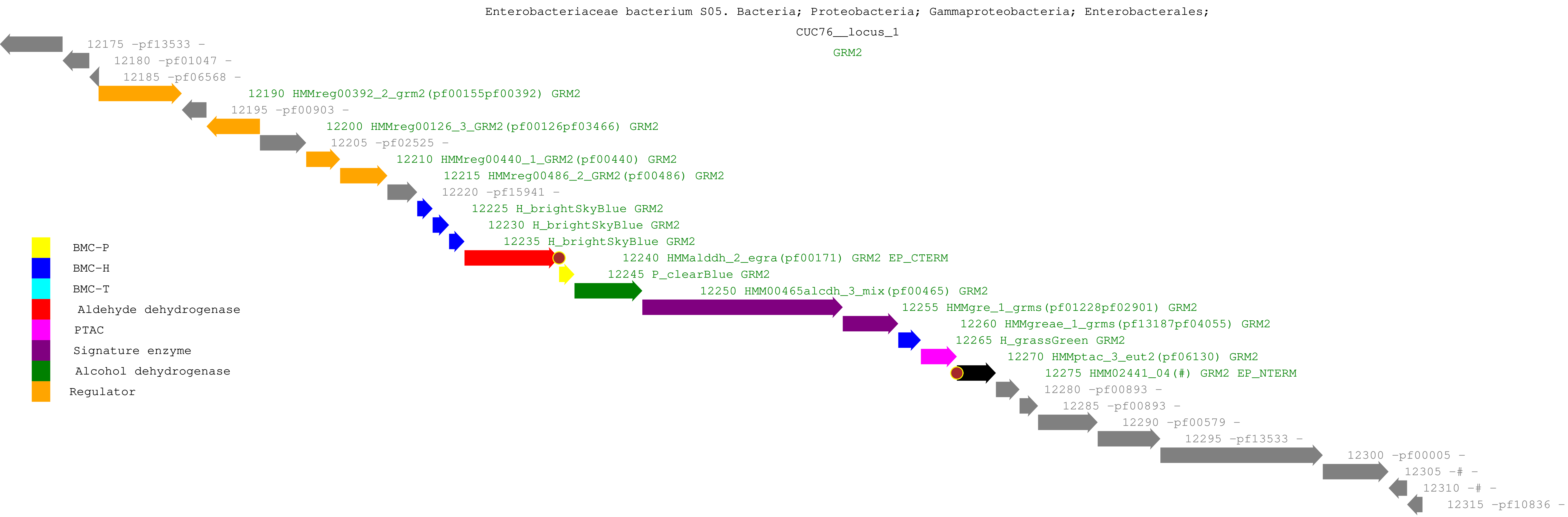
Enterobacteriaceae bacterium S05. Bacteria; Proteobacteria; Gammaproteobacteria; Enterobacterales;
CUC76__locus_1
GRM2
12175 -pf13533 -
12180 -pf01047 -
12185 -pf06568 -
12190 HMMreg00392_2_grm2(pf00155pf00392) GRM2
12195 -pf00903 -
12200 HMMreg00126_3_GRM2(pf00126pf03466) GRM2
12205 -pf02525 -
12210 HMMreg00440_1_GRM2(pf00440) GRM2
12215 HMMreg00486_2_GRM2(pf00486) GRM2
12220 -pf15941 -
12225 H_brightSkyBlue GRM2
12230 H_brightSkyBlue GRM2
12235 H_brightSkyBlue GRM2
12240 HMMaldhh_2_egra(pf00171) GRM2 EP_CTERM
12245 P_clearBlue GRM2
12250 HMM00465alcdh_3_mix(pf00465) GRM2
12255 HMMgre_1_grms(pf01228pf02901) GRM2
12260 HMMgreae_1_grms(pf13187pf04055) GRM2
12265 H_grassGreen GRM2
12270 HMMptac_3_eut2(pf06130) GRM2
12275 HMM02441_04(#) GRM2 EP_NTERM
12280 -pf00893 -
12285 -pf00893 -
12290 -pf00579 -
12295 -pf13533 -
12300 -pf00005 -
12305 -# -
12310 -# -
12315 -pf10836 -
BMC-P
BMC-H
BMC-T
Aldehyde dehydrogenase
PTAC
Signature enzyme
Alcohol dehydrogenase
Regulator

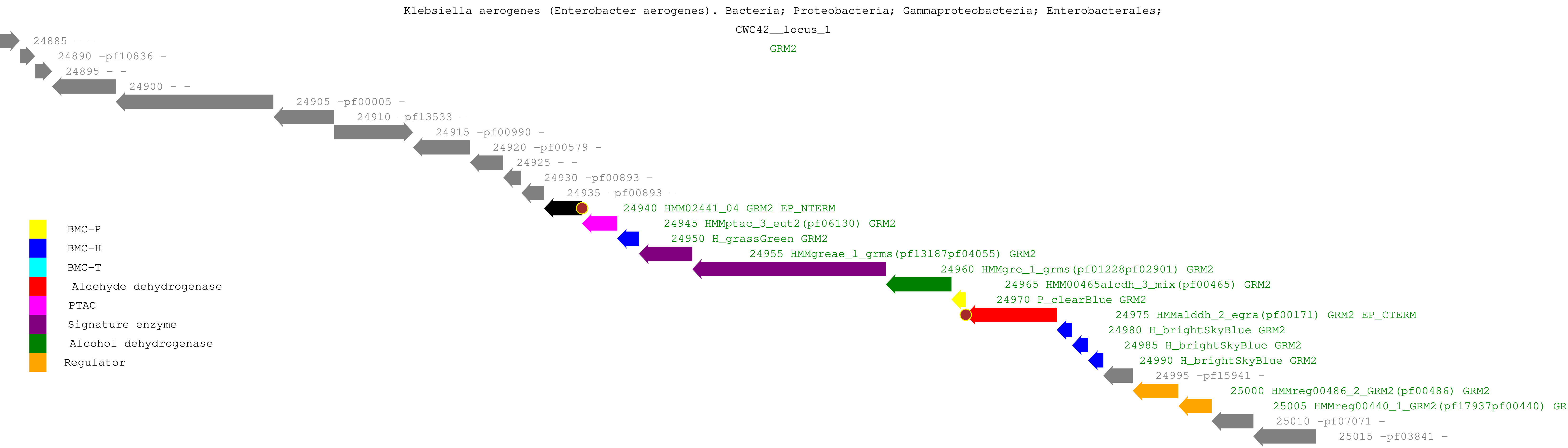
Klebsiella aerogenes (Enterobacter aerogenes). Bacteria; Proteobacteria; Gammaproteobacteria; Enterobacterales;
CWC42__locus_1
GRM2
24885 - -
24890 -pf10836 -
24895 - -
24900 - -
24905 -pf00005 -
24910 -pf13533 -
24915 -pf00990 -
24920 -pf00579 -
24925 - -
24930 -pf00893 -
24935 -pf00893 -
24940 HMM02441_04 GRM2 EP_NTERM
24945 HMMptac_3_eut2(pf06130) GRM2
24950 H_grassGreen GRM2
24955 HMMgreae_1_grms(pf13187pf04055) GRM2
24960 HMMgre_1_grms(pf01228pf02901) GRM2
24965 HMM00465alcdh_3_mix(pf00465) GRM2
24970 P_clearBlue GRM2
24975 HMMalddh_2_egra(pf00171) GRM2 EP_CTERM
24980 H_brightSkyBlue GRM2
24985 H_brightSkyBlue GRM2
24990 H_brightSkyBlue GRM2
24995 -pf15941 -
25000 HMMreg00486_2_GRM2(pf00486) GRM2
25005 HMMreg00440_1_GRM2(pf17937pf00440) GR
25010 -pf07071 -
25015 -pf03841 -
BMC-P
BMC-H
BMC-T
Aldehyde dehydrogenase
PTAC
Signature enzyme
Alcohol dehydrogenase
Regulator

Raoultella sp. T31. Bacteria; Proteobacteria; Gammaproteobacteria; Enterobacterales;

CWM52__locus_2

GRM2

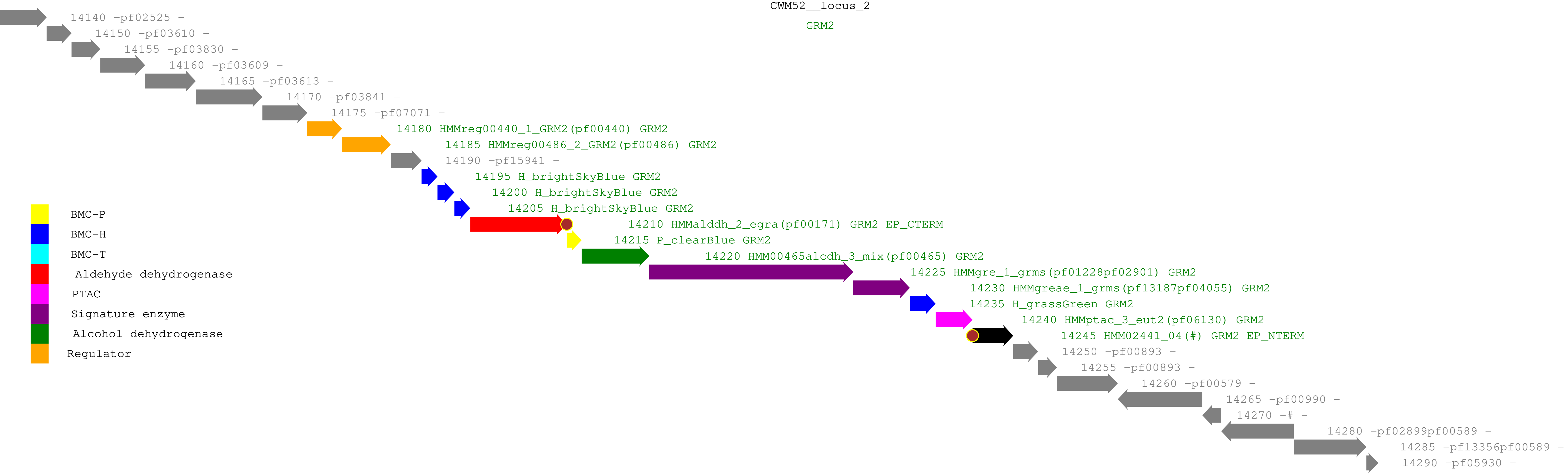

Klebsiella michiganensis. Bacteria; Proteobacteria; Gammaproteobacteria; Enterobacterales;

CWN04__locus_1

GRM2

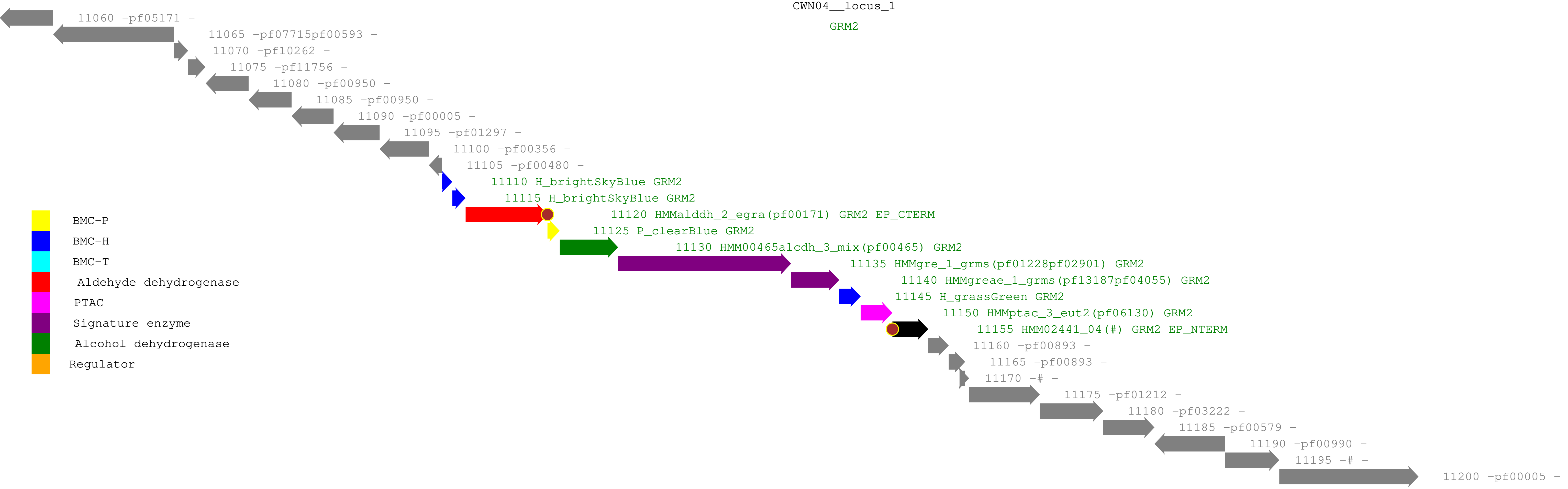

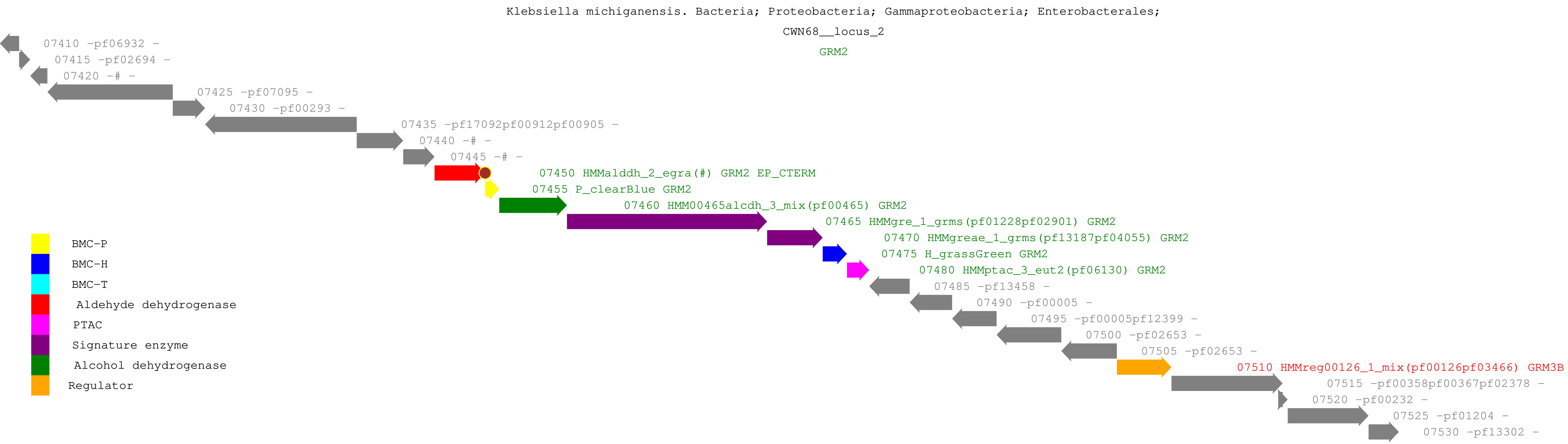
Klebsiella michiganensis. Bacteria; Proteobacteria; Gammaproteobacteria; Enterobacterales;
CWN68__locus_2
GRM2
07410 -pf06932 -
07415 -pf02694 -
07420 -# -
07425 -pf07095 -
07430 -pf00293 -
07435 -pf17092pf00912pf00905 -
07440 -# -
07445 -# -
07450 HMMaldh_2_egra(#) GRM2 EP_CTERM
07455 P_clearBlue GRM2
07460 HMM00465alcdh_3_mix(pf00465) GRM2
07465 HMMgre_1_grms(pf01228pf02901) GRM2
07470 HMMgreae_1_grms(pf13187pf04055) GRM2
07475 H_grassGreen GRM2
07480 HMMptac_3_eut2(pf06130) GRM2
07485 -pf13458 -
07490 -pf00005 -
07495 -pf00005pf12399 -
07500 -pf02653 -
07505 -pf02653 -
07510 HMMreg00126_1_mix(pf00126pf03466) GRM3B
07515 -pf00358pf00367pf02378 -
07520 -pf00232 -
07525 -pf01204 -
07530 -pf13302 -
BMC-P
BMC-H
BMC-T
Aldehyde dehydrogenase
PTAC
Signature enzyme
Alcohol dehydrogenase
Regulator

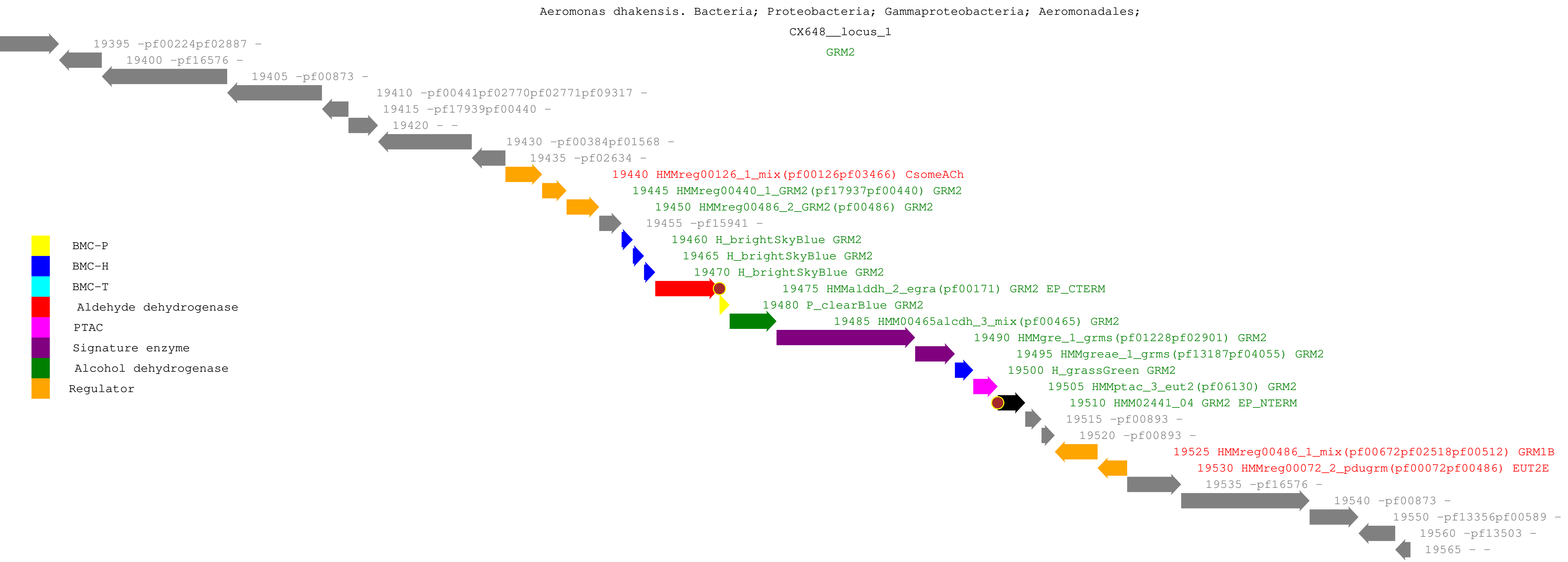
Aeromonas dhakensis. Bacteria; Proteobacteria; Gammaproteobacteria; Aeromonadales;
CX648__locus_1
GRM2
19395 -pf00224pf02887 -
19400 -pf16576 -
19405 -pf00873 -
19410 -pf00441pf02770pf02771pf09317 -
19415 -pf17939pf00440 -
19420 - -
19430 -pf00384pf01568 -
19435 -pf02634 -
19440 HMMreg00126_1_mix(pf00126pf03466) CsomeACh
19445 HMMreg00440_1_GRM2(pf17937pf00440) GRM2
19450 HMMreg00486_2_GRM2(pf00486) GRM2
19455 -pf15941 -
19460 H_brightSkyBlue GRM2
19465 H_brightSkyBlue GRM2
19470 H_brightSkyBlue GRM2
19475 HMMaldh_2_egra(pf00171) GRM2 EP_CTERM
19480 P_clearBlue GRM2
19485 HMM00465alcdh_3_mix(pf00465) GRM2
19490 HMMgre_1_grms(pf01228pf02901) GRM2
19495 HMMgreae_1_grms(pf13187pf04055) GRM2
19500 H_grassGreen GRM2
19505 HMMptac_3_eut2(pf06130) GRM2
19510 HMM02441_04 GRM2 EP_NTERM
19515 -pf00893 -
19520 -pf00893 -
19525 HMMreg00486_1_mix(pf00672pf02518pf00512) GRM1B
19530 HMMreg00072_2_pdugrm(pf00072pf00486) EUT2E
19535 -pf16576 -
19540 -pf00873 -
19550 -pf13356pf00589 -
19560 -pf13503 -
19565 - -
BMC-P
BMC-H
BMC-T
Aldehyde dehydrogenase
PTAC
Signature enzyme
Alcohol dehydrogenase
Regulator

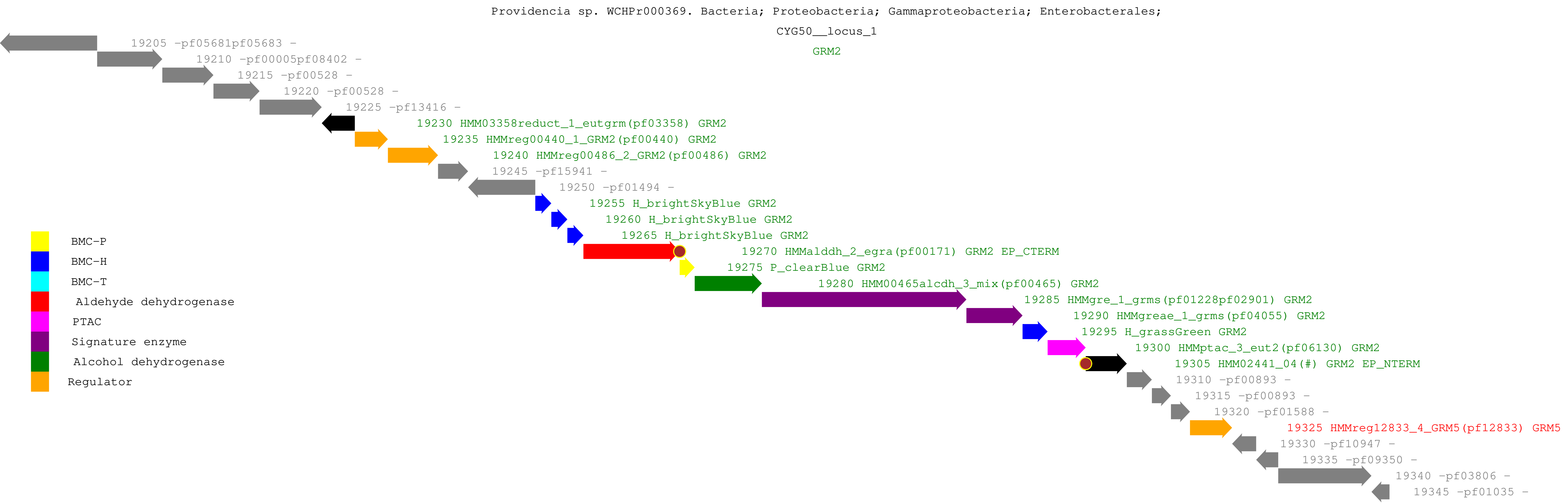
Providencia sp. WCHPr000369. Bacteria; Proteobacteria; Gammaproteobacteria; Enterobacterales;
CYG50__locus_1
GRM2
19205 -pf05681pf05683 -
19210 -pf00005pf08402 -
19215 -pf00528 -
19220 -pf00528 -
19225 -pf13416 -
19230 HMM03358reduct_1_eutgrm(pf03358) GRM2
19235 HMMreg00440_1_GRM2(pf00440) GRM2
19240 HMMreg00486_2_GRM2(pf00486) GRM2
19245 -pf15941 -
19250 -pf01494 -
19255 H_brightSkyBlue GRM2
19260 H_brightSkyBlue GRM2
19265 H_brightSkyBlue GRM2
19270 HMMalddh_2_egra(pf00171) GRM2 EP_CTERM
19275 P_clearBlue GRM2
19280 HMM00465alcdh_3_mix(pf00465) GRM2
19285 HMMgre_1_grms(pf01228pf02901) GRM2
19290 HMMgreae_1_grms(pf04055) GRM2
19295 H_grassGreen GRM2
19300 HMMptac_3_eut2(pf06130) GRM2
19305 HMM02441_04(#) GRM2 EP_NTERM
19310 -pf00893 -
19315 -pf00893 -
19320 -pf01588 -
19325 HMMreg12833_4_GRM5(pf12833) GRM5
19330 -pf10947 -
19335 -pf09350 -
19340 -pf03806 -
19345 -pf01035 -
BMC-P
BMC-H
BMC-T
Aldehyde dehydrogenase
PTAC
Signature enzyme
Alcohol dehydrogenase
Regulator

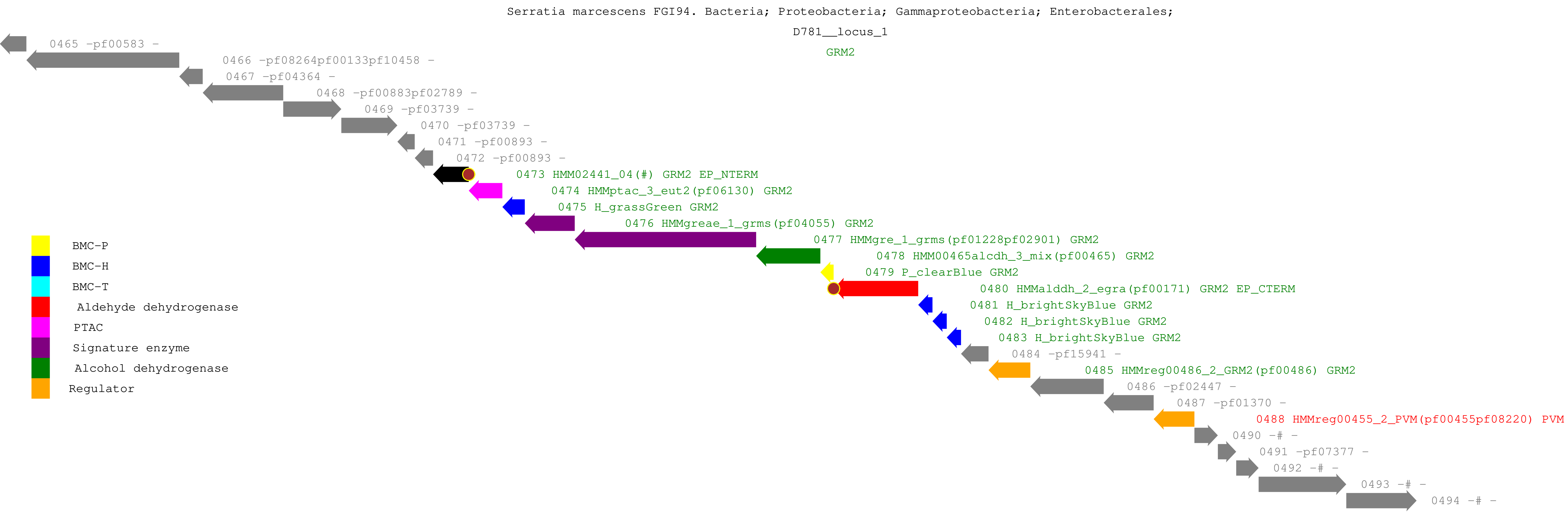
Serratia marcescens FGI94. Bacteria; Proteobacteria; Gammaproteobacteria; Enterobacterales;
D781__locus_1
GRM2
0465 -pf00583 -
0466 -pf08264pf00133pf10458 -
0467 -pf04364 -
0468 -pf00883pf02789 -
0469 -pf03739 -
0470 -pf03739 -
0471 -pf00893 -
0472 -pf00893 -
0473 HMM02441_04(#) GRM2 EP_NTERM
0474 HMMptac_3_eut2(pf06130) GRM2
0475 H_grassGreen GRM2
0476 HMMgreae_1_grms(pf04055) GRM2
0477 HMMgre_1_grms(pf01228pf02901) GRM2
0478 HMM00465alcdh_3_mix(pf00465) GRM2
0479 P_clearBlue GRM2
0480 HMMaldd h_2_egra(pf00171) GRM2 EP_CTERM
0481 H_brightSkyBlue GRM2
0482 H_brightSkyBlue GRM2
0483 H_brightSkyBlue GRM2
0484 -pf15941 -
0485 HMMreg00486_2_GRM2(pf00486) GRM2
0486 -pf02447 -
0487 -pf01370 -
0488 HMMreg00455_2_PVM(pf00455pf08220) PVM
0490 -# -
0491 -pf07377 -
0492 -# -
0493 -# -
0494 -# -
BMC-P
BMC-H
BMC-T
Aldehyde dehydrogenase
PTAC
Signature enzyme
Alcohol dehydrogenase
Regulator

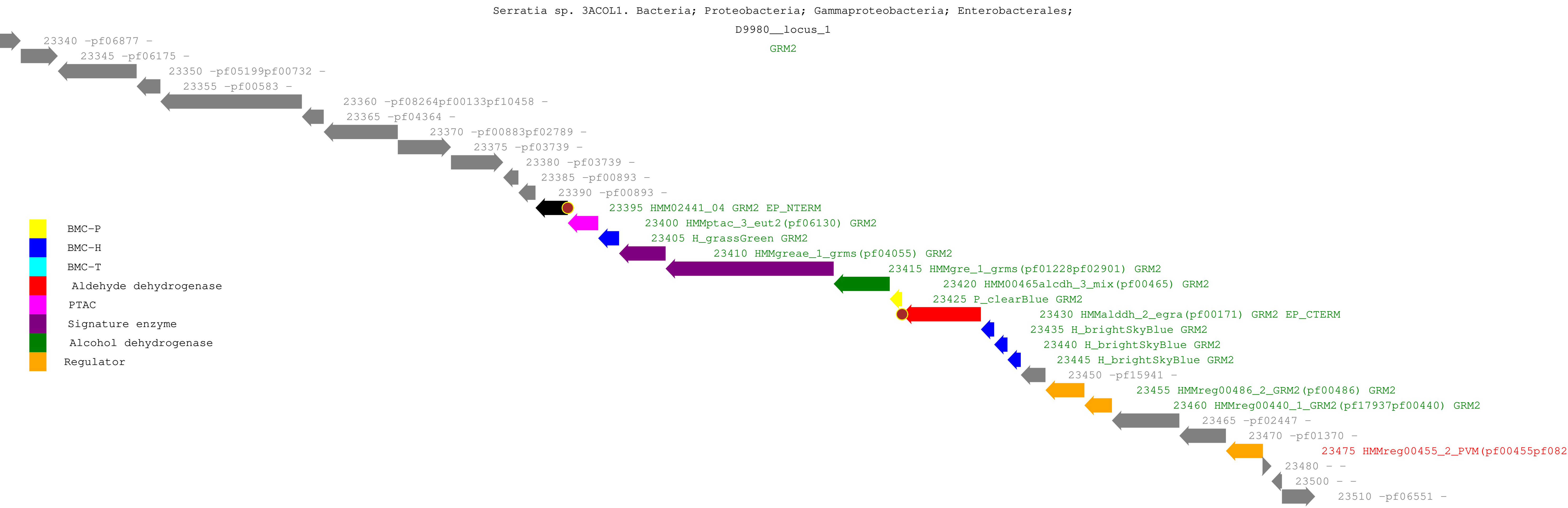
Serratia sp. 3ACOL1. Bacteria; Proteobacteria; Gammaproteobacteria; Enterobacterales;
D9980__locus_1
GRM2
23340 -pf06877 -
23345 -pf06175 -
23350 -pf05199pf00732 -
23355 -pf00583 -
23360 -pf08264pf00133pf10458 -
23365 -pf04364 -
23370 -pf00883pf02789 -
23375 -pf03739 -
23380 -pf03739 -
23385 -pf00893 -
23390 -pf00893 -
23395 HMM02441_04 GRM2 EP_NTERM
23400 HMMptac_3_eut2(pf06130) GRM2
23405 H_grassGreen GRM2
23410 HMMgreae_1_grms(pf04055) GRM2
23415 HMMgre_1_grms(pf01228pf02901) GRM2
23420 HMM00465alcdh_3_mix(pf00465) GRM2
23425 P_clearBlue GRM2
23430 HMMaldh_2_egra(pf00171) GRM2 EP_CTERM
23435 H_brightSkyBlue GRM2
23440 H_brightSkyBlue GRM2
23445 H_brightSkyBlue GRM2
23450 -pf15941 -
23455 HMMreg00486_2_GRM2(pf00486) GRM2
23460 HMMreg00440_1_GRM2(pf17937pf00440) GRM2
23465 -pf02447 -
23470 -pf01370 -
23475 HMMreg00455_2_PVM(pf00455pf082
23480 - -
23500 - -
23510 -pf06551 -
BMC-P
BMC-H
BMC-T
Aldehyde dehydrogenase
PTAC
Signature enzyme
Alcohol dehydrogenase
Regulator

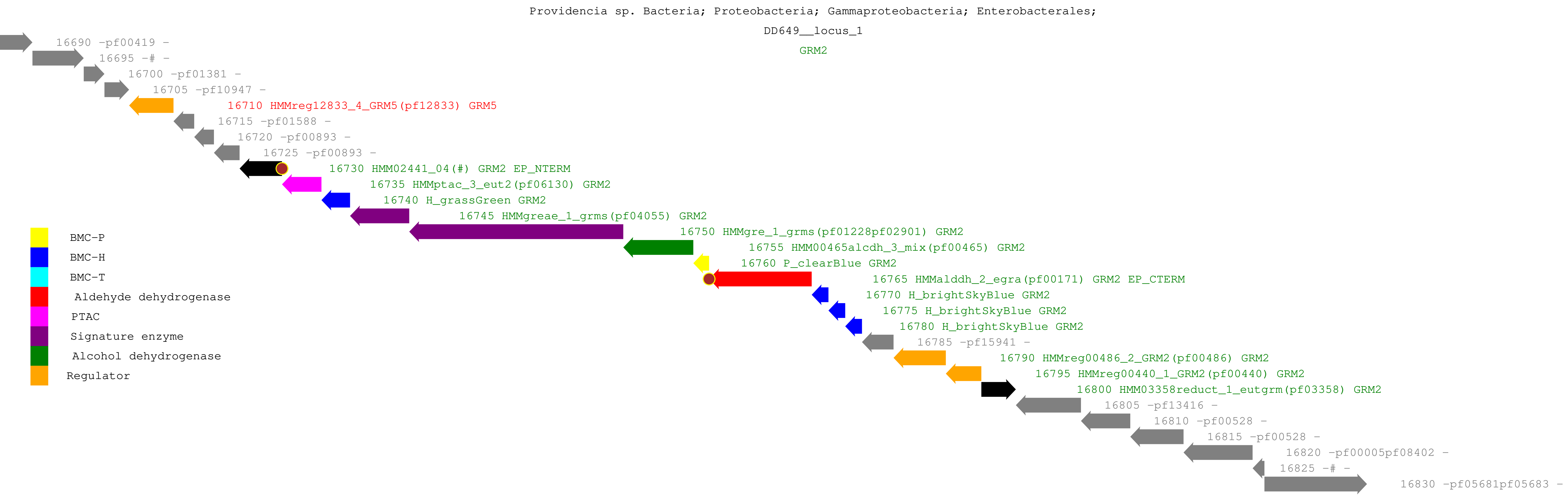
Providencia sp. Bacteria; Proteobacteria; Gammaproteobacteria; Enterobacterales;
DD649__locus_1
GRM2
16690 -pf00419 -
16695 -# -
16700 -pf01381 -
16705 -pf10947 -
16710 HMMreg12833_4_GRM5(pf12833) GRM5
16715 -pf01588 -
16720 -pf00893 -
16725 -pf00893 -
16730 HMM02441_04(#) GRM2 EP_NTERM
16735 HMMptac_3_eut2(pf06130) GRM2
16740 H_grassGreen GRM2
16745 HMMgreae_1_grms(pf04055) GRM2
16750 HMMgre_1_grms(pf01228pf02901) GRM2
16755 HMM00465alcdh_3_mix(pf00465) GRM2
16760 P_clearBlue GRM2
16765 HMMalddh_2_egra(pf00171) GRM2 EP_CTERM
16770 H_brightSkyBlue GRM2
16775 H_brightSkyBlue GRM2
16780 H_brightSkyBlue GRM2
16785 -pf15941 -
16790 HMMreg00486_2_GRM2(pf00486) GRM2
16795 HMMreg00440_1_GRM2(pf00440) GRM2
16800 HMM03358reduct_1_eutgrm(pf03358) GRM2
16805 -pf13416 -
16810 -pf00528 -
16815 -pf00528 -
16820 -pf00005pf08402 -
16825 -# -
16830 -pf05681pf05683 -
BMC-P
BMC-H
BMC-T
Aldehyde dehydrogenase
PTAC
Signature enzyme
Alcohol dehydrogenase
Regulator

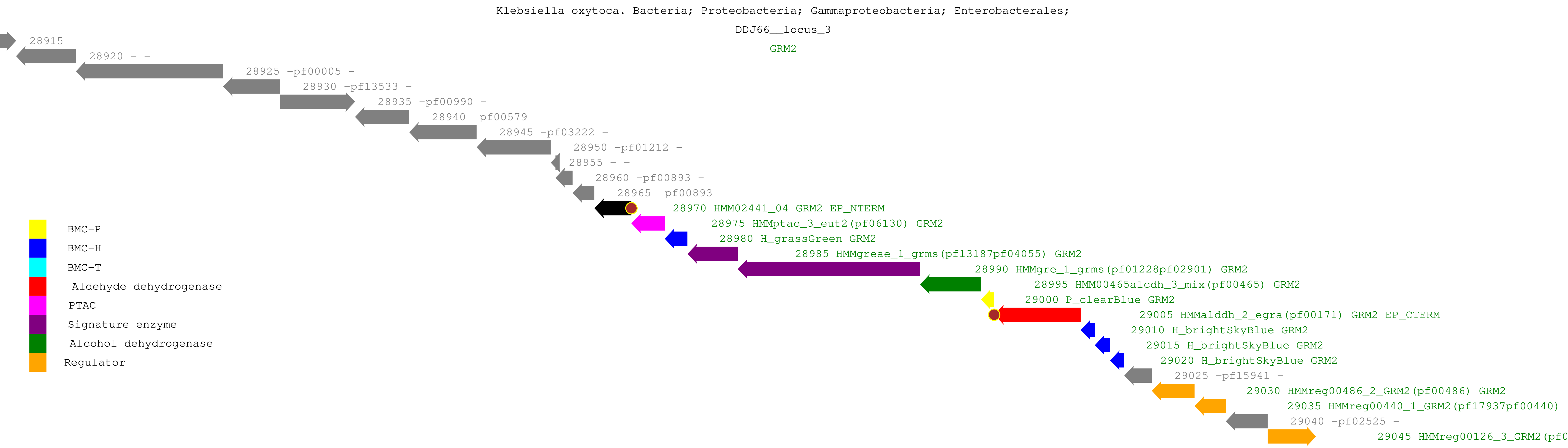

Klebsiella oxytoca. Bacteria; Proteobacteria; Gammaproteobacteria; Enterobacterales;
DDJ66__locus_3
GRM2
28915 - -
28920 - -
28925 -pf00005 -
28930 -pf13533 -
28935 -pf00990 -
28940 -pf00579 -
28945 -pf03222 -
28950 -pf01212 -
28955 - -
28960 -pf00893 -
28965 -pf00893 -
28970 HMM02441_04 GRM2 EP_NTERM
28975 HMMptac_3_eut2(pf06130) GRM2
28980 H_grassGreen GRM2
28985 HMMgreae_1_grms(pf13187pf04055) GRM2
28990 HMMgre_1_grms(pf01228pf02901) GRM2
28995 HMM00465alcdh_3_mix(pf00465) GRM2
29000 P_clearBlue GRM2
29005 HMMaldh_2_egra(pf00171) GRM2 EP_CTERM
29010 H_brightSkyBlue GRM2
29015 H_brightSkyBlue GRM2
29020 H_brightSkyBlue GRM2
29025 -pf15941 -
29030 HMMreg00486_2_GRM2(pf00486) GRM2
29035 HMMreg00440_1_GRM2(pf17937pf00440)
29040 -pf02525 -
29045 HMMreg00126_3_GRM2(pf0
BMC-P
BMC-H
BMC-T
Aldehyde dehydrogenase
PTAC
Signature enzyme
Alcohol dehydrogenase
Regulator

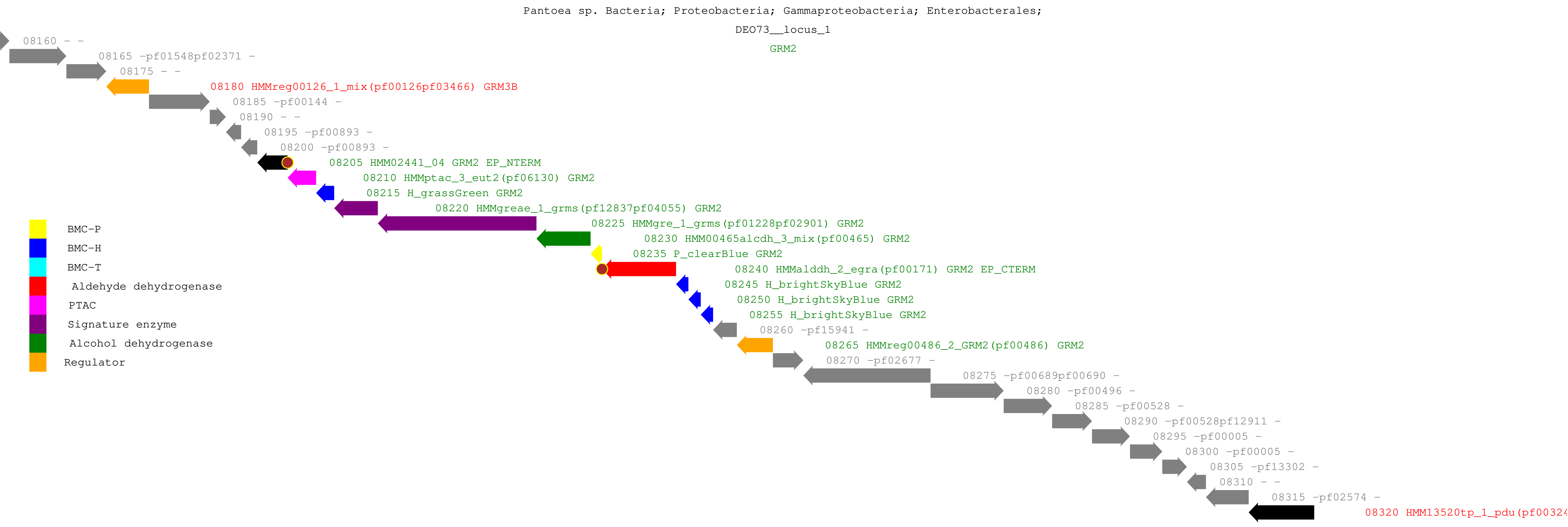
Pantoea sp. Bacteria; Proteobacteria; Gammaproteobacteria; Enterobacterales;
DEO73__locus_1
GRM2
08160 - -
08165 -pf01548pf02371 -
08175 - -
08180 HMMreg00126_1_mix(pf00126pf03466) GRM3B
08185 -pf00144 -
08190 - -
08195 -pf00893 -
08200 -pf00893 -
08205 HMM02441_04 GRM2 EP_NTERM
08210 HMMptac_3_eut2(pf06130) GRM2
08215 H_grassGreen GRM2
08220 HMMgreae_1_grms(pf12837pf04055) GRM2
08225 HMMgre_1_grms(pf01228pf02901) GRM2
08230 HMM00465alcdh_3_mix(pf00465) GRM2
08235 P_clearBlue GRM2
08240 HMMalddh_2_egra(pf00171) GRM2 EP_CTERM
08245 H_brightSkyBlue GRM2
08250 H_brightSkyBlue GRM2
08255 H_brightSkyBlue GRM2
08260 -pf15941 -
08265 HMMreg00486_2_GRM2(pf00486) GRM2
08270 -pf02677 -
08275 -pf00689pf00690 -
08280 -pf00496 -
08285 -pf00528 -
08290 -pf00528pf12911 -
08295 -pf00005 -
08300 -pf00005 -
08305 -pf13302 -
08310 - -
08315 -pf02574 -
08320 HMM13520tp_1_pdu(pf0032
BMC-P
BMC-H
BMC-T
Aldehyde dehydrogenase
PTAC
Signature enzyme
Alcohol dehydrogenase
Regulator

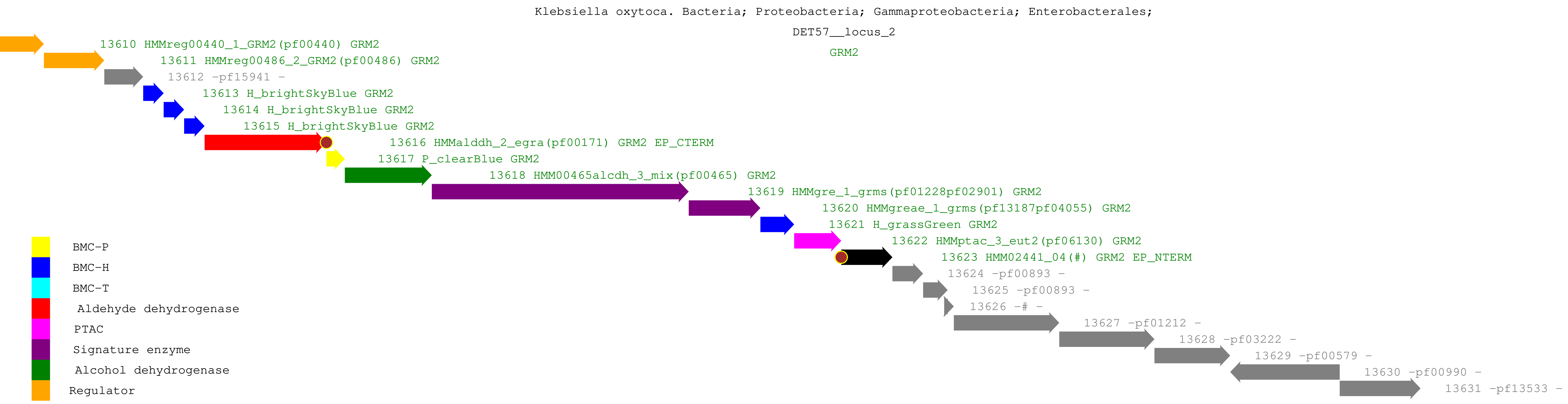
Klebsiella oxytoca. Bacteria; Proteobacteria; Gammaproteobacteria; Enterobacterales;
DET57__locus_2
GRM2
13610 HMMreg00440_1_GRM2(pf00440) GRM2
13611 HMMreg00486_2_GRM2(pf00486) GRM2
13612 -pf15941 -
13613 H_brightSkyBlue GRM2
13614 H_brightSkyBlue GRM2
13615 H_brightSkyBlue GRM2
13616 HMMalddh_2_egra(pf00171) GRM2 EP_CTERM
13617 P_clearBlue GRM2
13618 HMM00465alcdh_3_mix(pf00465) GRM2
13619 HMMgre_1_grms(pf01228pf02901) GRM2
13620 HMMgreae_1_grms(pf13187pf04055) GRM2
13621 H_grassGreen GRM2
13622 HMMptac_3_eut2(pf06130) GRM2
13623 HMM02441_04(#) GRM2 EP_NTERM
13624 -pf00893 -
13625 -pf00893 -
13626 -# -
13627 -pf01212 -
13628 -pf03222 -
13629 -pf00579 -
13630 -pf00990 -
13631 -pf13533 -
BMC-P
BMC-H
BMC-T
Aldehyde dehydrogenase
PTAC
Signature enzyme
Alcohol dehydrogenase
Regulator

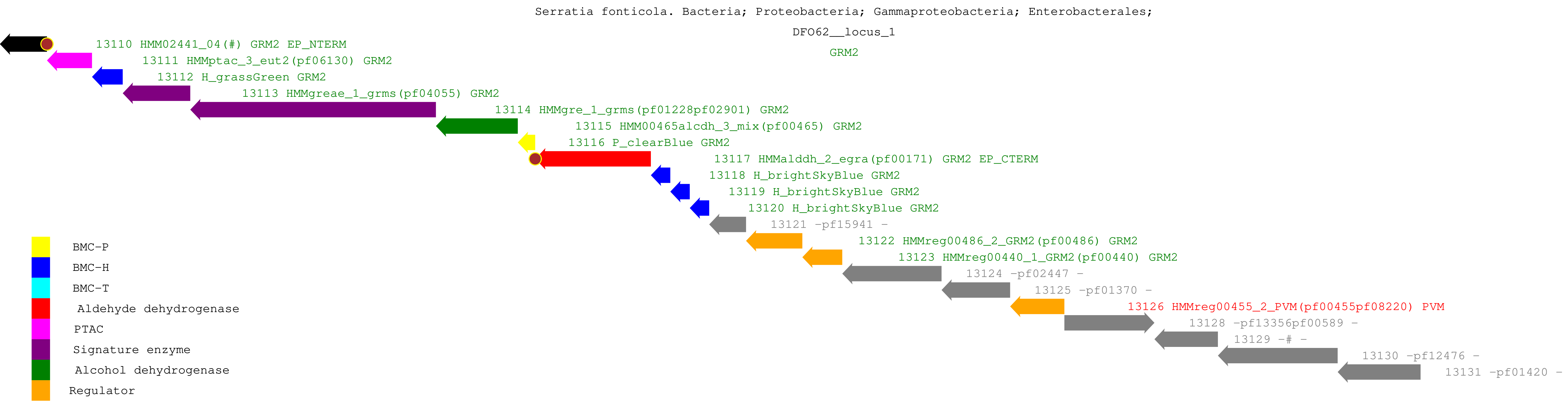
Serratia fonticola. Bacteria; Proteobacteria; Gammaproteobacteria; Enterobacterales;
DFO62__locus_1
GRM2
13110 HMM02441_04(#) GRM2 EP_NTERM
13111 HMMptac_3_eut2(pf06130) GRM2
13112 H_grassGreen GRM2
13113 HMMgreae_1_grms(pf04055) GRM2
13114 HMMgre_1_grms(pf01228pf02901) GRM2
13115 HMM00465alcdh_3_mix(pf00465) GRM2
13116 P_clearBlue GRM2
13117 HMMalddh_2_egra(pf00171) GRM2 EP_CTERM
13118 H_brightSkyBlue GRM2
13119 H_brightSkyBlue GRM2
13120 H_brightSkyBlue GRM2
13121 -pf15941 -
13122 HMMreg00486_2_GRM2(pf00486) GRM2
13123 HMMreg00440_1_GRM2(pf00440) GRM2
13124 -pf02447 -
13125 -pf01370 -
13126 HMMreg00455_2_PVM(pf00455pf08220) PVM
13128 -pf13356pf00589 -
13129 -# -
13130 -pf12476 -
13131 -pf01420 -
BMC-P
BMC-H
BMC-T
Aldehyde dehydrogenase
PTAC
Signature enzyme
Alcohol dehydrogenase
Regulator

Klebsiella michiganensis. Bacteria; Proteobacteria; Gammaproteobacteria; Enterobacterales;

DJ519__locus_2

GRM2

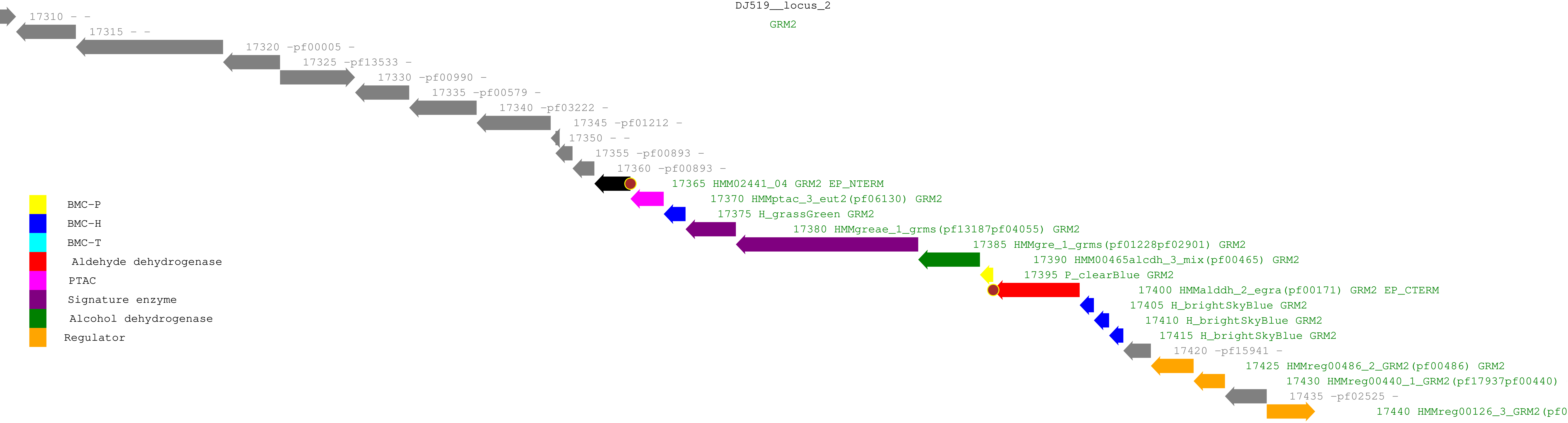

Klebsiella grimontii. Bacteria; Proteobacteria; Gammaproteobacteria; Enterobacterales;

DJ542__locus_1

GRM2

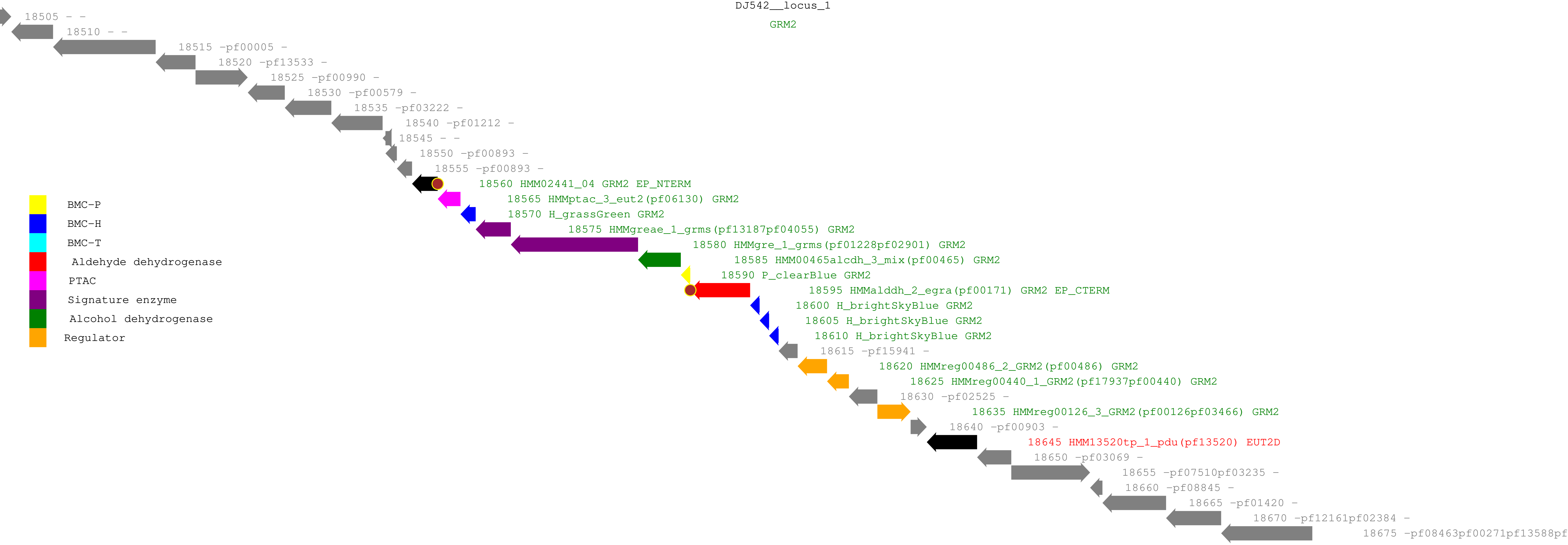

Klebsiella variicola. Bacteria; Proteobacteria; Gammaproteobacteria; Enterobacterales;

DMS35__locus_1

GRM2

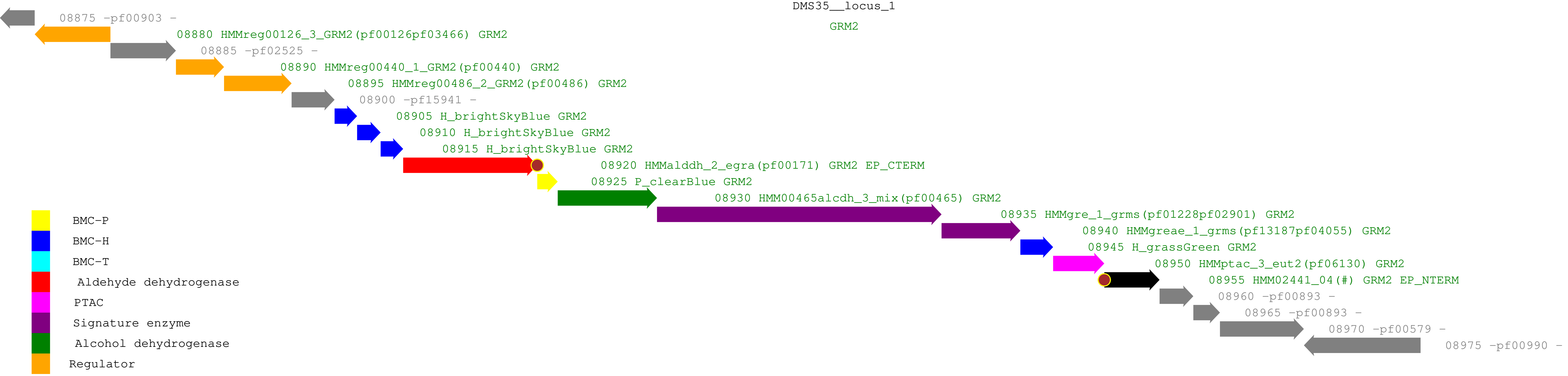

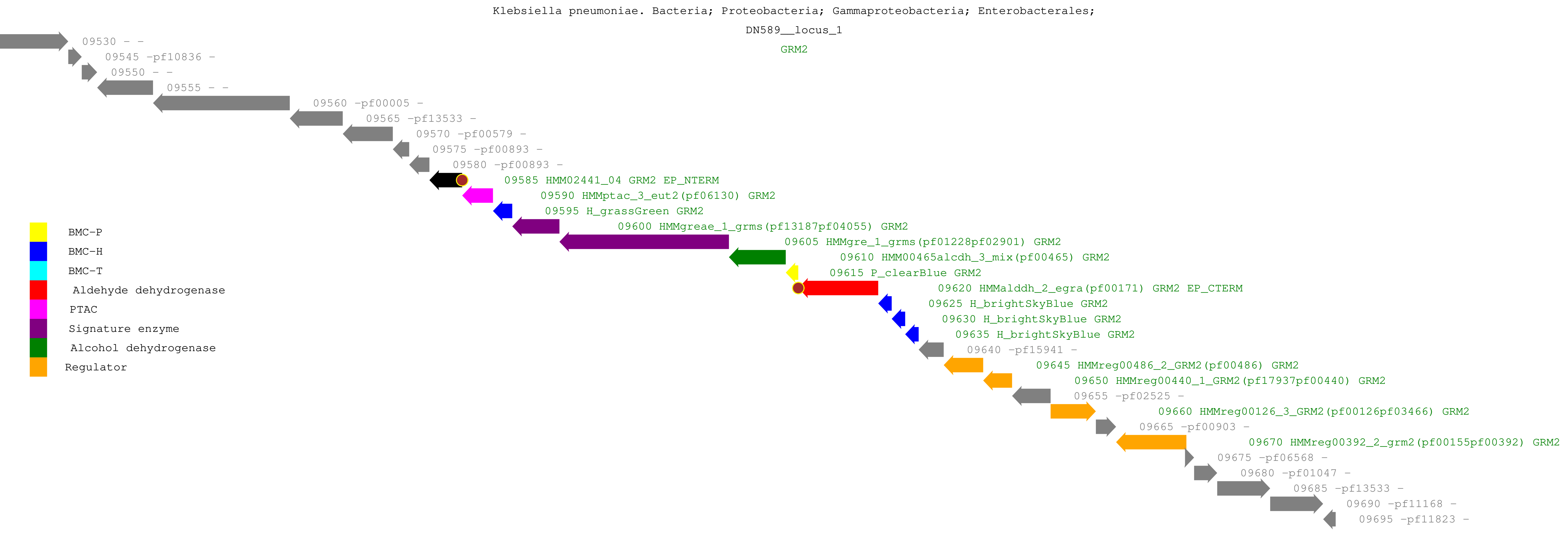
Klebsiella pneumoniae. Bacteria; Proteobacteria; Gammaproteobacteria; Enterobacterales;
DN589__locus_1
GRM2
09530 - -
09545 -pf10836 -
09550 - -
09555 - -
09560 -pf00005 -
09565 -pf13533 -
09570 -pf00579 -
09575 -pf00893 -
09580 -pf00893 -
09585 HMM02441_04 GRM2 EP_NTERM
09590 HMMptac_3_eut2(pf06130) GRM2
09595 H_grassGreen GRM2
09600 HMMgreae_1_grms(pf13187pf04055) GRM2
09605 HMMgre_1_grms(pf01228pf02901) GRM2
09610 HMM00465alcdh_3_mix(pf00465) GRM2
09615 P_clearBlue GRM2
09620 HMMalddh_2_egra(pf00171) GRM2 EP_CTERM
09625 H_brightSkyBlue GRM2
09630 H_brightSkyBlue GRM2
09635 H_brightSkyBlue GRM2
09640 -pf15941 -
09645 HMMreg00486_2_GRM2(pf00486) GRM2
09650 HMMreg00440_1_GRM2(pf17937pf00440) GRM2
09655 -pf02525 -
09660 HMMreg00126_3_GRM2(pf00126pf03466) GRM2
09665 -pf00903 -
09670 HMMreg00392_2_grm2(pf00155pf00392) GRM2
09675 -pf06568 -
09680 -pf01047 -
09685 -pf13533 -
09690 -pf11168 -
09695 -pf11823 -
BMC-P
BMC-H
BMC-T
Aldehyde dehydrogenase
PTAC
Signature enzyme
Alcohol dehydrogenase
Regulator

Raoultella planticola (Klebsiella planticola). Bacteria; Proteobacteria; Gammaproteobacteria; Enterobacterales;

DN603__locus_1

GRM2

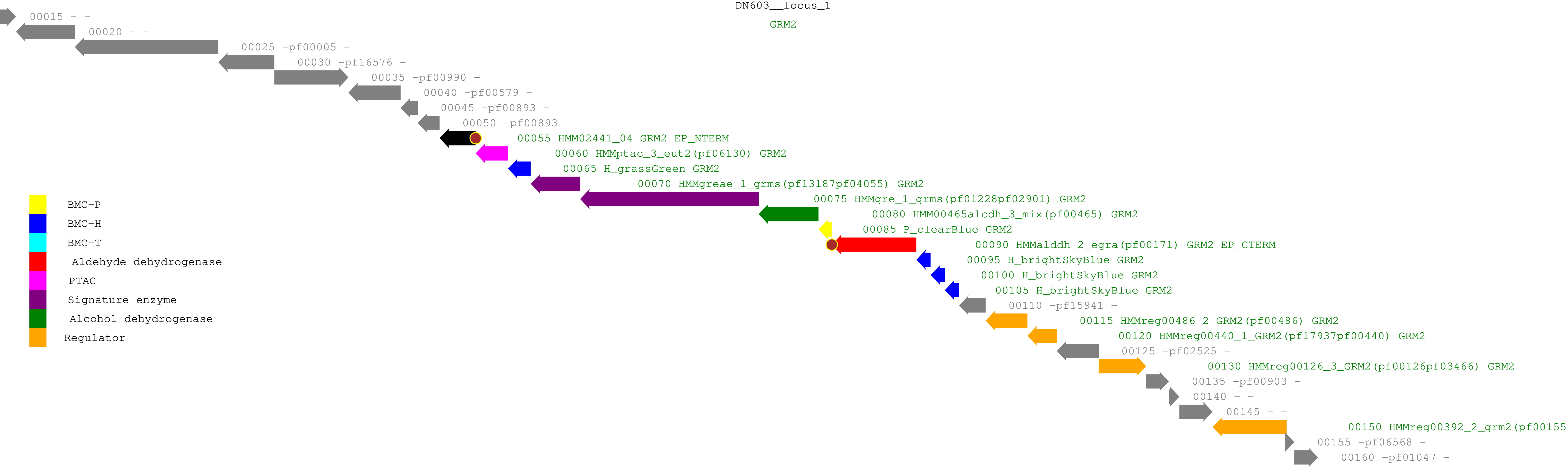

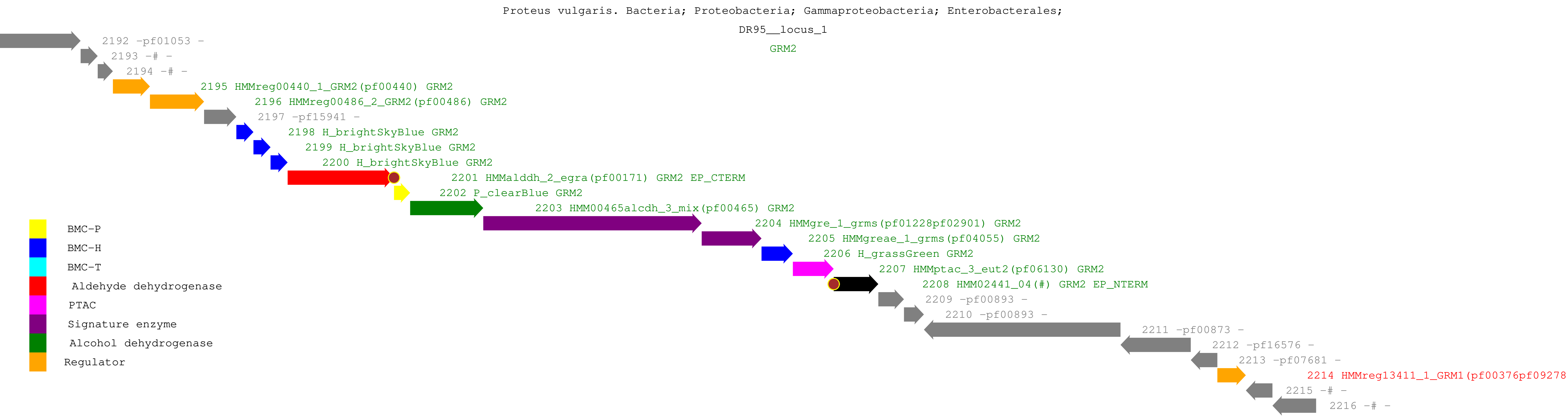

Proteus vulgaris. Bacteria; Proteobacteria; Gammaproteobacteria; Enterobacterales;
DR95__locus_1
GRM2
2192 -pf01053 -
2193 -# -
2194 -# -
2195 HMMreg00440_1_GRM2(pf00440) GRM2
2196 HMMreg00486_2_GRM2(pf00486) GRM2
2197 -pf15941 -
2198 H_brightSkyBlue GRM2
2199 H_brightSkyBlue GRM2
2200 H_brightSkyBlue GRM2
2201 HMMalddh_2_egra(pf00171) GRM2 EP_CTERM
2202 P_clearBlue GRM2
2203 HMM00465alcdh_3_mix(pf00465) GRM2
2204 HMMgre_1_grms(pf01228pf02901) GRM2
2205 HMMgreae_1_grms(pf04055) GRM2
2206 H_grassGreen GRM2
2207 HMMptac_3_eut2(pf06130) GRM2
2208 HMM02441_04(#) GRM2 EP_NTERM
2209 -pf00893 -
2210 -pf00893 -
2211 -pf00873 -
2212 -pf16576 -
2213 -pf07681 -
2214 HMMreg13411_1_GRM1(pf00376pf09278
2215 -# -
2216 -# -
BMC-P
BMC-H
BMC-T
Aldehyde dehydrogenase
PTAC
Signature enzyme
Alcohol dehydrogenase
Regulator

Klebsiella sp. P1CD1. Bacteria; Proteobacteria; Gammaproteobacteria; Enterobacterales;

DTA24__locus_3

GRM2

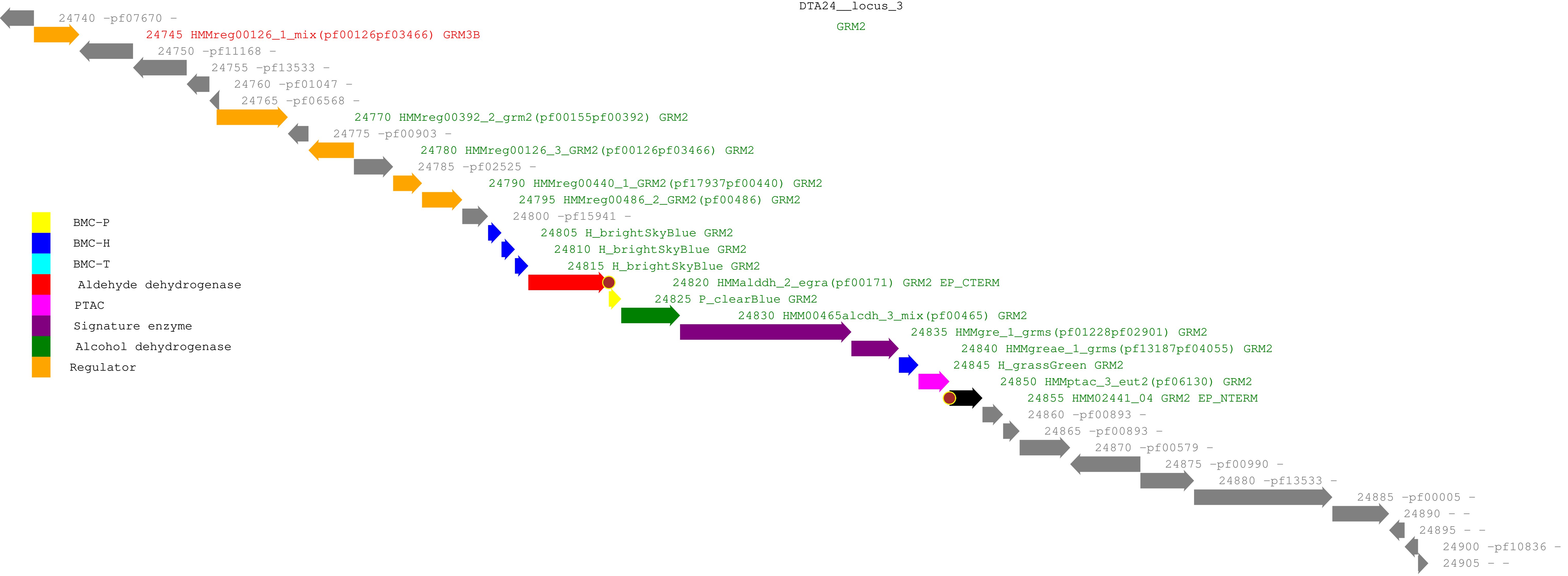

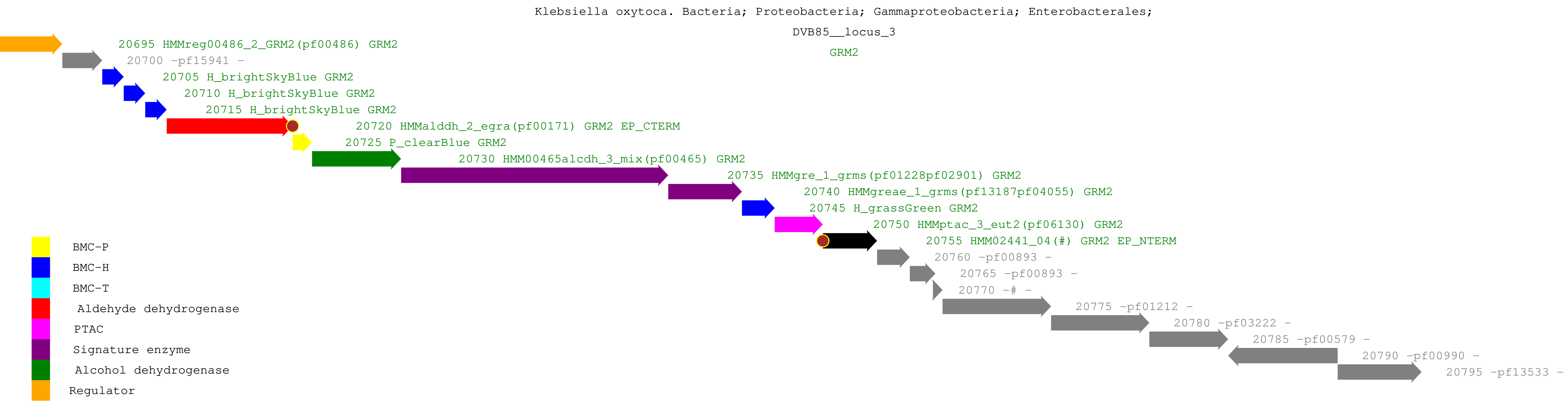
Klebsiella oxytoca. Bacteria; Proteobacteria; Gammaproteobacteria; Enterobacterales;
DVB85__locus_3
GRM2
20695 HMMreg00486_2_GRM2(pf00486) GRM2
20700 -pf15941 -
20705 H_brightSkyBlue GRM2
20710 H_brightSkyBlue GRM2
20715 H_brightSkyBlue GRM2
20720 HMMaldh_2_egra(pf00171) GRM2 EP_CTERM
20725 P_clearBlue GRM2
20730 HMM00465alcdh_3_mix(pf00465) GRM2
20735 HMMgre_1_grms(pf01228pf02901) GRM2
20740 HMMgreae_1_grms(pf13187pf04055) GRM2
20745 H_grassGreen GRM2
20750 HMMptac_3_eut2(pf06130) GRM2
20755 HMM02441_04(#) GRM2 EP_NTERM
20760 -pf00893 -
20765 -pf00893 -
20770 -# -
20775 -pf01212 -
20780 -pf03222 -
20785 -pf00579 -
20790 -pf00990 -
20795 -pf13533 -
BMC-P
BMC-H
BMC-T
Aldehyde dehydrogenase
PTAC
Signature enzyme
Alcohol dehydrogenase
Regulator

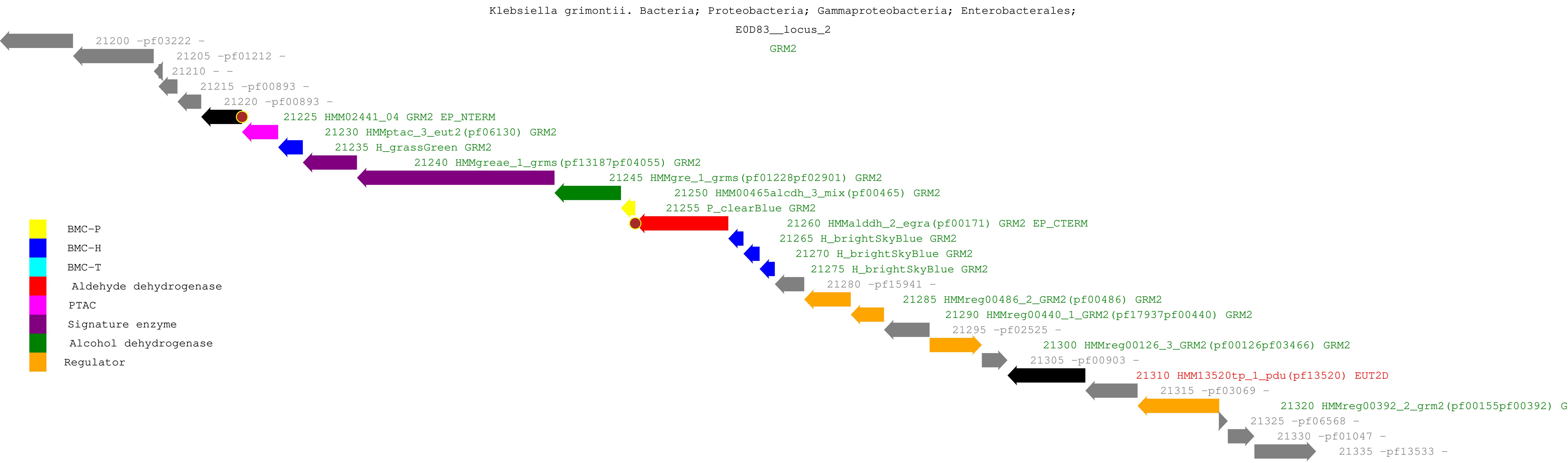
Klebsiella grimontii. Bacteria; Proteobacteria; Gammaproteobacteria; Enterobacterales;
E0D83__locus_2
GRM2
21200 -pf03222 -
21205 -pf01212 -
21210 - -
21215 -pf00893 -
21220 -pf00893 -
21225 HMM02441_04 GRM2 EP_NTERM
21230 HMMptac_3_eut2(pf06130) GRM2
21235 H_grassGreen GRM2
21240 HMMgreae_1_grms(pf13187pf04055) GRM2
21245 HMMgre_1_grms(pf01228pf02901) GRM2
21250 HMM00465alcdh_3_mix(pf00465) GRM2
21255 P_clearBlue GRM2
21260 HMMalddh_2_egra(pf00171) GRM2 EP_CTERM
21265 H_brightSkyBlue GRM2
21270 H_brightSkyBlue GRM2
21275 H_brightSkyBlue GRM2
21280 -pf15941 -
21285 HMMreg00486_2_GRM2(pf00486) GRM2
21290 HMMreg00440_1_GRM2(pf17937pf00440) GRM2
21295 -pf02525 -
21300 HMMreg00126_3_GRM2(pf00126pf03466) GRM2
21305 -pf00903 -
21310 HMM13520tp_1_pdu(pf13520) EUT2D
21315 -pf03069 -
21320 HMMreg00392_2_grm2(pf00155pf00392) G
21325 -pf06568 -
21330 -pf01047 -
21335 -pf13533 -
BMC-P
BMC-H
BMC-T
Aldehyde dehydrogenase
PTAC
Signature enzyme
Alcohol dehydrogenase
Regulator

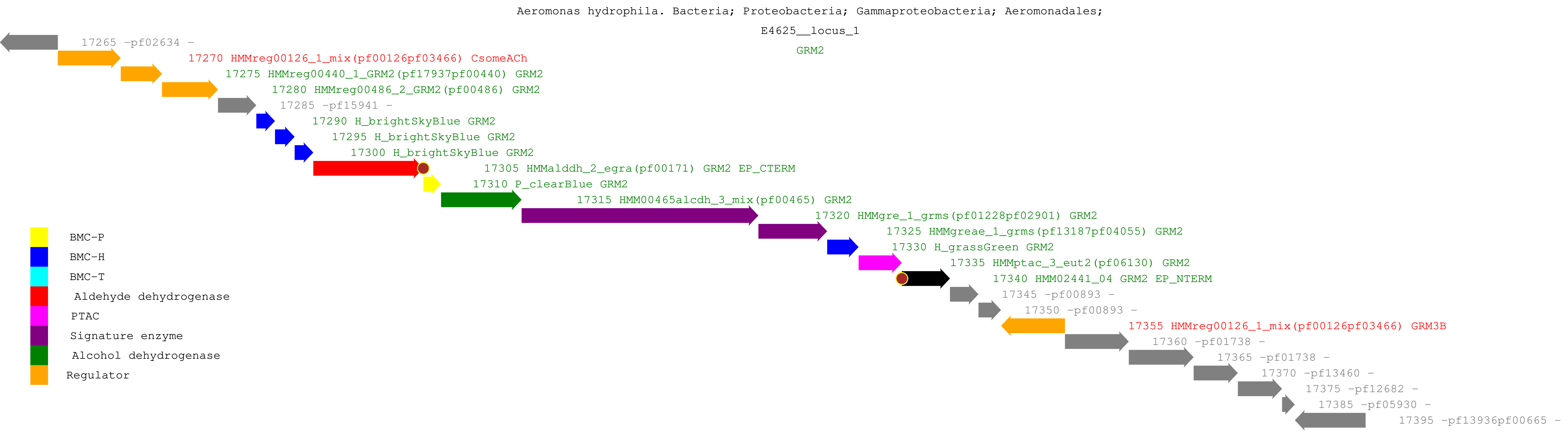

Aeromonas hydrophila. Bacteria; Proteobacteria; Gammaproteobacteria; Aeromonadales;
E4625__locus_1
GRM2
17265 -pf02634 -
17270 HMMreg00126_1_mix(pf00126pf03466) CsomeACh
17275 HMMreg00440_1_GRM2(pf17937pf00440) GRM2
17280 HMMreg00486_2_GRM2(pf00486) GRM2
17285 -pf15941 -
17290 H_brightSkyBlue GRM2
17295 H_brightSkyBlue GRM2
17300 H_brightSkyBlue GRM2
17305 HMMalddh_2_egra(pf00171) GRM2 EP_CTERM
17310 P_clearBlue GRM2
17315 HMM00465alcdh_3_mix(pf00465) GRM2
17320 HMMgre_1_grms(pf01228pf02901) GRM2
17325 HMMgreae_1_grms(pf13187pf04055) GRM2
17330 H_grassGreen GRM2
17335 HMMptac_3_eut2(pf06130) GRM2
17340 HMM02441_04 GRM2 EP_NTERM
17345 -pf00893 -
17350 -pf00893 -
17355 HMMreg00126_1_mix(pf00126pf03466) GRM3B
17360 -pf01738 -
17365 -pf01738 -
17370 -pf13460 -
17375 -pf12682 -
17385 -pf05930 -
17395 -pf13936pf00665 -
BMC-P
BMC-H
BMC-T
Aldehyde dehydrogenase
PTAC
Signature enzyme
Alcohol dehydrogenase
Regulator

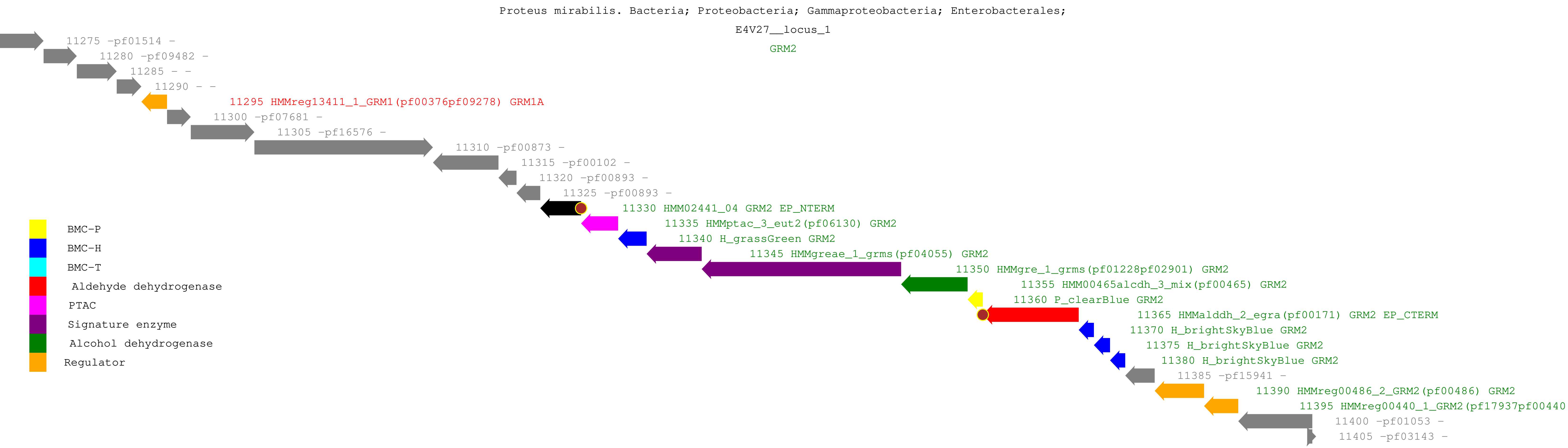

Proteus mirabilis. Bacteria; Proteobacteria; Gammaproteobacteria; Enterobacterales;
E4V27__locus_1
GRM2
11275 -pf01514 -
11280 -pf09482 -
11285 - -
11290 - -
11295 HMMreg13411_1_GRM1(pf00376pf09278) GRM1A
11300 -pf07681 -
11305 -pf16576 -
11310 -pf00873 -
11315 -pf00102 -
11320 -pf00893 -
11325 -pf00893 -
11330 HMM02441_04 GRM2 EP_NTERM
11335 HMMptac_3_eut2(pf06130) GRM2
11340 H_grassGreen GRM2
11345 HMMgreae_1_grms(pf04055) GRM2
11350 HMMgre_1_grms(pf01228pf02901) GRM2
11355 HMM00465alcdh_3_mix(pf00465) GRM2
11360 P_clearBlue GRM2
11365 HMMalddh_2_egra(pf00171) GRM2 EP_CTERM
11370 H_brightSkyBlue GRM2
11375 H_brightSkyBlue GRM2
11380 H_brightSkyBlue GRM2
11385 -pf15941 -
11390 HMMreg00486_2_GRM2(pf00486) GRM2
11395 HMMreg00440_1_GRM2(pf17937pf00440
11400 -pf01053 -
11405 -pf03143 -
BMC-P
BMC-H
BMC-T
Aldehyde dehydrogenase
PTAC
Signature enzyme
Alcohol dehydrogenase
Regulator

Proteus mirabilis. Bacteria; Proteobacteria; Gammaproteobacteria; Enterobacterales;

E4V37__locus_1

GRM2

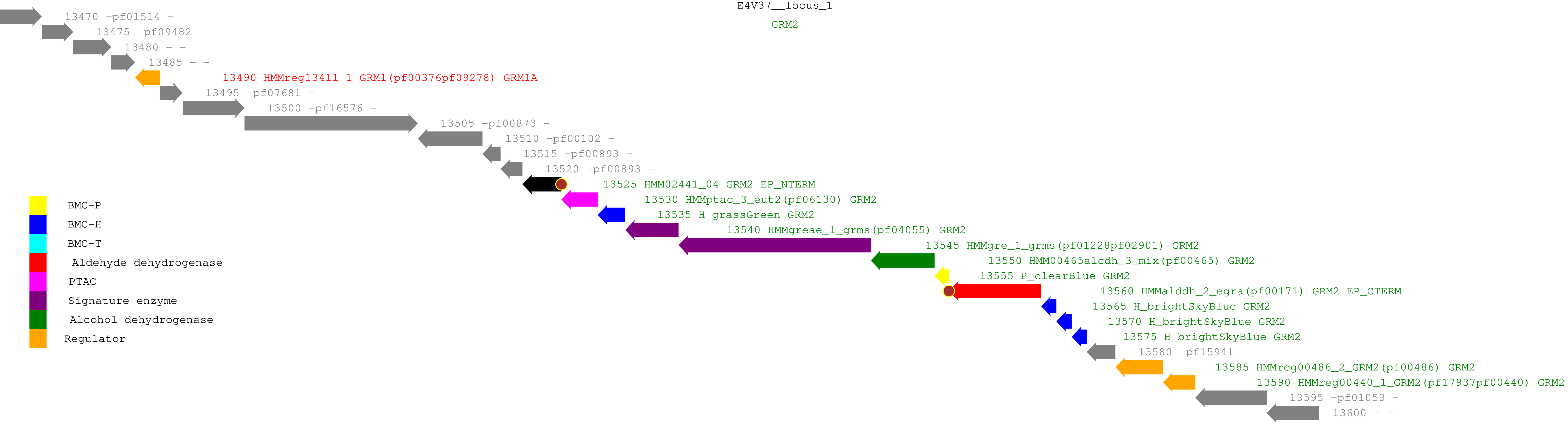

Proteus mirabilis. Bacteria; Proteobacteria; Gammaproteobacteria; Enterobacterales;

E4V48__locus_1

GRM2

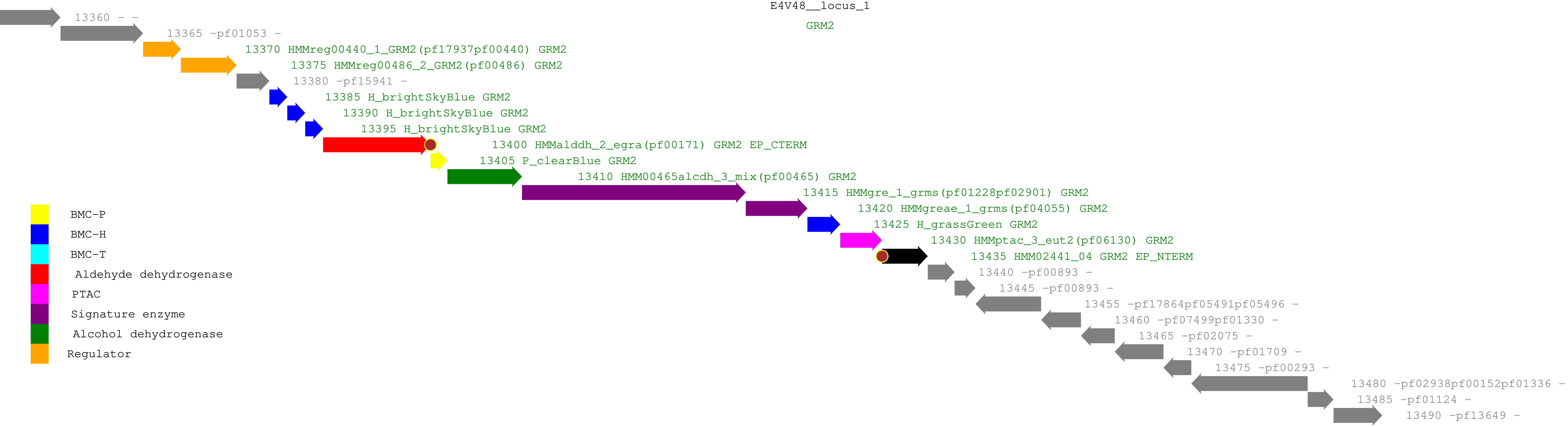

Providencia sp. MGF014. Bacteria; Proteobacteria; Gammaproteobacteria; Enterobacterales;

E6R27__locus_1

GRM2

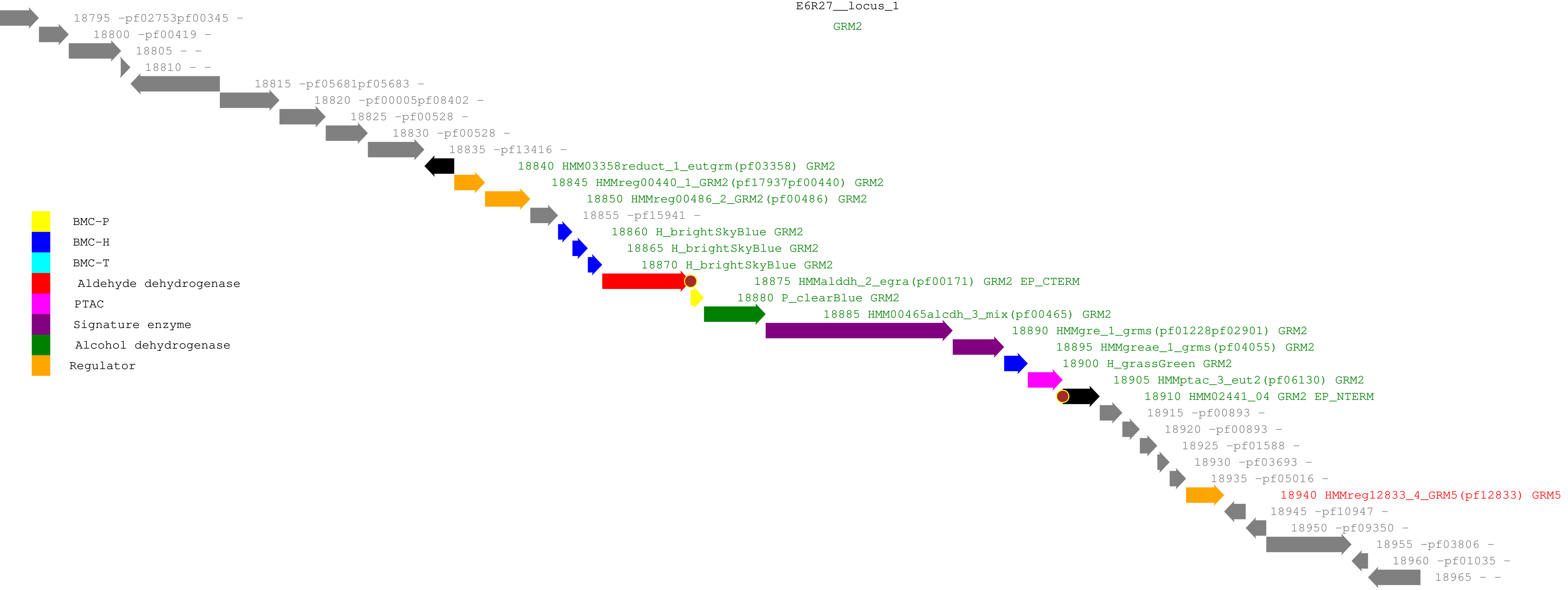

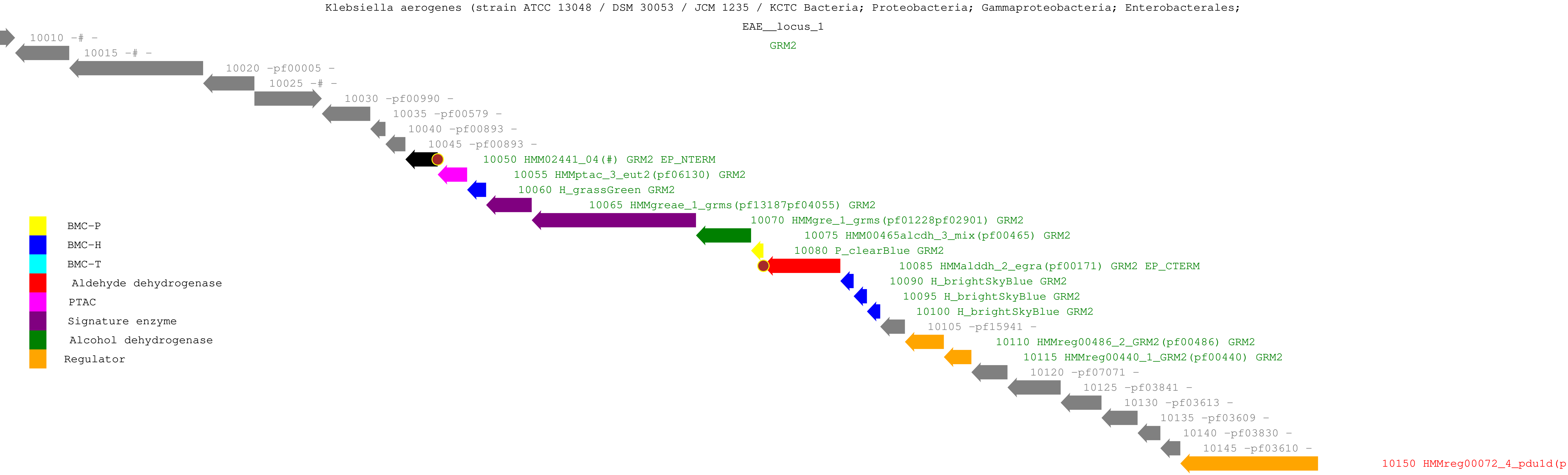
Klebsiella aerogenes (strain ATCC 13048 / DSM 30053 / JCM 1235 / KCTC Bacteria; Proteobacteria; Gammaproteobacteria; Enterobacterales;
EAE__locus_1
GRM2
10010 -# -
10015 -# -
10020 -pf00005 -
10025 -# -
10030 -pf00990 -
10035 -pf00579 -
10040 -pf00893 -
10045 -pf00893 -
10050 HMM02441_04(#) GRM2 EP_NTERM
10055 HMMptac_3_eut2(pf06130) GRM2
10060 H_grassGreen GRM2
10065 HMMgreae_1_grms(pf13187pf04055) GRM2
10070 HMMgre_1_grms(pf01228pf02901) GRM2
10075 HMM00465alcdh_3_mix(pf00465) GRM2
10080 P_clearBlue GRM2
10085 HMMalddh_2_egra(pf00171) GRM2 EP_CTERM
10090 H_brightSkyBlue GRM2
10095 H_brightSkyBlue GRM2
10100 H_brightSkyBlue GRM2
10105 -pf15941 -
10110 HMMreg00486_2_GRM2(pf00486) GRM2
10115 HMMreg00440_1_GRM2(pf00440) GRM2
10120 -pf07071 -
10125 -pf03841 -
10130 -pf03613 -
10135 -pf03609 -
10140 -pf03830 -
10145 -pf03610 -
10150 HMMreg00072_4_pdu1d(p
BMC-P
BMC-H
BMC-T
Aldehyde dehydrogenase
PTAC
Signature enzyme
Alcohol dehydrogenase
Regulator

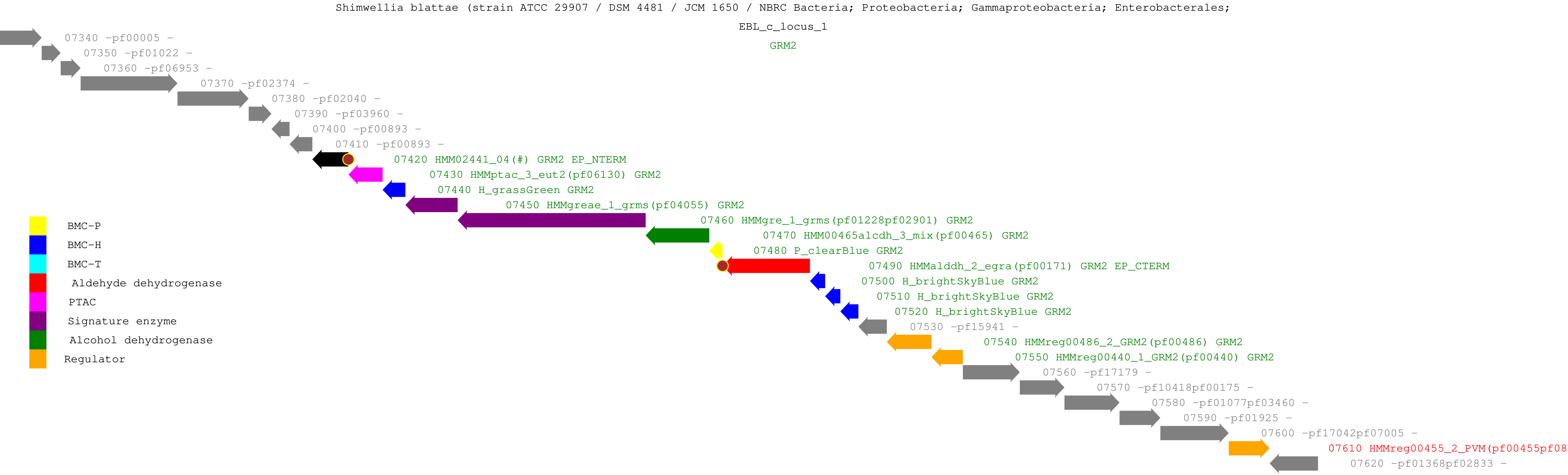
Shimwellia blattae (strain ATCC 29907 / DSM 4481 / JCM 1650 / NBRC Bacteria; Proteobacteria; Gammaproteobacteria; Enterobacterales;
EBL_c_locus_1
GRM2
07340 -pf00005 -
07350 -pf01022 -
07360 -pf06953 -
07370 -pf02374 -
07380 -pf02040 -
07390 -pf03960 -
07400 -pf00893 -
07410 -pf00893 -
07420 HMM02441_04(#) GRM2 EP_NTERM
07430 HMMptac_3_eut2(pf06130) GRM2
07440 H_grassGreen GRM2
07450 HMMgreae_1_grms(pf04055) GRM2
07460 HMMgre_1_grms(pf01228pf02901) GRM2
07470 HMM00465alcdh_3_mix(pf00465) GRM2
07480 P_clearBlue GRM2
07490 HMMalddh_2_egra(pf00171) GRM2 EP_CTERM
07500 H_brightSkyBlue GRM2
07510 H_brightSkyBlue GRM2
07520 H_brightSkyBlue GRM2
07530 -pf15941 -
07540 HMMreg00486_2_GRM2(pf00486) GRM2
07550 HMMreg00440_1_GRM2(pf00440) GRM2
07560 -pf17179 -
07570 -pf10418pf00175 -
07580 -pf01077pf03460 -
07590 -pf01925 -
07600 -pf17042pf07005 -
07610 HMMreg00455_2_PVM(pf00455pf08
07620 -pf01368pf02833 -
BMC-P
BMC-H
BMC-T
Aldehyde dehydrogenase
PTAC
Signature enzyme
Alcohol dehydrogenase
Regulator

Providencia alcalifaciens. Bacteria; Proteobacteria; Gammaproteobacteria; Enterobacterales;

EC835__locus_1

GRM2

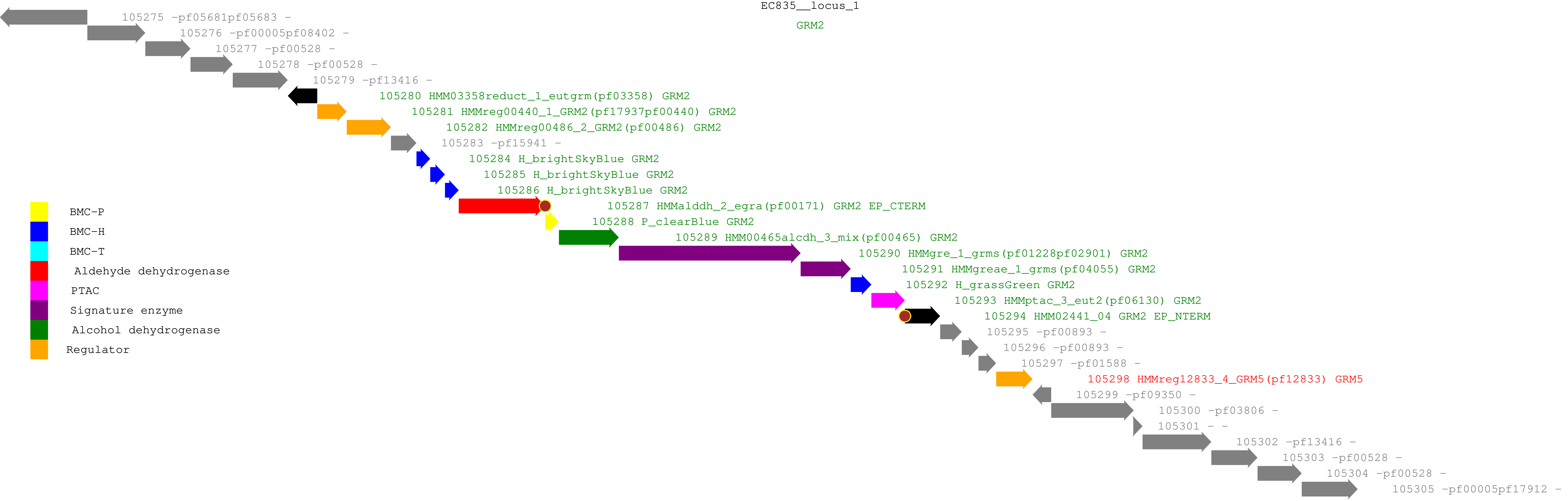

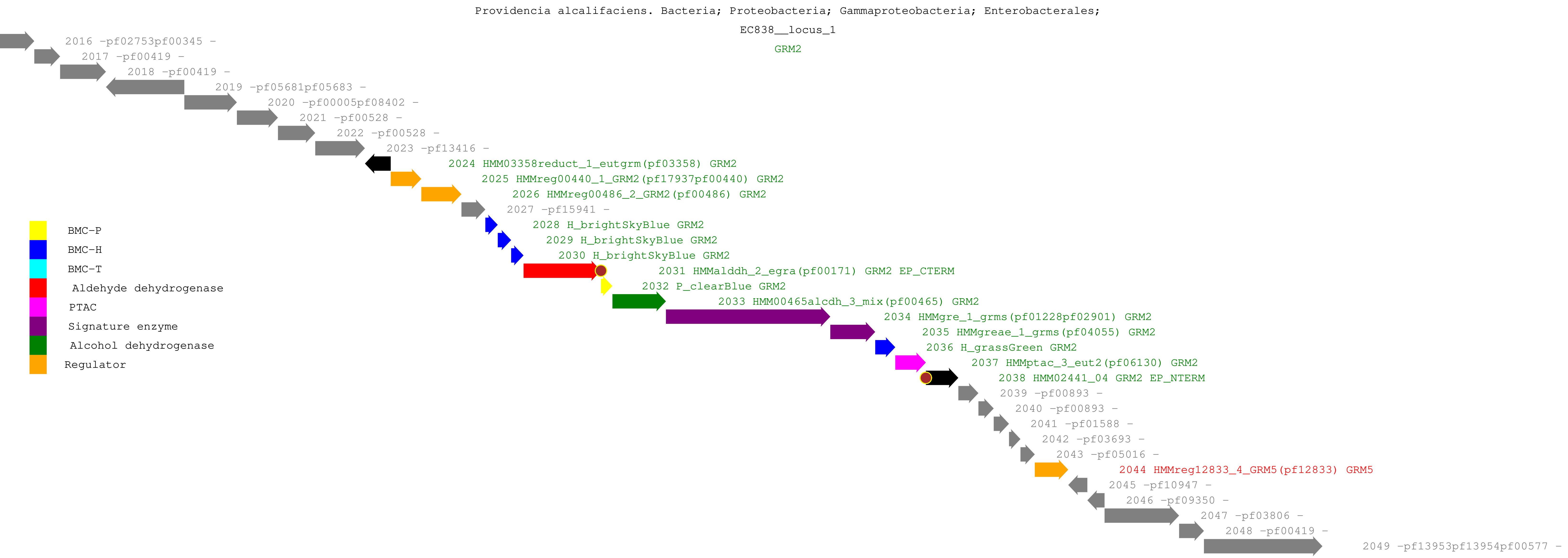
Providencia alcalifaciens. Bacteria; Proteobacteria; Gammaproteobacteria; Enterobacterales;
EC838__locus_1
GRM2
2016 -pf02753pf00345 -
2017 -pf00419 -
2018 -pf00419 -
2019 -pf05681pf05683 -
2020 -pf00005pf08402 -
2021 -pf00528 -
2022 -pf00528 -
2023 -pf13416 -
2024 HMM03358reduct_1_eutgrm(pf03358) GRM2
2025 HMMreg00440_1_GRM2(pf17937pf00440) GRM2
2026 HMMreg00486_2_GRM2(pf00486) GRM2
2027 -pf15941 -
2028 H_brightSkyBlue GRM2
2029 H_brightSkyBlue GRM2
2030 H_brightSkyBlue GRM2
2031 HMMalddh_2_egra(pf00171) GRM2 EP_CTERM
2032 P_clearBlue GRM2
2033 HMM00465alcdh_3_mix(pf00465) GRM2
2034 HMMgre_1_grms(pf01228pf02901) GRM2
2035 HMMgreae_1_grms(pf04055) GRM2
2036 H_grassGreen GRM2
2037 HMMptac_3_eut2(pf06130) GRM2
2038 HMM02441_04 GRM2 EP_NTERM
2039 -pf00893 -
2040 -pf00893 -
2041 -pf01588 -
2042 -pf03693 -
2043 -pf05016 -
2044 HMMreg12833_4_GRM5(pf12833) GRM5
2045 -pf10947 -
2046 -pf09350 -
2047 -pf03806 -
2048 -pf00419 -
2049 -pf13953pf13954pf00577 -
BMC-P
BMC-H
BMC-T
Aldehyde dehydrogenase
PTAC
Signature enzyme
Alcohol dehydrogenase
Regulator

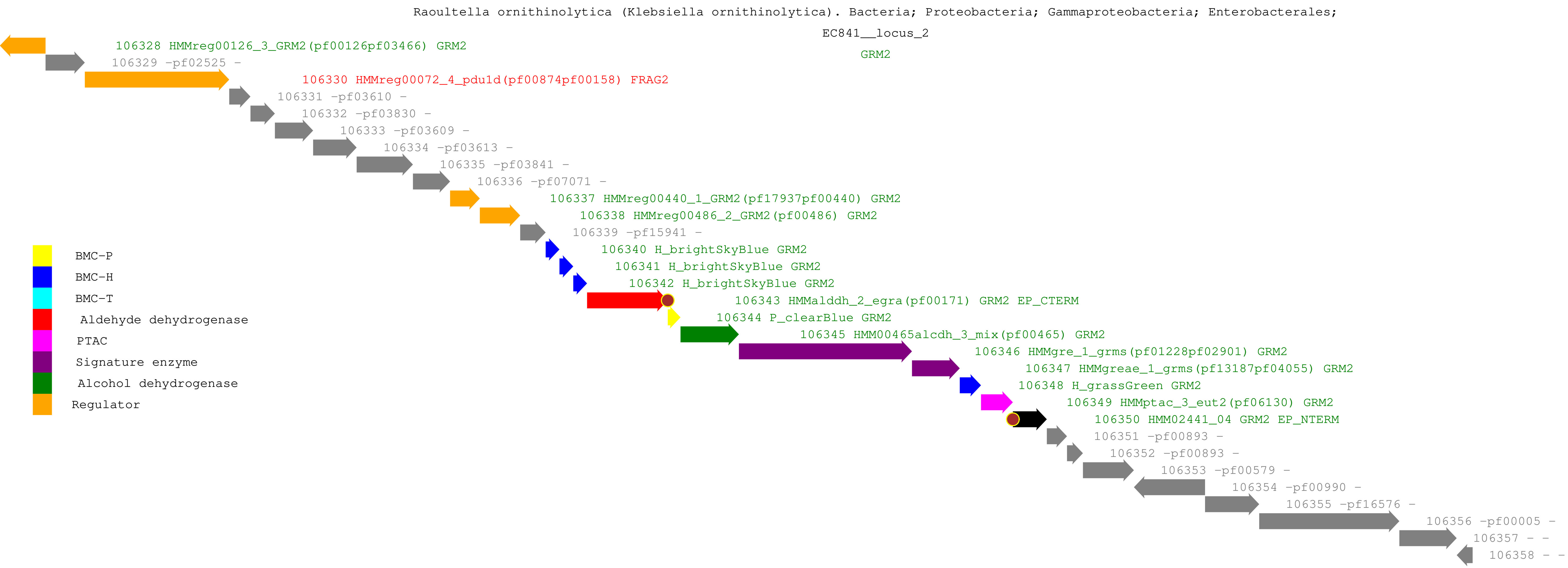

Raoultella ornithinolytica (Klebsiella ornithinolytica). Bacteria; Proteobacteria; Gammaproteobacteria; Enterobacterales;
EC841__locus_2
GRM2
106328 HMMreg00126_3_GRM2(pf00126pf03466) GRM2
106329 -pf02525 -
106330 HMMreg00072_4_pdu1d(pf00874pf00158) FRAG2
106331 -pf03610 -
106332 -pf03830 -
106333 -pf03609 -
106334 -pf03613 -
106335 -pf03841 -
106336 -pf07071 -
106337 HMMreg00440_1_GRM2(pf17937pf00440) GRM2
106338 HMMreg00486_2_GRM2(pf00486) GRM2
106339 -pf15941 -
106340 H_brightSkyBlue GRM2
106341 H_brightSkyBlue GRM2
106342 H_brightSkyBlue GRM2
106343 HMMaldh_2_egra(pf00171) GRM2 EP_CTERM
106344 P_clearBlue GRM2
106345 HMM00465alcdh_3_mix(pf00465) GRM2
106346 HMMgre_1_grms(pf01228pf02901) GRM2
106347 HMMgreae_1_grms(pf13187pf04055) GRM2
106348 H_grassGreen GRM2
106349 HMMptac_3_eut2(pf06130) GRM2
106350 HMM02441_04 GRM2 EP_NTERM
106351 -pf00893 -
106352 -pf00893 -
106353 -pf00579 -
106354 -pf00990 -
106355 -pf16576 -
106356 -pf00005 -
106357 - -
106358 - -
BMC-P
BMC-H
BMC-T
Aldehyde dehydrogenase
PTAC
Signature enzyme
Alcohol dehydrogenase
Regulator

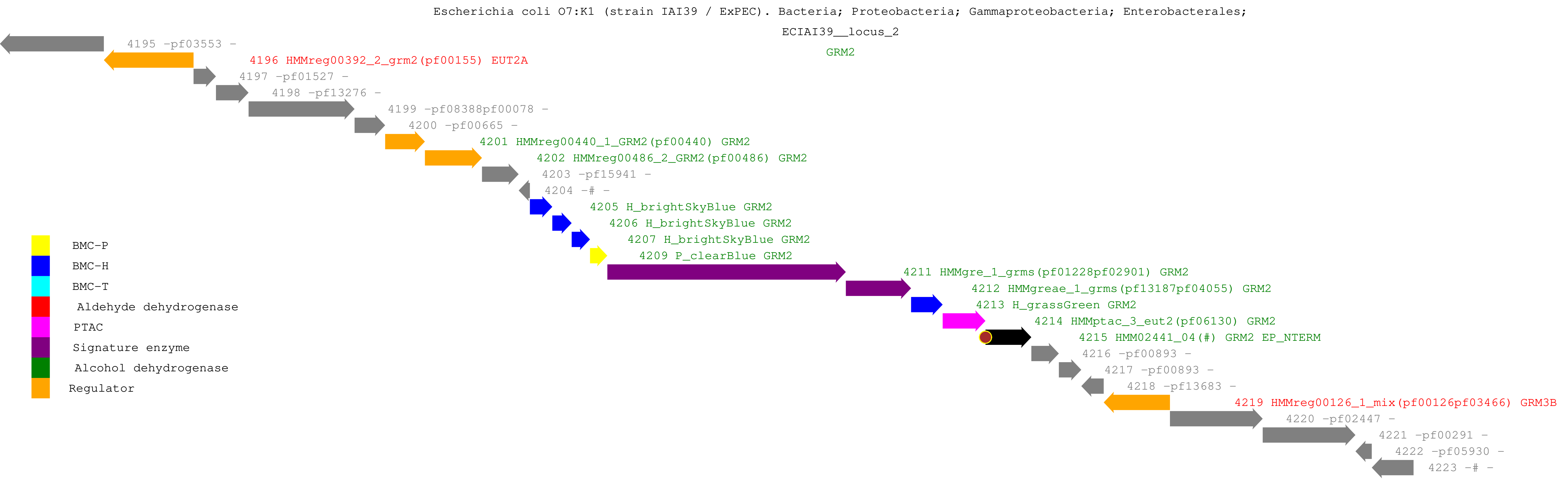
Escherichia coli O7:K1 (strain IAI39 / ExPEC). Bacteria; Proteobacteria; Gammaproteobacteria; Enterobacterales;
ECIAI39__locus_2
GRM2
4195 -pf03553 -
4196 HMMreg00392_2_grm2(pf00155) EUT2A
4197 -pf01527 -
4198 -pf13276 -
4199 -pf08388pf00078 -
4200 -pf00665 -
4201 HMMreg00440_1_GRM2(pf00440) GRM2
4202 HMMreg00486_2_GRM2(pf00486) GRM2
4203 -pf15941 -
4204 -# -
4205 H_brightSkyBlue GRM2
4206 H_brightSkyBlue GRM2
4207 H_brightSkyBlue GRM2
4209 P_clearBlue GRM2
4211 HMMgre_1_grms(pf01228pf02901) GRM2
4212 HMMgreae_1_grms(pf13187pf04055) GRM2
4213 H_grassGreen GRM2
4214 HMMptac_3_eut2(pf06130) GRM2
4215 HMM02441_04(#) GRM2 EP_NTERM
4216 -pf00893 -
4217 -pf00893 -
4218 -pf13683 -
4219 HMMreg00126_1_mix(pf00126pf03466) GRM3B
4220 -pf02447 -
4221 -pf00291 -
4222 -pf05930 -
4223 -# -
BMC-P
BMC-H
BMC-T
Aldehyde dehydrogenase
PTAC
Signature enzyme
Alcohol dehydrogenase
Regulator

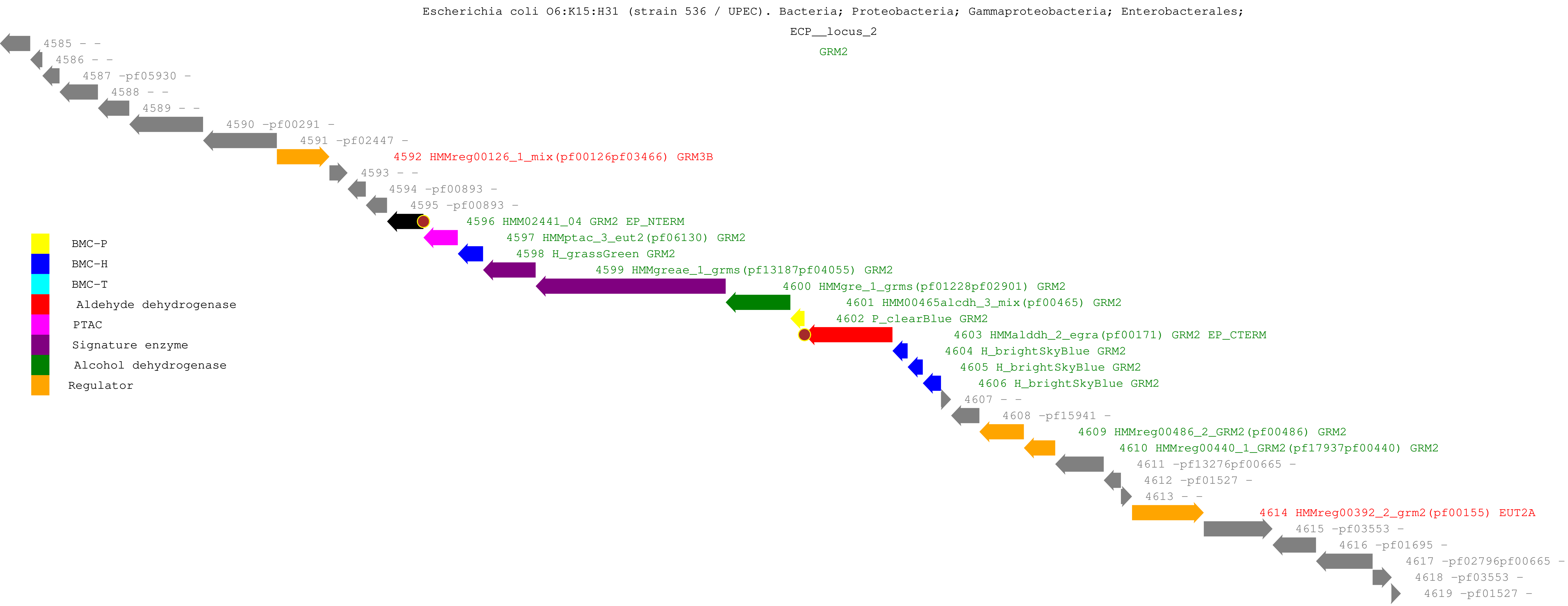

Escherichia coli O6:K15:H31 (strain 536 / UPEC). Bacteria; Proteobacteria; Gammaproteobacteria; Enterobacterales;
ECP__locus_2
GRM2
4585 - -
4586 - -
4587 -pf05930 -
4588 - -
4589 - -
4590 -pf00291 -
4591 -pf02447 -
4592 HMMreg00126_1_mix(pf00126pf03466) GRM3B
4593 - -
4594 -pf00893 -
4595 -pf00893 -
4596 HMM02441_04 GRM2 EP_NTERM
4597 HMMptac_3_eut2(pf06130) GRM2
4598 H_grassGreen GRM2
4599 HMMgreae_1_grms(pf13187pf04055) GRM2
4600 HMMgre_1_grms(pf01228pf02901) GRM2
4601 HMM00465alcdh_3_mix(pf00465) GRM2
4602 P_clearBlue GRM2
4603 HMMaldddh_2_egra(pf00171) GRM2 EP_CTERM
4604 H_brightSkyBlue GRM2
4605 H_brightSkyBlue GRM2
4606 H_brightSkyBlue GRM2
4607 - -
4608 -pf15941 -
4609 HMMreg00486_2_GRM2(pf00486) GRM2
4610 HMMreg00440_1_GRM2(pf17937pf00440) GRM2
4611 -pf13276pf00665 -
4612 -pf01527 -
4613 - -
4614 HMMreg00392_2_grm2(pf00155) EUT2A
4615 -pf03553 -
4616 -pf01695 -
4617 -pf02796pf00665 -
4618 -pf03553 -
4619 -pf01527 -
BMC-P
BMC-H
BMC-T
Aldehyde dehydrogenase
PTAC
Signature enzyme
Alcohol dehydrogenase
Regulator

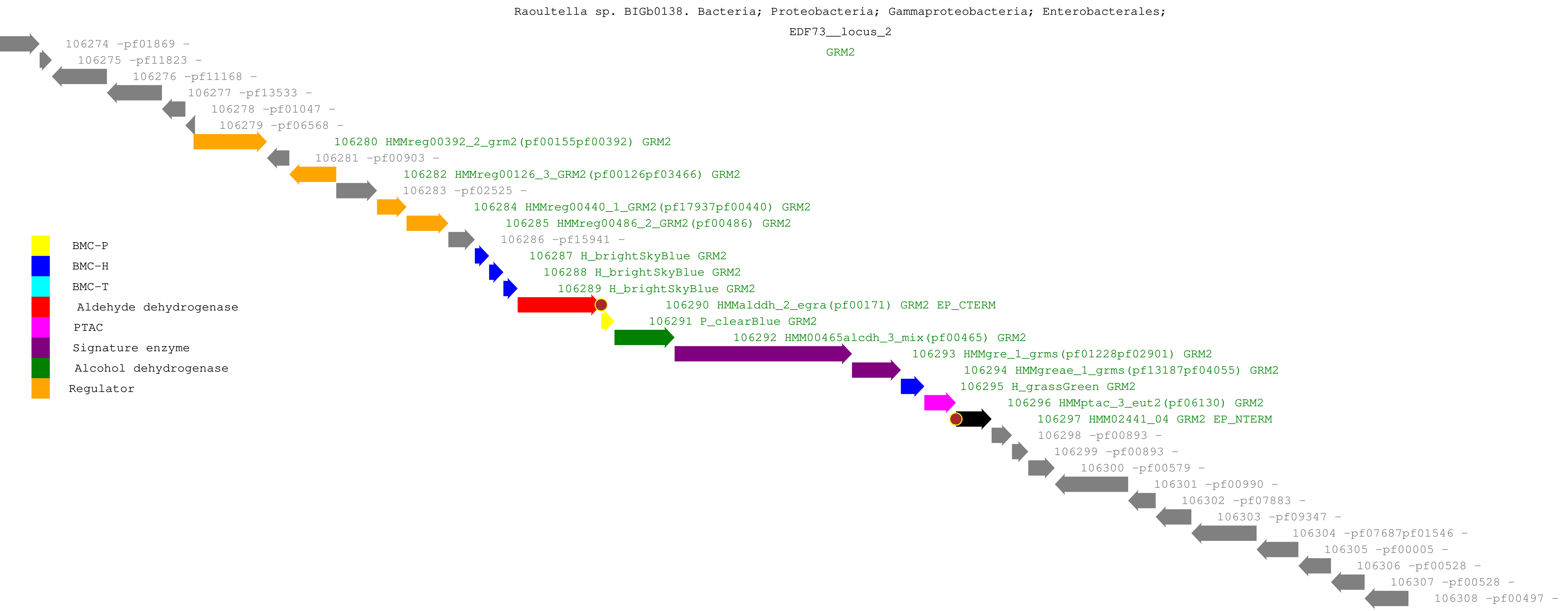
Raoultella sp. BIGb0138. Bacteria; Proteobacteria; Gammaproteobacteria; Enterobacterales;
EDF73__locus_2
GRM2
106274 -pf01869 -
106275 -pf11823 -
106276 -pf11168 -
106277 -pf13533 -
106278 -pf01047 -
106279 -pf06568 -
106280 HMMreg00392_2_grm2(pf00155pf00392) GRM2
106281 -pf00903 -
106282 HMMreg00126_3_GRM2(pf00126pf03466) GRM2
106283 -pf02525 -
106284 HMMreg00440_1_GRM2(pf17937pf00440) GRM2
106285 HMMreg00486_2_GRM2(pf00486) GRM2
106286 -pf15941 -
106287 H_brightSkyBlue GRM2
106288 H_brightSkyBlue GRM2
106289 H_brightSkyBlue GRM2
106290 HMMaldh_2_egra(pf00171) GRM2 EP_CTERM
106291 P_clearBlue GRM2
106292 HMM00465alcdh_3_mix(pf00465) GRM2
106293 HMMgre_1_grms(pf01228pf02901) GRM2
106294 HMMgreae_1_grms(pf13187pf04055) GRM2
106295 H_grassGreen GRM2
106296 HMMptac_3_eut2(pf06130) GRM2
106297 HMM02441_04 GRM2 EP_NTERM
106298 -pf00893 -
106299 -pf00893 -
106300 -pf00579 -
106301 -pf00990 -
106302 -pf07883 -
106303 -pf09347 -
106304 -pf07687pf01546 -
106305 -pf00005 -
106306 -pf00528 -
106307 -pf00528 -
106308 -pf00497 -
BMC-P
BMC-H
BMC-T
Aldehyde dehydrogenase
PTAC
Signature enzyme
Alcohol dehydrogenase
Regulator

Raoultella terrigena (Klebsiella terrigena). Bacteria; Proteobacteria; Gammaproteobacteria; Enterobacterales;

EDF76__locus_2

GRM2

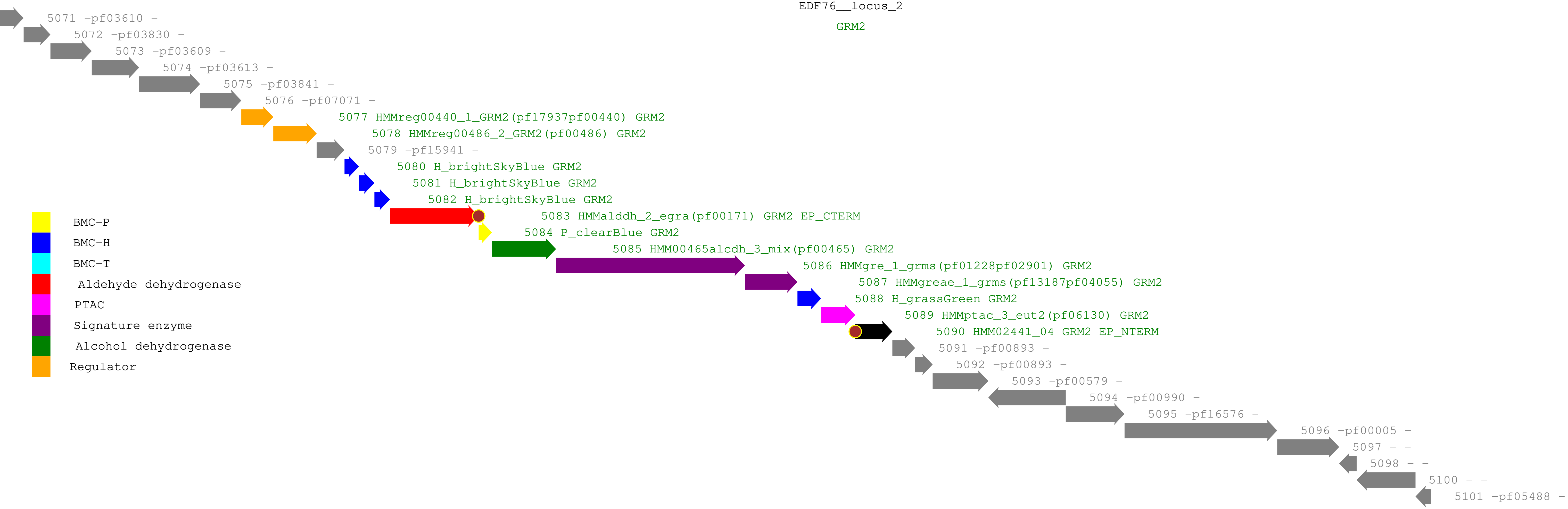

Raoultella sp. BIGb0399. Bacteria; Proteobacteria; Gammaproteobacteria; Enterobacterales;

EDF82__locus_2

GRM2

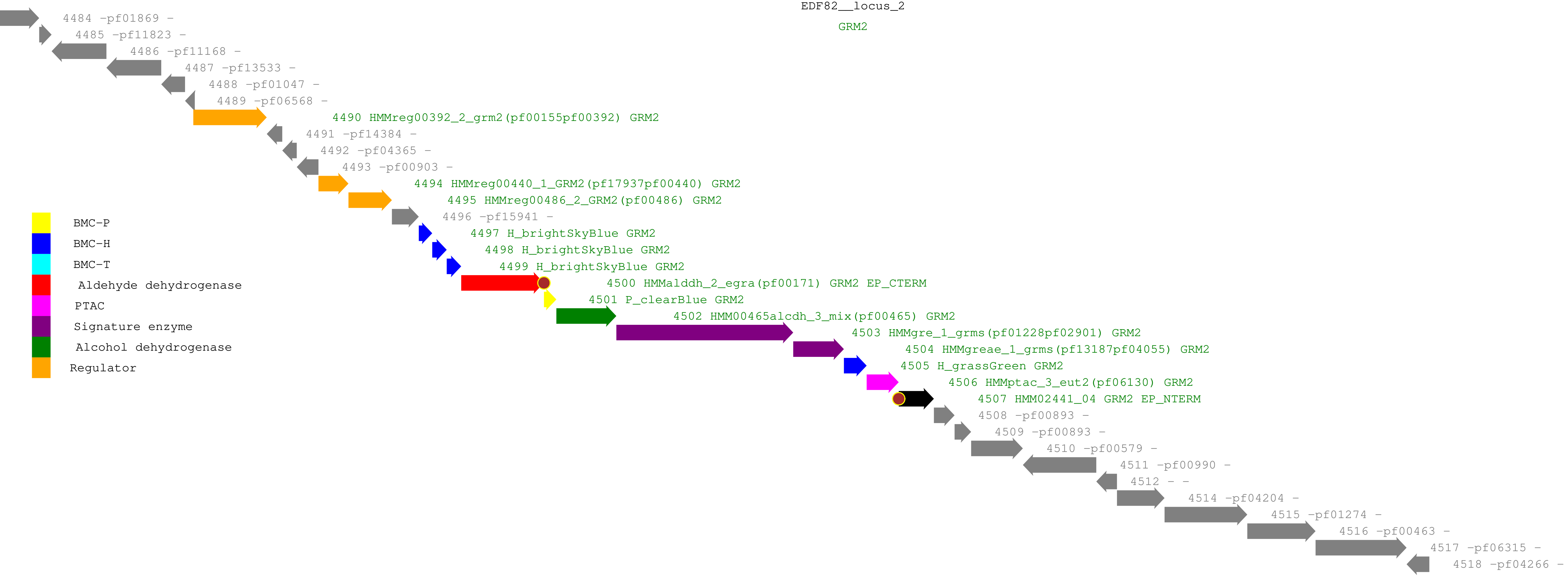

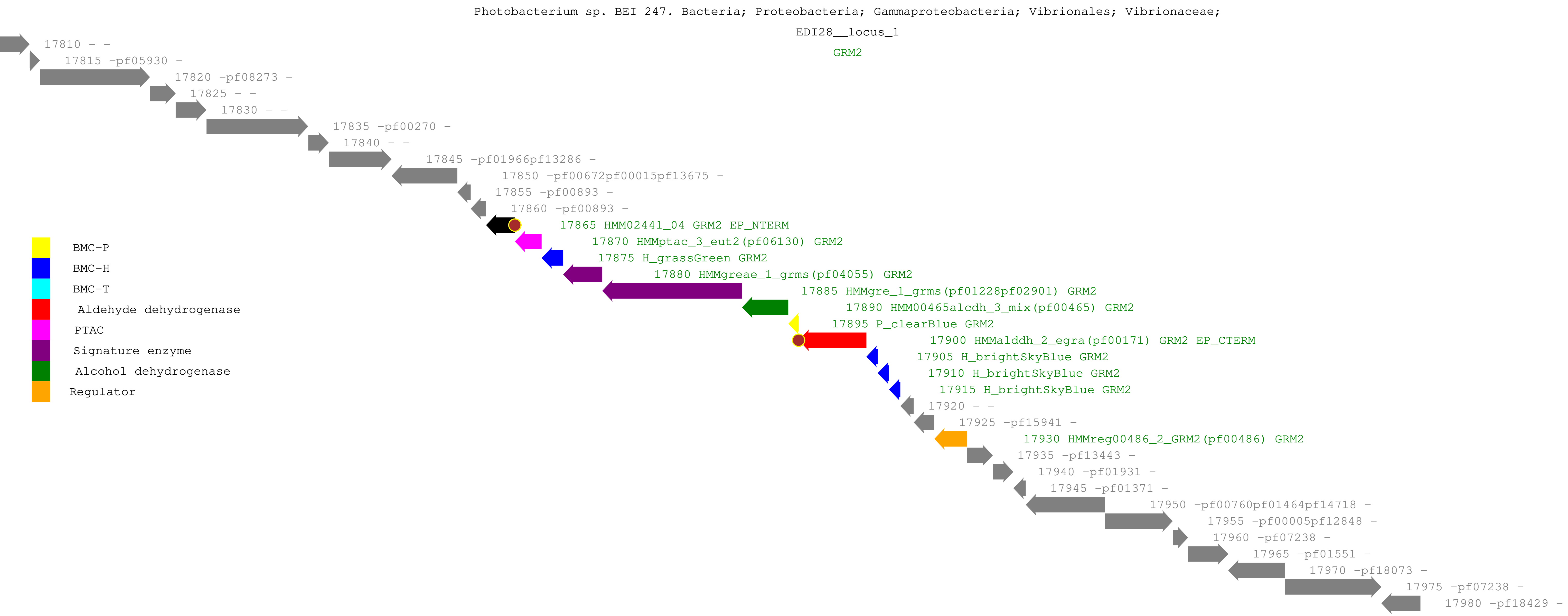

Photobacterium sp. BEI 247. Bacteria; Proteobacteria; Gammaproteobacteria; Vibrionales; Vibrionaceae;
EDI28__locus_1
GRM2
17810 - -
17815 -pf05930 -
17820 -pf08273 -
17825 - -
17830 - -
17835 -pf00270 -
17840 - -
17845 -pf01966pf13286 -
17850 -pf00672pf00015pf13675 -
17855 -pf00893 -
17860 -pf00893 -
17865 HMM02441_04 GRM2 EP_NTERM
17870 HMMptac_3_eut2(pf06130) GRM2
17875 H_grassGreen GRM2
17880 HMMgreae_1_grms(pf04055) GRM2
17885 HMMgre_1_grms(pf01228pf02901) GRM2
17890 HMM00465alcdh_3_mix(pf00465) GRM2
17895 P_clearBlue GRM2
17900 HMMalddh_2_egra(pf00171) GRM2 EP_CTERM
17905 H_brightSkyBlue GRM2
17910 H_brightSkyBlue GRM2
17915 H_brightSkyBlue GRM2
17920 - -
17925 -pf15941 -
17930 HMMreg00486_2_GRM2(pf00486) GRM2
17935 -pf13443 -
17940 -pf01931 -
17945 -pf01371 -
17950 -pf00760pf01464pf14718 -
17955 -pf00005pf12848 -
17960 -pf07238 -
17965 -pf01551 -
17970 -pf18073 -
17975 -pf07238 -
17980 -pf18429 -
BMC-P
BMC-H
BMC-T
Aldehyde dehydrogenase
PTAC
Signature enzyme
Alcohol dehydrogenase
Regulator

Providencia rettgeri. Bacteria; Proteobacteria; Gammaproteobacteria; Enterobacterales;

EGJ46__locus_1

GRM2

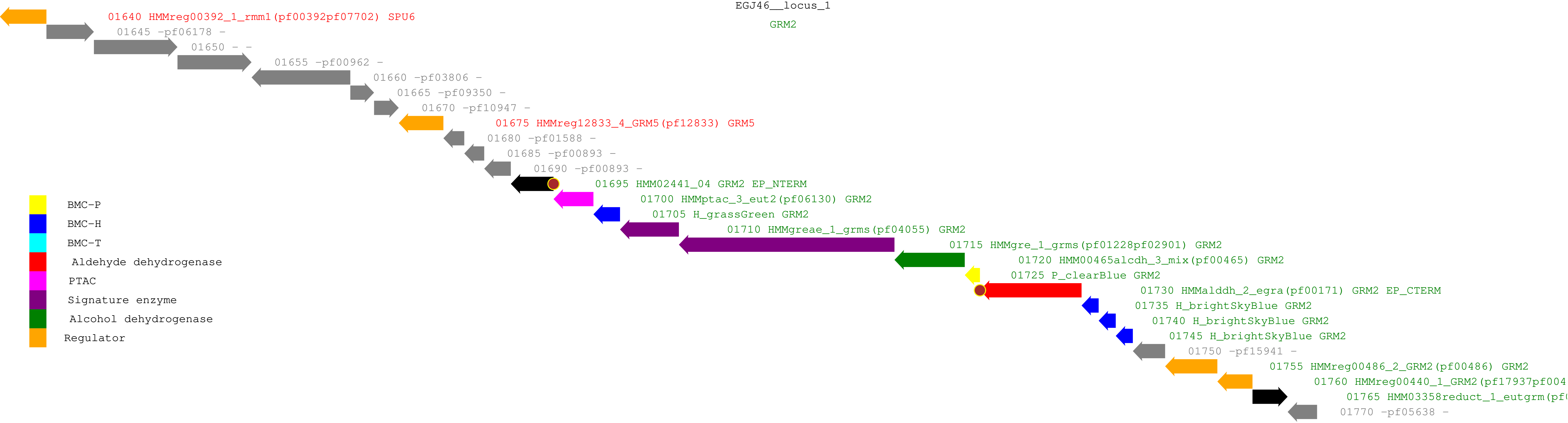

Pantoea dispersa. Bacteria; Proteobacteria; Gammaproteobacteria; Enterobacterales;

EKH82__locus_1

GRM2

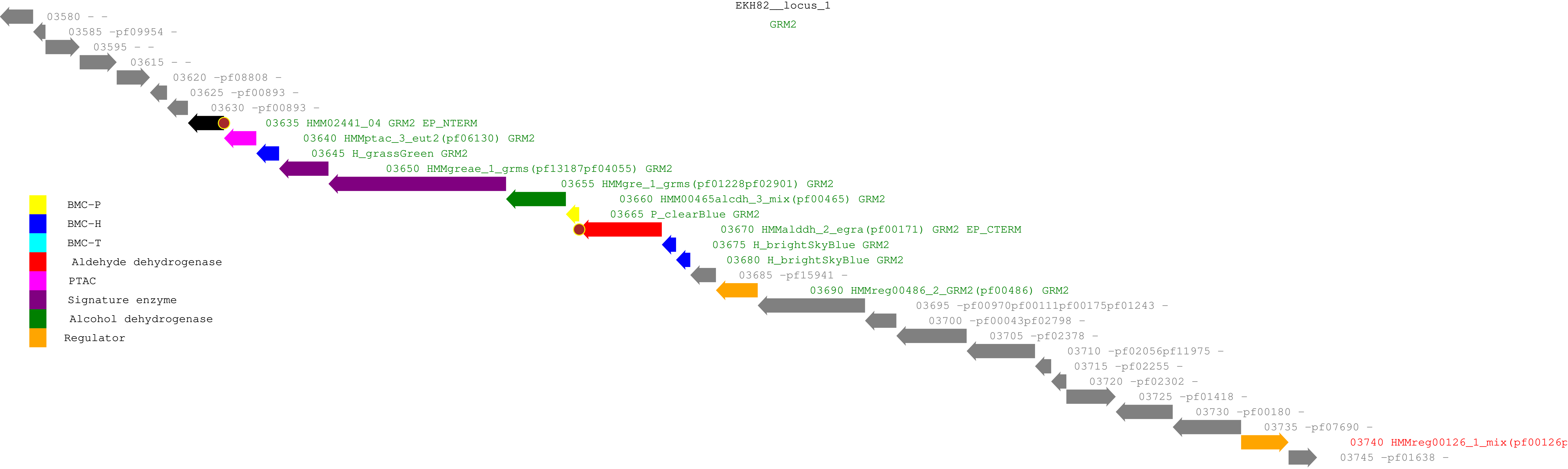

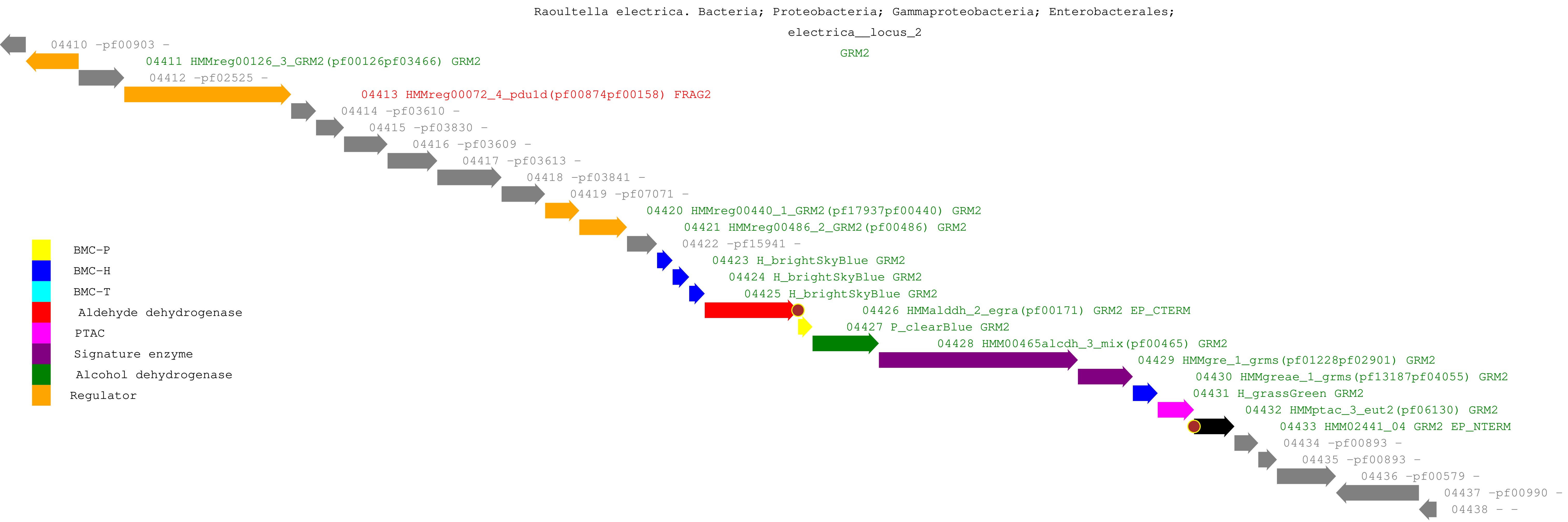

Raoultella electrica. Bacteria; Proteobacteria; Gammaproteobacteria; Enterobacterales;
electrica__locus_2
GRM2
04410 -pf00903 -
04411 HMMreg00126_3_GRM2(pf00126pf03466) GRM2
04412 -pf02525 -
04413 HMMreg00072_4_pdu1d(pf00874pf00158) FRAG2
04414 -pf03610 -
04415 -pf03830 -
04416 -pf03609 -
04417 -pf03613 -
04418 -pf03841 -
04419 -pf07071 -
04420 HMMreg00440_1_GRM2(pf17937pf00440) GRM2
04421 HMMreg00486_2_GRM2(pf00486) GRM2
04422 -pf15941 -
04423 H_brightSkyBlue GRM2
04424 H_brightSkyBlue GRM2
04425 H_brightSkyBlue GRM2
04426 HMMalddh_2_egra(pf00171) GRM2 EP_CTERM
04427 P_clearBlue GRM2
04428 HMM00465alcdh_3_mix(pf00465) GRM2
04429 HMMgre_1_grms(pf01228pf02901) GRM2
04430 HMMgreae_1_grms(pf13187pf04055) GRM2
04431 H_grassGreen GRM2
04432 HMMptac_3_eut2(pf06130) GRM2
04433 HMM02441_04 GRM2 EP_NTERM
04434 -pf00893 -
04435 -pf00893 -
04436 -pf00579 -
04437 -pf00990 -
04438 - -
BMC-P
BMC-H
BMC-T
Aldehyde dehydrogenase
PTAC
Signature enzyme
Alcohol dehydrogenase
Regulator

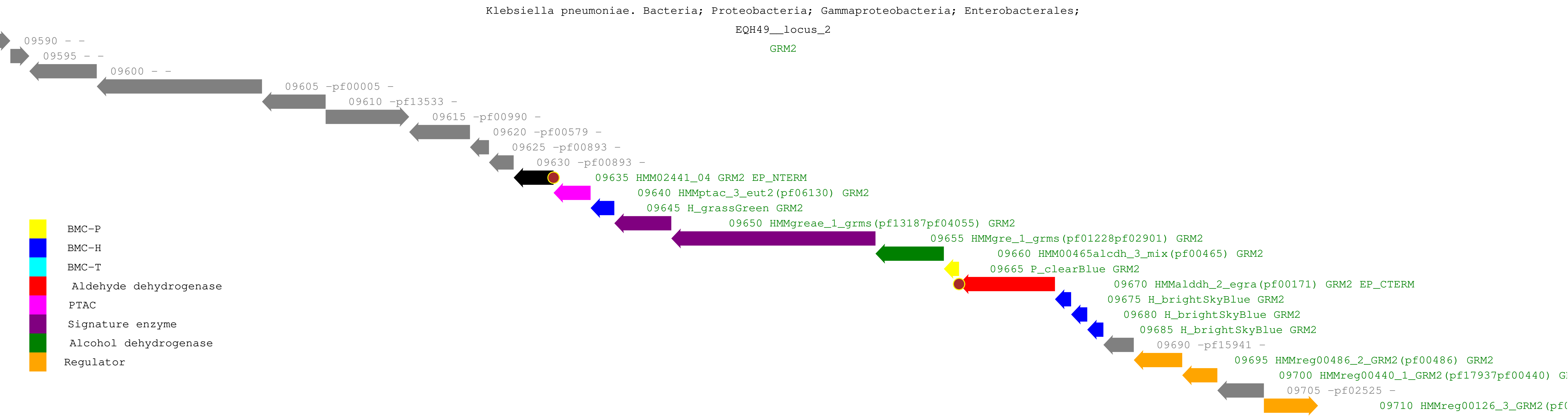
Klebsiella pneumoniae. Bacteria; Proteobacteria; Gammaproteobacteria; Enterobacterales;
EQH49__locus_2
GRM2
09590 - -
09595 - -
09600 - -
09605 -pf00005 -
09610 -pf13533 -
09615 -pf00990 -
09620 -pf00579 -
09625 -pf00893 -
09630 -pf00893 -
09635 HMM02441_04 GRM2 EP_NTERM
09640 HMMptac_3_eut2(pf06130) GRM2
09645 H_grassGreen GRM2
09650 HMMgreae_1_grms(pf13187pf04055) GRM2
09655 HMMgre_1_grms(pf01228pf02901) GRM2
09660 HMM00465alcdh_3_mix(pf00465) GRM2
09665 P_clearBlue GRM2
09670 HMMalddh_2_egra(pf00171) GRM2 EP_CTERM
09675 H_brightSkyBlue GRM2
09680 H_brightSkyBlue GRM2
09685 H_brightSkyBlue GRM2
09690 -pf15941 -
09695 HMMreg00486_2_GRM2(pf00486) GRM2
09700 HMMreg00440_1_GRM2(pf17937pf00440)
09705 -pf02525 -
09710 HMMreg00126_3_GRM2
BMC-P
BMC-H
BMC-T
Aldehyde dehydrogenase
PTAC
Signature enzyme
Alcohol dehydrogenase
Regulator

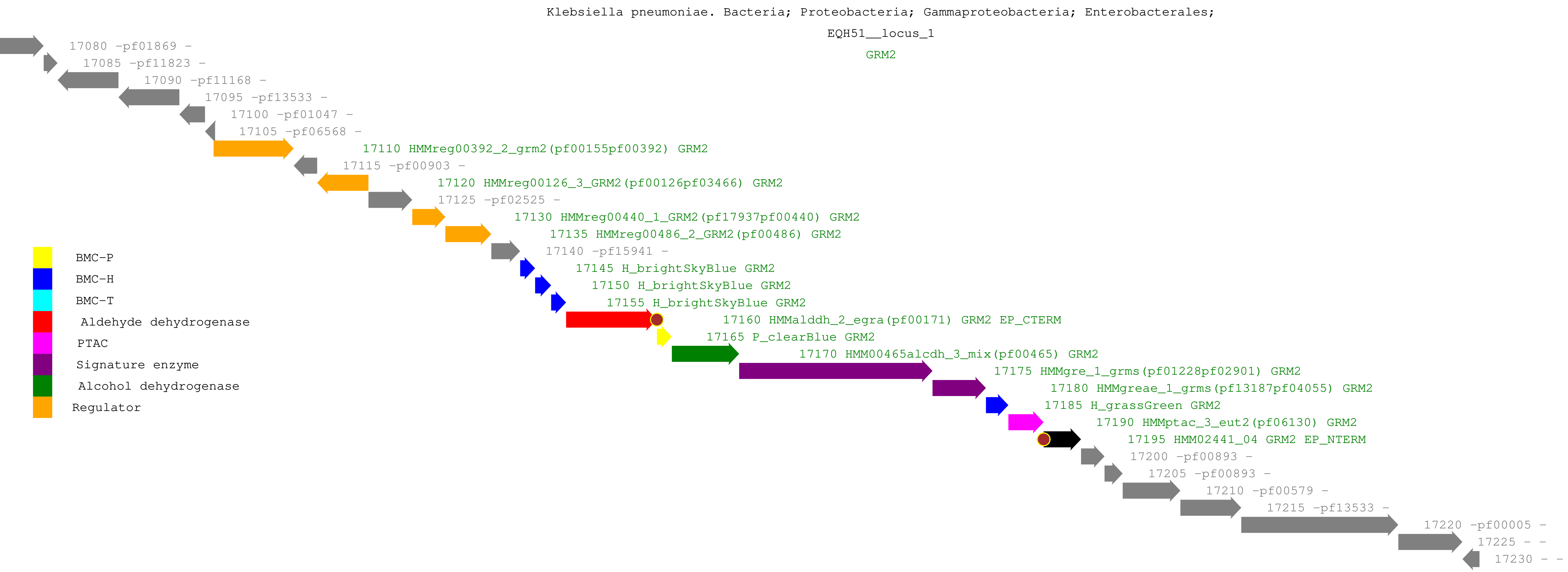
Klebsiella pneumoniae. Bacteria; Proteobacteria; Gammaproteobacteria; Enterobacterales;
EQH51__locus_1
GRM2
17080 -pf01869 -
17085 -pf11823 -
17090 -pf11168 -
17095 -pf13533 -
17100 -pf01047 -
17105 -pf06568 -
17110 HMMreg00392_2_grm2(pf00155pf00392) GRM2
17115 -pf00903 -
17120 HMMreg00126_3_GRM2(pf00126pf03466) GRM2
17125 -pf02525 -
17130 HMMreg00440_1_GRM2(pf17937pf00440) GRM2
17135 HMMreg00486_2_GRM2(pf00486) GRM2
17140 -pf15941 -
17145 H_brightSkyBlue GRM2
17150 H_brightSkyBlue GRM2
17155 H_brightSkyBlue GRM2
17160 HMMalddh_2_egra(pf00171) GRM2 EP_CTERM
17165 P_clearBlue GRM2
17170 HMM00465alcdh_3_mix(pf00465) GRM2
17175 HMMgre_1_grms(pf01228pf02901) GRM2
17180 HMMgreae_1_grms(pf13187pf04055) GRM2
17185 H_grassGreen GRM2
17190 HMMptac_3_eut2(pf06130) GRM2
17195 HMM02441_04 GRM2 EP_NTERM
17200 -pf00893 -
17205 -pf00893 -
17210 -pf00579 -
17215 -pf13533 -
17220 -pf00005 -
17225 - -
17230 - -
BMC-P
BMC-H
BMC-T
Aldehyde dehydrogenase
PTAC
Signature enzyme
Alcohol dehydrogenase
Regulator

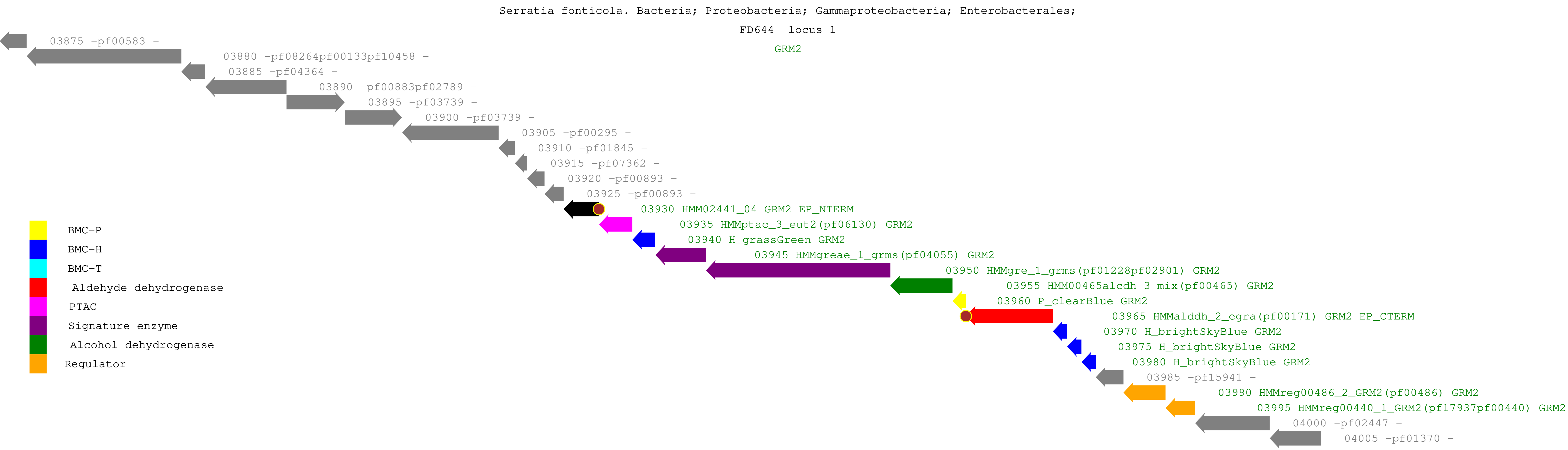
Serratia fonticola. Bacteria; Proteobacteria; Gammaproteobacteria; Enterobacterales;
FD644__locus_1
GRM2
03875 -pf00583 -
03880 -pf08264pf00133pf10458 -
03885 -pf04364 -
03890 -pf00883pf02789 -
03895 -pf03739 -
03900 -pf03739 -
03905 -pf00295 -
03910 -pf01845 -
03915 -pf07362 -
03920 -pf00893 -
03925 -pf00893 -
03930 HMM02441_04 GRM2 EP_NTERM
03935 HMMptac_3_eut2(pf06130) GRM2
03940 H_grassGreen GRM2
03945 HMMgreae_1_grms(pf04055) GRM2
03950 HMMgre_1_grms(pf01228pf02901) GRM2
03955 HMM00465alcdh_3_mix(pf00465) GRM2
03960 P_clearBlue GRM2
03965 HMMalddh_2_egra(pf00171) GRM2 EP_CTERM
03970 H_brightSkyBlue GRM2
03975 H_brightSkyBlue GRM2
03980 H_brightSkyBlue GRM2
03985 -pf15941 -
03990 HMMreg00486_2_GRM2(pf00486) GRM2
03995 HMMreg00440_1_GRM2(pf17937pf00440) GRM2
04000 -pf02447 -
04005 -pf01370 -
BMC-P
BMC-H
BMC-T
Aldehyde dehydrogenase
PTAC
Signature enzyme
Alcohol dehydrogenase
Regulator

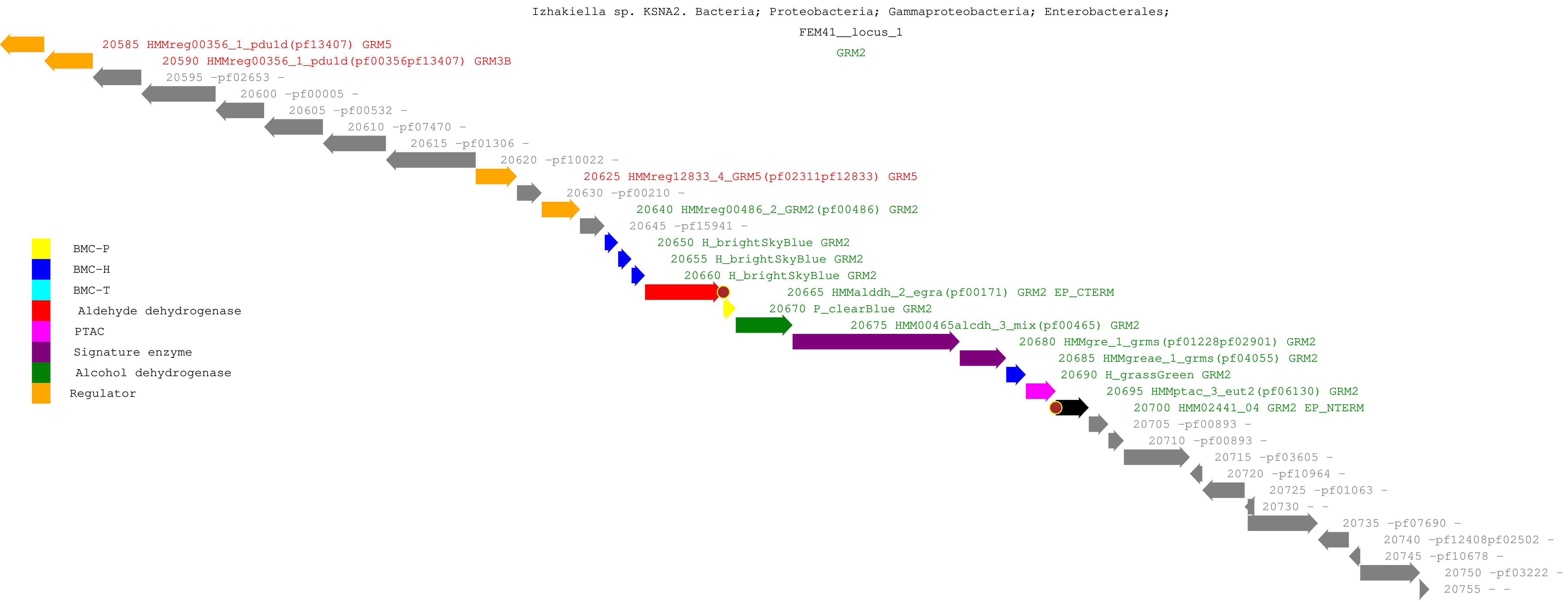
Izhakiella sp. KSNA2. Bacteria; Proteobacteria; Gammaproteobacteria; Enterobacterales;
FEM41__locus_1
GRM2
20585 HMMreg00356_1_pdu1d(pf13407) GRM5
20590 HMMreg00356_1_pdu1d(pf00356pf13407) GRM3B
20595 -pf02653 -
20600 -pf00005 -
20605 -pf00532 -
20610 -pf07470 -
20615 -pf01306 -
20620 -pf10022 -
20625 HMMreg12833_4_GRM5(pf02311pf12833) GRM5
20630 -pf00210 -
20640 HMMreg00486_2_GRM2(pf00486) GRM2
20645 -pf15941 -
20650 H_brightSkyBlue GRM2
20655 H_brightSkyBlue GRM2
20660 H_brightSkyBlue GRM2
20665 HMMaldh_2_egra(pf00171) GRM2 EP_CTERM
20670 P_clearBlue GRM2
20675 HMM00465alcdh_3_mix(pf00465) GRM2
20680 HMMgre_1_grms(pf01228pf02901) GRM2
20685 HMMgreae_1_grms(pf04055) GRM2
20690 H_grassGreen GRM2
20695 HMMptac_3_eut2(pf06130) GRM2
20700 HMM02441_04 GRM2 EP_NTERM
20705 -pf00893 -
20710 -pf00893 -
20715 -pf03605 -
20720 -pf10964 -
20725 -pf01063 -
20730 - -
20735 -pf07690 -
20740 -pf12408pf02502 -
20745 -pf10678 -
20750 -pf03222 -
20755 - -
BMC-P
BMC-H
BMC-T
Aldehyde dehydrogenase
PTAC
Signature enzyme
Alcohol dehydrogenase
Regulator

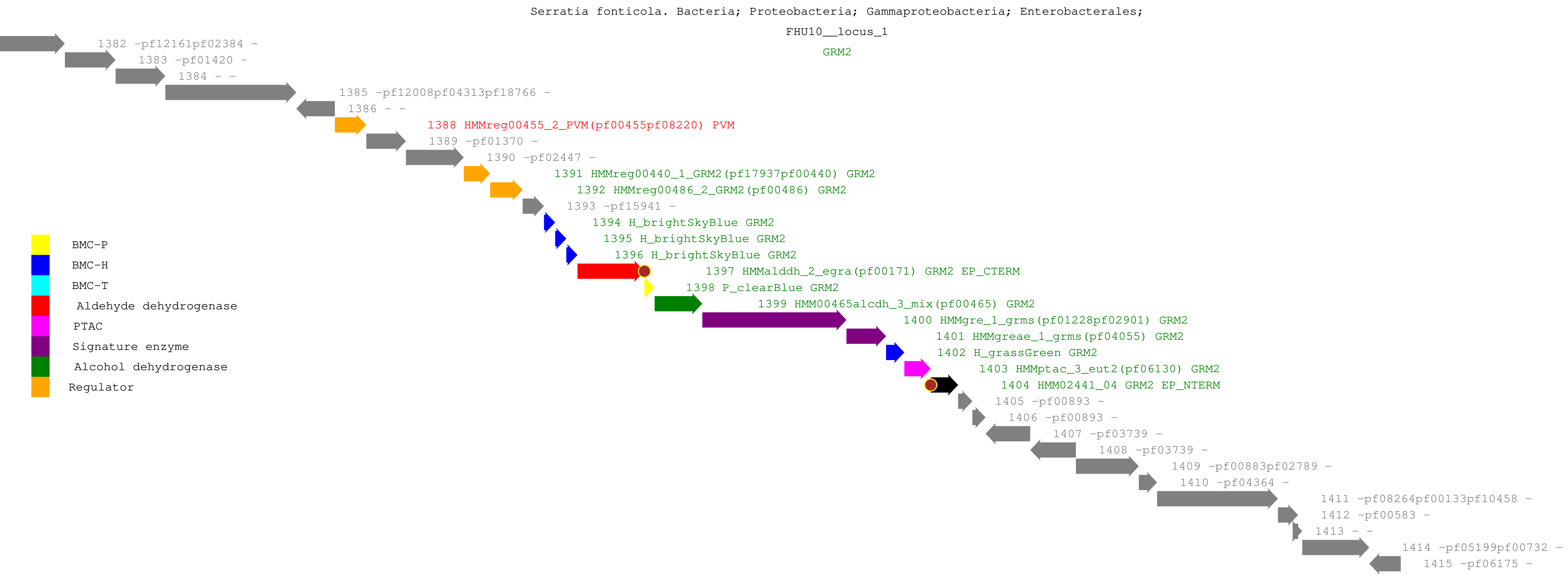

Serratia fonticola. Bacteria; Proteobacteria; Gammaproteobacteria; Enterobacterales;
FHU10__locus_1
GRM2
1382 -pf12161pf02384 -
1383 -pf01420 -
1384 - -
1385 -pf12008pf04313pf18766 -
1386 - -
1388 HMMreg00455_2_PVM(pf00455pf08220) PVM
1389 -pf01370 -
1390 -pf02447 -
1391 HMMreg00440_1_GRM2(pf17937pf00440) GRM2
1392 HMMreg00486_2_GRM2(pf00486) GRM2
1393 -pf15941 -
1394 H_brightSkyBlue GRM2
1395 H_brightSkyBlue GRM2
1396 H_brightSkyBlue GRM2
1397 HMMaldh_2_egra(pf00171) GRM2 EP_CTERM
1398 P_clearBlue GRM2
1399 HMM00465alcdh_3_mix(pf00465) GRM2
1400 HMMgre_1_grms(pf01228pf02901) GRM2
1401 HMMgreae_1_grms(pf04055) GRM2
1402 H_grassGreen GRM2
1403 HMMptac_3_eut2(pf06130) GRM2
1404 HMM02441_04 GRM2 EP_NTERM
1405 -pf00893 -
1406 -pf00893 -
1407 -pf03739 -
1408 -pf03739 -
1409 -pf00883pf02789 -
1410 -pf04364 -
1411 -pf08264pf00133pf10458 -
1412 -pf00583 -
1413 - -
1414 -pf05199pf00732 -
1415 -pf06175 -
BMC-P
BMC-H
BMC-T
Aldehyde dehydrogenase
PTAC
Signature enzyme
Alcohol dehydrogenase
Regulator

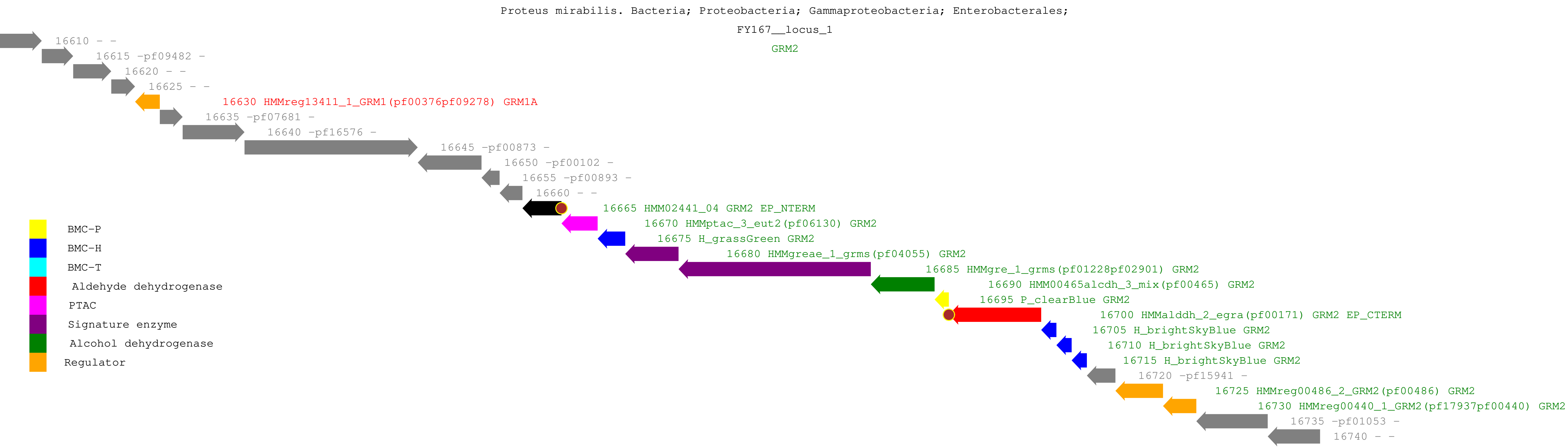
Proteus mirabilis. Bacteria; Proteobacteria; Gammaproteobacteria; Enterobacterales;
FY167__locus_1
GRM2
16610 - -
16615 -pf09482 -
16620 - -
16625 - -
16630 HMMreg13411_1_GRM1(pf00376pf09278) GRM1A
16635 -pf07681 -
16640 -pf16576 -
16645 -pf00873 -
16650 -pf00102 -
16655 -pf00893 -
16660 - -
16665 HMM02441_04 GRM2 EP_NTERM
16670 HMMptac_3_eut2(pf06130) GRM2
16675 H_grassGreen GRM2
16680 HMMgreae_1_grms(pf04055) GRM2
16685 HMMgre_1_grms(pf01228pf02901) GRM2
16690 HMM00465alcdh_3_mix(pf00465) GRM2
16695 P_clearBlue GRM2
16700 HMMalddh_2_egra(pf00171) GRM2 EP_CTERM
16705 H_brightSkyBlue GRM2
16710 H_brightSkyBlue GRM2
16715 H_brightSkyBlue GRM2
16720 -pf15941 -
16725 HMMreg00486_2_GRM2(pf00486) GRM2
16730 HMMreg00440_1_GRM2(pf17937pf00440) GRM2
16735 -pf01053 -
16740 - -
BMC-P
BMC-H
BMC-T
Aldehyde dehydrogenase
PTAC
Signature enzyme
Alcohol dehydrogenase
Regulator

Escherichia coli HVH 70 (4-2963531). Bacteria; Proteobacteria; Gammaproteobacteria; Enterobacterales;

G736__locus_3

GRM2

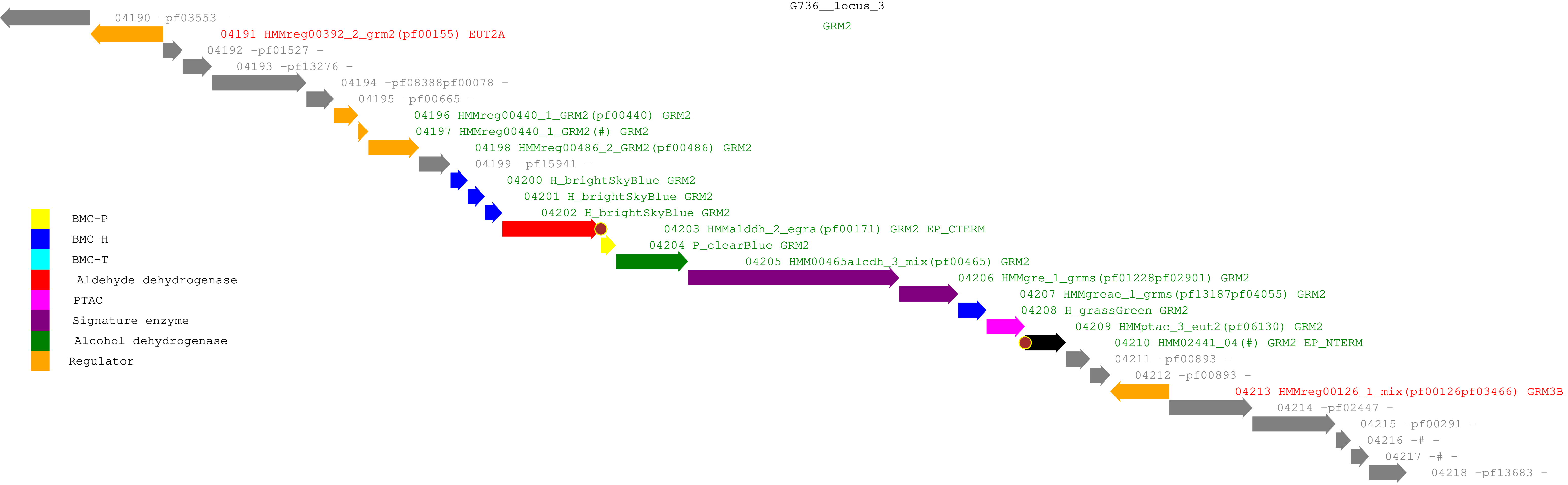

Escherichia coli UMEA 3212-1. Bacteria; Proteobacteria; Gammaproteobacteria; Enterobacterales;

G943__locus_2

GRM2

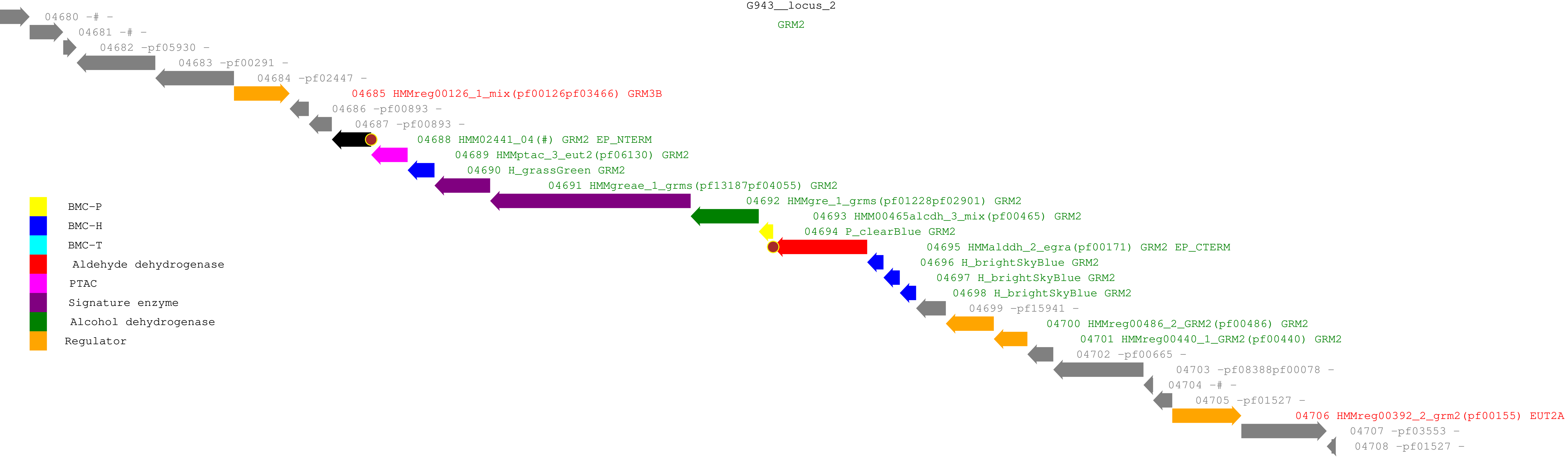

Kluyvera ascorbata ATCC 33433. Bacteria; Proteobacteria; Gammaproteobacteria; Enterobacterales;

GKAS__locus_1

GRM2

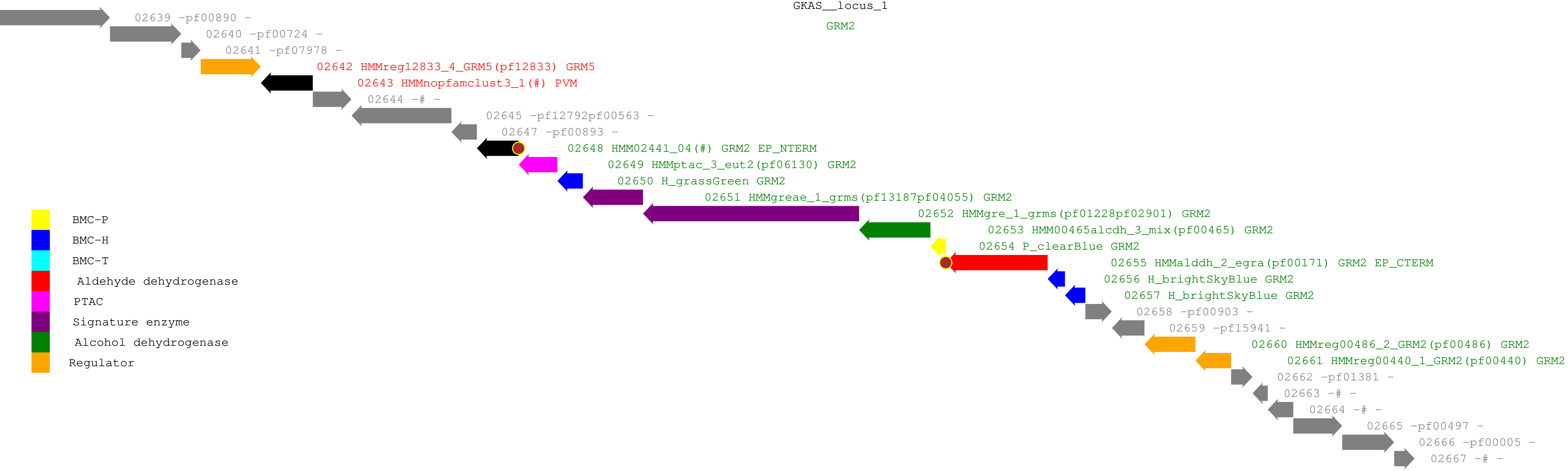

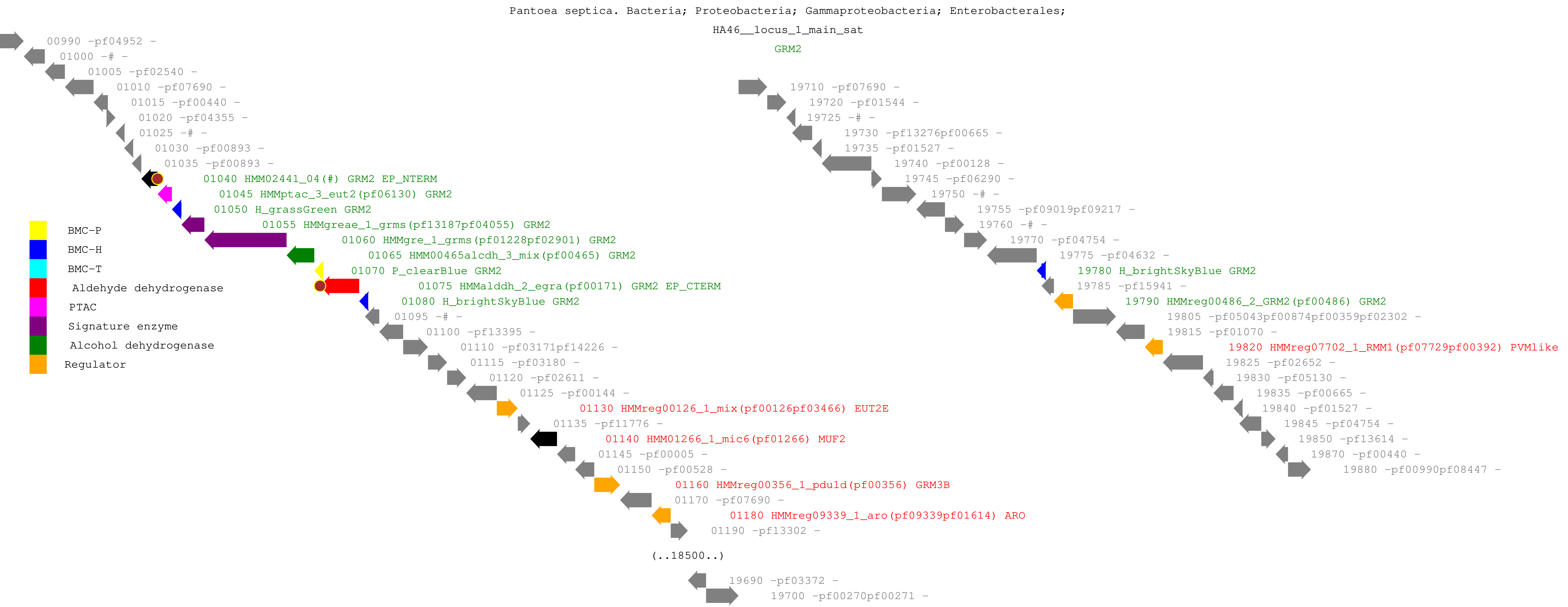
Pantoea septica. Bacteria; Proteobacteria; Gammaproteobacteria; Enterobacterales;
HA46__locus_1_main_sat
GRM2
00990 -pf04952 -
01000 -# -
01005 -pf02540 -
01010 -pf07690 -
01015 -pf00440 -
01020 -pf04355 -
01025 -# -
01030 -pf00893 -
01035 -pf00893 -
01040 HMM02441_04(#) GRM2 EP_NTERM
01045 HMMptac_3_eut2(pf06130) GRM2
01050 H_grassGreen GRM2
01055 HMMgreae_1_grms(pf13187pf04055) GRM2
01060 HMMgre_1_grms(pf01228pf02901) GRM2
01065 HMM00465alcdh_3_mix(pf00465) GRM2
01070 P_clearBlue GRM2
01075 HMMalddh_2_egra(pf00171) GRM2 EP_CTERM
01080 H_brightSkyBlue GRM2
01095 -# -
01100 -pf13395 -
01110 -pf03171pf14226 -
01115 -pf03180 -
01120 -pf02611 -
01125 -pf00144 -
01130 HMMreg00126_1_mix(pf00126pf03466) EUT2E
01135 -pf11776 -
01140 HMM01266_1_mic6(pf01266) MUF2
01145 -pf00005 -
01150 -pf00528 -
01160 HMMreg00356_1_pdu1d(pf00356) GRM3B
01170 -pf07690 -
01180 HMMreg09339_1_aro(pf09339pf01614) ARO
01190 -pf13302 -
(..18500..)
19690 -pf03372 -
19700 -pf00270pf00271 -
19710 -pf07690 -
19720 -pf01544 -
19725 -# -
19730 -pf13276pf00665 -
19735 -pf01527 -
19740 -pf00128 -
19745 -pf06290 -
19750 -# -
19755 -pf09019pf09217 -
19760 -# -
19770 -pf04754 -
19775 -pf04632 -
19780 H_brightSkyBlue GRM2
19785 -pf15941 -
19790 HMMreg00486_2_GRM2(pf00486) GRM2
19805 -pf05043pf00874pf00359pf02302 -
19815 -pf01070 -
19820 HMMreg07702_1_RMM1(pf07729pf00392) PVMlike
19825 -pf02652 -
19830 -pf05130 -
19835 -pf00665 -
19840 -pf01527 -
19845 -pf04754 -
19850 -pf13614 -
19870 -pf00440 -
19880 -pf00990pf08447 -
BMC-P
BMC-H
BMC-T
Aldehyde dehydrogenase
PTAC
Signature enzyme
Alcohol dehydrogenase
Regulator

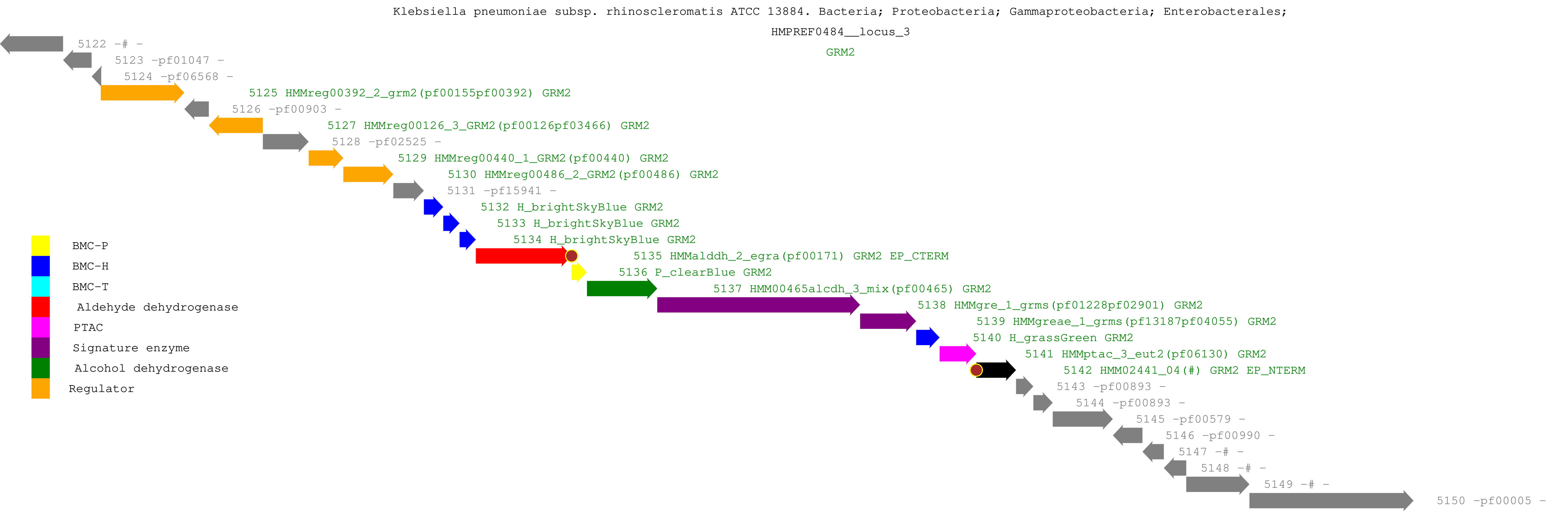
Klebsiella pneumoniae subsp. rhinoscleromatis ATCC 13884. Bacteria; Proteobacteria; Gammaproteobacteria; Enterobacterales;
HMPREF0484__locus_3
GRM2
5122 -# -
5123 -pf01047 -
5124 -pf06568 -
5125 HMMreg00392_2_grm2(pf00155pf00392) GRM2
5126 -pf00903 -
5127 HMMreg00126_3_GRM2(pf00126pf03466) GRM2
5128 -pf02525 -
5129 HMMreg00440_1_GRM2(pf00440) GRM2
5130 HMMreg00486_2_GRM2(pf00486) GRM2
5131 -pf15941 -
5132 H_brightSkyBlue GRM2
5133 H_brightSkyBlue GRM2
5134 H_brightSkyBlue GRM2
5135 HMMalddh_2_egra(pf00171) GRM2 EP_CTERM
5136 P_clearBlue GRM2
5137 HMM00465alcdh_3_mix(pf00465) GRM2
5138 HMMgre_1_grms(pf01228pf02901) GRM2
5139 HMMgreae_1_grms(pf13187pf04055) GRM2
5140 H_grassGreen GRM2
5141 HMMptac_3_eut2(pf06130) GRM2
5142 HMM02441_04(#) GRM2 EP_NTERM
5143 -pf00893 -
5144 -pf00893 -
5145 -pf00579 -
5146 -pf00990 -
5147 -# -
5148 -# -
5149 -# -
5150 -pf00005 -
BMC-P
BMC-H
BMC-T
Aldehyde dehydrogenase
PTAC
Signature enzyme
Alcohol dehydrogenase
Regulator

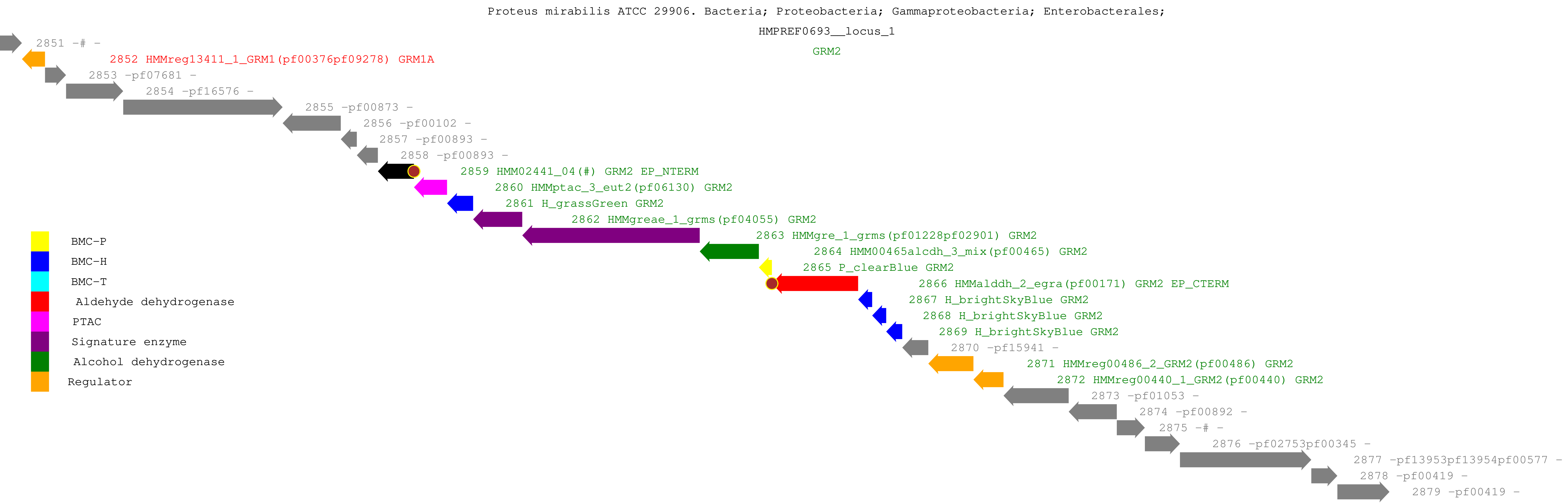
Proteus mirabilis ATCC 29906. Bacteria; Proteobacteria; Gammaproteobacteria; Enterobacterales;
HMPREF0693__locus_1
GRM2
2851 -# -
2852 HMMreg13411_1_GRM1(pf00376pf09278) GRM1A
2853 -pf07681 -
2854 -pf16576 -
2855 -pf00873 -
2856 -pf00102 -
2857 -pf00893 -
2858 -pf00893 -
2859 HMM02441_04(#) GRM2 EP_NTERM
2860 HMMptac_3_eut2(pf06130) GRM2
2861 H_grassGreen GRM2
2862 HMMgreae_1_grms(pf04055) GRM2
2863 HMMgre_1_grms(pf01228pf02901) GRM2
2864 HMM00465alcdh_3_mix(pf00465) GRM2
2865 P_clearBlue GRM2
2866 HMMalddh_2_egra(pf00171) GRM2 EP_CTERM
2867 H_brightSkyBlue GRM2
2868 H_brightSkyBlue GRM2
2869 H_brightSkyBlue GRM2
2870 -pf15941 -
2871 HMMreg00486_2_GRM2(pf00486) GRM2
2872 HMMreg00440_1_GRM2(pf00440) GRM2
2873 -pf01053 -
2874 -pf00892 -
2875 -# -
2876 -pf02753pf00345 -
2877 -pf13953pf13954pf00577 -
2878 -pf00419 -
2879 -pf00419 -
BMC-P
BMC-H
BMC-T
Aldehyde dehydrogenase
PTAC
Signature enzyme
Alcohol dehydrogenase
Regulator

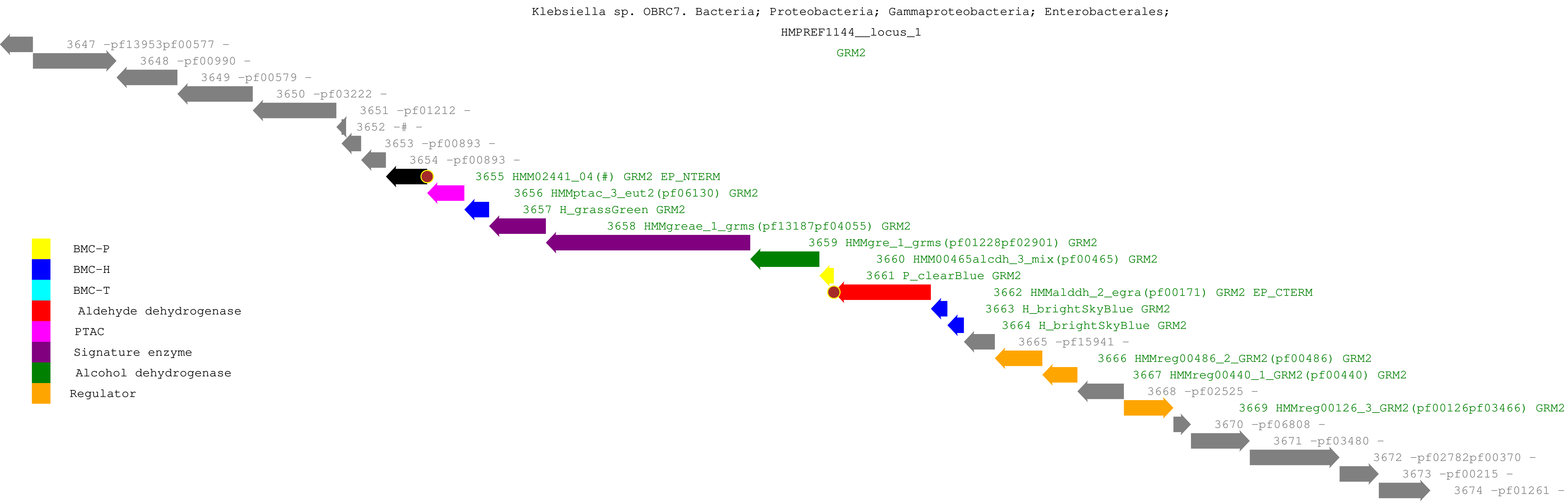

Klebsiella sp. OBRC7. Bacteria; Proteobacteria; Gammaproteobacteria; Enterobacterales;
HMPREF1144__locus_1
GRM2
3647 -pf13953pf00577 -
3648 -pf00990 -
3649 -pf00579 -
3650 -pf03222 -
3651 -pf01212 -
3652 -# -
3653 -pf00893 -
3654 -pf00893 -
3655 HMM02441_04(#) GRM2 EP_NTERM
3656 HMMptac_3_eut2(pf06130) GRM2
3657 H_grassGreen GRM2
3658 HMMgreae_1_grms(pf13187pf04055) GRM2
3659 HMMgre_1_grms(pf01228pf02901) GRM2
3660 HMM00465alcdh_3_mix(pf00465) GRM2
3661 P_clearBlue GRM2
3662 HMMaldh_2_egra(pf00171) GRM2 EP_CTERM
3663 H_brightSkyBlue GRM2
3664 H_brightSkyBlue GRM2
3665 -pf15941 -
3666 HMMreg00486_2_GRM2(pf00486) GRM2
3667 HMMreg00440_1_GRM2(pf00440) GRM2
3668 -pf02525 -
3669 HMMreg00126_3_GRM2(pf00126pf03466) GRM2
3670 -pf06808 -
3671 -pf03480 -
3672 -pf02782pf00370 -
3673 -pf00215 -
3674 -pf01261 -
BMC-P
BMC-H
BMC-T
Aldehyde dehydrogenase
PTAC
Signature enzyme
Alcohol dehydrogenase
Regulator

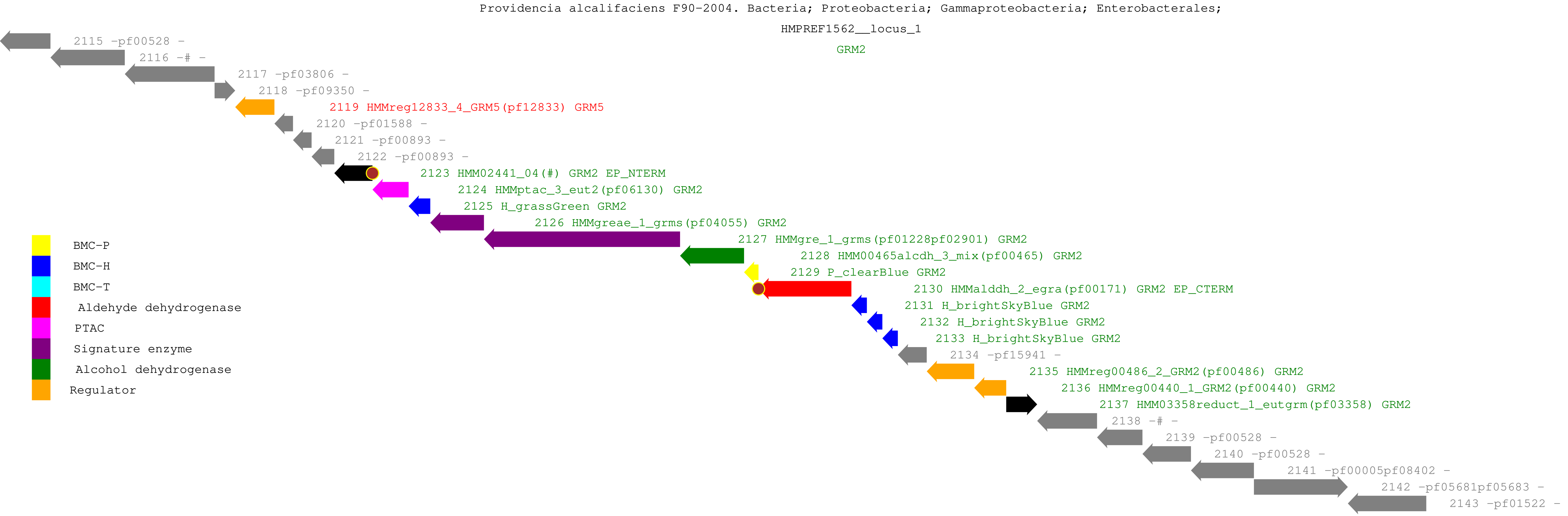
Providencia alcalifaciens F90-2004. Bacteria; Proteobacteria; Gammaproteobacteria; Enterobacterales;
HMPREF1562__locus_1
GRM2
2115 -pf00528 -
2116 -# -
2117 -pf03806 -
2118 -pf09350 -
2119 HMMreg12833_4_GRM5(pf12833) GRM5
2120 -pf01588 -
2121 -pf00893 -
2122 -pf00893 -
2123 HMM02441_04(#) GRM2 EP_NTERM
2124 HMMptac_3_eut2(pf06130) GRM2
2125 H_grassGreen GRM2
2126 HMMgreae_1_grms(pf04055) GRM2
2127 HMMgre_1_grms(pf01228pf02901) GRM2
2128 HMM00465alcdh_3_mix(pf00465) GRM2
2129 P_clearBlue GRM2
2130 HMMalddh_2_egra(pf00171) GRM2 EP_CTERM
2131 H_brightSkyBlue GRM2
2132 H_brightSkyBlue GRM2
2133 H_brightSkyBlue GRM2
2134 -pf15941 -
2135 HMMreg00486_2_GRM2(pf00486) GRM2
2136 HMMreg00440_1_GRM2(pf00440) GRM2
2137 HMM03358reduct_1_eutgrm(pf03358) GRM2
2138 -# -
2139 -pf00528 -
2140 -pf00528 -
2141 -pf00005pf08402 -
2142 -pf05681pf05683 -
2143 -pf01522 -
BMC-P
BMC-H
BMC-T
Aldehyde dehydrogenase
PTAC
Signature enzyme
Alcohol dehydrogenase
Regulator

Providencia alcalifaciens R90-1475. Bacteria; Proteobacteria; Gammaproteobacteria; Enterobacterales;

HMPREF1564__locus_1

GRM2

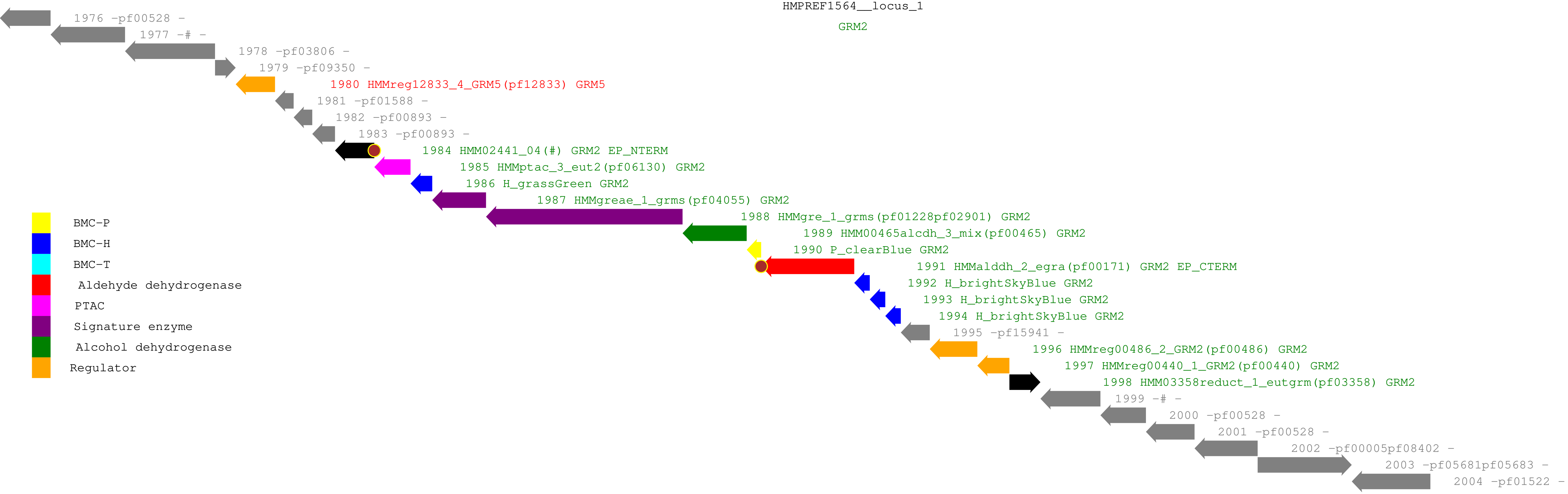

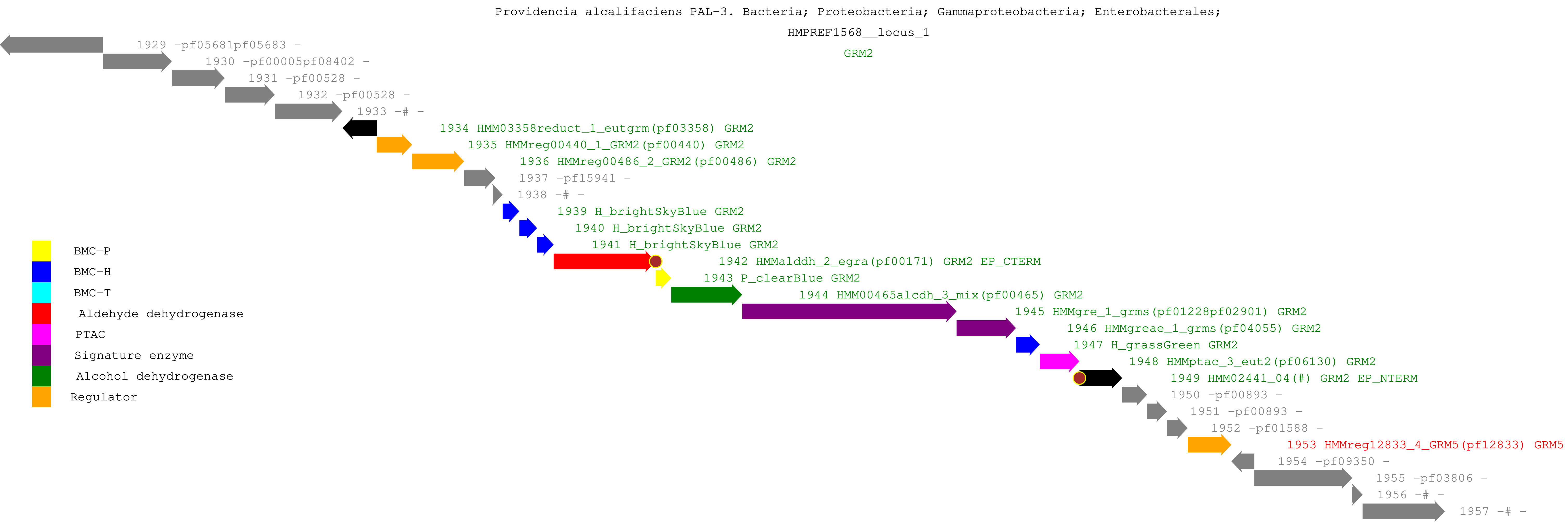
Providencia alcalifaciens PAL-3. Bacteria; Proteobacteria; Gammaproteobacteria; Enterobacterales;
HMPREF1568__locus_1
GRM2
1929 -pf05681pf05683 -
1930 -pf00005pf08402 -
1931 -pf00528 -
1932 -pf00528 -
1933 -# -
1934 HMM03358reduct_1_eutgrm(pf03358) GRM2
1935 HMMreg00440_1_GRM2(pf00440) GRM2
1936 HMMreg00486_2_GRM2(pf00486) GRM2
1937 -pf15941 -
1938 -# -
1939 H_brightSkyBlue GRM2
1940 H_brightSkyBlue GRM2
1941 H_brightSkyBlue GRM2
1942 HMMalddh_2_egra(pf00171) GRM2 EP_CTERM
1943 P_clearBlue GRM2
1944 HMM00465alcdh_3_mix(pf00465) GRM2
1945 HMMgre_1_grms(pf01228pf02901) GRM2
1946 HMMgreae_1_grms(pf04055) GRM2
1947 H_grassGreen GRM2
1948 HMMptac_3_eut2(pf06130) GRM2
1949 HMM02441_04(#) GRM2 EP_NTERM
1950 -pf00893 -
1951 -pf00893 -
1952 -pf01588 -
1953 HMMreg12833_4_GRM5(pf12833) GRM5
1954 -pf09350 -
1955 -pf03806 -
1956 -# -
1957 -# -
BMC-P
BMC-H
BMC-T
Aldehyde dehydrogenase
PTAC
Signature enzyme
Alcohol dehydrogenase
Regulator

Enterobacter sp. HMSC055A11. Bacteria; Proteobacteria; Gammaproteobacteria; Enterobacterales;

HMPREF2540__locus_1

GRM2

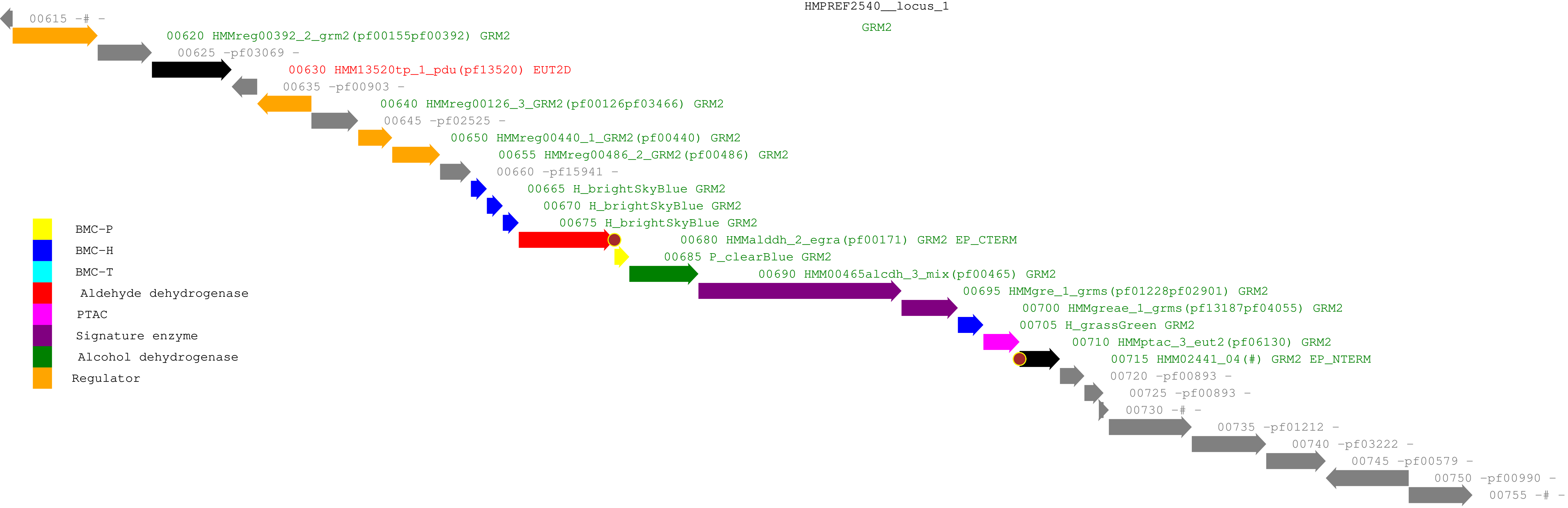

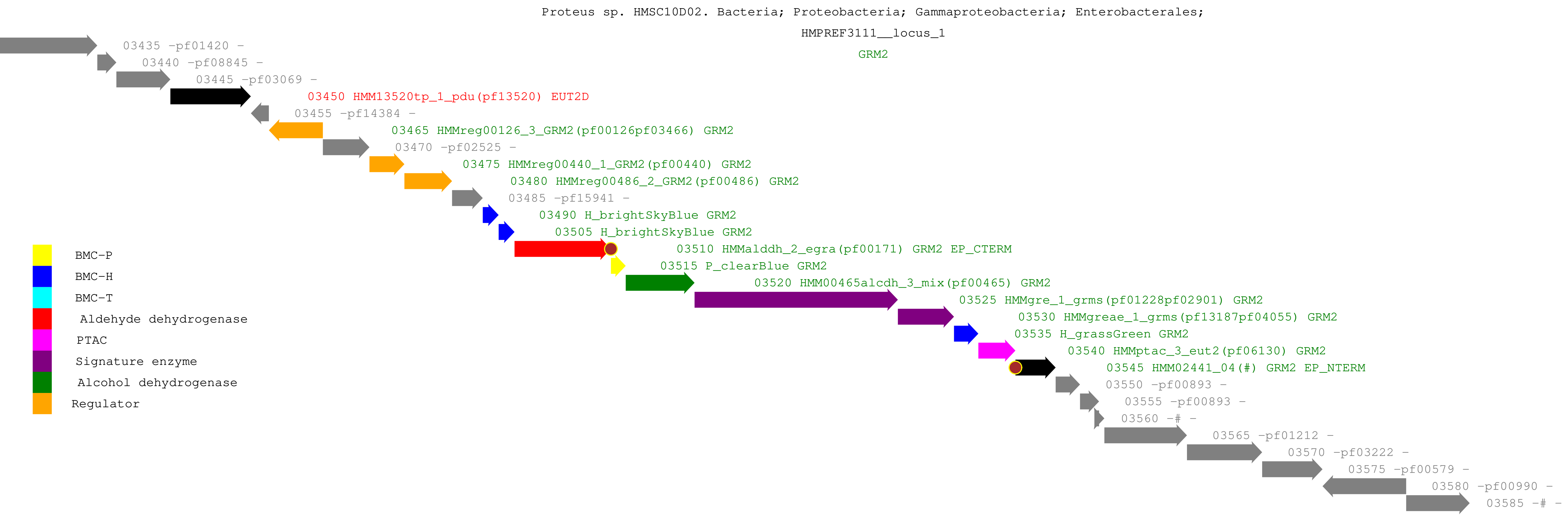
Proteus sp. HMSC10D02. Bacteria; Proteobacteria; Gammaproteobacteria; Enterobacterales;
HMPREF3111__locus_1
GRM2
03435 -pf01420 -
03440 -pf08845 -
03445 -pf03069 -
03450 HMM13520tp_1_pdu(pf13520) EUT2D
03455 -pf14384 -
03465 HMMreg00126_3_GRM2(pf00126pf03466) GRM2
03470 -pf02525 -
03475 HMMreg00440_1_GRM2(pf00440) GRM2
03480 HMMreg00486_2_GRM2(pf00486) GRM2
03485 -pf15941 -
03490 H_brightSkyBlue GRM2
03505 H_brightSkyBlue GRM2
03510 HMMaldh_2_egra(pf00171) GRM2 EP_CTERM
03515 P_clearBlue GRM2
03520 HMM00465alcdh_3_mix(pf00465) GRM2
03525 HMMgre_1_grms(pf01228pf02901) GRM2
03530 HMMgreae_1_grms(pf13187pf04055) GRM2
03535 H_grassGreen GRM2
03540 HMMptac_3_eut2(pf06130) GRM2
03545 HMM02441_04(#) GRM2 EP_NTERM
03550 -pf00893 -
03555 -pf00893 -
03560 -# -
03565 -pf01212 -
03570 -pf03222 -
03575 -pf00579 -
03580 -pf00990 -
03585 -# -
BMC-P
BMC-H
BMC-T
Aldehyde dehydrogenase
PTAC
Signature enzyme
Alcohol dehydrogenase
Regulator

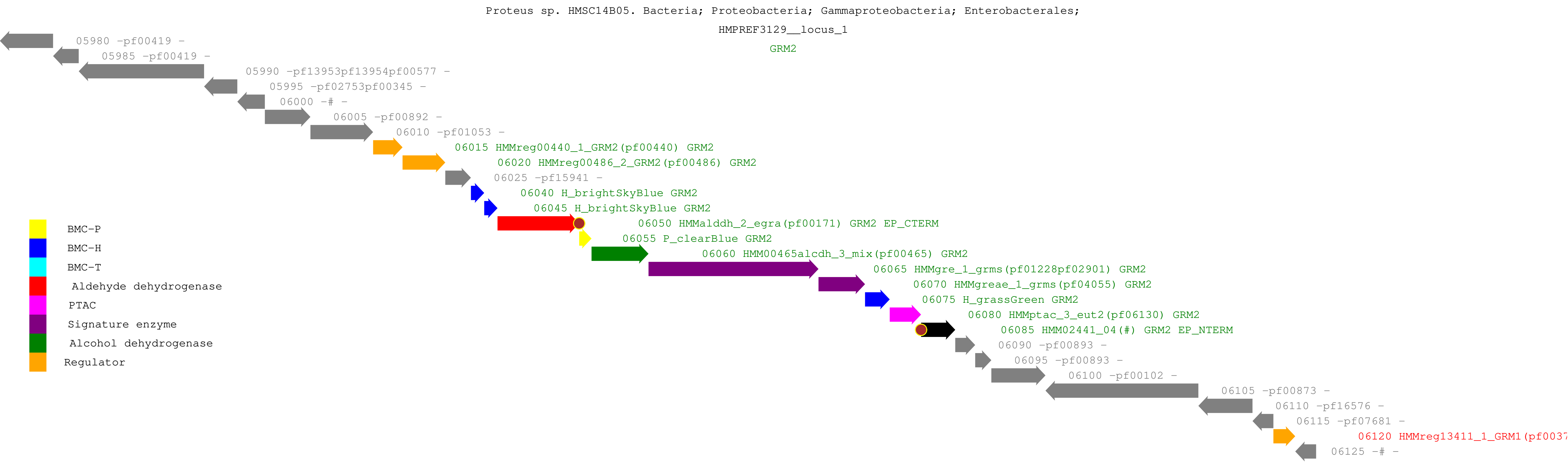
Proteus sp. HMSC14B05. Bacteria; Proteobacteria; Gammaproteobacteria; Enterobacterales;
HMPREF3129__locus_1
GRM2
05980 -pf00419 -
05985 -pf00419 -
05990 -pf13953pf13954pf00577 -
05995 -pf02753pf00345 -
06000 -# -
06005 -pf00892 -
06010 -pf01053 -
06015 HMMreg00440_1_GRM2(pf00440) GRM2
06020 HMMreg00486_2_GRM2(pf00486) GRM2
06025 -pf15941 -
06040 H_brightSkyBlue GRM2
06045 H_brightSkyBlue GRM2
06050 HMMalddh_2_egra(pf00171) GRM2 EP_CTERM
06055 P_clearBlue GRM2
06060 HMM00465alcdh_3_mix(pf00465) GRM2
06065 HMMgre_1_grms(pf01228pf02901) GRM2
06070 HMMgreae_1_grms(pf04055) GRM2
06075 H_grassGreen GRM2
06080 HMMptac_3_eut2(pf06130) GRM2
06085 HMM02441_04(#) GRM2 EP_NTERM
06090 -pf00893 -
06095 -pf00893 -
06100 -pf00102 -
06105 -pf00873 -
06110 -pf16576 -
06115 -pf07681 -
06120 HMMreg13411_1_GRM1(pf0037
06125 -# -
BMC-P
BMC-H
BMC-T
Aldehyde dehydrogenase
PTAC
Signature enzyme
Alcohol dehydrogenase
Regulator

Klebsiella sp. HMSC16C06. Bacteria; Proteobacteria; Gammaproteobacteria; Enterobacterales;

HMPREF3142__locus_1

GRM2

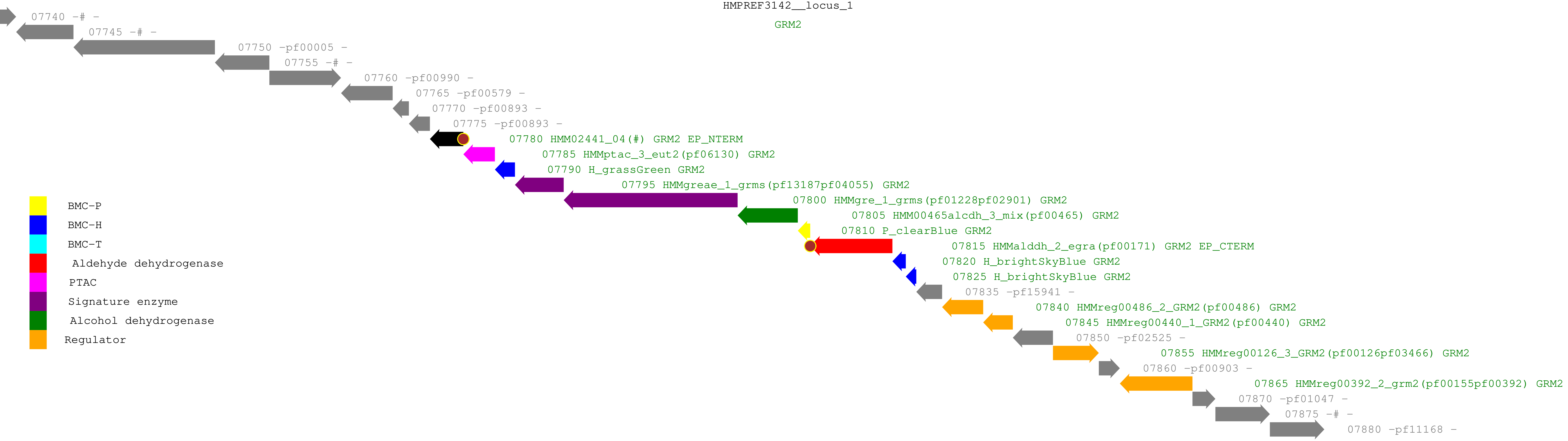

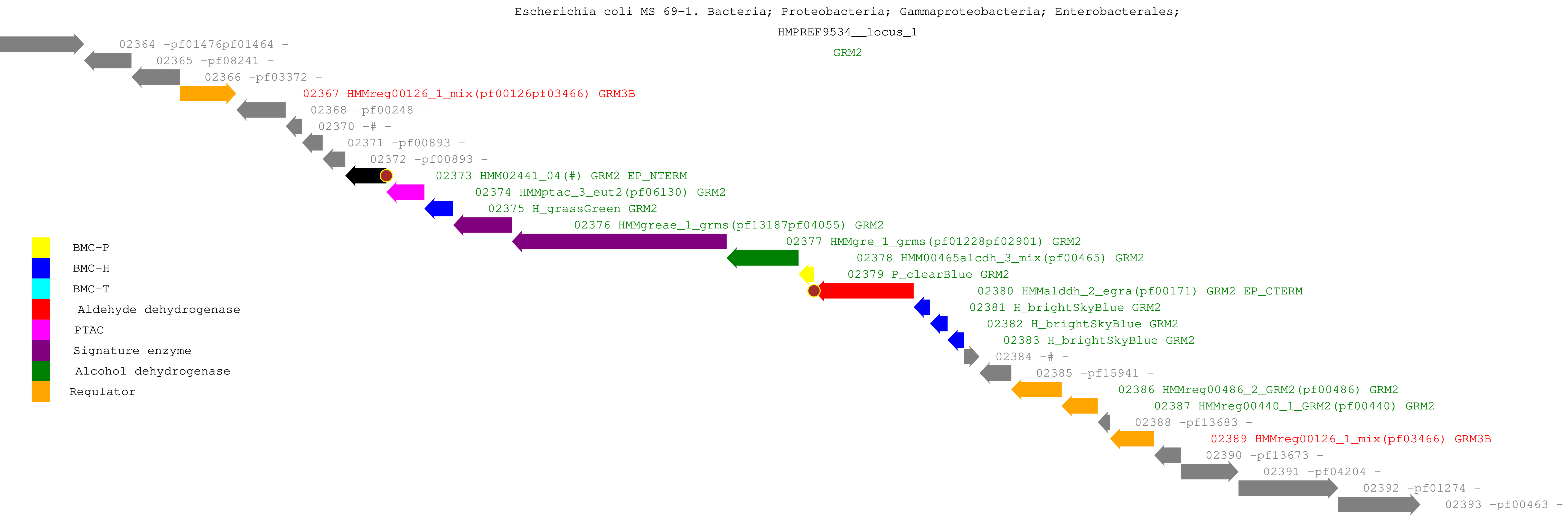
Escherichia coli MS 69-1. Bacteria; Proteobacteria; Gammaproteobacteria; Enterobacterales;
HMPREF9534__locus_1
GRM2
02364 -pf01476pf01464 -
02365 -pf08241 -
02366 -pf03372 -
02367 HMMreg00126_1_mix(pf00126pf03466) GRM3B
02368 -pf00248 -
02370 -# -
02371 -pf00893 -
02372 -pf00893 -
02373 HMM02441_04(#) GRM2 EP_NTERM
02374 HMMptac_3_eut2(pf06130) GRM2
02375 H_grassGreen GRM2
02376 HMMgreae_1_grms(pf13187pf04055) GRM2
02377 HMMgre_1_grms(pf01228pf02901) GRM2
02378 HMM00465alcdh_3_mix(pf00465) GRM2
02379 P_clearBlue GRM2
02380 HMMalddh_2_egra(pf00171) GRM2 EP_CTERM
02381 H_brightSkyBlue GRM2
02382 H_brightSkyBlue GRM2
02383 H_brightSkyBlue GRM2
02384 -# -
02385 -pf15941 -
02386 HMMreg00486_2_GRM2(pf00486) GRM2
02387 HMMreg00440_1_GRM2(pf00440) GRM2
02388 -pf13683 -
02389 HMMreg00126_1_mix(pf03466) GRM3B
02390 -pf13673 -
02391 -pf04204 -
02392 -pf01274 -
02393 -pf00463 -
BMC-P
BMC-H
BMC-T
Aldehyde dehydrogenase
PTAC
Signature enzyme
Alcohol dehydrogenase
Regulator

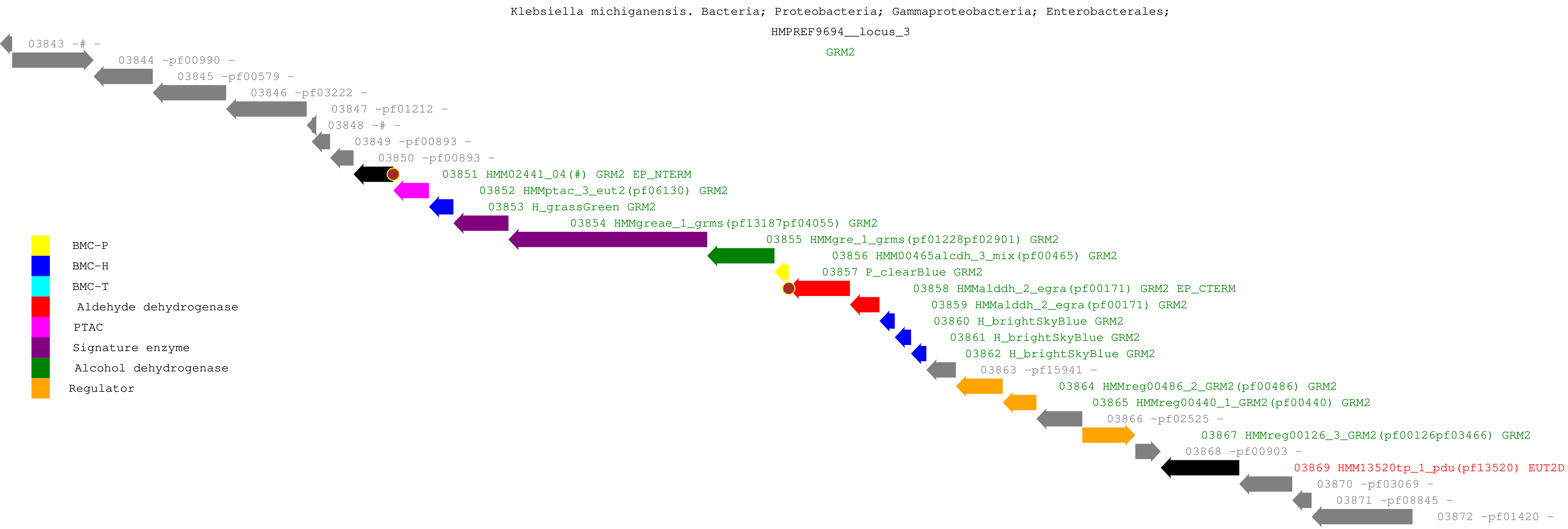
Klebsiella michiganensis. Bacteria; Proteobacteria; Gammaproteobacteria; Enterobacterales;
HMPREF9694__locus_3
GRM2
03843 -# -
03844 -pf00990 -
03845 -pf00579 -
03846 -pf03222 -
03847 -pf01212 -
03848 -# -
03849 -pf00893 -
03850 -pf00893 -
03851 HMM02441_04(#) GRM2 EP_NTERM
03852 HMMptac_3_eut2(pf06130) GRM2
03853 H_grassGreen GRM2
03854 HMMgreae_1_grms(pf13187pf04055) GRM2
03855 HMMgre_1_grms(pf01228pf02901) GRM2
03856 HMM00465alcdh_3_mix(pf00465) GRM2
03857 P_clearBlue GRM2
03858 HMMaldddh_2_egra(pf00171) GRM2 EP_CTERM
03859 HMMaldddh_2_egra(pf00171) GRM2
03860 H_brightSkyBlue GRM2
03861 H_brightSkyBlue GRM2
03862 H_brightSkyBlue GRM2
03863 -pf15941 -
03864 HMMreg00486_2_GRM2(pf00486) GRM2
03865 HMMreg00440_1_GRM2(pf00440) GRM2
03866 -pf02525 -
03867 HMMreg00126_3_GRM2(pf00126pf03466) GRM2
03868 -pf00903 -
03869 HMM13520tp_1_pdu(pf13520) EUT2D
03870 -pf03069 -
03871 -pf08845 -
03872 -pf01420 -
BMC-P
BMC-H
BMC-T
Aldehyde dehydrogenase
PTAC
Signature enzyme
Alcohol dehydrogenase
Regulator

Photobacterium aphoticum. Bacteria; Proteobacteria; Gammaproteobacteria; Vibrionales;

JCM19237__locus_1

GRM2

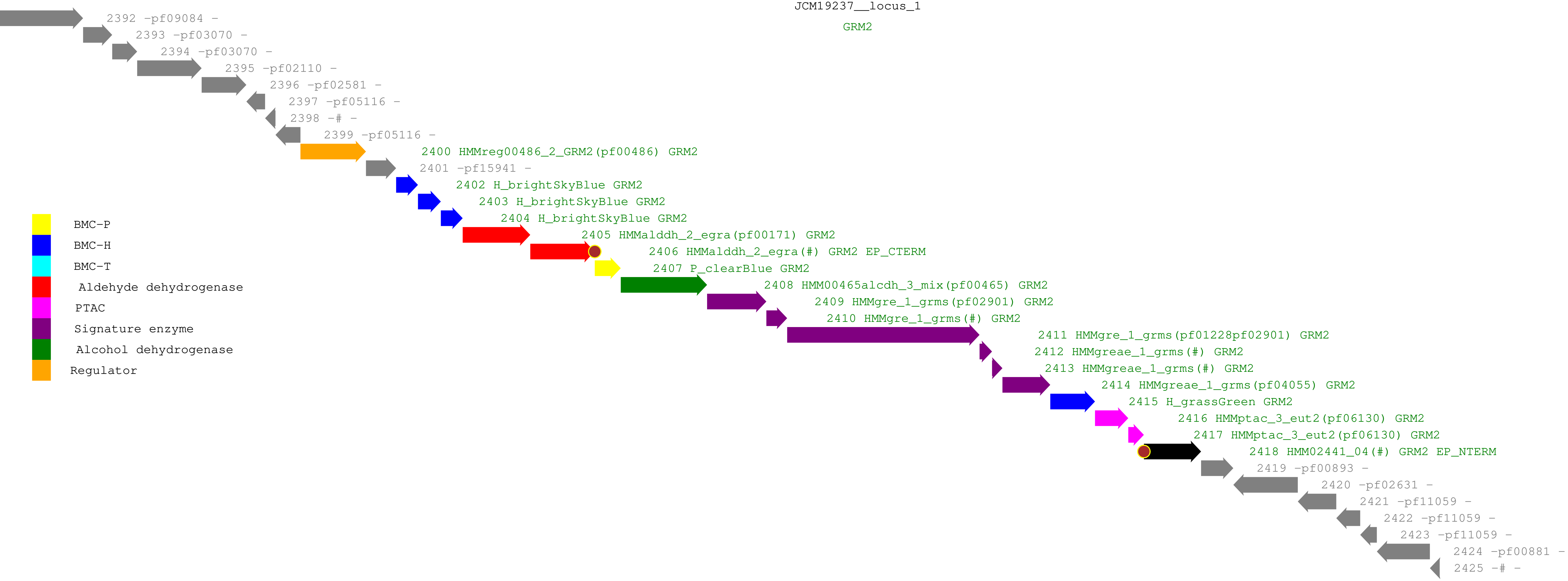

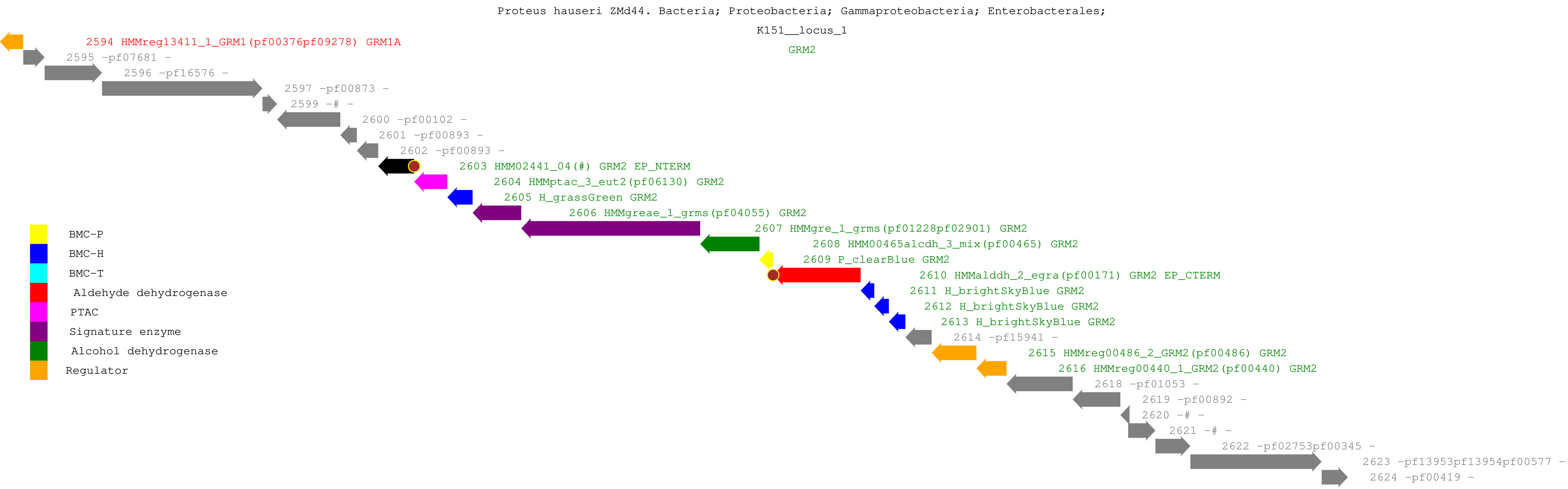
Proteus hauseri ZMd44. Bacteria; Proteobacteria; Gammaproteobacteria; Enterobacterales;
K151__locus_1
GRM2
2594 HMMreg13411_1_GRM1(pf00376pf09278) GRM1A
2595 -pf07681 -
2596 -pf16576 -
2597 -pf00873 -
2599 -# -
2600 -pf00102 -
2601 -pf00893 -
2602 -pf00893 -
2603 HMM02441_04(#) GRM2 EP_NTERM
2604 HMMptac_3_eut2(pf06130) GRM2
2605 H_grassGreen GRM2
2606 HMMgreae_1_grms(pf04055) GRM2
2607 HMMgre_1_grms(pf01228pf02901) GRM2
2608 HMM00465alcdh_3_mix(pf00465) GRM2
2609 P_clearBlue GRM2
2610 HMMalddh_2_egra(pf00171) GRM2 EP_CTERM
2611 H_brightSkyBlue GRM2
2612 H_brightSkyBlue GRM2
2613 H_brightSkyBlue GRM2
2614 -pf15941 -
2615 HMMreg00486_2_GRM2(pf00486) GRM2
2616 HMMreg00440_1_GRM2(pf00440) GRM2
2618 -pf01053 -
2619 -pf00892 -
2620 -# -
2621 -# -
2622 -pf02753pf00345 -
2623 -pf13953pf13954pf00577 -
2624 -pf00419 -
BMC-P
BMC-H
BMC-T
Aldehyde dehydrogenase
PTAC
Signature enzyme
Alcohol dehydrogenase
Regulator

Klebsiella oxytoca KONIH1. Bacteria; Proteobacteria; Gammaproteobacteria; Enterobacterales;

KONIH1__locus_1

GRM2

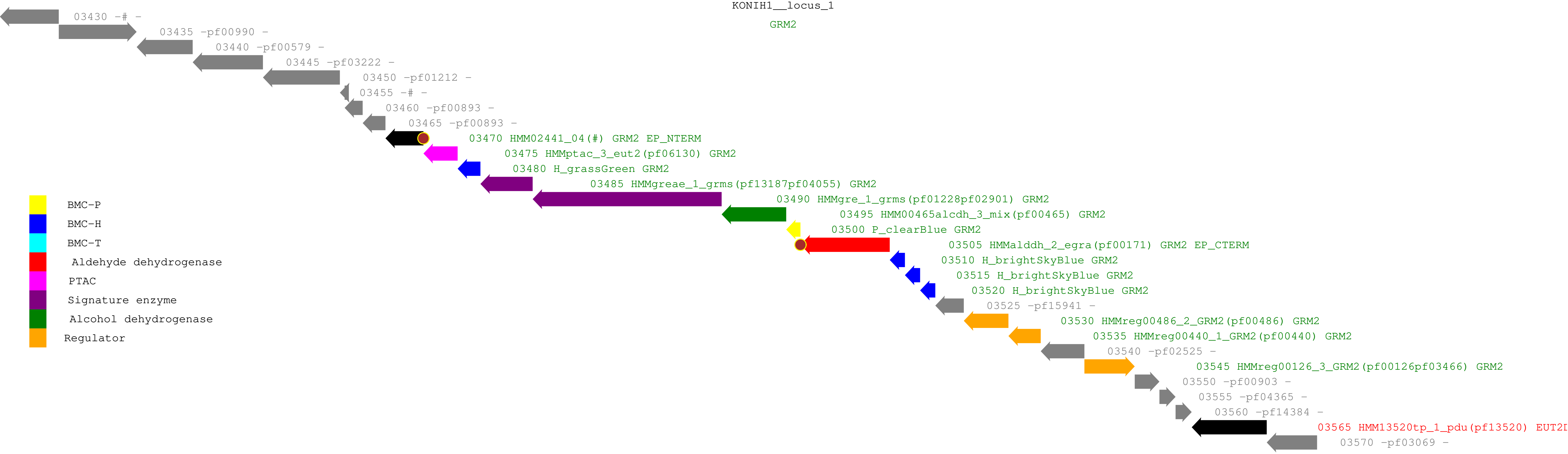

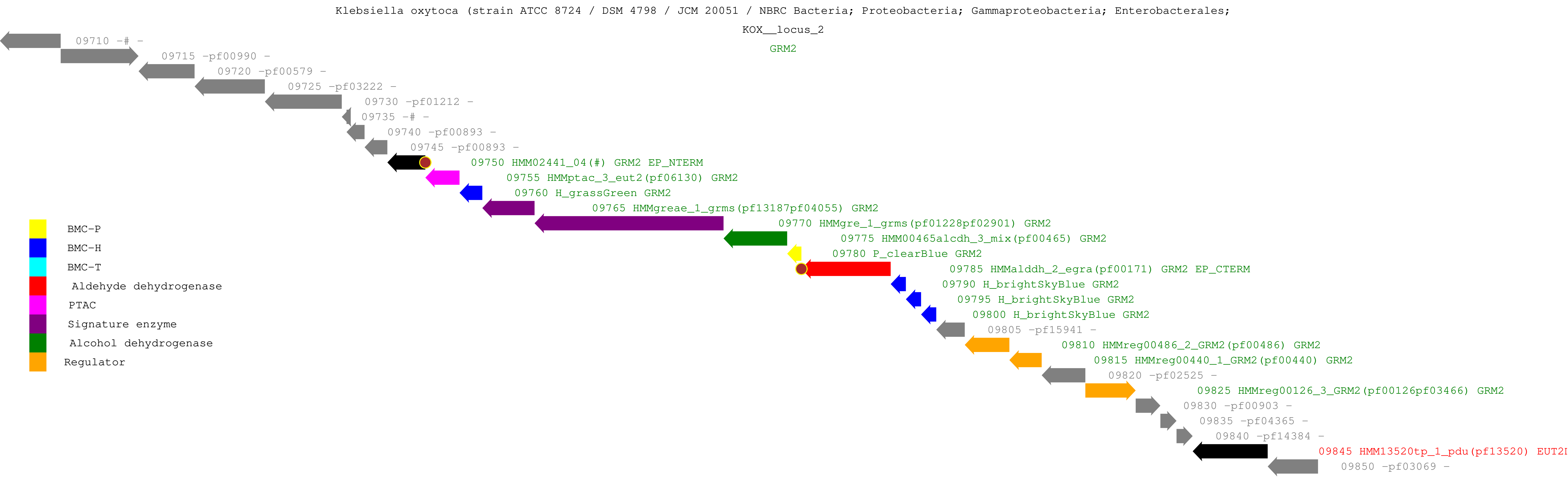
Klebsiella oxytoca (strain ATCC 8724 / DSM 4798 / JCM 20051 / NBRC Bacteria; Proteobacteria; Gammaproteobacteria; Enterobacterales;
KOX__locus_2
GRM2
09710 -# -
09715 -pf00990 -
09720 -pf00579 -
09725 -pf03222 -
09730 -pf01212 -
09735 -# -
09740 -pf00893 -
09745 -pf00893 -
09750 HMM02441_04(#) GRM2 EP_NTERM
09755 HMMptac_3_eut2(pf06130) GRM2
09760 H_grassGreen GRM2
09765 HMMgreae_1_grms(pf13187pf04055) GRM2
09770 HMMgre_1_grms(pf01228pf02901) GRM2
09775 HMM00465alcdh_3_mix(pf00465) GRM2
09780 P_clearBlue GRM2
09785 HMMaldh_2_egra(pf00171) GRM2 EP_CTERM
09790 H_brightSkyBlue GRM2
09795 H_brightSkyBlue GRM2
09800 H_brightSkyBlue GRM2
09805 -pf15941 -
09810 HMMreg00486_2_GRM2(pf00486) GRM2
09815 HMMreg00440_1_GRM2(pf00440) GRM2
09820 -pf02525 -
09825 HMMreg00126_3_GRM2(pf00126pf03466) GRM2
09830 -pf00903 -
09835 -pf04365 -
09840 -pf14384 -
09845 HMM13520tp_1_pdu(pf13520) EUT2I
09850 -pf03069 -
BMC-P
BMC-H
BMC-T
Aldehyde dehydrogenase
PTAC
Signature enzyme
Alcohol dehydrogenase
Regulator

Klebsiella pneumoniae (strain 342). Bacteria; Proteobacteria; Gammaproteobacteria; Enterobacterales;

KPK__locus_3

GRM2

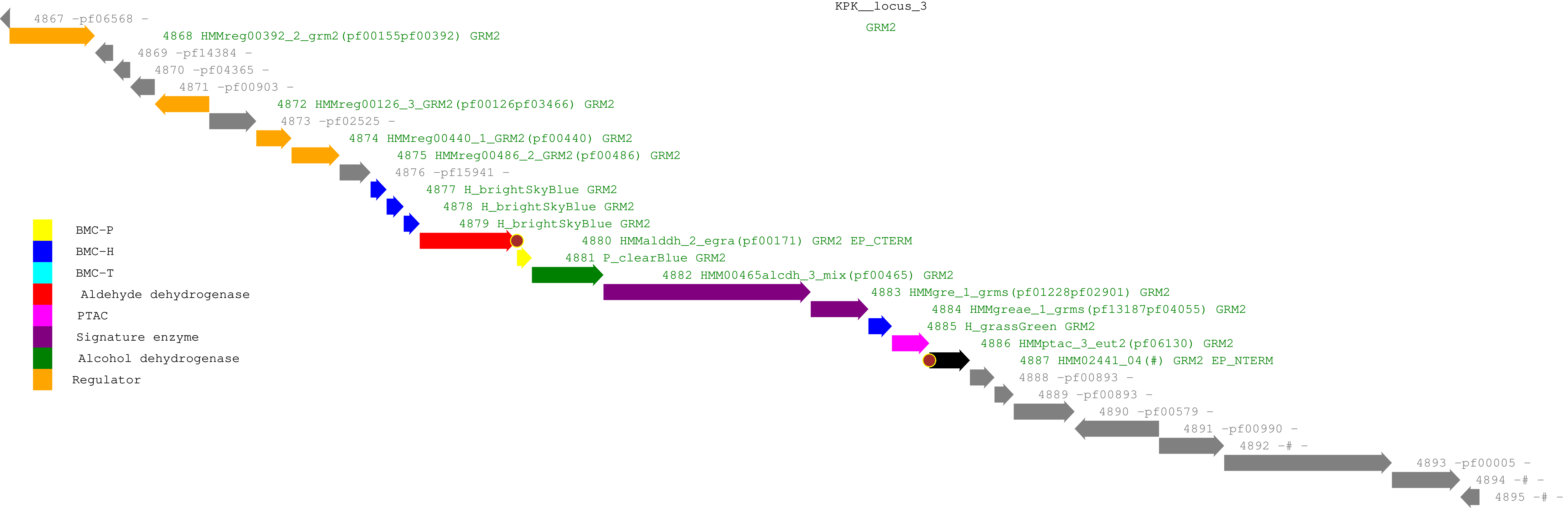

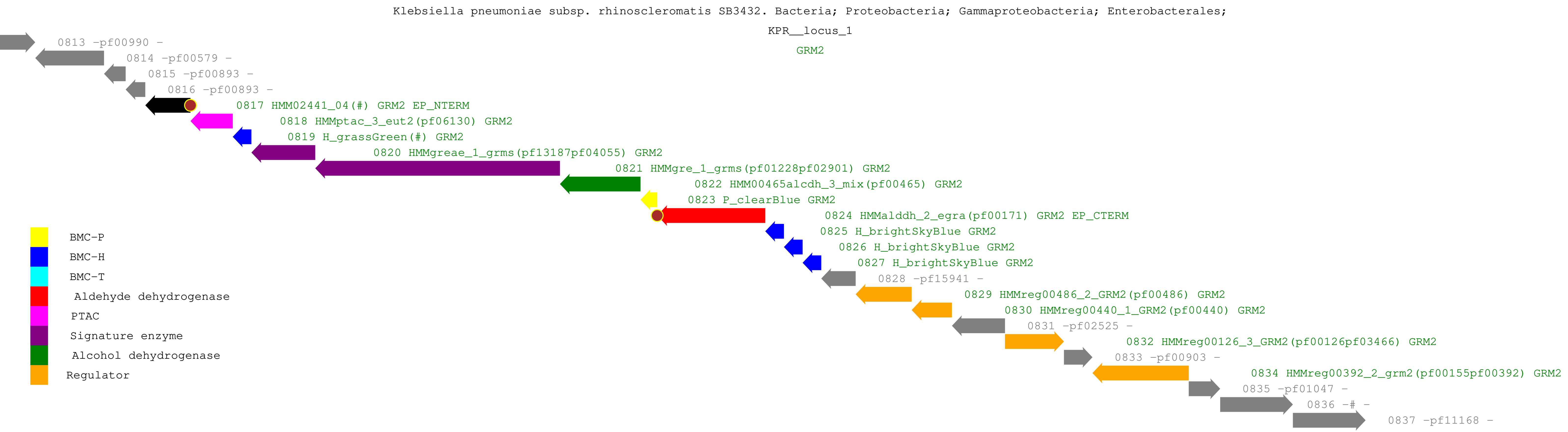

Klebsiella pneumoniae subsp. rhinoscleromatis SB3432. Bacteria; Proteobacteria; Gammaproteobacteria; Enterobacterales;
KPR__locus_1
GRM2
0813 -pf00990 -
0814 -pf00579 -
0815 -pf00893 -
0816 -pf00893 -
0817 HMM02441_04(#) GRM2 EP_NTERM
0818 HMMptac_3_eut2(pf06130) GRM2
0819 H_grassGreen(#) GRM2
0820 HMMgreae_1_grms(pf13187pf04055) GRM2
0821 HMMgre_1_grms(pf01228pf02901) GRM2
0822 HMM00465alcdh_3_mix(pf00465) GRM2
0823 P_clearBlue GRM2
0824 HMMaldh_2_egra(pf00171) GRM2 EP_CTERM
0825 H_brightSkyBlue GRM2
0826 H_brightSkyBlue GRM2
0827 H_brightSkyBlue GRM2
0828 -pf15941 -
0829 HMMreg00486_2_GRM2(pf00486) GRM2
0830 HMMreg00440_1_GRM2(pf00440) GRM2
0831 -pf02525 -
0832 HMMreg00126_3_GRM2(pf00126pf03466) GRM2
0833 -pf00903 -
0834 HMMreg00392_2_grm2(pf00155pf00392) GRM2
0835 -pf01047 -
0836 -# -
0837 -pf11168 -
BMC-P
BMC-H
BMC-T
Aldehyde dehydrogenase
PTAC
Signature enzyme
Alcohol dehydrogenase
Regulator

Serratia fonticola AU-P3(3). Bacteria; Proteobacteria; Gammaproteobacteria; Enterobacterales;

L580__locus_1

GRM2

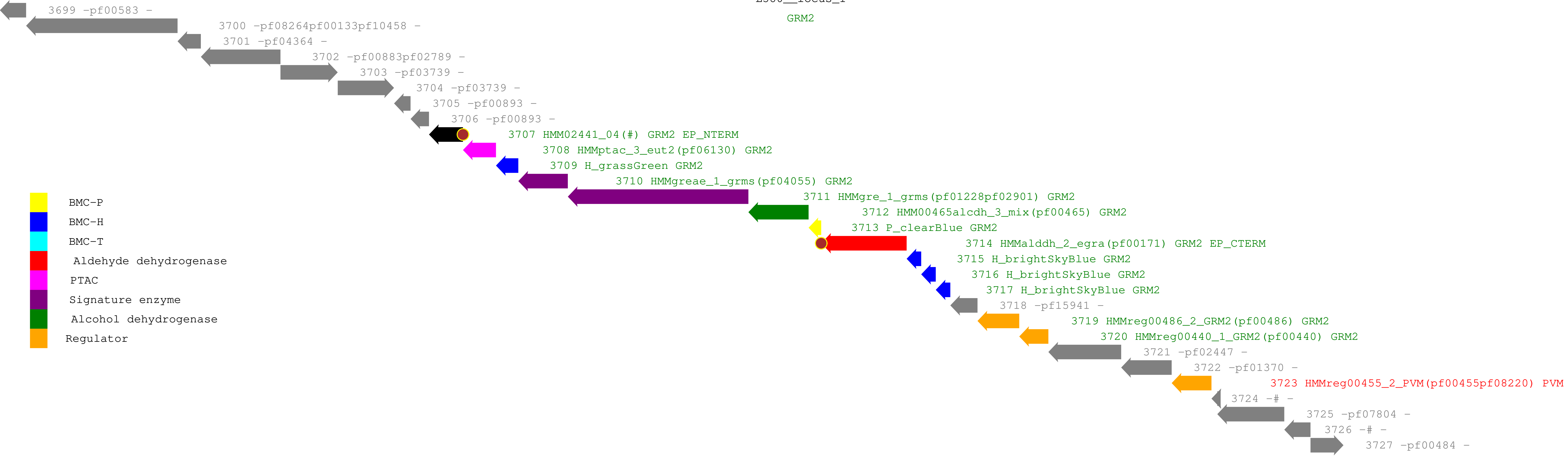

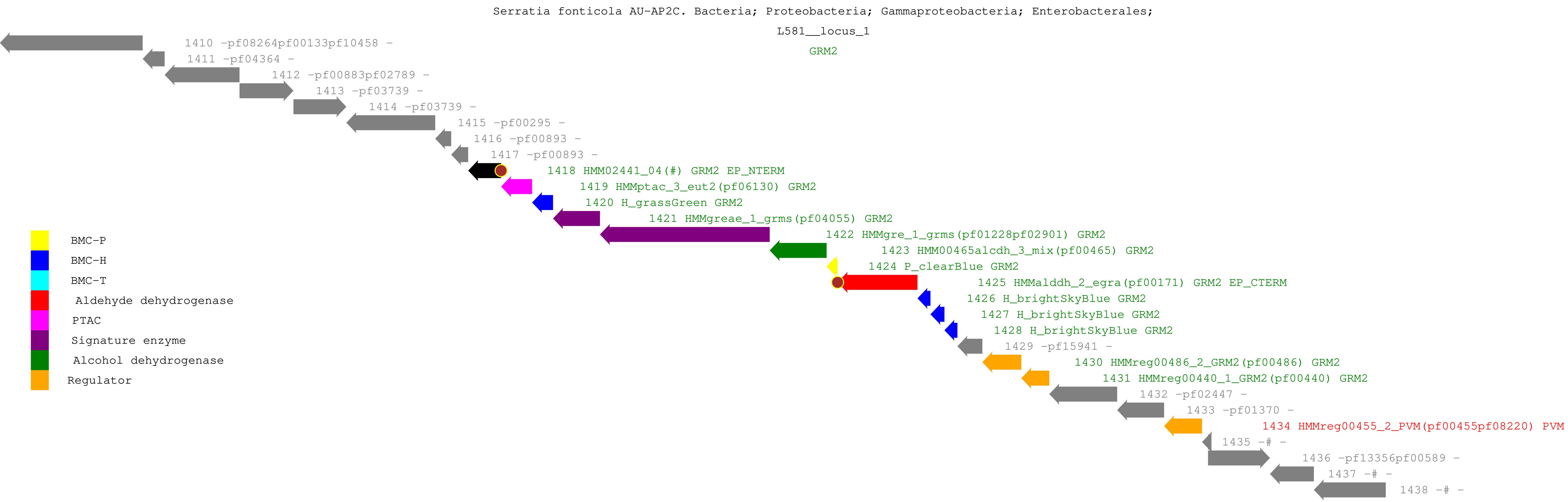
Serratia fonticola AU-AP2C. Bacteria; Proteobacteria; Gammaproteobacteria; Enterobacterales;
L581__locus_1
GRM2
1410 -pf08264pf00133pf10458 -
1411 -pf04364 -
1412 -pf00883pf02789 -
1413 -pf03739 -
1414 -pf03739 -
1415 -pf00295 -
1416 -pf00893 -
1417 -pf00893 -
1418 HMM02441_04(#) GRM2 EP_NTERM
1419 HMMptac_3_eut2(pf06130) GRM2
1420 H_grassGreen GRM2
1421 HMMgreae_1_grms(pf04055) GRM2
1422 HMMgre_1_grms(pf01228pf02901) GRM2
1423 HMM00465alcdh_3_mix(pf00465) GRM2
1424 P_clearBlue GRM2
1425 HMMalddh_2_egra(pf00171) GRM2 EP_CTERM
1426 H_brightSkyBlue GRM2
1427 H_brightSkyBlue GRM2
1428 H_brightSkyBlue GRM2
1429 -pf15941 -
1430 HMMreg00486_2_GRM2(pf00486) GRM2
1431 HMMreg00440_1_GRM2(pf00440) GRM2
1432 -pf02447 -
1433 -pf01370 -
1434 HMMreg00455_2_PVM(pf00455pf08220) PVM
1435 -# -
1436 -pf13356pf00589 -
1437 -# -
1438 -# -
BMC-P
BMC-H
BMC-T
Aldehyde dehydrogenase
PTAC
Signature enzyme
Alcohol dehydrogenase
Regulator

Moellerella wisconsensis ATCC 35017. Bacteria; Proteobacteria; Gammaproteobacteria; Enterobacterales;

M992__locus_1

GRM2

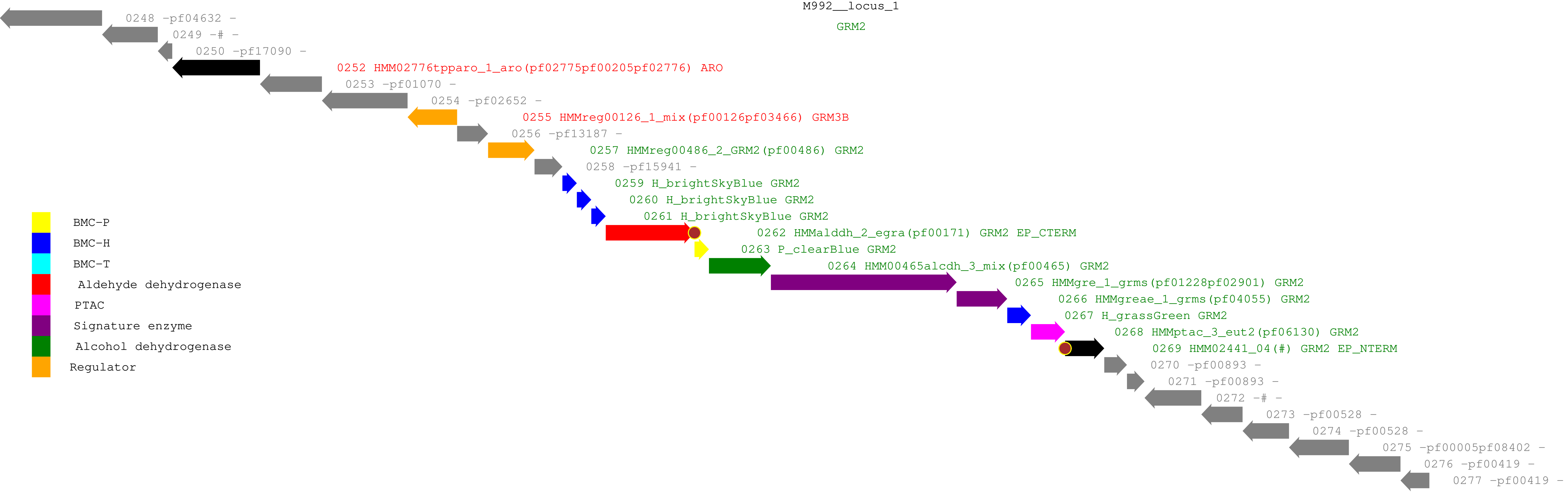

Proteus hauseri ATCC 700826. Bacteria; Proteobacteria; Gammaproteobacteria; Enterobacterales;

M997__locus_1

GRM2

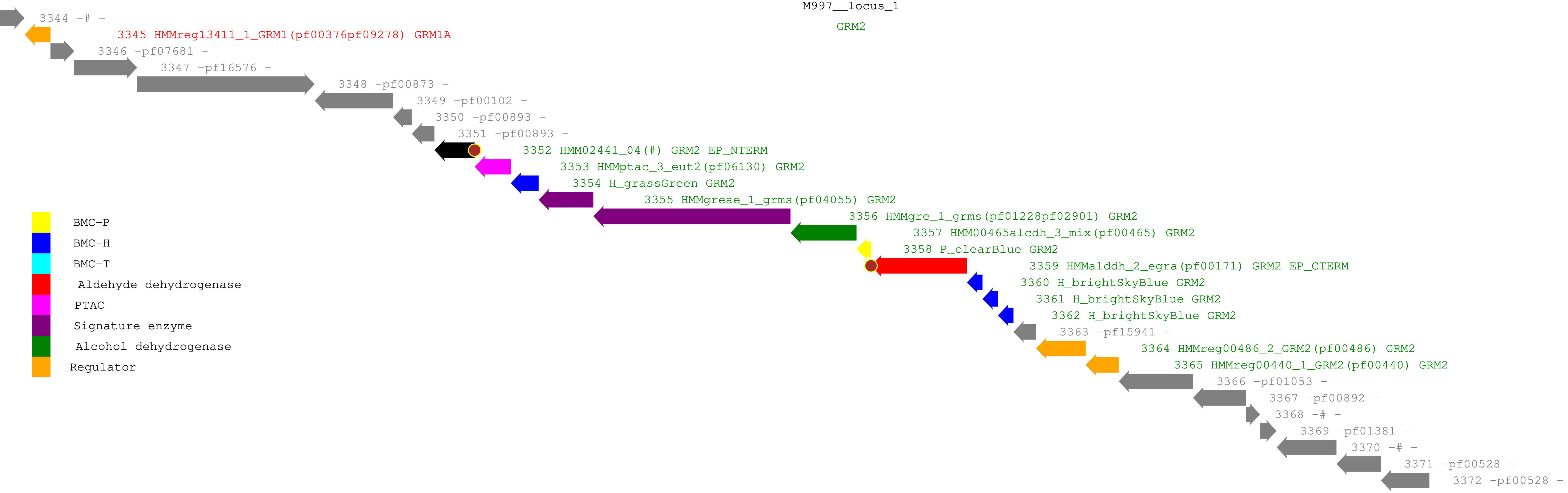

Providencia heimbachae ATCC 35613. Bacteria; Proteobacteria; Gammaproteobacteria; Enterobacterales;

M998__locus_1

GRM2

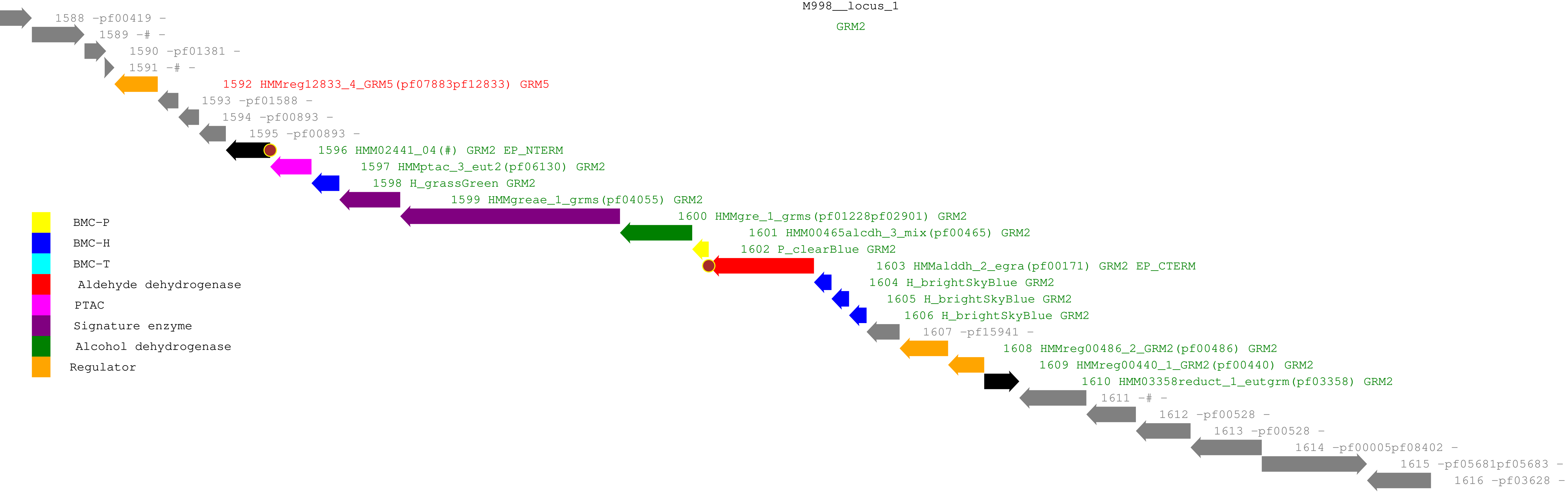

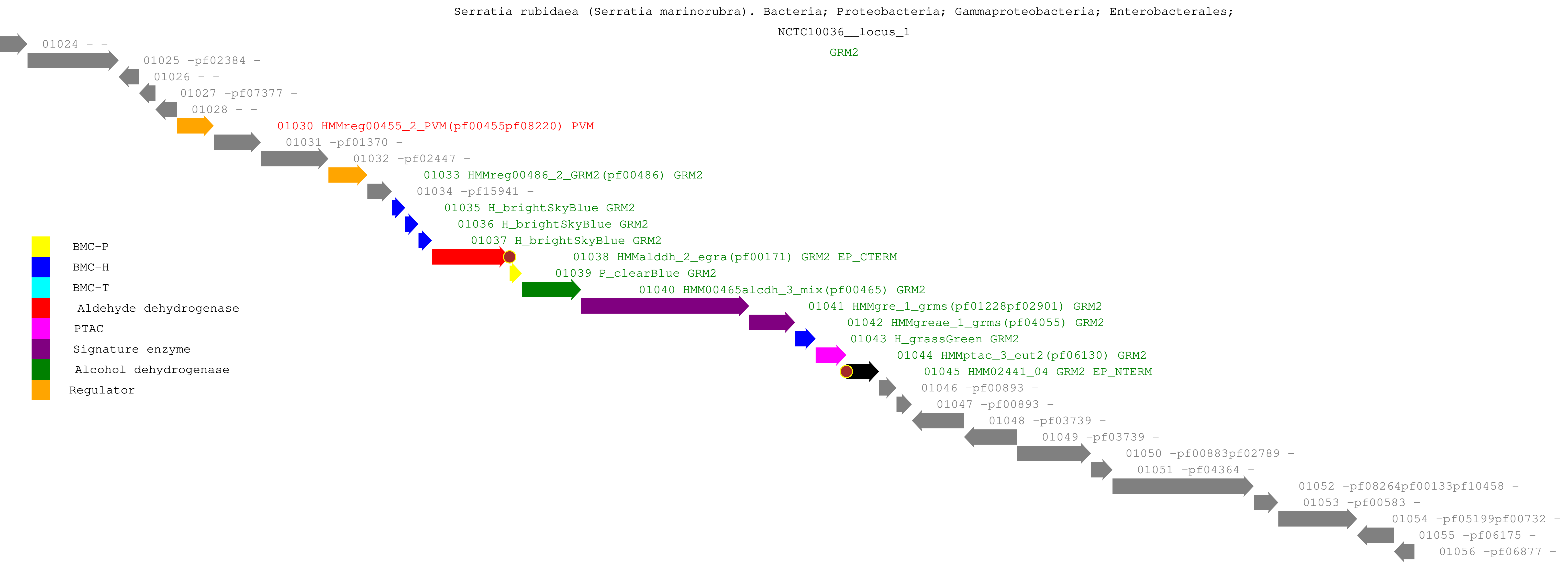
Serratia rubidaea (Serratia marinorubra). Bacteria; Proteobacteria; Gammaproteobacteria; Enterobacterales;
NCTC10036__locus_1
GRM2
01024 - -
01025 -pf02384 -
01026 - -
01027 -pf07377 -
01028 - -
01030 HMMreg00455_2_PVM(pf00455pf08220) PVM
01031 -pf01370 -
01032 -pf02447 -
01033 HMMreg00486_2_GRM2(pf00486) GRM2
01034 -pf15941 -
01035 H_brightSkyBlue GRM2
01036 H_brightSkyBlue GRM2
01037 H_brightSkyBlue GRM2
01038 HMMalddh_2_egra(pf00171) GRM2 EP_CTERM
01039 P_clearBlue GRM2
01040 HMM00465alcdh_3_mix(pf00465) GRM2
01041 HMMgre_1_grms(pf01228pf02901) GRM2
01042 HMMgreae_1_grms(pf04055) GRM2
01043 H_grassGreen GRM2
01044 HMMptac_3_eut2(pf06130) GRM2
01045 HMM02441_04 GRM2 EP_NTERM
01046 -pf00893 -
01047 -pf00893 -
01048 -pf03739 -
01049 -pf03739 -
01050 -pf00883pf02789 -
01051 -pf04364 -
01052 -pf08264pf00133pf10458 -
01053 -pf00583 -
01054 -pf05199pf00732 -
01055 -pf06175 -
01056 -pf06877 -
BMC-P
BMC-H
BMC-T
Aldehyde dehydrogenase
PTAC
Signature enzyme
Alcohol dehydrogenase
Regulator

Klebsiella pneumoniae. Bacteria; Proteobacteria; Gammaproteobacteria; Enterobacterales;

NCTC10262__locus_3

GRM2

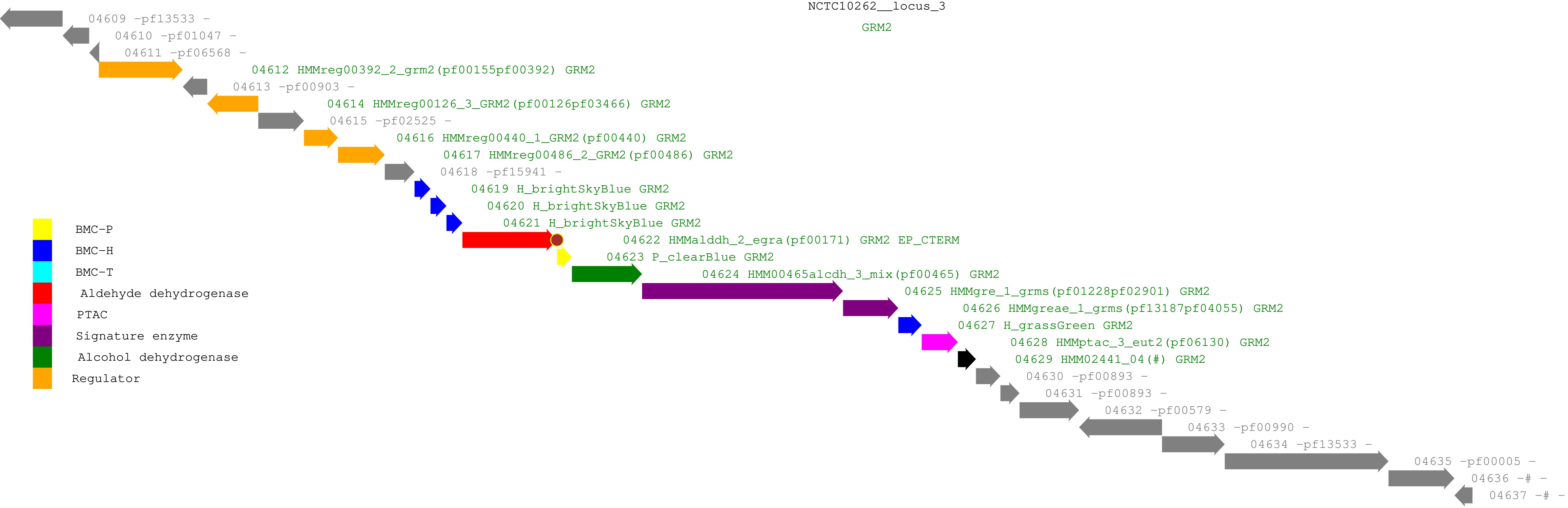

Proteus vulgaris. Bacteria; Proteobacteria; Gammaproteobacteria; Enterobacterales;

NCTC10376__locus_1

GRM2

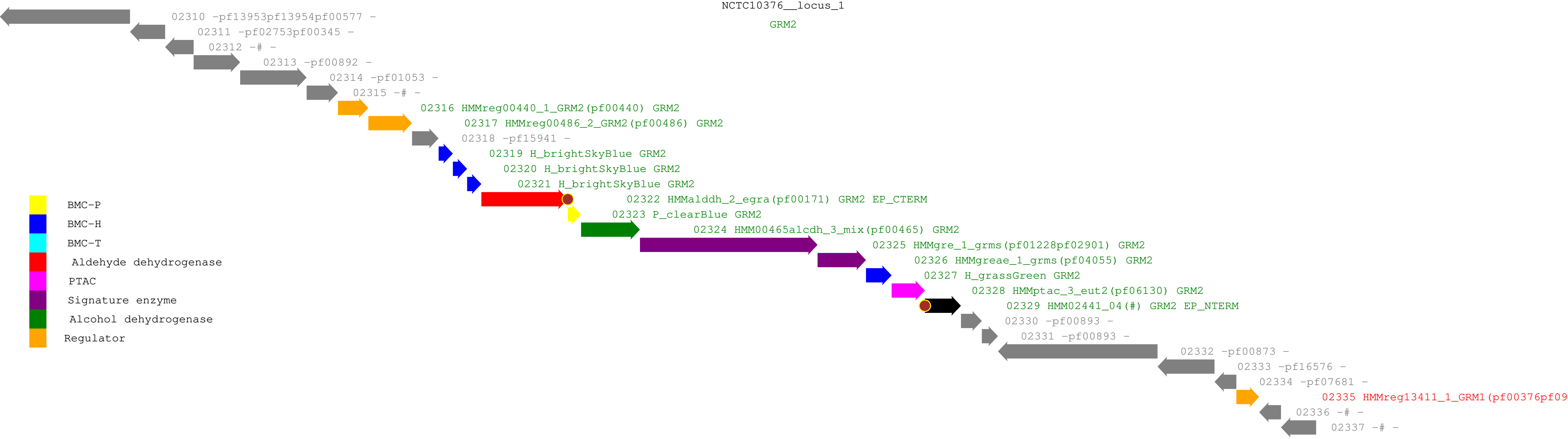

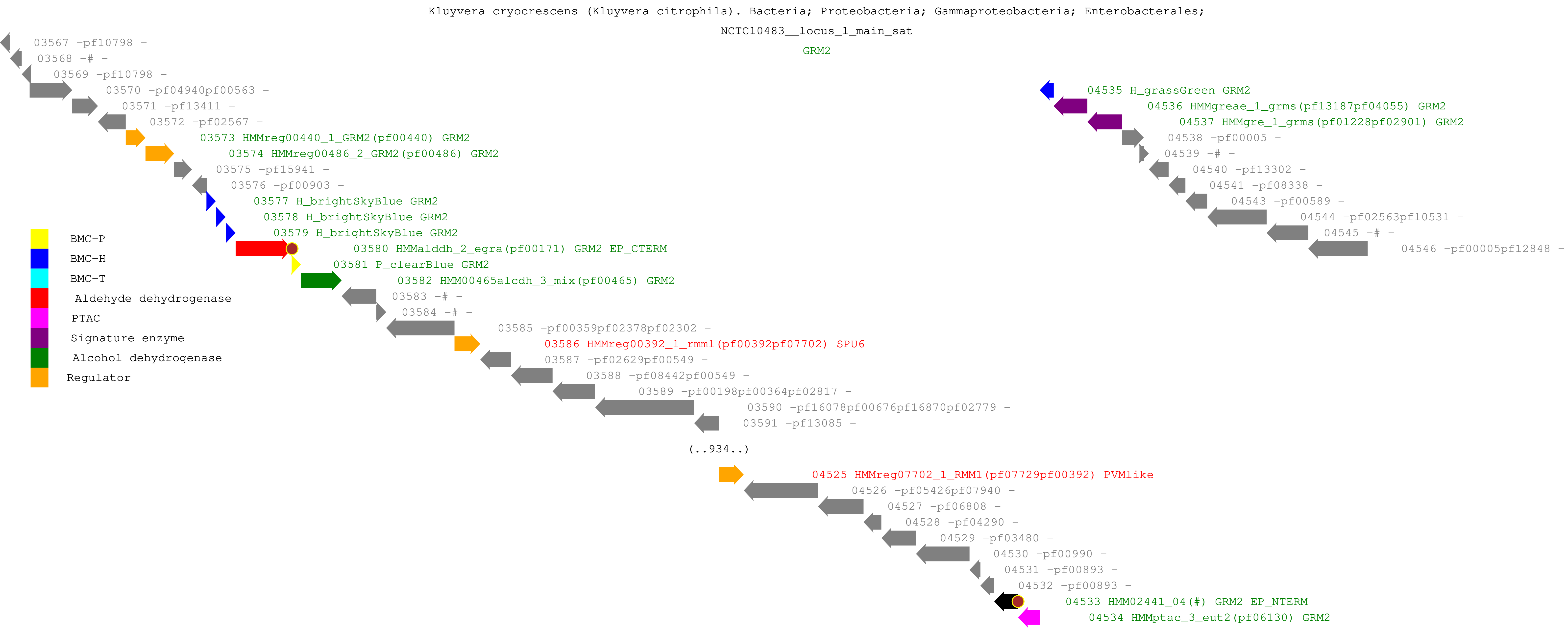

Kluyvera cryocrescens (Kluyvera citrophila). Bacteria; Proteobacteria; Gammaproteobacteria; Enterobacterales;
NCTC10483__locus_1_main_sat
GRM2
03567 -pf10798 -
03568 -# -
03569 -pf10798 -
03570 -pf04940pf00563 -
03571 -pf13411 -
03572 -pf02567 -
03573 HMMreg00440_1_GRM2(pf00440) GRM2
03574 HMMreg00486_2_GRM2(pf00486) GRM2
03575 -pf15941 -
03576 -pf00903 -
03577 H_brightSkyBlue GRM2
03578 H_brightSkyBlue GRM2
03579 H_brightSkyBlue GRM2
03580 HMMaldh_2_egra(pf00171) GRM2 EP_CTERM
03581 P_clearBlue GRM2
03582 HMM00465alcdh_3_mix(pf00465) GRM2
03583 -# -
03584 -# -
03585 -pf00359pf02378pf02302 -
03586 HMMreg00392_1_rmm1(pf00392pf07702) SPU6
03587 -pf02629pf00549 -
03588 -pf08442pf00549 -
03589 -pf00198pf00364pf02817 -
03590 -pf16078pf00676pf16870pf02779 -
03591 -pf13085 -
(..934..)
04525 HMMreg07702_1_RMM1(pf07729pf00392) PVMlike
04526 -pf05426pf07940 -
04527 -pf06808 -
04528 -pf04290 -
04529 -pf03480 -
04530 -pf00990 -
04531 -pf00893 -
04532 -pf00893 -
04533 HMM02441_04(#) GRM2 EP_NTERM
04534 HMMptac_3_eut2(pf06130) GRM2
04535 H_grassGreen GRM2
04536 HMMgreae_1_grms(pf13187pf04055) GRM2
04537 HMMgre_1_grms(pf01228pf02901) GRM2
04538 -pf00005 -
04539 -# -
04540 -pf13302 -
04541 -pf08338 -
04543 -pf00589 -
04544 -pf02563pf10531 -
04545 -# -
04546 -pf00005pf12848 -
BMC-P
BMC-H
BMC-T
Aldehyde dehydrogenase
PTAC
Signature enzyme
Alcohol dehydrogenase
Regulator

Serratia rubidaea (Serratia marinorubra). Bacteria; Proteobacteria; Gammaproteobacteria; Enterobacterales;

NCTC10848__locus_1

GRM2

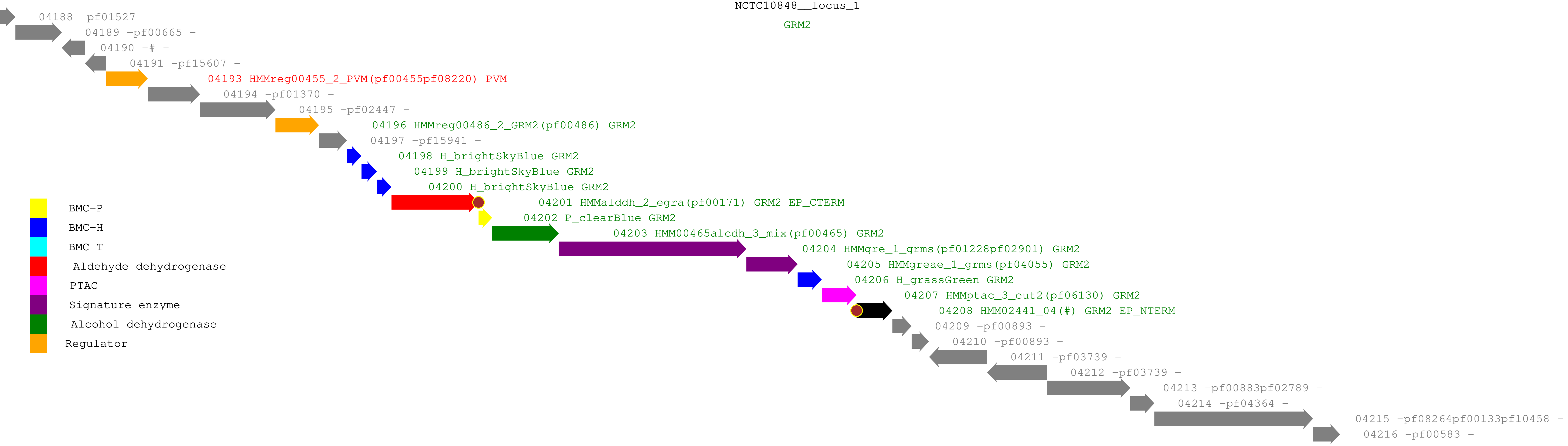

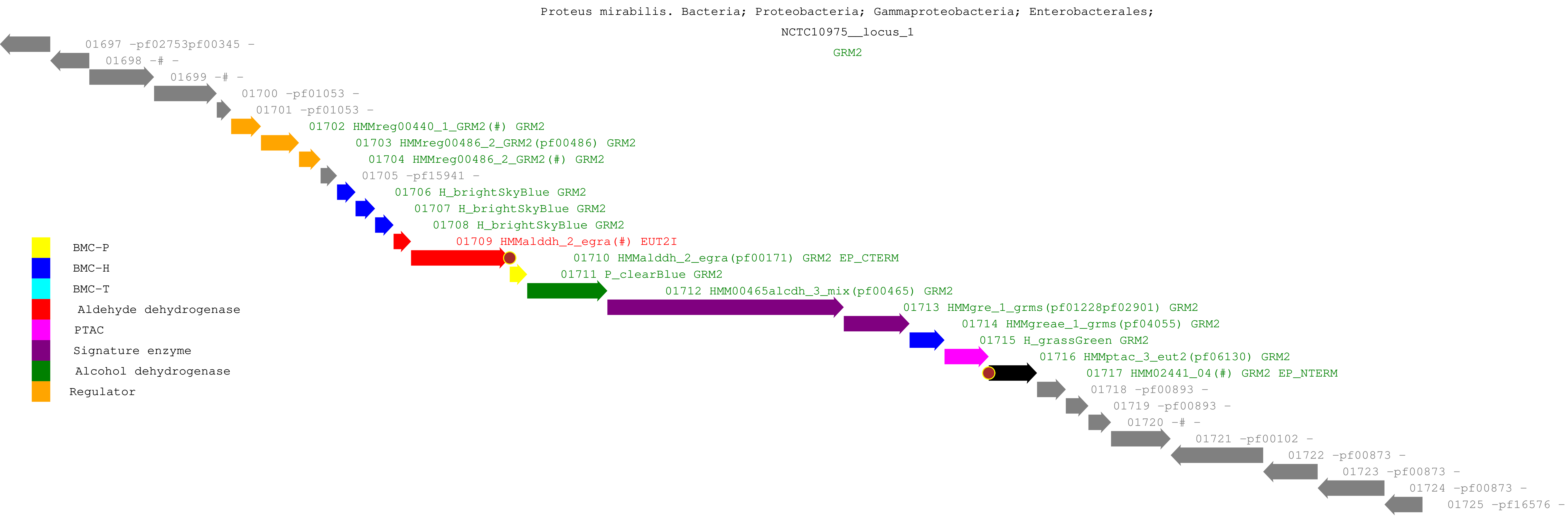

Proteus mirabilis. Bacteria; Proteobacteria; Gammaproteobacteria; Enterobacterales;
NCTC10975__locus_1
GRM2
01697 -pf02753pf00345 -
01698 -# -
01699 -# -
01700 -pf01053 -
01701 -pf01053 -
01702 HMMreg00440_1_GRM2(#) GRM2
01703 HMMreg00486_2_GRM2(pf00486) GRM2
01704 HMMreg00486_2_GRM2(#) GRM2
01705 -pf15941 -
01706 H_brightSkyBlue GRM2
01707 H_brightSkyBlue GRM2
01708 H_brightSkyBlue GRM2
01709 HMMalddh_2_egra(#) EUT2I
01710 HMMalddh_2_egra(pf00171) GRM2 EP_CTERM
01711 P_clearBlue GRM2
01712 HMM00465alcdh_3_mix(pf00465) GRM2
01713 HMMgre_1_grms(pf01228pf02901) GRM2
01714 HMMgreae_1_grms(pf04055) GRM2
01715 H_grassGreen GRM2
01716 HMMptac_3_eut2(pf06130) GRM2
01717 HMM02441_04(#) GRM2 EP_NTERM
01718 -pf00893 -
01719 -pf00893 -
01720 -# -
01721 -pf00102 -
01722 -pf00873 -
01723 -pf00873 -
01724 -pf00873 -
01725 -pf16576 -
BMC-P
BMC-H
BMC-T
Aldehyde dehydrogenase
PTAC
Signature enzyme
Alcohol dehydrogenase
Regulator

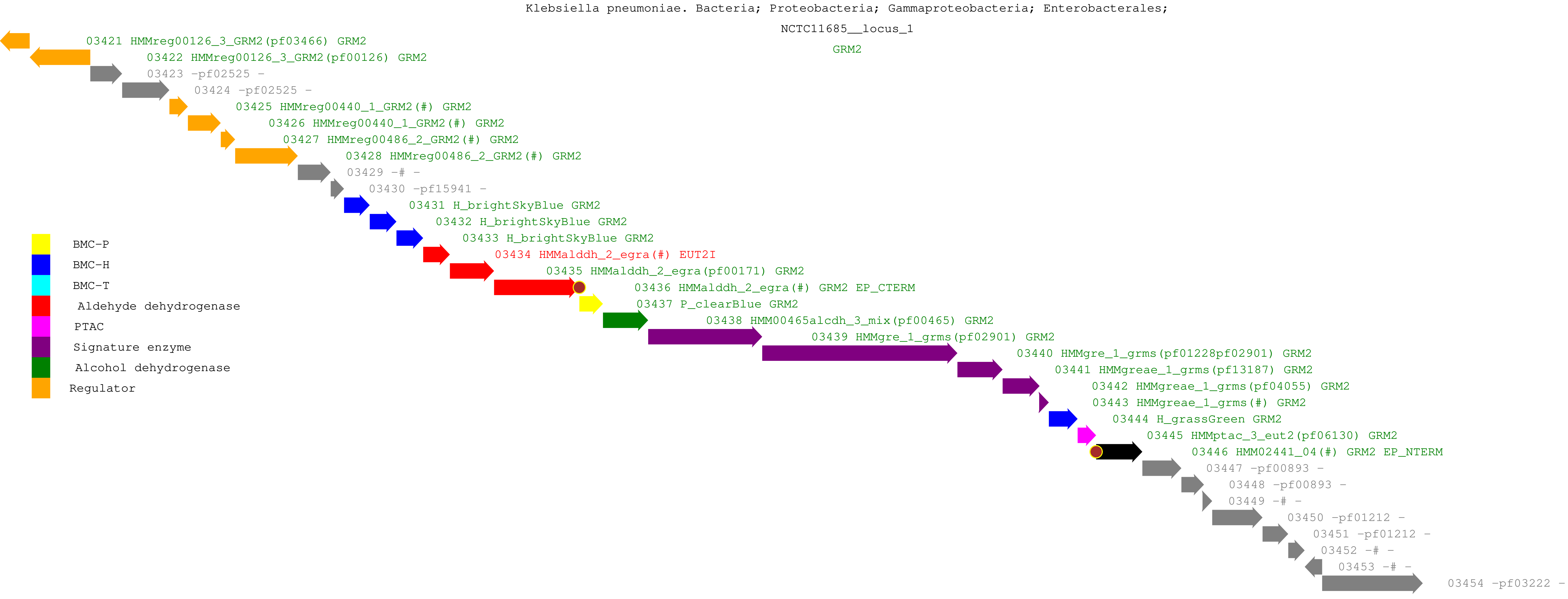

Klebsiella pneumoniae. Bacteria; Proteobacteria; Gammaproteobacteria; Enterobacterales;
NCTC11685__locus_1
GRM2
03421 HMMreg00126_3_GRM2(pf03466) GRM2
03422 HMMreg00126_3_GRM2(pf00126) GRM2
03423 -pf02525 -
03424 -pf02525 -
03425 HMMreg00440_1_GRM2(#) GRM2
03426 HMMreg00440_1_GRM2(#) GRM2
03427 HMMreg00486_2_GRM2(#) GRM2
03428 HMMreg00486_2_GRM2(#) GRM2
03429 -# -
03430 -pf15941 -
03431 H_brightSkyBlue GRM2
03432 H_brightSkyBlue GRM2
03433 H_brightSkyBlue GRM2
03434 HMMalddh_2_egra(#) EUT2I
03435 HMMalddh_2_egra(pf00171) GRM2
03436 HMMalddh_2_egra(#) GRM2 EP_CTERM
03437 P_clearBlue GRM2
03438 HMM00465alcdh_3_mix(pf00465) GRM2
03439 HMMgre_1_grms(pf02901) GRM2
03440 HMMgre_1_grms(pf01228pf02901) GRM2
03441 HMMgreae_1_grms(pf13187) GRM2
03442 HMMgreae_1_grms(pf04055) GRM2
03443 HMMgreae_1_grms(#) GRM2
03444 H_grassGreen GRM2
03445 HMMptac_3_eut2(pf06130) GRM2
03446 HMM02441_04(#) GRM2 EP_NTERM
03447 -pf00893 -
03448 -pf00893 -
03449 -# -
03450 -pf01212 -
03451 -pf01212 -
03452 -# -
03453 -# -
03454 -pf03222 -
BMC-P
BMC-H
BMC-T
Aldehyde dehydrogenase
PTAC
Signature enzyme
Alcohol dehydrogenase
Regulator

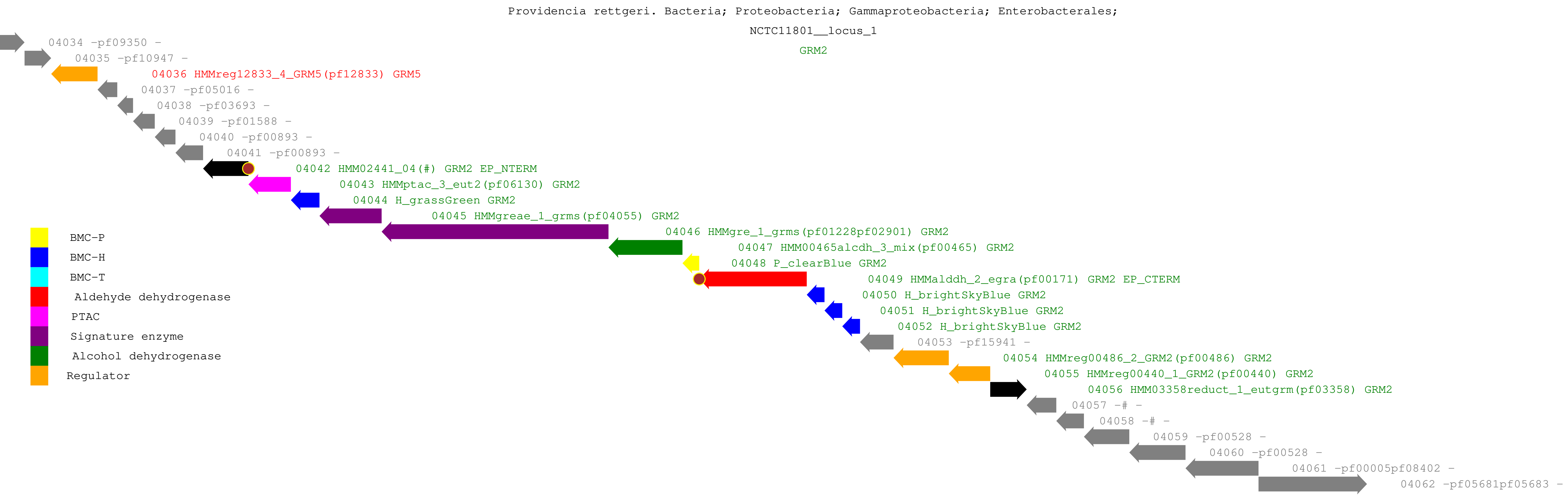

Providencia rettgeri. Bacteria; Proteobacteria; Gammaproteobacteria; Enterobacterales;
NCTC11801__locus_1
GRM2
04034 -pf09350 -
04035 -pf10947 -
04036 HMMreg12833_4_GRM5(pf12833) GRM5
04037 -pf05016 -
04038 -pf03693 -
04039 -pf01588 -
04040 -pf00893 -
04041 -pf00893 -
04042 HMM02441_04(#) GRM2 EP_NTERM
04043 HMMptac_3_eut2(pf06130) GRM2
04044 H_grassGreen GRM2
04045 HMMgreae_1_grms(pf04055) GRM2
04046 HMMgre_1_grms(pf01228pf02901) GRM2
04047 HMM00465alcdh_3_mix(pf00465) GRM2
04048 P_clearBlue GRM2
04049 HMMalddh_2_egra(pf00171) GRM2 EP_CTERM
04050 H_brightSkyBlue GRM2
04051 H_brightSkyBlue GRM2
04052 H_brightSkyBlue GRM2
04053 -pf15941 -
04054 HMMreg00486_2_GRM2(pf00486) GRM2
04055 HMMreg00440_1_GRM2(pf00440) GRM2
04056 HMM03358reduct_1_eutgrm(pf03358) GRM2
04057 -# -
04058 -# -
04059 -pf00528 -
04060 -pf00528 -
04061 -pf00005pf08402 -
04062 -pf05681pf05683 -
BMC-P
BMC-H
BMC-T
Aldehyde dehydrogenase
PTAC
Signature enzyme
Alcohol dehydrogenase
Regulator

Proteus mirabilis. Bacteria; Proteobacteria; Gammaproteobacteria; Enterobacterales;

NCTC11938__locus_1

GRM2

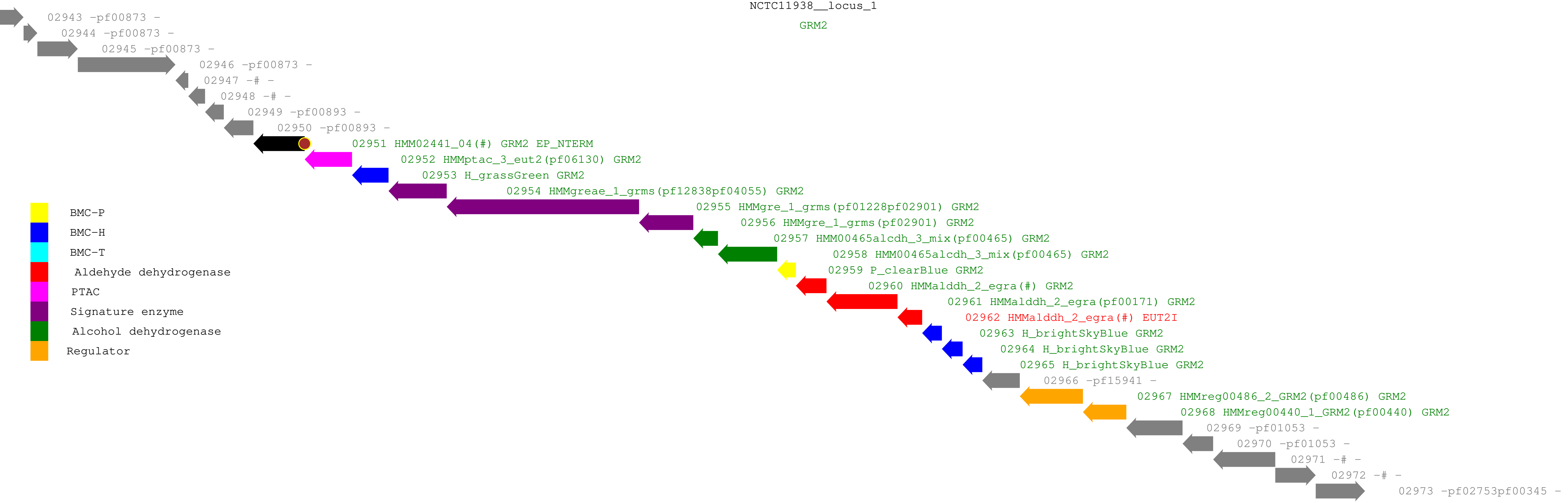

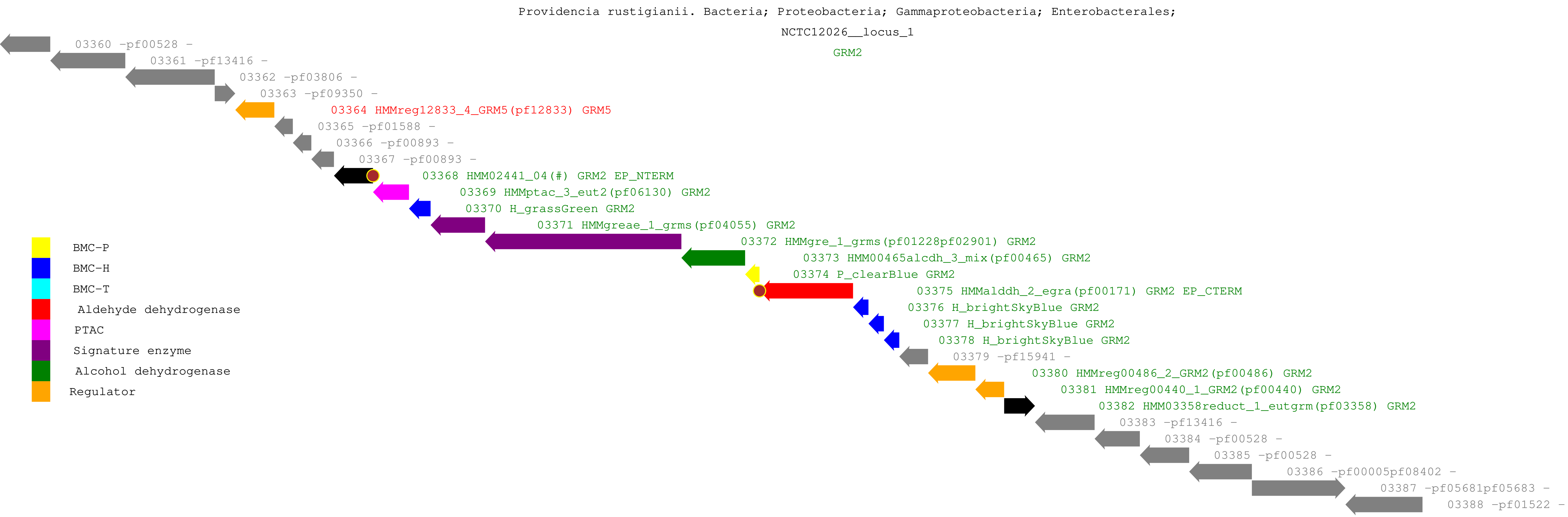
Providencia rustigianii. Bacteria; Proteobacteria; Gammaproteobacteria; Enterobacterales;
NCTC12026__locus_1
GRM2
03360 -pf00528 -
03361 -pf13416 -
03362 -pf03806 -
03363 -pf09350 -
03364 HMMreg12833_4_GRM5(pf12833) GRM5
03365 -pf01588 -
03366 -pf00893 -
03367 -pf00893 -
03368 HMM02441_04(#) GRM2 EP_NTERM
03369 HMMptac_3_eut2(pf06130) GRM2
03370 H_grassGreen GRM2
03371 HMMgreae_1_grms(pf04055) GRM2
03372 HMMgre_1_grms(pf01228pf02901) GRM2
03373 HMM00465alcdh_3_mix(pf00465) GRM2
03374 P_clearBlue GRM2
03375 HMMalddh_2_egra(pf00171) GRM2 EP_CTERM
03376 H_brightSkyBlue GRM2
03377 H_brightSkyBlue GRM2
03378 H_brightSkyBlue GRM2
03379 -pf15941 -
03380 HMMreg00486_2_GRM2(pf00486) GRM2
03381 HMMreg00440_1_GRM2(pf00440) GRM2
03382 HMM03358reduct_1_eutgrm(pf03358) GRM2
03383 -pf13416 -
03384 -pf00528 -
03385 -pf00528 -
03386 -pf00005pf08402 -
03387 -pf05681pf05683 -
03388 -pf01522 -
BMC-P
BMC-H
BMC-T
Aldehyde dehydrogenase
PTAC
Signature enzyme
Alcohol dehydrogenase
Regulator

Kluyvera intermedia (Enterobacter intermedius). Bacteria; Proteobacteria; Gammaproteobacteria; Enterobacterales;
NCTC12125__locus_1
GRM2

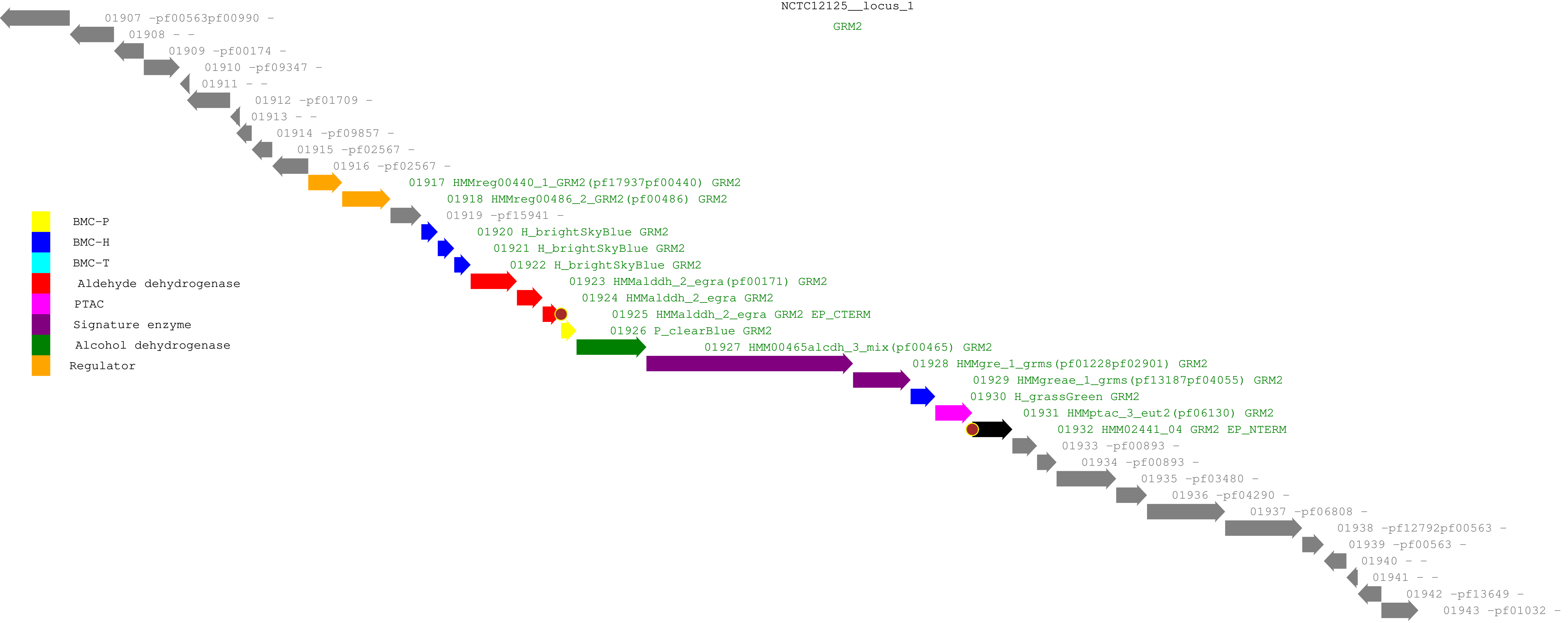

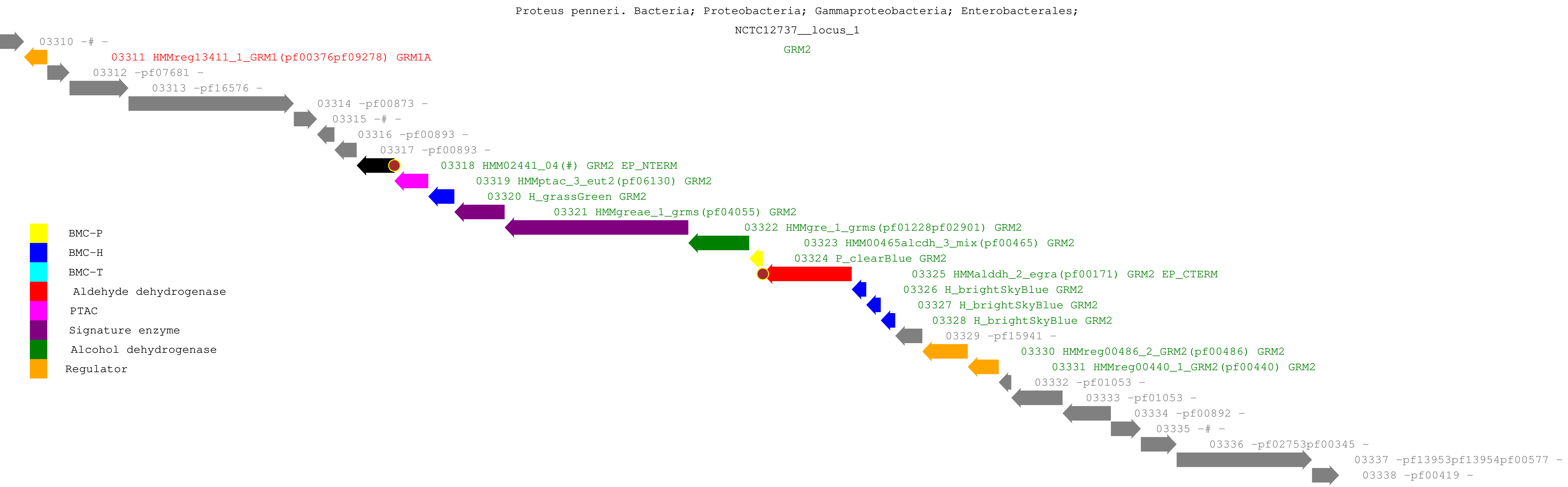

Proteus penneri. Bacteria; Proteobacteria; Gammaproteobacteria; Enterobacterales;
NCTC12737__locus_1
GRM2
03310 -# -
03311 HMMreg13411_1_GRM1(pf00376pf09278) GRM1A
03312 -pf07681 -
03313 -pf16576 -
03314 -pf00873 -
03315 -# -
03316 -pf00893 -
03317 -pf00893 -
03318 HMM02441_04(#) GRM2 EP_NTERM
03319 HMMptac_3_eut2(pf06130) GRM2
03320 H_grassGreen GRM2
03321 HMMgreae_1_grms(pf04055) GRM2
03322 HMMgre_1_grms(pf01228pf02901) GRM2
03323 HMM00465alcdh_3_mix(pf00465) GRM2
03324 P_clearBlue GRM2
03325 HMMaldh_2_egra(pf00171) GRM2 EP_CTERM
03326 H_brightSkyBlue GRM2
03327 H_brightSkyBlue GRM2
03328 H_brightSkyBlue GRM2
03329 -pf15941 -
03330 HMMreg00486_2_GRM2(pf00486) GRM2
03331 HMMreg00440_1_GRM2(pf00440) GRM2
03332 -pf01053 -
03333 -pf01053 -
03334 -pf00892 -
03335 -# -
03336 -pf02753pf00345 -
03337 -pf13953pf13954pf00577 -
03338 -pf00419 -
BMC-P
BMC-H
BMC-T
Aldehyde dehydrogenase
PTAC
Signature enzyme
Alcohol dehydrogenase
Regulator

Serratia rubidaea (Serratia marinorubra). Bacteria; Proteobacteria; Gammaproteobacteria; Enterobacterales;

NCTC12971__locus_1

GRM2

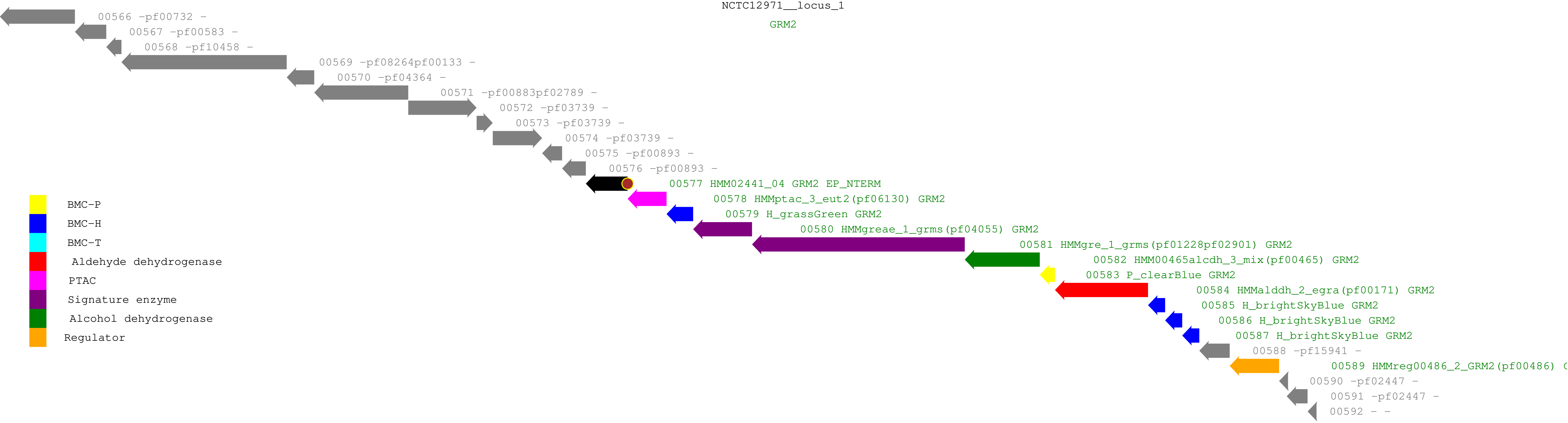

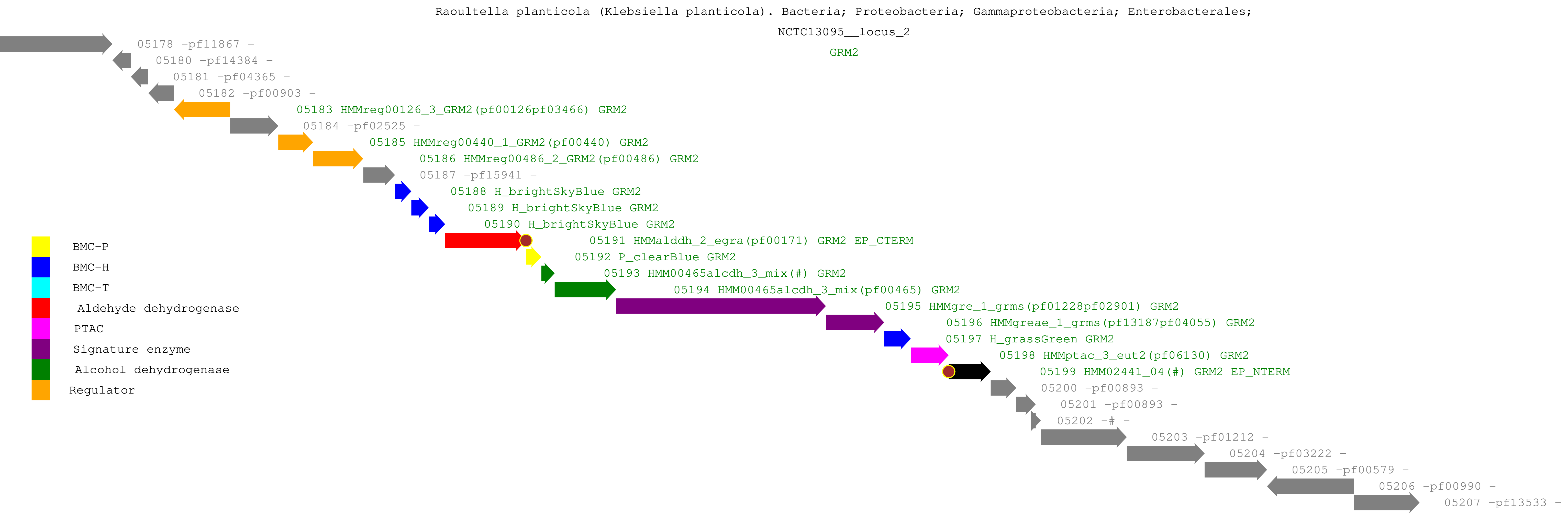

Raoultella planticola (Klebsiella planticola). Bacteria; Proteobacteria; Gammaproteobacteria; Enterobacterales;
NCTC13095__locus_2
GRM2
05178 -pf11867 -
05180 -pf14384 -
05181 -pf04365 -
05182 -pf00903 -
05183 HMMreg00126_3_GRM2(pf00126pf03466) GRM2
05184 -pf02525 -
05185 HMMreg00440_1_GRM2(pf00440) GRM2
05186 HMMreg00486_2_GRM2(pf00486) GRM2
05187 -pf15941 -
05188 H_brightSkyBlue GRM2
05189 H_brightSkyBlue GRM2
05190 H_brightSkyBlue GRM2
05191 HMMalddh_2_egra(pf00171) GRM2 EP_CTERM
05192 P_clearBlue GRM2
05193 HMM00465alcdh_3_mix(#) GRM2
05194 HMM00465alcdh_3_mix(pf00465) GRM2
05195 HMMgre_1_grms(pf01228pf02901) GRM2
05196 HMMgreae_1_grms(pf13187pf04055) GRM2
05197 H_grassGreen GRM2
05198 HMMptac_3_eut2(pf06130) GRM2
05199 HMM02441_04(#) GRM2 EP_NTERM
05200 -pf00893 -
05201 -pf00893 -
05202 -# -
05203 -pf01212 -
05204 -pf03222 -
05205 -pf00579 -
05206 -pf00990 -
05207 -pf13533 -
BMC-P
BMC-H
BMC-T
Aldehyde dehydrogenase
PTAC
Signature enzyme
Alcohol dehydrogenase
Regulator

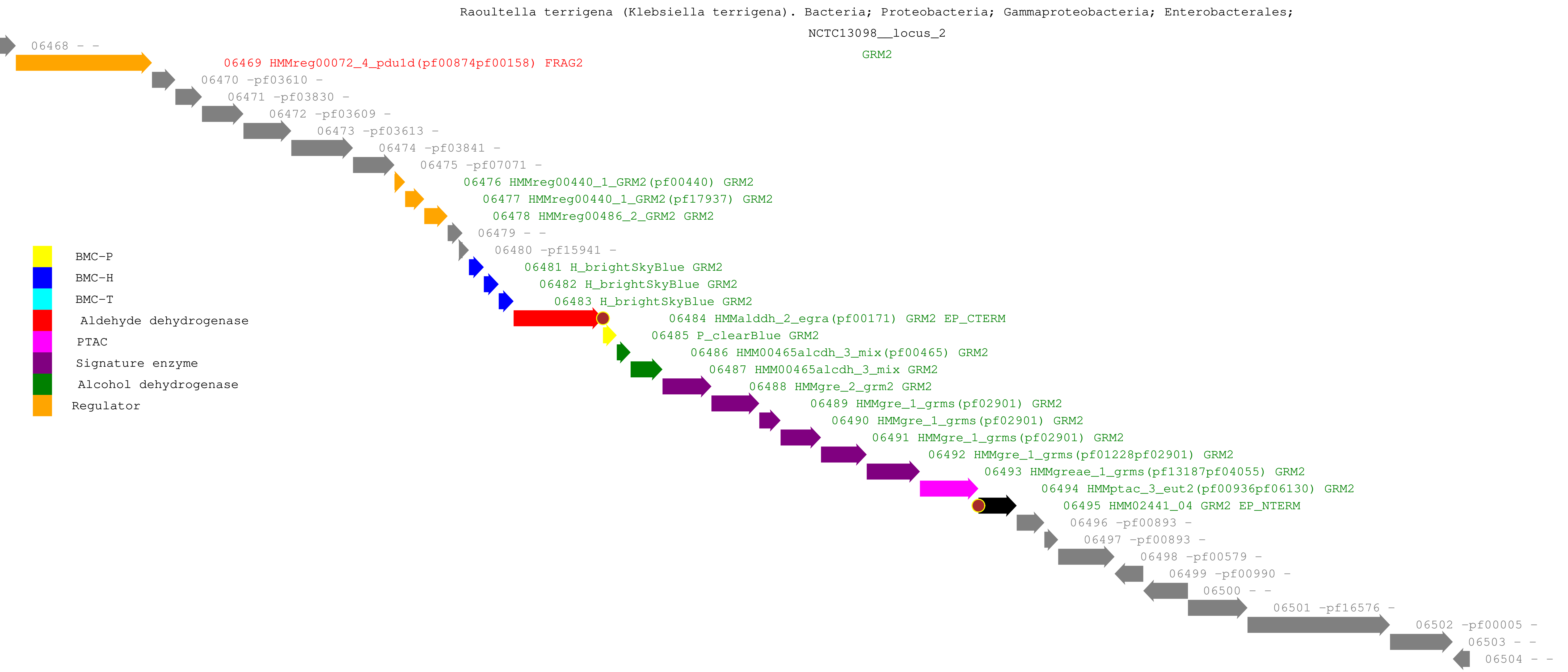
Raoultella terrigena (Klebsiella terrigena). Bacteria; Proteobacteria; Gammaproteobacteria; Enterobacterales;
NCTC13098__locus_2
GRM2
06468 – –
06469 HMMreg00072_4_pdu1d(pf00874pf00158) FRAG2
06470 –pf03610 –
06471 –pf03830 –
06472 –pf03609 –
06473 –pf03613 –
06474 –pf03841 –
06475 –pf07071 –
06476 HMMreg00440_1_GRM2(pf00440) GRM2
06477 HMMreg00440_1_GRM2(pf17937) GRM2
06478 HMMreg00486_2_GRM2 GRM2
06479 – –
06480 –pf15941 –
06481 H_brightSkyBlue GRM2
06482 H_brightSkyBlue GRM2
06483 H_brightSkyBlue GRM2
06484 HMMaldh_2_egra(pf00171) GRM2 EP_CTERM
06485 P_clearBlue GRM2
06486 HMM00465alcdh_3_mix(pf00465) GRM2
06487 HMM00465alcdh_3_mix GRM2
06488 HMMgre_2_grm2 GRM2
06489 HMMgre_1_grms(pf02901) GRM2
06490 HMMgre_1_grms(pf02901) GRM2
06491 HMMgre_1_grms(pf02901) GRM2
06492 HMMgre_1_grms(pf01228pf02901) GRM2
06493 HMMgreae_1_grms(pf13187pf04055) GRM2
06494 HMMptac_3_eut2(pf00936pf06130) GRM2
06495 HMM02441_04 GRM2 EP_NTERM
06496 –pf00893 –
06497 –pf00893 –
06498 –pf00579 –
06499 –pf00990 –
06500 – –
06501 –pf16576 –
06502 –pf00005 –
06503 – –
06504 – –
BMC-P
BMC-H
BMC-T
Aldehyde dehydrogenase
PTAC
Signature enzyme
Alcohol dehydrogenase
Regulator

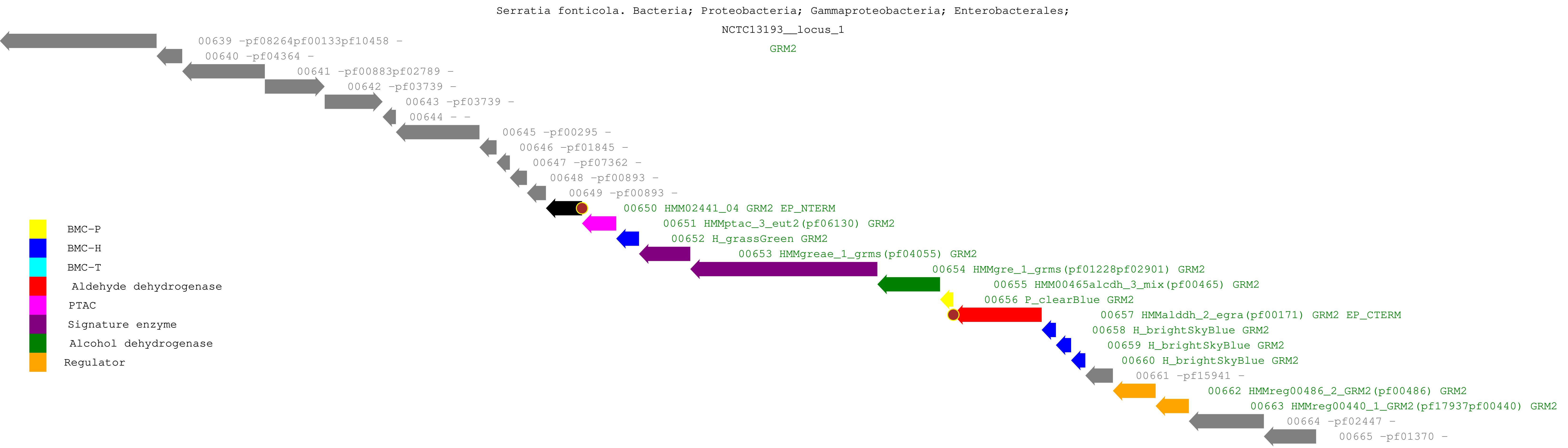

Serratia fonticola. Bacteria; Proteobacteria; Gammaproteobacteria; Enterobacterales;
NCTC13193__locus_1
GRM2
00639 -pf08264pf00133pf10458 -
00640 -pf04364 -
00641 -pf00883pf02789 -
00642 -pf03739 -
00643 -pf03739 -
00644 - -
00645 -pf00295 -
00646 -pf01845 -
00647 -pf07362 -
00648 -pf00893 -
00649 -pf00893 -
00650 HMM02441_04 GRM2 EP_NTERM
00651 HMMptac_3_eut2(pf06130) GRM2
00652 H_grassGreen GRM2
00653 HMMgreae_1_grms(pf04055) GRM2
00654 HMMgre_1_grms(pf01228pf02901) GRM2
00655 HMM00465alcdh_3_mix(pf00465) GRM2
00656 P_clearBlue GRM2
00657 HMMalddh_2_egra(pf00171) GRM2 EP_CTERM
00658 H_brightSkyBlue GRM2
00659 H_brightSkyBlue GRM2
00660 H_brightSkyBlue GRM2
00661 -pf15941 -
00662 HMMreg00486_2_GRM2(pf00486) GRM2
00663 HMMreg00440_1_GRM2(pf17937pf00440) GRM2
00664 -pf02447 -
00665 -pf01370 -
BMC-P
BMC-H
BMC-T
Aldehyde dehydrogenase
PTAC
Signature enzyme
Alcohol dehydrogenase
Regulator

Klebsiella pneumoniae subsp. rhinoscleromatis. Bacteria; Proteobacteria; Gammaproteobacteria; Enterobacterales;

NCTC5046__locus_1

GRM2

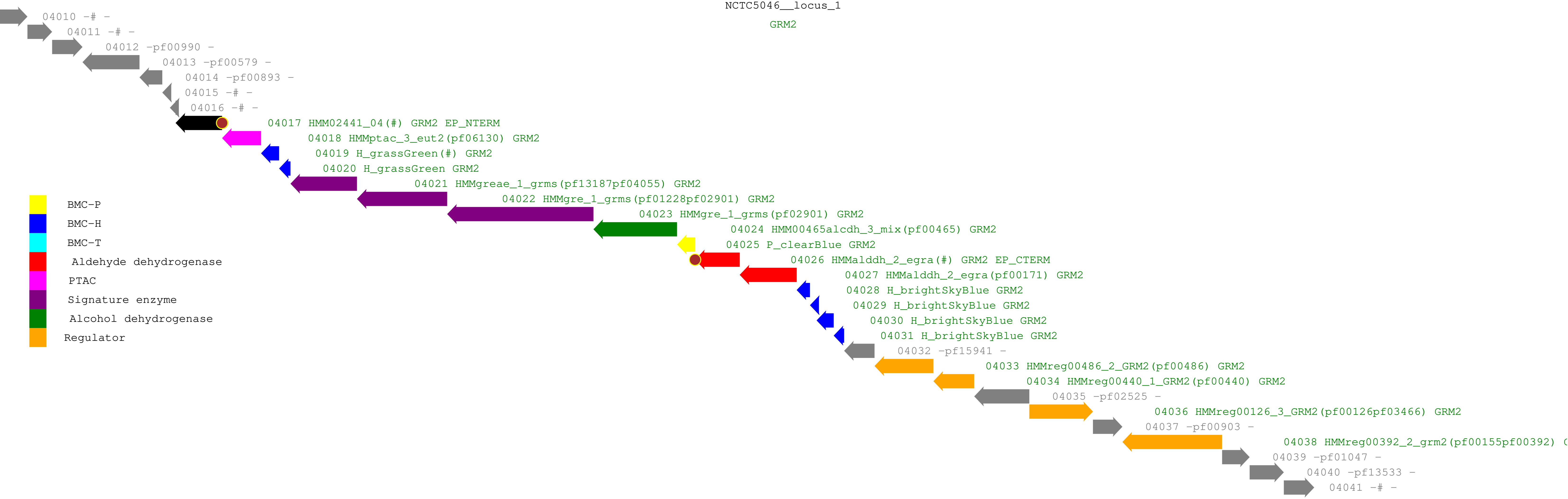

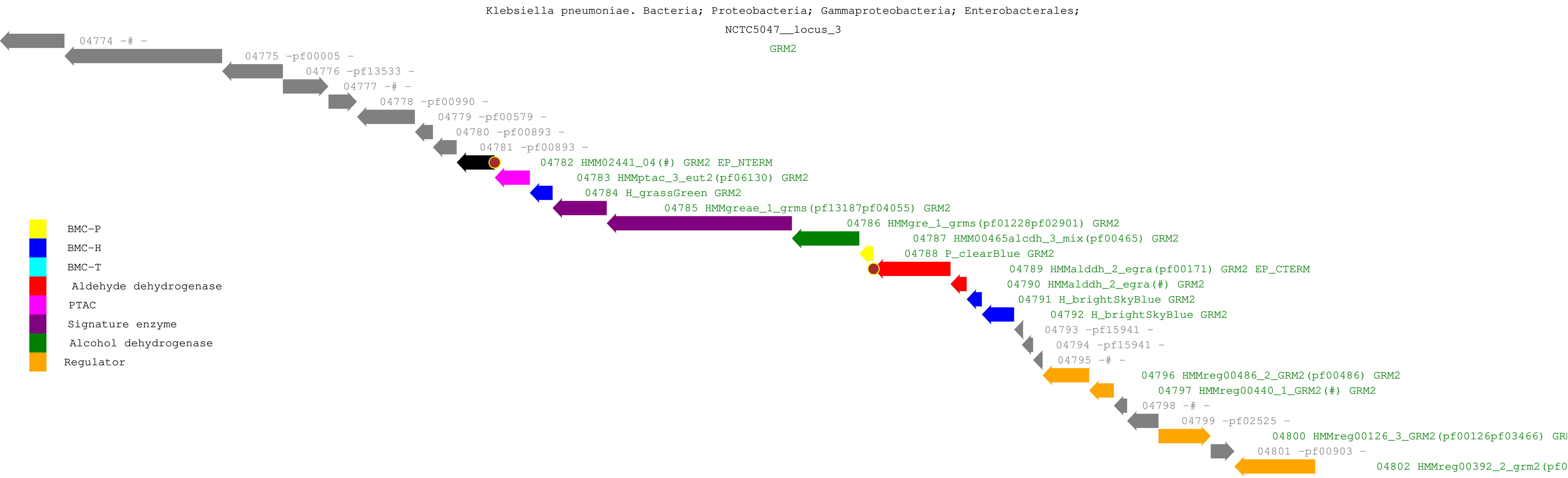
Klebsiella pneumoniae. Bacteria; Proteobacteria; Gammaproteobacteria; Enterobacterales;
NCTC5047__locus_3
GRM2
04774 -# -
04775 -pf00005 -
04776 -pf13533 -
04777 -# -
04778 -pf00990 -
04779 -pf00579 -
04780 -pf00893 -
04781 -pf00893 -
04782 HMM02441_04(#) GRM2 EP_NTERM
04783 HMMptac_3_eut2(pf06130) GRM2
04784 H_grassGreen GRM2
04785 HMMgreae_1_grms(pf13187pf04055) GRM2
04786 HMMgre_1_grms(pf01228pf02901) GRM2
04787 HMM00465alcdh_3_mix(pf00465) GRM2
04788 P_clearBlue GRM2
04789 HMMaldh_2_egra(pf00171) GRM2 EP_CTERM
04790 HMMaldh_2_egra(#) GRM2
04791 H_brightSkyBlue GRM2
04792 H_brightSkyBlue GRM2
04793 -pf15941 -
04794 -pf15941 -
04795 -# -
04796 HMMreg00486_2_GRM2(pf00486) GRM2
04797 HMMreg00440_1_GRM2(#) GRM2
04798 -# -
04799 -pf02525 -
04800 HMMreg00126_3_GRM2(pf00126pf03466) GR
04801 -pf00903 -
04802 HMMreg00392_2_grm2(pf0
BMC-P
BMC-H
BMC-T
Aldehyde dehydrogenase
PTAC
Signature enzyme
Alcohol dehydrogenase
Regulator

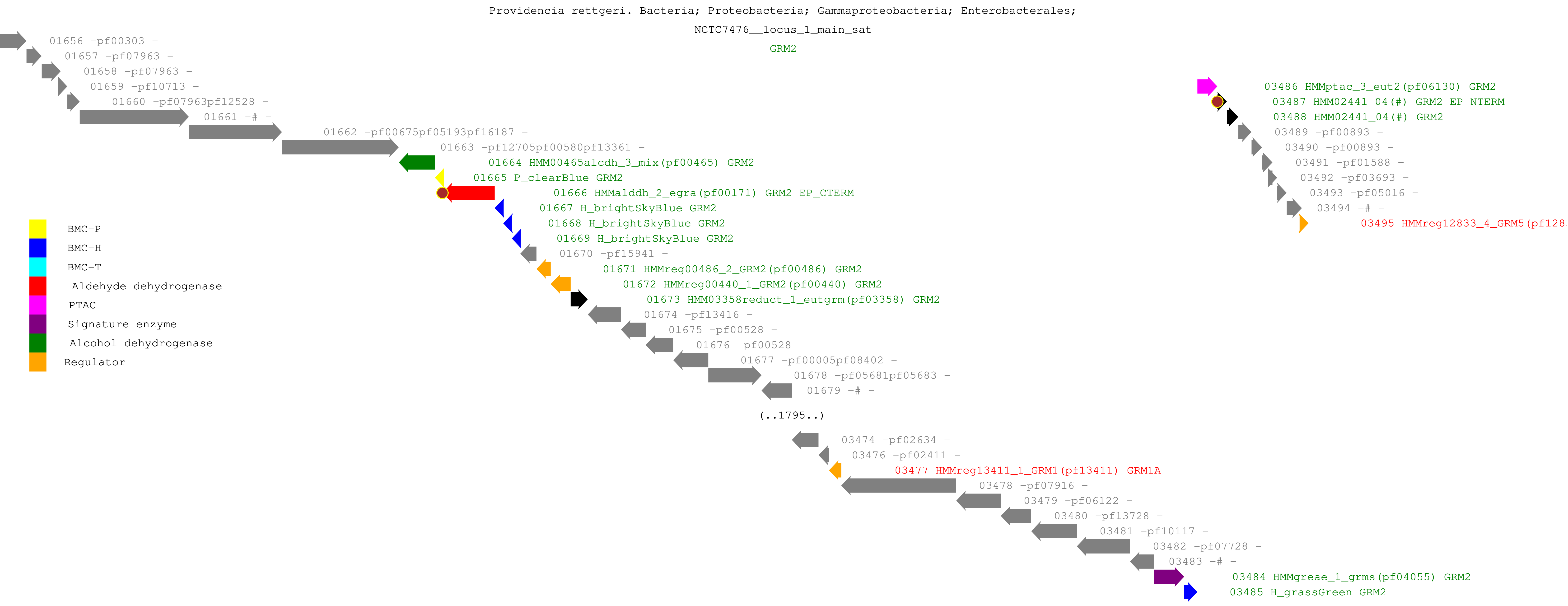
Providencia rettgeri. Bacteria; Proteobacteria; Gammaproteobacteria; Enterobacterales;
NCTC7476__locus_1_main_sat
GRM2
01656 -pf00303 -
01657 -pf07963 -
01658 -pf07963 -
01659 -pf10713 -
01660 -pf07963pf12528 -
01661 -# -
01662 -pf00675pf05193pf16187 -
01663 -pf12705pf00580pf13361 -
01664 HMM00465alcdh_3_mix(pf00465) GRM2
01665 P_clearBlue GRM2
01666 HMMalddh_2_egra(pf00171) GRM2 EP_CTERM
01667 H_brightSkyBlue GRM2
01668 H_brightSkyBlue GRM2
01669 H_brightSkyBlue GRM2
01670 -pf15941 -
01671 HMMreg00486_2_GRM2(pf00486) GRM2
01672 HMMreg00440_1_GRM2(pf00440) GRM2
01673 HMM03358reduct_1_eutgrm(pf03358) GRM2
01674 -pf13416 -
01675 -pf00528 -
01676 -pf00528 -
01677 -pf00005pf08402 -
01678 -pf05681pf05683 -
01679 -# -
(..1795..)
03474 -pf02634 -
03476 -pf02411 -
03477 HMMreg13411_1_GRM1(pf13411) GRM1A
03478 -pf07916 -
03479 -pf06122 -
03480 -pf13728 -
03481 -pf10117 -
03482 -pf07728 -
03483 -# -
03484 HMMgreae_1_grms(pf04055) GRM2
03485 H_grassGreen GRM2
03486 HMMptac_3_eut2(pf06130) GRM2
03487 HMM02441_04(#) GRM2 EP_NTERM
03488 HMM02441_04(#) GRM2
03489 -pf00893 -
03490 -pf00893 -
03491 -pf01588 -
03492 -pf03693 -
03493 -pf05016 -
03494 -# -
03495 HMMreg12833_4_GRM5(pf128
BMC-P
BMC-H
BMC-T
Aldehyde dehydrogenase
PTAC
Signature enzyme
Alcohol dehydrogenase
Regulator

Escherichia coli. Bacteria; Proteobacteria; Gammaproteobacteria; Enterobacterales;

NCTC7928__locus_2

GRM2

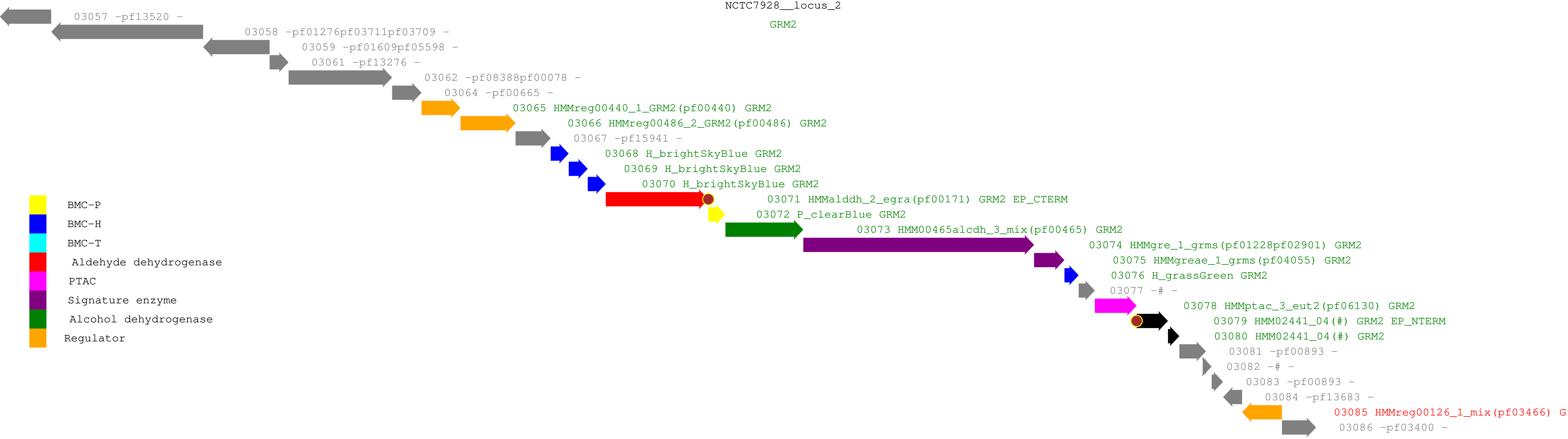

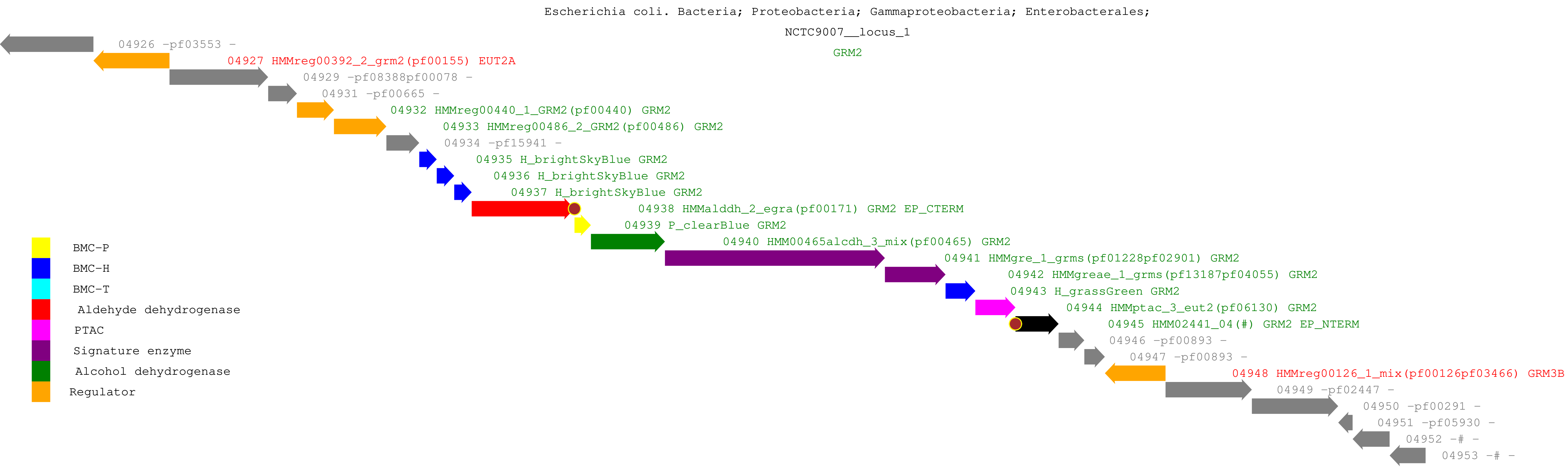

Escherichia coli. Bacteria; Proteobacteria; Gammaproteobacteria; Enterobacterales;
NCTC9007__locus_1
GRM2
04926 -pf03553 -
04927 HMMreg00392_2_grm2(pf00155) EUT2A
04929 -pf08388pf00078 -
04931 -pf00665 -
04932 HMMreg00440_1_GRM2(pf00440) GRM2
04933 HMMreg00486_2_GRM2(pf00486) GRM2
04934 -pf15941 -
04935 H_brightSkyBlue GRM2
04936 H_brightSkyBlue GRM2
04937 H_brightSkyBlue GRM2
04938 HMMalddh_2_egra(pf00171) GRM2 EP_CTERM
04939 P_clearBlue GRM2
04940 HMM00465alcdh_3_mix(pf00465) GRM2
04941 HMMgre_1_grms(pf01228pf02901) GRM2
04942 HMMgreae_1_grms(pf13187pf04055) GRM2
04943 H_grassGreen GRM2
04944 HMMptac_3_eut2(pf06130) GRM2
04945 HMM02441_04(#) GRM2 EP_NTERM
04946 -pf00893 -
04947 -pf00893 -
04948 HMMreg00126_1_mix(pf00126pf03466) GRM3B
04949 -pf02447 -
04950 -pf00291 -
04951 -pf05930 -
04952 -# -
04953 -# -
BMC-P
BMC-H
BMC-T
Aldehyde dehydrogenase
PTAC
Signature enzyme
Alcohol dehydrogenase
Regulator

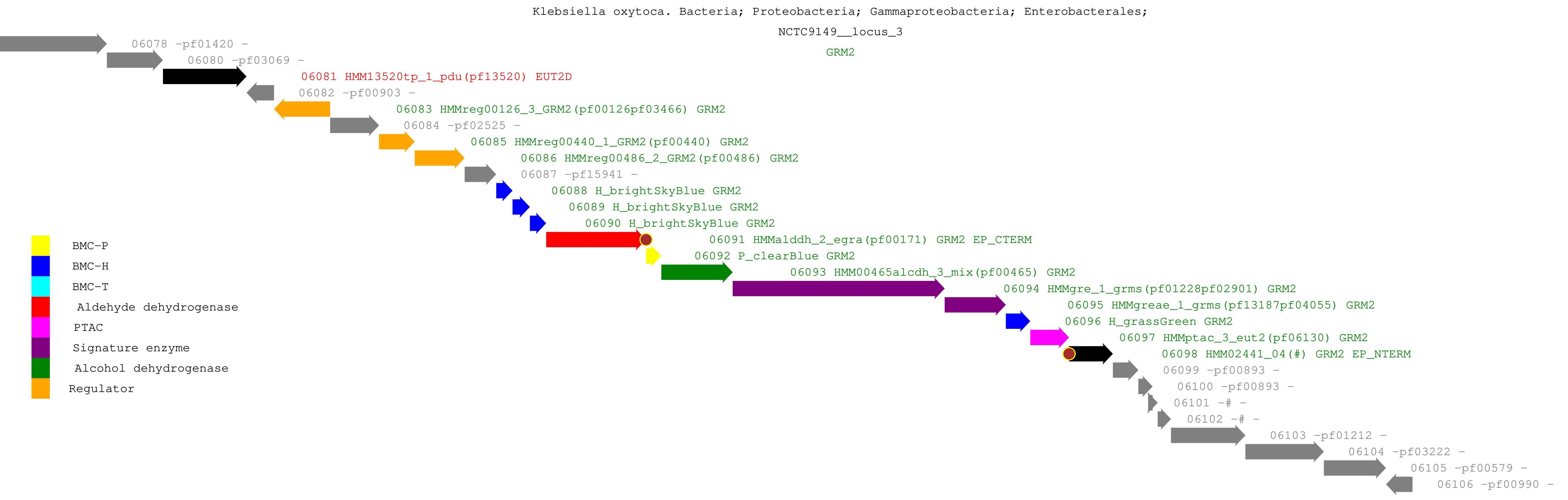
Klebsiella oxytoca. Bacteria; Proteobacteria; Gammaproteobacteria; Enterobacterales;
NCTC9149__locus_3
GRM2
06078 -pf01420 -
06080 -pf03069 -
06081 HMM13520tp_1_pdu(pf13520) EUT2D
06082 -pf00903 -
06083 HMMreg00126_3_GRM2(pf00126pf03466) GRM2
06084 -pf02525 -
06085 HMMreg00440_1_GRM2(pf00440) GRM2
06086 HMMreg00486_2_GRM2(pf00486) GRM2
06087 -pf15941 -
06088 H_brightSkyBlue GRM2
06089 H_brightSkyBlue GRM2
06090 H_brightSkyBlue GRM2
06091 HMMaldd h_2_egra(pf00171) GRM2 EP_CTERM
06092 P_clearBlue GRM2
06093 HMM00465alcdh_3_mix(pf00465) GRM2
06094 HMMgre_1_grms(pf01228pf02901) GRM2
06095 HMMgreae_1_grms(pf13187pf04055) GRM2
06096 H_grassGreen GRM2
06097 HMMptac_3_eut2(pf06130) GRM2
06098 HMM02441_04(#) GRM2 EP_NTERM
06099 -pf00893 -
06100 -pf00893 -
06101 -# -
06102 -# -
06103 -pf01212 -
06104 -pf03222 -
06105 -pf00579 -
06106 -pf00990 -
BMC-P
BMC-H
BMC-T
Aldehyde dehydrogenase
PTAC
Signature enzyme
Alcohol dehydrogenase
Regulator

Klebsiella pneumoniae. Bacteria; Proteobacteria; Gammaproteobacteria; Enterobacterales;

NCTC9177__locus_1

GRM2

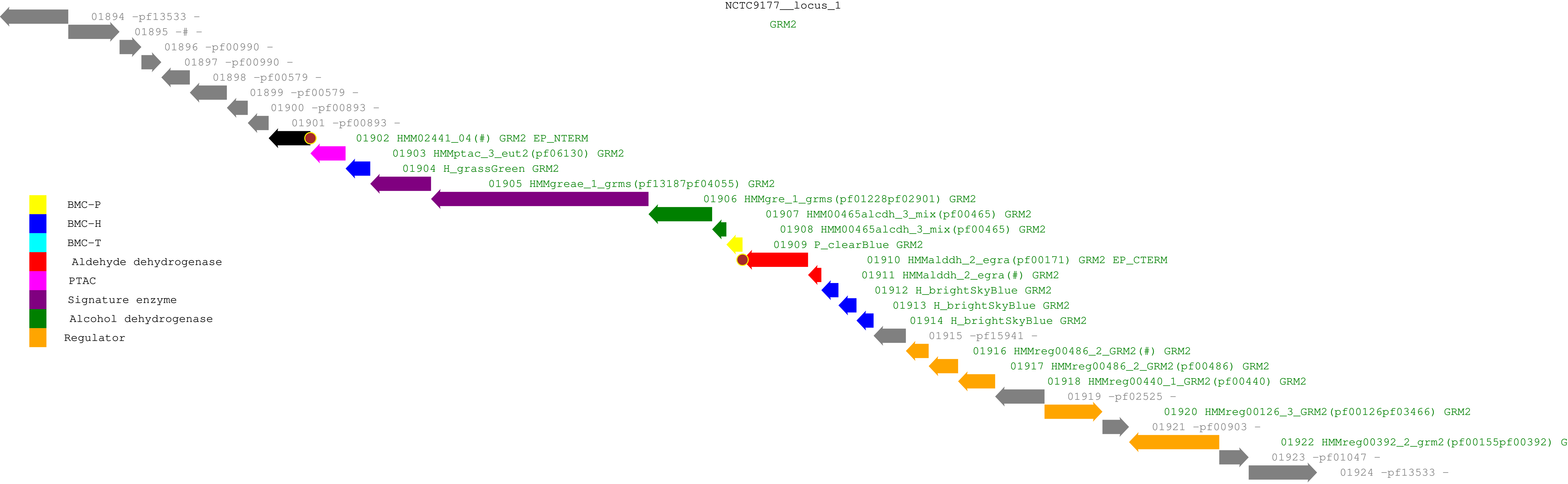

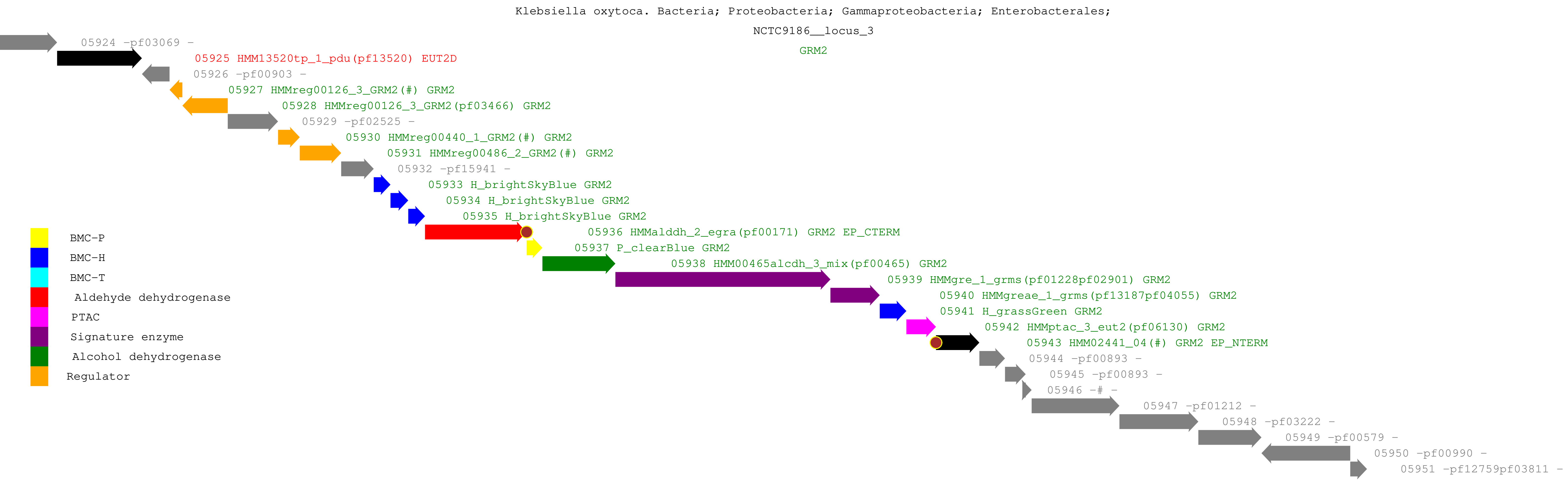
Klebsiella oxytoca. Bacteria; Proteobacteria; Gammaproteobacteria; Enterobacterales;
NCTC9186__locus_3
GRM2
05924 -pf03069 -
05925 HMM13520tp_1_pdu(pf13520) EUT2D
05926 -pf00903 -
05927 HMMreg00126_3_GRM2(#) GRM2
05928 HMMreg00126_3_GRM2(pf03466) GRM2
05929 -pf02525 -
05930 HMMreg00440_1_GRM2(#) GRM2
05931 HMMreg00486_2_GRM2(#) GRM2
05932 -pf15941 -
05933 H_brightSkyBlue GRM2
05934 H_brightSkyBlue GRM2
05935 H_brightSkyBlue GRM2
05936 HMMalddh_2_egra(pf00171) GRM2 EP_CTERM
05937 P_clearBlue GRM2
05938 HMM00465alcdh_3_mix(pf00465) GRM2
05939 HMMgre_1_grms(pf01228pf02901) GRM2
05940 HMMgreae_1_grms(pf13187pf04055) GRM2
05941 H_grassGreen GRM2
05942 HMMptac_3_eut2(pf06130) GRM2
05943 HMM02441_04(#) GRM2 EP_NTERM
05944 -pf00893 -
05945 -pf00893 -
05946 -# -
05947 -pf01212 -
05948 -pf03222 -
05949 -pf00579 -
05950 -pf00990 -
05951 -pf12759pf03811 -
BMC-P
BMC-H
BMC-T
Aldehyde dehydrogenase
PTAC
Signature enzyme
Alcohol dehydrogenase
Regulator

Raoultella sp. NCTC 9189. Bacteria; Proteobacteria; Gammaproteobacteria; Enterobacterales;

NCTC9189__locus_1

GRM2

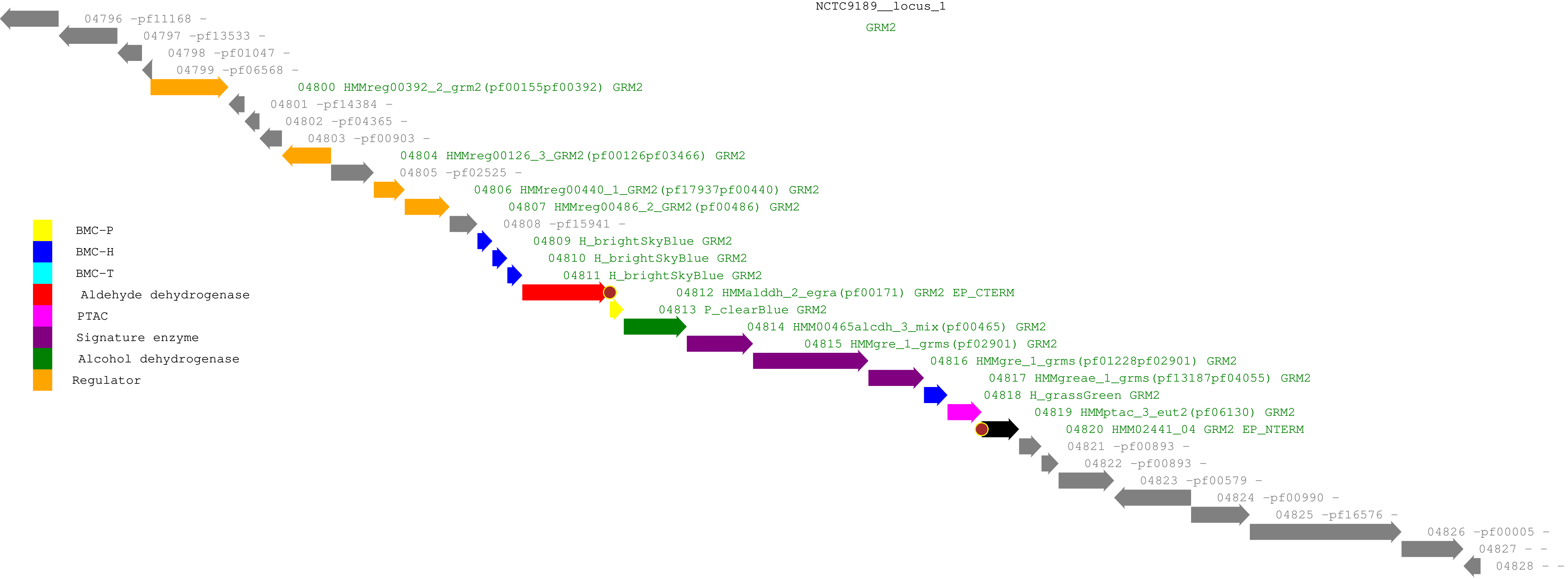

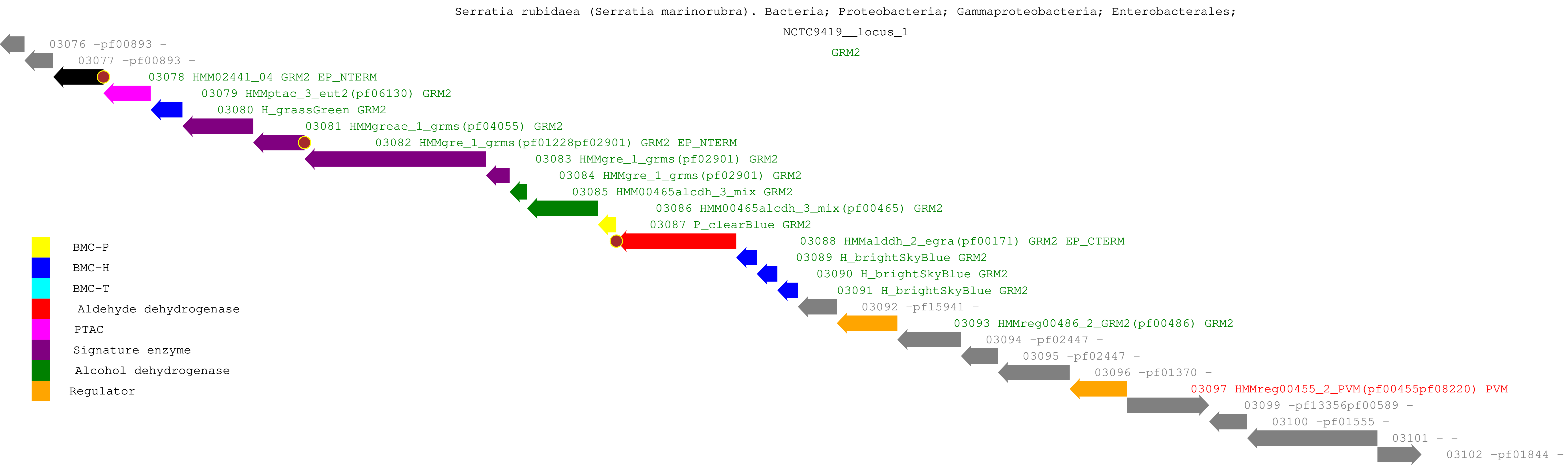
Serratia rubidaea (Serratia marinorubra). Bacteria; Proteobacteria; Gammaproteobacteria; Enterobacterales;
NCTC9419__locus_1
GRM2
03076 -pf00893 -
03077 -pf00893 -
03078 HMM02441_04 GRM2 EP_NTERM
03079 HMMptac_3_eut2(pf06130) GRM2
03080 H_grassGreen GRM2
03081 HMMgreae_1_grms(pf04055) GRM2
03082 HMMgre_1_grms(pf01228pf02901) GRM2 EP_NTERM
03083 HMMgre_1_grms(pf02901) GRM2
03084 HMMgre_1_grms(pf02901) GRM2
03085 HMM00465alcdh_3_mix GRM2
03086 HMM00465alcdh_3_mix(pf00465) GRM2
03087 P_clearBlue GRM2
03088 HMMalddh_2_egra(pf00171) GRM2 EP_CTERM
03089 H_brightSkyBlue GRM2
03090 H_brightSkyBlue GRM2
03091 H_brightSkyBlue GRM2
03092 -pf15941 -
03093 HMMreg00486_2_GRM2(pf00486) GRM2
03094 -pf02447 -
03095 -pf02447 -
03096 -pf01370 -
03097 HMMreg00455_2_PVM(pf00455pf08220) PVM
03099 -pf13356pf00589 -
03100 -pf01555 -
03101 - -
03102 -pf01844 -
BMC-P
BMC-H
BMC-T
Aldehyde dehydrogenase
PTAC
Signature enzyme
Alcohol dehydrogenase
Regulator

Klebsiella aerogenes (Enterobacter aerogenes). Bacteria; Proteobacteria; Gammaproteobacteria; Enterobacterales;

NCTC9499__locus_1

GRM2

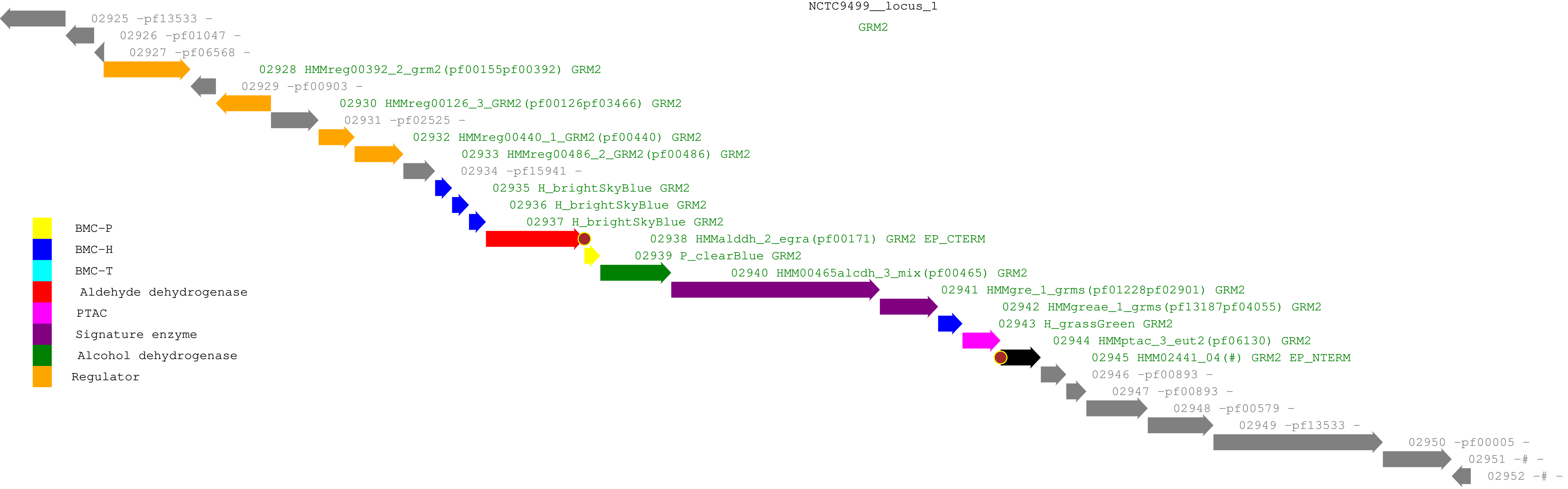

Klebsiella pneumoniae subsp. pneumoniae. Bacteria; Proteobacteria; Gammaproteobacteria; Enterobacterales;

NCTC9504__locus_3

GRM2

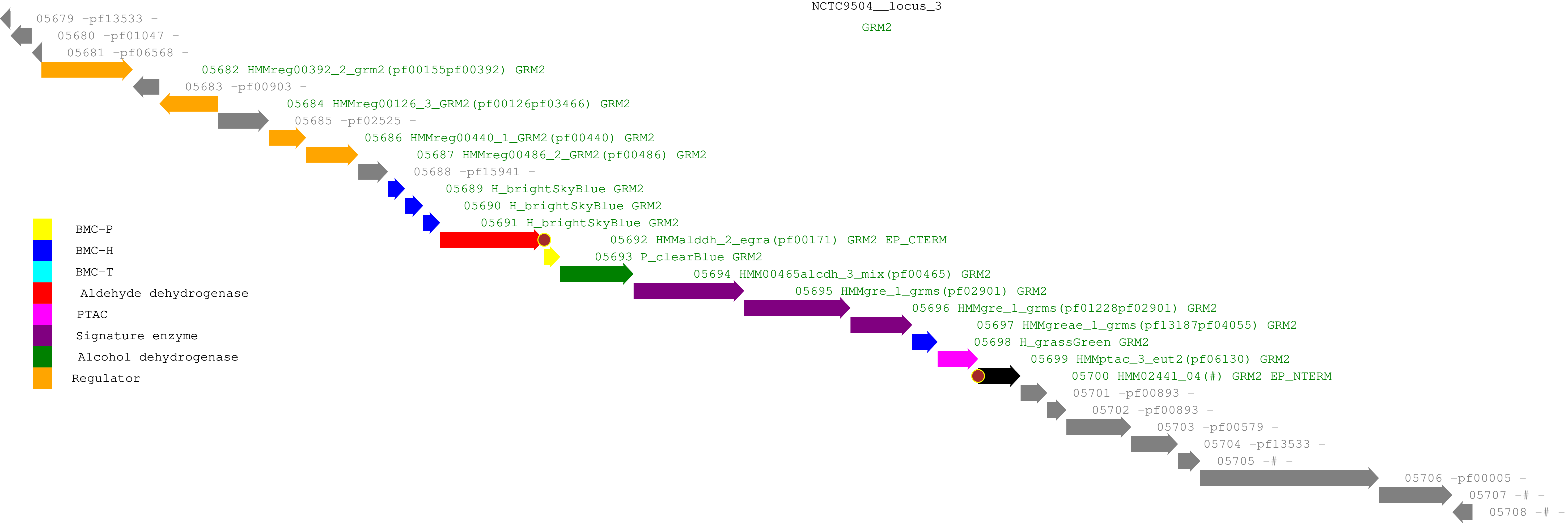

Raoultella planticola (Klebsiella planticola). Bacteria; Proteobacteria; Gammaproteobacteria; Enterobacterales;

NCTC9528__locus_1

GRM2

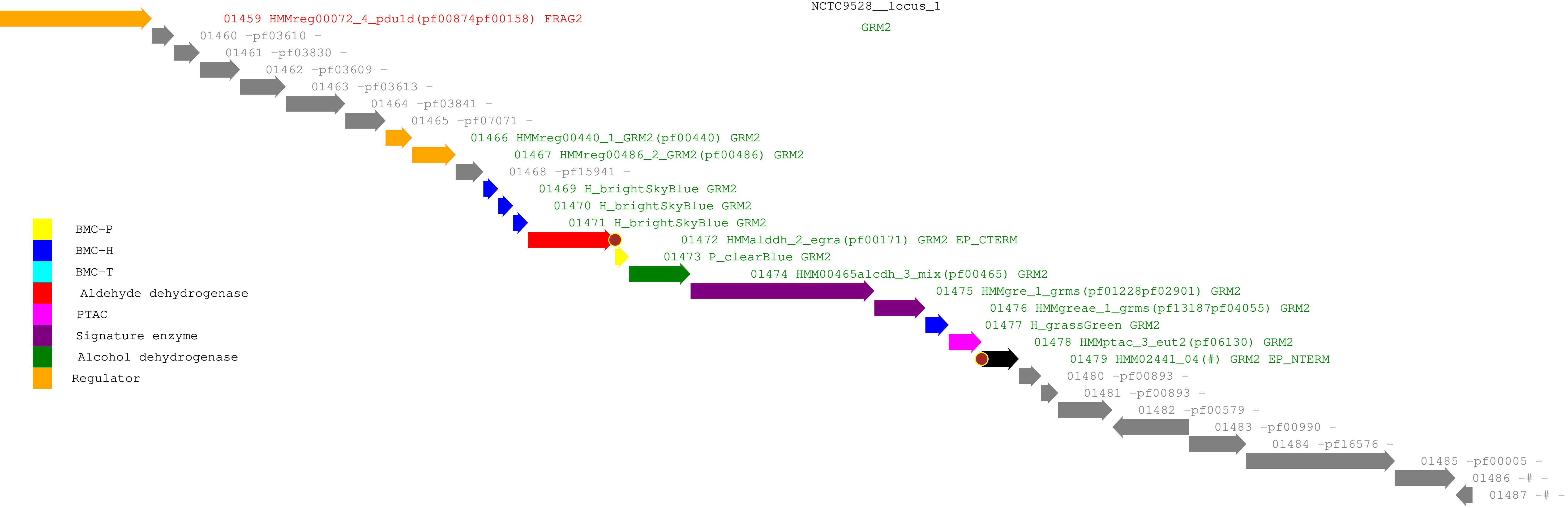

Kluyvera ascorbata. Bacteria; Proteobacteria; Gammaproteobacteria; Enterobacterales;

NCTC9737__locus_1

GRM2

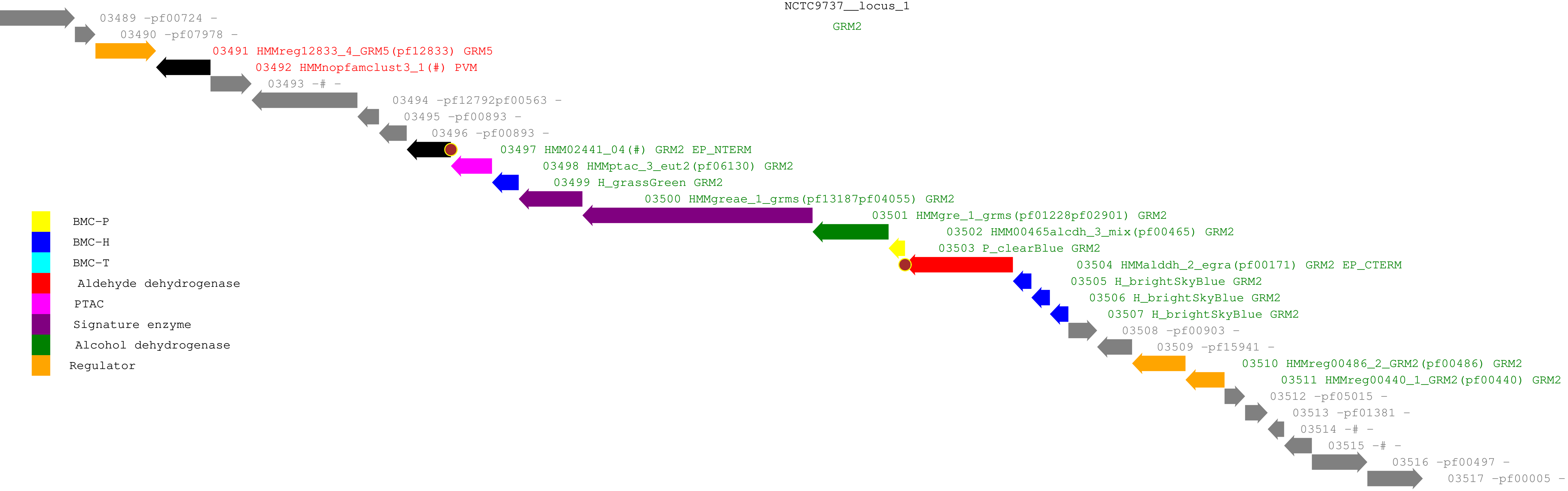

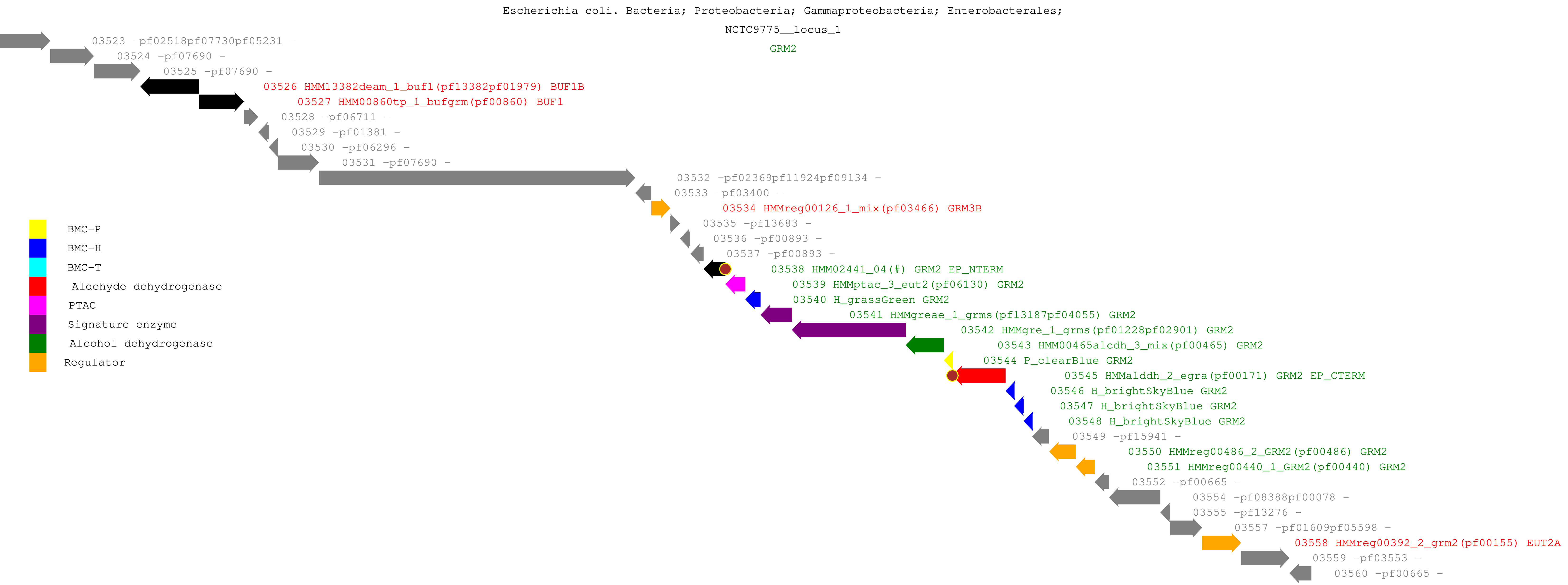

Escherichia coli. Bacteria; Proteobacteria; Gammaproteobacteria; Enterobacterales;
NCTC9775__locus_1
GRM2
03523 -pf02518pf07730pf05231 -
03524 -pf07690 -
03525 -pf07690 -
03526 HMM13382deam_1_buf1(pf13382pf01979) BUF1B
03527 HMM00860tp_1_bufgrm(pf00860) BUF1
03528 -pf06711 -
03529 -pf01381 -
03530 -pf06296 -
03531 -pf07690 -
03532 -pf02369pf11924pf09134 -
03533 -pf03400 -
03534 HMMreg00126_1_mix(pf03466) GRM3B
03535 -pf13683 -
03536 -pf00893 -
03537 -pf00893 -
03538 HMM02441_04(#) GRM2 EP_NTERM
03539 HMMptac_3_eut2(pf06130) GRM2
03540 H_grassGreen GRM2
03541 HMMgreae_1_grms(pf13187pf04055) GRM2
03542 HMMgre_1_grms(pf01228pf02901) GRM2
03543 HMM00465alcdh_3_mix(pf00465) GRM2
03544 P_clearBlue GRM2
03545 HMMalddh_2_egra(pf00171) GRM2 EP_CTERM
03546 H_brightSkyBlue GRM2
03547 H_brightSkyBlue GRM2
03548 H_brightSkyBlue GRM2
03549 -pf15941 -
03550 HMMreg00486_2_GRM2(pf00486) GRM2
03551 HMMreg00440_1_GRM2(pf00440) GRM2
03552 -pf00665 -
03554 -pf08388pf00078 -
03555 -pf13276 -
03557 -pf01609pf05598 -
03558 HMMreg00392_2_grm2(pf00155) EUT2A
03559 -pf03553 -
03560 -pf00665 -
BMC-P
BMC-H
BMC-T
Aldehyde dehydrogenase
PTAC
Signature enzyme
Alcohol dehydrogenase
Regulator

Providencia rettgeri Dmel1. Bacteria; Proteobacteria; Gammaproteobacteria; Enterobacterales;

OOC__locus_1

GRM2

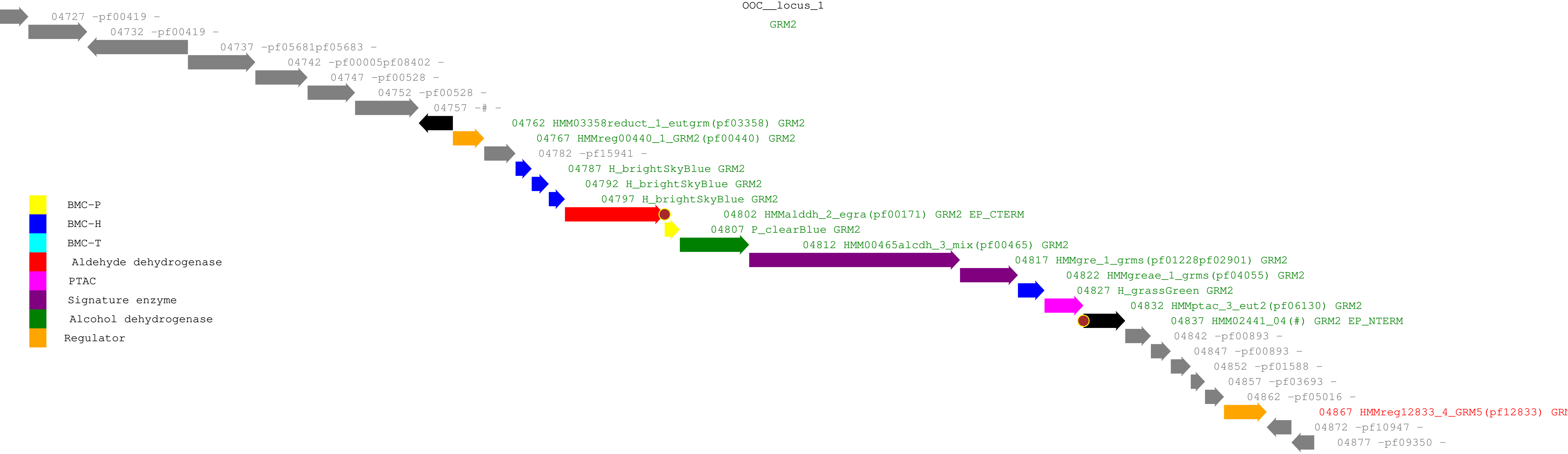

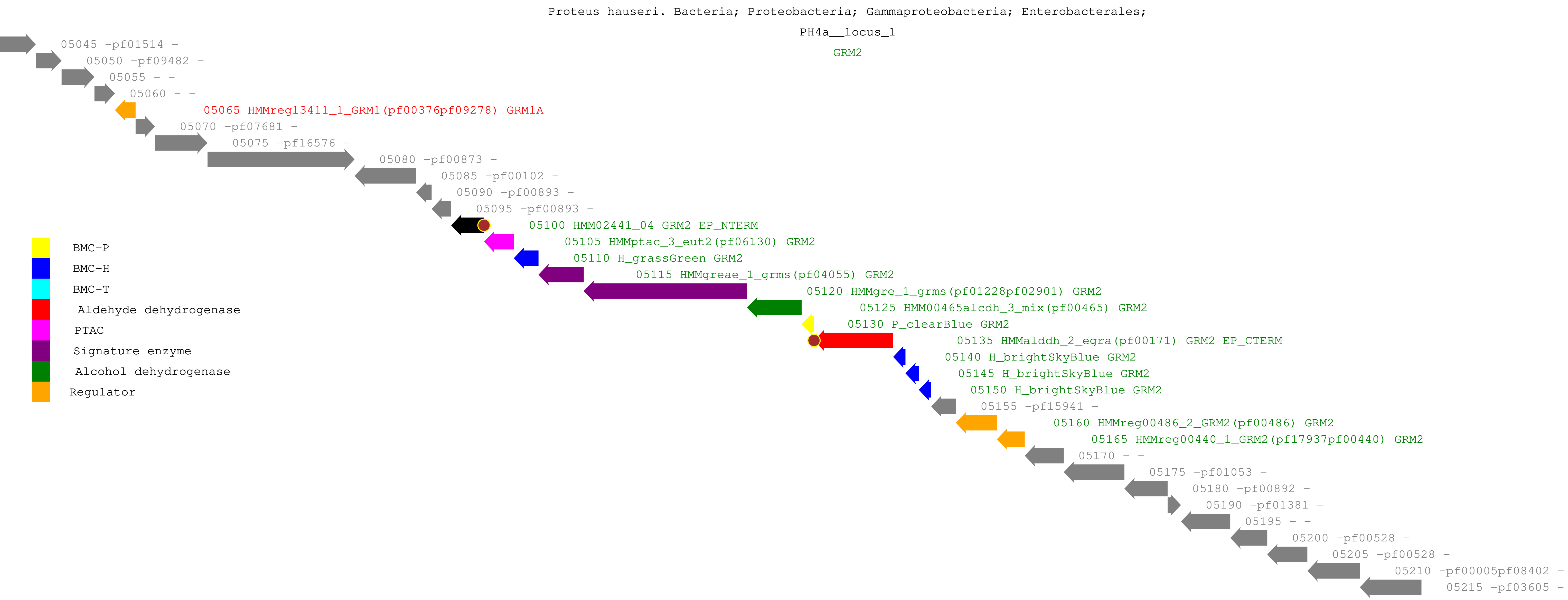
Proteus hauseri. Bacteria; Proteobacteria; Gammaproteobacteria; Enterobacterales;
PH4a__locus_1
GRM2
05045 -pf01514 -
05050 -pf09482 -
05055 - -
05060 - -
05065 HMMreg13411_1_GRM1(pf00376pf09278) GRM1A
05070 -pf07681 -
05075 -pf16576 -
05080 -pf00873 -
05085 -pf00102 -
05090 -pf00893 -
05095 -pf00893 -
05100 HMM02441_04 GRM2 EP_NTERM
05105 HMMptac_3_eut2(pf06130) GRM2
05110 H_grassGreen GRM2
05115 HMMgreae_1_grms(pf04055) GRM2
05120 HMMgre_1_grms(pf01228pf02901) GRM2
05125 HMM00465alcdh_3_mix(pf00465) GRM2
05130 P_clearBlue GRM2
05135 HMMalddh_2_egra(pf00171) GRM2 EP_CTERM
05140 H_brightSkyBlue GRM2
05145 H_brightSkyBlue GRM2
05150 H_brightSkyBlue GRM2
05155 -pf15941 -
05160 HMMreg00486_2_GRM2(pf00486) GRM2
05165 HMMreg00440_1_GRM2(pf17937pf00440) GRM2
05170 - -
05175 -pf01053 -
05180 -pf00892 -
05190 -pf01381 -
05195 - -
05200 -pf00528 -
05205 -pf00528 -
05210 -pf00005pf08402 -
05215 -pf03605 -
BMC-P
BMC-H
BMC-T
Aldehyde dehydrogenase
PTAC
Signature enzyme
Alcohol dehydrogenase
Regulator

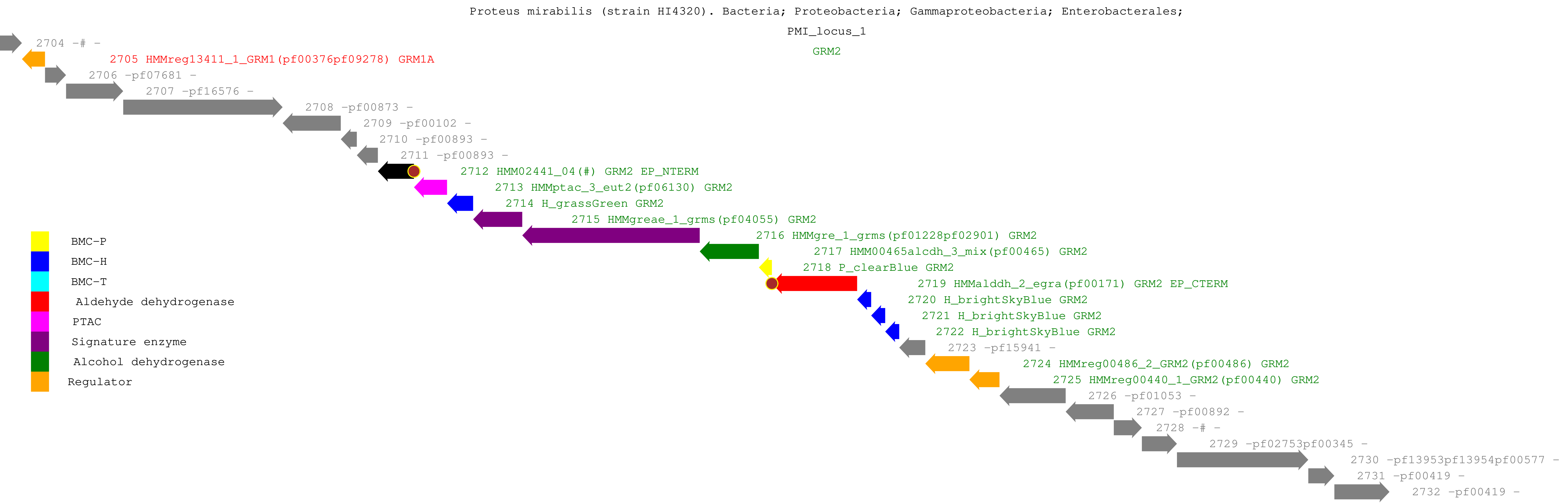
Proteus mirabilis (strain HI4320). Bacteria; Proteobacteria; Gammaproteobacteria; Enterobacterales;
PMI_locus_1
GRM2
2704 -# -
2705 HMMreg13411_1_GRM1(pf00376pf09278) GRM1A
2706 -pf07681 -
2707 -pf16576 -
2708 -pf00873 -
2709 -pf00102 -
2710 -pf00893 -
2711 -pf00893 -
2712 HMM02441_04(#) GRM2 EP_NTERM
2713 HMMptac_3_eut2(pf06130) GRM2
2714 H_grassGreen GRM2
2715 HMMgreae_1_grms(pf04055) GRM2
2716 HMMgre_1_grms(pf01228pf02901) GRM2
2717 HMM00465alcdh_3_mix(pf00465) GRM2
2718 P_clearBlue GRM2
2719 HMMalddh_2_egra(pf00171) GRM2 EP_CTERM
2720 H_brightSkyBlue GRM2
2721 H_brightSkyBlue GRM2
2722 H_brightSkyBlue GRM2
2723 -pf15941 -
2724 HMMreg00486_2_GRM2(pf00486) GRM2
2725 HMMreg00440_1_GRM2(pf00440) GRM2
2726 -pf01053 -
2727 -pf00892 -
2728 -# -
2729 -pf02753pf00345 -
2730 -pf13953pf13954pf00577 -
2731 -pf00419 -
2732 -pf00419 -
BMC-P
BMC-H
BMC-T
Aldehyde dehydrogenase
PTAC
Signature enzyme
Alcohol dehydrogenase
Regulator

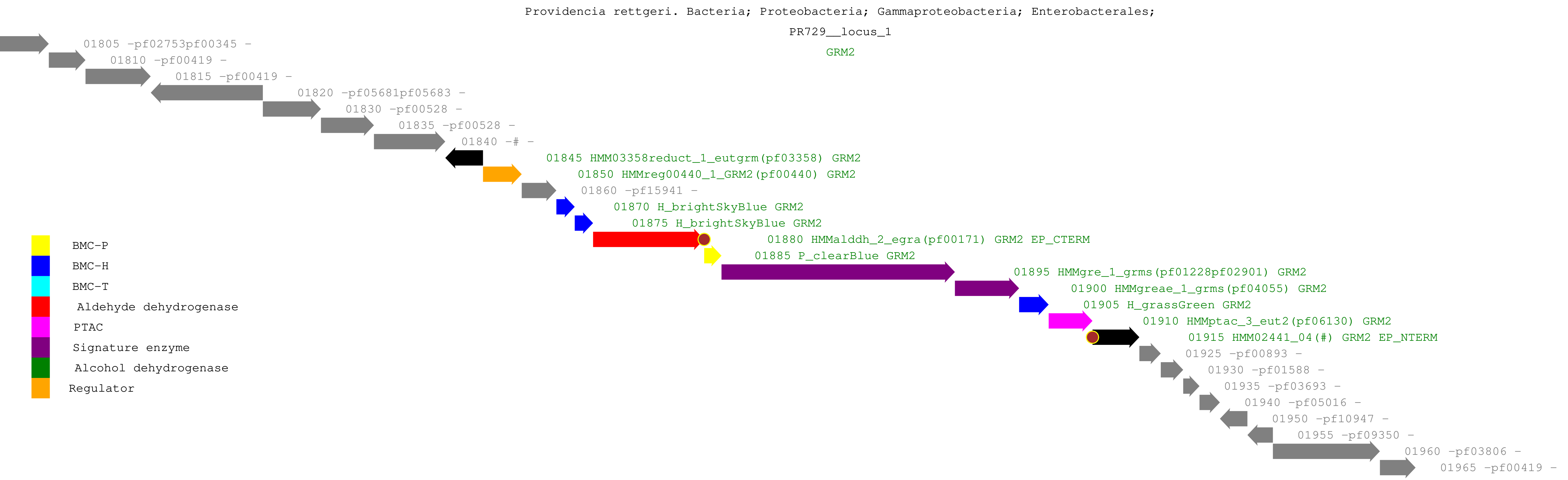

Providencia rettgeri. Bacteria; Proteobacteria; Gammaproteobacteria; Enterobacterales;
PR729__locus_1
GRM2
01805 -pf02753pf00345 -
01810 -pf00419 -
01815 -pf00419 -
01820 -pf05681pf05683 -
01830 -pf00528 -
01835 -pf00528 -
01840 -# -
01845 HMM03358reduct_1_eutgrm(pf03358) GRM2
01850 HMMreg00440_1_GRM2(pf00440) GRM2
01860 -pf15941 -
01870 H_brightSkyBlue GRM2
01875 H_brightSkyBlue GRM2
01880 HMMalddh_2_egra(pf00171) GRM2 EP_CTERM
01885 P_clearBlue GRM2
01895 HMMgre_1_grms(pf01228pf02901) GRM2
01900 HMMgreae_1_grms(pf04055) GRM2
01905 H_grassGreen GRM2
01910 HMMptac_3_eut2(pf06130) GRM2
01915 HMM02441_04(#) GRM2 EP_NTERM
01925 -pf00893 -
01930 -pf01588 -
01935 -pf03693 -
01940 -pf05016 -
01950 -pf10947 -
01955 -pf09350 -
01960 -pf03806 -
01965 -pf00419 -
BMC-P
BMC-H
BMC-T
Aldehyde dehydrogenase
PTAC
Signature enzyme
Alcohol dehydrogenase
Regulator

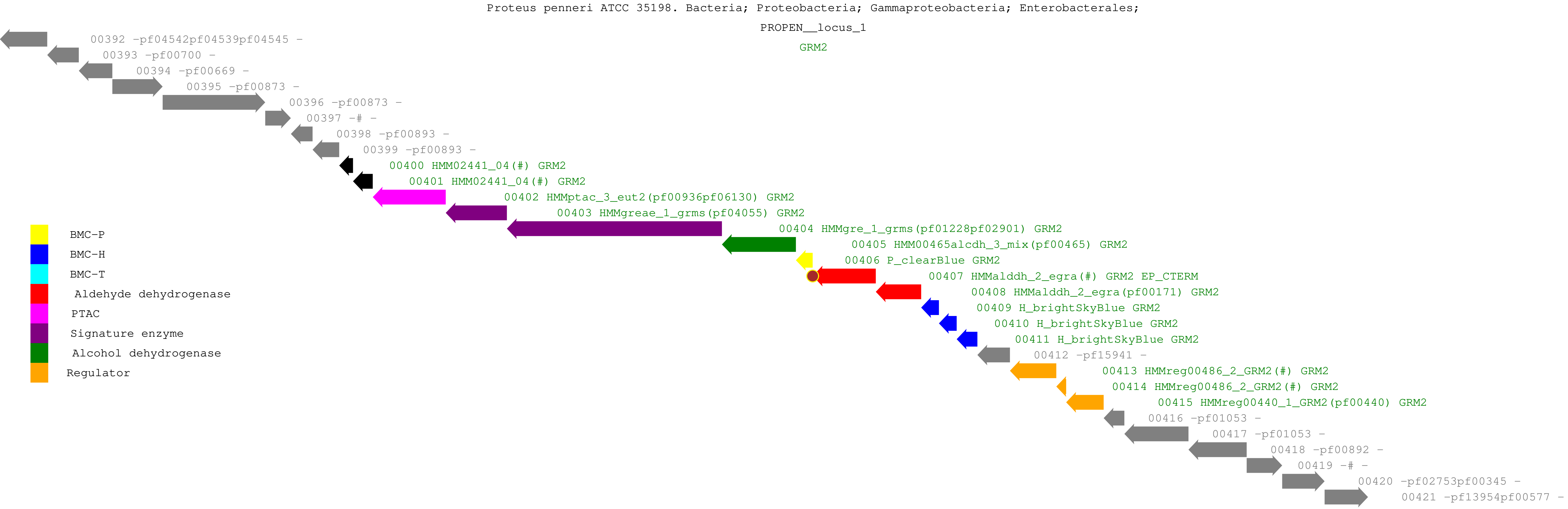

Proteus penneri ATCC 35198. Bacteria; Proteobacteria; Gammaproteobacteria; Enterobacterales;
PROPEN__locus_1
GRM2
00392 -pf04542pf04539pf04545 -
00393 -pf00700 -
00394 -pf00669 -
00395 -pf00873 -
00396 -pf00873 -
00397 -# -
00398 -pf00893 -
00399 -pf00893 -
00400 HMM02441_04(#) GRM2
00401 HMM02441_04(#) GRM2
00402 HMMptac_3_eut2(pf00936pf06130) GRM2
00403 HMMgreae_1_grms(pf04055) GRM2
00404 HMMgre_1_grms(pf01228pf02901) GRM2
00405 HMM00465alcdh_3_mix(pf00465) GRM2
00406 P_clearBlue GRM2
00407 HMMalddh_2_egra(#) GRM2 EP_CTERM
00408 HMMalddh_2_egra(pf00171) GRM2
00409 H_brightSkyBlue GRM2
00410 H_brightSkyBlue GRM2
00411 H_brightSkyBlue GRM2
00412 -pf15941 -
00413 HMMreg00486_2_GRM2(#) GRM2
00414 HMMreg00486_2_GRM2(#) GRM2
00415 HMMreg00440_1_GRM2(pf00440) GRM2
00416 -pf01053 -
00417 -pf01053 -
00418 -pf00892 -
00419 -# -
00420 -pf02753pf00345 -
00421 -pf13954pf00577 -
BMC-P
BMC-H
BMC-T
Aldehyde dehydrogenase
PTAC
Signature enzyme
Alcohol dehydrogenase
Regulator

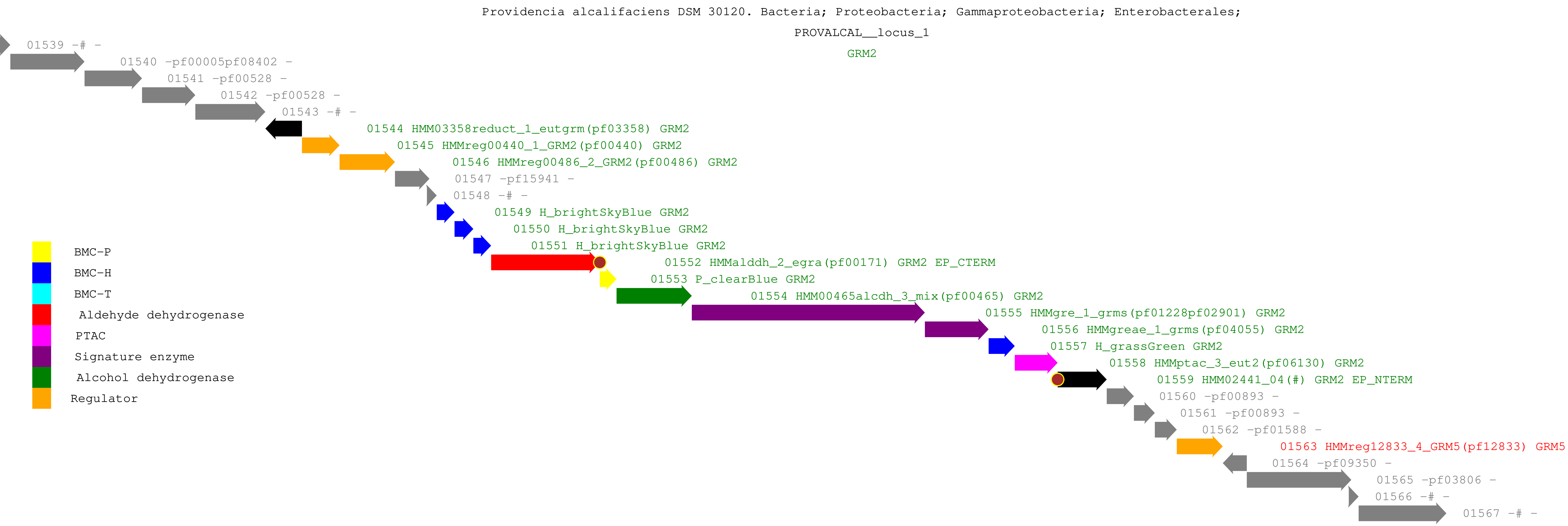
Providencia alcalifaciens DSM 30120. Bacteria; Proteobacteria; Gammaproteobacteria; Enterobacterales;
PROVALCAL__locus_1
GRM2
01539 -# -
01540 -pf00005pf08402 -
01541 -pf00528 -
01542 -pf00528 -
01543 -# -
01544 HMM03358reduct_1_eutgrm(pf03358) GRM2
01545 HMMreg00440_1_GRM2(pf00440) GRM2
01546 HMMreg00486_2_GRM2(pf00486) GRM2
01547 -pf15941 -
01548 -# -
01549 H_brightSkyBlue GRM2
01550 H_brightSkyBlue GRM2
01551 H_brightSkyBlue GRM2
01552 HMMalddh_2_egra(pf00171) GRM2 EP_CTERM
01553 P_clearBlue GRM2
01554 HMM00465alcdh_3_mix(pf00465) GRM2
01555 HMMgre_1_grms(pf01228pf02901) GRM2
01556 HMMgreae_1_grms(pf04055) GRM2
01557 H_grassGreen GRM2
01558 HMMptac_3_eut2(pf06130) GRM2
01559 HMM02441_04(#) GRM2 EP_NTERM
01560 -pf00893 -
01561 -pf00893 -
01562 -pf01588 -
01563 HMMreg12833_4_GRM5(pf12833) GRM5
01564 -pf09350 -
01565 -pf03806 -
01566 -# -
01567 -# -
BMC-P
BMC-H
BMC-T
Aldehyde dehydrogenase
PTAC
Signature enzyme
Alcohol dehydrogenase
Regulator

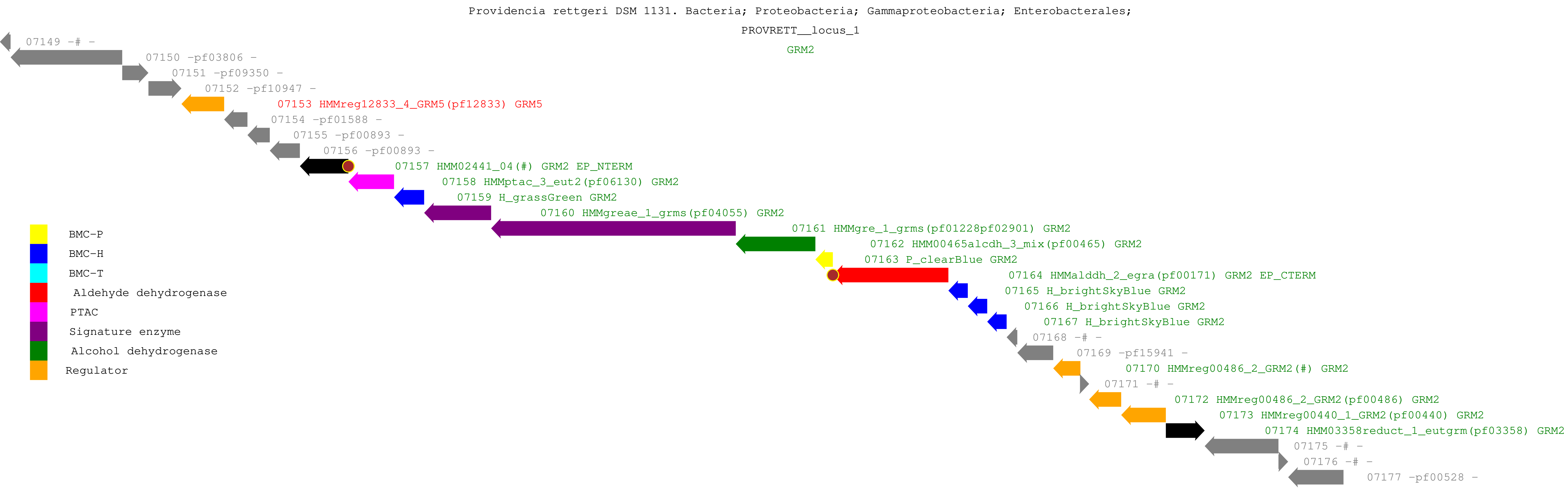
Providencia rettgeri DSM 1131. Bacteria; Proteobacteria; Gammaproteobacteria; Enterobacterales;
PROVRETT__locus_1
GRM2
07149 -# -
07150 -pf03806 -
07151 -pf09350 -
07152 -pf10947 -
07153 HMMreg12833_4_GRM5(pf12833) GRM5
07154 -pf01588 -
07155 -pf00893 -
07156 -pf00893 -
07157 HMM02441_04(#) GRM2 EP_NTERM
07158 HMMptac_3_eut2(pf06130) GRM2
07159 H_grassGreen GRM2
07160 HMMgreae_1_grms(pf04055) GRM2
07161 HMMgre_1_grms(pf01228pf02901) GRM2
07162 HMM00465alcdh_3_mix(pf00465) GRM2
07163 P_clearBlue GRM2
07164 HMMaldh_2_egra(pf00171) GRM2 EP_CTERM
07165 H_brightSkyBlue GRM2
07166 H_brightSkyBlue GRM2
07167 H_brightSkyBlue GRM2
07168 -# -
07169 -pf15941 -
07170 HMMreg00486_2_GRM2(#) GRM2
07171 -# -
07172 HMMreg00486_2_GRM2(pf00486) GRM2
07173 HMMreg00440_1_GRM2(pf00440) GRM2
07174 HMM03358reduct_1_eutgrm(pf03358) GRM2
07175 -# -
07176 -# -
07177 -pf00528 -
BMC-P
BMC-H
BMC-T
Aldehyde dehydrogenase
PTAC
Signature enzyme
Alcohol dehydrogenase
Regulator

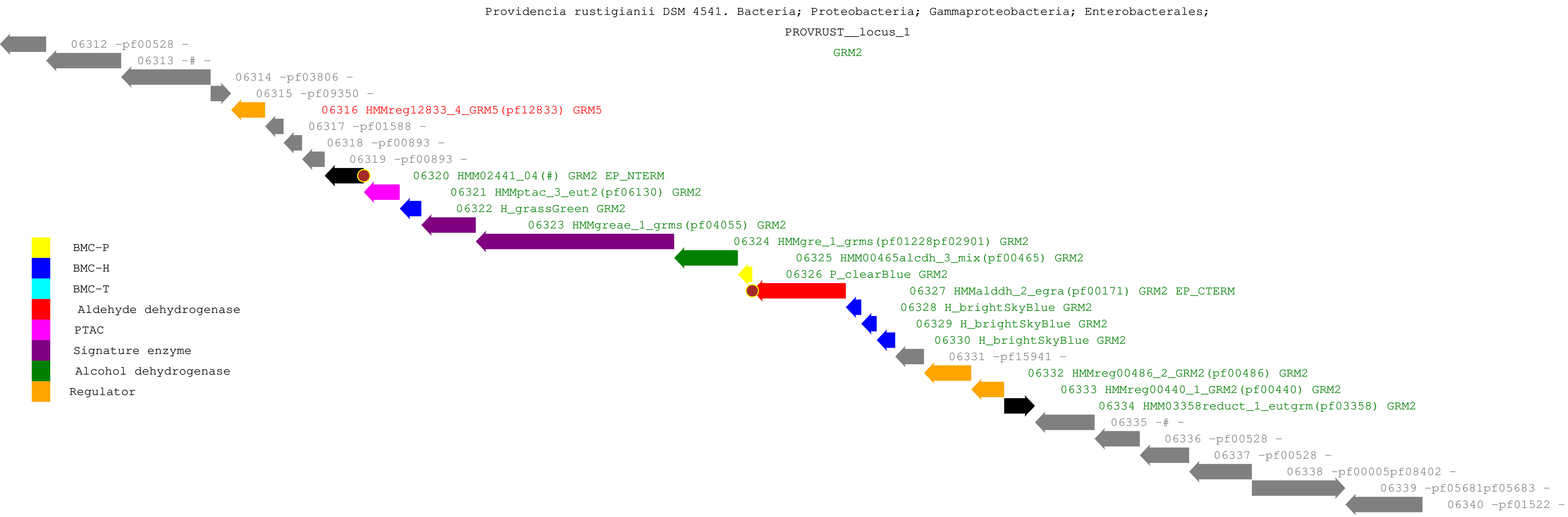

Providencia rustigianii DSM 4541. Bacteria; Proteobacteria; Gammaproteobacteria; Enterobacterales;
PROVRUST__locus_1
GRM2
06312 -pf00528 -
06313 -# -
06314 -pf03806 -
06315 -pf09350 -
06316 HMMreg12833_4_GRM5(pf12833) GRM5
06317 -pf01588 -
06318 -pf00893 -
06319 -pf00893 -
06320 HMM02441_04(#) GRM2 EP_NTERM
06321 HMMptac_3_eut2(pf06130) GRM2
06322 H_grassGreen GRM2
06323 HMMgreae_1_grms(pf04055) GRM2
06324 HMMgre_1_grms(pf01228pf02901) GRM2
06325 HMM00465alcdh_3_mix(pf00465) GRM2
06326 P_clearBlue GRM2
06327 HMMalddh_2_egra(pf00171) GRM2 EP_CTERM
06328 H_brightSkyBlue GRM2
06329 H_brightSkyBlue GRM2
06330 H_brightSkyBlue GRM2
06331 -pf15941 -
06332 HMMreg00486_2_GRM2(pf00486) GRM2
06333 HMMreg00440_1_GRM2(pf00440) GRM2
06334 HMM03358reduct_1_eutgrm(pf03358) GRM2
06335 -# -
06336 -pf00528 -
06337 -pf00528 -
06338 -pf00005pf08402 -
06339 -pf05681pf05683 -
06340 -pf01522 -
BMC-P
BMC-H
BMC-T
Aldehyde dehydrogenase
PTAC
Signature enzyme
Alcohol dehydrogenase
Regulator

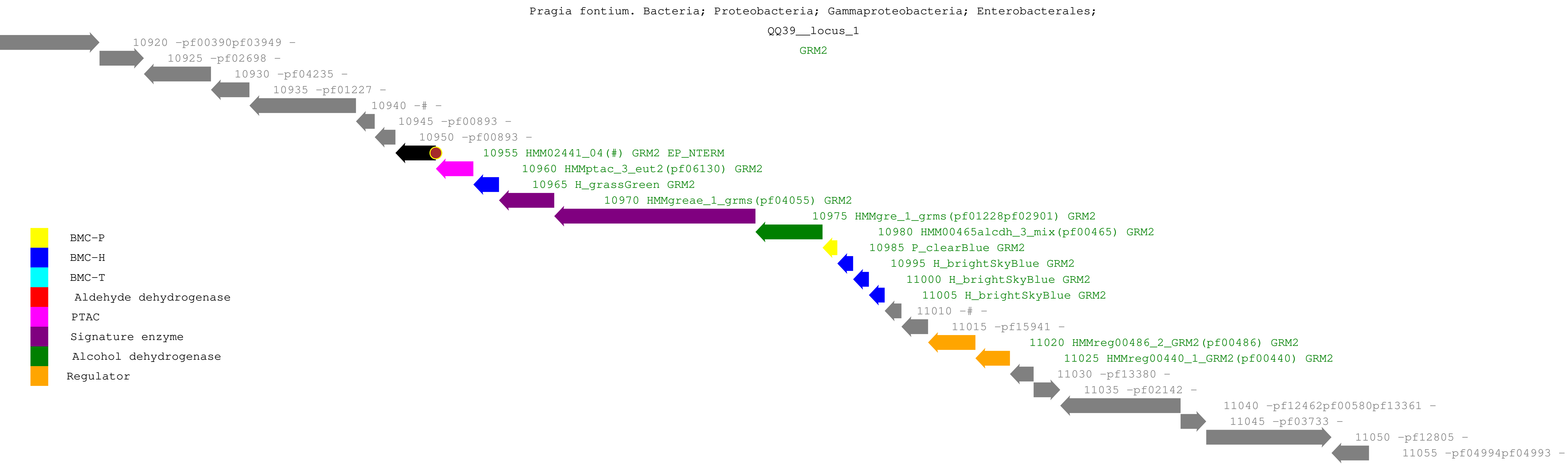
Pragia fontium. Bacteria; Proteobacteria; Gammaproteobacteria; Enterobacterales;
QQ39__locus_1
GRM2
10920 -pf00390pf03949 -
10925 -pf02698 -
10930 -pf04235 -
10935 -pf01227 -
10940 -# -
10945 -pf00893 -
10950 -pf00893 -
10955 HMM02441_04(#) GRM2 EP_NTERM
10960 HMMptac_3_eut2(pf06130) GRM2
10965 H_grassGreen GRM2
10970 HMMgreae_1_grms(pf04055) GRM2
10975 HMMgre_1_grms(pf01228pf02901) GRM2
10980 HMM00465alcdh_3_mix(pf00465) GRM2
10985 P_clearBlue GRM2
10995 H_brightSkyBlue GRM2
11000 H_brightSkyBlue GRM2
11005 H_brightSkyBlue GRM2
11010 -# -
11015 -pf15941 -
11020 HMMreg00486_2_GRM2(pf00486) GRM2
11025 HMMreg00440_1_GRM2(pf00440) GRM2
11030 -pf13380 -
11035 -pf02142 -
11040 -pf12462pf00580pf13361 -
11045 -pf03733 -
11050 -pf12805 -
11055 -pf04994pf04993 -
BMC-P
BMC-H
BMC-T
Aldehyde dehydrogenase
PTAC
Signature enzyme
Alcohol dehydrogenase
Regulator

Providencia rettgeri. Bacteria; Proteobacteria; Gammaproteobacteria; Enterobacterales;

RB151__locus_1

GRM2

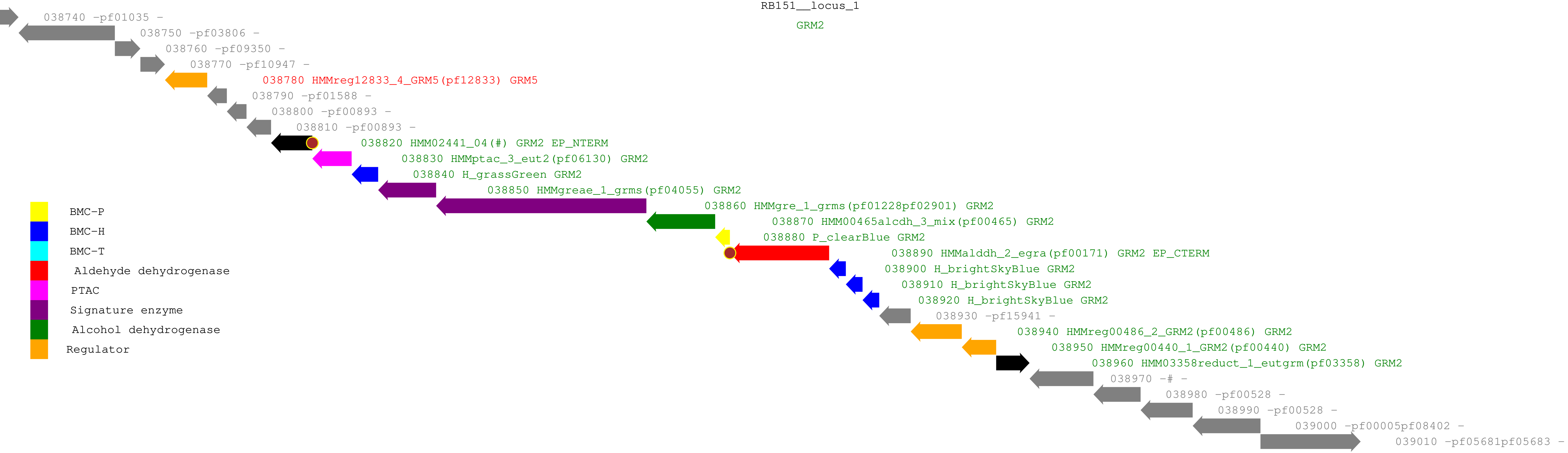

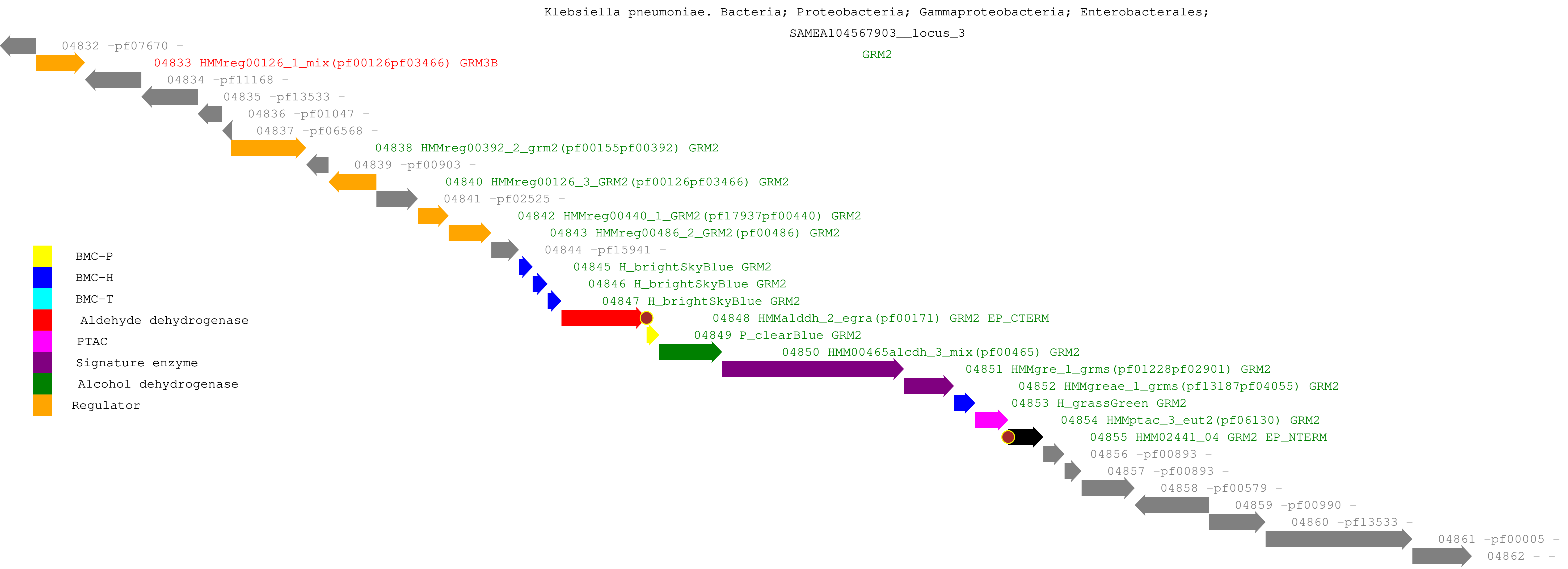
Klebsiella pneumoniae. Bacteria; Proteobacteria; Gammaproteobacteria; Enterobacterales;
SAMEA104567903__locus_3
GRM2
04832 -pf07670 -
04833 HMMreg00126_1_mix(pf00126pf03466) GRM3B
04834 -pf11168 -
04835 -pf13533 -
04836 -pf01047 -
04837 -pf06568 -
04838 HMMreg00392_2_grm2(pf00155pf00392) GRM2
04839 -pf00903 -
04840 HMMreg00126_3_GRM2(pf00126pf03466) GRM2
04841 -pf02525 -
04842 HMMreg00440_1_GRM2(pf17937pf00440) GRM2
04843 HMMreg00486_2_GRM2(pf00486) GRM2
04844 -pf15941 -
04845 H_brightSkyBlue GRM2
04846 H_brightSkyBlue GRM2
04847 H_brightSkyBlue GRM2
04848 HMMaldDh_2_egra(pf00171) GRM2 EP_CTERM
04849 P_clearBlue GRM2
04850 HMM00465alcdh_3_mix(pf00465) GRM2
04851 HMMgre_1_grms(pf01228pf02901) GRM2
04852 HMMgreae_1_grms(pf13187pf04055) GRM2
04853 H_grassGreen GRM2
04854 HMMptac_3_eut2(pf06130) GRM2
04855 HMM02441_04 GRM2 EP_NTERM
04856 -pf00893 -
04857 -pf00893 -
04858 -pf00579 -
04859 -pf00990 -
04860 -pf13533 -
04861 -pf00005 -
04862 - -
BMC-P
BMC-H
BMC-T
Aldehyde dehydrogenase
PTAC
Signature enzyme
Alcohol dehydrogenase
Regulator

Klebsiella pneumoniae. Bacteria; Proteobacteria; Gammaproteobacteria; Enterobacterales;

SAMEA2054088__locus_1

GRM2

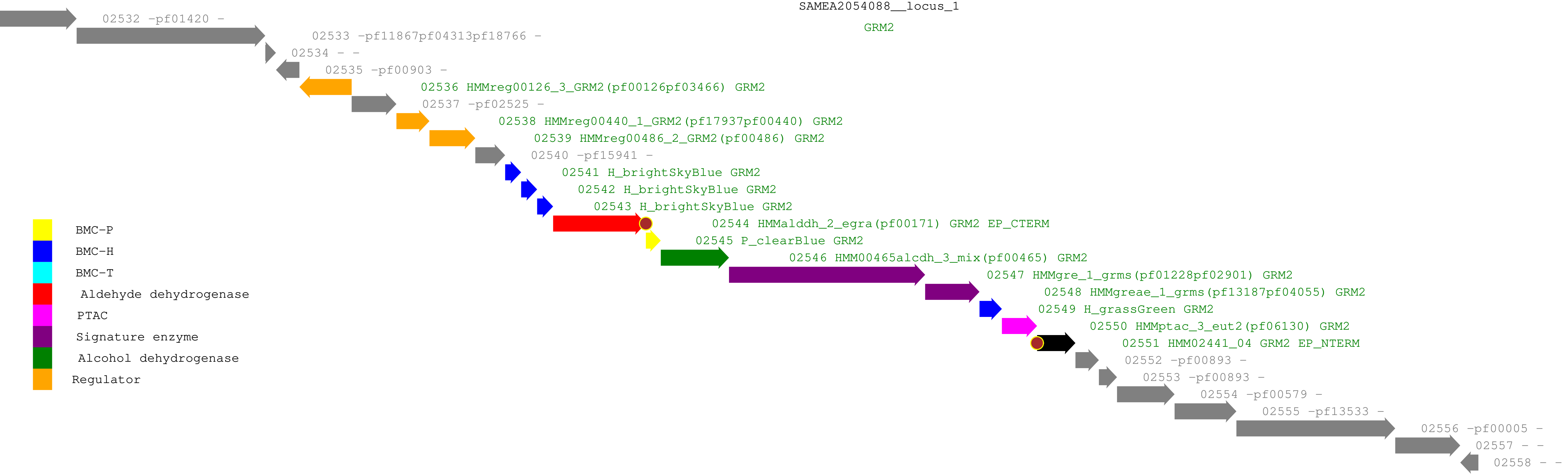

Klebsiella pneumoniae. Bacteria; Proteobacteria; Gammaproteobacteria; Enterobacterales;

SAMEA2055014__locus_2

GRM2

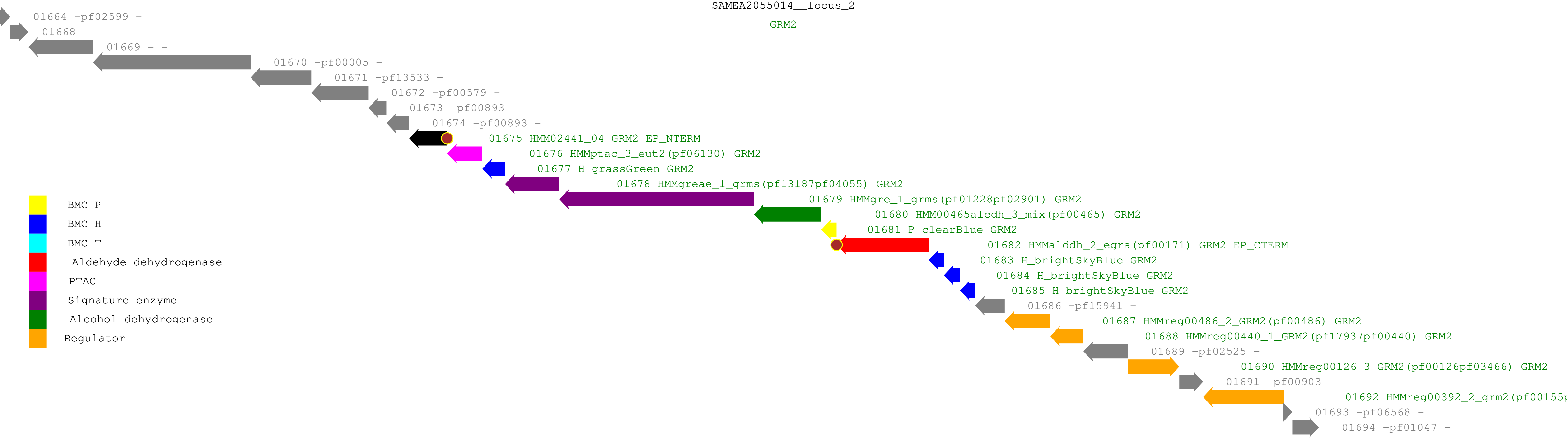

Enterobacter cloacae. Bacteria; Proteobacteria; Gammaproteobacteria; Enterobacterales;

SAMEA2273283__locus_1

GRM2

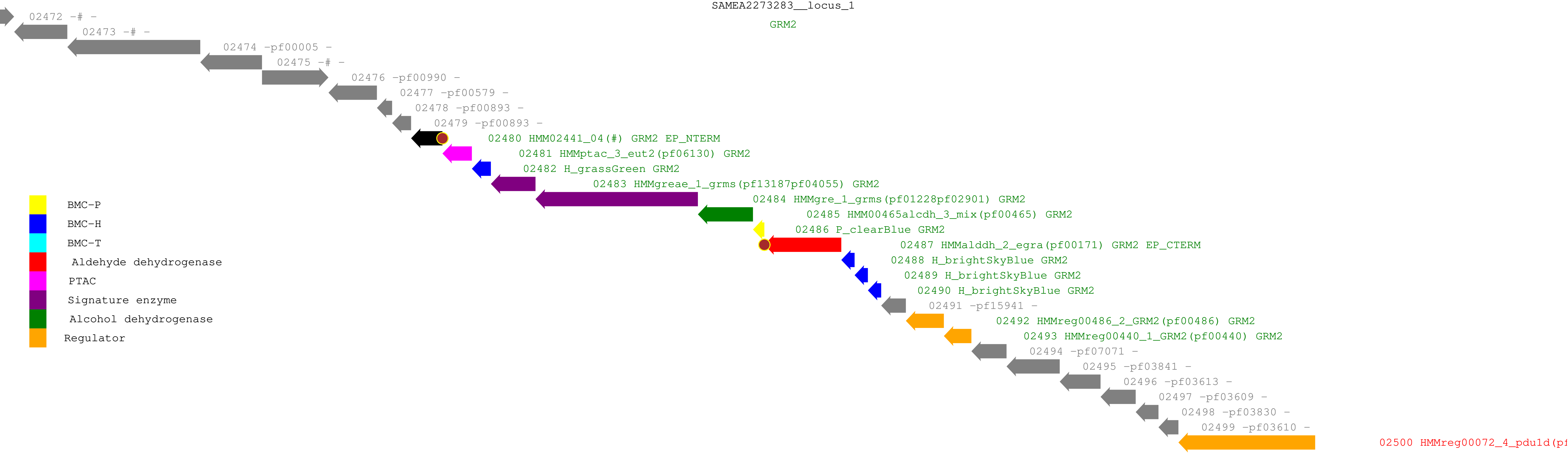

Klebsiella oxytoca. Bacteria; Proteobacteria; Gammaproteobacteria; Enterobacterales;

SAMEA2273573__locus_2

GRM2

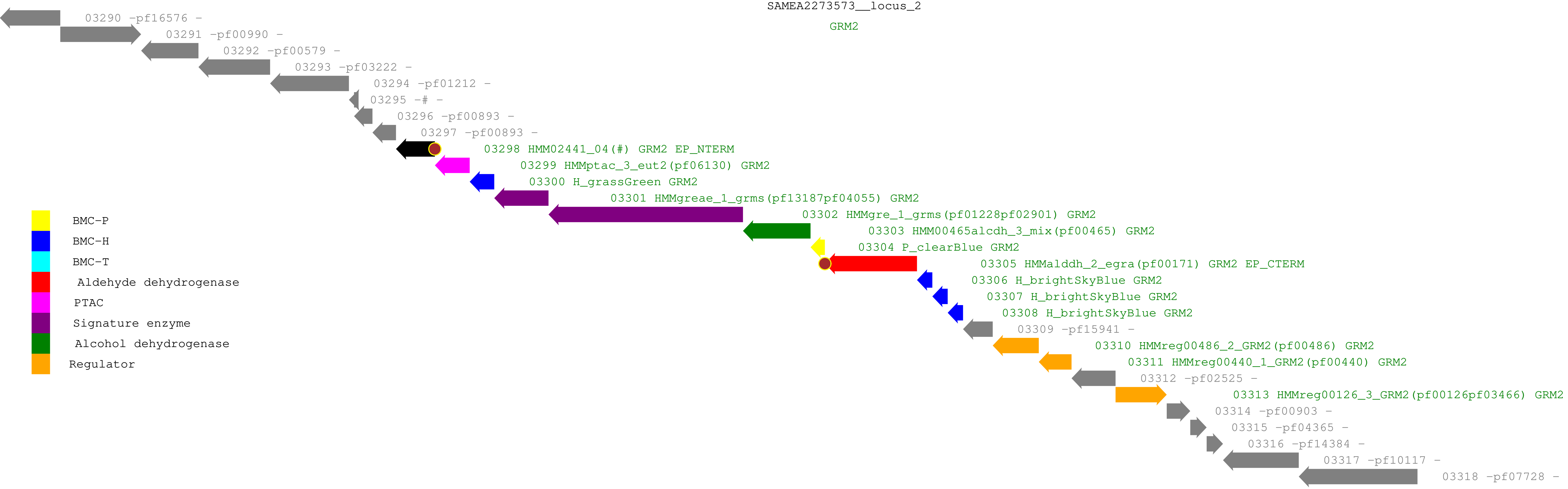

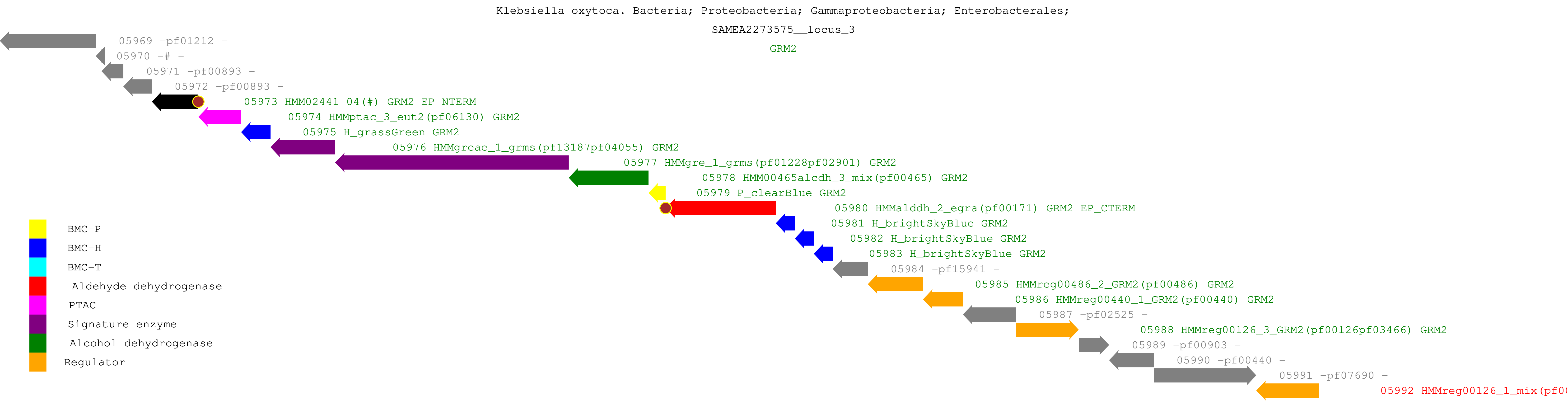

Klebsiella oxytoca. Bacteria; Proteobacteria; Gammaproteobacteria; Enterobacterales;
SAMEA2273575__locus_3
GRM2
05969 -pf01212 -
05970 -# -
05971 -pf00893 -
05972 -pf00893 -
05973 HMM02441_04(#) GRM2 EP_NTERM
05974 HMMptac_3_eut2(pf06130) GRM2
05975 H_grassGreen GRM2
05976 HMMgreae_1_grms(pf13187pf04055) GRM2
05977 HMMgre_1_grms(pf01228pf02901) GRM2
05978 HMM00465alcdh_3_mix(pf00465) GRM2
05979 P_clearBlue GRM2
05980 HMMalddh_2_egra(pf00171) GRM2 EP_CTERM
05981 H_brightSkyBlue GRM2
05982 H_brightSkyBlue GRM2
05983 H_brightSkyBlue GRM2
05984 -pf15941 -
05985 HMMreg00486_2_GRM2(pf00486) GRM2
05986 HMMreg00440_1_GRM2(pf00440) GRM2
05987 -pf02525 -
05988 HMMreg00126_3_GRM2(pf00126pf03466) GRM2
05989 -pf00903 -
05990 -pf00440 -
05991 -pf07690 -
05992 HMMreg00126_1_mix(pf0
BMC-P
BMC-H
BMC-T
Aldehyde dehydrogenase
PTAC
Signature enzyme
Alcohol dehydrogenase
Regulator

Klebsiella oxytoca. Bacteria; Proteobacteria; Gammaproteobacteria; Enterobacterales;

SAMEA2273639__locus_3

GRM2

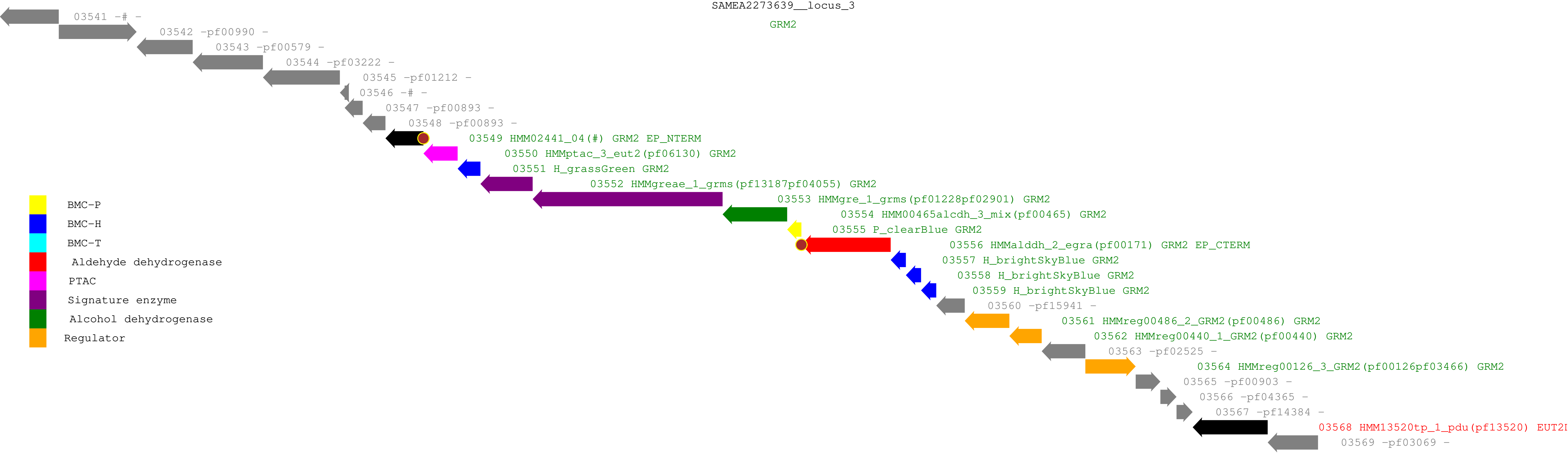

Klebsiella oxytoca. Bacteria; Proteobacteria; Gammaproteobacteria; Enterobacterales;

SAMEA2273697__locus_3

GRM2

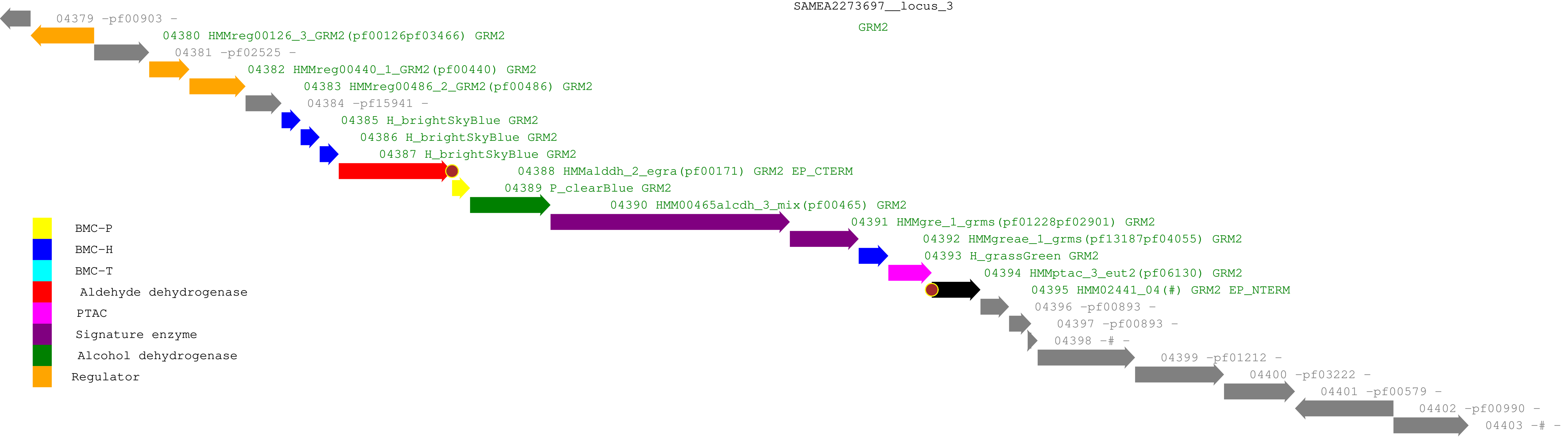

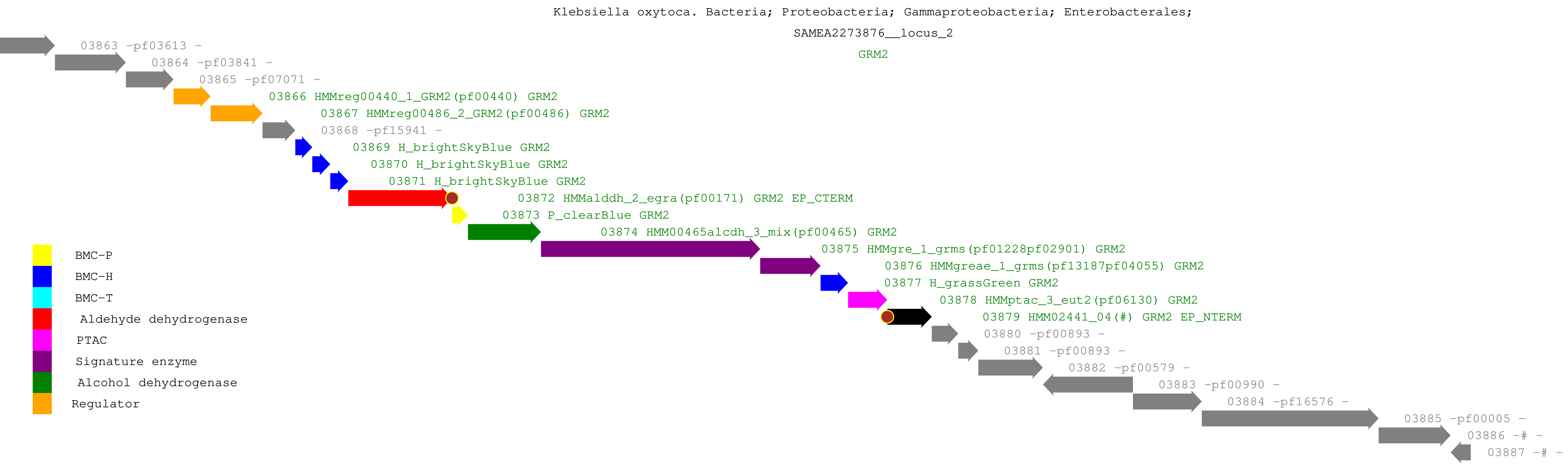

Klebsiella oxytoca. Bacteria; Proteobacteria; Gammaproteobacteria; Enterobacterales;
SAMEA2273876__locus_2
GRM2
03863 -pf03613 -
03864 -pf03841 -
03865 -pf07071 -
03866 HMMreg00440_1_GRM2(pf00440) GRM2
03867 HMMreg00486_2_GRM2(pf00486) GRM2
03868 -pf15941 -
03869 H_brightSkyBlue GRM2
03870 H_brightSkyBlue GRM2
03871 H_brightSkyBlue GRM2
03872 HMMalddh_2_egra(pf00171) GRM2 EP_CTERM
03873 P_clearBlue GRM2
03874 HMM00465alcdh_3_mix(pf00465) GRM2
03875 HMMgre_1_grms(pf01228pf02901) GRM2
03876 HMMgreae_1_grms(pf13187pf04055) GRM2
03877 H_grassGreen GRM2
03878 HMMptac_3_eut2(pf06130) GRM2
03879 HMM02441_04(#) GRM2 EP_NTERM
03880 -pf00893 -
03881 -pf00893 -
03882 -pf00579 -
03883 -pf00990 -
03884 -pf16576 -
03885 -pf00005 -
03886 -# -
03887 -# -
BMC-P
BMC-H
BMC-T
Aldehyde dehydrogenase
PTAC
Signature enzyme
Alcohol dehydrogenase
Regulator

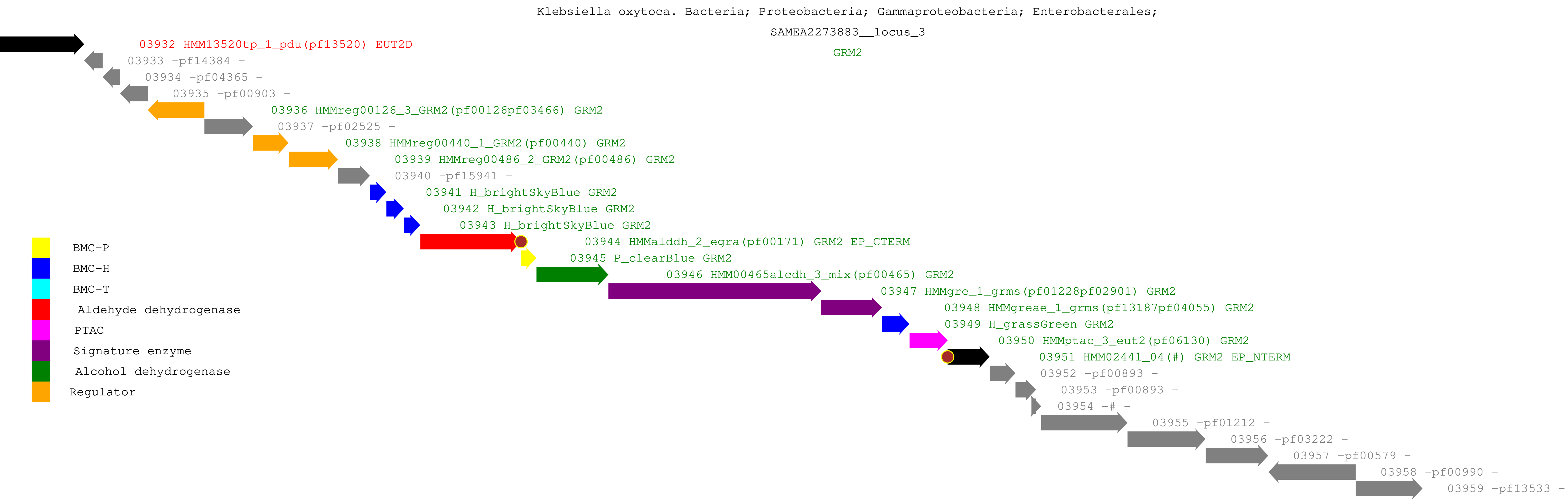
Klebsiella oxytoca. Bacteria; Proteobacteria; Gammaproteobacteria; Enterobacterales;
SAMEA2273883__locus_3
GRM2
03932 HMM13520tp_1_pdu(pf13520) EUT2D
03933 -pf14384 -
03934 -pf04365 -
03935 -pf00903 -
03936 HMMreg00126_3_GRM2(pf00126pf03466) GRM2
03937 -pf02525 -
03938 HMMreg00440_1_GRM2(pf00440) GRM2
03939 HMMreg00486_2_GRM2(pf00486) GRM2
03940 -pf15941 -
03941 H_brightSkyBlue GRM2
03942 H_brightSkyBlue GRM2
03943 H_brightSkyBlue GRM2
03944 HMMalddh_2_egra(pf00171) GRM2 EP_CTERM
03945 P_clearBlue GRM2
03946 HMM00465alcdh_3_mix(pf00465) GRM2
03947 HMMgre_1_grms(pf01228pf02901) GRM2
03948 HMMgreae_1_grms(pf13187pf04055) GRM2
03949 H_grassGreen GRM2
03950 HMMptac_3_eut2(pf06130) GRM2
03951 HMM02441_04(#) GRM2 EP_NTERM
03952 -pf00893 -
03953 -pf00893 -
03954 -# -
03955 -pf01212 -
03956 -pf03222 -
03957 -pf00579 -
03958 -pf00990 -
03959 -pf13533 -
BMC-P
BMC-H
BMC-T
Aldehyde dehydrogenase
PTAC
Signature enzyme
Alcohol dehydrogenase
Regulator

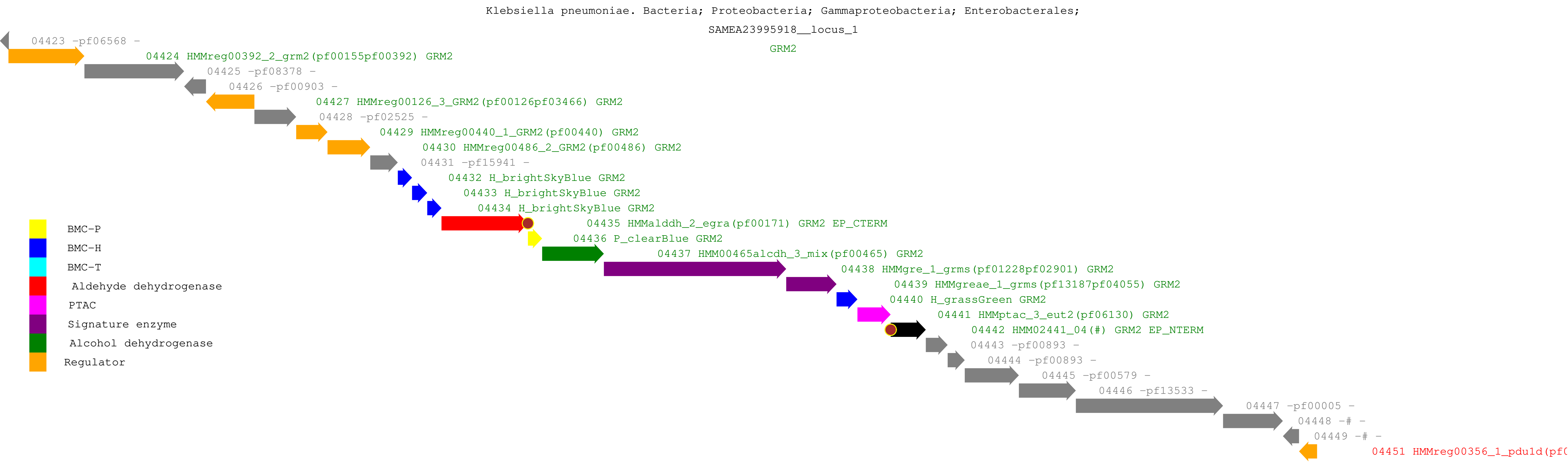
Klebsiella pneumoniae. Bacteria; Proteobacteria; Gammaproteobacteria; Enterobacterales;
SAMEA23995918__locus_1
GRM2
04423 -pf06568 -
04424 HMMreg00392_2_grm2(pf00155pf00392) GRM2
04425 -pf08378 -
04426 -pf00903 -
04427 HMMreg00126_3_GRM2(pf00126pf03466) GRM2
04428 -pf02525 -
04429 HMMreg00440_1_GRM2(pf00440) GRM2
04430 HMMreg00486_2_GRM2(pf00486) GRM2
04431 -pf15941 -
04432 H_brightSkyBlue GRM2
04433 H_brightSkyBlue GRM2
04434 H_brightSkyBlue GRM2
04435 HMMalddh_2_egra(pf00171) GRM2 EP_CTERM
04436 P_clearBlue GRM2
04437 HMM00465alcdh_3_mix(pf00465) GRM2
04438 HMMgre_1_grms(pf01228pf02901) GRM2
04439 HMMgreae_1_grms(pf13187pf04055) GRM2
04440 H_grassGreen GRM2
04441 HMMptac_3_eut2(pf06130) GRM2
04442 HMM02441_04(#) GRM2 EP_NTERM
04443 -pf00893 -
04444 -pf00893 -
04445 -pf00579 -
04446 -pf13533 -
04447 -pf00005 -
04448 -# -
04449 -# -
04451 HMMreg00356_1_pdu1d(pf0
BMC-P
BMC-H
BMC-T
Aldehyde dehydrogenase
PTAC
Signature enzyme
Alcohol dehydrogenase
Regulator

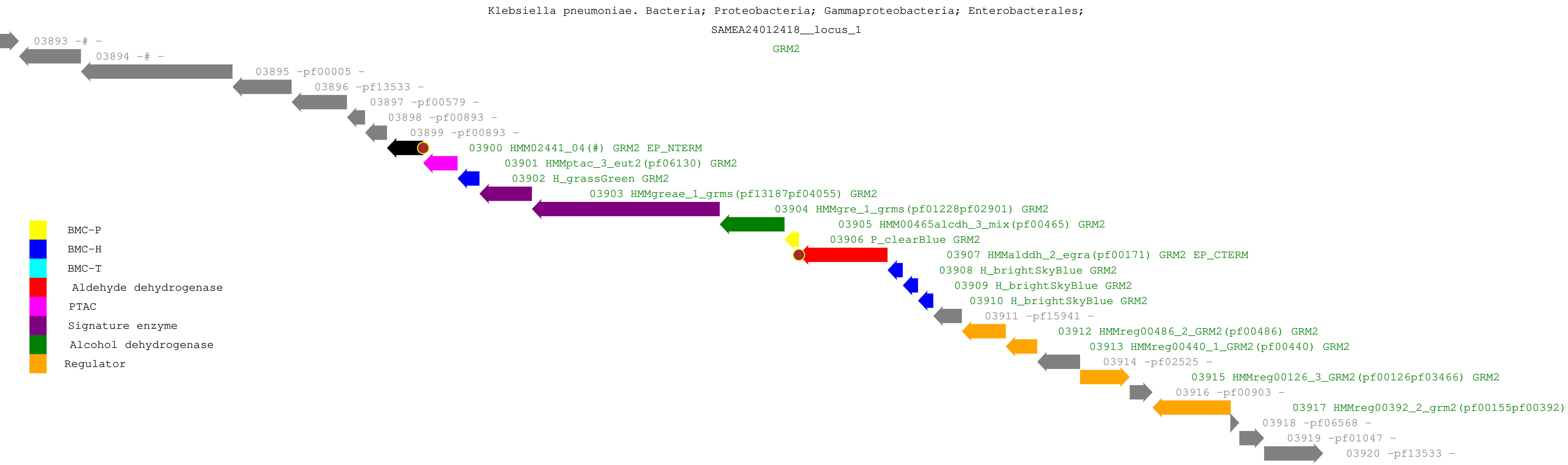
Klebsiella pneumoniae. Bacteria; Proteobacteria; Gammaproteobacteria; Enterobacterales;
SAMEA24012418__locus_1
GRM2
03893 -# -
03894 -# -
03895 -pf00005 -
03896 -pf13533 -
03897 -pf00579 -
03898 -pf00893 -
03899 -pf00893 -
03900 HMM02441_04(#) GRM2 EP_NTERM
03901 HMMptac_3_eut2(pf06130) GRM2
03902 H_grassGreen GRM2
03903 HMMgreae_1_grms(pf13187pf04055) GRM2
03904 HMMgre_1_grms(pf01228pf02901) GRM2
03905 HMM00465alcdh_3_mix(pf00465) GRM2
03906 P_clearBlue GRM2
03907 HMMalddh_2_egra(pf00171) GRM2 EP_CTERM
03908 H_brightSkyBlue GRM2
03909 H_brightSkyBlue GRM2
03910 H_brightSkyBlue GRM2
03911 -pf15941 -
03912 HMMreg00486_2_GRM2(pf00486) GRM2
03913 HMMreg00440_1_GRM2(pf00440) GRM2
03914 -pf02525 -
03915 HMMreg00126_3_GRM2(pf00126pf03466) GRM2
03916 -pf00903 -
03917 HMMreg00392_2_grm2(pf00155pf00392)
03918 -pf06568 -
03919 -pf01047 -
03920 -pf13533 -
BMC-P
BMC-H
BMC-T
Aldehyde dehydrogenase
PTAC
Signature enzyme
Alcohol dehydrogenase
Regulator

Raoultella ornithinolytica (Klebsiella ornithinolytica). Bacteria; Proteobacteria; Gammaproteobacteria; Enterobacterales;

SAMEA3181514__locus_2

GRM2

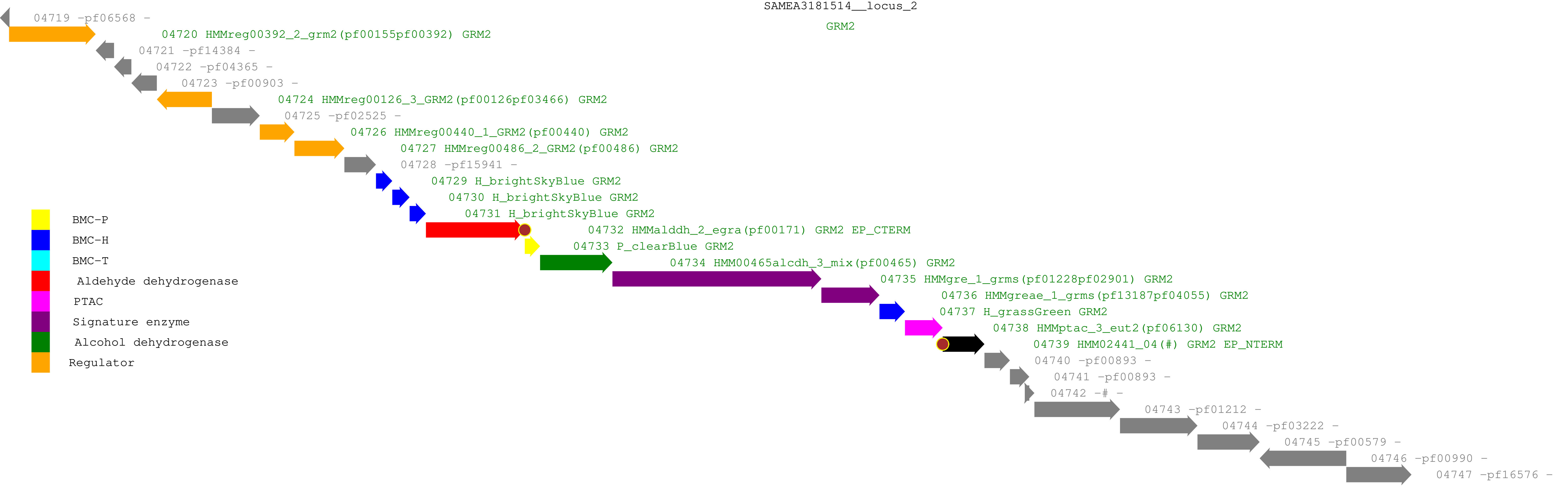

Raoultella terrigena (Klebsiella terrigena). Bacteria; Proteobacteria; Gammaproteobacteria; Enterobacterales;

SAMEA3368263__locus_1

GRM2

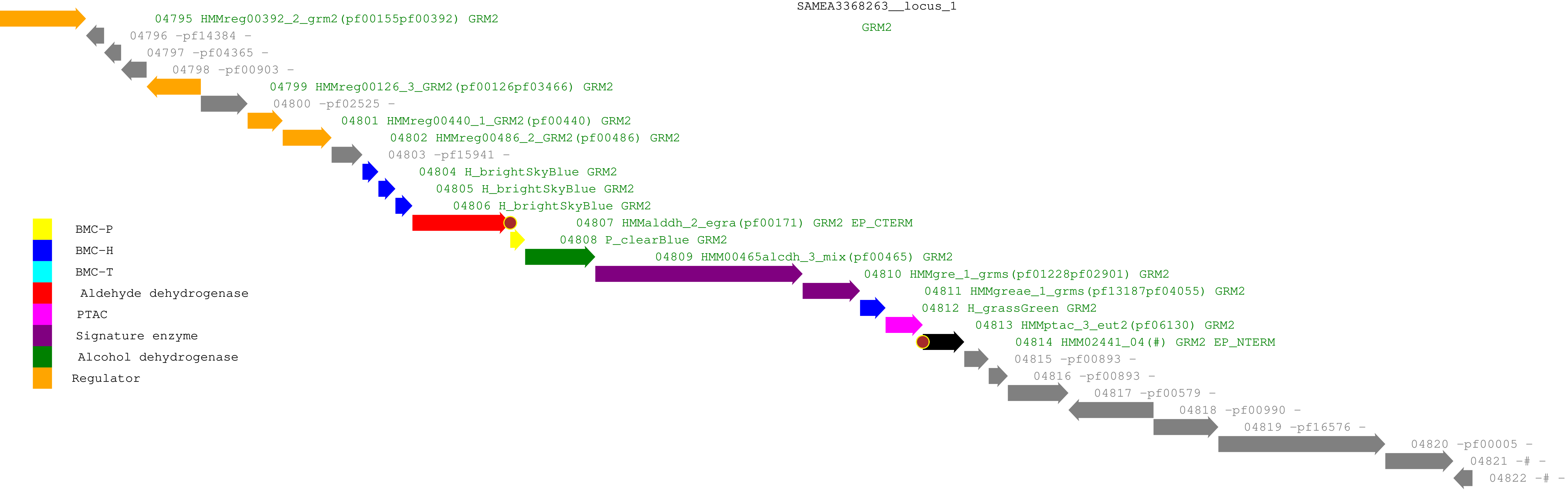

Klebsiella variicola. Bacteria; Proteobacteria; Gammaproteobacteria; Enterobacterales;

SAMEA3515120__locus_3

GRM2

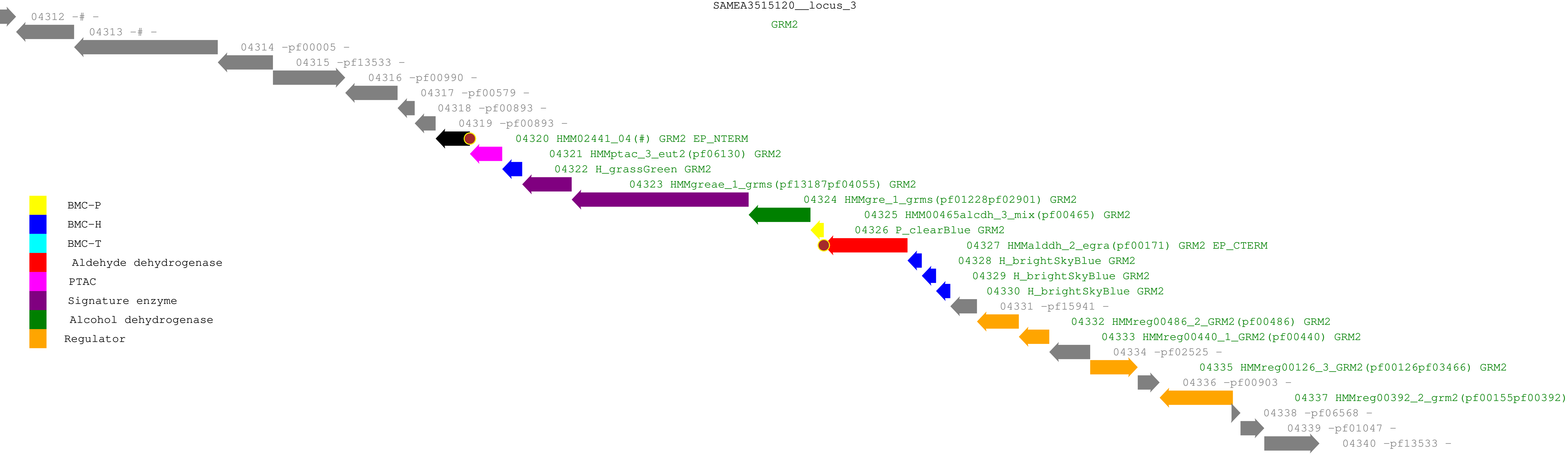

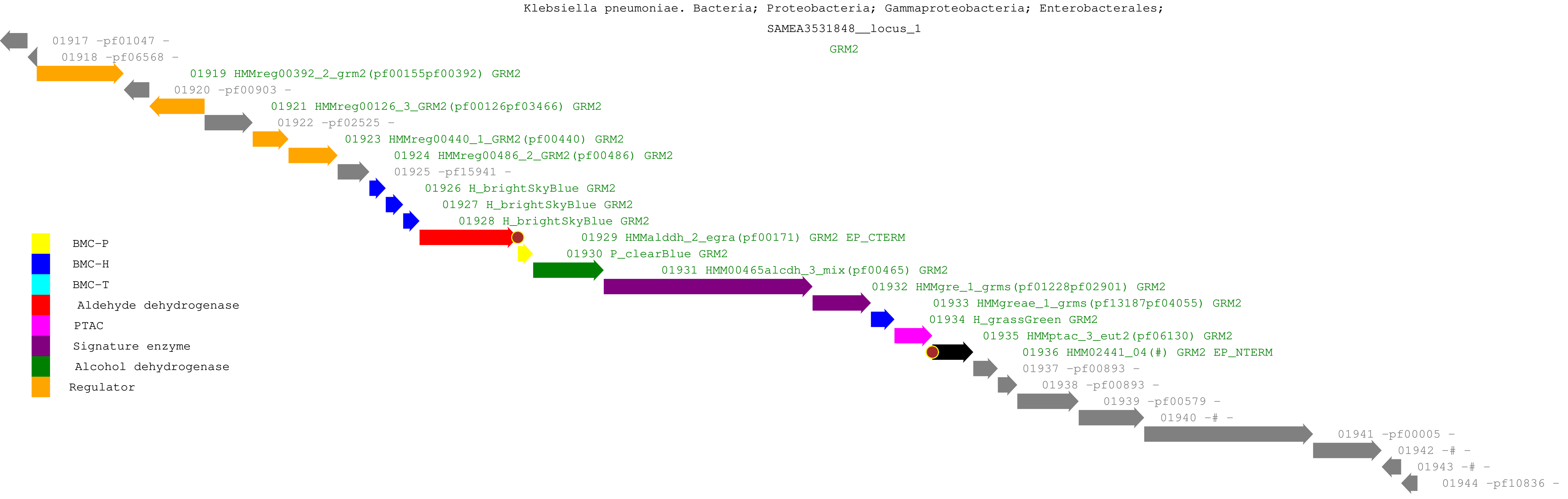
Klebsiella pneumoniae. Bacteria; Proteobacteria; Gammaproteobacteria; Enterobacterales;
SAMEA3531848__locus_1
GRM2
01917 -pf01047 -
01918 -pf06568 -
01919 HMMreg00392_2_grm2(pf00155pf00392) GRM2
01920 -pf00903 -
01921 HMMreg00126_3_GRM2(pf00126pf03466) GRM2
01922 -pf02525 -
01923 HMMreg00440_1_GRM2(pf00440) GRM2
01924 HMMreg00486_2_GRM2(pf00486) GRM2
01925 -pf15941 -
01926 H_brightSkyBlue GRM2
01927 H_brightSkyBlue GRM2
01928 H_brightSkyBlue GRM2
01929 HMMaldh_2_egra(pf00171) GRM2 EP_CTERM
01930 P_clearBlue GRM2
01931 HMM00465alcdh_3_mix(pf00465) GRM2
01932 HMMgre_1_grms(pf01228pf02901) GRM2
01933 HMMgreae_1_grms(pf13187pf04055) GRM2
01934 H_grassGreen GRM2
01935 HMMptac_3_eut2(pf06130) GRM2
01936 HMM02441_04(#) GRM2 EP_NTERM
01937 -pf00893 -
01938 -pf00893 -
01939 -pf00579 -
01940 -# -
01941 -pf00005 -
01942 -# -
01943 -# -
01944 -pf10836 -
BMC-P
BMC-H
BMC-T
Aldehyde dehydrogenase
PTAC
Signature enzyme
Alcohol dehydrogenase
Regulator

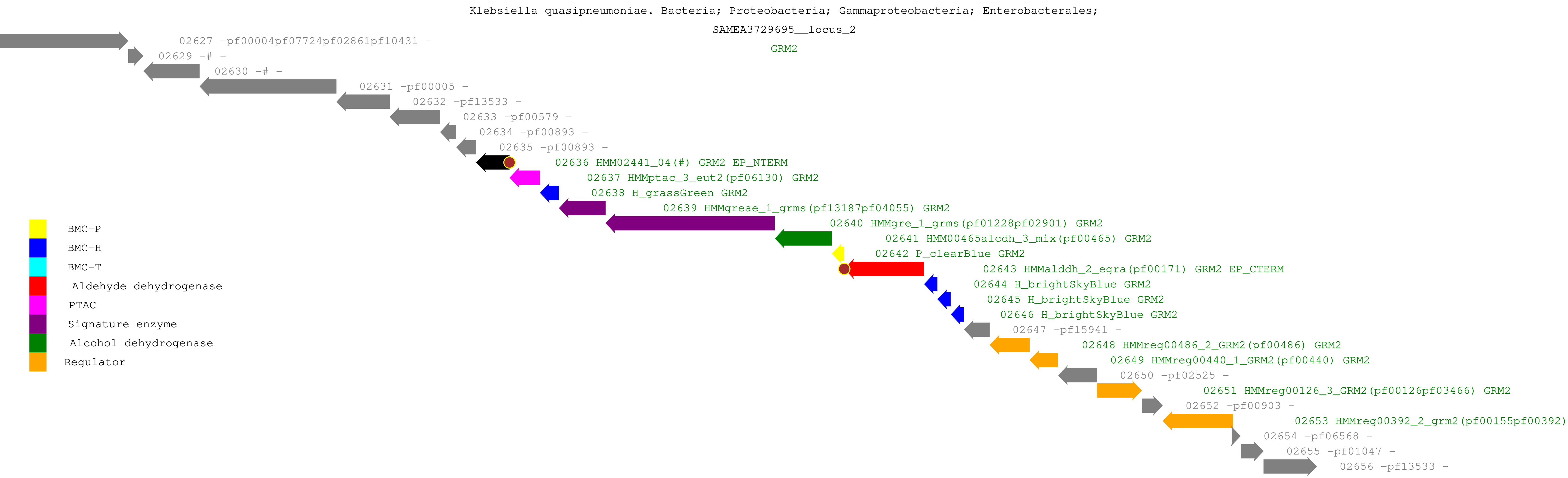

Klebsiella quasipneumoniae. Bacteria; Proteobacteria; Gammaproteobacteria; Enterobacterales;
SAMEA3729695__locus_2
GRM2
02627 -pf00004pf07724pf02861pf10431 -
02629 -# -
02630 -# -
02631 -pf00005 -
02632 -pf13533 -
02633 -pf00579 -
02634 -pf00893 -
02635 -pf00893 -
02636 HMM02441_04(#) GRM2 EP_NTERM
02637 HMMptac_3_eut2(pf06130) GRM2
02638 H_grassGreen GRM2
02639 HMMgreae_1_grms(pf13187pf04055) GRM2
02640 HMMgre_1_grms(pf01228pf02901) GRM2
02641 HMM00465alcdh_3_mix(pf00465) GRM2
02642 P_clearBlue GRM2
02643 HMMalddh_2_egra(pf00171) GRM2 EP_CTERM
02644 H_brightSkyBlue GRM2
02645 H_brightSkyBlue GRM2
02646 H_brightSkyBlue GRM2
02647 -pf15941 -
02648 HMMreg00486_2_GRM2(pf00486) GRM2
02649 HMMreg00440_1_GRM2(pf00440) GRM2
02650 -pf02525 -
02651 HMMreg00126_3_GRM2(pf00126pf03466) GRM2
02652 -pf00903 -
02653 HMMreg00392_2_grm2(pf00155pf00392)
02654 -pf06568 -
02655 -pf01047 -
02656 -pf13533 -
BMC-P
BMC-H
BMC-T
Aldehyde dehydrogenase
PTAC
Signature enzyme
Alcohol dehydrogenase
Regulator

Klebsiella pneumoniae. Bacteria; Proteobacteria; Gammaproteobacteria; Enterobacterales;
SAMEA4362693__locus_3
GRM2

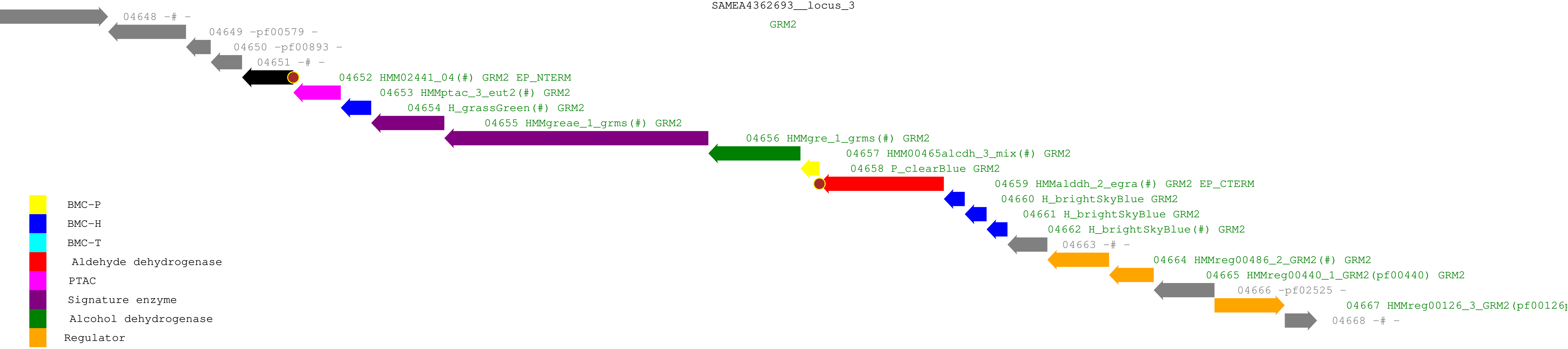

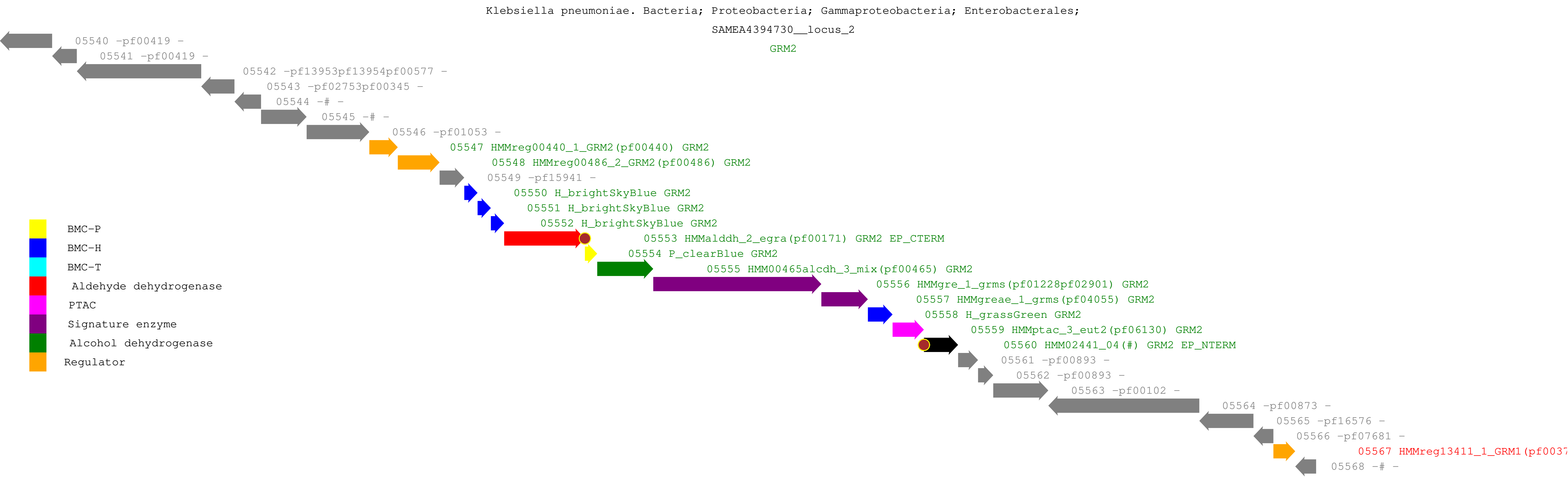
Klebsiella pneumoniae. Bacteria; Proteobacteria; Gammaproteobacteria; Enterobacterales;
SAMEA4394730__locus_2
GRM2
05540 -pf00419 -
05541 -pf00419 -
05542 -pf13953pf13954pf00577 -
05543 -pf02753pf00345 -
05544 -# -
05545 -# -
05546 -pf01053 -
05547 HMMreg00440_1_GRM2(pf00440) GRM2
05548 HMMreg00486_2_GRM2(pf00486) GRM2
05549 -pf15941 -
05550 H_brightSkyBlue GRM2
05551 H_brightSkyBlue GRM2
05552 H_brightSkyBlue GRM2
05553 HMMalddh_2_egra(pf00171) GRM2 EP_CTERM
05554 P_clearBlue GRM2
05555 HMM00465alcdh_3_mix(pf00465) GRM2
05556 HMMgre_1_grms(pf01228pf02901) GRM2
05557 HMMgreae_1_grms(pf04055) GRM2
05558 H_grassGreen GRM2
05559 HMMptac_3_eut2(pf06130) GRM2
05560 HMM02441_04(#) GRM2 EP_NTERM
05561 -pf00893 -
05562 -pf00893 -
05563 -pf00102 -
05564 -pf00873 -
05565 -pf16576 -
05566 -pf07681 -
05567 HMMreg13411_1_GRM1(pf0037
05568 -# -
BMC-P
BMC-H
BMC-T
Aldehyde dehydrogenase
PTAC
Signature enzyme
Alcohol dehydrogenase
Regulator

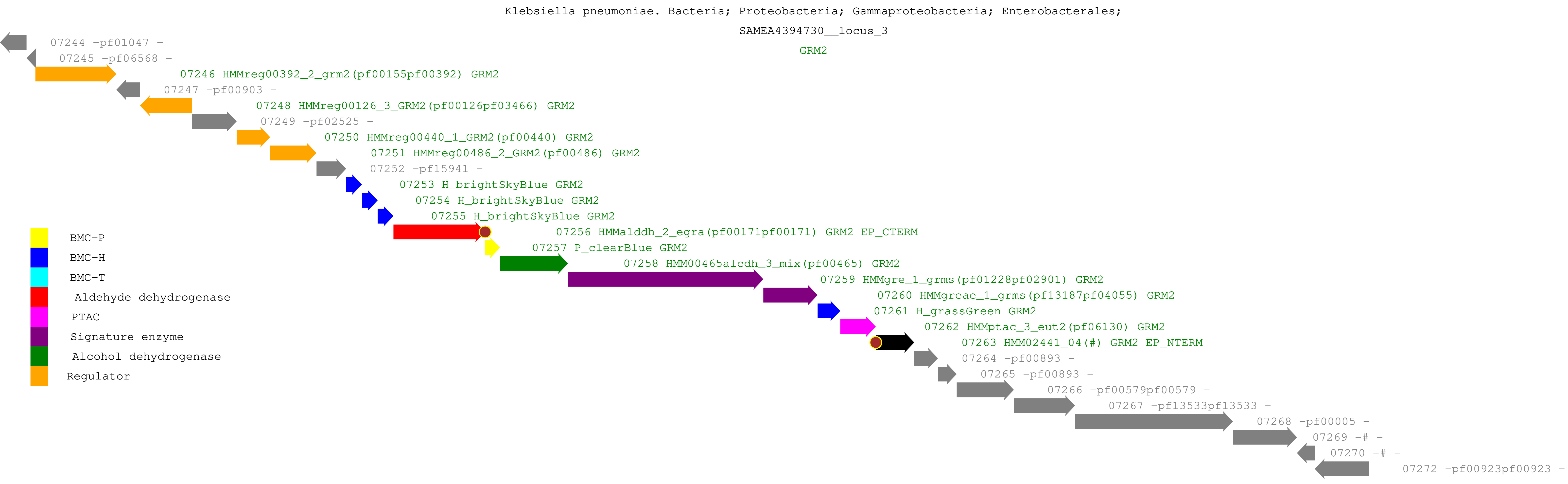
Klebsiella pneumoniae. Bacteria; Proteobacteria; Gammaproteobacteria; Enterobacterales;
SAMEA4394730__locus_3
GRM2
07244 -pf01047 -
07245 -pf06568 -
07246 HMMreg00392_2_grm2(pf00155pf00392) GRM2
07247 -pf00903 -
07248 HMMreg00126_3_GRM2(pf00126pf03466) GRM2
07249 -pf02525 -
07250 HMMreg00440_1_GRM2(pf00440) GRM2
07251 HMMreg00486_2_GRM2(pf00486) GRM2
07252 -pf15941 -
07253 H_brightSkyBlue GRM2
07254 H_brightSkyBlue GRM2
07255 H_brightSkyBlue GRM2
07256 HMMaldh_2_egra(pf00171pf00171) GRM2 EP_CTERM
07257 P_clearBlue GRM2
07258 HMM00465alcdh_3_mix(pf00465) GRM2
07259 HMMgre_1_grms(pf01228pf02901) GRM2
07260 HMMgreae_1_grms(pf13187pf04055) GRM2
07261 H_grassGreen GRM2
07262 HMMptac_3_eut2(pf06130) GRM2
07263 HMM02441_04(#) GRM2 EP_NTERM
07264 -pf00893 -
07265 -pf00893 -
07266 -pf00579pf00579 -
07267 -pf13533pf13533 -
07268 -pf00005 -
07269 -# -
07270 -# -
07272 -pf00923pf00923 -
BMC-P
BMC-H
BMC-T
Aldehyde dehydrogenase
PTAC
Signature enzyme
Alcohol dehydrogenase
Regulator

Klebsiella pneumoniae. Bacteria; Proteobacteria; Gammaproteobacteria; Enterobacterales;

SAMEA4873561__locus_1

GRM2

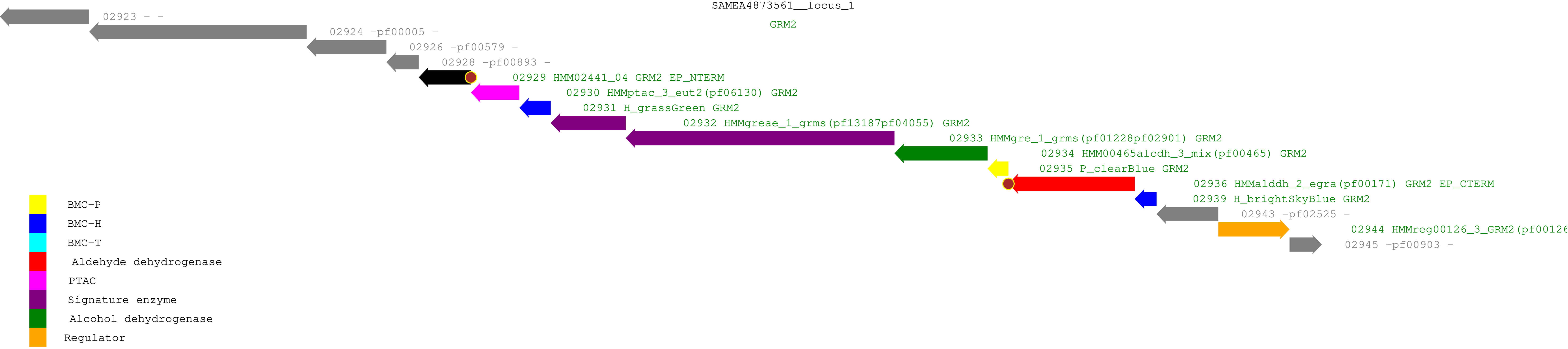

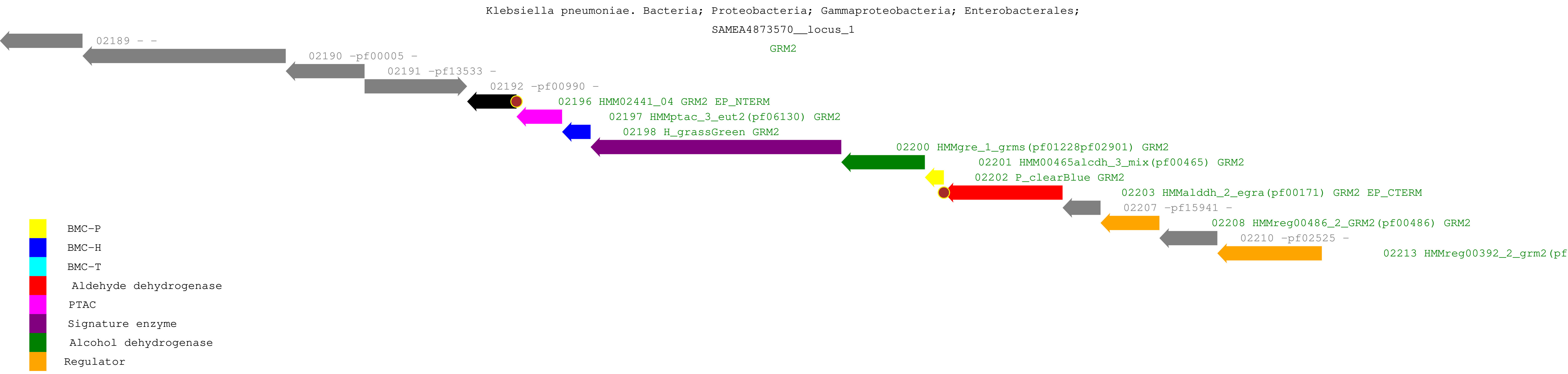
Klebsiella pneumoniae. Bacteria; Proteobacteria; Gammaproteobacteria; Enterobacterales;
SAMEA4873570__locus_1
GRM2
02189 - -
02190 -pf00005 -
02191 -pf13533 -
02192 -pf00990 -
02196 HMM02441_04 GRM2 EP_NTERM
02197 HMMptac_3_eut2(pf06130) GRM2
02198 H_grassGreen GRM2
02200 HMMgre_1_grms(pf01228pf02901) GRM2
02201 HMM00465alcdh_3_mix(pf00465) GRM2
02202 P_clearBlue GRM2
02203 HMMaldh_2_egra(pf00171) GRM2 EP_CTERM
02207 -pf15941 -
02208 HMMreg00486_2_GRM2(pf00486) GRM2
02210 -pf02525 -
02213 HMMreg00392_2_grm2(pf
BMC-P
BMC-H
BMC-T
Aldehyde dehydrogenase
PTAC
Signature enzyme
Alcohol dehydrogenase
Regulator

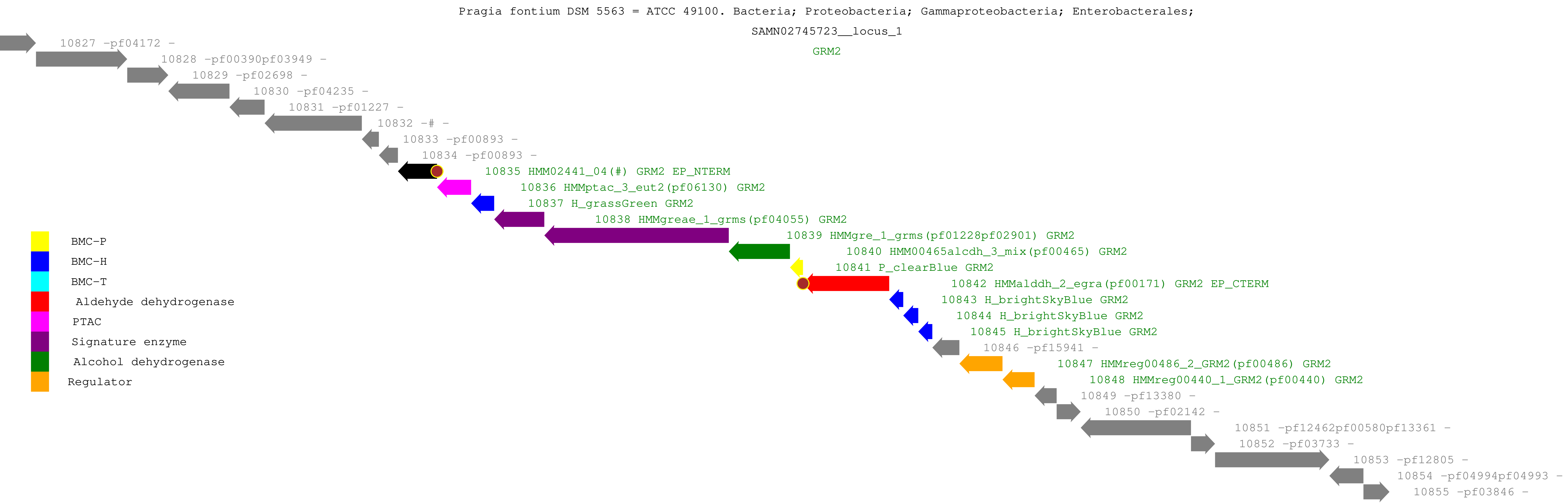
Pragia fontium DSM 5563 = ATCC 49100. Bacteria; Proteobacteria; Gammaproteobacteria; Enterobacterales;
SAMN02745723__locus_1
GRM2
10827 -pf04172 -
10828 -pf00390pf03949 -
10829 -pf02698 -
10830 -pf04235 -
10831 -pf01227 -
10832 -# -
10833 -pf00893 -
10834 -pf00893 -
10835 HMM02441_04(#) GRM2 EP_NTERM
10836 HMMptac_3_eut2(pf06130) GRM2
10837 H_grassGreen GRM2
10838 HMMgreae_1_grms(pf04055) GRM2
10839 HMMgre_1_grms(pf01228pf02901) GRM2
10840 HMM00465alcdh_3_mix(pf00465) GRM2
10841 P_clearBlue GRM2
10842 HMMalddh_2_egra(pf00171) GRM2 EP_CTERM
10843 H_brightSkyBlue GRM2
10844 H_brightSkyBlue GRM2
10845 H_brightSkyBlue GRM2
10846 -pf15941 -
10847 HMMreg00486_2_GRM2(pf00486) GRM2
10848 HMMreg00440_1_GRM2(pf00440) GRM2
10849 -pf13380 -
10850 -pf02142 -
10851 -pf12462pf00580pf13361 -
10852 -pf03733 -
10853 -pf12805 -
10854 -pf04994pf04993 -
10855 -pf03846 -
BMC-P
BMC-H
BMC-T
Aldehyde dehydrogenase
PTAC
Signature enzyme
Alcohol dehydrogenase
Regulator

Klebsiella sp. NFIX38. Bacteria; Proteobacteria; Gammaproteobacteria; Enterobacterales;

SAMN03159330__locus_2

GRM2

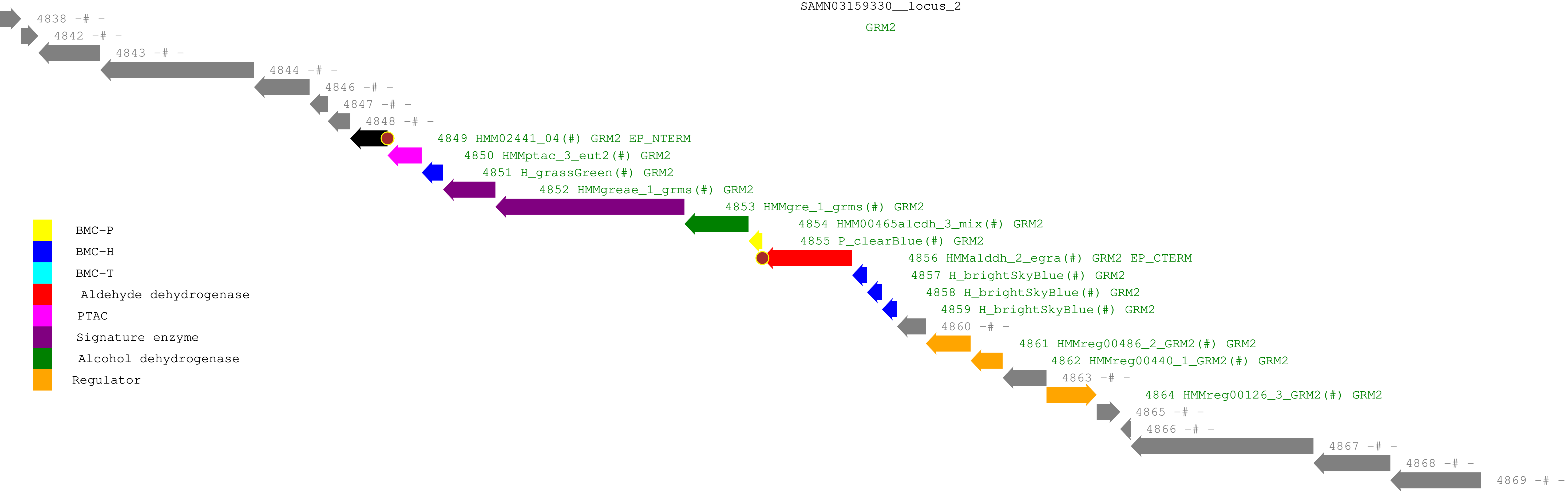

Klebsiella sp. NFIX14. Bacteria; Proteobacteria; Gammaproteobacteria; Enterobacterales;
SAMN03159388__locus_2
GRM2

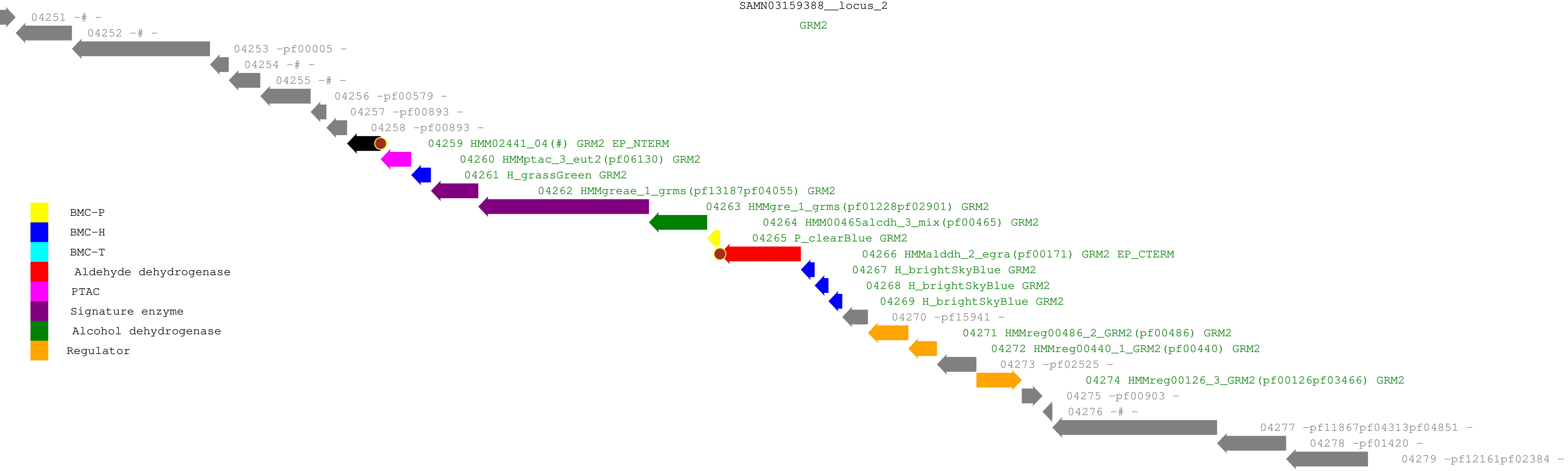

Aeromonas hydrophila. Bacteria; Proteobacteria; Gammaproteobacteria; Aeromonadales;

SAMN05878295__locus_1

GRM2

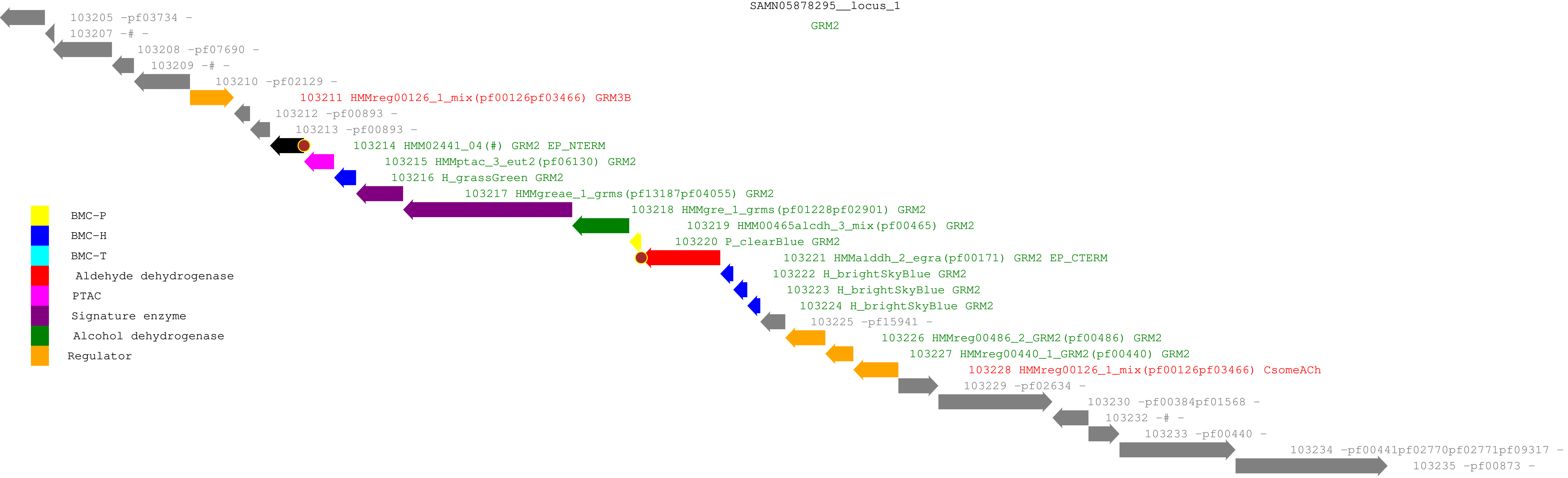

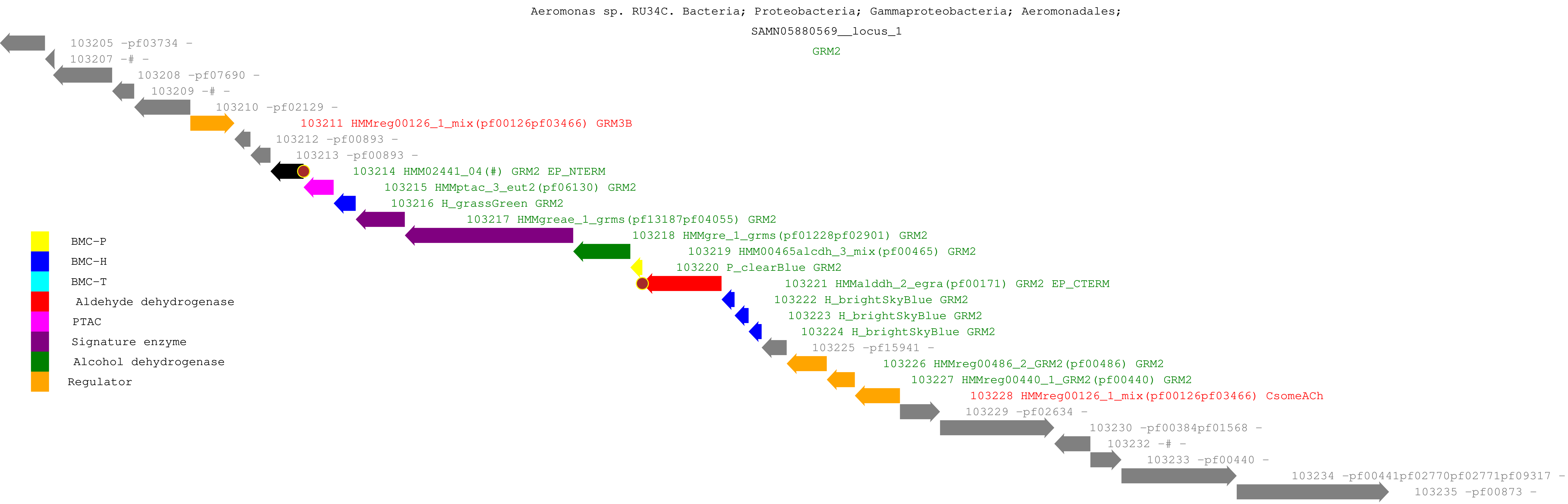
Aeromonas sp. RU34C. Bacteria; Proteobacteria; Gammaproteobacteria; Aeromonadales;
SAMN05880569__locus_1
GRM2
103205 -pf03734 -
103207 -# -
103208 -pf07690 -
103209 -# -
103210 -pf02129 -
103211 HMMreg00126_1_mix(pf00126pf03466) GRM3B
103212 -pf00893 -
103213 -pf00893 -
103214 HMM02441_04(#) GRM2 EP_NTERM
103215 HMMptac_3_eut2(pf06130) GRM2
103216 H_grassGreen GRM2
103217 HMMgreae_1_grms(pf13187pf04055) GRM2
103218 HMMgre_1_grms(pf01228pf02901) GRM2
103219 HMM00465alcdh_3_mix(pf00465) GRM2
103220 P_clearBlue GRM2
103221 HMMalddh_2_egra(pf00171) GRM2 EP_CTERM
103222 H_brightSkyBlue GRM2
103223 H_brightSkyBlue GRM2
103224 H_brightSkyBlue GRM2
103225 -pf15941 -
103226 HMMreg00486_2_GRM2(pf00486) GRM2
103227 HMMreg00440_1_GRM2(pf00440) GRM2
103228 HMMreg00126_1_mix(pf00126pf03466) CsomeACh
103229 -pf02634 -
103230 -pf00384pf01568 -
103232 -# -
103233 -pf00440 -
103234 -pf00441pf02770pf02771pf09317 -
103235 -pf00873 -
BMC-P
BMC-H
BMC-T
Aldehyde dehydrogenase
PTAC
Signature enzyme
Alcohol dehydrogenase
Regulator

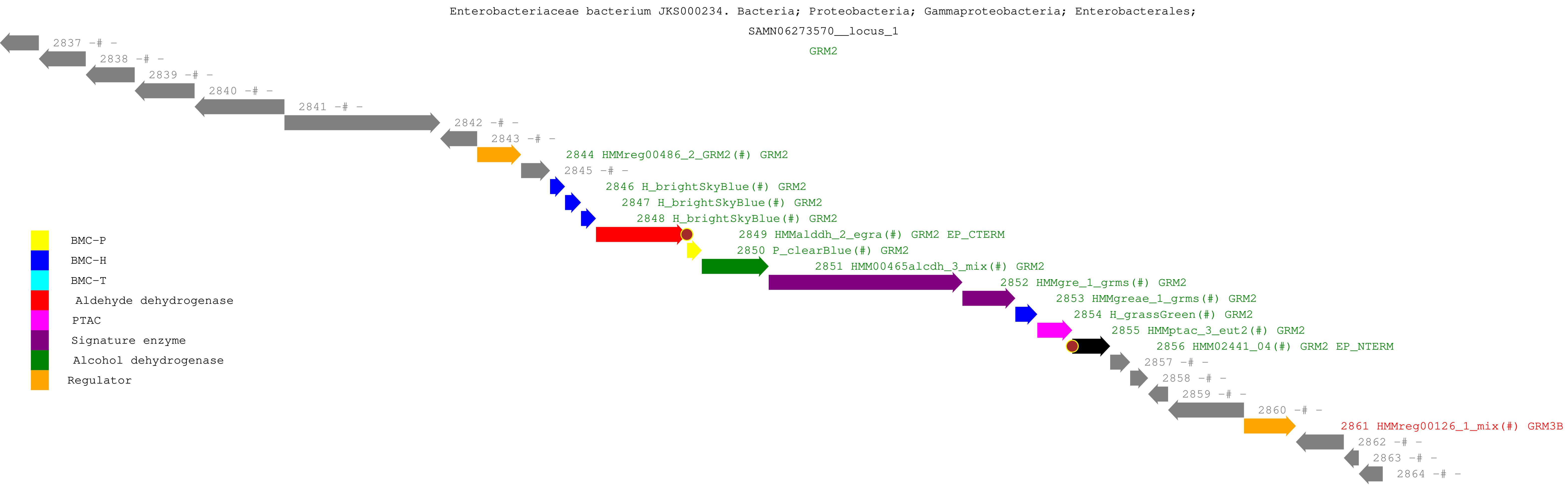
Enterobacteriaceae bacterium JKS000234. Bacteria; Proteobacteria; Gammaproteobacteria; Enterobacterales;
SAMN06273570__locus_1
GRM2
2837 -# -
2838 -# -
2839 -# -
2840 -# -
2841 -# -
2842 -# -
2843 -# -
2844 HMMreg00486_2_GRM2(#) GRM2
2845 -# -
2846 H_brightSkyBlue(#) GRM2
2847 H_brightSkyBlue(#) GRM2
2848 H_brightSkyBlue(#) GRM2
2849 HMMalddh_2_egra(#) GRM2 EP_CTERM
2850 P_clearBlue(#) GRM2
2851 HMM00465alcdh_3_mix(#) GRM2
2852 HMMgre_1_grms(#) GRM2
2853 HMMgreae_1_grms(#) GRM2
2854 H_grassGreen(#) GRM2
2855 HMMptac_3_eut2(#) GRM2
2856 HMM02441_04(#) GRM2 EP_NTERM
2857 -# -
2858 -# -
2859 -# -
2860 -# -
2861 HMMreg00126_1_mix(#) GRM3B
2862 -# -
2863 -# -
2864 -# -
BMC-P
BMC-H
BMC-T
Aldehyde dehydrogenase
PTAC
Signature enzyme
Alcohol dehydrogenase
Regulator

Klebsiella quasipneumoniae subsp. similipneumoniae. Bacteria; Proteobacteria; Gammaproteobacteria; Enterobacterales;

SB30__locus_1

GRM2

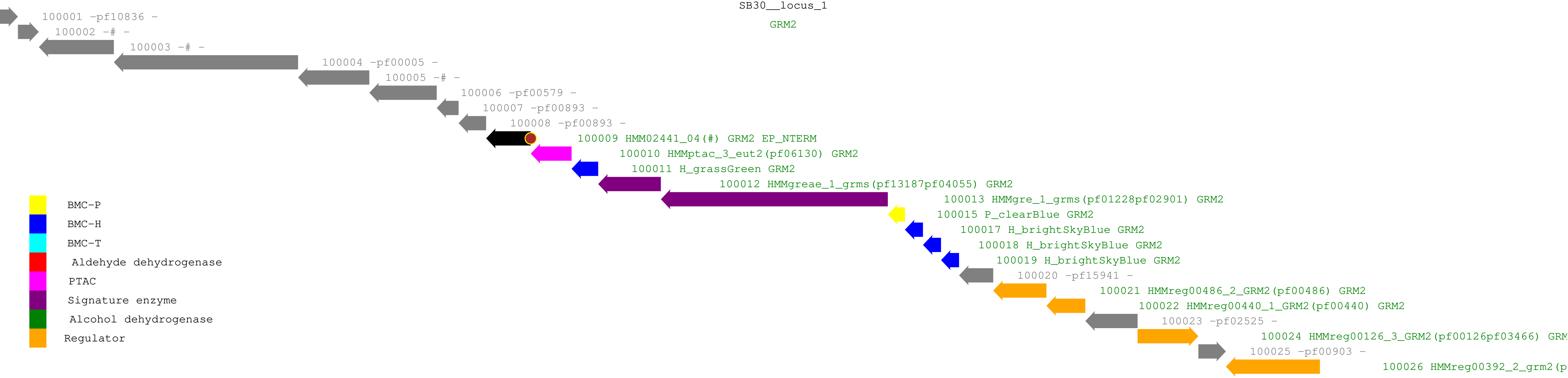

Klebsiella michiganensis. Bacteria; Proteobacteria; Gammaproteobacteria; Enterobacterales;

SB6408__locus_2

GRM2

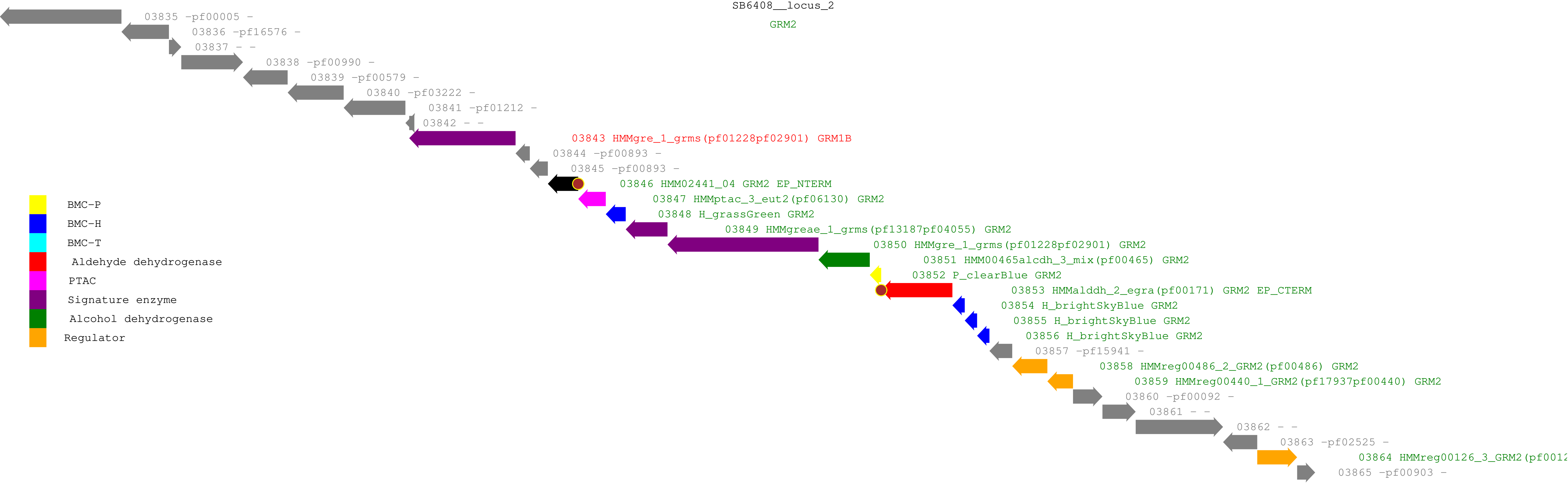

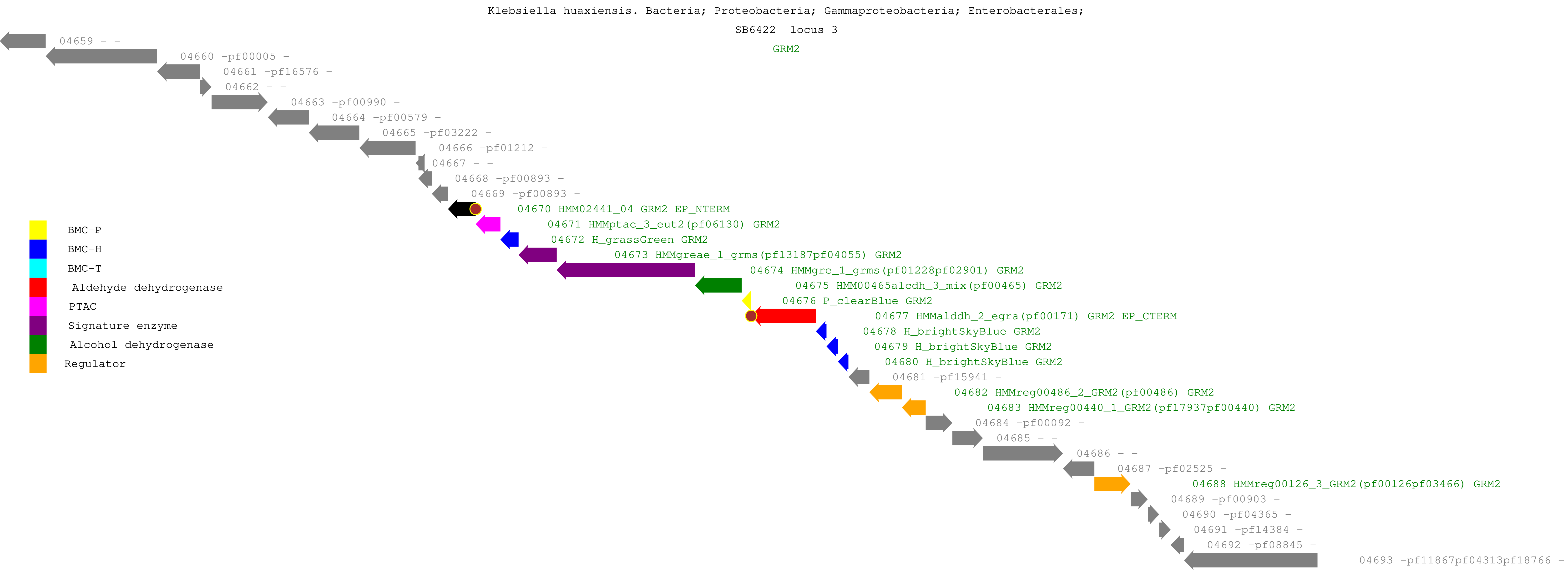
Klebsiella huaxiensis. Bacteria; Proteobacteria; Gammaproteobacteria; Enterobacterales;
SB6422__locus_3
GRM2
04659 - -
04660 -pf00005 -
04661 -pf16576 -
04662 - -
04663 -pf00990 -
04664 -pf00579 -
04665 -pf03222 -
04666 -pf01212 -
04667 - -
04668 -pf00893 -
04669 -pf00893 -
04670 HMM02441_04 GRM2 EP_NTERM
04671 HMMptac_3_eut2(pf06130) GRM2
04672 H_grassGreen GRM2
04673 HMMgreae_1_grms(pf13187pf04055) GRM2
04674 HMMgre_1_grms(pf01228pf02901) GRM2
04675 HMM00465alcdh_3_mix(pf00465) GRM2
04676 P_clearBlue GRM2
04677 HMMalddh_2_egra(pf00171) GRM2 EP_CTERM
04678 H_brightSkyBlue GRM2
04679 H_brightSkyBlue GRM2
04680 H_brightSkyBlue GRM2
04681 -pf15941 -
04682 HMMreg00486_2_GRM2(pf00486) GRM2
04683 HMMreg00440_1_GRM2(pf17937pf00440) GRM2
04684 -pf00092 -
04685 - -
04686 - -
04687 -pf02525 -
04688 HMMreg00126_3_GRM2(pf00126pf03466) GRM2
04689 -pf00903 -
04690 -pf04365 -
04691 -pf14384 -
04692 -pf08845 -
04693 -pf11867pf04313pf18766 -
BMC-P
BMC-H
BMC-T
Aldehyde dehydrogenase
PTAC
Signature enzyme
Alcohol dehydrogenase
Regulator

Klebsiella michiganensis. Bacteria; Proteobacteria; Gammaproteobacteria; Enterobacterales;

SK91__locus_2

GRM2

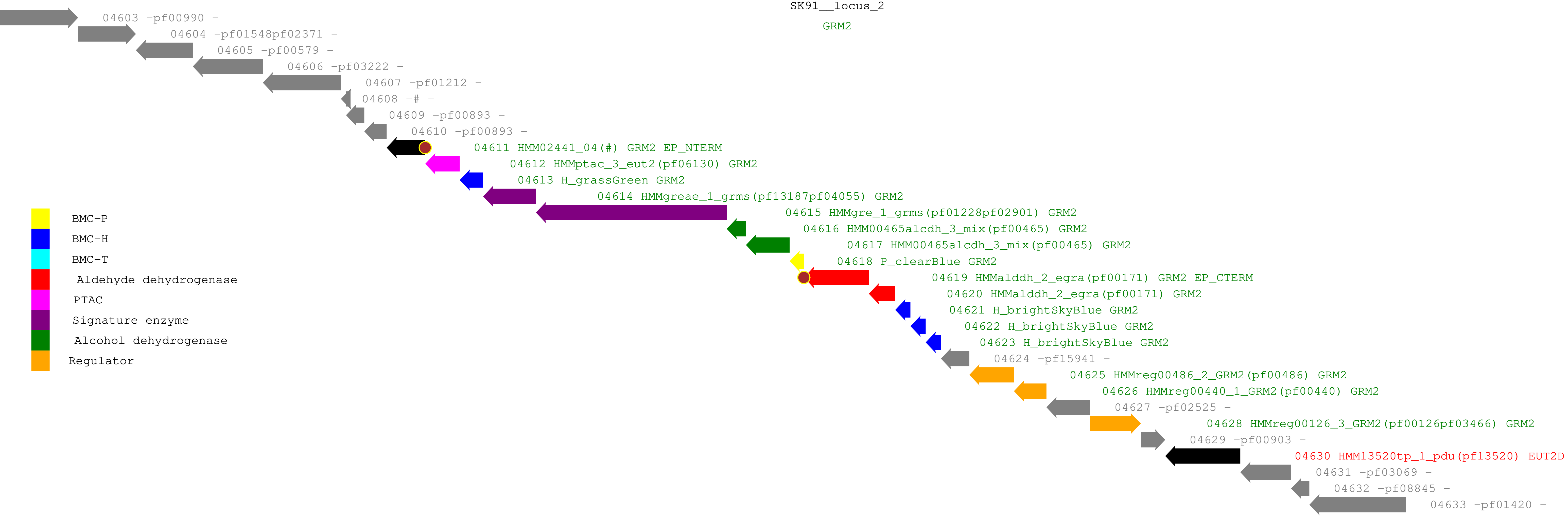

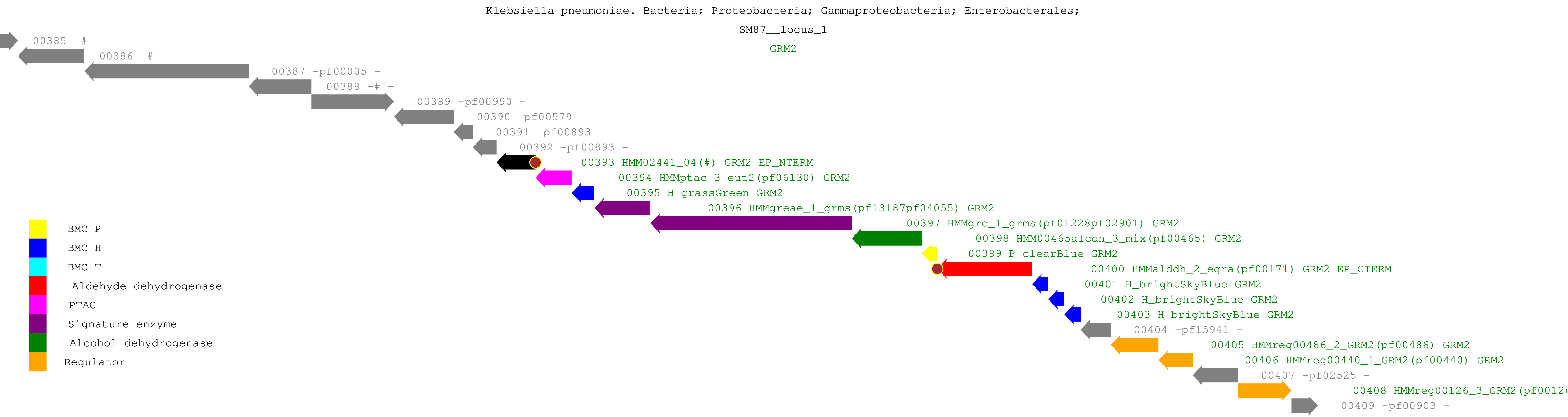
Klebsiella pneumoniae. Bacteria; Proteobacteria; Gammaproteobacteria; Enterobacterales;
SM87__locus_1
GRM2
00385 -# -
00386 -# -
00387 -pf00005 -
00388 -# -
00389 -pf00990 -
00390 -pf00579 -
00391 -pf00893 -
00392 -pf00893 -
00393 HMM02441_04(#) GRM2 EP_NTERM
00394 HMMptac_3_eut2(pf06130) GRM2
00395 H_grassGreen GRM2
00396 HMMgreae_1_grms(pf13187pf04055) GRM2
00397 HMMgre_1_grms(pf01228pf02901) GRM2
00398 HMM00465alcdh_3_mix(pf00465) GRM2
00399 P_clearBlue GRM2
00400 HMMaldh_2_egra(pf00171) GRM2 EP_CTERM
00401 H_brightSkyBlue GRM2
00402 H_brightSkyBlue GRM2
00403 H_brightSkyBlue GRM2
00404 -pf15941 -
00405 HMMreg00486_2_GRM2(pf00486) GRM2
00406 HMMreg00440_1_GRM2(pf00440) GRM2
00407 -pf02525 -
00408 HMMreg00126_3_GRM2(pf0012
00409 -pf00903 -
BMC-P
BMC-H
BMC-T
Aldehyde dehydrogenase
PTAC
Signature enzyme
Alcohol dehydrogenase
Regulator

Raoultella ornithinolytica (Klebsiella ornithinolytica). Bacteria; Proteobacteria; Gammaproteobacteria; Enterobacterales;

TE10__locus_1

GRM2

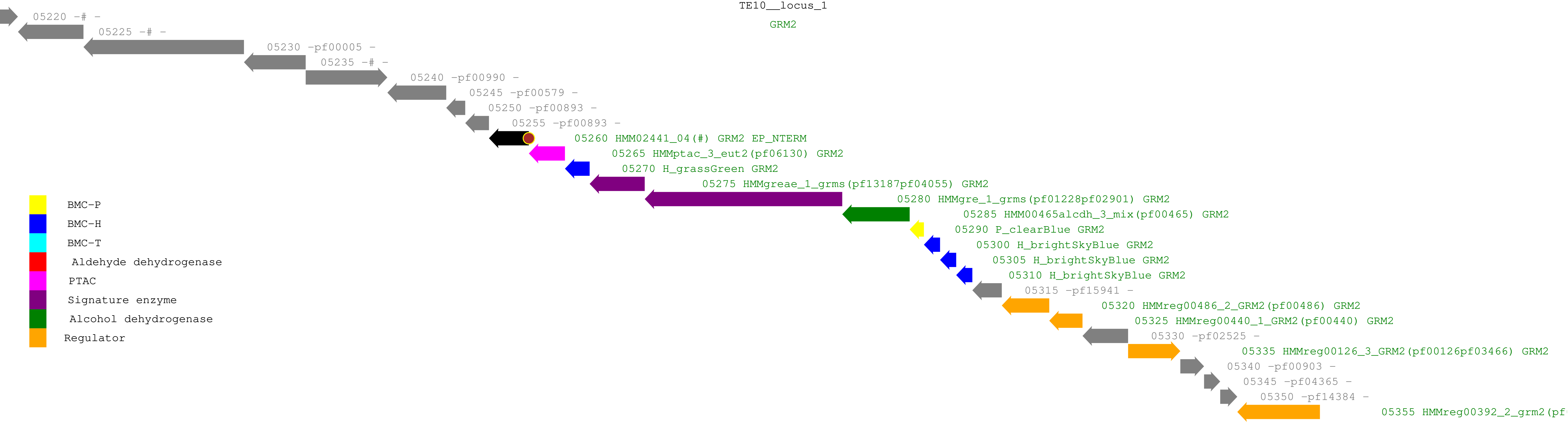

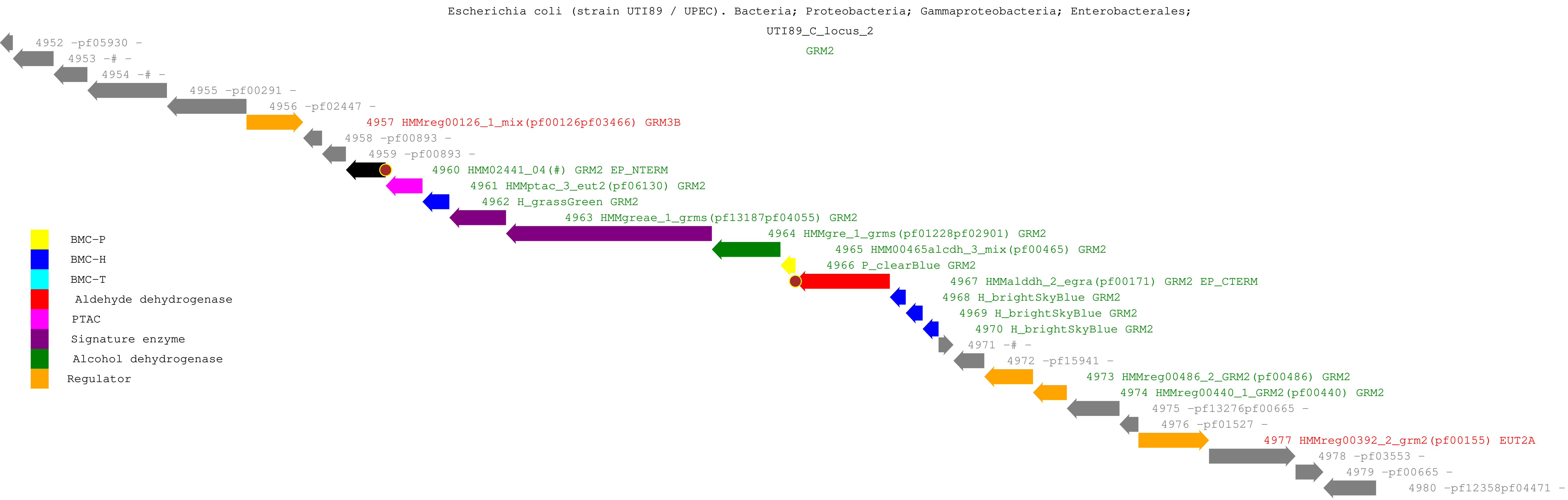
Escherichia coli (strain UTI89 / UPEC). Bacteria; Proteobacteria; Gammaproteobacteria; Enterobacterales;
UTI89_C_locus_2
GRM2
4952 -pf05930 -
4953 -# -
4954 -# -
4955 -pf00291 -
4956 -pf02447 -
4957 HMMreg00126_1_mix(pf00126pf03466) GRM3B
4958 -pf00893 -
4959 -pf00893 -
4960 HMM02441_04(#) GRM2 EP_NTERM
4961 HMMptac_3_eut2(pf06130) GRM2
4962 H_grassGreen GRM2
4963 HMMgreae_1_grms(pf13187pf04055) GRM2
4964 HMMgre_1_grms(pf01228pf02901) GRM2
4965 HMM00465alcdh_3_mix(pf00465) GRM2
4966 P_clearBlue GRM2
4967 HMMaldh_2_egra(pf00171) GRM2 EP_CTERM
4968 H_brightSkyBlue GRM2
4969 H_brightSkyBlue GRM2
4970 H_brightSkyBlue GRM2
4971 -# -
4972 -pf15941 -
4973 HMMreg00486_2_GRM2(pf00486) GRM2
4974 HMMreg00440_1_GRM2(pf00440) GRM2
4975 -pf13276pf00665 -
4976 -pf01527 -
4977 HMMreg00392_2_grm2(pf00155) EUT2A
4978 -pf03553 -
4979 -pf00665 -
4980 -pf12358pf04471 -
BMC-P
BMC-H
BMC-T
Aldehyde dehydrogenase
PTAC
Signature enzyme
Alcohol dehydrogenase
Regulator

Escherichia coli LAU-EC10. Bacteria; Proteobacteria; Gammaproteobacteria; Enterobacterales;

V415__locus_1

GRM2

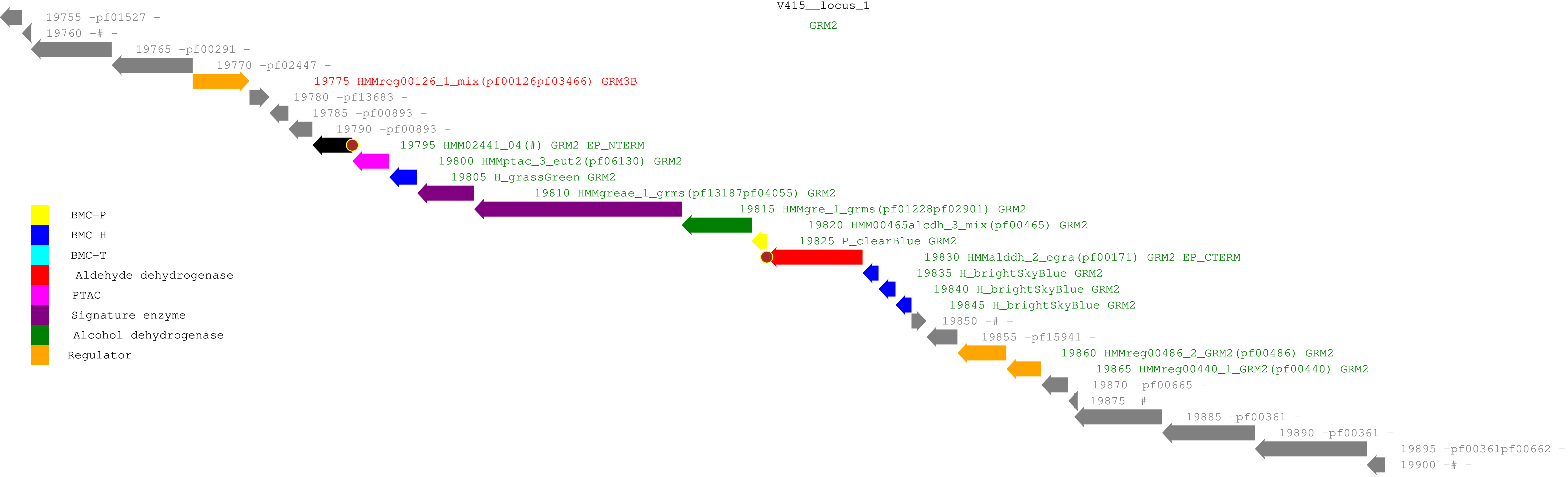

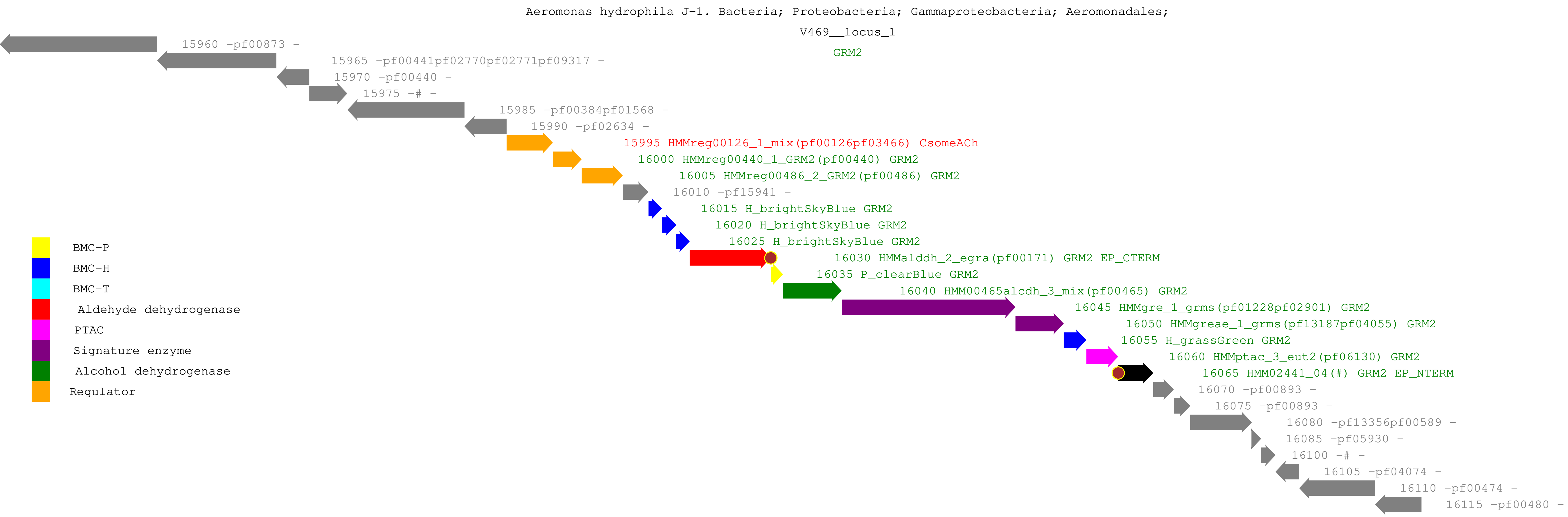

Aeromonas hydrophila J-1. Bacteria; Proteobacteria; Gammaproteobacteria; Aeromonadales;
V469__locus_1
GRM2
15960 -pf00873 -
15965 -pf00441pf02770pf02771pf09317 -
15970 -pf00440 -
15975 -# -
15985 -pf00384pf01568 -
15990 -pf02634 -
15995 HMMreg00126_1_mix(pf00126pf03466) CsomeACh
16000 HMMreg00440_1_GRM2(pf00440) GRM2
16005 HMMreg00486_2_GRM2(pf00486) GRM2
16010 -pf15941 -
16015 H_brightSkyBlue GRM2
16020 H_brightSkyBlue GRM2
16025 H_brightSkyBlue GRM2
16030 HMMaldh_2_egra(pf00171) GRM2 EP_CTERM
16035 P_clearBlue GRM2
16040 HMM00465alcdh_3_mix(pf00465) GRM2
16045 HMMgre_1_grms(pf01228pf02901) GRM2
16050 HMMgreae_1_grms(pf13187pf04055) GRM2
16055 H_grassGreen GRM2
16060 HMMptac_3_eut2(pf06130) GRM2
16065 HMM02441_04(#) GRM2 EP_NTERM
16070 -pf00893 -
16075 -pf00893 -
16080 -pf13356pf00589 -
16085 -pf05930 -
16100 -# -
16105 -pf04074 -
16110 -pf00474 -
16115 -pf00480 -
BMC-P
BMC-H
BMC-T
Aldehyde dehydrogenase
PTAC
Signature enzyme
Alcohol dehydrogenase
Regulator

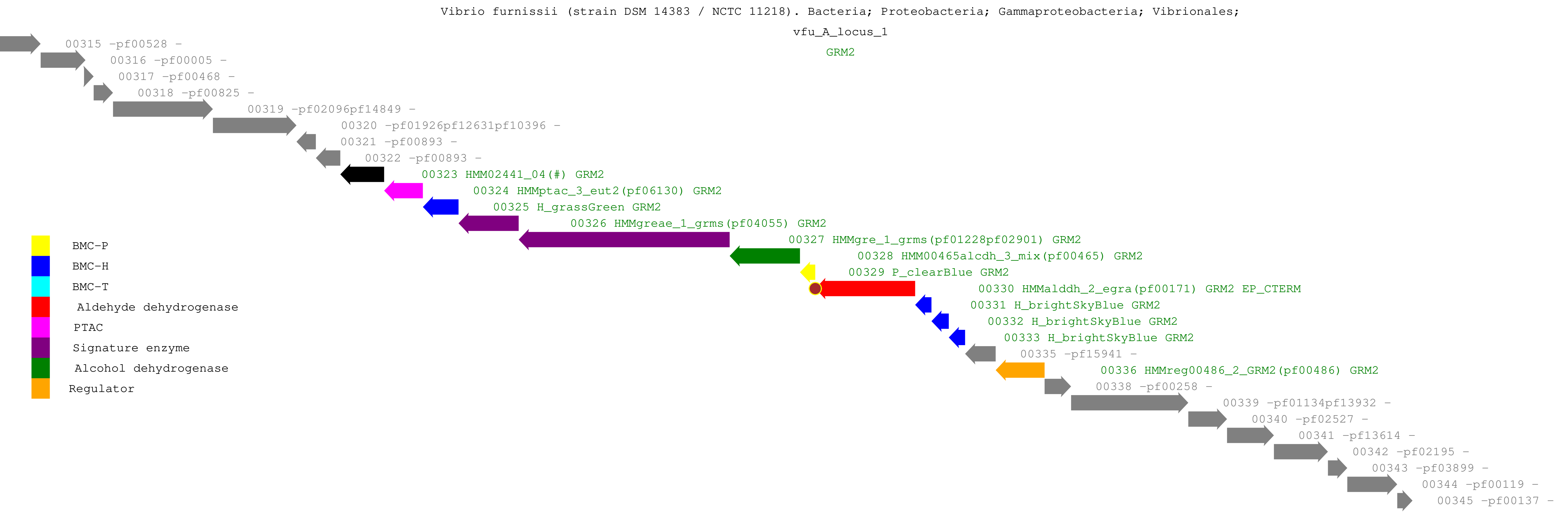
Vibrio furnissii (strain DSM 14383 / NCTC 11218). Bacteria; Proteobacteria; Gammaproteobacteria; Vibrionales;
vfu_A_locus_1
GRM2
00315 -pf00528 -
00316 -pf00005 -
00317 -pf00468 -
00318 -pf00825 -
00319 -pf02096pf14849 -
00320 -pf01926pf12631pf10396 -
00321 -pf00893 -
00322 -pf00893 -
00323 HMM02441_04(#) GRM2
00324 HMMptac_3_eut2(pf06130) GRM2
00325 H_grassGreen GRM2
00326 HMMgreae_1_grms(pf04055) GRM2
00327 HMMgre_1_grms(pf01228pf02901) GRM2
00328 HMM00465alcdh_3_mix(pf00465) GRM2
00329 P_clearBlue GRM2
00330 HMMaldh_2_egra(pf00171) GRM2 EP_CTERM
00331 H_brightSkyBlue GRM2
00332 H_brightSkyBlue GRM2
00333 H_brightSkyBlue GRM2
00335 -pf15941 -
00336 HMMreg00486_2_GRM2(pf00486) GRM2
00338 -pf00258 -
00339 -pf01134pf13932 -
00340 -pf02527 -
00341 -pf13614 -
00342 -pf02195 -
00343 -pf03899 -
00344 -pf00119 -
00345 -pf00137 -
BMC-P
BMC-H
BMC-T
Aldehyde dehydrogenase
PTAC
Signature enzyme
Alcohol dehydrogenase
Regulator

Moellerella wisconsensis. Bacteria; Proteobacteria; Gammaproteobacteria; Enterobacterales;

VK86__locus_1

GRM2

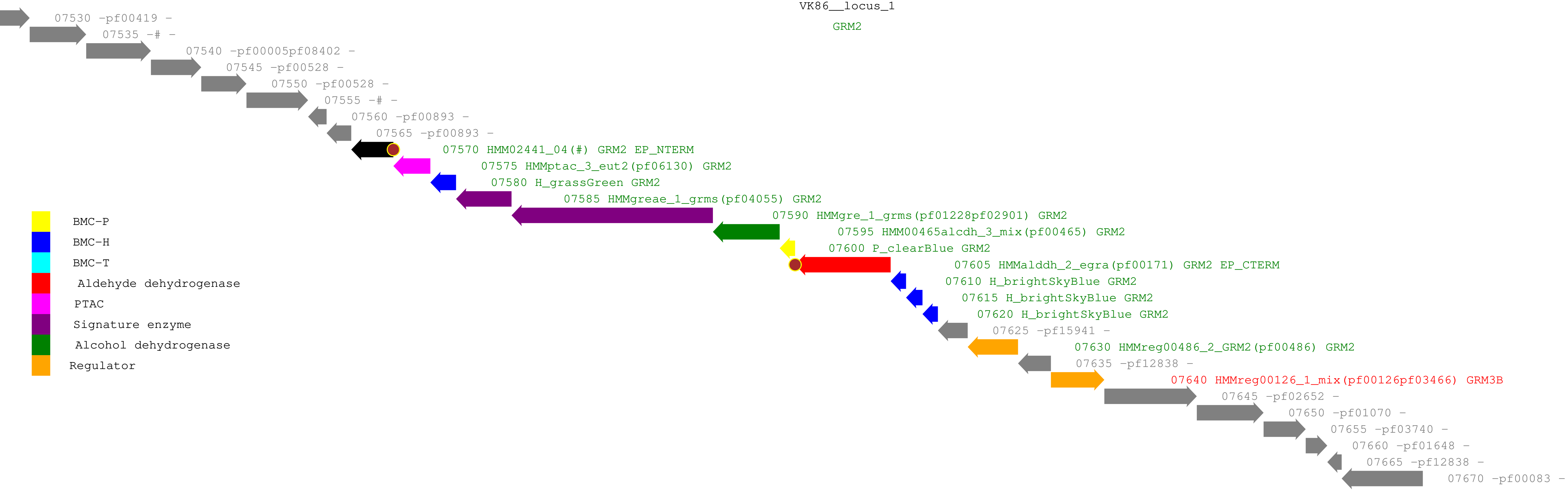

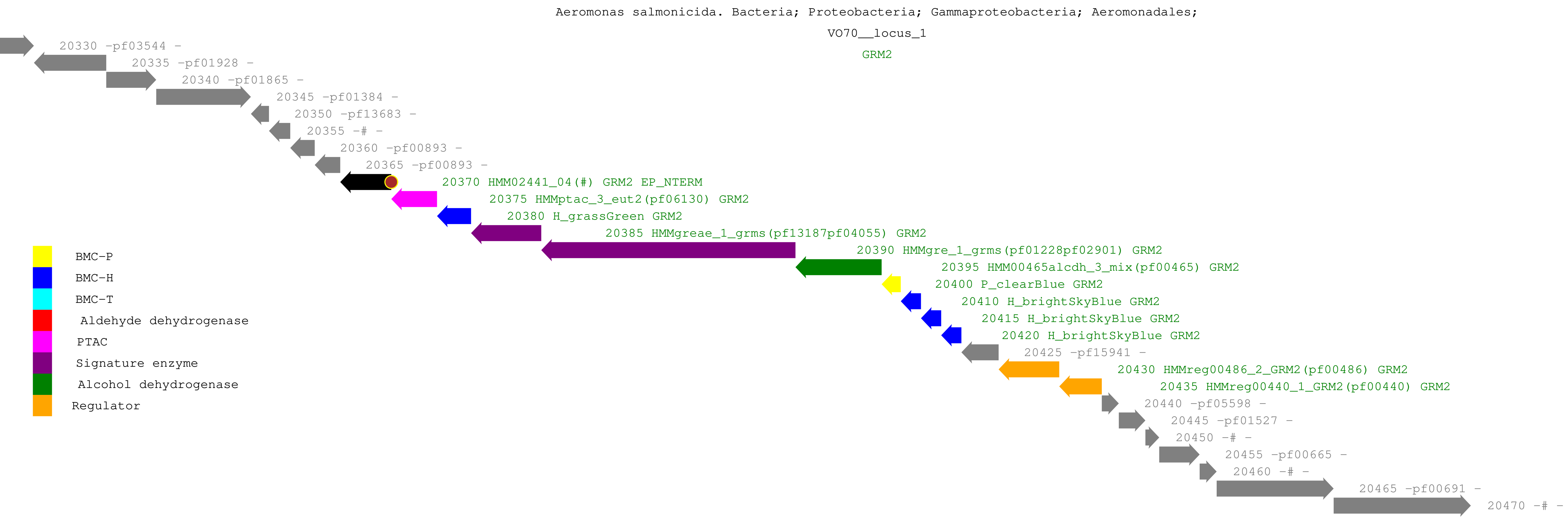

Aeromonas salmonicida. Bacteria; Proteobacteria; Gammaproteobacteria; Aeromonadales;
VO70__locus_1
GRM2
20330 -pf03544 -
20335 -pf01928 -
20340 -pf01865 -
20345 -pf01384 -
20350 -pf13683 -
20355 -# -
20360 -pf00893 -
20365 -pf00893 -
20370 HMM02441_04(#) GRM2 EP_NTERM
20375 HMMptac_3_eut2(pf06130) GRM2
20380 H_grassGreen GRM2
20385 HMMgreae_1_grms(pf13187pf04055) GRM2
20390 HMMgre_1_grms(pf01228pf02901) GRM2
20395 HMM00465alcdh_3_mix(pf00465) GRM2
20400 P_clearBlue GRM2
20410 H_brightSkyBlue GRM2
20415 H_brightSkyBlue GRM2
20420 H_brightSkyBlue GRM2
20425 -pf15941 -
20430 HMMreg00486_2_GRM2(pf00486) GRM2
20435 HMMreg00440_1_GRM2(pf00440) GRM2
20440 -pf05598 -
20445 -pf01527 -
20450 -# -
20455 -pf00665 -
20460 -# -
20465 -pf00691 -
20470 -# -
BMC-P
BMC-H
BMC-T
Aldehyde dehydrogenase
PTAC
Signature enzyme
Alcohol dehydrogenase
Regulator

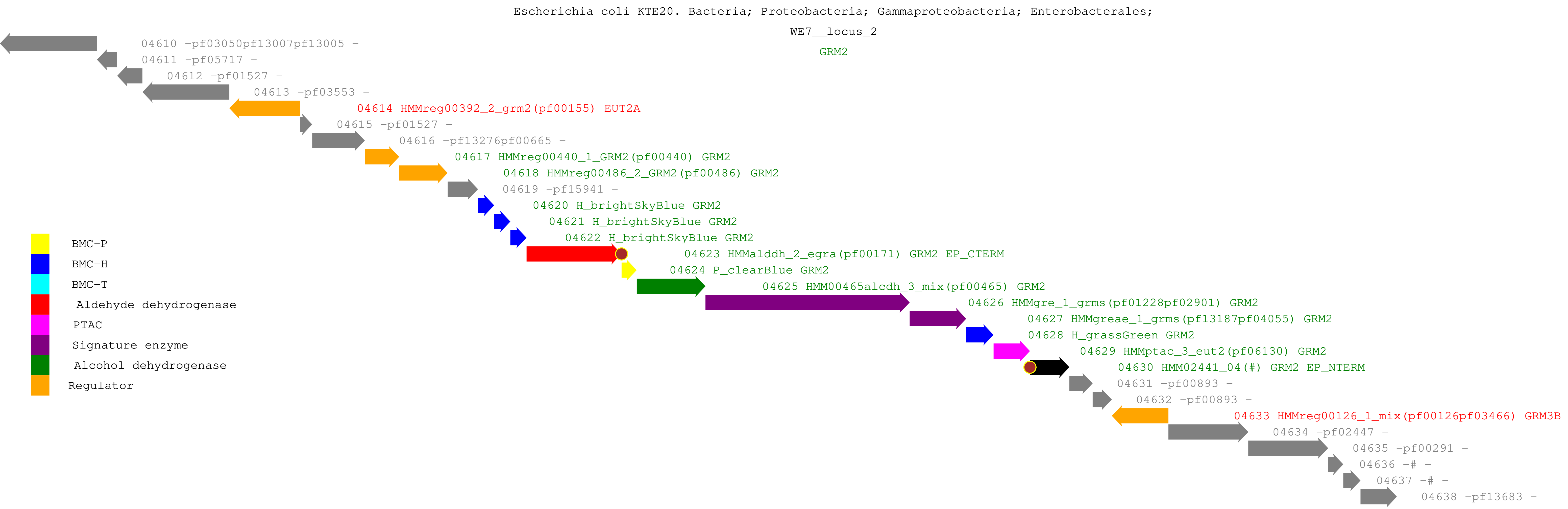
Escherichia coli KTE20. Bacteria; Proteobacteria; Gammaproteobacteria; Enterobacterales;
WE7__locus_2
GRM2
04610 -pf03050pf13007pf13005 -
04611 -pf05717 -
04612 -pf01527 -
04613 -pf03553 -
04614 HMMreg00392_2_grm2(pf00155) EUT2A
04615 -pf01527 -
04616 -pf13276pf00665 -
04617 HMMreg00440_1_GRM2(pf00440) GRM2
04618 HMMreg00486_2_GRM2(pf00486) GRM2
04619 -pf15941 -
04620 H_brightSkyBlue GRM2
04621 H_brightSkyBlue GRM2
04622 H_brightSkyBlue GRM2
04623 HMMalddh_2_egra(pf00171) GRM2 EP_CTERM
04624 P_clearBlue GRM2
04625 HMM00465alcdh_3_mix(pf00465) GRM2
04626 HMMgre_1_grms(pf01228pf02901) GRM2
04627 HMMgreae_1_grms(pf13187pf04055) GRM2
04628 H_grassGreen GRM2
04629 HMMptac_3_eut2(pf06130) GRM2
04630 HMM02441_04(#) GRM2 EP_NTERM
04631 -pf00893 -
04632 -pf00893 -
04633 HMMreg00126_1_mix(pf00126pf03466) GRM3B
04634 -pf02447 -
04635 -pf00291 -
04636 -# -
04637 -# -
04638 -pf13683 -
BMC-P
BMC-H
BMC-T
Aldehyde dehydrogenase
PTAC
Signature enzyme
Alcohol dehydrogenase
Regulator

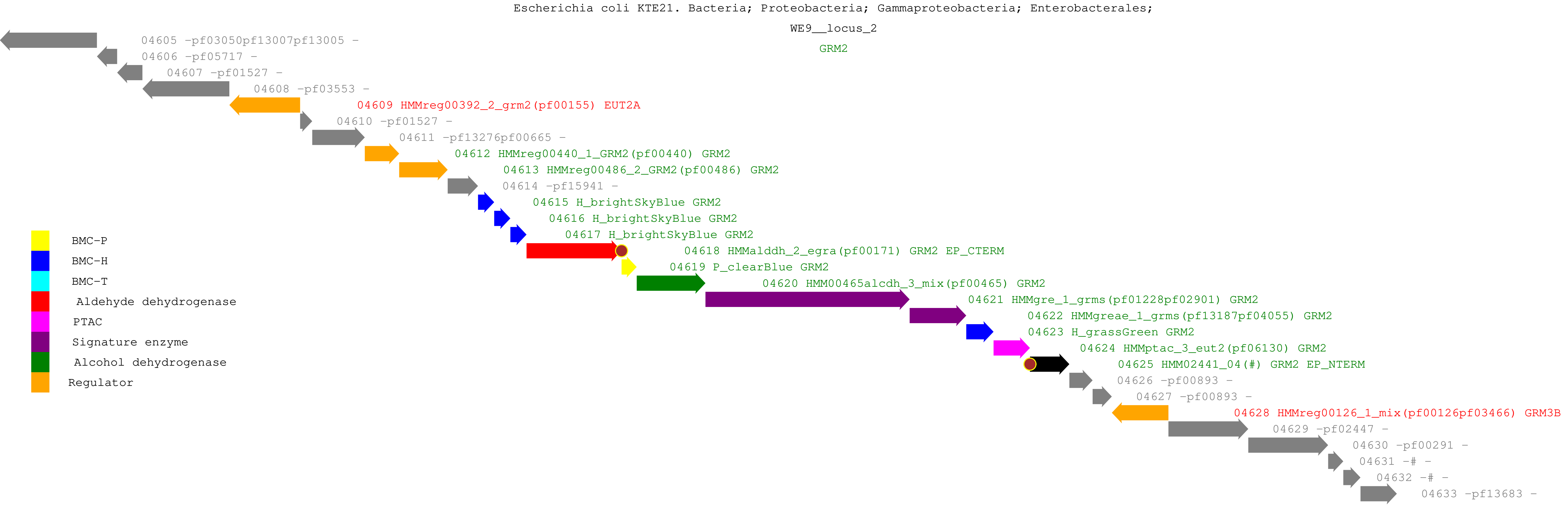

Escherichia coli KTE21. Bacteria; Proteobacteria; Gammaproteobacteria; Enterobacterales;
WE9__locus_2
GRM2
04605 -pf03050pf13007pf13005 -
04606 -pf05717 -
04607 -pf01527 -
04608 -pf03553 -
04609 HMMreg00392_2_grm2(pf00155) EUT2A
04610 -pf01527 -
04611 -pf13276pf00665 -
04612 HMMreg00440_1_GRM2(pf00440) GRM2
04613 HMMreg00486_2_GRM2(pf00486) GRM2
04614 -pf15941 -
04615 H_brightSkyBlue GRM2
04616 H_brightSkyBlue GRM2
04617 H_brightSkyBlue GRM2
04618 HMMaldddh_2_egra(pf00171) GRM2 EP_CTERM
04619 P_clearBlue GRM2
04620 HMM00465alcdh_3_mix(pf00465) GRM2
04621 HMMgre_1_grms(pf01228pf02901) GRM2
04622 HMMgreae_1_grms(pf13187pf04055) GRM2
04623 H_grassGreen GRM2
04624 HMMptac_3_eut2(pf06130) GRM2
04625 HMM02441_04(#) GRM2 EP_NTERM
04626 -pf00893 -
04627 -pf00893 -
04628 HMMreg00126_1_mix(pf00126pf03466) GRM3B
04629 -pf02447 -
04630 -pf00291 -
04631 -# -
04632 -# -
04633 -pf13683 -
BMC-P
BMC-H
BMC-T
Aldehyde dehydrogenase
PTAC
Signature enzyme
Alcohol dehydrogenase
Regulator

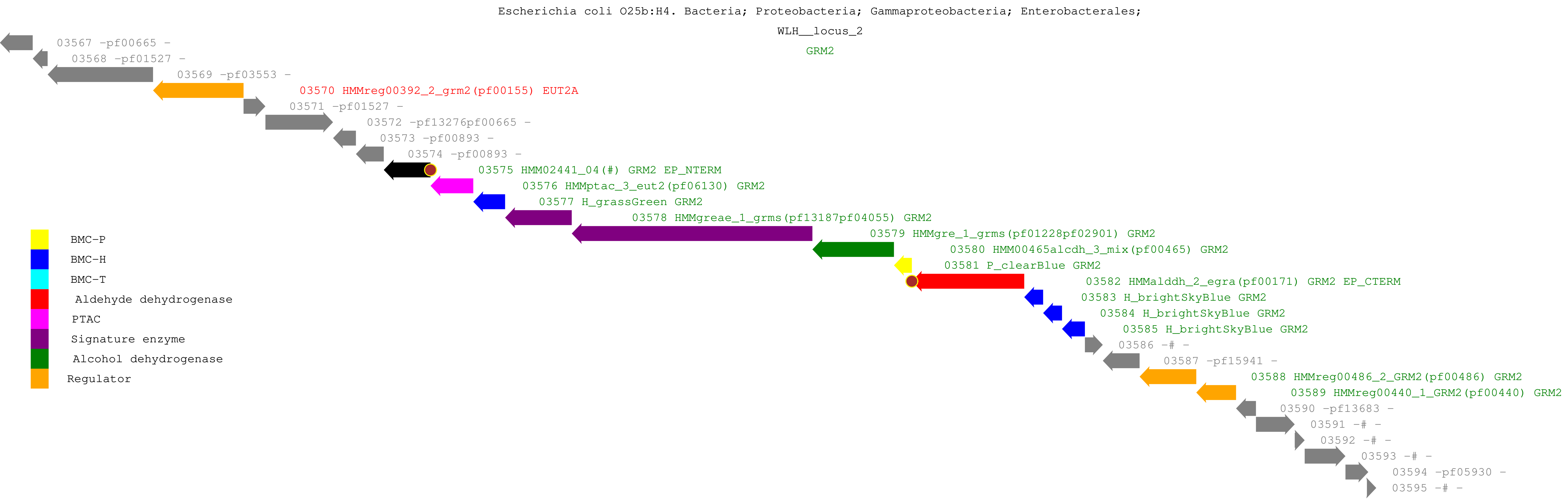
Escherichia coli O25b:H4. Bacteria; Proteobacteria; Gammaproteobacteria; Enterobacterales;
WLH__locus_2
GRM2
03567 -pf00665 -
03568 -pf01527 -
03569 -pf03553 -
03570 HMMreg00392_2_grm2(pf00155) EUT2A
03571 -pf01527 -
03572 -pf13276pf00665 -
03573 -pf00893 -
03574 -pf00893 -
03575 HMM02441_04(#) GRM2 EP_NTERM
03576 HMMptac_3_eut2(pf06130) GRM2
03577 H_grassGreen GRM2
03578 HMMgreae_1_grms(pf13187pf04055) GRM2
03579 HMMgre_1_grms(pf01228pf02901) GRM2
03580 HMM00465alcdh_3_mix(pf00465) GRM2
03581 P_clearBlue GRM2
03582 HMMaldh_2_egra(pf00171) GRM2 EP_CTERM
03583 H_brightSkyBlue GRM2
03584 H_brightSkyBlue GRM2
03585 H_brightSkyBlue GRM2
03586 -# -
03587 -pf15941 -
03588 HMMreg00486_2_GRM2(pf00486) GRM2
03589 HMMreg00440_1_GRM2(pf00440) GRM2
03590 -pf13683 -
03591 -# -
03592 -# -
03593 -# -
03594 -pf05930 -
03595 -# -
BMC-P
BMC-H
BMC-T
Aldehyde dehydrogenase
PTAC
Signature enzyme
Alcohol dehydrogenase
Regulator

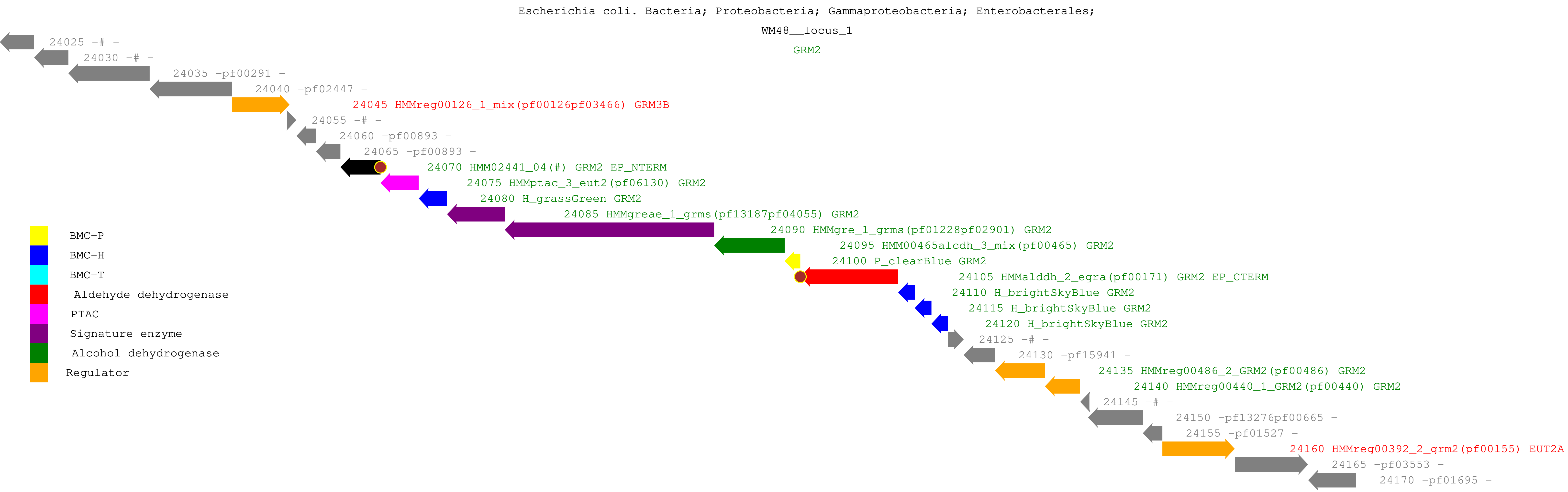
Escherichia coli. Bacteria; Proteobacteria; Gammaproteobacteria; Enterobacterales;
WM48__locus_1
GRM2
24025 -# -
24030 -# -
24035 -pf00291 -
24040 -pf02447 -
24045 HMMreg00126_1_mix(pf00126pf03466) GRM3B
24055 -# -
24060 -pf00893 -
24065 -pf00893 -
24070 HMM02441_04(#) GRM2 EP_NTERM
24075 HMMptac_3_eut2(pf06130) GRM2
24080 H_grassGreen GRM2
24085 HMMgreae_1_grms(pf13187pf04055) GRM2
24090 HMMgre_1_grms(pf01228pf02901) GRM2
24095 HMM00465alcdh_3_mix(pf00465) GRM2
24100 P_clearBlue GRM2
24105 HMMaldh_2_egra(pf00171) GRM2 EP_CTERM
24110 H_brightSkyBlue GRM2
24115 H_brightSkyBlue GRM2
24120 H_brightSkyBlue GRM2
24125 -# -
24130 -pf15941 -
24135 HMMreg00486_2_GRM2(pf00486) GRM2
24140 HMMreg00440_1_GRM2(pf00440) GRM2
24145 -# -
24150 -pf13276pf00665 -
24155 -pf01527 -
24160 HMMreg00392_2_grm2(pf00155) EUT2A
24165 -pf03553 -
24170 -pf01695 -
BMC-P
BMC-H
BMC-T
Aldehyde dehydrogenase
PTAC
Signature enzyme
Alcohol dehydrogenase
Regulator

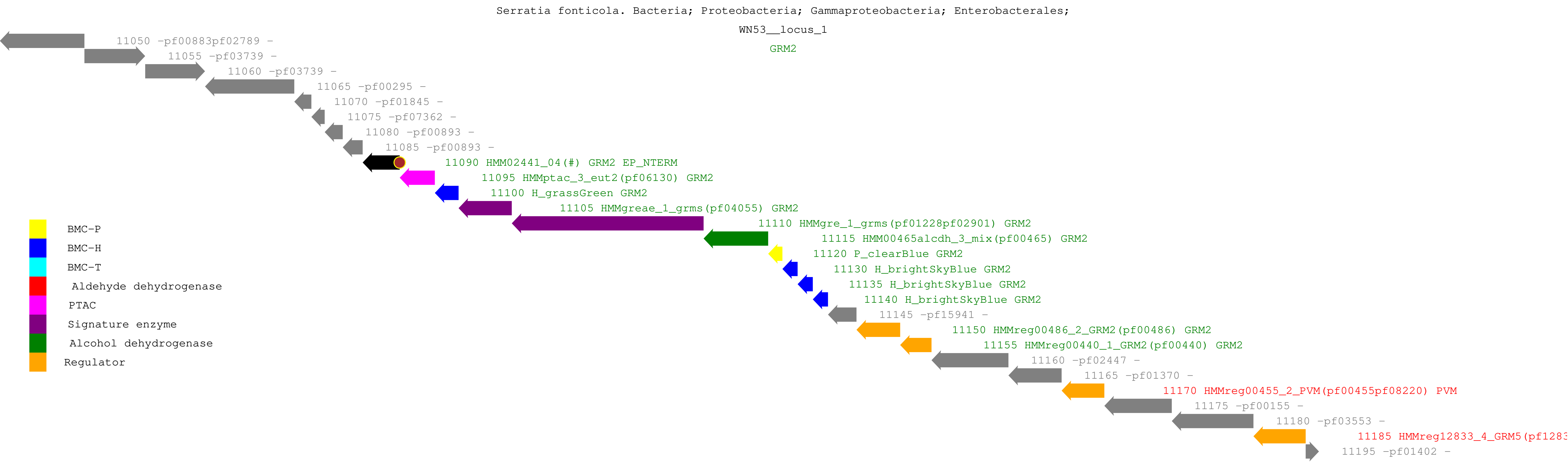
Serratia fonticola. Bacteria; Proteobacteria; Gammaproteobacteria; Enterobacterales;
WN53__locus_1
GRM2
11050 -pf00883pf02789 -
11055 -pf03739 -
11060 -pf03739 -
11065 -pf00295 -
11070 -pf01845 -
11075 -pf07362 -
11080 -pf00893 -
11085 -pf00893 -
11090 HMM02441_04(#) GRM2 EP_NTERM
11095 HMMptac_3_eut2(pf06130) GRM2
11100 H_grassGreen GRM2
11105 HMMgreae_1_grms(pf04055) GRM2
11110 HMMgre_1_grms(pf01228pf02901) GRM2
11115 HMM00465alcdh_3_mix(pf00465) GRM2
11120 P_clearBlue GRM2
11130 H_brightSkyBlue GRM2
11135 H_brightSkyBlue GRM2
11140 H_brightSkyBlue GRM2
11145 -pf15941 -
11150 HMMreg00486_2_GRM2(pf00486) GRM2
11155 HMMreg00440_1_GRM2(pf00440) GRM2
11160 -pf02447 -
11165 -pf01370 -
11170 HMMreg00455_2_PVM(pf00455pf08220) PVM
11175 -pf00155 -
11180 -pf03553 -
11185 HMMreg12833_4_GRM5(pf1283
11195 -pf01402 -
BMC-P
BMC-H
BMC-T
Aldehyde dehydrogenase
PTAC
Signature enzyme
Alcohol dehydrogenase
Regulator

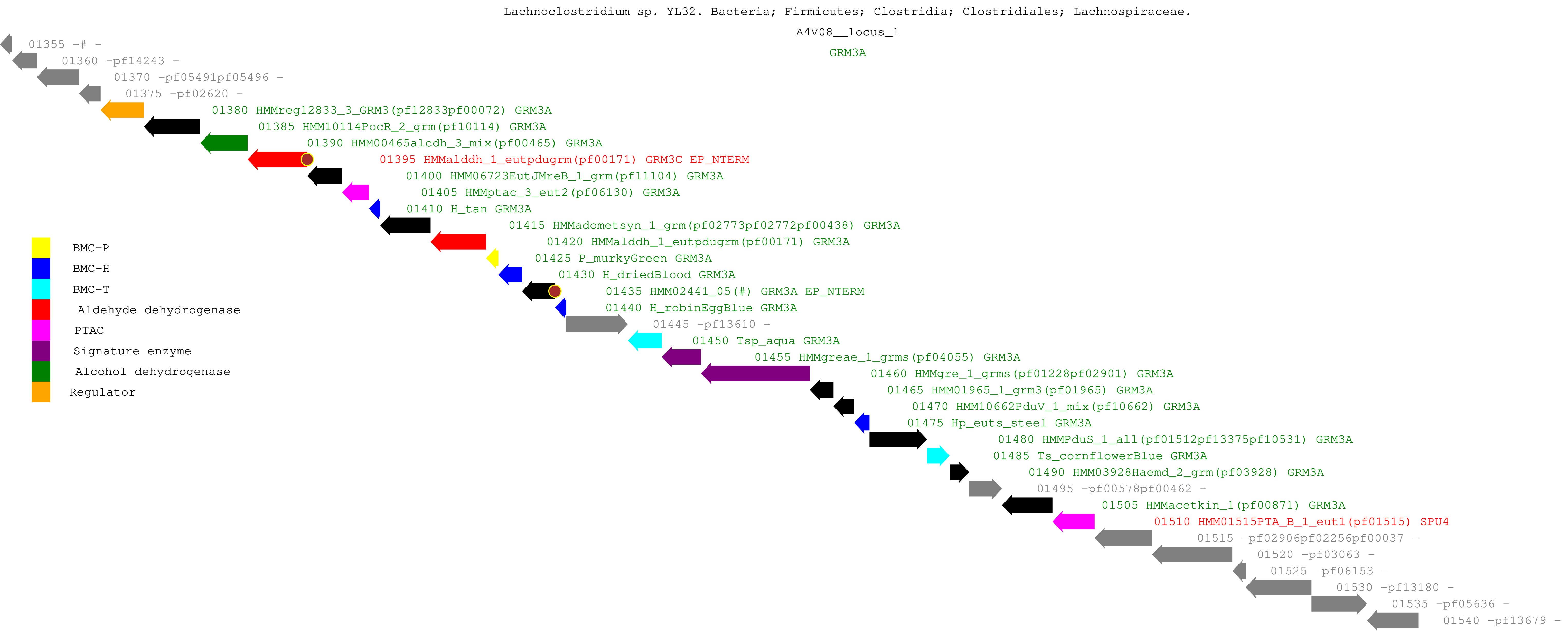
Lachnoclostridium sp. YL32. Bacteria; Firmicutes; Clostridia; Clostridiales; Lachnospiraceae.
A4V08__locus_1
GRM3A
01355 -# -
01360 -pf14243 -
01370 -pf05491pf05496 -
01375 -pf02620 -
01380 HMMreg12833_3_GRM3(pf12833pf00072) GRM3A
01385 HMM10114PocR_2_grm(pf10114) GRM3A
01390 HMM00465alcdh_3_mix(pf00465) GRM3A
01395 HMMaldDh_1_eutpdugrm(pf00171) GRM3C EP_NTERM
01400 HMM06723EutJMreB_1_grm(pf11104) GRM3A
01405 HMMptac_3_eut2(pf06130) GRM3A
01410 H_tan GRM3A
01415 HMMadometsyn_1_grm(pf02773pf02772pf00438) GRM3A
01420 HMMaldDh_1_eutpdugrm(pf00171) GRM3A
01425 P_murkyGreen GRM3A
01430 H_driedBlood GRM3A
01435 HMM02441_05(#) GRM3A EP_NTERM
01440 H_robinEggBlue GRM3A
01445 -pf13610 -
01450 Tsp_aqua GRM3A
01455 HMMgreae_1_grms(pf04055) GRM3A
01460 HMMgre_1_grms(pf01228pf02901) GRM3A
01465 HMM01965_1_grm3(pf01965) GRM3A
01470 HMM10662PduV_1_mix(pf10662) GRM3A
01475 Hp_euts_steel GRM3A
01480 HMMPduS_1_all(pf01512pf13375pf10531) GRM3A
01485 Ts_cornflowerBlue GRM3A
01490 HMM03928Haemd_2_grm(pf03928) GRM3A
01495 -pf00578pf00462 -
01505 HMMacetkin_1(pf00871) GRM3A
01510 HMM01515PTA_B_1_eut1(pf01515) SPU4
01515 -pf02906pf02256pf00037 -
01520 -pf03063 -
01525 -pf06153 -
01530 -pf13180 -
01535 -pf05636 -
01540 -pf13679 -
BMC-P
BMC-H
BMC-T
Aldehyde dehydrogenase
PTAC
Signature enzyme
Alcohol dehydrogenase
Regulator

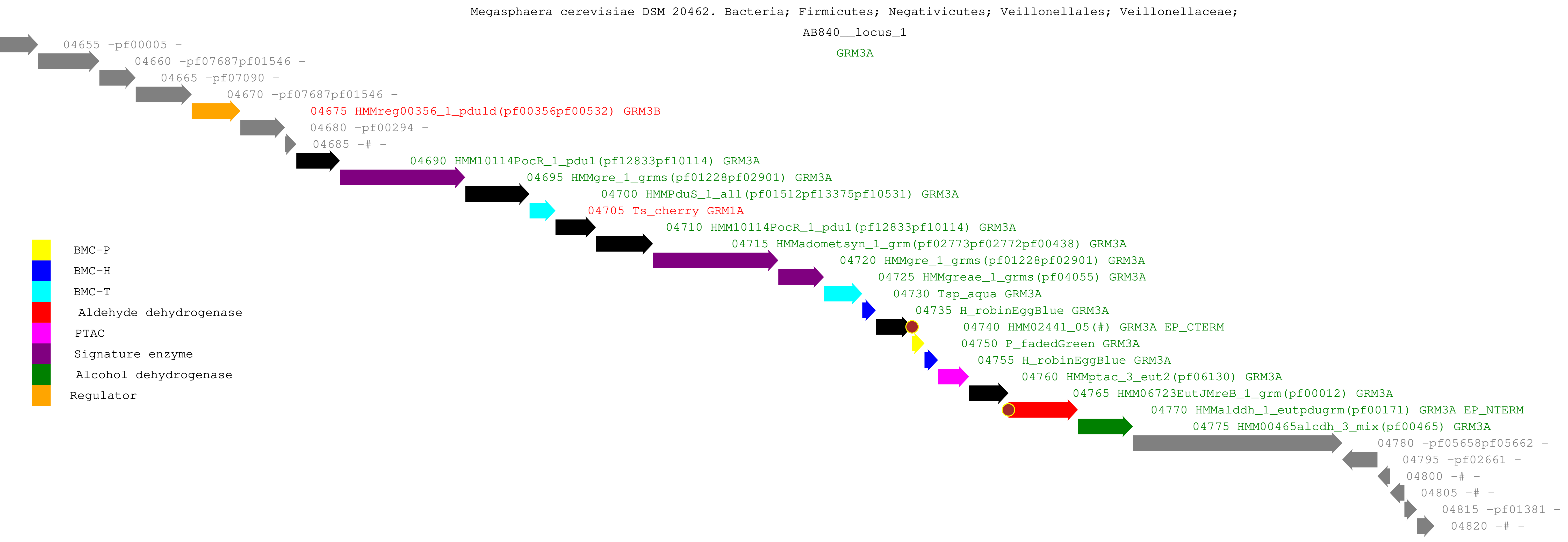
Megasphaera cerevisiae DSM 20462. Bacteria; Firmicutes; Negativicutes; Veillonellales; Veillonellaceae;
AB840__locus_1
GRM3A
04655 -pf00005 -
04660 -pf07687pf01546 -
04665 -pf07090 -
04670 -pf07687pf01546 -
04675 HMMreg00356_1_pdu1d(pf00356pf00532) GRM3B
04680 -pf00294 -
04685 -# -
04690 HMM10114PocR_1_pdu1(pf12833pf10114) GRM3A
04695 HMMgre_1_grms(pf01228pf02901) GRM3A
04700 HMMPduS_1_all(pf01512pf13375pf10531) GRM3A
04705 Ts_cherry GRM1A
04710 HMM10114PocR_1_pdu1(pf12833pf10114) GRM3A
04715 HMMadometsyn_1_grm(pf02773pf02772pf00438) GRM3A
04720 HMMgre_1_grms(pf01228pf02901) GRM3A
04725 HMMgreae_1_grms(pf04055) GRM3A
04730 Tsp_aqua GRM3A
04735 H_robinEggBlue GRM3A
04740 HMM02441_05(#) GRM3A EP_CTERM
04750 P_fadedGreen GRM3A
04755 H_robinEggBlue GRM3A
04760 HMMptac_3_eut2(pf06130) GRM3A
04765 HMM06723EutJMreB_1_grm(pf00012) GRM3A
04770 HMMalddh_1_eutpdugrm(pf00171) GRM3A EP_NTERM
04775 HMM00465alcdh_3_mix(pf00465) GRM3A
04780 -pf05658pf05662 -
04795 -pf02661 -
04800 -# -
04805 -# -
04815 -pf01381 -
04820 -# -
BMC-P
BMC-H
BMC-T
Aldehyde dehydrogenase
PTAC
Signature enzyme
Alcohol dehydrogenase
Regulator

Clostridium clostridioforme. Bacteria; Firmicutes; Clostridia; Clostridiales; Lachnospiraceae.

ADH76__locus_1

GRM3A

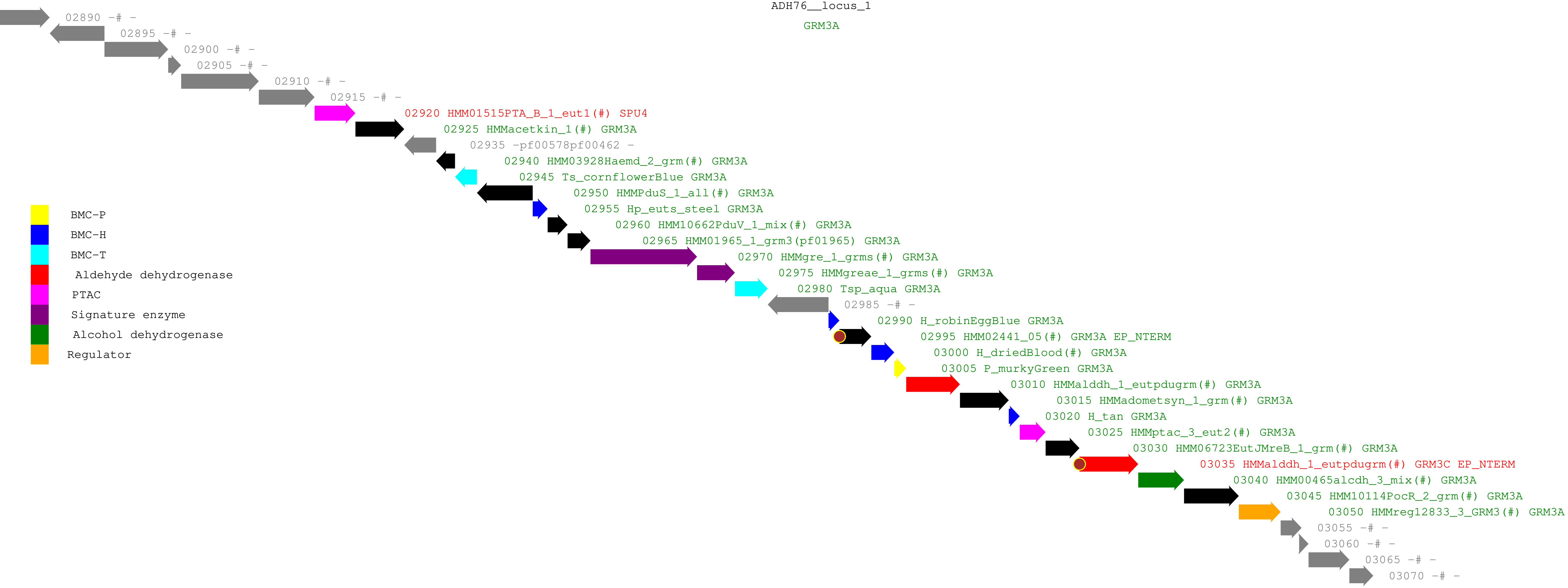

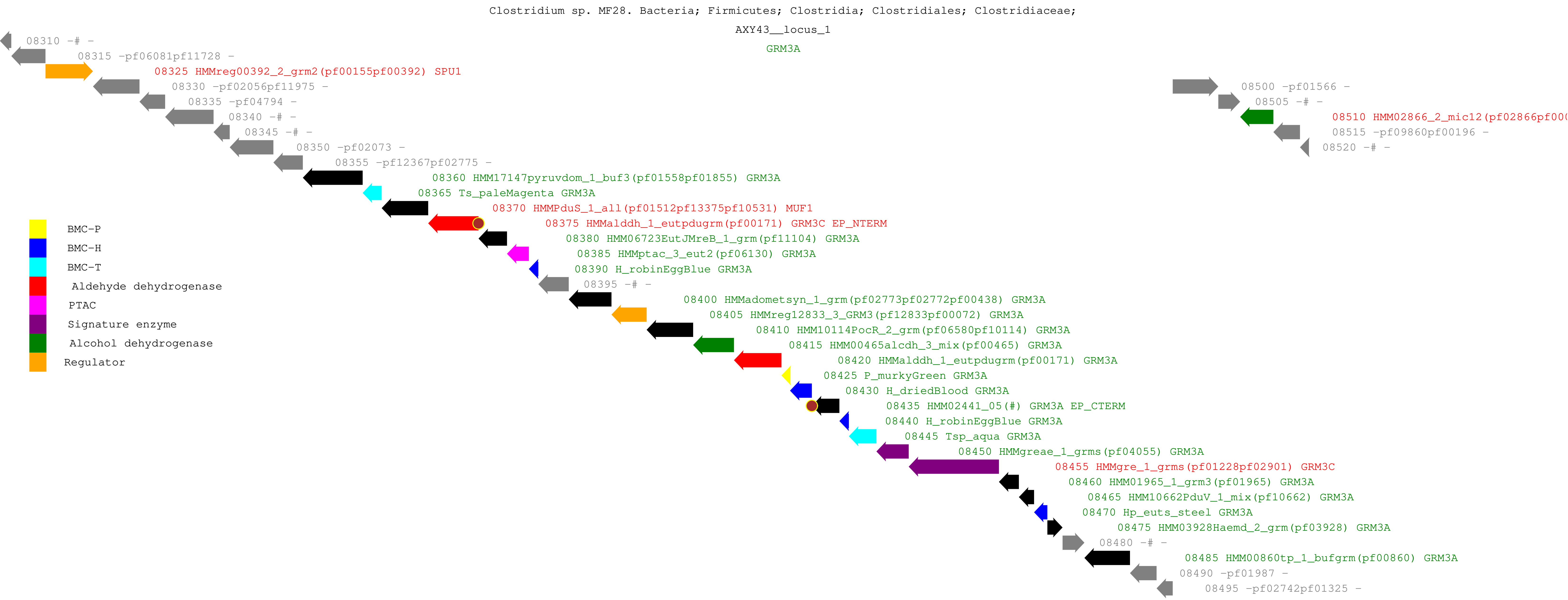
Clostridium sp. MF28. Bacteria; Firmicutes; Clostridia; Clostridiales; Clostridiaceae;
AXY43__locus_1
GRM3A
08310 -# -
08315 -pf06081pf11728 -
08325 HMMreg00392_2_grm2(pf00155pf00392) SPU1
08330 -pf02056pf11975 -
08335 -pf04794 -
08340 -# -
08345 -# -
08350 -pf02073 -
08355 -pf12367pf02775 -
08360 HMM17147pyruvdom_1_buf3(pf01558pf01855) GRM3A
08365 Ts_paleMagenta GRM3A
08370 HMMPduS_1_all(pf01512pf13375pf10531) MUF1
08375 HMMaldh_1_eutpdugrm(pf00171) GRM3C EP_NTERM
08380 HMM06723EutJMreB_1_grm(pf11104) GRM3A
08385 HMMptac_3_eut2(pf06130) GRM3A
08390 H_robinEggBlue GRM3A
08395 -# -
08400 HMMadometsyn_1_grm(pf02773pf02772pf00438) GRM3A
08405 HMMreg12833_3_GRM3(pf12833pf00072) GRM3A
08410 HMM10114PocR_2_grm(pf06580pf10114) GRM3A
08415 HMM00465alcdh_3_mix(pf00465) GRM3A
08420 HMMaldh_1_eutpdugrm(pf00171) GRM3A
08425 P_murkyGreen GRM3A
08430 H_driedBlood GRM3A
08435 HMM02441_05(#) GRM3A EP_CTERM
08440 H_robinEggBlue GRM3A
08445 Tsp_aqua GRM3A
08450 HMMgreae_1_grms(pf04055) GRM3A
08455 HMMgre_1_grms(pf01228pf02901) GRM3C
08460 HMM01965_1_grm3(pf01965) GRM3A
08465 HMM10662PduV_1_mix(pf10662) GRM3A
08470 Hp_euts_steel GRM3A
08475 HMM03928Haemd_2_grm(pf03928) GRM3A
08480 -# -
08485 HMM00860tp_1_bufgrm(pf00860) GRM3A
08490 -pf01987 -
08495 -pf02742pf01325 -
08500 -pf01566 -
08505 -# -
08510 HMM02866_2_mic12(pf02866pf00
08515 -pf09860pf00196 -
08520 -# -
BMC-P
BMC-H
BMC-T
Aldehyde dehydrogenase
PTAC
Signature enzyme
Alcohol dehydrogenase
Regulator

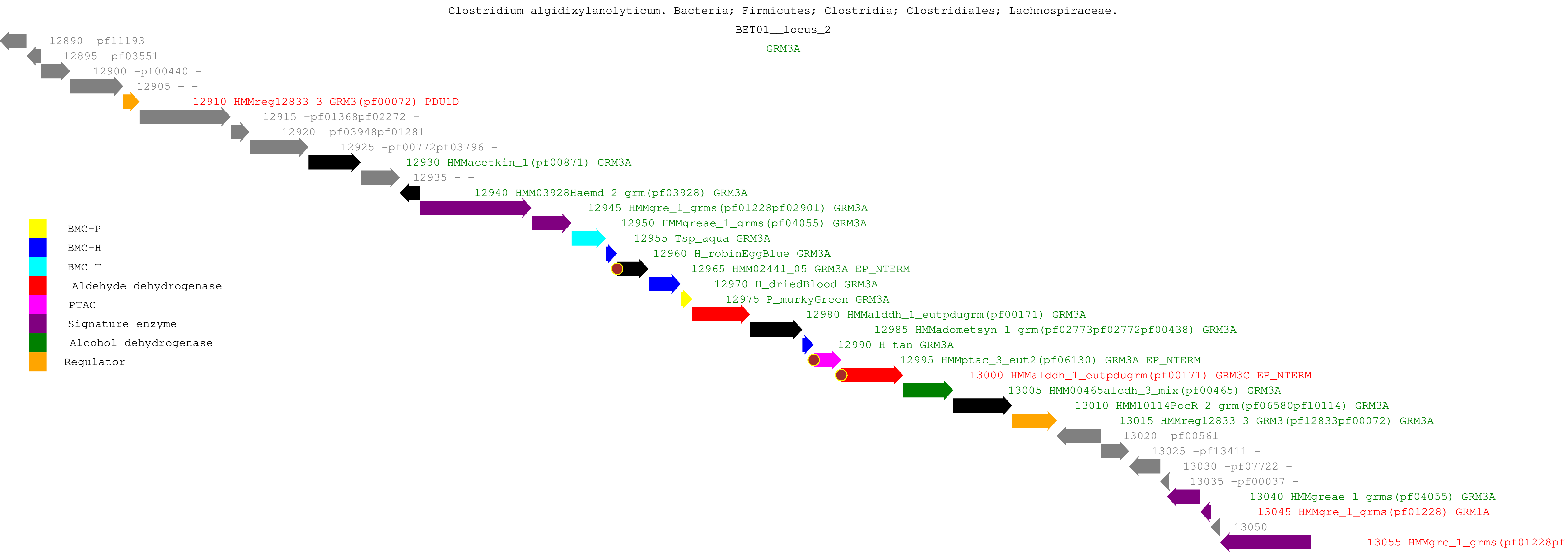
Clostridium algidixylanolyticum. Bacteria; Firmicutes; Clostridia; Clostridiales; Lachnospiraceae.
BET01__locus_2
GRM3A
12890 -pf11193 -
12895 -pf03551 -
12900 -pf00440 -
12905 - -
12910 HMMreg12833_3_GRM3(pf00072) PDU1D
12915 -pf01368pf02272 -
12920 -pf03948pf01281 -
12925 -pf00772pf03796 -
12930 HMMacetkin_1(pf00871) GRM3A
12935 - -
12940 HMM03928Haemd_2_grm(pf03928) GRM3A
12945 HMMgre_1_grms(pf01228pf02901) GRM3A
12950 HMMgreae_1_grms(pf04055) GRM3A
12955 Tsp_aqua GRM3A
12960 H_robinEggBlue GRM3A
12965 HMM02441_05 GRM3A EP_NTERM
12970 H_driedBlood GRM3A
12975 P_murkyGreen GRM3A
12980 HMMalddh_1_eutpdugrm(pf00171) GRM3A
12985 HMMadometsyn_1_grm(pf02773pf02772pf00438) GRM3A
12990 H_tan GRM3A
12995 HMMptac_3_eut2(pf06130) GRM3A EP_NTERM
13000 HMMalddh_1_eutpdugrm(pf00171) GRM3C EP_NTERM
13005 HMM00465alcdh_3_mix(pf00465) GRM3A
13010 HMM10114PocR_2_grm(pf06580pf10114) GRM3A
13015 HMMreg12833_3_GRM3(pf12833pf00072) GRM3A
13020 -pf00561 -
13025 -pf13411 -
13030 -pf07722 -
13035 -pf00037 -
13040 HMMgreae_1_grms(pf04055) GRM3A
13045 HMMgre_1_grms(pf01228) GRM1A
13050 - -
13055 HMMgre_1_grms(pf01228pf
BMC-P
BMC-H
BMC-T
Aldehyde dehydrogenase
PTAC
Signature enzyme
Alcohol dehydrogenase
Regulator

Clostridium beijerinckii (Clostridium MP). Bacteria; Firmicutes; Clostridia; Clostridiales; Clostridiaceae;

BGS1__locus_1

GRM3A

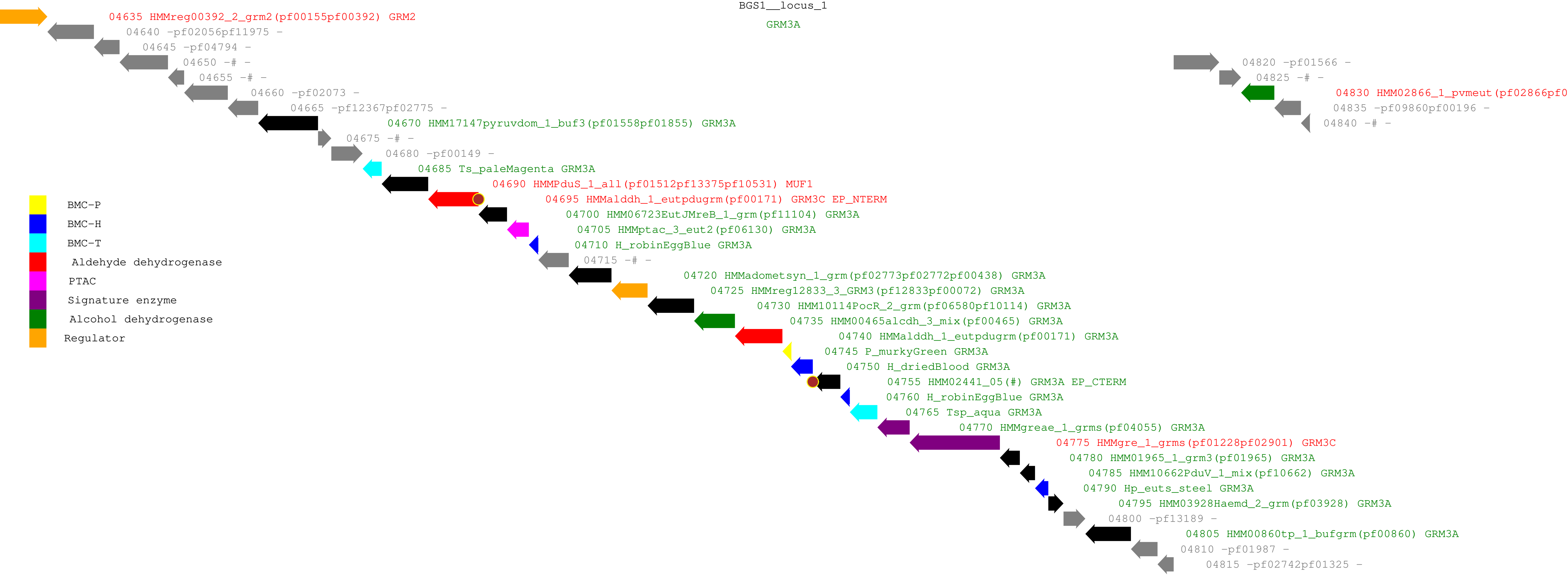

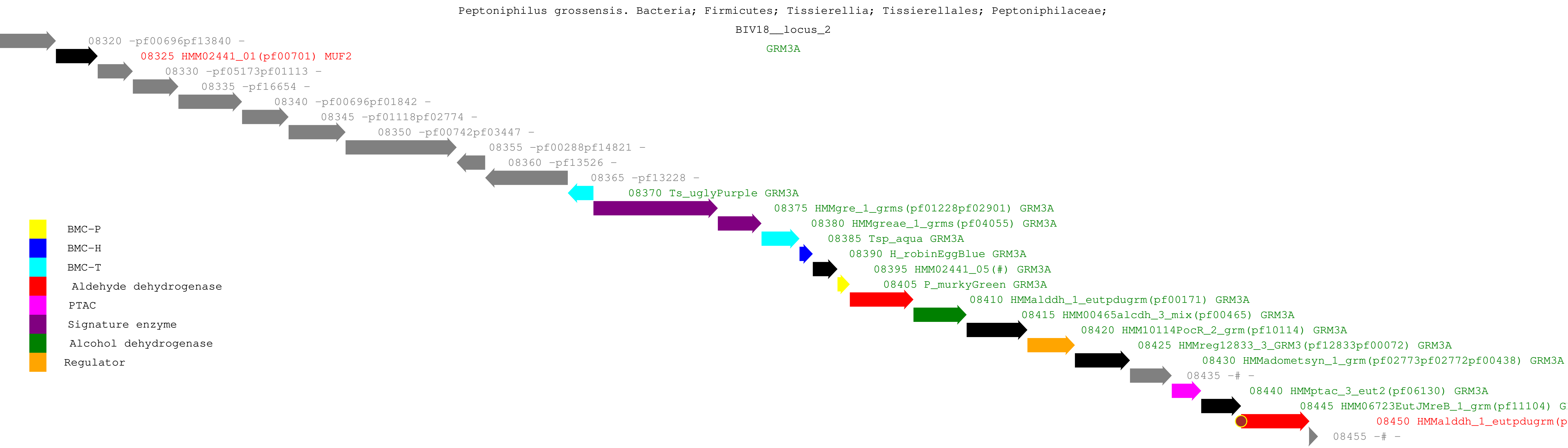
Peptoniphilus grossensis. Bacteria; Firmicutes; Tissierellia; Tissierellales; Peptoniphilaceae;
BIV18__locus_2
GRM3A
08320 -pf00696pf13840 -
08325 HMM02441_01(pf00701) MUF2
08330 -pf05173pf01113 -
08335 -pf16654 -
08340 -pf00696pf01842 -
08345 -pf01118pf02774 -
08350 -pf00742pf03447 -
08355 -pf00288pf14821 -
08360 -pf13526 -
08365 -pf13228 -
08370 Ts_uglyPurple GRM3A
08375 HMMgre_1_grms(pf01228pf02901) GRM3A
08380 HMMgreae_1_grms(pf04055) GRM3A
08385 Tsp_aqua GRM3A
08390 H_robinEggBlue GRM3A
08395 HMM02441_05(#) GRM3A
08405 P_murkyGreen GRM3A
08410 HMMaldh_1_eutpdugrm(pf00171) GRM3A
08415 HMM00465alcdh_3_mix(pf00465) GRM3A
08420 HMM10114PocR_2_grm(pf10114) GRM3A
08425 HMMreg12833_3_GRM3(pf12833pf00072) GRM3A
08430 HMMadometsyn_1_grm(pf02773pf02772pf00438) GRM3A
08435 -# -
08440 HMMptac_3_eut2(pf06130) GRM3A
08445 HMM06723EutJMreB_1_grm(pf11104) G
08450 HMMaldh_1_eutpdugrm(p
08455 -# -
BMC-P
BMC-H
BMC-T
Aldehyde dehydrogenase
PTAC
Signature enzyme
Alcohol dehydrogenase
Regulator

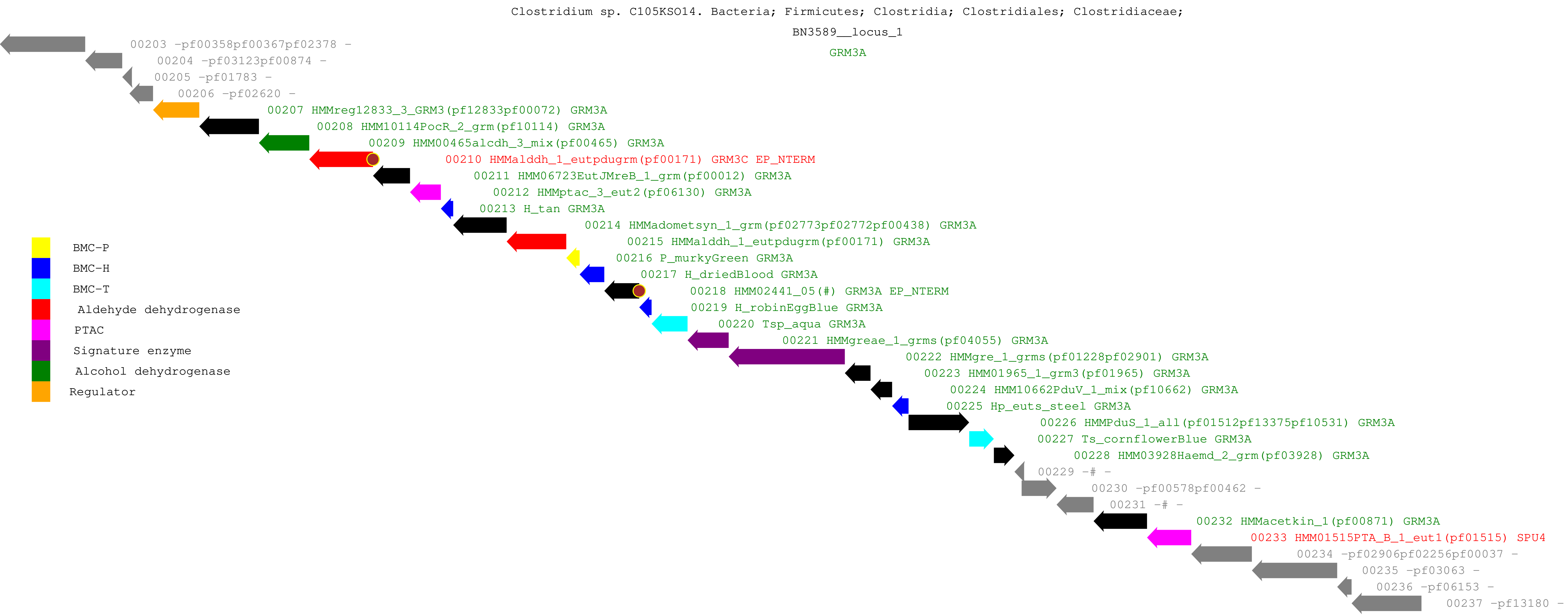

Clostridium sp. C105KSO14. Bacteria; Firmicutes; Clostridia; Clostridiales; Clostridiaceae;
BN3589__locus_1
GRM3A
00203 -pf00358pf00367pf02378 -
00204 -pf03123pf00874 -
00205 -pf01783 -
00206 -pf02620 -
00207 HMMreg12833_3_GRM3(pf12833pf00072) GRM3A
00208 HMM10114PocR_2_grm(pf10114) GRM3A
00209 HMM00465alcdh_3_mix(pf00465) GRM3A
00210 HMMaldh_1_eutpdugrm(pf00171) GRM3C EP_NTERM
00211 HMM06723EutJMreB_1_grm(pf00012) GRM3A
00212 HMMptac_3_eut2(pf06130) GRM3A
00213 H_tan GRM3A
00214 HMMadometsyn_1_grm(pf02773pf02772pf00438) GRM3A
00215 HMMaldh_1_eutpdugrm(pf00171) GRM3A
00216 P_murkyGreen GRM3A
00217 H_driedBlood GRM3A
00218 HMM02441_05(#) GRM3A EP_NTERM
00219 H_robinEggBlue GRM3A
00220 Tsp_aqua GRM3A
00221 HMMgreae_1_grms(pf04055) GRM3A
00222 HMMgre_1_grms(pf01228pf02901) GRM3A
00223 HMM01965_1_grm3(pf01965) GRM3A
00224 HMM10662PduV_1_mix(pf10662) GRM3A
00225 Hp_euts_steel GRM3A
00226 HMMPduS_1_all(pf01512pf13375pf10531) GRM3A
00227 Ts_cornflowerBlue GRM3A
00228 HMM03928Haemd_2_grm(pf03928) GRM3A
00229 -# -
00230 -pf00578pf00462 -
00231 -# -
00232 HMMacetkin_1(pf00871) GRM3A
00233 HMM01515PTA_B_1_eut1(pf01515) SPU4
00234 -pf02906pf02256pf00037 -
00235 -pf03063 -
00236 -pf06153 -
00237 -pf13180 -
BMC-P
BMC-H
BMC-T
Aldehyde dehydrogenase
PTAC
Signature enzyme
Alcohol dehydrogenase
Regulator

Clostridium clostridioforme CAG:132. Bacteria; Firmicutes; Clostridia; Clostridiales; Clostridiaceae;

BN486__locus_1

GRM3A

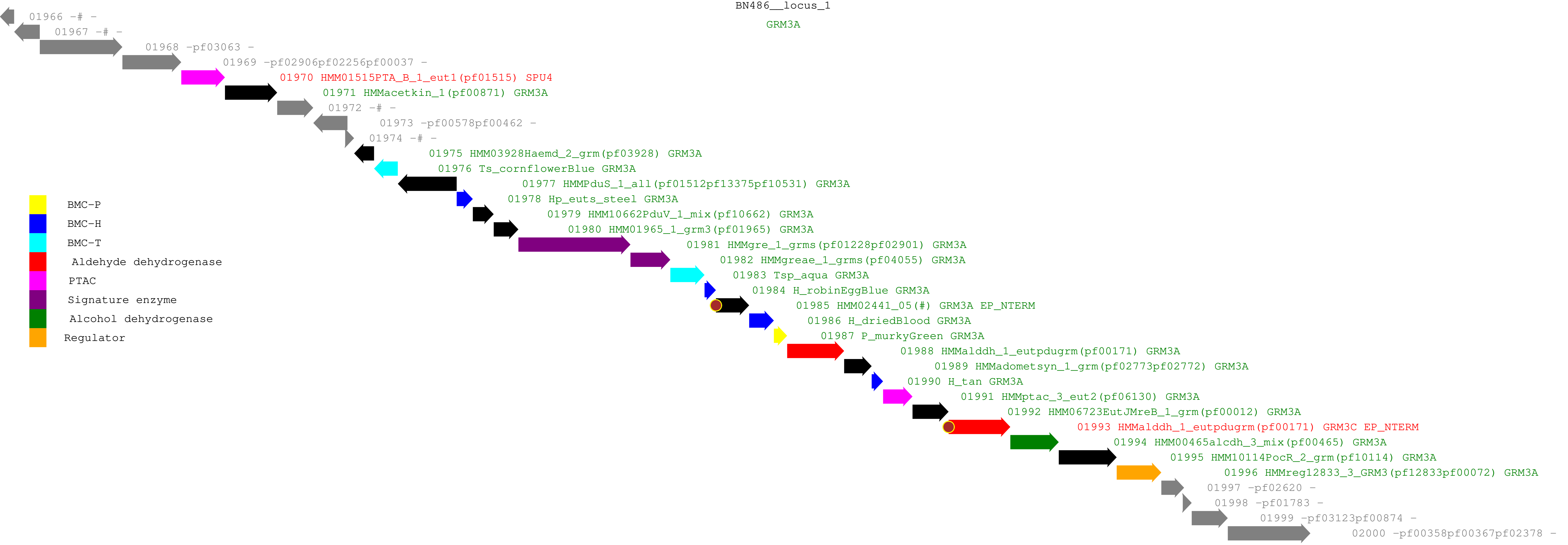

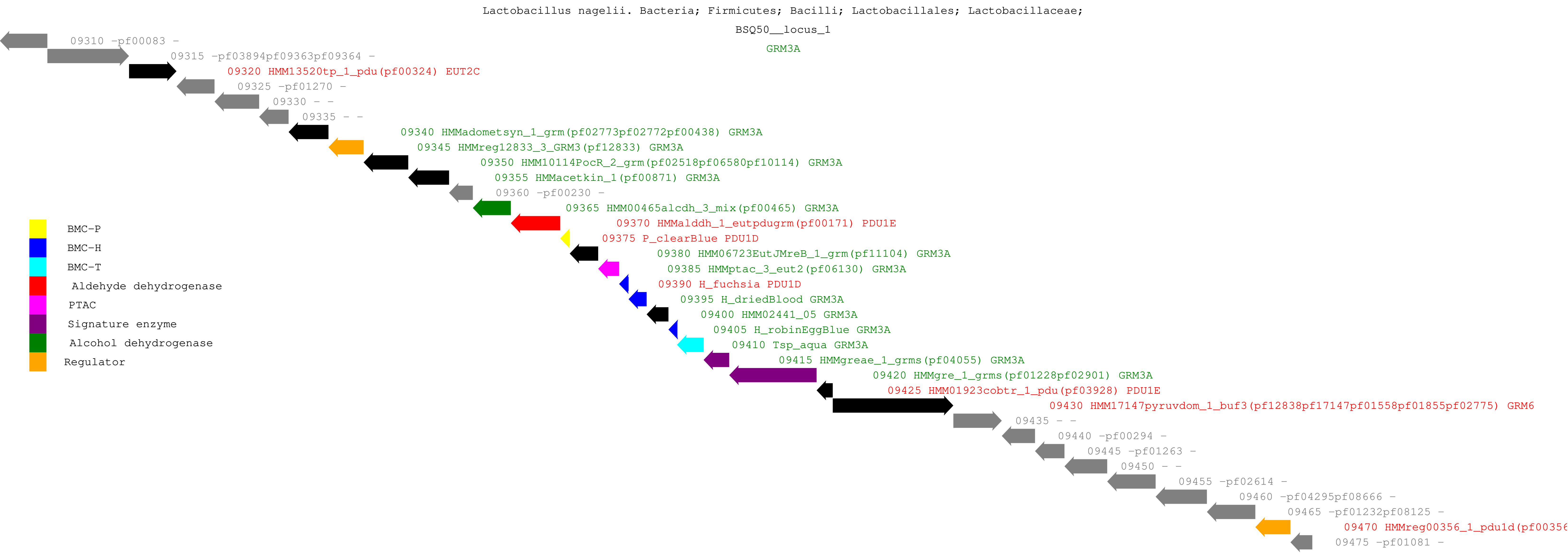
Lactobacillus nagelii. Bacteria; Firmicutes; Bacilli; Lactobacillales; Lactobacillaceae;
BSQ50__locus_1
GRM3A
09310 -pf00083 -
09315 -pf03894pf09363pf09364 -
09320 HMM13520tp_1_pdu(pf00324) EUT2C
09325 -pf01270 -
09330 - -
09335 - -
09340 HMMadometsyn_1_grm(pf02773pf02772pf00438) GRM3A
09345 HMMreg12833_3_GRM3(pf12833) GRM3A
09350 HMM10114PocR_2_grm(pf02518pf06580pf10114) GRM3A
09355 HMMacetkin_1(pf00871) GRM3A
09360 -pf00230 -
09365 HMM00465alcdh_3_mix(pf00465) GRM3A
09370 HMMalddh_1_eutpdugrm(pf00171) PDU1E
09375 P_clearBlue PDU1D
09380 HMM06723EutJMreB_1_grm(pf11104) GRM3A
09385 HMMptac_3_eut2(pf06130) GRM3A
09390 H_fuchsia PDU1D
09395 H_driedBlood GRM3A
09400 HMM02441_05 GRM3A
09405 H_robinEggBlue GRM3A
09410 Tsp_aqua GRM3A
09415 HMMgreae_1_grms(pf04055) GRM3A
09420 HMMgre_1_grms(pf01228pf02901) GRM3A
09425 HMM01923cobtr_1_pdu(pf03928) PDU1E
09430 HMM17147pyruvdom_1_buf3(pf12838pf17147pf01558pf01855pf02775) GRM6
09435 - -
09440 -pf00294 -
09445 -pf01263 -
09450 - -
09455 -pf02614 -
09460 -pf04295pf08666 -
09465 -pf01232pf08125 -
09470 HMMreg00356_1_pdu1d(pf0035
09475 -pf01081 -
BMC-P
BMC-H
BMC-T
Aldehyde dehydrogenase
PTAC
Signature enzyme
Alcohol dehydrogenase
Regulator

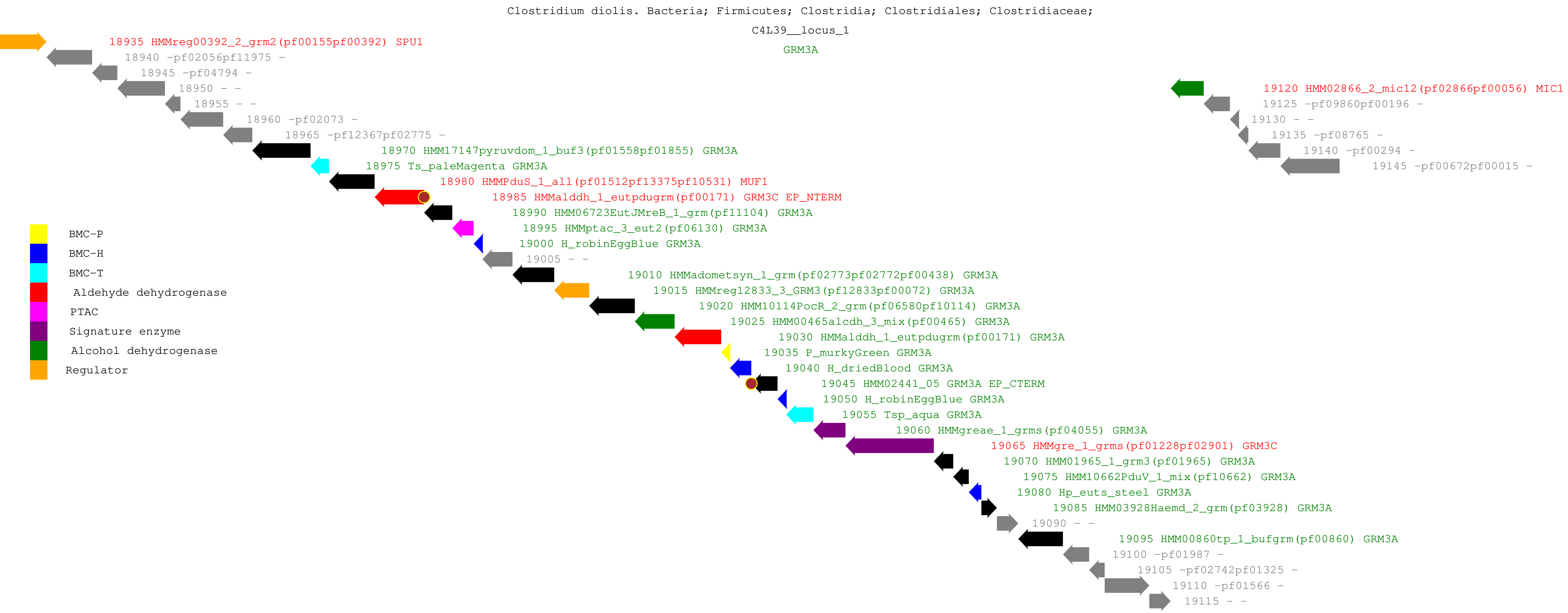

Clostridium diolis. Bacteria; Firmicutes; Clostridia; Clostridiales; Clostridiaceae;
C4L39__locus_1
GRM3A
18935 HMMreg00392_2_grm2(pf00155pf00392) SPU1
18940 -pf02056pf11975 -
18945 -pf04794 -
18950 - -
18955 - -
18960 -pf02073 -
18965 -pf12367pf02775 -
18970 HMM17147pyruvdom_1_buf3(pf01558pf01855) GRM3A
18975 Ts_paleMagenta GRM3A
18980 HMMPduS_1_all(pf01512pf13375pf10531) MUF1
18985 HMMalddh_1_eutpdugrm(pf00171) GRM3C EP_NTERM
18990 HMM06723EutJMreB_1_grm(pf11104) GRM3A
18995 HMMptac_3_eut2(pf06130) GRM3A
19000 H_robinEggBlue GRM3A
19005 - -
19010 HMMadometsyn_1_grm(pf02773pf02772pf00438) GRM3A
19015 HMMreg12833_3_GRM3(pf12833pf00072) GRM3A
19020 HMM10114PocR_2_grm(pf06580pf10114) GRM3A
19025 HMM00465alcdh_3_mix(pf00465) GRM3A
19030 HMMalddh_1_eutpdugrm(pf00171) GRM3A
19035 P_murkyGreen GRM3A
19040 H_driedBlood GRM3A
19045 HMM02441_05 GRM3A EP_CTERM
19050 H_robinEggBlue GRM3A
19055 Tsp_aqua GRM3A
19060 HMMgreae_1_grms(pf04055) GRM3A
19065 HMMgre_1_grms(pf01228pf02901) GRM3C
19070 HMM01965_1_grm3(pf01965) GRM3A
19075 HMM10662PduV_1_mix(pf10662) GRM3A
19080 Hp_euts_steel GRM3A
19085 HMM03928Haemd_2_grm(pf03928) GRM3A
19090 - -
19095 HMM00860tp_1_bufgrm(pf00860) GRM3A
19100 -pf01987 -
19105 -pf02742pf01325 -
19110 -pf01566 -
19115 - -
19120 HMM02866_2_mic12(pf02866pf00056) MIC1
19125 -pf09860pf00196 -
19130 - -
19135 -pf08765 -
19140 -pf00294 -
19145 -pf00672pf00015 -
BMC-P
BMC-H
BMC-T
Aldehyde dehydrogenase
PTAC
Signature enzyme
Alcohol dehydrogenase
Regulator

Megasphaera elsdenii. Bacteria; Firmicutes; Negativicutes; Veillonellales; Veillonellaceae;

C6Y28__locus_1

GRM3A

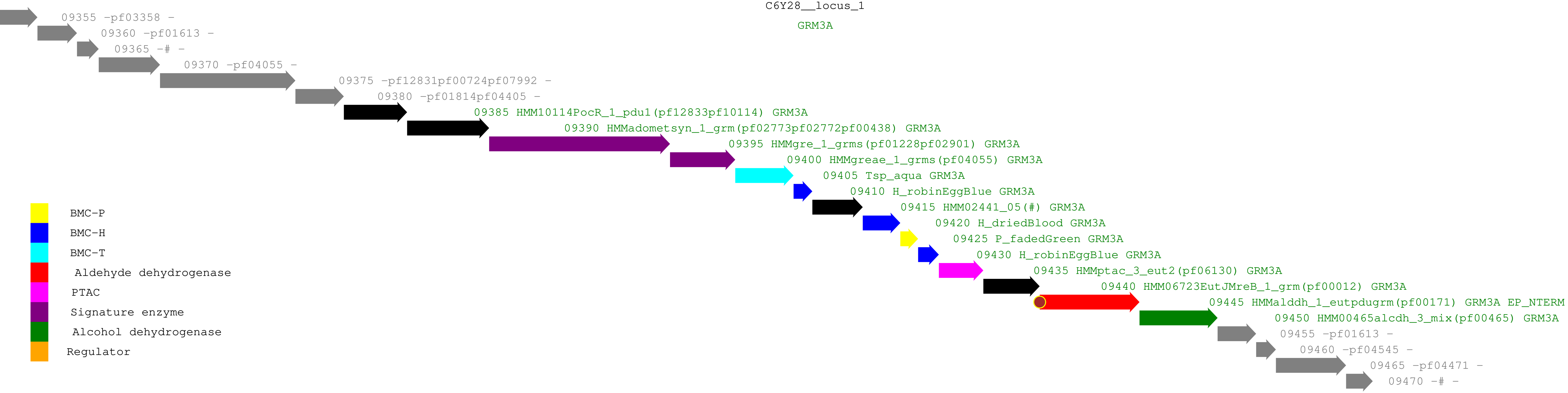

Megasphaera genomosp. type_2. Bacteria; Firmicutes; Negativicutes; Veillonellales; Veillonellaceae;

CAL30__locus_1

GRM3A

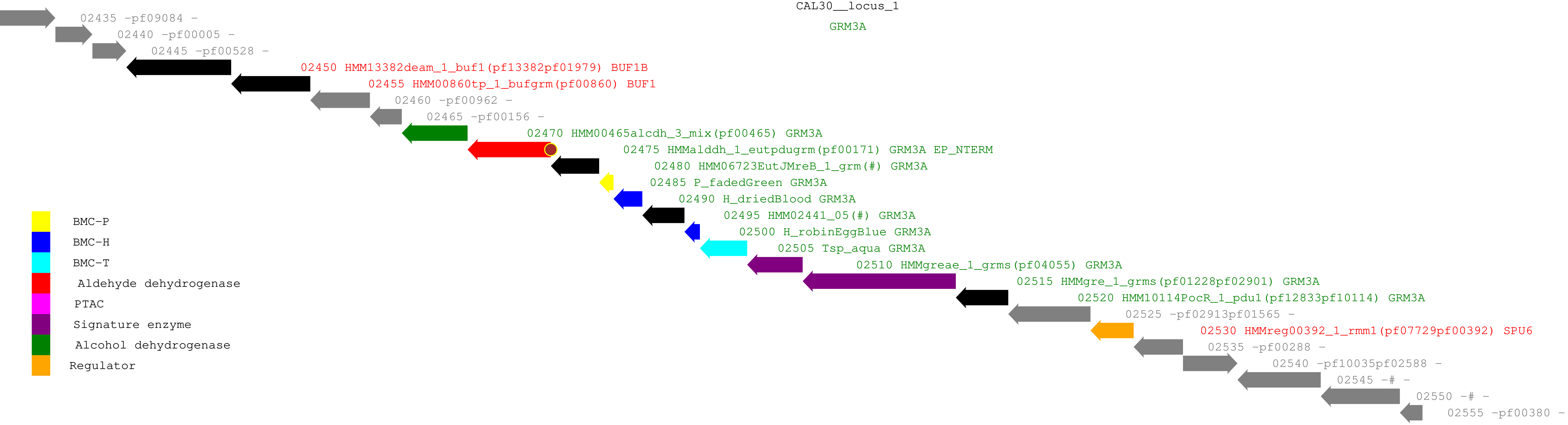

Clostridium beijerinckii (Clostridium MP). Bacteria; Firmicutes; Clostridia; Clostridiales; Clostridiaceae;

CBEIBR21__locus_1

GRM3A

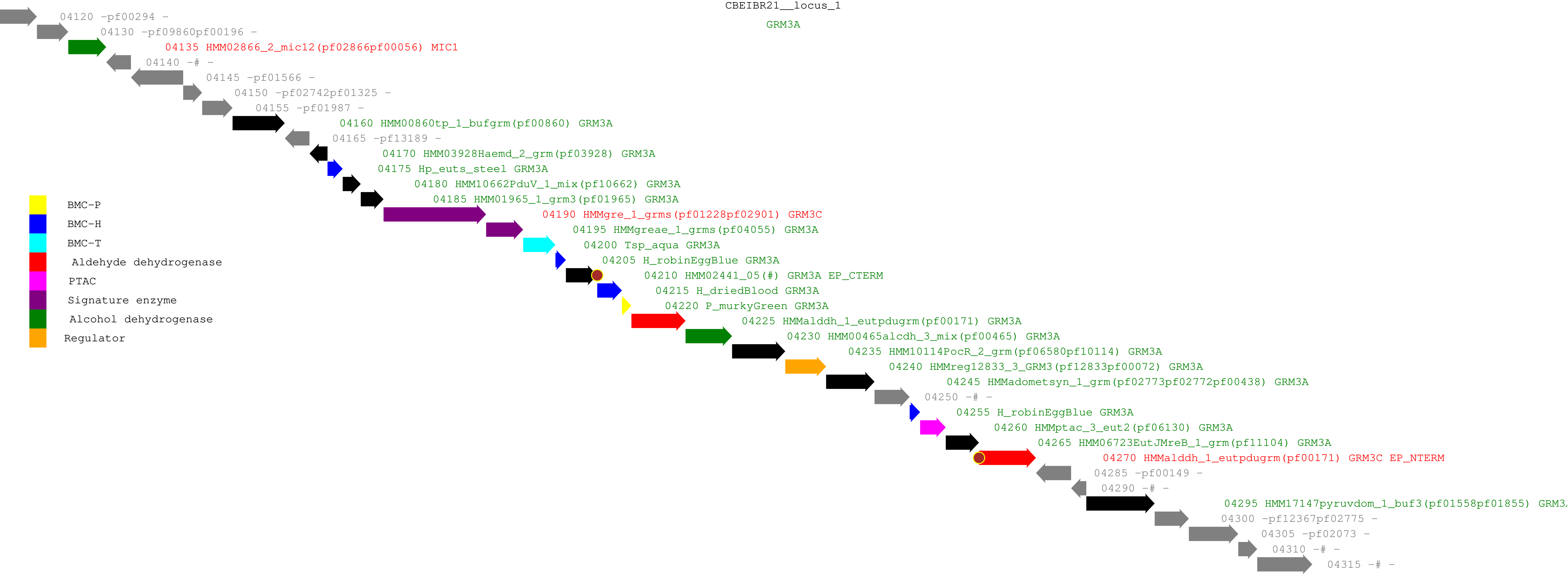

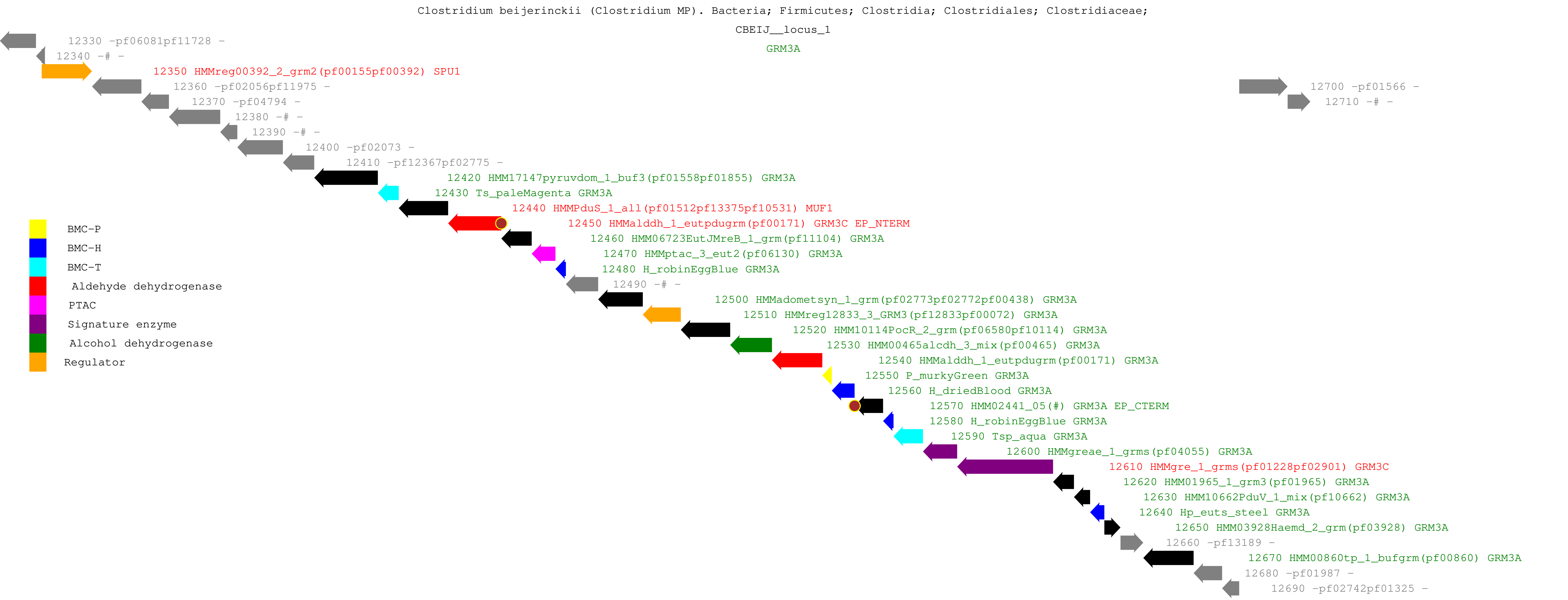
Clostridium beijerinckii (Clostridium MP). Bacteria; Firmicutes; Clostridia; Clostridiales; Clostridiaceae;
CBEIJ__locus_1
GRM3A
12330 -pf06081pf11728 -
12340 -# -
12350 HMMreg00392_2_grm2(pf00155pf00392) SPU1
12360 -pf02056pf11975 -
12370 -pf04794 -
12380 -# -
12390 -# -
12400 -pf02073 -
12410 -pf12367pf02775 -
12420 HMM17147pyruvdom_1_buf3(pf01558pf01855) GRM3A
12430 Ts_paleMagenta GRM3A
12440 HMMPduS_1_all(pf01512pf13375pf10531) MUF1
12450 HMMalddh_1_eutpdugrm(pf00171) GRM3C EP_NTERM
12460 HMM06723EutJMreB_1_grm(pf11104) GRM3A
12470 HMMptac_3_eut2(pf06130) GRM3A
12480 H_robinEggBlue GRM3A
12490 -# -
12500 HMMadometsyn_1_grm(pf02773pf02772pf00438) GRM3A
12510 HMMreg12833_3_GRM3(pf12833pf00072) GRM3A
12520 HMM10114PocR_2_grm(pf06580pf10114) GRM3A
12530 HMM00465alcdh_3_mix(pf00465) GRM3A
12540 HMMalddh_1_eutpdugrm(pf00171) GRM3A
12550 P_murkyGreen GRM3A
12560 H_driedBlood GRM3A
12570 HMM02441_05(#) GRM3A EP_CTERM
12580 H_robinEggBlue GRM3A
12590 Tsp_aqua GRM3A
12600 HMMgreae_1_grms(pf04055) GRM3A
12610 HMMgre_1_grms(pf01228pf02901) GRM3C
12620 HMM01965_1_grm3(pf01965) GRM3A
12630 HMM10662PduV_1_mix(pf10662) GRM3A
12640 Hp_euts_steel GRM3A
12650 HMM03928Haemd_2_grm(pf03928) GRM3A
12660 -pf13189 -
12670 HMM00860tp_1_bufgrm(pf00860) GRM3A
12680 -pf01987 -
12690 -pf02742pf01325 -
12700 -pf01566 -
12710 -# -
BMC-P
BMC-H
BMC-T
Aldehyde dehydrogenase
PTAC
Signature enzyme
Alcohol dehydrogenase
Regulator

Clostridium beijerinckii (strain ATCC 51743 / NCIMB 8052) (Clostridium Bacteria; Firmicutes; Clostridia; Clostridiales; Clostridiaceae;

Cbei__locus_1

GRM3A

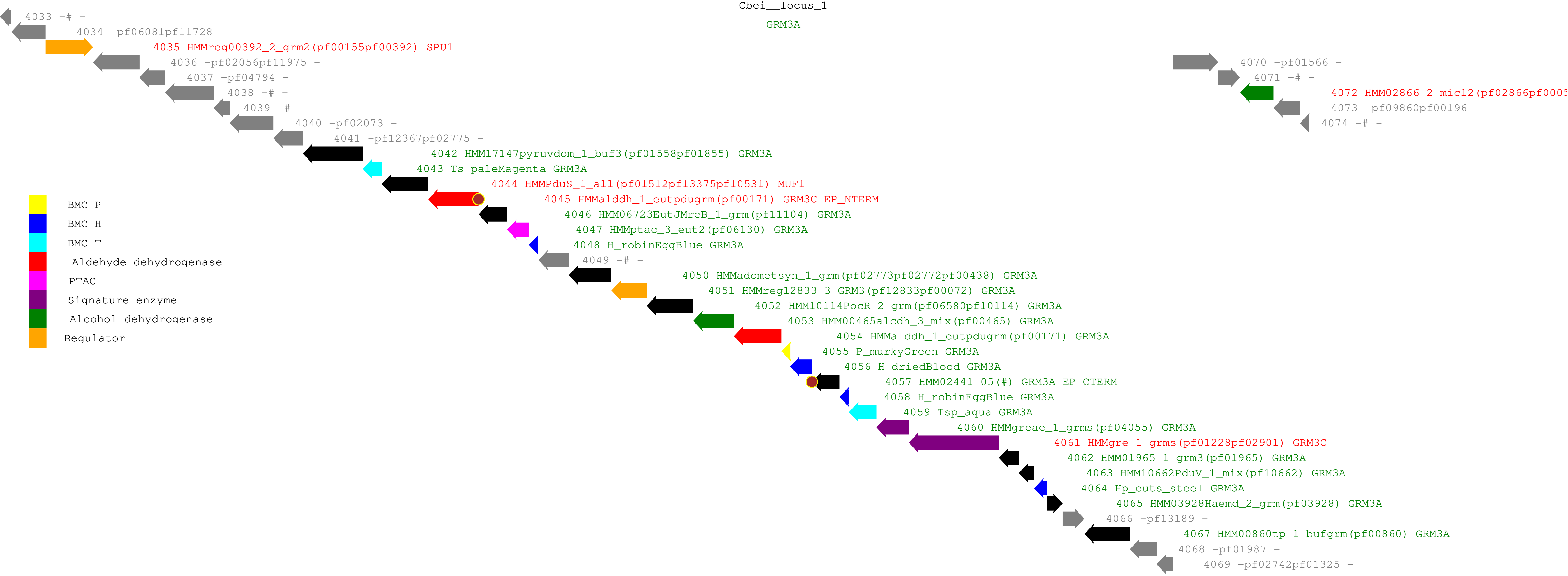

Clostridium diolis. Bacteria; Firmicutes; Clostridia; Clostridiales; Clostridiaceae;

CCS79__locus_1

GRM3A

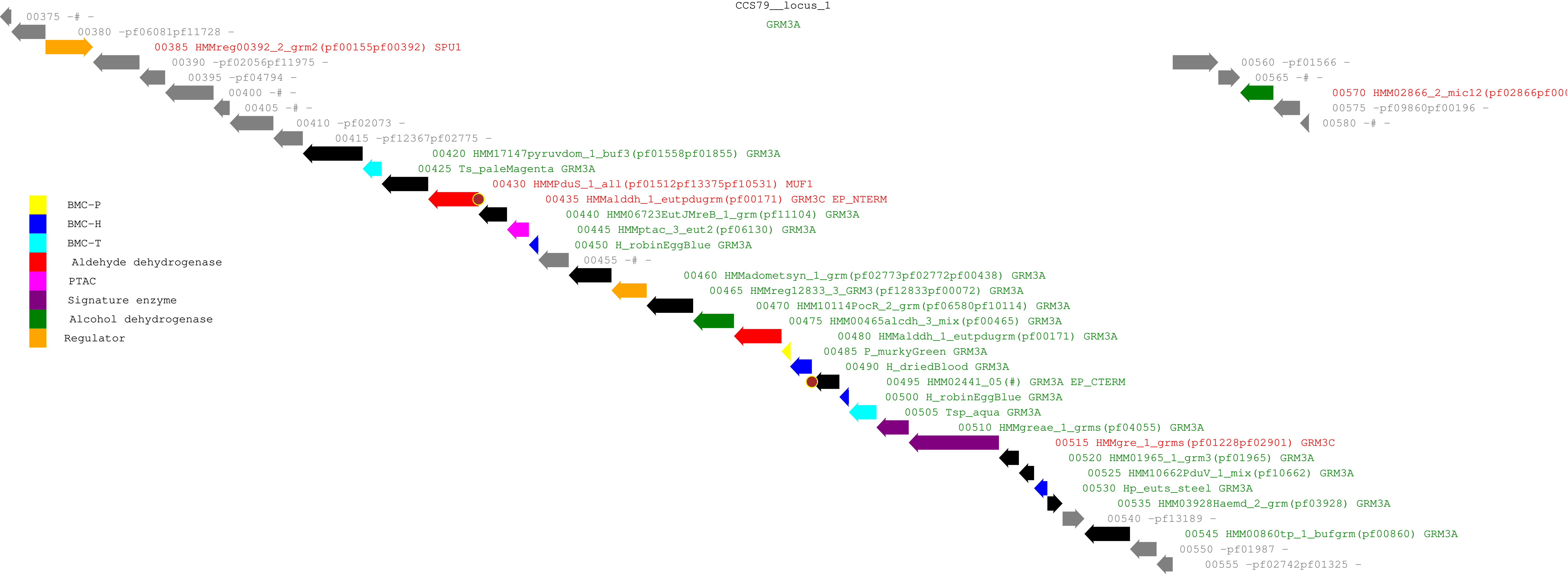

Clostridium bolteae. Bacteria; Firmicutes; Clostridia; Clostridiales; Lachnospiraceae.

CGC65__locus_1

GRM3A

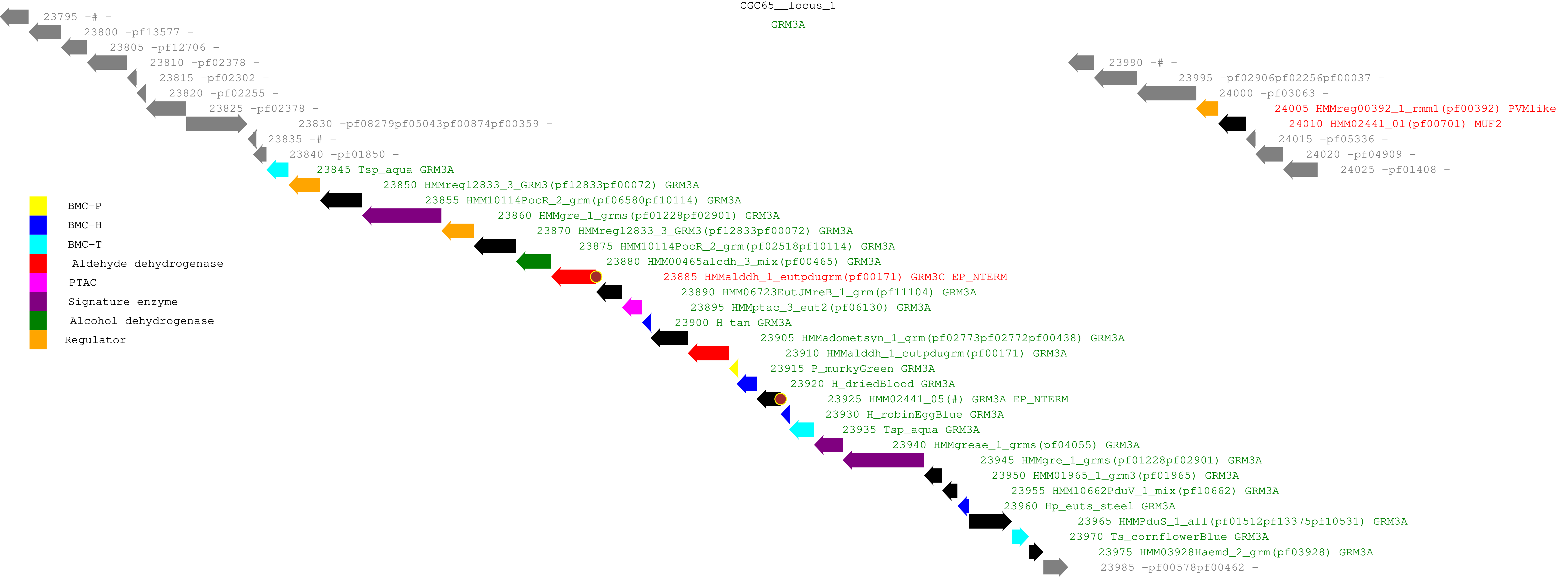

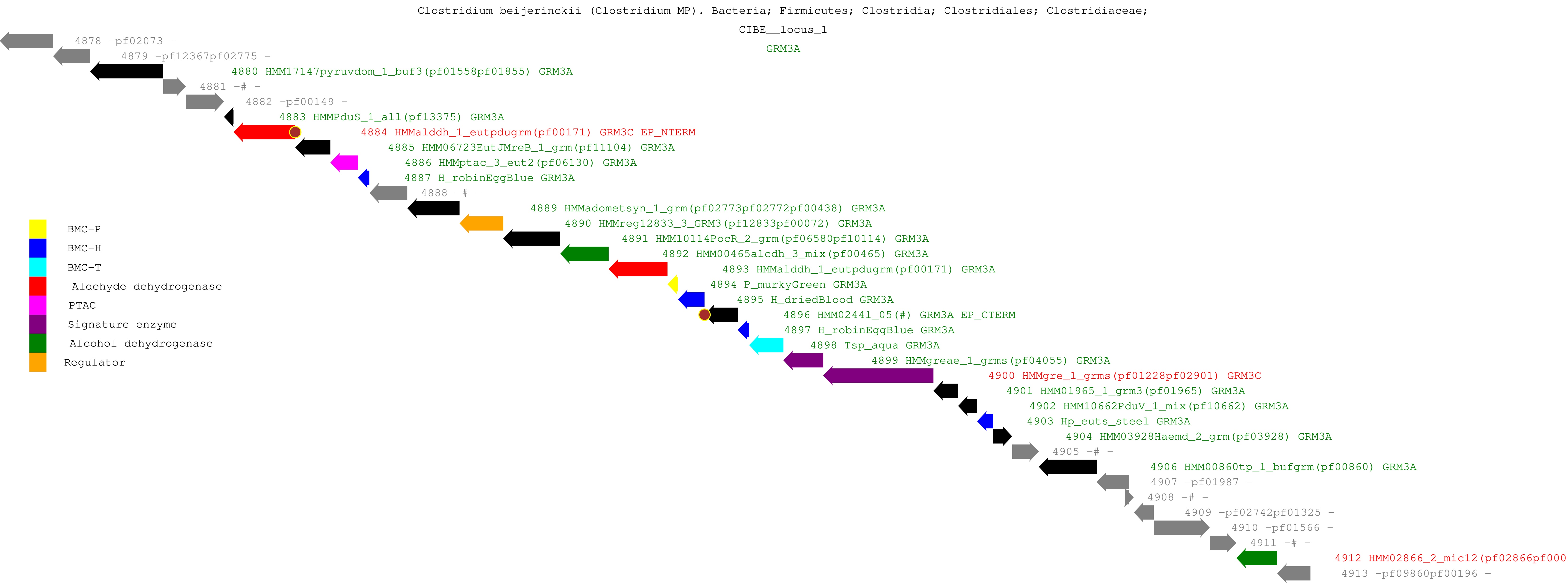
Clostridium beijerinckii (Clostridium MP). Bacteria; Firmicutes; Clostridia; Clostridiales; Clostridiaceae;
CIBE__locus_1
GRM3A
4878 -pf02073 -
4879 -pf12367pf02775 -
4880 HMM17147pyruvdom_1_buf3(pf01558pf01855) GRM3A
4881 -# -
4882 -pf00149 -
4883 HMMPduS_1_all(pf13375) GRM3A
4884 HMMaldh_1_eutpdugrm(pf00171) GRM3C EP_NTERM
4885 HMM06723EutJMreB_1_grm(pf11104) GRM3A
4886 HMMptac_3_eut2(pf06130) GRM3A
4887 H_robinEggBlue GRM3A
4888 -# -
4889 HMMadometsyn_1_grm(pf02773pf02772pf00438) GRM3A
4890 HMMreg12833_3_GRM3(pf12833pf00072) GRM3A
4891 HMM10114PocR_2_grm(pf06580pf10114) GRM3A
4892 HMM00465alcdh_3_mix(pf00465) GRM3A
4893 HMMaldh_1_eutpdugrm(pf00171) GRM3A
4894 P_murkyGreen GRM3A
4895 H_driedBlood GRM3A
4896 HMM02441_05(#) GRM3A EP_CTERM
4897 H_robinEggBlue GRM3A
4898 Tsp_aqua GRM3A
4899 HMMgreae_1_grms(pf04055) GRM3A
4900 HMMgre_1_grms(pf01228pf02901) GRM3C
4901 HMM01965_1_grm3(pf01965) GRM3A
4902 HMM10662PduV_1_mix(pf10662) GRM3A
4903 Hp_euts_steel GRM3A
4904 HMM03928Haemd_2_grm(pf03928) GRM3A
4905 -# -
4906 HMM00860tp_1_bufgrm(pf00860) GRM3A
4907 -pf01987 -
4908 -# -
4909 -pf02742pf01325 -
4910 -pf01566 -
4911 -# -
4912 HMM02866_2_mic12(pf02866pf000
4913 -pf09860pf00196 -
BMC-P
BMC-H
BMC-T
Aldehyde dehydrogenase
PTAC
Signature enzyme
Alcohol dehydrogenase
Regulator

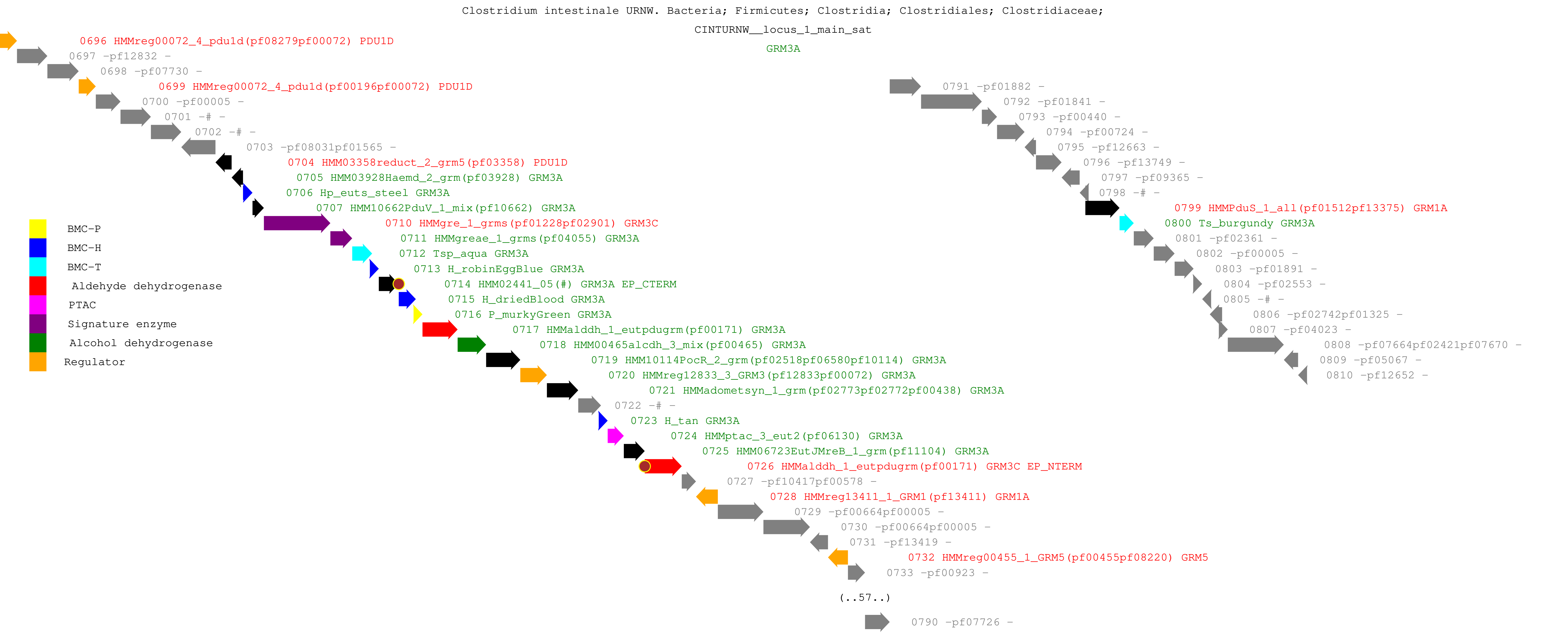

Clostridium intestinale URNW. Bacteria; Firmicutes; Clostridia; Clostridiales; Clostridiaceae;
CINTURNW__locus_1_main_sat
GRM3A
0696 HMMreg00072_4_pdu1d(pf08279pf00072) PDU1D
0697 -pf12832 -
0698 -pf07730 -
0699 HMMreg00072_4_pdu1d(pf00196pf00072) PDU1D
0700 -pf00005 -
0701 -# -
0702 -# -
0703 -pf08031pf01565 -
0704 HMM03358reduct_2_grm5(pf03358) PDU1D
0705 HMM03928Haemd_2_grm(pf03928) GRM3A
0706 Hp_euts_steel GRM3A
0707 HMM10662PduV_1_mix(pf10662) GRM3A
0710 HMMgre_1_grms(pf01228pf02901) GRM3C
0711 HMMgreae_1_grms(pf04055) GRM3A
0712 Tsp_aqua GRM3A
0713 H_robinEggBlue GRM3A
0714 HMM02441_05(#) GRM3A EP_CTERM
0715 H_driedBlood GRM3A
0716 P_murkyGreen GRM3A
0717 HMMalddh_1_eutpdugrm(pf00171) GRM3A
0718 HMM00465alcdh_3_mix(pf00465) GRM3A
0719 HMM10114PocR_2_grm(pf02518pf06580pf10114) GRM3A
0720 HMMreg12833_3_GRM3(pf12833pf00072) GRM3A
0721 HMMadometsyn_1_grm(pf02773pf02772pf00438) GRM3A
0722 -# -
0723 H_tan GRM3A
0724 HMMptac_3_eut2(pf06130) GRM3A
0725 HMM06723EutJMreB_1_grm(pf11104) GRM3A
0726 HMMalddh_1_eutpdugrm(pf00171) GRM3C EP_NTERM
0727 -pf10417pf00578 -
0728 HMMreg13411_1_GRM1(pf13411) GRM1A
0729 -pf00664pf00005 -
0730 -pf00664pf00005 -
0731 -pf13419 -
0732 HMMreg00455_1_GRM5(pf00455pf08220) GRM5
0733 -pf00923 -
(..57..)
0790 -pf07726 -
0791 -pf01882 -
0792 -pf01841 -
0793 -pf00440 -
0794 -pf00724 -
0795 -pf12663 -
0796 -pf13749 -
0797 -pf09365 -
0798 -# -
0799 HMMPduS_1_all(pf01512pf13375) GRM1A
0800 Ts_burgundy GRM3A
0801 -pf02361 -
0802 -pf00005 -
0803 -pf01891 -
0804 -pf02553 -
0805 -# -
0806 -pf02742pf01325 -
0807 -pf04023 -
0808 -pf07664pf02421pf07670 -
0809 -pf05067 -
0810 -pf12652 -
BMC-P
BMC-H
BMC-T
Aldehyde dehydrogenase
PTAC
Signature enzyme
Alcohol dehydrogenase
Regulator

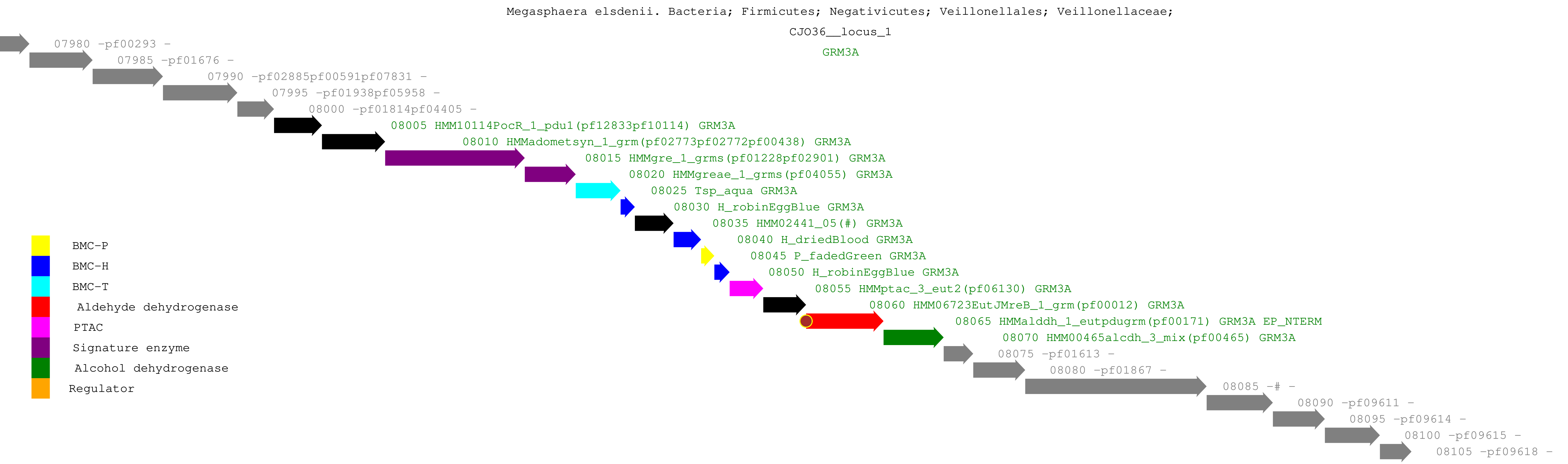

Megasphaera elsdenii. Bacteria; Firmicutes; Negativicutes; Veillonellales; Veillonellaceae;
CJO36__locus_1
GRM3A
07980 -pf00293 -
07985 -pf01676 -
07990 -pf02885pf00591pf07831 -
07995 -pf01938pf05958 -
08000 -pf01814pf04405 -
08005 HMM10114PocR_1_pdu1(pf12833pf10114) GRM3A
08010 HMMadometsyn_1_grm(pf02773pf02772pf00438) GRM3A
08015 HMMgre_1_grms(pf01228pf02901) GRM3A
08020 HMMgreae_1_grms(pf04055) GRM3A
08025 Tsp_aqua GRM3A
08030 H_robinEggBlue GRM3A
08035 HMM02441_05(#) GRM3A
08040 H_driedBlood GRM3A
08045 P_fadedGreen GRM3A
08050 H_robinEggBlue GRM3A
08055 HMMptac_3_eut2(pf06130) GRM3A
08060 HMM06723EutJMreB_1_grm(pf00012) GRM3A
08065 HMMaldh_1_eutpdugrm(pf00171) GRM3A EP_NTERM
08070 HMM00465alcdh_3_mix(pf00465) GRM3A
08075 -pf01613 -
08080 -pf01867 -
08085 -# -
08090 -pf09611 -
08095 -pf09614 -
08100 -pf09615 -
08105 -pf09618 -
BMC-P
BMC-H
BMC-T
Aldehyde dehydrogenase
PTAC
Signature enzyme
Alcohol dehydrogenase
Regulator

Clostridium beijerinckii (Clostridium MP). Bacteria; Firmicutes; Clostridia; Clostridiales; Clostridiaceae;

CLBEIC__locus_1

GRM3A

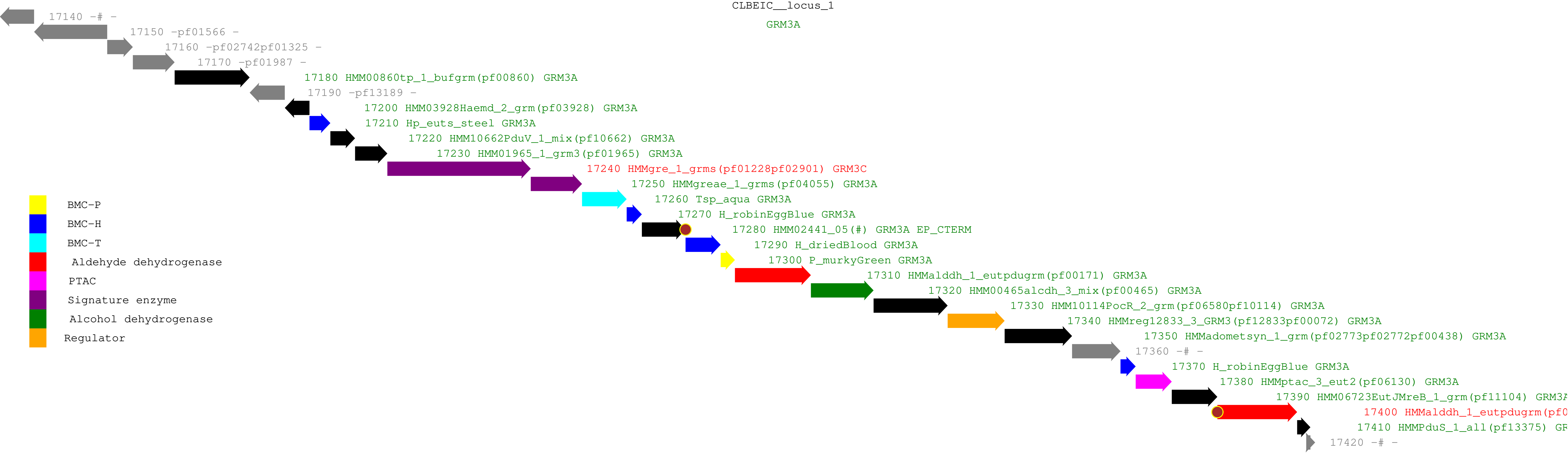

Clostridium beijerinckii (Clostridium MP). Bacteria; Firmicutes; Clostridia; Clostridiales; Clostridiaceae;

CLBIJ__locus_1

GRM3A

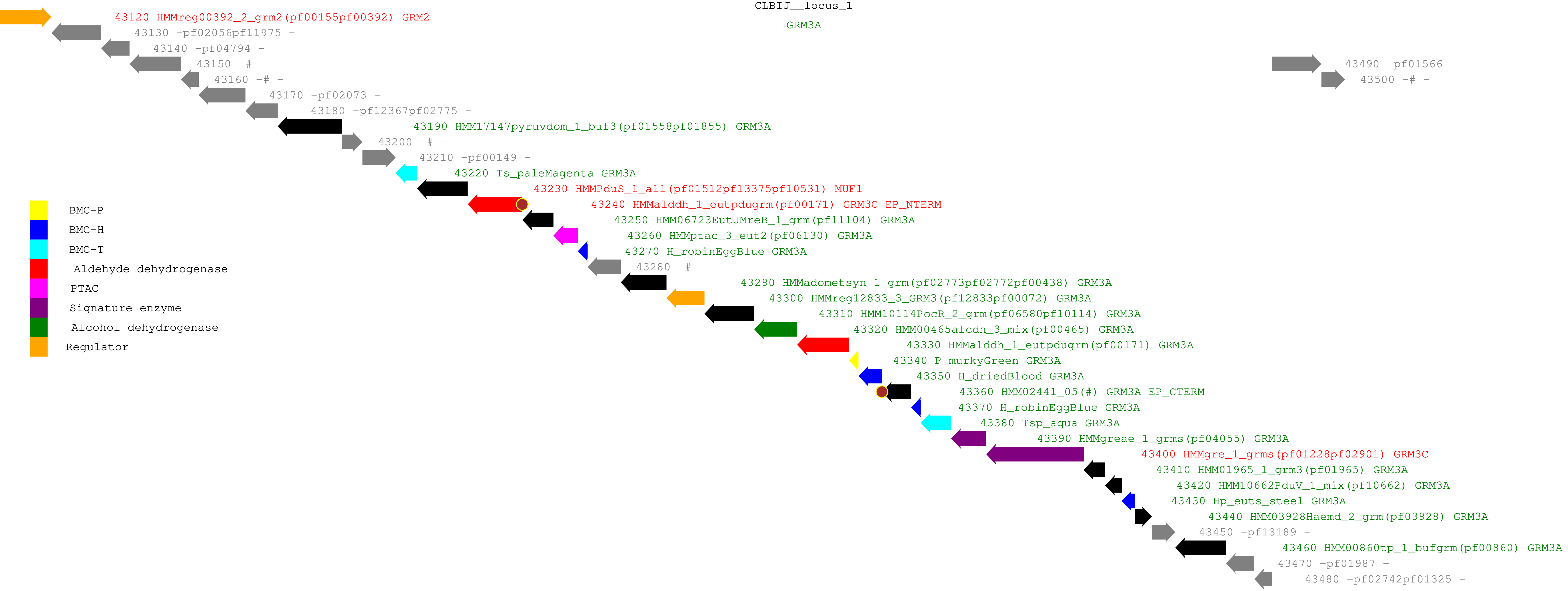

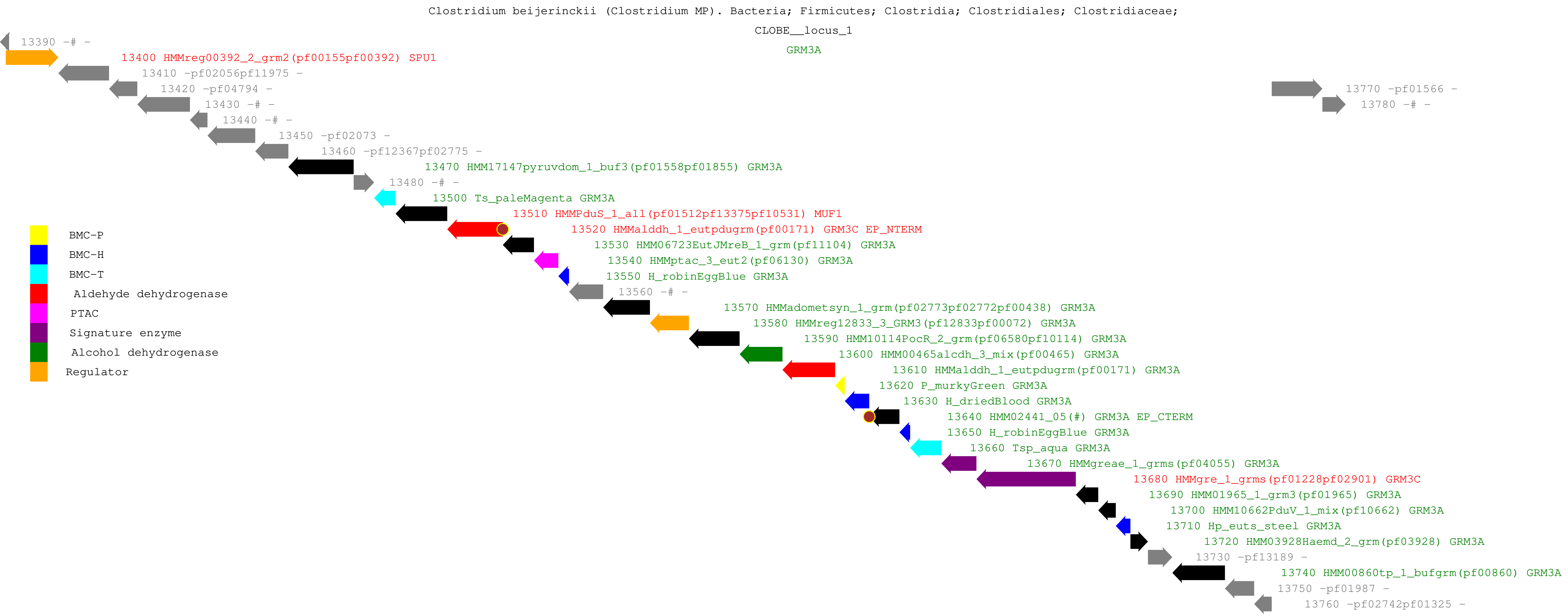
Clostridium beijerinckii (Clostridium MP). Bacteria; Firmicutes; Clostridia; Clostridiales; Clostridiaceae;
CLOBE__locus_1
GRM3A
13390 -# -
13400 HMMreg00392_2_grm2(pf00155pf00392) SPU1
13410 -pf02056pf11975 -
13420 -pf04794 -
13430 -# -
13440 -# -
13450 -pf02073 -
13460 -pf12367pf02775 -
13470 HMM17147pyruvdom_1_buf3(pf01558pf01855) GRM3A
13480 -# -
13500 Ts_paleMagenta GRM3A
13510 HMMPduS_1_all(pf01512pf13375pf10531) MUF1
13520 HMMaldh_1_eutpdugrm(pf00171) GRM3C EP_NTERM
13530 HMM06723EutJMreB_1_grm(pf11104) GRM3A
13540 HMMptac_3_eut2(pf06130) GRM3A
13550 H_robinEggBlue GRM3A
13560 -# -
13570 HMMadometsyn_1_grm(pf02773pf02772pf00438) GRM3A
13580 HMMreg12833_3_GRM3(pf12833pf00072) GRM3A
13590 HMM10114PocR_2_grm(pf06580pf10114) GRM3A
13600 HMM00465alcdh_3_mix(pf00465) GRM3A
13610 HMMaldh_1_eutpdugrm(pf00171) GRM3A
13620 P_murkyGreen GRM3A
13630 H_driedBlood GRM3A
13640 HMM02441_05(#) GRM3A EP_CTERM
13650 H_robinEggBlue GRM3A
13660 Tsp_aqua GRM3A
13670 HMMgreae_1_grms(pf04055) GRM3A
13680 HMMgre_1_grms(pf01228pf02901) GRM3C
13690 HMM01965_1_grm3(pf01965) GRM3A
13700 HMM10662PduV_1_mix(pf10662) GRM3A
13710 Hp_euts_steel GRM3A
13720 HMM03928Haemd_2_grm(pf03928) GRM3A
13730 -pf13189 -
13740 HMM00860tp_1_bufgrm(pf00860) GRM3A
13750 -pf01987 -
13760 -pf02742pf01325 -
13770 -pf01566 -
13780 -# -
BMC-P
BMC-H
BMC-T
Aldehyde dehydrogenase
PTAC
Signature enzyme
Alcohol dehydrogenase
Regulator

Clostridium sp. BL-8. Bacteria; Firmicutes; Clostridia; Clostridiales; Clostridiaceae;

CLOBL__locus_1

GRM3A

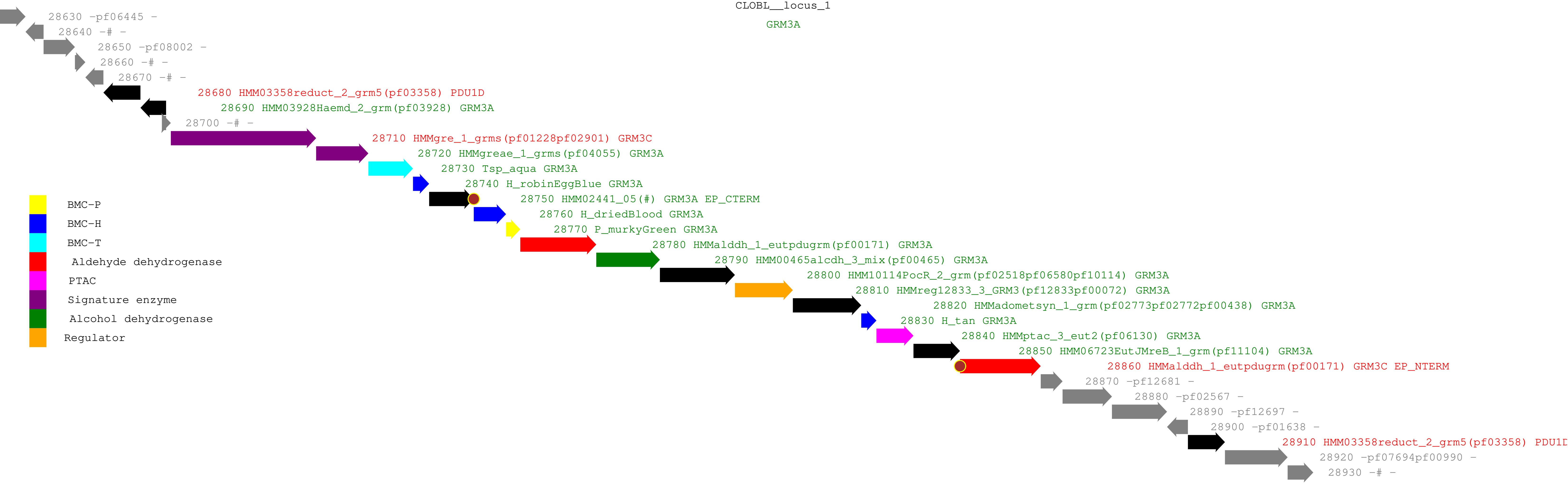

Clostridium bolteae (strain ATCC BAA-613 / WAL 16351). Bacteria; Firmicutes; Clostridia; Clostridiales; Lachnospiraceae.

CLOBOL__locus_1

GRM3A

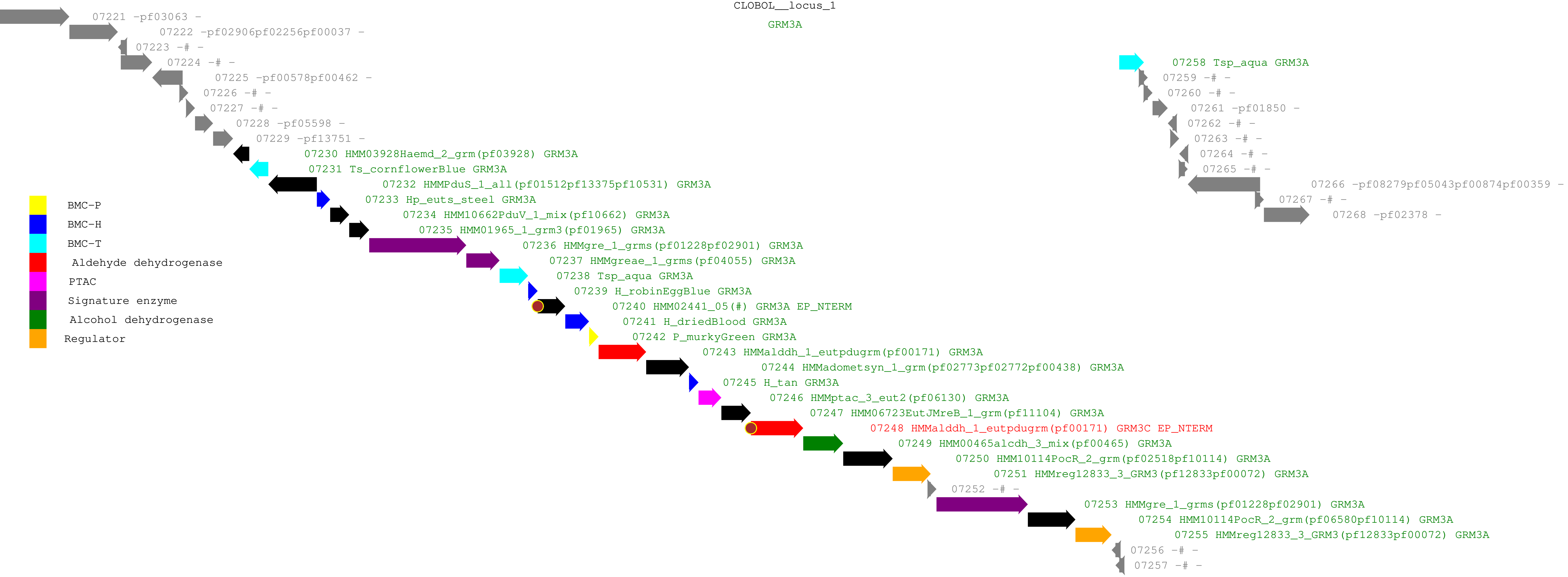

Clostridium hiranonis DSM 13275. Bacteria; Firmicutes; Clostridia; Clostridiales;

CLOHIR__locus_1

GRM3A

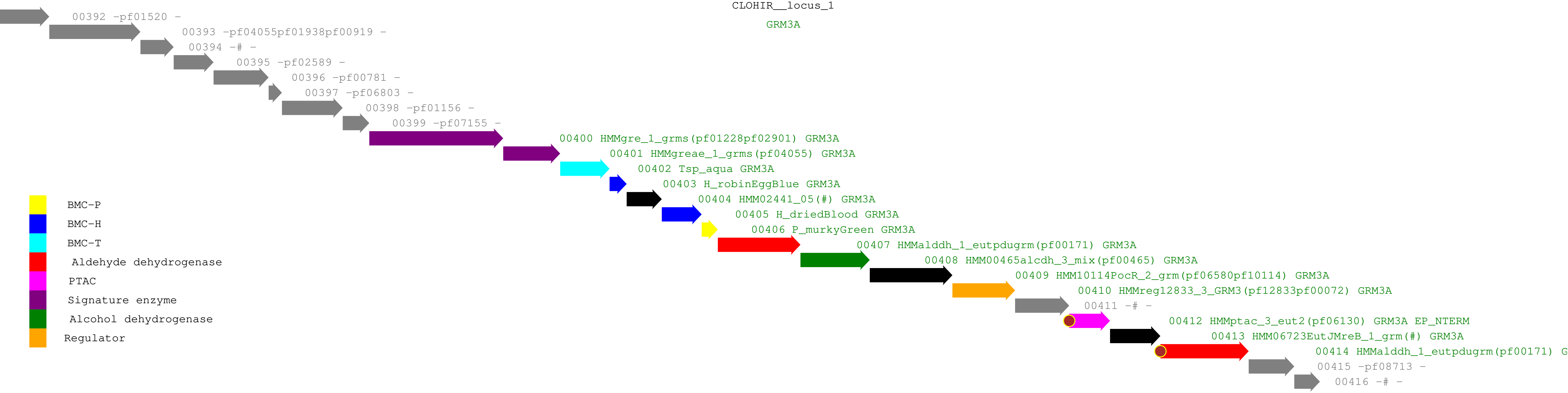

Clostridium cf. saccharolyticum K10. Bacteria; Firmicutes; Clostridia; Clostridiales; Lachnospiraceae.

CLS__locus_1

GRM3A

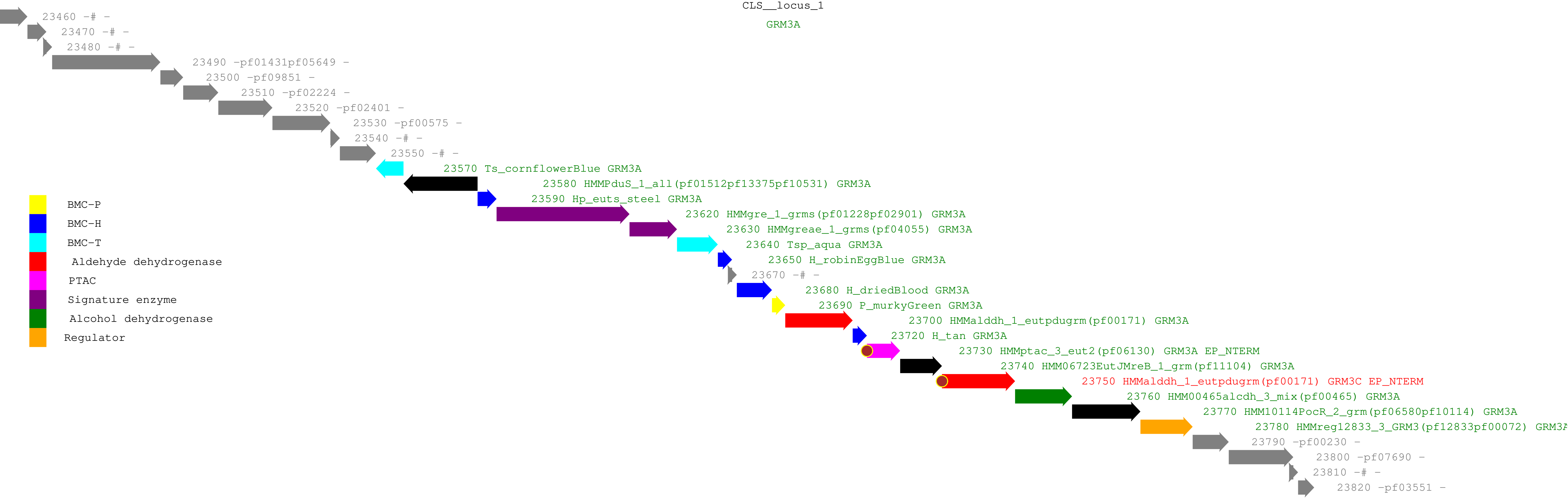

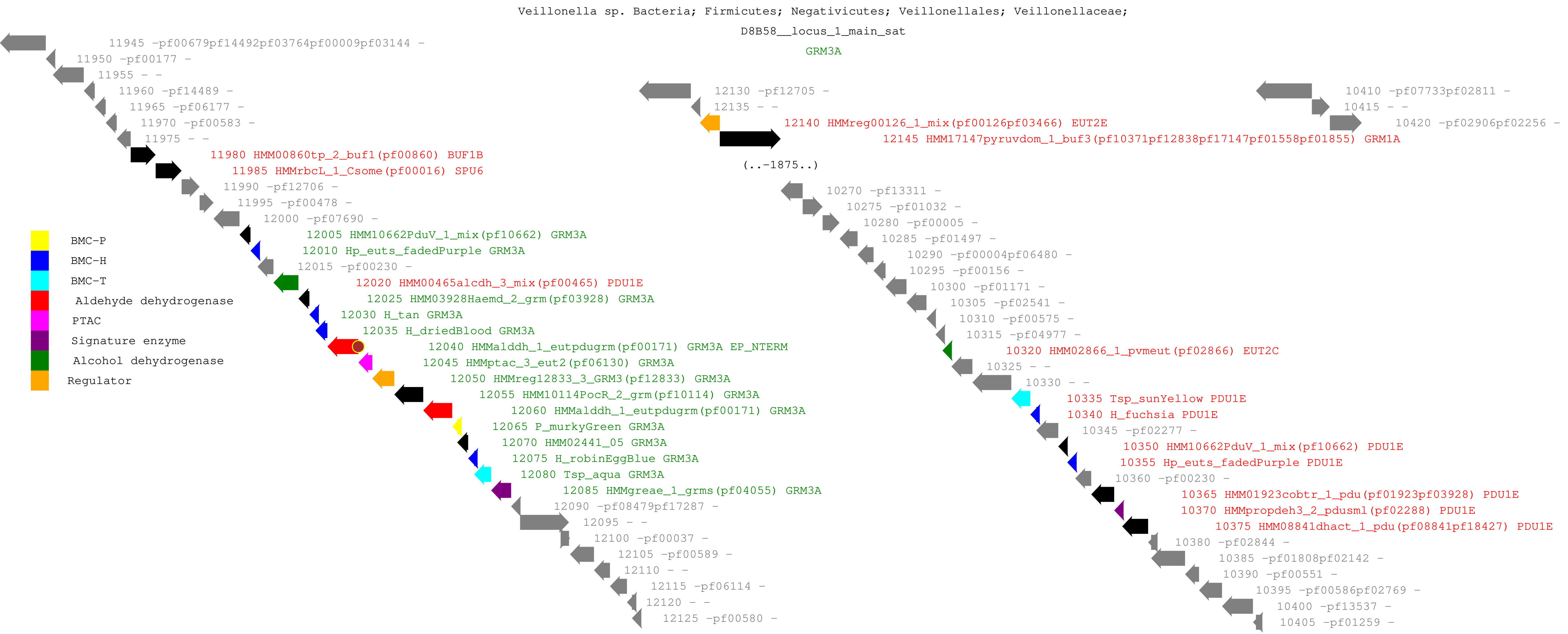

Veillonella sp. Bacteria; Firmicutes; Negativicutes; Veillonellales; Veillonellaceae;
D8B58__locus_1_main_sat
GRM3A
11945 -pf00679pf14492pf03764pf00009pf03144 -
11950 -pf00177 -
11955 - -
11960 -pf14489 -
11965 -pf06177 -
11970 -pf00583 -
11975 - -
11980 HMM00860tp_2_buf1(pf00860) BUF1B
11985 HMMrbcL_1_Csome(pf00016) SPU6
11990 -pf12706 -
11995 -pf00478 -
12000 -pf07690 -
12005 HMM10662PduV_1_mix(pf10662) GRM3A
12010 Hp_euts_fadedPurple GRM3A
12015 -pf00230 -
12020 HMM00465alcdh_3_mix(pf00465) PDU1E
12025 HMM03928Haemd_2_grm(pf03928) GRM3A
12030 H_tan GRM3A
12035 H_driedBlood GRM3A
12040 HMMaldh_1_eutpdugrm(pf00171) GRM3A EP_NTERM
12045 HMMptac_3_eut2(pf06130) GRM3A
12050 HMMreg12833_3_GRM3(pf12833) GRM3A
12055 HMM10114PocR_2_grm(pf10114) GRM3A
12060 HMMaldh_1_eutpdugrm(pf00171) GRM3A
12065 P_murkyGreen GRM3A
12070 HMM02441_05 GRM3A
12075 H_robinEggBlue GRM3A
12080 Tsp_aqua GRM3A
12085 HMMgreae_1_grms(pf04055) GRM3A
12090 -pf08479pf17287 -
12095 - -
12100 -pf00037 -
12105 -pf00589 -
12110 - -
12115 -pf06114 -
12120 - -
12125 -pf00580 -
12130 -pf12705 -
12135 - -
12140 HMMreg00126_1_mix(pf00126pf03466) EUT2E
12145 HMM17147pyruvdom_1_buf3(pf10371pf12838pf17147pf01558pf01855) GRM1A
(..-1875..)
10270 -pf13311 -
10275 -pf01032 -
10280 -pf00005 -
10285 -pf01497 -
10290 -pf00004pf06480 -
10295 -pf00156 -
10300 -pf01171 -
10305 -pf02541 -
10310 -pf00575 -
10315 -pf04977 -
10320 HMM02866_1_pvmeut(pf02866) EUT2C
10325 - -
10330 - -
10335 Tsp_sunYellow PDU1E
10340 H_fuchsia PDU1E
10345 -pf02277 -
10350 HMM10662PduV_1_mix(pf10662) PDU1E
10355 Hp_euts_fadedPurple PDU1E
10360 -pf00230 -
10365 HMM01923cobtr_1_pdu(pf01923pf03928) PDU1E
10370 HMMpropdeh3_2_pdusml(pf02288) PDU1E
10375 HMM08841dhact_1_pdu(pf08841pf18427) PDU1E
10380 -pf02844 -
10385 -pf01808pf02142 -
10390 -pf00551 -
10395 -pf00586pf02769 -
10400 -pf13537 -
10405 -pf01259 -
10410 -pf07733pf02811 -
10415 - -
10420 -pf02906pf02256 -
BMC-P
BMC-H
BMC-T
Aldehyde dehydrogenase
PTAC
Signature enzyme
Alcohol dehydrogenase
Regulator

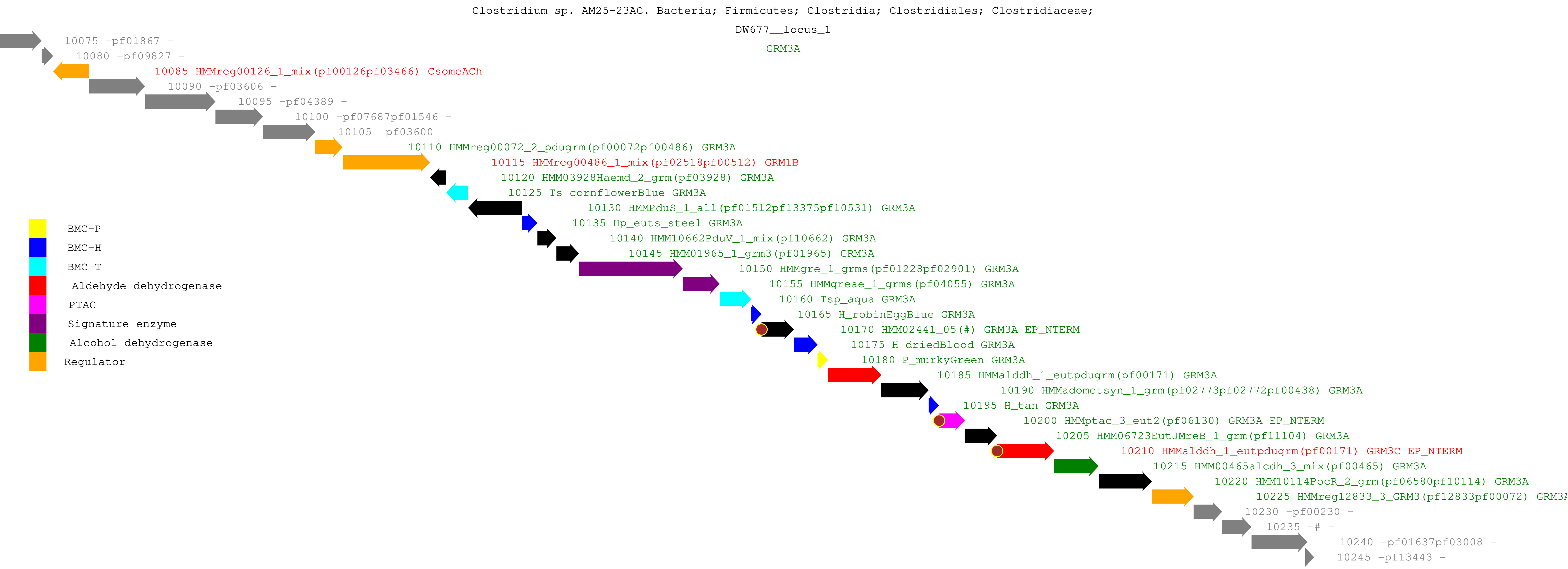

Clostridium sp. AM25-23AC. Bacteria; Firmicutes; Clostridia; Clostridiales; Clostridiaceae;
DW677__locus_1
GRM3A
10075 -pf01867 -
10080 -pf09827 -
10085 HMMreg00126_1_mix(pf00126pf03466) CsomeACh
10090 -pf03606 -
10095 -pf04389 -
10100 -pf07687pf01546 -
10105 -pf03600 -
10110 HMMreg00072_2_pdugrm(pf00072pf00486) GRM3A
10115 HMMreg00486_1_mix(pf02518pf00512) GRM1B
10120 HMM03928Haemd_2_grm(pf03928) GRM3A
10125 Ts_cornflowerBlue GRM3A
10130 HMMPduS_1_all(pf01512pf13375pf10531) GRM3A
10135 Hp_euts_steel GRM3A
10140 HMM10662PduV_1_mix(pf10662) GRM3A
10145 HMM01965_1_grm3(pf01965) GRM3A
10150 HMMgre_1_grms(pf01228pf02901) GRM3A
10155 HMMgreae_1_grms(pf04055) GRM3A
10160 Tsp_aqua GRM3A
10165 H_robinEggBlue GRM3A
10170 HMM02441_05(#) GRM3A EP_NTERM
10175 H_driedBlood GRM3A
10180 P_murkyGreen GRM3A
10185 HMMalddh_1_eutpdugrm(pf00171) GRM3A
10190 HMMadometsyn_1_grm(pf02773pf02772pf00438) GRM3A
10195 H_tan GRM3A
10200 HMMptac_3_eut2(pf06130) GRM3A EP_NTERM
10205 HMM06723EutJMreB_1_grm(pf11104) GRM3A
10210 HMMalddh_1_eutpdugrm(pf00171) GRM3C EP_NTERM
10215 HMM00465alcdh_3_mix(pf00465) GRM3A
10220 HMM10114PocR_2_grm(pf06580pf10114) GRM3A
10225 HMMreg12833_3_GRM3(pf12833pf00072) GRM3A
10230 -pf00230 -
10235 -# -
10240 -pf01637pf03008 -
10245 -pf13443 -
BMC-P
BMC-H
BMC-T
Aldehyde dehydrogenase
PTAC
Signature enzyme
Alcohol dehydrogenase
Regulator

Clostridium sp. AM29-11AC. Bacteria; Firmicutes; Clostridia; Clostridiales; Clostridiaceae;

DW757__locus_1

GRM3A

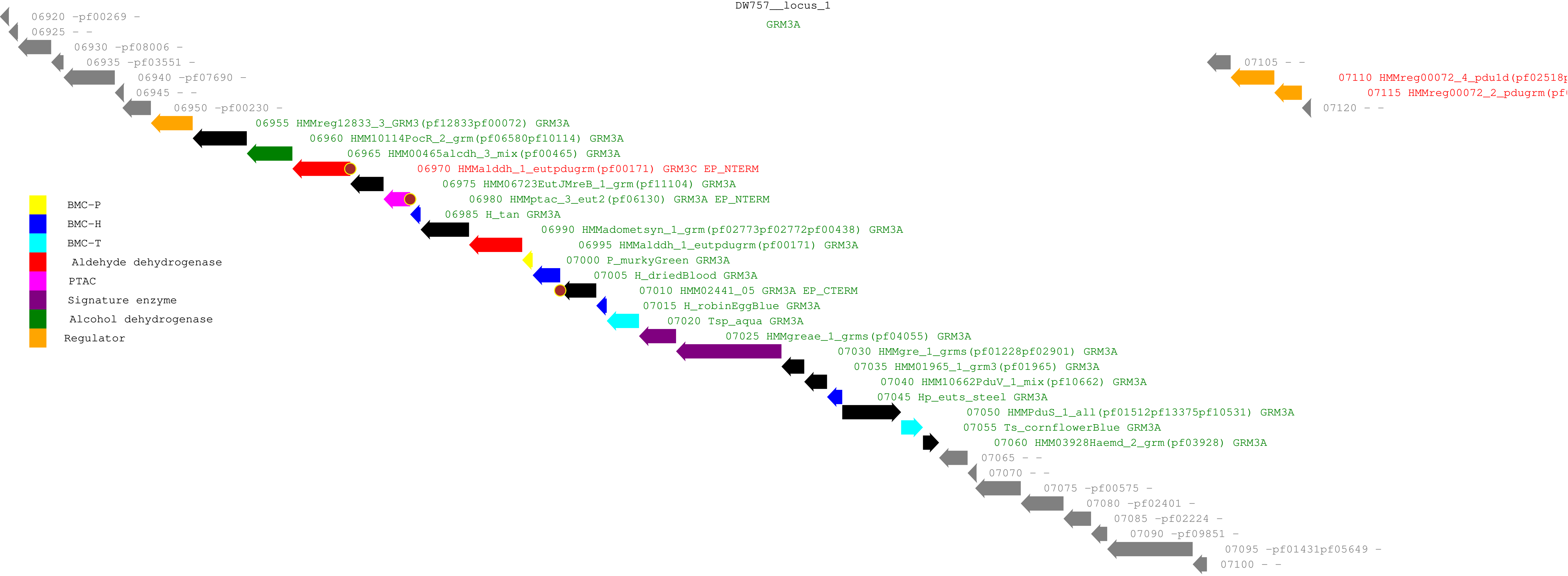

Clostridium bolteae. Bacteria; Firmicutes; Clostridia; Clostridiales; Lachnospiraceae.

DW839__locus_1

GRM3A

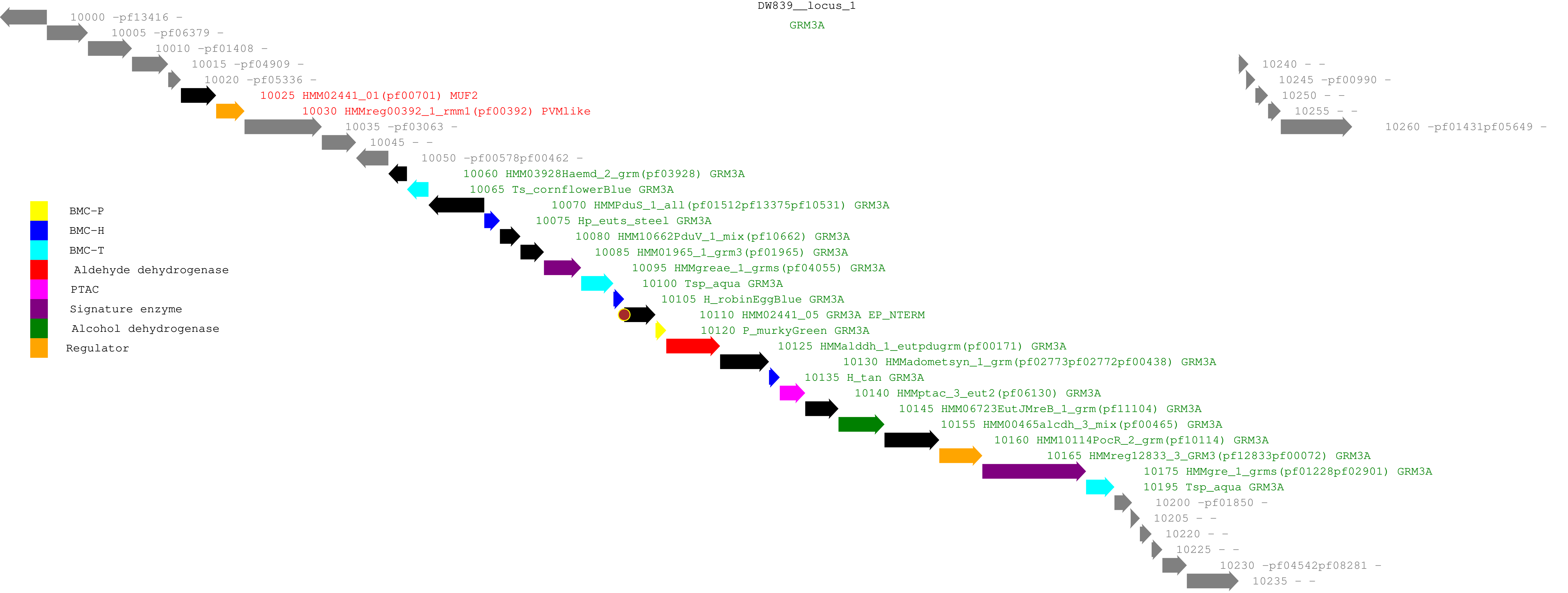

Clostridiaceae bacterium AF02-42. Bacteria; Firmicutes; Clostridia; Clostridiales; Clostridiaceae.

DWV19__locus_1

GRM3A

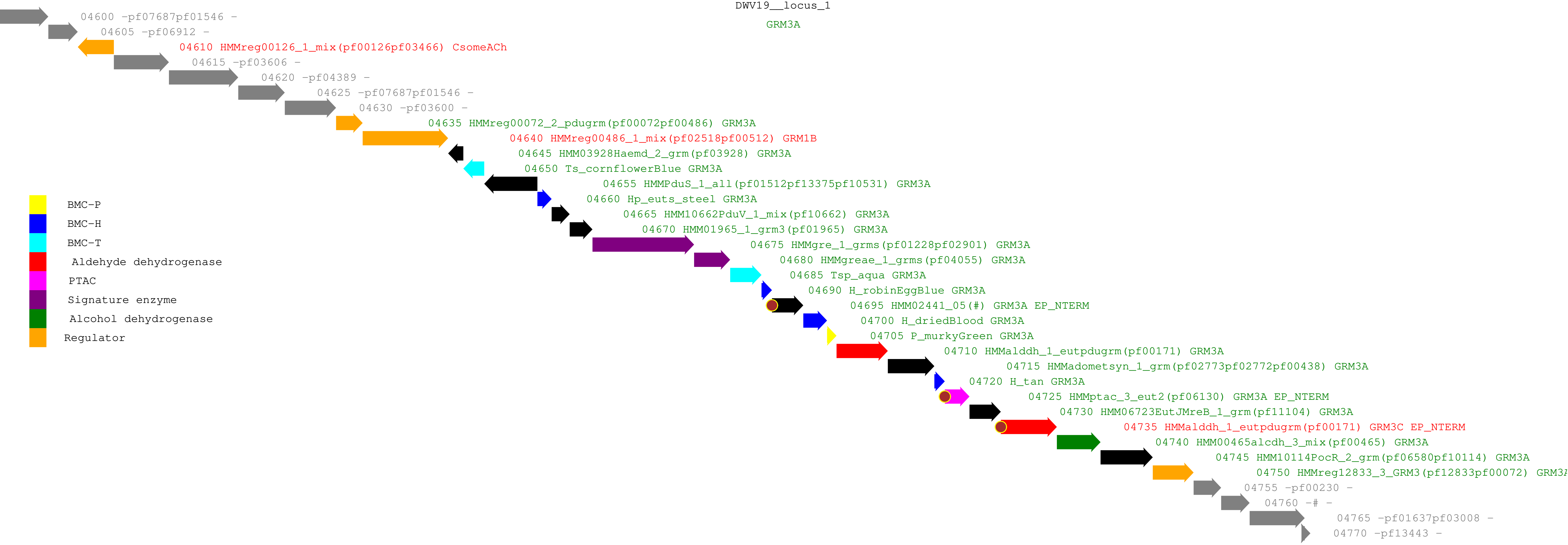

Veillonella parvula. Bacteria; Firmicutes; Negativicutes; Veillonellales; Veillonellaceae;

DWV36__locus_1

GRM3A

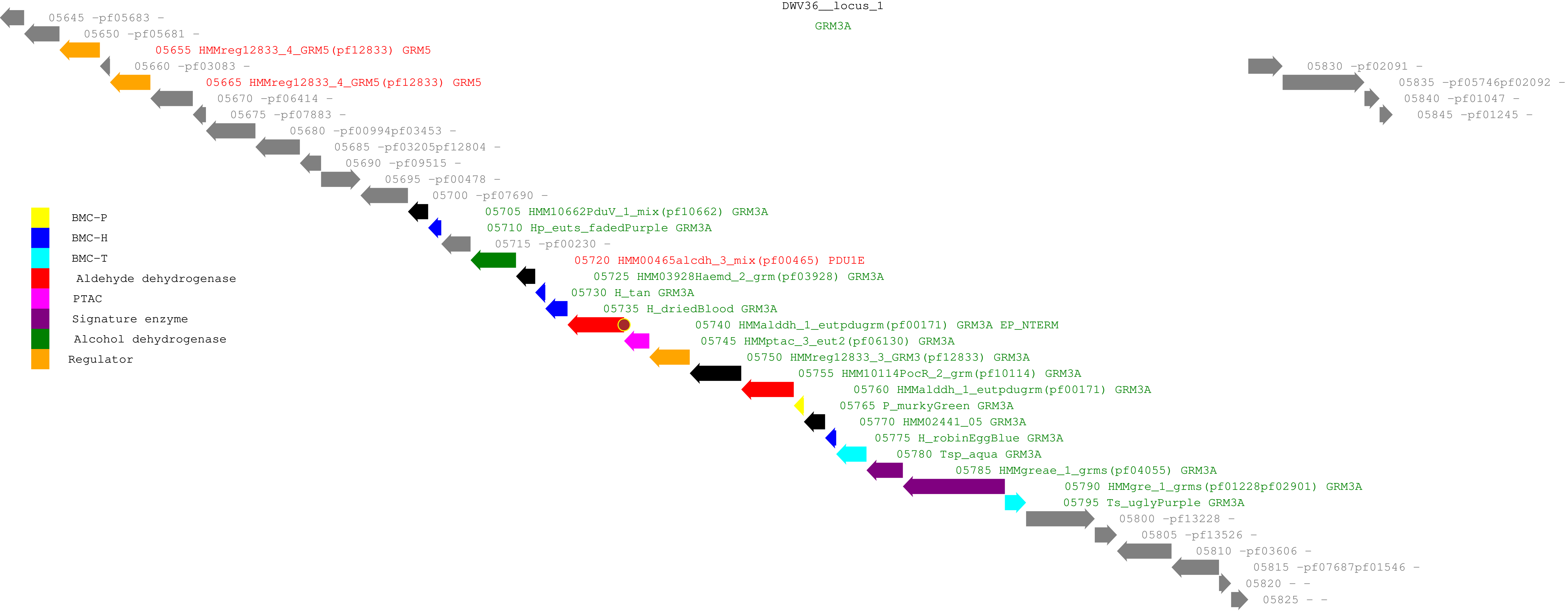

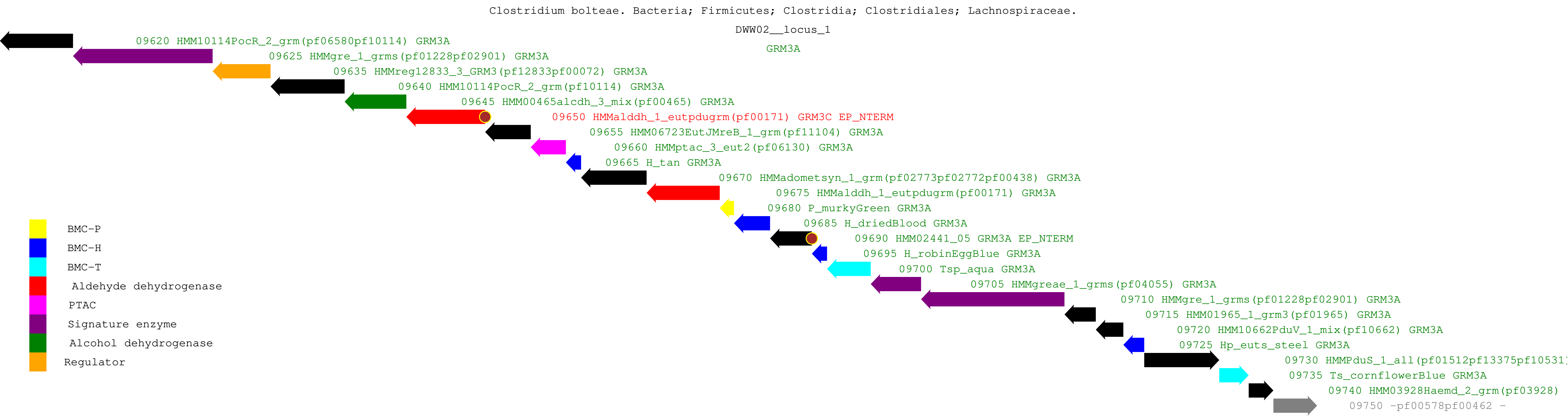

Clostridium bolteae. Bacteria; Firmicutes; Clostridia; Clostridiales; Lachnospiraceae.
DWW02__locus_1
GRM3A
09620 HMM10114PocR_2_grm(pf06580pf10114) GRM3A
09625 HMMgre_1_grms(pf01228pf02901) GRM3A
09635 HMMreg12833_3_GRM3(pf12833pf00072) GRM3A
09640 HMM10114PocR_2_grm(pf10114) GRM3A
09645 HMM00465alcdh_3_mix(pf00465) GRM3A
09650 HMMalddh_1_eutpdugrm(pf00171) GRM3C EP_NTERM
09655 HMM06723EutJMreB_1_grm(pf11104) GRM3A
09660 HMMptac_3_eut2(pf06130) GRM3A
09665 H_tan GRM3A
09670 HMMadometsyn_1_grm(pf02773pf02772pf00438) GRM3A
09675 HMMalddh_1_eutpdugrm(pf00171) GRM3A
09680 P_murkyGreen GRM3A
09685 H_driedBlood GRM3A
09690 HMM02441_05 GRM3A EP_NTERM
09695 H_robinEggBlue GRM3A
09700 Tsp_aqua GRM3A
09705 HMMgreae_1_grms(pf04055) GRM3A
09710 HMMgre_1_grms(pf01228pf02901) GRM3A
09715 HMM01965_1_grm3(pf01965) GRM3A
09720 HMM10662PduV_1_mix(pf10662) GRM3A
09725 Hp_euts_steel GRM3A
09730 HMMPduS_1_all(pf01512pf13375pf10531
09735 Ts_cornflowerBlue GRM3A
09740 HMM03928Haemd_2_grm(pf03928)
09750 -pf00578pf00462 -
BMC-P
BMC-H
BMC-T
Aldehyde dehydrogenase
PTAC
Signature enzyme
Alcohol dehydrogenase
Regulator

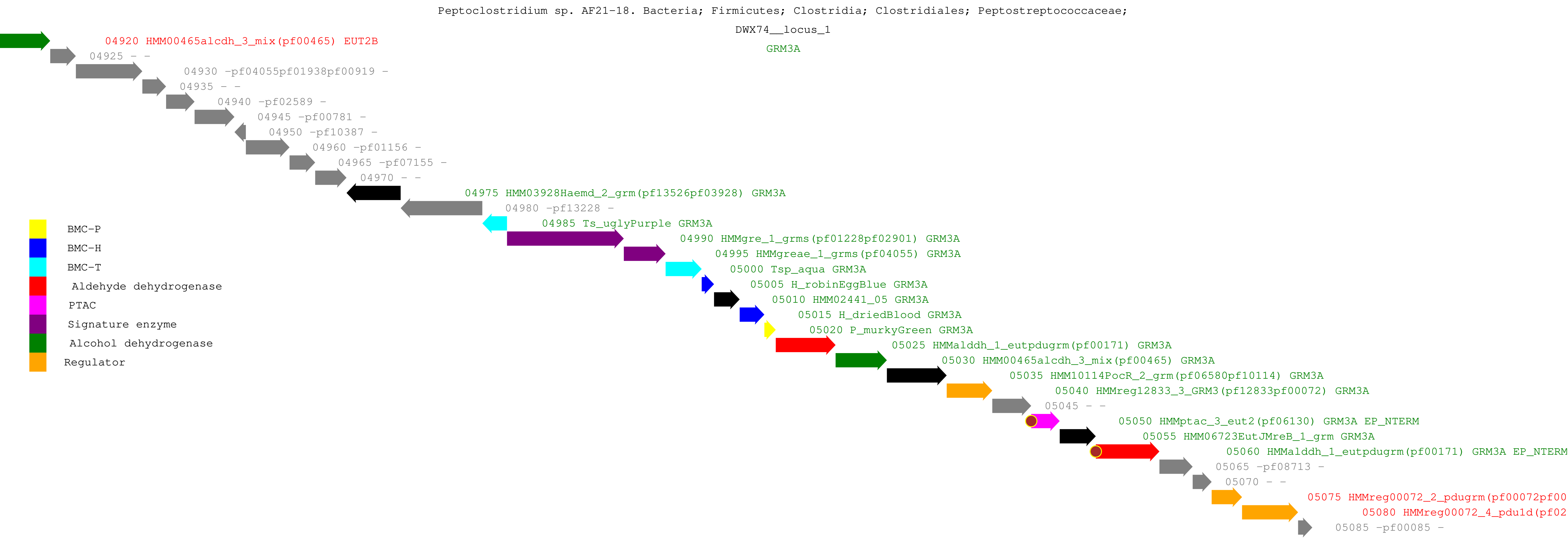
Peptoclostridium sp. AF21-18. Bacteria; Firmicutes; Clostridia; Clostridiales; Peptostreptococcaceae;
DWX74__locus_1
GRM3A
04920 HMM00465alcdh_3_mix(pf00465) EUT2B
04925 - -
04930 -pf04055pf01938pf00919 -
04935 - -
04940 -pf02589 -
04945 -pf00781 -
04950 -pf10387 -
04960 -pf01156 -
04965 -pf07155 -
04970 - -
04975 HMM03928Haemd_2_grm(pf13526pf03928) GRM3A
04980 -pf13228 -
04985 Ts_uglyPurple GRM3A
04990 HMMgre_1_grms(pf01228pf02901) GRM3A
04995 HMMgreae_1_grms(pf04055) GRM3A
05000 Tsp_aqua GRM3A
05005 H_robinEggBlue GRM3A
05010 HMM02441_05 GRM3A
05015 H_driedBlood GRM3A
05020 P_murkyGreen GRM3A
05025 HMMalddh_1_eutpdugrm(pf00171) GRM3A
05030 HMM00465alcdh_3_mix(pf00465) GRM3A
05035 HMM10114PocR_2_grm(pf06580pf10114) GRM3A
05040 HMMreg12833_3_GRM3(pf12833pf00072) GRM3A
05045 - -
05050 HMMptac_3_eut2(pf06130) GRM3A EP_NTERM
05055 HMM06723EutJMreB_1_grm GRM3A
05060 HMMalddh_1_eutpdugrm(pf00171) GRM3A EP_NTERM
05065 -pf08713 -
05070 - -
05075 HMMreg00072_2_pdugrm(pf00072pf00
05080 HMMreg00072_4_pdu1d(pf02
05085 -pf00085 -
BMC-P
BMC-H
BMC-T
Aldehyde dehydrogenase
PTAC
Signature enzyme
Alcohol dehydrogenase
Regulator

Clostridium sp. AF28-12. Bacteria; Firmicutes; Clostridia; Clostridiales; Clostridiaceae;

DWY93__locus_1

GRM3A

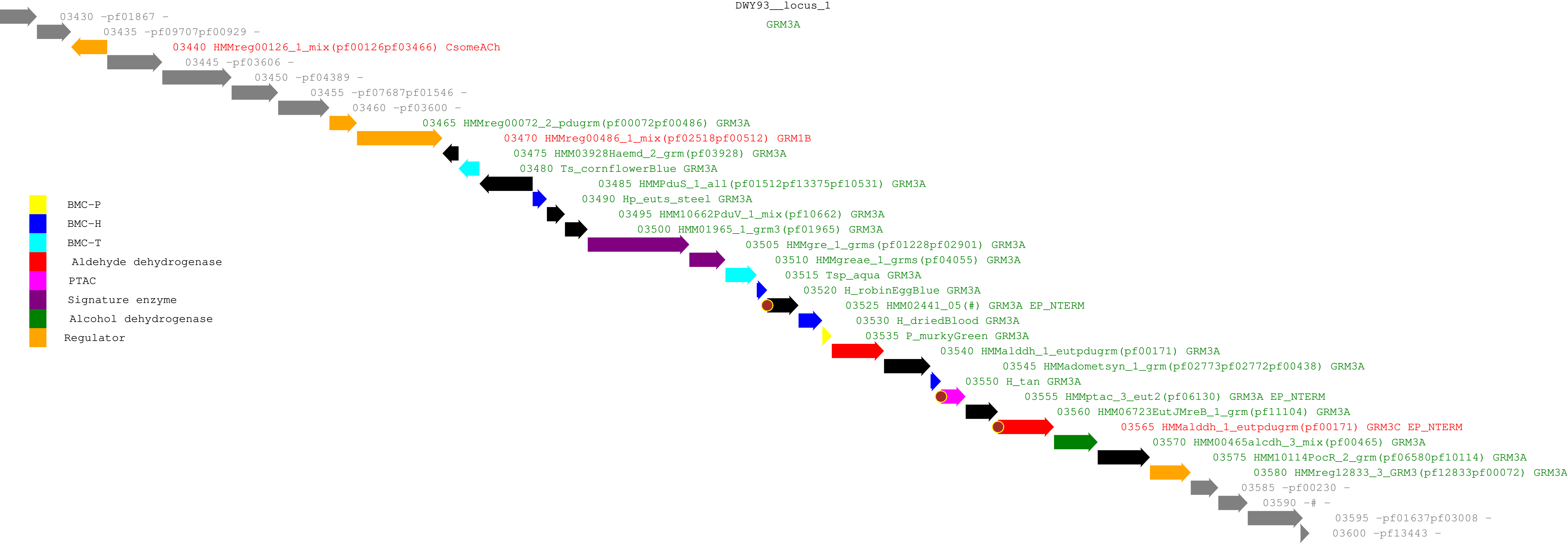

Clostridium citroniae. Bacteria; Firmicutes; Clostridia; Clostridiales; Lachnospiraceae.

DWZ14__locus_2

GRM3A

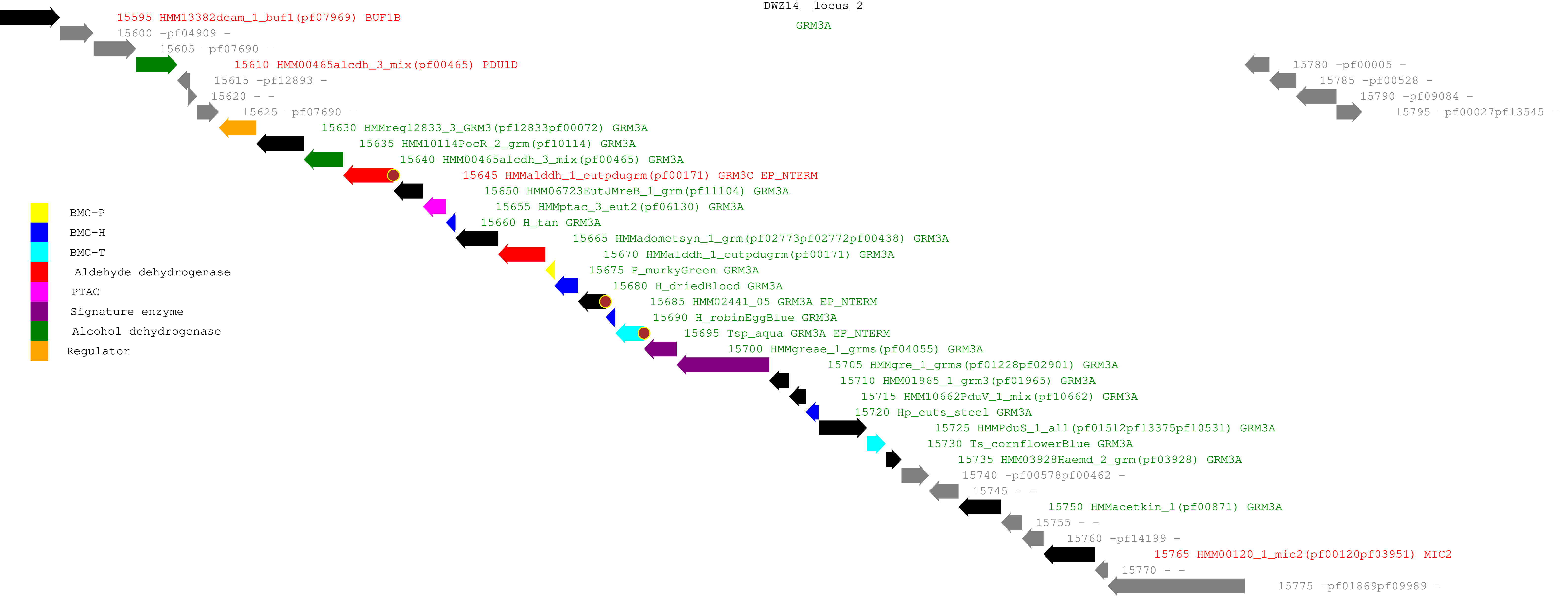

Clostridiales bacterium AF36-10. Bacteria; Firmicutes; Clostridia; Clostridiales.

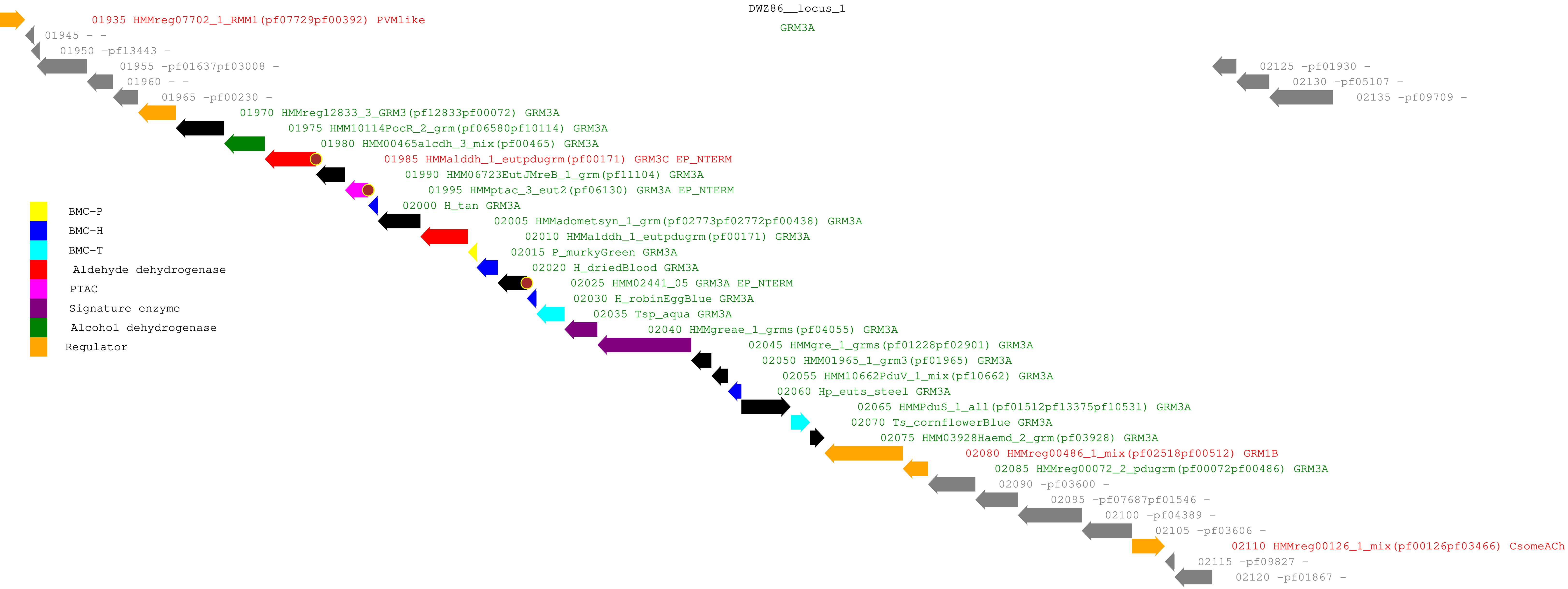

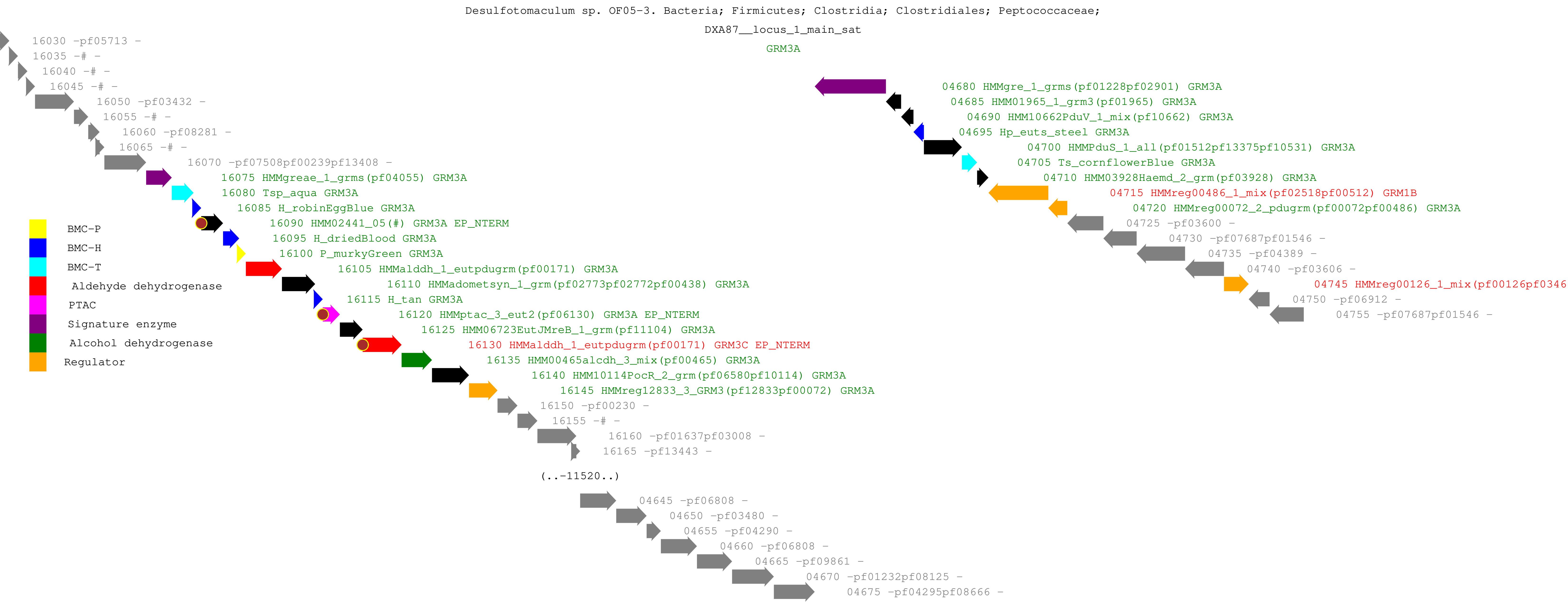

Desulfotomaculum sp. OF05-3. Bacteria; Firmicutes; Clostridia; Clostridiales; Peptococcaceae;
DXA87__locus_1_main_sat
GRM3A
16030 -pf05713 -
16035 -# -
16040 -# -
16045 -# -
16050 -pf03432 -
16055 -# -
16060 -pf08281 -
16065 -# -
16070 -pf07508pf00239pf13408 -
16075 HMMgreae_1_grms(pf04055) GRM3A
16080 Tsp_aqua GRM3A
16085 H_robinEggBlue GRM3A
16090 HMM02441_05(#) GRM3A EP_NTERM
16095 H_driedBlood GRM3A
16100 P_murkyGreen GRM3A
16105 HMMaldh_1_eutpdugrm(pf00171) GRM3A
16110 HMMadometsyn_1_grm(pf02773pf02772pf00438) GRM3A
16115 H_tan GRM3A
16120 HMMptac_3_eut2(pf06130) GRM3A EP_NTERM
16125 HMM06723EutJMreB_1_grm(pf11104) GRM3A
16130 HMMaldh_1_eutpdugrm(pf00171) GRM3C EP_NTERM
16135 HMM00465alcdh_3_mix(pf00465) GRM3A
16140 HMM10114PocR_2_grm(pf06580pf10114) GRM3A
16145 HMMreg12833_3_GRM3(pf12833pf00072) GRM3A
16150 -pf00230 -
16155 -# -
16160 -pf01637pf03008 -
16165 -pf13443 -
(..-11520..)
04645 -pf06808 -
04650 -pf03480 -
04655 -pf04290 -
04660 -pf06808 -
04665 -pf09861 -
04670 -pf01232pf08125 -
04675 -pf04295pf08666 -
04680 HMMgre_1_grms(pf01228pf02901) GRM3A
04685 HMM01965_1_grm3(pf01965) GRM3A
04690 HMM10662PduV_1_mix(pf10662) GRM3A
04695 Hp_euts_steel GRM3A
04700 HMMPduS_1_all(pf01512pf13375pf10531) GRM3A
04705 Ts_cornflowerBlue GRM3A
04710 HMM03928Haemd_2_grm(pf03928) GRM3A
04715 HMMreg00486_1_mix(pf02518pf00512) GRM1B
04720 HMMreg00072_2_pdugrm(pf00072pf00486) GRM3A
04725 -pf03600 -
04730 -pf07687pf01546 -
04735 -pf04389 -
04740 -pf03606 -
04745 HMMreg00126_1_mix(pf00126pf0346
04750 -pf06912 -
04755 -pf07687pf01546 -
BMC-P
BMC-H
BMC-T
Aldehyde dehydrogenase
PTAC
Signature enzyme
Alcohol dehydrogenase
Regulator

Clostridiaceae bacterium TF01-6. Bacteria; Firmicutes; Clostridia; Clostridiales; Clostridiaceae.

DXC33__locus_1

GRM3A

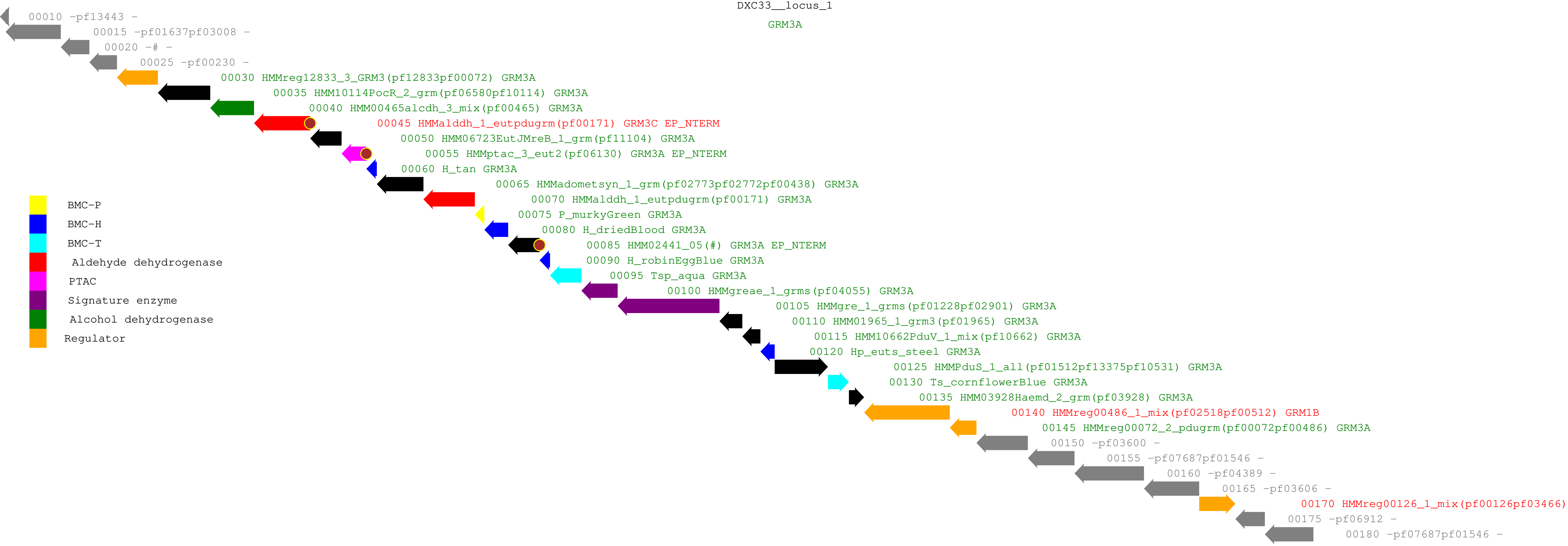

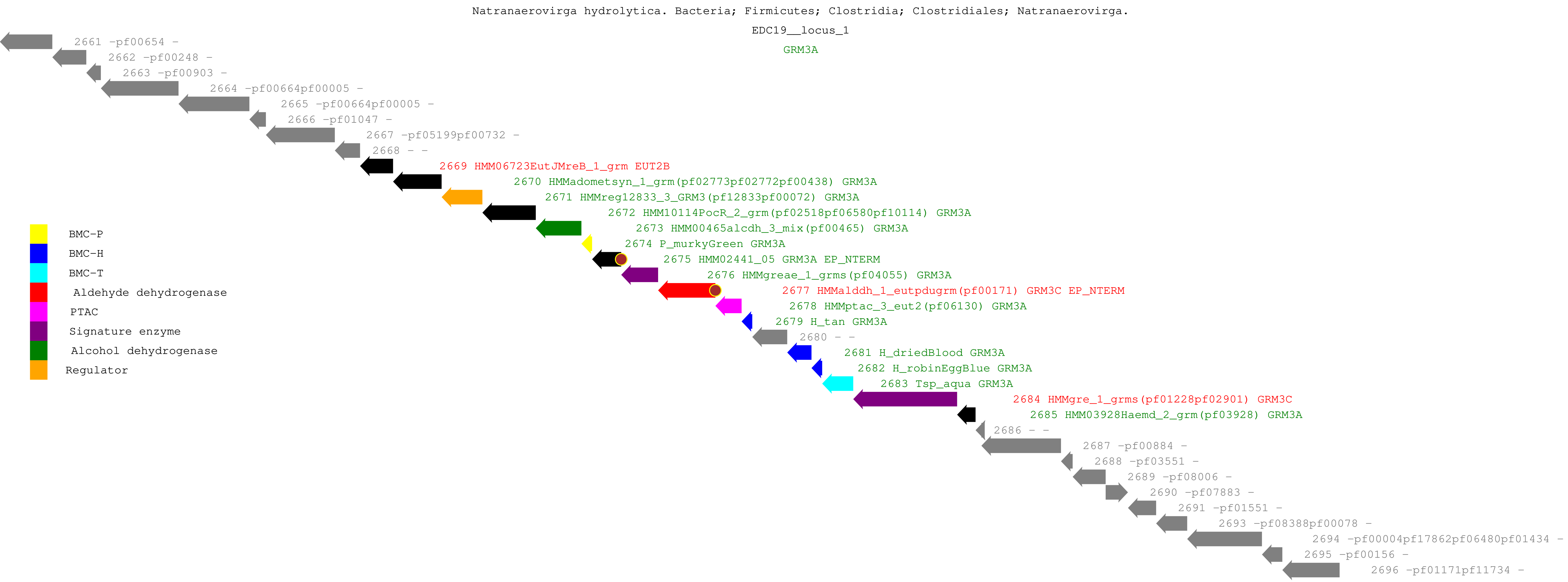
Natranaerovirga hydrolytica. Bacteria; Firmicutes; Clostridia; Clostridiales; Natranaerovirga.
EDC19__locus_1
GRM3A
2661 -pf00654 -
2662 -pf00248 -
2663 -pf00903 -
2664 -pf00664pf00005 -
2665 -pf00664pf00005 -
2666 -pf01047 -
2667 -pf05199pf00732 -
2668 - -
2669 HMM06723EutJMreB_1_grm EUT2B
2670 HMMadometsyn_1_grm(pf02773pf02772pf00438) GRM3A
2671 HMMreg12833_3_GRM3(pf12833pf00072) GRM3A
2672 HMM10114PocR_2_grm(pf02518pf06580pf10114) GRM3A
2673 HMM00465alcdh_3_mix(pf00465) GRM3A
2674 P_murkyGreen GRM3A
2675 HMM02441_05 GRM3A EP_NTERM
2676 HMMgreae_1_grms(pf04055) GRM3A
2677 HMMalddh_1_eutpdugrm(pf00171) GRM3C EP_NTERM
2678 HMMptac_3_eut2(pf06130) GRM3A
2679 H_tan GRM3A
2680 - -
2681 H_driedBlood GRM3A
2682 H_robinEggBlue GRM3A
2683 Tsp_aqua GRM3A
2684 HMMgre_1_grms(pf01228pf02901) GRM3C
2685 HMM03928Haemd_2_grm(pf03928) GRM3A
2686 - -
2687 -pf00884 -
2688 -pf03551 -
2689 -pf08006 -
2690 -pf07883 -
2691 -pf01551 -
2693 -pf08388pf00078 -
2694 -pf00004pf17862pf06480pf01434 -
2695 -pf00156 -
2696 -pf01171pf11734 -
BMC-P
BMC-H
BMC-T
Aldehyde dehydrogenase
PTAC
Signature enzyme
Alcohol dehydrogenase
Regulator

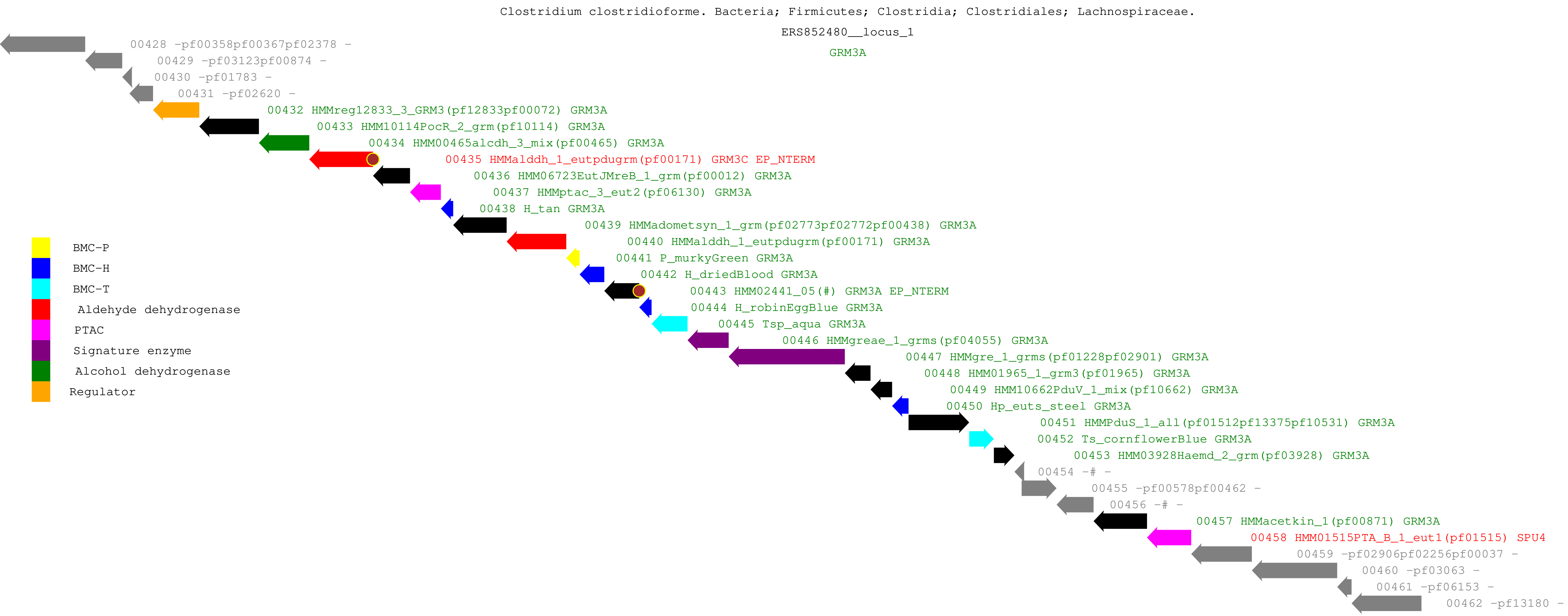

Clostridium clostridioforme. Bacteria; Firmicutes; Clostridia; Clostridiales; Lachnospiraceae.
ERS852480__locus_1
GRM3A
00428 -pf00358pf00367pf02378 -
00429 -pf03123pf00874 -
00430 -pf01783 -
00431 -pf02620 -
00432 HMMreg12833_3_GRM3(pf12833pf00072) GRM3A
00433 HMM10114PocR_2_grm(pf10114) GRM3A
00434 HMM00465alcdh_3_mix(pf00465) GRM3A
00435 HMMaldh_1_eutpdugrm(pf00171) GRM3C EP_NTERM
00436 HMM06723EutJMreB_1_grm(pf00012) GRM3A
00437 HMMptac_3_eut2(pf06130) GRM3A
00438 H_tan GRM3A
00439 HMMadometsyn_1_grm(pf02773pf02772pf00438) GRM3A
00440 HMMaldh_1_eutpdugrm(pf00171) GRM3A
00441 P_murkyGreen GRM3A
00442 H_driedBlood GRM3A
00443 HMM02441_05(#) GRM3A EP_NTERM
00444 H_robinEggBlue GRM3A
00445 Tsp_aqua GRM3A
00446 HMMgreae_1_grms(pf04055) GRM3A
00447 HMMgre_1_grms(pf01228pf02901) GRM3A
00448 HMM01965_1_grm3(pf01965) GRM3A
00449 HMM10662PduV_1_mix(pf10662) GRM3A
00450 Hp_euts_steel GRM3A
00451 HMMPduS_1_all(pf01512pf13375pf10531) GRM3A
00452 Ts_cornflowerBlue GRM3A
00453 HMM03928Haemd_2_grm(pf03928) GRM3A
00454 -# -
00455 -pf00578pf00462 -
00456 -# -
00457 HMMacetkin_1(pf00871) GRM3A
00458 HMM01515PTA_B_1_eut1(pf01515) SPU4
00459 -pf02906pf02256pf00037 -
00460 -pf03063 -
00461 -pf06153 -
00462 -pf13180 -
BMC-P
BMC-H
BMC-T
Aldehyde dehydrogenase
PTAC
Signature enzyme
Alcohol dehydrogenase
Regulator

Lactobacillus ghanensis DSM 18630. Bacteria; Firmicutes; Bacilli; Lactobacillales; Lactobacillaceae;

FC89_GL_locus_1

GRM3A

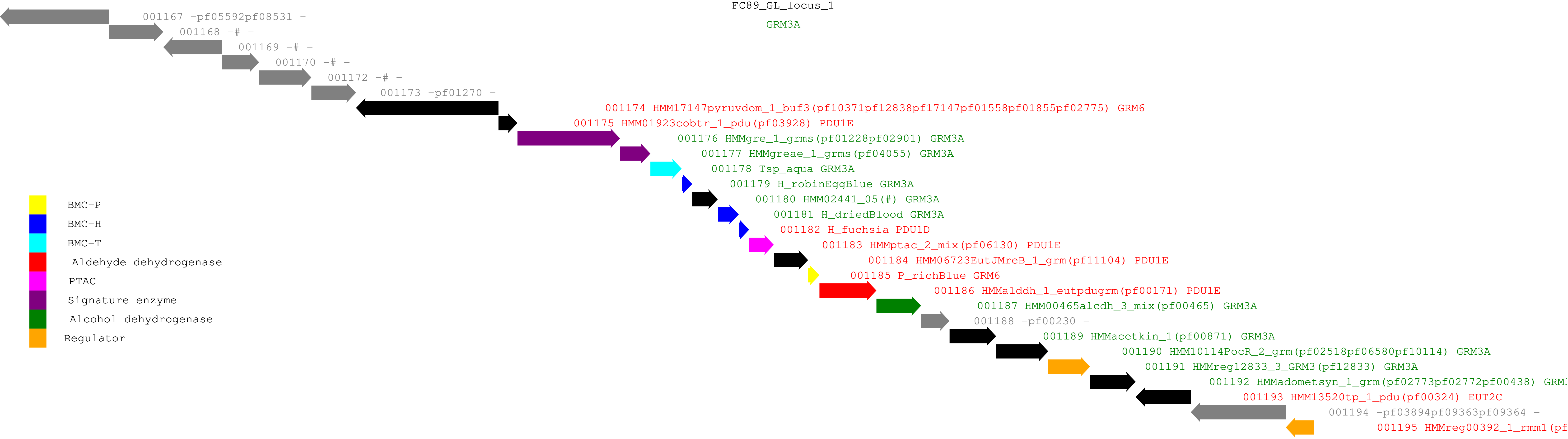

Eubacterium yurii subsp. margaretiae ATCC 43715. Bacteria; Firmicutes; Clostridia; Clostridiales;

HMPREF0379__locus_1

GRM3A

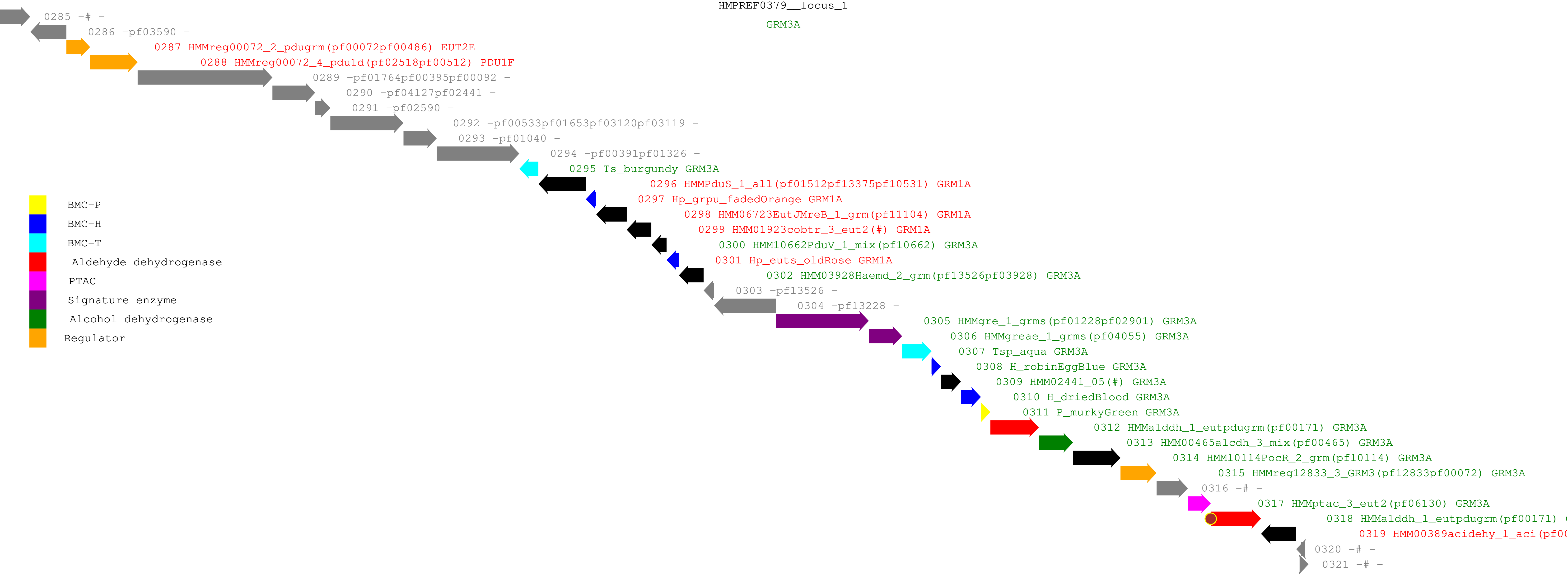

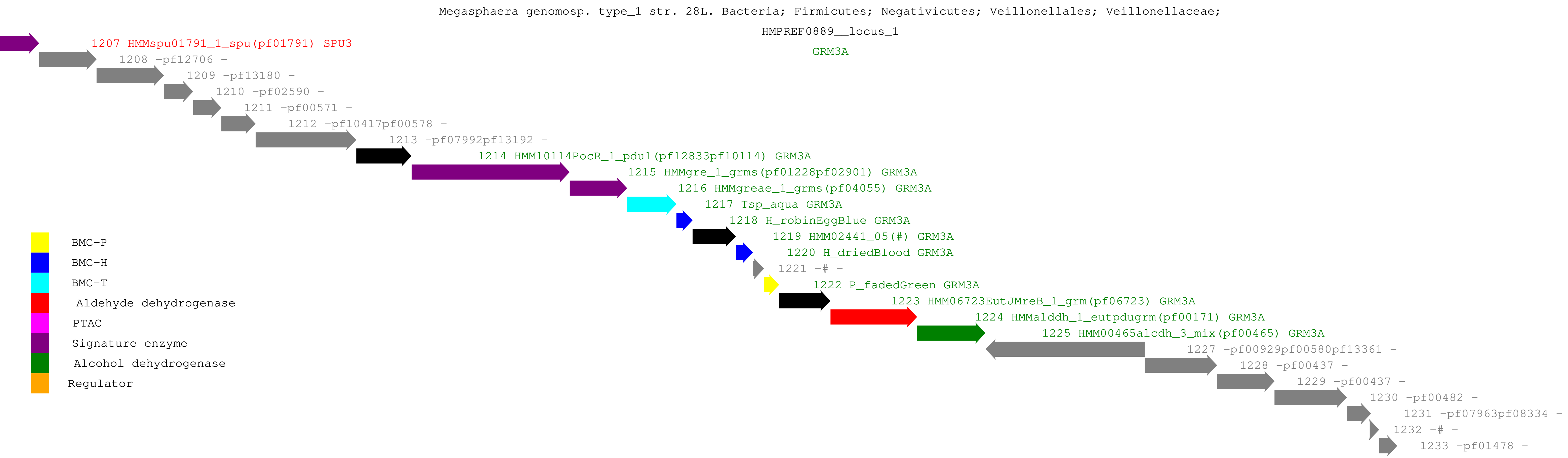

Megasphaera genomosp. type_1 str. 28L. Bacteria; Firmicutes; Negativicutes; Veillonellales; Veillonellaceae;
HMPREF0889__locus_1
GRM3A
1207 HMMspu01791_1_spu(pf01791) SPU3
1208 -pf12706 -
1209 -pf13180 -
1210 -pf02590 -
1211 -pf00571 -
1212 -pf10417pf00578 -
1213 -pf07992pf13192 -
1214 HMM10114PocR_1_pdu1(pf12833pf10114) GRM3A
1215 HMMgre_1_grms(pf01228pf02901) GRM3A
1216 HMMgreae_1_grms(pf04055) GRM3A
1217 Tsp_aqua GRM3A
1218 H_robinEggBlue GRM3A
1219 HMM02441_05(#) GRM3A
1220 H_driedBlood GRM3A
1221 -# -
1222 P_fadedGreen GRM3A
1223 HMM06723EutJMreB_1_grm(pf06723) GRM3A
1224 HMMaldh_1_eutpdugrm(pf00171) GRM3A
1225 HMM00465alcdh_3_mix(pf00465) GRM3A
1227 -pf00929pf00580pf13361 -
1228 -pf00437 -
1229 -pf00437 -
1230 -pf00482 -
1231 -pf07963pf08334 -
1232 -# -
1233 -pf01478 -
BMC-P
BMC-H
BMC-T
Aldehyde dehydrogenase
PTAC
Signature enzyme
Alcohol dehydrogenase
Regulator

Megasphaera sp. UPII 135-E. Bacteria; Firmicutes; Negativicutes; Veillonellales; Veillonellaceae;

HMPREF1040__locus_1

GRM3A

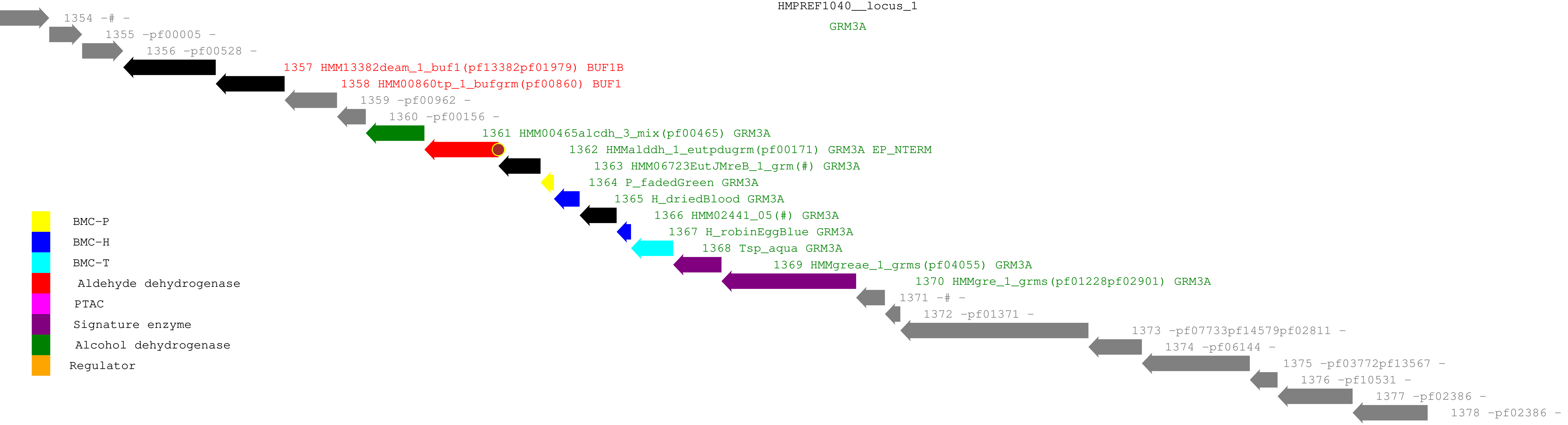

Clostridium clostridioforme 90A6. Bacteria; Firmicutes; Clostridia; Clostridiales; Lachnospiraceae.

HMPREF1083__locus_1

GRM3A

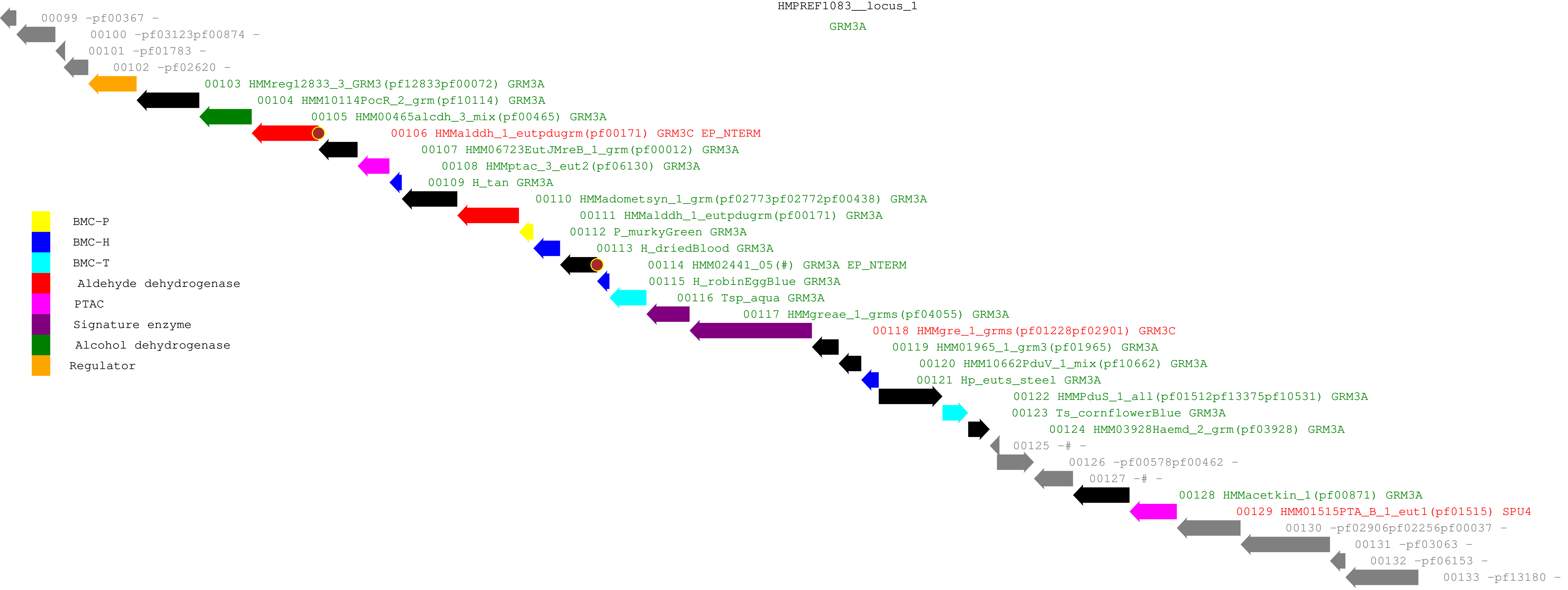

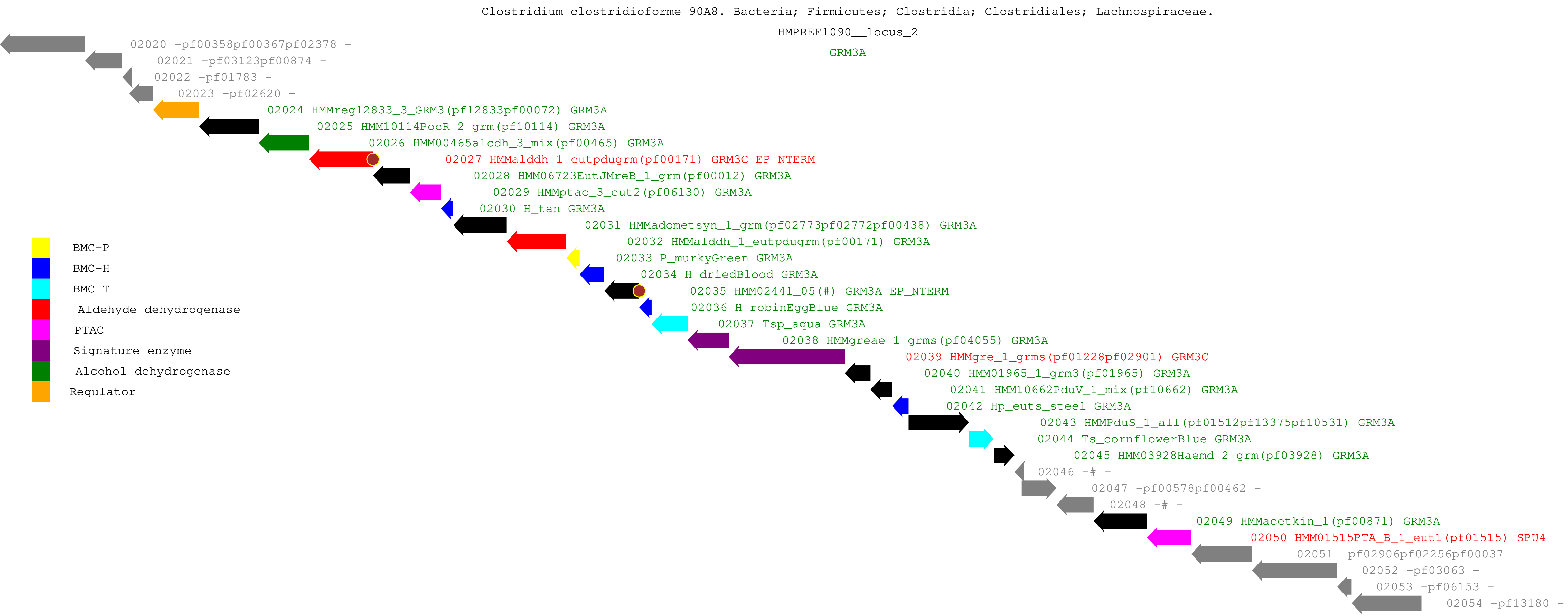
Clostridium clostridioforme 90A8. Bacteria; Firmicutes; Clostridia; Clostridiales; Lachnospiraceae.
HMPREF1090__locus_2
GRM3A
02020 -pf00358pf00367pf02378 -
02021 -pf03123pf00874 -
02022 -pf01783 -
02023 -pf02620 -
02024 HMMreg12833_3_GRM3(pf12833pf00072) GRM3A
02025 HMM10114PocR_2_grm(pf10114) GRM3A
02026 HMM00465alcdh_3_mix(pf00465) GRM3A
02027 HMMaldd h_1_eutpdugrm(pf00171) GRM3C EP_NTERM
02028 HMM06723EutJMreB_1_grm(pf00012) GRM3A
02029 HMMptac_3_eut2(pf06130) GRM3A
02030 H_tan GRM3A
02031 HMMadometsyn_1_grm(pf02773pf02772pf00438) GRM3A
02032 HMMaldd h_1_eutpdugrm(pf00171) GRM3A
02033 P_murkyGreen GRM3A
02034 H_driedBlood GRM3A
02035 HMM02441_05(#) GRM3A EP_NTERM
02036 H_robinEggBlue GRM3A
02037 Tsp_aqua GRM3A
02038 HMMgreae_1_grms(pf04055) GRM3A
02039 HMMgre_1_grms(pf01228pf02901) GRM3C
02040 HMM01965_1_grm3(pf01965) GRM3A
02041 HMM10662PduV_1_mix(pf10662) GRM3A
02042 Hp_euts_steel GRM3A
02043 HMMPduS_1_all(pf01512pf13375pf10531) GRM3A
02044 Ts_cornflowerBlue GRM3A
02045 HMM03928Haemd_2_grm(pf03928) GRM3A
02046 -# -
02047 -pf00578pf00462 -
02048 -# -
02049 HMMacetkin_1(pf00871) GRM3A
02050 HMM01515PTA_B_1_eut1(pf01515) SPU4
02051 -pf02906pf02256pf00037 -
02052 -pf03063 -
02053 -pf06153 -
02054 -pf13180 -
BMC-P
BMC-H
BMC-T
Aldehyde dehydrogenase
PTAC
Signature enzyme
Alcohol dehydrogenase
Regulator

Clostridium bolteae 90B8. Bacteria; Firmicutes; Clostridia; Clostridiales; Lachnospiraceae.

HMPREF1097__locus_1

GRM3A

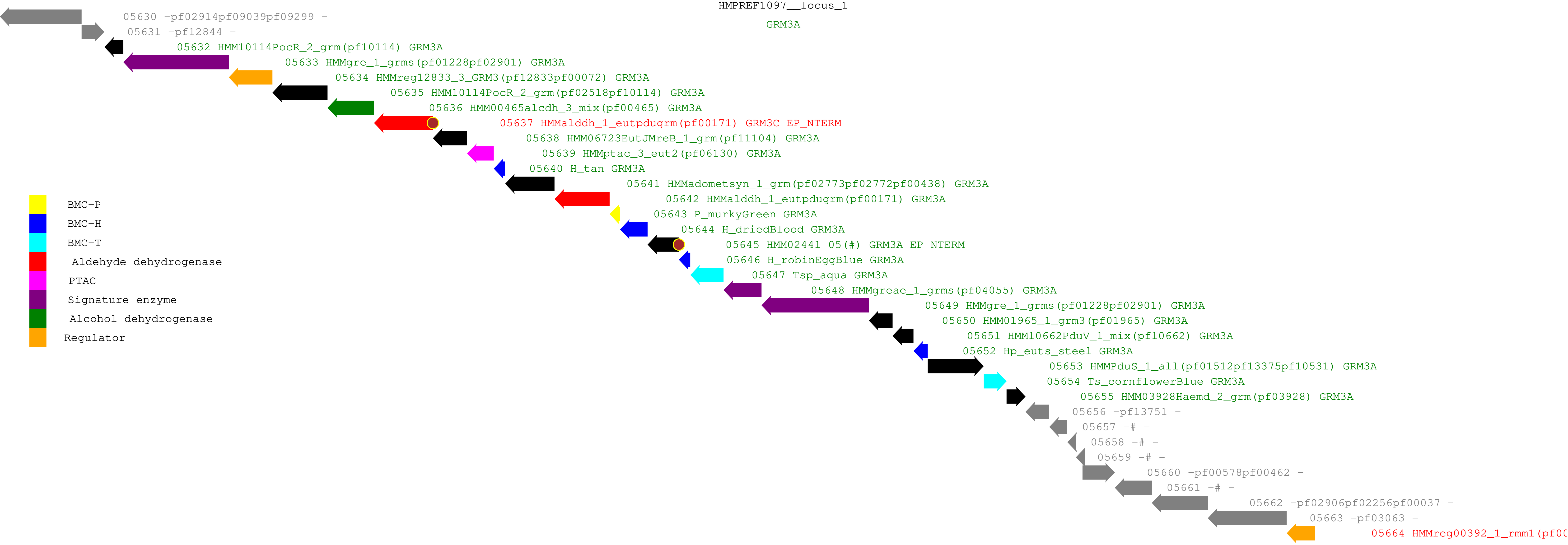

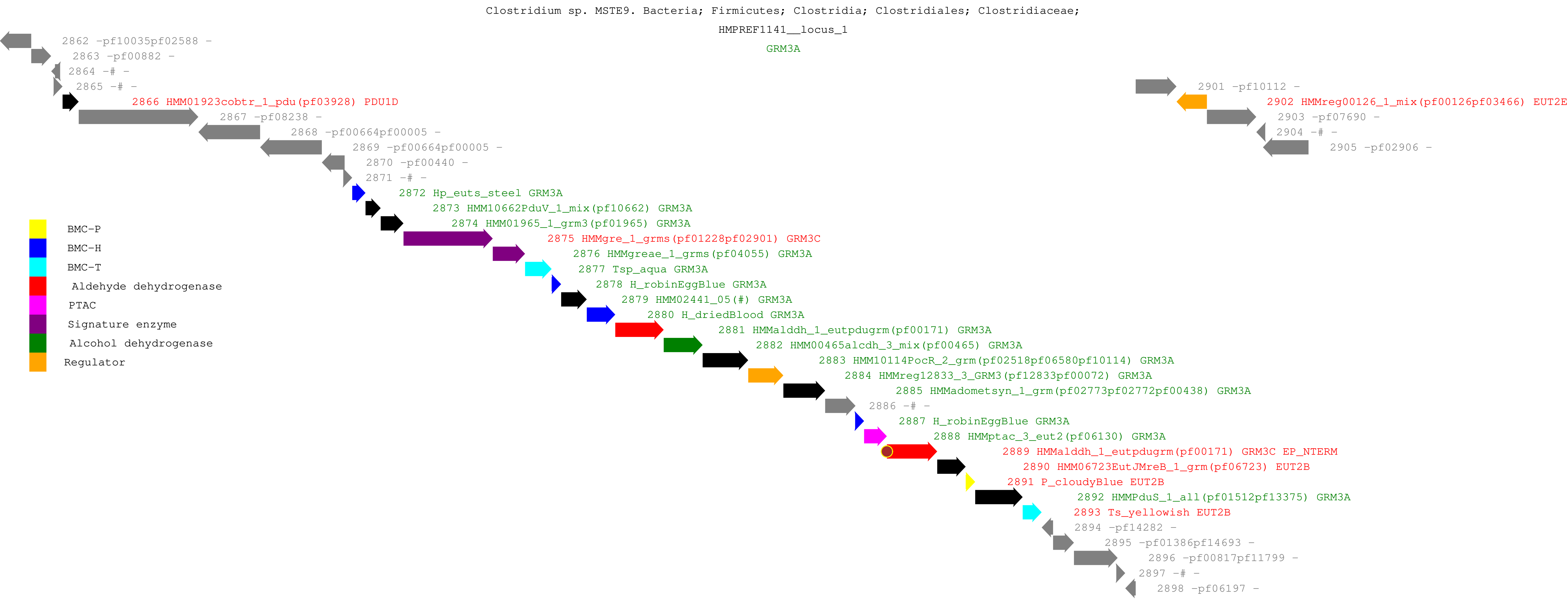
Clostridium sp. MSTE9. Bacteria; Firmicutes; Clostridia; Clostridiales; Clostridiaceae;
HMPREF1141__locus_1
GRM3A
2862 -pf10035pf02588 -
2863 -pf00882 -
2864 -# -
2865 -# -
2866 HMM01923cobtr_1_pdu(pf03928) PDU1D
2867 -pf08238 -
2868 -pf00664pf00005 -
2869 -pf00664pf00005 -
2870 -pf00440 -
2871 -# -
2872 Hp_euts_steel GRM3A
2873 HMM10662PduV_1_mix(pf10662) GRM3A
2874 HMM01965_1_grm3(pf01965) GRM3A
2875 HMMgre_1_grms(pf01228pf02901) GRM3C
2876 HMMgreae_1_grms(pf04055) GRM3A
2877 Tsp_aqua GRM3A
2878 H_robinEggBlue GRM3A
2879 HMM02441_05(#) GRM3A
2880 H_driedBlood GRM3A
2881 HMMalddh_1_eutpdugrm(pf00171) GRM3A
2882 HMM00465alcdh_3_mix(pf00465) GRM3A
2883 HMM10114PocR_2_grm(pf02518pf06580pf10114) GRM3A
2884 HMMreg12833_3_GRM3(pf12833pf00072) GRM3A
2885 HMMadometsyn_1_grm(pf02773pf02772pf00438) GRM3A
2886 -# -
2887 H_robinEggBlue GRM3A
2888 HMMptac_3_eut2(pf06130) GRM3A
2889 HMMalddh_1_eutpdugrm(pf00171) GRM3C EP_NTERM
2890 HMM06723EutJMreB_1_grm(pf06723) EUT2B
2891 P_cloudyBlue EUT2B
2892 HMMPduS_1_all(pf01512pf13375) GRM3A
2893 Ts_yellowish EUT2B
2894 -pf14282 -
2895 -pf01386pf14693 -
2896 -pf00817pf11799 -
2897 -# -
2898 -pf06197 -
2901 -pf10112 -
2902 HMMreg00126_1_mix(pf00126pf03466) EUT2E
2903 -pf07690 -
2904 -# -
2905 -pf02906 -
BMC-P
BMC-H
BMC-T
Aldehyde dehydrogenase
PTAC
Signature enzyme
Alcohol dehydrogenase
Regulator

Peptostreptococcaceae bacterium AS15. Bacteria; Firmicutes; Clostridia; Clostridiales;

HMPREF1142__locus_1

GRM3A

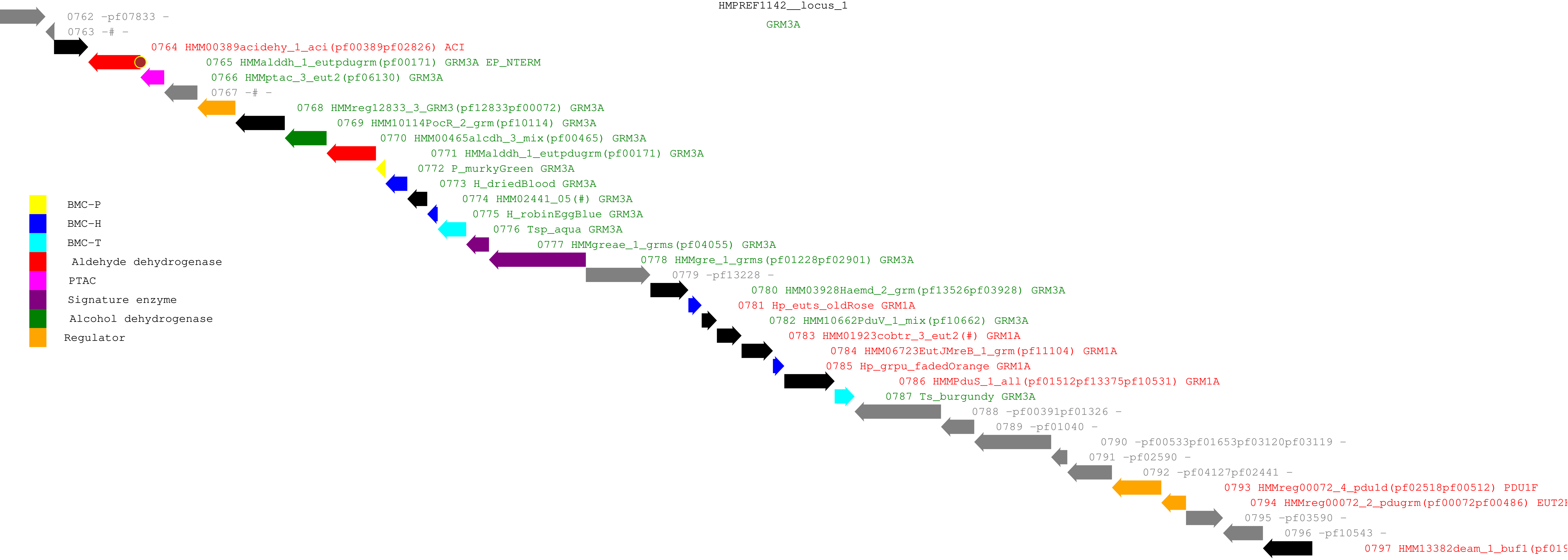

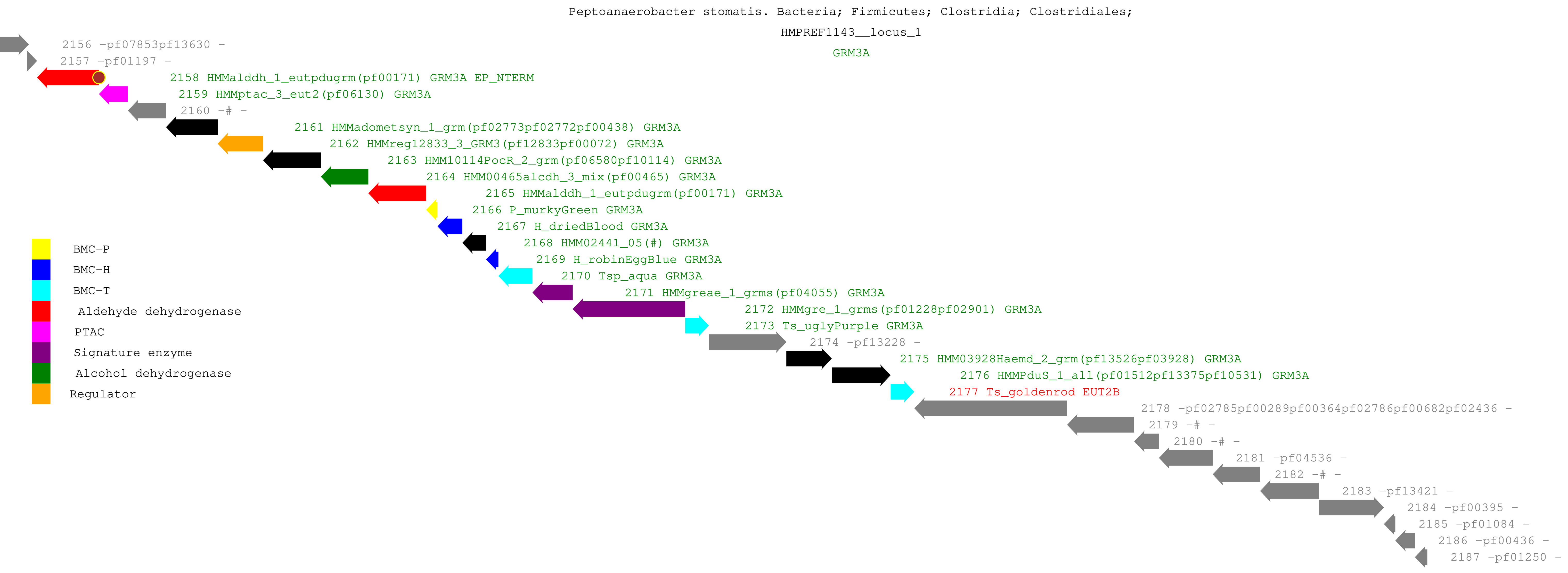
Peptoanaerobacter stomatis. Bacteria; Firmicutes; Clostridia; Clostridiales;
HMPREF1143__locus_1
GRM3A
2156 -pf07853pf13630 -
2157 -pf01197 -
2158 HMMalddh_1_eutpdugrm(pf00171) GRM3A EP_NTERM
2159 HMMptac_3_eut2(pf06130) GRM3A
2160 -# -
2161 HMMadometsyn_1_grm(pf02773pf02772pf00438) GRM3A
2162 HMMreg12833_3_GRM3(pf12833pf00072) GRM3A
2163 HMM10114PocR_2_grm(pf06580pf10114) GRM3A
2164 HMM00465alcdh_3_mix(pf00465) GRM3A
2165 HMMalddh_1_eutpdugrm(pf00171) GRM3A
2166 P_murkyGreen GRM3A
2167 H_driedBlood GRM3A
2168 HMM02441_05(#) GRM3A
2169 H_robinEggBlue GRM3A
2170 Tsp_aqua GRM3A
2171 HMMgreae_1_grms(pf04055) GRM3A
2172 HMMgre_1_grms(pf01228pf02901) GRM3A
2173 Ts_uglyPurple GRM3A
2174 -pf13228 -
2175 HMM03928Haemd_2_grm(pf13526pf03928) GRM3A
2176 HMMPduS_1_all(pf01512pf13375pf10531) GRM3A
2177 Ts_goldenrod EUT2B
2178 -pf02785pf00289pf00364pf02786pf00682pf02436 -
2179 -# -
2180 -# -
2181 -pf04536 -
2182 -# -
2183 -pf13421 -
2184 -pf00395 -
2185 -pf01084 -
2186 -pf00436 -
2187 -pf01250 -
BMC-P
BMC-H
BMC-T
Aldehyde dehydrogenase
PTAC
Signature enzyme
Alcohol dehydrogenase
Regulator

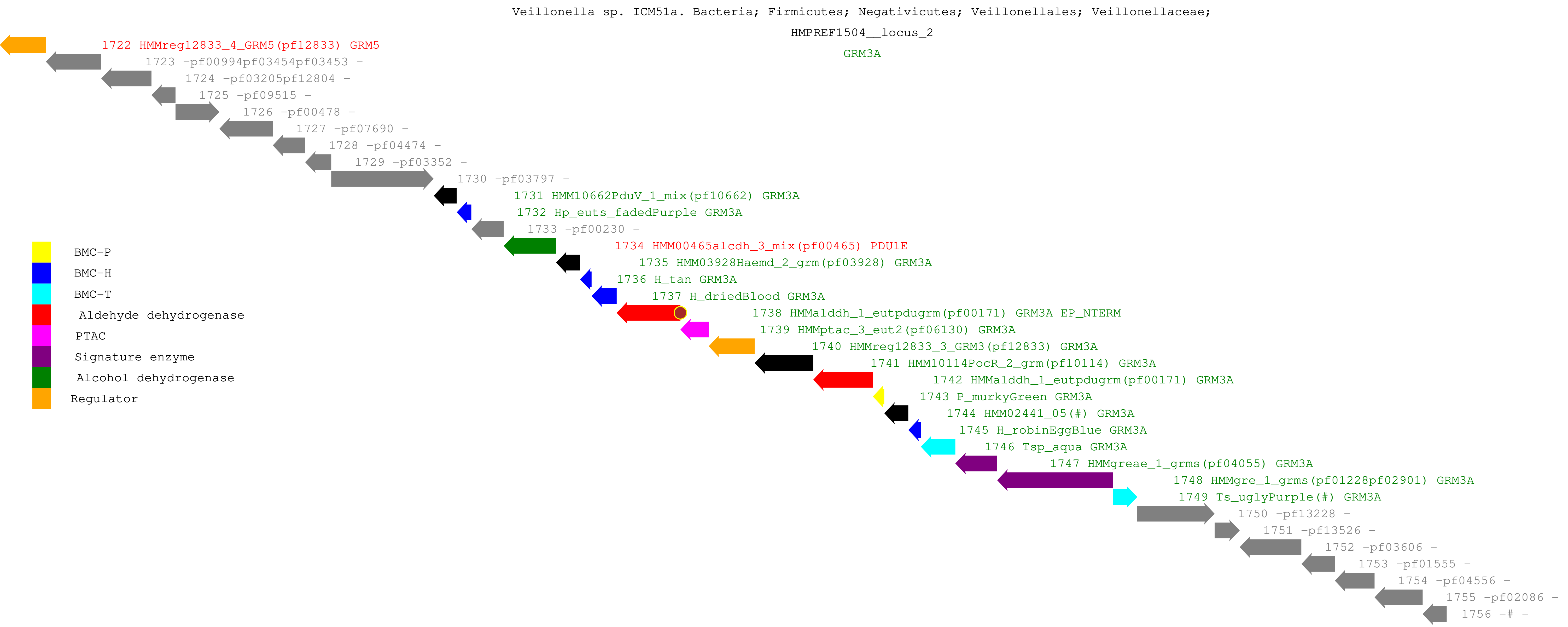
Veillonella sp. ICM51a. Bacteria; Firmicutes; Negativicutes; Veillonellales; Veillonellaceae;
HMPREF1504__locus_2
GRM3A
1722 HMMreg12833_4_GRM5(pf12833) GRM5
1723 -pf00994pf03454pf03453 -
1724 -pf03205pf12804 -
1725 -pf09515 -
1726 -pf00478 -
1727 -pf07690 -
1728 -pf04474 -
1729 -pf03352 -
1730 -pf03797 -
1731 HMM10662PduV_1_mix(pf10662) GRM3A
1732 Hp_euts_fadedPurple GRM3A
1733 -pf00230 -
1734 HMM00465alcdh_3_mix(pf00465) PDU1E
1735 HMM03928Haemd_2_grm(pf03928) GRM3A
1736 H_tan GRM3A
1737 H_driedBlood GRM3A
1738 HMMaldh_1_eutpdugrm(pf00171) GRM3A EP_NTERM
1739 HMMptac_3_eut2(pf06130) GRM3A
1740 HMMreg12833_3_GRM3(pf12833) GRM3A
1741 HMM10114PocR_2_grm(pf10114) GRM3A
1742 HMMaldh_1_eutpdugrm(pf00171) GRM3A
1743 P_murkyGreen GRM3A
1744 HMM02441_05(#) GRM3A
1745 H_robinEggBlue GRM3A
1746 Tsp_aqua GRM3A
1747 HMMgreae_1_grms(pf04055) GRM3A
1748 HMMgre_1_grms(pf01228pf02901) GRM3A
1749 Ts_uglyPurple(#) GRM3A
1750 -pf13228 -
1751 -pf13526 -
1752 -pf03606 -
1753 -pf01555 -
1754 -pf04556 -
1755 -pf02086 -
1756 -# -
BMC-P
BMC-H
BMC-T
Aldehyde dehydrogenase
PTAC
Signature enzyme
Alcohol dehydrogenase
Regulator

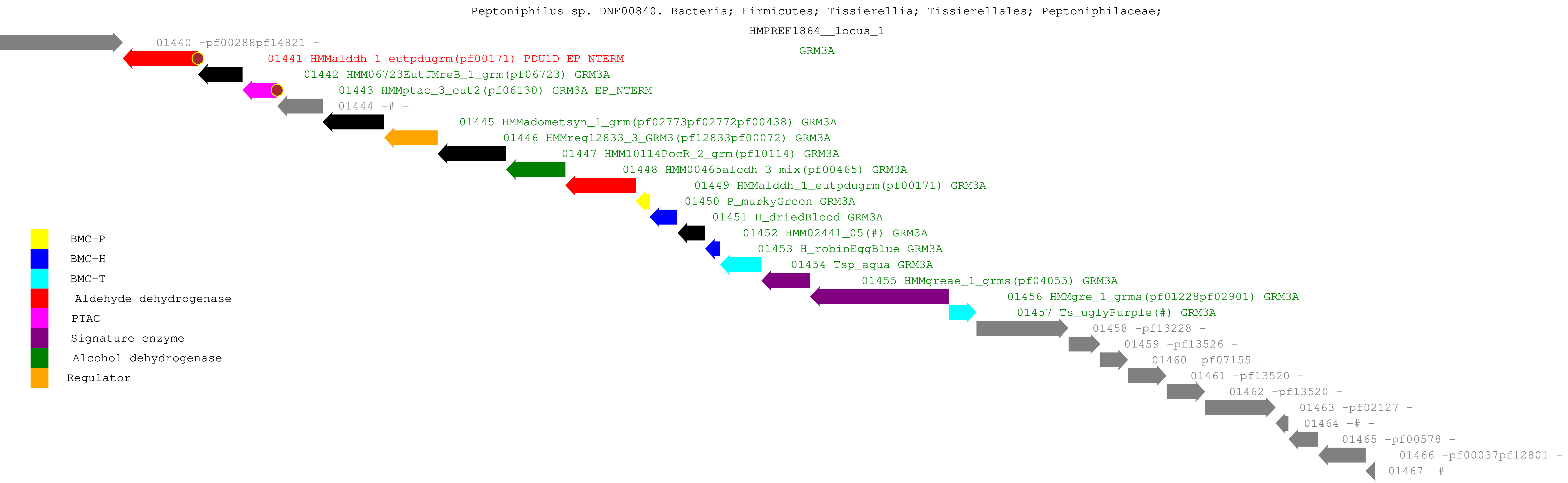
Peptoniphilus sp. DNF00840. Bacteria; Firmicutes; Tissierellia; Tissierellales; Peptoniphilaceae;
HMPREF1864__locus_1
GRM3A
01440 -pf00288pf14821 -
01441 HMMalddh_1_eutpdugrm(pf00171) PDU1D EP_NTERM
01442 HMM06723EutJMreB_1_grm(pf06723) GRM3A
01443 HMMptac_3_eut2(pf06130) GRM3A EP_NTERM
01444 -# -
01445 HMMadometsyn_1_grm(pf02773pf02772pf00438) GRM3A
01446 HMMreg12833_3_GRM3(pf12833pf00072) GRM3A
01447 HMM10114PocR_2_grm(pf10114) GRM3A
01448 HMM00465alcdh_3_mix(pf00465) GRM3A
01449 HMMalddh_1_eutpdugrm(pf00171) GRM3A
01450 P_murkyGreen GRM3A
01451 H_driedBlood GRM3A
01452 HMM02441_05(#) GRM3A
01453 H_robinEggBlue GRM3A
01454 Tsp_aqua GRM3A
01455 HMMgreae_1_grms(pf04055) GRM3A
01456 HMMgre_1_grms(pf01228pf02901) GRM3A
01457 Ts_uglyPurple(#) GRM3A
01458 -pf13228 -
01459 -pf13526 -
01460 -pf07155 -
01461 -pf13520 -
01462 -pf13520 -
01463 -pf02127 -
01464 -# -
01465 -pf00578 -
01466 -pf00037pf12801 -
01467 -# -
BMC-P
BMC-H
BMC-T
Aldehyde dehydrogenase
PTAC
Signature enzyme
Alcohol dehydrogenase
Regulator

Veillonellaceae bacterium KA00182. Bacteria; Firmicutes; Negativicutes; Veillonellales; Veillonellaceae.

HMPREF3182__locus_1

GRM3A

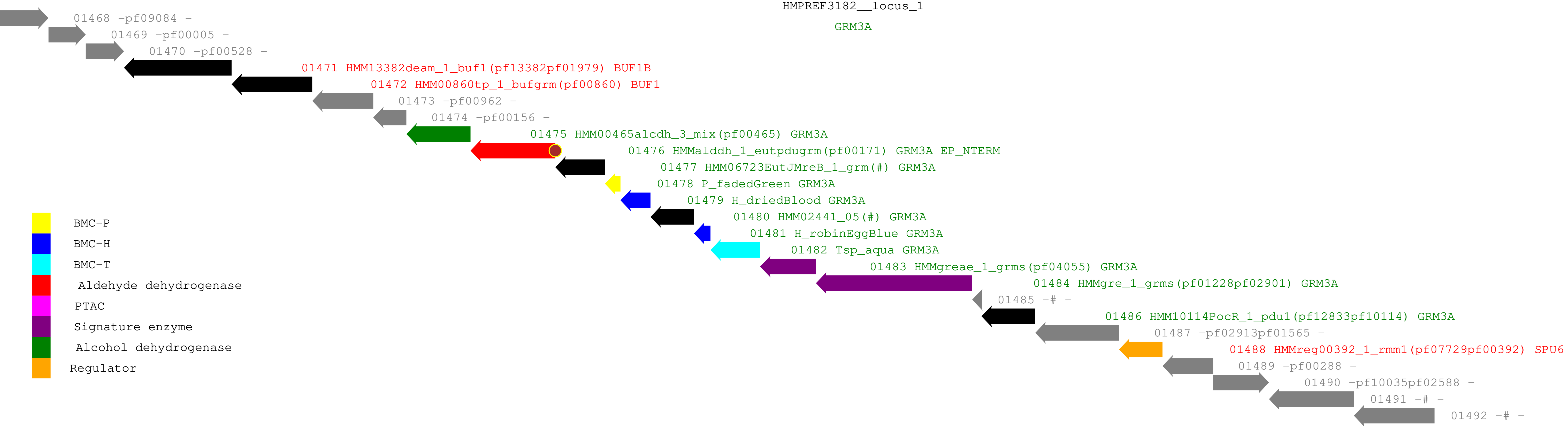

Megasphaera sp. MJR8396C. Bacteria; Firmicutes; Negativicutes; Veillonellales; Veillonellaceae;

HMPREF3201__locus_1

GRM3A

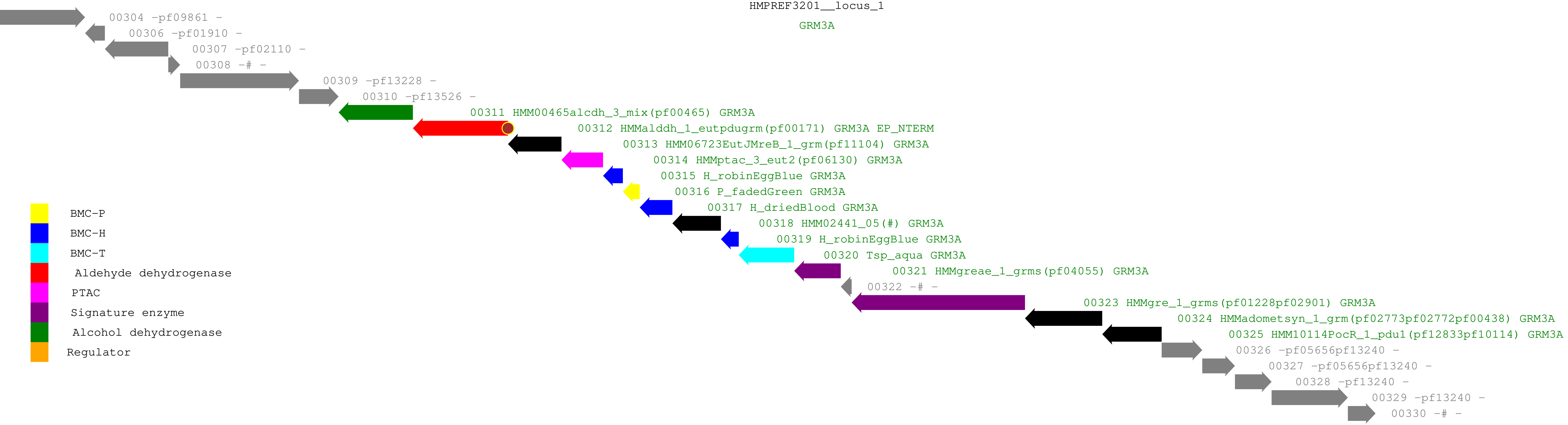

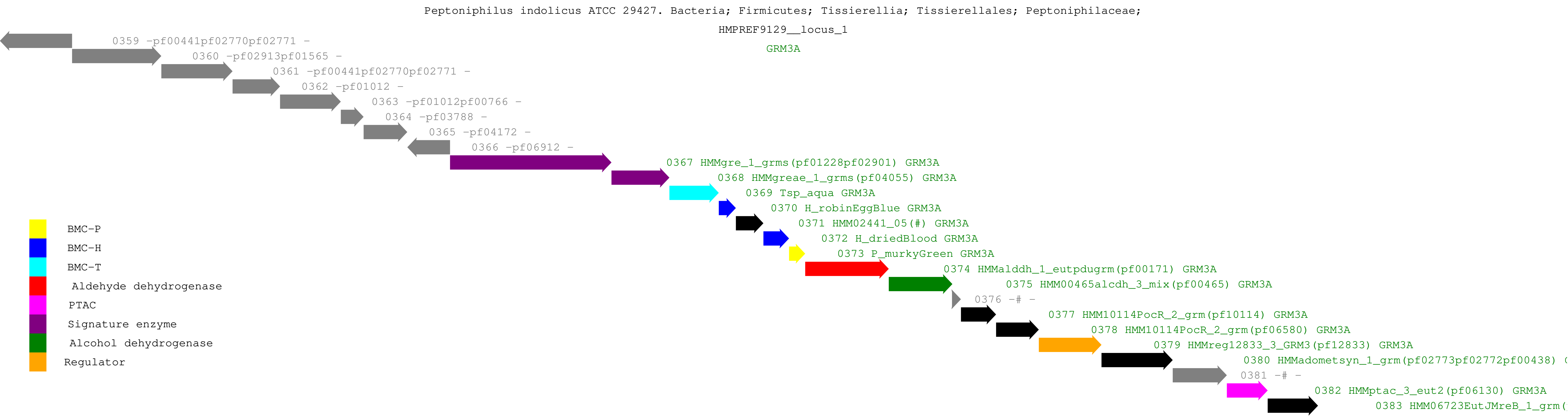
Peptoniphilus indolicus ATCC 29427. Bacteria; Firmicutes; Tissierellia; Tissierellales; Peptoniphilaceae;
HMPREF9129__locus_1
GRM3A
0359 -pf00441pf02770pf02771 -
0360 -pf02913pf01565 -
0361 -pf00441pf02770pf02771 -
0362 -pf01012 -
0363 -pf01012pf00766 -
0364 -pf03788 -
0365 -pf04172 -
0366 -pf06912 -
0367 HMMgre_1_grms(pf01228pf02901) GRM3A
0368 HMMgreae_1_grms(pf04055) GRM3A
0369 Tsp_aqua GRM3A
0370 H_robinEggBlue GRM3A
0371 HMM02441_05(#) GRM3A
0372 H_driedBlood GRM3A
0373 P_murkyGreen GRM3A
0374 HMMalddh_1_eutpdugrm(pf00171) GRM3A
0375 HMM00465alcdh_3_mix(pf00465) GRM3A
0376 -# -
0377 HMM10114PocR_2_grm(pf10114) GRM3A
0378 HMM10114PocR_2_grm(pf06580) GRM3A
0379 HMMreg12833_3_GRM3(pf12833) GRM3A
0380 HMMadometsyn_1_grm(pf02773pf02772pf00438)
0381 -# -
0382 HMMptac_3_eut2(pf06130) GRM3A
0383 HMM06723EutJMreB_1_grm(
BMC-P
BMC-H
BMC-T
Aldehyde dehydrogenase
PTAC
Signature enzyme
Alcohol dehydrogenase
Regulator

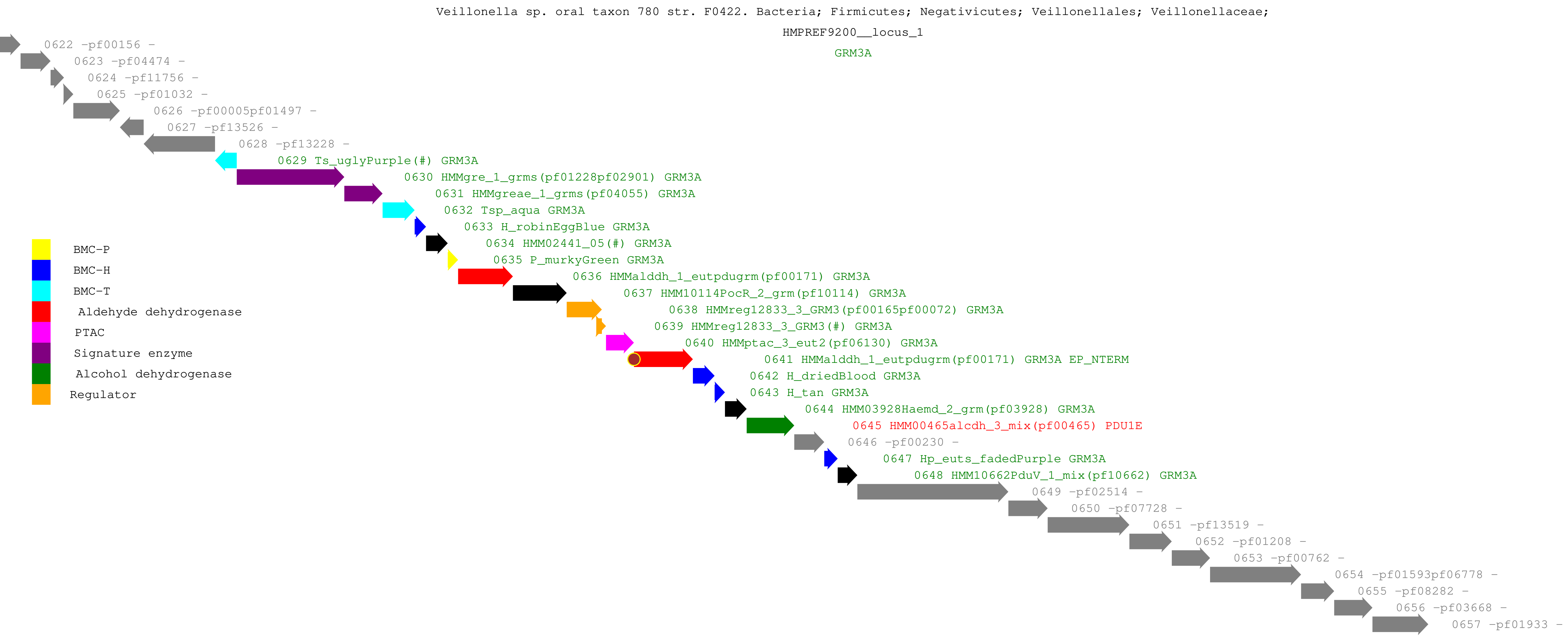
Veillonella sp. oral taxon 780 str. F0422. Bacteria; Firmicutes; Negativicutes; Veillonellales; Veillonellaceae;
HMPREF9200__locus_1
GRM3A
0622 -pf00156 -
0623 -pf04474 -
0624 -pf11756 -
0625 -pf01032 -
0626 -pf00005pf01497 -
0627 -pf13526 -
0628 -pf13228 -
0629 Ts_uglyPurple(#) GRM3A
0630 HMMgre_1_grms(pf01228pf02901) GRM3A
0631 HMMgreae_1_grms(pf04055) GRM3A
0632 Tsp_aqua GRM3A
0633 H_robinEggBlue GRM3A
0634 HMM02441_05(#) GRM3A
0635 P_murkyGreen GRM3A
0636 HMMalddh_1_eutpdugrm(pf00171) GRM3A
0637 HMM10114PocR_2_grm(pf10114) GRM3A
0638 HMMreg12833_3_GRM3(pf00165pf00072) GRM3A
0639 HMMreg12833_3_GRM3(#) GRM3A
0640 HMMptac_3_eut2(pf06130) GRM3A
0641 HMMalddh_1_eutpdugrm(pf00171) GRM3A EP_NTERM
0642 H_driedBlood GRM3A
0643 H_tan GRM3A
0644 HMM03928Haemd_2_grm(pf03928) GRM3A
0645 HMM00465alcdh_3_mix(pf00465) PDU1E
0646 -pf00230 -
0647 Hp_euts_fadedPurple GRM3A
0648 HMM10662PduV_1_mix(pf10662) GRM3A
0649 -pf02514 -
0650 -pf07728 -
0651 -pf13519 -
0652 -pf01208 -
0653 -pf00762 -
0654 -pf01593pf06778 -
0655 -pf08282 -
0656 -pf03668 -
0657 -pf01933 -
BMC-P
BMC-H
BMC-T
Aldehyde dehydrogenase
PTAC
Signature enzyme
Alcohol dehydrogenase
Regulator

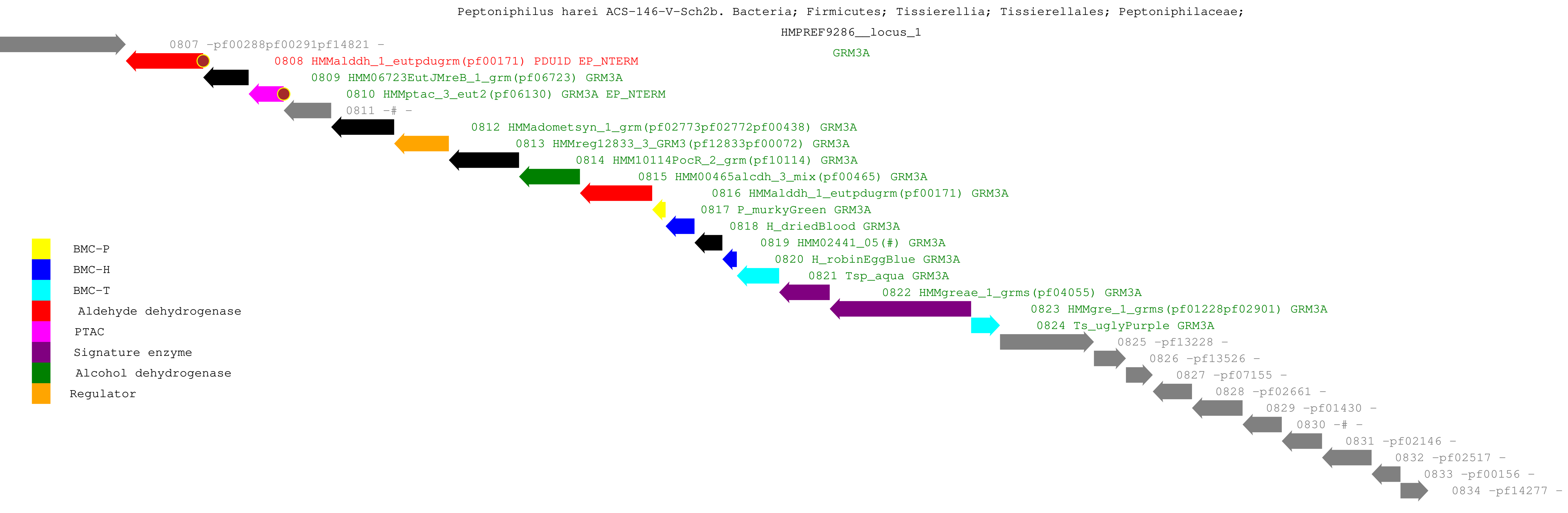
Peptoniphilus harei ACS-146-V-Sch2b. Bacteria; Firmicutes; Tissierellia; Tissierellales; Peptoniphilaceae;
HMPREF9286__locus_1
GRM3A
0807 -pf00288pf00291pf14821 -
0808 HMMalddh_1_eutpdugrm(pf00171) PDU1D EP_NTERM
0809 HMM06723EutJMreB_1_grm(pf06723) GRM3A
0810 HMMptac_3_eut2(pf06130) GRM3A EP_NTERM
0811 -# -
0812 HMMadometsyn_1_grm(pf02773pf02772pf00438) GRM3A
0813 HMMreg12833_3_GRM3(pf12833pf00072) GRM3A
0814 HMM10114PocR_2_grm(pf10114) GRM3A
0815 HMM00465alcdh_3_mix(pf00465) GRM3A
0816 HMMalddh_1_eutpdugrm(pf00171) GRM3A
0817 P_murkyGreen GRM3A
0818 H_driedBlood GRM3A
0819 HMM02441_05(#) GRM3A
0820 H_robinEggBlue GRM3A
0821 Tsp_aqua GRM3A
0822 HMMgreae_1_grms(pf04055) GRM3A
0823 HMMgre_1_grms(pf01228pf02901) GRM3A
0824 Ts_uglyPurple GRM3A
0825 -pf13228 -
0826 -pf13526 -
0827 -pf07155 -
0828 -pf02661 -
0829 -pf01430 -
0830 -# -
0831 -pf02146 -
0832 -pf02517 -
0833 -pf00156 -
0834 -pf14277 -
BMC-P
BMC-H
BMC-T
Aldehyde dehydrogenase
PTAC
Signature enzyme
Alcohol dehydrogenase
Regulator

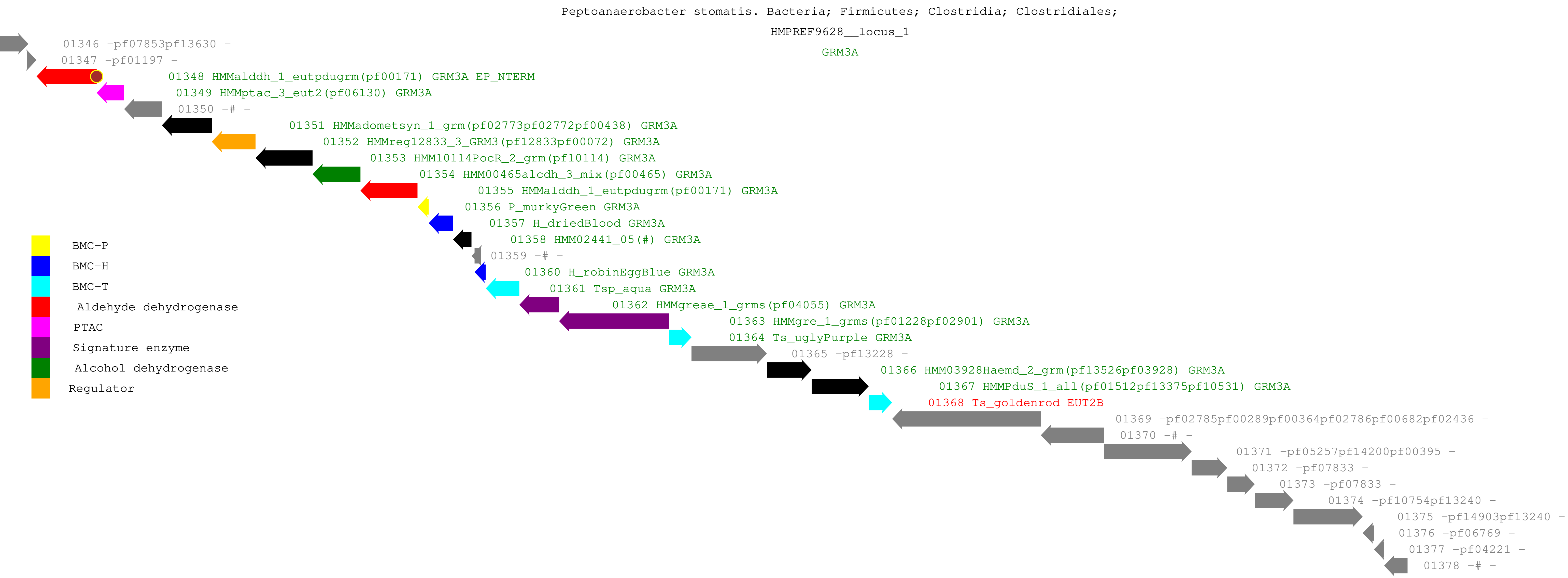

Peptoanaerobacter stomatis. Bacteria; Firmicutes; Clostridia; Clostridiales;
HMPREF9628__locus_1
GRM3A
01346 -pf07853pf13630 -
01347 -pf01197 -
01348 HMMalddh_1_eutpdugrm(pf00171) GRM3A EP_NTERM
01349 HMMptac_3_eut2(pf06130) GRM3A
01350 -# -
01351 HMMadometsyn_1_grm(pf02773pf02772pf00438) GRM3A
01352 HMMreg12833_3_GRM3(pf12833pf00072) GRM3A
01353 HMM10114PocR_2_grm(pf10114) GRM3A
01354 HMM00465alcdh_3_mix(pf00465) GRM3A
01355 HMMalddh_1_eutpdugrm(pf00171) GRM3A
01356 P_murkyGreen GRM3A
01357 H_driedBlood GRM3A
01358 HMM02441_05(#) GRM3A
01359 -# -
01360 H_robinEggBlue GRM3A
01361 Tsp_aqua GRM3A
01362 HMMgreae_1_grms(pf04055) GRM3A
01363 HMMgre_1_grms(pf01228pf02901) GRM3A
01364 Ts_uglyPurple GRM3A
01365 -pf13228 -
01366 HMM03928Haemd_2_grm(pf13526pf03928) GRM3A
01367 HMMPduS_1_all(pf01512pf13375pf10531) GRM3A
01368 Ts_goldenrod EUT2B
01369 -pf02785pf00289pf00364pf02786pf00682pf02436 -
01370 -# -
01371 -pf05257pf14200pf00395 -
01372 -pf07833 -
01373 -pf07833 -
01374 -pf10754pf13240 -
01375 -pf14903pf13240 -
01376 -pf06769 -
01377 -pf04221 -
01378 -# -
BMC-P
BMC-H
BMC-T
Aldehyde dehydrogenase
PTAC
Signature enzyme
Alcohol dehydrogenase
Regulator

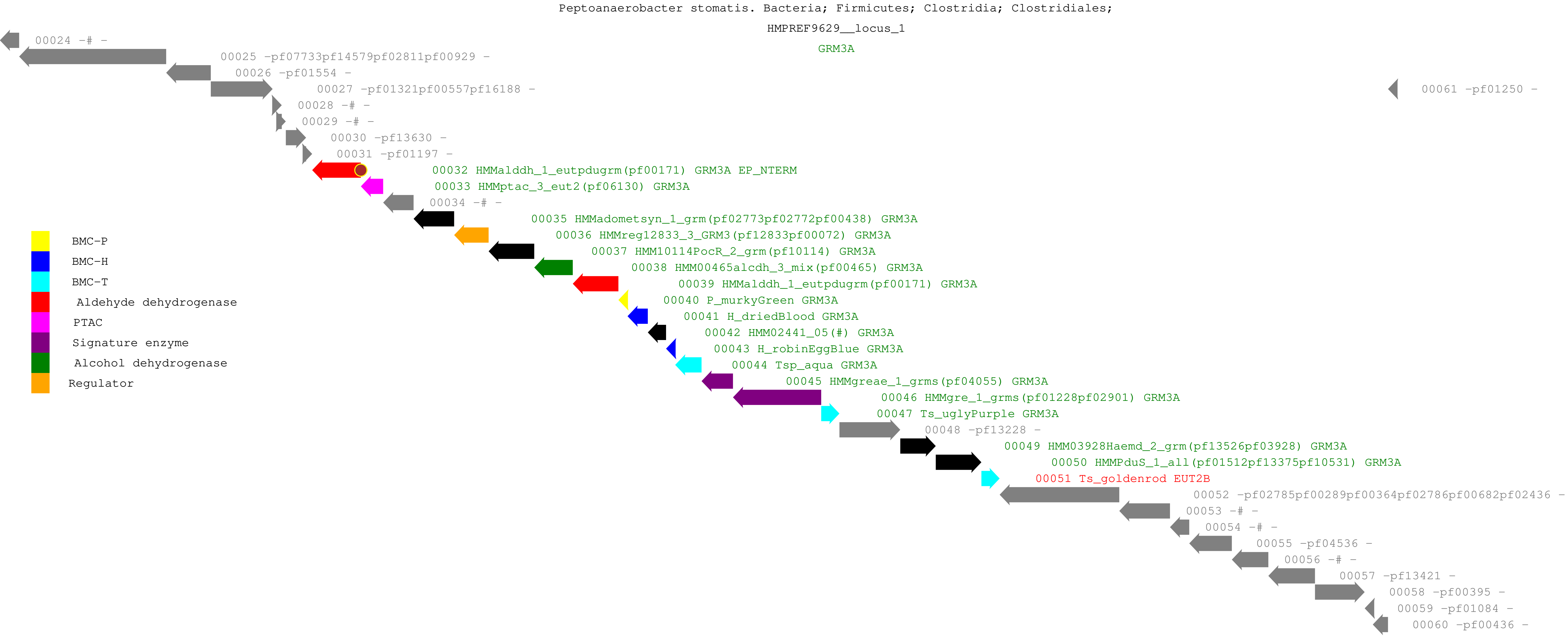
Peptoanaerobacter stomatis. Bacteria; Firmicutes; Clostridia; Clostridiales;
HMPREF9629__locus_1
GRM3A
00024 -# -
00025 -pf07733pf14579pf02811pf00929 -
00026 -pf01554 -
00027 -pf01321pf00557pf16188 -
00028 -# -
00029 -# -
00030 -pf13630 -
00031 -pf01197 -
00032 HMMalddh_1_eutpdugrm(pf00171) GRM3A EP_NTERM
00033 HMMptac_3_eut2(pf06130) GRM3A
00034 -# -
00035 HMMadometsyn_1_grm(pf02773pf02772pf00438) GRM3A
00036 HMMreg12833_3_GRM3(pf12833pf00072) GRM3A
00037 HMM10114PocR_2_grm(pf10114) GRM3A
00038 HMM00465alcdh_3_mix(pf00465) GRM3A
00039 HMMalddh_1_eutpdugrm(pf00171) GRM3A
00040 P_murkyGreen GRM3A
00041 H_driedBlood GRM3A
00042 HMM02441_05(#) GRM3A
00043 H_robinEggBlue GRM3A
00044 Tsp_aqua GRM3A
00045 HMMgreae_1_grms(pf04055) GRM3A
00046 HMMgre_1_grms(pf01228pf02901) GRM3A
00047 Ts_uglyPurple GRM3A
00048 -pf13228 -
00049 HMM03928Haemd_2_grm(pf13526pf03928) GRM3A
00050 HMMPduS_1_all(pf01512pf13375pf10531) GRM3A
00051 Ts_goldenrod EUT2B
00052 -pf02785pf00289pf00364pf02786pf00682pf02436 -
00053 -# -
00054 -# -
00055 -pf04536 -
00056 -# -
00057 -pf13421 -
00058 -pf00395 -
00059 -pf01084 -
00060 -pf00436 -
00061 -pf01250 -
BMC-P
BMC-H
BMC-T
Aldehyde dehydrogenase
PTAC
Signature enzyme
Alcohol dehydrogenase
Regulator

Peptoanaerobacter stomatis. Bacteria; Firmicutes; Clostridia; Clostridiales;

HMPREF9630__locus_1

GRM3A

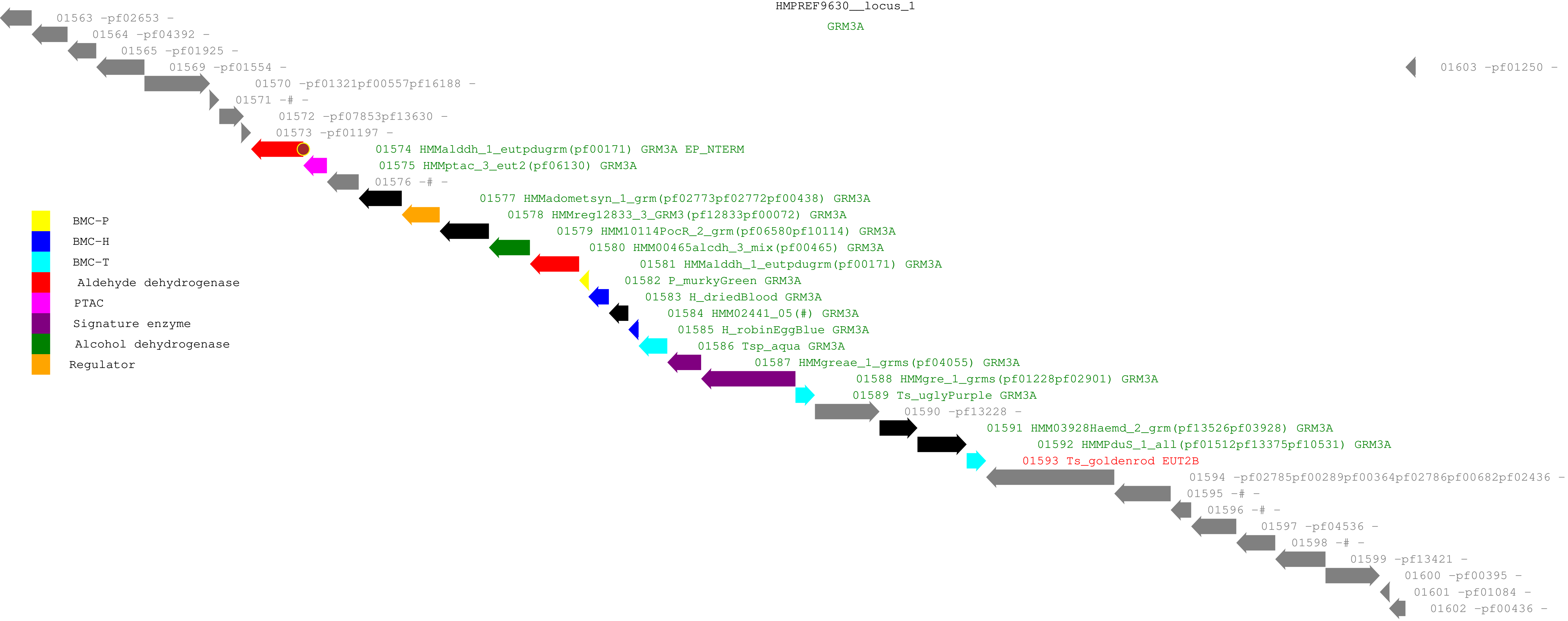

Clostridium beijerinckii (Clostridium MP). Bacteria; Firmicutes; Clostridia; Clostridiales; Clostridiaceae;

LF65__locus_1

GRM3A

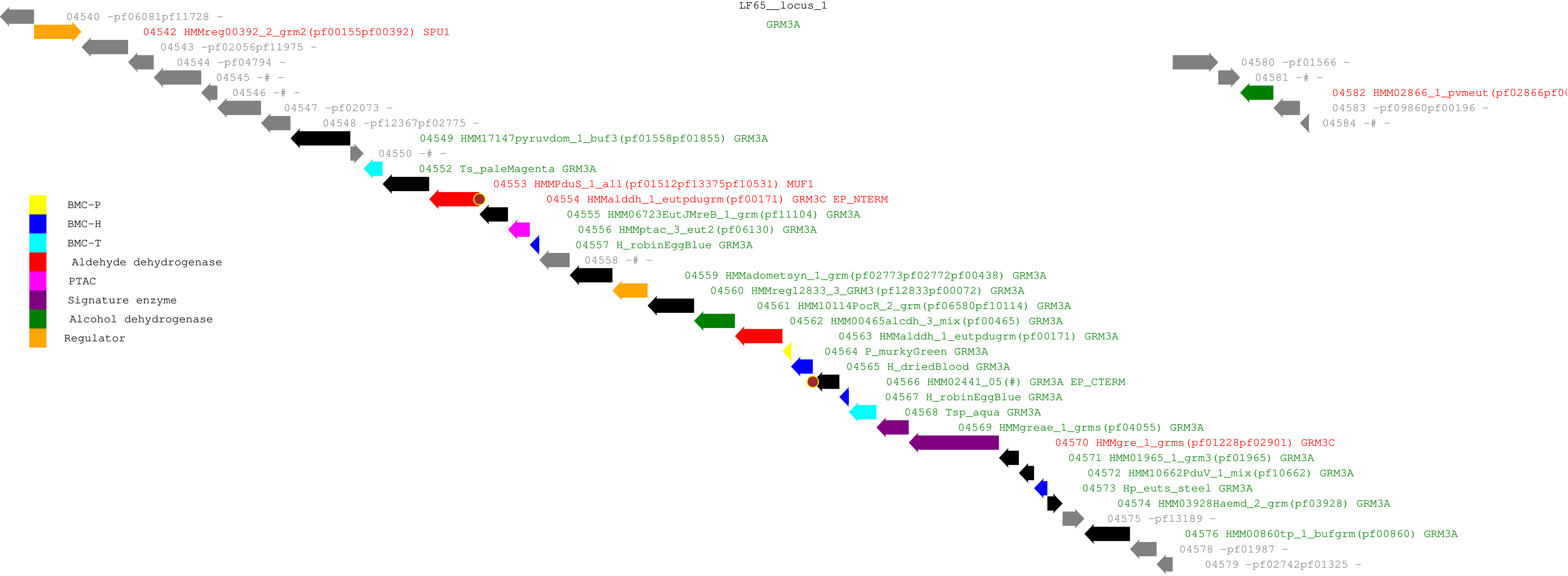

Clostridium sp. KNHs216. Bacteria; Firmicutes; Clostridia; Clostridiales; Clostridiaceae;

LY85__locus_1

GRM3A

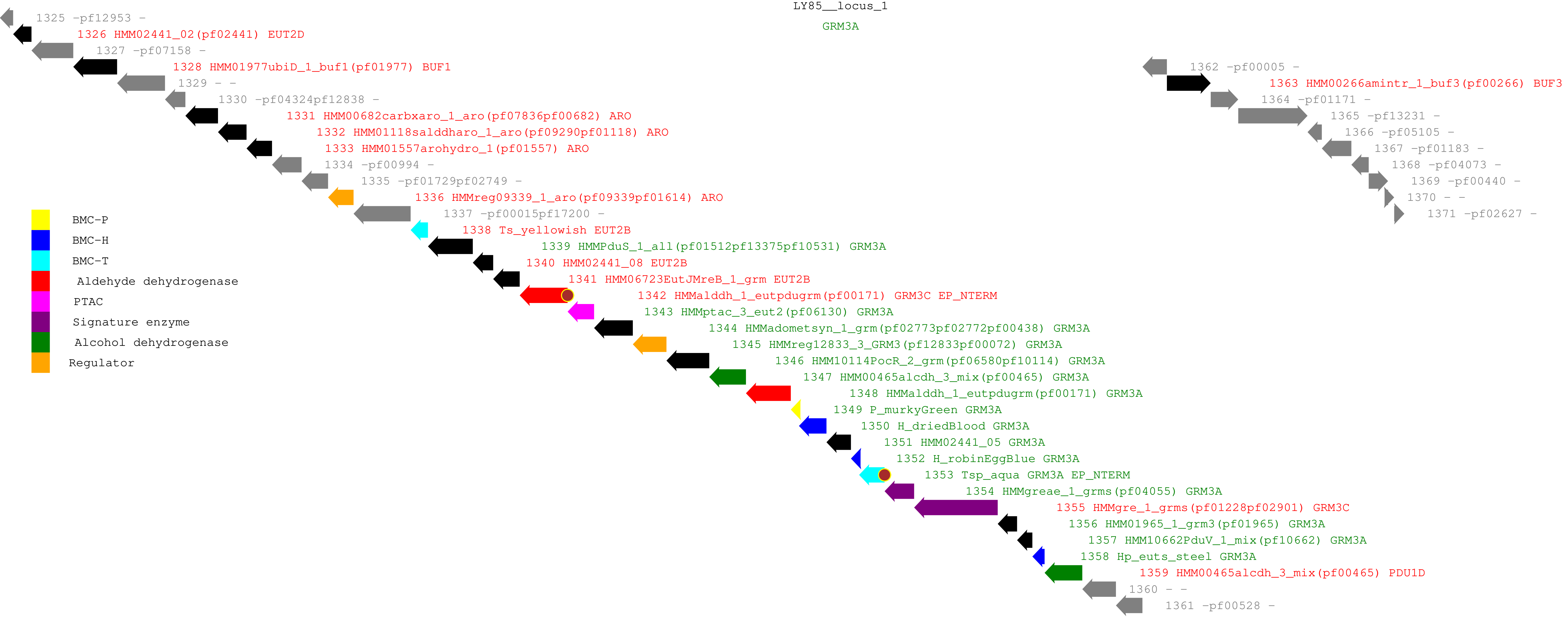

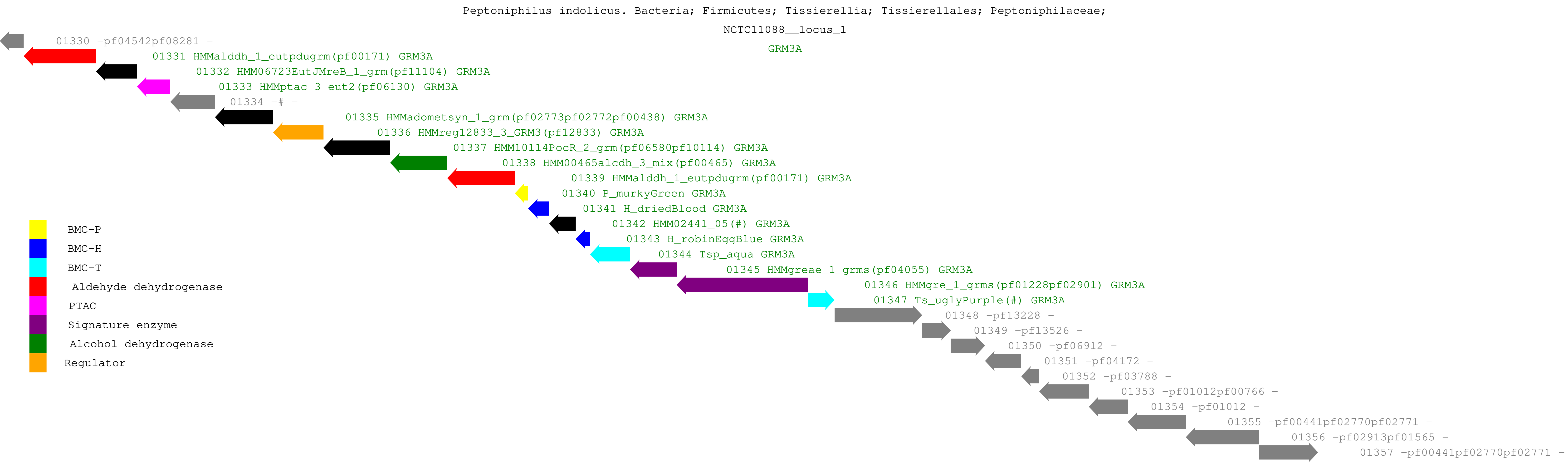
Peptoniphilus indolicus. Bacteria; Firmicutes; Tissierellia; Tissierellales; Peptoniphilaceae;
NCTC11088__locus_1
GRM3A
01330 -pf04542pf08281 -
01331 HMMalddh_1_eutpdugrm(pf00171) GRM3A
01332 HMM06723EutJMreB_1_grm(pf11104) GRM3A
01333 HMMptac_3_eut2(pf06130) GRM3A
01334 -# -
01335 HMMadometsyn_1_grm(pf02773pf02772pf00438) GRM3A
01336 HMMreg12833_3_GRM3(pf12833) GRM3A
01337 HMM10114PocR_2_grm(pf06580pf10114) GRM3A
01338 HMM00465alcdh_3_mix(pf00465) GRM3A
01339 HMMalddh_1_eutpdugrm(pf00171) GRM3A
01340 P_murkyGreen GRM3A
01341 H_driedBlood GRM3A
01342 HMM02441_05(#) GRM3A
01343 H_robinEggBlue GRM3A
01344 Tsp_aqua GRM3A
01345 HMMgreae_1_grms(pf04055) GRM3A
01346 HMMgre_1_grms(pf01228pf02901) GRM3A
01347 Ts_uglyPurple(#) GRM3A
01348 -pf13228 -
01349 -pf13526 -
01350 -pf06912 -
01351 -pf04172 -
01352 -pf03788 -
01353 -pf01012pf00766 -
01354 -pf01012 -
01355 -pf00441pf02770pf02771 -
01356 -pf02913pf01565 -
01357 -pf00441pf02770pf02771 -
BMC-P
BMC-H
BMC-T
Aldehyde dehydrogenase
PTAC
Signature enzyme
Alcohol dehydrogenase
Regulator

Clostridium clostridioforme. Bacteria; Firmicutes; Clostridia; Clostridiales; Lachnospiraceae.

NCTC11224__locus_1

GRM3A

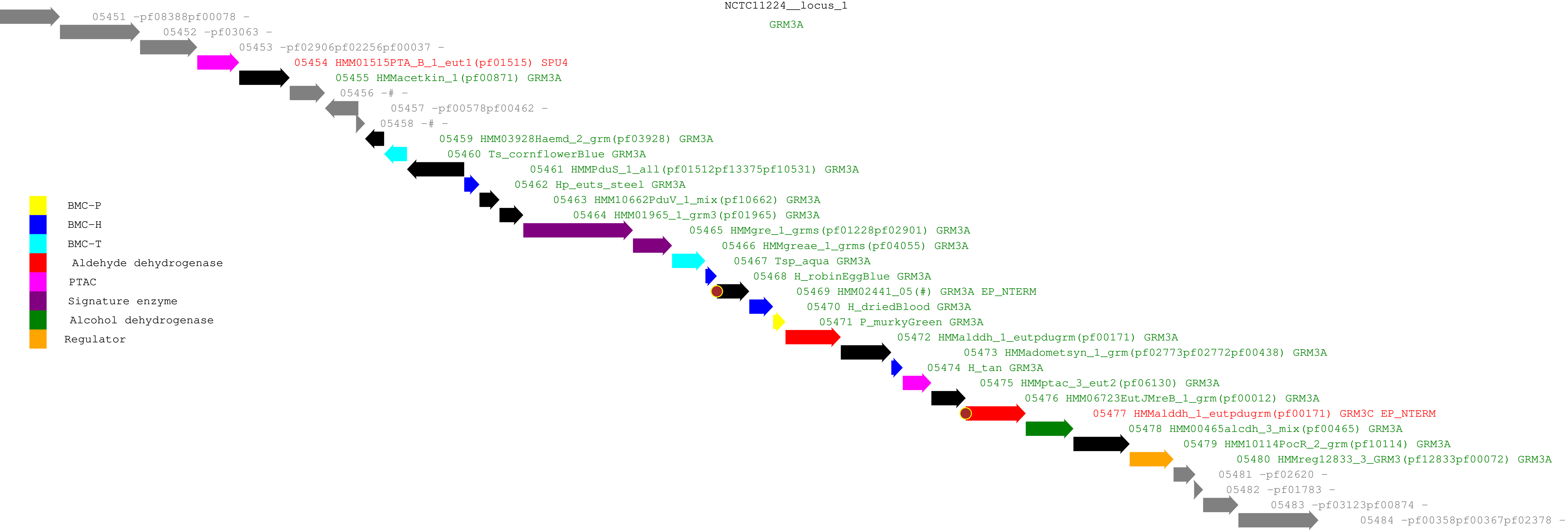

Streptococcus massiliensis. Bacteria; Firmicutes; Bacilli; Lactobacillales; Streptococcaceae;

NCTC13765__locus_1

GRM3A

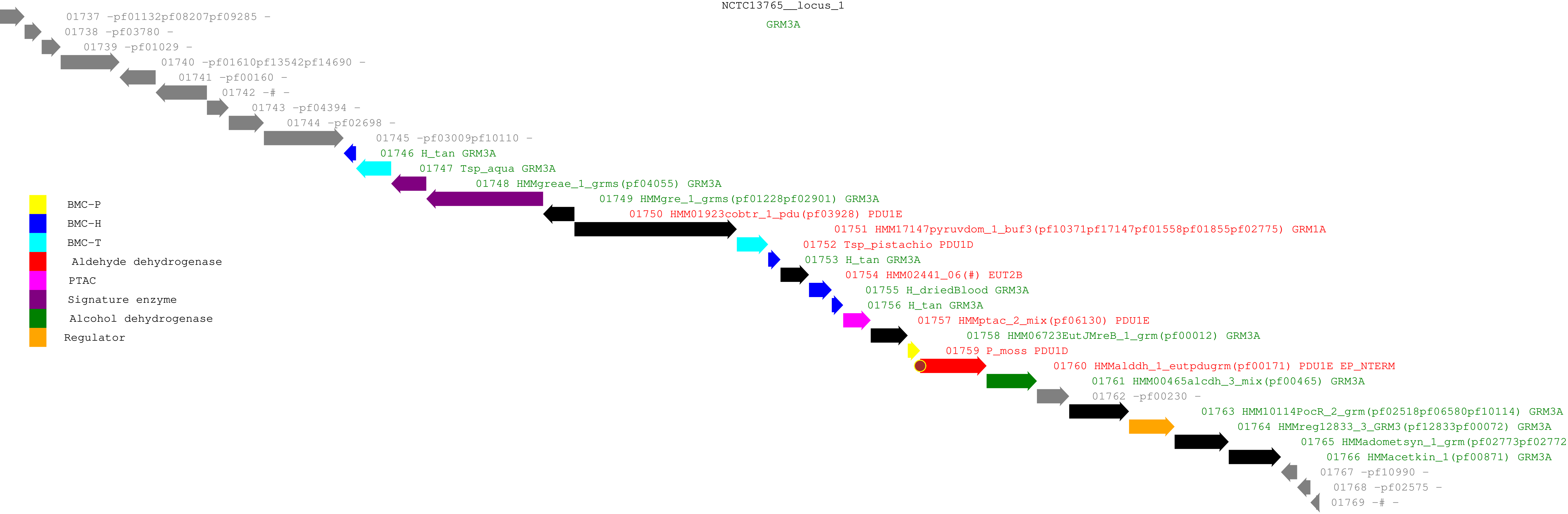

Oscillibacter valericigenes (strain DSM 18026 / NBRC 101213 / Bacteria; Firmicutes; Clostridia; Clostridiales; Oscillospiraceae;

OBV__locus_1

GRM3A

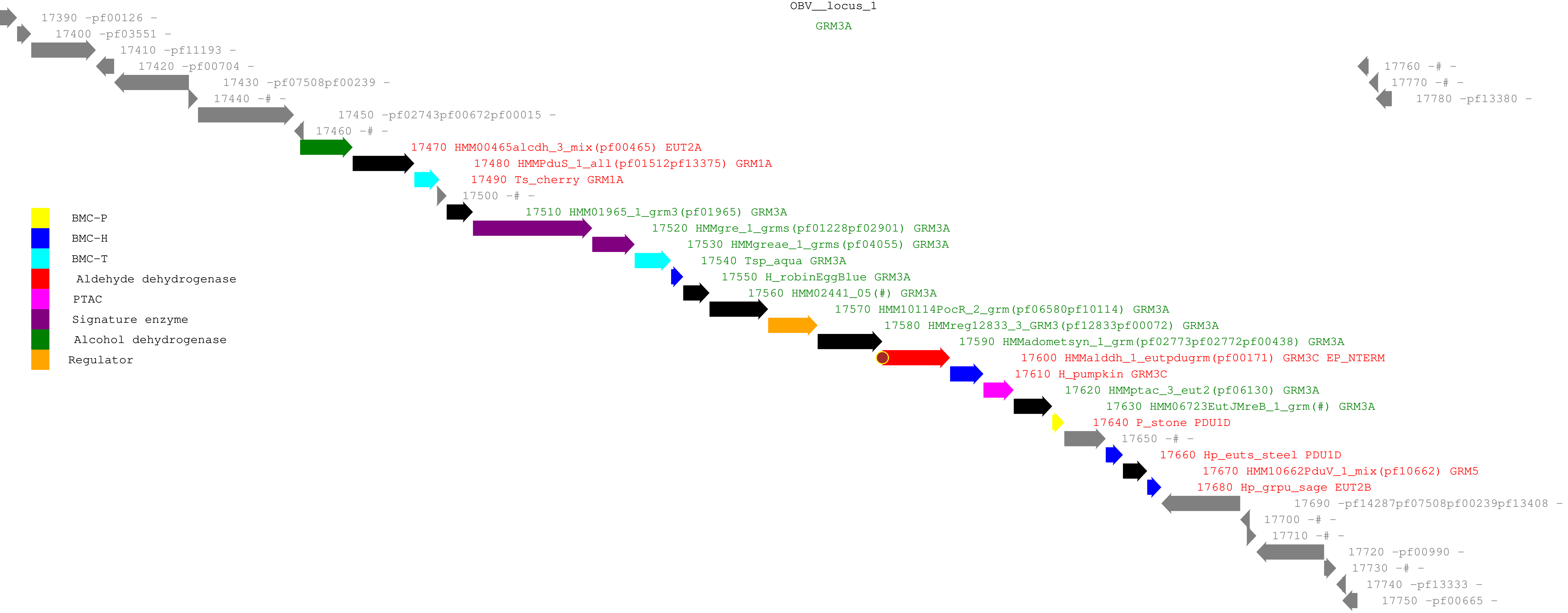

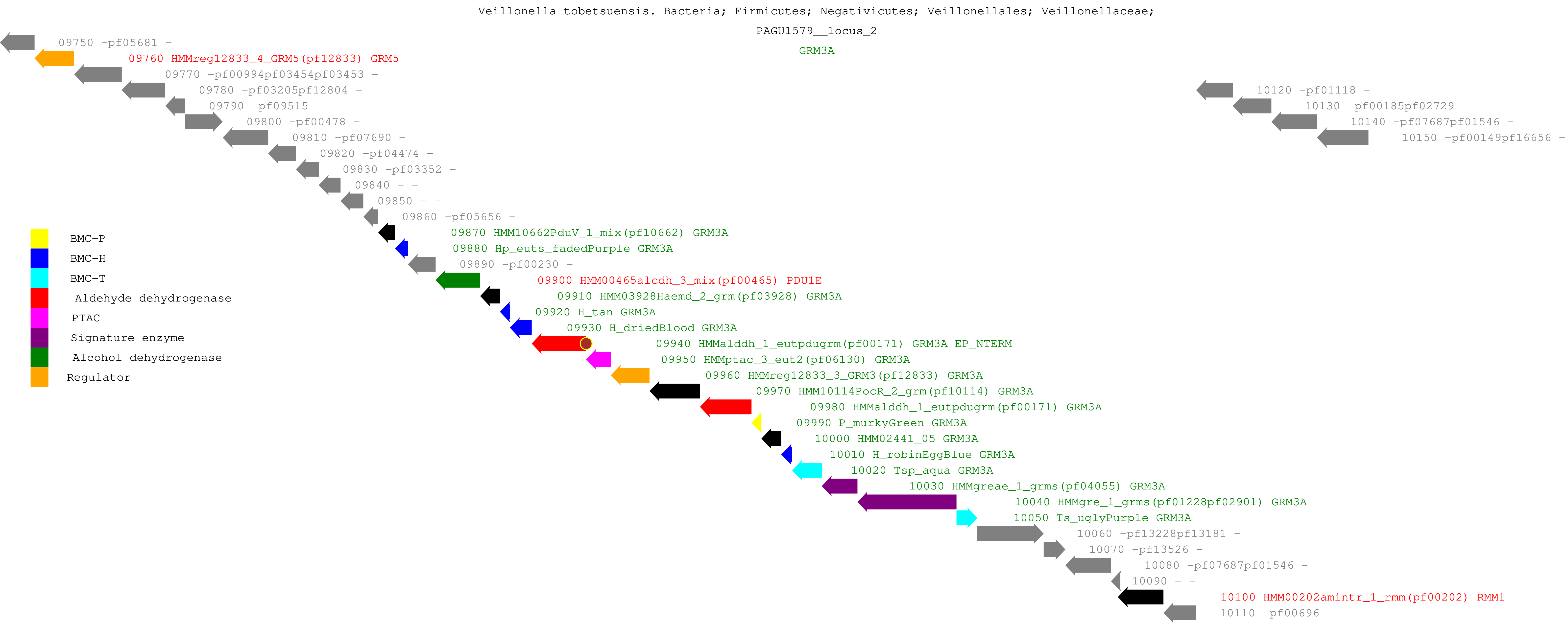

Veillonella tobetsuensis. Bacteria; Firmicutes; Negativicutes; Veillonellales; Veillonellaceae;
PAGU1579__locus_2
GRM3A
09750 -pf05681 -
09760 HMMreg12833_4_GRM5(pf12833) GRM5
09770 -pf00994pf03454pf03453 -
09780 -pf03205pf12804 -
09790 -pf09515 -
09800 -pf00478 -
09810 -pf07690 -
09820 -pf04474 -
09830 -pf03352 -
09840 - -
09850 - -
09860 -pf05656 -
09870 HMM10662PduV_1_mix(pf10662) GRM3A
09880 Hp_euts_fadedPurple GRM3A
09890 -pf00230 -
09900 HMM00465alcdh_3_mix(pf00465) PDU1E
09910 HMM03928Haemd_2_grm(pf03928) GRM3A
09920 H_tan GRM3A
09930 H_driedBlood GRM3A
09940 HMMaldh_1_eutpdugrm(pf00171) GRM3A EP_NTERM
09950 HMMptac_3_eut2(pf06130) GRM3A
09960 HMMreg12833_3_GRM3(pf12833) GRM3A
09970 HMM10114PocR_2_grm(pf10114) GRM3A
09980 HMMaldh_1_eutpdugrm(pf00171) GRM3A
09990 P_murkyGreen GRM3A
10000 HMM02441_05 GRM3A
10010 H_robinEggBlue GRM3A
10020 Tsp_aqua GRM3A
10030 HMMgreae_1_grms(pf04055) GRM3A
10040 HMMgre_1_grms(pf01228pf02901) GRM3A
10050 Ts_uglyPurple GRM3A
10060 -pf13228pf13181 -
10070 -pf13526 -
10080 -pf07687pf01546 -
10090 - -
10100 HMM00202amintr_1_rmm(pf00202) RMM1
10110 -pf00696 -
10120 -pf01118 -
10130 -pf00185pf02729 -
10140 -pf07687pf01546 -
10150 -pf00149pf16656 -
BMC-P
BMC-H
BMC-T
Aldehyde dehydrogenase
PTAC
Signature enzyme
Alcohol dehydrogenase
Regulator

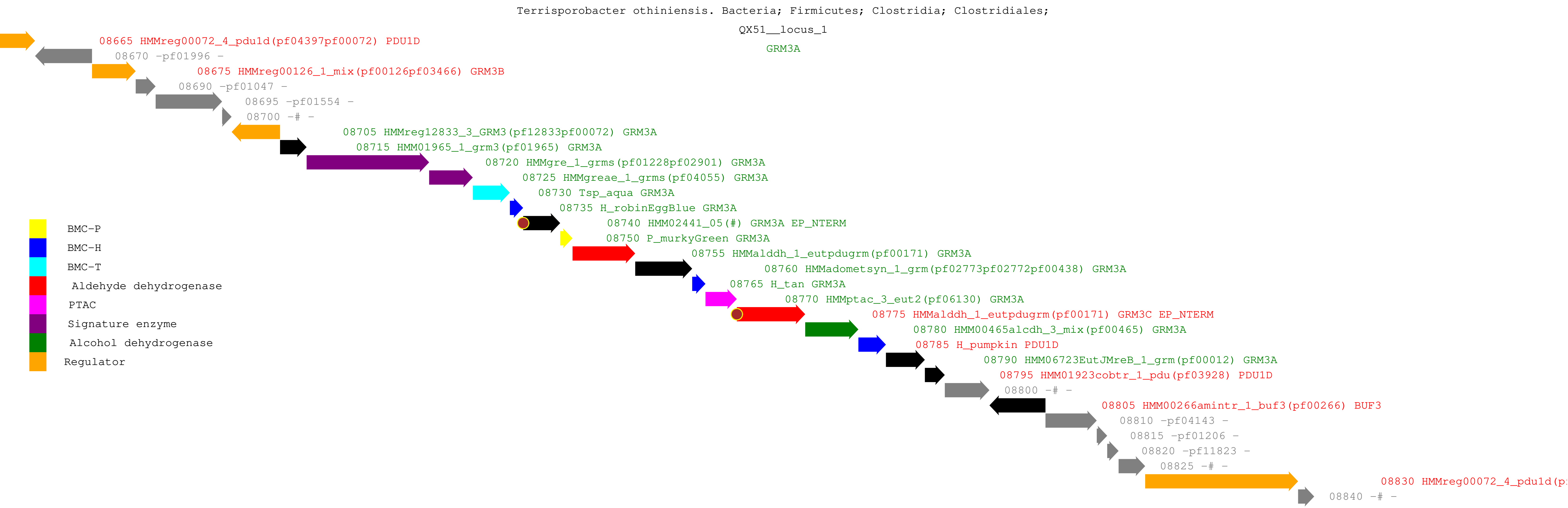

Terrisporobacter othiniensis. Bacteria; Firmicutes; Clostridia; Clostridiales;
QX51__locus_1
GRM3A
08665 HMMreg00072_4_pdu1d(pf04397pf00072) PDU1D
08670 -pf01996 -
08675 HMMreg00126_1_mix(pf00126pf03466) GRM3B
08690 -pf01047 -
08695 -pf01554 -
08700 -# -
08705 HMMreg12833_3_GRM3(pf12833pf00072) GRM3A
08715 HMM01965_1_grm3(pf01965) GRM3A
08720 HMMgre_1_grms(pf01228pf02901) GRM3A
08725 HMMgreae_1_grms(pf04055) GRM3A
08730 Tsp_aqua GRM3A
08735 H_robinEggBlue GRM3A
08740 HMM02441_05(#) GRM3A EP_NTERM
08750 P_murkyGreen GRM3A
08755 HMMaldh_1_eutpdugrm(pf00171) GRM3A
08760 HMMadometsyn_1_grm(pf02773pf02772pf00438) GRM3A
08765 H_tan GRM3A
08770 HMMptac_3_eut2(pf06130) GRM3A
08775 HMMaldh_1_eutpdugrm(pf00171) GRM3C EP_NTERM
08780 HMM00465alcdh_3_mix(pf00465) GRM3A
08785 H_pumpkin PDU1D
08790 HMM06723EutJMreB_1_grm(pf00012) GRM3A
08795 HMM01923cobtr_1_pdu(pf03928) PDU1D
08800 -# -
08805 HMM00266amintr_1_buf3(pf00266) BUF3
08810 -pf04143 -
08815 -pf01206 -
08820 -pf11823 -
08825 -# -
08830 HMMreg00072_4_pdu1d(p
08840 -# -
BMC-P
BMC-H
BMC-T
Aldehyde dehydrogenase
PTAC
Signature enzyme
Alcohol dehydrogenase
Regulator

uncultured Clostridium sp. Bacteria; Firmicutes; Clostridia; Clostridiales; Clostridiaceae;

SAMEA3545303__locus_1

GRM3A

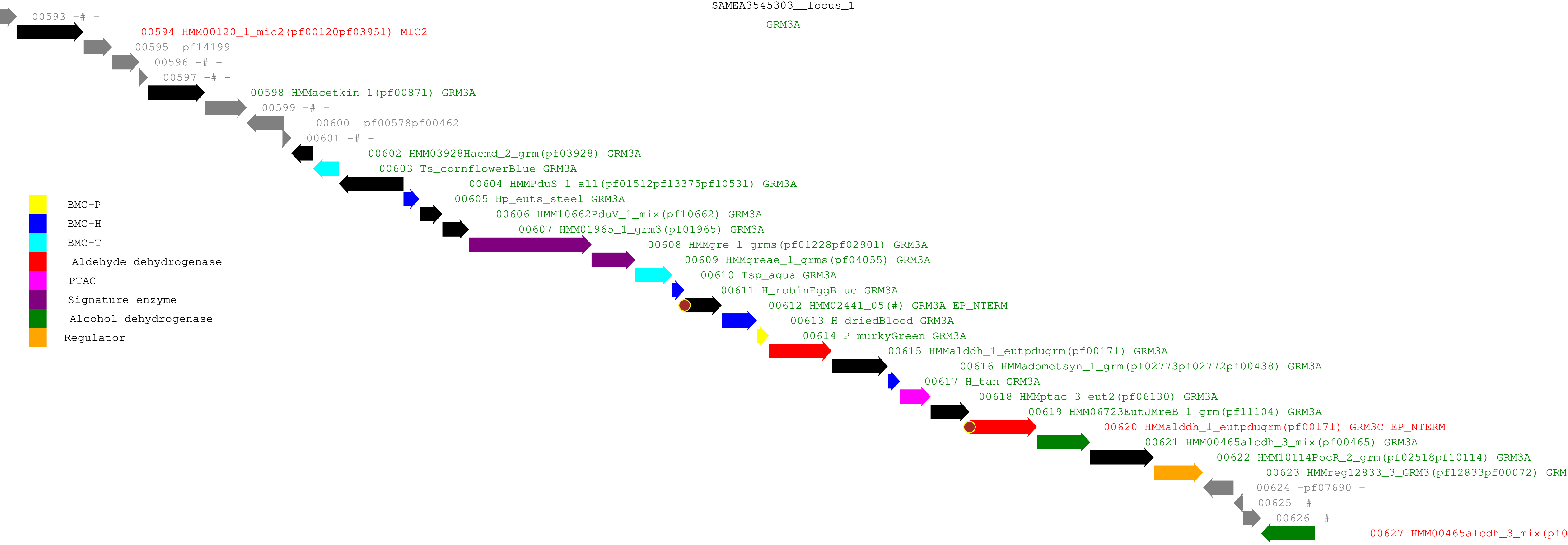

uncultured Clostridium sp. Bacteria; Firmicutes; Clostridia; Clostridiales; Clostridiaceae;

SAMEA3545303__locus_3

GRM3A

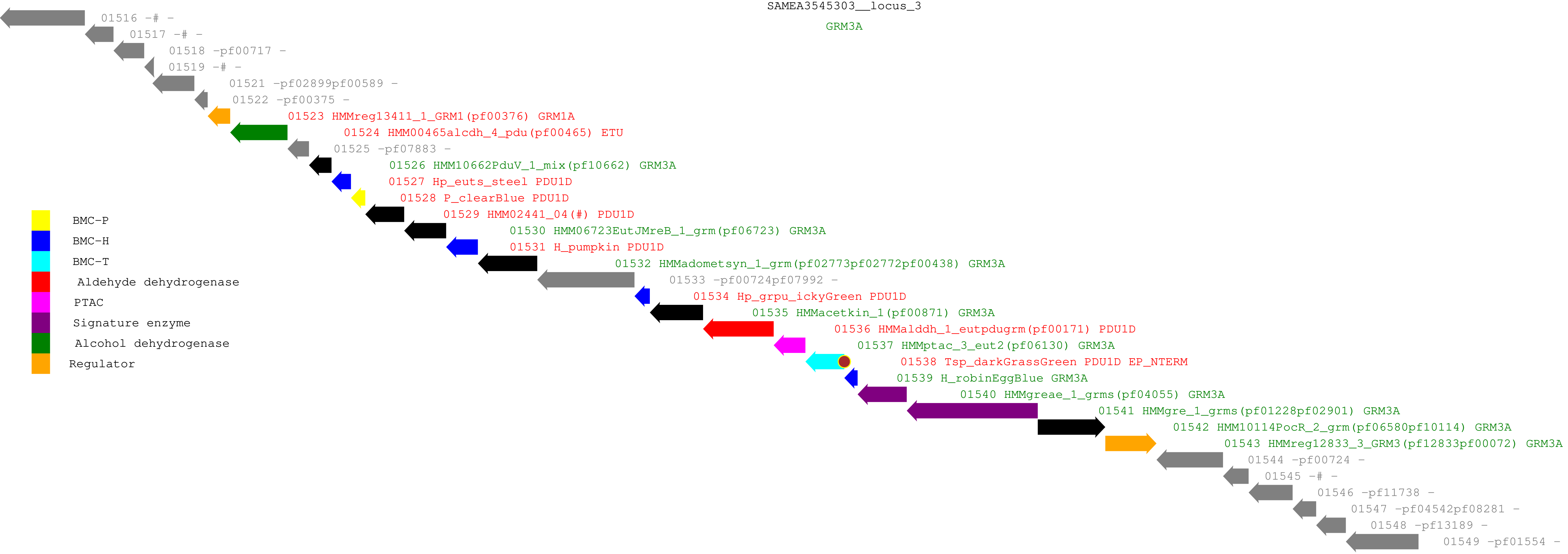

Peptoniphilus asaccharolyticus DSM 20463. Bacteria; Firmicutes; Tissierellia; Tissierellales; Peptoniphilaceae;

SAMN00017477__locus_1

GRM3A

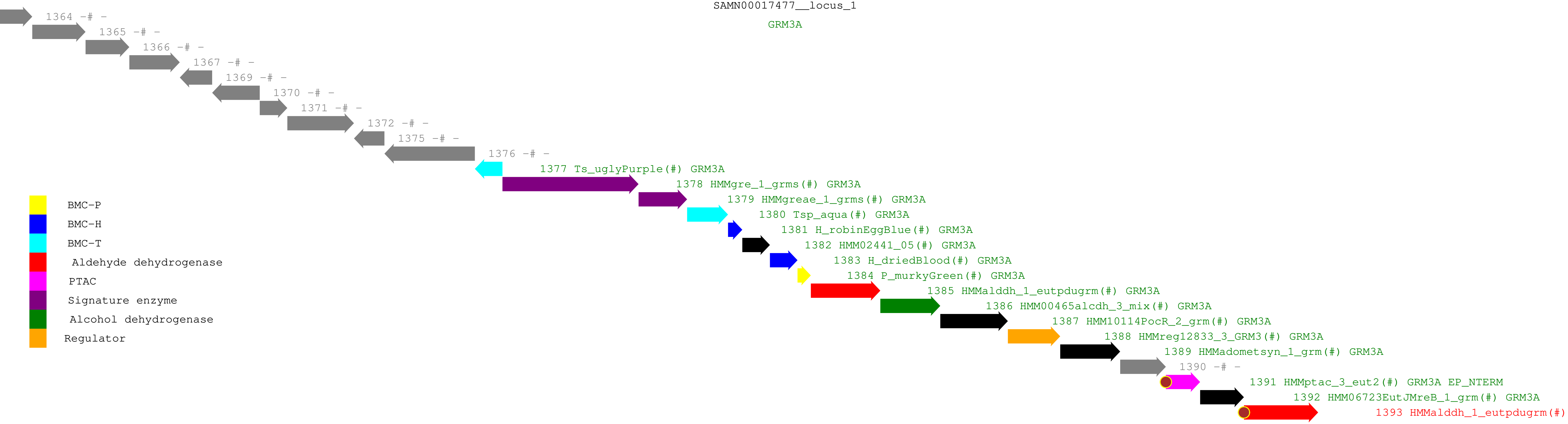

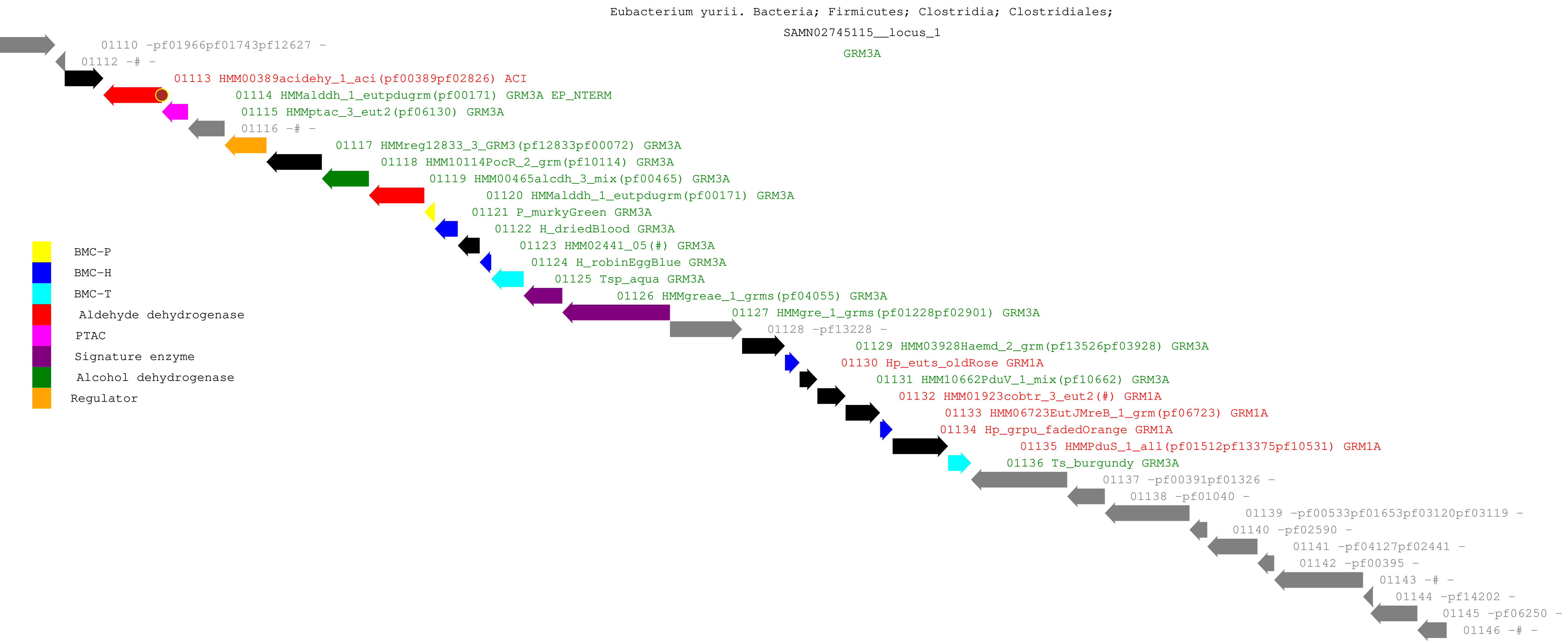
Eubacterium yurii. Bacteria; Firmicutes; Clostridia; Clostridiales;
SAMN02745115__locus_1
GRM3A
01110 -pf01966pf01743pf12627 -
01112 -# -
01113 HMM00389acidehy_1_aci(pf00389pf02826) ACI
01114 HMMalddh_1_eutpdugrm(pf00171) GRM3A EP_NTERM
01115 HMMptac_3_eut2(pf06130) GRM3A
01116 -# -
01117 HMMreg12833_3_GRM3(pf12833pf00072) GRM3A
01118 HMM10114PocR_2_grm(pf10114) GRM3A
01119 HMM00465alcdh_3_mix(pf00465) GRM3A
01120 HMMalddh_1_eutpdugrm(pf00171) GRM3A
01121 P_murkyGreen GRM3A
01122 H_driedBlood GRM3A
01123 HMM02441_05(#) GRM3A
01124 H_robinEggBlue GRM3A
01125 Tsp_aqua GRM3A
01126 HMMgreae_1_grms(pf04055) GRM3A
01127 HMMgre_1_grms(pf01228pf02901) GRM3A
01128 -pf13228 -
01129 HMM03928Haemd_2_grm(pf13526pf03928) GRM3A
01130 Hp_euts_oldRose GRM1A
01131 HMM10662PduV_1_mix(pf10662) GRM3A
01132 HMM01923cobtr_3_eut2(#) GRM1A
01133 HMM06723EutJMreB_1_grm(pf06723) GRM1A
01134 Hp_grpu_fadedOrange GRM1A
01135 HMMPduS_1_all(pf01512pf13375pf10531) GRM1A
01136 Ts_burgundy GRM3A
01137 -pf00391pf01326 -
01138 -pf01040 -
01139 -pf00533pf01653pf03120pf03119 -
01140 -pf02590 -
01141 -pf04127pf02441 -
01142 -pf00395 -
01143 -# -
01144 -pf14202 -
01145 -pf06250 -
01146 -# -
BMC-P
BMC-H
BMC-T
Aldehyde dehydrogenase
PTAC
Signature enzyme
Alcohol dehydrogenase
Regulator

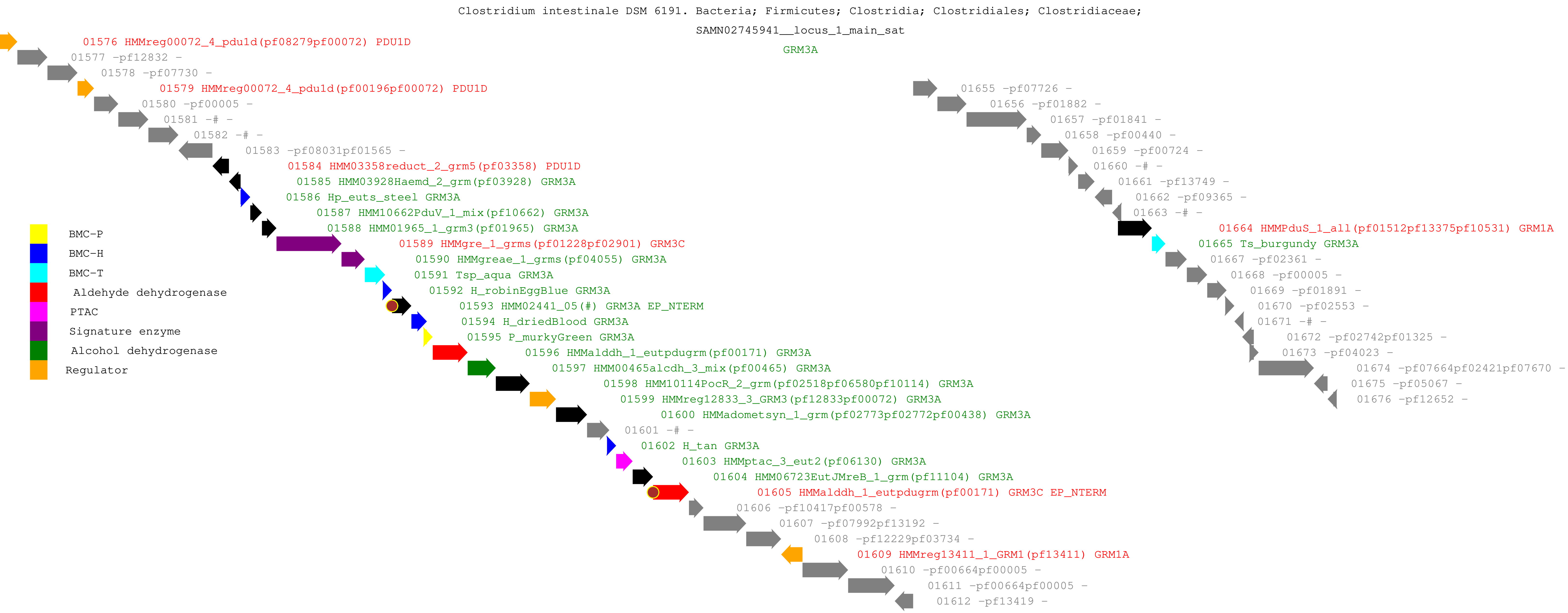

Clostridium intestinale DSM 6191. Bacteria; Firmicutes; Clostridia; Clostridiales; Clostridiaceae;
SAMN02745941__locus_1_main_sat
GRM3A
01576 HMMreg00072_4_pdu1d(pf08279pf00072) PDU1D
01577 -pf12832 -
01578 -pf07730 -
01579 HMMreg00072_4_pdu1d(pf00196pf00072) PDU1D
01580 -pf00005 -
01581 -# -
01582 -# -
01583 -pf08031pf01565 -
01584 HMM03358reduct_2_grm5(pf03358) PDU1D
01585 HMM03928Haemd_2_grm(pf03928) GRM3A
01586 Hp_euts_steel GRM3A
01587 HMM10662PduV_1_mix(pf10662) GRM3A
01588 HMM01965_1_grm3(pf01965) GRM3A
01589 HMMgre_1_grms(pf01228pf02901) GRM3C
01590 HMMgreae_1_grms(pf04055) GRM3A
01591 Tsp_aqua GRM3A
01592 H_robinEggBlue GRM3A
01593 HMM02441_05(#) GRM3A EP_NTERM
01594 H_driedBlood GRM3A
01595 P_murkyGreen GRM3A
01596 HMMalddh_1_eutpdugrm(pf00171) GRM3A
01597 HMM00465alcdh_3_mix(pf00465) GRM3A
01598 HMM10114PocR_2_grm(pf02518pf06580pf10114) GRM3A
01599 HMMreg12833_3_GRM3(pf12833pf00072) GRM3A
01600 HMMadometsyn_1_grm(pf02773pf02772pf00438) GRM3A
01601 -# -
01602 H_tan GRM3A
01603 HMMptac_3_eut2(pf06130) GRM3A
01604 HMM06723EutJMreB_1_grm(pf11104) GRM3A
01605 HMMaldddh_1_eutpdugrm(pf00171) GRM3C EP_NTERM
01606 -pf10417pf00578 -
01607 -pf07992pf13192 -
01608 -pf12229pf03734 -
01609 HMMreg13411_1_GRM1(pf13411) GRM1A
01610 -pf00664pf00005 -
01611 -pf00664pf00005 -
01612 -pf13419 -
01655 -pf07726 -
01656 -pf01882 -
01657 -pf01841 -
01658 -pf00440 -
01659 -pf00724 -
01660 -# -
01661 -pf13749 -
01662 -pf09365 -
01663 -# -
01664 HMMPduS_1_all(pf01512pf13375pf10531) GRM1A
01665 Ts_burgundy GRM3A
01667 -pf02361 -
01668 -pf00005 -
01669 -pf01891 -
01670 -pf02553 -
01671 -# -
01672 -pf02742pf01325 -
01673 -pf04023 -
01674 -pf07664pf02421pf07670 -
01675 -pf05067 -
01676 -pf12652 -
BMC-P
BMC-H
BMC-T
Aldehyde dehydrogenase
PTAC
Signature enzyme
Alcohol dehydrogenase
Regulator

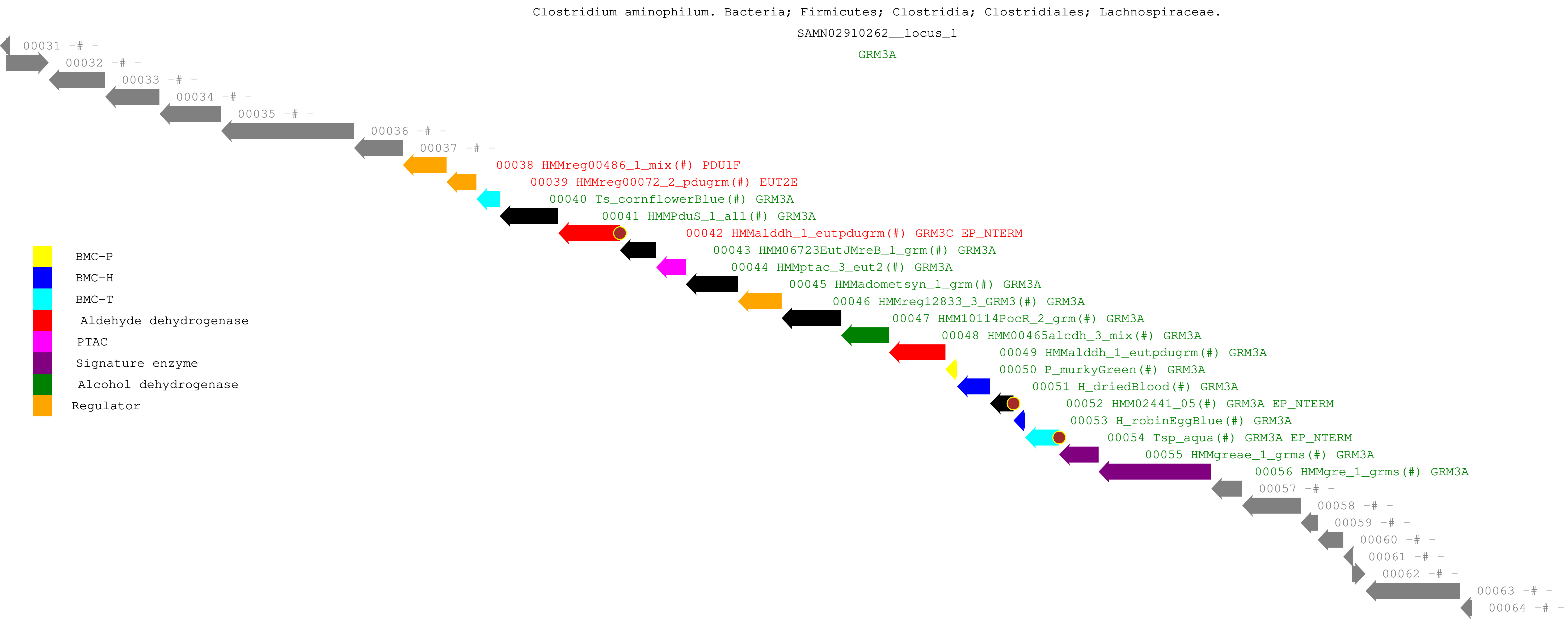

Clostridium aminophilum. Bacteria; Firmicutes; Clostridia; Clostridiales; Lachnospiraceae.
SAMN02910262__locus_1
GRM3A
00031 -# -
00032 -# -
00033 -# -
00034 -# -
00035 -# -
00036 -# -
00037 -# -
00038 HMMreg00486_1_mix(#) PDU1F
00039 HMMreg00072_2_pdugrm(#) EUT2E
00040 Ts_cornflowerBlue(#) GRM3A
00041 HMMPduS_1_all(#) GRM3A
00042 HMMalddh_1_eutpdugrm(#) GRM3C EP_NTERM
00043 HMM06723EutJMreB_1_grm(#) GRM3A
00044 HMMptac_3_eut2(#) GRM3A
00045 HMMadometsyn_1_grm(#) GRM3A
00046 HMMreg12833_3_GRM3(#) GRM3A
00047 HMM10114PocR_2_grm(#) GRM3A
00048 HMM00465alcdh_3_mix(#) GRM3A
00049 HMMalddh_1_eutpdugrm(#) GRM3A
00050 P_murkyGreen(#) GRM3A
00051 H_driedBlood(#) GRM3A
00052 HMM02441_05(#) GRM3A EP_NTERM
00053 H_robinEggBlue(#) GRM3A
00054 Tsp_aqua(#) GRM3A EP_NTERM
00055 HMMgreae_1_grms(#) GRM3A
00056 HMMgre_1_grms(#) GRM3A
00057 -# -
00058 -# -
00059 -# -
00060 -# -
00061 -# -
00062 -# -
00063 -# -
00064 -# -
BMC-P
BMC-H
BMC-T
Aldehyde dehydrogenase
PTAC
Signature enzyme
Alcohol dehydrogenase
Regulator

Pseudobutyrivibrio sp. ACV-2. Bacteria; Firmicutes; Clostridia; Clostridiales; Lachnospiraceae;

SAMN02910384__locus_1

GRM3A

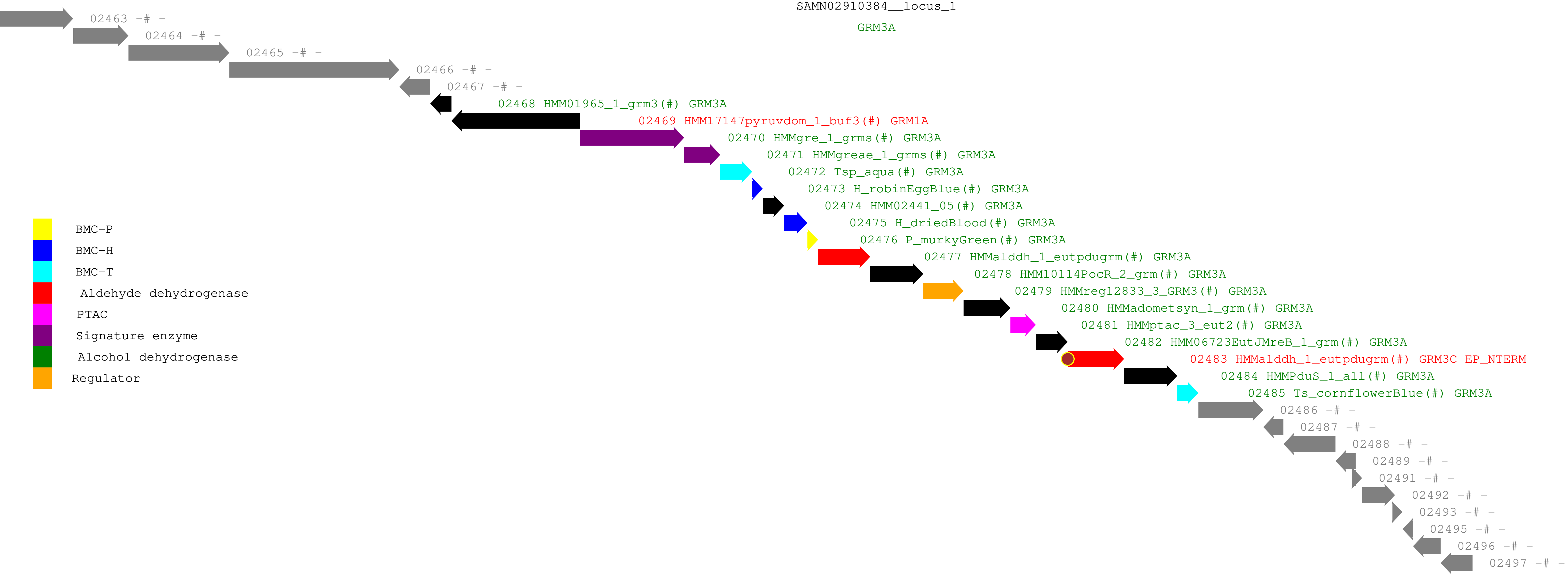

Eubacterium oxidoreducens. Bacteria; Firmicutes; Clostridia; Clostridiales; Eubacteriaceae;

SAMN02910417__locus_1

GRM3A

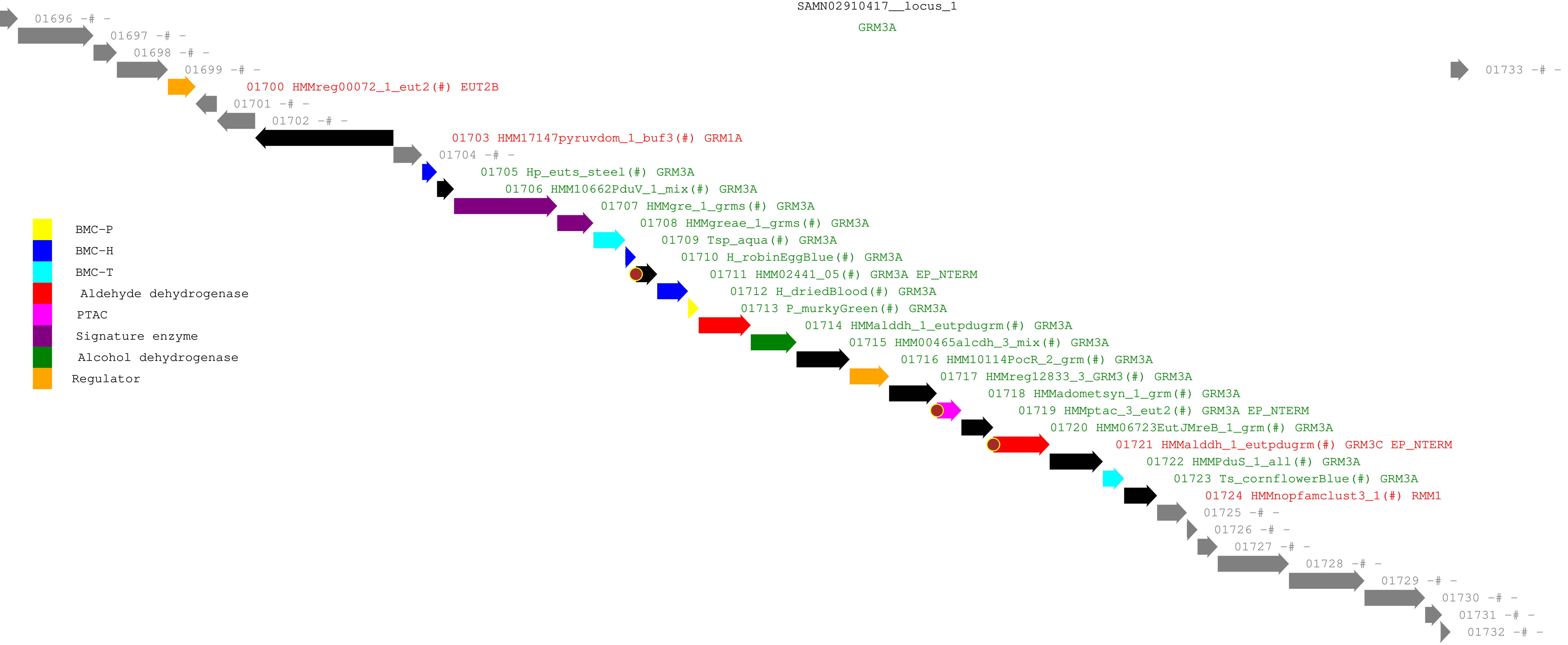

Clostridium aminophilum. Bacteria; Firmicutes; Clostridia; Clostridiales; Lachnospiraceae.

SAMN04487771__locus_1

GRM3A

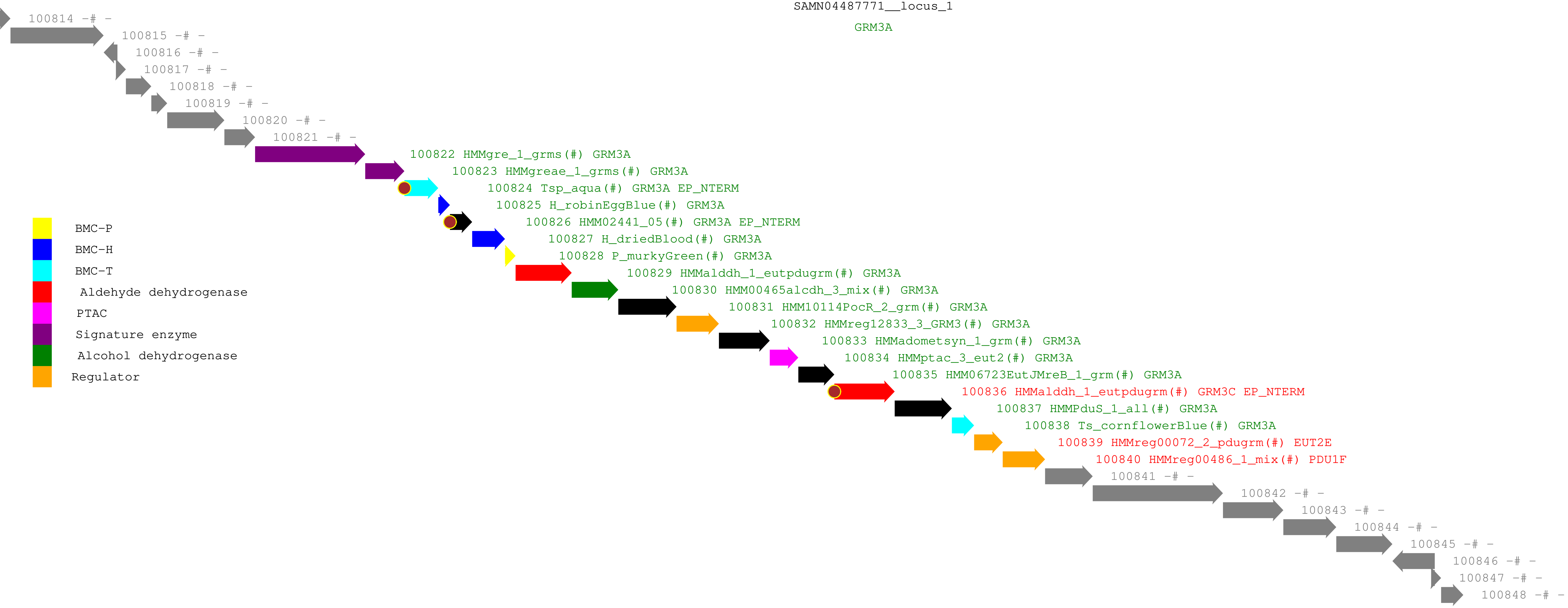

Clostridium cadaveris. Bacteria; Firmicutes; Clostridia; Clostridiales; Clostridiaceae;
SAMN04487885__locus_3
GRM3A

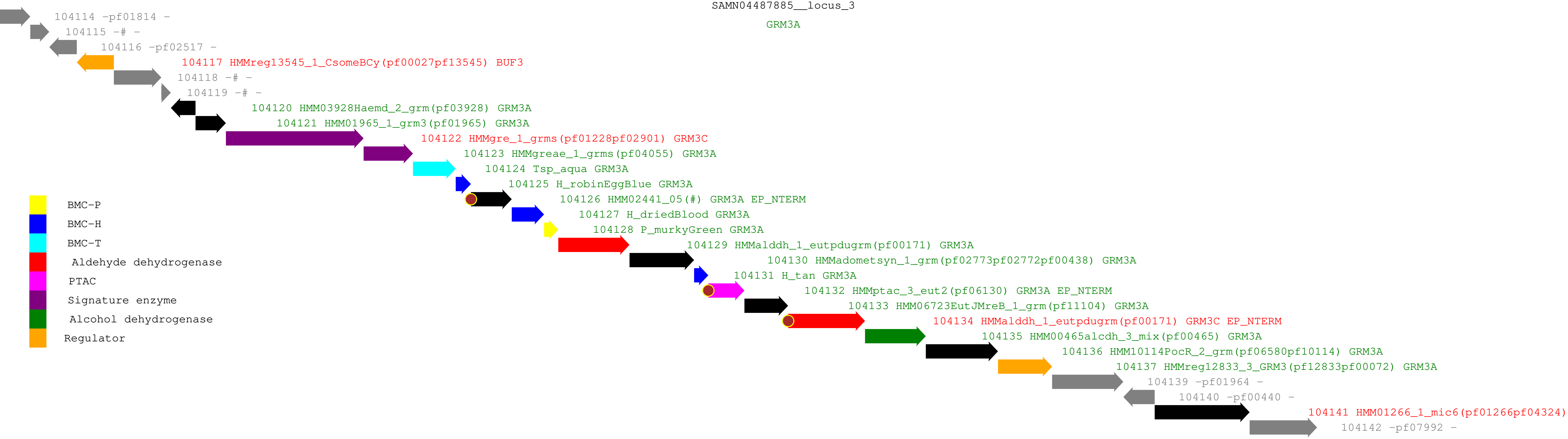

Megasphaera paucivorans. Bacteria; Firmicutes; Negativicutes; Veillonellales; Veillonellaceae;

SAMN05660299__locus_1

GRM3A

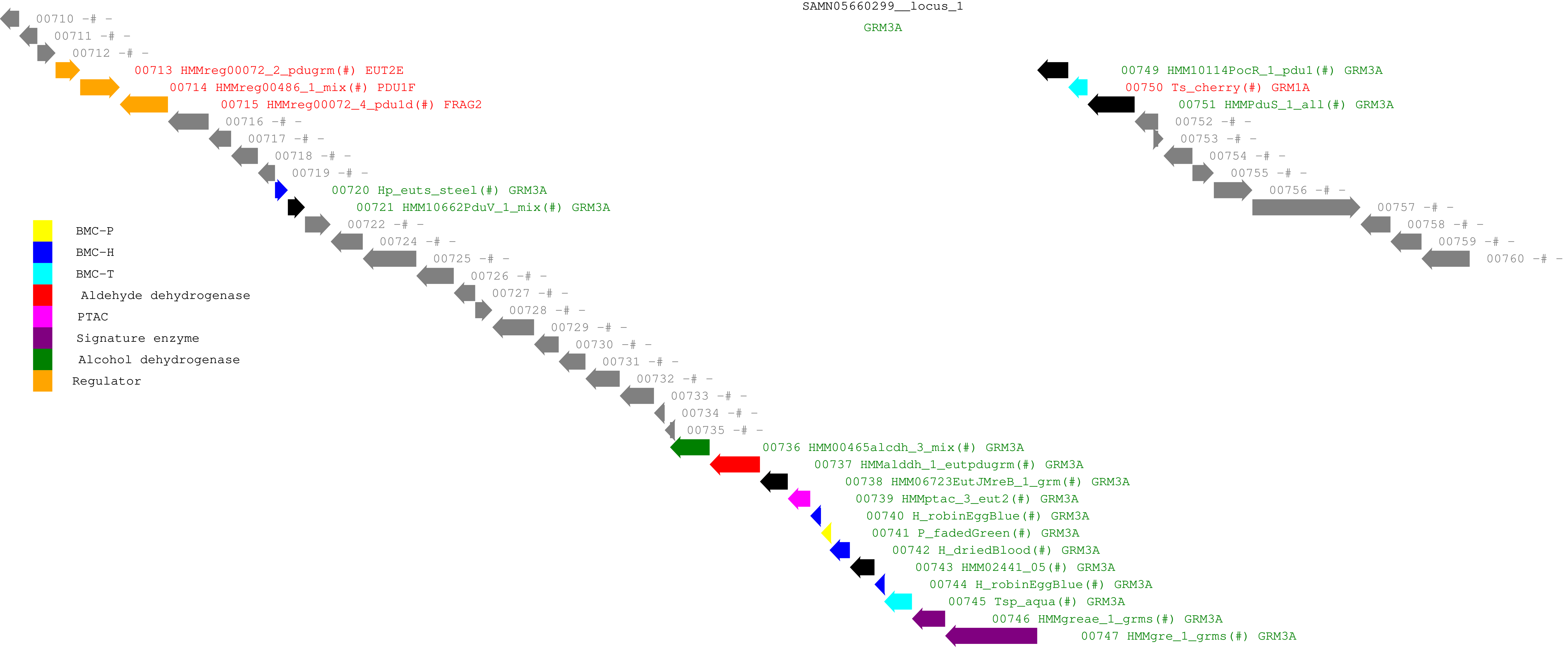

Megasphaera cerevisiae DSM 20462. Bacteria; Firmicutes; Negativicutes; Veillonellales; Veillonellaceae;

SAMN05660900__locus_1_main_sat

GRM3A

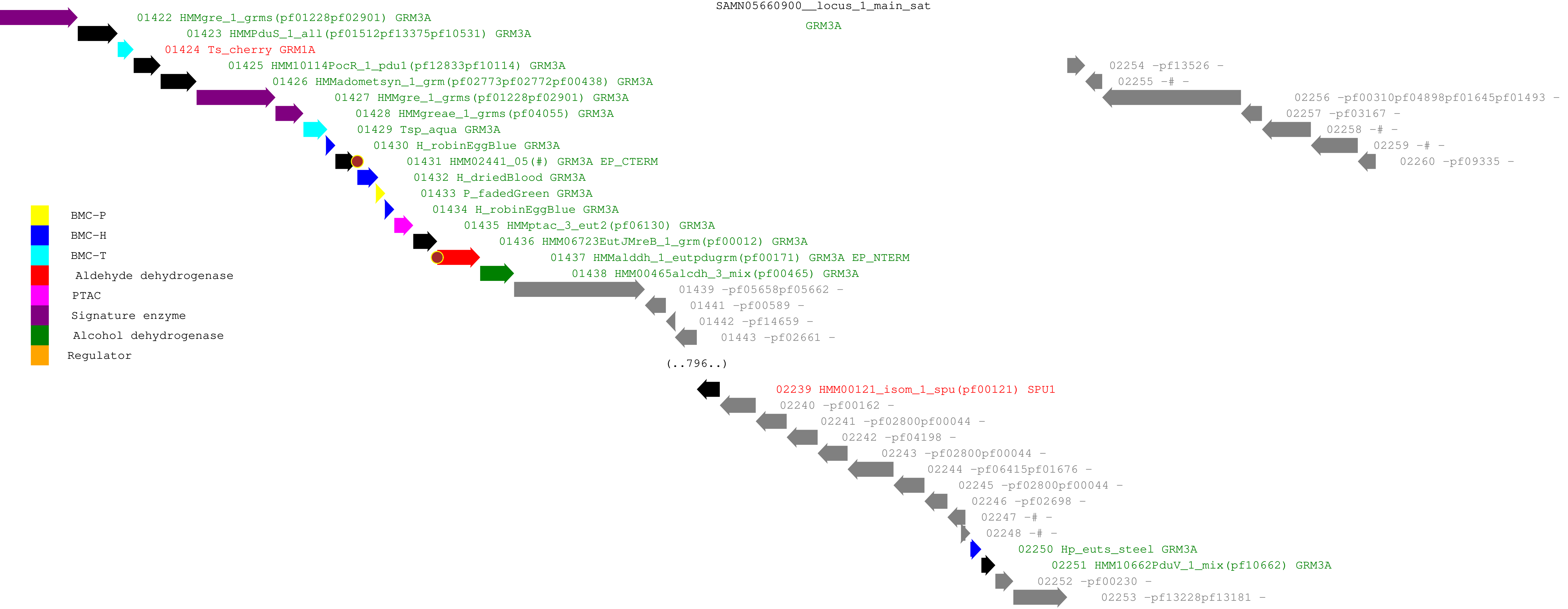

Clostridium beijerinckii NRRL B-598. Bacteria; Firmicutes; Clostridia; Clostridiales; Clostridiaceae;

X276__locus_1

GRM3A

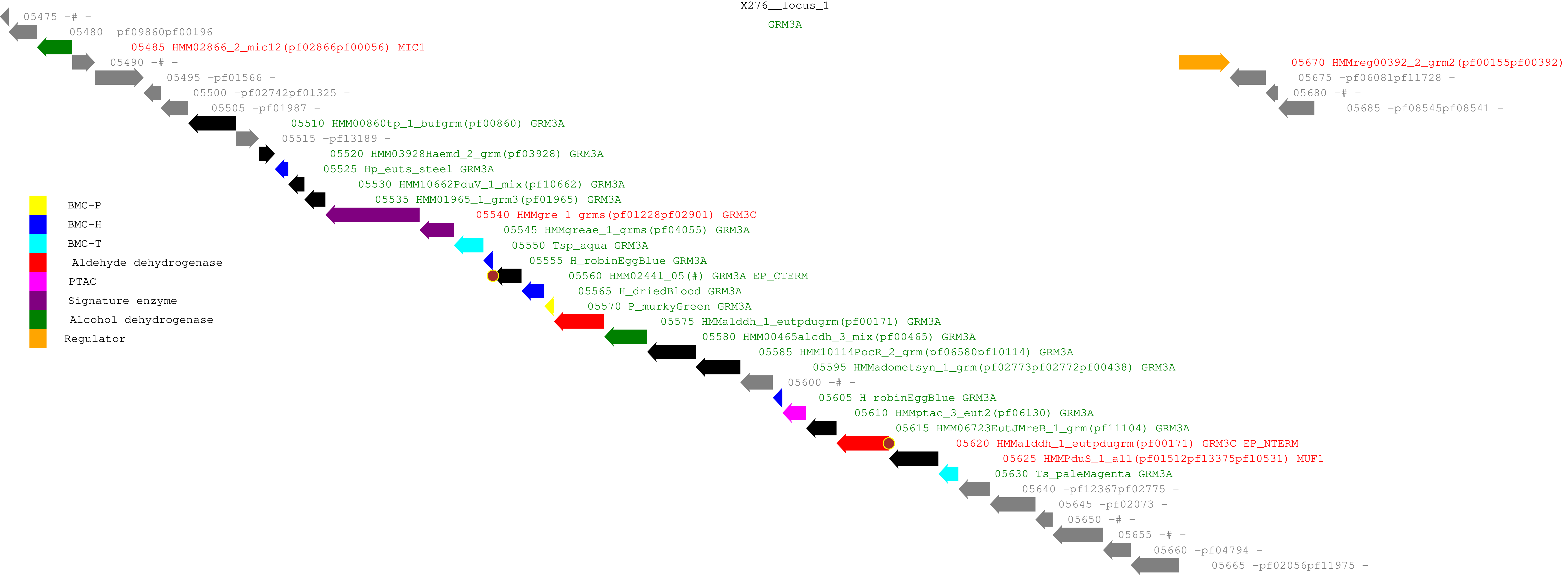

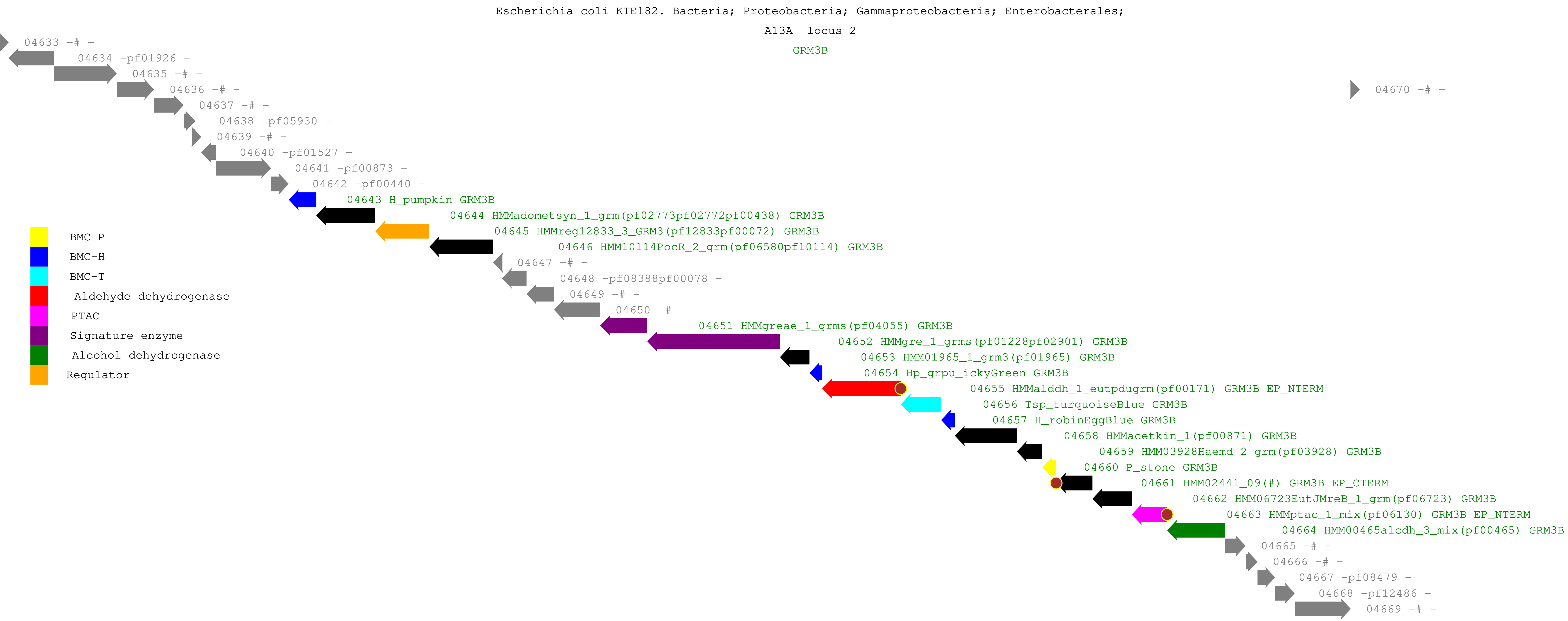
Escherichia coli KTE182. Bacteria; Proteobacteria; Gammaproteobacteria; Enterobacterales;
A13A__locus_2
GRM3B
04633 -# -
04634 -pf01926 -
04635 -# -
04636 -# -
04637 -# -
04638 -pf05930 -
04639 -# -
04640 -pf01527 -
04641 -pf00873 -
04642 -pf00440 -
04643 H_pumpkin GRM3B
04644 HMMadometsyn_1_grm(pf02773pf02772pf00438) GRM3B
04645 HMMreg12833_3_GRM3(pf12833pf00072) GRM3B
04646 HMM10114PocR_2_grm(pf06580pf10114) GRM3B
04647 -# -
04648 -pf08388pf00078 -
04649 -# -
04650 -# -
04651 HMMgreae_1_grms(pf04055) GRM3B
04652 HMMgre_1_grms(pf01228pf02901) GRM3B
04653 HMM01965_1_grm3(pf01965) GRM3B
04654 Hp_grpu_ickyGreen GRM3B
04655 HMMalddh_1_eutpdugrm(pf00171) GRM3B EP_NTERM
04656 Tsp_turquoiseBlue GRM3B
04657 H_robinEggBlue GRM3B
04658 HMMacetkin_1(pf00871) GRM3B
04659 HMM03928Haemd_2_grm(pf03928) GRM3B
04660 P_stone GRM3B
04661 HMM02441_09(#) GRM3B EP_CTERM
04662 HMM06723EutJMreB_1_grm(pf06723) GRM3B
04663 HMMptac_1_mix(pf06130) GRM3B EP_NTERM
04664 HMM00465alcdh_3_mix(pf00465) GRM3B
04665 -# -
04666 -# -
04667 -pf08479 -
04668 -pf12486 -
04669 -# -
04670 -# -
BMC-P
BMC-H
BMC-T
Aldehyde dehydrogenase
PTAC
Signature enzyme
Alcohol dehydrogenase
Regulator

Escherichia coli KTE76. Bacteria; Proteobacteria; Gammaproteobacteria; Enterobacterales;

A1UO__locus_2

GRM3B

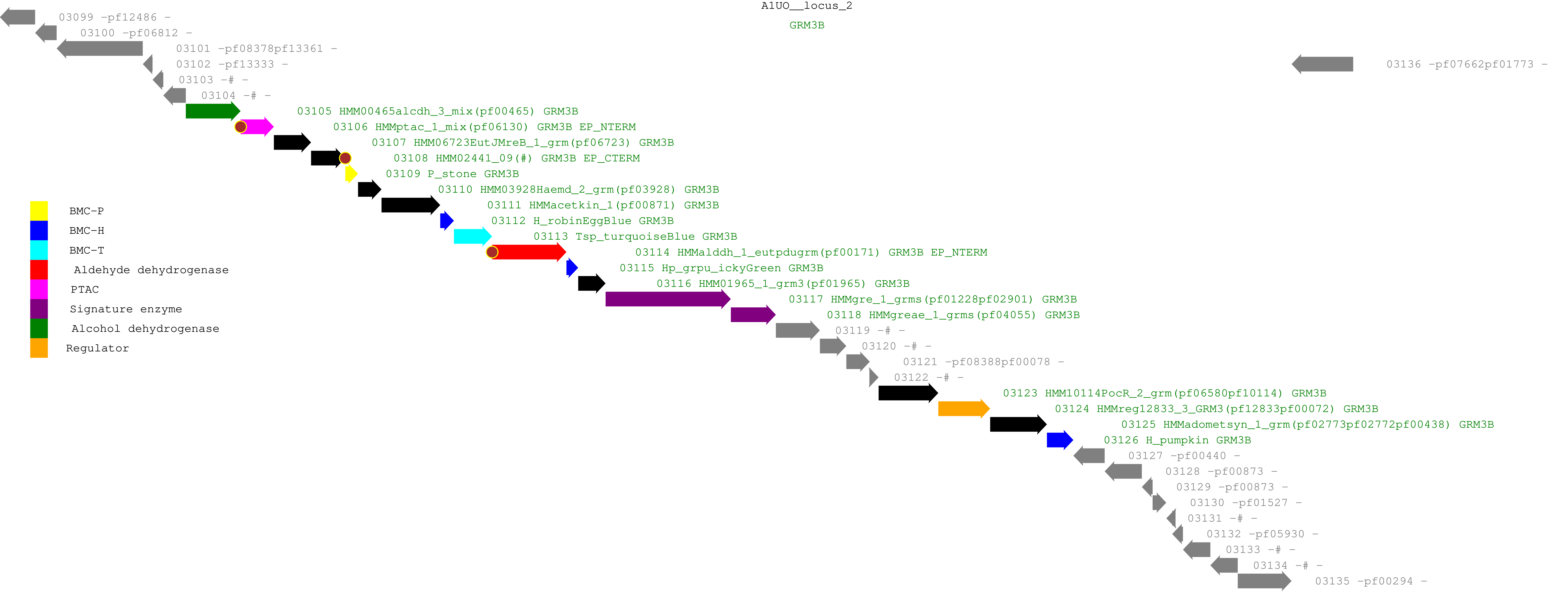

Pectobacterium wasabiae CFBP 3304. Bacteria; Proteobacteria; Gammaproteobacteria; Enterobacterales;

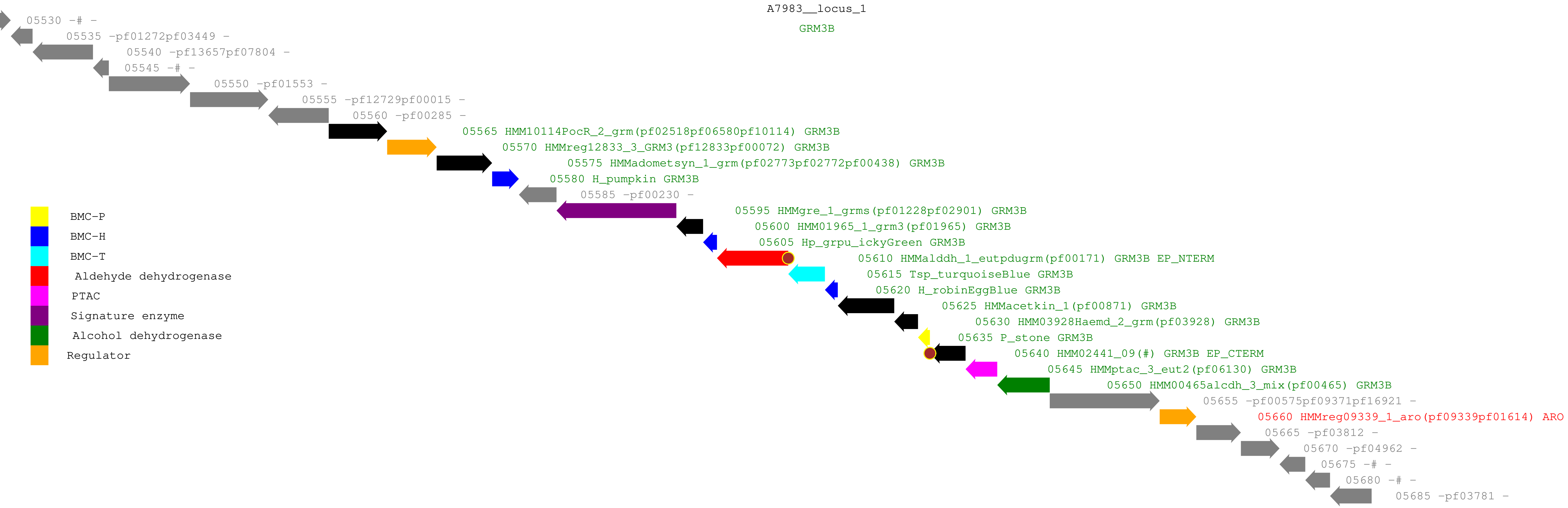

Escherichia coli. Bacteria; Proteobacteria; Gammaproteobacteria; Enterobacterales;

A8M81__locus_2

GRM3B

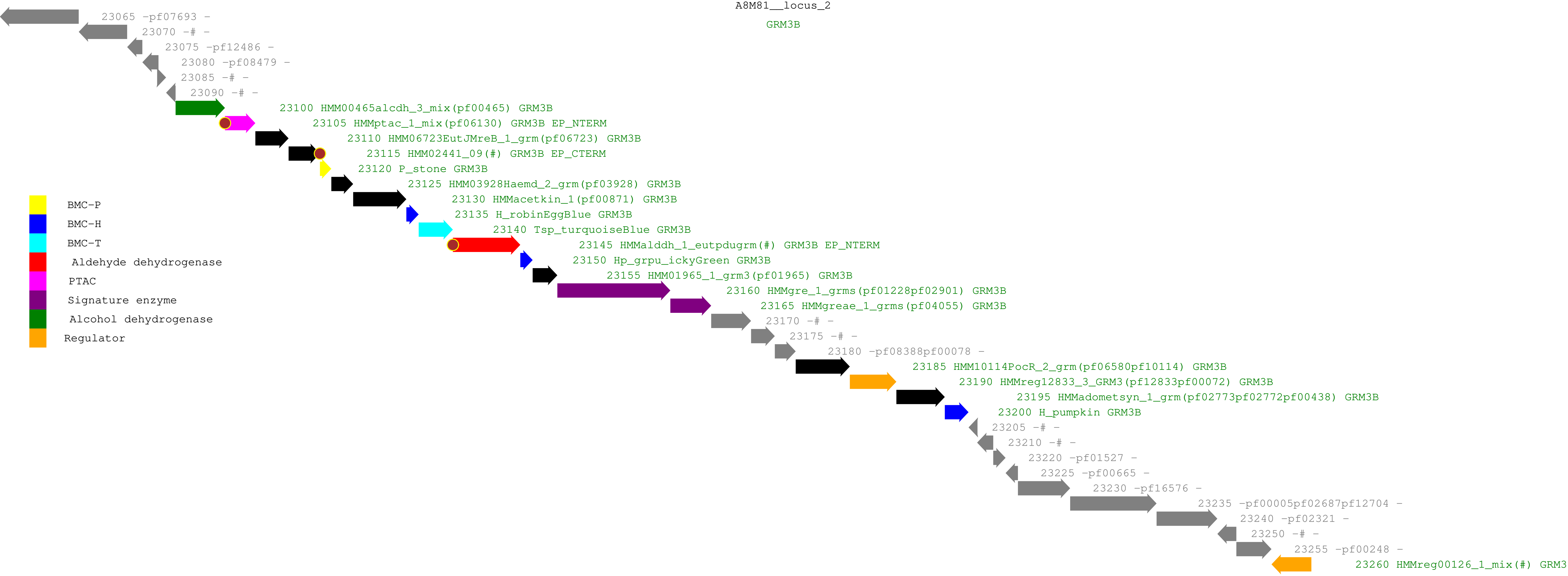

Rahnella victoriana. Bacteria; Proteobacteria; Gammaproteobacteria; Enterobacterales;

A9993__locus_1

GRM3B

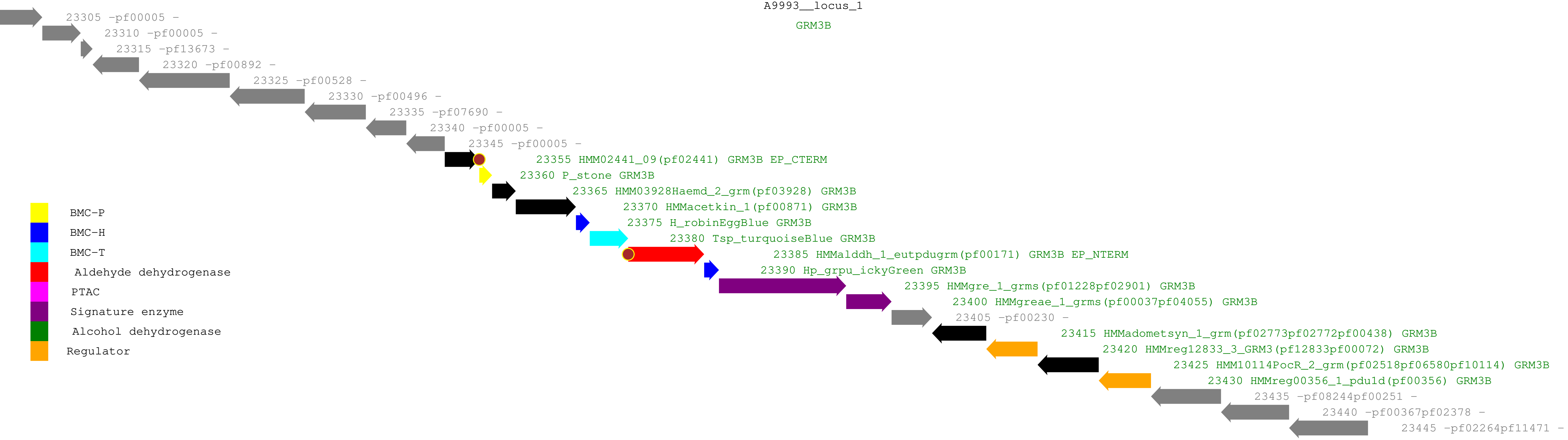

Escherichia coli 2-005-03_S4_C2. Bacteria; Proteobacteria; Gammaproteobacteria; Enterobacterales;

AC93__locus_2

GRM3B

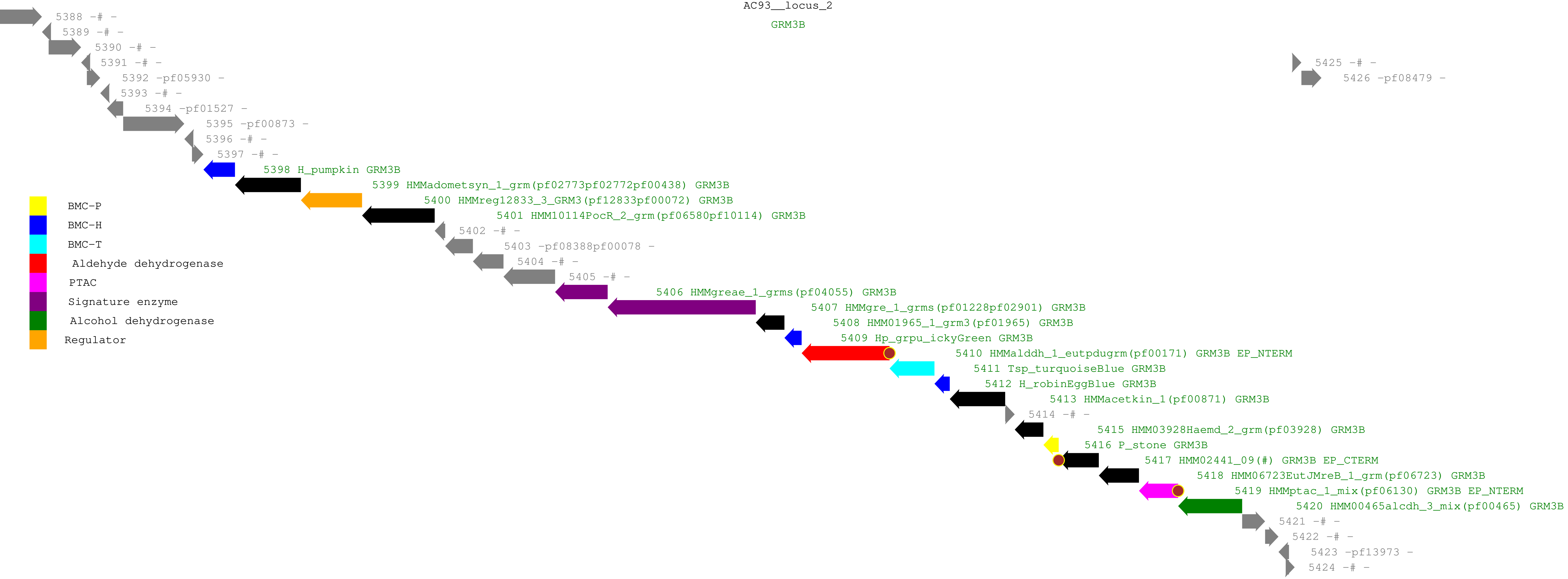

Escherichia coli 2-005-03_S4_C3. Bacteria; Proteobacteria; Gammaproteobacteria; Enterobacterales;

AD23__locus_2

GRM3B

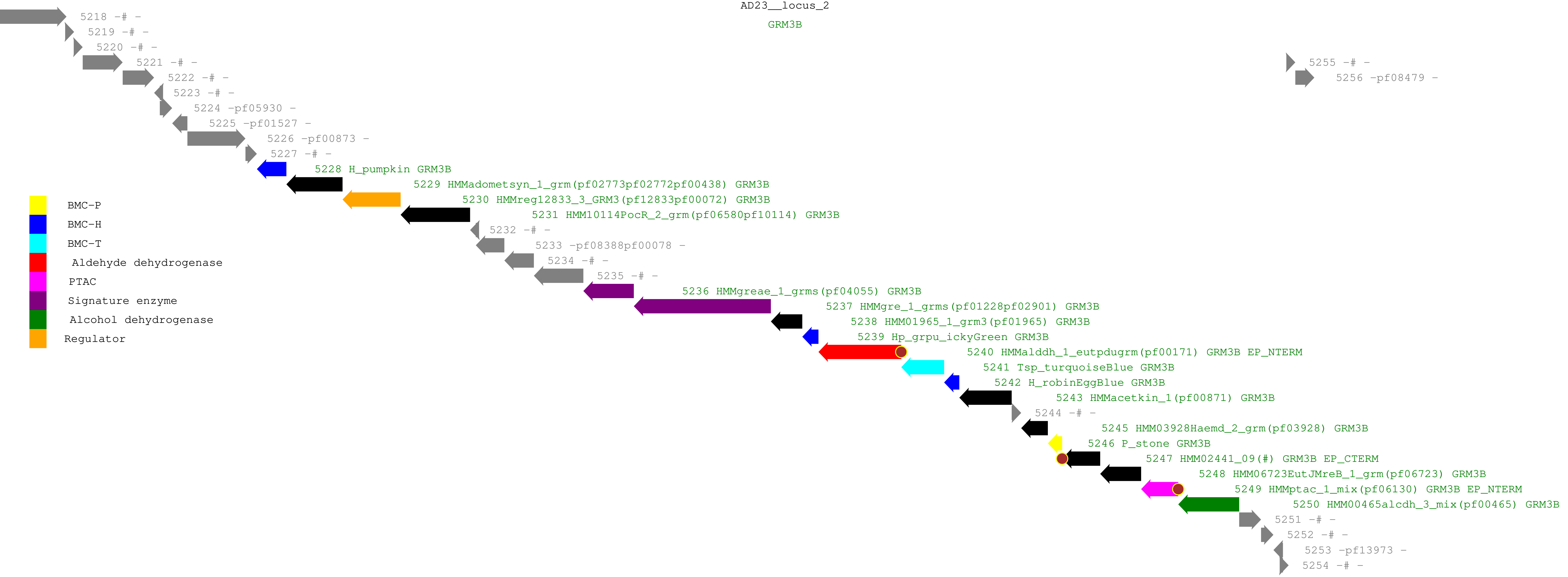

Escherichia coli UCI 65. Bacteria; Proteobacteria; Gammaproteobacteria; Enterobacterales;

AE24__locus_2

GRM3B

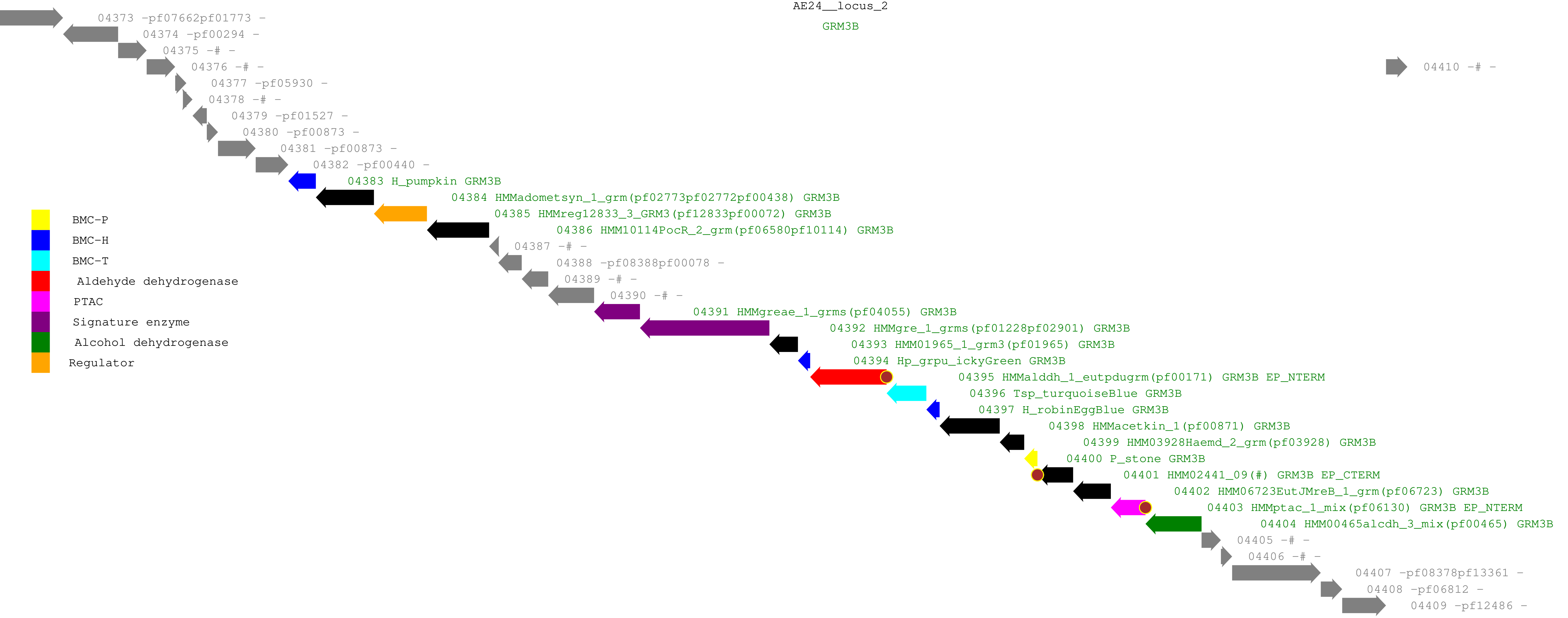

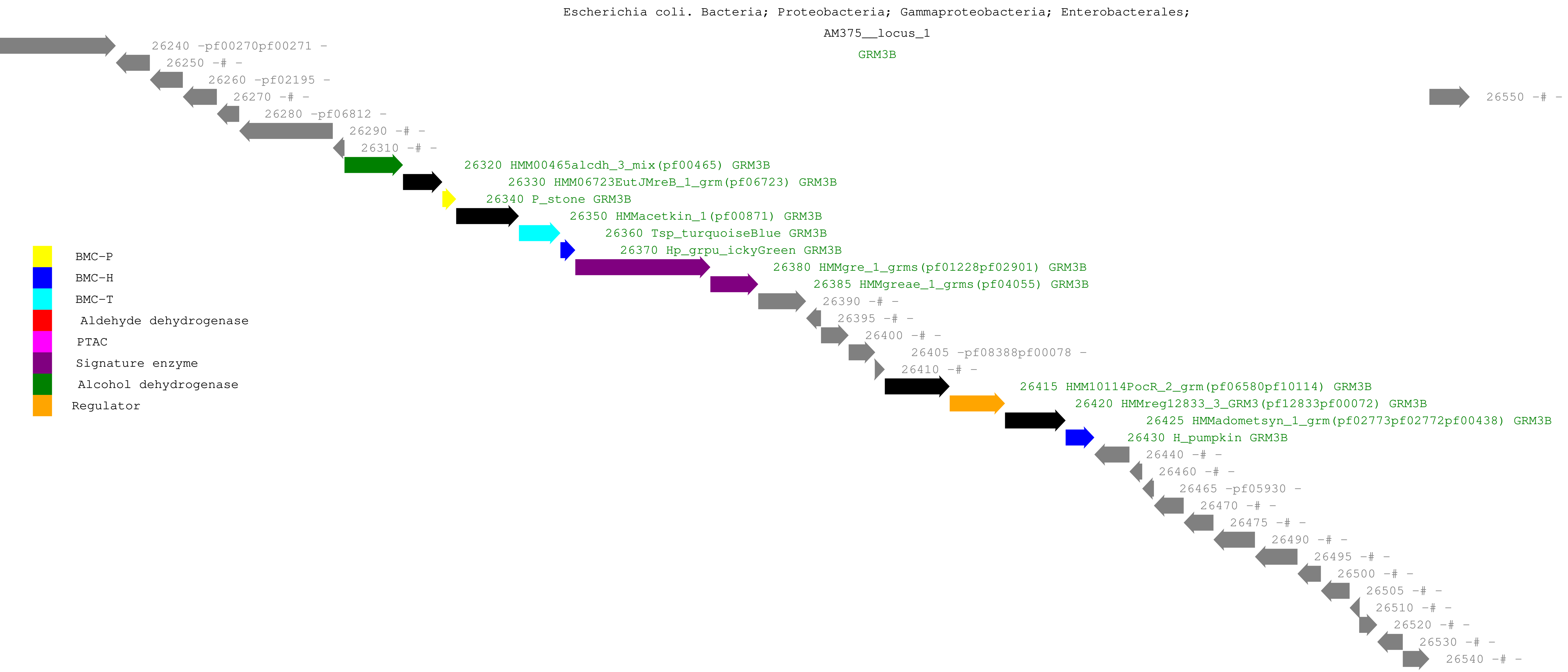

Escherichia coli. Bacteria; Proteobacteria; Gammaproteobacteria; Enterobacterales;
AM375__locus_1
GRM3B
26240 -pf00270pf00271 -
26250 -# -
26260 -pf02195 -
26270 -# -
26280 -pf06812 -
26290 -# -
26310 -# -
26320 HMM00465alcdh_3_mix(pf00465) GRM3B
26330 HMM06723EutJMreB_1_grm(pf06723) GRM3B
26340 P_stone GRM3B
26350 HMMacetkin_1(pf00871) GRM3B
26360 Tsp_turquoiseBlue GRM3B
26370 Hp_grpu_ickyGreen GRM3B
26380 HMMgre_1_grms(pf01228pf02901) GRM3B
26385 HMMgreae_1_grms(pf04055) GRM3B
26390 -# -
26395 -# -
26400 -# -
26405 -pf08388pf00078 -
26410 -# -
26415 HMM10114PocR_2_grm(pf06580pf10114) GRM3B
26420 HMMreg12833_3_GRM3(pf12833pf00072) GRM3B
26425 HMMadometsyn_1_grm(pf02773pf02772pf00438) GRM3B
26430 H_pumpkin GRM3B
26440 -# -
26460 -# -
26465 -pf05930 -
26470 -# -
26475 -# -
26490 -# -
26495 -# -
26500 -# -
26505 -# -
26510 -# -
26520 -# -
26530 -# -
26540 -# -
26550 -# -
BMC-P
BMC-H
BMC-T
Aldehyde dehydrogenase
PTAC
Signature enzyme
Alcohol dehydrogenase
Regulator

Escherichia coli O1:K1 / APEC. Bacteria; Proteobacteria; Gammaproteobacteria; Enterobacterales;

APECO1__locus_1

GRM3B

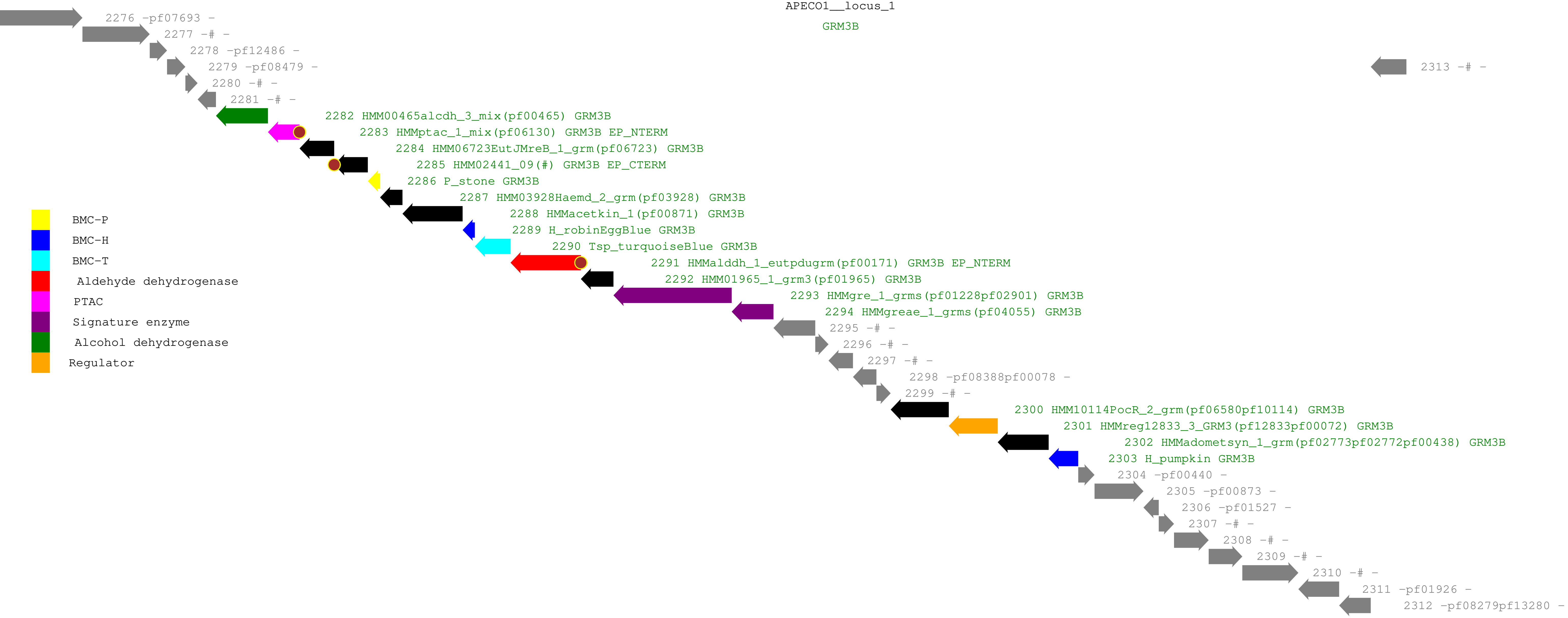

Escherichia coli. Bacteria; Proteobacteria; Gammaproteobacteria; Enterobacterales;

APT94__locus_2

GRM3B

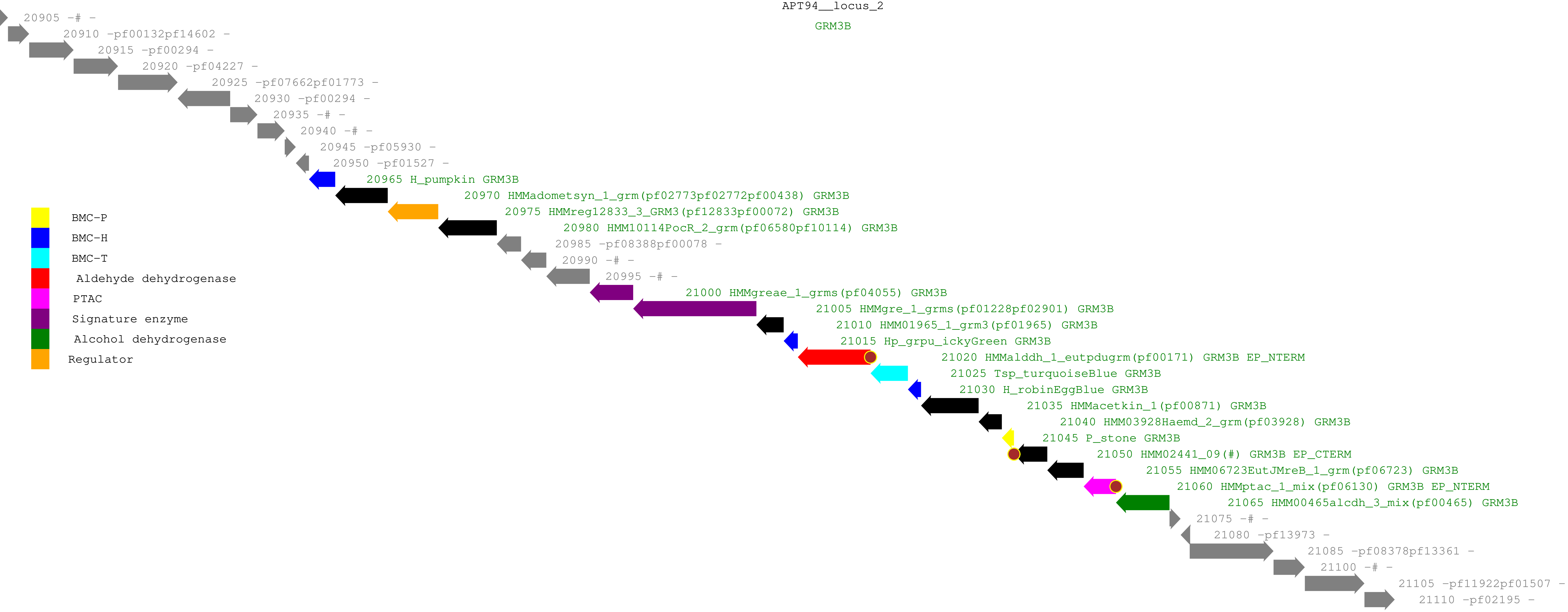

Gibbsiella quercinecans. Bacteria; Proteobacteria; Gammaproteobacteria; Enterobacterales;

AWC35__locus_1

GRM3B

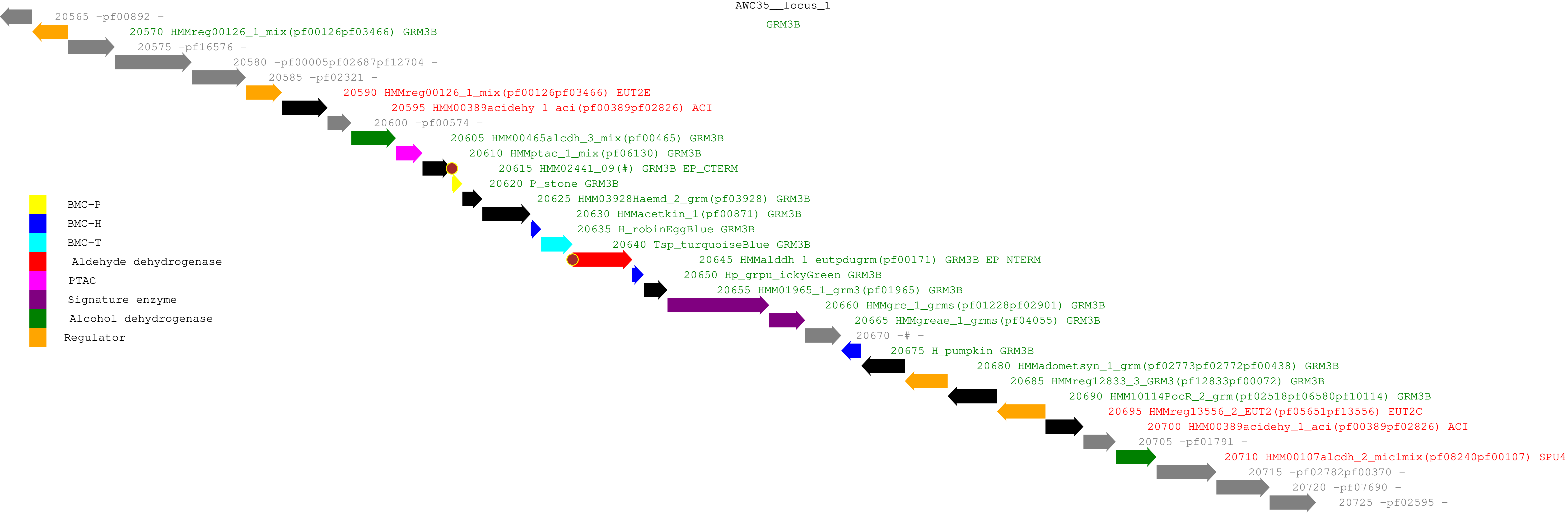

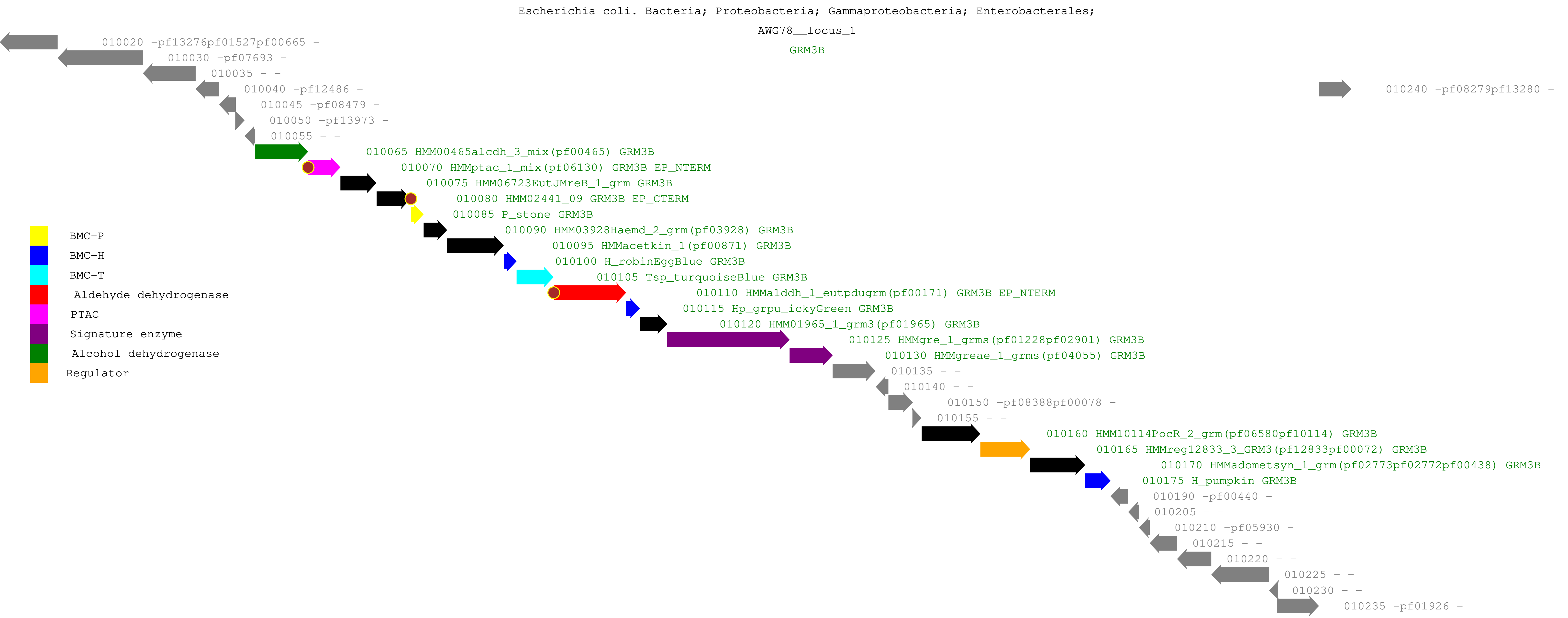

Escherichia coli. Bacteria; Proteobacteria; Gammaproteobacteria; Enterobacterales;
AWG78__locus_1
GRM3B
010020 -pf13276pf01527pf00665 -
010030 -pf07693 -
010035 - -
010040 -pf12486 -
010045 -pf08479 -
010050 -pf13973 -
010055 - -
010065 HMM00465alcdh_3_mix(pf00465) GRM3B
010070 HMMptac_1_mix(pf06130) GRM3B EP_NTERM
010075 HMM06723EutJMreB_1_grm GRM3B
010080 HMM02441_09 GRM3B EP_CTERM
010085 P_stone GRM3B
010090 HMM03928Haemd_2_grm(pf03928) GRM3B
010095 HMMacetkin_1(pf00871) GRM3B
010100 H_robinEggBlue GRM3B
010105 Tsp_turquoiseBlue GRM3B
010110 HMMalddh_1_eutpdugrm(pf00171) GRM3B EP_NTERM
010115 Hp_grpu_ickyGreen GRM3B
010120 HMM01965_1_grm3(pf01965) GRM3B
010125 HMMgre_1_grms(pf01228pf02901) GRM3B
010130 HMMgreae_1_grms(pf04055) GRM3B
010135 - -
010140 - -
010150 -pf08388pf00078 -
010155 - -
010160 HMM10114PocR_2_grm(pf06580pf10114) GRM3B
010165 HMMreg12833_3_GRM3(pf12833pf00072) GRM3B
010170 HMMadometsyn_1_grm(pf02773pf02772pf00438) GRM3B
010175 H_pumpkin GRM3B
010190 -pf00440 -
010205 - -
010210 -pf05930 -
010215 - -
010220 - -
010225 - -
010230 - -
010235 -pf01926 -
010240 -pf08279pf13280 -
BMC-P
BMC-H
BMC-T
Aldehyde dehydrogenase
PTAC
Signature enzyme
Alcohol dehydrogenase
Regulator

Raoultella planticola (Klebsiella planticola). Bacteria; Proteobacteria; Gammaproteobacteria; Enterobacterales;

AYO04__locus_2

GRM3B

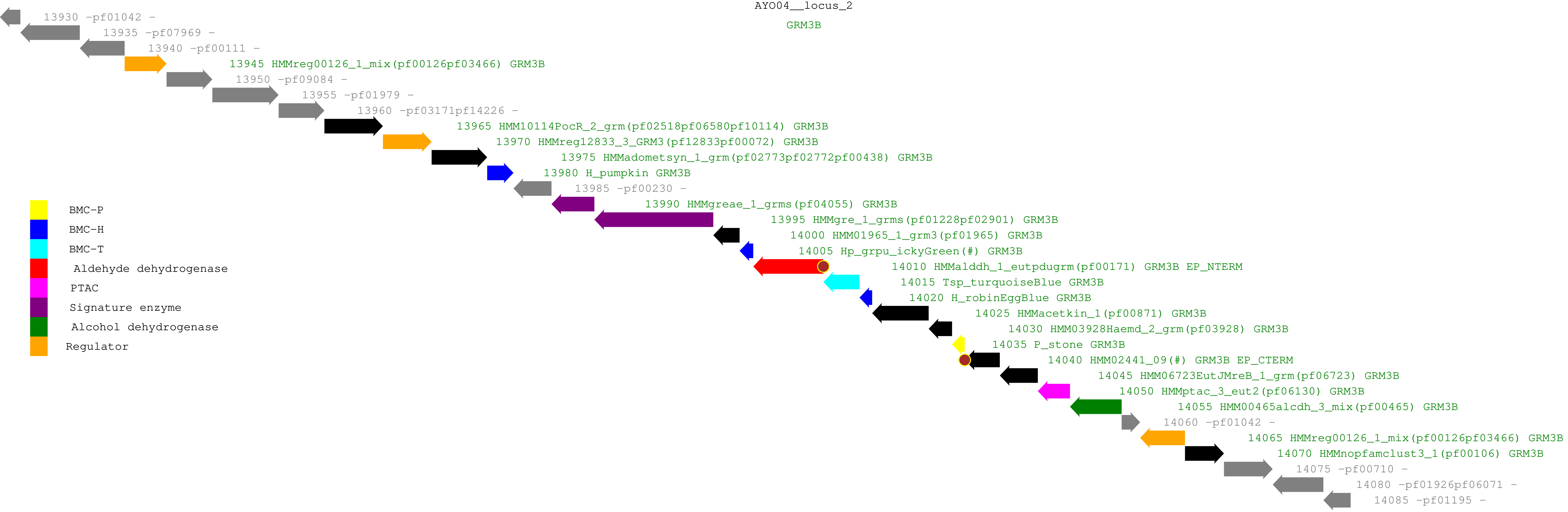

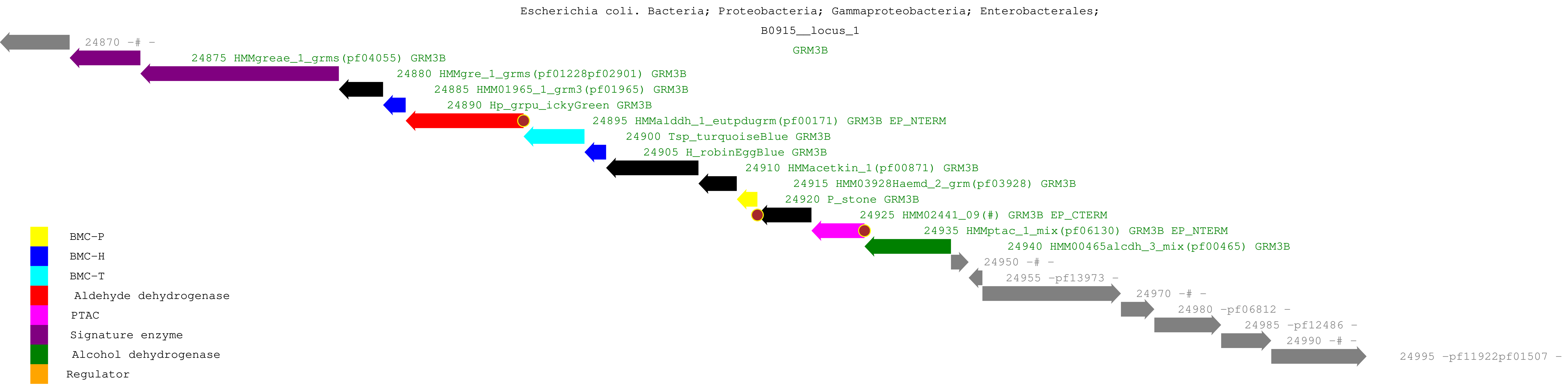
Escherichia coli. Bacteria; Proteobacteria; Gammaproteobacteria; Enterobacterales;
B0915__locus_1
GRM3B
24870 -# -
24875 HMMgreae_1_grms(pf04055) GRM3B
24880 HMMgre_1_grms(pf01228pf02901) GRM3B
24885 HMM01965_1_grm3(pf01965) GRM3B
24890 Hp_grpu_ickyGreen GRM3B
24895 HMMalddh_1_eutpdugrm(pf00171) GRM3B EP_NTERM
24900 Tsp_turquoiseBlue GRM3B
24905 H_robinEggBlue GRM3B
24910 HMMacetkin_1(pf00871) GRM3B
24915 HMM03928Haemd_2_grm(pf03928) GRM3B
24920 P_stone GRM3B
24925 HMM02441_09(#) GRM3B EP_CTERM
24935 HMMptac_1_mix(pf06130) GRM3B EP_NTERM
24940 HMM00465alcdh_3_mix(pf00465) GRM3B
24950 -# -
24955 -pf13973 -
24970 -# -
24980 -pf06812 -
24985 -pf12486 -
24990 -# -
24995 -pf11922pf01507 -
BMC-P
BMC-H
BMC-T
Aldehyde dehydrogenase
PTAC
Signature enzyme
Alcohol dehydrogenase
Regulator

Vibrio natriegens. Bacteria; Proteobacteria; Gammaproteobacteria; Vibrionales;

BA894__locus_1

GRM3B

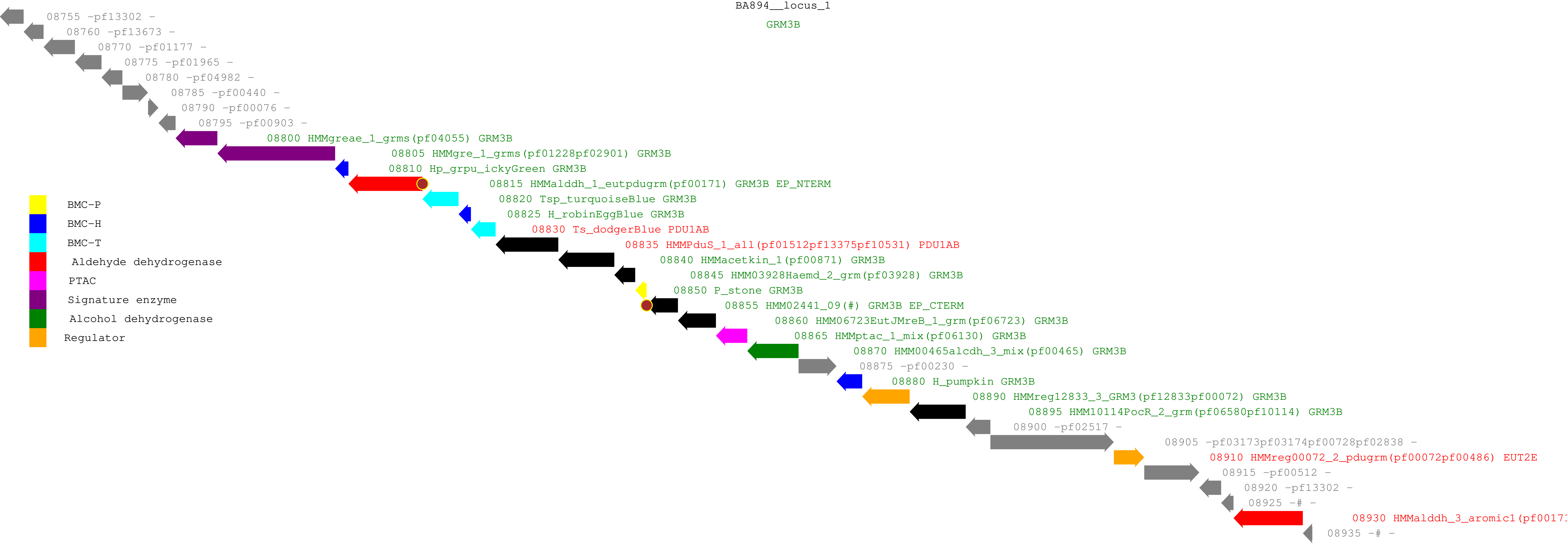

Escherichia coli. Bacteria; Proteobacteria; Gammaproteobacteria; Enterobacterales;

BET08__locus_1_main_sat

GRM3B

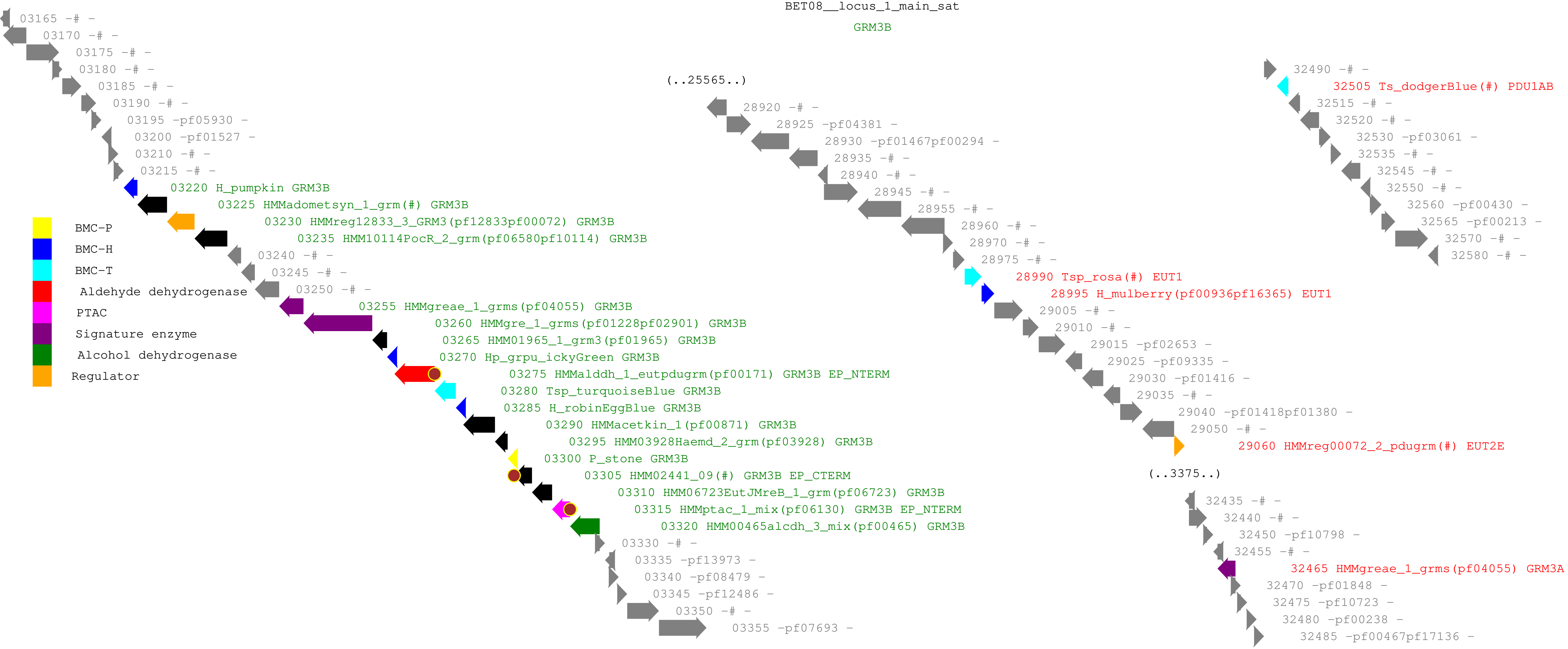

Escherichia coli. Bacteria; Proteobacteria; Gammaproteobacteria; Enterobacterales;

BK337__locus_1

GRM3B

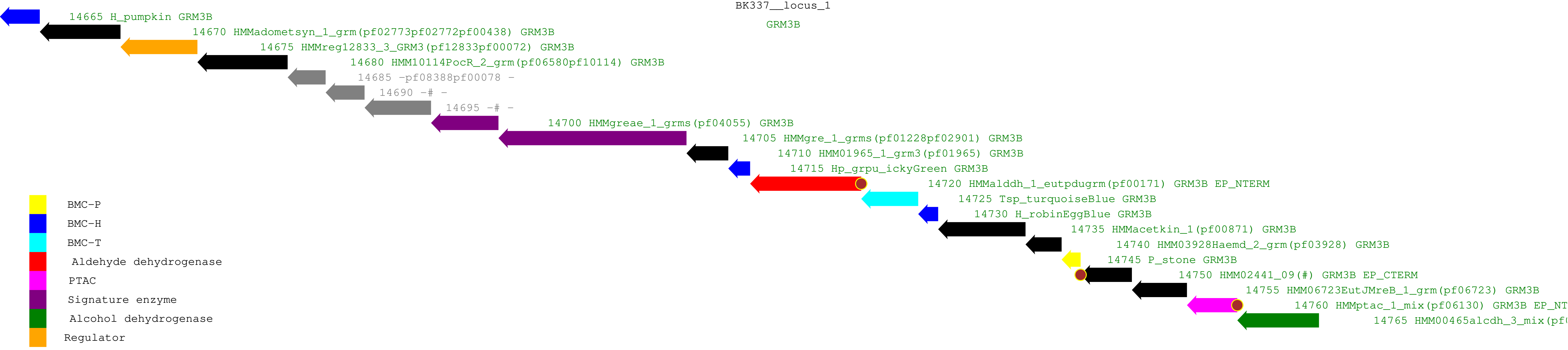

Raoultella ornithinolytica (Klebsiella ornithinolytica). Bacteria; Proteobacteria; Gammaproteobacteria; Enterobacterales;

BK817__locus_2

GRM3B

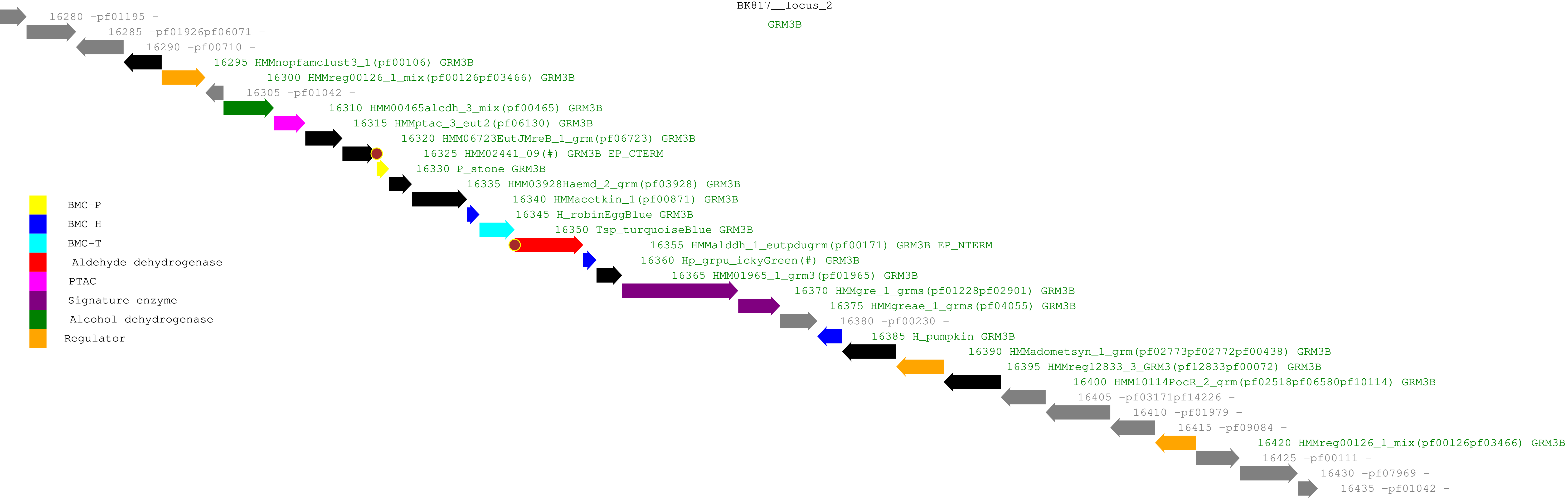

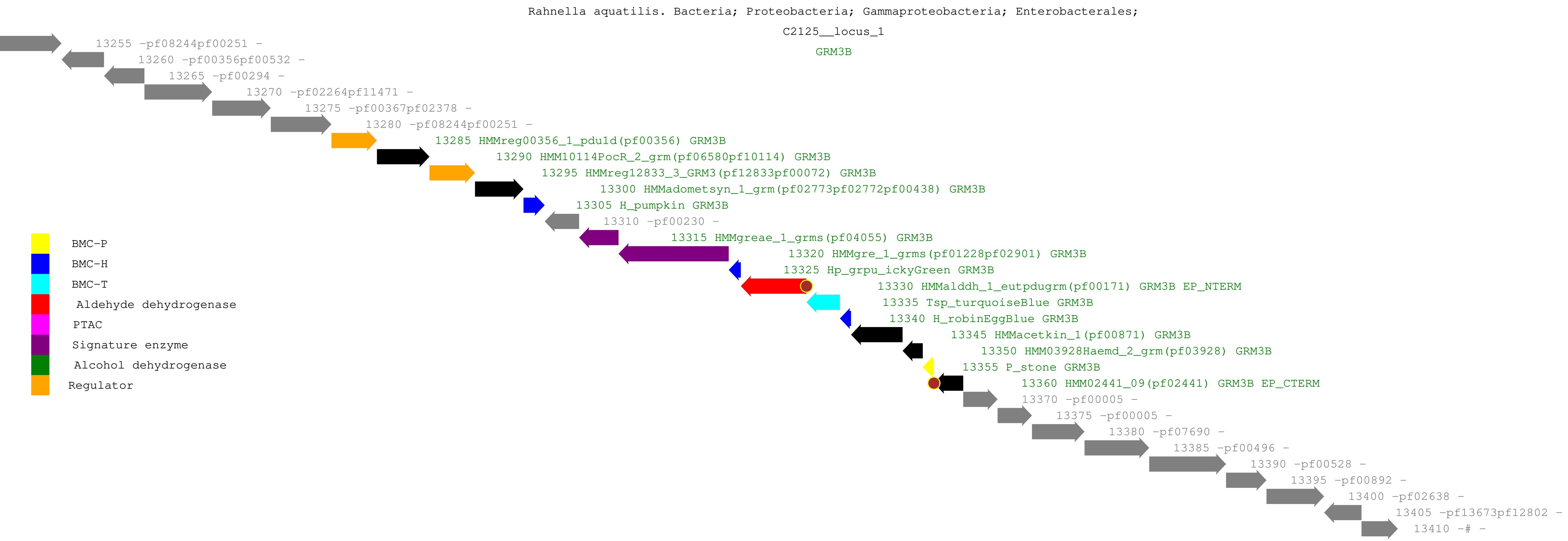
Rahnella aquatilis. Bacteria; Proteobacteria; Gammaproteobacteria; Enterobacterales;
C2125__locus_1
GRM3B
13255 -pf08244pf00251 -
13260 -pf00356pf00532 -
13265 -pf00294 -
13270 -pf02264pf11471 -
13275 -pf00367pf02378 -
13280 -pf08244pf00251 -
13285 HMMreg00356_1_pdu1d(pf00356) GRM3B
13290 HMM10114PocR_2_grm(pf06580pf10114) GRM3B
13295 HMMreg12833_3_GRM3(pf12833pf00072) GRM3B
13300 HMMadometsyn_1_grm(pf02773pf02772pf00438) GRM3B
13305 H_pumpkin GRM3B
13310 -pf00230 -
13315 HMMgreae_1_grms(pf04055) GRM3B
13320 HMMgre_1_grms(pf01228pf02901) GRM3B
13325 Hp_grpu_ickyGreen GRM3B
13330 HMMalddh_1_eutpdugrm(pf00171) GRM3B EP_NTERM
13335 Tsp_turquoiseBlue GRM3B
13340 H_robinEggBlue GRM3B
13345 HMMacetkin_1(pf00871) GRM3B
13350 HMM03928Haemd_2_grm(pf03928) GRM3B
13355 P_stone GRM3B
13360 HMM02441_09(pf02441) GRM3B EP_CTERM
13370 -pf00005 -
13375 -pf00005 -
13380 -pf07690 -
13385 -pf00496 -
13390 -pf00528 -
13395 -pf00892 -
13400 -pf02638 -
13405 -pf13673pf12802 -
13410 -# -
BMC-P
BMC-H
BMC-T
Aldehyde dehydrogenase
PTAC
Signature enzyme
Alcohol dehydrogenase
Regulator

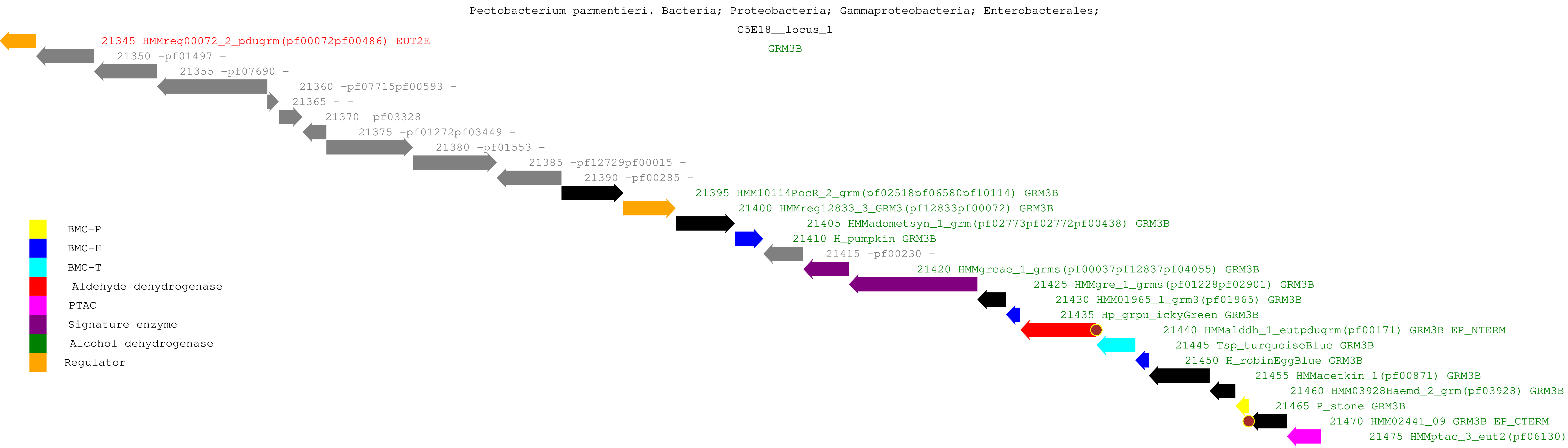

Pectobacterium parmentieri. Bacteria; Proteobacteria; Gammaproteobacteria; Enterobacterales;
C5E18__locus_1
GRM3B
21345 HMMreg00072_2_pdugrm(pf00072pf00486) EUT2E
21350 -pf01497 -
21355 -pf07690 -
21360 -pf07715pf00593 -
21365 - -
21370 -pf03328 -
21375 -pf01272pf03449 -
21380 -pf01553 -
21385 -pf12729pf00015 -
21390 -pf00285 -
21395 HMM10114PocR_2_grm(pf02518pf06580pf10114) GRM3B
21400 HMMreg12833_3_GRM3(pf12833pf00072) GRM3B
21405 HMMadometsyn_1_grm(pf02773pf02772pf00438) GRM3B
21410 H_pumpkin GRM3B
21415 -pf00230 -
21420 HMMgreae_1_grms(pf00037pf12837pf04055) GRM3B
21425 HMMgre_1_grms(pf01228pf02901) GRM3B
21430 HMM01965_1_grm3(pf01965) GRM3B
21435 Hp_grpu_ickyGreen GRM3B
21440 HMMalddh_1_eutpdugrm(pf00171) GRM3B EP_NTERM
21445 Tsp_turquoiseBlue GRM3B
21450 H_robinEggBlue GRM3B
21455 HMMacetkin_1(pf00871) GRM3B
21460 HMM03928Haemd_2_grm(pf03928) GRM3B
21465 P_stone GRM3B
21470 HMM02441_09 GRM3B EP_CTERM
21475 HMMptac_3_eut2(pf06130)
BMC-P
BMC-H
BMC-T
Aldehyde dehydrogenase
PTAC
Signature enzyme
Alcohol dehydrogenase
Regulator

Raoultella ornithinolytica (Klebsiella ornithinolytica). Bacteria; Proteobacteria; Gammaproteobacteria; Enterobacterales;

CAC00__locus_1

GRM3B

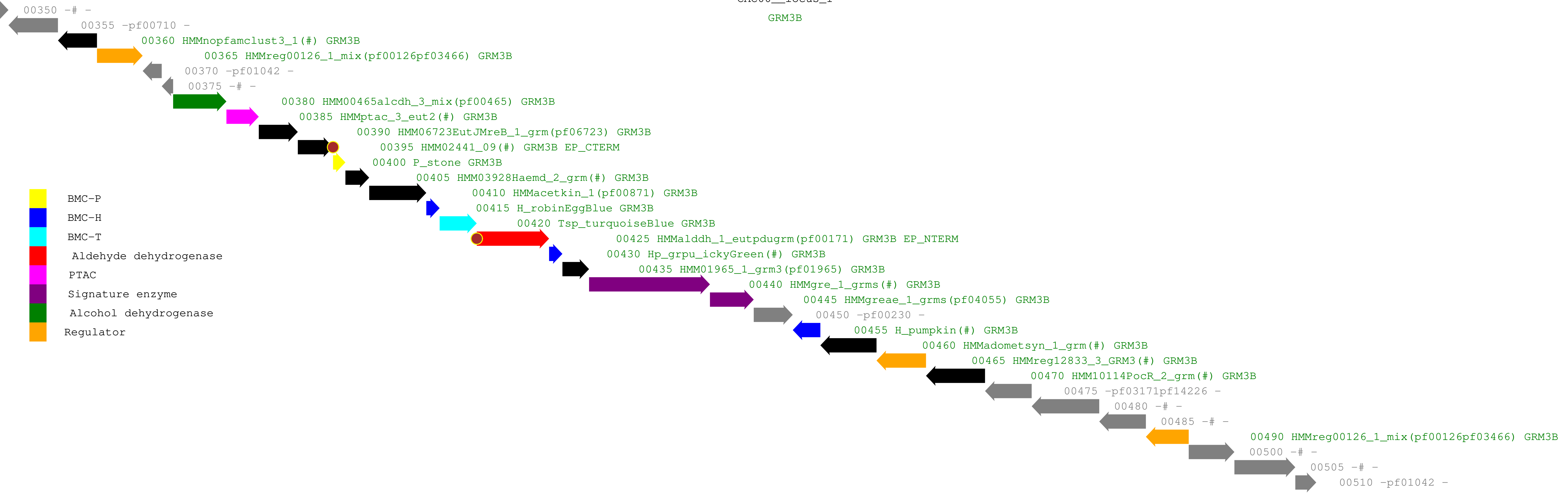

Enterobacteriaceae bacterium BP-1. Bacteria; Proteobacteria; Gammaproteobacteria; Enterobacterales;

CHU32__locus_1

GRM3B

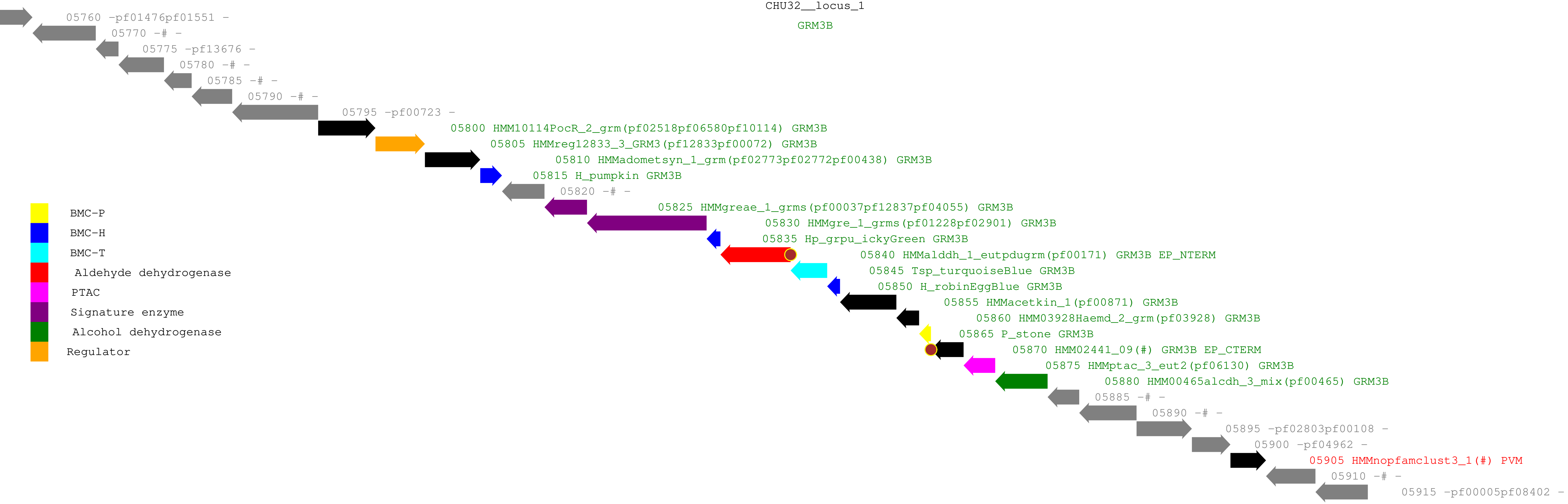

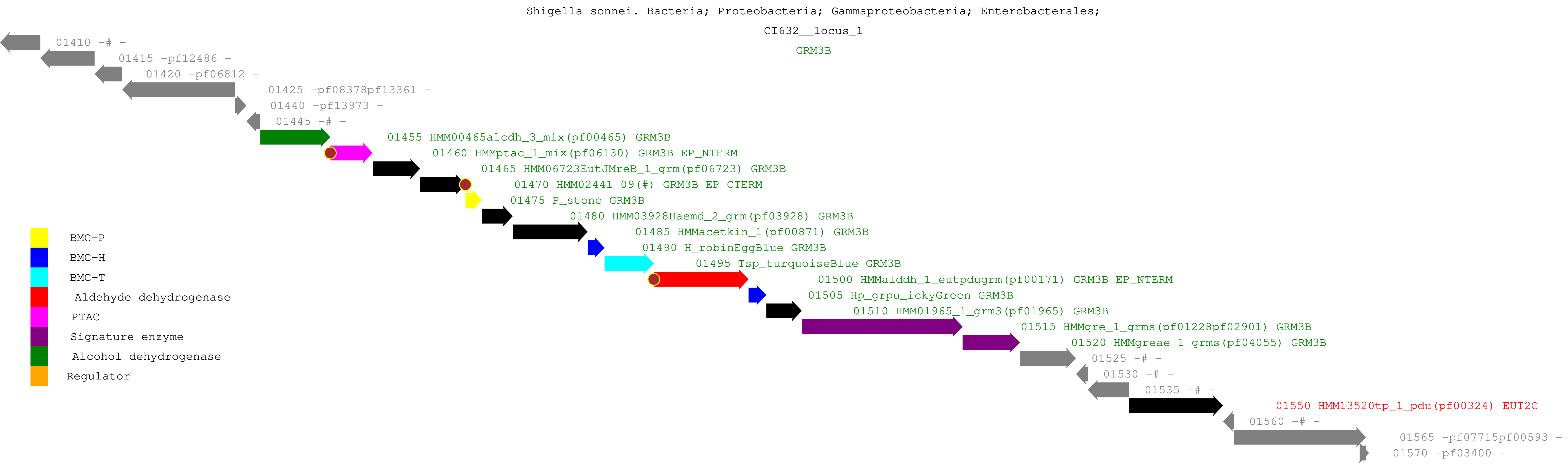
Shigella sonnei. Bacteria; Proteobacteria; Gammaproteobacteria; Enterobacterales;
CI632__locus_1
GRM3B
01410 -# -
01415 -pf12486 -
01420 -pf06812 -
01425 -pf08378pf13361 -
01440 -pf13973 -
01445 -# -
01455 HMM00465alcdh_3_mix(pf00465) GRM3B
01460 HMMptac_1_mix(pf06130) GRM3B EP_NTERM
01465 HMM06723EutJMreB_1_grm(pf06723) GRM3B
01470 HMM02441_09(#) GRM3B EP_CTERM
01475 P_stone GRM3B
01480 HMM03928Haemd_2_grm(pf03928) GRM3B
01485 HMMacetkin_1(pf00871) GRM3B
01490 H_robinEggBlue GRM3B
01495 Tsp_turquoiseBlue GRM3B
01500 HMMalddh_1_eutpdugrm(pf00171) GRM3B EP_NTERM
01505 Hp_grpu_ickyGreen GRM3B
01510 HMM01965_1_grm3(pf01965) GRM3B
01515 HMMgre_1_grms(pf01228pf02901) GRM3B
01520 HMMgreae_1_grms(pf04055) GRM3B
01525 -# -
01530 -# -
01535 -# -
01550 HMM13520tp_1_pdu(pf00324) EUT2C
01560 -# -
01565 -pf07715pf00593 -
01570 -pf03400 -
BMC-P
BMC-H
BMC-T
Aldehyde dehydrogenase
PTAC
Signature enzyme
Alcohol dehydrogenase
Regulator

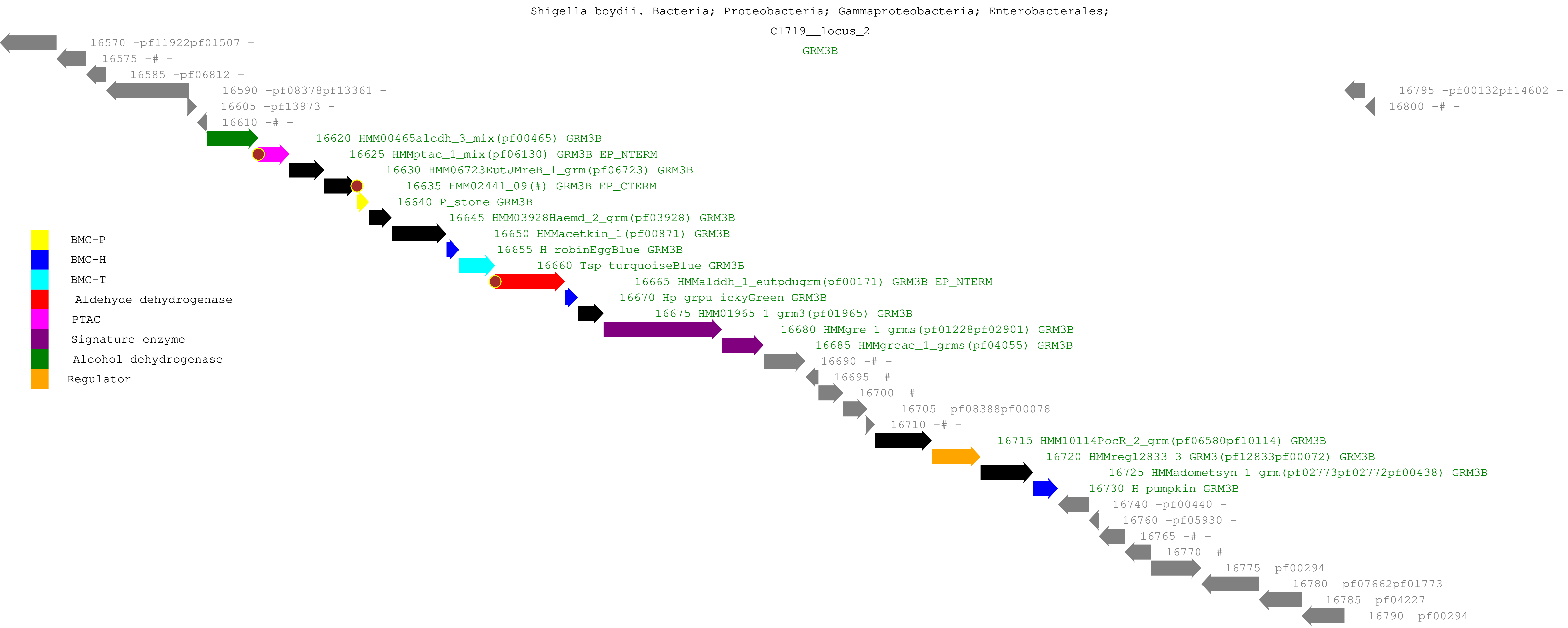
Shigella boydii. Bacteria; Proteobacteria; Gammaproteobacteria; Enterobacterales;
CI719__locus_2
GRM3B
16570 -pf11922pf01507 -
16575 -# -
16585 -pf06812 -
16590 -pf08378pf13361 -
16605 -pf13973 -
16610 -# -
16620 HMM00465alcdh_3_mix(pf00465) GRM3B
16625 HMMptac_1_mix(pf06130) GRM3B EP_NTERM
16630 HMM06723EutJMreB_1_grm(pf06723) GRM3B
16635 HMM02441_09(#) GRM3B EP_CTERM
16640 P_stone GRM3B
16645 HMM03928Haemd_2_grm(pf03928) GRM3B
16650 HMMacetkin_1(pf00871) GRM3B
16655 H_robinEggBlue GRM3B
16660 Tsp_turquoiseBlue GRM3B
16665 HMMalddh_1_eutpdugrm(pf00171) GRM3B EP_NTERM
16670 Hp_grpu_ickyGreen GRM3B
16675 HMM01965_1_grm3(pf01965) GRM3B
16680 HMMgre_1_grms(pf01228pf02901) GRM3B
16685 HMMgreae_1_grms(pf04055) GRM3B
16690 -# -
16695 -# -
16700 -# -
16705 -pf08388pf00078 -
16710 -# -
16715 HMM10114PocR_2_grm(pf06580pf10114) GRM3B
16720 HMMreg12833_3_GRM3(pf12833pf00072) GRM3B
16725 HMMadometsyn_1_grm(pf02773pf02772pf00438) GRM3B
16730 H_pumpkin GRM3B
16740 -pf00440 -
16760 -pf05930 -
16765 -# -
16770 -# -
16775 -pf00294 -
16780 -pf07662pf01773 -
16785 -pf04227 -
16790 -pf00294 -
16795 -pf00132pf14602 -
16800 -# -
BMC-P
BMC-H
BMC-T
Aldehyde dehydrogenase
PTAC
Signature enzyme
Alcohol dehydrogenase
Regulator

Shigella boydii. Bacteria; Proteobacteria; Gammaproteobacteria; Enterobacterales;

CI767__locus_1

GRM3B

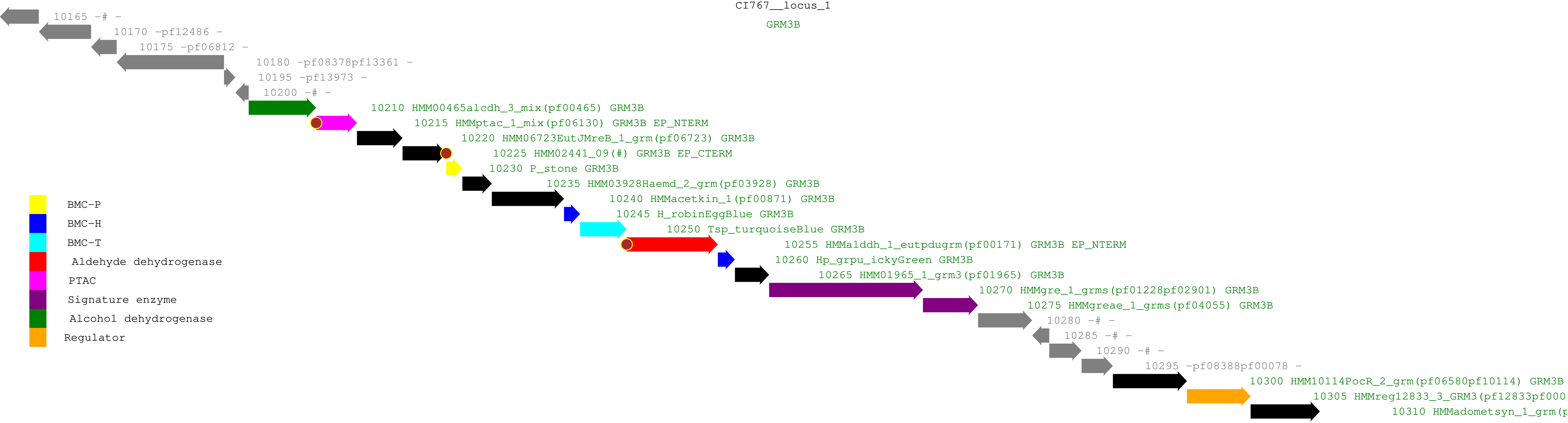

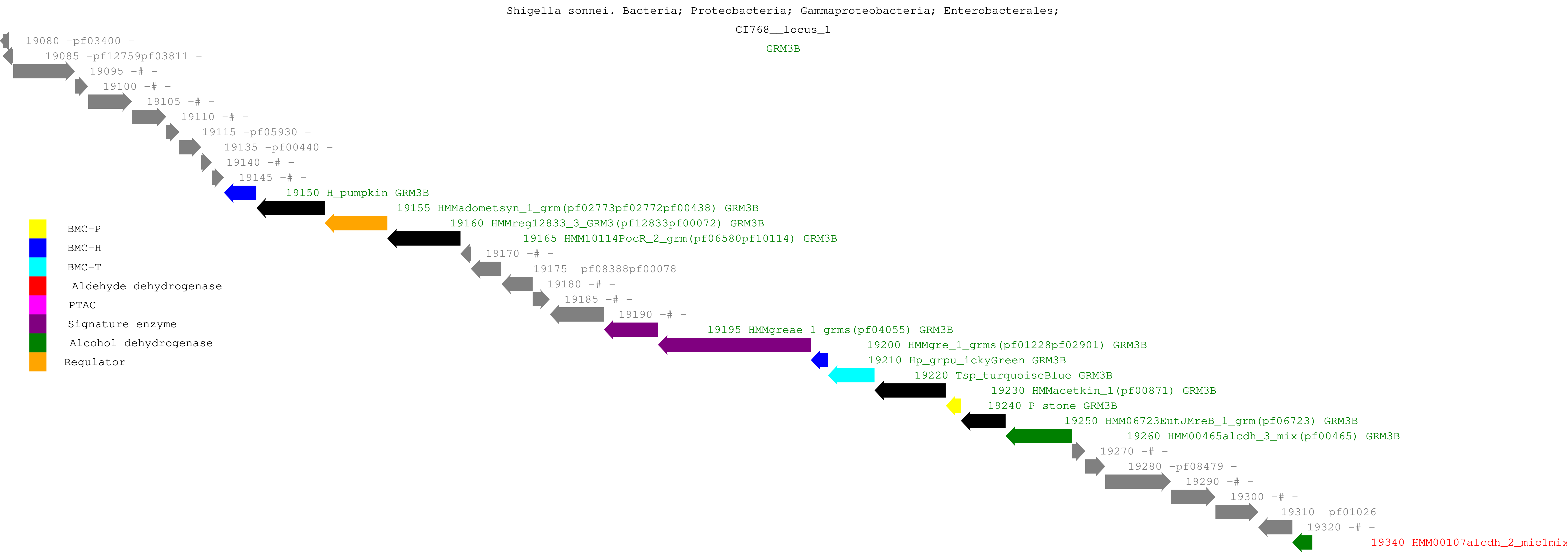

Shigella sonnei. Bacteria; Proteobacteria; Gammaproteobacteria; Enterobacterales;
CI768__locus_1
GRM3B
19080 -pf03400 -
19085 -pf12759pf03811 -
19095 -# -
19100 -# -
19105 -# -
19110 -# -
19115 -pf05930 -
19135 -pf00440 -
19140 -# -
19145 -# -
19150 H_pumpkin GRM3B
19155 HMMadometsyn_1_grm(pf02773pf02772pf00438) GRM3B
19160 HMMreg12833_3_GRM3(pf12833pf00072) GRM3B
19165 HMM10114PocR_2_grm(pf06580pf10114) GRM3B
19170 -# -
19175 -pf08388pf00078 -
19180 -# -
19185 -# -
19190 -# -
19195 HMMgreae_1_grms(pf04055) GRM3B
19200 HMMgre_1_grms(pf01228pf02901) GRM3B
19210 Hp_grpu_ickyGreen GRM3B
19220 Tsp_turquoiseBlue GRM3B
19230 HMMacetkin_1(pf00871) GRM3B
19240 P_stone GRM3B
19250 HMM06723EutJMreB_1_grm(pf06723) GRM3B
19260 HMM00465alcdh_3_mix(pf00465) GRM3B
19270 -# -
19280 -pf08479 -
19290 -# -
19300 -# -
19310 -pf01026 -
19320 -# -
19340 HMM00107alcdh_2_mic1mix
BMC-P
BMC-H
BMC-T
Aldehyde dehydrogenase
PTAC
Signature enzyme
Alcohol dehydrogenase
Regulator

Raoultella planticola (Klebsiella planticola). Bacteria; Proteobacteria; Gammaproteobacteria; Enterobacterales;

CIG23__locus_1

GRM3B

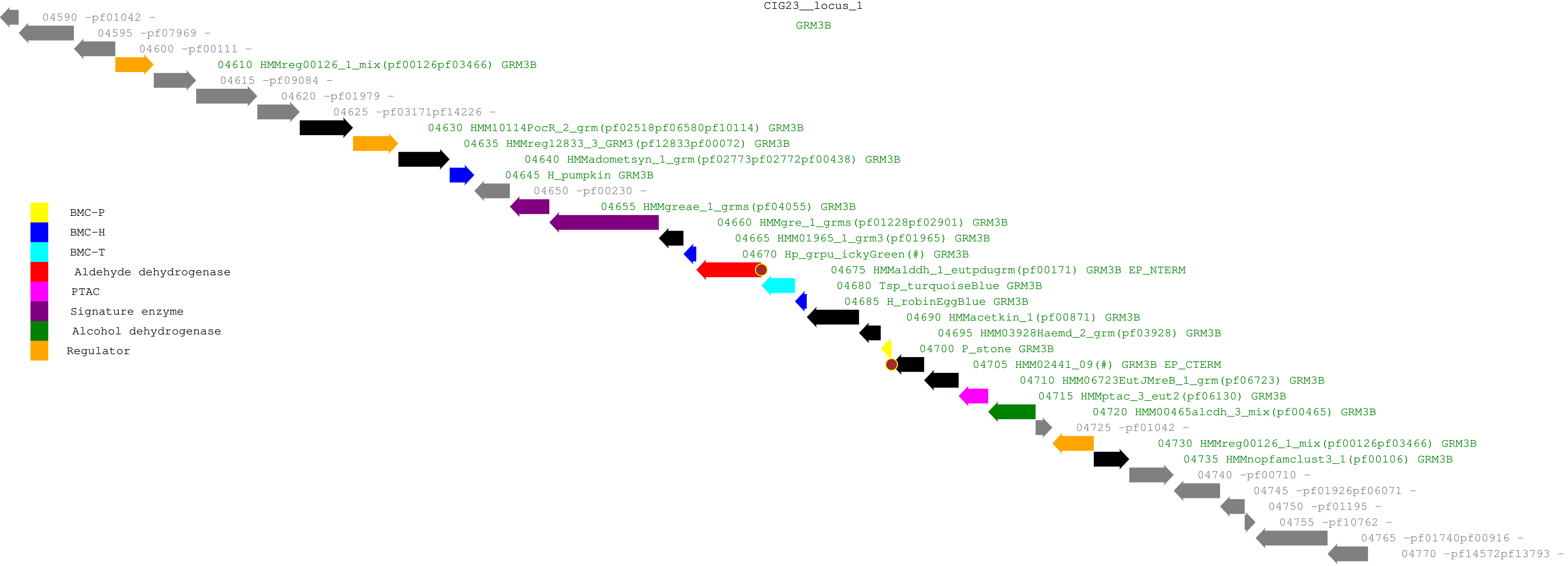

Escherichia coli O6:H1 (strain CFT073 / ATCC 700928 / UPEC). Bacteria; Proteobacteria; Gammaproteobacteria; Enterobacterales;

c_locus_2

GRM3B

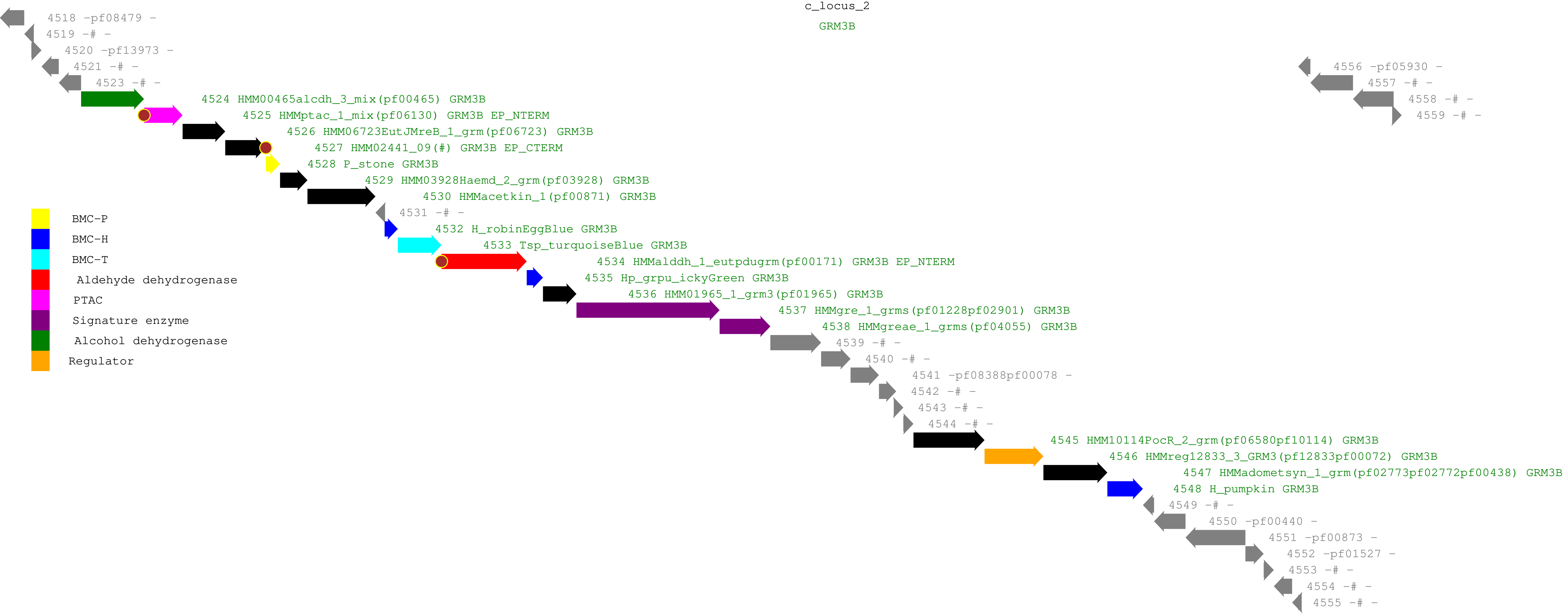

Budvicia aquatica. Bacteria; Proteobacteria; Gammaproteobacteria; Enterobacterales;

CRN84__locus_1

GRM3B

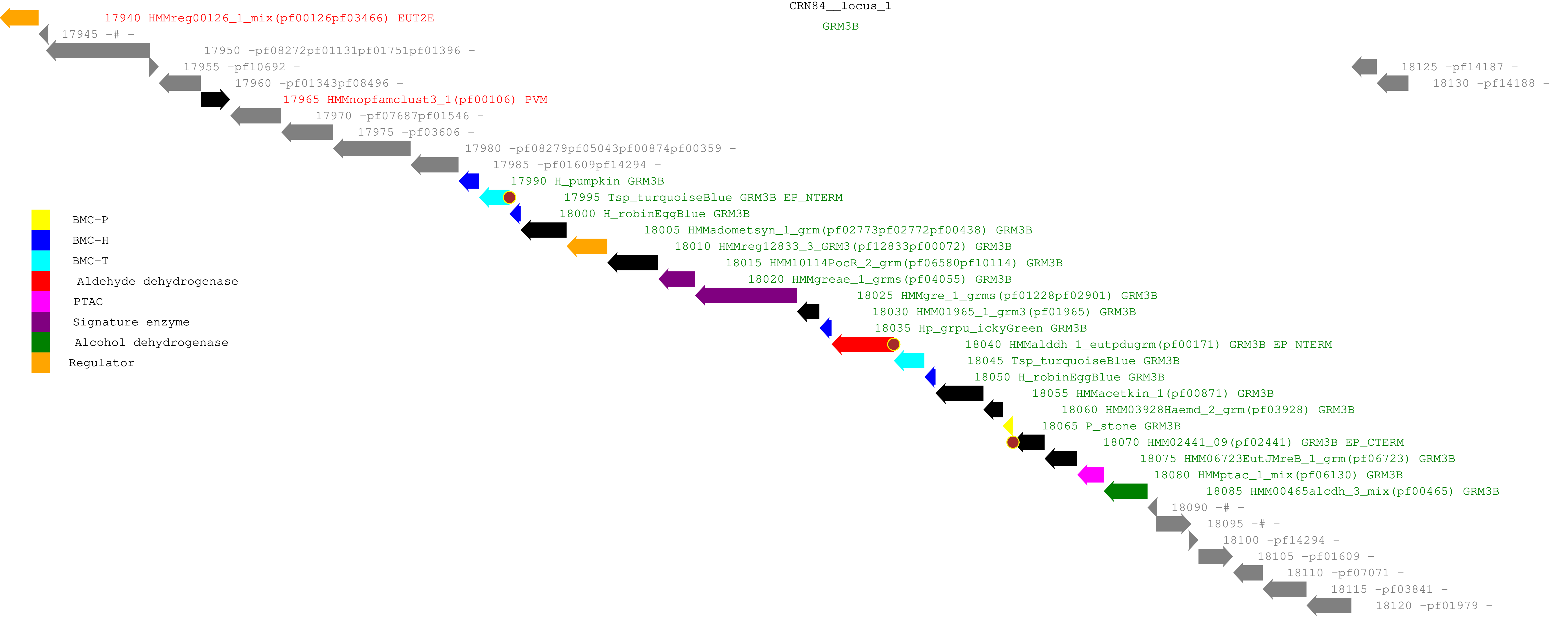

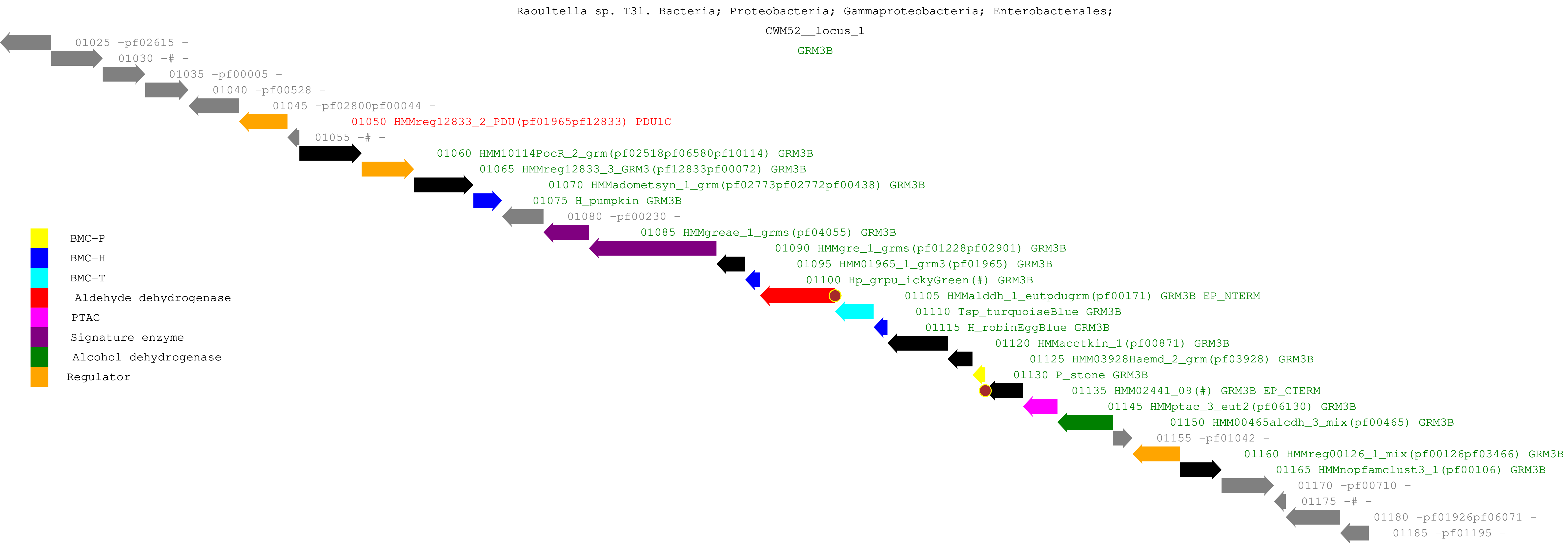
Raoultella sp. T31. Bacteria; Proteobacteria; Gammaproteobacteria; Enterobacterales;
CWM52__locus_1
GRM3B
01025 -pf02615 -
01030 -# -
01035 -pf00005 -
01040 -pf00528 -
01045 -pf02800pf00044 -
01050 HMMreg12833_2_PDU(pf01965pf12833) PDU1C
01055 -# -
01060 HMM10114PocR_2_grm(pf02518pf06580pf10114) GRM3B
01065 HMMreg12833_3_GRM3(pf12833pf00072) GRM3B
01070 HMMadometsyn_1_grm(pf02773pf02772pf00438) GRM3B
01075 H_pumpkin GRM3B
01080 -pf00230 -
01085 HMMgreae_1_grms(pf04055) GRM3B
01090 HMMgre_1_grms(pf01228pf02901) GRM3B
01095 HMM01965_1_grm3(pf01965) GRM3B
01100 Hp_grpu_ickyGreen(#) GRM3B
01105 HMMalddh_1_eutpdugrm(pf00171) GRM3B EP_NTERM
01110 Tsp_turquoiseBlue GRM3B
01115 H_robinEggBlue GRM3B
01120 HMMacetkin_1(pf00871) GRM3B
01125 HMM03928Haemd_2_grm(pf03928) GRM3B
01130 P_stone GRM3B
01135 HMM02441_09(#) GRM3B EP_CTERM
01145 HMMptac_3_eut2(pf06130) GRM3B
01150 HMM00465alcdh_3_mix(pf00465) GRM3B
01155 -pf01042 -
01160 HMMreg00126_1_mix(pf00126pf03466) GRM3B
01165 HMMnopfamclust3_1(pf00106) GRM3B
01170 -pf00710 -
01175 -# -
01180 -pf01926pf06071 -
01185 -pf01195 -
BMC-P
BMC-H
BMC-T
Aldehyde dehydrogenase
PTAC
Signature enzyme
Alcohol dehydrogenase
Regulator

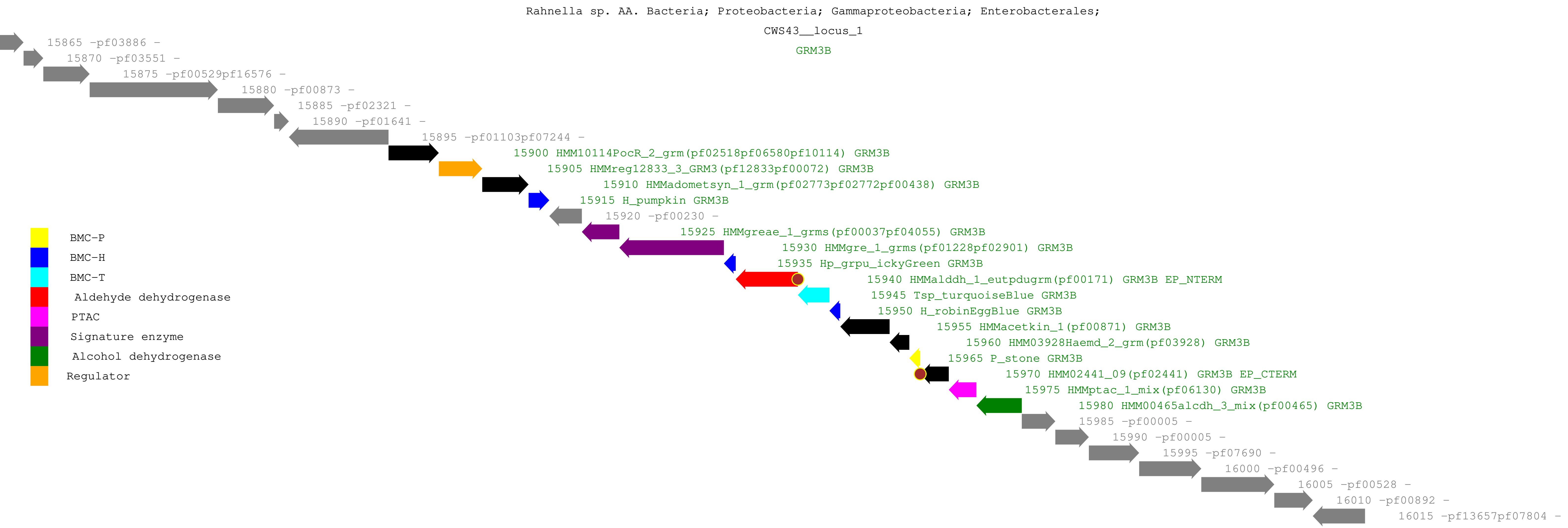
Rahnella sp. AA. Bacteria; Proteobacteria; Gammaproteobacteria; Enterobacterales;
CWS43__locus_1
GRM3B
15865 -pf03886 -
15870 -pf03551 -
15875 -pf00529pf16576 -
15880 -pf00873 -
15885 -pf02321 -
15890 -pf01641 -
15895 -pf01103pf07244 -
15900 HMM10114PocR_2_grm(pf02518pf06580pf10114) GRM3B
15905 HMMreg12833_3_GRM3(pf12833pf00072) GRM3B
15910 HMMadometsyn_1_grm(pf02773pf02772pf00438) GRM3B
15915 H_pumpkin GRM3B
15920 -pf00230 -
15925 HMMgreae_1_grms(pf00037pf04055) GRM3B
15930 HMMgre_1_grms(pf01228pf02901) GRM3B
15935 Hp_grpu_ickyGreen GRM3B
15940 HMMalddh_1_eutpdugrm(pf00171) GRM3B EP_NTERM
15945 Tsp_turquoiseBlue GRM3B
15950 H_robinEggBlue GRM3B
15955 HMMacetkin_1(pf00871) GRM3B
15960 HMM03928Haemd_2_grm(pf03928) GRM3B
15965 P_stone GRM3B
15970 HMM02441_09(pf02441) GRM3B EP_CTERM
15975 HMMptac_1_mix(pf06130) GRM3B
15980 HMM00465alcdh_3_mix(pf00465) GRM3B
15985 -pf00005 -
15990 -pf00005 -
15995 -pf07690 -
16000 -pf00496 -
16005 -pf00528 -
16010 -pf00892 -
16015 -pf13657pf07804 -
BMC-P
BMC-H
BMC-T
Aldehyde dehydrogenase
PTAC
Signature enzyme
Alcohol dehydrogenase
Regulator

Buttiauxella sp. 3AFRM03. Bacteria; Proteobacteria; Gammaproteobacteria; Enterobacterales;

D8682__locus_1

GRM3B

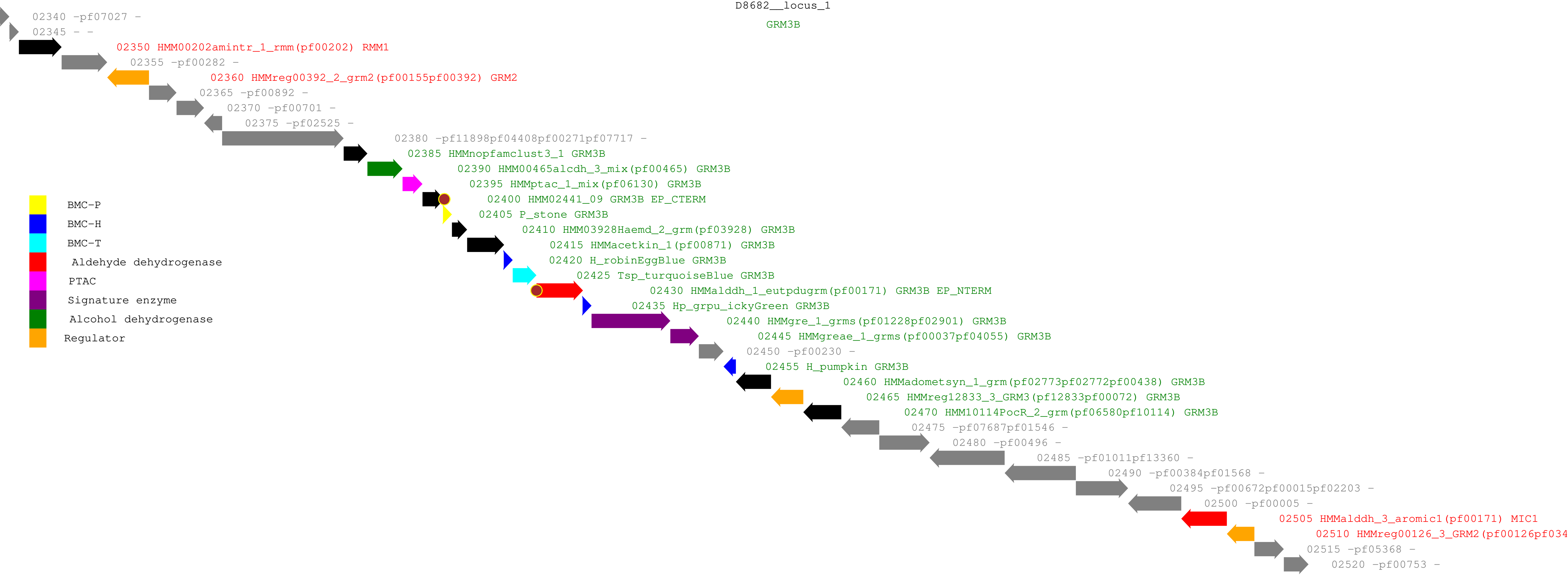

Enterobacteriaceae bacterium. Bacteria; Proteobacteria; Gammaproteobacteria; Enterobacterales;

DCM53__locus_1

GRM3B

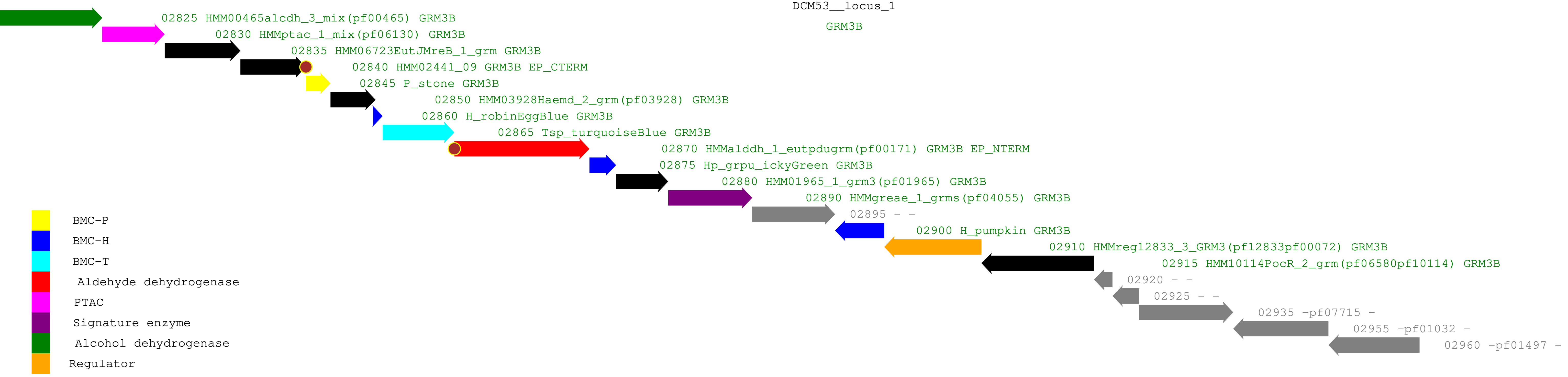

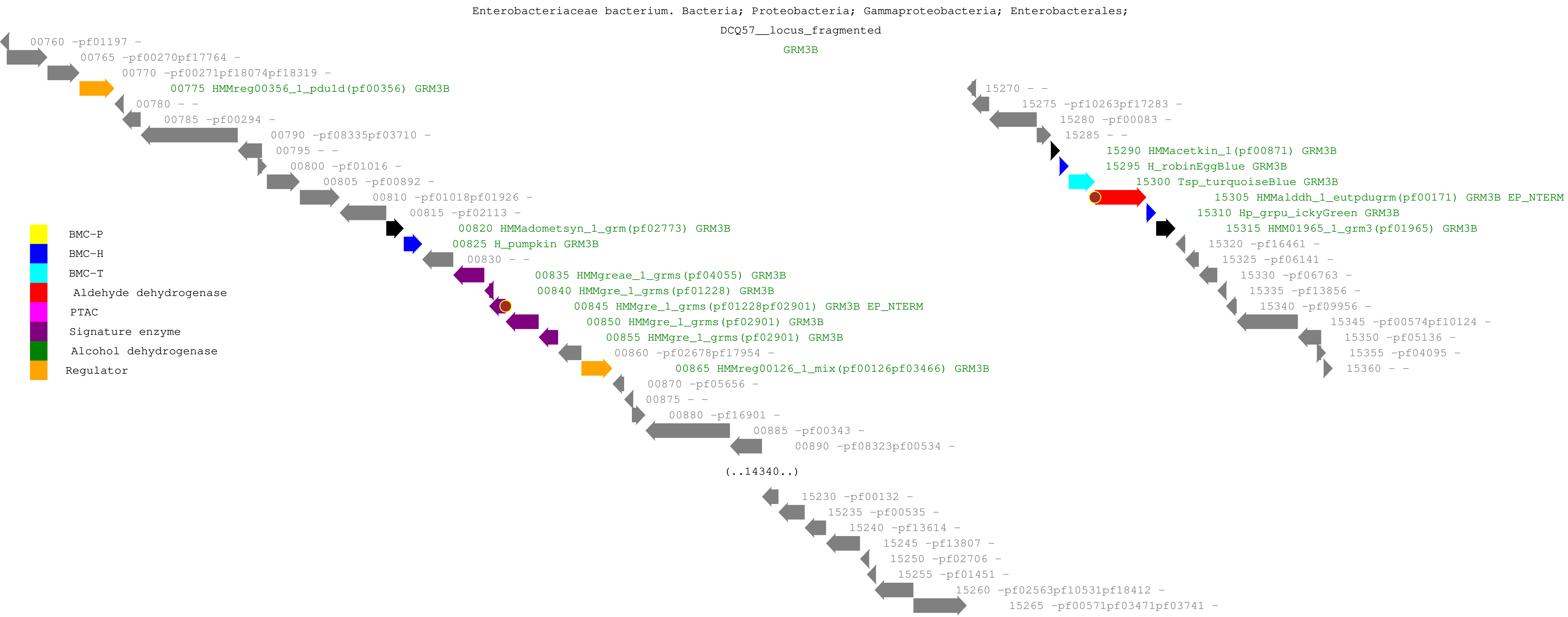

Enterobacteriaceae bacterium. Bacteria; Proteobacteria; Gammaproteobacteria; Enterobacterales;
DCQ57__locus_fragmented
GRM3B
00760 -pf01197 -
00765 -pf00270pf17764 -
00770 -pf00271pf18074pf18319 -
00775 HMMreg00356_1_pdu1d(pf00356) GRM3B
00780 - -
00785 -pf00294 -
00790 -pf08335pf03710 -
00795 - -
00800 -pf01016 -
00805 -pf00892 -
00810 -pf01018pf01926 -
00815 -pf02113 -
00820 HMMadometsyn_1_grm(pf02773) GRM3B
00825 H_pumpkin GRM3B
00830 - -
00835 HMMgreae_1_grms(pf04055) GRM3B
00840 HMMgre_1_grms(pf01228) GRM3B
00845 HMMgre_1_grms(pf01228pf02901) GRM3B EP_NTERM
00850 HMMgre_1_grms(pf02901) GRM3B
00855 HMMgre_1_grms(pf02901) GRM3B
00860 -pf02678pf17954 -
00865 HMMreg00126_1_mix(pf00126pf03466) GRM3B
00870 -pf05656 -
00875 - -
00880 -pf16901 -
00885 -pf00343 -
00890 -pf08323pf00534 -
(..14340..)
15230 -pf00132 -
15235 -pf00535 -
15240 -pf13614 -
15245 -pf13807 -
15250 -pf02706 -
15255 -pf01451 -
15260 -pf02563pf10531pf18412 -
15265 -pf00571pf03471pf03741 -
15270 - -
15275 -pf10263pf17283 -
15280 -pf00083 -
15285 - -
15290 HMMacetkin_1(pf00871) GRM3B
15295 H_robinEggBlue GRM3B
15300 Tsp_turquoiseBlue GRM3B
15305 HMMalddh_1_eutpdugrm(pf00171) GRM3B EP_NTERM
15310 Hp_grpu_ickyGreen GRM3B
15315 HMM01965_1_grm3(pf01965) GRM3B
15320 -pf16461 -
15325 -pf06141 -
15330 -pf06763 -
15335 -pf13856 -
15340 -pf09956 -
15345 -pf00574pf10124 -
15350 -pf05136 -
15355 -pf04095 -
15360 - -
BMC-P
BMC-H
BMC-T
Aldehyde dehydrogenase
PTAC
Signature enzyme
Alcohol dehydrogenase
Regulator

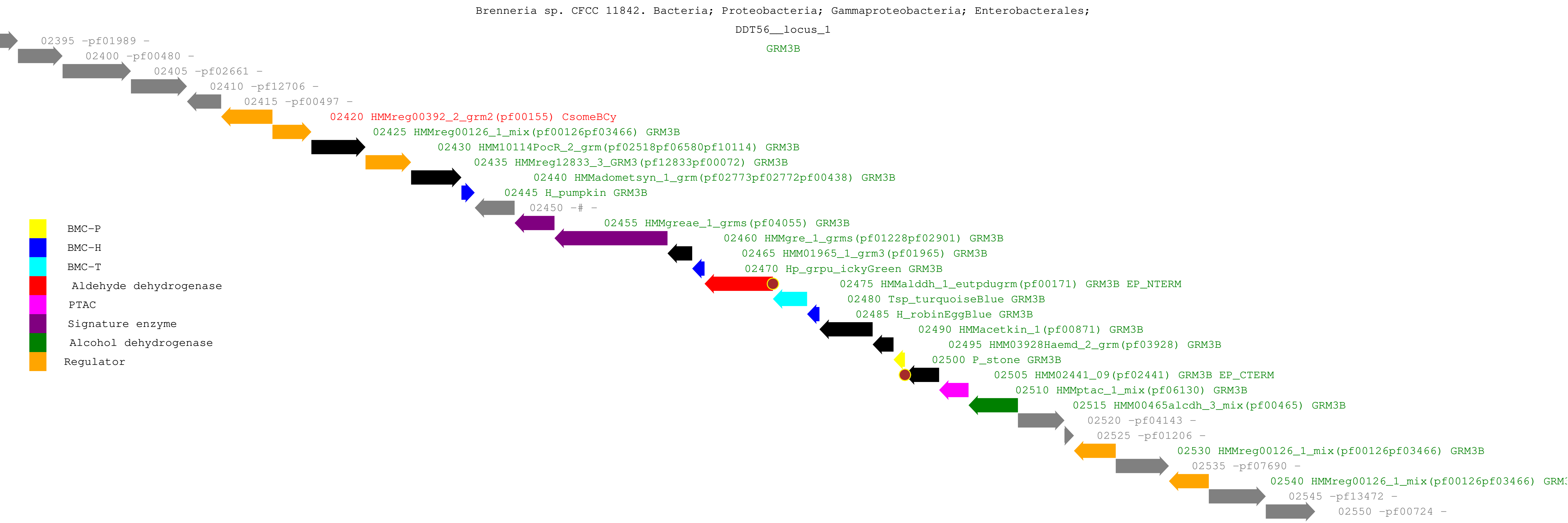

Brenneria sp. CFCC 11842. Bacteria; Proteobacteria; Gammaproteobacteria; Enterobacterales;
DDT56__locus_1
GRM3B
02395 -pf01989 -
02400 -pf00480 -
02405 -pf02661 -
02410 -pf12706 -
02415 -pf00497 -
02420 HMMreg00392_2_grm2(pf00155) CsomeBCy
02425 HMMreg00126_1_mix(pf00126pf03466) GRM3B
02430 HMM10114PocR_2_grm(pf02518pf06580pf10114) GRM3B
02435 HMMreg12833_3_GRM3(pf12833pf00072) GRM3B
02440 HMMadometsyn_1_grm(pf02773pf02772pf00438) GRM3B
02445 H_pumpkin GRM3B
02450 -# -
02455 HMMgreae_1_grms(pf04055) GRM3B
02460 HMMgre_1_grms(pf01228pf02901) GRM3B
02465 HMM01965_1_grm3(pf01965) GRM3B
02470 Hp_grpu_ickyGreen GRM3B
02475 HMMalddh_1_eutpdugrm(pf00171) GRM3B EP_NTERM
02480 Tsp_turquoiseBlue GRM3B
02485 H_robinEggBlue GRM3B
02490 HMMacetkin_1(pf00871) GRM3B
02495 HMM03928Haemd_2_grm(pf03928) GRM3B
02500 P_stone GRM3B
02505 HMM02441_09(pf02441) GRM3B EP_CTERM
02510 HMMptac_1_mix(pf06130) GRM3B
02515 HMM00465alcdh_3_mix(pf00465) GRM3B
02520 -pf04143 -
02525 -pf01206 -
02530 HMMreg00126_1_mix(pf00126pf03466) GRM3B
02535 -pf07690 -
02540 HMMreg00126_1_mix(pf00126pf03466) GRM3
02545 -pf13472 -
02550 -pf00724 -
BMC-P
BMC-H
BMC-T
Aldehyde dehydrogenase
PTAC
Signature enzyme
Alcohol dehydrogenase
Regulator

Raoultella planticola (Klebsiella planticola). Bacteria; Proteobacteria; Gammaproteobacteria; Enterobacterales;

DN603__locus_2

GRM3B

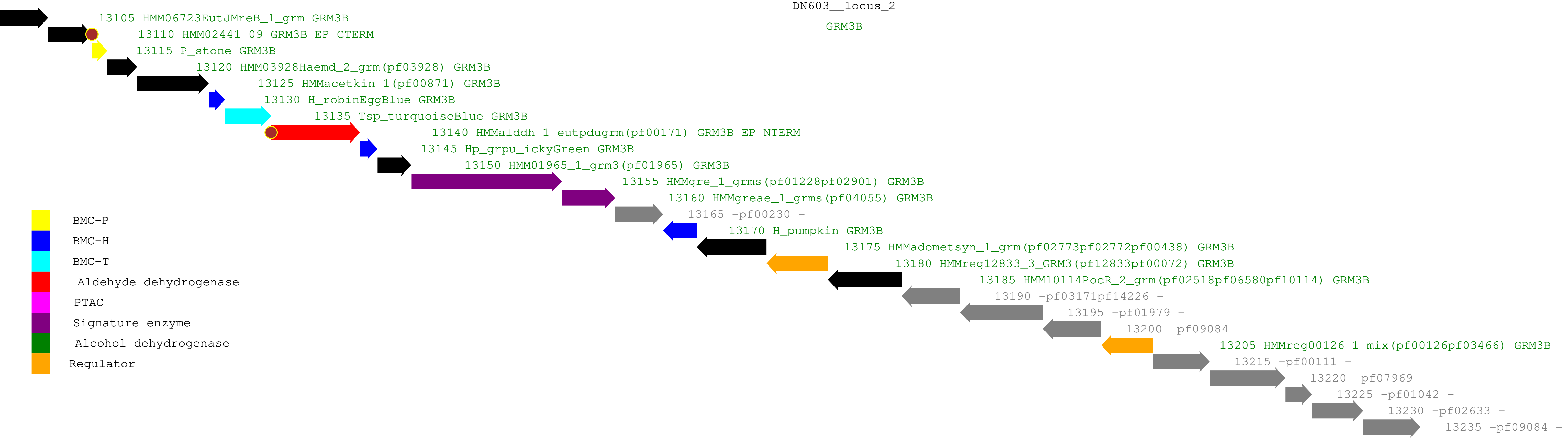

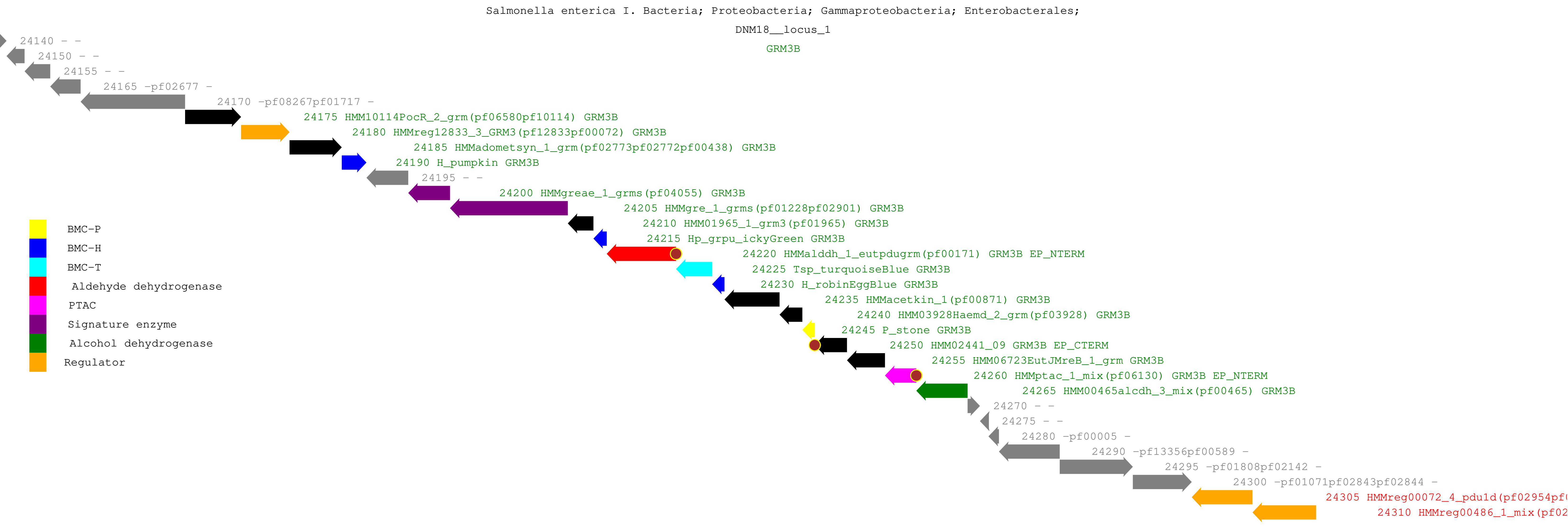
Salmonella enterica I. Bacteria; Proteobacteria; Gammaproteobacteria; Enterobacterales;
DNM18__locus_1
GRM3B
24140 - -
24150 - -
24155 - -
24165 -pf02677 -
24170 -pf08267pf01717 -
24175 HMM10114PocR_2_grm(pf06580pf10114) GRM3B
24180 HMMreg12833_3_GRM3(pf12833pf00072) GRM3B
24185 HMMadometsyn_1_grm(pf02773pf02772pf00438) GRM3B
24190 H_pumpkin GRM3B
24195 - -
24200 HMMgreae_1_grms(pf04055) GRM3B
24205 HMMgre_1_grms(pf01228pf02901) GRM3B
24210 HMM01965_1_grm3(pf01965) GRM3B
24215 Hp_grpu_ickyGreen GRM3B
24220 HMMalddh_1_eutpdugrm(pf00171) GRM3B EP_NTERM
24225 Tsp_turquoiseBlue GRM3B
24230 H_robinEggBlue GRM3B
24235 HMMacetkin_1(pf00871) GRM3B
24240 HMM03928Haemd_2_grm(pf03928) GRM3B
24245 P_stone GRM3B
24250 HMM02441_09 GRM3B EP_CTERM
24255 HMM06723EutJMreB_1_grm GRM3B
24260 HMMptac_1_mix(pf06130) GRM3B EP_NTERM
24265 HMM00465alcdh_3_mix(pf00465) GRM3B
24270 - -
24275 - -
24280 -pf00005 -
24290 -pf13356pf00589 -
24295 -pf01808pf02142 -
24300 -pf01071pf02843pf02844 -
24305 HMMreg00072_4_pdu1d(pf02954pf
24310 HMMreg00486_1_mix(pf02
BMC-P
BMC-H
BMC-T
Aldehyde dehydrogenase
PTAC
Signature enzyme
Alcohol dehydrogenase
Regulator

Escherichia coli. Bacteria; Proteobacteria; Gammaproteobacteria; Enterobacterales;

DNQ41__locus_1

GRM3B

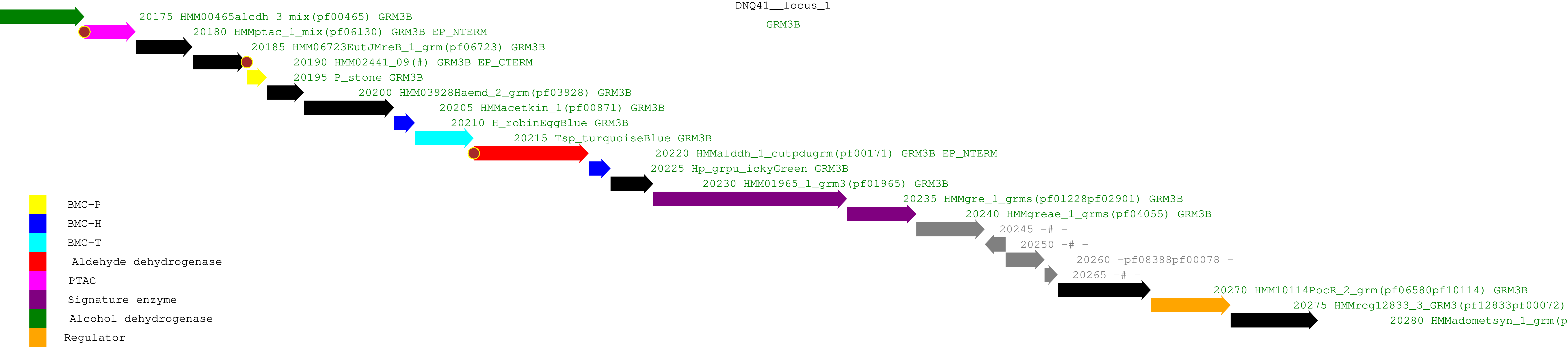

Rahnella sp. JUb53. Bacteria; Proteobacteria; Gammaproteobacteria; Enterobacterales;

EC840__locus_1

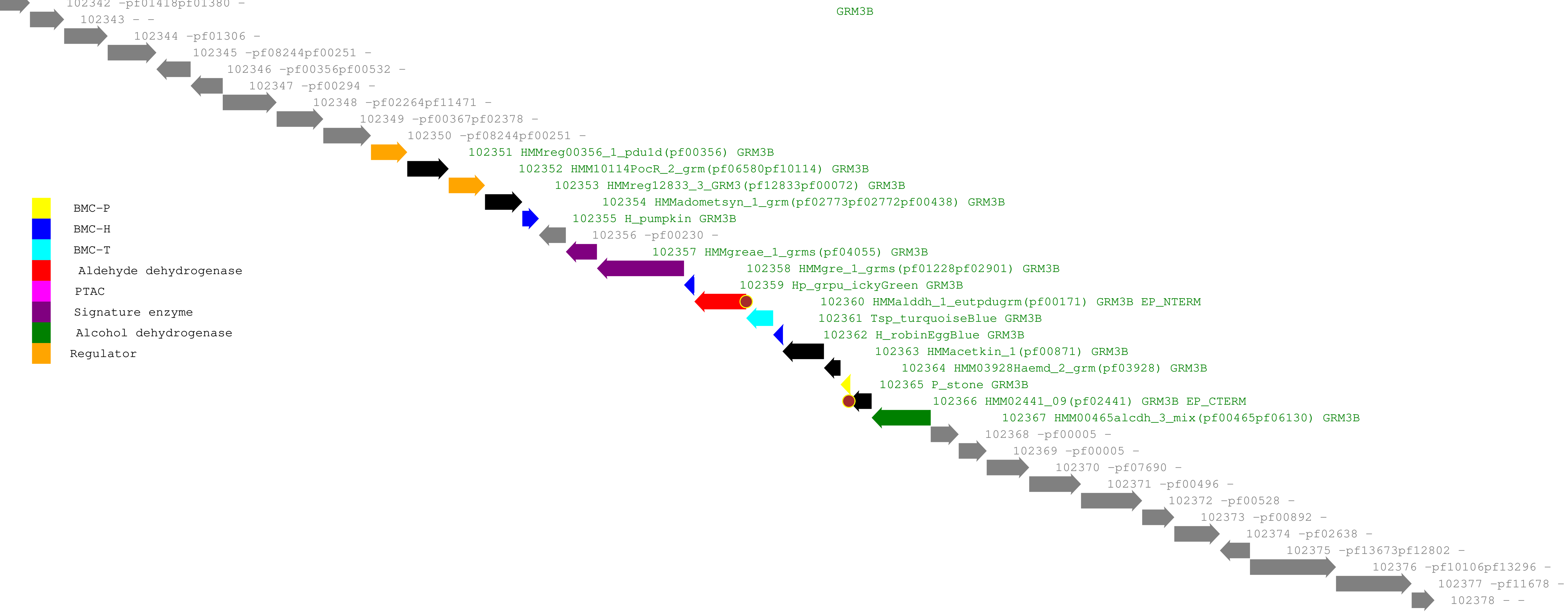

Raoultella ornithinolytica (Klebsiella ornithinolytica). Bacteria; Proteobacteria; Gammaproteobacteria; Enterobacterales;

EC841__locus_1

GRM3B

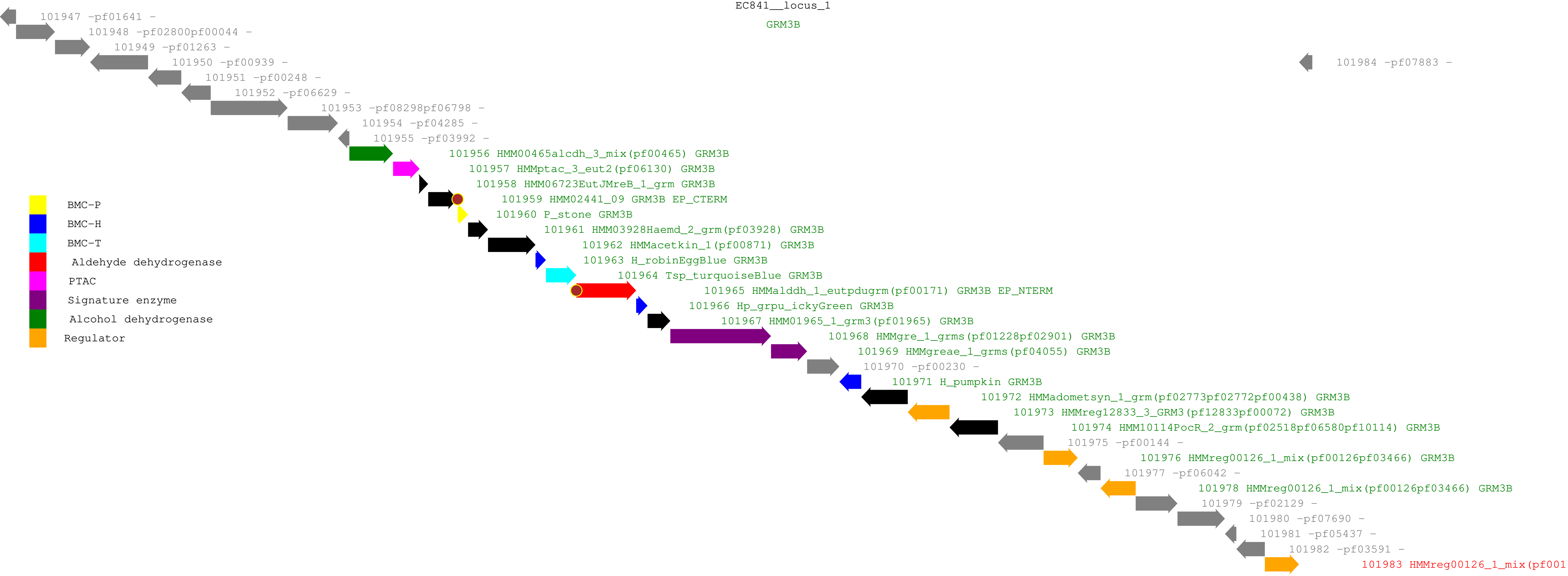

Buttiauxella sp. JUb87. Bacteria; Proteobacteria; Gammaproteobacteria; Enterobacterales;

EC843__locus_2

GRM3B

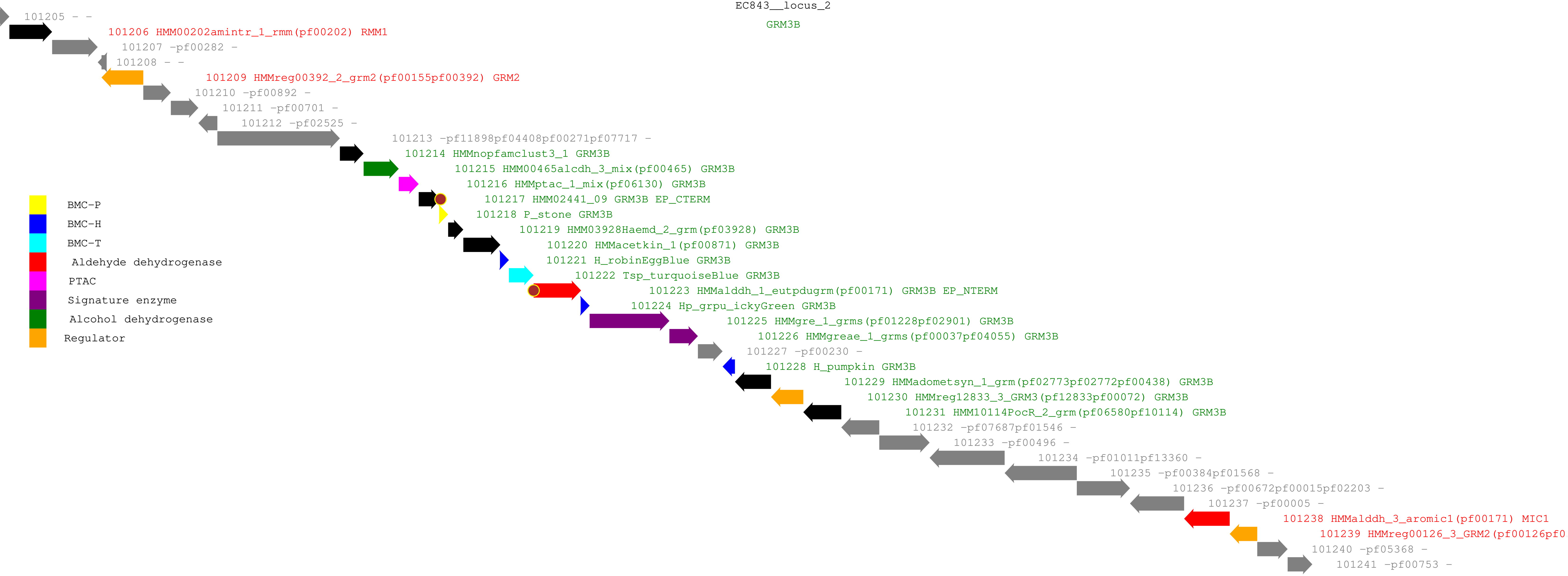

Enterobacter sp. BIGb0359. Bacteria; Proteobacteria; Gammaproteobacteria; Enterobacterales;

EC848__locus_1

GRM3B

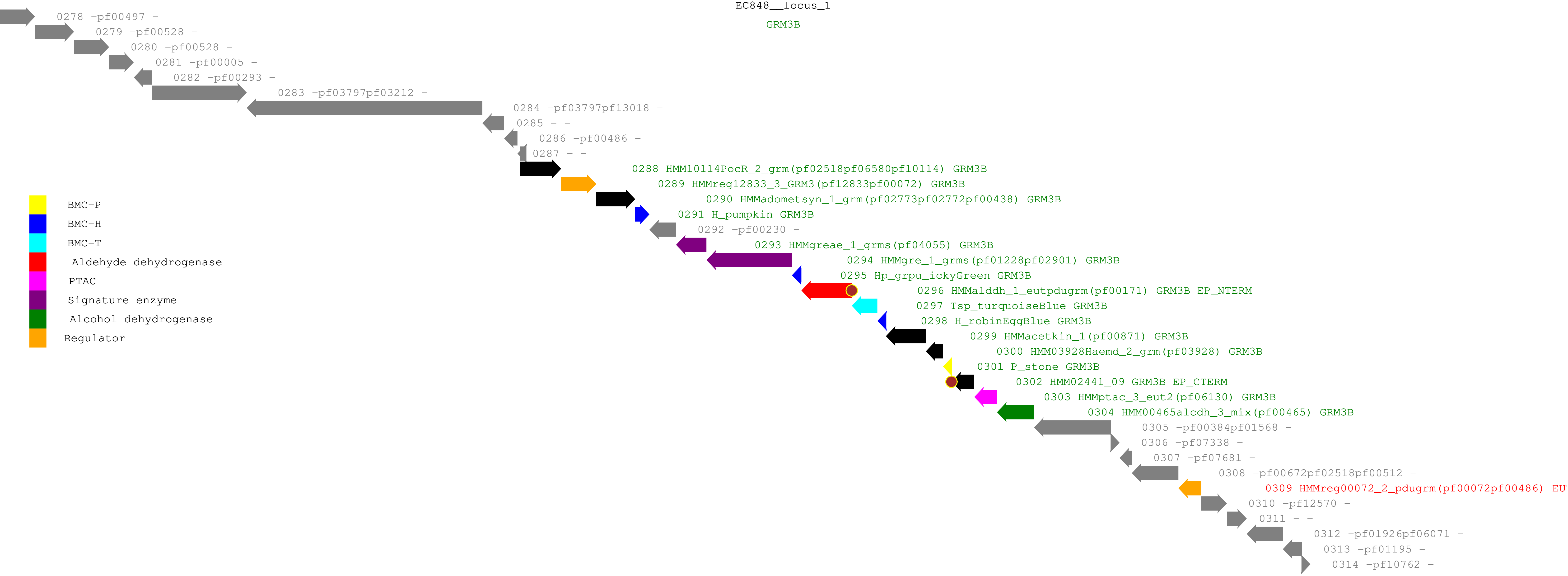

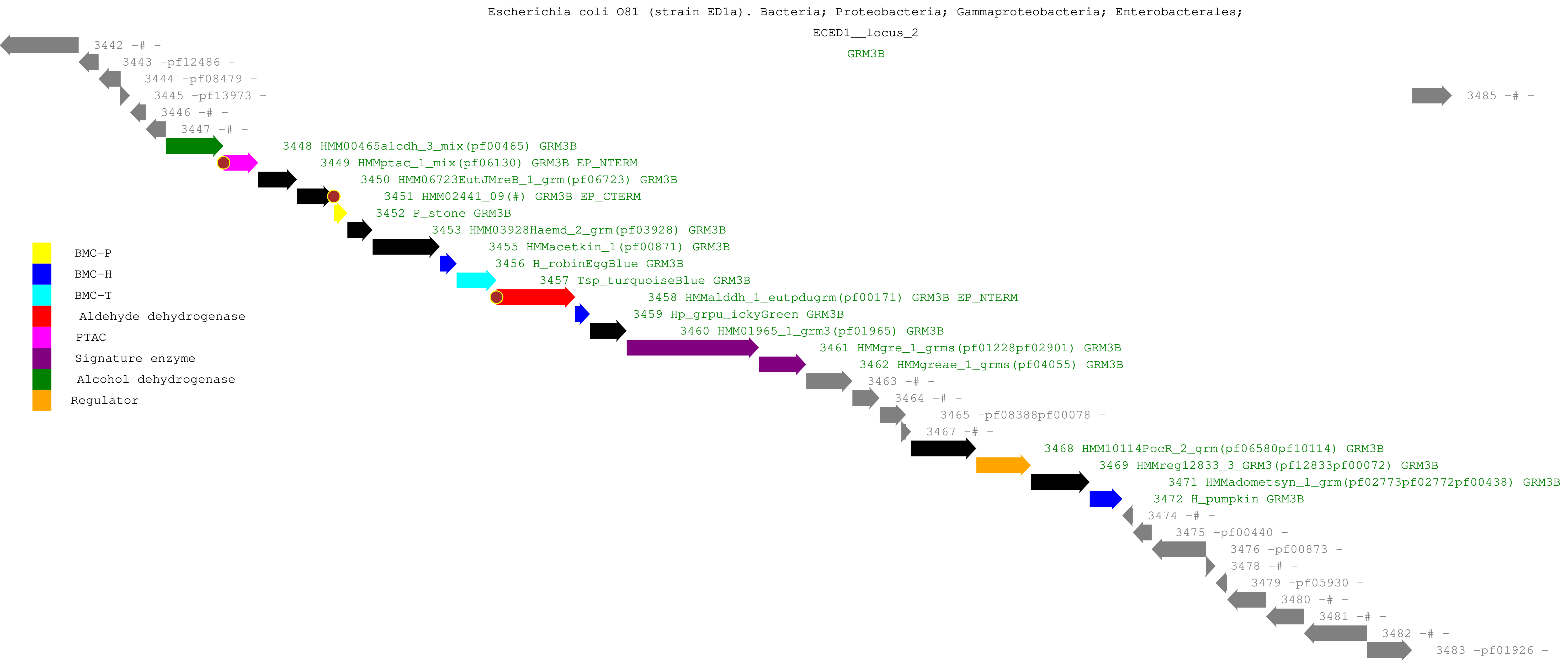

Escherichia coli O81 (strain ED1a). Bacteria; Proteobacteria; Gammaproteobacteria; Enterobacterales;
ECED1__locus_2
GRM3B
3442 -# -
3443 -pf12486 -
3444 -pf08479 -
3445 -pf13973 -
3446 -# -
3447 -# -
3448 HMM00465alcdh_3_mix(pf00465) GRM3B
3449 HMMptac_1_mix(pf06130) GRM3B EP_NTERM
3450 HMM06723EutJMreB_1_grm(pf06723) GRM3B
3451 HMM02441_09(#) GRM3B EP_CTERM
3452 P_stone GRM3B
3453 HMM03928Haemd_2_grm(pf03928) GRM3B
3455 HMMacetkin_1(pf00871) GRM3B
3456 H_robinEggBlue GRM3B
3457 Tsp_turquoiseBlue GRM3B
3458 HMMalddh_1_eutpdugrm(pf00171) GRM3B EP_NTERM
3459 Hp_grpu_ickyGreen GRM3B
3460 HMM01965_1_grm3(pf01965) GRM3B
3461 HMMgre_1_grms(pf01228pf02901) GRM3B
3462 HMMgreae_1_grms(pf04055) GRM3B
3463 -# -
3464 -# -
3465 -pf08388pf00078 -
3467 -# -
3468 HMM10114PocR_2_grm(pf06580pf10114) GRM3B
3469 HMMreg12833_3_GRM3(pf12833pf00072) GRM3B
3471 HMMadometsyn_1_grm(pf02773pf02772pf00438) GRM3B
3472 H_pumpkin GRM3B
3474 -# -
3475 -pf00440 -
3476 -pf00873 -
3478 -# -
3479 -pf05930 -
3480 -# -
3481 -# -
3482 -# -
3483 -pf01926 -
3485 -# -
BMC-P
BMC-H
BMC-T
Aldehyde dehydrogenase
PTAC
Signature enzyme
Alcohol dehydrogenase
Regulator

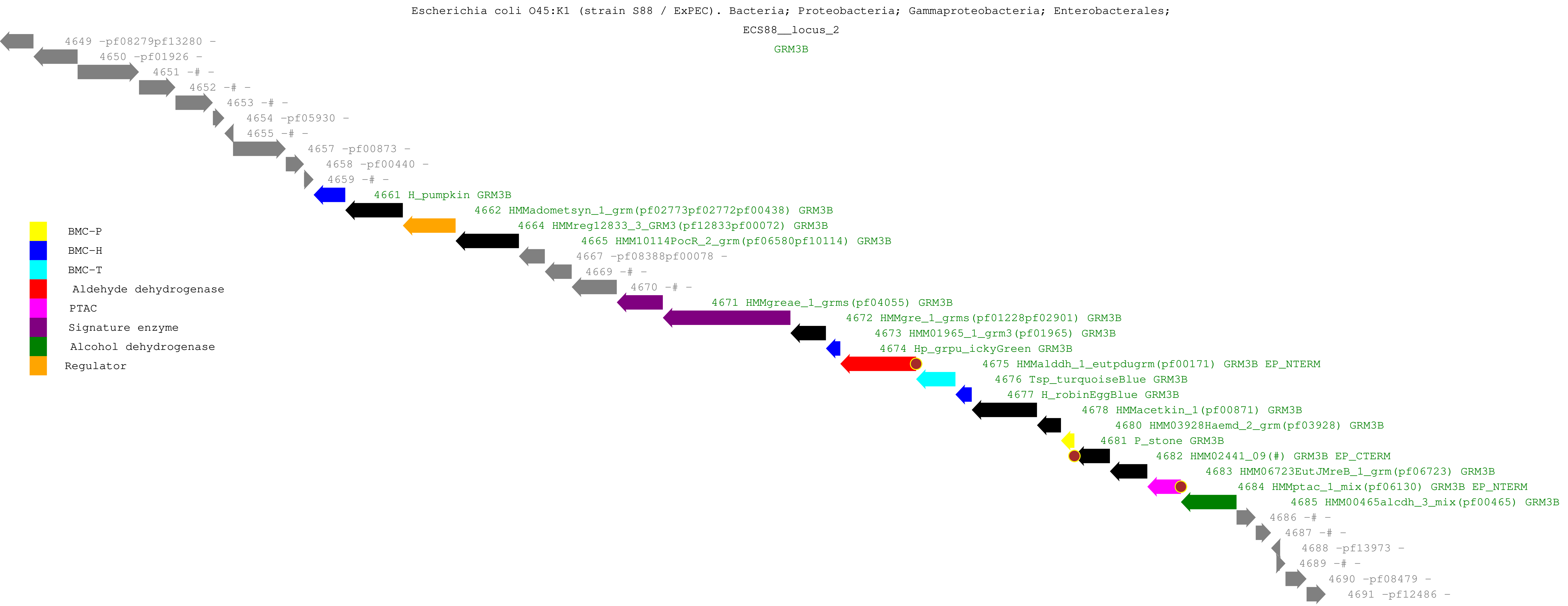
Escherichia coli O45:K1 (strain S88 / ExPEC). Bacteria; Proteobacteria; Gammaproteobacteria; Enterobacterales;
ECS88__locus_2
GRM3B
4649 -pf08279pf13280 -
4650 -pf01926 -
4651 -# -
4652 -# -
4653 -# -
4654 -pf05930 -
4655 -# -
4657 -pf00873 -
4658 -pf00440 -
4659 -# -
4661 H_pumpkin GRM3B
4662 HMMadometsyn_1_grm(pf02773pf02772pf00438) GRM3B
4664 HMMreg12833_3_GRM3(pf12833pf00072) GRM3B
4665 HMM10114PocR_2_grm(pf06580pf10114) GRM3B
4667 -pf08388pf00078 -
4669 -# -
4670 -# -
4671 HMMgreae_1_grms(pf04055) GRM3B
4672 HMMgre_1_grms(pf01228pf02901) GRM3B
4673 HMM01965_1_grm3(pf01965) GRM3B
4674 Hp_grpu_ickyGreen GRM3B
4675 HMMalddh_1_eutpdugrm(pf00171) GRM3B EP_NTERM
4676 Tsp_turquoiseBlue GRM3B
4677 H_robinEggBlue GRM3B
4678 HMMacetkin_1(pf00871) GRM3B
4680 HMM03928Haemd_2_grm(pf03928) GRM3B
4681 P_stone GRM3B
4682 HMM02441_09(#) GRM3B EP_CTERM
4683 HMM06723EutJMreB_1_grm(pf06723) GRM3B
4684 HMMptac_1_mix(pf06130) GRM3B EP_NTERM
4685 HMM00465alcdh_3_mix(pf00465) GRM3B
4686 -# -
4687 -# -
4688 -pf13973 -
4689 -# -
4690 -pf08479 -
4691 -pf12486 -
BMC-P
BMC-H
BMC-T
Aldehyde dehydrogenase
PTAC
Signature enzyme
Alcohol dehydrogenase
Regulator

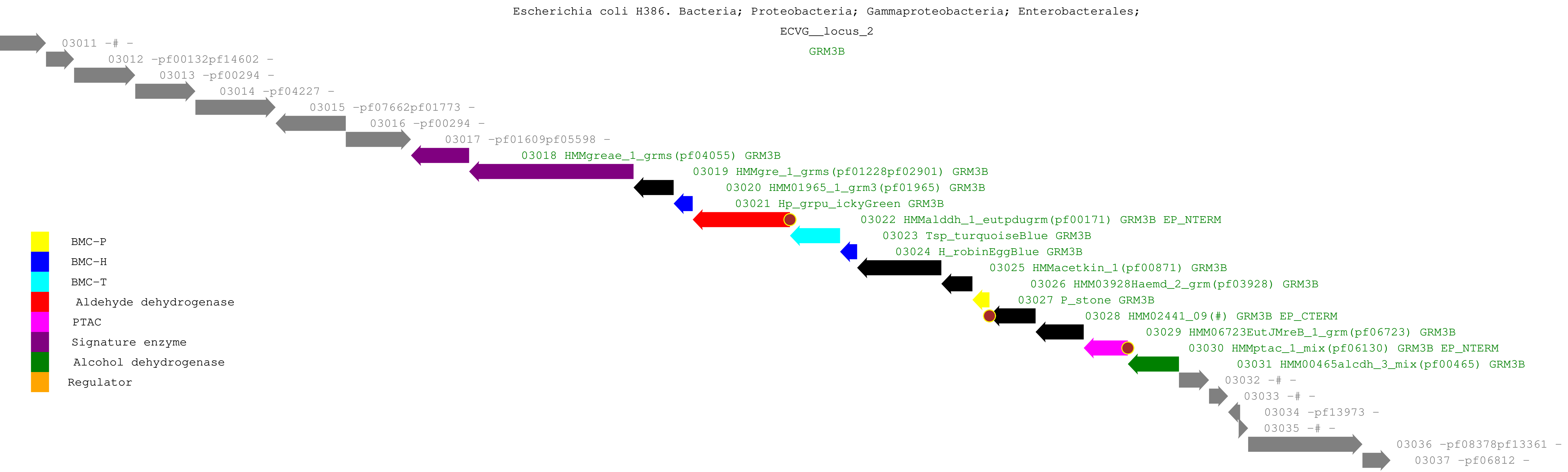
Escherichia coli H386. Bacteria; Proteobacteria; Gammaproteobacteria; Enterobacterales;
ECVG__locus_2
GRM3B
03011 -# -
03012 -pf00132pf14602 -
03013 -pf00294 -
03014 -pf04227 -
03015 -pf07662pf01773 -
03016 -pf00294 -
03017 -pf01609pf05598 -
03018 HMMgreae_1_grms(pf04055) GRM3B
03019 HMMgre_1_grms(pf01228pf02901) GRM3B
03020 HMM01965_1_grm3(pf01965) GRM3B
03021 Hp_grpu_ickyGreen GRM3B
03022 HMMalddh_1_eutpdugrm(pf00171) GRM3B EP_NTERM
03023 Tsp_turquoiseBlue GRM3B
03024 H_robinEggBlue GRM3B
03025 HMMacetkin_1(pf00871) GRM3B
03026 HMM03928Haemd_2_grm(pf03928) GRM3B
03027 P_stone GRM3B
03028 HMM02441_09(#) GRM3B EP_CTERM
03029 HMM06723EutJMreB_1_grm(pf06723) GRM3B
03030 HMMptac_1_mix(pf06130) GRM3B EP_NTERM
03031 HMM00465alcdh_3_mix(pf00465) GRM3B
03032 -# -
03033 -# -
03034 -pf13973 -
03035 -# -
03036 -pf08378pf13361 -
03037 -pf06812 -
BMC-P
BMC-H
BMC-T
Aldehyde dehydrogenase
PTAC
Signature enzyme
Alcohol dehydrogenase
Regulator

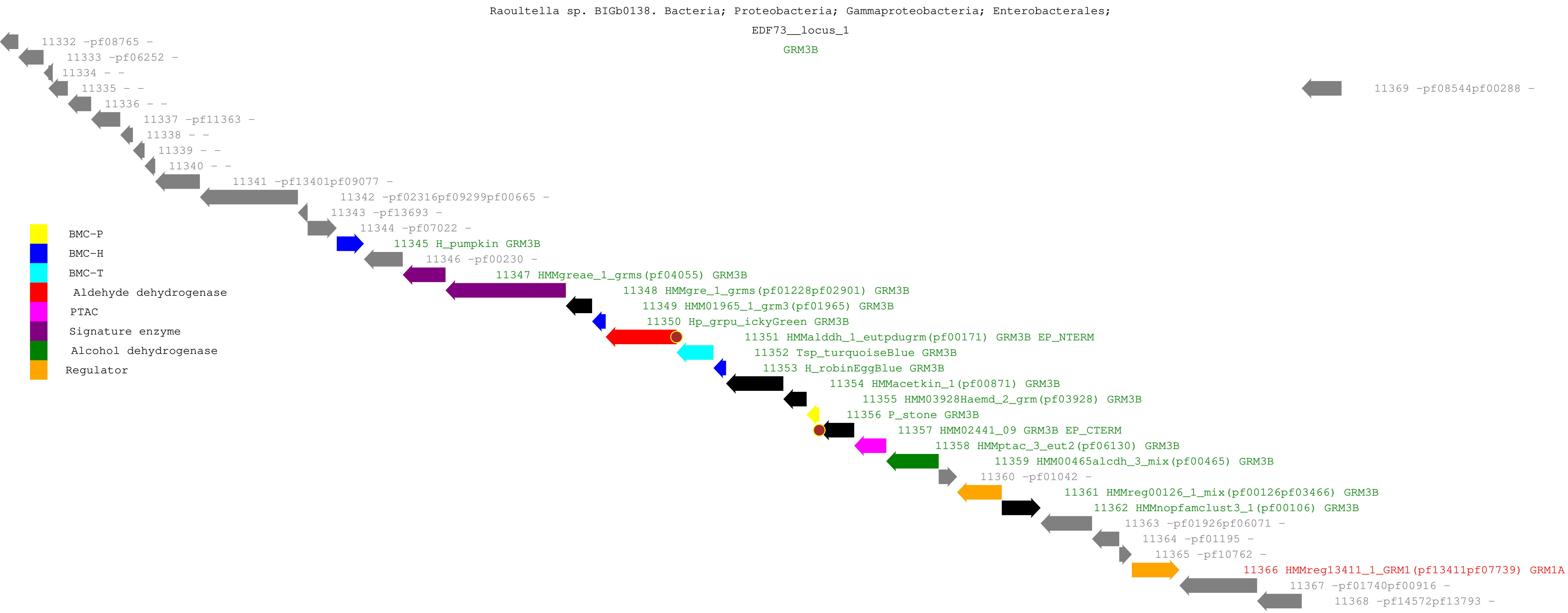
Raoultella sp. BIGb0138. Bacteria; Proteobacteria; Gammaproteobacteria; Enterobacterales;
EDF73__locus_1
GRM3B
11332 -pf08765 -
11333 -pf06252 -
11334 - -
11335 - -
11336 - -
11337 -pf11363 -
11338 - -
11339 - -
11340 - -
11341 -pf13401pf09077 -
11342 -pf02316pf09299pf00665 -
11343 -pf13693 -
11344 -pf07022 -
11345 H_pumpkin GRM3B
11346 -pf00230 -
11347 HMMgreae_1_grms(pf04055) GRM3B
11348 HMMgre_1_grms(pf01228pf02901) GRM3B
11349 HMM01965_1_grm3(pf01965) GRM3B
11350 Hp_grpu_ickyGreen GRM3B
11351 HMMalddh_1_eutpdugrm(pf00171) GRM3B EP_NTERM
11352 Tsp_turquoiseBlue GRM3B
11353 H_robinEggBlue GRM3B
11354 HMMacetkin_1(pf00871) GRM3B
11355 HMM03928Haemd_2_grm(pf03928) GRM3B
11356 P_stone GRM3B
11357 HMM02441_09 GRM3B EP_CTERM
11358 HMMptac_3_eut2(pf06130) GRM3B
11359 HMM00465alcdh_3_mix(pf00465) GRM3B
11360 -pf01042 -
11361 HMMreg00126_1_mix(pf00126pf03466) GRM3B
11362 HMMnopfamclust3_1(pf00106) GRM3B
11363 -pf01926pf06071 -
11364 -pf01195 -
11365 -pf10762 -
11366 HMMreg13411_1_GRM1(pf13411pf07739) GRM1A
11367 -pf01740pf00916 -
11368 -pf14572pf13793 -
11369 -pf08544pf00288 -
BMC-P
BMC-H
BMC-T
Aldehyde dehydrogenase
PTAC
Signature enzyme
Alcohol dehydrogenase
Regulator

Raoultella terrigena (Klebsiella terrigena). Bacteria; Proteobacteria; Gammaproteobacteria; Enterobacterales;

EDF76__locus_1

GRM3B

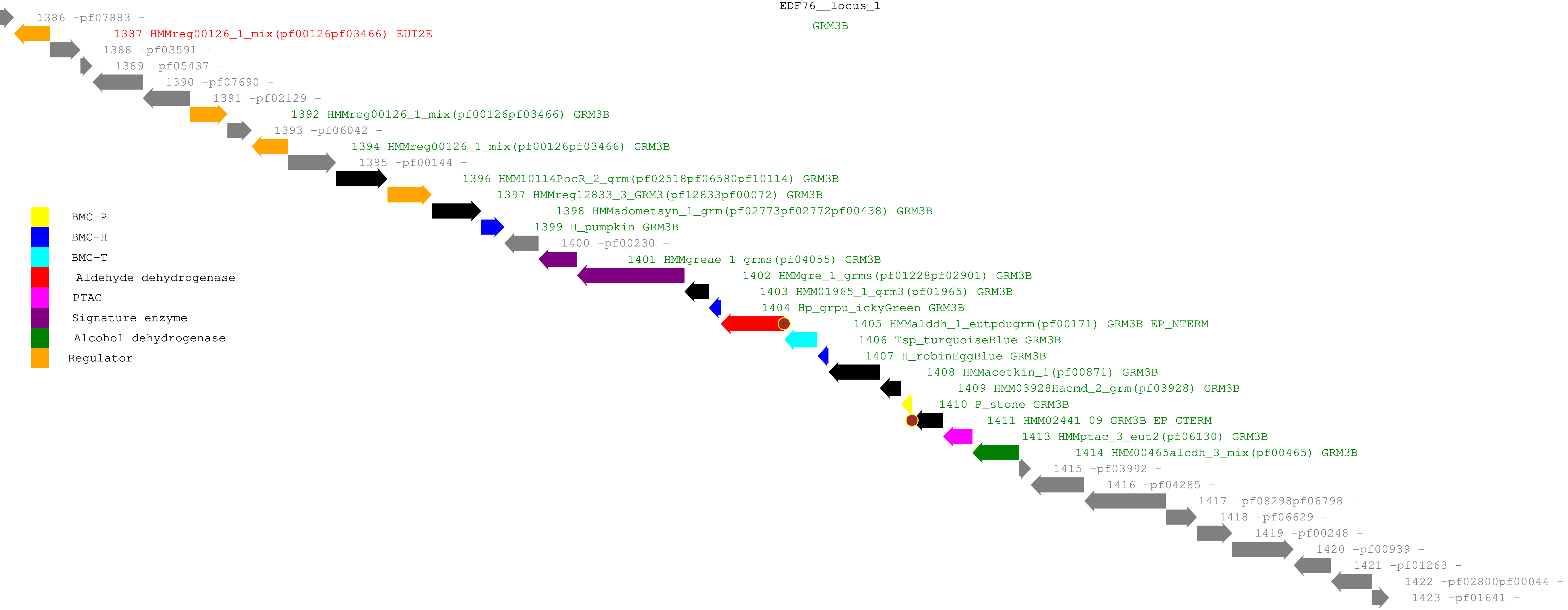

Rahnella sp. BIGb0236. Bacteria; Proteobacteria; Gammaproteobacteria; Enterobacterales;

EDF78__locus_1

GRM3B

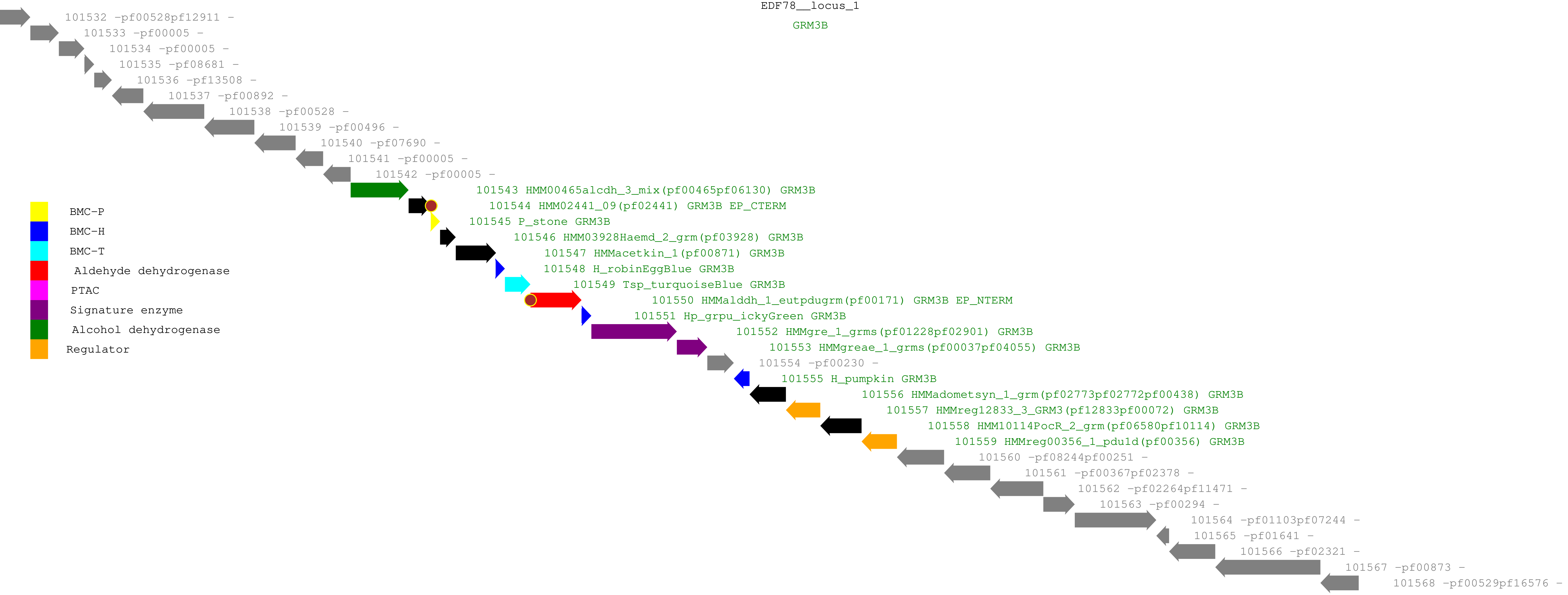

Enterobacter sp. BIGb0383. Bacteria; Proteobacteria; Gammaproteobacteria; Enterobacterales;

EDF81__locus_1

GRM3B

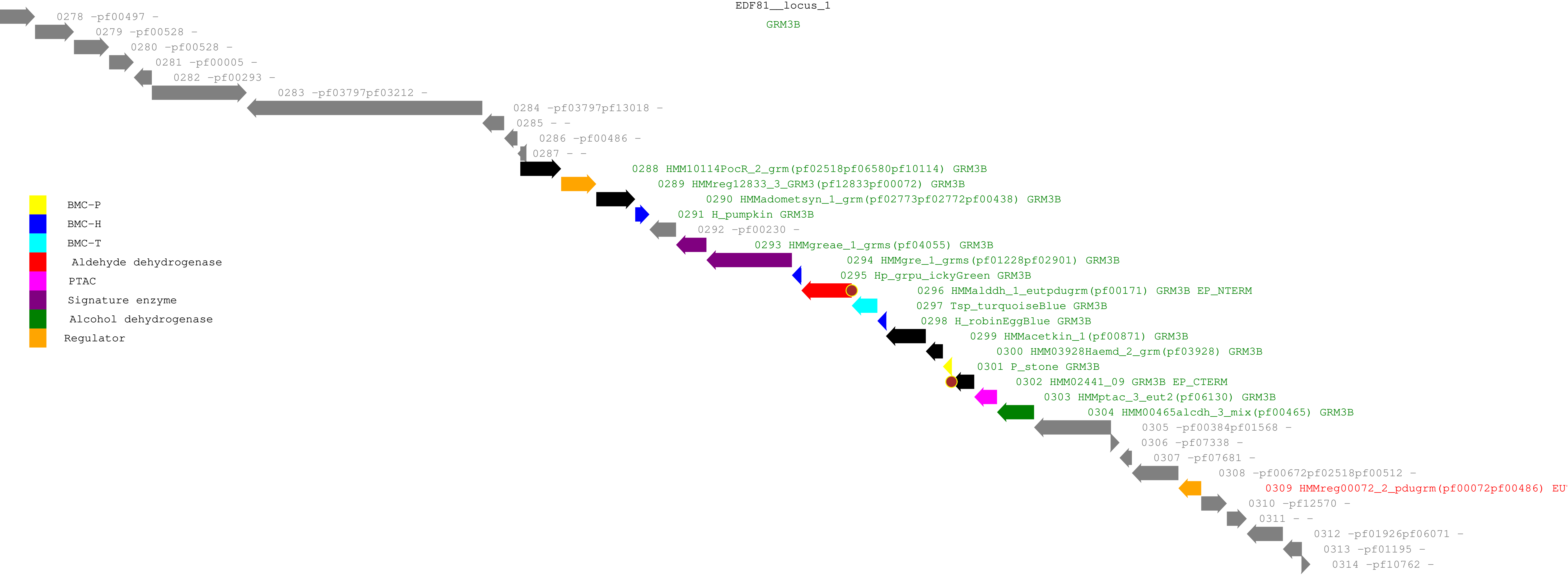

Raoultella sp. BIGb0399. Bacteria; Proteobacteria; Gammaproteobacteria; Enterobacterales;

EDF82__locus_1

GRM3B

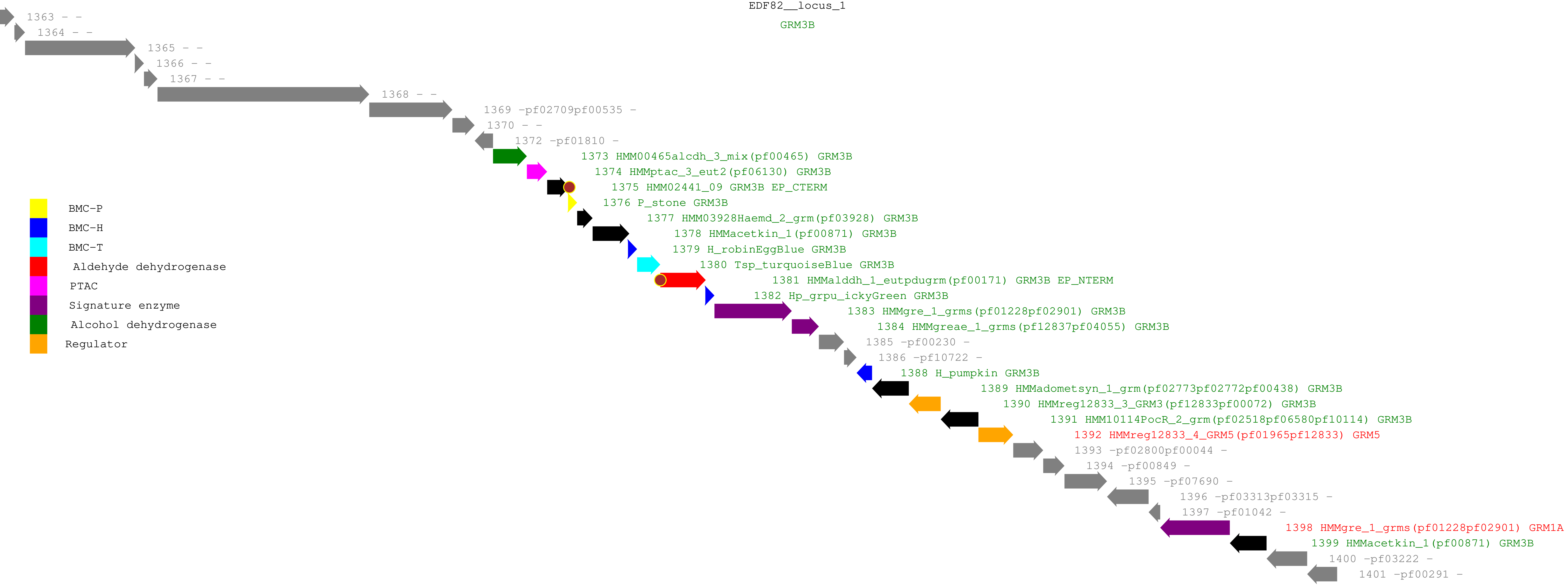

Atlantibacter subterranea. Bacteria; Proteobacteria; Gammaproteobacteria; Enterobacterales;

EGT71__locus_1

GRM3B

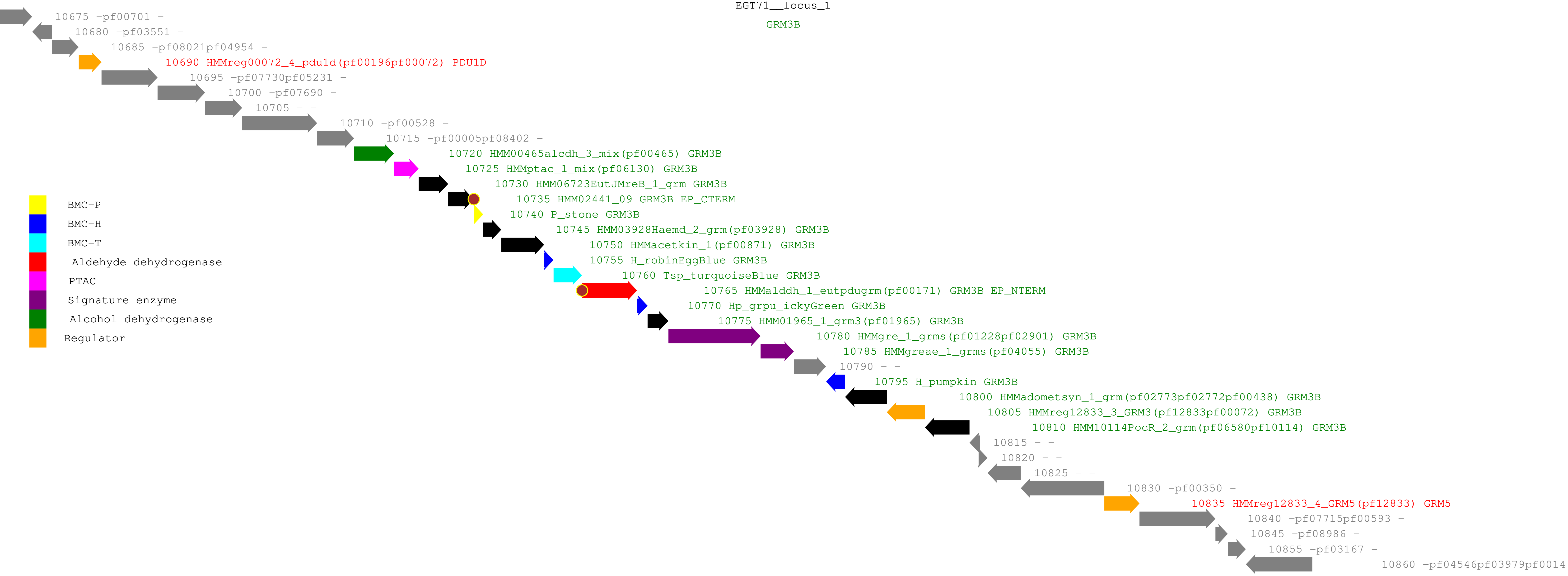

Atlantibacter hermannii NBRC 105704. Bacteria; Proteobacteria; Gammaproteobacteria; Enterobacterales;

EH105704_20__locus_1

GRM3B

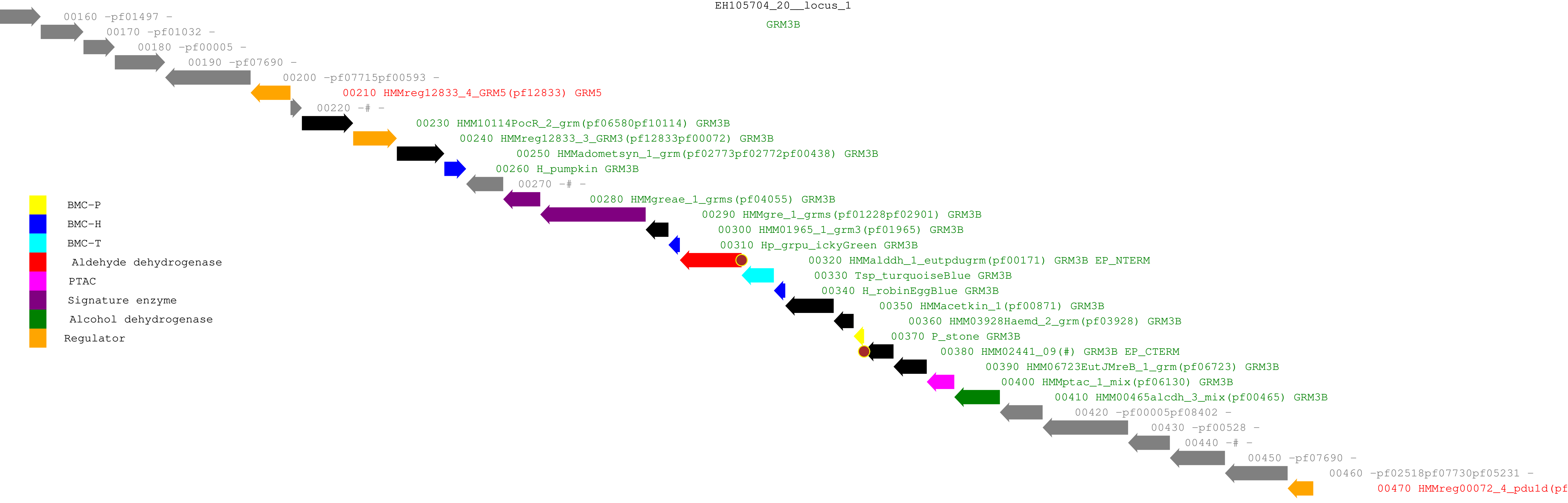

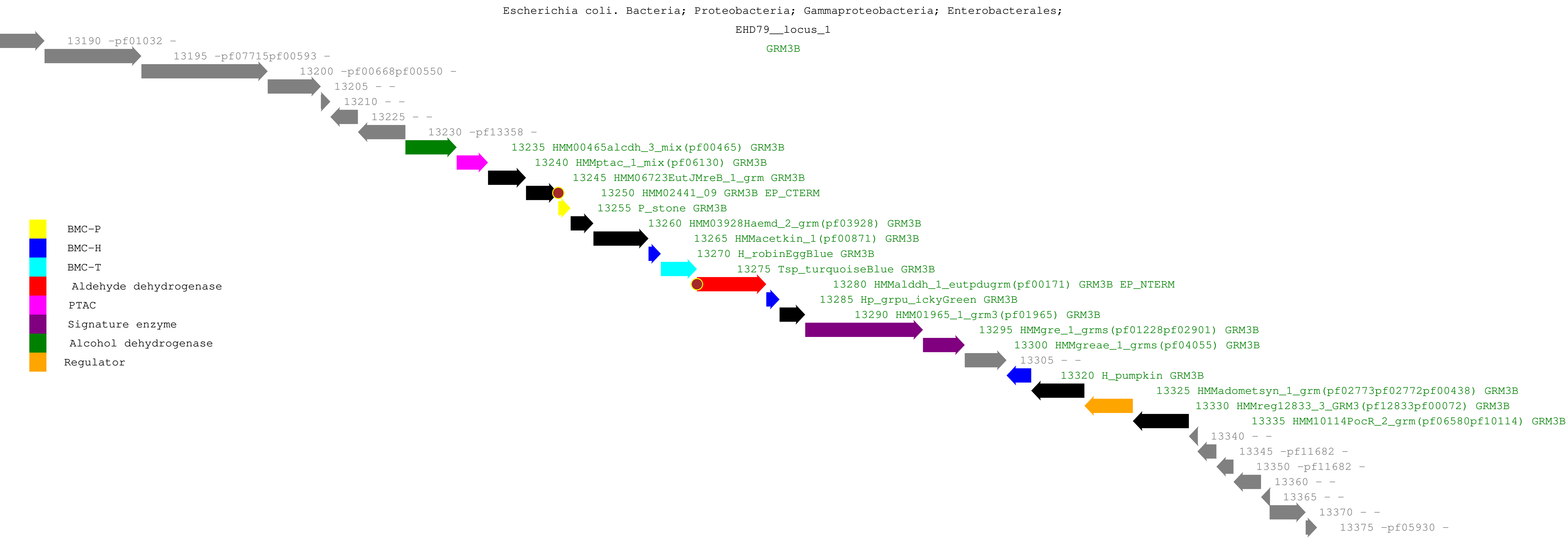
Escherichia coli. Bacteria; Proteobacteria; Gammaproteobacteria; Enterobacterales;
EHD79__locus_1
GRM3B
13190 -pf01032 -
13195 -pf07715pf00593 -
13200 -pf00668pf00550 -
13205 - -
13210 - -
13225 - -
13230 -pf13358 -
13235 HMM00465alcdh_3_mix(pf00465) GRM3B
13240 HMMptac_1_mix(pf06130) GRM3B
13245 HMM06723EutJMreB_1_grm GRM3B
13250 HMM02441_09 GRM3B EP_CTERM
13255 P_stone GRM3B
13260 HMM03928Haemd_2_grm(pf03928) GRM3B
13265 HMMacetkin_1(pf00871) GRM3B
13270 H_robinEggBlue GRM3B
13275 Tsp_turquoiseBlue GRM3B
13280 HMMalddh_1_eutpdugrm(pf00171) GRM3B EP_NTERM
13285 Hp_grpu_ickyGreen GRM3B
13290 HMM01965_1_grm3(pf01965) GRM3B
13295 HMMgre_1_grms(pf01228pf02901) GRM3B
13300 HMMgreae_1_grms(pf04055) GRM3B
13305 - -
13320 H_pumpkin GRM3B
13325 HMMadometsyn_1_grm(pf02773pf02772pf00438) GRM3B
13330 HMMreg12833_3_GRM3(pf12833pf00072) GRM3B
13335 HMM10114PocR_2_grm(pf06580pf10114) GRM3B
13340 - -
13345 -pf11682 -
13350 -pf11682 -
13360 - -
13365 - -
13370 - -
13375 -pf05930 -
BMC-P
BMC-H
BMC-T
Aldehyde dehydrogenase
PTAC
Signature enzyme
Alcohol dehydrogenase
Regulator

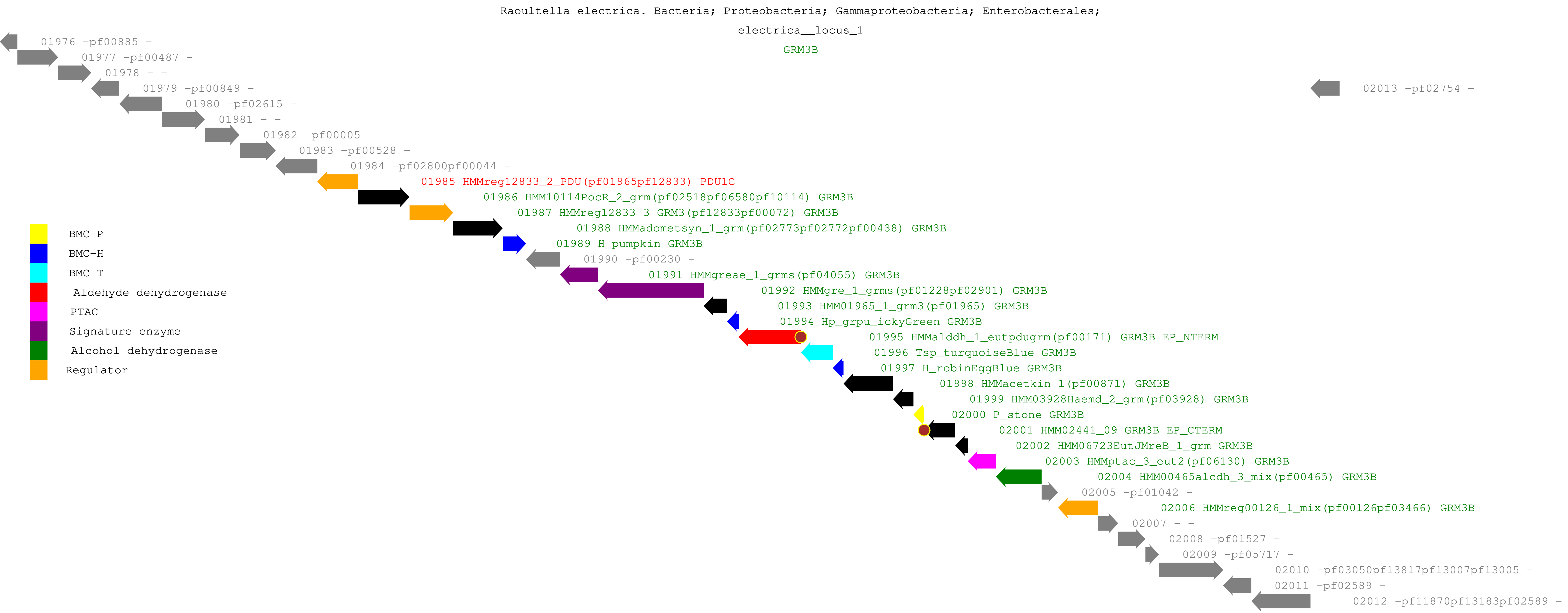
Raoultella electrica. Bacteria; Proteobacteria; Gammaproteobacteria; Enterobacterales;
electrica__locus_1
GRM3B
01976 -pf00885 -
01977 -pf00487 -
01978 - -
01979 -pf00849 -
01980 -pf02615 -
01981 - -
01982 -pf00005 -
01983 -pf00528 -
01984 -pf02800pf00044 -
01985 HMMreg12833_2_PDU(pf01965pf12833) PDU1C
01986 HMM10114PocR_2_grm(pf02518pf06580pf10114) GRM3B
01987 HMMreg12833_3_GRM3(pf12833pf00072) GRM3B
01988 HMMadometsyn_1_grm(pf02773pf02772pf00438) GRM3B
01989 H_pumpkin GRM3B
01990 -pf00230 -
01991 HMMgreae_1_grms(pf04055) GRM3B
01992 HMMgre_1_grms(pf01228pf02901) GRM3B
01993 HMM01965_1_grm3(pf01965) GRM3B
01994 Hp_grpu_ickyGreen GRM3B
01995 HMMalddh_1_eutpdugrm(pf00171) GRM3B EP_NTERM
01996 Tsp_turquoiseBlue GRM3B
01997 H_robinEggBlue GRM3B
01998 HMMacetkin_1(pf00871) GRM3B
01999 HMM03928Haemd_2_grm(pf03928) GRM3B
02000 P_stone GRM3B
02001 HMM02441_09 GRM3B EP_CTERM
02002 HMM06723EutJMreB_1_grm GRM3B
02003 HMMptac_3_eut2(pf06130) GRM3B
02004 HMM00465alcdh_3_mix(pf00465) GRM3B
02005 -pf01042 -
02006 HMMreg00126_1_mix(pf00126pf03466) GRM3B
02007 - -
02008 -pf01527 -
02009 -pf05717 -
02010 -pf03050pf13817pf13007pf13005 -
02011 -pf02589 -
02012 -pf11870pf13183pf02589 -
02013 -pf02754 -
BMC-P
BMC-H
BMC-T
Aldehyde dehydrogenase
PTAC
Signature enzyme
Alcohol dehydrogenase
Regulator

Escherichia coli. Bacteria; Proteobacteria; Gammaproteobacteria; Enterobacterales;

EPT01__locus_2

GRM3B

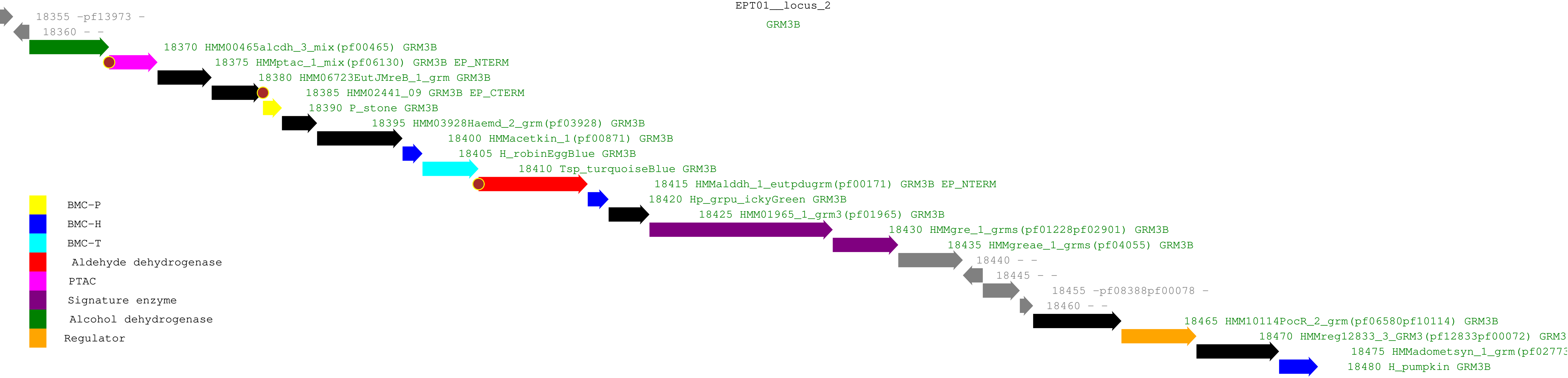

Escherichia coli. Bacteria; Proteobacteria; Gammaproteobacteria; Enterobacterales;

ERS085374__locus_1

GRM3B

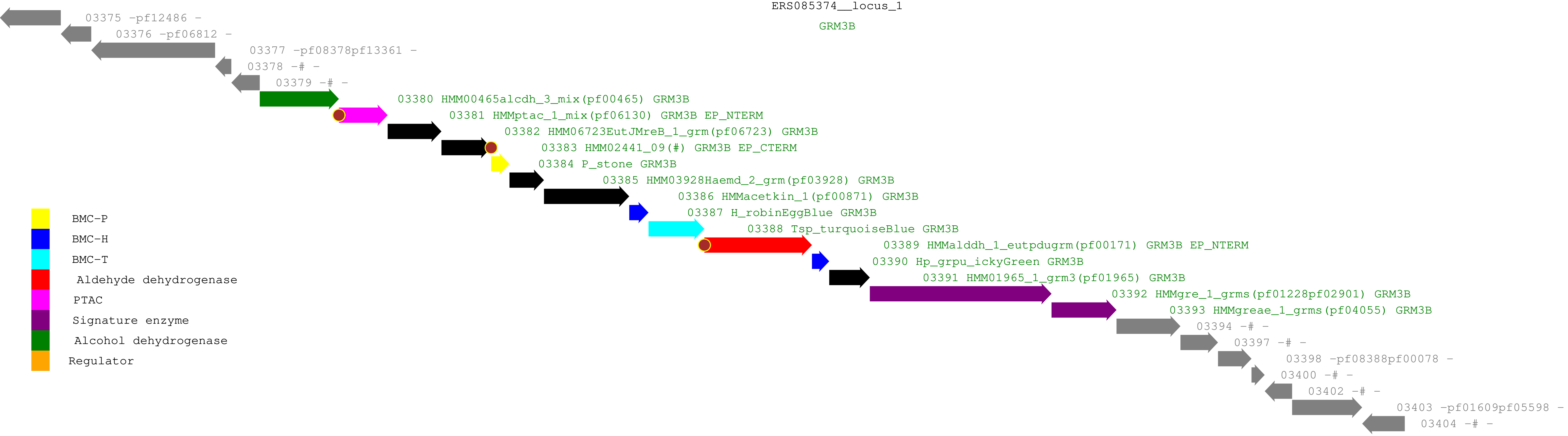

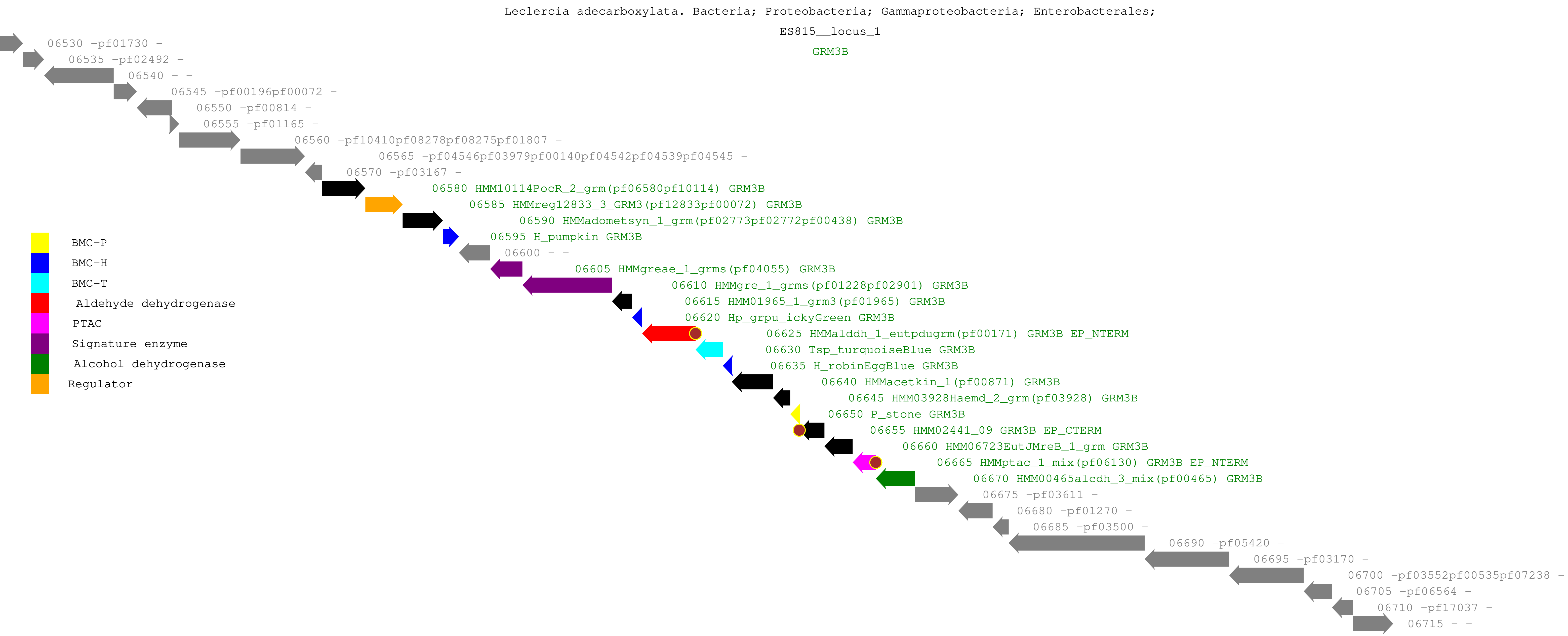

Leclercia adecarboxylata. Bacteria; Proteobacteria; Gammaproteobacteria; Enterobacterales;
ES815__locus_1
GRM3B
06530 -pf01730 -
06535 -pf02492 -
06540 - -
06545 -pf00196pf00072 -
06550 -pf00814 -
06555 -pf01165 -
06560 -pf10410pf08278pf08275pf01807 -
06565 -pf04546pf03979pf00140pf04542pf04539pf04545 -
06570 -pf03167 -
06580 HMM10114PocR_2_grm(pf06580pf10114) GRM3B
06585 HMMreg12833_3_GRM3(pf12833pf00072) GRM3B
06590 HMMadometsyn_1_grm(pf02773pf02772pf00438) GRM3B
06595 H_pumpkin GRM3B
06600 - -
06605 HMMgreae_1_grms(pf04055) GRM3B
06610 HMMgre_1_grms(pf01228pf02901) GRM3B
06615 HMM01965_1_grm3(pf01965) GRM3B
06620 Hp_grpu_ickyGreen GRM3B
06625 HMMalddh_1_eutpdugrm(pf00171) GRM3B EP_NTERM
06630 Tsp_turquoiseBlue GRM3B
06635 H_robinEggBlue GRM3B
06640 HMMacetkin_1(pf00871) GRM3B
06645 HMM03928Haemd_2_grm(pf03928) GRM3B
06650 P_stone GRM3B
06655 HMM02441_09 GRM3B EP_CTERM
06660 HMM06723EutJMreB_1_grm GRM3B
06665 HMMptac_1_mix(pf06130) GRM3B EP_NTERM
06670 HMM00465alcdh_3_mix(pf00465) GRM3B
06675 -pf03611 -
06680 -pf01270 -
06685 -pf03500 -
06690 -pf05420 -
06695 -pf03170 -
06700 -pf03552pf00535pf07238 -
06705 -pf06564 -
06710 -pf17037 -
06715 - -
BMC-P
BMC-H
BMC-T
Aldehyde dehydrogenase
PTAC
Signature enzyme
Alcohol dehydrogenase
Regulator

Lonepinella koalarum. Bacteria; Proteobacteria; Gammaproteobacteria; Pasteurellales;

EV692__locus_1

GRM3B

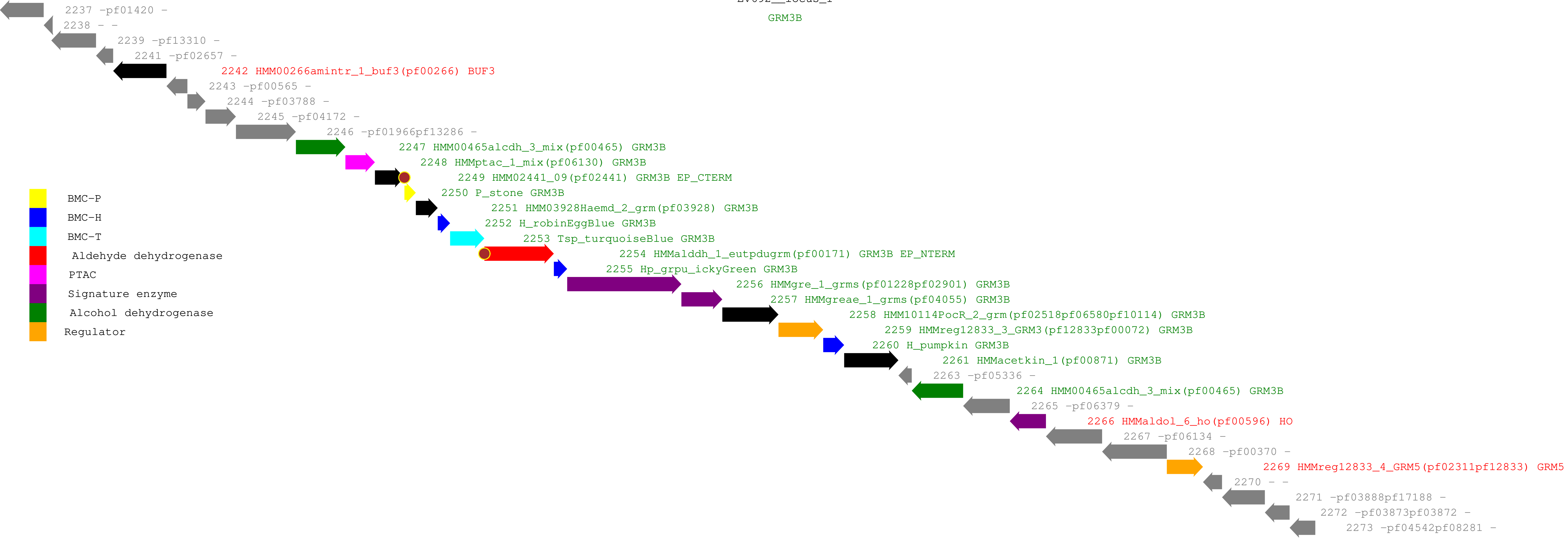

Salmonella enterica subsp. enterica serovar Adelaide. Bacteria; Proteobacteria; Gammaproteobacteria; Enterobacterales;

EZ821__locus_1

GRM3B

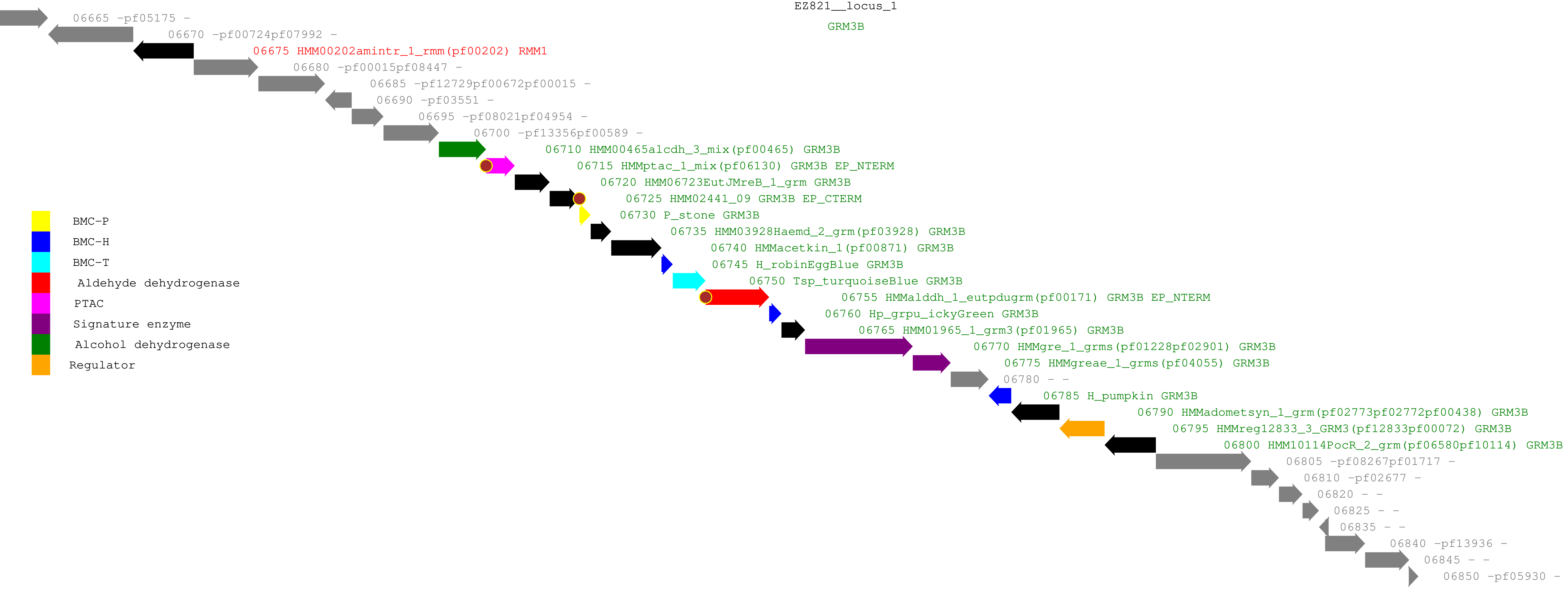

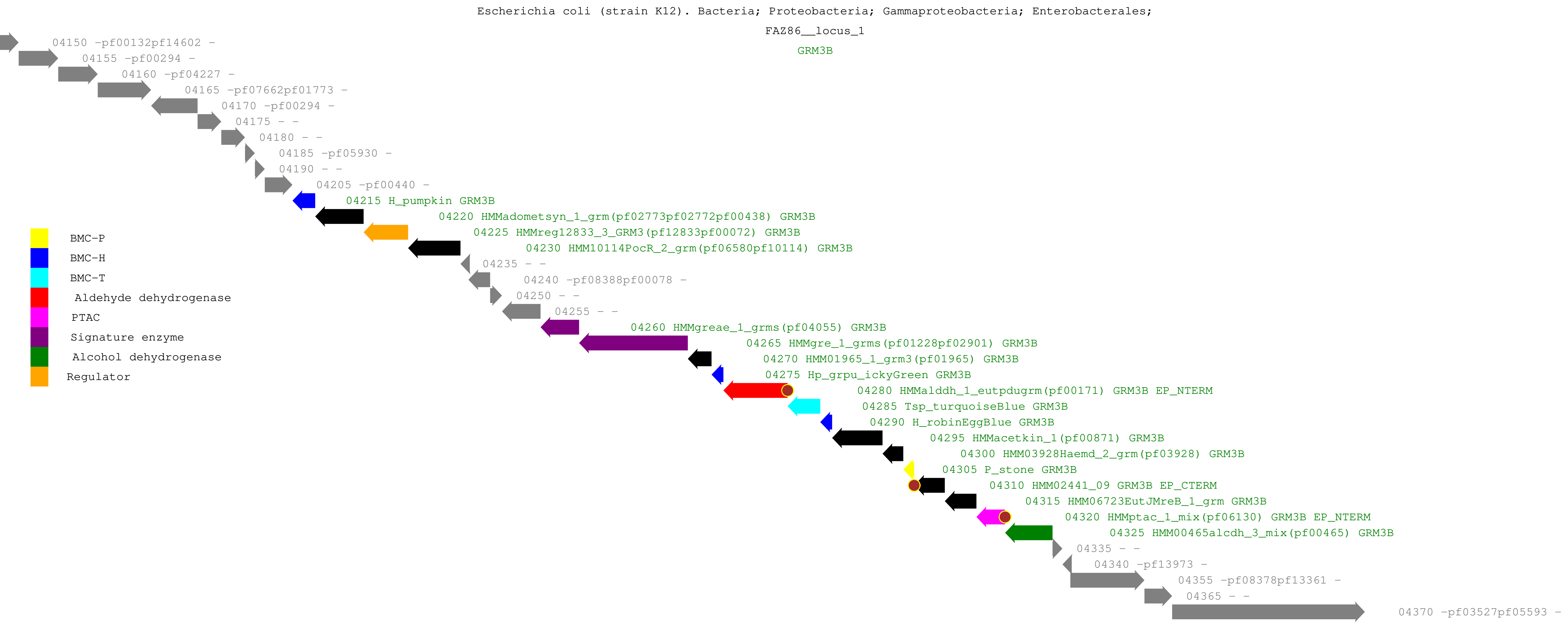
Escherichia coli (strain K12). Bacteria; Proteobacteria; Gammaproteobacteria; Enterobacterales;
FAZ86__locus_1
GRM3B
04150 -pf00132pf14602 -
04155 -pf00294 -
04160 -pf04227 -
04165 -pf07662pf01773 -
04170 -pf00294 -
04175 - -
04180 - -
04185 -pf05930 -
04190 - -
04205 -pf00440 -
04215 H_pumpkin GRM3B
04220 HMMadometsyn_1_grm(pf02773pf02772pf00438) GRM3B
04225 HMMreg12833_3_GRM3(pf12833pf00072) GRM3B
04230 HMM10114PocR_2_grm(pf06580pf10114) GRM3B
04235 - -
04240 -pf08388pf00078 -
04250 - -
04255 - -
04260 HMMgreae_1_grms(pf04055) GRM3B
04265 HMMgre_1_grms(pf01228pf02901) GRM3B
04270 HMM01965_1_grm3(pf01965) GRM3B
04275 Hp_grpu_ickyGreen GRM3B
04280 HMMalddh_1_eutpdugrm(pf00171) GRM3B EP_NTERM
04285 Tsp_turquoiseBlue GRM3B
04290 H_robinEggBlue GRM3B
04295 HMMacetkin_1(pf00871) GRM3B
04300 HMM03928Haemd_2_grm(pf03928) GRM3B
04305 P_stone GRM3B
04310 HMM02441_09 GRM3B EP_CTERM
04315 HMM06723EutJMreB_1_grm GRM3B
04320 HMMptac_1_mix(pf06130) GRM3B EP_NTERM
04325 HMM00465alcdh_3_mix(pf00465) GRM3B
04335 - -
04340 -pf13973 -
04355 -pf08378pf13361 -
04365 - -
04370 -pf03527pf05593 -
BMC-P
BMC-H
BMC-T
Aldehyde dehydrogenase
PTAC
Signature enzyme
Alcohol dehydrogenase
Regulator

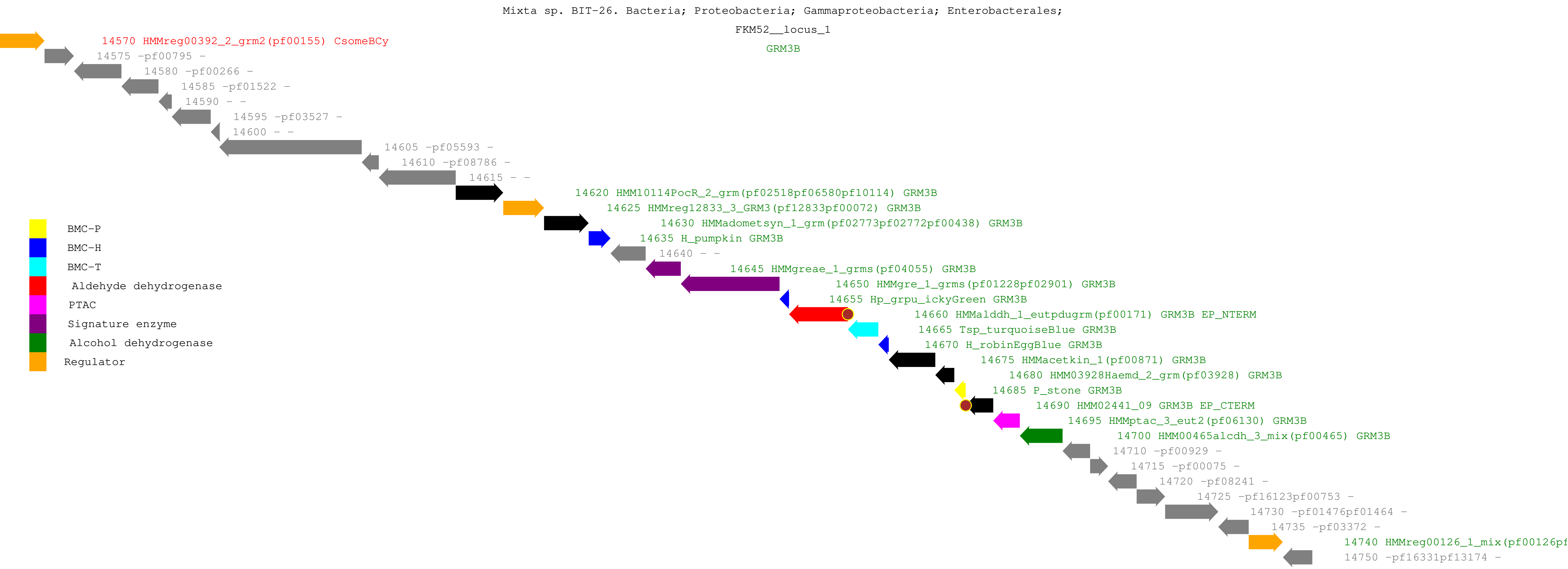
Mixta sp. BIT-26. Bacteria; Proteobacteria; Gammaproteobacteria; Enterobacterales;
FKM52__locus_1
GRM3B
14570 HMMreg00392_2_grm2(pf00155) CsomeBCy
14575 -pf00795 -
14580 -pf00266 -
14585 -pf01522 -
14590 - -
14595 -pf03527 -
14600 - -
14605 -pf05593 -
14610 -pf08786 -
14615 - -
14620 HMM10114PocR_2_grm(pf02518pf06580pf10114) GRM3B
14625 HMMreg12833_3_GRM3(pf12833pf00072) GRM3B
14630 HMMadometsyn_1_grm(pf02773pf02772pf00438) GRM3B
14635 H_pumpkin GRM3B
14640 - -
14645 HMMgreae_1_grms(pf04055) GRM3B
14650 HMMgre_1_grms(pf01228pf02901) GRM3B
14655 Hp_grpu_ickyGreen GRM3B
14660 HMMaldh_1_eutpdugrm(pf00171) GRM3B EP_NTERM
14665 Tsp_turquoiseBlue GRM3B
14670 H_robinEggBlue GRM3B
14675 HMMacetkin_1(pf00871) GRM3B
14680 HMM03928Haemd_2_grm(pf03928) GRM3B
14685 P_stone GRM3B
14690 HMM02441_09 GRM3B EP_CTERM
14695 HMMptac_3_eut2(pf06130) GRM3B
14700 HMM00465alcdh_3_mix(pf00465) GRM3B
14710 -pf00929 -
14715 -pf00075 -
14720 -pf08241 -
14725 -pf16123pf00753 -
14730 -pf01476pf01464 -
14735 -pf03372 -
14740 HMMreg00126_1_mix(pf00126pf
14750 -pf16331pf13174 -
BMC-P
BMC-H
BMC-T
Aldehyde dehydrogenase
PTAC
Signature enzyme
Alcohol dehydrogenase
Regulator

Enterobacter sp. E76. Bacteria; Proteobacteria; Gammaproteobacteria; Enterobacterales;

FR762__locus_1

GRM3B

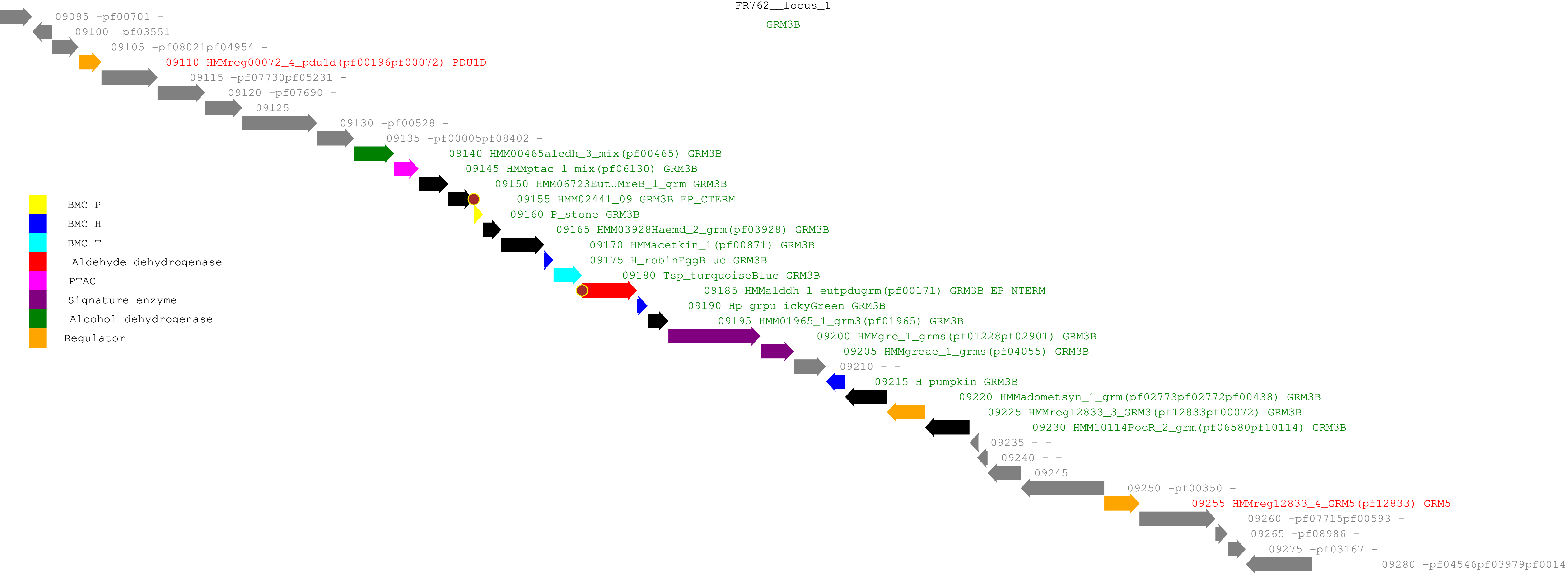

Escherichia coli HVH 25 (4-5851939). Bacteria; Proteobacteria; Gammaproteobacteria; Enterobacterales;

G701__locus_2

GRM3B

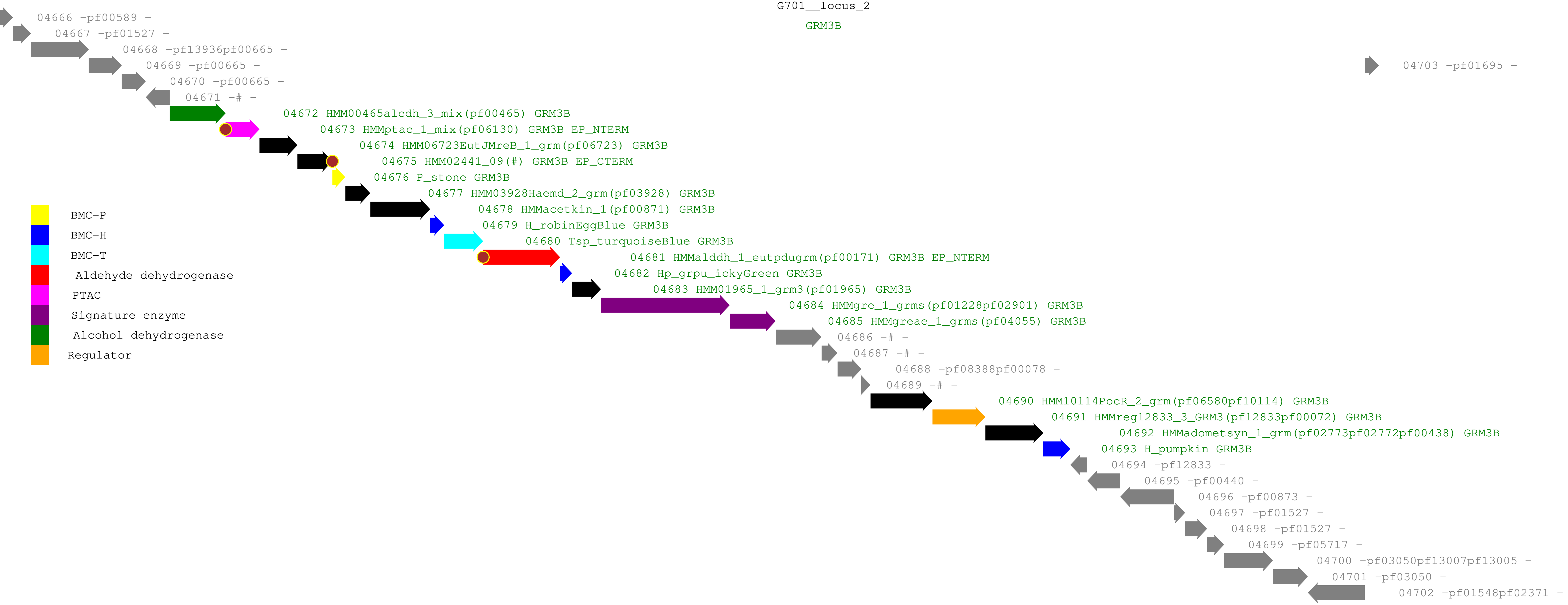

Escherichia coli HVH 87 (4-5977630). Bacteria; Proteobacteria; Gammaproteobacteria; Enterobacterales;

G749__locus_1_main_sat

GRM3B

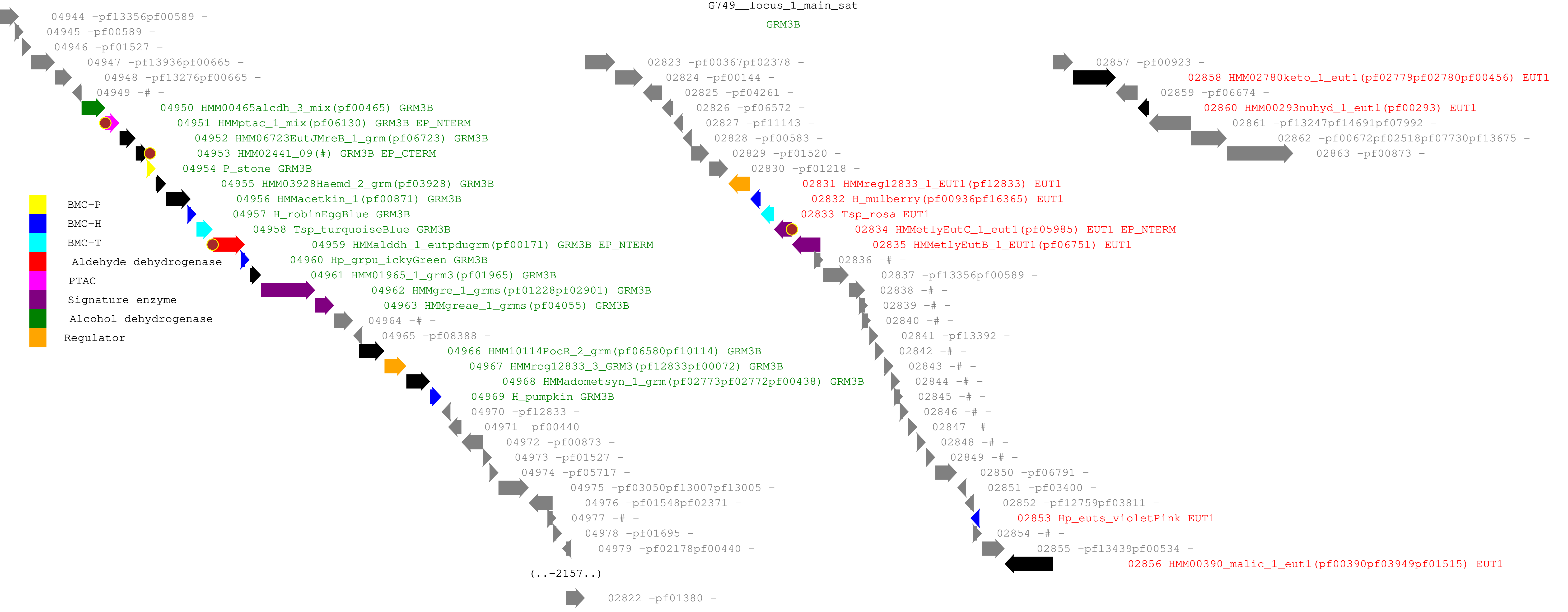

Escherichia coli (strain UMEA 3162-1). Bacteria; Proteobacteria; Gammaproteobacteria; Enterobacterales;

G925__locus_2

GRM3B

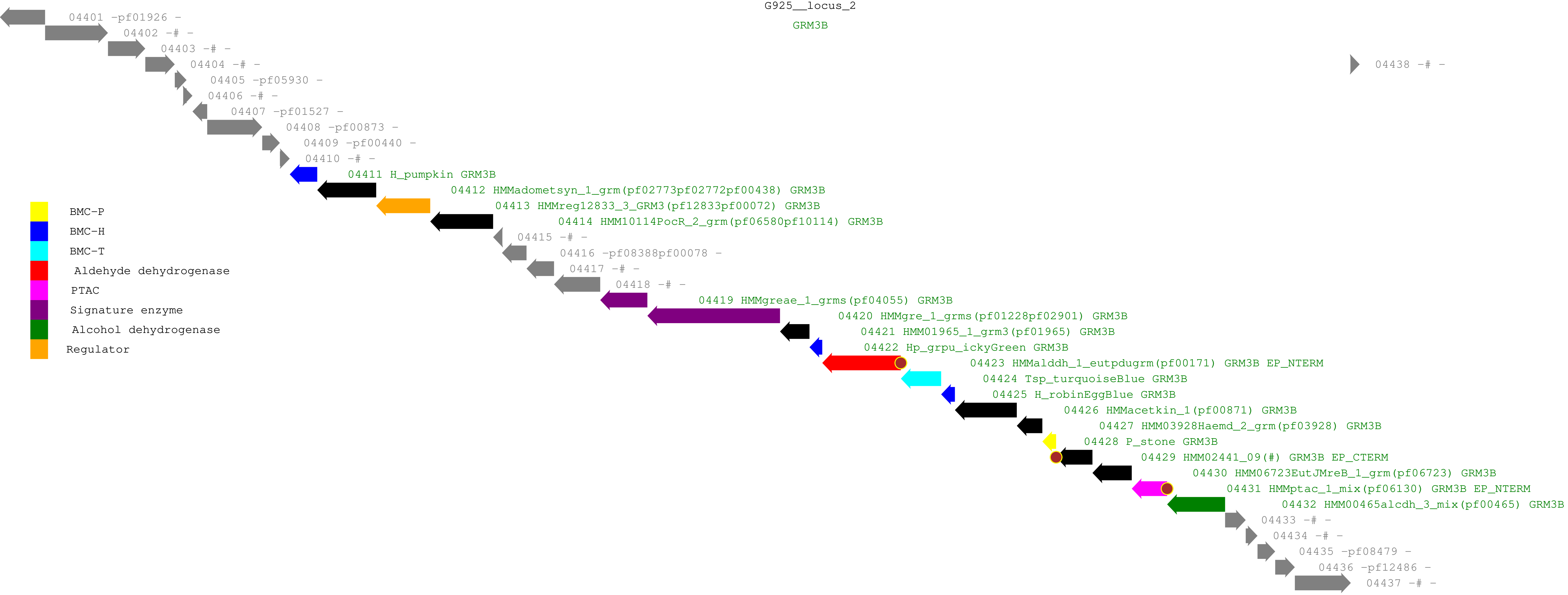

Escherichia coli UMEA 3323-1. Bacteria; Proteobacteria; Gammaproteobacteria; Enterobacterales;

G966__locus_2

GRM3B

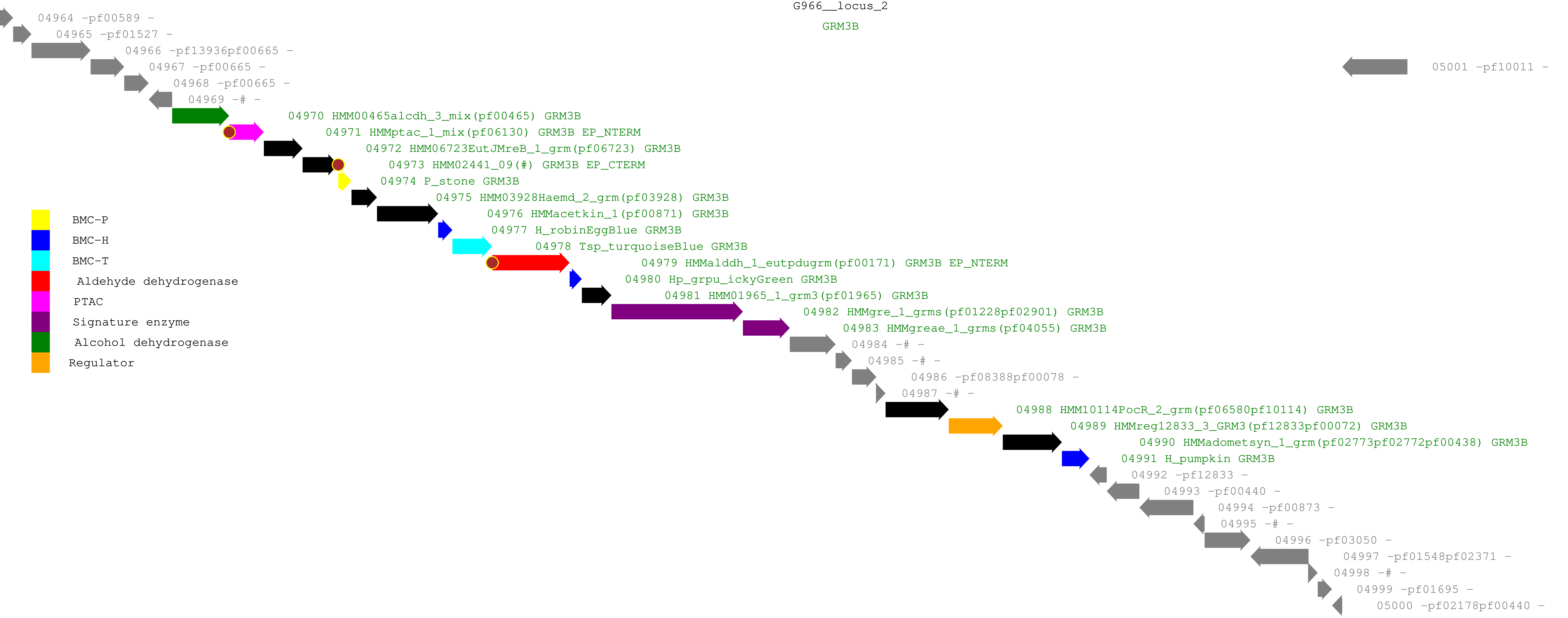

Escherichia coli (strain MS 21-1). Bacteria; Proteobacteria; Gammaproteobacteria; Enterobacterales;

HMPREF9530__locus_1

GRM3B

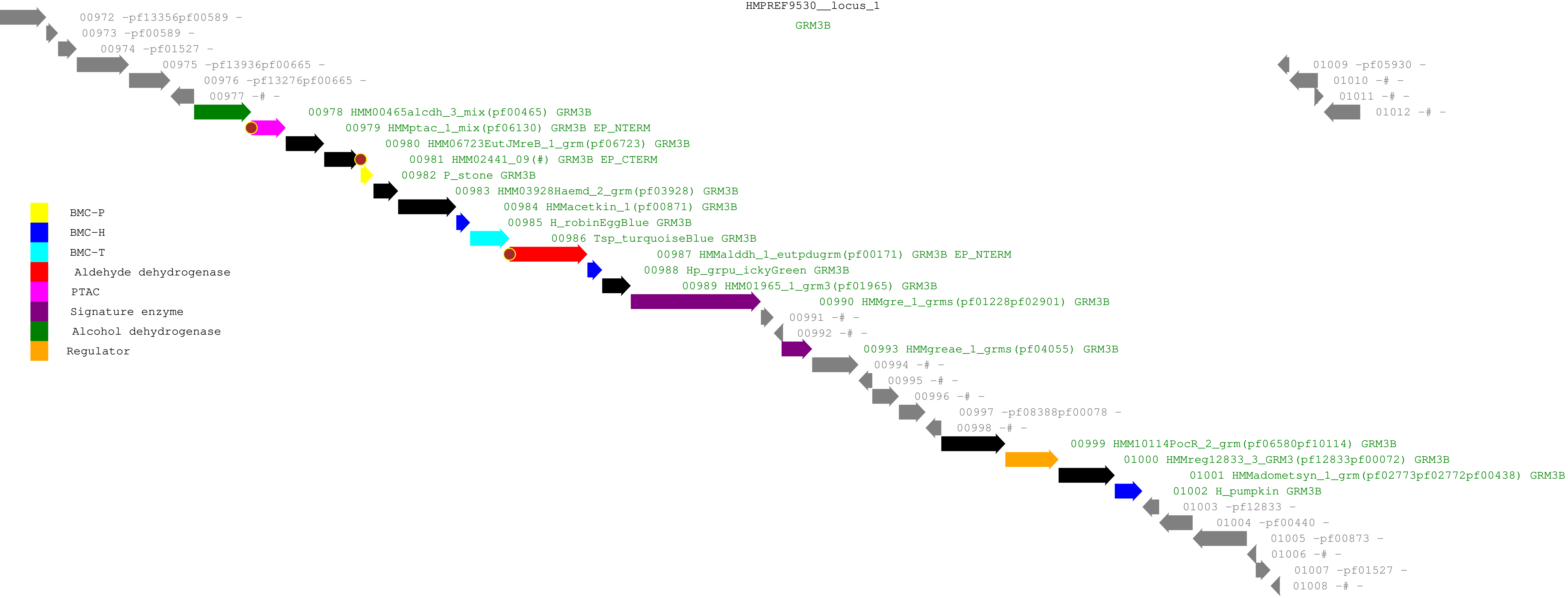

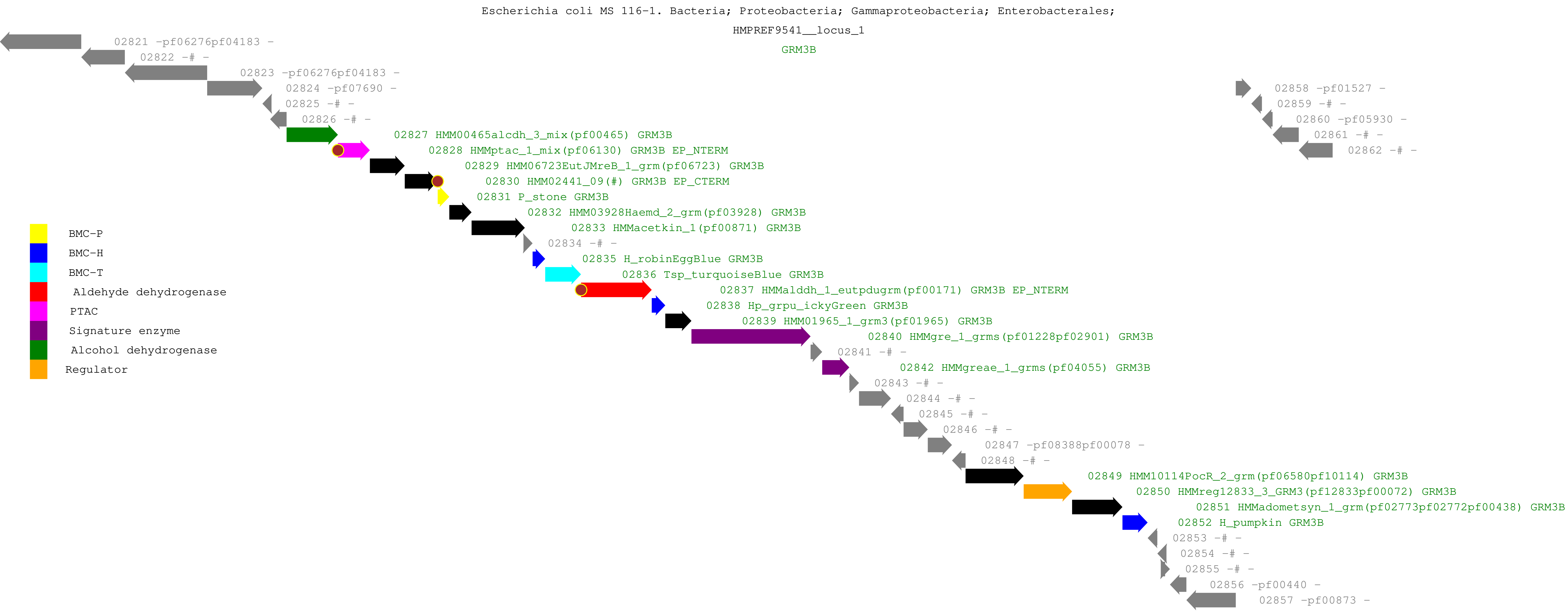
Escherichia coli MS 116-1. Bacteria; Proteobacteria; Gammaproteobacteria; Enterobacterales;
HMPREF9541__locus_1
GRM3B
02821 -pf06276pf04183 -
02822 -# -
02823 -pf06276pf04183 -
02824 -pf07690 -
02825 -# -
02826 -# -
02827 HMM00465alcdh_3_mix(pf00465) GRM3B
02828 HMMptac_1_mix(pf06130) GRM3B EP_NTERM
02829 HMM06723EutJMreB_1_grm(pf06723) GRM3B
02830 HMM02441_09(#) GRM3B EP_CTERM
02831 P_stone GRM3B
02832 HMM03928Haemd_2_grm(pf03928) GRM3B
02833 HMMacetkin_1(pf00871) GRM3B
02834 -# -
02835 H_robinEggBlue GRM3B
02836 Tsp_turquoiseBlue GRM3B
02837 HMMalddh_1_eutpdugrm(pf00171) GRM3B EP_NTERM
02838 Hp_grpu_ickyGreen GRM3B
02839 HMM01965_1_grm3(pf01965) GRM3B
02840 HMMgre_1_grms(pf01228pf02901) GRM3B
02841 -# -
02842 HMMgreae_1_grms(pf04055) GRM3B
02843 -# -
02844 -# -
02845 -# -
02846 -# -
02847 -pf08388pf00078 -
02848 -# -
02849 HMM10114PocR_2_grm(pf06580pf10114) GRM3B
02850 HMMreg12833_3_GRM3(pf12833pf00072) GRM3B
02851 HMMadometsyn_1_grm(pf02773pf02772pf00438) GRM3B
02852 H_pumpkin GRM3B
02853 -# -
02854 -# -
02855 -# -
02856 -pf00440 -
02857 -pf00873 -
02858 -pf01527 -
02859 -# -
02860 -pf05930 -
02861 -# -
02862 -# -
BMC-P
BMC-H
BMC-T
Aldehyde dehydrogenase
PTAC
Signature enzyme
Alcohol dehydrogenase
Regulator

Buttiauxella ferragutiae ATCC 51602. Bacteria; Proteobacteria; Gammaproteobacteria; Enterobacterales;

M976__locus_1

GRM3B

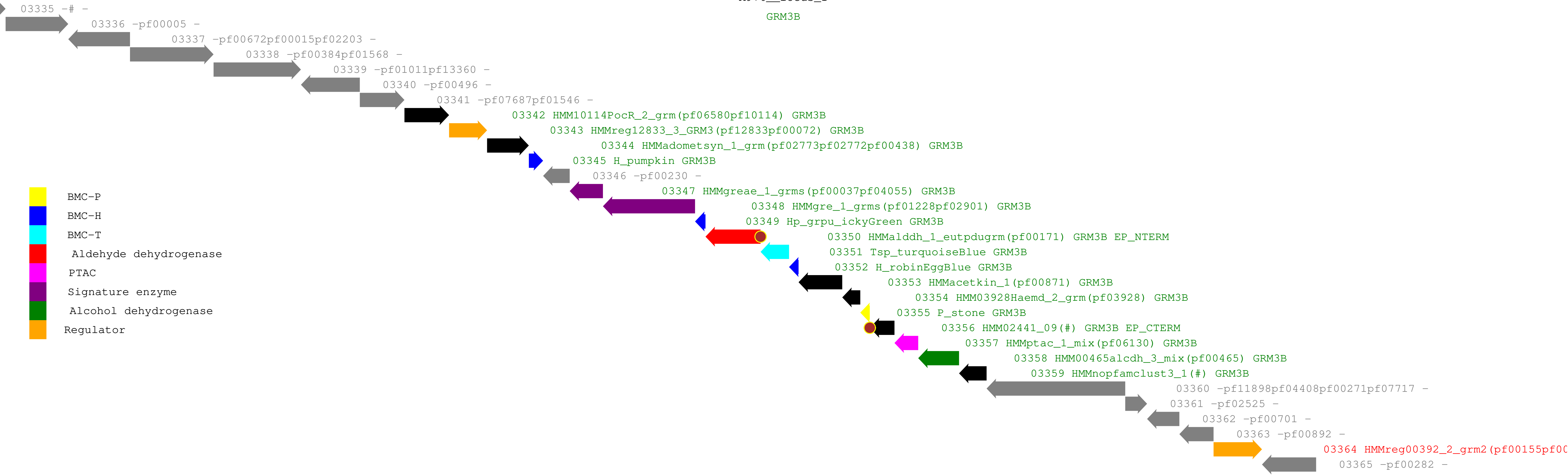

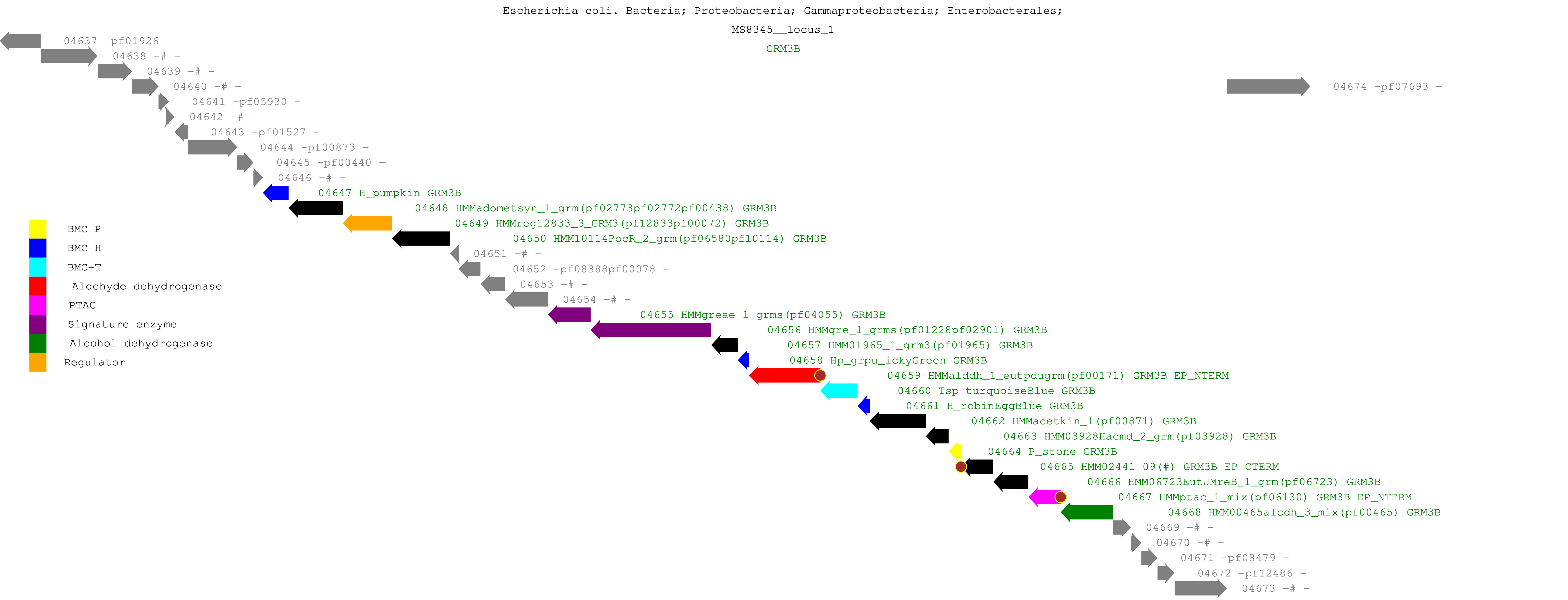

Escherichia coli. Bacteria; Proteobacteria; Gammaproteobacteria; Enterobacterales;
MS8345__locus_1
GRM3B
04637 -pf01926 -
04638 -# -
04639 -# -
04640 -# -
04641 -pf05930 -
04642 -# -
04643 -pf01527 -
04644 -pf00873 -
04645 -pf00440 -
04646 -# -
04647 H_pumpkin GRM3B
04648 HMMadometsyn_1_grm(pf02773pf02772pf00438) GRM3B
04649 HMMreg12833_3_GRM3(pf12833pf00072) GRM3B
04650 HMM10114PocR_2_grm(pf06580pf10114) GRM3B
04651 -# -
04652 -pf08388pf00078 -
04653 -# -
04654 -# -
04655 HMMgreae_1_grms(pf04055) GRM3B
04656 HMMgre_1_grms(pf01228pf02901) GRM3B
04657 HMM01965_1_grm3(pf01965) GRM3B
04658 Hp_grpu_ickyGreen GRM3B
04659 HMMalddh_1_eutpdugrm(pf00171) GRM3B EP_NTERM
04660 Tsp_turquoiseBlue GRM3B
04661 H_robinEggBlue GRM3B
04662 HMMacetkin_1(pf00871) GRM3B
04663 HMM03928Haemd_2_grm(pf03928) GRM3B
04664 P_stone GRM3B
04665 HMM02441_09(#) GRM3B EP_CTERM
04666 HMM06723EutJMreB_1_grm(pf06723) GRM3B
04667 HMMptac_1_mix(pf06130) GRM3B EP_NTERM
04668 HMM00465alcdh_3_mix(pf00465) GRM3B
04669 -# -
04670 -# -
04671 -pf08479 -
04672 -pf12486 -
04673 -# -
04674 -pf07693 -
BMC-P
BMC-H
BMC-T
Aldehyde dehydrogenase
PTAC
Signature enzyme
Alcohol dehydrogenase
Regulator

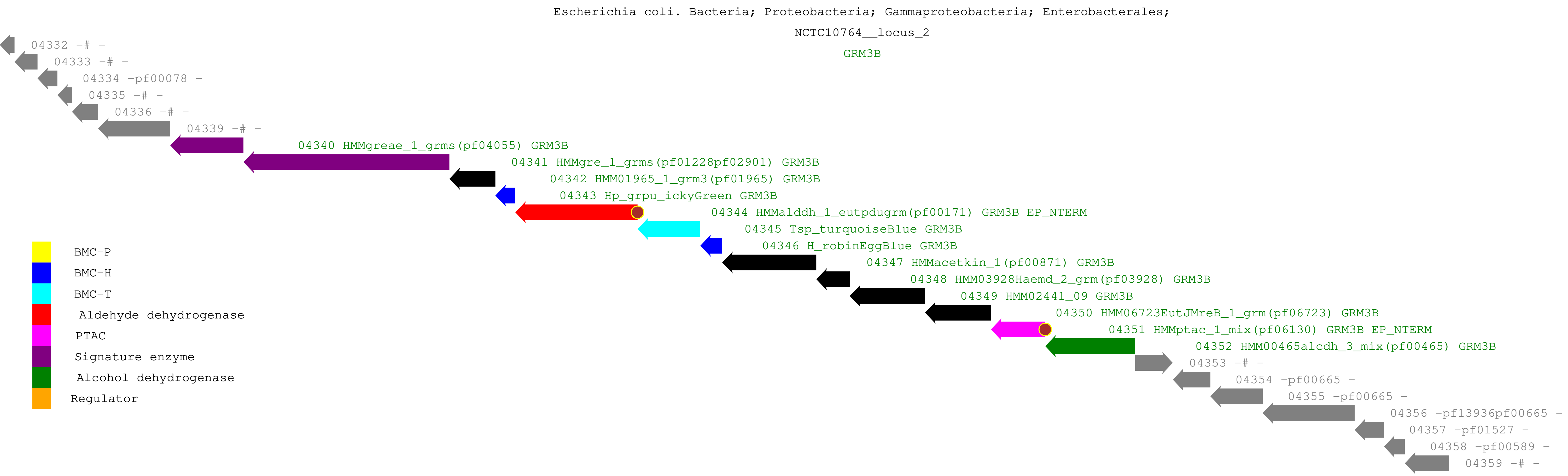

Escherichia coli. Bacteria; Proteobacteria; Gammaproteobacteria; Enterobacterales;
NCTC10764__locus_2
GRM3B
04332 -# -
04333 -# -
04334 -pf00078 -
04335 -# -
04336 -# -
04339 -# -
04340 HMMgreae_1_grms(pf04055) GRM3B
04341 HMMgre_1_grms(pf01228pf02901) GRM3B
04342 HMM01965_1_grm3(pf01965) GRM3B
04343 Hp_grpu_ickyGreen GRM3B
04344 HMMaldh_1_eutpdugrm(pf00171) GRM3B EP_NTERM
04345 Tsp_turquoiseBlue GRM3B
04346 H_robinEggBlue GRM3B
04347 HMMacetkin_1(pf00871) GRM3B
04348 HMM03928Haemd_2_grm(pf03928) GRM3B
04349 HMM02441_09 GRM3B
04350 HMM06723EutJMreB_1_grm(pf06723) GRM3B
04351 HMMptac_1_mix(pf06130) GRM3B EP_NTERM
04352 HMM00465alcdh_3_mix(pf00465) GRM3B
04353 -# -
04354 -pf00665 -
04355 -pf00665 -
04356 -pf13936pf00665 -
04357 -pf01527 -
04358 -pf00589 -
04359 -# -
BMC-P
BMC-H
BMC-T
Aldehyde dehydrogenase
PTAC
Signature enzyme
Alcohol dehydrogenase
Regulator

Atlantibacter hermannii (Escherichia hermannii). Bacteria; Proteobacteria; Gammaproteobacteria; Enterobacterales;

NCTC12129__locus_1

GRM3B

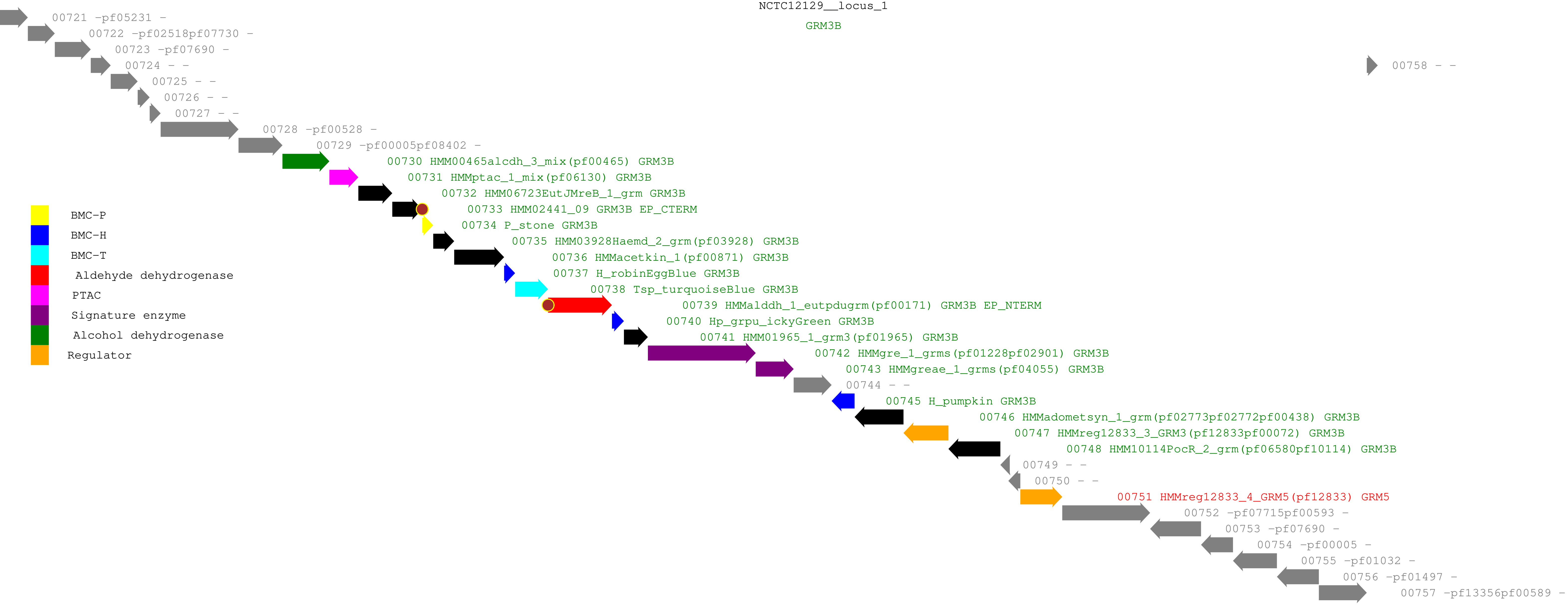

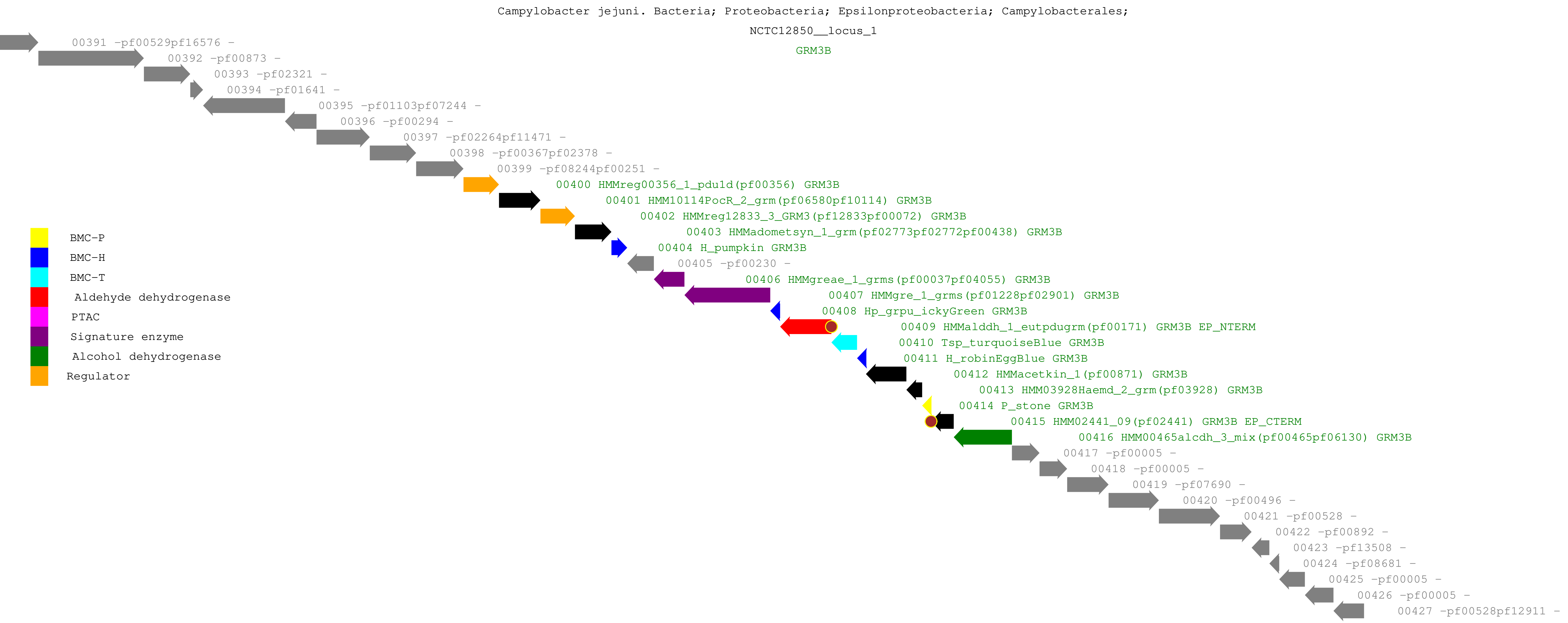
Campylobacter jejuni. Bacteria; Proteobacteria; Epsilonproteobacteria; Campylobacterales;
NCTC12850__locus_1
GRM3B
00391 -pf00529pf16576 -
00392 -pf00873 -
00393 -pf02321 -
00394 -pf01641 -
00395 -pf01103pf07244 -
00396 -pf00294 -
00397 -pf02264pf11471 -
00398 -pf00367pf02378 -
00399 -pf08244pf00251 -
00400 HMMreg00356_1_pdu1d(pf00356) GRM3B
00401 HMM10114PocR_2_grm(pf06580pf10114) GRM3B
00402 HMMreg12833_3_GRM3(pf12833pf00072) GRM3B
00403 HMMadometsyn_1_grm(pf02773pf02772pf00438) GRM3B
00404 H_pumpkin GRM3B
00405 -pf00230 -
00406 HMMgreae_1_grms(pf00037pf04055) GRM3B
00407 HMMgre_1_grms(pf01228pf02901) GRM3B
00408 Hp_grpu_ickyGreen GRM3B
00409 HMMalddh_1_eutpdugrm(pf00171) GRM3B EP_NTERM
00410 Tsp_turquoiseBlue GRM3B
00411 H_robinEggBlue GRM3B
00412 HMMacetkin_1(pf00871) GRM3B
00413 HMM03928Haemd_2_grm(pf03928) GRM3B
00414 P_stone GRM3B
00415 HMM02441_09(pf02441) GRM3B EP_CTERM
00416 HMM00465alcdh_3_mix(pf00465pf06130) GRM3B
00417 -pf00005 -
00418 -pf00005 -
00419 -pf07690 -
00420 -pf00496 -
00421 -pf00528 -
00422 -pf00892 -
00423 -pf13508 -
00424 -pf08681 -
00425 -pf00005 -
00426 -pf00005 -
00427 -pf00528pf12911 -
BMC-P
BMC-H
BMC-T
Aldehyde dehydrogenase
PTAC
Signature enzyme
Alcohol dehydrogenase
Regulator

Raoultella planticola (Klebsiella planticola). Bacteria; Proteobacteria; Gammaproteobacteria; Enterobacterales;

NCTC13095__locus_1

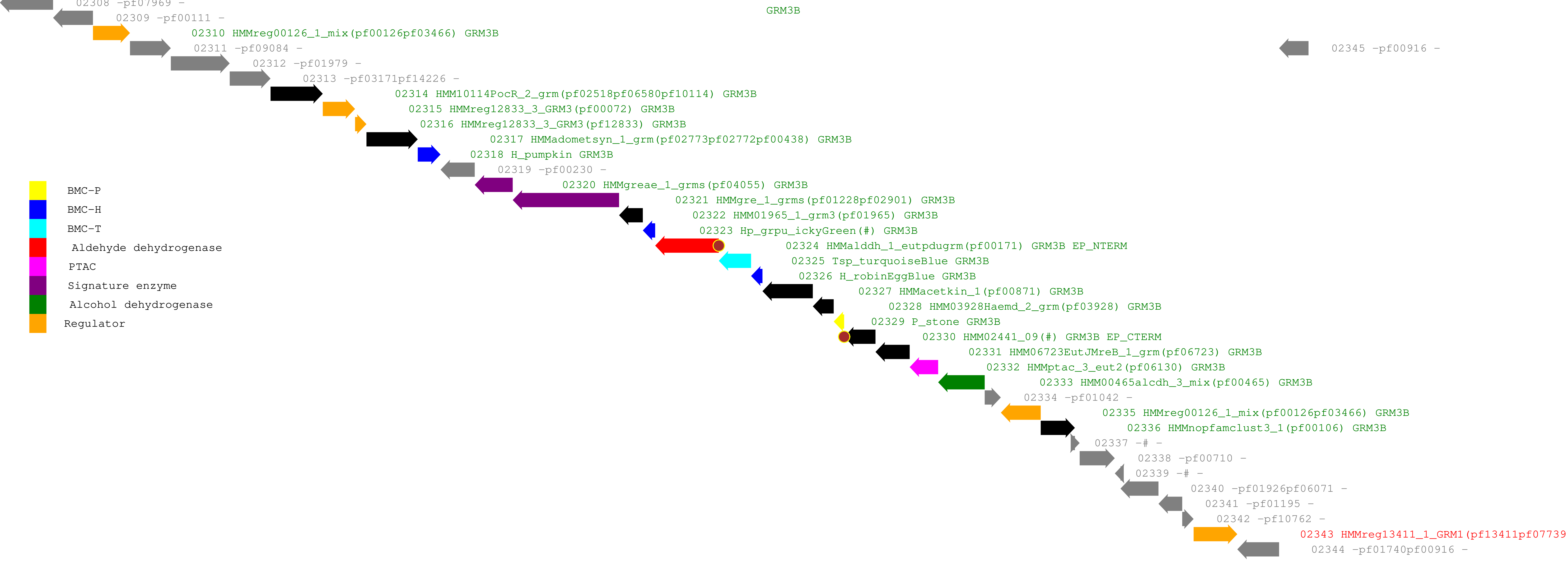

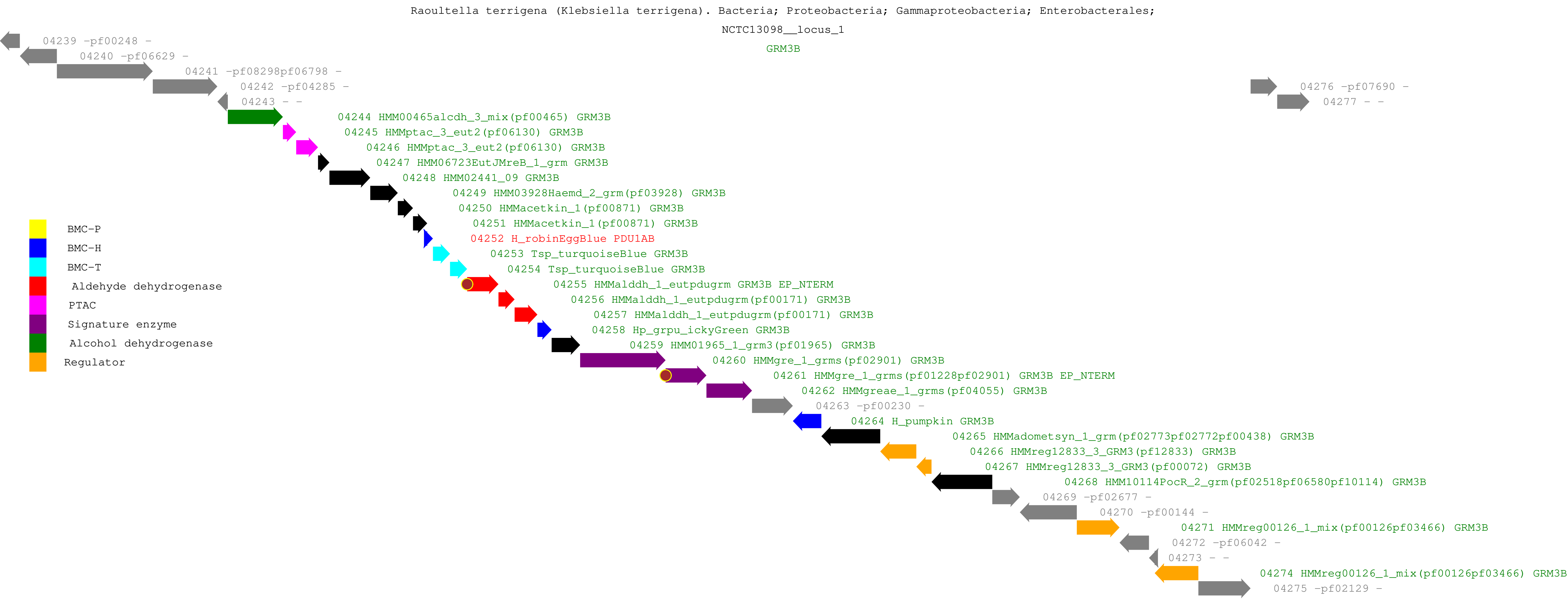
Raoultella terrigena (Klebsiella terrigena). Bacteria; Proteobacteria; Gammaproteobacteria; Enterobacterales;
NCTC13098__locus_1
GRM3B
04239 -pf00248 -
04240 -pf06629 -
04241 -pf08298pf06798 -
04242 -pf04285 -
04243 - -
04244 HMM00465alcdh_3_mix(pf00465) GRM3B
04245 HMMptac_3_eut2(pf06130) GRM3B
04246 HMMptac_3_eut2(pf06130) GRM3B
04247 HMM06723EutJMreB_1_grm GRM3B
04248 HMM02441_09 GRM3B
04249 HMM03928Haemd_2_grm(pf03928) GRM3B
04250 HMMacetkin_1(pf00871) GRM3B
04251 HMMacetkin_1(pf00871) GRM3B
04252 H_robinEggBlue PDU1AB
04253 Tsp_turquoiseBlue GRM3B
04254 Tsp_turquoiseBlue GRM3B
04255 HMMaldh_1_eutpdugrm GRM3B EP_NTERM
04256 HMMaldh_1_eutpdugrm(pf00171) GRM3B
04257 HMMaldh_1_eutpdugrm(pf00171) GRM3B
04258 Hp_grpu_ickyGreen GRM3B
04259 HMM01965_1_grm3(pf01965) GRM3B
04260 HMMgre_1_grms(pf02901) GRM3B
04261 HMMgre_1_grms(pf01228pf02901) GRM3B EP_NTERM
04262 HMMgreae_1_grms(pf04055) GRM3B
04263 -pf00230 -
04264 H_pumpkin GRM3B
04265 HMMadometsyn_1_grm(pf02773pf02772pf00438) GRM3B
04266 HMMreg12833_3_GRM3(pf12833) GRM3B
04267 HMMreg12833_3_GRM3(pf00072) GRM3B
04268 HMM10114PocR_2_grm(pf02518pf06580pf10114) GRM3B
04269 -pf02677 -
04270 -pf00144 -
04271 HMMreg00126_1_mix(pf00126pf03466) GRM3B
04272 -pf06042 -
04273 - -
04274 HMMreg00126_1_mix(pf00126pf03466) GRM3B
04275 -pf02129 -
04276 -pf07690 -
04277 - -
BMC-P
BMC-H
BMC-T
Aldehyde dehydrogenase
PTAC
Signature enzyme
Alcohol dehydrogenase
Regulator

Escherichia coli. Bacteria; Proteobacteria; Gammaproteobacteria; Enterobacterales;

NCTC7927__locus_2

GRM3B

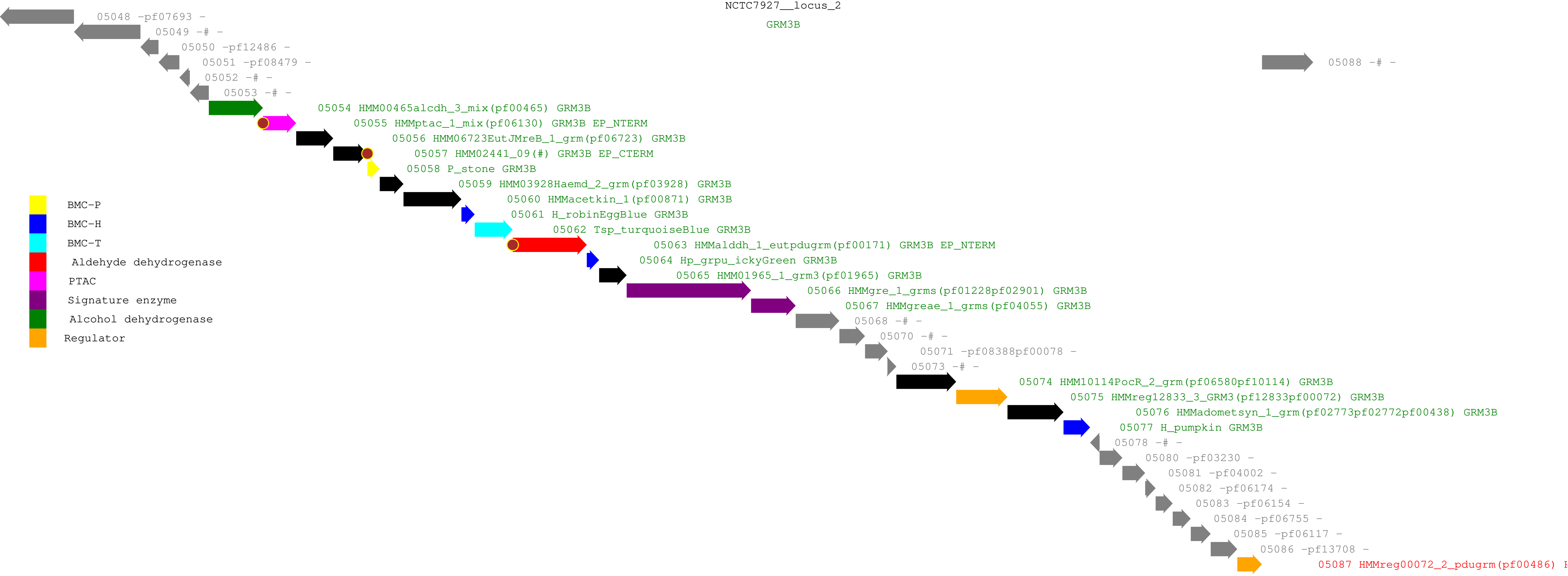

Escherichia coli. Bacteria; Proteobacteria; Gammaproteobacteria; Enterobacterales;

NCTC9050__locus_1

GRM3B

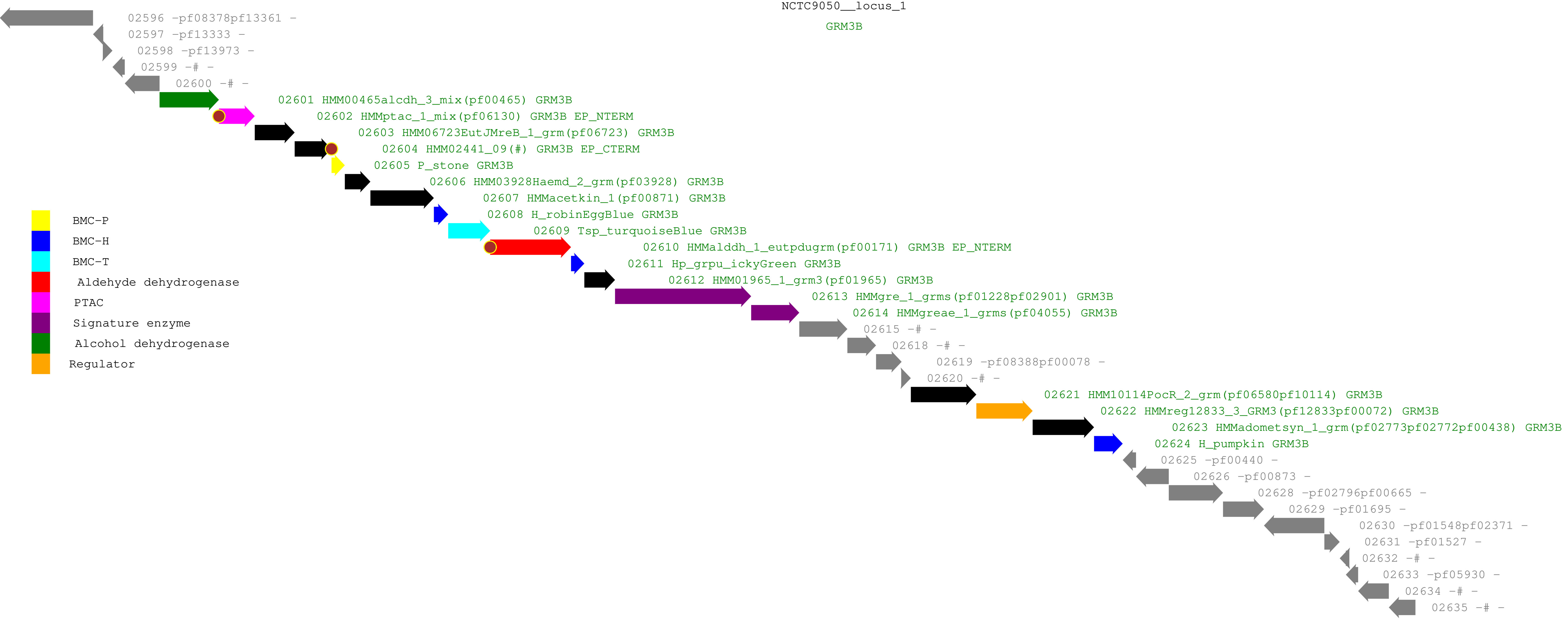

Raoultella planticola (Klebsiella planticola). Bacteria; Proteobacteria; Gammaproteobacteria; Enterobacterales;

NCTC9528__locus_2

GRM3B

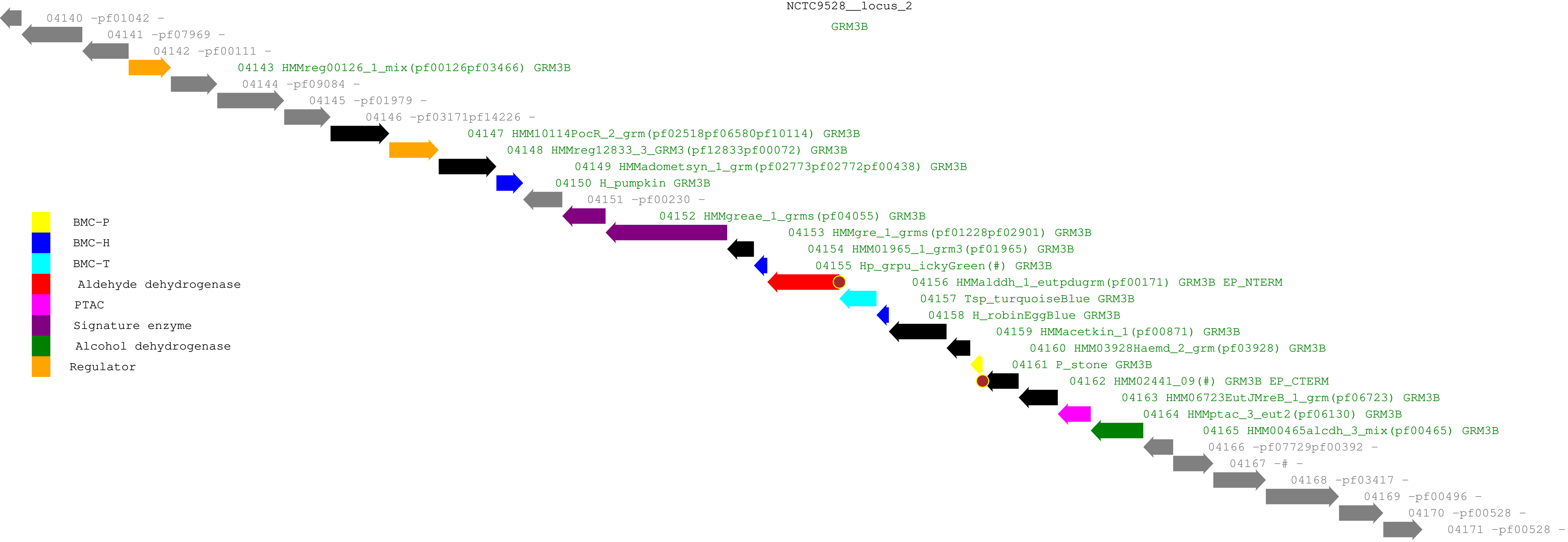

Pectobacterium parmentieri (strain WPP163) (Pectobacterium wasabiae Bacteria; Proteobacteria; Gammaproteobacteria; Enterobacterales;

Pecwa__locus_1

GRM3B

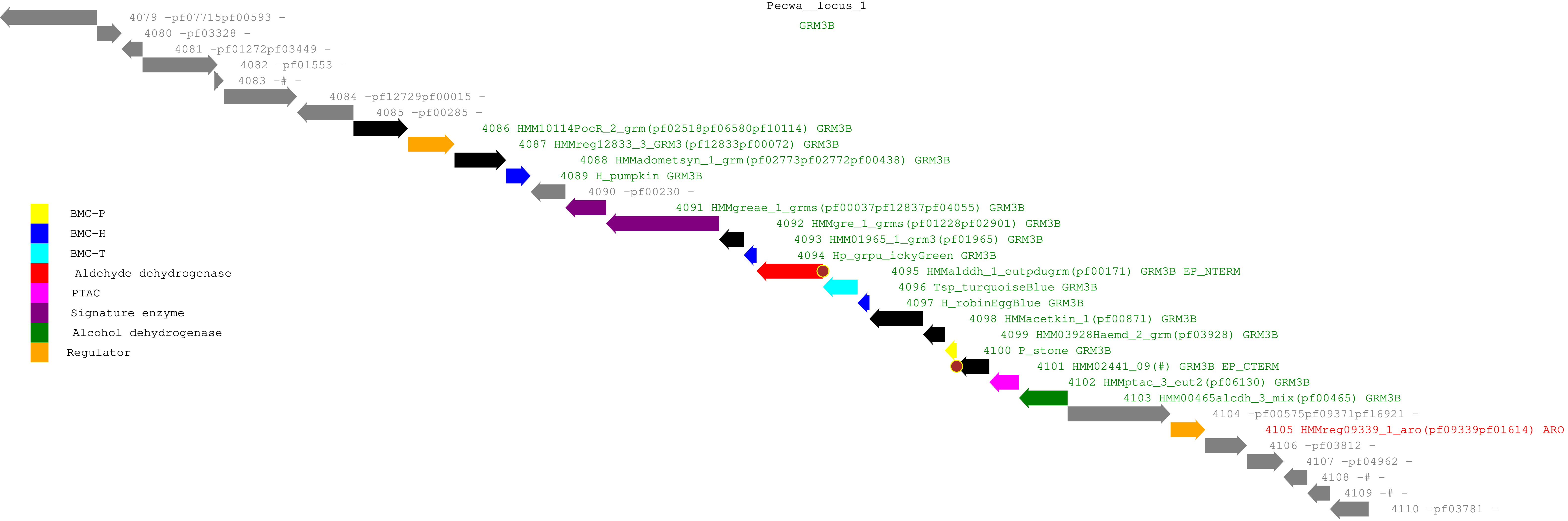

Klebsiella oxytoca. Bacteria; Proteobacteria; Gammaproteobacteria; Enterobacterales;

SAMEA2273573__locus_1

GRM3B

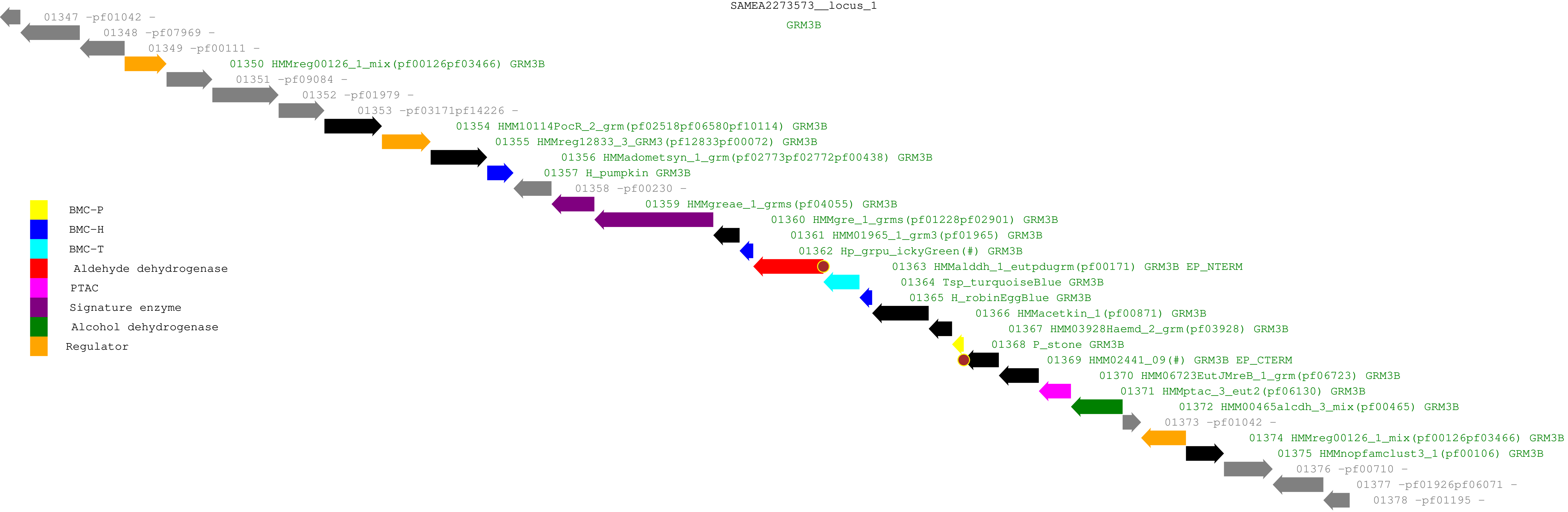

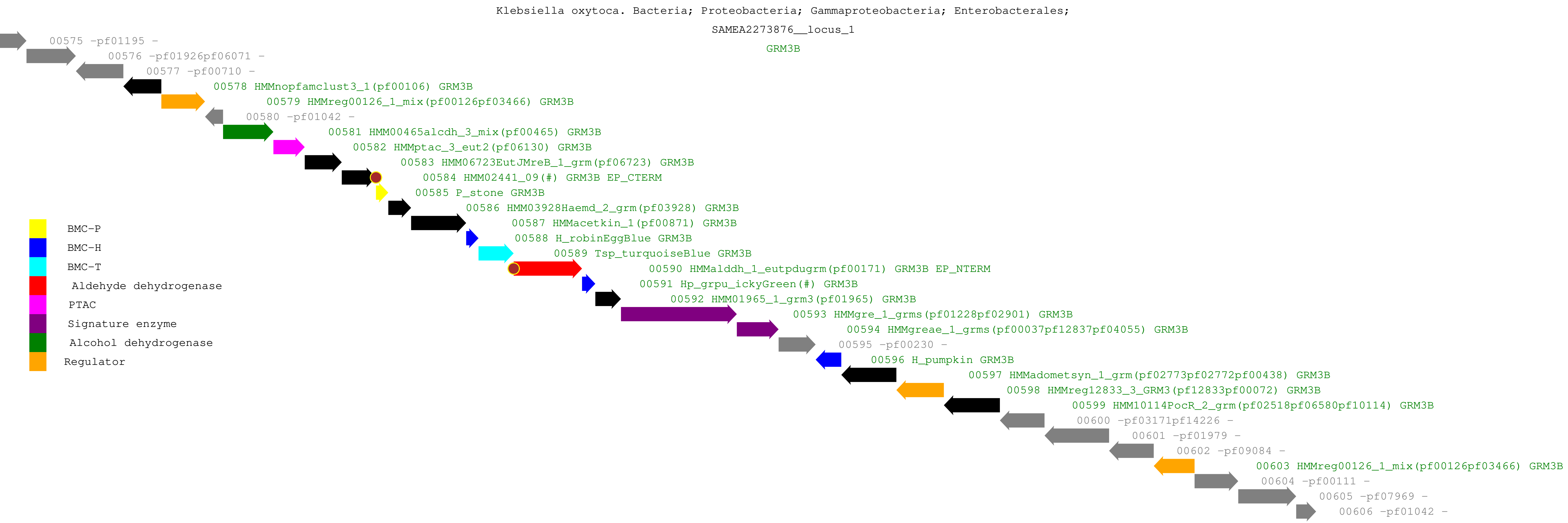
Klebsiella oxytoca. Bacteria; Proteobacteria; Gammaproteobacteria; Enterobacterales;
SAMEA2273876__locus_1
GRM3B
00575 -pf01195 -
00576 -pf01926pf06071 -
00577 -pf00710 -
00578 HMMnopfamclust3_1(pf00106) GRM3B
00579 HMMreg00126_1_mix(pf00126pf03466) GRM3B
00580 -pf01042 -
00581 HMM00465alcdh_3_mix(pf00465) GRM3B
00582 HMMptac_3_eut2(pf06130) GRM3B
00583 HMM06723EutJMreB_1_grm(pf06723) GRM3B
00584 HMM02441_09(#) GRM3B EP_CTERM
00585 P_stone GRM3B
00586 HMM03928Haemd_2_grm(pf03928) GRM3B
00587 HMMacetkin_1(pf00871) GRM3B
00588 H_robinEggBlue GRM3B
00589 Tsp_turquoiseBlue GRM3B
00590 HMMaldh_1_eutpdugrm(pf00171) GRM3B EP_NTERM
00591 Hp_grpu_ickyGreen(#) GRM3B
00592 HMM01965_1_grm3(pf01965) GRM3B
00593 HMMgre_1_grms(pf01228pf02901) GRM3B
00594 HMMgreae_1_grms(pf00037pf12837pf04055) GRM3B
00595 -pf00230 -
00596 H_pumpkin GRM3B
00597 HMMadometsyn_1_grm(pf02773pf02772pf00438) GRM3B
00598 HMMreg12833_3_GRM3(pf12833pf00072) GRM3B
00599 HMM10114PocR_2_grm(pf02518pf06580pf10114) GRM3B
00600 -pf03171pf14226 -
00601 -pf01979 -
00602 -pf09084 -
00603 HMMreg00126_1_mix(pf00126pf03466) GRM3B
00604 -pf00111 -
00605 -pf07969 -
00606 -pf01042 -
BMC-P
BMC-H
BMC-T
Aldehyde dehydrogenase
PTAC
Signature enzyme
Alcohol dehydrogenase
Regulator

Raoultella ornithinolytica (Klebsiella ornithinolytica). Bacteria; Proteobacteria; Gammaproteobacteria; Enterobacterales;

SAMEA3181514__locus_1

GRM3B

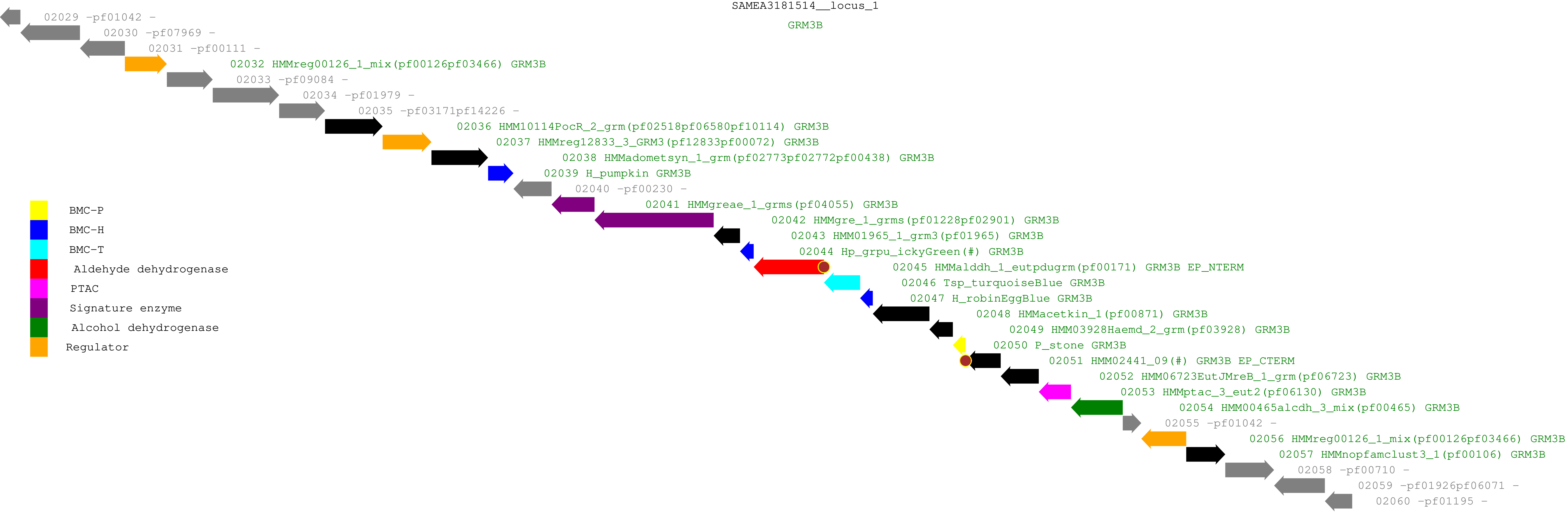

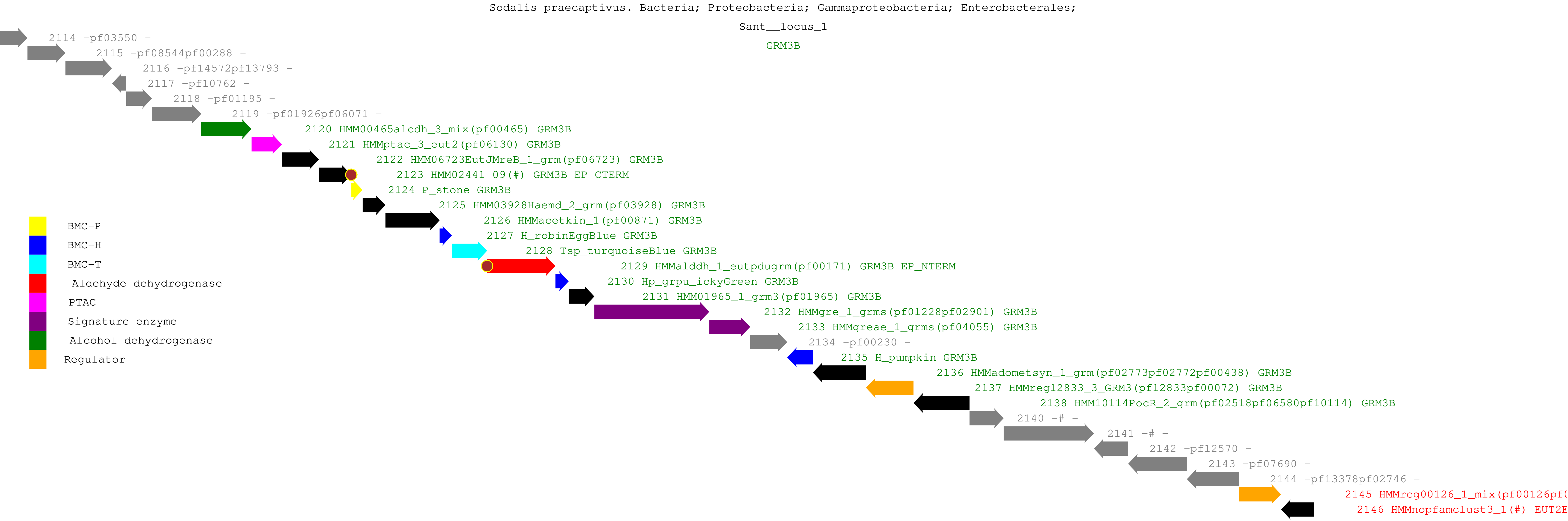
Sodalis praecaptivus. Bacteria; Proteobacteria; Gammaproteobacteria; Enterobacterales;
Sant__locus_1
GRM3B
2114 -pf03550 -
2115 -pf08544pf00288 -
2116 -pf14572pf13793 -
2117 -pf10762 -
2118 -pf01195 -
2119 -pf01926pf06071 -
2120 HMM00465alcdh_3_mix(pf00465) GRM3B
2121 HMMptac_3_eut2(pf06130) GRM3B
2122 HMM06723EutJMreB_1_grm(pf06723) GRM3B
2123 HMM02441_09(#) GRM3B EP_CTERM
2124 P_stone GRM3B
2125 HMM03928Haemd_2_grm(pf03928) GRM3B
2126 HMMacetkin_1(pf00871) GRM3B
2127 H_robinEggBlue GRM3B
2128 Tsp_turquoiseBlue GRM3B
2129 HMMalddh_1_eutpdugrm(pf00171) GRM3B EP_NTERM
2130 Hp_grpu_ickyGreen GRM3B
2131 HMM01965_1_grm3(pf01965) GRM3B
2132 HMMgre_1_grms(pf01228pf02901) GRM3B
2133 HMMgreae_1_grms(pf04055) GRM3B
2134 -pf00230 -
2135 H_pumpkin GRM3B
2136 HMMadometsyn_1_grm(pf02773pf02772pf00438) GRM3B
2137 HMMreg12833_3_GRM3(pf12833pf00072) GRM3B
2138 HMM10114PocR_2_grm(pf02518pf06580pf10114) GRM3B
2140 -# -
2141 -# -
2142 -pf12570 -
2143 -pf07690 -
2144 -pf13378pf02746 -
2145 HMMreg00126_1_mix(pf00126pf
2146 HMMnopfamclust3_1(#) EUT2E
BMC-P
BMC-H
BMC-T
Aldehyde dehydrogenase
PTAC
Signature enzyme
Alcohol dehydrogenase
Regulator

Snodgrassella alvi SCGC AB-598-002. Bacteria; Proteobacteria; Betaproteobacteria; Neisseriales;

SASC598002__locus_1_main_sat

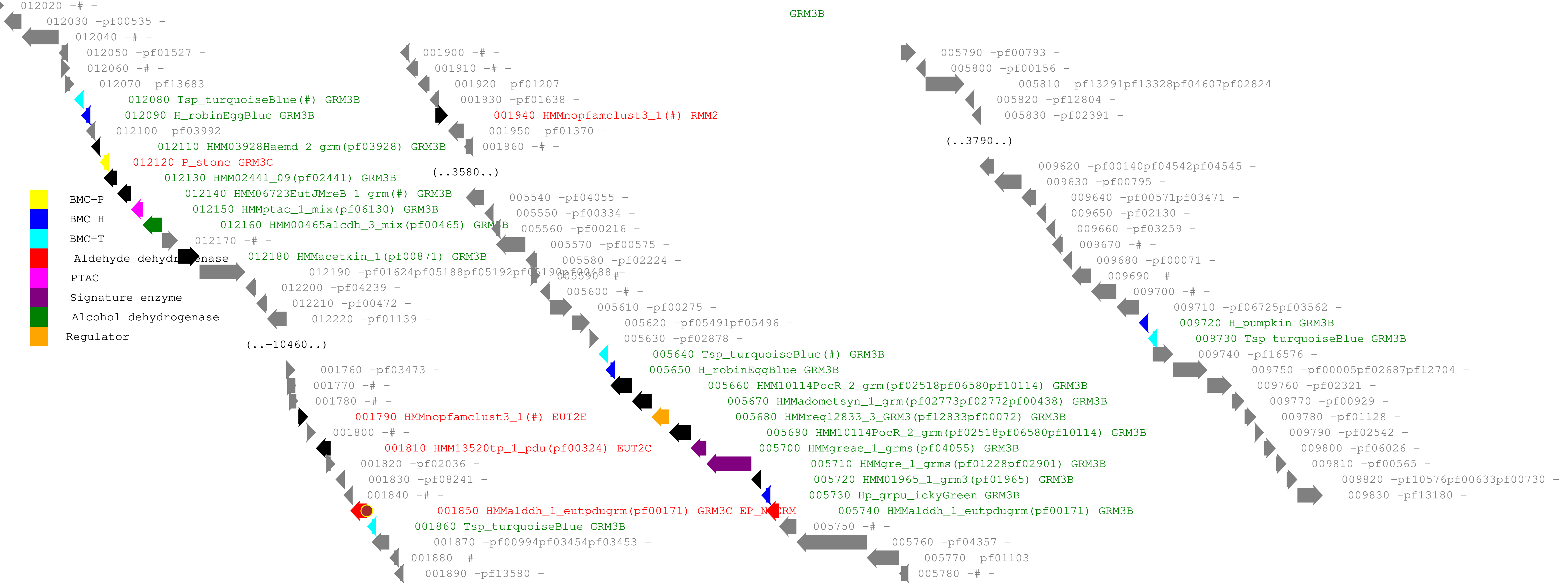

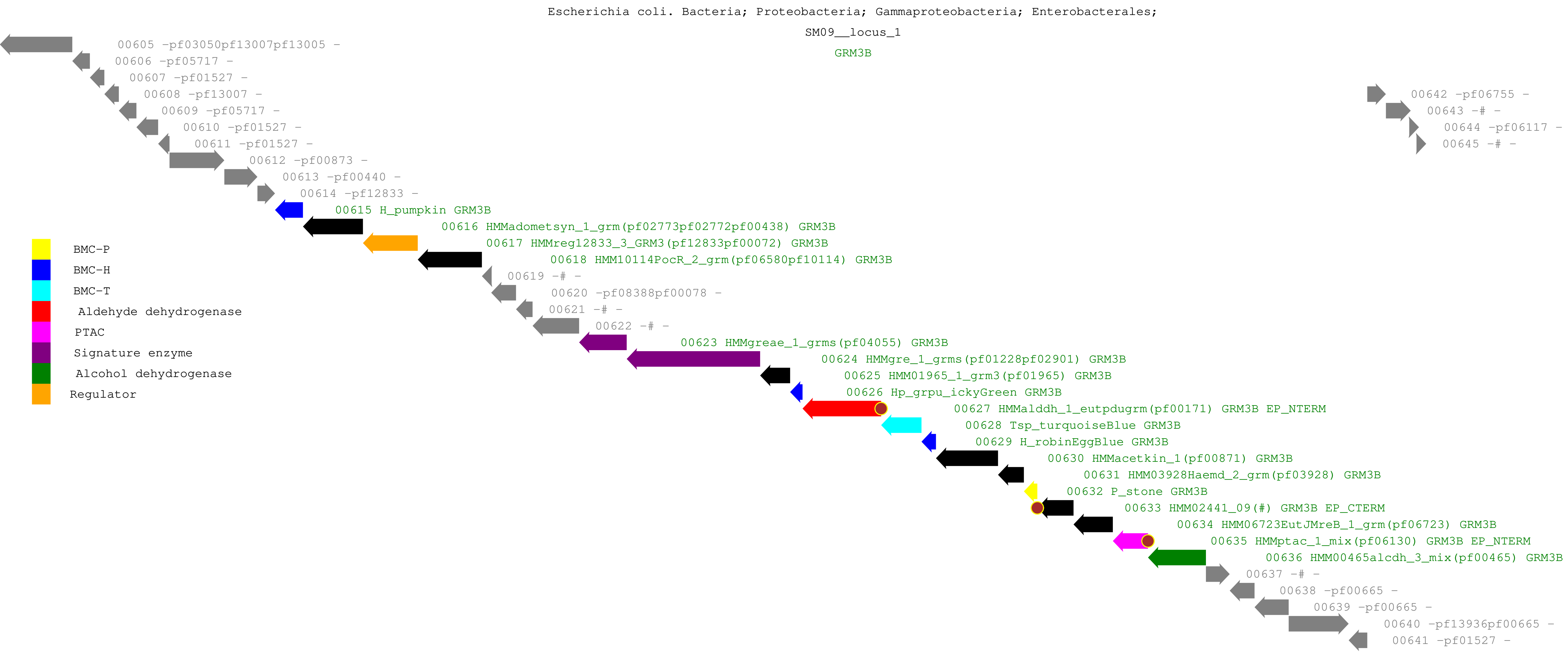
Escherichia coli. Bacteria; Proteobacteria; Gammaproteobacteria; Enterobacterales;
SM09__locus_1
GRM3B
00605 -pf03050pf13007pf13005 -
00606 -pf05717 -
00607 -pf01527 -
00608 -pf13007 -
00609 -pf05717 -
00610 -pf01527 -
00611 -pf01527 -
00612 -pf00873 -
00613 -pf00440 -
00614 -pf12833 -
00615 H_pumpkin GRM3B
00616 HMMadometsyn_1_grm(pf02773pf02772pf00438) GRM3B
00617 HMMreg12833_3_GRM3(pf12833pf00072) GRM3B
00618 HMM10114PocR_2_grm(pf06580pf10114) GRM3B
00619 -# -
00620 -pf08388pf00078 -
00621 -# -
00622 -# -
00623 HMMgreae_1_grms(pf04055) GRM3B
00624 HMMgre_1_grms(pf01228pf02901) GRM3B
00625 HMM01965_1_grm3(pf01965) GRM3B
00626 Hp_grpu_ickyGreen GRM3B
00627 HMMalddh_1_eutpdugrm(pf00171) GRM3B EP_NTERM
00628 Tsp_turquoiseBlue GRM3B
00629 H_robinEggBlue GRM3B
00630 HMMacetkin_1(pf00871) GRM3B
00631 HMM03928Haemd_2_grm(pf03928) GRM3B
00632 P_stone GRM3B
00633 HMM02441_09(#) GRM3B EP_CTERM
00634 HMM06723EutJMreB_1_grm(pf06723) GRM3B
00635 HMMptac_1_mix(pf06130) GRM3B EP_NTERM
00636 HMM00465alcdh_3_mix(pf00465) GRM3B
00637 -# -
00638 -pf00665 -
00639 -pf00665 -
00640 -pf13936pf00665 -
00641 -pf01527 -
00642 -pf06755 -
00643 -# -
00644 -pf06117 -
00645 -# -
BMC-P
BMC-H
BMC-T
Aldehyde dehydrogenase
PTAC
Signature enzyme
Alcohol dehydrogenase
Regulator

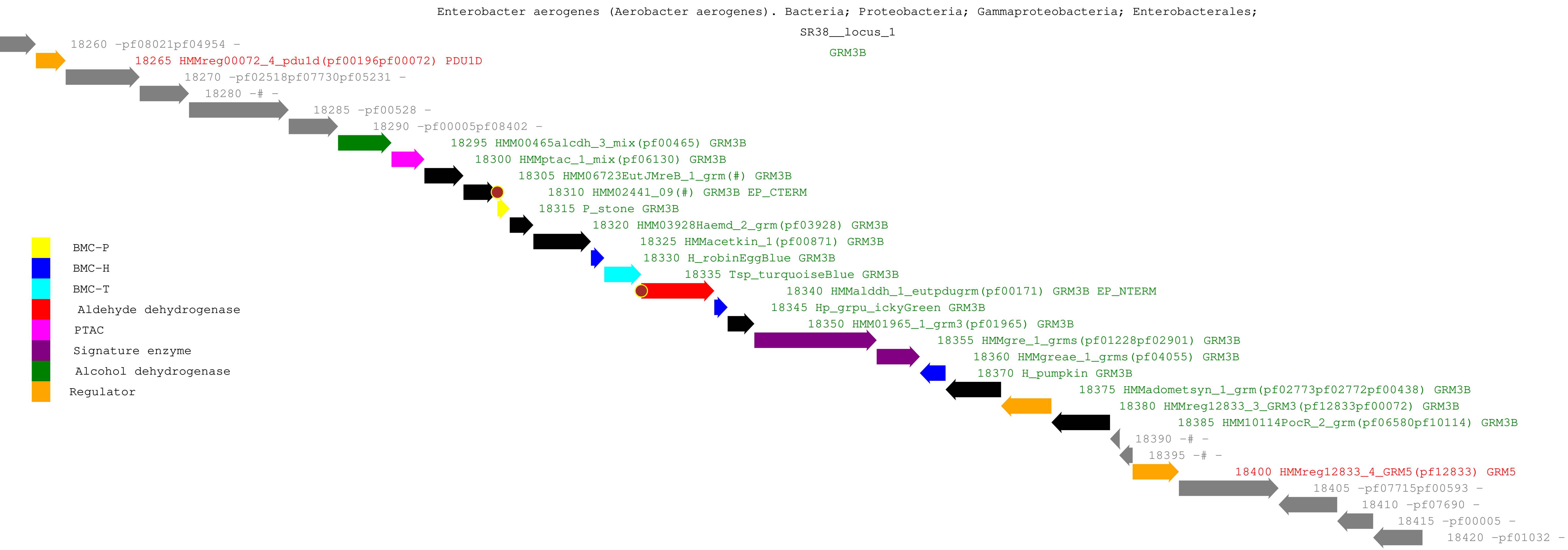
Enterobacter aerogenes (Aerobacter aerogenes). Bacteria; Proteobacteria; Gammaproteobacteria; Enterobacterales;
SR38__locus_1
GRM3B
18260 -pf08021pf04954 -
18265 HMMreg00072_4_pdu1d(pf00196pf00072) PDU1D
18270 -pf02518pf07730pf05231 -
18280 -# -
18285 -pf00528 -
18290 -pf00005pf08402 -
18295 HMM00465alcdh_3_mix(pf00465) GRM3B
18300 HMMptac_1_mix(pf06130) GRM3B
18305 HMM06723EutJMreB_1_grm(#) GRM3B
18310 HMM02441_09(#) GRM3B EP_CTERM
18315 P_stone GRM3B
18320 HMM03928Haemd_2_grm(pf03928) GRM3B
18325 HMMacetkin_1(pf00871) GRM3B
18330 H_robinEggBlue GRM3B
18335 Tsp_turquoiseBlue GRM3B
18340 HMMalddh_1_eutpdugrm(pf00171) GRM3B EP_NTERM
18345 Hp_grpu_ickyGreen GRM3B
18350 HMM01965_1_grm3(pf01965) GRM3B
18355 HMMgre_1_grms(pf01228pf02901) GRM3B
18360 HMMgreae_1_grms(pf04055) GRM3B
18370 H_pumpkin GRM3B
18375 HMMadometsyn_1_grm(pf02773pf02772pf00438) GRM3B
18380 HMMreg12833_3_GRM3(pf12833pf00072) GRM3B
18385 HMM10114PocR_2_grm(pf06580pf10114) GRM3B
18390 -# -
18395 -# -
18400 HMMreg12833_4_GRM5(pf12833) GRM5
18405 -pf07715pf00593 -
18410 -pf07690 -
18415 -pf00005 -
18420 -pf01032 -
BMC-P
BMC-H
BMC-T
Aldehyde dehydrogenase
PTAC
Signature enzyme
Alcohol dehydrogenase
Regulator

Vibrio celticus. Bacteria; Proteobacteria; Gammaproteobacteria; Vibrionales;

VCE7224__locus_1

GRM3B

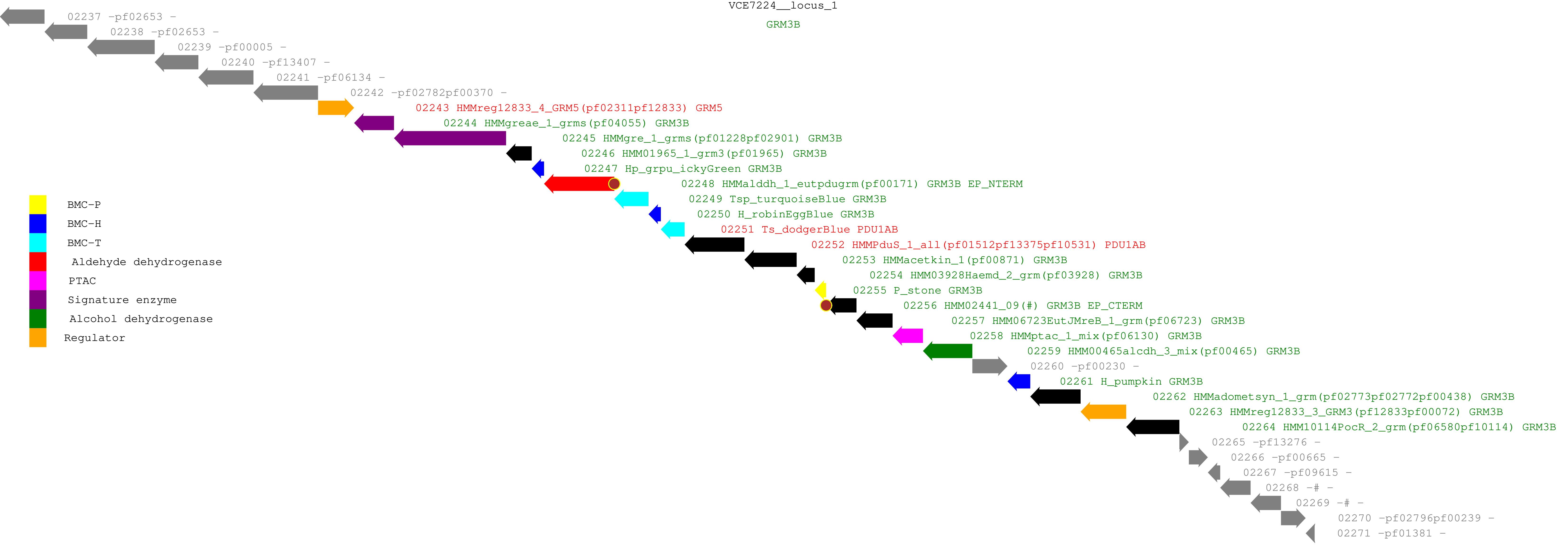

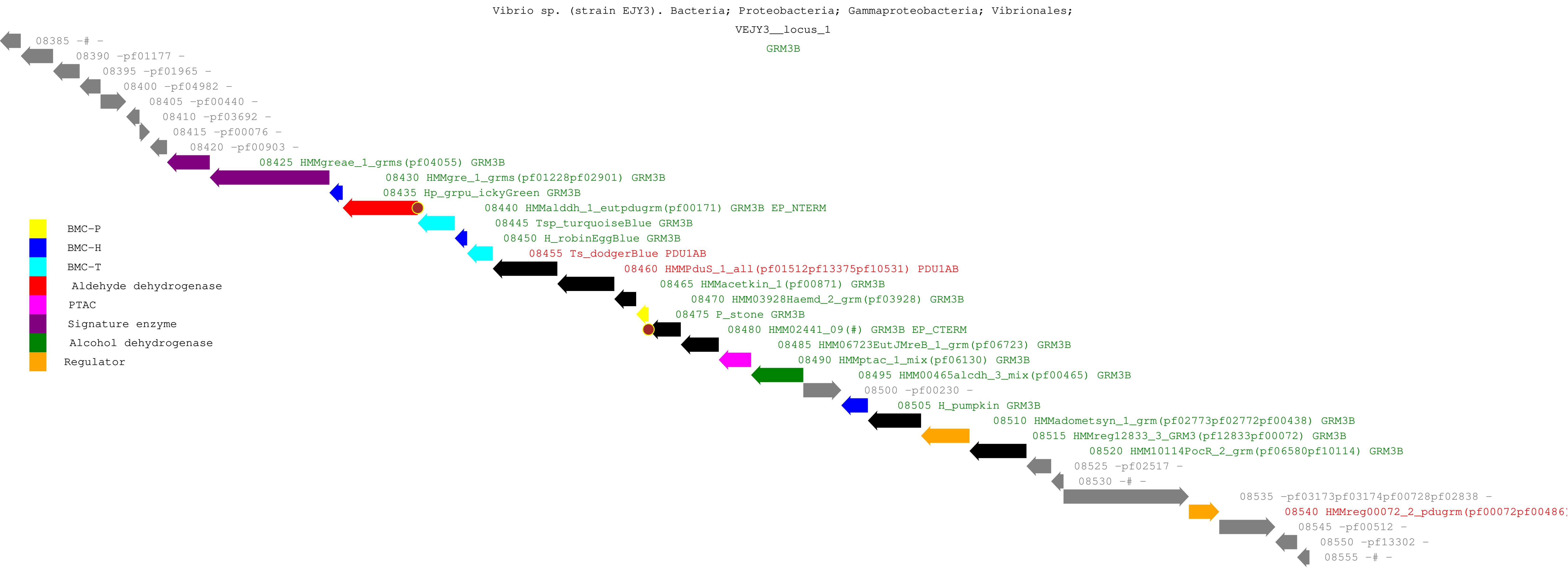

Vibrio sp. (strain EJY3). Bacteria; Proteobacteria; Gammaproteobacteria; Vibrionales;
VEJY3__locus_1
GRM3B
08385 -# -
08390 -pf01177 -
08395 -pf01965 -
08400 -pf04982 -
08405 -pf00440 -
08410 -pf03692 -
08415 -pf00076 -
08420 -pf00903 -
08425 HMMgreae_1_grms(pf04055) GRM3B
08430 HMMgre_1_grms(pf01228pf02901) GRM3B
08435 Hp_grpu_ickyGreen GRM3B
08440 HMMalddh_1_eutpdugrm(pf00171) GRM3B EP_NTERM
08445 Tsp_turquoiseBlue GRM3B
08450 H_robinEggBlue GRM3B
08455 Ts_dodgerBlue PDU1AB
08460 HMMPduS_1_all(pf01512pf13375pf10531) PDU1AB
08465 HMMacetkin_1(pf00871) GRM3B
08470 HMM03928Haemd_2_grm(pf03928) GRM3B
08475 P_stone GRM3B
08480 HMM02441_09(#) GRM3B EP_CTERM
08485 HMM06723EutJMreB_1_grm(pf06723) GRM3B
08490 HMMptac_1_mix(pf06130) GRM3B
08495 HMM00465alcdh_3_mix(pf00465) GRM3B
08500 -pf00230 -
08505 H_pumpkin GRM3B
08510 HMMadometsyn_1_grm(pf02773pf02772pf00438) GRM3B
08515 HMMreg12833_3_GRM3(pf12833pf00072) GRM3B
08520 HMM10114PocR_2_grm(pf06580pf10114) GRM3B
08525 -pf02517 -
08530 -# -
08535 -pf03173pf03174pf00728pf02838 -
08540 HMMreg00072_2_pdugrm(pf00072pf00486)
08545 -pf00512 -
08550 -pf13302 -
08555 -# -
BMC-P
BMC-H
BMC-T
Aldehyde dehydrogenase
PTAC
Signature enzyme
Alcohol dehydrogenase
Regulator

Pectobacterium sp. (strain SCC3193). Bacteria; Proteobacteria; Gammaproteobacteria; Enterobacterales;

W5S__locus_1

GRM3B

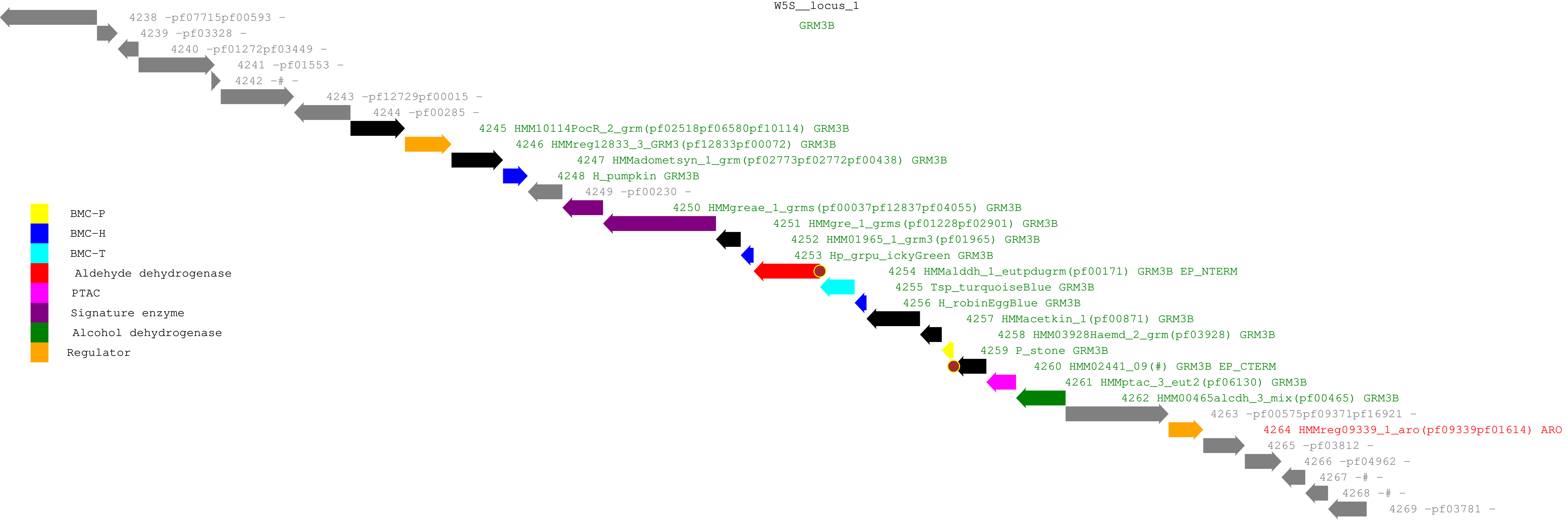

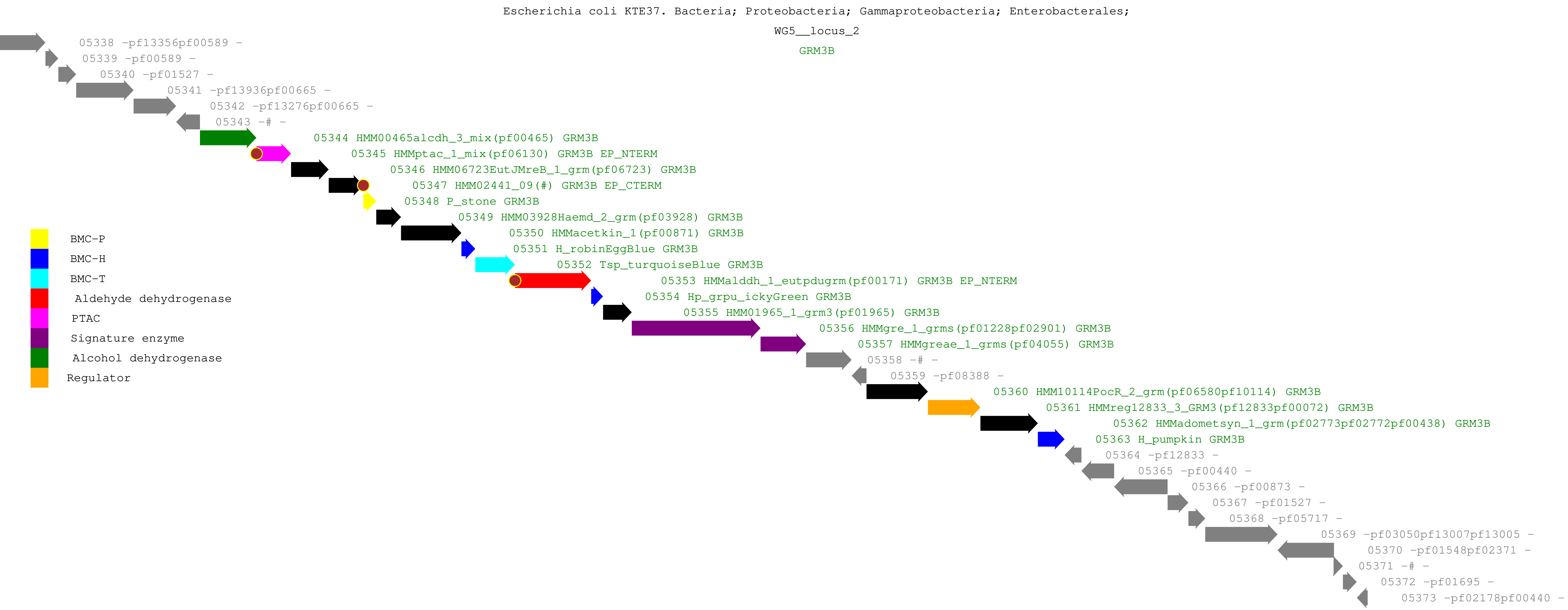

Escherichia coli KTE37. Bacteria; Proteobacteria; Gammaproteobacteria; Enterobacterales;
WG5__locus_2
GRM3B
05338 -pf13356pf00589 -
05339 -pf00589 -
05340 -pf01527 -
05341 -pf13936pf00665 -
05342 -pf13276pf00665 -
05343 -# -
05344 HMM00465alcdh_3_mix(pf00465) GRM3B
05345 HMMptac_1_mix(pf06130) GRM3B EP_NTERM
05346 HMM06723EutJMreB_1_grm(pf06723) GRM3B
05347 HMM02441_09(#) GRM3B EP_CTERM
05348 P_stone GRM3B
05349 HMM03928Haemd_2_grm(pf03928) GRM3B
05350 HMMacetkin_1(pf00871) GRM3B
05351 H_robinEggBlue GRM3B
05352 Tsp_turquoiseBlue GRM3B
05353 HMMalddh_1_eutpdugrm(pf00171) GRM3B EP_NTERM
05354 Hp_grpu_ickyGreen GRM3B
05355 HMM01965_1_grm3(pf01965) GRM3B
05356 HMMgre_1_grms(pf01228pf02901) GRM3B
05357 HMMgreae_1_grms(pf04055) GRM3B
05358 -# -
05359 -pf08388 -
05360 HMM10114PocR_2_grm(pf06580pf10114) GRM3B
05361 HMMreg12833_3_GRM3(pf12833pf00072) GRM3B
05362 HMMadometsyn_1_grm(pf02773pf02772pf00438) GRM3B
05363 H_pumpkin GRM3B
05364 -pf12833 -
05365 -pf00440 -
05366 -pf00873 -
05367 -pf01527 -
05368 -pf05717 -
05369 -pf03050pf13007pf13005 -
05370 -pf01548pf02371 -
05371 -# -
05372 -pf01695 -
05373 -pf02178pf00440 -
BMC-P
BMC-H
BMC-T
Aldehyde dehydrogenase
PTAC
Signature enzyme
Alcohol dehydrogenase
Regulator

Escherichia coli KTE38. Bacteria; Proteobacteria; Gammaproteobacteria; Enterobacterales;

WG7__locus_2

GRM3B

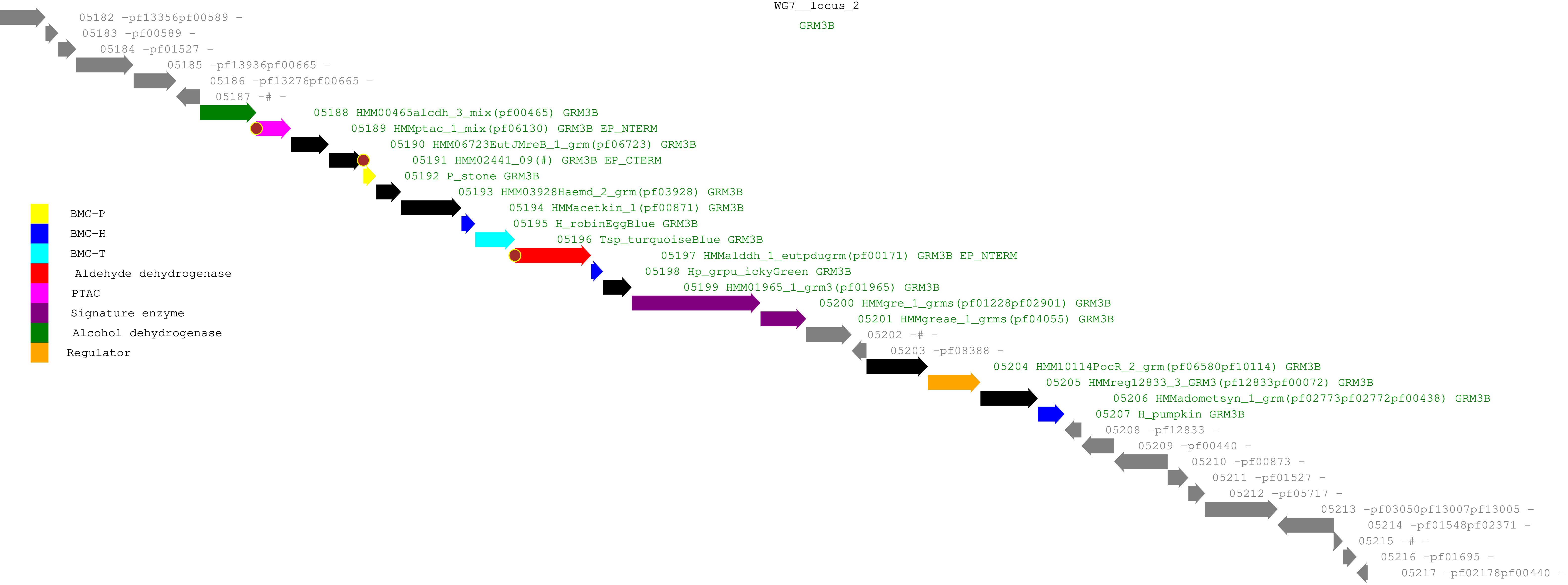

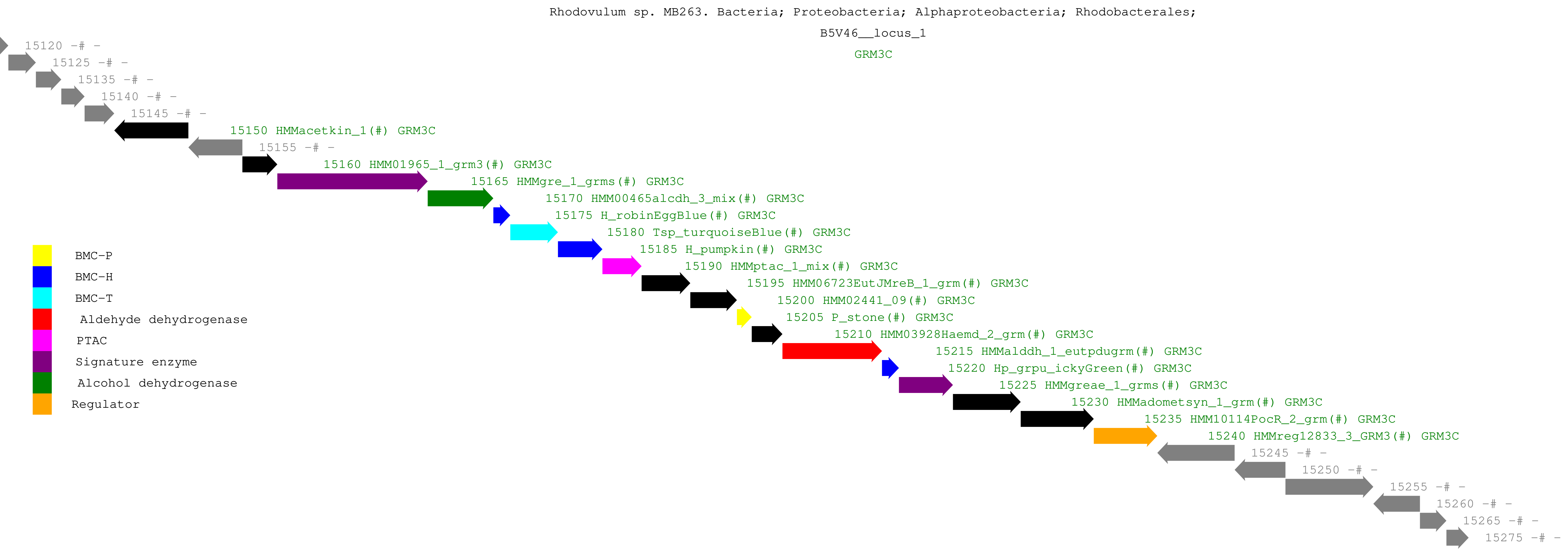
Rhodovulum sp. MB263. Bacteria; Proteobacteria; Alphaproteobacteria; Rhodobacterales;
B5V46__locus_1
GRM3C
15120 -# -
15125 -# -
15135 -# -
15140 -# -
15145 -# -
15150 HMMacetkin_1(#) GRM3C
15155 -# -
15160 HMM01965_1_grm3(#) GRM3C
15165 HMMgre_1_grms(#) GRM3C
15170 HMM00465alcdh_3_mix(#) GRM3C
15175 H_robinEggBlue(#) GRM3C
15180 Tsp_turquoiseBlue(#) GRM3C
15185 H_pumpkin(#) GRM3C
15190 HMMptac_1_mix(#) GRM3C
15195 HMM06723EutJMreB_1_grm(#) GRM3C
15200 HMM02441_09(#) GRM3C
15205 P_stone(#) GRM3C
15210 HMM03928Haemd_2_grm(#) GRM3C
15215 HMMalddh_1_eutpdugrm(#) GRM3C
15220 Hp_grpu_ickyGreen(#) GRM3C
15225 HMMgreae_1_grms(#) GRM3C
15230 HMMadometsyn_1_grm(#) GRM3C
15235 HMM10114PocR_2_grm(#) GRM3C
15240 HMMreg12833_3_GRM3(#) GRM3C
15245 -# -
15250 -# -
15255 -# -
15260 -# -
15265 -# -
15275 -# -
BMC-P
BMC-H
BMC-T
Aldehyde dehydrogenase
PTAC
Signature enzyme
Alcohol dehydrogenase
Regulator

Rhodovulum sulfidophilum (Rhodobacter sulfidophilus). Bacteria; Proteobacteria; Alphaproteobacteria; Rhodobacterales;

BV392__locus_1

GRM3C

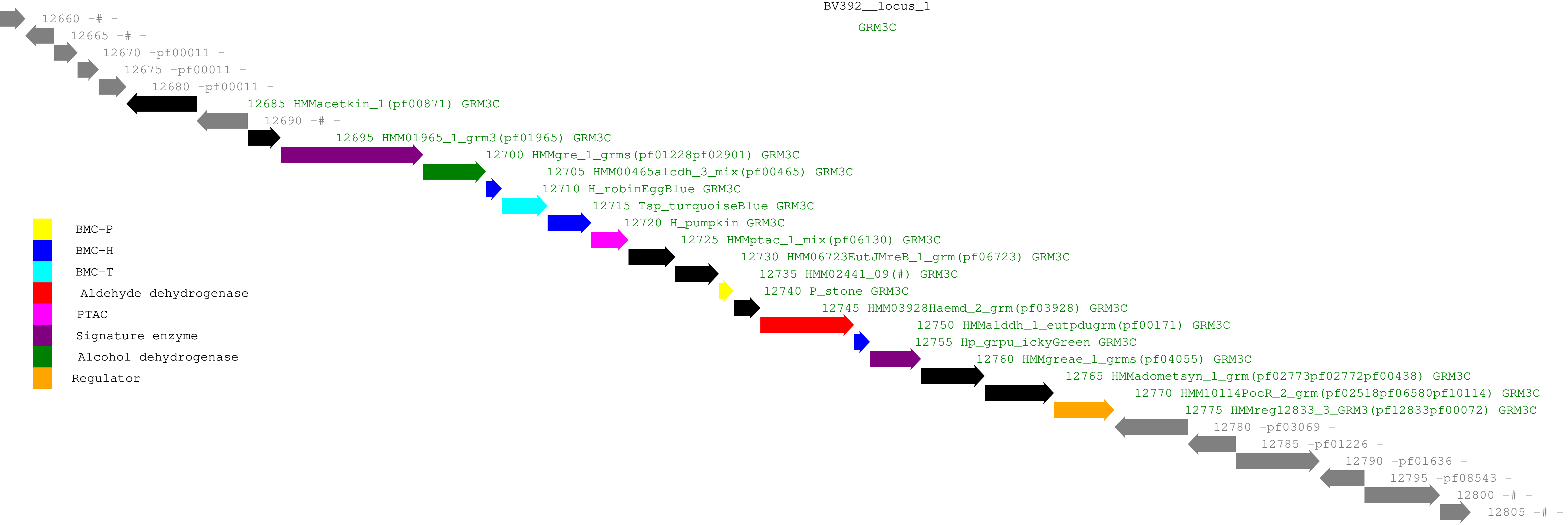

Rhodovulum sulfidophilum (Rhodobacter sulfidophilus). Bacteria; Proteobacteria; Alphaproteobacteria; Rhodobacterales;

BV509__locus_1

GRM3C

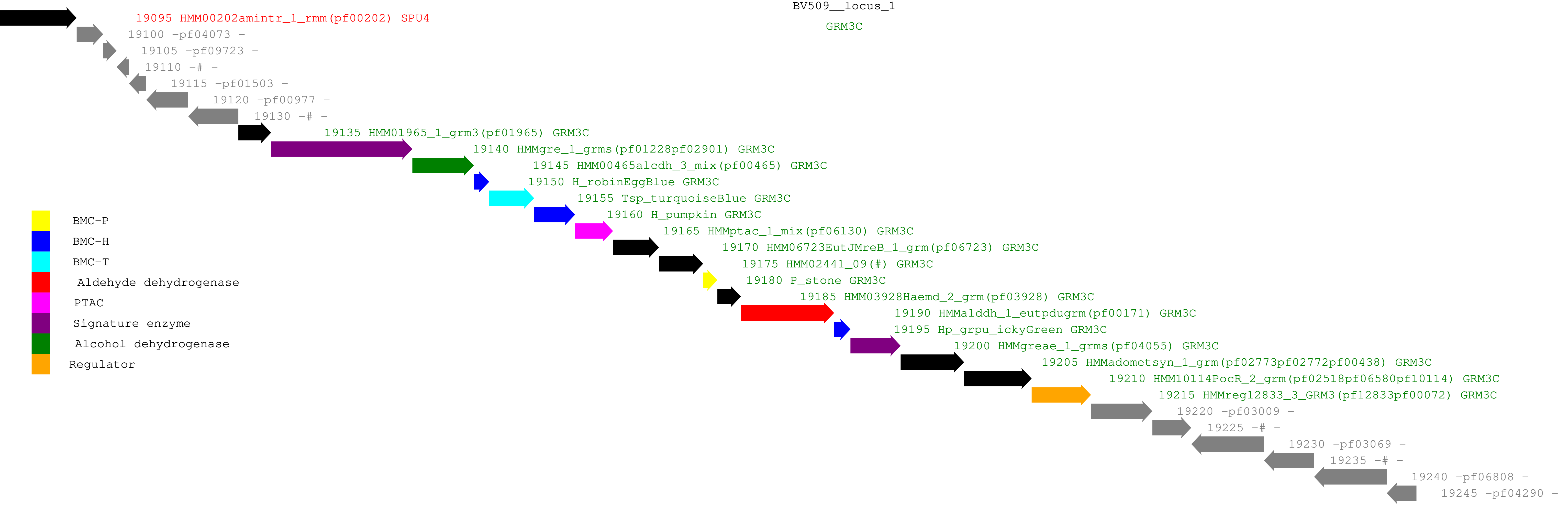

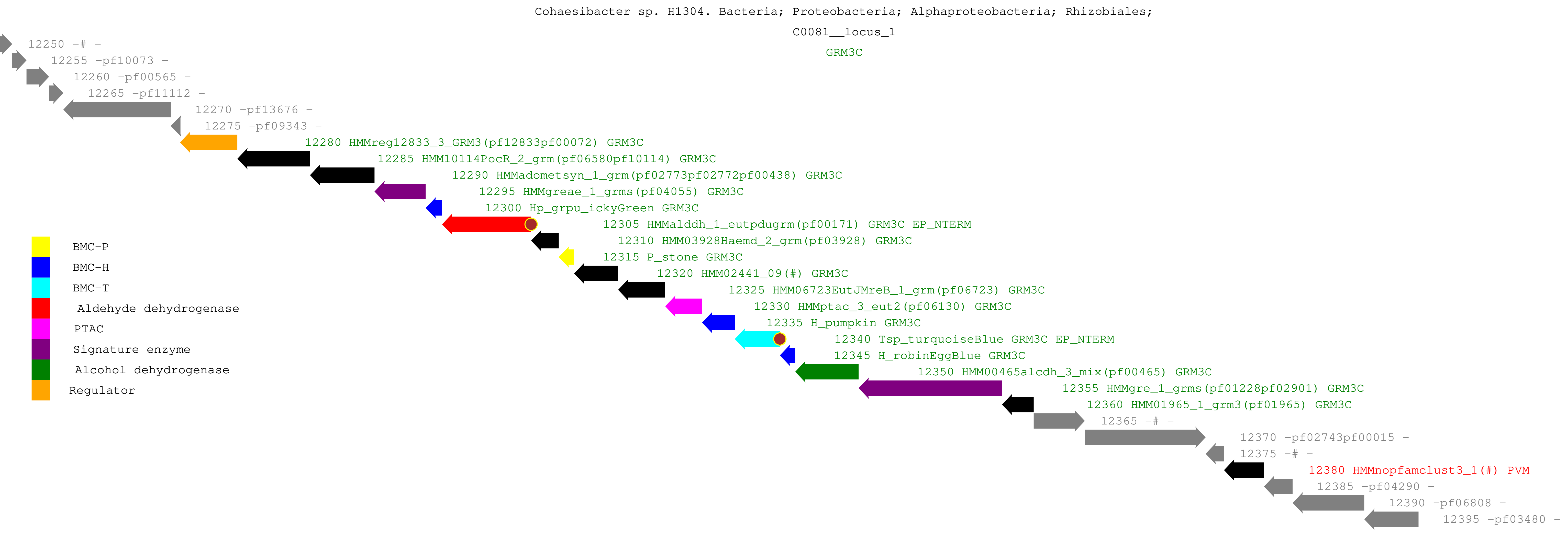
Cohaesibacter sp. H1304. Bacteria; Proteobacteria; Alphaproteobacteria; Rhizobiales;
C0081__locus_1
GRM3C
12250 -# -
12255 -pf10073 -
12260 -pf00565 -
12265 -pf11112 -
12270 -pf13676 -
12275 -pf09343 -
12280 HMMreg12833_3_GRM3(pf12833pf00072) GRM3C
12285 HMM10114PocR_2_grm(pf06580pf10114) GRM3C
12290 HMMadometsyn_1_grm(pf02773pf02772pf00438) GRM3C
12295 HMMgreae_1_grms(pf04055) GRM3C
12300 Hp_grpu_ickyGreen GRM3C
12305 HMMalddh_1_eutpdugrm(pf00171) GRM3C EP_NTERM
12310 HMM03928Haemd_2_grm(pf03928) GRM3C
12315 P_stone GRM3C
12320 HMM02441_09(#) GRM3C
12325 HMM06723EutJMreB_1_grm(pf06723) GRM3C
12330 HMMptac_3_eut2(pf06130) GRM3C
12335 H_pumpkin GRM3C
12340 Tsp_turquoiseBlue GRM3C EP_NTERM
12345 H_robinEggBlue GRM3C
12350 HMM00465alcdh_3_mix(pf00465) GRM3C
12355 HMMgre_1_grms(pf01228pf02901) GRM3C
12360 HMM01965_1_grm3(pf01965) GRM3C
12365 -# -
12370 -pf02743pf00015 -
12375 -# -
12380 HMMnopfamclust3_1(#) PVM
12385 -pf04290 -
12390 -pf06808 -
12395 -pf03480 -
BMC-P
BMC-H
BMC-T
Aldehyde dehydrogenase
PTAC
Signature enzyme
Alcohol dehydrogenase
Regulator

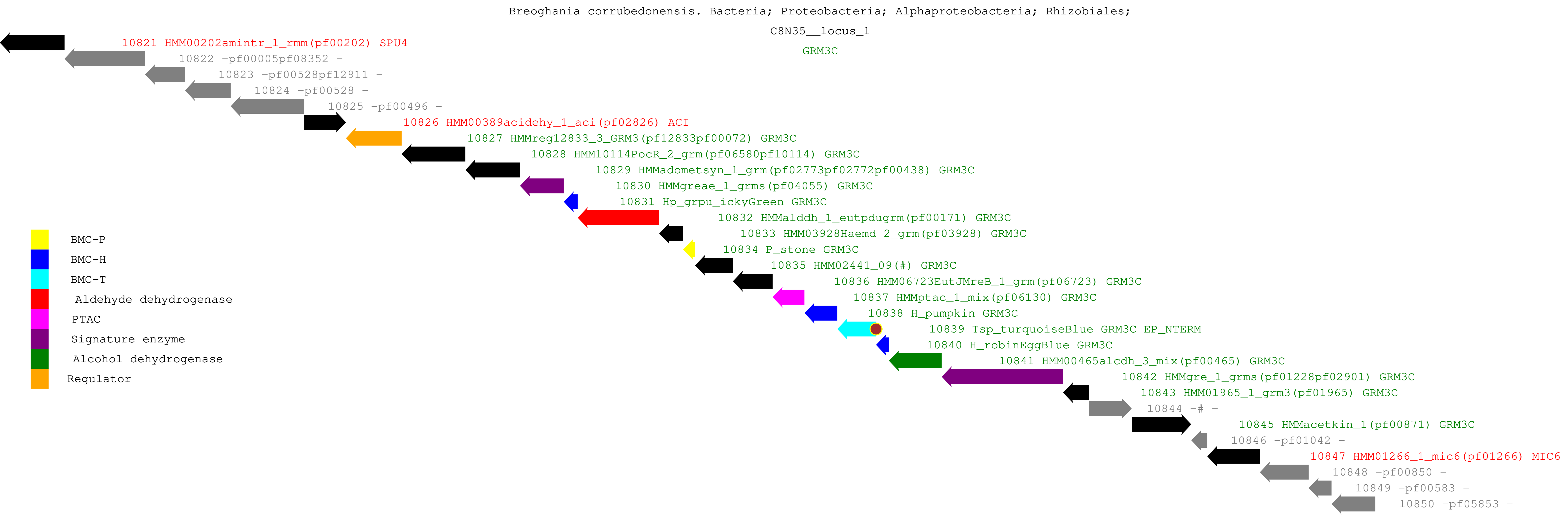
Breoghania corrubedonensis. Bacteria; Proteobacteria; Alphaproteobacteria; Rhizobiales;
C8N35__locus_1
GRM3C
10821 HMM00202amintr_1_rmm(pf00202) SPU4
10822 -pf00005pf08352 -
10823 -pf00528pf12911 -
10824 -pf00528 -
10825 -pf00496 -
10826 HMM00389acidehy_1_aci(pf02826) ACI
10827 HMMreg12833_3_GRM3(pf12833pf00072) GRM3C
10828 HMM10114PocR_2_grm(pf06580pf10114) GRM3C
10829 HMMadometsyn_1_grm(pf02773pf02772pf00438) GRM3C
10830 HMMgreae_1_grms(pf04055) GRM3C
10831 Hp_grpu_ickyGreen GRM3C
10832 HMMalddh_1_eutpdugrm(pf00171) GRM3C
10833 HMM03928Haemd_2_grm(pf03928) GRM3C
10834 P_stone GRM3C
10835 HMM02441_09(#) GRM3C
10836 HMM06723EutJMreB_1_grm(pf06723) GRM3C
10837 HMMptac_1_mix(pf06130) GRM3C
10838 H_pumpkin GRM3C
10839 Tsp_turquoiseBlue GRM3C EP_NTERM
10840 H_robinEggBlue GRM3C
10841 HMM00465alcdh_3_mix(pf00465) GRM3C
10842 HMMgre_1_grms(pf01228pf02901) GRM3C
10843 HMM01965_1_grm3(pf01965) GRM3C
10844 -# -
10845 HMMacetkin_1(pf00871) GRM3C
10846 -pf01042 -
10847 HMM01266_1_mic6(pf01266) MIC6
10848 -pf00850 -
10849 -pf00583 -
10850 -pf05853 -
BMC-P
BMC-H
BMC-T
Aldehyde dehydrogenase
PTAC
Signature enzyme
Alcohol dehydrogenase
Regulator

Rhodovulum kholense. Bacteria; Proteobacteria; Alphaproteobacteria; Rhodobacterales;

C8N38__locus_1

GRM3C

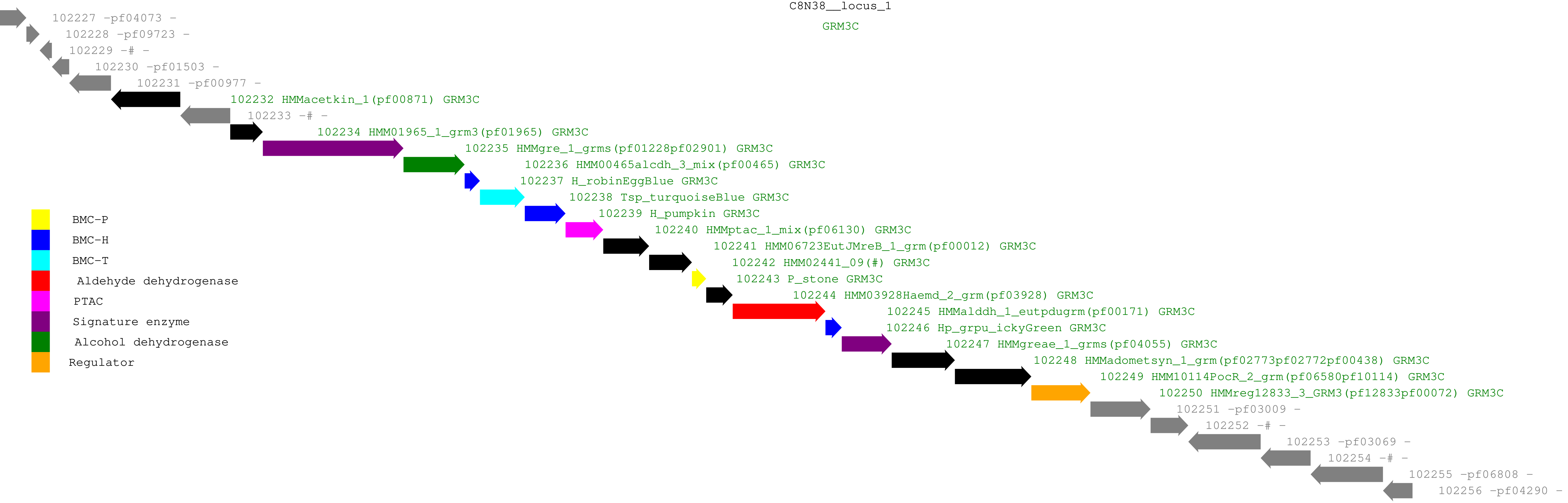

Rhodobium orientis. Bacteria; Proteobacteria; Alphaproteobacteria; Rhizobiales;

CH339__locus_1

GRM3C

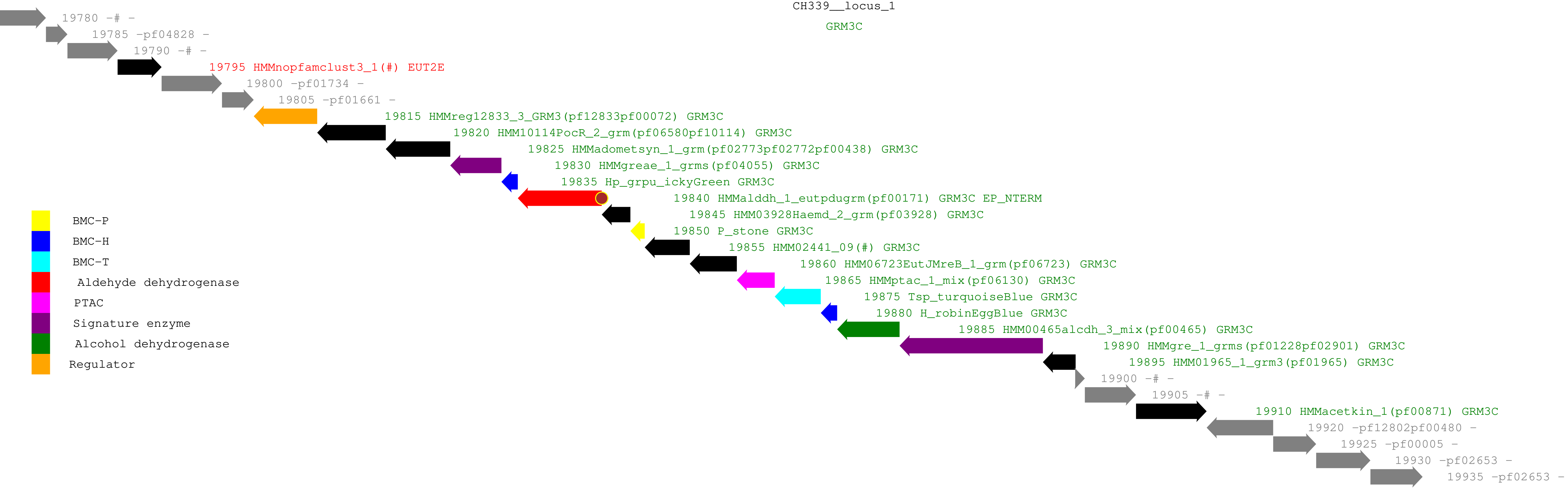

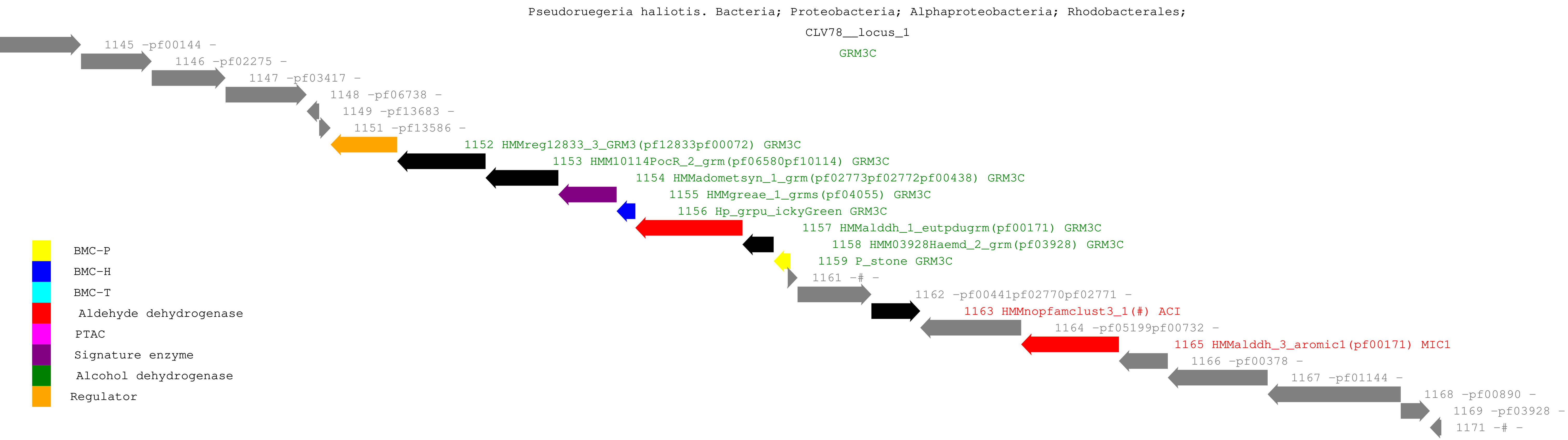
Pseudoruegeria haliotis. Bacteria; Proteobacteria; Alphaproteobacteria; Rhodobacterales;
CLV78__locus_1
GRM3C
1145 -pf00144 -
1146 -pf02275 -
1147 -pf03417 -
1148 -pf06738 -
1149 -pf13683 -
1151 -pf13586 -
1152 HMMreg12833_3_GRM3(pf12833pf00072) GRM3C
1153 HMM10114PocR_2_grm(pf06580pf10114) GRM3C
1154 HMMadometsyn_1_grm(pf02773pf02772pf00438) GRM3C
1155 HMMgreae_1_grms(pf04055) GRM3C
1156 Hp_grpu_ickyGreen GRM3C
1157 HMMalddh_1_eutpdugrm(pf00171) GRM3C
1158 HMM03928Haemd_2_grm(pf03928) GRM3C
1159 P_stone GRM3C
1161 -# -
1162 -pf00441pf02770pf02771 -
1163 HMMnopfamclust3_1(#) ACI
1164 -pf05199pf00732 -
1165 HMMalddh_3_aromic1(pf00171) MIC1
1166 -pf00378 -
1167 -pf01144 -
1168 -pf00890 -
1169 -pf03928 -
1171 -# -
BMC-P
BMC-H
BMC-T
Aldehyde dehydrogenase
PTAC
Signature enzyme
Alcohol dehydrogenase
Regulator

Rhodovulum sp. BSW8. Bacteria; Proteobacteria; Alphaproteobacteria; Rhodobacterales;

DSD19__locus_1

GRM3C

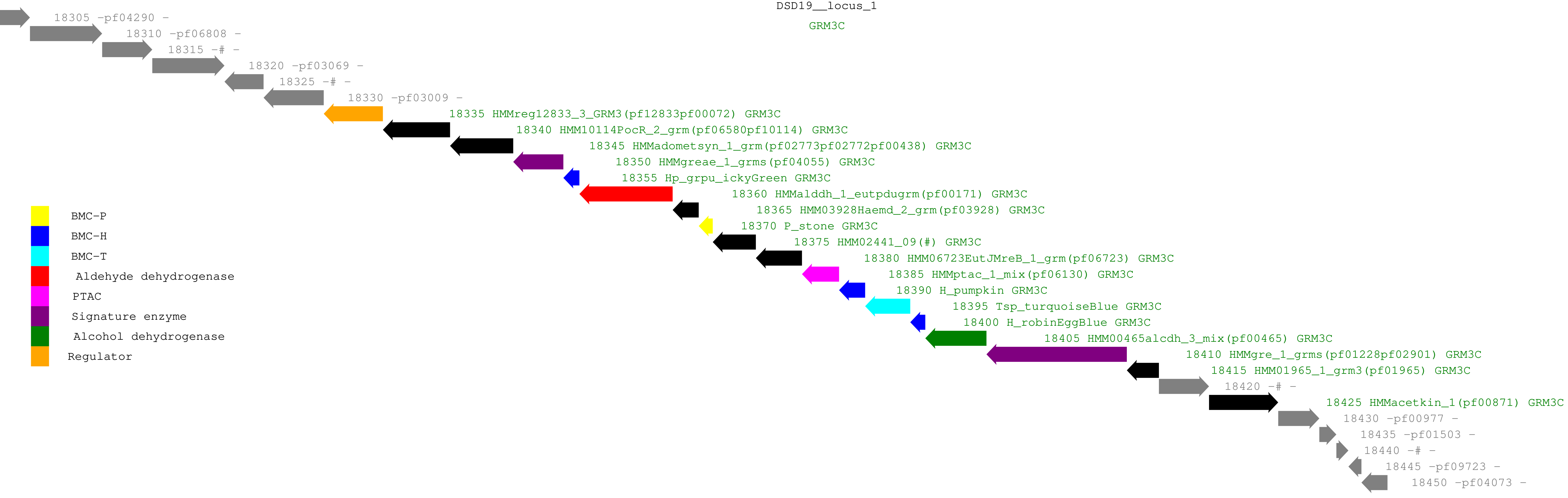

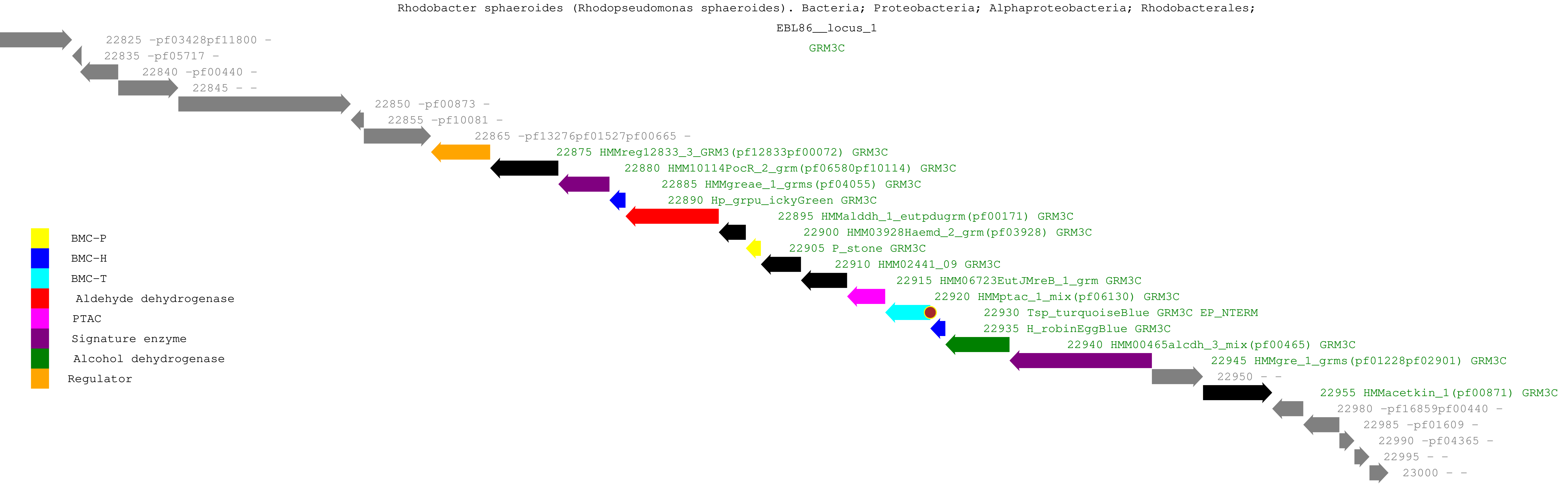
Rhodobacter sphaeroides (Rhodopseudomonas sphaeroides). Bacteria; Proteobacteria; Alphaproteobacteria; Rhodobacterales;
EBL86__locus_1
GRM3C
22825 -pf03428pf11800 -
22835 -pf05717 -
22840 -pf00440 -
22845 - -
22850 -pf00873 -
22855 -pf10081 -
22865 -pf13276pf01527pf00665 -
22875 HMMreg12833_3_GRM3(pf12833pf00072) GRM3C
22880 HMM10114PocR_2_grm(pf06580pf10114) GRM3C
22885 HMMgreae_1_grms(pf04055) GRM3C
22890 Hp_grpu_ickyGreen GRM3C
22895 HMMalddh_1_eutpdugrm(pf00171) GRM3C
22900 HMM03928Haemd_2_grm(pf03928) GRM3C
22905 P_stone GRM3C
22910 HMM02441_09 GRM3C
22915 HMM06723EutJMreB_1_grm GRM3C
22920 HMMptac_1_mix(pf06130) GRM3C
22930 Tsp_turquoiseBlue GRM3C EP_NTERM
22935 H_robinEggBlue GRM3C
22940 HMM00465alcdh_3_mix(pf00465) GRM3C
22945 HMMgre_1_grms(pf01228pf02901) GRM3C
22950 - -
22955 HMMacetkin_1(pf00871) GRM3C
22980 -pf16859pf00440 -
22985 -pf01609 -
22990 -pf04365 -
22995 - -
23000 - -
BMC-P
BMC-H
BMC-T
Aldehyde dehydrogenase
PTAC
Signature enzyme
Alcohol dehydrogenase
Regulator

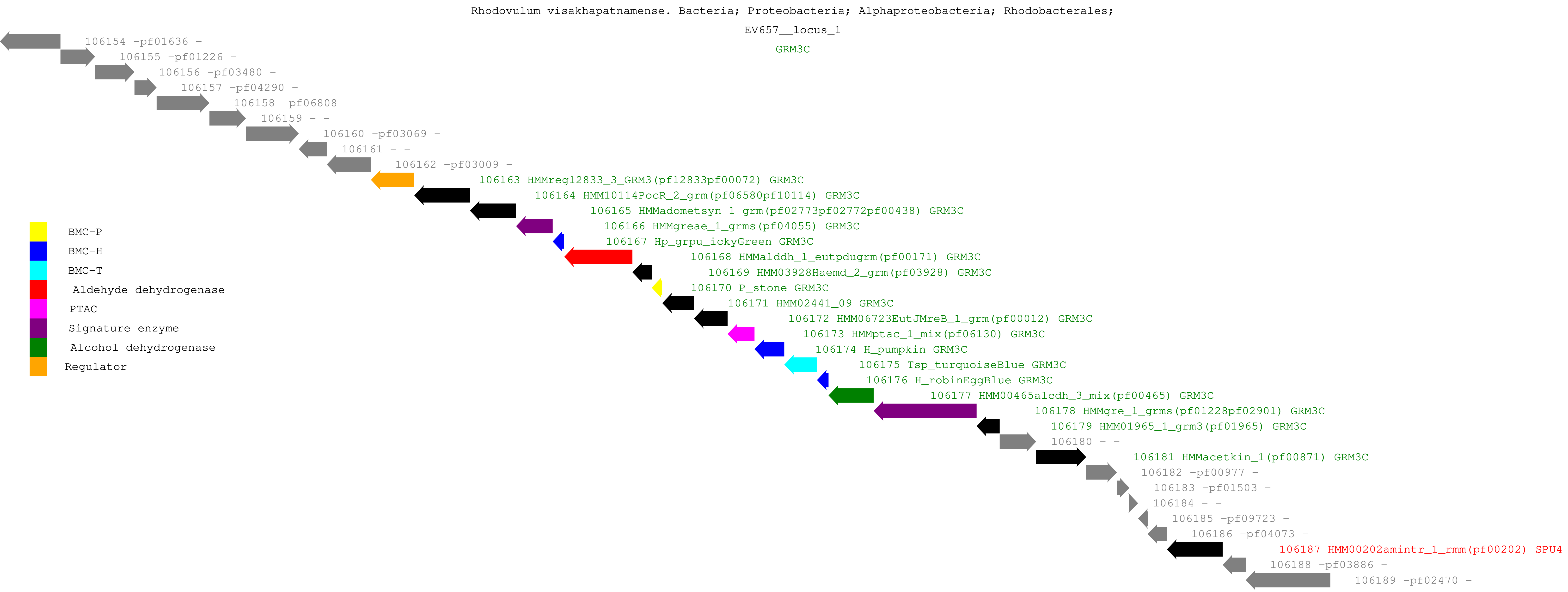
Rhodovulum visakhapatnamense. Bacteria; Proteobacteria; Alphaproteobacteria; Rhodobacterales;
EV657__locus_1
GRM3C
106154 -pf01636 -
106155 -pf01226 -
106156 -pf03480 -
106157 -pf04290 -
106158 -pf06808 -
106159 - -
106160 -pf03069 -
106161 - -
106162 -pf03009 -
106163 HMMreg12833_3_GRM3(pf12833pf00072) GRM3C
106164 HMM10114PocR_2_grm(pf06580pf10114) GRM3C
106165 HMMadometsyn_1_grm(pf02773pf02772pf00438) GRM3C
106166 HMMgreae_1_grms(pf04055) GRM3C
106167 Hp_grpu_ickyGreen GRM3C
106168 HMMaldh_1_eutpdugrm(pf00171) GRM3C
106169 HMM03928Haemd_2_grm(pf03928) GRM3C
106170 P_stone GRM3C
106171 HMM02441_09 GRM3C
106172 HMM06723EutJMreB_1_grm(pf00012) GRM3C
106173 HMMptac_1_mix(pf06130) GRM3C
106174 H_pumpkin GRM3C
106175 Tsp_turquoiseBlue GRM3C
106176 H_robinEggBlue GRM3C
106177 HMM00465alcdh_3_mix(pf00465) GRM3C
106178 HMMgre_1_grms(pf01228pf02901) GRM3C
106179 HMM01965_1_grm3(pf01965) GRM3C
106180 - -
106181 HMMacetkin_1(pf00871) GRM3C
106182 -pf00977 -
106183 -pf01503 -
106184 - -
106185 -pf09723 -
106186 -pf04073 -
106187 HMM00202amintr_1_rmm(pf00202) SPU4
106188 -pf03886 -
106189 -pf02470 -
BMC-P
BMC-H
BMC-T
Aldehyde dehydrogenase
PTAC
Signature enzyme
Alcohol dehydrogenase
Regulator

Rhodobacter capsulatus (Rhodopseudomonas capsulata). Bacteria; Proteobacteria; Alphaproteobacteria; Rhodobacterales;

FBT96__locus_1

GRM3C

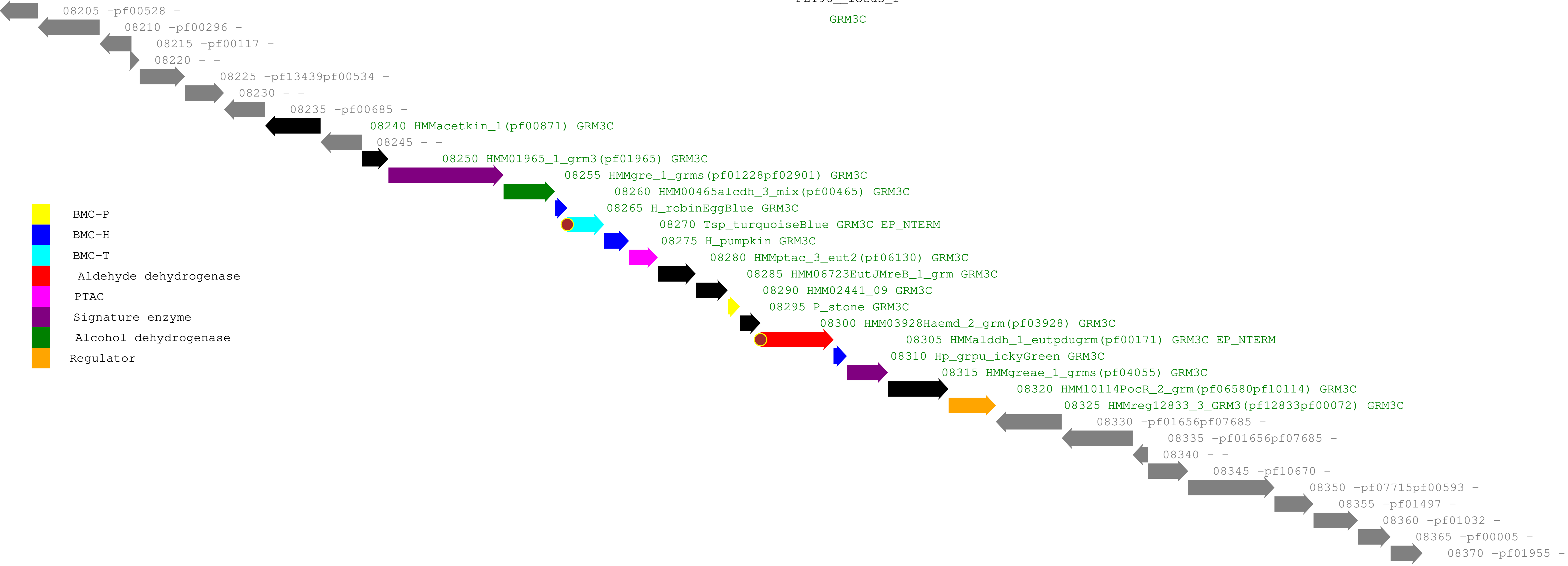

Rhizobium smilacinae. Bacteria; Proteobacteria; Alphaproteobacteria; Rhizobiales; Rhizobiaceae;

FHP24__locus_1

GRM3C

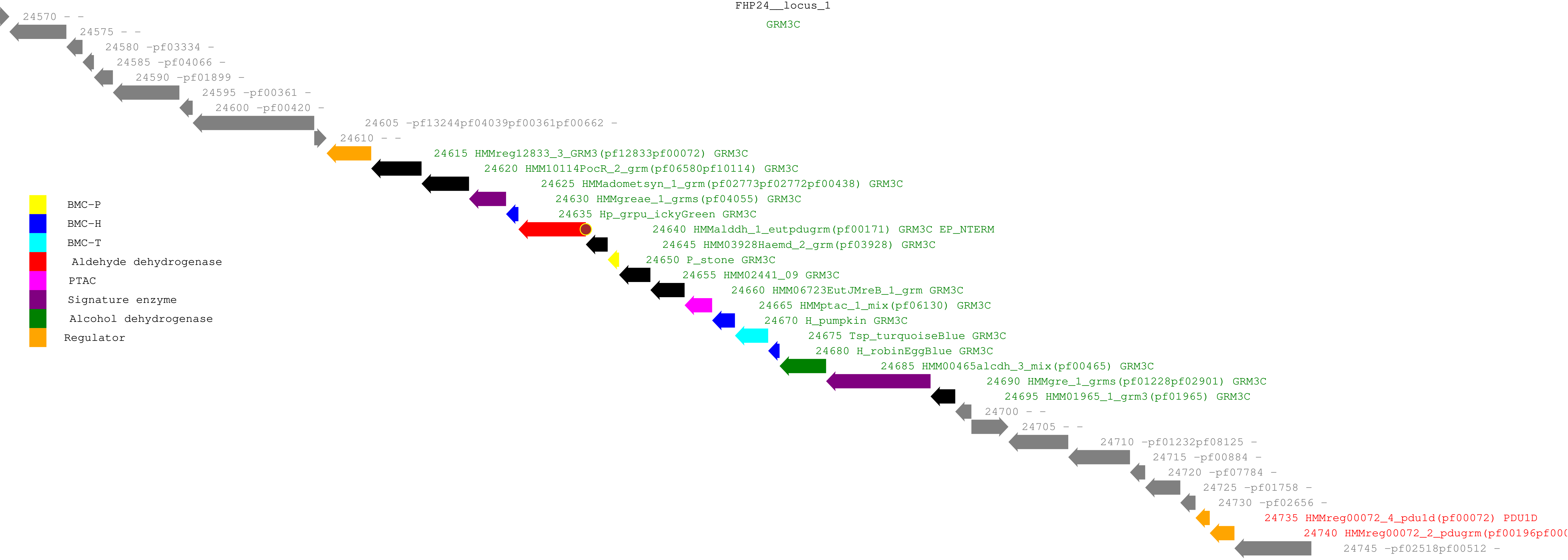

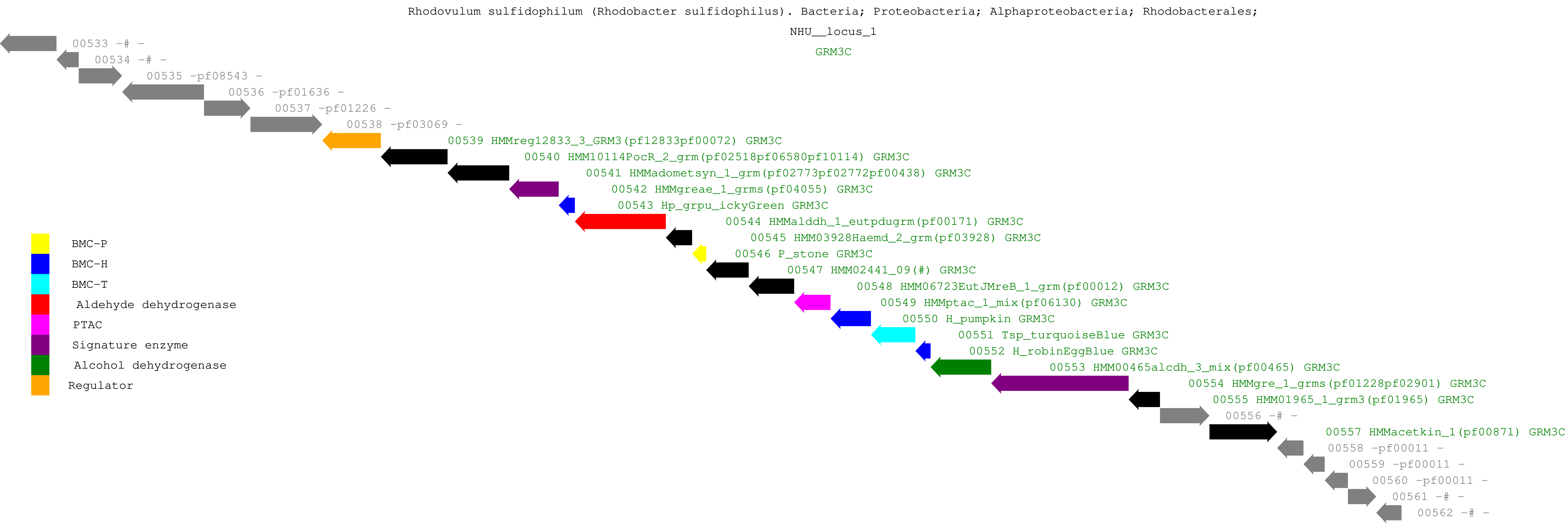
Rhodovulum sulfidophilum (Rhodobacter sulfidophilus). Bacteria; Proteobacteria; Alphaproteobacteria; Rhodobacterales;
NHU__locus_1
GRM3C
00533 -# -
00534 -# -
00535 -pf08543 -
00536 -pf01636 -
00537 -pf01226 -
00538 -pf03069 -
00539 HMMreg12833_3_GRM3(pf12833pf00072) GRM3C
00540 HMM10114PocR_2_grm(pf02518pf06580pf10114) GRM3C
00541 HMMadometsyn_1_grm(pf02773pf02772pf00438) GRM3C
00542 HMMgreae_1_grms(pf04055) GRM3C
00543 Hp_grpu_ickyGreen GRM3C
00544 HMMalddh_1_eutpdugrm(pf00171) GRM3C
00545 HMM03928Haemd_2_grm(pf03928) GRM3C
00546 P_stone GRM3C
00547 HMM02441_09(#) GRM3C
00548 HMM06723EutJMreB_1_grm(pf00012) GRM3C
00549 HMMptac_1_mix(pf06130) GRM3C
00550 H_pumpkin GRM3C
00551 Tsp_turquoiseBlue GRM3C
00552 H_robinEggBlue GRM3C
00553 HMM00465alcdh_3_mix(pf00465) GRM3C
00554 HMMgre_1_grms(pf01228pf02901) GRM3C
00555 HMM01965_1_grm3(pf01965) GRM3C
00556 -# -
00557 HMMacetkin_1(pf00871) GRM3C
00558 -pf00011 -
00559 -pf00011 -
00560 -pf00011 -
00561 -# -
00562 -# -
BMC-P
BMC-H
BMC-T
Aldehyde dehydrogenase
PTAC
Signature enzyme
Alcohol dehydrogenase
Regulator

Rhodobacter capsulatus (strain ATCC BAA-309 / NBRC 16581 / SB1003). Bacteria; Proteobacteria; Alphaproteobacteria; Rhodobacterales;

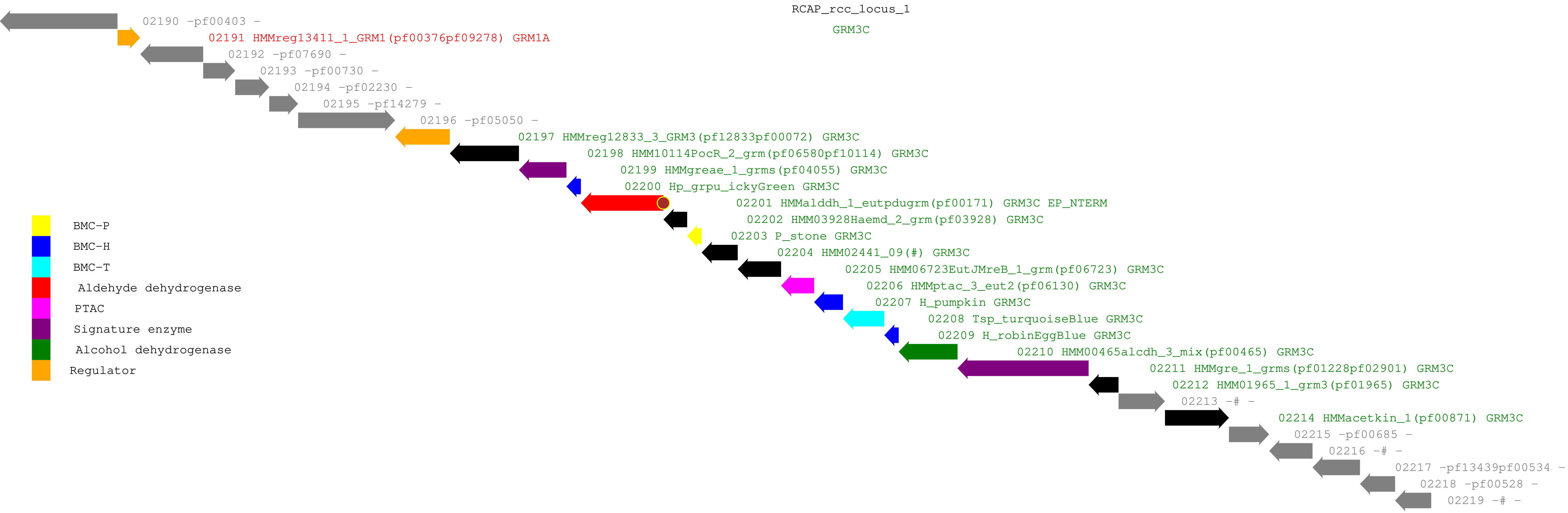

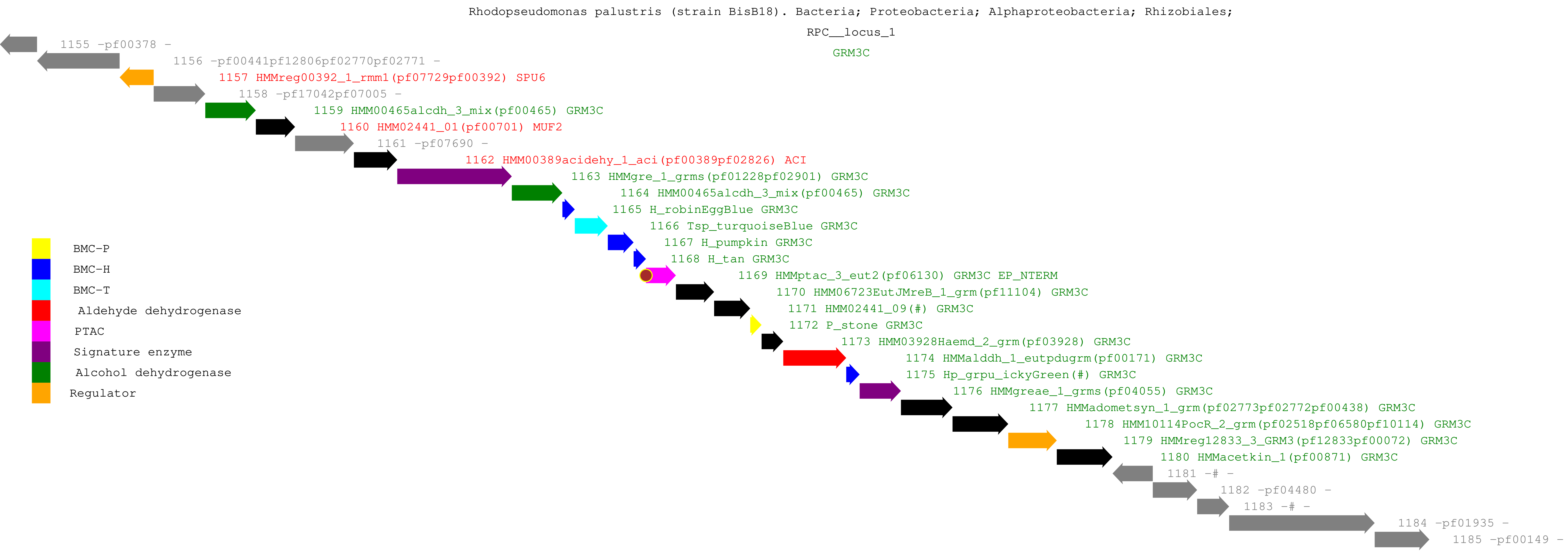
Rhodopseudomonas palustris (strain BisB18). Bacteria; Proteobacteria; Alphaproteobacteria; Rhizobiales;
RPC__locus_1
GRM3C
1155 -pf00378 -
1156 -pf00441pf12806pf02770pf02771 -
1157 HMMreg00392_1_rmm1(pf07729pf00392) SPU6
1158 -pf17042pf07005 -
1159 HMM00465alcdh_3_mix(pf00465) GRM3C
1160 HMM02441_01(pf00701) MUF2
1161 -pf07690 -
1162 HMM00389acidehy_1_aci(pf00389pf02826) ACI
1163 HMMgre_1_grms(pf01228pf02901) GRM3C
1164 HMM00465alcdh_3_mix(pf00465) GRM3C
1165 H_robinEggBlue GRM3C
1166 Tsp_turquoiseBlue GRM3C
1167 H_pumpkin GRM3C
1168 H_tan GRM3C
1169 HMMptac_3_eut2(pf06130) GRM3C EP_NTERM
1170 HMM06723EutJMreB_1_grm(pf11104) GRM3C
1171 HMM02441_09(#) GRM3C
1172 P_stone GRM3C
1173 HMM03928Haemd_2_grm(pf03928) GRM3C
1174 HMMalddh_1_eutpdugrm(pf00171) GRM3C
1175 Hp_grpu_ickyGreen(#) GRM3C
1176 HMMgreae_1_grms(pf04055) GRM3C
1177 HMMadometsyn_1_grm(pf02773pf02772pf00438) GRM3C
1178 HMM10114PocR_2_grm(pf02518pf06580pf10114) GRM3C
1179 HMMreg12833_3_GRM3(pf12833pf00072) GRM3C
1180 HMMacetkin_1(pf00871) GRM3C
1181 -# -
1182 -pf04480 -
1183 -# -
1184 -pf01935 -
1185 -pf00149 -
BMC-P
BMC-H
BMC-T
Aldehyde dehydrogenase
PTAC
Signature enzyme
Alcohol dehydrogenase
Regulator

Rhodospirillum rubrum (strain ATCC 11170 / ATH 1.1.1 / DSM 467 / LMG Bacteria; Proteobacteria; Alphaproteobacteria; Rhodospirillales;

Rru_A_locus_1

GRM3C

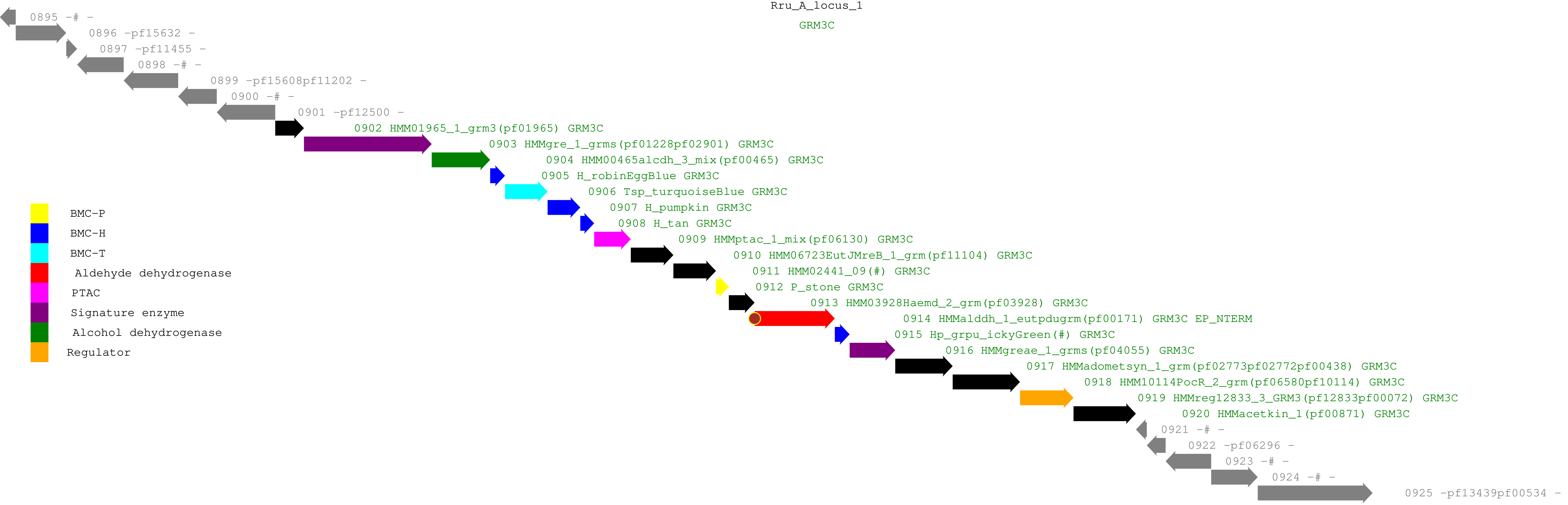

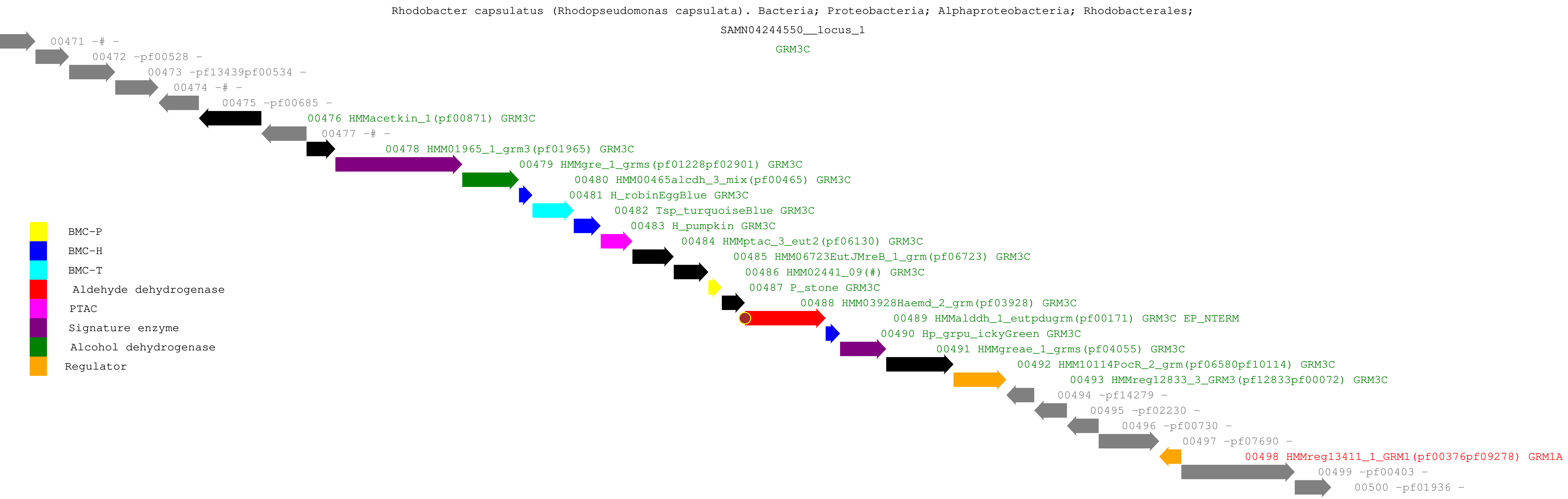

Rhodobacter capsulatus (Rhodopseudomonas capsulata). Bacteria; Proteobacteria; Alphaproteobacteria; Rhodobacterales;
SAMN04244550__locus_1
GRM3C
00471 -# -
00472 -pf00528 -
00473 -pf13439pf00534 -
00474 -# -
00475 -pf00685 -
00476 HMMacetkin_1(pf00871) GRM3C
00477 -# -
00478 HMM01965_1_grm3(pf01965) GRM3C
00479 HMMgre_1_grms(pf01228pf02901) GRM3C
00480 HMM00465alcdh_3_mix(pf00465) GRM3C
00481 H_robinEggBlue GRM3C
00482 Tsp_turquoiseBlue GRM3C
00483 H_pumpkin GRM3C
00484 HMMptac_3_eut2(pf06130) GRM3C
00485 HMM06723EutJMreB_1_grm(pf06723) GRM3C
00486 HMM02441_09(#) GRM3C
00487 P_stone GRM3C
00488 HMM03928Haemd_2_grm(pf03928) GRM3C
00489 HMMalddh_1_eutpdugrm(pf00171) GRM3C EP_NTERM
00490 Hp_grpu_ickyGreen GRM3C
00491 HMMgreae_1_grms(pf04055) GRM3C
00492 HMM10114PocR_2_grm(pf06580pf10114) GRM3C
00493 HMMreg12833_3_GRM3(pf12833pf00072) GRM3C
00494 -pf14279 -
00495 -pf02230 -
00496 -pf00730 -
00497 -pf07690 -
00498 HMMreg13411_1_GRM1(pf00376pf09278) GRM1A
00499 -pf00403 -
00500 -pf01936 -
BMC-P
BMC-H
BMC-T
Aldehyde dehydrogenase
PTAC
Signature enzyme
Alcohol dehydrogenase
Regulator

Cohaesibacter marisflavi. Bacteria; Proteobacteria; Alphaproteobacteria; Rhizobiales;

SAMN04488056__locus_1_main_sat

GRM3C

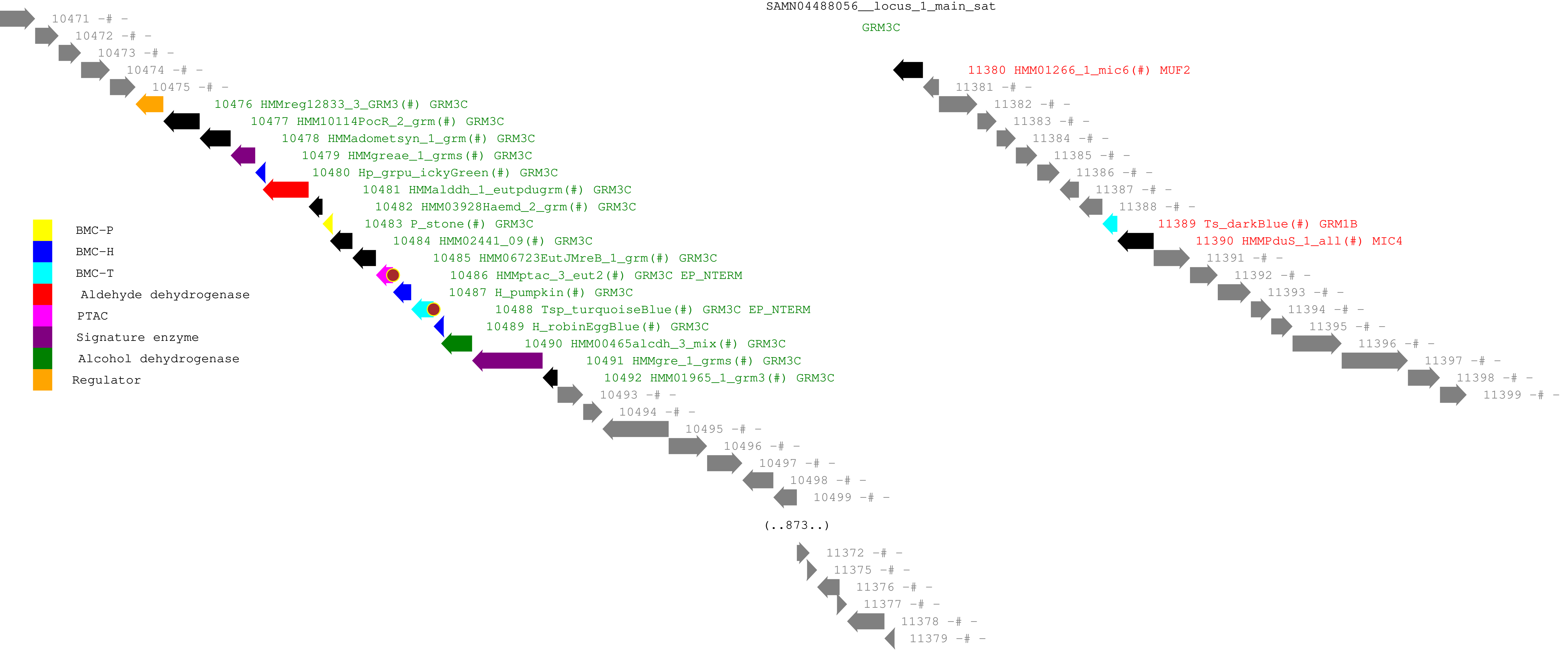

Rhodobacter aestuarii. Bacteria; Proteobacteria; Alphaproteobacteria; Rhodobacterales;

SAMN05421580__locus_1

GRM3C

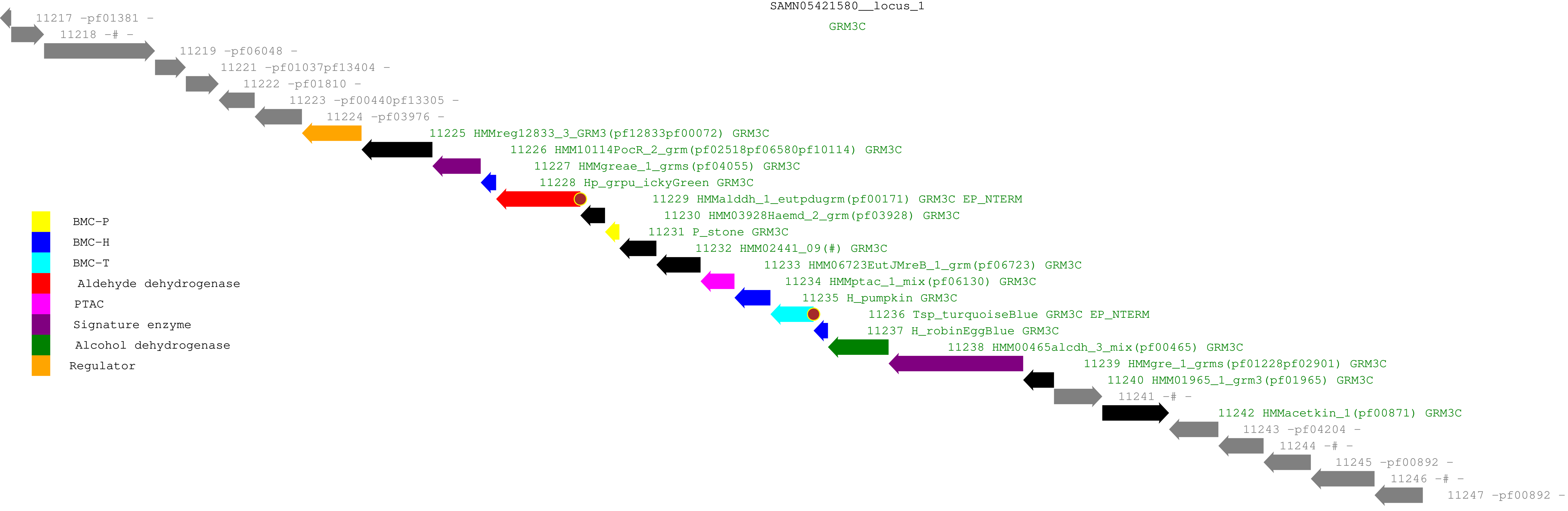

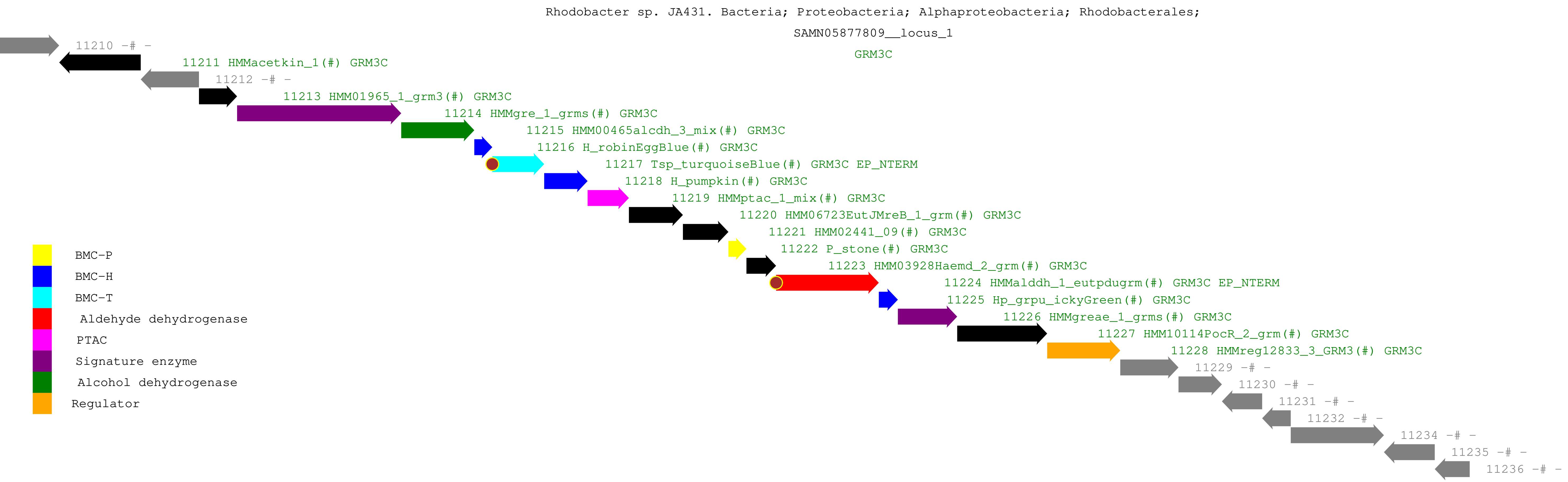
Rhodobacter sp. JA431. Bacteria; Proteobacteria; Alphaproteobacteria; Rhodobacterales;
SAMN05877809__locus_1
GRM3C
11210 -# -
11211 HMMacetkin_1(#) GRM3C
11212 -# -
11213 HMM01965_1_grm3(#) GRM3C
11214 HMMgre_1_grms(#) GRM3C
11215 HMM00465alcdh_3_mix(#) GRM3C
11216 H_robinEggBlue(#) GRM3C
11217 Tsp_turquoiseBlue(#) GRM3C EP_NTERM
11218 H_pumpkin(#) GRM3C
11219 HMMptac_1_mix(#) GRM3C
11220 HMM06723EutJMreB_1_grm(#) GRM3C
11221 HMM02441_09(#) GRM3C
11222 P_stone(#) GRM3C
11223 HMM03928Haemd_2_grm(#) GRM3C
11224 HMMalddh_1_eutpdugrm(#) GRM3C EP_NTERM
11225 Hp_grpu_ickyGreen(#) GRM3C
11226 HMMgreae_1_grms(#) GRM3C
11227 HMM10114PocR_2_grm(#) GRM3C
11228 HMMreg12833_3_GRM3(#) GRM3C
11229 -# -
11230 -# -
11231 -# -
11232 -# -
11234 -# -
11235 -# -
11236 -# -
BMC-P
BMC-H
BMC-T
Aldehyde dehydrogenase
PTAC
Signature enzyme
Alcohol dehydrogenase
Regulator

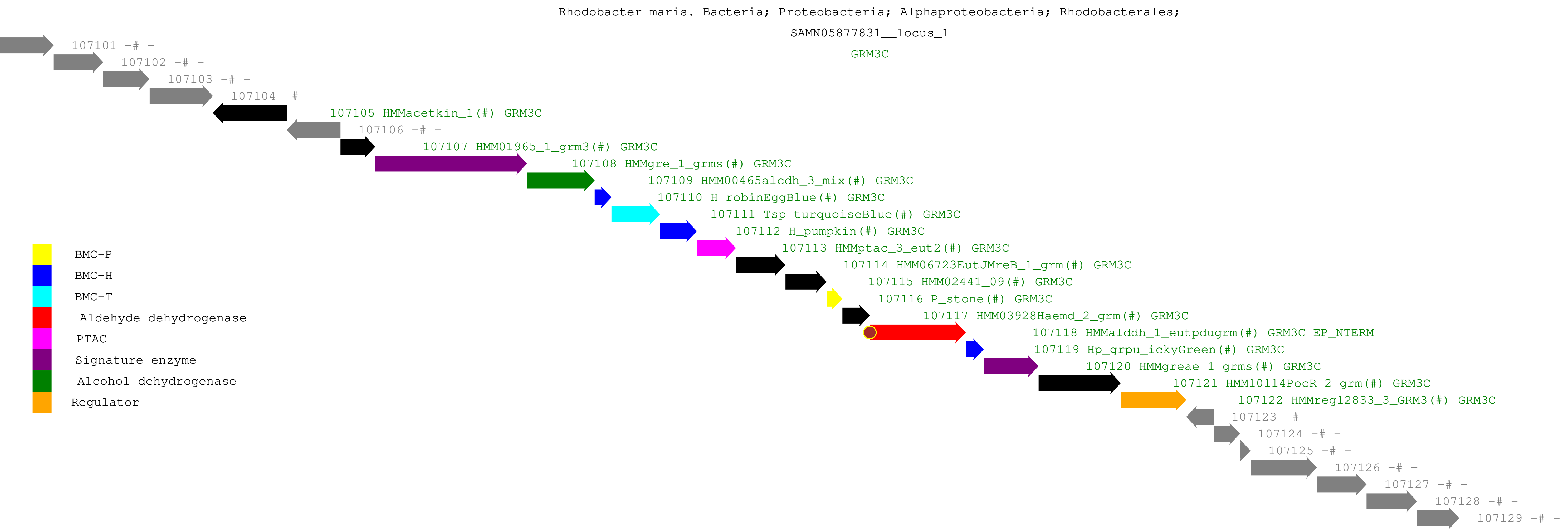
Rhodobacter maris. Bacteria; Proteobacteria; Alphaproteobacteria; Rhodobacterales;
SAMN05877831__locus_1
GRM3C
107101 -# -
107102 -# -
107103 -# -
107104 -# -
107105 HMMacetkin_1(#) GRM3C
107106 -# -
107107 HMM01965_1_grm3(#) GRM3C
107108 HMMgre_1_grms(#) GRM3C
107109 HMM00465alcdh_3_mix(#) GRM3C
107110 H_robinEggBlue(#) GRM3C
107111 Tsp_turquoiseBlue(#) GRM3C
107112 H_pumpkin(#) GRM3C
107113 HMMptac_3_eut2(#) GRM3C
107114 HMM06723EutJMreB_1_grm(#) GRM3C
107115 HMM02441_09(#) GRM3C
107116 P_stone(#) GRM3C
107117 HMM03928Haemd_2_grm(#) GRM3C
107118 HMMaldh_1_eutpdugrm(#) GRM3C EP_NTERM
107119 Hp_grpu_ickyGreen(#) GRM3C
107120 HMMgreae_1_grms(#) GRM3C
107121 HMM10114PocR_2_grm(#) GRM3C
107122 HMMreg12833_3_GRM3(#) GRM3C
107123 -# -
107124 -# -
107125 -# -
107126 -# -
107127 -# -
107128 -# -
107129 -# -
BMC-P
BMC-H
BMC-T
Aldehyde dehydrogenase
PTAC
Signature enzyme
Alcohol dehydrogenase
Regulator

Azospirillum sp. TSO35-2. Bacteria; Proteobacteria; Alphaproteobacteria; Rhodospirillales;

TSO352__locus_1

GRM3C

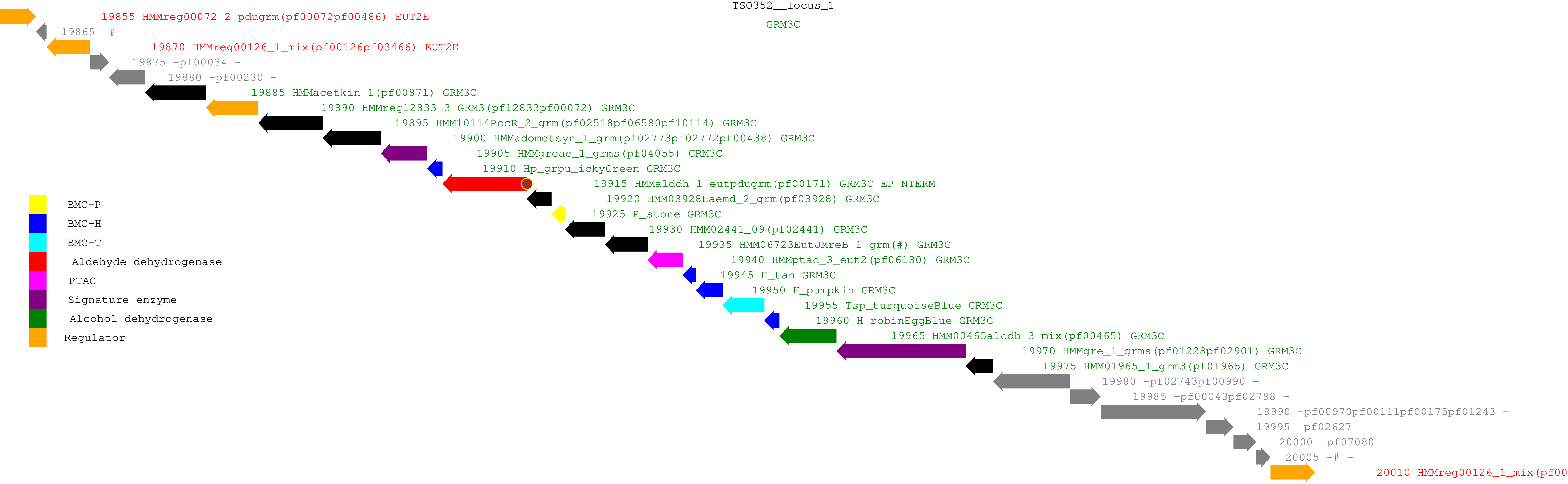

Rhodobacter capsulatus Y262. Bacteria; Proteobacteria; Alphaproteobacteria; Rhodobacterales;

U715__locus_1

GRM3C

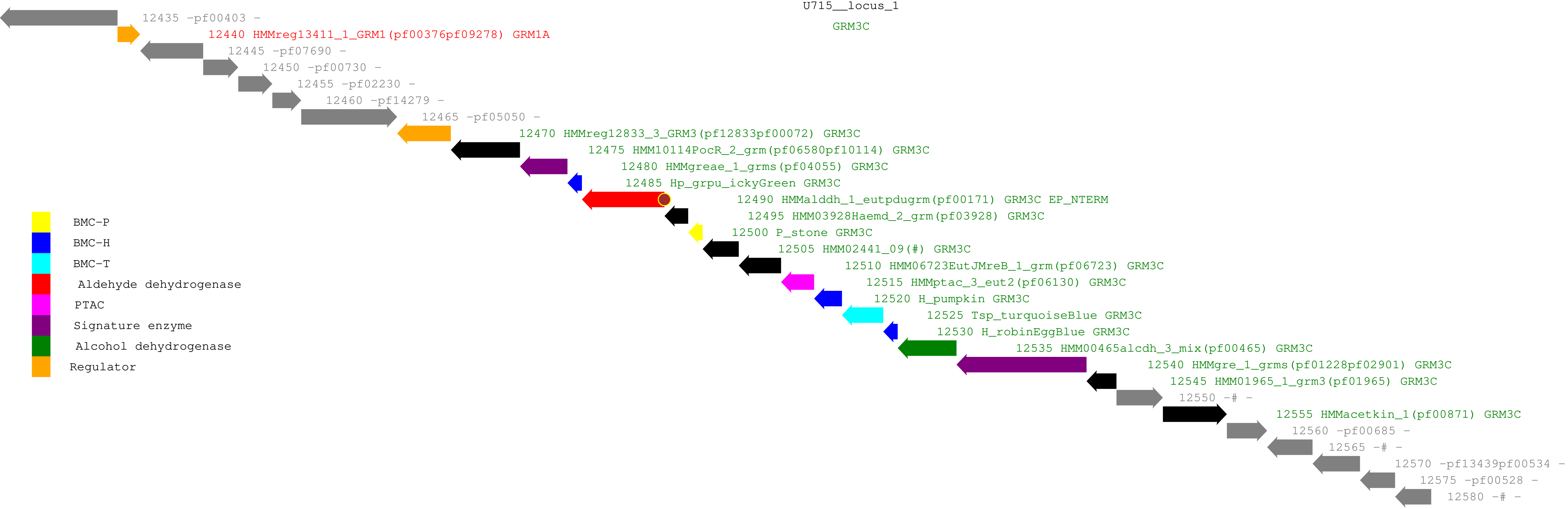

Salmonella enterica subsp. enterica serovar Senftenberg str. Bacteria; Proteobacteria; Gammaproteobacteria; Enterobacterales;

AV984__locus_3

GRM4

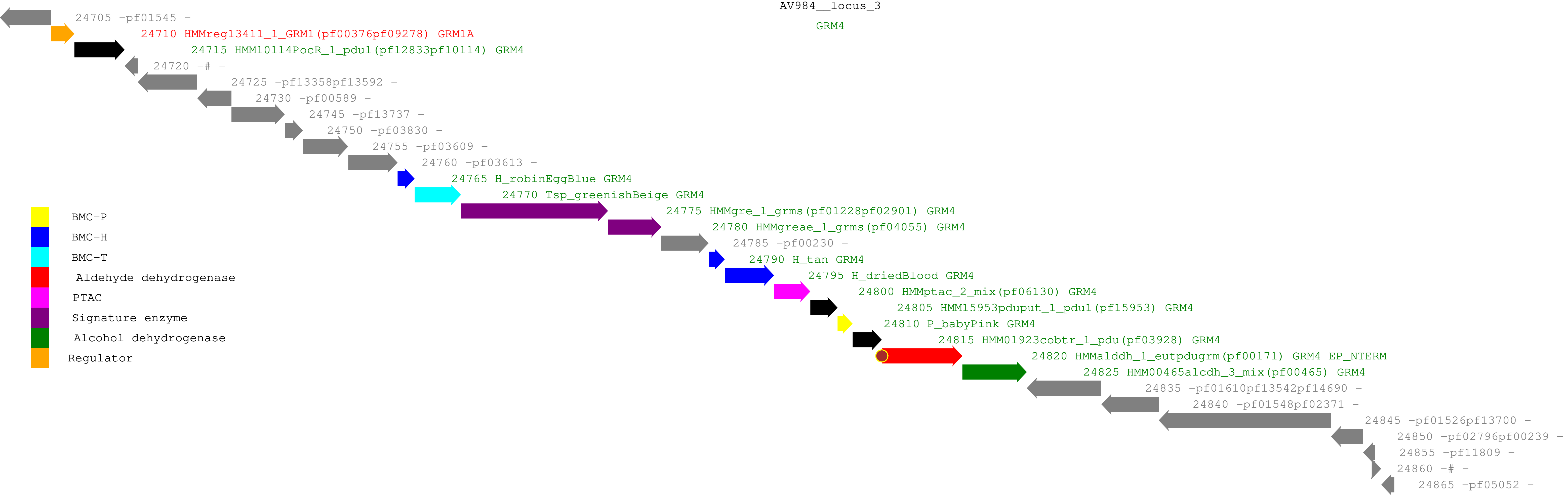

Escherichia coli. Bacteria; Proteobacteria; Gammaproteobacteria; Enterobacterales;

C6985__locus_3

GRM4

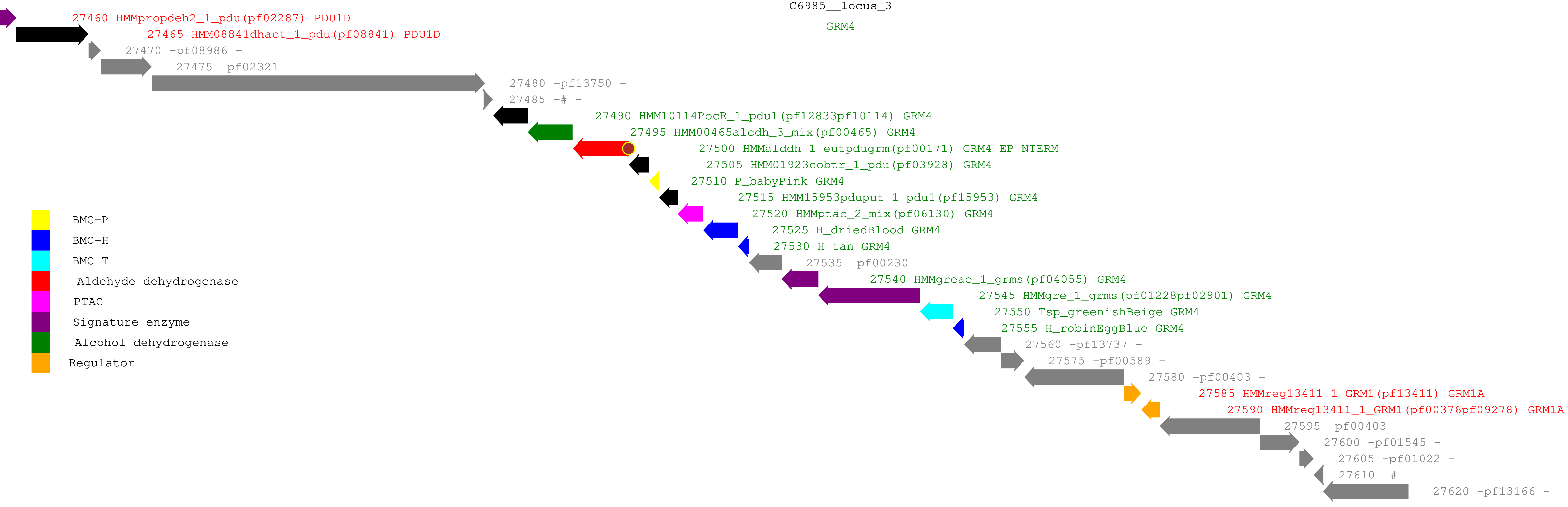

Klebsiella oxytoca. Bacteria; Proteobacteria; Gammaproteobacteria; Enterobacterales;

DET57__locus_1

GRM4

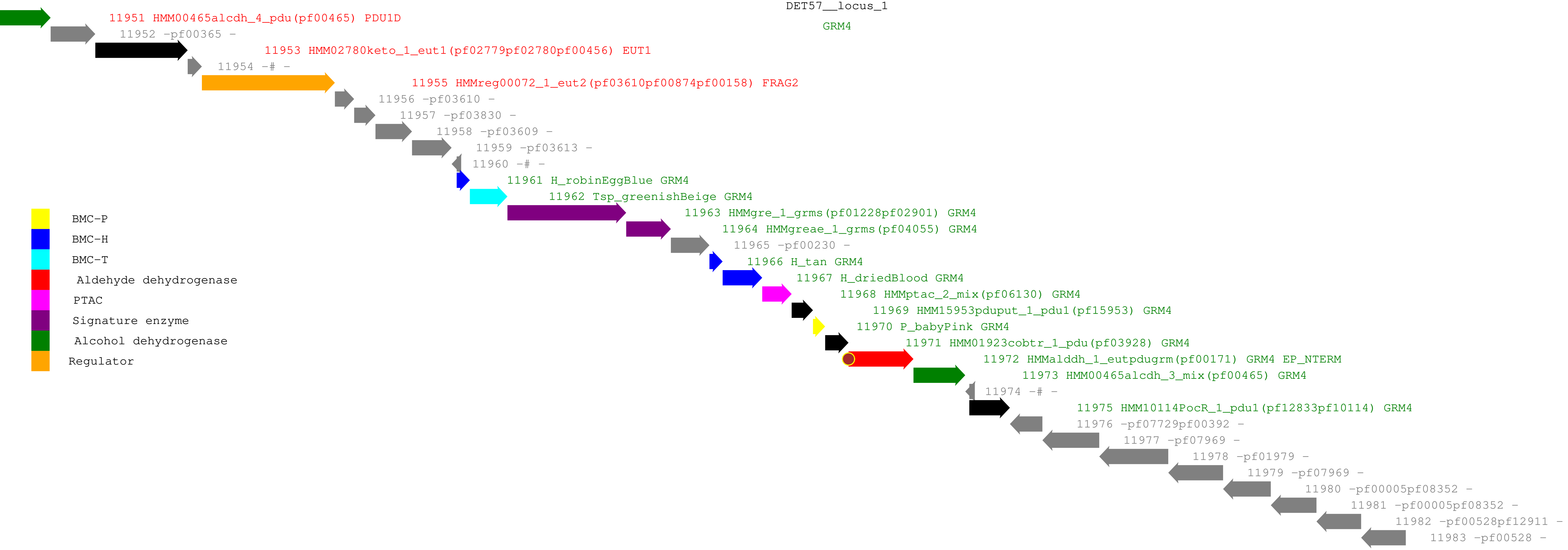

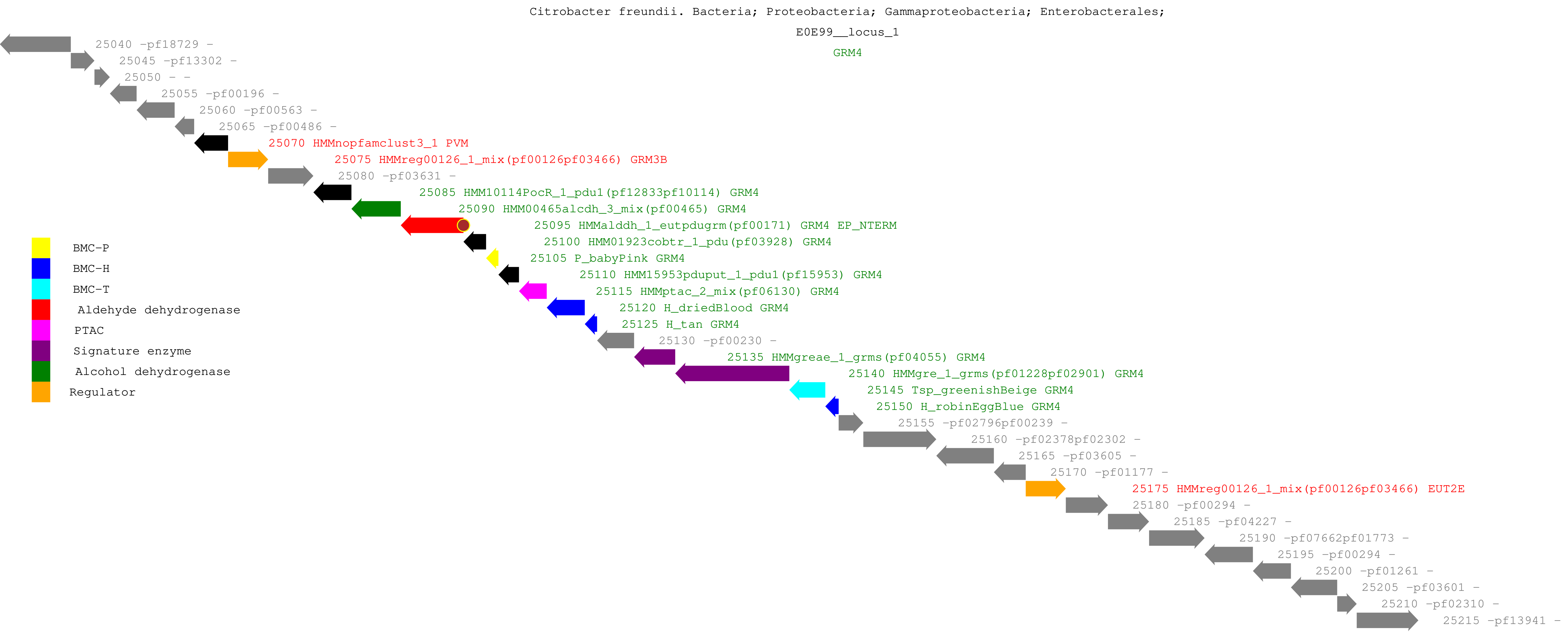
Citrobacter freundii. Bacteria; Proteobacteria; Gammaproteobacteria; Enterobacterales;
E0E99__locus_1
GRM4
25040 -pf18729 -
25045 -pf13302 -
25050 - -
25055 -pf00196 -
25060 -pf00563 -
25065 -pf00486 -
25070 HMMnopfamclust3_1 PVM
25075 HMMreg00126_1_mix(pf00126pf03466) GRM3B
25080 -pf03631 -
25085 HMM10114PocR_1_pdu1(pf12833pf10114) GRM4
25090 HMM00465alcdh_3_mix(pf00465) GRM4
25095 HMMalddh_1_eutpdugrm(pf00171) GRM4 EP_NTERM
25100 HMM01923cobtr_1_pdu(pf03928) GRM4
25105 P_babyPink GRM4
25110 HMM15953pduput_1_pdu1(pf15953) GRM4
25115 HMMptac_2_mix(pf06130) GRM4
25120 H_driedBlood GRM4
25125 H_tan GRM4
25130 -pf00230 -
25135 HMMgreae_1_grms(pf04055) GRM4
25140 HMMgre_1_grms(pf01228pf02901) GRM4
25145 Tsp_greenishBeige GRM4
25150 H_robinEggBlue GRM4
25155 -pf02796pf00239 -
25160 -pf02378pf02302 -
25165 -pf03605 -
25170 -pf01177 -
25175 HMMreg00126_1_mix(pf00126pf03466) EUT2E
25180 -pf00294 -
25185 -pf04227 -
25190 -pf07662pf01773 -
25195 -pf00294 -
25200 -pf01261 -
25205 -pf03601 -
25210 -pf02310 -
25215 -pf13941 -
BMC-P
BMC-H
BMC-T
Aldehyde dehydrogenase
PTAC
Signature enzyme
Alcohol dehydrogenase
Regulator

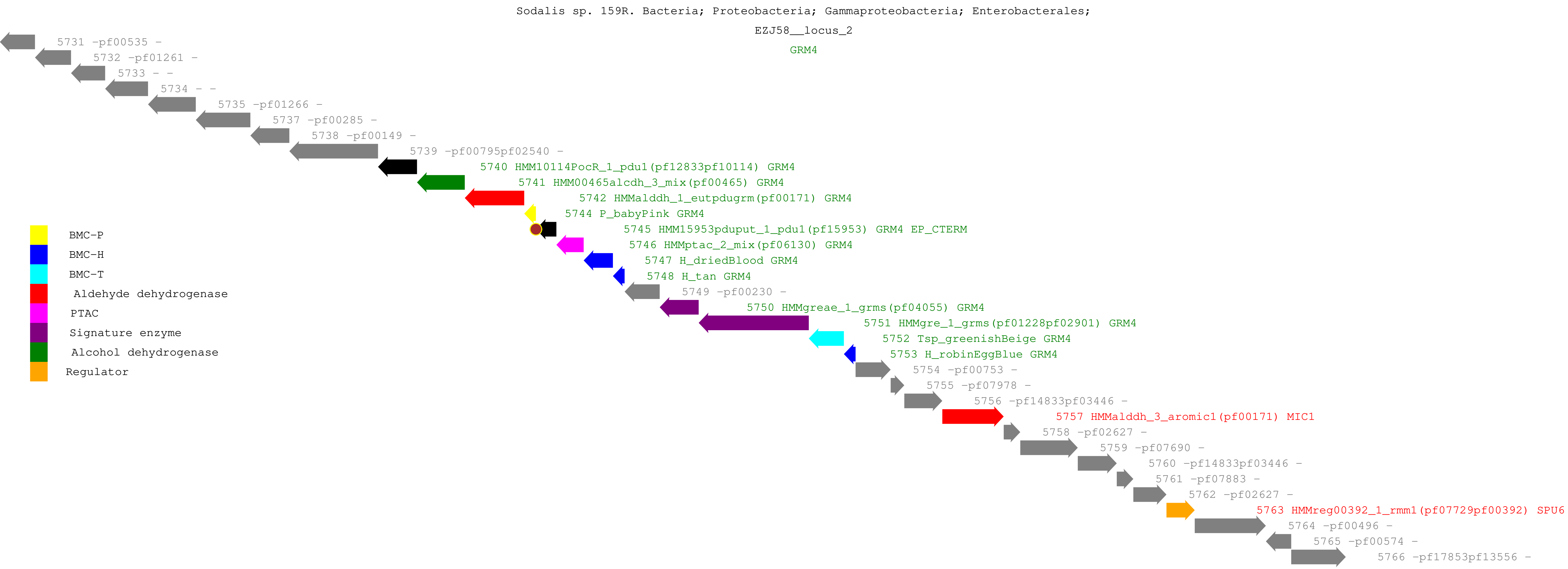

Sodalis sp. 159R. Bacteria; Proteobacteria; Gammaproteobacteria; Enterobacterales;
EZJ58__locus_2
GRM4
5731 -pf00535 -
5732 -pf01261 -
5733 - -
5734 - -
5735 -pf01266 -
5737 -pf00285 -
5738 -pf00149 -
5739 -pf00795pf02540 -
5740 HMM10114PocR_1_pdu1(pf12833pf10114) GRM4
5741 HMM00465alcdh_3_mix(pf00465) GRM4
5742 HMMalddh_1_eutpdugrm(pf00171) GRM4
5744 P_babyPink GRM4
5745 HMM15953pduput_1_pdu1(pf15953) GRM4 EP_CTERM
5746 HMMptac_2_mix(pf06130) GRM4
5747 H_driedBlood GRM4
5748 H_tan GRM4
5749 -pf00230 -
5750 HMMgreae_1_grms(pf04055) GRM4
5751 HMMgre_1_grms(pf01228pf02901) GRM4
5752 Tsp_greenishBeige GRM4
5753 H_robinEggBlue GRM4
5754 -pf00753 -
5755 -pf07978 -
5756 -pf14833pf03446 -
5757 HMMalddh_3_aromic1(pf00171) MIC1
5758 -pf02627 -
5759 -pf07690 -
5760 -pf14833pf03446 -
5761 -pf07883 -
5762 -pf02627 -
5763 HMMreg00392_1_rmm1(pf07729pf00392) SPU6
5764 -pf00496 -
5765 -pf00574 -
5766 -pf17853pf13556 -
BMC-P
BMC-H
BMC-T
Aldehyde dehydrogenase
PTAC
Signature enzyme
Alcohol dehydrogenase
Regulator

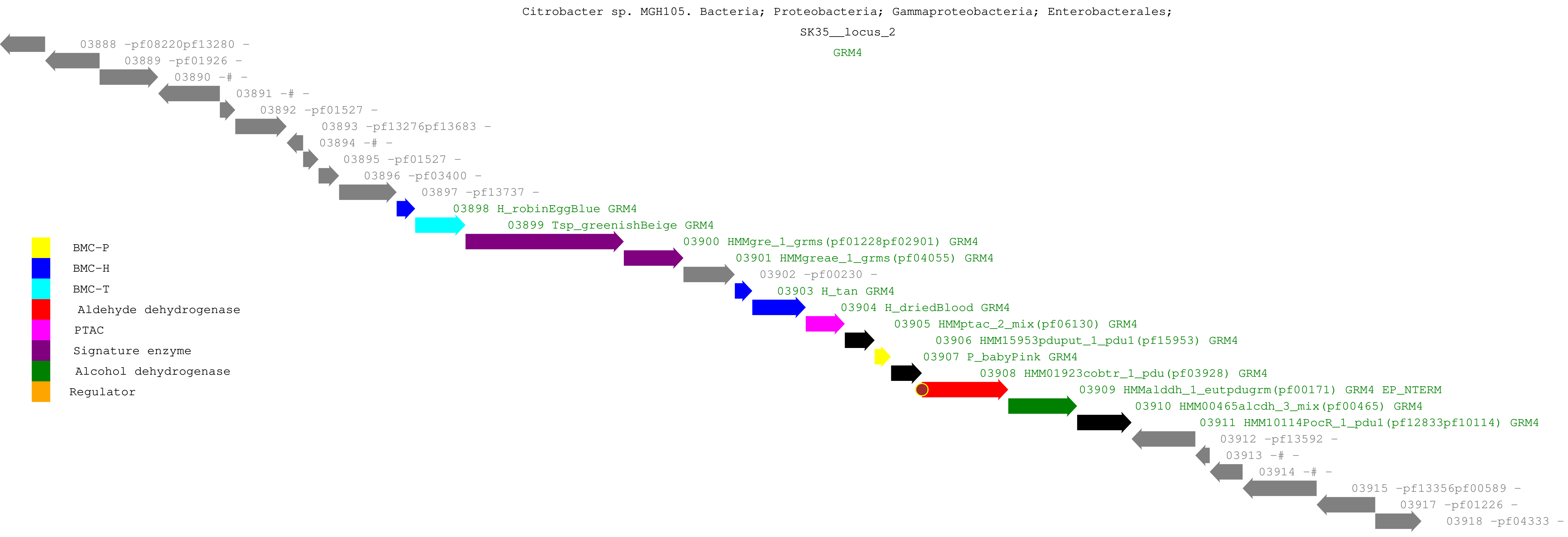
Citrobacter sp. MGH105. Bacteria; Proteobacteria; Gammaproteobacteria; Enterobacterales;
SK35__locus_2
GRM4
03888 -pf08220pf13280 -
03889 -pf01926 -
03890 -# -
03891 -# -
03892 -pf01527 -
03893 -pf13276pf13683 -
03894 -# -
03895 -pf01527 -
03896 -pf03400 -
03897 -pf13737 -
03898 H_robinEggBlue GRM4
03899 Tsp_greenishBeige GRM4
03900 HMMgre_1_grms(pf01228pf02901) GRM4
03901 HMMgreae_1_grms(pf04055) GRM4
03902 -pf00230 -
03903 H_tan GRM4
03904 H_driedBlood GRM4
03905 HMMptac_2_mix(pf06130) GRM4
03906 HMM15953pduput_1_pdu1(pf15953) GRM4
03907 P_babyPink GRM4
03908 HMM01923cobtr_1_pdu(pf03928) GRM4
03909 HMMalddh_1_eutpdugrm(pf00171) GRM4 EP_NTERM
03910 HMM00465alcdh_3_mix(pf00465) GRM4
03911 HMM10114PocR_1_pdu1(pf12833pf10114) GRM4
03912 -pf13592 -
03913 -# -
03914 -# -
03915 -pf13356pf00589 -
03917 -pf01226 -
03918 -pf04333 -
BMC-P
BMC-H
BMC-T
Aldehyde dehydrogenase
PTAC
Signature enzyme
Alcohol dehydrogenase
Regulator

Shewanella putrefaciens (strain CN-32 / ATCC BAA-453). Bacteria; Proteobacteria; Gammaproteobacteria; Alteromonadales;

Sputcn32__locus_1

GRM4

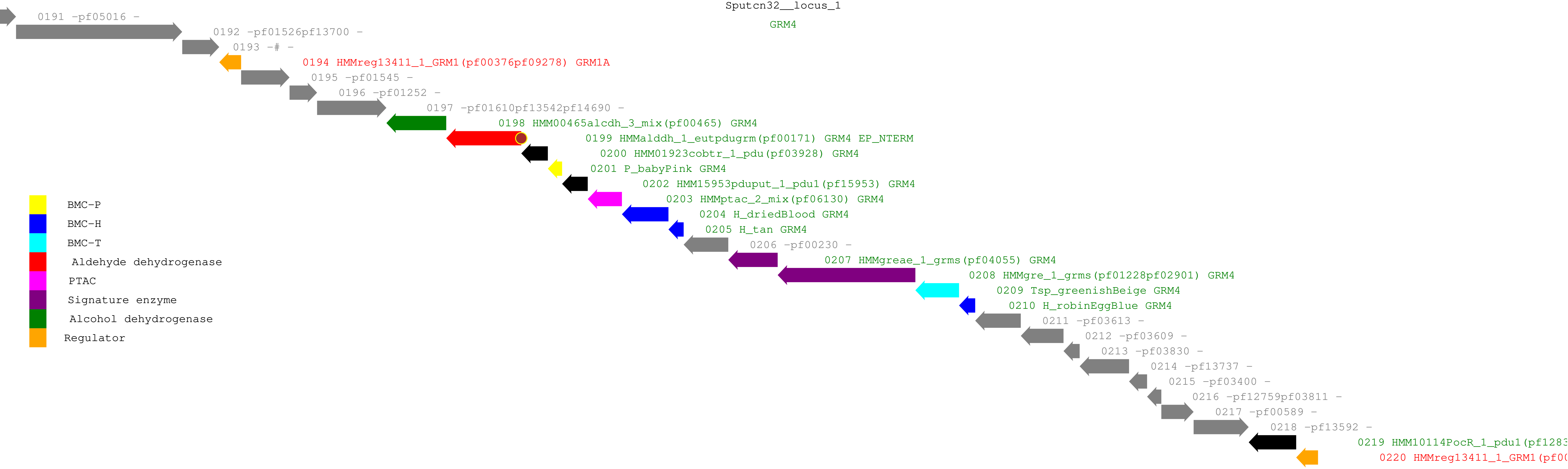

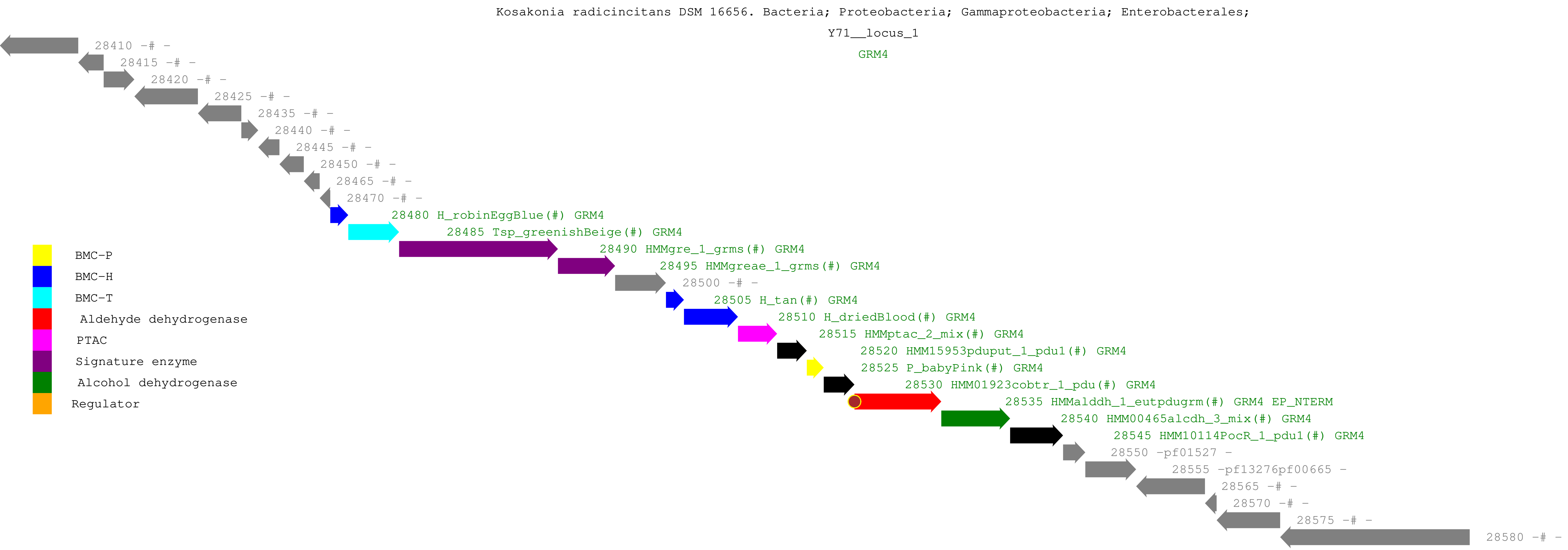
Kosakonia radicincitans DSM 16656. Bacteria; Proteobacteria; Gammaproteobacteria; Enterobacterales;
Y71__locus_1
GRM4
28410 -# -
28415 -# -
28420 -# -
28425 -# -
28435 -# -
28440 -# -
28445 -# -
28450 -# -
28465 -# -
28470 -# -
28480 H_robinEggBlue(#) GRM4
28485 Tsp_greenishBeige(#) GRM4
28490 HMMgre_1_grms(#) GRM4
28495 HMMgreae_1_grms(#) GRM4
28500 -# -
28505 H_tan(#) GRM4
28510 H_driedBlood(#) GRM4
28515 HMMptac_2_mix(#) GRM4
28520 HMM15953pduput_1_pdu1(#) GRM4
28525 P_babyPink(#) GRM4
28530 HMM01923cobtr_1_pdu(#) GRM4
28535 HMMalddh_1_eutpdugrm(#) GRM4 EP_NTERM
28540 HMM00465alcdh_3_mix(#) GRM4
28545 HMM10114PocR_1_pdu1(#) GRM4
28550 -pf01527 -
28555 -pf13276pf00665 -
28565 -# -
28570 -# -
28575 -# -
28580 -# -
BMC-P
BMC-H
BMC-T
Aldehyde dehydrogenase
PTAC
Signature enzyme
Alcohol dehydrogenase
Regulator

Acetatifactor muris. Bacteria; Firmicutes; Clostridia; Clostridiales; Lachnospiraceae;

AMURIS__locus_1

GRM5

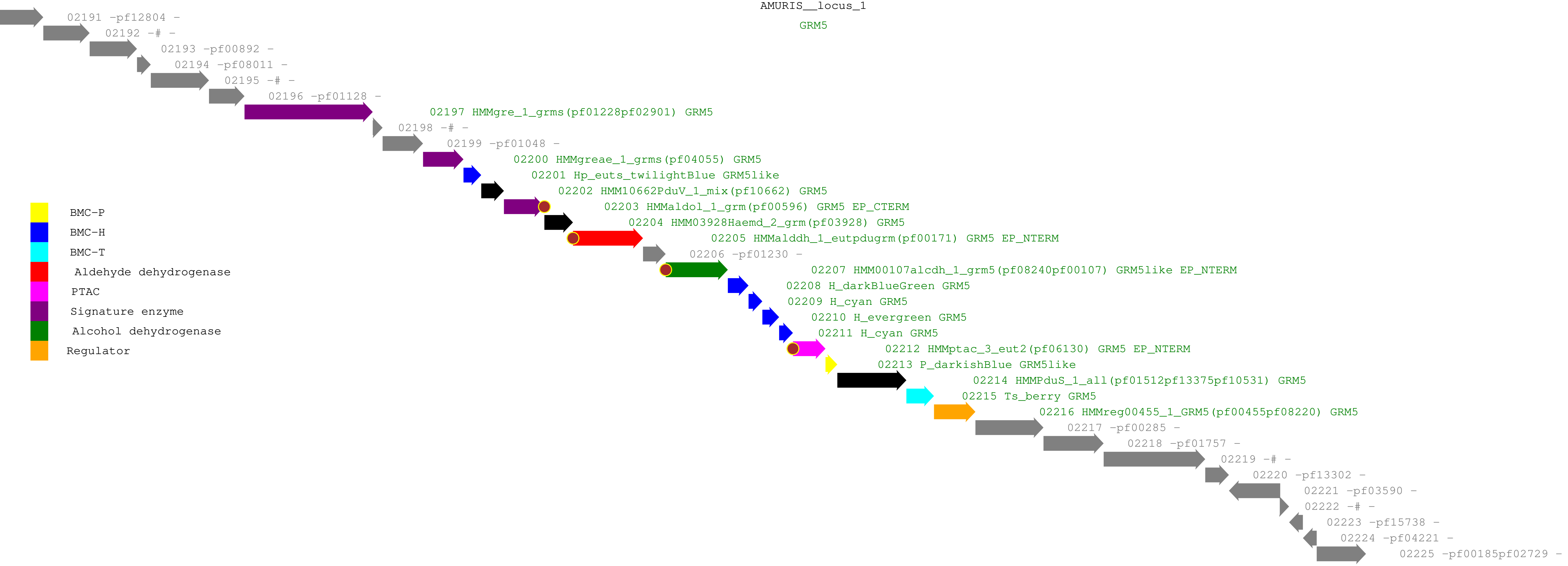

Eisenbergiella tayi. Bacteria; Firmicutes; Clostridia; Clostridiales; Lachnospiraceae;

BEH84__locus_1

GRM5

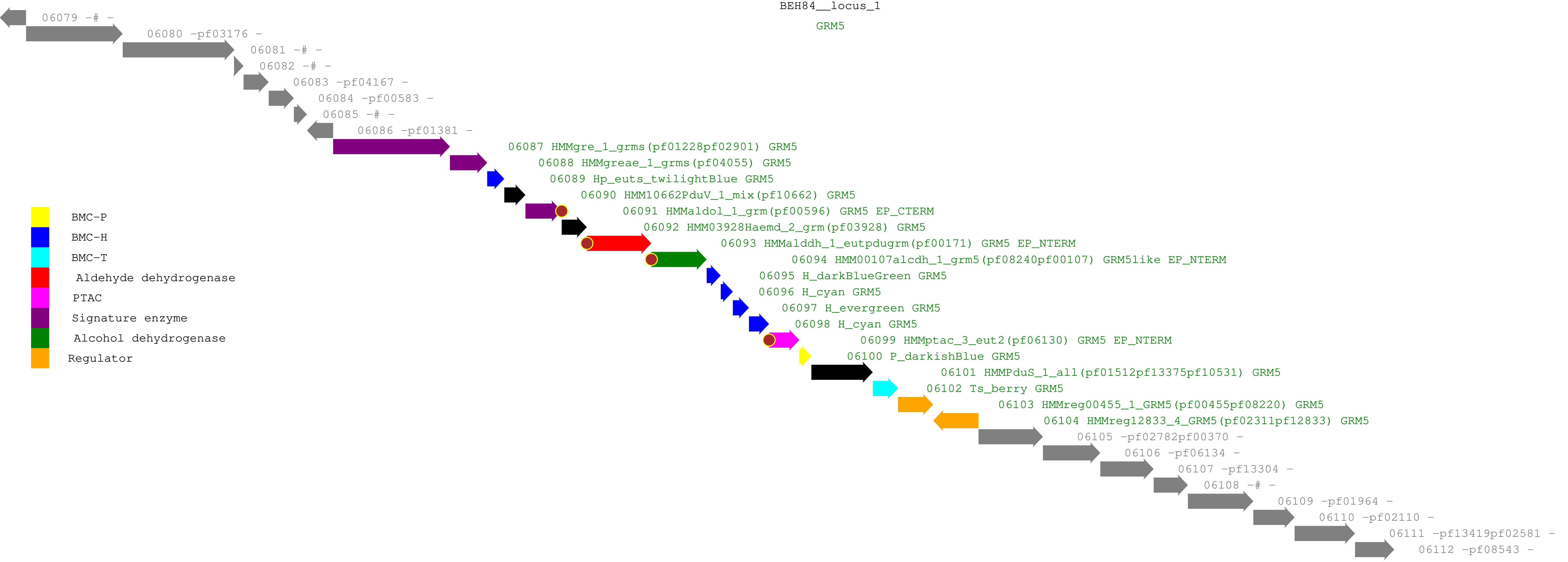

Eisenbergiella tayi. Bacteria; Firmicutes; Clostridia; Clostridiales; Lachnospiraceae;

BEI59__locus_2

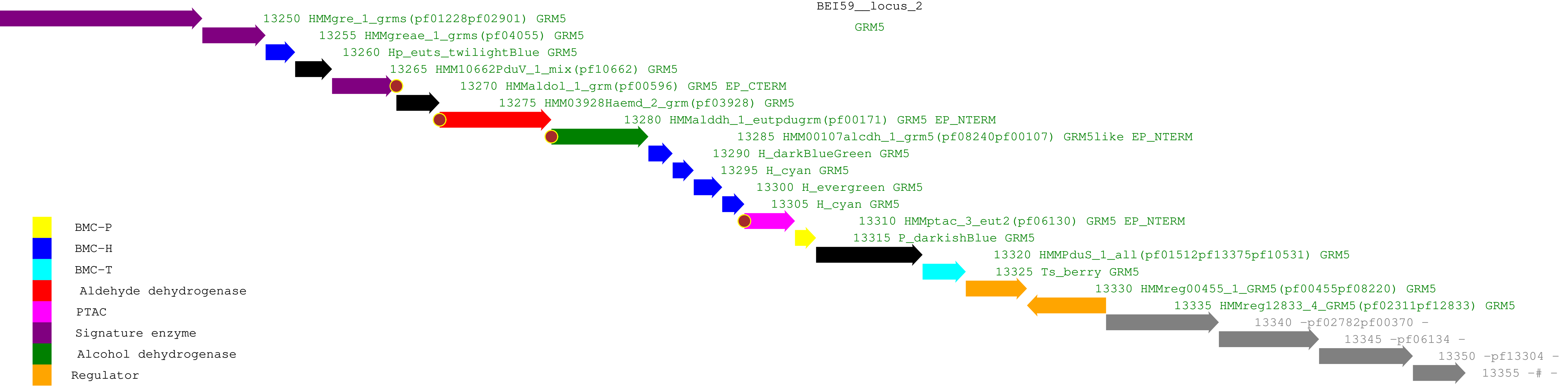

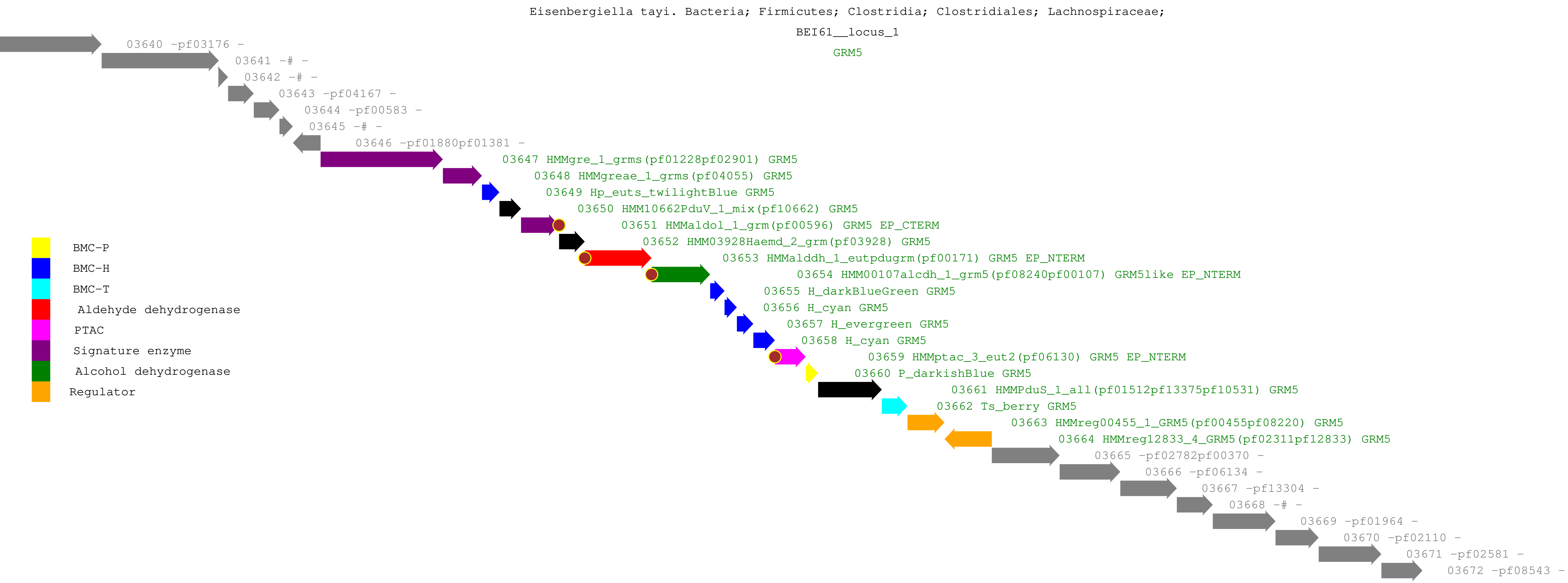
Eisenbergiella tayi. Bacteria; Firmicutes; Clostridia; Clostridiales; Lachnospiraceae;
BEI61__locus_1
GRM5
03640 -pf03176 -
03641 -# -
03642 -# -
03643 -pf04167 -
03644 -pf00583 -
03645 -# -
03646 -pf01880pf01381 -
03647 HMMgre_1_grms(pf01228pf02901) GRM5
03648 HMMgreae_1_grms(pf04055) GRM5
03649 Hp_euts_twilightBlue GRM5
03650 HMM10662PduV_1_mix(pf10662) GRM5
03651 HMMaldol_1_grm(pf00596) GRM5 EP_CTERM
03652 HMM03928Haemd_2_grm(pf03928) GRM5
03653 HMMalddh_1_eutpdugrm(pf00171) GRM5 EP_NTERM
03654 HMM00107alcdh_1_grm5(pf08240pf00107) GRM5like EP_NTERM
03655 H_darkBlueGreen GRM5
03656 H_cyan GRM5
03657 H_evergreen GRM5
03658 H_cyan GRM5
03659 HMMptac_3_eut2(pf06130) GRM5 EP_NTERM
03660 P_darkishBlue GRM5
03661 HMMPduS_1_all(pf01512pf13375pf10531) GRM5
03662 Ts_berry GRM5
03663 HMMreg00455_1_GRM5(pf00455pf08220) GRM5
03664 HMMreg12833_4_GRM5(pf02311pf12833) GRM5
03665 -pf02782pf00370 -
03666 -pf06134 -
03667 -pf13304 -
03668 -# -
03669 -pf01964 -
03670 -pf02110 -
03671 -pf02581 -
03672 -pf08543 -
BMC-P
BMC-H
BMC-T
Aldehyde dehydrogenase
PTAC
Signature enzyme
Alcohol dehydrogenase
Regulator

Clostridiales bacterium 42_27. Bacteria; Firmicutes; Clostridia; Clostridiales.

BHV90__locus_1_main_sat

GRM5

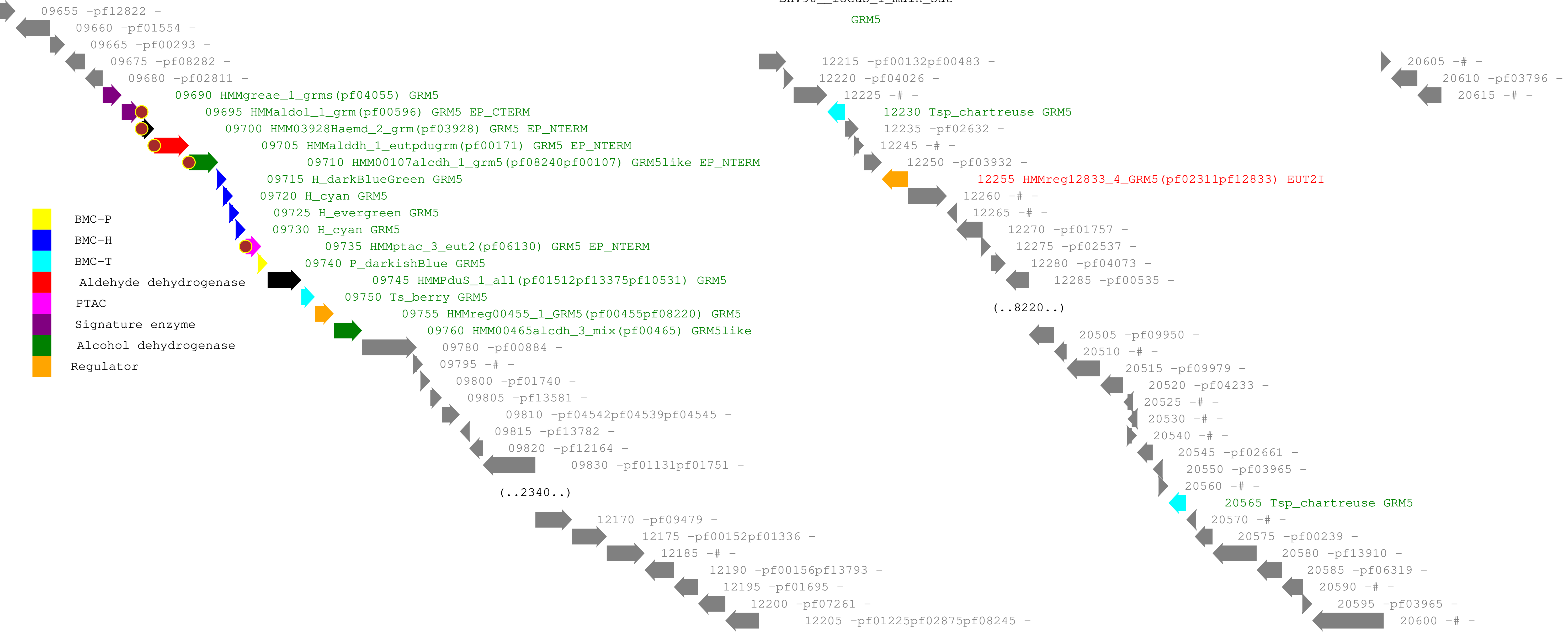

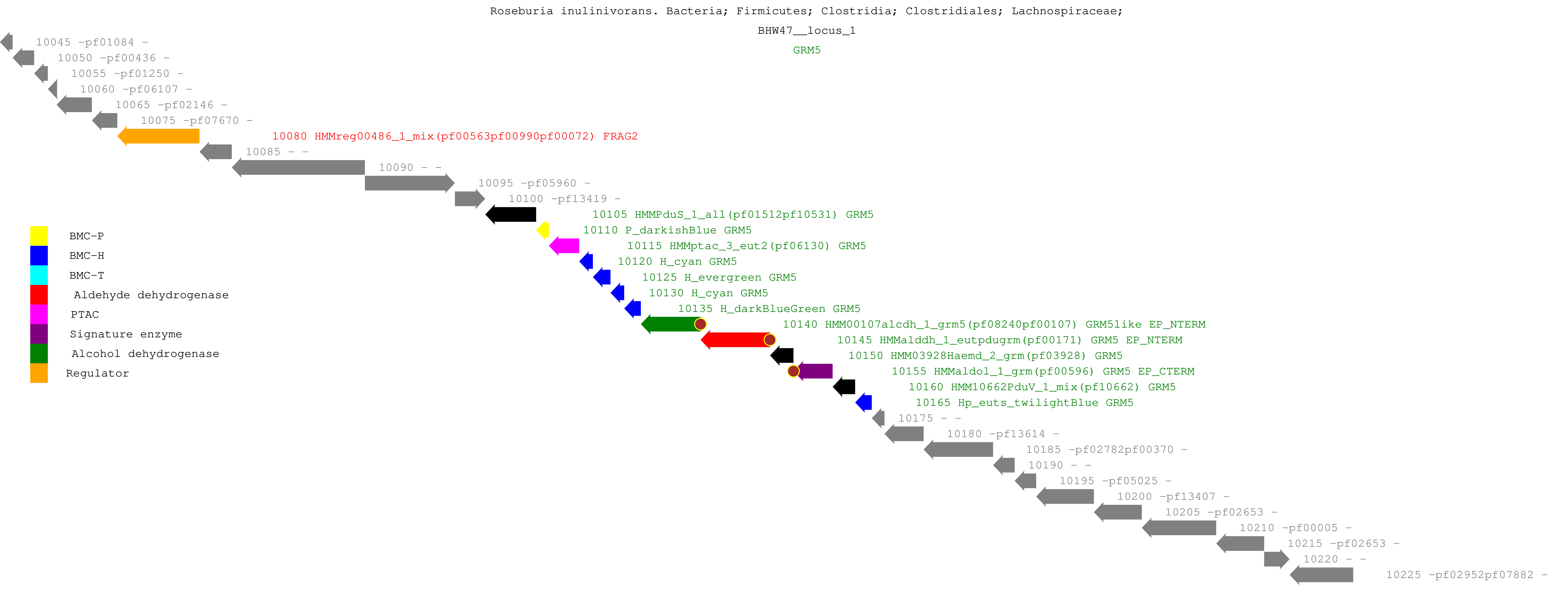
Roseburia inulinivorans. Bacteria; Firmicutes; Clostridia; Clostridiales; Lachnospiraceae;
BHW47__locus_1
GRM5
10045 -pf01084 -
10050 -pf00436 -
10055 -pf01250 -
10060 -pf06107 -
10065 -pf02146 -
10075 -pf07670 -
10080 HMMreg00486_1_mix(pf00563pf00990pf00072) FRAG2
10085 - -
10090 - -
10095 -pf05960 -
10100 -pf13419 -
10105 HMMPduS_1_all(pf01512pf10531) GRM5
10110 P_darkishBlue GRM5
10115 HMMptac_3_eut2(pf06130) GRM5
10120 H_cyan GRM5
10125 H_evergreen GRM5
10130 H_cyan GRM5
10135 H_darkBlueGreen GRM5
10140 HMM00107alcdh_1_grm5(pf08240pf00107) GRM5like EP_NTERM
10145 HMMaldh_1_eutpdugrm(pf00171) GRM5 EP_NTERM
10150 HMM03928Haemd_2_grm(pf03928) GRM5
10155 HMMaldol_1_grm(pf00596) GRM5 EP_CTERM
10160 HMM10662PduV_1_mix(pf10662) GRM5
10165 Hp_euts_twilightBlue GRM5
10175 - -
10180 -pf13614 -
10185 -pf02782pf00370 -
10190 - -
10195 -pf05025 -
10200 -pf13407 -
10205 -pf02653 -
10210 -pf00005 -
10215 -pf02653 -
10220 - -
10225 -pf02952pf07882 -
BMC-P
BMC-H
BMC-T
Aldehyde dehydrogenase
PTAC
Signature enzyme
Alcohol dehydrogenase
Regulator

Clostridium sp. C105KSO13. Bacteria; Firmicutes; Clostridia; Clostridiales; Clostridiaceae;

BN3456__locus_1

GRM5

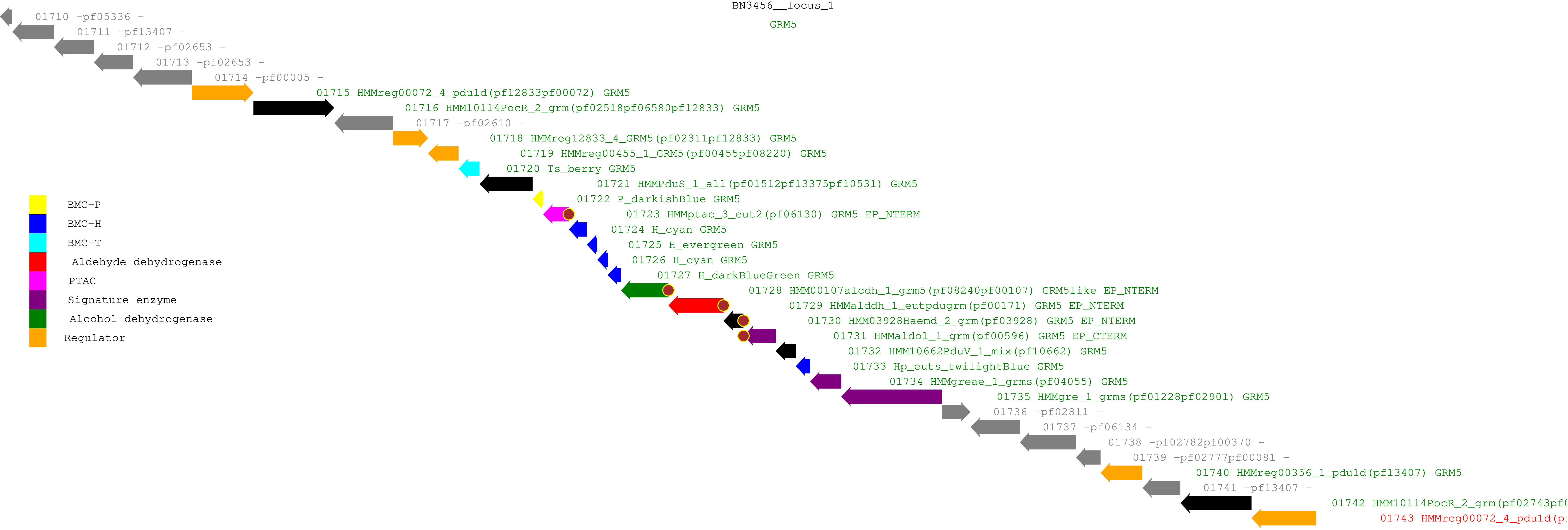

Ruminococcus gnavus CAG:126. Bacteria; Firmicutes; Clostridia; Clostridiales; Ruminococcaceae;

BN481__locus_1_main_sat

GRM5

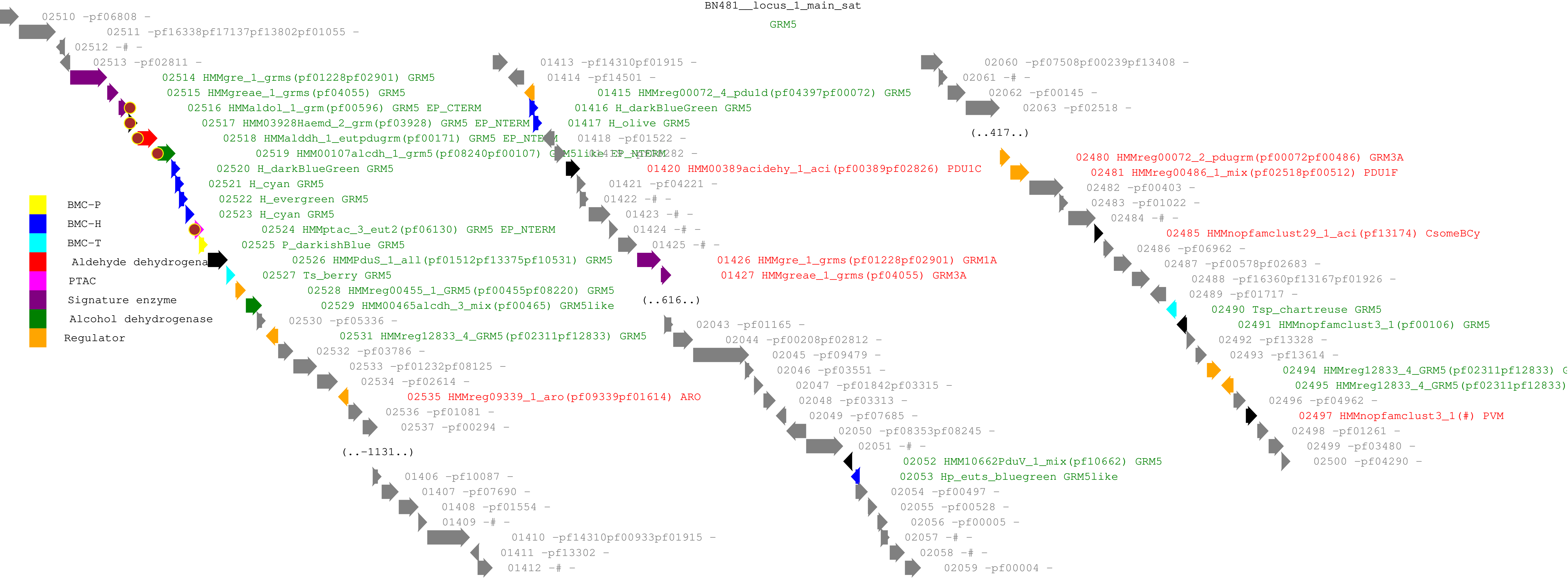

Ruminococcus sp. CAG:17. Bacteria; Firmicutes; Clostridia; Clostridiales; Ruminococcaceae;

BN514__locus_1_main_sat

GRM5

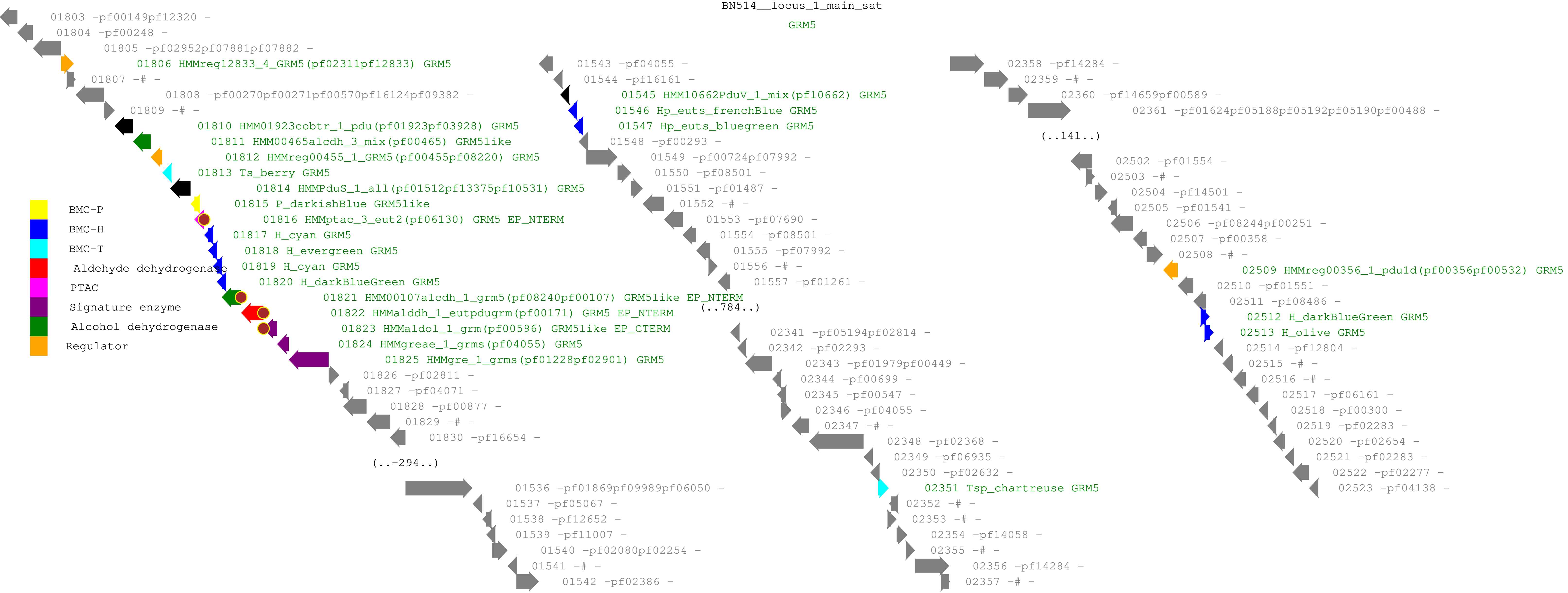

Firmicutes bacterium CAG:212. Bacteria; Firmicutes; environmental samples.

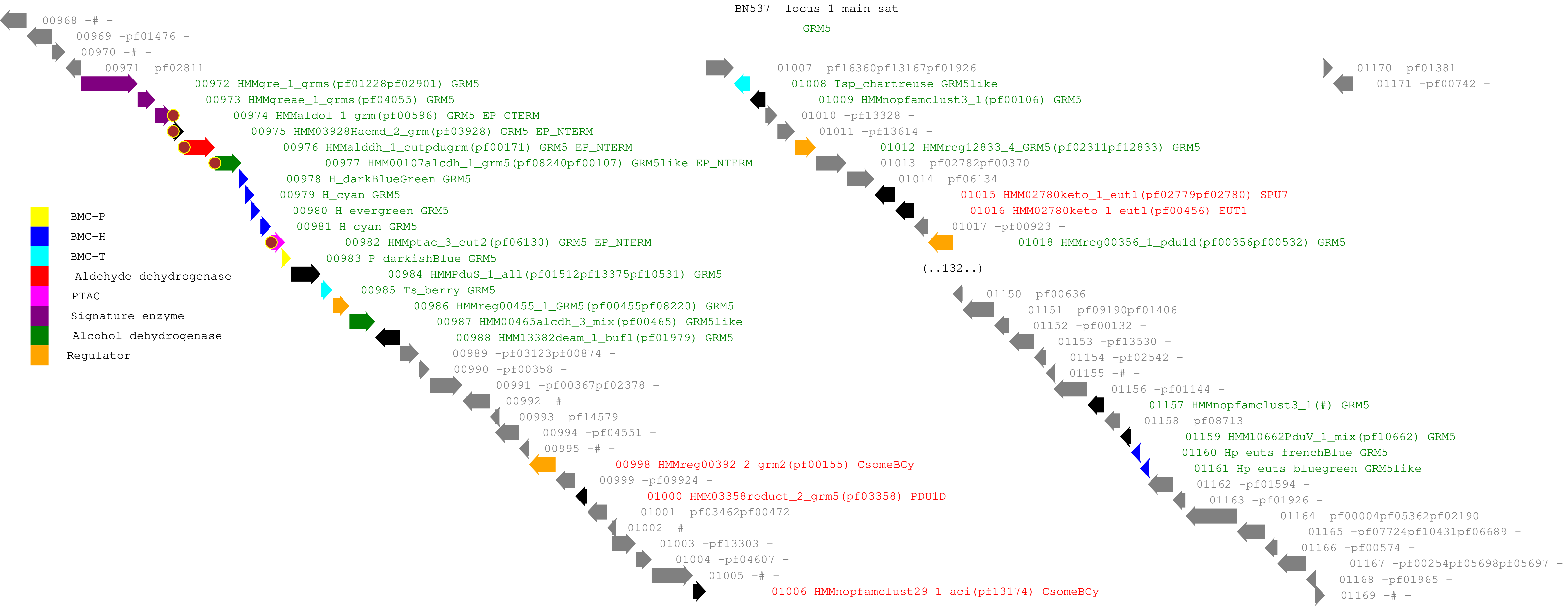

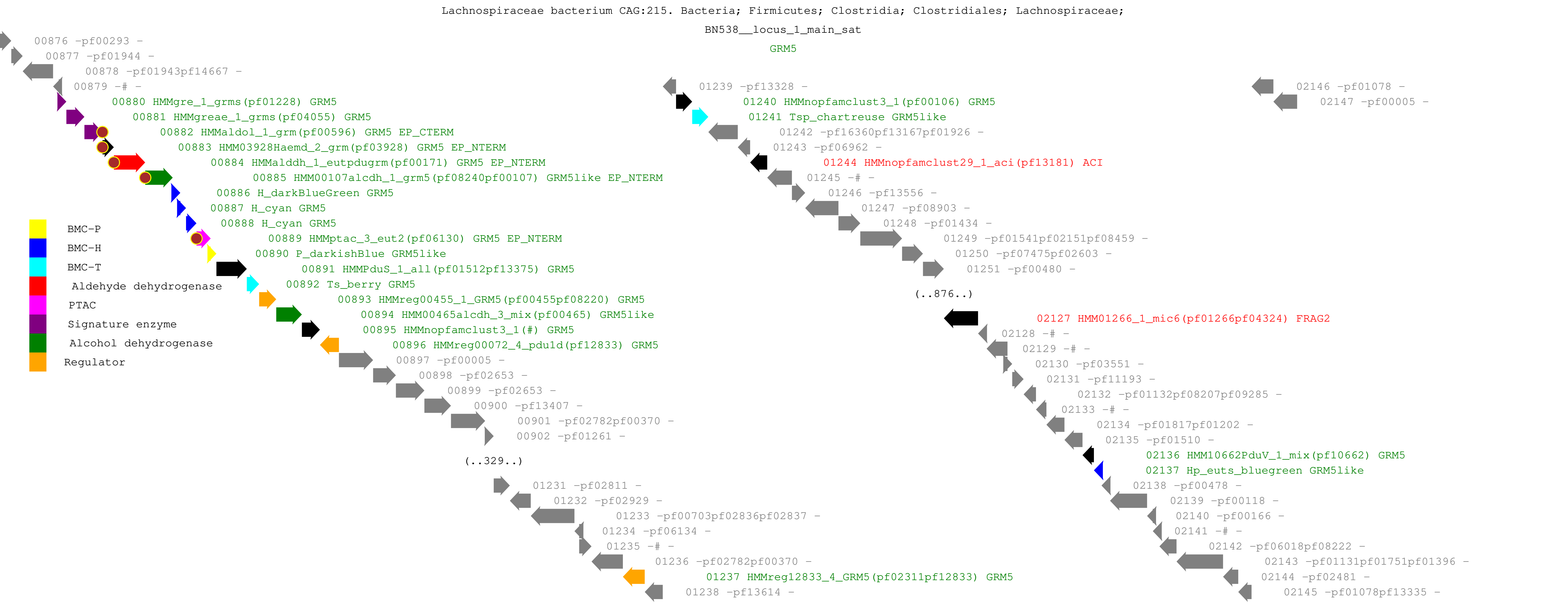
Lachnospiraceae bacterium CAG:215. Bacteria; Firmicutes; Clostridia; Clostridiales; Lachnospiraceae;
BN538__locus_1_main_sat
GRM5
00876 -pf00293 -
00877 -pf01944 -
00878 -pf01943pf14667 -
00879 -# -
00880 HMMgre_1_grms(pf01228) GRM5
00881 HMMgreae_1_grms(pf04055) GRM5
00882 HMMaldol_1_grm(pf00596) GRM5 EP_CTERM
00883 HMM03928Haemd_2_grm(pf03928) GRM5 EP_NTERM
00884 HMMalddh_1_eutpdugrm(pf00171) GRM5 EP_NTERM
00885 HMM00107alcdh_1_grm5(pf08240pf00107) GRM5like EP_NTERM
00886 H_darkBlueGreen GRM5
00887 H_cyan GRM5
00888 H_cyan GRM5
00889 HMMptac_3_eut2(pf06130) GRM5 EP_NTERM
00890 P_darkishBlue GRM5like
00891 HMMPduS_1_all(pf01512pf13375) GRM5
00892 Ts_berry GRM5
00893 HMMreg00455_1_GRM5(pf00455pf08220) GRM5
00894 HMM00465alcdh_3_mix(pf00465) GRM5like
00895 HMMnopfamclust3_1(#) GRM5
00896 HMMreg00072_4_pduld(pf12833) GRM5
00897 -pf00005 -
00898 -pf02653 -
00899 -pf02653 -
00900 -pf13407 -
00901 -pf02782pf00370 -
00902 -pf01261 -
(..329..)
01231 -pf02811 -
01232 -pf02929 -
01233 -pf00703pf02836pf02837 -
01234 -pf06134 -
01235 -# -
01236 -pf02782pf00370 -
01237 HMMreg12833_4_GRM5(pf02311pf12833) GRM5
01238 -pf13614 -
BMC-P
BMC-H
BMC-T
Aldehyde dehydrogenase
PTAC
Signature enzyme
Alcohol dehydrogenase
Regulator
01239 -pf13328 -
01240 HMMnopfamclust3_1(pf00106) GRM5
01241 Tsp_chartreuse GRM5like
01242 -pf16360pf13167pf01926 -
01243 -pf06962 -
01244 HMMnopfamclust29_1_aci(pf13181) ACI
01245 -# -
01246 -pf13556 -
01247 -pf08903 -
01248 -pf01434 -
01249 -pf01541pf02151pf08459 -
01250 -pf07475pf02603 -
01251 -pf00480 -
(..876..)
02127 HMM01266_1_mic6(pf01266pf04324) FRAG2
02128 -# -
02129 -# -
02130 -pf03551 -
02131 -pf11193 -
02132 -pf01132pf08207pf09285 -
02133 -# -
02134 -pf01817pf01202 -
02135 -pf01510 -
02136 HMM10662PduV_1_mix(pf10662) GRM5
02137 Hp_euts_bluegreen GRM5like
02138 -pf00478 -
02139 -pf00118 -
02140 -pf00166 -
02141 -# -
02142 -pf06018pf08222 -
02143 -pf01131pf01751pf01396 -
02144 -pf02481 -
02145 -pf01078pf13335 -
02146 -pf01078 -
02147 -pf00005 -

Blautia sp. CAG:237. Bacteria; Firmicutes; Clostridia; Clostridiales; Lachnospiraceae;

BN552__locus_1_main_sat

GRM5

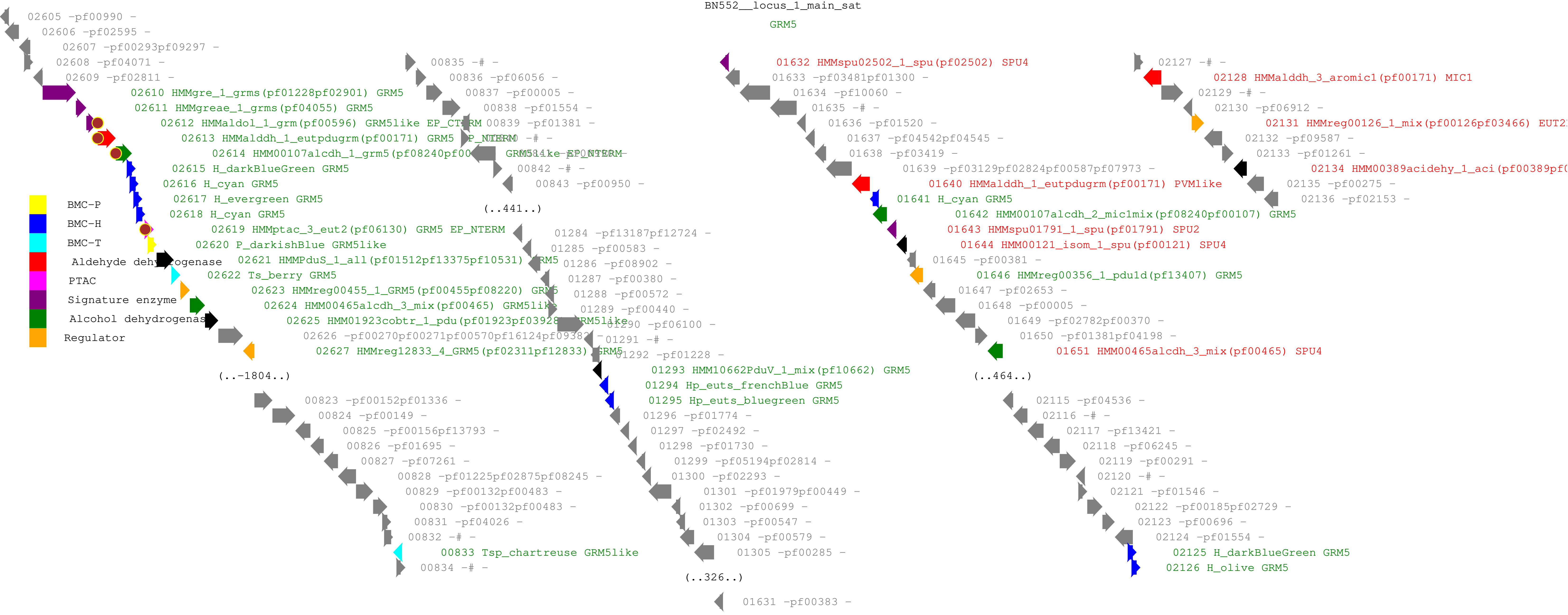

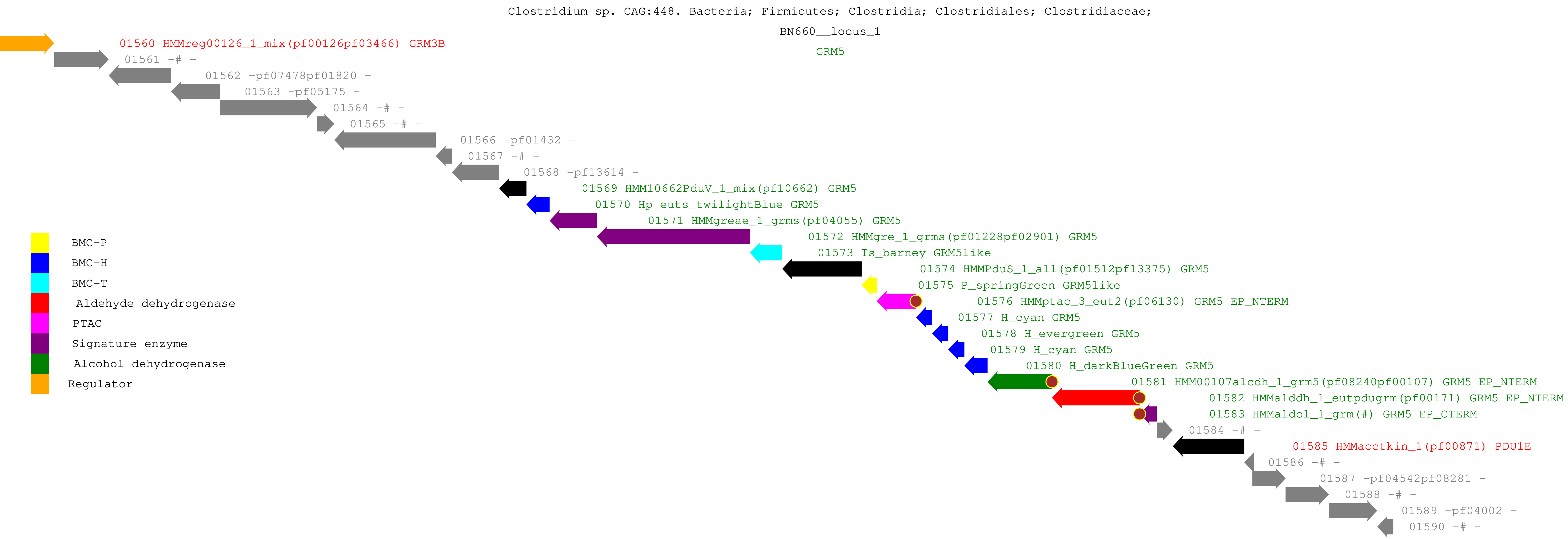
Clostridium sp. CAG:448. Bacteria; Firmicutes; Clostridia; Clostridiales; Clostridiaceae;
BN660__locus_1
GRM5
01560 HMMreg00126_1_mix(pf00126pf03466) GRM3B
01561 -# -
01562 -pf07478pf01820 -
01563 -pf05175 -
01564 -# -
01565 -# -
01566 -pf01432 -
01567 -# -
01568 -pf13614 -
01569 HMM10662PduV_1_mix(pf10662) GRM5
01570 Hp_euts_twilightBlue GRM5
01571 HMMgreae_1_grms(pf04055) GRM5
01572 HMMgre_1_grms(pf01228pf02901) GRM5
01573 Ts_barney GRM5like
01574 HMMPduS_1_all(pf01512pf13375) GRM5
01575 P_springGreen GRM5like
01576 HMMptac_3_eut2(pf06130) GRM5 EP_NTERM
01577 H_cyan GRM5
01578 H_evergreen GRM5
01579 H_cyan GRM5
01580 H_darkBlueGreen GRM5
01581 HMM00107alcdh_1_grm5(pf08240pf00107) GRM5 EP_NTERM
01582 HMMaldh_1_eutpdugrm(pf00171) GRM5 EP_NTERM
01583 HMMaldol_1_grm(#) GRM5 EP_CTERM
01584 -# -
01585 HMMacetkin_1(pf00871) PDU1E
01586 -# -
01587 -pf04542pf08281 -
01588 -# -
01589 -pf04002 -
01590 -# -
BMC-P
BMC-H
BMC-T
Aldehyde dehydrogenase
PTAC
Signature enzyme
Alcohol dehydrogenase
Regulator

Ruminococcus sp. CAG:55. Bacteria; Firmicutes; Clostridia; Clostridiales; Ruminococcaceae;

BN703__locus_1_main_sat

GRM5

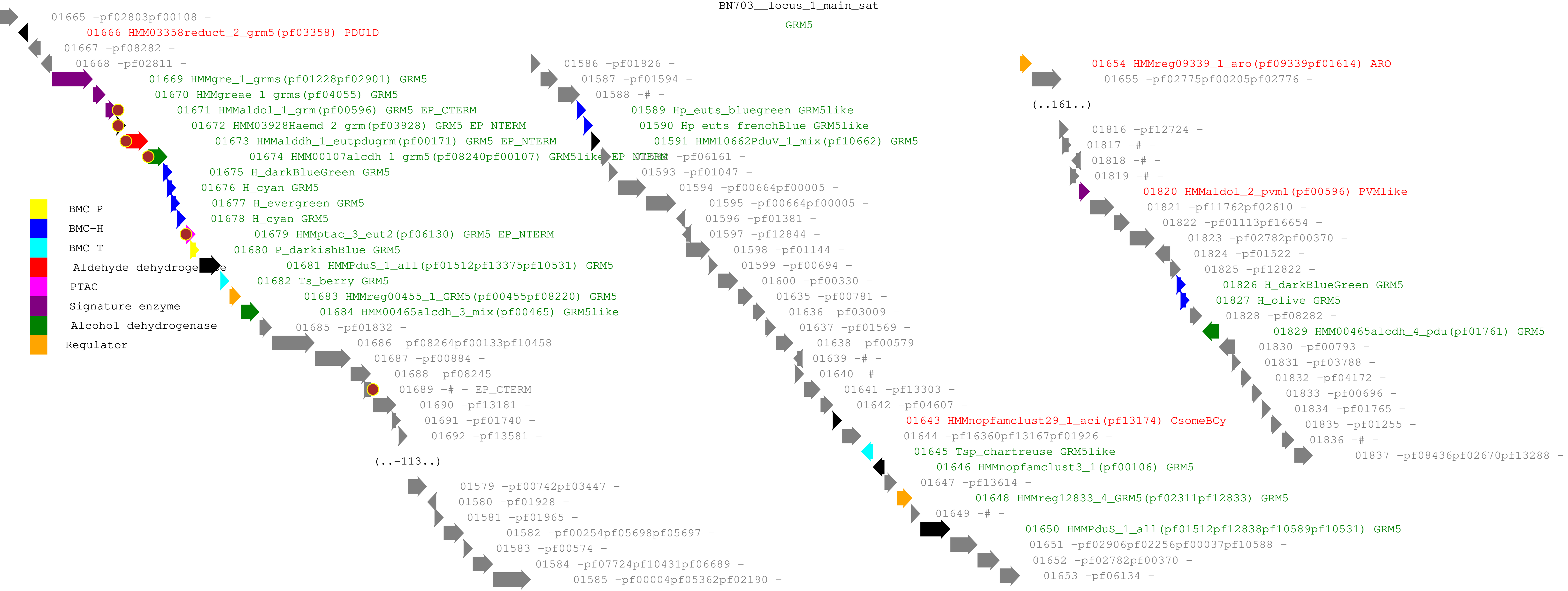

Firmicutes bacterium CAG:882. Bacteria; Firmicutes; environmental samples.

BN803__locus_1

GRM5

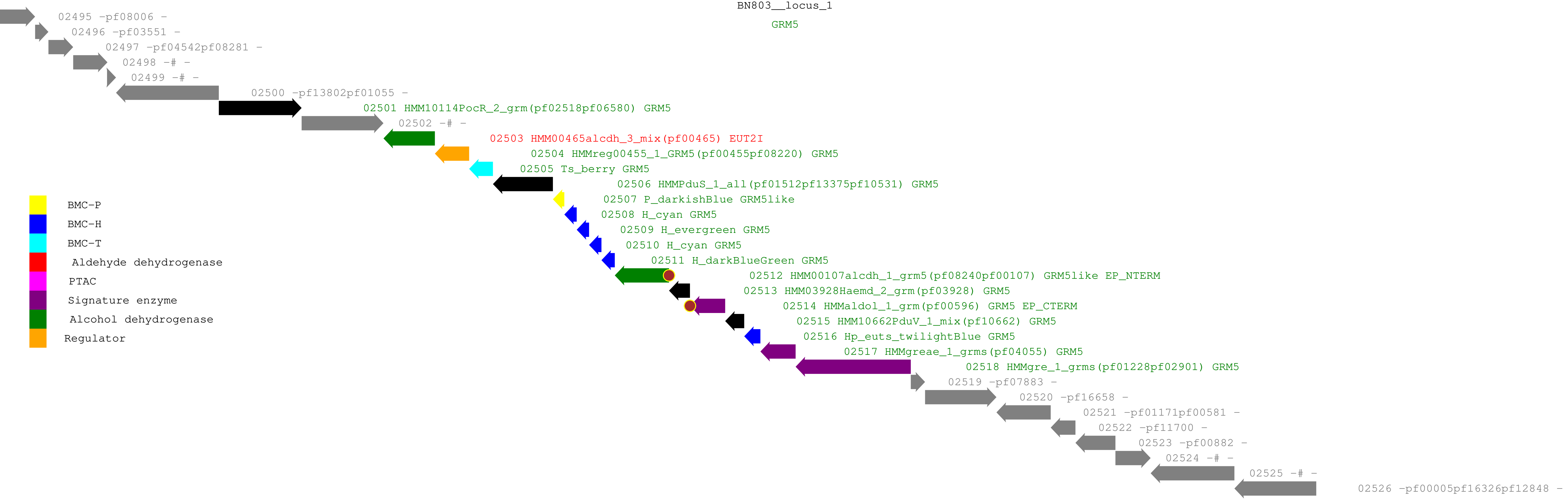

Blautia obeum. Bacteria; Firmicutes; Clostridia; Clostridiales; Lachnospiraceae;

C4886__locus_1_main_sat

GRM5

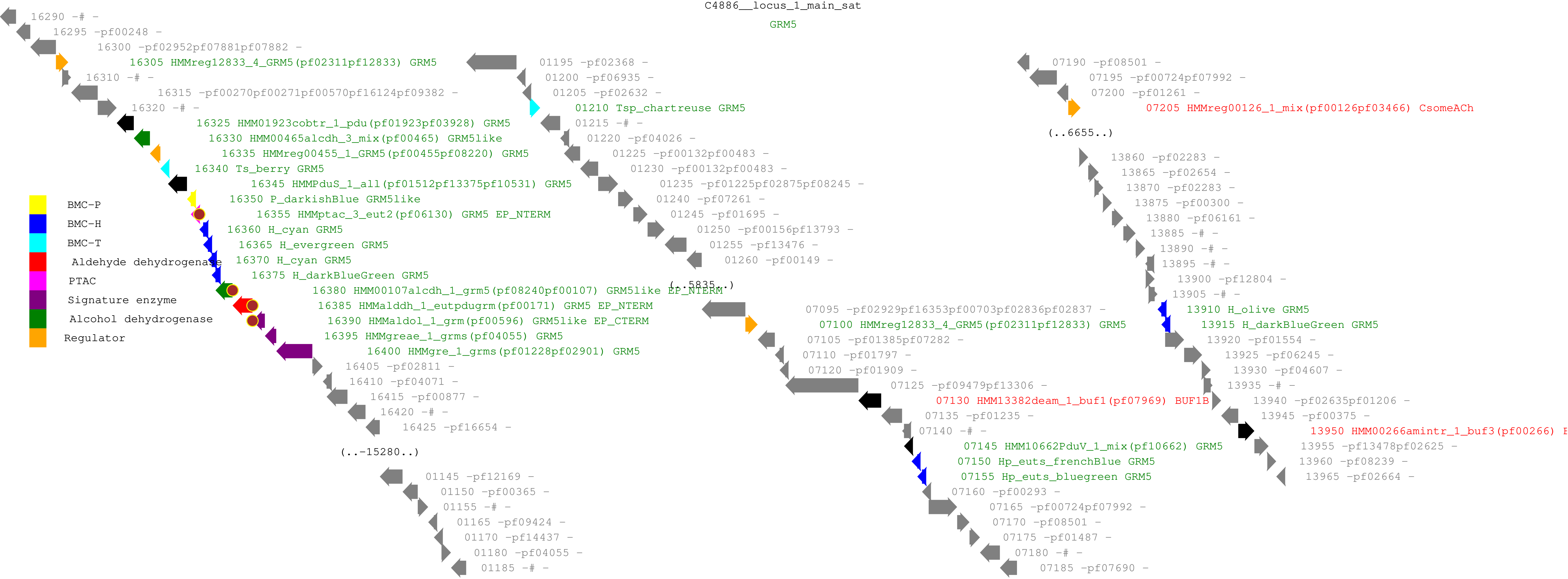

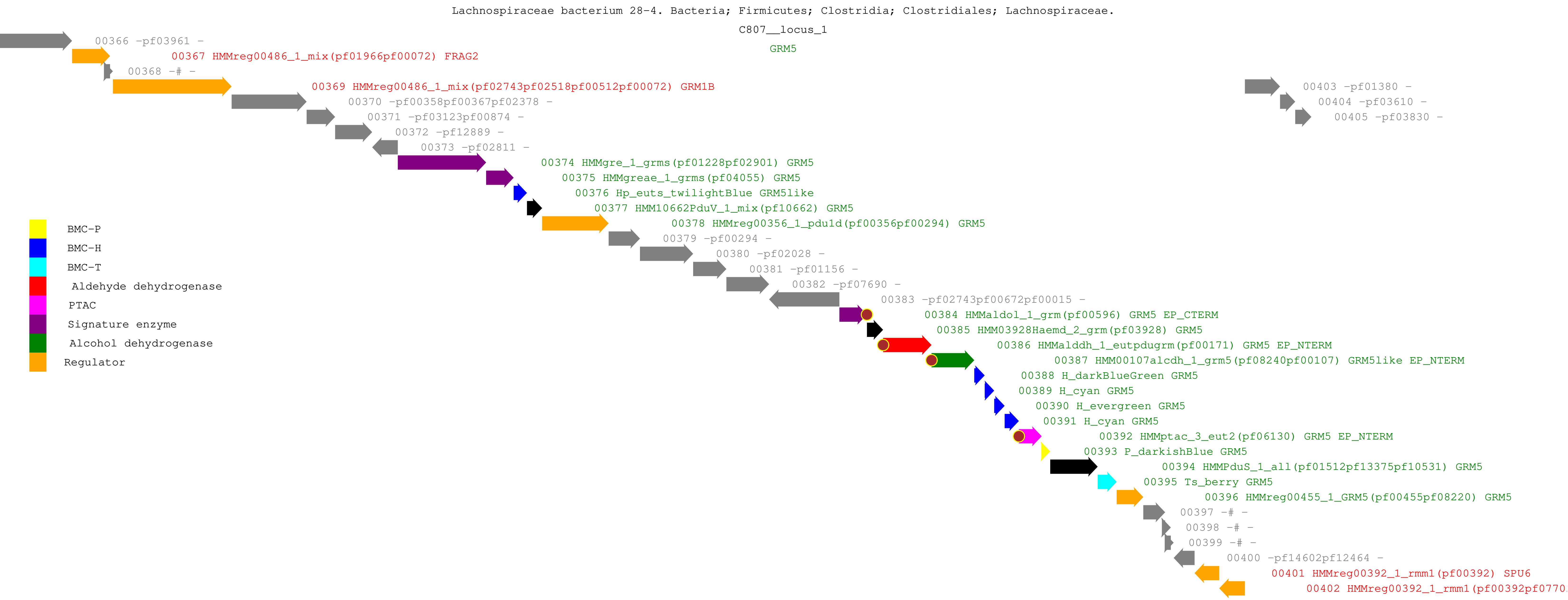
Lachnospiraceae bacterium 28-4. Bacteria; Firmicutes; Clostridia; Clostridiales; Lachnospiraceae.
C807__locus_1
GRM5
00366 -pf03961 -
00367 HMMreg00486_1_mix(pf01966pf00072) FRAG2
00368 -# -
00369 HMMreg00486_1_mix(pf02743pf02518pf00512pf00072) GRM1B
00370 -pf00358pf00367pf02378 -
00371 -pf03123pf00874 -
00372 -pf12889 -
00373 -pf02811 -
00374 HMMgre_1_grms(pf01228pf02901) GRM5
00375 HMMgreae_1_grms(pf04055) GRM5
00376 Hp_euts_twilightBlue GRM5like
00377 HMM10662PduV_1_mix(pf10662) GRM5
00378 HMMreg00356_1_pdu1d(pf00356pf00294) GRM5
00379 -pf00294 -
00380 -pf02028 -
00381 -pf01156 -
00382 -pf07690 -
00383 -pf02743pf00672pf00015 -
00384 HMMaldol_1_grm(pf00596) GRM5 EP_CTERM
00385 HMM03928Haemd_2_grm(pf03928) GRM5
00386 HMMalddh_1_eutpdugrm(pf00171) GRM5 EP_NTERM
00387 HMM00107alcdh_1_grm5(pf08240pf00107) GRM5like EP_NTERM
00388 H_darkBlueGreen GRM5
00389 H_cyan GRM5
00390 H_evergreen GRM5
00391 H_cyan GRM5
00392 HMMptac_3_eut2(pf06130) GRM5 EP_NTERM
00393 P_darkishBlue GRM5
00394 HMMPduS_1_all(pf01512pf13375pf10531) GRM5
00395 Ts_berry GRM5
00396 HMMreg00455_1_GRM5(pf00455pf08220) GRM5
00397 -# -
00398 -# -
00399 -# -
00400 -pf14602pf12464 -
00401 HMMreg00392_1_rmm1(pf00392) SPU6
00402 HMMreg00392_1_rmm1(pf00392pf0770
00403 -pf01380 -
00404 -pf03610 -
00405 -pf03830 -
BMC-P
BMC-H
BMC-T
Aldehyde dehydrogenase
PTAC
Signature enzyme
Alcohol dehydrogenase
Regulator

Lachnospiraceae bacterium M18-1. Bacteria; Firmicutes; Clostridia; Clostridiales; Lachnospiraceae.

C808__locus_1

GRM5

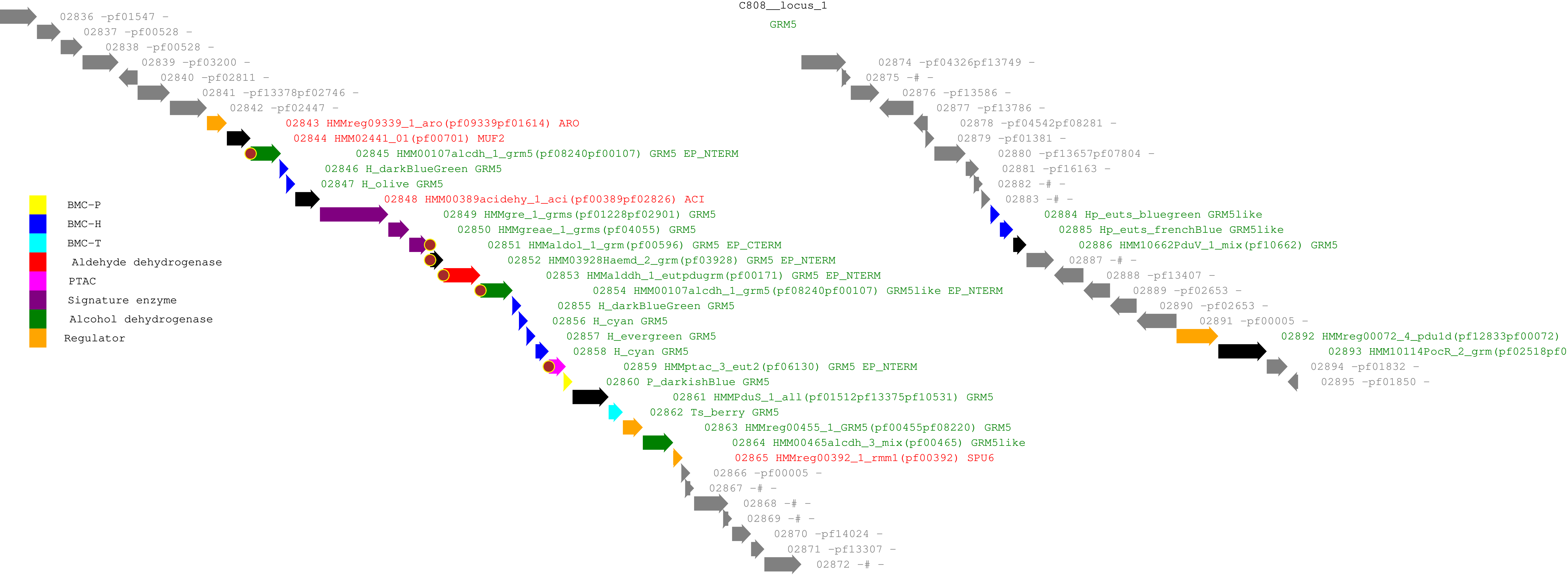

Lachnospiraceae bacterium 3-2. Bacteria; Firmicutes; Clostridia; Clostridiales; Lachnospiraceae.

C818__locus_1

GRM5

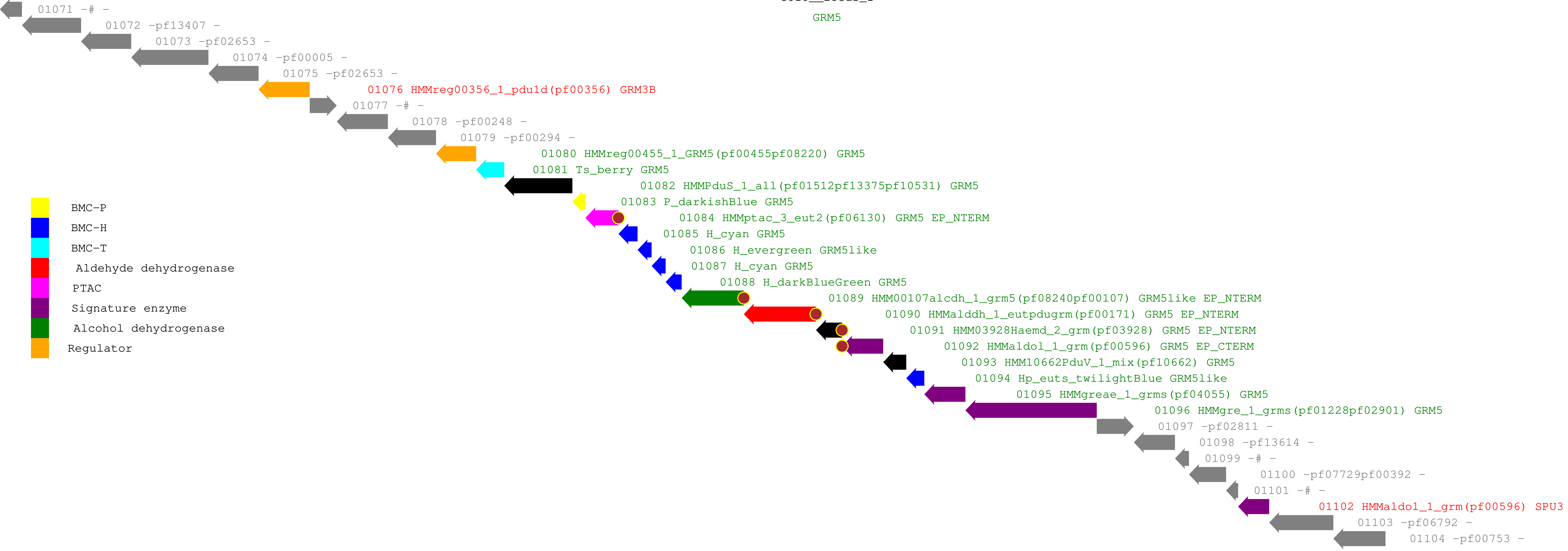

Eubacterium plexicaudatum ASF492. Bacteria; Firmicutes; Clostridia; Clostridiales; Eubacteriaceae;

C823__locus_1

GRM5

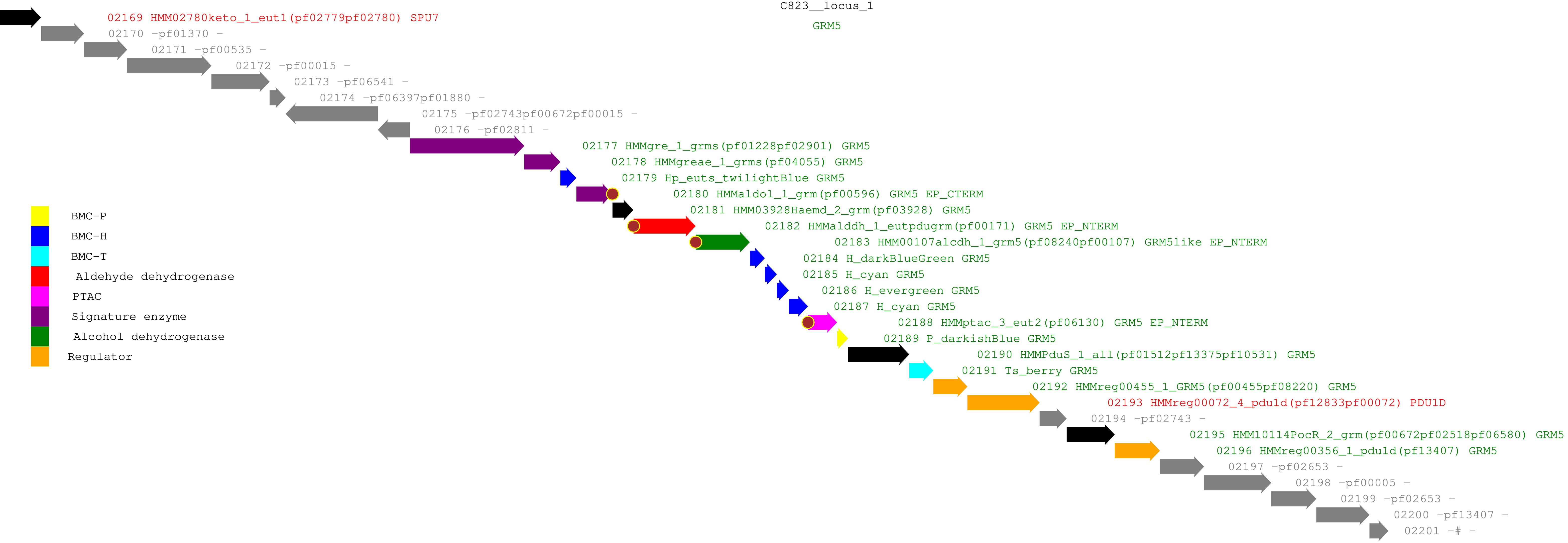

Clostridium sp. ASF502. Bacteria; Firmicutes; Clostridia; Clostridiales; Clostridiaceae;

C824__locus_1

GRM5

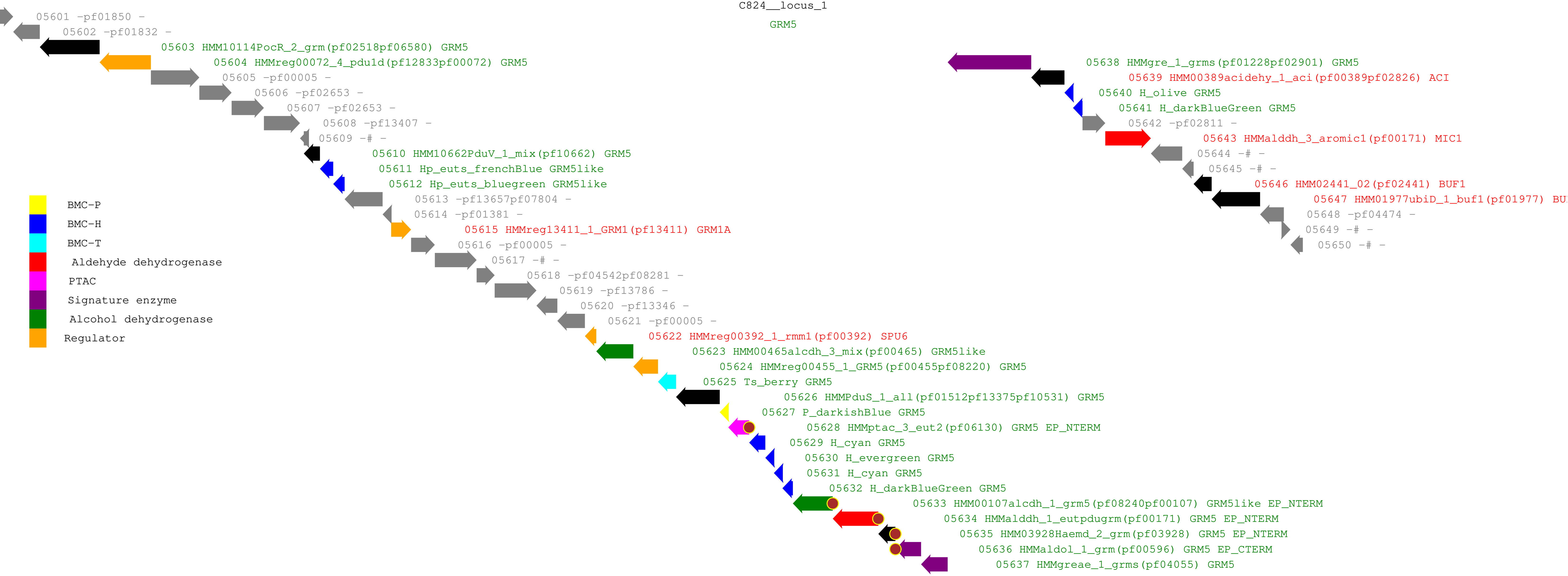

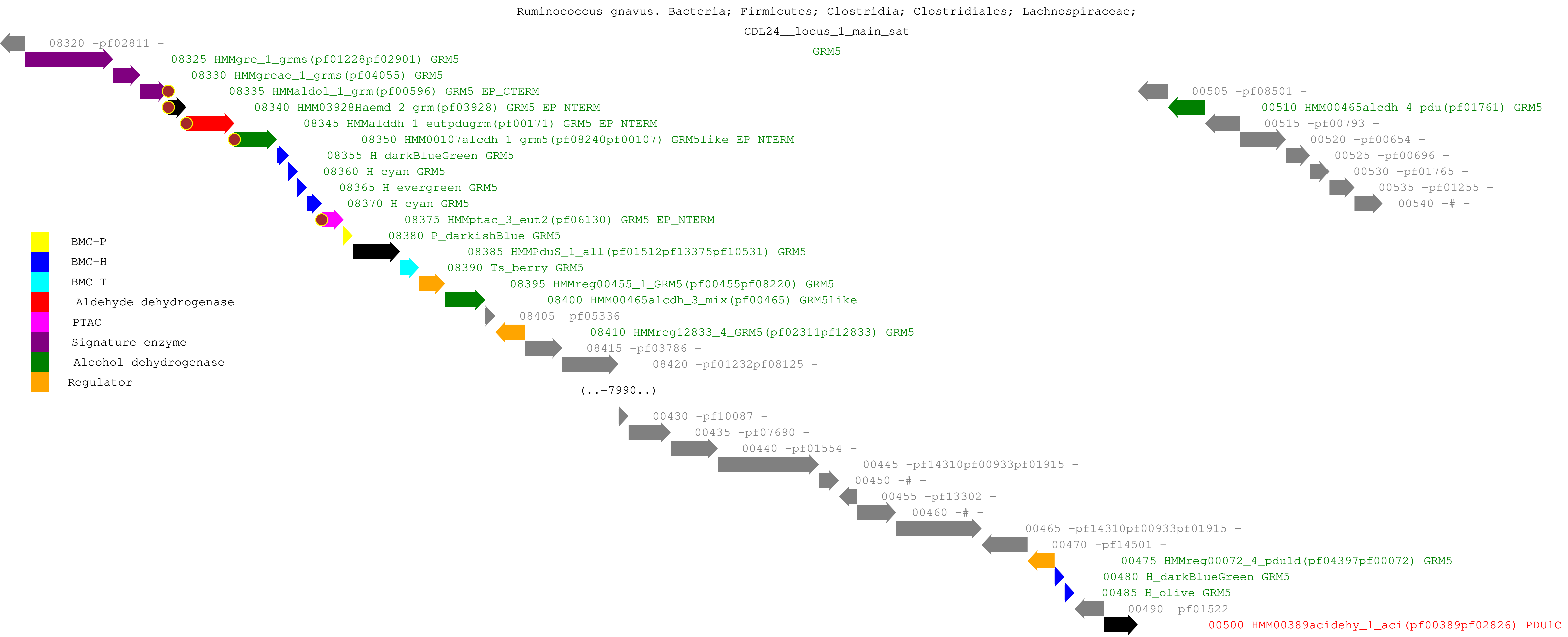
Ruminococcus gnavus. Bacteria; Firmicutes; Clostridia; Clostridiales; Lachnospiraceae;
CDL24__locus_1_main_sat
GRM5
08320 -pf02811 -
08325 HMMgre_1_grms(pf01228pf02901) GRM5
08330 HMMgreae_1_grms(pf04055) GRM5
08335 HMMaldol_1_grm(pf00596) GRM5 EP_CTERM
08340 HMM03928Haemd_2_grm(pf03928) GRM5 EP_NTERM
08345 HMMalddh_1_eutpdugrm(pf00171) GRM5 EP_NTERM
08350 HMM00107alcdh_1_grm5(pf08240pf00107) GRM5like EP_NTERM
08355 H_darkBlueGreen GRM5
08360 H_cyan GRM5
08365 H_evergreen GRM5
08370 H_cyan GRM5
08375 HMMptac_3_eut2(pf06130) GRM5 EP_NTERM
08380 P_darkishBlue GRM5
08385 HMMPduS_1_all(pf01512pf13375pf10531) GRM5
08390 Ts_berry GRM5
08395 HMMreg00455_1_GRM5(pf00455pf08220) GRM5
08400 HMM00465alcdh_3_mix(pf00465) GRM5like
08405 -pf05336 -
08410 HMMreg12833_4_GRM5(pf02311pf12833) GRM5
08415 -pf03786 -
08420 -pf01232pf08125 -
(..-7990..)
00430 -pf10087 -
00435 -pf07690 -
00440 -pf01554 -
00445 -pf14310pf00933pf01915 -
00450 -# -
00455 -pf13302 -
00460 -# -
00465 -pf14310pf00933pf01915 -
00470 -pf14501 -
00475 HMMreg00072_4_pdu1d(pf04397pf00072) GRM5
00480 H_darkBlueGreen GRM5
00485 H_olive GRM5
00490 -pf01522 -
00500 HMM00389acidehy_1_aci(pf00389pf02826) PDU1C
00505 -pf08501 -
00510 HMM00465alcdh_4_pdu(pf01761) GRM5
00515 -pf00793 -
00520 -pf00654 -
00525 -pf00696 -
00530 -pf01765 -
00535 -pf01255 -
00540 -# -
BMC-P
BMC-H
BMC-T
Aldehyde dehydrogenase
PTAC
Signature enzyme
Alcohol dehydrogenase
Regulator

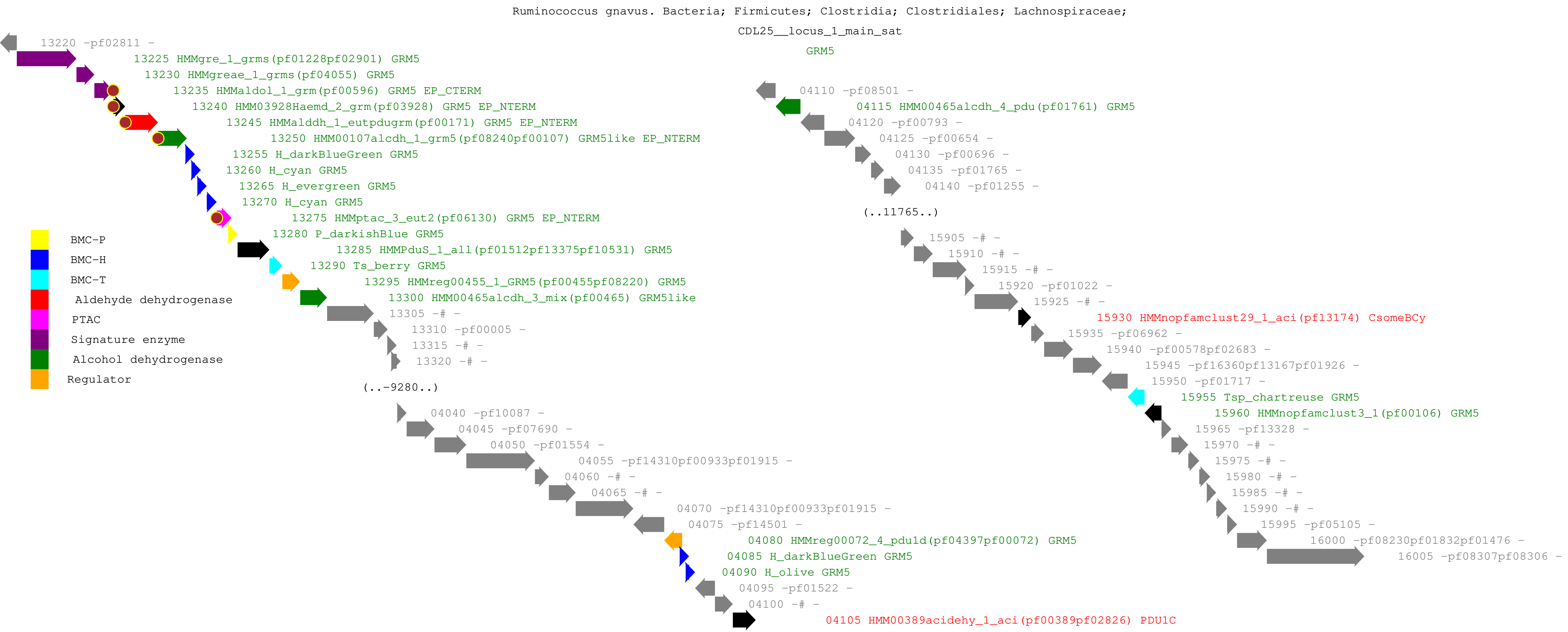
Ruminococcus gnavus. Bacteria; Firmicutes; Clostridia; Clostridiales; Lachnospiraceae;
CDL25__locus_1_main_sat
GRM5
13220 -pf02811 -
13225 HMMgre_1_grms(pf01228pf02901) GRM5
13230 HMMgreae_1_grms(pf04055) GRM5
13235 HMMaldol_1_grm(pf00596) GRM5 EP_CTERM
13240 HMM03928Haemd_2_grm(pf03928) GRM5 EP_NTERM
13245 HMMalddh_1_eutpdugrm(pf00171) GRM5 EP_NTERM
13250 HMM00107alcdh_1_grm5(pf08240pf00107) GRM5like EP_NTERM
13255 H_darkBlueGreen GRM5
13260 H_cyan GRM5
13265 H_evergreen GRM5
13270 H_cyan GRM5
13275 HMMptac_3_eut2(pf06130) GRM5 EP_NTERM
13280 P_darkishBlue GRM5
13285 HMMPduS_1_all(pf01512pf13375pf10531) GRM5
13290 Ts_berry GRM5
13295 HMMreg00455_1_GRM5(pf00455pf08220) GRM5
13300 HMM00465alcdh_3_mix(pf00465) GRM5like
13305 -# -
13310 -pf00005 -
13315 -# -
13320 -# -
(..-9280..)
04040 -pf10087 -
04045 -pf07690 -
04050 -pf01554 -
04055 -pf14310pf00933pf01915 -
04060 -# -
04065 -# -
04070 -pf14310pf00933pf01915 -
04075 -pf14501 -
04080 HMMreg00072_4_pdu1d(pf04397pf00072) GRM5
04085 H_darkBlueGreen GRM5
04090 H_olive GRM5
04095 -pf01522 -
04100 -# -
04105 HMM00389acidehy_1_aci(pf00389pf02826) PDU1C
04110 -pf08501 -
04115 HMM00465alcdh_4_pdu(pf01761) GRM5
04120 -pf00793 -
04125 -pf00654 -
04130 -pf00696 -
04135 -pf01765 -
04140 -pf01255 -
(..11765..)
15905 -# -
15910 -# -
15915 -# -
15920 -pf01022 -
15925 -# -
15930 HMMnopfamclust29_1_aci(pf13174) CsomeBCy
15935 -pf06962 -
15940 -pf00578pf02683 -
15945 -pf16360pf13167pf01926 -
15950 -pf01717 -
15955 Tsp_chartreuse GRM5
15960 HMMnopfamclust3_1(pf00106) GRM5
15965 -pf13328 -
15970 -# -
15975 -# -
15980 -# -
15985 -# -
15990 -# -
15995 -pf05105 -
16000 -pf08230pf01832pf01476 -
16005 -pf08307pf08306 -
BMC-P
BMC-H
BMC-T
Aldehyde dehydrogenase
PTAC
Signature enzyme
Alcohol dehydrogenase
Regulator

Ruminococcus gnavus. Bacteria; Firmicutes; Clostridia; Clostridiales; Lachnospiraceae;

CDL27__locus_1_main_sat

GRM5

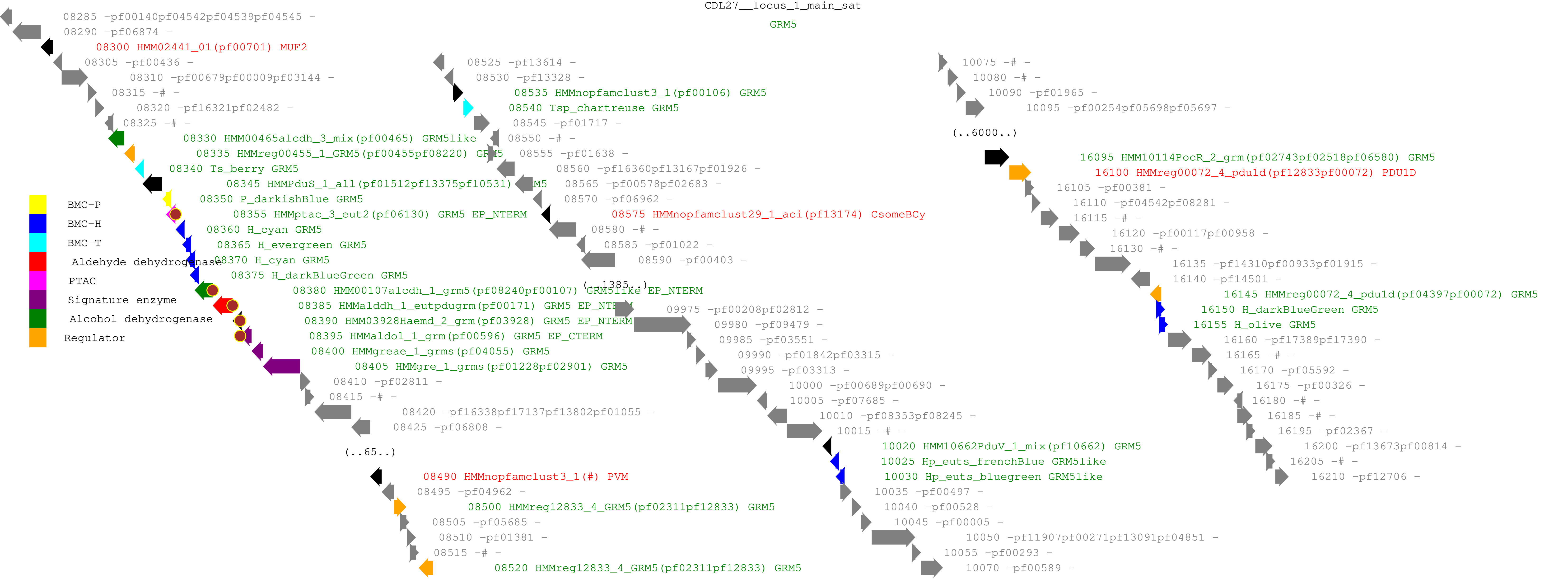

Ruminococcus sp. SR1/5. Bacteria; Firmicutes; Clostridia; Clostridiales; Ruminococcaceae;

CK1__locus_1_main_sat

GRM5

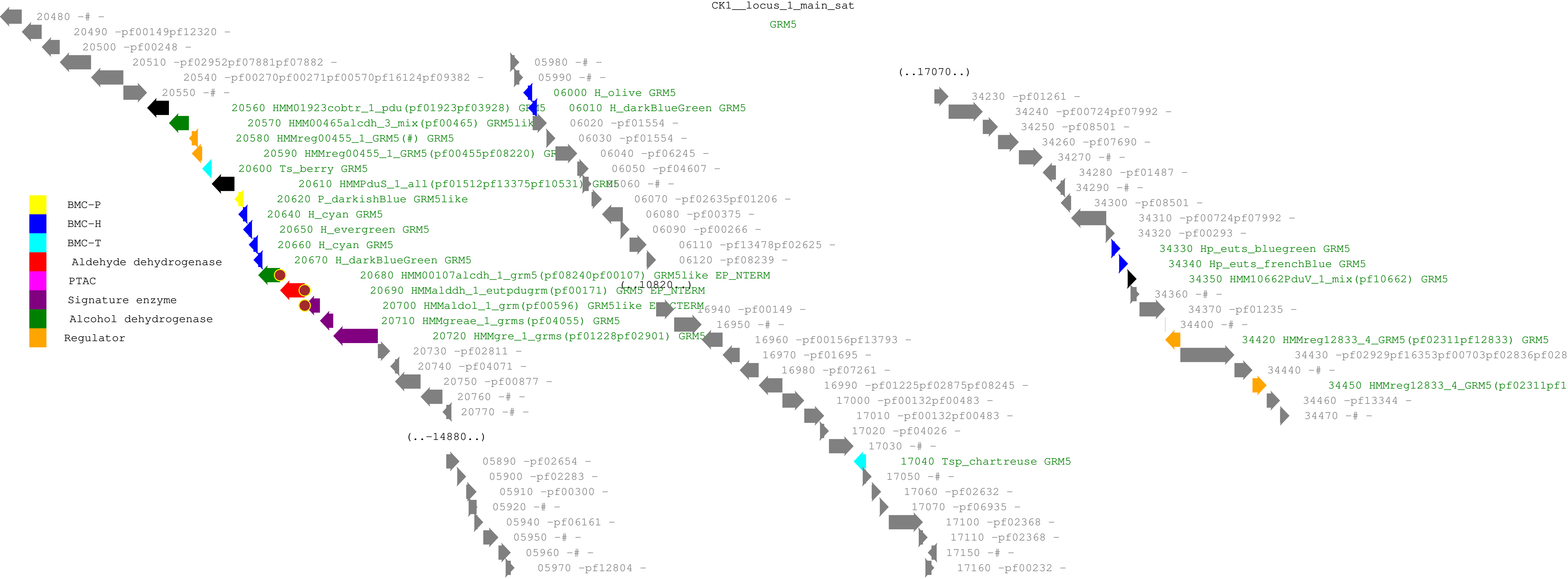

Blautia obeum A2-162. Bacteria; Firmicutes; Clostridia; Clostridiales; Lachnospiraceae;

CK5__locus_1_main_sat

GRM5

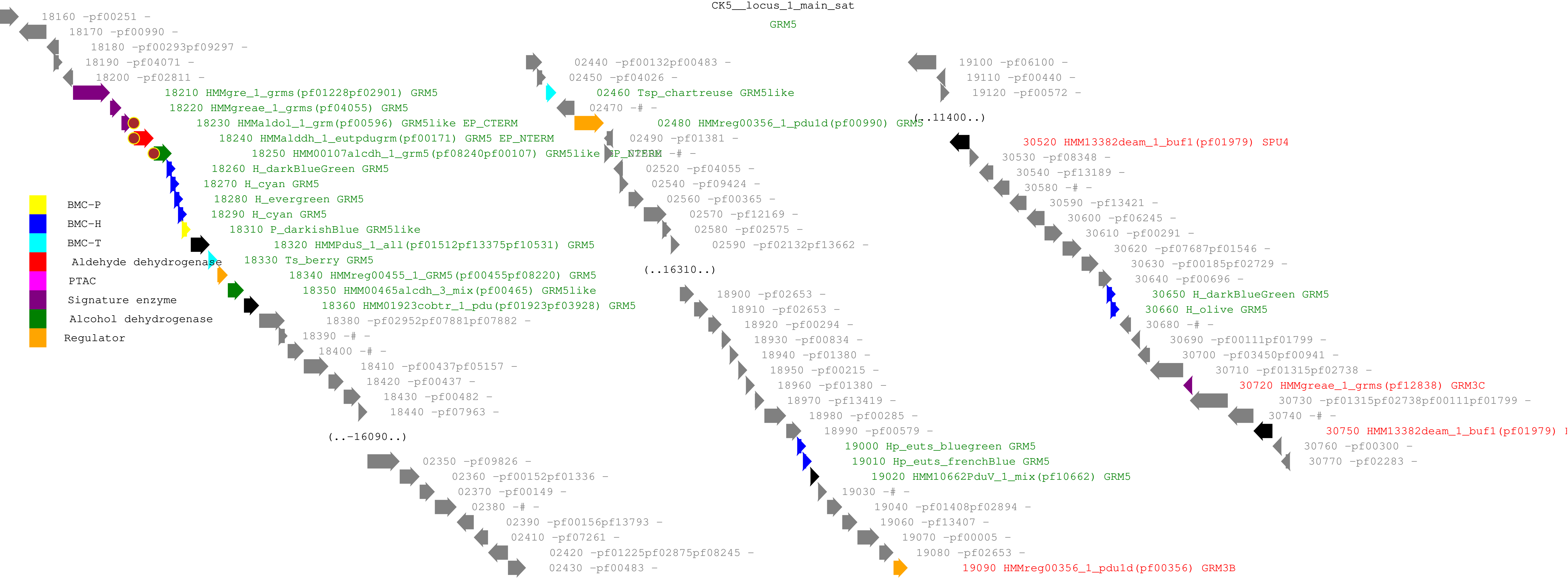

Lachnoclostridium phytofermentans (strain ATCC 700394 / DSM 18823 / Bacteria; Firmicutes; Clostridia; Clostridiales; Lachnospiraceae.

Cphy__locus_1

GRM5

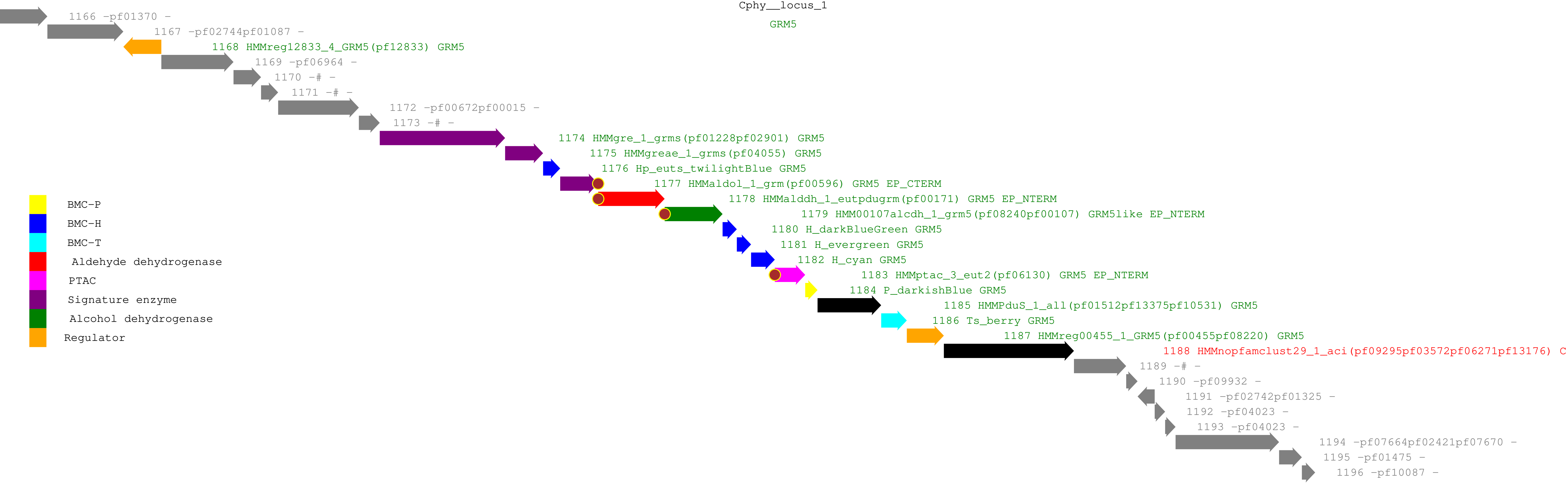

Clostridium neonatale. Bacteria; Firmicutes; Clostridia; Clostridiales; Clostridiaceae;

CQ394__locus_1

GRM5

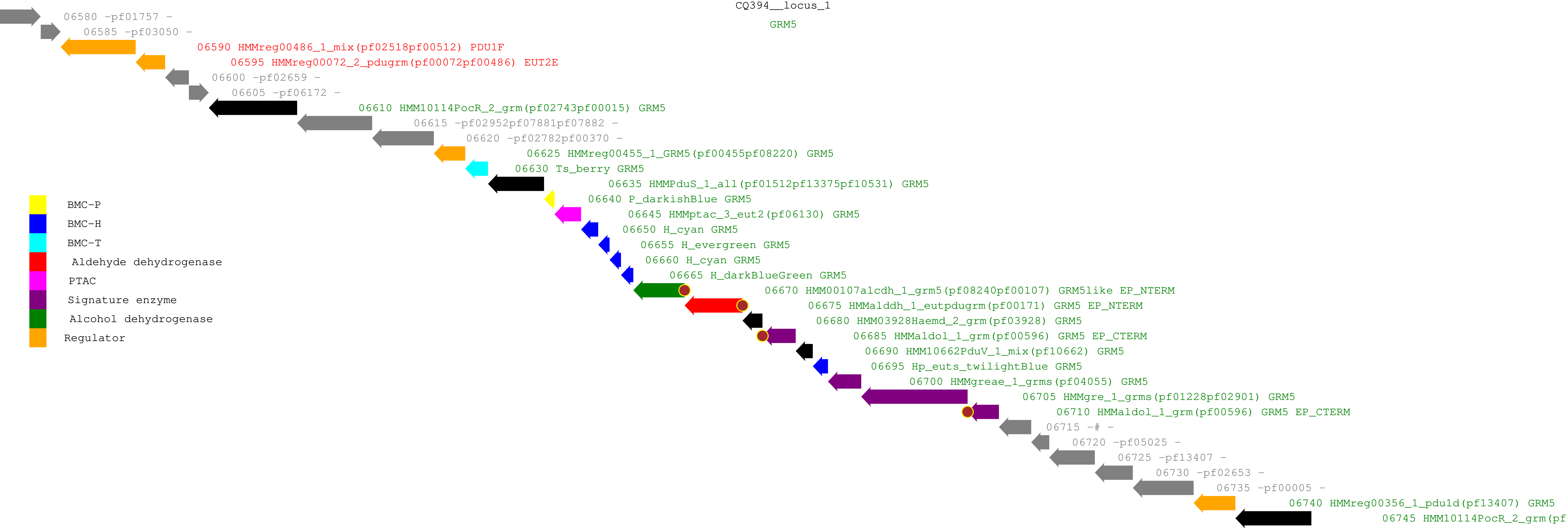

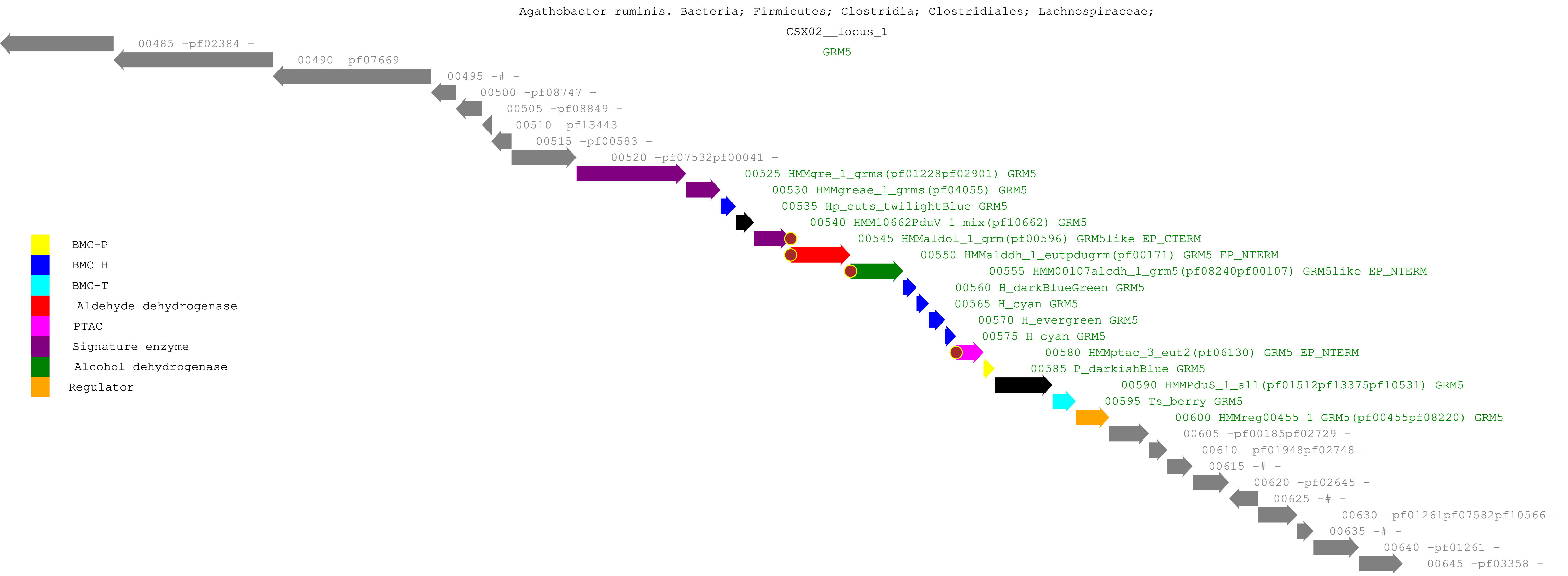
Agathobacter ruminis. Bacteria; Firmicutes; Clostridia; Clostridiales; Lachnospiraceae;
CSX02__locus_1
GRM5
00485 -pf02384 -
00490 -pf07669 -
00495 -# -
00500 -pf08747 -
00505 -pf08849 -
00510 -pf13443 -
00515 -pf00583 -
00520 -pf07532pf00041 -
00525 HMMgre_1_grms(pf01228pf02901) GRM5
00530 HMMgreae_1_grms(pf04055) GRM5
00535 Hp_euts_twilightBlue GRM5
00540 HMM10662PduV_1_mix(pf10662) GRM5
00545 HMMaldol_1_grm(pf00596) GRM5like EP_CTERM
00550 HMMalddh_1_eutpdugrm(pf00171) GRM5 EP_NTERM
00555 HMM00107alcdh_1_grm5(pf08240pf00107) GRM5like EP_NTERM
00560 H_darkBlueGreen GRM5
00565 H_cyan GRM5
00570 H_evergreen GRM5
00575 H_cyan GRM5
00580 HMMptac_3_eut2(pf06130) GRM5 EP_NTERM
00585 P_darkishBlue GRM5
00590 HMMPduS_1_all(pf01512pf13375pf10531) GRM5
00595 Ts_berry GRM5
00600 HMMreg00455_1_GRM5(pf00455pf08220) GRM5
00605 -pf00185pf02729 -
00610 -pf01948pf02748 -
00615 -# -
00620 -pf02645 -
00625 -# -
00630 -pf01261pf07582pf10566 -
00635 -# -
00640 -pf01261 -
00645 -pf03358 -
BMC-P
BMC-H
BMC-T
Aldehyde dehydrogenase
PTAC
Signature enzyme
Alcohol dehydrogenase
Regulator

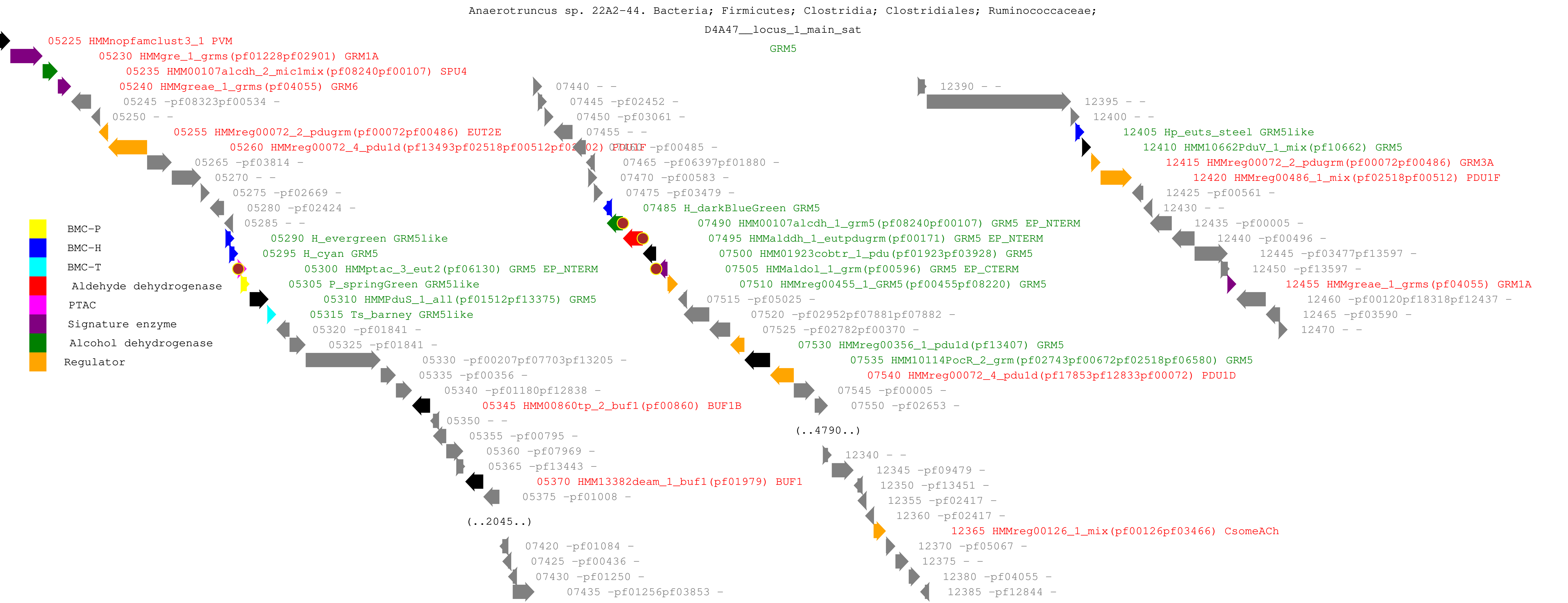
Anaerotruncus sp. 22A2-44. Bacteria; Firmicutes; Clostridia; Clostridiales; Ruminococcaceae;
D4A47__locus_1_main_sat
GRM5
05225 HMMnopfamclust3_1 PVM
05230 HMMgre_1_grms(pf01228pf02901) GRM1A
05235 HMM00107alcdh_2_mic1mix(pf08240pf00107) SPU4
05240 HMMgreae_1_grms(pf04055) GRM6
05245 -pf08323pf00534 -
05250 - -
05255 HMMreg00072_2_pdugrm(pf00072pf00486) EUT2E
05260 HMMreg00072_4_pdu1d(pf13493pf02518pf00512pf02[illegible]02) PDU1F
05265 -pf03814 -
05270 - -
05275 -pf02669 -
05280 -pf02424 -
05285 - -
05290 H_evergreen GRM5like
05295 H_cyan GRM5
05300 HMMptac_3_eut2(pf06130) GRM5 EP_NTERM
05305 P_springGreen GRM5like
05310 HMMPduS_1_all(pf01512pf13375) GRM5
05315 Ts_barney GRM5like
05320 -pf01841 -
05325 -pf01841 -
05330 -pf00207pf07703pf13205 -
05335 -pf00356 -
05340 -pf01180pf12838 -
05345 HMM00860tp_2_buf1(pf00860) BUF1B
05350 - -
05355 -pf00795 -
05360 -pf07969 -
05365 -pf13443 -
05370 HMM13382deam_1_buf1(pf01979) BUF1
05375 -pf01008 -
(..2045..)
07420 -pf01084 -
07425 -pf00436 -
07430 -pf01250 -
07435 -pf01256pf03853 -
07440 - -
07445 -pf02452 -
07450 -pf03061 -
07455 - -
07460 -pf00485 -
07465 -pf06397pf01880 -
07470 -pf00583 -
07475 -pf03479 -
07485 H_darkBlueGreen GRM5
07490 HMM00107alcdh_1_grm5(pf08240pf00107) GRM5 EP_NTERM
07495 HMMalddh_1_eutpdugrm(pf00171) GRM5 EP_NTERM
07500 HMM01923cobtr_1_pdu(pf01923pf03928) GRM5
07505 HMMaldol_1_grm(pf00596) GRM5 EP_CTERM
07510 HMMreg00455_1_GRM5(pf00455pf08220) GRM5
07515 -pf05025 -
07520 -pf02952pf07881pf07882 -
07525 -pf02782pf00370 -
07530 HMMreg00356_1_pdu1d(pf13407) GRM5
07535 HMM10114PocR_2_grm(pf02743pf00672pf02518pf06580) GRM5
07540 HMMreg00072_4_pdu1d(pf17853pf12833pf00072) PDU1D
07545 -pf00005 -
07550 -pf02653 -
(..4790..)
12340 - -
12345 -pf09479 -
12350 -pf13451 -
12355 -pf02417 -
12360 -pf02417 -
12365 HMMreg00126_1_mix(pf00126pf03466) CsomeACh
12370 -pf05067 -
12375 - -
12380 -pf04055 -
12385 -pf12844 -
12390 - -
12395 - -
12400 - -
12405 Hp_euts_steel GRM5like
12410 HMM10662PduV_1_mix(pf10662) GRM5
12415 HMMreg00072_2_pdugrm(pf00072pf00486) GRM3A
12420 HMMreg00486_1_mix(pf02518pf00512) PDU1F
12425 -pf00561 -
12430 - -
12435 -pf00005 -
12440 -pf00496 -
12445 -pf03477pf13597 -
12450 -pf13597 -
12455 HMMgreae_1_grms(pf04055) GRM1A
12460 -pf00120pf18318pf12437 -
12465 -pf03590 -
12470 - -
BMC-P
BMC-H
BMC-T
Aldehyde dehydrogenase
PTAC
Signature enzyme
Alcohol dehydrogenase
Regulator

bacterium 1XD8-92. Bacteria.

D6D47__locus_1

GRM5

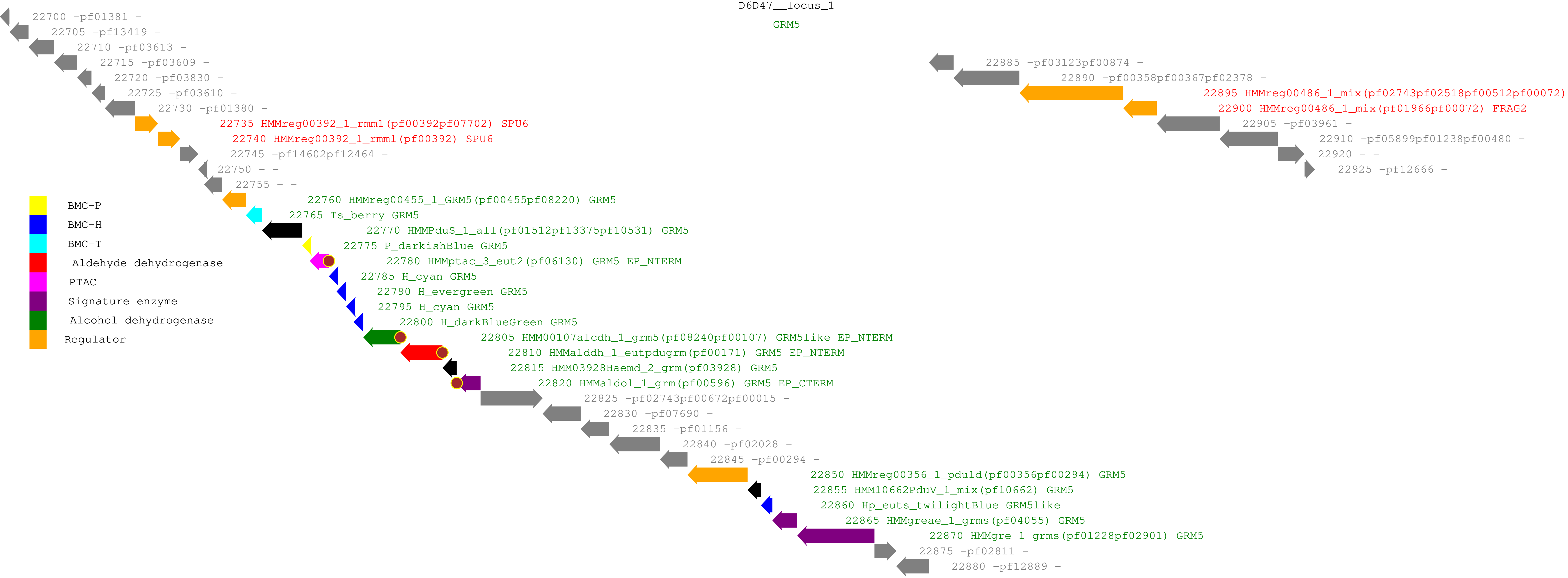

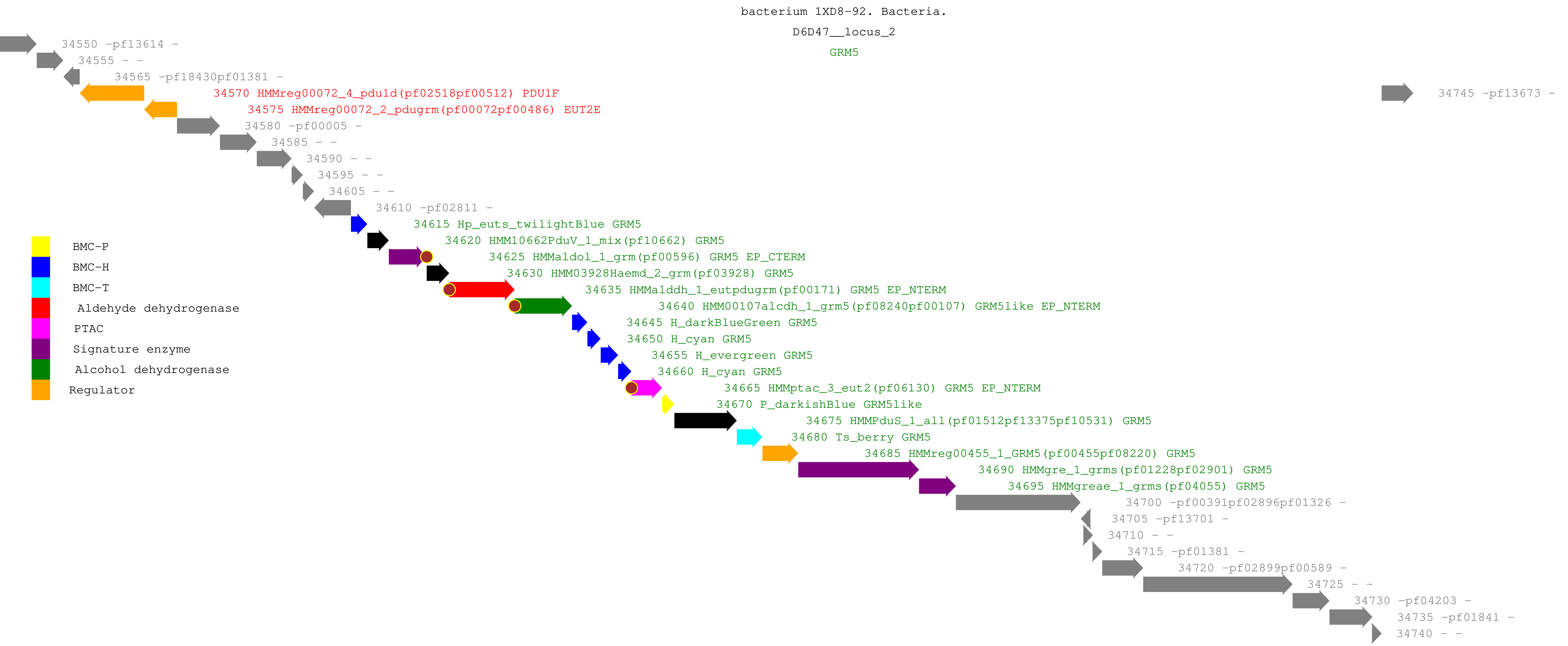
bacterium 1XD8-92. Bacteria.
D6D47__locus_2
GRM5
34550 -pf13614 -
34555 - -
34565 -pf18430pf01381 -
34570 HMMreg00072_4_pdu1d(pf02518pf00512) PDU1F
34575 HMMreg00072_2_pdugrm(pf00072pf00486) EUT2E
34580 -pf00005 -
34585 - -
34590 - -
34595 - -
34605 - -
34610 -pf02811 -
34615 Hp_euts_twilightBlue GRM5
34620 HMM10662PduV_1_mix(pf10662) GRM5
34625 HMMaldol_1_grm(pf00596) GRM5 EP_CTERM
34630 HMM03928Haemd_2_grm(pf03928) GRM5
34635 HMMalddh_1_eutpdugrm(pf00171) GRM5 EP_NTERM
34640 HMM00107alcdh_1_grm5(pf08240pf00107) GRM5like EP_NTERM
34645 H_darkBlueGreen GRM5
34650 H_cyan GRM5
34655 H_evergreen GRM5
34660 H_cyan GRM5
34665 HMMptac_3_eut2(pf06130) GRM5 EP_NTERM
34670 P_darkishBlue GRM5like
34675 HMMPduS_1_all(pf01512pf13375pf10531) GRM5
34680 Ts_berry GRM5
34685 HMMreg00455_1_GRM5(pf00455pf08220) GRM5
34690 HMMgre_1_grms(pf01228pf02901) GRM5
34695 HMMgreae_1_grms(pf04055) GRM5
34700 -pf00391pf02896pf01326 -
34705 -pf13701 -
34710 - -
34715 -pf01381 -
34720 -pf02899pf00589 -
34725 - -
34730 -pf04203 -
34735 -pf01841 -
34740 - -
34745 -pf13673 -
BMC-P
BMC-H
BMC-T
Aldehyde dehydrogenase
PTAC
Signature enzyme
Alcohol dehydrogenase
Regulator

bacterium 1XD42-8. Bacteria.

D7X25__locus_2

GRM5

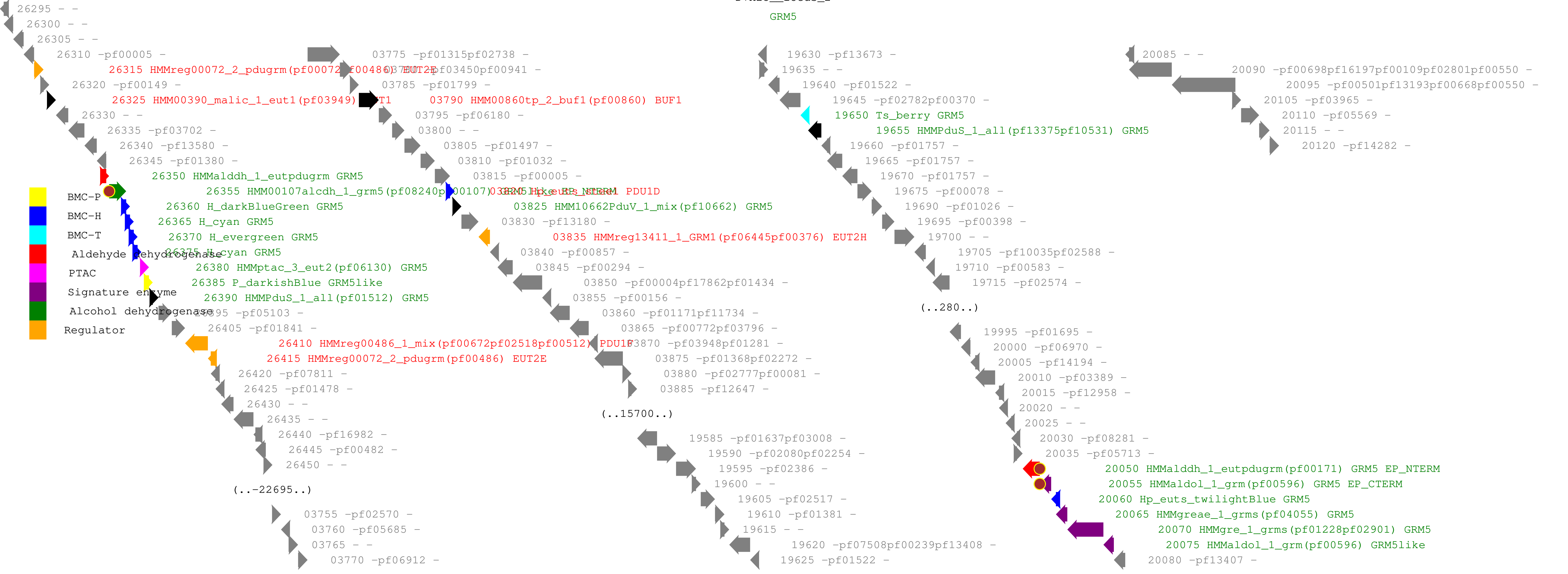

bacterium D16-50. Bacteria.

D7X48__locus_1

GRM5

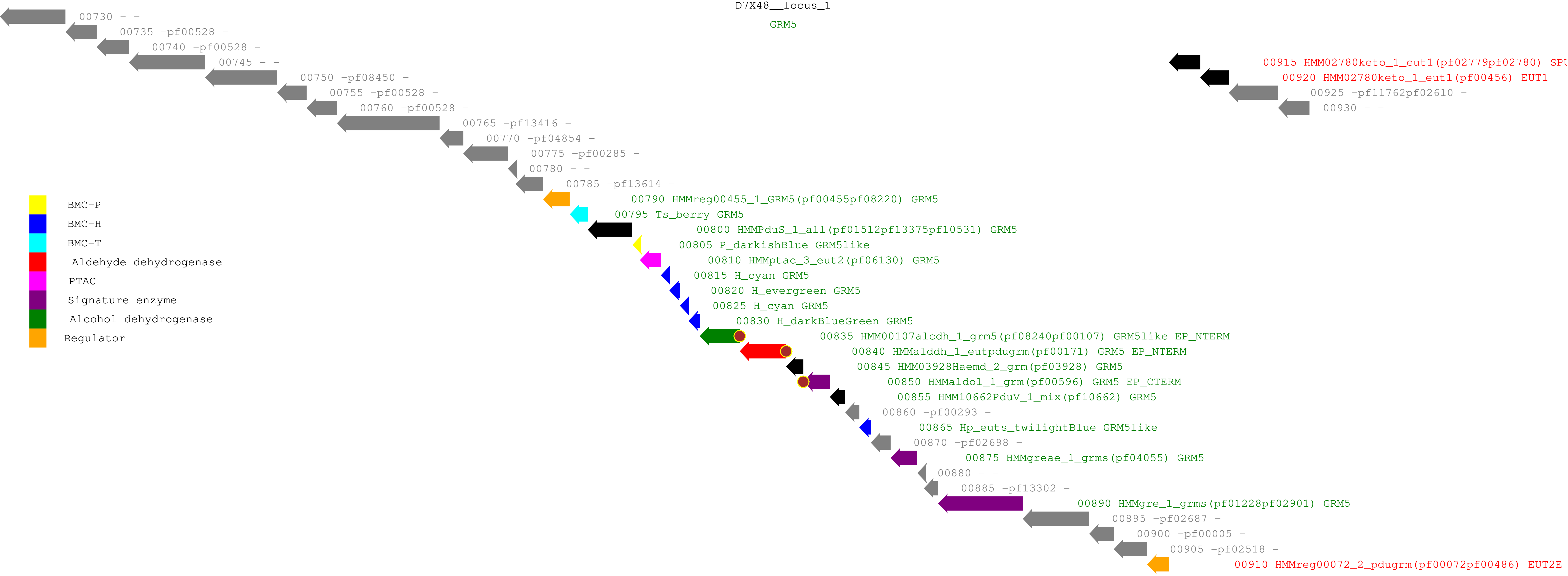

bacterium C-53. Bacteria.

D7X88__locus_1

GRM5

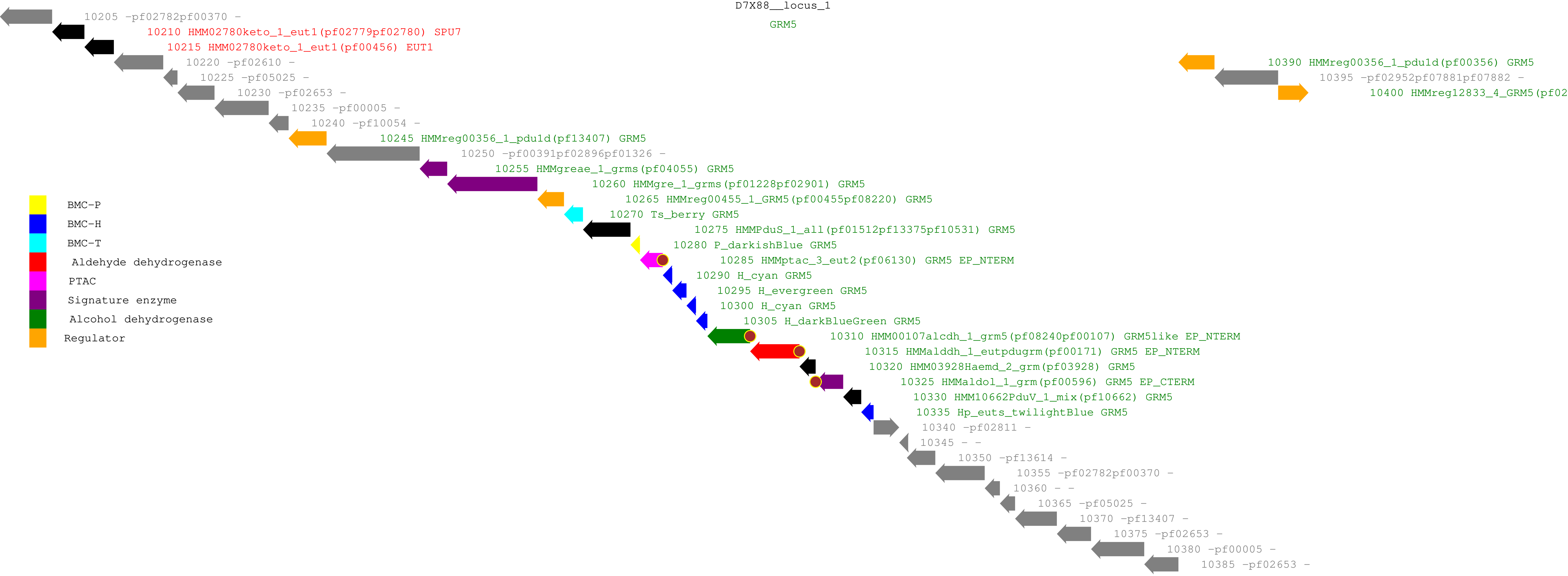

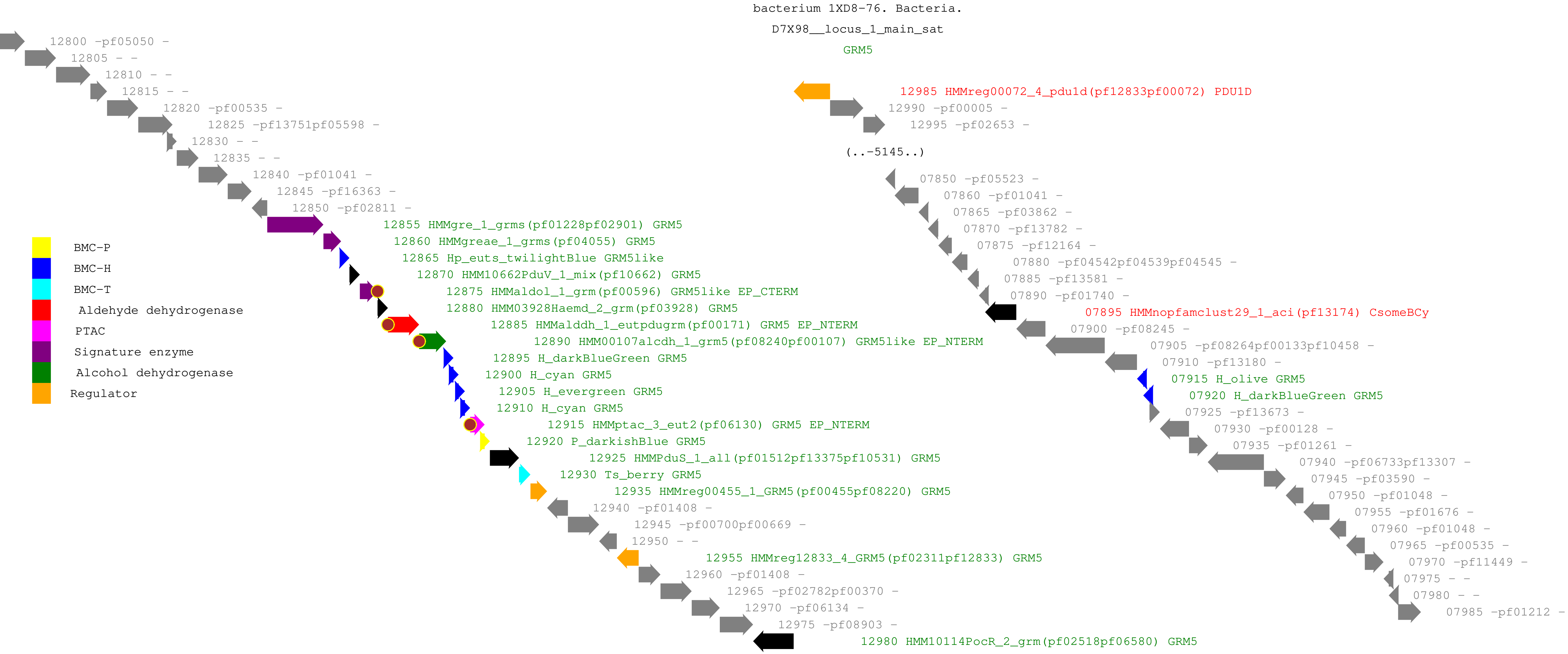
bacterium 1XD8-76. Bacteria.
D7X98__locus_1_main_sat
GRM5
12800 -pf05050 -
12805 - -
12810 - -
12815 - -
12820 -pf00535 -
12825 -pf13751pf05598 -
12830 - -
12835 - -
12840 -pf01041 -
12845 -pf16363 -
12850 -pf02811 -
12855 HMMgre_1_grms(pf01228pf02901) GRM5
12860 HMMgreae_1_grms(pf04055) GRM5
12865 Hp_euts_twilightBlue GRM5like
12870 HMM10662PduV_1_mix(pf10662) GRM5
12875 HMMaldol_1_grm(pf00596) GRM5like EP_CTERM
12880 HMM03928Haemd_2_grm(pf03928) GRM5
12885 HMMaldh_1_eutpdugrm(pf00171) GRM5 EP_NTERM
12890 HMM00107alcdh_1_grm5(pf08240pf00107) GRM5like EP_NTERM
12895 H_darkBlueGreen GRM5
12900 H_cyan GRM5
12905 H_evergreen GRM5
12910 H_cyan GRM5
12915 HMMptac_3_eut2(pf06130) GRM5 EP_NTERM
12920 P_darkishBlue GRM5
12925 HMMPduS_1_all(pf01512pf13375pf10531) GRM5
12930 Ts_berry GRM5
12935 HMMreg00455_1_GRM5(pf00455pf08220) GRM5
12940 -pf01408 -
12945 -pf00700pf00669 -
12950 - -
12955 HMMreg12833_4_GRM5(pf02311pf12833) GRM5
12960 -pf01408 -
12965 -pf02782pf00370 -
12970 -pf06134 -
12975 -pf08903 -
12980 HMM10114PocR_2_grm(pf02518pf06580) GRM5
12985 HMMreg00072_4_pdu1d(pf12833pf00072) PDU1D
12990 -pf00005 -
12995 -pf02653 -
(..-5145..)
07850 -pf05523 -
07860 -pf01041 -
07865 -pf03862 -
07870 -pf13782 -
07875 -pf12164 -
07880 -pf04542pf04539pf04545 -
07885 -pf13581 -
07890 -pf01740 -
07895 HMMnopfamclust29_1_aci(pf13174) CsomeBCy
07900 -pf08245 -
07905 -pf08264pf00133pf10458 -
07910 -pf13180 -
07915 H_olive GRM5
07920 H_darkBlueGreen GRM5
07925 -pf13673 -
07930 -pf00128 -
07935 -pf01261 -
07940 -pf06733pf13307 -
07945 -pf03590 -
07950 -pf01048 -
07955 -pf01676 -
07960 -pf01048 -
07965 -pf00535 -
07970 -pf11449 -
07975 - -
07980 - -
07985 -pf01212 -
BMC-P
BMC-H
BMC-T
Aldehyde dehydrogenase
PTAC
Signature enzyme
Alcohol dehydrogenase
Regulator

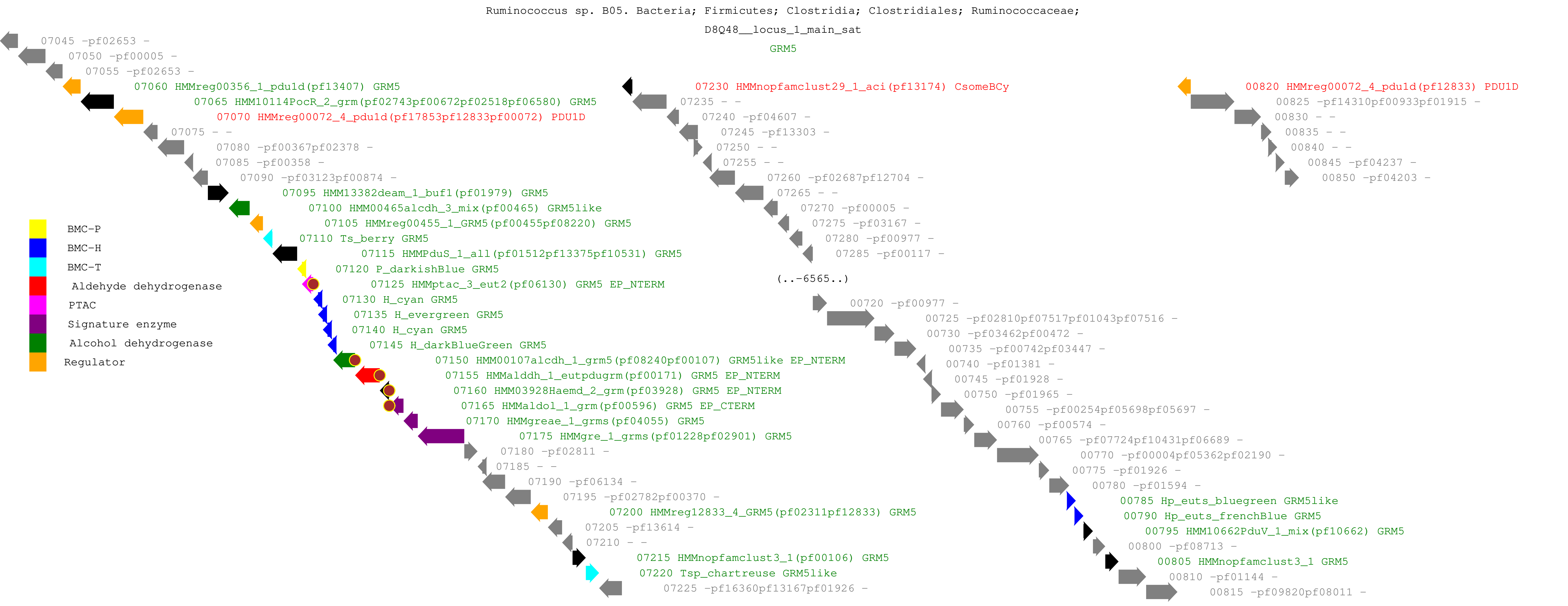

Ruminococcus sp. B05. Bacteria; Firmicutes; Clostridia; Clostridiales; Ruminococcaceae;
D8Q48__locus_1_main_sat
GRM5
07045 -pf02653 -
07050 -pf00005 -
07055 -pf02653 -
07060 HMMreg00356_1_pdu1d(pf13407) GRM5
07065 HMM10114PocR_2_grm(pf02743pf00672pf02518pf06580) GRM5
07070 HMMreg00072_4_pdu1d(pf17853pf12833pf00072) PDU1D
07075 - -
07080 -pf00367pf02378 -
07085 -pf00358 -
07090 -pf03123pf00874 -
07095 HMM13382deam_1_buf1(pf01979) GRM5
07100 HMM00465alcdh_3_mix(pf00465) GRM5like
07105 HMMreg00455_1_GRM5(pf00455pf08220) GRM5
07110 Ts_berry GRM5
07115 HMMPduS_1_all(pf01512pf13375pf10531) GRM5
07120 P_darkishBlue GRM5
07125 HMMptac_3_eut2(pf06130) GRM5 EP_NTERM
07130 H_cyan GRM5
07135 H_evergreen GRM5
07140 H_cyan GRM5
07145 H_darkBlueGreen GRM5
07150 HMM00107alcdh_1_grm5(pf08240pf00107) GRM5like EP_NTERM
07155 HMMalddh_1_eutpdugrm(pf00171) GRM5 EP_NTERM
07160 HMM03928Haemd_2_grm(pf03928) GRM5 EP_NTERM
07165 HMMaldol_1_grm(pf00596) GRM5 EP_CTERM
07170 HMMgreae_1_grms(pf04055) GRM5
07175 HMMgre_1_grms(pf01228pf02901) GRM5
07180 -pf02811 -
07185 - -
07190 -pf06134 -
07195 -pf02782pf00370 -
07200 HMMreg12833_4_GRM5(pf02311pf12833) GRM5
07205 -pf13614 -
07210 - -
07215 HMMnopfamclust3_1(pf00106) GRM5
07220 Tsp_chartreuse GRM5like
07225 -pf16360pf13167pf01926 -
07230 HMMnopfamclust29_1_aci(pf13174) CsomeBCy
07235 - -
07240 -pf04607 -
07245 -pf13303 -
07250 - -
07255 - -
07260 -pf02687pf12704 -
07265 - -
07270 -pf00005 -
07275 -pf03167 -
07280 -pf00977 -
07285 -pf00117 -
(..-6565..)
00720 -pf00977 -
00725 -pf02810pf07517pf01043pf07516 -
00730 -pf03462pf00472 -
00735 -pf00742pf03447 -
00740 -pf01381 -
00745 -pf01928 -
00750 -pf01965 -
00755 -pf00254pf05698pf05697 -
00760 -pf00574 -
00765 -pf07724pf10431pf06689 -
00770 -pf00004pf05362pf02190 -
00775 -pf01926 -
00780 -pf01594 -
00785 Hp_euts_bluegreen GRM5like
00790 Hp_euts_frenchBlue GRM5
00795 HMM10662PduV_1_mix(pf10662) GRM5
00800 -pf08713 -
00805 HMMnopfamclust3_1 GRM5
00810 -pf01144 -
00815 -pf09820pf08011 -
00820 HMMreg00072_4_pdu1d(pf12833) PDU1D
00825 -pf14310pf00933pf01915 -
00830 - -
00835 - -
00840 - -
00845 -pf04237 -
00850 -pf04203 -
BMC-P
BMC-H
BMC-T
Aldehyde dehydrogenase
PTAC
Signature enzyme
Alcohol dehydrogenase
Regulator

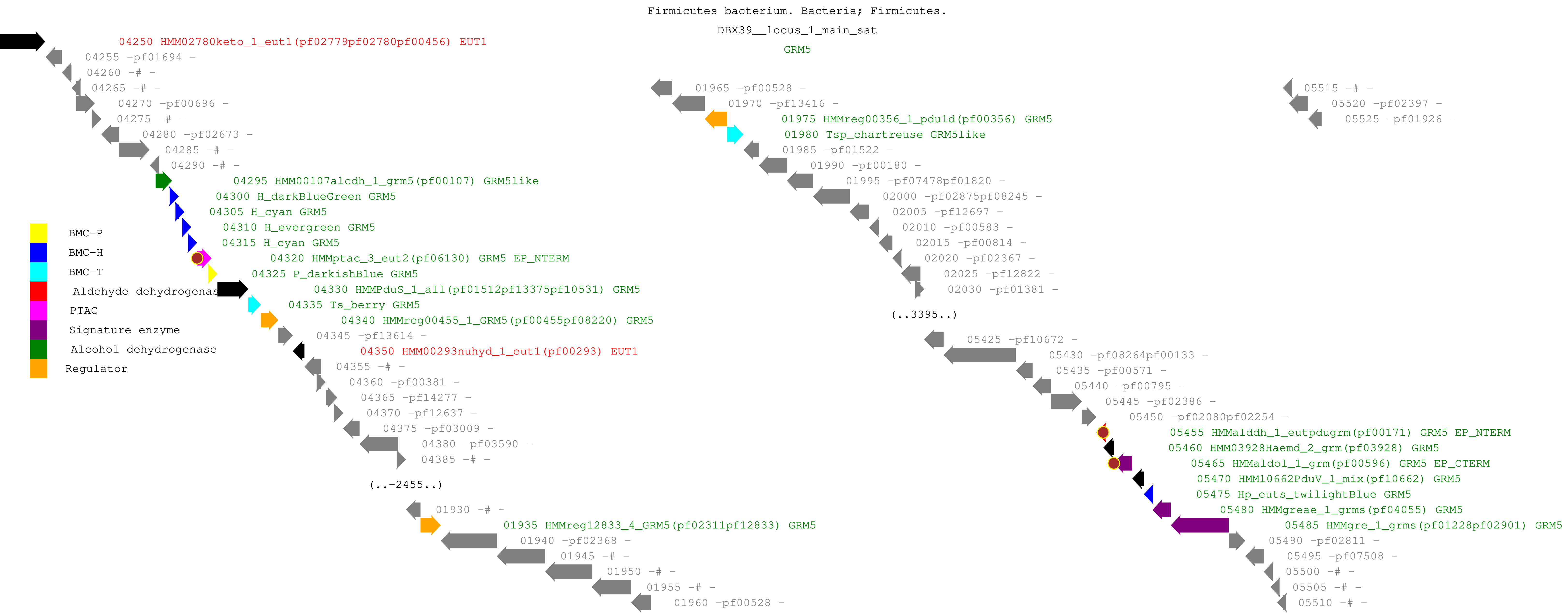
Firmicutes bacterium. Bacteria; Firmicutes.
DBX39__locus_1_main_sat
GRM5
04250 HMM02780keto_1_eut1(pf02779pf02780pf00456) EUT1
04255 -pf01694 -
04260 -# -
04265 -# -
04270 -pf00696 -
04275 -# -
04280 -pf02673 -
04285 -# -
04290 -# -
04295 HMM00107alcdh_1_grm5(pf00107) GRM5like
04300 H_darkBlueGreen GRM5
04305 H_cyan GRM5
04310 H_evergreen GRM5
04315 H_cyan GRM5
04320 HMMptac_3_eut2(pf06130) GRM5 EP_NTERM
04325 P_darkishBlue GRM5
04330 HMMPduS_1_all(pf01512pf13375pf10531) GRM5
04335 Ts_berry GRM5
04340 HMMreg00455_1_GRM5(pf00455pf08220) GRM5
04345 -pf13614 -
04350 HMM00293nuhyd_1_eut1(pf00293) EUT1
04355 -# -
04360 -pf00381 -
04365 -pf14277 -
04370 -pf12637 -
04375 -pf03009 -
04380 -pf03590 -
04385 -# -
(..-2455..)
01930 -# -
01935 HMMreg12833_4_GRM5(pf02311pf12833) GRM5
01940 -pf02368 -
01945 -# -
01950 -# -
01955 -# -
01960 -pf00528 -
BMC-P
BMC-H
BMC-T
Aldehyde dehydrogenas
PTAC
Signature enzyme
Alcohol dehydrogenase
Regulator
01965 -pf00528 -
01970 -pf13416 -
01975 HMMreg00356_1_pduld(pf00356) GRM5
01980 Tsp_chartreuse GRM5like
01985 -pf01522 -
01990 -pf00180 -
01995 -pf07478pf01820 -
02000 -pf02875pf08245 -
02005 -pf12697 -
02010 -pf00583 -
02015 -pf00814 -
02020 -pf02367 -
02025 -pf12822 -
02030 -pf01381 -
(..3395..)
05425 -pf10672 -
05430 -pf08264pf00133 -
05435 -pf00571 -
05440 -pf00795 -
05445 -pf02386 -
05450 -pf02080pf02254 -
05455 HMMaldDh_1_eutpdugrm(pf00171) GRM5 EP_NTERM
05460 HMM03928Haemd_2_grm(pf03928) GRM5
05465 HMMaldol_1_grm(pf00596) GRM5 EP_CTERM
05470 HMM10662PduV_1_mix(pf10662) GRM5
05475 Hp_euts_twilightBlue GRM5
05480 HMMgreae_1_grms(pf04055) GRM5
05485 HMMgre_1_grms(pf01228pf02901) GRM5
05490 -pf02811 -
05495 -pf07508 -
05500 -# -
05505 -# -
05510 -# -
05515 -# -
05520 -pf02397 -
05525 -pf01926 -

Clostridiales bacterium. Bacteria; Firmicutes; Clostridia; Clostridiales.

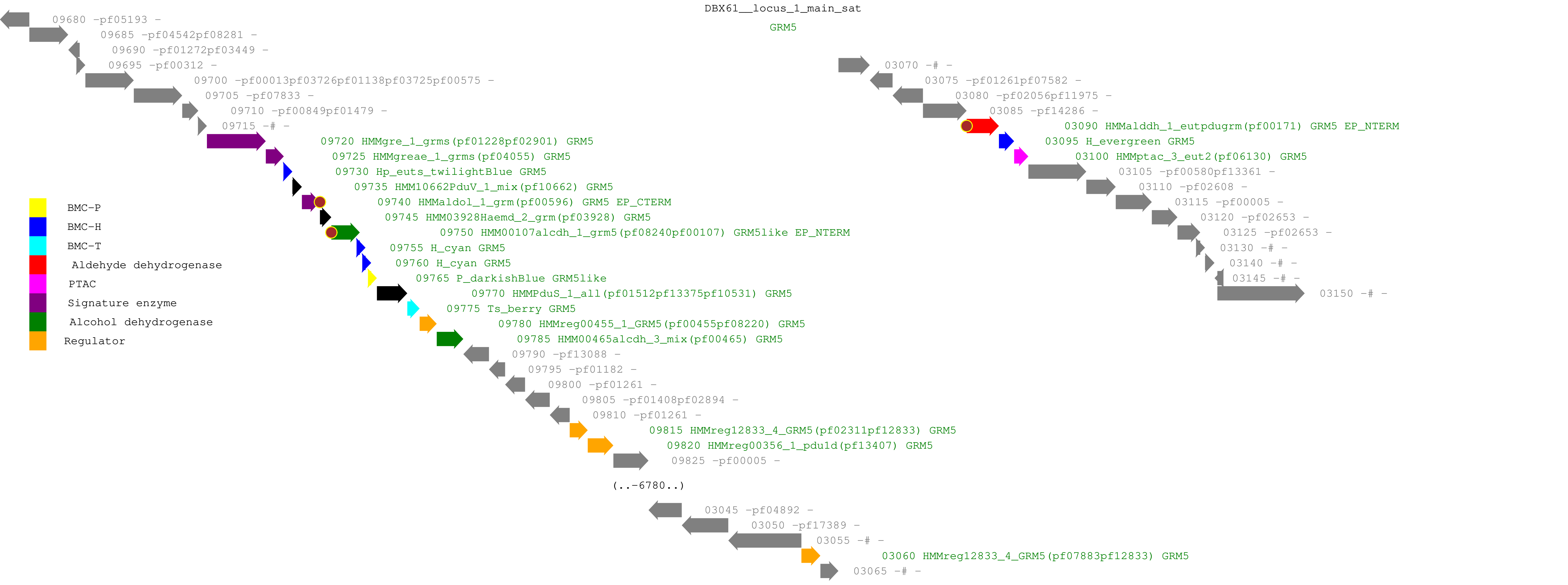

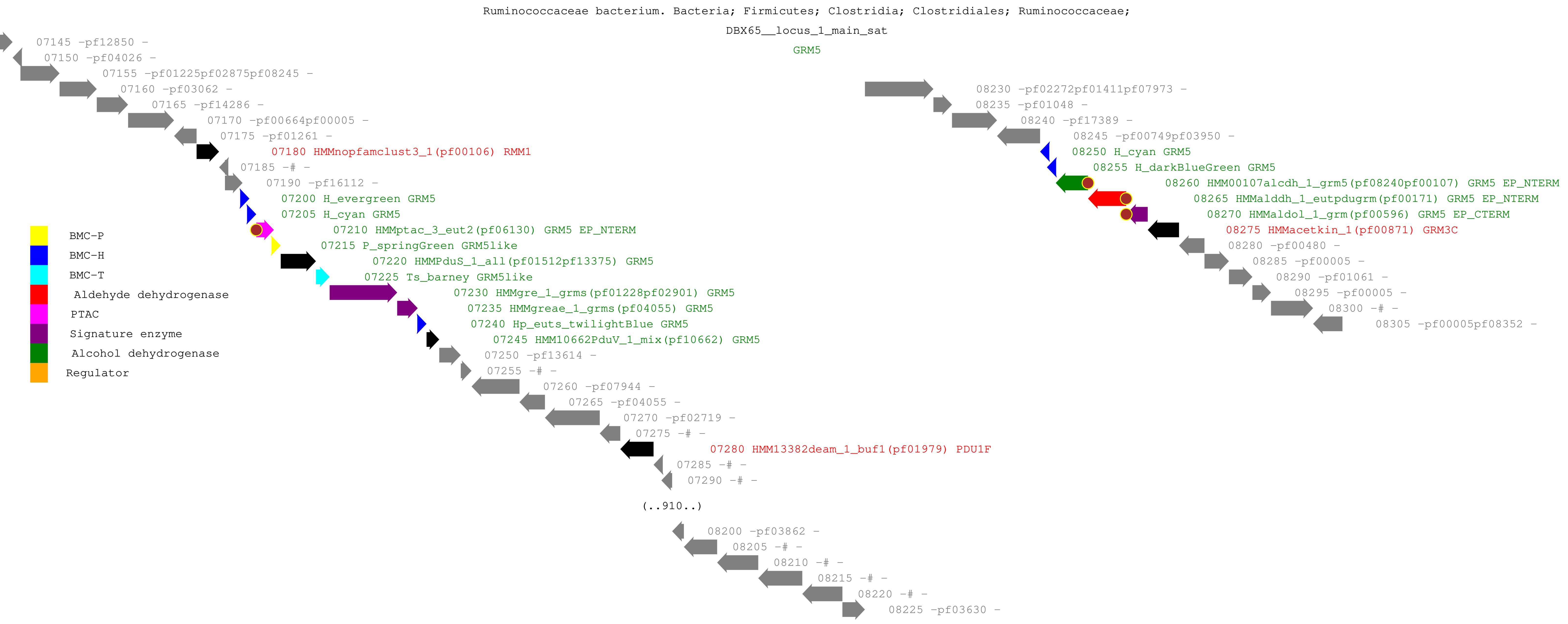

Ruminococcaceae bacterium. Bacteria; Firmicutes; Clostridia; Clostridiales; Ruminococcaceae;
DBX65__locus_1_main_sat
GRM5
07145 -pf12850 -
07150 -pf04026 -
07155 -pf01225pf02875pf08245 -
07160 -pf03062 -
07165 -pf14286 -
07170 -pf00664pf00005 -
07175 -pf01261 -
07180 HMMnopfamclust3_1(pf00106) RMM1
07185 -# -
07190 -pf16112 -
07200 H_evergreen GRM5
07205 H_cyan GRM5
07210 HMMptac_3_eut2(pf06130) GRM5 EP_NTERM
07215 P_springGreen GRM5like
07220 HMMPduS_1_all(pf01512pf13375) GRM5
07225 Ts_barney GRM5like
07230 HMMgre_1_grms(pf01228pf02901) GRM5
07235 HMMgreae_1_grms(pf04055) GRM5
07240 Hp_euts_twilightBlue GRM5
07245 HMM10662PduV_1_mix(pf10662) GRM5
07250 -pf13614 -
07255 -# -
07260 -pf07944 -
07265 -pf04055 -
07270 -pf02719 -
07275 -# -
07280 HMM13382deam_1_buf1(pf01979) PDU1F
07285 -# -
07290 -# -
(..910..)
08200 -pf03862 -
08205 -# -
08210 -# -
08215 -# -
08220 -# -
08225 -pf03630 -
08230 -pf02272pf01411pf07973 -
08235 -pf01048 -
08240 -pf17389 -
08245 -pf00749pf03950 -
08250 H_cyan GRM5
08255 H_darkBlueGreen GRM5
08260 HMM00107alcdh_1_grm5(pf08240pf00107) GRM5 EP_NTERM
08265 HMMalddh_1_eutpdugrm(pf00171) GRM5 EP_NTERM
08270 HMMaldol_1_grm(pf00596) GRM5 EP_CTERM
08275 HMMacetkin_1(pf00871) GRM3C
08280 -pf00480 -
08285 -pf00005 -
08290 -pf01061 -
08295 -pf00005 -
08300 -# -
08305 -pf00005pf08352 -
BMC-P
BMC-H
BMC-T
Aldehyde dehydrogenase
PTAC
Signature enzyme
Alcohol dehydrogenase
Regulator

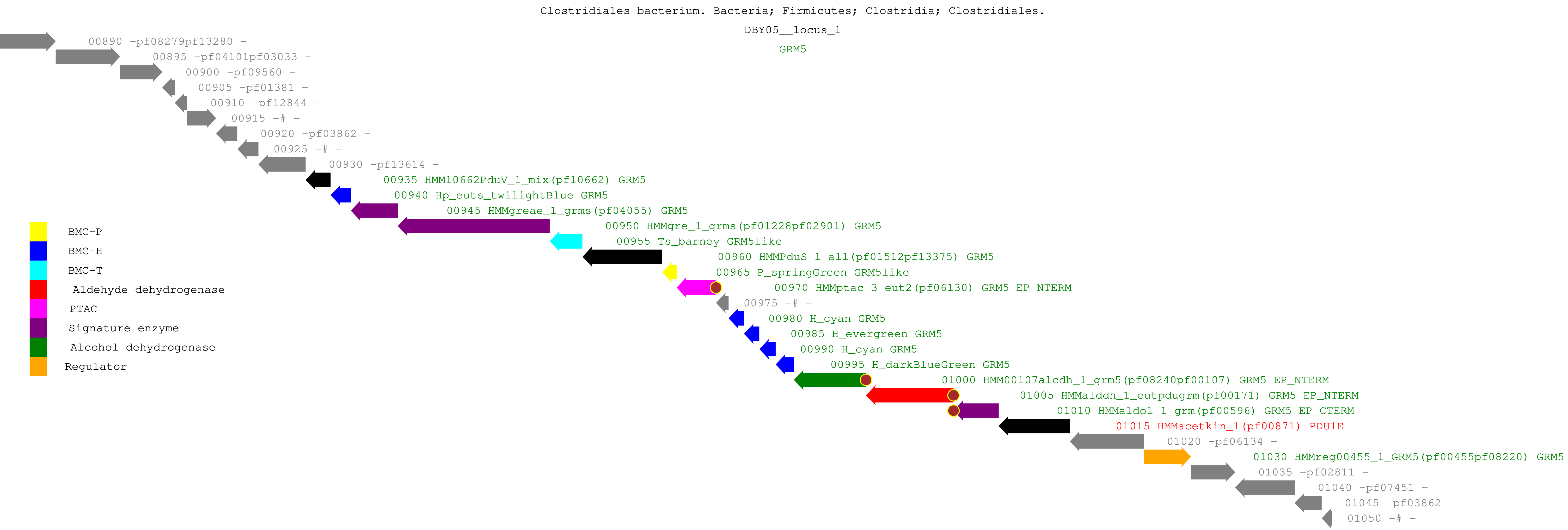
Clostridiales bacterium. Bacteria; Firmicutes; Clostridia; Clostridiales.
DBY05__locus_1
GRM5
00890 -pf08279pf13280 -
00895 -pf04101pf03033 -
00900 -pf09560 -
00905 -pf01381 -
00910 -pf12844 -
00915 -# -
00920 -pf03862 -
00925 -# -
00930 -pf13614 -
00935 HMM10662PduV_1_mix(pf10662) GRM5
00940 Hp_euts_twilightBlue GRM5
00945 HMMgreae_1_grms(pf04055) GRM5
00950 HMMgre_1_grms(pf01228pf02901) GRM5
00955 Ts_barney GRM5like
00960 HMMPduS_1_all(pf01512pf13375) GRM5
00965 P_springGreen GRM5like
00970 HMMptac_3_eut2(pf06130) GRM5 EP_NTERM
00975 -# -
00980 H_cyan GRM5
00985 H_evergreen GRM5
00990 H_cyan GRM5
00995 H_darkBlueGreen GRM5
01000 HMM00107alcdh_1_grm5(pf08240pf00107) GRM5 EP_NTERM
01005 HMMalddh_1_eutpdugrm(pf00171) GRM5 EP_NTERM
01010 HMMaldol_1_grm(pf00596) GRM5 EP_CTERM
01015 HMMacetkin_1(pf00871) PDU1E
01020 -pf06134 -
01030 HMMreg00455_1_GRM5(pf00455pf08220) GRM5
01035 -pf02811 -
01040 -pf07451 -
01045 -pf03862 -
01050 -# -
BMC-P
BMC-H
BMC-T
Aldehyde dehydrogenase
PTAC
Signature enzyme
Alcohol dehydrogenase
Regulator

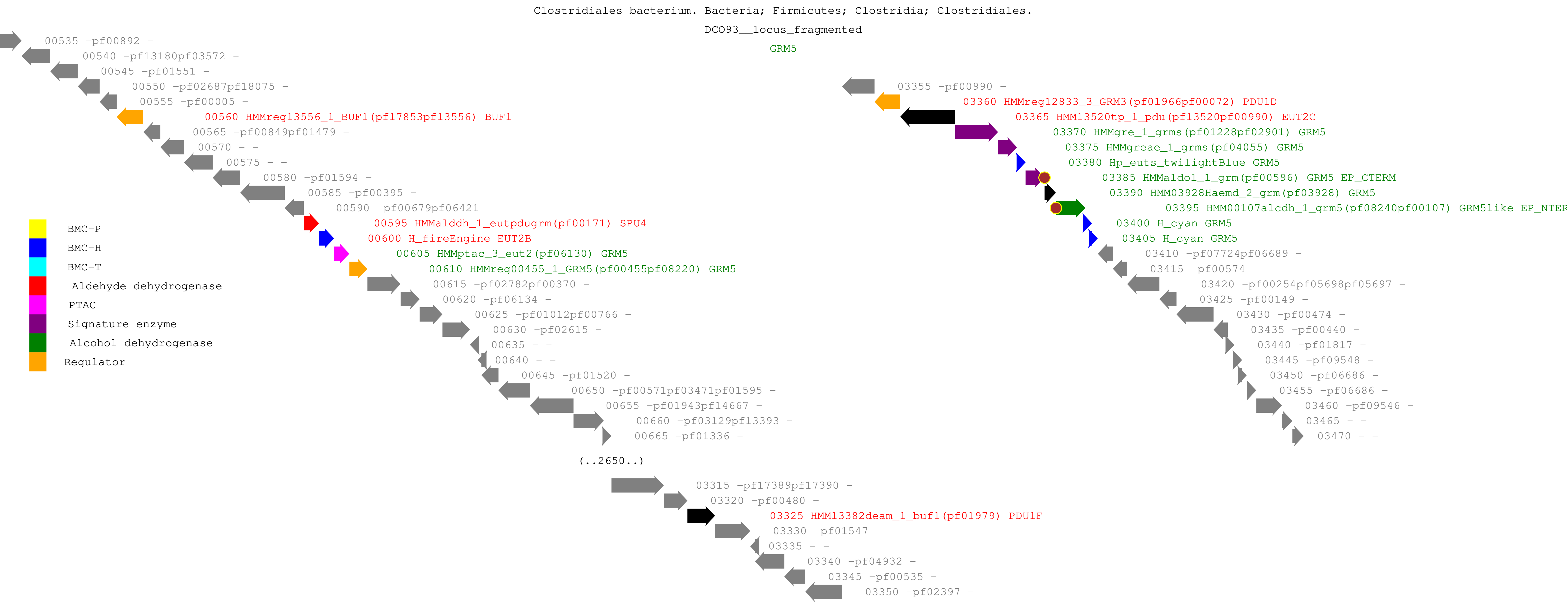

Clostridiales bacterium. Bacteria; Firmicutes; Clostridia; Clostridiales.
DCO93__locus_fragmented
GRM5
00535 -pf00892 -
00540 -pf13180pf03572 -
00545 -pf01551 -
00550 -pf02687pf18075 -
00555 -pf00005 -
00560 HMMreg13556_1_BUF1(pf17853pf13556) BUF1
00565 -pf00849pf01479 -
00570 - -
00575 - -
00580 -pf01594 -
00585 -pf00395 -
00590 -pf00679pf06421 -
00595 HMMalddh_1_eutpdugrm(pf00171) SPU4
00600 H_fireEngine EUT2B
00605 HMMptac_3_eut2(pf06130) GRM5
00610 HMMreg00455_1_GRM5(pf00455pf08220) GRM5
00615 -pf02782pf00370 -
00620 -pf06134 -
00625 -pf01012pf00766 -
00630 -pf02615 -
00635 - -
00640 - -
00645 -pf01520 -
00650 -pf00571pf03471pf01595 -
00655 -pf01943pf14667 -
00660 -pf03129pf13393 -
00665 -pf01336 -
(..2650..)
03315 -pf17389pf17390 -
03320 -pf00480 -
03325 HMM13382deam_1_buf1(pf01979) PDU1F
03330 -pf01547 -
03335 - -
03340 -pf04932 -
03345 -pf00535 -
03350 -pf02397 -
03355 -pf00990 -
03360 HMMreg12833_3_GRM3(pf01966pf00072) PDU1D
03365 HMM13520tp_1_pdu(pf13520pf00990) EUT2C
03370 HMMgre_1_grms(pf01228pf02901) GRM5
03375 HMMgreae_1_grms(pf04055) GRM5
03380 Hp_euts_twilightBlue GRM5
03385 HMMaldol_1_grm(pf00596) GRM5 EP_CTERM
03390 HMM03928Haemd_2_grm(pf03928) GRM5
03395 HMM00107alcdh_1_grm5(pf08240pf00107) GRM5like EP_NTER
03400 H_cyan GRM5
03405 H_cyan GRM5
03410 -pf07724pf06689 -
03415 -pf00574 -
03420 -pf00254pf05698pf05697 -
03425 -pf00149 -
03430 -pf00474 -
03435 -pf00440 -
03440 -pf01817 -
03445 -pf09548 -
03450 -pf06686 -
03455 -pf06686 -
03460 -pf09546 -
03465 - -
03470 - -
BMC-P
BMC-H
BMC-T
Aldehyde dehydrogenase
PTAC
Signature enzyme
Alcohol dehydrogenase
Regulator

Lachnospiraceae bacterium. Bacteria; Firmicutes; Clostridia; Clostridiales; Lachnospiraceae.

DCP46__locus_1

GRM5

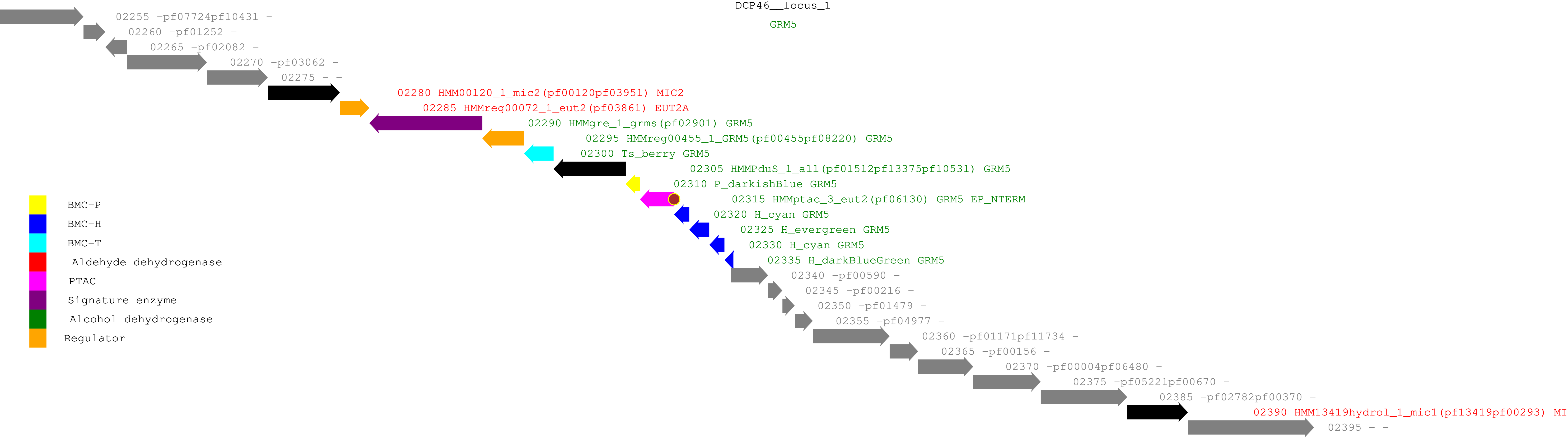

Lachnospiraceae bacterium. Bacteria; Firmicutes; Clostridia; Clostridiales; Lachnospiraceae.

DCX82__locus_1

GRM5

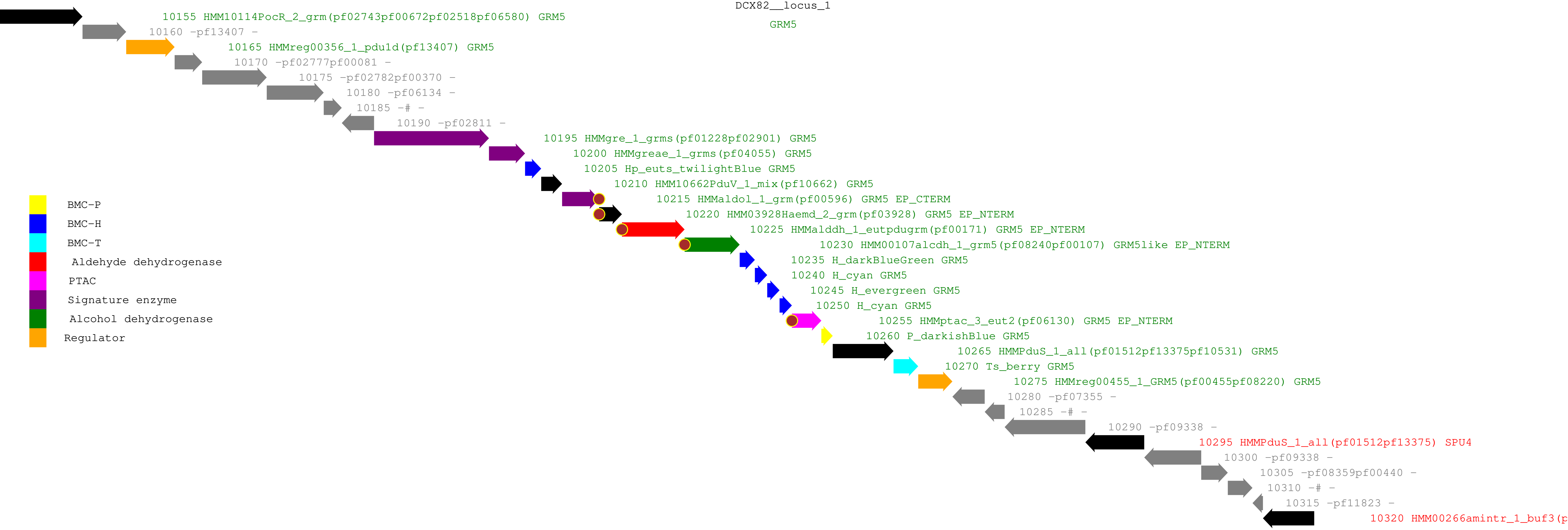

Lachnospiraceae bacterium. Bacteria; Firmicutes; Clostridia; Clostridiales; Lachnospiraceae.

DCZ40__locus_1

GRM5

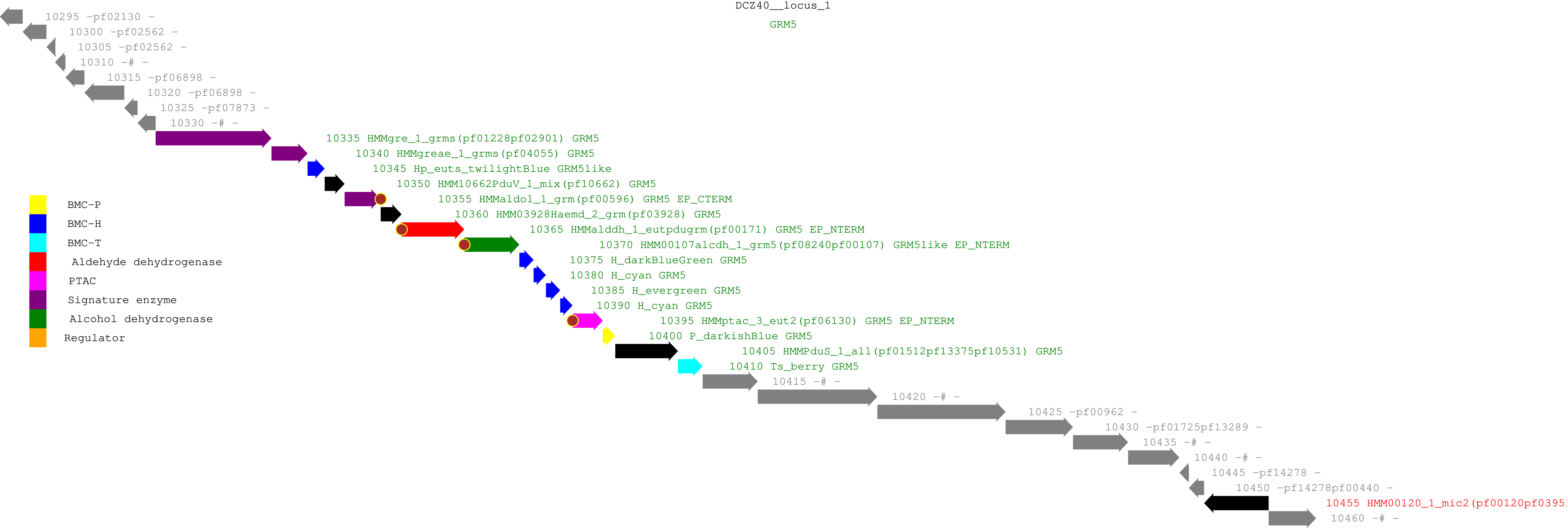

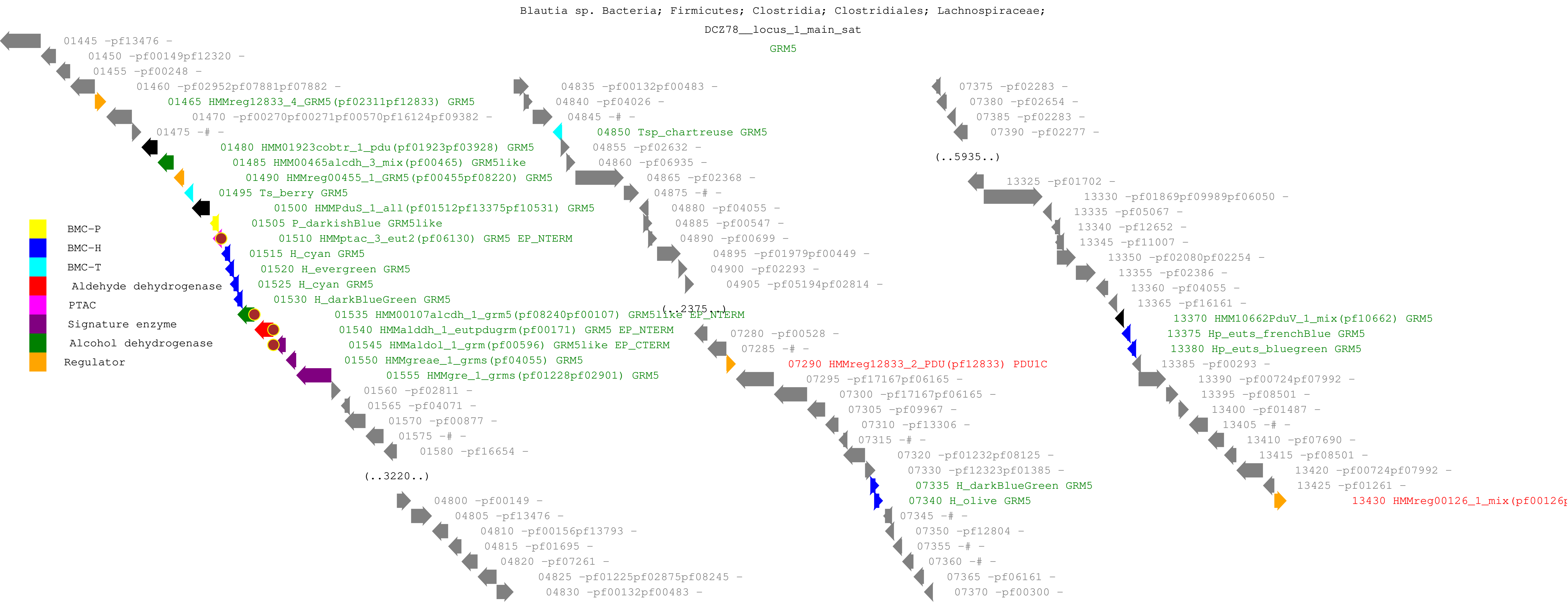
Blautia sp. Bacteria; Firmicutes; Clostridia; Clostridiales; Lachnospiraceae;
DCZ78__locus_1_main_sat
GRM5
01445 -pf13476 -
01450 -pf00149pf12320 -
01455 -pf00248 -
01460 -pf02952pf07881pf07882 -
01465 HMMreg12833_4_GRM5(pf02311pf12833) GRM5
01470 -pf00270pf00271pf00570pf16124pf09382 -
01475 -# -
01480 HMM01923cobtr_1_pdu(pf01923pf03928) GRM5
01485 HMM00465alcdh_3_mix(pf00465) GRM5like
01490 HMMreg00455_1_GRM5(pf00455pf08220) GRM5
01495 Ts_berry GRM5
01500 HMMPduS_1_all(pf01512pf13375pf10531) GRM5
01505 P_darkishBlue GRM5like
01510 HMMptac_3_eut2(pf06130) GRM5 EP_NTERM
01515 H_cyan GRM5
01520 H_evergreen GRM5
01525 H_cyan GRM5
01530 H_darkBlueGreen GRM5
01535 HMM00107alcdh_1_grm5(pf08240pf00107) GRM5like EP_NTERM
01540 HMMalddh_1_eutpdugrm(pf00171) GRM5 EP_NTERM
01545 HMMaldol_1_grm(pf00596) GRM5like EP_CTERM
01550 HMMgreae_1_grms(pf04055) GRM5
01555 HMMgre_1_grms(pf01228pf02901) GRM5
01560 -pf02811 -
01565 -pf04071 -
01570 -pf00877 -
01575 -# -
01580 -pf16654 -
(..3220..)
04800 -pf00149 -
04805 -pf13476 -
04810 -pf00156pf13793 -
04815 -pf01695 -
04820 -pf07261 -
04825 -pf01225pf02875pf08245 -
04830 -pf00132pf00483 -
04835 -pf00132pf00483 -
04840 -pf04026 -
04845 -# -
04850 Tsp_chartreuse GRM5
04855 -pf02632 -
04860 -pf06935 -
04865 -pf02368 -
04875 -# -
04880 -pf04055 -
04885 -pf00547 -
04890 -pf00699 -
04895 -pf01979pf00449 -
04900 -pf02293 -
04905 -pf05194pf02814 -
(..2375..)
07280 -pf00528 -
07285 -# -
07290 HMMreg12833_2_PDU(pf12833) PDU1C
07295 -pf17167pf06165 -
07300 -pf17167pf06165 -
07305 -pf09967 -
07310 -pf13306 -
07315 -# -
07320 -pf01232pf08125 -
07330 -pf12323pf01385 -
07335 H_darkBlueGreen GRM5
07340 H_olive GRM5
07345 -# -
07350 -pf12804 -
07355 -# -
07360 -# -
07365 -pf06161 -
07370 -pf00300 -
07375 -pf02283 -
07380 -pf02654 -
07385 -pf02283 -
07390 -pf02277 -
(..5935..)
13325 -pf01702 -
13330 -pf01869pf09989pf06050 -
13335 -pf05067 -
13340 -pf12652 -
13345 -pf11007 -
13350 -pf02080pf02254 -
13355 -pf02386 -
13360 -pf04055 -
13365 -pf16161 -
13370 HMM10662PduV_1_mix(pf10662) GRM5
13375 Hp_euts_frenchBlue GRM5
13380 Hp_euts_bluegreen GRM5
13385 -pf00293 -
13390 -pf00724pf07992 -
13395 -pf08501 -
13400 -pf01487 -
13405 -# -
13410 -pf07690 -
13415 -pf08501 -
13420 -pf00724pf07992 -
13425 -pf01261 -
13430 HMMreg00126_1_mix(pf00126
BMC-P
BMC-H
BMC-T
Aldehyde dehydrogenase
PTAC
Signature enzyme
Alcohol dehydrogenase
Regulator

Clostridiales bacterium. Bacteria; Firmicutes; Clostridia; Clostridiales.

DD391__locus_1

GRM5

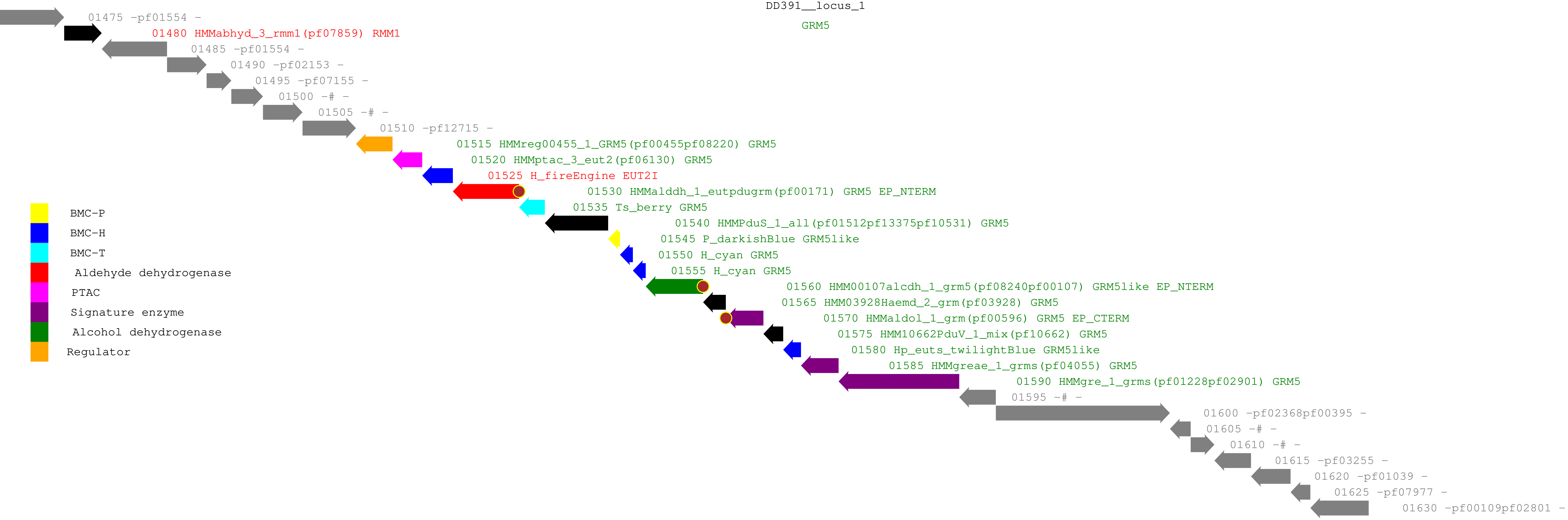

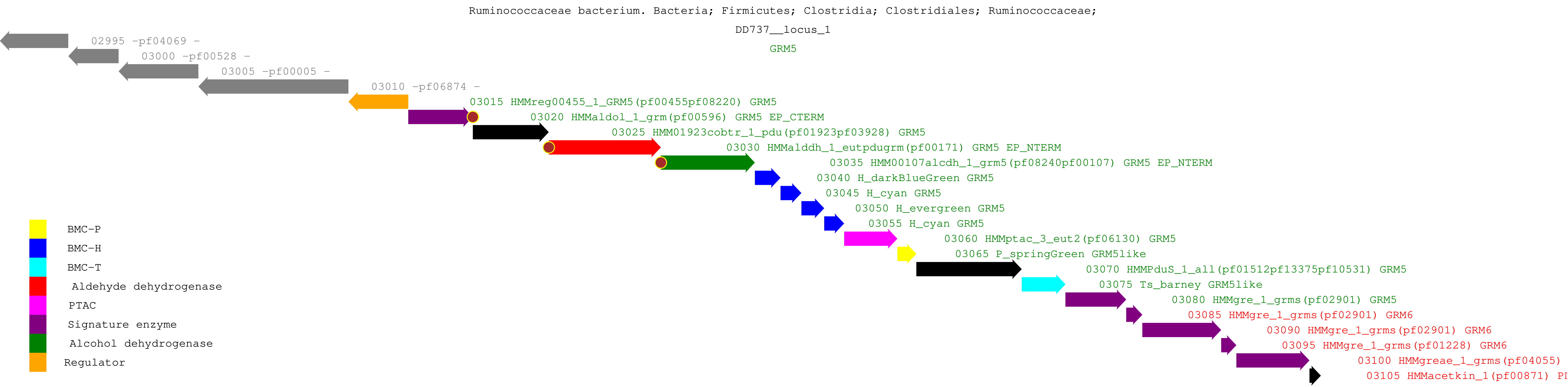
Ruminococcaceae bacterium. Bacteria; Firmicutes; Clostridia; Clostridiales; Ruminococcaceae;
DD737__locus_1
GRM5
02995 -pf04069 -
03000 -pf00528 -
03005 -pf00005 -
03010 -pf06874 -
03015 HMMreg00455_1_GRM5(pf00455pf08220) GRM5
03020 HMMaldol_1_grm(pf00596) GRM5 EP_CTERM
03025 HMM01923cobtr_1_pdu(pf01923pf03928) GRM5
03030 HMMaldh_1_eutpdugrm(pf00171) GRM5 EP_NTERM
03035 HMM00107alcdh_1_grm5(pf08240pf00107) GRM5 EP_NTERM
03040 H_darkBlueGreen GRM5
03045 H_cyan GRM5
03050 H_evergreen GRM5
03055 H_cyan GRM5
03060 HMMptac_3_eut2(pf06130) GRM5
03065 P_springGreen GRM5like
03070 HMMPduS_1_all(pf01512pf13375pf10531) GRM5
03075 Ts_barney GRM5like
03080 HMMgre_1_grms(pf02901) GRM5
03085 HMMgre_1_grms(pf02901) GRM6
03090 HMMgre_1_grms(pf02901) GRM6
03095 HMMgre_1_grms(pf01228) GRM6
03100 HMMgreae_1_grms(pf04055)
03105 HMMacetkin_1(pf00871)
BMC-P
BMC-H
BMC-T
Aldehyde dehydrogenase
PTAC
Signature enzyme
Alcohol dehydrogenase
Regulator

Ruminococcaceae bacterium. Bacteria; Firmicutes; Clostridia; Clostridiales; Ruminococcaceae;

DDW99__locus_1

GRM5

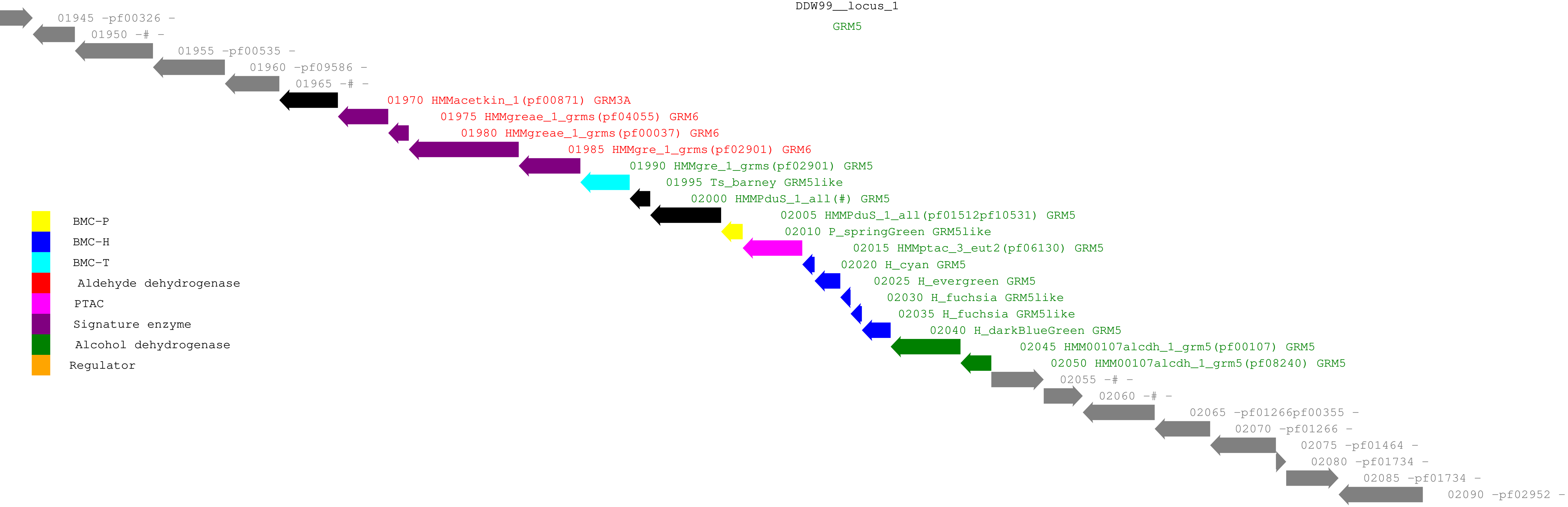

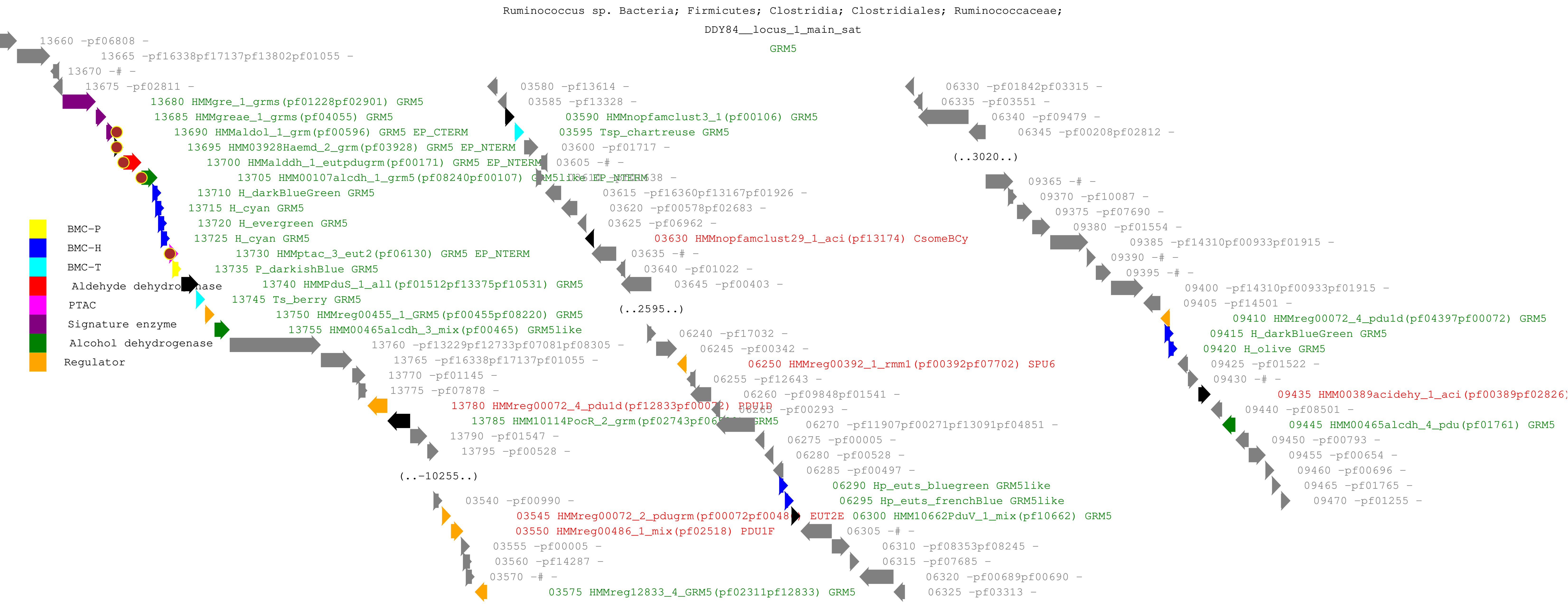

Ruminococcus sp. Bacteria; Firmicutes; Clostridia; Clostridiales; Ruminococcaceae;
DDY84__locus_1_main_sat
GRM5
13660 -pf06808 -
13665 -pf16338pf17137pf13802pf01055 -
13670 -# -
13675 -pf02811 -
13680 HMMgre_1_grms(pf01228pf02901) GRM5
13685 HMMgreae_1_grms(pf04055) GRM5
13690 HMMaldol_1_grm(pf00596) GRM5 EP_CTERM
13695 HMM03928Haemd_2_grm(pf03928) GRM5 EP_NTERM
13700 HMMalddh_1_eutpdugrm(pf00171) GRM5 EP_NTERM
13705 HMM00107alcdh_1_grm5(pf08240pf00107) GRM5like EP_NTERM
13710 H_darkBlueGreen GRM5
13715 H_cyan GRM5
13720 H_evergreen GRM5
13725 H_cyan GRM5
13730 HMMptac_3_eut2(pf06130) GRM5 EP_NTERM
13735 P_darkishBlue GRM5
13740 HMMPduS_1_all(pf01512pf13375pf10531) GRM5
13745 Ts_berry GRM5
13750 HMMreg00455_1_GRM5(pf00455pf08220) GRM5
13755 HMM00465alcdh_3_mix(pf00465) GRM5like
13760 -pf13229pf12733pf07081pf08305 -
13765 -pf16338pf17137pf01055 -
13770 -pf01145 -
13775 -pf07878 -
13780 HMMreg00072_4_pdu1d(pf12833pf00072) PDU1D
13785 HMM10114PocR_2_grm GRM5
13790 -pf01547 -
13795 -pf00528 -
(..-10255..)
03540 -pf00990 -
03545 HMMreg00072_2_pdugrm EUT2E
03550 HMMreg00486_1_mix(pf02518) PDU1F
03555 -pf00005 -
03560 -pf14287 -
03570 -# -
03575 HMMreg12833_4_GRM5(pf02311pf12833) GRM5
03580 -pf13614 -
03585 -pf13328 -
03590 HMMnopfamclust3_1(pf00106) GRM5
03595 Tsp_chartreuse GRM5
03600 -pf01717 -
03605 -# -
03615 -pf16360pf13167pf01926 -
03620 -pf00578pf02683 -
03625 -pf06962 -
03630 HMMnopfamclust29_1_aci(pf13174) CsomeBCy
03635 -# -
03640 -pf01022 -
03645 -pf00403 -
(..2595..)
06240 -pf17032 -
06245 -pf00342 -
06250 HMMreg00392_1_rmm1(pf00392pf07702) SPU6
06255 -pf12643 -
06260 -pf09848pf01541 -
06265 -pf00293 -
06270 -pf11907pf00271pf13091pf04851 -
06275 -pf00005 -
06280 -pf00528 -
06285 -pf00497 -
06290 Hp_euts_bluegreen GRM5like
06295 Hp_euts_frenchBlue GRM5like
06300 HMM10662PduV_1_mix(pf10662) GRM5
06305 -# -
06310 -pf08353pf08245 -
06315 -pf07685 -
06320 -pf00689pf00690 -
06325 -pf03313 -
06330 -pf01842pf03315 -
06335 -pf03551 -
06340 -pf09479 -
06345 -pf00208pf02812 -
(..3020..)
09365 -# -
09370 -pf10087 -
09375 -pf07690 -
09380 -pf01554 -
09385 -pf14310pf00933pf01915 -
09390 -# -
09395 -# -
09400 -pf14310pf00933pf01915 -
09405 -pf14501 -
09410 HMMreg00072_4_pdu1d(pf04397pf00072) GRM5
09415 H_darkBlueGreen GRM5
09420 H_olive GRM5
09425 -pf01522 -
09430 -# -
09435 HMM00389acidehy_1_aci(pf00389pf02826)
09440 -pf08501 -
09445 HMM00465alcdh_4_pdu(pf01761) GRM5
09450 -pf00793 -
09455 -pf00654 -
09460 -pf00696 -
09465 -pf01765 -
09470 -pf01255 -
BMC-P
BMC-H
BMC-T
Aldehyde dehydrogenase
PTAC
Signature enzyme
Alcohol dehydrogenase
Regulator

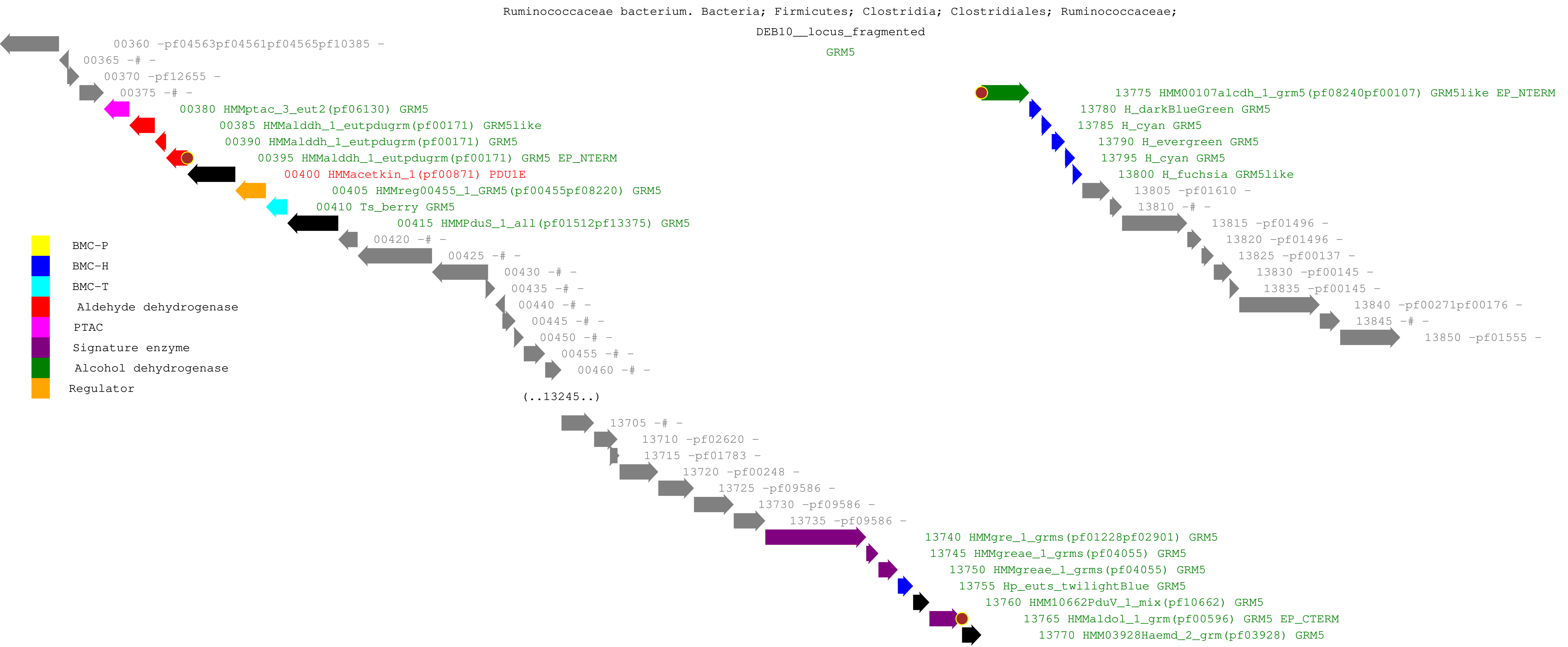

Ruminococcaceae bacterium. Bacteria; Firmicutes; Clostridia; Clostridiales; Ruminococcaceae;
DEB10__locus_fragmented
GRM5
00360 -pf04563pf04561pf04565pf10385 -
00365 -# -
00370 -pf12655 -
00375 -# -
00380 HMMptac_3_eut2(pf06130) GRM5
00385 HMMalddh_1_eutpdugrm(pf00171) GRM5like
00390 HMMalddh_1_eutpdugrm(pf00171) GRM5
00395 HMMalddh_1_eutpdugrm(pf00171) GRM5 EP_NTERM
00400 HMMacetkin_1(pf00871) PDU1E
00405 HMMreg00455_1_GRM5(pf00455pf08220) GRM5
00410 Ts_berry GRM5
00415 HMMPduS_1_all(pf01512pf13375) GRM5
00420 -# -
00425 -# -
00430 -# -
00435 -# -
00440 -# -
00445 -# -
00450 -# -
00455 -# -
00460 -# -
(..13245..)
13705 -# -
13710 -pf02620 -
13715 -pf01783 -
13720 -pf00248 -
13725 -pf09586 -
13730 -pf09586 -
13735 -pf09586 -
13740 HMMgre_1_grms(pf01228pf02901) GRM5
13745 HMMgreae_1_grms(pf04055) GRM5
13750 HMMgreae_1_grms(pf04055) GRM5
13755 Hp_euts_twilightBlue GRM5
13760 HMM10662PduV_1_mix(pf10662) GRM5
13765 HMMaldol_1_grm(pf00596) GRM5 EP_CTERM
13770 HMM03928Haemd_2_grm(pf03928) GRM5
13775 HMM00107alcdh_1_grm5(pf08240pf00107) GRM5like EP_NTERM
13780 H_darkBlueGreen GRM5
13785 H_cyan GRM5
13790 H_evergreen GRM5
13795 H_cyan GRM5
13800 H_fuchsia GRM5like
13805 -pf01610 -
13810 -# -
13815 -pf01496 -
13820 -pf01496 -
13825 -pf00137 -
13830 -pf00145 -
13835 -pf00145 -
13840 -pf00271pf00176 -
13845 -# -
13850 -pf01555 -
BMC-P
BMC-H
BMC-T
Aldehyde dehydrogenase
PTAC
Signature enzyme
Alcohol dehydrogenase
Regulator

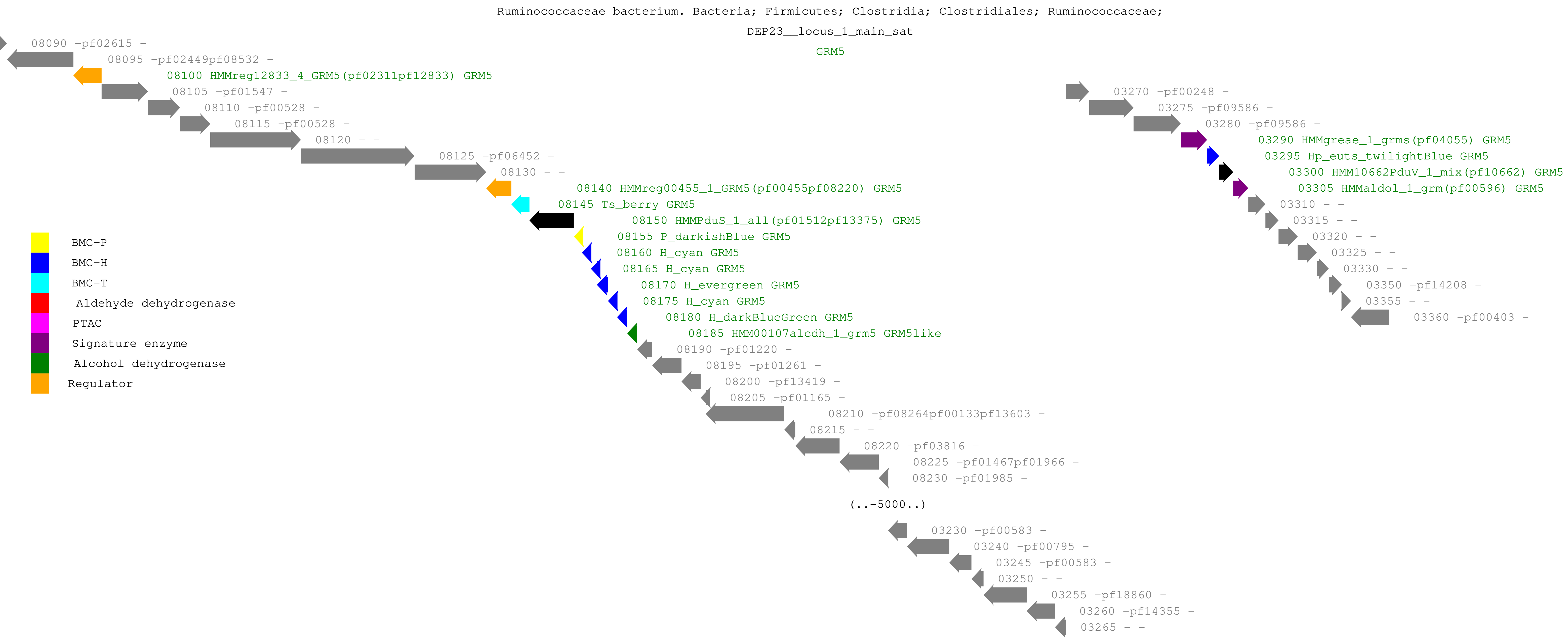
Ruminococcaceae bacterium. Bacteria; Firmicutes; Clostridia; Clostridiales; Ruminococcaceae;
DEP23__locus_1_main_sat
GRM5
08090 -pf02615 -
08095 -pf02449pf08532 -
08100 HMMreg12833_4_GRM5(pf02311pf12833) GRM5
08105 -pf01547 -
08110 -pf00528 -
08115 -pf00528 -
08120 - -
08125 -pf06452 -
08130 - -
08140 HMMreg00455_1_GRM5(pf00455pf08220) GRM5
08145 Ts_berry GRM5
08150 HMMPduS_1_all(pf01512pf13375) GRM5
08155 P_darkishBlue GRM5
08160 H_cyan GRM5
08165 H_cyan GRM5
08170 H_evergreen GRM5
08175 H_cyan GRM5
08180 H_darkBlueGreen GRM5
08185 HMM00107alcdh_1_grm5 GRM5like
08190 -pf01220 -
08195 -pf01261 -
08200 -pf13419 -
08205 -pf01165 -
08210 -pf08264pf00133pf13603 -
08215 - -
08220 -pf03816 -
08225 -pf01467pf01966 -
08230 -pf01985 -
(..-5000..)
03230 -pf00583 -
03240 -pf00795 -
03245 -pf00583 -
03250 - -
03255 -pf18860 -
03260 -pf14355 -
03265 - -
03270 -pf00248 -
03275 -pf09586 -
03280 -pf09586 -
03290 HMMgreae_1_grms(pf04055) GRM5
03295 Hp_euts_twilightBlue GRM5
03300 HMM10662PduV_1_mix(pf10662) GRM5
03305 HMMaldol_1_grm(pf00596) GRM5
03310 - -
03315 - -
03320 - -
03325 - -
03330 - -
03350 -pf14208 -
03355 - -
03360 -pf00403 -
BMC-P
BMC-H
BMC-T
Aldehyde dehydrogenase
PTAC
Signature enzyme
Alcohol dehydrogenase
Regulator

Lachnospiraceae bacterium. Bacteria; Firmicutes; Clostridia; Clostridiales; Lachnospiraceae.

DF613__locus_1_main_sat

GRM5

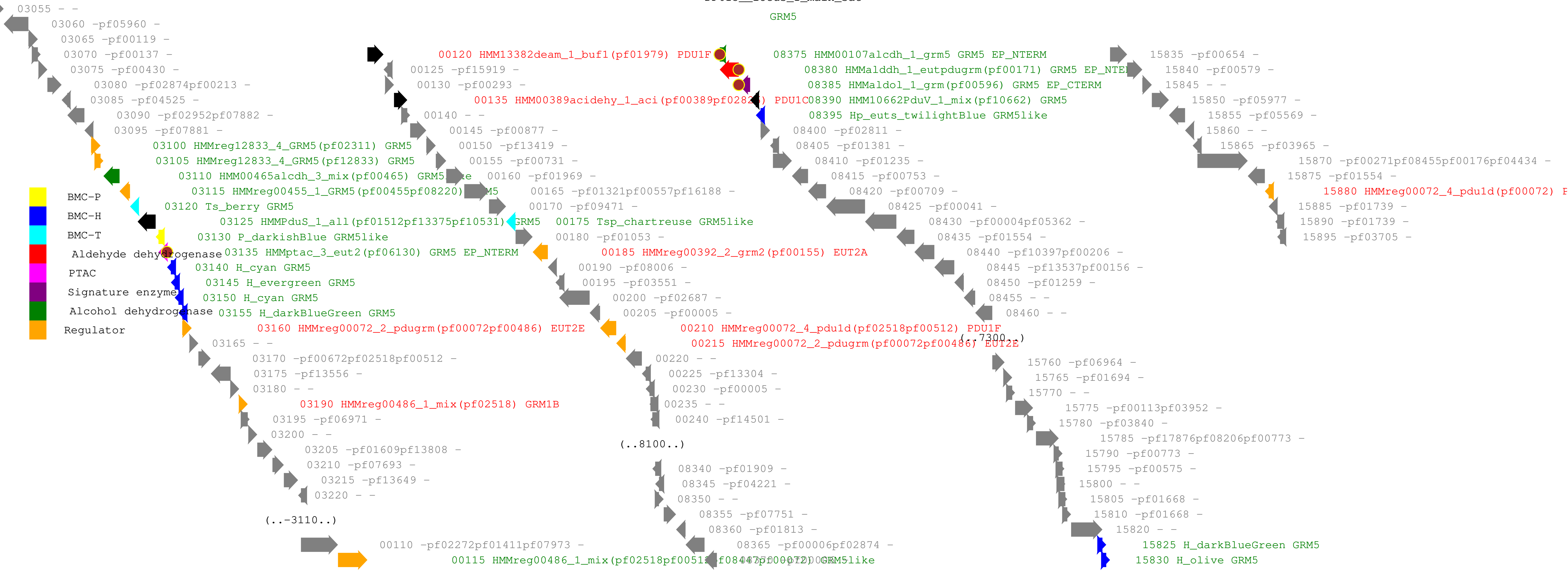

Roseburia sp. Bacteria; Firmicutes; Clostridia; Clostridiales; Lachnospiraceae;

DGH14__locus_1

GRM5

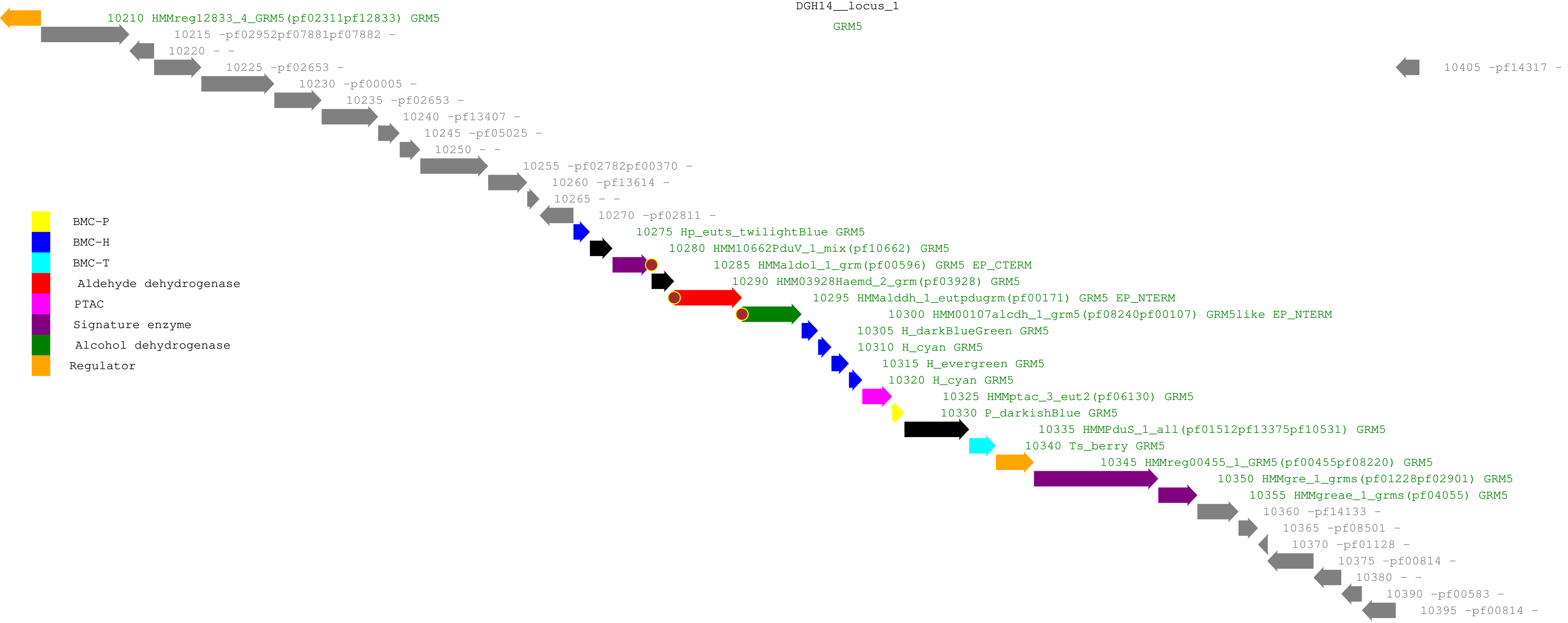

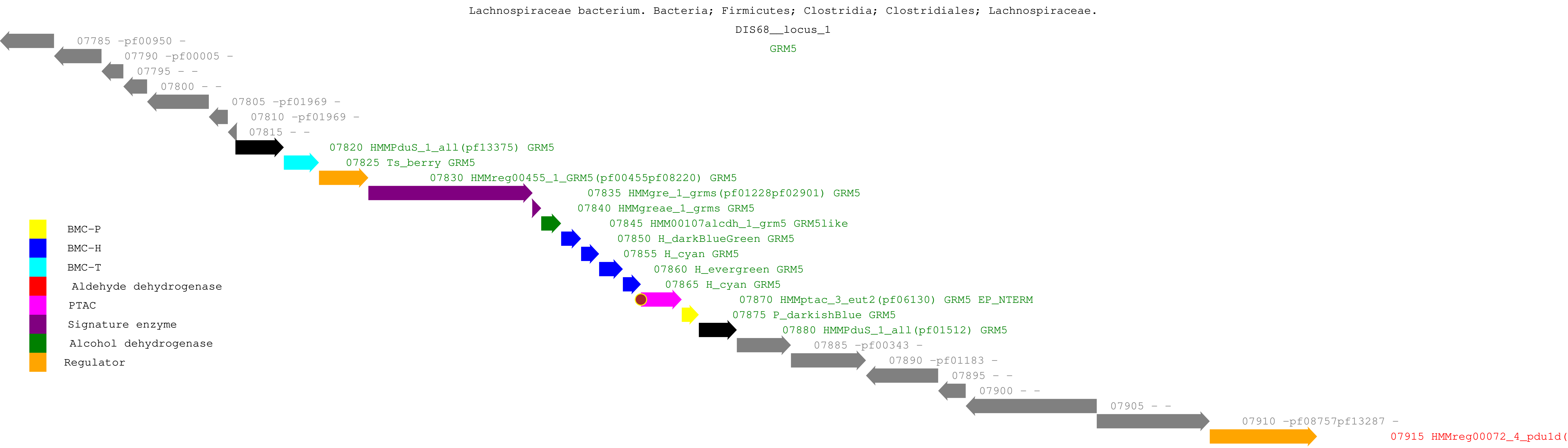

Lachnospiraceae bacterium. Bacteria; Firmicutes; Clostridia; Clostridiales; Lachnospiraceae.
DIS68__locus_1
GRM5
07785 -pf00950 -
07790 -pf00005 -
07795 - -
07800 - -
07805 -pf01969 -
07810 -pf01969 -
07815 - -
07820 HMMPduS_1_all(pf13375) GRM5
07825 Ts_berry GRM5
07830 HMMreg00455_1_GRM5(pf00455pf08220) GRM5
07835 HMMgre_1_grms(pf01228pf02901) GRM5
07840 HMMgreae_1_grms GRM5
07845 HMM00107alcdh_1_grm5 GRM5like
07850 H_darkBlueGreen GRM5
07855 H_cyan GRM5
07860 H_evergreen GRM5
07865 H_cyan GRM5
07870 HMMptac_3_eut2(pf06130) GRM5 EP_NTERM
07875 P_darkishBlue GRM5
07880 HMMPduS_1_all(pf01512) GRM5
07885 -pf00343 -
07890 -pf01183 -
07895 - -
07900 - -
07905 - -
07910 -pf08757pf13287 -
07915 HMMreg00072_4_pdu1d(
BMC-P
BMC-H
BMC-T
Aldehyde dehydrogenase
PTAC
Signature enzyme
Alcohol dehydrogenase
Regulator

Blautia sp. BCRC 81119. Bacteria; Firmicutes; Clostridia; Clostridiales; Lachnospiraceae;

DMI82__locus_1_main_sat

GRM5

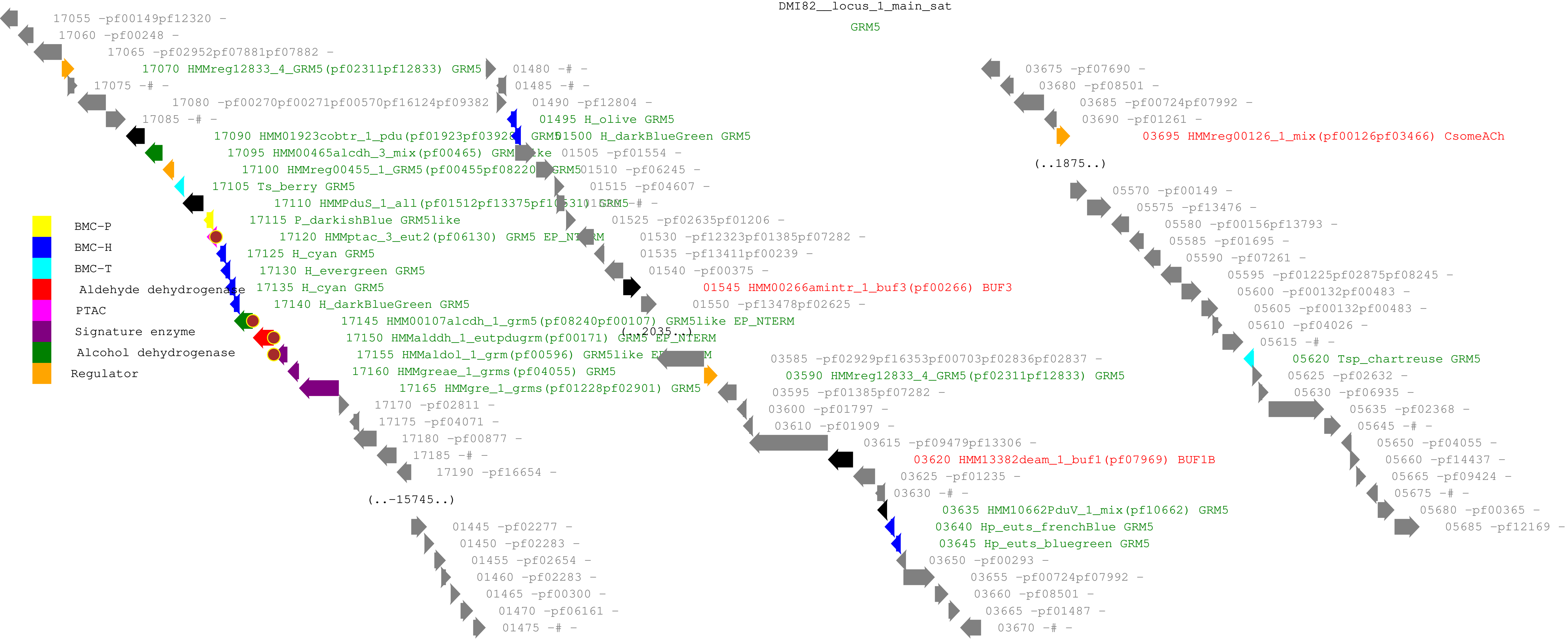

Ruminococcus sp. AF37-6AT. Bacteria; Firmicutes; Clostridia; Clostridiales; Ruminococcaceae;

DW022__locus_1_main_sat

GRM5

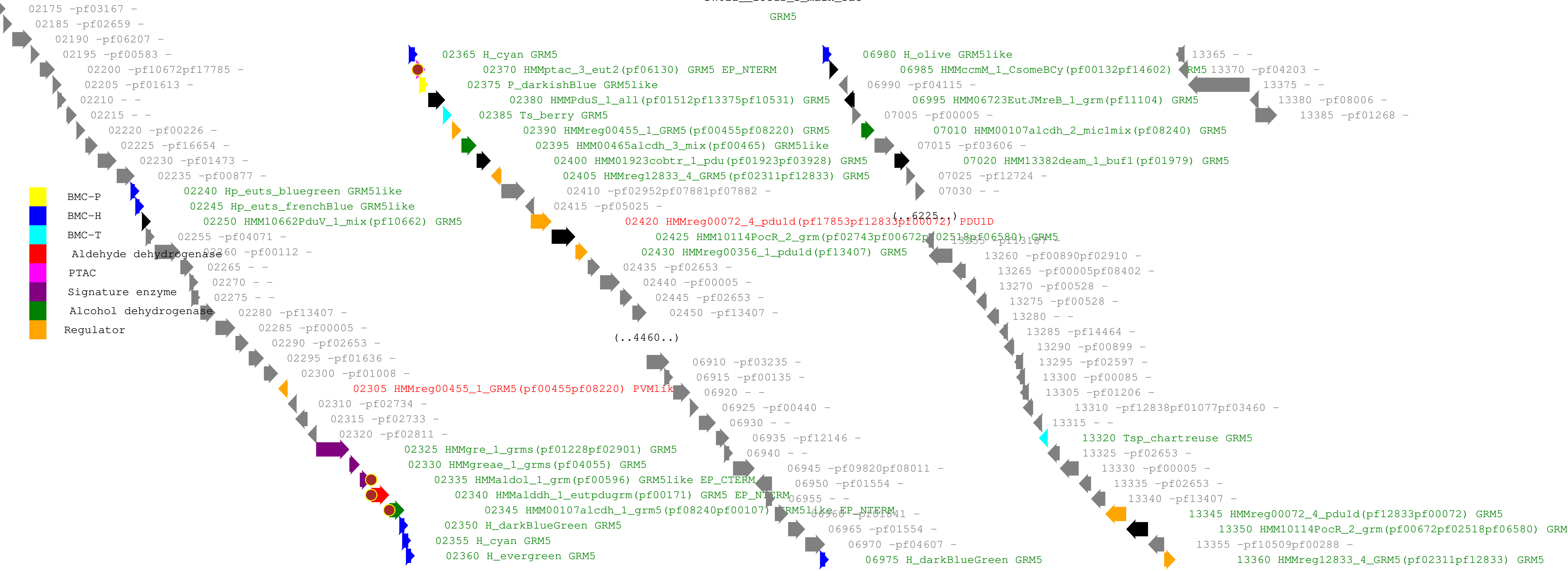

Ruminococcus sp. AF42-9BH. Bacteria; Firmicutes; Clostridia; Clostridiales; Ruminococcaceae;

DW061__locus_1_main_sat

GRM5

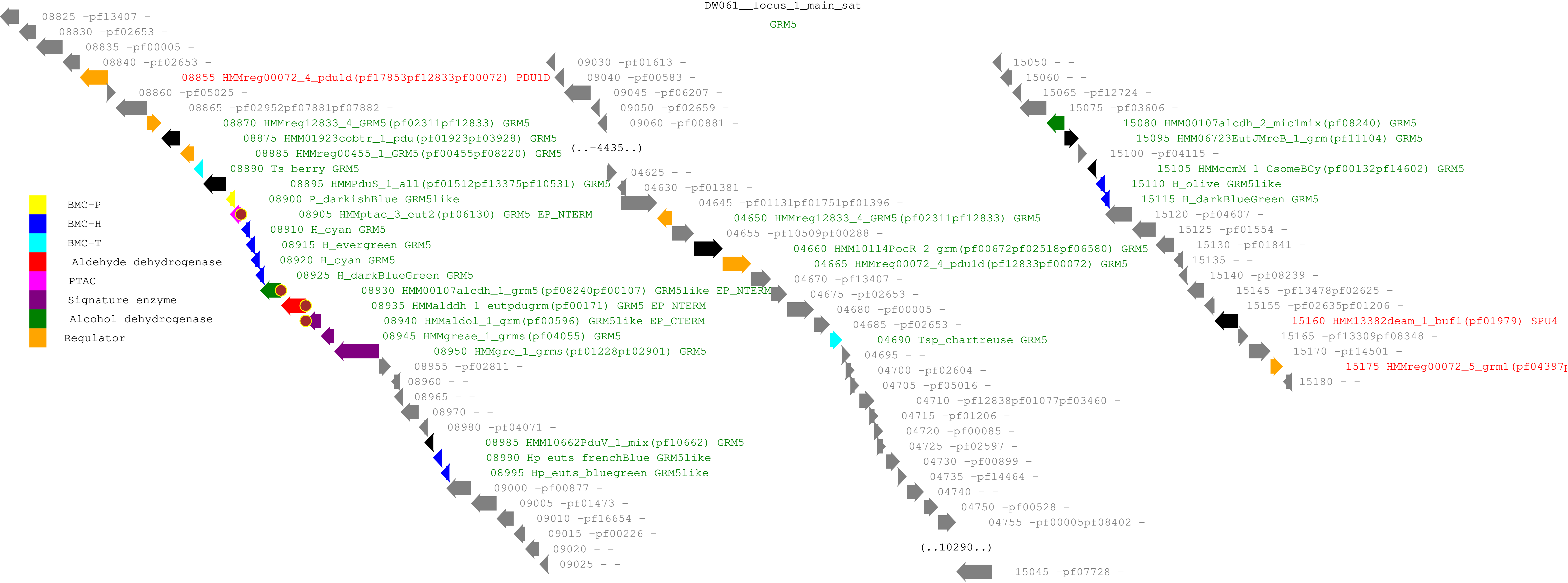

Ruminococcus sp. AF45-4BH. Bacteria; Firmicutes; Clostridia; Clostridiales; Ruminococcaceae;

DW073__locus_1_main_sat

GRM5

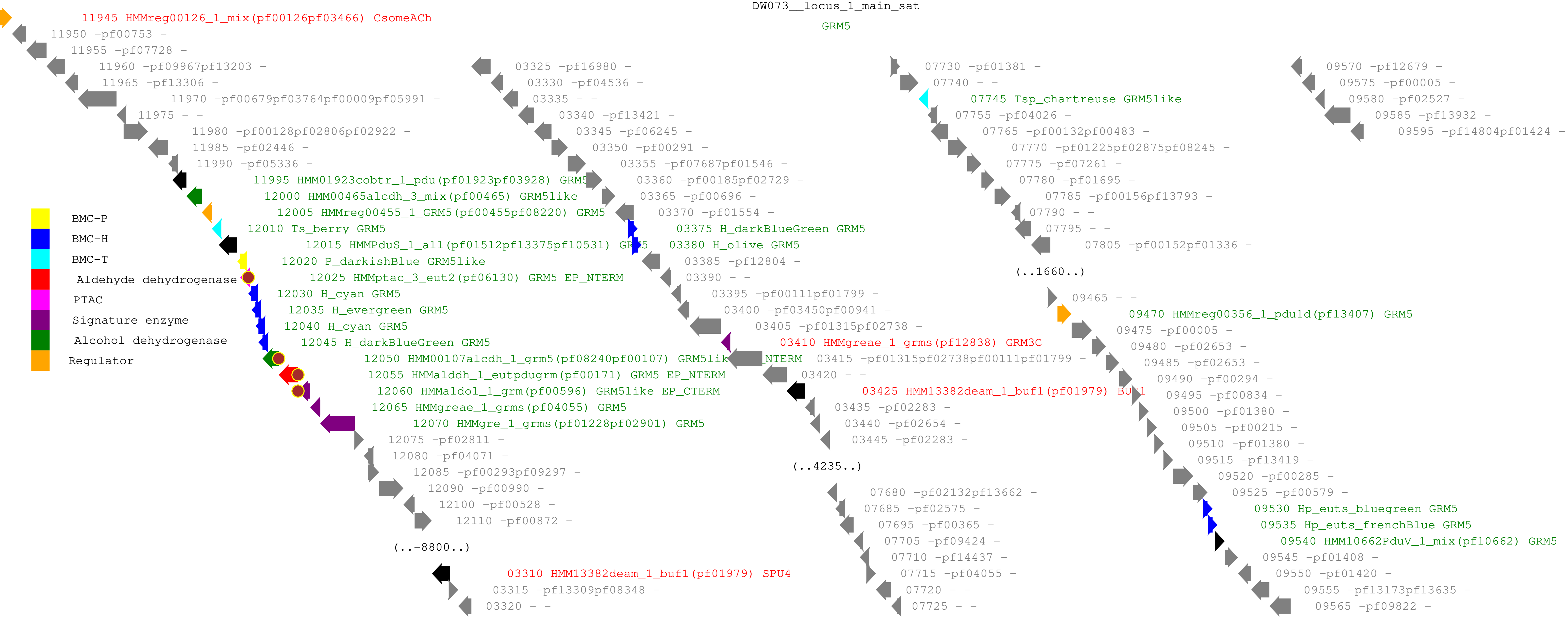

Ruminococcus sp. AF46-10NS. Bacteria; Firmicutes; Clostridia; Clostridiales; Ruminococcaceae;

DW074__locus_1_main_sat

GRM5

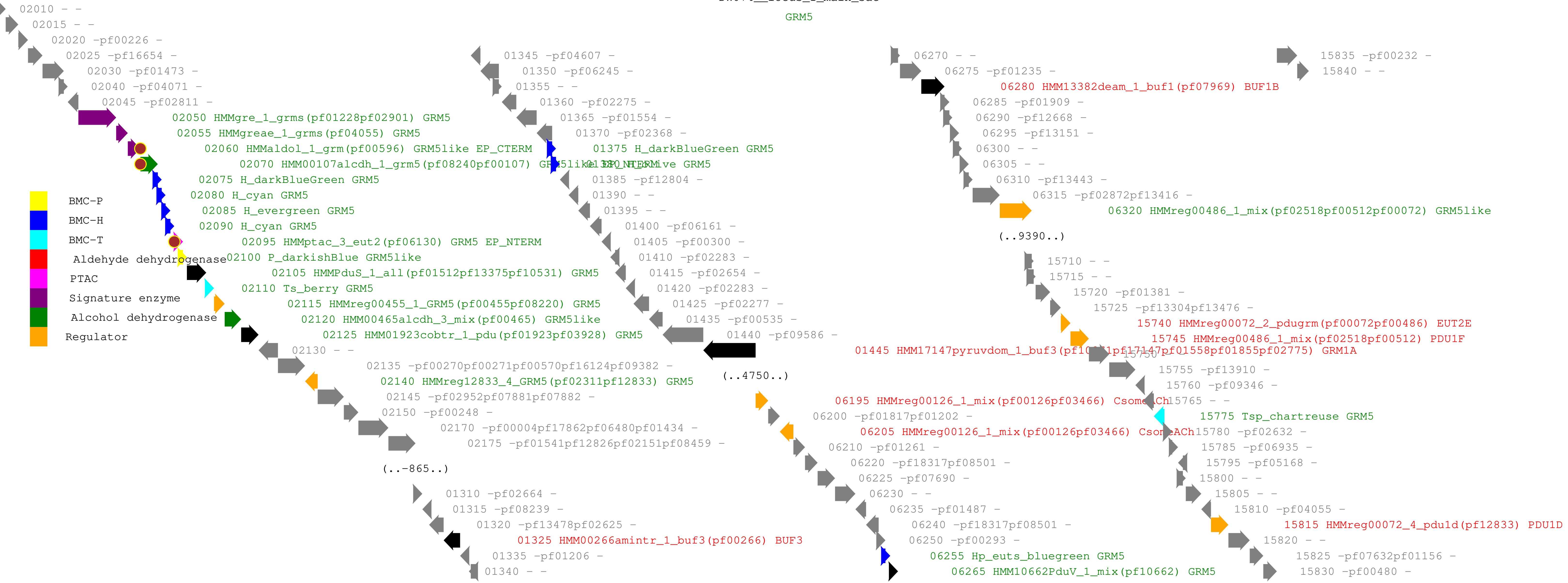

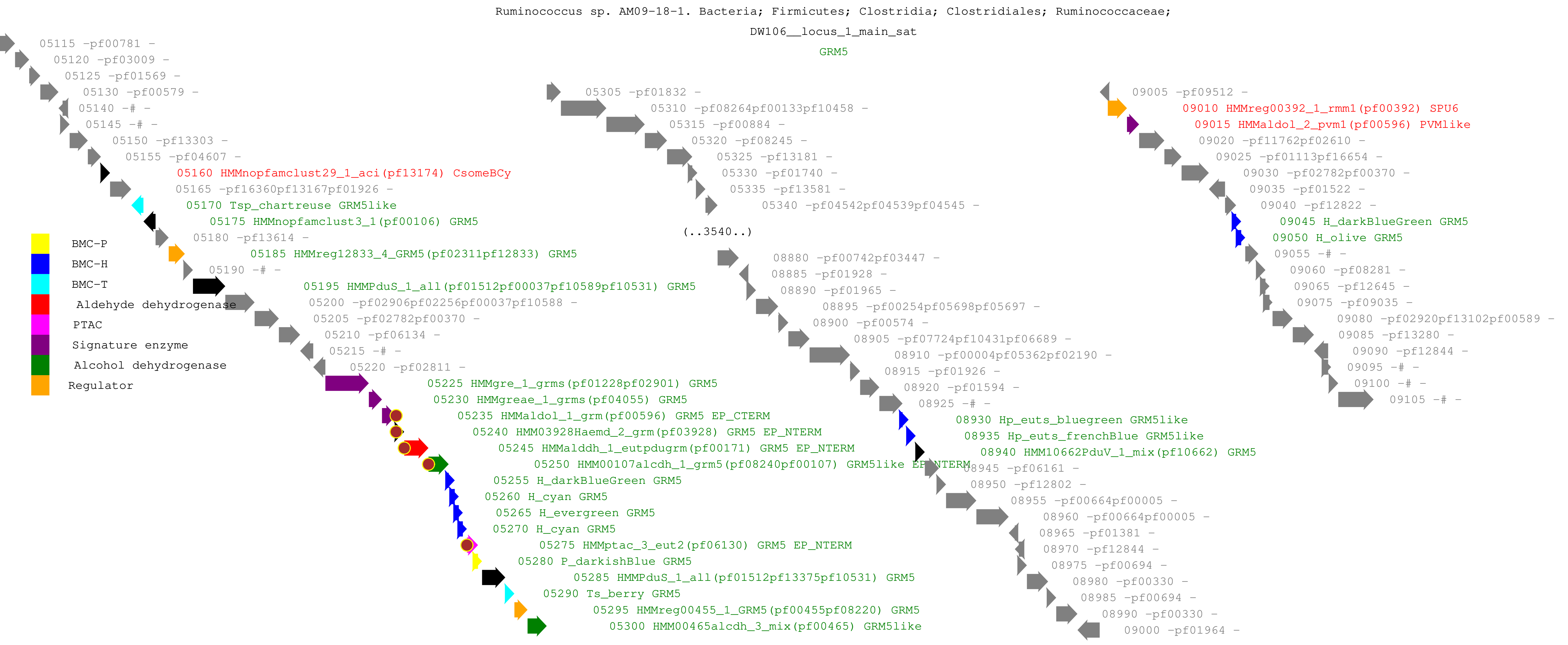
Ruminococcus sp. AM09-18-1. Bacteria; Firmicutes; Clostridia; Clostridiales; Ruminococcaceae;
DW106__locus_1_main_sat
GRM5
05115 -pf00781 -
05120 -pf03009 -
05125 -pf01569 -
05130 -pf00579 -
05140 -# -
05145 -# -
05150 -pf13303 -
05155 -pf04607 -
05160 HMMnopfamclust29_1_aci(pf13174) CsomeBCy
05165 -pf16360pf13167pf01926 -
05170 Tsp_chartreuse GRM5like
05175 HMMnopfamclust3_1(pf00106) GRM5
05180 -pf13614 -
05185 HMMreg12833_4_GRM5(pf02311pf12833) GRM5
05190 -# -
05195 HMMPduS_1_all(pf01512pf00037pf10589pf10531) GRM5
05200 -pf02906pf02256pf00037pf10588 -
05205 -pf02782pf00370 -
05210 -pf06134 -
05215 -# -
05220 -pf02811 -
05225 HMMgre_1_grms(pf01228pf02901) GRM5
05230 HMMgreae_1_grms(pf04055) GRM5
05235 HMMaldol_1_grm(pf00596) GRM5 EP_CTERM
05240 HMM03928Haemd_2_grm(pf03928) GRM5 EP_NTERM
05245 HMMalddh_1_eutpdugrm(pf00171) GRM5 EP_NTERM
05250 HMM00107alcdh_1_grm5(pf08240pf00107) GRM5like EP_NTERM
05255 H_darkBlueGreen GRM5
05260 H_cyan GRM5
05265 H_evergreen GRM5
05270 H_cyan GRM5
05275 HMMptac_3_eut2(pf06130) GRM5 EP_NTERM
05280 P_darkishBlue GRM5
05285 HMMPduS_1_all(pf01512pf13375pf10531) GRM5
05290 Ts_berry GRM5
05295 HMMreg00455_1_GRM5(pf00455pf08220) GRM5
05300 HMM00465alcdh_3_mix(pf00465) GRM5like
BMC-P
BMC-H
BMC-T
Aldehyde dehydrogenase
PTAC
Signature enzyme
Alcohol dehydrogenase
Regulator
05305 -pf01832 -
05310 -pf08264pf00133pf10458 -
05315 -pf00884 -
05320 -pf08245 -
05325 -pf13181 -
05330 -pf01740 -
05335 -pf13581 -
05340 -pf04542pf04539pf04545 -
(..3540..)
08880 -pf00742pf03447 -
08885 -pf01928 -
08890 -pf01965 -
08895 -pf00254pf05698pf05697 -
08900 -pf00574 -
08905 -pf07724pf10431pf06689 -
08910 -pf00004pf05362pf02190 -
08915 -pf01926 -
08920 -pf01594 -
08925 -# -
08930 Hp_euts_bluegreen GRM5like
08935 Hp_euts_frenchBlue GRM5like
08940 HMM10662PduV_1_mix(pf10662) GRM5
08945 -pf06161 -
08950 -pf12802 -
08955 -pf00664pf00005 -
08960 -pf00664pf00005 -
08965 -pf01381 -
08970 -pf12844 -
08975 -pf00694 -
08980 -pf00330 -
08985 -pf00694 -
08990 -pf00330 -
09000 -pf01964 -
09005 -pf09512 -
09010 HMMreg00392_1_rmm1(pf00392) SPU6
09015 HMMaldol_2_pvm1(pf00596) PVMlike
09020 -pf11762pf02610 -
09025 -pf01113pf16654 -
09030 -pf02782pf00370 -
09035 -pf01522 -
09040 -pf12822 -
09045 H_darkBlueGreen GRM5
09050 H_olive GRM5
09055 -# -
09060 -pf08281 -
09065 -pf12645 -
09075 -pf09035 -
09080 -pf02920pf13102pf00589 -
09085 -pf13280 -
09090 -pf12844 -
09095 -# -
09100 -# -
09105 -# -

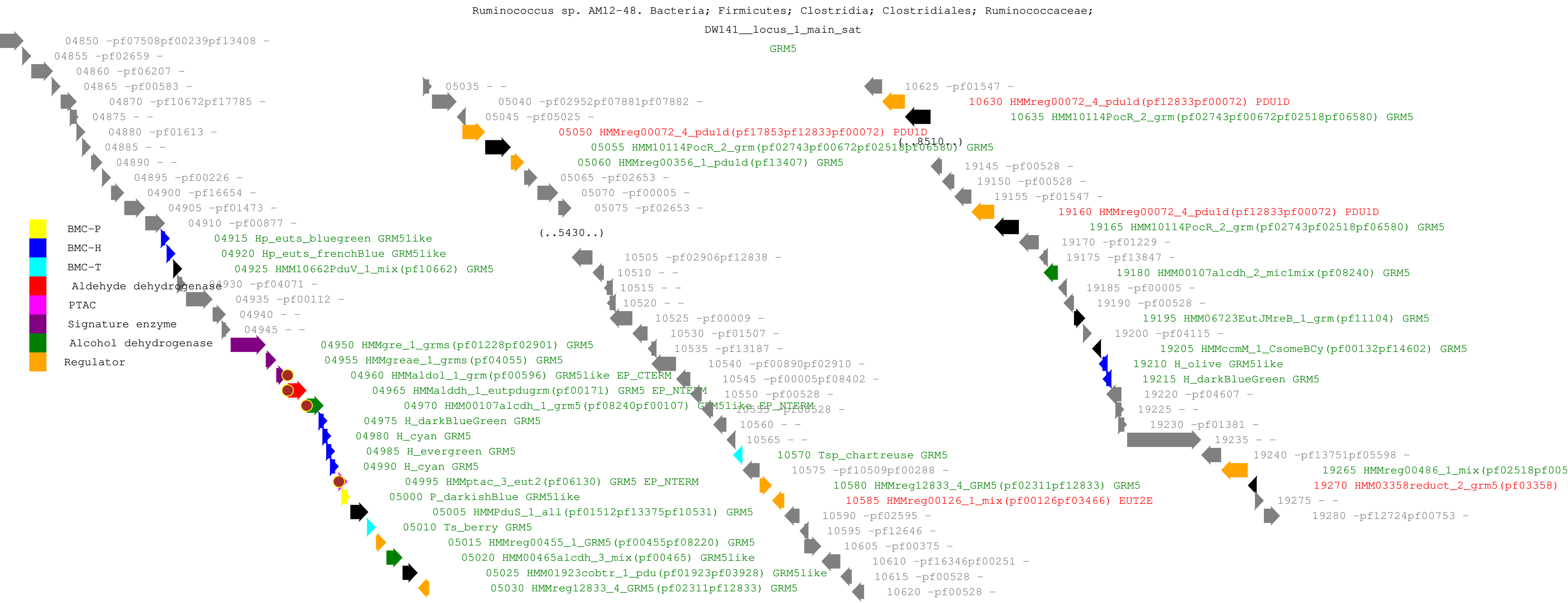
Ruminococcus sp. AM12-48. Bacteria; Firmicutes; Clostridia; Clostridiales; Ruminococcaceae;
DW141__locus_1_main_sat
GRM5
BMC-P
BMC-H
BMC-T
Aldehyde dehydrogenase
PTAC
Signature enzyme
Alcohol dehydrogenase
Regulator
04850 -pf07508pf00239pf13408 -
04855 -pf02659 -
04860 -pf06207 -
04865 -pf00583 -
04870 -pf10672pf17785 -
04875 - -
04880 -pf01613 -
04885 - -
04890 - -
04895 -pf00226 -
04900 -pf16654 -
04905 -pf01473 -
04910 -pf00877 -
04915 Hp_euts_bluegreen GRM5like
04920 Hp_euts_frenchBlue GRM5like
04925 HMM10662PduV_1_mix(pf10662) GRM5
04930 -pf04071 -
04935 -pf00112 -
04940 - -
04945 - -
04950 HMMgre_1_grms(pf01228pf02901) GRM5
04955 HMMgreae_1_grms(pf04055) GRM5
04960 HMMaldol_1_grm(pf00596) GRM5like EP_CTERM
04965 HMMalddh_1_eutpdugrm(pf00171) GRM5 EP_NTERM
04970 HMM00107alcdh_1_grm5(pf08240pf00107) GRM5like EP_NTERM
04975 H_darkBlueGreen GRM5
04980 H_cyan GRM5
04985 H_evergreen GRM5
04990 H_cyan GRM5
04995 HMMptac_3_eut2(pf06130) GRM5 EP_NTERM
05000 P_darkishBlue GRM5like
05005 HMMPduS_1_all(pf01512pf13375pf10531) GRM5
05010 Ts_berry GRM5
05015 HMMreg00455_1_GRM5(pf00455pf08220) GRM5
05020 HMM00465alcdh_3_mix(pf00465) GRM5like
05025 HMM01923cobtr_1_pdu(pf01923pf03928) GRM5like
05030 HMMreg12833_4_GRM5(pf02311pf12833) GRM5
05035 - -
05040 -pf02952pf07881pf07882 -
05045 -pf05025 -
05050 HMMreg00072_4_pdu1d(pf17853pf12833pf00072) PDU1D
05055 HMM10114PocR_2_grm(pf02743pf00672pf02518pf06580) GRM5
05060 HMMreg00356_1_pdu1d(pf13407) GRM5
05065 -pf02653 -
05070 -pf00005 -
05075 -pf02653 -
(..5430..)
10505 -pf02906pf12838 -
10510 - -
10515 - -
10520 - -
10525 -pf00009 -
10530 -pf01507 -
10535 -pf13187 -
10540 -pf00890pf02910 -
10545 -pf00005pf08402 -
10550 -pf00528 -
10555 -pf00528 -
10560 - -
10565 - -
10570 Tsp_chartreuse GRM5
10575 -pf10509pf00288 -
10580 HMMreg12833_4_GRM5(pf02311pf12833) GRM5
10585 HMMreg00126_1_mix(pf00126pf03466) EUT2E
10590 -pf02595 -
10595 -pf12646 -
10605 -pf00375 -
10610 -pf16346pf00251 -
10615 -pf00528 -
10620 -pf00528 -
10625 -pf01547 -
10630 HMMreg00072_4_pdu1d(pf12833pf00072) PDU1D
10635 HMM10114PocR_2_grm(pf02743pf00672pf02518pf06580) GRM5
(..8510..)
19145 -pf00528 -
19150 -pf00528 -
19155 -pf01547 -
19160 HMMreg00072_4_pdu1d(pf12833pf00072) PDU1D
19165 HMM10114PocR_2_grm(pf02743pf02518pf06580) GRM5
19170 -pf01229 -
19175 -pf13847 -
19180 HMM00107alcdh_2_mic1mix(pf08240) GRM5
19185 -pf00005 -
19190 -pf00528 -
19195 HMM06723EutJMreB_1_grm(pf11104) GRM5
19200 -pf04115 -
19205 HMMccmM_1_CsomeBCy(pf00132pf14602) GRM5
19210 H_olive GRM5like
19215 H_darkBlueGreen GRM5
19220 -pf04607 -
19225 - -
19230 -pf01381 -
19235 - -
19240 -pf13751pf05598 -
19265 HMMreg00486_1_mix(pf02518pf005
19270 HMM03358reduct_2_grm5(pf03358)
19275 - -
19280 -pf12724pf00753 -

Ruminococcus sp. AM16-34. Bacteria; Firmicutes; Clostridia; Clostridiales; Ruminococcaceae;

DW187__locus_1_main_sat

GRM5

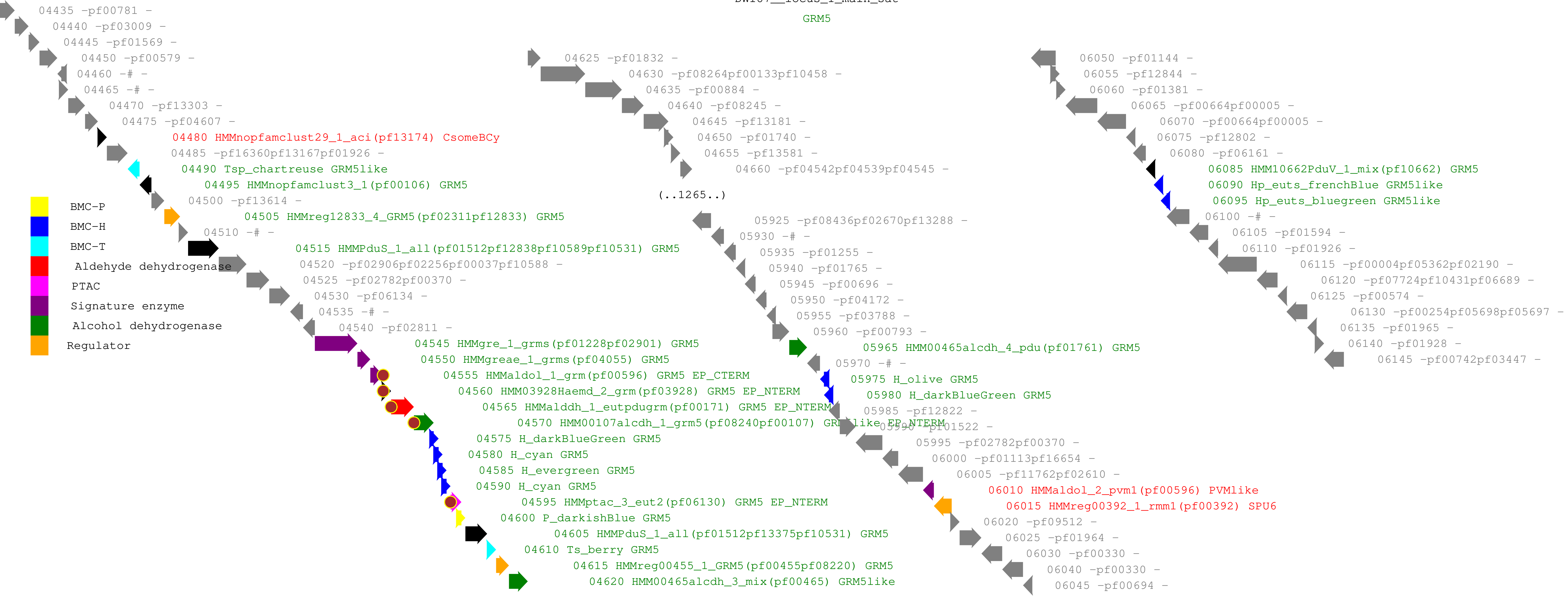

Ruminococcus sp. AM18-44. Bacteria; Firmicutes; Clostridia; Clostridiales; Ruminococcaceae;

DW225__locus_1_main_sat

GRM5

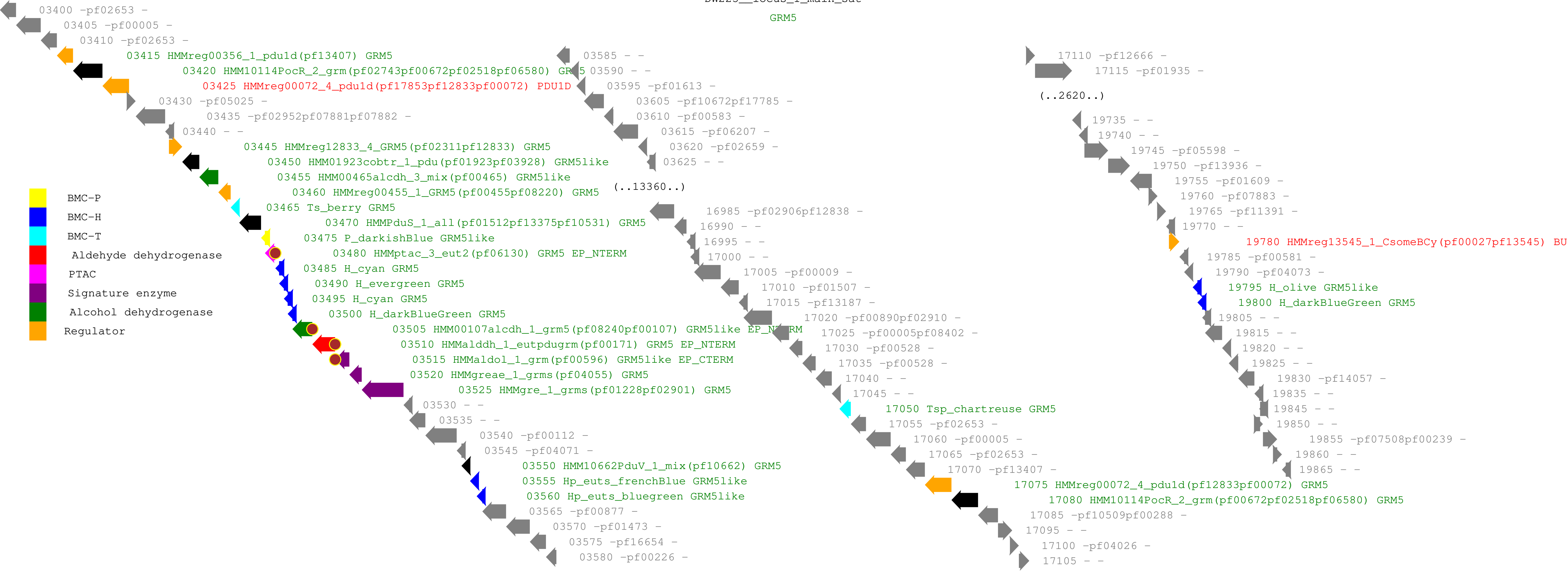

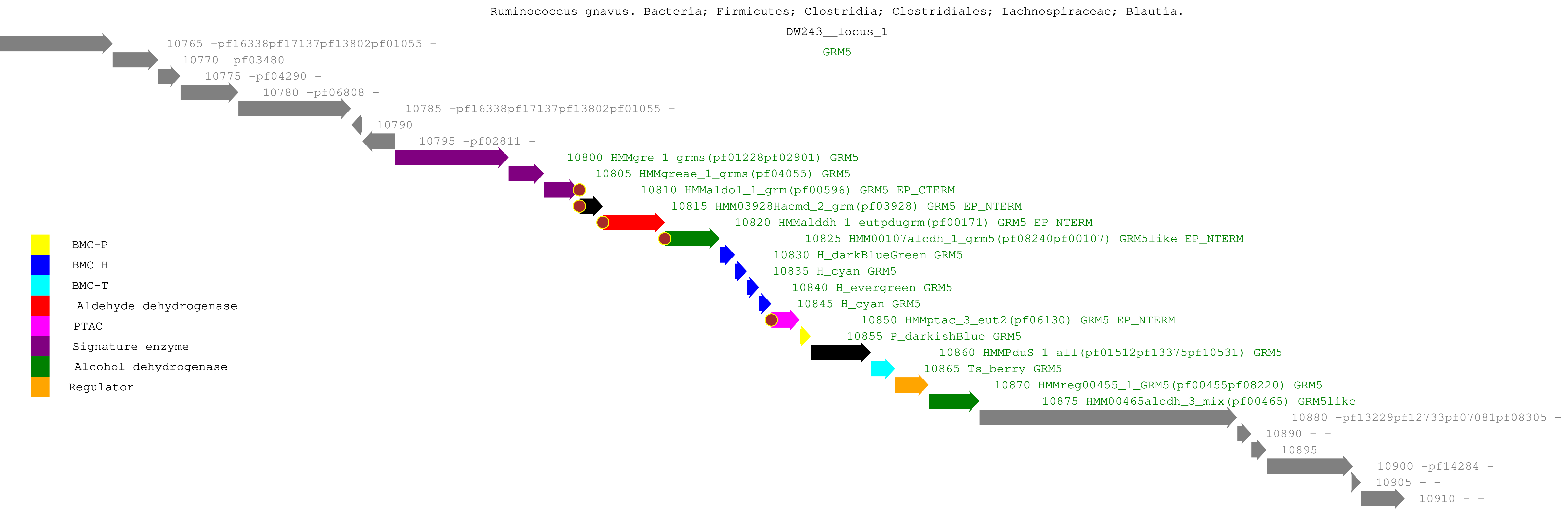
Ruminococcus gnavus. Bacteria; Firmicutes; Clostridia; Clostridiales; Lachnospiraceae; Blautia.
DW243__locus_1
GRM5
10765 -pf16338pf17137pf13802pf01055 -
10770 -pf03480 -
10775 -pf04290 -
10780 -pf06808 -
10785 -pf16338pf17137pf13802pf01055 -
10790 - -
10795 -pf02811 -
10800 HMMgre_1_grms(pf01228pf02901) GRM5
10805 HMMgreae_1_grms(pf04055) GRM5
10810 HMMaldol_1_grm(pf00596) GRM5 EP_CTERM
10815 HMM03928Haemd_2_grm(pf03928) GRM5 EP_NTERM
10820 HMMalddh_1_eutpdugrm(pf00171) GRM5 EP_NTERM
10825 HMM00107alcdh_1_grm5(pf08240pf00107) GRM5like EP_NTERM
10830 H_darkBlueGreen GRM5
10835 H_cyan GRM5
10840 H_evergreen GRM5
10845 H_cyan GRM5
10850 HMMptac_3_eut2(pf06130) GRM5 EP_NTERM
10855 P_darkishBlue GRM5
10860 HMMPduS_1_all(pf01512pf13375pf10531) GRM5
10865 Ts_berry GRM5
10870 HMMreg00455_1_GRM5(pf00455pf08220) GRM5
10875 HMM00465alcdh_3_mix(pf00465) GRM5like
10880 -pf13229pf12733pf07081pf08305 -
10890 - -
10895 - -
10900 -pf14284 -
10905 - -
10910 - -
BMC-P
BMC-H
BMC-T
Aldehyde dehydrogenase
PTAC
Signature enzyme
Alcohol dehydrogenase
Regulator

Ruminococcus sp. AM22-14LB. Bacteria; Firmicutes; Clostridia; Clostridiales; Ruminococcaceae;

DW256__locus_1_main_sat

GRM5

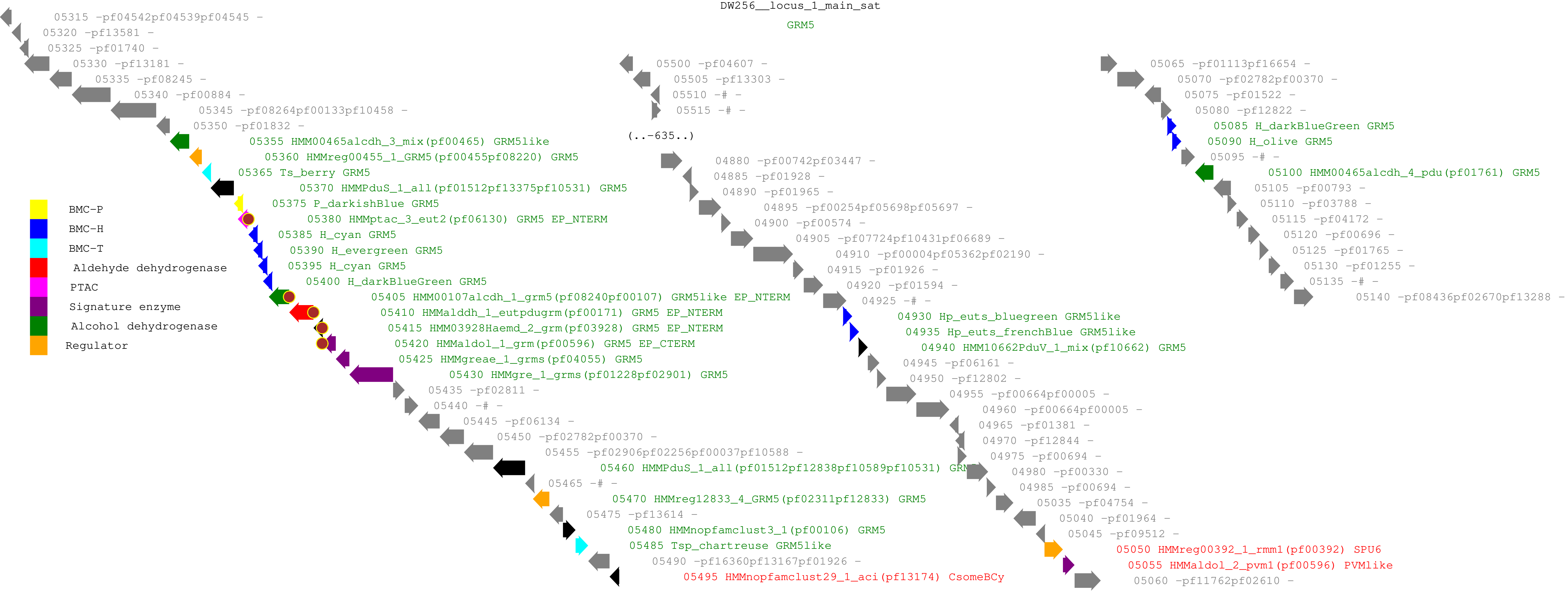

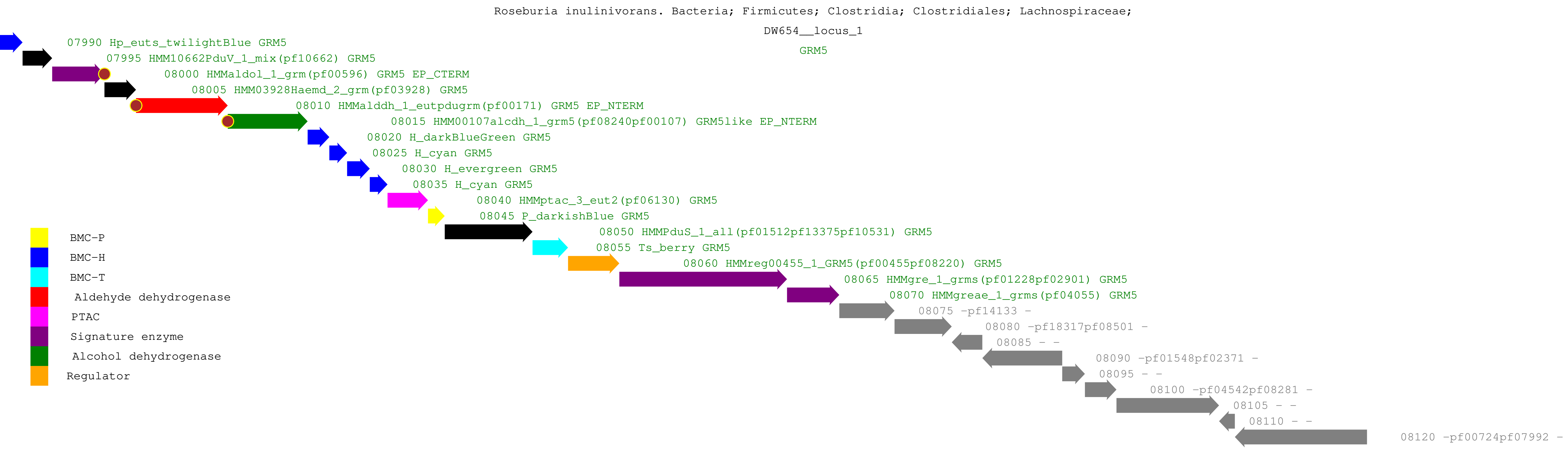
Roseburia inulinivorans. Bacteria; Firmicutes; Clostridia; Clostridiales; Lachnospiraceae;
DW654__locus_1
GRM5
07990 Hp_euts_twilightBlue GRM5
07995 HMM10662PduV_1_mix(pf10662) GRM5
08000 HMMaldol_1_grm(pf00596) GRM5 EP_CTERM
08005 HMM03928Haemd_2_grm(pf03928) GRM5
08010 HMMalddh_1_eutpdugrm(pf00171) GRM5 EP_NTERM
08015 HMM00107alcdh_1_grm5(pf08240pf00107) GRM5like EP_NTERM
08020 H_darkBlueGreen GRM5
08025 H_cyan GRM5
08030 H_evergreen GRM5
08035 H_cyan GRM5
08040 HMMptac_3_eut2(pf06130) GRM5
08045 P_darkishBlue GRM5
08050 HMMPduS_1_all(pf01512pf13375pf10531) GRM5
08055 Ts_berry GRM5
08060 HMMreg00455_1_GRM5(pf00455pf08220) GRM5
08065 HMMgre_1_grms(pf01228pf02901) GRM5
08070 HMMgreae_1_grms(pf04055) GRM5
08075 -pf14133 -
08080 -pf18317pf08501 -
08085 - -
08090 -pf01548pf02371 -
08095 - -
08100 -pf04542pf08281 -
08105 - -
08110 - -
08120 -pf00724pf07992 -
BMC-P
BMC-H
BMC-T
Aldehyde dehydrogenase
PTAC
Signature enzyme
Alcohol dehydrogenase
Regulator

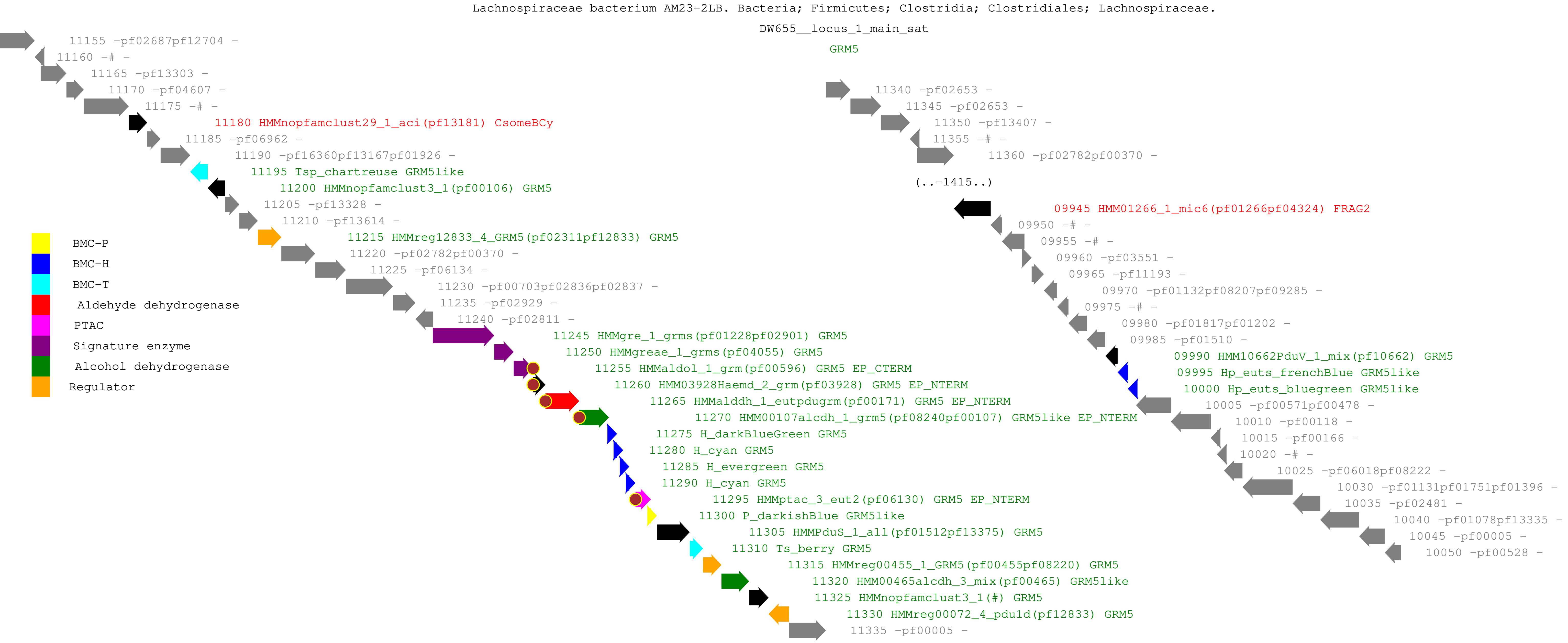
Lachnospiraceae bacterium AM23-2LB. Bacteria; Firmicutes; Clostridia; Clostridiales; Lachnospiraceae.
DW655__locus_1_main_sat
GRM5
11155 -pf02687pf12704 -
11160 -# -
11165 -pf13303 -
11170 -pf04607 -
11175 -# -
11180 HMMnopfamclust29_1_aci(pf13181) CsomeBCy
11185 -pf06962 -
11190 -pf16360pf13167pf01926 -
11195 Tsp_chartreuse GRM5like
11200 HMMnopfamclust3_1(pf00106) GRM5
11205 -pf13328 -
11210 -pf13614 -
11215 HMMreg12833_4_GRM5(pf02311pf12833) GRM5
11220 -pf02782pf00370 -
11225 -pf06134 -
11230 -pf00703pf02836pf02837 -
11235 -pf02929 -
11240 -pf02811 -
11245 HMMgre_1_grms(pf01228pf02901) GRM5
11250 HMMgreae_1_grms(pf04055) GRM5
11255 HMMaldol_1_grm(pf00596) GRM5 EP_CTERM
11260 HMM03928Haemd_2_grm(pf03928) GRM5 EP_NTERM
11265 HMMalddh_1_eutpdugrm(pf00171) GRM5 EP_NTERM
11270 HMM00107alcdh_1_grm5(pf08240pf00107) GRM5like EP_NTERM
11275 H_darkBlueGreen GRM5
11280 H_cyan GRM5
11285 H_evergreen GRM5
11290 H_cyan GRM5
11295 HMMptac_3_eut2(pf06130) GRM5 EP_NTERM
11300 P_darkishBlue GRM5like
11305 HMMPduS_1_all(pf01512pf13375) GRM5
11310 Ts_berry GRM5
11315 HMMreg00455_1_GRM5(pf00455pf08220) GRM5
11320 HMM00465alcdh_3_mix(pf00465) GRM5like
11325 HMMnopfamclust3_1(#) GRM5
11330 HMMreg00072_4_pdu1d(pf12833) GRM5
11335 -pf00005 -
11340 -pf02653 -
11345 -pf02653 -
11350 -pf13407 -
11355 -# -
11360 -pf02782pf00370 -
(..-1415..)
09945 HMM01266_1_mic6(pf01266pf04324) FRAG2
09950 -# -
09955 -# -
09960 -pf03551 -
09965 -pf11193 -
09970 -pf01132pf08207pf09285 -
09975 -# -
09980 -pf01817pf01202 -
09985 -pf01510 -
09990 HMM10662PduV_1_mix(pf10662) GRM5
09995 Hp_euts_frenchBlue GRM5like
10000 Hp_euts_bluegreen GRM5like
10005 -pf00571pf00478 -
10010 -pf00118 -
10015 -pf00166 -
10020 -# -
10025 -pf06018pf08222 -
10030 -pf01131pf01751pf01396 -
10035 -pf02481 -
10040 -pf01078pf13335 -
10045 -pf00005 -
10050 -pf00528 -
BMC-P
BMC-H
BMC-T
Aldehyde dehydrogenase
PTAC
Signature enzyme
Alcohol dehydrogenase
Regulator

Ruminococcus sp. AM26-12LB. Bacteria; Firmicutes; Clostridia; Clostridiales; Ruminococcaceae;

DW694__locus_1_main_sat

GRM5

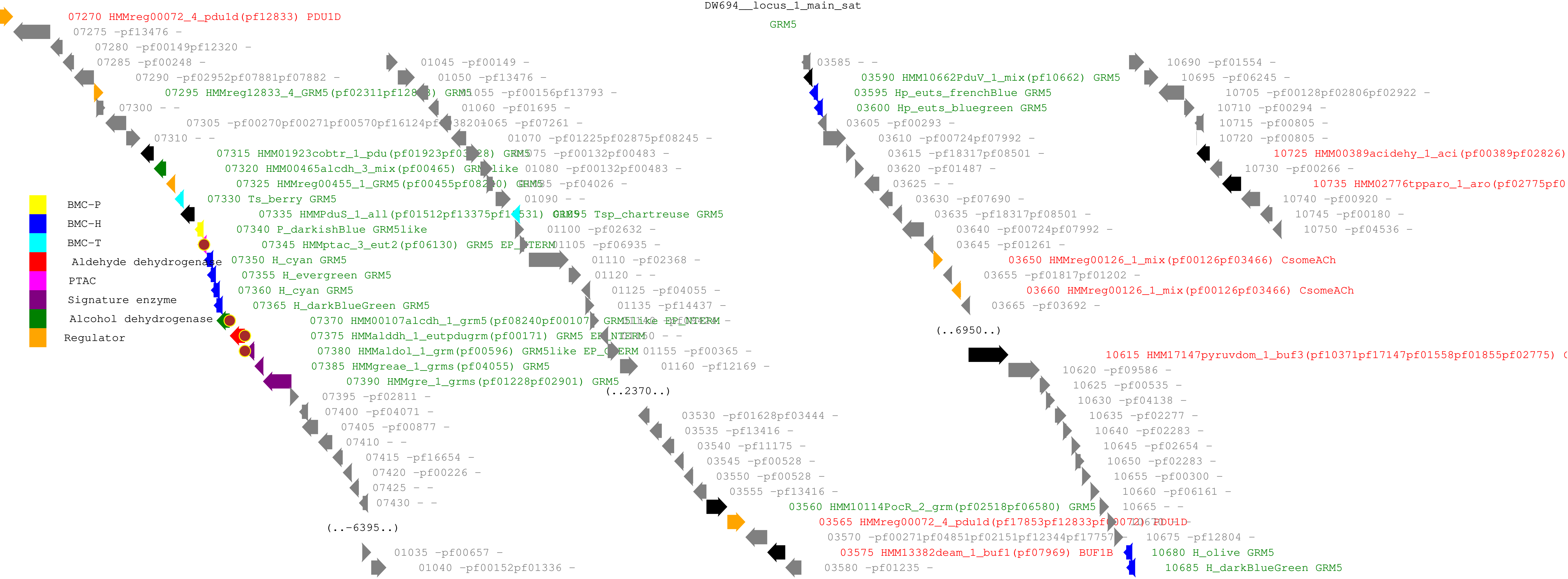

Ruminococcus sp. AM27-11LB. Bacteria; Firmicutes; Clostridia; Clostridiales; Ruminococcaceae;

DW708__locus_1_main_sat

GRM5

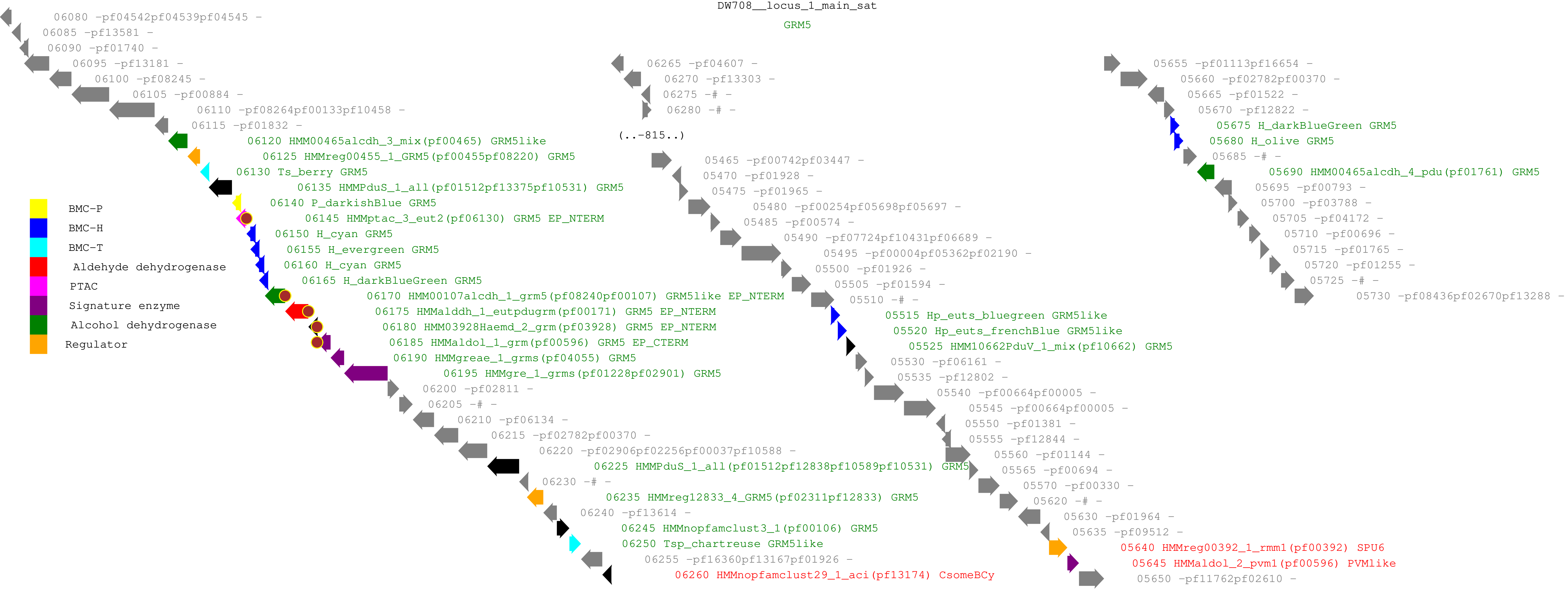

Ruminococcus sp. AM27-16. Bacteria; Firmicutes; Clostridia; Clostridiales; Ruminococcaceae;

DW711__locus_1_main_sat

GRM5

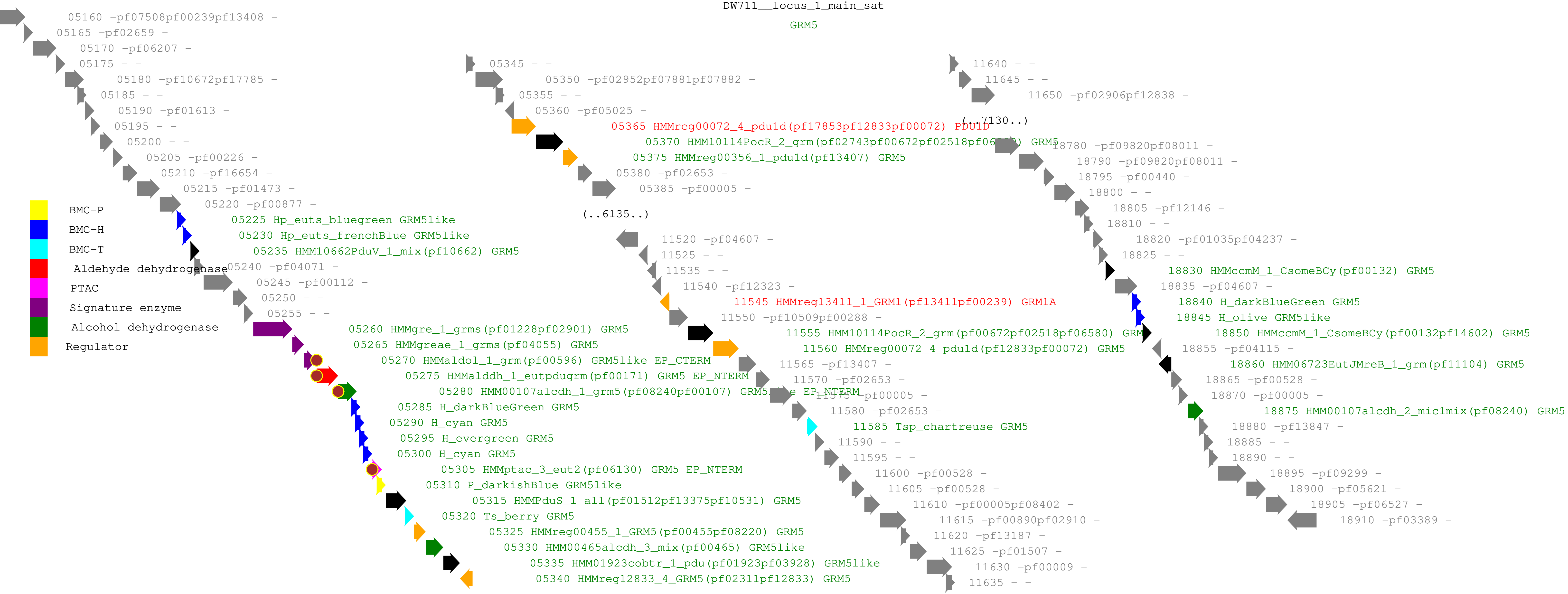

Ruminococcus sp. AM27-27. Bacteria; Firmicutes; Clostridia; Clostridiales; Ruminococcaceae;

DW719__locus_1_main_sat

GRM5

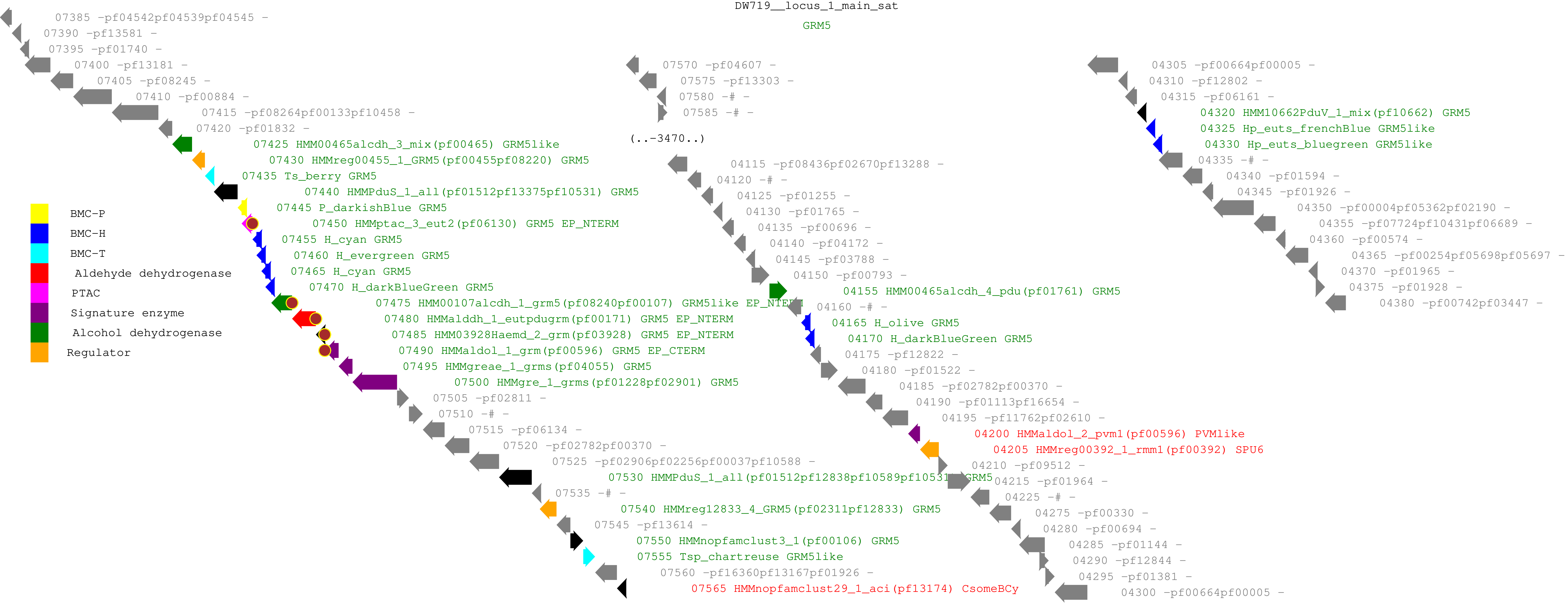

Blautia obeum. Bacteria; Firmicutes; Clostridia; Clostridiales; Lachnospiraceae; Blautia.

DW740__locus_1_main_sat

GRM5

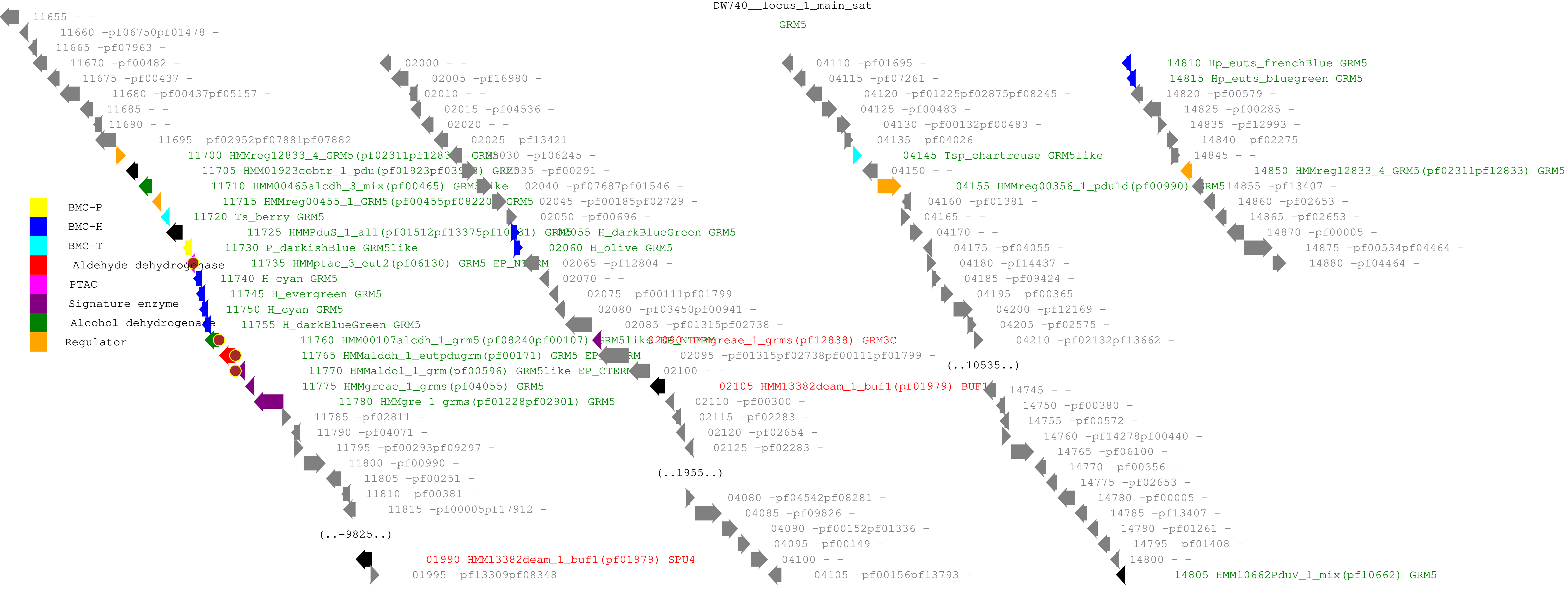

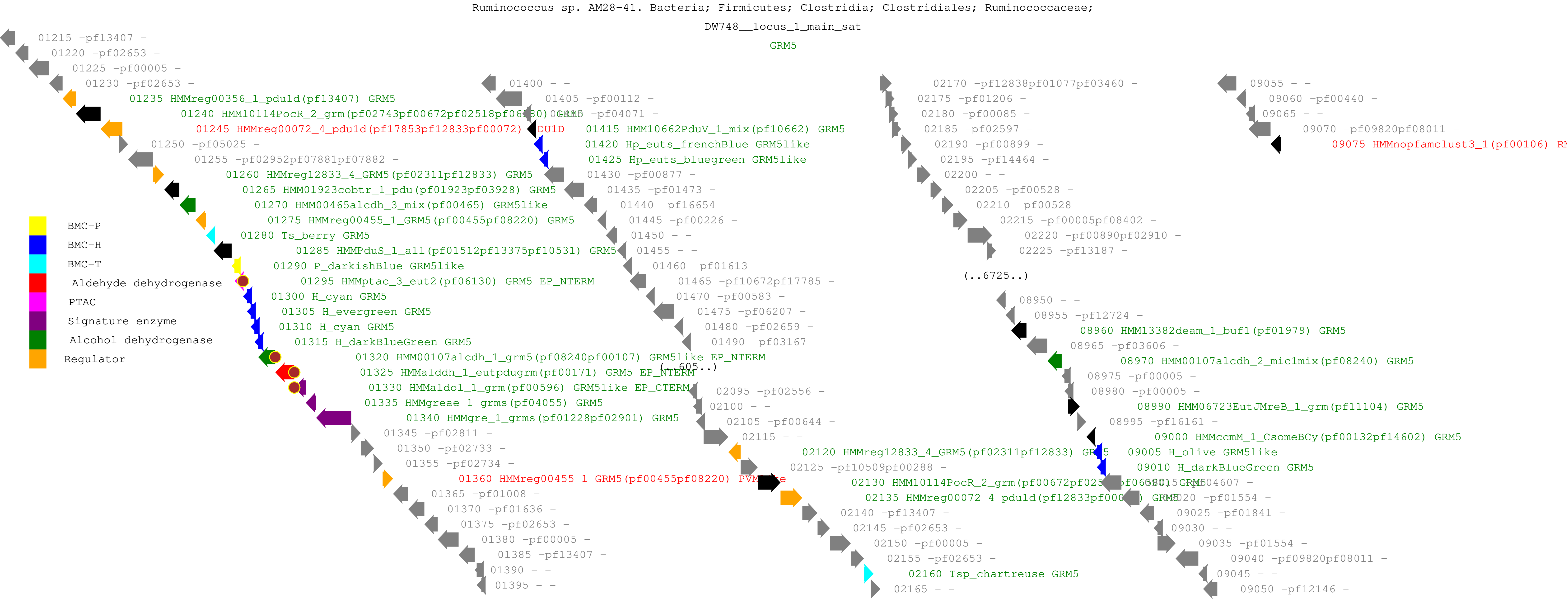
Ruminococcus sp. AM28-41. Bacteria; Firmicutes; Clostridia; Clostridiales; Ruminococcaceae;
DW748__locus_1_main_sat
GRM5
BMC-P
BMC-H
BMC-T
Aldehyde dehydrogenase
PTAC
Signature enzyme
Alcohol dehydrogenase
Regulator
01215 -pf13407 -
01220 -pf02653 -
01225 -pf00005 -
01230 -pf02653 -
01235 HMMreg00356_1_pduld(pf13407) GRM5
01245 HMMreg00072_4_pduld(pf17853pf12833pf00072)
01250 -pf05025 -
01255 -pf02952pf07881pf07882 -
01260 HMMreg12833_4_GRM5(pf02311pf12833) GRM5
01265 HMM01923cobtr_1_pdu(pf01923pf03928) GRM5
01270 HMM00465alcdh_3_mix(pf00465) GRM5like
01275 HMMreg00455_1_GRM5(pf00455pf08220) GRM5
01280 Ts_berry GRM5
01285 HMMPduS_1_all(pf01512pf13375pf10531) GRM5
01290 P_darkishBlue GRM5like
01295 HMMptac_3_eut2(pf06130) GRM5 EP_NTERM
01300 H_cyan GRM5
01305 H_evergreen GRM5
01310 H_cyan GRM5
01315 H_darkBlueGreen GRM5
01320 HMM00107alcdh_1_grm5(pf08240pf00107) GRM5like EP_NTERM
01325 HMMalddh_1_eutpdugrm(pf00171) GRM5 EP_NTERM
01330 HMMaldol_1_grm(pf00596) GRM5like EP_CTERM
01335 HMMgreae_1_grms(pf04055) GRM5
01340 HMMgre_1_grms(pf01228pf02901) GRM5
01345 -pf02811 -
01350 -pf02733 -
01355 -pf02734 -
01360 HMMreg00455_1_GRM5(pf00455pf08220)
01365 -pf01008 -
01370 -pf01636 -
01375 -pf02653 -
01380 -pf00005 -
01385 -pf13407 -
01390 - -
01395 - -
01400 - -
01405 -pf00112 -
01415 HMM10662PduV_1_mix(pf10662) GRM5
01420 Hp_euts_frenchBlue GRM5like
01425 Hp_euts_bluegreen GRM5like
01430 -pf00877 -
01435 -pf01473 -
01440 -pf16654 -
01445 -pf00226 -
01450 - -
01455 - -
01460 -pf01613 -
01465 -pf10672pf17785 -
01470 -pf00583 -
01475 -pf06207 -
01480 -pf02659 -
01490 -pf03167 -
(..605..)
02095 -pf02556 -
02100 - -
02105 -pf00644 -
02115 - -
02120 HMMreg12833_4_GRM5(pf02311pf12833)
02125 -pf10509pf00288 -
02140 -pf13407 -
02145 -pf02653 -
02150 -pf00005 -
02155 -pf02653 -
02160 Tsp_chartreuse GRM5
02165 - -
02170 -pf12838pf01077pf03460 -
02175 -pf01206 -
02180 -pf00085 -
02185 -pf02597 -
02190 -pf00899 -
02195 -pf14464 -
02200 - -
02205 -pf00528 -
02210 -pf00528 -
02215 -pf00005pf08402 -
02220 -pf00890pf02910 -
02225 -pf13187 -
(..6725..)
08950 - -
08955 -pf12724 -
08960 HMM13382deam_1_buf1(pf01979) GRM5
08965 -pf03606 -
08970 HMM00107alcdh_2_mic1mix(pf08240) GRM5
08975 -pf00005 -
08980 -pf00005 -
08990 HMM06723EutJMreB_1_grm(pf11104) GRM5
08995 -pf16161 -
09000 HMMccmM_1_CsomeBCy(pf00132pf14602) GRM5
09005 H_olive GRM5like
09010 H_darkBlueGreen GRM5
09020 -pf01554 -
09025 -pf01841 -
09030 - -
09035 -pf01554 -
09040 -pf09820pf08011 -
09045 - -
09050 -pf12146 -
09055 - -
09060 -pf00440 -
09065 - -
09070 -pf09820pf08011 -
09075 HMMnopfamclust3_1(pf00106)

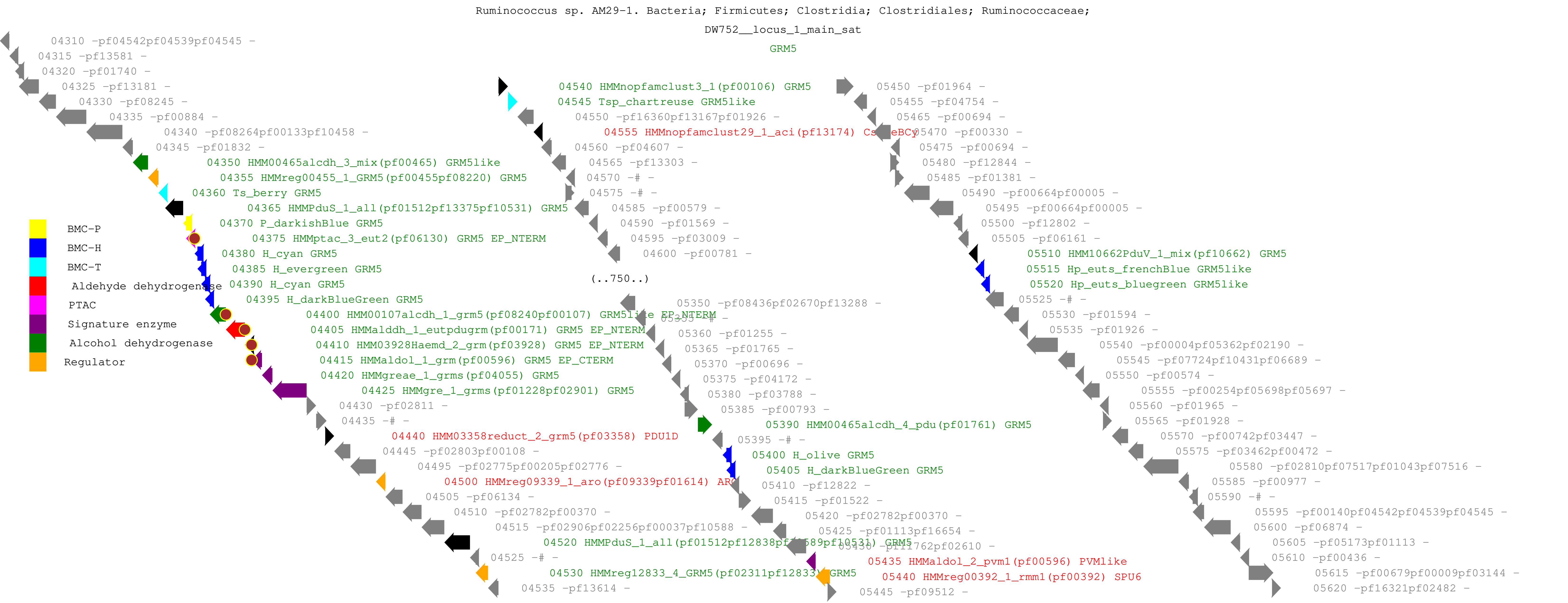

Ruminococcus sp. AM29-1. Bacteria; Firmicutes; Clostridia; Clostridiales; Ruminococcaceae;
DW752__locus_1_main_sat
GRM5
BMC-P
BMC-H
BMC-T
Aldehyde dehydrogenase
PTAC
Signature enzyme
Alcohol dehydrogenase
Regulator
04310 -pf04542pf04539pf04545 -
04315 -pf13581 -
04320 -pf01740 -
04325 -pf13181 -
04330 -pf08245 -
04335 -pf00884 -
04340 -pf08264pf00133pf10458 -
04345 -pf01832 -
04350 HMM00465alcdh_3_mix(pf00465) GRM5like
04355 HMMreg00455_1_GRM5(pf00455pf08220) GRM5
04360 Ts_berry GRM5
04365 HMMPduS_1_all(pf01512pf13375pf10531) GRM5
04370 P_darkishBlue GRM5
04375 HMMptac_3_eut2(pf06130) GRM5 EP_NTERM
04380 H_cyan GRM5
04385 H_evergreen GRM5
04390 H_cyan GRM5
04395 H_darkBlueGreen GRM5
04400 HMM00107alcdh_1_grm5(pf08240pf00107) GRM5like EP_NTERM
04405 HMMaldh_1_eutpdugrm(pf00171) GRM5 EP_NTERM
04410 HMM03928Haemd_2_grm(pf03928) GRM5 EP_NTERM
04415 HMMaldol_1_grm(pf00596) GRM5 EP_CTERM
04420 HMMgreae_1_grms(pf04055) GRM5
04425 HMMgre_1_grms(pf01228pf02901) GRM5
04430 -pf02811 -
04435 -# -
04440 HMM03358reduct_2_grm5(pf03358) PDU1D
04445 -pf02803pf00108 -
04495 -pf02775pf00205pf02776 -
04500 HMMreg09339_1_aro(pf09339pf01614) ARO
04505 -pf06134 -
04510 -pf02782pf00370 -
04515 -pf02906pf02256pf00037pf10588 -
04520 HMMPduS_1_all(pf01512pf12838pf13589pf10531) GRM5
04525 -# -
04530 HMMreg12833_4_GRM5(pf02311pf12833) GRM5
04535 -pf13614 -
04540 HMMnopfamclust3_1(pf00106) GRM5
04545 Tsp_chartreuse GRM5like
04550 -pf16360pf13167pf01926 -
04555 HMMnopfamclust29_1_aci(pf13174) CsoeBCy
04560 -pf04607 -
04565 -pf13303 -
04570 -# -
04575 -# -
04585 -pf00579 -
04590 -pf01569 -
04595 -pf03009 -
04600 -pf00781 -
(..750..)
05350 -pf08436pf02670pf13288 -
05355 -# -
05360 -pf01255 -
05365 -pf01765 -
05370 -pf00696 -
05375 -pf04172 -
05380 -pf03788 -
05385 -pf00793 -
05390 HMM00465alcdh_4_pdu(pf01761) GRM5
05395 -# -
05400 H_olive GRM5
05405 H_darkBlueGreen GRM5
05410 -pf12822 -
05415 -pf01522 -
05420 -pf02782pf00370 -
05425 -pf01113pf16654 -
05430 -pf11762pf02610 -
05435 HMMaldol_2_pvm1(pf00596) PVMlike
05440 HMMreg00392_1_rmm1(pf00392) SPU6
05445 -pf09512 -
05450 -pf01964 -
05455 -pf04754 -
05465 -pf00694 -
05470 -pf00330 -
05475 -pf00694 -
05480 -pf12844 -
05485 -pf01381 -
05490 -pf00664pf00005 -
05495 -pf00664pf00005 -
05500 -pf12802 -
05505 -pf06161 -
05510 HMM10662PduV_1_mix(pf10662) GRM5
05515 Hp_euts_frenchBlue GRM5like
05520 Hp_euts_bluegreen GRM5like
05525 -# -
05530 -pf01594 -
05535 -pf01926 -
05540 -pf00004pf05362pf02190 -
05545 -pf07724pf10431pf06689 -
05550 -pf00574 -
05555 -pf00254pf05698pf05697 -
05560 -pf01965 -
05565 -pf01928 -
05570 -pf00742pf03447 -
05575 -pf03462pf00472 -
05580 -pf02810pf07517pf01043pf07516 -
05585 -pf00977 -
05590 -# -
05595 -pf00140pf04542pf04539pf04545 -
05600 -pf06874 -
05605 -pf05173pf01113 -
05610 -pf00436 -
05615 -pf00679pf00009pf03144 -
05620 -pf16321pf02482 -

Ruminococcus sp. AM29-10LB. Bacteria; Firmicutes; Clostridia; Clostridiales; Ruminococcaceae;

DW755__locus_1_main_sat

GRM5

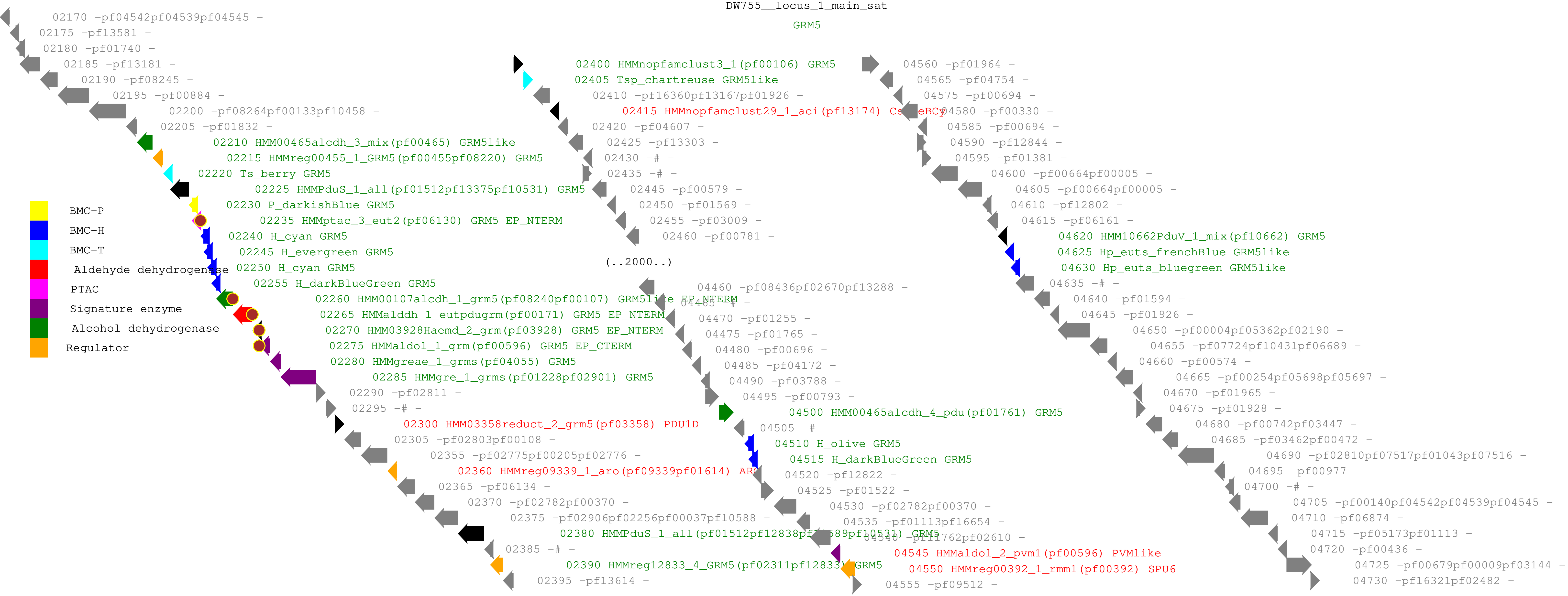

Ruminococcus sp. AM29-12LB. Bacteria; Firmicutes; Clostridia; Clostridiales; Ruminococcaceae;

DW759__locus_1_main_sat

GRM5

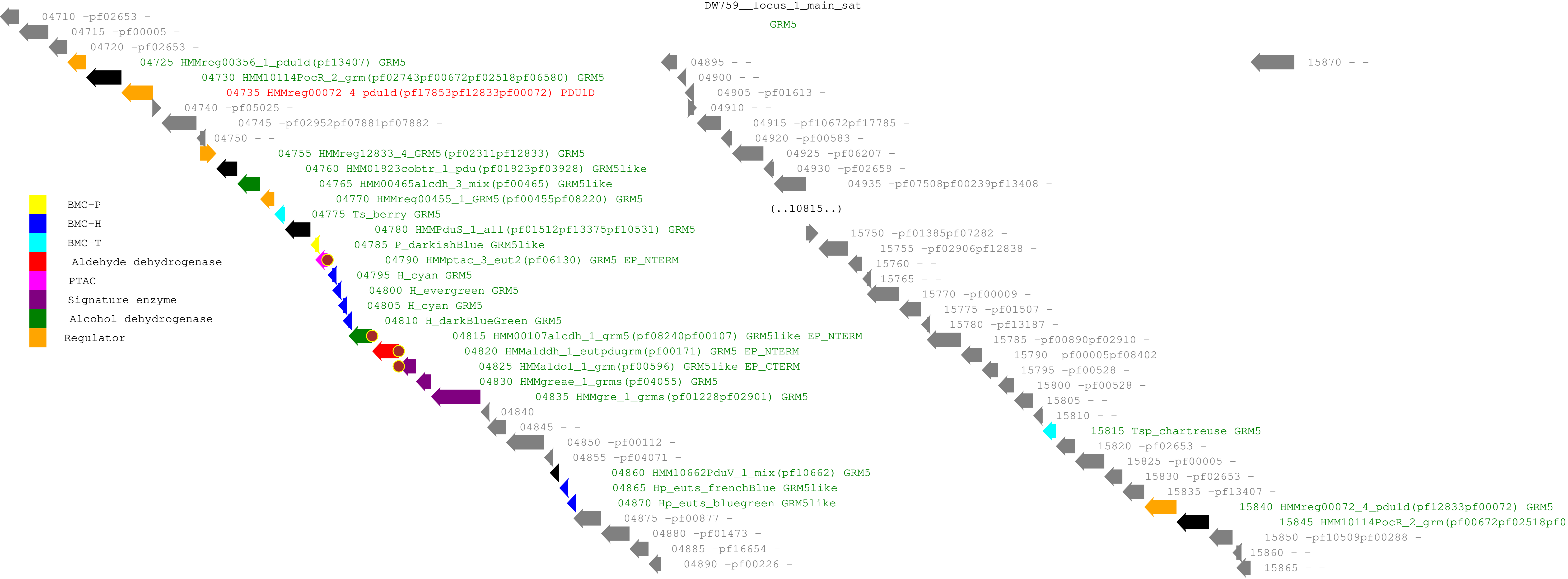

Ruminococcus sp. AM29-19LB. Bacteria; Firmicutes; Clostridia; Clostridiales; Ruminococcaceae;

DW762__locus_1_main_sat

GRM5

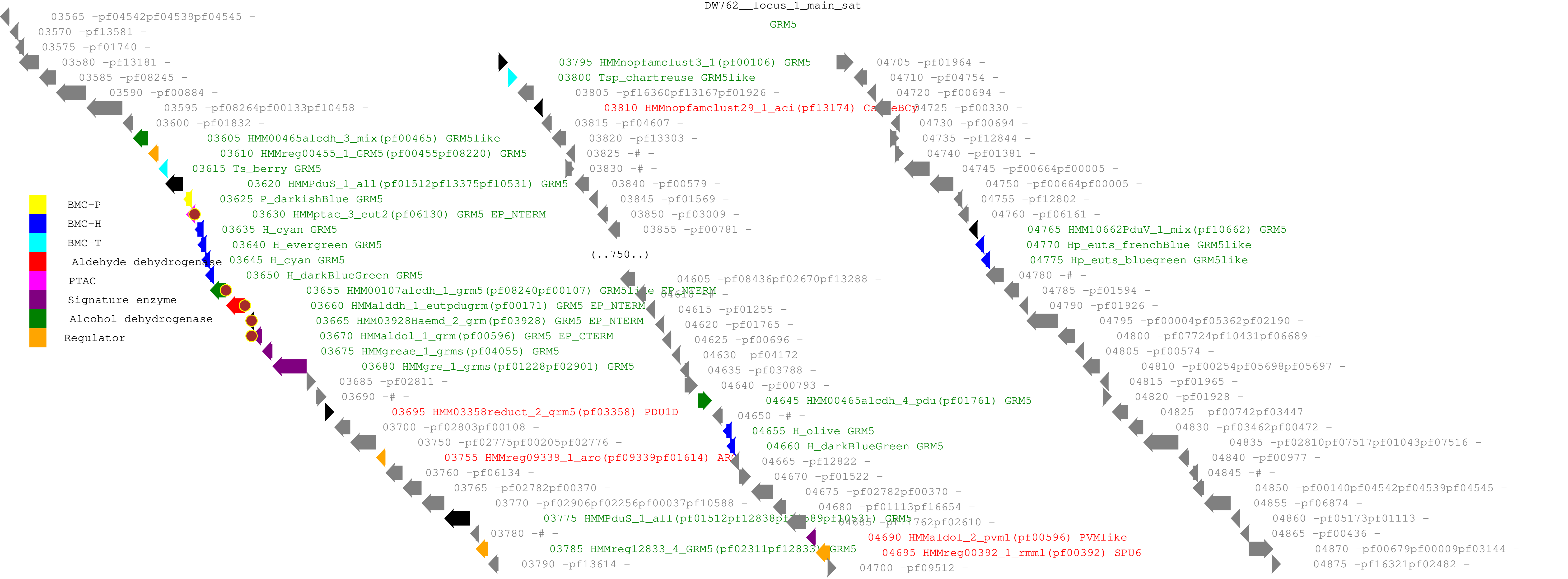

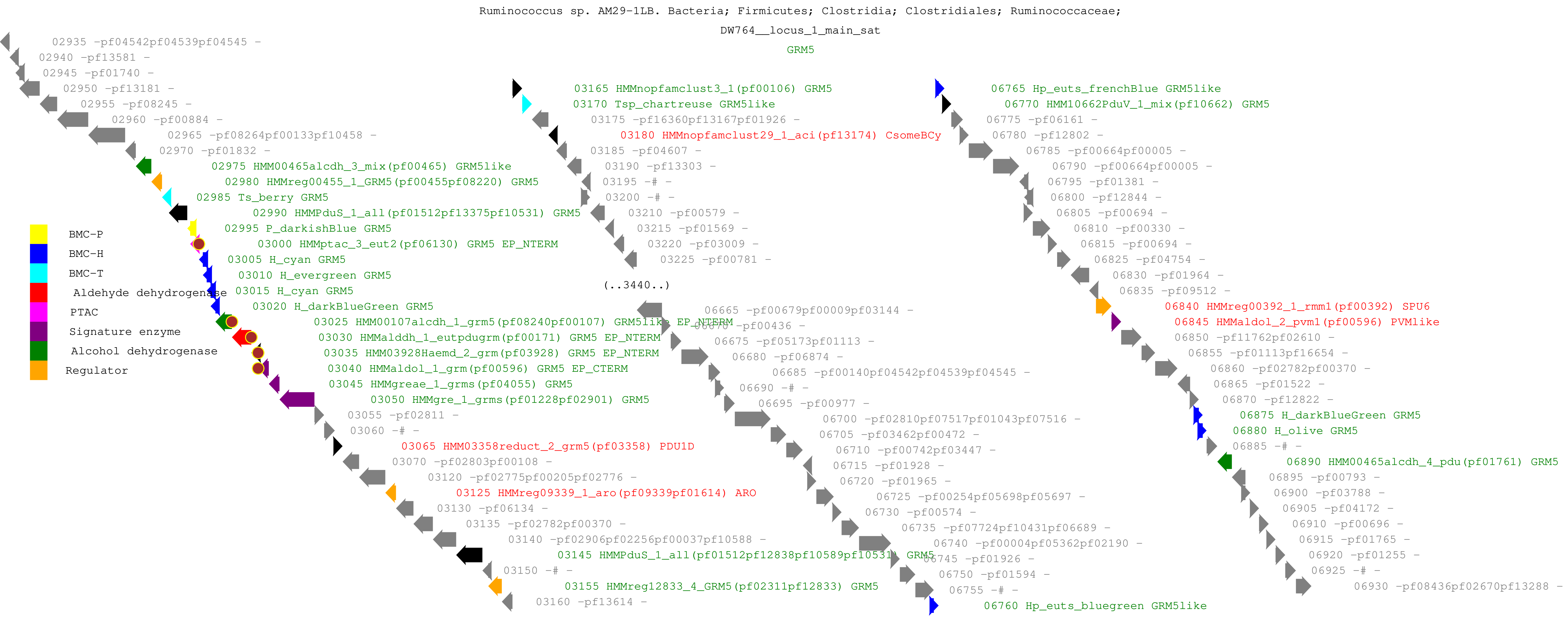
Ruminococcus sp. AM29-1LB. Bacteria; Firmicutes; Clostridia; Clostridiales; Ruminococcaceae;
DW764__locus_1_main_sat
GRM5
BMC-P
BMC-H
BMC-T
Aldehyde dehydrogenase
PTAC
Signature enzyme
Alcohol dehydrogenase
Regulator
02935 -pf04542pf04539pf04545 -
02940 -pf13581 -
02945 -pf01740 -
02950 -pf13181 -
02955 -pf08245 -
02960 -pf00884 -
02965 -pf08264pf00133pf10458 -
02970 -pf01832 -
02975 HMM00465alcdh_3_mix(pf00465) GRM5like
02980 HMMreg00455_1_GRM5(pf00455pf08220) GRM5
02985 Ts_berry GRM5
02990 HMMPduS_1_all(pf01512pf13375pf10531) GRM5
02995 P_darkishBlue GRM5
03000 HMMptac_3_eut2(pf06130) GRM5 EP_NTERM
03005 H_cyan GRM5
03010 H_evergreen GRM5
03015 H_cyan GRM5
03020 H_darkBlueGreen GRM5
03025 HMM00107alcdh_1_grm5(pf08240pf00107) GRM5like EP_NTERM
03030 HMMaldh_1_eutpdugrm(pf00171) GRM5 EP_NTERM
03035 HMM03928Haemd_2_grm(pf03928) GRM5 EP_NTERM
03040 HMMaldol_1_grm(pf00596) GRM5 EP_CTERM
03045 HMMgreae_1_grms(pf04055) GRM5
03050 HMMgre_1_grms(pf01228pf02901) GRM5
03055 -pf02811 -
03060 -# -
03065 HMM03358reduct_2_grm5(pf03358) PDU1D
03070 -pf02803pf00108 -
03120 -pf02775pf00205pf02776 -
03125 HMMreg09339_1_aro(pf09339pf01614) ARO
03130 -pf06134 -
03135 -pf02782pf00370 -
03140 -pf02906pf02256pf00037pf10588 -
03145 HMMPduS_1_all(pf01512pf12838pf10589pf10531) GRM5
03150 -# -
03155 HMMreg12833_4_GRM5(pf02311pf12833) GRM5
03160 -pf13614 -
03165 HMMnopfamclust3_1(pf00106) GRM5
03170 Tsp_chartreuse GRM5like
03175 -pf16360pf13167pf01926 -
03180 HMMnopfamclust29_1_aci(pf13174) CsomeBCy
03185 -pf04607 -
03190 -pf13303 -
03195 -# -
03200 -# -
03210 -pf00579 -
03215 -pf01569 -
03220 -pf03009 -
03225 -pf00781 -
(..3440..)
06665 -pf00679pf00009pf03144 -
06670 -pf00436 -
06675 -pf05173pf01113 -
06680 -pf06874 -
06685 -pf00140pf04542pf04539pf04545 -
06690 -# -
06695 -pf00977 -
06700 -pf02810pf07517pf01043pf07516 -
06705 -pf03462pf00472 -
06710 -pf00742pf03447 -
06715 -pf01928 -
06720 -pf01965 -
06725 -pf00254pf05698pf05697 -
06730 -pf00574 -
06735 -pf07724pf10431pf06689 -
06740 -pf00004pf05362pf02190 -
06745 -pf01926 -
06750 -pf01594 -
06755 -# -
06760 Hp_euts_bluegreen GRM5like
06765 Hp_euts_frenchBlue GRM5like
06770 HMM10662PduV_1_mix(pf10662) GRM5
06775 -pf06161 -
06780 -pf12802 -
06785 -pf00664pf00005 -
06790 -pf00664pf00005 -
06795 -pf01381 -
06800 -pf12844 -
06805 -pf00694 -
06810 -pf00330 -
06815 -pf00694 -
06825 -pf04754 -
06830 -pf01964 -
06835 -pf09512 -
06840 HMMreg00392_1_rmm1(pf00392) SPU6
06845 HMMaldol_2_pvm1(pf00596) PVMlike
06850 -pf11762pf02610 -
06855 -pf01113pf16654 -
06860 -pf02782pf00370 -
06865 -pf01522 -
06870 -pf12822 -
06875 H_darkBlueGreen GRM5
06880 H_olive GRM5
06885 -# -
06890 HMM00465alcdh_4_pdu(pf01761) GRM5
06895 -pf00793 -
06900 -pf03788 -
06905 -pf04172 -
06910 -pf00696 -
06915 -pf01765 -
06920 -pf01255 -
06925 -# -
06930 -pf08436pf02670pf13288 -

Ruminococcus sp. AM29-26. Bacteria; Firmicutes; Clostridia; Clostridiales; Ruminococcaceae;

DW768__locus_1_main_sat

GRM5

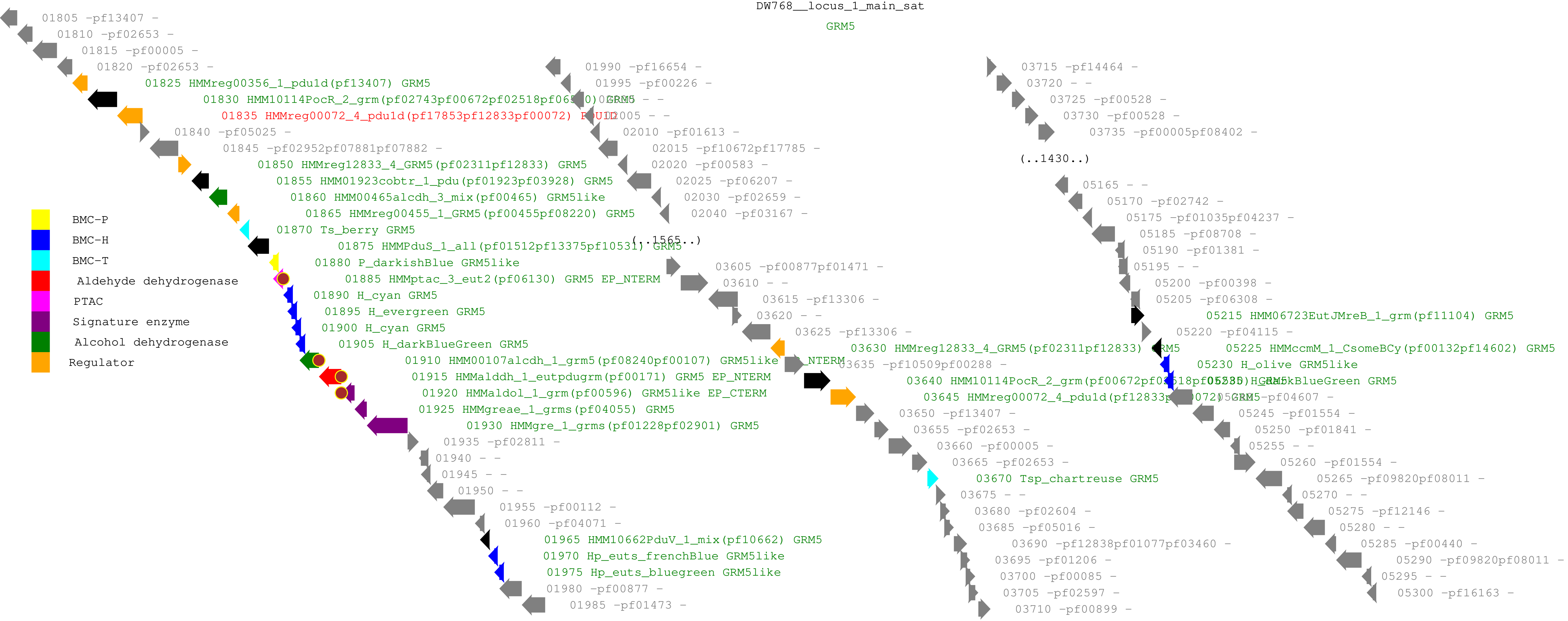

Ruminococcus sp. AM29-5AC. Bacteria; Firmicutes; Clostridia; Clostridiales; Ruminococcaceae;

DW772__locus_1_main_sat

GRM5

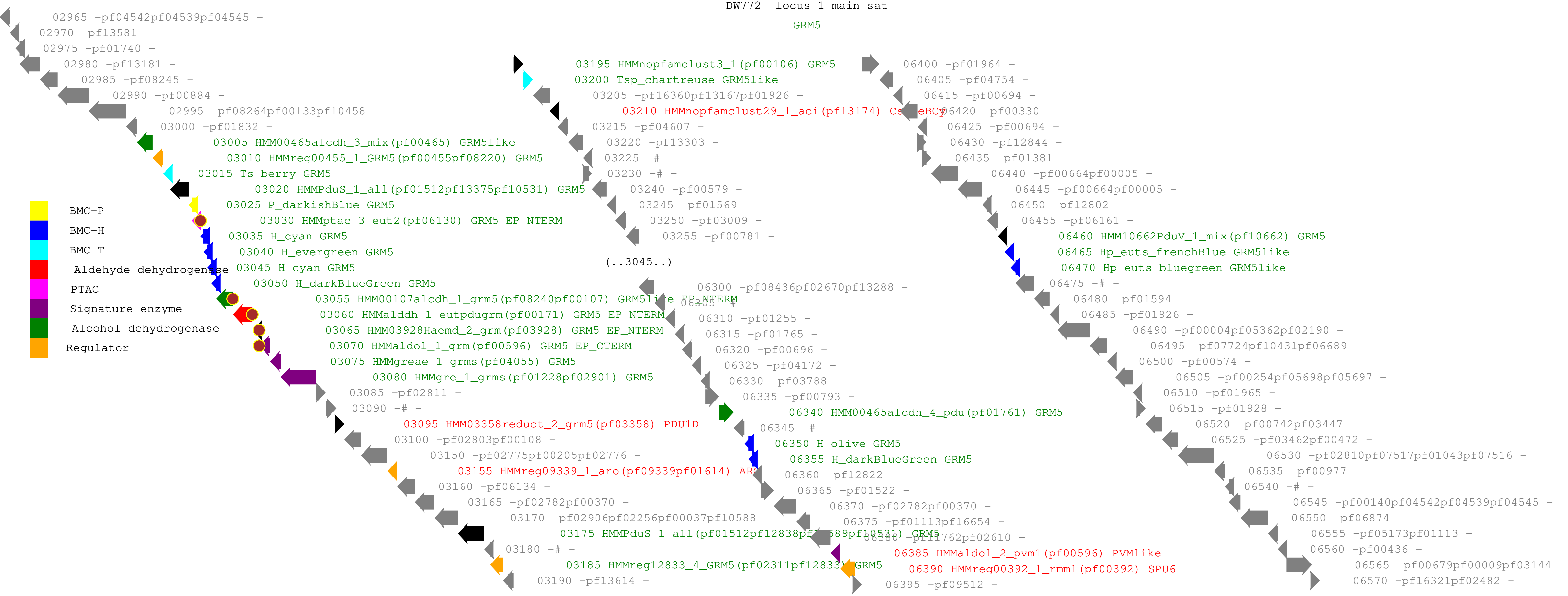

Firmicutes bacterium AM31-12AC. Bacteria; Firmicutes.

DW790__locus_1_main_sat

GRM5

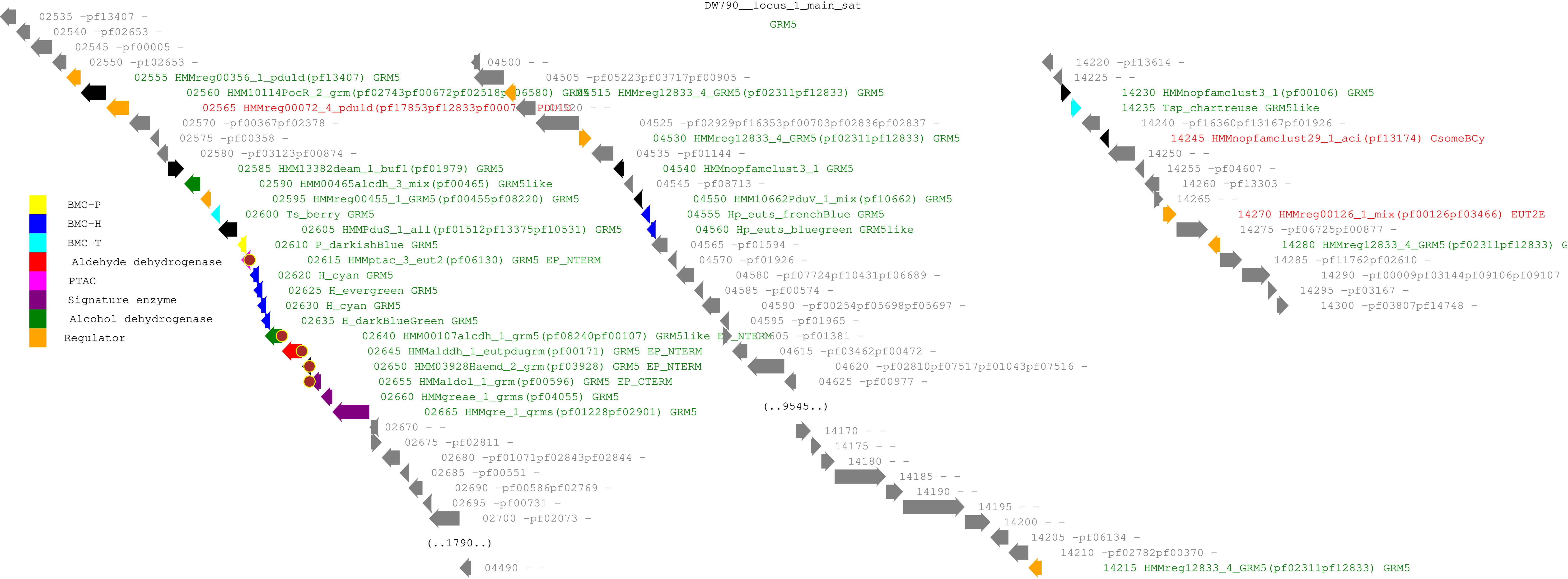

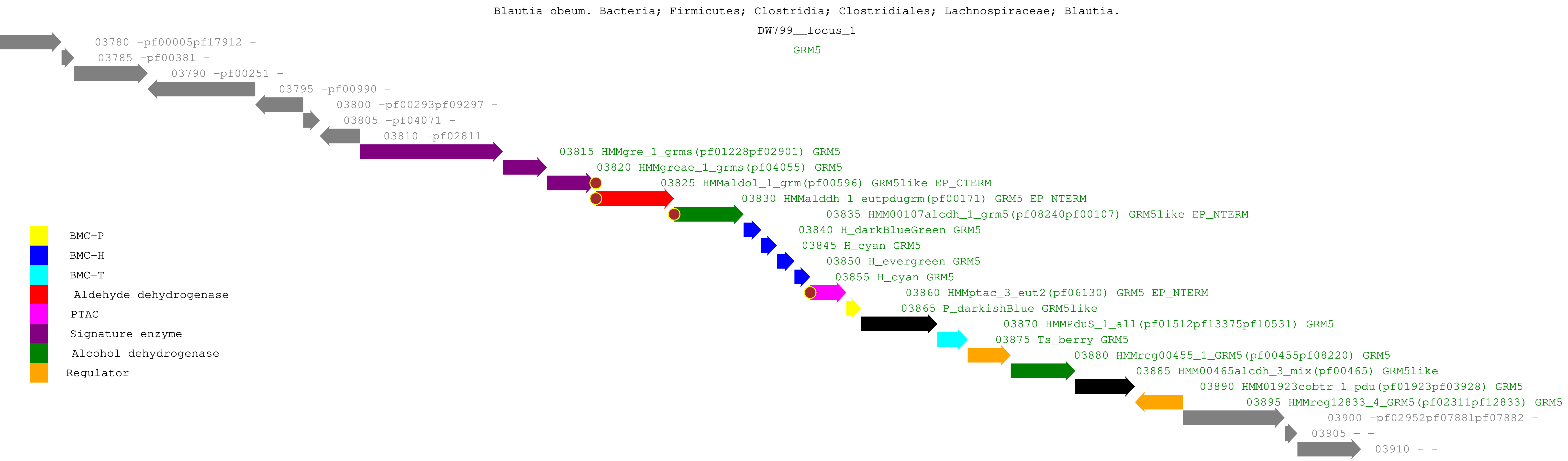

Blautia obeum. Bacteria; Firmicutes; Clostridia; Clostridiales; Lachnospiraceae; Blautia.
DW799__locus_1
GRM5
03780 -pf00005pf17912 -
03785 -pf00381 -
03790 -pf00251 -
03795 -pf00990 -
03800 -pf00293pf09297 -
03805 -pf04071 -
03810 -pf02811 -
03815 HMMgre_1_grms(pf01228pf02901) GRM5
03820 HMMgreae_1_grms(pf04055) GRM5
03825 HMMaldol_1_grm(pf00596) GRM5like EP_CTERM
03830 HMMaldh_1_eutpdugrm(pf00171) GRM5 EP_NTERM
03835 HMM00107alcdh_1_grm5(pf08240pf00107) GRM5like EP_NTERM
03840 H_darkBlueGreen GRM5
03845 H_cyan GRM5
03850 H_evergreen GRM5
03855 H_cyan GRM5
03860 HMMptac_3_eut2(pf06130) GRM5 EP_NTERM
03865 P_darkishBlue GRM5like
03870 HMMPduS_1_all(pf01512pf13375pf10531) GRM5
03875 Ts_berry GRM5
03880 HMMreg00455_1_GRM5(pf00455pf08220) GRM5
03885 HMM00465alcdh_3_mix(pf00465) GRM5like
03890 HMM01923cobtr_1_pdu(pf01923pf03928) GRM5
03895 HMMreg12833_4_GRM5(pf02311pf12833) GRM5
03900 -pf02952pf07881pf07882 -
03905 - -
03910 - -
BMC-P
BMC-H
BMC-T
Aldehyde dehydrogenase
PTAC
Signature enzyme
Alcohol dehydrogenase
Regulator

Ruminococcus sp. AM32-17LB. Bacteria; Firmicutes; Clostridia; Clostridiales; Ruminococcaceae;

DW805__locus_1_main_sat

GRM5

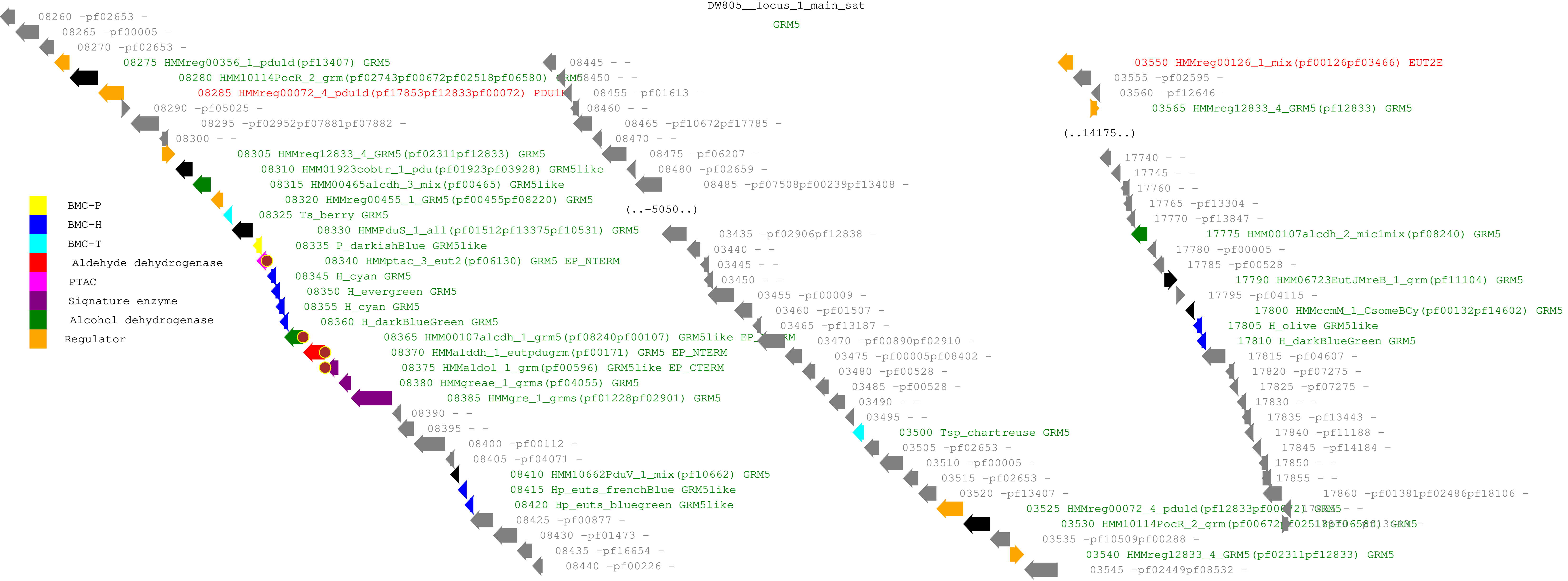

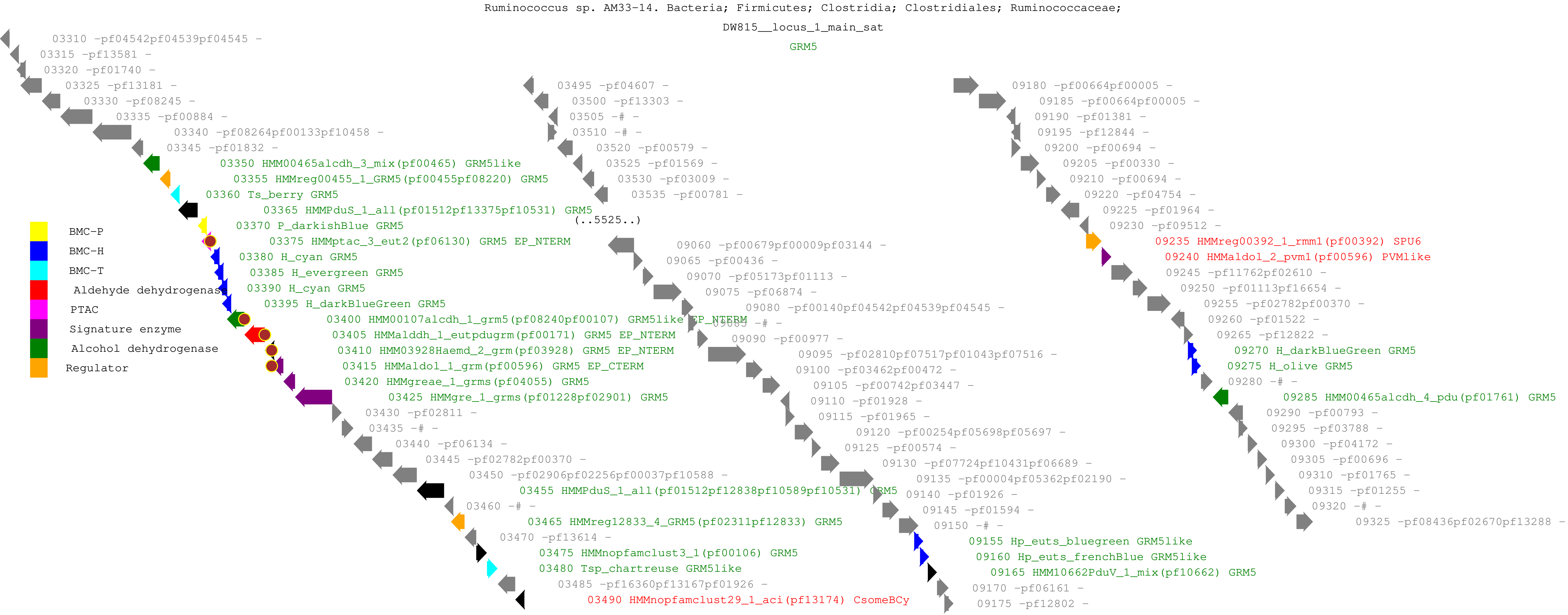
Ruminococcus sp. AM33-14. Bacteria; Firmicutes; Clostridia; Clostridiales; Ruminococcaceae;
DW815__locus_1_main_sat
GRM5
03310 -pf04542pf04539pf04545 -
03315 -pf13581 -
03320 -pf01740 -
03325 -pf13181 -
03330 -pf08245 -
03335 -pf00884 -
03340 -pf08264pf00133pf10458 -
03345 -pf01832 -
03350 HMM00465alcdh_3_mix(pf00465) GRM5like
03355 HMMreg00455_1_GRM5(pf00455pf08220) GRM5
03360 Ts_berry GRM5
03365 HMMPduS_1_all(pf01512pf13375pf10531) GRM5
03370 P_darkishBlue GRM5
03375 HMMptac_3_eut2(pf06130) GRM5 EP_NTERM
03380 H_cyan GRM5
03385 H_evergreen GRM5
03390 H_cyan GRM5
03395 H_darkBlueGreen GRM5
03400 HMM00107alcdh_1_grm5(pf08240pf00107) GRM5like EP_NTERM
03405 HMMaldh_1_eutpdugrm(pf00171) GRM5 EP_NTERM
03410 HMM03928Haemd_2_grm(pf03928) GRM5 EP_NTERM
03415 HMMaldol_1_grm(pf00596) GRM5 EP_CTERM
03420 HMMgreae_1_grms(pf04055) GRM5
03425 HMMgre_1_grms(pf01228pf02901) GRM5
03430 -pf02811 -
03435 -# -
03440 -pf06134 -
03445 -pf02782pf00370 -
03450 -pf02906pf02256pf00037pf10588 -
03455 HMMPduS_1_all(pf01512pf12838pf10589pf10531) GRM5
03460 -# -
03465 HMMreg12833_4_GRM5(pf02311pf12833) GRM5
03470 -pf13614 -
03475 HMMnopfamclust3_1(pf00106) GRM5
03480 Tsp_chartreuse GRM5like
03485 -pf16360pf13167pf01926 -
03490 HMMnopfamclust29_1_aci(pf13174) CsomeBCy
BMC-P
BMC-H
BMC-T
Aldehyde dehydrogenase
PTAC
Signature enzyme
Alcohol dehydrogenase
Regulator
03495 -pf04607 -
03500 -pf13303 -
03505 -# -
03510 -# -
03520 -pf00579 -
03525 -pf01569 -
03530 -pf03009 -
03535 -pf00781 -
(..5525..)
09060 -pf00679pf00009pf03144 -
09065 -pf00436 -
09070 -pf05173pf01113 -
09075 -pf06874 -
09080 -pf00140pf04542pf04539pf04545 -
09085 -# -
09090 -pf00977 -
09095 -pf02810pf07517pf01043pf07516 -
09100 -pf03462pf00472 -
09105 -pf00742pf03447 -
09110 -pf01928 -
09115 -pf01965 -
09120 -pf00254pf05698pf05697 -
09125 -pf00574 -
09130 -pf07724pf10431pf06689 -
09135 -pf00004pf05362pf02190 -
09140 -pf01926 -
09145 -pf01594 -
09150 -# -
09155 Hp_euts_bluegreen GRM5like
09160 Hp_euts_frenchBlue GRM5like
09165 HMM10662PduV_1_mix(pf10662) GRM5
09170 -pf06161 -
09175 -pf12802 -
09180 -pf00664pf00005 -
09185 -pf00664pf00005 -
09190 -pf01381 -
09195 -pf12844 -
09200 -pf00694 -
09205 -pf00330 -
09210 -pf00694 -
09220 -pf04754 -
09225 -pf01964 -
09230 -pf09512 -
09235 HMMreg00392_1_rmm1(pf00392) SPU6
09240 HMMaldol_2_pvm1(pf00596) PVMlike
09245 -pf11762pf02610 -
09250 -pf01113pf16654 -
09255 -pf02782pf00370 -
09260 -pf01522 -
09265 -pf12822 -
09270 H_darkBlueGreen GRM5
09275 H_olive GRM5
09280 -# -
09285 HMM00465alcdh_4_pdu(pf01761) GRM5
09290 -pf00793 -
09295 -pf03788 -
09300 -pf04172 -
09305 -pf00696 -
09310 -pf01765 -
09315 -pf01255 -
09320 -# -
09325 -pf08436pf02670pf13288 -

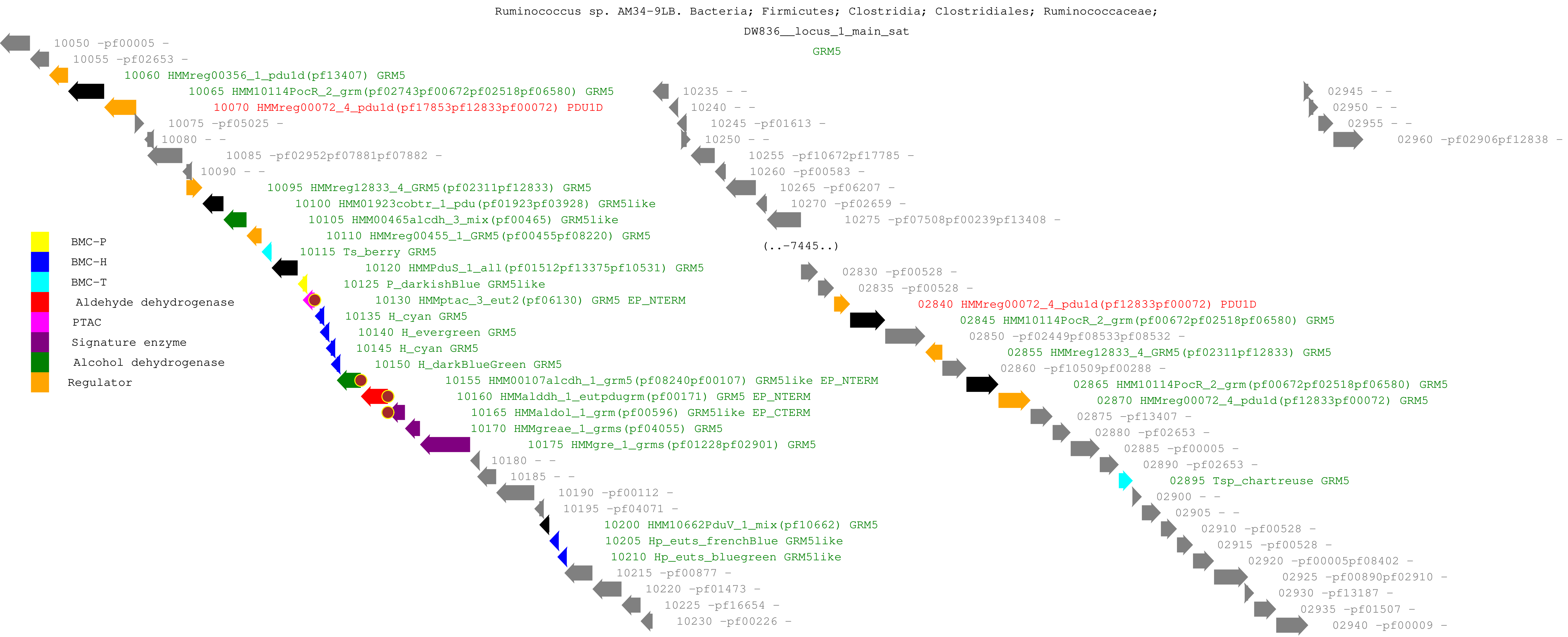
Ruminococcus sp. AM34-9LB. Bacteria; Firmicutes; Clostridia; Clostridiales; Ruminococcaceae;
DW836__locus_1_main_sat
GRM5
10050 -pf00005 -
10055 -pf02653 -
10060 HMMreg00356_1_pdu1d(pf13407) GRM5
10065 HMM10114PocR_2_grm(pf02743pf00672pf02518pf06580) GRM5
10070 HMMreg00072_4_pdu1d(pf17853pf12833pf00072) PDU1D
10075 -pf05025 -
10080 - -
10085 -pf02952pf07881pf07882 -
10090 - -
10095 HMMreg12833_4_GRM5(pf02311pf12833) GRM5
10100 HMM01923cobtr_1_pdu(pf01923pf03928) GRM5like
10105 HMM00465alcdh_3_mix(pf00465) GRM5like
10110 HMMreg00455_1_GRM5(pf00455pf08220) GRM5
10115 Ts_berry GRM5
10120 HMMPduS_1_all(pf01512pf13375pf10531) GRM5
10125 P_darkishBlue GRM5like
10130 HMMptac_3_eut2(pf06130) GRM5 EP_NTERM
10135 H_cyan GRM5
10140 H_evergreen GRM5
10145 H_cyan GRM5
10150 H_darkBlueGreen GRM5
10155 HMM00107alcdh_1_grm5(pf08240pf00107) GRM5like EP_NTERM
10160 HMMaldddh_1_eutpdugrm(pf00171) GRM5 EP_NTERM
10165 HMMaldol_1_grm(pf00596) GRM5like EP_CTERM
10170 HMMgreae_1_grms(pf04055) GRM5
10175 HMMgre_1_grms(pf01228pf02901) GRM5
10180 - -
10185 - -
10190 -pf00112 -
10195 -pf04071 -
10200 HMM10662PduV_1_mix(pf10662) GRM5
10205 Hp_euts_frenchBlue GRM5like
10210 Hp_euts_bluegreen GRM5like
10215 -pf00877 -
10220 -pf01473 -
10225 -pf16654 -
10230 -pf00226 -
10235 - -
10240 - -
10245 -pf01613 -
10250 - -
10255 -pf10672pf17785 -
10260 -pf00583 -
10265 -pf06207 -
10270 -pf02659 -
10275 -pf07508pf00239pf13408 -
(..-7445..)
02830 -pf00528 -
02835 -pf00528 -
02840 HMMreg00072_4_pdu1d(pf12833pf00072) PDU1D
02845 HMM10114PocR_2_grm(pf00672pf02518pf06580) GRM5
02850 -pf02449pf08533pf08532 -
02855 HMMreg12833_4_GRM5(pf02311pf12833) GRM5
02860 -pf10509pf00288 -
02865 HMM10114PocR_2_grm(pf00672pf02518pf06580) GRM5
02870 HMMreg00072_4_pdu1d(pf12833pf00072) GRM5
02875 -pf13407 -
02880 -pf02653 -
02885 -pf00005 -
02890 -pf02653 -
02895 Tsp_chartreuse GRM5
02900 - -
02905 - -
02910 -pf00528 -
02915 -pf00528 -
02920 -pf00005pf08402 -
02925 -pf00890pf02910 -
02930 -pf13187 -
02935 -pf01507 -
02940 -pf00009 -
02945 - -
02950 - -
02955 - -
02960 -pf02906pf12838 -
BMC-P
BMC-H
BMC-T
Aldehyde dehydrogenase
PTAC
Signature enzyme
Alcohol dehydrogenase
Regulator

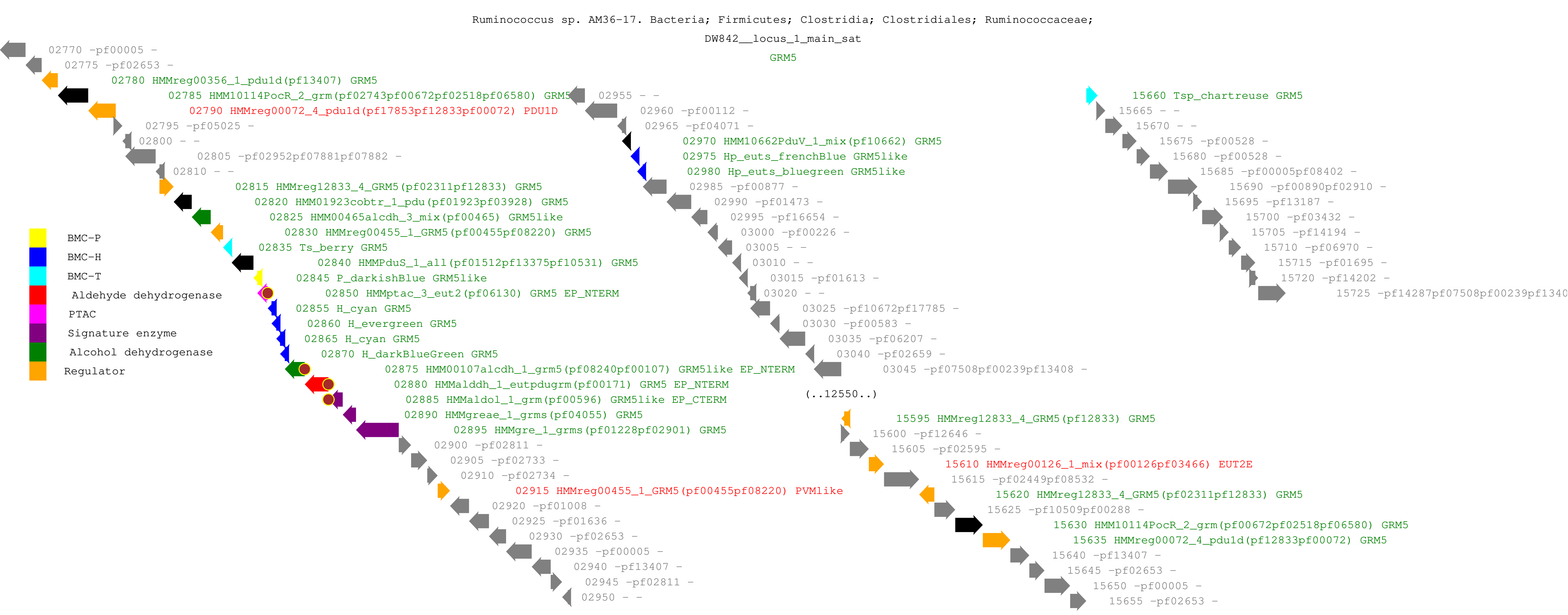
Ruminococcus sp. AM36-17. Bacteria; Firmicutes; Clostridia; Clostridiales; Ruminococcaceae;
DW842__locus_1_main_sat
GRM5
02770 -pf00005 -
02775 -pf02653 -
02780 HMMreg00356_1_pdu1d(pf13407) GRM5
02785 HMM10114PocR_2_grm(pf02743pf00672pf02518pf06580) GRM5
02790 HMMreg00072_4_pdu1d(pf17853pf12833pf00072) PDU1D
02795 -pf05025 -
02800 - -
02805 -pf02952pf07881pf07882 -
02810 - -
02815 HMMreg12833_4_GRM5(pf02311pf12833) GRM5
02820 HMM01923cobtr_1_pdu(pf01923pf03928) GRM5
02825 HMM00465alcdh_3_mix(pf00465) GRM5like
02830 HMMreg00455_1_GRM5(pf00455pf08220) GRM5
02835 Ts_berry GRM5
02840 HMMPduS_1_all(pf01512pf13375pf10531) GRM5
02845 P_darkishBlue GRM5like
02850 HMMptac_3_eut2(pf06130) GRM5 EP_NTERM
02855 H_cyan GRM5
02860 H_evergreen GRM5
02865 H_cyan GRM5
02870 H_darkBlueGreen GRM5
02875 HMM00107alcdh_1_grm5(pf08240pf00107) GRM5like EP_NTERM
02880 HMMaldh_1_eutpdugrm(pf00171) GRM5 EP_NTERM
02885 HMMaldol_1_grm(pf00596) GRM5like EP_CTERM
02890 HMMgreae_1_grms(pf04055) GRM5
02895 HMMgre_1_grms(pf01228pf02901) GRM5
02900 -pf02811 -
02905 -pf02733 -
02910 -pf02734 -
02915 HMMreg00455_1_GRM5(pf00455pf08220) PVMlike
02920 -pf01008 -
02925 -pf01636 -
02930 -pf02653 -
02935 -pf00005 -
02940 -pf13407 -
02945 -pf02811 -
02950 - -
02955 - -
02960 -pf00112 -
02965 -pf04071 -
02970 HMM10662PduV_1_mix(pf10662) GRM5
02975 Hp_euts_frenchBlue GRM5like
02980 Hp_euts_bluegreen GRM5like
02985 -pf00877 -
02990 -pf01473 -
02995 -pf16654 -
03000 -pf00226 -
03005 - -
03010 - -
03015 -pf01613 -
03020 - -
03025 -pf10672pf17785 -
03030 -pf00583 -
03035 -pf06207 -
03040 -pf02659 -
03045 -pf07508pf00239pf13408 -
(..12550..)
15595 HMMreg12833_4_GRM5(pf12833) GRM5
15600 -pf12646 -
15605 -pf02595 -
15610 HMMreg00126_1_mix(pf00126pf03466) EUT2E
15615 -pf02449pf08532 -
15620 HMMreg12833_4_GRM5(pf02311pf12833) GRM5
15625 -pf10509pf00288 -
15630 HMM10114PocR_2_grm(pf00672pf02518pf06580) GRM5
15635 HMMreg00072_4_pdu1d(pf12833pf00072) GRM5
15640 -pf13407 -
15645 -pf02653 -
15650 -pf00005 -
15655 -pf02653 -
15660 Tsp_chartreuse GRM5
15665 - -
15670 - -
15675 -pf00528 -
15680 -pf00528 -
15685 -pf00005pf08402 -
15690 -pf00890pf02910 -
15695 -pf13187 -
15700 -pf03432 -
15705 -pf14194 -
15710 -pf06970 -
15715 -pf01695 -
15720 -pf14202 -
15725 -pf14287pf07508pf00239pf1340
BMC-P
BMC-H
BMC-T
Aldehyde dehydrogenase
PTAC
Signature enzyme
Alcohol dehydrogenase
Regulator

Ruminococcus sp. AM36-2AA. Bacteria; Firmicutes; Clostridia; Clostridiales; Ruminococcaceae;

DW846__locus_1_main_sat

GRM5

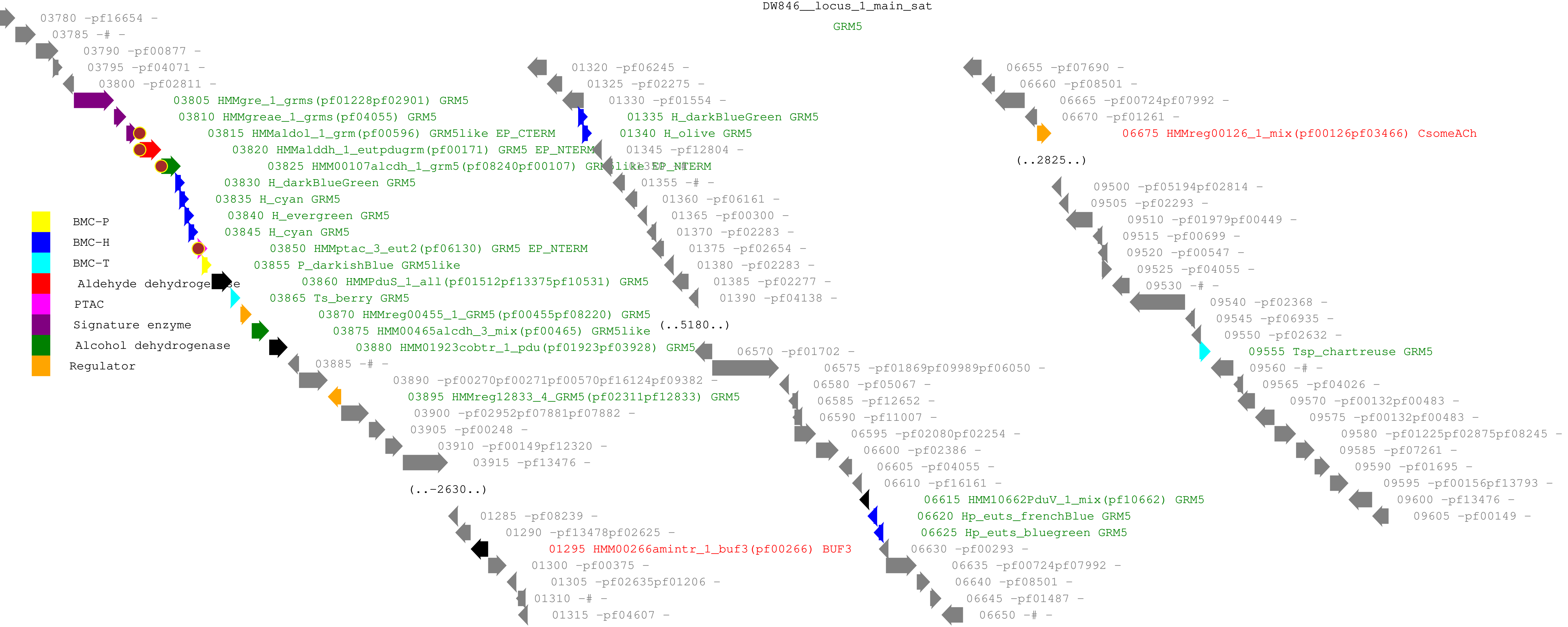

Ruminococcus sp. AM36-5. Bacteria; Firmicutes; Clostridia; Clostridiales; Ruminococcaceae;

DW851__locus_1_main_sat

GRM5

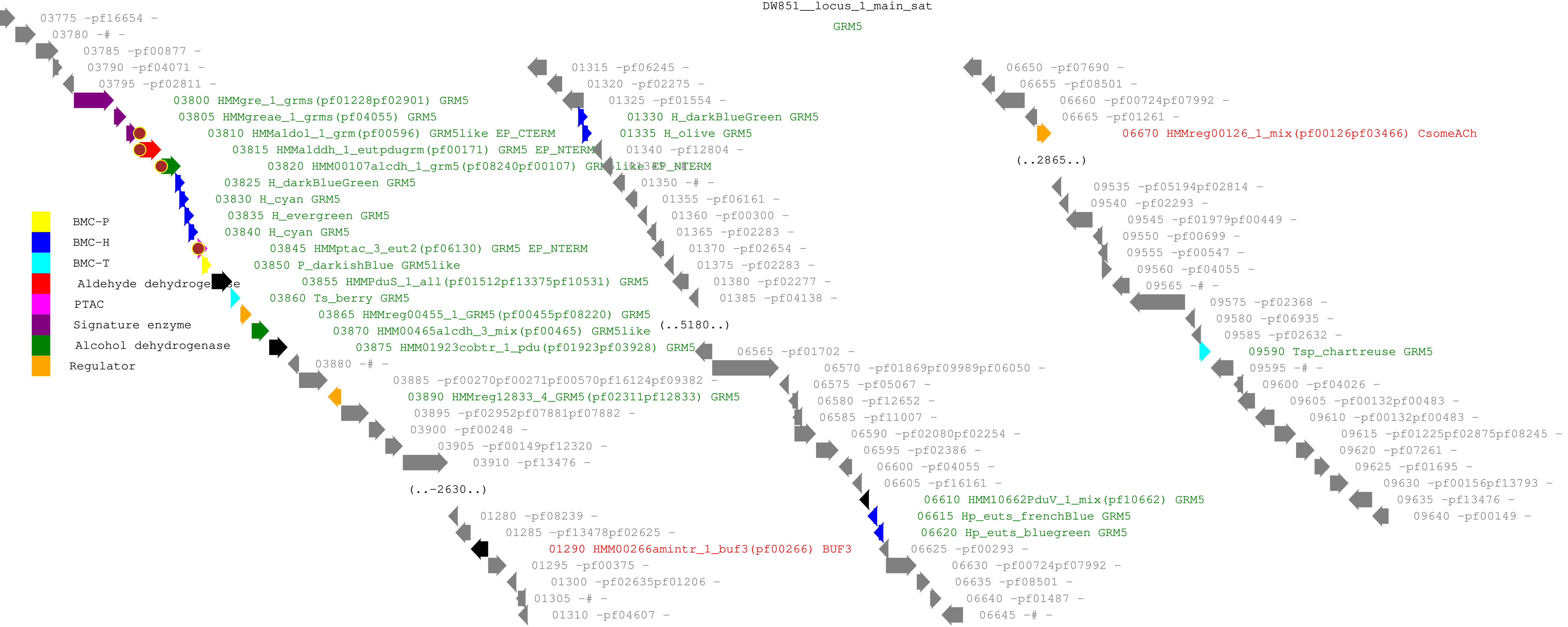

Ruminococcus sp. AM40-10AC. Bacteria; Firmicutes; Clostridia; Clostridiales; Ruminococcaceae;

DW884__locus_1_main_sat

GRM5

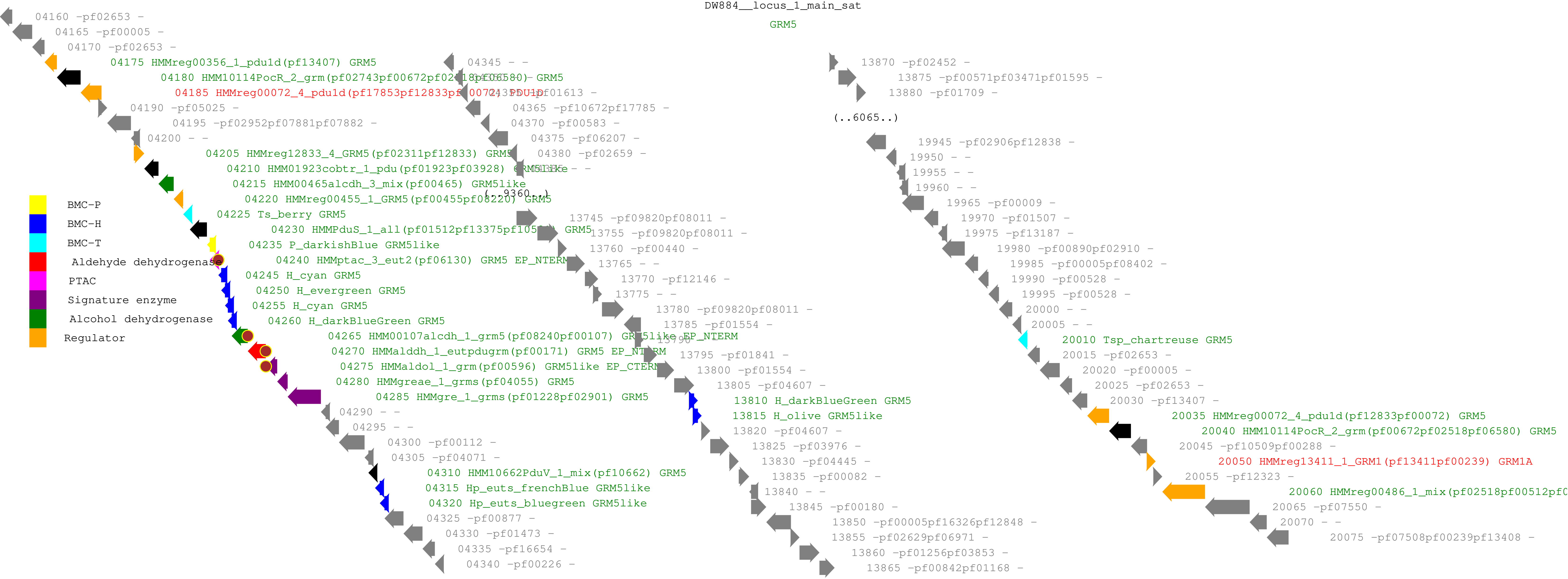

Lachnospiraceae bacterium AM40-2BH. Bacteria; Firmicutes; Clostridia; Clostridiales; Lachnospiraceae.

DW887__locus_1_main_sat

GRM5

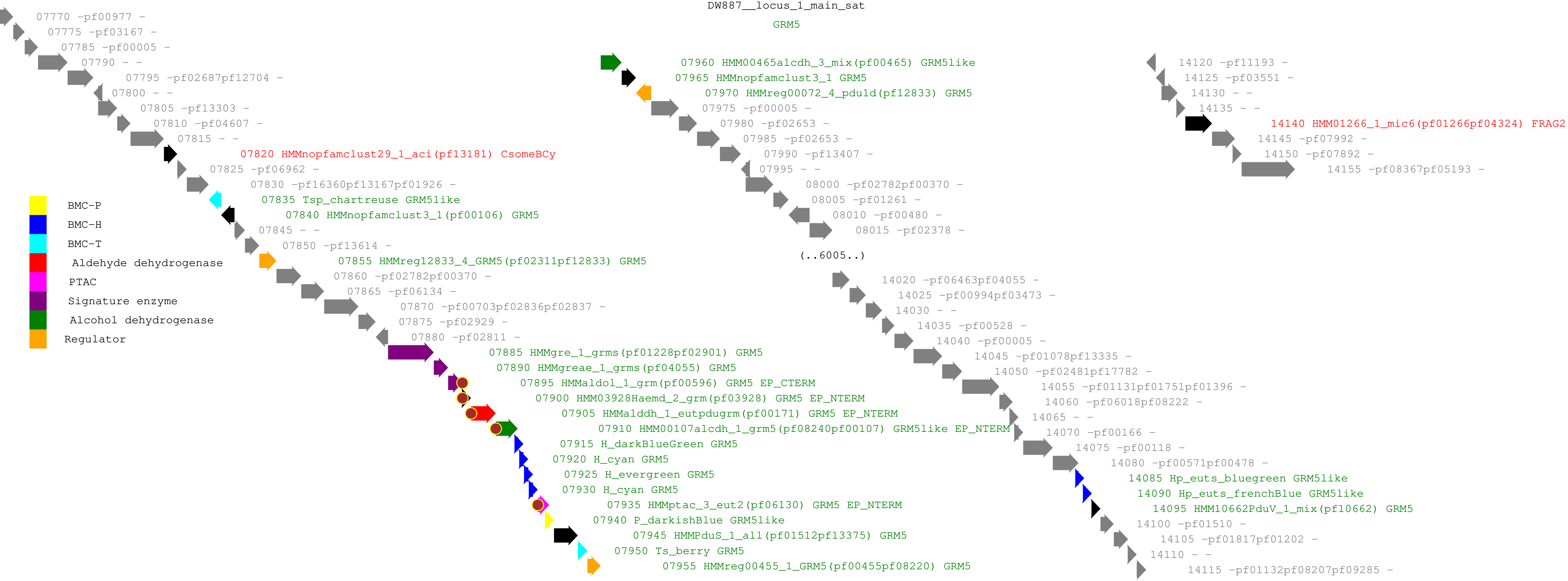

Ruminococcus sp. AM41-10BH. Bacteria; Firmicutes; Clostridia; Clostridiales; Ruminococcaceae;

DW894__locus_1_main_sat

GRM5

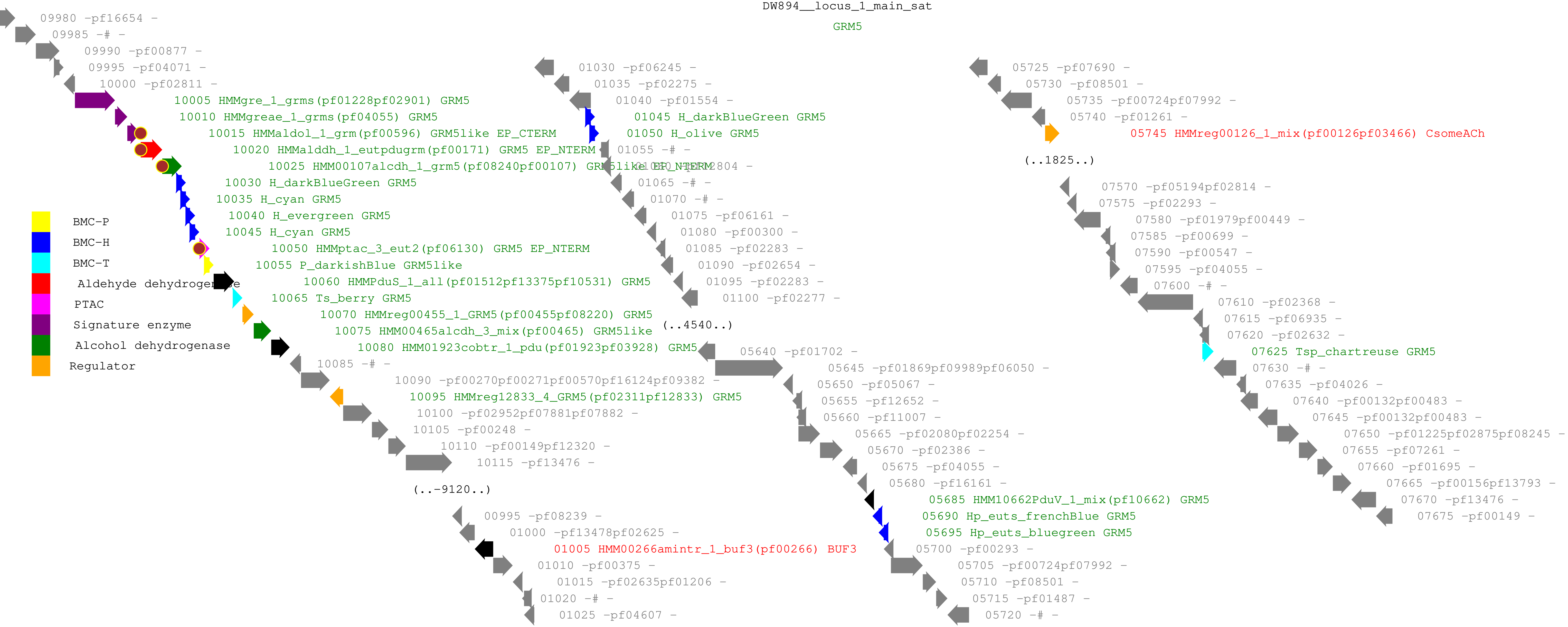

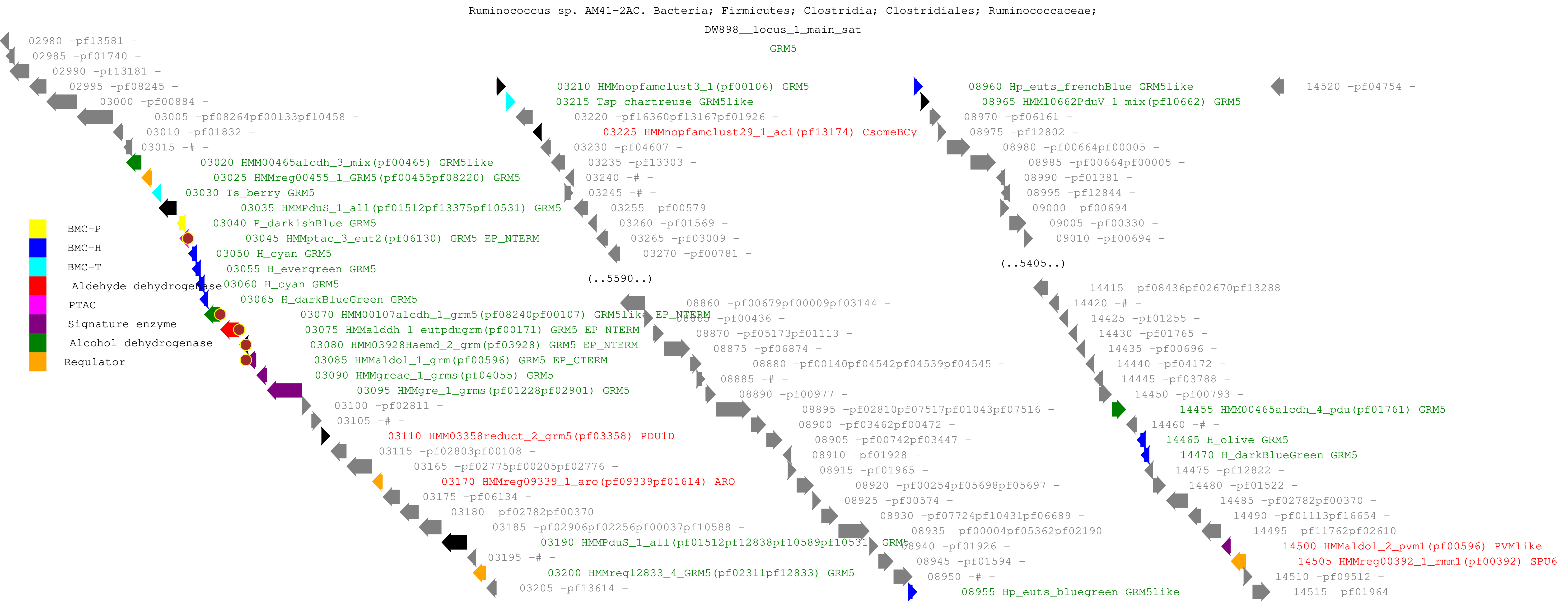

Ruminococcus sp. AM41-2AC. Bacteria; Firmicutes; Clostridia; Clostridiales; Ruminococcaceae;
DW898__locus_1_main_sat
GRM5
02980 -pf13581 -
02985 -pf01740 -
02990 -pf13181 -
02995 -pf08245 -
03000 -pf00884 -
03005 -pf08264pf00133pf10458 -
03010 -pf01832 -
03015 -# -
03020 HMM00465alcdh_3_mix(pf00465) GRM5like
03025 HMMreg00455_1_GRM5(pf00455pf08220) GRM5
03030 Ts_berry GRM5
03035 HMMPduS_1_all(pf01512pf13375pf10531) GRM5
03040 P_darkishBlue GRM5
03045 HMMptac_3_eut2(pf06130) GRM5 EP_NTERM
03050 H_cyan GRM5
03055 H_evergreen GRM5
03060 H_cyan GRM5
03065 H_darkBlueGreen GRM5
03070 HMM00107alcdh_1_grm5(pf08240pf00107) GRM5like EP_NTERM
03075 HMMalddh_1_eutpdugrm(pf00171) GRM5 EP_NTERM
03080 HMM03928Haemd_2_grm(pf03928) GRM5 EP_NTERM
03085 HMMaldol_1_grm(pf00596) GRM5 EP_CTERM
03090 HMMgreae_1_grms(pf04055) GRM5
03095 HMMgre_1_grms(pf01228pf02901) GRM5
03100 -pf02811 -
03105 -# -
03110 HMM03358reduct_2_grm5(pf03358) PDU1D
03115 -pf02803pf00108 -
03165 -pf02775pf00205pf02776 -
03170 HMMreg09339_1_aro(pf09339pf01614) ARO
03175 -pf06134 -
03180 -pf02782pf00370 -
03185 -pf02906pf02256pf00037pf10588 -
03190 HMMPduS_1_all(pf01512pf12838pf10589pf10531) GRM5
03195 -# -
03200 HMMreg12833_4_GRM5(pf02311pf12833) GRM5
03205 -pf13614 -
BMC-P
BMC-H
BMC-T
Aldehyde dehydrogenase
PTAC
Signature enzyme
Alcohol dehydrogenase
Regulator
03210 HMMnopfamclust3_1(pf00106) GRM5
03215 Tsp_chartreuse GRM5like
03220 -pf16360pf13167pf01926 -
03225 HMMnopfamclust29_1_aci(pf13174) CsomeBCy
03230 -pf04607 -
03235 -pf13303 -
03240 -# -
03245 -# -
03255 -pf00579 -
03260 -pf01569 -
03265 -pf03009 -
03270 -pf00781 -
(..5590..)
08860 -pf00679pf00009pf03144 -
08865 -pf00436 -
08870 -pf05173pf01113 -
08875 -pf06874 -
08880 -pf00140pf04542pf04539pf04545 -
08885 -# -
08890 -pf00977 -
08895 -pf02810pf07517pf01043pf07516 -
08900 -pf03462pf00472 -
08905 -pf00742pf03447 -
08910 -pf01928 -
08915 -pf01965 -
08920 -pf00254pf05698pf05697 -
08925 -pf00574 -
08930 -pf07724pf10431pf06689 -
08935 -pf00004pf05362pf02190 -
08940 -pf01926 -
08945 -pf01594 -
08950 -# -
08955 Hp_euts_bluegreen GRM5like
08960 Hp_euts_frenchBlue GRM5like
08965 HMM10662PduV_1_mix(pf10662) GRM5
08970 -pf06161 -
08975 -pf12802 -
08980 -pf00664pf00005 -
08985 -pf00664pf00005 -
08990 -pf01381 -
08995 -pf12844 -
09000 -pf00694 -
09005 -pf00330 -
09010 -pf00694 -
(..5405..)
14415 -pf08436pf02670pf13288 -
14420 -# -
14425 -pf01255 -
14430 -pf01765 -
14435 -pf00696 -
14440 -pf04172 -
14445 -pf03788 -
14450 -pf00793 -
14455 HMM00465alcdh_4_pdu(pf01761) GRM5
14460 -# -
14465 H_olive GRM5
14470 H_darkBlueGreen GRM5
14475 -pf12822 -
14480 -pf01522 -
14485 -pf02782pf00370 -
14490 -pf01113pf16654 -
14495 -pf11762pf02610 -
14500 HMMaldol_2_pvm1(pf00596) PVMlike
14505 HMMreg00392_1_rmm1(pf00392) SPU6
14510 -pf09512 -
14515 -pf01964 -
14520 -pf04754 -

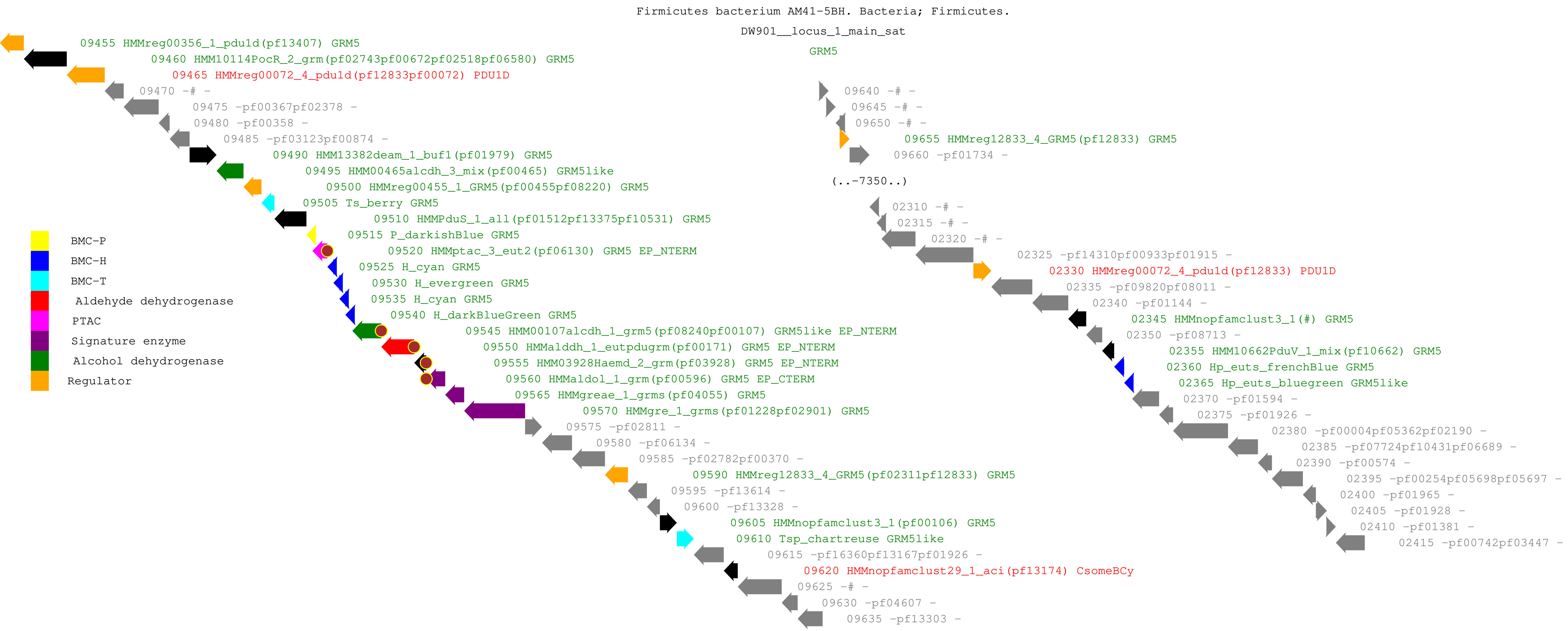
Firmicutes bacterium AM41-5BH. Bacteria; Firmicutes.
DW901__locus_1_main_sat
GRM5
09455 HMMreg00356_1_pdu1d(pf13407) GRM5
09460 HMM10114PocR_2_grm(pf02743pf00672pf02518pf06580) GRM5
09465 HMMreg00072_4_pdu1d(pf12833pf00072) PDU1D
09470 -# -
09475 -pf00367pf02378 -
09480 -pf00358 -
09485 -pf03123pf00874 -
09490 HMM13382deam_1_buf1(pf01979) GRM5
09495 HMM00465alcdh_3_mix(pf00465) GRM5like
09500 HMMreg00455_1_GRM5(pf00455pf08220) GRM5
09505 Ts_berry GRM5
09510 HMMPduS_1_all(pf01512pf13375pf10531) GRM5
09515 P_darkishBlue GRM5
09520 HMMptac_3_eut2(pf06130) GRM5 EP_NTERM
09525 H_cyan GRM5
09530 H_evergreen GRM5
09535 H_cyan GRM5
09540 H_darkBlueGreen GRM5
09545 HMM00107alcdh_1_grm5(pf08240pf00107) GRM5like EP_NTERM
09550 HMMalddh_1_eutpdugrm(pf00171) GRM5 EP_NTERM
09555 HMM03928Haemd_2_grm(pf03928) GRM5 EP_NTERM
09560 HMMaldol_1_grm(pf00596) GRM5 EP_CTERM
09565 HMMgreae_1_grms(pf04055) GRM5
09570 HMMgre_1_grms(pf01228pf02901) GRM5
09575 -pf02811 -
09580 -pf06134 -
09585 -pf02782pf00370 -
09590 HMMreg12833_4_GRM5(pf02311pf12833) GRM5
09595 -pf13614 -
09600 -pf13328 -
09605 HMMnopfamclust3_1(pf00106) GRM5
09610 Tsp_chartreuse GRM5like
09615 -pf16360pf13167pf01926 -
09620 HMMnopfamclust29_1_aci(pf13174) CsomeBCy
09625 -# -
09630 -pf04607 -
09635 -pf13303 -
BMC-P
BMC-H
BMC-T
Aldehyde dehydrogenase
PTAC
Signature enzyme
Alcohol dehydrogenase
Regulator
09640 -# -
09645 -# -
09650 -# -
09655 HMMreg12833_4_GRM5(pf12833) GRM5
09660 -pf01734 -
(..-7350..)
02310 -# -
02315 -# -
02320 -# -
02325 -pf14310pf00933pf01915 -
02330 HMMreg00072_4_pdu1d(pf12833) PDU1D
02335 -pf09820pf08011 -
02340 -pf01144 -
02345 HMMnopfamclust3_1(#) GRM5
02350 -pf08713 -
02355 HMM10662PduV_1_mix(pf10662) GRM5
02360 Hp_euts_frenchBlue GRM5
02365 Hp_euts_bluegreen GRM5like
02370 -pf01594 -
02375 -pf01926 -
02380 -pf00004pf05362pf02190 -
02385 -pf07724pf10431pf06689 -
02390 -pf00574 -
02395 -pf00254pf05698pf05697 -
02400 -pf01965 -
02405 -pf01928 -
02410 -pf01381 -
02415 -pf00742pf03447 -

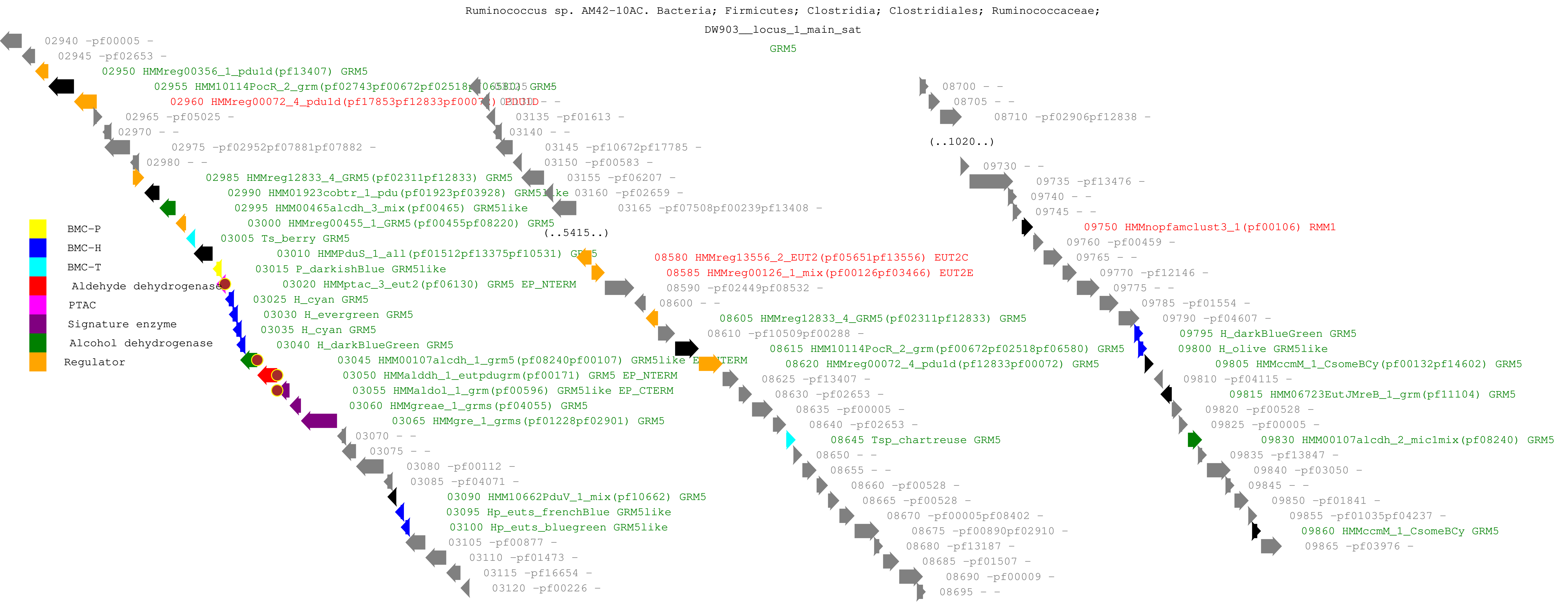

Ruminococcus sp. AM42-10AC. Bacteria; Firmicutes; Clostridia; Clostridiales; Ruminococcaceae;
DW903__locus_1_main_sat
GRM5
BMC-P
BMC-H
BMC-T
Aldehyde dehydrogenase
PTAC
Signature enzyme
Alcohol dehydrogenase
Regulator
02940 -pf00005 -
02945 -pf02653 -
02950 HMMreg00356_1_pduld(pf13407) GRM5
02955 HMM10114PocR_2_grm(pf02743pf00672pf02518pf06580) GRM5
02960 HMMreg00072_4_pduld(pf17853pf12833pf00072) PDUD
02965 -pf05025 -
02970 - -
02975 -pf02952pf07881pf07882 -
02980 - -
02985 HMMreg12833_4_GRM5(pf02311pf12833) GRM5
02990 HMM01923cobtr_1_pdu(pf01923pf03928) GRM5like
02995 HMM00465alcdh_3_mix(pf00465) GRM5like
03000 HMMreg00455_1_GRM5(pf00455pf08220) GRM5
03005 Ts_berry GRM5
03010 HMMPduS_1_all(pf01512pf13375pf10531) GRM5
03015 P_darkishBlue GRM5like
03020 HMMptac_3_eut2(pf06130) GRM5 EP_NTERM
03025 H_cyan GRM5
03030 H_evergreen GRM5
03035 H_cyan GRM5
03040 H_darkBlueGreen GRM5
03045 HMM00107alcdh_1_grm5(pf08240pf00107) GRM5like EP_NTERM
03050 HMMaldh_1_eutpdugrm(pf00171) GRM5 EP_NTERM
03055 HMMaldol_1_grm(pf00596) GRM5like EP_CTERM
03060 HMMgreae_1_grms(pf04055) GRM5
03065 HMMgre_1_grms(pf01228pf02901) GRM5
03070 - -
03075 - -
03080 -pf00112 -
03085 -pf04071 -
03090 HMM10662PduV_1_mix(pf10662) GRM5
03095 Hp_euts_frenchBlue GRM5like
03100 Hp_euts_bluegreen GRM5like
03105 -pf00877 -
03110 -pf01473 -
03115 -pf16654 -
03120 -pf00226 -
03125 GRM5
03130 - -
03135 -pf01613 -
03140 - -
03145 -pf10672pf17785 -
03150 -pf00583 -
03155 -pf06207 -
03160 -pf02659 -
03165 -pf07508pf00239pf13408 -
(..5415..)
08580 HMMreg13556_2_EUT2(pf05651pf13556) EUT2C
08585 HMMreg00126_1_mix(pf00126pf03466) EUT2E
08590 -pf02449pf08532 -
08600 - -
08605 HMMreg12833_4_GRM5(pf02311pf12833) GRM5
08610 -pf10509pf00288 -
08615 HMM10114PocR_2_grm(pf00672pf02518pf06580) GRM5
08620 HMMreg00072_4_pduld(pf12833pf00072) GRM5
08625 -pf13407 -
08630 -pf02653 -
08635 -pf00005 -
08640 -pf02653 -
08645 Tsp_chartreuse GRM5
08650 - -
08655 - -
08660 -pf00528 -
08665 -pf00528 -
08670 -pf00005pf08402 -
08675 -pf00890pf02910 -
08680 -pf13187 -
08685 -pf01507 -
08690 -pf00009 -
08695 - -
08700 - -
08705 - -
08710 -pf02906pf12838 -
(..1020..)
09730 - -
09735 -pf13476 -
09740 - -
09745 - -
09750 HMMnopfamclust3_1(pf00106) RMM1
09760 -pf00459 -
09765 - -
09770 -pf12146 -
09775 - -
09785 -pf01554 -
09790 -pf04607 -
09795 H_darkBlueGreen GRM5
09800 H_olive GRM5like
09805 HMMccmM_1_CsomeBCy(pf00132pf14602) GRM5
09810 -pf04115 -
09815 HMM06723EutJMreB_1_grm(pf11104) GRM5
09820 -pf00528 -
09825 -pf00005 -
09830 HMM00107alcdh_2_mic1mix(pf08240) GRM5
09835 -pf13847 -
09840 -pf03050 -
09845 - -
09850 -pf01841 -
09855 -pf01035pf04237 -
09860 HMMccmM_1_CsomeBCy GRM5
09865 -pf03976 -

Firmicutes bacterium AM43-11BH. Bacteria; Firmicutes.

DW928__locus_1_main_sat

GRM5

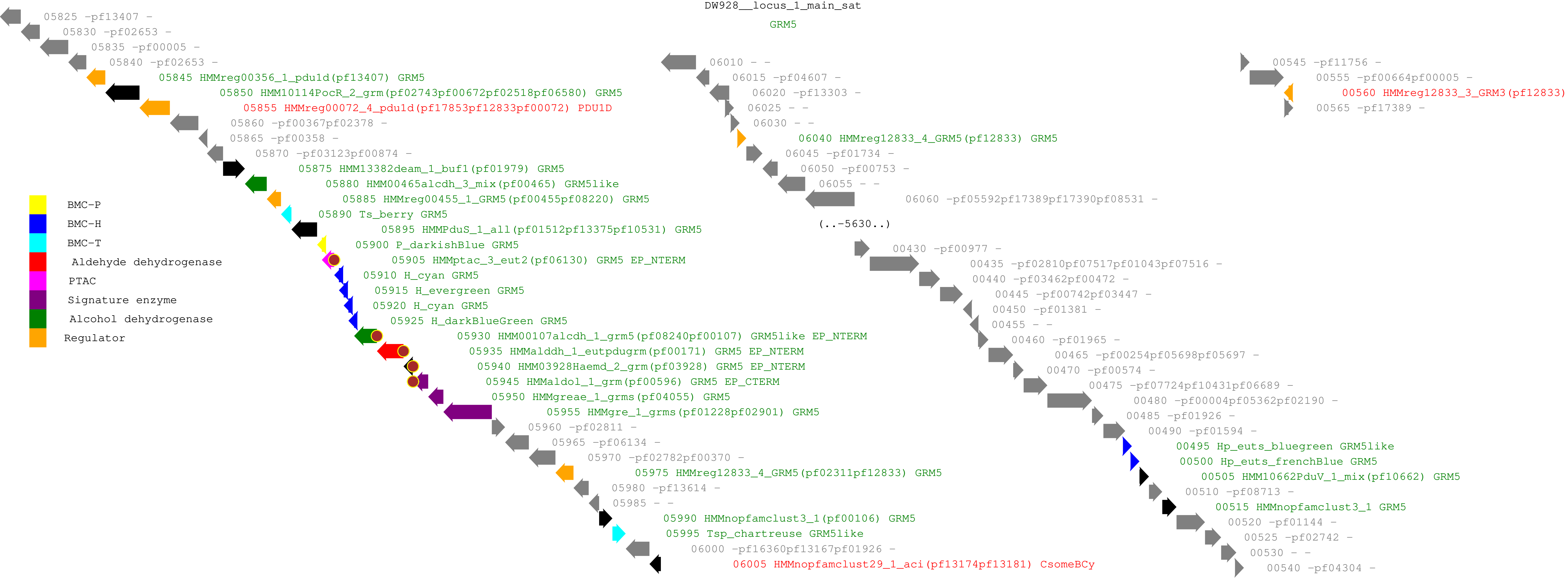

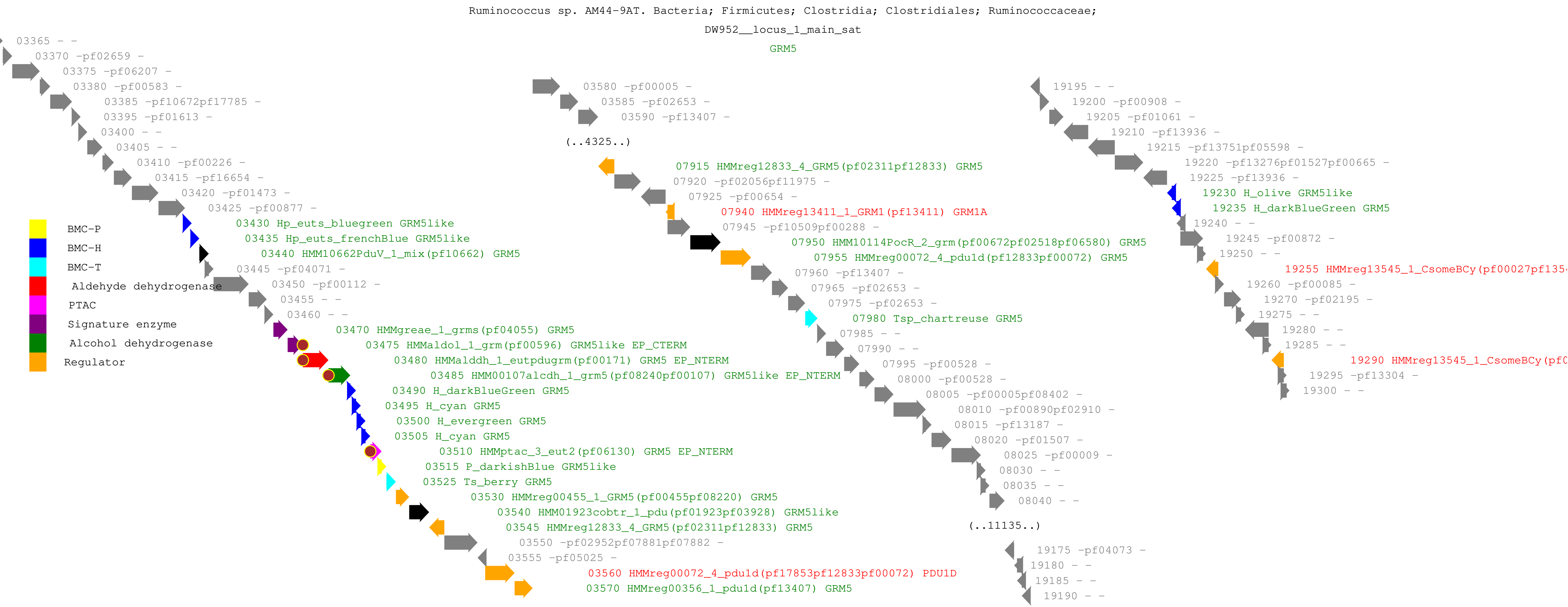
Ruminococcus sp. AM44-9AT. Bacteria; Firmicutes; Clostridia; Clostridiales; Ruminococcaceae;
DW952__locus_1_main_sat
GRM5
03365 - -
03370 -pf02659 -
03375 -pf06207 -
03380 -pf00583 -
03385 -pf10672pf17785 -
03395 -pf01613 -
03400 - -
03405 - -
03410 -pf00226 -
03415 -pf16654 -
03420 -pf01473 -
03425 -pf00877 -
03430 Hp_euts_bluegreen GRM5like
03435 Hp_euts_frenchBlue GRM5like
03440 HMM10662PduV_1_mix(pf10662) GRM5
03445 -pf04071 -
03450 -pf00112 -
03455 - -
03460 - -
03470 HMMgreae_1_grms(pf04055) GRM5
03475 HMMaldol_1_grm(pf00596) GRM5like EP_CTERM
03480 HMMaldh_1_eutpdugrm(pf00171) GRM5 EP_NTERM
03485 HMM00107alcdh_1_grm5(pf08240pf00107) GRM5like EP_NTERM
03490 H_darkBlueGreen GRM5
03495 H_cyan GRM5
03500 H_evergreen GRM5
03505 H_cyan GRM5
03510 HMMptac_3_eut2(pf06130) GRM5 EP_NTERM
03515 P_darkishBlue GRM5like
03525 Ts_berry GRM5
03530 HMMreg00455_1_GRM5(pf00455pf08220) GRM5
03540 HMM01923cobtr_1_pdu(pf01923pf03928) GRM5like
03545 HMMreg12833_4_GRM5(pf02311pf12833) GRM5
03550 -pf02952pf07881pf07882 -
03555 -pf05025 -
03560 HMMreg00072_4_pdu1d(pf17853pf12833pf00072) PDU1D
03570 HMMreg00356_1_pdu1d(pf13407) GRM5
BMC-P
BMC-H
BMC-T
Aldehyde dehydrogenase
PTAC
Signature enzyme
Alcohol dehydrogenase
Regulator
03580 -pf00005 -
03585 -pf02653 -
03590 -pf13407 -
(..4325..)
07915 HMMreg12833_4_GRM5(pf02311pf12833) GRM5
07920 -pf02056pf11975 -
07925 -pf00654 -
07940 HMMreg13411_1_GRM1(pf13411) GRM1A
07945 -pf10509pf00288 -
07950 HMM10114PocR_2_grm(pf00672pf02518pf06580) GRM5
07955 HMMreg00072_4_pdu1d(pf12833pf00072) GRM5
07960 -pf13407 -
07965 -pf02653 -
07975 -pf02653 -
07980 Tsp_chartreuse GRM5
07985 - -
07990 - -
07995 -pf00528 -
08000 -pf00528 -
08005 -pf00005pf08402 -
08010 -pf00890pf02910 -
08015 -pf13187 -
08020 -pf01507 -
08025 -pf00009 -
08030 - -
08035 - -
08040 - -
(..11135..)
19175 -pf04073 -
19180 - -
19185 - -
19190 - -
19195 - -
19200 -pf00908 -
19205 -pf01061 -
19210 -pf13936 -
19215 -pf13751pf05598 -
19220 -pf13276pf01527pf00665 -
19225 -pf13936 -
19230 H_olive GRM5like
19235 H_darkBlueGreen GRM5
19240 - -
19245 -pf00872 -
19250 - -
19255 HMMreg13545_1_CsomeBCy(pf00027pf135
19260 -pf00085 -
19270 -pf02195 -
19275 - -
19280 - -
19285 - -
19290 HMMreg13545_1_CsomeBCy(pf0
19295 -pf13304 -
19300 - -

Ruminococcus sp. AM45-2. Bacteria; Firmicutes; Clostridia; Clostridiales; Ruminococcaceae;

DW953__locus_1_main_sat

GRM5

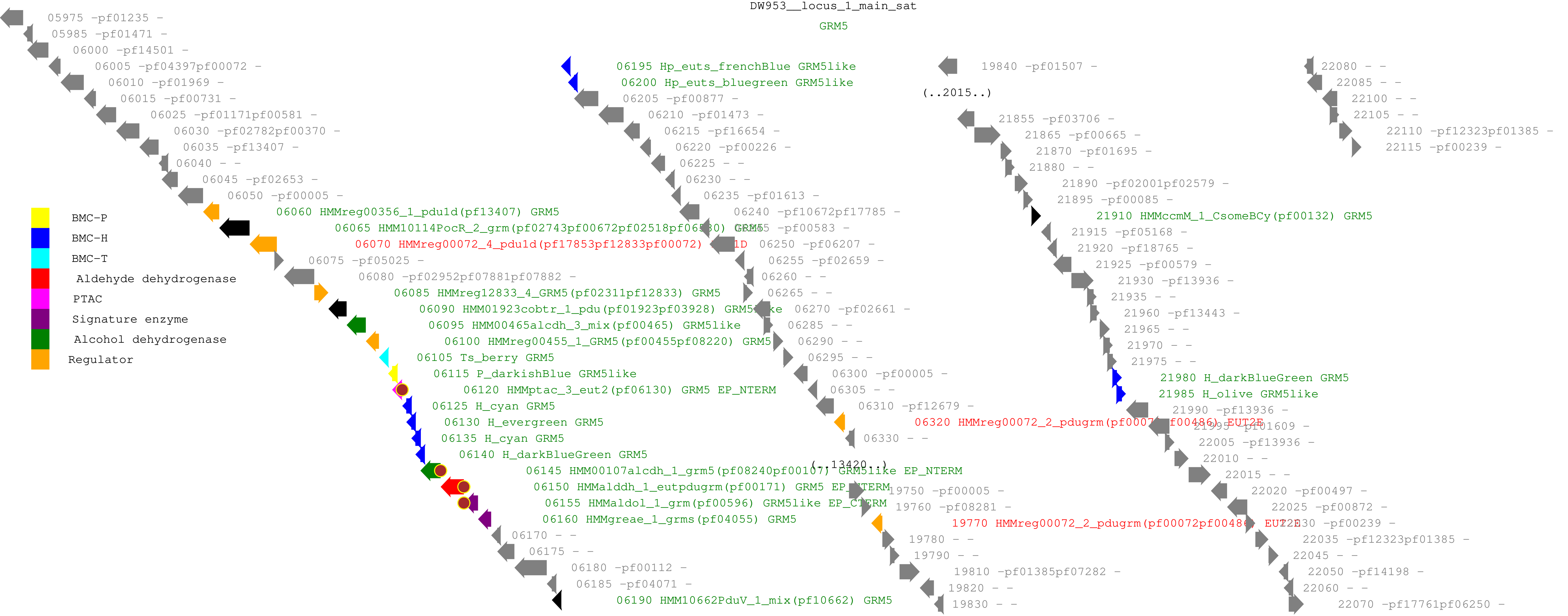

Ruminococcus sp. AM46-18. Bacteria; Firmicutes; Clostridia; Clostridiales; Ruminococcaceae;

DW958__locus_1

GRM5

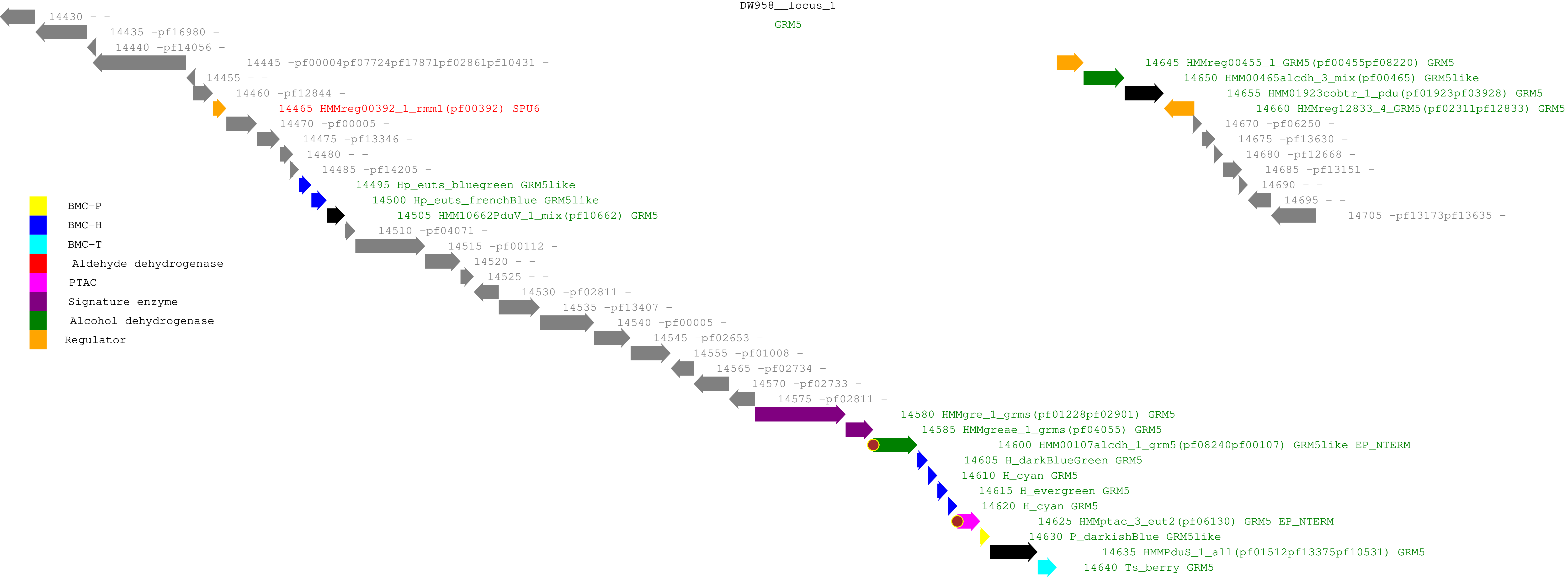

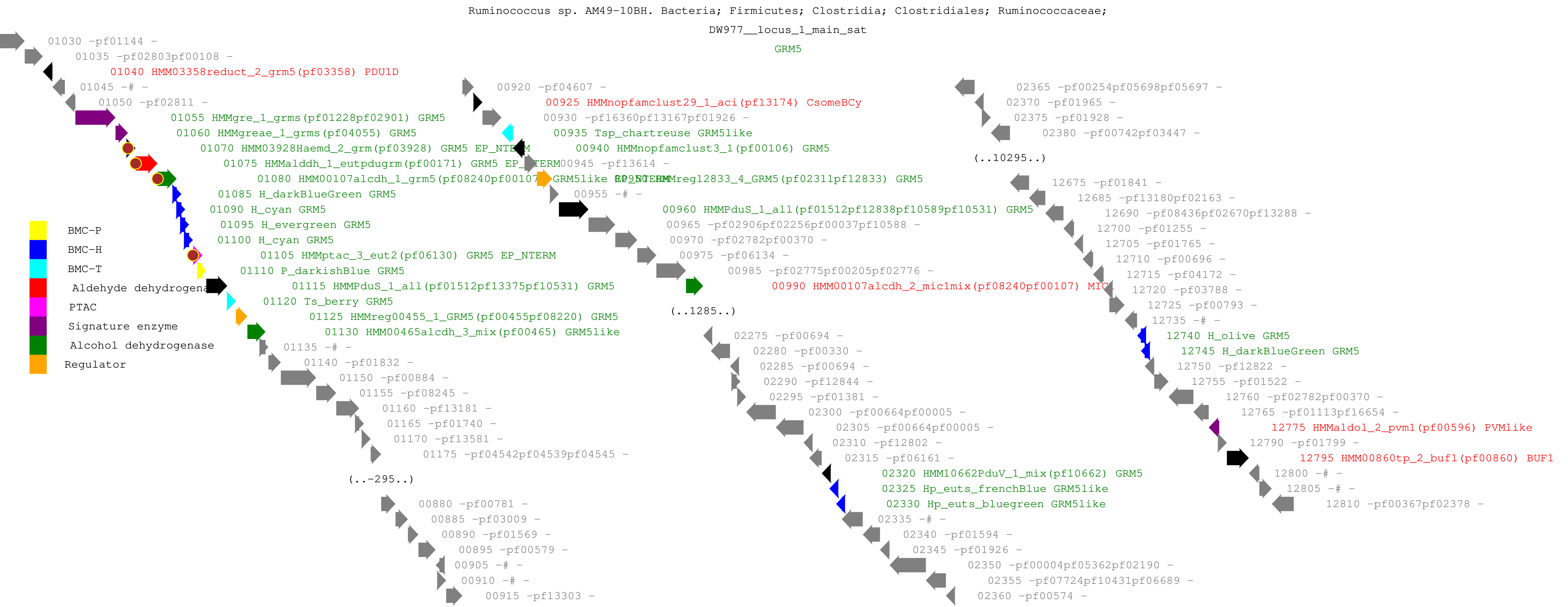

Ruminococcus sp. AM49-10BH. Bacteria; Firmicutes; Clostridia; Clostridiales; Ruminococcaceae;
DW977__locus_1_main_sat
GRM5
01030 -pf01144 -
01035 -pf02803pf00108 -
01040 HMM03358reduct_2_grm5(pf03358) PDU1D
01045 -# -
01050 -pf02811 -
01055 HMMgre_1_grms(pf01228pf02901) GRM5
01060 HMMgreae_1_grms(pf04055) GRM5
01070 HMM03928Haemd_2_grm(pf03928) GRM5 EP_NTERM
01075 HMMalddh_1_eutpdugrm(pf00171) GRM5 EP_NTERM
01080 HMM00107alcdh_1_grm5(pf08240pf00107) GRM5like EP_NTERM
01085 H_darkBlueGreen GRM5
01090 H_cyan GRM5
01095 H_evergreen GRM5
01100 H_cyan GRM5
01105 HMMptac_3_eut2(pf06130) GRM5 EP_NTERM
01110 P_darkishBlue GRM5
01115 HMMPduS_1_all(pf01512pf13375pf10531) GRM5
01120 Ts_berry GRM5
01125 HMMreg00455_1_GRM5(pf00455pf08220) GRM5
01130 HMM00465alcdh_3_mix(pf00465) GRM5like
01135 -# -
01140 -pf01832 -
01150 -pf00884 -
01155 -pf08245 -
01160 -pf13181 -
01165 -pf01740 -
01170 -pf13581 -
01175 -pf04542pf04539pf04545 -
(..-295..)
00880 -pf00781 -
00885 -pf03009 -
00890 -pf01569 -
00895 -pf00579 -
00905 -# -
00910 -# -
00915 -pf13303 -
00920 -pf04607 -
00925 HMMnopfamclust29_1_aci(pf13174) CsomeBCy
00930 -pf16360pf13167pf01926 -
00935 Tsp_chartreuse GRM5like
00940 HMMnopfamclust3_1(pf00106) GRM5
00945 -pf13614 -
00950 HMMreg12833_4_GRM5(pf02311pf12833) GRM5
00955 -# -
00960 HMMPduS_1_all(pf01512pf12838pf10589pf10531) GRM5
00965 -pf02906pf02256pf00037pf10588 -
00970 -pf02782pf00370 -
00975 -pf06134 -
00985 -pf02775pf00205pf02776 -
00990 HMM00107alcdh_2_mic1mix(pf08240pf00107) MIC
(..1285..)
02275 -pf00694 -
02280 -pf00330 -
02285 -pf00694 -
02290 -pf12844 -
02295 -pf01381 -
02300 -pf00664pf00005 -
02305 -pf00664pf00005 -
02310 -pf12802 -
02315 -pf06161 -
02320 HMM10662PduV_1_mix(pf10662) GRM5
02325 Hp_euts_frenchBlue GRM5like
02330 Hp_euts_bluegreen GRM5like
02335 -# -
02340 -pf01594 -
02345 -pf01926 -
02350 -pf00004pf05362pf02190 -
02355 -pf07724pf10431pf06689 -
02360 -pf00574 -
02365 -pf00254pf05698pf05697 -
02370 -pf01965 -
02375 -pf01928 -
02380 -pf00742pf03447 -
(..10295..)
12675 -pf01841 -
12685 -pf13180pf02163 -
12690 -pf08436pf02670pf13288 -
12700 -pf01255 -
12705 -pf01765 -
12710 -pf00696 -
12715 -pf04172 -
12720 -pf03788 -
12725 -pf00793 -
12735 -# -
12740 H_olive GRM5
12745 H_darkBlueGreen GRM5
12750 -pf12822 -
12755 -pf01522 -
12760 -pf02782pf00370 -
12765 -pf01113pf16654 -
12775 HMMaldol_2_pvm1(pf00596) PVMlike
12790 -pf01799 -
12795 HMM00860tp_2_buf1(pf00860) BUF1
12800 -# -
12805 -# -
12810 -pf00367pf02378 -
BMC-P
BMC-H
BMC-T
Aldehyde dehydrogena
PTAC
Signature enzyme
Alcohol dehydrogenase
Regulator

Ruminococcus sp. AM49-8. Bacteria; Firmicutes; Clostridia; Clostridiales; Ruminococcaceae;

DW983__locus_1_main_sat

GRM5

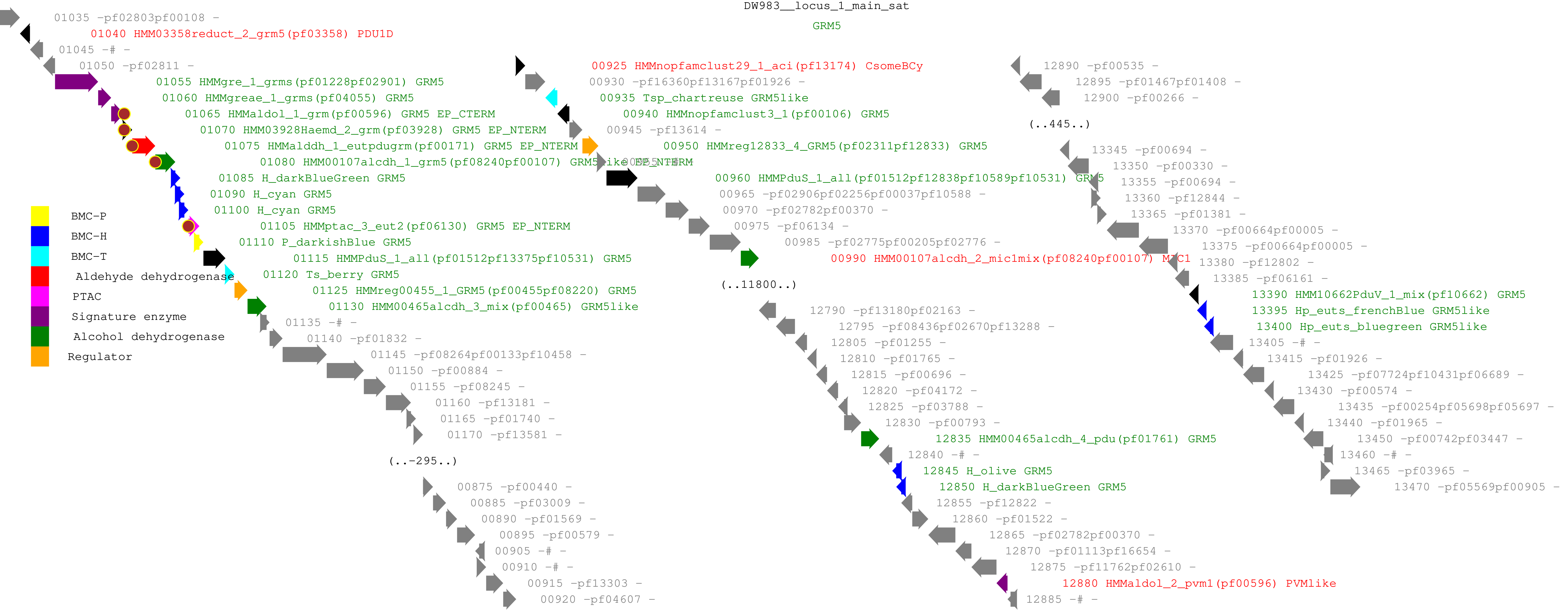

Ruminococcus sp. AM54-14NS. Bacteria; Firmicutes; Clostridia; Clostridiales; Ruminococcaceae;

DW999__locus_1_main_sat

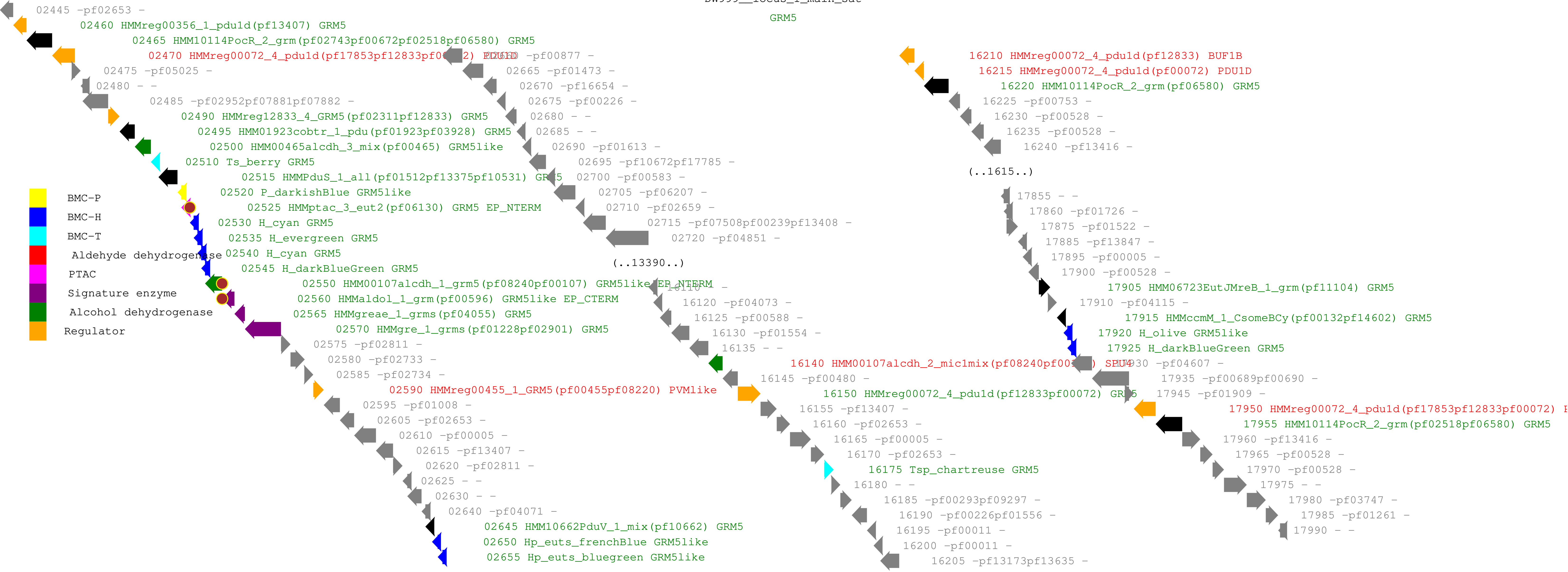

Ruminococcus sp. AF12-5. Bacteria; Firmicutes; Clostridia; Clostridiales; Ruminococcaceae;

DWV75__locus_1_main_sat

GRM5

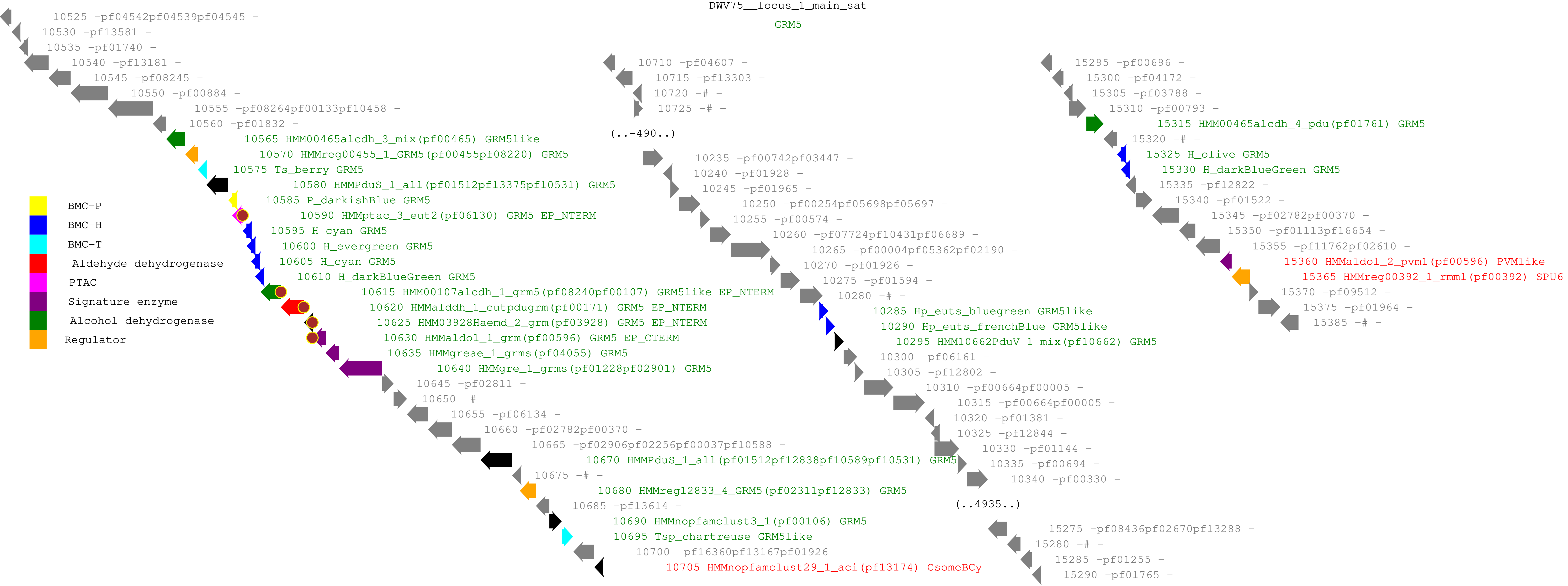

Ruminococcus sp. AF13-37. Bacteria; Firmicutes; Clostridia; Clostridiales; Ruminococcaceae;

DWV90__locus_1_main_sat

GRM5

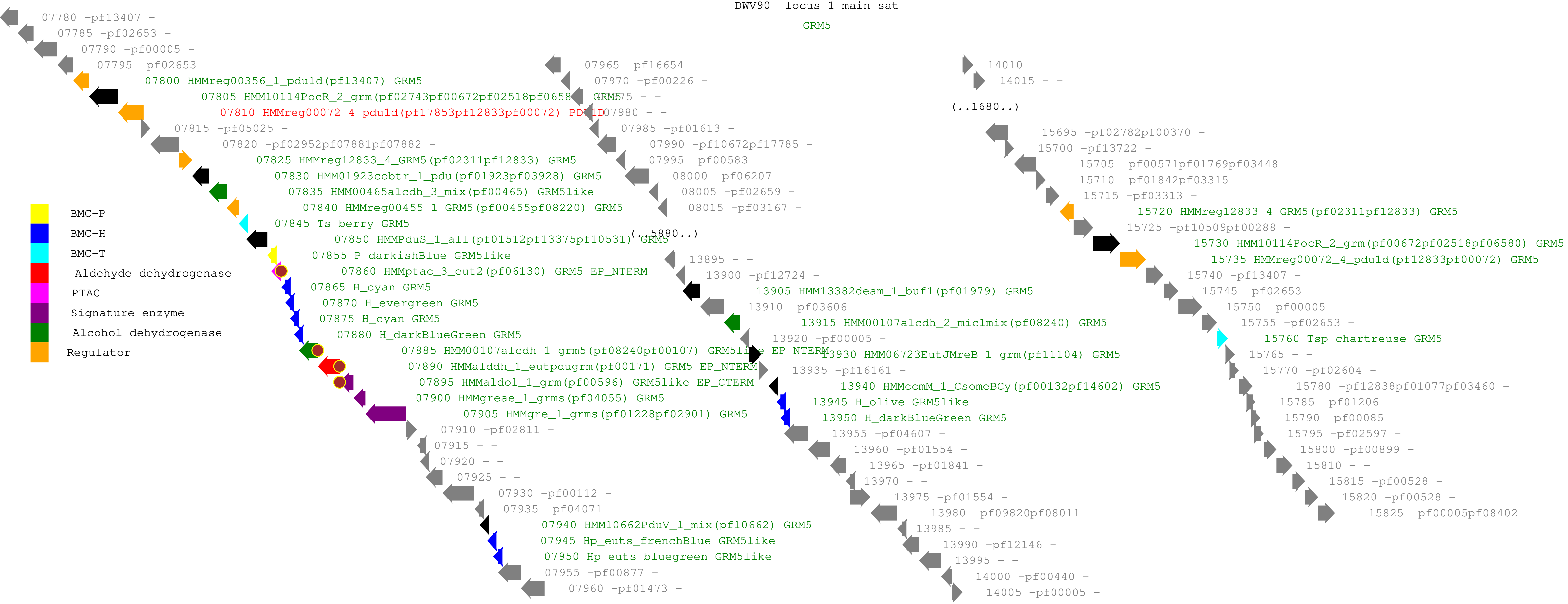

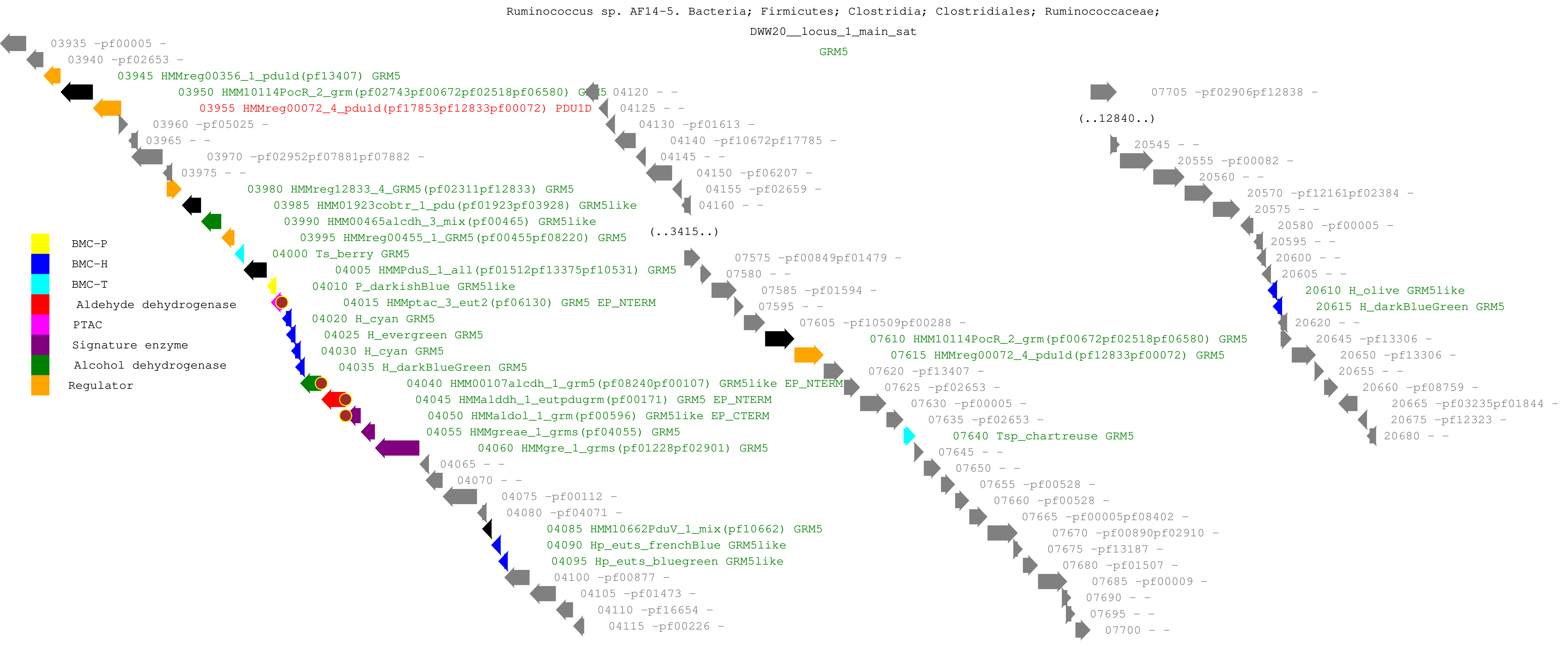

Ruminococcus sp. AF14-5. Bacteria; Firmicutes; Clostridia; Clostridiales; Ruminococcaceae;
DWW20__locus_1_main_sat
GRM5
03935 -pf00005 -
03940 -pf02653 -
03945 HMMreg00356_1_pdu1d(pf13407) GRM5
03950 HMM10114PocR_2_grm(pf02743pf00672pf02518pf06580) GRM5
03955 HMMreg00072_4_pdu1d(pf17853pf12833pf00072) PDU1D
03960 -pf05025 -
03965 - -
03970 -pf02952pf07881pf07882 -
03975 - -
03980 HMMreg12833_4_GRM5(pf02311pf12833) GRM5
03985 HMM01923cobtr_1_pdu(pf01923pf03928) GRM5like
03990 HMM00465alcdh_3_mix(pf00465) GRM5like
03995 HMMreg00455_1_GRM5(pf00455pf08220) GRM5
04000 Ts_berry GRM5
04005 HMMPduS_1_all(pf01512pf13375pf10531) GRM5
04010 P_darkishBlue GRM5like
04015 HMMptac_3_eut2(pf06130) GRM5 EP_NTERM
04020 H_cyan GRM5
04025 H_evergreen GRM5
04030 H_cyan GRM5
04035 H_darkBlueGreen GRM5
04040 HMM00107alcdh_1_grm5(pf08240pf00107) GRM5like EP_NTERM
04045 HMMalddh_1_eutpdugrm(pf00171) GRM5 EP_NTERM
04050 HMMaldol_1_grm(pf00596) GRM5like EP_CTERM
04055 HMMgreae_1_grms(pf04055) GRM5
04060 HMMgre_1_grms(pf01228pf02901) GRM5
04065 - -
04070 - -
04075 -pf00112 -
04080 -pf04071 -
04085 HMM10662PduV_1_mix(pf10662) GRM5
04090 Hp_euts_frenchBlue GRM5like
04095 Hp_euts_bluegreen GRM5like
04100 -pf00877 -
04105 -pf01473 -
04110 -pf16654 -
04115 -pf00226 -
04120 - -
04125 - -
04130 -pf01613 -
04140 -pf10672pf17785 -
04145 - -
04150 -pf06207 -
04155 -pf02659 -
04160 - -
(..3415..)
07575 -pf00849pf01479 -
07580 - -
07585 -pf01594 -
07595 - -
07605 -pf10509pf00288 -
07610 HMM10114PocR_2_grm(pf00672pf02518pf06580) GRM5
07615 HMMreg00072_4_pdu1d(pf12833pf00072) GRM5
07620 -pf13407 -
07625 -pf02653 -
07630 -pf00005 -
07635 -pf02653 -
07640 Tsp_chartreuse GRM5
07645 - -
07650 - -
07655 -pf00528 -
07660 -pf00528 -
07665 -pf00005pf08402 -
07670 -pf00890pf02910 -
07675 -pf13187 -
07680 -pf01507 -
07685 -pf00009 -
07690 - -
07695 - -
07700 - -
07705 -pf02906pf12838 -
(..12840..)
20545 - -
20555 -pf00082 -
20560 - -
20570 -pf12161pf02384 -
20575 - -
20580 -pf00005 -
20595 - -
20600 - -
20605 - -
20610 H_olive GRM5like
20615 H_darkBlueGreen GRM5
20620 - -
20645 -pf13306 -
20650 -pf13306 -
20655 - -
20660 -pf08759 -
20665 -pf03235pf01844 -
20675 -pf12323 -
20680 - -
BMC-P
BMC-H
BMC-T
Aldehyde dehydrogenase
PTAC
Signature enzyme
Alcohol dehydrogenase
Regulator

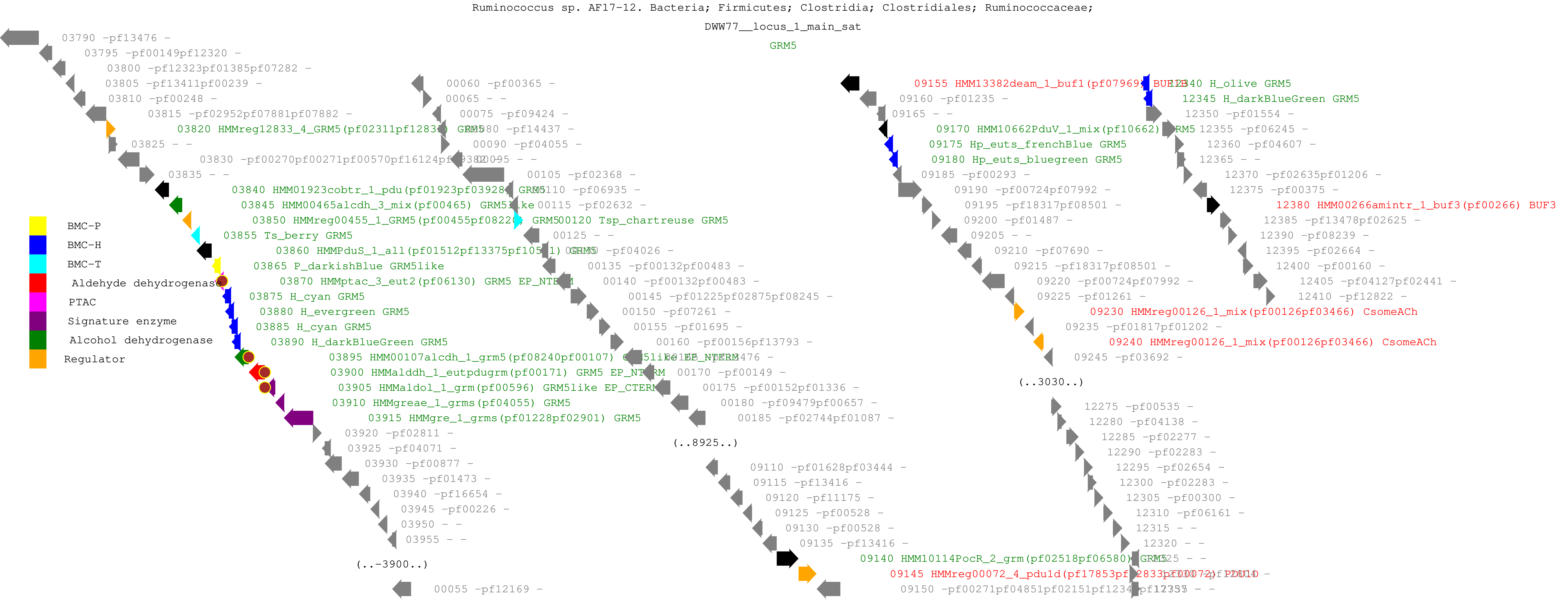
Ruminococcus sp. AF17-12. Bacteria; Firmicutes; Clostridia; Clostridiales; Ruminococcaceae;
DWW77__locus_1_main_sat
GRM5
BMC-P
BMC-H
BMC-T
Aldehyde dehydrogenase
PTAC
Signature enzyme
Alcohol dehydrogenase
Regulator
03790 -pf13476 -
03795 -pf00149pf12320 -
03800 -pf12323pf01385pf07282 -
03805 -pf13411pf00239 -
03810 -pf00248 -
03815 -pf02952pf07881pf07882 -
03820 HMMreg12833_4_GRM5(pf02311pf12833) GRM5
03825 - -
03830 -pf00270pf00271pf00570pf16124pf09382
03835 - -
03840 HMM01923cobtr_1_pdu(pf01923pf03928) GRM5
03845 HMM00465alcdh_3_mix(pf00465) GRM5like
03850 HMMreg00455_1_GRM5(pf00455pf08220) GRM5
03855 Ts_berry GRM5
03860 HMMPduS_1_all(pf01512pf13375pf10531) GRM5
03865 P_darkishBlue GRM5like
03870 HMMptac_3_eut2(pf06130) GRM5 EP_NTERM
03875 H_cyan GRM5
03880 H_evergreen GRM5
03885 H_cyan GRM5
03890 H_darkBlueGreen GRM5
03895 HMM00107alcdh_1_grm5(pf08240pf00107) GRM5like EP_NTERM
03900 HMMalddh_1_eutpdugrm(pf00171) GRM5 EP_NTERM
03905 HMMaldol_1_grm(pf00596) GRM5like EP_CTERM
03910 HMMgreae_1_grms(pf04055) GRM5
03915 HMMgre_1_grms(pf01228pf02901) GRM5
03920 -pf02811 -
03925 -pf04071 -
03930 -pf00877 -
03935 -pf01473 -
03940 -pf16654 -
03945 -pf00226 -
03950 - -
03955 - -
(..-3900..)
00055 -pf12169 -
00060 -pf00365 -
00065 - -
00075 -pf09424 -
00080 -pf14437 -
00090 -pf04055 -
00095 - -
00105 -pf02368 -
00110 -pf06935 -
00115 -pf02632 -
00120 Tsp_chartreuse GRM5
00125 - -
00130 -pf04026 -
00135 -pf00132pf00483 -
00140 -pf00132pf00483 -
00145 -pf01225pf02875pf08245 -
00150 -pf07261 -
00155 -pf01695 -
00160 -pf00156pf13793 -
00165 -pf13476 -
00170 -pf00149 -
00175 -pf00152pf01336 -
00180 -pf09479pf00657 -
00185 -pf02744pf01087 -
(..8925..)
09110 -pf01628pf03444 -
09115 -pf13416 -
09120 -pf11175 -
09125 -pf00528 -
09130 -pf00528 -
09135 -pf13416 -
09140 HMM10114PocR_2_grm(pf02518pf06580) GRM5
09145 HMMreg00072_4_pdu1d(pf17853pf13283pf00072) PDU1D
09150 -pf00271pf04851pf02151pf12344
09155 HMM13382deam_1_buf1(pf07969) BUF1B
09160 -pf01235 -
09165 - -
09170 HMM10662PduV_1_mix(pf10662) GRM5
09175 Hp_euts_frenchBlue GRM5
09180 Hp_euts_bluegreen GRM5
09185 -pf00293 -
09190 -pf00724pf07992 -
09195 -pf18317pf08501 -
09200 -pf01487 -
09205 - -
09210 -pf07690 -
09215 -pf18317pf08501 -
09220 -pf00724pf07992 -
09225 -pf01261 -
09230 HMMreg00126_1_mix(pf00126pf03466) CsomeACh
09235 -pf01817pf01202 -
09240 HMMreg00126_1_mix(pf00126pf03466) CsomeACh
09245 -pf03692 -
(..3030..)
12275 -pf00535 -
12280 -pf04138 -
12285 -pf02277 -
12290 -pf02283 -
12295 -pf02654 -
12300 -pf02283 -
12305 -pf00300 -
12310 -pf06161 -
12315 - -
12320 - -
12325 - -
12330 -pf12804 -
12335 - -
12340 H_olive GRM5
12345 H_darkBlueGreen GRM5
12350 -pf01554 -
12355 -pf06245 -
12360 -pf04607 -
12365 - -
12370 -pf02635pf01206 -
12375 -pf00375 -
12380 HMM00266amintr_1_buf3(pf00266) BUF3
12385 -pf13478pf02625 -
12390 -pf08239 -
12395 -pf02664 -
12400 -pf00160 -
12405 -pf04127pf02441 -
12410 -pf12822 -

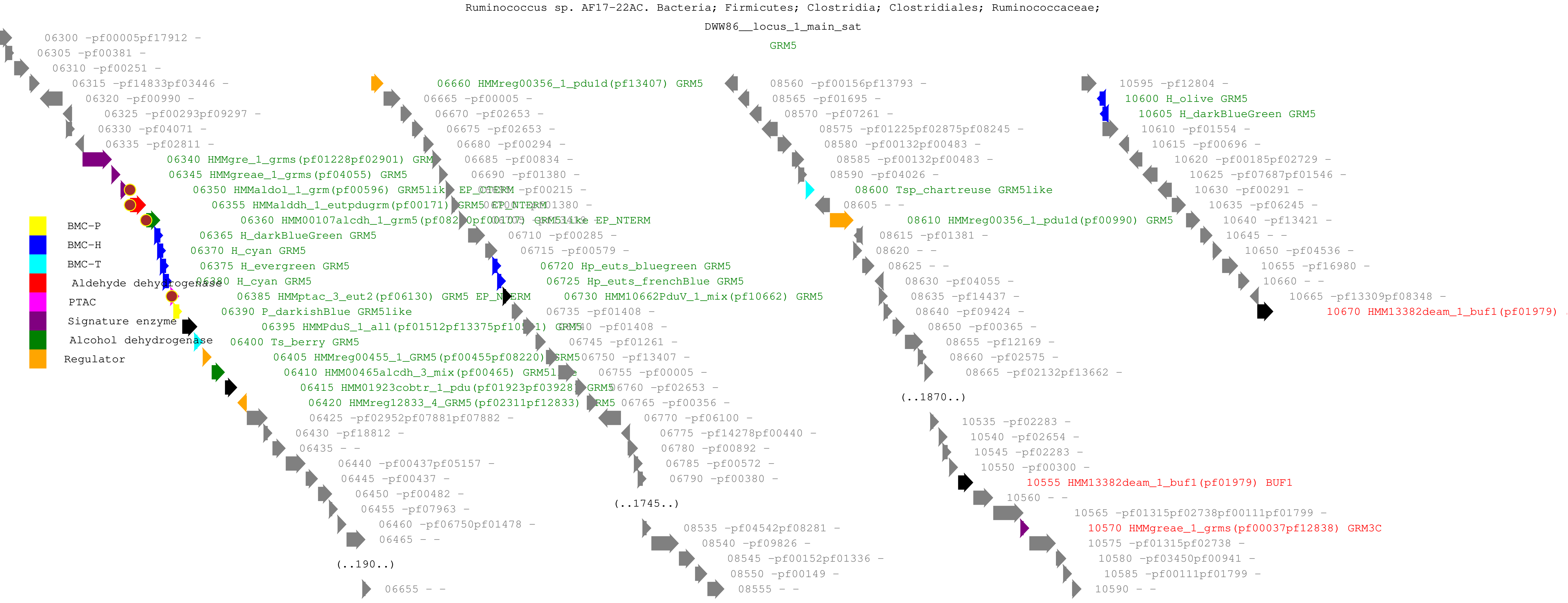
Ruminococcus sp. AF17-22AC. Bacteria; Firmicutes; Clostridia; Clostridiales; Ruminococcaceae;
DWW86__locus_1_main_sat
GRM5
BMC-P
BMC-H
BMC-T
Aldehyde dehydrogenase
PTAC
Signature enzyme
Alcohol dehydrogenase
Regulator
06300 -pf00005pf17912 -
06305 -pf00381 -
06310 -pf00251 -
06315 -pf14833pf03446 -
06320 -pf00990 -
06325 -pf00293pf09297 -
06330 -pf04071 -
06335 -pf02811 -
06340 HMMgre_1_grms(pf01228pf02901) GRM
06345 HMMgreae_1_grms(pf04055) GRM5
06350 HMMaldol_1_grm(pf00596) GRM5lik
06355 HMMaldh_1_eutpdugrm(pf00171) GRM5
06360 HMM00107alcdh_1_grm5(pf082
06365 H_darkBlueGreen GRM5
06370 H_cyan GRM5
06375 H_evergreen GRM5
06380 H_cyan GRM5
06385 HMMptac_3_eut2(pf06130) GRM5 EP_N
06390 P_darkishBlue GRM5like
06395 HMMPduS_1_all(pf01512pf13375pf10
06400 Ts_berry GRM5
06405 HMMreg00455_1_GRM5(pf00455pf08220)
06410 HMM00465alcdh_3_mix(pf00465) GRM5l
06415 HMM01923cobtr_1_pdu(pf01923pf03928
06420 HMMreg12833_4_GRM5(pf02311pf12833)
06425 -pf02952pf07881pf07882 -
06430 -pf18812 -
06435 - -
06440 -pf00437pf05157 -
06445 -pf00437 -
06450 -pf00482 -
06455 -pf07963 -
06460 -pf06750pf01478 -
06465 - -
(..190..)
06655 - -
06660 HMMreg00356_1_pdu1d(pf13407) GRM5
06665 -pf00005 -
06670 -pf02653 -
06675 -pf02653 -
06680 -pf00294 -
06685 -pf00834 -
06690 -pf01380 -
EP_CTERM -pf00215 -
EP_NTERM
06710 -pf00285 -
06715 -pf00579 -
06720 Hp_euts_bluegreen GRM5
06725 Hp_euts_frenchBlue GRM5
06730 HMM10662PduV_1_mix(pf10662) GRM5
06735 -pf01408 -
06740 -pf01408 -
06745 -pf01261 -
06750 -pf13407 -
06755 -pf00005 -
06760 -pf02653 -
06765 -pf00356 -
06770 -pf06100 -
06775 -pf14278pf00440 -
06780 -pf00892 -
06785 -pf00572 -
06790 -pf00380 -
(..1745..)
08535 -pf04542pf08281 -
08540 -pf09826 -
08545 -pf00152pf01336 -
08550 -pf00149 -
08555 - -
08560 -pf00156pf13793 -
08565 -pf01695 -
08570 -pf07261 -
08575 -pf01225pf02875pf08245 -
08580 -pf00132pf00483 -
08585 -pf00132pf00483 -
08590 -pf04026 -
08600 Tsp_chartreuse GRM5like
08605 - -
08610 HMMreg00356_1_pdu1d(pf00990) GRM5
08615 -pf01381 -
08620 - -
08625 - -
08630 -pf04055 -
08635 -pf14437 -
08640 -pf09424 -
08650 -pf00365 -
08655 -pf12169 -
08660 -pf02575 -
08665 -pf02132pf13662 -
(..1870..)
10535 -pf02283 -
10540 -pf02654 -
10545 -pf02283 -
10550 -pf00300 -
10555 HMM13382deam_1_buf1(pf01979) BUF1
10560 - -
10565 -pf01315pf02738pf00111pf01799 -
10570 HMMgreae_1_grms(pf00037pf12838) GRM3C
10575 -pf01315pf02738 -
10580 -pf03450pf00941 -
10585 -pf00111pf01799 -
10590 - -
10595 -pf12804 -
10600 H_olive GRM5
10605 H_darkBlueGreen GRM5
10610 -pf01554 -
10615 -pf00696 -
10620 -pf00185pf02729 -
10625 -pf07687pf01546 -
10630 -pf00291 -
10635 -pf06245 -
10640 -pf13421 -
10645 - -
10650 -pf04536 -
10655 -pf16980 -
10660 - -
10665 -pf13309pf08348 -
10670 HMM13382deam_1_buf1(pf01979)

Blautia sp. AF19-10LB. Bacteria; Firmicutes; Clostridia; Clostridiales; Lachnospiraceae;

DWX28__locus_1_main_sat

GRM5

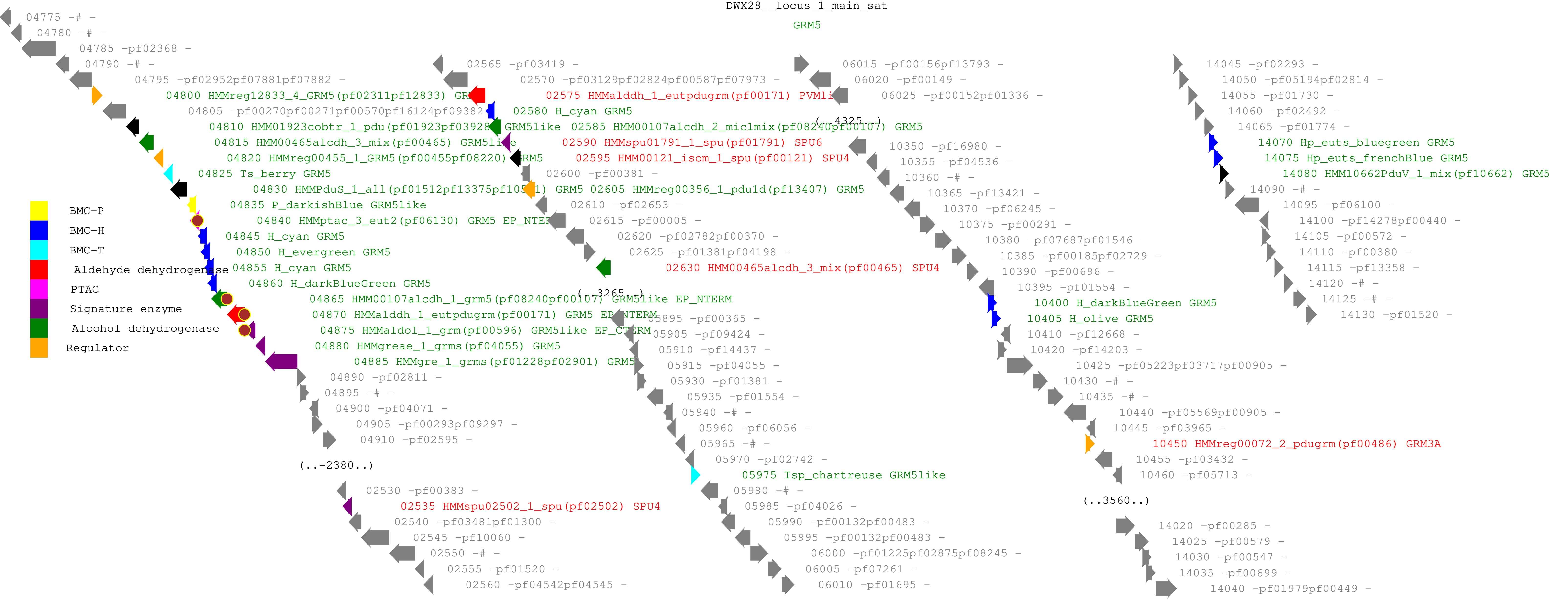

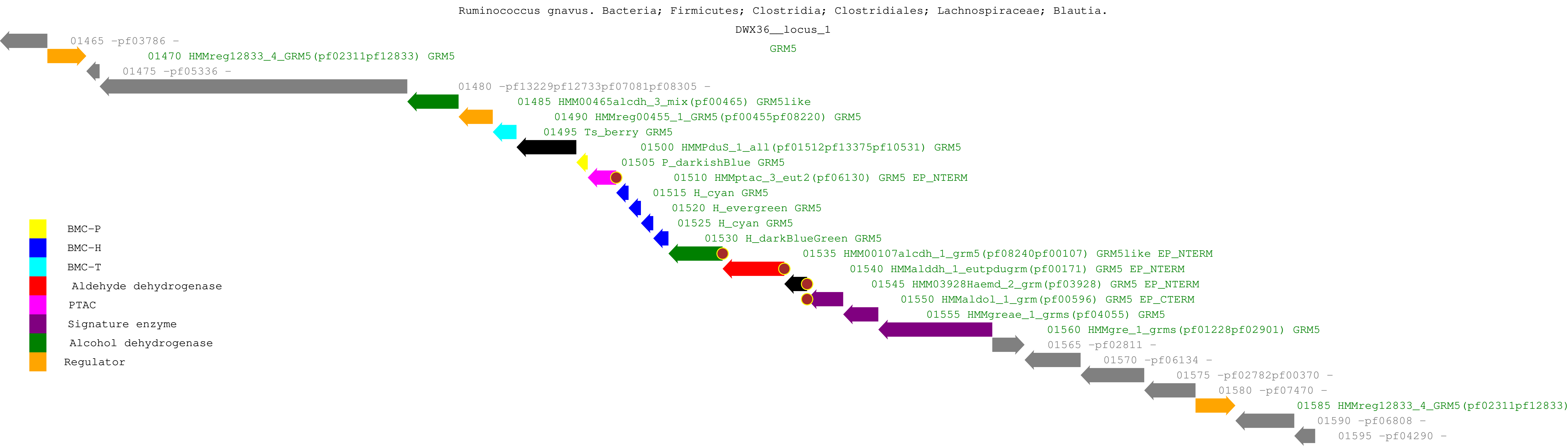
Ruminococcus gnavus. Bacteria; Firmicutes; Clostridia; Clostridiales; Lachnospiraceae; Blautia.
DWX36__locus_1
GRM5
01465 -pf03786 -
01470 HMMreg12833_4_GRM5(pf02311pf12833) GRM5
01475 -pf05336 -
01480 -pf13229pf12733pf07081pf08305 -
01485 HMM00465alcdh_3_mix(pf00465) GRM5like
01490 HMMreg00455_1_GRM5(pf00455pf08220) GRM5
01495 Ts_berry GRM5
01500 HMMPduS_1_all(pf01512pf13375pf10531) GRM5
01505 P_darkishBlue GRM5
01510 HMMptac_3_eut2(pf06130) GRM5 EP_NTERM
01515 H_cyan GRM5
01520 H_evergreen GRM5
01525 H_cyan GRM5
01530 H_darkBlueGreen GRM5
01535 HMM00107alcdh_1_grm5(pf08240pf00107) GRM5like EP_NTERM
01540 HMMalddh_1_eutpdugrm(pf00171) GRM5 EP_NTERM
01545 HMM03928Haemd_2_grm(pf03928) GRM5 EP_NTERM
01550 HMMaldol_1_grm(pf00596) GRM5 EP_CTERM
01555 HMMgreae_1_grms(pf04055) GRM5
01560 HMMgre_1_grms(pf01228pf02901) GRM5
01565 -pf02811 -
01570 -pf06134 -
01575 -pf02782pf00370 -
01580 -pf07470 -
01585 HMMreg12833_4_GRM5(pf02311pf12833)
01590 -pf06808 -
01595 -pf04290 -
BMC-P
BMC-H
BMC-T
Aldehyde dehydrogenase
PTAC
Signature enzyme
Alcohol dehydrogenase
Regulator

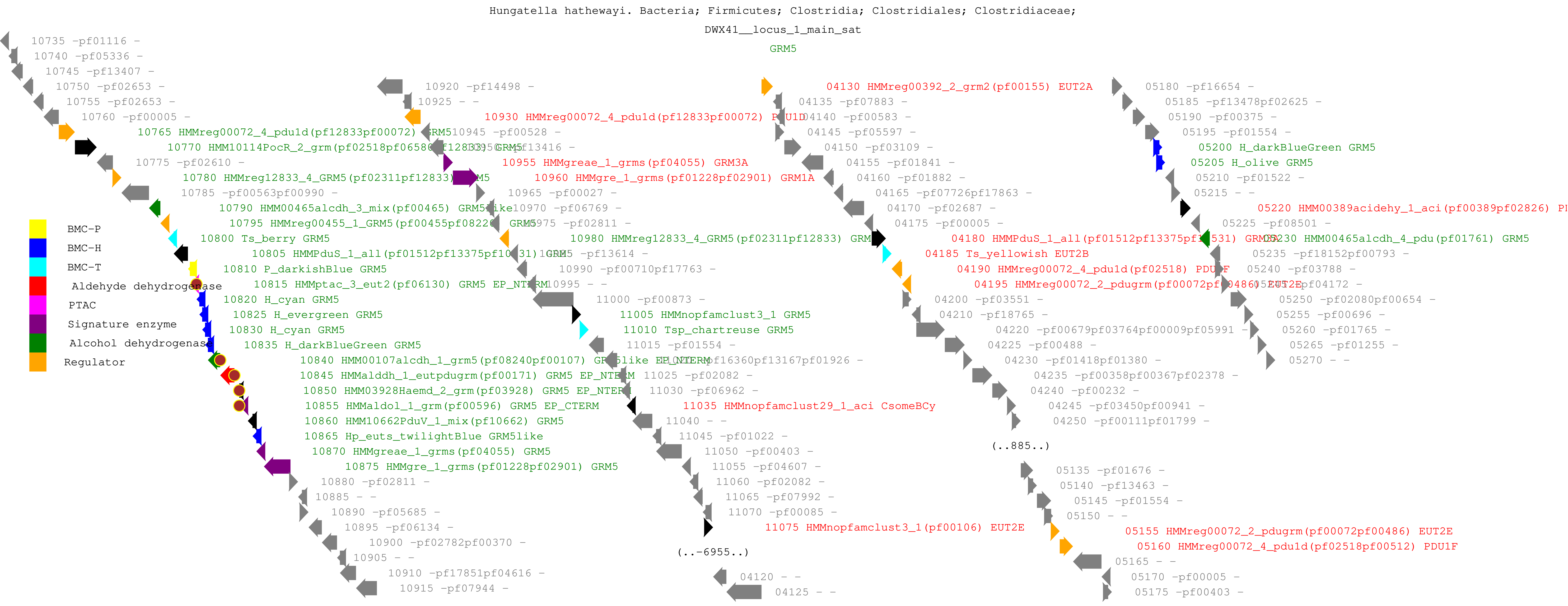
Hungatella hathewayi. Bacteria; Firmicutes; Clostridia; Clostridiales; Clostridiaceae;
DWX41__locus_1_main_sat
GRM5
BMC-P
BMC-H
BMC-T
Aldehyde dehydrogenase
PTAC
Signature enzyme
Alcohol dehydrogenase
Regulator
10735 -pf01116 -
10740 -pf05336 -
10745 -pf13407 -
10750 -pf02653 -
10755 -pf02653 -
10760 -pf00005 -
10765 HMMreg00072_4_pdu1d(pf12833pf00072)
10770 HMM10114PocR_2_grm(pf02518pf06580)
10775 -pf02610 -
10780 HMMreg12833_4_GRM5(pf02311pf12833)
10785 -pf00563pf00990 -
10790 HMM00465alcdh_3_mix(pf00465) GRM5like
10795 HMMreg00455_1_GRM5(pf00455pf08220) GRM5
10800 Ts_berry GRM5
10805 HMMPduS_1_all(pf01512pf13375pf10531)
10810 P_darkishBlue GRM5
10815 HMMptac_3_eut2(pf06130) GRM5 EP_NTERM
10820 H_cyan GRM5
10825 H_evergreen GRM5
10830 H_cyan GRM5
10835 H_darkBlueGreen GRM5
10840 HMM00107alcdh_1_grm5(pf08240pf00107) GRM5like EP_NTERM
10845 HMMaldдh_1_eutpdugrm(pf00171) GRM5 EP_NTERM
10850 HMM03928Haemd_2_grm(pf03928) GRM5 EP_NTERM
10855 HMMaldol_1_grm(pf00596) GRM5 EP_CTERM
10860 HMM10662PduV_1_mix(pf10662) GRM5
10865 Hp_euts_twilightBlue GRM5like
10870 HMMgreae_1_grms(pf04055) GRM5
10875 HMMgre_1_grms(pf01228pf02901) GRM5
10880 -pf02811 -
10885 - -
10890 -pf05685 -
10895 -pf06134 -
10900 -pf02782pf00370 -
10905 - -
10910 -pf17851pf04616 -
10915 -pf07944 -
10920 -pf14498 -
10925 - -
10930 HMMreg00072_4_pdu1d(pf12833pf00072) PDU1D
10945 -pf00528 -
10950 -pf13416 -
10955 HMMgreae_1_grms(pf04055) GRM3A
10960 HMMgre_1_grms(pf01228pf02901) GRM1A
10965 -pf00027 -
10970 -pf06769 -
10975 -pf02811 -
10980 HMMreg12833_4_GRM5(pf02311pf12833) GRM5
10985 -pf13614 -
10990 -pf00710pf17763 -
10995 - -
11000 -pf00873 -
11005 HMMnopfamclust3_1 GRM5
11010 Tsp_chartreuse GRM5
11015 -pf01554 -
11020 -pf16360pf13167pf01926 -
11025 -pf02082 -
11030 -pf06962 -
11035 HMMnopfamclust29_1_aci CsomeBCy
11040 - -
11045 -pf01022 -
11050 -pf00403 -
11055 -pf04607 -
11060 -pf02082 -
11065 -pf07992 -
11070 -pf00085 -
11075 HMMnopfamclust3_1(pf00106) EUT2E
(..-6955..)
04120 - -
04125 - -
04130 HMMreg00392_2_grm2(pf00155) EUT2A
04135 -pf07883 -
04140 -pf00583 -
04145 -pf05597 -
04150 -pf03109 -
04155 -pf01841 -
04160 -pf01882 -
04165 -pf07726pf17863 -
04170 -pf02687 -
04175 -pf00005 -
04180 HMMPduS_1_all(pf01512pf13375pf10531) GRM3
04185 Ts_yellowish EUT2B
04190 HMMreg00072_4_pdu1d(pf02518) PDU1F
04195 HMMreg00072_2_pdugrm(pf00072pf00486) EUT2E
04200 -pf03551 -
04210 -pf18765 -
04220 -pf00679pf03764pf00009pf05991 -
04225 -pf00488 -
04230 -pf01418pf01380 -
04235 -pf00358pf00367pf02378 -
04240 -pf00232 -
04245 -pf03450pf00941 -
04250 -pf00111pf01799 -
(..885..)
05135 -pf01676 -
05140 -pf13463 -
05145 -pf01554 -
05150 - -
05155 HMMreg00072_2_pdugrm(pf00072pf00486) EUT2E
05160 HMMreg00072_4_pdu1d(pf02518pf00512) PDU1F
05165 - -
05170 -pf00005 -
05175 -pf00403 -
05180 -pf16654 -
05185 -pf13478pf02625 -
05190 -pf00375 -
05195 -pf01554 -
05200 H_darkBlueGreen GRM5
05205 H_olive GRM5
05210 -pf01522 -
05215 - -
05220 HMM00389acidehy_1_aci(pf00389pf02826) PI
05225 -pf08501 -
05230 HMM00465alcdh_4_pdu(pf01761) GRM5
05235 -pf18152pf00793 -
05240 -pf03788 -
05245 -pf04172 -
05250 -pf02080pf00654 -
05255 -pf00696 -
05260 -pf01765 -
05265 -pf01255 -
05270 - -

Clostridium sp. AF19-22AC. Bacteria; Firmicutes; Clostridia; Clostridiales; Clostridiaceae;

DWX43__locus_1_main_sat

GRM5

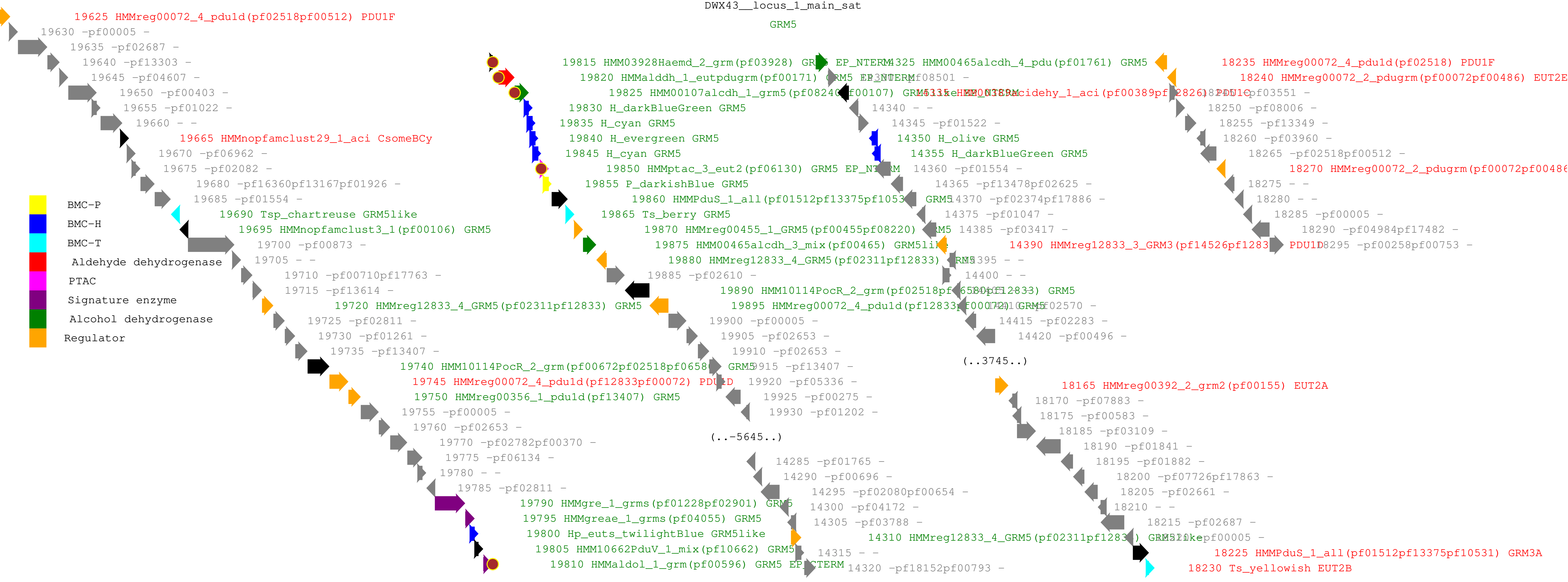

Ruminococcus sp. AF19-29. Bacteria; Firmicutes; Clostridia; Clostridiales; Ruminococcaceae;

DWX46__locus_1_main_sat

GRM5

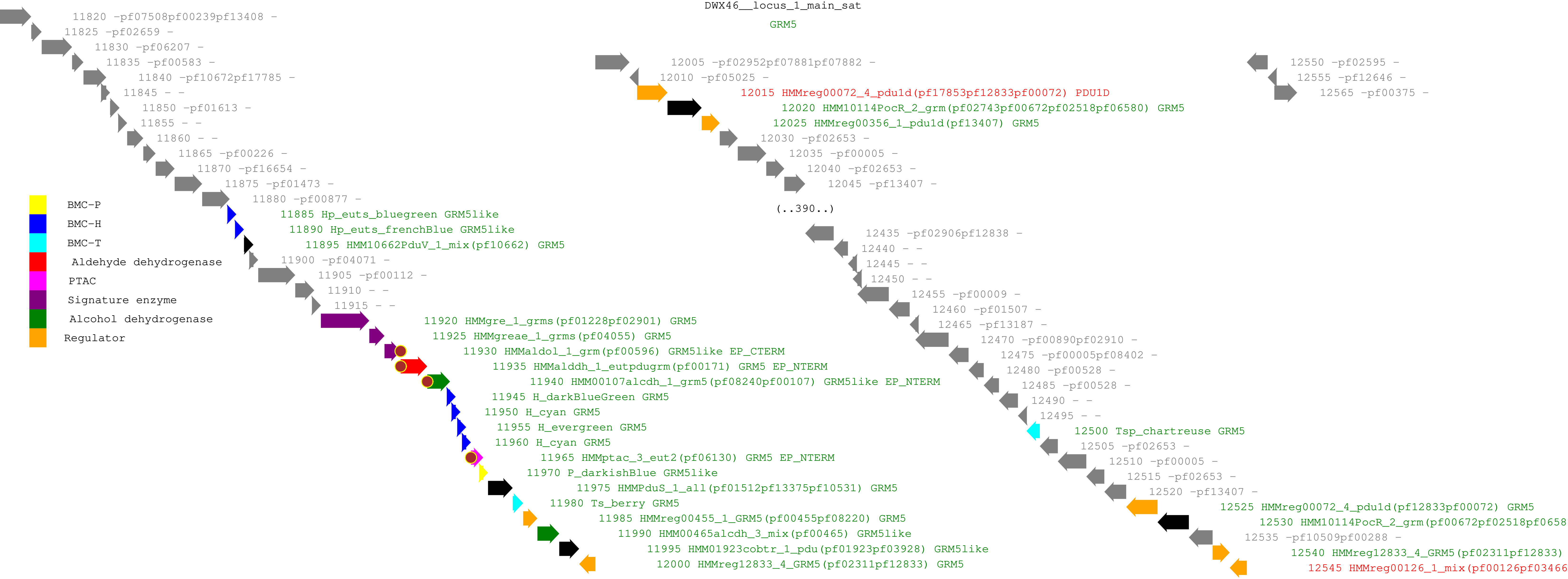

Ruminococcus sp. AF19-4LB. Bacteria; Firmicutes; Clostridia; Clostridiales; Ruminococcaceae;

DWX54__locus_1_main_sat

GRM5

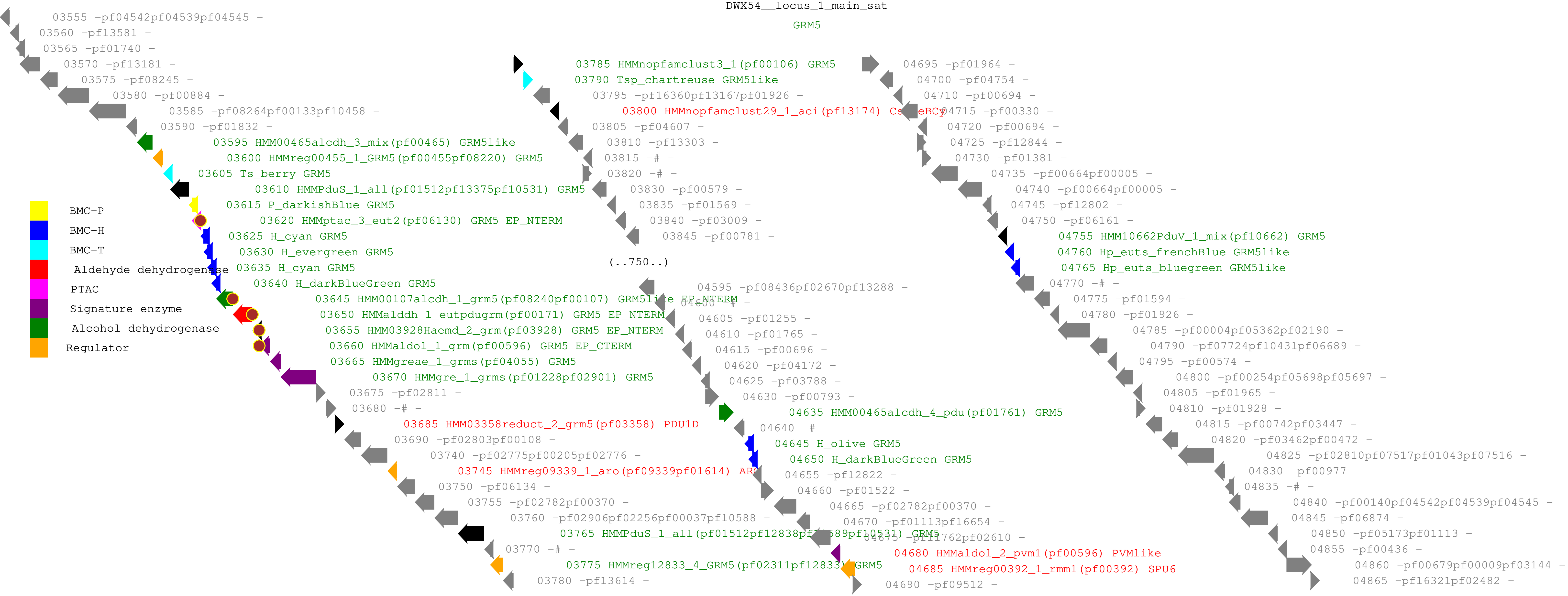

Ruminococcus sp. AF20-12LB. Bacteria; Firmicutes; Clostridia; Clostridiales; Ruminococcaceae;

DWX61__locus_1_main_sat

GRM5

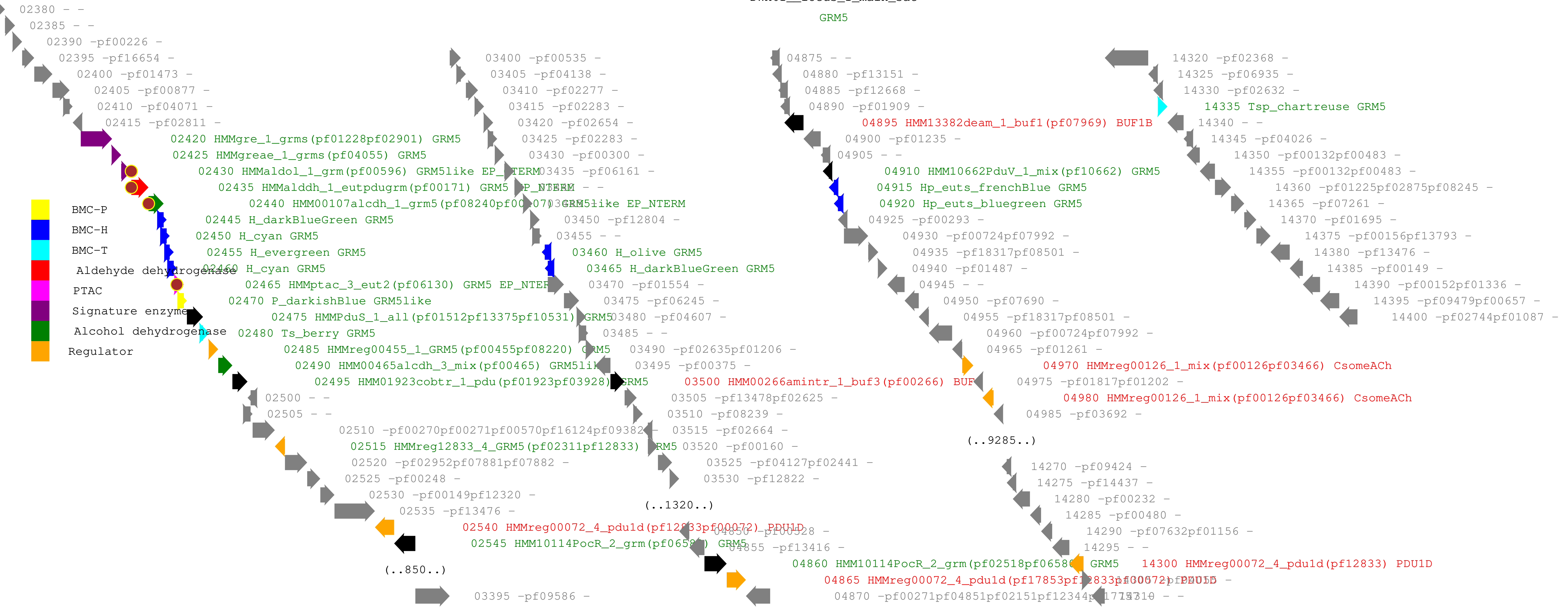

Ruminococcus sp. AF21-3. Bacteria; Firmicutes; Clostridia; Clostridiales; Ruminococcaceae;

DWX80__locus_1_main_sat

GRM5

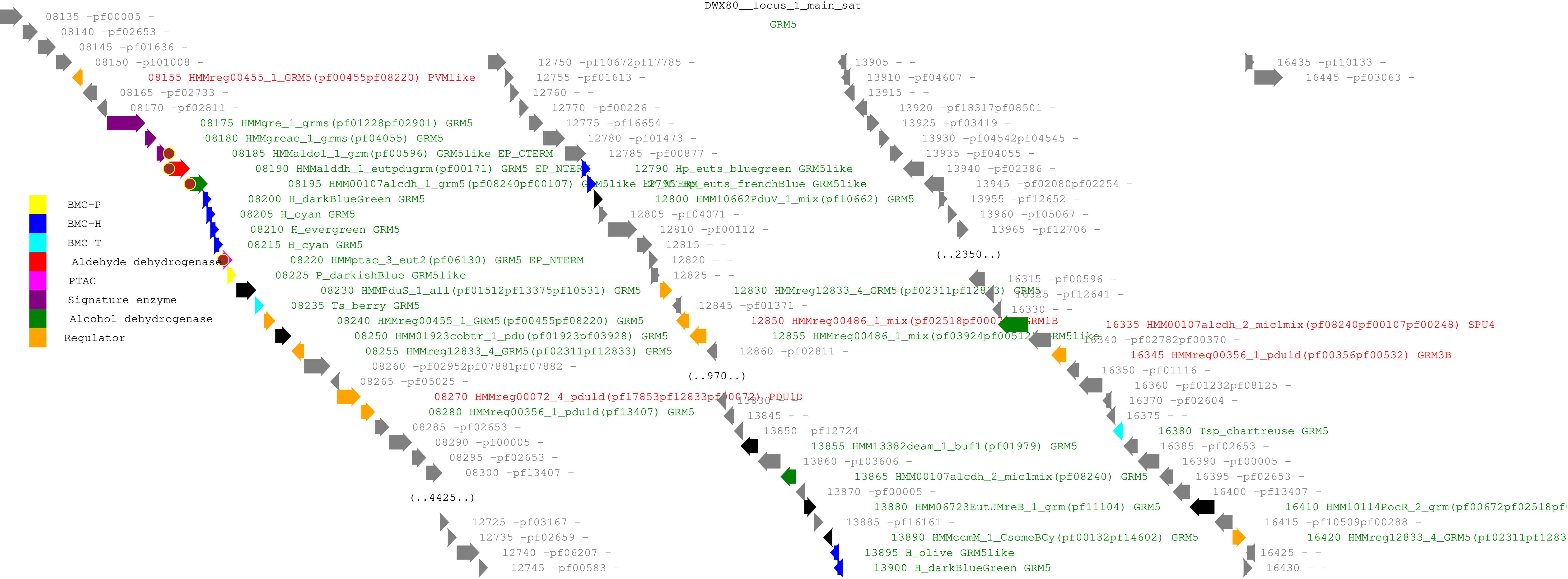

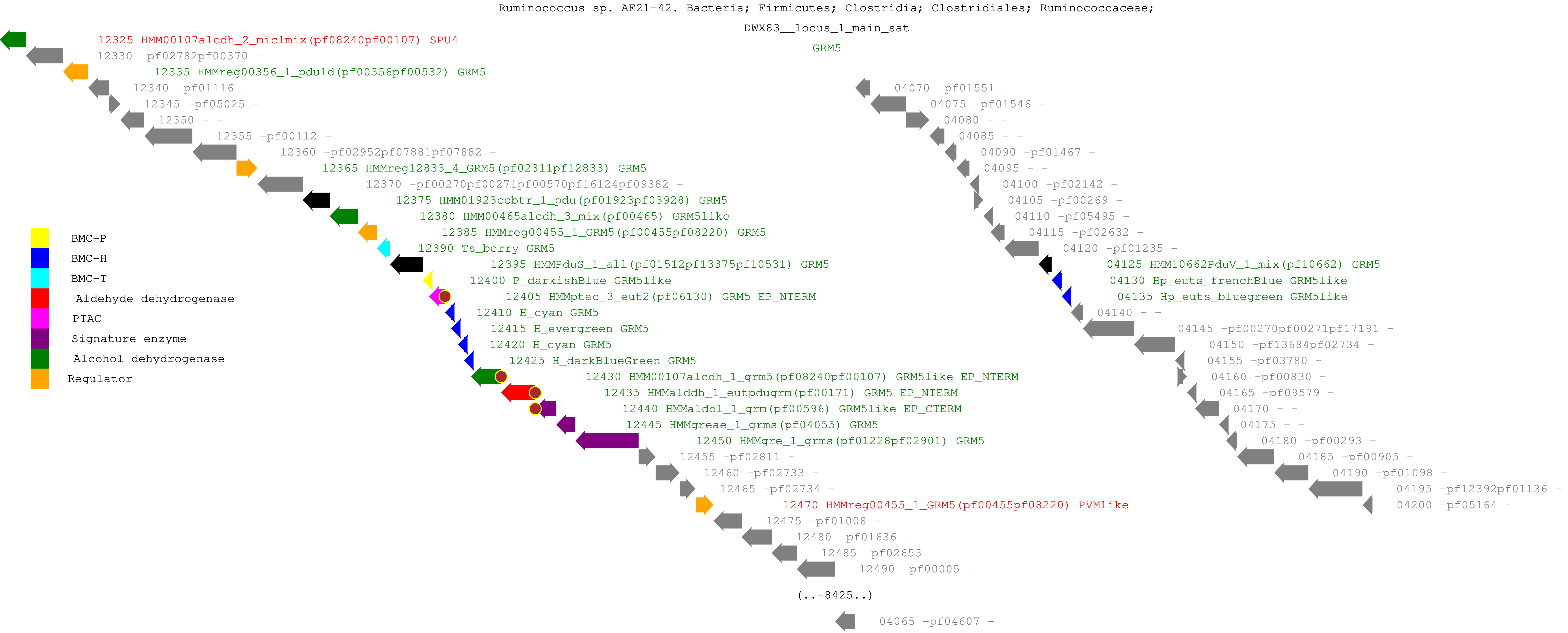
Ruminococcus sp. AF21-42. Bacteria; Firmicutes; Clostridia; Clostridiales; Ruminococcaceae;
DWX83__locus_1_main_sat
GRM5
12325 HMM00107alcdh_2_mic1mix(pf08240pf00107) SPU4
12330 -pf02782pf00370 -
12335 HMMreg00356_1_pdu1d(pf00356pf00532) GRM5
12340 -pf01116 -
12345 -pf05025 -
12350 - -
12355 -pf00112 -
12360 -pf02952pf07881pf07882 -
12365 HMMreg12833_4_GRM5(pf02311pf12833) GRM5
12370 -pf00270pf00271pf00570pf16124pf09382 -
12375 HMM01923cobtr_1_pdu(pf01923pf03928) GRM5
12380 HMM00465alcdh_3_mix(pf00465) GRM5like
12385 HMMreg00455_1_GRM5(pf00455pf08220) GRM5
12390 Ts_berry GRM5
12395 HMMPduS_1_all(pf01512pf13375pf10531) GRM5
12400 P_darkishBlue GRM5like
12405 HMMptac_3_eut2(pf06130) GRM5 EP_NTERM
12410 H_cyan GRM5
12415 H_evergreen GRM5
12420 H_cyan GRM5
12425 H_darkBlueGreen GRM5
12430 HMM00107alcdh_1_grm5(pf08240pf00107) GRM5like EP_NTERM
12435 HMMaldh_1_eutpdugrm(pf00171) GRM5 EP_NTERM
12440 HMMaldol_1_grm(pf00596) GRM5like EP_CTERM
12445 HMMgreae_1_grms(pf04055) GRM5
12450 HMMgre_1_grms(pf01228pf02901) GRM5
12455 -pf02811 -
12460 -pf02733 -
12465 -pf02734 -
12470 HMMreg00455_1_GRM5(pf00455pf08220) PVMlike
12475 -pf01008 -
12480 -pf01636 -
12485 -pf02653 -
12490 -pf00005 -
(..-8425..)
04065 -pf04607 -
04070 -pf01551 -
04075 -pf01546 -
04080 - -
04085 - -
04090 -pf01467 -
04095 - -
04100 -pf02142 -
04105 -pf00269 -
04110 -pf05495 -
04115 -pf02632 -
04120 -pf01235 -
04125 HMM10662PduV_1_mix(pf10662) GRM5
04130 Hp_euts_frenchBlue GRM5like
04135 Hp_euts_bluegreen GRM5like
04140 - -
04145 -pf00270pf00271pf17191 -
04150 -pf13684pf02734 -
04155 -pf03780 -
04160 -pf00830 -
04165 -pf09579 -
04170 - -
04175 - -
04180 -pf00293 -
04185 -pf00905 -
04190 -pf01098 -
04195 -pf12392pf01136 -
04200 -pf05164 -
BMC-P
BMC-H
BMC-T
Aldehyde dehydrogenase
PTAC
Signature enzyme
Alcohol dehydrogenase
Regulator

Ruminococcus sp. AF24-16. Bacteria; Firmicutes; Clostridia; Clostridiales; Ruminococcaceae;

DWY13__locus_1_main_sat

GRM5

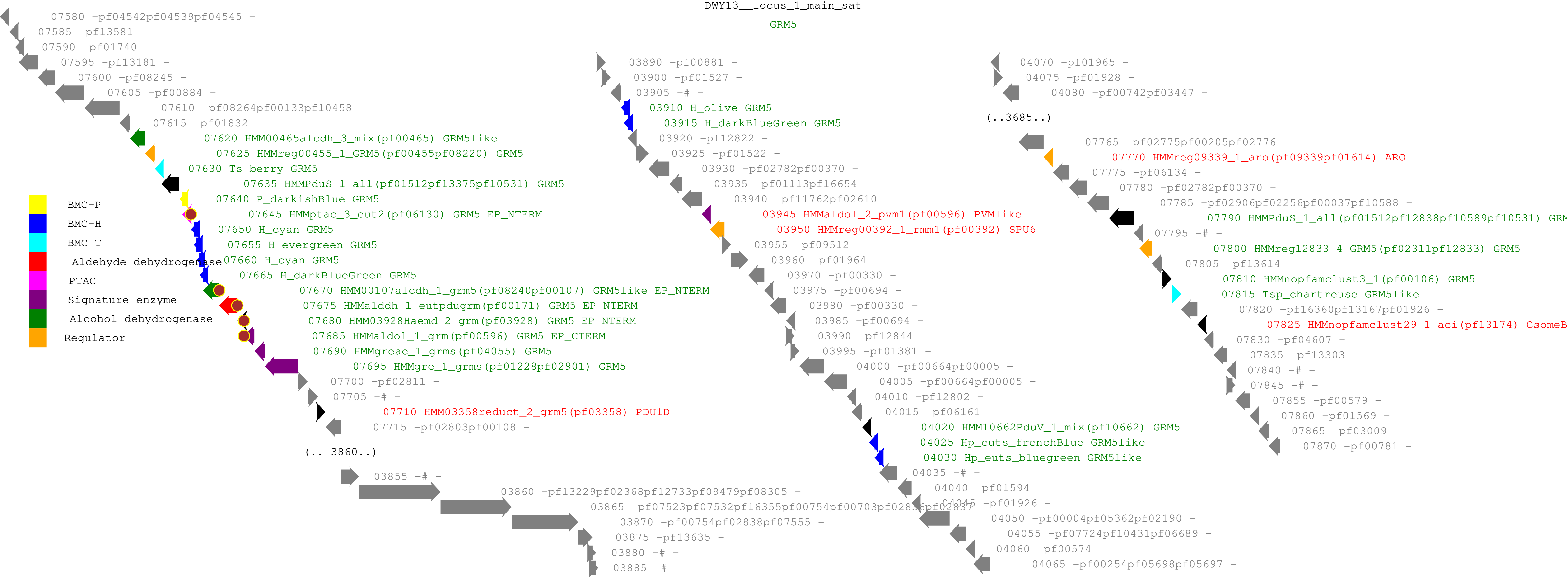

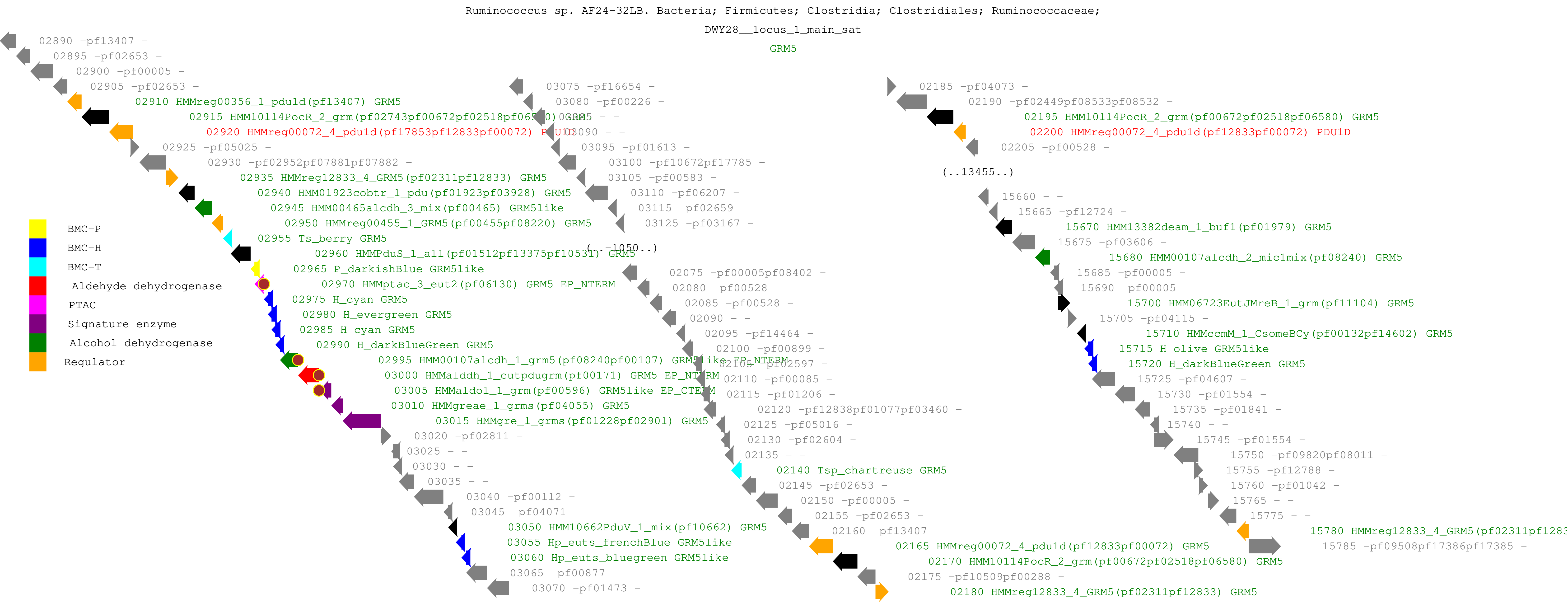

Ruminococcus sp. AF24-32LB. Bacteria; Firmicutes; Clostridia; Clostridiales; Ruminococcaceae;
DWY28__locus_1_main_sat
GRM5
BMC-P
BMC-H
BMC-T
Aldehyde dehydrogenase
PTAC
Signature enzyme
Alcohol dehydrogenase
Regulator
02890 -pf13407 -
02895 -pf02653 -
02900 -pf00005 -
02905 -pf02653 -
02910 HMMreg00356_1_pdu1d(pf13407) GRM5
02915 HMM10114PocR_2_grm(pf02743pf00672pf02518pf06580) GRM5
02920 HMMreg00072_4_pdu1d(pf17853pf12833pf00072) PDU1D
02925 -pf05025 -
02930 -pf02952pf07881pf07882 -
02935 HMMreg12833_4_GRM5(pf02311pf12833) GRM5
02940 HMM01923cobtr_1_pdu(pf01923pf03928) GRM5
02945 HMM00465alcdh_3_mix(pf00465) GRM5like
02950 HMMreg00455_1_GRM5(pf00455pf08220) GRM5
02955 Ts_berry GRM5
02960 HMMPduS_1_all(pf01512pf13375pf10531) GRM5
02965 P_darkishBlue GRM5like
02970 HMMptac_3_eut2(pf06130) GRM5 EP_NTERM
02975 H_cyan GRM5
02980 H_evergreen GRM5
02985 H_cyan GRM5
02990 H_darkBlueGreen GRM5
02995 HMM00107alcdh_1_grm5(pf08240pf00107) GRM5like EP_NTERM
03000 HMMaldh_1_eutpdugrm(pf00171) GRM5 EP_NTERM
03005 HMMaldol_1_grm(pf00596) GRM5like EP_CTERM
03010 HMMgreae_1_grms(pf04055) GRM5
03015 HMMgre_1_grms(pf01228pf02901) GRM5
03020 -pf02811 -
03025 - -
03030 - -
03035 - -
03040 -pf00112 -
03045 -pf04071 -
03050 HMM10662PduV_1_mix(pf10662) GRM5
03055 Hp_euts_frenchBlue GRM5like
03060 Hp_euts_bluegreen GRM5like
03065 -pf00877 -
03070 -pf01473 -
03075 -pf16654 -
03080 -pf00226 -
03085 - -
03090 - -
03095 -pf01613 -
03100 -pf10672pf17785 -
03105 -pf00583 -
03110 -pf06207 -
03115 -pf02659 -
03125 -pf03167 -
(..-1050..)
02075 -pf00005pf08402 -
02080 -pf00528 -
02085 -pf00528 -
02090 - -
02095 -pf14464 -
02100 -pf00899 -
02105 -pf02597 -
02110 -pf00085 -
02115 -pf01206 -
02120 -pf12838pf01077pf03460 -
02125 -pf05016 -
02130 -pf02604 -
02135 - -
02140 Tsp_chartreuse GRM5
02145 -pf02653 -
02150 -pf00005 -
02155 -pf02653 -
02160 -pf13407 -
02165 HMMreg00072_4_pdu1d(pf12833pf00072) GRM5
02170 HMM10114PocR_2_grm(pf00672pf02518pf06580) GRM5
02175 -pf10509pf00288 -
02180 HMMreg12833_4_GRM5(pf02311pf12833) GRM5
02185 -pf04073 -
02190 -pf02449pf08533pf08532 -
02195 HMM10114PocR_2_grm(pf00672pf02518pf06580) GRM5
02200 HMMreg00072_4_pdu1d(pf12833pf00072) PDU1D
02205 -pf00528 -
(..13455..)
15660 - -
15665 -pf12724 -
15670 HMM13382deam_1_buf1(pf01979) GRM5
15675 -pf03606 -
15680 HMM00107alcdh_2_mic1mix(pf08240) GRM5
15685 -pf00005 -
15690 -pf00005 -
15700 HMM06723EutJMreB_1_grm(pf11104) GRM5
15705 -pf04115 -
15710 HMMccmM_1_CsomeBCy(pf00132pf14602) GRM5
15715 H_olive GRM5like
15720 H_darkBlueGreen GRM5
15725 -pf04607 -
15730 -pf01554 -
15735 -pf01841 -
15740 - -
15745 -pf01554 -
15750 -pf09820pf08011 -
15755 -pf12788 -
15760 -pf01042 -
15765 - -
15775 - -
15780 HMMreg12833_4_GRM5(pf02311pf1283
15785 -pf09508pf17386pf17385 -

Ruminococcus sp. AF25-13. Bacteria; Firmicutes; Clostridia; Clostridiales; Ruminococcaceae;

DWY35__locus_1_main_sat

GRM5

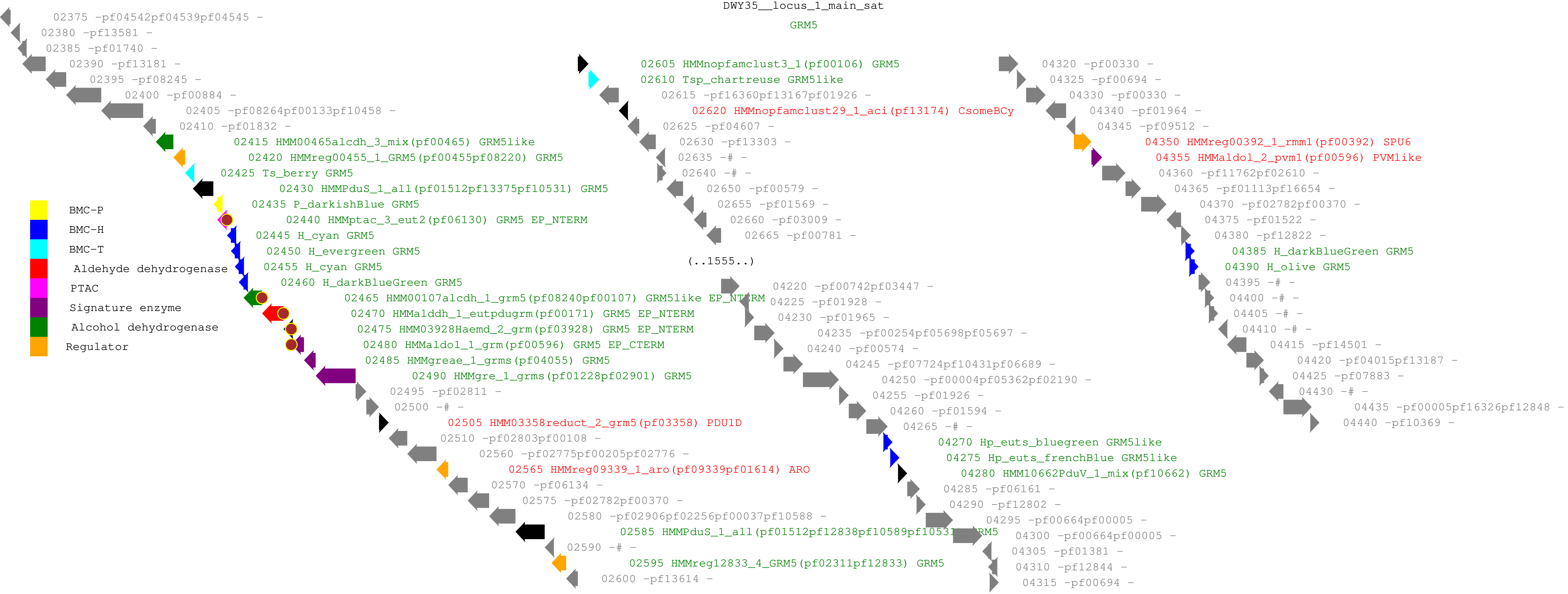

Ruminococcus sp. AF25-17. Bacteria; Firmicutes; Clostridia; Clostridiales; Ruminococcaceae;

DWY40__locus_1_main_sat

GRM5

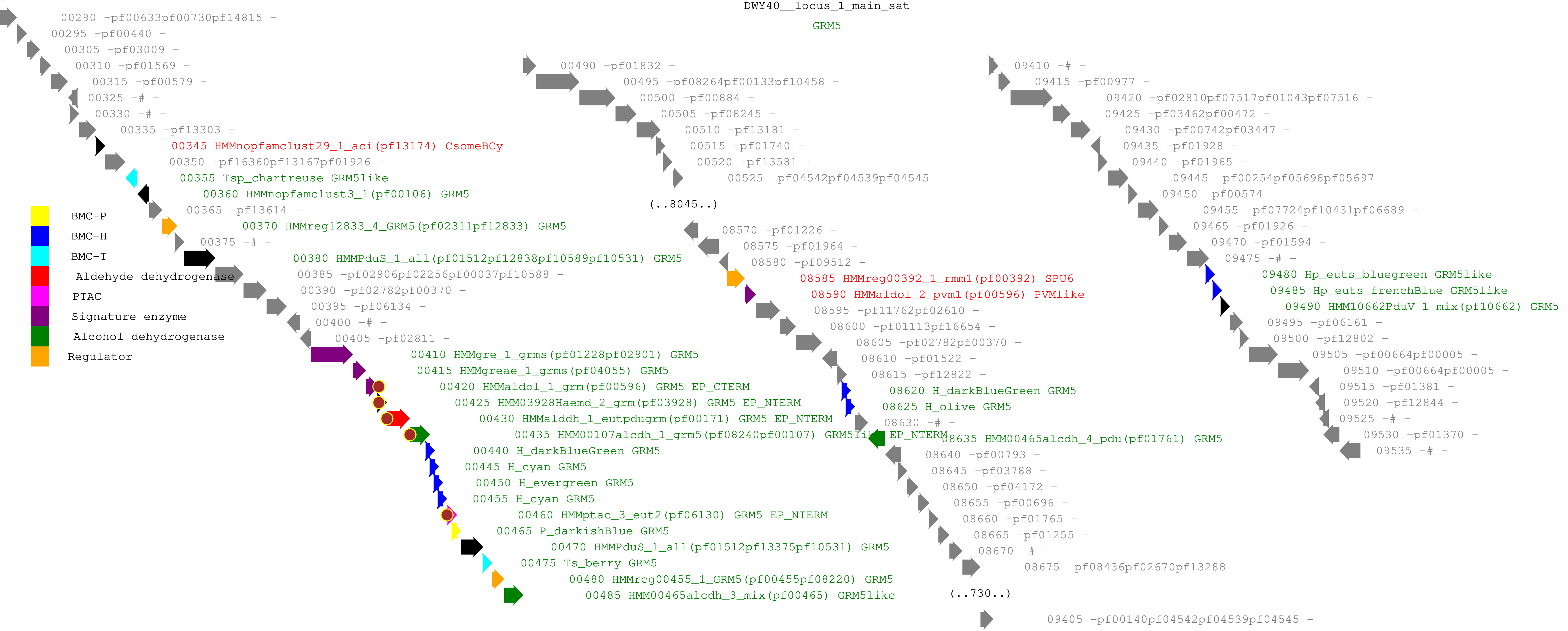

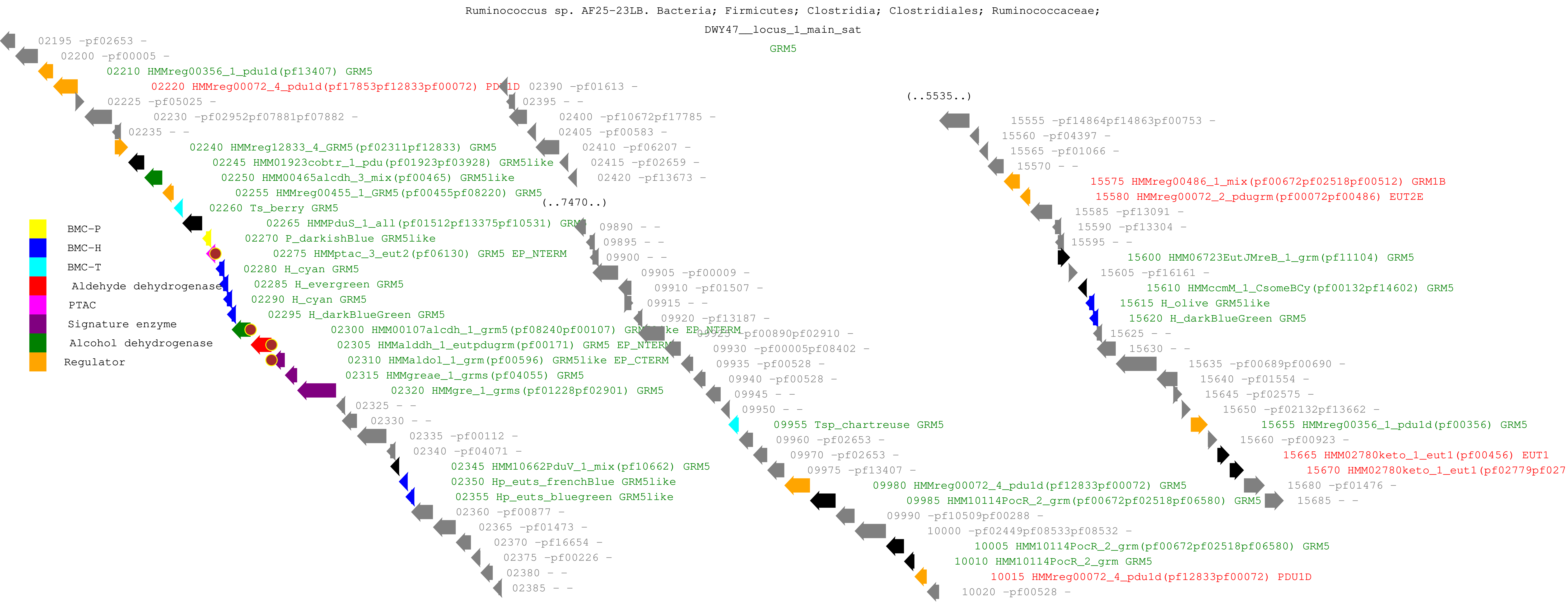
Ruminococcus sp. AF25-23LB. Bacteria; Firmicutes; Clostridia; Clostridiales; Ruminococcaceae;
DWY47__locus_1_main_sat
GRM5
02195 -pf02653 -
02200 -pf00005 -
02210 HMMreg00356_1_pdu1d(pf13407) GRM5
02220 HMMreg00072_4_pdu1d(pf17853pf12833pf00072) PDU1D
02225 -pf05025 -
02230 -pf02952pf07881pf07882 -
02235 - -
02240 HMMreg12833_4_GRM5(pf02311pf12833) GRM5
02245 HMM01923cobtr_1_pdu(pf01923pf03928) GRM5like
02250 HMM00465alcdh_3_mix(pf00465) GRM5like
02255 HMMreg00455_1_GRM5(pf00455pf08220) GRM5
02260 Ts_berry GRM5
02265 HMMPduS_1_all(pf01512pf13375pf10531) GRM5
02270 P_darkishBlue GRM5like
02275 HMMptac_3_eut2(pf06130) GRM5 EP_NTERM
02280 H_cyan GRM5
02285 H_evergreen GRM5
02290 H_cyan GRM5
02295 H_darkBlueGreen GRM5
02300 HMM00107alcdh_1_grm5(pf08240pf00107) GRM5like EP_NTERM
02305 HMMaldh_1_eutpdugrm(pf00171) GRM5 EP_NTERM
02310 HMMaldol_1_grm(pf00596) GRM5like EP_CTERM
02315 HMMgreae_1_grms(pf04055) GRM5
02320 HMMgre_1_grms(pf01228pf02901) GRM5
02325 - -
02330 - -
02335 -pf00112 -
02340 -pf04071 -
02345 HMM10662PduV_1_mix(pf10662) GRM5
02350 Hp_euts_frenchBlue GRM5like
02355 Hp_euts_bluegreen GRM5like
02360 -pf00877 -
02365 -pf01473 -
02370 -pf16654 -
02375 -pf00226 -
02380 - -
02385 - -
02390 -pf01613 -
02395 - -
02400 -pf10672pf17785 -
02405 -pf00583 -
02410 -pf06207 -
02415 -pf02659 -
02420 -pf13673 -
(..7470..)
09890 - -
09895 - -
09900 - -
09905 -pf00009 -
09910 -pf01507 -
09915 - -
09920 -pf13187 -
09925 -pf00890pf02910 -
09930 -pf00005pf08402 -
09935 -pf00528 -
09940 -pf00528 -
09945 - -
09950 - -
09955 Tsp_chartreuse GRM5
09960 -pf02653 -
09970 -pf02653 -
09975 -pf13407 -
09980 HMMreg00072_4_pdu1d(pf12833pf00072) GRM5
09985 HMM10114PocR_2_grm(pf00672pf02518pf06580) GRM5
09990 -pf10509pf00288 -
10000 -pf02449pf08533pf08532 -
10005 HMM10114PocR_2_grm(pf00672pf02518pf06580) GRM5
10010 HMM10114PocR_2_grm GRM5
10015 HMMreg00072_4_pdu1d(pf12833pf00072) PDU1D
10020 -pf00528 -
(..5535..)
15555 -pf14864pf14863pf00753 -
15560 -pf04397 -
15565 -pf01066 -
15570 - -
15575 HMMreg00486_1_mix(pf00672pf02518pf00512) GRM1B
15580 HMMreg00072_2_pdugrm(pf00072pf00486) EUT2E
15585 -pf13091 -
15590 -pf13304 -
15595 - -
15600 HMM06723EutJMreB_1_grm(pf11104) GRM5
15605 -pf16161 -
15610 HMMccmM_1_CsomeBCy(pf00132pf14602) GRM5
15615 H_olive GRM5like
15620 H_darkBlueGreen GRM5
15625 - -
15630 - -
15635 -pf00689pf00690 -
15640 -pf01554 -
15645 -pf02575 -
15650 -pf02132pf13662 -
15655 HMMreg00356_1_pdu1d(pf00356) GRM5
15660 -pf00923 -
15665 HMM02780keto_1_eut1(pf00456) EUT1
15670 HMM02780keto_1_eut1(pf02779pf027
15680 -pf01476 -
15685 - -
BMC-P
BMC-H
BMC-T
Aldehyde dehydrogenase
PTAC
Signature enzyme
Alcohol dehydrogenase
Regulator

Ruminococcus sp. AF25-28AC. Bacteria; Firmicutes; Clostridia; Clostridiales; Ruminococcaceae;

DWY50__locus_1_main_sat

GRM5

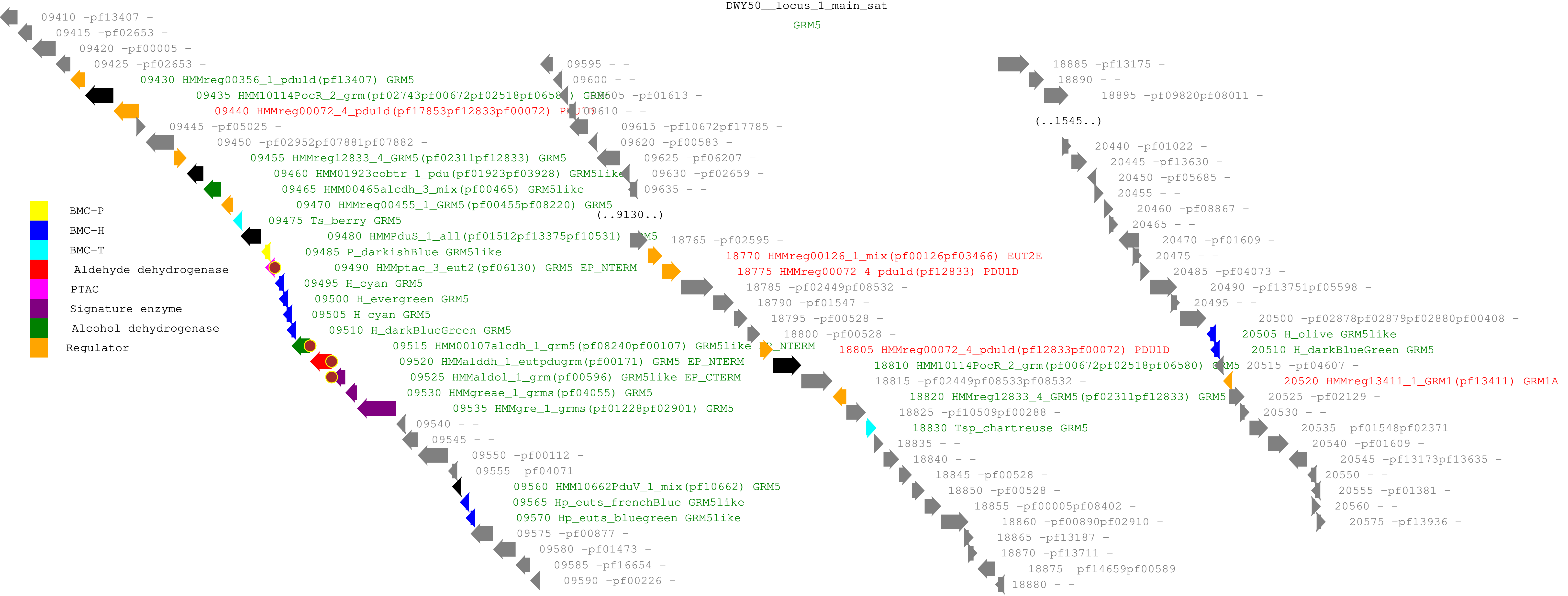

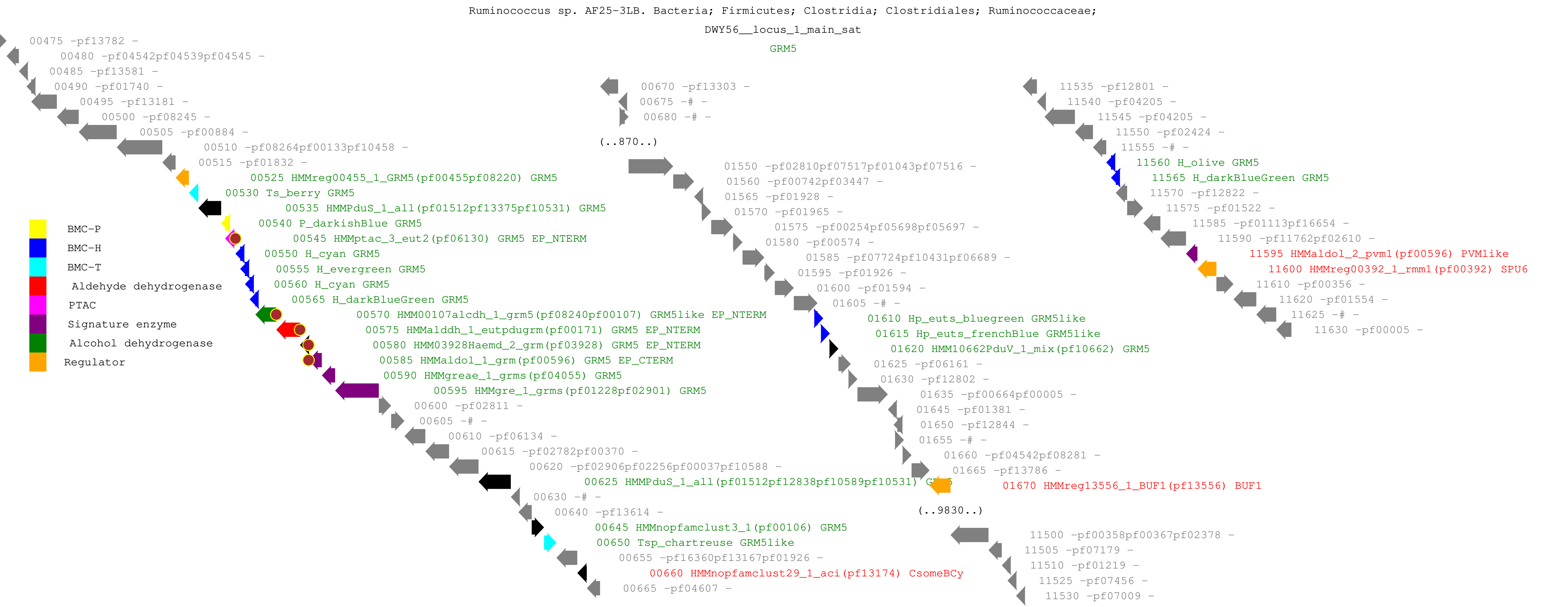

Ruminococcus sp. AF25-3LB. Bacteria; Firmicutes; Clostridia; Clostridiales; Ruminococcaceae;
DWY56__locus_1_main_sat
GRM5
BMC-P
BMC-H
BMC-T
Aldehyde dehydrogenase
PTAC
Signature enzyme
Alcohol dehydrogenase
Regulator
00475 -pf13782 -
00480 -pf04542pf04539pf04545 -
00485 -pf13581 -
00490 -pf01740 -
00495 -pf13181 -
00500 -pf08245 -
00505 -pf00884 -
00510 -pf08264pf00133pf10458 -
00515 -pf01832 -
00525 HMMreg00455_1_GRM5(pf00455pf08220) GRM5
00530 Ts_berry GRM5
00535 HMMPduS_1_all(pf01512pf13375pf10531) GRM5
00540 P_darkishBlue GRM5
00545 HMMptac_3_eut2(pf06130) GRM5 EP_NTERM
00550 H_cyan GRM5
00555 H_evergreen GRM5
00560 H_cyan GRM5
00565 H_darkBlueGreen GRM5
00570 HMM00107alcdh_1_grm5(pf08240pf00107) GRM5like EP_NTERM
00575 HMMaldddh_1_eutpdugrm(pf00171) GRM5 EP_NTERM
00580 HMM03928Haemd_2_grm(pf03928) GRM5 EP_NTERM
00585 HMMaldol_1_grm(pf00596) GRM5 EP_CTERM
00590 HMMgreae_1_grms(pf04055) GRM5
00595 HMMgre_1_grms(pf01228pf02901) GRM5
00600 -pf02811 -
00605 -# -
00610 -pf06134 -
00615 -pf02782pf00370 -
00620 -pf02906pf02256pf00037pf10588 -
00625 HMMPduS_1_all(pf01512pf12838pf10589pf10531) GRM5
00630 -# -
00640 -pf13614 -
00645 HMMnopfamclust3_1(pf00106) GRM5
00650 Tsp_chartreuse GRM5like
00655 -pf16360pf13167pf01926 -
00660 HMMnopfamclust29_1_aci(pf13174) CsomeBCy
00665 -pf04607 -
00670 -pf13303 -
00675 -# -
00680 -# -
(..870..)
01550 -pf02810pf07517pf01043pf07516 -
01560 -pf00742pf03447 -
01565 -pf01928 -
01570 -pf01965 -
01575 -pf00254pf05698pf05697 -
01580 -pf00574 -
01585 -pf07724pf10431pf06689 -
01595 -pf01926 -
01600 -pf01594 -
01605 -# -
01610 Hp_euts_bluegreen GRM5like
01615 Hp_euts_frenchBlue GRM5like
01620 HMM10662PduV_1_mix(pf10662) GRM5
01625 -pf06161 -
01630 -pf12802 -
01635 -pf00664pf00005 -
01645 -pf01381 -
01650 -pf12844 -
01655 -# -
01660 -pf04542pf08281 -
01665 -pf13786 -
01670 HMMreg13556_1_BUF1(pf13556) BUF1
(..9830..)
11500 -pf00358pf00367pf02378 -
11505 -pf07179 -
11510 -pf01219 -
11525 -pf07456 -
11530 -pf07009 -
11535 -pf12801 -
11540 -pf04205 -
11545 -pf04205 -
11550 -pf02424 -
11555 -# -
11560 H_olive GRM5
11565 H_darkBlueGreen GRM5
11570 -pf12822 -
11575 -pf01522 -
11585 -pf01113pf16654 -
11590 -pf11762pf02610 -
11595 HMMaldol_2_pvm1(pf00596) PVMlike
11600 HMMreg00392_1_rmm1(pf00392) SPU6
11610 -pf00356 -
11620 -pf01554 -
11625 -# -
11630 -pf00005 -

Eisenbergiella massiliensis. Bacteria; Firmicutes; Clostridia; Clostridiales; Lachnospiraceae;

DWY69__locus_1

GRM5

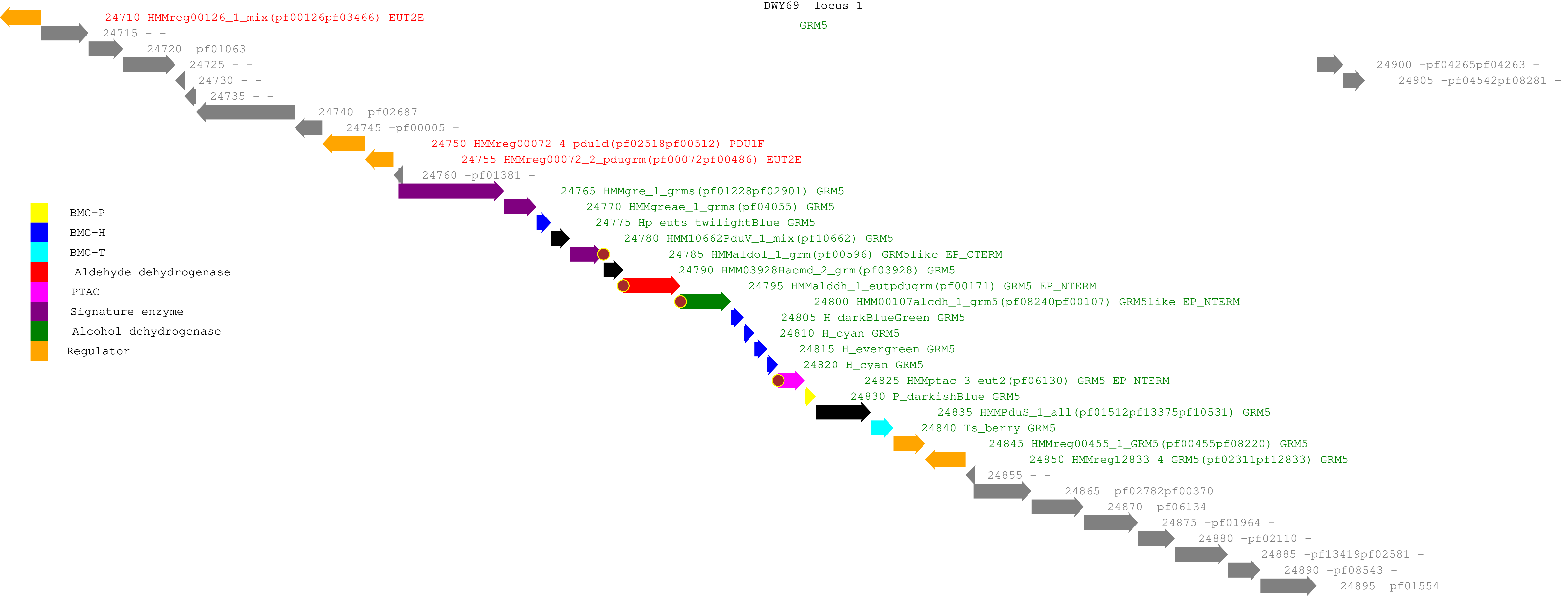

Ruminococcus sp. AF27-11AA. Bacteria; Firmicutes; Clostridia; Clostridiales; Ruminococcaceae;

DWY75__locus_1_main_sat

GRM5

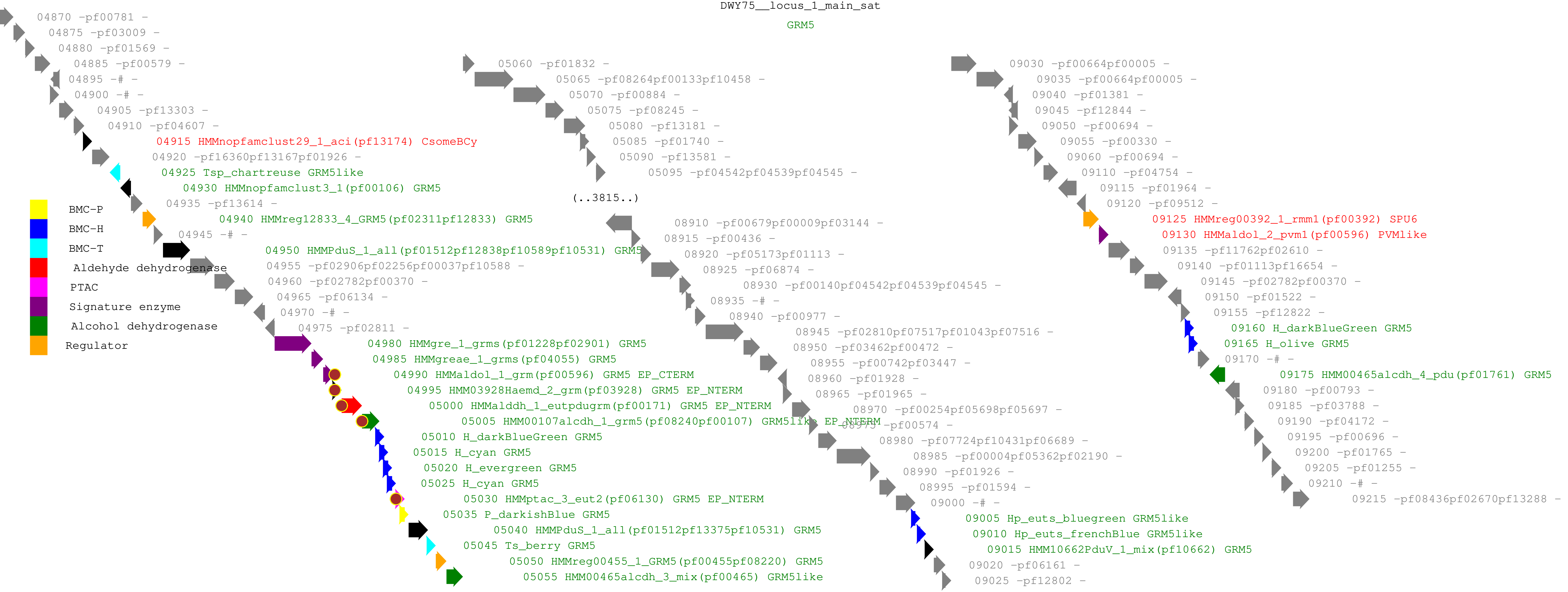

Ruminococcus sp. AF27-12AA. Bacteria; Firmicutes; Clostridia; Clostridiales; Ruminococcaceae;

DWY78__locus_1_main_sat

GRM5

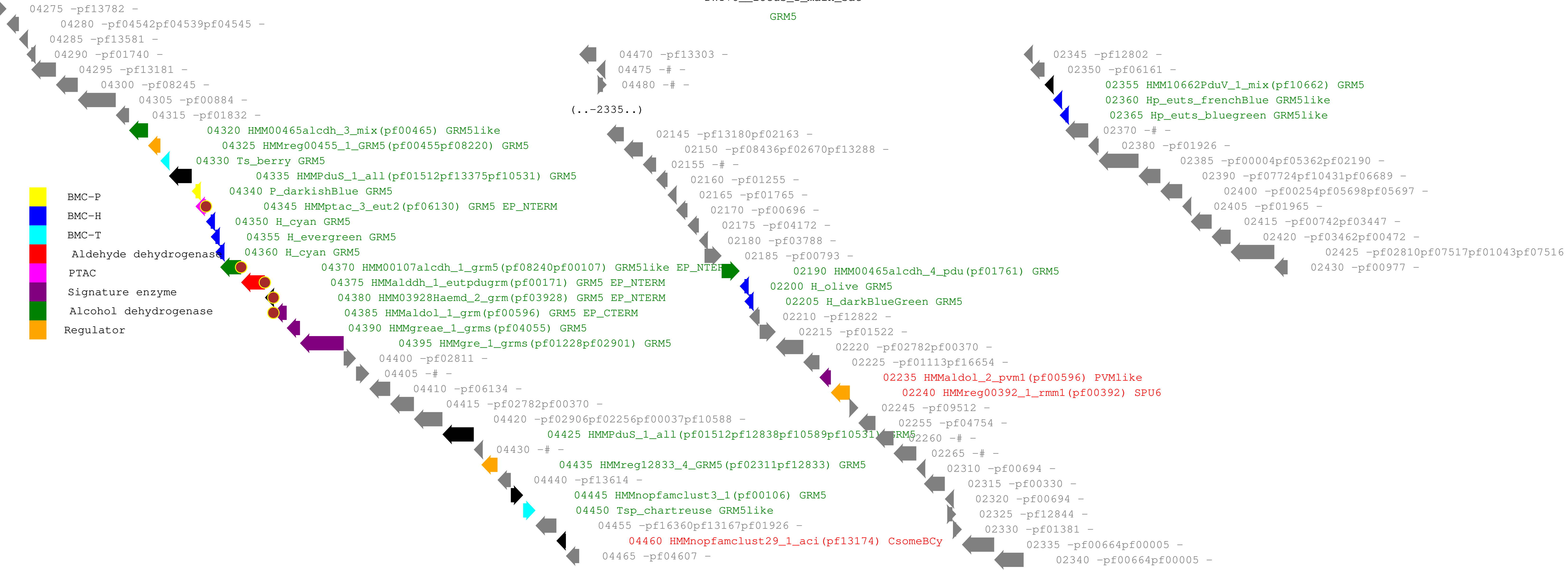

Ruminococcus sp. AF27-3. Bacteria; Firmicutes; Clostridia; Clostridiales; Ruminococcaceae;

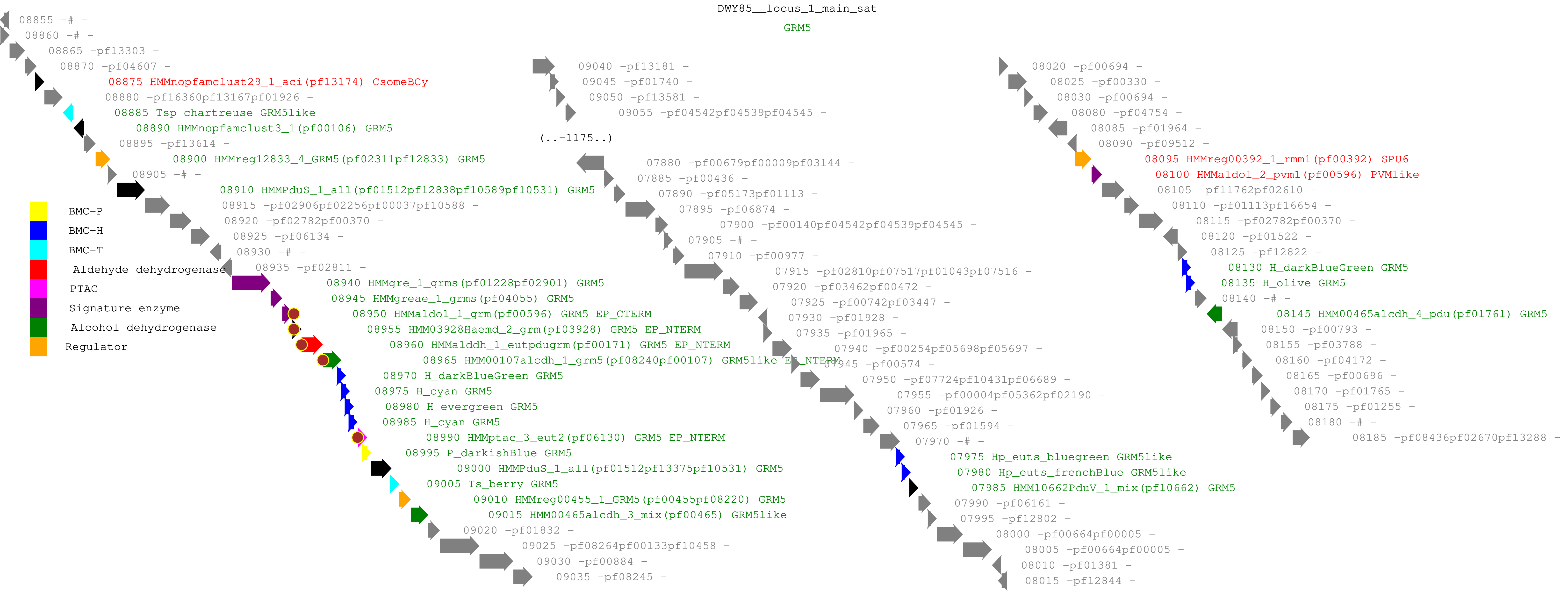

Roseburia inulinivorans. Bacteria; Firmicutes; Clostridia; Clostridiales; Lachnospiraceae;

DWY96__locus_1

GRM5

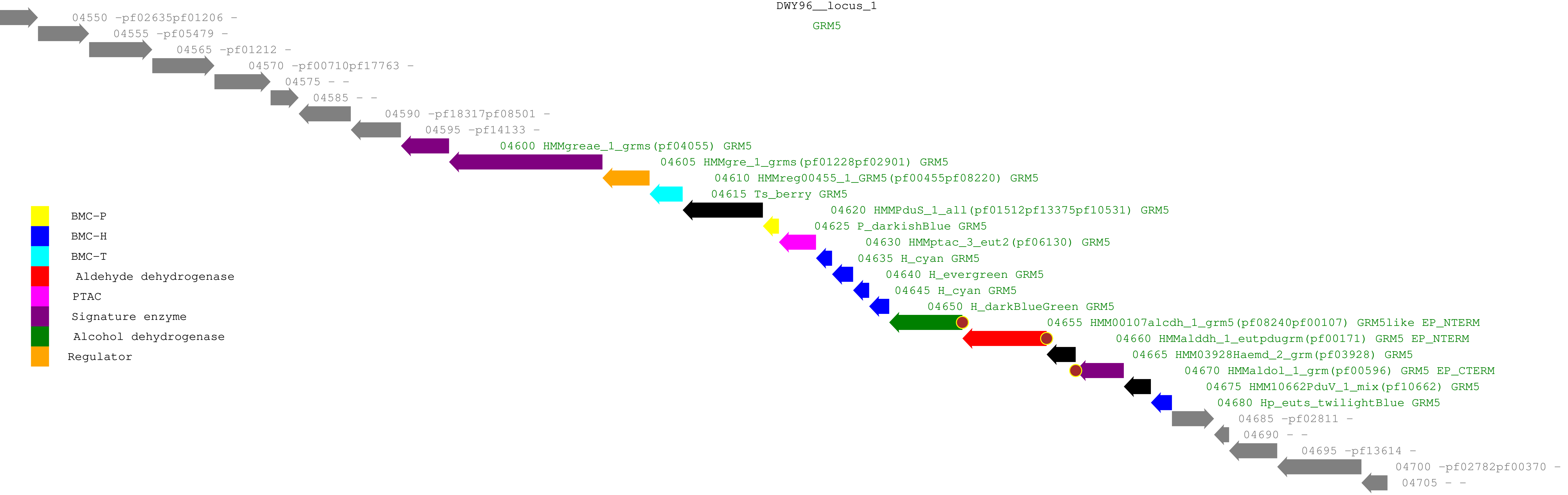

Ruminococcus sp. AF31-14BH. Bacteria; Firmicutes; Clostridia; Clostridiales; Ruminococcaceae;

DWZ26__locus_1_main_sat

GRM5

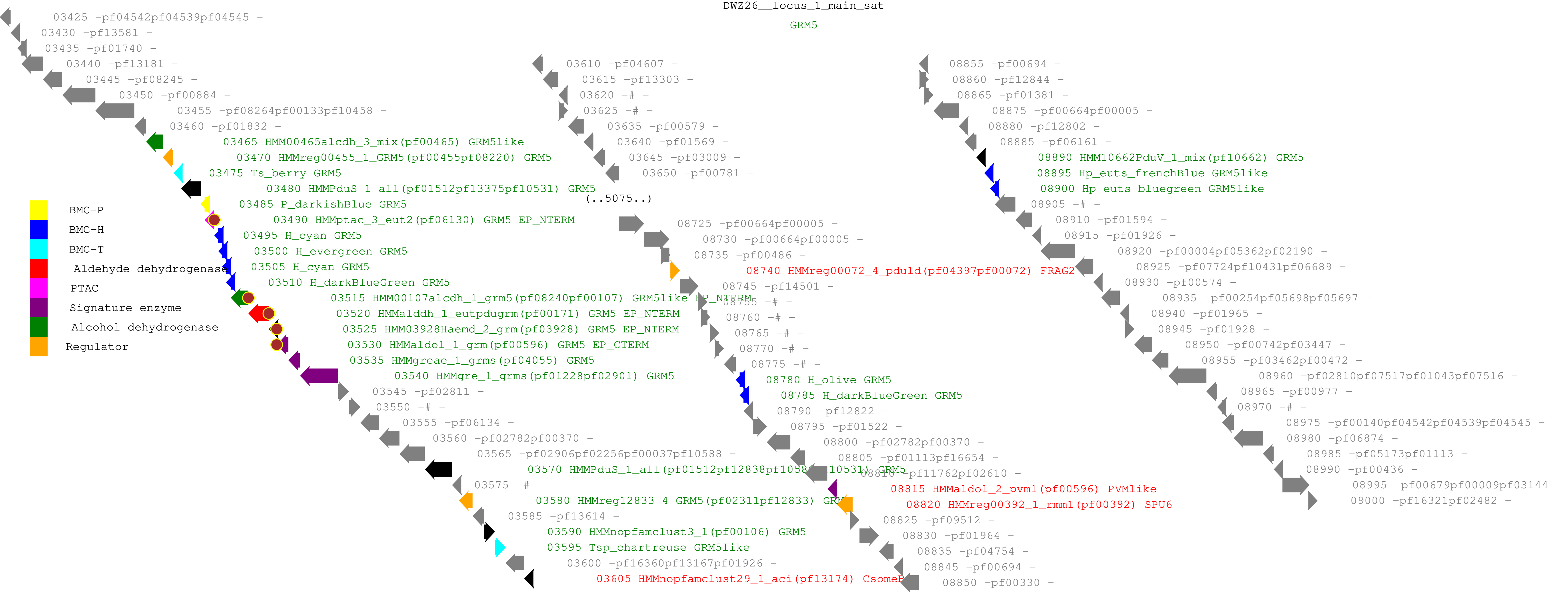

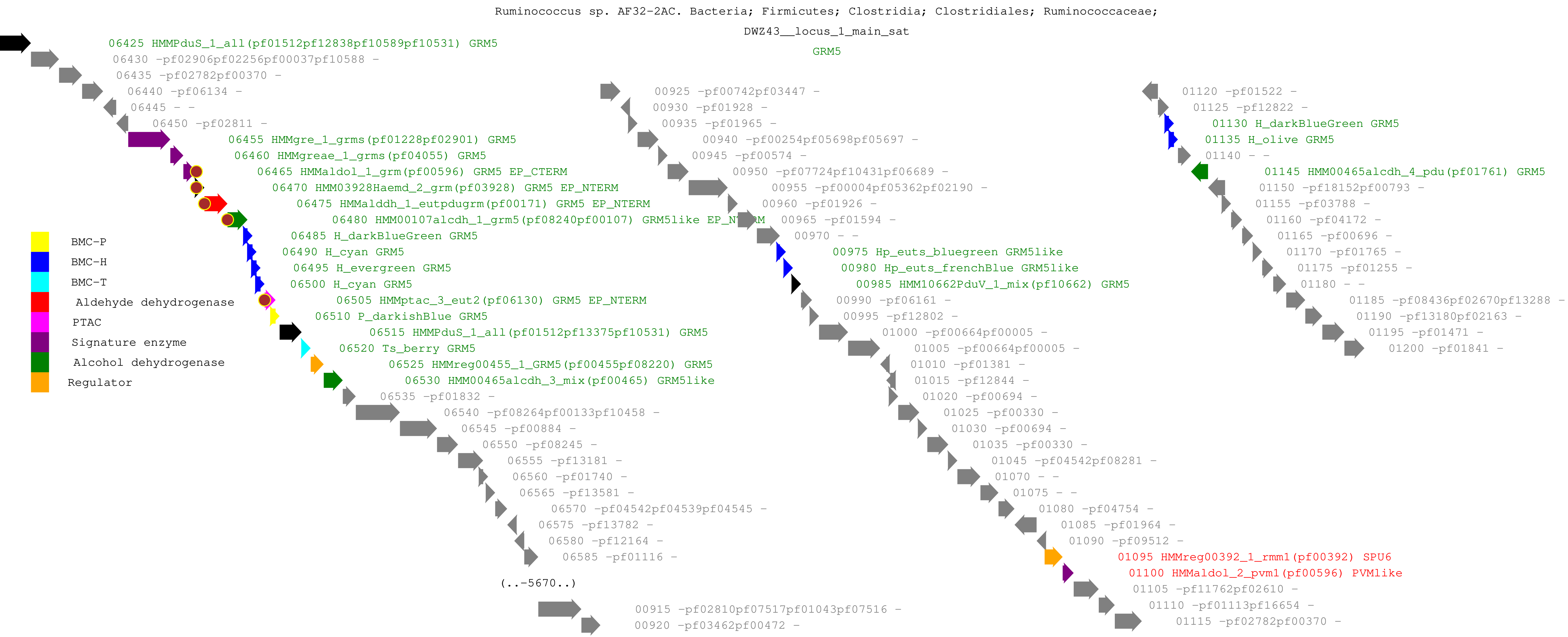
Ruminococcus sp. AF32-2AC. Bacteria; Firmicutes; Clostridia; Clostridiales; Ruminococcaceae;
DWZ43__locus_1_main_sat
GRM5
06425 HMMPduS_1_all(pf01512pf12838pf10589pf10531) GRM5
06430 -pf02906pf02256pf00037pf10588 -
06435 -pf02782pf00370 -
06440 -pf06134 -
06445 - -
06450 -pf02811 -
06455 HMMgre_1_grms(pf01228pf02901) GRM5
06460 HMMgreae_1_grms(pf04055) GRM5
06465 HMMaldol_1_grm(pf00596) GRM5 EP_CTERM
06470 HMM03928Haemd_2_grm(pf03928) GRM5 EP_NTERM
06475 HMMaldddh_1_eutpdugrm(pf00171) GRM5 EP_NTERM
06480 HMM00107alcdh_1_grm5(pf08240pf00107) GRM5like EP_N
06485 H_darkBlueGreen GRM5
06490 H_cyan GRM5
06495 H_evergreen GRM5
06500 H_cyan GRM5
06505 HMMptac_3_eut2(pf06130) GRM5 EP_NTERM
06510 P_darkishBlue GRM5
06515 HMMPduS_1_all(pf01512pf13375pf10531) GRM5
06520 Ts_berry GRM5
06525 HMMreg00455_1_GRM5(pf00455pf08220) GRM5
06530 HMM00465alcdh_3_mix(pf00465) GRM5like
06535 -pf01832 -
06540 -pf08264pf00133pf10458 -
06545 -pf00884 -
06550 -pf08245 -
06555 -pf13181 -
06560 -pf01740 -
06565 -pf13581 -
06570 -pf04542pf04539pf04545 -
06575 -pf13782 -
06580 -pf12164 -
06585 -pf01116 -
(..-5670..)
00915 -pf02810pf07517pf01043pf07516 -
00920 -pf03462pf00472 -
00925 -pf00742pf03447 -
00930 -pf01928 -
00935 -pf01965 -
00940 -pf00254pf05698pf05697 -
00945 -pf00574 -
00950 -pf07724pf10431pf06689 -
00955 -pf00004pf05362pf02190 -
00960 -pf01926 -
00965 -pf01594 -
00970 - -
00975 Hp_euts_bluegreen GRM5like
00980 Hp_euts_frenchBlue GRM5like
00985 HMM10662PduV_1_mix(pf10662) GRM5
00990 -pf06161 -
00995 -pf12802 -
01000 -pf00664pf00005 -
01005 -pf00664pf00005 -
01010 -pf01381 -
01015 -pf12844 -
01020 -pf00694 -
01025 -pf00330 -
01030 -pf00694 -
01035 -pf00330 -
01045 -pf04542pf08281 -
01070 - -
01075 - -
01080 -pf04754 -
01085 -pf01964 -
01090 -pf09512 -
01095 HMMreg00392_1_rmm1(pf00392) SPU6
01100 HMMaldol_2_pvm1(pf00596) PVMlike
01105 -pf11762pf02610 -
01110 -pf01113pf16654 -
01115 -pf02782pf00370 -
01120 -pf01522 -
01125 -pf12822 -
01130 H_darkBlueGreen GRM5
01135 H_olive GRM5
01140 - -
01145 HMM00465alcdh_4_pdu(pf01761) GRM5
01150 -pf18152pf00793 -
01155 -pf03788 -
01160 -pf04172 -
01165 -pf00696 -
01170 -pf01765 -
01175 -pf01255 -
01180 - -
01185 -pf08436pf02670pf13288 -
01190 -pf13180pf02163 -
01195 -pf01471 -
01200 -pf01841 -
BMC-P
BMC-H
BMC-T
Aldehyde dehydrogenase
PTAC
Signature enzyme
Alcohol dehydrogenase
Regulator

Ruminococcus gnavus. Bacteria; Firmicutes; Clostridia; Clostridiales; Lachnospiraceae; Blautia.

DWZ50__locus_1

GRM5

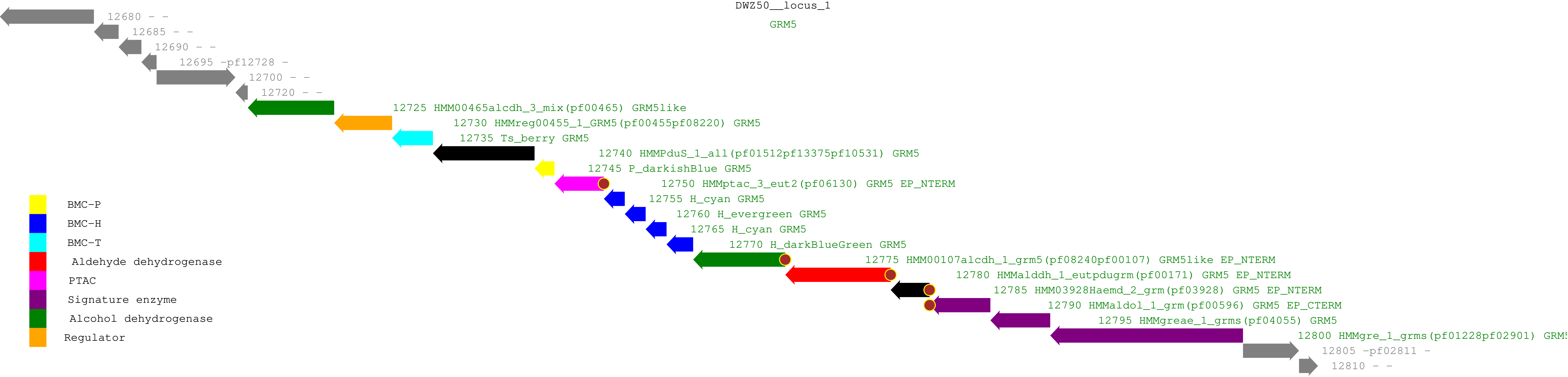

Lachnotalea sp. AF33-28. Bacteria; Firmicutes; Clostridia; Clostridiales; Lachnospiraceae;

DWZ56__locus_1_main_sat

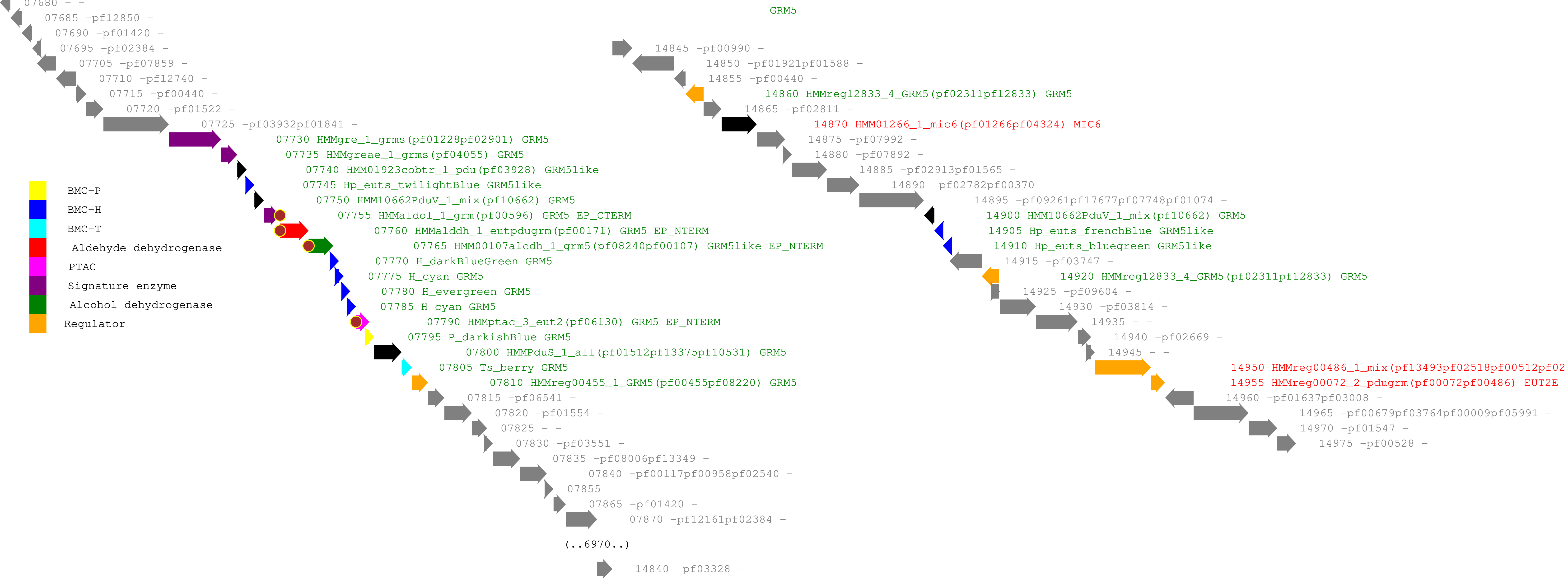

Ruminococcus sp. AM57-5. Bacteria; Firmicutes; Clostridia; Clostridiales; Ruminococcaceae;

DXA12__locus_1_main_sat

GRM5

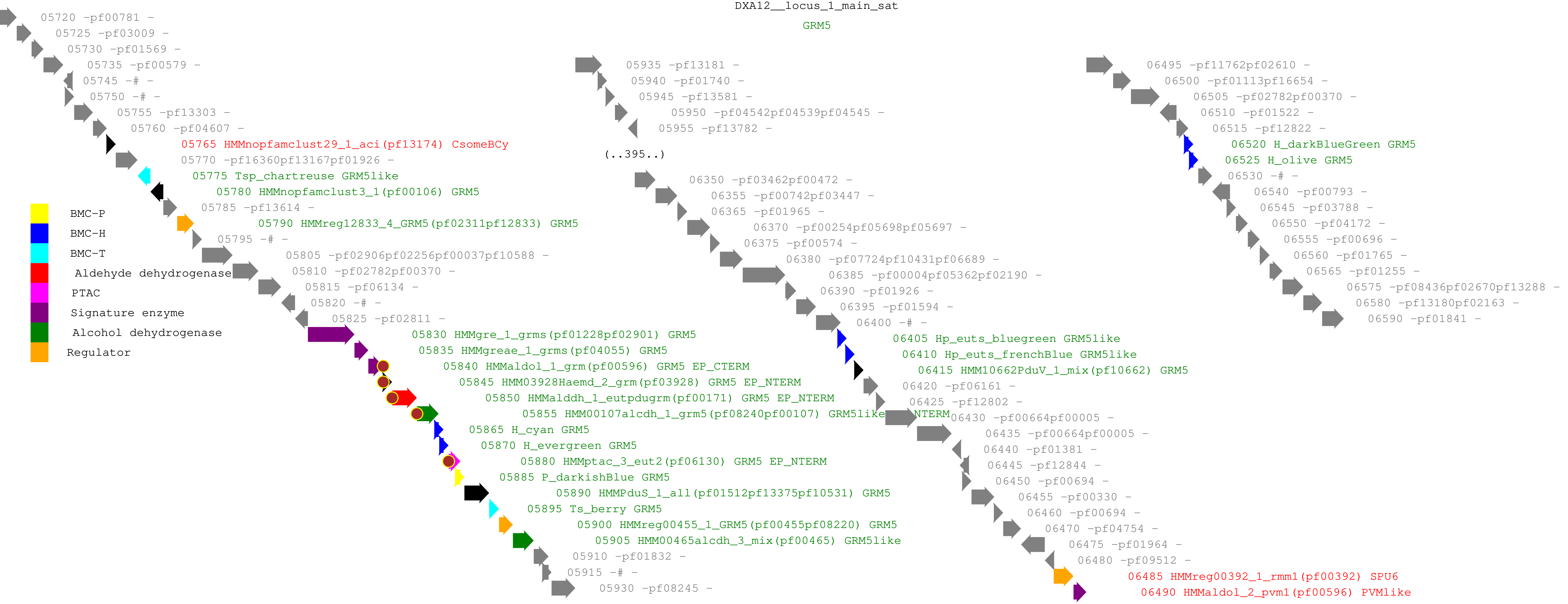

Ruminococcus sp. AM58-7XD. Bacteria; Firmicutes; Clostridia; Clostridiales; Ruminococcaceae;

DXA17__locus_1_main_sat

GRM5

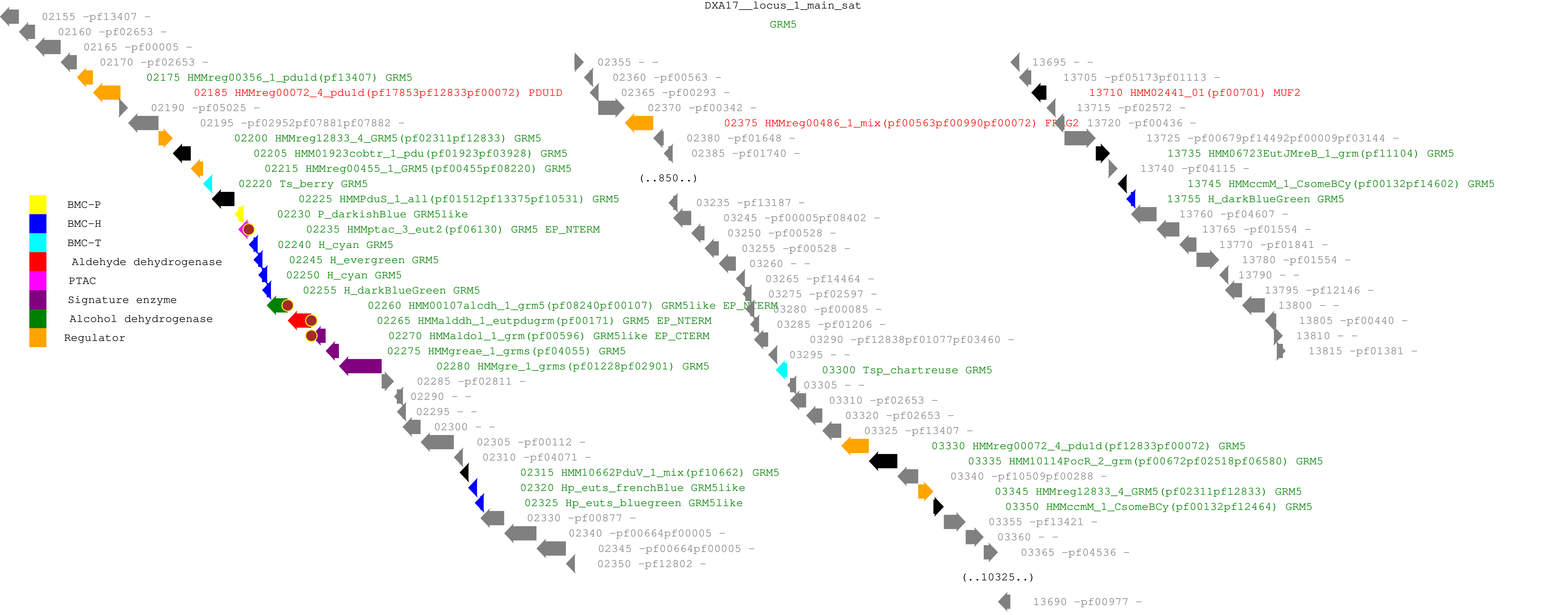

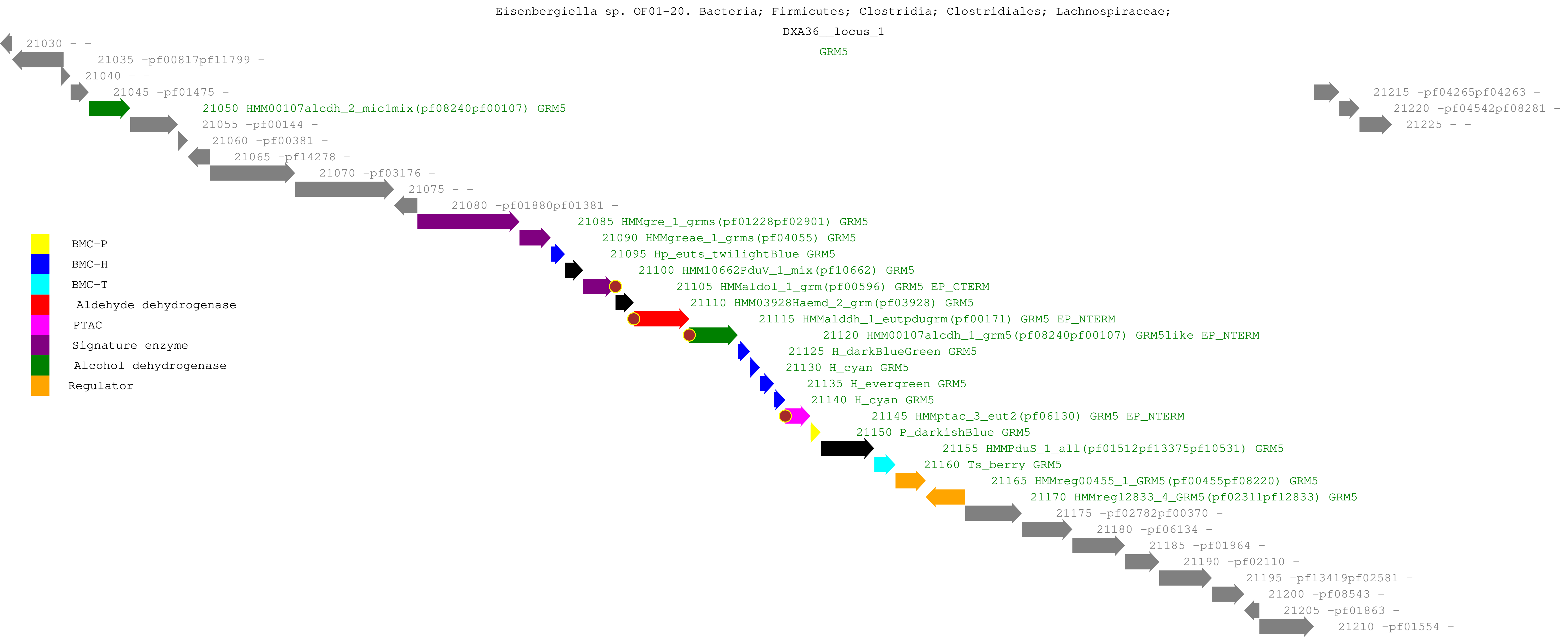
Eisenbergiella sp. OF01-20. Bacteria; Firmicutes; Clostridia; Clostridiales; Lachnospiraceae;
DXA36__locus_1
GRM5
21030 - -
21035 -pf00817pf11799 -
21040 - -
21045 -pf01475 -
21050 HMM00107alcdh_2_mic1mix(pf08240pf00107) GRM5
21055 -pf00144 -
21060 -pf00381 -
21065 -pf14278 -
21070 -pf03176 -
21075 - -
21080 -pf01880pf01381 -
21085 HMMgre_1_grms(pf01228pf02901) GRM5
21090 HMMgreae_1_grms(pf04055) GRM5
21095 Hp_euts_twilightBlue GRM5
21100 HMM10662PduV_1_mix(pf10662) GRM5
21105 HMMaldol_1_grm(pf00596) GRM5 EP_CTERM
21110 HMM03928Haemd_2_grm(pf03928) GRM5
21115 HMMalddh_1_eutpdugrm(pf00171) GRM5 EP_NTERM
21120 HMM00107alcdh_1_grm5(pf08240pf00107) GRM5like EP_NTERM
21125 H_darkBlueGreen GRM5
21130 H_cyan GRM5
21135 H_evergreen GRM5
21140 H_cyan GRM5
21145 HMMptac_3_eut2(pf06130) GRM5 EP_NTERM
21150 P_darkishBlue GRM5
21155 HMMPduS_1_all(pf01512pf13375pf10531) GRM5
21160 Ts_berry GRM5
21165 HMMreg00455_1_GRM5(pf00455pf08220) GRM5
21170 HMMreg12833_4_GRM5(pf02311pf12833) GRM5
21175 -pf02782pf00370 -
21180 -pf06134 -
21185 -pf01964 -
21190 -pf02110 -
21195 -pf13419pf02581 -
21200 -pf08543 -
21205 -pf01863 -
21210 -pf01554 -
21215 -pf04265pf04263 -
21220 -pf04542pf08281 -
21225 - -
BMC-P
BMC-H
BMC-T
Aldehyde dehydrogenase
PTAC
Signature enzyme
Alcohol dehydrogenase
Regulator

Ruminococcus sp. OF02-6. Bacteria; Firmicutes; Clostridia; Clostridiales; Ruminococcaceae;

DXA48__locus_1_main_sat

GRM5

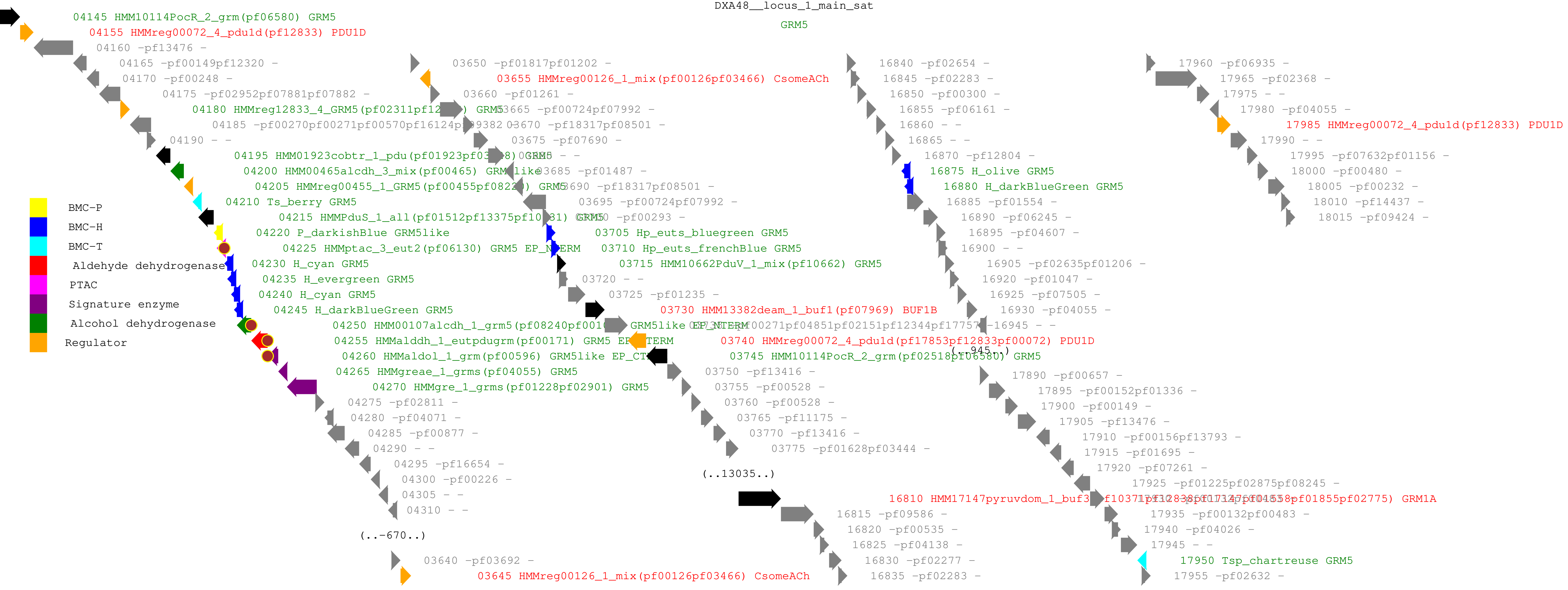

Ruminococcus sp. OF03-6AA. Bacteria; Firmicutes; Clostridia; Clostridiales; Ruminococcaceae;

DXA65__locus_1_main_sat

GRM5

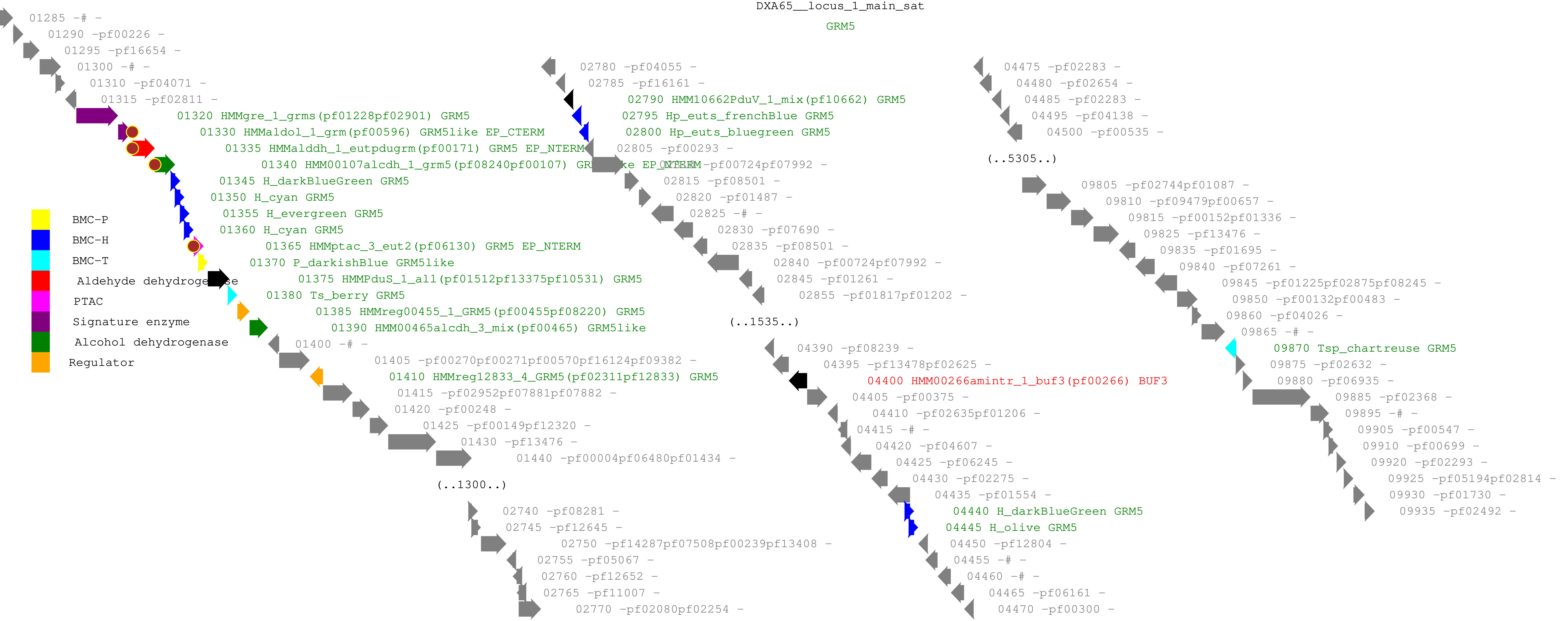

Ruminococcus sp. OF05-2BH. Bacteria; Firmicutes; Clostridia; Clostridiales; Ruminococcaceae;

DXA86__locus_1_main_sat

GRM5

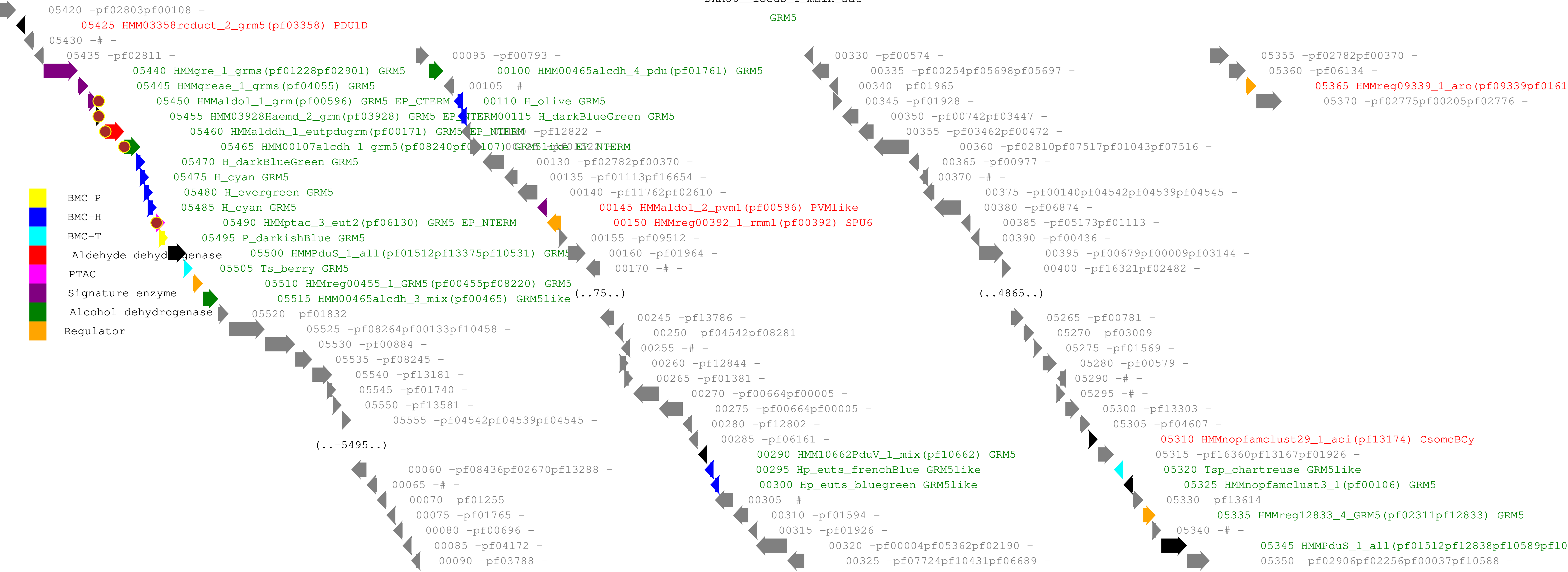

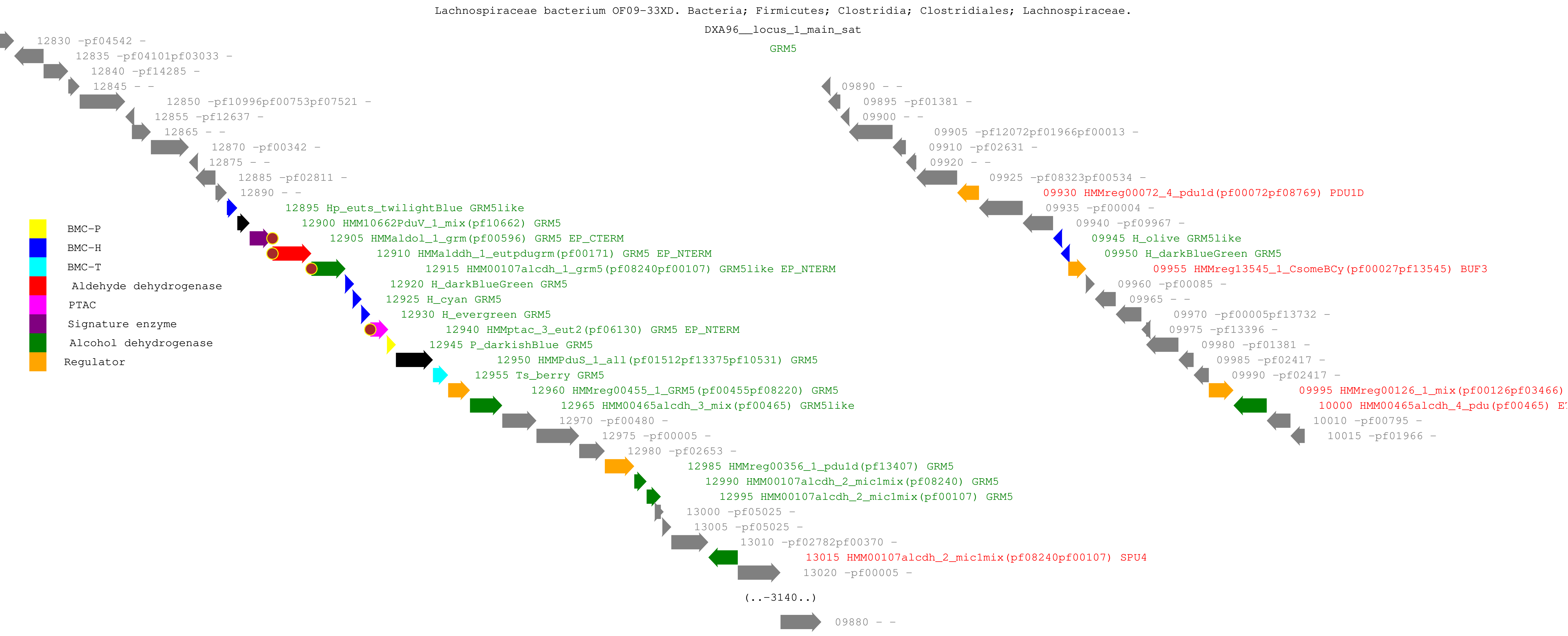

Lachnospiraceae bacterium OF09-33XD. Bacteria; Firmicutes; Clostridia; Clostridiales; Lachnospiraceae.
DXA96__locus_1_main_sat
GRM5
12830 -pf04542 -
12835 -pf04101pf03033 -
12840 -pf14285 -
12845 - -
12850 -pf10996pf00753pf07521 -
12855 -pf12637 -
12865 - -
12870 -pf00342 -
12875 - -
12885 -pf02811 -
12890 - -
12895 Hp_euts_twilightBlue GRM5like
12900 HMM10662PduV_1_mix(pf10662) GRM5
12905 HMMaldol_1_grm(pf00596) GRM5 EP_CTERM
12910 HMMalddh_1_eutpdugrm(pf00171) GRM5 EP_NTERM
12915 HMM00107alcdh_1_grm5(pf08240pf00107) GRM5like EP_NTERM
12920 H_darkBlueGreen GRM5
12925 H_cyan GRM5
12930 H_evergreen GRM5
12940 HMMptac_3_eut2(pf06130) GRM5 EP_NTERM
12945 P_darkishBlue GRM5
12950 HMMPduS_1_all(pf01512pf13375pf10531) GRM5
12955 Ts_berry GRM5
12960 HMMreg00455_1_GRM5(pf00455pf08220) GRM5
12965 HMM00465alcdh_3_mix(pf00465) GRM5like
12970 -pf00480 -
12975 -pf00005 -
12980 -pf02653 -
12985 HMMreg00356_1_pdu1d(pf13407) GRM5
12990 HMM00107alcdh_2_mic1mix(pf08240) GRM5
12995 HMM00107alcdh_2_mic1mix(pf00107) GRM5
13000 -pf05025 -
13005 -pf05025 -
13010 -pf02782pf00370 -
13015 HMM00107alcdh_2_mic1mix(pf08240pf00107) SPU4
13020 -pf00005 -
(..-3140..)
09880 - -
BMC-P
BMC-H
BMC-T
Aldehyde dehydrogenase
PTAC
Signature enzyme
Alcohol dehydrogenase
Regulator
09890 - -
09895 -pf01381 -
09900 - -
09905 -pf12072pf01966pf00013 -
09910 -pf02631 -
09920 - -
09925 -pf08323pf00534 -
09930 HMMreg00072_4_pdu1d(pf00072pf08769) PDU1D
09935 -pf00004 -
09940 -pf09967 -
09945 H_olive GRM5like
09950 H_darkBlueGreen GRM5
09955 HMMreg13545_1_CsomeBCy(pf00027pf13545) BUF3
09960 -pf00085 -
09965 - -
09970 -pf00005pf13732 -
09975 -pf13396 -
09980 -pf01381 -
09985 -pf02417 -
09990 -pf02417 -
09995 HMMreg00126_1_mix(pf00126pf03466)
10000 HMM00465alcdh_4_pdu(pf00465)
10010 -pf00795 -
10015 -pf01966 -

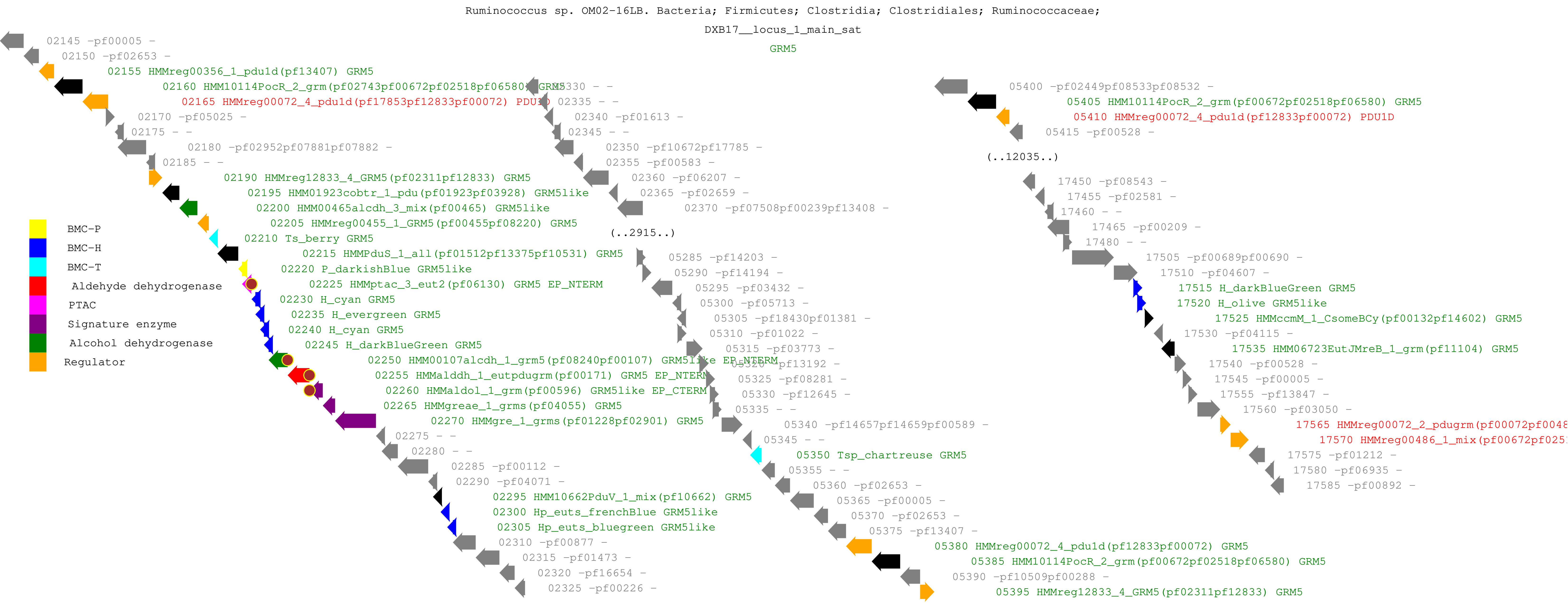

Ruminococcus sp. OM02-16LB. Bacteria; Firmicutes; Clostridia; Clostridiales; Ruminococcaceae;
DXB17__locus_1_main_sat
GRM5
BMC-P
BMC-H
BMC-T
Aldehyde dehydrogenase
PTAC
Signature enzyme
Alcohol dehydrogenase
Regulator
02145 -pf00005 -
02150 -pf02653 -
02155 HMMreg00356_1_pdu1d(pf13407) GRM5
02160 HMM10114PocR_2_grm(pf02743pf00672pf02518pf06580) GRM5
02165 HMMreg00072_4_pdu1d(pf17853pf12833pf00072) PDU1D
02170 -pf05025 -
02175 - -
02180 -pf02952pf07881pf07882 -
02185 - -
02190 HMMreg12833_4_GRM5(pf02311pf12833) GRM5
02195 HMM01923cobtr_1_pdu(pf01923pf03928) GRM5like
02200 HMM00465alcdh_3_mix(pf00465) GRM5like
02205 HMMreg00455_1_GRM5(pf00455pf08220) GRM5
02210 Ts_berry GRM5
02215 HMMPduS_1_all(pf01512pf13375pf10531) GRM5
02220 P_darkishBlue GRM5like
02225 HMMptac_3_eut2(pf06130) GRM5 EP_NTERM
02230 H_cyan GRM5
02235 H_evergreen GRM5
02240 H_cyan GRM5
02245 H_darkBlueGreen GRM5
02250 HMM00107alcdh_1_grm5(pf08240pf00107) GRM5like EP_NTERM
02255 HMMalddh_1_eutpdugrm(pf00171) GRM5 EP_NTERM
02260 HMMaldol_1_grm(pf00596) GRM5like EP_CTERM
02265 HMMgreae_1_grms(pf04055) GRM5
02270 HMMgre_1_grms(pf01228pf02901) GRM5
02275 - -
02280 - -
02285 -pf00112 -
02290 -pf04071 -
02295 HMM10662PduV_1_mix(pf10662) GRM5
02300 Hp_euts_frenchBlue GRM5like
02305 Hp_euts_bluegreen GRM5like
02310 -pf00877 -
02315 -pf01473 -
02320 -pf16654 -
02325 -pf00226 -
02330 - -
02335 - -
02340 -pf01613 -
02345 - -
02350 -pf10672pf17785 -
02355 -pf00583 -
02360 -pf06207 -
02365 -pf02659 -
02370 -pf07508pf00239pf13408 -
(..2915..)
05285 -pf14203 -
05290 -pf14194 -
05295 -pf03432 -
05300 -pf05713 -
05305 -pf18430pf01381 -
05310 -pf01022 -
05315 -pf03773 -
05320 -pf13192 -
05325 -pf08281 -
05330 -pf12645 -
05335 - -
05340 -pf14657pf14659pf00589 -
05345 - -
05350 Tsp_chartreuse GRM5
05355 - -
05360 -pf02653 -
05365 -pf00005 -
05370 -pf02653 -
05375 -pf13407 -
05380 HMMreg00072_4_pdu1d(pf12833pf00072) GRM5
05385 HMM10114PocR_2_grm(pf00672pf02518pf06580) GRM5
05390 -pf10509pf00288 -
05395 HMMreg12833_4_GRM5(pf02311pf12833) GRM5
05400 -pf02449pf08533pf08532 -
05405 HMM10114PocR_2_grm(pf00672pf02518pf06580) GRM5
05410 HMMreg00072_4_pdu1d(pf12833pf00072) PDU1D
05415 -pf00528 -
(..12035..)
17450 -pf08543 -
17455 -pf02581 -
17460 - -
17465 -pf00209 -
17480 - -
17505 -pf00689pf00690 -
17510 -pf04607 -
17515 H_darkBlueGreen GRM5
17520 H_olive GRM5like
17525 HMMccmM_1_CsomeBCy(pf00132pf14602) GRM5
17530 -pf04115 -
17535 HMM06723EutJMreB_1_grm(pf11104) GRM5
17540 -pf00528 -
17545 -pf00005 -
17555 -pf13847 -
17560 -pf03050 -
17565 HMMreg00072_2_pdugrm(pf00072pf004
17570 HMMreg00486_1_mix(pf00672pf025
17575 -pf01212 -
17580 -pf06935 -
17585 -pf00892 -

Ruminococcus sp. OM04-4AA. Bacteria; Firmicutes; Clostridia; Clostridiales; Ruminococcaceae;

DXB52__locus_1_main_sat

GRM5

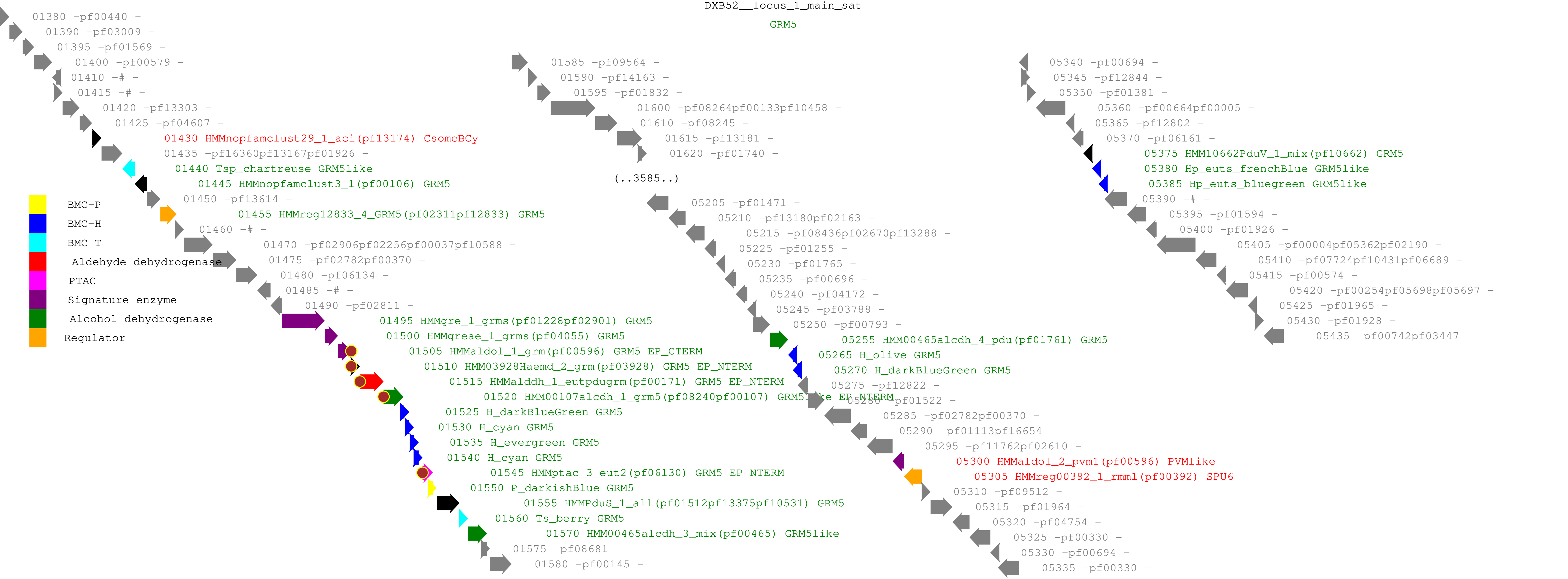

Ruminococcus sp. OM05-7. Bacteria; Firmicutes; Clostridia; Clostridiales; Ruminococcaceae;

DXB74__locus_1_main_sat

GRM5

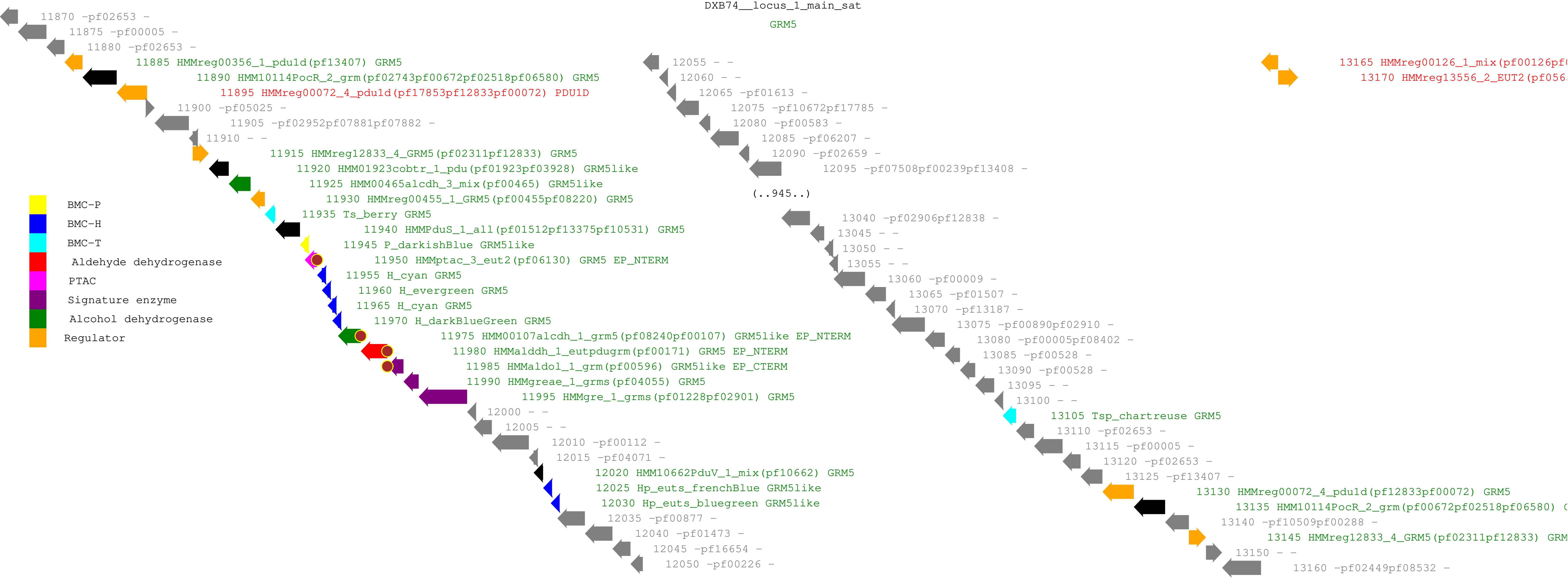

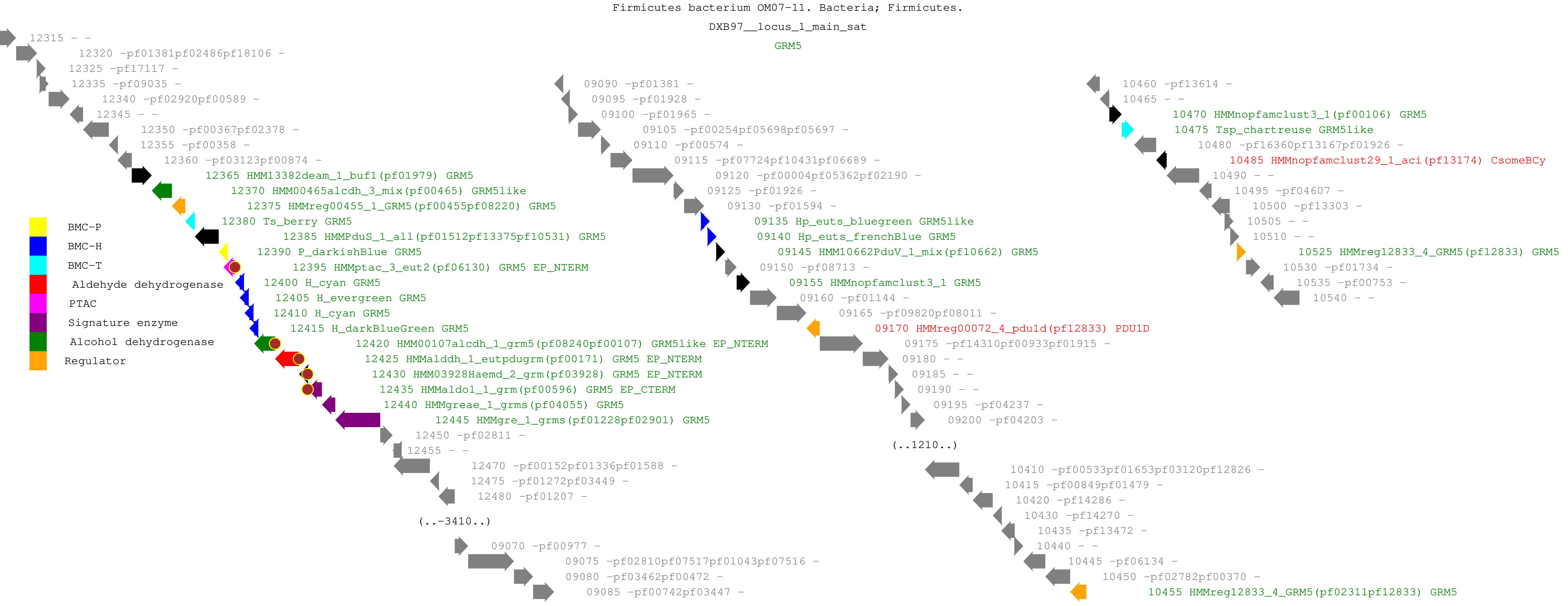
Firmicutes bacterium OM07-11. Bacteria; Firmicutes.
DXB97__locus_1_main_sat
GRM5
12315 - -
12320 -pf01381pf02486pf18106 -
12325 -pf17117 -
12335 -pf09035 -
12340 -pf02920pf00589 -
12345 - -
12350 -pf00367pf02378 -
12355 -pf00358 -
12360 -pf03123pf00874 -
12365 HMM13382deam_1_buf1(pf01979) GRM5
12370 HMM00465alcdh_3_mix(pf00465) GRM5like
12375 HMMreg00455_1_GRM5(pf00455pf08220) GRM5
12380 Ts_berry GRM5
12385 HMMPduS_1_all(pf01512pf13375pf10531) GRM5
12390 P_darkishBlue GRM5
12395 HMMptac_3_eut2(pf06130) GRM5 EP_NTERM
12400 H_cyan GRM5
12405 H_evergreen GRM5
12410 H_cyan GRM5
12415 H_darkBlueGreen GRM5
12420 HMM00107alcdh_1_grm5(pf08240pf00107) GRM5like EP_NTERM
12425 HMMalddh_1_eutpdugrm(pf00171) GRM5 EP_NTERM
12430 HMM03928Haemd_2_grm(pf03928) GRM5 EP_NTERM
12435 HMMaldol_1_grm(pf00596) GRM5 EP_CTERM
12440 HMMgreae_1_grms(pf04055) GRM5
12445 HMMgre_1_grms(pf01228pf02901) GRM5
12450 -pf02811 -
12455 - -
12470 -pf00152pf01336pf01588 -
12475 -pf01272pf03449 -
12480 -pf01207 -
(..-3410..)
09070 -pf00977 -
09075 -pf02810pf07517pf01043pf07516 -
09080 -pf03462pf00472 -
09085 -pf00742pf03447 -
BMC-P
BMC-H
BMC-T
Aldehyde dehydrogenase
PTAC
Signature enzyme
Alcohol dehydrogenase
Regulator
09090 -pf01381 -
09095 -pf01928 -
09100 -pf01965 -
09105 -pf00254pf05698pf05697 -
09110 -pf00574 -
09115 -pf07724pf10431pf06689 -
09120 -pf00004pf05362pf02190 -
09125 -pf01926 -
09130 -pf01594 -
09135 Hp_euts_bluegreen GRM5like
09140 Hp_euts_frenchBlue GRM5
09145 HMM10662PduV_1_mix(pf10662) GRM5
09150 -pf08713 -
09155 HMMnopfamclust3_1 GRM5
09160 -pf01144 -
09165 -pf09820pf08011 -
09170 HMMreg00072_4_pdu1d(pf12833) PDU1D
09175 -pf14310pf00933pf01915 -
09180 - -
09185 - -
09190 - -
09195 -pf04237 -
09200 -pf04203 -
(..1210..)
10410 -pf00533pf01653pf03120pf12826 -
10415 -pf00849pf01479 -
10420 -pf14286 -
10430 -pf14270 -
10435 -pf13472 -
10440 - -
10445 -pf06134 -
10450 -pf02782pf00370 -
10455 HMMreg12833_4_GRM5(pf02311pf12833) GRM5
10460 -pf13614 -
10465 - -
10470 HMMnopfamclust3_1(pf00106) GRM5
10475 Tsp_chartreuse GRM5like
10480 -pf16360pf13167pf01926 -
10485 HMMnopfamclust29_1_aci(pf13174) CsomeBCy
10490 - -
10495 -pf04607 -
10500 -pf13303 -
10505 - -
10510 - -
10525 HMMreg12833_4_GRM5(pf12833) GRM5
10530 -pf01734 -
10535 -pf00753 -
10540 - -

Blautia sp. OM07-19. Bacteria; Firmicutes; Clostridia; Clostridiales; Lachnospiraceae; Blautia;

DXC01__locus_1_main_sat

GRM5

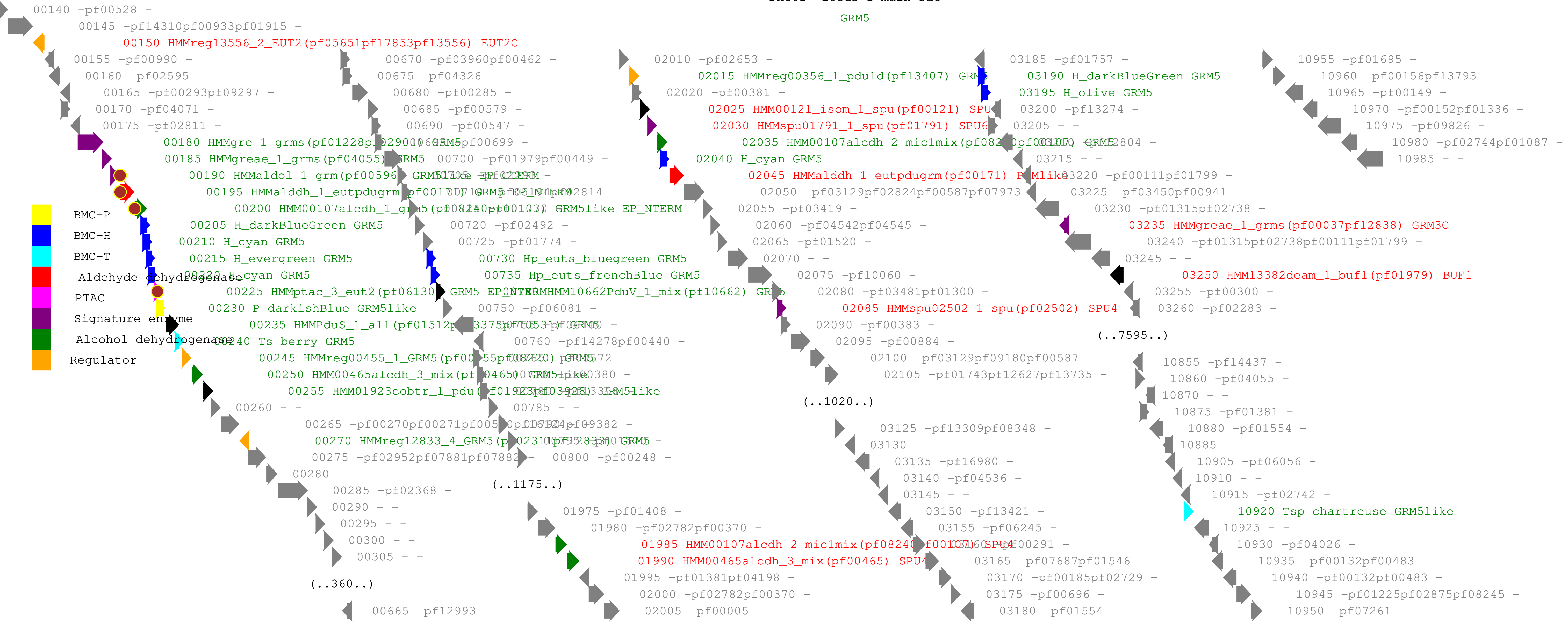

Ruminococcus sp. OM07-7. Bacteria; Firmicutes; Clostridia; Clostridiales; Ruminococcaceae;

DXC06__locus_1_main_sat

GRM5

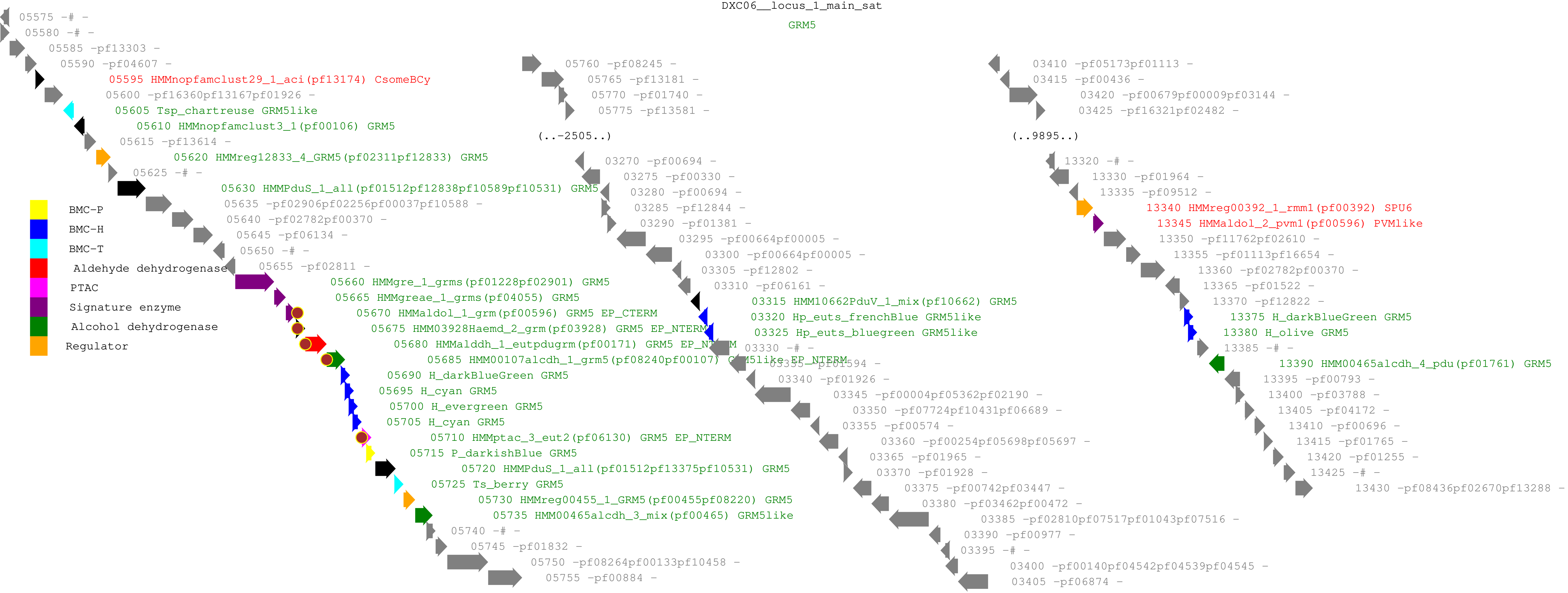

Ruminococcus sp. OM08-13AT. Bacteria; Firmicutes; Clostridia; Clostridiales; Ruminococcaceae;

DXC15__locus_1_main_sat

GRM5

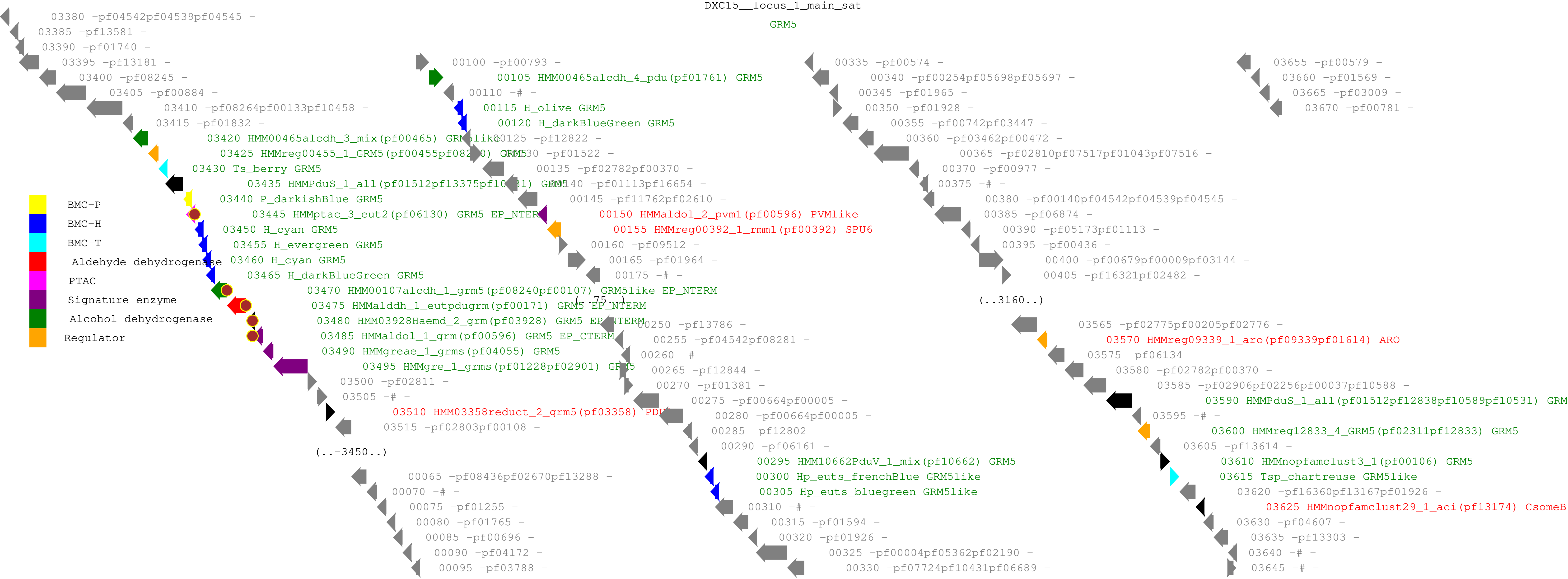

Ruminococcus sp. OM08-7. Bacteria; Firmicutes; Clostridia; Clostridiales; Ruminococcaceae;

DXC27__locus_1_main_sat

GRM5

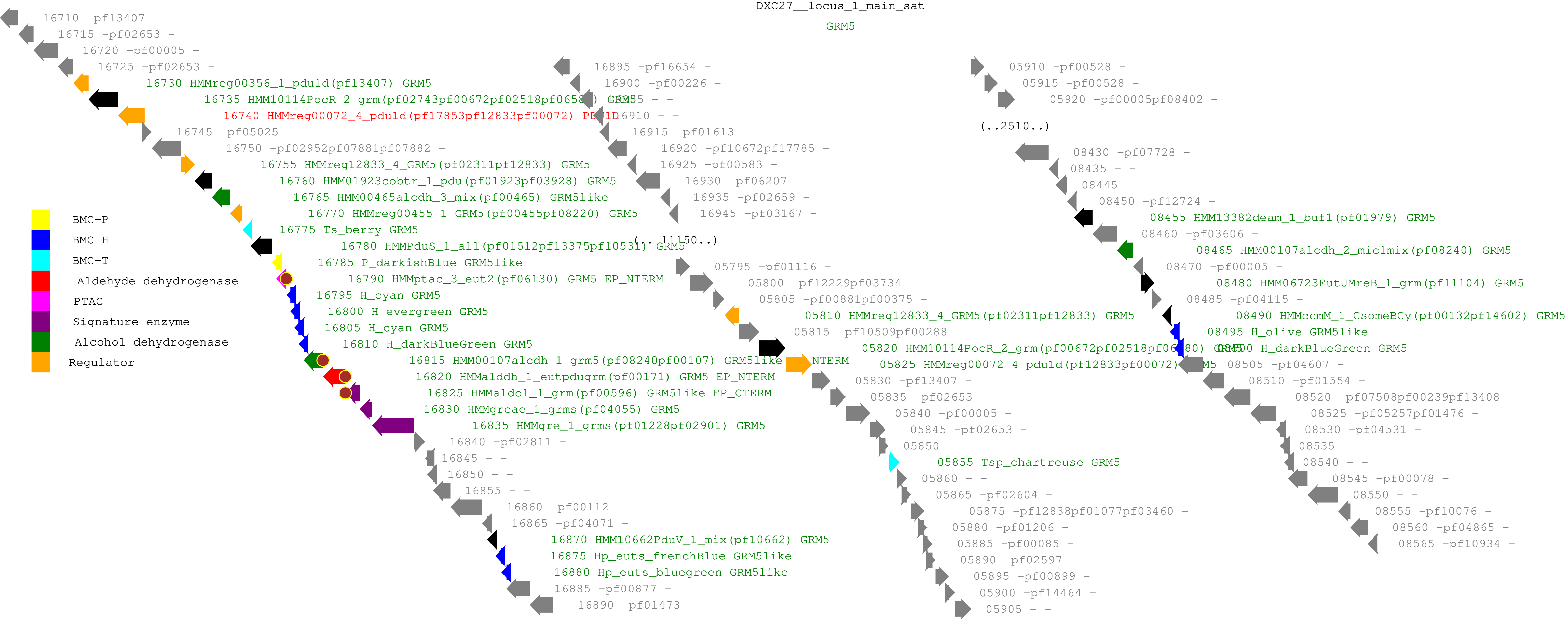

Ruminococcus sp. OM08-9BH. Bacteria; Firmicutes; Clostridia; Clostridiales; Ruminococcaceae;

DXC28__locus_1_main_sat

GRM5

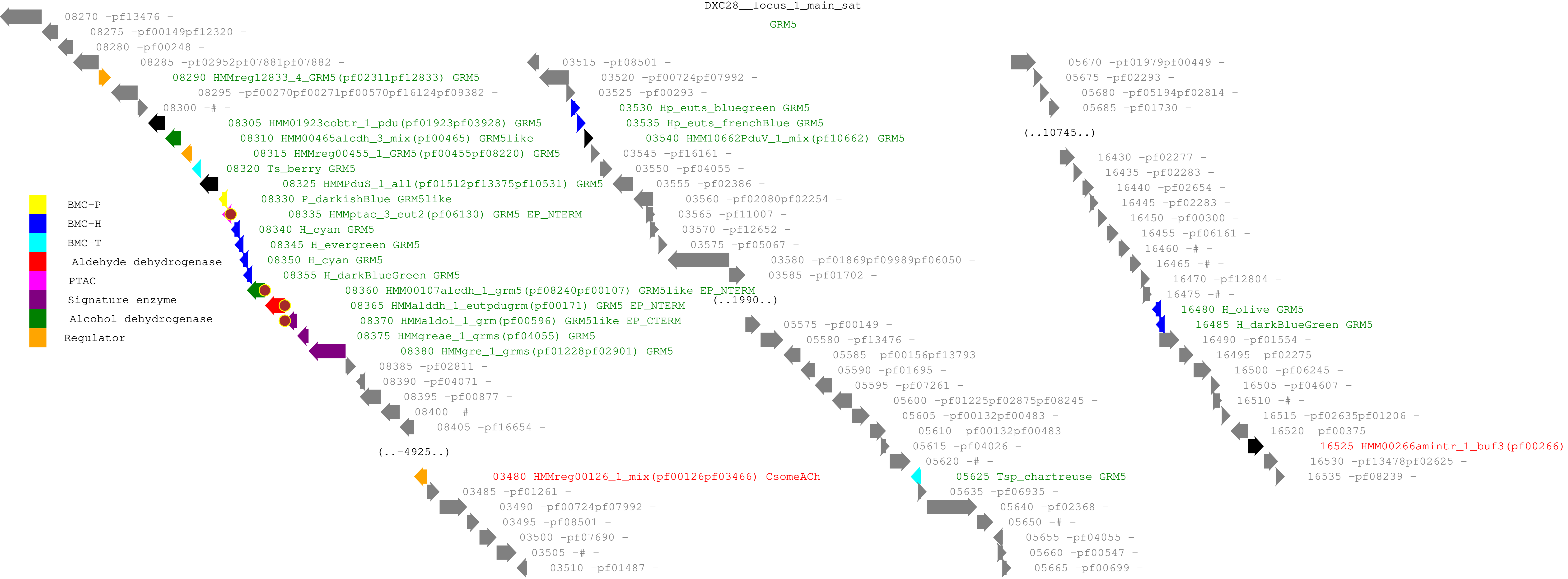

Eisenbergiella massiliensis. Bacteria; Firmicutes; Clostridia; Clostridiales; Lachnospiraceae;

DXC51__locus_1

GRM5

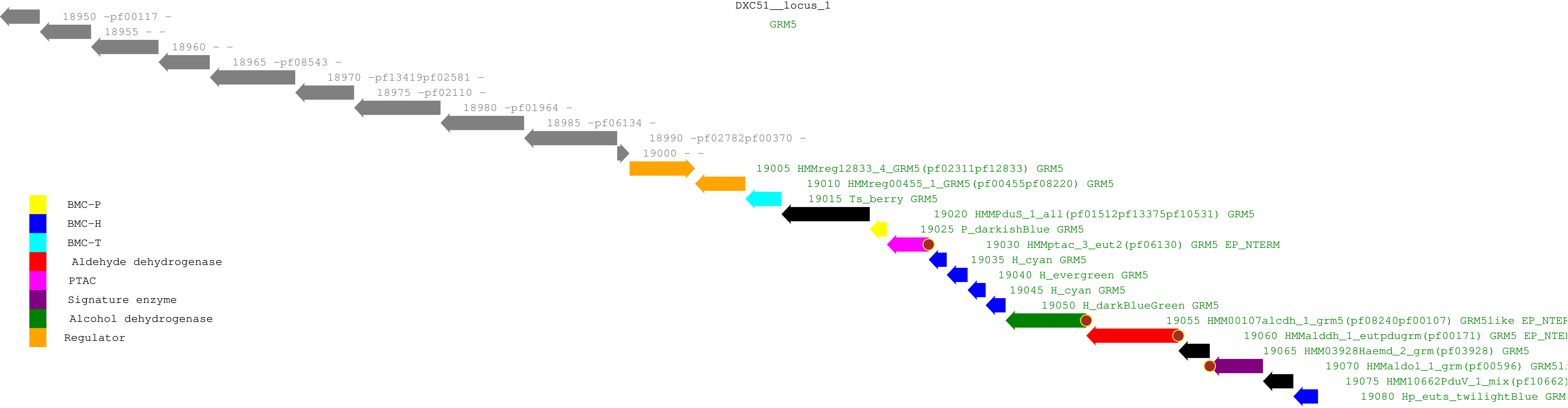

Ruminococcus sp. TF06-23. Bacteria; Firmicutes; Clostridia; Clostridiales; Ruminococcaceae;

DXC58__locus_1_main_sat

GRM5

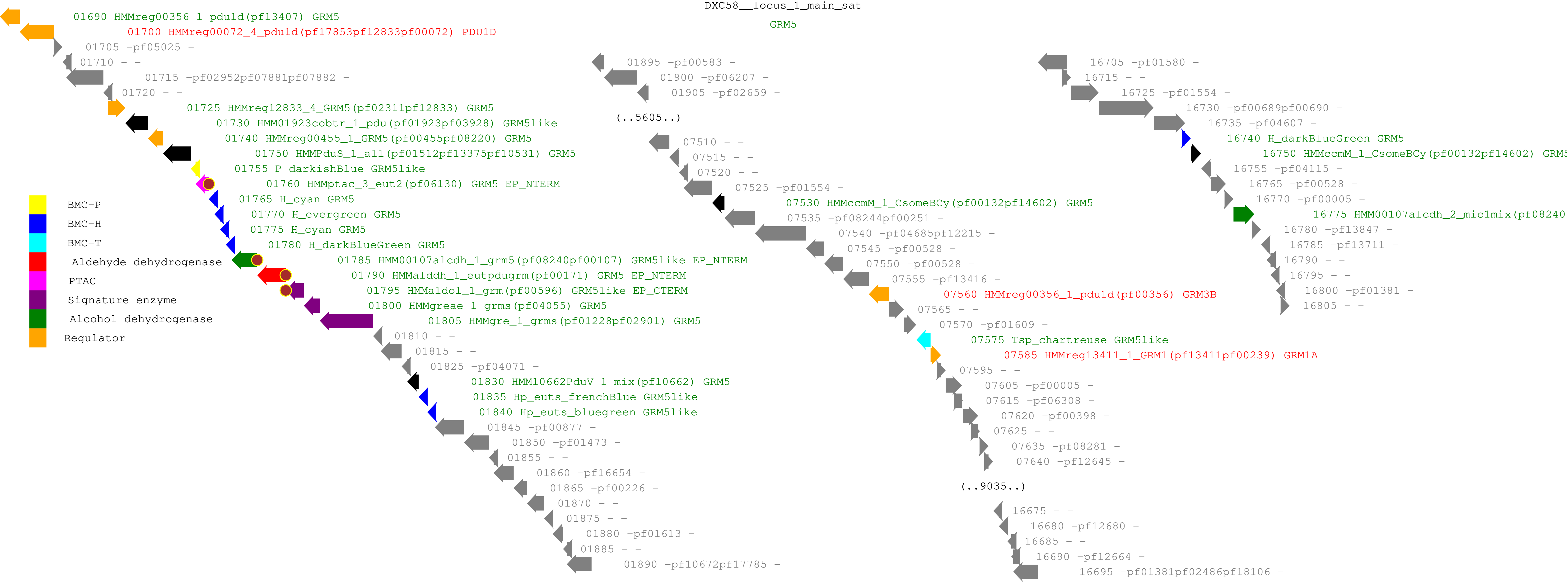

Ruminococcus sp. TF08-4. Bacteria; Firmicutes; Clostridia; Clostridiales; Ruminococcaceae;

DXC84__locus_1_main_sat

GRM5

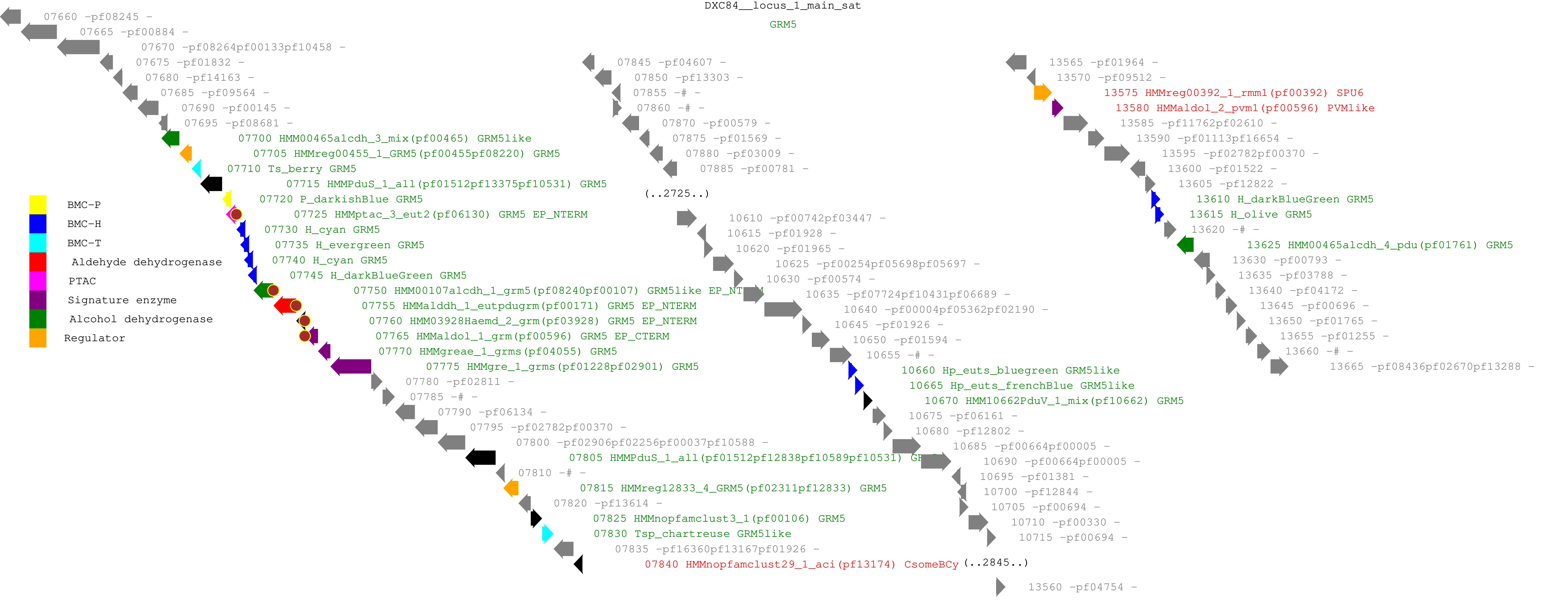

Lachnospiraceae bacterium TF09-5. Bacteria; Firmicutes; Clostridia; Clostridiales; Lachnospiraceae.

DXC97__locus_1

GRM5

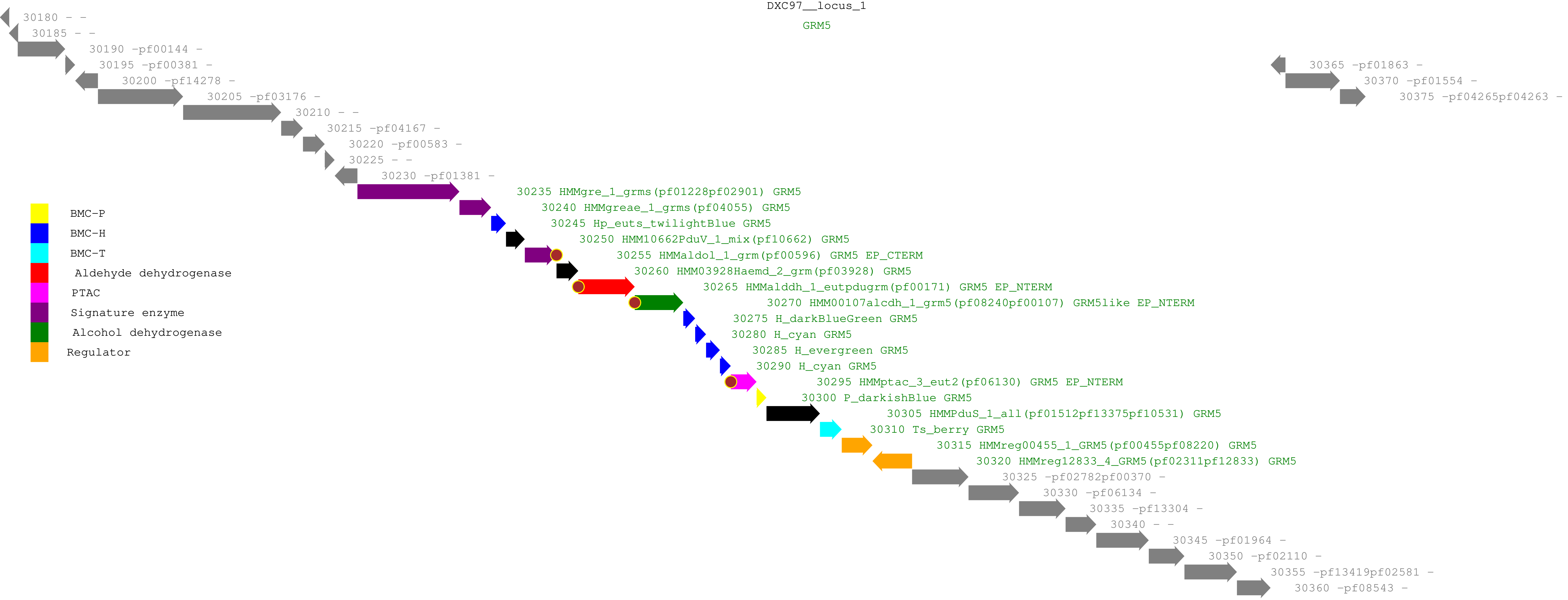

Ruminococcus sp. TF10-12AC. Bacteria; Firmicutes; Clostridia; Clostridiales; Ruminococcaceae;

DXD00__locus_1_main_sat

GRM5

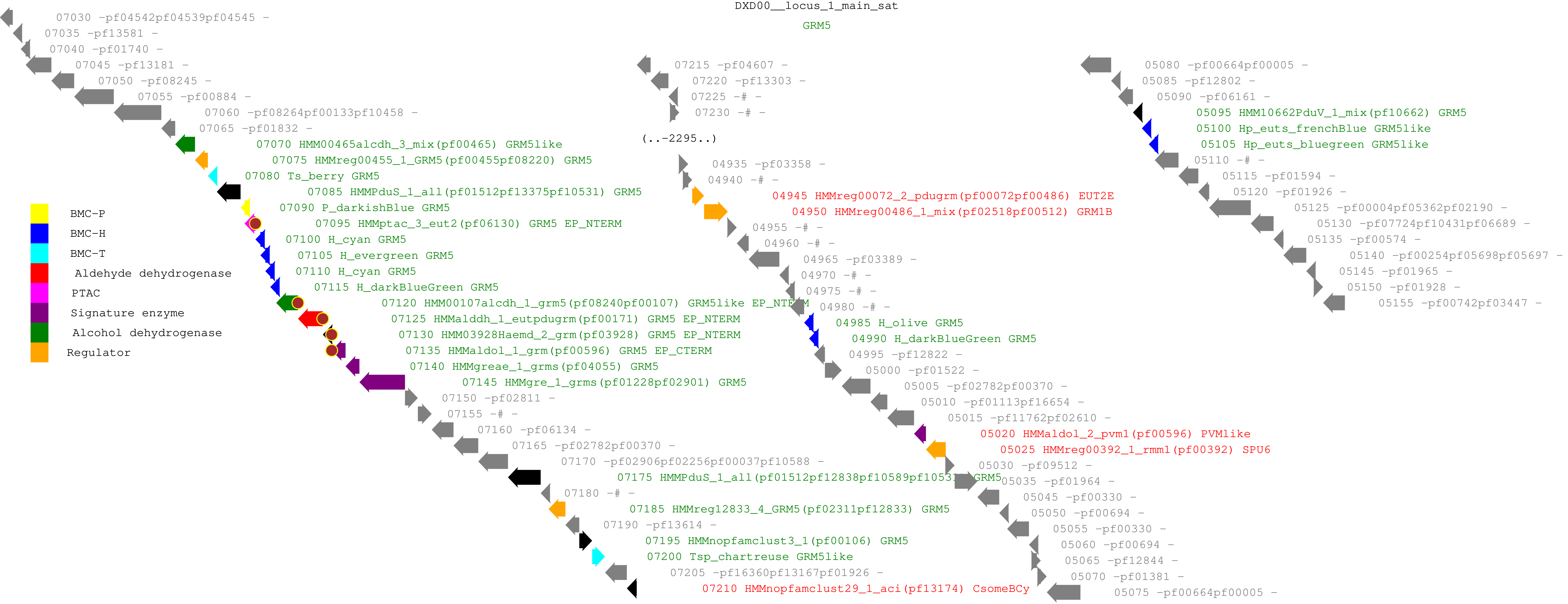

Ruminococcus sp. TF10-6. Bacteria; Firmicutes; Clostridia; Clostridiales; Ruminococcaceae;

DXD07__locus_1_main_sat

GRM5

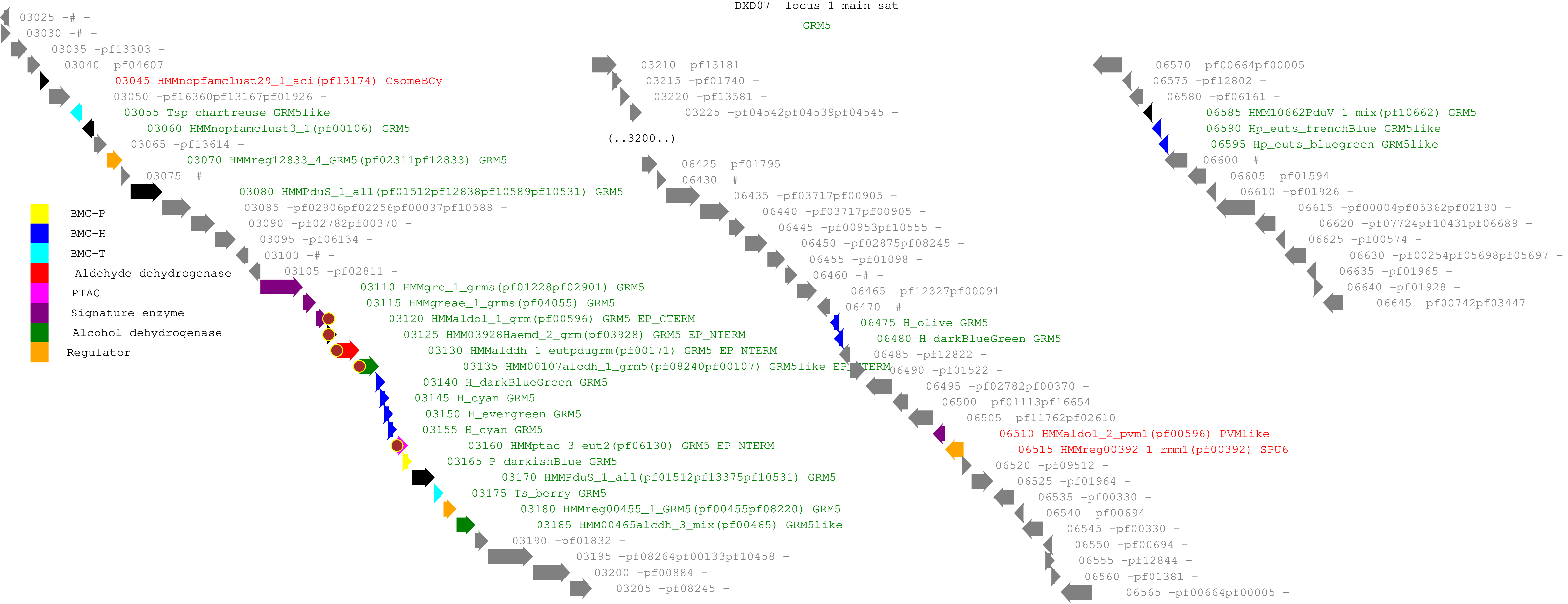

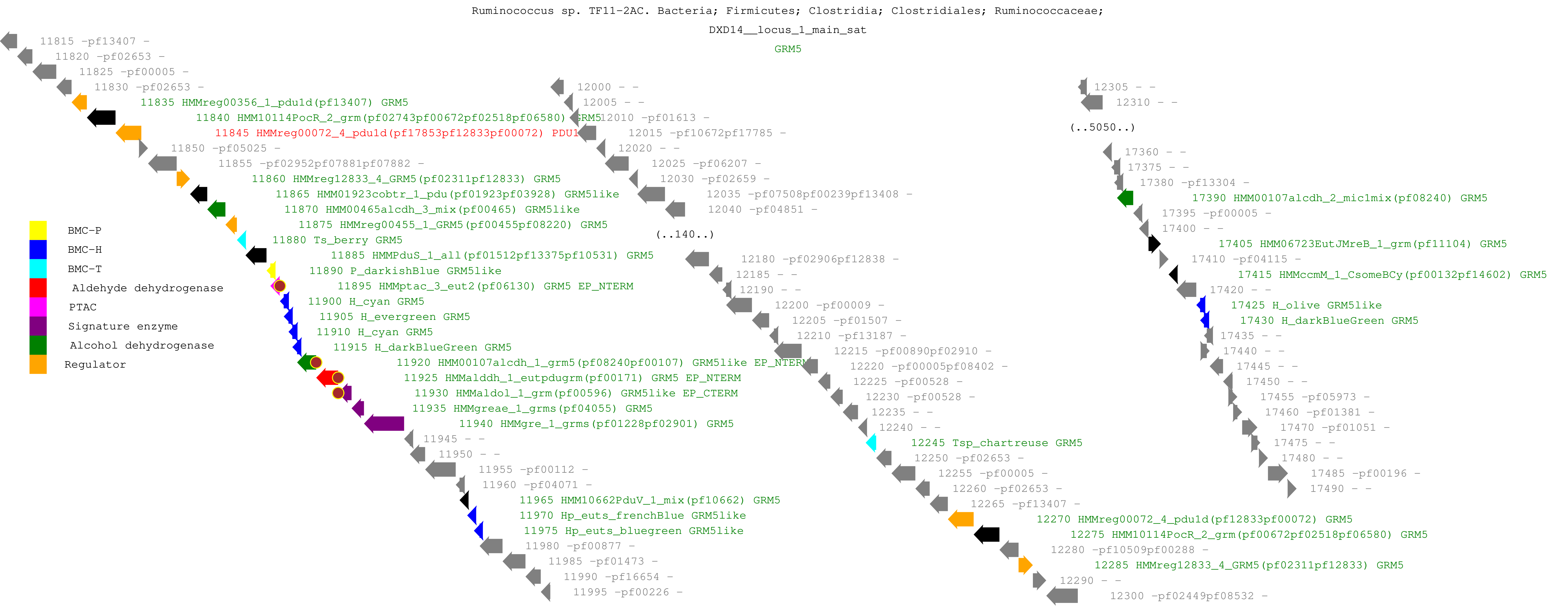

Ruminococcus sp. TF11-2AC. Bacteria; Firmicutes; Clostridia; Clostridiales; Ruminococcaceae;
DXD14__locus_1_main_sat
GRM5
11815 -pf13407 -
11820 -pf02653 -
11825 -pf00005 -
11830 -pf02653 -
11835 HMMreg00356_1_pdu1d(pf13407) GRM5
11840 HMM10114PocR_2_grm(pf02743pf00672pf02518pf06580) GRM5
11845 HMMreg00072_4_pdu1d(pf17853pf12833pf00072) PDU1
11850 -pf05025 -
11855 -pf02952pf07881pf07882 -
11860 HMMreg12833_4_GRM5(pf02311pf12833) GRM5
11865 HMM01923cobtr_1_pdu(pf01923pf03928) GRM5like
11870 HMM00465alcdh_3_mix(pf00465) GRM5like
11875 HMMreg00455_1_GRM5(pf00455pf08220) GRM5
11880 Ts_berry GRM5
11885 HMMPduS_1_all(pf01512pf13375pf10531) GRM5
11890 P_darkishBlue GRM5like
11895 HMMptac_3_eut2(pf06130) GRM5 EP_NTERM
11900 H_cyan GRM5
11905 H_evergreen GRM5
11910 H_cyan GRM5
11915 H_darkBlueGreen GRM5
11920 HMM00107alcdh_1_grm5(pf08240pf00107) GRM5like EP_NTERM
11925 HMMalddh_1_eutpdugrm(pf00171) GRM5 EP_NTERM
11930 HMMaldol_1_grm(pf00596) GRM5like EP_CTERM
11935 HMMgreae_1_grms(pf04055) GRM5
11940 HMMgre_1_grms(pf01228pf02901) GRM5
11945 - -
11950 - -
11955 -pf00112 -
11960 -pf04071 -
11965 HMM10662PduV_1_mix(pf10662) GRM5
11970 Hp_euts_frenchBlue GRM5like
11975 Hp_euts_bluegreen GRM5like
11980 -pf00877 -
11985 -pf01473 -
11990 -pf16654 -
11995 -pf00226 -
12000 - -
12005 - -
12010 -pf01613 -
12015 -pf10672pf17785 -
12020 - -
12025 -pf06207 -
12030 -pf02659 -
12035 -pf07508pf00239pf13408 -
12040 -pf04851 -
(..140..)
12180 -pf02906pf12838 -
12185 - -
12190 - -
12200 -pf00009 -
12205 -pf01507 -
12210 -pf13187 -
12215 -pf00890pf02910 -
12220 -pf00005pf08402 -
12225 -pf00528 -
12230 -pf00528 -
12235 - -
12240 - -
12245 Tsp_chartreuse GRM5
12250 -pf02653 -
12255 -pf00005 -
12260 -pf02653 -
12265 -pf13407 -
12270 HMMreg00072_4_pdu1d(pf12833pf00072) GRM5
12275 HMM10114PocR_2_grm(pf00672pf02518pf06580) GRM5
12280 -pf10509pf00288 -
12285 HMMreg12833_4_GRM5(pf02311pf12833) GRM5
12290 - -
12300 -pf02449pf08532 -
12305 - -
12310 - -
(..5050..)
17360 - -
17375 - -
17380 -pf13304 -
17390 HMM00107alcdh_2_mic1mix(pf08240) GRM5
17395 -pf00005 -
17400 - -
17405 HMM06723EutJMreB_1_grm(pf11104) GRM5
17410 -pf04115 -
17415 HMMccmM_1_CsomeBCy(pf00132pf14602) GRM5
17420 - -
17425 H_olive GRM5like
17430 H_darkBlueGreen GRM5
17435 - -
17440 - -
17445 - -
17450 - -
17455 -pf05973 -
17460 -pf01381 -
17470 -pf01051 -
17475 - -
17480 - -
17485 -pf00196 -
17490 - -
BMC-P
BMC-H
BMC-T
Aldehyde dehydrogenase
PTAC
Signature enzyme
Alcohol dehydrogenase
Regulator

Lachnospiraceae bacterium TM07-2AC. Bacteria; Firmicutes; Clostridia; Clostridiales; Lachnospiraceae.

DXD70__locus_1_main_sat

GRM5

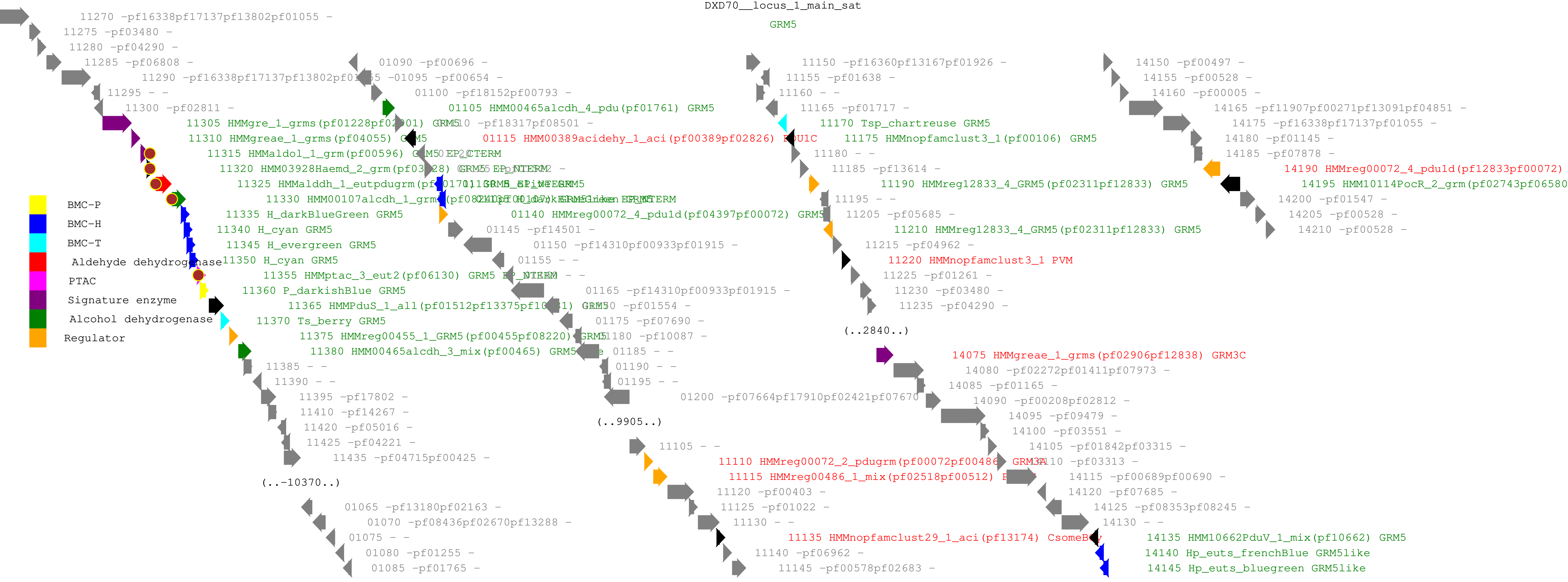

Ruminococcus sp. TM09-4. Bacteria; Firmicutes; Clostridia; Clostridiales; Ruminococcaceae;

DXD85__locus_1_main_sat

GRM5

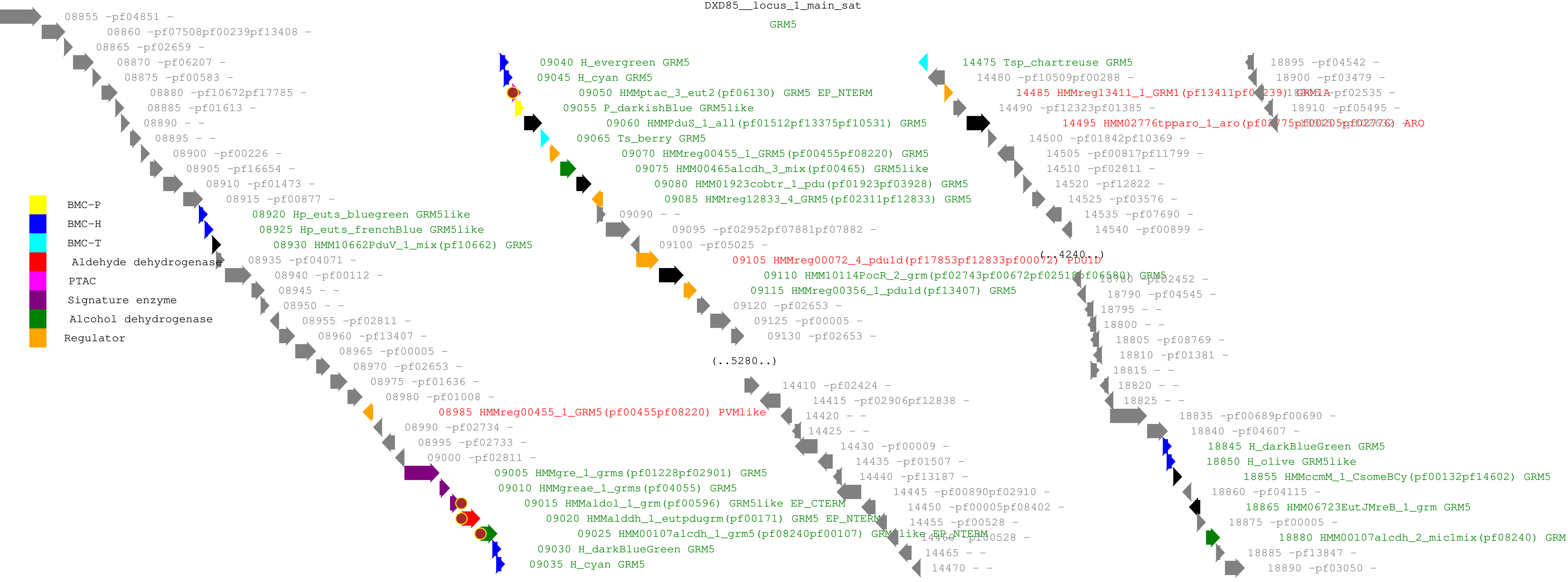

Ruminococcus sp. TM10-9AT. Bacteria; Firmicutes; Clostridia; Clostridiales; Ruminococcaceae;

DXD97__locus_1_main_sat

GRM5

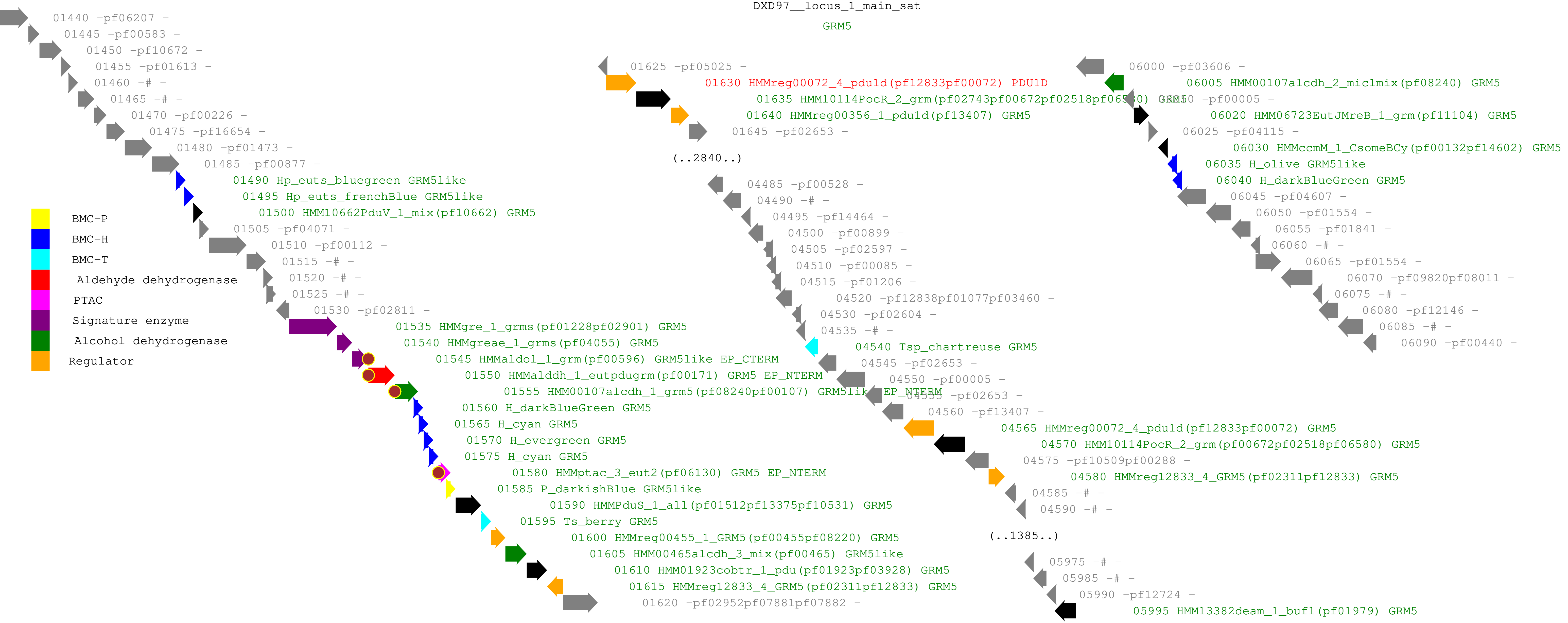

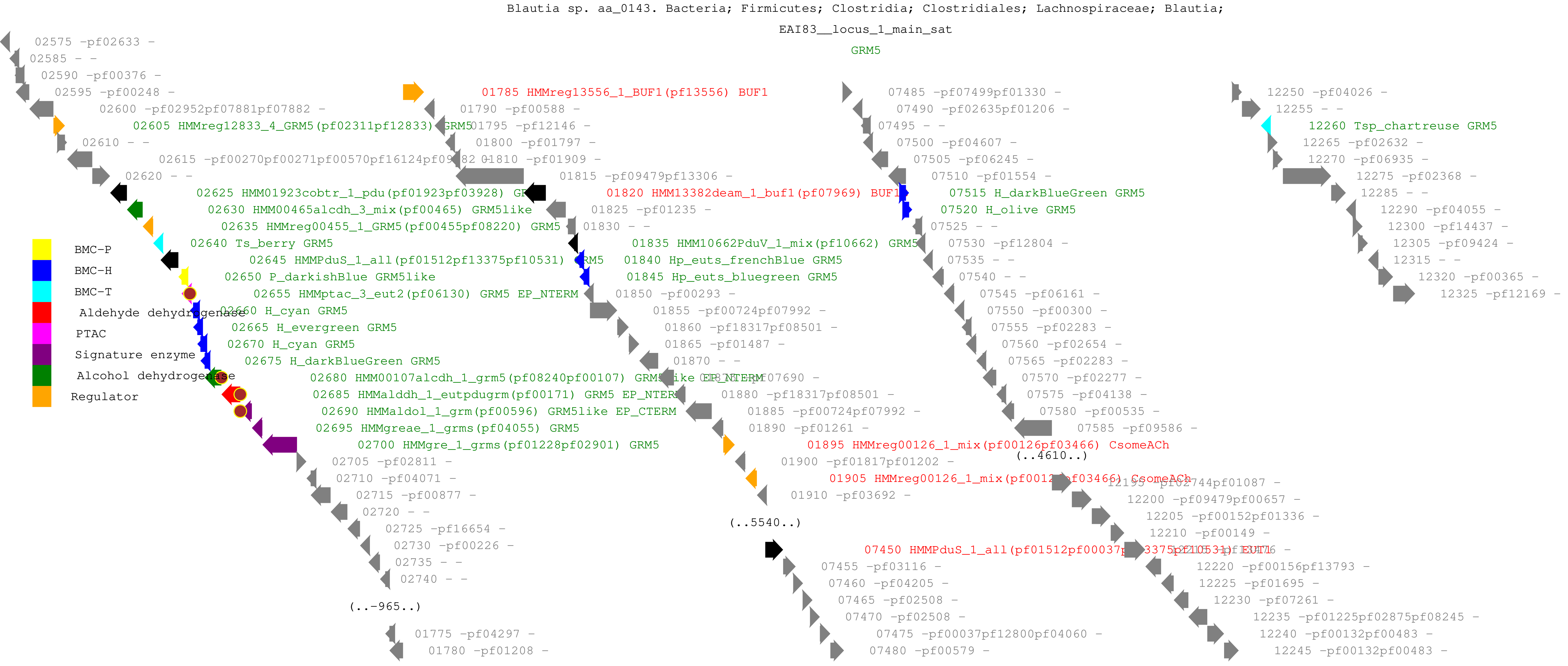

Blautia sp. aa_0143. Bacteria; Firmicutes; Clostridia; Clostridiales; Lachnospiraceae; Blautia;
EAI83__locus_1_main_sat
GRM5
BMC-P
BMC-H
BMC-T
Aldehyde dehydrogenase
PTAC
Signature enzyme
Alcohol dehydrogenase
Regulator
02575 -pf02633 -
02585 - -
02590 -pf00376 -
02595 -pf00248 -
02600 -pf02952pf07881pf07882 -
02605 HMMreg12833_4_GRM5(pf02311pf12833) GRM5
02610 - -
02615 -pf00270pf00271pf00570pf16124pf09382 -
02620 - -
02625 HMM01923cobtr_1_pdu(pf01923pf03928) GRM5
02630 HMM00465alcdh_3_mix(pf00465) GRM5like
02635 HMMreg00455_1_GRM5(pf00455pf08220) GRM5
02640 Ts_berry GRM5
02645 HMMPduS_1_all(pf01512pf13375pf10531) GRM5
02650 P_darkishBlue GRM5like
02655 HMMptac_3_eut2(pf06130) GRM5 EP_NTERM
02660 H_cyan GRM5
02665 H_evergreen GRM5
02670 H_cyan GRM5
02675 H_darkBlueGreen GRM5
02680 HMM00107alcdh_1_grm5(pf08240pf00107) GRM5like EP_NTERM
02685 HMMaldh_1_eutpdugrm(pf00171) GRM5 EP_NTERM
02690 HMMaldol_1_grm(pf00596) GRM5like EP_CTERM
02695 HMMgreae_1_grms(pf04055) GRM5
02700 HMMgre_1_grms(pf01228pf02901) GRM5
02705 -pf02811 -
02710 -pf04071 -
02715 -pf00877 -
02720 - -
02725 -pf16654 -
02730 -pf00226 -
02735 - -
02740 - -
(..-965..)
01775 -pf04297 -
01780 -pf01208 -
01785 HMMreg13556_1_BUF1(pf13556) BUF1
01790 -pf00588 -
01795 -pf12146 -
01800 -pf01797 -
01810 -pf01909 -
01815 -pf09479pf13306 -
01820 HMM13382deam_1_buf1(pf07969) BUF1
01825 -pf01235 -
01830 - -
01835 HMM10662PduV_1_mix(pf10662) GRM5
01840 Hp_euts_frenchBlue GRM5
01845 Hp_euts_bluegreen GRM5
01850 -pf00293 -
01855 -pf00724pf07992 -
01860 -pf18317pf08501 -
01865 -pf01487 -
01870 - -
01875 -pf07690 -
01880 -pf18317pf08501 -
01885 -pf00724pf07992 -
01890 -pf01261 -
01895 HMMreg00126_1_mix(pf00126pf03466) CsomeACh
01900 -pf01817pf01202 -
01905 HMMreg00126_1_mix(pf00126pf03466) CsomeACh
01910 -pf03692 -
(..5540..)
07450 HMMPduS_1_all(pf01512pf00037pf13375pf10531) EUT1
07455 -pf03116 -
07460 -pf04205 -
07465 -pf02508 -
07470 -pf02508 -
07475 -pf00037pf12800pf04060 -
07480 -pf00579 -
07485 -pf07499pf01330 -
07490 -pf02635pf01206 -
07495 - -
07500 -pf04607 -
07505 -pf06245 -
07510 -pf01554 -
07515 H_darkBlueGreen GRM5
07520 H_olive GRM5
07525 - -
07530 -pf12804 -
07535 - -
07540 - -
07545 -pf06161 -
07550 -pf00300 -
07555 -pf02283 -
07560 -pf02654 -
07565 -pf02283 -
07570 -pf02277 -
07575 -pf04138 -
07580 -pf00535 -
07585 -pf09586 -
(..4610..)
12195 -pf02744pf01087 -
12200 -pf09479pf00657 -
12205 -pf00152pf01336 -
12210 -pf00149 -
12215 -pf13416 -
12220 -pf00156pf13793 -
12225 -pf01695 -
12230 -pf07261 -
12235 -pf01225pf02875pf08245 -
12240 -pf00132pf00483 -
12245 -pf00132pf00483 -
12250 -pf04026 -
12255 - -
12260 Tsp_chartreuse GRM5
12265 -pf02632 -
12270 -pf06935 -
12275 -pf02368 -
12285 - -
12290 -pf04055 -
12300 -pf14437 -
12305 -pf09424 -
12315 - -
12320 -pf00365 -
12325 -pf12169 -

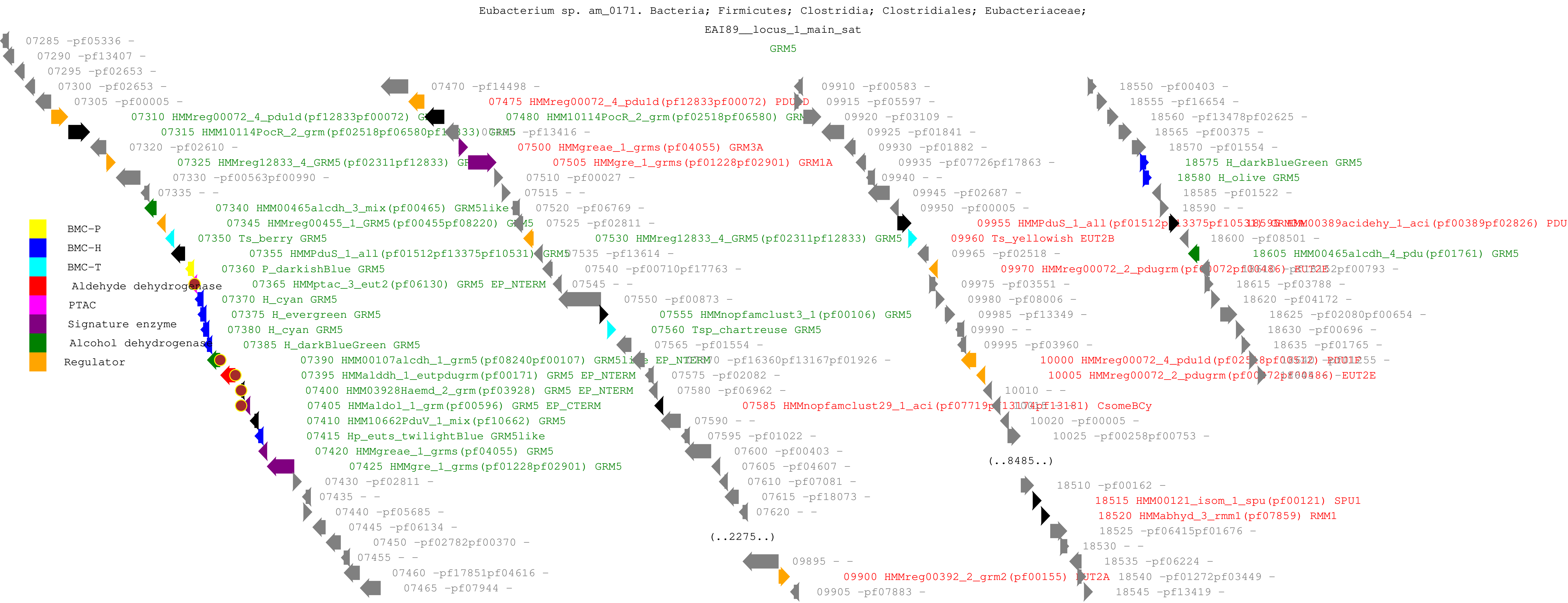
Eubacterium sp. am_0171. Bacteria; Firmicutes; Clostridia; Clostridiales; Eubacteriaceae;
EAI89__locus_1_main_sat
GRM5
BMC-P
BMC-H
BMC-T
Aldehyde dehydrogenase
PTAC
Signature enzyme
Alcohol dehydrogenase
Regulator
07285 -pf05336 -
07290 -pf13407 -
07295 -pf02653 -
07300 -pf02653 -
07305 -pf00005 -
07310 HMMreg00072_4_pdu1d(pf12833pf00072) GRM5
07315 HMM10114PocR_2_grm(pf02518pf06580pf12833) GRM5
07320 -pf02610 -
07325 HMMreg12833_4_GRM5(pf02311pf12833) GRM5
07330 -pf00563pf00990 -
07335 - -
07340 HMM00465alcdh_3_mix(pf00465) GRM5like
07345 HMMreg00455_1_GRM5(pf00455pf08220) GRM5
07350 Ts_berry GRM5
07355 HMMPduS_1_all(pf01512pf13375pf10531) GRM5
07360 P_darkishBlue GRM5
07365 HMMptac_3_eut2(pf06130) GRM5 EP_NTERM
07370 H_cyan GRM5
07375 H_evergreen GRM5
07380 H_cyan GRM5
07385 H_darkBlueGreen GRM5
07390 HMM00107alcdh_1_grm5(pf08240pf00107) GRM5like EP_NTERM
07395 HMMalddh_1_eutpdugrm(pf00171) GRM5 EP_NTERM
07400 HMM03928Haemd_2_grm(pf03928) GRM5 EP_NTERM
07405 HMMaldol_1_grm(pf00596) GRM5 EP_CTERM
07410 HMM10662PduV_1_mix(pf10662) GRM5
07415 Hp_euts_twilightBlue GRM5like
07420 HMMgreae_1_grms(pf04055) GRM5
07425 HMMgre_1_grms(pf01228pf02901) GRM5
07430 -pf02811 -
07435 - -
07440 -pf05685 -
07445 -pf06134 -
07450 -pf02782pf00370 -
07455 - -
07460 -pf17851pf04616 -
07465 -pf07944 -
07470 -pf14498 -
07475 HMMreg00072_4_pdu1d(pf12833pf00072) PDU1D
07480 HMM10114PocR_2_grm(pf02518pf06580) GRM5
07495 -pf13416 -
07500 HMMgreae_1_grms(pf04055) GRM3A
07505 HMMgre_1_grms(pf01228pf02901) GRM1A
07510 -pf00027 -
07515 - -
07520 -pf06769 -
07525 -pf02811 -
07530 HMMreg12833_4_GRM5(pf02311pf12833) GRM5
07535 -pf13614 -
07540 -pf00710pf17763 -
07545 - -
07550 -pf00873 -
07555 HMMnopfamclust3_1(pf00106) GRM5
07560 Tsp_chartreuse GRM5
07565 -pf01554 -
07570 -pf16360pf13167pf01926 -
07575 -pf02082 -
07580 -pf06962 -
07585 HMMnopfamclust29_1_aci(pf07719pf13174pf13181) CsomeBCy
07590 - -
07595 -pf01022 -
07600 -pf00403 -
07605 -pf04607 -
07610 -pf07081 -
07615 -pf18073 -
07620 - -
(..2275..)
09895 - -
09900 HMMreg00392_2_grm2(pf00155) EUT2A
09905 -pf07883 -
09910 -pf00583 -
09915 -pf05597 -
09920 -pf03109 -
09925 -pf01841 -
09930 -pf01882 -
09935 -pf07726pf17863 -
09940 - -
09945 -pf02687 -
09950 -pf00005 -
09955 HMMPduS_1_all(pf01512pf13375pf10531) GRM5
09960 Ts_yellowish EUT2B
09965 -pf02518 -
09970 HMMreg00072_2_pdugrm(pf00072pf00486) EUT2E
09975 -pf03551 -
09980 -pf08006 -
09985 -pf13349 -
09990 - -
09995 -pf03960 -
10000 HMMreg00072_4_pdu1d(pf02518pf00512) PDU1E
10005 HMMreg00072_2_pdugrm(pf00072pf00486) EUT2E
10010 - -
10015 - -
10020 -pf00005 -
10025 -pf00258pf00753 -
(..8485..)
18510 -pf00162 -
18515 HMM00121_isom_1_spu(pf00121) SPU1
18520 HMMabhyd_3_rmm1(pf07859) RMM1
18525 -pf06415pf01676 -
18530 - -
18535 -pf06224 -
18540 -pf01272pf03449 -
18545 -pf13419 -
18550 -pf00403 -
18555 -pf16654 -
18560 -pf13478pf02625 -
18565 -pf00375 -
18570 -pf01554 -
18575 H_darkBlueGreen GRM5
18580 H_olive GRM5
18585 -pf01522 -
18590 - -
18595 HMM00389acidehy_1_aci(pf00389pf02826) PDU
18600 -pf08501 -
18605 HMM00465alcdh_4_pdu(pf01761) GRM5
18610 -pf02652pf00793 -
18615 -pf03788 -
18620 -pf04172 -
18625 -pf02080pf00654 -
18630 -pf00696 -
18635 -pf01765 -
18640 -pf02255 -

Schaedlerella arabinosiphila. Bacteria; Firmicutes; Clostridia; Clostridiales; Lachnospiraceae;

EBB54__locus_1

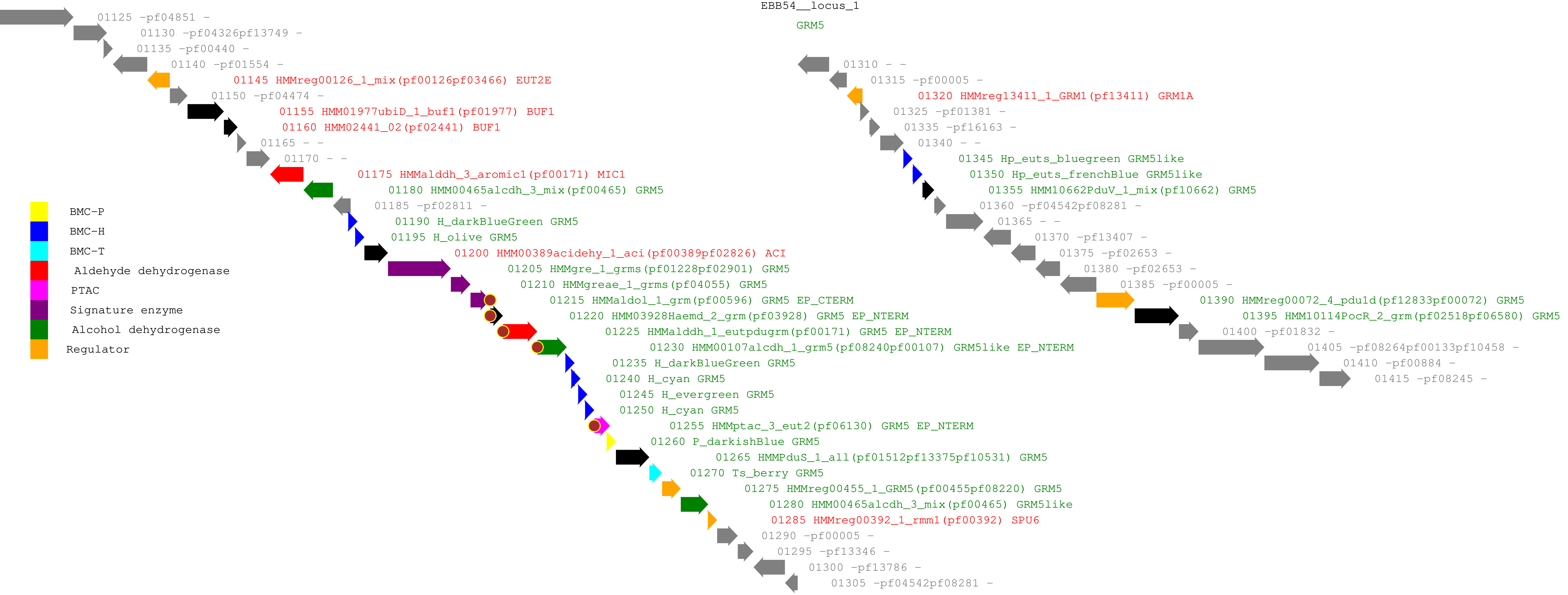

Muricomes intestini. Bacteria; Firmicutes; Clostridia; Clostridiales; Lachnospiraceae;

EDD59__locus_1

GRM5

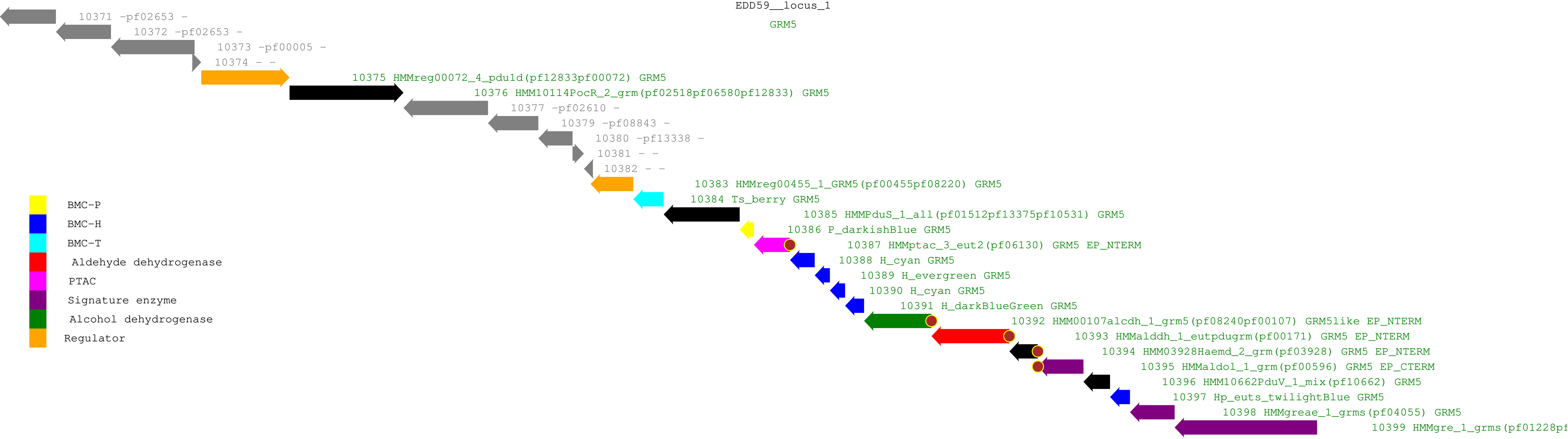

Blautia obeum. Bacteria; Firmicutes; Clostridia; Clostridiales; Lachnospiraceae;

ERS852395__locus_1_main_sat

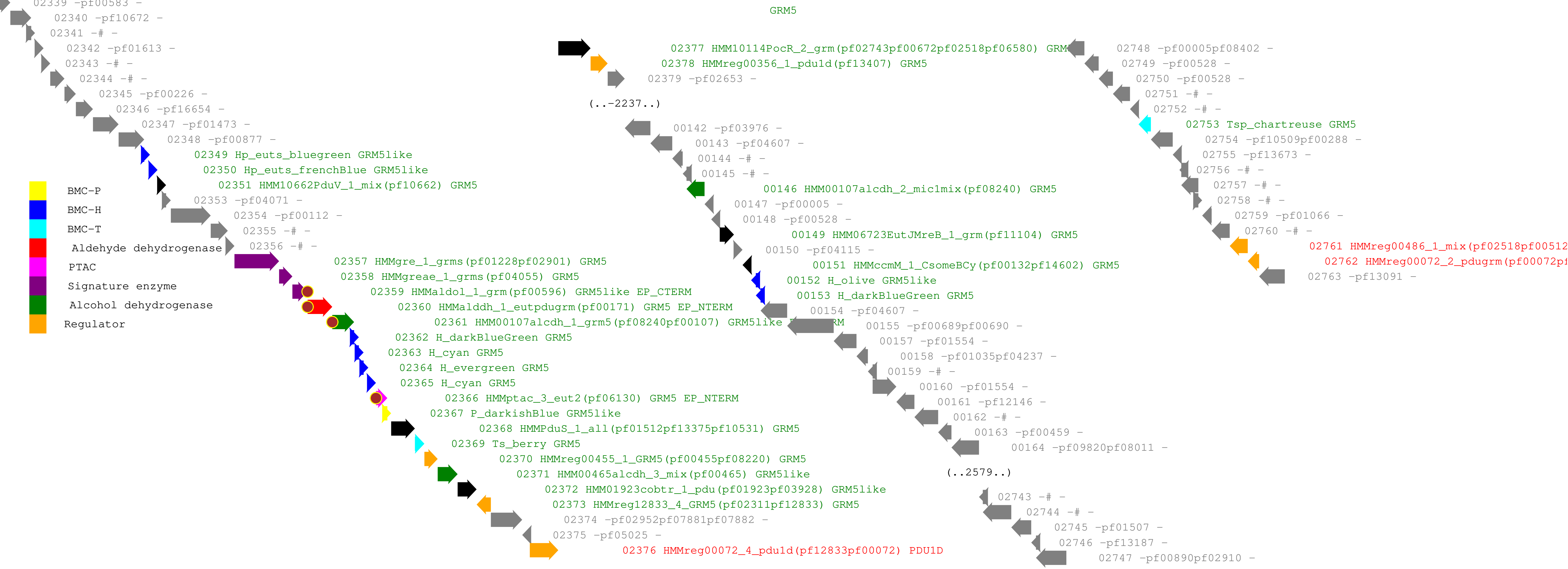

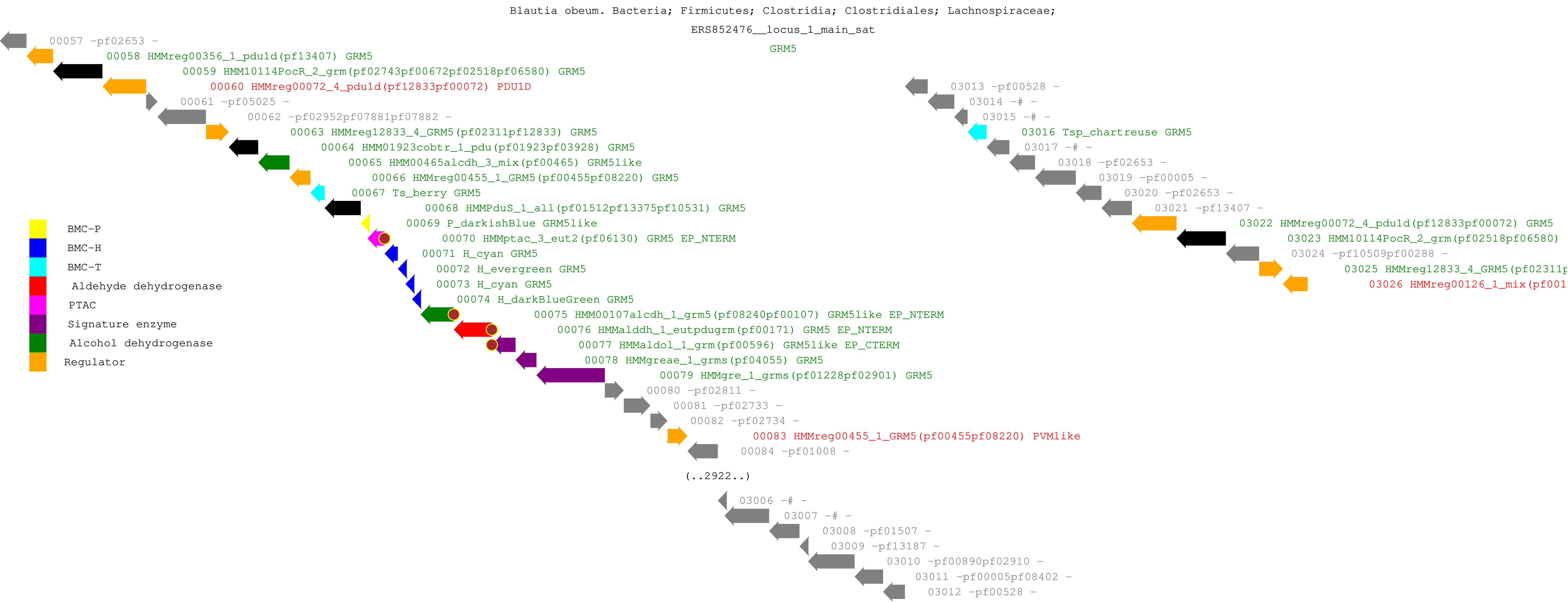
Blautia obeum. Bacteria; Firmicutes; Clostridia; Clostridiales; Lachnospiraceae;
ERS852476__locus_1_main_sat
GRM5
00057 -pf02653 -
00058 HMMreg00356_1_pdu1d(pf13407) GRM5
00059 HMM10114PocR_2_grm(pf02743pf00672pf02518pf06580) GRM5
00060 HMMreg00072_4_pdu1d(pf12833pf00072) PDU1D
00061 -pf05025 -
00062 -pf02952pf07881pf07882 -
00063 HMMreg12833_4_GRM5(pf02311pf12833) GRM5
00064 HMM01923cobtr_1_pdu(pf01923pf03928) GRM5
00065 HMM00465alcdh_3_mix(pf00465) GRM5like
00066 HMMreg00455_1_GRM5(pf00455pf08220) GRM5
00067 Ts_berry GRM5
00068 HMMPduS_1_all(pf01512pf13375pf10531) GRM5
00069 P_darkishBlue GRM5like
00070 HMMptac_3_eut2(pf06130) GRM5 EP_NTERM
00071 H_cyan GRM5
00072 H_evergreen GRM5
00073 H_cyan GRM5
00074 H_darkBlueGreen GRM5
00075 HMM00107alcdh_1_grm5(pf08240pf00107) GRM5like EP_NTERM
00076 HMMalddh_1_eutpdugrm(pf00171) GRM5 EP_NTERM
00077 HMMaldol_1_grm(pf00596) GRM5like EP_CTERM
00078 HMMgreae_1_grms(pf04055) GRM5
00079 HMMgre_1_grms(pf01228pf02901) GRM5
00080 -pf02811 -
00081 -pf02733 -
00082 -pf02734 -
00083 HMMreg00455_1_GRM5(pf00455pf08220) PVMlike
00084 -pf01008 -
(..2922..)
03006 -# -
03007 -# -
03008 -pf01507 -
03009 -pf13187 -
03010 -pf00890pf02910 -
03011 -pf00005pf08402 -
03012 -pf00528 -
03013 -pf00528 -
03014 -# -
03015 -# -
03016 Tsp_chartreuse GRM5
03017 -# -
03018 -pf02653 -
03019 -pf00005 -
03020 -pf02653 -
03021 -pf13407 -
03022 HMMreg00072_4_pdu1d(pf12833pf00072) GRM5
03023 HMM10114PocR_2_grm(pf02518pf06580)
03024 -pf10509pf00288 -
03025 HMMreg12833_4_GRM5(pf02311p
03026 HMMreg00126_1_mix(pf001
BMC-P
BMC-H
BMC-T
Aldehyde dehydrogenase
PTAC
Signature enzyme
Alcohol dehydrogenase
Regulator

Blautia wexlerae. Bacteria; Firmicutes; Clostridia; Clostridiales; Lachnospiraceae;

ERS852478__locus_1_main_sat

GRM5

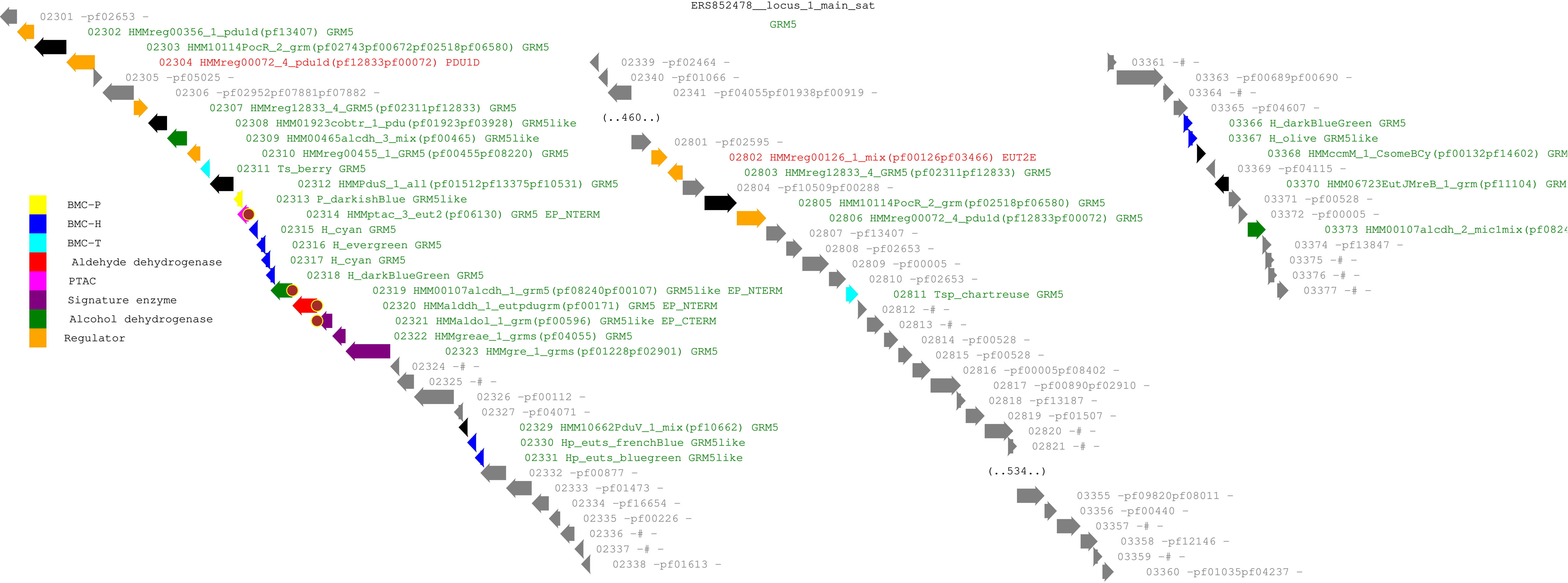

Eubacterium contortum. Bacteria; Firmicutes; Clostridia; Clostridiales; Lachnospiraceae.

ERS852491__locus_1_main_sat

GRM5

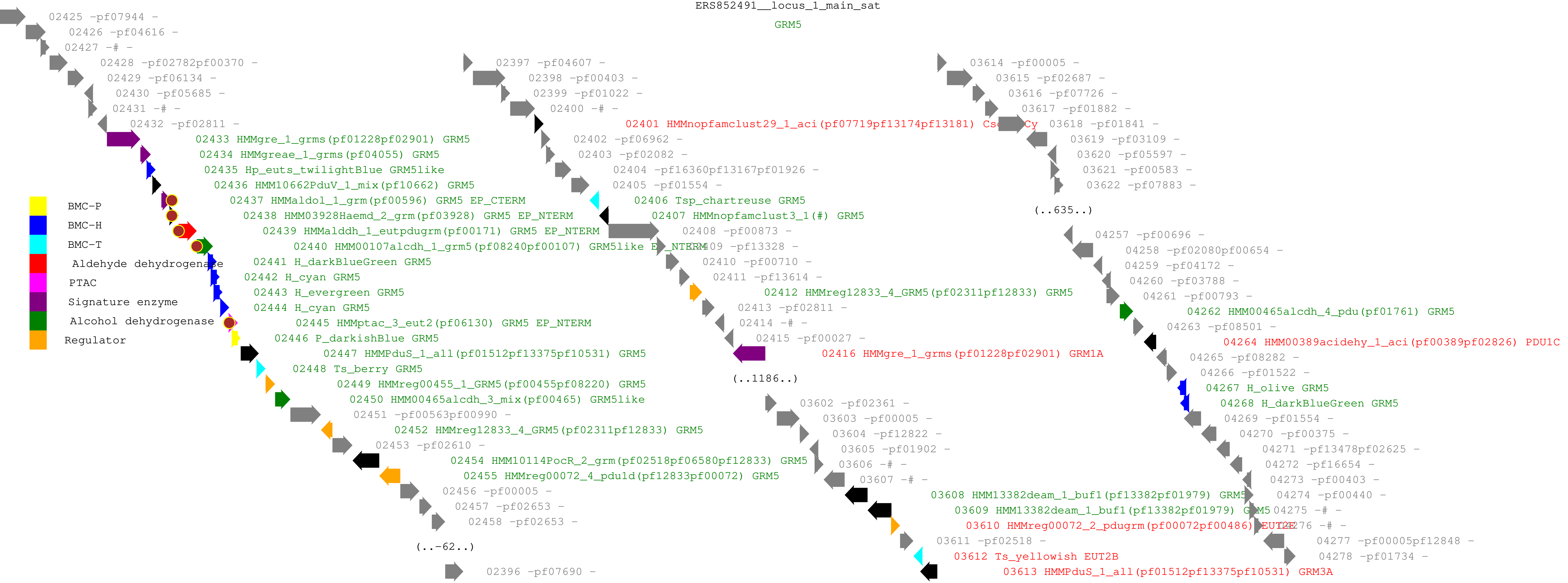

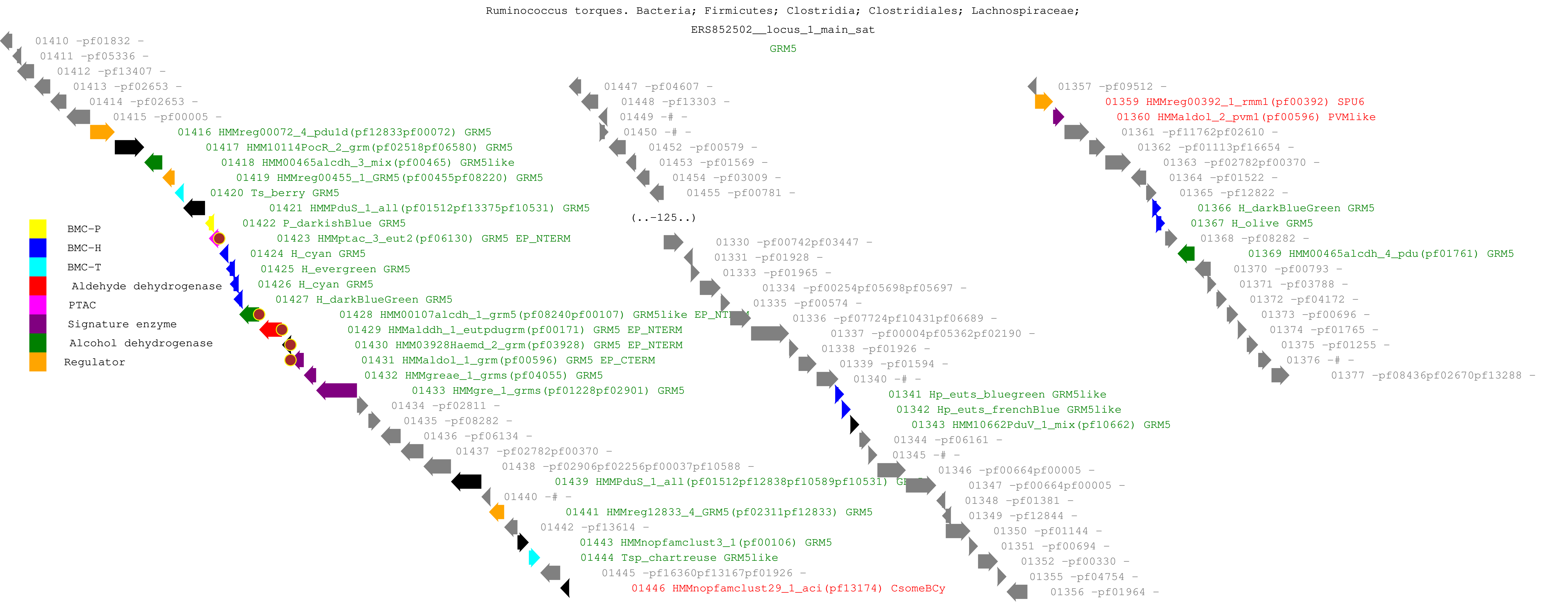

Ruminococcus torques. Bacteria; Firmicutes; Clostridia; Clostridiales; Lachnospiraceae;
ERS852502__locus_1_main_sat
GRM5
01410 -pf01832 -
01411 -pf05336 -
01412 -pf13407 -
01413 -pf02653 -
01414 -pf02653 -
01415 -pf00005 -
01416 HMMreg00072_4_pdu1d(pf12833pf00072) GRM5
01417 HMM10114PocR_2_grm(pf02518pf06580) GRM5
01418 HMM00465alcdh_3_mix(pf00465) GRM5like
01419 HMMreg00455_1_GRM5(pf00455pf08220) GRM5
01420 Ts_berry GRM5
01421 HMMPduS_1_all(pf01512pf13375pf10531) GRM5
01422 P_darkishBlue GRM5
01423 HMMptac_3_eut2(pf06130) GRM5 EP_NTERM
01424 H_cyan GRM5
01425 H_evergreen GRM5
01426 H_cyan GRM5
01427 H_darkBlueGreen GRM5
01428 HMM00107alcdh_1_grm5(pf08240pf00107) GRM5like EP_NT
01429 HMMalddh_1_eutpdugrm(pf00171) GRM5 EP_NTERM
01430 HMM03928Haemd_2_grm(pf03928) GRM5 EP_NTERM
01431 HMMaldol_1_grm(pf00596) GRM5 EP_CTERM
01432 HMMgreae_1_grms(pf04055) GRM5
01433 HMMgre_1_grms(pf01228pf02901) GRM5
01434 -pf02811 -
01435 -pf08282 -
01436 -pf06134 -
01437 -pf02782pf00370 -
01438 -pf02906pf02256pf00037pf10588 -
01439 HMMPduS_1_all(pf01512pf12838pf10589pf10531)
01440 -# -
01441 HMMreg12833_4_GRM5(pf02311pf12833) GRM5
01442 -pf13614 -
01443 HMMnopfamclust3_1(pf00106) GRM5
01444 Tsp_chartreuse GRM5like
01445 -pf16360pf13167pf01926 -
01446 HMMnopfamclust29_1_aci(pf13174) CsomeBCy
BMC-P
BMC-H
BMC-T
Aldehyde dehydrogenase
PTAC
Signature enzyme
Alcohol dehydrogenase
Regulator
01447 -pf04607 -
01448 -pf13303 -
01449 -# -
01450 -# -
01452 -pf00579 -
01453 -pf01569 -
01454 -pf03009 -
01455 -pf00781 -
(..-125..)
01330 -pf00742pf03447 -
01331 -pf01928 -
01333 -pf01965 -
01334 -pf00254pf05698pf05697 -
01335 -pf00574 -
01336 -pf07724pf10431pf06689 -
01337 -pf00004pf05362pf02190 -
01338 -pf01926 -
01339 -pf01594 -
01340 -# -
01341 Hp_euts_bluegreen GRM5like
01342 Hp_euts_frenchBlue GRM5like
01343 HMM10662PduV_1_mix(pf10662) GRM5
01344 -pf06161 -
01345 -# -
01346 -pf00664pf00005 -
01347 -pf00664pf00005 -
01348 -pf01381 -
01349 -pf12844 -
01350 -pf01144 -
01351 -pf00694 -
01352 -pf00330 -
01355 -pf04754 -
01356 -pf01964 -
01357 -pf09512 -
01359 HMMreg00392_1_rmm1(pf00392) SPU6
01360 HMMaldol_2_pvm1(pf00596) PVMlike
01361 -pf11762pf02610 -
01362 -pf01113pf16654 -
01363 -pf02782pf00370 -
01364 -pf01522 -
01365 -pf12822 -
01366 H_darkBlueGreen GRM5
01367 H_olive GRM5
01368 -pf08282 -
01369 HMM00465alcdh_4_pdu(pf01761) GRM5
01370 -pf00793 -
01371 -pf03788 -
01372 -pf04172 -
01373 -pf00696 -
01374 -pf01765 -
01375 -pf01255 -
01376 -# -
01377 -pf08436pf02670pf13288 -

Blautia wexlerae. Bacteria; Firmicutes; Clostridia; Clostridiales; Lachnospiraceae;

ERS852523__locus_1_main_sat

GRM5

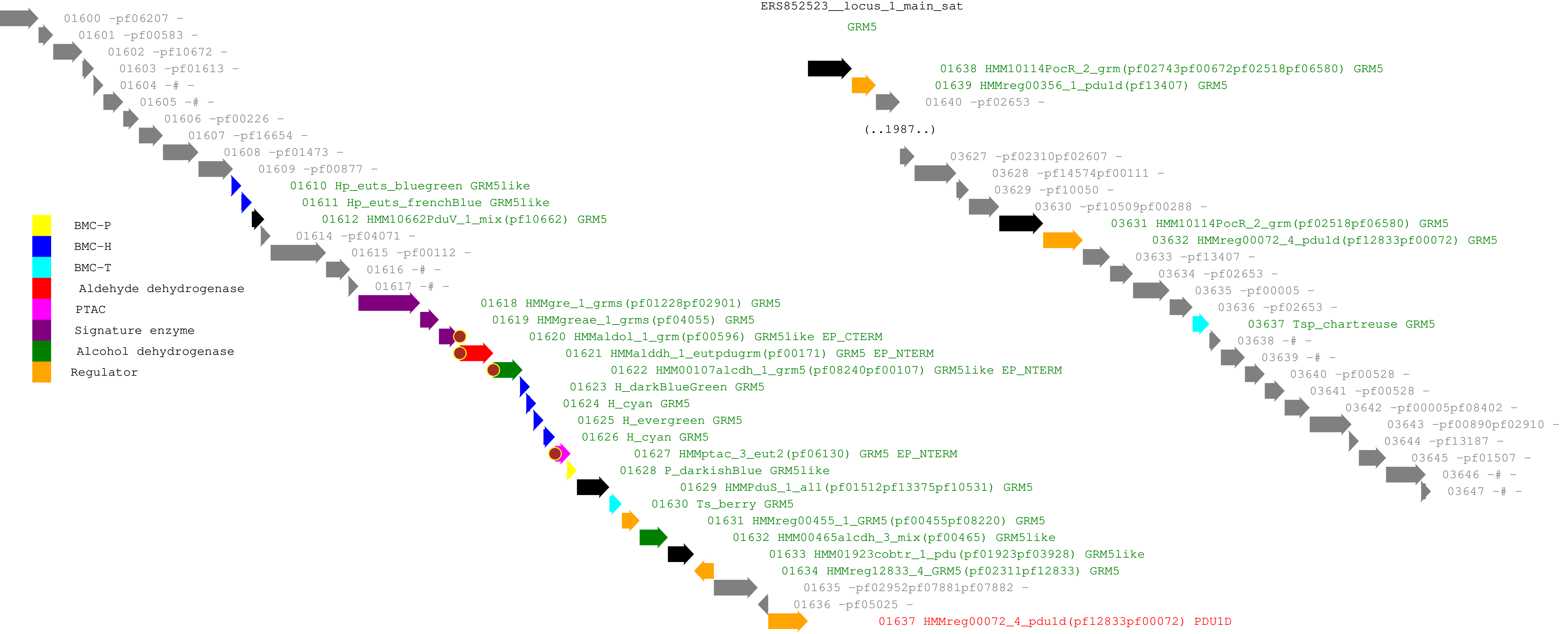

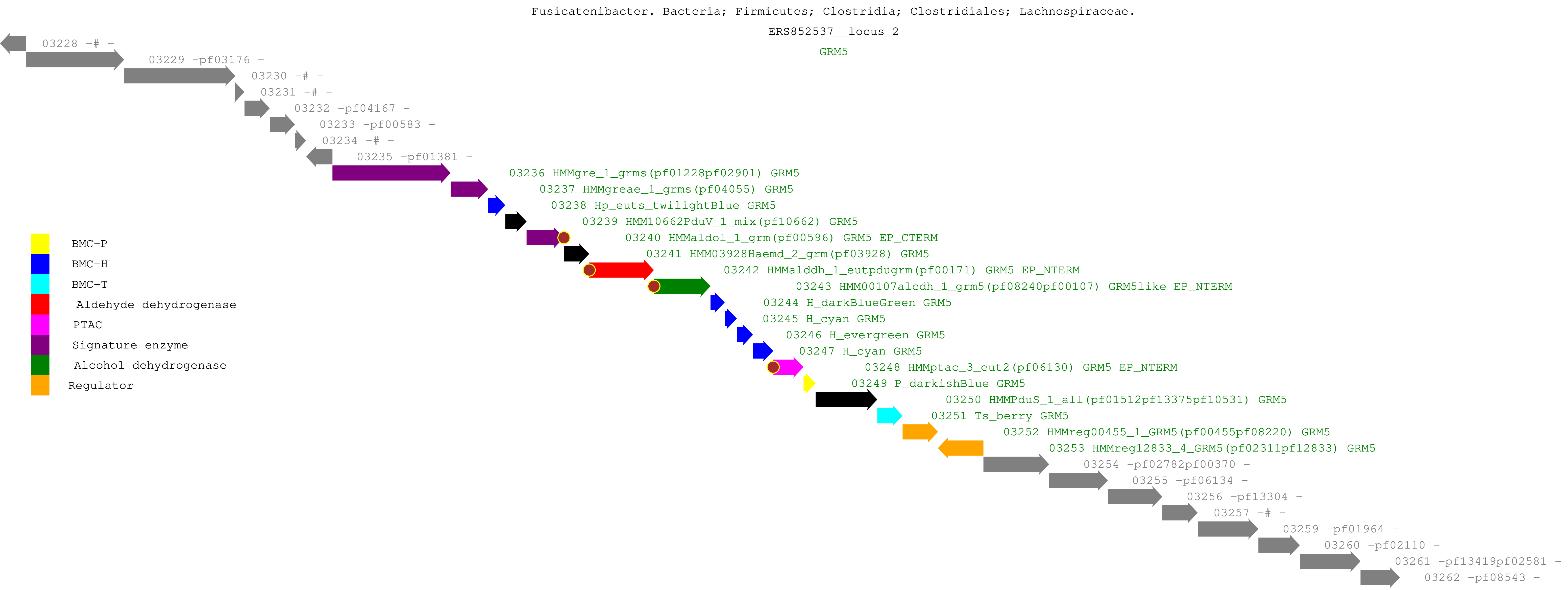
Fusicatenibacter. Bacteria; Firmicutes; Clostridia; Clostridiales; Lachnospiraceae.
ERS852537__locus_2
GRM5
03228 -# -
03229 -pf03176 -
03230 -# -
03231 -# -
03232 -pf04167 -
03233 -pf00583 -
03234 -# -
03235 -pf01381 -
03236 HMMgre_1_grms(pf01228pf02901) GRM5
03237 HMMgreae_1_grms(pf04055) GRM5
03238 Hp_euts_twilightBlue GRM5
03239 HMM10662PduV_1_mix(pf10662) GRM5
03240 HMMaldol_1_grm(pf00596) GRM5 EP_CTERM
03241 HMM03928Haemd_2_grm(pf03928) GRM5
03242 HMMalddh_1_eutpdugrm(pf00171) GRM5 EP_NTERM
03243 HMM00107alcdh_1_grm5(pf08240pf00107) GRM5like EP_NTERM
03244 H_darkBlueGreen GRM5
03245 H_cyan GRM5
03246 H_evergreen GRM5
03247 H_cyan GRM5
03248 HMMptac_3_eut2(pf06130) GRM5 EP_NTERM
03249 P_darkishBlue GRM5
03250 HMMPduS_1_all(pf01512pf13375pf10531) GRM5
03251 Ts_berry GRM5
03252 HMMreg00455_1_GRM5(pf00455pf08220) GRM5
03253 HMMreg12833_4_GRM5(pf02311pf12833) GRM5
03254 -pf02782pf00370 -
03255 -pf06134 -
03256 -pf13304 -
03257 -# -
03259 -pf01964 -
03260 -pf02110 -
03261 -pf13419pf02581 -
03262 -pf08543 -
BMC-P
BMC-H
BMC-T
Aldehyde dehydrogenase
PTAC
Signature enzyme
Alcohol dehydrogenase
Regulator

Ruminococcus torques. Bacteria; Firmicutes; Clostridia; Clostridiales; Lachnospiraceae;

ERS852577__locus_1

GRM5

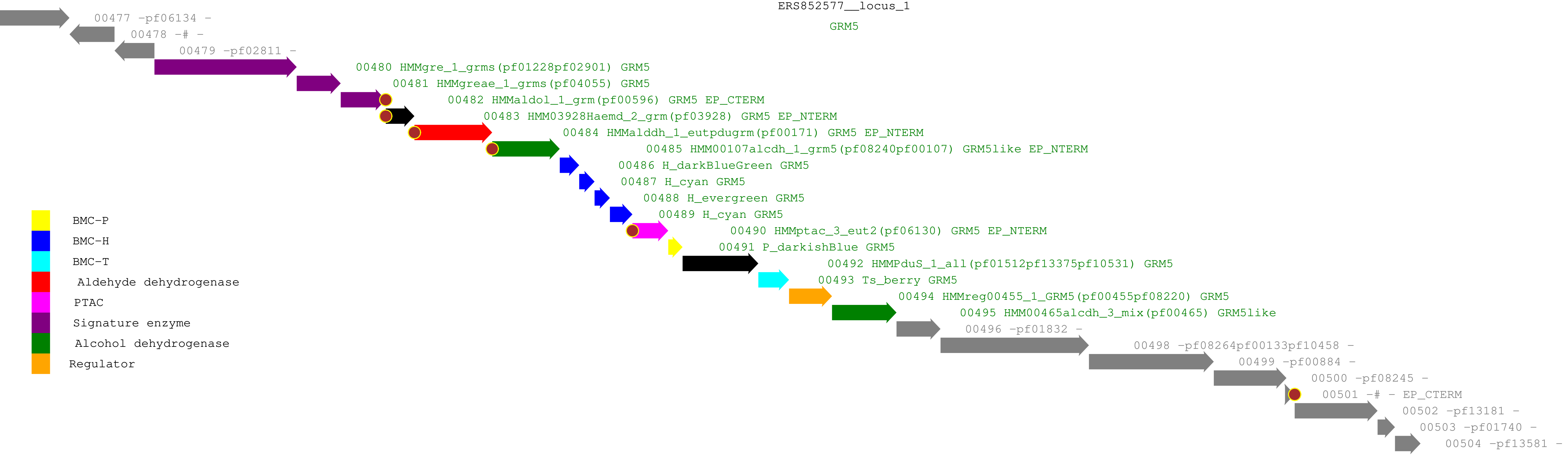

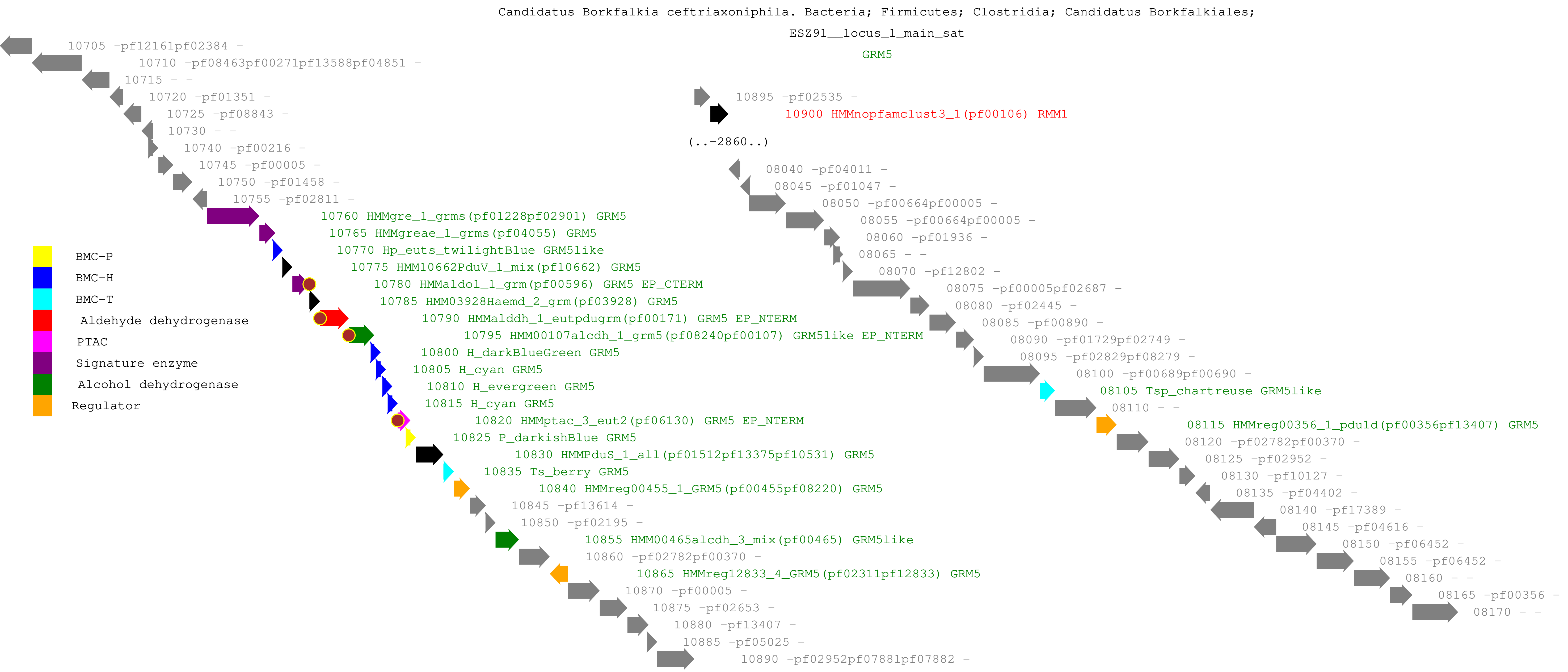

Candidatus Borkfalkia ceftriaxoniphila. Bacteria; Firmicutes; Clostridia; Candidatus Borkfalkiales;
ESZ91__locus_1_main_sat
GRM5
10705 -pf12161pf02384 -
10710 -pf08463pf00271pf13588pf04851 -
10715 - -
10720 -pf01351 -
10725 -pf08843 -
10730 - -
10740 -pf00216 -
10745 -pf00005 -
10750 -pf01458 -
10755 -pf02811 -
10760 HMMgre_1_grms(pf01228pf02901) GRM5
10765 HMMgreae_1_grms(pf04055) GRM5
10770 Hp_euts_twilightBlue GRM5like
10775 HMM10662PduV_1_mix(pf10662) GRM5
10780 HMMaldol_1_grm(pf00596) GRM5 EP_CTERM
10785 HMM03928Haemd_2_grm(pf03928) GRM5
10790 HMMalddh_1_eutpdugrm(pf00171) GRM5 EP_NTERM
10795 HMM00107alcdh_1_grm5(pf08240pf00107) GRM5like EP_NTERM
10800 H_darkBlueGreen GRM5
10805 H_cyan GRM5
10810 H_evergreen GRM5
10815 H_cyan GRM5
10820 HMMptac_3_eut2(pf06130) GRM5 EP_NTERM
10825 P_darkishBlue GRM5
10830 HMMPduS_1_all(pf01512pf13375pf10531) GRM5
10835 Ts_berry GRM5
10840 HMMreg00455_1_GRM5(pf00455pf08220) GRM5
10845 -pf13614 -
10850 -pf02195 -
10855 HMM00465alcdh_3_mix(pf00465) GRM5like
10860 -pf02782pf00370 -
10865 HMMreg12833_4_GRM5(pf02311pf12833) GRM5
10870 -pf00005 -
10875 -pf02653 -
10880 -pf13407 -
10885 -pf05025 -
10890 -pf02952pf07881pf07882 -
10895 -pf02535 -
10900 HMMnopfamclust3_1(pf00106) RMM1
(..-2860..)
08040 -pf04011 -
08045 -pf01047 -
08050 -pf00664pf00005 -
08055 -pf00664pf00005 -
08060 -pf01936 -
08065 - -
08070 -pf12802 -
08075 -pf00005pf02687 -
08080 -pf02445 -
08085 -pf00890 -
08090 -pf01729pf02749 -
08095 -pf02829pf08279 -
08100 -pf00689pf00690 -
08105 Tsp_chartreuse GRM5like
08110 - -
08115 HMMreg00356_1_pdu1d(pf00356pf13407) GRM5
08120 -pf02782pf00370 -
08125 -pf02952 -
08130 -pf10127 -
08135 -pf04402 -
08140 -pf17389 -
08145 -pf04616 -
08150 -pf06452 -
08155 -pf06452 -
08160 - -
08165 -pf00356 -
08170 - -
BMC-P
BMC-H
BMC-T
Aldehyde dehydrogenase
PTAC
Signature enzyme
Alcohol dehydrogenase
Regulator

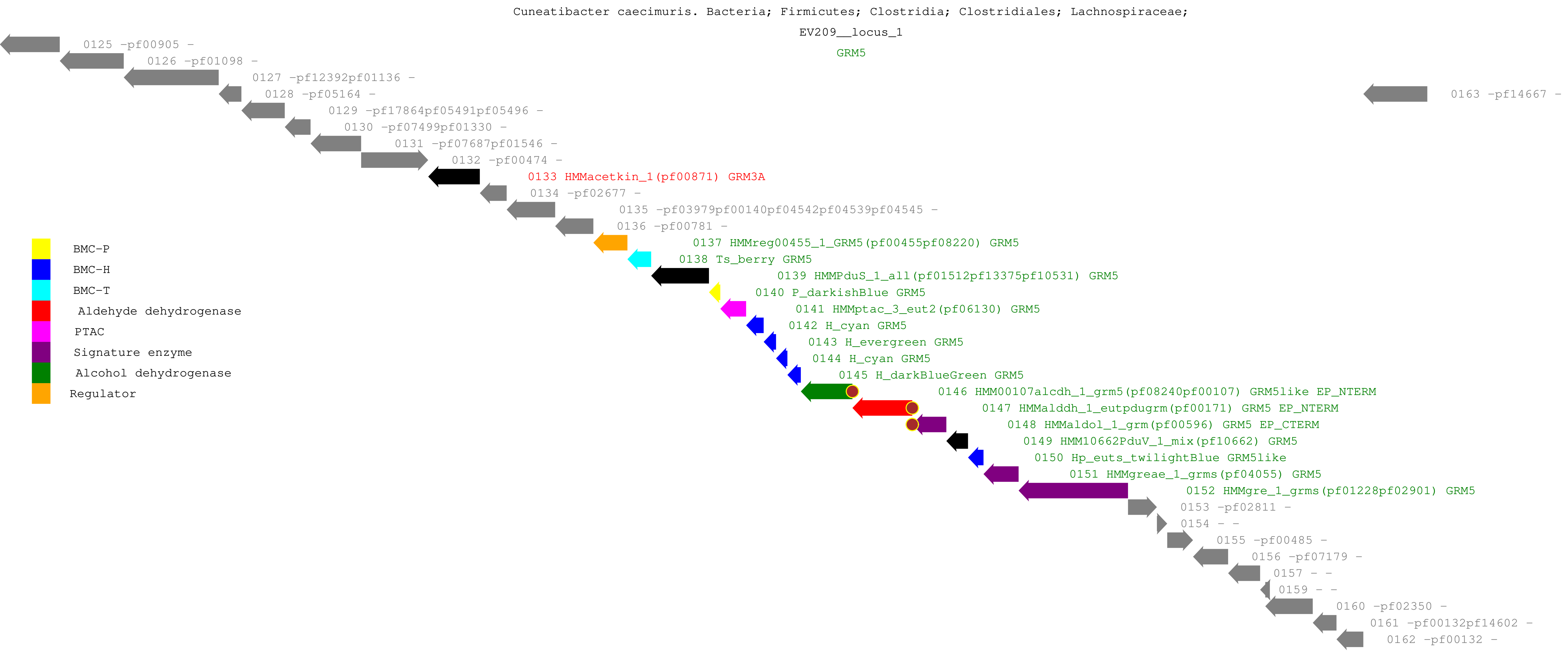
Cuneatibacter caecimuris. Bacteria; Firmicutes; Clostridia; Clostridiales; Lachnospiraceae;
EV209__locus_1
GRM5
0125 -pf00905 -
0126 -pf01098 -
0127 -pf12392pf01136 -
0128 -pf05164 -
0129 -pf17864pf05491pf05496 -
0130 -pf07499pf01330 -
0131 -pf07687pf01546 -
0132 -pf00474 -
0133 HMMacetkin_1(pf00871) GRM3A
0134 -pf02677 -
0135 -pf03979pf00140pf04542pf04539pf04545 -
0136 -pf00781 -
0137 HMMreg00455_1_GRM5(pf00455pf08220) GRM5
0138 Ts_berry GRM5
0139 HMMPduS_1_all(pf01512pf13375pf10531) GRM5
0140 P_darkishBlue GRM5
0141 HMMptac_3_eut2(pf06130) GRM5
0142 H_cyan GRM5
0143 H_evergreen GRM5
0144 H_cyan GRM5
0145 H_darkBlueGreen GRM5
0146 HMM00107alcdh_1_grm5(pf08240pf00107) GRM5like EP_NTERM
0147 HMMaldh_1_eutpdugrm(pf00171) GRM5 EP_NTERM
0148 HMMaldol_1_grm(pf00596) GRM5 EP_CTERM
0149 HMM10662PduV_1_mix(pf10662) GRM5
0150 Hp_euts_twilightBlue GRM5like
0151 HMMgreae_1_grms(pf04055) GRM5
0152 HMMgre_1_grms(pf01228pf02901) GRM5
0153 -pf02811 -
0154 - -
0155 -pf00485 -
0156 -pf07179 -
0157 - -
0159 - -
0160 -pf02350 -
0161 -pf00132pf14602 -
0162 -pf00132 -
0163 -pf14667 -
BMC-P
BMC-H
BMC-T
Aldehyde dehydrogenase
PTAC
Signature enzyme
Alcohol dehydrogenase
Regulator

Mediterraneibacter sp. gm002. Bacteria; Firmicutes; Clostridia; Clostridiales; Lachnospiraceae;

EYA86__locus_1_main_sat

GRM5

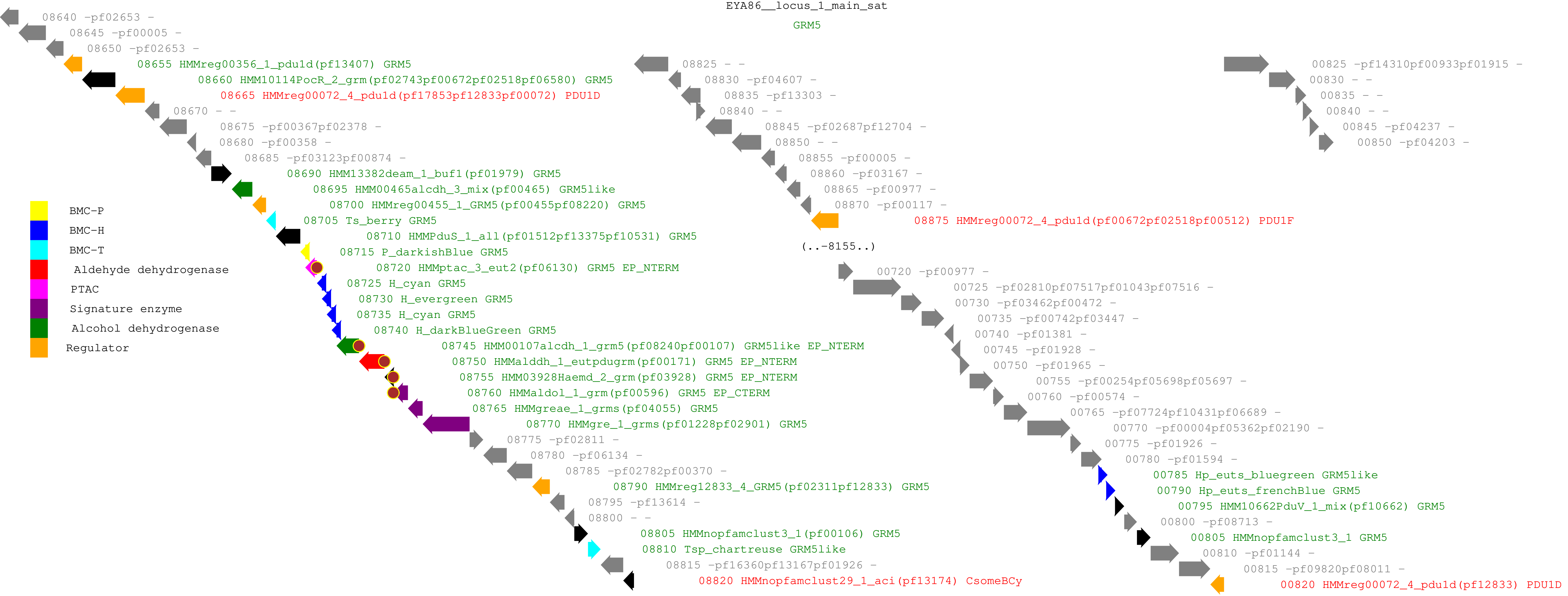

Blautia sp. SC05B48. Bacteria; Firmicutes; Clostridia; Clostridiales; Lachnospiraceae; Blautia;

EYS05__locus_1_main_sat

GRM5

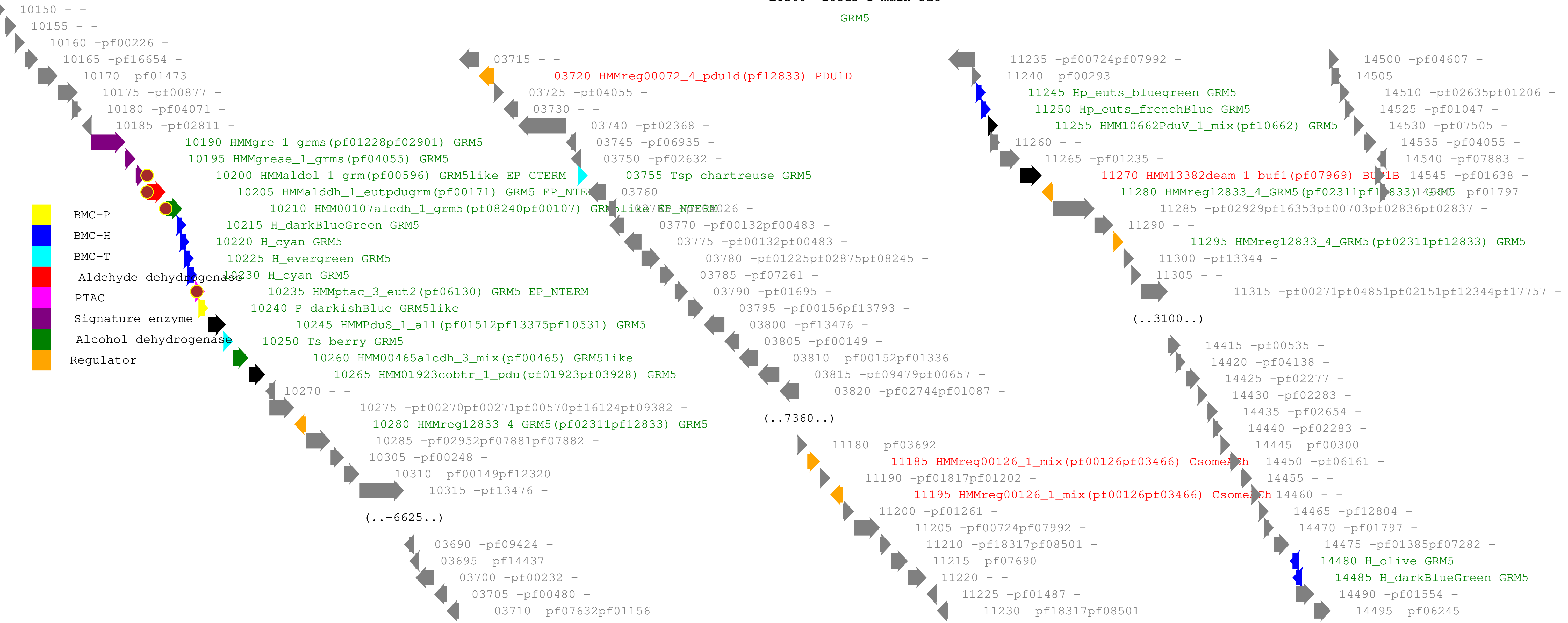

Ruminococcus gnavus (strain ATCC 29149 / VPI C7-9). Bacteria; Firmicutes; Clostridia; Clostridiales; Lachnospiraceae; Blautia.

FXV78__locus_1_main_sat

GRM5

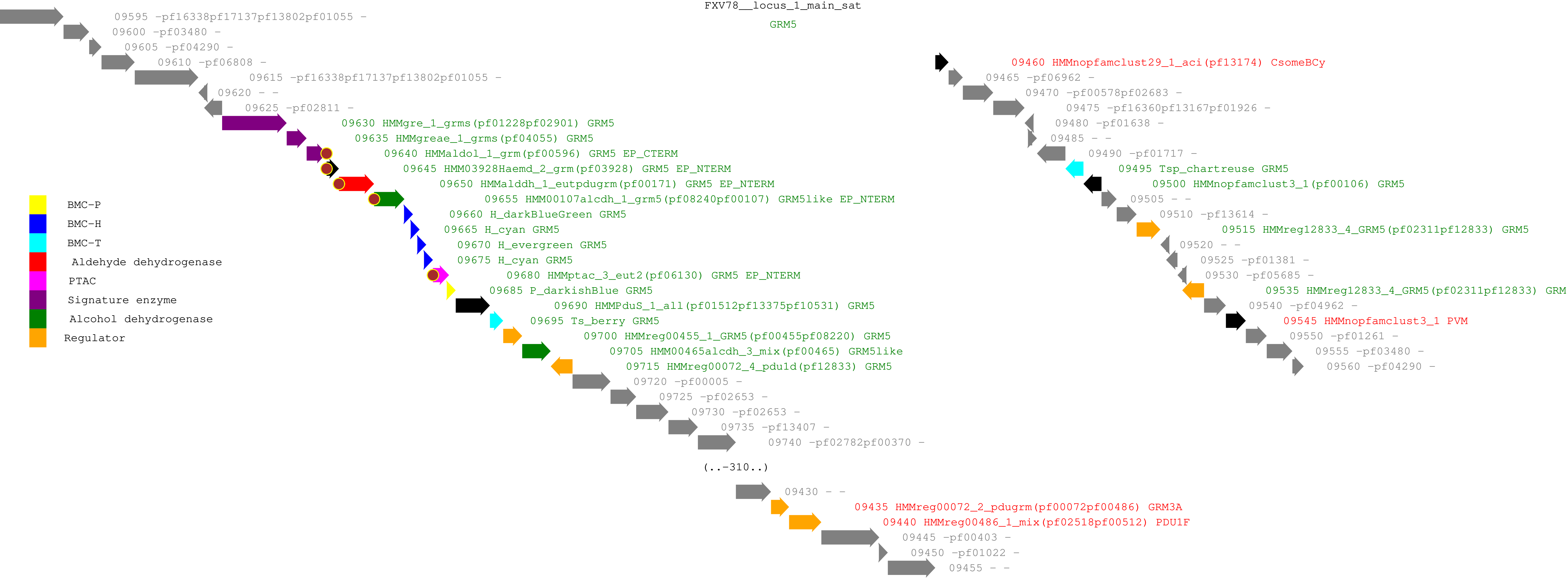

Shuttleworthia satelles DSM 14600. Bacteria; Firmicutes; Clostridia; Clostridiales; Lachnospiraceae;

GCWU000342__locus_1

GRM5

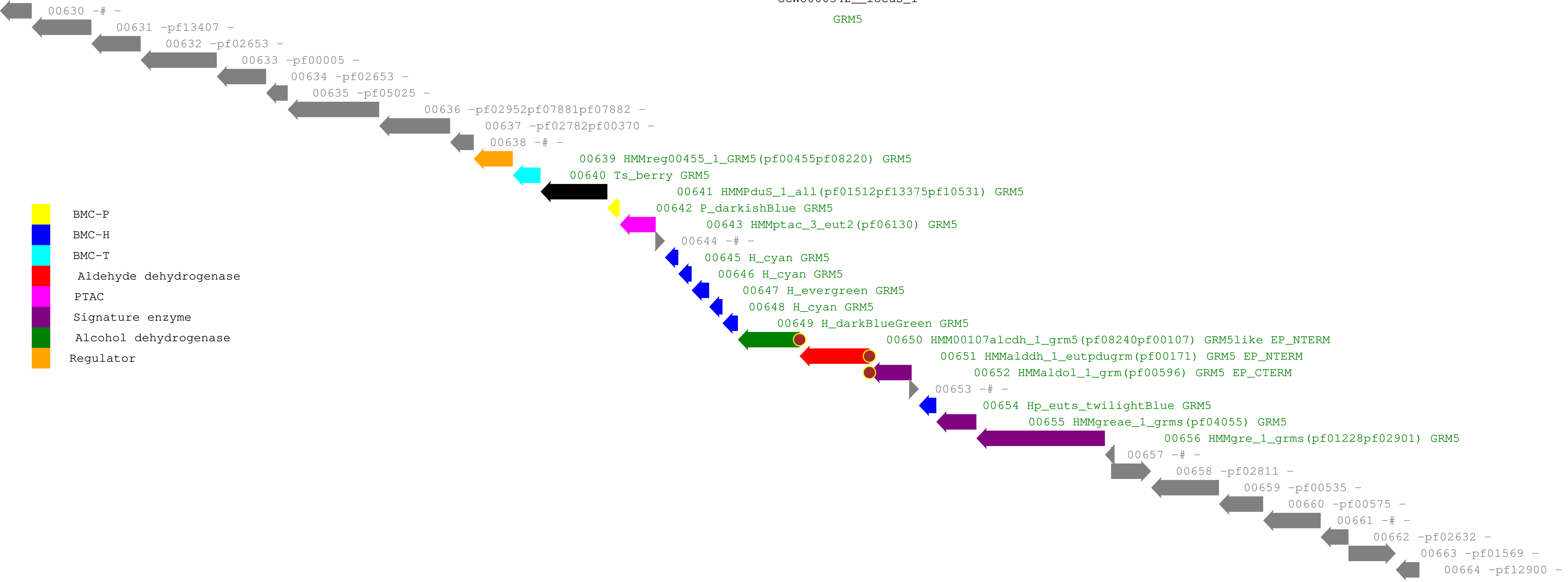

Clostridium sp. D5. Bacteria; Firmicutes; Clostridia; Clostridiales; Clostridiaceae;

HMPREF0240__locus_1_main_sat

GRM5

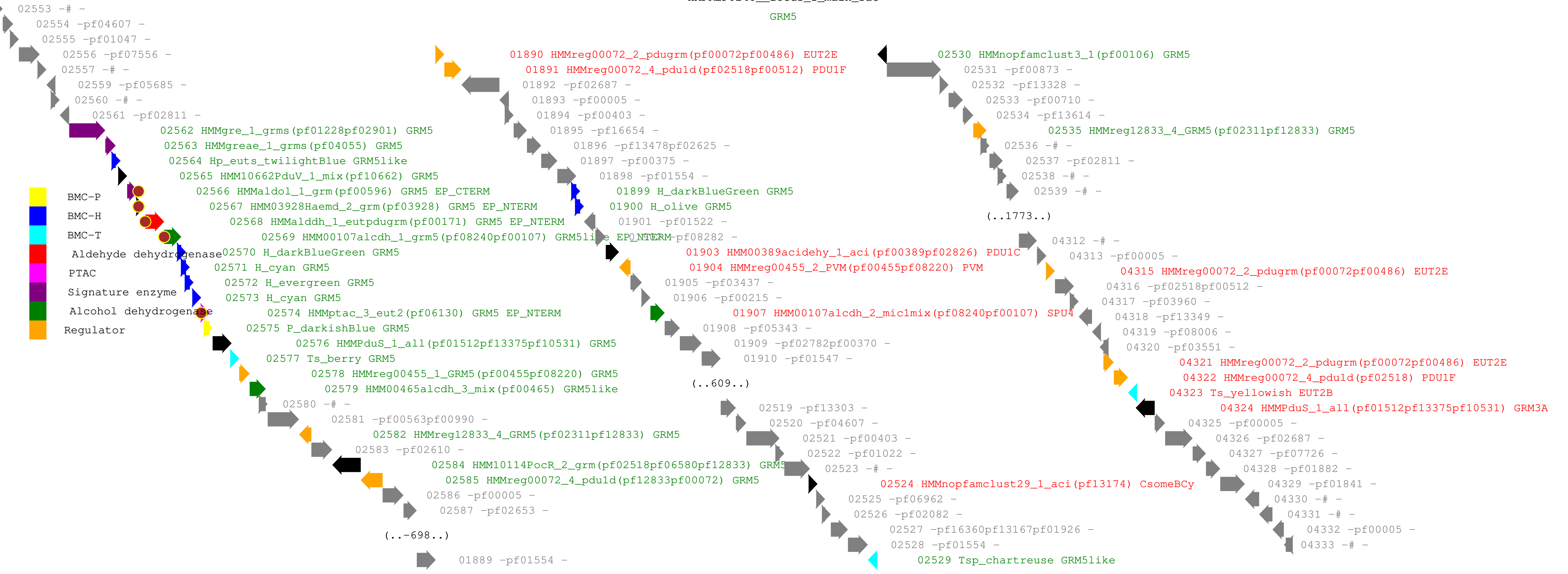

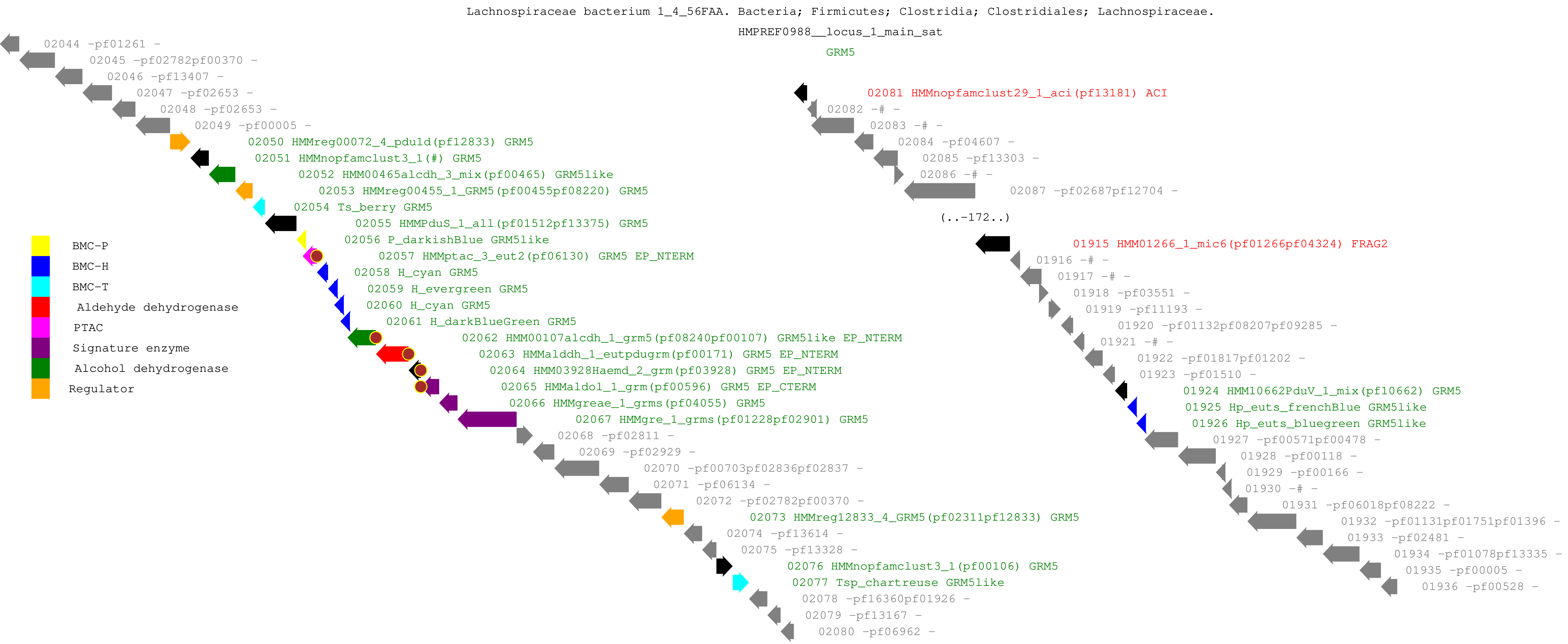
Lachnospiraceae bacterium 1_4_56FAA. Bacteria; Firmicutes; Clostridia; Clostridiales; Lachnospiraceae.
HMPREF0988__locus_1_main_sat
GRM5
02044 -pf01261 -
02045 -pf02782pf00370 -
02046 -pf13407 -
02047 -pf02653 -
02048 -pf02653 -
02049 -pf00005 -
02050 HMMreg00072_4_pduld(pf12833) GRM5
02051 HMMnopfamclust3_1(#) GRM5
02052 HMM00465alcdh_3_mix(pf00465) GRM5like
02053 HMMreg00455_1_GRM5(pf00455pf08220) GRM5
02054 Ts_berry GRM5
02055 HMMPduS_1_all(pf01512pf13375) GRM5
02056 P_darkishBlue GRM5like
02057 HMMptac_3_eut2(pf06130) GRM5 EP_NTERM
02058 H_cyan GRM5
02059 H_evergreen GRM5
02060 H_cyan GRM5
02061 H_darkBlueGreen GRM5
02062 HMM00107alcdh_1_grm5(pf08240pf00107) GRM5like EP_NTERM
02063 HMMaldh_1_eutpdugrm(pf00171) GRM5 EP_NTERM
02064 HMM03928Haemd_2_grm(pf03928) GRM5 EP_NTERM
02065 HMMaldol_1_grm(pf00596) GRM5 EP_CTERM
02066 HMMgreae_1_grms(pf04055) GRM5
02067 HMMgre_1_grms(pf01228pf02901) GRM5
02068 -pf02811 -
02069 -pf02929 -
02070 -pf00703pf02836pf02837 -
02071 -pf06134 -
02072 -pf02782pf00370 -
02073 HMMreg12833_4_GRM5(pf02311pf12833) GRM5
02074 -pf13614 -
02075 -pf13328 -
02076 HMMnopfamclust3_1(pf00106) GRM5
02077 Tsp_chartreuse GRM5like
02078 -pf16360pf01926 -
02079 -pf13167 -
02080 -pf06962 -
BMC-P
BMC-H
BMC-T
Aldehyde dehydrogenase
PTAC
Signature enzyme
Alcohol dehydrogenase
Regulator
02081 HMMnopfamclust29_1_aci(pf13181) ACI
02082 -# -
02083 -# -
02084 -pf04607 -
02085 -pf13303 -
02086 -# -
02087 -pf02687pf12704 -
(..-172..)
01915 HMM01266_1_mic6(pf01266pf04324) FRAG2
01916 -# -
01917 -# -
01918 -pf03551 -
01919 -pf11193 -
01920 -pf01132pf08207pf09285 -
01921 -# -
01922 -pf01817pf01202 -
01923 -pf01510 -
01924 HMM10662PduV_1_mix(pf10662) GRM5
01925 Hp_euts_frenchBlue GRM5like
01926 Hp_euts_bluegreen GRM5like
01927 -pf00571pf00478 -
01928 -pf00118 -
01929 -pf00166 -
01930 -# -
01931 -pf06018pf08222 -
01932 -pf01131pf01751pf01396 -
01933 -pf02481 -
01934 -pf01078pf13335 -
01935 -pf00005 -
01936 -pf00528 -

Lachnospiraceae bacterium 2_1_58FAA. Bacteria; Firmicutes; Clostridia; Clostridiales; Lachnospiraceae.

HMPREF0991__locus_1_main_sat

GRM5

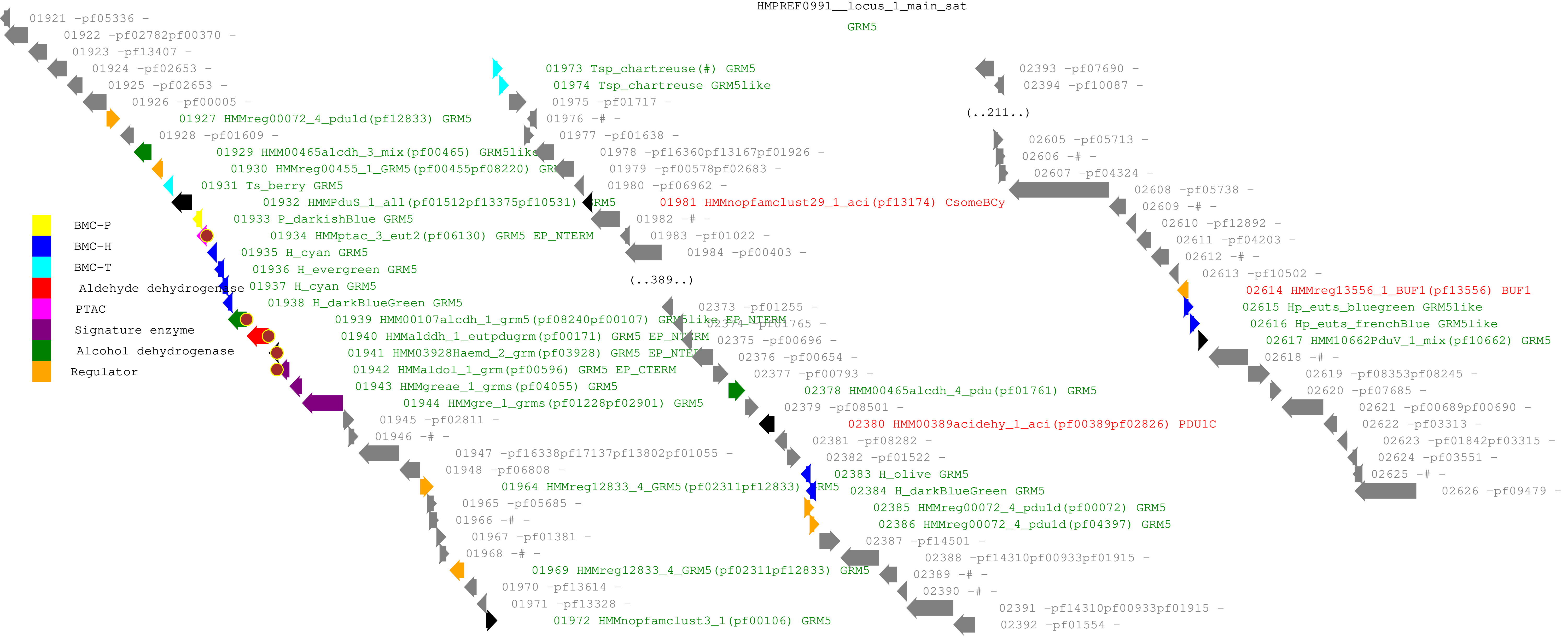

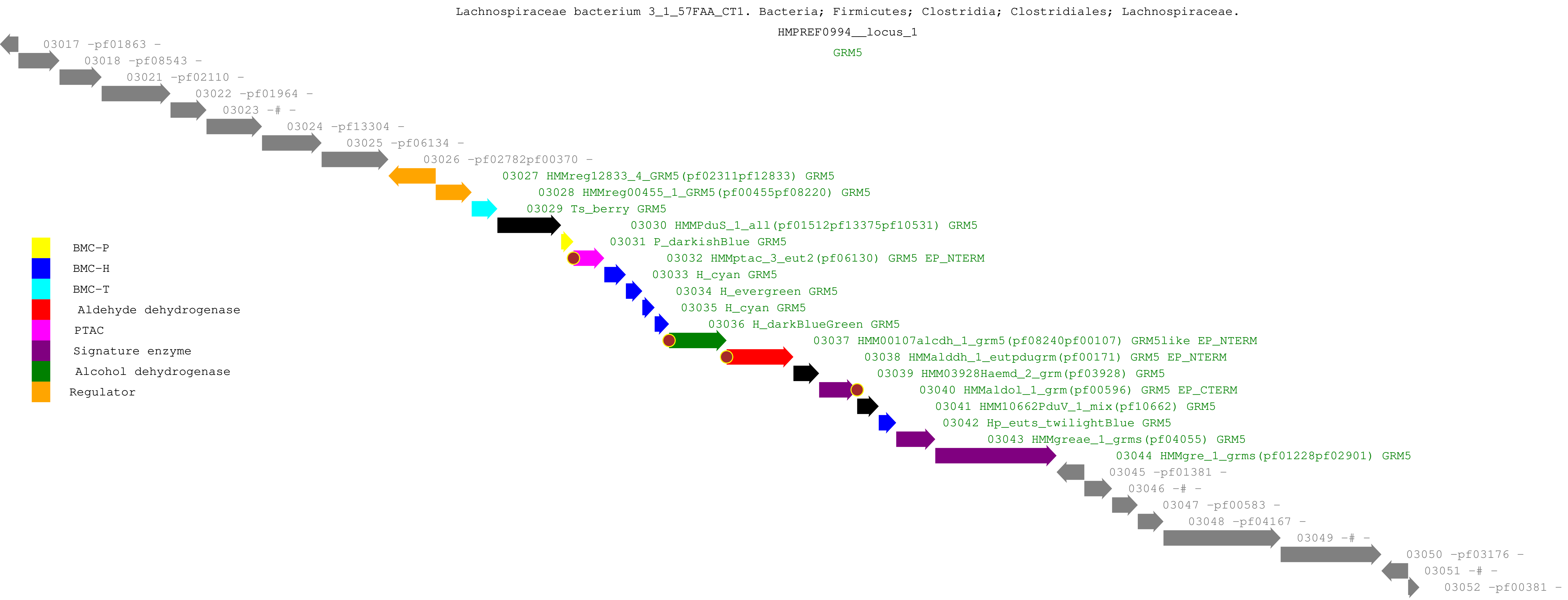
Lachnospiraceae bacterium 3_1_57FAA_CT1. Bacteria; Firmicutes; Clostridia; Clostridiales; Lachnospiraceae.
HMPREF0994__locus_1
GRM5
03017 -pf01863 -
03018 -pf08543 -
03021 -pf02110 -
03022 -pf01964 -
03023 -# -
03024 -pf13304 -
03025 -pf06134 -
03026 -pf02782pf00370 -
03027 HMMreg12833_4_GRM5(pf02311pf12833) GRM5
03028 HMMreg00455_1_GRM5(pf00455pf08220) GRM5
03029 Ts_berry GRM5
03030 HMMPduS_1_all(pf01512pf13375pf10531) GRM5
03031 P_darkishBlue GRM5
03032 HMMptac_3_eut2(pf06130) GRM5 EP_NTERM
03033 H_cyan GRM5
03034 H_evergreen GRM5
03035 H_cyan GRM5
03036 H_darkBlueGreen GRM5
03037 HMM00107alcdh_1_grm5(pf08240pf00107) GRM5like EP_NTERM
03038 HMMalddh_1_eutpdugrm(pf00171) GRM5 EP_NTERM
03039 HMM03928Haemd_2_grm(pf03928) GRM5
03040 HMMaldol_1_grm(pf00596) GRM5 EP_CTERM
03041 HMM10662PduV_1_mix(pf10662) GRM5
03042 Hp_euts_twilightBlue GRM5
03043 HMMgreae_1_grms(pf04055) GRM5
03044 HMMgre_1_grms(pf01228pf02901) GRM5
03045 -pf01381 -
03046 -# -
03047 -pf00583 -
03048 -pf04167 -
03049 -# -
03050 -pf03176 -
03051 -# -
03052 -pf00381 -
BMC-P
BMC-H
BMC-T
Aldehyde dehydrogenase
PTAC
Signature enzyme
Alcohol dehydrogenase
Regulator

Ruminococcus gnavus CC55_001C. Bacteria; Firmicutes; Clostridia; Clostridiales; Lachnospiraceae;

HMPREF1201__locus_1_main_sat

GRM5

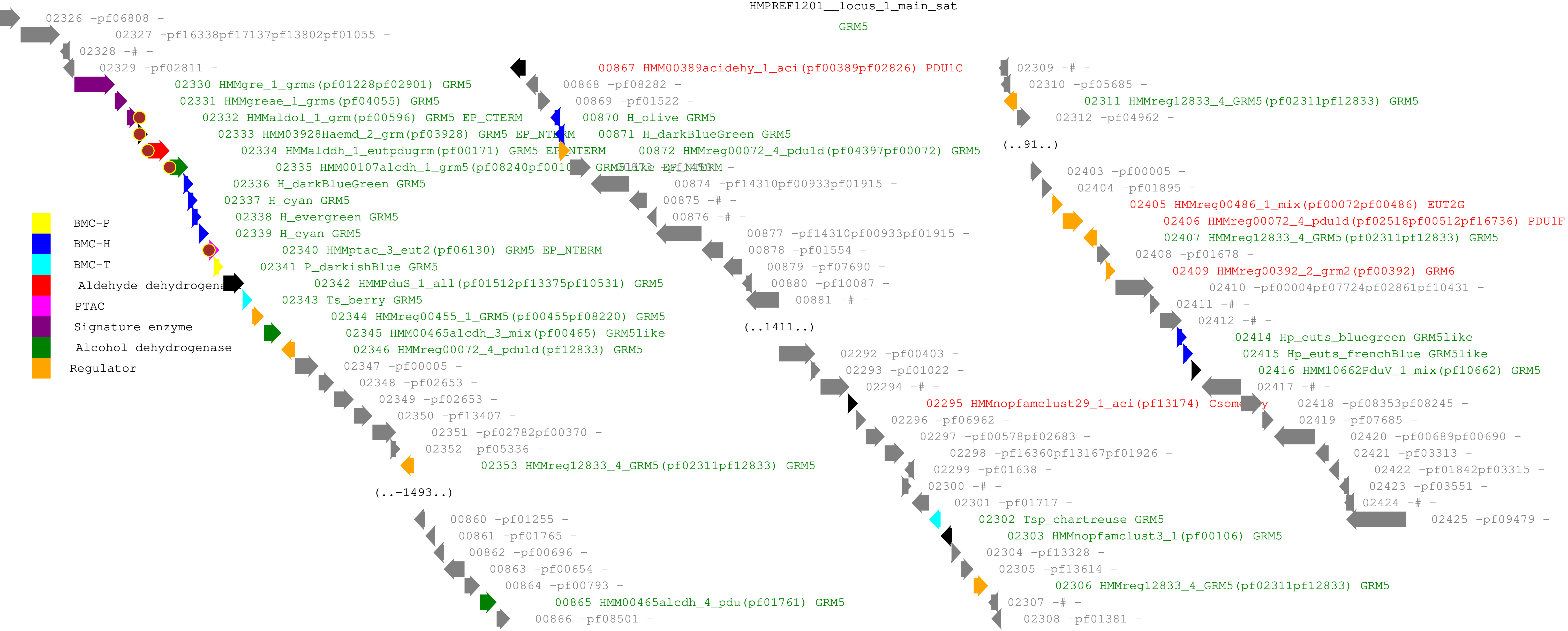

Shuttleworthia sp. MSX8B. Bacteria; Firmicutes; Clostridia; Clostridiales; Lachnospiraceae;

HMPREF1508__locus_1_main_sat

GRM5

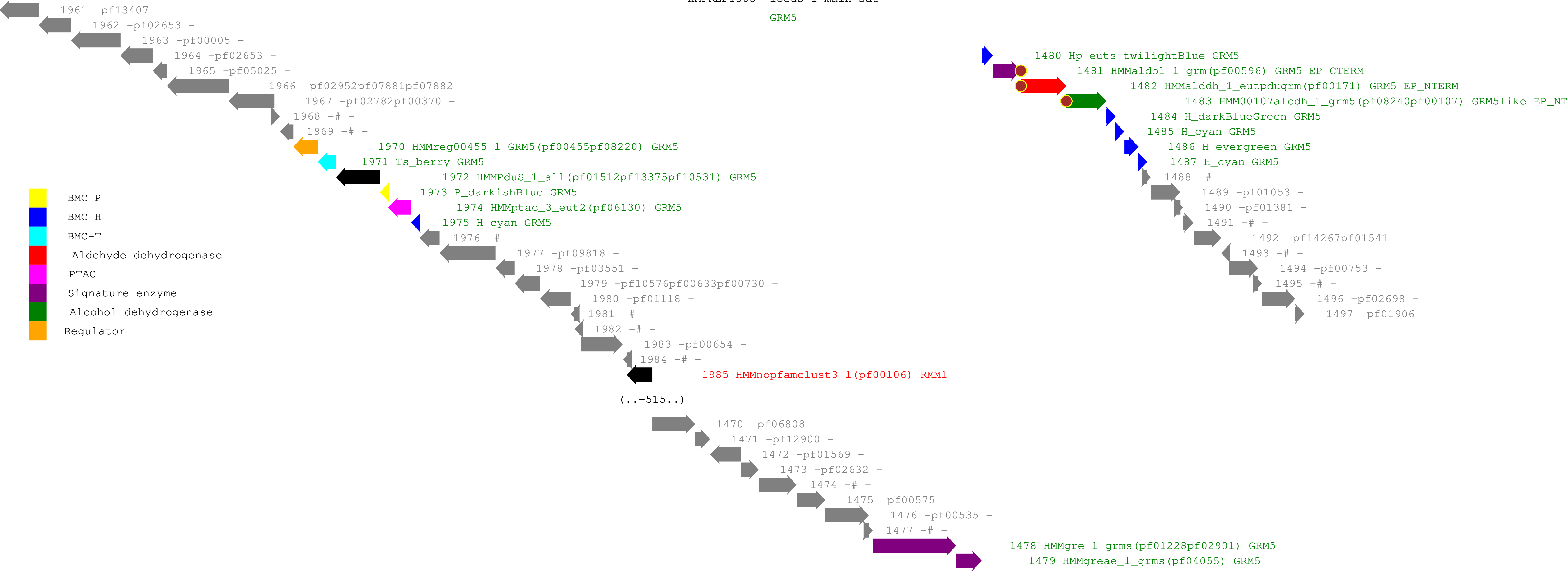

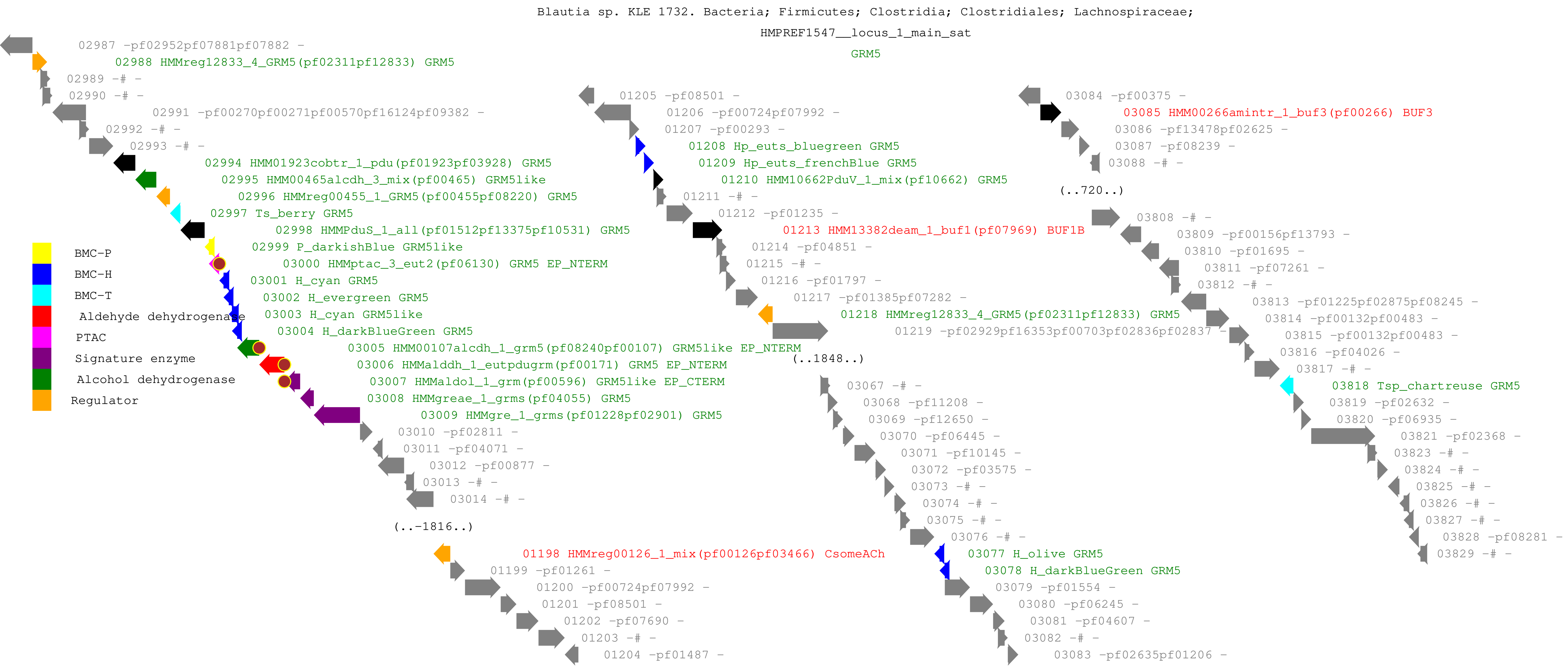
Blautia sp. KLE 1732. Bacteria; Firmicutes; Clostridia; Clostridiales; Lachnospiraceae;
HMPREF1547__locus_1_main_sat
GRM5
BMC-P
BMC-H
BMC-T
Aldehyde dehydrogenase
PTAC
Signature enzyme
Alcohol dehydrogenase
Regulator
02987 -pf02952pf07881pf07882 -
02988 HMMreg12833_4_GRM5(pf02311pf12833) GRM5
02989 -# -
02990 -# -
02991 -pf00270pf00271pf00570pf16124pf09382 -
02992 -# -
02993 -# -
02994 HMM01923cobtr_1_pdu(pf01923pf03928) GRM5
02995 HMM00465alcdh_3_mix(pf00465) GRM5like
02996 HMMreg00455_1_GRM5(pf00455pf08220) GRM5
02997 Ts_berry GRM5
02998 HMMPduS_1_all(pf01512pf13375pf10531) GRM5
02999 P_darkishBlue GRM5like
03000 HMMptac_3_eut2(pf06130) GRM5 EP_NTERM
03001 H_cyan GRM5
03002 H_evergreen GRM5
03003 H_cyan GRM5like
03004 H_darkBlueGreen GRM5
03005 HMM00107alcdh_1_grm5(pf08240pf00107) GRM5like EP_NTERM
03006 HMMaldh_1_eutpdugrm(pf00171) GRM5 EP_NTERM
03007 HMMaldol_1_grm(pf00596) GRM5like EP_CTERM
03008 HMMgreae_1_grms(pf04055) GRM5
03009 HMMgre_1_grms(pf01228pf02901) GRM5
03010 -pf02811 -
03011 -pf04071 -
03012 -pf00877 -
03013 -# -
03014 -# -
(..-1816..)
01198 HMMreg00126_1_mix(pf00126pf03466) CsomeACh
01199 -pf01261 -
01200 -pf00724pf07992 -
01201 -pf08501 -
01202 -pf07690 -
01203 -# -
01204 -pf01487 -
01205 -pf08501 -
01206 -pf00724pf07992 -
01207 -pf00293 -
01208 Hp_euts_bluegreen GRM5
01209 Hp_euts_frenchBlue GRM5
01210 HMM10662PduV_1_mix(pf10662) GRM5
01211 -# -
01212 -pf01235 -
01213 HMM13382deam_1_buf1(pf07969) BUF1B
01214 -pf04851 -
01215 -# -
01216 -pf01797 -
01217 -pf01385pf07282 -
01218 HMMreg12833_4_GRM5(pf02311pf12833) GRM5
01219 -pf02929pf16353pf00703pf02836pf02837 -
(..1848..)
03067 -# -
03068 -pf11208 -
03069 -pf12650 -
03070 -pf06445 -
03071 -pf10145 -
03072 -pf03575 -
03073 -# -
03074 -# -
03075 -# -
03076 -# -
03077 H_olive GRM5
03078 H_darkBlueGreen GRM5
03079 -pf01554 -
03080 -pf06245 -
03081 -pf04607 -
03082 -# -
03083 -pf02635pf01206 -
03084 -pf00375 -
03085 HMM00266amintr_1_buf3(pf00266) BUF3
03086 -pf13478pf02625 -
03087 -pf08239 -
03088 -# -
(..720..)
03808 -# -
03809 -pf00156pf13793 -
03810 -pf01695 -
03811 -pf07261 -
03812 -# -
03813 -pf01225pf02875pf08245 -
03814 -pf00132pf00483 -
03815 -pf00132pf00483 -
03816 -pf04026 -
03817 -# -
03818 Tsp_chartreuse GRM5
03819 -pf02632 -
03820 -pf06935 -
03821 -pf02368 -
03823 -# -
03824 -# -
03825 -# -
03826 -# -
03827 -# -
03828 -pf08281 -
03829 -# -

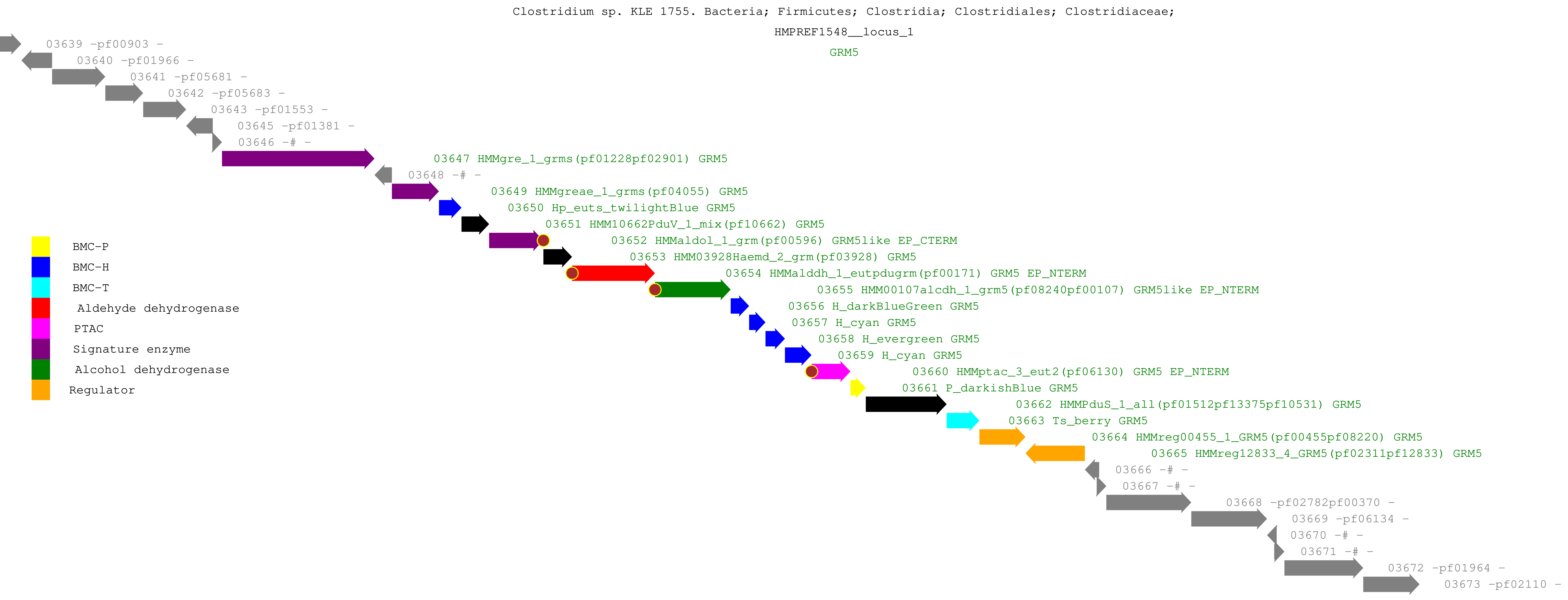
Clostridium sp. KLE 1755. Bacteria; Firmicutes; Clostridia; Clostridiales; Clostridiaceae;
HMPREF1548__locus_1
GRM5
03639 -pf00903 -
03640 -pf01966 -
03641 -pf05681 -
03642 -pf05683 -
03643 -pf01553 -
03645 -pf01381 -
03646 -# -
03647 HMMgre_1_grms(pf01228pf02901) GRM5
03648 -# -
03649 HMMgreae_1_grms(pf04055) GRM5
03650 Hp_euts_twilightBlue GRM5
03651 HMM10662PduV_1_mix(pf10662) GRM5
03652 HMMaldol_1_grm(pf00596) GRM5like EP_CTERM
03653 HMM03928Haemd_2_grm(pf03928) GRM5
03654 HMMalddh_1_eutpdugrm(pf00171) GRM5 EP_NTERM
03655 HMM00107alcdh_1_grm5(pf08240pf00107) GRM5like EP_NTERM
03656 H_darkBlueGreen GRM5
03657 H_cyan GRM5
03658 H_evergreen GRM5
03659 H_cyan GRM5
03660 HMMptac_3_eut2(pf06130) GRM5 EP_NTERM
03661 P_darkishBlue GRM5
03662 HMMPduS_1_all(pf01512pf13375pf10531) GRM5
03663 Ts_berry GRM5
03664 HMMreg00455_1_GRM5(pf00455pf08220) GRM5
03665 HMMreg12833_4_GRM5(pf02311pf12833) GRM5
03666 -# -
03667 -# -
03668 -pf02782pf00370 -
03669 -pf06134 -
03670 -# -
03671 -# -
03672 -pf01964 -
03673 -pf02110 -
BMC-P
BMC-H
BMC-T
Aldehyde dehydrogenase
PTAC
Signature enzyme
Alcohol dehydrogenase
Regulator

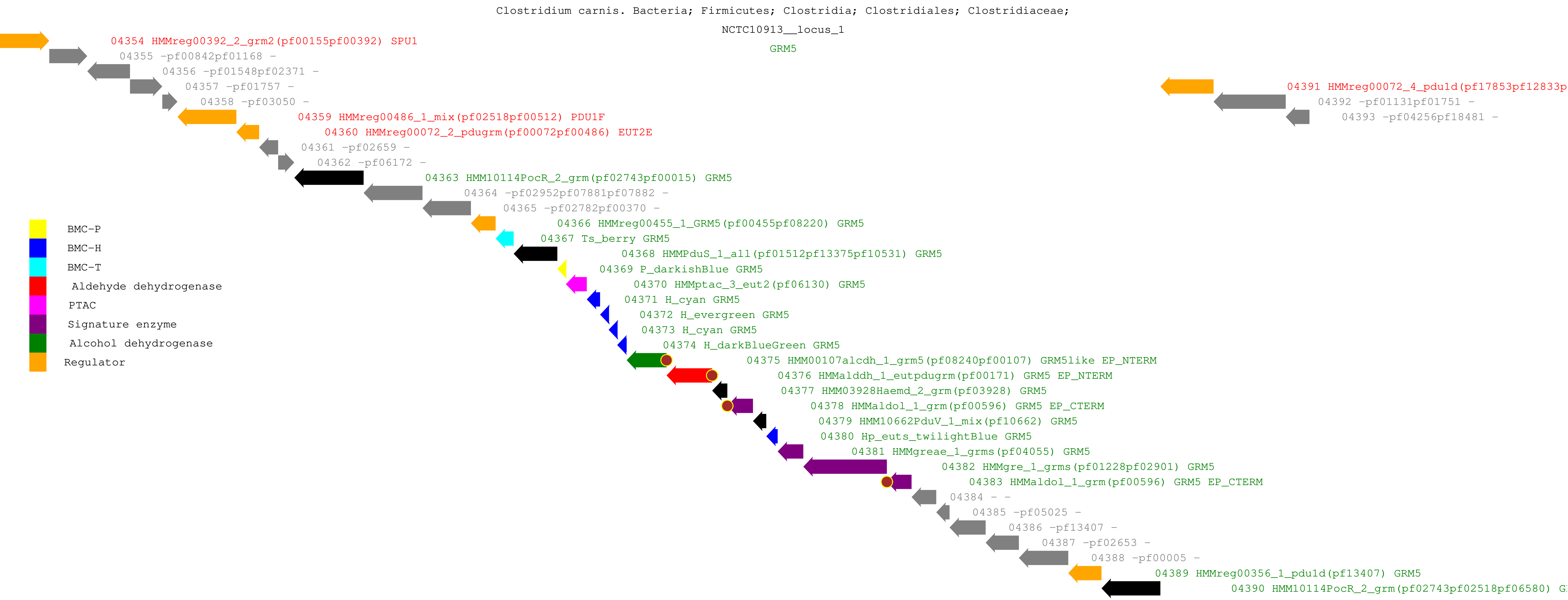
Clostridium carnis. Bacteria; Firmicutes; Clostridia; Clostridiales; Clostridiaceae;
NCTC10913__locus_1
GRM5
04354 HMMreg00392_2_grm2(pf00155pf00392) SPU1
04355 -pf00842pf01168 -
04356 -pf01548pf02371 -
04357 -pf01757 -
04358 -pf03050 -
04359 HMMreg00486_1_mix(pf02518pf00512) PDU1F
04360 HMMreg00072_2_pdugrm(pf00072pf00486) EUT2E
04361 -pf02659 -
04362 -pf06172 -
04363 HMM10114PocR_2_grm(pf02743pf00015) GRM5
04364 -pf02952pf07881pf07882 -
04365 -pf02782pf00370 -
04366 HMMreg00455_1_GRM5(pf00455pf08220) GRM5
04367 Ts_berry GRM5
04368 HMMPduS_1_all(pf01512pf13375pf10531) GRM5
04369 P_darkishBlue GRM5
04370 HMMptac_3_eut2(pf06130) GRM5
04371 H_cyan GRM5
04372 H_evergreen GRM5
04373 H_cyan GRM5
04374 H_darkBlueGreen GRM5
04375 HMM00107alcdh_1_grm5(pf08240pf00107) GRM5like EP_NTERM
04376 HMMaldh_1_eutpdugrm(pf00171) GRM5 EP_NTERM
04377 HMM03928Haemd_2_grm(pf03928) GRM5
04378 HMMaldol_1_grm(pf00596) GRM5 EP_CTERM
04379 HMM10662PduV_1_mix(pf10662) GRM5
04380 Hp_euts_twilightBlue GRM5
04381 HMMgreae_1_grms(pf04055) GRM5
04382 HMMgre_1_grms(pf01228pf02901) GRM5
04383 HMMaldol_1_grm(pf00596) GRM5 EP_CTERM
04384 - -
04385 -pf05025 -
04386 -pf13407 -
04387 -pf02653 -
04388 -pf00005 -
04389 HMMreg00356_1_pdu1d(pf13407) GRM5
04390 HMM10114PocR_2_grm(pf02743pf02518pf06580) G
04391 HMMreg00072_4_pdu1d(pf17853pf12833p
04392 -pf01131pf01751 -
04393 -pf04256pf18481 -
BMC-P
BMC-H
BMC-T
Aldehyde dehydrogenase
PTAC
Signature enzyme
Alcohol dehydrogenase
Regulator

Roseburia inulinivorans DSM 16841. Bacteria; Firmicutes; Clostridia; Clostridiales; Lachnospiraceae;

ROSEINA2194__locus_1

GRM5

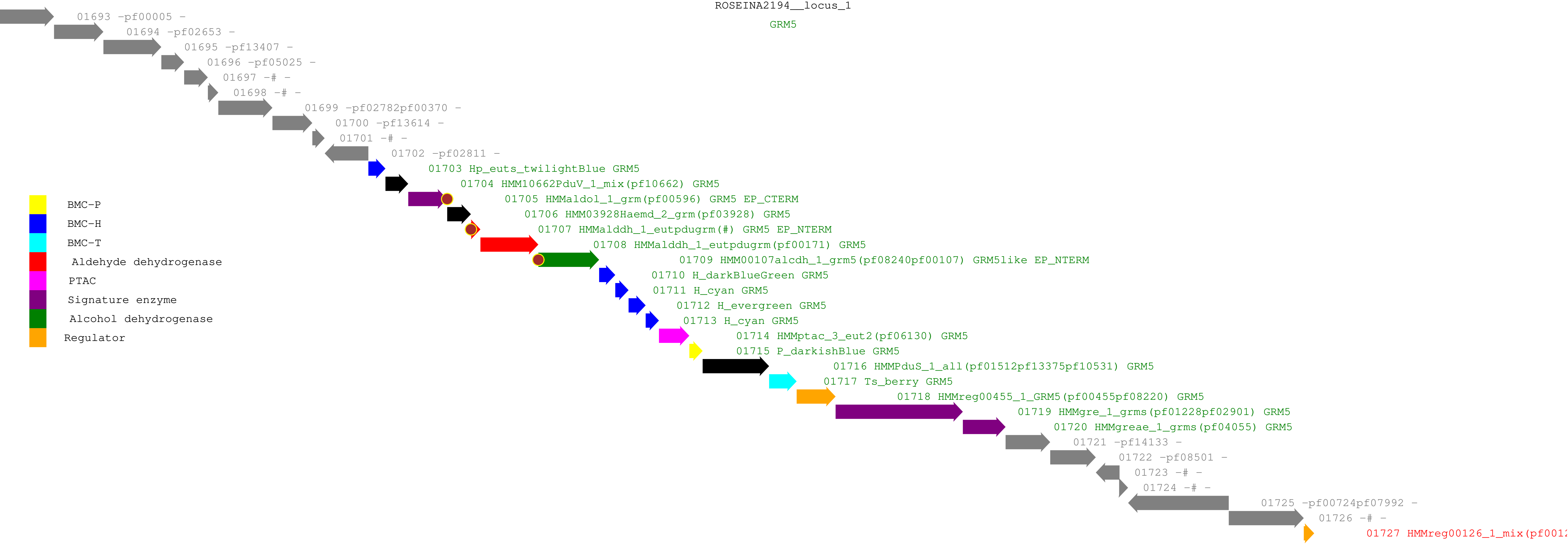

Ruminococcus sp. 5_1_39BFAA. Bacteria; Firmicutes; Clostridia; Clostridiales; Ruminococcaceae;

RSAG__locus_1_main_sat

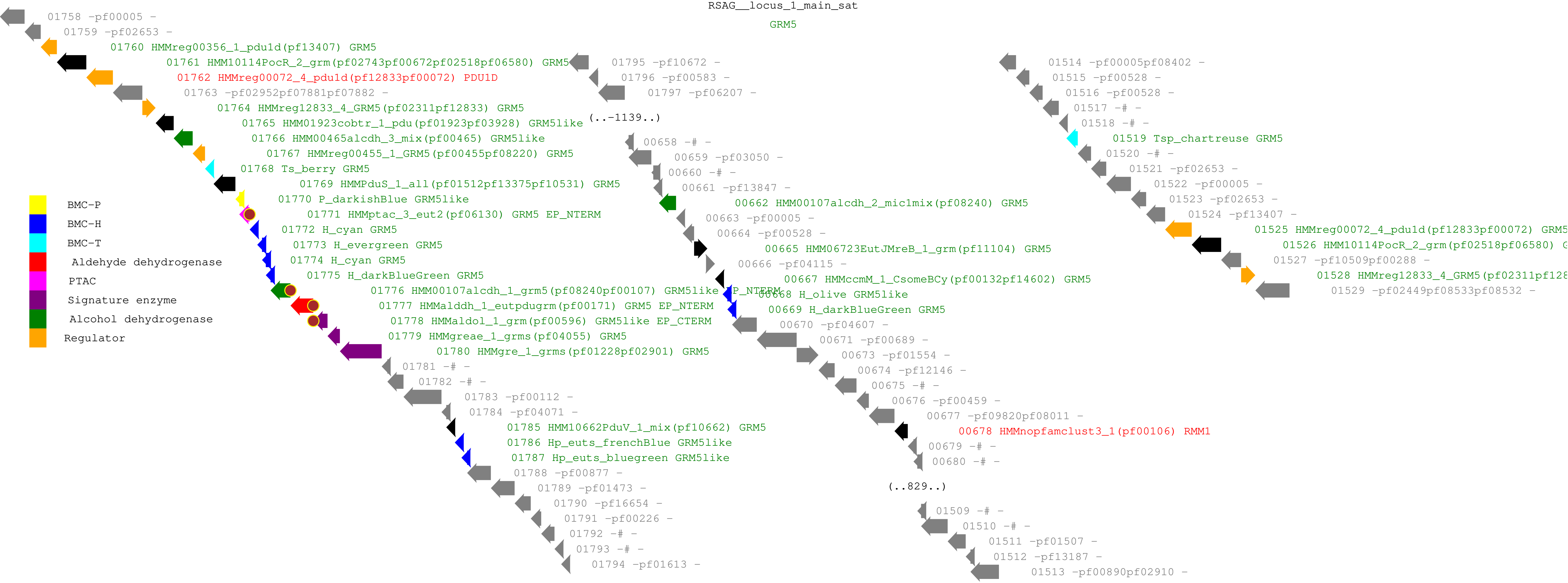

Blautia luti. Bacteria; Firmicutes; Clostridia; Clostridiales; Lachnospiraceae; Blautia.

RSSSTS7063__locus_1_main_sat

GRM5

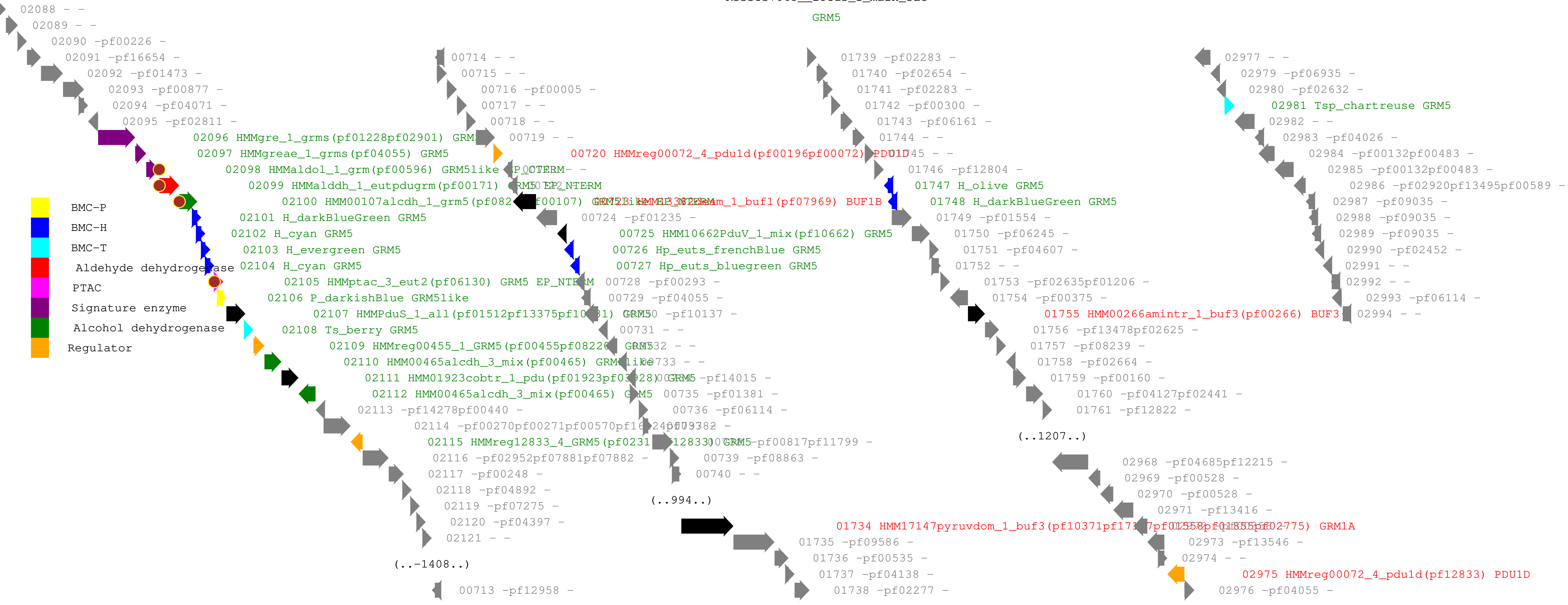

Ruminococcus torques L2-14. Bacteria; Firmicutes; Clostridia; Clostridiales; Lachnospiraceae;

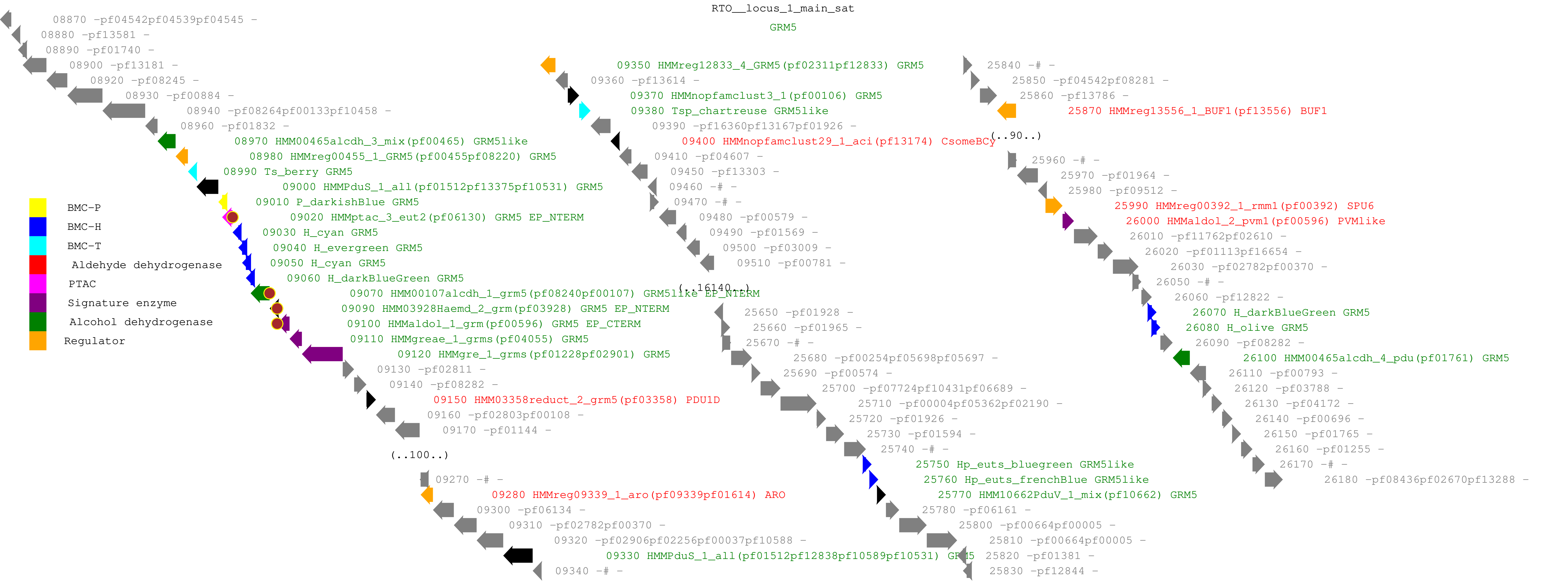

Ruminococcus gnavus ATCC 29149. Bacteria; Firmicutes; Clostridia; Clostridiales; Lachnospiraceae;

RUMGNA__locus_1_main_sat

GRM5

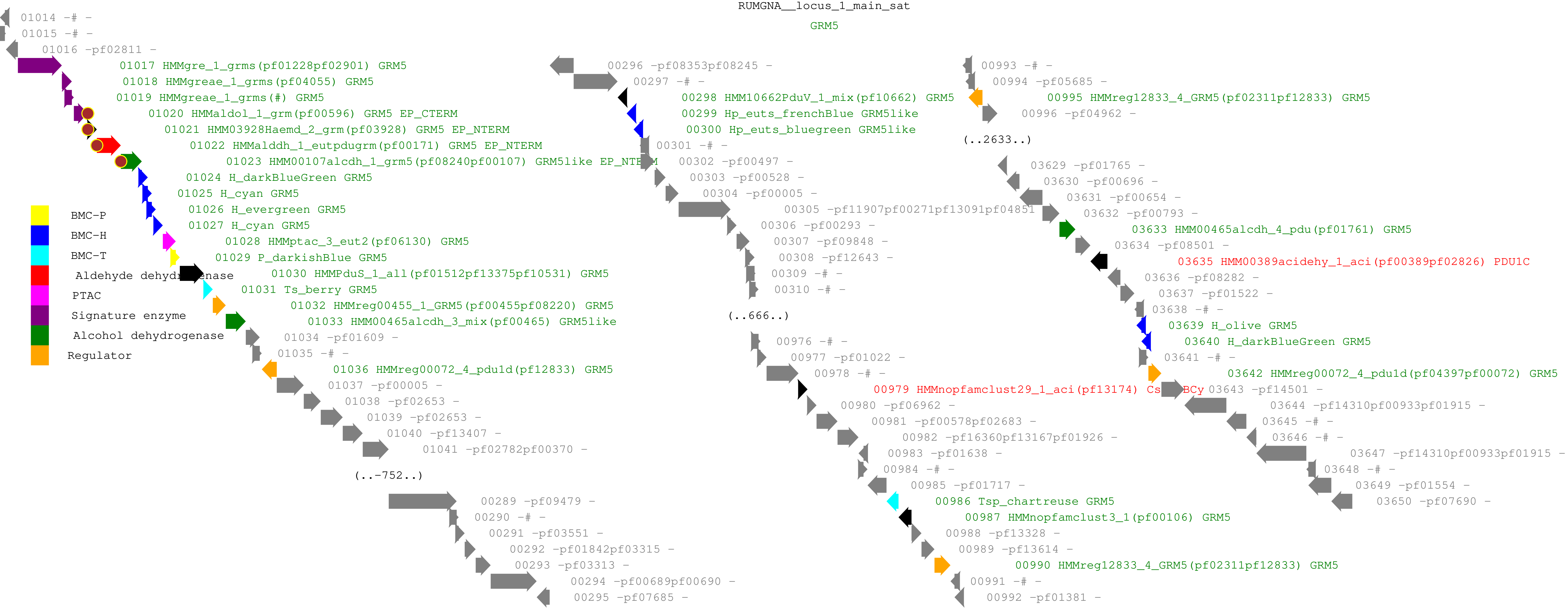

uncultured Blautia sp. Bacteria; Firmicutes; Clostridia; Clostridiales; Lachnospiraceae;

SAMEA2393092__locus_1_main_sat

GRM5

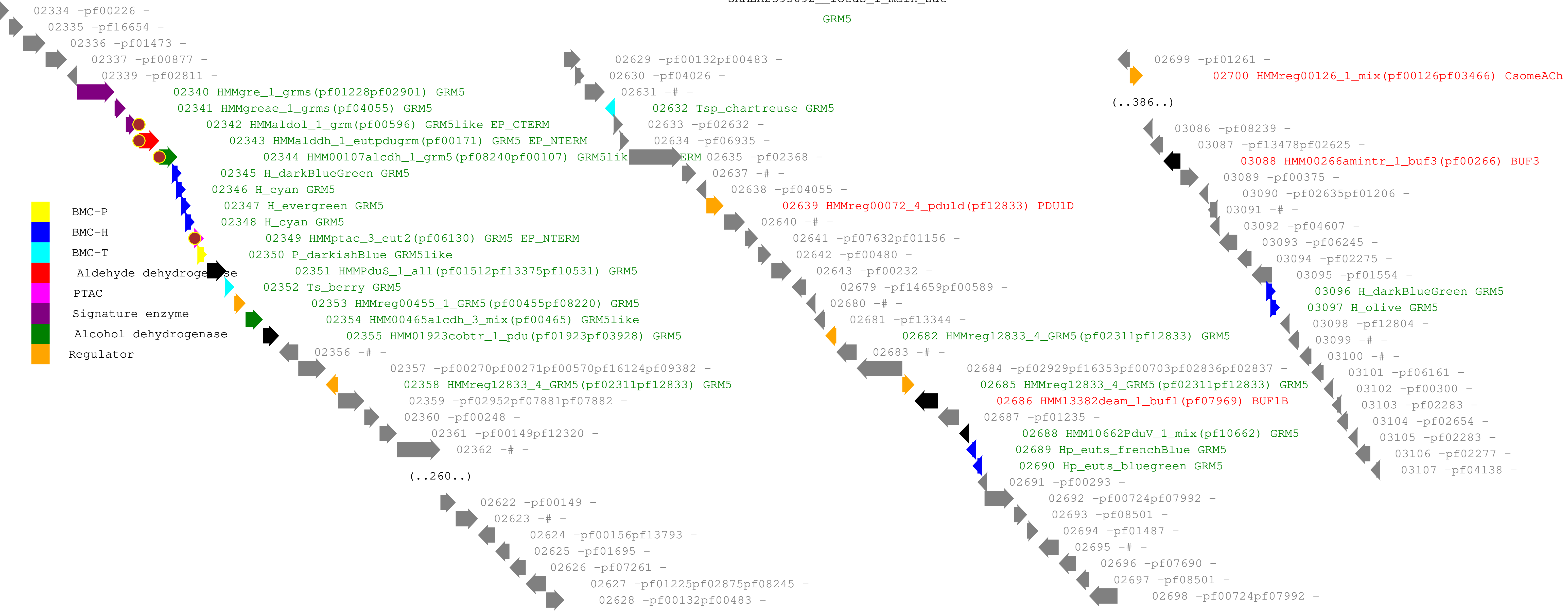

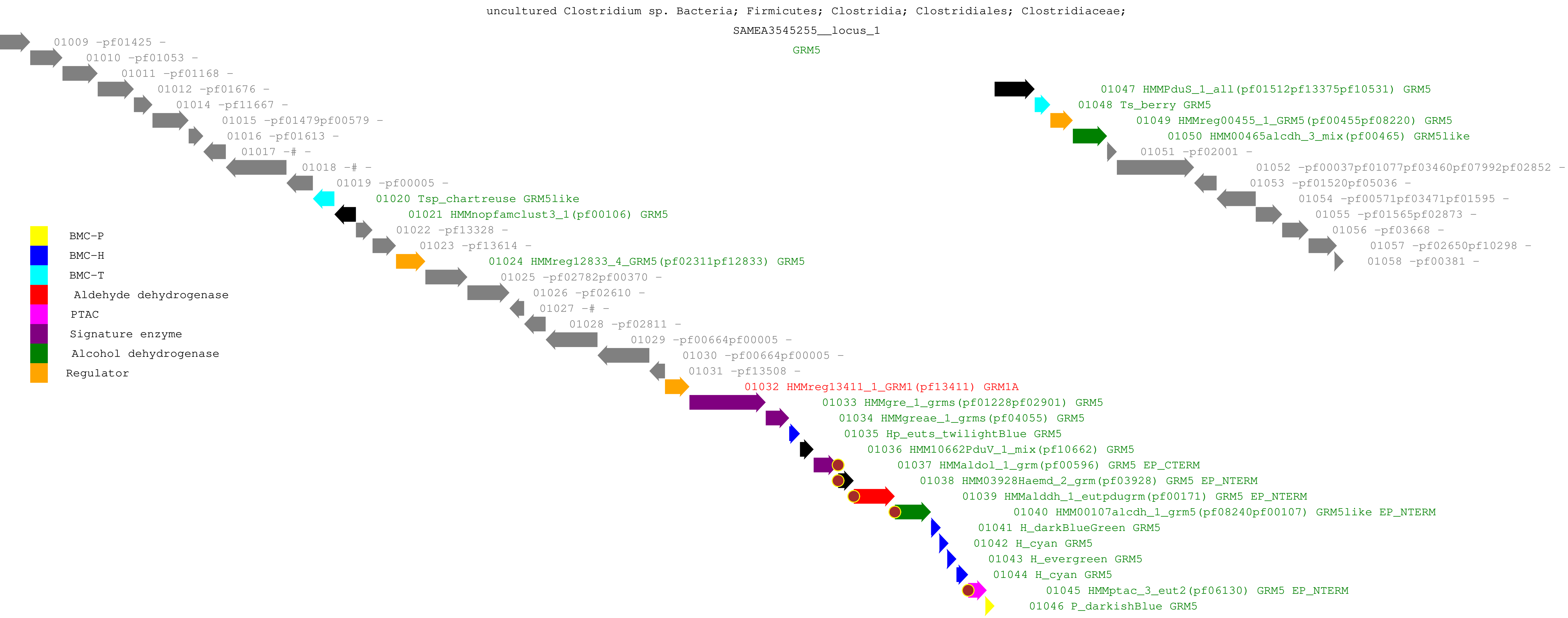

uncultured Clostridium sp. Bacteria; Firmicutes; Clostridia; Clostridiales; Clostridiaceae;
SAMEA3545255__locus_1
GRM5
01009 -pf01425 -
01010 -pf01053 -
01011 -pf01168 -
01012 -pf01676 -
01014 -pf11667 -
01015 -pf01479pf00579 -
01016 -pf01613 -
01017 -# -
01018 -# -
01019 -pf00005 -
01020 Tsp_chartreuse GRM5like
01021 HMMnopfamclust3_1(pf00106) GRM5
01022 -pf13328 -
01023 -pf13614 -
01024 HMMreg12833_4_GRM5(pf02311pf12833) GRM5
01025 -pf02782pf00370 -
01026 -pf02610 -
01027 -# -
01028 -pf02811 -
01029 -pf00664pf00005 -
01030 -pf00664pf00005 -
01031 -pf13508 -
01032 HMMreg13411_1_GRM1(pf13411) GRM1A
01033 HMMgre_1_grms(pf01228pf02901) GRM5
01034 HMMgreae_1_grms(pf04055) GRM5
01035 Hp_euts_twilightBlue GRM5
01036 HMM10662PduV_1_mix(pf10662) GRM5
01037 HMMaldol_1_grm(pf00596) GRM5 EP_CTERM
01038 HMM03928Haemd_2_grm(pf03928) GRM5 EP_NTERM
01039 HMMalddh_1_eutpdugrm(pf00171) GRM5 EP_NTERM
01040 HMM00107alcdh_1_grm5(pf08240pf00107) GRM5like EP_NTERM
01041 H_darkBlueGreen GRM5
01042 H_cyan GRM5
01043 H_evergreen GRM5
01044 H_cyan GRM5
01045 HMMptac_3_eut2(pf06130) GRM5 EP_NTERM
01046 P_darkishBlue GRM5
01047 HMMPduS_1_all(pf01512pf13375pf10531) GRM5
01048 Ts_berry GRM5
01049 HMMreg00455_1_GRM5(pf00455pf08220) GRM5
01050 HMM00465alcdh_3_mix(pf00465) GRM5like
01051 -pf02001 -
01052 -pf00037pf01077pf03460pf07992pf02852 -
01053 -pf01520pf05036 -
01054 -pf00571pf03471pf01595 -
01055 -pf01565pf02873 -
01056 -pf03668 -
01057 -pf02650pf10298 -
01058 -pf00381 -
BMC-P
BMC-H
BMC-T
Aldehyde dehydrogenase
PTAC
Signature enzyme
Alcohol dehydrogenase
Regulator

uncultured Blautia sp. Bacteria; Firmicutes; Clostridia; Clostridiales; Lachnospiraceae;

SAMEA3545287__locus_1_main_sat

GRM5

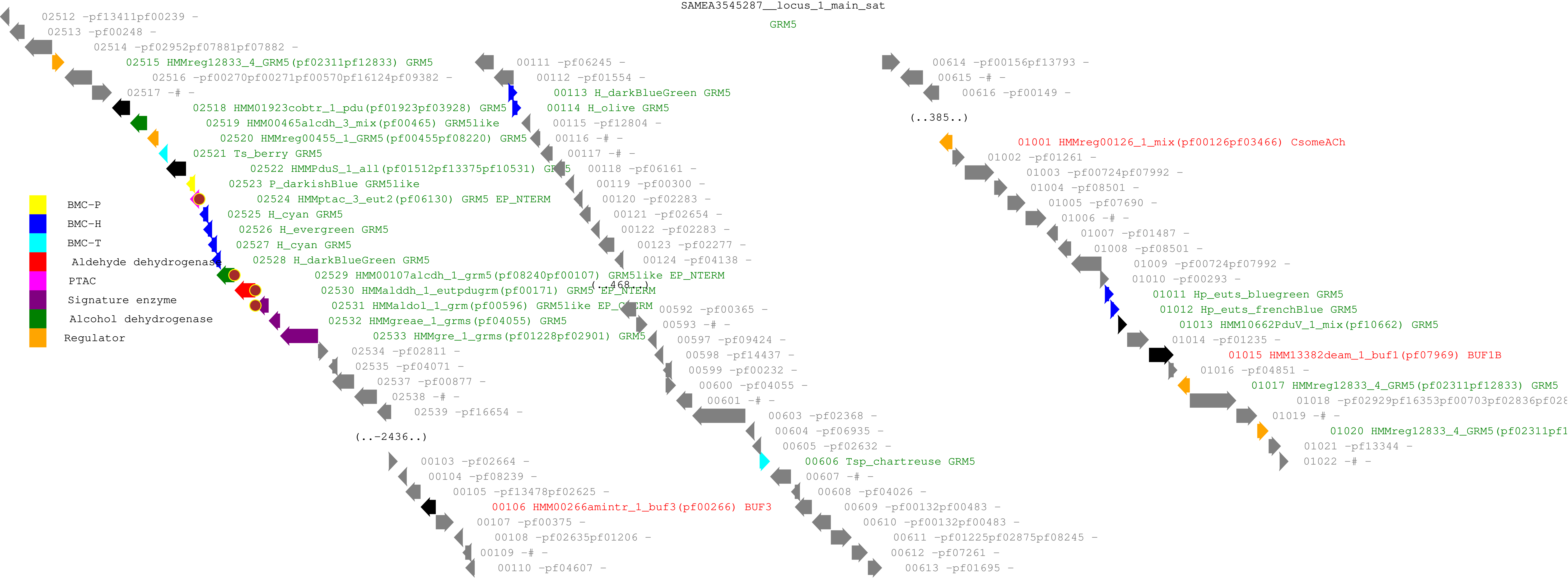

uncultured Ruminococcus sp. Bacteria; Firmicutes; Clostridia; Clostridiales; Ruminococcaceae;

SAMEA3545290__locus_1_main_sat

GRM5

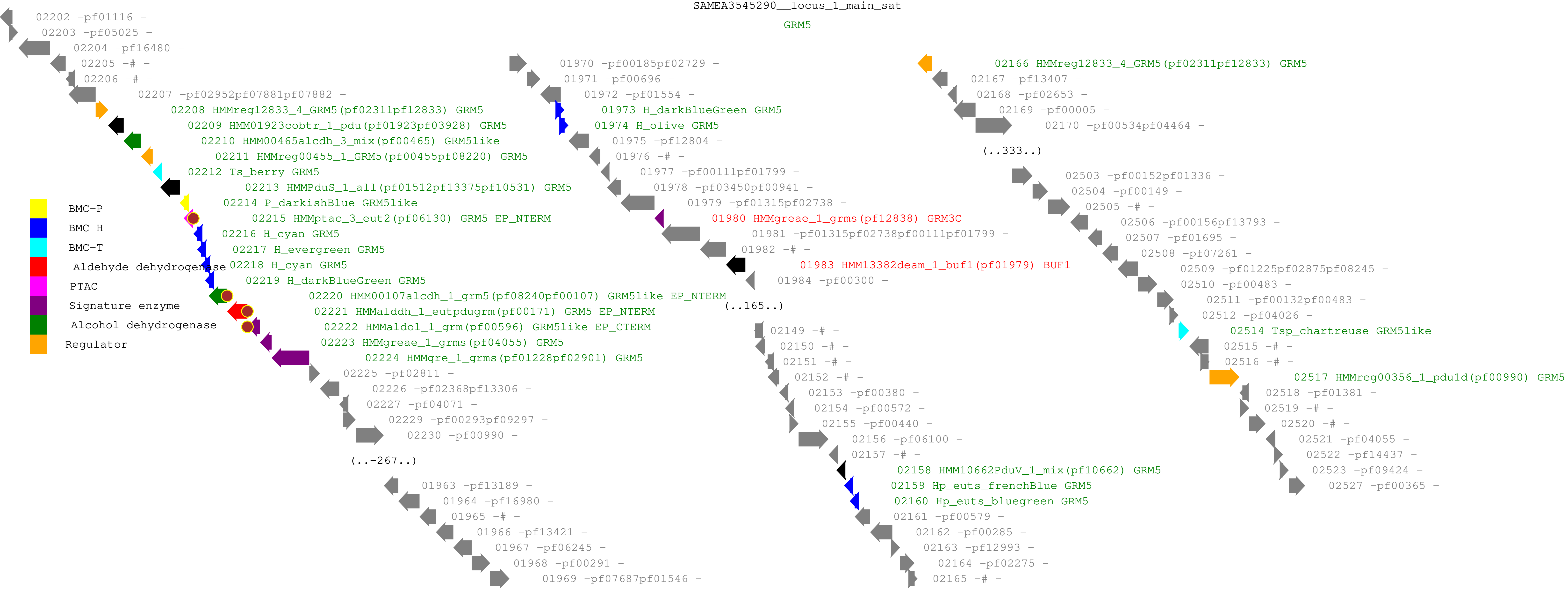

uncultured Clostridium sp. Bacteria; Firmicutes; Clostridia; Clostridiales; Clostridiaceae;

SAMEA3545337__locus_1_main_sat

GRM5

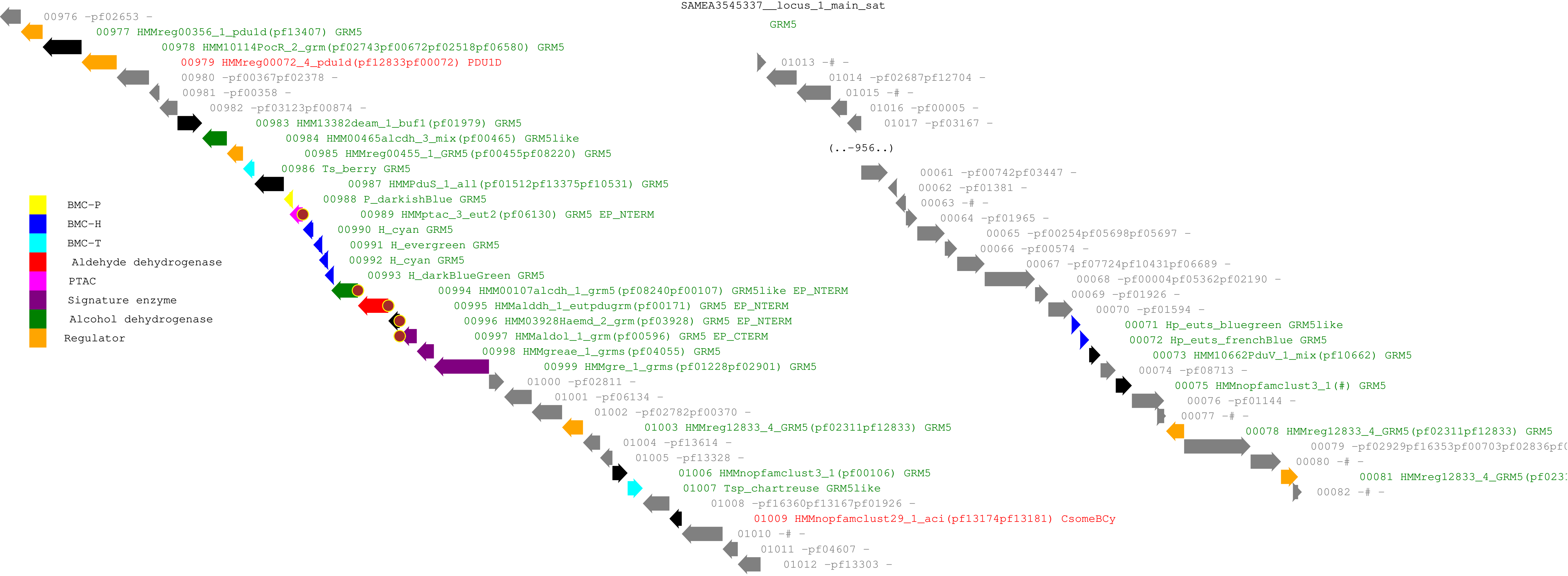

uncultured Ruminococcus sp. Bacteria; Firmicutes; Clostridia; Clostridiales; Ruminococcaceae;

SAMEA3545352__locus_1_main_sat

GRM5

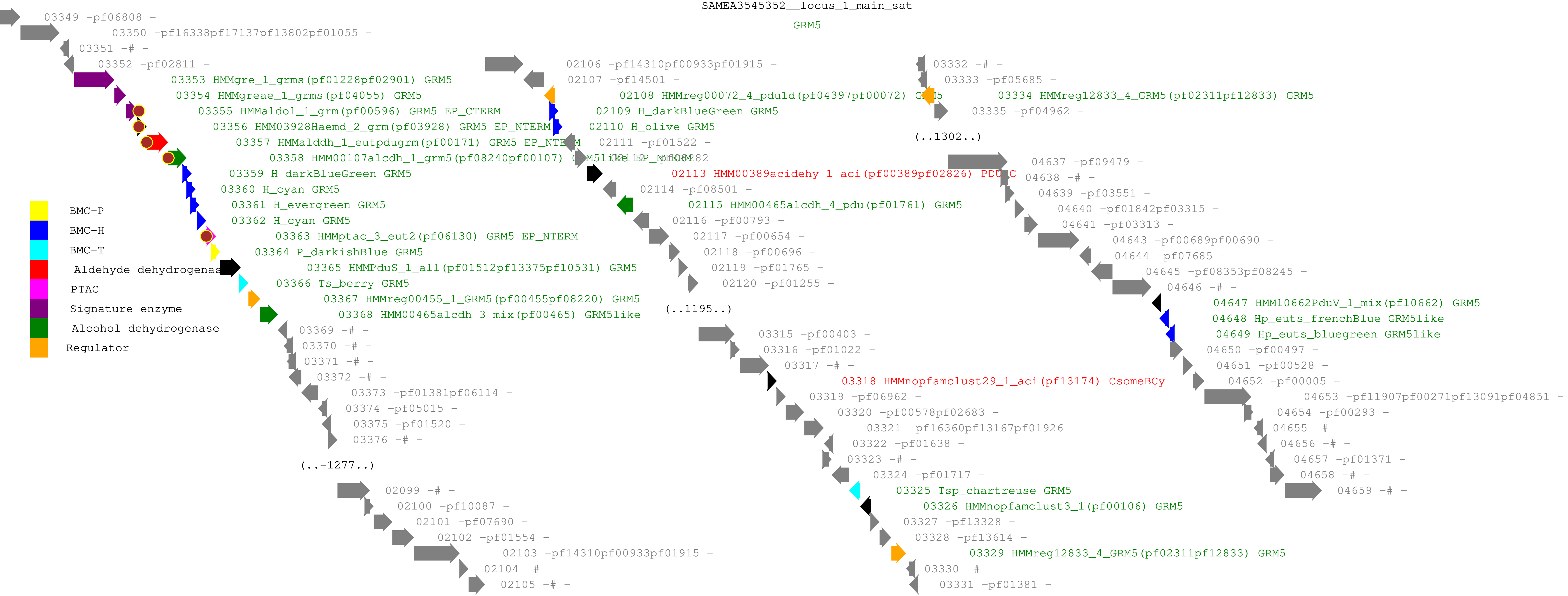

Anaerocolumna jejuensis DSM 15929. Bacteria; Firmicutes; Clostridia; Clostridiales; Lachnospiraceae;

SAMN02745136__locus_1

GRM5

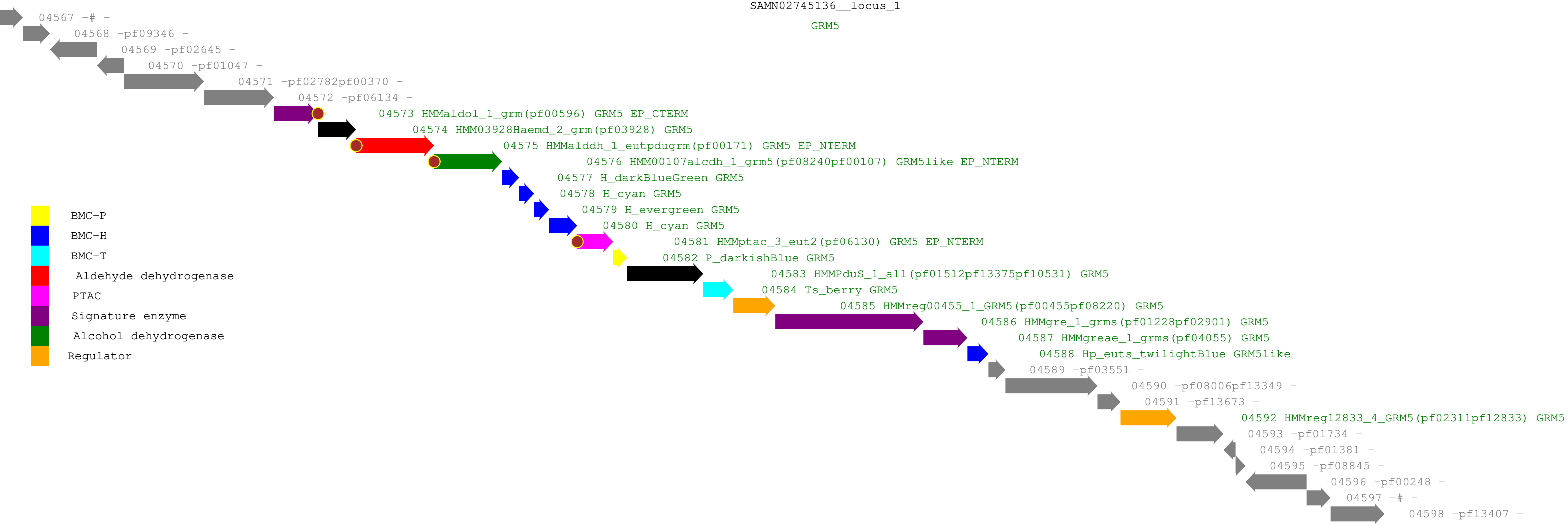

Blautia sp. SF-50. Bacteria; Firmicutes; Clostridia; Clostridiales; Lachnospiraceae;

SAMN02910433__locus_1_main_sat

GRM5

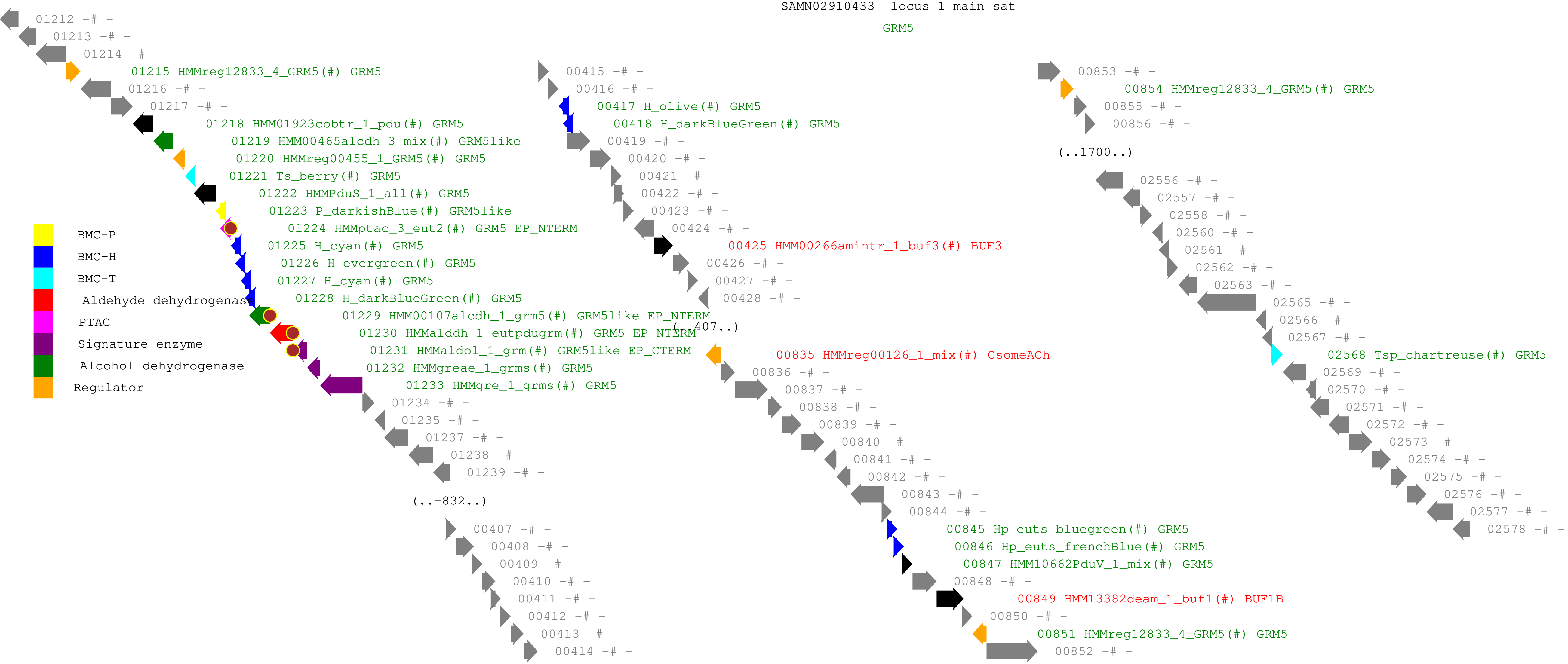

Lachnospiraceae bacterium NLAE-zl-G231. Bacteria; Firmicutes; Clostridia; Clostridiales; Lachnospiraceae.
SAMN05216405__locus_1
GRM5

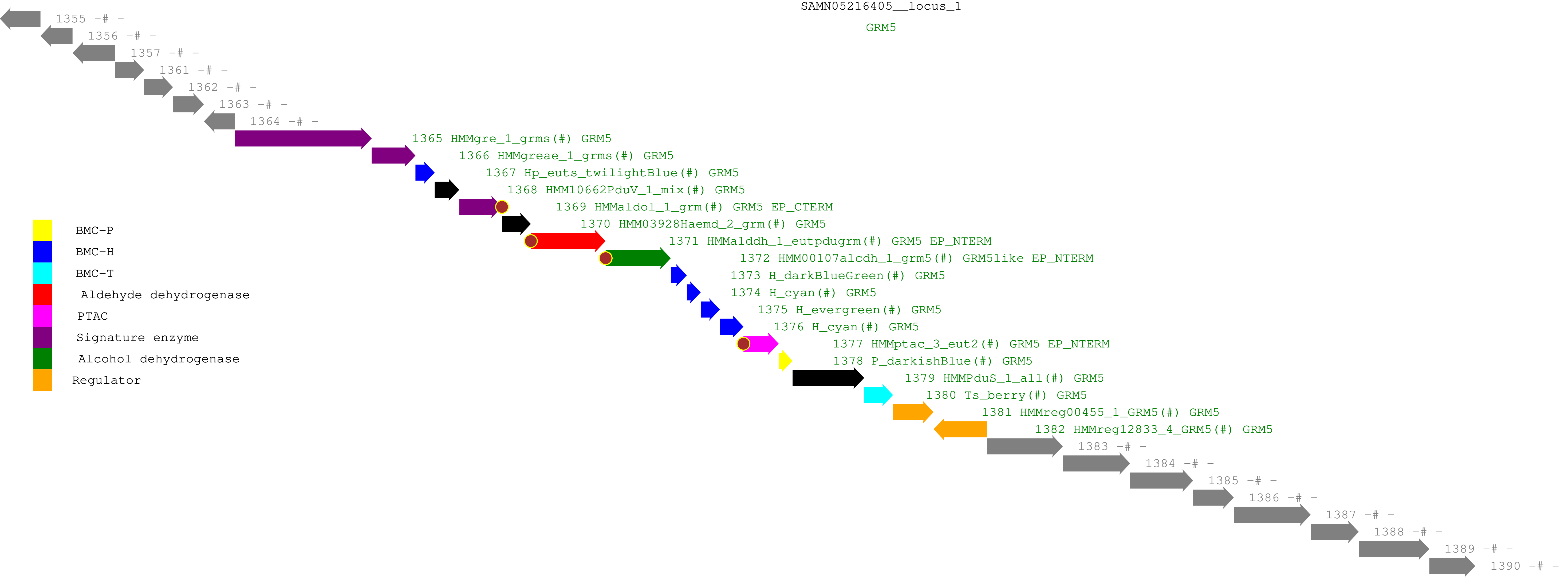

Faecalicatena contorta. Bacteria; Firmicutes; Clostridia; Clostridiales; Lachnospiraceae;

SAMN05216529__locus_1_main_sat

GRM5

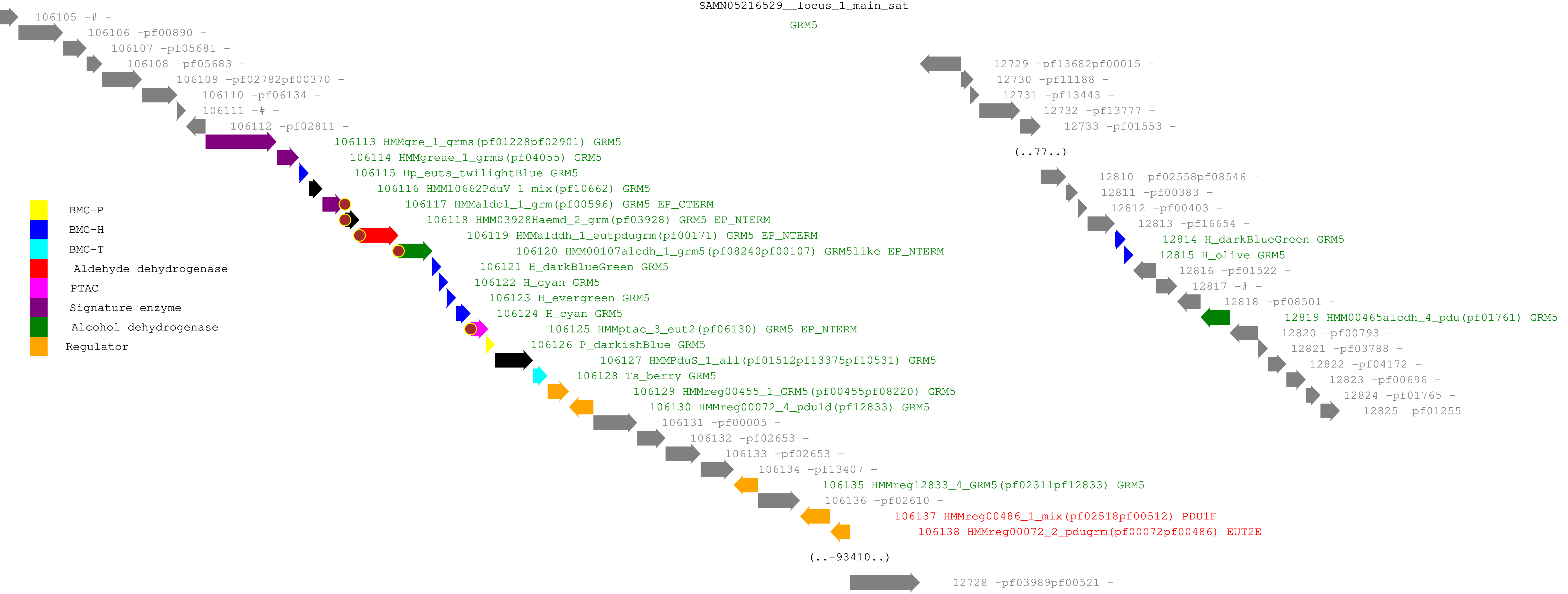

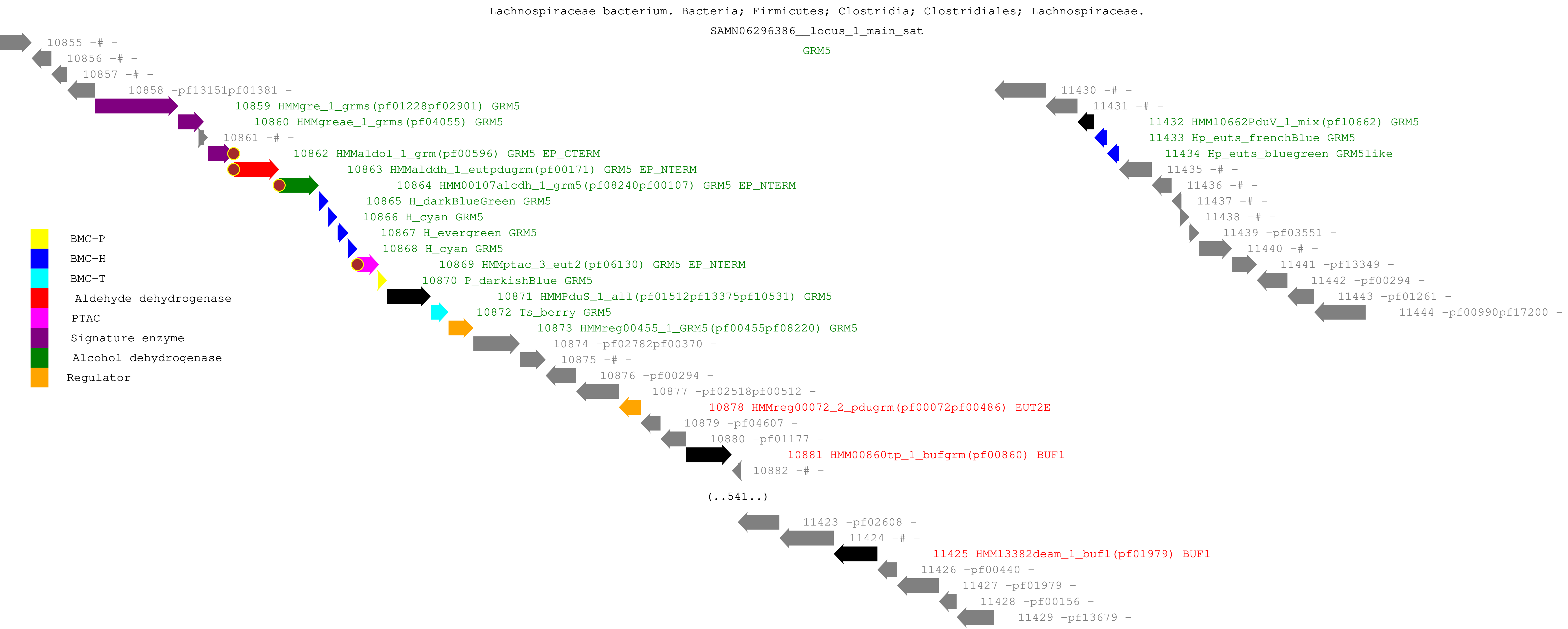
Lachnospiraceae bacterium. Bacteria; Firmicutes; Clostridia; Clostridiales; Lachnospiraceae.
SAMN06296386__locus_1_main_sat
GRM5
10855 -# -
10856 -# -
10857 -# -
10858 -pf13151pf01381 -
10859 HMMgre_1_grms(pf01228pf02901) GRM5
10860 HMMgreae_1_grms(pf04055) GRM5
10861 -# -
10862 HMMaldol_1_grm(pf00596) GRM5 EP_CTERM
10863 HMMalddh_1_eutpdugrm(pf00171) GRM5 EP_NTERM
10864 HMM00107alcdh_1_grm5(pf08240pf00107) GRM5 EP_NTERM
10865 H_darkBlueGreen GRM5
10866 H_cyan GRM5
10867 H_evergreen GRM5
10868 H_cyan GRM5
10869 HMMptac_3_eut2(pf06130) GRM5 EP_NTERM
10870 P_darkishBlue GRM5
10871 HMMPduS_1_all(pf01512pf13375pf10531) GRM5
10872 Ts_berry GRM5
10873 HMMreg00455_1_GRM5(pf00455pf08220) GRM5
10874 -pf02782pf00370 -
10875 -# -
10876 -pf00294 -
10877 -pf02518pf00512 -
10878 HMMreg00072_2_pdugrm(pf00072pf00486) EUT2E
10879 -pf04607 -
10880 -pf01177 -
10881 HMM00860tp_1_bufgrm(pf00860) BUF1
10882 -# -
(..541..)
11423 -pf02608 -
11424 -# -
11425 HMM13382deam_1_buf1(pf01979) BUF1
11426 -pf00440 -
11427 -pf01979 -
11428 -pf00156 -
11429 -pf13679 -
11430 -# -
11431 -# -
11432 HMM10662PduV_1_mix(pf10662) GRM5
11433 Hp_euts_frenchBlue GRM5
11434 Hp_euts_bluegreen GRM5like
11435 -# -
11436 -# -
11437 -# -
11438 -# -
11439 -pf03551 -
11440 -# -
11441 -pf13349 -
11442 -pf00294 -
11443 -pf01261 -
11444 -pf00990pf17200 -
BMC-P
BMC-H
BMC-T
Aldehyde dehydrogenase
PTAC
Signature enzyme
Alcohol dehydrogenase
Regulator

Lachnoclostridium sp. An169. Bacteria; Firmicutes; Clostridia; Clostridiales; Lachnospiraceae.

B5F07__locus_1_main_sat

GRM5like

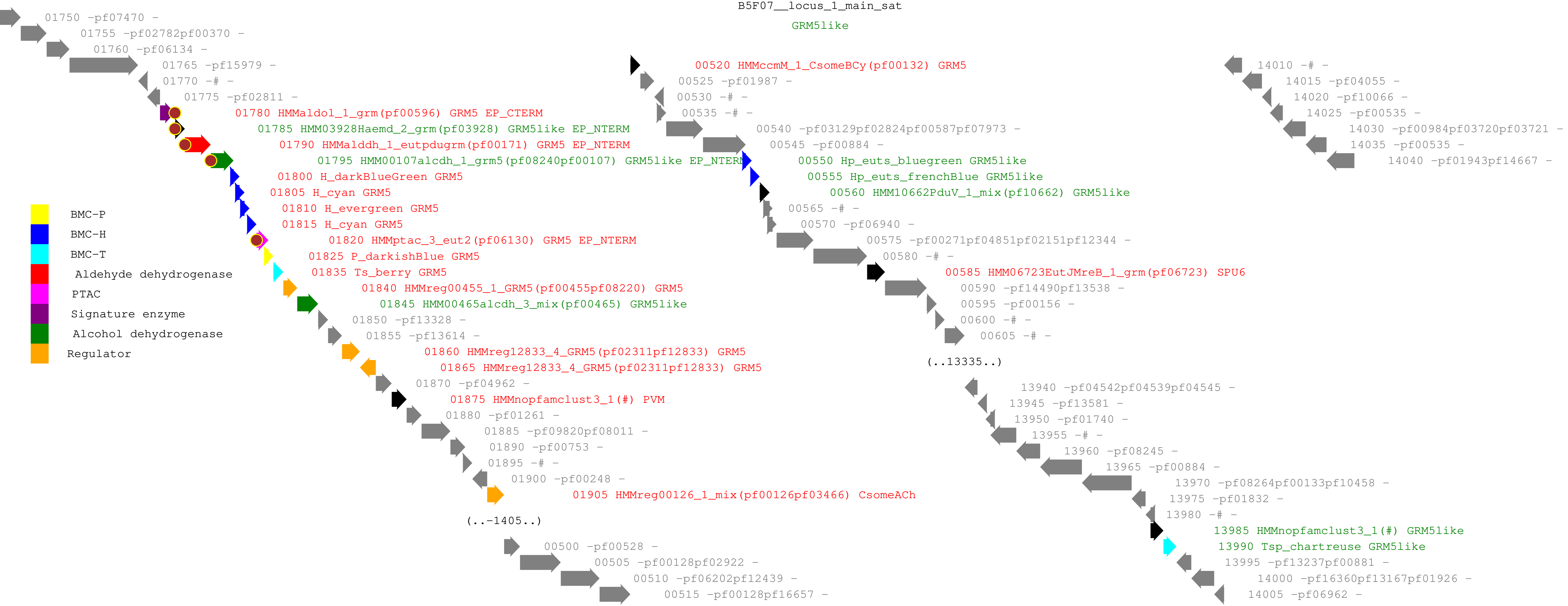

Lachnoclostridium sp. An298. Bacteria; Firmicutes; Clostridia; Clostridiales; Lachnospiraceae.

B5F86__locus_1_main_sat

GRM5like

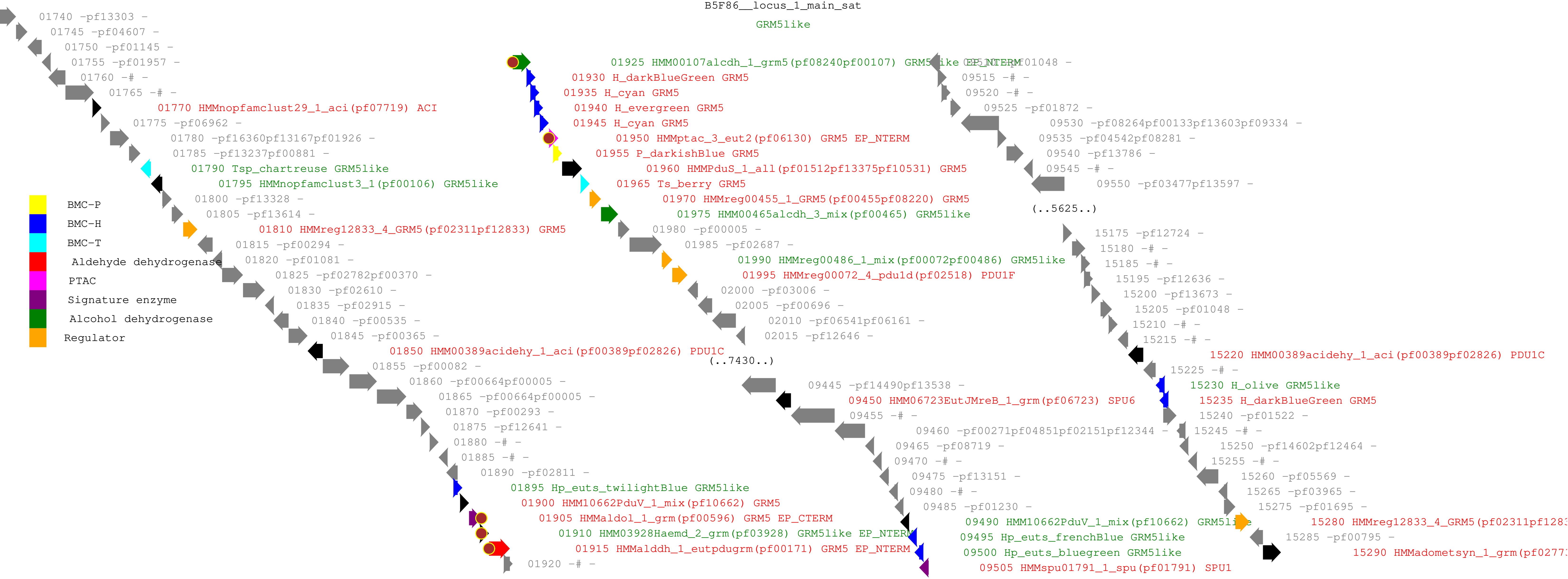

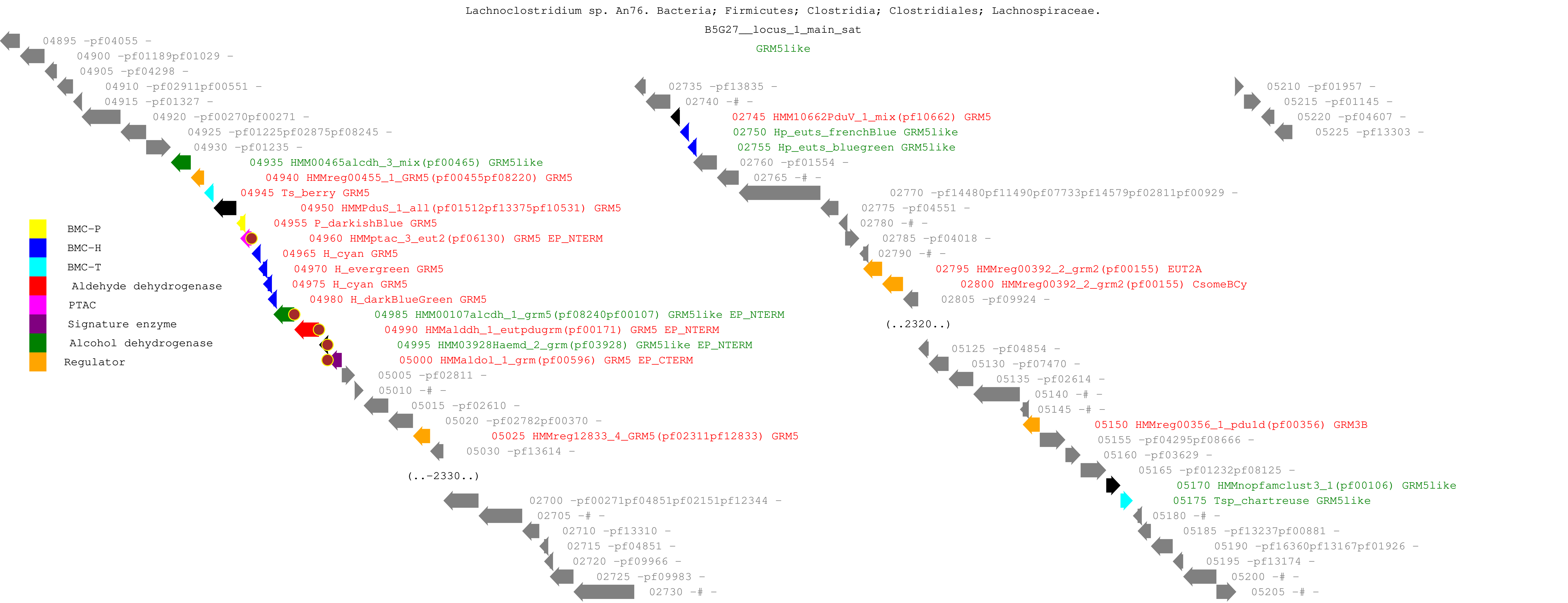
Lachnoclostridium sp. An76. Bacteria; Firmicutes; Clostridia; Clostridiales; Lachnospiraceae.
B5G27__locus_1_main_sat
GRM5like
04895 -pf04055 -
04900 -pf01189pf01029 -
04905 -pf04298 -
04910 -pf02911pf00551 -
04915 -pf01327 -
04920 -pf00270pf00271 -
04925 -pf01225pf02875pf08245 -
04930 -pf01235 -
04935 HMM00465alcdh_3_mix(pf00465) GRM5like
04940 HMMreg00455_1_GRM5(pf00455pf08220) GRM5
04945 Ts_berry GRM5
04950 HMMPduS_1_all(pf01512pf13375pf10531) GRM5
04955 P_darkishBlue GRM5
04960 HMMptac_3_eut2(pf06130) GRM5 EP_NTERM
04965 H_cyan GRM5
04970 H_evergreen GRM5
04975 H_cyan GRM5
04980 H_darkBlueGreen GRM5
04985 HMM00107alcdh_1_grm5(pf08240pf00107) GRM5like EP_NTERM
04990 HMMaldd h_1_eutpdugrm(pf00171) GRM5 EP_NTERM
04995 HMM03928Haemd_2_grm(pf03928) GRM5like EP_NTERM
05000 HMMaldol_1_grm(pf00596) GRM5 EP_CTERM
05005 -pf02811 -
05010 -# -
05015 -pf02610 -
05020 -pf02782pf00370 -
05025 HMMreg12833_4_GRM5(pf02311pf12833) GRM5
05030 -pf13614 -
(..-2330..)
02700 -pf00271pf04851pf02151pf12344 -
02705 -# -
02710 -pf13310 -
02715 -pf04851 -
02720 -pf09966 -
02725 -pf09983 -
02730 -# -
BMC-P
BMC-H
BMC-T
Aldehyde dehydrogenase
PTAC
Signature enzyme
Alcohol dehydrogenase
Regulator
02735 -pf13835 -
02740 -# -
02745 HMM10662PduV_1_mix(pf10662) GRM5
02750 Hp_euts_frenchBlue GRM5like
02755 Hp_euts_bluegreen GRM5like
02760 -pf01554 -
02765 -# -
02770 -pf14480pf11490pf07733pf14579pf02811pf00929 -
02775 -pf04551 -
02780 -# -
02785 -pf04018 -
02790 -# -
02795 HMMreg00392_2_grm2(pf00155) EUT2A
02800 HMMreg00392_2_grm2(pf00155) CsomeBCy
02805 -pf09924 -
(..2320..)
05125 -pf04854 -
05130 -pf07470 -
05135 -pf02614 -
05140 -# -
05145 -# -
05150 HMMreg00356_1_pdu1d(pf00356) GRM3B
05155 -pf04295pf08666 -
05160 -pf03629 -
05165 -pf01232pf08125 -
05170 HMMnopfamclust3_1(pf00106) GRM5like
05175 Tsp_chartreuse GRM5like
05180 -# -
05185 -pf13237pf00881 -
05190 -pf16360pf13167pf01926 -
05195 -pf13174 -
05200 -# -
05205 -# -
05210 -pf01957 -
05215 -pf01145 -
05220 -pf04607 -
05225 -pf13303 -

Ruminococcus sp. CAG:9-related_41_34. Bacteria; Firmicutes; Clostridia; Clostridiales; Ruminococcaceae;

BHW51__locus_1_main_sat

GRM5like

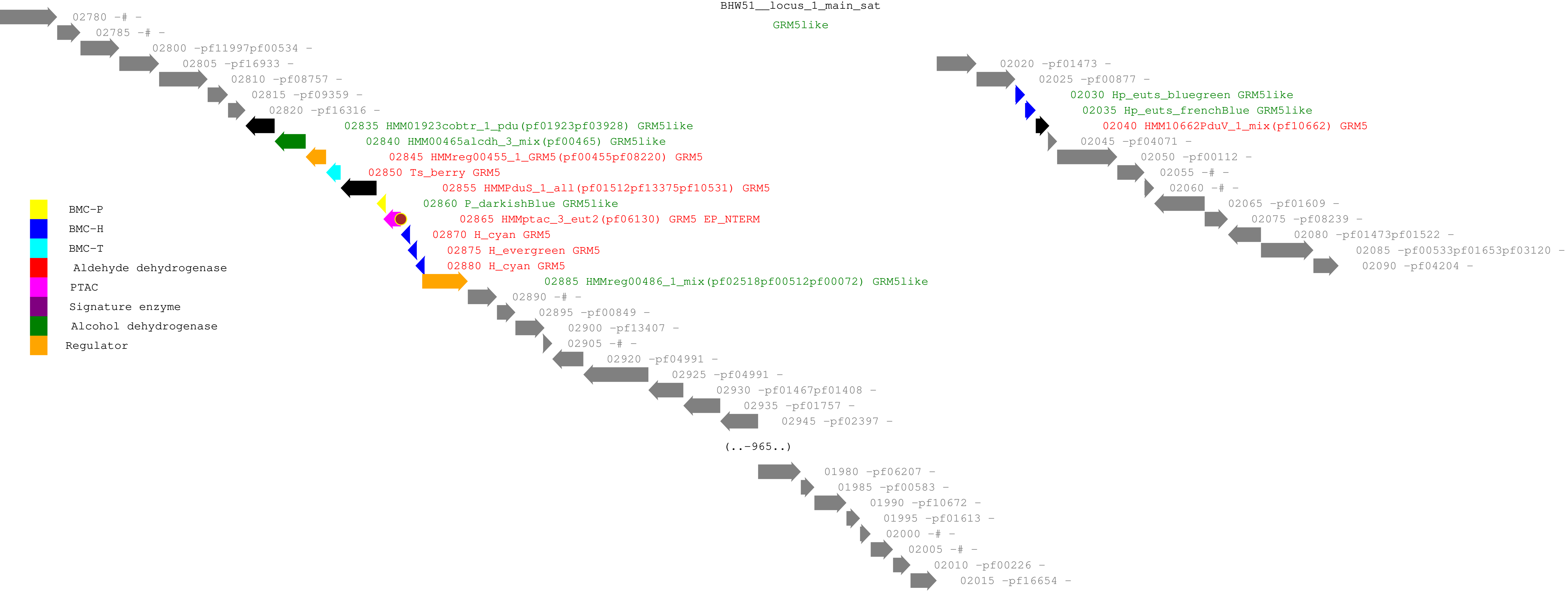

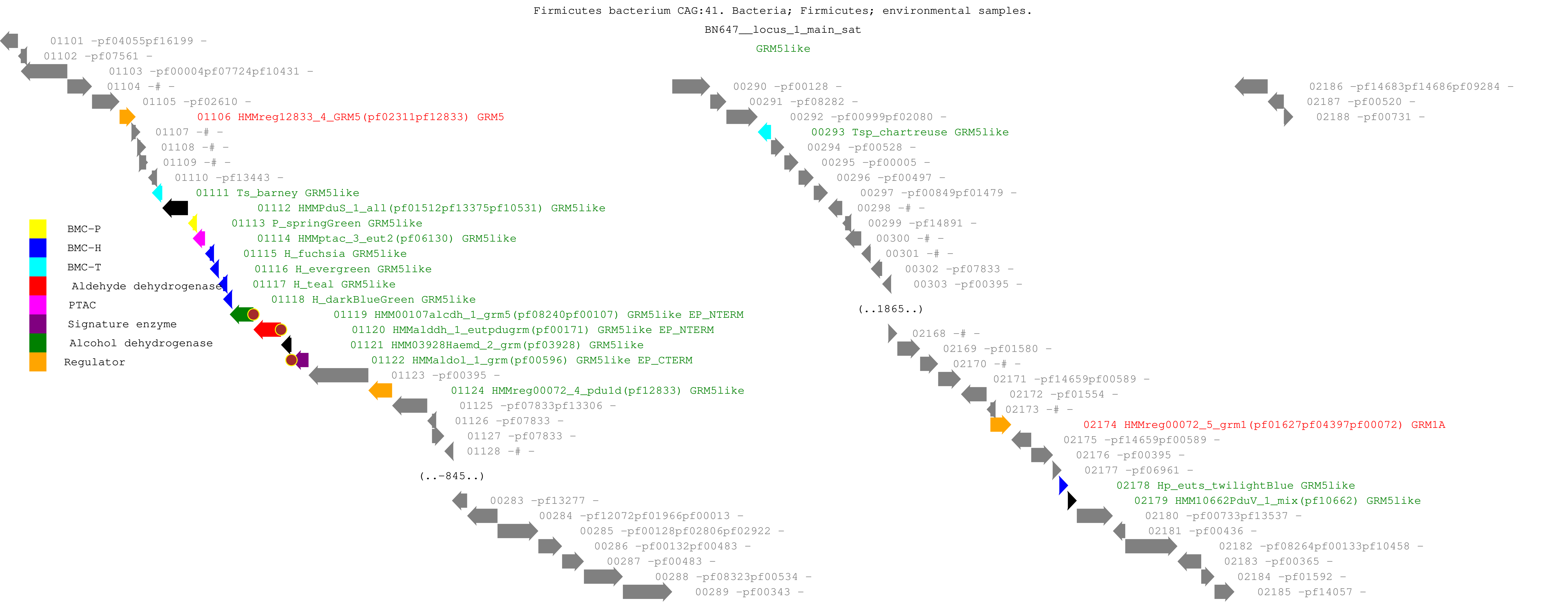

Firmicutes bacterium CAG:41. Bacteria; Firmicutes; environmental samples.
BN647__locus_1_main_sat
GRM5like
01101 -pf04055pf16199 -
01102 -pf07561 -
01103 -pf00004pf07724pf10431 -
01104 -# -
01105 -pf02610 -
01106 HMMreg12833_4_GRM5(pf02311pf12833) GRM5
01107 -# -
01108 -# -
01109 -# -
01110 -pf13443 -
01111 Ts_barney GRM5like
01112 HMMPduS_1_all(pf01512pf13375pf10531) GRM5like
01113 P_springGreen GRM5like
01114 HMMptac_3_eut2(pf06130) GRM5like
01115 H_fuchsia GRM5like
01116 H_evergreen GRM5like
01117 H_teal GRM5like
01118 H_darkBlueGreen GRM5like
01119 HMM00107alcdh_1_grm5(pf08240pf00107) GRM5like EP_NTERM
01120 HMMaldh_1_eutpdugrm(pf00171) GRM5like EP_NTERM
01121 HMM03928Haemd_2_grm(pf03928) GRM5like
01122 HMMaldol_1_grm(pf00596) GRM5like EP_CTERM
01123 -pf00395 -
01124 HMMreg00072_4_pdu1d(pf12833) GRM5like
01125 -pf07833pf13306 -
01126 -pf07833 -
01127 -pf07833 -
01128 -# -
(..-845..)
00283 -pf13277 -
00284 -pf12072pf01966pf00013 -
00285 -pf00128pf02806pf02922 -
00286 -pf00132pf00483 -
00287 -pf00483 -
00288 -pf08323pf00534 -
00289 -pf00343 -
BMC-P
BMC-H
BMC-T
Aldehyde dehydrogenase
PTAC
Signature enzyme
Alcohol dehydrogenase
Regulator
00290 -pf00128 -
00291 -pf08282 -
00292 -pf00999pf02080 -
00293 Tsp_chartreuse GRM5like
00294 -pf00528 -
00295 -pf00005 -
00296 -pf00497 -
00297 -pf00849pf01479 -
00298 -# -
00299 -pf14891 -
00300 -# -
00301 -# -
00302 -pf07833 -
00303 -pf00395 -
(..1865..)
02168 -# -
02169 -pf01580 -
02170 -# -
02171 -pf14659pf00589 -
02172 -pf01554 -
02173 -# -
02174 HMMreg00072_5_grm1(pf01627pf04397pf00072) GRM1A
02175 -pf14659pf00589 -
02176 -pf00395 -
02177 -pf06961 -
02178 Hp_euts_twilightBlue GRM5like
02179 HMM10662PduV_1_mix(pf10662) GRM5like
02180 -pf00733pf13537 -
02181 -pf00436 -
02182 -pf08264pf00133pf10458 -
02183 -pf00365 -
02184 -pf01592 -
02185 -pf14057 -
02186 -pf14683pf14686pf09284 -
02187 -pf00520 -
02188 -pf00731 -

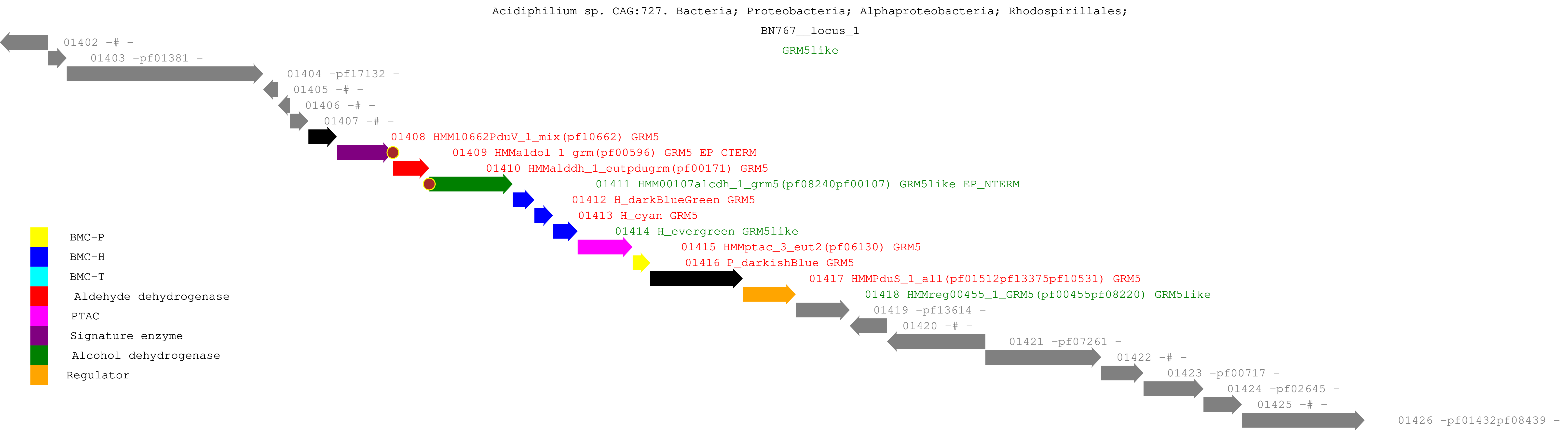

Acidiphilium sp. CAG:727. Bacteria; Proteobacteria; Alphaproteobacteria; Rhodospirillales;
BN767__locus_1
GRM5like
01402 -# -
01403 -pf01381 -
01404 -pf17132 -
01405 -# -
01406 -# -
01407 -# -
01408 HMM10662PduV_1_mix(pf10662) GRM5
01409 HMMaldol_1_grm(pf00596) GRM5 EP_CTERM
01410 HMMalddh_1_eutpdugrm(pf00171) GRM5
01411 HMM00107alcdh_1_grm5(pf08240pf00107) GRM5like EP_NTERM
01412 H_darkBlueGreen GRM5
01413 H_cyan GRM5
01414 H_evergreen GRM5like
01415 HMMptac_3_eut2(pf06130) GRM5
01416 P_darkishBlue GRM5
01417 HMMPduS_1_all(pf01512pf13375pf10531) GRM5
01418 HMMreg00455_1_GRM5(pf00455pf08220) GRM5like
01419 -pf13614 -
01420 -# -
01421 -pf07261 -
01422 -# -
01423 -pf00717 -
01424 -pf02645 -
01425 -# -
01426 -pf01432pf08439 -
BMC-P
BMC-H
BMC-T
Aldehyde dehydrogenase
PTAC
Signature enzyme
Alcohol dehydrogenase
Regulator

Ruminococcus sp. CAG:9. Bacteria; Firmicutes; Clostridia; Clostridiales; Ruminococcaceae;

BN806__locus_1_main_sat

GRM5like

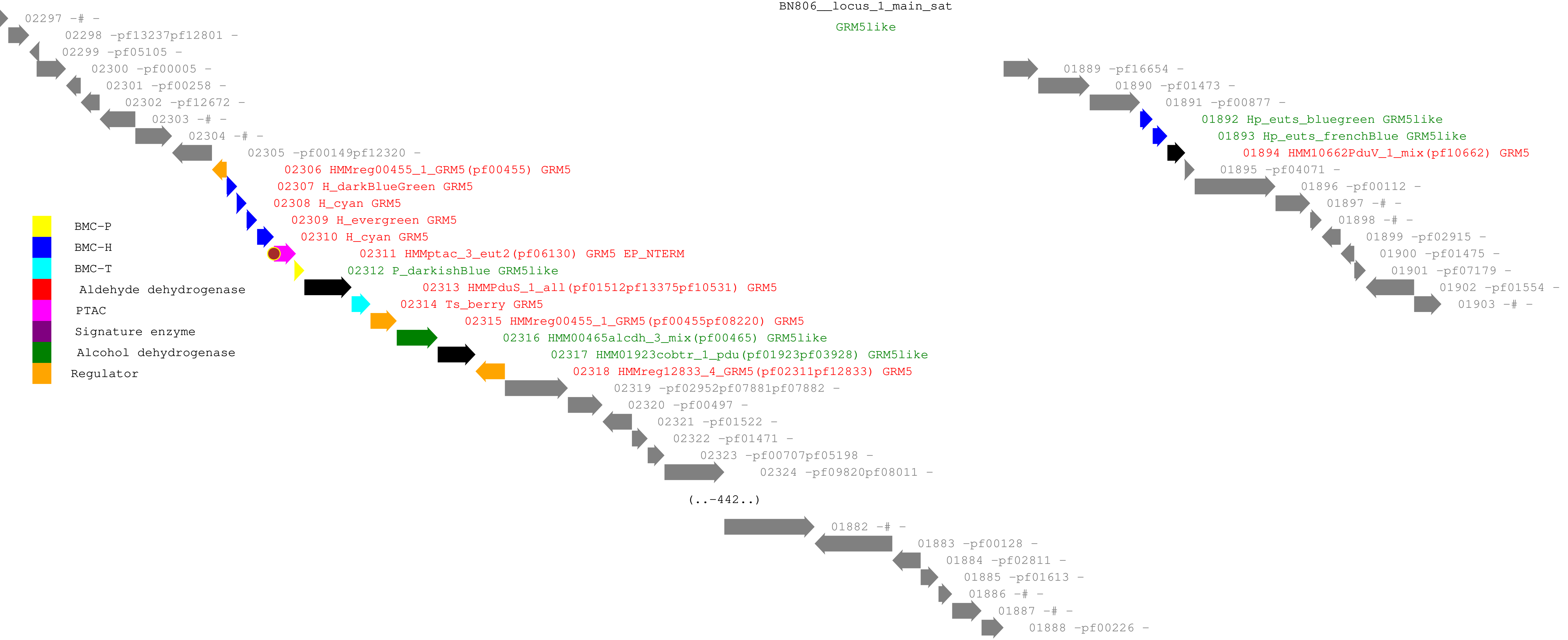

Murimonas intestini. Bacteria; Firmicutes; Clostridia; Clostridiales; Lachnospiraceae;

C7383__locus_1_main_sat

GRM5like

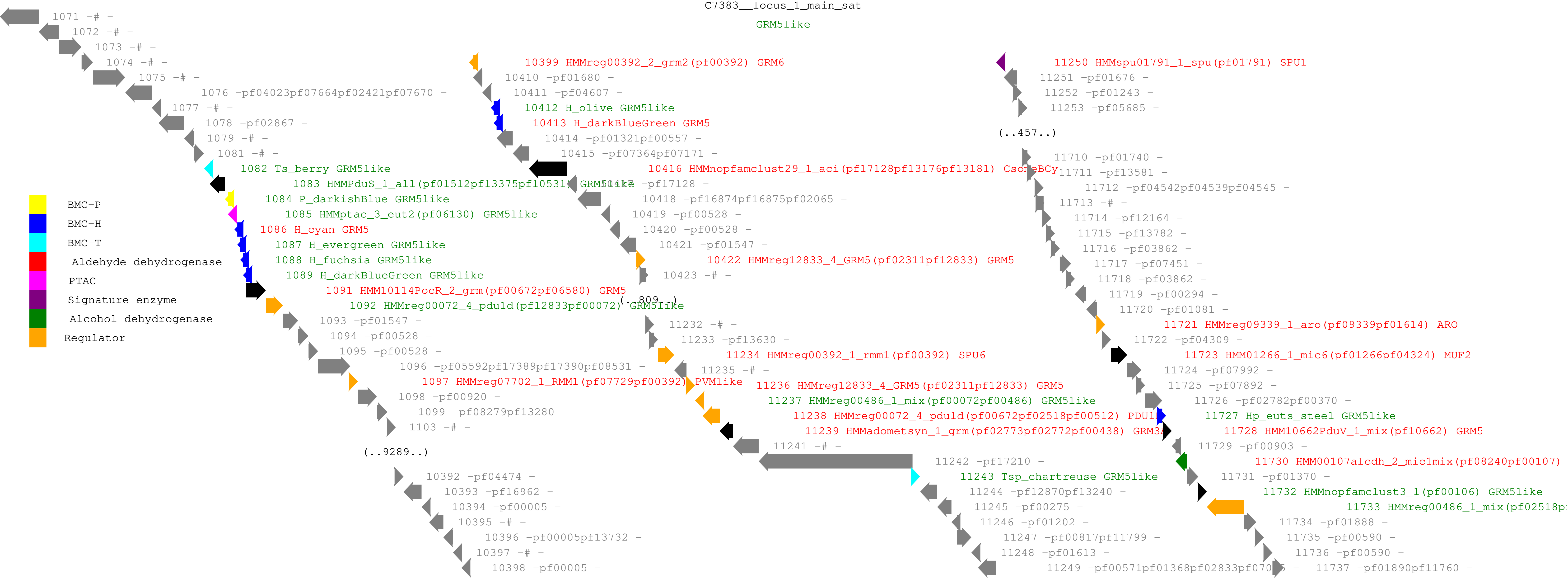

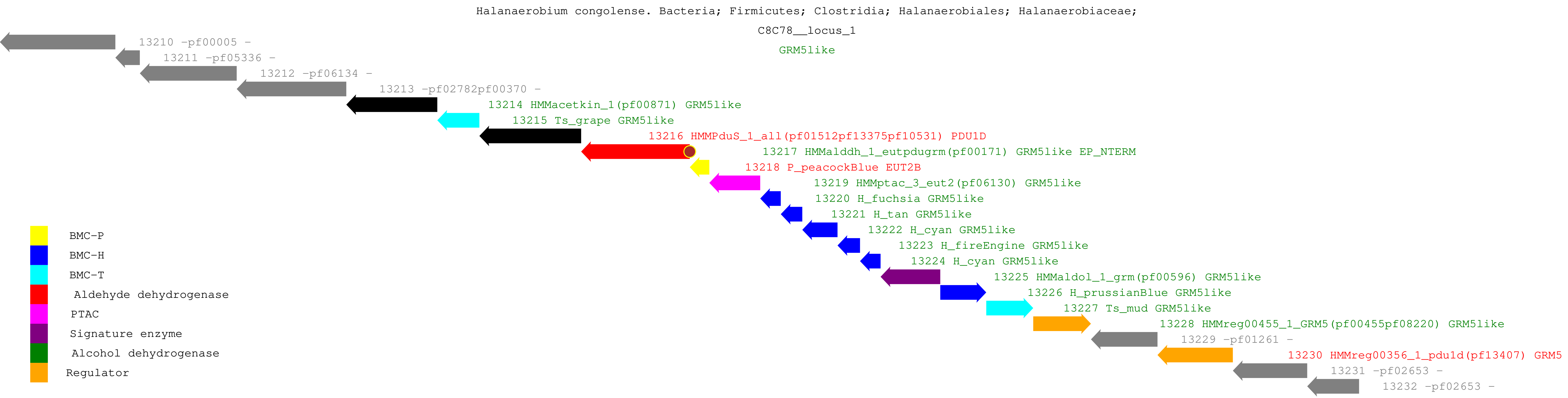

Halanaerobium congolense. Bacteria; Firmicutes; Clostridia; Halanaerobiales; Halanaerobiaceae;
C8C78__locus_1
GRM5like
13210 -pf00005 -
13211 -pf05336 -
13212 -pf06134 -
13213 -pf02782pf00370 -
13214 HMMacetkin_1(pf00871) GRM5like
13215 Ts_grape GRM5like
13216 HMMPduS_1_all(pf01512pf13375pf10531) PDU1D
13217 HMMalddh_1_eutpdugrm(pf00171) GRM5like EP_NTERM
13218 P_peacockBlue EUT2B
13219 HMMptac_3_eut2(pf06130) GRM5like
13220 H_fuchsia GRM5like
13221 H_tan GRM5like
13222 H_cyan GRM5like
13223 H_fireEngine GRM5like
13224 H_cyan GRM5like
13225 HMMaldol_1_grm(pf00596) GRM5like
13226 H_prussianBlue GRM5like
13227 Ts_mud GRM5like
13228 HMMreg00455_1_GRM5(pf00455pf08220) GRM5like
13229 -pf01261 -
13230 HMMreg00356_1_pdu1d(pf13407) GRM5
13231 -pf02653 -
13232 -pf02653 -
BMC-P
BMC-H
BMC-T
Aldehyde dehydrogenase
PTAC
Signature enzyme
Alcohol dehydrogenase
Regulator

Lachnotalea glycerini. Bacteria; Firmicutes; Clostridia; Clostridiales; Lachnospiraceae;

C8E03__locus_2

GRM5like

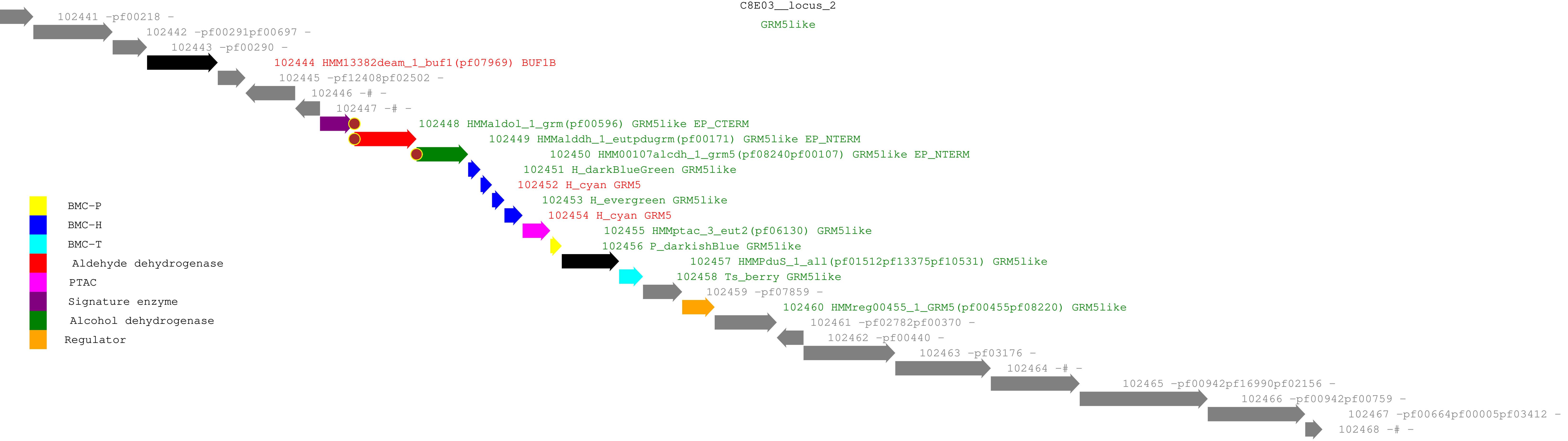

Clostridium chauvoei JF4335. Bacteria; Firmicutes; Clostridia; Clostridiales; Clostridiaceae;

CCH01__locus_1

GRM5like

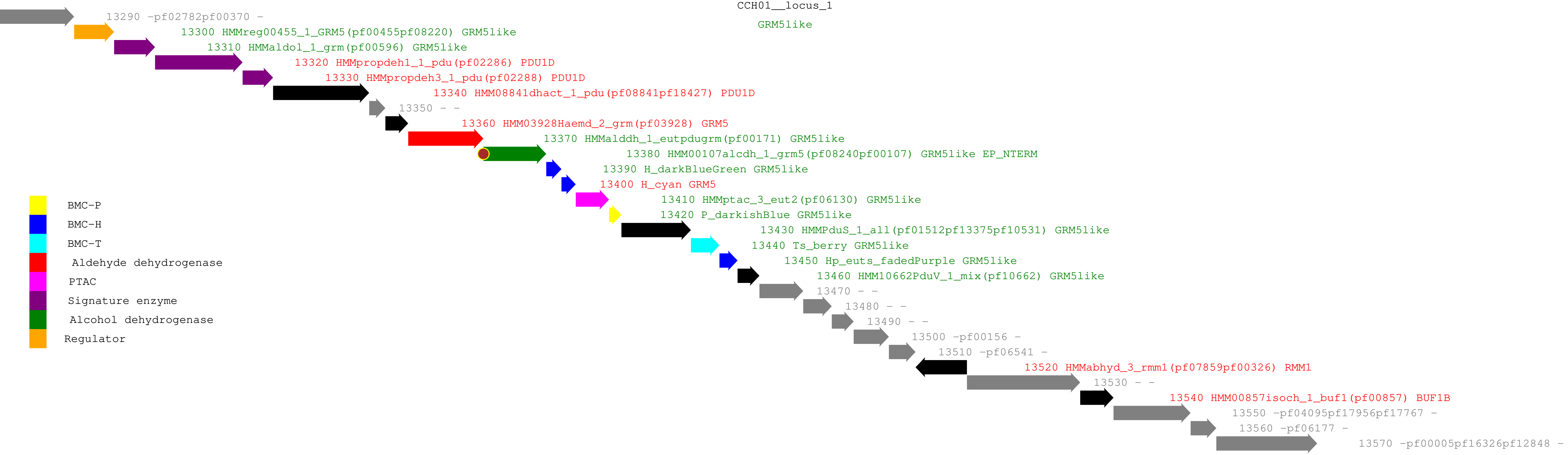

Lachnotalea glycerini. Bacteria; Firmicutes; Clostridia; Clostridiales; Lachnospiraceae;

CG710__locus_1

GRM5like

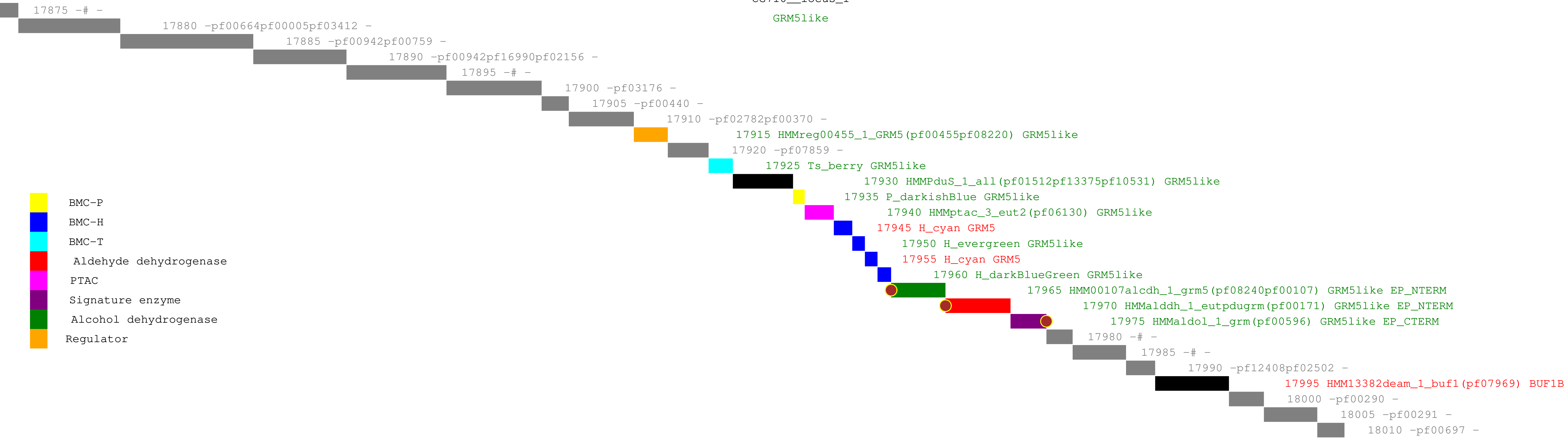

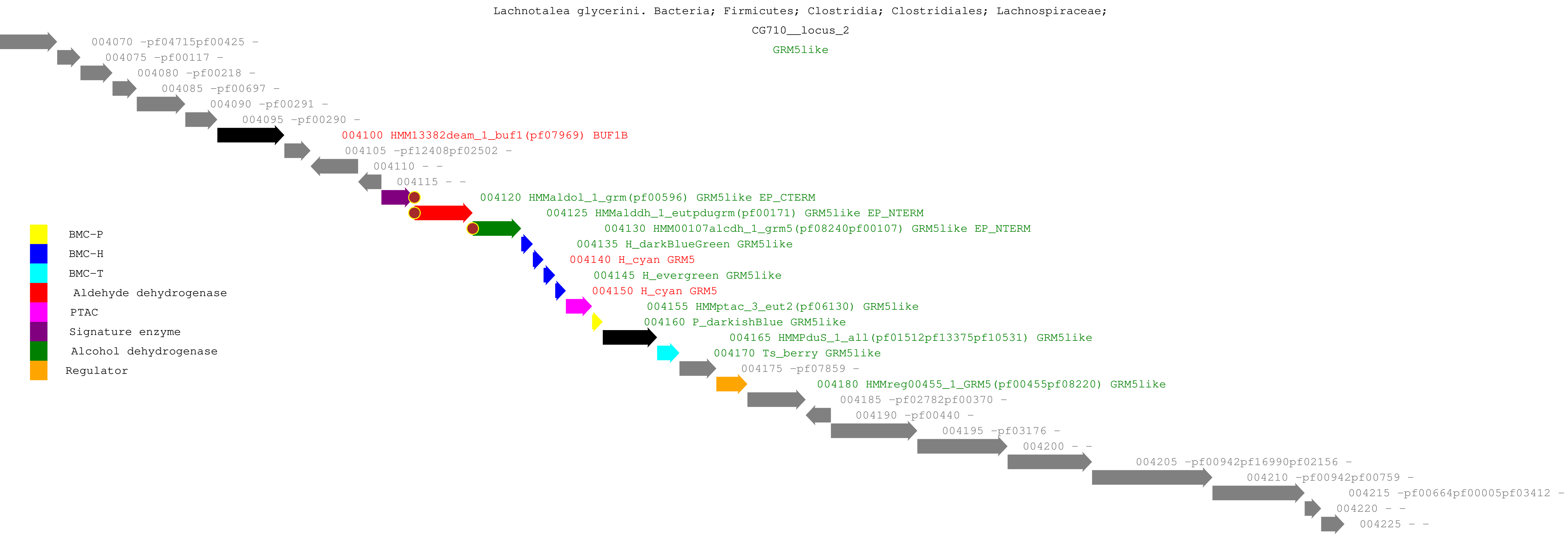
Lachnotalea glycerini. Bacteria; Firmicutes; Clostridia; Clostridiales; Lachnospiraceae;
CG710__locus_2
GRM5like
004070 -pf04715pf00425 -
004075 -pf00117 -
004080 -pf00218 -
004085 -pf00697 -
004090 -pf00291 -
004095 -pf00290 -
004100 HMM13382deam_1_buf1(pf07969) BUF1B
004105 -pf12408pf02502 -
004110 - -
004115 - -
004120 HMMaldol_1_grm(pf00596) GRM5like EP_CTERM
004125 HMMalddh_1_eutpdugrm(pf00171) GRM5like EP_NTERM
004130 HMM00107alcdh_1_grm5(pf08240pf00107) GRM5like EP_NTERM
004135 H_darkBlueGreen GRM5like
004140 H_cyan GRM5
004145 H_evergreen GRM5like
004150 H_cyan GRM5
004155 HMMptac_3_eut2(pf06130) GRM5like
004160 P_darkishBlue GRM5like
004165 HMMPduS_1_all(pf01512pf13375pf10531) GRM5like
004170 Ts_berry GRM5like
004175 -pf07859 -
004180 HMMreg00455_1_GRM5(pf00455pf08220) GRM5like
004185 -pf02782pf00370 -
004190 -pf00440 -
004195 -pf03176 -
004200 - -
004205 -pf00942pf16990pf02156 -
004210 -pf00942pf00759 -
004215 -pf00664pf00005pf03412 -
004220 - -
004225 - -
BMC-P
BMC-H
BMC-T
Aldehyde dehydrogenase
PTAC
Signature enzyme
Alcohol dehydrogenase
Regulator

Lachnospiraceae bacterium. Bacteria; Firmicutes; Clostridia; Clostridiales; Lachnospiraceae.

CS063__locus_1

GRM5like

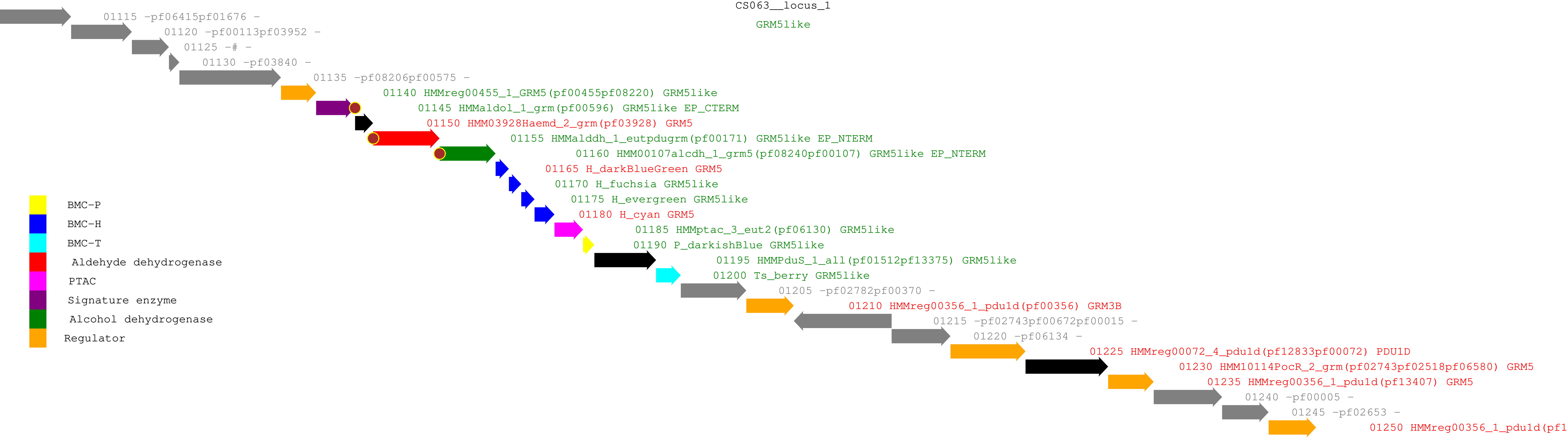

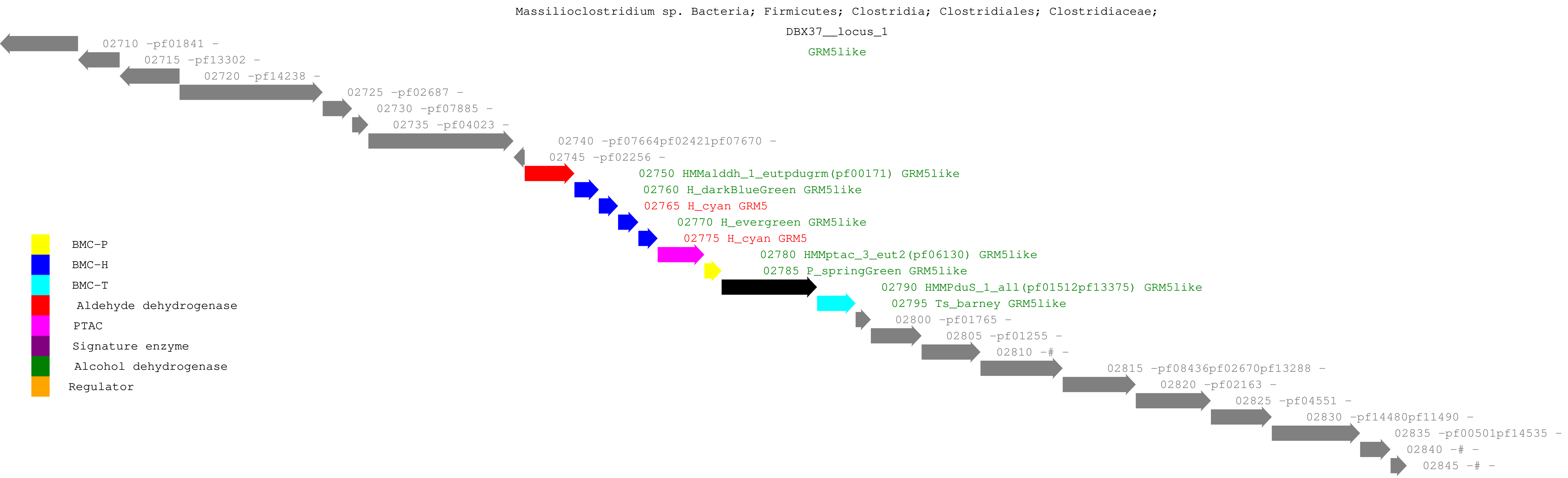
Massilioclostridium sp. Bacteria; Firmicutes; Clostridia; Clostridiales; Clostridiaceae;
DBX37__locus_1
GRM5like
02710 -pf01841 -
02715 -pf13302 -
02720 -pf14238 -
02725 -pf02687 -
02730 -pf07885 -
02735 -pf04023 -
02740 -pf07664pf02421pf07670 -
02745 -pf02256 -
02750 HMMaldh_1_eutpdugrm(pf00171) GRM5like
02760 H_darkBlueGreen GRM5like
02765 H_cyan GRM5
02770 H_evergreen GRM5like
02775 H_cyan GRM5
02780 HMMptac_3_eut2(pf06130) GRM5like
02785 P_springGreen GRM5like
02790 HMMPduS_1_all(pf01512pf13375) GRM5like
02795 Ts_barney GRM5like
02800 -pf01765 -
02805 -pf01255 -
02810 -# -
02815 -pf08436pf02670pf13288 -
02820 -pf02163 -
02825 -pf04551 -
02830 -pf14480pf11490 -
02835 -pf00501pf14535 -
02840 -# -
02845 -# -
BMC-P
BMC-H
BMC-T
Aldehyde dehydrogenase
PTAC
Signature enzyme
Alcohol dehydrogenase
Regulator

Massilioclostridium sp. Bacteria; Firmicutes; Clostridia; Clostridiales; Clostridiaceae;

DBX37__locus_2

GRM5like

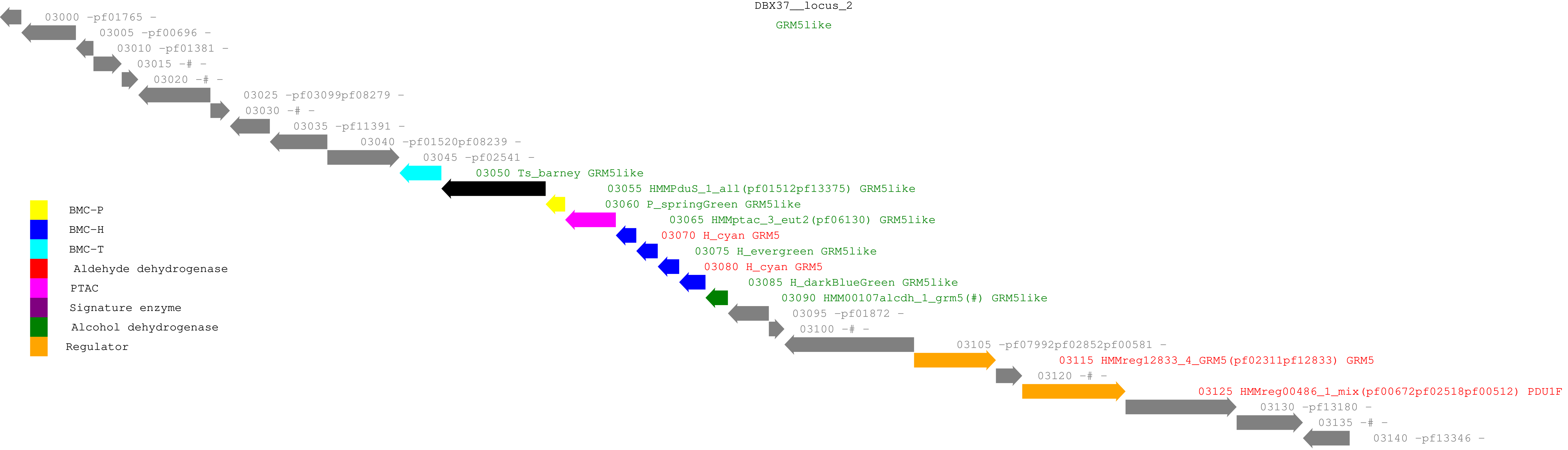

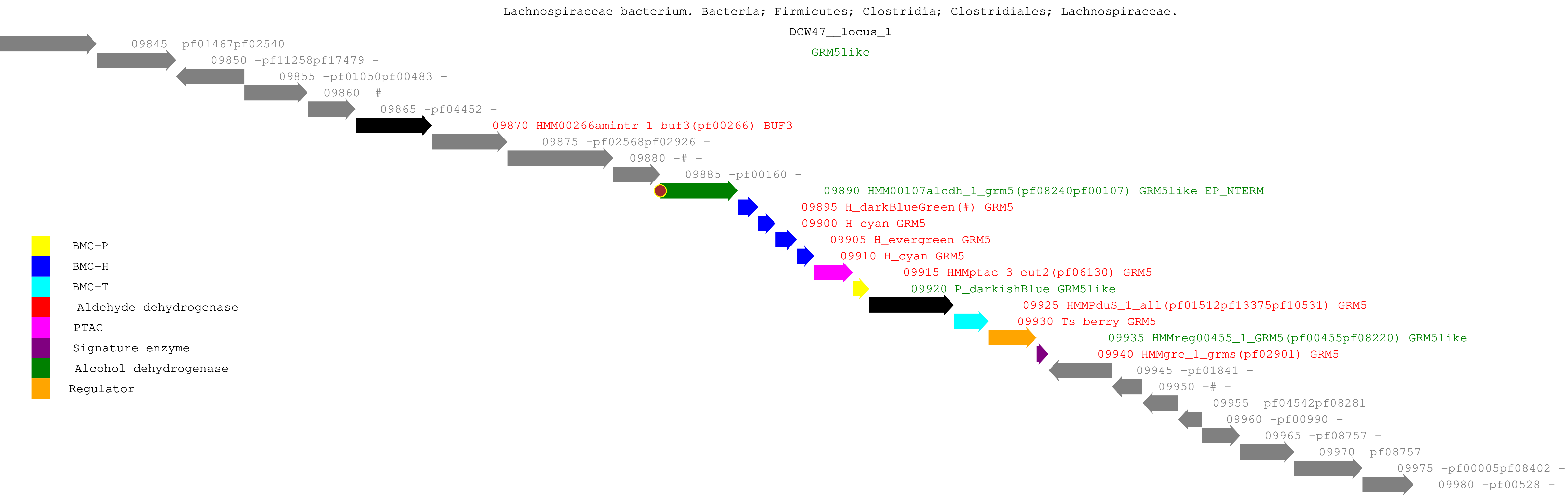
Lachnospiraceae bacterium. Bacteria; Firmicutes; Clostridia; Clostridiales; Lachnospiraceae.
DCW47__locus_1
GRM5like
09845 -pf01467pf02540 -
09850 -pf11258pf17479 -
09855 -pf01050pf00483 -
09860 -# -
09865 -pf04452 -
09870 HMM00266amintr_1_buf3(pf00266) BUF3
09875 -pf02568pf02926 -
09880 -# -
09885 -pf00160 -
09890 HMM00107alcdh_1_grm5(pf08240pf00107) GRM5like EP_NTERM
09895 H_darkBlueGreen(#) GRM5
09900 H_cyan GRM5
09905 H_evergreen GRM5
09910 H_cyan GRM5
09915 HMMptac_3_eut2(pf06130) GRM5
09920 P_darkishBlue GRM5like
09925 HMMPduS_1_all(pf01512pf13375pf10531) GRM5
09930 Ts_berry GRM5
09935 HMMreg00455_1_GRM5(pf00455pf08220) GRM5like
09940 HMMgre_1_grms(pf02901) GRM5
09945 -pf01841 -
09950 -# -
09955 -pf04542pf08281 -
09960 -pf00990 -
09965 -pf08757 -
09970 -pf08757 -
09975 -pf00005pf08402 -
09980 -pf00528 -
BMC-P
BMC-H
BMC-T
Aldehyde dehydrogenase
PTAC
Signature enzyme
Alcohol dehydrogenase
Regulator

Lachnospiraceae bacterium. Bacteria; Firmicutes; Clostridia; Clostridiales; Lachnospiraceae.

DCZ91__locus_1_main_sat

GRM5like

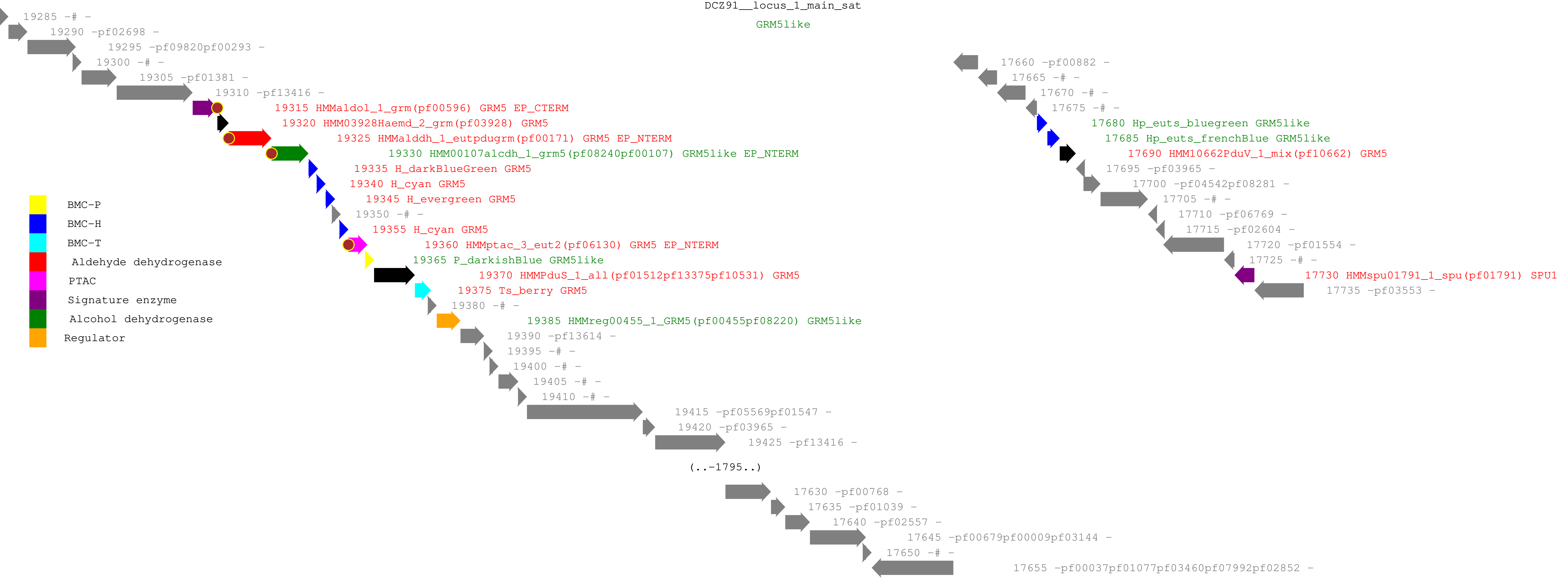

Clostridiales bacterium. Bacteria; Firmicutes; Clostridia; Clostridiales.

DEO51__locus_1_main_sat

GRM5like

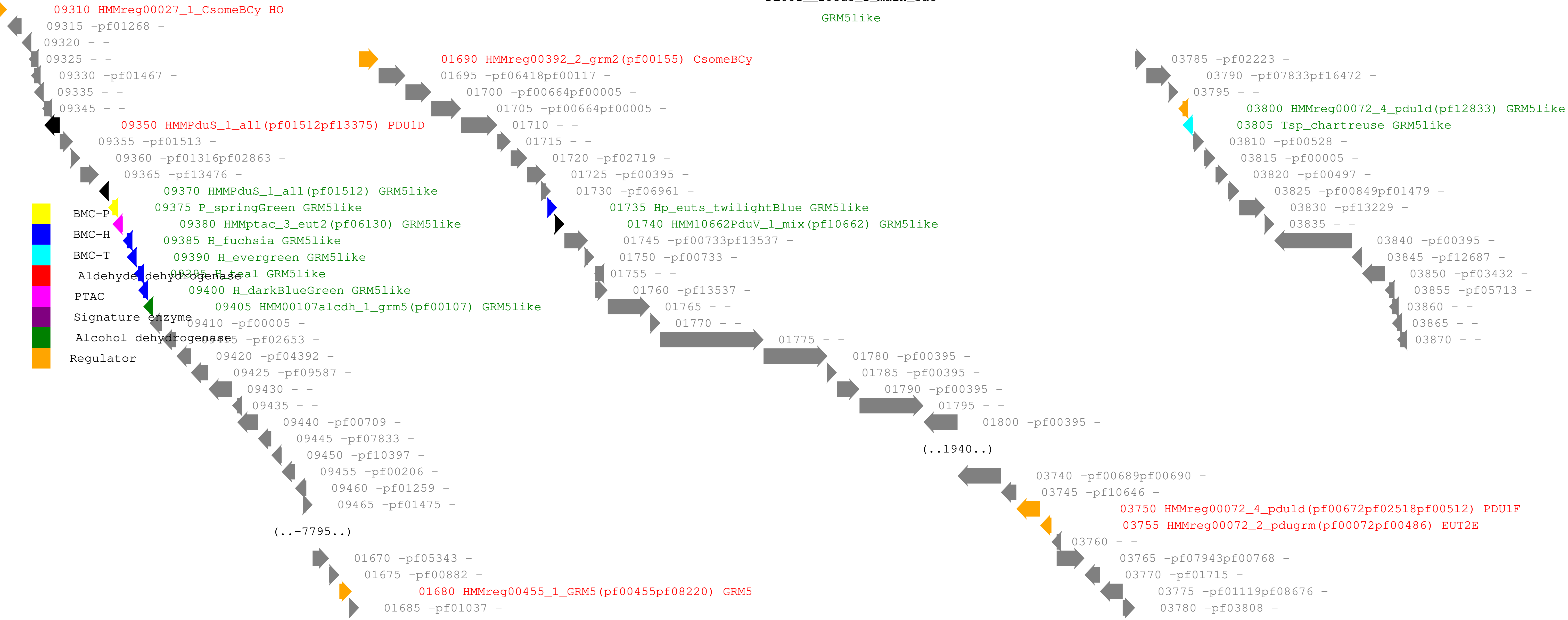

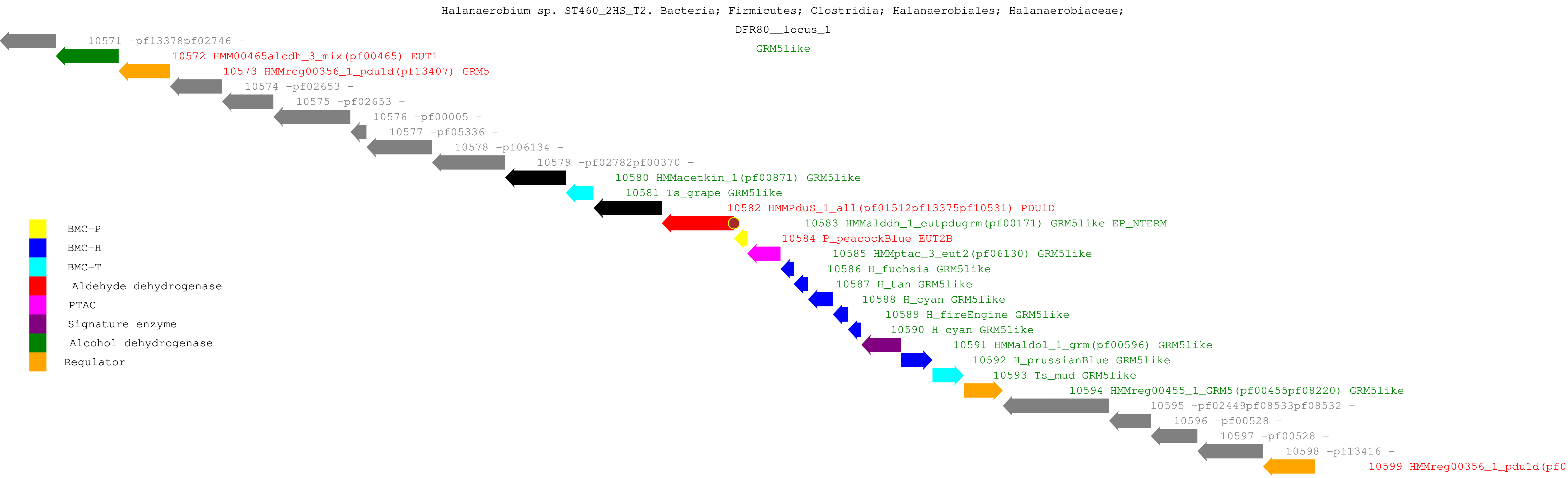
Halanaerobium sp. ST460_2HS_T2. Bacteria; Firmicutes; Clostridia; Halanaerobiales; Halanaerobiaceae;
DFR80__locus_1
GRM5like
10571 -pf13378pf02746 -
10572 HMM00465alcdh_3_mix(pf00465) EUT1
10573 HMMreg00356_1_pdu1d(pf13407) GRM5
10574 -pf02653 -
10575 -pf02653 -
10576 -pf00005 -
10577 -pf05336 -
10578 -pf06134 -
10579 -pf02782pf00370 -
10580 HMMacetkin_1(pf00871) GRM5like
10581 Ts_grape GRM5like
10582 HMMPduS_1_all(pf01512pf13375pf10531) PDU1D
10583 HMMalddh_1_eutpdugrm(pf00171) GRM5like EP_NTERM
10584 P_peacockBlue EUT2B
10585 HMMptac_3_eut2(pf06130) GRM5like
10586 H_fuchsia GRM5like
10587 H_tan GRM5like
10588 H_cyan GRM5like
10589 H_fireEngine GRM5like
10590 H_cyan GRM5like
10591 HMMaldol_1_grm(pf00596) GRM5like
10592 H_prussianBlue GRM5like
10593 Ts_mud GRM5like
10594 HMMreg00455_1_GRM5(pf00455pf08220) GRM5like
10595 -pf02449pf08533pf08532 -
10596 -pf00528 -
10597 -pf00528 -
10598 -pf13416 -
10599 HMMreg00356_1_pdu1d(pf0
BMC-P
BMC-H
BMC-T
Aldehyde dehydrogenase
PTAC
Signature enzyme
Alcohol dehydrogenase
Regulator

Blautia sp. Bacteria; Firmicutes; Clostridia; Clostridiales; Lachnospiraceae; Blautia;

DHW25__locus_1_main_sat

GRM5like

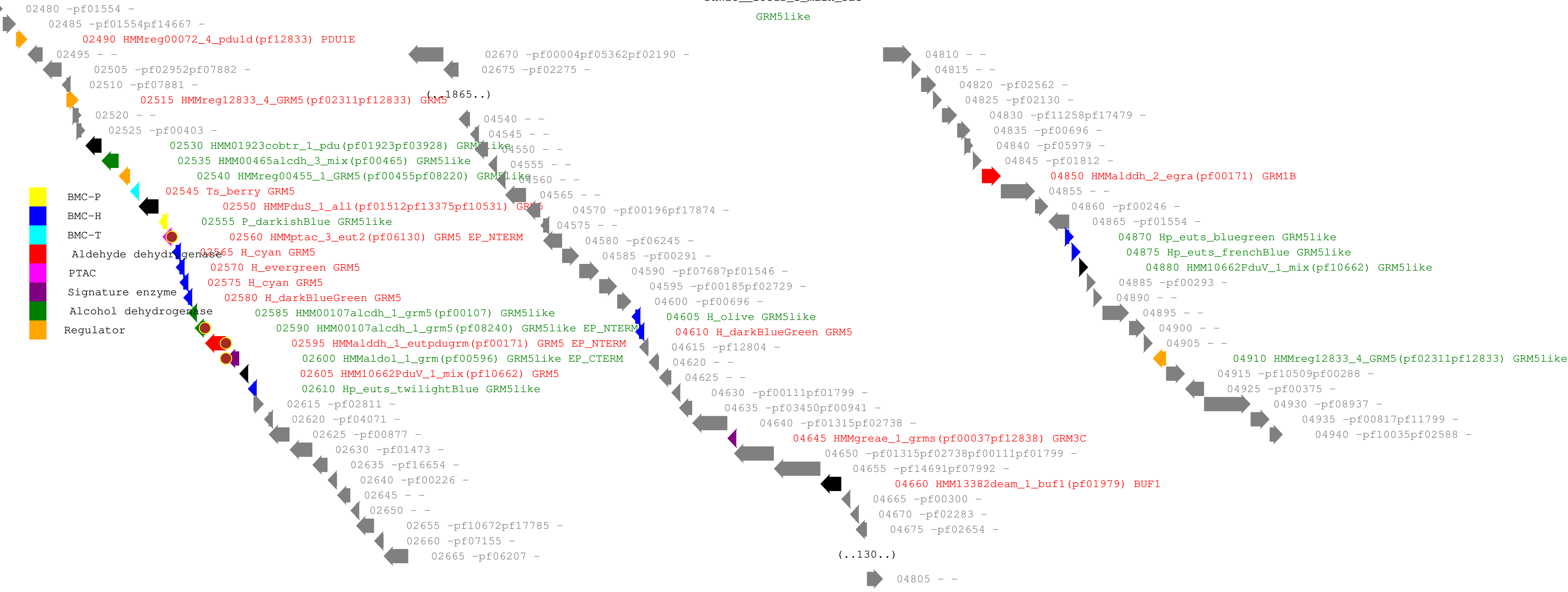

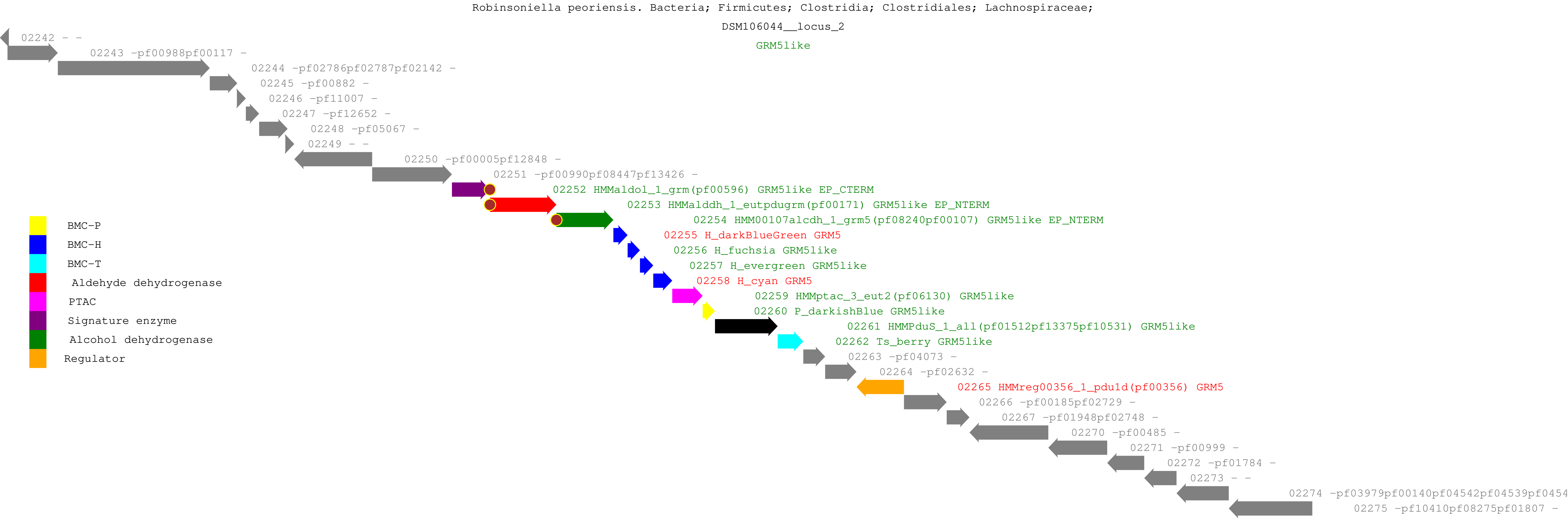

Robinsoniella peoriensis. Bacteria; Firmicutes; Clostridia; Clostridiales; Lachnospiraceae;
DSM106044__locus_2
GRM5like
02242 - -
02243 -pf00988pf00117 -
02244 -pf02786pf02787pf02142 -
02245 -pf00882 -
02246 -pf11007 -
02247 -pf12652 -
02248 -pf05067 -
02249 - -
02250 -pf00005pf12848 -
02251 -pf00990pf08447pf13426 -
02252 HMMaldol_1_grm(pf00596) GRM5like EP_CTERM
02253 HMMalddh_1_eutpdugrm(pf00171) GRM5like EP_NTERM
02254 HMM00107alcdh_1_grm5(pf08240pf00107) GRM5like EP_NTERM
02255 H_darkBlueGreen GRM5
02256 H_fuchsia GRM5like
02257 H_evergreen GRM5like
02258 H_cyan GRM5
02259 HMMptac_3_eut2(pf06130) GRM5like
02260 P_darkishBlue GRM5like
02261 HMMPduS_1_all(pf01512pf13375pf10531) GRM5like
02262 Ts_berry GRM5like
02263 -pf04073 -
02264 -pf02632 -
02265 HMMreg00356_1_pdu1d(pf00356) GRM5
02266 -pf00185pf02729 -
02267 -pf01948pf02748 -
02270 -pf00485 -
02271 -pf00999 -
02272 -pf01784 -
02273 - -
02274 -pf03979pf00140pf04542pf04539pf0454
02275 -pf10410pf08275pf01807 -
BMC-P
BMC-H
BMC-T
Aldehyde dehydrogenase
PTAC
Signature enzyme
Alcohol dehydrogenase
Regulator

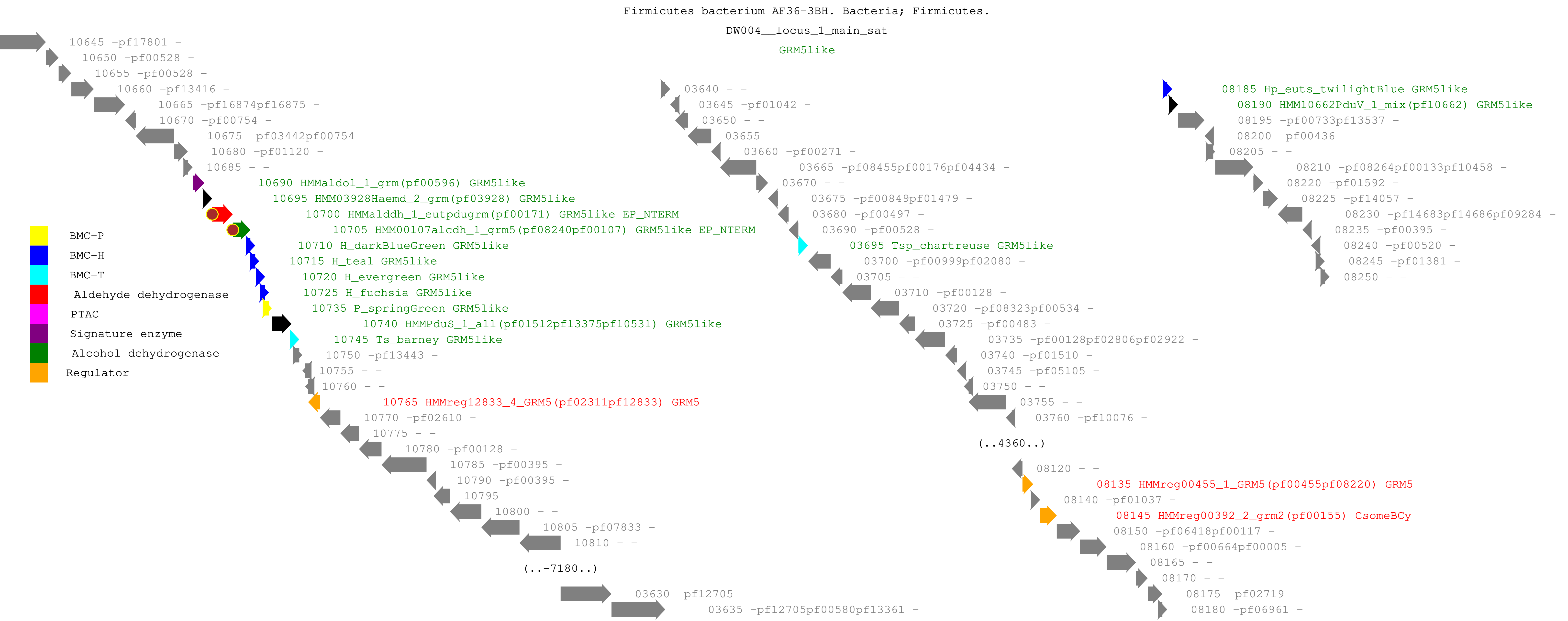
Firmicutes bacterium AF36-3BH. Bacteria; Firmicutes.
DW004__locus_1_main_sat
GRM5like
10645 -pf17801 -
10650 -pf00528 -
10655 -pf00528 -
10660 -pf13416 -
10665 -pf16874pf16875 -
10670 -pf00754 -
10675 -pf03442pf00754 -
10680 -pf01120 -
10685 - -
10690 HMMaldol_1_grm(pf00596) GRM5like
10695 HMM03928Haemd_2_grm(pf03928) GRM5like
10700 HMMalddh_1_eutpdugrm(pf00171) GRM5like EP_NTERM
10705 HMM00107alcdh_1_grm5(pf08240pf00107) GRM5like EP_NTERM
10710 H_darkBlueGreen GRM5like
10715 H_teal GRM5like
10720 H_evergreen GRM5like
10725 H_fuchsia GRM5like
10735 P_springGreen GRM5like
10740 HMMPduS_1_all(pf01512pf13375pf10531) GRM5like
10745 Ts_barney GRM5like
10750 -pf13443 -
10755 - -
10760 - -
10765 HMMreg12833_4_GRM5(pf02311pf12833) GRM5
10770 -pf02610 -
10775 - -
10780 -pf00128 -
10785 -pf00395 -
10790 -pf00395 -
10795 - -
10800 - -
10805 -pf07833 -
10810 - -
(..-7180..)
03630 -pf12705 -
03635 -pf12705pf00580pf13361 -
BMC-P
BMC-H
BMC-T
Aldehyde dehydrogenase
PTAC
Signature enzyme
Alcohol dehydrogenase
Regulator
03640 - -
03645 -pf01042 -
03650 - -
03655 - -
03660 -pf00271 -
03665 -pf08455pf00176pf04434 -
03670 - -
03675 -pf00849pf01479 -
03680 -pf00497 -
03690 -pf00528 -
03695 Tsp_chartreuse GRM5like
03700 -pf00999pf02080 -
03705 - -
03710 -pf00128 -
03720 -pf08323pf00534 -
03725 -pf00483 -
03735 -pf00128pf02806pf02922 -
03740 -pf01510 -
03745 -pf05105 -
03750 - -
03755 - -
03760 -pf10076 -
(..4360..)
08120 - -
08135 HMMreg00455_1_GRM5(pf00455pf08220) GRM5
08140 -pf01037 -
08145 HMMreg00392_2_grm2(pf00155) CsomeBCy
08150 -pf06418pf00117 -
08160 -pf00664pf00005 -
08165 - -
08170 - -
08175 -pf02719 -
08180 -pf06961 -
08185 Hp_euts_twilightBlue GRM5like
08190 HMM10662PduV_1_mix(pf10662) GRM5like
08195 -pf00733pf13537 -
08200 -pf00436 -
08205 - -
08210 -pf08264pf00133pf10458 -
08220 -pf01592 -
08225 -pf14057 -
08230 -pf14683pf14686pf09284 -
08235 -pf00395 -
08240 -pf00520 -
08245 -pf01381 -
08250 - -

Ruminococcus sp. AM42-11. Bacteria; Firmicutes; Clostridia; Clostridiales; Ruminococcaceae;

DW904__locus_1_main_sat

GRM5like

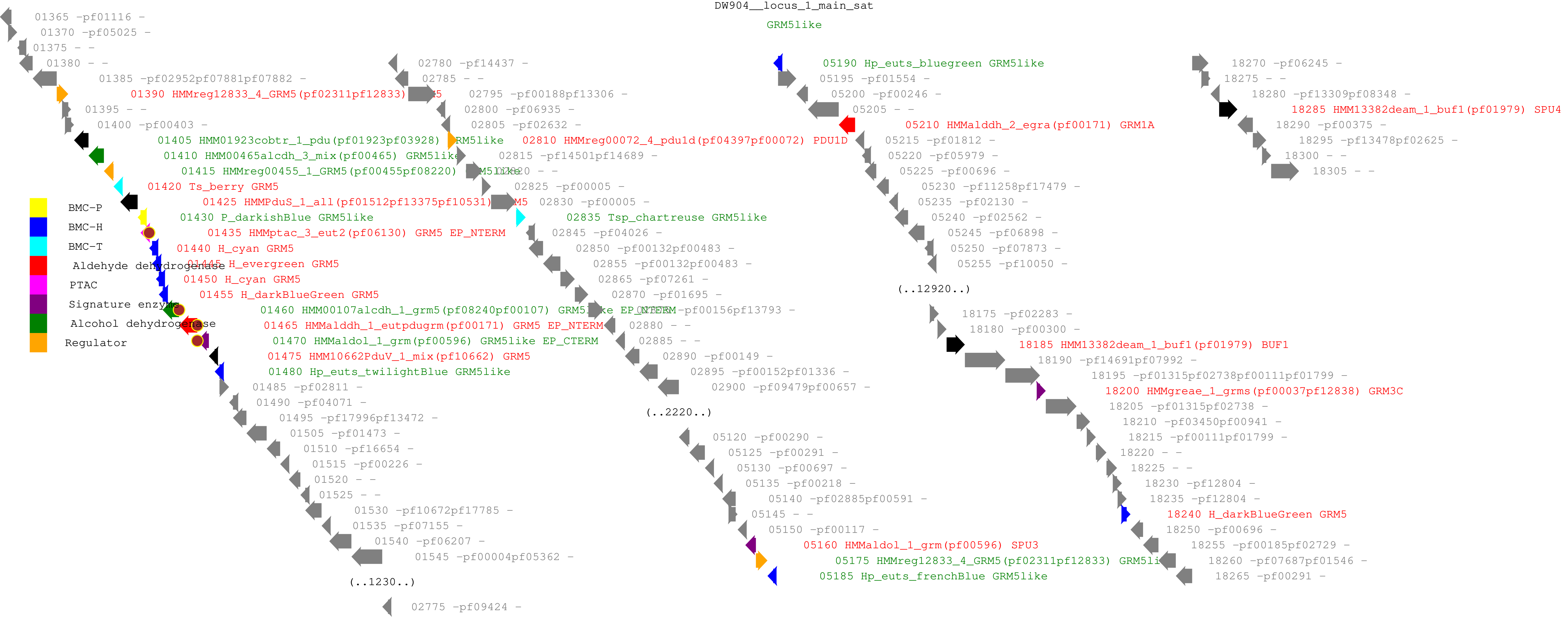

Ruminococcus sp. AF31-8BH. Bacteria; Firmicutes; Clostridia; Clostridiales; Ruminococcaceae;
DWZ38__locus_1_main_sat
GRM5like

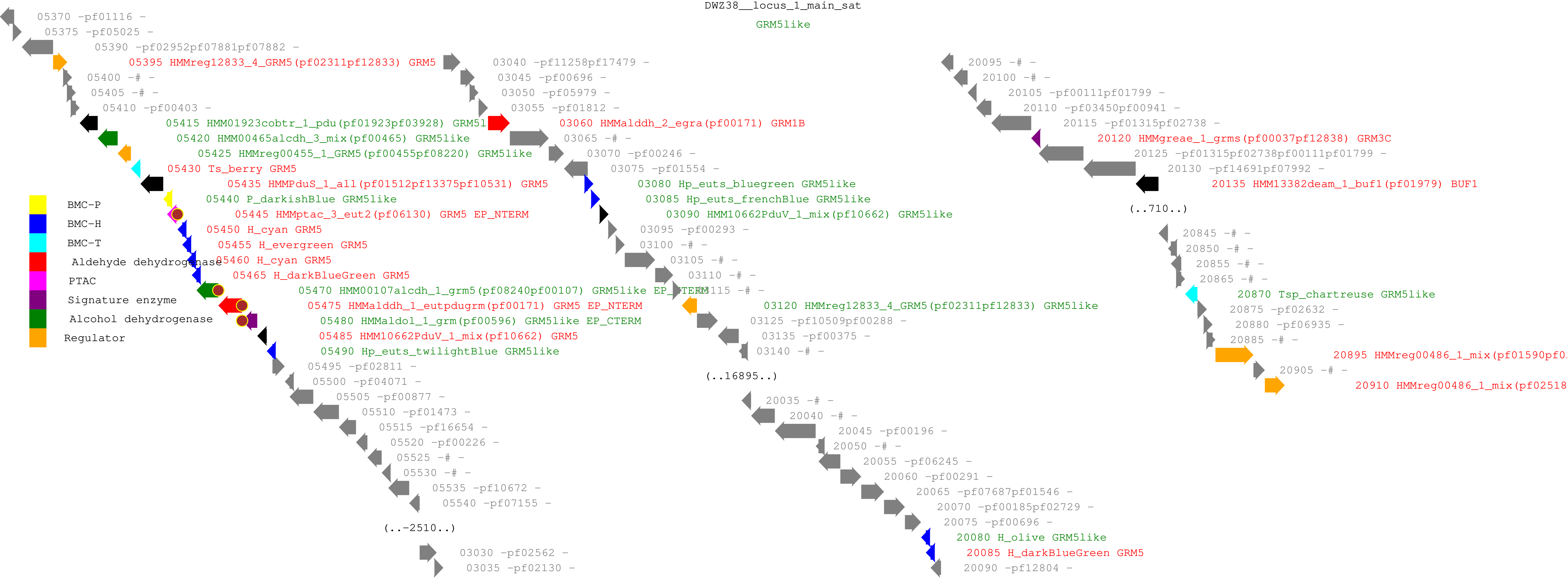

Firmicutes bacterium AM55-24TS. Bacteria; Firmicutes.

DXA10__locus_1_main_sat

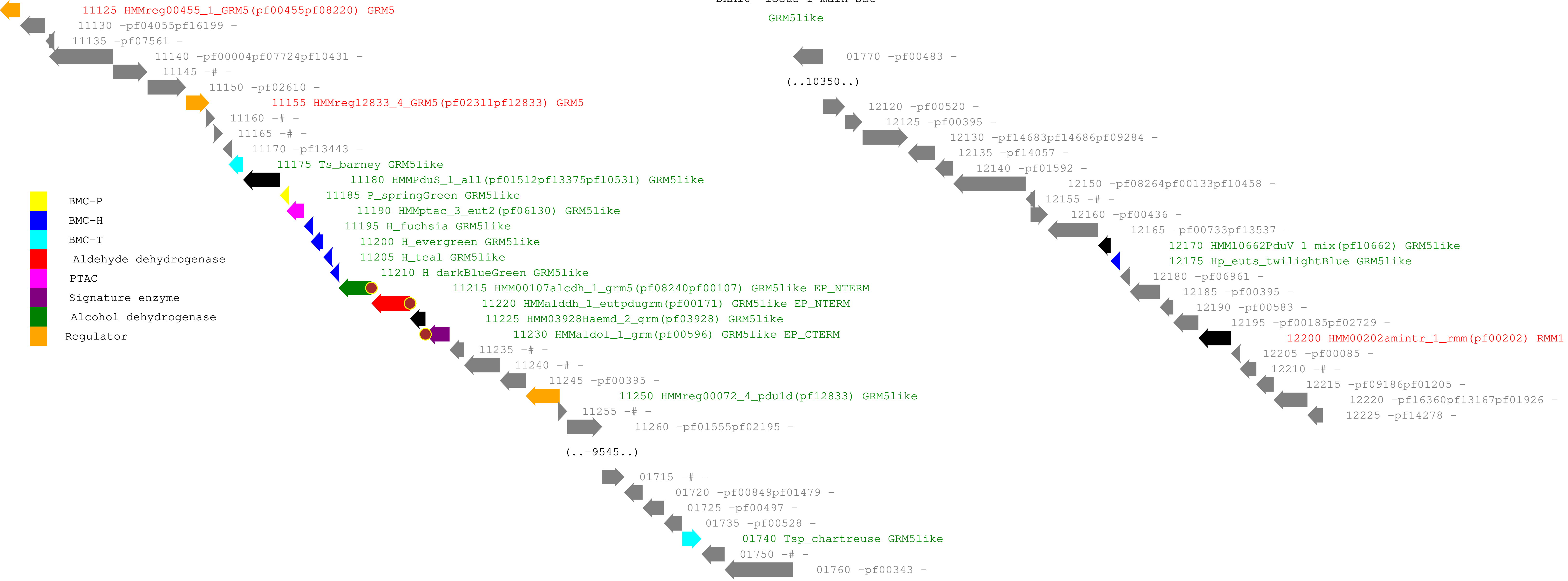

Blautia sp. OF03-15BH. Bacteria; Firmicutes; Clostridia; Clostridiales; Lachnospiraceae; Blautia;

DXA57__locus_1_main_sat

GRM5like

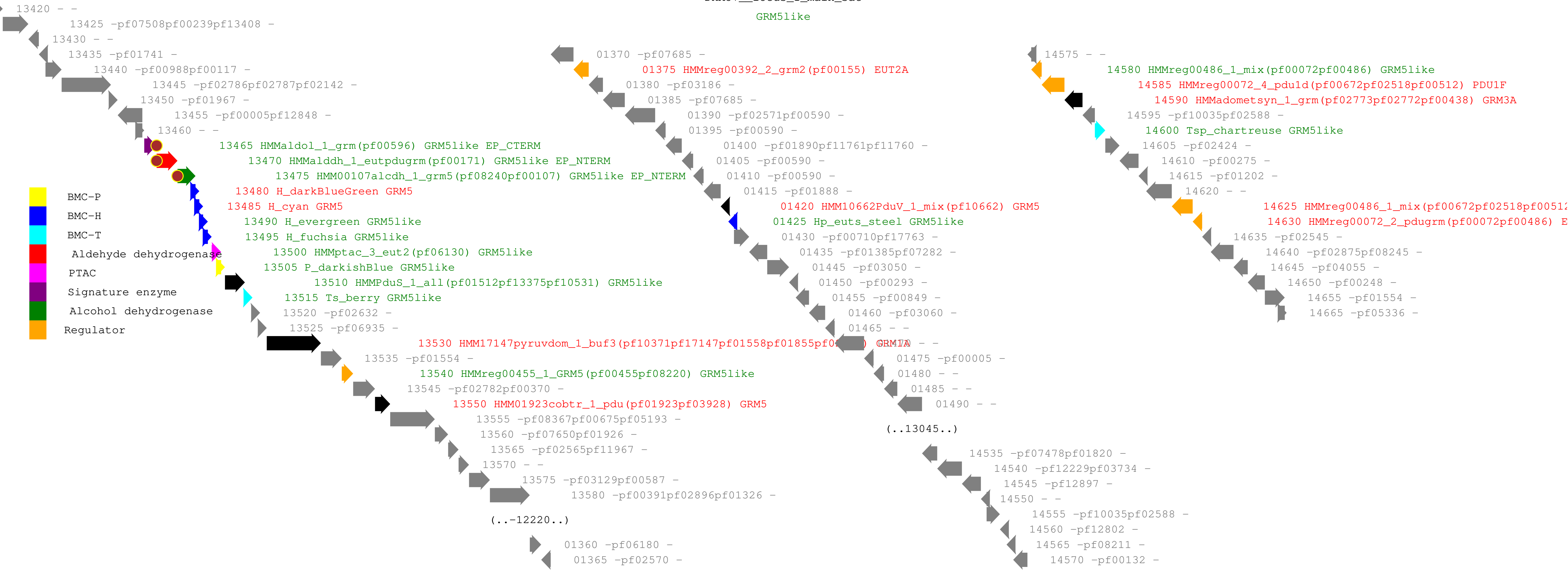

Clostridium disporicum. Bacteria; Firmicutes; Clostridia; Clostridiales; Clostridiaceae;

ERS852471__locus_1

GRM5like

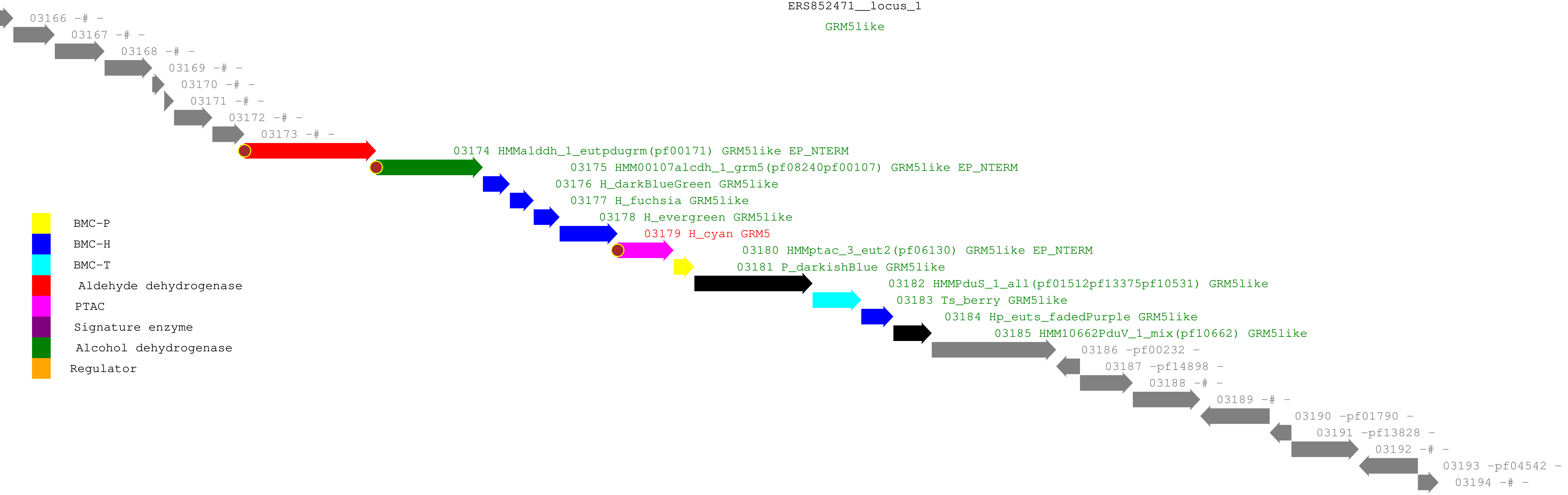

Ruminococcus torques. Bacteria; Firmicutes; Clostridia; Clostridiales; Lachnospiraceae;

ERS852530__locus_1_main_sat

GRM5like

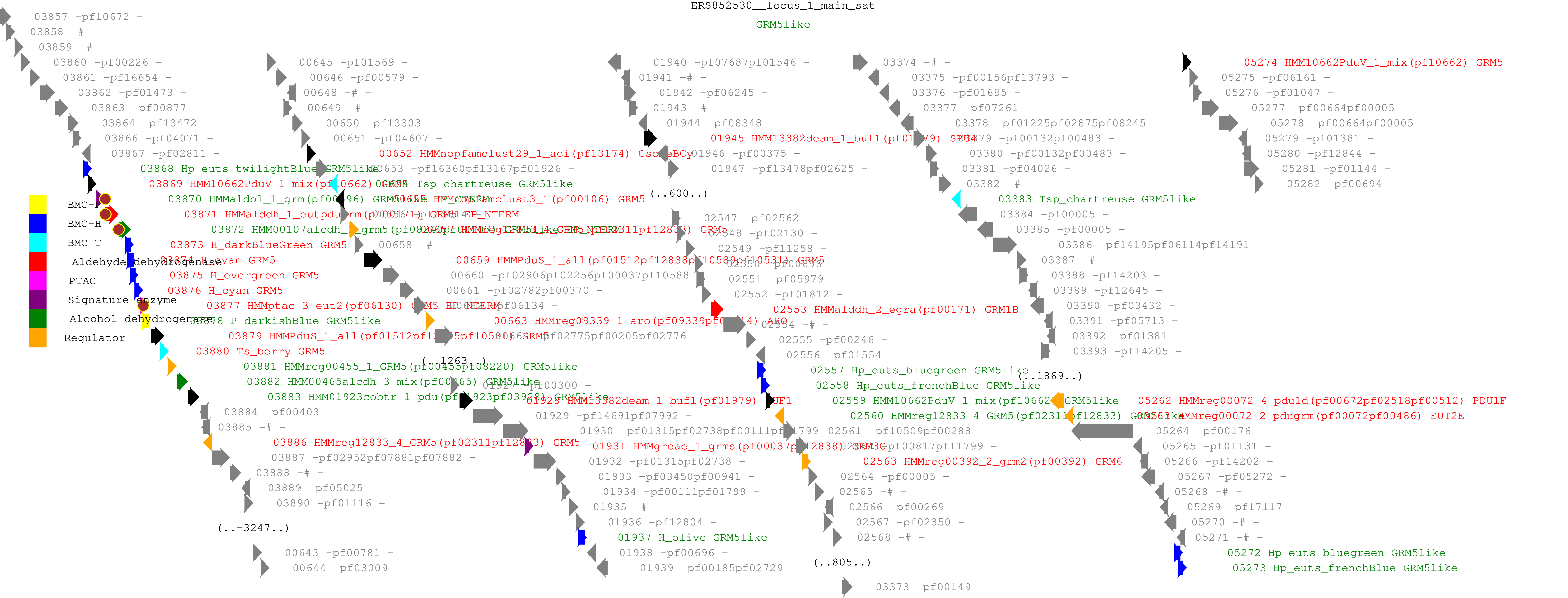

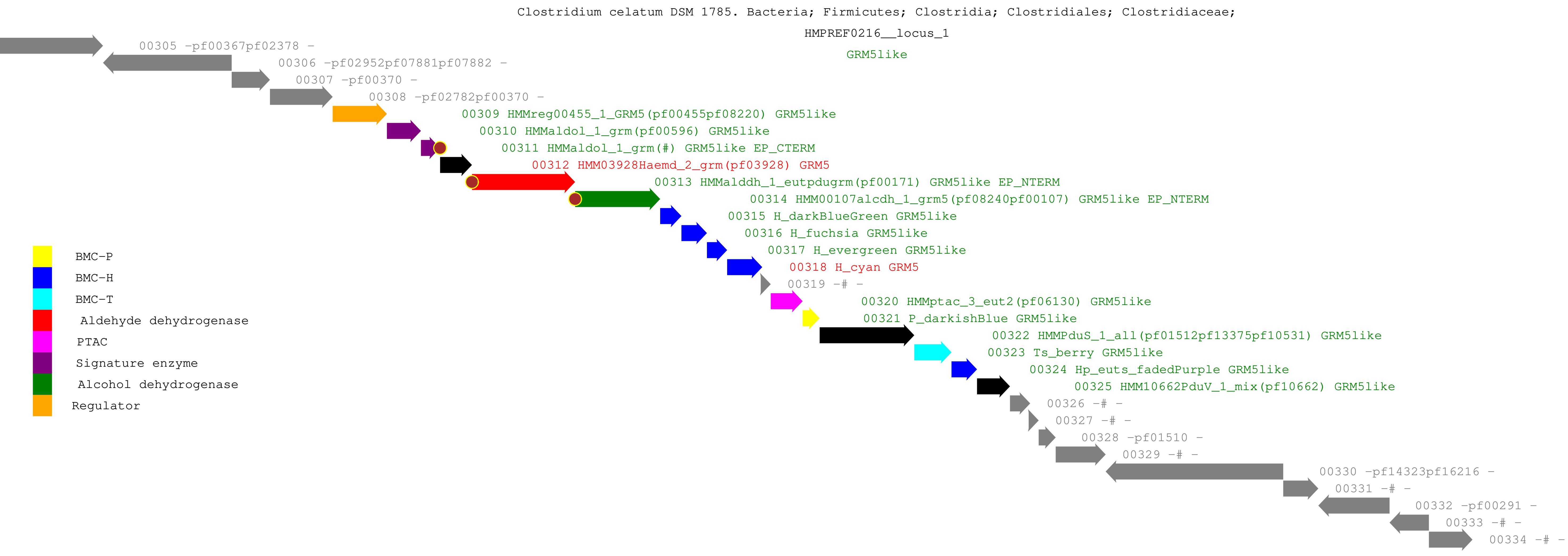
Clostridium celatum DSM 1785. Bacteria; Firmicutes; Clostridia; Clostridiales; Clostridiaceae;
HMPREF0216__locus_1
GRM5like
00305 -pf00367pf02378 -
00306 -pf02952pf07881pf07882 -
00307 -pf00370 -
00308 -pf02782pf00370 -
00309 HMMreg00455_1_GRM5(pf00455pf08220) GRM5like
00310 HMMaldol_1_grm(pf00596) GRM5like
00311 HMMaldol_1_grm(#) GRM5like EP_CTERM
00312 HMM03928Haemd_2_grm(pf03928) GRM5
00313 HMMalddh_1_eutpdugrm(pf00171) GRM5like EP_NTERM
00314 HMM00107alcdh_1_grm5(pf08240pf00107) GRM5like EP_NTERM
00315 H_darkBlueGreen GRM5like
00316 H_fuchsia GRM5like
00317 H_evergreen GRM5like
00318 H_cyan GRM5
00319 -# -
00320 HMMptac_3_eut2(pf06130) GRM5like
00321 P_darkishBlue GRM5like
00322 HMMPduS_1_all(pf01512pf13375pf10531) GRM5like
00323 Ts_berry GRM5like
00324 Hp_euts_fadedPurple GRM5like
00325 HMM10662PduV_1_mix(pf10662) GRM5like
00326 -# -
00327 -# -
00328 -pf01510 -
00329 -# -
00330 -pf14323pf16216 -
00331 -# -
00332 -pf00291 -
00333 -# -
00334 -# -
BMC-P
BMC-H
BMC-T
Aldehyde dehydrogenase
PTAC
Signature enzyme
Alcohol dehydrogenase
Regulator

Christensenella minuta. Bacteria; Firmicutes; Clostridia; Clostridiales; Christensenellaceae;

HMPREF3293__locus_1_main_sat

GRM5like

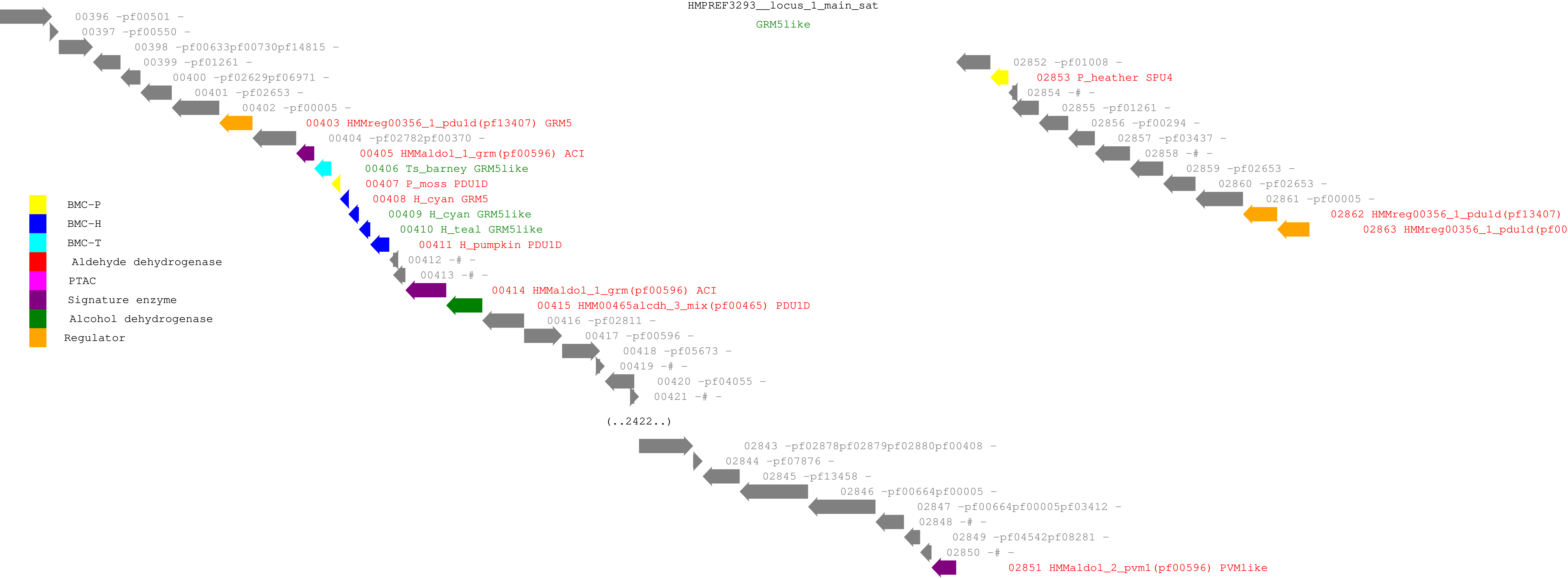

Robinsoniella sp. RHS. Bacteria; Firmicutes; Clostridia; Clostridiales; Lachnospiraceae;

RHS__locus_1

GRM5like

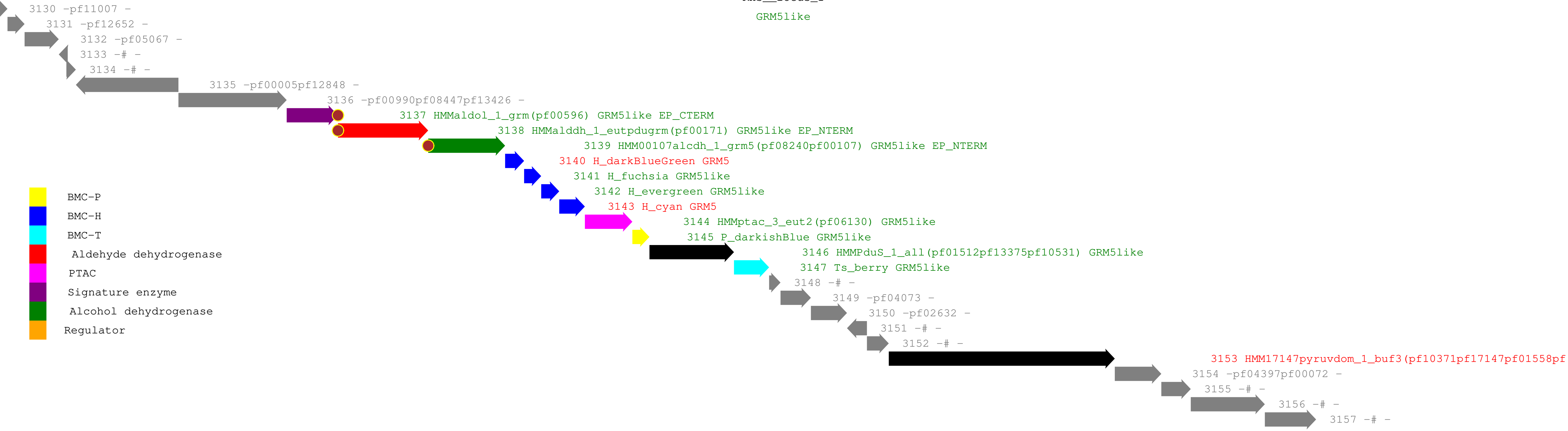

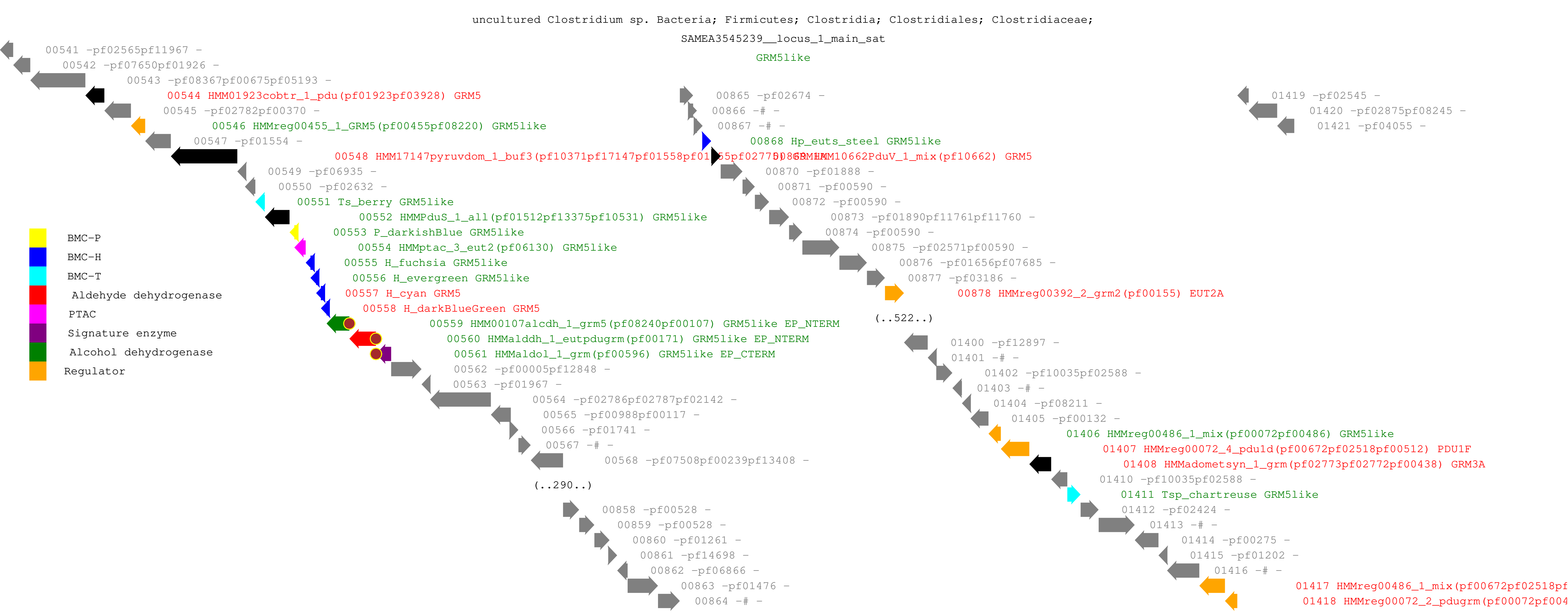

uncultured Clostridium sp. Bacteria; Firmicutes; Clostridia; Clostridiales; Clostridiaceae;
SAMEA3545239__locus_1_main_sat
GRM5like
00541 -pf02565pf11967 -
00542 -pf07650pf01926 -
00543 -pf08367pf00675pf05193 -
00544 HMM01923cobtr_1_pdu(pf01923pf03928) GRM5
00545 -pf02782pf00370 -
00546 HMMreg00455_1_GRM5(pf00455pf08220) GRM5like
00547 -pf01554 -
00549 -pf06935 -
00550 -pf02632 -
00551 Ts_berry GRM5like
00552 HMMPduS_1_all(pf01512pf13375pf10531) GRM5like
00553 P_darkishBlue GRM5like
00554 HMMptac_3_eut2(pf06130) GRM5like
00555 H_fuchsia GRM5like
00556 H_evergreen GRM5like
00557 H_cyan GRM5
00558 H_darkBlueGreen GRM5
00559 HMM00107alcdh_1_grm5(pf08240pf00107) GRM5like EP_NTERM
00560 HMMaldh_1_eutpdugrm(pf00171) GRM5like EP_NTERM
00561 HMMaldol_1_grm(pf00596) GRM5like EP_CTERM
00562 -pf00005pf12848 -
00563 -pf01967 -
00564 -pf02786pf02787pf02142 -
00565 -pf00988pf00117 -
00566 -pf01741 -
00567 -# -
00568 -pf07508pf00239pf13408 -
(..290..)
00858 -pf00528 -
00859 -pf00528 -
00860 -pf01261 -
00861 -pf14698 -
00862 -pf06866 -
00863 -pf01476 -
00864 -# -
00865 -pf02674 -
00866 -# -
00867 -# -
00868 Hp_euts_steel GRM5like
00870 -pf01888 -
00871 -pf00590 -
00872 -pf00590 -
00873 -pf01890pf11761pf11760 -
00874 -pf00590 -
00875 -pf02571pf00590 -
00876 -pf01656pf07685 -
00877 -pf03186 -
00878 HMMreg00392_2_grm2(pf00155) EUT2A
(..522..)
01400 -pf12897 -
01401 -# -
01402 -pf10035pf02588 -
01403 -# -
01404 -pf08211 -
01405 -pf00132 -
01406 HMMreg00486_1_mix(pf00072pf00486) GRM5like
01407 HMMreg00072_4_pdu1d(pf00672pf02518pf00512) PDU1F
01408 HMMadometsyn_1_grm(pf02773pf02772pf00438) GRM3A
01410 -pf10035pf02588 -
01411 Tsp_chartreuse GRM5like
01412 -pf02424 -
01413 -# -
01414 -pf00275 -
01415 -pf01202 -
01416 -# -
01417 HMMreg00486_1_mix(pf00672pf02518pf
01418 HMMreg00072_2_pdugrm(pf00072pf004
01419 -pf02545 -
01420 -pf02875pf08245 -
01421 -pf04055 -
BMC-P
BMC-H
BMC-T
Aldehyde dehydrogenase
PTAC
Signature enzyme
Alcohol dehydrogenase
Regulator

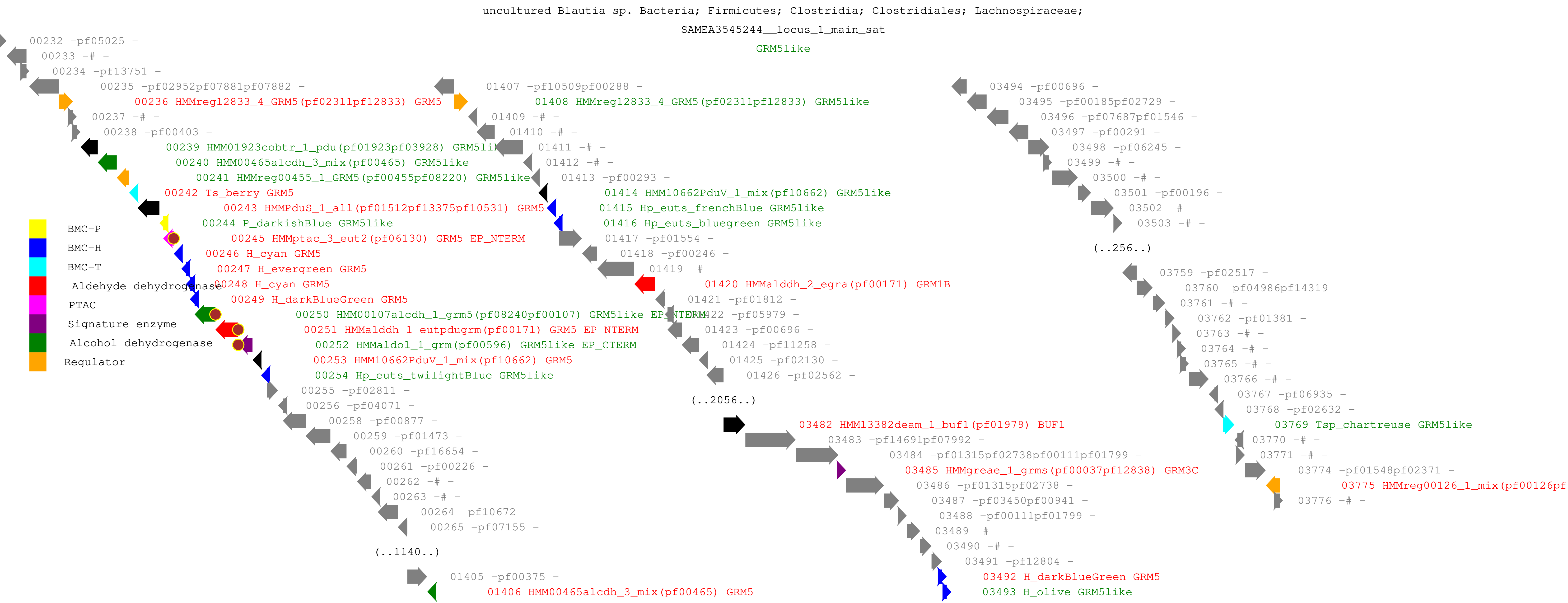
uncultured Blautia sp. Bacteria; Firmicutes; Clostridia; Clostridiales; Lachnospiraceae;
SAMEA3545244__locus_1_main_sat
GRM5like
BMC-P
BMC-H
BMC-T
Aldehyde dehydrogenase
PTAC
Signature enzyme
Alcohol dehydrogenase
Regulator
00232 -pf05025 -
00233 -# -
00234 -pf13751 -
00235 -pf02952pf07881pf07882 -
00236 HMMreg12833_4_GRM5(pf02311pf12833) GRM5
00237 -# -
00238 -pf00403 -
00239 HMM01923cobtr_1_pdu(pf01923pf03928) GRM5like
00240 HMM00465alcdh_3_mix(pf00465) GRM5like
00241 HMMreg00455_1_GRM5(pf00455pf08220) GRM5like
00242 Ts_berry GRM5
00243 HMMPduS_1_all(pf01512pf13375pf10531) GRM5
00244 P_darkishBlue GRM5like
00245 HMMptac_3_eut2(pf06130) GRM5 EP_NTERM
00246 H_cyan GRM5
00247 H_evergreen GRM5
00248 H_cyan GRM5
00249 H_darkBlueGreen GRM5
00250 HMM00107alcdh_1_grm5(pf08240pf00107) GRM5like EP_NTERM
00251 HMMalddh_1_eutpdugrm(pf00171) GRM5 EP_NTERM
00252 HMMaldol_1_grm(pf00596) GRM5like EP_CTERM
00253 HMM10662PduV_1_mix(pf10662) GRM5
00254 Hp_euts_twilightBlue GRM5like
00255 -pf02811 -
00256 -pf04071 -
00258 -pf00877 -
00259 -pf01473 -
00260 -pf16654 -
00261 -pf00226 -
00262 -# -
00263 -# -
00264 -pf10672 -
00265 -pf07155 -
(..1140..)
01405 -pf00375 -
01406 HMM00465alcdh_3_mix(pf00465) GRM5
01407 -pf10509pf00288 -
01408 HMMreg12833_4_GRM5(pf02311pf12833) GRM5like
01409 -# -
01410 -# -
01411 -# -
01412 -# -
01413 -pf00293 -
01414 HMM10662PduV_1_mix(pf10662) GRM5like
01415 Hp_euts_frenchBlue GRM5like
01416 Hp_euts_bluegreen GRM5like
01417 -pf01554 -
01418 -pf00246 -
01419 -# -
01420 HMMalddh_2_egra(pf00171) GRM1B
01421 -pf01812 -
01422 -pf05979 -
01423 -pf00696 -
01424 -pf11258 -
01425 -pf02130 -
01426 -pf02562 -
(..2056..)
03482 HMM13382deam_1_buf1(pf01979) BUF1
03483 -pf14691pf07992 -
03484 -pf01315pf02738pf00111pf01799 -
03485 HMMgreae_1_grms(pf00037pf12838) GRM3C
03486 -pf01315pf02738 -
03487 -pf03450pf00941 -
03488 -pf00111pf01799 -
03489 -# -
03490 -# -
03491 -pf12804 -
03492 H_darkBlueGreen GRM5
03493 H_olive GRM5like
03494 -pf00696 -
03495 -pf00185pf02729 -
03496 -pf07687pf01546 -
03497 -pf00291 -
03498 -pf06245 -
03499 -# -
03500 -# -
03501 -pf00196 -
03502 -# -
03503 -# -
(..256..)
03759 -pf02517 -
03760 -pf04986pf14319 -
03761 -# -
03762 -pf01381 -
03763 -# -
03764 -# -
03765 -# -
03766 -# -
03767 -pf06935 -
03768 -pf02632 -
03769 Tsp_chartreuse GRM5like
03770 -# -
03771 -# -
03774 -pf01548pf02371 -
03775 HMMreg00126_1_mix(pf00126pf
03776 -# -

uncultured Blautia sp. Bacteria; Firmicutes; Clostridia; Clostridiales; Lachnospiraceae;

SAMEA3545252__locus_1_main_sat

GRM5like

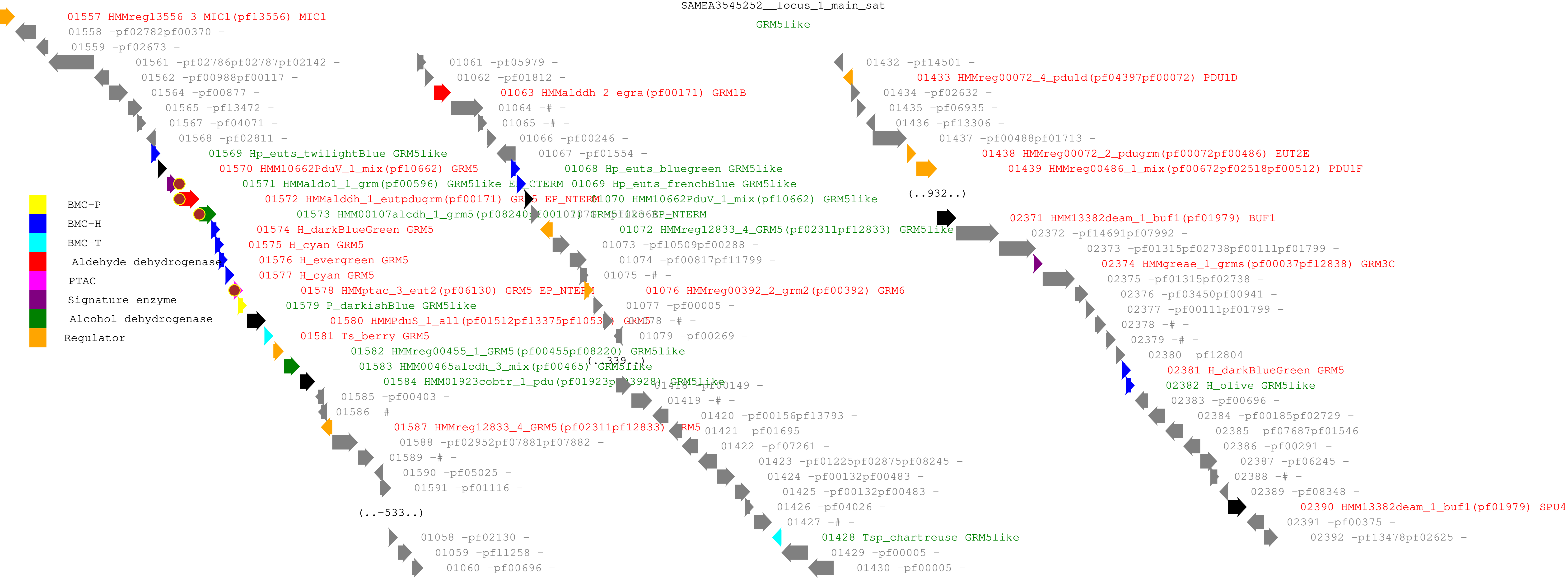

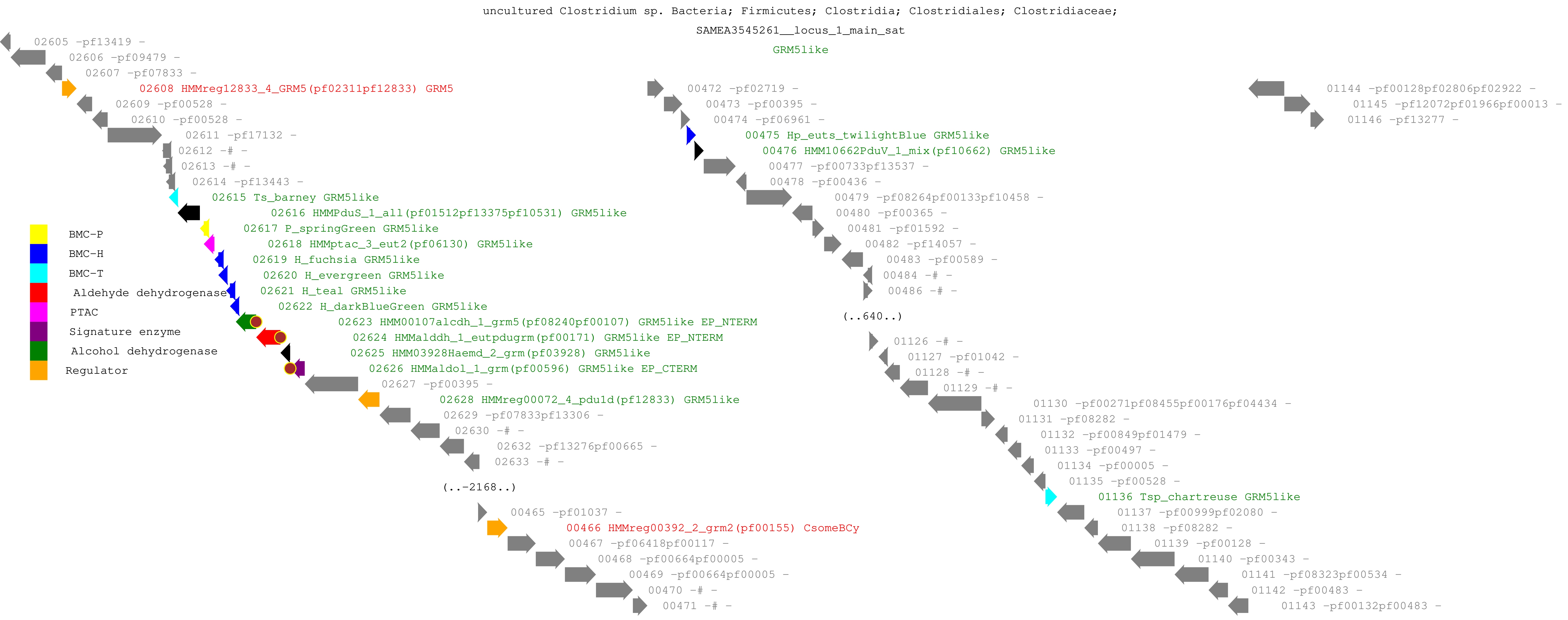
uncultured Clostridium sp. Bacteria; Firmicutes; Clostridia; Clostridiales; Clostridiaceae;
SAMEA3545261__locus_1_main_sat
GRM5like
02605 -pf13419 -
02606 -pf09479 -
02607 -pf07833 -
02608 HMMreg12833_4_GRM5(pf02311pf12833) GRM5
02609 -pf00528 -
02610 -pf00528 -
02611 -pf17132 -
02612 -# -
02613 -# -
02614 -pf13443 -
02615 Ts_barney GRM5like
02616 HMMPduS_1_all(pf01512pf13375pf10531) GRM5like
02617 P_springGreen GRM5like
02618 HMMptac_3_eut2(pf06130) GRM5like
02619 H_fuchsia GRM5like
02620 H_evergreen GRM5like
02621 H_teal GRM5like
02622 H_darkBlueGreen GRM5like
02623 HMM00107alcdh_1_grm5(pf08240pf00107) GRM5like EP_NTERM
02624 HMMaldd h_1_eutpdugrm(pf00171) GRM5like EP_NTERM
02625 HMM03928Haemd_2_grm(pf03928) GRM5like
02626 HMMaldol_1_grm(pf00596) GRM5like EP_CTERM
02627 -pf00395 -
02628 HMMreg00072_4_pdu1d(pf12833) GRM5like
02629 -pf07833pf13306 -
02630 -# -
02632 -pf13276pf00665 -
02633 -# -
(..-2168..)
00465 -pf01037 -
00466 HMMreg00392_2_grm2(pf00155) CsomeBCy
00467 -pf06418pf00117 -
00468 -pf00664pf00005 -
00469 -pf00664pf00005 -
00470 -# -
00471 -# -
BMC-P
BMC-H
BMC-T
Aldehyde dehydrogenase
PTAC
Signature enzyme
Alcohol dehydrogenase
Regulator
00472 -pf02719 -
00473 -pf00395 -
00474 -pf06961 -
00475 Hp_euts_twilightBlue GRM5like
00476 HMM10662PduV_1_mix(pf10662) GRM5like
00477 -pf00733pf13537 -
00478 -pf00436 -
00479 -pf08264pf00133pf10458 -
00480 -pf00365 -
00481 -pf01592 -
00482 -pf14057 -
00483 -pf00589 -
00484 -# -
00486 -# -
(..640..)
01126 -# -
01127 -pf01042 -
01128 -# -
01129 -# -
01130 -pf00271pf08455pf00176pf04434 -
01131 -pf08282 -
01132 -pf00849pf01479 -
01133 -pf00497 -
01134 -pf00005 -
01135 -pf00528 -
01136 Tsp_chartreuse GRM5like
01137 -pf00999pf02080 -
01138 -pf08282 -
01139 -pf00128 -
01140 -pf00343 -
01141 -pf08323pf00534 -
01142 -pf00483 -
01143 -pf00132pf00483 -
01144 -pf00128pf02806pf02922 -
01145 -pf12072pf01966pf00013 -
01146 -pf13277 -

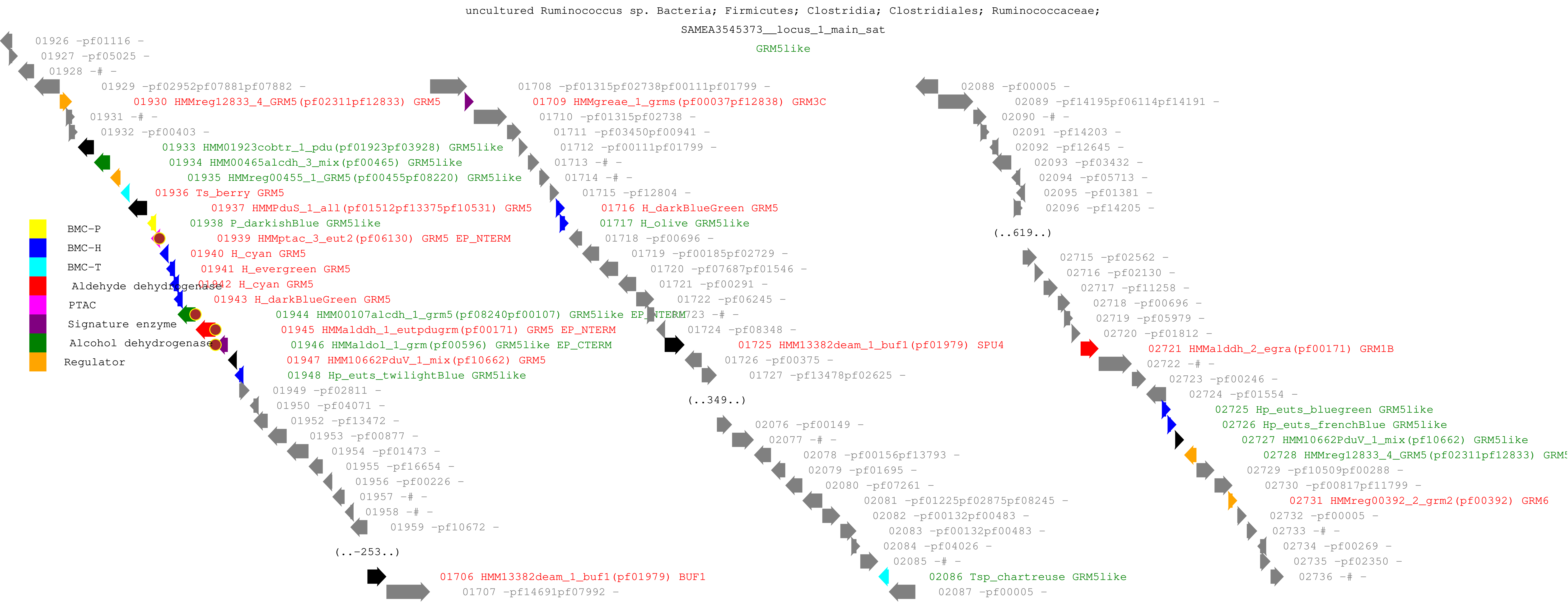
uncultured Ruminococcus sp. Bacteria; Firmicutes; Clostridia; Clostridiales; Ruminococcaceae;
SAMEA3545373__locus_1_main_sat
GRM5like
BMC-P
BMC-H
BMC-T
Aldehyde dehydrogenase
PTAC
Signature enzyme
Alcohol dehydrogenase
Regulator
01926 -pf01116 -
01927 -pf05025 -
01928 -# -
01929 -pf02952pf07881pf07882 -
01930 HMMreg12833_4_GRM5(pf02311pf12833) GRM5
01931 -# -
01932 -pf00403 -
01933 HMM01923cobtr_1_pdu(pf01923pf03928) GRM5like
01934 HMM00465alcdh_3_mix(pf00465) GRM5like
01935 HMMreg00455_1_GRM5(pf00455pf08220) GRM5like
01936 Ts_berry GRM5
01937 HMMPduS_1_all(pf01512pf13375pf10531) GRM5
01938 P_darkishBlue GRM5like
01939 HMMptac_3_eut2(pf06130) GRM5 EP_NTERM
01940 H_cyan GRM5
01941 H_evergreen GRM5
01942 H_cyan GRM5
01943 H_darkBlueGreen GRM5
01944 HMM00107alcdh_1_grm5(pf08240pf00107) GRM5like EP_NTERM
01945 HMMalddh_1_eutpdugrm(pf00171) GRM5 EP_NTERM
01946 HMMaldol_1_grm(pf00596) GRM5like EP_CTERM
01947 HMM10662PduV_1_mix(pf10662) GRM5
01948 Hp_euts_twilightBlue GRM5like
01949 -pf02811 -
01950 -pf04071 -
01952 -pf13472 -
01953 -pf00877 -
01954 -pf01473 -
01955 -pf16654 -
01956 -pf00226 -
01957 -# -
01958 -# -
01959 -pf10672 -
(..-253..)
01706 HMM13382deam_1_buf1(pf01979) BUF1
01707 -pf14691pf07992 -
01708 -pf01315pf02738pf00111pf01799 -
01709 HMMgreae_1_grms(pf00037pf12838) GRM3C
01710 -pf01315pf02738 -
01711 -pf03450pf00941 -
01712 -pf00111pf01799 -
01713 -# -
01714 -# -
01715 -pf12804 -
01716 H_darkBlueGreen GRM5
01717 H_olive GRM5like
01718 -pf00696 -
01719 -pf00185pf02729 -
01720 -pf07687pf01546 -
01721 -pf00291 -
01722 -pf06245 -
01723 -# -
01724 -pf08348 -
01725 HMM13382deam_1_buf1(pf01979) SPU4
01726 -pf00375 -
01727 -pf13478pf02625 -
(..349..)
02076 -pf00149 -
02077 -# -
02078 -pf00156pf13793 -
02079 -pf01695 -
02080 -pf07261 -
02081 -pf01225pf02875pf08245 -
02082 -pf00132pf00483 -
02083 -pf00132pf00483 -
02084 -pf04026 -
02085 -# -
02086 Tsp_chartreuse GRM5like
02087 -pf00005 -
02088 -pf00005 -
02089 -pf14195pf06114pf14191 -
02090 -# -
02091 -pf14203 -
02092 -pf12645 -
02093 -pf03432 -
02094 -pf05713 -
02095 -pf01381 -
02096 -pf14205 -
(..619..)
02715 -pf02562 -
02716 -pf02130 -
02717 -pf11258 -
02718 -pf00696 -
02719 -pf05979 -
02720 -pf01812 -
02721 HMMalddh_2_egra(pf00171) GRM1B
02722 -# -
02723 -pf00246 -
02724 -pf01554 -
02725 Hp_euts_bluegreen GRM5like
02726 Hp_euts_frenchBlue GRM5like
02727 HMM10662PduV_1_mix(pf10662) GRM5like
02728 HMMreg12833_4_GRM5(pf02311pf12833) GRM
02729 -pf10509pf00288 -
02730 -pf00817pf11799 -
02731 HMMreg00392_2_grm2(pf00392) GRM6
02732 -pf00005 -
02733 -# -
02734 -pf00269 -
02735 -pf02350 -
02736 -# -

uncultured Clostridium sp. Bacteria; Firmicutes; Clostridia; Clostridiales; Clostridiaceae;

SAMEA3545382__locus_1_main_sat

GRM5like

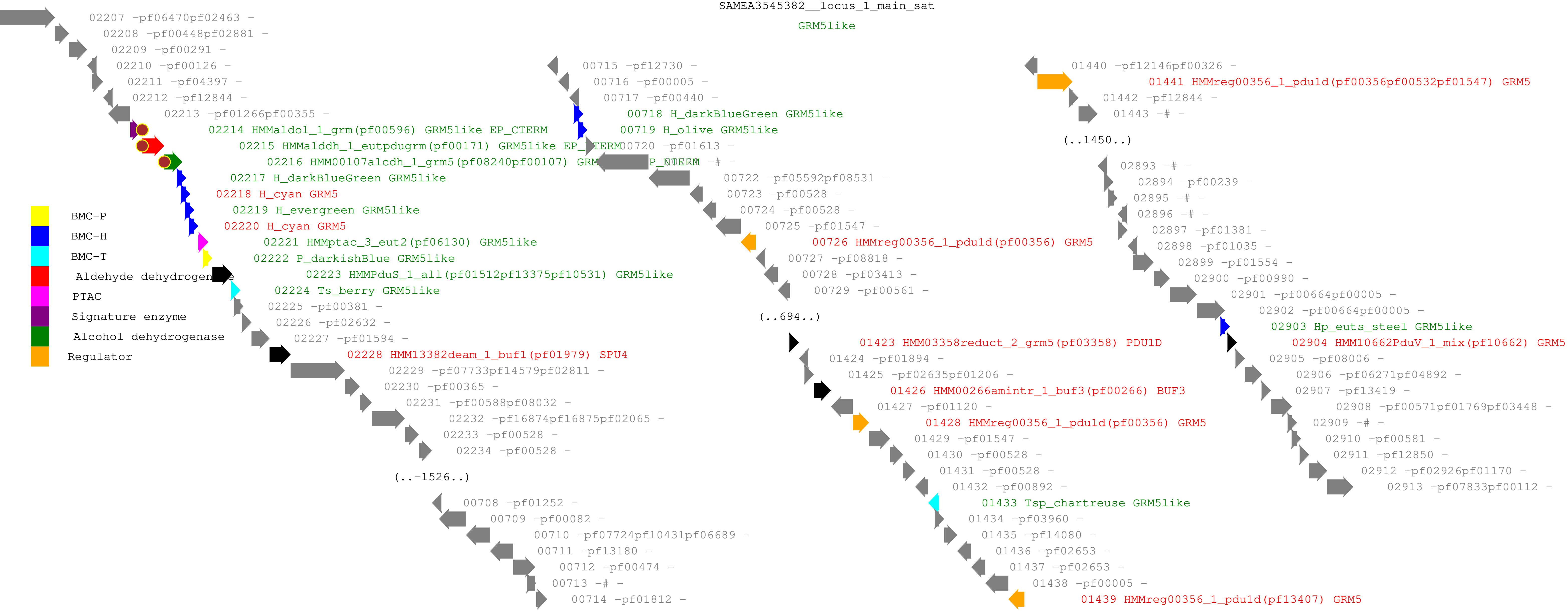

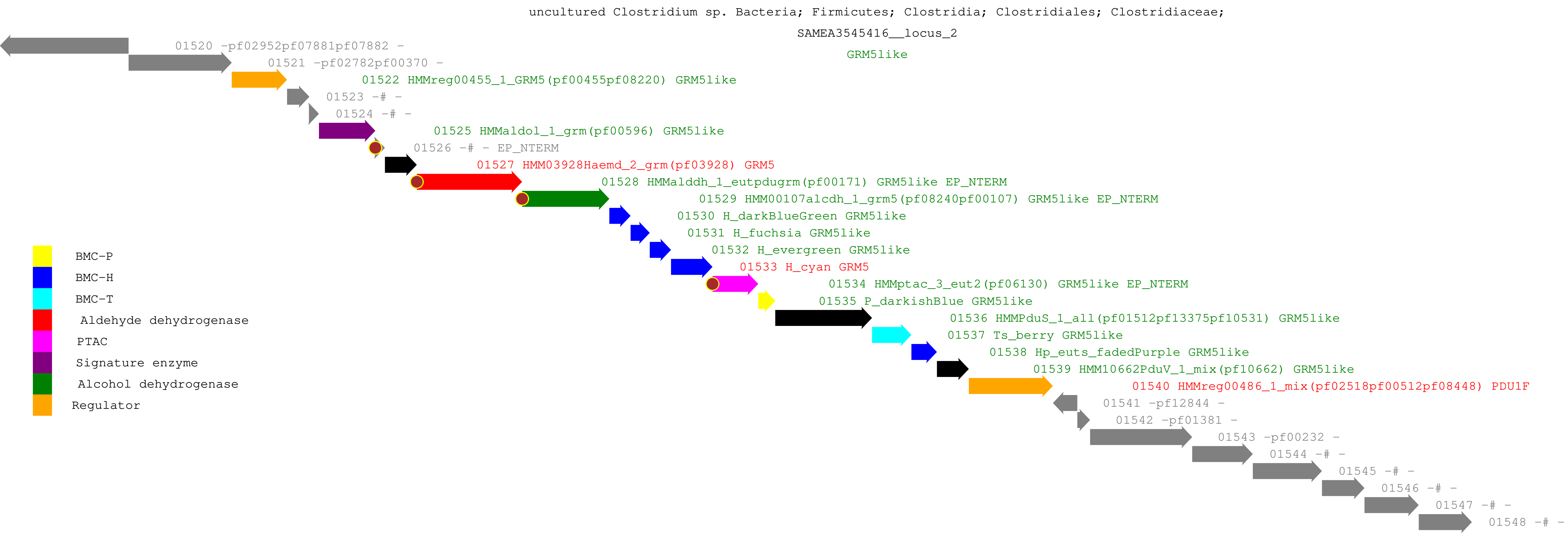
uncultured Clostridium sp. Bacteria; Firmicutes; Clostridia; Clostridiales; Clostridiaceae;
SAMEA3545416__locus_2
GRM5like
01520 -pf02952pf07881pf07882 -
01521 -pf02782pf00370 -
01522 HMMreg00455_1_GRM5(pf00455pf08220) GRM5like
01523 -# -
01524 -# -
01525 HMMaldol_1_grm(pf00596) GRM5like
01526 -# - EP_NTERM
01527 HMM03928Haemd_2_grm(pf03928) GRM5
01528 HMMalddh_1_eutpdugrm(pf00171) GRM5like EP_NTERM
01529 HMM00107alcdh_1_grm5(pf08240pf00107) GRM5like EP_NTERM
01530 H_darkBlueGreen GRM5like
01531 H_fuchsia GRM5like
01532 H_evergreen GRM5like
01533 H_cyan GRM5
01534 HMMptac_3_eut2(pf06130) GRM5like EP_NTERM
01535 P_darkishBlue GRM5like
01536 HMMPduS_1_all(pf01512pf13375pf10531) GRM5like
01537 Ts_berry GRM5like
01538 Hp_euts_fadedPurple GRM5like
01539 HMM10662PduV_1_mix(pf10662) GRM5like
01540 HMMreg00486_1_mix(pf02518pf00512pf08448) PDU1F
01541 -pf12844 -
01542 -pf01381 -
01543 -pf00232 -
01544 -# -
01545 -# -
01546 -# -
01547 -# -
01548 -# -
BMC-P
BMC-H
BMC-T
Aldehyde dehydrogenase
PTAC
Signature enzyme
Alcohol dehydrogenase
Regulator

uncultured Clostridium sp. Bacteria; Firmicutes; Clostridia; Clostridiales; Clostridiaceae;

SAMEA3545416__locus_3

GRM5like

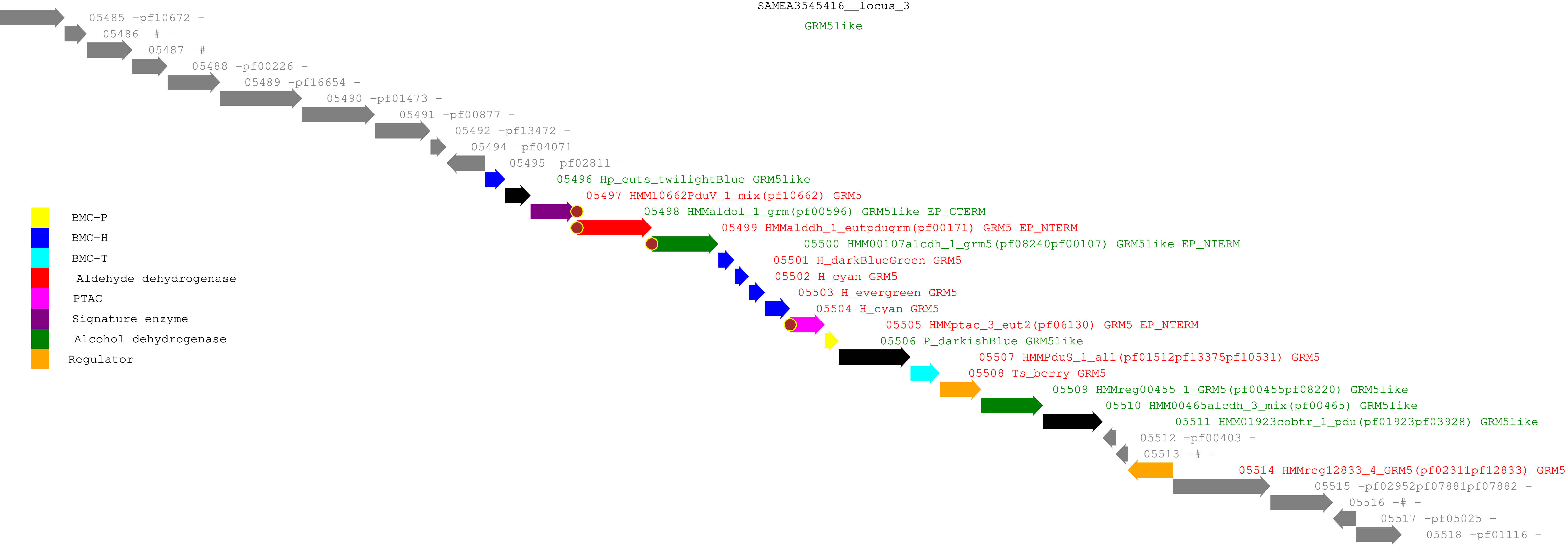

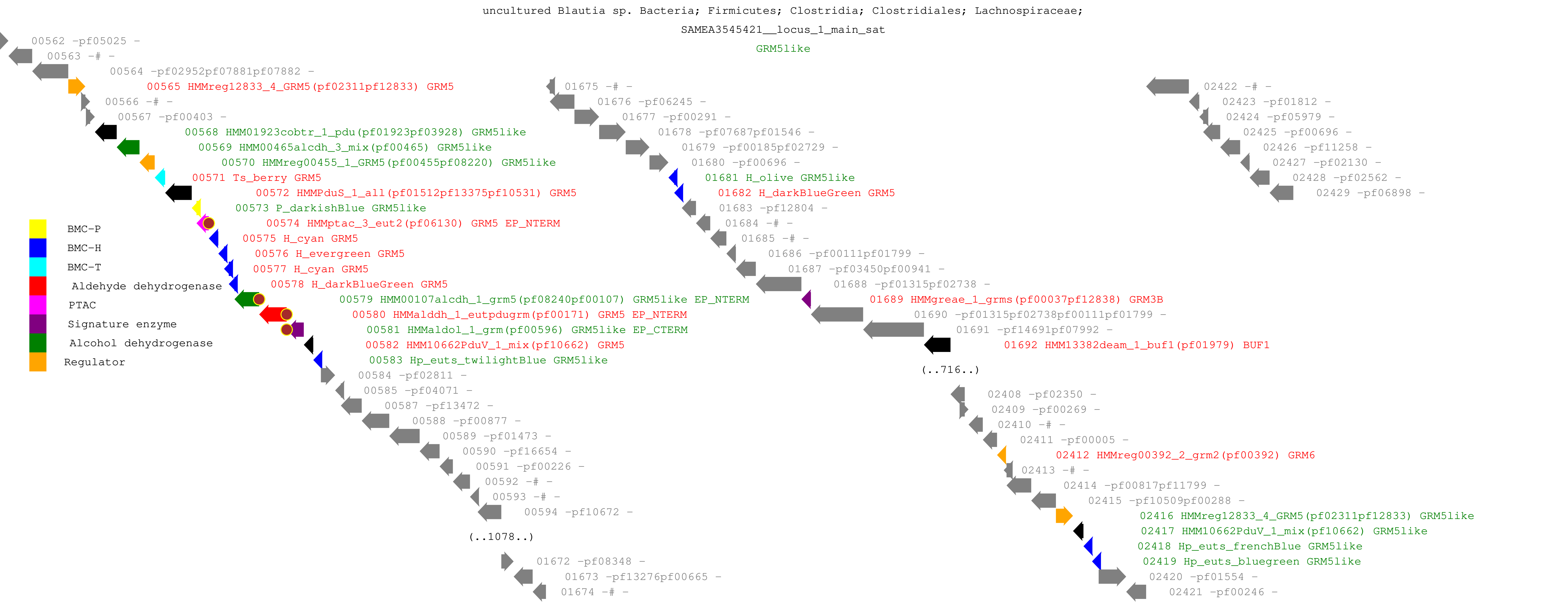

uncultured Blautia sp. Bacteria; Firmicutes; Clostridia; Clostridiales; Lachnospiraceae;
SAMEA3545421__locus_1_main_sat
GRM5like
00562 -pf05025 -
00563 -# -
00564 -pf02952pf07881pf07882 -
00565 HMMreg12833_4_GRM5(pf02311pf12833) GRM5
00566 -# -
00567 -pf00403 -
00568 HMM01923cobtr_1_pdu(pf01923pf03928) GRM5like
00569 HMM00465alcdh_3_mix(pf00465) GRM5like
00570 HMMreg00455_1_GRM5(pf00455pf08220) GRM5like
00571 Ts_berry GRM5
00572 HMMPduS_1_all(pf01512pf13375pf10531) GRM5
00573 P_darkishBlue GRM5like
00574 HMMptac_3_eut2(pf06130) GRM5 EP_NTERM
00575 H_cyan GRM5
00576 H_evergreen GRM5
00577 H_cyan GRM5
00578 H_darkBlueGreen GRM5
00579 HMM00107alcdh_1_grm5(pf08240pf00107) GRM5like EP_NTERM
00580 HMMalddh_1_eutpdugrm(pf00171) GRM5 EP_NTERM
00581 HMMaldol_1_grm(pf00596) GRM5like EP_CTERM
00582 HMM10662PduV_1_mix(pf10662) GRM5
00583 Hp_euts_twilightBlue GRM5like
00584 -pf02811 -
00585 -pf04071 -
00587 -pf13472 -
00588 -pf00877 -
00589 -pf01473 -
00590 -pf16654 -
00591 -pf00226 -
00592 -# -
00593 -# -
00594 -pf10672 -
(..1078..)
01672 -pf08348 -
01673 -pf13276pf00665 -
01674 -# -
01675 -# -
01676 -pf06245 -
01677 -pf00291 -
01678 -pf07687pf01546 -
01679 -pf00185pf02729 -
01680 -pf00696 -
01681 H_olive GRM5like
01682 H_darkBlueGreen GRM5
01683 -pf12804 -
01684 -# -
01685 -# -
01686 -pf00111pf01799 -
01687 -pf03450pf00941 -
01688 -pf01315pf02738 -
01689 HMMgreae_1_grms(pf00037pf12838) GRM3B
01690 -pf01315pf02738pf00111pf01799 -
01691 -pf14691pf07992 -
01692 HMM13382deam_1_buf1(pf01979) BUF1
(..716..)
02408 -pf02350 -
02409 -pf00269 -
02410 -# -
02411 -pf00005 -
02412 HMMreg00392_2_grm2(pf00392) GRM6
02413 -# -
02414 -pf00817pf11799 -
02415 -pf10509pf00288 -
02416 HMMreg12833_4_GRM5(pf02311pf12833) GRM5like
02417 HMM10662PduV_1_mix(pf10662) GRM5like
02418 Hp_euts_frenchBlue GRM5like
02419 Hp_euts_bluegreen GRM5like
02420 -pf01554 -
02421 -pf00246 -
02422 -# -
02423 -pf01812 -
02424 -pf05979 -
02425 -pf00696 -
02426 -pf11258 -
02427 -pf02130 -
02428 -pf02562 -
02429 -pf06898 -
BMC-P
BMC-H
BMC-T
Aldehyde dehydrogenase
PTAC
Signature enzyme
Alcohol dehydrogenase
Regulator

Halanaerobium congolense. Bacteria; Firmicutes; Clostridia; Halanaerobiales; Halanaerobiaceae;

SAMN04488597__locus_1

GRM5like

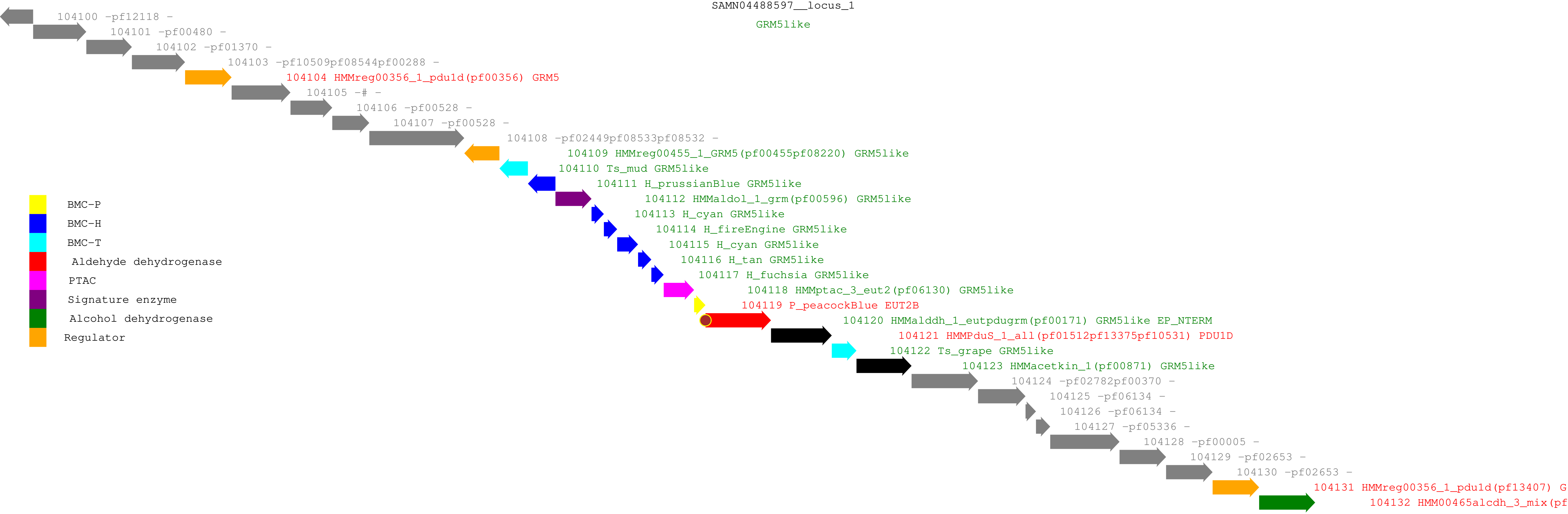

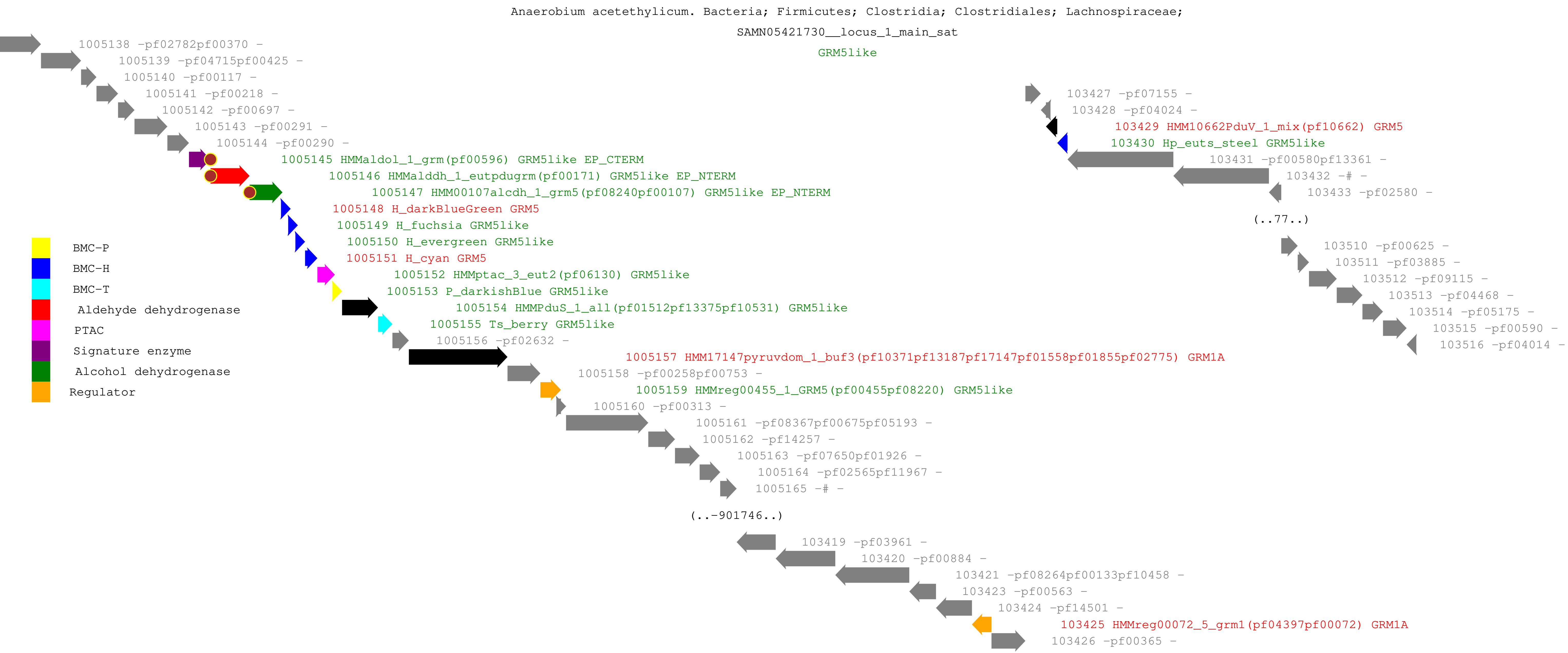
Anaerobium acetethylicum. Bacteria; Firmicutes; Clostridia; Clostridiales; Lachnospiraceae;
SAMN05421730__locus_1_main_sat
GRM5like
1005138 -pf02782pf00370 -
1005139 -pf04715pf00425 -
1005140 -pf00117 -
1005141 -pf00218 -
1005142 -pf00697 -
1005143 -pf00291 -
1005144 -pf00290 -
1005145 HMMaldol_1_grm(pf00596) GRM5like EP_CTERM
1005146 HMMalddh_1_eutpdugrm(pf00171) GRM5like EP_NTERM
1005147 HMM00107alcdh_1_grm5(pf08240pf00107) GRM5like EP_NTERM
1005148 H_darkBlueGreen GRM5
1005149 H_fuchsia GRM5like
1005150 H_evergreen GRM5like
1005151 H_cyan GRM5
1005152 HMMptac_3_eut2(pf06130) GRM5like
1005153 P_darkishBlue GRM5like
1005154 HMMPduS_1_all(pf01512pf13375pf10531) GRM5like
1005155 Ts_berry GRM5like
1005156 -pf02632 -
1005157 HMM17147pyruvdom_1_buf3(pf10371pf13187pf17147pf01558pf01855pf02775) GRM1A
1005158 -pf00258pf00753 -
1005159 HMMreg00455_1_GRM5(pf00455pf08220) GRM5like
1005160 -pf00313 -
1005161 -pf08367pf00675pf05193 -
1005162 -pf14257 -
1005163 -pf07650pf01926 -
1005164 -pf02565pf11967 -
1005165 -# -
(..-901746..)
103419 -pf03961 -
103420 -pf00884 -
103421 -pf08264pf00133pf10458 -
103423 -pf00563 -
103424 -pf14501 -
103425 HMMreg00072_5_grm1(pf04397pf00072) GRM1A
103426 -pf00365 -
103427 -pf07155 -
103428 -pf04024 -
103429 HMM10662PduV_1_mix(pf10662) GRM5
103430 Hp_euts_steel GRM5like
103431 -pf00580pf13361 -
103432 -# -
103433 -pf02580 -
(..77..)
103510 -pf00625 -
103511 -pf03885 -
103512 -pf09115 -
103513 -pf04468 -
103514 -pf05175 -
103515 -pf00590 -
103516 -pf04014 -
BMC-P
BMC-H
BMC-T
Aldehyde dehydrogenase
PTAC
Signature enzyme
Alcohol dehydrogenase
Regulator

Halanaerobium kushneri. Bacteria; Firmicutes; Clostridia; Halanaerobiales; Halanaerobiaceae;

SAMN05421834__locus_1

GRM5like

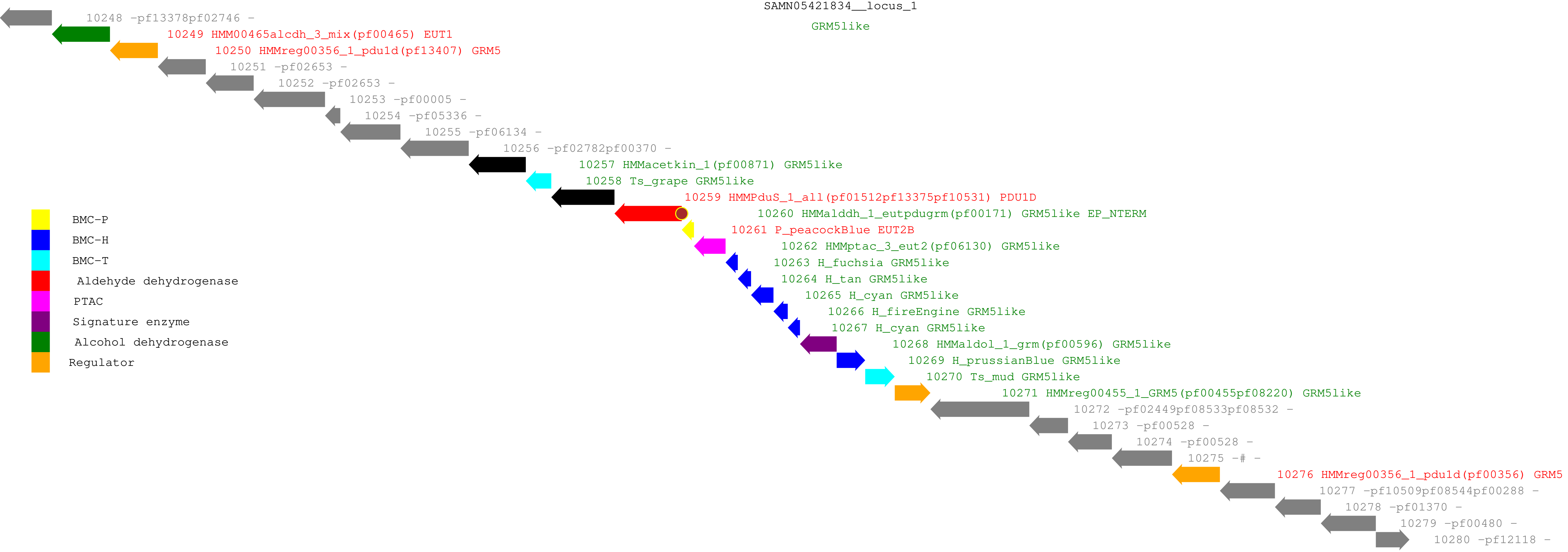

Virgibacillus pantothenticus. Bacteria; Firmicutes; Bacilli; Bacillales; Bacillaceae; Virgibacillus.

AFK71__locus_1

GRM6

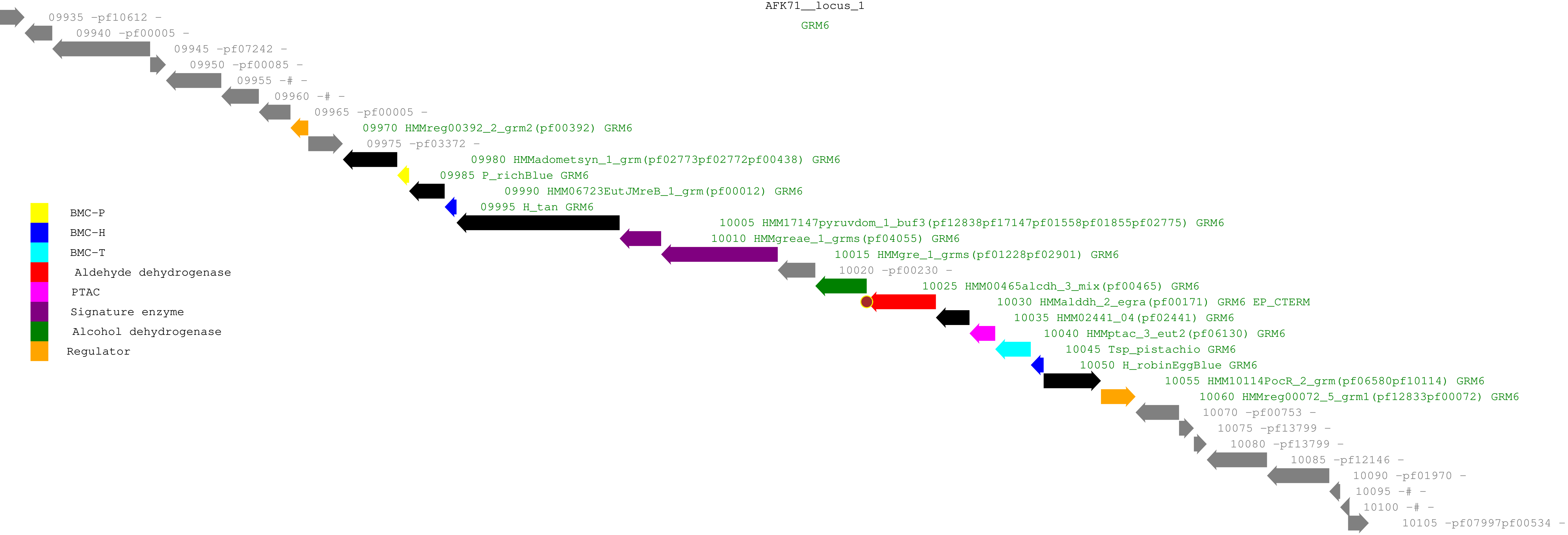

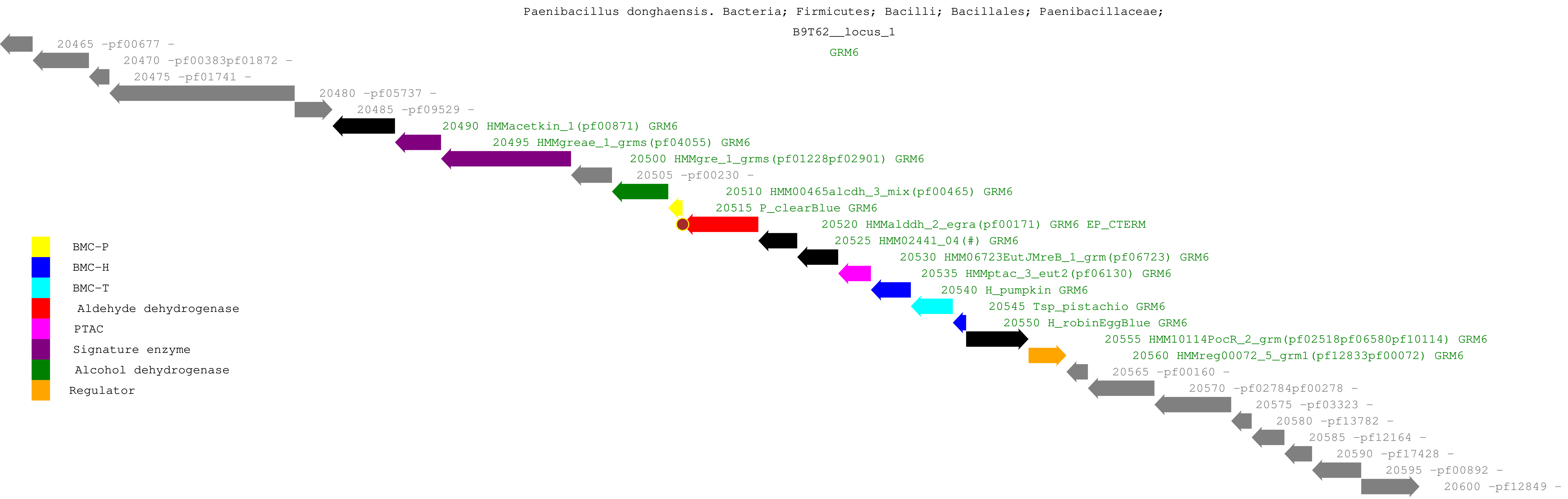
Paenibacillus donghaensis. Bacteria; Firmicutes; Bacilli; Bacillales; Paenibacillaceae;
B9T62__locus_1
GRM6
20465 -pf00677 -
20470 -pf00383pf01872 -
20475 -pf01741 -
20480 -pf05737 -
20485 -pf09529 -
20490 HMMacetkin_1(pf00871) GRM6
20495 HMMgreae_1_grms(pf04055) GRM6
20500 HMMgre_1_grms(pf01228pf02901) GRM6
20505 -pf00230 -
20510 HMM00465alcdh_3_mix(pf00465) GRM6
20515 P_clearBlue GRM6
20520 HMMalddh_2_egra(pf00171) GRM6 EP_CTERM
20525 HMM02441_04(#) GRM6
20530 HMM06723EutJMreB_1_grm(pf06723) GRM6
20535 HMMptac_3_eut2(pf06130) GRM6
20540 H_pumpkin GRM6
20545 Tsp_pistachio GRM6
20550 H_robinEggBlue GRM6
20555 HMM10114PocR_2_grm(pf02518pf06580pf10114) GRM6
20560 HMMreg00072_5_grm1(pf12833pf00072) GRM6
20565 -pf00160 -
20570 -pf02784pf00278 -
20575 -pf03323 -
20580 -pf13782 -
20585 -pf12164 -
20590 -pf17428 -
20595 -pf00892 -
20600 -pf12849 -
BMC-P
BMC-H
BMC-T
Aldehyde dehydrogenase
PTAC
Signature enzyme
Alcohol dehydrogenase
Regulator

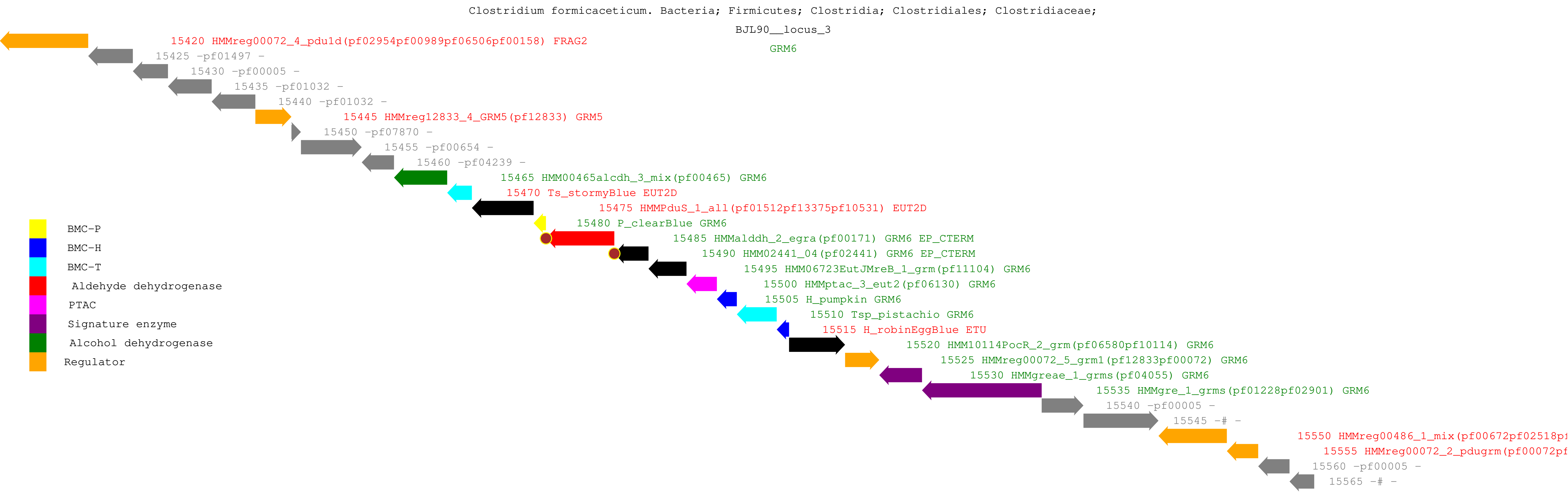
Clostridium formicaceticum. Bacteria; Firmicutes; Clostridia; Clostridiales; Clostridiaceae;
BJL90__locus_3
GRM6
15420 HMMreg00072_4_pdu1d(pf02954pf00989pf06506pf00158) FRAG2
15425 -pf01497 -
15430 -pf00005 -
15435 -pf01032 -
15440 -pf01032 -
15445 HMMreg12833_4_GRM5(pf12833) GRM5
15450 -pf07870 -
15455 -pf00654 -
15460 -pf04239 -
15465 HMM00465alcdh_3_mix(pf00465) GRM6
15470 Ts_stormyBlue EUT2D
15475 HMMPduS_1_all(pf01512pf13375pf10531) EUT2D
15480 P_clearBlue GRM6
15485 HMMalddh_2_egra(pf00171) GRM6 EP_CTERM
15490 HMM02441_04(pf02441) GRM6 EP_CTERM
15495 HMM06723EutJMreB_1_grm(pf11104) GRM6
15500 HMMptac_3_eut2(pf06130) GRM6
15505 H_pumpkin GRM6
15510 Tsp_pistachio GRM6
15515 H_robinEggBlue ETU
15520 HMM10114PocR_2_grm(pf06580pf10114) GRM6
15525 HMMreg00072_5_grm1(pf12833pf00072) GRM6
15530 HMMgreae_1_grms(pf04055) GRM6
15535 HMMgre_1_grms(pf01228pf02901) GRM6
15540 -pf00005 -
15545 -# -
15550 HMMreg00486_1_mix(pf00672pf02518pf
15555 HMMreg00072_2_pdugrm(pf00072pf
15560 -pf00005 -
15565 -# -
BMC-P
BMC-H
BMC-T
Aldehyde dehydrogenase
PTAC
Signature enzyme
Alcohol dehydrogenase
Regulator

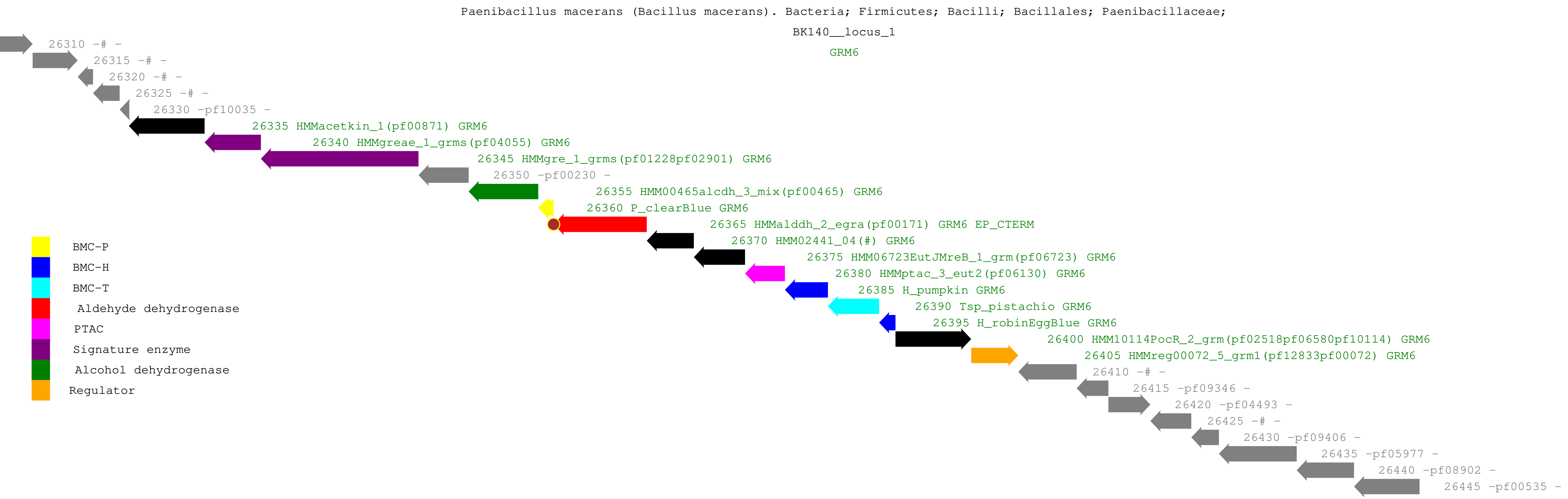
Paenibacillus macerans (Bacillus macerans). Bacteria; Firmicutes; Bacilli; Bacillales; Paenibacillaceae;
BK140__locus_1
GRM6
26310 -# -
26315 -# -
26320 -# -
26325 -# -
26330 -pf10035 -
26335 HMMacetkin_1(pf00871) GRM6
26340 HMMgreae_1_grms(pf04055) GRM6
26345 HMMgre_1_grms(pf01228pf02901) GRM6
26350 -pf00230 -
26355 HMM00465alcdh_3_mix(pf00465) GRM6
26360 P_clearBlue GRM6
26365 HMMalddh_2_egra(pf00171) GRM6 EP_CTERM
26370 HMM02441_04(#) GRM6
26375 HMM06723EutJMreB_1_grm(pf06723) GRM6
26380 HMMptac_3_eut2(pf06130) GRM6
26385 H_pumpkin GRM6
26390 Tsp_pistachio GRM6
26395 H_robinEggBlue GRM6
26400 HMM10114PocR_2_grm(pf02518pf06580pf10114) GRM6
26405 HMMreg00072_5_grm1(pf12833pf00072) GRM6
26410 -# -
26415 -pf09346 -
26420 -pf04493 -
26425 -# -
26430 -pf09406 -
26435 -pf05977 -
26440 -pf08902 -
26445 -pf00535 -
BMC-P
BMC-H
BMC-T
Aldehyde dehydrogenase
PTAC
Signature enzyme
Alcohol dehydrogenase
Regulator

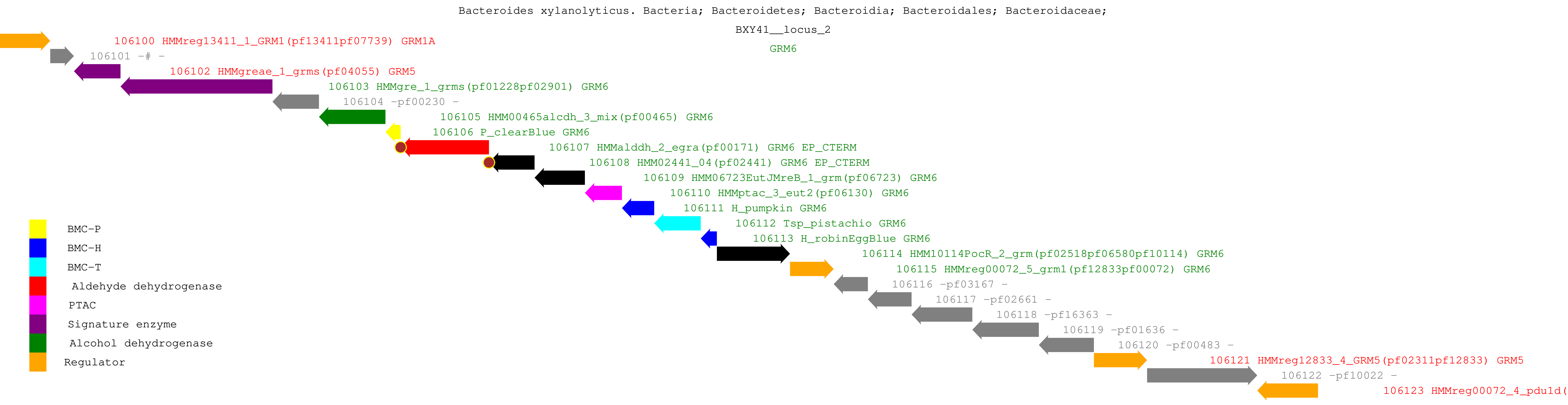

Bacteroides xylanolyticus. Bacteria; Bacteroidetes; Bacteroidia; Bacteroidales; Bacteroidaceae;
BXY41__locus_2
GRM6
106100 HMMreg13411_1_GRM1(pf13411pf07739) GRM1A
106101 -# -
106102 HMMgreae_1_grms(pf04055) GRM5
106103 HMMgre_1_grms(pf01228pf02901) GRM6
106104 -pf00230 -
106105 HMM00465alcdh_3_mix(pf00465) GRM6
106106 P_clearBlue GRM6
106107 HMMalddh_2_egra(pf00171) GRM6 EP_CTERM
106108 HMM02441_04(pf02441) GRM6 EP_CTERM
106109 HMM06723EutJMreB_1_grm(pf06723) GRM6
106110 HMMptac_3_eut2(pf06130) GRM6
106111 H_pumpkin GRM6
106112 Tsp_pistachio GRM6
106113 H_robinEggBlue GRM6
106114 HMM10114PocR_2_grm(pf02518pf06580pf10114) GRM6
106115 HMMreg00072_5_grm1(pf12833pf00072) GRM6
106116 -pf03167 -
106117 -pf02661 -
106118 -pf16363 -
106119 -pf01636 -
106120 -pf00483 -
106121 HMMreg12833_4_GRM5(pf02311pf12833) GRM5
106122 -pf10022 -
106123 HMMreg00072_4_pdu1d(
BMC-P
BMC-H
BMC-T
Aldehyde dehydrogenase
PTAC
Signature enzyme
Alcohol dehydrogenase
Regulator

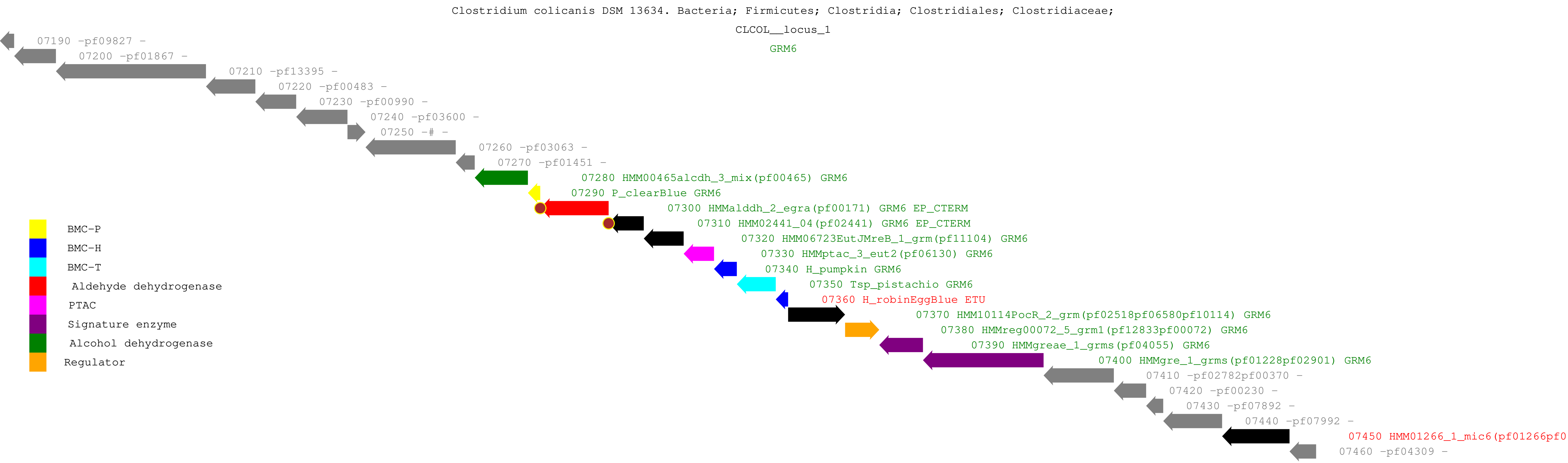
Clostridium colicanis DSM 13634. Bacteria; Firmicutes; Clostridia; Clostridiales; Clostridiaceae;
CLCOL__locus_1
GRM6
07190 -pf09827 -
07200 -pf01867 -
07210 -pf13395 -
07220 -pf00483 -
07230 -pf00990 -
07240 -pf03600 -
07250 -# -
07260 -pf03063 -
07270 -pf01451 -
07280 HMM00465alcdh_3_mix(pf00465) GRM6
07290 P_clearBlue GRM6
07300 HMMalddh_2_egra(pf00171) GRM6 EP_CTERM
07310 HMM02441_04(pf02441) GRM6 EP_CTERM
07320 HMM06723EutJMreB_1_grm(pf11104) GRM6
07330 HMMptac_3_eut2(pf06130) GRM6
07340 H_pumpkin GRM6
07350 Tsp_pistachio GRM6
07360 H_robinEggBlue ETU
07370 HMM10114PocR_2_grm(pf02518pf06580pf10114) GRM6
07380 HMMreg00072_5_grm1(pf12833pf00072) GRM6
07390 HMMgreae_1_grms(pf04055) GRM6
07400 HMMgre_1_grms(pf01228pf02901) GRM6
07410 -pf02782pf00370 -
07420 -pf00230 -
07430 -pf07892 -
07440 -pf07992 -
07450 HMM01266_1_mic6(pf01266pf0
07460 -pf04309 -
BMC-P
BMC-H
BMC-T
Aldehyde dehydrogenase
PTAC
Signature enzyme
Alcohol dehydrogenase
Regulator

Clostridium magnum DSM 2767. Bacteria; Firmicutes; Clostridia; Clostridiales; Clostridiaceae;

CLMAG__locus_3

GRM6

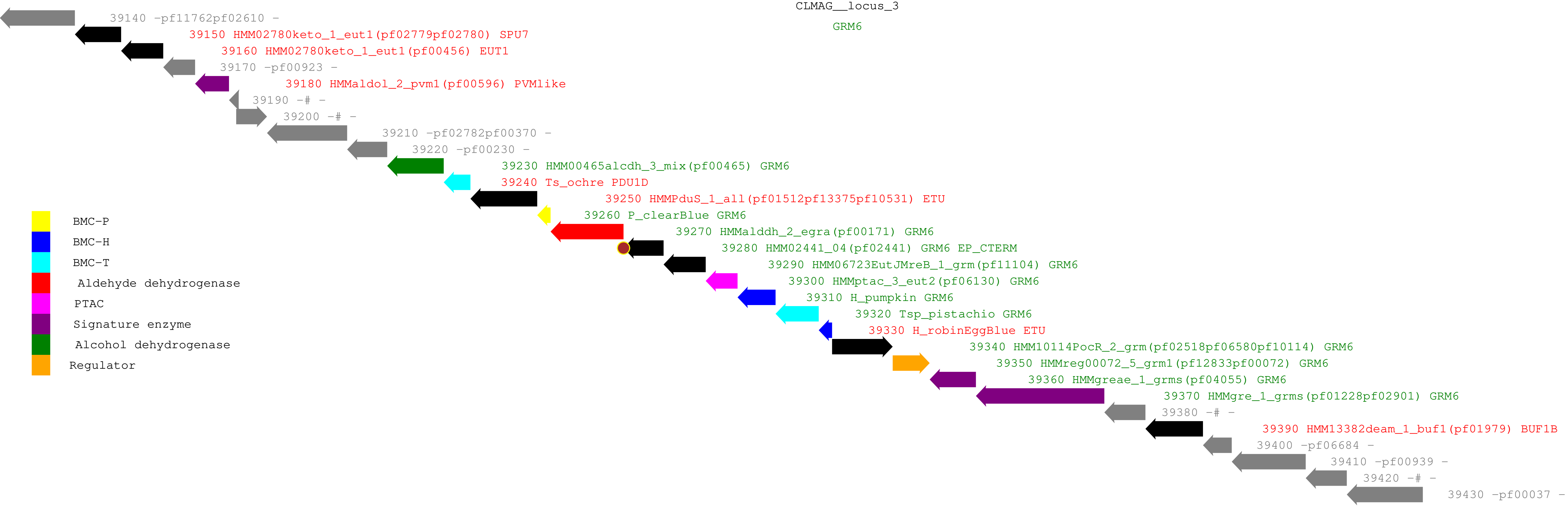

Clostridium thermopalmarium DSM 5974. Bacteria; Firmicutes; Clostridia; Clostridiales; Clostridiaceae;

CPAL__locus_1

GRM6

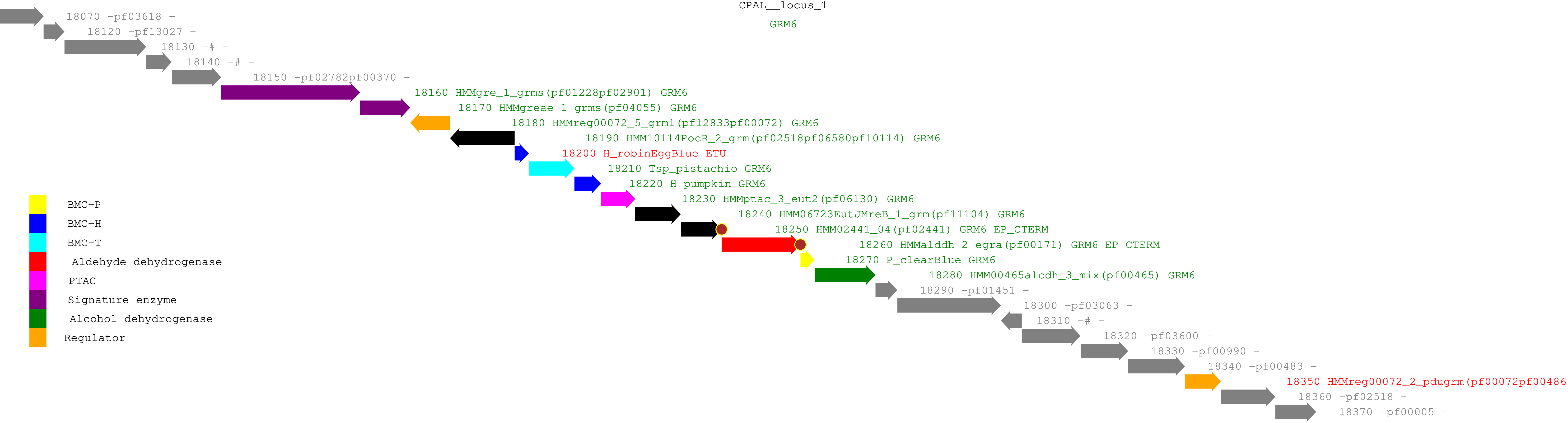

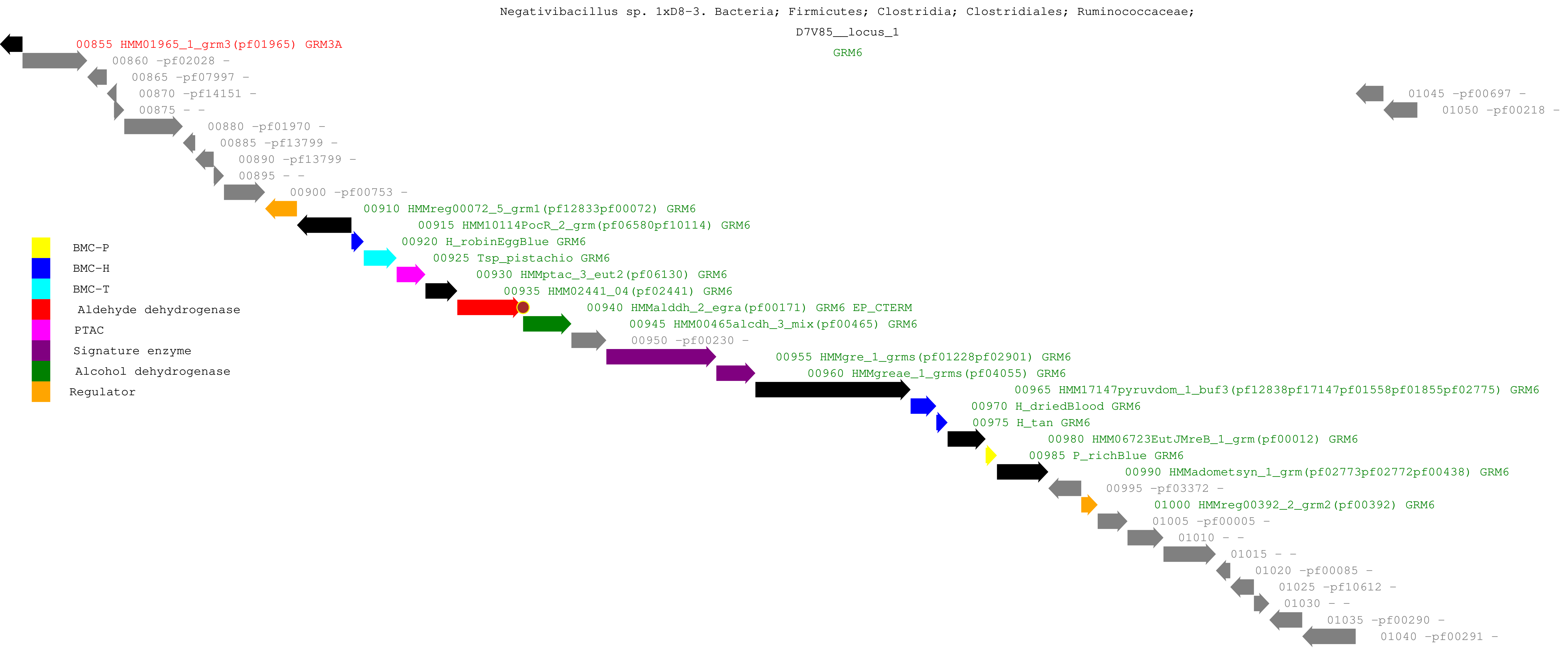
Negativibacillus sp. 1xD8-3. Bacteria; Firmicutes; Clostridia; Clostridiales; Ruminococcaceae;
D7V85__locus_1
GRM6
00855 HMM01965_1_grm3(pf01965) GRM3A
00860 -pf02028 -
00865 -pf07997 -
00870 -pf14151 -
00875 - -
00880 -pf01970 -
00885 -pf13799 -
00890 -pf13799 -
00895 - -
00900 -pf00753 -
00910 HMMreg00072_5_grm1(pf12833pf00072) GRM6
00915 HMM10114PocR_2_grm(pf06580pf10114) GRM6
00920 H_robinEggBlue GRM6
00925 Tsp_pistachio GRM6
00930 HMMptac_3_eut2(pf06130) GRM6
00935 HMM02441_04(pf02441) GRM6
00940 HMMalddh_2_egra(pf00171) GRM6 EP_CTERM
00945 HMM00465alcdh_3_mix(pf00465) GRM6
00950 -pf00230 -
00955 HMMgre_1_grms(pf01228pf02901) GRM6
00960 HMMgreae_1_grms(pf04055) GRM6
00965 HMM17147pyruvdom_1_buf3(pf12838pf17147pf01558pf01855pf02775) GRM6
00970 H_driedBlood GRM6
00975 H_tan GRM6
00980 HMM06723EutJMreB_1_grm(pf00012) GRM6
00985 P_richBlue GRM6
00990 HMMadometsyn_1_grm(pf02773pf02772pf00438) GRM6
00995 -pf03372 -
01000 HMMreg00392_2_grm2(pf00392) GRM6
01005 -pf00005 -
01010 - -
01015 - -
01020 -pf00085 -
01025 -pf10612 -
01030 - -
01035 -pf00290 -
01040 -pf00291 -
01045 -pf00697 -
01050 -pf00218 -
BMC-P
BMC-H
BMC-T
Aldehyde dehydrogenase
PTAC
Signature enzyme
Alcohol dehydrogenase
Regulator

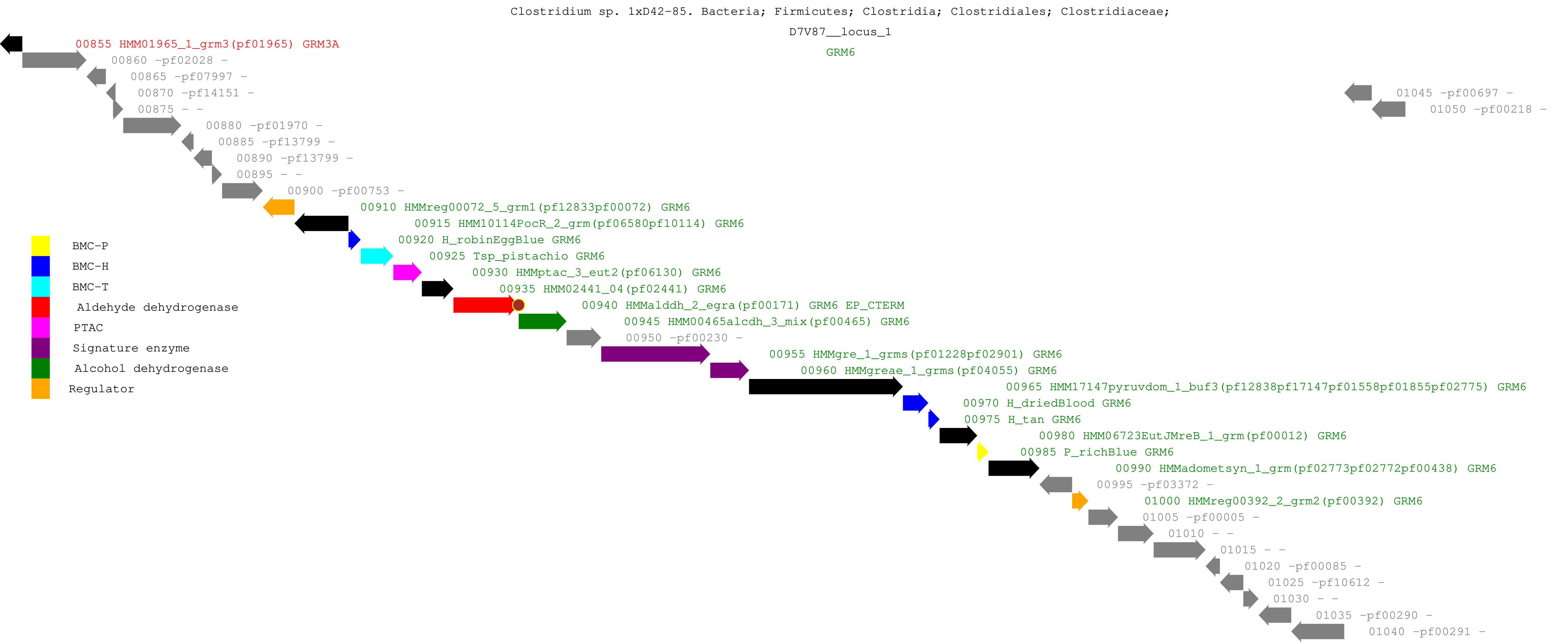
Clostridium sp. 1xD42-85. Bacteria; Firmicutes; Clostridia; Clostridiales; Clostridiaceae;
D7V87__locus_1
GRM6
00855 HMM01965_1_grm3(pf01965) GRM3A
00860 -pf02028 -
00865 -pf07997 -
00870 -pf14151 -
00875 - -
00880 -pf01970 -
00885 -pf13799 -
00890 -pf13799 -
00895 - -
00900 -pf00753 -
00910 HMMreg00072_5_grm1(pf12833pf00072) GRM6
00915 HMM10114PocR_2_grm(pf06580pf10114) GRM6
00920 H_robinEggBlue GRM6
00925 Tsp_pistachio GRM6
00930 HMMptac_3_eut2(pf06130) GRM6
00935 HMM02441_04(pf02441) GRM6
00940 HMMalddh_2_egra(pf00171) GRM6 EP_CTERM
00945 HMM00465alcdh_3_mix(pf00465) GRM6
00950 -pf00230 -
00955 HMMgre_1_grms(pf01228pf02901) GRM6
00960 HMMgreae_1_grms(pf04055) GRM6
00965 HMM17147pyruvdom_1_buf3(pf12838pf17147pf01558pf01855pf02775) GRM6
00970 H_driedBlood GRM6
00975 H_tan GRM6
00980 HMM06723EutJMreB_1_grm(pf00012) GRM6
00985 P_richBlue GRM6
00990 HMMadometsyn_1_grm(pf02773pf02772pf00438) GRM6
00995 -pf03372 -
01000 HMMreg00392_2_grm2(pf00392) GRM6
01005 -pf00005 -
01010 - -
01015 - -
01020 -pf00085 -
01025 -pf10612 -
01030 - -
01035 -pf00290 -
01040 -pf00291 -
01045 -pf00697 -
01050 -pf00218 -
BMC-P
BMC-H
BMC-T
Aldehyde dehydrogenase
PTAC
Signature enzyme
Alcohol dehydrogenase
Regulator

bacterium 1XD42-44. Bacteria.

D7Y14__locus_1

GRM6

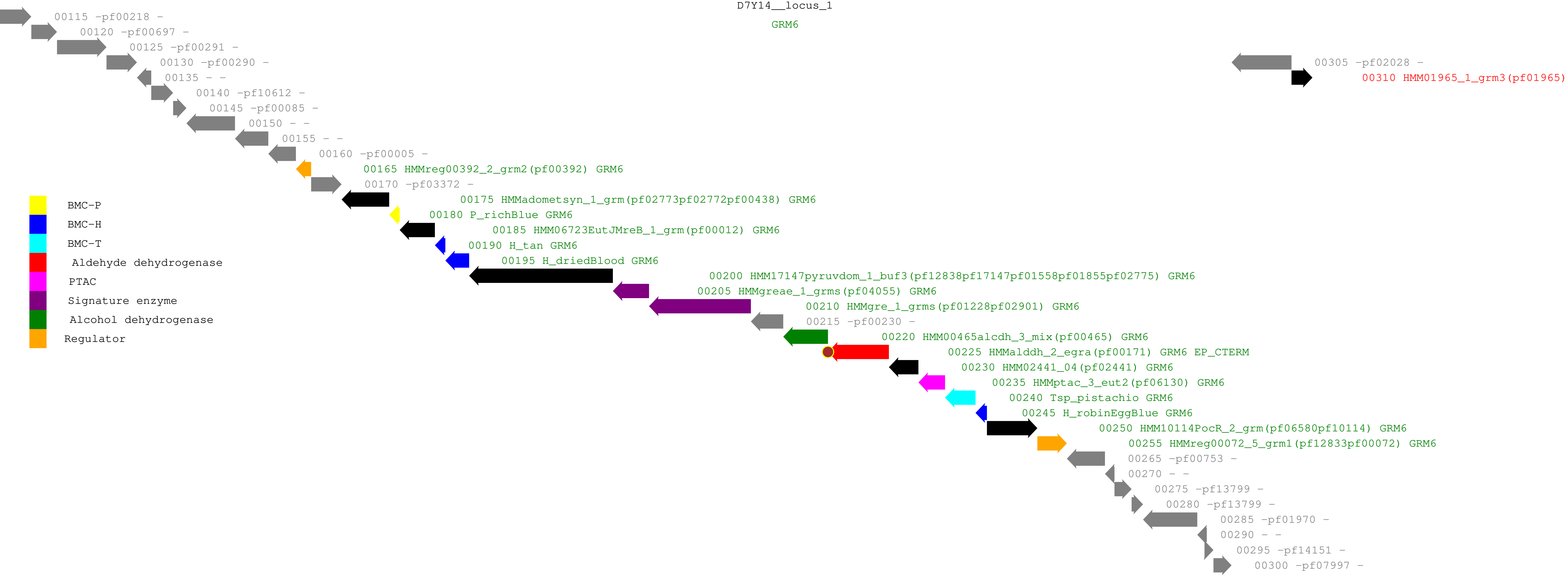

Paenibacillus macerans (Bacillus macerans). Bacteria; Firmicutes; Bacilli; Bacillales; Paenibacillaceae;

DJ90__locus_1

GRM6

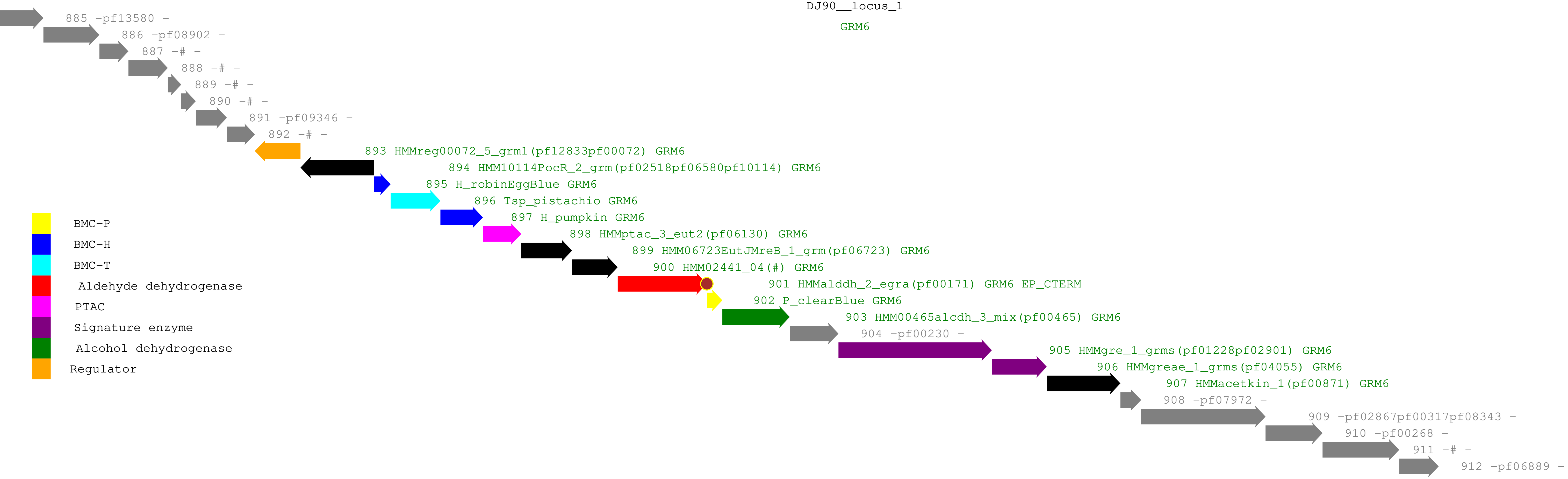

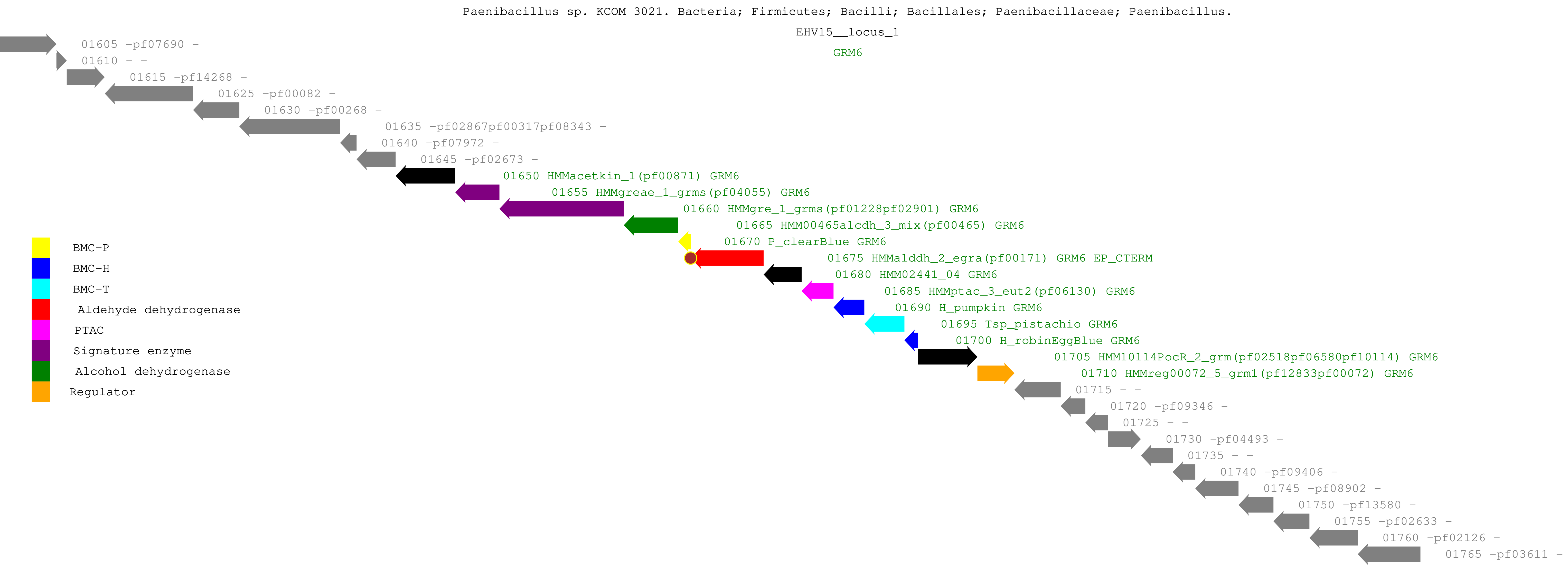
Paenibacillus sp. KCOM 3021. Bacteria; Firmicutes; Bacilli; Bacillales; Paenibacillaceae; Paenibacillus.
EHV15__locus_1
GRM6
01605 -pf07690 -
01610 - -
01615 -pf14268 -
01625 -pf00082 -
01630 -pf00268 -
01635 -pf02867pf00317pf08343 -
01640 -pf07972 -
01645 -pf02673 -
01650 HMMacetkin_1(pf00871) GRM6
01655 HMMgreae_1_grms(pf04055) GRM6
01660 HMMgre_1_grms(pf01228pf02901) GRM6
01665 HMM00465alcdh_3_mix(pf00465) GRM6
01670 P_clearBlue GRM6
01675 HMMalddh_2_egra(pf00171) GRM6 EP_CTERM
01680 HMM02441_04 GRM6
01685 HMMptac_3_eut2(pf06130) GRM6
01690 H_pumpkin GRM6
01695 Tsp_pistachio GRM6
01700 H_robinEggBlue GRM6
01705 HMM10114PocR_2_grm(pf02518pf06580pf10114) GRM6
01710 HMMreg00072_5_grm1(pf12833pf00072) GRM6
01715 - -
01720 -pf09346 -
01725 - -
01730 -pf04493 -
01735 - -
01740 -pf09406 -
01745 -pf08902 -
01750 -pf13580 -
01755 -pf02633 -
01760 -pf02126 -
01765 -pf03611 -
BMC-P
BMC-H
BMC-T
Aldehyde dehydrogenase
PTAC
Signature enzyme
Alcohol dehydrogenase
Regulator

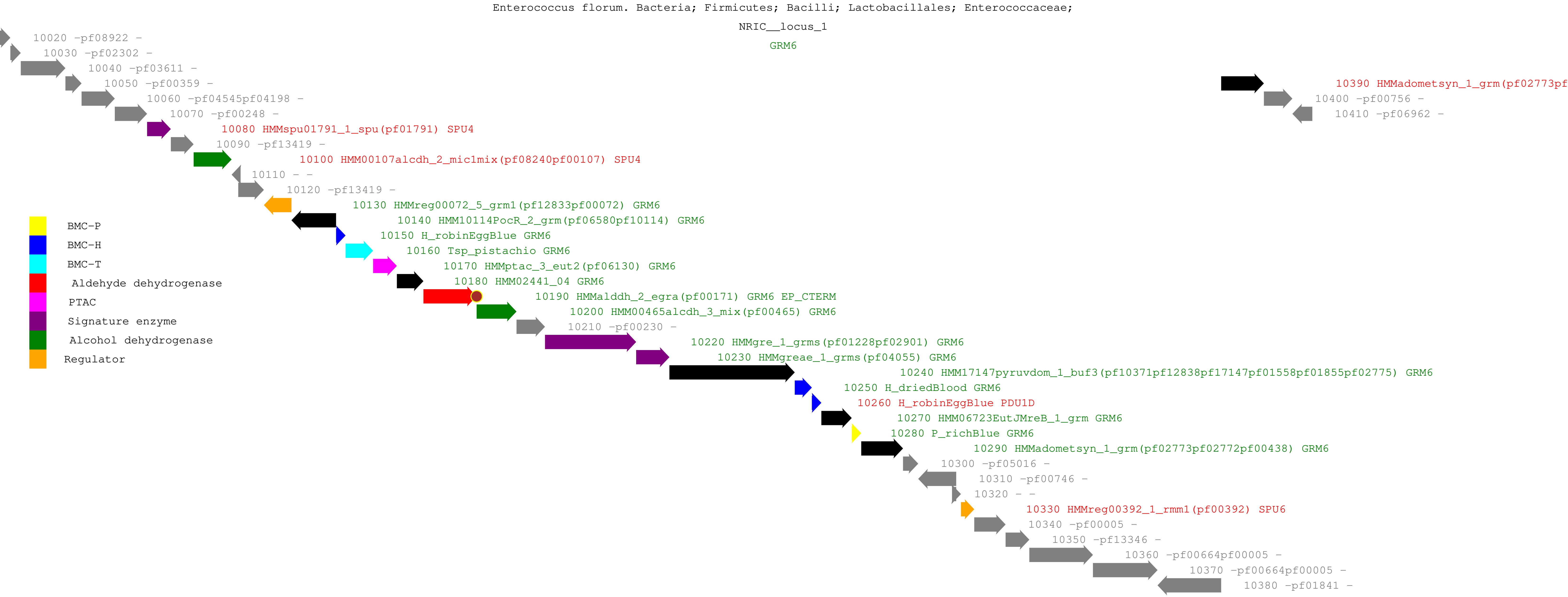

Enterococcus florum. Bacteria; Firmicutes; Bacilli; Lactobacillales; Enterococcaceae;
NRIC__locus_1
GRM6
10020 -pf08922 -
10030 -pf02302 -
10040 -pf03611 -
10050 -pf00359 -
10060 -pf04545pf04198 -
10070 -pf00248 -
10080 HMMspu01791_1_spu(pf01791) SPU4
10090 -pf13419 -
10100 HMM00107alcdh_2_mic1mix(pf08240pf00107) SPU4
10110 - -
10120 -pf13419 -
10130 HMMreg00072_5_grm1(pf12833pf00072) GRM6
10140 HMM10114PocR_2_grm(pf06580pf10114) GRM6
10150 H_robinEggBlue GRM6
10160 Tsp_pistachio GRM6
10170 HMMptac_3_eut2(pf06130) GRM6
10180 HMM02441_04 GRM6
10190 HMMalddh_2_egra(pf00171) GRM6 EP_CTERM
10200 HMM00465alcdh_3_mix(pf00465) GRM6
10210 -pf00230 -
10220 HMMgre_1_grms(pf01228pf02901) GRM6
10230 HMMgreae_1_grms(pf04055) GRM6
10240 HMM17147pyruvdom_1_buf3(pf10371pf12838pf17147pf01558pf01855pf02775) GRM6
10250 H_driedBlood GRM6
10260 H_robinEggBlue PDU1D
10270 HMM06723EutJMreB_1_grm GRM6
10280 P_richBlue GRM6
10290 HMMadometsyn_1_grm(pf02773pf02772pf00438) GRM6
10300 -pf05016 -
10310 -pf00746 -
10320 - -
10330 HMMreg00392_1_rmm1(pf00392) SPU6
10340 -pf00005 -
10350 -pf13346 -
10360 -pf00664pf00005 -
10370 -pf00664pf00005 -
10380 -pf01841 -
10390 HMMadometsyn_1_grm(pf02773pf
10400 -pf00756 -
10410 -pf06962 -
BMC-P
BMC-H
BMC-T
Aldehyde dehydrogenase
PTAC
Signature enzyme
Alcohol dehydrogenase
Regulator

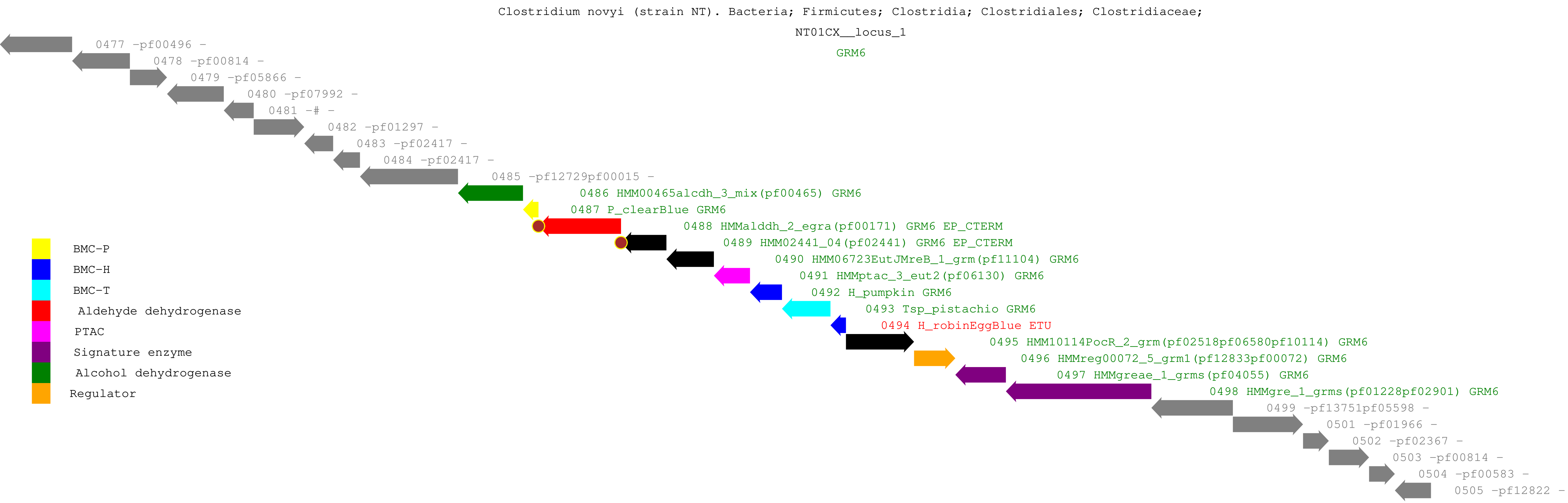
Clostridium novyi (strain NT). Bacteria; Firmicutes; Clostridia; Clostridiales; Clostridiaceae;
NT01CX__locus_1
GRM6
0477 -pf00496 -
0478 -pf00814 -
0479 -pf05866 -
0480 -pf07992 -
0481 -# -
0482 -pf01297 -
0483 -pf02417 -
0484 -pf02417 -
0485 -pf12729pf00015 -
0486 HMM00465alcdh_3_mix(pf00465) GRM6
0487 P_clearBlue GRM6
0488 HMMalddh_2_egra(pf00171) GRM6 EP_CTERM
0489 HMM02441_04(pf02441) GRM6 EP_CTERM
0490 HMM06723EutJMreB_1_grm(pf11104) GRM6
0491 HMMptac_3_eut2(pf06130) GRM6
0492 H_pumpkin GRM6
0493 Tsp_pistachio GRM6
0494 H_robinEggBlue ETU
0495 HMM10114PocR_2_grm(pf02518pf06580pf10114) GRM6
0496 HMMreg00072_5_grm1(pf12833pf00072) GRM6
0497 HMMgreae_1_grms(pf04055) GRM6
0498 HMMgre_1_grms(pf01228pf02901) GRM6
0499 -pf13751pf05598 -
0501 -pf01966 -
0502 -pf02367 -
0503 -pf00814 -
0504 -pf00583 -
0505 -pf12822 -
BMC-P
BMC-H
BMC-T
Aldehyde dehydrogenase
PTAC
Signature enzyme
Alcohol dehydrogenase
Regulator

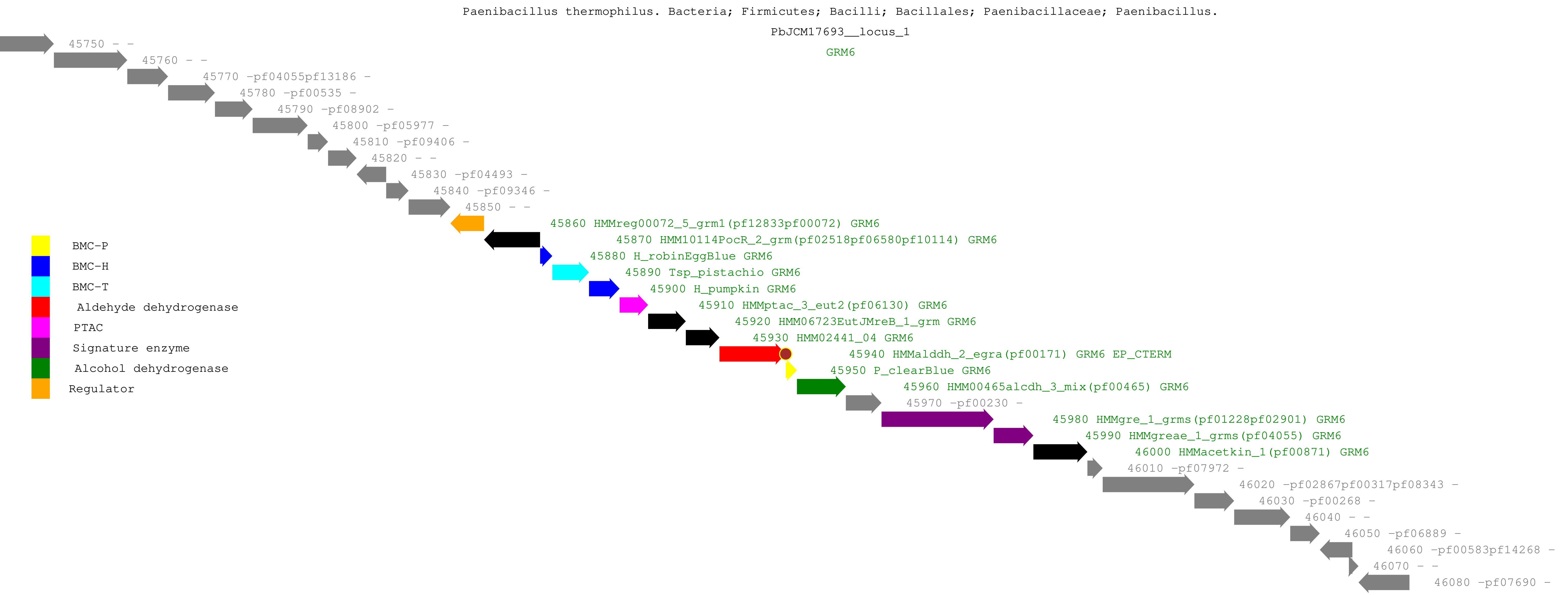
Paenibacillus thermophilus. Bacteria; Firmicutes; Bacilli; Bacillales; Paenibacillaceae; Paenibacillus.
PbJCM17693__locus_1
GRM6
45750 - -
45760 - -
45770 -pf04055pf13186 -
45780 -pf00535 -
45790 -pf08902 -
45800 -pf05977 -
45810 -pf09406 -
45820 - -
45830 -pf04493 -
45840 -pf09346 -
45850 - -
45860 HMMreg00072_5_grm1(pf12833pf00072) GRM6
45870 HMM10114PocR_2_grm(pf02518pf06580pf10114) GRM6
45880 H_robinEggBlue GRM6
45890 Tsp_pistachio GRM6
45900 H_pumpkin GRM6
45910 HMMptac_3_eut2(pf06130) GRM6
45920 HMM06723EutJMreB_1_grm GRM6
45930 HMM02441_04 GRM6
45940 HMMalddh_2_egra(pf00171) GRM6 EP_CTERM
45950 P_clearBlue GRM6
45960 HMM00465alcdh_3_mix(pf00465) GRM6
45970 -pf00230 -
45980 HMMgre_1_grms(pf01228pf02901) GRM6
45990 HMMgreae_1_grms(pf04055) GRM6
46000 HMMacetkin_1(pf00871) GRM6
46010 -pf07972 -
46020 -pf02867pf00317pf08343 -
46030 -pf00268 -
46040 - -
46050 -pf06889 -
46060 -pf00583pf14268 -
46070 - -
46080 -pf07690 -
BMC-P
BMC-H
BMC-T
Aldehyde dehydrogenase
PTAC
Signature enzyme
Alcohol dehydrogenase
Regulator

Desulfotomaculum putei DSM 12395. Bacteria; Firmicutes; Clostridia; Clostridiales; Peptococcaceae;

SAMN02745133__locus_1

GRM6

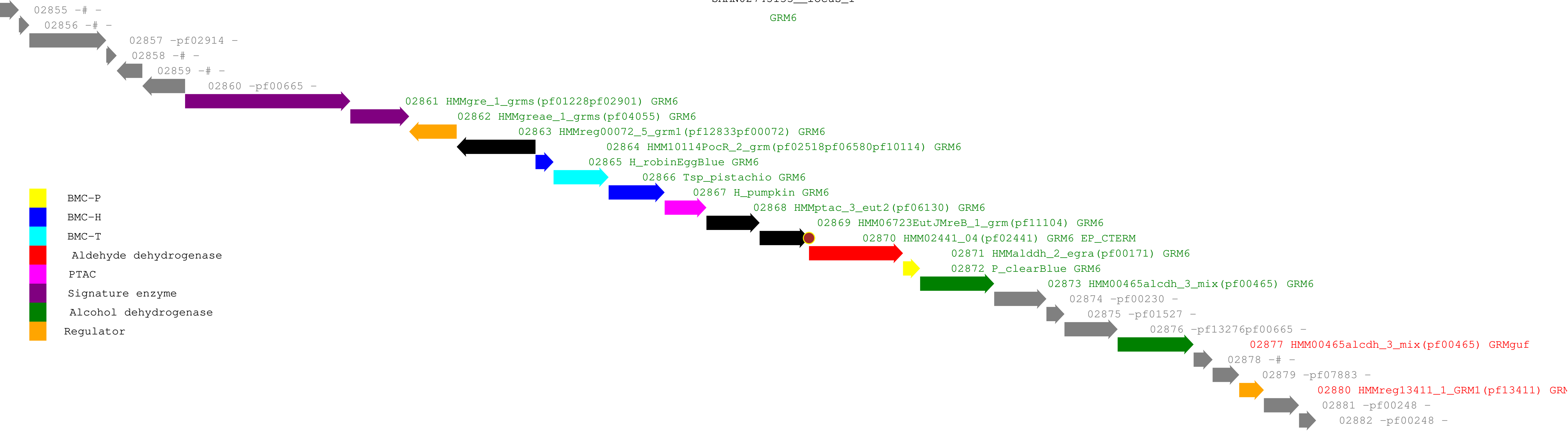

Propionispora hippei DSM 15287. Bacteria; Firmicutes; Negativicutes; Selenomonadales; Sporomusaceae;

SAMN02745170__locus_1

GRM6

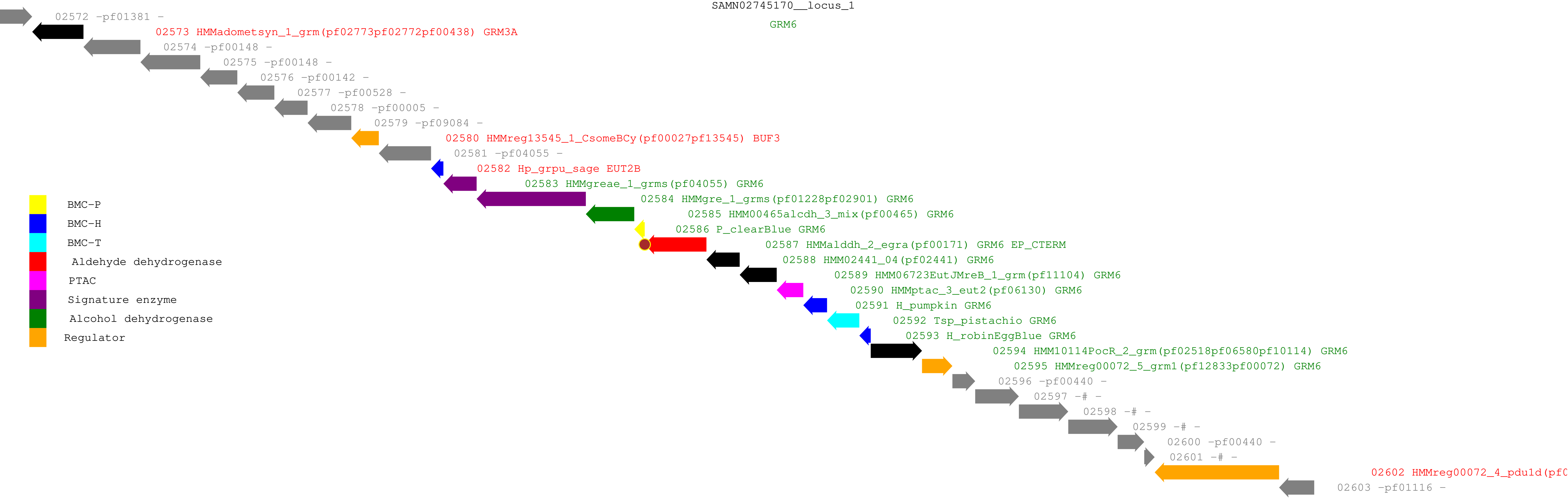

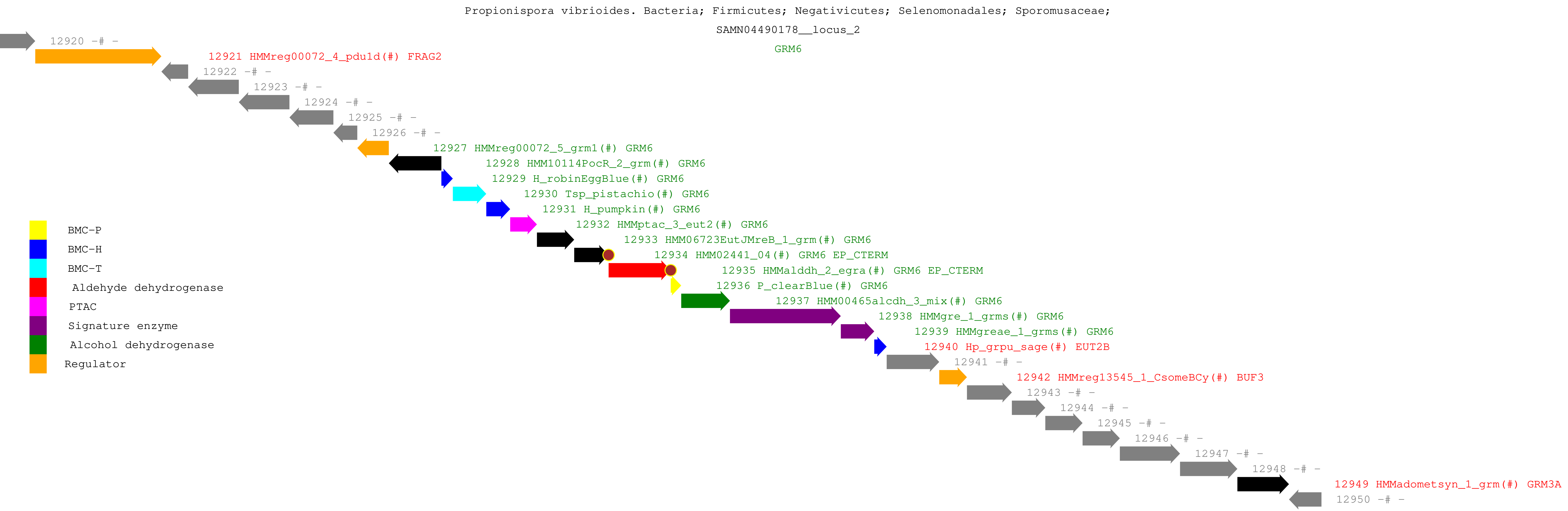
Propionispora vibrioides. Bacteria; Firmicutes; Negativicutes; Selenomonadales; Sporomusaceae;
SAMN04490178__locus_2
GRM6
12920 -# -
12921 HMMreg00072_4_pdu1d(#) FRAG2
12922 -# -
12923 -# -
12924 -# -
12925 -# -
12926 -# -
12927 HMMreg00072_5_grm1(#) GRM6
12928 HMM10114PocR_2_grm(#) GRM6
12929 H_robinEggBlue(#) GRM6
12930 Tsp_pistachio(#) GRM6
12931 H_pumpkin(#) GRM6
12932 HMMptac_3_eut2(#) GRM6
12933 HMM06723EutJMreB_1_grm(#) GRM6
12934 HMM02441_04(#) GRM6 EP_CTERM
12935 HMMalddh_2_egra(#) GRM6 EP_CTERM
12936 P_clearBlue(#) GRM6
12937 HMM00465alcdh_3_mix(#) GRM6
12938 HMMgre_1_grms(#) GRM6
12939 HMMgreae_1_grms(#) GRM6
12940 Hp_grpu_sage(#) EUT2B
12941 -# -
12942 HMMreg13545_1_CsomeBCy(#) BUF3
12943 -# -
12944 -# -
12945 -# -
12946 -# -
12947 -# -
12948 -# -
12949 HMMadometsyn_1_grm(#) GRM3A
12950 -# -
BMC-P
BMC-H
BMC-T
Aldehyde dehydrogenase
PTAC
Signature enzyme
Alcohol dehydrogenase
Regulator

Desulfitobacterium hafniense (Desulfitobacterium frappieri). Bacteria; Firmicutes; Clostridia; Clostridiales; Peptococcaceae;

AT727__locus_3

GRMguf

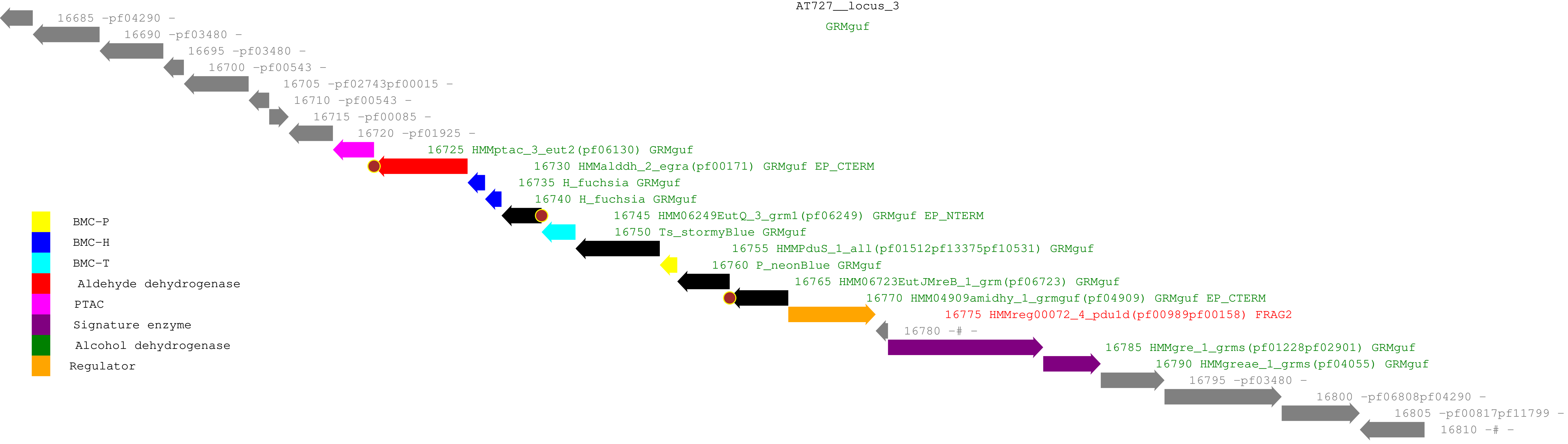

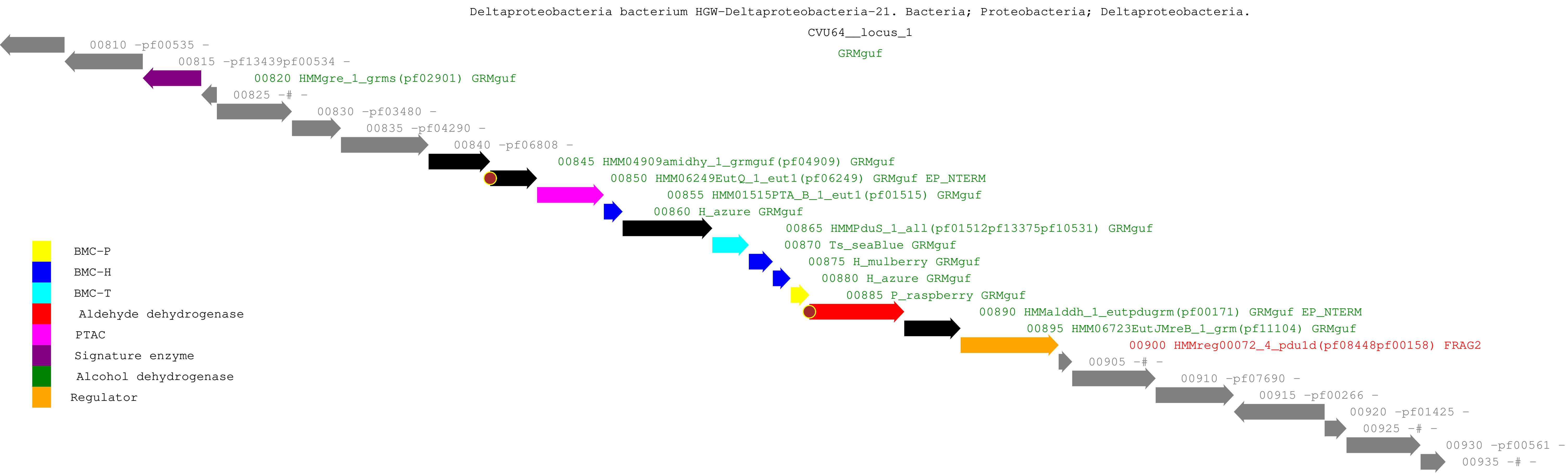
Deltaproteobacteria bacterium HGW-Deltaproteobacteria-21. Bacteria; Proteobacteria; Deltaproteobacteria.
CVU64__locus_1
GRMguf
00810 -pf00535 -
00815 -pf13439pf00534 -
00820 HMMgre_1_grms(pf02901) GRMguf
00825 -# -
00830 -pf03480 -
00835 -pf04290 -
00840 -pf06808 -
00845 HMM04909amidhy_1_grmguf(pf04909) GRMguf
00850 HMM06249EutQ_1_eut1(pf06249) GRMguf EP_NTERM
00855 HMM01515PTA_B_1_eut1(pf01515) GRMguf
00860 H_azure GRMguf
00865 HMMPduS_1_all(pf01512pf13375pf10531) GRMguf
00870 Ts_seaBlue GRMguf
00875 H_mulberry GRMguf
00880 H_azure GRMguf
00885 P_raspberry GRMguf
00890 HMMalddh_1_eutpdugrm(pf00171) GRMguf EP_NTERM
00895 HMM06723EutJMreB_1_grm(pf11104) GRMguf
00900 HMMreg00072_4_pdu1d(pf08448pf00158) FRAG2
00905 -# -
00910 -pf07690 -
00915 -pf00266 -
00920 -pf01425 -
00925 -# -
00930 -pf00561 -
00935 -# -
BMC-P
BMC-H
BMC-T
Aldehyde dehydrogenase
PTAC
Signature enzyme
Alcohol dehydrogenase
Regulator

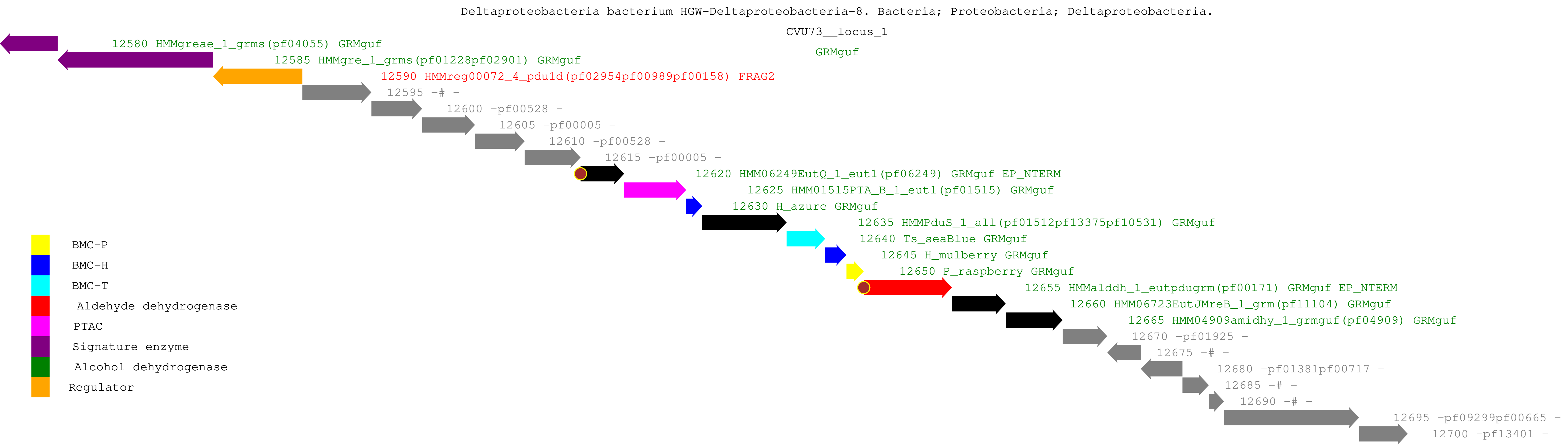

Deltaproteobacteria bacterium HGW-Deltaproteobacteria-8. Bacteria; Proteobacteria; Deltaproteobacteria.
CVU73__locus_1
GRMguf
12580 HMMgreae_1_grms(pf04055) GRMguf
12585 HMMgre_1_grms(pf01228pf02901) GRMguf
12590 HMMreg00072_4_pdu1d(pf02954pf00989pf00158) FRAG2
12595 -# -
12600 -pf00528 -
12605 -pf00005 -
12610 -pf00528 -
12615 -pf00005 -
12620 HMM06249EutQ_1_eut1(pf06249) GRMguf EP_NTERM
12625 HMM01515PTA_B_1_eut1(pf01515) GRMguf
12630 H_azure GRMguf
12635 HMMPduS_1_all(pf01512pf13375pf10531) GRMguf
12640 Ts_seaBlue GRMguf
12645 H_mulberry GRMguf
12650 P_raspberry GRMguf
12655 HMMalddh_1_eutpdugrm(pf00171) GRMguf EP_NTERM
12660 HMM06723EutJMreB_1_grm(pf11104) GRMguf
12665 HMM04909amidhy_1_grmguf(pf04909) GRMguf
12670 -pf01925 -
12675 -# -
12680 -pf01381pf00717 -
12685 -# -
12690 -# -
12695 -pf09299pf00665 -
12700 -pf13401 -
BMC-P
BMC-H
BMC-T
Aldehyde dehydrogenase
PTAC
Signature enzyme
Alcohol dehydrogenase
Regulator

Desulfosporosinus sp. Bacteria; Firmicutes; Clostridia; Clostridiales; Peptococcaceae;

DEF42__locus_2

GRMguf

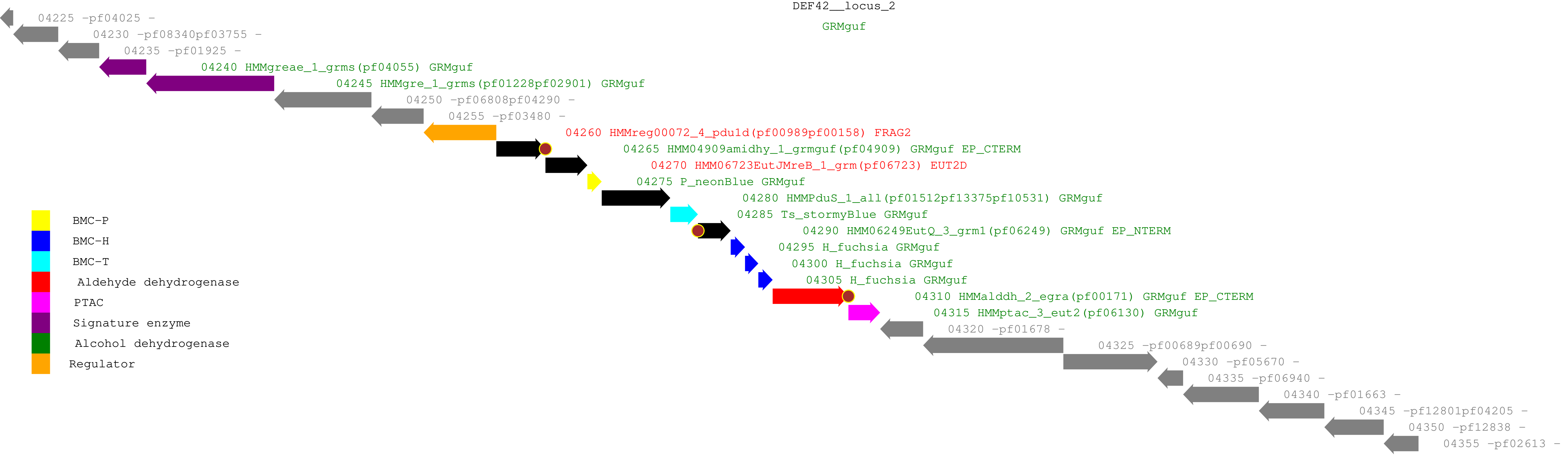

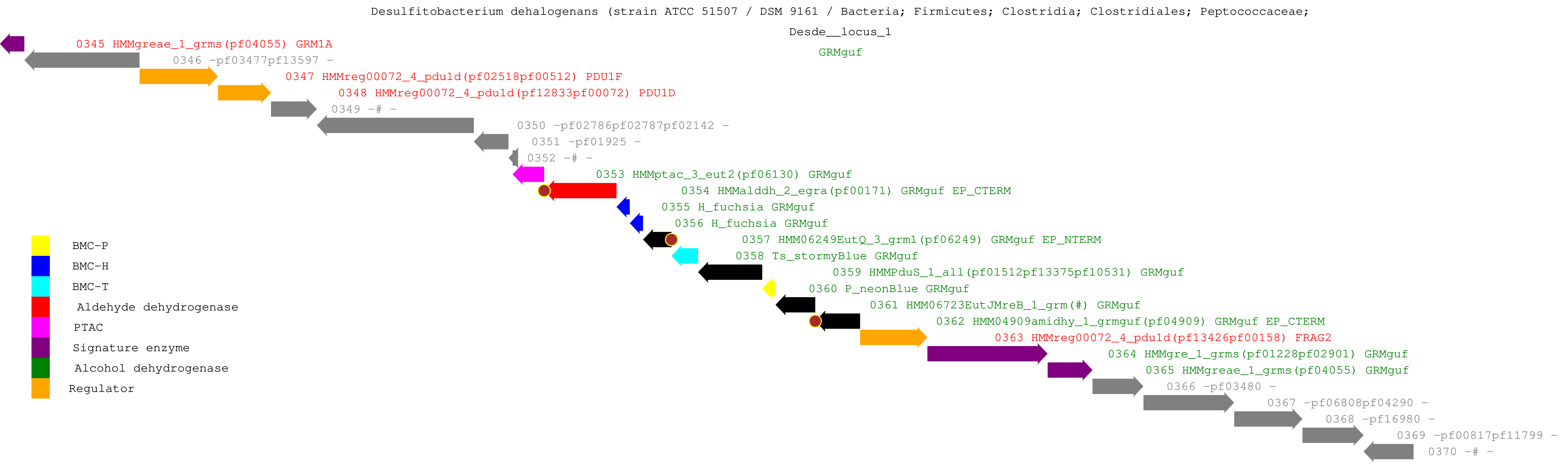
Desulfitobacterium dehalogenans (strain ATCC 51507 / DSM 9161 / Bacteria; Firmicutes; Clostridia; Clostridiales; Peptococcaceae;
Desde__locus_1
GRMguf
0345 HMMgreae_1_grms(pf04055) GRM1A
0346 -pf03477pf13597 -
0347 HMMreg00072_4_pdu1d(pf02518pf00512) PDU1F
0348 HMMreg00072_4_pdu1d(pf12833pf00072) PDU1D
0349 -# -
0350 -pf02786pf02787pf02142 -
0351 -pf01925 -
0352 -# -
0353 HMMptac_3_eut2(pf06130) GRMguf
0354 HMMalddh_2_egra(pf00171) GRMguf EP_CTERM
0355 H_fuchsia GRMguf
0356 H_fuchsia GRMguf
0357 HMM06249EutQ_3_grm1(pf06249) GRMguf EP_NTERM
0358 Ts_stormyBlue GRMguf
0359 HMMPduS_1_all(pf01512pf13375pf10531) GRMguf
0360 P_neonBlue GRMguf
0361 HMM06723EutJMreB_1_grm(#) GRMguf
0362 HMM04909amidhy_1_grmguf(pf04909) GRMguf EP_CTERM
0363 HMMreg00072_4_pdu1d(pf13426pf00158) FRAG2
0364 HMMgre_1_grms(pf01228pf02901) GRMguf
0365 HMMgreae_1_grms(pf04055) GRMguf
0366 -pf03480 -
0367 -pf06808pf04290 -
0368 -pf16980 -
0369 -pf00817pf11799 -
0370 -# -
BMC-P
BMC-H
BMC-T
Aldehyde dehydrogenase
PTAC
Signature enzyme
Alcohol dehydrogenase
Regulator

Desulfitobacterium sp. LBE. Bacteria; Firmicutes; Clostridia; Clostridiales; Peptococcaceae;

DesLBE__locus_1

GRMguf

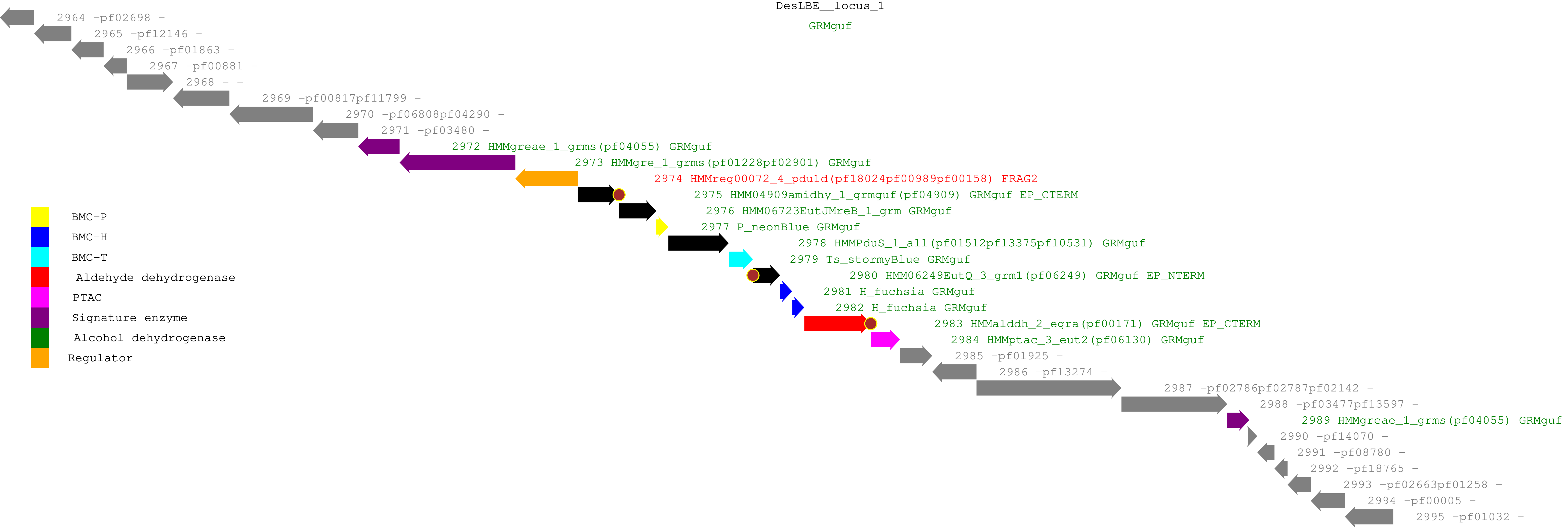

Desulfosporosinus youngiae DSM 17734. Bacteria; Firmicutes; Clostridia; Clostridiales; Peptococcaceae;

DesyoDRAFT__locus_2

GRMguf

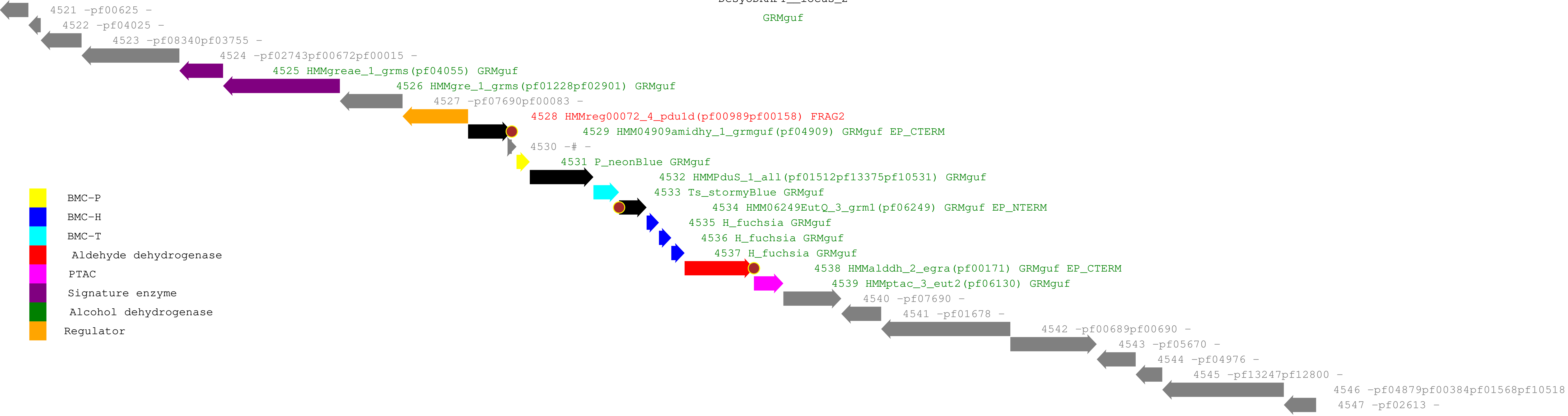

Desulfitobacterium hafniense (strain DCB-2 / DSM 10664). Bacteria; Firmicutes; Clostridia; Clostridiales; Peptococcaceae;

Dhaf__locus_1

GRMguf

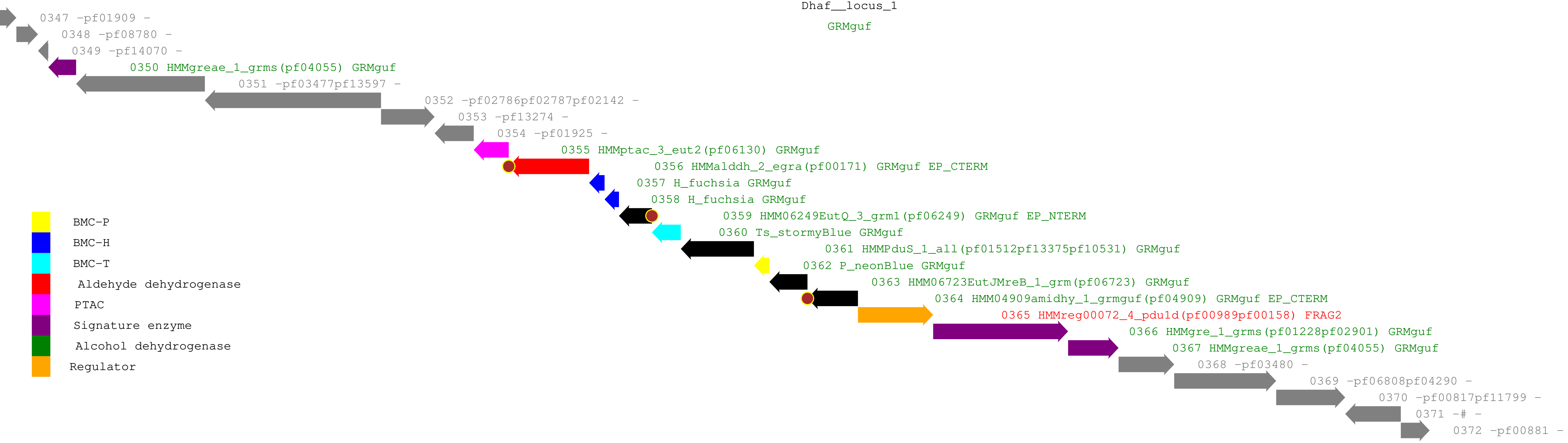

Desulfotalea psychrophila (strain LSv54 / DSM 12343). Bacteria; Proteobacteria; Deltaproteobacteria; Desulfobacterales;

DP_locus_1_main_sat

GRMguf

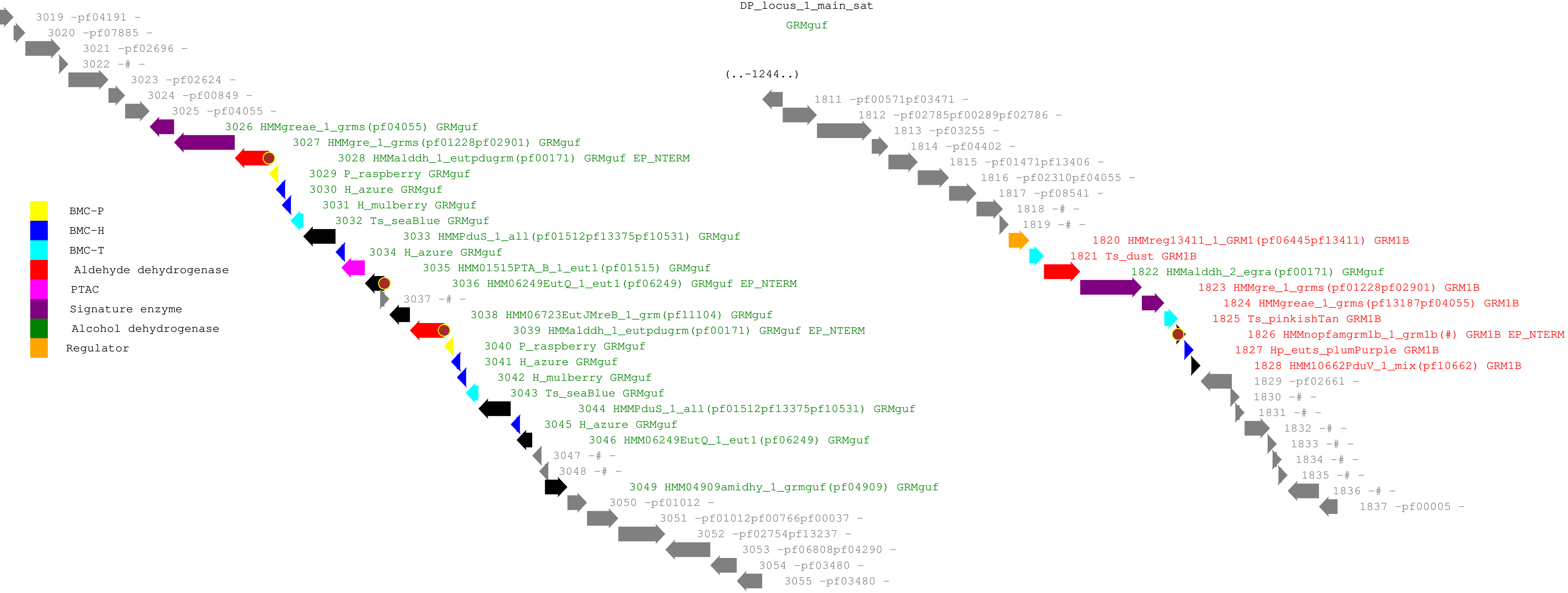

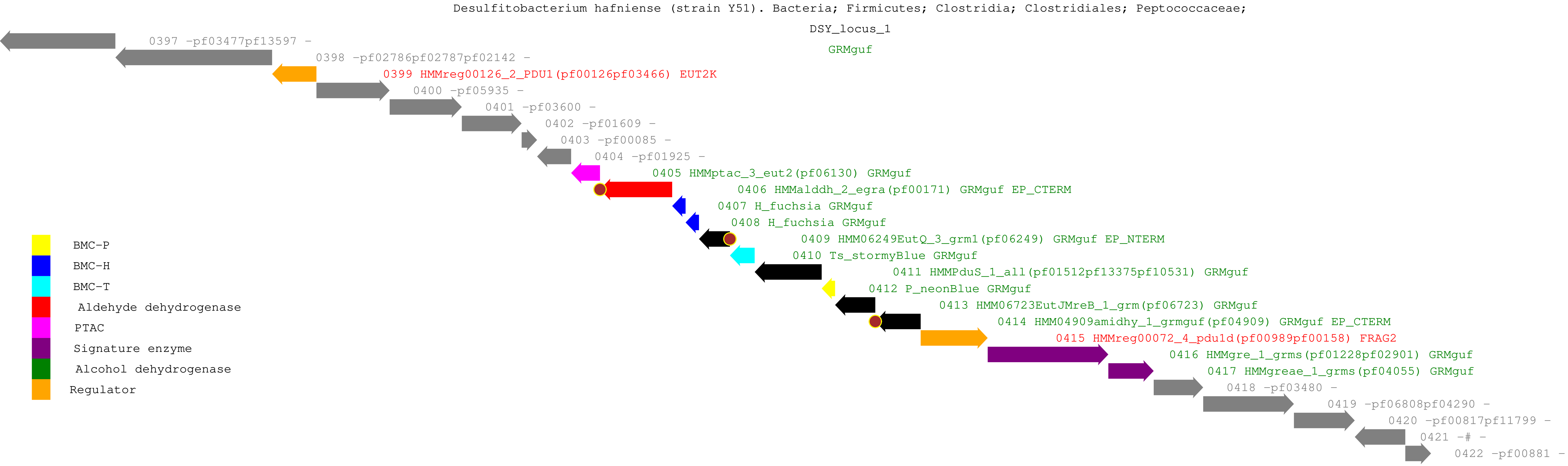
Desulfitobacterium hafniense (strain Y51). Bacteria; Firmicutes; Clostridia; Clostridiales; Peptococcaceae;
DSY_locus_1
GRMguf
0397 -pf03477pf13597 -
0398 -pf02786pf02787pf02142 -
0399 HMMreg00126_2_PDU1(pf00126pf03466) EUT2K
0400 -pf05935 -
0401 -pf03600 -
0402 -pf01609 -
0403 -pf00085 -
0404 -pf01925 -
0405 HMMptac_3_eut2(pf06130) GRMguf
0406 HMMaldh_2_egra(pf00171) GRMguf EP_CTERM
0407 H_fuchsia GRMguf
0408 H_fuchsia GRMguf
0409 HMM06249EutQ_3_grm1(pf06249) GRMguf EP_NTERM
0410 Ts_stormyBlue GRMguf
0411 HMMPduS_1_all(pf01512pf13375pf10531) GRMguf
0412 P_neonBlue GRMguf
0413 HMM06723EutJMreB_1_grm(pf06723) GRMguf
0414 HMM04909amidhy_1_grmguf(pf04909) GRMguf EP_CTERM
0415 HMMreg00072_4_pdu1d(pf00989pf00158) FRAG2
0416 HMMgre_1_grms(pf01228pf02901) GRMguf
0417 HMMgreae_1_grms(pf04055) GRMguf
0418 -pf03480 -
0419 -pf06808pf04290 -
0420 -pf00817pf11799 -
0421 -# -
0422 -pf00881 -
BMC-P
BMC-H
BMC-T
Aldehyde dehydrogenase
PTAC
Signature enzyme
Alcohol dehydrogenase
Regulator

Bilophila sp. 4_1_30. Bacteria; Proteobacteria; Deltaproteobacteria; Desulfovibrionales;

HMPREF0178__locus_1_main_sat

GRMguf

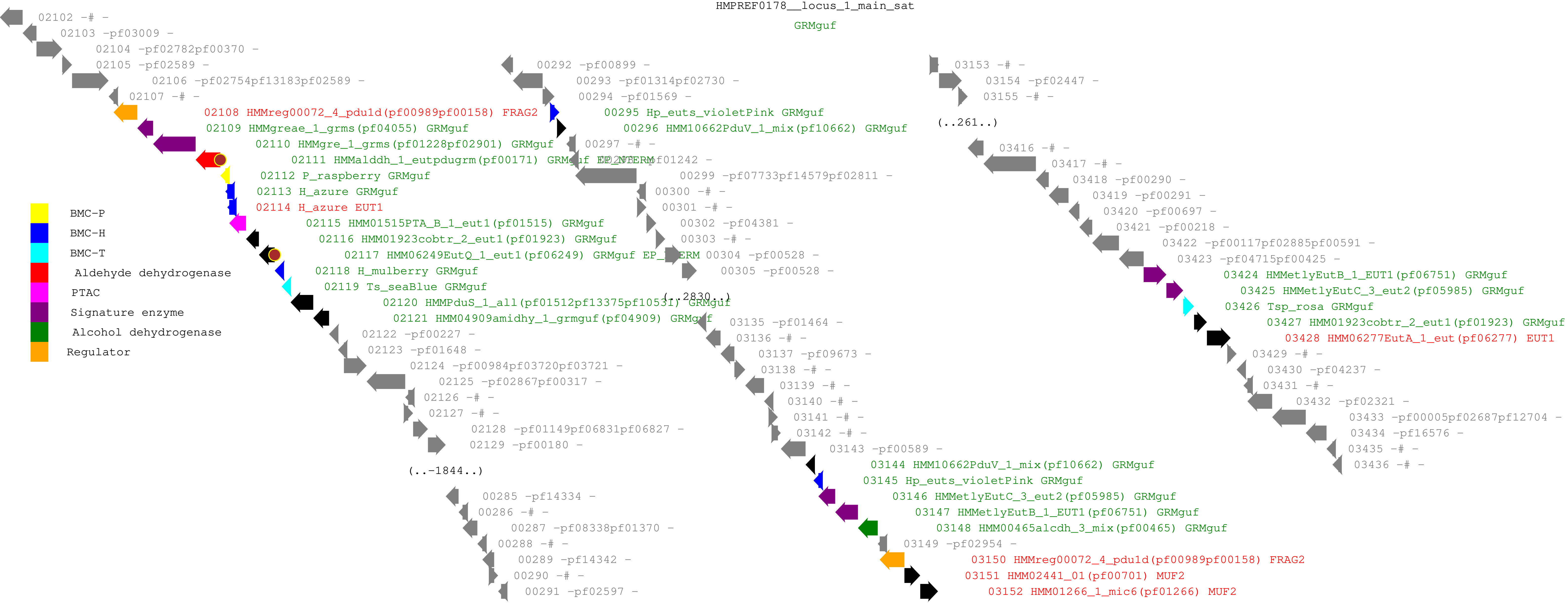

Bilophila wadsworthia 3_1_6. Bacteria; Proteobacteria; Deltaproteobacteria; Desulfovibrionales;

HMPREF0179__locus_1_main_sat

GRMguf

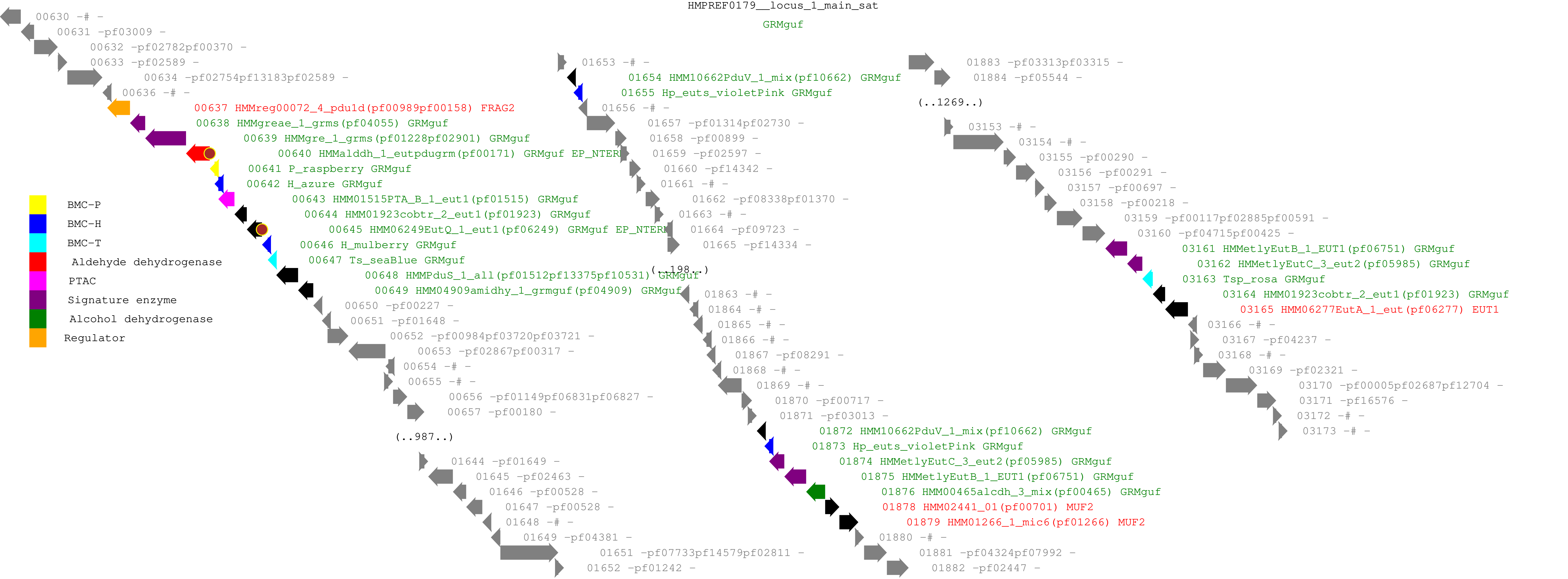

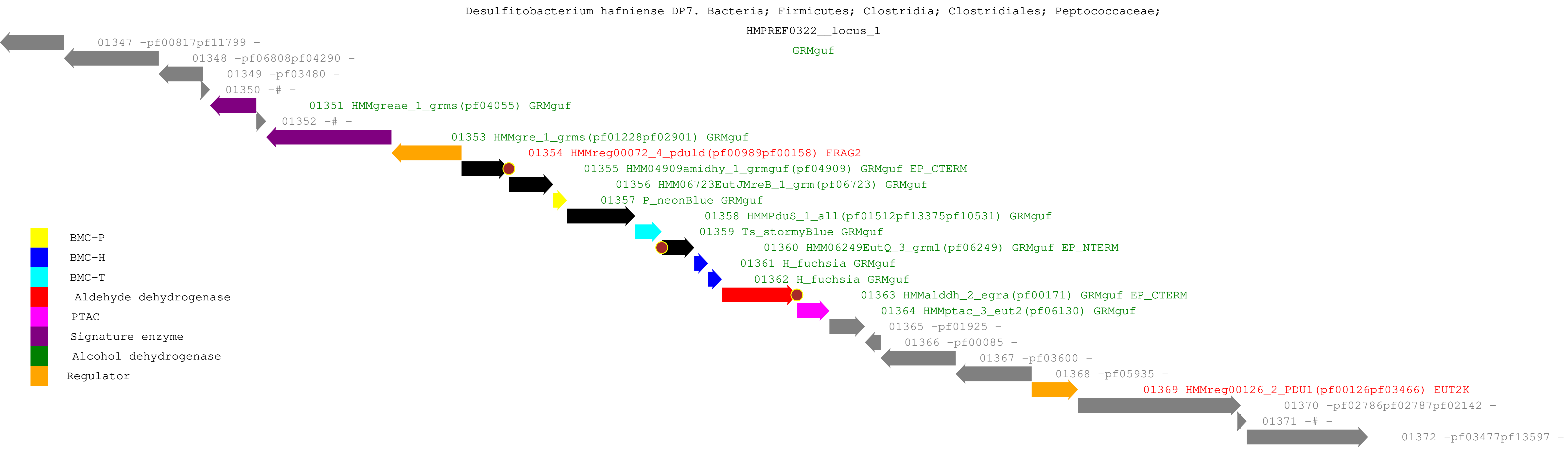
Desulfitobacterium hafniense DP7. Bacteria; Firmicutes; Clostridia; Clostridiales; Peptococcaceae;
HMPREF0322__locus_1
GRMguf
01347 -pf00817pf11799 -
01348 -pf06808pf04290 -
01349 -pf03480 -
01350 -# -
01351 HMMgreae_1_grms(pf04055) GRMguf
01352 -# -
01353 HMMgre_1_grms(pf01228pf02901) GRMguf
01354 HMMreg00072_4_pdu1d(pf00989pf00158) FRAG2
01355 HMM04909amidhy_1_grmguf(pf04909) GRMguf EP_CTERM
01356 HMM06723EutJMreB_1_grm(pf06723) GRMguf
01357 P_neonBlue GRMguf
01358 HMMPduS_1_all(pf01512pf13375pf10531) GRMguf
01359 Ts_stormyBlue GRMguf
01360 HMM06249EutQ_3_grm1(pf06249) GRMguf EP_NTERM
01361 H_fuchsia GRMguf
01362 H_fuchsia GRMguf
01363 HMMalddh_2_egra(pf00171) GRMguf EP_CTERM
01364 HMMptac_3_eut2(pf06130) GRMguf
01365 -pf01925 -
01366 -pf00085 -
01367 -pf03600 -
01368 -pf05935 -
01369 HMMreg00126_2_PDU1(pf00126pf03466) EUT2K
01370 -pf02786pf02787pf02142 -
01371 -# -
01372 -pf03477pf13597 -
BMC-P
BMC-H
BMC-T
Aldehyde dehydrogenase
PTAC
Signature enzyme
Alcohol dehydrogenase
Regulator

Desulfotomaculum aquiferis. Bacteria; Firmicutes; Clostridia; Clostridiales; Peptococcaceae;

N752__locus_2

GRMguf

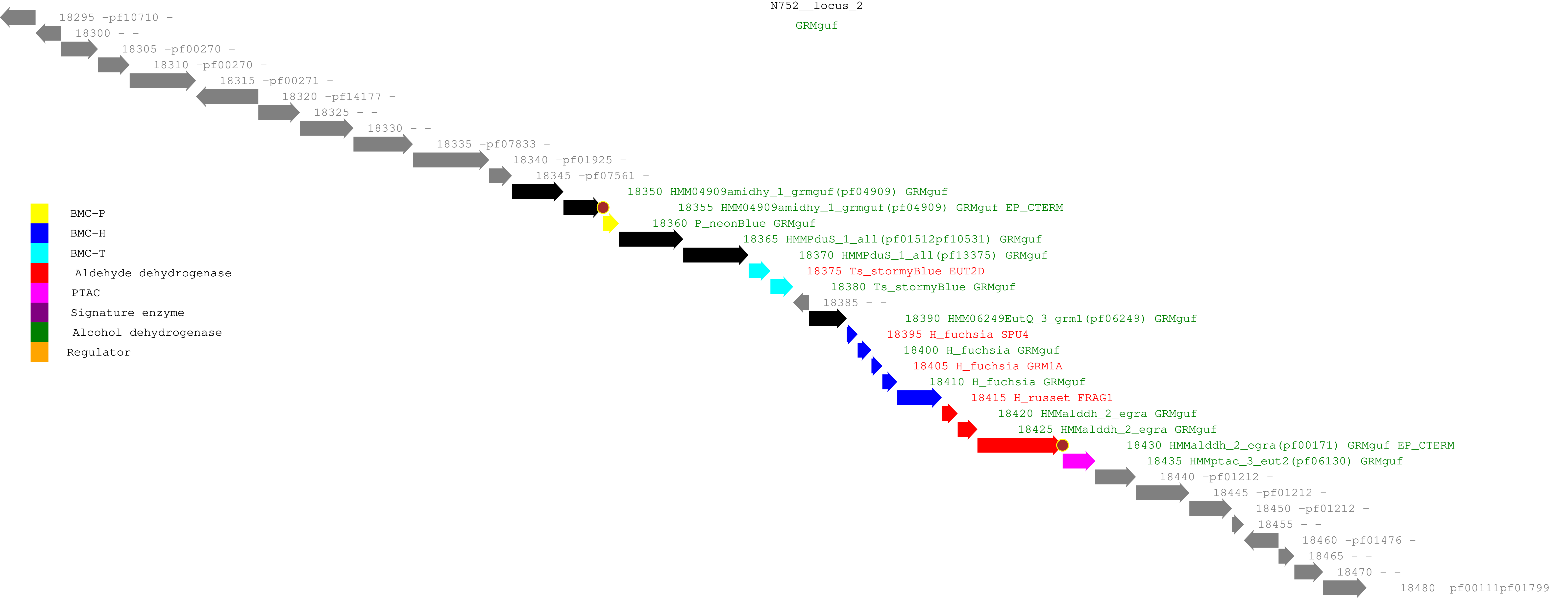

Desulfacinum infernum DSM 9756. Bacteria; Proteobacteria; Deltaproteobacteria; Syntrophobacterales;

SAMN02745206__locus_1

GRMguf

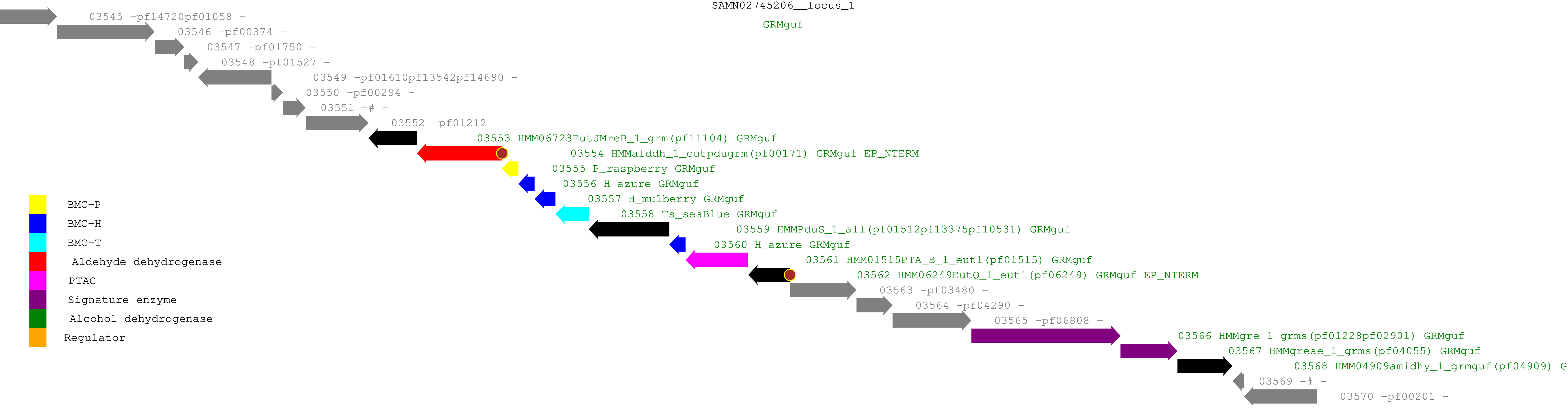

Desulfacinum hydrothermale DSM 13146. Bacteria; Proteobacteria; Deltaproteobacteria; Syntrophobacterales;

SAMN02746041__locus_1

GRMguf

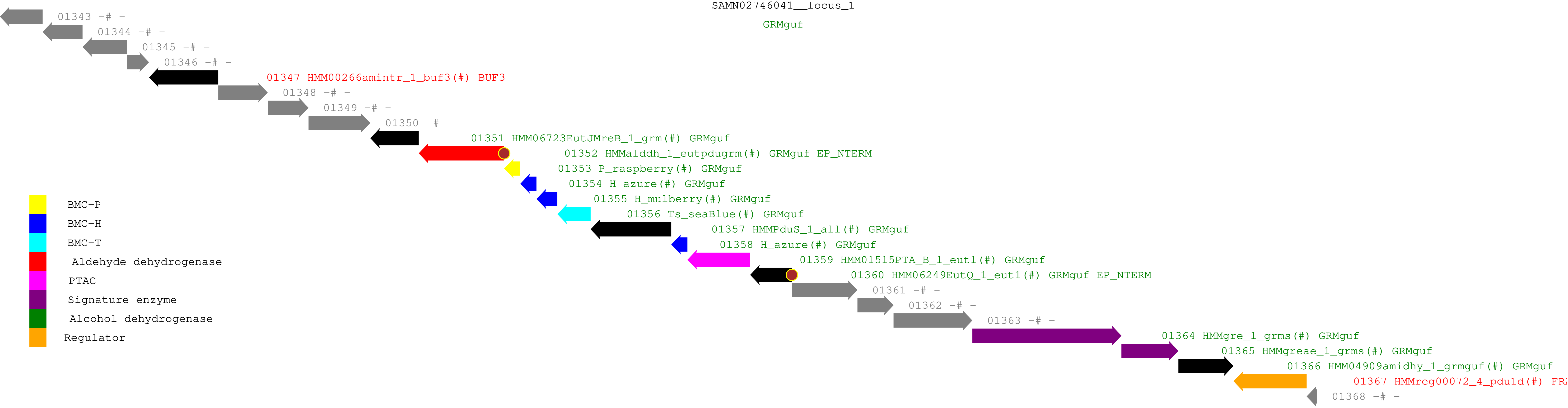

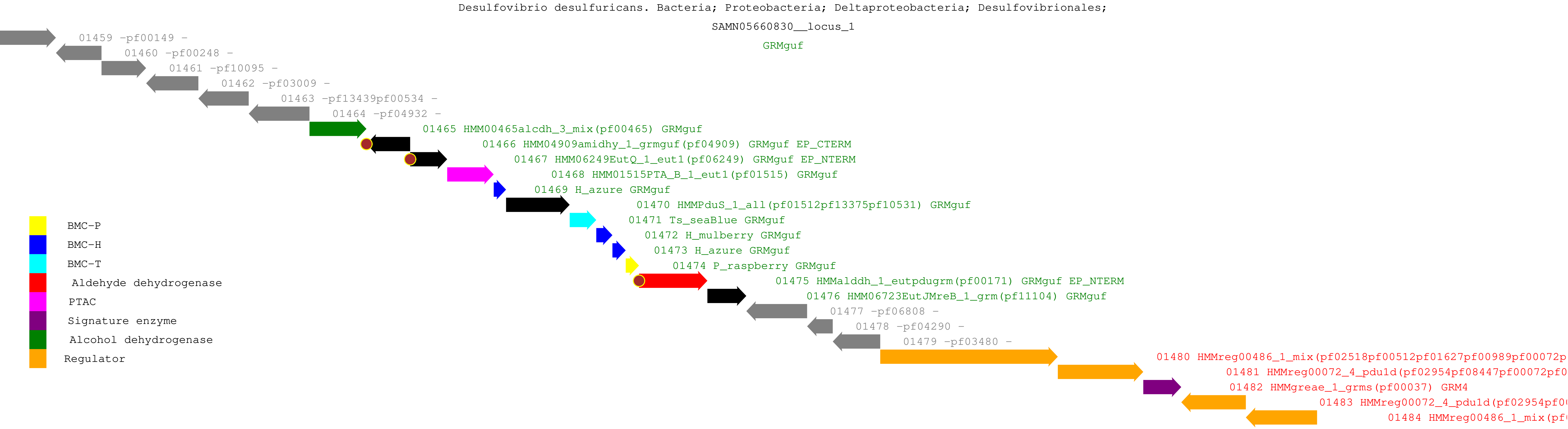
Desulfovibrio desulfuricans. Bacteria; Proteobacteria; Deltaproteobacteria; Desulfovibrionales;
SAMN05660830__locus_1
GRMguf
01459 -pf00149 -
01460 -pf00248 -
01461 -pf10095 -
01462 -pf03009 -
01463 -pf13439pf00534 -
01464 -pf04932 -
01465 HMM00465alcdh_3_mix(pf00465) GRMguf
01466 HMM04909amidhy_1_grmguf(pf04909) GRMguf EP_CTERM
01467 HMM06249EutQ_1_eut1(pf06249) GRMguf EP_NTERM
01468 HMM01515PTA_B_1_eut1(pf01515) GRMguf
01469 H_azure GRMguf
01470 HMMPduS_1_all(pf01512pf13375pf10531) GRMguf
01471 Ts_seaBlue GRMguf
01472 H_mulberry GRMguf
01473 H_azure GRMguf
01474 P_raspberry GRMguf
01475 HMMalddh_1_eutpdugrm(pf00171) GRMguf EP_NTERM
01476 HMM06723EutJMreB_1_grm(pf11104) GRMguf
01477 -pf06808 -
01478 -pf04290 -
01479 -pf03480 -
01480 HMMreg00486_1_mix(pf02518pf00512pf01627pf00989pf00072p
01481 HMMreg00072_4_pdu1d(pf02954pf08447pf00072pf0
01482 HMMgreae_1_grms(pf00037) GRM4
01483 HMMreg00072_4_pdu1d(pf02954pf0
01484 HMMreg00486_1_mix(pf
BMC-P
BMC-H
BMC-T
Aldehyde dehydrogenase
PTAC
Signature enzyme
Alcohol dehydrogenase
Regulator

Halodesulfovibrio spirochaetisodalis. Bacteria; Proteobacteria; Deltaproteobacteria; Desulfovibrionales;

SP90__locus_1

GRMguf

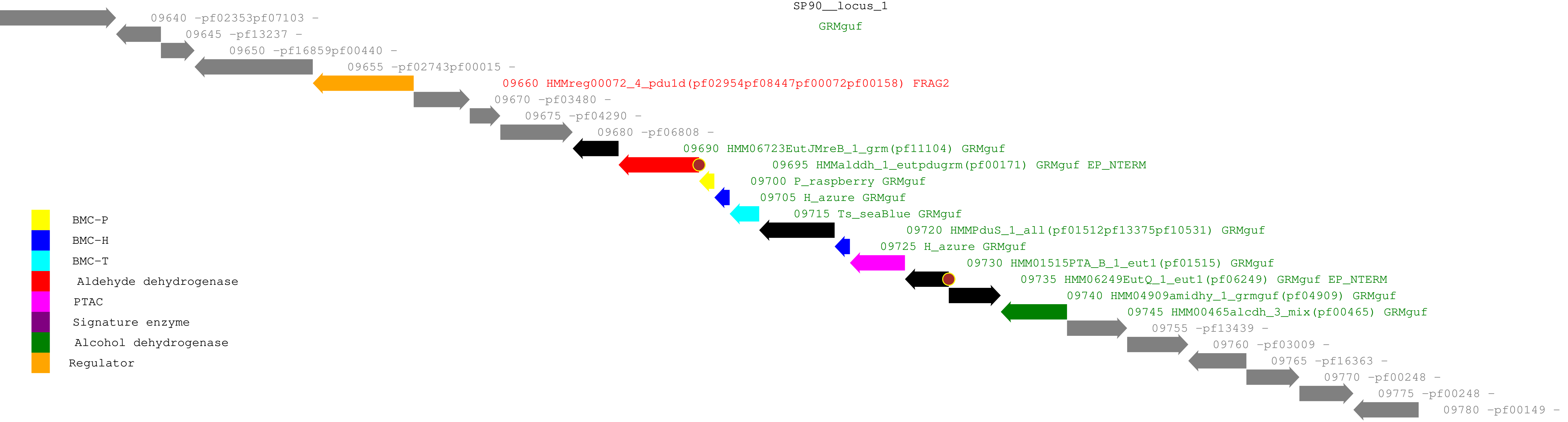

Deltaproteobacteria bacterium RIFOXYA12_FULL_61_11. Bacteria; Proteobacteria; Deltaproteobacteria.

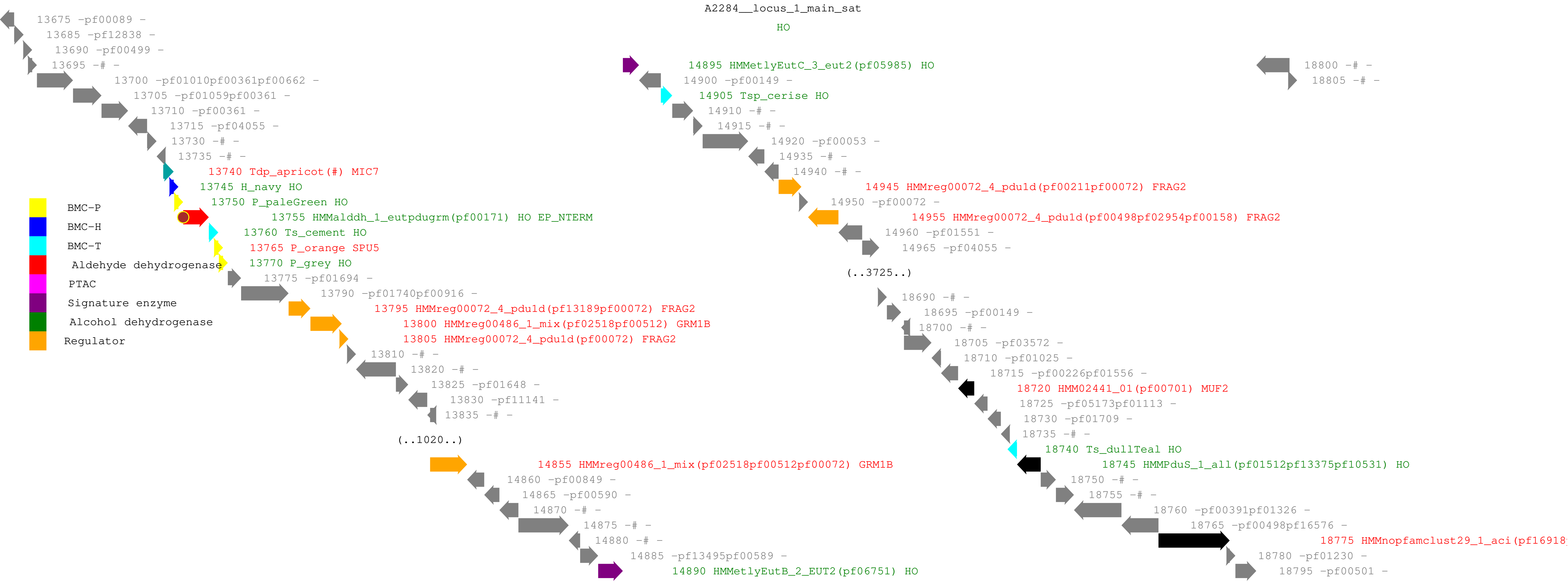

Deltaproteobacteria bacterium RIFOXYA12_FULL_58_15. Bacteria; Proteobacteria; Deltaproteobacteria.

A2289__locus_1_main_sat

HO

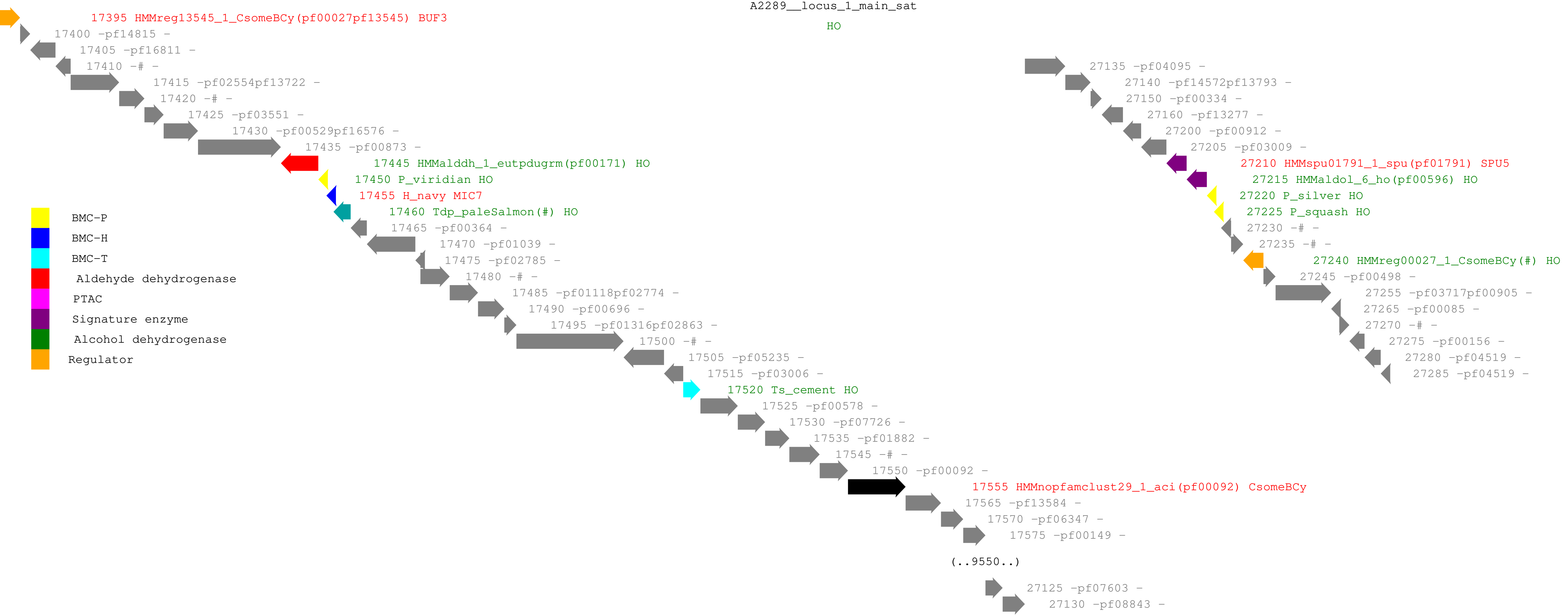

Deltaproteobacteria bacterium RIFOXYB12_FULL_58_9. Bacteria; Proteobacteria; Deltaproteobacteria.

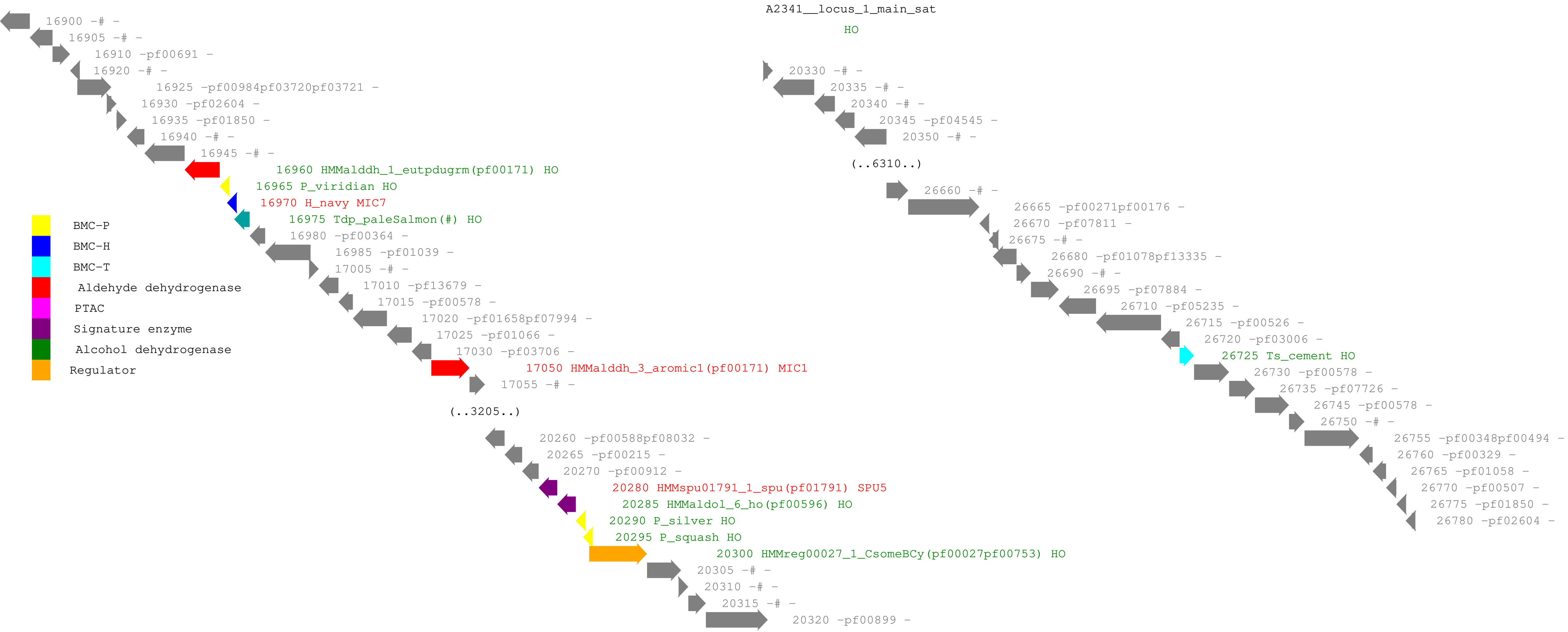

Deltaproteobacteria bacterium RIFCSPHIGHO2_02_FULL_40_11. Bacteria; Proteobacteria; Deltaproteobacteria.

A3B70__locus_1

HO

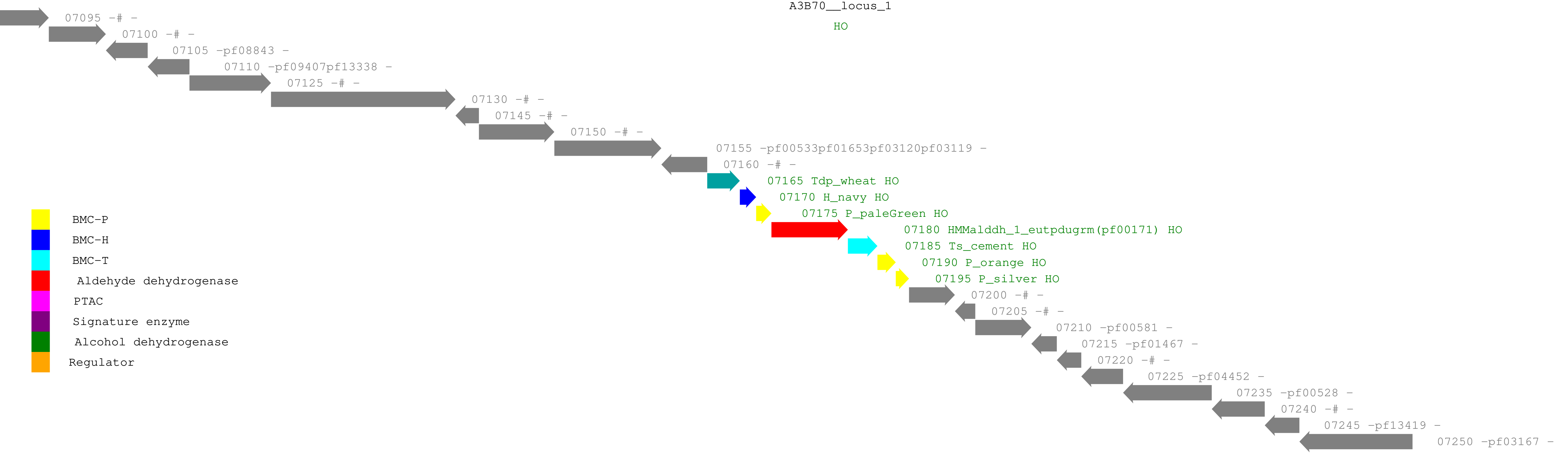

Deltaproteobacteria bacterium RIFCSPHIGHO2_02_FULL_38_15. Bacteria; Proteobacteria; Deltaproteobacteria.

A3D19__locus_1

HO

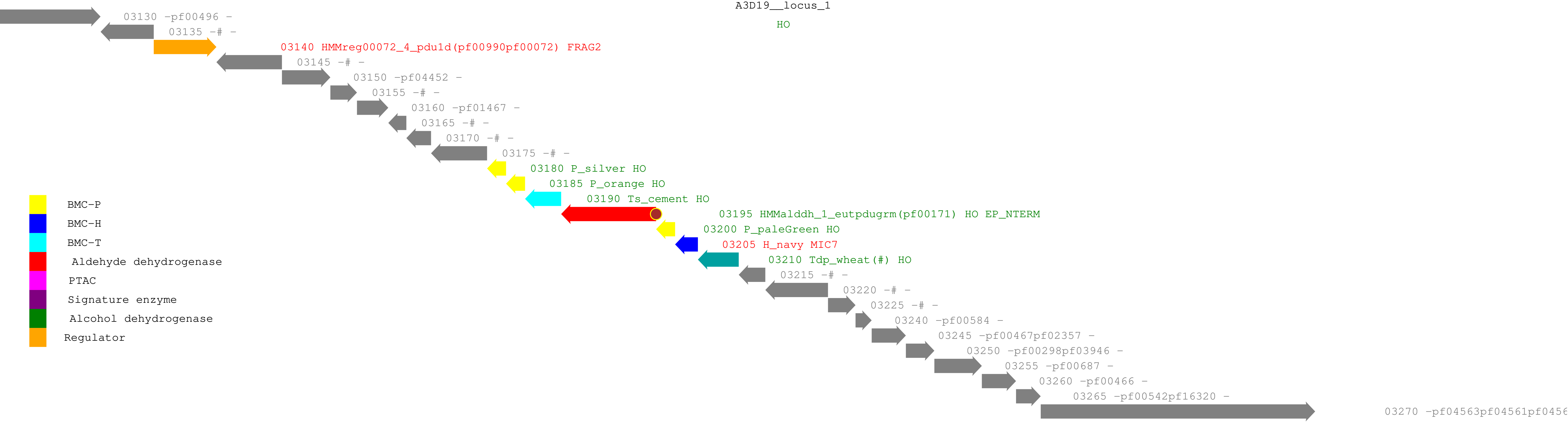

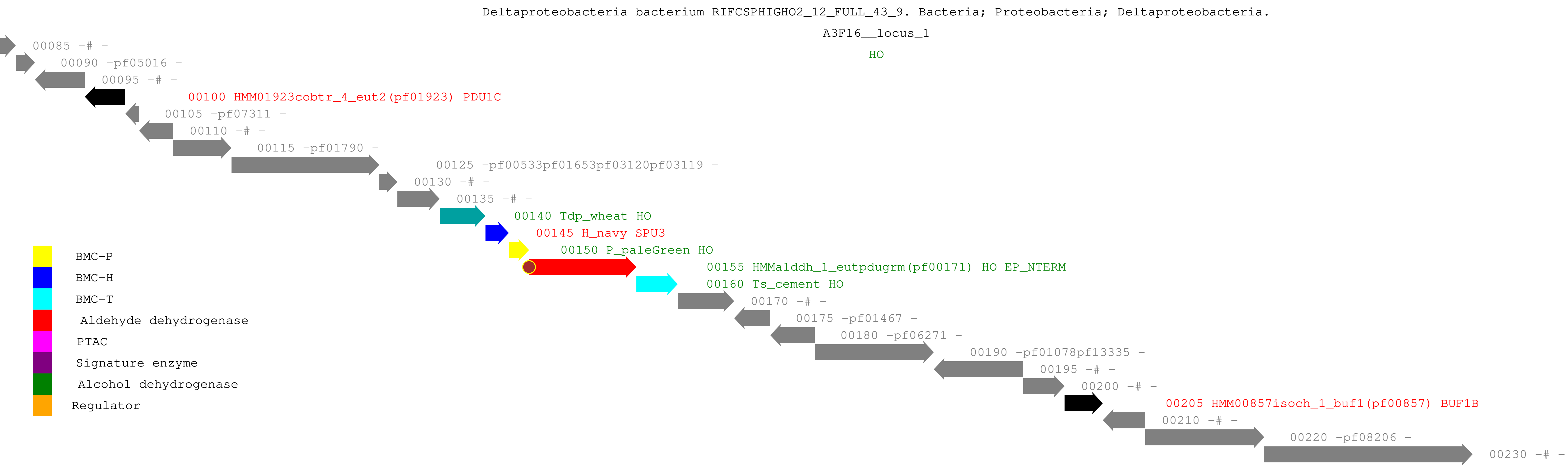
Deltaproteobacteria bacterium RIFCSPHIGHO2_12_FULL_43_9. Bacteria; Proteobacteria; Deltaproteobacteria.
A3F16__locus_1
HO
00085 -# -
00090 -pf05016 -
00095 -# -
00100 HMM01923cobtr_4_eut2(pf01923) PDU1C
00105 -pf07311 -
00110 -# -
00115 -pf01790 -
00125 -pf00533pf01653pf03120pf03119 -
00130 -# -
00135 -# -
00140 Tdp_wheat HO
00145 H_navy SPU3
00150 P_paleGreen HO
00155 HMMalddh_1_eutpdugrm(pf00171) HO EP_NTERM
00160 Ts_cement HO
00170 -# -
00175 -pf01467 -
00180 -pf06271 -
00190 -pf01078pf13335 -
00195 -# -
00200 -# -
00205 HMM00857isoch_1_buf1(pf00857) BUF1B
00210 -# -
00220 -pf08206 -
00230 -# -
BMC-P
BMC-H
BMC-T
Aldehyde dehydrogenase
PTAC
Signature enzyme
Alcohol dehydrogenase
Regulator

Bdellovibrionales bacterium RIFOXYD1_FULL_44_7. Bacteria; Proteobacteria; Deltaproteobacteria; Bdellovibrionales.

A3K03__locus_1

HO

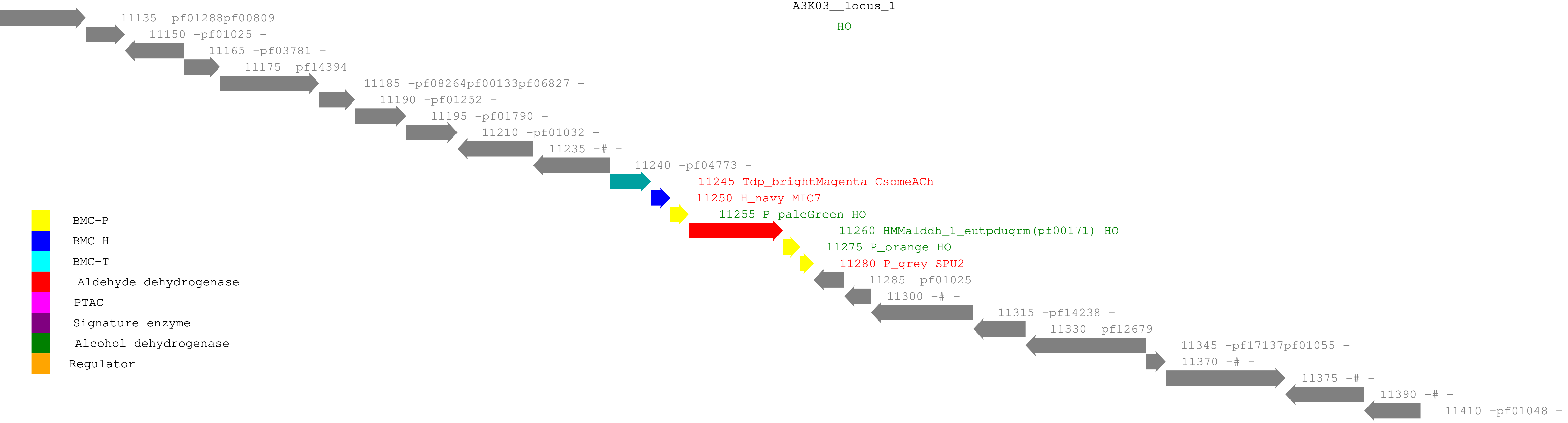

Deltaproteobacteria bacterium CG2_30_63_29. Bacteria; Proteobacteria; Deltaproteobacteria.

AUK47__locus_1_main_sat

HO

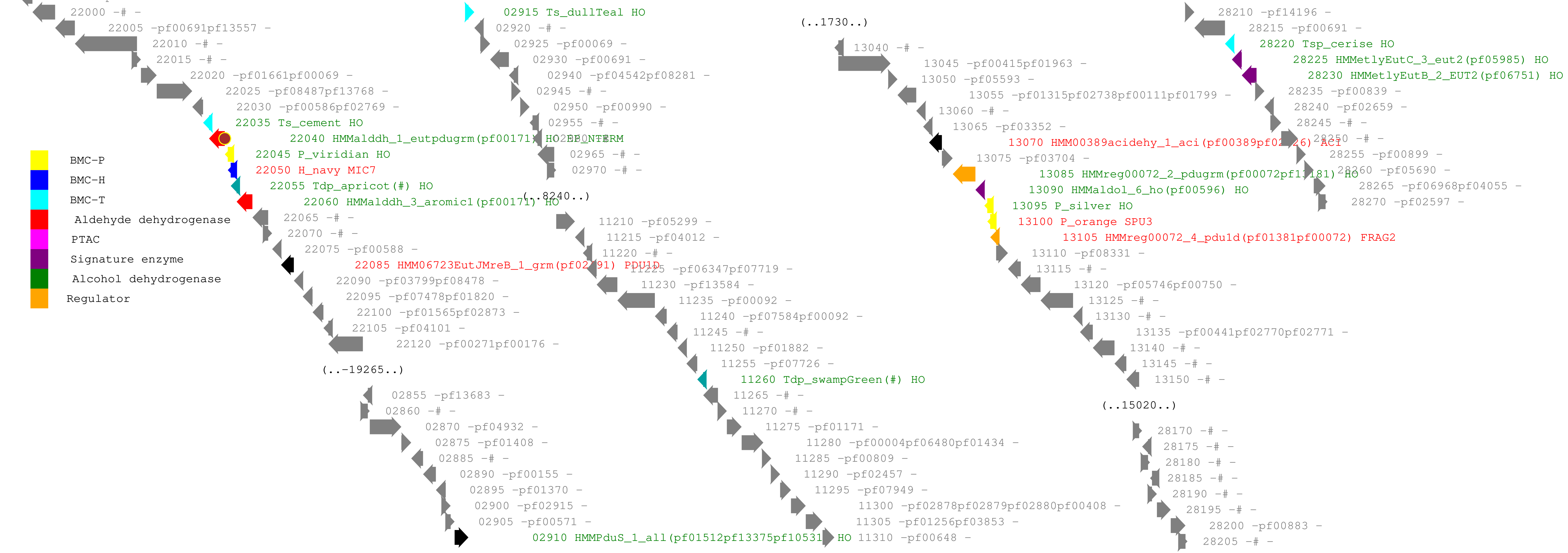

Deltaproteobacteria bacterium ADurb.Bin058. Bacteria; Proteobacteria; Deltaproteobacteria.

BWX66__locus_1_main_sat

HO

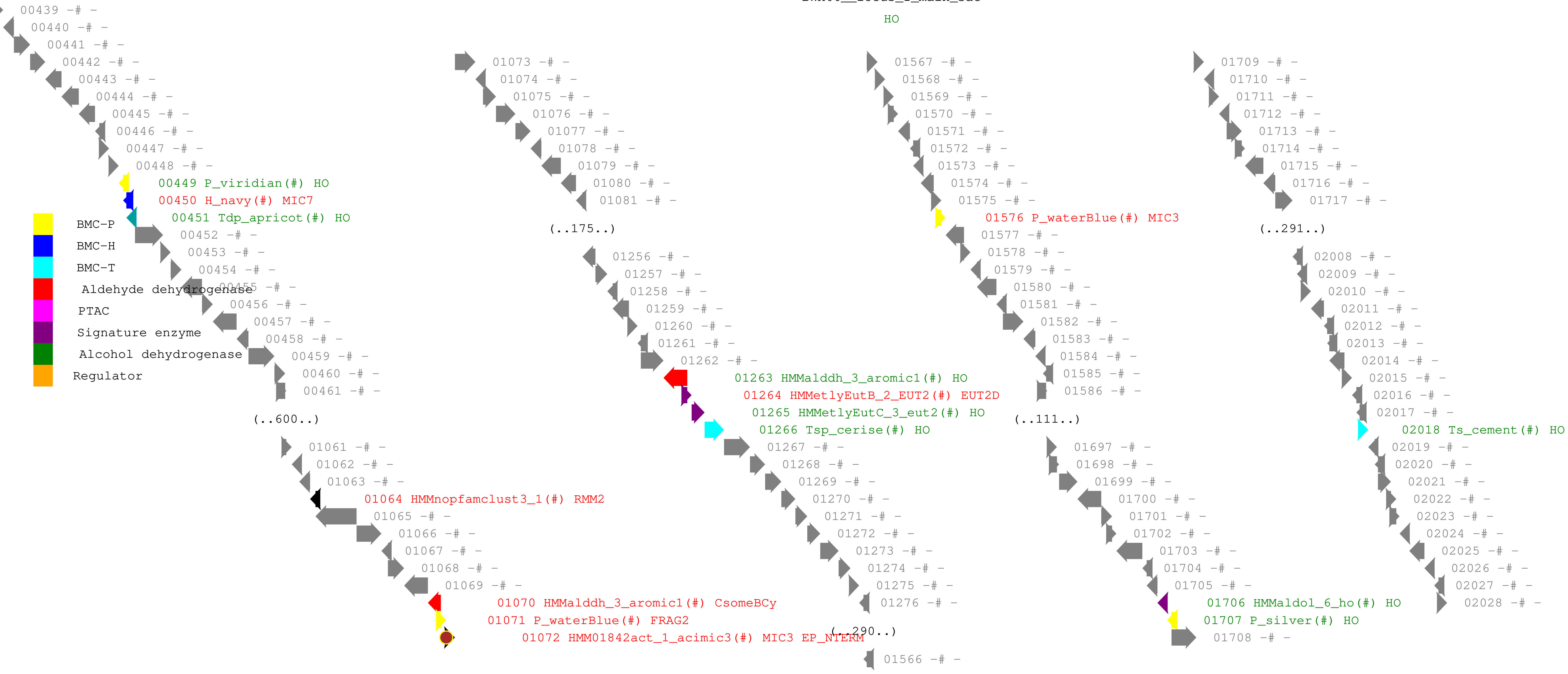

Deltaproteobacteria bacterium. Bacteria; Proteobacteria; Deltaproteobacteria.

CL916__locus_1_main_sat

HO

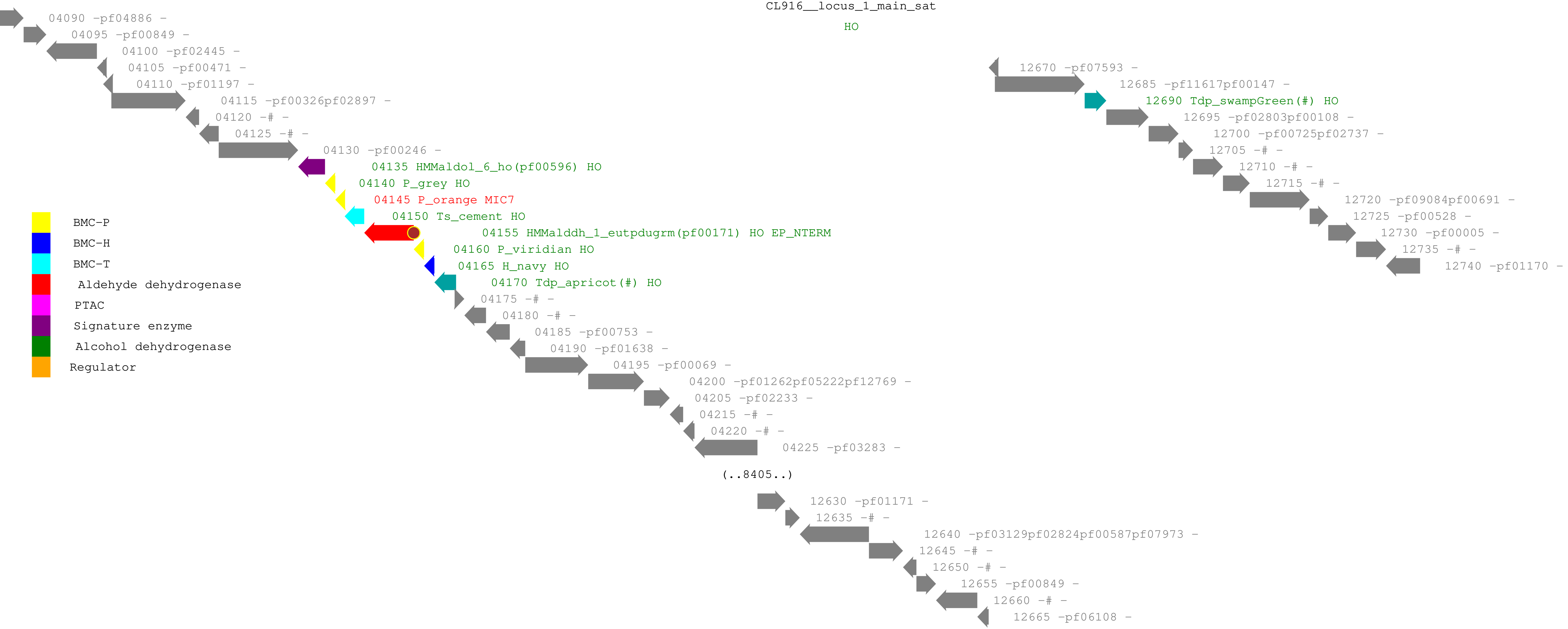

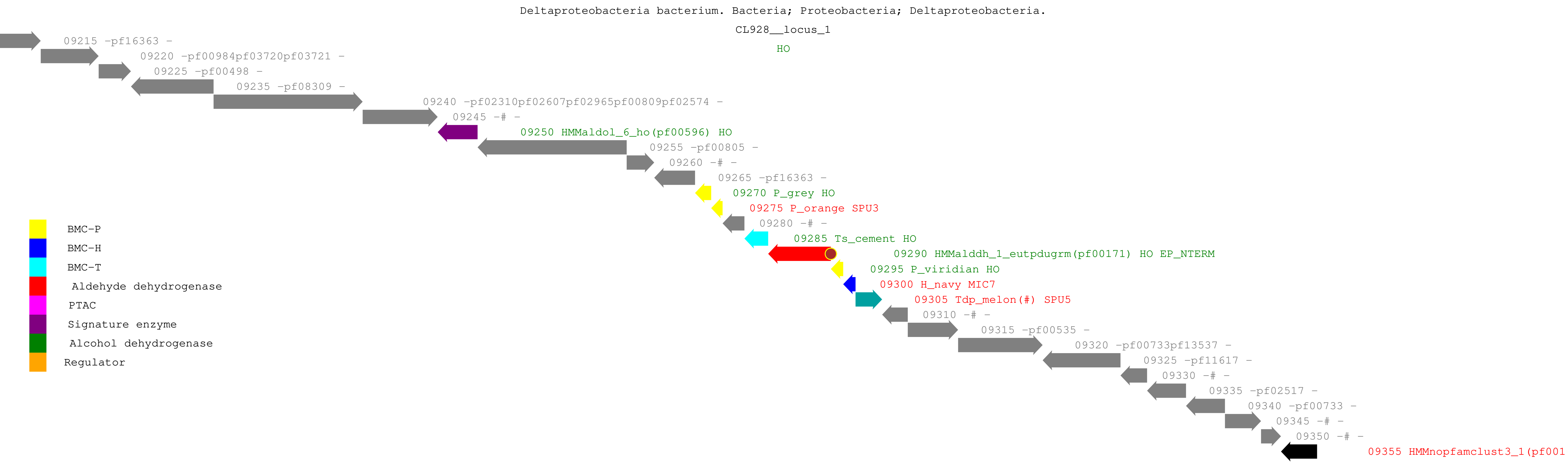
Deltaproteobacteria bacterium. Bacteria; Proteobacteria; Deltaproteobacteria.
CL928__locus_1
HO
09215 -pf16363 -
09220 -pf00984pf03720pf03721 -
09225 -pf00498 -
09235 -pf08309 -
09240 -pf02310pf02607pf02965pf00809pf02574 -
09245 -# -
09250 HMMaldol_6_ho(pf00596) HO
09255 -pf00805 -
09260 -# -
09265 -pf16363 -
09270 P_grey HO
09275 P_orange SPU3
09280 -# -
09285 Ts_cement HO
09290 HMMaldh_1_eutpdugrm(pf00171) HO EP_NTERM
09295 P_viridian HO
09300 H_navy MIC7
09305 Tdp_melon(#) SPU5
09310 -# -
09315 -pf00535 -
09320 -pf00733pf13537 -
09325 -pf11617 -
09330 -# -
09335 -pf02517 -
09340 -pf00733 -
09345 -# -
09350 -# -
09355 HMMnopfamclust3_1(pf001
BMC-P
BMC-H
BMC-T
Aldehyde dehydrogenase
PTAC
Signature enzyme
Alcohol dehydrogenase
Regulator

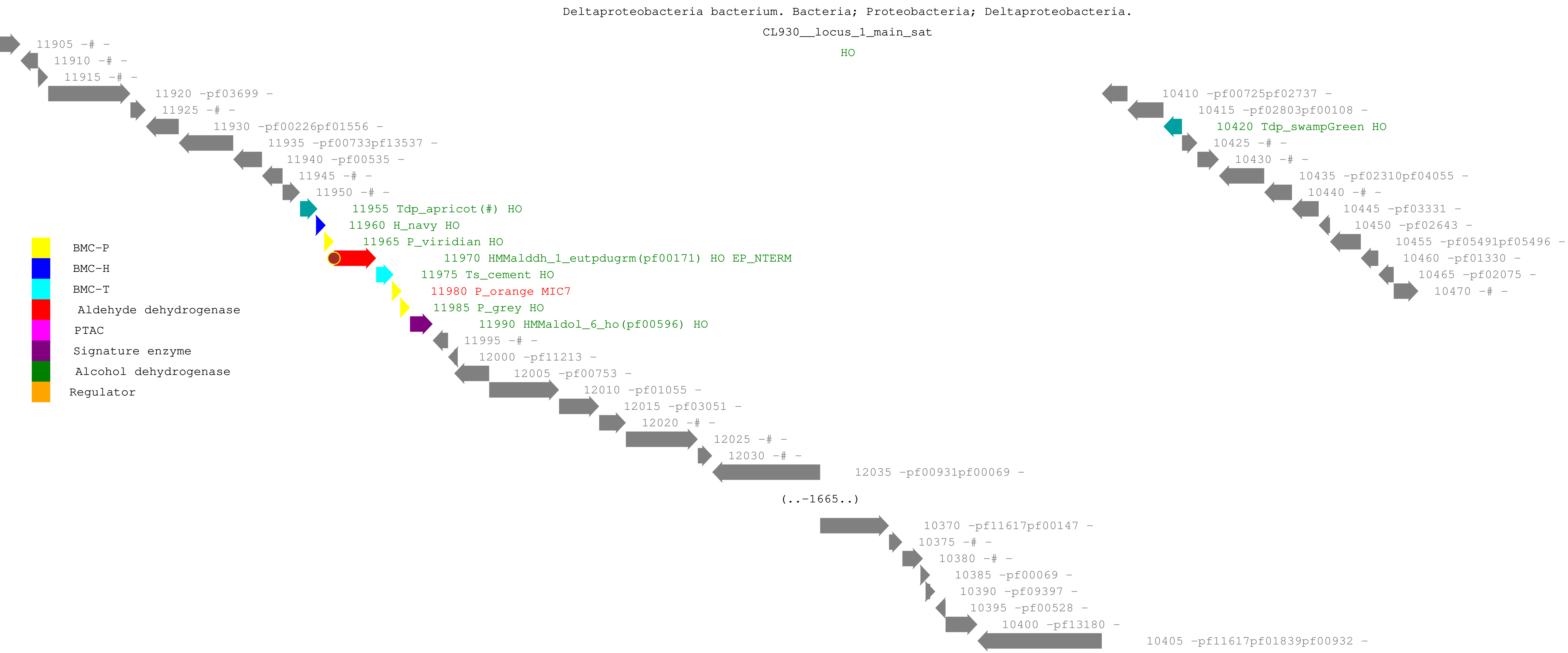
Deltaproteobacteria bacterium. Bacteria; Proteobacteria; Deltaproteobacteria.
CL930__locus_1_main_sat
HO
11905 -# -
11910 -# -
11915 -# -
11920 -pf03699 -
11925 -# -
11930 -pf00226pf01556 -
11935 -pf00733pf13537 -
11940 -pf00535 -
11945 -# -
11950 -# -
11955 Tdp_apricot(#) HO
11960 H_navy HO
11965 P_viridian HO
11970 HMMalddh_1_eutpdugrm(pf00171) HO EP_NTERM
11975 Ts_cement HO
11980 P_orange MIC7
11985 P_grey HO
11990 HMMaldol_6_ho(pf00596) HO
11995 -# -
12000 -pf11213 -
12005 -pf00753 -
12010 -pf01055 -
12015 -pf03051 -
12020 -# -
12025 -# -
12030 -# -
12035 -pf00931pf00069 -
(..-1665..)
10370 -pf11617pf00147 -
10375 -# -
10380 -# -
10385 -pf00069 -
10390 -pf09397 -
10395 -pf00528 -
10400 -pf13180 -
10405 -pf11617pf01839pf00932 -
10410 -pf00725pf02737 -
10415 -pf02803pf00108 -
10420 Tdp_swampGreen HO
10425 -# -
10430 -# -
10435 -pf02310pf04055 -
10440 -# -
10445 -pf03331 -
10450 -pf02643 -
10455 -pf05491pf05496 -
10460 -pf01330 -
10465 -pf02075 -
10470 -# -
BMC-P
BMC-H
BMC-T
Aldehyde dehydrogenase
PTAC
Signature enzyme
Alcohol dehydrogenase
Regulator

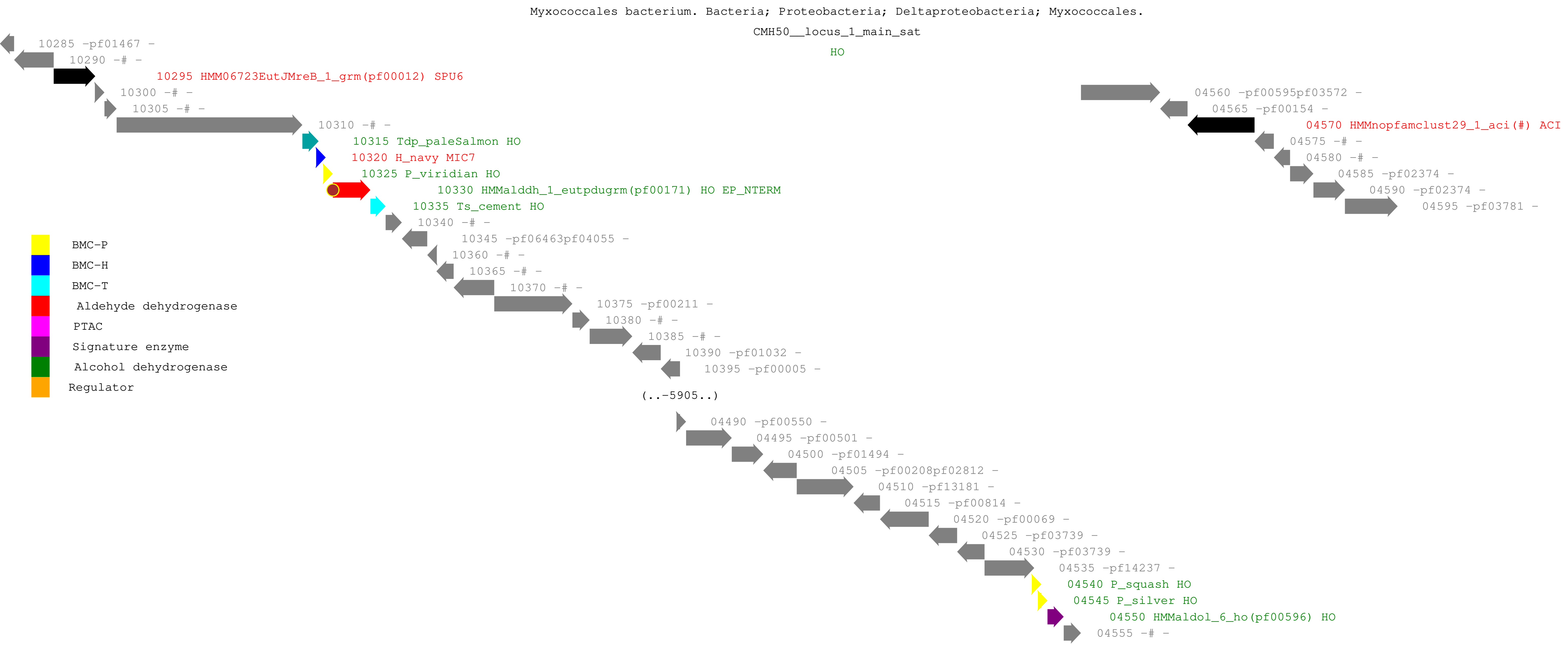
Myxococcales bacterium. Bacteria; Proteobacteria; Deltaproteobacteria; Myxococcales.
CMH50__locus_1_main_sat
HO
10285 -pf01467 -
10290 -# -
10295 HMM06723EutJMreB_1_grm(pf00012) SPU6
10300 -# -
10305 -# -
10310 -# -
10315 Tdp_paleSalmon HO
10320 H_navy MIC7
10325 P_viridian HO
10330 HMMaldh_1_eutpdugrm(pf00171) HO EP_NTERM
10335 Ts_cement HO
10340 -# -
10345 -pf06463pf04055 -
10360 -# -
10365 -# -
10370 -# -
10375 -pf00211 -
10380 -# -
10385 -# -
10390 -pf01032 -
10395 -pf00005 -
(..-5905..)
04490 -pf00550 -
04495 -pf00501 -
04500 -pf01494 -
04505 -pf00208pf02812 -
04510 -pf13181 -
04515 -pf00814 -
04520 -pf00069 -
04525 -pf03739 -
04530 -pf03739 -
04535 -pf14237 -
04540 P_squash HO
04545 P_silver HO
04550 HMMaldol_6_ho(pf00596) HO
04555 -# -
04560 -pf00595pf03572 -
04565 -pf00154 -
04570 HMMnopfamclust29_1_aci(#) ACI
04575 -# -
04580 -# -
04585 -pf02374 -
04590 -pf02374 -
04595 -pf03781 -
BMC-P
BMC-H
BMC-T
Aldehyde dehydrogenase
PTAC
Signature enzyme
Alcohol dehydrogenase
Regulator

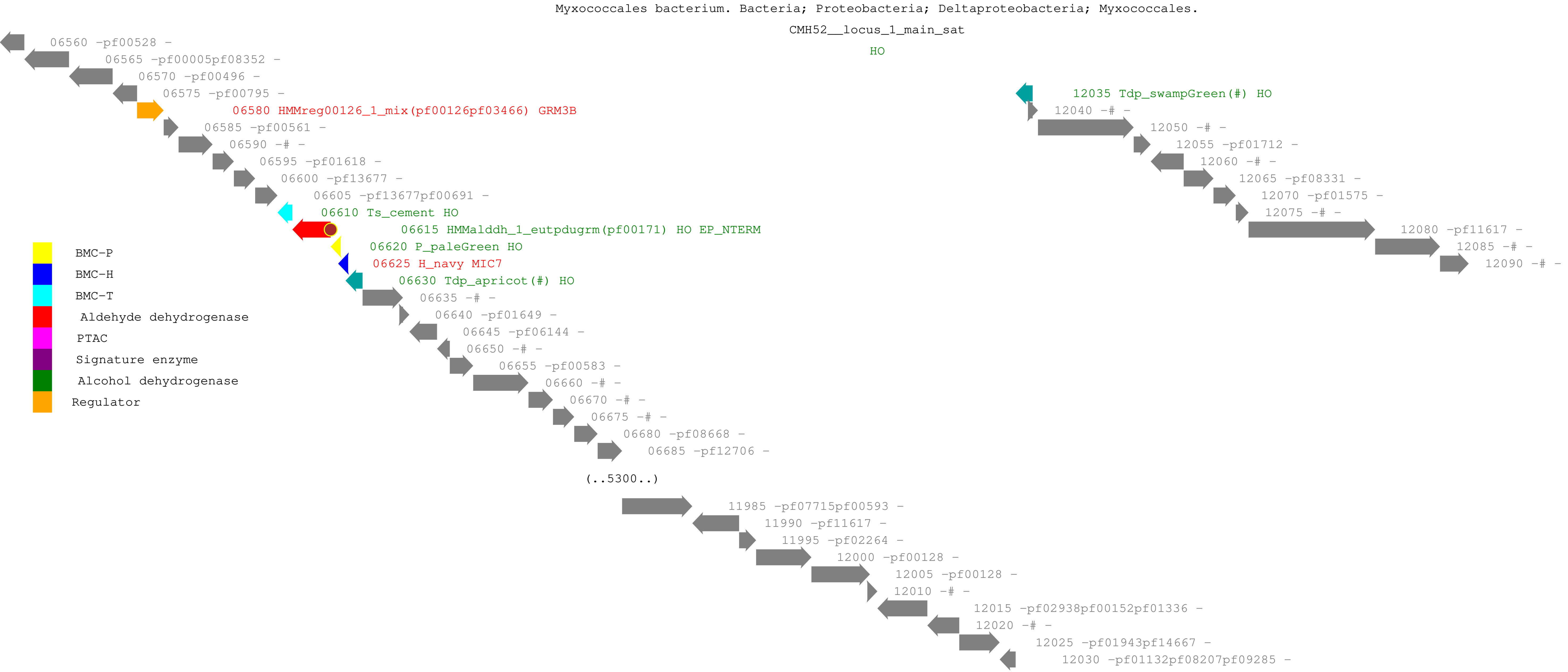
Myxococcales bacterium. Bacteria; Proteobacteria; Deltaproteobacteria; Myxococcales.
CMH52__locus_1_main_sat
HO
06560 -pf00528 -
06565 -pf00005pf08352 -
06570 -pf00496 -
06575 -pf00795 -
06580 HMMreg00126_1_mix(pf00126pf03466) GRM3B
06585 -pf00561 -
06590 -# -
06595 -pf01618 -
06600 -pf13677 -
06605 -pf13677pf00691 -
06610 Ts_cement HO
06615 HMMalddh_1_eutpdugrm(pf00171) HO EP_NTERM
06620 P_paleGreen HO
06625 H_navy MIC7
06630 Tdp_apricot(#) HO
06635 -# -
06640 -pf01649 -
06645 -pf06144 -
06650 -# -
06655 -pf00583 -
06660 -# -
06670 -# -
06675 -# -
06680 -pf08668 -
06685 -pf12706 -
(..5300..)
11985 -pf07715pf00593 -
11990 -pf11617 -
11995 -pf02264 -
12000 -pf00128 -
12005 -pf00128 -
12010 -# -
12015 -pf02938pf00152pf01336 -
12020 -# -
12025 -pf01943pf14667 -
12030 -pf01132pf08207pf09285 -
12035 Tdp_swampGreen(#) HO
12040 -# -
12050 -# -
12055 -pf01712 -
12060 -# -
12065 -pf08331 -
12070 -pf01575 -
12075 -# -
12080 -pf11617 -
12085 -# -
12090 -# -
BMC-P
BMC-H
BMC-T
Aldehyde dehydrogenase
PTAC
Signature enzyme
Alcohol dehydrogenase
Regulator

Myxococcales bacterium. Bacteria; Proteobacteria; Deltaproteobacteria; Myxococcales.

CMH53__locus_fragmented

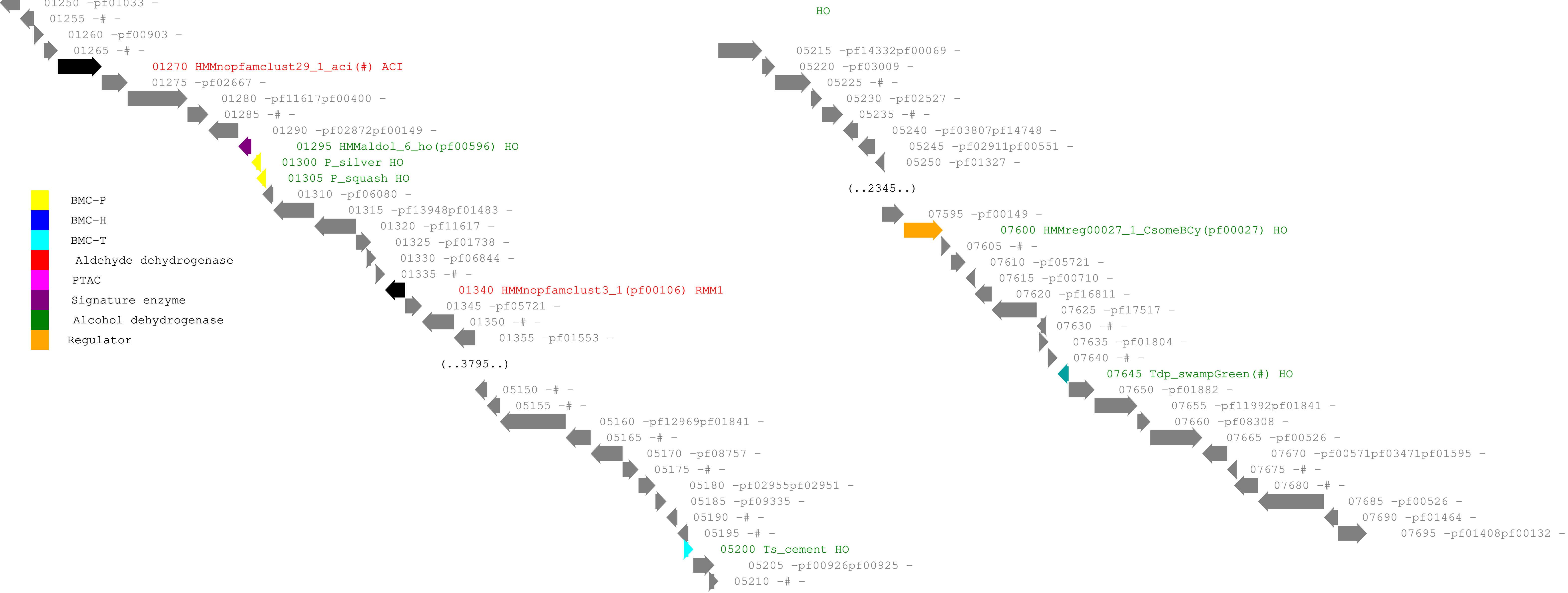

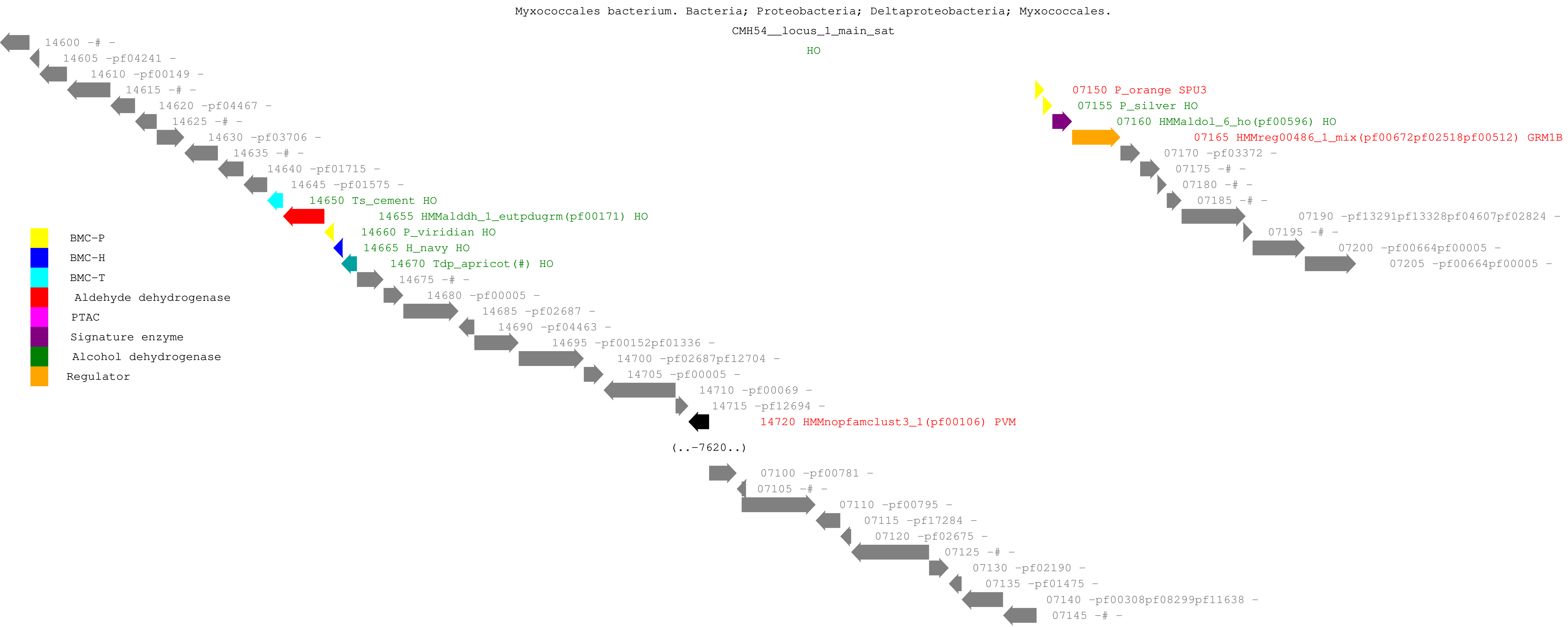

Myxococcales bacterium. Bacteria; Proteobacteria; Deltaproteobacteria; Myxococcales.
CMH54__locus_1_main_sat
HO
14600 -# -
14605 -pf04241 -
14610 -pf00149 -
14615 -# -
14620 -pf04467 -
14625 -# -
14630 -pf03706 -
14635 -# -
14640 -pf01715 -
14645 -pf01575 -
14650 Ts_cement HO
14655 HMMalddh_1_eutpdugrm(pf00171) HO
14660 P_viridian HO
14665 H_navy HO
14670 Tdp_apricot(#) HO
14675 -# -
14680 -pf00005 -
14685 -pf02687 -
14690 -pf04463 -
14695 -pf00152pf01336 -
14700 -pf02687pf12704 -
14705 -pf00005 -
14710 -pf00069 -
14715 -pf12694 -
14720 HMMnopfamclust3_1(pf00106) PVM
(..-7620..)
07100 -pf00781 -
07105 -# -
07110 -pf00795 -
07115 -pf17284 -
07120 -pf02675 -
07125 -# -
07130 -pf02190 -
07135 -pf01475 -
07140 -pf00308pf08299pf11638 -
07145 -# -
07150 P_orange SPU3
07155 P_silver HO
07160 HMMaldol_6_ho(pf00596) HO
07165 HMMreg00486_1_mix(pf00672pf02518pf00512) GRM1B
07170 -pf03372 -
07175 -# -
07180 -# -
07185 -# -
07190 -pf13291pf13328pf04607pf02824 -
07195 -# -
07200 -pf00664pf00005 -
07205 -pf00664pf00005 -
BMC-P
BMC-H
BMC-T
Aldehyde dehydrogenase
PTAC
Signature enzyme
Alcohol dehydrogenase
Regulator

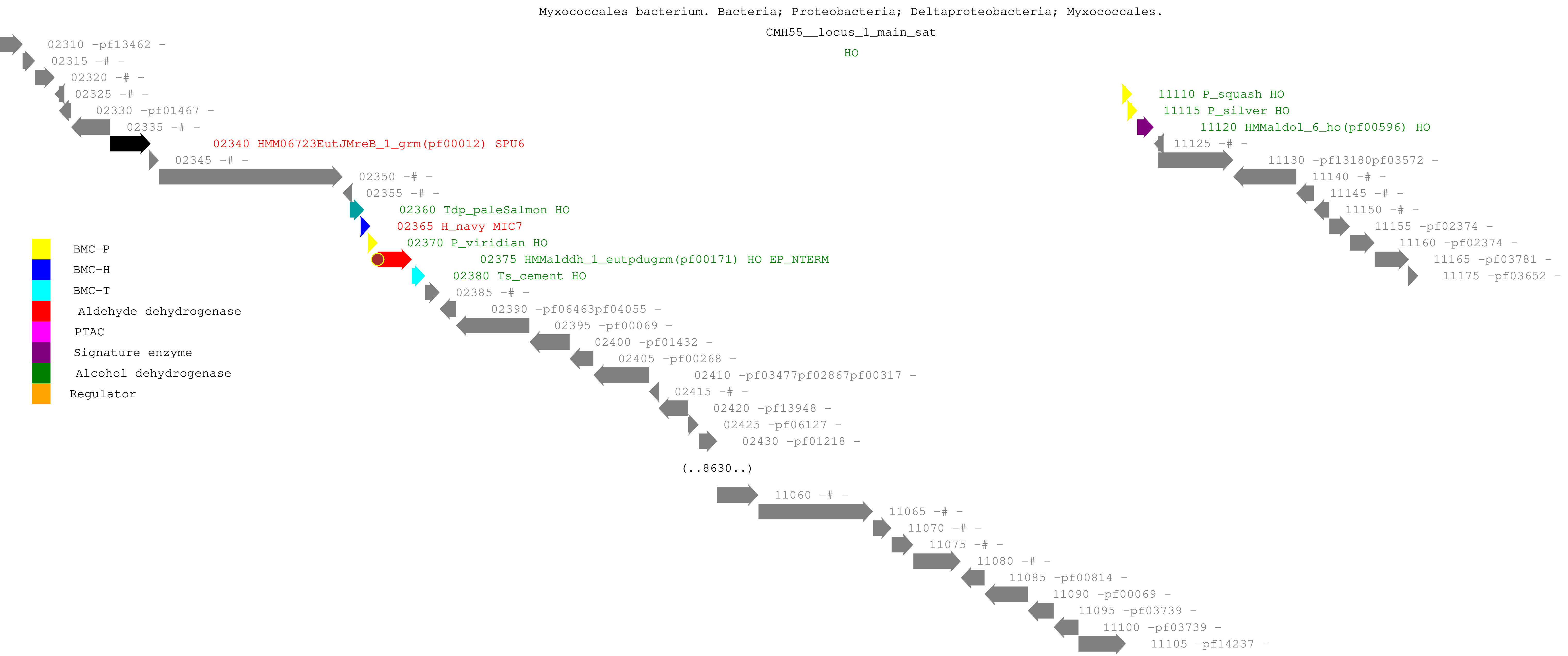
Myxococcales bacterium. Bacteria; Proteobacteria; Deltaproteobacteria; Myxococcales.
CMH55__locus_1_main_sat
HO
02310 -pf13462 -
02315 -# -
02320 -# -
02325 -# -
02330 -pf01467 -
02335 -# -
02340 HMM06723EutJMreB_1_grm(pf00012) SPU6
02345 -# -
02350 -# -
02355 -# -
02360 Tdp_paleSalmon HO
02365 H_navy MIC7
02370 P_viridian HO
02375 HMMaldddh_1_eutpdugrm(pf00171) HO EP_NTERM
02380 Ts_cement HO
02385 -# -
02390 -pf06463pf04055 -
02395 -pf00069 -
02400 -pf01432 -
02405 -pf00268 -
02410 -pf03477pf02867pf00317 -
02415 -# -
02420 -pf13948 -
02425 -pf06127 -
02430 -pf01218 -
(..8630..)
11060 -# -
11065 -# -
11070 -# -
11075 -# -
11080 -# -
11085 -pf00814 -
11090 -pf00069 -
11095 -pf03739 -
11100 -pf03739 -
11105 -pf14237 -
11110 P_squash HO
11115 P_silver HO
11120 HMMaldol_6_ho(pf00596) HO
11125 -# -
11130 -pf13180pf03572 -
11140 -# -
11145 -# -
11150 -# -
11155 -pf02374 -
11160 -pf02374 -
11165 -pf03781 -
11175 -pf03652 -
BMC-P
BMC-H
BMC-T
Aldehyde dehydrogenase
PTAC
Signature enzyme
Alcohol dehydrogenase
Regulator

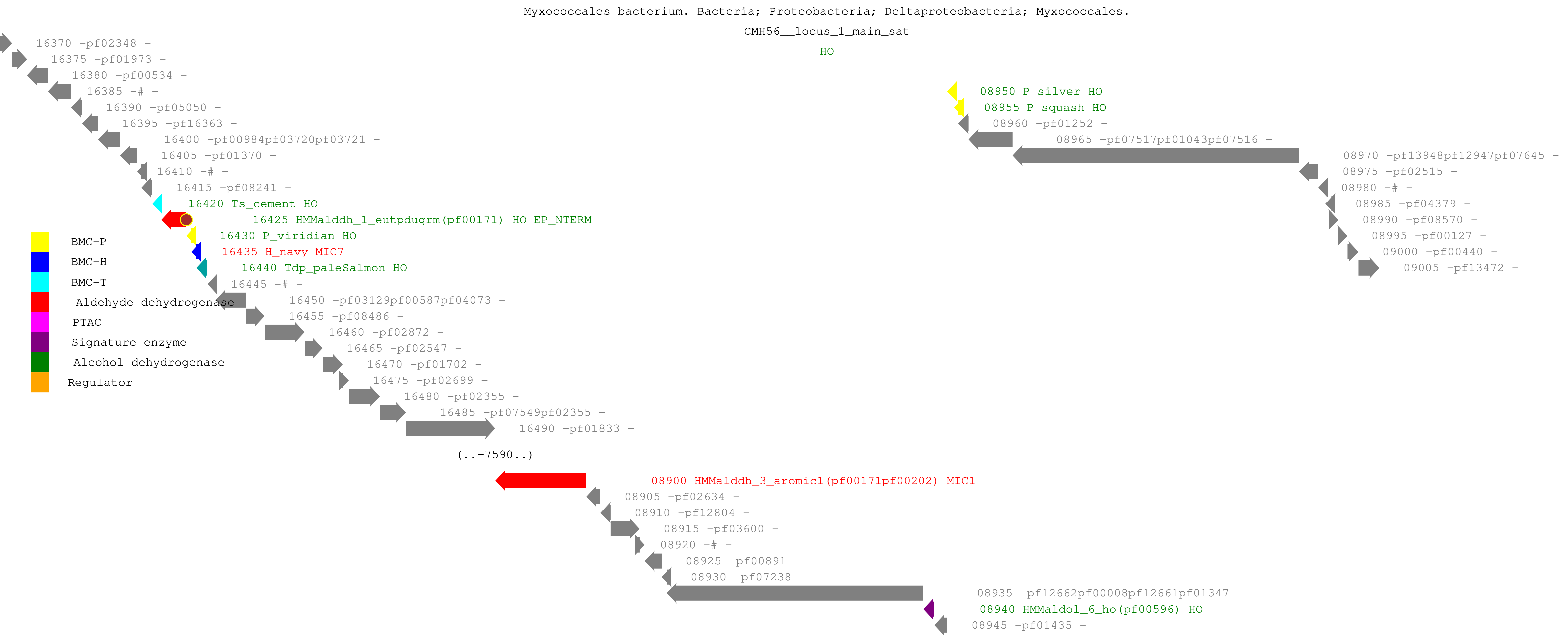
Myxococcales bacterium. Bacteria; Proteobacteria; Deltaproteobacteria; Myxococcales.
CMH56__locus_1_main_sat
HO
16370 -pf02348 -
16375 -pf01973 -
16380 -pf00534 -
16385 -# -
16390 -pf05050 -
16395 -pf16363 -
16400 -pf00984pf03720pf03721 -
16405 -pf01370 -
16410 -# -
16415 -pf08241 -
16420 Ts_cement HO
16425 HMMalddh_1_eutpdugrm(pf00171) HO EP_NTERM
16430 P_viridian HO
16435 H_navy MIC7
16440 Tdp_paleSalmon HO
16445 -# -
16450 -pf03129pf00587pf04073 -
16455 -pf08486 -
16460 -pf02872 -
16465 -pf02547 -
16470 -pf01702 -
16475 -pf02699 -
16480 -pf02355 -
16485 -pf07549pf02355 -
16490 -pf01833 -
(..-7590..)
08900 HMMalddh_3_aromic1(pf00171pf00202) MIC1
08905 -pf02634 -
08910 -pf12804 -
08915 -pf03600 -
08920 -# -
08925 -pf00891 -
08930 -pf07238 -
08935 -pf12662pf00008pf12661pf01347 -
08940 HMMaldol_6_ho(pf00596) HO
08945 -pf01435 -
08950 P_silver HO
08955 P_squash HO
08960 -pf01252 -
08965 -pf07517pf01043pf07516 -
08970 -pf13948pf12947pf07645 -
08975 -pf02515 -
08980 -# -
08985 -pf04379 -
08990 -pf08570 -
08995 -pf00127 -
09000 -pf00440 -
09005 -pf13472 -
BMC-P
BMC-H
BMC-T
Aldehyde dehydrogenase
PTAC
Signature enzyme
Alcohol dehydrogenase
Regulator

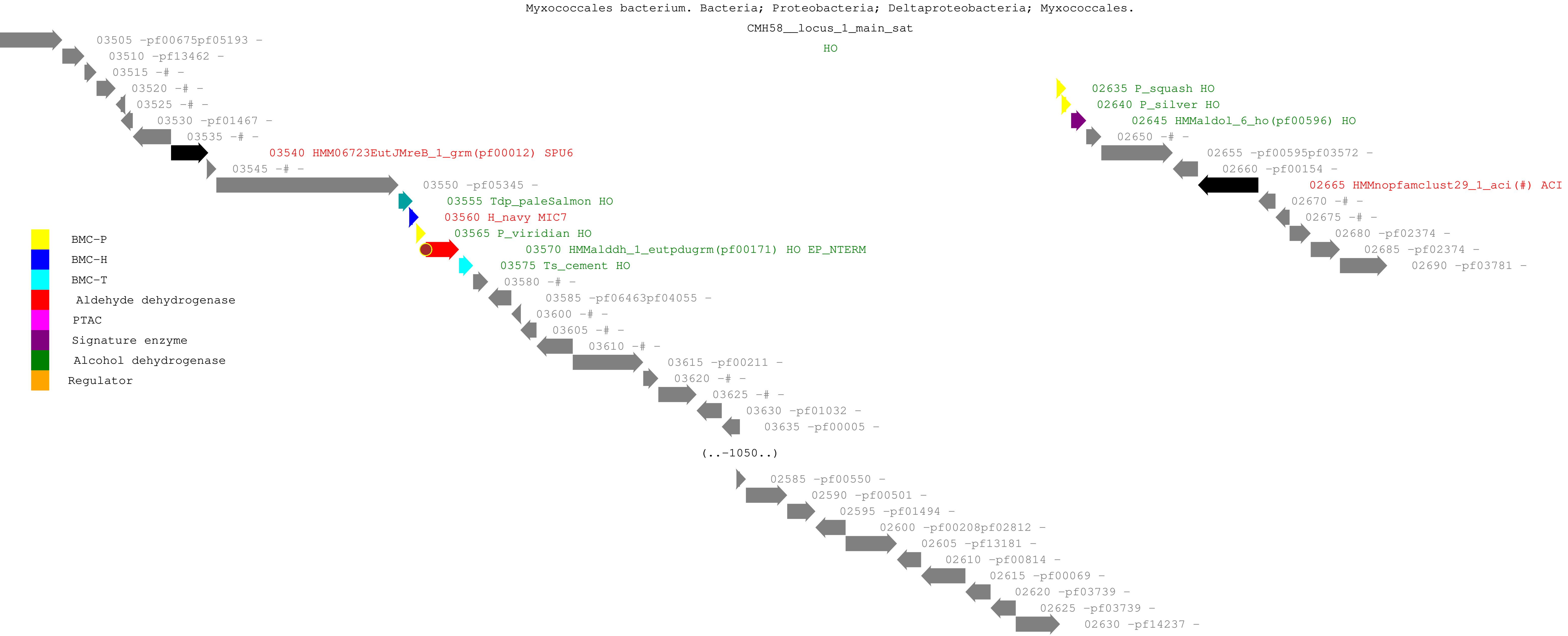

Myxococcales bacterium. Bacteria; Proteobacteria; Deltaproteobacteria; Myxococcales.
CMH58__locus_1_main_sat
HO
03505 -pf00675pf05193 -
03510 -pf13462 -
03515 -# -
03520 -# -
03525 -# -
03530 -pf01467 -
03535 -# -
03540 HMM06723EutJMreB_1_grm(pf00012) SPU6
03545 -# -
03550 -pf05345 -
03555 Tdp_paleSalmon HO
03560 H_navy MIC7
03565 P_viridian HO
03570 HMMalddh_1_eutpdugrm(pf00171) HO EP_NTERM
03575 Ts_cement HO
03580 -# -
03585 -pf06463pf04055 -
03600 -# -
03605 -# -
03610 -# -
03615 -pf00211 -
03620 -# -
03625 -# -
03630 -pf01032 -
03635 -pf00005 -
(..-1050..)
02585 -pf00550 -
02590 -pf00501 -
02595 -pf01494 -
02600 -pf00208pf02812 -
02605 -pf13181 -
02610 -pf00814 -
02615 -pf00069 -
02620 -pf03739 -
02625 -pf03739 -
02630 -pf14237 -
02635 P_squash HO
02640 P_silver HO
02645 HMMaldol_6_ho(pf00596) HO
02650 -# -
02655 -pf00595pf03572 -
02660 -pf00154 -
02665 HMMnopfamclust29_1_aci(#) ACI
02670 -# -
02675 -# -
02680 -pf02374 -
02685 -pf02374 -
02690 -pf03781 -
BMC-P
BMC-H
BMC-T
Aldehyde dehydrogenase
PTAC
Signature enzyme
Alcohol dehydrogenase
Regulator

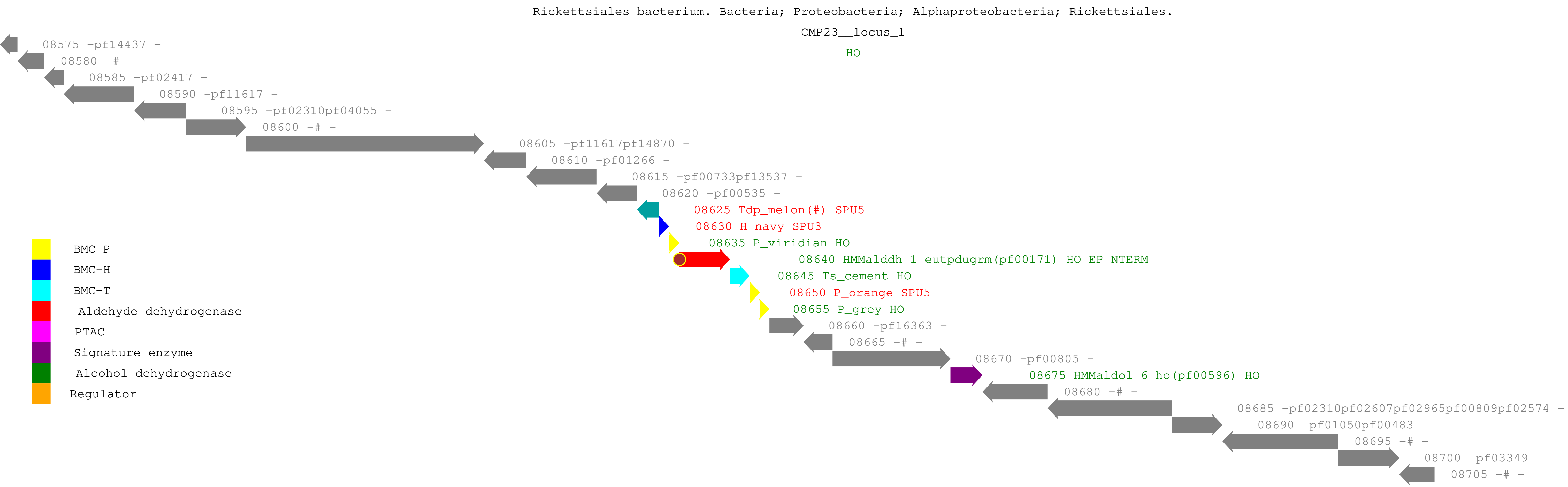
Rickettsiales bacterium. Bacteria; Proteobacteria; Alphaproteobacteria; Rickettsiales.
CMP23__locus_1
HO
08575 -pf14437 -
08580 -# -
08585 -pf02417 -
08590 -pf11617 -
08595 -pf02310pf04055 -
08600 -# -
08605 -pf11617pf14870 -
08610 -pf01266 -
08615 -pf00733pf13537 -
08620 -pf00535 -
08625 Tdp_melon(#) SPU5
08630 H_navy SPU3
08635 P_viridian HO
08640 HMMalddh_1_eutpdugrm(pf00171) HO EP_NTERM
08645 Ts_cement HO
08650 P_orange SPU5
08655 P_grey HO
08660 -pf16363 -
08665 -# -
08670 -pf00805 -
08675 HMMaldol_6_ho(pf00596) HO
08680 -# -
08685 -pf02310pf02607pf02965pf00809pf02574 -
08690 -pf01050pf00483 -
08695 -# -
08700 -pf03349 -
08705 -# -
BMC-P
BMC-H
BMC-T
Aldehyde dehydrogenase
PTAC
Signature enzyme
Alcohol dehydrogenase
Regulator

Deltaproteobacteria bacterium CG_4_9_14_3_um_filter_63_12. Bacteria; Proteobacteria; Deltaproteobacteria.

CO108__locus_1_main_sat

HO

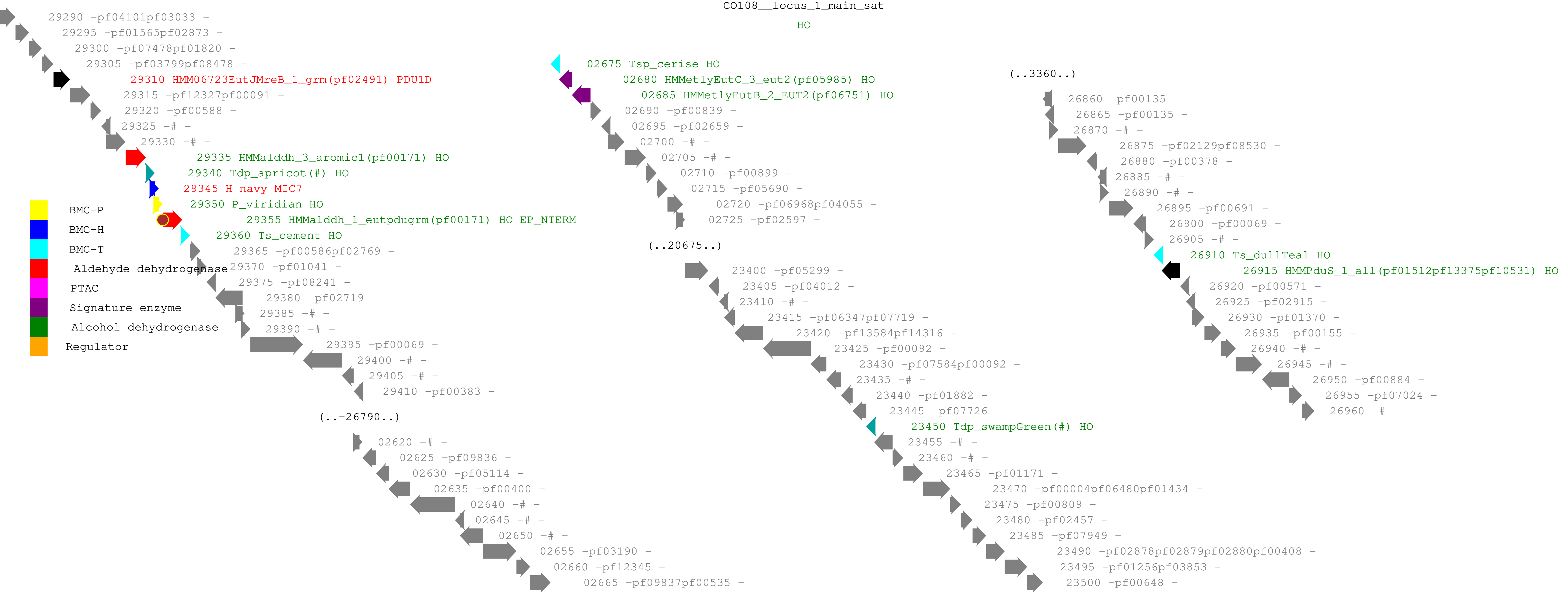

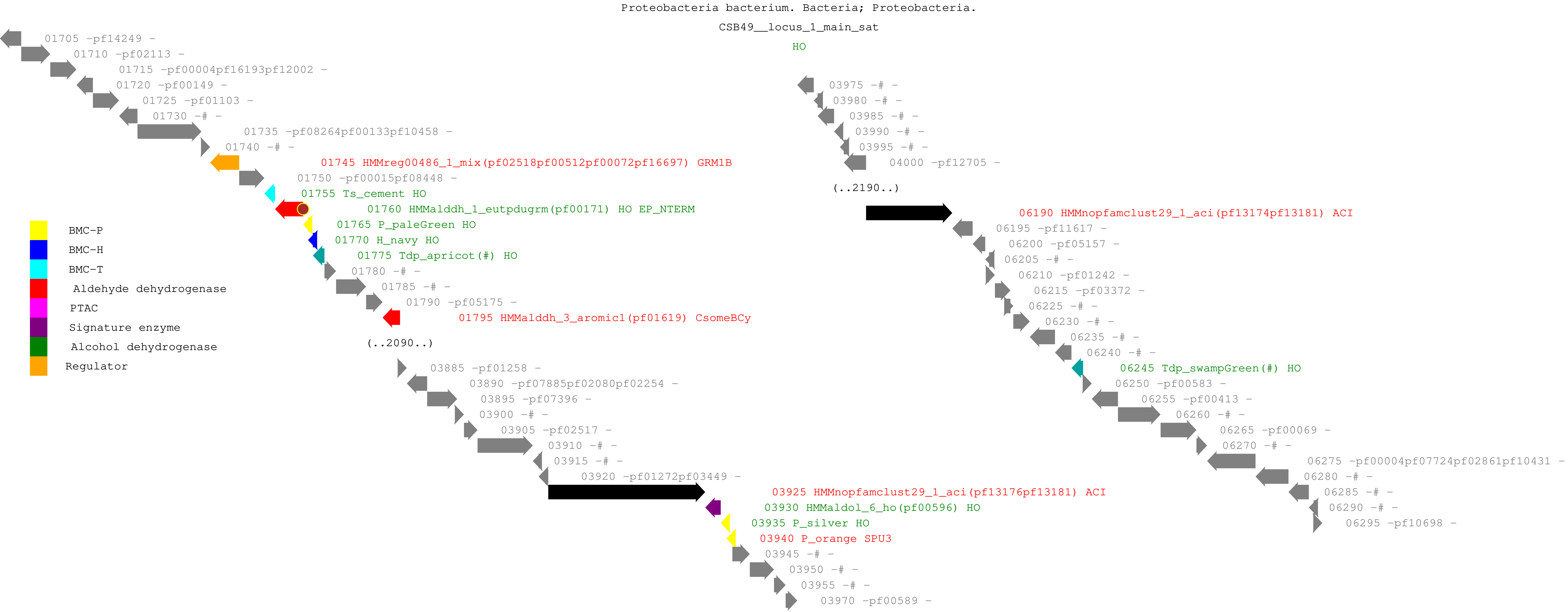
Proteobacteria bacterium. Bacteria; Proteobacteria.
CSB49__locus_1_main_sat
HO
01705 -pf14249 -
01710 -pf02113 -
01715 -pf00004pf16193pf12002 -
01720 -pf00149 -
01725 -pf01103 -
01730 -# -
01735 -pf08264pf00133pf10458 -
01740 -# -
01745 HMMreg00486_1_mix(pf02518pf00512pf00072pf16697) GRM1B
01750 -pf00015pf08448 -
01755 Ts_cement HO
01760 HMMalddh_1_eutpdugrm(pf00171) HO EP_NTERM
01765 P_paleGreen HO
01770 H_navy HO
01775 Tdp_apricot(#) HO
01780 -# -
01785 -# -
01790 -pf05175 -
01795 HMMalddh_3_aromic1(pf01619) CsomeBCy
(..2090..)
03885 -pf01258 -
03890 -pf07885pf02080pf02254 -
03895 -pf07396 -
03900 -# -
03905 -pf02517 -
03910 -# -
03915 -# -
03920 -pf01272pf03449 -
03925 HMMnopfamclust29_1_aci(pf13176pf13181) ACI
03930 HMMaldol_6_ho(pf00596) HO
03935 P_silver HO
03940 P_orange SPU3
03945 -# -
03950 -# -
03955 -# -
03970 -pf00589 -
03975 -# -
03980 -# -
03985 -# -
03990 -# -
03995 -# -
04000 -pf12705 -
(..2190..)
06190 HMMnopfamclust29_1_aci(pf13174pf13181) ACI
06195 -pf11617 -
06200 -pf05157 -
06205 -# -
06210 -pf01242 -
06215 -pf03372 -
06225 -# -
06230 -# -
06235 -# -
06240 -# -
06245 Tdp_swampGreen(#) HO
06250 -pf00583 -
06255 -pf00413 -
06260 -# -
06265 -pf00069 -
06270 -# -
06275 -pf00004pf07724pf02861pf10431 -
06280 -# -
06285 -# -
06290 -# -
06295 -pf10698 -
BMC-P
BMC-H
BMC-T
Aldehyde dehydrogenase
PTAC
Signature enzyme
Alcohol dehydrogenase
Regulator

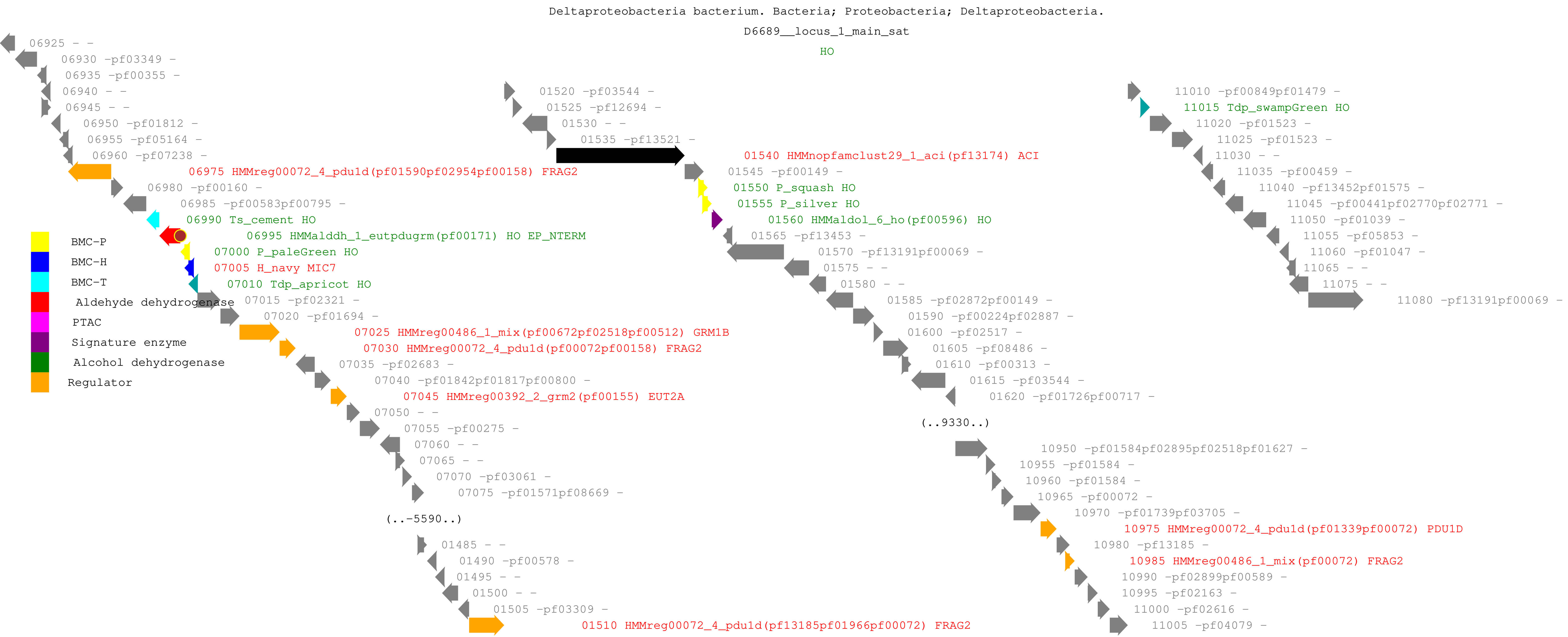
Deltaproteobacteria bacterium. Bacteria; Proteobacteria; Deltaproteobacteria.
D6689__locus_1_main_sat
HO
06925 - -
06930 -pf03349 -
06935 -pf00355 -
06940 - -
06945 - -
06950 -pf01812 -
06955 -pf05164 -
06960 -pf07238 -
06975 HMMreg00072_4_pdu1d(pf01590pf02954pf00158) FRAG2
06980 -pf00160 -
06985 -pf00583pf00795 -
06990 Ts_cement HO
06995 HMMalddh_1_eutpdugrm(pf00171) HO EP_NTERM
07000 P_paleGreen HO
07005 H_navy MIC7
07010 Tdp_apricot HO
07015 -pf02321 -
07020 -pf01694 -
07025 HMMreg00486_1_mix(pf00672pf02518pf00512) GRM1B
07030 HMMreg00072_4_pdu1d(pf00072pf00158) FRAG2
07035 -pf02683 -
07040 -pf01842pf01817pf00800 -
07045 HMMreg00392_2_grm2(pf00155) EUT2A
07050 - -
07055 -pf00275 -
07060 - -
07065 - -
07070 -pf03061 -
07075 -pf01571pf08669 -
(..-5590..)
01485 - -
01490 -pf00578 -
01495 - -
01500 - -
01505 -pf03309 -
01510 HMMreg00072_4_pdu1d(pf13185pf01966pf00072) FRAG2
01520 -pf03544 -
01525 -pf12694 -
01530 - -
01535 -pf13521 -
01540 HMMnopfamclust29_1_aci(pf13174) ACI
01545 -pf00149 -
01550 P_squash HO
01555 P_silver HO
01560 HMMaldol_6_ho(pf00596) HO
01565 -pf13453 -
01570 -pf13191pf00069 -
01575 - -
01580 - -
01585 -pf02872pf00149 -
01590 -pf00224pf02887 -
01600 -pf02517 -
01605 -pf08486 -
01610 -pf00313 -
01615 -pf03544 -
01620 -pf01726pf00717 -
(..9330..)
10950 -pf01584pf02895pf02518pf01627 -
10955 -pf01584 -
10960 -pf01584 -
10965 -pf00072 -
10970 -pf01739pf03705 -
10975 HMMreg00072_4_pdu1d(pf01339pf00072) PDU1D
10980 -pf13185 -
10985 HMMreg00486_1_mix(pf00072) FRAG2
10990 -pf02899pf00589 -
10995 -pf02163 -
11000 -pf02616 -
11005 -pf04079 -
11010 -pf00849pf01479 -
11015 Tdp_swampGreen HO
11020 -pf01523 -
11025 -pf01523 -
11030 - -
11035 -pf00459 -
11040 -pf13452pf01575 -
11045 -pf00441pf02770pf02771 -
11050 -pf01039 -
11055 -pf05853 -
11060 -pf01047 -
11065 - -
11075 - -
11080 -pf13191pf00069 -
BMC-P
BMC-H
BMC-T
Aldehyde dehydrogenase
PTAC
Signature enzyme
Alcohol dehydrogenase
Regulator

Deltaproteobacteria bacterium. Bacteria; Proteobacteria; Deltaproteobacteria.

D6729__locus_1_main_sat

HO

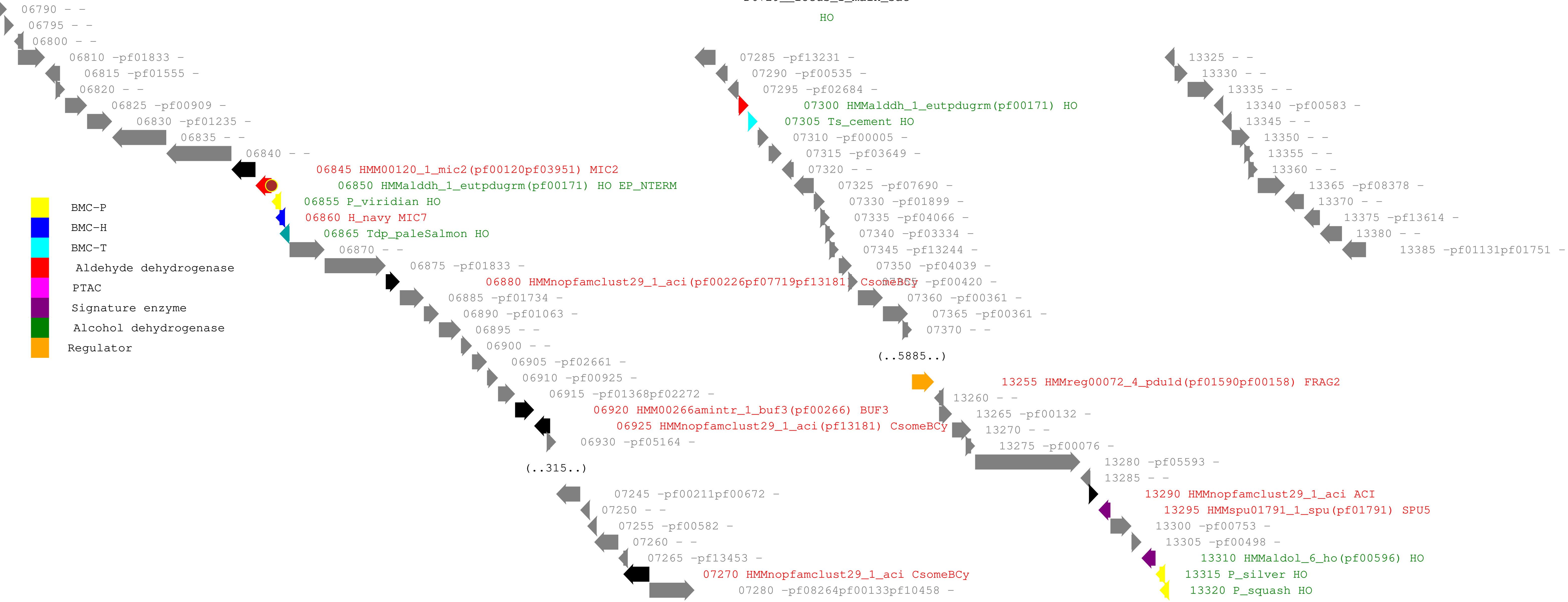

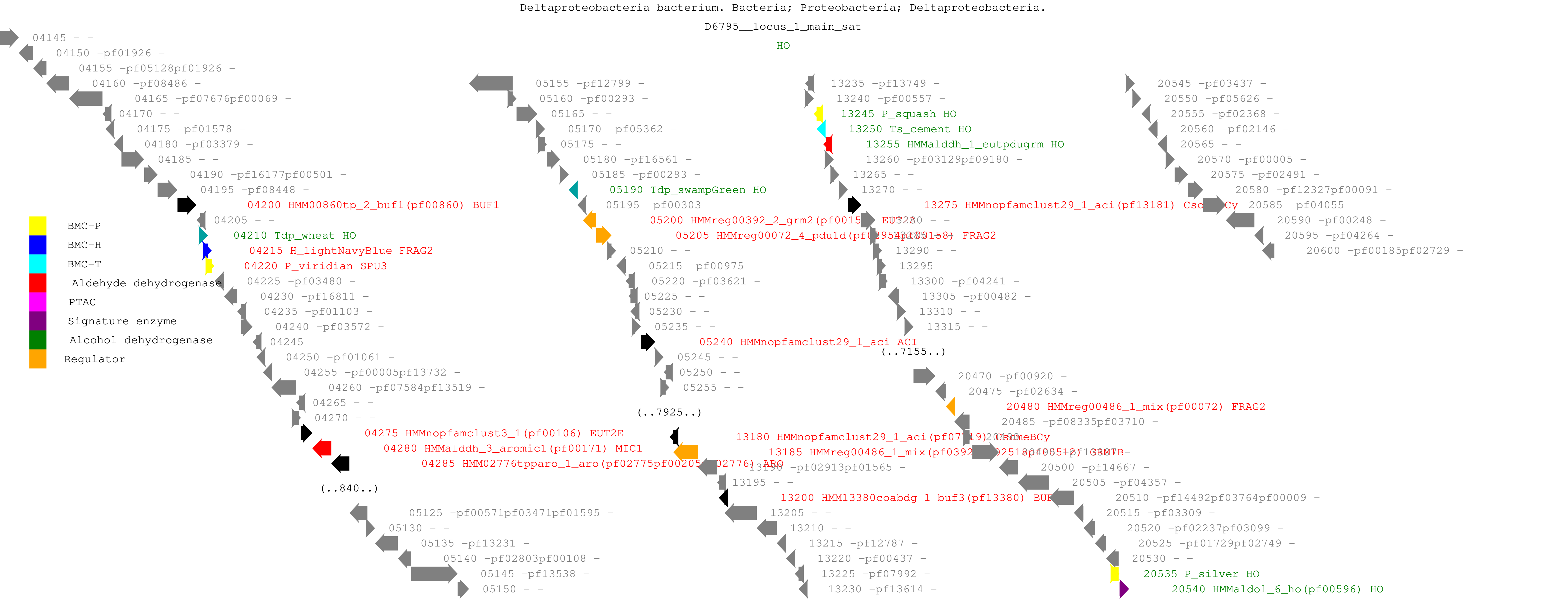

Deltaproteobacteria bacterium. Bacteria; Proteobacteria; Deltaproteobacteria.
D6795__locus_1_main_sat
HO
BMC-P
BMC-H
BMC-T
Aldehyde dehydrogenase
PTAC
Signature enzyme
Alcohol dehydrogenase
Regulator
04145 - -
04150 -pf01926 -
04155 -pf05128pf01926 -
04160 -pf08486 -
04165 -pf07676pf00069 -
04170 - -
04175 -pf01578 -
04180 -pf03379 -
04185 - -
04190 -pf16177pf00501 -
04195 -pf08448 -
04200 HMM00860tp_2_buf1(pf00860) BUF1
04205 - -
04210 Tdp_wheat HO
04215 H_lightNavyBlue FRAG2
04220 P_viridian SPU3
04225 -pf03480 -
04230 -pf16811 -
04235 -pf01103 -
04240 -pf03572 -
04245 - -
04250 -pf01061 -
04255 -pf00005pf13732 -
04260 -pf07584pf13519 -
04265 - -
04270 - -
04275 HMMnopfamclust3_1(pf00106) EUT2E
04280 HMMaldd h_3_aromic1(pf00171) MIC1
04285 HMM02776tpparo_1_aro(pf02775pf00205pf02776) ARO
(..840..)
05125 -pf00571pf03471pf01595 -
05130 - -
05135 -pf13231 -
05140 -pf02803pf00108 -
05145 -pf13538 -
05150 - -
05155 -pf12799 -
05160 -pf00293 -
05165 - -
05170 -pf05362 -
05175 - -
05180 -pf16561 -
05185 -pf00293 -
05190 Tdp_swampGreen HO
05195 -pf00303 -
05200 HMMreg00392_2_grm2(pf0015 EUT2E
05205 HMMreg00072_4_pdu1d(pf0 pf00158) FRAG2
05210 - -
05215 -pf00975 -
05220 -pf03621 -
05225 - -
05230 - -
05235 - -
05240 HMMnopfamclust29_1_aci ACI
05245 - -
05250 - -
05255 - -
(..7925..)
13180 HMMnopfamclust29_1_aci(pf07 9) CsomeBCy
13185 HMMreg00486_1_mix(pf0392 GRM1B
13190 -pf02913pf01565 -
13195 - -
13200 HMM13380coabdg_1_buf3(pf13380) BUF
13205 - -
13210 - -
13215 -pf12787 -
13220 -pf00437 -
13225 -pf07992 -
13230 -pf13614 -
13235 -pf13749 -
13240 -pf00557 -
13245 P_squash HO
13250 Ts_cement HO
13255 HMMaldd h_1_eutpdugrm HO
13260 -pf03129pf09180 -
13265 - -
13270 - -
13275 HMMnopfamclust29_1_aci(pf13181) CsomeBCy
13290 - -
13295 - -
13300 -pf04241 -
13305 -pf00482 -
13310 - -
13315 - -
(..7155..)
20470 -pf00920 -
20475 -pf02634 -
20480 HMMreg00486_1_mix(pf00072) FRAG2
20485 -pf08335pf03710 -
20500 -pf14667 -
20505 -pf04357 -
20510 -pf14492pf03764pf00009 -
20515 -pf03309 -
20520 -pf02237pf03099 -
20525 -pf01729pf02749 -
20530 - -
20535 P_silver HO
20540 HMMaldol_6_ho(pf00596) HO
20545 -pf03437 -
20550 -pf05626 -
20555 -pf02368 -
20560 -pf02146 -
20565 - -
20570 -pf00005 -
20575 -pf02491 -
20580 -pf12327pf00091 -
20585 -pf04055 -
20590 -pf00248 -
20595 -pf04264 -
20600 -pf00185pf02729 -

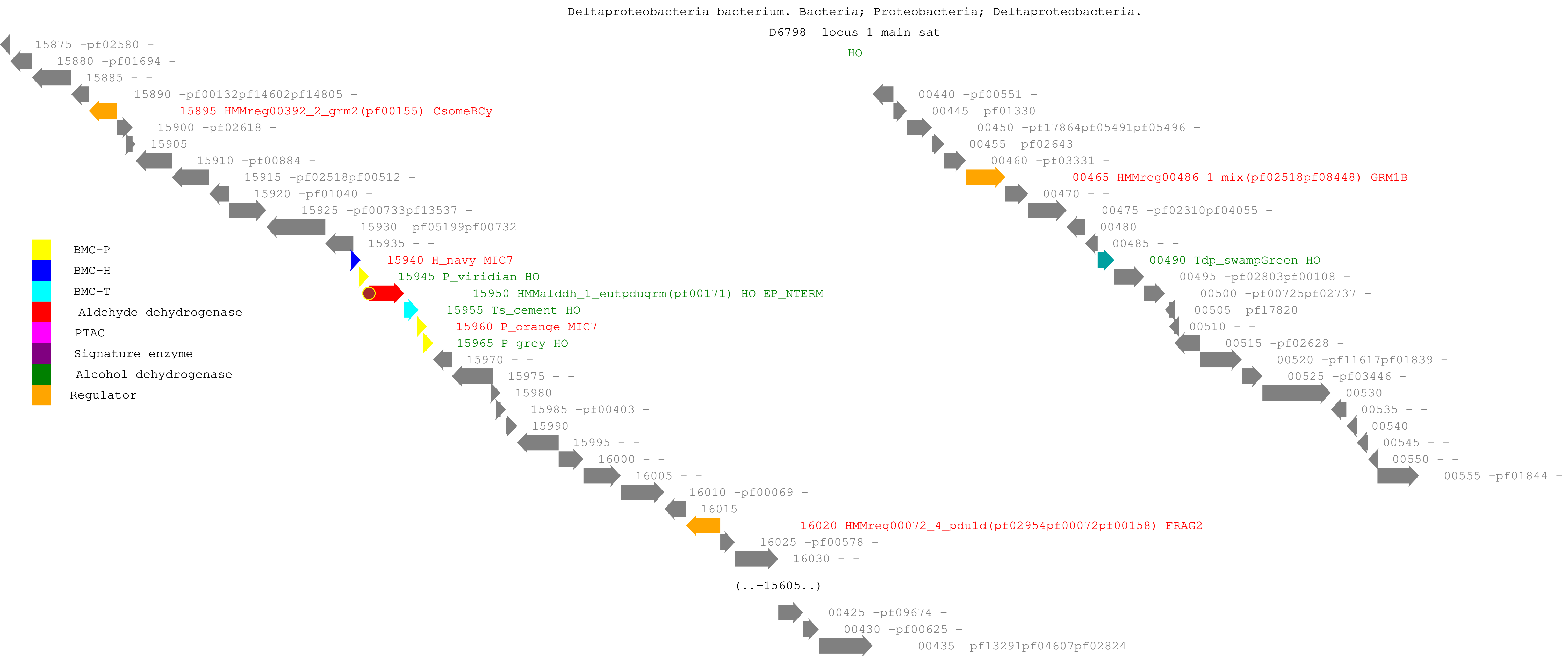

Deltaproteobacteria bacterium. Bacteria; Proteobacteria; Deltaproteobacteria.
D6798__locus_1_main_sat
HO
15875 -pf02580 -
15880 -pf01694 -
15885 - -
15890 -pf00132pf14602pf14805 -
15895 HMMreg00392_2_grm2(pf00155) CsomeBCy
15900 -pf02618 -
15905 - -
15910 -pf00884 -
15915 -pf02518pf00512 -
15920 -pf01040 -
15925 -pf00733pf13537 -
15930 -pf05199pf00732 -
15935 - -
15940 H_navy MIC7
15945 P_viridian HO
15950 HMMalddh_1_eutpdugrm(pf00171) HO EP_NTERM
15955 Ts_cement HO
15960 P_orange MIC7
15965 P_grey HO
15970 - -
15975 - -
15980 - -
15985 -pf00403 -
15990 - -
15995 - -
16000 - -
16005 - -
16010 -pf00069 -
16015 - -
16020 HMMreg00072_4_pdu1d(pf02954pf00072pf00158) FRAG2
16025 -pf00578 -
16030 - -
(..-15605..)
00425 -pf09674 -
00430 -pf00625 -
00435 -pf13291pf04607pf02824 -
00440 -pf00551 -
00445 -pf01330 -
00450 -pf17864pf05491pf05496 -
00455 -pf02643 -
00460 -pf03331 -
00465 HMMreg00486_1_mix(pf02518pf08448) GRM1B
00470 - -
00475 -pf02310pf04055 -
00480 - -
00485 - -
00490 Tdp_swampGreen HO
00495 -pf02803pf00108 -
00500 -pf00725pf02737 -
00505 -pf17820 -
00510 - -
00515 -pf02628 -
00520 -pf11617pf01839 -
00525 -pf03446 -
00530 - -
00535 - -
00540 - -
00545 - -
00550 - -
00555 -pf01844 -
BMC-P
BMC-H
BMC-T
Aldehyde dehydrogenase
PTAC
Signature enzyme
Alcohol dehydrogenase
Regulator

Hyalangium minutum. Bacteria; Proteobacteria; Deltaproteobacteria; Myxococcales;

DB31__locus_1_main_sat

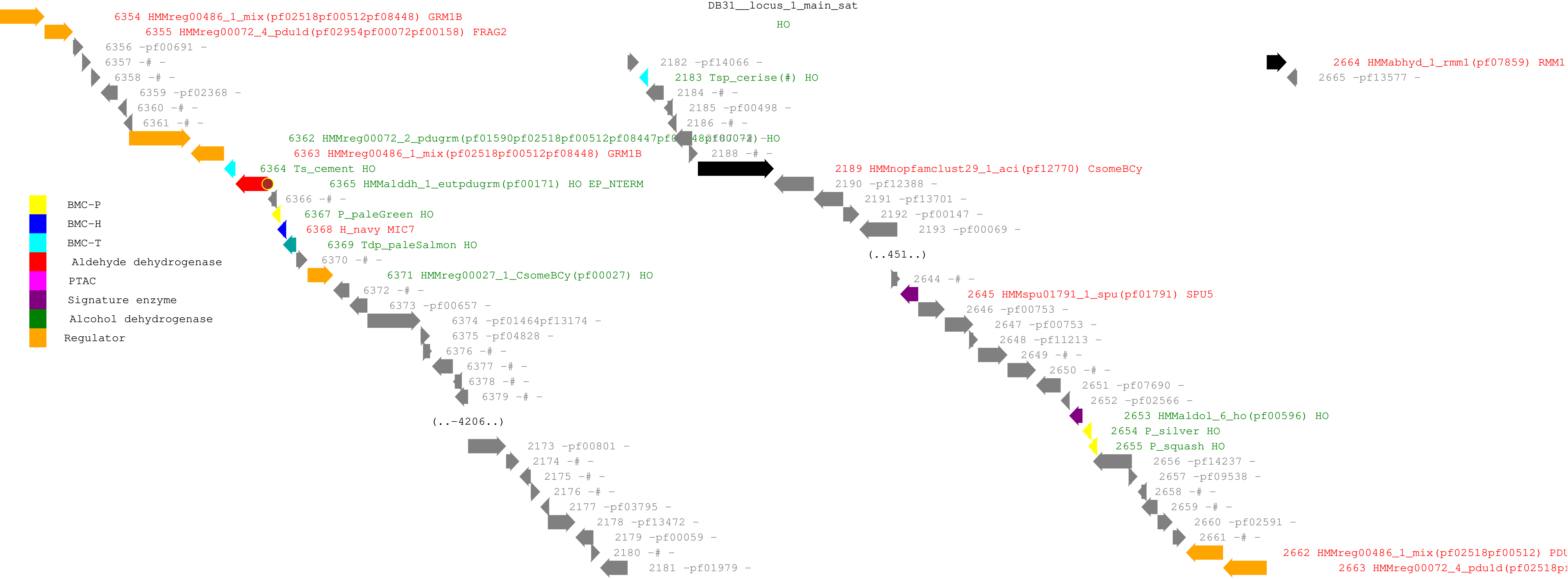

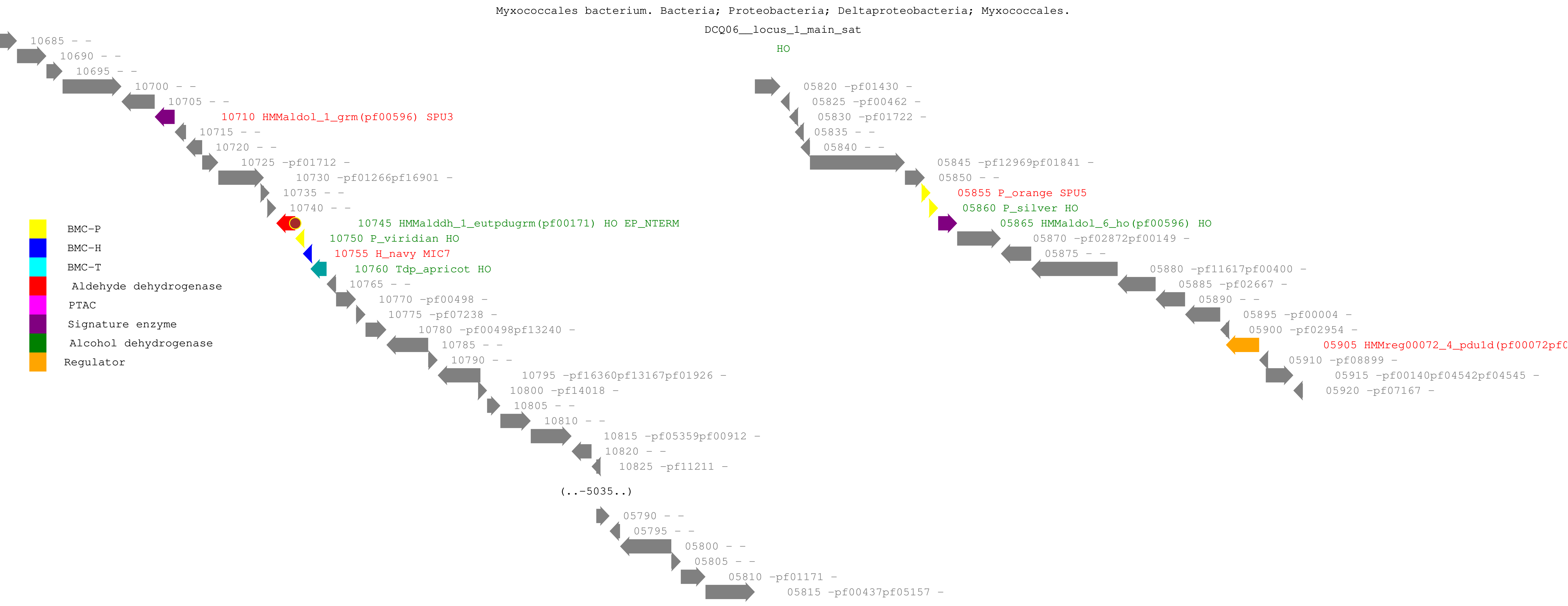

Myxococcales bacterium. Bacteria; Proteobacteria; Deltaproteobacteria; Myxococcales.
DCQ06__locus_1_main_sat
HO
10685 - -
10690 - -
10695 - -
10700 - -
10705 - -
10710 HMMaldol_1_grm(pf00596) SPU3
10715 - -
10720 - -
10725 -pf01712 -
10730 -pf01266pf16901 -
10735 - -
10740 - -
10745 HMMalddh_1_eutpdugrm(pf00171) HO EP_NTERM
10750 P_viridian HO
10755 H_navy MIC7
10760 Tdp_apricot HO
10765 - -
10770 -pf00498 -
10775 -pf07238 -
10780 -pf00498pf13240 -
10785 - -
10790 - -
10795 -pf16360pf13167pf01926 -
10800 -pf14018 -
10805 - -
10810 - -
10815 -pf05359pf00912 -
10820 - -
10825 -pf11211 -
(..-5035..)
05790 - -
05795 - -
05800 - -
05805 - -
05810 -pf01171 -
05815 -pf00437pf05157 -
05820 -pf01430 -
05825 -pf00462 -
05830 -pf01722 -
05835 - -
05840 - -
05845 -pf12969pf01841 -
05850 - -
05855 P_orange SPU5
05860 P_silver HO
05865 HMMaldol_6_ho(pf00596) HO
05870 -pf02872pf00149 -
05875 - -
05880 -pf11617pf00400 -
05885 -pf02667 -
05890 - -
05895 -pf00004 -
05900 -pf02954 -
05905 HMMreg00072_4_pdu1d(pf00072pf0
05910 -pf08899 -
05915 -pf00140pf04542pf04545 -
05920 -pf07167 -
BMC-P
BMC-H
BMC-T
Aldehyde dehydrogenase
PTAC
Signature enzyme
Alcohol dehydrogenase
Regulator

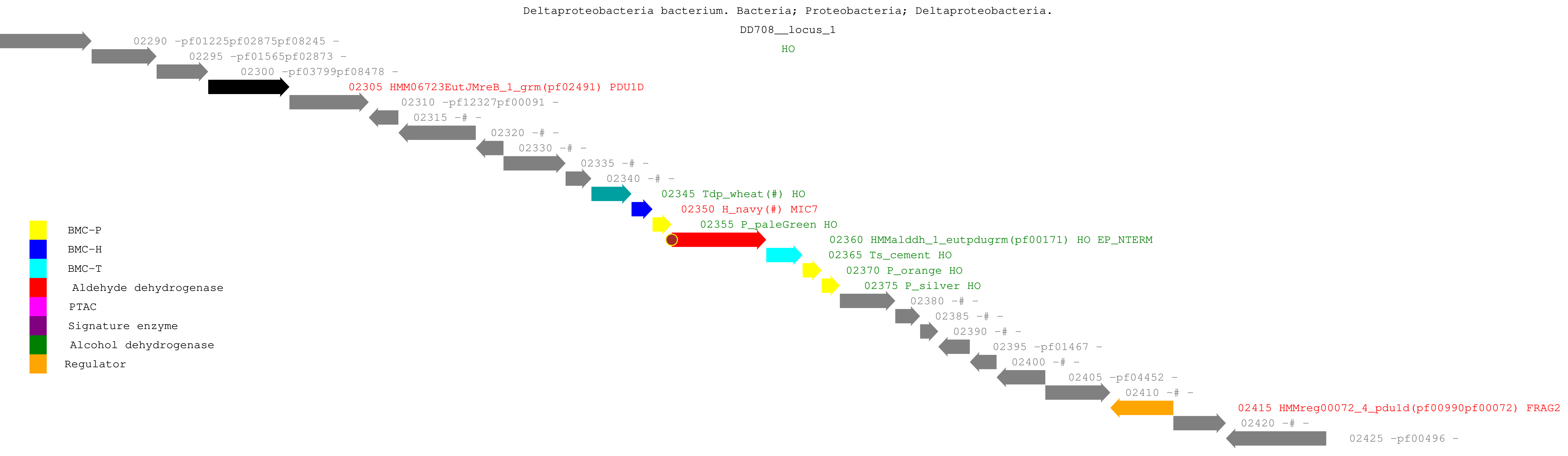
Deltaproteobacteria bacterium. Bacteria; Proteobacteria; Deltaproteobacteria.
DD708__locus_1
HO
02290 -pf01225pf02875pf08245 -
02295 -pf01565pf02873 -
02300 -pf03799pf08478 -
02305 HMM06723EutJMreB_1_grm(pf02491) PDU1D
02310 -pf12327pf00091 -
02315 -# -
02320 -# -
02330 -# -
02335 -# -
02340 -# -
02345 Tdp_wheat(#) HO
02350 H_navy(#) MIC7
02355 P_paleGreen HO
02360 HMMalddh_1_eutpdugrm(pf00171) HO EP_NTERM
02365 Ts_cement HO
02370 P_orange HO
02375 P_silver HO
02380 -# -
02385 -# -
02390 -# -
02395 -pf01467 -
02400 -# -
02405 -pf04452 -
02410 -# -
02415 HMMreg00072_4_pdu1d(pf00990pf00072) FRAG2
02420 -# -
02425 -pf00496 -
BMC-P
BMC-H
BMC-T
Aldehyde dehydrogenase
PTAC
Signature enzyme
Alcohol dehydrogenase
Regulator

Myxococcales bacterium. Bacteria; Proteobacteria; Deltaproteobacteria; Myxococcales.

DEB46__locus_fragmented

HO

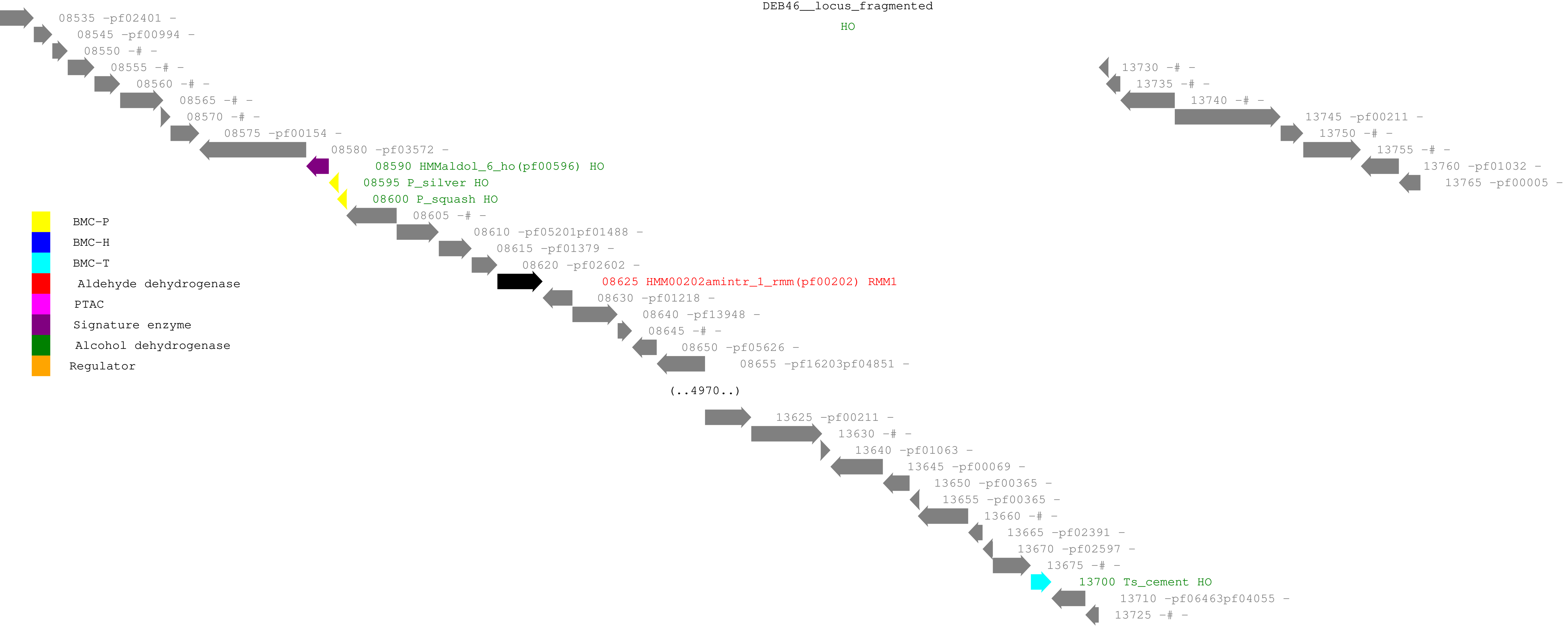

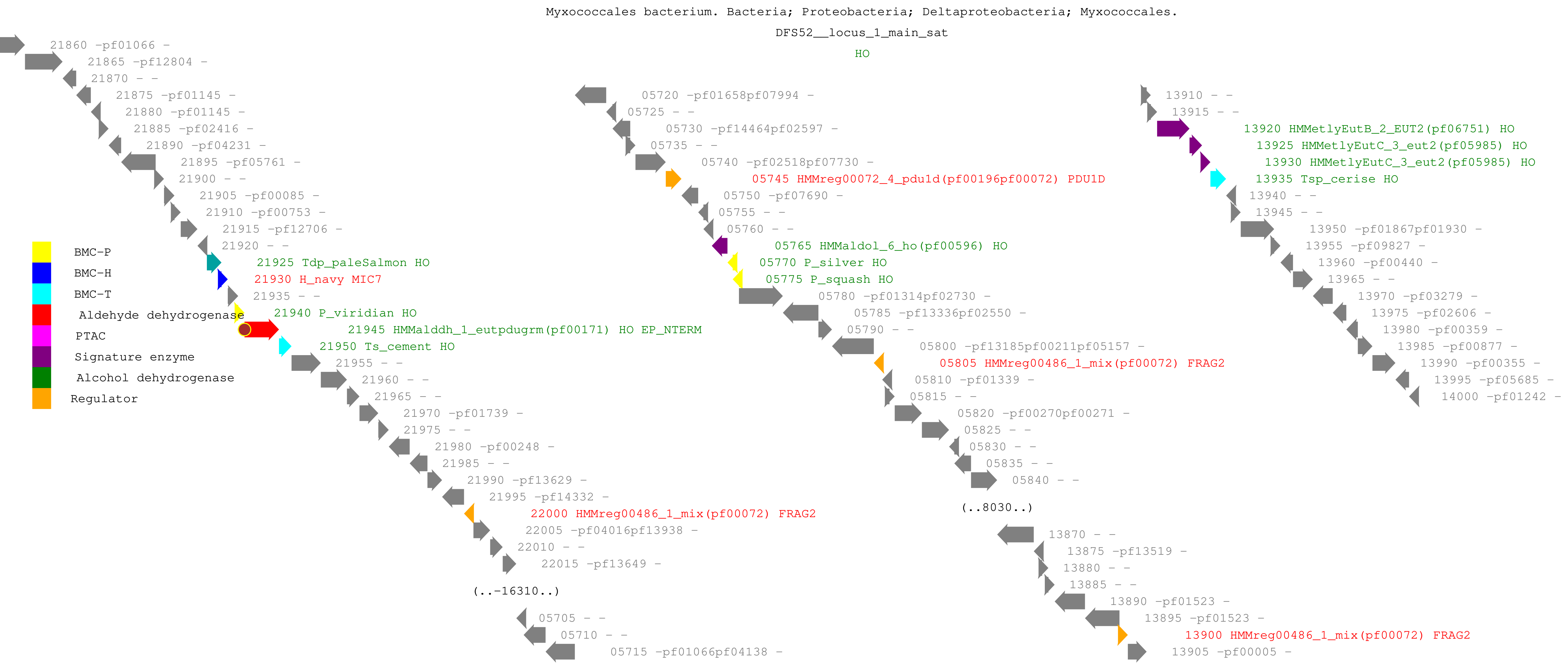
Myxococcales bacterium. Bacteria; Proteobacteria; Deltaproteobacteria; Myxococcales.
DFS52__locus_1_main_sat
HO
21860 -pf01066 -
21865 -pf12804 -
21870 - -
21875 -pf01145 -
21880 -pf01145 -
21885 -pf02416 -
21890 -pf04231 -
21895 -pf05761 -
21900 - -
21905 -pf00085 -
21910 -pf00753 -
21915 -pf12706 -
21920 - -
21925 Tdp_paleSalmon HO
21930 H_navy MIC7
21935 - -
21940 P_viridian HO
21945 HMMalddh_1_eutpdugrm(pf00171) HO EP_NTERM
21950 Ts_cement HO
21955 - -
21960 - -
21965 - -
21970 -pf01739 -
21975 - -
21980 -pf00248 -
21985 - -
21990 -pf13629 -
21995 -pf14332 -
22000 HMMreg00486_1_mix(pf00072) FRAG2
22005 -pf04016pf13938 -
22010 - -
22015 -pf13649 -
(..-16310..)
05705 - -
05710 - -
05715 -pf01066pf04138 -
05720 -pf01658pf07994 -
05725 - -
05730 -pf14464pf02597 -
05735 - -
05740 -pf02518pf07730 -
05745 HMMreg00072_4_pdu1d(pf00196pf00072) PDU1D
05750 -pf07690 -
05755 - -
05760 - -
05765 HMMaldol_6_ho(pf00596) HO
05770 P_silver HO
05775 P_squash HO
05780 -pf01314pf02730 -
05785 -pf13336pf02550 -
05790 - -
05800 -pf13185pf00211pf05157 -
05805 HMMreg00486_1_mix(pf00072) FRAG2
05810 -pf01339 -
05815 - -
05820 -pf00270pf00271 -
05825 - -
05830 - -
05835 - -
05840 - -
(..8030..)
13870 - -
13875 -pf13519 -
13880 - -
13885 - -
13890 -pf01523 -
13895 -pf01523 -
13900 HMMreg00486_1_mix(pf00072) FRAG2
13905 -pf00005 -
13910 - -
13915 - -
13920 HMMetlyEutB_2_EUT2(pf06751) HO
13925 HMMetlyEutC_3_eut2(pf05985) HO
13930 HMMetlyEutC_3_eut2(pf05985) HO
13935 Tsp_cerise HO
13940 - -
13945 - -
13950 -pf01867pf01930 -
13955 -pf09827 -
13960 -pf00440 -
13965 - -
13970 -pf03279 -
13975 -pf02606 -
13980 -pf00359 -
13985 -pf00877 -
13990 -pf00355 -
13995 -pf05685 -
14000 -pf01242 -
BMC-P
BMC-H
BMC-T
Aldehyde dehydrogenase
PTAC
Signature enzyme
Alcohol dehydrogenase
Regulator

Archangium gephyra. Bacteria; Proteobacteria; Deltaproteobacteria; Myxococcales;

DI536__locus_1_main_sat

HO

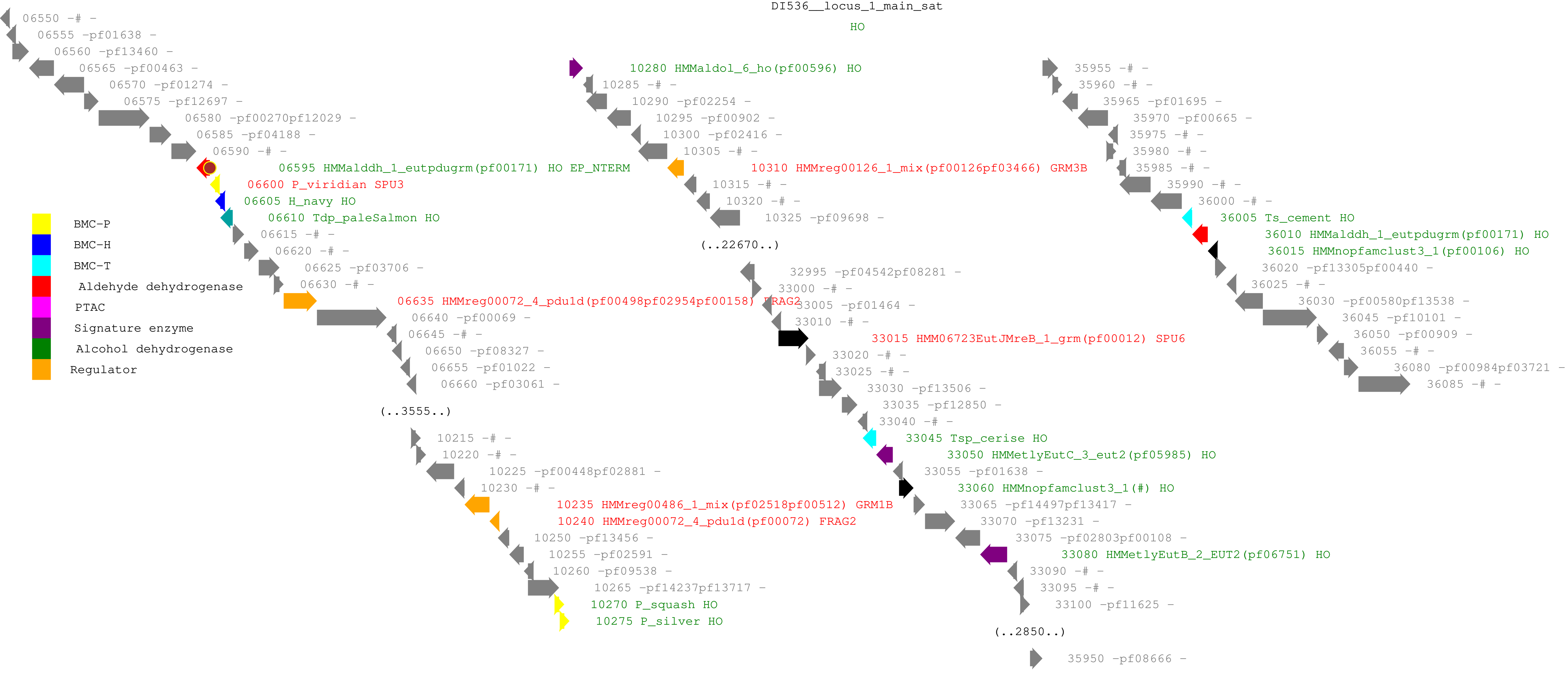

Deltaproteobacteria bacterium. Bacteria; Proteobacteria; Deltaproteobacteria.

E6J61__locus_1_main_sat

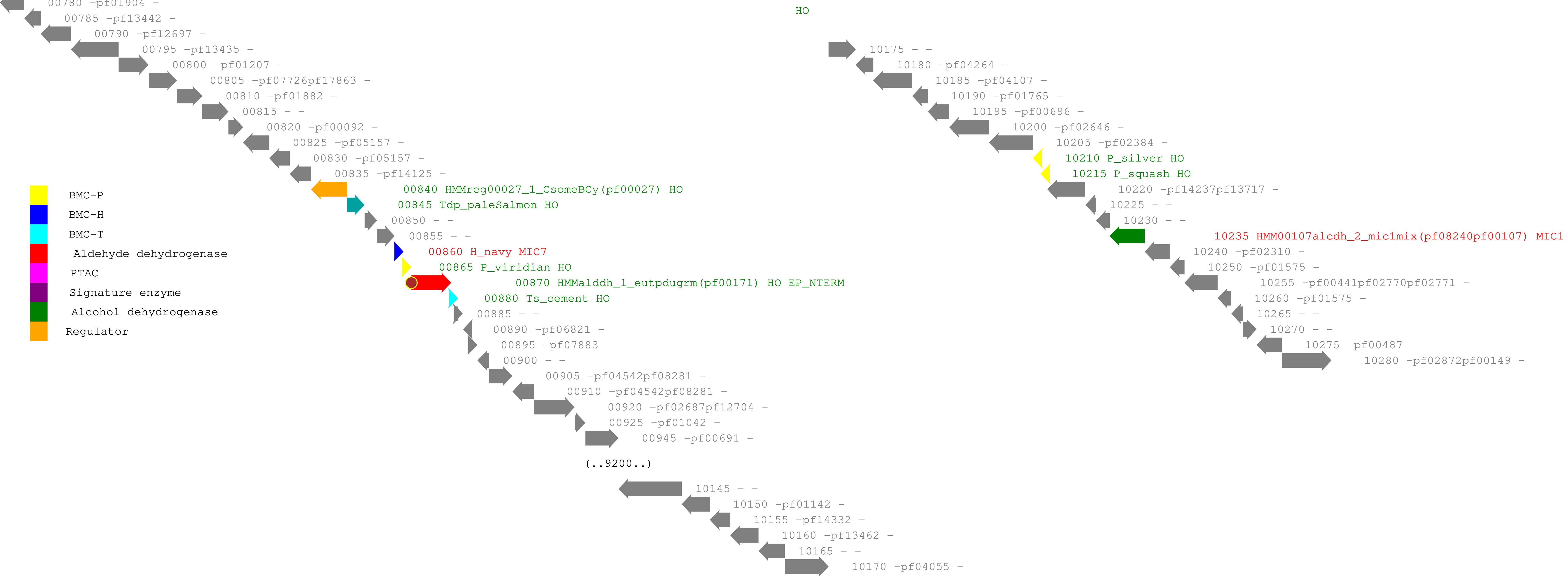

Deltaproteobacteria bacterium. Bacteria; Proteobacteria; Deltaproteobacteria.

E6J78__locus_1_main_sat

HO

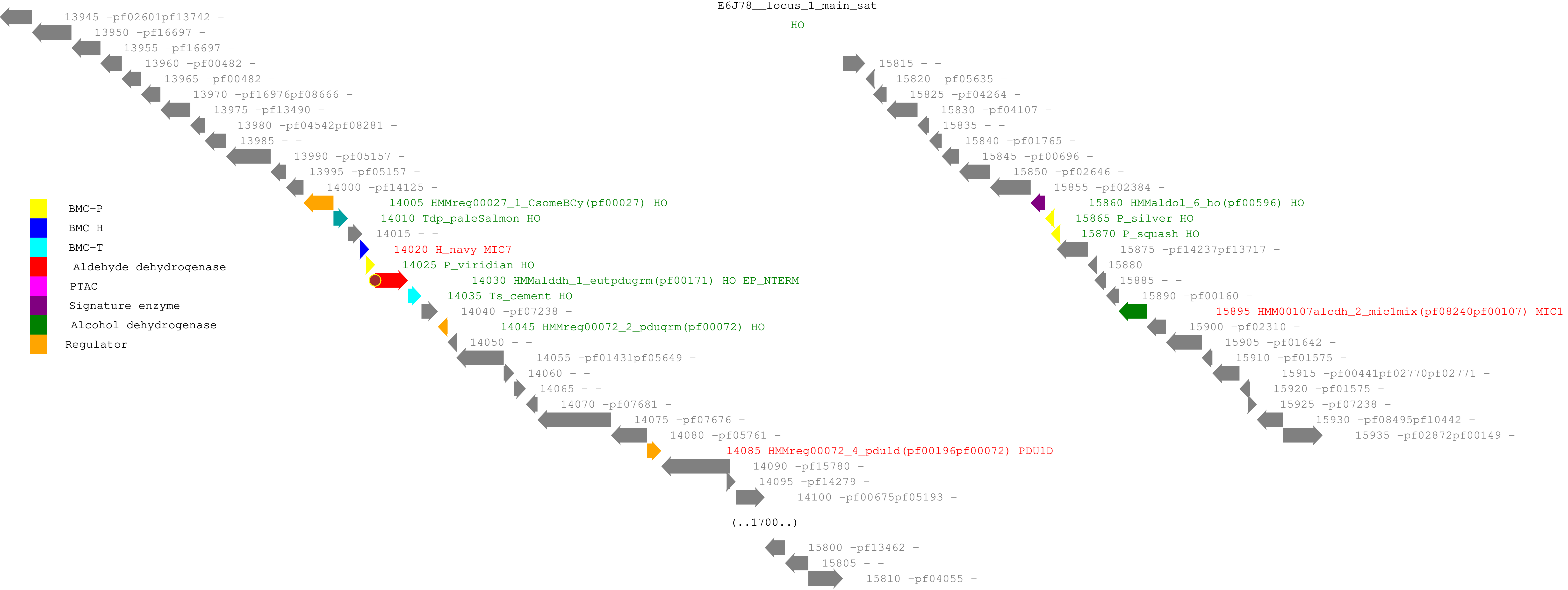

Deltaproteobacteria bacterium. Bacteria; Proteobacteria; Deltaproteobacteria.

E6J88__locus_1_main_sat

HO

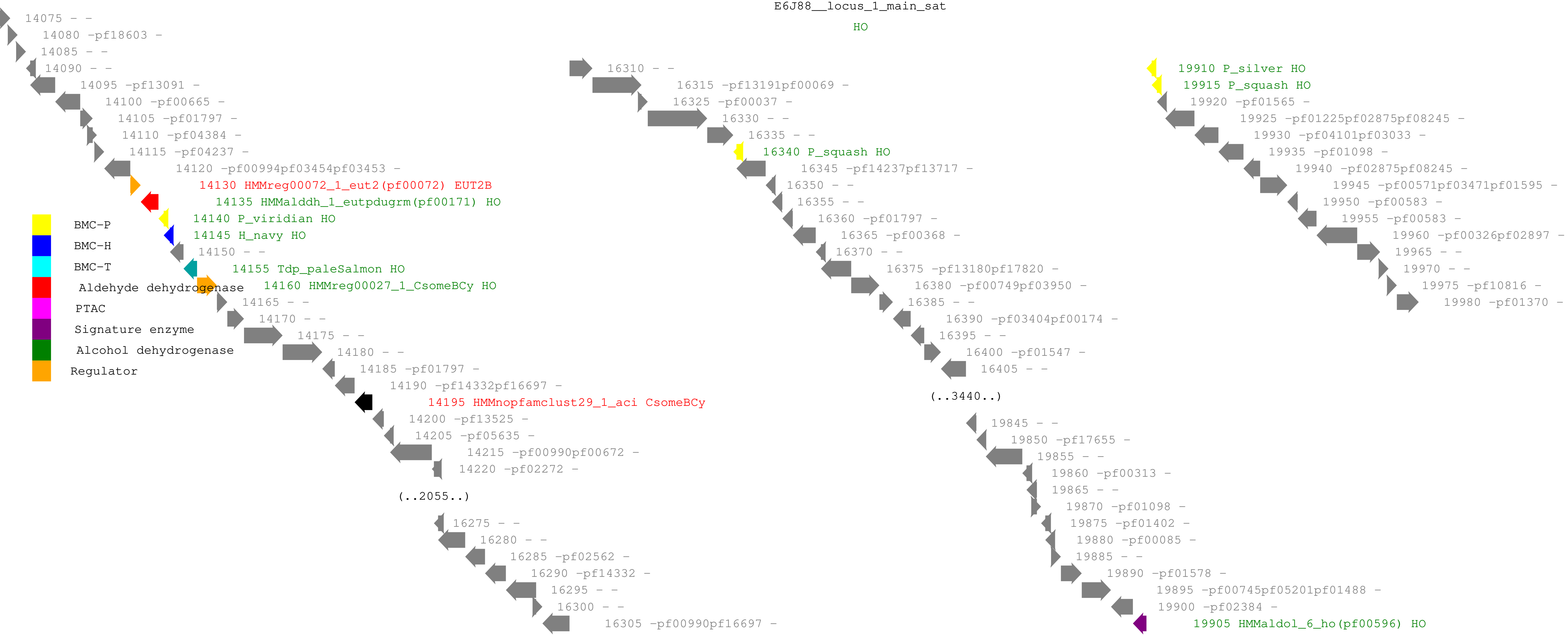

Deltaproteobacteria bacterium. Bacteria; Proteobacteria; Deltaproteobacteria.

E6J90__locus_1_main_sat

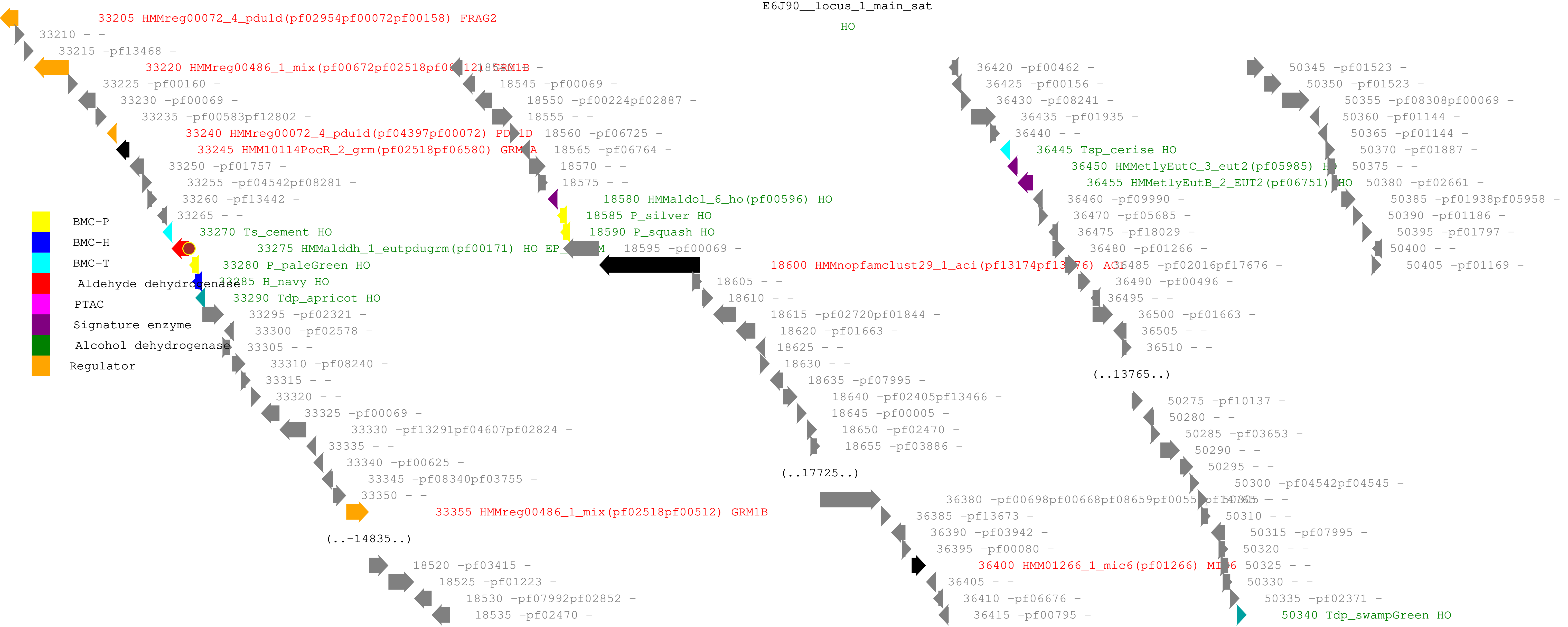

Candidatus Latescibacteria bacterium. Bacteria; Candidatus Latescibacteria.

EHM19__locus_1_main_sat

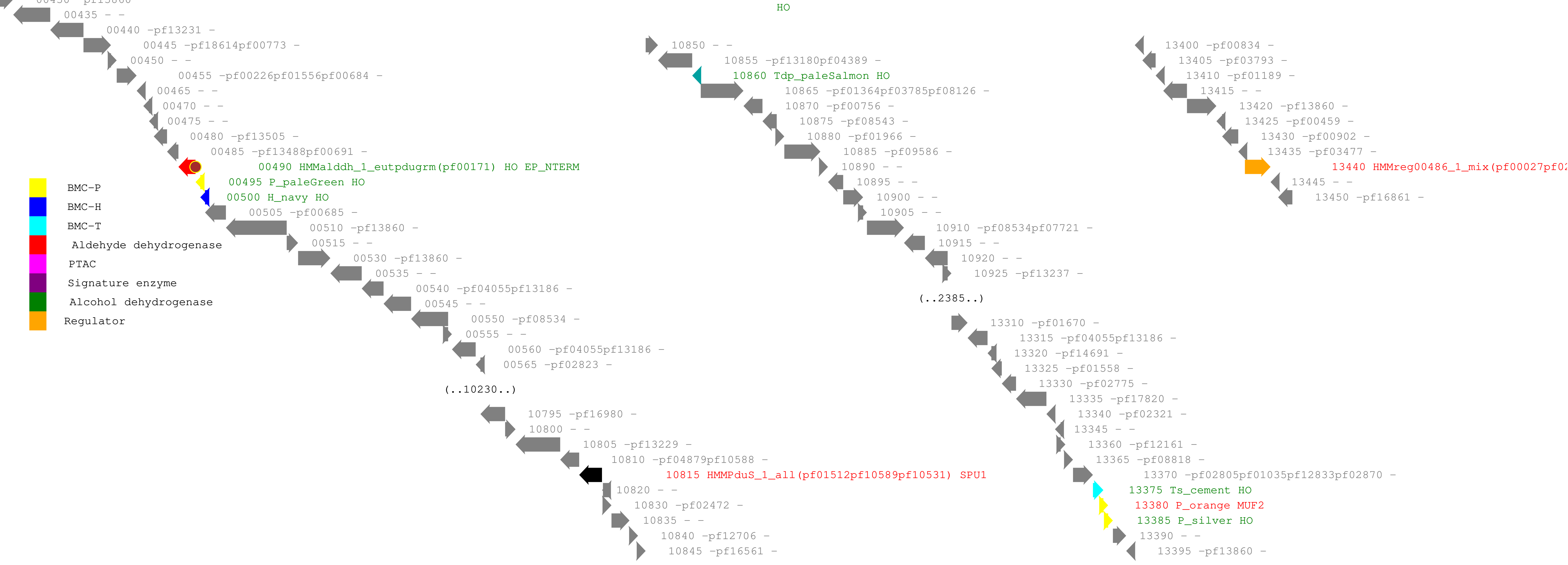

Myxococcaceae bacterium. Bacteria; Proteobacteria; Deltaproteobacteria; Myxococcales;

EPO40__locus_1_main_sat

HO

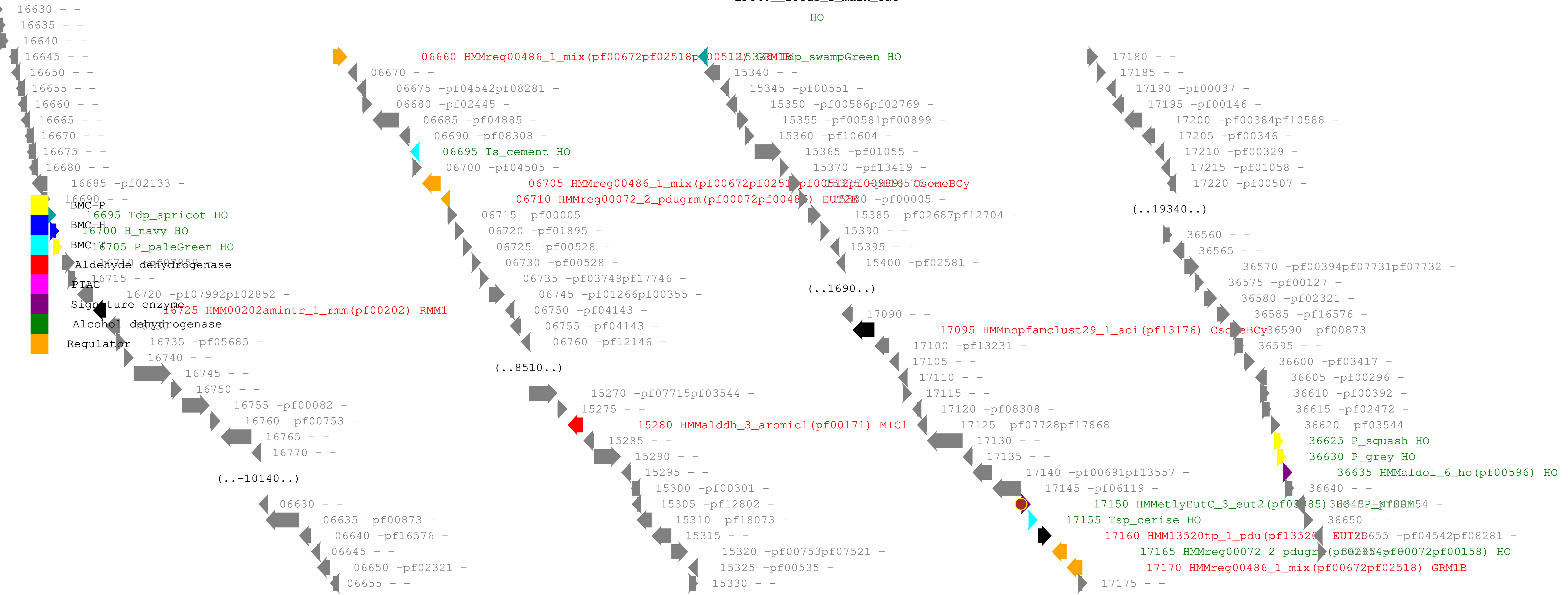

Bradymonadales bacterium V1718. Bacteria; Proteobacteria; Deltaproteobacteria; Bradymonadales;

FRD01__locus_1_main_sat

HO

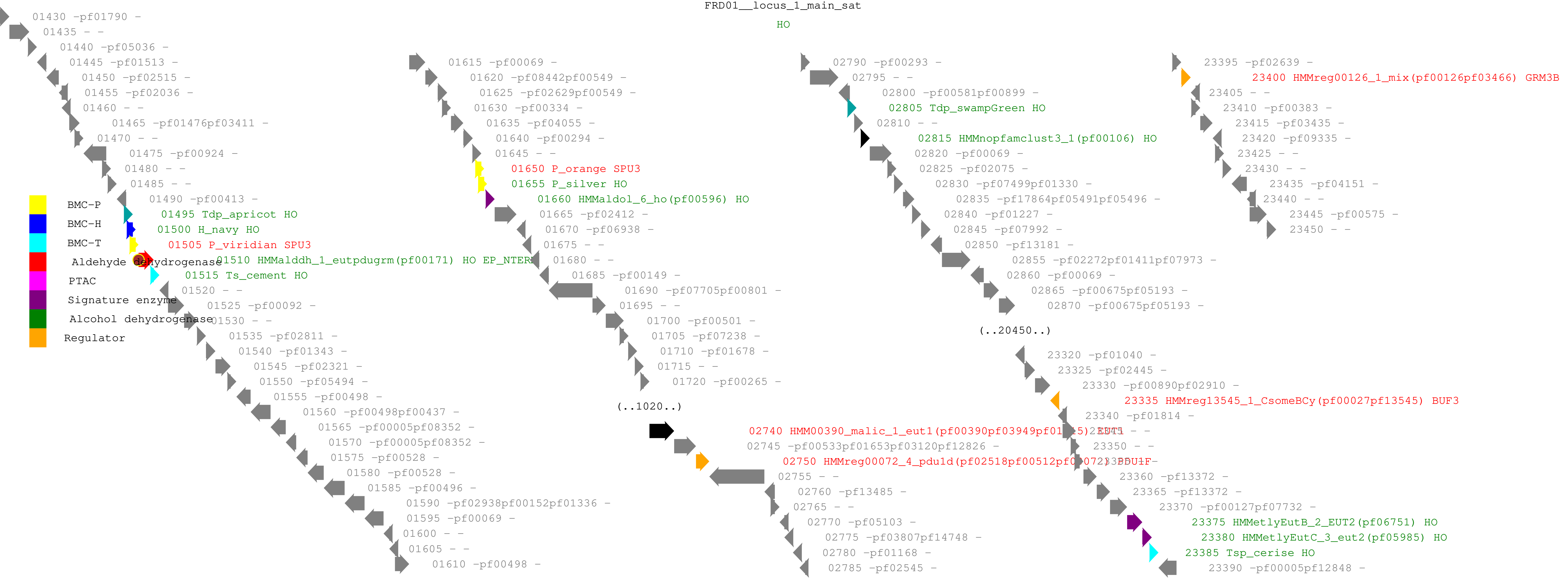

Haliangium ochraceum (strain DSM 14365 / JCM 11303 / SMP-2). Bacteria; Proteobacteria; Deltaproteobacteria; Myxococcales;

Hoch__locus_1_main_sat

HO

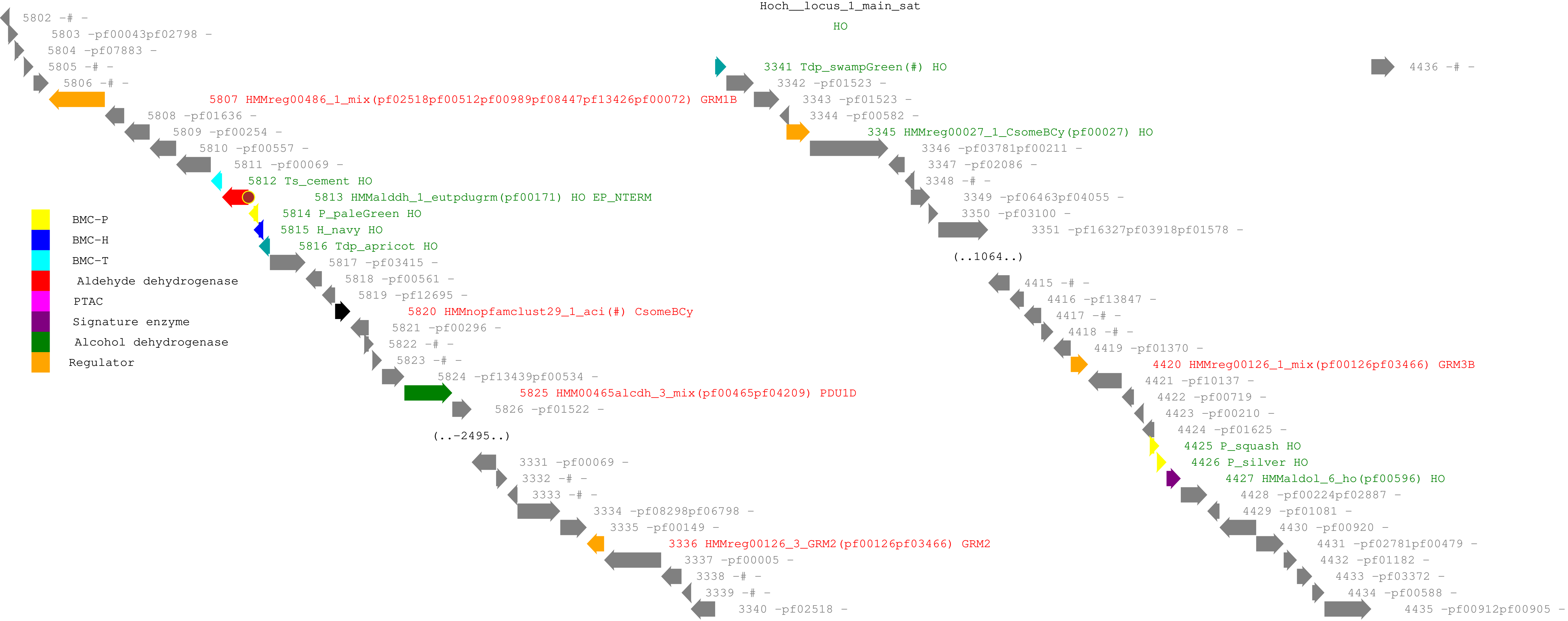

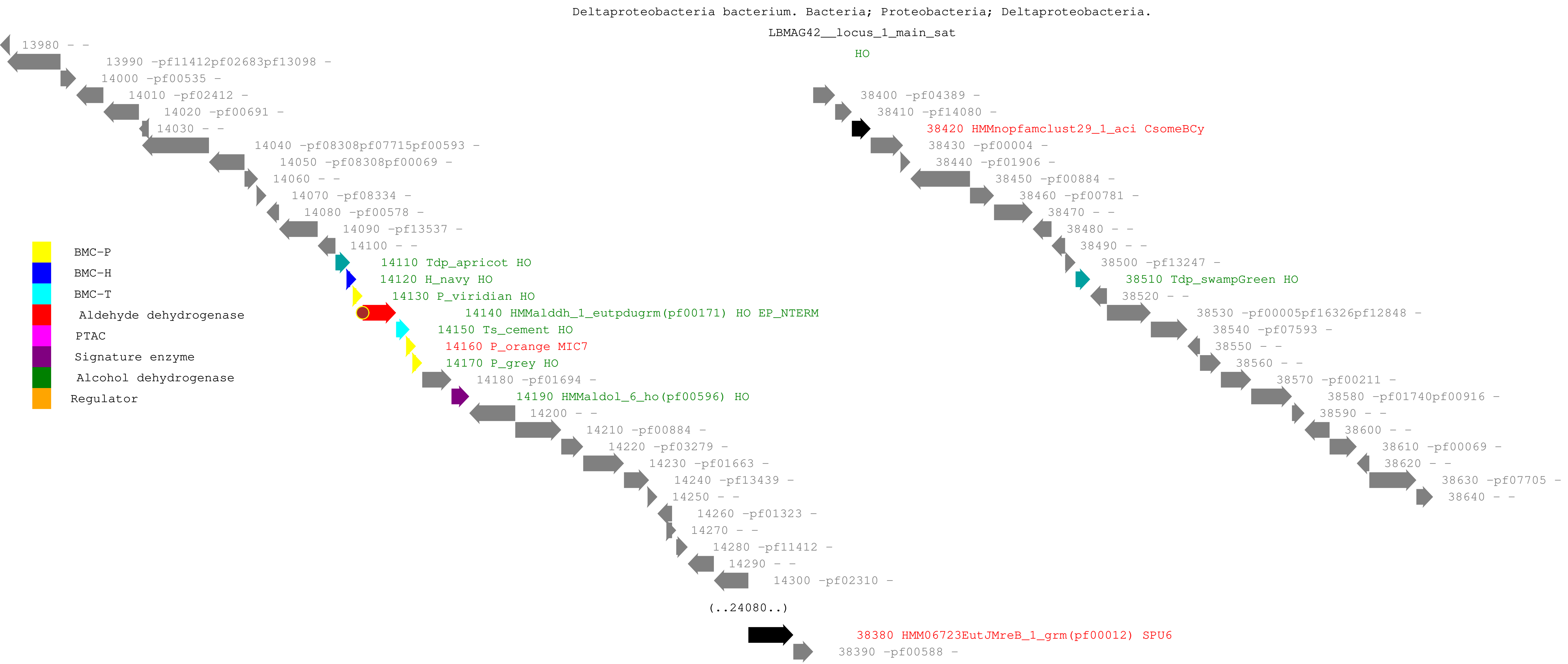

Deltaproteobacteria bacterium. Bacteria; Proteobacteria; Deltaproteobacteria.
LBMAG42__locus_1_main_sat
HO
13980 - -
13990 -pf11412pf02683pf13098 -
14000 -pf00535 -
14010 -pf02412 -
14020 -pf00691 -
14030 - -
14040 -pf08308pf07715pf00593 -
14050 -pf08308pf00069 -
14060 - -
14070 -pf08334 -
14080 -pf00578 -
14090 -pf13537 -
14100 - -
14110 Tdp_apricot HO
14120 H_navy HO
14130 P_viridian HO
14140 HMMalddh_1_eutpdugrm(pf00171) HO EP_NTERM
14150 Ts_cement HO
14160 P_orange MIC7
14170 P_grey HO
14180 -pf01694 -
14190 HMMaldol_6_ho(pf00596) HO
14200 - -
14210 -pf00884 -
14220 -pf03279 -
14230 -pf01663 -
14240 -pf13439 -
14250 - -
14260 -pf01323 -
14270 - -
14280 -pf11412 -
14290 - -
14300 -pf02310 -
(..24080..)
38380 HMM06723EutJMreB_1_grm(pf00012) SPU6
38390 -pf00588 -
38400 -pf04389 -
38410 -pf14080 -
38420 HMMnopfamclust29_1_aci CsomeBCy
38430 -pf00004 -
38440 -pf01906 -
38450 -pf00884 -
38460 -pf00781 -
38470 - -
38480 - -
38490 - -
38500 -pf13247 -
38510 Tdp_swampGreen HO
38520 - -
38530 -pf00005pf16326pf12848 -
38540 -pf07593 -
38550 - -
38560 - -
38570 -pf00211 -
38580 -pf01740pf00916 -
38590 - -
38600 - -
38610 -pf00069 -
38620 - -
38630 -pf07705 -
38640 - -
BMC-P
BMC-H
BMC-T
Aldehyde dehydrogenase
PTAC
Signature enzyme
Alcohol dehydrogenase
Regulator

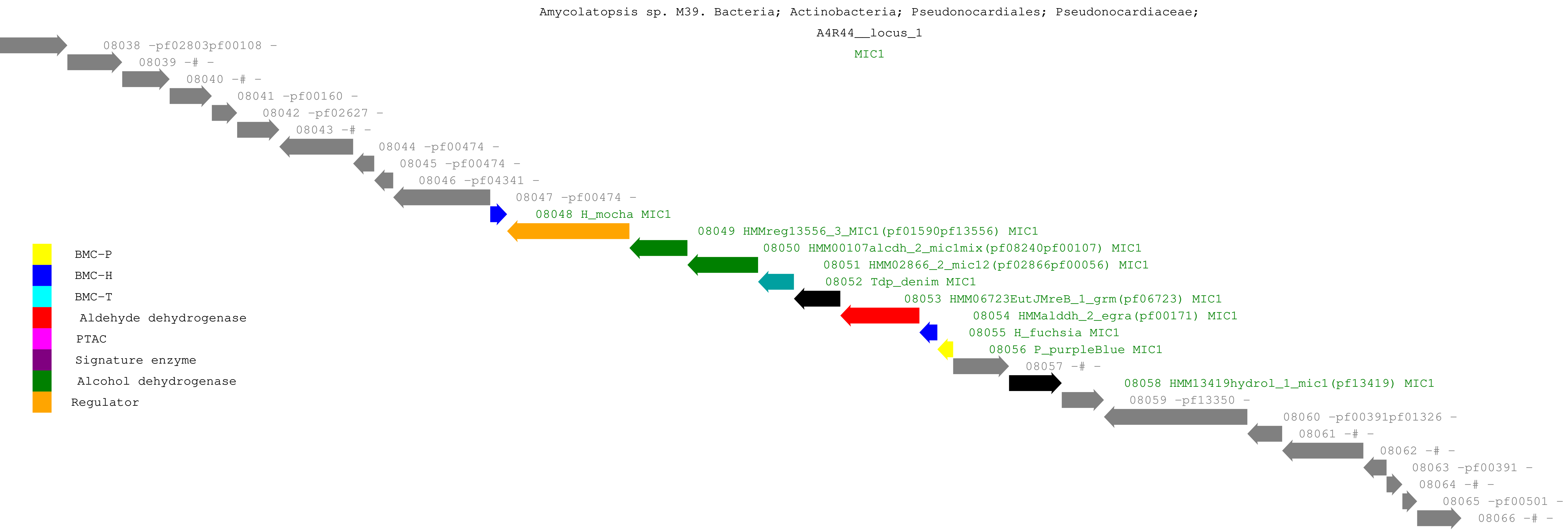
Amycolatopsis sp. M39. Bacteria; Actinobacteria; Pseudonocardiales; Pseudonocardiaceae;
A4R44__locus_1
MIC1
08038 -pf02803pf00108 -
08039 -# -
08040 -# -
08041 -pf00160 -
08042 -pf02627 -
08043 -# -
08044 -pf00474 -
08045 -pf00474 -
08046 -pf04341 -
08047 -pf00474 -
08048 H_mocha MIC1
08049 HMMreg13556_3_MIC1(pf01590pf13556) MIC1
08050 HMM00107alcdh_2_mic1mix(pf08240pf00107) MIC1
08051 HMM02866_2_mic12(pf02866pf00056) MIC1
08052 Tdp_denim MIC1
08053 HMM06723EutJMreB_1_grm(pf06723) MIC1
08054 HMMalddh_2_egra(pf00171) MIC1
08055 H_fuchsia MIC1
08056 P_purpleBlue MIC1
08057 -# -
08058 HMM13419hydrol_1_mic1(pf13419) MIC1
08059 -pf13350 -
08060 -pf00391pf01326 -
08061 -# -
08062 -# -
08063 -pf00391 -
08064 -# -
08065 -pf00501 -
08066 -# -
BMC-P
BMC-H
BMC-T
Aldehyde dehydrogenase
PTAC
Signature enzyme
Alcohol dehydrogenase
Regulator

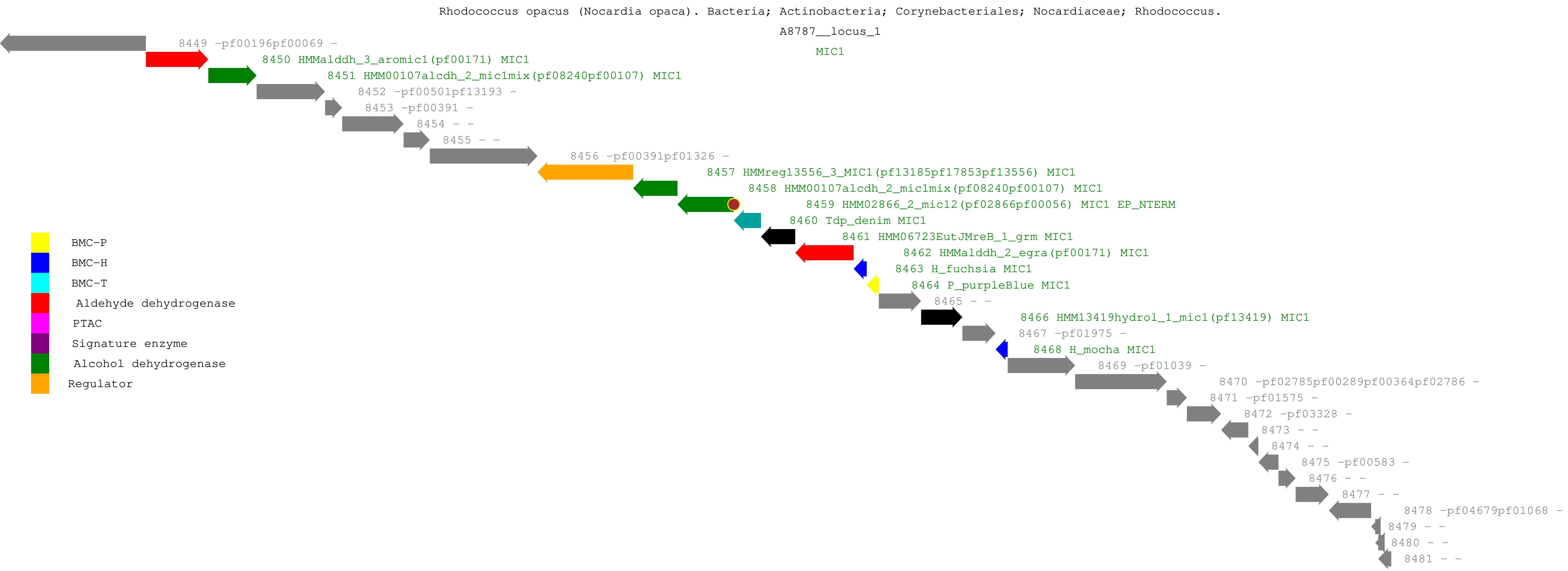
Rhodococcus opacus (Nocardia opaca). Bacteria; Actinobacteria; Corynebacteriales; Nocardiaceae; Rhodococcus.
A8787__locus_1
MIC1
8449 -pf00196pf00069 -
8450 HMMalddh_3_aromic1(pf00171) MIC1
8451 HMM00107alcdh_2_mic1mix(pf08240pf00107) MIC1
8452 -pf00501pf13193 -
8453 -pf00391 -
8454 - -
8455 - -
8456 -pf00391pf01326 -
8457 HMMreg13556_3_MIC1(pf13185pf17853pf13556) MIC1
8458 HMM00107alcdh_2_mic1mix(pf08240pf00107) MIC1
8459 HMM02866_2_mic12(pf02866pf00056) MIC1 EP_NTERM
8460 Tdp_denim MIC1
8461 HMM06723EutJMreB_1_grm MIC1
8462 HMMalddh_2_egra(pf00171) MIC1
8463 H_fuchsia MIC1
8464 P_purpleBlue MIC1
8465 - -
8466 HMM13419hydrol_1_mic1(pf13419) MIC1
8467 -pf01975 -
8468 H_mocha MIC1
8469 -pf01039 -
8470 -pf02785pf00289pf00364pf02786 -
8471 -pf01575 -
8472 -pf03328 -
8473 - -
8474 - -
8475 -pf00583 -
8476 - -
8477 - -
8478 -pf04679pf01068 -
8479 - -
8480 - -
8481 - -
BMC-P
BMC-H
BMC-T
Aldehyde dehydrogenase
PTAC
Signature enzyme
Alcohol dehydrogenase
Regulator

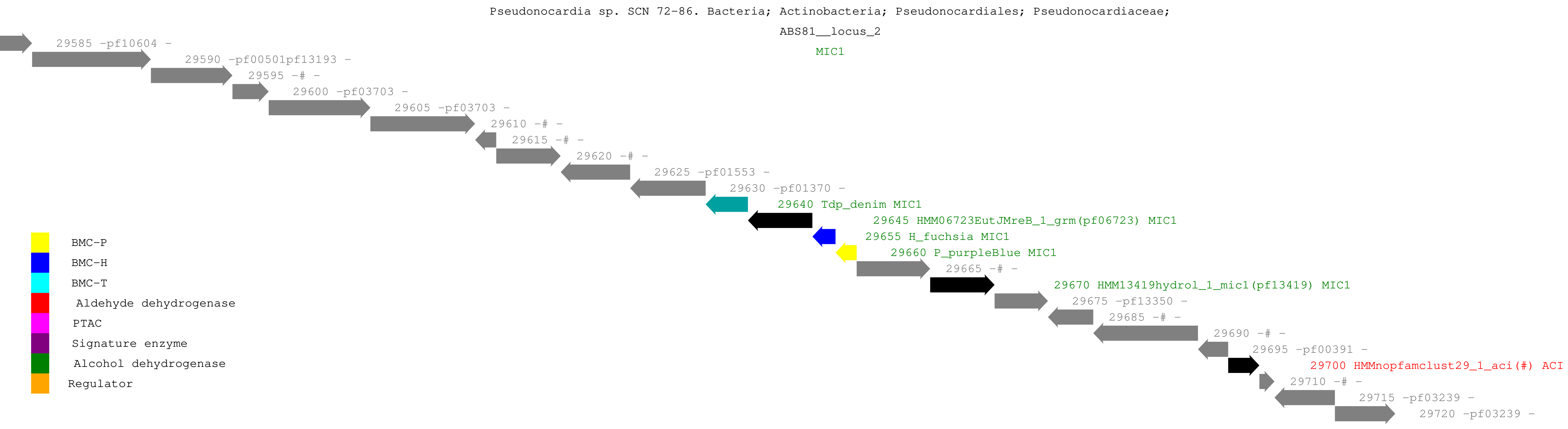

Pseudonocardia sp. SCN 72-86. Bacteria; Actinobacteria; Pseudonocardiales; Pseudonocardiaceae;
ABS81__locus_2
MIC1
29585 -pf10604 -
29590 -pf00501pf13193 -
29595 -# -
29600 -pf03703 -
29605 -pf03703 -
29610 -# -
29615 -# -
29620 -# -
29625 -pf01553 -
29630 -pf01370 -
29640 Tdp_denim MIC1
29645 HMM06723EutJMreB_1_grm(pf06723) MIC1
29655 H_fuchsia MIC1
29660 P_purpleBlue MIC1
29665 -# -
29670 HMM13419hydrol_1_mic1(pf13419) MIC1
29675 -pf13350 -
29685 -# -
29690 -# -
29695 -pf00391 -
29700 HMMnopfamclust29_1_aci(#) ACI
29710 -# -
29715 -pf03239 -
29720 -pf03239 -
BMC-P
BMC-H
BMC-T
Aldehyde dehydrogenase
PTAC
Signature enzyme
Alcohol dehydrogenase
Regulator

Pseudonocardia sp. SCN 73-27. Bacteria; Actinobacteria; Pseudonocardiales; Pseudonocardiaceae;

ABT15__locus_2

MIC1

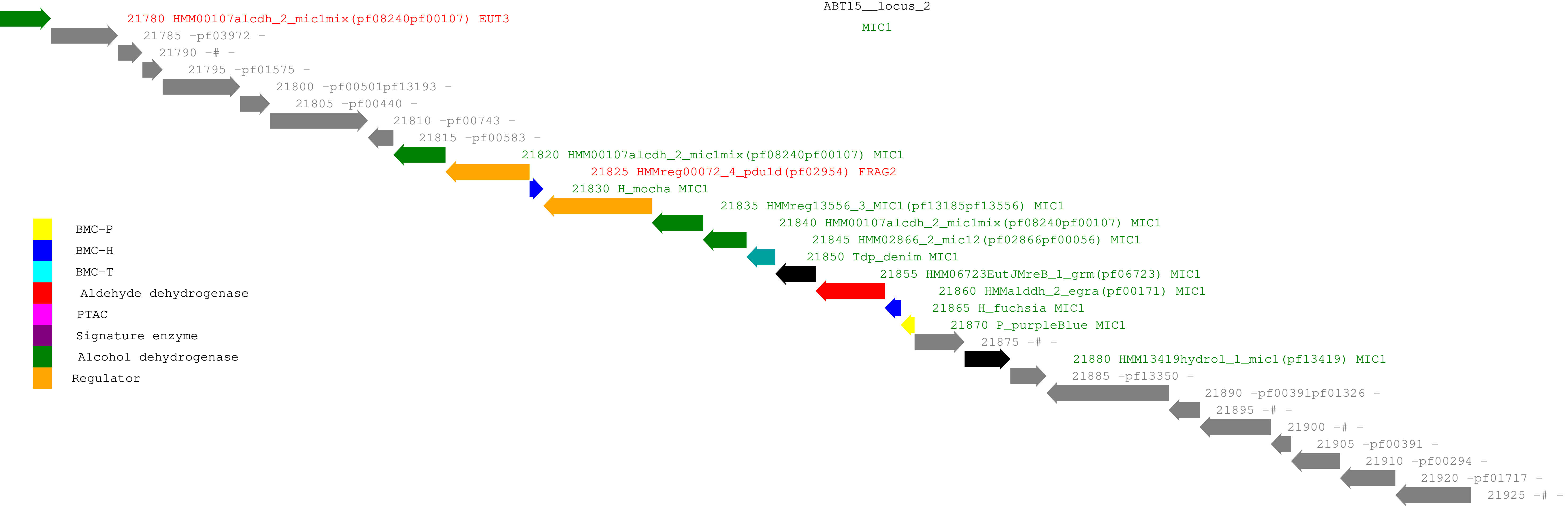

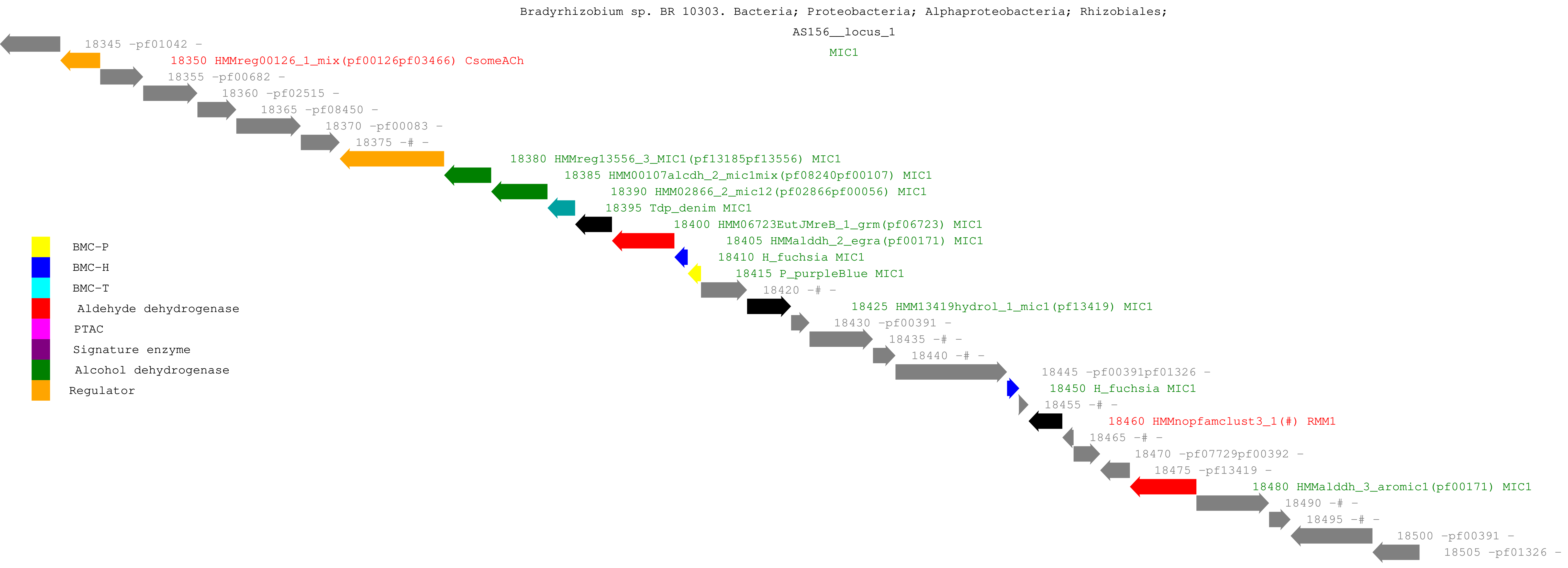
Bradyrhizobium sp. BR 10303. Bacteria; Proteobacteria; Alphaproteobacteria; Rhizobiales;
AS156__locus_1
MIC1
18345 -pf01042 -
18350 HMMreg00126_1_mix(pf00126pf03466) CsomeACh
18355 -pf00682 -
18360 -pf02515 -
18365 -pf08450 -
18370 -pf00083 -
18375 -# -
18380 HMMreg13556_3_MIC1(pf13185pf13556) MIC1
18385 HMM00107alcdh_2_mic1mix(pf08240pf00107) MIC1
18390 HMM02866_2_mic12(pf02866pf00056) MIC1
18395 Tdp_denim MIC1
18400 HMM06723EutJMreB_1_grm(pf06723) MIC1
18405 HMMalddh_2_egra(pf00171) MIC1
18410 H_fuchsia MIC1
18415 P_purpleBlue MIC1
18420 -# -
18425 HMM13419hydrol_1_mic1(pf13419) MIC1
18430 -pf00391 -
18435 -# -
18440 -# -
18445 -pf00391pf01326 -
18450 H_fuchsia MIC1
18455 -# -
18460 HMMnopfamclust3_1(#) RMM1
18465 -# -
18470 -pf07729pf00392 -
18475 -pf13419 -
18480 HMMalddh_3_aromic1(pf00171) MIC1
18490 -# -
18495 -# -
18500 -pf00391 -
18505 -pf01326 -
BMC-P
BMC-H
BMC-T
Aldehyde dehydrogenase
PTAC
Signature enzyme
Alcohol dehydrogenase
Regulator

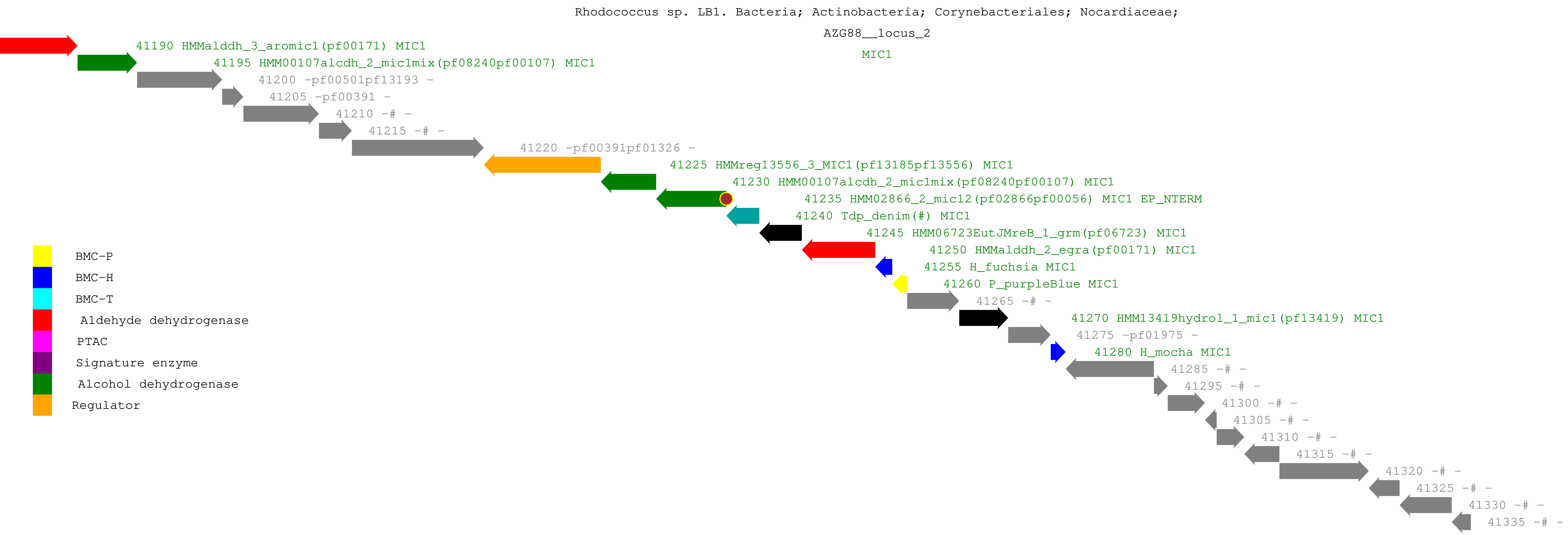
Rhodococcus sp. LB1. Bacteria; Actinobacteria; Corynebacteriales; Nocardiaceae;
AZG88__locus_2
MIC1
41190 HMMalddh_3_aromic1(pf00171) MIC1
41195 HMM00107alcdh_2_mic1mix(pf08240pf00107) MIC1
41200 -pf00501pf13193 -
41205 -pf00391 -
41210 -# -
41215 -# -
41220 -pf00391pf01326 -
41225 HMMreg13556_3_MIC1(pf13185pf13556) MIC1
41230 HMM00107alcdh_2_mic1mix(pf08240pf00107) MIC1
41235 HMM02866_2_mic12(pf02866pf00056) MIC1 EP_NTERM
41240 Tdp_denim(#) MIC1
41245 HMM06723EutJMreB_1_grm(pf06723) MIC1
41250 HMMalddh_2_egra(pf00171) MIC1
41255 H_fuchsia MIC1
41260 P_purpleBlue MIC1
41265 -# -
41270 HMM13419hydrol_1_mic1(pf13419) MIC1
41275 -pf01975 -
41280 H_mocha MIC1
41285 -# -
41295 -# -
41300 -# -
41305 -# -
41310 -# -
41315 -# -
41320 -# -
41325 -# -
41330 -# -
41335 -# -
BMC-P
BMC-H
BMC-T
Aldehyde dehydrogenase
PTAC
Signature enzyme
Alcohol dehydrogenase
Regulator

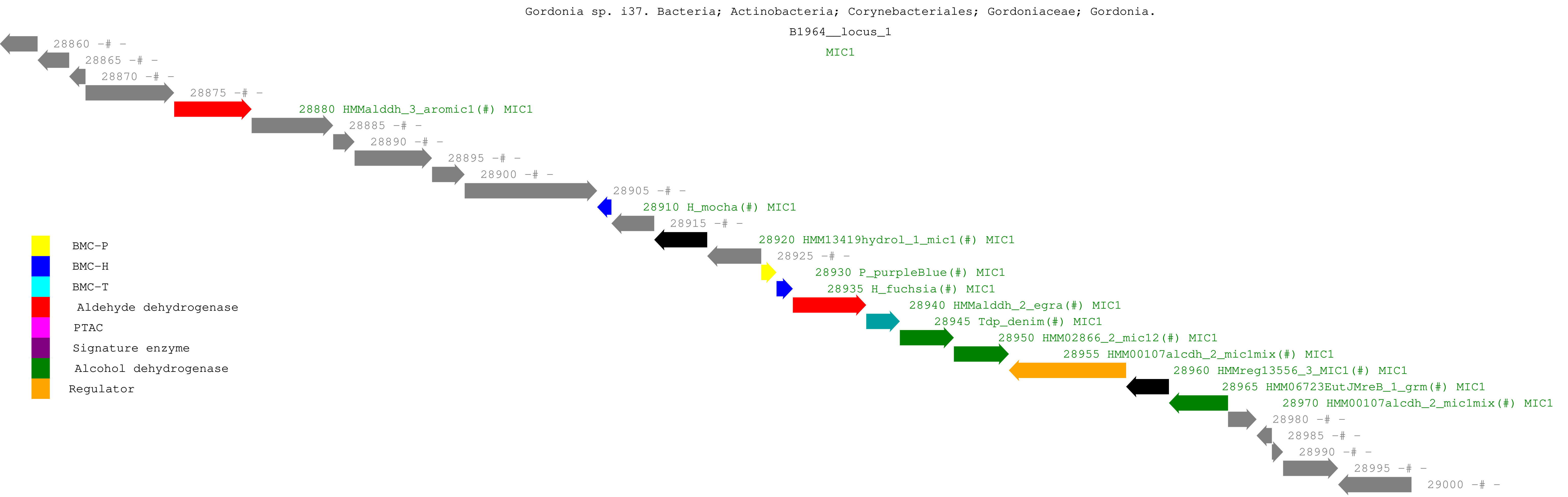
Gordonia sp. i37. Bacteria; Actinobacteria; Corynebacteriales; Gordoniaceae; Gordonia.
B1964__locus_1
MIC1
28860 -# -
28865 -# -
28870 -# -
28875 -# -
28880 HMMalddh_3_aromic1(#) MIC1
28885 -# -
28890 -# -
28895 -# -
28900 -# -
28905 -# -
28910 H_mocha(#) MIC1
28915 -# -
28920 HMM13419hydrol_1_mic1(#) MIC1
28925 -# -
28930 P_purpleBlue(#) MIC1
28935 H_fuchsia(#) MIC1
28940 HMMalddh_2_egra(#) MIC1
28945 Tdp_denim(#) MIC1
28950 HMM02866_2_mic12(#) MIC1
28955 HMM00107alcdh_2_mic1mix(#) MIC1
28960 HMMreg13556_3_MIC1(#) MIC1
28965 HMM06723EutJMreB_1_grm(#) MIC1
28970 HMM00107alcdh_2_mic1mix(#) MIC1
28980 -# -
28985 -# -
28990 -# -
28995 -# -
29000 -# -
BMC-P
BMC-H
BMC-T
Aldehyde dehydrogenase
PTAC
Signature enzyme
Alcohol dehydrogenase
Regulator

Pseudonocardia autotrophica (Amycolata autotrophica) (Nocardia Bacteria; Actinobacteria; Pseudonocardiales; Pseudonocardiaceae;

BG845__locus_1_main_sat

MIC1

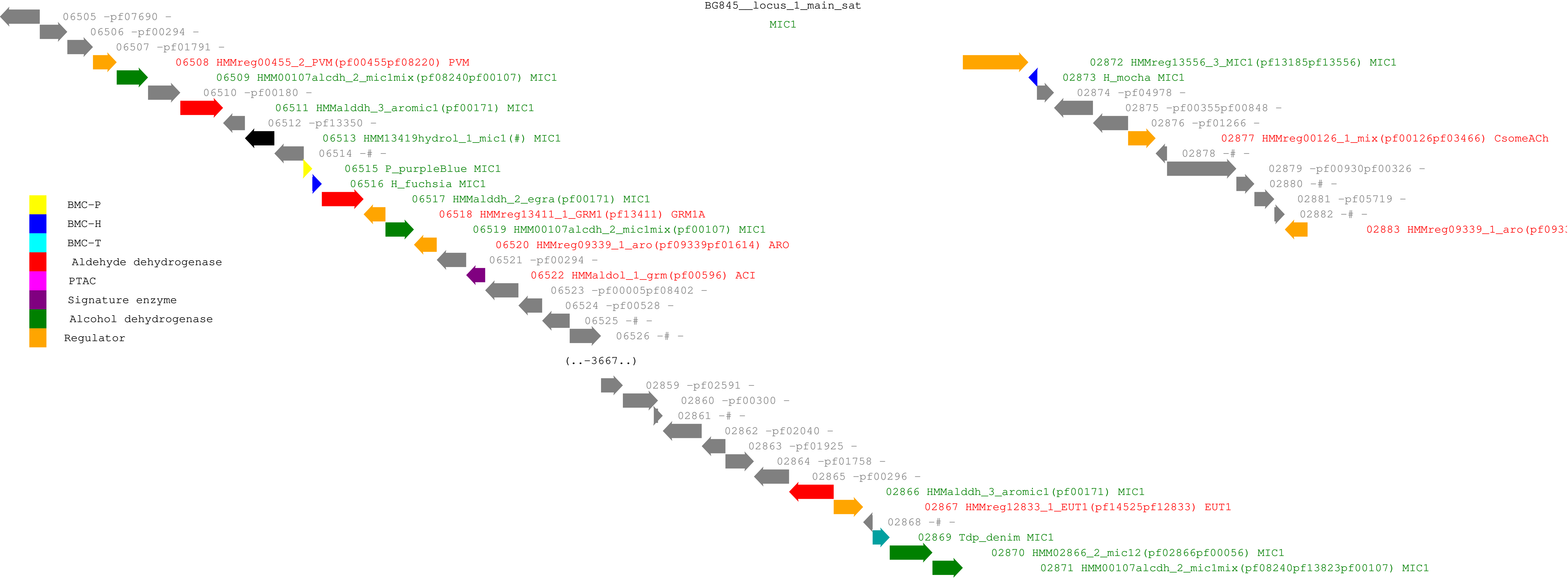

Burkholderiales bacterium 66-5. Bacteria; Proteobacteria; Betaproteobacteria; Burkholderiales.

BGO13__locus_1_main_sat

MIC1

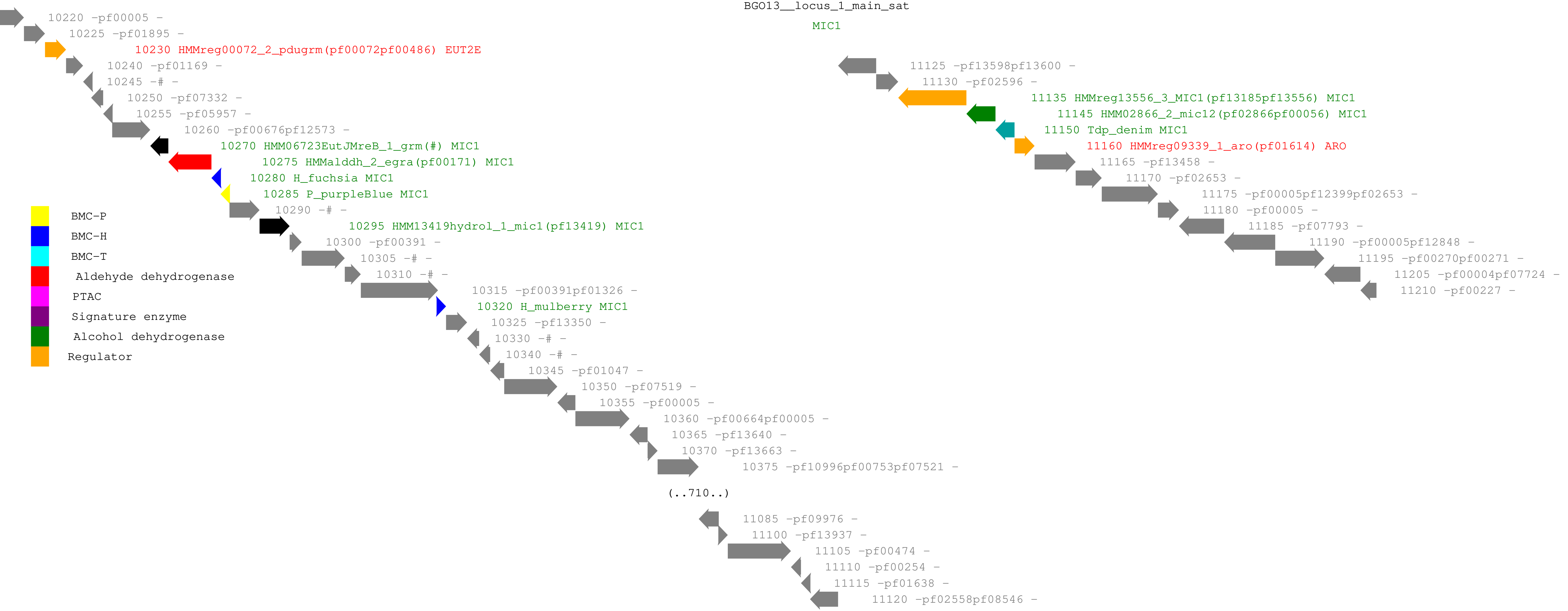

Pseudonocardia sp. 73-21. Bacteria; Actinobacteria; Pseudonocardiales; Pseudonocardiaceae;

BGP03__locus_1

MIC1

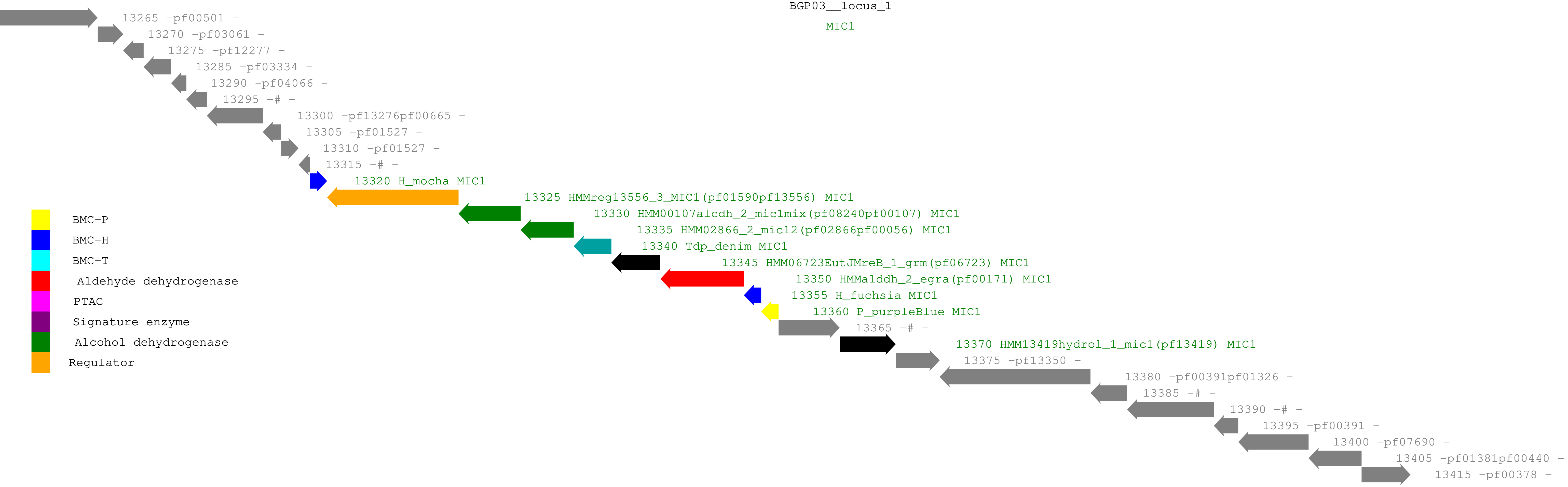

Actinomycetospora cinnamomea. Bacteria; Actinobacteria; Pseudonocardiales; Pseudonocardiaceae;

C8D89__locus_1

MIC1

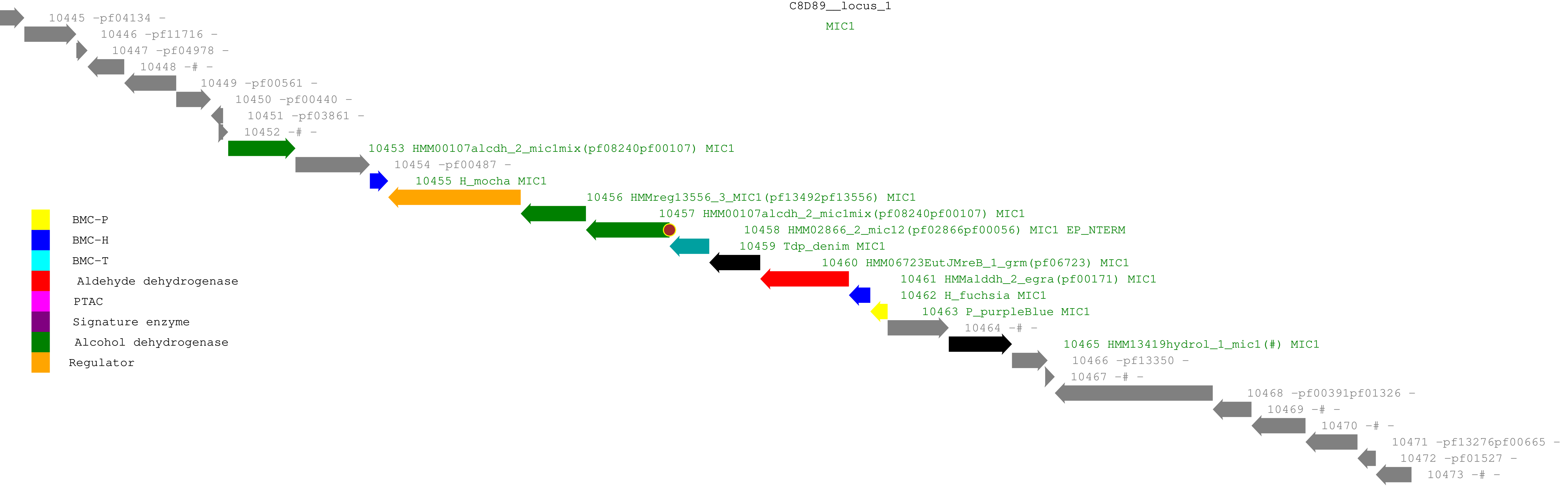

Rhodococcus sp. NCIMB 12038. Bacteria; Actinobacteria; Corynebacteriales; Nocardiaceae;

CA951__locus_1

MIC1

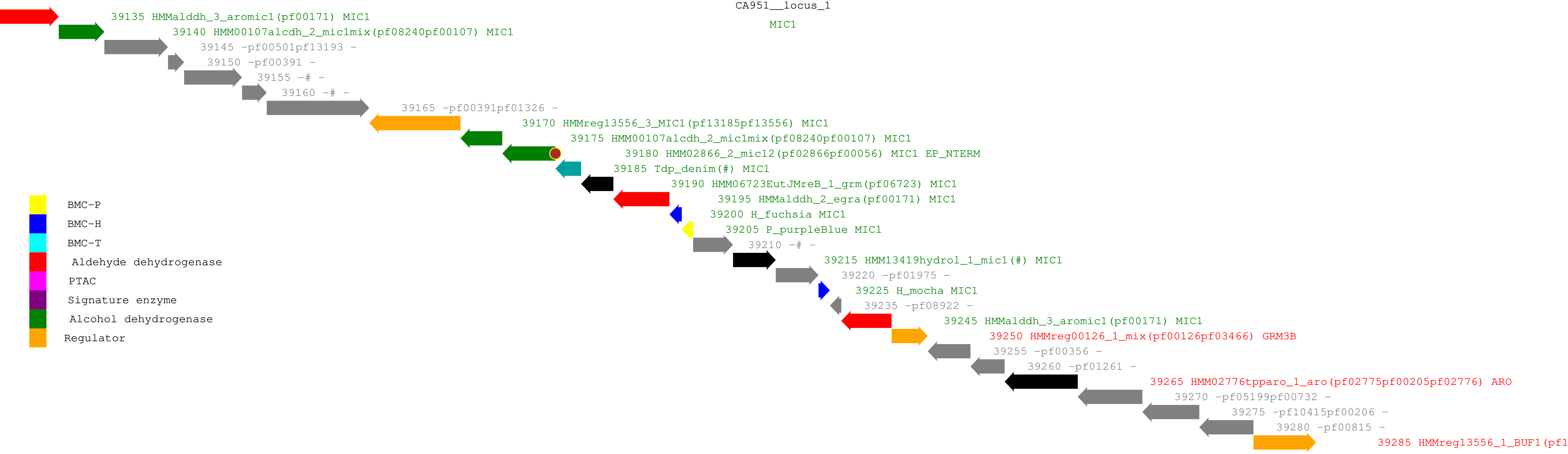

Rhodococcus sp. S2-17. Bacteria; Actinobacteria; Corynebacteriales; Nocardiaceae;

CBI38__locus_1

MIC1

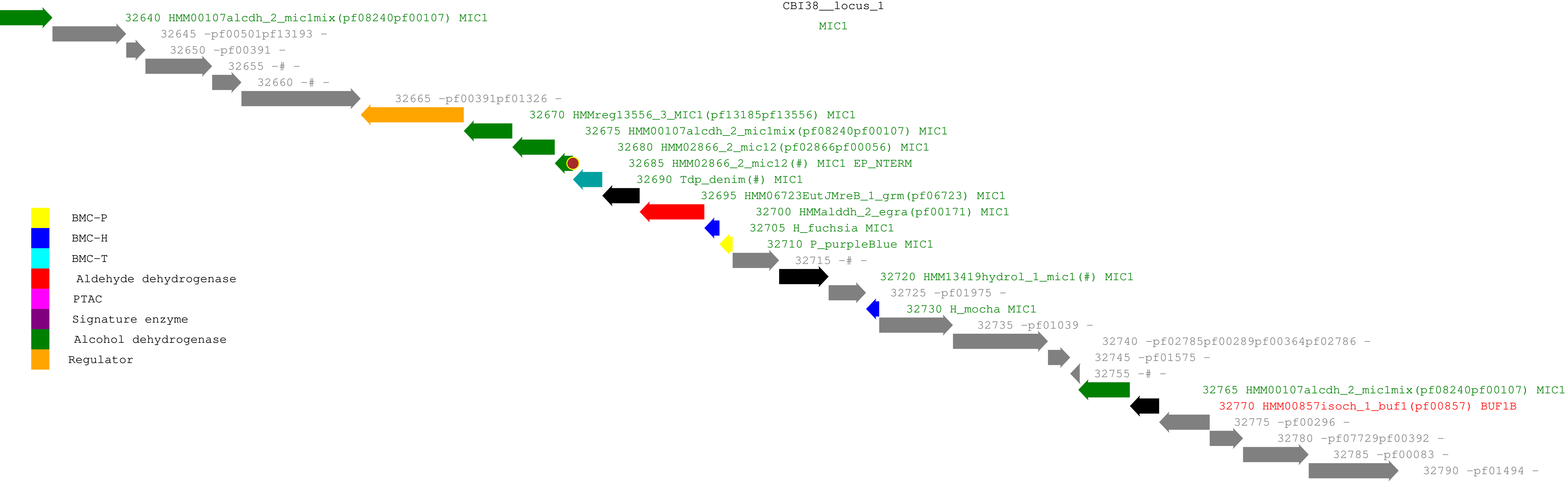

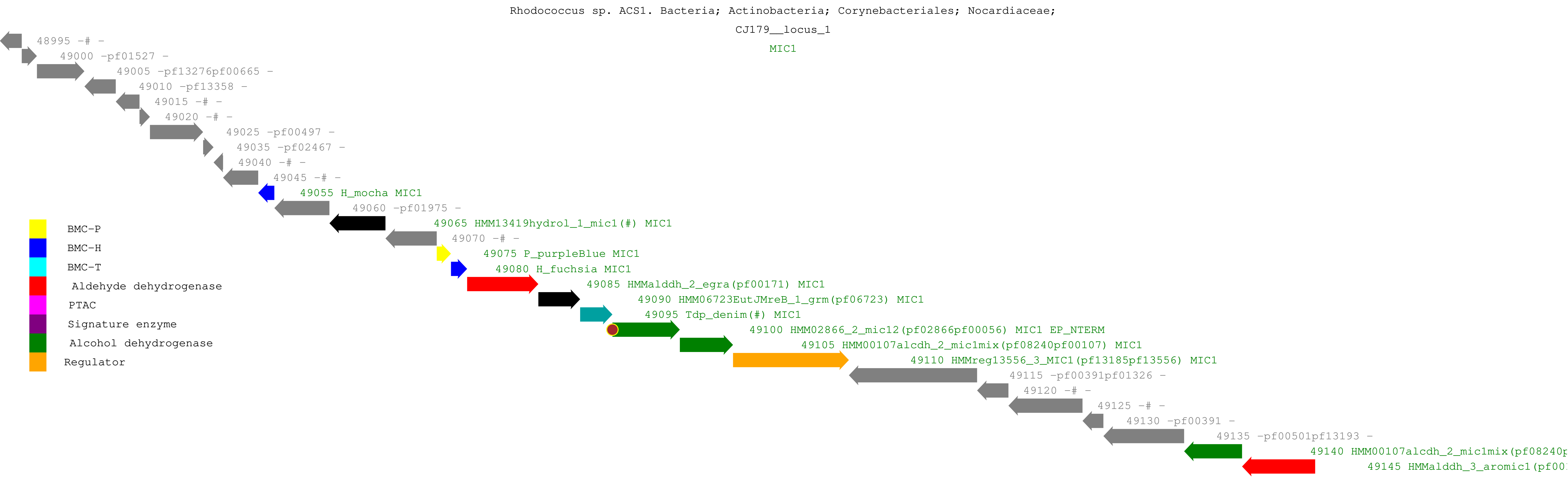
Rhodococcus sp. ACS1. Bacteria; Actinobacteria; Corynebacteriales; Nocardiaceae;
CJ179__locus_1
MIC1
48995 -# -
49000 -pf01527 -
49005 -pf13276pf00665 -
49010 -pf13358 -
49015 -# -
49020 -# -
49025 -pf00497 -
49035 -pf02467 -
49040 -# -
49045 -# -
49055 H_mocha MIC1
49060 -pf01975 -
49065 HMM13419hydrol_1_mic1(#) MIC1
49070 -# -
49075 P_purpleBlue MIC1
49080 H_fuchsia MIC1
49085 HMMalddh_2_egra(pf00171) MIC1
49090 HMM06723EutJMreB_1_grm(pf06723) MIC1
49095 Tdp_denim(#) MIC1
49100 HMM02866_2_mic12(pf02866pf00056) MIC1 EP_NTERM
49105 HMM00107alcdh_2_mic1mix(pf08240pf00107) MIC1
49110 HMMreg13556_3_MIC1(pf13185pf13556) MIC1
49115 -pf00391pf01326 -
49120 -# -
49125 -# -
49130 -pf00391 -
49135 -pf00501pf13193 -
49140 HMM00107alcdh_2_mic1mix(pf08240p
49145 HMMalddh_3_aromic1(pf00
BMC-P
BMC-H
BMC-T
Aldehyde dehydrogenase
PTAC
Signature enzyme
Alcohol dehydrogenase
Regulator

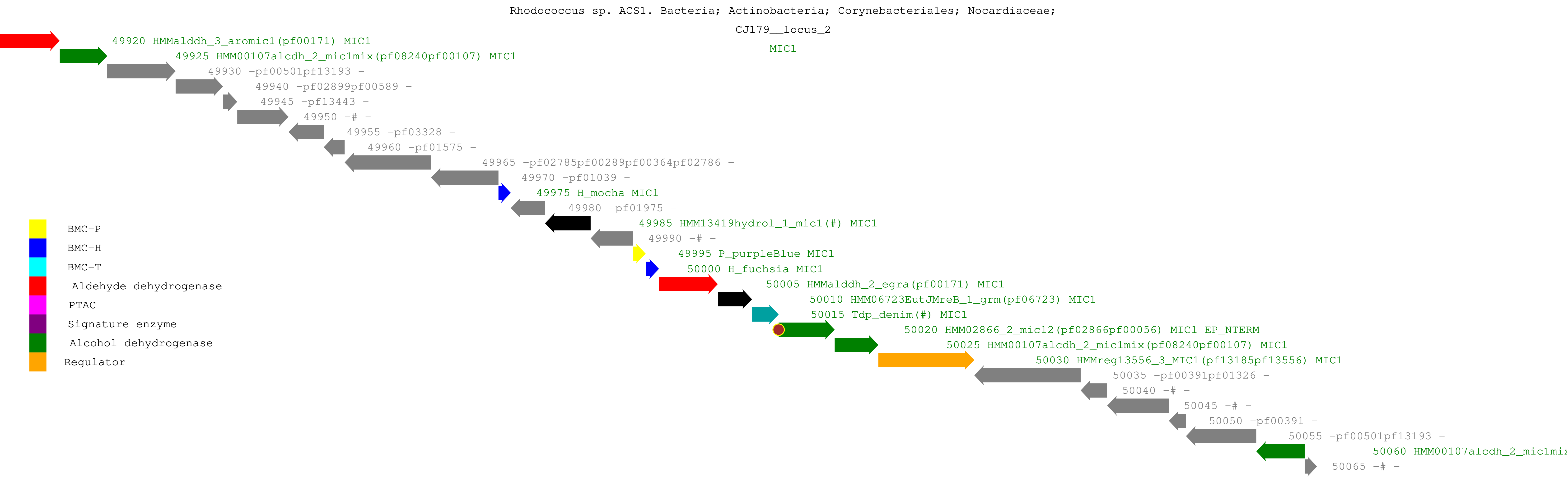
Rhodococcus sp. ACS1. Bacteria; Actinobacteria; Corynebacteriales; Nocardiaceae;
CJ179__locus_2
MIC1
49920 HMMalddh_3_aromic1(pf00171) MIC1
49925 HMM00107alcdh_2_mic1mix(pf08240pf00107) MIC1
49930 -pf00501pf13193 -
49940 -pf02899pf00589 -
49945 -pf13443 -
49950 -# -
49955 -pf03328 -
49960 -pf01575 -
49965 -pf02785pf00289pf00364pf02786 -
49970 -pf01039 -
49975 H_mocha MIC1
49980 -pf01975 -
49985 HMM13419hydrol_1_mic1(#) MIC1
49990 -# -
49995 P_purpleBlue MIC1
50000 H_fuchsia MIC1
50005 HMMalddh_2_egra(pf00171) MIC1
50010 HMM06723EutJMreB_1_grm(pf06723) MIC1
50015 Tdp_denim(#) MIC1
50020 HMM02866_2_mic12(pf02866pf00056) MIC1 EP_NTERM
50025 HMM00107alcdh_2_mic1mix(pf08240pf00107) MIC1
50030 HMMreg13556_3_MIC1(pf13185pf13556) MIC1
50035 -pf00391pf01326 -
50040 -# -
50045 -# -
50050 -pf00391 -
50055 -pf00501pf13193 -
50060 HMM00107alcdh_2_mic1mi
50065 -# -
BMC-P
BMC-H
BMC-T
Aldehyde dehydrogenase
PTAC
Signature enzyme
Alcohol dehydrogenase
Regulator

Bradyrhizobium sp. Leo170. Bacteria; Proteobacteria; Alphaproteobacteria; Rhizobiales;

CWO89__locus_1_main_sat

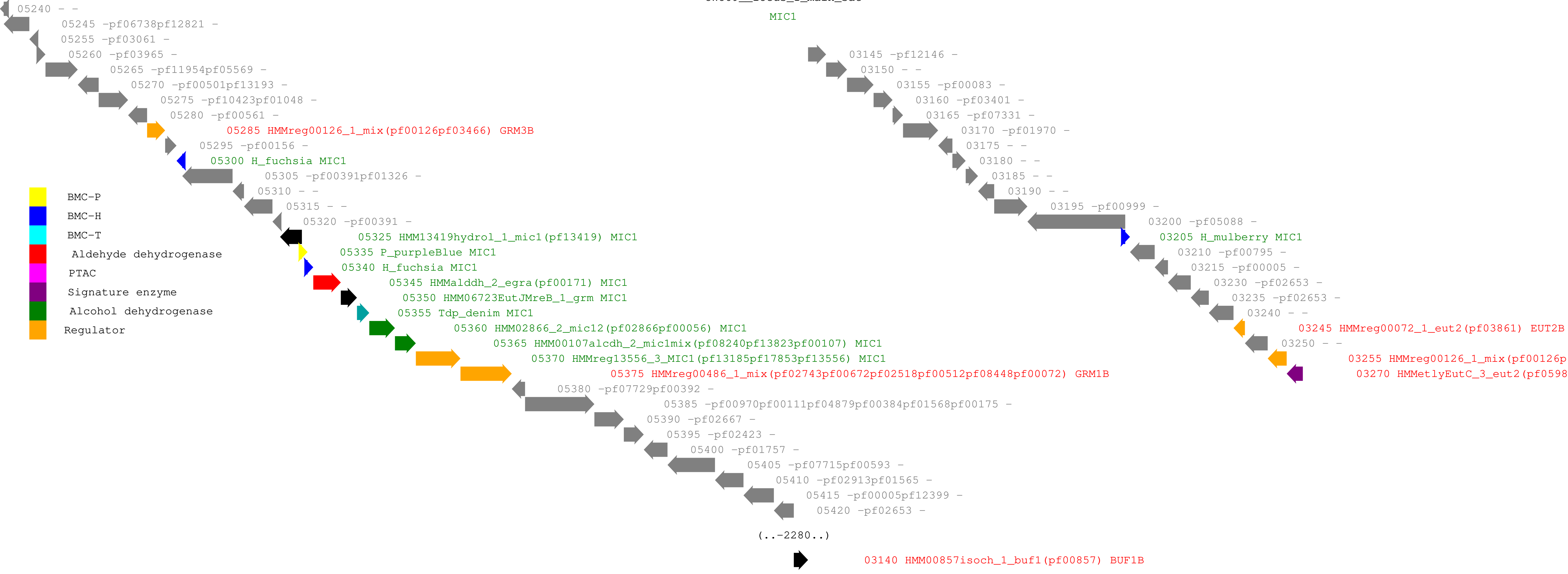

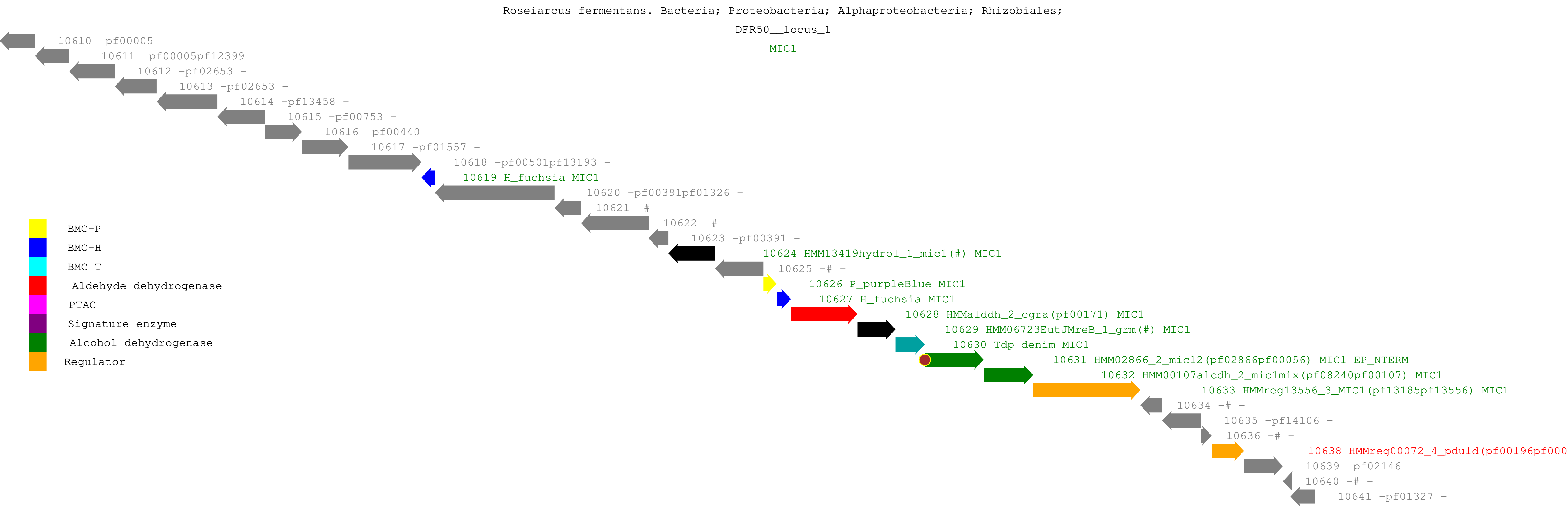
Roseiarcus fermentans. Bacteria; Proteobacteria; Alphaproteobacteria; Rhizobiales;
DFR50__locus_1
MIC1
10610 -pf00005 -
10611 -pf00005pf12399 -
10612 -pf02653 -
10613 -pf02653 -
10614 -pf13458 -
10615 -pf00753 -
10616 -pf00440 -
10617 -pf01557 -
10618 -pf00501pf13193 -
10619 H_fuchsia MIC1
10620 -pf00391pf01326 -
10621 -# -
10622 -# -
10623 -pf00391 -
10624 HMM13419hydrol_1_mic1(#) MIC1
10625 -# -
10626 P_purpleBlue MIC1
10627 H_fuchsia MIC1
10628 HMMaldh_2_egra(pf00171) MIC1
10629 HMM06723EutJMreB_1_grm(#) MIC1
10630 Tdp_denim MIC1
10631 HMM02866_2_mic12(pf02866pf00056) MIC1 EP_NTERM
10632 HMM00107alcdh_2_mic1mix(pf08240pf00107) MIC1
10633 HMMreg13556_3_MIC1(pf13185pf13556) MIC1
10634 -# -
10635 -pf14106 -
10636 -# -
10638 HMMreg00072_4_pdu1d(pf00196pf000
10639 -pf02146 -
10640 -# -
10641 -pf01327 -
BMC-P
BMC-H
BMC-T
Aldehyde dehydrogenase
PTAC
Signature enzyme
Alcohol dehydrogenase
Regulator

Cryobacterium sp. Hh14. Bacteria; Actinobacteria; Micrococcales; Microbacteriaceae; Cryobacterium;

E3T55__locus_1

MIC1

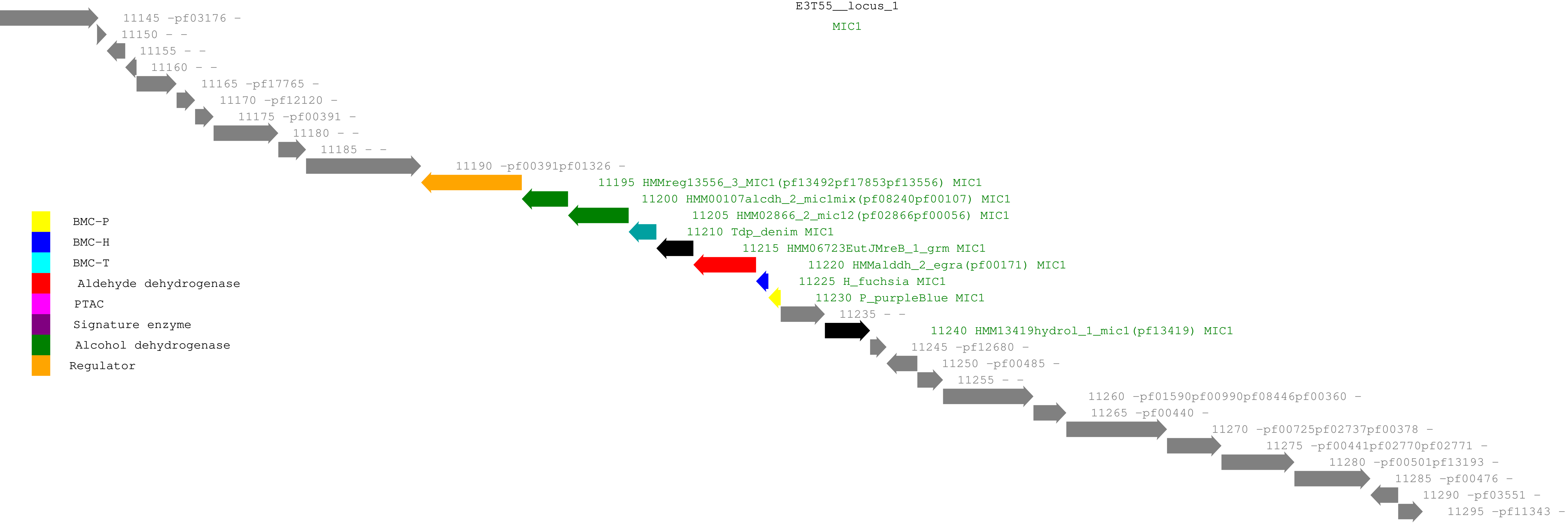

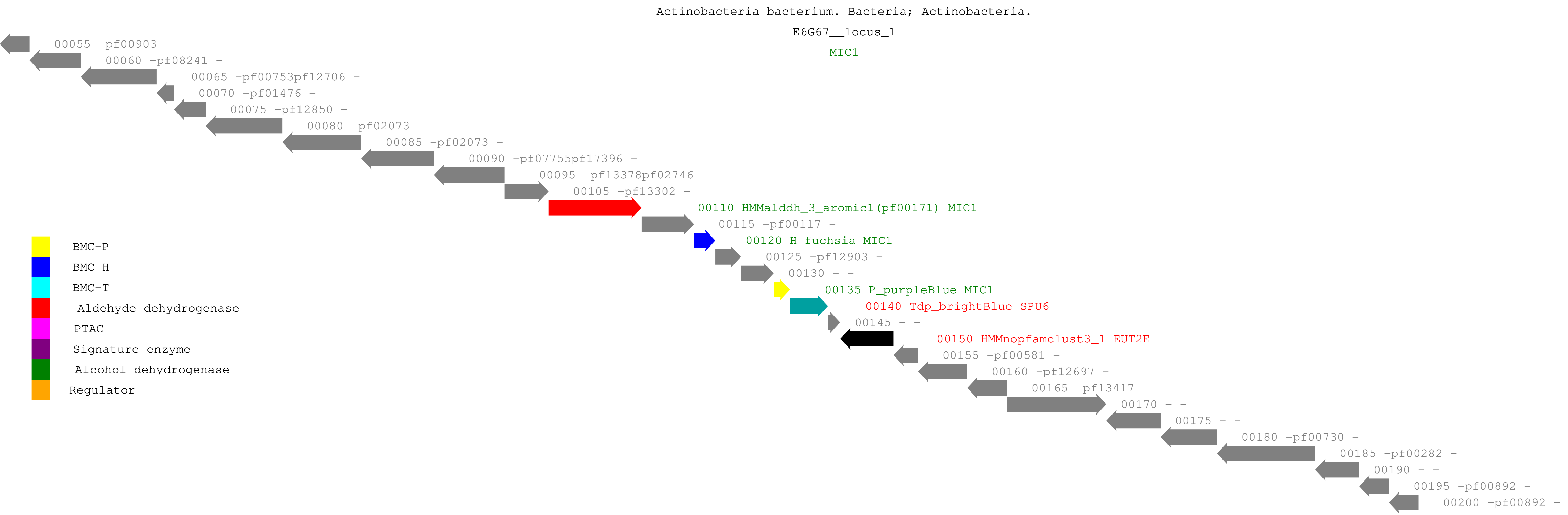

Actinobacteria bacterium. Bacteria; Actinobacteria.
E6G67__locus_1
MIC1
00055 -pf00903 -
00060 -pf08241 -
00065 -pf00753pf12706 -
00070 -pf01476 -
00075 -pf12850 -
00080 -pf02073 -
00085 -pf02073 -
00090 -pf07755pf17396 -
00095 -pf13378pf02746 -
00105 -pf13302 -
00110 HMMalddh_3_aromic1(pf00171) MIC1
00115 -pf00117 -
00120 H_fuchsia MIC1
00125 -pf12903 -
00130 - -
00135 P_purpleBlue MIC1
00140 Tdp_brightBlue SPU6
00145 - -
00150 HMMnopfamclust3_1 EUT2E
00155 -pf00581 -
00160 -pf12697 -
00165 -pf13417 -
00170 - -
00175 - -
00180 -pf00730 -
00185 -pf00282 -
00190 - -
00195 -pf00892 -
00200 -pf00892 -
BMC-P
BMC-H
BMC-T
Aldehyde dehydrogenase
PTAC
Signature enzyme
Alcohol dehydrogenase
Regulator

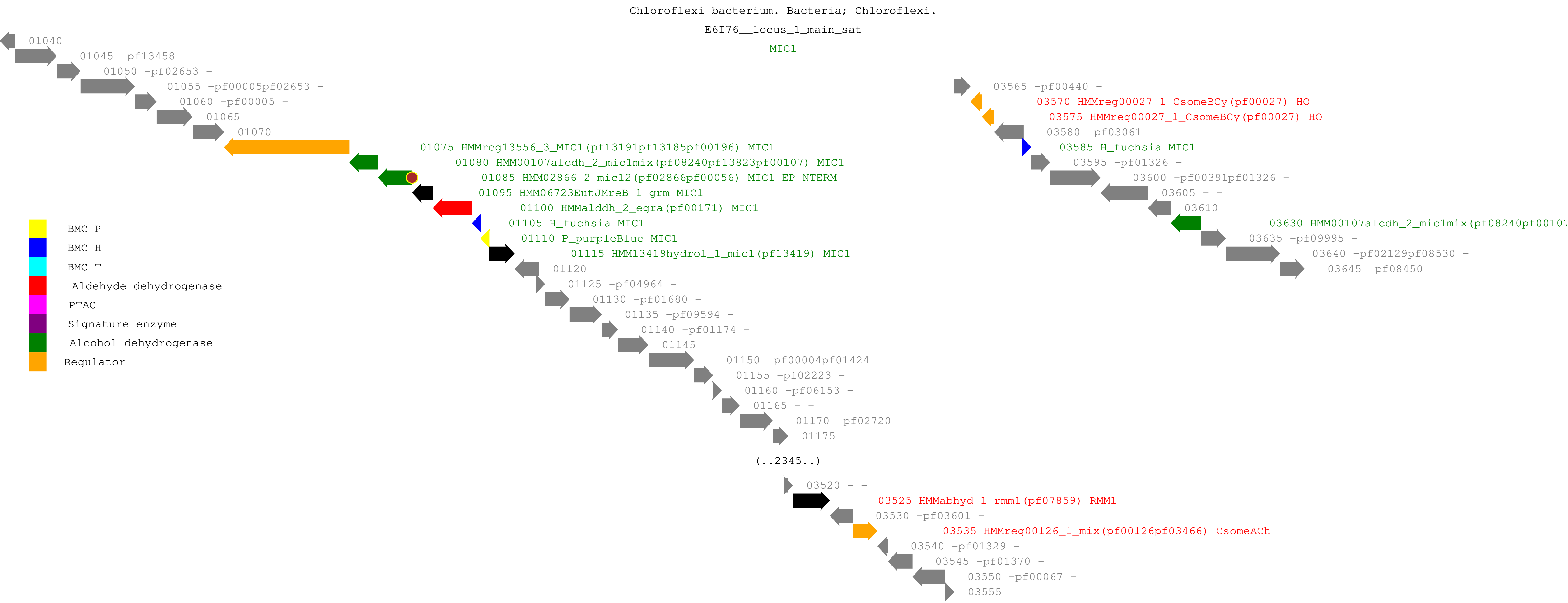

Chloroflexi bacterium. Bacteria; Chloroflexi.
E6I76__locus_1_main_sat
MIC1
01040 - -
01045 -pf13458 -
01050 -pf02653 -
01055 -pf00005pf02653 -
01060 -pf00005 -
01065 - -
01070 - -
01075 HMMreg13556_3_MIC1(pf13191pf13185pf00196) MIC1
01080 HMM00107alcdh_2_mic1mix(pf08240pf13823pf00107) MIC1
01085 HMM02866_2_mic12(pf02866pf00056) MIC1 EP_NTERM
01095 HMM06723EutJMreB_1_grm MIC1
01100 HMMalddh_2_egra(pf00171) MIC1
01105 H_fuchsia MIC1
01110 P_purpleBlue MIC1
01115 HMM13419hydrol_1_mic1(pf13419) MIC1
01120 - -
01125 -pf04964 -
01130 -pf01680 -
01135 -pf09594 -
01140 -pf01174 -
01145 - -
01150 -pf00004pf01424 -
01155 -pf02223 -
01160 -pf06153 -
01165 - -
01170 -pf02720 -
01175 - -
(..2345..)
03520 - -
03525 HMMabhyd_1_rmm1(pf07859) RMM1
03530 -pf03601 -
03535 HMMreg00126_1_mix(pf00126pf03466) CsomeACh
03540 -pf01329 -
03545 -pf01370 -
03550 -pf00067 -
03555 - -
03565 -pf00440 -
03570 HMMreg00027_1_CsomeBCy(pf00027) HO
03575 HMMreg00027_1_CsomeBCy(pf00027) HO
03580 -pf03061 -
03585 H_fuchsia MIC1
03595 -pf01326 -
03600 -pf00391pf01326 -
03605 - -
03610 - -
03630 HMM00107alcdh_2_mic1mix(pf08240pf0010
03635 -pf09995 -
03640 -pf02129pf08530 -
03645 -pf08450 -
BMC-P
BMC-H
BMC-T
Aldehyde dehydrogenase
PTAC
Signature enzyme
Alcohol dehydrogenase
Regulator

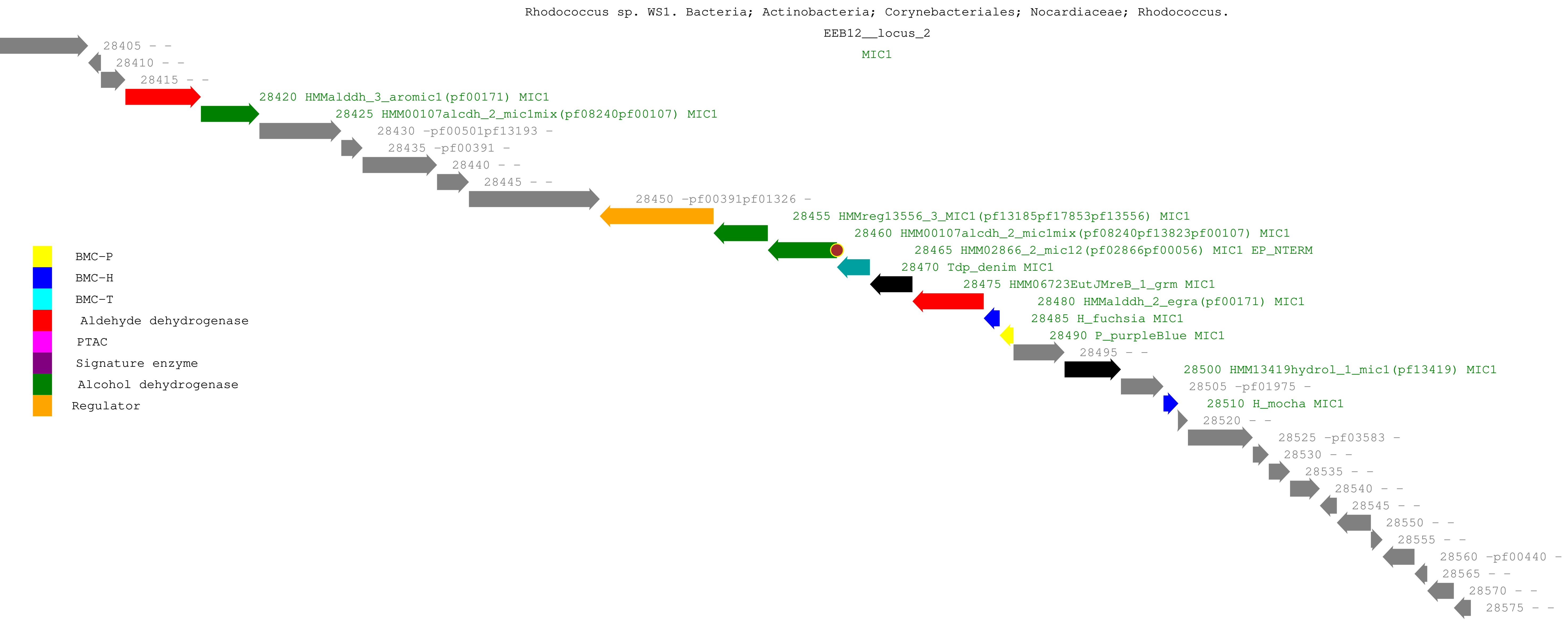
Rhodococcus sp. WS1. Bacteria; Actinobacteria; Corynebacteriales; Nocardiaceae; Rhodococcus.
EEB12__locus_2
MIC1
28405 - -
28410 - -
28415 - -
28420 HMMaldh_3_aromic1(pf00171) MIC1
28425 HMM00107alcdh_2_mic1mix(pf08240pf00107) MIC1
28430 -pf00501pf13193 -
28435 -pf00391 -
28440 - -
28445 - -
28450 -pf00391pf01326 -
28455 HMMreg13556_3_MIC1(pf13185pf17853pf13556) MIC1
28460 HMM00107alcdh_2_mic1mix(pf08240pf13823pf00107) MIC1
28465 HMM02866_2_mic12(pf02866pf00056) MIC1 EP_NTERM
28470 Tdp_denim MIC1
28475 HMM06723EutJMreB_1_grm MIC1
28480 HMMaldh_2_egra(pf00171) MIC1
28485 H_fuchsia MIC1
28490 P_purpleBlue MIC1
28495 - -
28500 HMM13419hydrol_1_mic1(pf13419) MIC1
28505 -pf01975 -
28510 H_mocha MIC1
28520 - -
28525 -pf03583 -
28530 - -
28535 - -
28540 - -
28545 - -
28550 - -
28555 - -
28560 -pf00440 -
28565 - -
28570 - -
28575 - -
BMC-P
BMC-H
BMC-T
Aldehyde dehydrogenase
PTAC
Signature enzyme
Alcohol dehydrogenase
Regulator

Pseudonocardiaceae bacterium. Bacteria; Actinobacteria; Pseudonocardiales; Pseudonocardiaceae.

EKK42__locus_1

MIC1

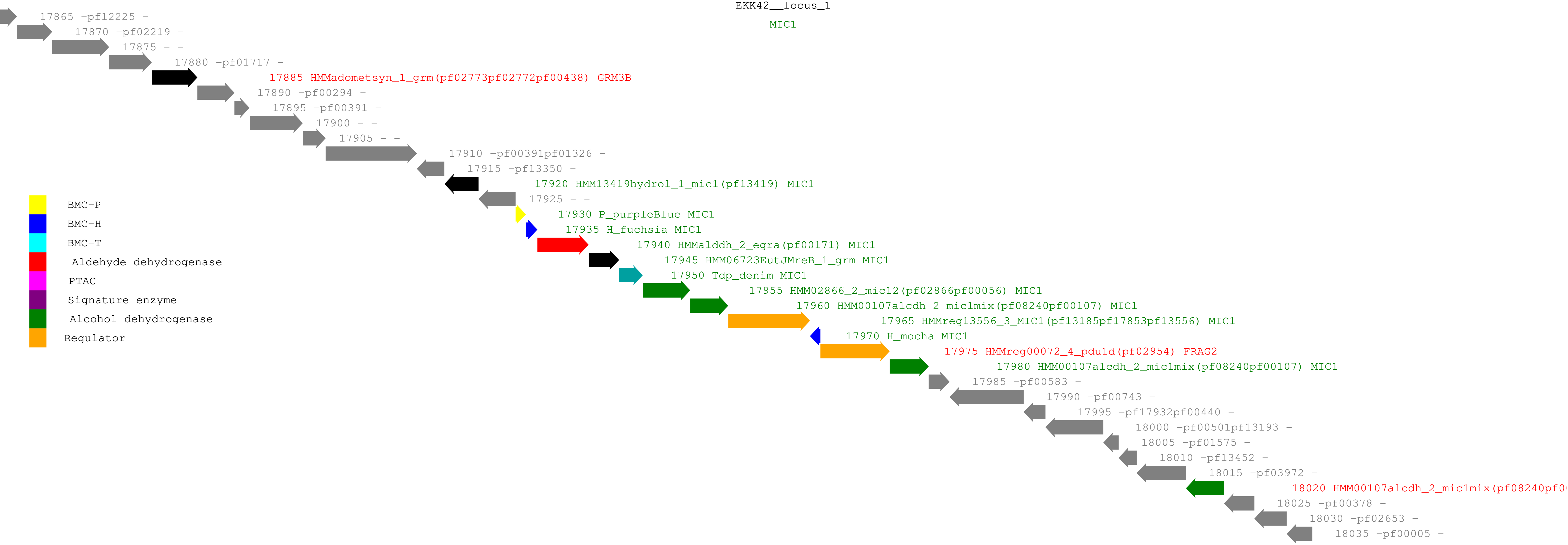

Rhizobiales bacterium. Bacteria; Proteobacteria; Alphaproteobacteria; Rhizobiales.

EOP24__locus_1

MIC1

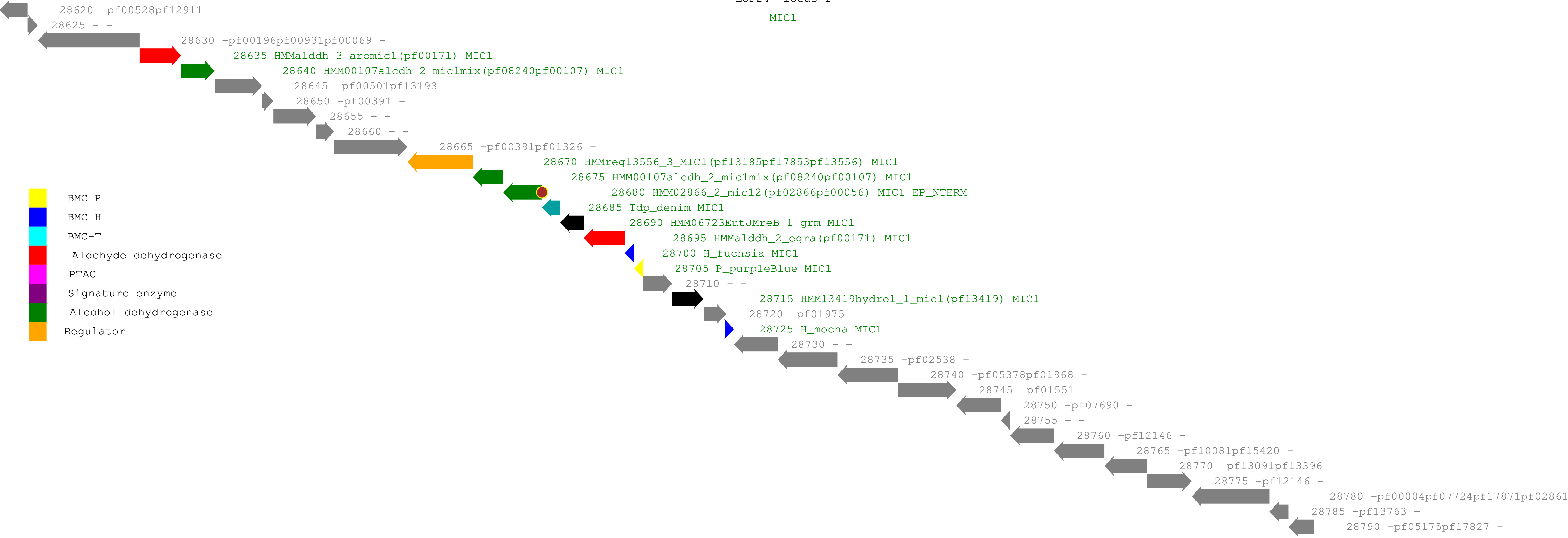

Rhodococcus opacus (Nocardia opaca). Bacteria; Actinobacteria; Corynebacteriales; Nocardiaceae;

EP51__locus_1

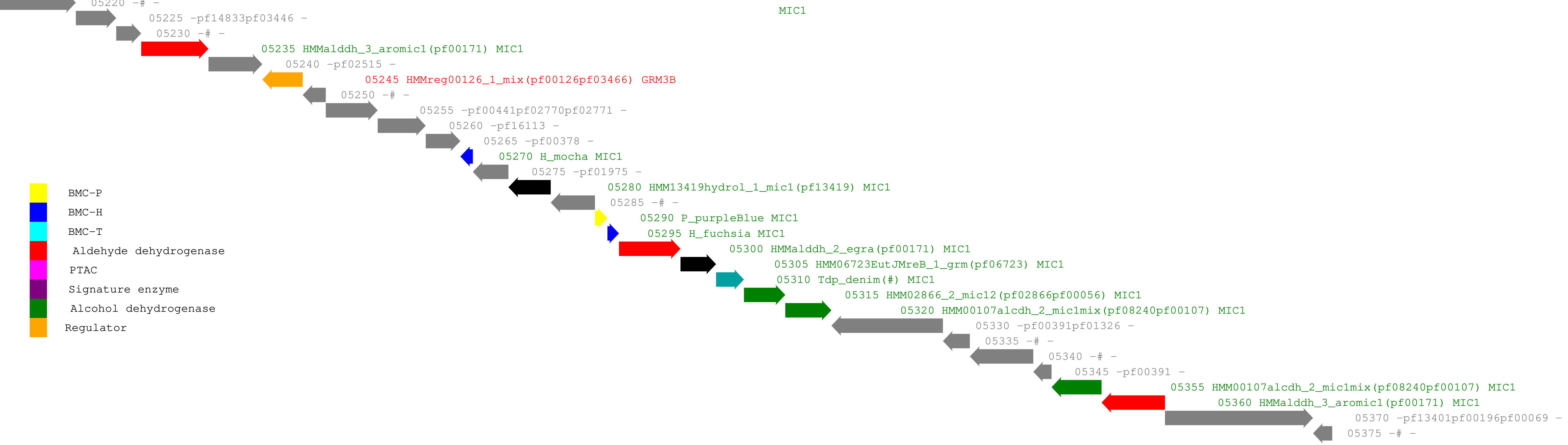

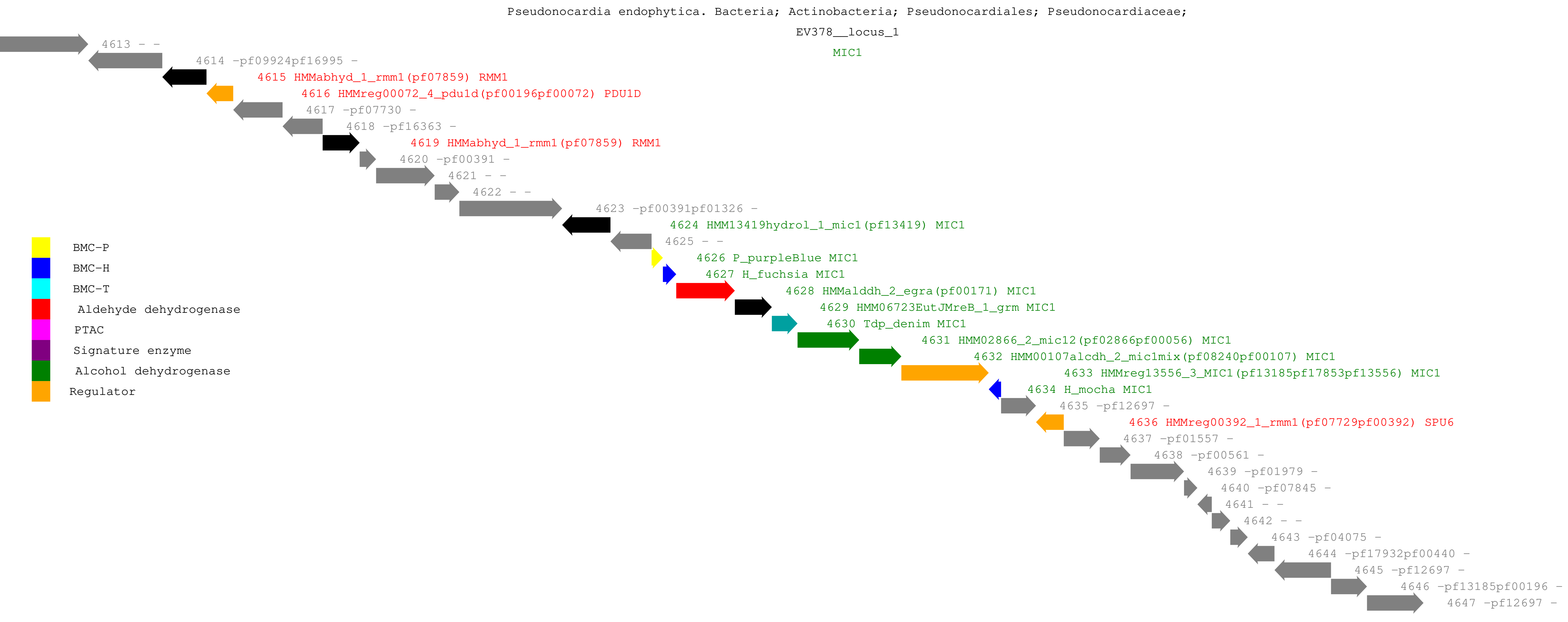
Pseudonocardia endophytica. Bacteria; Actinobacteria; Pseudonocardiales; Pseudonocardiaceae;
EV378__locus_1
MIC1
4613 - -
4614 -pf09924pf16995 -
4615 HMMabhyd_1_rmm1(pf07859) RMM1
4616 HMMreg00072_4_pdu1d(pf00196pf00072) PDU1D
4617 -pf07730 -
4618 -pf16363 -
4619 HMMabhyd_1_rmm1(pf07859) RMM1
4620 -pf00391 -
4621 - -
4622 - -
4623 -pf00391pf01326 -
4624 HMM13419hydrol_1_mic1(pf13419) MIC1
4625 - -
4626 P_purpleBlue MIC1
4627 H_fuchsia MIC1
4628 HMMaldh_2_egra(pf00171) MIC1
4629 HMM06723EutJMreB_1_grm MIC1
4630 Tdp_denim MIC1
4631 HMM02866_2_mic12(pf02866pf00056) MIC1
4632 HMM00107alcdh_2_mic1mix(pf08240pf00107) MIC1
4633 HMMreg13556_3_MIC1(pf13185pf17853pf13556) MIC1
4634 H_mocha MIC1
4635 -pf12697 -
4636 HMMreg00392_1_rmm1(pf07729pf00392) SPU6
4637 -pf01557 -
4638 -pf00561 -
4639 -pf01979 -
4640 -pf07845 -
4641 - -
4642 - -
4643 -pf04075 -
4644 -pf17932pf00440 -
4645 -pf12697 -
4646 -pf13185pf00196 -
4647 -pf12697 -
BMC-P
BMC-H
BMC-T
Aldehyde dehydrogenase
PTAC
Signature enzyme
Alcohol dehydrogenase
Regulator

Pseudonocardia endophytica. Bacteria; Actinobacteria; Pseudonocardiales; Pseudonocardiaceae;

EV378__locus_2

MIC1

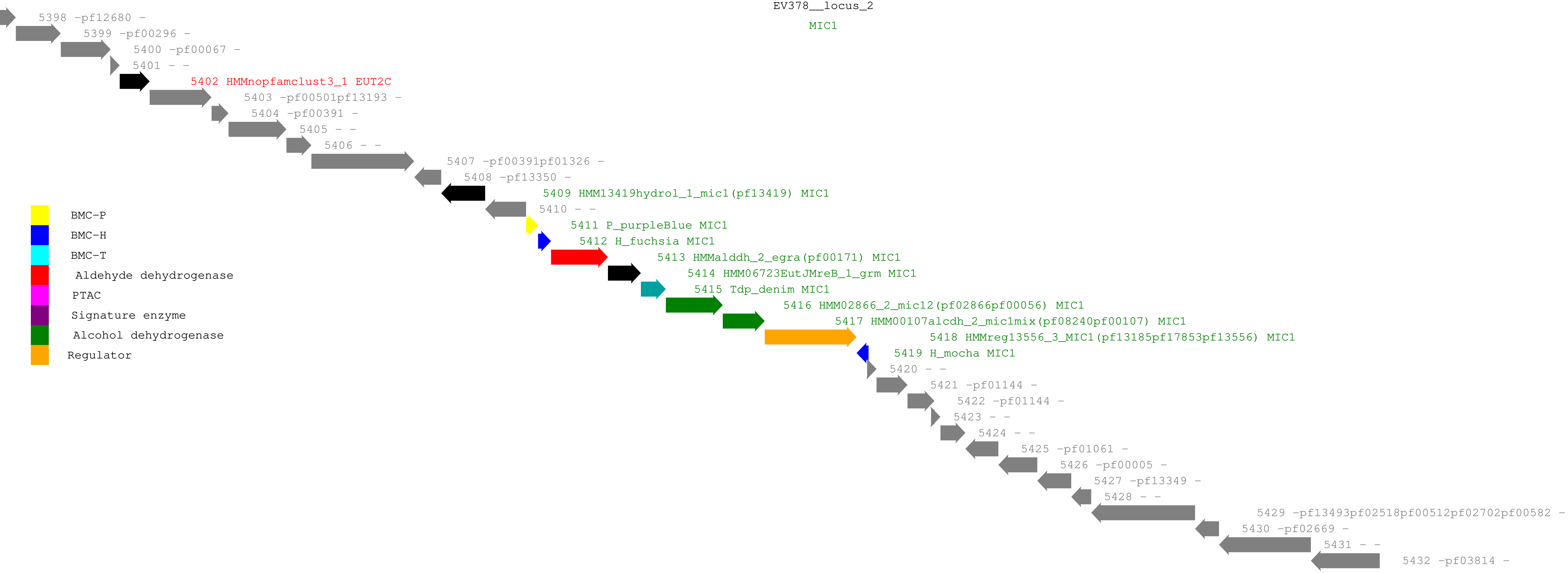

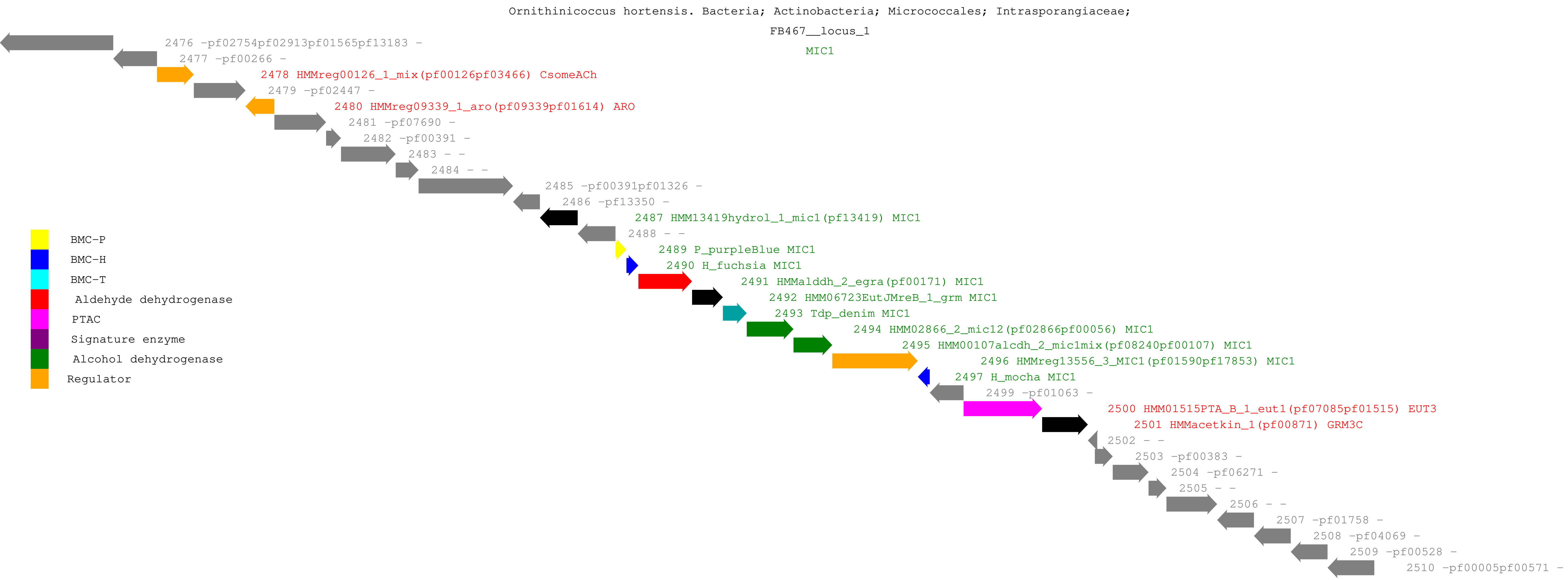
Ornithinicoccus hortensis. Bacteria; Actinobacteria; Micrococcales; Intrasporangiaceae;
FB467__locus_1
MIC1
2476 -pf02754pf02913pf01565pf13183 -
2477 -pf00266 -
2478 HMMreg00126_1_mix(pf00126pf03466) CsomeACh
2479 -pf02447 -
2480 HMMreg09339_1_aro(pf09339pf01614) ARO
2481 -pf07690 -
2482 -pf00391 -
2483 - -
2484 - -
2485 -pf00391pf01326 -
2486 -pf13350 -
2487 HMM13419hydrol_1_mic1(pf13419) MIC1
2488 - -
2489 P_purpleBlue MIC1
2490 H_fuchsia MIC1
2491 HMMaldh_2_egra(pf00171) MIC1
2492 HMM06723EutJMreB_1_grm MIC1
2493 Tdp_denim MIC1
2494 HMM02866_2_mic12(pf02866pf00056) MIC1
2495 HMM00107alcdh_2_mic1mix(pf08240pf00107) MIC1
2496 HMMreg13556_3_MIC1(pf01590pf17853) MIC1
2497 H_mocha MIC1
2499 -pf01063 -
2500 HMM01515PTA_B_1_eut1(pf07085pf01515) EUT3
2501 HMMacetkin_1(pf00871) GRM3C
2502 - -
2503 -pf00383 -
2504 -pf06271 -
2505 - -
2506 - -
2507 -pf01758 -
2508 -pf04069 -
2509 -pf00528 -
2510 -pf00005pf00571 -
BMC-P
BMC-H
BMC-T
Aldehyde dehydrogenase
PTAC
Signature enzyme
Alcohol dehydrogenase
Regulator

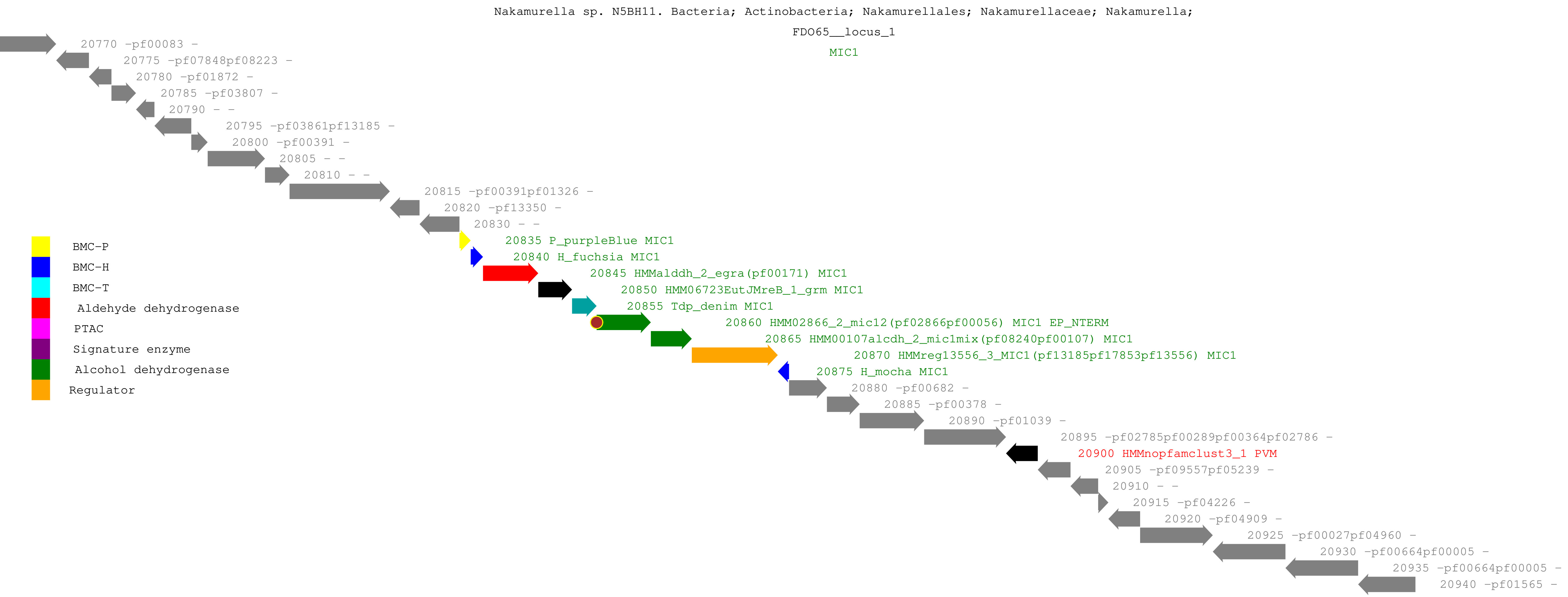
Nakamurella sp. N5BH11. Bacteria; Actinobacteria; Nakamurellales; Nakamurellaceae; Nakamurella;
FDO65__locus_1
MIC1
20770 -pf00083 -
20775 -pf07848pf08223 -
20780 -pf01872 -
20785 -pf03807 -
20790 - -
20795 -pf03861pf13185 -
20800 -pf00391 -
20805 - -
20810 - -
20815 -pf00391pf01326 -
20820 -pf13350 -
20830 - -
20835 P_purpleBlue MIC1
20840 H_fuchsia MIC1
20845 HMMalddh_2_egra(pf00171) MIC1
20850 HMM06723EutJMreB_1_grm MIC1
20855 Tdp_denim MIC1
20860 HMM02866_2_mic12(pf02866pf00056) MIC1 EP_NTERM
20865 HMM00107alcdh_2_mic1mix(pf08240pf00107) MIC1
20870 HMMreg13556_3_MIC1(pf13185pf17853pf13556) MIC1
20875 H_mocha MIC1
20880 -pf00682 -
20885 -pf00378 -
20890 -pf01039 -
20895 -pf02785pf00289pf00364pf02786 -
20900 HMMnopfamclust3_1 PVM
20905 -pf09557pf05239 -
20910 - -
20915 -pf04226 -
20920 -pf04909 -
20925 -pf00027pf04960 -
20930 -pf00664pf00005 -
20935 -pf00664pf00005 -
20940 -pf01565 -
BMC-P
BMC-H
BMC-T
Aldehyde dehydrogenase
PTAC
Signature enzyme
Alcohol dehydrogenase
Regulator

Amycolatopsis sp. SYSUP0005. Bacteria; Actinobacteria; Pseudonocardiales; Pseudonocardiaceae;

FG385__locus_1_main_sat

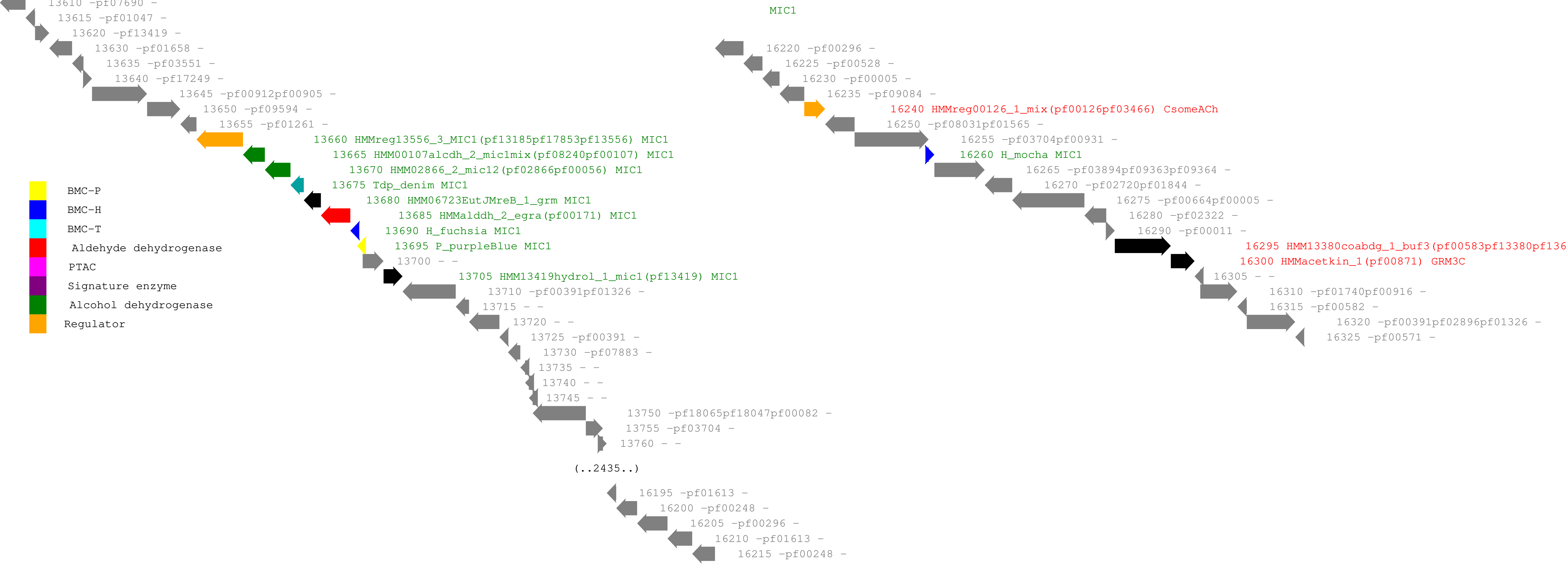

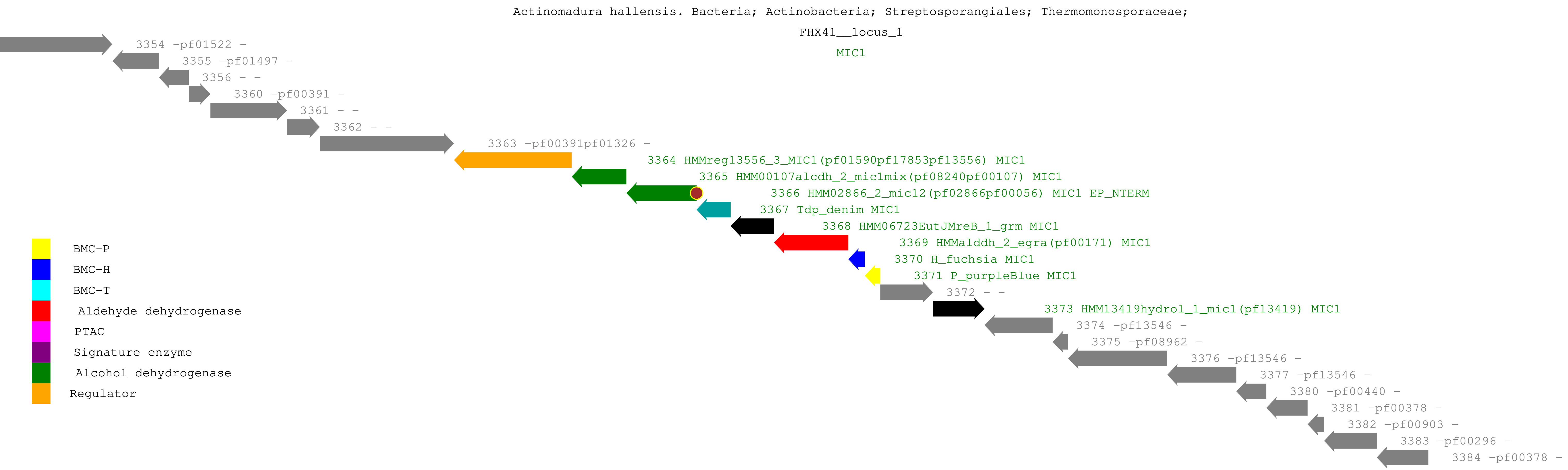
Actinomadura hallensis. Bacteria; Actinobacteria; Streptosporangiales; Thermomonosporaceae;
FHX41__locus_1
MIC1
3354 -pf01522 -
3355 -pf01497 -
3356 - -
3360 -pf00391 -
3361 - -
3362 - -
3363 -pf00391pf01326 -
3364 HMMreg13556_3_MIC1(pf01590pf17853pf13556) MIC1
3365 HMM00107alcdh_2_mic1mix(pf08240pf00107) MIC1
3366 HMM02866_2_mic12(pf02866pf00056) MIC1 EP_NTERM
3367 Tdp_denim MIC1
3368 HMM06723EutJMreB_1_grm MIC1
3369 HMMalddh_2_egra(pf00171) MIC1
3370 H_fuchsia MIC1
3371 P_purpleBlue MIC1
3372 - -
3373 HMM13419hydrol_1_mic1(pf13419) MIC1
3374 -pf13546 -
3375 -pf08962 -
3376 -pf13546 -
3377 -pf13546 -
3380 -pf00440 -
3381 -pf00378 -
3382 -pf00903 -
3383 -pf00296 -
3384 -pf00378 -
BMC-P
BMC-H
BMC-T
Aldehyde dehydrogenase
PTAC
Signature enzyme
Alcohol dehydrogenase
Regulator

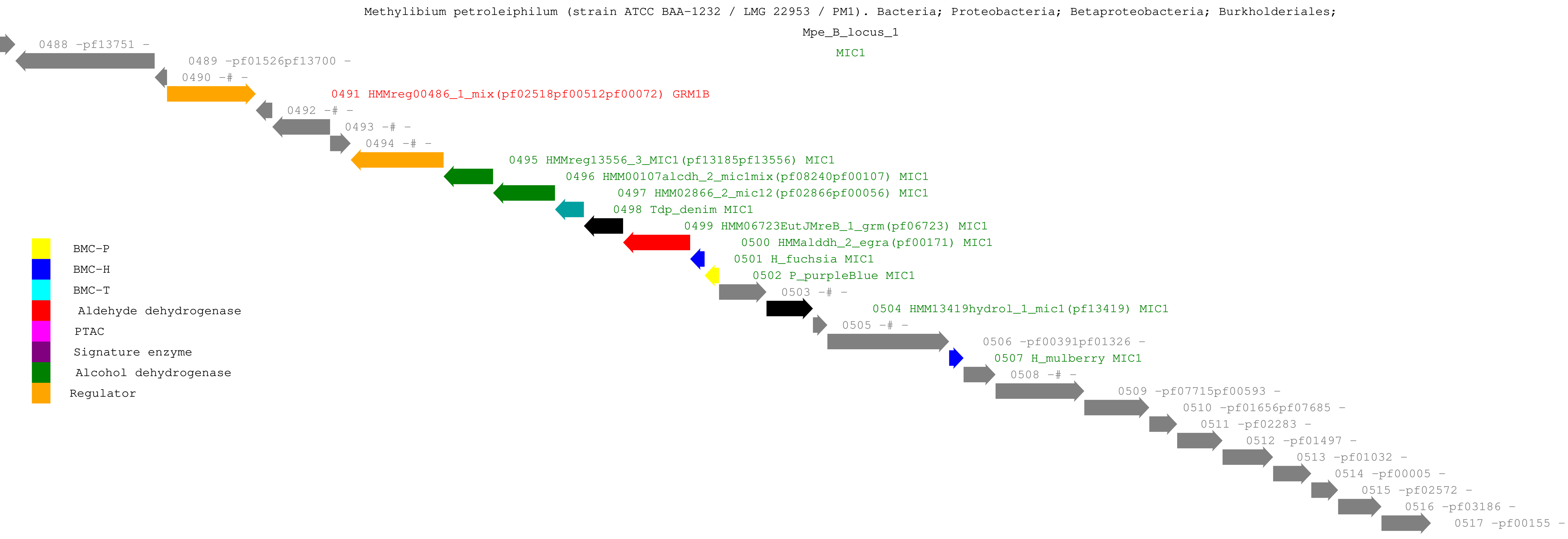
Methylibium petroleiphilum (strain ATCC BAA-1232 / LMG 22953 / PM1). Bacteria; Proteobacteria; Betaproteobacteria; Burkholderiales;
Mpe_B_locus_1
MIC1
0488 -pf13751 -
0489 -pf01526pf13700 -
0490 -# -
0491 HMMreg00486_1_mix(pf02518pf00512pf00072) GRM1B
0492 -# -
0493 -# -
0494 -# -
0495 HMMreg13556_3_MIC1(pf13185pf13556) MIC1
0496 HMM00107alcdh_2_mic1mix(pf08240pf00107) MIC1
0497 HMM02866_2_mic12(pf02866pf00056) MIC1
0498 Tdp_denim MIC1
0499 HMM06723EutJMreB_1_grm(pf06723) MIC1
0500 HMMaldh_2_egra(pf00171) MIC1
0501 H_fuchsia MIC1
0502 P_purpleBlue MIC1
0503 -# -
0504 HMM13419hydrol_1_mic1(pf13419) MIC1
0505 -# -
0506 -pf00391pf01326 -
0507 H_mulberry MIC1
0508 -# -
0509 -pf07715pf00593 -
0510 -pf01656pf07685 -
0511 -pf02283 -
0512 -pf01497 -
0513 -pf01032 -
0514 -pf00005 -
0515 -pf02572 -
0516 -pf03186 -
0517 -pf00155 -
BMC-P
BMC-H
BMC-T
Aldehyde dehydrogenase
PTAC
Signature enzyme
Alcohol dehydrogenase
Regulator

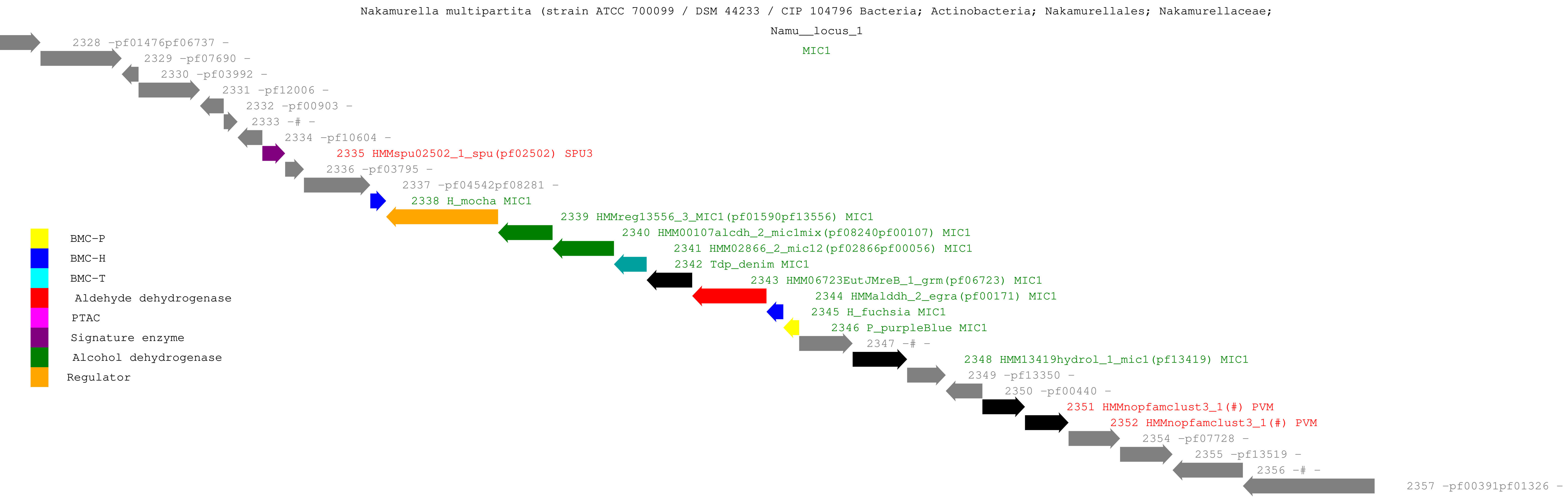

Nakamurella multipartita (strain ATCC 700099 / DSM 44233 / CIP 104796 Bacteria; Actinobacteria; Nakamurellales; Nakamurellaceae;
Namu__locus_1
MIC1
2328 -pf01476pf06737 -
2329 -pf07690 -
2330 -pf03992 -
2331 -pf12006 -
2332 -pf00903 -
2333 -# -
2334 -pf10604 -
2335 HMMspu02502_1_spu(pf02502) SPU3
2336 -pf03795 -
2337 -pf04542pf08281 -
2338 H_mocha MIC1
2339 HMMreg13556_3_MIC1(pf01590pf13556) MIC1
2340 HMM00107alcdh_2_mic1mix(pf08240pf00107) MIC1
2341 HMM02866_2_mic12(pf02866pf00056) MIC1
2342 Tdp_denim MIC1
2343 HMM06723EutJMreB_1_grm(pf06723) MIC1
2344 HMMalddh_2_egra(pf00171) MIC1
2345 H_fuchsia MIC1
2346 P_purpleBlue MIC1
2347 -# -
2348 HMM13419hydrol_1_mic1(pf13419) MIC1
2349 -pf13350 -
2350 -pf00440 -
2351 HMMnopfamclust3_1(#) PVM
2352 HMMnopfamclust3_1(#) PVM
2354 -pf07728 -
2355 -pf13519 -
2356 -# -
2357 -pf00391pf01326 -
BMC-P
BMC-H
BMC-T
Aldehyde dehydrogenase
PTAC
Signature enzyme
Alcohol dehydrogenase
Regulator

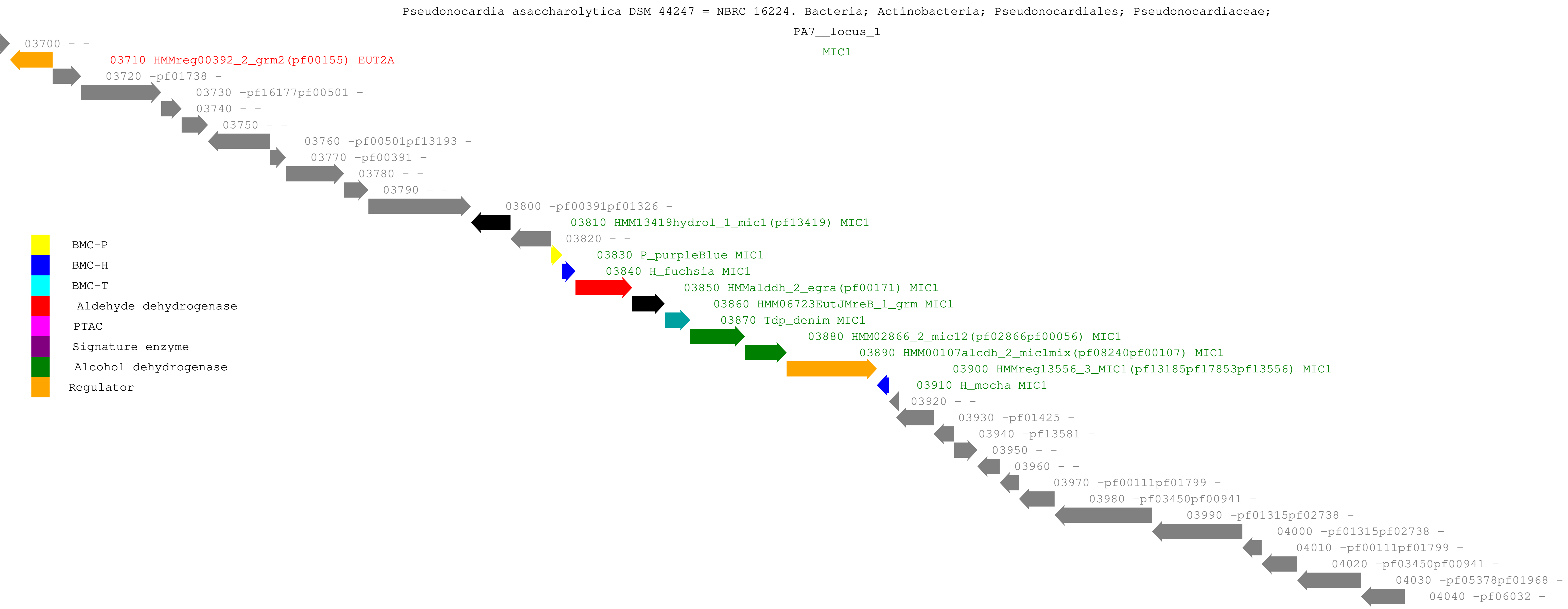

Pseudonocardia asaccharolytica DSM 44247 = NBRC 16224. Bacteria; Actinobacteria; Pseudonocardiales; Pseudonocardiaceae;
PA7__locus_1
MIC1
03700 - -
03710 HMMreg00392_2_grm2(pf00155) EUT2A
03720 -pf01738 -
03730 -pf16177pf00501 -
03740 - -
03750 - -
03760 -pf00501pf13193 -
03770 -pf00391 -
03780 - -
03790 - -
03800 -pf00391pf01326 -
03810 HMM13419hydrol_1_mic1(pf13419) MIC1
03820 - -
03830 P_purpleBlue MIC1
03840 H_fuchsia MIC1
03850 HMMaldh_2_egra(pf00171) MIC1
03860 HMM06723EutJMreB_1_grm MIC1
03870 Tdp_denim MIC1
03880 HMM02866_2_mic12(pf02866pf00056) MIC1
03890 HMM00107alcdh_2_mic1mix(pf08240pf00107) MIC1
03900 HMMreg13556_3_MIC1(pf13185pf17853pf13556) MIC1
03910 H_mocha MIC1
03920 - -
03930 -pf01425 -
03940 -pf13581 -
03950 - -
03960 - -
03970 -pf00111pf01799 -
03980 -pf03450pf00941 -
03990 -pf01315pf02738 -
04000 -pf01315pf02738 -
04010 -pf00111pf01799 -
04020 -pf03450pf00941 -
04030 -pf05378pf01968 -
04040 -pf06032 -
BMC-P
BMC-H
BMC-T
Aldehyde dehydrogenase
PTAC
Signature enzyme
Alcohol dehydrogenase
Regulator

Pseudonocardia hydrocarbonoxydans. Bacteria; Actinobacteria; Pseudonocardiales; Pseudonocardiaceae;

PHY01__locus_1

MIC1

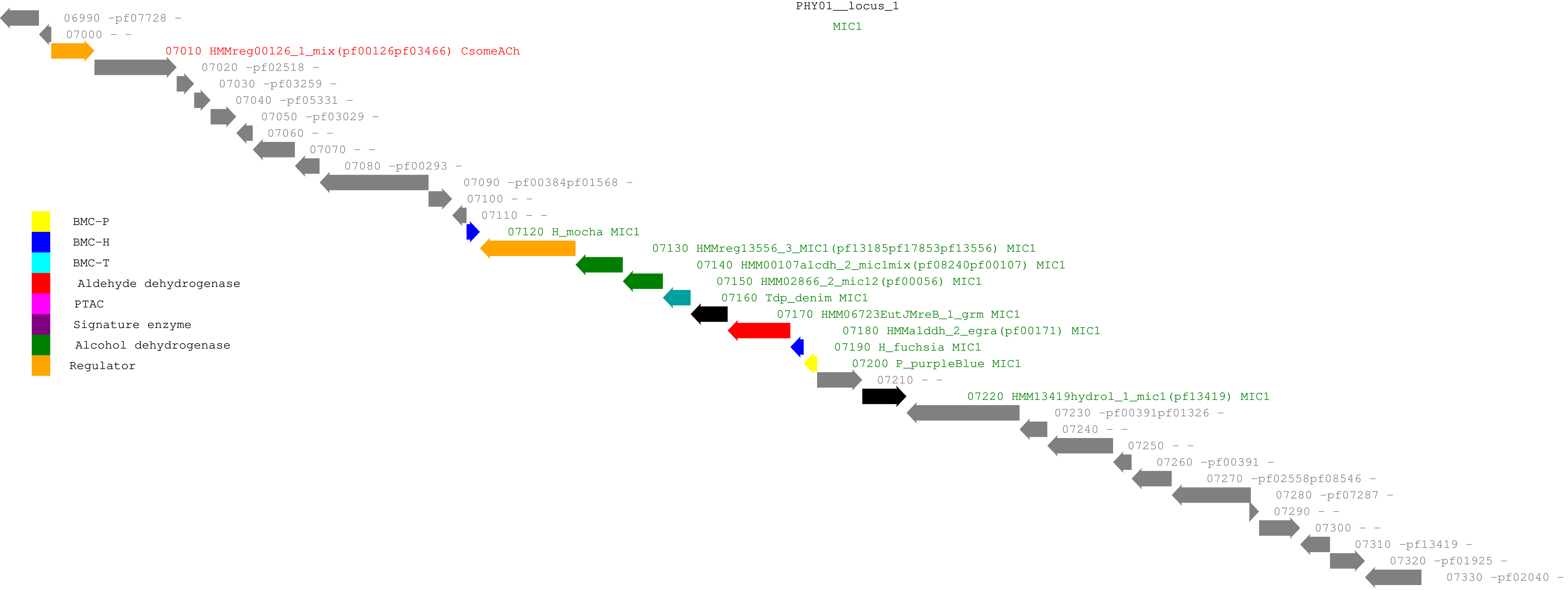

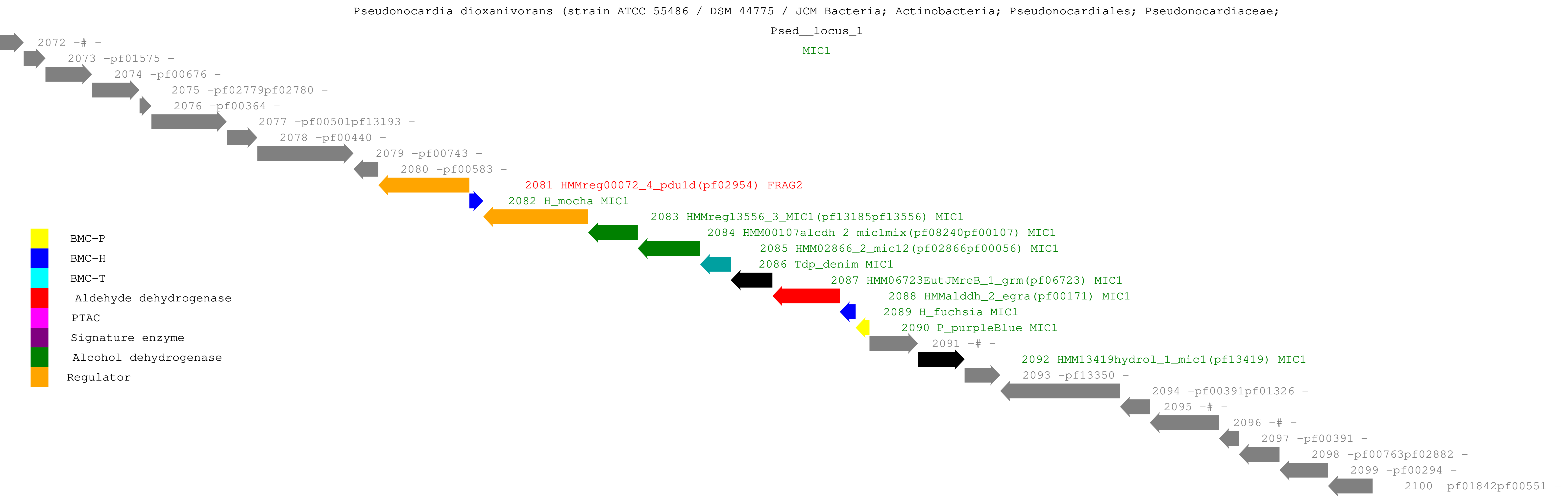
Pseudonocardia dioxanivorans (strain ATCC 55486 / DSM 44775 / JCM Bacteria; Actinobacteria; Pseudonocardiales; Pseudonocardiaceae;
Psed__locus_1
MIC1
2072 -# -
2073 -pf01575 -
2074 -pf00676 -
2075 -pf02779pf02780 -
2076 -pf00364 -
2077 -pf00501pf13193 -
2078 -pf00440 -
2079 -pf00743 -
2080 -pf00583 -
2081 HMMreg00072_4_pdu1d(pf02954) FRAG2
2082 H_mocha MIC1
2083 HMMreg13556_3_MIC1(pf13185pf13556) MIC1
2084 HMM00107alcdh_2_mic1mix(pf08240pf00107) MIC1
2085 HMM02866_2_mic12(pf02866pf00056) MIC1
2086 Tdp_denim MIC1
2087 HMM06723EutJMreB_1_grm(pf06723) MIC1
2088 HMMalddh_2_egra(pf00171) MIC1
2089 H_fuchsia MIC1
2090 P_purpleBlue MIC1
2091 -# -
2092 HMM13419hydrol_1_mic1(pf13419) MIC1
2093 -pf13350 -
2094 -pf00391pf01326 -
2095 -# -
2096 -# -
2097 -pf00391 -
2098 -pf00763pf02882 -
2099 -pf00294 -
2100 -pf01842pf00551 -
BMC-P
BMC-H
BMC-T
Aldehyde dehydrogenase
PTAC
Signature enzyme
Alcohol dehydrogenase
Regulator

Pseudonocardia sulfidoxydans NBRC 16205. Bacteria; Actinobacteria; Pseudonocardiales; Pseudonocardiaceae;

PSU4__locus_1

MIC1

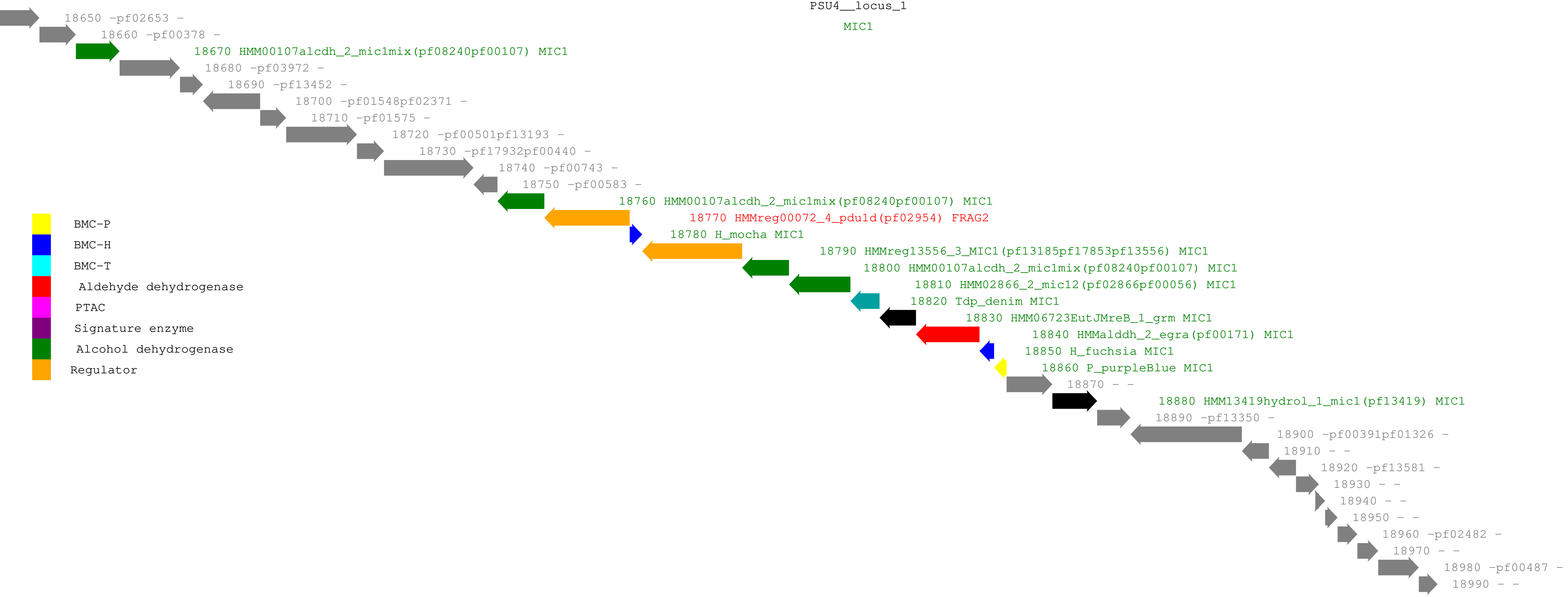

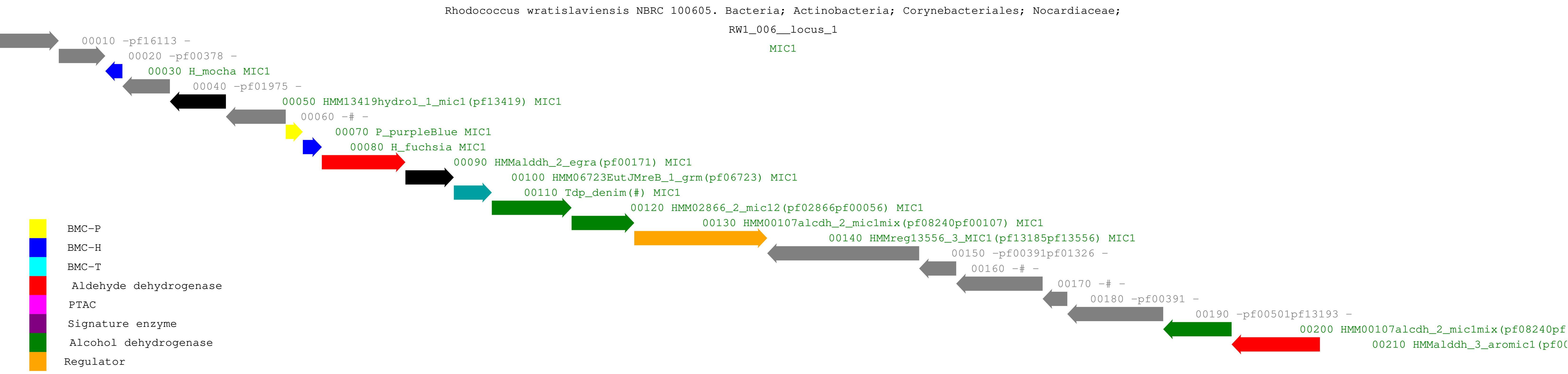
Rhodococcus wratislaviensis NBRC 100605. Bacteria; Actinobacteria; Corynebacteriales; Nocardiaceae;
RW1_006__locus_1
MIC1
00010 -pf16113 -
00020 -pf00378 -
00030 H_mocha MIC1
00040 -pf01975 -
00050 HMM13419hydrol_1_mic1(pf13419) MIC1
00060 -# -
00070 P_purpleBlue MIC1
00080 H_fuchsia MIC1
00090 HMMalddh_2_egra(pf00171) MIC1
00100 HMM06723EutJMreB_1_grm(pf06723) MIC1
00110 Tdp_denim(#) MIC1
00120 HMM02866_2_mic12(pf02866pf00056) MIC1
00130 HMM00107alcdh_2_mic1mix(pf08240pf00107) MIC1
00140 HMMreg13556_3_MIC1(pf13185pf13556) MIC1
00150 -pf00391pf01326 -
00160 -# -
00170 -# -
00180 -pf00391 -
00190 -pf00501pf13193 -
00200 HMM00107alcdh_2_mic1mix(pf08240pf
00210 HMMalddh_3_aromic1(pf00
BMC-P
BMC-H
BMC-T
Aldehyde dehydrogenase
PTAC
Signature enzyme
Alcohol dehydrogenase
Regulator

Rhodococcus jostii. Bacteria; Actinobacteria; Corynebacteriales; Nocardiaceae;

SAMN04490220__locus_2

MIC1

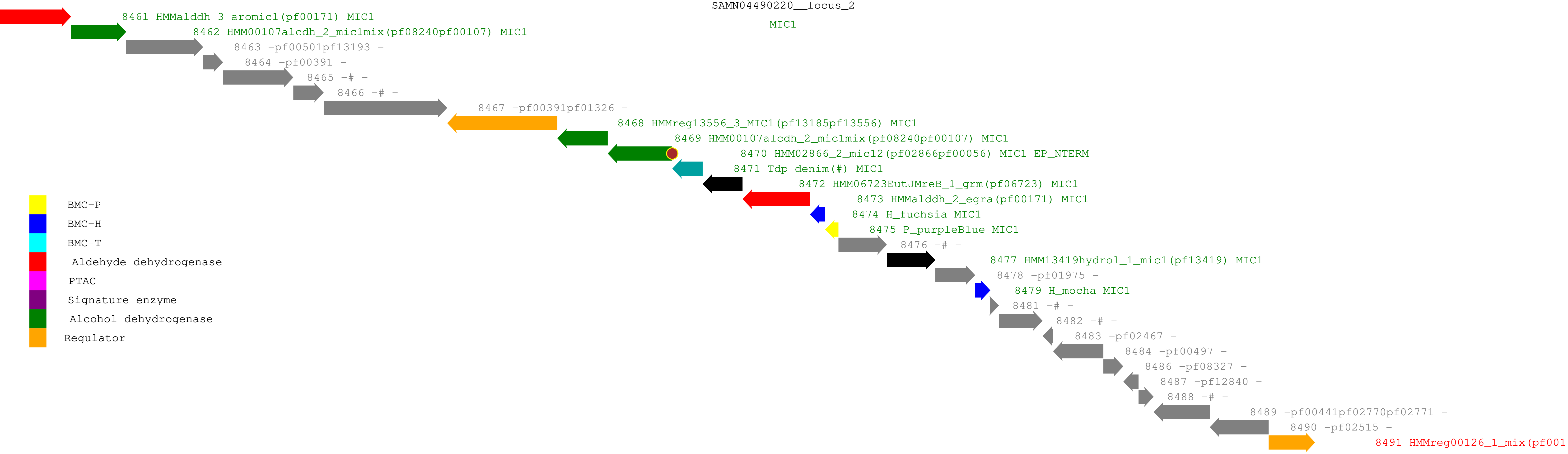

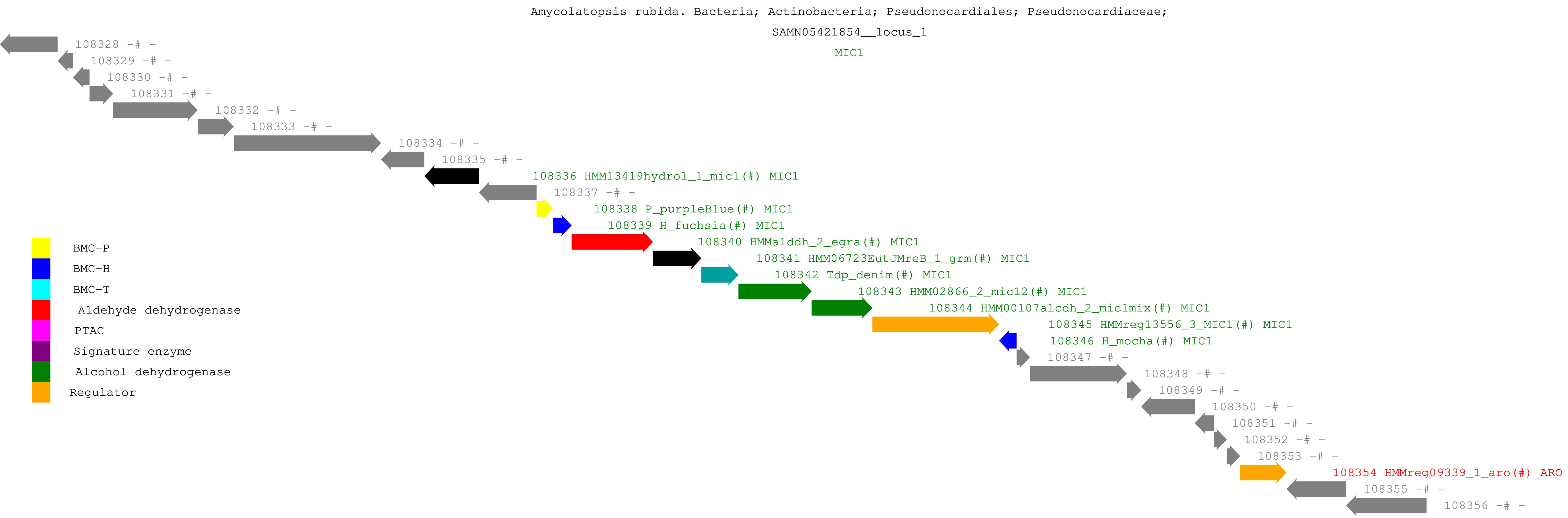
Amycolatopsis rubida. Bacteria; Actinobacteria; Pseudonocardiales; Pseudonocardiaceae;
SAMN05421854__locus_1
MIC1
108328 -# -
108329 -# -
108330 -# -
108331 -# -
108332 -# -
108333 -# -
108334 -# -
108335 -# -
108336 HMM13419hydrol_1_mic1(#) MIC1
108337 -# -
108338 P_purpleBlue(#) MIC1
108339 H_fuchsia(#) MIC1
108340 HMMalddh_2_egra(#) MIC1
108341 HMM06723EutJMreB_1_grm(#) MIC1
108342 Tdp_denim(#) MIC1
108343 HMM02866_2_mic12(#) MIC1
108344 HMM00107alcdh_2_mic1mix(#) MIC1
108345 HMMreg13556_3_MIC1(#) MIC1
108346 H_mocha(#) MIC1
108347 -# -
108348 -# -
108349 -# -
108350 -# -
108351 -# -
108352 -# -
108353 -# -
108354 HMMreg09339_1_aro(#) ARO
108355 -# -
108356 -# -
BMC-P
BMC-H
BMC-T
Aldehyde dehydrogenase
PTAC
Signature enzyme
Alcohol dehydrogenase
Regulator

Pseudonocardia thermophila. Bacteria; Actinobacteria; Pseudonocardiales; Pseudonocardiaceae;

SAMN05443637__locus_1

MIC1

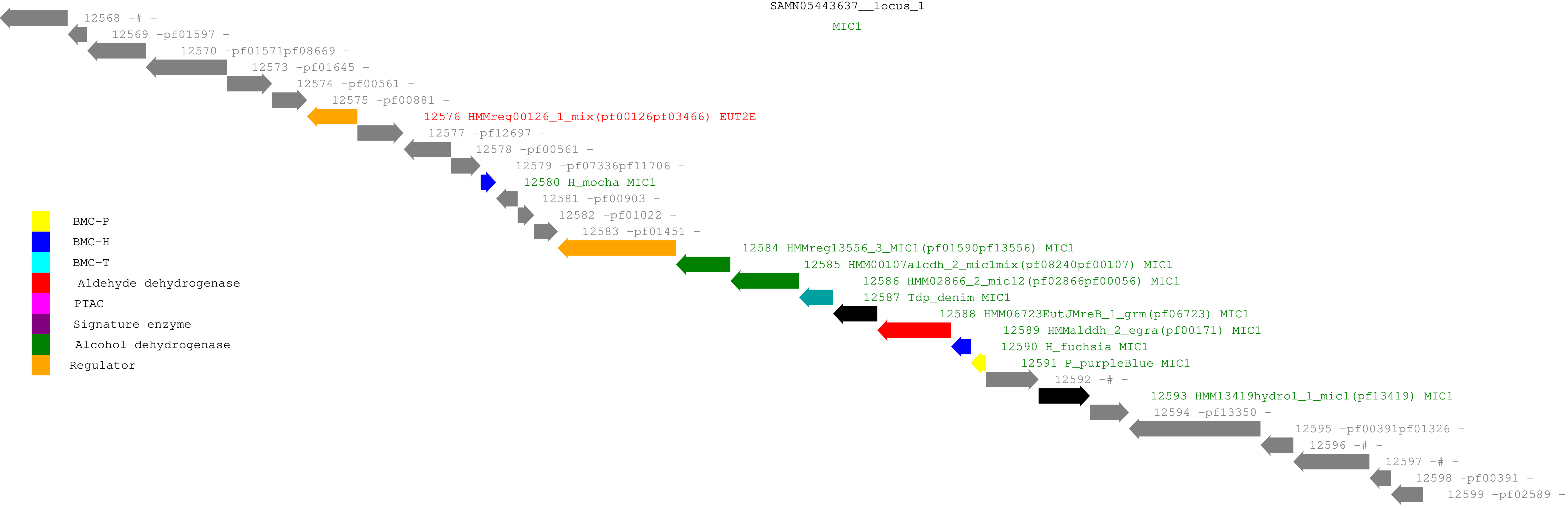

Williamsia sterculiae. Bacteria; Actinobacteria; Corynebacteriales; Williamsiaceae;

SAMN05445060__locus_1

MIC1

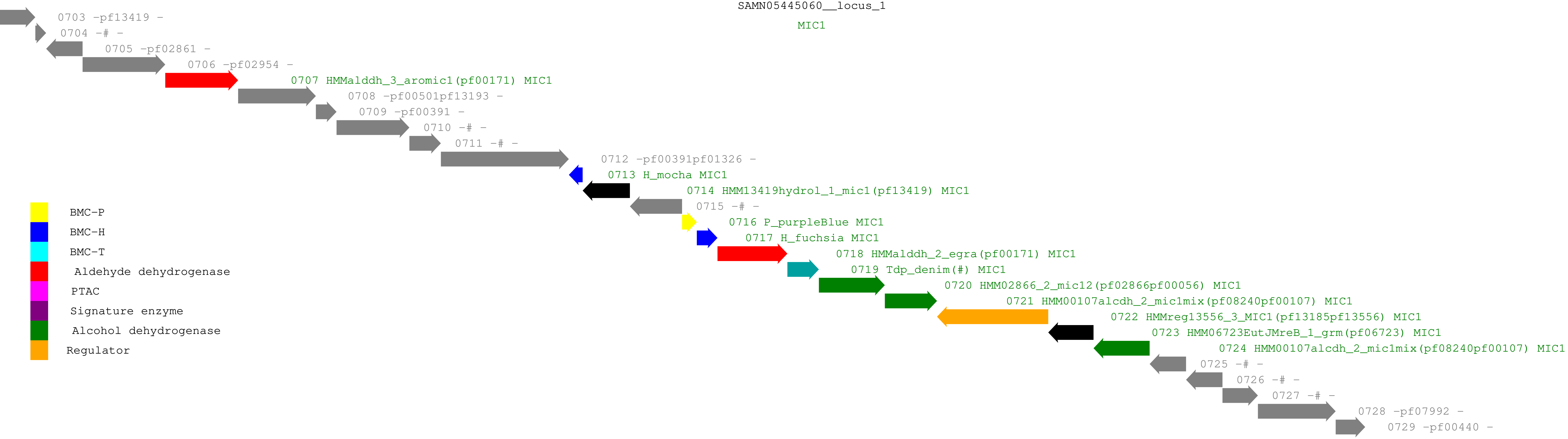

Pseudonocardia sp. N23. Bacteria; Actinobacteria; Pseudonocardiales; Pseudonocardiaceae;

TOK__locus_1

MIC1

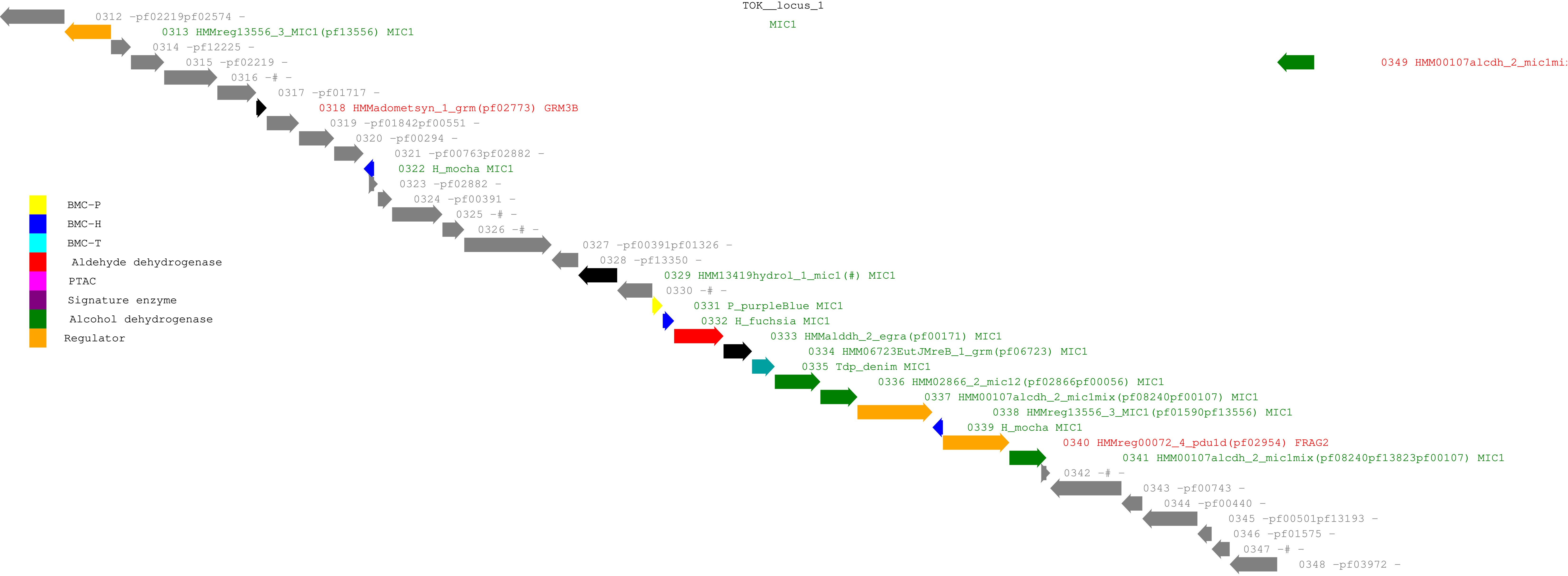

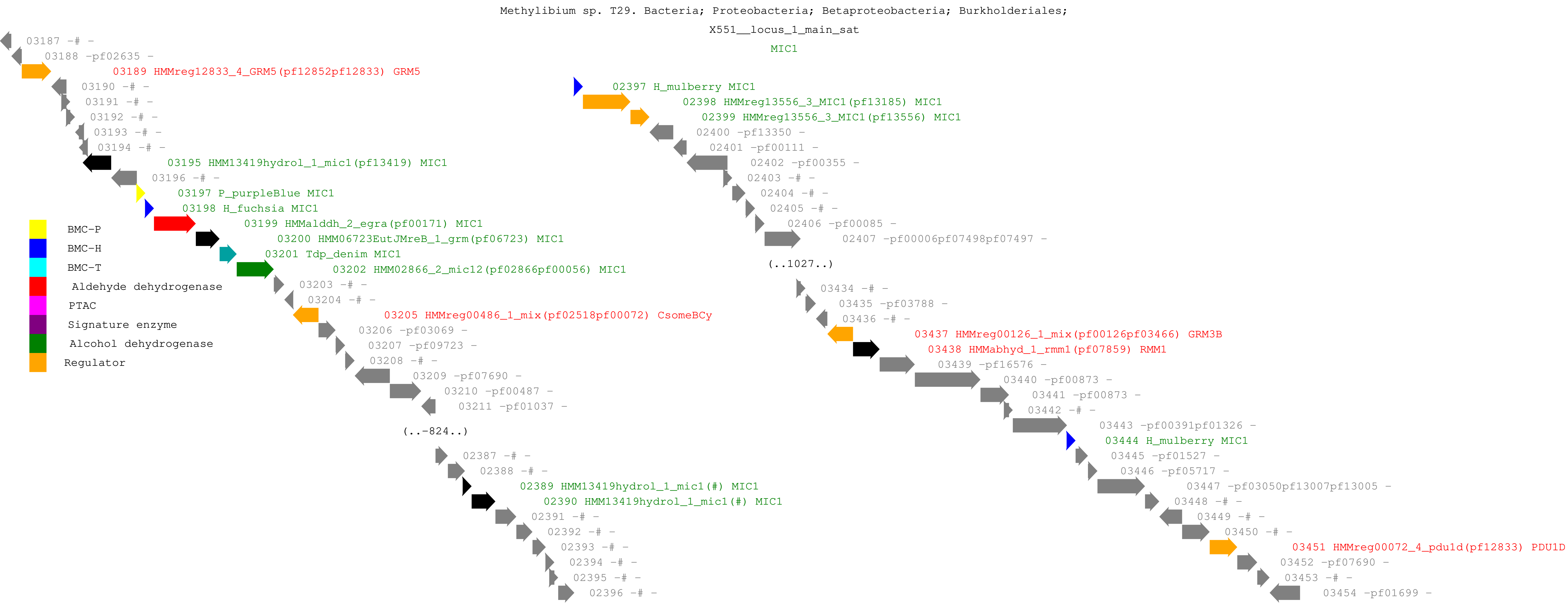
Methylibium sp. T29. Bacteria; Proteobacteria; Betaproteobacteria; Burkholderiales;
X551__locus_1_main_sat
MIC1
03187 -# -
03188 -pf02635 -
03189 HMMreg12833_4_GRM5(pf12852pf12833) GRM5
03190 -# -
03191 -# -
03192 -# -
03193 -# -
03194 -# -
03195 HMM13419hydrol_1_mic1(pf13419) MIC1
03196 -# -
03197 P_purpleBlue MIC1
03198 H_fuchsia MIC1
03199 HMMalddh_2_egra(pf00171) MIC1
03200 HMM06723EutJMreB_1_grm(pf06723) MIC1
03201 Tdp_denim MIC1
03202 HMM02866_2_mic12(pf02866pf00056) MIC1
03203 -# -
03204 -# -
03205 HMMreg00486_1_mix(pf02518pf00072) CsomeBCy
03206 -pf03069 -
03207 -pf09723 -
03208 -# -
03209 -pf07690 -
03210 -pf00487 -
03211 -pf01037 -
(..-824..)
02387 -# -
02388 -# -
02389 HMM13419hydrol_1_mic1(#) MIC1
02390 HMM13419hydrol_1_mic1(#) MIC1
02391 -# -
02392 -# -
02393 -# -
02394 -# -
02395 -# -
02396 -# -
02397 H_mulberry MIC1
02398 HMMreg13556_3_MIC1(pf13185) MIC1
02399 HMMreg13556_3_MIC1(pf13556) MIC1
02400 -pf13350 -
02401 -pf00111 -
02402 -pf00355 -
02403 -# -
02404 -# -
02405 -# -
02406 -pf00085 -
02407 -pf00006pf07498pf07497 -
(..1027..)
03434 -# -
03435 -pf03788 -
03436 -# -
03437 HMMreg00126_1_mix(pf00126pf03466) GRM3B
03438 HMMabhyd_1_rmm1(pf07859) RMM1
03439 -pf16576 -
03440 -pf00873 -
03441 -pf00873 -
03442 -# -
03443 -pf00391pf01326 -
03444 H_mulberry MIC1
03445 -pf01527 -
03446 -pf05717 -
03447 -pf03050pf13007pf13005 -
03448 -# -
03449 -# -
03450 -# -
03451 HMMreg00072_4_pdu1d(pf12833) PDU1D
03452 -pf07690 -
03453 -# -
03454 -pf01699 -
BMC-P
BMC-H
BMC-T
Aldehyde dehydrogenase
PTAC
Signature enzyme
Alcohol dehydrogenase
Regulator

Methylibium sp. T29-B. Bacteria; Proteobacteria; Betaproteobacteria; Burkholderiales;

Y694__locus_1_main_sat

MIC1

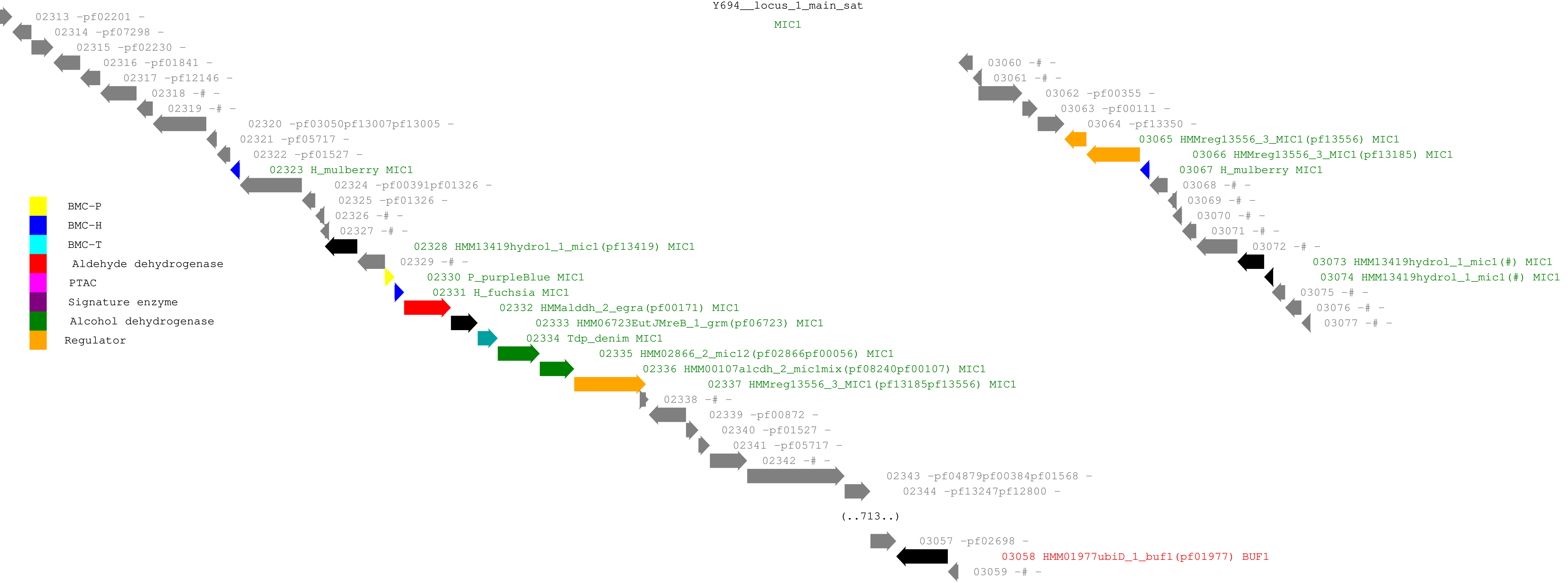

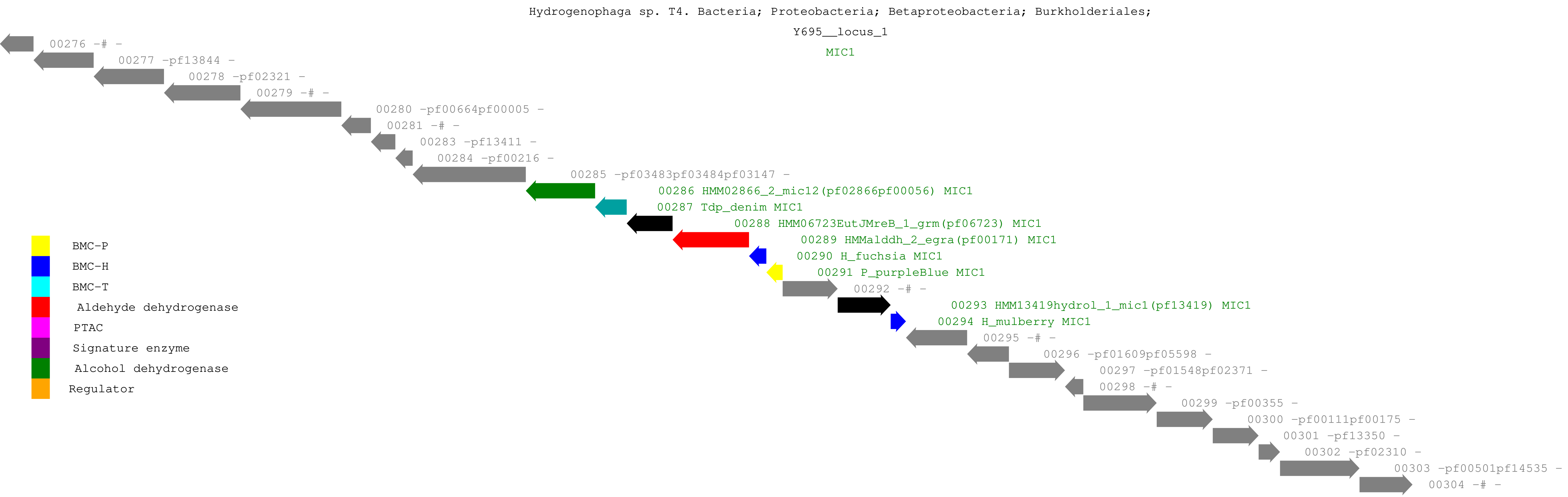
Hydrogenophaga sp. T4. Bacteria; Proteobacteria; Betaproteobacteria; Burkholderiales;
Y695__locus_1
MIC1
00276 -# -
00277 -pf13844 -
00278 -pf02321 -
00279 -# -
00280 -pf00664pf00005 -
00281 -# -
00283 -pf13411 -
00284 -pf00216 -
00285 -pf03483pf03484pf03147 -
00286 HMM02866_2_mic12(pf02866pf00056) MIC1
00287 Tdp_denim MIC1
00288 HMM06723EutJMreB_1_grm(pf06723) MIC1
00289 HMMaldh_2_egra(pf00171) MIC1
00290 H_fuchsia MIC1
00291 P_purpleBlue MIC1
00292 -# -
00293 HMM13419hydrol_1_mic1(pf13419) MIC1
00294 H_mulberry MIC1
00295 -# -
00296 -pf01609pf05598 -
00297 -pf01548pf02371 -
00298 -# -
00299 -pf00355 -
00300 -pf00111pf00175 -
00301 -pf13350 -
00302 -pf02310 -
00303 -pf00501pf14535 -
00304 -# -
BMC-P
BMC-H
BMC-T
Aldehyde dehydrogenase
PTAC
Signature enzyme
Alcohol dehydrogenase
Regulator

Chromatiales bacterium (ex Bugula neritina AB1). Bacteria; Proteobacteria; Gammaproteobacteria; Chromatiales;

AB833__locus_1

MIC2

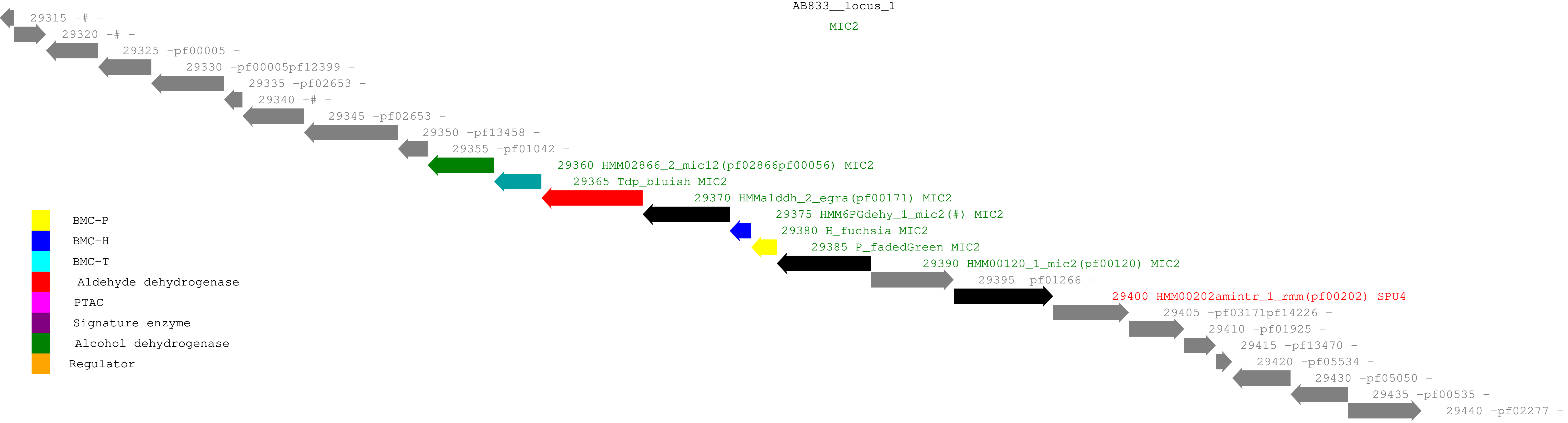

Candidatus Rhodobacter lobularis. Bacteria; Proteobacteria; Alphaproteobacteria; Rhodobacterales;

AIOL__locus_1

MIC2

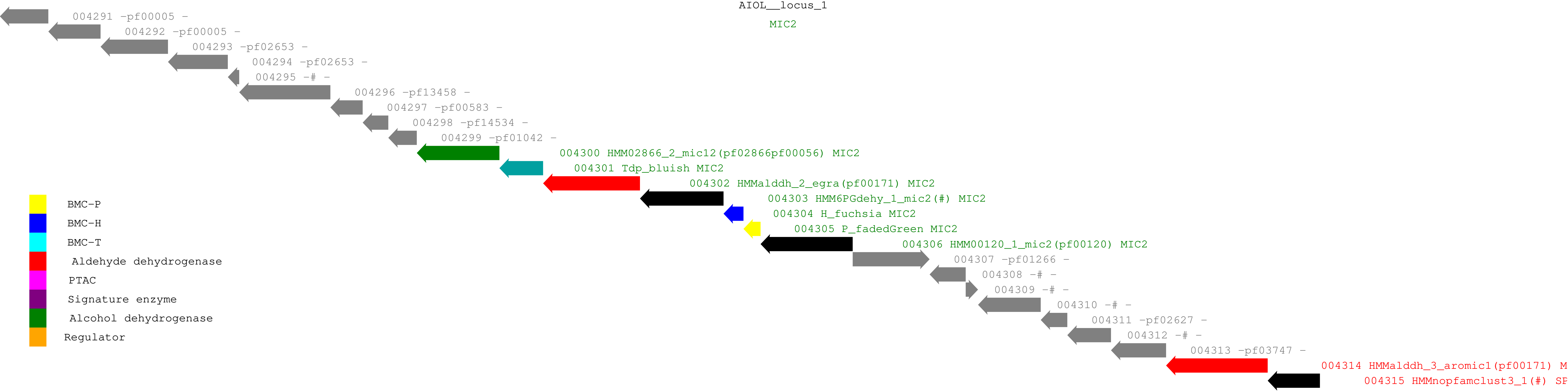

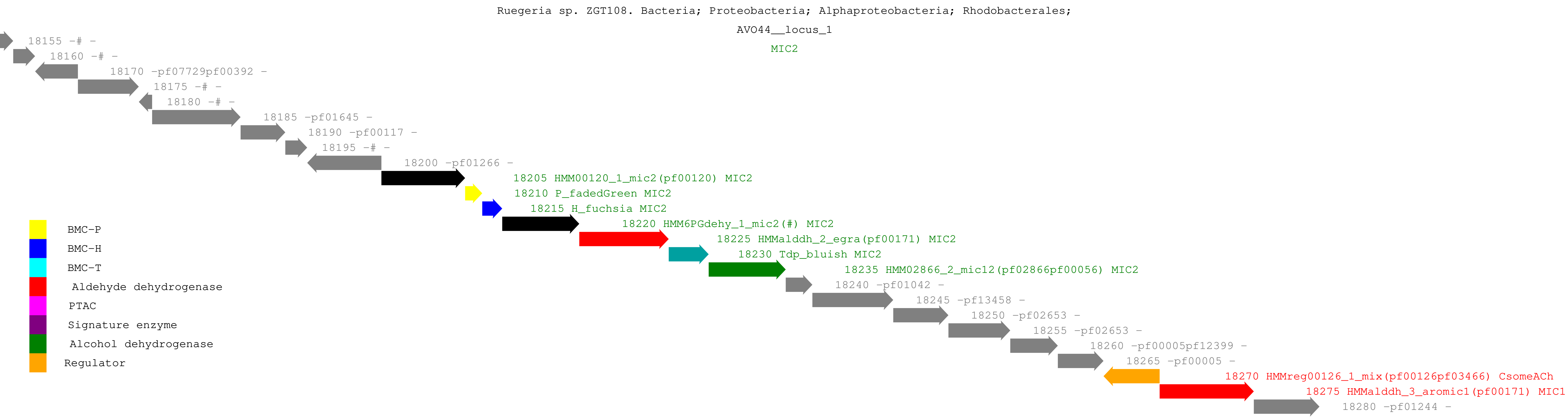

Ruegeria sp. ZGT108. Bacteria; Proteobacteria; Alphaproteobacteria; Rhodobacterales;
AVO44__locus_1
MIC2
18155 -# -
18160 -# -
18170 -pf07729pf00392 -
18175 -# -
18180 -# -
18185 -pf01645 -
18190 -pf00117 -
18195 -# -
18200 -pf01266 -
18205 HMM00120_1_mic2(pf00120) MIC2
18210 P_fadedGreen MIC2
18215 H_fuchsia MIC2
18220 HMM6PGdehy_1_mic2(#) MIC2
18225 HMMalddh_2_egra(pf00171) MIC2
18230 Tdp_bluish MIC2
18235 HMM02866_2_mic12(pf02866pf00056) MIC2
18240 -pf01042 -
18245 -pf13458 -
18250 -pf02653 -
18255 -pf02653 -
18260 -pf00005pf12399 -
18265 -pf00005 -
18270 HMMreg00126_1_mix(pf00126pf03466) CsomeACh
18275 HMMalddh_3_aromic1(pf00171) MIC1
18280 -pf01244 -
BMC-P
BMC-H
BMC-T
Aldehyde dehydrogenase
PTAC
Signature enzyme
Alcohol dehydrogenase
Regulator

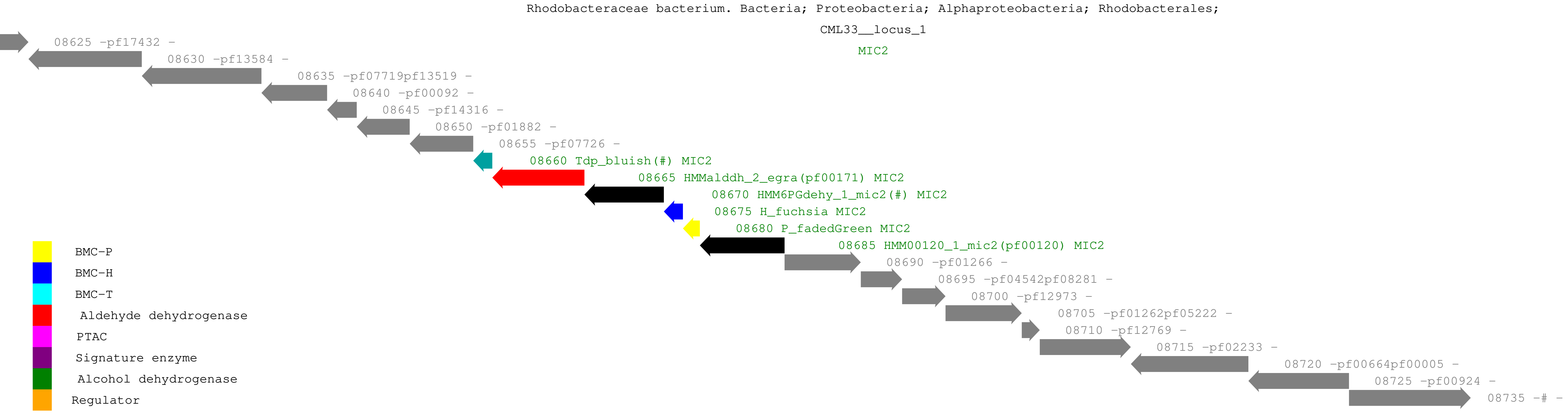

Rhodobacteraceae bacterium. Bacteria; Proteobacteria; Alphaproteobacteria; Rhodobacterales;
CML33__locus_1
MIC2
08625 -pf17432 -
08630 -pf13584 -
08635 -pf07719pf13519 -
08640 -pf00092 -
08645 -pf14316 -
08650 -pf01882 -
08655 -pf07726 -
08660 Tdp_bluish(#) MIC2
08665 HMMaldh_2_egra(pf00171) MIC2
08670 HMM6PGdehy_1_mic2(#) MIC2
08675 H_fuchsia MIC2
08680 P_fadedGreen MIC2
08685 HMM00120_1_mic2(pf00120) MIC2
08690 -pf01266 -
08695 -pf04542pf08281 -
08700 -pf12973 -
08705 -pf01262pf05222 -
08710 -pf12769 -
08715 -pf02233 -
08720 -pf00664pf00005 -
08725 -pf00924 -
08735 -# -
BMC-P
BMC-H
BMC-T
Aldehyde dehydrogenase
PTAC
Signature enzyme
Alcohol dehydrogenase
Regulator

Rhodobacteraceae bacterium. Bacteria; Proteobacteria; Alphaproteobacteria; Rhodobacterales;

CML41__locus_1

MIC2

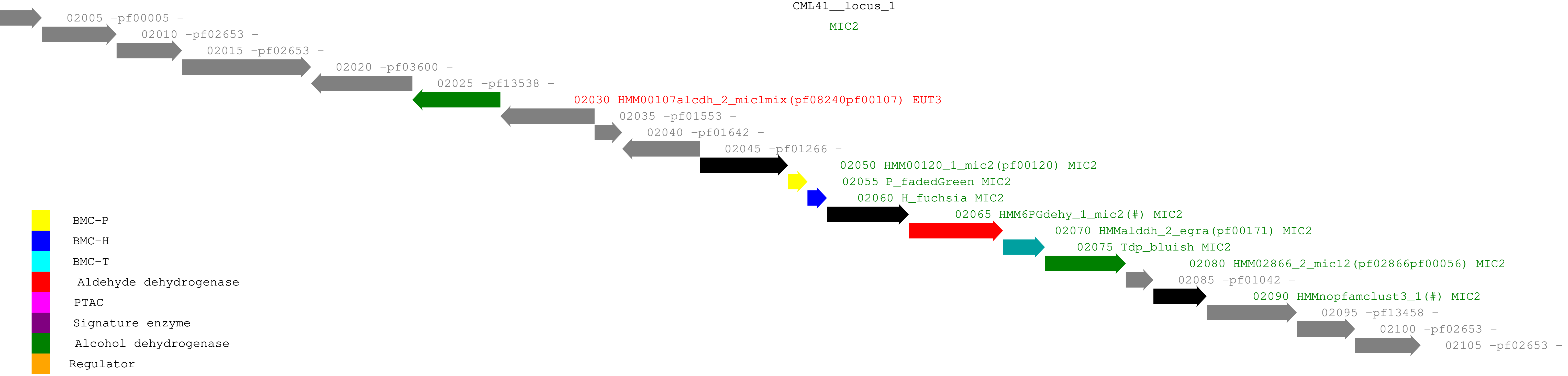

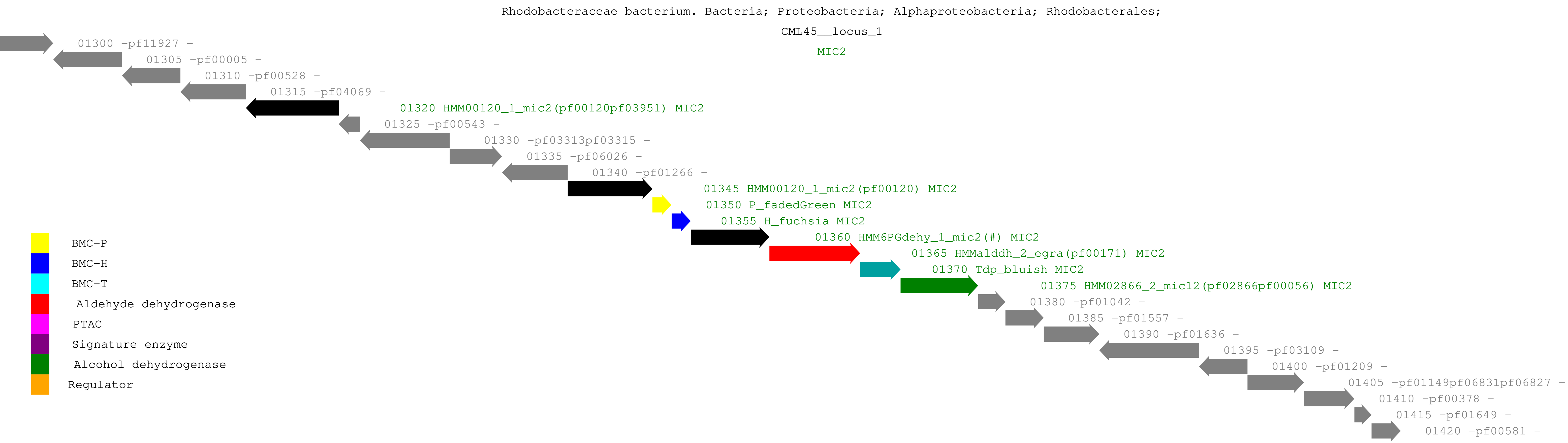
Rhodobacteraceae bacterium. Bacteria; Proteobacteria; Alphaproteobacteria; Rhodobacterales;
CML45__locus_1
MIC2
01300 -pf11927 -
01305 -pf00005 -
01310 -pf00528 -
01315 -pf04069 -
01320 HMM00120_1_mic2(pf00120pf03951) MIC2
01325 -pf00543 -
01330 -pf03313pf03315 -
01335 -pf06026 -
01340 -pf01266 -
01345 HMM00120_1_mic2(pf00120) MIC2
01350 P_fadedGreen MIC2
01355 H_fuchsia MIC2
01360 HMM6PGdehy_1_mic2(#) MIC2
01365 HMMalddh_2_egra(pf00171) MIC2
01370 Tdp_bluish MIC2
01375 HMM02866_2_mic12(pf02866pf00056) MIC2
01380 -pf01042 -
01385 -pf01557 -
01390 -pf01636 -
01395 -pf03109 -
01400 -pf01209 -
01405 -pf01149pf06831pf06827 -
01410 -pf00378 -
01415 -pf01649 -
01420 -pf00581 -
BMC-P
BMC-H
BMC-T
Aldehyde dehydrogenase
PTAC
Signature enzyme
Alcohol dehydrogenase
Regulator

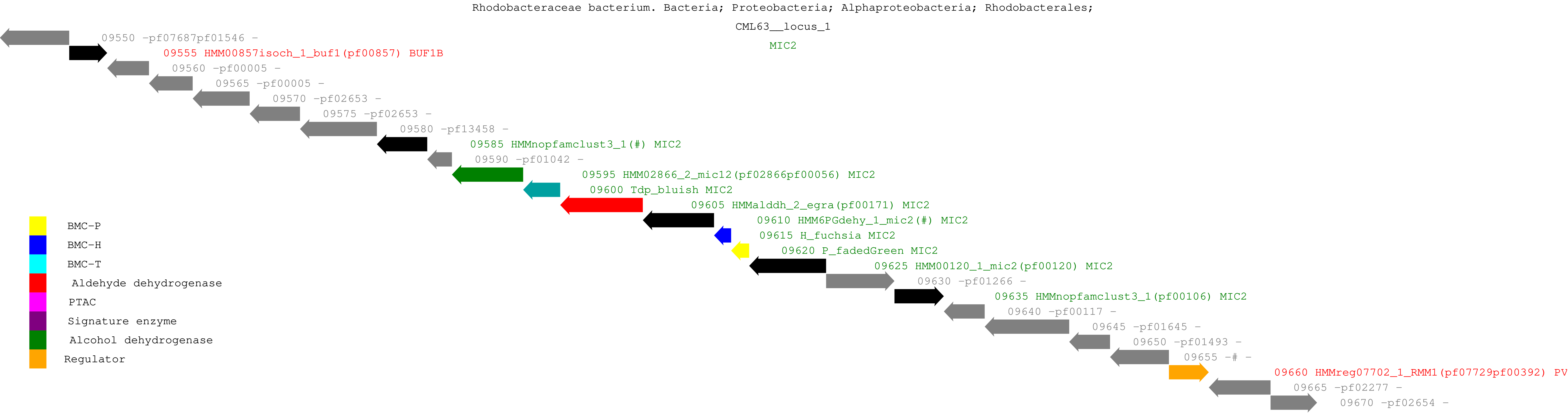
Rhodobacteraceae bacterium. Bacteria; Proteobacteria; Alphaproteobacteria; Rhodobacterales;
CML63__locus_1
MIC2
09550 -pf07687pf01546 -
09555 HMM00857isoch_1_buf1(pf00857) BUF1B
09560 -pf00005 -
09565 -pf00005 -
09570 -pf02653 -
09575 -pf02653 -
09580 -pf13458 -
09585 HMMnopfamclust3_1(#) MIC2
09590 -pf01042 -
09595 HMM02866_2_mic12(pf02866pf00056) MIC2
09600 Tdp_bluish MIC2
09605 HMMalddh_2_egra(pf00171) MIC2
09610 HMM6PGdehy_1_mic2(#) MIC2
09615 H_fuchsia MIC2
09620 P_fadedGreen MIC2
09625 HMM00120_1_mic2(pf00120) MIC2
09630 -pf01266 -
09635 HMMnopfamclust3_1(pf00106) MIC2
09640 -pf00117 -
09645 -pf01645 -
09650 -pf01493 -
09655 -# -
09660 HMMreg07702_1_RMM1(pf07729pf00392) PV
09665 -pf02277 -
09670 -pf02654 -
BMC-P
BMC-H
BMC-T
Aldehyde dehydrogenase
PTAC
Signature enzyme
Alcohol dehydrogenase
Regulator

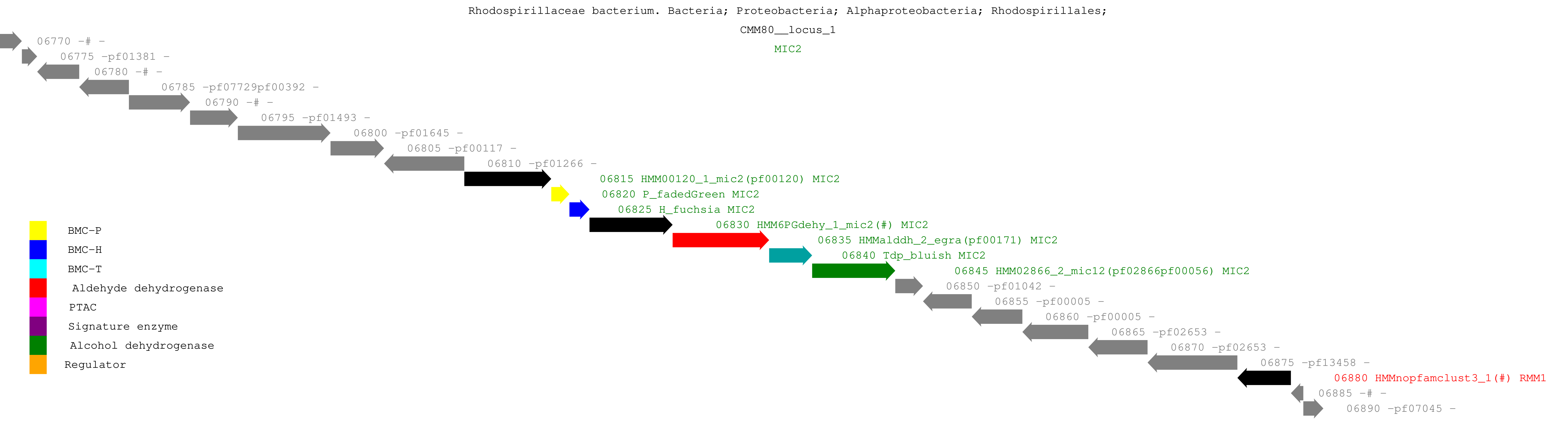
Rhodospirillaceae bacterium. Bacteria; Proteobacteria; Alphaproteobacteria; Rhodospirillales;
CMM80__locus_1
MIC2
06770 -# -
06775 -pf01381 -
06780 -# -
06785 -pf07729pf00392 -
06790 -# -
06795 -pf01493 -
06800 -pf01645 -
06805 -pf00117 -
06810 -pf01266 -
06815 HMM00120_1_mic2(pf00120) MIC2
06820 P_fadedGreen MIC2
06825 H_fuchsia MIC2
06830 HMM6PGdehy_1_mic2(#) MIC2
06835 HMMalddh_2_egra(pf00171) MIC2
06840 Tdp_bluish MIC2
06845 HMM02866_2_mic12(pf02866pf00056) MIC2
06850 -pf01042 -
06855 -pf00005 -
06860 -pf00005 -
06865 -pf02653 -
06870 -pf02653 -
06875 -pf13458 -
06880 HMMnopfamclust3_1(#) RMM1
06885 -# -
06890 -pf07045 -
BMC-P
BMC-H
BMC-T
Aldehyde dehydrogenase
PTAC
Signature enzyme
Alcohol dehydrogenase
Regulator

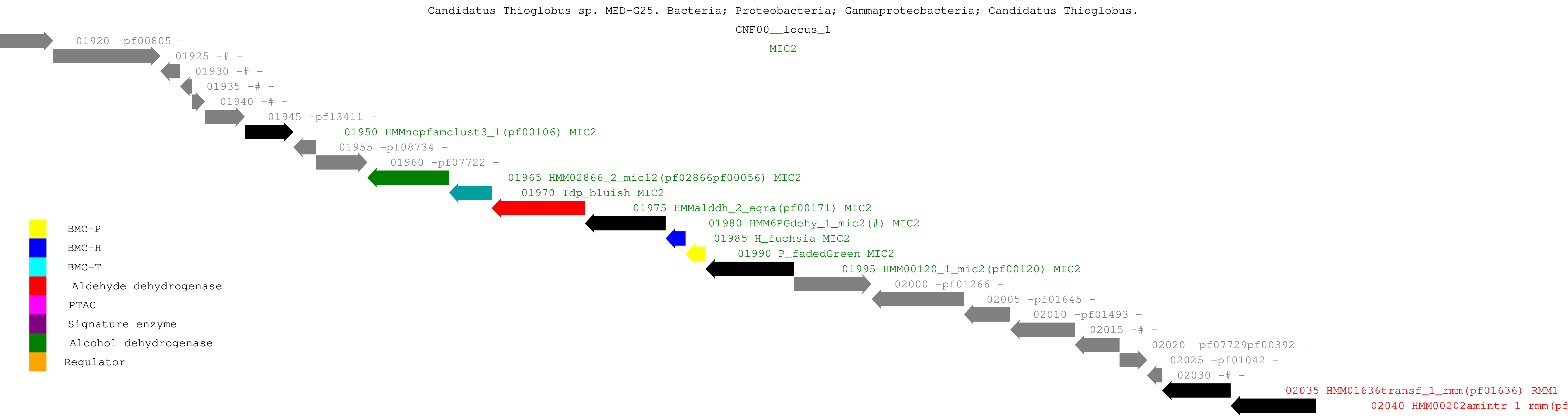
Candidatus Thioglobus sp. MED-G25. Bacteria; Proteobacteria; Gammaproteobacteria; Candidatus Thioglobus.
CNF00__locus_1
MIC2
01920 -pf00805 -
01925 -# -
01930 -# -
01935 -# -
01940 -# -
01945 -pf13411 -
01950 HMMnopfamclust3_1(pf00106) MIC2
01955 -pf08734 -
01960 -pf07722 -
01965 HMM02866_2_mic12(pf02866pf00056) MIC2
01970 Tdp_bluish MIC2
01975 HMMalddh_2_egra(pf00171) MIC2
01980 HMM6PGdehy_1_mic2(#) MIC2
01985 H_fuchsia MIC2
01990 P_fadedGreen MIC2
01995 HMM00120_1_mic2(pf00120) MIC2
02000 -pf01266 -
02005 -pf01645 -
02010 -pf01493 -
02015 -# -
02020 -pf07729pf00392 -
02025 -pf01042 -
02030 -# -
02035 HMM01636transf_1_rmm(pf01636) RMM1
02040 HMM00202amintr_1_rmm(pf
BMC-P
BMC-H
BMC-T
Aldehyde dehydrogenase
PTAC
Signature enzyme
Alcohol dehydrogenase
Regulator

Acidithiobacillus sp. Bacteria; Proteobacteria; Acidithiobacillia; Acidithiobacillales;

COA89__locus_1

MIC2

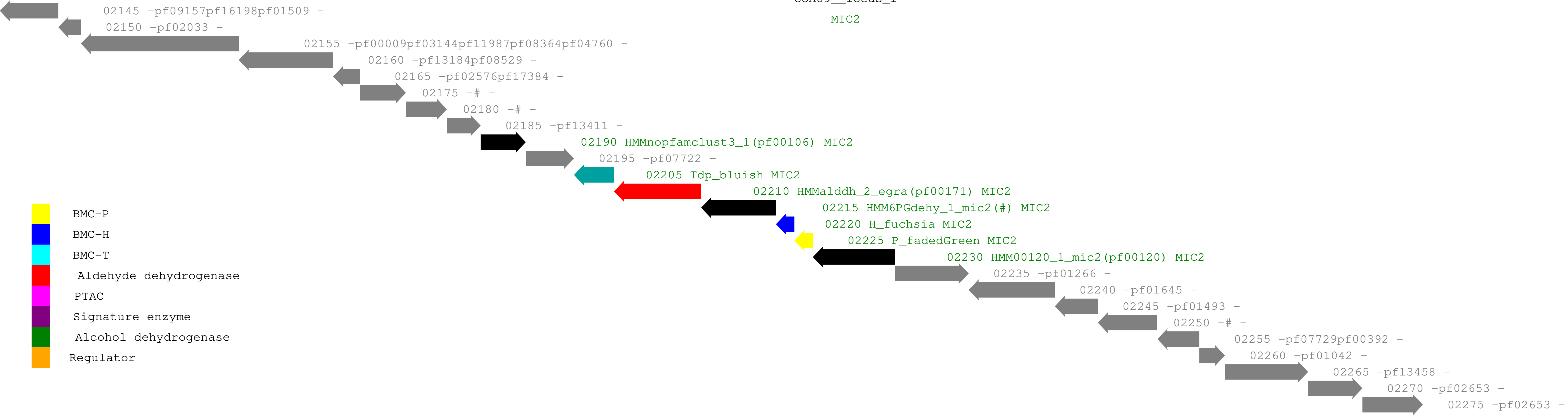

Ruegeria sp. AD91A. Bacteria; Proteobacteria; Alphaproteobacteria; Rhodobacterales;

D1823__locus_1

MIC2

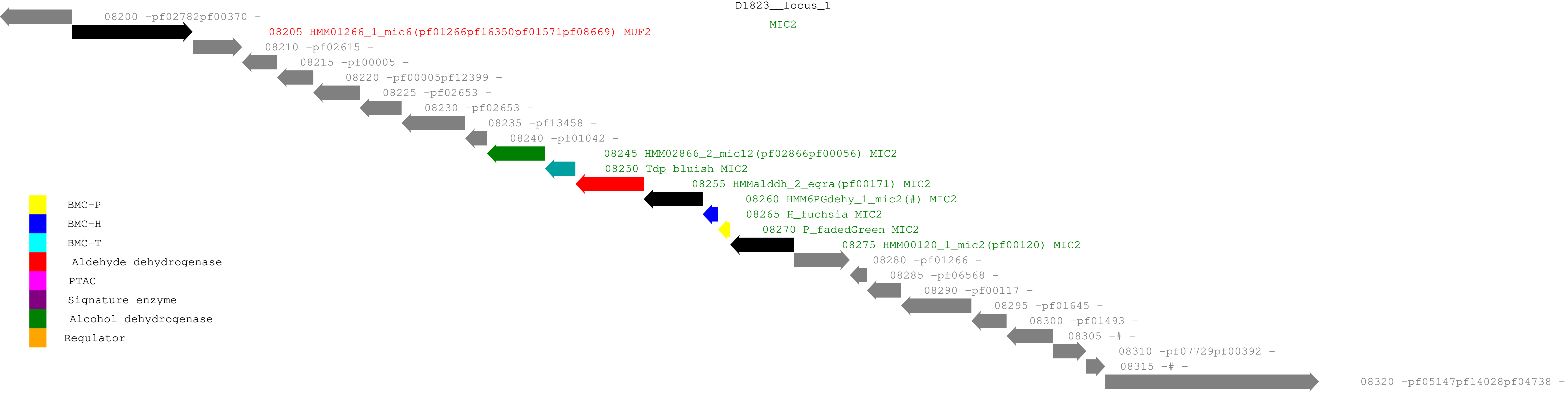

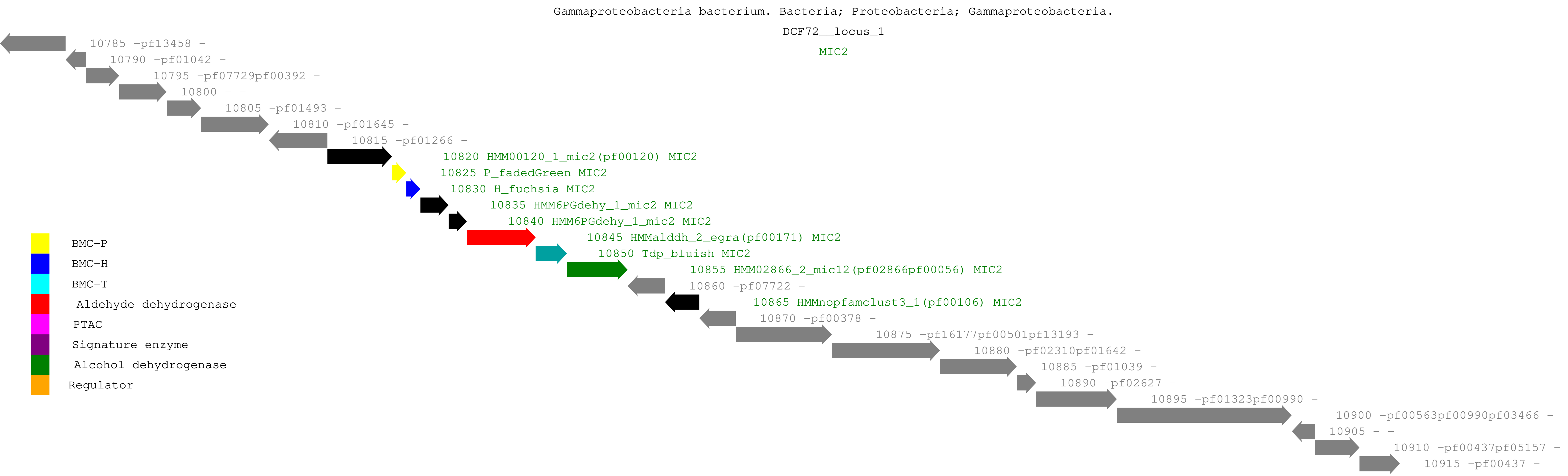
Gammaproteobacteria bacterium. Bacteria; Proteobacteria; Gammaproteobacteria.
DCF72__locus_1
MIC2
10785 -pf13458 -
10790 -pf01042 -
10795 -pf07729pf00392 -
10800 - -
10805 -pf01493 -
10810 -pf01645 -
10815 -pf01266 -
10820 HMM00120_1_mic2(pf00120) MIC2
10825 P_fadedGreen MIC2
10830 H_fuchsia MIC2
10835 HMM6PGdehy_1_mic2 MIC2
10840 HMM6PGdehy_1_mic2 MIC2
10845 HMMalddh_2_egra(pf00171) MIC2
10850 Tdp_bluish MIC2
10855 HMM02866_2_mic12(pf02866pf00056) MIC2
10860 -pf07722 -
10865 HMMnopfamclust3_1(pf00106) MIC2
10870 -pf00378 -
10875 -pf16177pf00501pf13193 -
10880 -pf02310pf01642 -
10885 -pf01039 -
10890 -pf02627 -
10895 -pf01323pf00990 -
10900 -pf00563pf00990pf03466 -
10905 - -
10910 -pf00437pf05157 -
10915 -pf00437 -
BMC-P
BMC-H
BMC-T
Aldehyde dehydrogenase
PTAC
Signature enzyme
Alcohol dehydrogenase
Regulator

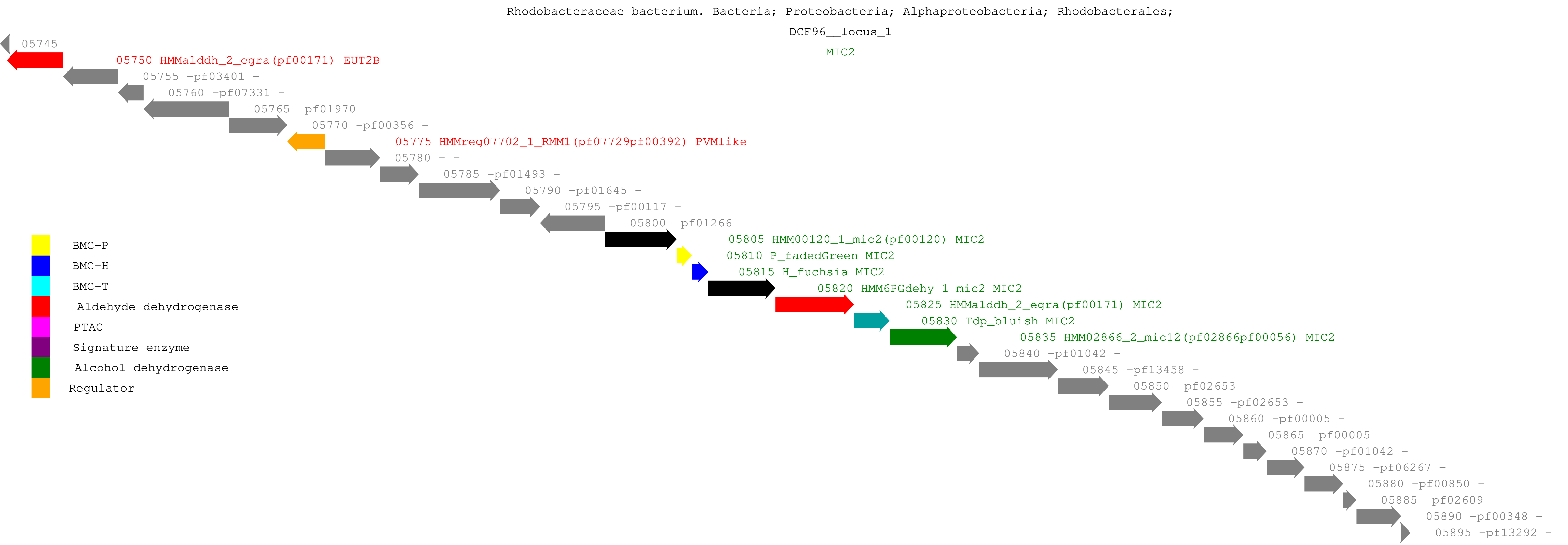

Rhodobacteraceae bacterium. Bacteria; Proteobacteria; Alphaproteobacteria; Rhodobacterales;
DCF96__locus_1
MIC2
05745 - -
05750 HMMalddh_2_egra(pf00171) EUT2B
05755 -pf03401 -
05760 -pf07331 -
05765 -pf01970 -
05770 -pf00356 -
05775 HMMreg07702_1_RMM1(pf07729pf00392) PVMlike
05780 - -
05785 -pf01493 -
05790 -pf01645 -
05795 -pf00117 -
05800 -pf01266 -
05805 HMM00120_1_mic2(pf00120) MIC2
05810 P_fadedGreen MIC2
05815 H_fuchsia MIC2
05820 HMM6PGdehy_1_mic2 MIC2
05825 HMMalddh_2_egra(pf00171) MIC2
05830 Tdp_bluish MIC2
05835 HMM02866_2_mic12(pf02866pf00056) MIC2
05840 -pf01042 -
05845 -pf13458 -
05850 -pf02653 -
05855 -pf02653 -
05860 -pf00005 -
05865 -pf00005 -
05870 -pf01042 -
05875 -pf06267 -
05880 -pf00850 -
05885 -pf02609 -
05890 -pf00348 -
05895 -pf13292 -
BMC-P
BMC-H
BMC-T
Aldehyde dehydrogenase
PTAC
Signature enzyme
Alcohol dehydrogenase
Regulator

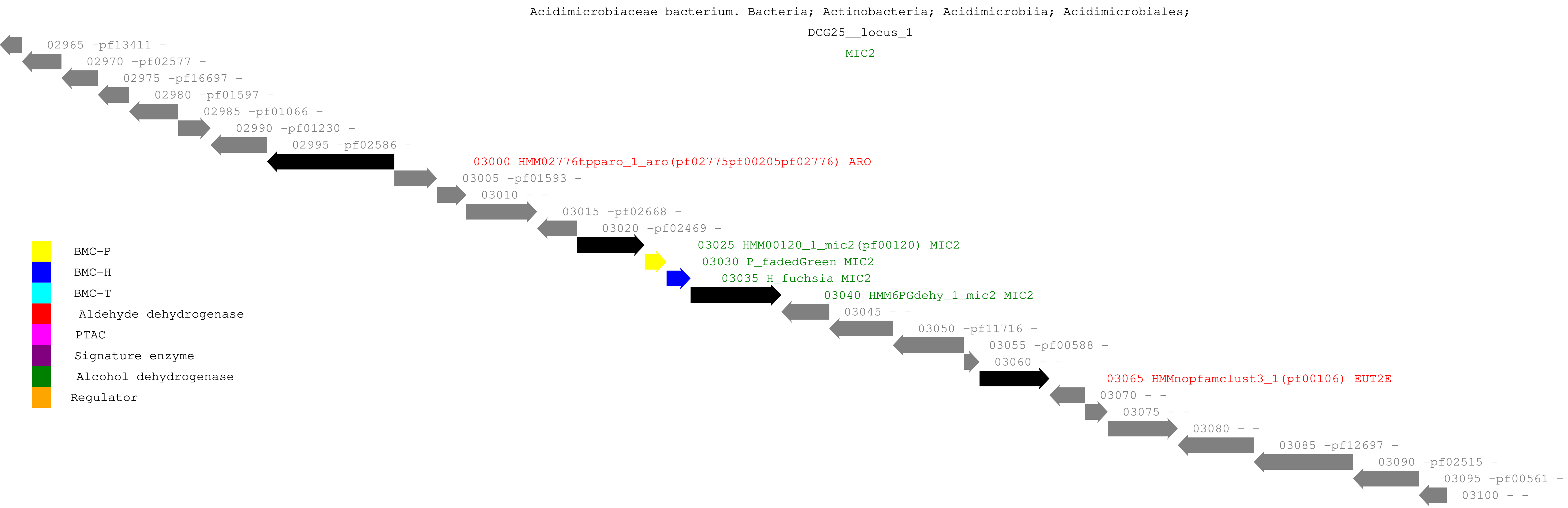

Acidimicrobiaceae bacterium. Bacteria; Actinobacteria; Acidimicrobiia; Acidimicrobiales;
DCG25__locus_1
MIC2
02965 -pf13411 -
02970 -pf02577 -
02975 -pf16697 -
02980 -pf01597 -
02985 -pf01066 -
02990 -pf01230 -
02995 -pf02586 -
03000 HMM02776tpparo_1_aro(pf02775pf00205pf02776) ARO
03005 -pf01593 -
03010 - -
03015 -pf02668 -
03020 -pf02469 -
03025 HMM00120_1_mic2(pf00120) MIC2
03030 P_fadedGreen MIC2
03035 H_fuchsia MIC2
03040 HMM6PGdehy_1_mic2 MIC2
03045 - -
03050 -pf11716 -
03055 -pf00588 -
03060 - -
03065 HMMnopfamclust3_1(pf00106) EUT2E
03070 - -
03075 - -
03080 - -
03085 -pf12697 -
03090 -pf02515 -
03095 -pf00561 -
03100 - -
BMC-P
BMC-H
BMC-T
Aldehyde dehydrogenase
PTAC
Signature enzyme
Alcohol dehydrogenase
Regulator

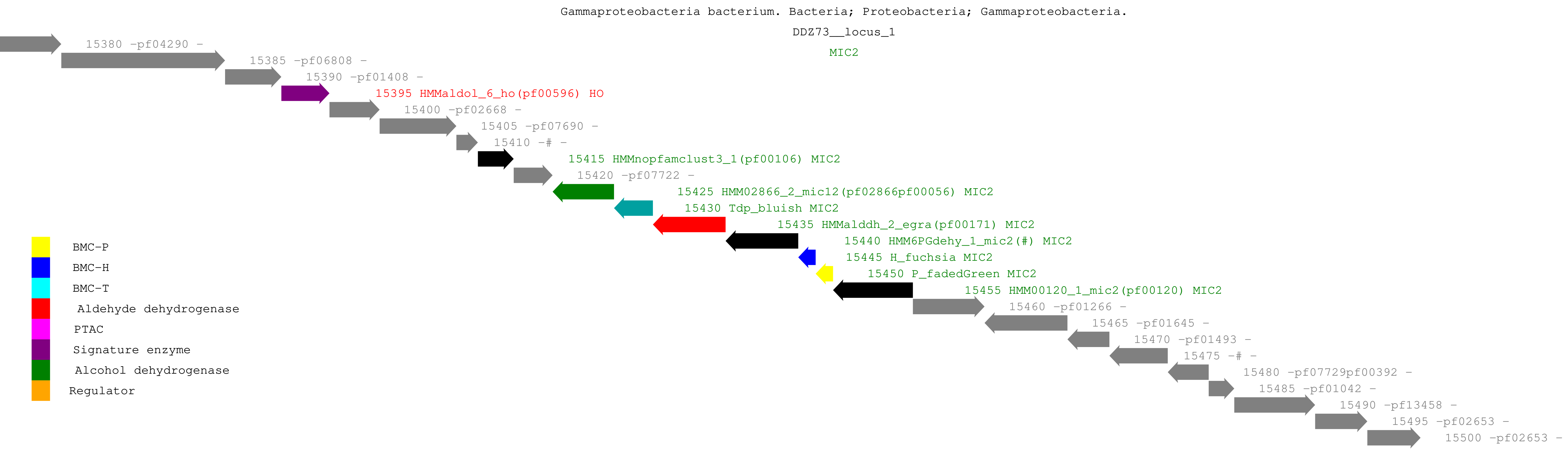
Gammaproteobacteria bacterium. Bacteria; Proteobacteria; Gammaproteobacteria.
DDZ73__locus_1
MIC2
15380 -pf04290 -
15385 -pf06808 -
15390 -pf01408 -
15395 HMMaldol_6_ho(pf00596) HO
15400 -pf02668 -
15405 -pf07690 -
15410 -# -
15415 HMMnopfamclust3_1(pf00106) MIC2
15420 -pf07722 -
15425 HMM02866_2_mic12(pf02866pf00056) MIC2
15430 Tdp_bluish MIC2
15435 HMMalddh_2_egra(pf00171) MIC2
15440 HMM6PGdehy_1_mic2(#) MIC2
15445 H_fuchsia MIC2
15450 P_fadedGreen MIC2
15455 HMM00120_1_mic2(pf00120) MIC2
15460 -pf01266 -
15465 -pf01645 -
15470 -pf01493 -
15475 -# -
15480 -pf07729pf00392 -
15485 -pf01042 -
15490 -pf13458 -
15495 -pf02653 -
15500 -pf02653 -
BMC-P
BMC-H
BMC-T
Aldehyde dehydrogenase
PTAC
Signature enzyme
Alcohol dehydrogenase
Regulator

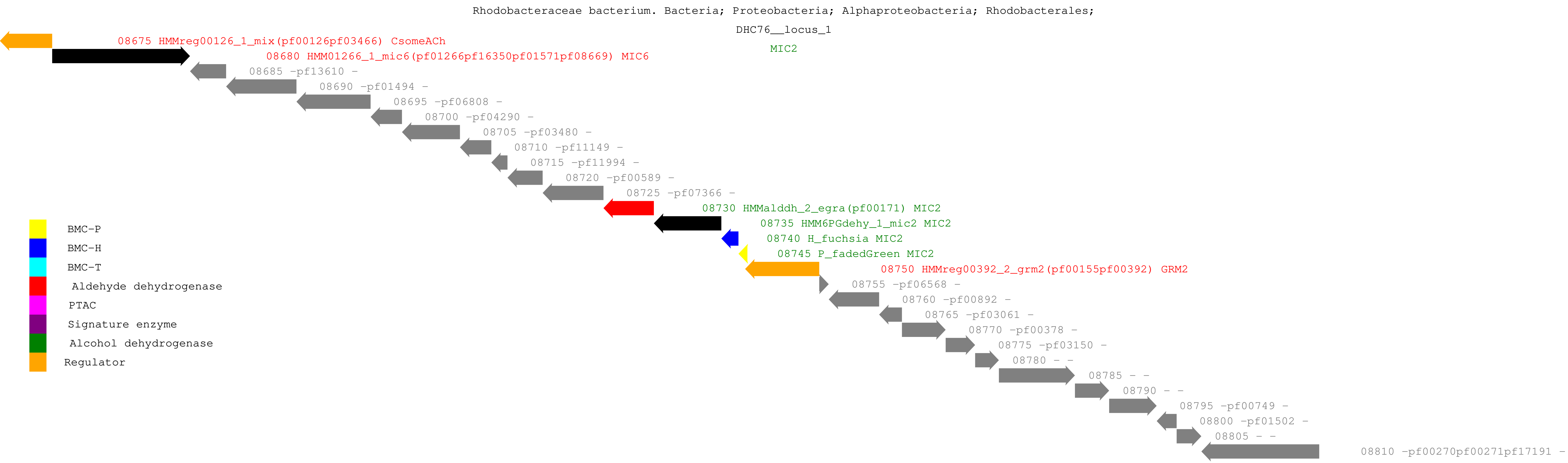
Rhodobacteraceae bacterium. Bacteria; Proteobacteria; Alphaproteobacteria; Rhodobacterales;
DHC76__locus_1
MIC2
08675 HMMreg00126_1_mix(pf00126pf03466) CsomeACh
08680 HMM01266_1_mic6(pf01266pf16350pf01571pf08669) MIC6
08685 -pf13610 -
08690 -pf01494 -
08695 -pf06808 -
08700 -pf04290 -
08705 -pf03480 -
08710 -pf11149 -
08715 -pf11994 -
08720 -pf00589 -
08725 -pf07366 -
08730 HMMaldh_2_egra(pf00171) MIC2
08735 HMM6PGdehy_1_mic2 MIC2
08740 H_fuchsia MIC2
08745 P_fadedGreen MIC2
08750 HMMreg00392_2_grm2(pf00155pf00392) GRM2
08755 -pf06568 -
08760 -pf00892 -
08765 -pf03061 -
08770 -pf00378 -
08775 -pf03150 -
08780 - -
08785 - -
08790 - -
08795 -pf00749 -
08800 -pf01502 -
08805 - -
08810 -pf00270pf00271pf17191 -
BMC-P
BMC-H
BMC-T
Aldehyde dehydrogenase
PTAC
Signature enzyme
Alcohol dehydrogenase
Regulator

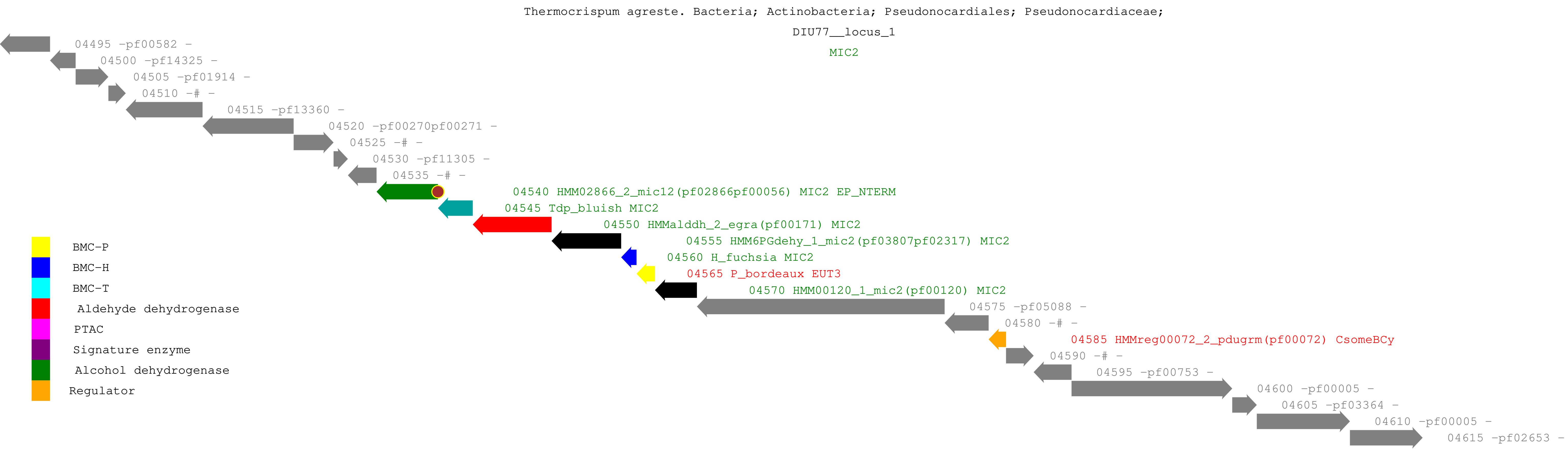
Thermocrispum agreste. Bacteria; Actinobacteria; Pseudonocardiales; Pseudonocardiaceae;
DIU77__locus_1
MIC2
04495 -pf00582 -
04500 -pf14325 -
04505 -pf01914 -
04510 -# -
04515 -pf13360 -
04520 -pf00270pf00271 -
04525 -# -
04530 -pf11305 -
04535 -# -
04540 HMM02866_2_mic12(pf02866pf00056) MIC2 EP_NTERM
04545 Tdp_bluish MIC2
04550 HMMaldh_2_egra(pf00171) MIC2
04555 HMM6PGdehy_1_mic2(pf03807pf02317) MIC2
04560 H_fuchsia MIC2
04565 P_bordeaux EUT3
04570 HMM00120_1_mic2(pf00120) MIC2
04575 -pf05088 -
04580 -# -
04585 HMMreg00072_2_pdugrm(pf00072) CsomeBCy
04590 -# -
04595 -pf00753 -
04600 -pf00005 -
04605 -pf03364 -
04610 -pf00005 -
04615 -pf02653 -
BMC-P
BMC-H
BMC-T
Aldehyde dehydrogenase
PTAC
Signature enzyme
Alcohol dehydrogenase
Regulator

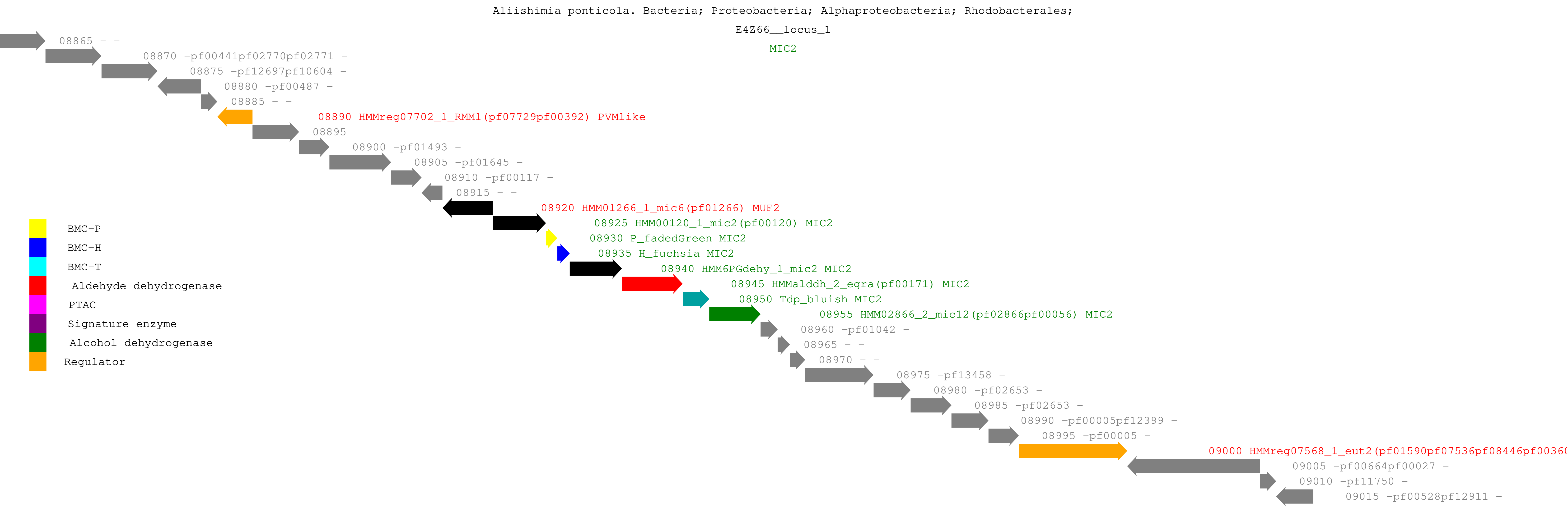
Aliishimia ponticola. Bacteria; Proteobacteria; Alphaproteobacteria; Rhodobacterales;
E4Z66__locus_1
MIC2
08865 - -
08870 -pf00441pf02770pf02771 -
08875 -pf12697pf10604 -
08880 -pf00487 -
08885 - -
08890 HMMreg07702_1_RMM1(pf07729pf00392) PVMlike
08895 - -
08900 -pf01493 -
08905 -pf01645 -
08910 -pf00117 -
08915 - -
08920 HMM01266_1_mic6(pf01266) MUF2
08925 HMM00120_1_mic2(pf00120) MIC2
08930 P_fadedGreen MIC2
08935 H_fuchsia MIC2
08940 HMM6PGdehy_1_mic2 MIC2
08945 HMMalddh_2_egra(pf00171) MIC2
08950 Tdp_bluish MIC2
08955 HMM02866_2_mic12(pf02866pf00056) MIC2
08960 -pf01042 -
08965 - -
08970 - -
08975 -pf13458 -
08980 -pf02653 -
08985 -pf02653 -
08990 -pf00005pf12399 -
08995 -pf00005 -
09000 HMMreg07568_1_eut2(pf01590pf07536pf08446pf0036
09005 -pf00664pf00027 -
09010 -pf11750 -
09015 -pf00528pf12911 -
BMC-P
BMC-H
BMC-T
Aldehyde dehydrogenase
PTAC
Signature enzyme
Alcohol dehydrogenase
Regulator

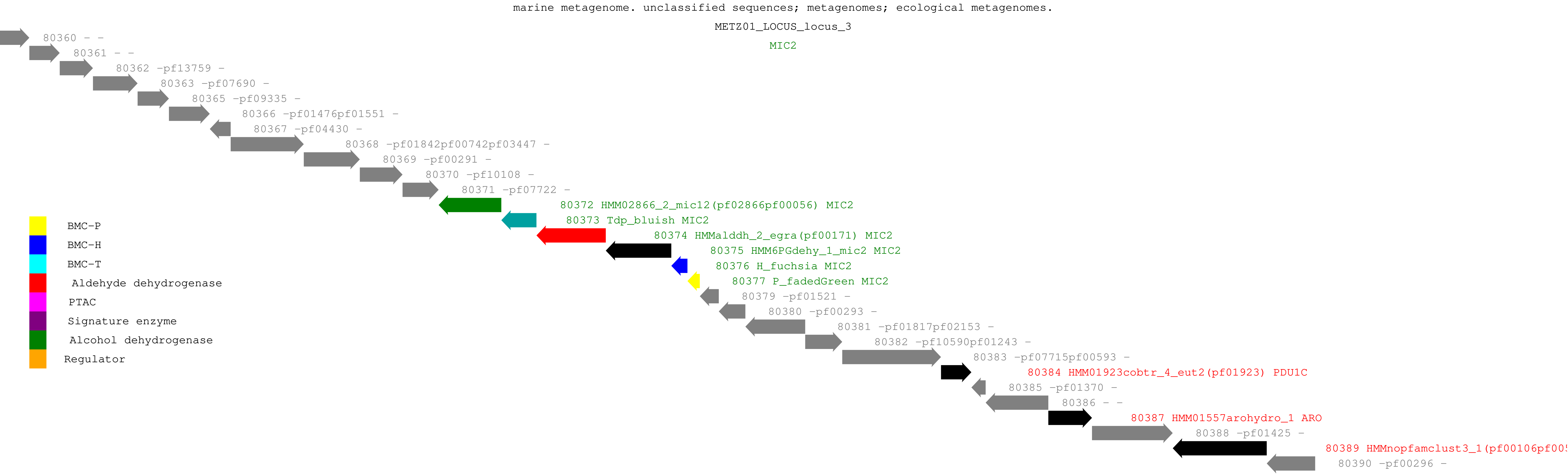
marine metagenome. unclassified sequences; metagenomes; ecological metagenomes.
METZ01_LOCUS_locus_3
MIC2
80360 - -
80361 - -
80362 -pf13759 -
80363 -pf07690 -
80365 -pf09335 -
80366 -pf01476pf01551 -
80367 -pf04430 -
80368 -pf01842pf00742pf03447 -
80369 -pf00291 -
80370 -pf10108 -
80371 -pf07722 -
80372 HMM02866_2_mic12(pf02866pf00056) MIC2
80373 Tdp_bluish MIC2
80374 HMMalddh_2_egra(pf00171) MIC2
80375 HMM6PGdehy_1_mic2 MIC2
80376 H_fuchsia MIC2
80377 P_fadedGreen MIC2
80379 -pf01521 -
80380 -pf00293 -
80381 -pf01817pf02153 -
80382 -pf10590pf01243 -
80383 -pf07715pf00593 -
80384 HMM01923cobtr_4_eut2(pf01923) PDU1C
80385 -pf01370 -
80386 - -
80387 HMM01557arohydro_1 ARO
80388 -pf01425 -
80389 HMMnopfamclust3_1(pf00106pf00
80390 -pf00296 -
BMC-P
BMC-H
BMC-T
Aldehyde dehydrogenase
PTAC
Signature enzyme
Alcohol dehydrogenase
Regulator

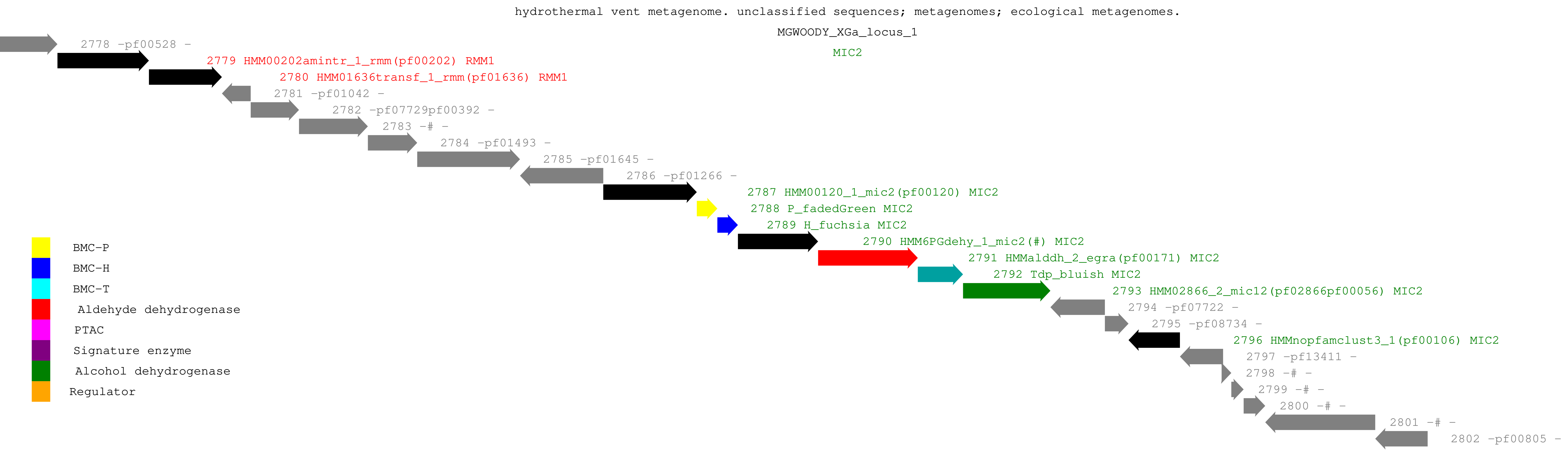
hydrothermal vent metagenome. unclassified sequences; metagenomes; ecological metagenomes.
MGWOODY_XGa_locus_1
MIC2
2778 -pf00528 -
2779 HMM00202amintr_1_rmm(pf00202) RMM1
2780 HMM01636transf_1_rmm(pf01636) RMM1
2781 -pf01042 -
2782 -pf07729pf00392 -
2783 -# -
2784 -pf01493 -
2785 -pf01645 -
2786 -pf01266 -
2787 HMM00120_1_mic2(pf00120) MIC2
2788 P_fadedGreen MIC2
2789 H_fuchsia MIC2
2790 HMM6PGdehy_1_mic2(#) MIC2
2791 HMMalddh_2_egra(pf00171) MIC2
2792 Tdp_bluish MIC2
2793 HMM02866_2_mic12(pf02866pf00056) MIC2
2794 -pf07722 -
2795 -pf08734 -
2796 HMMnopfamclust3_1(pf00106) MIC2
2797 -pf13411 -
2798 -# -
2799 -# -
2800 -# -
2801 -# -
2802 -pf00805 -
BMC-P
BMC-H
BMC-T
Aldehyde dehydrogenase
PTAC
Signature enzyme
Alcohol dehydrogenase
Regulator

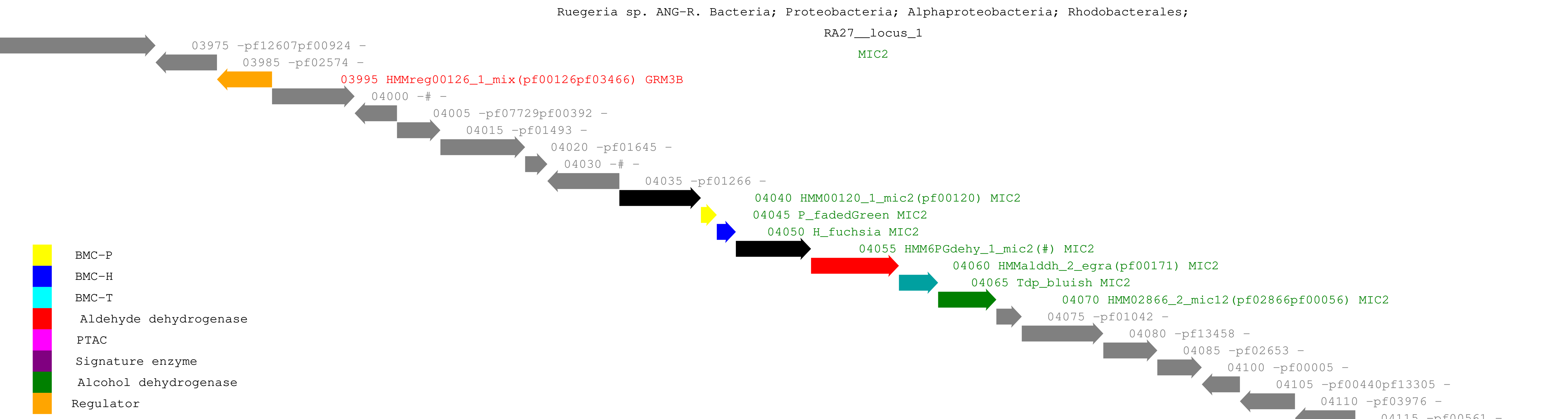

Ruegeria sp. ANG-R. Bacteria; Proteobacteria; Alphaproteobacteria; Rhodobacterales;
RA27__locus_1
MIC2
03975 -pf12607pf00924 -
03985 -pf02574 -
03995 HMMreg00126_1_mix(pf00126pf03466) GRM3B
04000 -# -
04005 -pf07729pf00392 -
04015 -pf01493 -
04020 -pf01645 -
04030 -# -
04035 -pf01266 -
04040 HMM00120_1_mic2(pf00120) MIC2
04045 P_fadedGreen MIC2
04050 H_fuchsia MIC2
04055 HMM6PGdehy_1_mic2(#) MIC2
04060 HMMalddh_2_egra(pf00171) MIC2
04065 Tdp_bluish MIC2
04070 HMM02866_2_mic12(pf02866pf00056) MIC2
04075 -pf01042 -
04080 -pf13458 -
04085 -pf02653 -
04100 -pf00005 -
04105 -pf00440pf13305 -
04110 -pf03976 -
04115 -pf00561 -
04120 -pf04204 -
04125 -# -
BMC-P
BMC-H
BMC-T
Aldehyde dehydrogenase
PTAC
Signature enzyme
Alcohol dehydrogenase
Regulator

Ruegeria mediterranea M15O_3. Bacteria; Proteobacteria; Alphaproteobacteria; Rhodobacterales;

TRM7615__locus_1

MIC2

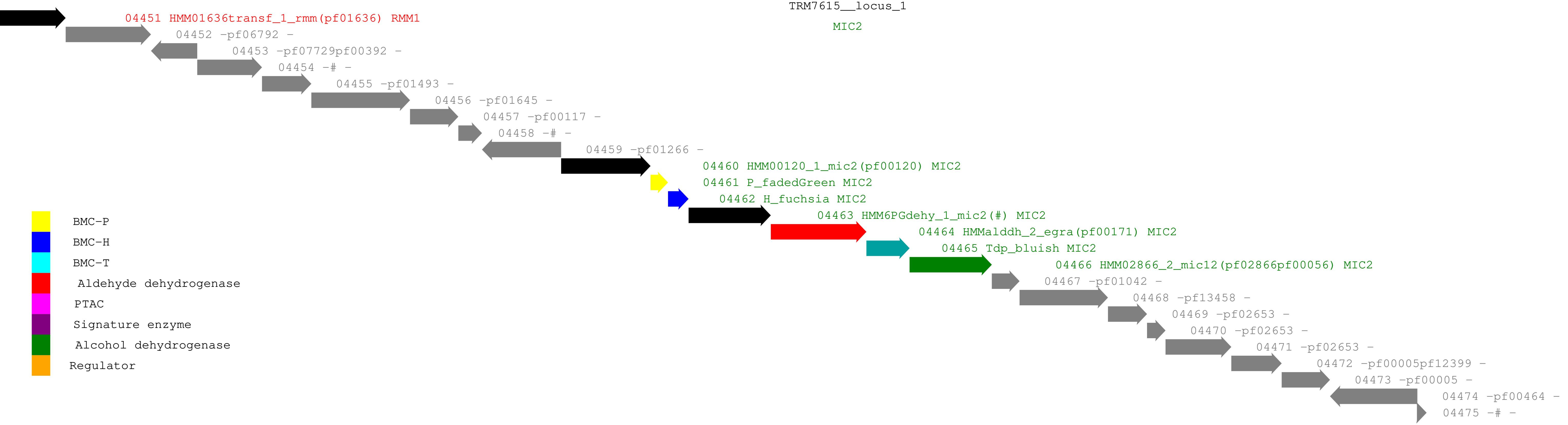

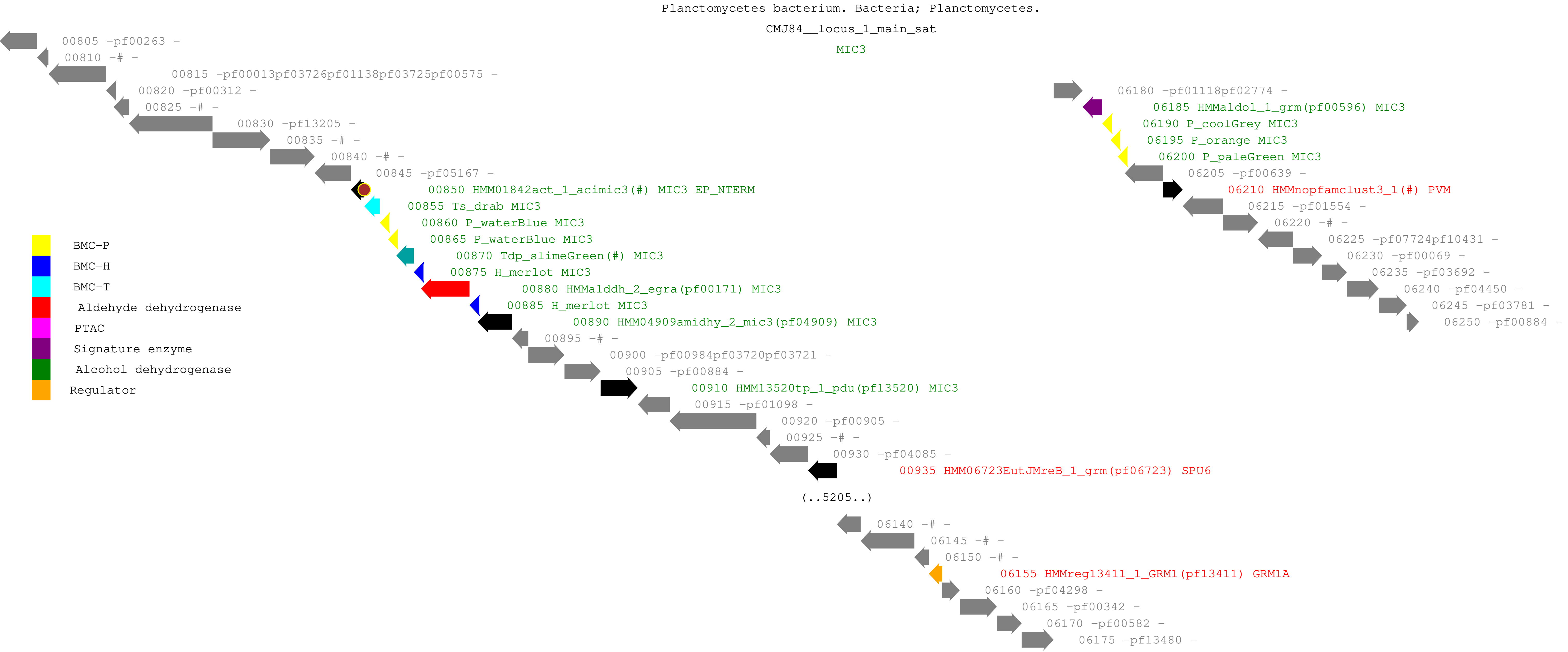

Planctomycetes bacterium. Bacteria; Planctomycetes.
CMJ84__locus_1_main_sat
MIC3
00805 -pf00263 -
00810 -# -
00815 -pf00013pf03726pf01138pf03725pf00575 -
00820 -pf00312 -
00825 -# -
00830 -pf13205 -
00835 -# -
00840 -# -
00845 -pf05167 -
00850 HMM01842act_1_acimic3(#) MIC3 EP_NTERM
00855 Ts_drab MIC3
00860 P_waterBlue MIC3
00865 P_waterBlue MIC3
00870 Tdp_slimeGreen(#) MIC3
00875 H_merlot MIC3
00880 HMMaldDh_2_egra(pf00171) MIC3
00885 H_merlot MIC3
00890 HMM04909amidhy_2_mic3(pf04909) MIC3
00895 -# -
00900 -pf00984pf03720pf03721 -
00905 -pf00884 -
00910 HMM13520tp_1_pdu(pf13520) MIC3
00915 -pf01098 -
00920 -pf00905 -
00925 -# -
00930 -pf04085 -
00935 HMM06723EutJMreB_1_grm(pf06723) SPU6
(..5205..)
06140 -# -
06145 -# -
06150 -# -
06155 HMMreg13411_1_GRM1(pf13411) GRM1A
06160 -pf04298 -
06165 -pf00342 -
06170 -pf00582 -
06175 -pf13480 -
06180 -pf01118pf02774 -
06185 HMMaldol_1_grm(pf00596) MIC3
06190 P_coolGrey MIC3
06195 P_orange MIC3
06200 P_paleGreen MIC3
06205 -pf00639 -
06210 HMMnopfamclust3_1(#) PVM
06215 -pf01554 -
06220 -# -
06225 -pf07724pf10431 -
06230 -pf00069 -
06235 -pf03692 -
06240 -pf04450 -
06245 -pf03781 -
06250 -pf00884 -
BMC-P
BMC-H
BMC-T
Aldehyde dehydrogenase
PTAC
Signature enzyme
Alcohol dehydrogenase
Regulator

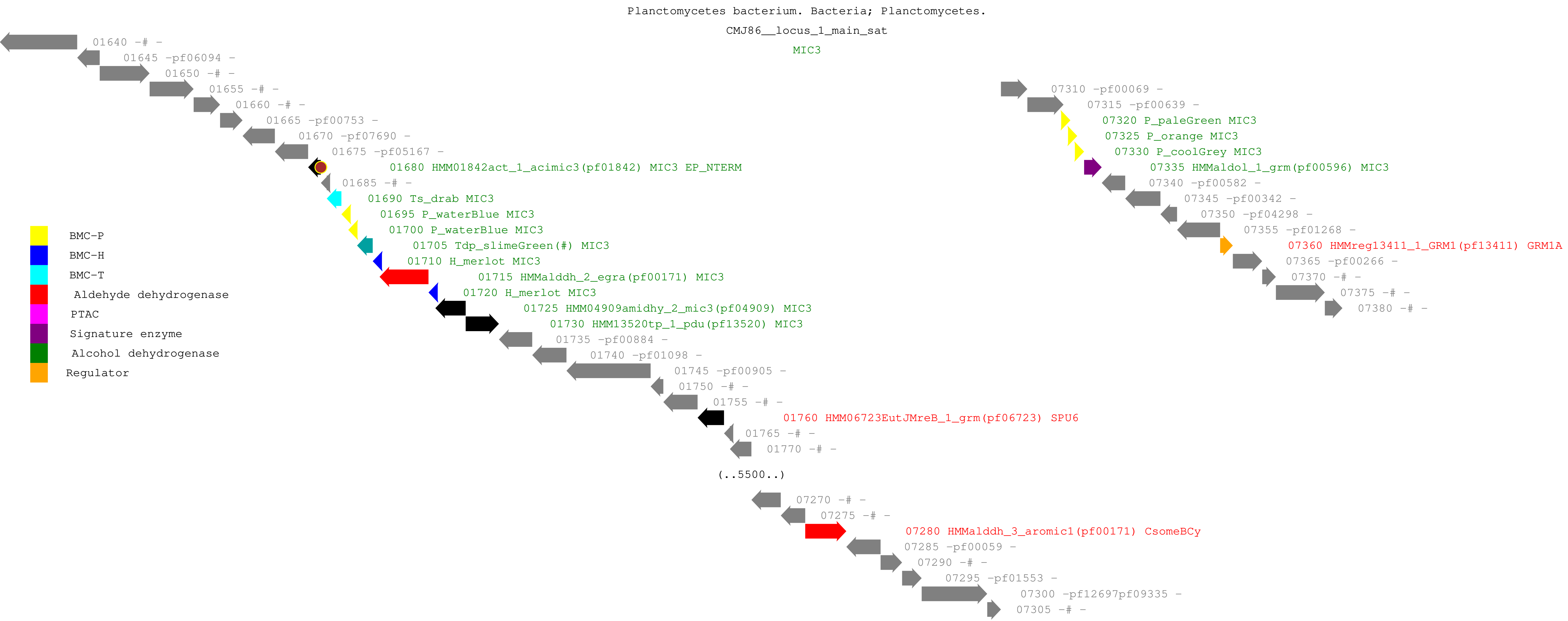
Planctomycetes bacterium. Bacteria; Planctomycetes.
CMJ86__locus_1_main_sat
MIC3
01640 -# -
01645 -pf06094 -
01650 -# -
01655 -# -
01660 -# -
01665 -pf00753 -
01670 -pf07690 -
01675 -pf05167 -
01680 HMM01842act_1_acimic3(pf01842) MIC3 EP_NTERM
01685 -# -
01690 Ts_drab MIC3
01695 P_waterBlue MIC3
01700 P_waterBlue MIC3
01705 Tdp_slimeGreen(#) MIC3
01710 H_merlot MIC3
01715 HMMaldh_2_egra(pf00171) MIC3
01720 H_merlot MIC3
01725 HMM04909amidhy_2_mic3(pf04909) MIC3
01730 HMM13520tp_1_pdu(pf13520) MIC3
01735 -pf00884 -
01740 -pf01098 -
01745 -pf00905 -
01750 -# -
01755 -# -
01760 HMM06723EutJMreB_1_grm(pf06723) SPU6
01765 -# -
01770 -# -
(..5500..)
07270 -# -
07275 -# -
07280 HMMaldh_3_aromic1(pf00171) CsomeBCy
07285 -pf00059 -
07290 -# -
07295 -pf01553 -
07300 -pf12697pf09335 -
07305 -# -
07310 -pf00069 -
07315 -pf00639 -
07320 P_paleGreen MIC3
07325 P_orange MIC3
07330 P_coolGrey MIC3
07335 HMMaldol_1_grm(pf00596) MIC3
07340 -pf00582 -
07345 -pf00342 -
07350 -pf04298 -
07355 -pf01268 -
07360 HMMreg13411_1_GRM1(pf13411) GRM1A
07365 -pf00266 -
07370 -# -
07375 -# -
07380 -# -
BMC-P
BMC-H
BMC-T
Aldehyde dehydrogenase
PTAC
Signature enzyme
Alcohol dehydrogenase
Regulator

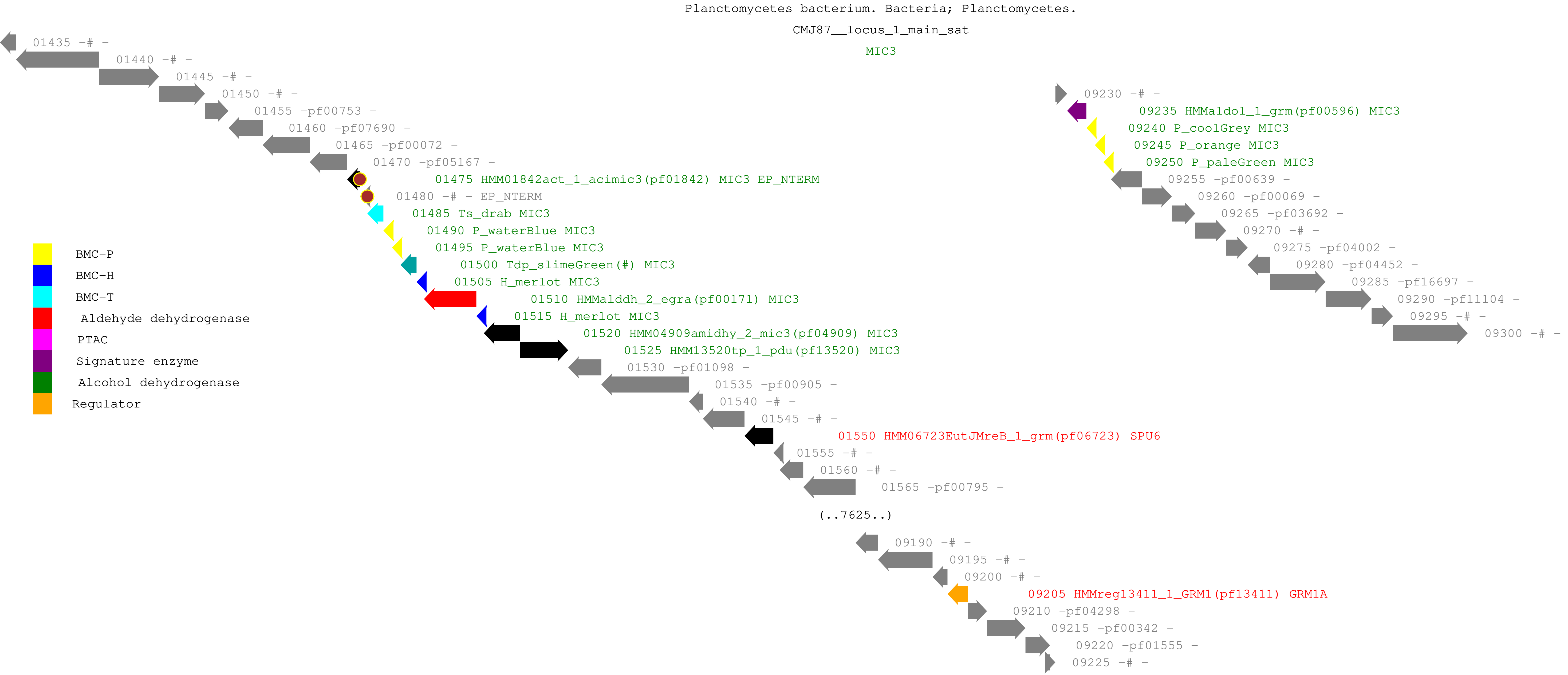
Planctomycetes bacterium. Bacteria; Planctomycetes.
CMJ87__locus_1_main_sat
MIC3
01435 -# -
01440 -# -
01445 -# -
01450 -# -
01455 -pf00753 -
01460 -pf07690 -
01465 -pf00072 -
01470 -pf05167 -
01475 HMM01842act_1_acimic3(pf01842) MIC3 EP_NTERM
01480 -# - EP_NTERM
01485 Ts_drab MIC3
01490 P_waterBlue MIC3
01495 P_waterBlue MIC3
01500 Tdp_slimeGreen(#) MIC3
01505 H_merlot MIC3
01510 HMMalddh_2_egra(pf00171) MIC3
01515 H_merlot MIC3
01520 HMM04909amidhy_2_mic3(pf04909) MIC3
01525 HMM13520tp_1_pdu(pf13520) MIC3
01530 -pf01098 -
01535 -pf00905 -
01540 -# -
01545 -# -
01550 HMM06723EutJMreB_1_grm(pf06723) SPU6
01555 -# -
01560 -# -
01565 -pf00795 -
(..7625..)
09190 -# -
09195 -# -
09200 -# -
09205 HMMreg13411_1_GRM1(pf13411) GRM1A
09210 -pf04298 -
09215 -pf00342 -
09220 -pf01555 -
09225 -# -
09230 -# -
09235 HMMaldol_1_grm(pf00596) MIC3
09240 P_coolGrey MIC3
09245 P_orange MIC3
09250 P_paleGreen MIC3
09255 -pf00639 -
09260 -pf00069 -
09265 -pf03692 -
09270 -# -
09275 -pf04002 -
09280 -pf04452 -
09285 -pf16697 -
09290 -pf11104 -
09295 -# -
09300 -# -
BMC-P
BMC-H
BMC-T
Aldehyde dehydrogenase
PTAC
Signature enzyme
Alcohol dehydrogenase
Regulator

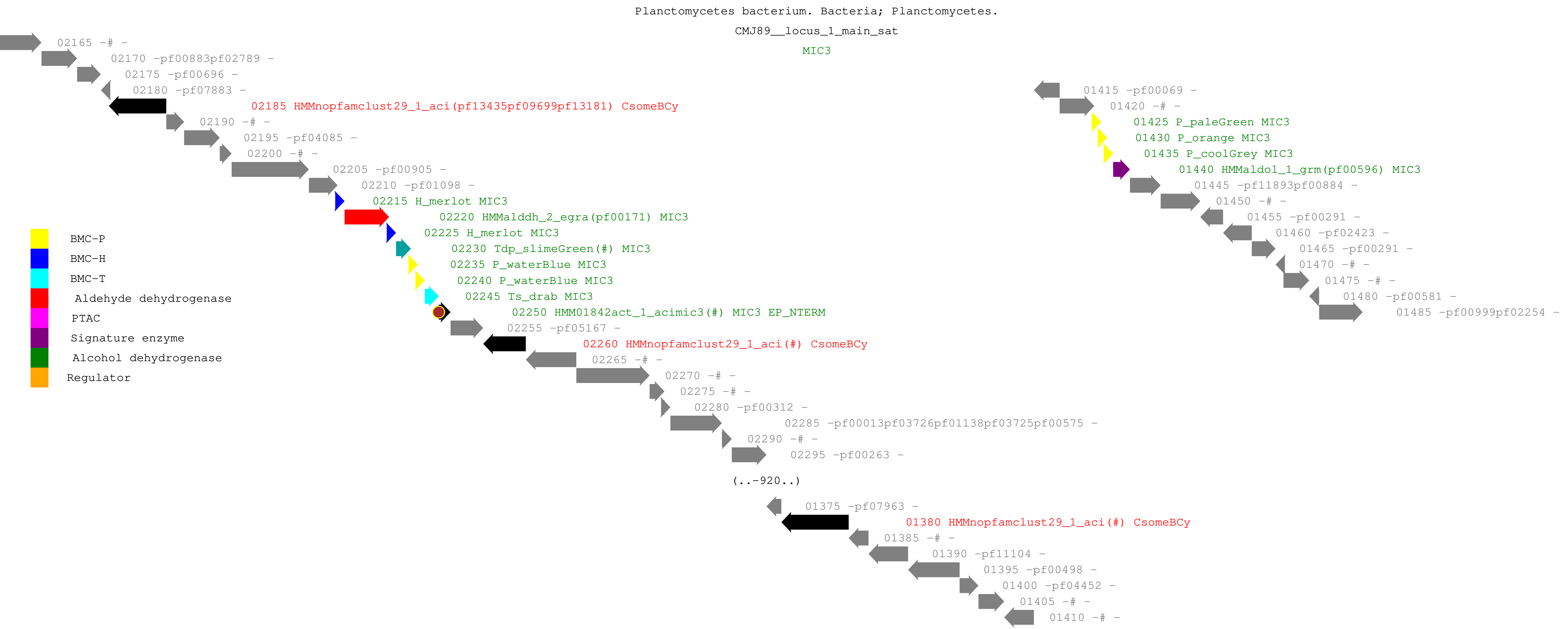
Planctomycetes bacterium. Bacteria; Planctomycetes.
CMJ89__locus_1_main_sat
MIC3
02165 -# -
02170 -pf00883pf02789 -
02175 -pf00696 -
02180 -pf07883 -
02185 HMMnopfamclust29_1_aci(pf13435pf09699pf13181) CsomeBCy
02190 -# -
02195 -pf04085 -
02200 -# -
02205 -pf00905 -
02210 -pf01098 -
02215 H_merlot MIC3
02220 HMMalddh_2_egra(pf00171) MIC3
02225 H_merlot MIC3
02230 Tdp_slimeGreen(#) MIC3
02235 P_waterBlue MIC3
02240 P_waterBlue MIC3
02245 Ts_drab MIC3
02250 HMM01842act_1_acimic3(#) MIC3 EP_NTERM
02255 -pf05167 -
02260 HMMnopfamclust29_1_aci(#) CsomeBCy
02265 -# -
02270 -# -
02275 -# -
02280 -pf00312 -
02285 -pf00013pf03726pf01138pf03725pf00575 -
02290 -# -
02295 -pf00263 -
(..-920..)
01375 -pf07963 -
01380 HMMnopfamclust29_1_aci(#) CsomeBCy
01385 -# -
01390 -pf11104 -
01395 -pf00498 -
01400 -pf04452 -
01405 -# -
01410 -# -
01415 -pf00069 -
01420 -# -
01425 P_paleGreen MIC3
01430 P_orange MIC3
01435 P_coolGrey MIC3
01440 HMMaldol_1_grm(pf00596) MIC3
01445 -pf11893pf00884 -
01450 -# -
01455 -pf00291 -
01460 -pf02423 -
01465 -pf00291 -
01470 -# -
01475 -# -
01480 -pf00581 -
01485 -pf00999pf02254 -
BMC-P
BMC-H
BMC-T
Aldehyde dehydrogenase
PTAC
Signature enzyme
Alcohol dehydrogenase
Regulator

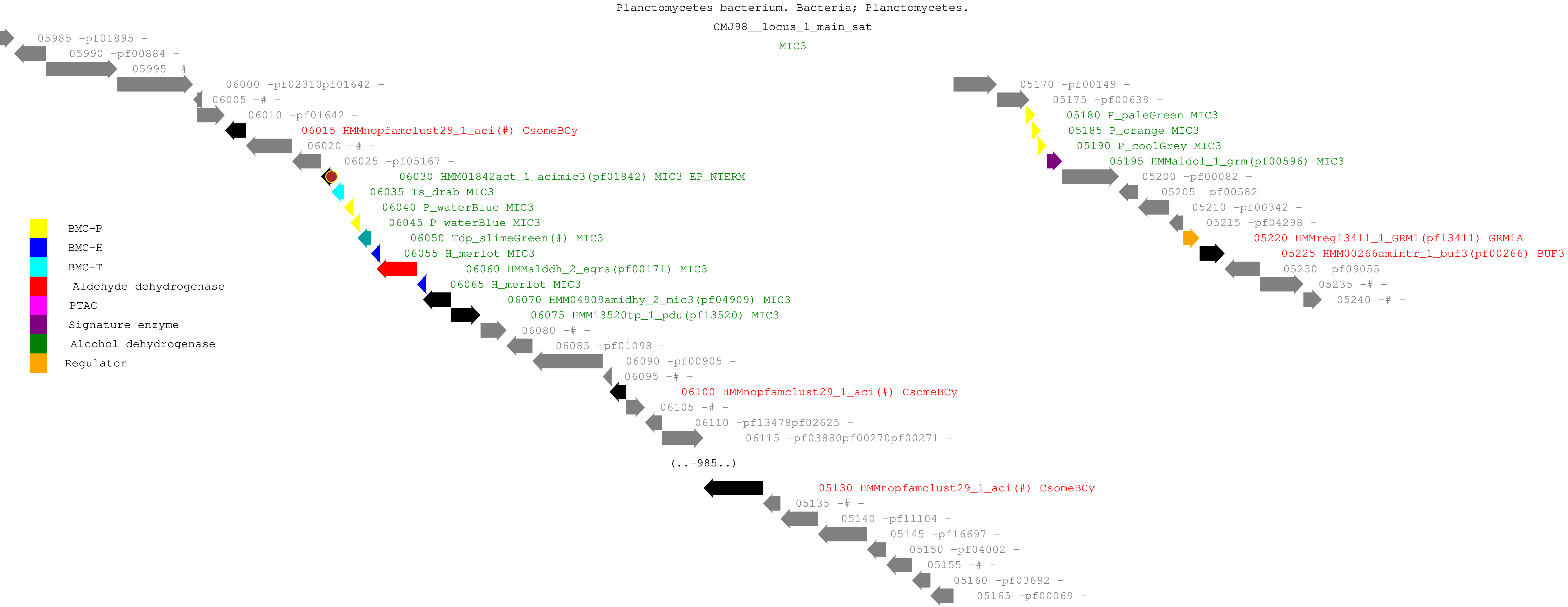
Planctomycetes bacterium. Bacteria; Planctomycetes.
CMJ98__locus_1_main_sat
MIC3
05985 -pf01895 -
05990 -pf00884 -
05995 -# -
06000 -pf02310pf01642 -
06005 -# -
06010 -pf01642 -
06015 HMMnopfamclust29_1_aci(#) CsomeBCy
06020 -# -
06025 -pf05167 -
06030 HMM01842act_1_acimic3(pf01842) MIC3 EP_NTERM
06035 Ts_drab MIC3
06040 P_waterBlue MIC3
06045 P_waterBlue MIC3
06050 Tdp_slimeGreen(#) MIC3
06055 H_merlot MIC3
06060 HMMalddh_2_egra(pf00171) MIC3
06065 H_merlot MIC3
06070 HMM04909amidhy_2_mic3(pf04909) MIC3
06075 HMM13520tp_1_pdu(pf13520) MIC3
06080 -# -
06085 -pf01098 -
06090 -pf00905 -
06095 -# -
06100 HMMnopfamclust29_1_aci(#) CsomeBCy
06105 -# -
06110 -pf13478pf02625 -
06115 -pf03880pf00270pf00271 -
(..-985..)
05130 HMMnopfamclust29_1_aci(#) CsomeBCy
05135 -# -
05140 -pf11104 -
05145 -pf16697 -
05150 -pf04002 -
05155 -# -
05160 -pf03692 -
05165 -pf00069 -
05170 -pf00149 -
05175 -pf00639 -
05180 P_paleGreen MIC3
05185 P_orange MIC3
05190 P_coolGrey MIC3
05195 HMMaldol_1_grm(pf00596) MIC3
05200 -pf00082 -
05205 -pf00582 -
05210 -pf00342 -
05215 -pf04298 -
05220 HMMreg13411_1_GRM1(pf13411) GRM1A
05225 HMM00266amintr_1_buf3(pf00266) BUF3
05230 -pf09055 -
05235 -# -
05240 -# -
BMC-P
BMC-H
BMC-T
Aldehyde dehydrogenase
PTAC
Signature enzyme
Alcohol dehydrogenase
Regulator

Planctomycetes bacterium. Bacteria; Planctomycetes.

CMK00__locus_1_main_sat

MIC3

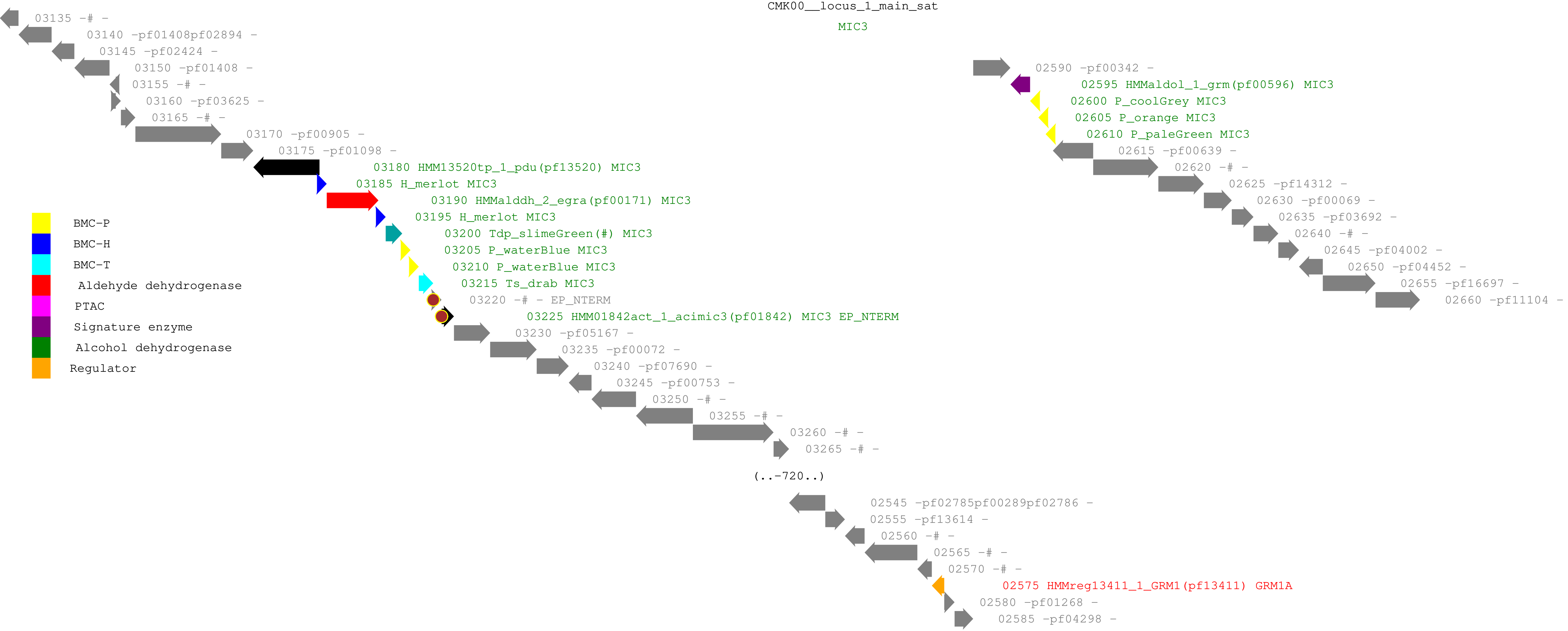

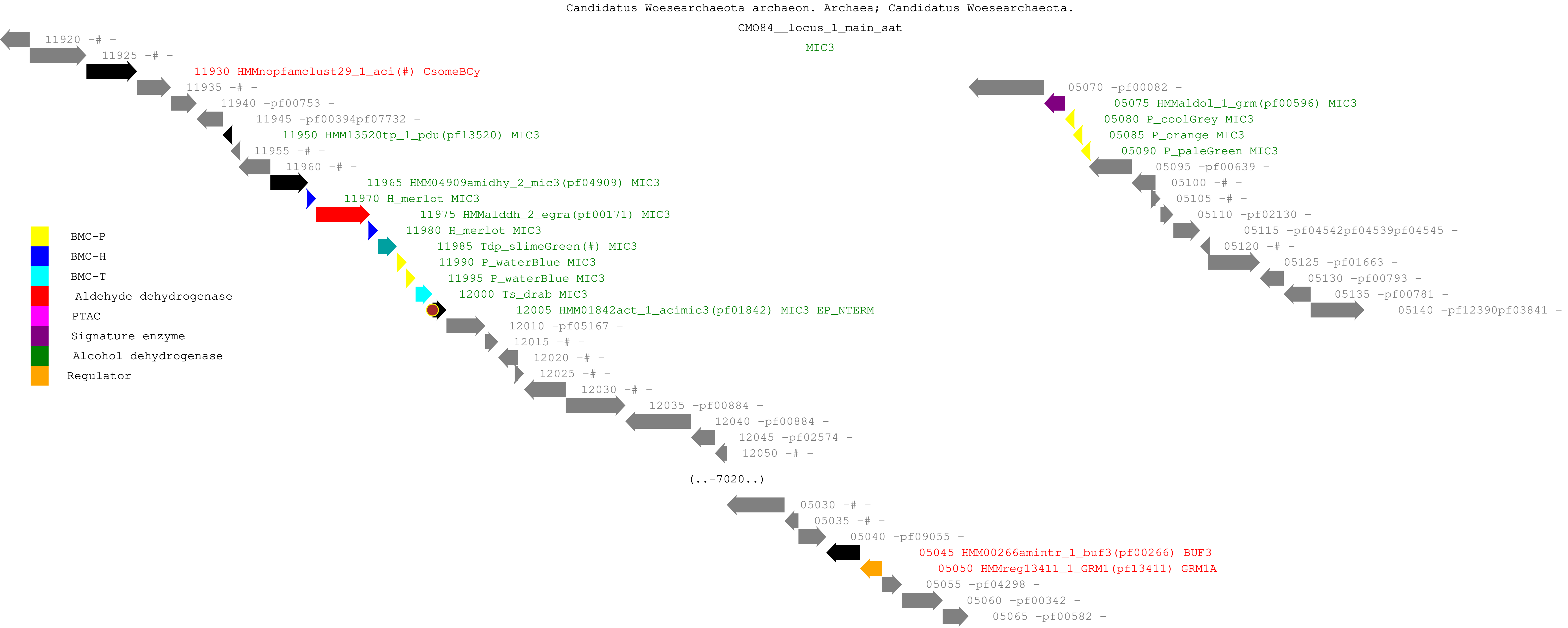

Candidatus Woesearchaeota archaeon. Archaea; Candidatus Woesearchaeota.
CMO84__locus_1_main_sat
MIC3
11920 -# -
11925 -# -
11930 HMMnopfamclust29_1_aci(#) CsomeBCy
11935 -# -
11940 -pf00753 -
11945 -pf00394pf07732 -
11950 HMM13520tp_1_pdu(pf13520) MIC3
11955 -# -
11960 -# -
11965 HMM04909amidhy_2_mic3(pf04909) MIC3
11970 H_merlot MIC3
11975 HMMalddh_2_egra(pf00171) MIC3
11980 H_merlot MIC3
11985 Tdp_slimeGreen(#) MIC3
11990 P_waterBlue MIC3
11995 P_waterBlue MIC3
12000 Ts_drab MIC3
12005 HMM01842act_1_acimic3(pf01842) MIC3 EP_NTERM
12010 -pf05167 -
12015 -# -
12020 -# -
12025 -# -
12030 -# -
12035 -pf00884 -
12040 -pf00884 -
12045 -pf02574 -
12050 -# -
(..-7020..)
05030 -# -
05035 -# -
05040 -pf09055 -
05045 HMM00266amintr_1_buf3(pf00266) BUF3
05050 HMMreg13411_1_GRM1(pf13411) GRM1A
05055 -pf04298 -
05060 -pf00342 -
05065 -pf00582 -
05070 -pf00082 -
05075 HMMaldol_1_grm(pf00596) MIC3
05080 P_coolGrey MIC3
05085 P_orange MIC3
05090 P_paleGreen MIC3
05095 -pf00639 -
05100 -# -
05105 -# -
05110 -pf02130 -
05115 -pf04542pf04539pf04545 -
05120 -# -
05125 -pf01663 -
05130 -pf00793 -
05135 -pf00781 -
05140 -pf12390pf03841 -
BMC-P
BMC-H
BMC-T
Aldehyde dehydrogenase
PTAC
Signature enzyme
Alcohol dehydrogenase
Regulator

Planctomycetes bacterium. Bacteria; Planctomycetes.

E2039__locus_1_main_sat

MIC3

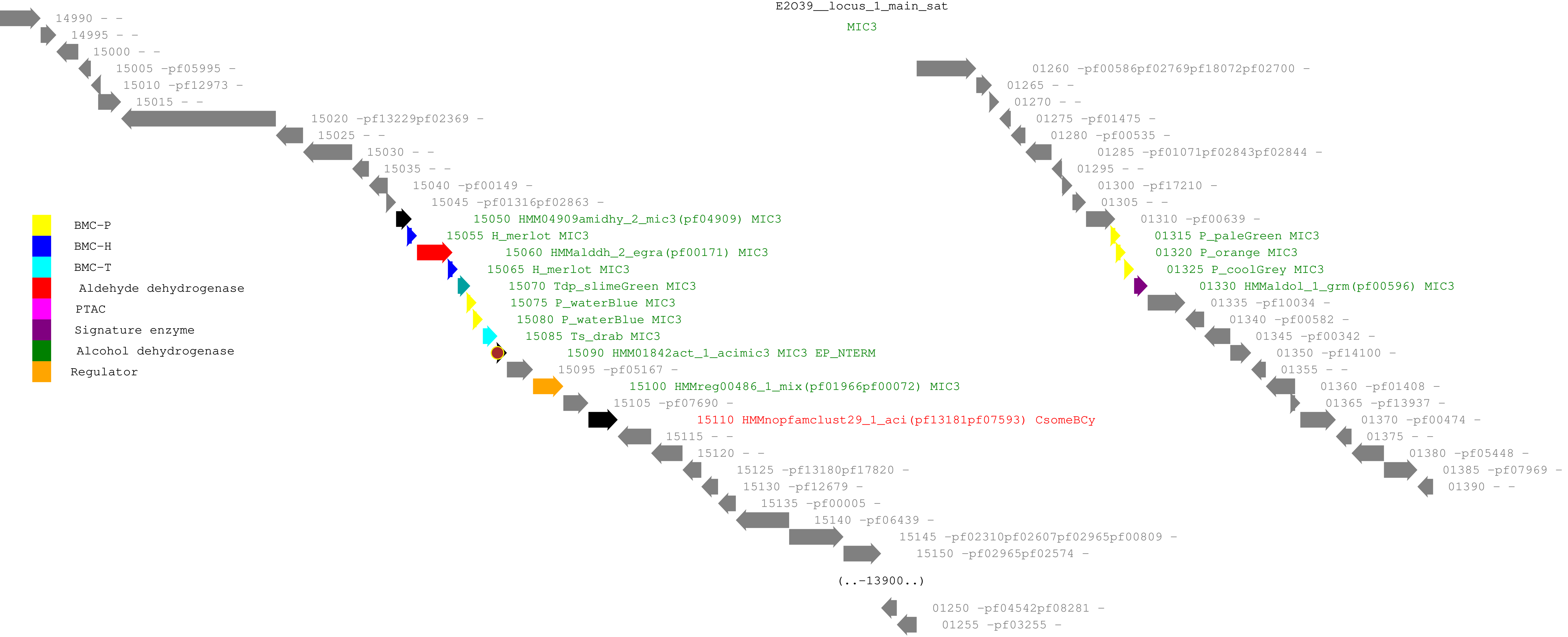

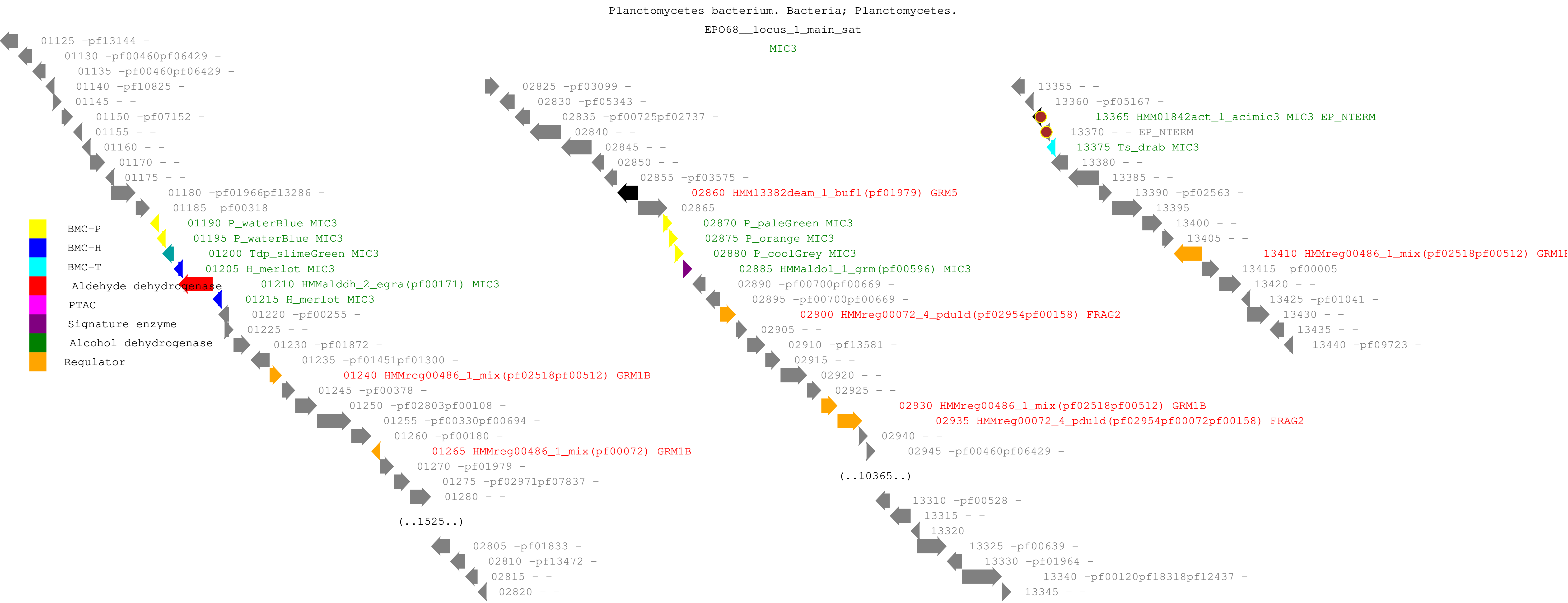

Planctomycetes bacterium. Bacteria; Planctomycetes.
EPO68__locus_1_main_sat
MIC3
BMC-P
BMC-H
BMC-T
Aldehyde dehydrogenase
PTAC
Signature enzyme
Alcohol dehydrogenase
Regulator
01125 -pf13144 -
01130 -pf00460pf06429 -
01135 -pf00460pf06429 -
01140 -pf10825 -
01145 - -
01150 -pf07152 -
01155 - -
01160 - -
01170 - -
01175 - -
01180 -pf01966pf13286 -
01185 -pf00318 -
01190 P_waterBlue MIC3
01195 P_waterBlue MIC3
01200 Tdp_slimeGreen MIC3
01205 H_merlot MIC3
01210 HMMalddh_2_egra(pf00171) MIC3
01215 H_merlot MIC3
01220 -pf00255 -
01225 - -
01230 -pf01872 -
01235 -pf01451pf01300 -
01240 HMMreg00486_1_mix(pf02518pf00512) GRM1B
01245 -pf00378 -
01250 -pf02803pf00108 -
01255 -pf00330pf00694 -
01260 -pf00180 -
01265 HMMreg00486_1_mix(pf00072) GRM1B
01270 -pf01979 -
01275 -pf02971pf07837 -
01280 - -
(..1525..)
02805 -pf01833 -
02810 -pf13472 -
02815 - -
02820 - -
02825 -pf03099 -
02830 -pf05343 -
02835 -pf00725pf02737 -
02840 - -
02845 - -
02850 - -
02855 -pf03575 -
02860 HMM13382deam_1_buf1(pf01979) GRM5
02865 - -
02870 P_paleGreen MIC3
02875 P_orange MIC3
02880 P_coolGrey MIC3
02885 HMMaldol_1_grm(pf00596) MIC3
02890 -pf00700pf00669 -
02895 -pf00700pf00669 -
02900 HMMreg00072_4_pdu1d(pf02954pf00158) FRAG2
02905 - -
02910 -pf13581 -
02915 - -
02920 - -
02925 - -
02930 HMMreg00486_1_mix(pf02518pf00512) GRM1B
02935 HMMreg00072_4_pdu1d(pf02954pf00072pf00158) FRAG2
02940 - -
02945 -pf00460pf06429 -
(..10365..)
13310 -pf00528 -
13315 - -
13320 - -
13325 -pf00639 -
13330 -pf01964 -
13340 -pf00120pf18318pf12437 -
13345 - -
13355 - -
13360 -pf05167 -
13365 HMM01842act_1_acimic3 MIC3 EP_NTERM
13370 - - EP_NTERM
13375 Ts_drab MIC3
13380 - -
13385 - -
13390 -pf02563 -
13395 - -
13400 - -
13405 - -
13410 HMMreg00486_1_mix(pf02518pf00512) GRM1
13415 -pf00005 -
13420 - -
13425 -pf01041 -
13430 - -
13435 - -
13440 -pf09723 -

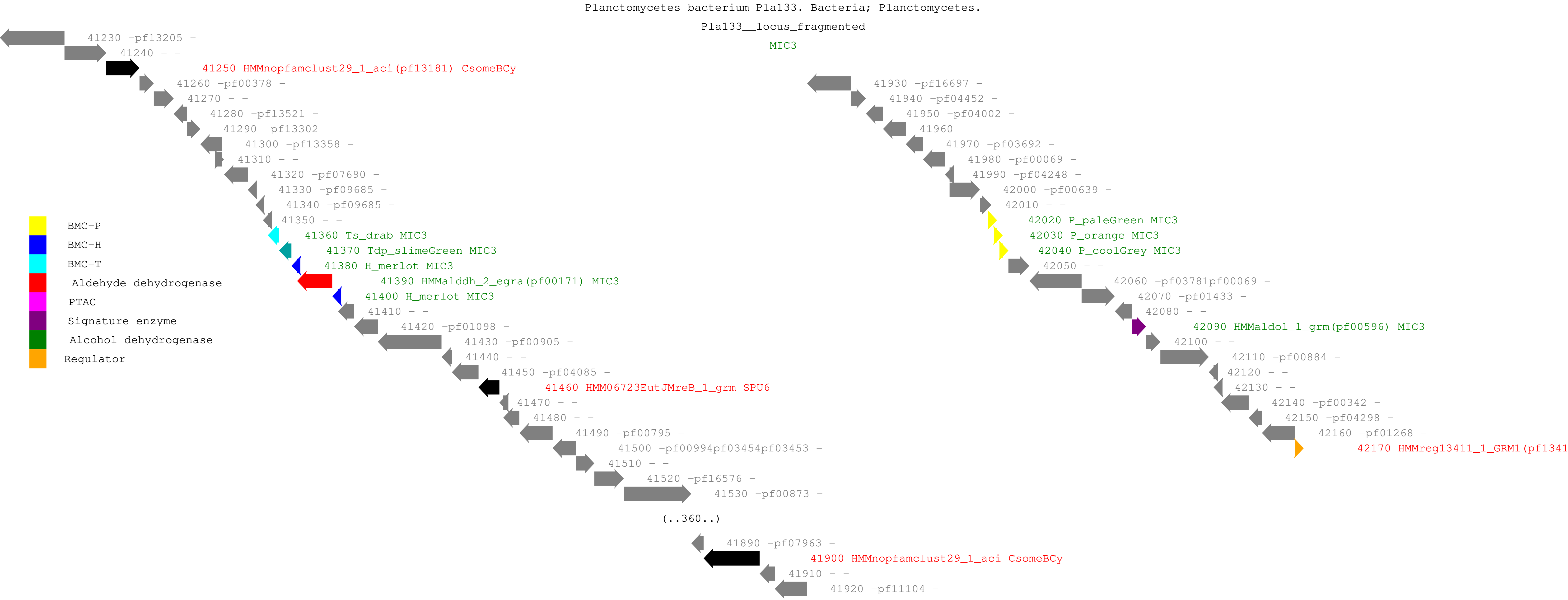
Planctomycetes bacterium Pla133. Bacteria; Planctomycetes.
Pla133__locus_fragmented
MIC3
41230 -pf13205 -
41240 - -
41250 HMMnopfamclust29_1_aci(pf13181) CsomeBCy
41260 -pf00378 -
41270 - -
41280 -pf13521 -
41290 -pf13302 -
41300 -pf13358 -
41310 - -
41320 -pf07690 -
41330 -pf09685 -
41340 -pf09685 -
41350 - -
41360 Ts_drab MIC3
41370 Tdp_slimeGreen MIC3
41380 H_merlot MIC3
41390 HMMalddh_2_egra(pf00171) MIC3
41400 H_merlot MIC3
41410 - -
41420 -pf01098 -
41430 -pf00905 -
41440 - -
41450 -pf04085 -
41460 HMM06723EutJMreB_1_grm SPU6
41470 - -
41480 - -
41490 -pf00795 -
41500 -pf00994pf03454pf03453 -
41510 - -
41520 -pf16576 -
41530 -pf00873 -
(..360..)
41890 -pf07963 -
41900 HMMnopfamclust29_1_aci CsomeBCy
41910 - -
41920 -pf11104 -
41930 -pf16697 -
41940 -pf04452 -
41950 -pf04002 -
41960 - -
41970 -pf03692 -
41980 -pf00069 -
41990 -pf04248 -
42000 -pf00639 -
42010 - -
42020 P_paleGreen MIC3
42030 P_orange MIC3
42040 P_coolGrey MIC3
42050 - -
42060 -pf03781pf00069 -
42070 -pf01433 -
42080 - -
42090 HMMaldol_1_grm(pf00596) MIC3
42100 - -
42110 -pf00884 -
42120 - -
42130 - -
42140 -pf00342 -
42150 -pf04298 -
42160 -pf01268 -
42170 HMMreg13411_1_GRM1(pf1341
BMC-P
BMC-H
BMC-T
Aldehyde dehydrogenase
PTAC
Signature enzyme
Alcohol dehydrogenase
Regulator

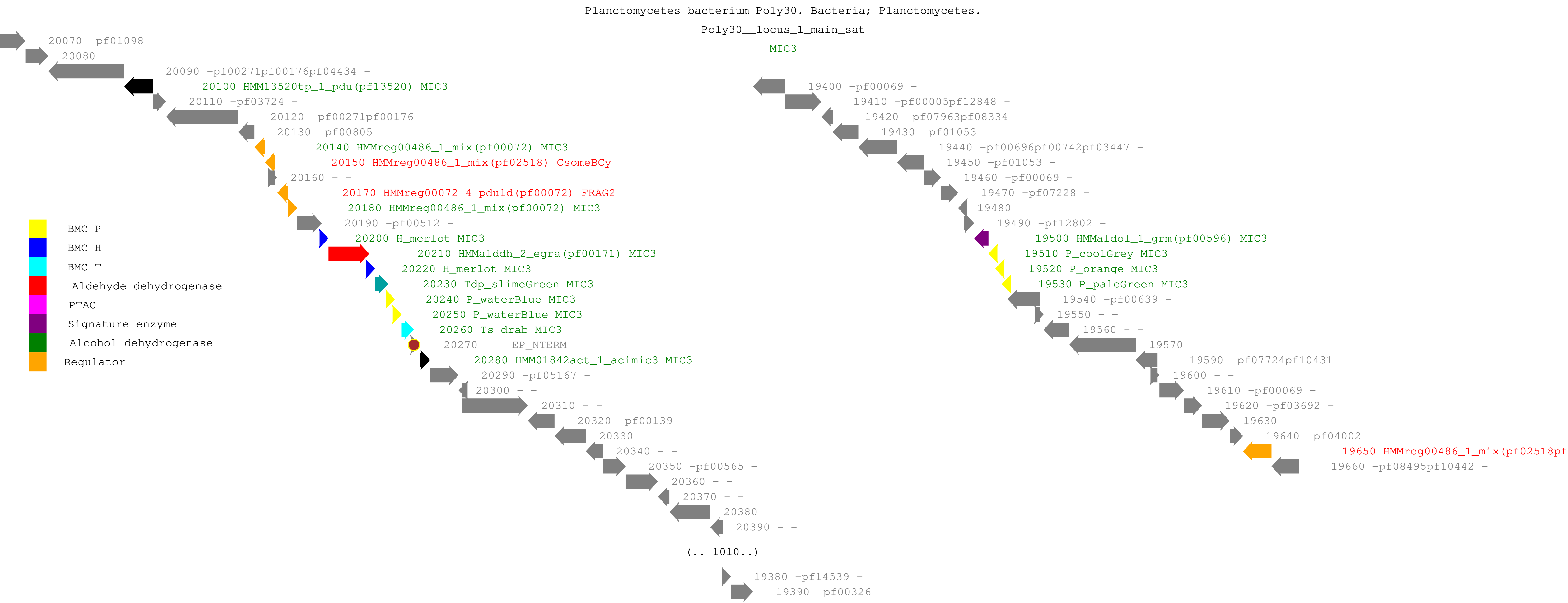
Planctomycetes bacterium Poly30. Bacteria; Planctomycetes.
Poly30__locus_1_main_sat
MIC3
20070 -pf01098 -
20080 - -
20090 -pf00271pf00176pf04434 -
20100 HMM13520tp_1_pdu(pf13520) MIC3
20110 -pf03724 -
20120 -pf00271pf00176 -
20130 -pf00805 -
20140 HMMreg00486_1_mix(pf00072) MIC3
20150 HMMreg00486_1_mix(pf02518) CsomeBCy
20160 - -
20170 HMMreg00072_4_pdu1d(pf00072) FRAG2
20180 HMMreg00486_1_mix(pf00072) MIC3
20190 -pf00512 -
20200 H_merlot MIC3
20210 HMMaldh_2_egra(pf00171) MIC3
20220 H_merlot MIC3
20230 Tdp_slimeGreen MIC3
20240 P_waterBlue MIC3
20250 P_waterBlue MIC3
20260 Ts_drab MIC3
20270 - - EP_NTERM
20280 HMM01842act_1_acimic3 MIC3
20290 -pf05167 -
20300 - -
20310 - -
20320 -pf00139 -
20330 - -
20340 - -
20350 -pf00565 -
20360 - -
20370 - -
20380 - -
20390 - -
(..-1010..)
19380 -pf14539 -
19390 -pf00326 -
19400 -pf00069 -
19410 -pf00005pf12848 -
19420 -pf07963pf08334 -
19430 -pf01053 -
19440 -pf00696pf00742pf03447 -
19450 -pf01053 -
19460 -pf00069 -
19470 -pf07228 -
19480 - -
19490 -pf12802 -
19500 HMMaldol_1_grm(pf00596) MIC3
19510 P_coolGrey MIC3
19520 P_orange MIC3
19530 P_paleGreen MIC3
19540 -pf00639 -
19550 - -
19560 - -
19570 - -
19590 -pf07724pf10431 -
19600 - -
19610 -pf00069 -
19620 -pf03692 -
19630 - -
19640 -pf04002 -
19650 HMMreg00486_1_mix(pf02518pf
19660 -pf08495pf10442 -
BMC-P
BMC-H
BMC-T
Aldehyde dehydrogenase
PTAC
Signature enzyme
Alcohol dehydrogenase
Regulator

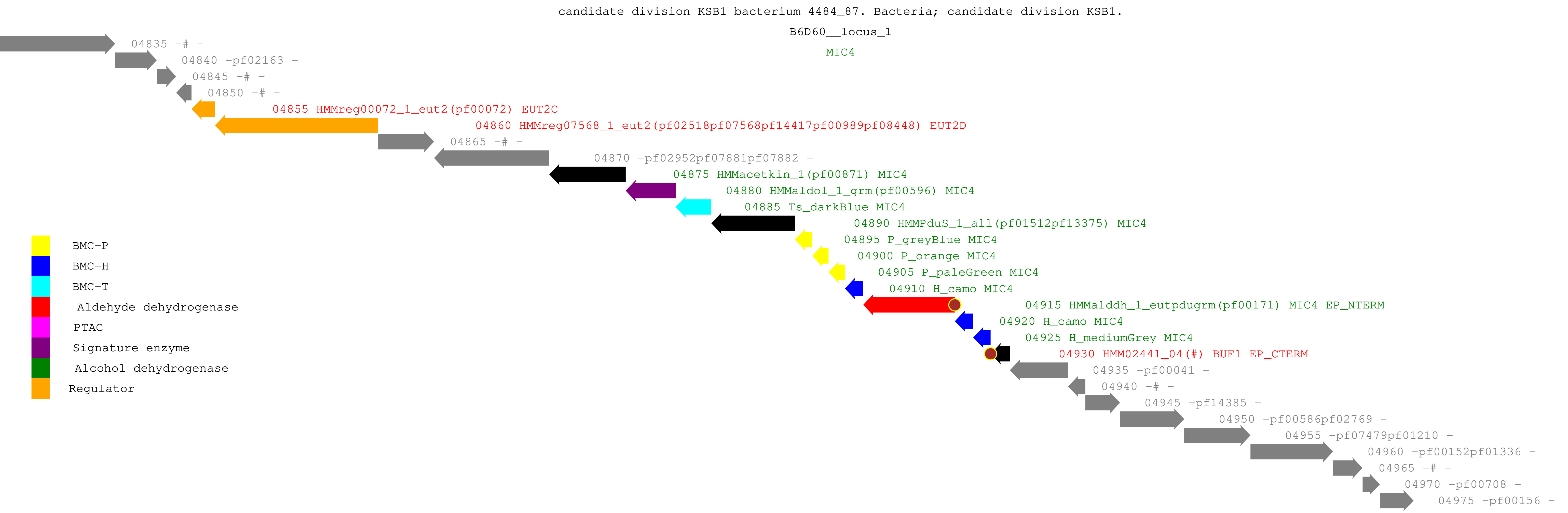
candidate division KSB1 bacterium 4484_87. Bacteria; candidate division KSB1.
B6D60__locus_1
MIC4
04835 -# -
04840 -pf02163 -
04845 -# -
04850 -# -
04855 HMMreg00072_1_eut2(pf00072) EUT2C
04860 HMMreg07568_1_eut2(pf02518pf07568pf14417pf00989pf08448) EUT2D
04865 -# -
04870 -pf02952pf07881pf07882 -
04875 HMMacetkin_1(pf00871) MIC4
04880 HMMaldol_1_grm(pf00596) MIC4
04885 Ts_darkBlue MIC4
04890 HMMPduS_1_all(pf01512pf13375) MIC4
04895 P_greyBlue MIC4
04900 P_orange MIC4
04905 P_paleGreen MIC4
04910 H_camo MIC4
04915 HMMaldh_1_eutpdugrm(pf00171) MIC4 EP_NTERM
04920 H_camo MIC4
04925 H_mediumGrey MIC4
04930 HMM02441_04(#) BUF1 EP_CTERM
04935 -pf00041 -
04940 -# -
04945 -pf14385 -
04950 -pf00586pf02769 -
04955 -pf07479pf01210 -
04960 -pf00152pf01336 -
04965 -# -
04970 -pf00708 -
04975 -pf00156 -
BMC-P
BMC-H
BMC-T
Aldehyde dehydrogenase
PTAC
Signature enzyme
Alcohol dehydrogenase
Regulator

Actinobacteria bacterium ADurb.Bin346. Bacteria; Actinobacteria.

BWY60__locus_1_main_sat

MIC4

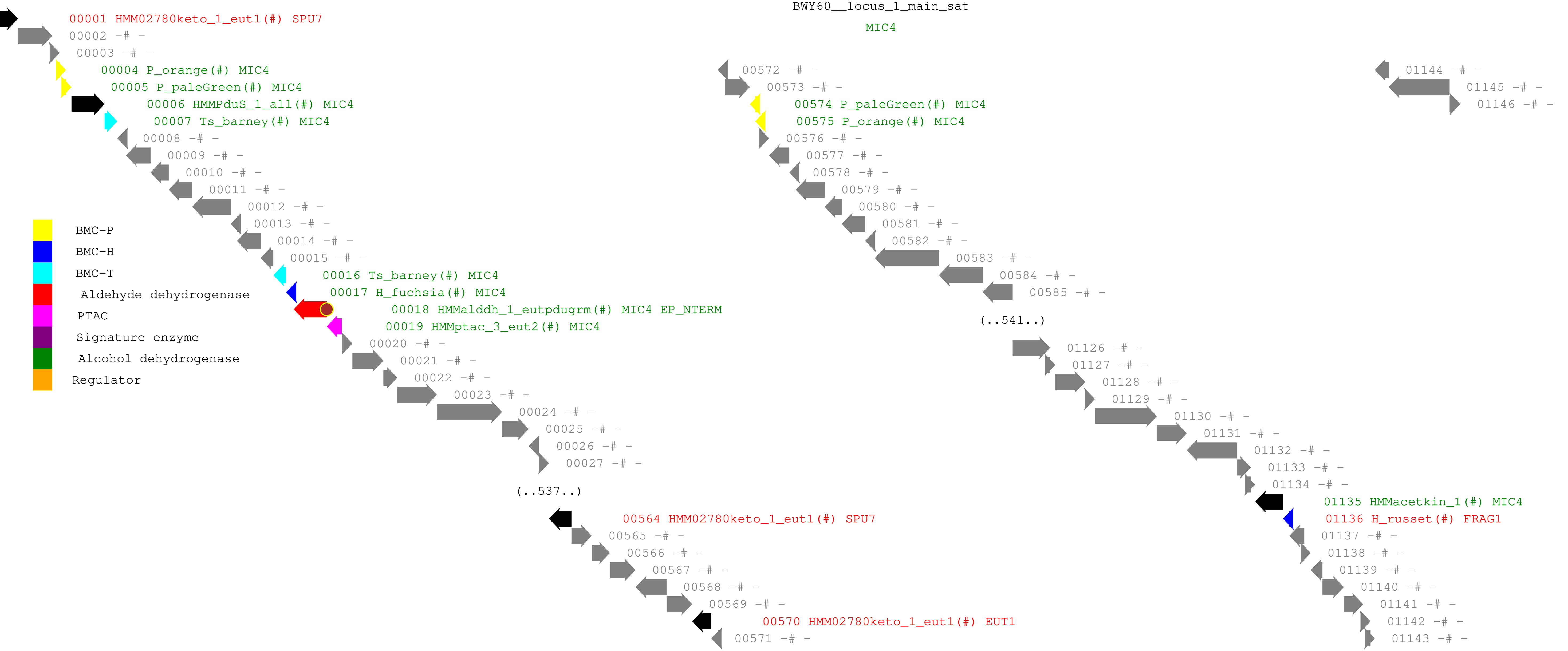

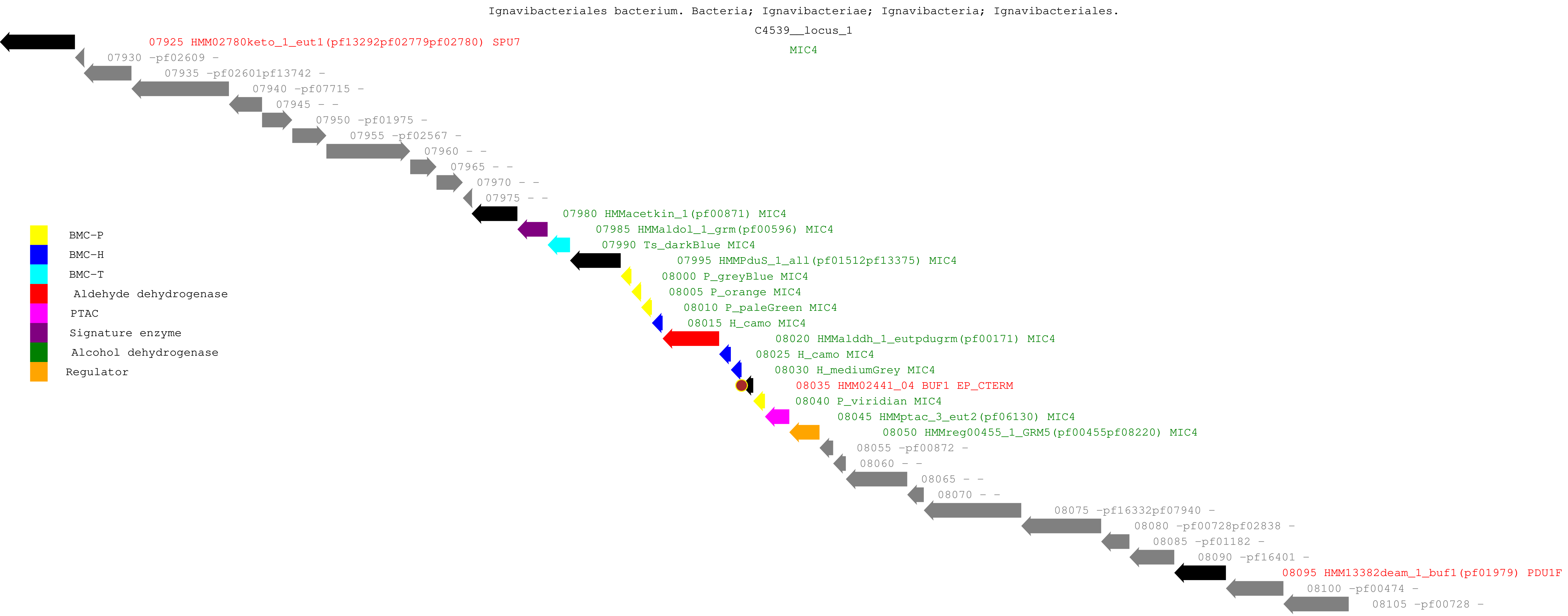
Ignavibacteriales bacterium. Bacteria; Ignavibacteriae; Ignavibacteria; Ignavibacteriales.
C4539__locus_1
MIC4
07925 HMM02780keto_1_eut1(pf13292pf02779pf02780) SPU7
07930 -pf02609 -
07935 -pf02601pf13742 -
07940 -pf07715 -
07945 - -
07950 -pf01975 -
07955 -pf02567 -
07960 - -
07965 - -
07970 - -
07975 - -
07980 HMMacetkin_1(pf00871) MIC4
07985 HMMaldol_1_grm(pf00596) MIC4
07990 Ts_darkBlue MIC4
07995 HMMPduS_1_all(pf01512pf13375) MIC4
08000 P_greyBlue MIC4
08005 P_orange MIC4
08010 P_paleGreen MIC4
08015 H_camo MIC4
08020 HMMalddh_1_eutpdugrm(pf00171) MIC4
08025 H_camo MIC4
08030 H_mediumGrey MIC4
08035 HMM02441_04 BUF1 EP_CTERM
08040 P_viridian MIC4
08045 HMMptac_3_eut2(pf06130) MIC4
08050 HMMreg00455_1_GRM5(pf00455pf08220) MIC4
08055 -pf00872 -
08060 - -
08065 - -
08070 - -
08075 -pf16332pf07940 -
08080 -pf00728pf02838 -
08085 -pf01182 -
08090 -pf16401 -
08095 HMM13382deam_1_buf1(pf01979) PDU1F
08100 -pf00474 -
08105 -pf00728 -
BMC-P
BMC-H
BMC-T
Aldehyde dehydrogenase
PTAC
Signature enzyme
Alcohol dehydrogenase
Regulator

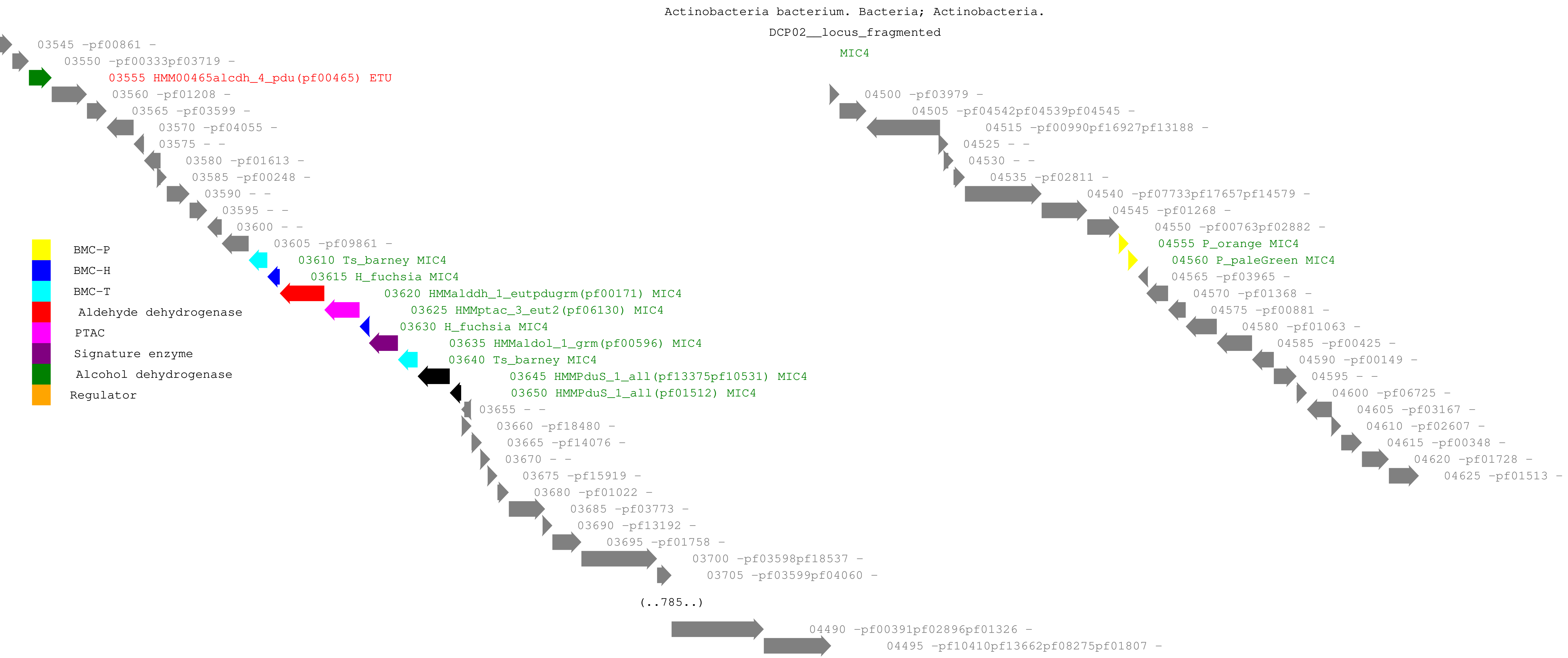

Actinobacteria bacterium. Bacteria; Actinobacteria.
DCP02__locus_fragmented
MIC4
03545 -pf00861 -
03550 -pf00333pf03719 -
03555 HMM00465alcdh_4_pdu(pf00465) ETU
03560 -pf01208 -
03565 -pf03599 -
03570 -pf04055 -
03575 - -
03580 -pf01613 -
03585 -pf00248 -
03590 - -
03595 - -
03600 - -
03605 -pf09861 -
03610 Ts_barney MIC4
03615 H_fuchsia MIC4
03620 HMMaldh_1_eutpdugrm(pf00171) MIC4
03625 HMMptac_3_eut2(pf06130) MIC4
03630 H_fuchsia MIC4
03635 HMMaldol_1_grm(pf00596) MIC4
03640 Ts_barney MIC4
03645 HMMPduS_1_all(pf13375pf10531) MIC4
03650 HMMPduS_1_all(pf01512) MIC4
03655 - -
03660 -pf18480 -
03665 -pf14076 -
03670 - -
03675 -pf15919 -
03680 -pf01022 -
03685 -pf03773 -
03690 -pf13192 -
03695 -pf01758 -
03700 -pf03598pf18537 -
03705 -pf03599pf04060 -
(..785..)
04490 -pf00391pf02896pf01326 -
04495 -pf10410pf13662pf08275pf01807 -
04500 -pf03979 -
04505 -pf04542pf04539pf04545 -
04515 -pf00990pf16927pf13188 -
04525 - -
04530 - -
04535 -pf02811 -
04540 -pf07733pf17657pf14579 -
04545 -pf01268 -
04550 -pf00763pf02882 -
04555 P_orange MIC4
04560 P_paleGreen MIC4
04565 -pf03965 -
04570 -pf01368 -
04575 -pf00881 -
04580 -pf01063 -
04585 -pf00425 -
04590 -pf00149 -
04595 - -
04600 -pf06725 -
04605 -pf03167 -
04610 -pf02607 -
04615 -pf00348 -
04620 -pf01728 -
04625 -pf01513 -
BMC-P
BMC-H
BMC-T
Aldehyde dehydrogenase
PTAC
Signature enzyme
Alcohol dehydrogenase
Regulator

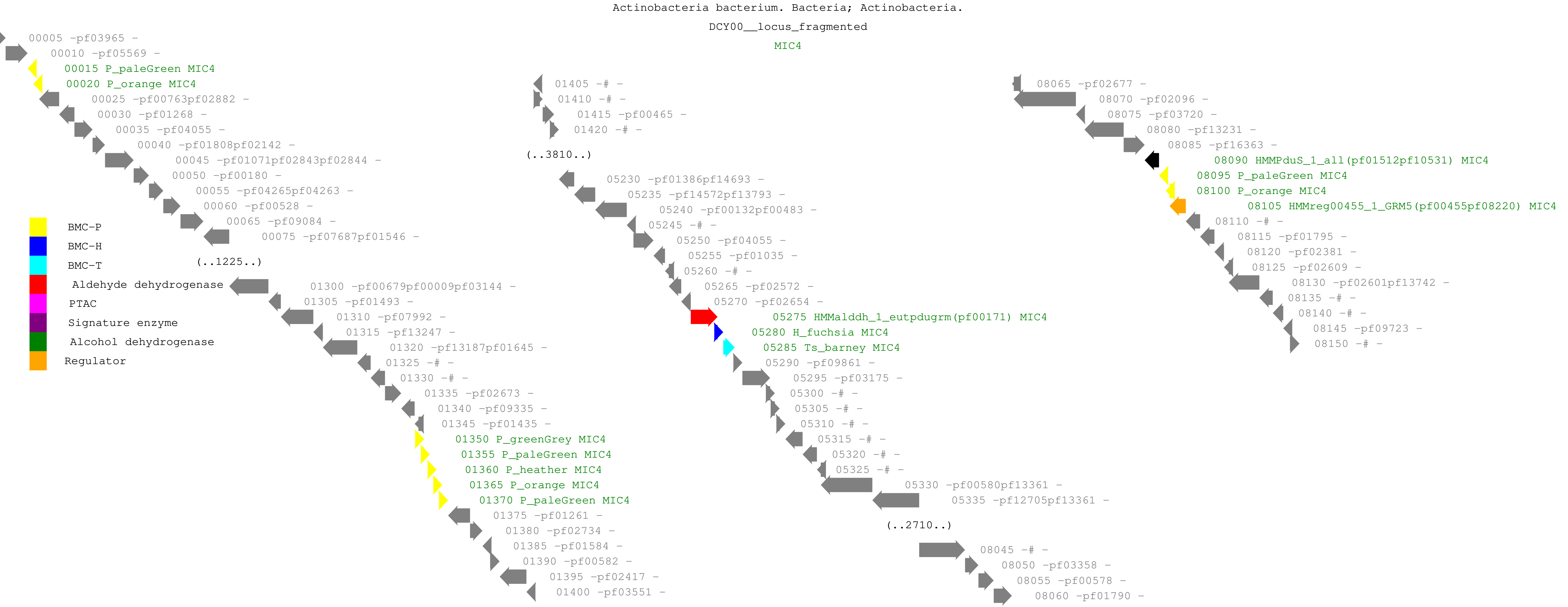

Actinobacteria bacterium. Bacteria; Actinobacteria.
DCY00__locus_fragmented
MIC4
00005 -pf03965 -
00010 -pf05569 -
00015 P_paleGreen MIC4
00020 P_orange MIC4
00025 -pf00763pf02882 -
00030 -pf01268 -
00035 -pf04055 -
00040 -pf01808pf02142 -
00045 -pf01071pf02843pf02844 -
00050 -pf00180 -
00055 -pf04265pf04263 -
00060 -pf00528 -
00065 -pf09084 -
00075 -pf07687pf01546 -
(..1225..)
01300 -pf00679pf00009pf03144 -
01305 -pf01493 -
01310 -pf07992 -
01315 -pf13247 -
01320 -pf13187pf01645 -
01325 -# -
01330 -# -
01335 -pf02673 -
01340 -pf09335 -
01345 -pf01435 -
01350 P_greenGrey MIC4
01355 P_paleGreen MIC4
01360 P_heather MIC4
01365 P_orange MIC4
01370 P_paleGreen MIC4
01375 -pf01261 -
01380 -pf02734 -
01385 -pf01584 -
01390 -pf00582 -
01395 -pf02417 -
01400 -pf03551 -
01405 -# -
01410 -# -
01415 -pf00465 -
01420 -# -
(..3810..)
05230 -pf01386pf14693 -
05235 -pf14572pf13793 -
05240 -pf00132pf00483 -
05245 -# -
05250 -pf04055 -
05255 -pf01035 -
05260 -# -
05265 -pf02572 -
05270 -pf02654 -
05275 HMMalddh_1_eutpdugrm(pf00171) MIC4
05280 H_fuchsia MIC4
05285 Ts_barney MIC4
05290 -pf09861 -
05295 -pf03175 -
05300 -# -
05305 -# -
05310 -# -
05315 -# -
05320 -# -
05325 -# -
05330 -pf00580pf13361 -
05335 -pf12705pf13361 -
(..2710..)
08045 -# -
08050 -pf03358 -
08055 -pf00578 -
08060 -pf01790 -
08065 -pf02677 -
08070 -pf02096 -
08075 -pf03720 -
08080 -pf13231 -
08085 -pf16363 -
08090 HMMPduS_1_all(pf01512pf10531) MIC4
08095 P_paleGreen MIC4
08100 P_orange MIC4
08105 HMMreg00455_1_GRM5(pf00455pf08220) MIC4
08110 -# -
08115 -pf01795 -
08120 -pf02381 -
08125 -pf02609 -
08130 -pf02601pf13742 -
08135 -# -
08140 -# -
08145 -pf09723 -
08150 -# -
BMC-P
BMC-H
BMC-T
Aldehyde dehydrogenase
PTAC
Signature enzyme
Alcohol dehydrogenase
Regulator

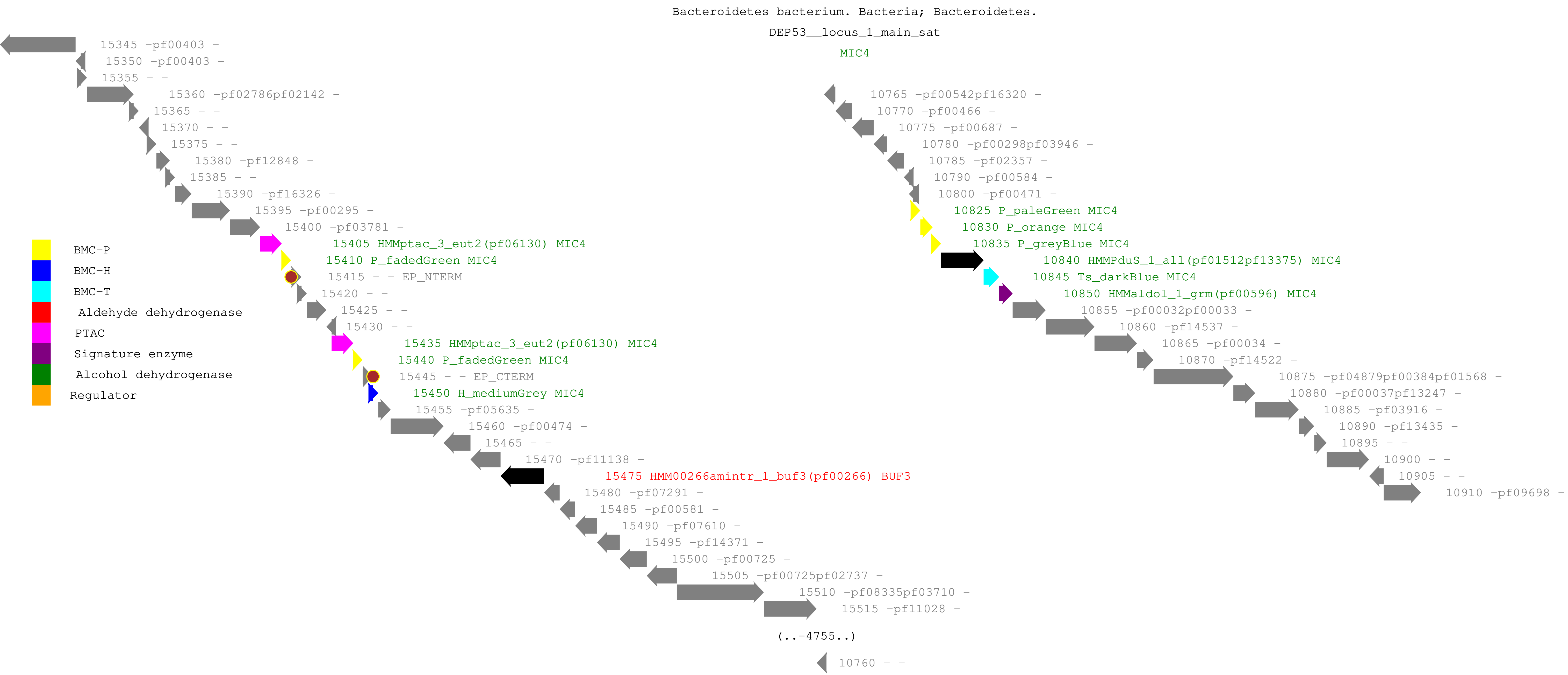

Bacteroidetes bacterium. Bacteria; Bacteroidetes.
DEP53__locus_1_main_sat
MIC4
15345 -pf00403 -
15350 -pf00403 -
15355 - -
15360 -pf02786pf02142 -
15365 - -
15370 - -
15375 - -
15380 -pf12848 -
15385 - -
15390 -pf16326 -
15395 -pf00295 -
15400 -pf03781 -
15405 HMMptac_3_eut2(pf06130) MIC4
15410 P_fadedGreen MIC4
15415 - - EP_NTERM
15420 - -
15425 - -
15430 - -
15435 HMMptac_3_eut2(pf06130) MIC4
15440 P_fadedGreen MIC4
15445 - - EP_CTERM
15450 H_mediumGrey MIC4
15455 -pf05635 -
15460 -pf00474 -
15465 - -
15470 -pf11138 -
15475 HMM00266amintr_1_buf3(pf00266) BUF3
15480 -pf07291 -
15485 -pf00581 -
15490 -pf07610 -
15495 -pf14371 -
15500 -pf00725 -
15505 -pf00725pf02737 -
15510 -pf08335pf03710 -
15515 -pf11028 -
(..-4755..)
10760 - -
10765 -pf00542pf16320 -
10770 -pf00466 -
10775 -pf00687 -
10780 -pf00298pf03946 -
10785 -pf02357 -
10790 -pf00584 -
10800 -pf00471 -
10825 P_paleGreen MIC4
10830 P_orange MIC4
10835 P_greyBlue MIC4
10840 HMMPduS_1_all(pf01512pf13375) MIC4
10845 Ts_darkBlue MIC4
10850 HMMaldol_1_grm(pf00596) MIC4
10855 -pf00032pf00033 -
10860 -pf14537 -
10865 -pf00034 -
10870 -pf14522 -
10875 -pf04879pf00384pf01568 -
10880 -pf00037pf13247 -
10885 -pf03916 -
10890 -pf13435 -
10895 - -
10900 - -
10905 - -
10910 -pf09698 -
BMC-P
BMC-H
BMC-T
Aldehyde dehydrogenase
PTAC
Signature enzyme
Alcohol dehydrogenase
Regulator

Actinobacteria bacterium. Bacteria; Actinobacteria.

E3J58__locus_1_main_sat

MIC4

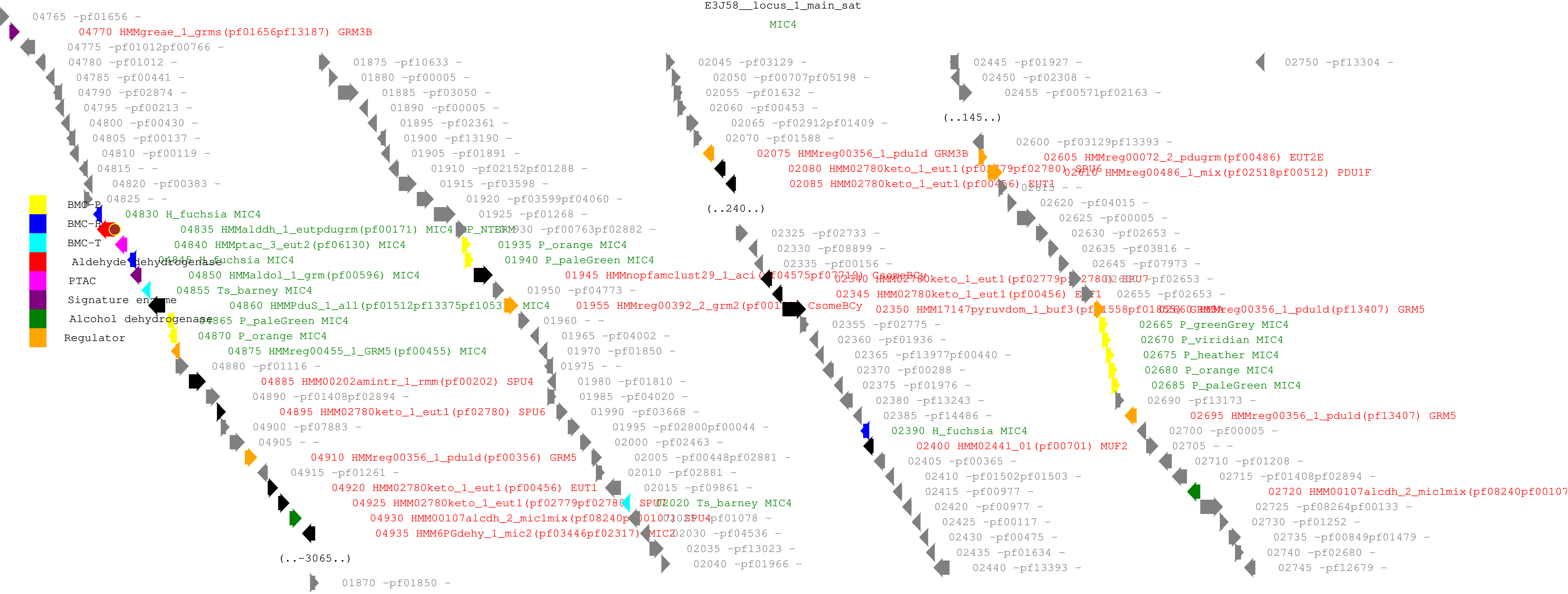

Hydrogenispora ethanolica. Bacteria; Firmicutes; Hydrogenispora.

EDC14__locus_1

MIC4

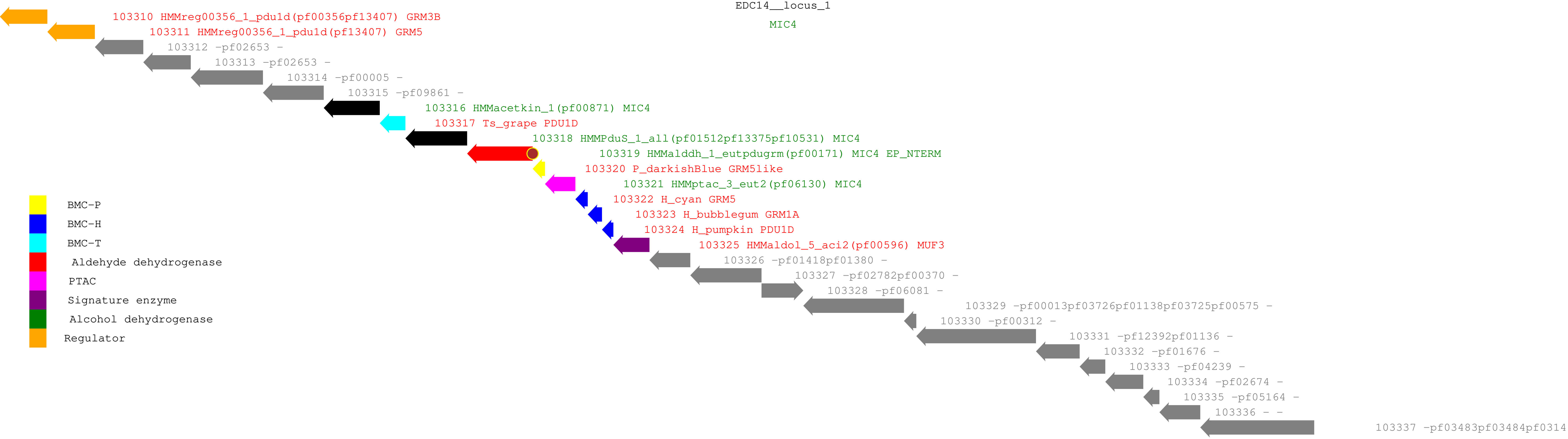

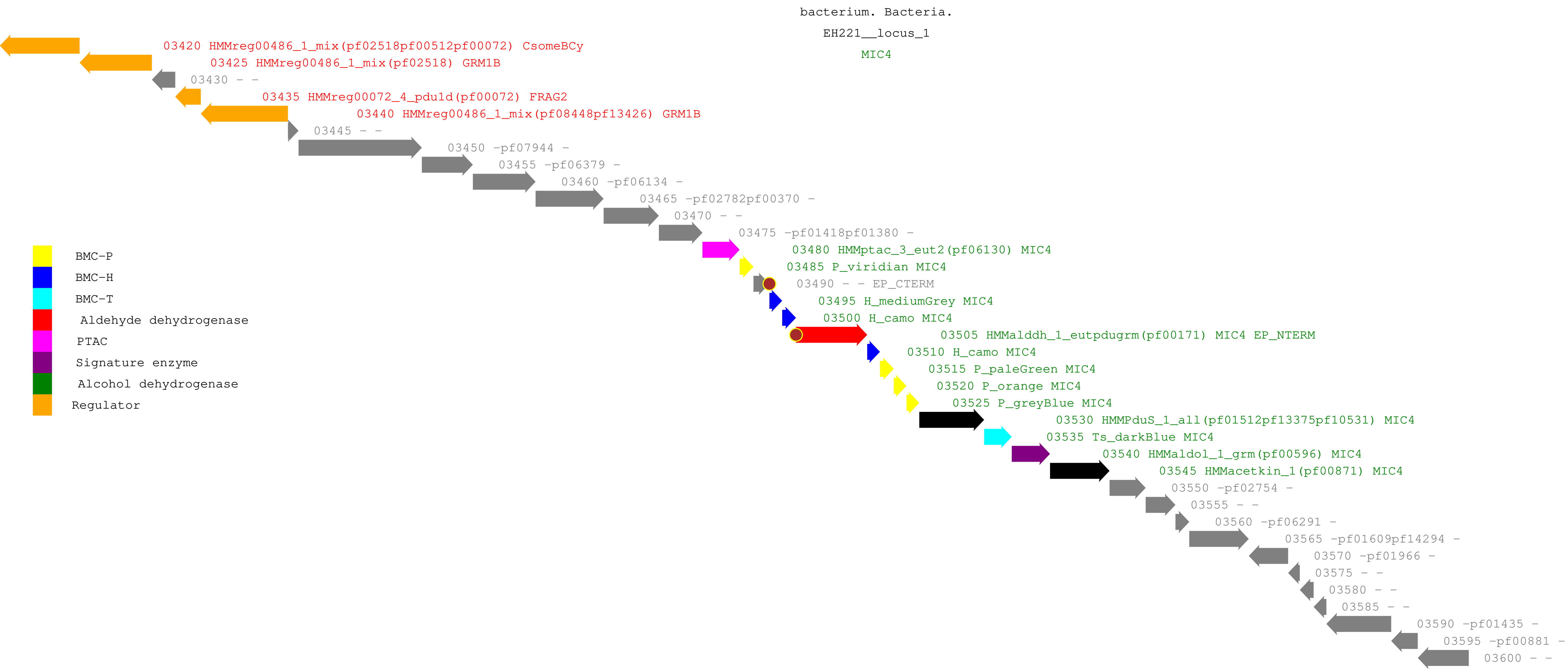
bacterium. Bacteria.
EH221__locus_1
MIC4
03420 HMMreg00486_1_mix(pf02518pf00512pf00072) CsomeBCy
03425 HMMreg00486_1_mix(pf02518) GRM1B
03430 - -
03435 HMMreg00072_4_pdu1d(pf00072) FRAG2
03440 HMMreg00486_1_mix(pf08448pf13426) GRM1B
03445 - -
03450 -pf07944 -
03455 -pf06379 -
03460 -pf06134 -
03465 -pf02782pf00370 -
03470 - -
03475 -pf01418pf01380 -
03480 HMMptac_3_eut2(pf06130) MIC4
03485 P_viridian MIC4
03490 - - EP_CTERM
03495 H_mediumGrey MIC4
03500 H_camo MIC4
03505 HMMaldh_1_eutpdugrm(pf00171) MIC4 EP_NTERM
03510 H_camo MIC4
03515 P_paleGreen MIC4
03520 P_orange MIC4
03525 P_greyBlue MIC4
03530 HMMPduS_1_all(pf01512pf13375pf10531) MIC4
03535 Ts_darkBlue MIC4
03540 HMMaldol_1_grm(pf00596) MIC4
03545 HMMacetkin_1(pf00871) MIC4
03550 -pf02754 -
03555 - -
03560 -pf06291 -
03565 -pf01609pf14294 -
03570 -pf01966 -
03575 - -
03580 - -
03585 - -
03590 -pf01435 -
03595 -pf00881 -
03600 - -
BMC-P
BMC-H
BMC-T
Aldehyde dehydrogenase
PTAC
Signature enzyme
Alcohol dehydrogenase
Regulator

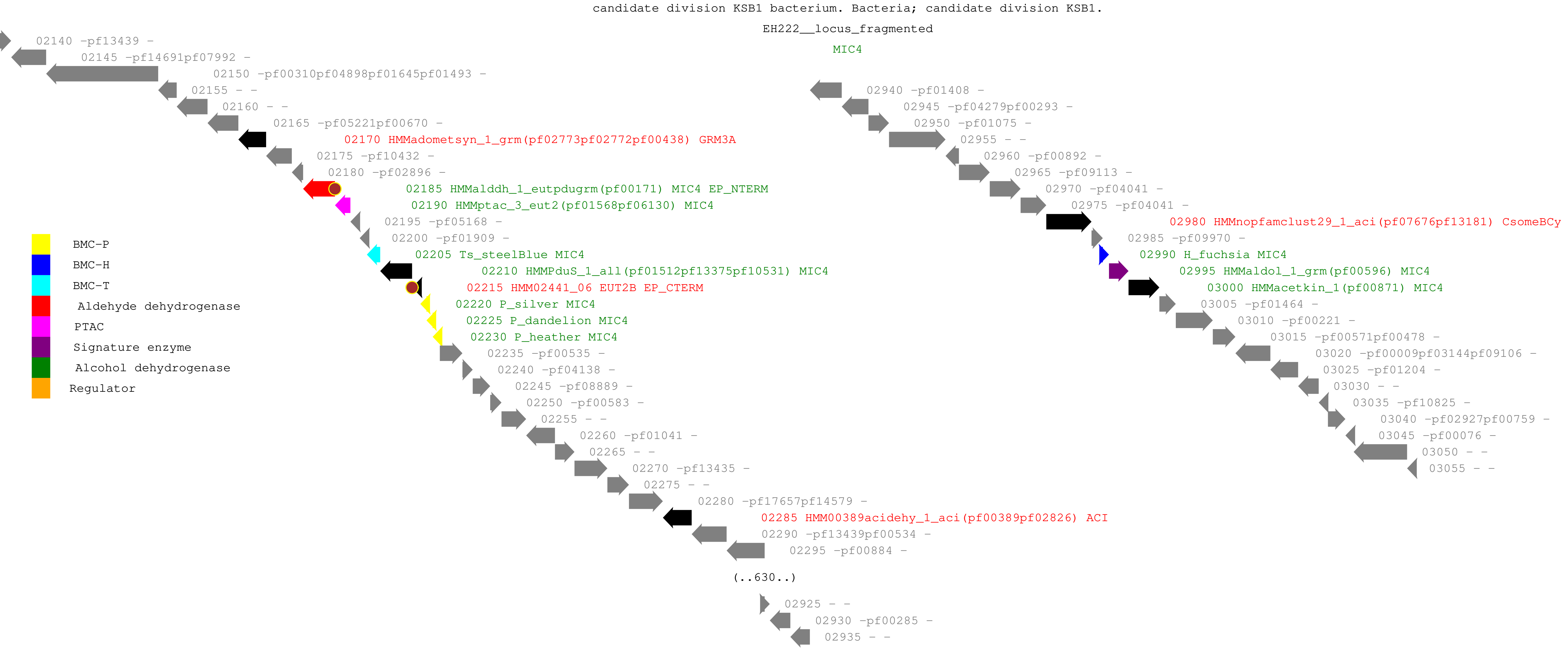
candidate division KSB1 bacterium. Bacteria; candidate division KSB1.
EH222__locus_fragmented
MIC4
02140 -pf13439 -
02145 -pf14691pf07992 -
02150 -pf00310pf04898pf01645pf01493 -
02155 - -
02160 - -
02165 -pf05221pf00670 -
02170 HMMadometsyn_1_grm(pf02773pf02772pf00438) GRM3A
02175 -pf10432 -
02180 -pf02896 -
02185 HMMalddh_1_eutpdugrm(pf00171) MIC4 EP_NTERM
02190 HMMptac_3_eut2(pf01568pf06130) MIC4
02195 -pf05168 -
02200 -pf01909 -
02205 Ts_steelBlue MIC4
02210 HMMPduS_1_all(pf01512pf13375pf10531) MIC4
02215 HMM02441_06 EUT2B EP_CTERM
02220 P_silver MIC4
02225 P_dandelion MIC4
02230 P_heather MIC4
02235 -pf00535 -
02240 -pf04138 -
02245 -pf08889 -
02250 -pf00583 -
02255 - -
02260 -pf01041 -
02265 - -
02270 -pf13435 -
02275 - -
02280 -pf17657pf14579 -
02285 HMM00389acidehy_1_aci(pf00389pf02826) ACI
02290 -pf13439pf00534 -
02295 -pf00884 -
(..630..)
02925 - -
02930 -pf00285 -
02935 - -
02940 -pf01408 -
02945 -pf04279pf00293 -
02950 -pf01075 -
02955 - -
02960 -pf00892 -
02965 -pf09113 -
02970 -pf04041 -
02975 -pf04041 -
02980 HMMnopfamclust29_1_aci(pf07676pf13181) CsomeBCy
02985 -pf09970 -
02990 H_fuchsia MIC4
02995 HMMaldol_1_grm(pf00596) MIC4
03000 HMMacetkin_1(pf00871) MIC4
03005 -pf01464 -
03010 -pf00221 -
03015 -pf00571pf00478 -
03020 -pf00009pf03144pf09106 -
03025 -pf01204 -
03030 - -
03035 -pf10825 -
03040 -pf02927pf00759 -
03045 -pf00076 -
03050 - -
03055 - -
BMC-P
BMC-H
BMC-T
Aldehyde dehydrogenase
PTAC
Signature enzyme
Alcohol dehydrogenase
Regulator

candidate division KSB1 bacterium. Bacteria; candidate division KSB1.

EH223__locus_1_main_sat

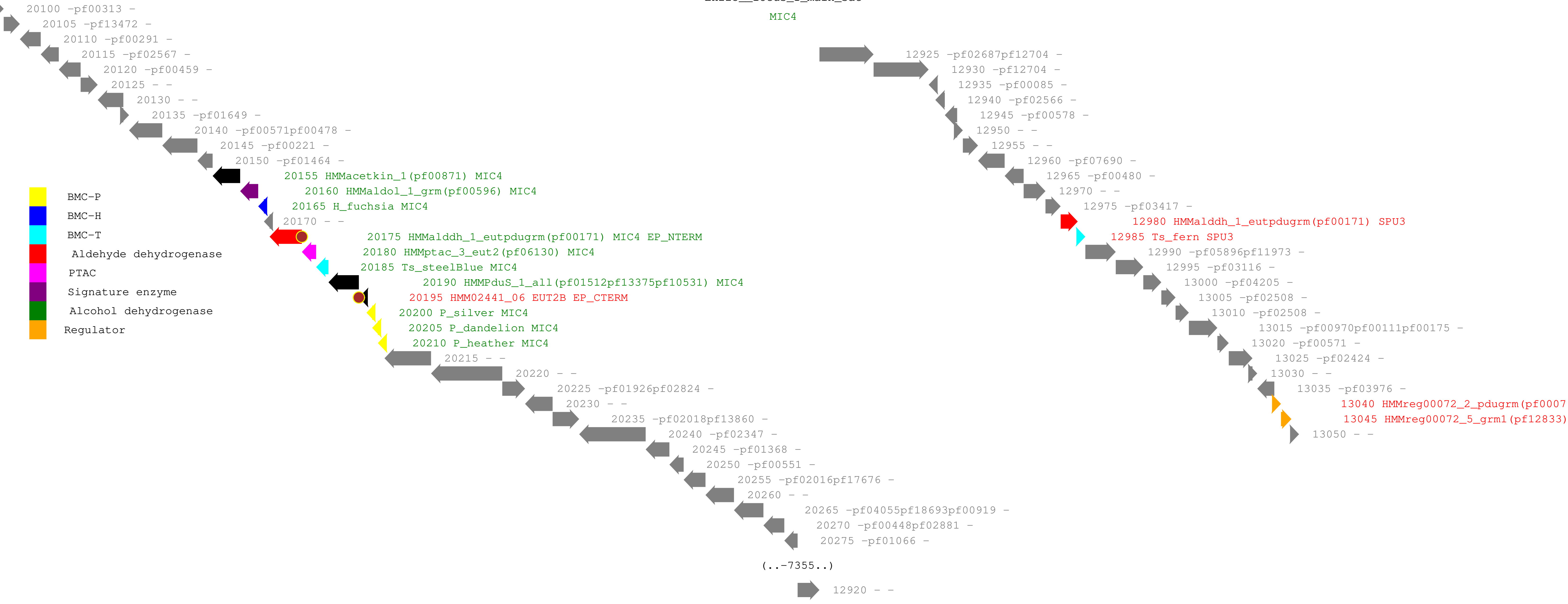

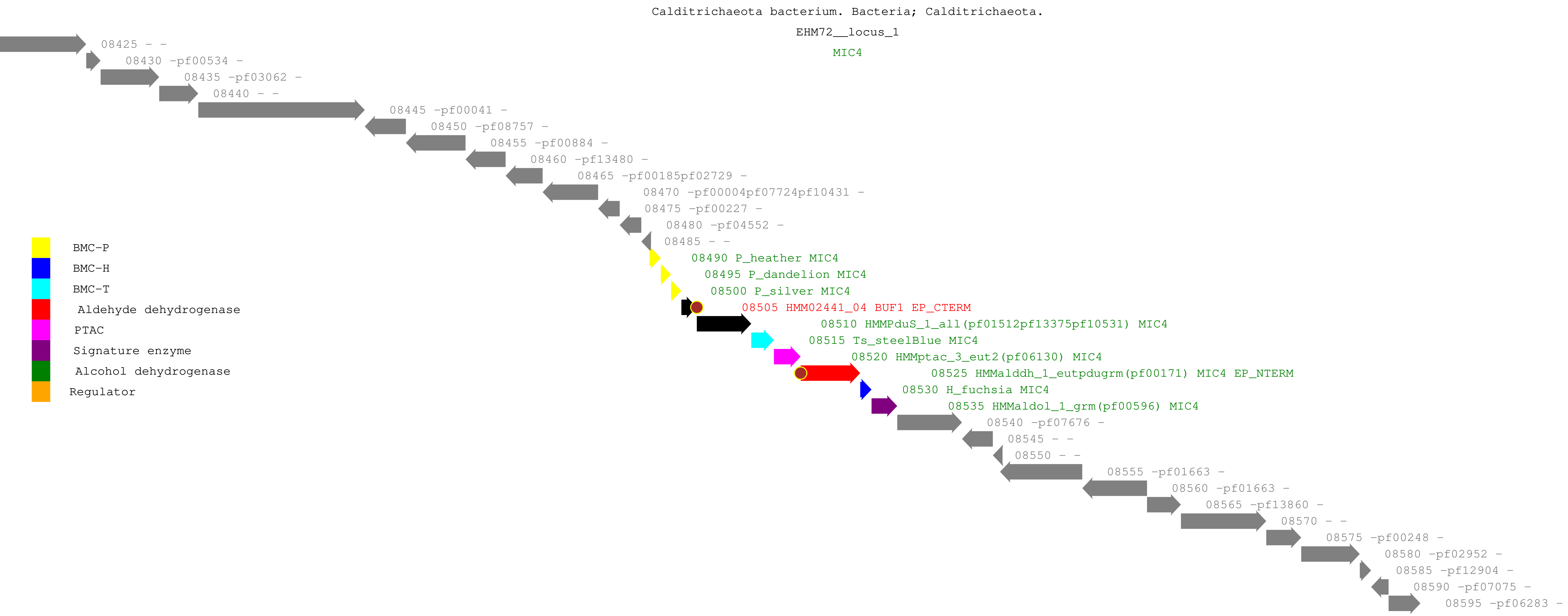
Calditrichaeota bacterium. Bacteria; Calditrichaeota.
EHM72__locus_1
MIC4
08425 - -
08430 -pf00534 -
08435 -pf03062 -
08440 - -
08445 -pf00041 -
08450 -pf08757 -
08455 -pf00884 -
08460 -pf13480 -
08465 -pf00185pf02729 -
08470 -pf00004pf07724pf10431 -
08475 -pf00227 -
08480 -pf04552 -
08485 - -
08490 P_heather MIC4
08495 P_dandelion MIC4
08500 P_silver MIC4
08505 HMM02441_04 BUF1 EP_CTERM
08510 HMMPduS_1_all(pf01512pf13375pf10531) MIC4
08515 Ts_steelBlue MIC4
08520 HMMptac_3_eut2(pf06130) MIC4
08525 HMMaldh_1_eutpdugrm(pf00171) MIC4 EP_NTERM
08530 H_fuchsia MIC4
08535 HMMaldol_1_grm(pf00596) MIC4
08540 -pf07676 -
08545 - -
08550 - -
08555 -pf01663 -
08560 -pf01663 -
08565 -pf13860 -
08570 - -
08575 -pf00248 -
08580 -pf02952 -
08585 -pf12904 -
08590 -pf07075 -
08595 -pf06283 -
BMC-P
BMC-H
BMC-T
Aldehyde dehydrogenase
PTAC
Signature enzyme
Alcohol dehydrogenase
Regulator

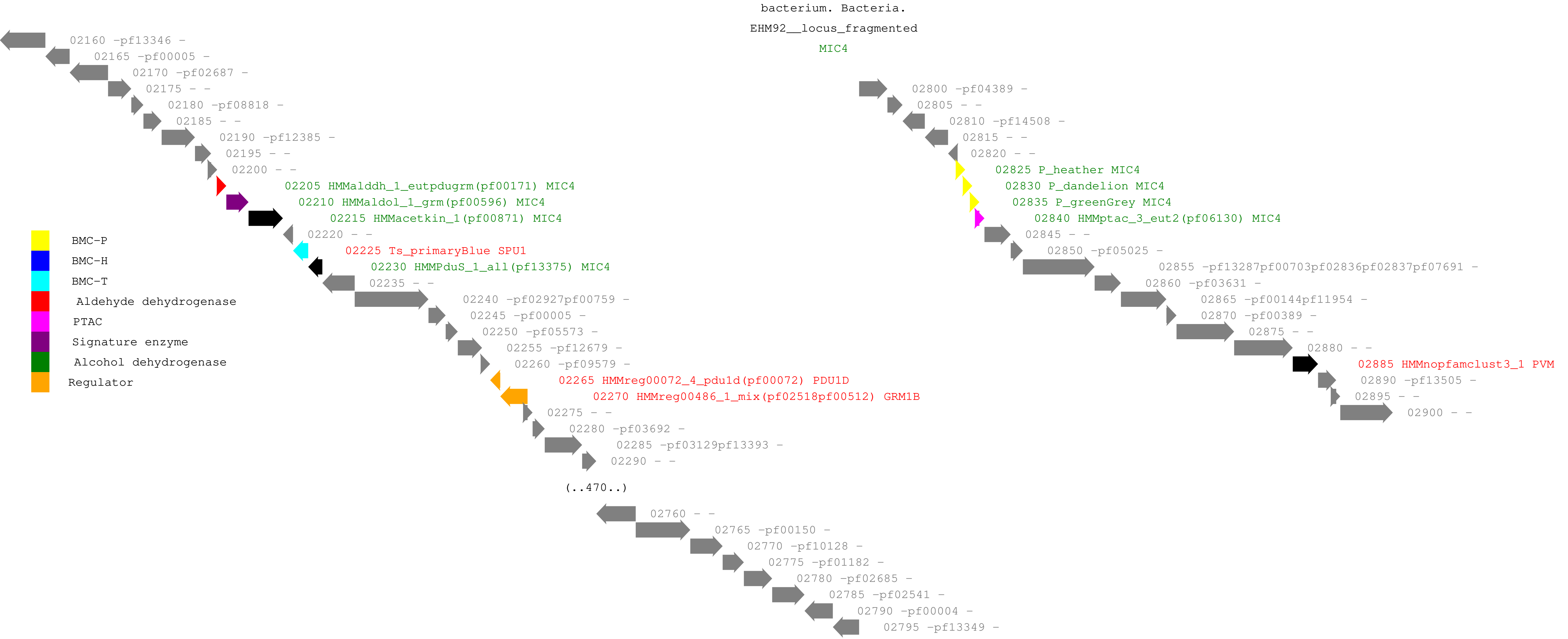
bacterium. Bacteria.
EHM92__locus_fragmented
MIC4
02160 -pf13346 -
02165 -pf00005 -
02170 -pf02687 -
02175 - -
02180 -pf08818 -
02185 - -
02190 -pf12385 -
02195 - -
02200 - -
02205 HMMalddh_1_eutpdugrm(pf00171) MIC4
02210 HMMaldol_1_grm(pf00596) MIC4
02215 HMMacetkin_1(pf00871) MIC4
02220 - -
02225 Ts_primaryBlue SPU1
02230 HMMPduS_1_all(pf13375) MIC4
02235 - -
02240 -pf02927pf00759 -
02245 -pf00005 -
02250 -pf05573 -
02255 -pf12679 -
02260 -pf09579 -
02265 HMMreg00072_4_pdu1d(pf00072) PDU1D
02270 HMMreg00486_1_mix(pf02518pf00512) GRM1B
02275 - -
02280 -pf03692 -
02285 -pf03129pf13393 -
02290 - -
(..470..)
02760 - -
02765 -pf00150 -
02770 -pf10128 -
02775 -pf01182 -
02780 -pf02685 -
02785 -pf02541 -
02790 -pf00004 -
02795 -pf13349 -
02800 -pf04389 -
02805 - -
02810 -pf14508 -
02815 - -
02820 - -
02825 P_heather MIC4
02830 P_dandelion MIC4
02835 P_greenGrey MIC4
02840 HMMptac_3_eut2(pf06130) MIC4
02845 - -
02850 -pf05025 -
02855 -pf13287pf00703pf02836pf02837pf07691 -
02860 -pf03631 -
02865 -pf00144pf11954 -
02870 -pf00389 -
02875 - -
02880 - -
02885 HMMnopfamclust3_1 PVM
02890 -pf13505 -
02895 - -
02900 - -
BMC-P
BMC-H
BMC-T
Aldehyde dehydrogenase
PTAC
Signature enzyme
Alcohol dehydrogenase
Regulator

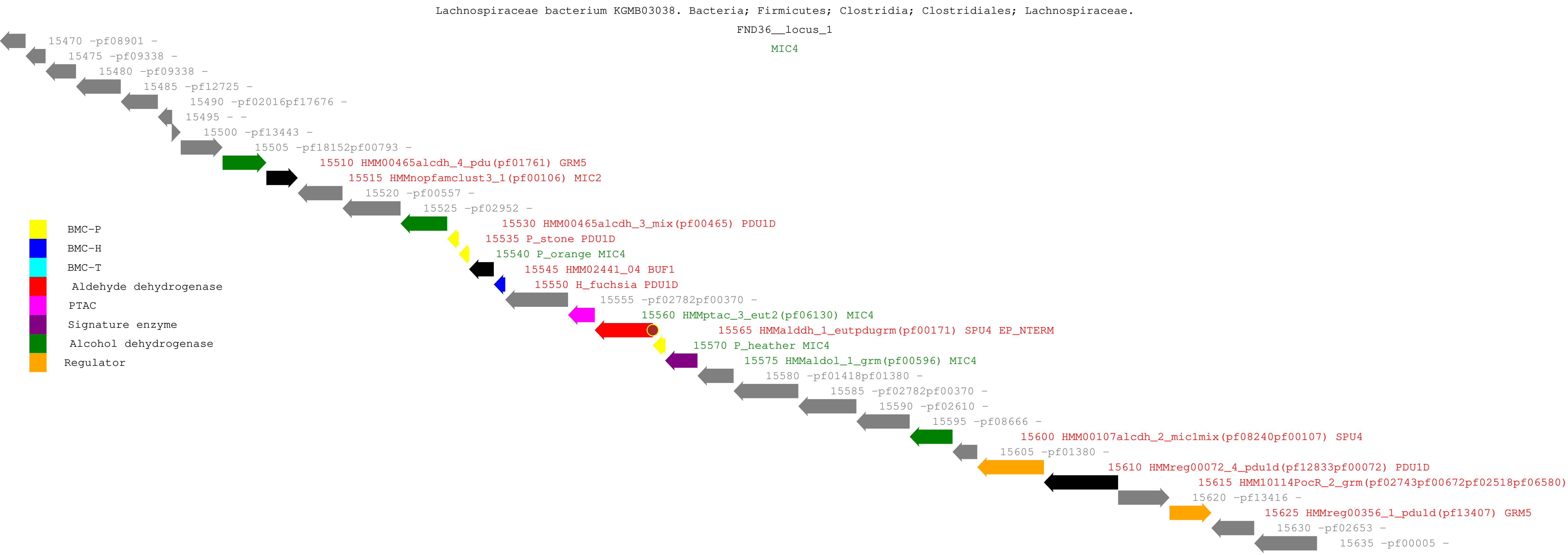

Lachnospiraceae bacterium KGMB03038. Bacteria; Firmicutes; Clostridia; Clostridiales; Lachnospiraceae.
FND36__locus_1
MIC4
15470 -pf08901 -
15475 -pf09338 -
15480 -pf09338 -
15485 -pf12725 -
15490 -pf02016pf17676 -
15495 - -
15500 -pf13443 -
15505 -pf18152pf00793 -
15510 HMM00465alcdh_4_pdu(pf01761) GRM5
15515 HMMnopfamclust3_1(pf00106) MIC2
15520 -pf00557 -
15525 -pf02952 -
15530 HMM00465alcdh_3_mix(pf00465) PDU1D
15535 P_stone PDU1D
15540 P_orange MIC4
15545 HMM02441_04 BUF1
15550 H_fuchsia PDU1D
15555 -pf02782pf00370 -
15560 HMMptac_3_eut2(pf06130) MIC4
15565 HMMalddh_1_eutpdugrm(pf00171) SPU4 EP_NTERM
15570 P_heather MIC4
15575 HMMaldol_1_grm(pf00596) MIC4
15580 -pf01418pf01380 -
15585 -pf02782pf00370 -
15590 -pf02610 -
15595 -pf08666 -
15600 HMM00107alcdh_2_mic1mix(pf08240pf00107) SPU4
15605 -pf01380 -
15610 HMMreg00072_4_pdu1d(pf12833pf00072) PDU1D
15615 HMM10114PocR_2_grm(pf02743pf00672pf02518pf06580)
15620 -pf13416 -
15625 HMMreg00356_1_pdu1d(pf13407) GRM5
15630 -pf02653 -
15635 -pf00005 -
BMC-P
BMC-H
BMC-T
Aldehyde dehydrogenase
PTAC
Signature enzyme
Alcohol dehydrogenase
Regulator

Melioribacter roseus (strain JCM 17771 / P3M-2). Bacteria; Ignavibacteriae; Ignavibacteria; Ignavibacteriales;

MROS__locus_1

MIC4

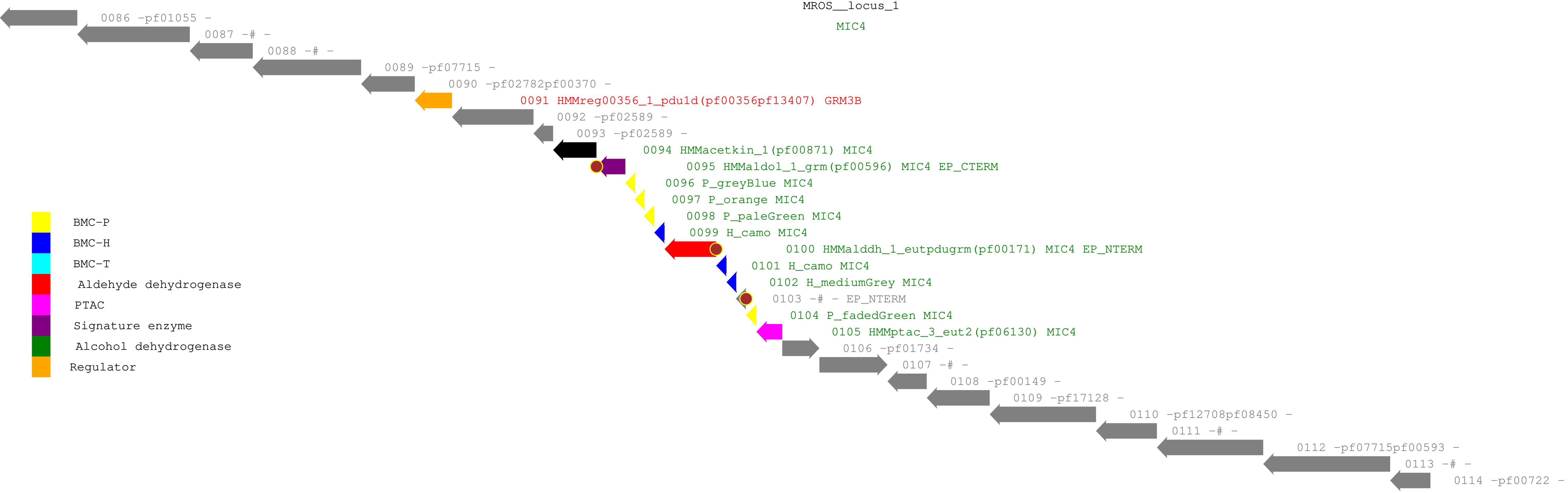

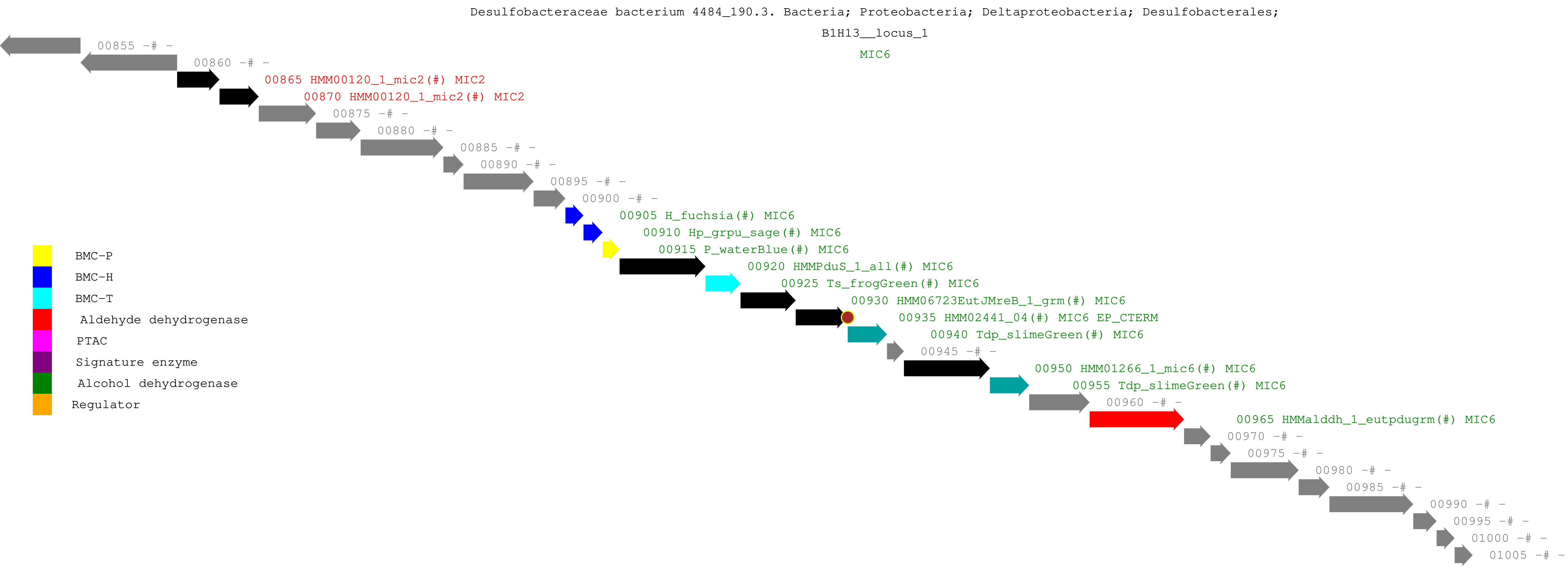
Desulfobacteraceae bacterium 4484_190.3. Bacteria; Proteobacteria; Deltaproteobacteria; Desulfobacterales;
B1H13__locus_1
MIC6
00855 -# -
00860 -# -
00865 HMM00120_1_mic2(#) MIC2
00870 HMM00120_1_mic2(#) MIC2
00875 -# -
00880 -# -
00885 -# -
00890 -# -
00895 -# -
00900 -# -
00905 H_fuchsia(#) MIC6
00910 Hp_grpu_sage(#) MIC6
00915 P_waterBlue(#) MIC6
00920 HMMPduS_1_all(#) MIC6
00925 Ts_frogGreen(#) MIC6
00930 HMM06723EutJMreB_1_grm(#) MIC6
00935 HMM02441_04(#) MIC6 EP_CTERM
00940 Tdp_slimeGreen(#) MIC6
00945 -# -
00950 HMM01266_1_mic6(#) MIC6
00955 Tdp_slimeGreen(#) MIC6
00960 -# -
00965 HMMaldDh_1_eutpdugrm(#) MIC6
00970 -# -
00975 -# -
00980 -# -
00985 -# -
00990 -# -
00995 -# -
01000 -# -
01005 -# -
BMC-P
BMC-H
BMC-T
Aldehyde dehydrogenase
PTAC
Signature enzyme
Alcohol dehydrogenase
Regulator

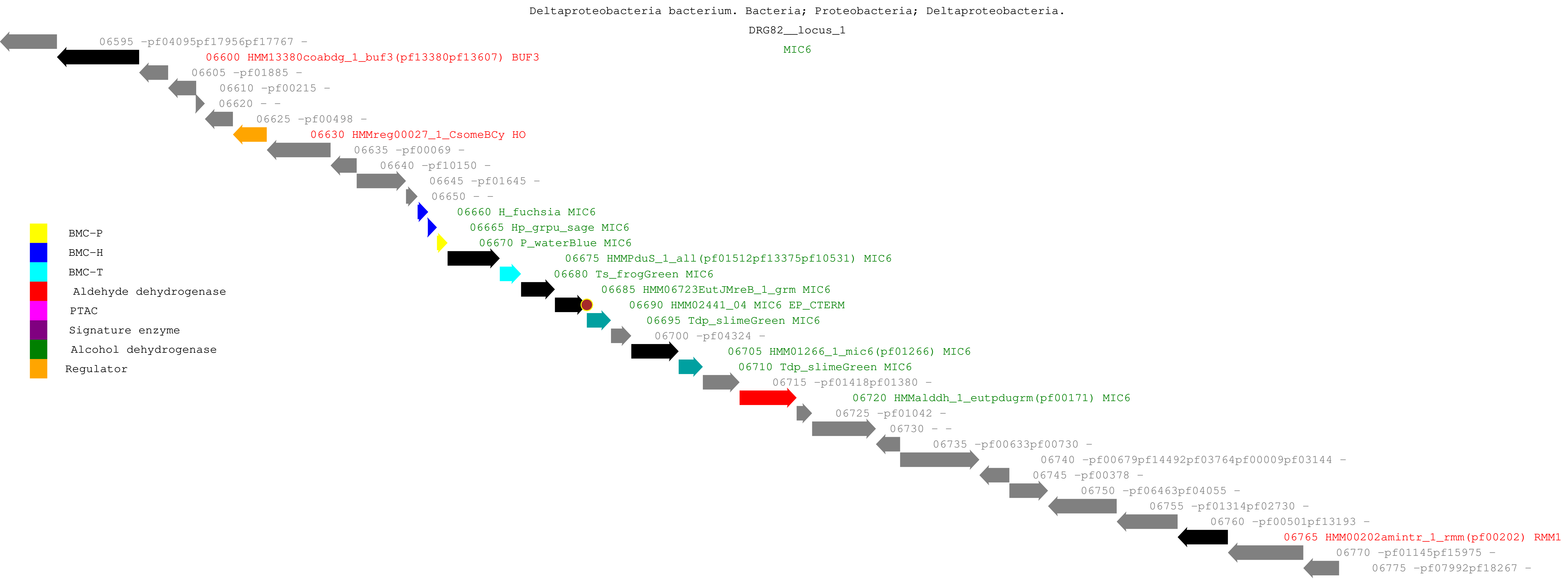
Deltaproteobacteria bacterium. Bacteria; Proteobacteria; Deltaproteobacteria.
DRG82__locus_1
MIC6
06595 -pf04095pf17956pf17767 -
06600 HMM13380coabdg_1_buf3(pf13380pf13607) BUF3
06605 -pf01885 -
06610 -pf00215 -
06620 - -
06625 -pf00498 -
06630 HMMreg00027_1_CsomeBCy HO
06635 -pf00069 -
06640 -pf10150 -
06645 -pf01645 -
06650 - -
06660 H_fuchsia MIC6
06665 Hp_grpu_sage MIC6
06670 P_waterBlue MIC6
06675 HMMPduS_1_all(pf01512pf13375pf10531) MIC6
06680 Ts_frogGreen MIC6
06685 HMM06723EutJMreB_1_grm MIC6
06690 HMM02441_04 MIC6 EP_CTERM
06695 Tdp_slimeGreen MIC6
06700 -pf04324 -
06705 HMM01266_1_mic6(pf01266) MIC6
06710 Tdp_slimeGreen MIC6
06715 -pf01418pf01380 -
06720 HMMaldh_1_eutpdugrm(pf00171) MIC6
06725 -pf01042 -
06730 - -
06735 -pf00633pf00730 -
06740 -pf00679pf14492pf03764pf00009pf03144 -
06745 -pf00378 -
06750 -pf06463pf04055 -
06755 -pf01314pf02730 -
06760 -pf00501pf13193 -
06765 HMM00202amintr_1_rmm(pf00202) RMM1
06770 -pf01145pf15975 -
06775 -pf07992pf18267 -
BMC-P
BMC-H
BMC-T
Aldehyde dehydrogenase
PTAC
Signature enzyme
Alcohol dehydrogenase
Regulator

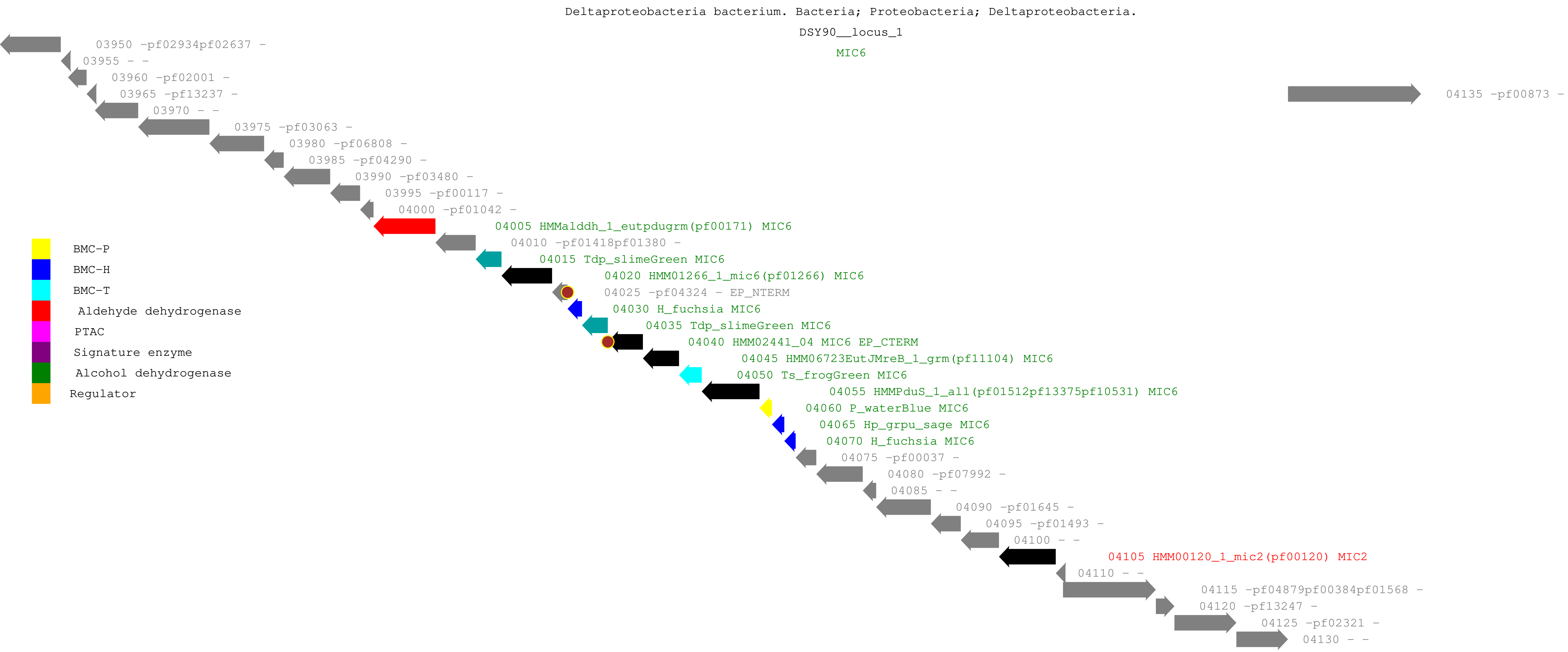

Deltaproteobacteria bacterium. Bacteria; Proteobacteria; Deltaproteobacteria.
DSY90__locus_1
MIC6
03950 -pf02934pf02637 -
03955 - -
03960 -pf02001 -
03965 -pf13237 -
03970 - -
03975 -pf03063 -
03980 -pf06808 -
03985 -pf04290 -
03990 -pf03480 -
03995 -pf00117 -
04000 -pf01042 -
04005 HMMalddh_1_eutpdugrm(pf00171) MIC6
04010 -pf01418pf01380 -
04015 Tdp_slimeGreen MIC6
04020 HMM01266_1_mic6(pf01266) MIC6
04025 -pf04324 - EP_NTERM
04030 H_fuchsia MIC6
04035 Tdp_slimeGreen MIC6
04040 HMM02441_04 MIC6 EP_CTERM
04045 HMM06723EutJMreB_1_grm(pf11104) MIC6
04050 Ts_frogGreen MIC6
04055 HMMPduS_1_all(pf01512pf13375pf10531) MIC6
04060 P_waterBlue MIC6
04065 Hp_grpu_sage MIC6
04070 H_fuchsia MIC6
04075 -pf00037 -
04080 -pf07992 -
04085 - -
04090 -pf01645 -
04095 -pf01493 -
04100 - -
04105 HMM00120_1_mic2(pf00120) MIC2
04110 - -
04115 -pf04879pf00384pf01568 -
04120 -pf13247 -
04125 -pf02321 -
04130 - -
04135 -pf00873 -
BMC-P
BMC-H
BMC-T
Aldehyde dehydrogenase
PTAC
Signature enzyme
Alcohol dehydrogenase
Regulator

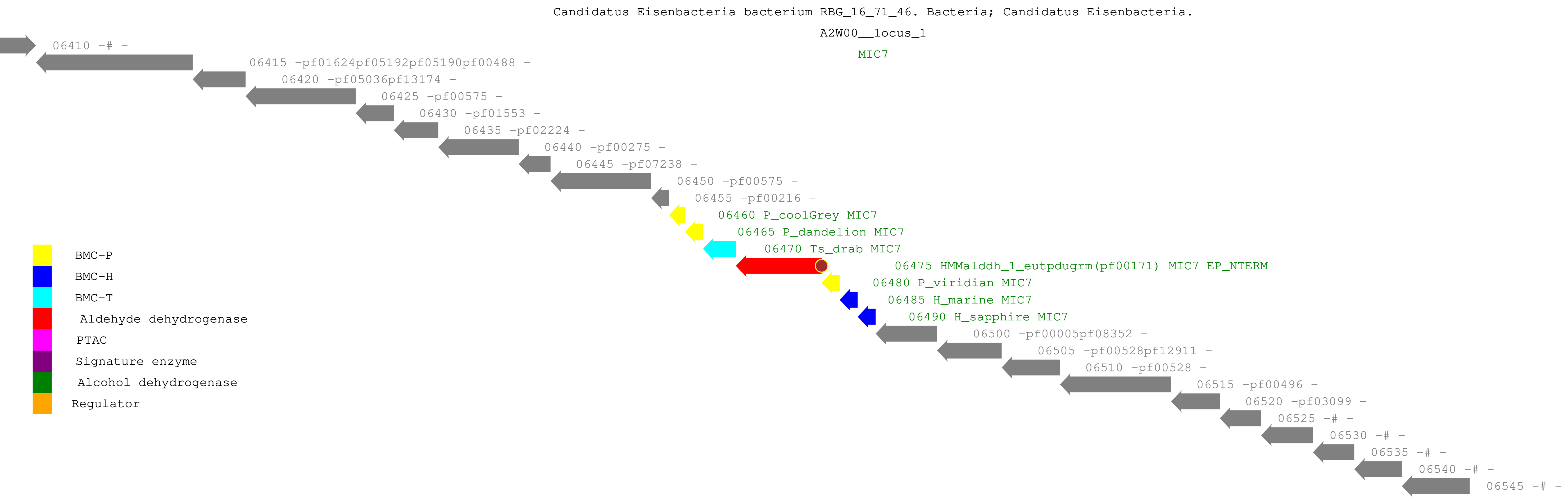
Candidatus Eisenbacteria bacterium RBG_16_71_46. Bacteria; Candidatus Eisenbacteria.
A2W00__locus_1
MIC7
06410 -# -
06415 -pf01624pf05192pf05190pf00488 -
06420 -pf05036pf13174 -
06425 -pf00575 -
06430 -pf01553 -
06435 -pf02224 -
06440 -pf00275 -
06445 -pf07238 -
06450 -pf00575 -
06455 -pf00216 -
06460 P_coolGrey MIC7
06465 P_dandelion MIC7
06470 Ts_drab MIC7
06475 HMMalddh_1_eutpdugrm(pf00171) MIC7 EP_NTERM
06480 P_viridian MIC7
06485 H_marine MIC7
06490 H_sapphire MIC7
06500 -pf00005pf08352 -
06505 -pf00528pf12911 -
06510 -pf00528 -
06515 -pf00496 -
06520 -pf03099 -
06525 -# -
06530 -# -
06535 -# -
06540 -# -
06545 -# -
BMC-P
BMC-H
BMC-T
Aldehyde dehydrogenase
PTAC
Signature enzyme
Alcohol dehydrogenase
Regulator

Latescibacteria bacterium DG_63. Bacteria; Candidatus Latescibacteria.

AMJ46__locus_1_main_sat

MIC7

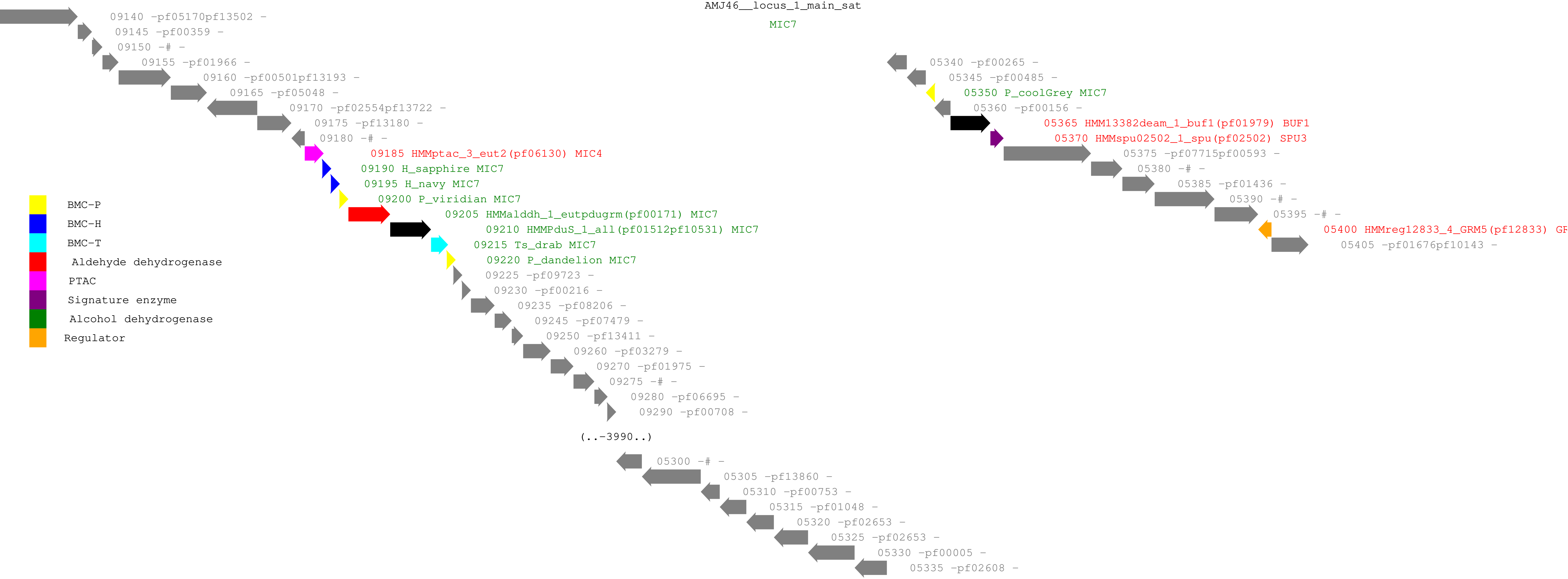

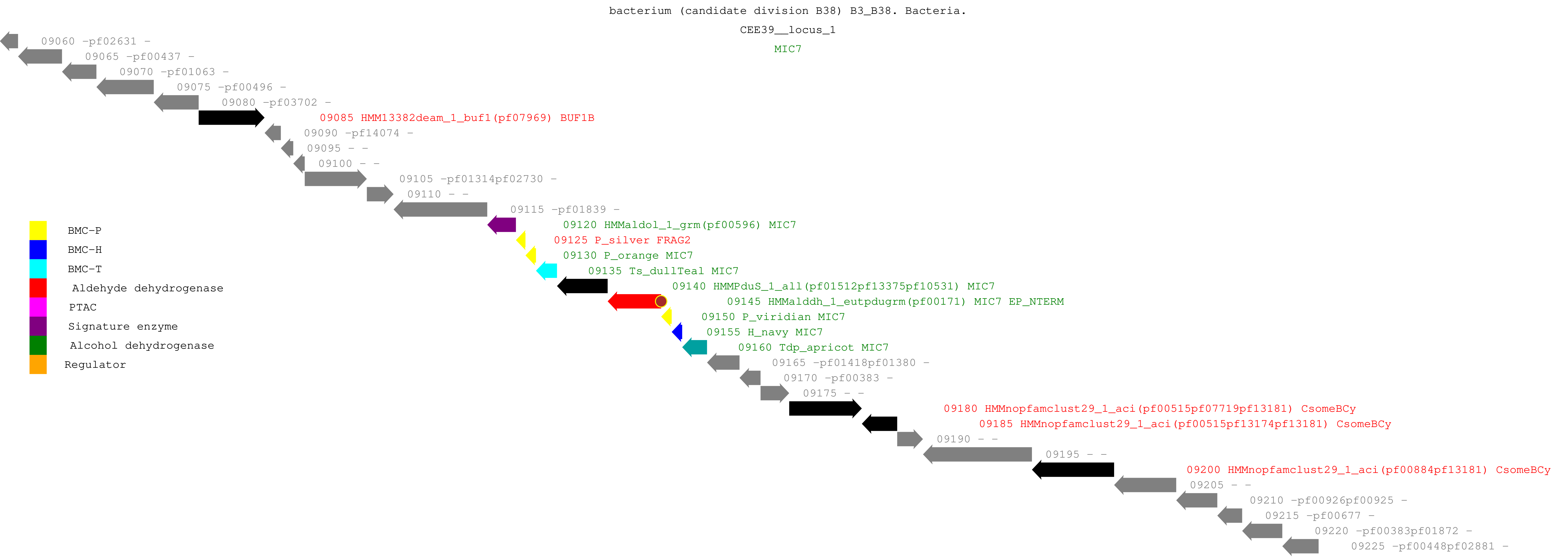
bacterium (candidate division B38) B3_B38. Bacteria.
CEE39__locus_1
MIC7
09060 -pf02631 -
09065 -pf00437 -
09070 -pf01063 -
09075 -pf00496 -
09080 -pf03702 -
09085 HMM13382deam_1_buf1(pf07969) BUF1B
09090 -pf14074 -
09095 - -
09100 - -
09105 -pf01314pf02730 -
09110 - -
09115 -pf01839 -
09120 HMMaldol_1_grm(pf00596) MIC7
09125 P_silver FRAG2
09130 P_orange MIC7
09135 Ts_dullTeal MIC7
09140 HMMPduS_1_all(pf01512pf13375pf10531) MIC7
09145 HMMalddh_1_eutpdugrm(pf00171) MIC7 EP_NTERM
09150 P_viridian MIC7
09155 H_navy MIC7
09160 Tdp_apricot MIC7
09165 -pf01418pf01380 -
09170 -pf00383 -
09175 - -
09180 HMMnopfamclust29_1_aci(pf00515pf07719pf13181) CsomeBCy
09185 HMMnopfamclust29_1_aci(pf00515pf13174pf13181) CsomeBCy
09190 - -
09195 - -
09200 HMMnopfamclust29_1_aci(pf00884pf13181) CsomeBCy
09205 - -
09210 -pf00926pf00925 -
09215 -pf00677 -
09220 -pf00383pf01872 -
09225 -pf00448pf02881 -
BMC-P
BMC-H
BMC-T
Aldehyde dehydrogenase
PTAC
Signature enzyme
Alcohol dehydrogenase
Regulator

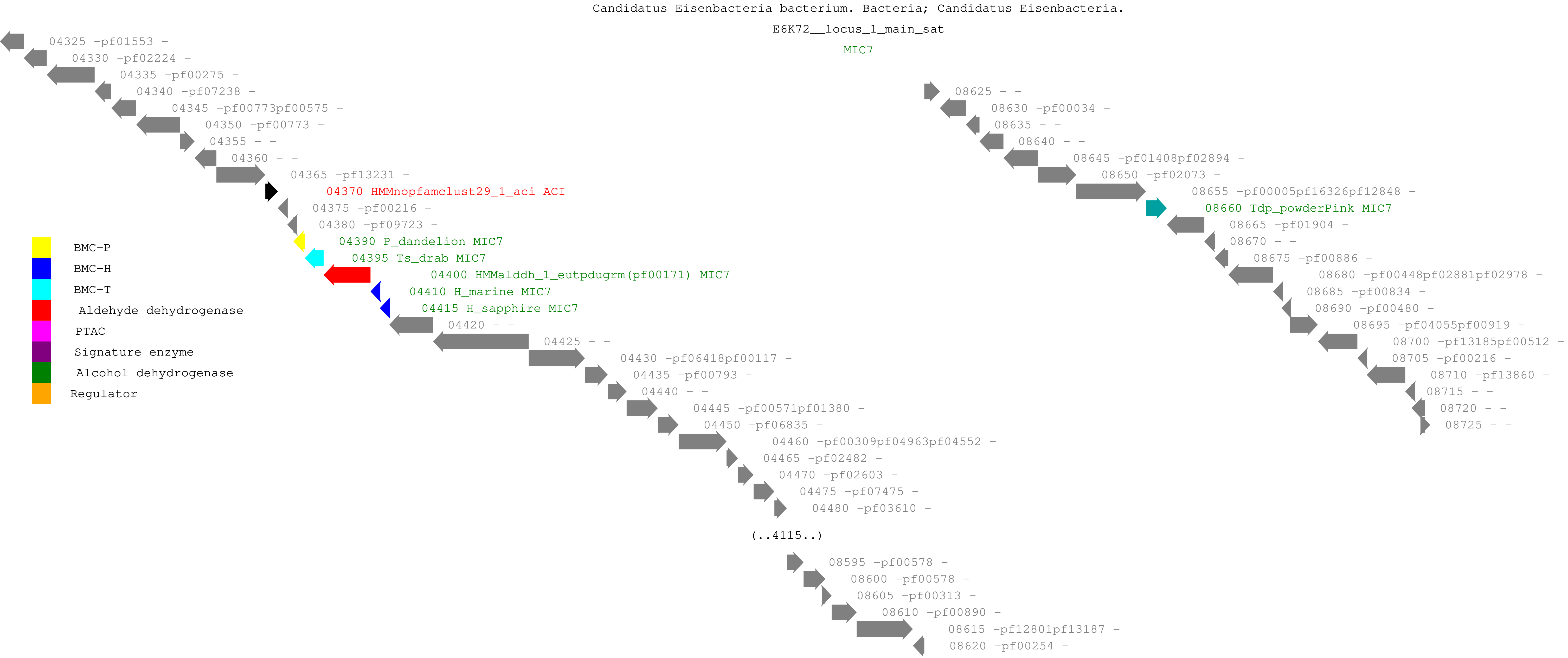
Candidatus Eisenbacteria bacterium. Bacteria; Candidatus Eisenbacteria.
E6K72__locus_1_main_sat
MIC7
04325 -pf01553 -
04330 -pf02224 -
04335 -pf00275 -
04340 -pf07238 -
04345 -pf00773pf00575 -
04350 -pf00773 -
04355 - -
04360 - -
04365 -pf13231 -
04370 HMMnopfamclust29_1_aci ACI
04375 -pf00216 -
04380 -pf09723 -
04390 P_dandelion MIC7
04395 Ts_drab MIC7
04400 HMMalddh_1_eutpdugrm(pf00171) MIC7
04410 H_marine MIC7
04415 H_sapphire MIC7
04420 - -
04425 - -
04430 -pf06418pf00117 -
04435 -pf00793 -
04440 - -
04445 -pf00571pf01380 -
04450 -pf06835 -
04460 -pf00309pf04963pf04552 -
04465 -pf02482 -
04470 -pf02603 -
04475 -pf07475 -
04480 -pf03610 -
(..4115..)
08595 -pf00578 -
08600 -pf00578 -
08605 -pf00313 -
08610 -pf00890 -
08615 -pf12801pf13187 -
08620 -pf00254 -
08625 - -
08630 -pf00034 -
08635 - -
08640 - -
08645 -pf01408pf02894 -
08650 -pf02073 -
08655 -pf00005pf16326pf12848 -
08660 Tdp_powderPink MIC7
08665 -pf01904 -
08670 - -
08675 -pf00886 -
08680 -pf00448pf02881pf02978 -
08685 -pf00834 -
08690 -pf00480 -
08695 -pf04055pf00919 -
08700 -pf13185pf00512 -
08705 -pf00216 -
08710 -pf13860 -
08715 - -
08720 - -
08725 - -
BMC-P
BMC-H
BMC-T
Aldehyde dehydrogenase
PTAC
Signature enzyme
Alcohol dehydrogenase
Regulator

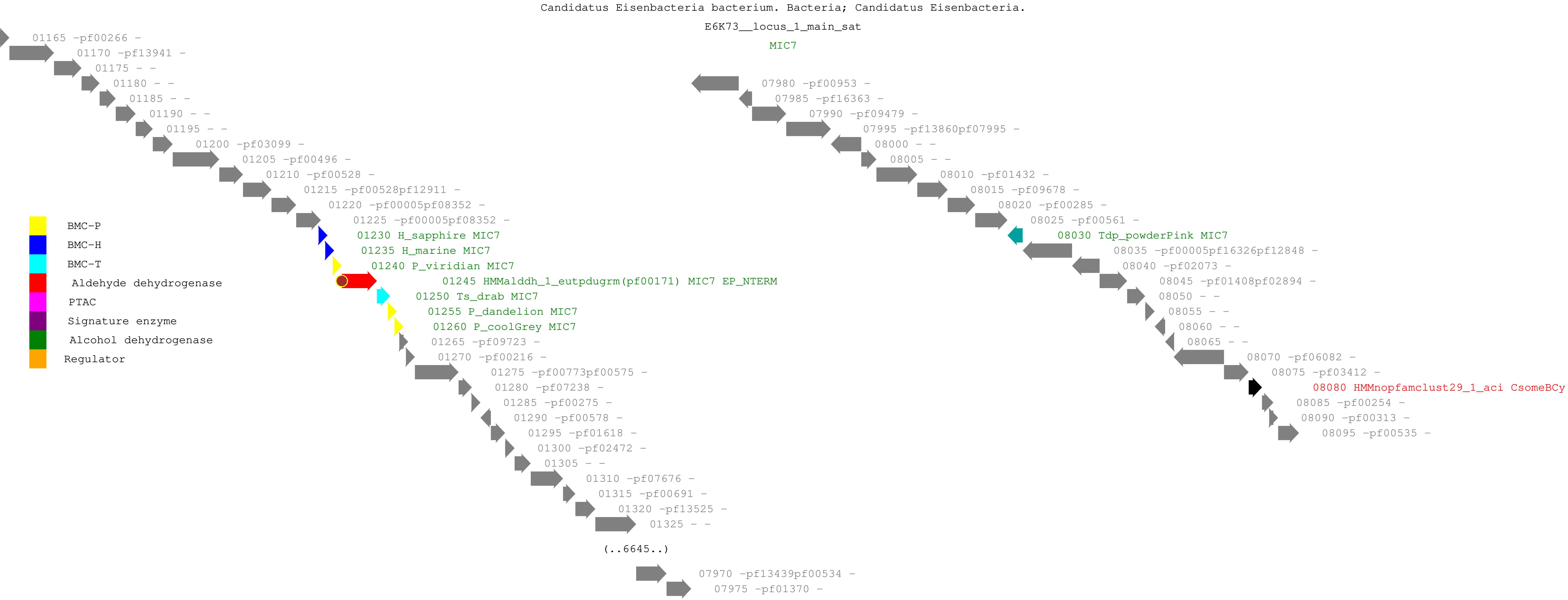

Candidatus Eisenbacteria bacterium. Bacteria; Candidatus Eisenbacteria.
E6K73__locus_1_main_sat
MIC7
01165 -pf00266 -
01170 -pf13941 -
01175 - -
01180 - -
01185 - -
01190 - -
01195 - -
01200 -pf03099 -
01205 -pf00496 -
01210 -pf00528 -
01215 -pf00528pf12911 -
01220 -pf00005pf08352 -
01225 -pf00005pf08352 -
01230 H_sapphire MIC7
01235 H_marine MIC7
01240 P_viridian MIC7
01245 HMMaldh_1_eutpdugrm(pf00171) MIC7 EP_NTERM
01250 Ts_drab MIC7
01255 P_dandelion MIC7
01260 P_coolGrey MIC7
01265 -pf09723 -
01270 -pf00216 -
01275 -pf00773pf00575 -
01280 -pf07238 -
01285 -pf00275 -
01290 -pf00578 -
01295 -pf01618 -
01300 -pf02472 -
01305 - -
01310 -pf07676 -
01315 -pf00691 -
01320 -pf13525 -
01325 - -
(..6645..)
07970 -pf13439pf00534 -
07975 -pf01370 -
07980 -pf00953 -
07985 -pf16363 -
07990 -pf09479 -
07995 -pf13860pf07995 -
08000 - -
08005 - -
08010 -pf01432 -
08015 -pf09678 -
08020 -pf00285 -
08025 -pf00561 -
08030 Tdp_powderPink MIC7
08035 -pf00005pf16326pf12848 -
08040 -pf02073 -
08045 -pf01408pf02894 -
08050 - -
08055 - -
08060 - -
08065 - -
08070 -pf06082 -
08075 -pf03412 -
08080 HMMnopfamclust29_1_aci CsomeBCy
08085 -pf00254 -
08090 -pf00313 -
08095 -pf00535 -
BMC-P
BMC-H
BMC-T
Aldehyde dehydrogenase
PTAC
Signature enzyme
Alcohol dehydrogenase
Regulator

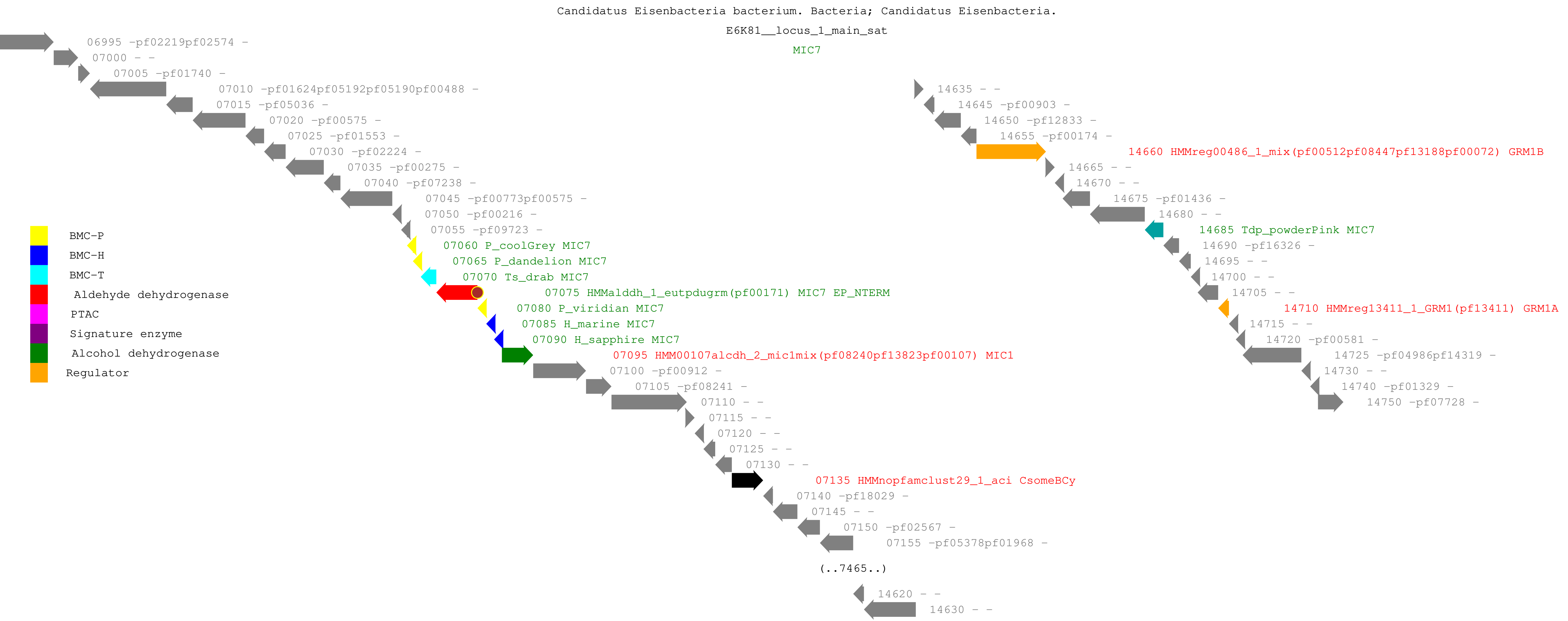
Candidatus Eisenbacteria bacterium. Bacteria; Candidatus Eisenbacteria.
E6K81__locus_1_main_sat
MIC7
06995 -pf02219pf02574 -
07000 - -
07005 -pf01740 -
07010 -pf01624pf05192pf05190pf00488 -
07015 -pf05036 -
07020 -pf00575 -
07025 -pf01553 -
07030 -pf02224 -
07035 -pf00275 -
07040 -pf07238 -
07045 -pf00773pf00575 -
07050 -pf00216 -
07055 -pf09723 -
07060 P_coolGrey MIC7
07065 P_dandelion MIC7
07070 Ts_drab MIC7
07075 HMMalddh_1_eutpdugrm(pf00171) MIC7 EP_NTERM
07080 P_viridian MIC7
07085 H_marine MIC7
07090 H_sapphire MIC7
07095 HMM00107alcdh_2_mic1mix(pf08240pf13823pf00107) MIC1
07100 -pf00912 -
07105 -pf08241 -
07110 - -
07115 - -
07120 - -
07125 - -
07130 - -
07135 HMMnopfamclust29_1_aci CsomeBCy
07140 -pf18029 -
07145 - -
07150 -pf02567 -
07155 -pf05378pf01968 -
(..7465..)
14620 - -
14630 - -
14635 - -
14645 -pf00903 -
14650 -pf12833 -
14655 -pf00174 -
14660 HMMreg00486_1_mix(pf00512pf08447pf13188pf00072) GRM1B
14665 - -
14670 - -
14675 -pf01436 -
14680 - -
14685 Tdp_powderPink MIC7
14690 -pf16326 -
14695 - -
14700 - -
14705 - -
14710 HMMreg13411_1_GRM1(pf13411) GRM1A
14715 - -
14720 -pf00581 -
14725 -pf04986pf14319 -
14730 - -
14740 -pf01329 -
14750 -pf07728 -
BMC-P
BMC-H
BMC-T
Aldehyde dehydrogenase
PTAC
Signature enzyme
Alcohol dehydrogenase
Regulator

hydrothermal vent metagenome. unclassified sequences; metagenomes; ecological metagenomes.

MNBD_NITROSPINAE01-_locus_1

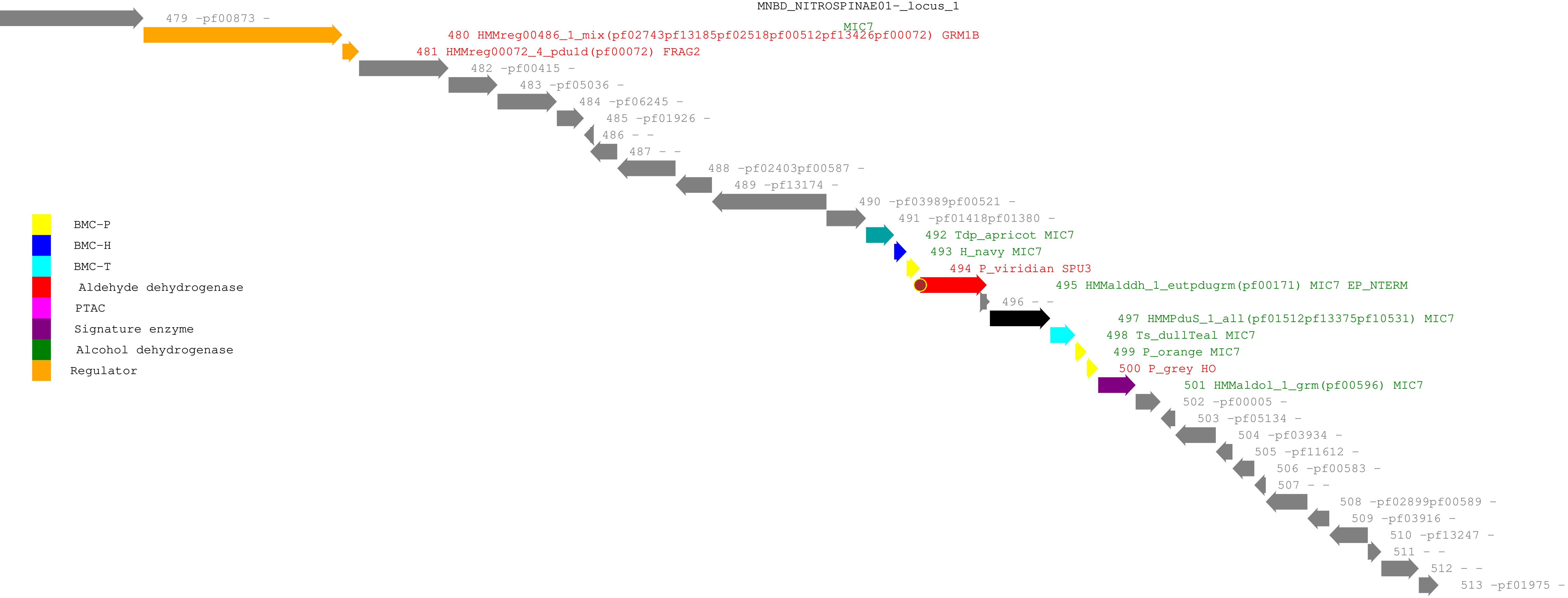

hydrothermal vent metagenome. unclassified sequences; metagenomes; ecological metagenomes.

MNBD_NITROSPINAE02-_locus_1

MIC7

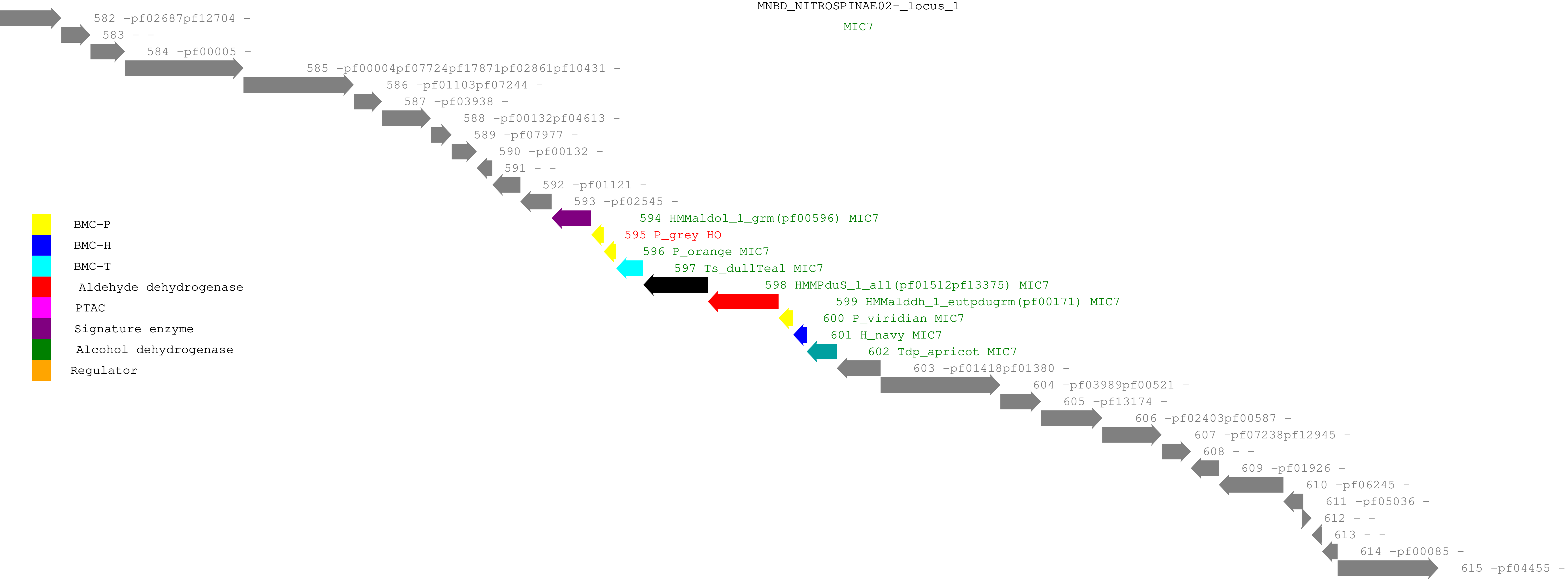

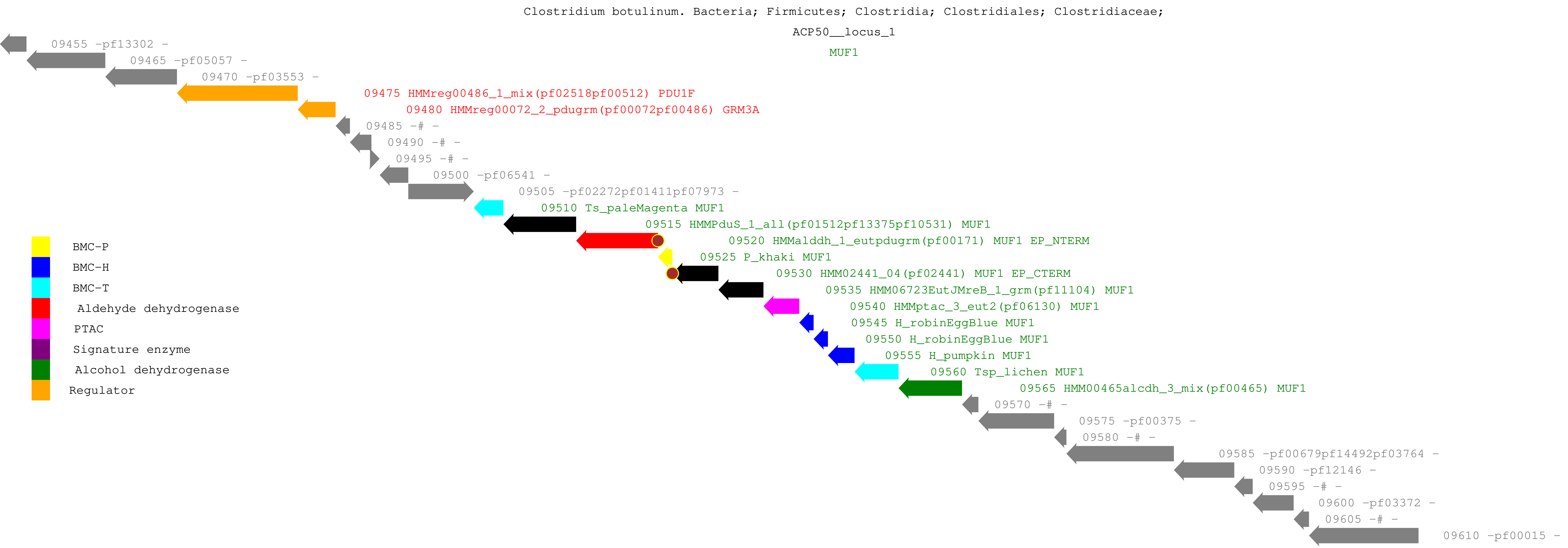

Clostridium botulinum. Bacteria; Firmicutes; Clostridia; Clostridiales; Clostridiaceae;
ACP50__locus_1
MUF1
09455 -pf13302 -
09465 -pf05057 -
09470 -pf03553 -
09475 HMMreg00486_1_mix(pf02518pf00512) PDU1F
09480 HMMreg00072_2_pdugrm(pf00072pf00486) GRM3A
09485 -# -
09490 -# -
09495 -# -
09500 -pf06541 -
09505 -pf02272pf01411pf07973 -
09510 Ts_paleMagenta MUF1
09515 HMMPduS_1_all(pf01512pf13375pf10531) MUF1
09520 HMMaldh_1_eutpdugrm(pf00171) MUF1 EP_NTERM
09525 P_khaki MUF1
09530 HMM02441_04(pf02441) MUF1 EP_CTERM
09535 HMM06723EutJMreB_1_grm(pf11104) MUF1
09540 HMMptac_3_eut2(pf06130) MUF1
09545 H_robinEggBlue MUF1
09550 H_robinEggBlue MUF1
09555 H_pumpkin MUF1
09560 Tsp_lichen MUF1
09565 HMM00465alcdh_3_mix(pf00465) MUF1
09570 -# -
09575 -pf00375 -
09580 -# -
09585 -pf00679pf14492pf03764 -
09590 -pf12146 -
09595 -# -
09600 -pf03372 -
09605 -# -
09610 -pf00015 -
BMC-P
BMC-H
BMC-T
Aldehyde dehydrogenase
PTAC
Signature enzyme
Alcohol dehydrogenase
Regulator

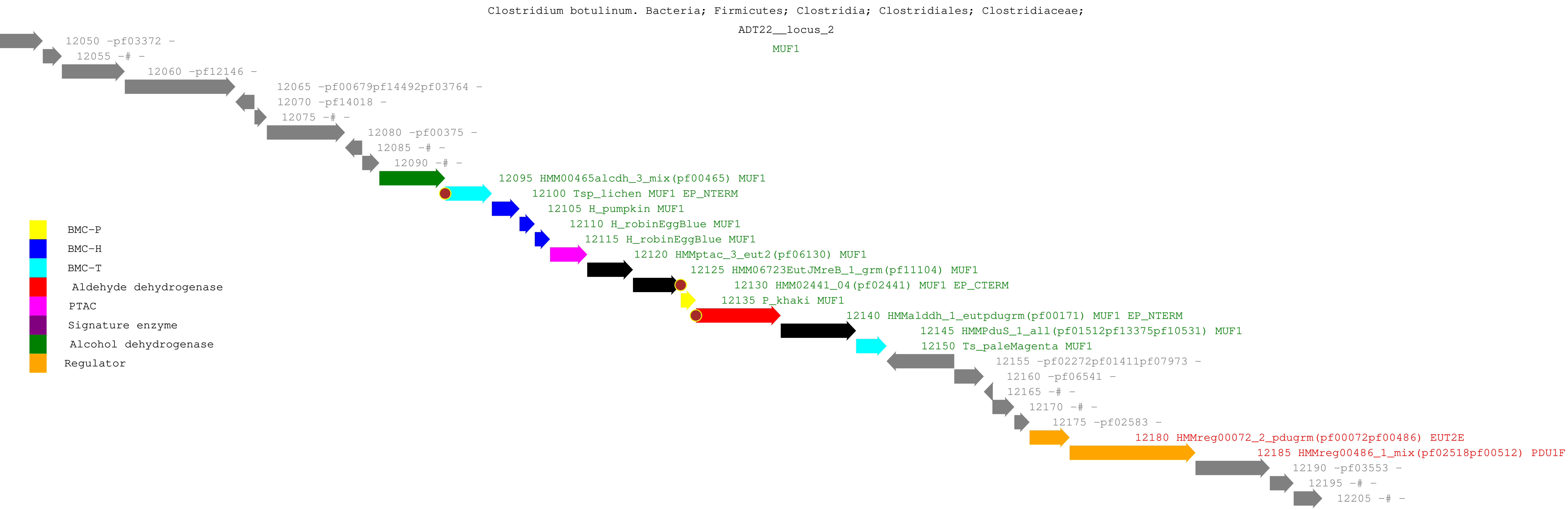

Clostridium botulinum. Bacteria; Firmicutes; Clostridia; Clostridiales; Clostridiaceae;
ADT22__locus_2
MUF1
12050 -pf03372 -
12055 -# -
12060 -pf12146 -
12065 -pf00679pf14492pf03764 -
12070 -pf14018 -
12075 -# -
12080 -pf00375 -
12085 -# -
12090 -# -
12095 HMM00465alcdh_3_mix(pf00465) MUF1
12100 Tsp_lichen MUF1 EP_NTERM
12105 H_pumpkin MUF1
12110 H_robinEggBlue MUF1
12115 H_robinEggBlue MUF1
12120 HMMptac_3_eut2(pf06130) MUF1
12125 HMM06723EutJMreB_1_grm(pf11104) MUF1
12130 HMM02441_04(pf02441) MUF1 EP_CTERM
12135 P_khaki MUF1
12140 HMMalddh_1_eutpdugrm(pf00171) MUF1 EP_NTERM
12145 HMMPduS_1_all(pf01512pf13375pf10531) MUF1
12150 Ts_paleMagenta MUF1
12155 -pf02272pf01411pf07973 -
12160 -pf06541 -
12165 -# -
12170 -# -
12175 -pf02583 -
12180 HMMreg00072_2_pdugrm(pf00072pf00486) EUT2E
12185 HMMreg00486_1_mix(pf02518pf00512) PDU1F
12190 -pf03553 -
12195 -# -
12205 -# -
BMC-P
BMC-H
BMC-T
Aldehyde dehydrogenase
PTAC
Signature enzyme
Alcohol dehydrogenase
Regulator

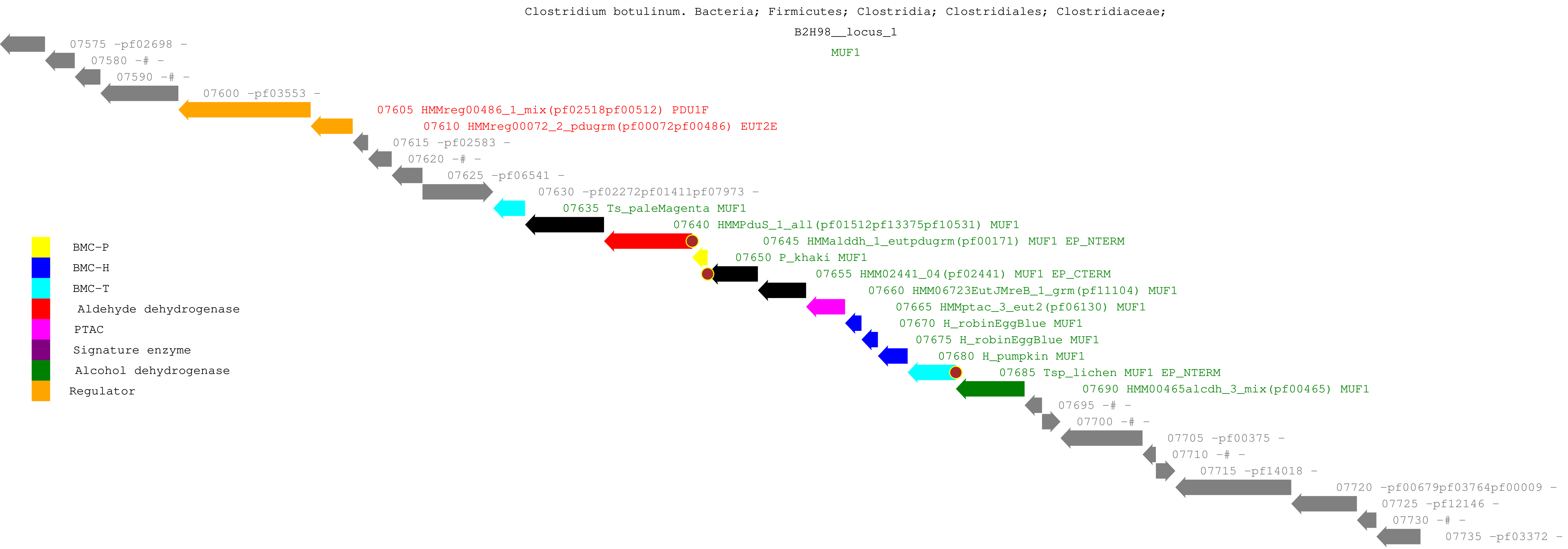

Clostridium botulinum. Bacteria; Firmicutes; Clostridia; Clostridiales; Clostridiaceae;
B2H98__locus_1
MUF1
07575 -pf02698 -
07580 -# -
07590 -# -
07600 -pf03553 -
07605 HMMreg00486_1_mix(pf02518pf00512) PDU1F
07610 HMMreg00072_2_pdugrm(pf00072pf00486) EUT2E
07615 -pf02583 -
07620 -# -
07625 -pf06541 -
07630 -pf02272pf01411pf07973 -
07635 Ts_paleMagenta MUF1
07640 HMMPduS_1_all(pf01512pf13375pf10531) MUF1
07645 HMMaldh_1_eutpdugrm(pf00171) MUF1 EP_NTERM
07650 P_khaki MUF1
07655 HMM02441_04(pf02441) MUF1 EP_CTERM
07660 HMM06723EutJMreB_1_grm(pf11104) MUF1
07665 HMMptac_3_eut2(pf06130) MUF1
07670 H_robinEggBlue MUF1
07675 H_robinEggBlue MUF1
07680 H_pumpkin MUF1
07685 Tsp_lichen MUF1 EP_NTERM
07690 HMM00465alcdh_3_mix(pf00465) MUF1
07695 -# -
07700 -# -
07705 -pf00375 -
07710 -# -
07715 -pf14018 -
07720 -pf00679pf03764pf00009 -
07725 -pf12146 -
07730 -# -
07735 -pf03372 -
BMC-P
BMC-H
BMC-T
Aldehyde dehydrogenase
PTAC
Signature enzyme
Alcohol dehydrogenase
Regulator

Clostridium taeniosporum. Bacteria; Firmicutes; Clostridia; Clostridiales; Clostridiaceae;

BGI42__locus_1

MUF1

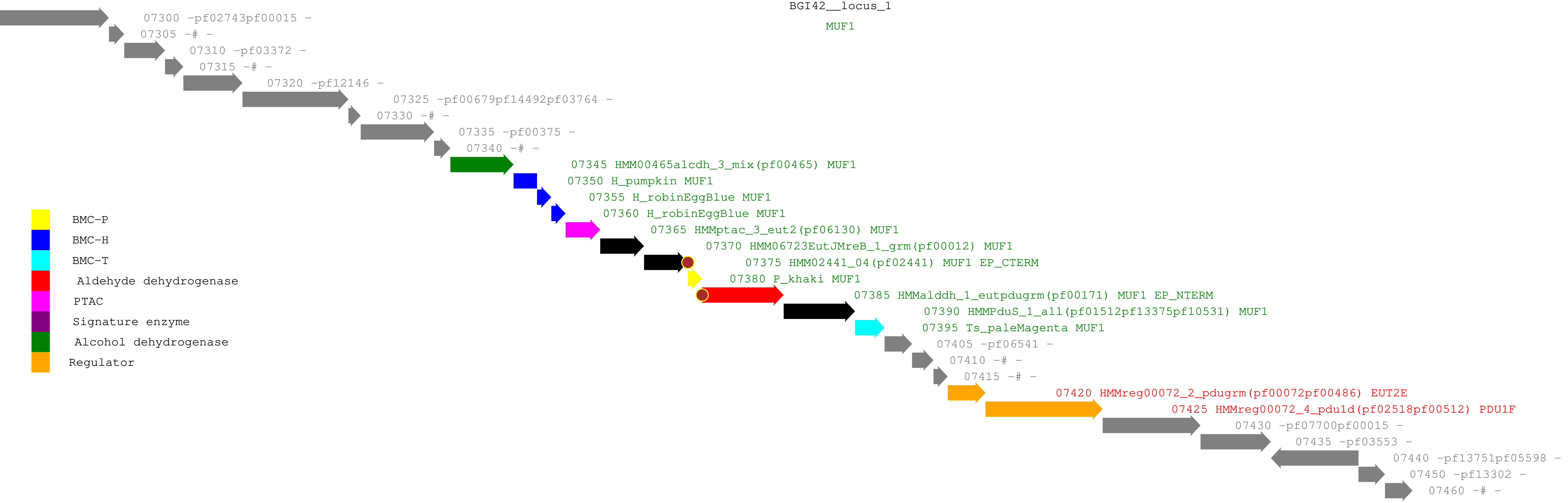

Clostridium botulinum (strain Eklund 17B / Type B). Bacteria; Firmicutes; Clostridia; Clostridiales; Clostridiaceae;

CLL_A_locus_2

MUF1

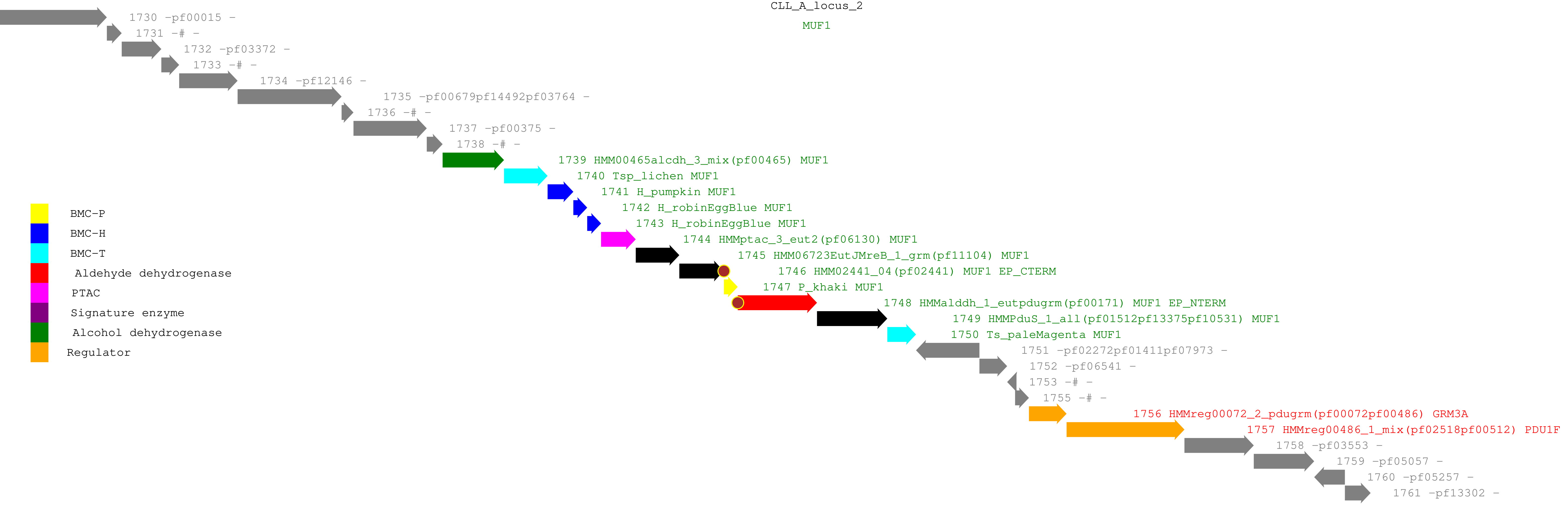

Clostridium botulinum E1 str. 'BoNT E Beluga'. Bacteria; Firmicutes; Clostridia; Clostridiales; Clostridiaceae;

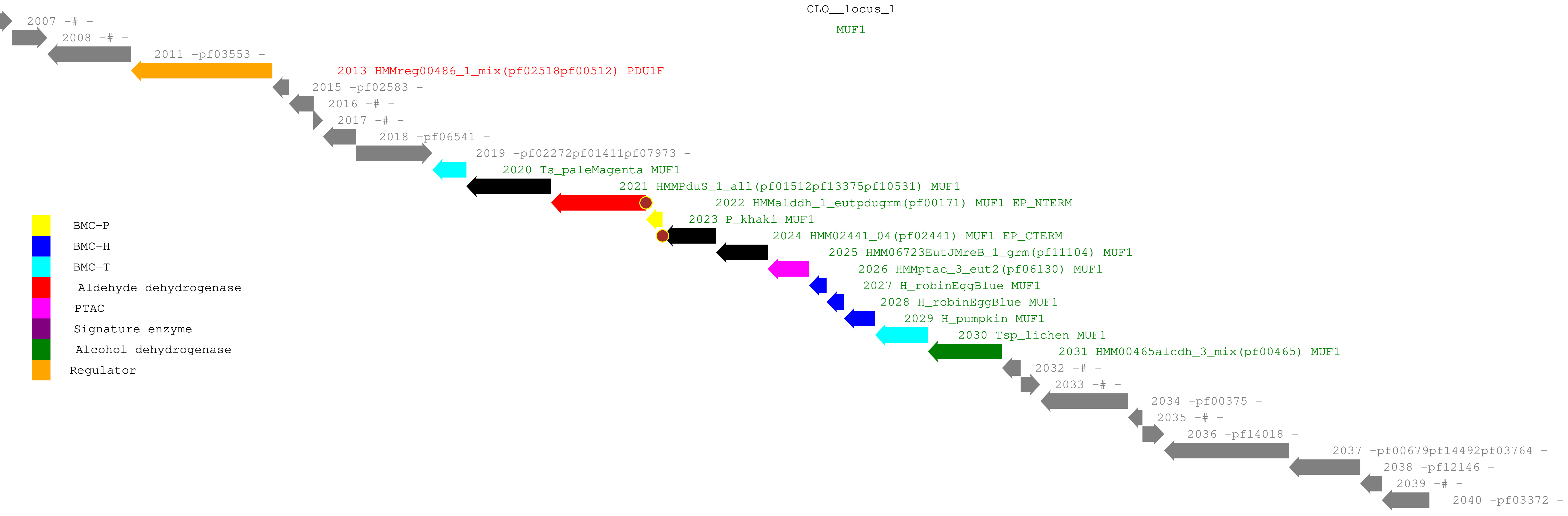

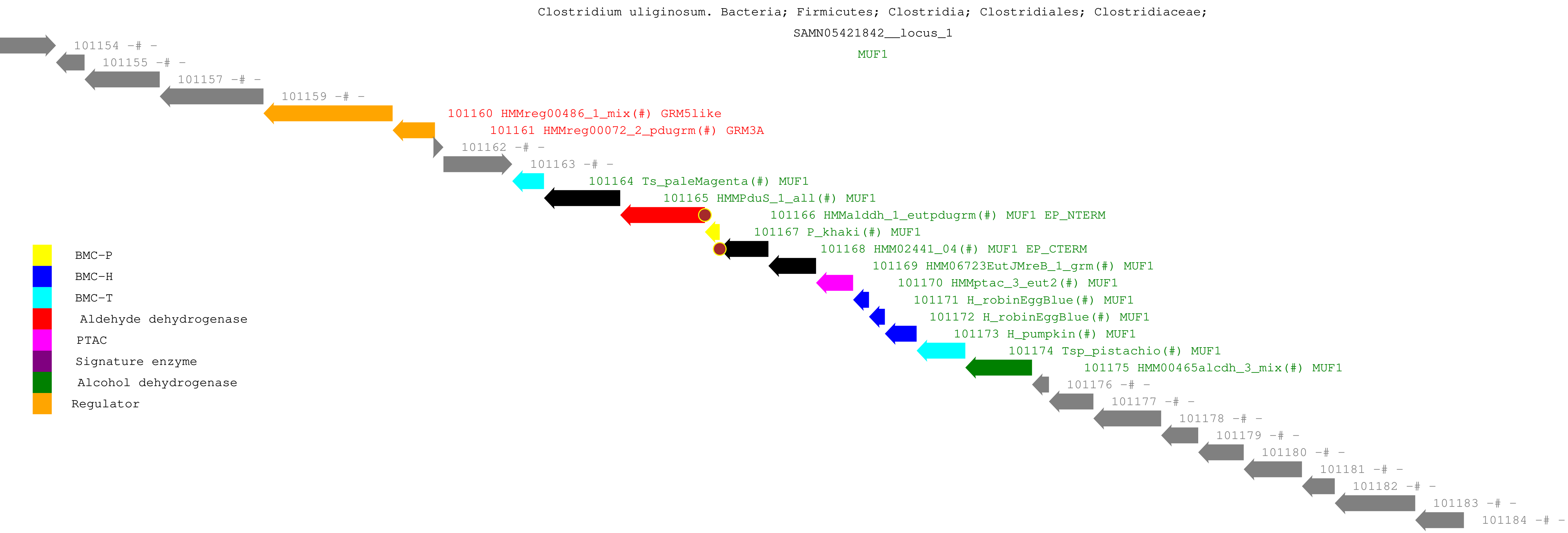
Clostridium uliginosum. Bacteria; Firmicutes; Clostridia; Clostridiales; Clostridiaceae;
SAMN05421842__locus_1
MUF1
101154 -# -
101155 -# -
101157 -# -
101159 -# -
101160 HMMreg00486_1_mix(#) GRM5like
101161 HMMreg00072_2_pdugrm(#) GRM3A
101162 -# -
101163 -# -
101164 Ts_paleMagenta(#) MUF1
101165 HMMPduS_1_all(#) MUF1
101166 HMMalddh_1_eutpdugrm(#) MUF1 EP_NTERM
101167 P_khaki(#) MUF1
101168 HMM02441_04(#) MUF1 EP_CTERM
101169 HMM06723EutJMreB_1_grm(#) MUF1
101170 HMMptac_3_eut2(#) MUF1
101171 H_robinEggBlue(#) MUF1
101172 H_robinEggBlue(#) MUF1
101173 H_pumpkin(#) MUF1
101174 Tsp_pistachio(#) MUF1
101175 HMM00465alcdh_3_mix(#) MUF1
101176 -# -
101177 -# -
101178 -# -
101179 -# -
101180 -# -
101181 -# -
101182 -# -
101183 -# -
101184 -# -
BMC-P
BMC-H
BMC-T
Aldehyde dehydrogenase
PTAC
Signature enzyme
Alcohol dehydrogenase
Regulator

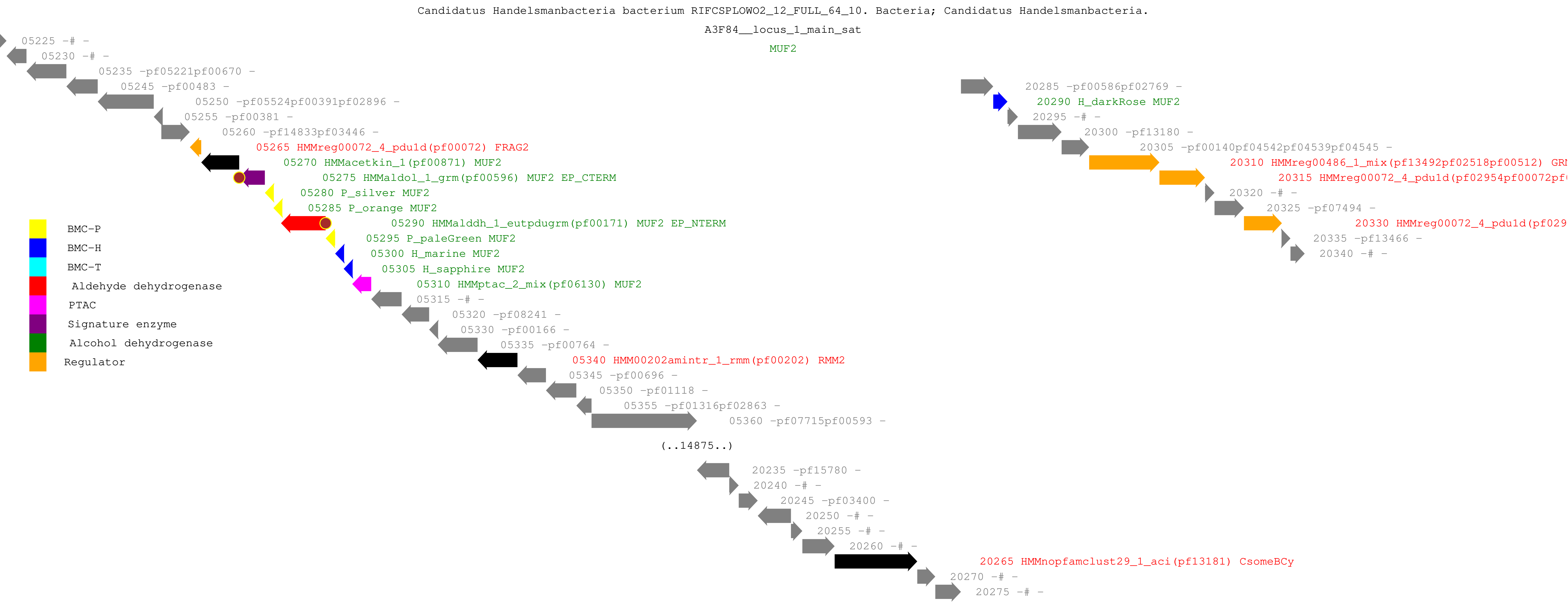
Candidatus Handelsmanbacteria bacterium RIFCSPLOWO2_12_FULL_64_10. Bacteria; Candidatus Handelsmanbacteria.
A3F84__locus_1_main_sat
MUF2
05225 -# -
05230 -# -
05235 -pf05221pf00670 -
05245 -pf00483 -
05250 -pf05524pf00391pf02896 -
05255 -pf00381 -
05260 -pf14833pf03446 -
05265 HMMreg00072_4_pdu1d(pf00072) FRAG2
05270 HMMacetkin_1(pf00871) MUF2
05275 HMMaldol_1_grm(pf00596) MUF2 EP_CTERM
05280 P_silver MUF2
05285 P_orange MUF2
05290 HMMalddh_1_eutpdugrm(pf00171) MUF2 EP_NTERM
05295 P_paleGreen MUF2
05300 H_marine MUF2
05305 H_sapphire MUF2
05310 HMMptac_2_mix(pf06130) MUF2
05315 -# -
05320 -pf08241 -
05330 -pf00166 -
05335 -pf00764 -
05340 HMM00202amintr_1_rmm(pf00202) RMM2
05345 -pf00696 -
05350 -pf01118 -
05355 -pf01316pf02863 -
05360 -pf07715pf00593 -
(..14875..)
20235 -pf15780 -
20240 -# -
20245 -pf03400 -
20250 -# -
20255 -# -
20260 -# -
20265 HMMnopfamclust29_1_aci(pf13181) CsomeBCy
20270 -# -
20275 -# -
20285 -pf00586pf02769 -
20290 H_darkRose MUF2
20295 -# -
20300 -pf13180 -
20305 -pf00140pf04542pf04539pf04545 -
20310 HMMreg00486_1_mix(pf13492pf02518pf00512) GR
20315 HMMreg00072_4_pdu1d(pf02954pf00072pf
20320 -# -
20325 -pf07494 -
20330 HMMreg00072_4_pdu1d(pf029
20335 -pf13466 -
20340 -# -
BMC-P
BMC-H
BMC-T
Aldehyde dehydrogenase
PTAC
Signature enzyme
Alcohol dehydrogenase
Regulator

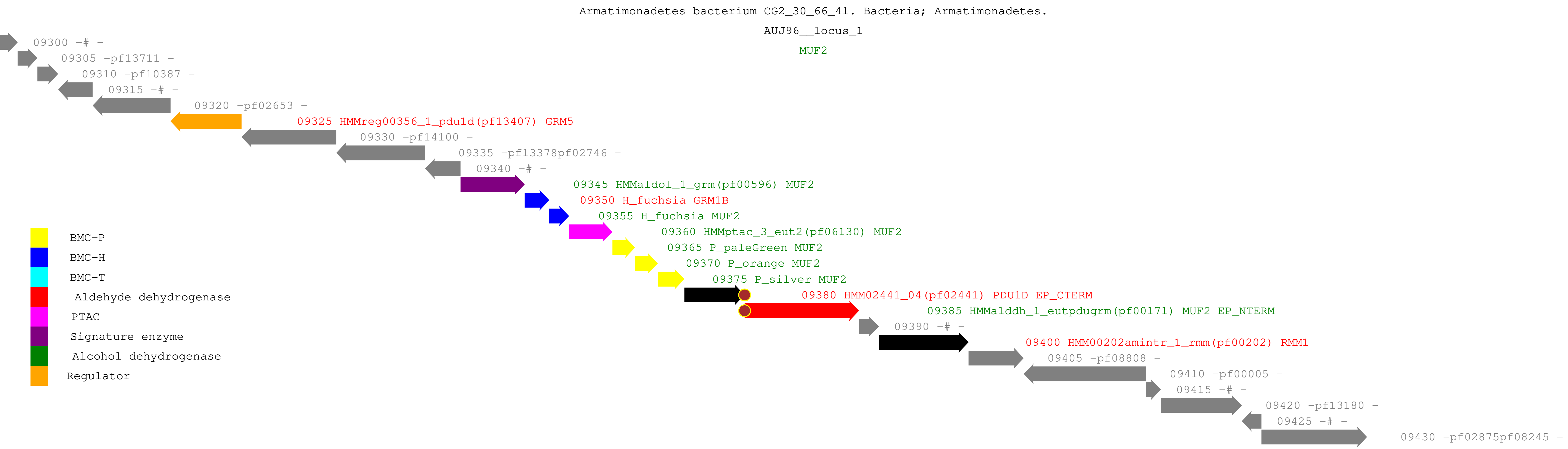
Armatimonadetes bacterium CG2_30_66_41. Bacteria; Armatimonadetes.
AUJ96__locus_1
MUF2
09300 -# -
09305 -pf13711 -
09310 -pf10387 -
09315 -# -
09320 -pf02653 -
09325 HMMreg00356_1_pdu1d(pf13407) GRM5
09330 -pf14100 -
09335 -pf13378pf02746 -
09340 -# -
09345 HMMaldol_1_grm(pf00596) MUF2
09350 H_fuchsia GRM1B
09355 H_fuchsia MUF2
09360 HMMptac_3_eut2(pf06130) MUF2
09365 P_paleGreen MUF2
09370 P_orange MUF2
09375 P_silver MUF2
09380 HMM02441_04(pf02441) PDU1D EP_CTERM
09385 HMMalddh_1_eutpdugrm(pf00171) MUF2 EP_NTERM
09390 -# -
09400 HMM00202amintr_1_rmm(pf00202) RMM1
09405 -pf08808 -
09410 -pf00005 -
09415 -# -
09420 -pf13180 -
09425 -# -
09430 -pf02875pf08245 -
BMC-P
BMC-H
BMC-T
Aldehyde dehydrogenase
PTAC
Signature enzyme
Alcohol dehydrogenase
Regulator

Gemmatimonadaceae bacterium. Bacteria; Gemmatimonadetes; Gemmatimonadales; Gemmatimonadaceae;

CME04__locus_1_main_sat

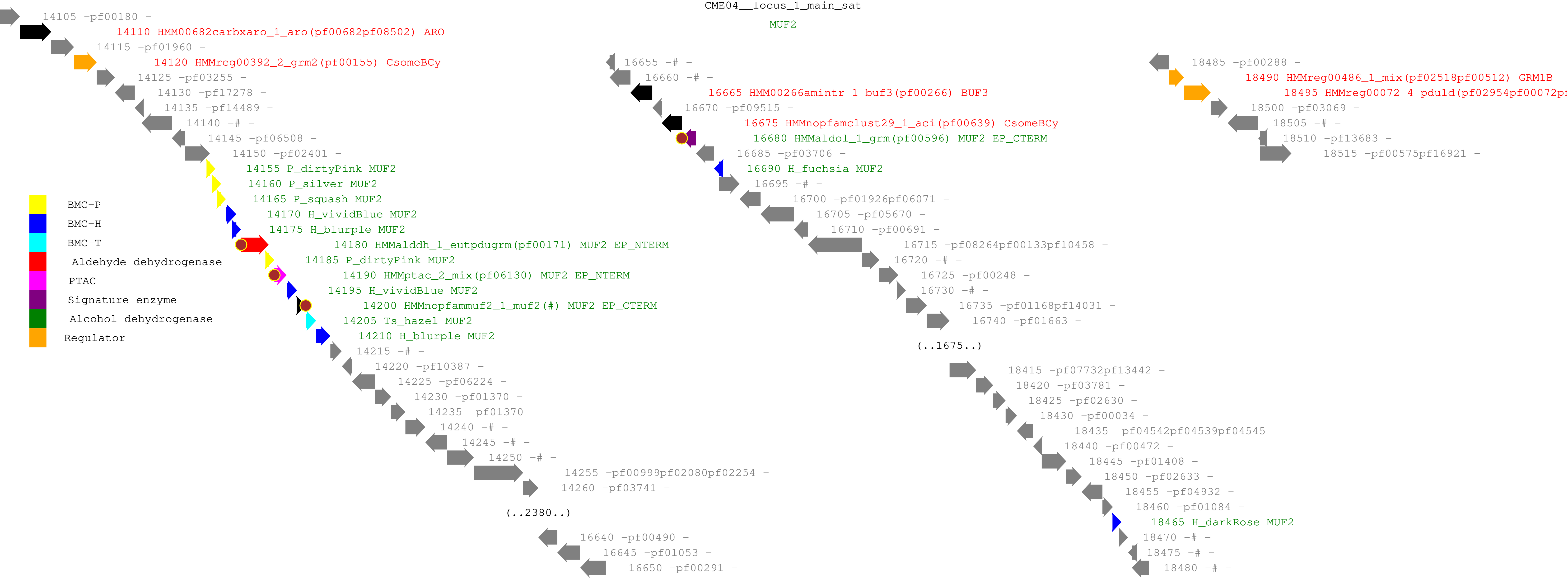

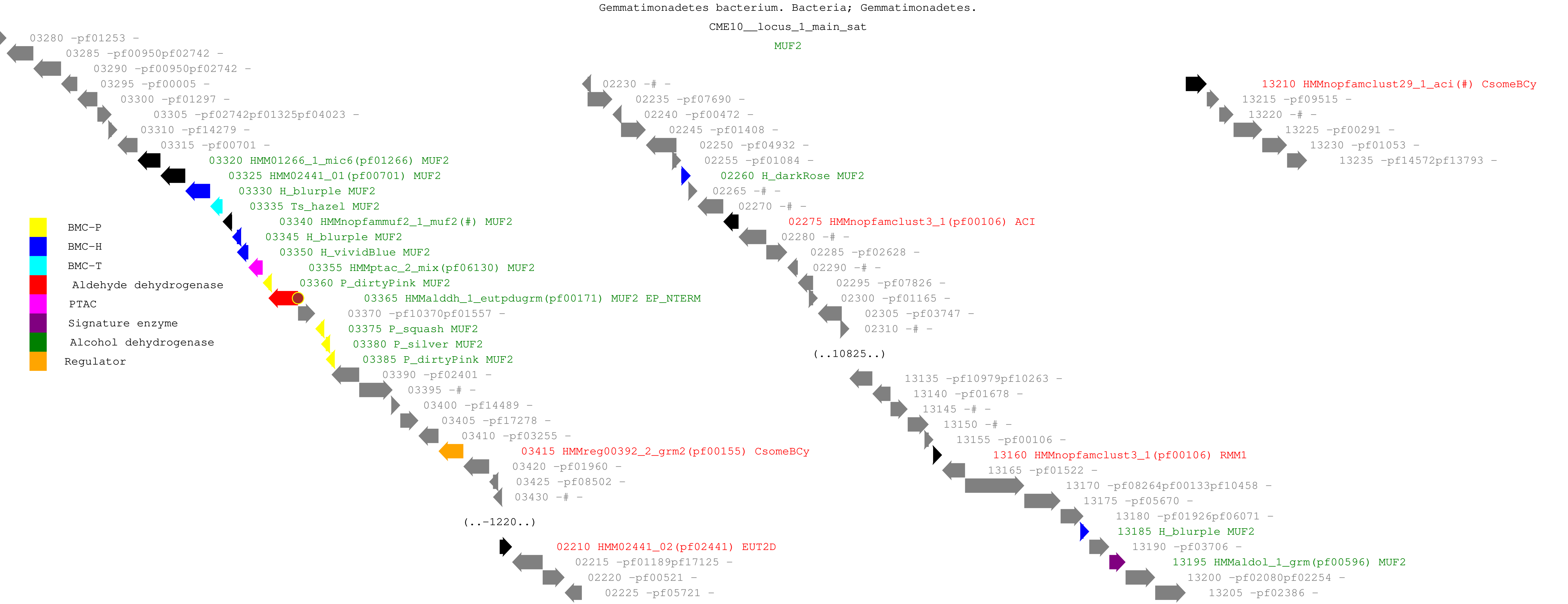
Gemmatimonadetes bacterium. Bacteria; Gemmatimonadetes.
CME10__locus_1_main_sat
MUF2
03280 -pf01253 -
03285 -pf00950pf02742 -
03290 -pf00950pf02742 -
03295 -pf00005 -
03300 -pf01297 -
03305 -pf02742pf01325pf04023 -
03310 -pf14279 -
03315 -pf00701 -
03320 HMM01266_1_mic6(pf01266) MUF2
03325 HMM02441_01(pf00701) MUF2
03330 H_blurple MUF2
03335 Ts_hazel MUF2
03340 HMMnopfammuf2_1_muf2(#) MUF2
03345 H_blurple MUF2
03350 H_vividBlue MUF2
03355 HMMptac_2_mix(pf06130) MUF2
03360 P_dirtyPink MUF2
03365 HMMalddh_1_eutpdugrm(pf00171) MUF2 EP_NTERM
03370 -pf10370pf01557 -
03375 P_squash MUF2
03380 P_silver MUF2
03385 P_dirtyPink MUF2
03390 -pf02401 -
03395 -# -
03400 -pf14489 -
03405 -pf17278 -
03410 -pf03255 -
03415 HMMreg00392_2_grm2(pf00155) CsomeBCy
03420 -pf01960 -
03425 -pf08502 -
03430 -# -
(..-1220..)
02210 HMM02441_02(pf02441) EUT2D
02215 -pf01189pf17125 -
02220 -pf00521 -
02225 -pf05721 -
BMC-P
BMC-H
BMC-T
Aldehyde dehydrogenase
PTAC
Signature enzyme
Alcohol dehydrogenase
Regulator
02230 -# -
02235 -pf07690 -
02240 -pf00472 -
02245 -pf01408 -
02250 -pf04932 -
02255 -pf01084 -
02260 H_darkRose MUF2
02265 -# -
02270 -# -
02275 HMMnopfamclust3_1(pf00106) ACI
02280 -# -
02285 -pf02628 -
02290 -# -
02295 -pf07826 -
02300 -pf01165 -
02305 -pf03747 -
02310 -# -
(..10825..)
13135 -pf10979pf10263 -
13140 -pf01678 -
13145 -# -
13150 -# -
13155 -pf00106 -
13160 HMMnopfamclust3_1(pf00106) RMM1
13165 -pf01522 -
13170 -pf08264pf00133pf10458 -
13175 -pf05670 -
13180 -pf01926pf06071 -
13185 H_blurple MUF2
13190 -pf03706 -
13195 HMMaldol_1_grm(pf00596) MUF2
13200 -pf02080pf02254 -
13205 -pf02386 -
13210 HMMnopfamclust29_1_aci(#) CsomeBCy
13215 -pf09515 -
13220 -# -
13225 -pf00291 -
13230 -pf01053 -
13235 -pf14572pf13793 -

Gemmatimonadetes bacterium. Bacteria; Gemmatimonadetes.

CME13__locus_1_main_sat

MUF2

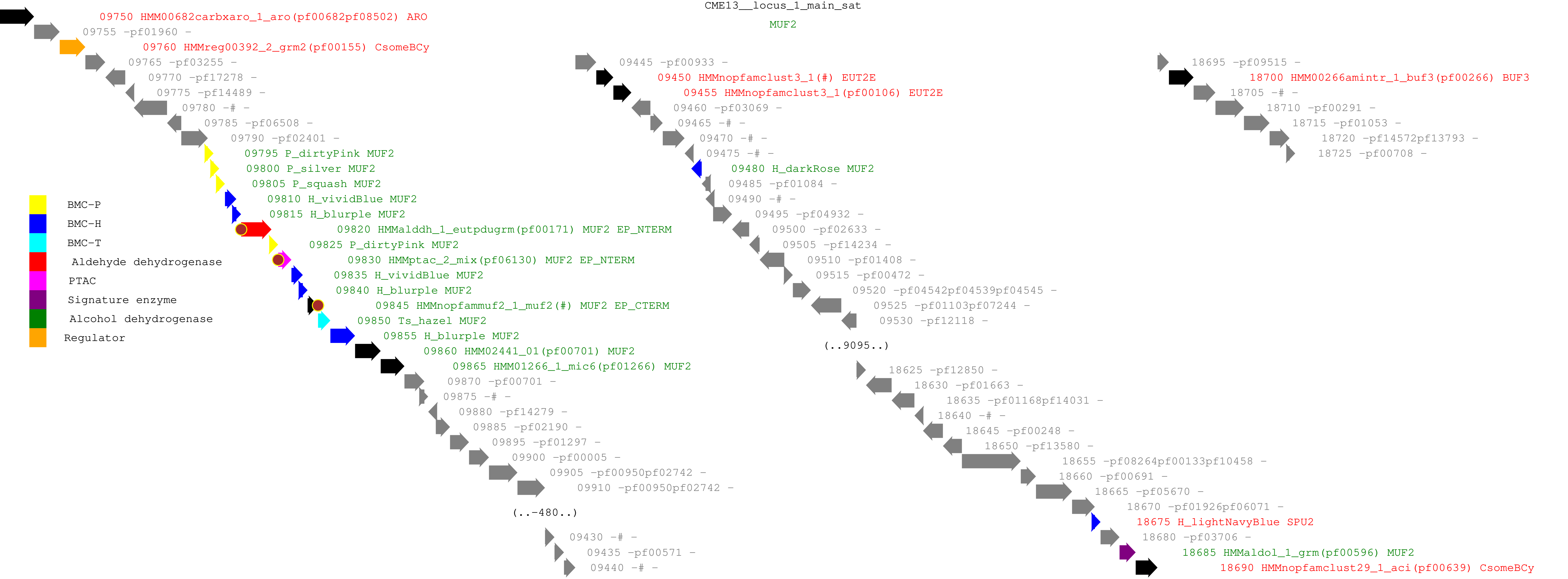

Gemmatimonadetes bacterium. Bacteria; Gemmatimonadetes.

CME19__locus_1_main_sat

MUF2

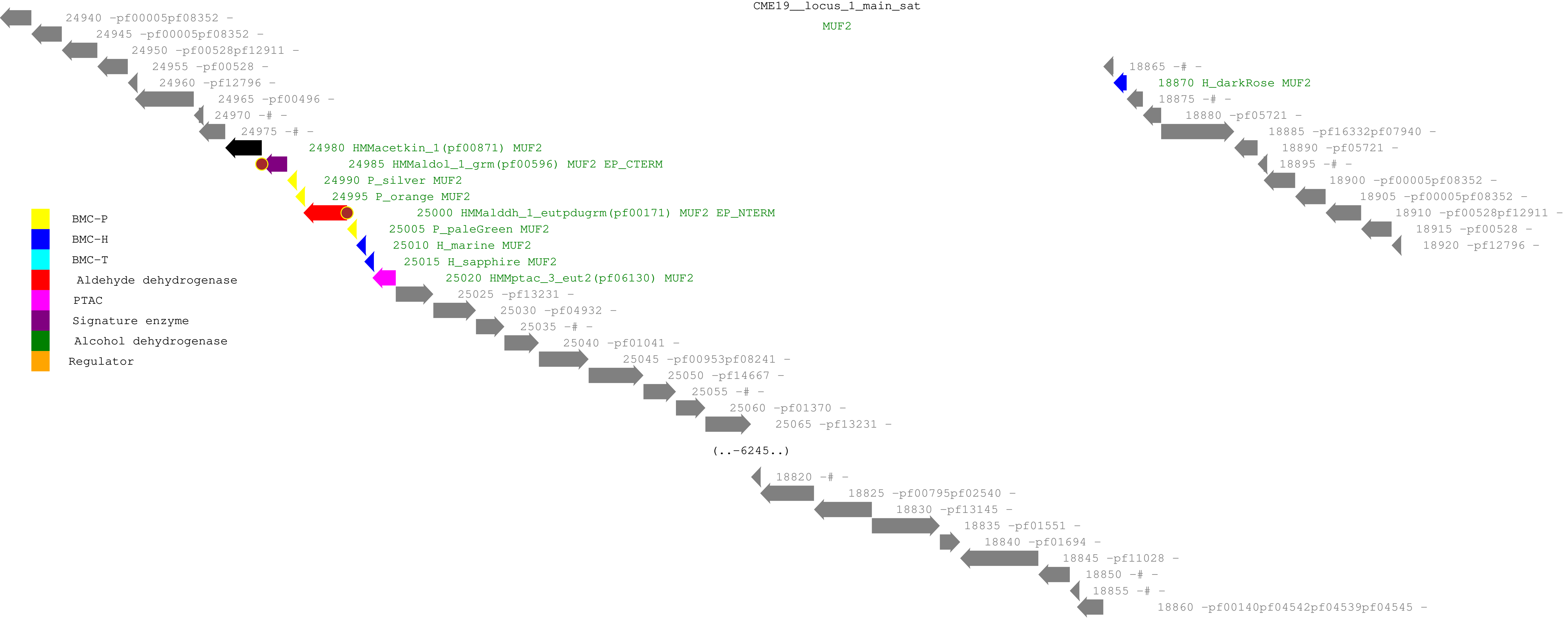

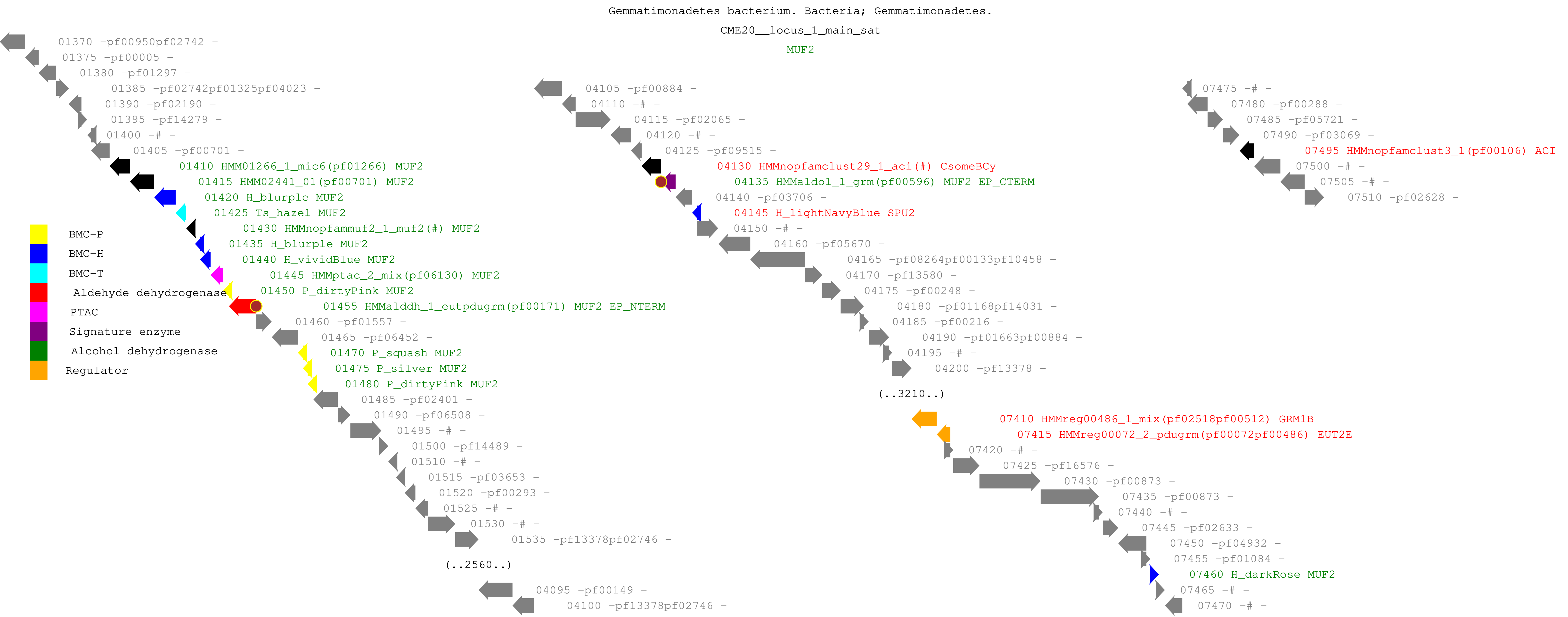
Gemmatimonadetes bacterium. Bacteria; Gemmatimonadetes.
CME20__locus_1_main_sat
MUF2
01370 -pf00950pf02742 -
01375 -pf00005 -
01380 -pf01297 -
01385 -pf02742pf01325pf04023 -
01390 -pf02190 -
01395 -pf14279 -
01400 -# -
01405 -pf00701 -
01410 HMM01266_1_mic6(pf01266) MUF2
01415 HMM02441_01(pf00701) MUF2
01420 H_blurple MUF2
01425 Ts_hazel MUF2
01430 HMMnopfammuf2_1_muf2(#) MUF2
01435 H_blurple MUF2
01440 H_vividBlue MUF2
01445 HMMptac_2_mix(pf06130) MUF2
01450 P_dirtyPink MUF2
01455 HMMalddh_1_eutpdugrm(pf00171) MUF2 EP_NTERM
01460 -pf01557 -
01465 -pf06452 -
01470 P_squash MUF2
01475 P_silver MUF2
01480 P_dirtyPink MUF2
01485 -pf02401 -
01490 -pf06508 -
01495 -# -
01500 -pf14489 -
01510 -# -
01515 -pf03653 -
01520 -pf00293 -
01525 -# -
01530 -# -
01535 -pf13378pf02746 -
(..2560..)
04095 -pf00149 -
04100 -pf13378pf02746 -
BMC-P
BMC-H
BMC-T
Aldehyde dehydrogenase
PTAC
Signature enzyme
Alcohol dehydrogenase
Regulator
04105 -pf00884 -
04110 -# -
04115 -pf02065 -
04120 -# -
04125 -pf09515 -
04130 HMMnopfamclust29_1_aci(#) CsomeBCy
04135 HMMaldol_1_grm(pf00596) MUF2 EP_CTERM
04140 -pf03706 -
04145 H_lightNavyBlue SPU2
04150 -# -
04160 -pf05670 -
04165 -pf08264pf00133pf10458 -
04170 -pf13580 -
04175 -pf00248 -
04180 -pf01168pf14031 -
04185 -pf00216 -
04190 -pf01663pf00884 -
04195 -# -
04200 -pf13378 -
(..3210..)
07410 HMMreg00486_1_mix(pf02518pf00512) GRM1B
07415 HMMreg00072_2_pdugrm(pf00072pf00486) EUT2E
07420 -# -
07425 -pf16576 -
07430 -pf00873 -
07435 -pf00873 -
07440 -# -
07445 -pf02633 -
07450 -pf04932 -
07455 -pf01084 -
07460 H_darkRose MUF2
07465 -# -
07470 -# -
07475 -# -
07480 -pf00288 -
07485 -pf05721 -
07490 -pf03069 -
07495 HMMnopfamclust3_1(pf00106) ACI
07500 -# -
07505 -# -
07510 -pf02628 -

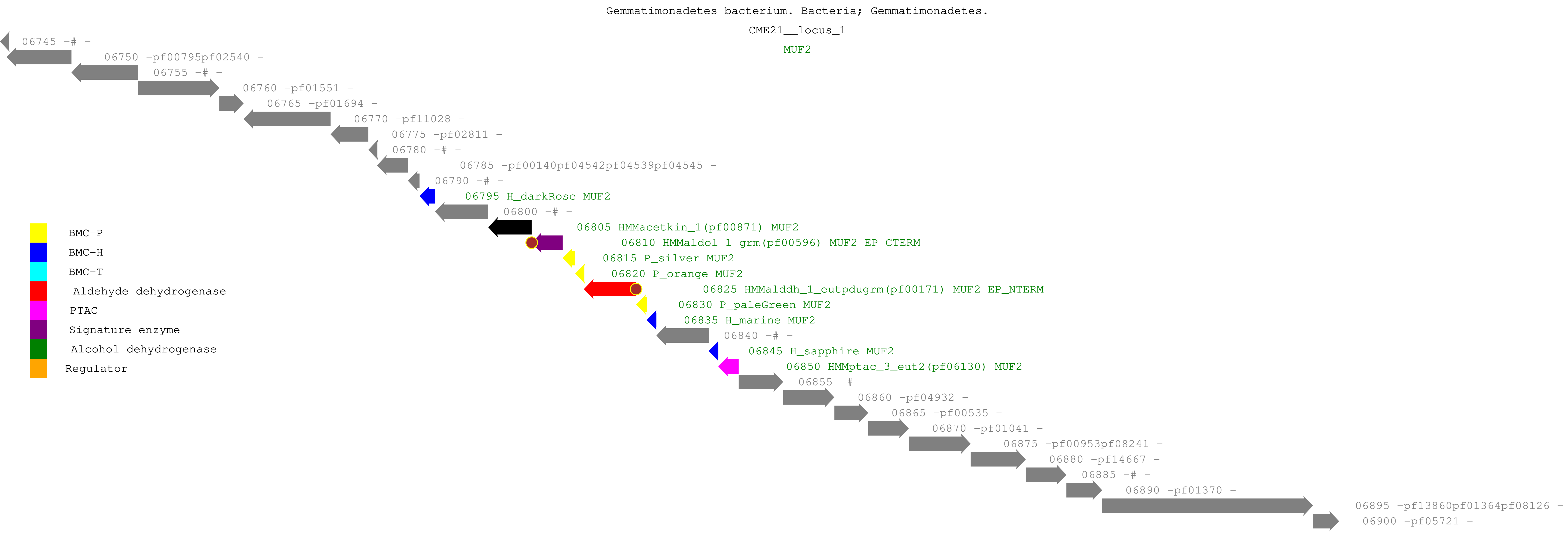
Gemmatimonadetes bacterium. Bacteria; Gemmatimonadetes.
CME21__locus_1
MUF2
06745 -# -
06750 -pf00795pf02540 -
06755 -# -
06760 -pf01551 -
06765 -pf01694 -
06770 -pf11028 -
06775 -pf02811 -
06780 -# -
06785 -pf00140pf04542pf04539pf04545 -
06790 -# -
06795 H_darkRose MUF2
06800 -# -
06805 HMMacetkin_1(pf00871) MUF2
06810 HMMaldol_1_grm(pf00596) MUF2 EP_CTERM
06815 P_silver MUF2
06820 P_orange MUF2
06825 HMMalddh_1_eutpdugrm(pf00171) MUF2 EP_NTERM
06830 P_paleGreen MUF2
06835 H_marine MUF2
06840 -# -
06845 H_sapphire MUF2
06850 HMMptac_3_eut2(pf06130) MUF2
06855 -# -
06860 -pf04932 -
06865 -pf00535 -
06870 -pf01041 -
06875 -pf00953pf08241 -
06880 -pf14667 -
06885 -# -
06890 -pf01370 -
06895 -pf13860pf01364pf08126 -
06900 -pf05721 -
BMC-P
BMC-H
BMC-T
Aldehyde dehydrogenase
PTAC
Signature enzyme
Alcohol dehydrogenase
Regulator

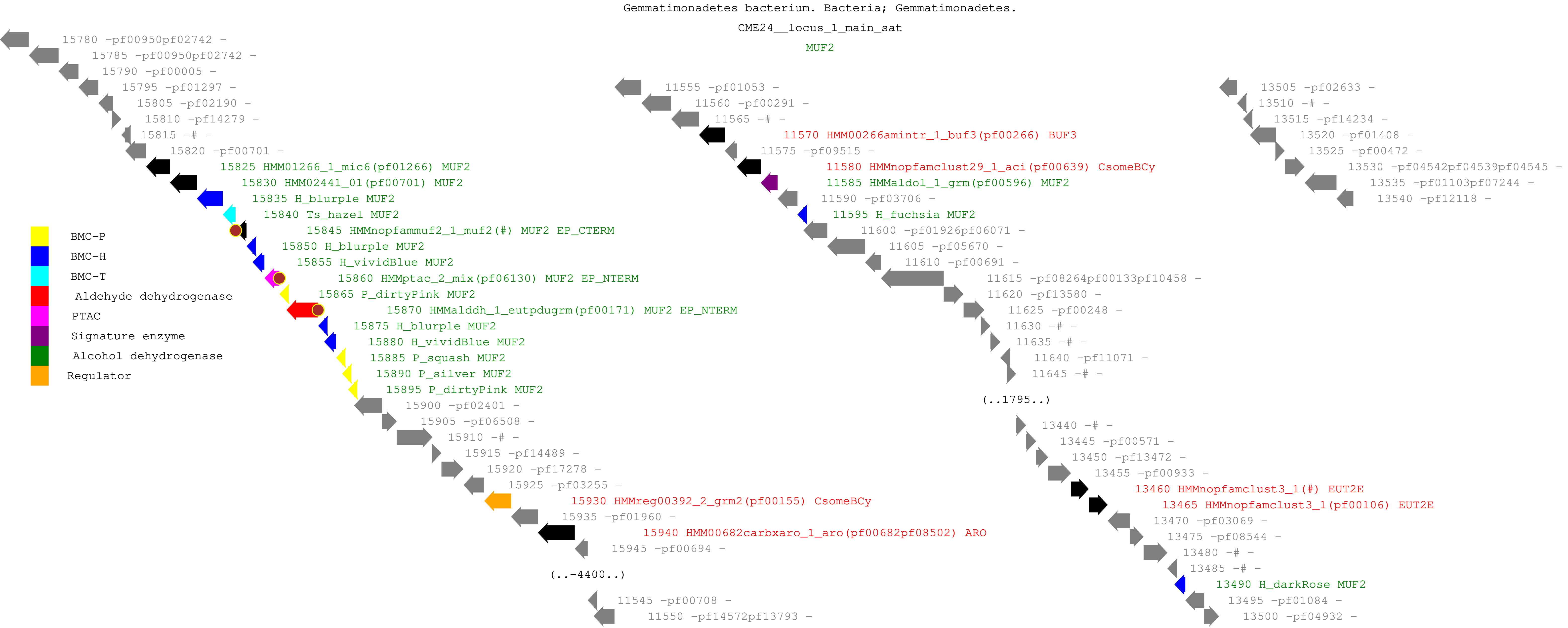
Gemmatimonadetes bacterium. Bacteria; Gemmatimonadetes.
CME24__locus_1_main_sat
MUF2
15780 -pf00950pf02742 -
15785 -pf00950pf02742 -
15790 -pf00005 -
15795 -pf01297 -
15805 -pf02190 -
15810 -pf14279 -
15815 -# -
15820 -pf00701 -
15825 HMM01266_1_mic6(pf01266) MUF2
15830 HMM02441_01(pf00701) MUF2
15835 H_blurple MUF2
15840 Ts_hazel MUF2
15845 HMMnopfammuf2_1_muf2(#) MUF2 EP_CTERM
15850 H_blurple MUF2
15855 H_vividBlue MUF2
15860 HMMptac_2_mix(pf06130) MUF2 EP_NTERM
15865 P_dirtyPink MUF2
15870 HMMalddh_1_eutpdugrm(pf00171) MUF2 EP_NTERM
15875 H_blurple MUF2
15880 H_vividBlue MUF2
15885 P_squash MUF2
15890 P_silver MUF2
15895 P_dirtyPink MUF2
15900 -pf02401 -
15905 -pf06508 -
15910 -# -
15915 -pf14489 -
15920 -pf17278 -
15925 -pf03255 -
15930 HMMreg00392_2_grm2(pf00155) CsomeBCy
15935 -pf01960 -
15940 HMM00682carbxaro_1_aro(pf00682pf08502) ARO
15945 -pf00694 -
(..-4400..)
11545 -pf00708 -
11550 -pf14572pf13793 -
11555 -pf01053 -
11560 -pf00291 -
11565 -# -
11570 HMM00266amintr_1_buf3(pf00266) BUF3
11575 -pf09515 -
11580 HMMnopfamclust29_1_aci(pf00639) CsomeBCy
11585 HMMaldol_1_grm(pf00596) MUF2
11590 -pf03706 -
11595 H_fuchsia MUF2
11600 -pf01926pf06071 -
11605 -pf05670 -
11610 -pf00691 -
11615 -pf08264pf00133pf10458 -
11620 -pf13580 -
11625 -pf00248 -
11630 -# -
11635 -# -
11640 -pf11071 -
11645 -# -
(..1795..)
13440 -# -
13445 -pf00571 -
13450 -pf13472 -
13455 -pf00933 -
13460 HMMnopfamclust3_1(#) EUT2E
13465 HMMnopfamclust3_1(pf00106) EUT2E
13470 -pf03069 -
13475 -pf08544 -
13480 -# -
13485 -# -
13490 H_darkRose MUF2
13495 -pf01084 -
13500 -pf04932 -
13505 -pf02633 -
13510 -# -
13515 -pf14234 -
13520 -pf01408 -
13525 -pf00472 -
13530 -pf04542pf04539pf04545 -
13535 -pf01103pf07244 -
13540 -pf12118 -
BMC-P
BMC-H
BMC-T
Aldehyde dehydrogenase
PTAC
Signature enzyme
Alcohol dehydrogenase
Regulator

Gemmatimonadetes bacterium. Bacteria; Gemmatimonadetes.

CME25__locus_1_main_sat

MUF2

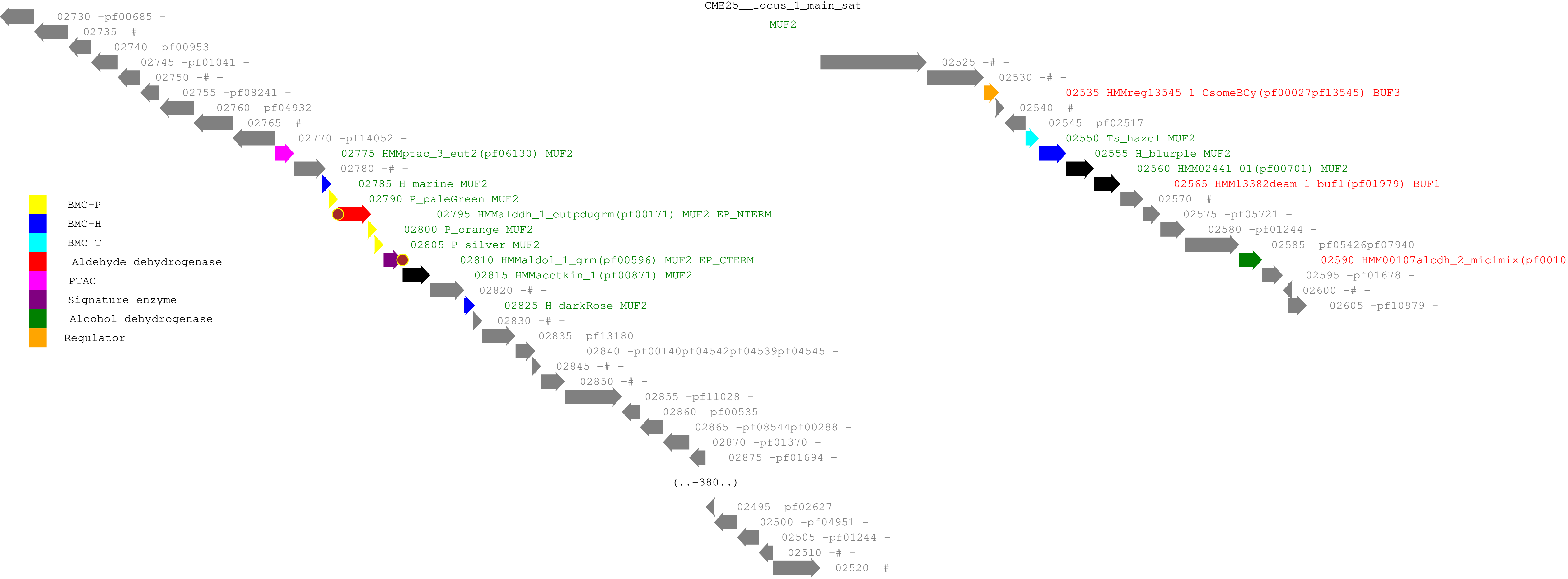

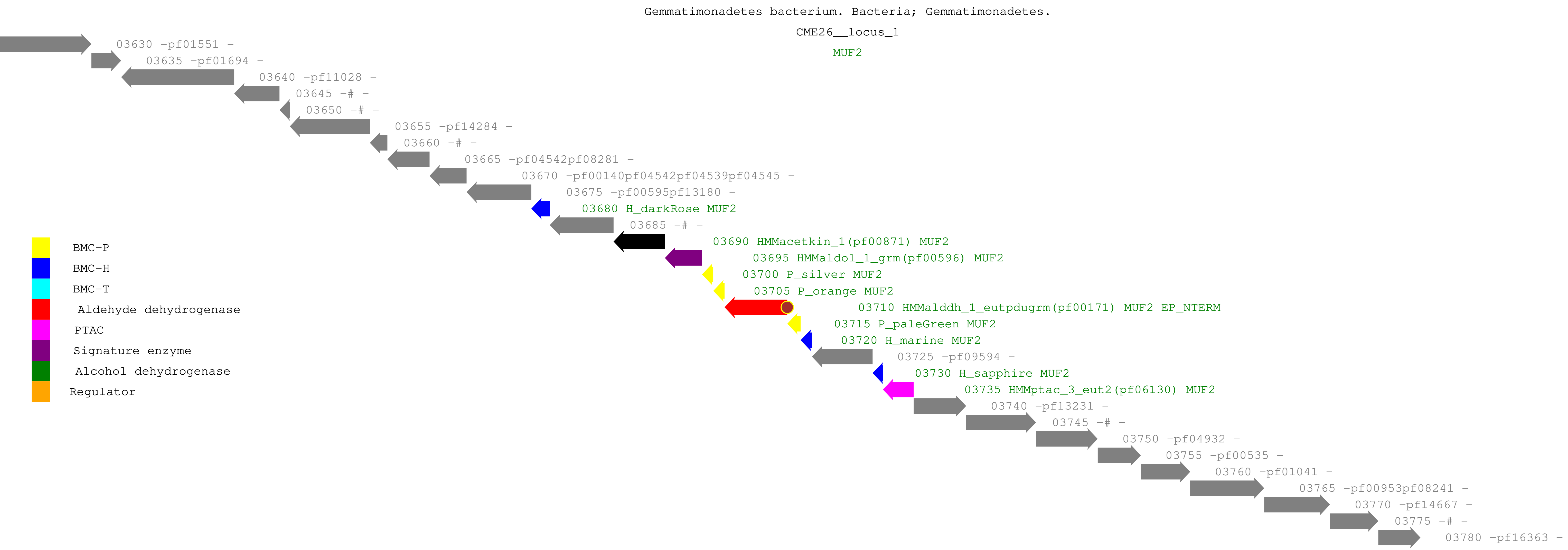
Gemmatimonadetes bacterium. Bacteria; Gemmatimonadetes.
CME26__locus_1
MUF2
03630 -pf01551 -
03635 -pf01694 -
03640 -pf11028 -
03645 -# -
03650 -# -
03655 -pf14284 -
03660 -# -
03665 -pf04542pf08281 -
03670 -pf00140pf04542pf04539pf04545 -
03675 -pf00595pf13180 -
03680 H_darkRose MUF2
03685 -# -
03690 HMMacetkin_1(pf00871) MUF2
03695 HMMaldol_1_grm(pf00596) MUF2
03700 P_silver MUF2
03705 P_orange MUF2
03710 HMMalddh_1_eutpdugrm(pf00171) MUF2 EP_NTERM
03715 P_paleGreen MUF2
03720 H_marine MUF2
03725 -pf09594 -
03730 H_sapphire MUF2
03735 HMMptac_3_eut2(pf06130) MUF2
03740 -pf13231 -
03745 -# -
03750 -pf04932 -
03755 -pf00535 -
03760 -pf01041 -
03765 -pf00953pf08241 -
03770 -pf14667 -
03775 -# -
03780 -pf16363 -
BMC-P
BMC-H
BMC-T
Aldehyde dehydrogenase
PTAC
Signature enzyme
Alcohol dehydrogenase
Regulator

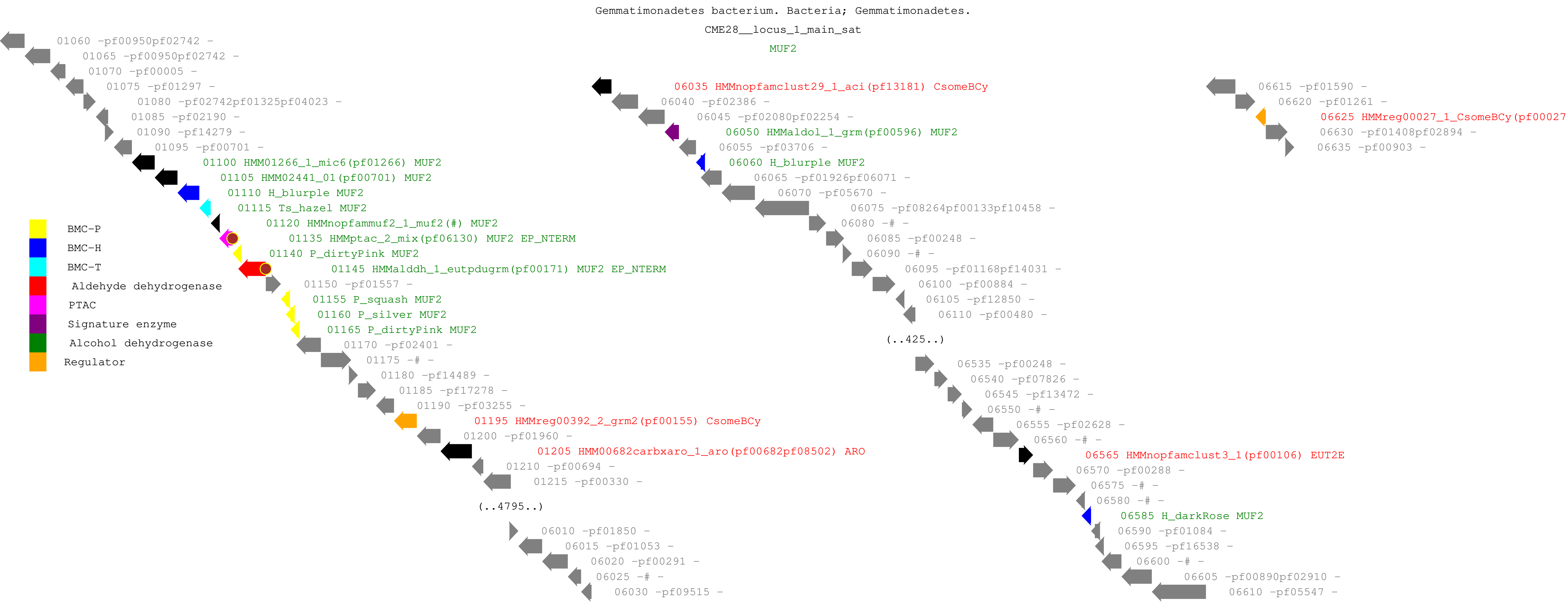
Gemmatimonadetes bacterium. Bacteria; Gemmatimonadetes.
CME28__locus_1_main_sat
MUF2
01060 -pf00950pf02742 -
01065 -pf00950pf02742 -
01070 -pf00005 -
01075 -pf01297 -
01080 -pf02742pf01325pf04023 -
01085 -pf02190 -
01090 -pf14279 -
01095 -pf00701 -
01100 HMM01266_1_mic6(pf01266) MUF2
01105 HMM02441_01(pf00701) MUF2
01110 H_blurple MUF2
01115 Ts_hazel MUF2
01120 HMMnopfammuf2_1_muf2(#) MUF2
01135 HMMptac_2_mix(pf06130) MUF2 EP_NTERM
01140 P_dirtyPink MUF2
01145 HMMaldddh_1_eutpdugrm(pf00171) MUF2 EP_NTERM
01150 -pf01557 -
01155 P_squash MUF2
01160 P_silver MUF2
01165 P_dirtyPink MUF2
01170 -pf02401 -
01175 -# -
01180 -pf14489 -
01185 -pf17278 -
01190 -pf03255 -
01195 HMMreg00392_2_grm2(pf00155) CsomeBCy
01200 -pf01960 -
01205 HMM00682carbxaro_1_aro(pf00682pf08502) ARO
01210 -pf00694 -
01215 -pf00330 -
(..4795..)
06010 -pf01850 -
06015 -pf01053 -
06020 -pf00291 -
06025 -# -
06030 -pf09515 -
06035 HMMnopfamclust29_1_aci(pf13181) CsomeBCy
06040 -pf02386 -
06045 -pf02080pf02254 -
06050 HMMaldol_1_grm(pf00596) MUF2
06055 -pf03706 -
06060 H_blurple MUF2
06065 -pf01926pf06071 -
06070 -pf05670 -
06075 -pf08264pf00133pf10458 -
06080 -# -
06085 -pf00248 -
06090 -# -
06095 -pf01168pf14031 -
06100 -pf00884 -
06105 -pf12850 -
06110 -pf00480 -
(..425..)
06535 -pf00248 -
06540 -pf07826 -
06545 -pf13472 -
06550 -# -
06555 -pf02628 -
06560 -# -
06565 HMMnopfamclust3_1(pf00106) EUT2E
06570 -pf00288 -
06575 -# -
06580 -# -
06585 H_darkRose MUF2
06590 -pf01084 -
06595 -pf16538 -
06600 -# -
06605 -pf00890pf02910 -
06610 -pf05547 -
06615 -pf01590 -
06620 -pf01261 -
06625 HMMreg00027_1_CsomeBCy(pf00027
06630 -pf01408pf02894 -
06635 -pf00903 -
BMC-P
BMC-H
BMC-T
Aldehyde dehydrogenase
PTAC
Signature enzyme
Alcohol dehydrogenase
Regulator

Armatimonadetes bacterium CG_4_8_14_3_um_filter_66_20. Bacteria; Armatimonadetes.

COZ57__locus_1_main_sat

MUF2

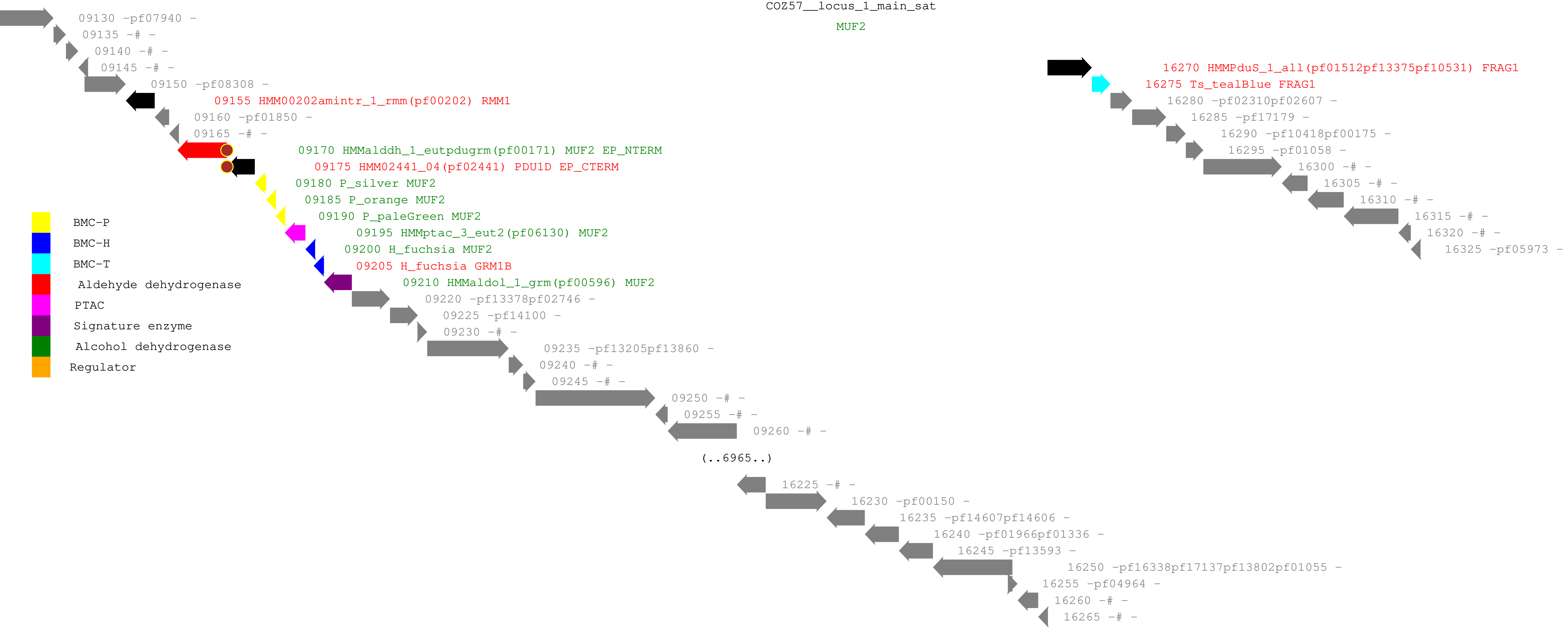

Candidatus Latescibacteria bacterium. Bacteria; Candidatus Latescibacteria.

DCE37__locus_1_main_sat

MUF2

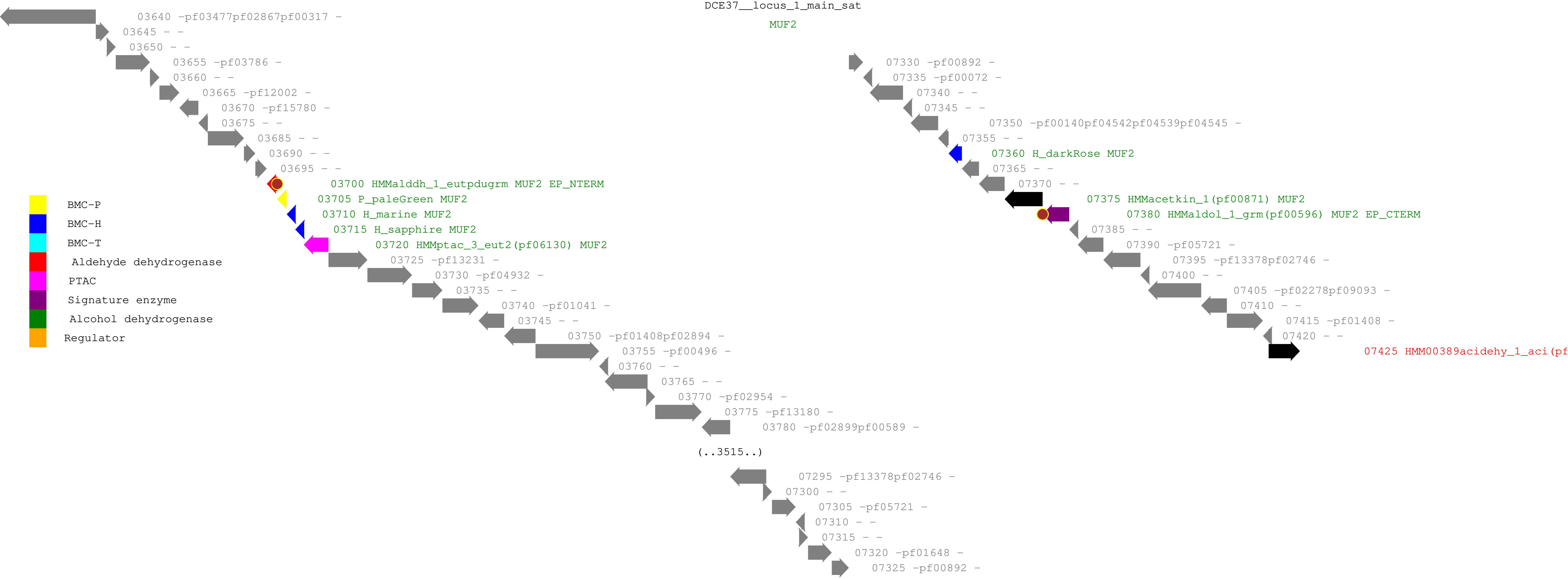

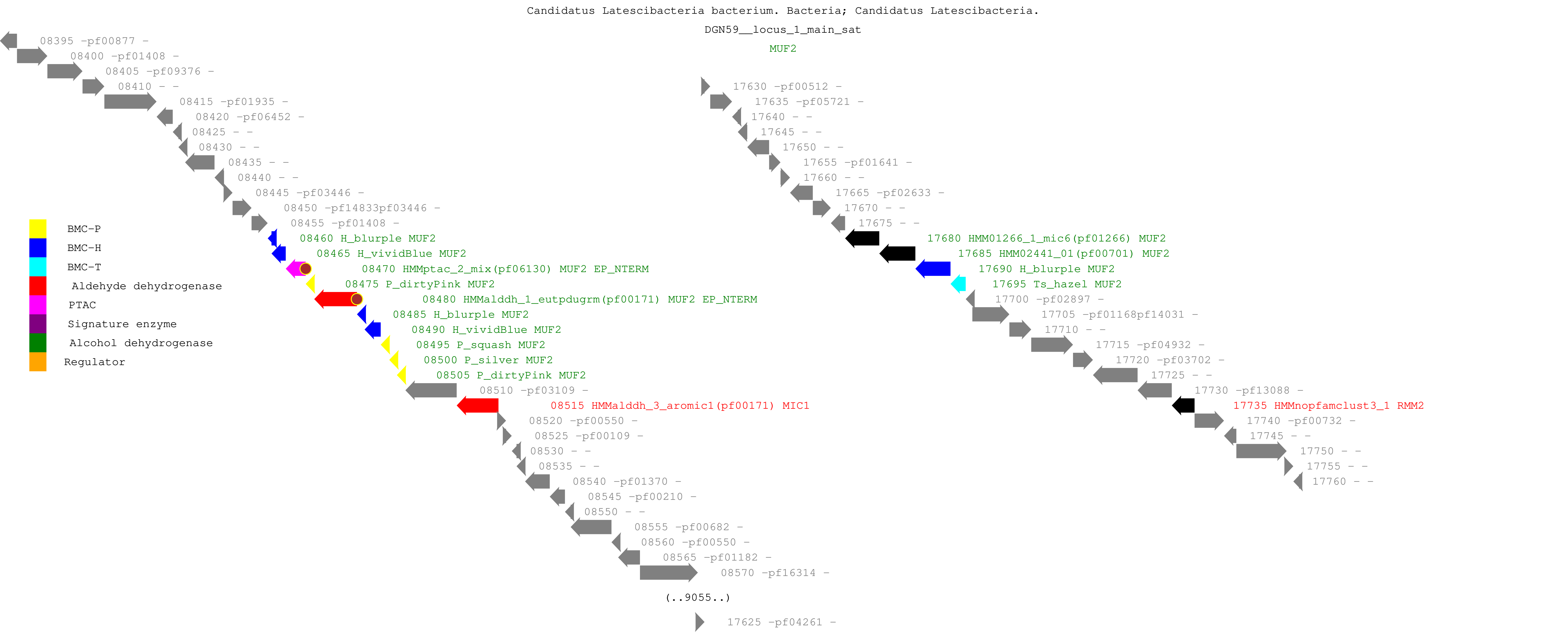

Candidatus Latescibacteria bacterium. Bacteria; Candidatus Latescibacteria.
DGN59__locus_1_main_sat
MUF2
08395 -pf00877 -
08400 -pf01408 -
08405 -pf09376 -
08410 - -
08415 -pf01935 -
08420 -pf06452 -
08425 - -
08430 - -
08435 - -
08440 - -
08445 -pf03446 -
08450 -pf14833pf03446 -
08455 -pf01408 -
08460 H_blurple MUF2
08465 H_vividBlue MUF2
08470 HMMptac_2_mix(pf06130) MUF2 EP_NTERM
08475 P_dirtyPink MUF2
08480 HMMalddh_1_eutpdugrm(pf00171) MUF2 EP_NTERM
08485 H_blurple MUF2
08490 H_vividBlue MUF2
08495 P_squash MUF2
08500 P_silver MUF2
08505 P_dirtyPink MUF2
08510 -pf03109 -
08515 HMMalddh_3_aromic1(pf00171) MIC1
08520 -pf00550 -
08525 -pf00109 -
08530 - -
08535 - -
08540 -pf01370 -
08545 -pf00210 -
08550 - -
08555 -pf00682 -
08560 -pf00550 -
08565 -pf01182 -
08570 -pf16314 -
(..9055..)
17625 -pf04261 -
17630 -pf00512 -
17635 -pf05721 -
17640 - -
17645 - -
17650 - -
17655 -pf01641 -
17660 - -
17665 -pf02633 -
17670 - -
17675 - -
17680 HMM01266_1_mic6(pf01266) MUF2
17685 HMM02441_01(pf00701) MUF2
17690 H_blurple MUF2
17695 Ts_hazel MUF2
17700 -pf02897 -
17705 -pf01168pf14031 -
17710 - -
17715 -pf04932 -
17720 -pf03702 -
17725 - -
17730 -pf13088 -
17735 HMMnopfamclust3_1 RMM2
17740 -pf00732 -
17745 - -
17750 - -
17755 - -
17760 - -
BMC-P
BMC-H
BMC-T
Aldehyde dehydrogenase
PTAC
Signature enzyme
Alcohol dehydrogenase
Regulator

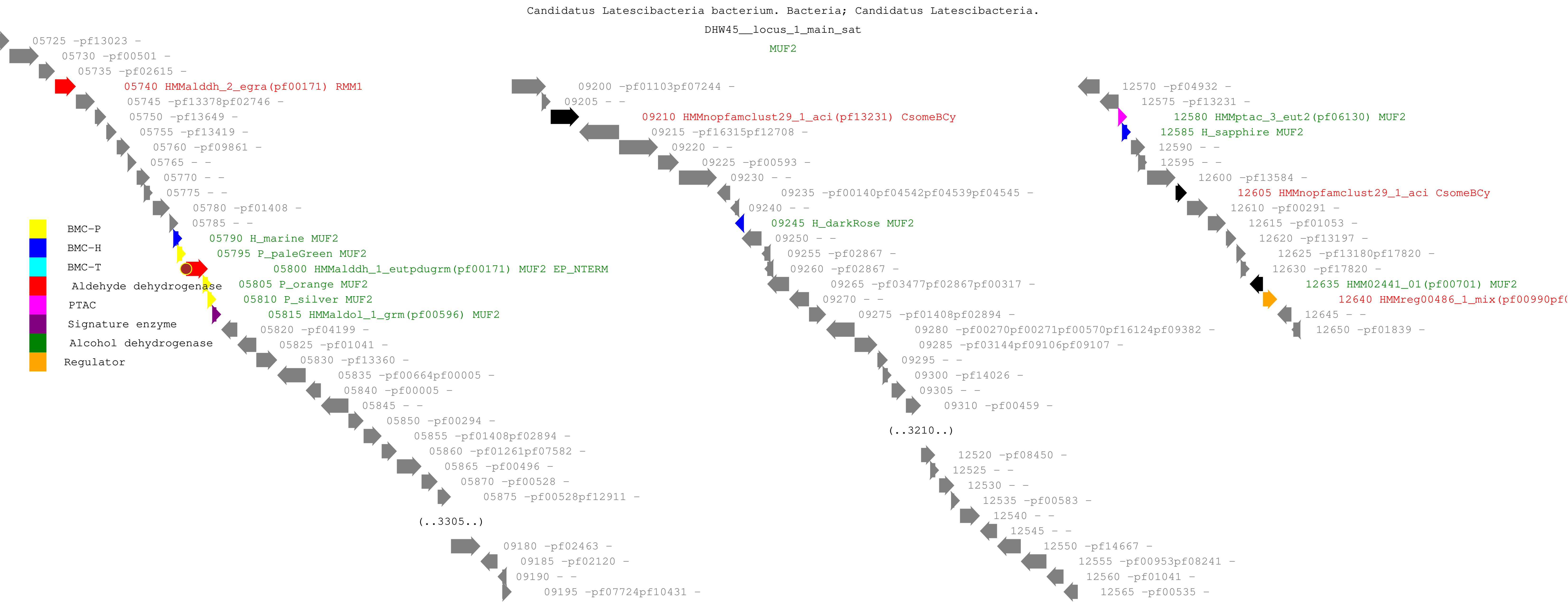

Candidatus Latescibacteria bacterium. Bacteria; Candidatus Latescibacteria.
DHW45__locus_1_main_sat
MUF2
05725 -pf13023 -
05730 -pf00501 -
05735 -pf02615 -
05740 HMMaldh_2_egra(pf00171) RMM1
05745 -pf13378pf02746 -
05750 -pf13649 -
05755 -pf13419 -
05760 -pf09861 -
05765 - -
05770 - -
05775 - -
05780 -pf01408 -
05785 - -
05790 H_marine MUF2
05795 P_paleGreen MUF2
05800 HMMaldh_1_eutpdugrm(pf00171) MUF2 EP_NTERM
05805 P_orange MUF2
05810 P_silver MUF2
05815 HMMaldol_1_grm(pf00596) MUF2
05820 -pf04199 -
05825 -pf01041 -
05830 -pf13360 -
05835 -pf00664pf00005 -
05840 -pf00005 -
05845 - -
05850 -pf00294 -
05855 -pf01408pf02894 -
05860 -pf01261pf07582 -
05865 -pf00496 -
05870 -pf00528 -
05875 -pf00528pf12911 -
(..3305..)
09180 -pf02463 -
09185 -pf02120 -
09190 - -
09195 -pf07724pf10431 -
BMC-P
BMC-H
BMC-T
Aldehyde dehydrogenase
PTAC
Signature enzyme
Alcohol dehydrogenase
Regulator
09200 -pf01103pf07244 -
09205 - -
09210 HMMnopfamclust29_1_aci(pf13231) CsomeBCy
09215 -pf16315pf12708 -
09220 - -
09225 -pf00593 -
09230 - -
09235 -pf00140pf04542pf04539pf04545 -
09240 - -
09245 H_darkRose MUF2
09250 - -
09255 -pf02867 -
09260 -pf02867 -
09265 -pf03477pf02867pf00317 -
09270 - -
09275 -pf01408pf02894 -
09280 -pf00270pf00271pf00570pf16124pf09382 -
09285 -pf03144pf09106pf09107 -
09295 - -
09300 -pf14026 -
09305 - -
09310 -pf00459 -
(..3210..)
12520 -pf08450 -
12525 - -
12530 - -
12535 -pf00583 -
12540 - -
12545 - -
12550 -pf14667 -
12555 -pf00953pf08241 -
12560 -pf01041 -
12565 -pf00535 -
12570 -pf04932 -
12575 -pf13231 -
12580 HMMptac_3_eut2(pf06130) MUF2
12585 H_sapphire MUF2
12590 - -
12595 - -
12600 -pf13584 -
12605 HMMnopfamclust29_1_aci CsomeBCy
12610 -pf00291 -
12615 -pf01053 -
12620 -pf13197 -
12625 -pf13180pf17820 -
12630 -pf17820 -
12635 HMM02441_01(pf00701) MUF2
12640 HMMreg00486_1_mix(pf00990pf0
12645 - -
12650 -pf01839 -

Candidatus Latescibacteria bacterium. Bacteria; Candidatus Latescibacteria.

DIC52__locus_1_main_sat

MUF2

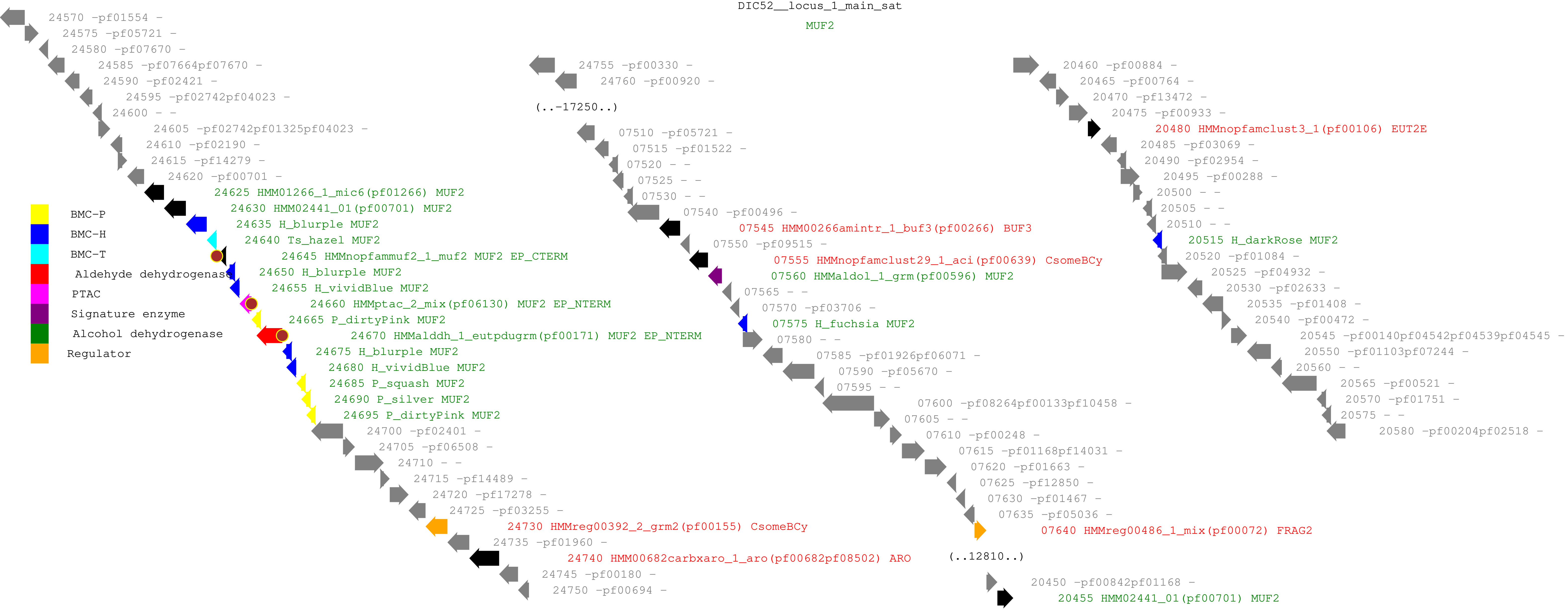

Candidatus Latescibacteria bacterium. Bacteria; Candidatus Latescibacteria.

DIU35__locus_1

MUF2

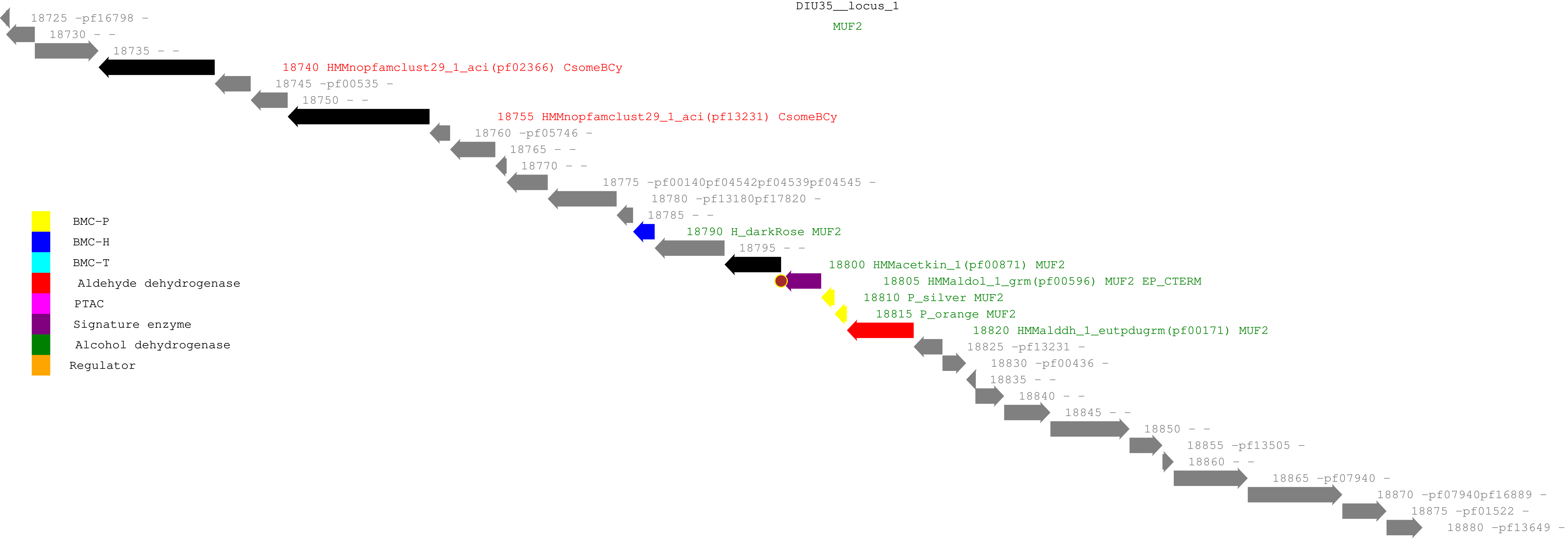

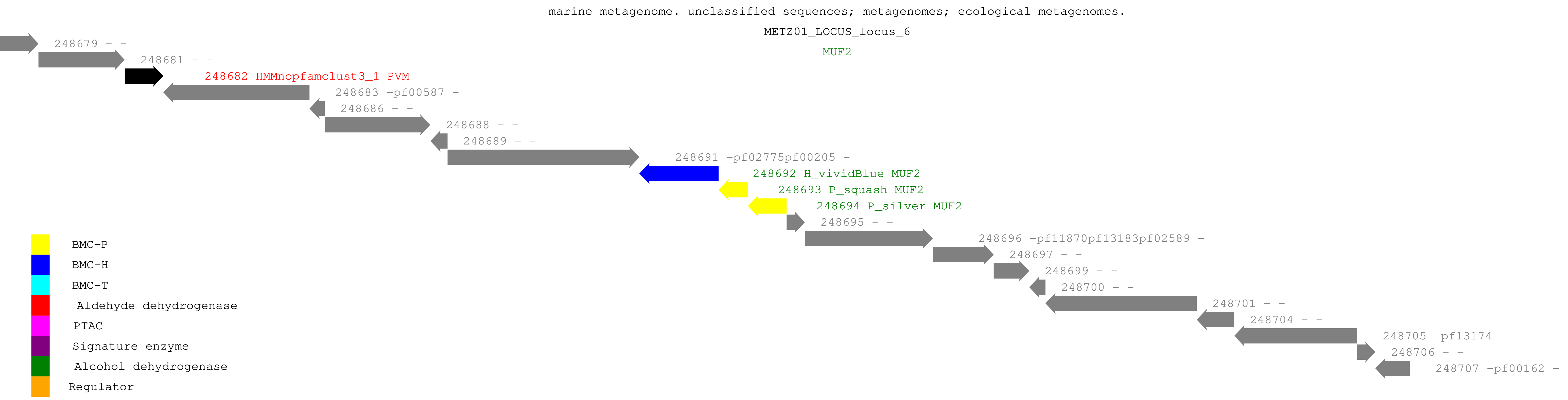
marine metagenome. unclassified sequences; metagenomes; ecological metagenomes.
METZ01_LOCUS_locus_6
MUF2
248679 - -
248681 - -
248682 HMMnopfamclust3_1 PVM
248683 -pf00587 -
248686 - -
248688 - -
248689 - -
248691 -pf02775pf00205 -
248692 H_vividBlue MUF2
248693 P_squash MUF2
248694 P_silver MUF2
248695 - -
248696 -pf11870pf13183pf02589 -
248697 - -
248699 - -
248700 - -
248701 - -
248704 - -
248705 -pf13174 -
248706 - -
248707 -pf00162 -
BMC-P
BMC-H
BMC-T
Aldehyde dehydrogenase
PTAC
Signature enzyme
Alcohol dehydrogenase
Regulator

Peptococcaceae bacterium CEB3. Bacteria; Firmicutes; Clostridia; Clostridiales; Peptococcaceae.

CEB3_c_locus_1

MUF3

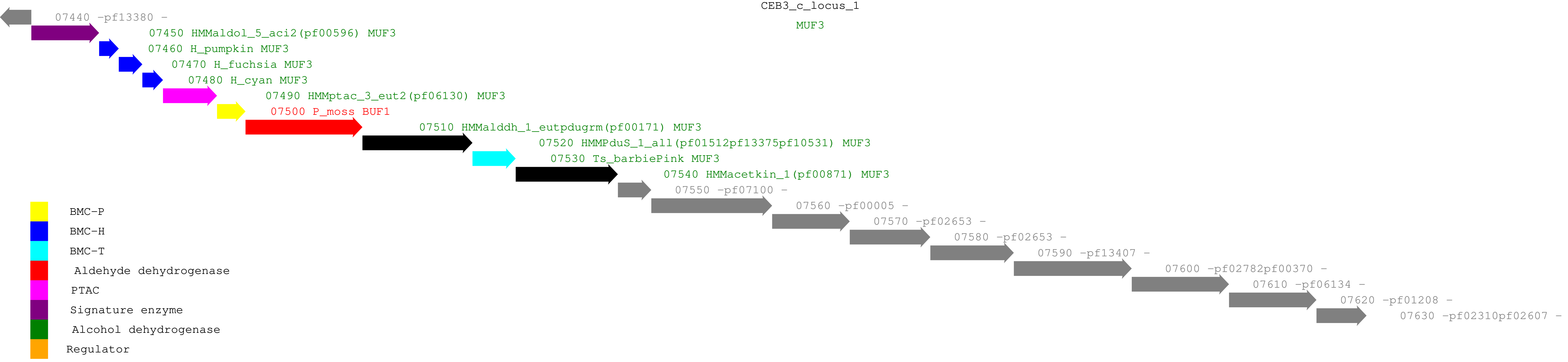

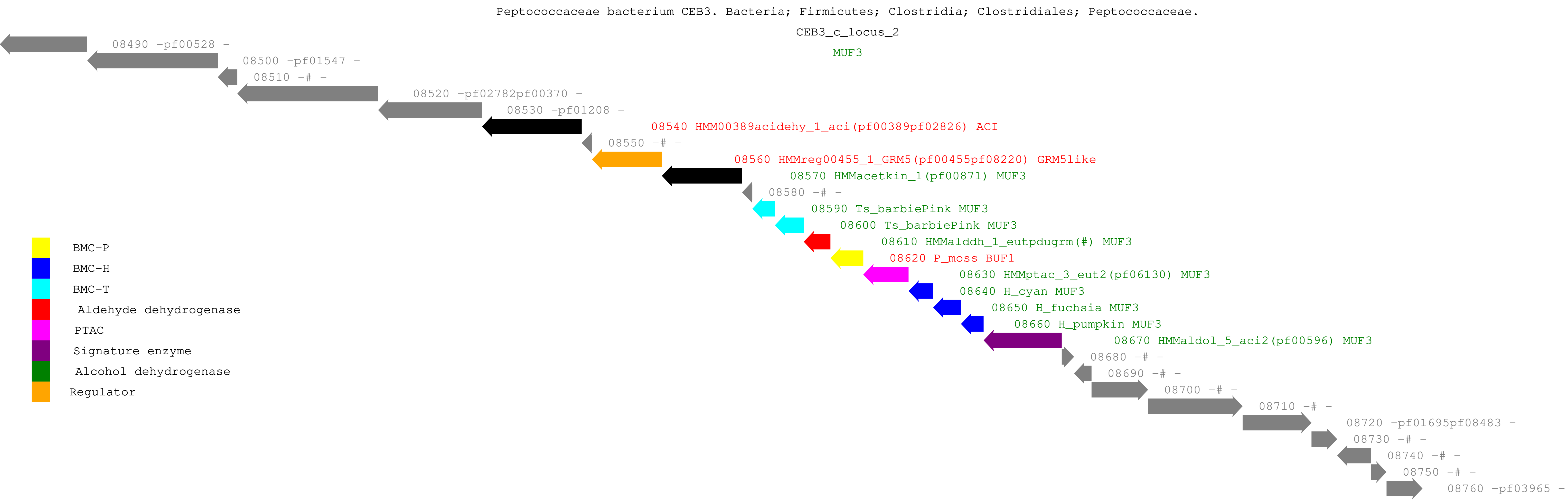
Peptococcaceae bacterium CEB3. Bacteria; Firmicutes; Clostridia; Clostridiales; Peptococcaceae.
CEB3_c_locus_2
MUF3
08490 -pf00528 -
08500 -pf01547 -
08510 -# -
08520 -pf02782pf00370 -
08530 -pf01208 -
08540 HMM00389acidehy_1_aci(pf00389pf02826) ACI
08550 -# -
08560 HMMreg00455_1_GRM5(pf00455pf08220) GRM5like
08570 HMMacetkin_1(pf00871) MUF3
08580 -# -
08590 Ts_barbiePink MUF3
08600 Ts_barbiePink MUF3
08610 HMMalddh_1_eutpdugrm(#) MUF3
08620 P_moss BUF1
08630 HMMptac_3_eut2(pf06130) MUF3
08640 H_cyan MUF3
08650 H_fuchsia MUF3
08660 H_pumpkin MUF3
08670 HMMaldol_5_aci2(pf00596) MUF3
08680 -# -
08690 -# -
08700 -# -
08710 -# -
08720 -pf01695pf08483 -
08730 -# -
08740 -# -
08750 -# -
08760 -pf03965 -
BMC-P
BMC-H
BMC-T
Aldehyde dehydrogenase
PTAC
Signature enzyme
Alcohol dehydrogenase
Regulator

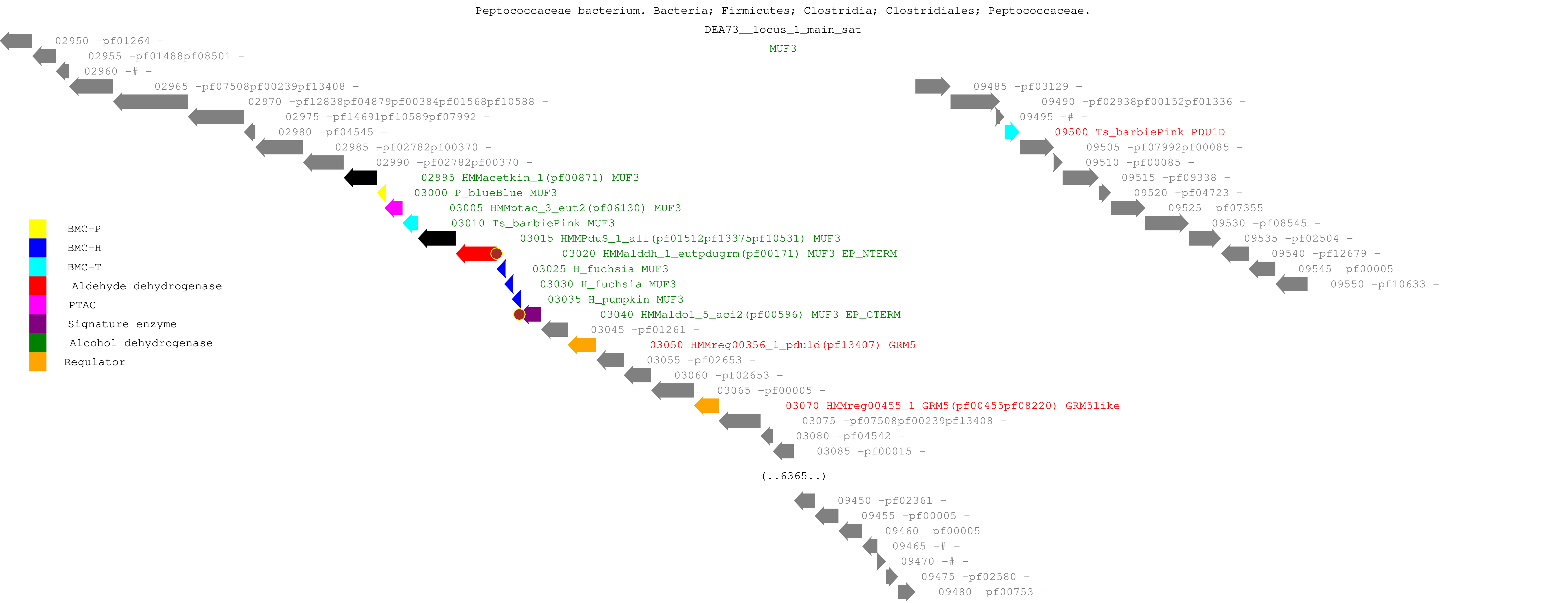

Peptococcaceae bacterium. Bacteria; Firmicutes; Clostridia; Clostridiales; Peptococcaceae.
DEA73__locus_1_main_sat
MUF3
02950 -pf01264 -
02955 -pf01488pf08501 -
02960 -# -
02965 -pf07508pf00239pf13408 -
02970 -pf12838pf04879pf00384pf01568pf10588 -
02975 -pf14691pf10589pf07992 -
02980 -pf04545 -
02985 -pf02782pf00370 -
02990 -pf02782pf00370 -
02995 HMMacetkin_1(pf00871) MUF3
03000 P_blueBlue MUF3
03005 HMMptac_3_eut2(pf06130) MUF3
03010 Ts_barbiePink MUF3
03015 HMMPduS_1_all(pf01512pf13375pf10531) MUF3
03020 HMMaldh_1_eutpdugrm(pf00171) MUF3 EP_NTERM
03025 H_fuchsia MUF3
03030 H_fuchsia MUF3
03035 H_pumpkin MUF3
03040 HMMaldol_5_aci2(pf00596) MUF3 EP_CTERM
03045 -pf01261 -
03050 HMMreg00356_1_pdu1d(pf13407) GRM5
03055 -pf02653 -
03060 -pf02653 -
03065 -pf00005 -
03070 HMMreg00455_1_GRM5(pf00455pf08220) GRM5like
03075 -pf07508pf00239pf13408 -
03080 -pf04542 -
03085 -pf00015 -
(..6365..)
09450 -pf02361 -
09455 -pf00005 -
09460 -pf00005 -
09465 -# -
09470 -# -
09475 -pf02580 -
09480 -pf00753 -
09485 -pf03129 -
09490 -pf02938pf00152pf01336 -
09495 -# -
09500 Ts_barbiePink PDU1D
09505 -pf07992pf00085 -
09510 -pf00085 -
09515 -pf09338 -
09520 -pf04723 -
09525 -pf07355 -
09530 -pf08545 -
09535 -pf02504 -
09540 -pf12679 -
09545 -pf00005 -
09550 -pf10633 -
BMC-P
BMC-H
BMC-T
Aldehyde dehydrogenase
PTAC
Signature enzyme
Alcohol dehydrogenase
Regulator

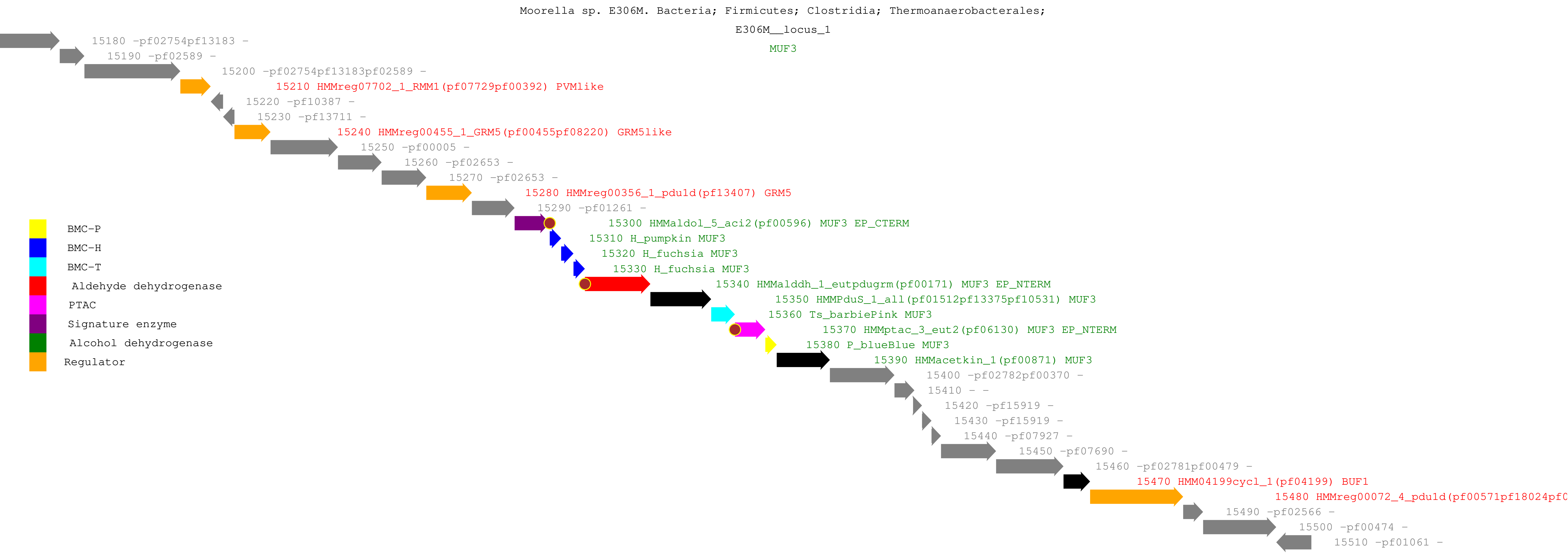

Moorella sp. E306M. Bacteria; Firmicutes; Clostridia; Thermoanaerobacterales;
E306M__locus_1
MUF3
15180 -pf02754pf13183 -
15190 -pf02589 -
15200 -pf02754pf13183pf02589 -
15210 HMMreg07702_1_RMM1(pf07729pf00392) PVMlike
15220 -pf10387 -
15230 -pf13711 -
15240 HMMreg00455_1_GRM5(pf00455pf08220) GRM5like
15250 -pf00005 -
15260 -pf02653 -
15270 -pf02653 -
15280 HMMreg00356_1_pdu1d(pf13407) GRM5
15290 -pf01261 -
15300 HMMaldol_5_aci2(pf00596) MUF3 EP_CTERM
15310 H_pumpkin MUF3
15320 H_fuchsia MUF3
15330 H_fuchsia MUF3
15340 HMMaldh_1_eutpdugrm(pf00171) MUF3 EP_NTERM
15350 HMMPduS_1_all(pf01512pf13375pf10531) MUF3
15360 Ts_barbiePink MUF3
15370 HMMptac_3_eut2(pf06130) MUF3 EP_NTERM
15380 P_blueBlue MUF3
15390 HMMacetkin_1(pf00871) MUF3
15400 -pf02782pf00370 -
15410 - -
15420 -pf15919 -
15430 -pf15919 -
15440 -pf07927 -
15450 -pf07690 -
15460 -pf02781pf00479 -
15470 HMM04199cycl_1(pf04199) BUF1
15480 HMMreg00072_4_pdu1d(pf00571pf18024pf0
15490 -pf02566 -
15500 -pf00474 -
15510 -pf01061 -
BMC-P
BMC-H
BMC-T
Aldehyde dehydrogenase
PTAC
Signature enzyme
Alcohol dehydrogenase
Regulator

Mahella australiensis (strain DSM 15567 / CIP 107919 / 50-1 BON). Bacteria; Firmicutes; Clostridia; Thermoanaerobacterales;

Mahau__locus_1

MUF3

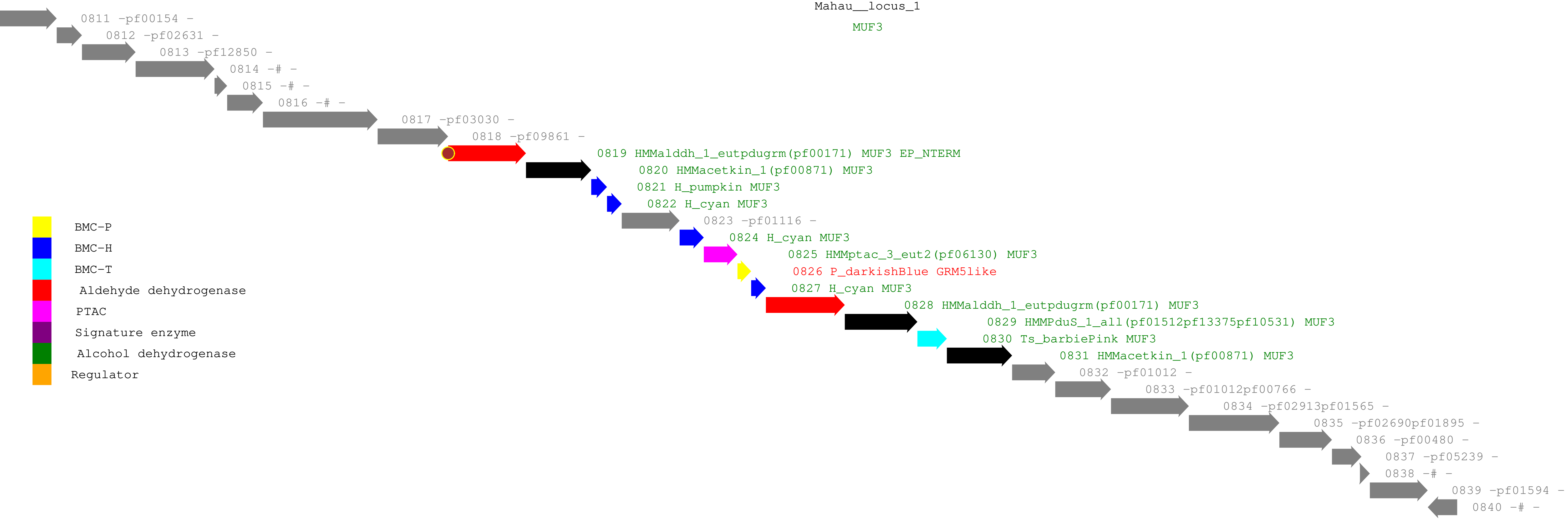

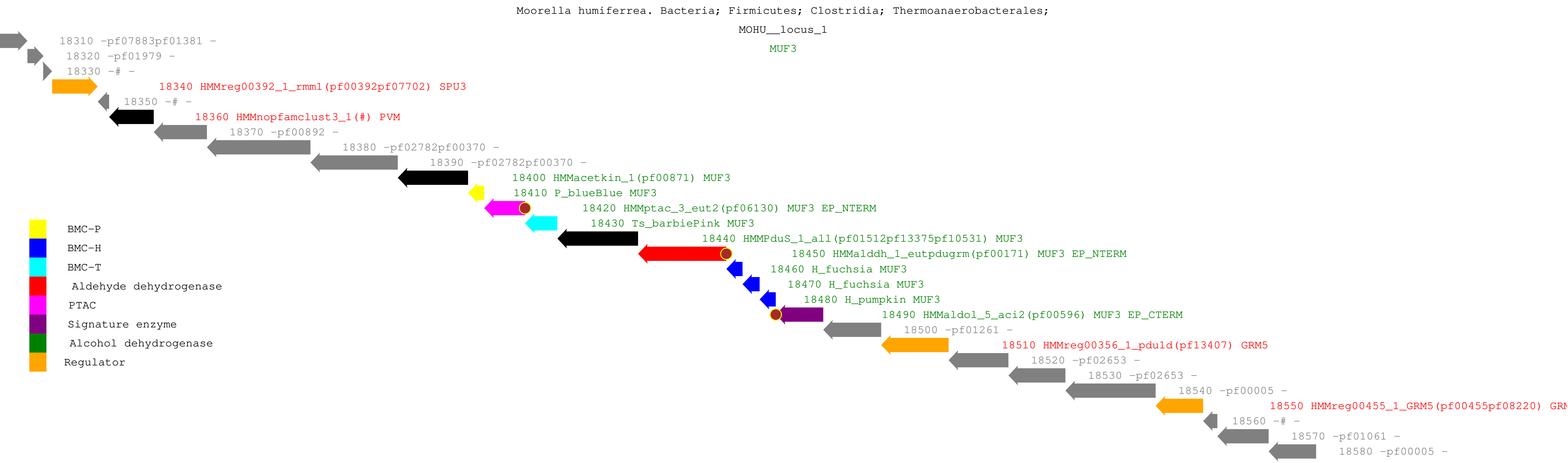

Moorella humiferrea. Bacteria; Firmicutes; Clostridia; Thermoanaerobacterales;
MOHU__locus_1
MUF3
18310 -pf07883pf01381 -
18320 -pf01979 -
18330 -# -
18340 HMMreg00392_1_rmm1(pf00392pf07702) SPU3
18350 -# -
18360 HMMnopfamclust3_1(#) PVM
18370 -pf00892 -
18380 -pf02782pf00370 -
18390 -pf02782pf00370 -
18400 HMMacetkin_1(pf00871) MUF3
18410 P_blueBlue MUF3
18420 HMMptac_3_eut2(pf06130) MUF3 EP_NTERM
18430 Ts_barbiePink MUF3
18440 HMMPduS_1_all(pf01512pf13375pf10531) MUF3
18450 HMMalddh_1_eutpdugrm(pf00171) MUF3 EP_NTERM
18460 H_fuchsia MUF3
18470 H_fuchsia MUF3
18480 H_pumpkin MUF3
18490 HMMaldol_5_aci2(pf00596) MUF3 EP_CTERM
18500 -pf01261 -
18510 HMMreg00356_1_pdu1d(pf13407) GRM5
18520 -pf02653 -
18530 -pf02653 -
18540 -pf00005 -
18550 HMMreg00455_1_GRM5(pf00455pf08220) GRM
18560 -# -
18570 -pf01061 -
18580 -pf00005 -
BMC-P
BMC-H
BMC-T
Aldehyde dehydrogenase
PTAC
Signature enzyme
Alcohol dehydrogenase
Regulator

Moorella mulderi DSM 14980. Bacteria; Firmicutes; Clostridia; Thermoanaerobacterales;

MOMUL__locus_1

MUF3

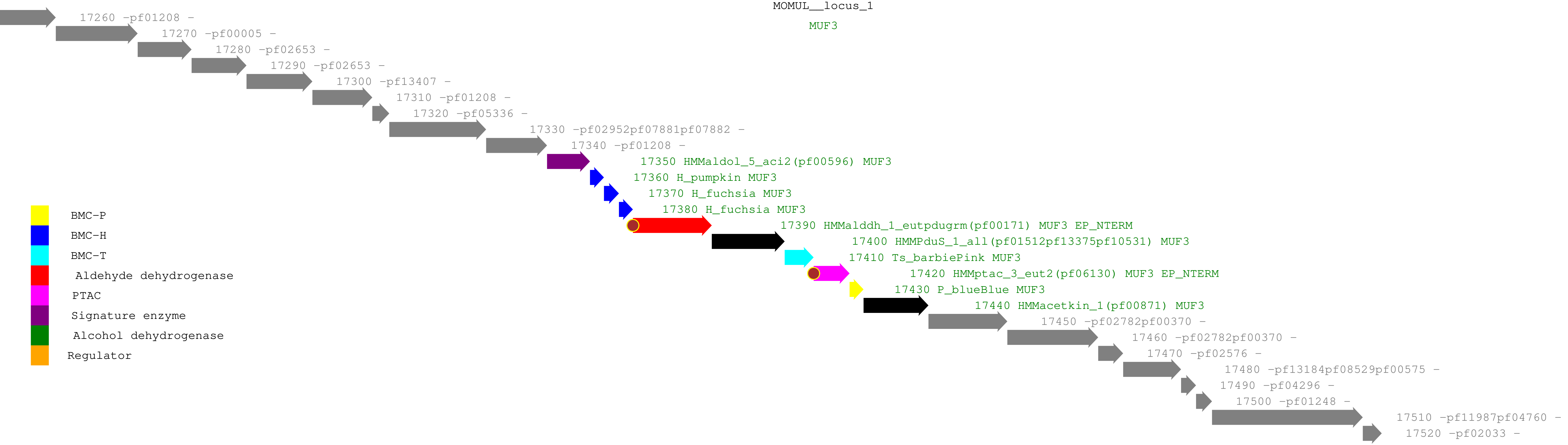

Thermanaeromonas toyohensis ToBE. Bacteria; Firmicutes; Clostridia; Thermoanaerobacterales;

SAMN00808754__locus_1

MUF3

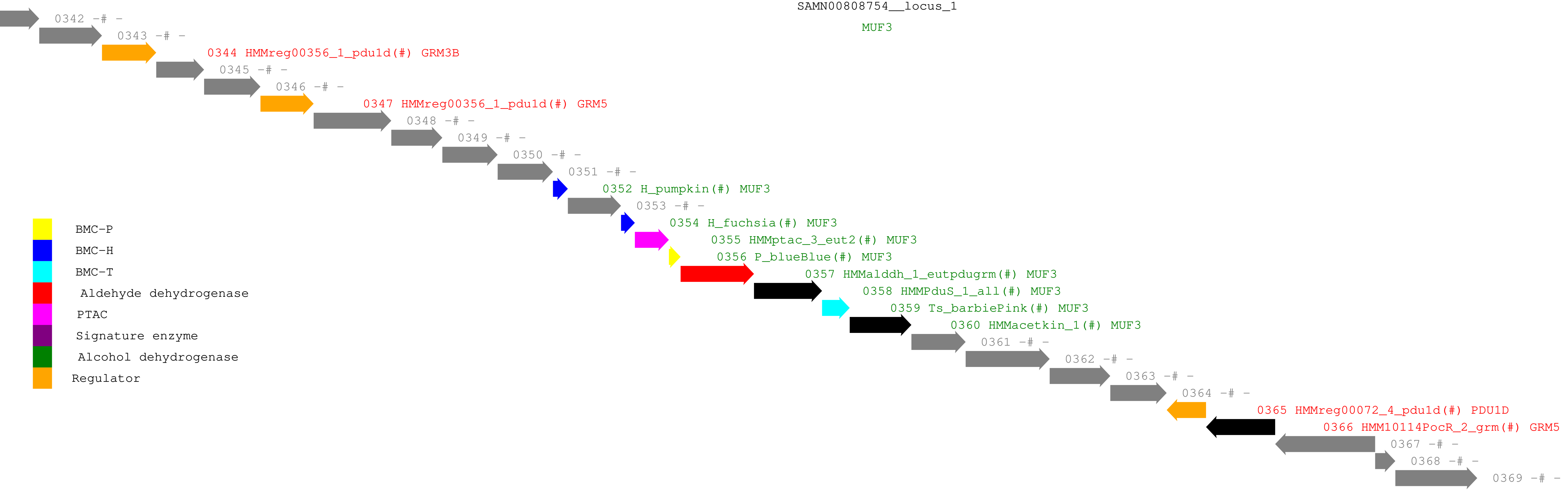

Moorella sp. 60_41. Bacteria; Firmicutes; Clostridia; Thermoanaerobacterales;

XD51__locus_1

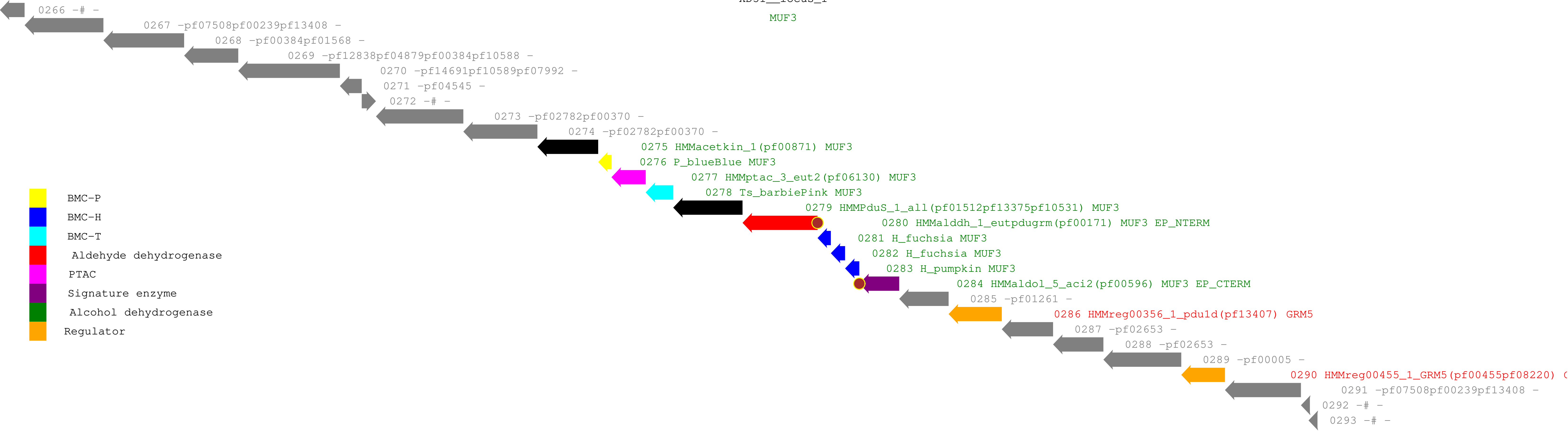

Salmonella muenster. Bacteria; Proteobacteria; Gammaproteobacteria; Enterobacterales;

A0F71__locus_2_main_sat

PDU1AB

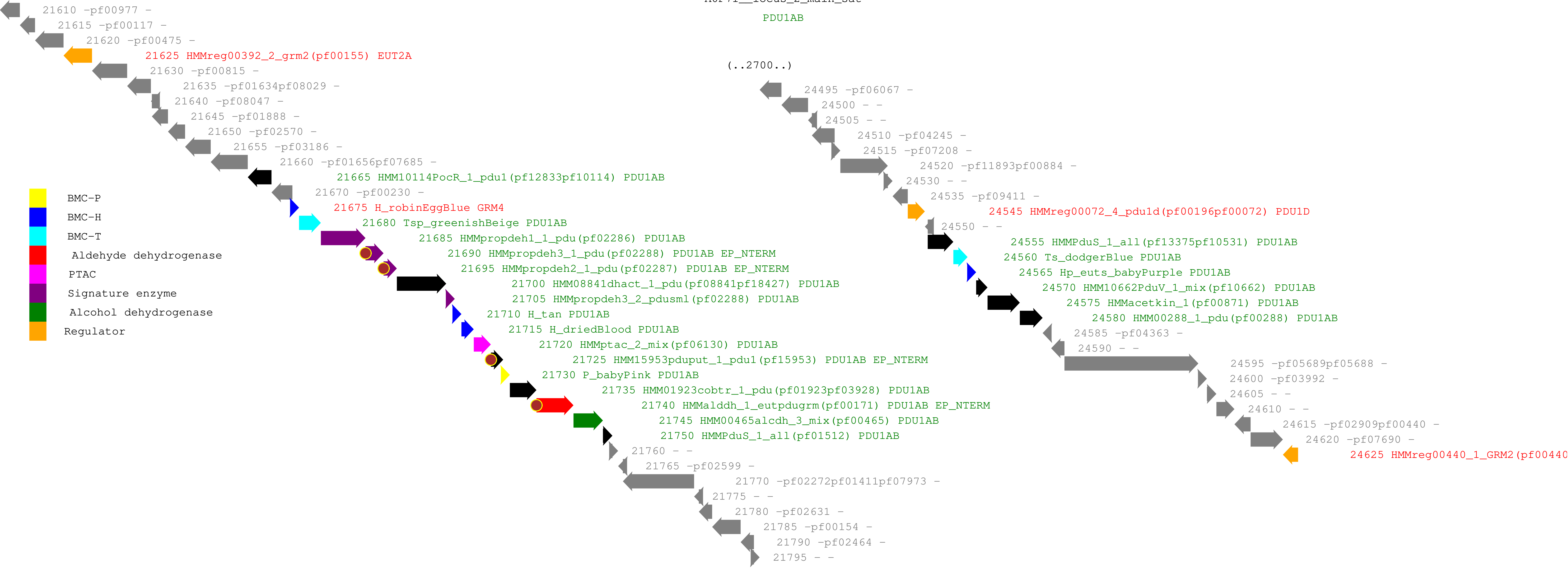

Salmonella muenster. Bacteria; Proteobacteria; Gammaproteobacteria; Enterobacterales;

A0F77__locus_1

PDU1AB

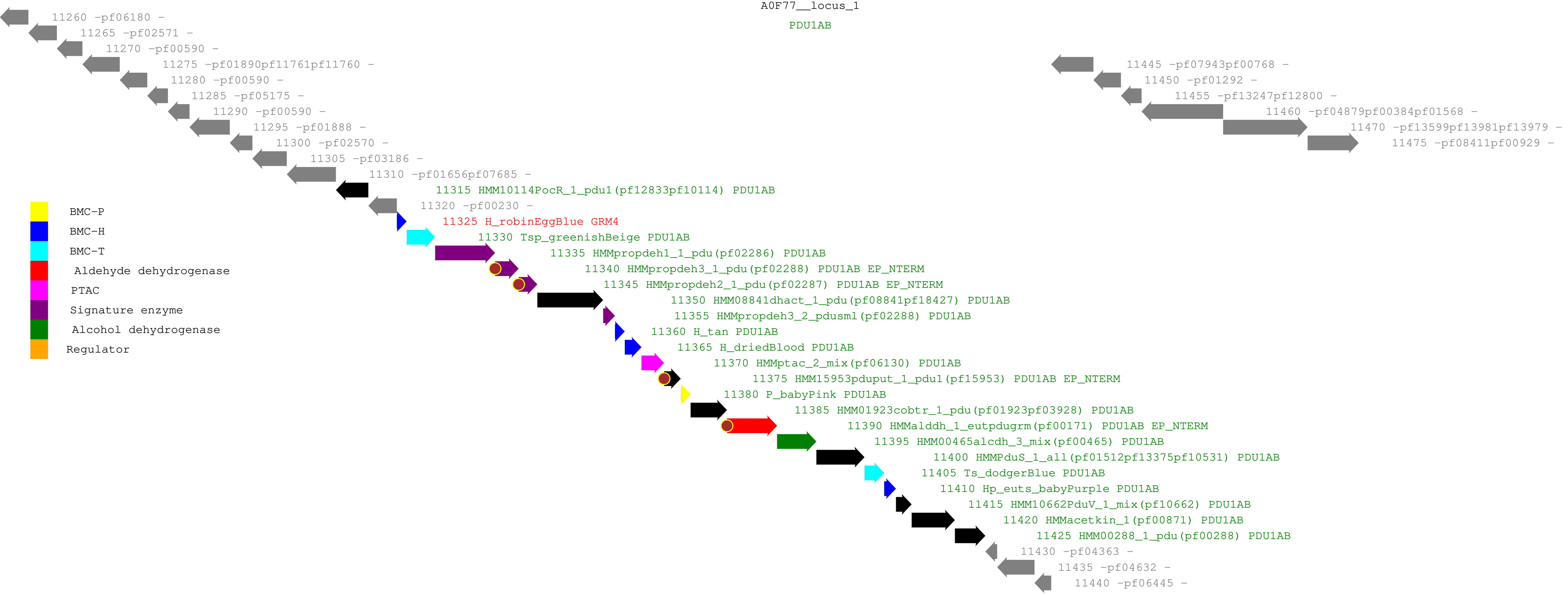

Escherichia sp. KTE52. Bacteria; Proteobacteria; Gammaproteobacteria; Enterobacterales;

A1SC__locus_1

PDU1AB

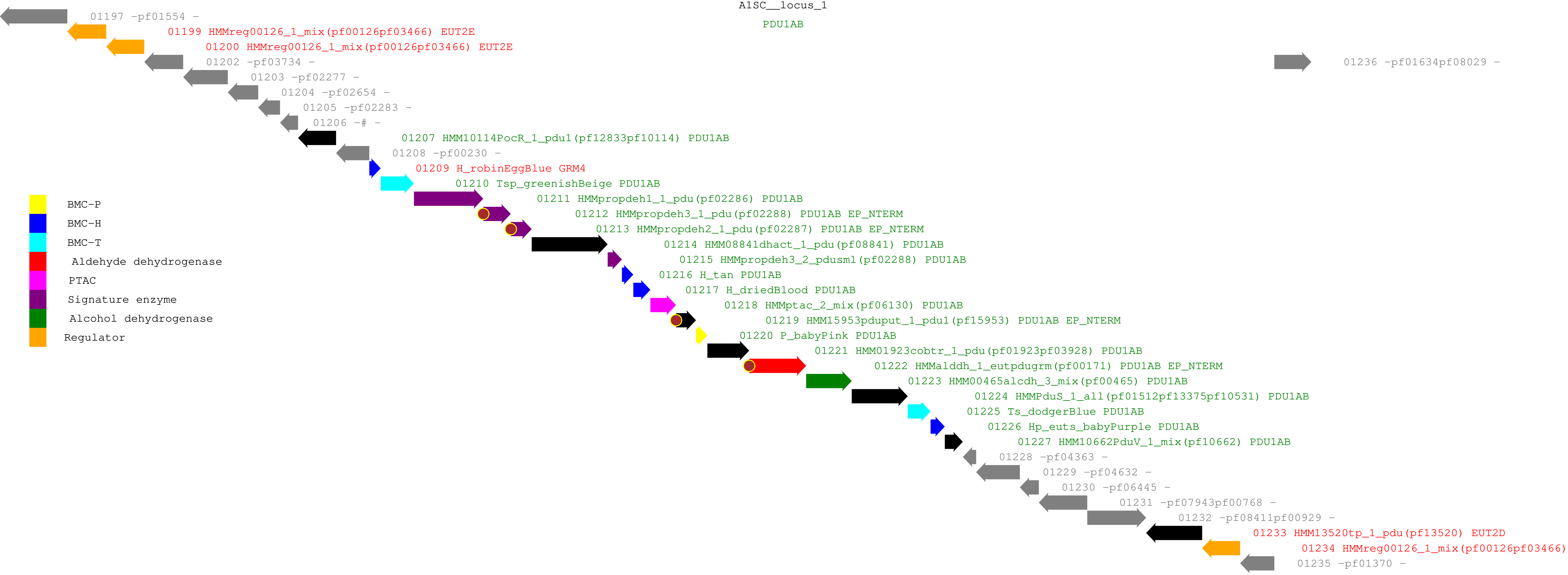

Escherichia coli KTE75. Bacteria; Proteobacteria; Gammaproteobacteria; Enterobacterales;

A1UM__locus_1

PDU1AB

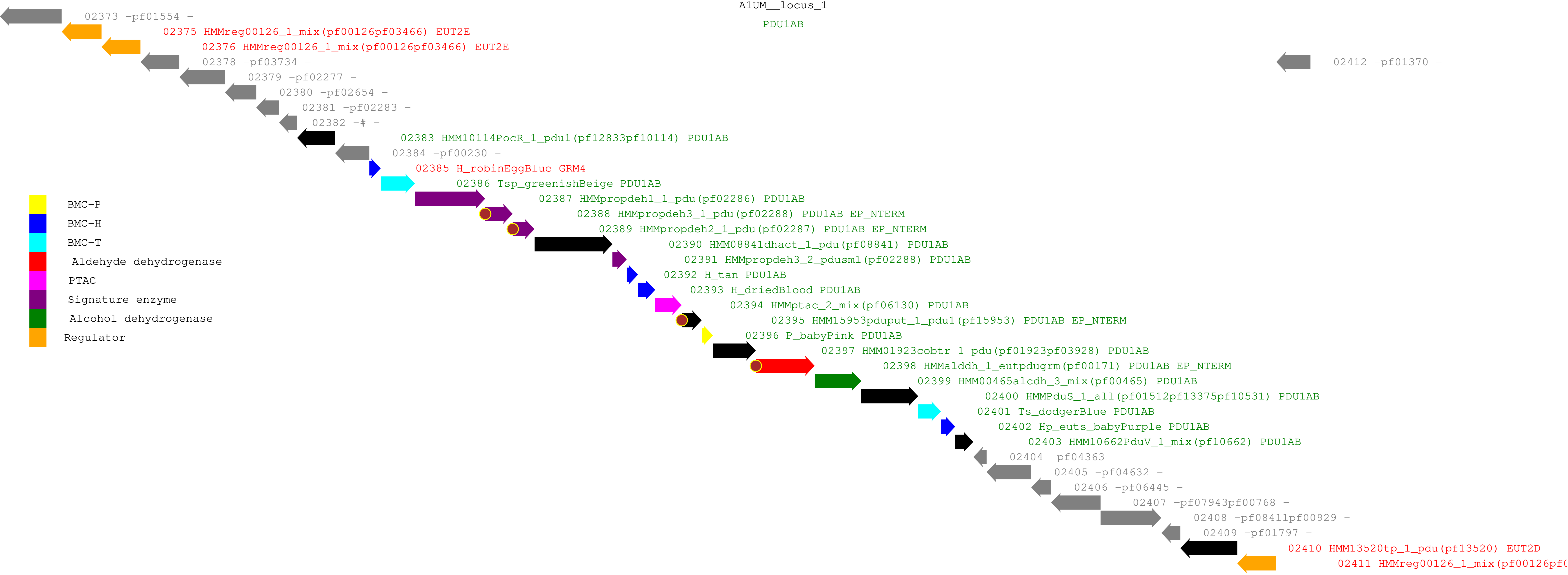

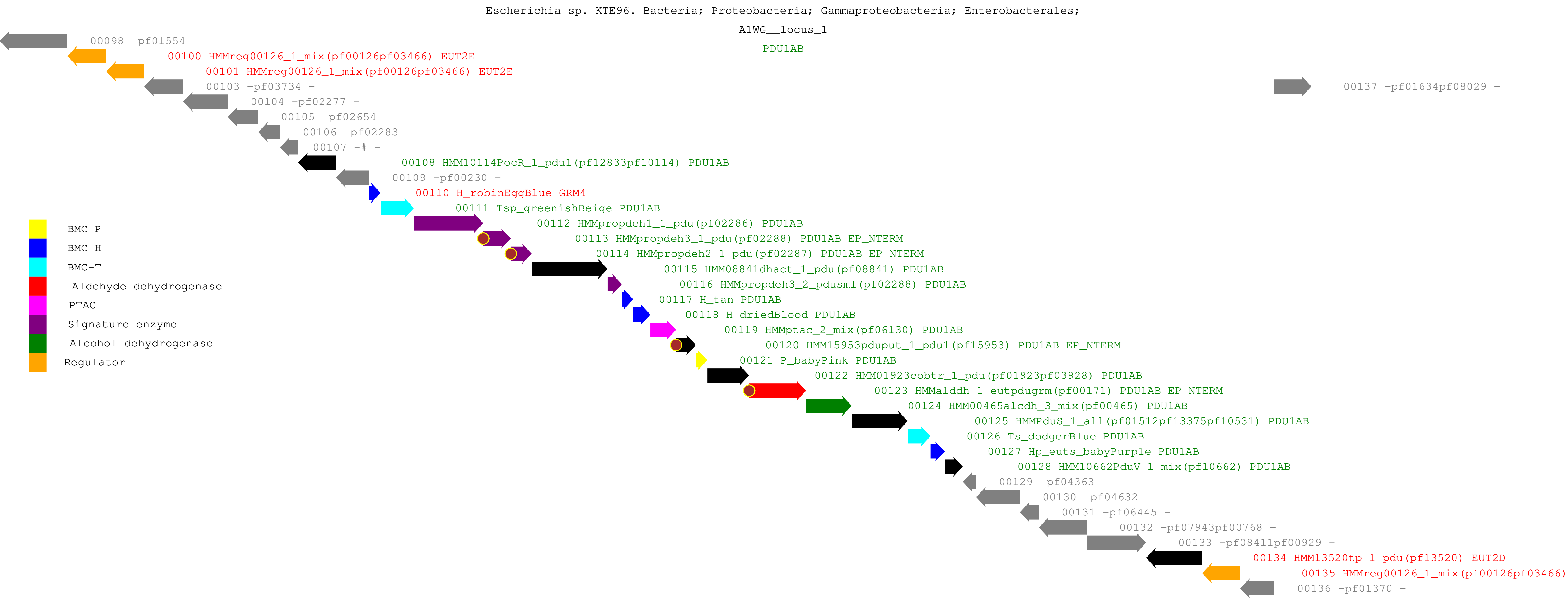
Escherichia sp. KTE96. Bacteria; Proteobacteria; Gammaproteobacteria; Enterobacterales;
A1WG__locus_1
PDU1AB
00098 -pf01554 -
00100 HMMreg00126_1_mix(pf00126pf03466) EUT2E
00101 HMMreg00126_1_mix(pf00126pf03466) EUT2E
00103 -pf03734 -
00104 -pf02277 -
00105 -pf02654 -
00106 -pf02283 -
00107 -# -
00108 HMM10114PocR_1_pdu1(pf12833pf10114) PDU1AB
00109 -pf00230 -
00110 H_robinEggBlue GRM4
00111 Tsp_greenishBeige PDU1AB
00112 HMMpropdeh1_1_pdu(pf02286) PDU1AB
00113 HMMpropdeh3_1_pdu(pf02288) PDU1AB EP_NTERM
00114 HMMpropdeh2_1_pdu(pf02287) PDU1AB EP_NTERM
00115 HMM08841dhact_1_pdu(pf08841) PDU1AB
00116 HMMpropdeh3_2_pdusml(pf02288) PDU1AB
00117 H_tan PDU1AB
00118 H_driedBlood PDU1AB
00119 HMMptac_2_mix(pf06130) PDU1AB
00120 HMM15953pduput_1_pdu1(pf15953) PDU1AB EP_NTERM
00121 P_babyPink PDU1AB
00122 HMM01923cobtr_1_pdu(pf01923pf03928) PDU1AB
00123 HMMalddh_1_eutpdugrm(pf00171) PDU1AB EP_NTERM
00124 HMM00465alcdh_3_mix(pf00465) PDU1AB
00125 HMMPduS_1_all(pf01512pf13375pf10531) PDU1AB
00126 Ts_dodgerBlue PDU1AB
00127 Hp_euts_babyPurple PDU1AB
00128 HMM10662PduV_1_mix(pf10662) PDU1AB
00129 -pf04363 -
00130 -pf04632 -
00131 -pf06445 -
00132 -pf07943pf00768 -
00133 -pf08411pf00929 -
00134 HMM13520tp_1_pdu(pf13520) EUT2D
00135 HMMreg00126_1_mix(pf00126pf03466)
00136 -pf01370 -
00137 -pf01634pf08029 -
BMC-P
BMC-H
BMC-T
Aldehyde dehydrogenase
PTAC
Signature enzyme
Alcohol dehydrogenase
Regulator

Klebsiella michiganensis E718. Bacteria; Proteobacteria; Gammaproteobacteria; Enterobacterales;

A225__locus_3

PDU1AB

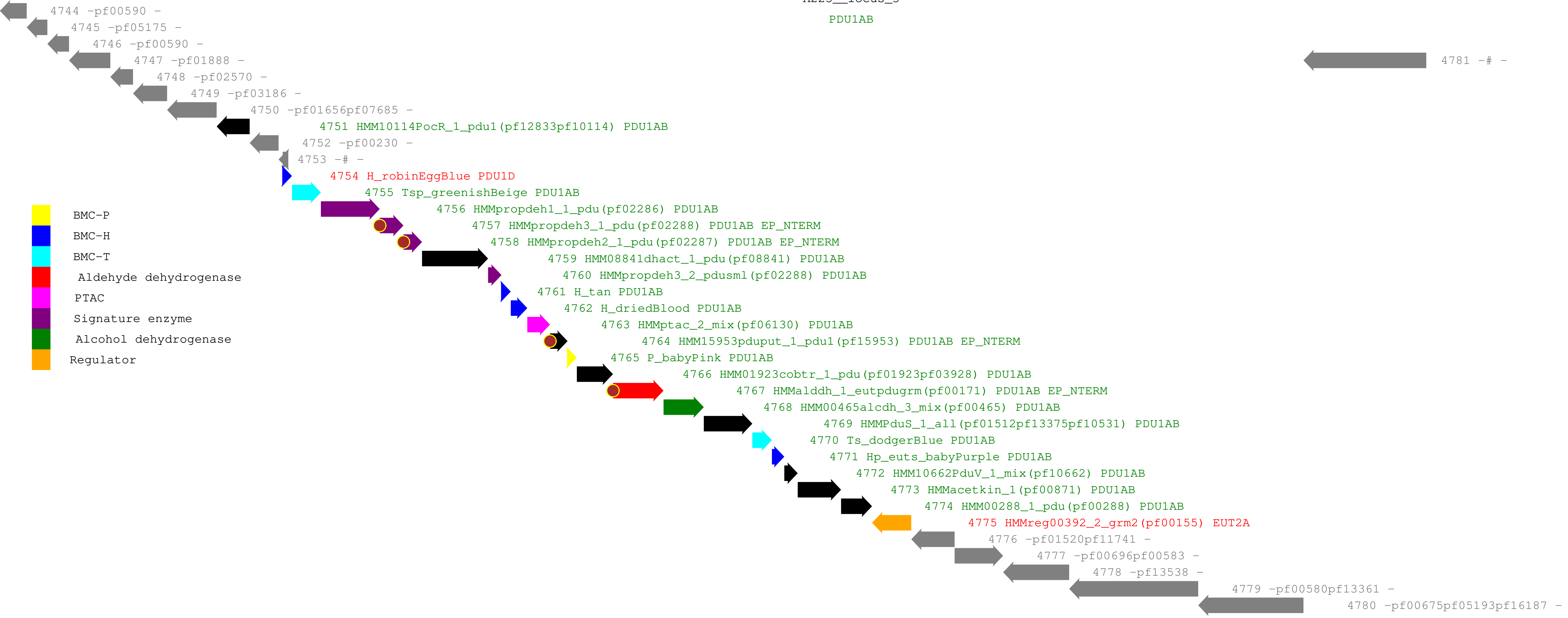

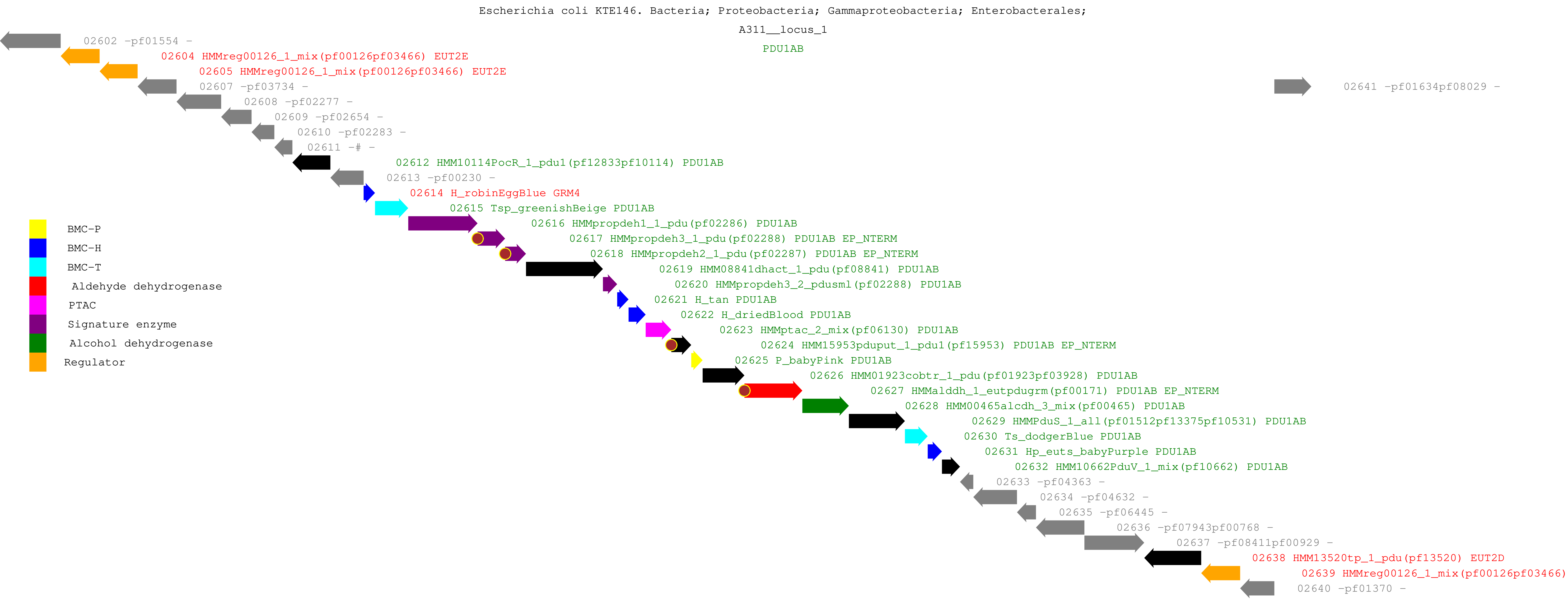
Escherichia coli KTE146. Bacteria; Proteobacteria; Gammaproteobacteria; Enterobacterales;
A311__locus_1
PDU1AB
02602 -pf01554 -
02604 HMMreg00126_1_mix(pf00126pf03466) EUT2E
02605 HMMreg00126_1_mix(pf00126pf03466) EUT2E
02607 -pf03734 -
02608 -pf02277 -
02609 -pf02654 -
02610 -pf02283 -
02611 -# -
02612 HMM10114PocR_1_pdu1(pf12833pf10114) PDU1AB
02613 -pf00230 -
02614 H_robinEggBlue GRM4
02615 Tsp_greenishBeige PDU1AB
02616 HMMpropdeh1_1_pdu(pf02286) PDU1AB
02617 HMMpropdeh3_1_pdu(pf02288) PDU1AB EP_NTERM
02618 HMMpropdeh2_1_pdu(pf02287) PDU1AB EP_NTERM
02619 HMM08841dhact_1_pdu(pf08841) PDU1AB
02620 HMMpropdeh3_2_pdusml(pf02288) PDU1AB
02621 H_tan PDU1AB
02622 H_driedBlood PDU1AB
02623 HMMptac_2_mix(pf06130) PDU1AB
02624 HMM15953pduput_1_pdu1(pf15953) PDU1AB EP_NTERM
02625 P_babyPink PDU1AB
02626 HMM01923cobtr_1_pdu(pf01923pf03928) PDU1AB
02627 HMMaldh_1_eutpdugrm(pf00171) PDU1AB EP_NTERM
02628 HMM00465alcdh_3_mix(pf00465) PDU1AB
02629 HMMPduS_1_all(pf01512pf13375pf10531) PDU1AB
02630 Ts_dodgerBlue PDU1AB
02631 Hp_euts_babyPurple PDU1AB
02632 HMM10662PduV_1_mix(pf10662) PDU1AB
02633 -pf04363 -
02634 -pf04632 -
02635 -pf06445 -
02636 -pf07943pf00768 -
02637 -pf08411pf00929 -
02638 HMM13520tp_1_pdu(pf13520) EUT2D
02639 HMMreg00126_1_mix(pf00126pf03466)
02640 -pf01370 -
02641 -pf01634pf08029 -
BMC-P
BMC-H
BMC-T
Aldehyde dehydrogenase
PTAC
Signature enzyme
Alcohol dehydrogenase
Regulator

Escherichia sp. KTE159. Bacteria; Proteobacteria; Gammaproteobacteria; Enterobacterales;

A31E__locus_1

PDU1AB

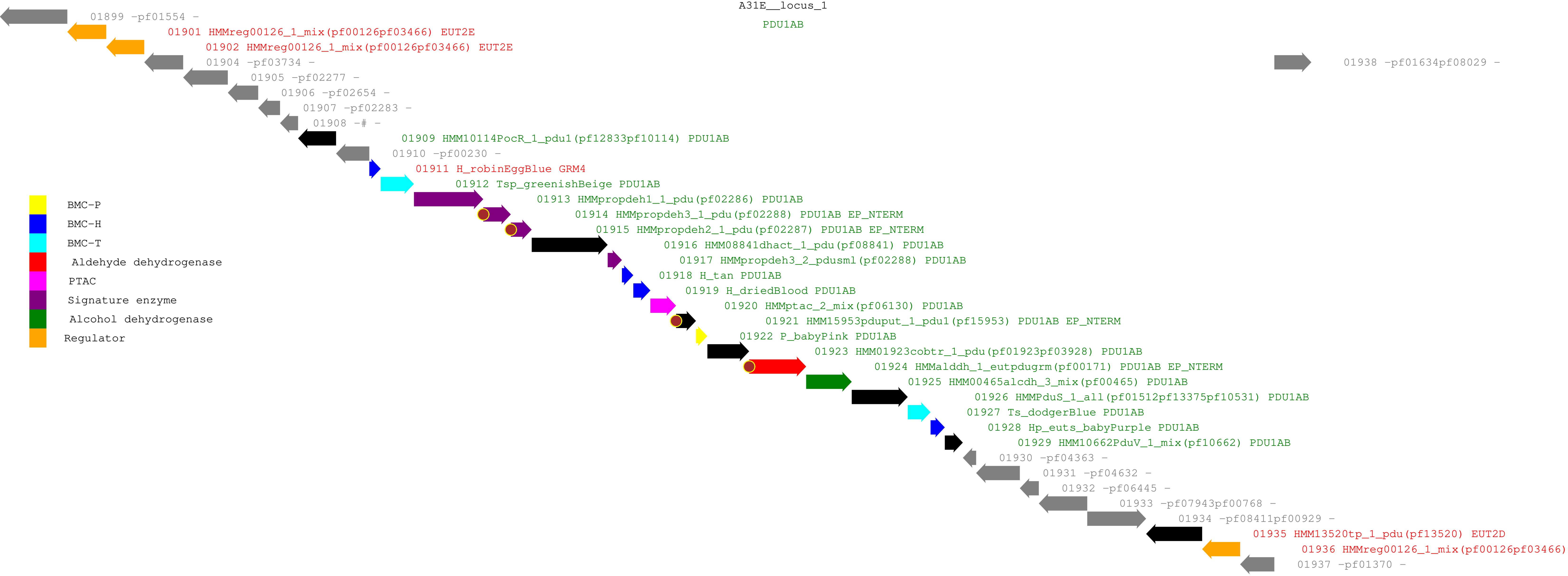

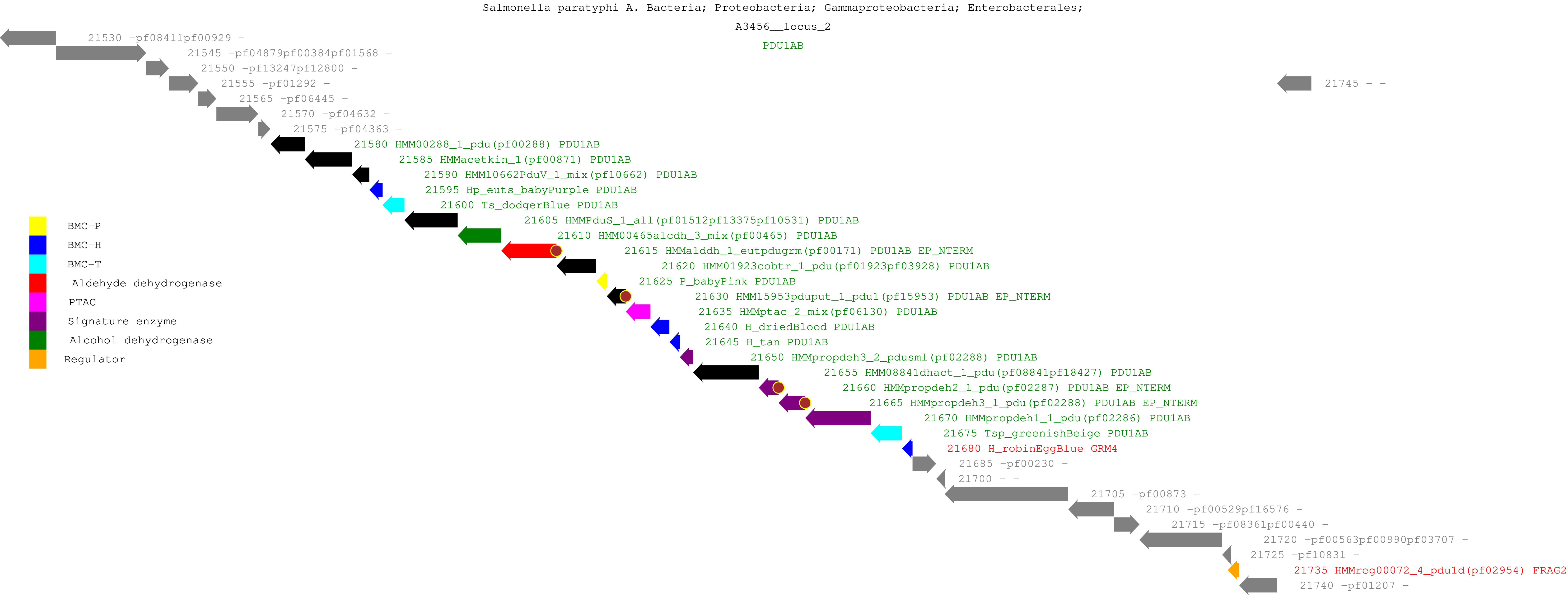
Salmonella paratyphi A. Bacteria; Proteobacteria; Gammaproteobacteria; Enterobacterales;
A3456__locus_2
PDU1AB
21530 -pf08411pf00929 -
21545 -pf04879pf00384pf01568 -
21550 -pf13247pf12800 -
21555 -pf01292 -
21565 -pf06445 -
21570 -pf04632 -
21575 -pf04363 -
21580 HMM00288_1_pdu(pf00288) PDU1AB
21585 HMMacetkin_1(pf00871) PDU1AB
21590 HMM10662PduV_1_mix(pf10662) PDU1AB
21595 Hp_euts_babyPurple PDU1AB
21600 Ts_dodgerBlue PDU1AB
21605 HMMPduS_1_all(pf01512pf13375pf10531) PDU1AB
21610 HMM00465alcdh_3_mix(pf00465) PDU1AB
21615 HMMalddh_1_eutpdugrm(pf00171) PDU1AB EP_NTERM
21620 HMM01923cobtr_1_pdu(pf01923pf03928) PDU1AB
21625 P_babyPink PDU1AB
21630 HMM15953pduput_1_pdu1(pf15953) PDU1AB EP_NTERM
21635 HMMptac_2_mix(pf06130) PDU1AB
21640 H_driedBlood PDU1AB
21645 H_tan PDU1AB
21650 HMMpropdeh3_2_pdusml(pf02288) PDU1AB
21655 HMM08841dhact_1_pdu(pf08841pf18427) PDU1AB
21660 HMMpropdeh2_1_pdu(pf02287) PDU1AB EP_NTERM
21665 HMMpropdeh3_1_pdu(pf02288) PDU1AB EP_NTERM
21670 HMMpropdeh1_1_pdu(pf02286) PDU1AB
21675 Tsp_greenishBeige PDU1AB
21680 H_robinEggBlue GRM4
21685 -pf00230 -
21700 - -
21705 -pf00873 -
21710 -pf00529pf16576 -
21715 -pf08361pf00440 -
21720 -pf00563pf00990pf03707 -
21725 -pf10831 -
21735 HMMreg00072_4_pdu1d(pf02954) FRAG2
21740 -pf01207 -
21745 - -
BMC-P
BMC-H
BMC-T
Aldehyde dehydrogenase
PTAC
Signature enzyme
Alcohol dehydrogenase
Regulator

Salmonella enterica I. Bacteria; Proteobacteria; Gammaproteobacteria; Enterobacterales;

A3T25__locus_2

PDU1AB

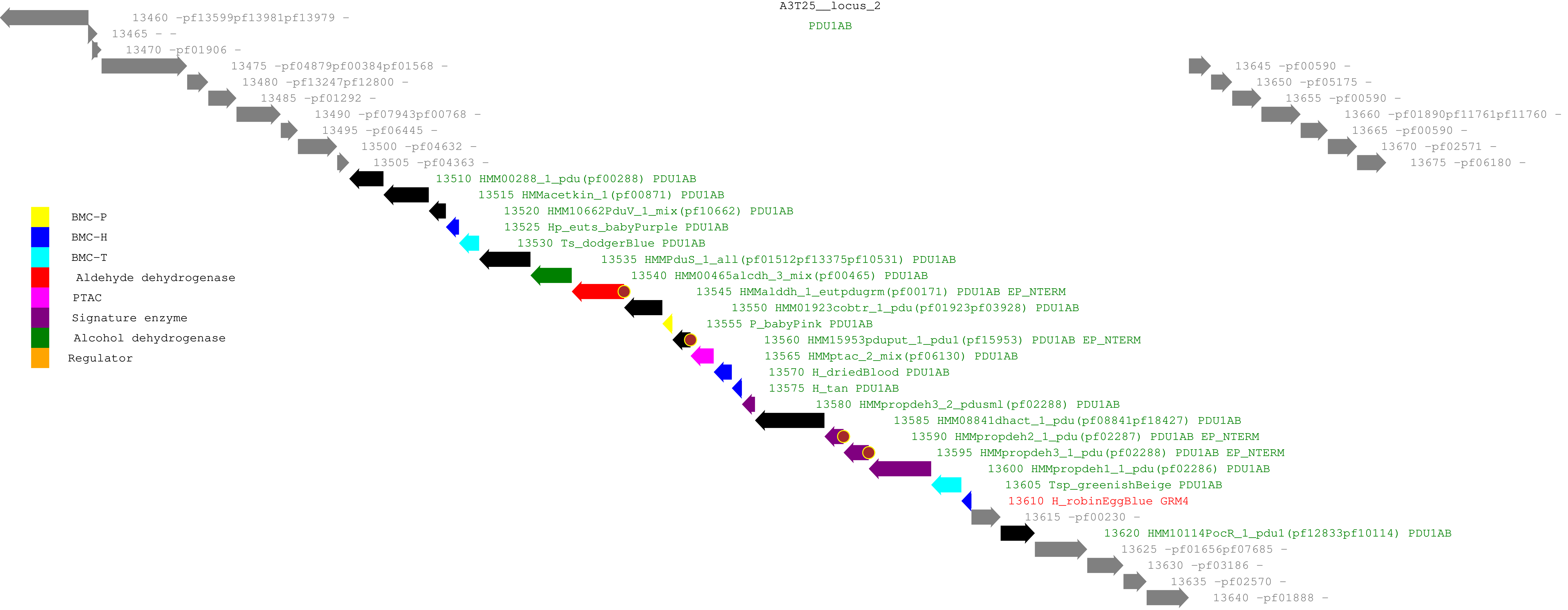

Salmonella enterica subsp. enterica serovar Heidelberg. Bacteria; Proteobacteria; Gammaproteobacteria; Enterobacterales;

A3X64__locus_1

PDU1AB

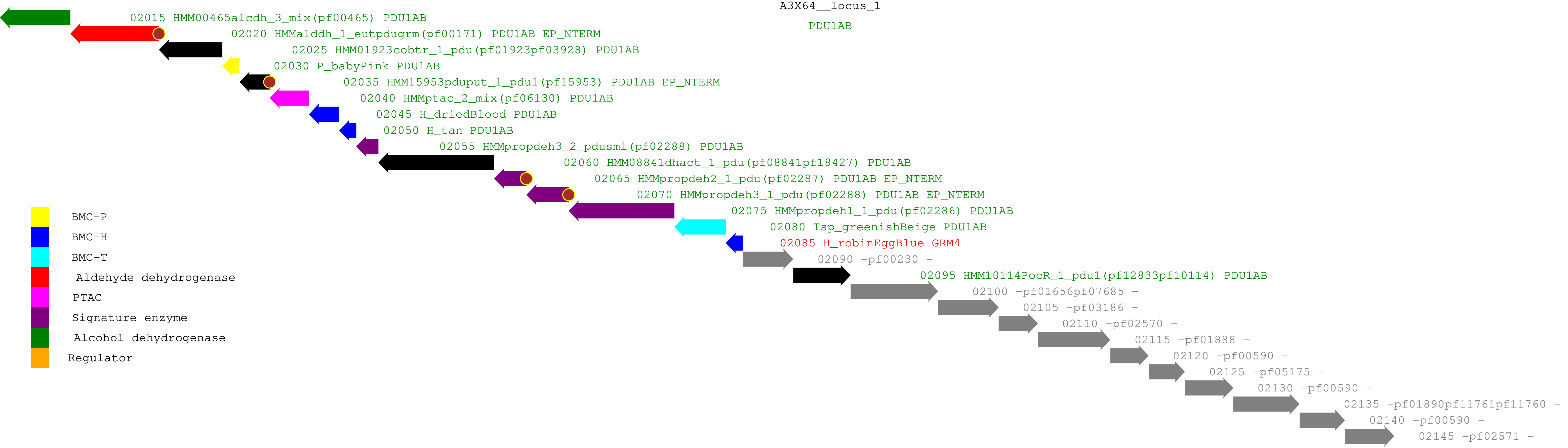

Salmonella enterica subsp. enterica serovar Bareilly str. CFSAN001114. Bacteria; Proteobacteria; Gammaproteobacteria; Enterobacterales;

A4266__locus_1

PDU1AB

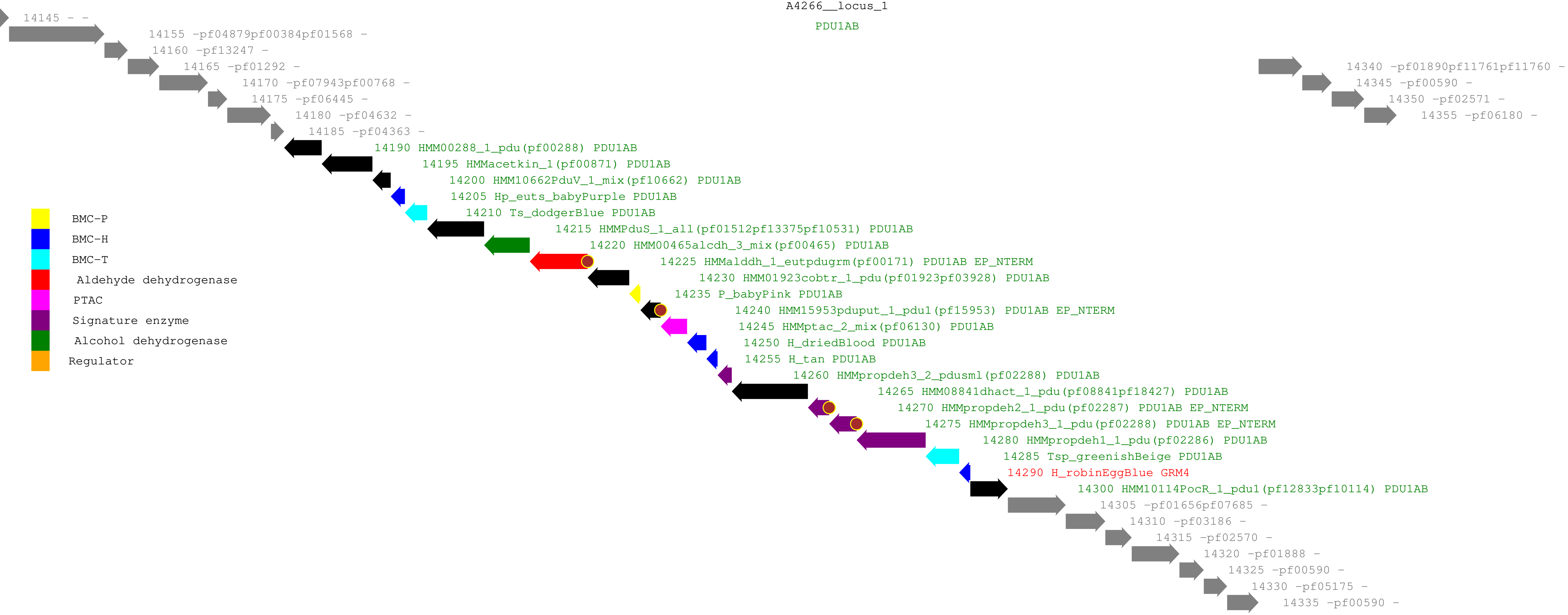

Salmonella enterica subsp. enterica serovar Choleraesuis. Bacteria; Proteobacteria; Gammaproteobacteria; Enterobacterales;

A4I48__locus_2

PDU1AB

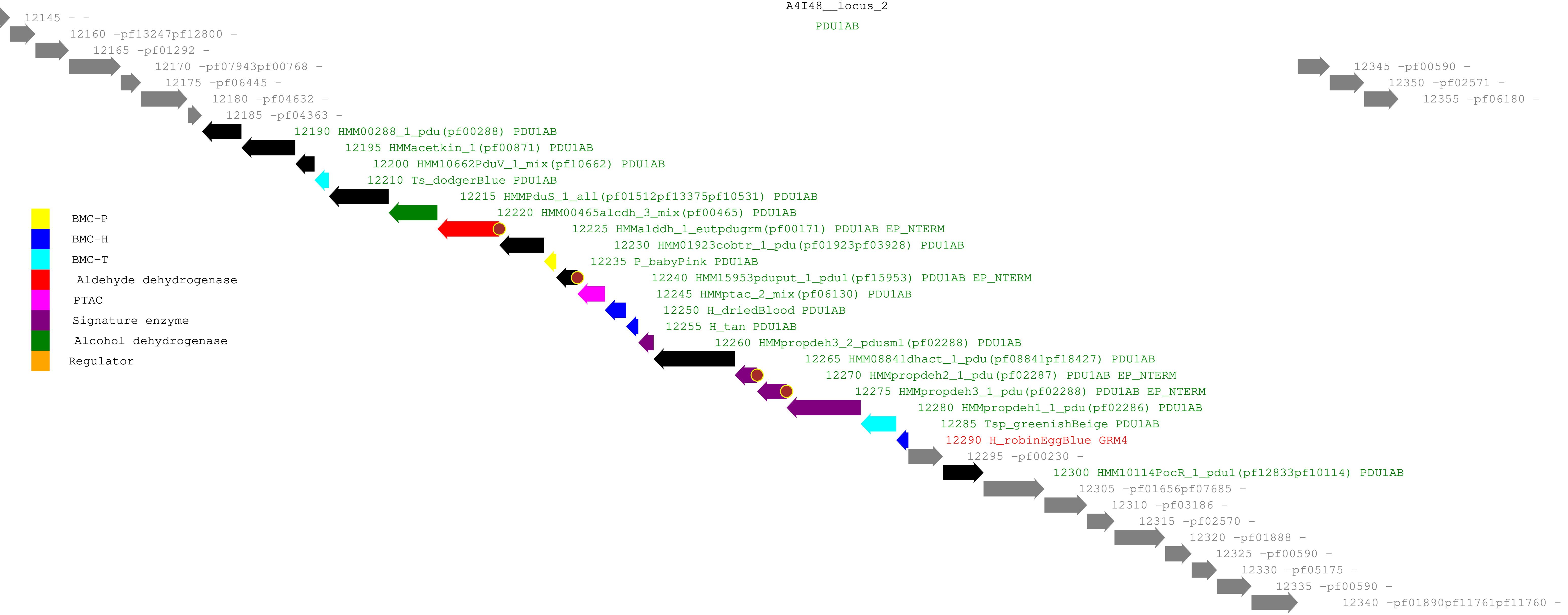

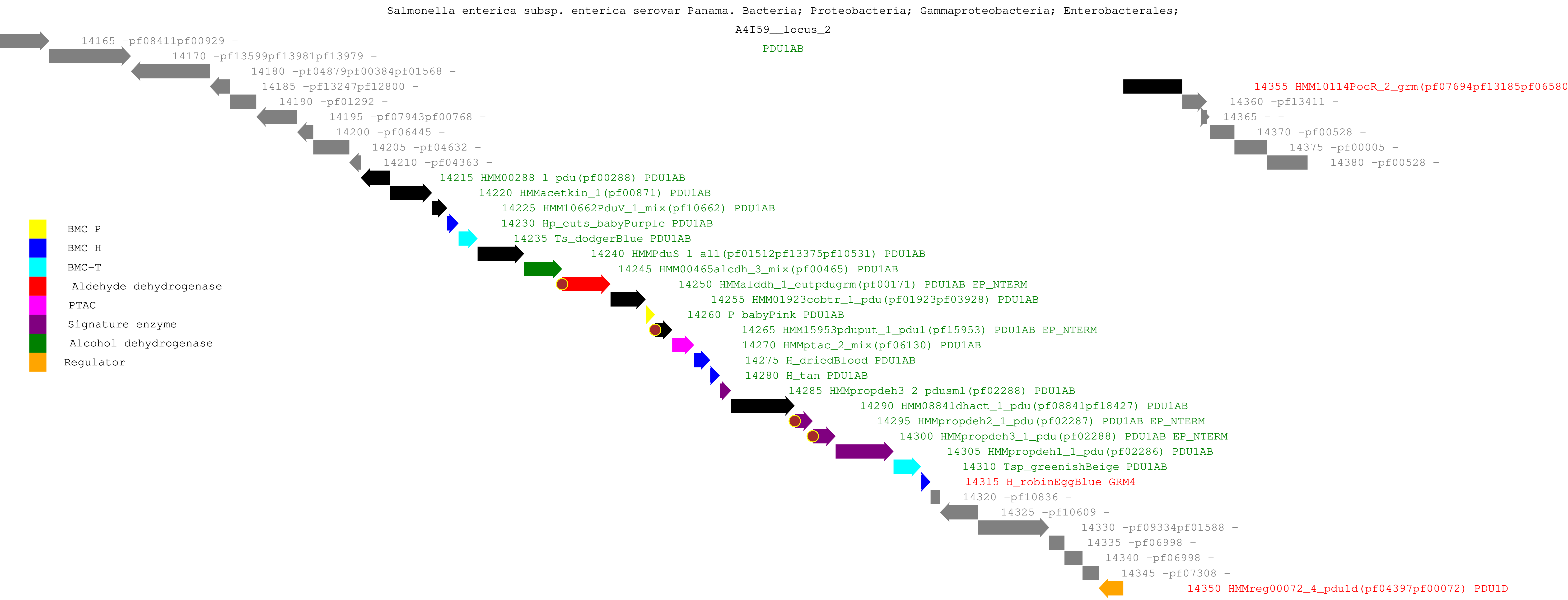
Salmonella enterica subsp. enterica serovar Panama. Bacteria; Proteobacteria; Gammaproteobacteria; Enterobacterales;
A4I59__locus_2
PDU1AB
14165 -pf08411pf00929 -
14170 -pf13599pf13981pf13979 -
14180 -pf04879pf00384pf01568 -
14185 -pf13247pf12800 -
14190 -pf01292 -
14195 -pf07943pf00768 -
14200 -pf06445 -
14205 -pf04632 -
14210 -pf04363 -
14215 HMM00288_1_pdu(pf00288) PDU1AB
14220 HMMacetkin_1(pf00871) PDU1AB
14225 HMM10662PduV_1_mix(pf10662) PDU1AB
14230 Hp_euts_babyPurple PDU1AB
14235 Ts_dodgerBlue PDU1AB
14240 HMMPduS_1_all(pf01512pf13375pf10531) PDU1AB
14245 HMM00465alcdh_3_mix(pf00465) PDU1AB
14250 HMMalddh_1_eutpdugrm(pf00171) PDU1AB EP_NTERM
14255 HMM01923cobtr_1_pdu(pf01923pf03928) PDU1AB
14260 P_babyPink PDU1AB
14265 HMM15953pduput_1_pdu1(pf15953) PDU1AB EP_NTERM
14270 HMMptac_2_mix(pf06130) PDU1AB
14275 H_driedBlood PDU1AB
14280 H_tan PDU1AB
14285 HMMpropdeh3_2_pdusml(pf02288) PDU1AB
14290 HMM08841dhact_1_pdu(pf08841pf18427) PDU1AB
14295 HMMpropdeh2_1_pdu(pf02287) PDU1AB EP_NTERM
14300 HMMpropdeh3_1_pdu(pf02288) PDU1AB EP_NTERM
14305 HMMpropdeh1_1_pdu(pf02286) PDU1AB
14310 Tsp_greenishBeige PDU1AB
14315 H_robinEggBlue GRM4
14320 -pf10836 -
14325 -pf10609 -
14330 -pf09334pf01588 -
14335 -pf06998 -
14340 -pf06998 -
14345 -pf07308 -
14350 HMMreg00072_4_pdu1d(pf04397pf00072) PDU1D
14355 HMM10114PocR_2_grm(pf07694pf13185pf06580
14360 -pf13411 -
14365 - -
14370 -pf00528 -
14375 -pf00005 -
14380 -pf00528 -
BMC-P
BMC-H
BMC-T
Aldehyde dehydrogenase
PTAC
Signature enzyme
Alcohol dehydrogenase
Regulator

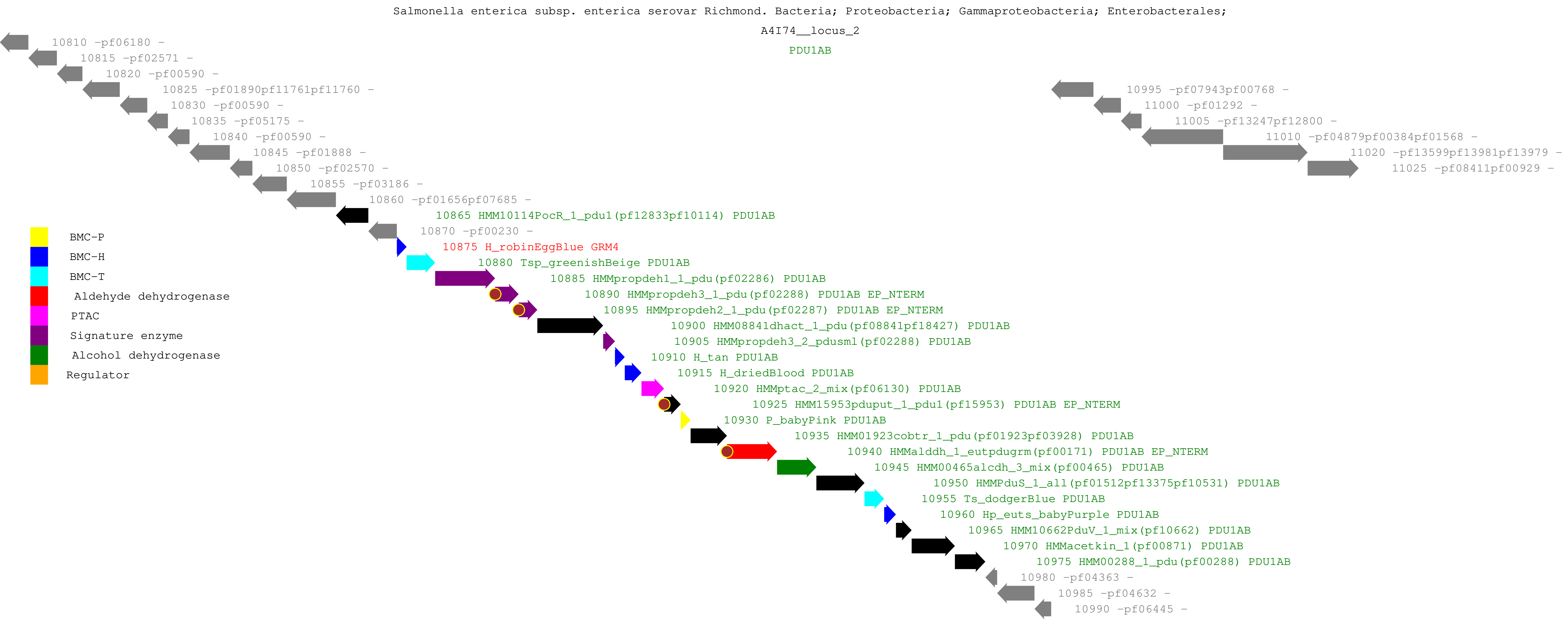

Salmonella enterica subsp. enterica serovar Richmond. Bacteria; Proteobacteria; Gammaproteobacteria; Enterobacterales;
A4I74__locus_2
PDU1AB
10810 -pf06180 -
10815 -pf02571 -
10820 -pf00590 -
10825 -pf01890pf11761pf11760 -
10830 -pf00590 -
10835 -pf05175 -
10840 -pf00590 -
10845 -pf01888 -
10850 -pf02570 -
10855 -pf03186 -
10860 -pf01656pf07685 -
10865 HMM10114PocR_1_pdu1(pf12833pf10114) PDU1AB
10870 -pf00230 -
10875 H_robinEggBlue GRM4
10880 Tsp_greenishBeige PDU1AB
10885 HMMpropdeh1_1_pdu(pf02286) PDU1AB
10890 HMMpropdeh3_1_pdu(pf02288) PDU1AB EP_NTERM
10895 HMMpropdeh2_1_pdu(pf02287) PDU1AB EP_NTERM
10900 HMM08841dhact_1_pdu(pf08841pf18427) PDU1AB
10905 HMMpropdeh3_2_pdusml(pf02288) PDU1AB
10910 H_tan PDU1AB
10915 H_driedBlood PDU1AB
10920 HMMptac_2_mix(pf06130) PDU1AB
10925 HMM15953pduput_1_pdu1(pf15953) PDU1AB EP_NTERM
10930 P_babyPink PDU1AB
10935 HMM01923cobtr_1_pdu(pf01923pf03928) PDU1AB
10940 HMMalddh_1_eutpdugrm(pf00171) PDU1AB EP_NTERM
10945 HMM00465alcdh_3_mix(pf00465) PDU1AB
10950 HMMPduS_1_all(pf01512pf13375pf10531) PDU1AB
10955 Ts_dodgerBlue PDU1AB
10960 Hp_euts_babyPurple PDU1AB
10965 HMM10662PduV_1_mix(pf10662) PDU1AB
10970 HMMacetkin_1(pf00871) PDU1AB
10975 HMM00288_1_pdu(pf00288) PDU1AB
10980 -pf04363 -
10985 -pf04632 -
10990 -pf06445 -
10995 -pf07943pf00768 -
11000 -pf01292 -
11005 -pf13247pf12800 -
11010 -pf04879pf00384pf01568 -
11020 -pf13599pf13981pf13979 -
11025 -pf08411pf00929 -
BMC-P
BMC-H
BMC-T
Aldehyde dehydrogenase
PTAC
Signature enzyme
Alcohol dehydrogenase
Regulator

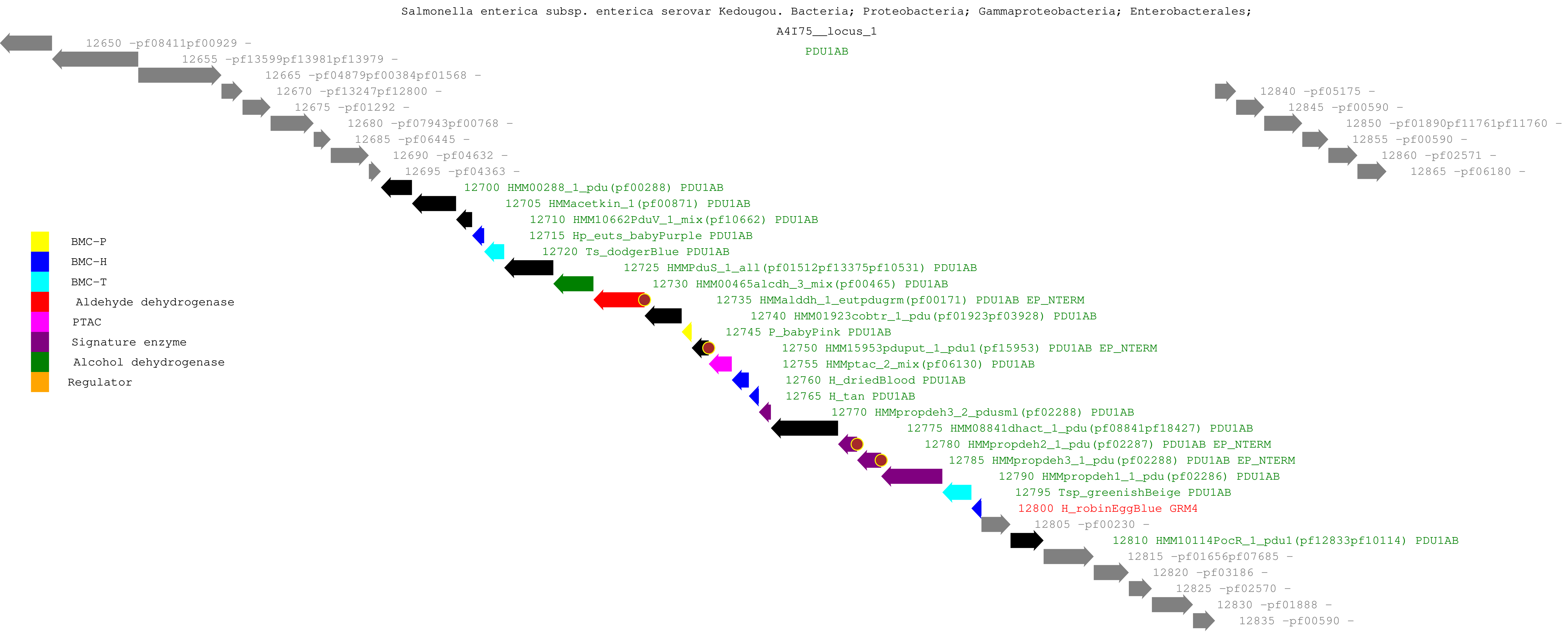
Salmonella enterica subsp. enterica serovar Kedougou. Bacteria; Proteobacteria; Gammaproteobacteria; Enterobacterales;
A4I75__locus_1
PDU1AB
12650 -pf08411pf00929 -
12655 -pf13599pf13981pf13979 -
12665 -pf04879pf00384pf01568 -
12670 -pf13247pf12800 -
12675 -pf01292 -
12680 -pf07943pf00768 -
12685 -pf06445 -
12690 -pf04632 -
12695 -pf04363 -
12700 HMM00288_1_pdu(pf00288) PDU1AB
12705 HMMacetkin_1(pf00871) PDU1AB
12710 HMM10662PduV_1_mix(pf10662) PDU1AB
12715 Hp_euts_babyPurple PDU1AB
12720 Ts_dodgerBlue PDU1AB
12725 HMMPduS_1_all(pf01512pf13375pf10531) PDU1AB
12730 HMM00465alcdh_3_mix(pf00465) PDU1AB
12735 HMMalddh_1_eutpdugrm(pf00171) PDU1AB EP_NTERM
12740 HMM01923cobtr_1_pdu(pf01923pf03928) PDU1AB
12745 P_babyPink PDU1AB
12750 HMM15953pduput_1_pdu1(pf15953) PDU1AB EP_NTERM
12755 HMMptac_2_mix(pf06130) PDU1AB
12760 H_driedBlood PDU1AB
12765 H_tan PDU1AB
12770 HMMpropdeh3_2_pdusml(pf02288) PDU1AB
12775 HMM08841dhact_1_pdu(pf08841pf18427) PDU1AB
12780 HMMpropdeh2_1_pdu(pf02287) PDU1AB EP_NTERM
12785 HMMpropdeh3_1_pdu(pf02288) PDU1AB EP_NTERM
12790 HMMpropdeh1_1_pdu(pf02286) PDU1AB
12795 Tsp_greenishBeige PDU1AB
12800 H_robinEggBlue GRM4
12805 -pf00230 -
12810 HMM10114PocR_1_pdu1(pf12833pf10114) PDU1AB
12815 -pf01656pf07685 -
12820 -pf03186 -
12825 -pf02570 -
12830 -pf01888 -
12835 -pf00590 -
12840 -pf05175 -
12845 -pf00590 -
12850 -pf01890pf11761pf11760 -
12855 -pf00590 -
12860 -pf02571 -
12865 -pf06180 -
BMC-P
BMC-H
BMC-T
Aldehyde dehydrogenase
PTAC
Signature enzyme
Alcohol dehydrogenase
Regulator

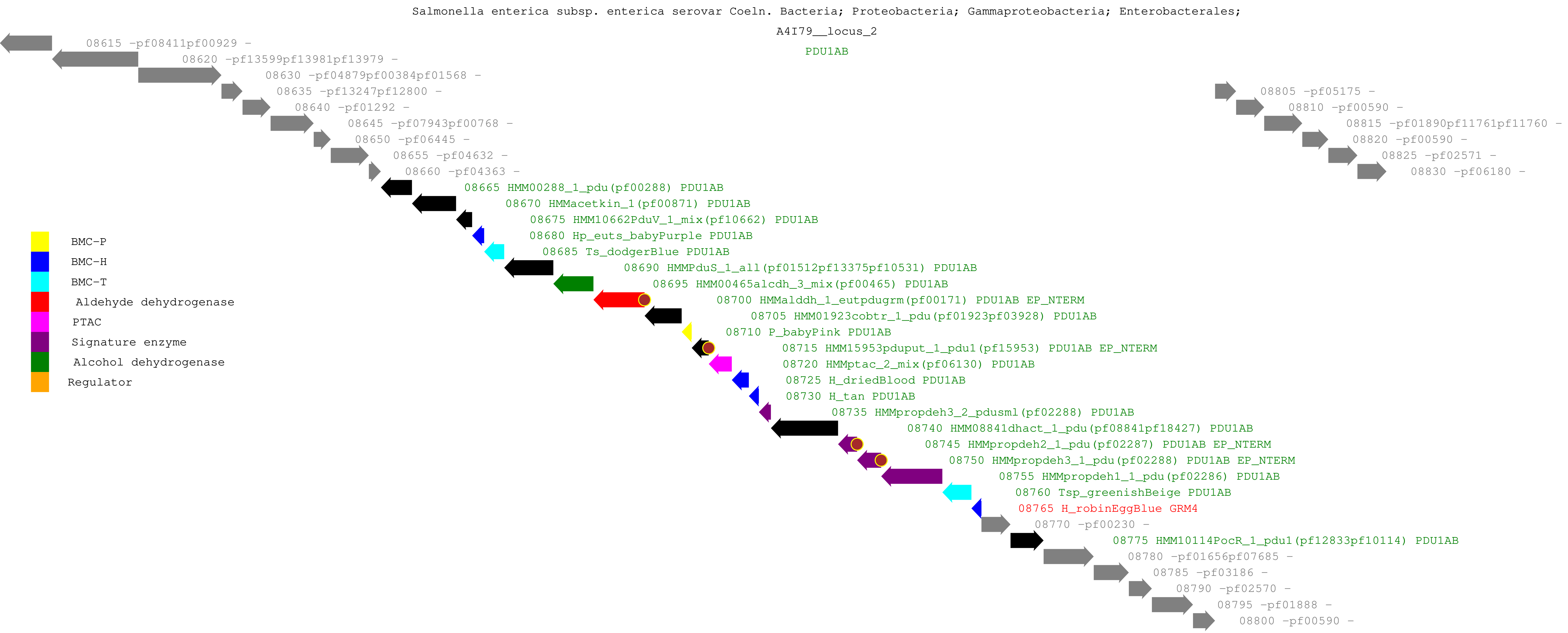

Salmonella enterica subsp. enterica serovar Coeln. Bacteria; Proteobacteria; Gammaproteobacteria; Enterobacterales;
A4I79__locus_2
PDU1AB
08615 -pf08411pf00929 -
08620 -pf13599pf13981pf13979 -
08630 -pf04879pf00384pf01568 -
08635 -pf13247pf12800 -
08640 -pf01292 -
08645 -pf07943pf00768 -
08650 -pf06445 -
08655 -pf04632 -
08660 -pf04363 -
08665 HMM00288_1_pdu(pf00288) PDU1AB
08670 HMMacetkin_1(pf00871) PDU1AB
08675 HMM10662PduV_1_mix(pf10662) PDU1AB
08680 Hp_euts_babyPurple PDU1AB
08685 Ts_dodgerBlue PDU1AB
08690 HMMPduS_1_all(pf01512pf13375pf10531) PDU1AB
08695 HMM00465alcdh_3_mix(pf00465) PDU1AB
08700 HMMalddh_1_eutpdugrm(pf00171) PDU1AB EP_NTERM
08705 HMM01923cobtr_1_pdu(pf01923pf03928) PDU1AB
08710 P_babyPink PDU1AB
08715 HMM15953pduput_1_pdu1(pf15953) PDU1AB EP_NTERM
08720 HMMptac_2_mix(pf06130) PDU1AB
08725 H_driedBlood PDU1AB
08730 H_tan PDU1AB
08735 HMMpropdeh3_2_pdusml(pf02288) PDU1AB
08740 HMM08841dhact_1_pdu(pf08841pf18427) PDU1AB
08745 HMMpropdeh2_1_pdu(pf02287) PDU1AB EP_NTERM
08750 HMMpropdeh3_1_pdu(pf02288) PDU1AB EP_NTERM
08755 HMMpropdeh1_1_pdu(pf02286) PDU1AB
08760 Tsp_greenishBeige PDU1AB
08765 H_robinEggBlue GRM4
08770 -pf00230 -
08775 HMM10114PocR_1_pdu1(pf12833pf10114) PDU1AB
08780 -pf01656pf07685 -
08785 -pf03186 -
08790 -pf02570 -
08795 -pf01888 -
08800 -pf00590 -
08805 -pf05175 -
08810 -pf00590 -
08815 -pf01890pf11761pf11760 -
08820 -pf00590 -
08825 -pf02571 -
08830 -pf06180 -
BMC-P
BMC-H
BMC-T
Aldehyde dehydrogenase
PTAC
Signature enzyme
Alcohol dehydrogenase
Regulator

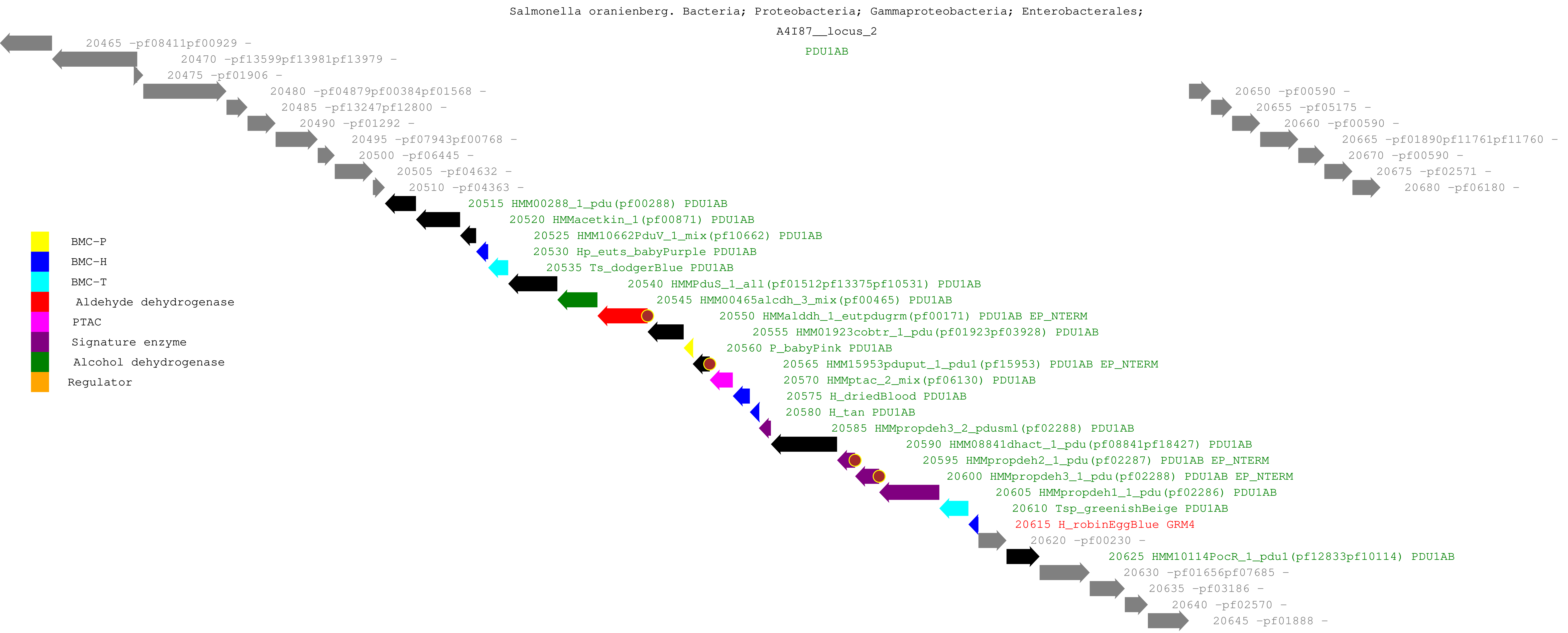
Salmonella oranienberg. Bacteria; Proteobacteria; Gammaproteobacteria; Enterobacterales;
A4I87__locus_2
PDU1AB
20465 -pf08411pf00929 -
20470 -pf13599pf13981pf13979 -
20475 -pf01906 -
20480 -pf04879pf00384pf01568 -
20485 -pf13247pf12800 -
20490 -pf01292 -
20495 -pf07943pf00768 -
20500 -pf06445 -
20505 -pf04632 -
20510 -pf04363 -
20515 HMM00288_1_pdu(pf00288) PDU1AB
20520 HMMacetkin_1(pf00871) PDU1AB
20525 HMM10662PduV_1_mix(pf10662) PDU1AB
20530 Hp_euts_babyPurple PDU1AB
20535 Ts_dodgerBlue PDU1AB
20540 HMMPduS_1_all(pf01512pf13375pf10531) PDU1AB
20545 HMM00465alcdh_3_mix(pf00465) PDU1AB
20550 HMMaldh_1_eutpdugrm(pf00171) PDU1AB EP_NTERM
20555 HMM01923cobtr_1_pdu(pf01923pf03928) PDU1AB
20560 P_babyPink PDU1AB
20565 HMM15953pduput_1_pdu1(pf15953) PDU1AB EP_NTERM
20570 HMMptac_2_mix(pf06130) PDU1AB
20575 H_driedBlood PDU1AB
20580 H_tan PDU1AB
20585 HMMpropdeh3_2_pdusml(pf02288) PDU1AB
20590 HMM08841dhact_1_pdu(pf08841pf18427) PDU1AB
20595 HMMpropdeh2_1_pdu(pf02287) PDU1AB EP_NTERM
20600 HMMpropdeh3_1_pdu(pf02288) PDU1AB EP_NTERM
20605 HMMpropdeh1_1_pdu(pf02286) PDU1AB
20610 Tsp_greenishBeige PDU1AB
20615 H_robinEggBlue GRM4
20620 -pf00230 -
20625 HMM10114PocR_1_pdu1(pf12833pf10114) PDU1AB
20630 -pf01656pf07685 -
20635 -pf03186 -
20640 -pf02570 -
20645 -pf01888 -
20650 -pf00590 -
20655 -pf05175 -
20660 -pf00590 -
20665 -pf01890pf11761pf11760 -
20670 -pf00590 -
20675 -pf02571 -
20680 -pf06180 -
BMC-P
BMC-H
BMC-T
Aldehyde dehydrogenase
PTAC
Signature enzyme
Alcohol dehydrogenase
Regulator

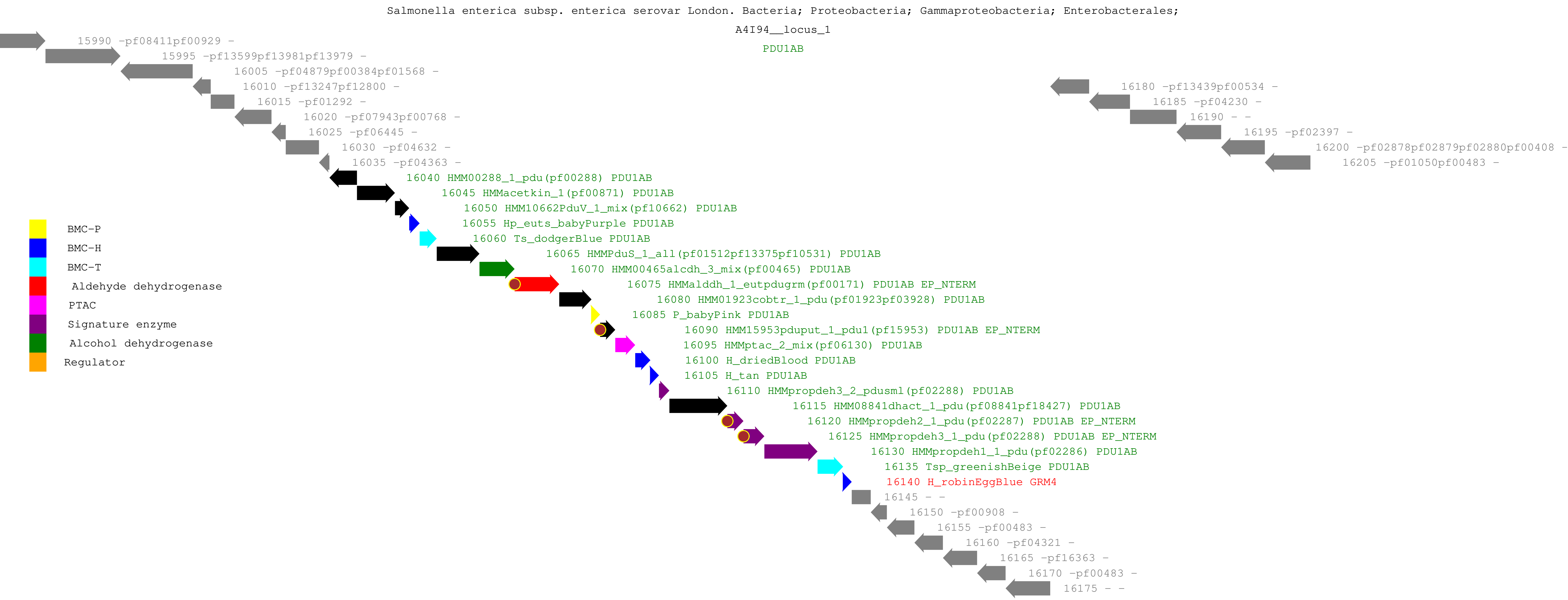
Salmonella enterica subsp. enterica serovar London. Bacteria; Proteobacteria; Gammaproteobacteria; Enterobacterales;
A4I94__locus_1
PDU1AB
15990 -pf08411pf00929 -
15995 -pf13599pf13981pf13979 -
16005 -pf04879pf00384pf01568 -
16010 -pf13247pf12800 -
16015 -pf01292 -
16020 -pf07943pf00768 -
16025 -pf06445 -
16030 -pf04632 -
16035 -pf04363 -
16040 HMM00288_1_pdu(pf00288) PDU1AB
16045 HMMacetkin_1(pf00871) PDU1AB
16050 HMM10662PduV_1_mix(pf10662) PDU1AB
16055 Hp_euts_babyPurple PDU1AB
16060 Ts_dodgerBlue PDU1AB
16065 HMMPduS_1_all(pf01512pf13375pf10531) PDU1AB
16070 HMM00465alcdh_3_mix(pf00465) PDU1AB
16075 HMMalddh_1_eutpdugrm(pf00171) PDU1AB EP_NTERM
16080 HMM01923cobtr_1_pdu(pf01923pf03928) PDU1AB
16085 P_babyPink PDU1AB
16090 HMM15953pduput_1_pdu1(pf15953) PDU1AB EP_NTERM
16095 HMMptac_2_mix(pf06130) PDU1AB
16100 H_driedBlood PDU1AB
16105 H_tan PDU1AB
16110 HMMpropdeh3_2_pdusml(pf02288) PDU1AB
16115 HMM08841dhact_1_pdu(pf08841pf18427) PDU1AB
16120 HMMpropdeh2_1_pdu(pf02287) PDU1AB EP_NTERM
16125 HMMpropdeh3_1_pdu(pf02288) PDU1AB EP_NTERM
16130 HMMpropdeh1_1_pdu(pf02286) PDU1AB
16135 Tsp_greenishBeige PDU1AB
16140 H_robinEggBlue GRM4
16145 - -
16150 -pf00908 -
16155 -pf00483 -
16160 -pf04321 -
16165 -pf16363 -
16170 -pf00483 -
16175 - -
16180 -pf13439pf00534 -
16185 -pf04230 -
16190 - -
16195 -pf02397 -
16200 -pf02878pf02879pf02880pf00408 -
16205 -pf01050pf00483 -
BMC-P
BMC-H
BMC-T
Aldehyde dehydrogenase
PTAC
Signature enzyme
Alcohol dehydrogenase
Regulator

Salmonella enterica subsp. enterica serovar Sandiego. Bacteria; Proteobacteria; Gammaproteobacteria; Enterobacterales;

A4J53__locus_1

PDU1AB

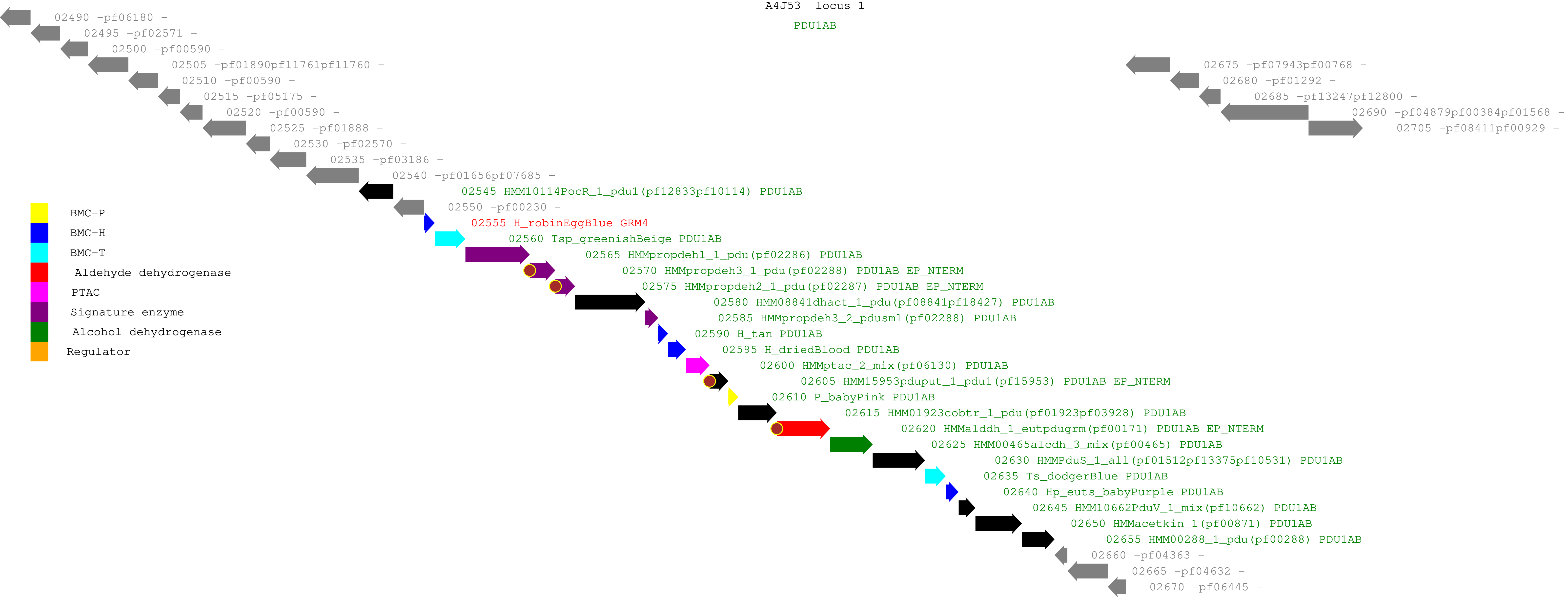

Salmonella enterica subsp. enterica serovar Joal. Bacteria; Proteobacteria; Gammaproteobacteria; Enterobacterales;

A4J61__locus_2

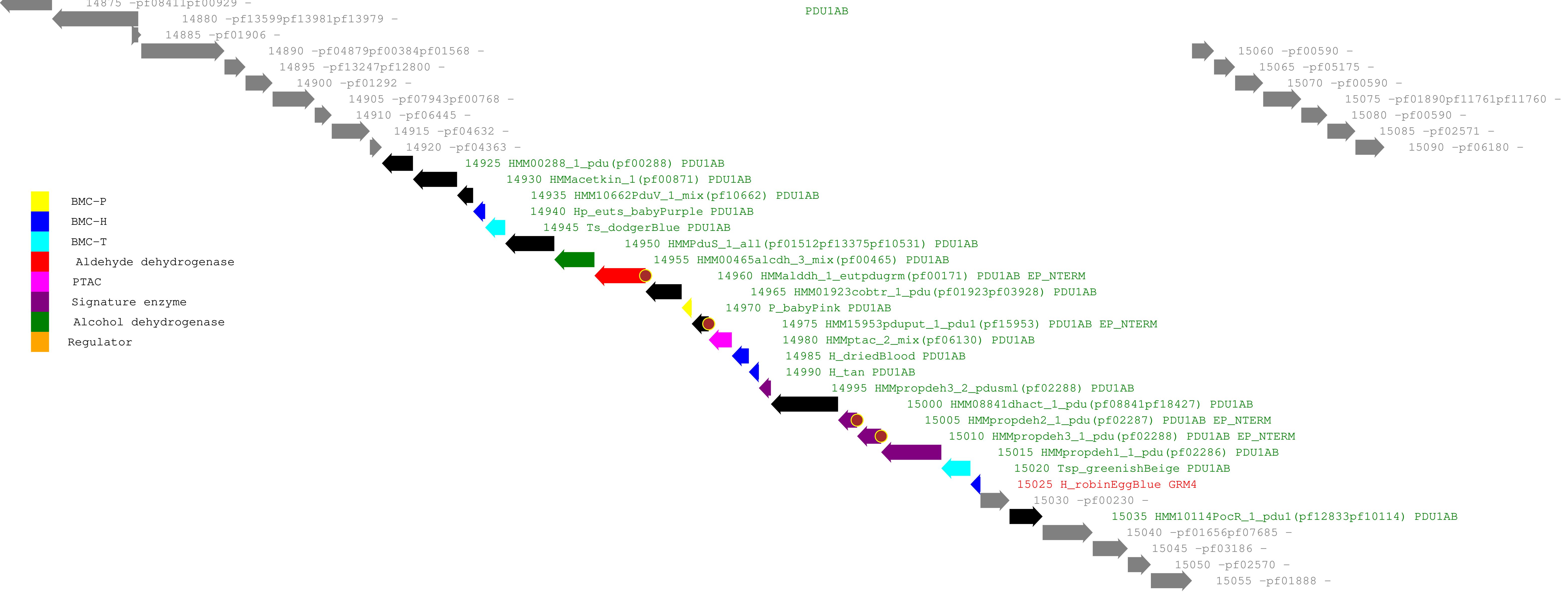

Salmonella enterica subsp. enterica serovar Chester. Bacteria; Proteobacteria; Gammaproteobacteria; Enterobacterales;

A4J74__locus_1

PDU1AB

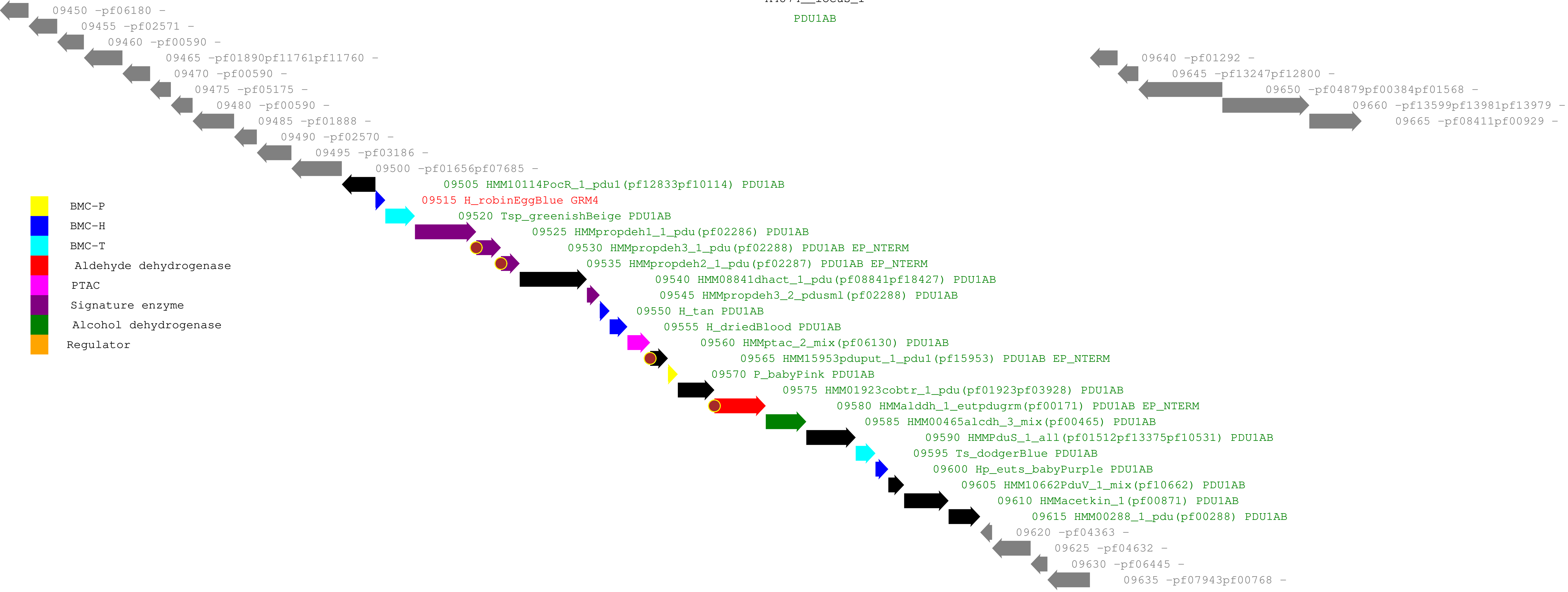

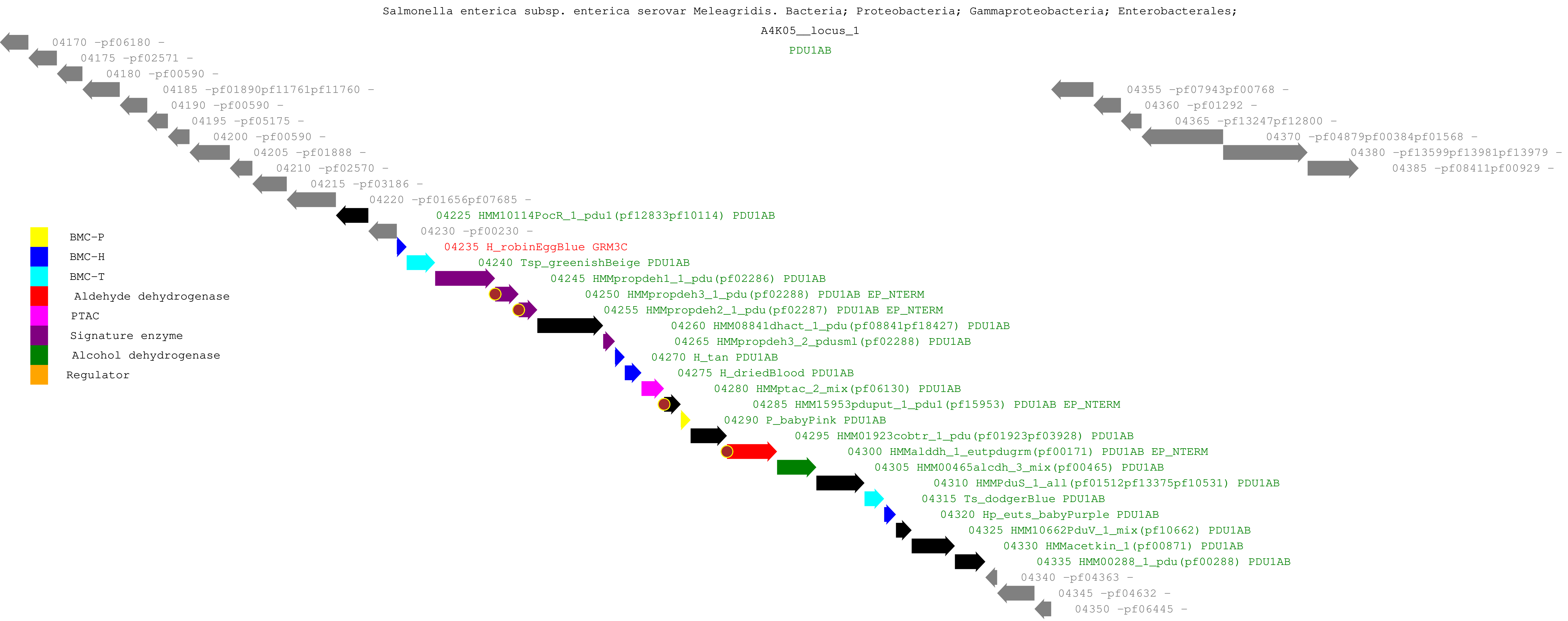
Salmonella enterica subsp. enterica serovar Meleagridis. Bacteria; Proteobacteria; Gammaproteobacteria; Enterobacterales;
A4K05__locus_1
PDU1AB
04170 -pf06180 -
04175 -pf02571 -
04180 -pf00590 -
04185 -pf01890pf11761pf11760 -
04190 -pf00590 -
04195 -pf05175 -
04200 -pf00590 -
04205 -pf01888 -
04210 -pf02570 -
04215 -pf03186 -
04220 -pf01656pf07685 -
04225 HMM10114PocR_1_pdu1(pf12833pf10114) PDU1AB
04230 -pf00230 -
04235 H_robinEggBlue GRM3C
04240 Tsp_greenishBeige PDU1AB
04245 HMMpropdeh1_1_pdu(pf02286) PDU1AB
04250 HMMpropdeh3_1_pdu(pf02288) PDU1AB EP_NTERM
04255 HMMpropdeh2_1_pdu(pf02287) PDU1AB EP_NTERM
04260 HMM08841dhact_1_pdu(pf08841pf18427) PDU1AB
04265 HMMpropdeh3_2_pdusml(pf02288) PDU1AB
04270 H_tan PDU1AB
04275 H_driedBlood PDU1AB
04280 HMMptac_2_mix(pf06130) PDU1AB
04285 HMM15953pduput_1_pdu1(pf15953) PDU1AB EP_NTERM
04290 P_babyPink PDU1AB
04295 HMM01923cobtr_1_pdu(pf01923pf03928) PDU1AB
04300 HMMalddh_1_eutpdugrm(pf00171) PDU1AB EP_NTERM
04305 HMM00465alcdh_3_mix(pf00465) PDU1AB
04310 HMMPduS_1_all(pf01512pf13375pf10531) PDU1AB
04315 Ts_dodgerBlue PDU1AB
04320 Hp_euts_babyPurple PDU1AB
04325 HMM10662PduV_1_mix(pf10662) PDU1AB
04330 HMMacetkin_1(pf00871) PDU1AB
04335 HMM00288_1_pdu(pf00288) PDU1AB
04340 -pf04363 -
04345 -pf04632 -
04350 -pf06445 -
04355 -pf07943pf00768 -
04360 -pf01292 -
04365 -pf13247pf12800 -
04370 -pf04879pf00384pf01568 -
04380 -pf13599pf13981pf13979 -
04385 -pf08411pf00929 -
BMC-P
BMC-H
BMC-T
Aldehyde dehydrogenase
PTAC
Signature enzyme
Alcohol dehydrogenase
Regulator

Salmonella enterica subsp. enterica serovar Bakau. Bacteria; Proteobacteria; Gammaproteobacteria; Enterobacterales;

A4K08__locus_1

PDU1AB

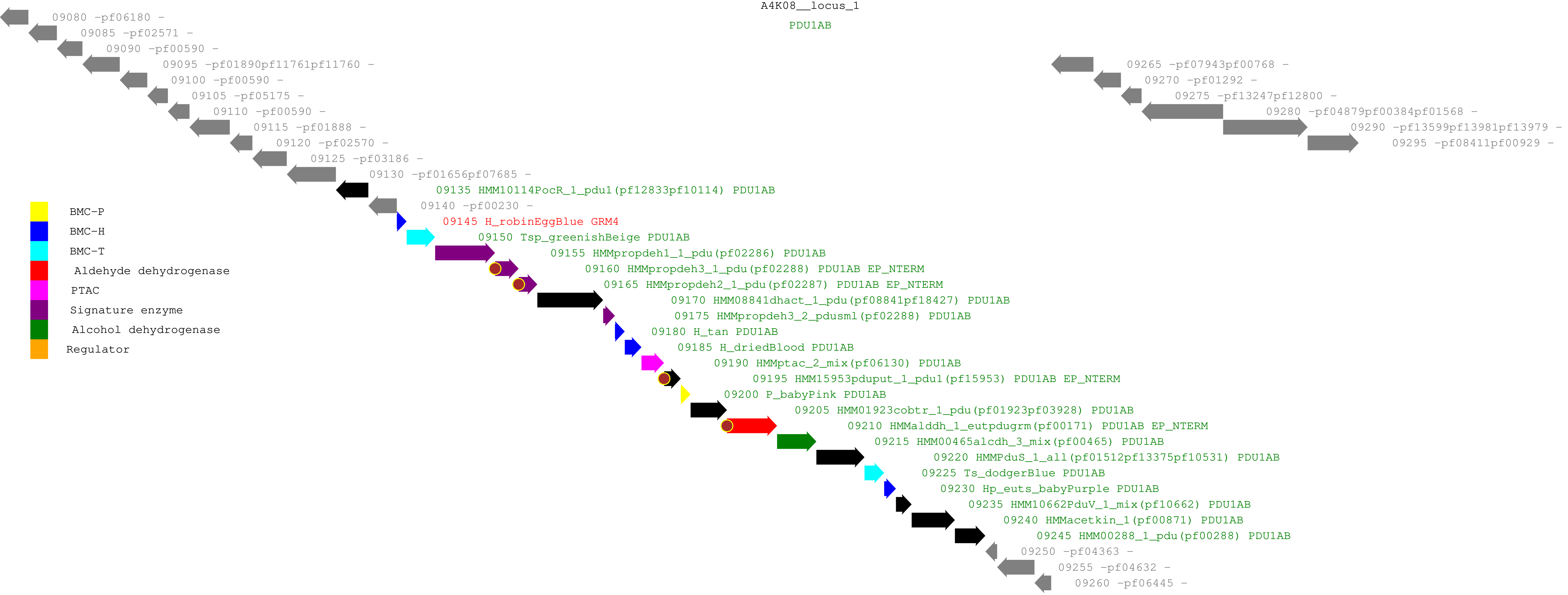

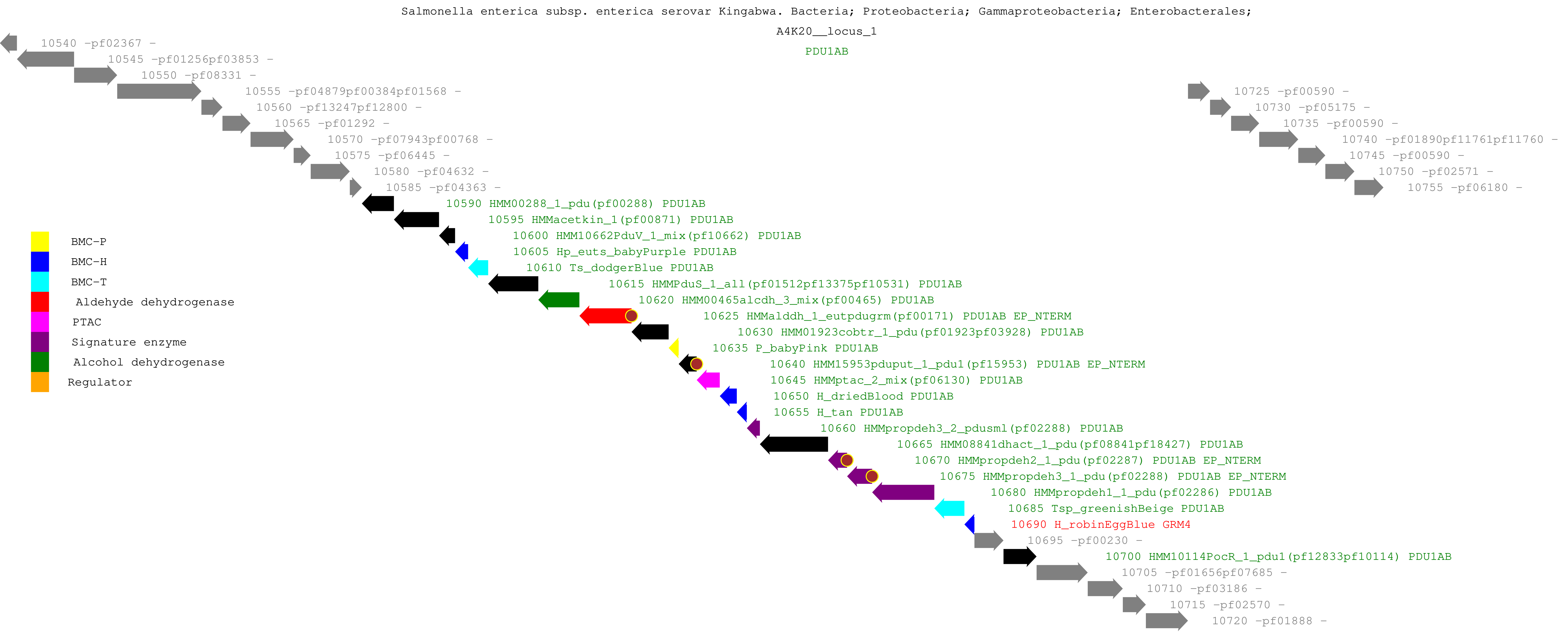

Salmonella enterica subsp. enterica serovar Kingabwa. Bacteria; Proteobacteria; Gammaproteobacteria; Enterobacterales;
A4K20__locus_1
PDU1AB
10540 -pf02367 -
10545 -pf01256pf03853 -
10550 -pf08331 -
10555 -pf04879pf00384pf01568 -
10560 -pf13247pf12800 -
10565 -pf01292 -
10570 -pf07943pf00768 -
10575 -pf06445 -
10580 -pf04632 -
10585 -pf04363 -
10590 HMM00288_1_pdu(pf00288) PDU1AB
10595 HMMacetkin_1(pf00871) PDU1AB
10600 HMM10662PduV_1_mix(pf10662) PDU1AB
10605 Hp_euts_babyPurple PDU1AB
10610 Ts_dodgerBlue PDU1AB
10615 HMMPduS_1_all(pf01512pf13375pf10531) PDU1AB
10620 HMM00465alcdh_3_mix(pf00465) PDU1AB
10625 HMMaldh_1_eutpdugrm(pf00171) PDU1AB EP_NTERM
10630 HMM01923cobtr_1_pdu(pf01923pf03928) PDU1AB
10635 P_babyPink PDU1AB
10640 HMM15953pduput_1_pdu1(pf15953) PDU1AB EP_NTERM
10645 HMMptac_2_mix(pf06130) PDU1AB
10650 H_driedBlood PDU1AB
10655 H_tan PDU1AB
10660 HMMpropdeh3_2_pdusml(pf02288) PDU1AB
10665 HMM08841dhact_1_pdu(pf08841pf18427) PDU1AB
10670 HMMpropdeh2_1_pdu(pf02287) PDU1AB EP_NTERM
10675 HMMpropdeh3_1_pdu(pf02288) PDU1AB EP_NTERM
10680 HMMpropdeh1_1_pdu(pf02286) PDU1AB
10685 Tsp_greenishBeige PDU1AB
10690 H_robinEggBlue GRM4
10695 -pf00230 -
10700 HMM10114PocR_1_pdu1(pf12833pf10114) PDU1AB
10705 -pf01656pf07685 -
10710 -pf03186 -
10715 -pf02570 -
10720 -pf01888 -
10725 -pf00590 -
10730 -pf05175 -
10735 -pf00590 -
10740 -pf01890pf11761pf11760 -
10745 -pf00590 -
10750 -pf02571 -
10755 -pf06180 -
BMC-P
BMC-H
BMC-T
Aldehyde dehydrogenase
PTAC
Signature enzyme
Alcohol dehydrogenase
Regulator

Salmonella enterica subsp. enterica serovar Singapore. Bacteria; Proteobacteria; Gammaproteobacteria; Enterobacterales;

A4K41__locus_2

PDU1AB

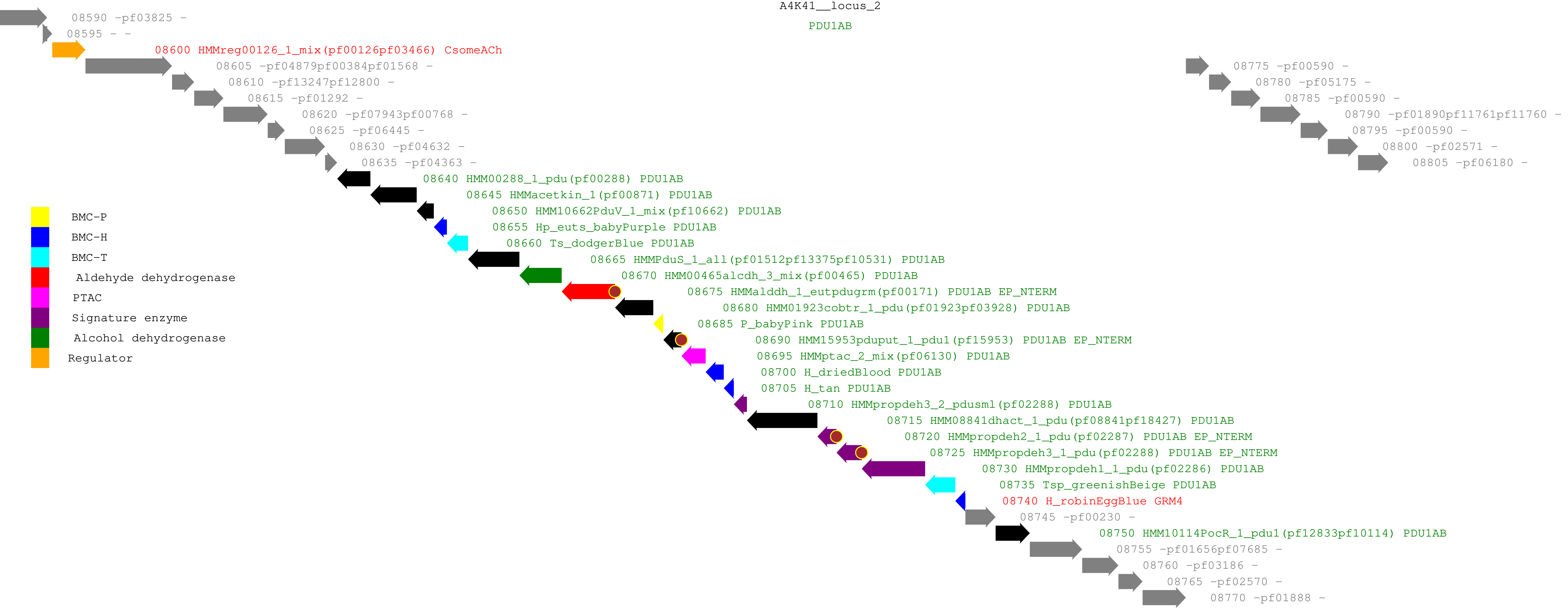

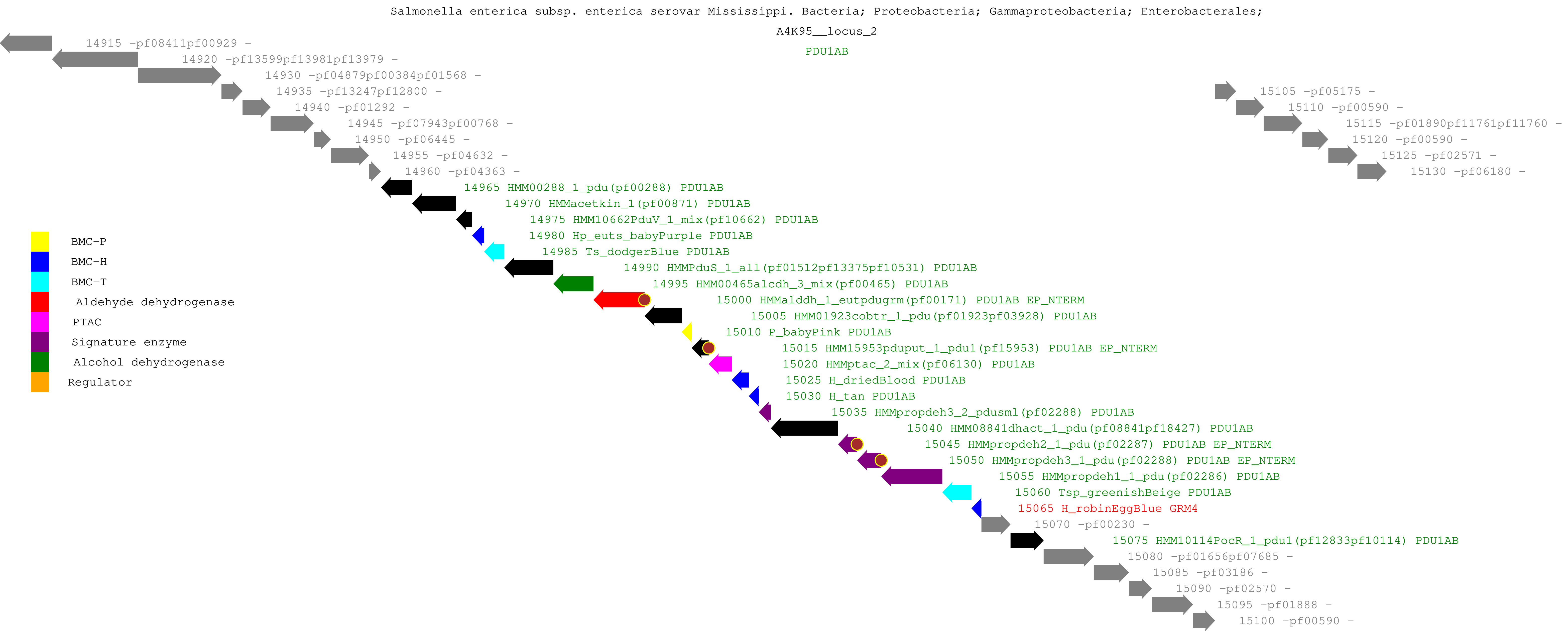

Salmonella enterica subsp. enterica serovar Mississippi. Bacteria; Proteobacteria; Gammaproteobacteria; Enterobacterales;
A4K95__locus_2
PDU1AB
14915 -pf08411pf00929 -
14920 -pf13599pf13981pf13979 -
14930 -pf04879pf00384pf01568 -
14935 -pf13247pf12800 -
14940 -pf01292 -
14945 -pf07943pf00768 -
14950 -pf06445 -
14955 -pf04632 -
14960 -pf04363 -
14965 HMM00288_1_pdu(pf00288) PDU1AB
14970 HMMacetkin_1(pf00871) PDU1AB
14975 HMM10662PduV_1_mix(pf10662) PDU1AB
14980 Hp_euts_babyPurple PDU1AB
14985 Ts_dodgerBlue PDU1AB
14990 HMMPduS_1_all(pf01512pf13375pf10531) PDU1AB
14995 HMM00465alcdh_3_mix(pf00465) PDU1AB
15000 HMMalddh_1_eutpdugrm(pf00171) PDU1AB EP_NTERM
15005 HMM01923cobtr_1_pdu(pf01923pf03928) PDU1AB
15010 P_babyPink PDU1AB
15015 HMM15953pduput_1_pdu1(pf15953) PDU1AB EP_NTERM
15020 HMMptac_2_mix(pf06130) PDU1AB
15025 H_driedBlood PDU1AB
15030 H_tan PDU1AB
15035 HMMpropdeh3_2_pdusml(pf02288) PDU1AB
15040 HMM08841dhact_1_pdu(pf08841pf18427) PDU1AB
15045 HMMpropdeh2_1_pdu(pf02287) PDU1AB EP_NTERM
15050 HMMpropdeh3_1_pdu(pf02288) PDU1AB EP_NTERM
15055 HMMpropdeh1_1_pdu(pf02286) PDU1AB
15060 Tsp_greenishBeige PDU1AB
15065 H_robinEggBlue GRM4
15070 -pf00230 -
15075 HMM10114PocR_1_pdu1(pf12833pf10114) PDU1AB
15080 -pf01656pf07685 -
15085 -pf03186 -
15090 -pf02570 -
15095 -pf01888 -
15100 -pf00590 -
15105 -pf05175 -
15110 -pf00590 -
15115 -pf01890pf11761pf11760 -
15120 -pf00590 -
15125 -pf02571 -
15130 -pf06180 -
BMC-P
BMC-H
BMC-T
Aldehyde dehydrogenase
PTAC
Signature enzyme
Alcohol dehydrogenase
Regulator

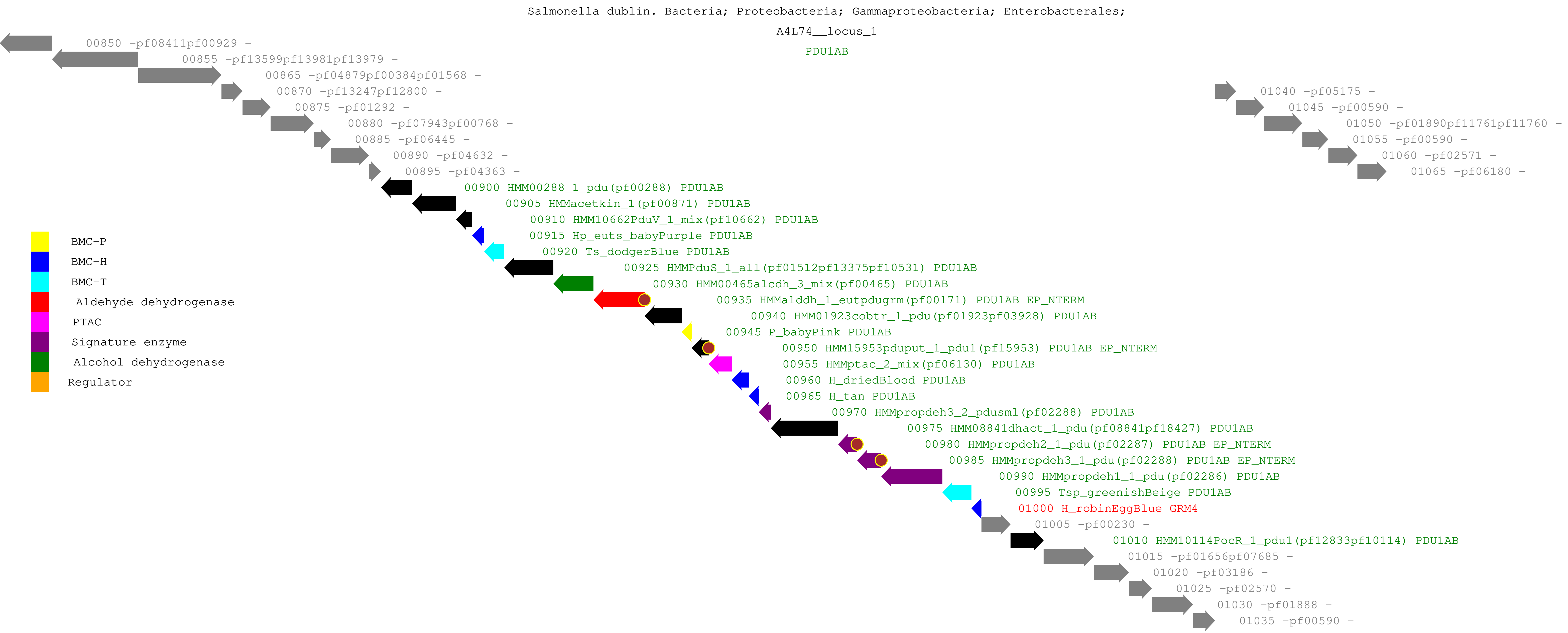
Salmonella dublin. Bacteria; Proteobacteria; Gammaproteobacteria; Enterobacterales;
A4L74__locus_1
PDU1AB
00850 -pf08411pf00929 -
00855 -pf13599pf13981pf13979 -
00865 -pf04879pf00384pf01568 -
00870 -pf13247pf12800 -
00875 -pf01292 -
00880 -pf07943pf00768 -
00885 -pf06445 -
00890 -pf04632 -
00895 -pf04363 -
00900 HMM00288_1_pdu(pf00288) PDU1AB
00905 HMMacetkin_1(pf00871) PDU1AB
00910 HMM10662PduV_1_mix(pf10662) PDU1AB
00915 Hp_euts_babyPurple PDU1AB
00920 Ts_dodgerBlue PDU1AB
00925 HMMPduS_1_all(pf01512pf13375pf10531) PDU1AB
00930 HMM00465alcdh_3_mix(pf00465) PDU1AB
00935 HMMalddh_1_eutpdugrm(pf00171) PDU1AB EP_NTERM
00940 HMM01923cobtr_1_pdu(pf01923pf03928) PDU1AB
00945 P_babyPink PDU1AB
00950 HMM15953pduput_1_pdu1(pf15953) PDU1AB EP_NTERM
00955 HMMptac_2_mix(pf06130) PDU1AB
00960 H_driedBlood PDU1AB
00965 H_tan PDU1AB
00970 HMMpropdeh3_2_pdusml(pf02288) PDU1AB
00975 HMM08841dhact_1_pdu(pf08841pf18427) PDU1AB
00980 HMMpropdeh2_1_pdu(pf02287) PDU1AB EP_NTERM
00985 HMMpropdeh3_1_pdu(pf02288) PDU1AB EP_NTERM
00990 HMMpropdeh1_1_pdu(pf02286) PDU1AB
00995 Tsp_greenishBeige PDU1AB
01000 H_robinEggBlue GRM4
01005 -pf00230 -
01010 HMM10114PocR_1_pdu1(pf12833pf10114) PDU1AB
01015 -pf01656pf07685 -
01020 -pf03186 -
01025 -pf02570 -
01030 -pf01888 -
01035 -pf00590 -
01040 -pf05175 -
01045 -pf00590 -
01050 -pf01890pf11761pf11760 -
01055 -pf00590 -
01060 -pf02571 -
01065 -pf06180 -
BMC-P
BMC-H
BMC-T
Aldehyde dehydrogenase
PTAC
Signature enzyme
Alcohol dehydrogenase
Regulator

Salmonella virchow. Bacteria; Proteobacteria; Gammaproteobacteria; Enterobacterales;

A4L79__locus_2

PDU1AB

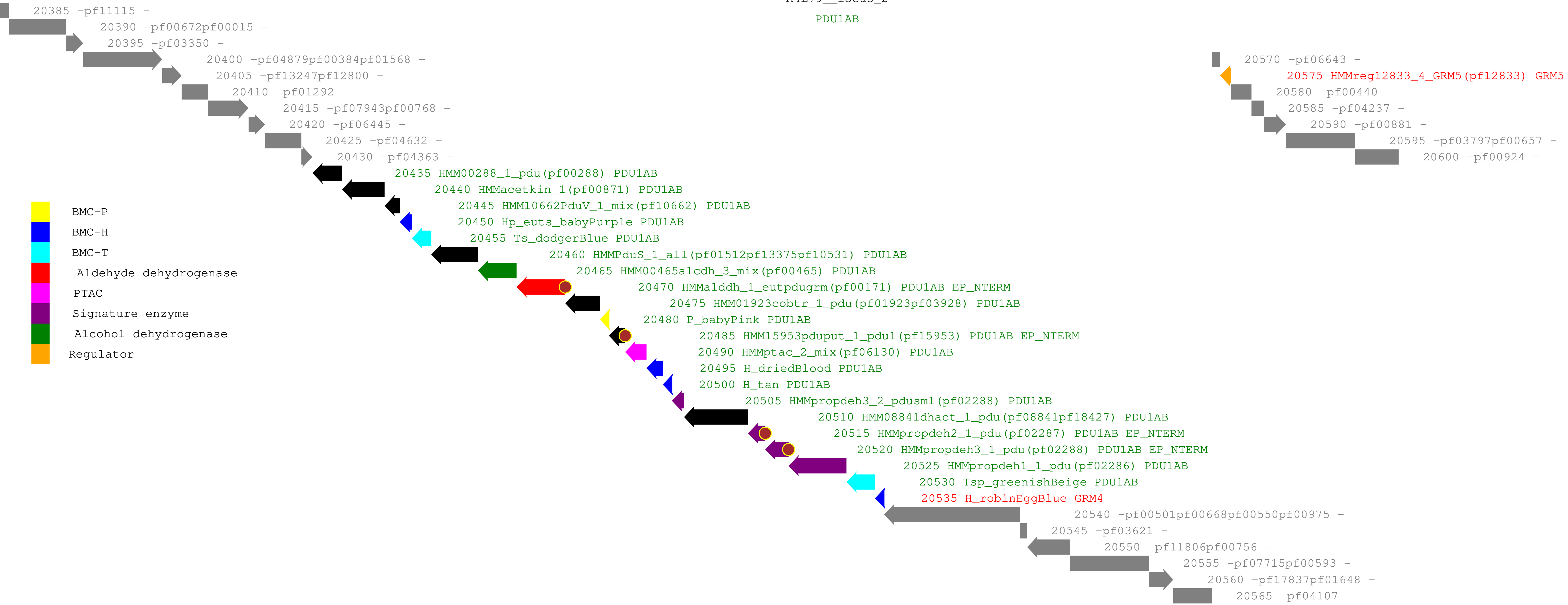

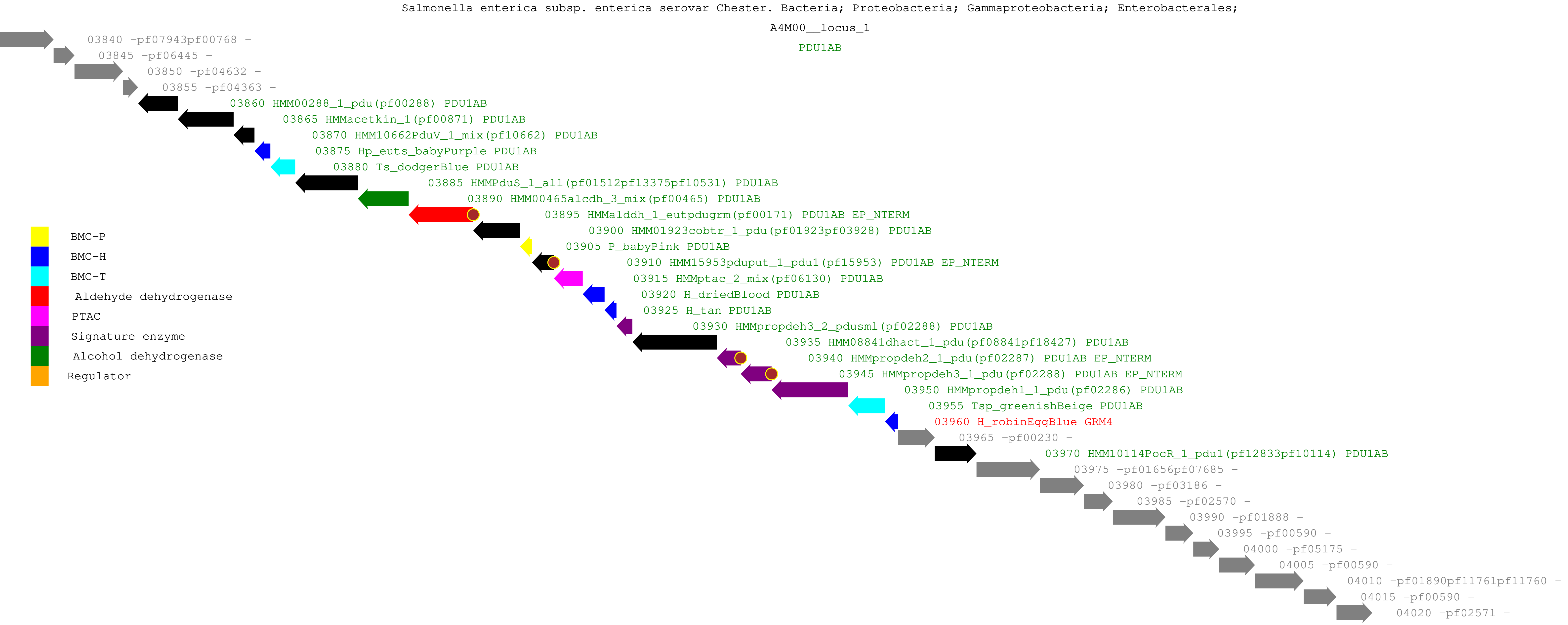

Salmonella enterica subsp. enterica serovar Chester. Bacteria; Proteobacteria; Gammaproteobacteria; Enterobacterales;
A4M00__locus_1
PDU1AB
03840 -pf07943pf00768 -
03845 -pf06445 -
03850 -pf04632 -
03855 -pf04363 -
03860 HMM00288_1_pdu(pf00288) PDU1AB
03865 HMMacetkin_1(pf00871) PDU1AB
03870 HMM10662PduV_1_mix(pf10662) PDU1AB
03875 Hp_euts_babyPurple PDU1AB
03880 Ts_dodgerBlue PDU1AB
03885 HMMPduS_1_all(pf01512pf13375pf10531) PDU1AB
03890 HMM00465alcdh_3_mix(pf00465) PDU1AB
03895 HMMaldh_1_eutpdugrm(pf00171) PDU1AB EP_NTERM
03900 HMM01923cobtr_1_pdu(pf01923pf03928) PDU1AB
03905 P_babyPink PDU1AB
03910 HMM15953pduput_1_pdu1(pf15953) PDU1AB EP_NTERM
03915 HMMptac_2_mix(pf06130) PDU1AB
03920 H_driedBlood PDU1AB
03925 H_tan PDU1AB
03930 HMMpropdeh3_2_pdusml(pf02288) PDU1AB
03935 HMM08841dhact_1_pdu(pf08841pf18427) PDU1AB
03940 HMMpropdeh2_1_pdu(pf02287) PDU1AB EP_NTERM
03945 HMMpropdeh3_1_pdu(pf02288) PDU1AB EP_NTERM
03950 HMMpropdeh1_1_pdu(pf02286) PDU1AB
03955 Tsp_greenishBeige PDU1AB
03960 H_robinEggBlue GRM4
03965 -pf00230 -
03970 HMM10114PocR_1_pdu1(pf12833pf10114) PDU1AB
03975 -pf01656pf07685 -
03980 -pf03186 -
03985 -pf02570 -
03990 -pf01888 -
03995 -pf00590 -
04000 -pf05175 -
04005 -pf00590 -
04010 -pf01890pf11761pf11760 -
04015 -pf00590 -
04020 -pf02571 -
BMC-P
BMC-H
BMC-T
Aldehyde dehydrogenase
PTAC
Signature enzyme
Alcohol dehydrogenase
Regulator

Salmonella enterica subsp. enterica serovar Haifa. Bacteria; Proteobacteria; Gammaproteobacteria; Enterobacterales;

A4M55__locus_1

PDU1AB

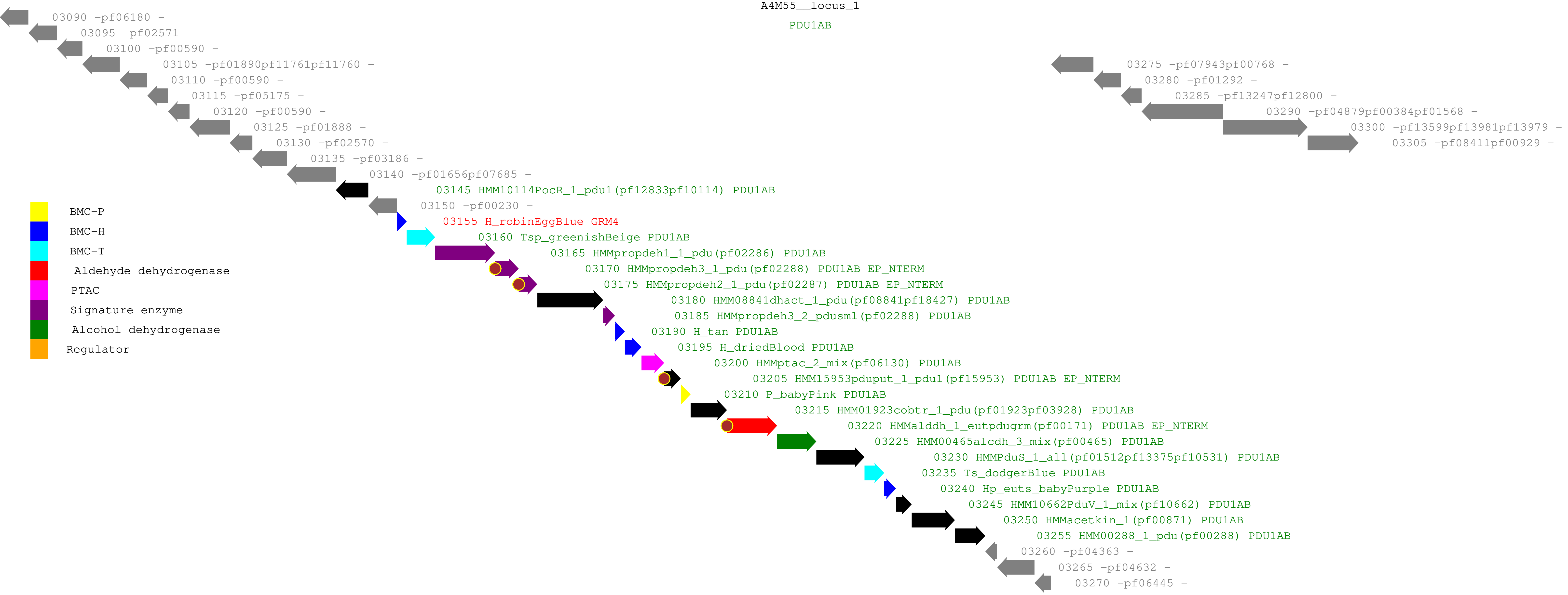

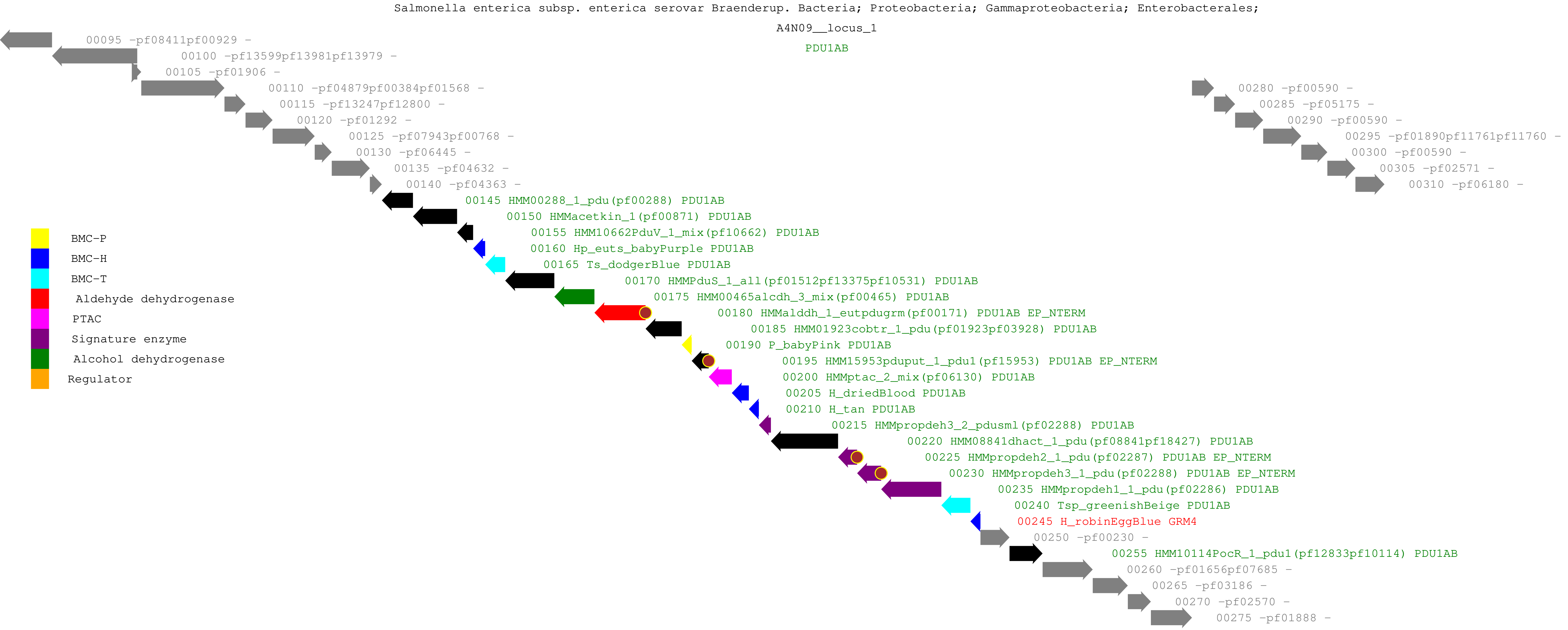
Salmonella enterica subsp. enterica serovar Braenderup. Bacteria; Proteobacteria; Gammaproteobacteria; Enterobacterales;
A4N09__locus_1
PDU1AB
00095 -pf08411pf00929 -
00100 -pf13599pf13981pf13979 -
00105 -pf01906 -
00110 -pf04879pf00384pf01568 -
00115 -pf13247pf12800 -
00120 -pf01292 -
00125 -pf07943pf00768 -
00130 -pf06445 -
00135 -pf04632 -
00140 -pf04363 -
00145 HMM00288_1_pdu(pf00288) PDU1AB
00150 HMMacetkin_1(pf00871) PDU1AB
00155 HMM10662PduV_1_mix(pf10662) PDU1AB
00160 Hp_euts_babyPurple PDU1AB
00165 Ts_dodgerBlue PDU1AB
00170 HMMPduS_1_all(pf01512pf13375pf10531) PDU1AB
00175 HMM00465alcdh_3_mix(pf00465) PDU1AB
00180 HMMalddh_1_eutpdugrm(pf00171) PDU1AB EP_NTERM
00185 HMM01923cobtr_1_pdu(pf01923pf03928) PDU1AB
00190 P_babyPink PDU1AB
00195 HMM15953pduput_1_pdu1(pf15953) PDU1AB EP_NTERM
00200 HMMptac_2_mix(pf06130) PDU1AB
00205 H_driedBlood PDU1AB
00210 H_tan PDU1AB
00215 HMMpropdeh3_2_pdusml(pf02288) PDU1AB
00220 HMM08841dhact_1_pdu(pf08841pf18427) PDU1AB
00225 HMMpropdeh2_1_pdu(pf02287) PDU1AB EP_NTERM
00230 HMMpropdeh3_1_pdu(pf02288) PDU1AB EP_NTERM
00235 HMMpropdeh1_1_pdu(pf02286) PDU1AB
00240 Tsp_greenishBeige PDU1AB
00245 H_robinEggBlue GRM4
00250 -pf00230 -
00255 HMM10114PocR_1_pdu1(pf12833pf10114) PDU1AB
00260 -pf01656pf07685 -
00265 -pf03186 -
00270 -pf02570 -
00275 -pf01888 -
00280 -pf00590 -
00285 -pf05175 -
00290 -pf00590 -
00295 -pf01890pf11761pf11760 -
00300 -pf00590 -
00305 -pf02571 -
00310 -pf06180 -
BMC-P
BMC-H
BMC-T
Aldehyde dehydrogenase
PTAC
Signature enzyme
Alcohol dehydrogenase
Regulator

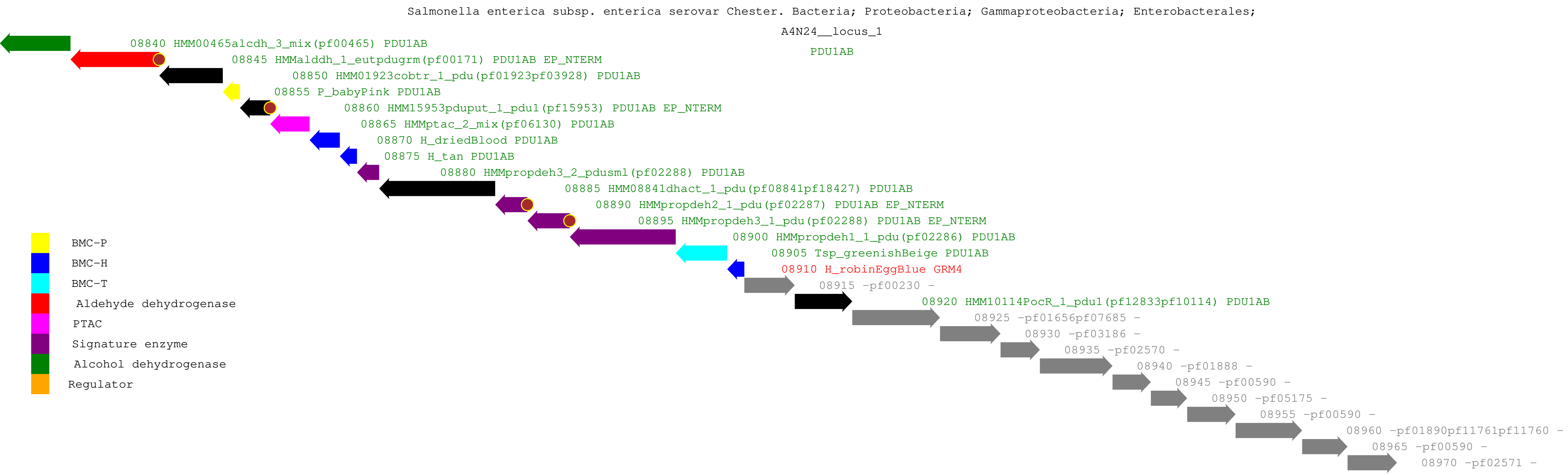
Salmonella enterica subsp. enterica serovar Chester. Bacteria; Proteobacteria; Gammaproteobacteria; Enterobacterales;
A4N24__locus_1
PDU1AB
08840 HMM00465alcdh_3_mix(pf00465) PDU1AB
08845 HMMalddh_1_eutpdugrm(pf00171) PDU1AB EP_NTERM
08850 HMM01923cobtr_1_pdu(pf01923pf03928) PDU1AB
08855 P_babyPink PDU1AB
08860 HMM15953pduput_1_pdu1(pf15953) PDU1AB EP_NTERM
08865 HMMptac_2_mix(pf06130) PDU1AB
08870 H_driedBlood PDU1AB
08875 H_tan PDU1AB
08880 HMMpropdeh3_2_pdusml(pf02288) PDU1AB
08885 HMM08841dhact_1_pdu(pf08841pf18427) PDU1AB
08890 HMMpropdeh2_1_pdu(pf02287) PDU1AB EP_NTERM
08895 HMMpropdeh3_1_pdu(pf02288) PDU1AB EP_NTERM
08900 HMMpropdeh1_1_pdu(pf02286) PDU1AB
08905 Tsp_greenishBeige PDU1AB
08910 H_robinEggBlue GRM4
08915 -pf00230 -
08920 HMM10114PocR_1_pdu1(pf12833pf10114) PDU1AB
08925 -pf01656pf07685 -
08930 -pf03186 -
08935 -pf02570 -
08940 -pf01888 -
08945 -pf00590 -
08950 -pf05175 -
08955 -pf00590 -
08960 -pf01890pf11761pf11760 -
08965 -pf00590 -
08970 -pf02571 -
BMC-P
BMC-H
BMC-T
Aldehyde dehydrogenase
PTAC
Signature enzyme
Alcohol dehydrogenase
Regulator

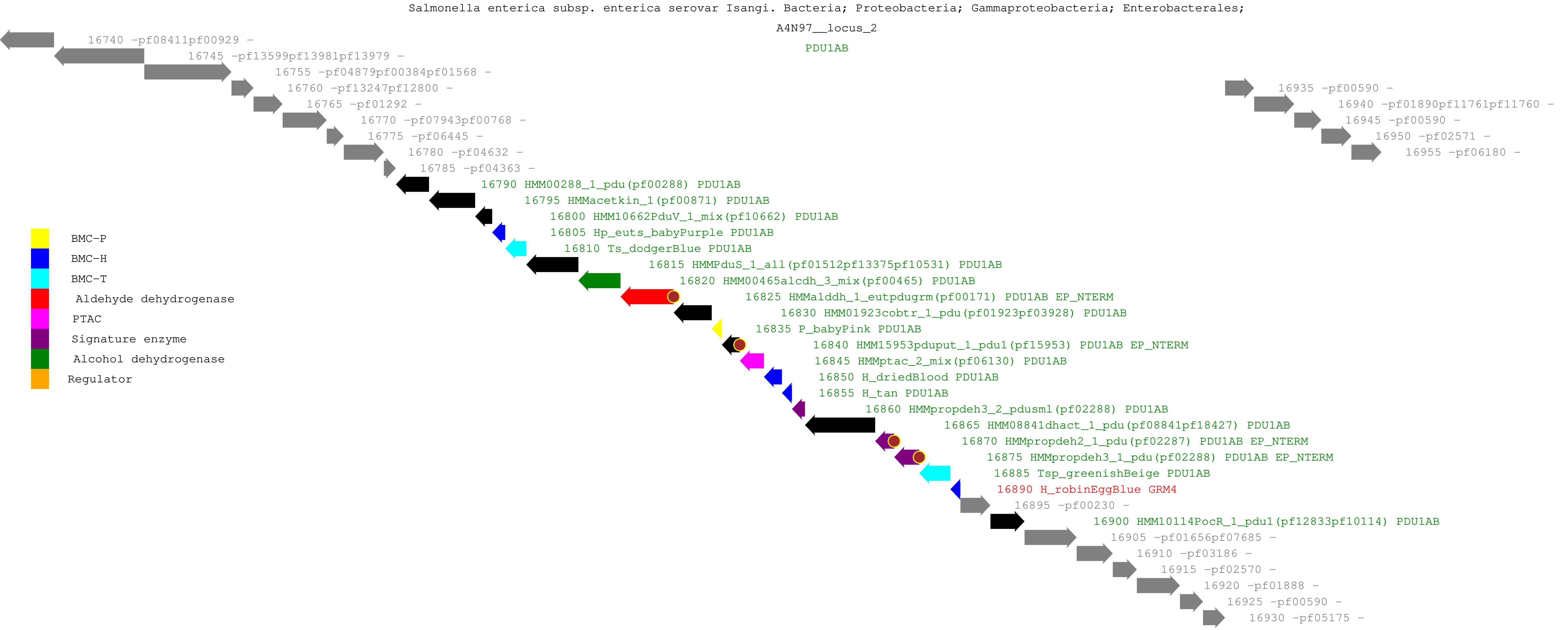

Salmonella enterica subsp. enterica serovar Isangi. Bacteria; Proteobacteria; Gammaproteobacteria; Enterobacterales;
A4N97__locus_2
PDU1AB
16740 -pf08411pf00929 -
16745 -pf13599pf13981pf13979 -
16755 -pf04879pf00384pf01568 -
16760 -pf13247pf12800 -
16765 -pf01292 -
16770 -pf07943pf00768 -
16775 -pf06445 -
16780 -pf04632 -
16785 -pf04363 -
16790 HMM00288_1_pdu(pf00288) PDU1AB
16795 HMMacetkin_1(pf00871) PDU1AB
16800 HMM10662PduV_1_mix(pf10662) PDU1AB
16805 Hp_euts_babyPurple PDU1AB
16810 Ts_dodgerBlue PDU1AB
16815 HMMPduS_1_all(pf01512pf13375pf10531) PDU1AB
16820 HMM00465alcdh_3_mix(pf00465) PDU1AB
16825 HMMalddh_1_eutpdugrm(pf00171) PDU1AB EP_NTERM
16830 HMM01923cobtr_1_pdu(pf01923pf03928) PDU1AB
16835 P_babyPink PDU1AB
16840 HMM15953pduput_1_pdu1(pf15953) PDU1AB EP_NTERM
16845 HMMptac_2_mix(pf06130) PDU1AB
16850 H_driedBlood PDU1AB
16855 H_tan PDU1AB
16860 HMMpropdeh3_2_pdusml(pf02288) PDU1AB
16865 HMM08841dhact_1_pdu(pf08841pf18427) PDU1AB
16870 HMMpropdeh2_1_pdu(pf02287) PDU1AB EP_NTERM
16875 HMMpropdeh3_1_pdu(pf02288) PDU1AB EP_NTERM
16885 Tsp_greenishBeige PDU1AB
16890 H_robinEggBlue GRM4
16895 -pf00230 -
16900 HMM10114PocR_1_pdu1(pf12833pf10114) PDU1AB
16905 -pf01656pf07685 -
16910 -pf03186 -
16915 -pf02570 -
16920 -pf01888 -
16925 -pf00590 -
16930 -pf05175 -
16935 -pf00590 -
16940 -pf01890pf11761pf11760 -
16945 -pf00590 -
16950 -pf02571 -
16955 -pf06180 -
BMC-P
BMC-H
BMC-T
Aldehyde dehydrogenase
PTAC
Signature enzyme
Alcohol dehydrogenase
Regulator

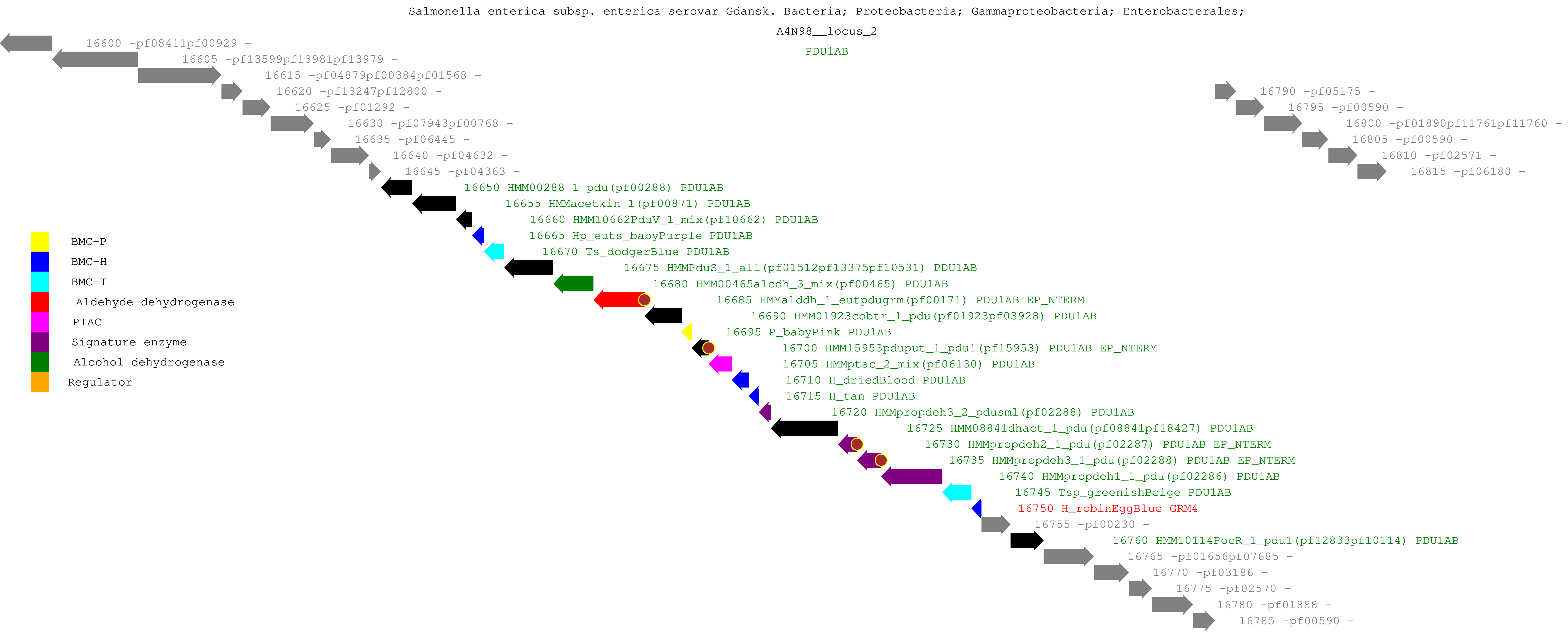
Salmonella enterica subsp. enterica serovar Gdansk. Bacteria; Proteobacteria; Gammaproteobacteria; Enterobacterales;
A4N98__locus_2
PDU1AB
16600 -pf08411pf00929 -
16605 -pf13599pf13981pf13979 -
16615 -pf04879pf00384pf01568 -
16620 -pf13247pf12800 -
16625 -pf01292 -
16630 -pf07943pf00768 -
16635 -pf06445 -
16640 -pf04632 -
16645 -pf04363 -
16650 HMM00288_1_pdu(pf00288) PDU1AB
16655 HMMacetkin_1(pf00871) PDU1AB
16660 HMM10662PduV_1_mix(pf10662) PDU1AB
16665 Hp_euts_babyPurple PDU1AB
16670 Ts_dodgerBlue PDU1AB
16675 HMMPduS_1_all(pf01512pf13375pf10531) PDU1AB
16680 HMM00465alcdh_3_mix(pf00465) PDU1AB
16685 HMMalddh_1_eutpdugrm(pf00171) PDU1AB EP_NTERM
16690 HMM01923cobtr_1_pdu(pf01923pf03928) PDU1AB
16695 P_babyPink PDU1AB
16700 HMM15953pduput_1_pdu1(pf15953) PDU1AB EP_NTERM
16705 HMMptac_2_mix(pf06130) PDU1AB
16710 H_driedBlood PDU1AB
16715 H_tan PDU1AB
16720 HMMpropdeh3_2_pdusml(pf02288) PDU1AB
16725 HMM08841dhact_1_pdu(pf08841pf18427) PDU1AB
16730 HMMpropdeh2_1_pdu(pf02287) PDU1AB EP_NTERM
16735 HMMpropdeh3_1_pdu(pf02288) PDU1AB EP_NTERM
16740 HMMpropdeh1_1_pdu(pf02286) PDU1AB
16745 Tsp_greenishBeige PDU1AB
16750 H_robinEggBlue GRM4
16755 -pf00230 -
16760 HMM10114PocR_1_pdu1(pf12833pf10114) PDU1AB
16765 -pf01656pf07685 -
16770 -pf03186 -
16775 -pf02570 -
16780 -pf01888 -
16785 -pf00590 -
16790 -pf05175 -
16795 -pf00590 -
16800 -pf01890pf11761pf11760 -
16805 -pf00590 -
16810 -pf02571 -
16815 -pf06180 -
BMC-P
BMC-H
BMC-T
Aldehyde dehydrogenase
PTAC
Signature enzyme
Alcohol dehydrogenase
Regulator

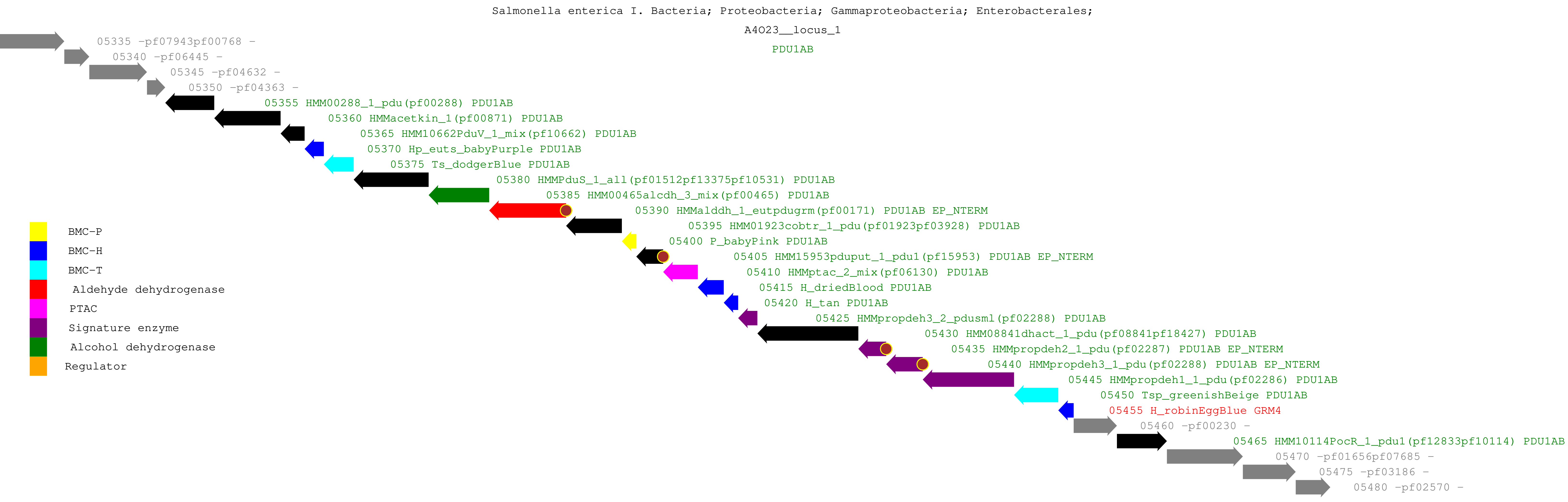
Salmonella enterica I. Bacteria; Proteobacteria; Gammaproteobacteria; Enterobacterales;
A4O23__locus_1
PDU1AB
05335 -pf07943pf00768 -
05340 -pf06445 -
05345 -pf04632 -
05350 -pf04363 -
05355 HMM00288_1_pdu(pf00288) PDU1AB
05360 HMMacetkin_1(pf00871) PDU1AB
05365 HMM10662PduV_1_mix(pf10662) PDU1AB
05370 Hp_euts_babyPurple PDU1AB
05375 Ts_dodgerBlue PDU1AB
05380 HMMPduS_1_all(pf01512pf13375pf10531) PDU1AB
05385 HMM00465alcdh_3_mix(pf00465) PDU1AB
05390 HMMalddh_1_eutpdugrm(pf00171) PDU1AB EP_NTERM
05395 HMM01923cobtr_1_pdu(pf01923pf03928) PDU1AB
05400 P_babyPink PDU1AB
05405 HMM15953pduput_1_pdu1(pf15953) PDU1AB EP_NTERM
05410 HMMptac_2_mix(pf06130) PDU1AB
05415 H_driedBlood PDU1AB
05420 H_tan PDU1AB
05425 HMMpropdeh3_2_pdusml(pf02288) PDU1AB
05430 HMM08841dhact_1_pdu(pf08841pf18427) PDU1AB
05435 HMMpropdeh2_1_pdu(pf02287) PDU1AB EP_NTERM
05440 HMMpropdeh3_1_pdu(pf02288) PDU1AB EP_NTERM
05445 HMMpropdeh1_1_pdu(pf02286) PDU1AB
05450 Tsp_greenishBeige PDU1AB
05455 H_robinEggBlue GRM4
05460 -pf00230 -
05465 HMM10114PocR_1_pdu1(pf12833pf10114) PDU1AB
05470 -pf01656pf07685 -
05475 -pf03186 -
05480 -pf02570 -
BMC-P
BMC-H
BMC-T
Aldehyde dehydrogenase
PTAC
Signature enzyme
Alcohol dehydrogenase
Regulator

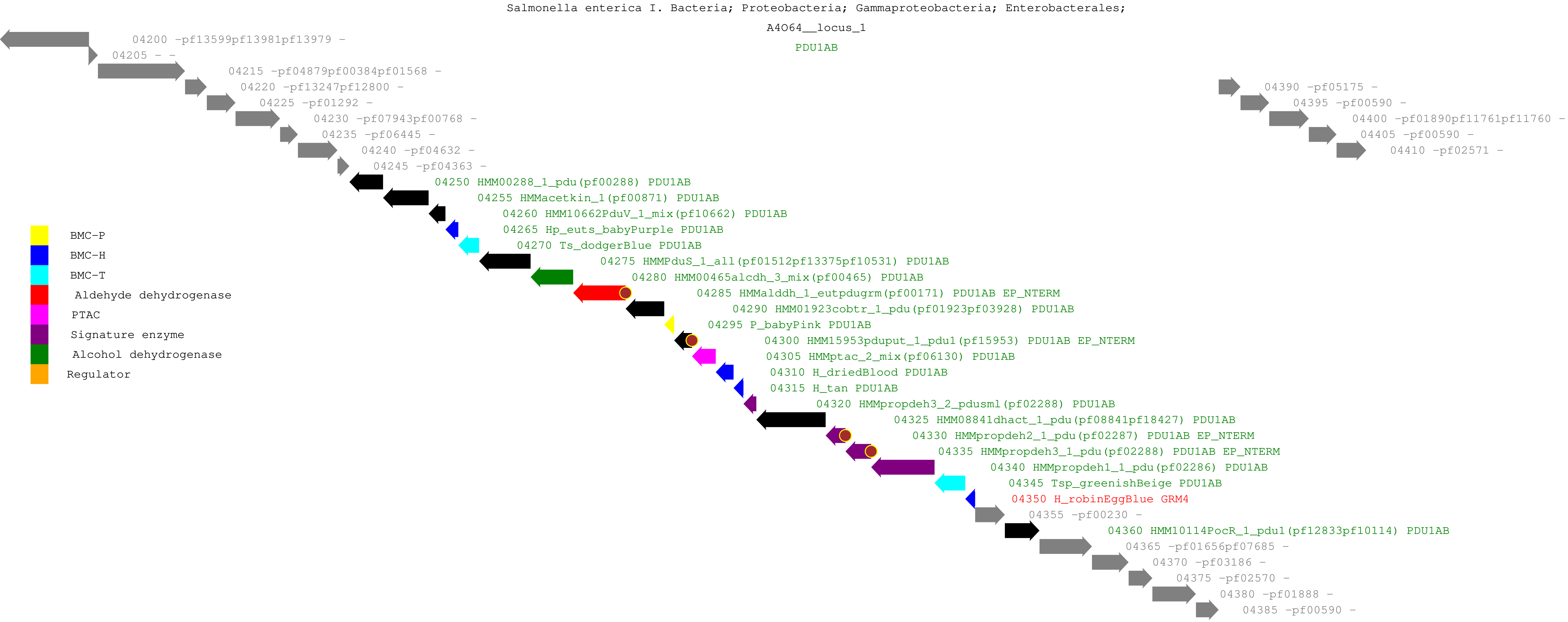

Salmonella enterica I. Bacteria; Proteobacteria; Gammaproteobacteria; Enterobacterales;
A4O64__locus_1
PDU1AB
04200 -pf13599pf13981pf13979 -
04205 - -
04215 -pf04879pf00384pf01568 -
04220 -pf13247pf12800 -
04225 -pf01292 -
04230 -pf07943pf00768 -
04235 -pf06445 -
04240 -pf04632 -
04245 -pf04363 -
04250 HMM00288_1_pdu(pf00288) PDU1AB
04255 HMMacetkin_1(pf00871) PDU1AB
04260 HMM10662PduV_1_mix(pf10662) PDU1AB
04265 Hp_euts_babyPurple PDU1AB
04270 Ts_dodgerBlue PDU1AB
04275 HMMPduS_1_all(pf01512pf13375pf10531) PDU1AB
04280 HMM00465alcdh_3_mix(pf00465) PDU1AB
04285 HMMalddh_1_eutpdugrm(pf00171) PDU1AB EP_NTERM
04290 HMM01923cobtr_1_pdu(pf01923pf03928) PDU1AB
04295 P_babyPink PDU1AB
04300 HMM15953pduput_1_pdu1(pf15953) PDU1AB EP_NTERM
04305 HMMptac_2_mix(pf06130) PDU1AB
04310 H_driedBlood PDU1AB
04315 H_tan PDU1AB
04320 HMMpropdeh3_2_pdusml(pf02288) PDU1AB
04325 HMM08841dhact_1_pdu(pf08841pf18427) PDU1AB
04330 HMMpropdeh2_1_pdu(pf02287) PDU1AB EP_NTERM
04335 HMMpropdeh3_1_pdu(pf02288) PDU1AB EP_NTERM
04340 HMMpropdeh1_1_pdu(pf02286) PDU1AB
04345 Tsp_greenishBeige PDU1AB
04350 H_robinEggBlue GRM4
04355 -pf00230 -
04360 HMM10114PocR_1_pdu1(pf12833pf10114) PDU1AB
04365 -pf01656pf07685 -
04370 -pf03186 -
04375 -pf02570 -
04380 -pf01888 -
04385 -pf00590 -
04390 -pf05175 -
04395 -pf00590 -
04400 -pf01890pf11761pf11760 -
04405 -pf00590 -
04410 -pf02571 -
BMC-P
BMC-H
BMC-T
Aldehyde dehydrogenase
PTAC
Signature enzyme
Alcohol dehydrogenase
Regulator

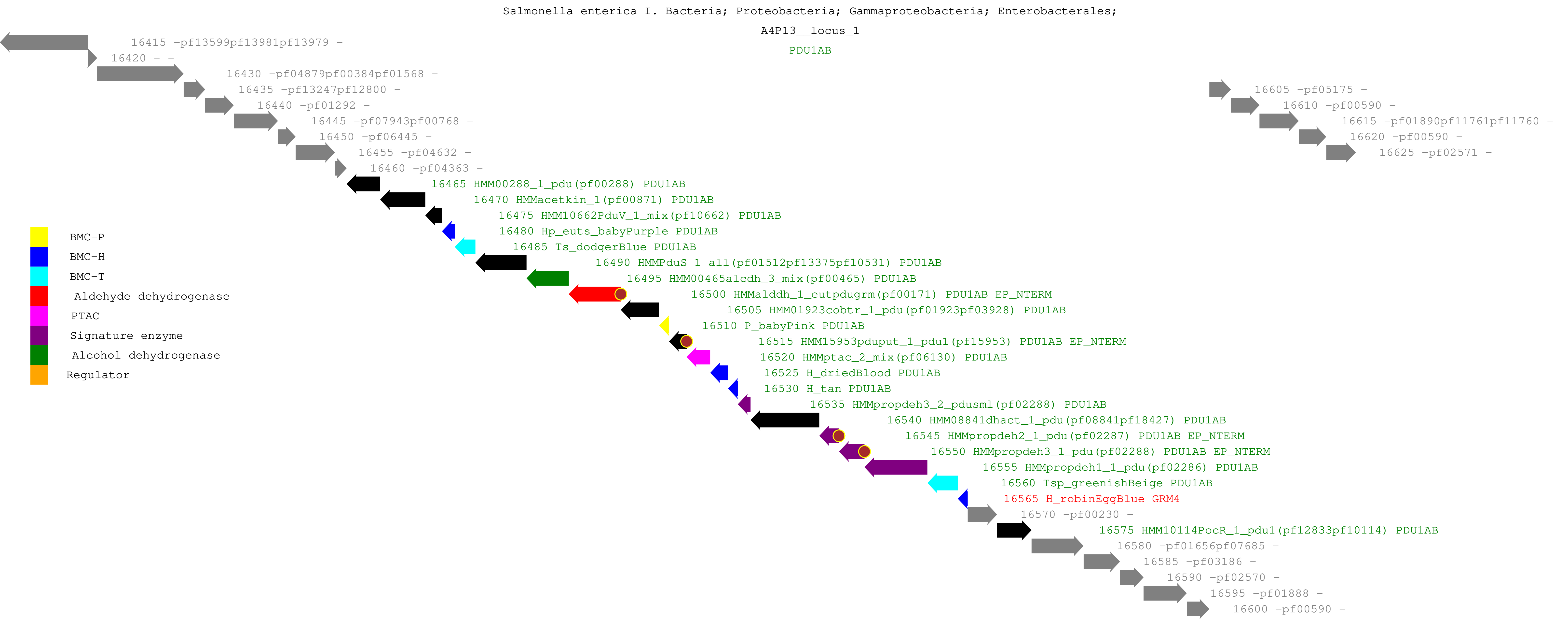

Salmonella enterica I. Bacteria; Proteobacteria; Gammaproteobacteria; Enterobacterales;
A4P13__locus_1
PDU1AB
16415 -pf13599pf13981pf13979 -
16420 - -
16430 -pf04879pf00384pf01568 -
16435 -pf13247pf12800 -
16440 -pf01292 -
16445 -pf07943pf00768 -
16450 -pf06445 -
16455 -pf04632 -
16460 -pf04363 -
16465 HMM00288_1_pdu(pf00288) PDU1AB
16470 HMMacetkin_1(pf00871) PDU1AB
16475 HMM10662PduV_1_mix(pf10662) PDU1AB
16480 Hp_euts_babyPurple PDU1AB
16485 Ts_dodgerBlue PDU1AB
16490 HMMPduS_1_all(pf01512pf13375pf10531) PDU1AB
16495 HMM00465alcdh_3_mix(pf00465) PDU1AB
16500 HMMalddh_1_eutpdugrm(pf00171) PDU1AB EP_NTERM
16505 HMM01923cobtr_1_pdu(pf01923pf03928) PDU1AB
16510 P_babyPink PDU1AB
16515 HMM15953pduput_1_pdu1(pf15953) PDU1AB EP_NTERM
16520 HMMptac_2_mix(pf06130) PDU1AB
16525 H_driedBlood PDU1AB
16530 H_tan PDU1AB
16535 HMMpropdeh3_2_pdusml(pf02288) PDU1AB
16540 HMM08841dhact_1_pdu(pf08841pf18427) PDU1AB
16545 HMMpropdeh2_1_pdu(pf02287) PDU1AB EP_NTERM
16550 HMMpropdeh3_1_pdu(pf02288) PDU1AB EP_NTERM
16555 HMMpropdeh1_1_pdu(pf02286) PDU1AB
16560 Tsp_greenishBeige PDU1AB
16565 H_robinEggBlue GRM4
16570 -pf00230 -
16575 HMM10114PocR_1_pdu1(pf12833pf10114) PDU1AB
16580 -pf01656pf07685 -
16585 -pf03186 -
16590 -pf02570 -
16595 -pf01888 -
16600 -pf00590 -
16605 -pf05175 -
16610 -pf00590 -
16615 -pf01890pf11761pf11760 -
16620 -pf00590 -
16625 -pf02571 -
BMC-P
BMC-H
BMC-T
Aldehyde dehydrogenase
PTAC
Signature enzyme
Alcohol dehydrogenase
Regulator

Salmonella rubislaw. Bacteria; Proteobacteria; Gammaproteobacteria; Enterobacterales;

A4T11__locus_2

PDU1AB

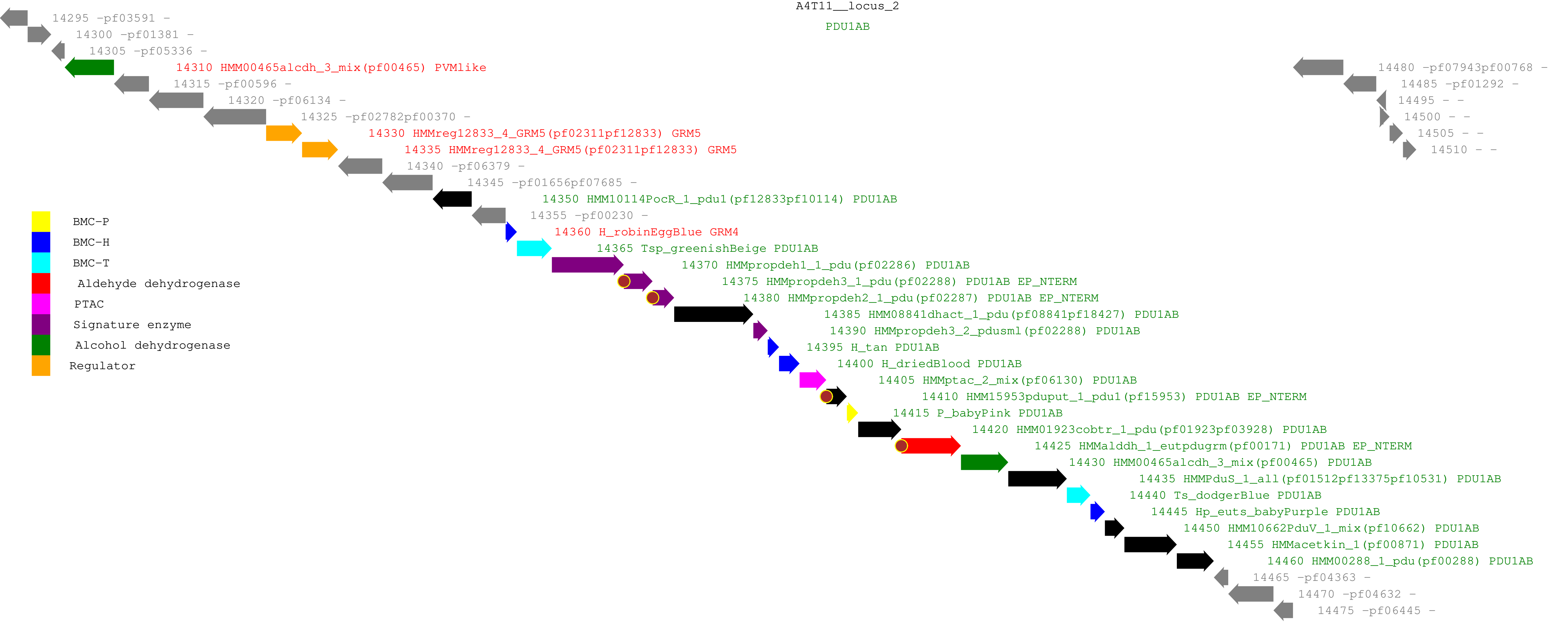

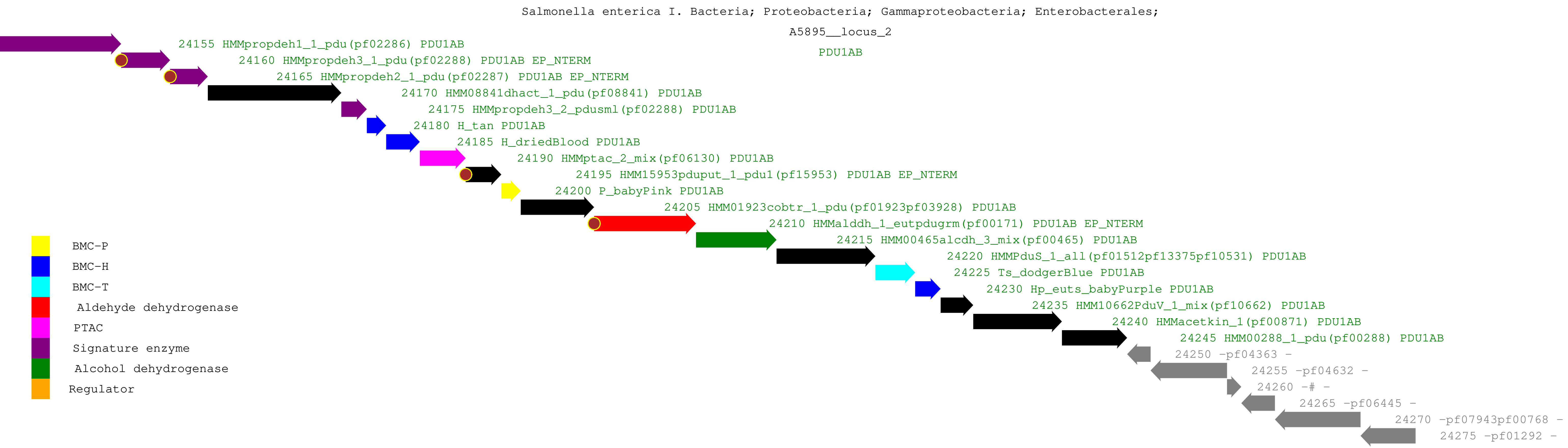
Salmonella enterica I. Bacteria; Proteobacteria; Gammaproteobacteria; Enterobacterales;
A5895__locus_2
PDU1AB
24155 HMMpropdeh1_1_pdu(pf02286) PDU1AB
24160 HMMpropdeh3_1_pdu(pf02288) PDU1AB EP_NTERM
24165 HMMpropdeh2_1_pdu(pf02287) PDU1AB EP_NTERM
24170 HMM08841dhact_1_pdu(pf08841) PDU1AB
24175 HMMpropdeh3_2_pdusml(pf02288) PDU1AB
24180 H_tan PDU1AB
24185 H_driedBlood PDU1AB
24190 HMMptac_2_mix(pf06130) PDU1AB
24195 HMM15953pduput_1_pdu1(pf15953) PDU1AB EP_NTERM
24200 P_babyPink PDU1AB
24205 HMM01923cobtr_1_pdu(pf01923pf03928) PDU1AB
24210 HMMaldh_1_eutpdugrm(pf00171) PDU1AB EP_NTERM
24215 HMM00465alcdh_3_mix(pf00465) PDU1AB
24220 HMMPduS_1_all(pf01512pf13375pf10531) PDU1AB
24225 Ts_dodgerBlue PDU1AB
24230 Hp_euts_babyPurple PDU1AB
24235 HMM10662PduV_1_mix(pf10662) PDU1AB
24240 HMMacetkin_1(pf00871) PDU1AB
24245 HMM00288_1_pdu(pf00288) PDU1AB
24250 -pf04363 -
24255 -pf04632 -
24260 -# -
24265 -pf06445 -
24270 -pf07943pf00768 -
24275 -pf01292 -
BMC-P
BMC-H
BMC-T
Aldehyde dehydrogenase
PTAC
Signature enzyme
Alcohol dehydrogenase
Regulator

Salmonella enterica subsp. enterica serovar Cubana str. 76814. Bacteria; Proteobacteria; Gammaproteobacteria; Enterobacterales;

A628__locus_2

PDU1AB

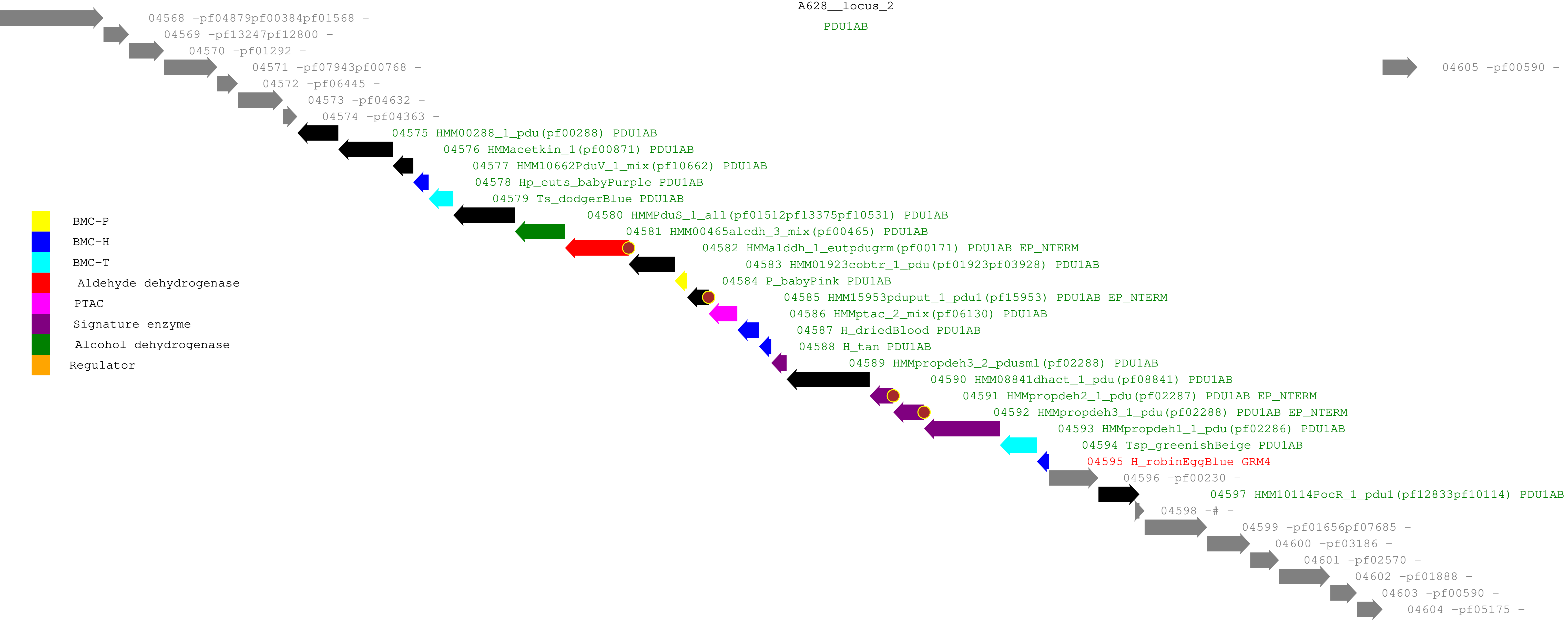

Salmonella enterica subsp. enterica serovar Dublin str. UC16. Bacteria; Proteobacteria; Gammaproteobacteria; Enterobacterales;

A670__locus_2

PDU1AB

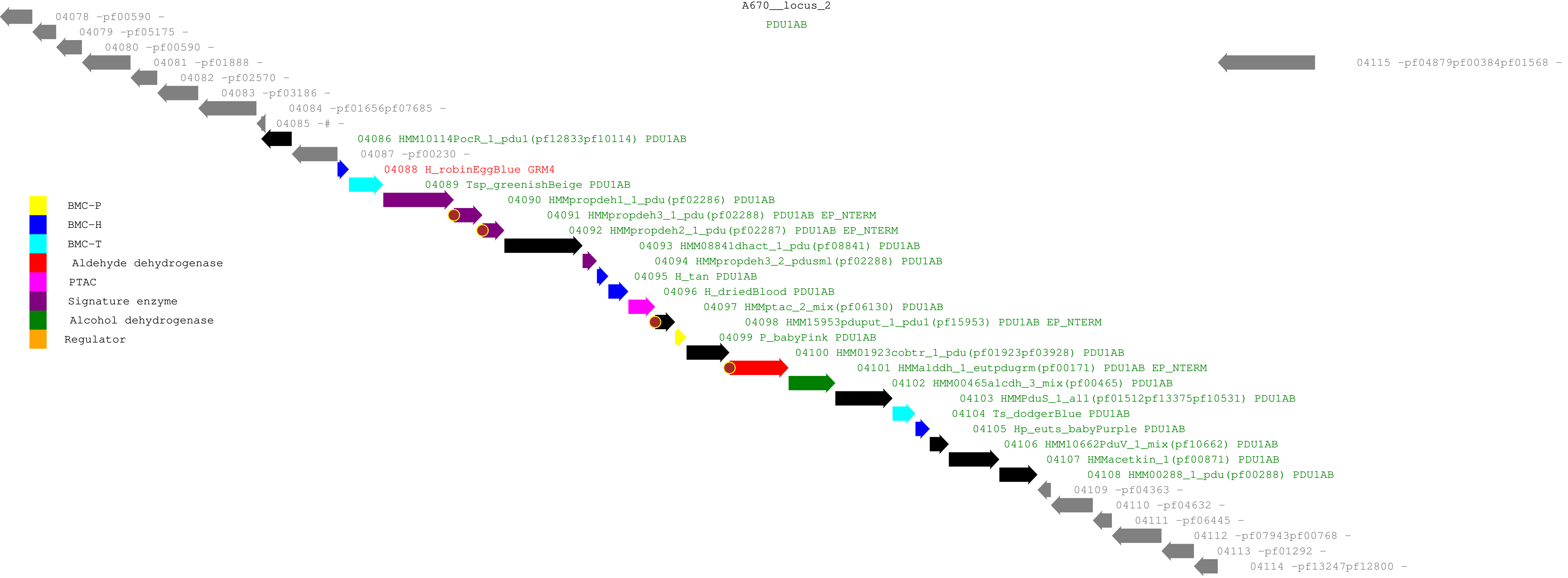

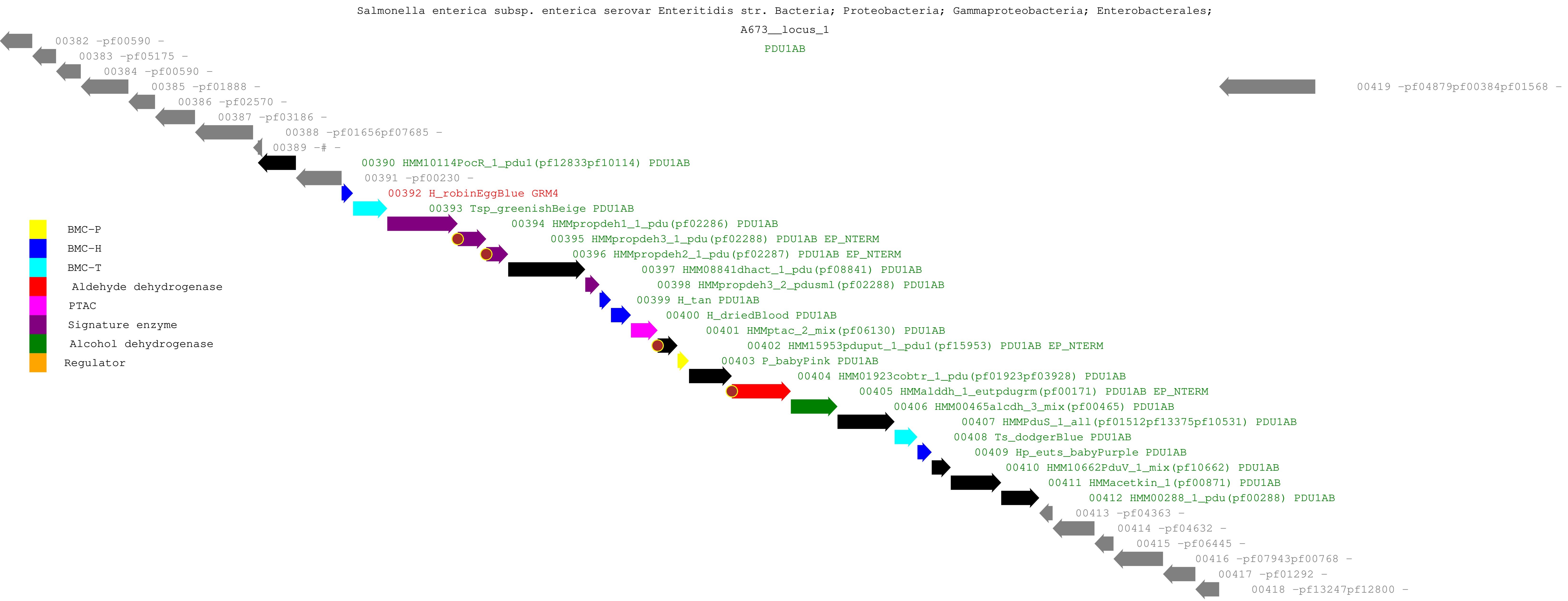
Salmonella enterica subsp. enterica serovar Enteritidis str. Bacteria; Proteobacteria; Gammaproteobacteria; Enterobacterales;
A673__locus_1
PDU1AB
00382 -pf00590 -
00383 -pf05175 -
00384 -pf00590 -
00385 -pf01888 -
00386 -pf02570 -
00387 -pf03186 -
00388 -pf01656pf07685 -
00389 -# -
00390 HMM10114PocR_1_pdu1(pf12833pf10114) PDU1AB
00391 -pf00230 -
00392 H_robinEggBlue GRM4
00393 Tsp_greenishBeige PDU1AB
00394 HMMpropdeh1_1_pdu(pf02286) PDU1AB
00395 HMMpropdeh3_1_pdu(pf02288) PDU1AB EP_NTERM
00396 HMMpropdeh2_1_pdu(pf02287) PDU1AB EP_NTERM
00397 HMM08841dhact_1_pdu(pf08841) PDU1AB
00398 HMMpropdeh3_2_pdusml(pf02288) PDU1AB
00399 H_tan PDU1AB
00400 H_driedBlood PDU1AB
00401 HMMptac_2_mix(pf06130) PDU1AB
00402 HMM15953pduput_1_pdu1(pf15953) PDU1AB EP_NTERM
00403 P_babyPink PDU1AB
00404 HMM01923cobtr_1_pdu(pf01923pf03928) PDU1AB
00405 HMMalddh_1_eutpdugrm(pf00171) PDU1AB EP_NTERM
00406 HMM00465alcdh_3_mix(pf00465) PDU1AB
00407 HMMPduS_1_all(pf01512pf13375pf10531) PDU1AB
00408 Ts_dodgerBlue PDU1AB
00409 Hp_euts_babyPurple PDU1AB
00410 HMM10662PduV_1_mix(pf10662) PDU1AB
00411 HMMacetkin_1(pf00871) PDU1AB
00412 HMM00288_1_pdu(pf00288) PDU1AB
00413 -pf04363 -
00414 -pf04632 -
00415 -pf06445 -
00416 -pf07943pf00768 -
00417 -pf01292 -
00418 -pf13247pf12800 -
00419 -pf04879pf00384pf01568 -
BMC-P
BMC-H
BMC-T
Aldehyde dehydrogenase
PTAC
Signature enzyme
Alcohol dehydrogenase
Regulator

Klebsiella michiganensis. Bacteria; Proteobacteria; Gammaproteobacteria; Enterobacterales;

A6A30__locus_3

PDU1AB

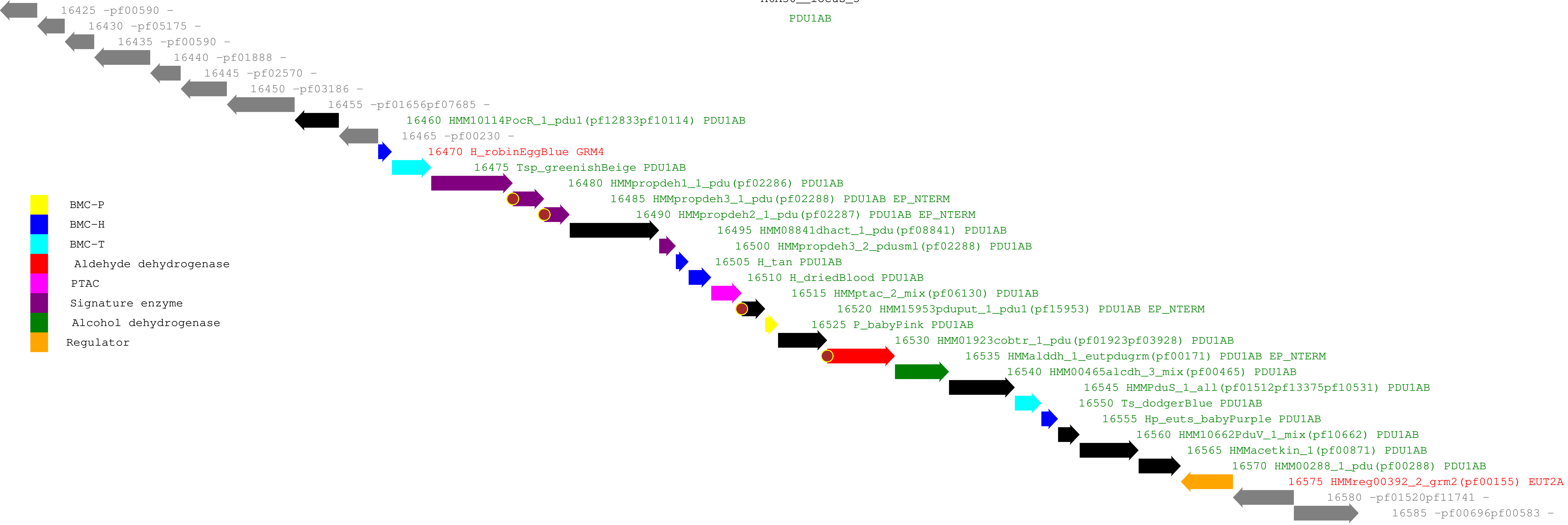

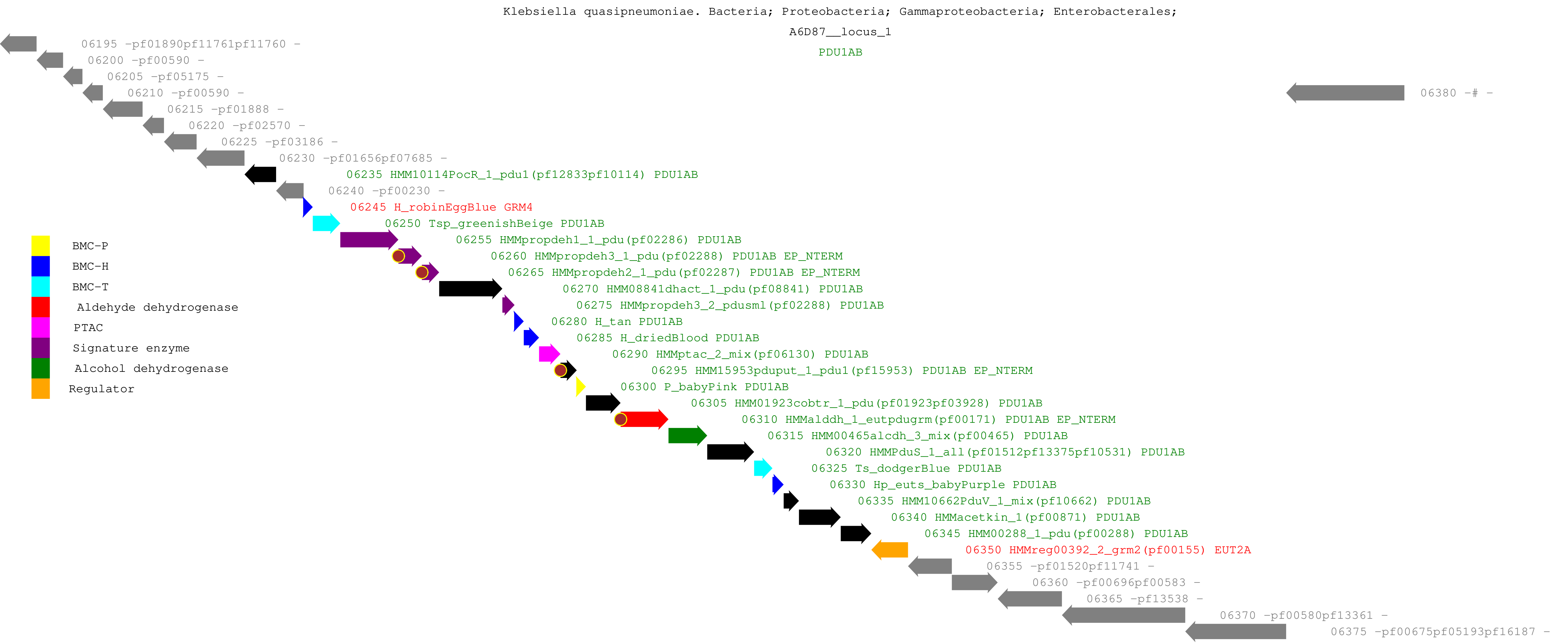
Klebsiella quasipneumoniae. Bacteria; Proteobacteria; Gammaproteobacteria; Enterobacterales;
A6D87__locus_1
PDU1AB
06195 -pf01890pf11761pf11760 -
06200 -pf00590 -
06205 -pf05175 -
06210 -pf00590 -
06215 -pf01888 -
06220 -pf02570 -
06225 -pf03186 -
06230 -pf01656pf07685 -
06235 HMM10114PocR_1_pdu1(pf12833pf10114) PDU1AB
06240 -pf00230 -
06245 H_robinEggBlue GRM4
06250 Tsp_greenishBeige PDU1AB
06255 HMMpropdeh1_1_pdu(pf02286) PDU1AB
06260 HMMpropdeh3_1_pdu(pf02288) PDU1AB EP_NTERM
06265 HMMpropdeh2_1_pdu(pf02287) PDU1AB EP_NTERM
06270 HMM08841dhact_1_pdu(pf08841) PDU1AB
06275 HMMpropdeh3_2_pdusml(pf02288) PDU1AB
06280 H_tan PDU1AB
06285 H_driedBlood PDU1AB
06290 HMMptac_2_mix(pf06130) PDU1AB
06295 HMM15953pduput_1_pdu1(pf15953) PDU1AB EP_NTERM
06300 P_babyPink PDU1AB
06305 HMM01923cobtr_1_pdu(pf01923pf03928) PDU1AB
06310 HMMalddh_1_eutpdugrm(pf00171) PDU1AB EP_NTERM
06315 HMM00465alcdh_3_mix(pf00465) PDU1AB
06320 HMMPduS_1_all(pf01512pf13375pf10531) PDU1AB
06325 Ts_dodgerBlue PDU1AB
06330 Hp_euts_babyPurple PDU1AB
06335 HMM10662PduV_1_mix(pf10662) PDU1AB
06340 HMMacetkin_1(pf00871) PDU1AB
06345 HMM00288_1_pdu(pf00288) PDU1AB
06350 HMMreg00392_2_grm2(pf00155) EUT2A
06355 -pf01520pf11741 -
06360 -pf00696pf00583 -
06365 -pf13538 -
06370 -pf00580pf13361 -
06375 -pf00675pf05193pf16187 -
06380 -# -
BMC-P
BMC-H
BMC-T
Aldehyde dehydrogenase
PTAC
Signature enzyme
Alcohol dehydrogenase
Regulator

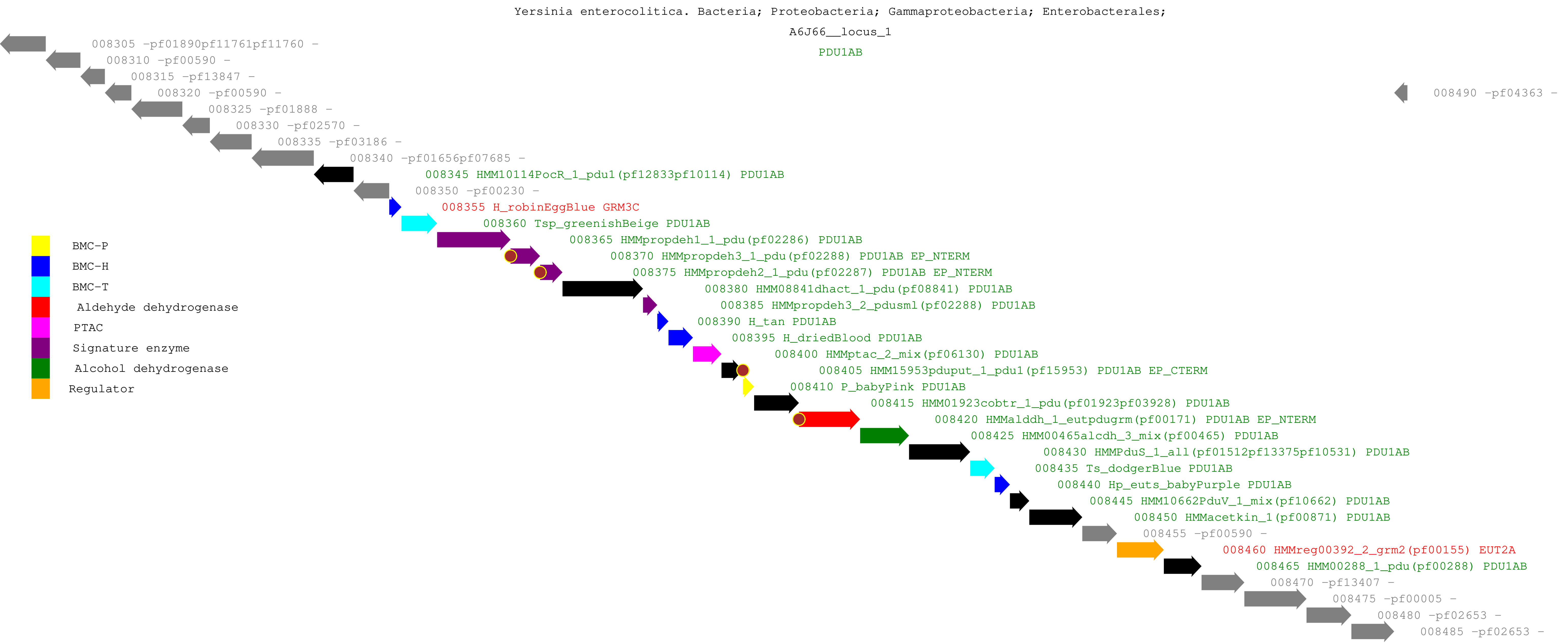
Yersinia enterocolitica. Bacteria; Proteobacteria; Gammaproteobacteria; Enterobacterales;
A6J66__locus_1
PDU1AB
008305 -pf01890pf11761pf11760 -
008310 -pf00590 -
008315 -pf13847 -
008320 -pf00590 -
008325 -pf01888 -
008330 -pf02570 -
008335 -pf03186 -
008340 -pf01656pf07685 -
008345 HMM10114PocR_1_pdu1(pf12833pf10114) PDU1AB
008350 -pf00230 -
008355 H_robinEggBlue GRM3C
008360 Tsp_greenishBeige PDU1AB
008365 HMMpropdeh1_1_pdu(pf02286) PDU1AB
008370 HMMpropdeh3_1_pdu(pf02288) PDU1AB EP_NTERM
008375 HMMpropdeh2_1_pdu(pf02287) PDU1AB EP_NTERM
008380 HMM08841dhact_1_pdu(pf08841) PDU1AB
008385 HMMpropdeh3_2_pdusml(pf02288) PDU1AB
008390 H_tan PDU1AB
008395 H_driedBlood PDU1AB
008400 HMMptac_2_mix(pf06130) PDU1AB
008405 HMM15953pduput_1_pdu1(pf15953) PDU1AB EP_CTERM
008410 P_babyPink PDU1AB
008415 HMM01923cobtr_1_pdu(pf01923pf03928) PDU1AB
008420 HMMalddh_1_eutpdugrm(pf00171) PDU1AB EP_NTERM
008425 HMM00465alcdh_3_mix(pf00465) PDU1AB
008430 HMMPduS_1_all(pf01512pf13375pf10531) PDU1AB
008435 Ts_dodgerBlue PDU1AB
008440 Hp_euts_babyPurple PDU1AB
008445 HMM10662PduV_1_mix(pf10662) PDU1AB
008450 HMMacetkin_1(pf00871) PDU1AB
008455 -pf00590 -
008460 HMMreg00392_2_grm2(pf00155) EUT2A
008465 HMM00288_1_pdu(pf00288) PDU1AB
008470 -pf13407 -
008475 -pf00005 -
008480 -pf02653 -
008485 -pf02653 -
008490 -pf04363 -
BMC-P
BMC-H
BMC-T
Aldehyde dehydrogenase
PTAC
Signature enzyme
Alcohol dehydrogenase
Regulator

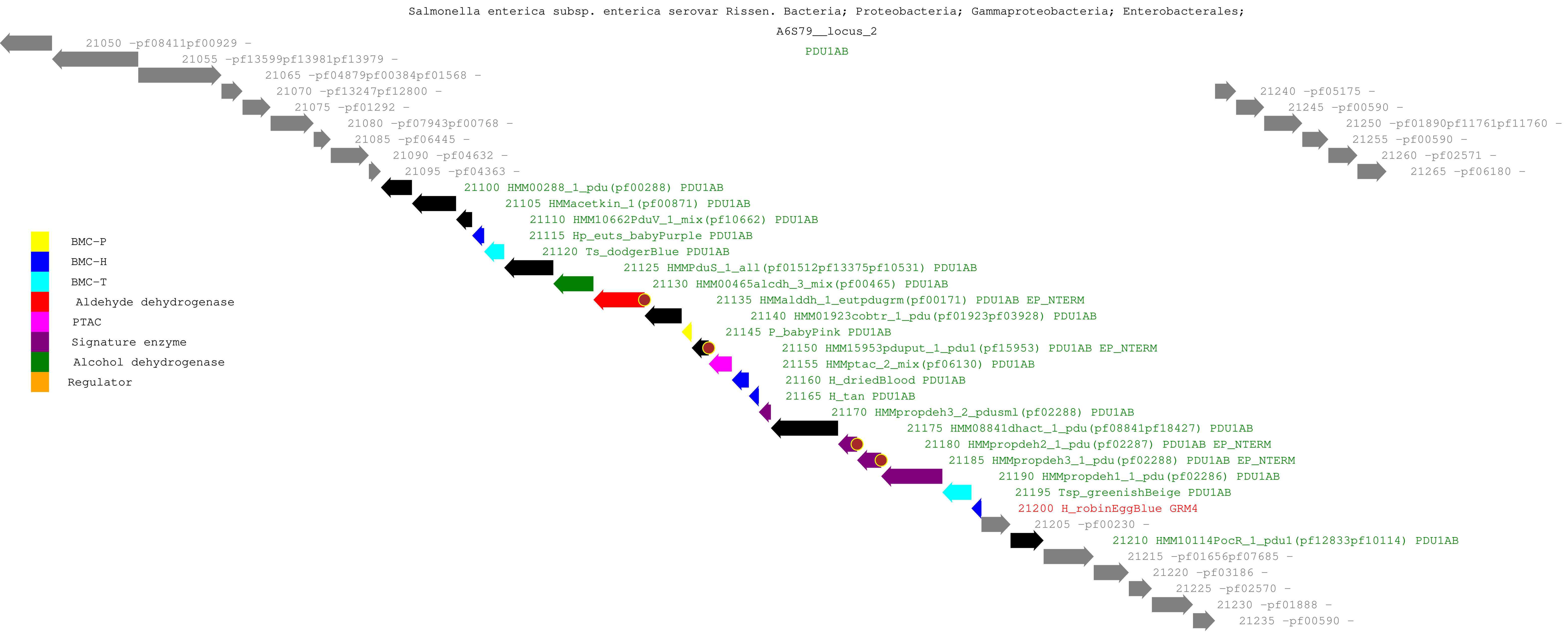

Salmonella enterica subsp. enterica serovar Rissen. Bacteria; Proteobacteria; Gammaproteobacteria; Enterobacterales;
A6S79__locus_2
PDU1AB
21050 -pf08411pf00929 -
21055 -pf13599pf13981pf13979 -
21065 -pf04879pf00384pf01568 -
21070 -pf13247pf12800 -
21075 -pf01292 -
21080 -pf07943pf00768 -
21085 -pf06445 -
21090 -pf04632 -
21095 -pf04363 -
21100 HMM00288_1_pdu(pf00288) PDU1AB
21105 HMMacetkin_1(pf00871) PDU1AB
21110 HMM10662PduV_1_mix(pf10662) PDU1AB
21115 Hp_euts_babyPurple PDU1AB
21120 Ts_dodgerBlue PDU1AB
21125 HMMPduS_1_all(pf01512pf13375pf10531) PDU1AB
21130 HMM00465alcdh_3_mix(pf00465) PDU1AB
21135 HMMalddh_1_eutpdugrm(pf00171) PDU1AB EP_NTERM
21140 HMM01923cobtr_1_pdu(pf01923pf03928) PDU1AB
21145 P_babyPink PDU1AB
21150 HMM15953pduput_1_pdu1(pf15953) PDU1AB EP_NTERM
21155 HMMptac_2_mix(pf06130) PDU1AB
21160 H_driedBlood PDU1AB
21165 H_tan PDU1AB
21170 HMMpropdeh3_2_pdusml(pf02288) PDU1AB
21175 HMM08841dhact_1_pdu(pf08841pf18427) PDU1AB
21180 HMMpropdeh2_1_pdu(pf02287) PDU1AB EP_NTERM
21185 HMMpropdeh3_1_pdu(pf02288) PDU1AB EP_NTERM
21190 HMMpropdeh1_1_pdu(pf02286) PDU1AB
21195 Tsp_greenishBeige PDU1AB
21200 H_robinEggBlue GRM4
21205 -pf00230 -
21210 HMM10114PocR_1_pdu1(pf12833pf10114) PDU1AB
21215 -pf01656pf07685 -
21220 -pf03186 -
21225 -pf02570 -
21230 -pf01888 -
21235 -pf00590 -
21240 -pf05175 -
21245 -pf00590 -
21250 -pf01890pf11761pf11760 -
21255 -pf00590 -
21260 -pf02571 -
21265 -pf06180 -
BMC-P
BMC-H
BMC-T
Aldehyde dehydrogenase
PTAC
Signature enzyme
Alcohol dehydrogenase
Regulator

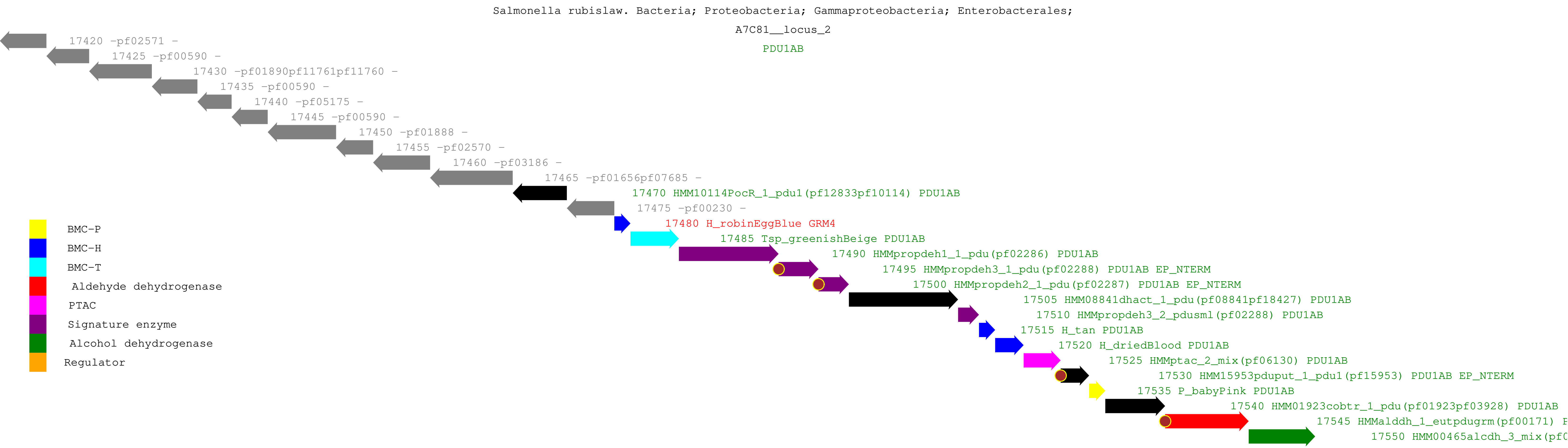
Salmonella rubislaw. Bacteria; Proteobacteria; Gammaproteobacteria; Enterobacterales;
A7C81__locus_2
PDU1AB
17420 -pf02571 -
17425 -pf00590 -
17430 -pf01890pf11761pf11760 -
17435 -pf00590 -
17440 -pf05175 -
17445 -pf00590 -
17450 -pf01888 -
17455 -pf02570 -
17460 -pf03186 -
17465 -pf01656pf07685 -
17470 HMM10114PocR_1_pdu1(pf12833pf10114) PDU1AB
17475 -pf00230 -
17480 H_robinEggBlue GRM4
17485 Tsp_greenishBeige PDU1AB
17490 HMMpropdeh1_1_pdu(pf02286) PDU1AB
17495 HMMpropdeh3_1_pdu(pf02288) PDU1AB EP_NTERM
17500 HMMpropdeh2_1_pdu(pf02287) PDU1AB EP_NTERM
17505 HMM08841dhact_1_pdu(pf08841pf18427) PDU1AB
17510 HMMpropdeh3_2_pdusml(pf02288) PDU1AB
17515 H_tan PDU1AB
17520 H_driedBlood PDU1AB
17525 HMMptac_2_mix(pf06130) PDU1AB
17530 HMM15953pduput_1_pdu1(pf15953) PDU1AB EP_NTERM
17535 P_babyPink PDU1AB
17540 HMM01923cobtr_1_pdu(pf01923pf03928) PDU1AB
17545 HMMalddh_1_eutpdugrm(pf00171) P
17550 HMM00465alcdh_3_mix(pf0
BMC-P
BMC-H
BMC-T
Aldehyde dehydrogenase
PTAC
Signature enzyme
Alcohol dehydrogenase
Regulator

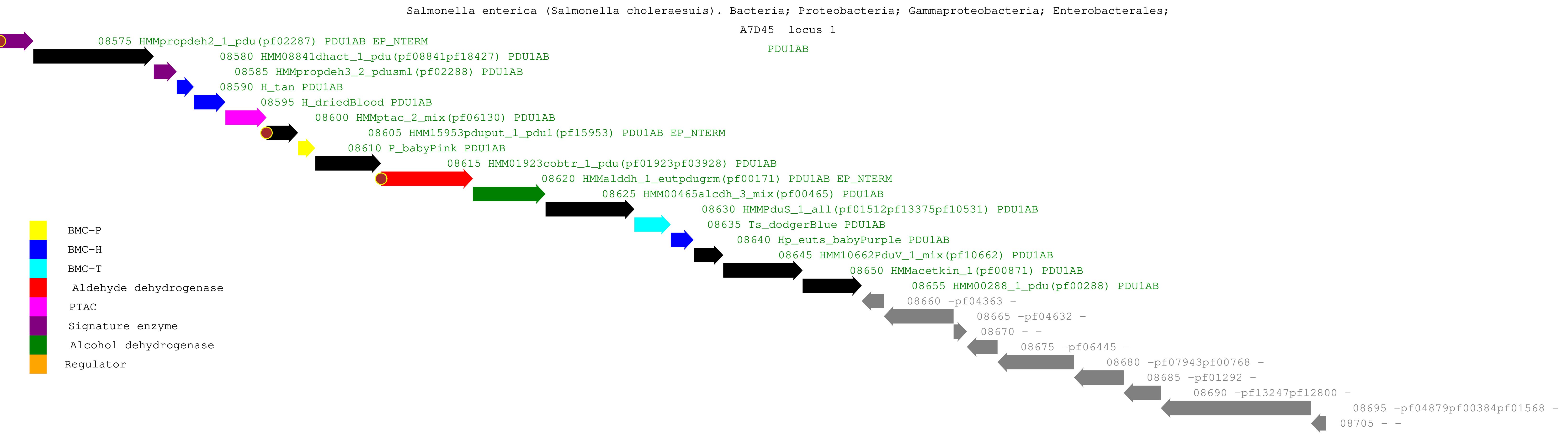

Salmonella enterica (Salmonella choleraesuis). Bacteria; Proteobacteria; Gammaproteobacteria; Enterobacterales;
A7D45__locus_1
PDU1AB
08575 HMMpropdeh2_1_pdu(pf02287) PDU1AB EP_NTERM
08580 HMM08841dhact_1_pdu(pf08841pf18427) PDU1AB
08585 HMMpropdeh3_2_pdusml(pf02288) PDU1AB
08590 H_tan PDU1AB
08595 H_driedBlood PDU1AB
08600 HMMptac_2_mix(pf06130) PDU1AB
08605 HMM15953pduput_1_pdu1(pf15953) PDU1AB EP_NTERM
08610 P_babyPink PDU1AB
08615 HMM01923cobtr_1_pdu(pf01923pf03928) PDU1AB
08620 HMMalddh_1_eutpdugrm(pf00171) PDU1AB EP_NTERM
08625 HMM00465alcdh_3_mix(pf00465) PDU1AB
08630 HMMPduS_1_all(pf01512pf13375pf10531) PDU1AB
08635 Ts_dodgerBlue PDU1AB
08640 Hp_euts_babyPurple PDU1AB
08645 HMM10662PduV_1_mix(pf10662) PDU1AB
08650 HMMacetkin_1(pf00871) PDU1AB
08655 HMM00288_1_pdu(pf00288) PDU1AB
08660 -pf04363 -
08665 -pf04632 -
08670 - -
08675 -pf06445 -
08680 -pf07943pf00768 -
08685 -pf01292 -
08690 -pf13247pf12800 -
08695 -pf04879pf00384pf01568 -
08705 - -
BMC-P
BMC-H
BMC-T
Aldehyde dehydrogenase
PTAC
Signature enzyme
Alcohol dehydrogenase
Regulator

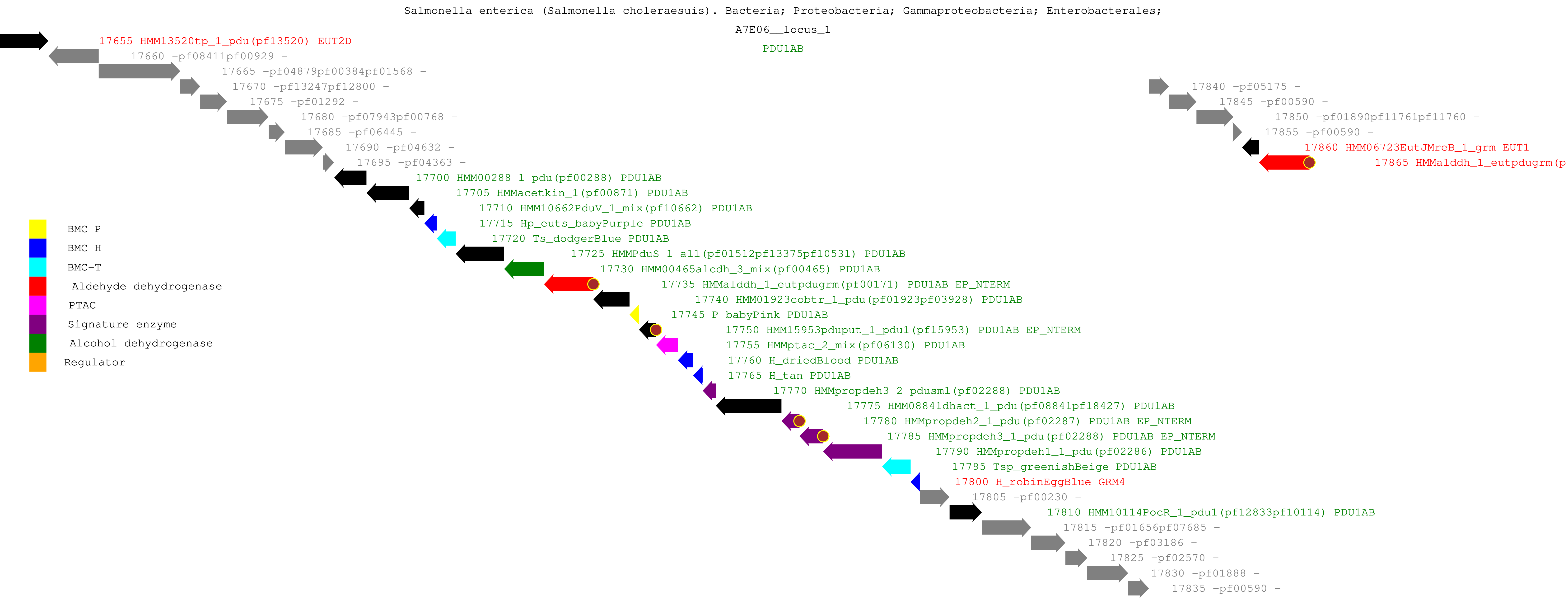
Salmonella enterica (Salmonella choleraesuis). Bacteria; Proteobacteria; Gammaproteobacteria; Enterobacterales;
A7E06__locus_1
PDU1AB
17655 HMM13520tp_1_pdu(pf13520) EUT2D
17660 -pf08411pf00929 -
17665 -pf04879pf00384pf01568 -
17670 -pf13247pf12800 -
17675 -pf01292 -
17680 -pf07943pf00768 -
17685 -pf06445 -
17690 -pf04632 -
17695 -pf04363 -
17700 HMM00288_1_pdu(pf00288) PDU1AB
17705 HMMacetkin_1(pf00871) PDU1AB
17710 HMM10662PduV_1_mix(pf10662) PDU1AB
17715 Hp_euts_babyPurple PDU1AB
17720 Ts_dodgerBlue PDU1AB
17725 HMMPduS_1_all(pf01512pf13375pf10531) PDU1AB
17730 HMM00465alcdh_3_mix(pf00465) PDU1AB
17735 HMMalddh_1_eutpdugrm(pf00171) PDU1AB EP_NTERM
17740 HMM01923cobtr_1_pdu(pf01923pf03928) PDU1AB
17745 P_babyPink PDU1AB
17750 HMM15953pduput_1_pdu1(pf15953) PDU1AB EP_NTERM
17755 HMMptac_2_mix(pf06130) PDU1AB
17760 H_driedBlood PDU1AB
17765 H_tan PDU1AB
17770 HMMpropdeh3_2_pdusml(pf02288) PDU1AB
17775 HMM08841dhact_1_pdu(pf08841pf18427) PDU1AB
17780 HMMpropdeh2_1_pdu(pf02287) PDU1AB EP_NTERM
17785 HMMpropdeh3_1_pdu(pf02288) PDU1AB EP_NTERM
17790 HMMpropdeh1_1_pdu(pf02286) PDU1AB
17795 Tsp_greenishBeige PDU1AB
17800 H_robinEggBlue GRM4
17805 -pf00230 -
17810 HMM10114PocR_1_pdu1(pf12833pf10114) PDU1AB
17815 -pf01656pf07685 -
17820 -pf03186 -
17825 -pf02570 -
17830 -pf01888 -
17835 -pf00590 -
17840 -pf05175 -
17845 -pf00590 -
17850 -pf01890pf11761pf11760 -
17855 -pf00590 -
17860 HMM06723EutJMreB_1_grm EUT1
17865 HMMalddh_1_eutpdugrm(p
BMC-P
BMC-H
BMC-T
Aldehyde dehydrogenase
PTAC
Signature enzyme
Alcohol dehydrogenase
Regulator

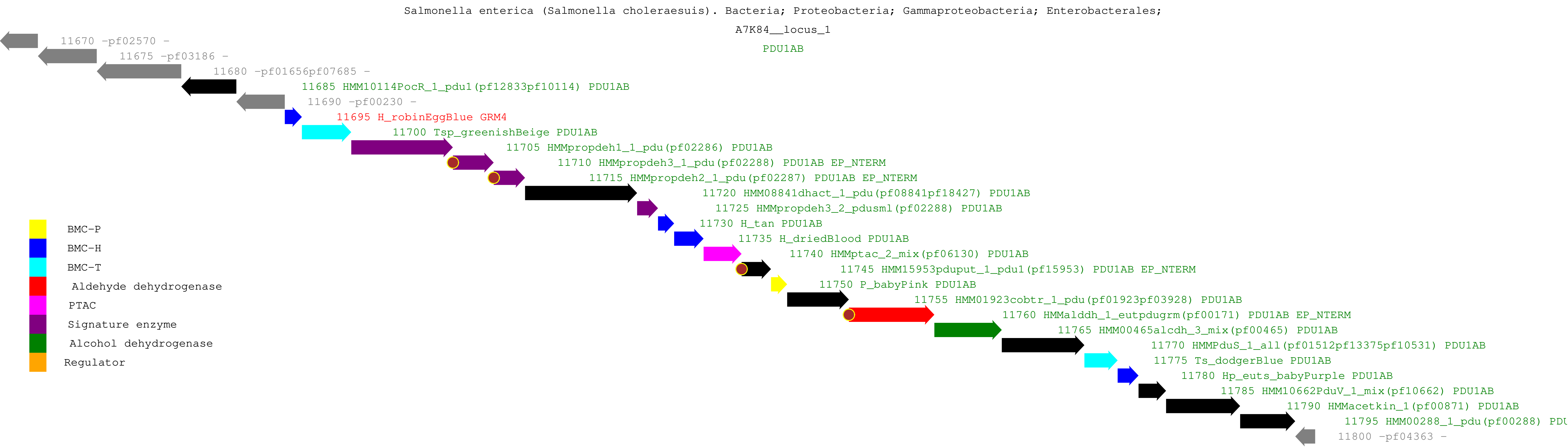
Salmonella enterica (Salmonella choleraesuis). Bacteria; Proteobacteria; Gammaproteobacteria; Enterobacterales;
A7K84__locus_1
PDU1AB
11670 -pf02570 -
11675 -pf03186 -
11680 -pf01656pf07685 -
11685 HMM10114PocR_1_pdu1(pf12833pf10114) PDU1AB
11690 -pf00230 -
11695 H_robinEggBlue GRM4
11700 Tsp_greenishBeige PDU1AB
11705 HMMpropdeh1_1_pdu(pf02286) PDU1AB
11710 HMMpropdeh3_1_pdu(pf02288) PDU1AB EP_NTERM
11715 HMMpropdeh2_1_pdu(pf02287) PDU1AB EP_NTERM
11720 HMM08841dhact_1_pdu(pf08841pf18427) PDU1AB
11725 HMMpropdeh3_2_pdusml(pf02288) PDU1AB
11730 H_tan PDU1AB
11735 H_driedBlood PDU1AB
11740 HMMptac_2_mix(pf06130) PDU1AB
11745 HMM15953pduput_1_pdu1(pf15953) PDU1AB EP_NTERM
11750 P_babyPink PDU1AB
11755 HMM01923cobtr_1_pdu(pf01923pf03928) PDU1AB
11760 HMMaldh_1_eutpdugrm(pf00171) PDU1AB EP_NTERM
11765 HMM00465alcdh_3_mix(pf00465) PDU1AB
11770 HMMPduS_1_all(pf01512pf13375pf10531) PDU1AB
11775 Ts_dodgerBlue PDU1AB
11780 Hp_euts_babyPurple PDU1AB
11785 HMM10662PduV_1_mix(pf10662) PDU1AB
11790 HMMacetkin_1(pf00871) PDU1AB
11795 HMM00288_1_pdu(pf00288) PDU
11800 -pf04363 -
BMC-P
BMC-H
BMC-T
Aldehyde dehydrogenase
PTAC
Signature enzyme
Alcohol dehydrogenase
Regulator

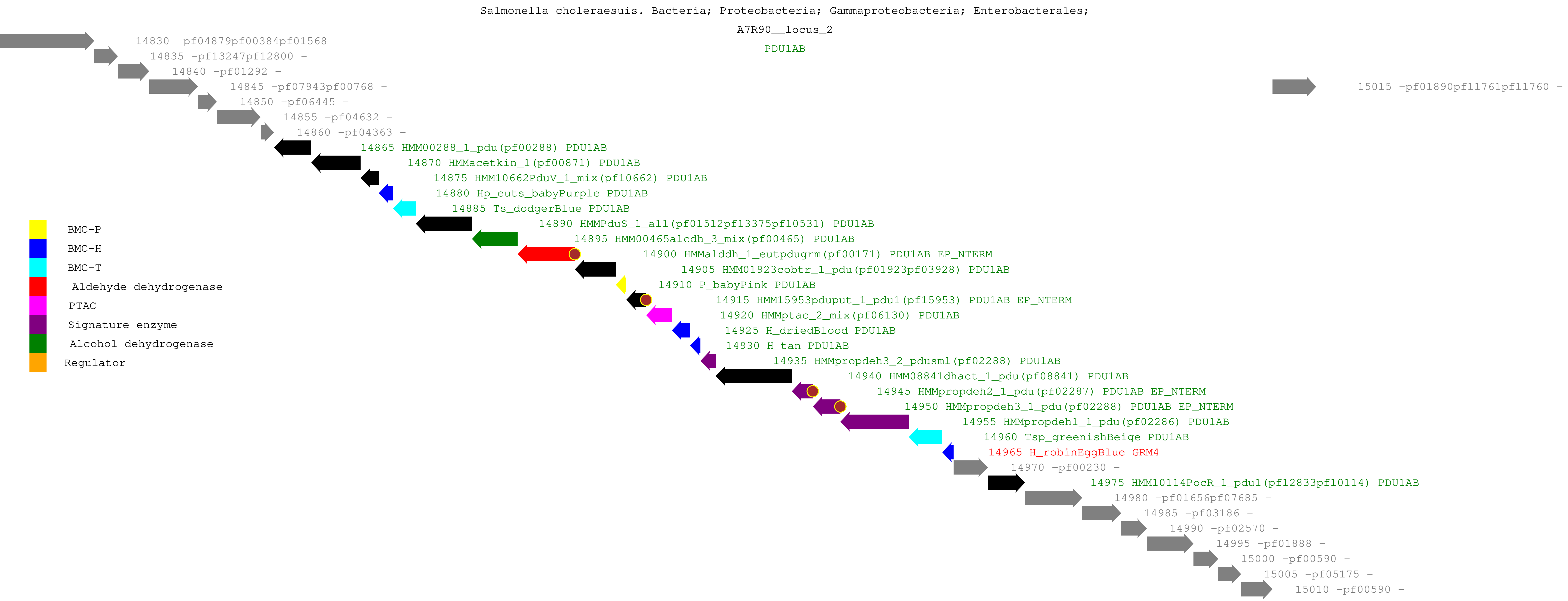
Salmonella choleraesuis. Bacteria; Proteobacteria; Gammaproteobacteria; Enterobacterales;
A7R90__locus_2
PDU1AB
14830 -pf04879pf00384pf01568 -
14835 -pf13247pf12800 -
14840 -pf01292 -
14845 -pf07943pf00768 -
14850 -pf06445 -
14855 -pf04632 -
14860 -pf04363 -
14865 HMM00288_1_pdu(pf00288) PDU1AB
14870 HMMacetkin_1(pf00871) PDU1AB
14875 HMM10662PduV_1_mix(pf10662) PDU1AB
14880 Hp_euts_babyPurple PDU1AB
14885 Ts_dodgerBlue PDU1AB
14890 HMMPduS_1_all(pf01512pf13375pf10531) PDU1AB
14895 HMM00465alcdh_3_mix(pf00465) PDU1AB
14900 HMMalddh_1_eutpdugrm(pf00171) PDU1AB EP_NTERM
14905 HMM01923cobtr_1_pdu(pf01923pf03928) PDU1AB
14910 P_babyPink PDU1AB
14915 HMM15953pduput_1_pdu1(pf15953) PDU1AB EP_NTERM
14920 HMMptac_2_mix(pf06130) PDU1AB
14925 H_driedBlood PDU1AB
14930 H_tan PDU1AB
14935 HMMpropdeh3_2_pdusml(pf02288) PDU1AB
14940 HMM08841dhact_1_pdu(pf08841) PDU1AB
14945 HMMpropdeh2_1_pdu(pf02287) PDU1AB EP_NTERM
14950 HMMpropdeh3_1_pdu(pf02288) PDU1AB EP_NTERM
14955 HMMpropdeh1_1_pdu(pf02286) PDU1AB
14960 Tsp_greenishBeige PDU1AB
14965 H_robinEggBlue GRM4
14970 -pf00230 -
14975 HMM10114PocR_1_pdu1(pf12833pf10114) PDU1AB
14980 -pf01656pf07685 -
14985 -pf03186 -
14990 -pf02570 -
14995 -pf01888 -
15000 -pf00590 -
15005 -pf05175 -
15010 -pf00590 -
15015 -pf01890pf11761pf11760 -
BMC-P
BMC-H
BMC-T
Aldehyde dehydrogenase
PTAC
Signature enzyme
Alcohol dehydrogenase
Regulator

Salmonella choleraesuis. Bacteria; Proteobacteria; Gammaproteobacteria; Enterobacterales;

A7S24__locus_1

PDU1AB

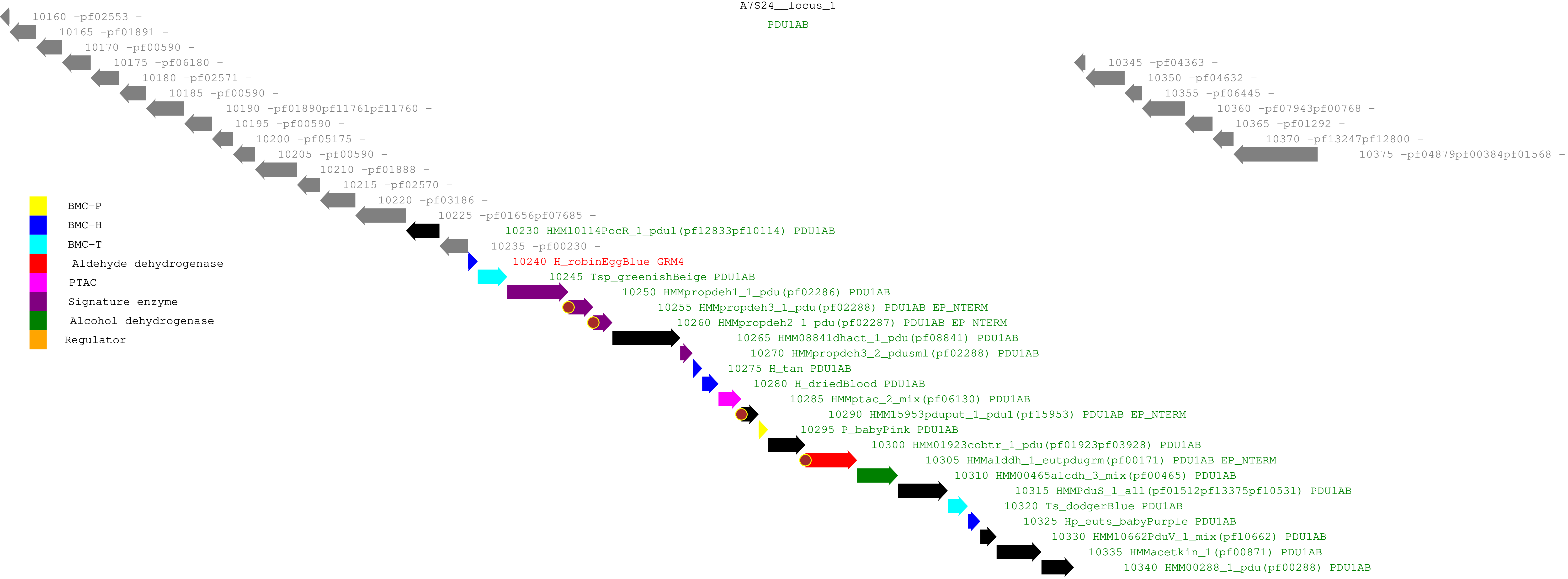

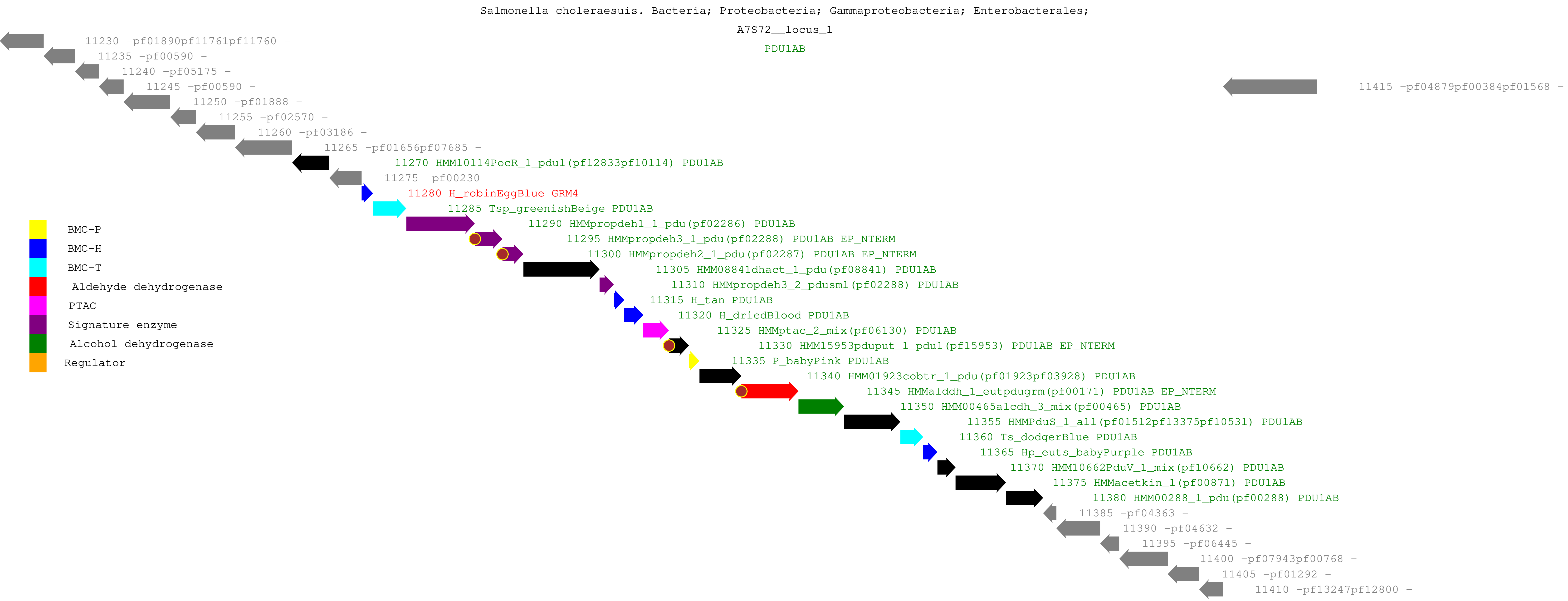
Salmonella choleraesuis. Bacteria; Proteobacteria; Gammaproteobacteria; Enterobacterales;
A7S72__locus_1
PDU1AB
11230 -pf01890pf11761pf11760 -
11235 -pf00590 -
11240 -pf05175 -
11245 -pf00590 -
11250 -pf01888 -
11255 -pf02570 -
11260 -pf03186 -
11265 -pf01656pf07685 -
11270 HMM10114PocR_1_pdu1(pf12833pf10114) PDU1AB
11275 -pf00230 -
11280 H_robinEggBlue GRM4
11285 Tsp_greenishBeige PDU1AB
11290 HMMpropdeh1_1_pdu(pf02286) PDU1AB
11295 HMMpropdeh3_1_pdu(pf02288) PDU1AB EP_NTERM
11300 HMMpropdeh2_1_pdu(pf02287) PDU1AB EP_NTERM
11305 HMM08841dhact_1_pdu(pf08841) PDU1AB
11310 HMMpropdeh3_2_pdusml(pf02288) PDU1AB
11315 H_tan PDU1AB
11320 H_driedBlood PDU1AB
11325 HMMptac_2_mix(pf06130) PDU1AB
11330 HMM15953pduput_1_pdu1(pf15953) PDU1AB EP_NTERM
11335 P_babyPink PDU1AB
11340 HMM01923cobtr_1_pdu(pf01923pf03928) PDU1AB
11345 HMMaldh_1_eutpdugrm(pf00171) PDU1AB EP_NTERM
11350 HMM00465alcdh_3_mix(pf00465) PDU1AB
11355 HMMPduS_1_all(pf01512pf13375pf10531) PDU1AB
11360 Ts_dodgerBlue PDU1AB
11365 Hp_euts_babyPurple PDU1AB
11370 HMM10662PduV_1_mix(pf10662) PDU1AB
11375 HMMacetkin_1(pf00871) PDU1AB
11380 HMM00288_1_pdu(pf00288) PDU1AB
11385 -pf04363 -
11390 -pf04632 -
11395 -pf06445 -
11400 -pf07943pf00768 -
11405 -pf01292 -
11410 -pf13247pf12800 -
11415 -pf04879pf00384pf01568 -
BMC-P
BMC-H
BMC-T
Aldehyde dehydrogenase
PTAC
Signature enzyme
Alcohol dehydrogenase
Regulator

Salmonella enterica subsp. enterica serovar Saintpaul. Bacteria; Proteobacteria; Gammaproteobacteria; Enterobacterales;

A7T00__locus_2

PDU1AB

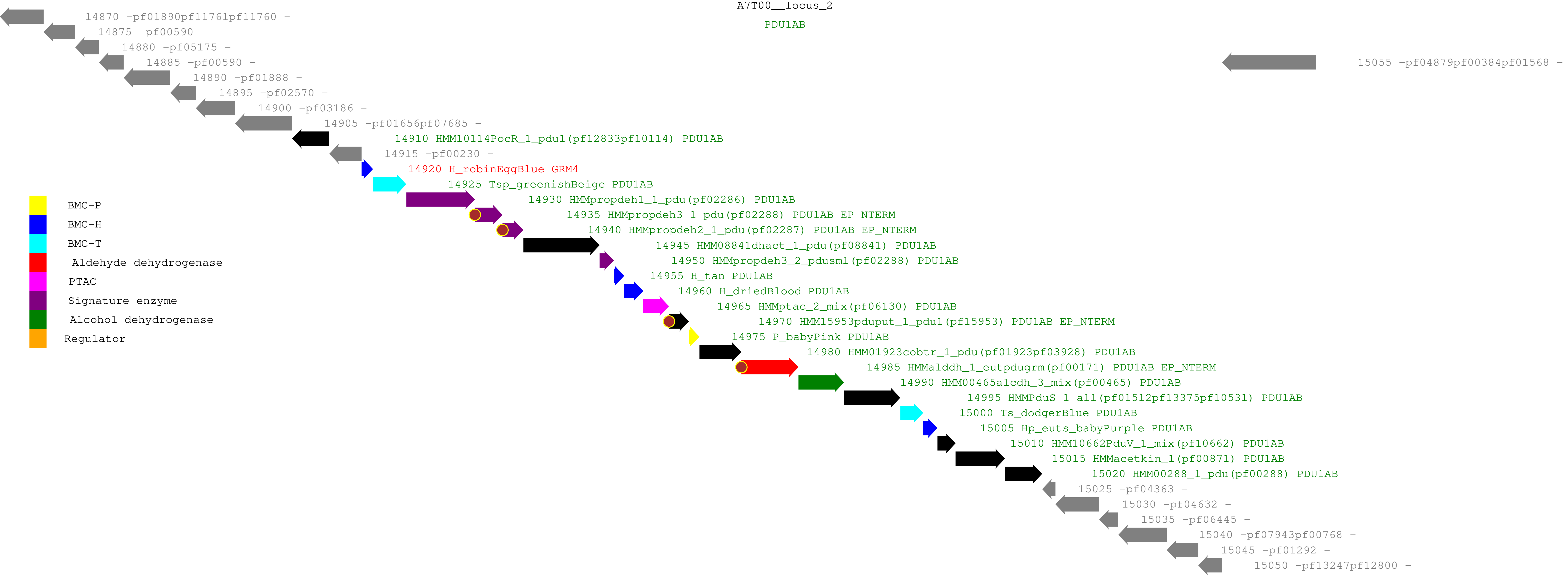

Salmonella enterica subsp. enterica serovar Inverness. Bacteria; Proteobacteria; Gammaproteobacteria; Enterobacterales;

A7T22__locus_2

PDU1AB

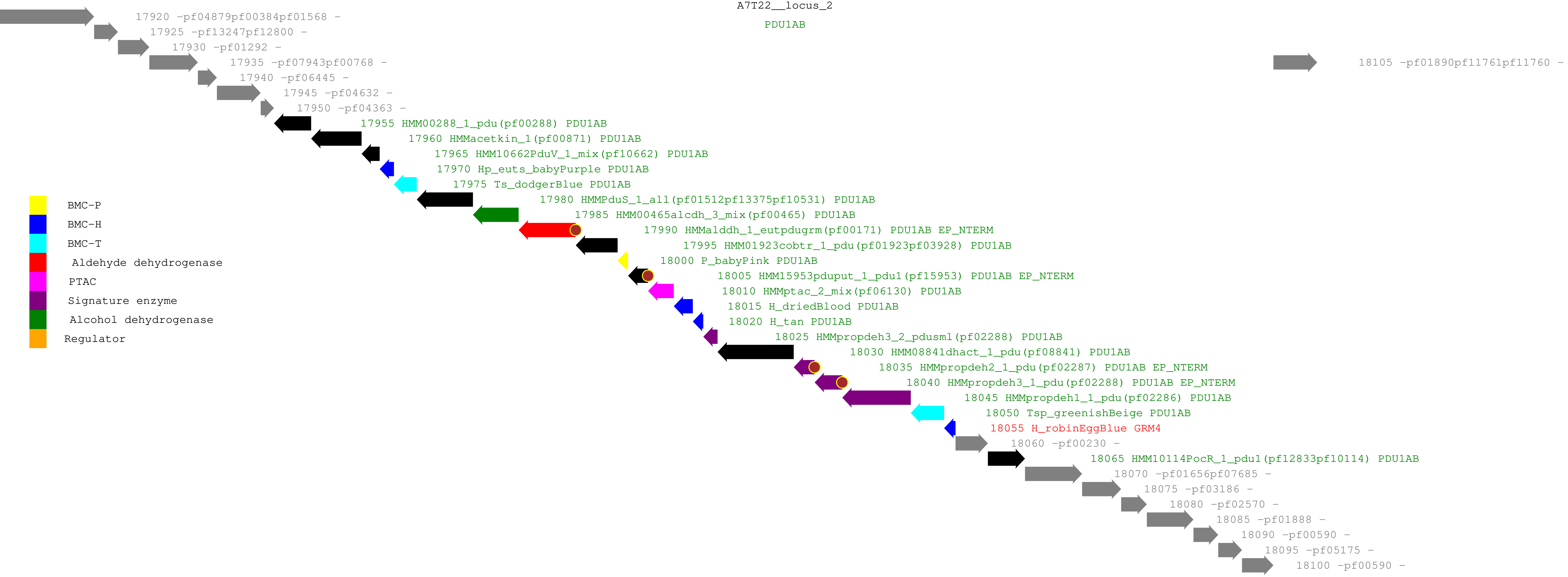

Salmonella enterica subsp. enterica serovar Bareilly. Bacteria; Proteobacteria; Gammaproteobacteria; Enterobacterales;

A7T25__locus_1

PDU1AB

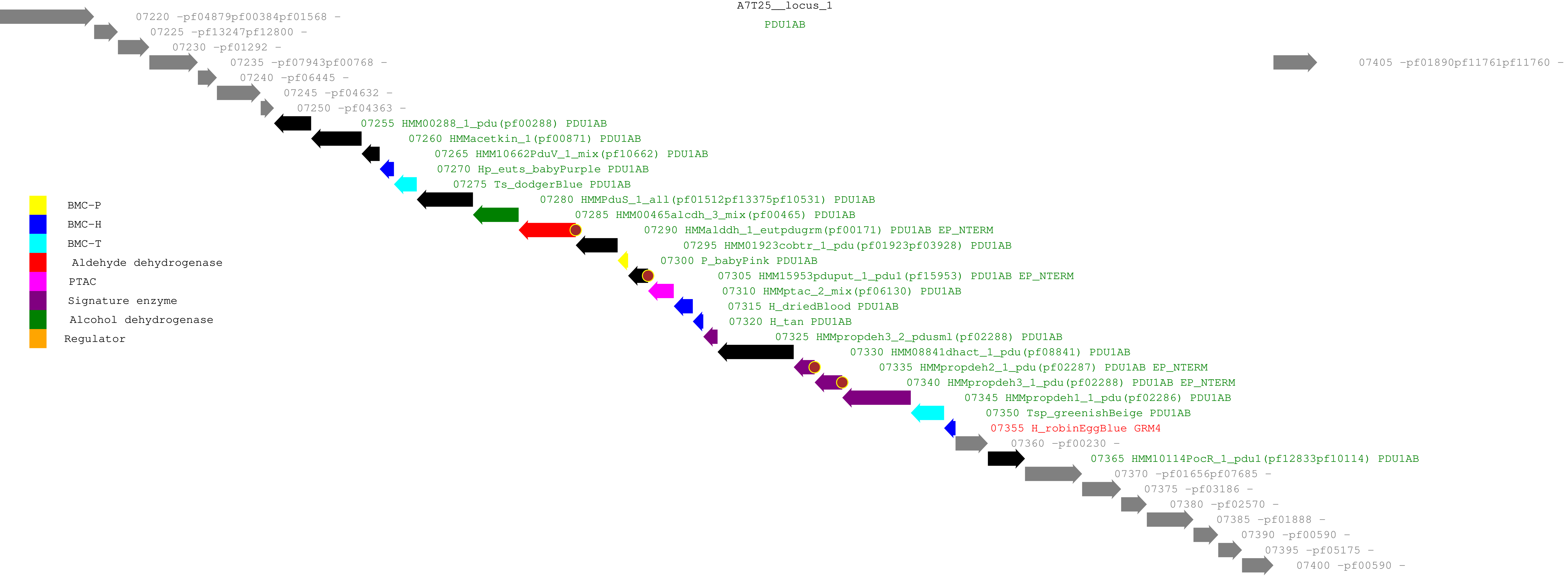

Salmonella anatum. Bacteria; Proteobacteria; Gammaproteobacteria; Enterobacterales;

A7T43__locus_1

PDU1AB

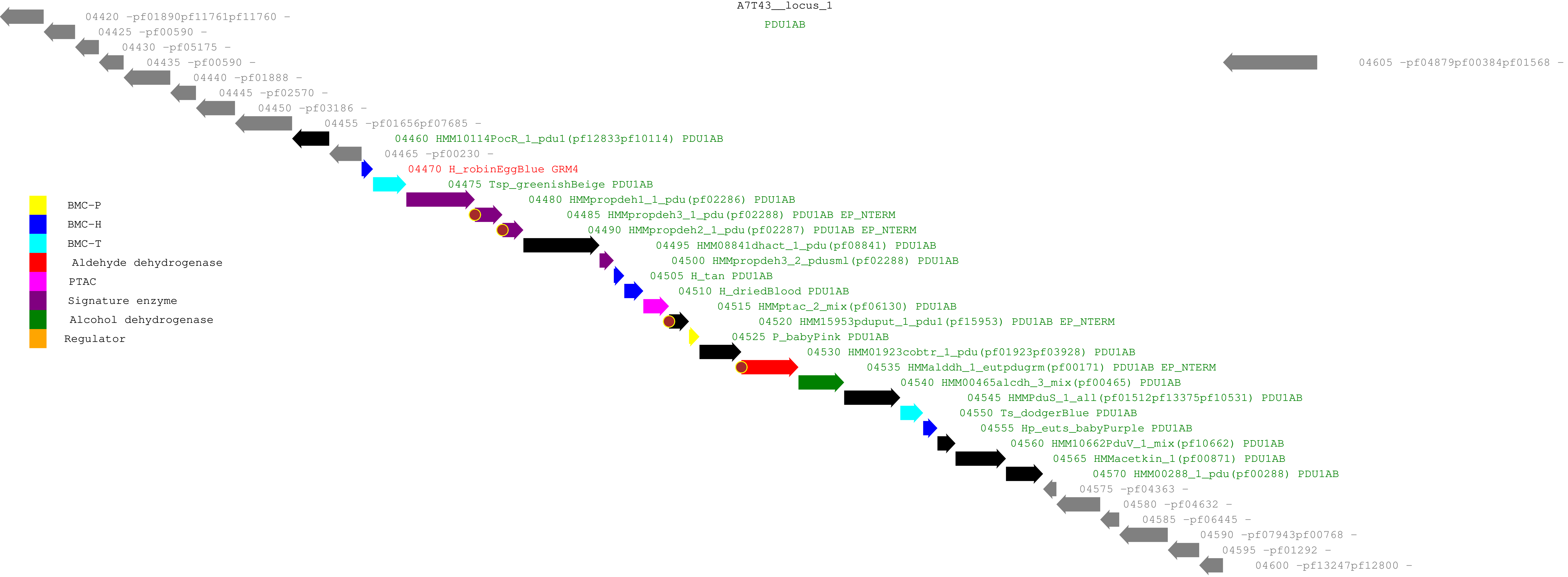

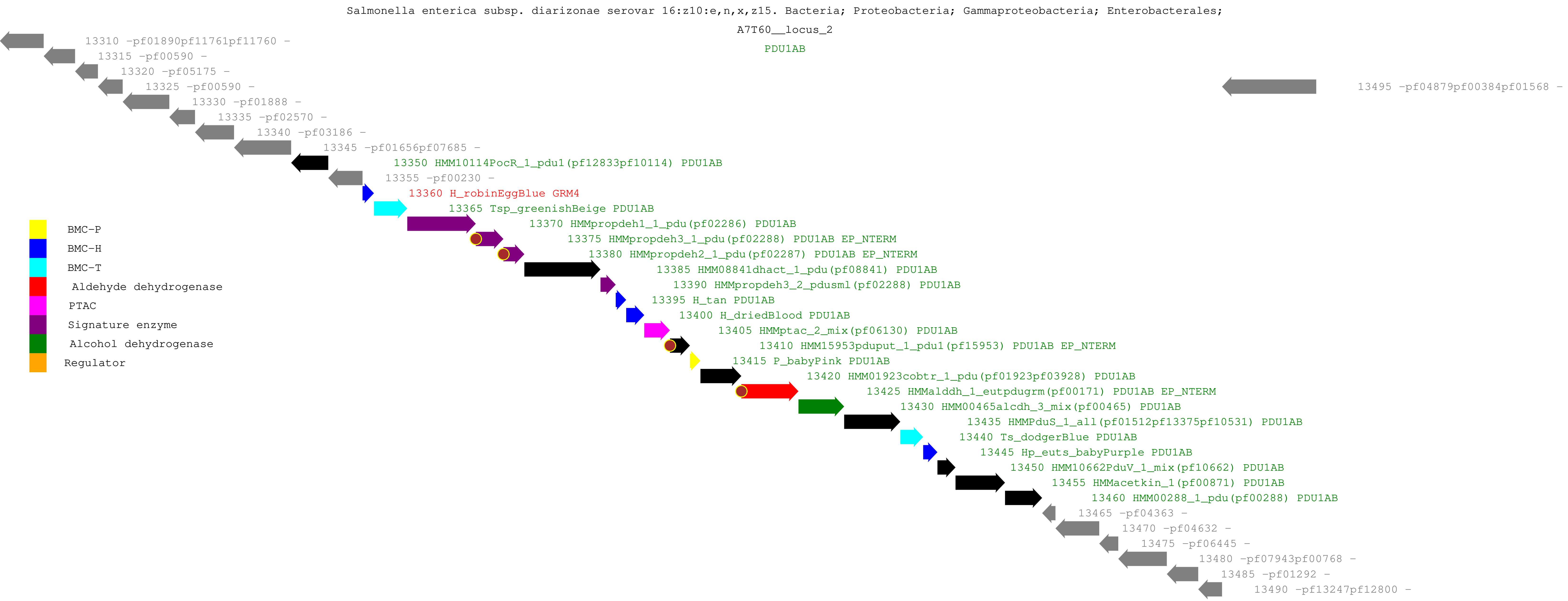
Salmonella enterica subsp. diarizonae serovar 16:z10:e,n,x,z15. Bacteria; Proteobacteria; Gammaproteobacteria; Enterobacterales;
A7T60__locus_2
PDU1AB
13310 -pf01890pf11761pf11760 -
13315 -pf00590 -
13320 -pf05175 -
13325 -pf00590 -
13330 -pf01888 -
13335 -pf02570 -
13340 -pf03186 -
13345 -pf01656pf07685 -
13350 HMM10114PocR_1_pdu1(pf12833pf10114) PDU1AB
13355 -pf00230 -
13360 H_robinEggBlue GRM4
13365 Tsp_greenishBeige PDU1AB
13370 HMMpropdeh1_1_pdu(pf02286) PDU1AB
13375 HMMpropdeh3_1_pdu(pf02288) PDU1AB EP_NTERM
13380 HMMpropdeh2_1_pdu(pf02287) PDU1AB EP_NTERM
13385 HMM08841dhact_1_pdu(pf08841) PDU1AB
13390 HMMpropdeh3_2_pdusml(pf02288) PDU1AB
13395 H_tan PDU1AB
13400 H_driedBlood PDU1AB
13405 HMMptac_2_mix(pf06130) PDU1AB
13410 HMM15953pduput_1_pdu1(pf15953) PDU1AB EP_NTERM
13415 P_babyPink PDU1AB
13420 HMM01923cobtr_1_pdu(pf01923pf03928) PDU1AB
13425 HMMaldh_1_eutpdugrm(pf00171) PDU1AB EP_NTERM
13430 HMM00465alcdh_3_mix(pf00465) PDU1AB
13435 HMMPduS_1_all(pf01512pf13375pf10531) PDU1AB
13440 Ts_dodgerBlue PDU1AB
13445 Hp_euts_babyPurple PDU1AB
13450 HMM10662PduV_1_mix(pf10662) PDU1AB
13455 HMMacetkin_1(pf00871) PDU1AB
13460 HMM00288_1_pdu(pf00288) PDU1AB
13465 -pf04363 -
13470 -pf04632 -
13475 -pf06445 -
13480 -pf07943pf00768 -
13485 -pf01292 -
13490 -pf13247pf12800 -
13495 -pf04879pf00384pf01568 -
BMC-P
BMC-H
BMC-T
Aldehyde dehydrogenase
PTAC
Signature enzyme
Alcohol dehydrogenase
Regulator

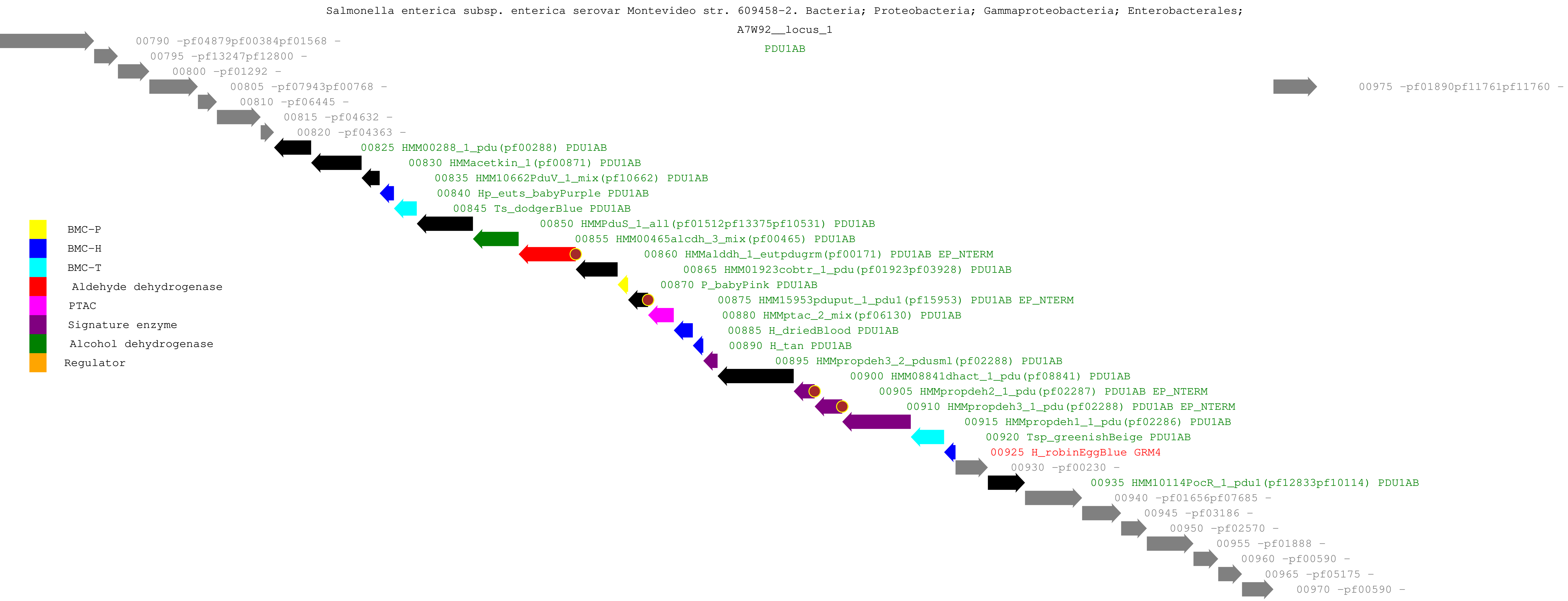
Salmonella enterica subsp. enterica serovar Montevideo str. 609458-2. Bacteria; Proteobacteria; Gammaproteobacteria; Enterobacterales;
A7W92__locus_1
PDU1AB
00790 -pf04879pf00384pf01568 -
00795 -pf13247pf12800 -
00800 -pf01292 -
00805 -pf07943pf00768 -
00975 -pf01890pf11761pf11760 -
00810 -pf06445 -
00815 -pf04632 -
00820 -pf04363 -
00825 HMM00288_1_pdu(pf00288) PDU1AB
00830 HMMacetkin_1(pf00871) PDU1AB
00835 HMM10662PduV_1_mix(pf10662) PDU1AB
00840 Hp_euts_babyPurple PDU1AB
00845 Ts_dodgerBlue PDU1AB
00850 HMMPduS_1_all(pf01512pf13375pf10531) PDU1AB
00855 HMM00465alcdh_3_mix(pf00465) PDU1AB
00860 HMMalddh_1_eutpdugrm(pf00171) PDU1AB EP_NTERM
00865 HMM01923cobtr_1_pdu(pf01923pf03928) PDU1AB
00870 P_babyPink PDU1AB
00875 HMM15953pduput_1_pdu1(pf15953) PDU1AB EP_NTERM
00880 HMMptac_2_mix(pf06130) PDU1AB
00885 H_driedBlood PDU1AB
00890 H_tan PDU1AB
00895 HMMpropdeh3_2_pdusml(pf02288) PDU1AB
00900 HMM08841dhact_1_pdu(pf08841) PDU1AB
00905 HMMpropdeh2_1_pdu(pf02287) PDU1AB EP_NTERM
00910 HMMpropdeh3_1_pdu(pf02288) PDU1AB EP_NTERM
00915 HMMpropdeh1_1_pdu(pf02286) PDU1AB
00920 Tsp_greenishBeige PDU1AB
00925 H_robinEggBlue GRM4
00930 -pf00230 -
00935 HMM10114PocR_1_pdu1(pf12833pf10114) PDU1AB
00940 -pf01656pf07685 -
00945 -pf03186 -
00950 -pf02570 -
00955 -pf01888 -
00960 -pf00590 -
00965 -pf05175 -
00970 -pf00590 -
BMC-P
BMC-H
BMC-T
Aldehyde dehydrogenase
PTAC
Signature enzyme
Alcohol dehydrogenase
Regulator

Escherichia coli. Bacteria; Proteobacteria; Gammaproteobacteria; Enterobacterales;

A8M44__locus_2

PDU1AB

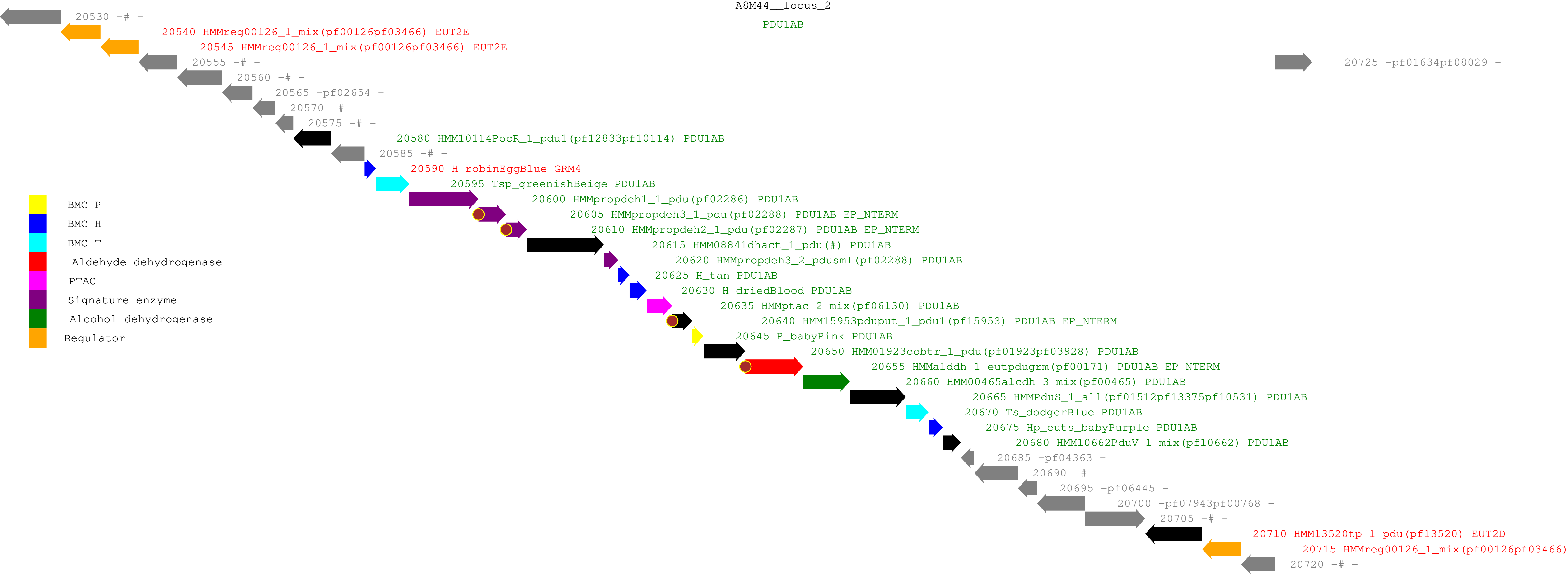

Salmonella enterica subsp. enterica serovar Napoli. Bacteria; Proteobacteria; Gammaproteobacteria; Enterobacterales;

A8N98__locus_1

PDU1AB

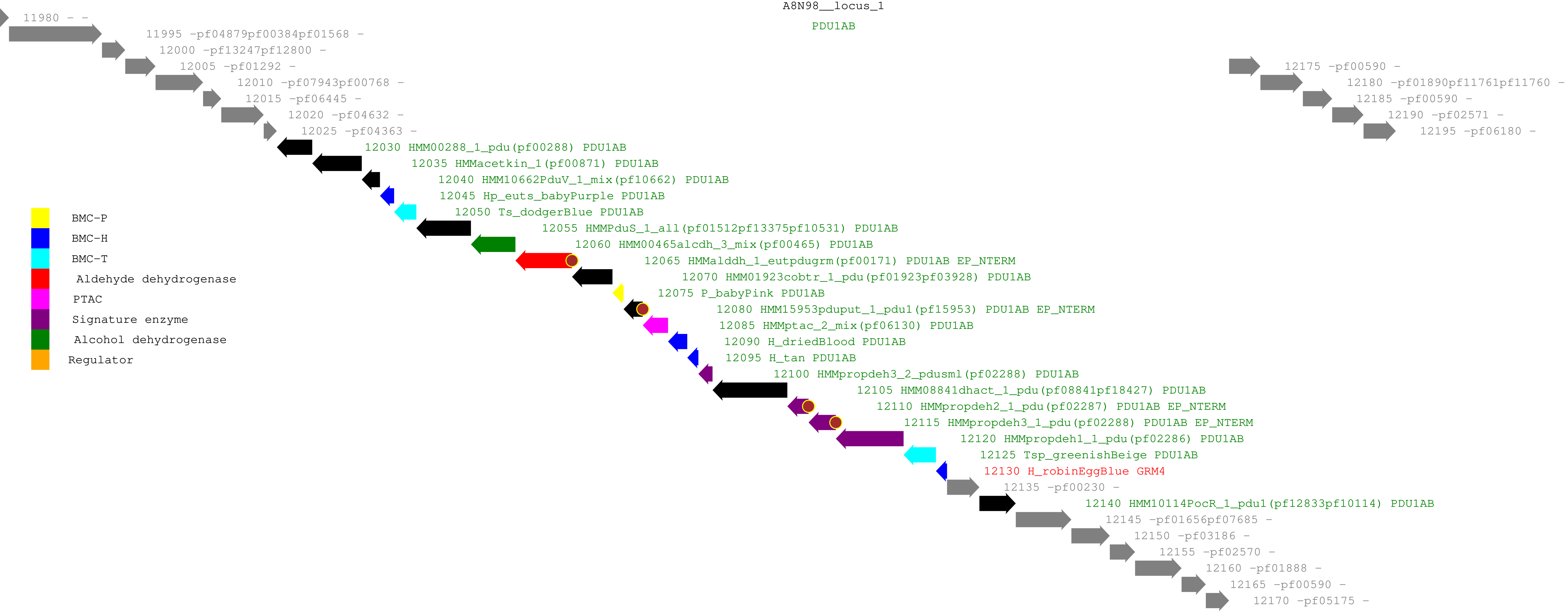

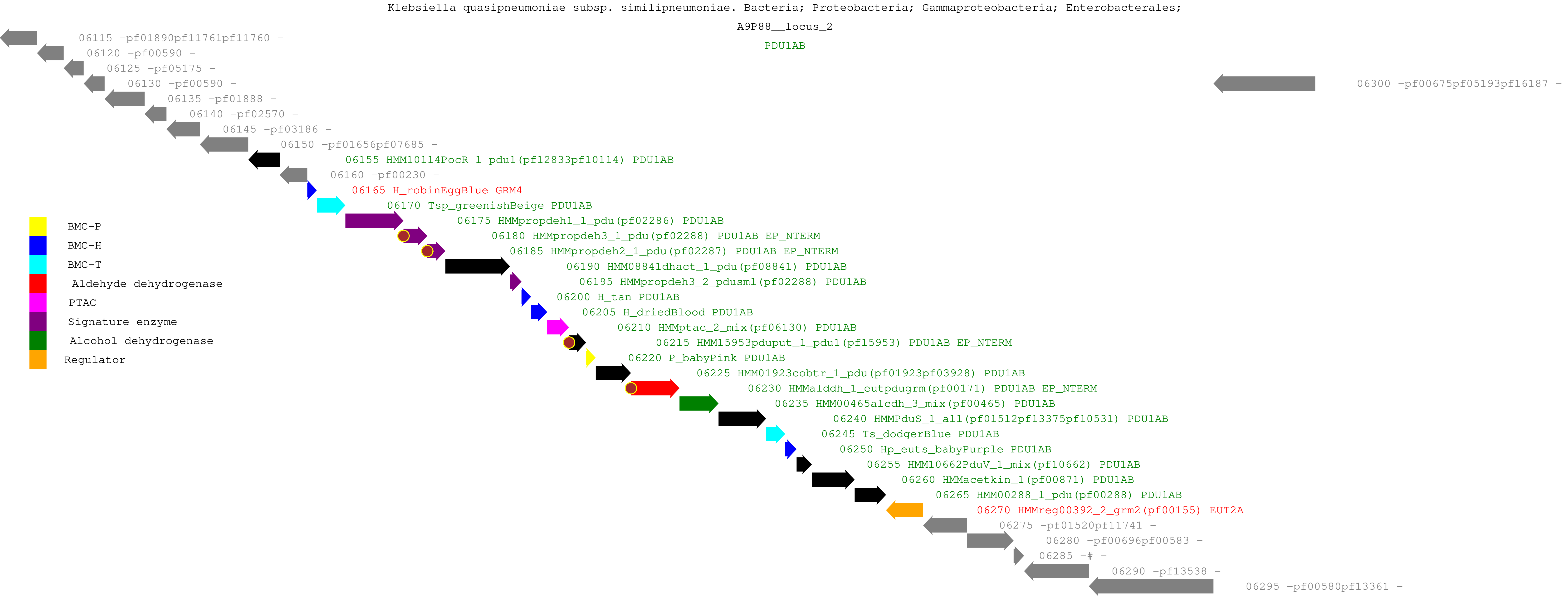
Klebsiella quasipneumoniae subsp. similipneumoniae. Bacteria; Proteobacteria; Gammaproteobacteria; Enterobacterales;
A9P88__locus_2
PDU1AB
06115 -pf01890pf11761pf11760 -
06120 -pf00590 -
06125 -pf05175 -
06130 -pf00590 -
06135 -pf01888 -
06140 -pf02570 -
06145 -pf03186 -
06150 -pf01656pf07685 -
06155 HMM10114PocR_1_pdu1(pf12833pf10114) PDU1AB
06160 -pf00230 -
06165 H_robinEggBlue GRM4
06170 Tsp_greenishBeige PDU1AB
06175 HMMpropdeh1_1_pdu(pf02286) PDU1AB
06180 HMMpropdeh3_1_pdu(pf02288) PDU1AB EP_NTERM
06185 HMMpropdeh2_1_pdu(pf02287) PDU1AB EP_NTERM
06190 HMM08841dhact_1_pdu(pf08841) PDU1AB
06195 HMMpropdeh3_2_pdusml(pf02288) PDU1AB
06200 H_tan PDU1AB
06205 H_driedBlood PDU1AB
06210 HMMptac_2_mix(pf06130) PDU1AB
06215 HMM15953pduput_1_pdu1(pf15953) PDU1AB EP_NTERM
06220 P_babyPink PDU1AB
06225 HMM01923cobtr_1_pdu(pf01923pf03928) PDU1AB
06230 HMMalddh_1_eutpdugrm(pf00171) PDU1AB EP_NTERM
06235 HMM00465alcdh_3_mix(pf00465) PDU1AB
06240 HMMPduS_1_all(pf01512pf13375pf10531) PDU1AB
06245 Ts_dodgerBlue PDU1AB
06250 Hp_euts_babyPurple PDU1AB
06255 HMM10662PduV_1_mix(pf10662) PDU1AB
06260 HMMacetkin_1(pf00871) PDU1AB
06265 HMM00288_1_pdu(pf00288) PDU1AB
06270 HMMreg00392_2_grm2(pf00155) EUT2A
06275 -pf01520pf11741 -
06280 -pf00696pf00583 -
06285 -# -
06290 -pf13538 -
06295 -pf00580pf13361 -
06300 -pf00675pf05193pf16187 -
BMC-P
BMC-H
BMC-T
Aldehyde dehydrogenase
PTAC
Signature enzyme
Alcohol dehydrogenase
Regulator

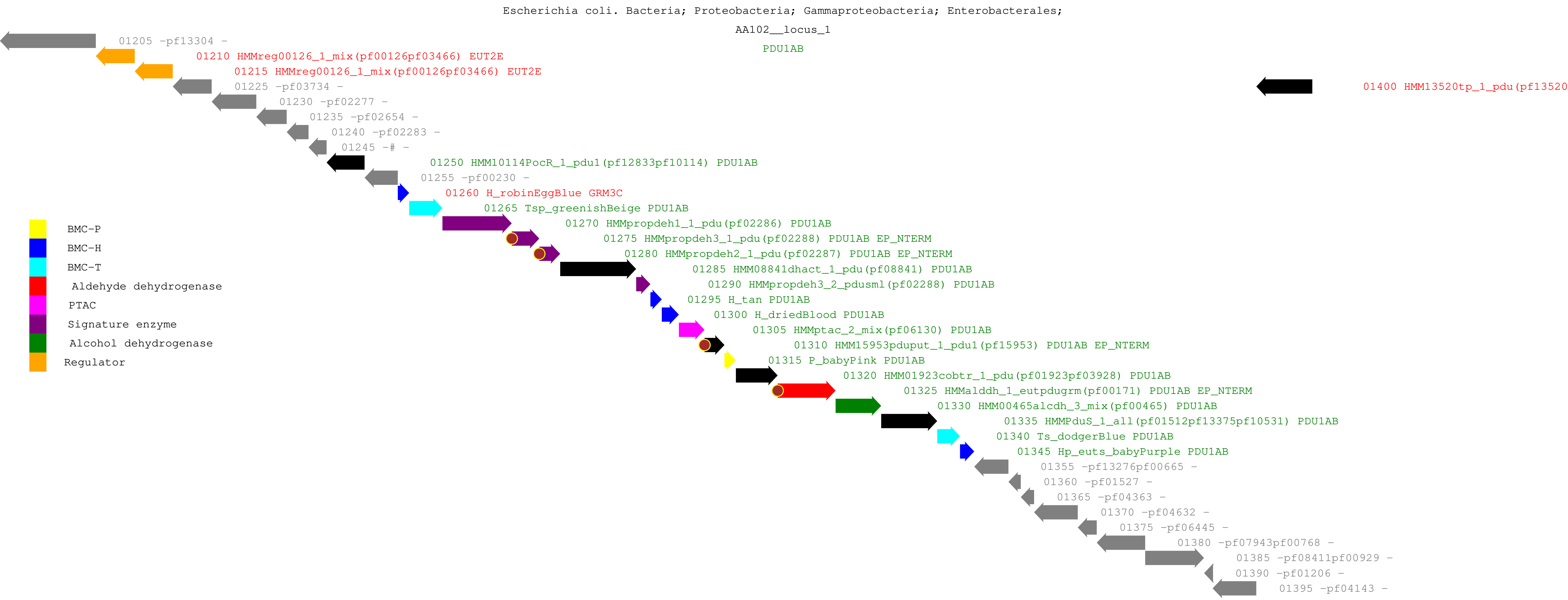
Escherichia coli. Bacteria; Proteobacteria; Gammaproteobacteria; Enterobacterales;
AA102__locus_1
PDU1AB
01205 -pf13304 -
01210 HMMreg00126_1_mix(pf00126pf03466) EUT2E
01215 HMMreg00126_1_mix(pf00126pf03466) EUT2E
01225 -pf03734 -
01230 -pf02277 -
01235 -pf02654 -
01240 -pf02283 -
01245 -# -
01250 HMM10114PocR_1_pdu1(pf12833pf10114) PDU1AB
01255 -pf00230 -
01260 H_robinEggBlue GRM3C
01265 Tsp_greenishBeige PDU1AB
01270 HMMpropdeh1_1_pdu(pf02286) PDU1AB
01275 HMMpropdeh3_1_pdu(pf02288) PDU1AB EP_NTERM
01280 HMMpropdeh2_1_pdu(pf02287) PDU1AB EP_NTERM
01285 HMM08841dhact_1_pdu(pf08841) PDU1AB
01290 HMMpropdeh3_2_pdusml(pf02288) PDU1AB
01295 H_tan PDU1AB
01300 H_driedBlood PDU1AB
01305 HMMptac_2_mix(pf06130) PDU1AB
01310 HMM15953pduput_1_pdu1(pf15953) PDU1AB EP_NTERM
01315 P_babyPink PDU1AB
01320 HMM01923cobtr_1_pdu(pf01923pf03928) PDU1AB
01325 HMMaldh_1_eutpdugrm(pf00171) PDU1AB EP_NTERM
01330 HMM00465alcdh_3_mix(pf00465) PDU1AB
01335 HMMPduS_1_all(pf01512pf13375pf10531) PDU1AB
01340 Ts_dodgerBlue PDU1AB
01345 Hp_euts_babyPurple PDU1AB
01355 -pf13276pf00665 -
01360 -pf01527 -
01365 -pf04363 -
01370 -pf04632 -
01375 -pf06445 -
01380 -pf07943pf00768 -
01385 -pf08411pf00929 -
01390 -pf01206 -
01395 -pf04143 -
01400 HMM13520tp_1_pdu(pf13520
BMC-P
BMC-H
BMC-T
Aldehyde dehydrogenase
PTAC
Signature enzyme
Alcohol dehydrogenase
Regulator

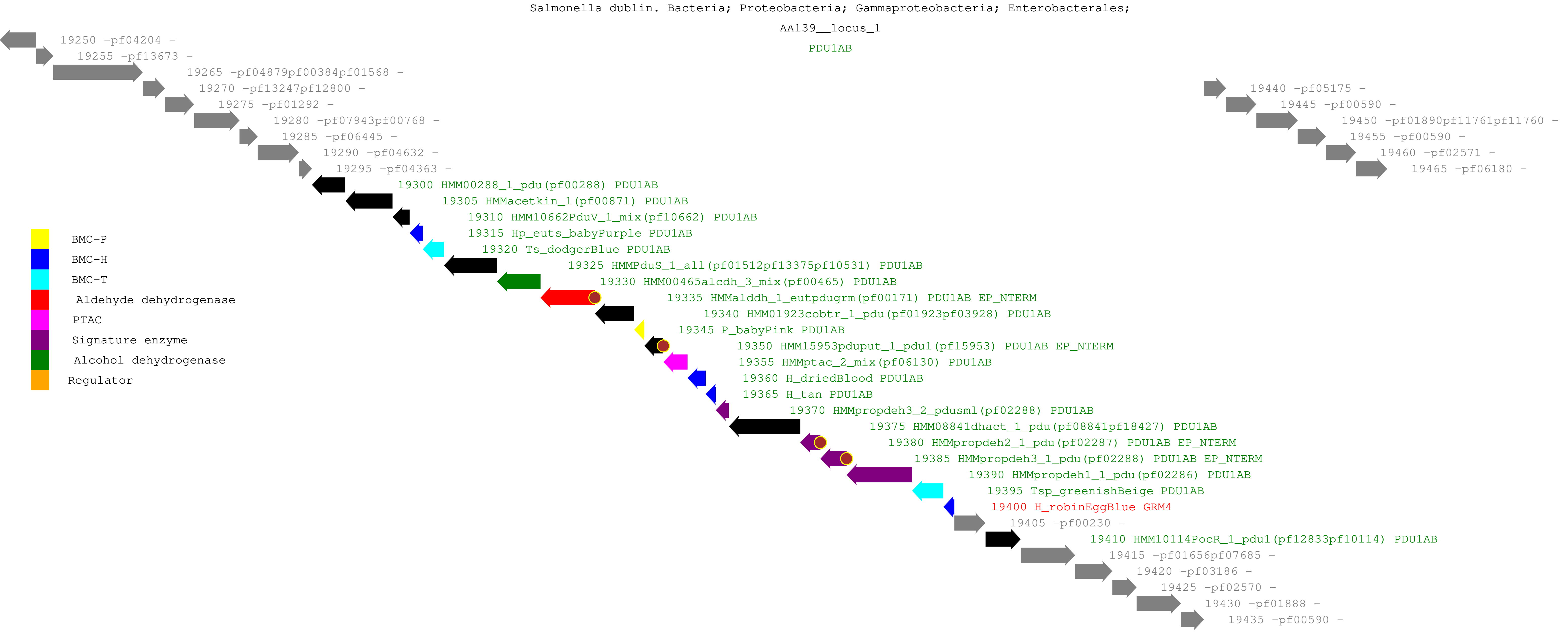

Salmonella dublin. Bacteria; Proteobacteria; Gammaproteobacteria; Enterobacterales;
AA139__locus_1
PDU1AB
19250 -pf04204 -
19255 -pf13673 -
19265 -pf04879pf00384pf01568 -
19270 -pf13247pf12800 -
19275 -pf01292 -
19280 -pf07943pf00768 -
19285 -pf06445 -
19290 -pf04632 -
19295 -pf04363 -
19300 HMM00288_1_pdu(pf00288) PDU1AB
19305 HMMacetkin_1(pf00871) PDU1AB
19310 HMM10662PduV_1_mix(pf10662) PDU1AB
19315 Hp_euts_babyPurple PDU1AB
19320 Ts_dodgerBlue PDU1AB
19325 HMMPduS_1_all(pf01512pf13375pf10531) PDU1AB
19330 HMM00465alcdh_3_mix(pf00465) PDU1AB
19335 HMMalddh_1_eutpdugrm(pf00171) PDU1AB EP_NTERM
19340 HMM01923cobtr_1_pdu(pf01923pf03928) PDU1AB
19345 P_babyPink PDU1AB
19350 HMM15953pduput_1_pdu1(pf15953) PDU1AB EP_NTERM
19355 HMMptac_2_mix(pf06130) PDU1AB
19360 H_driedBlood PDU1AB
19365 H_tan PDU1AB
19370 HMMpropdeh3_2_pdusml(pf02288) PDU1AB
19375 HMM08841dhact_1_pdu(pf08841pf18427) PDU1AB
19380 HMMpropdeh2_1_pdu(pf02287) PDU1AB EP_NTERM
19385 HMMpropdeh3_1_pdu(pf02288) PDU1AB EP_NTERM
19390 HMMpropdeh1_1_pdu(pf02286) PDU1AB
19395 Tsp_greenishBeige PDU1AB
19400 H_robinEggBlue GRM4
19405 -pf00230 -
19410 HMM10114PocR_1_pdu1(pf12833pf10114) PDU1AB
19415 -pf01656pf07685 -
19420 -pf03186 -
19425 -pf02570 -
19430 -pf01888 -
19435 -pf00590 -
19440 -pf05175 -
19445 -pf00590 -
19450 -pf01890pf11761pf11760 -
19455 -pf00590 -
19460 -pf02571 -
19465 -pf06180 -
BMC-P
BMC-H
BMC-T
Aldehyde dehydrogenase
PTAC
Signature enzyme
Alcohol dehydrogenase
Regulator

Salmonella anatum. Bacteria; Proteobacteria; Gammaproteobacteria; Enterobacterales;

AA325__locus_1

PDU1AB

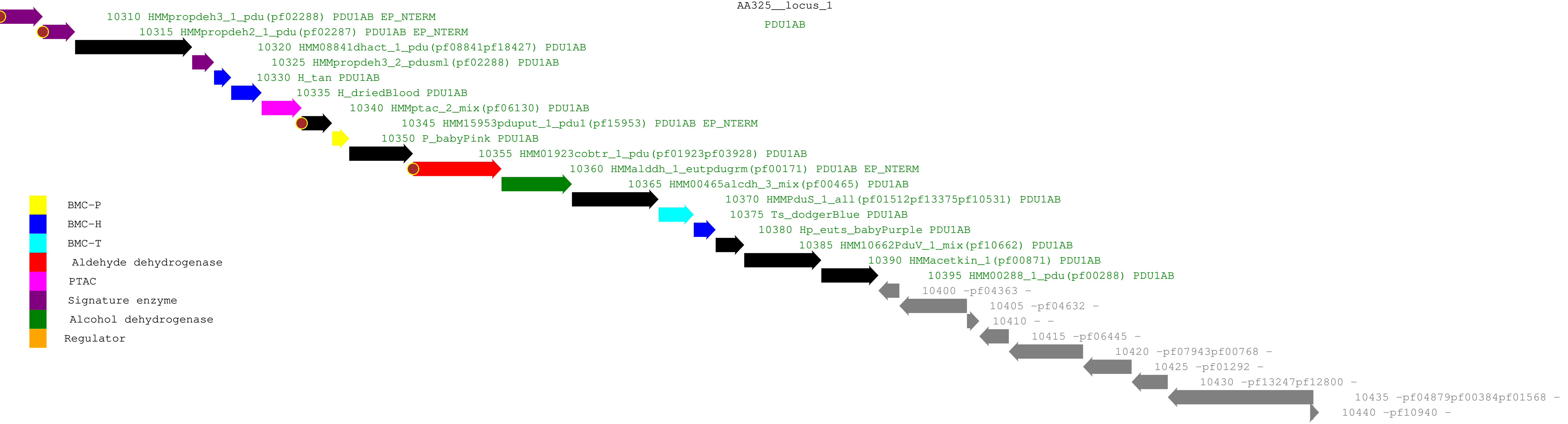

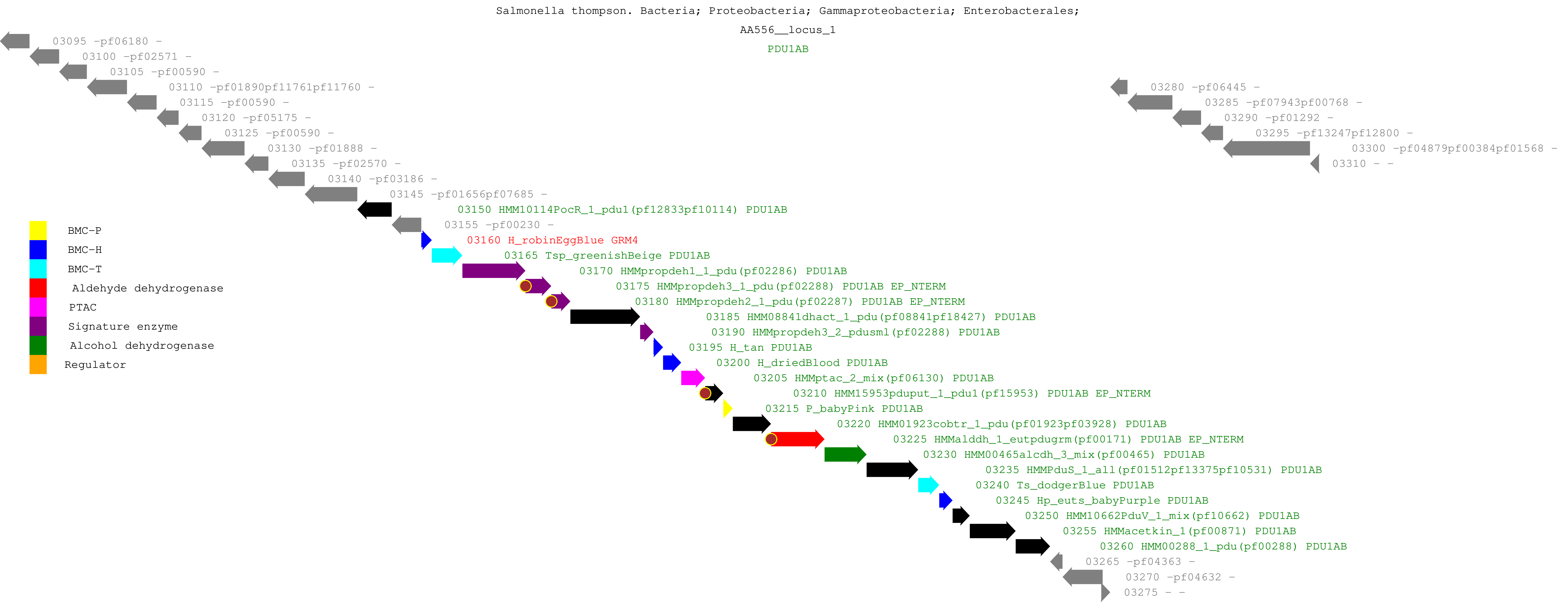
Salmonella thompson. Bacteria; Proteobacteria; Gammaproteobacteria; Enterobacterales;
AA556__locus_1
PDU1AB
03095 -pf06180 -
03100 -pf02571 -
03105 -pf00590 -
03110 -pf01890pf11761pf11760 -
03115 -pf00590 -
03120 -pf05175 -
03125 -pf00590 -
03130 -pf01888 -
03135 -pf02570 -
03140 -pf03186 -
03145 -pf01656pf07685 -
03150 HMM10114PocR_1_pdu1(pf12833pf10114) PDU1AB
03155 -pf00230 -
03160 H_robinEggBlue GRM4
03165 Tsp_greenishBeige PDU1AB
03170 HMMpropdeh1_1_pdu(pf02286) PDU1AB
03175 HMMpropdeh3_1_pdu(pf02288) PDU1AB EP_NTERM
03180 HMMpropdeh2_1_pdu(pf02287) PDU1AB EP_NTERM
03185 HMM08841dhact_1_pdu(pf08841pf18427) PDU1AB
03190 HMMpropdeh3_2_pdusml(pf02288) PDU1AB
03195 H_tan PDU1AB
03200 H_driedBlood PDU1AB
03205 HMMptac_2_mix(pf06130) PDU1AB
03210 HMM15953pduput_1_pdu1(pf15953) PDU1AB EP_NTERM
03215 P_babyPink PDU1AB
03220 HMM01923cobtr_1_pdu(pf01923pf03928) PDU1AB
03225 HMMaldh_1_eutpdugrm(pf00171) PDU1AB EP_NTERM
03230 HMM00465alcdh_3_mix(pf00465) PDU1AB
03235 HMMPduS_1_all(pf01512pf13375pf10531) PDU1AB
03240 Ts_dodgerBlue PDU1AB
03245 Hp_euts_babyPurple PDU1AB
03250 HMM10662PduV_1_mix(pf10662) PDU1AB
03255 HMMacetkin_1(pf00871) PDU1AB
03260 HMM00288_1_pdu(pf00288) PDU1AB
03265 -pf04363 -
03270 -pf04632 -
03275 - -
03280 -pf06445 -
03285 -pf07943pf00768 -
03290 -pf01292 -
03295 -pf13247pf12800 -
03300 -pf04879pf00384pf01568 -
03310 - -
BMC-P
BMC-H
BMC-T
Aldehyde dehydrogenase
PTAC
Signature enzyme
Alcohol dehydrogenase
Regulator

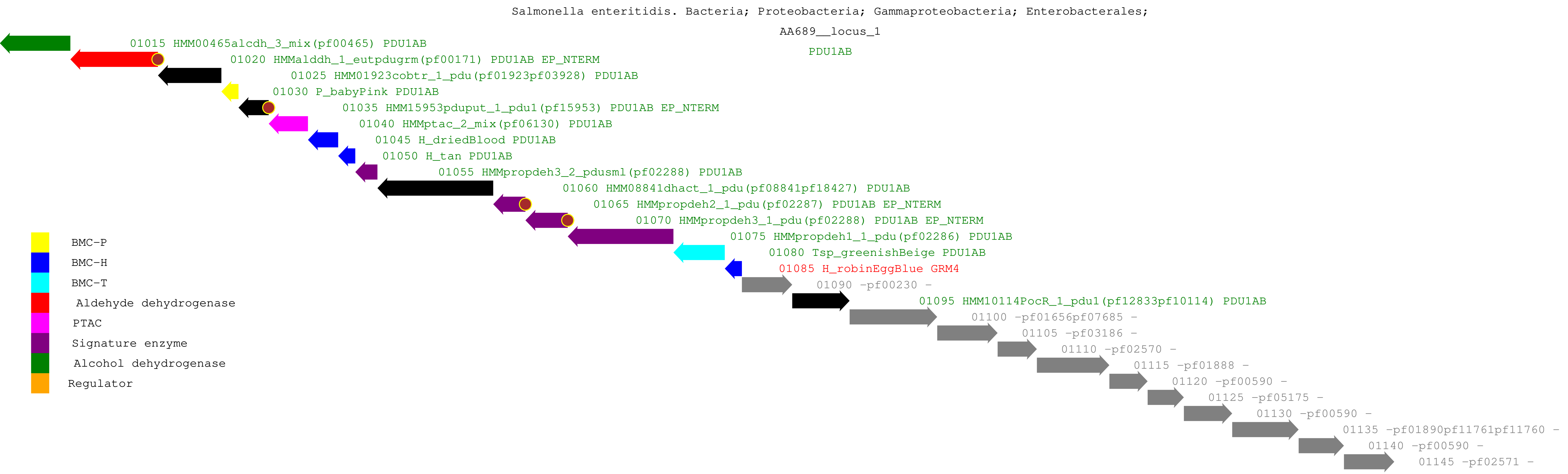

Salmonella enteritidis. Bacteria; Proteobacteria; Gammaproteobacteria; Enterobacterales;
AA689__locus_1
PDU1AB
01015 HMM00465alcdh_3_mix(pf00465) PDU1AB
01020 HMMalddh_1_eutpdugrm(pf00171) PDU1AB EP_NTERM
01025 HMM01923cobtr_1_pdu(pf01923pf03928) PDU1AB
01030 P_babyPink PDU1AB
01035 HMM15953pduput_1_pdu1(pf15953) PDU1AB EP_NTERM
01040 HMMptac_2_mix(pf06130) PDU1AB
01045 H_driedBlood PDU1AB
01050 H_tan PDU1AB
01055 HMMpropdeh3_2_pdusml(pf02288) PDU1AB
01060 HMM08841dhact_1_pdu(pf08841pf18427) PDU1AB
01065 HMMpropdeh2_1_pdu(pf02287) PDU1AB EP_NTERM
01070 HMMpropdeh3_1_pdu(pf02288) PDU1AB EP_NTERM
01075 HMMpropdeh1_1_pdu(pf02286) PDU1AB
01080 Tsp_greenishBeige PDU1AB
01085 H_robinEggBlue GRM4
01090 -pf00230 -
01095 HMM10114PocR_1_pdu1(pf12833pf10114) PDU1AB
01100 -pf01656pf07685 -
01105 -pf03186 -
01110 -pf02570 -
01115 -pf01888 -
01120 -pf00590 -
01125 -pf05175 -
01130 -pf00590 -
01135 -pf01890pf11761pf11760 -
01140 -pf00590 -
01145 -pf02571 -
BMC-P
BMC-H
BMC-T
Aldehyde dehydrogenase
PTAC
Signature enzyme
Alcohol dehydrogenase
Regulator

Salmonella enterica I. Bacteria; Proteobacteria; Gammaproteobacteria; Enterobacterales;

AAB19__locus_1

PDU1AB

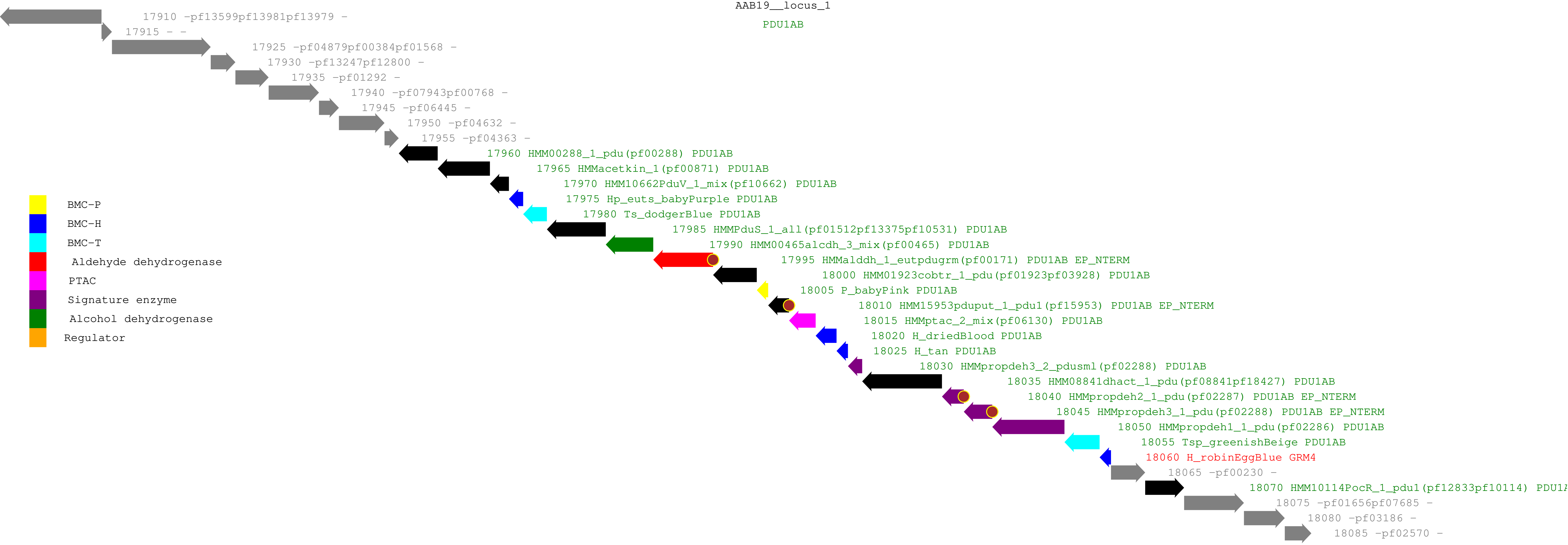

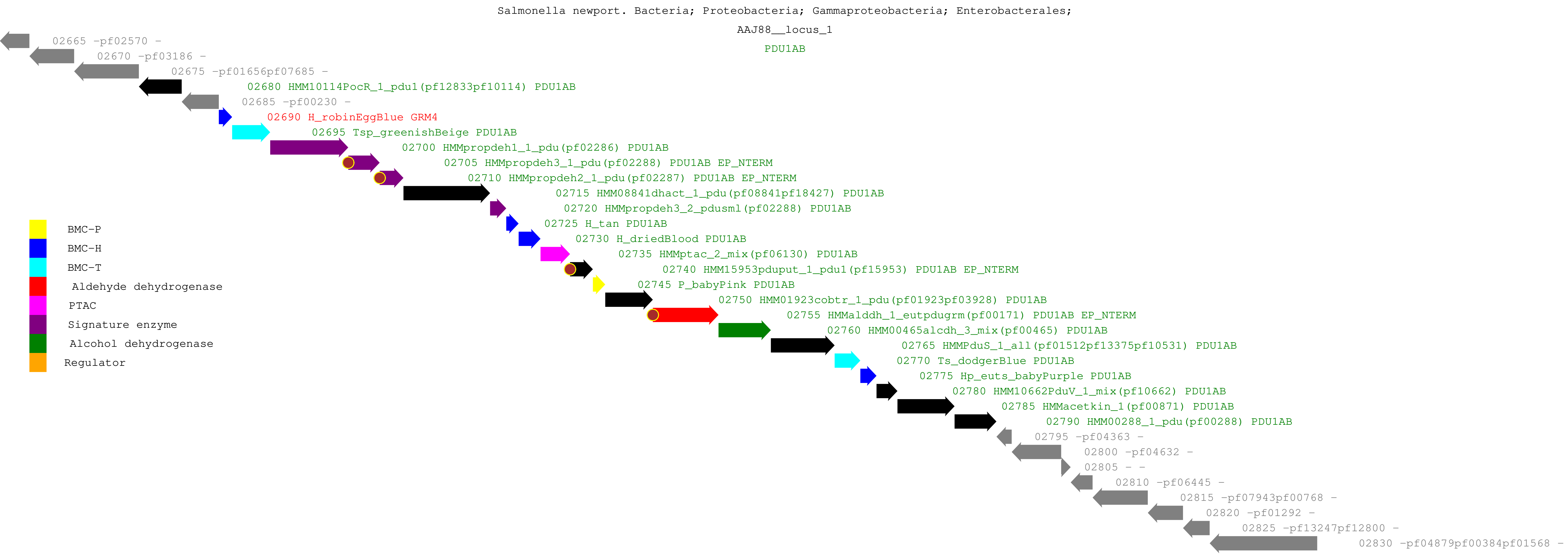

Salmonella newport. Bacteria; Proteobacteria; Gammaproteobacteria; Enterobacterales;
AAJ88__locus_1
PDU1AB
02665 -pf02570 -
02670 -pf03186 -
02675 -pf01656pf07685 -
02680 HMM10114PocR_1_pdu1(pf12833pf10114) PDU1AB
02685 -pf00230 -
02690 H_robinEggBlue GRM4
02695 Tsp_greenishBeige PDU1AB
02700 HMMpropdeh1_1_pdu(pf02286) PDU1AB
02705 HMMpropdeh3_1_pdu(pf02288) PDU1AB EP_NTERM
02710 HMMpropdeh2_1_pdu(pf02287) PDU1AB EP_NTERM
02715 HMM08841dhact_1_pdu(pf08841pf18427) PDU1AB
02720 HMMpropdeh3_2_pdusml(pf02288) PDU1AB
02725 H_tan PDU1AB
02730 H_driedBlood PDU1AB
02735 HMMptac_2_mix(pf06130) PDU1AB
02740 HMM15953pduput_1_pdu1(pf15953) PDU1AB EP_NTERM
02745 P_babyPink PDU1AB
02750 HMM01923cobtr_1_pdu(pf01923pf03928) PDU1AB
02755 HMMalddh_1_eutpdugrm(pf00171) PDU1AB EP_NTERM
02760 HMM00465alcdh_3_mix(pf00465) PDU1AB
02765 HMMPduS_1_all(pf01512pf13375pf10531) PDU1AB
02770 Ts_dodgerBlue PDU1AB
02775 Hp_euts_babyPurple PDU1AB
02780 HMM10662PduV_1_mix(pf10662) PDU1AB
02785 HMMacetkin_1(pf00871) PDU1AB
02790 HMM00288_1_pdu(pf00288) PDU1AB
02795 -pf04363 -
02800 -pf04632 -
02805 - -
02810 -pf06445 -
02815 -pf07943pf00768 -
02820 -pf01292 -
02825 -pf13247pf12800 -
02830 -pf04879pf00384pf01568 -
BMC-P
BMC-H
BMC-T
Aldehyde dehydrogenase
PTAC
Signature enzyme
Alcohol dehydrogenase
Regulator

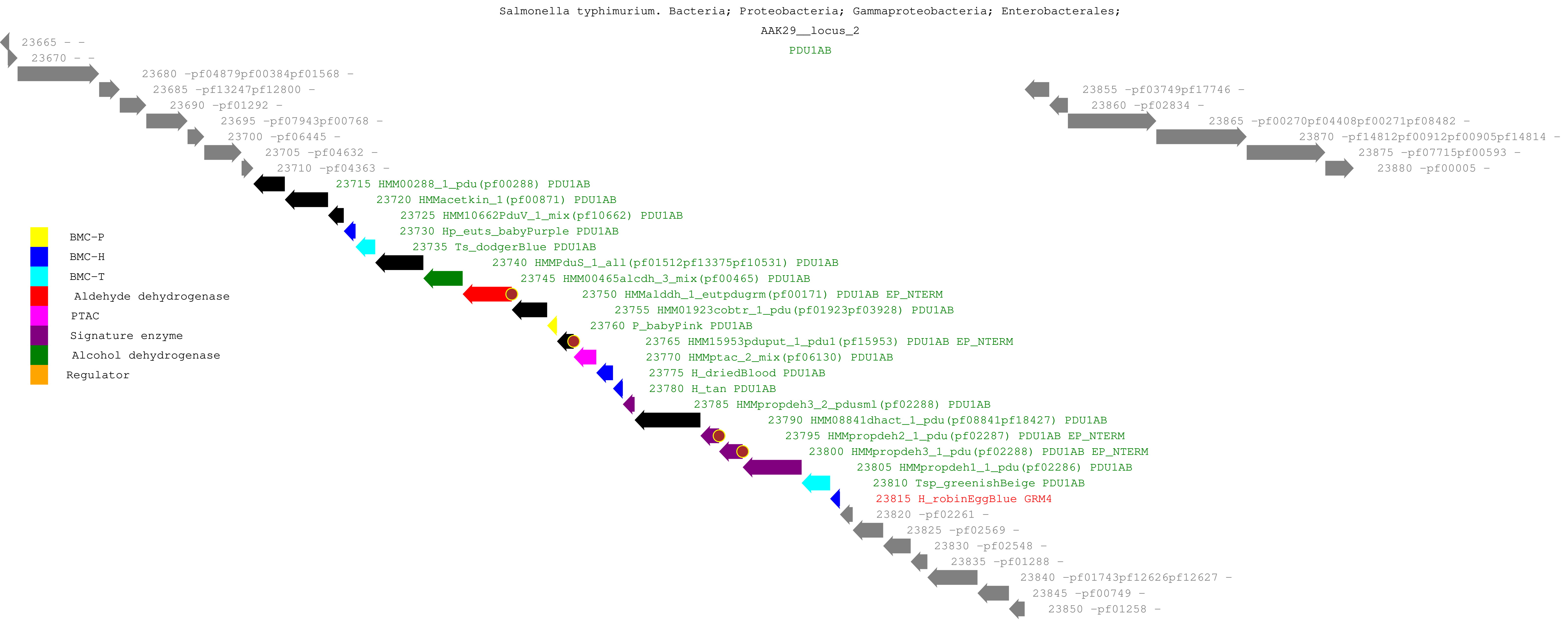
Salmonella typhimurium. Bacteria; Proteobacteria; Gammaproteobacteria; Enterobacterales;
AAK29__locus_2
PDU1AB
23665 - -
23670 - -
23680 -pf04879pf00384pf01568 -
23685 -pf13247pf12800 -
23690 -pf01292 -
23695 -pf07943pf00768 -
23700 -pf06445 -
23705 -pf04632 -
23710 -pf04363 -
23715 HMM00288_1_pdu(pf00288) PDU1AB
23720 HMMacetkin_1(pf00871) PDU1AB
23725 HMM10662PduV_1_mix(pf10662) PDU1AB
23730 Hp_euts_babyPurple PDU1AB
23735 Ts_dodgerBlue PDU1AB
23740 HMMPduS_1_all(pf01512pf13375pf10531) PDU1AB
23745 HMM00465alcdh_3_mix(pf00465) PDU1AB
23750 HMMalddh_1_eutpdugrm(pf00171) PDU1AB EP_NTERM
23755 HMM01923cobtr_1_pdu(pf01923pf03928) PDU1AB
23760 P_babyPink PDU1AB
23765 HMM15953pduput_1_pdu1(pf15953) PDU1AB EP_NTERM
23770 HMMptac_2_mix(pf06130) PDU1AB
23775 H_driedBlood PDU1AB
23780 H_tan PDU1AB
23785 HMMpropdeh3_2_pdusml(pf02288) PDU1AB
23790 HMM08841dhact_1_pdu(pf08841pf18427) PDU1AB
23795 HMMpropdeh2_1_pdu(pf02287) PDU1AB EP_NTERM
23800 HMMpropdeh3_1_pdu(pf02288) PDU1AB EP_NTERM
23805 HMMpropdeh1_1_pdu(pf02286) PDU1AB
23810 Tsp_greenishBeige PDU1AB
23815 H_robinEggBlue GRM4
23820 -pf02261 -
23825 -pf02569 -
23830 -pf02548 -
23835 -pf01288 -
23840 -pf01743pf12626pf12627 -
23845 -pf00749 -
23850 -pf01258 -
23855 -pf03749pf17746 -
23860 -pf02834 -
23865 -pf00270pf04408pf00271pf08482 -
23870 -pf14812pf00912pf00905pf14814 -
23875 -pf07715pf00593 -
23880 -pf00005 -
BMC-P
BMC-H
BMC-T
Aldehyde dehydrogenase
PTAC
Signature enzyme
Alcohol dehydrogenase
Regulator

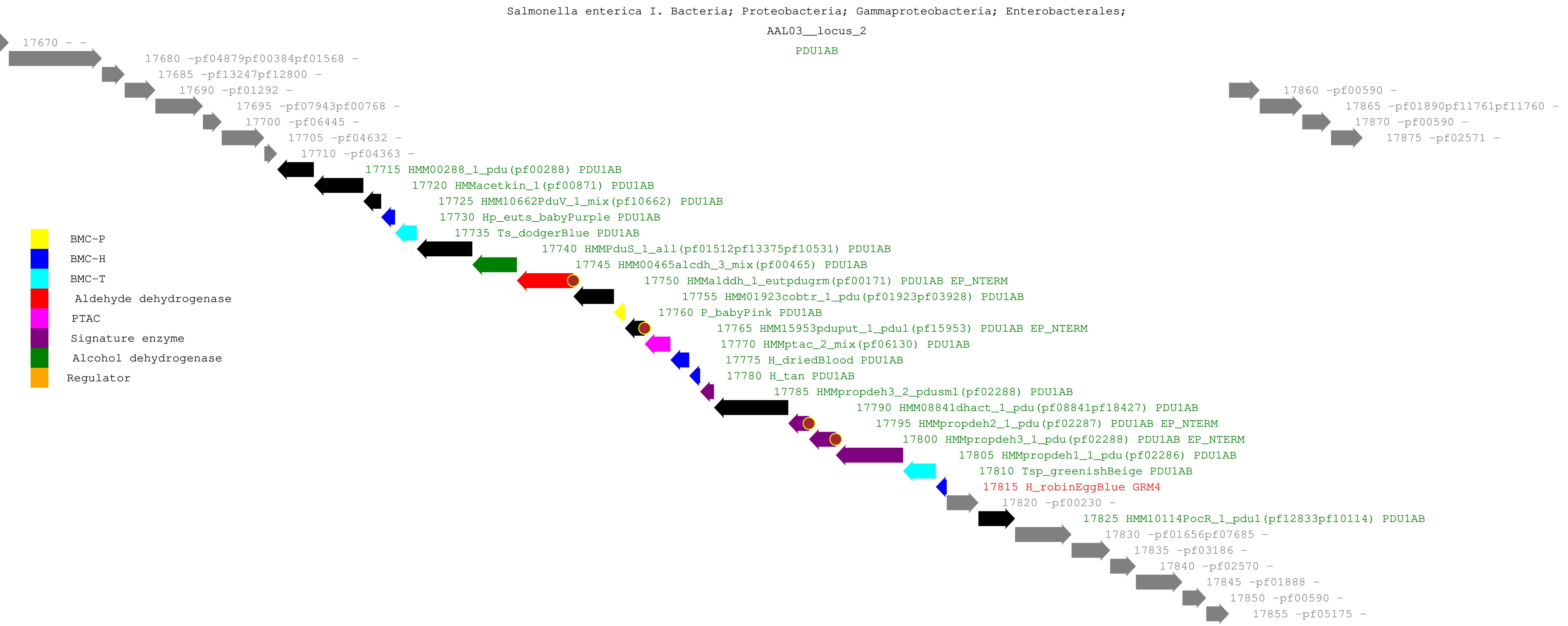

Salmonella enterica I. Bacteria; Proteobacteria; Gammaproteobacteria; Enterobacterales;
AAL03__locus_2
PDU1AB
17670 - -
17680 -pf04879pf00384pf01568 -
17685 -pf13247pf12800 -
17690 -pf01292 -
17695 -pf07943pf00768 -
17700 -pf06445 -
17705 -pf04632 -
17710 -pf04363 -
17715 HMM00288_1_pdu(pf00288) PDU1AB
17720 HMMacetkin_1(pf00871) PDU1AB
17725 HMM10662PduV_1_mix(pf10662) PDU1AB
17730 Hp_euts_babyPurple PDU1AB
17735 Ts_dodgerBlue PDU1AB
17740 HMMPduS_1_all(pf01512pf13375pf10531) PDU1AB
17745 HMM00465alcdh_3_mix(pf00465) PDU1AB
17750 HMMalddh_1_eutpdugrm(pf00171) PDU1AB EP_NTERM
17755 HMM01923cobtr_1_pdu(pf01923pf03928) PDU1AB
17760 P_babyPink PDU1AB
17765 HMM15953pduput_1_pdu1(pf15953) PDU1AB EP_NTERM
17770 HMMptac_2_mix(pf06130) PDU1AB
17775 H_driedBlood PDU1AB
17780 H_tan PDU1AB
17785 HMMpropdeh3_2_pdusml(pf02288) PDU1AB
17790 HMM08841dhact_1_pdu(pf08841pf18427) PDU1AB
17795 HMMpropdeh2_1_pdu(pf02287) PDU1AB EP_NTERM
17800 HMMpropdeh3_1_pdu(pf02288) PDU1AB EP_NTERM
17805 HMMpropdeh1_1_pdu(pf02286) PDU1AB
17810 Tsp_greenishBeige PDU1AB
17815 H_robinEggBlue GRM4
17820 -pf00230 -
17825 HMM10114PocR_1_pdu1(pf12833pf10114) PDU1AB
17830 -pf01656pf07685 -
17835 -pf03186 -
17840 -pf02570 -
17845 -pf01888 -
17850 -pf00590 -
17855 -pf05175 -
17860 -pf00590 -
17865 -pf01890pf11761pf11760 -
17870 -pf00590 -
17875 -pf02571 -
BMC-P
BMC-H
BMC-T
Aldehyde dehydrogenase
PTAC
Signature enzyme
Alcohol dehydrogenase
Regulator

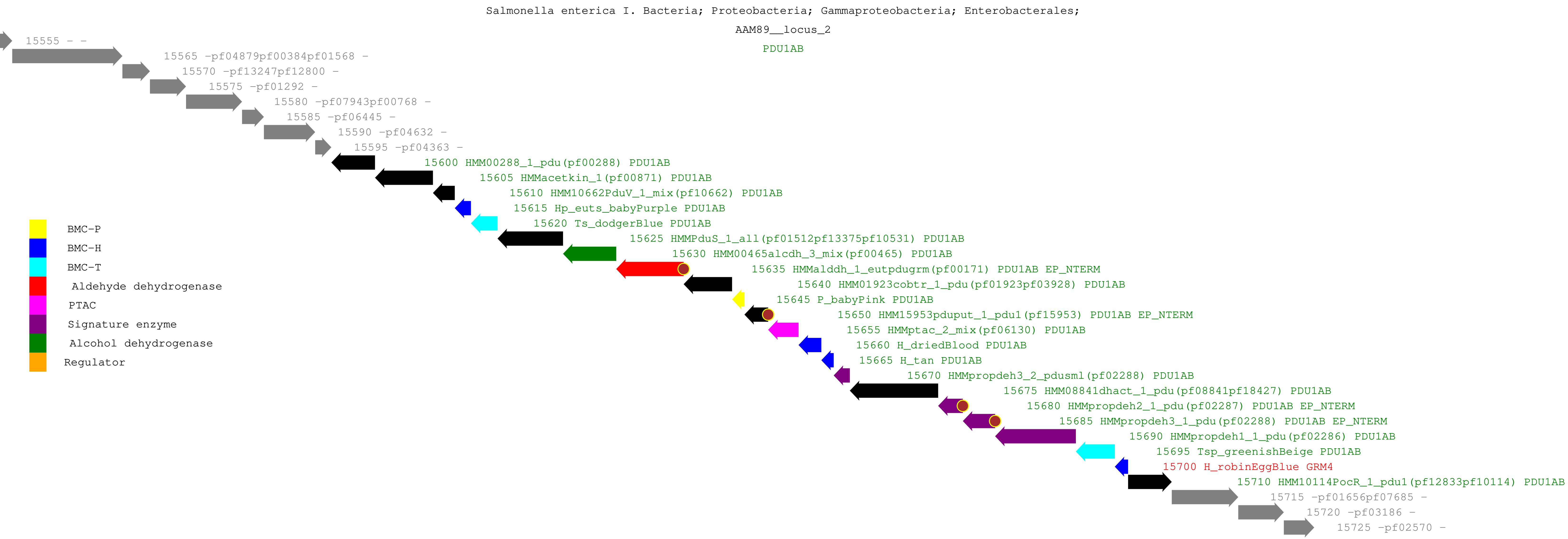
Salmonella enterica I. Bacteria; Proteobacteria; Gammaproteobacteria; Enterobacterales;
AAM89__locus_2
PDU1AB
15555 - -
15565 -pf04879pf00384pf01568 -
15570 -pf13247pf12800 -
15575 -pf01292 -
15580 -pf07943pf00768 -
15585 -pf06445 -
15590 -pf04632 -
15595 -pf04363 -
15600 HMM00288_1_pdu(pf00288) PDU1AB
15605 HMMacetkin_1(pf00871) PDU1AB
15610 HMM10662PduV_1_mix(pf10662) PDU1AB
15615 Hp_euts_babyPurple PDU1AB
15620 Ts_dodgerBlue PDU1AB
15625 HMMPduS_1_all(pf01512pf13375pf10531) PDU1AB
15630 HMM00465alcdh_3_mix(pf00465) PDU1AB
15635 HMMalddh_1_eutpdugrm(pf00171) PDU1AB EP_NTERM
15640 HMM01923cobtr_1_pdu(pf01923pf03928) PDU1AB
15645 P_babyPink PDU1AB
15650 HMM15953pduput_1_pdu1(pf15953) PDU1AB EP_NTERM
15655 HMMptac_2_mix(pf06130) PDU1AB
15660 H_driedBlood PDU1AB
15665 H_tan PDU1AB
15670 HMMpropdeh3_2_pdusml(pf02288) PDU1AB
15675 HMM08841dhact_1_pdu(pf08841pf18427) PDU1AB
15680 HMMpropdeh2_1_pdu(pf02287) PDU1AB EP_NTERM
15685 HMMpropdeh3_1_pdu(pf02288) PDU1AB EP_NTERM
15690 HMMpropdeh1_1_pdu(pf02286) PDU1AB
15695 Tsp_greenishBeige PDU1AB
15700 H_robinEggBlue GRM4
15710 HMM10114PocR_1_pdu1(pf12833pf10114) PDU1AB
15715 -pf01656pf07685 -
15720 -pf03186 -
15725 -pf02570 -
BMC-P
BMC-H
BMC-T
Aldehyde dehydrogenase
PTAC
Signature enzyme
Alcohol dehydrogenase
Regulator

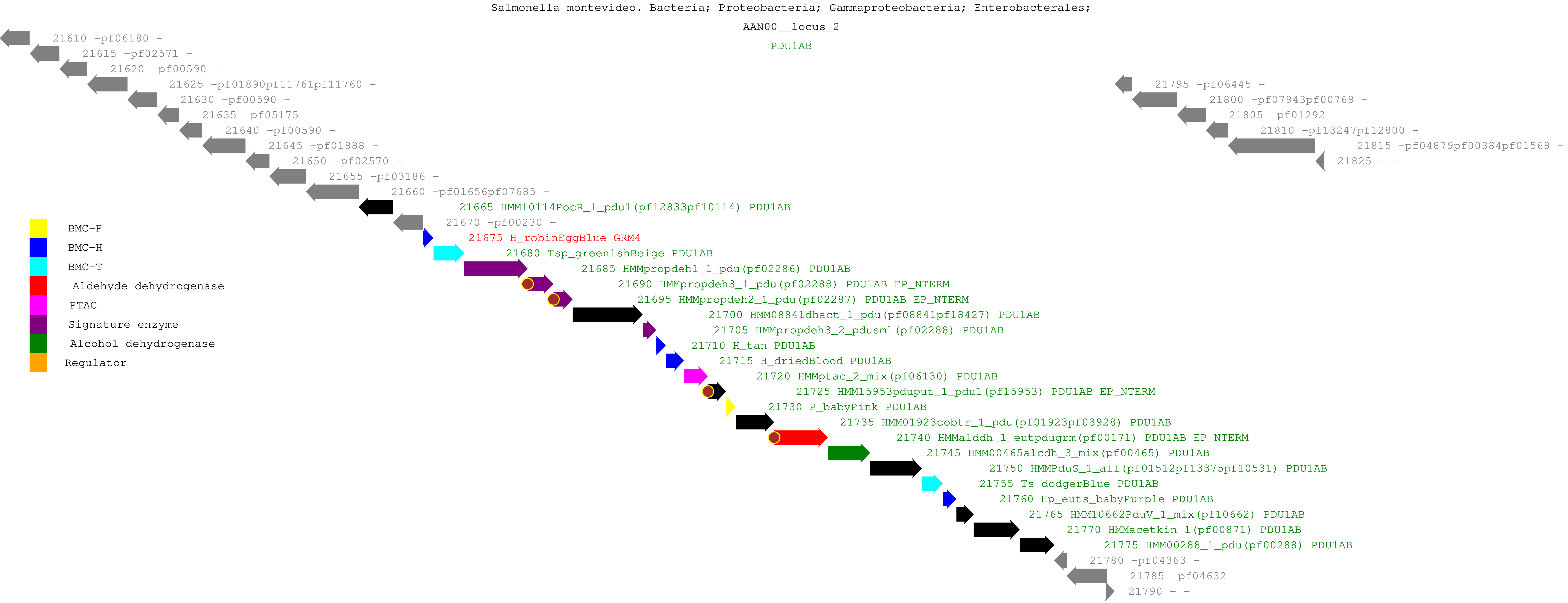

Salmonella montevideo. Bacteria; Proteobacteria; Gammaproteobacteria; Enterobacterales;
AAN00__locus_2
PDU1AB
21610 -pf06180 -
21615 -pf02571 -
21620 -pf00590 -
21625 -pf01890pf11761pf11760 -
21630 -pf00590 -
21635 -pf05175 -
21640 -pf00590 -
21645 -pf01888 -
21650 -pf02570 -
21655 -pf03186 -
21660 -pf01656pf07685 -
21665 HMM10114PocR_1_pdu1(pf12833pf10114) PDU1AB
21670 -pf00230 -
21675 H_robinEggBlue GRM4
21680 Tsp_greenishBeige PDU1AB
21685 HMMpropdeh1_1_pdu(pf02286) PDU1AB
21690 HMMpropdeh3_1_pdu(pf02288) PDU1AB EP_NTERM
21695 HMMpropdeh2_1_pdu(pf02287) PDU1AB EP_NTERM
21700 HMM08841dhact_1_pdu(pf08841pf18427) PDU1AB
21705 HMMpropdeh3_2_pdusml(pf02288) PDU1AB
21710 H_tan PDU1AB
21715 H_driedBlood PDU1AB
21720 HMMptac_2_mix(pf06130) PDU1AB
21725 HMM15953pduput_1_pdu1(pf15953) PDU1AB EP_NTERM
21730 P_babyPink PDU1AB
21735 HMM01923cobtr_1_pdu(pf01923pf03928) PDU1AB
21740 HMMalddh_1_eutpdugrm(pf00171) PDU1AB EP_NTERM
21745 HMM00465alcdh_3_mix(pf00465) PDU1AB
21750 HMMPduS_1_all(pf01512pf13375pf10531) PDU1AB
21755 Ts_dodgerBlue PDU1AB
21760 Hp_euts_babyPurple PDU1AB
21765 HMM10662PduV_1_mix(pf10662) PDU1AB
21770 HMMacetkin_1(pf00871) PDU1AB
21775 HMM00288_1_pdu(pf00288) PDU1AB
21780 -pf04363 -
21785 -pf04632 -
21790 - -
21795 -pf06445 -
21800 -pf07943pf00768 -
21805 -pf01292 -
21810 -pf13247pf12800 -
21815 -pf04879pf00384pf01568 -
21825 - -
BMC-P
BMC-H
BMC-T
Aldehyde dehydrogenase
PTAC
Signature enzyme
Alcohol dehydrogenase
Regulator

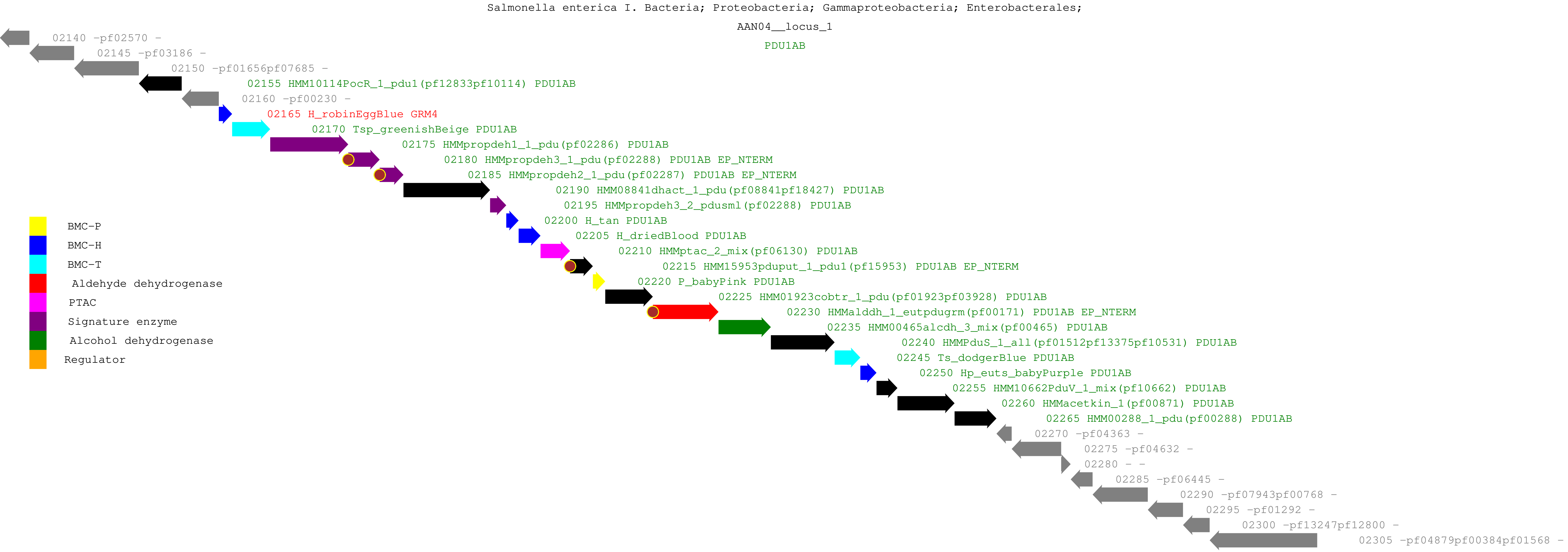

Salmonella enterica I. Bacteria; Proteobacteria; Gammaproteobacteria; Enterobacterales;
AAN04__locus_1
PDU1AB
02140 -pf02570 -
02145 -pf03186 -
02150 -pf01656pf07685 -
02155 HMM10114PocR_1_pdu1(pf12833pf10114) PDU1AB
02160 -pf00230 -
02165 H_robinEggBlue GRM4
02170 Tsp_greenishBeige PDU1AB
02175 HMMpropdeh1_1_pdu(pf02286) PDU1AB
02180 HMMpropdeh3_1_pdu(pf02288) PDU1AB EP_NTERM
02185 HMMpropdeh2_1_pdu(pf02287) PDU1AB EP_NTERM
02190 HMM08841dhact_1_pdu(pf08841pf18427) PDU1AB
02195 HMMpropdeh3_2_pdusml(pf02288) PDU1AB
02200 H_tan PDU1AB
02205 H_driedBlood PDU1AB
02210 HMMptac_2_mix(pf06130) PDU1AB
02215 HMM15953pduput_1_pdu1(pf15953) PDU1AB EP_NTERM
02220 P_babyPink PDU1AB
02225 HMM01923cobtr_1_pdu(pf01923pf03928) PDU1AB
02230 HMMalddh_1_eutpdugrm(pf00171) PDU1AB EP_NTERM
02235 HMM00465alcdh_3_mix(pf00465) PDU1AB
02240 HMMPduS_1_all(pf01512pf13375pf10531) PDU1AB
02245 Ts_dodgerBlue PDU1AB
02250 Hp_euts_babyPurple PDU1AB
02255 HMM10662PduV_1_mix(pf10662) PDU1AB
02260 HMMacetkin_1(pf00871) PDU1AB
02265 HMM00288_1_pdu(pf00288) PDU1AB
02270 -pf04363 -
02275 -pf04632 -
02280 - -
02285 -pf06445 -
02290 -pf07943pf00768 -
02295 -pf01292 -
02300 -pf13247pf12800 -
02305 -pf04879pf00384pf01568 -
BMC-P
BMC-H
BMC-T
Aldehyde dehydrogenase
PTAC
Signature enzyme
Alcohol dehydrogenase
Regulator

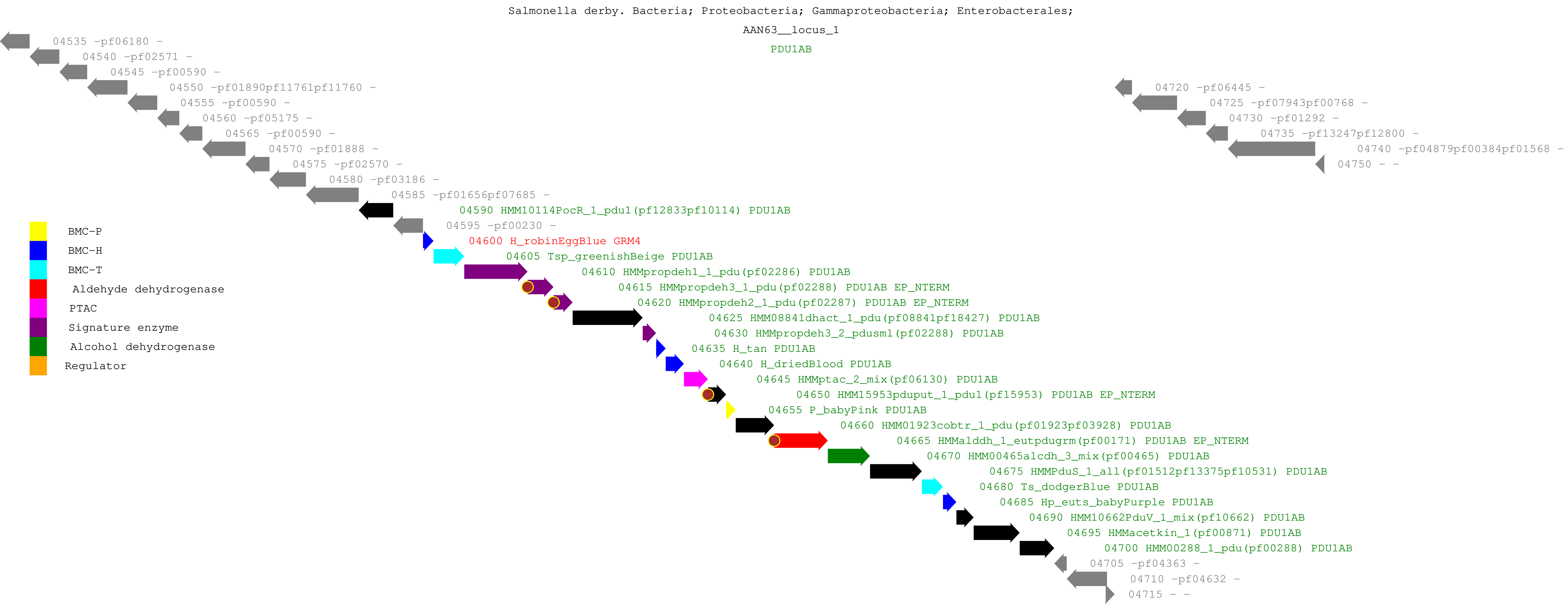

Salmonella derby. Bacteria; Proteobacteria; Gammaproteobacteria; Enterobacterales;
AAN63__locus_1
PDU1AB
04535 -pf06180 -
04540 -pf02571 -
04545 -pf00590 -
04550 -pf01890pf11761pf11760 -
04555 -pf00590 -
04560 -pf05175 -
04565 -pf00590 -
04570 -pf01888 -
04575 -pf02570 -
04580 -pf03186 -
04585 -pf01656pf07685 -
04590 HMM10114PocR_1_pdu1(pf12833pf10114) PDU1AB
04595 -pf00230 -
04600 H_robinEggBlue GRM4
04605 Tsp_greenishBeige PDU1AB
04610 HMMpropdeh1_1_pdu(pf02286) PDU1AB
04615 HMMpropdeh3_1_pdu(pf02288) PDU1AB EP_NTERM
04620 HMMpropdeh2_1_pdu(pf02287) PDU1AB EP_NTERM
04625 HMM08841dhact_1_pdu(pf08841pf18427) PDU1AB
04630 HMMpropdeh3_2_pdusml(pf02288) PDU1AB
04635 H_tan PDU1AB
04640 H_driedBlood PDU1AB
04645 HMMptac_2_mix(pf06130) PDU1AB
04650 HMM15953pduput_1_pdu1(pf15953) PDU1AB EP_NTERM
04655 P_babyPink PDU1AB
04660 HMM01923cobtr_1_pdu(pf01923pf03928) PDU1AB
04665 HMMalddh_1_eutpdugrm(pf00171) PDU1AB EP_NTERM
04670 HMM00465alcdh_3_mix(pf00465) PDU1AB
04675 HMMPduS_1_all(pf01512pf13375pf10531) PDU1AB
04680 Ts_dodgerBlue PDU1AB
04685 Hp_euts_babyPurple PDU1AB
04690 HMM10662PduV_1_mix(pf10662) PDU1AB
04695 HMMacetkin_1(pf00871) PDU1AB
04700 HMM00288_1_pdu(pf00288) PDU1AB
04705 -pf04363 -
04710 -pf04632 -
04715 - -
04720 -pf06445 -
04725 -pf07943pf00768 -
04730 -pf01292 -
04735 -pf13247pf12800 -
04740 -pf04879pf00384pf01568 -
04750 - -
BMC-P
BMC-H
BMC-T
Aldehyde dehydrogenase
PTAC
Signature enzyme
Alcohol dehydrogenase
Regulator

Salmonella enterica I. Bacteria; Proteobacteria; Gammaproteobacteria; Enterobacterales;

AAV76__locus_1

PDU1AB

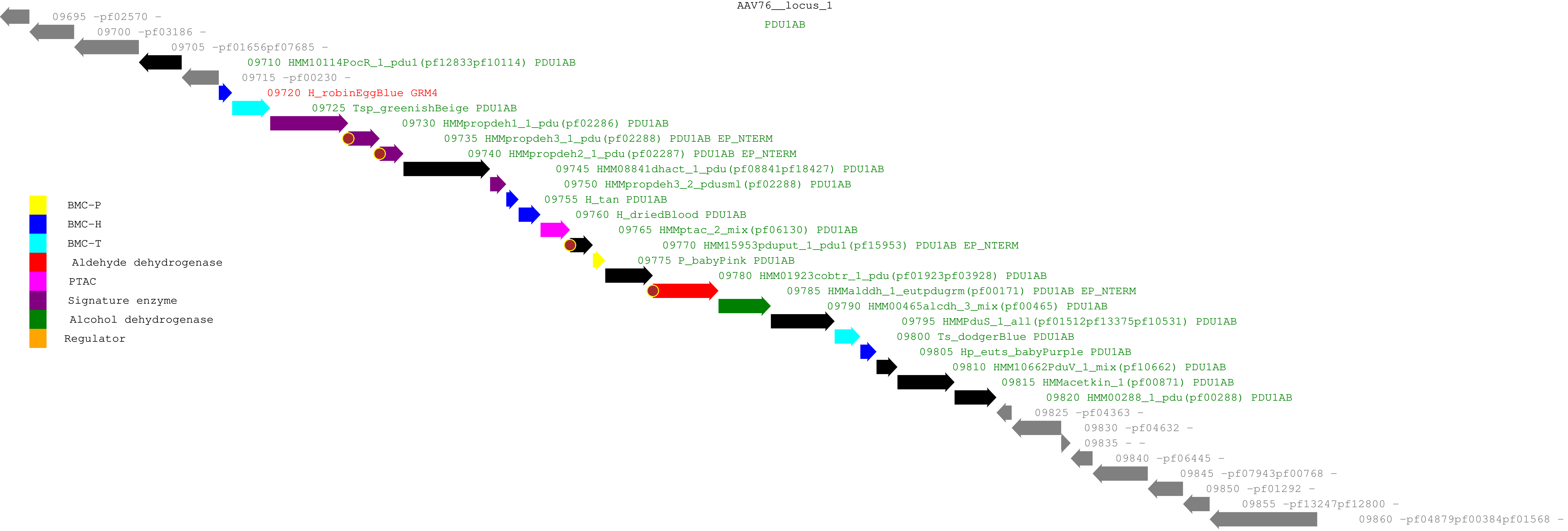

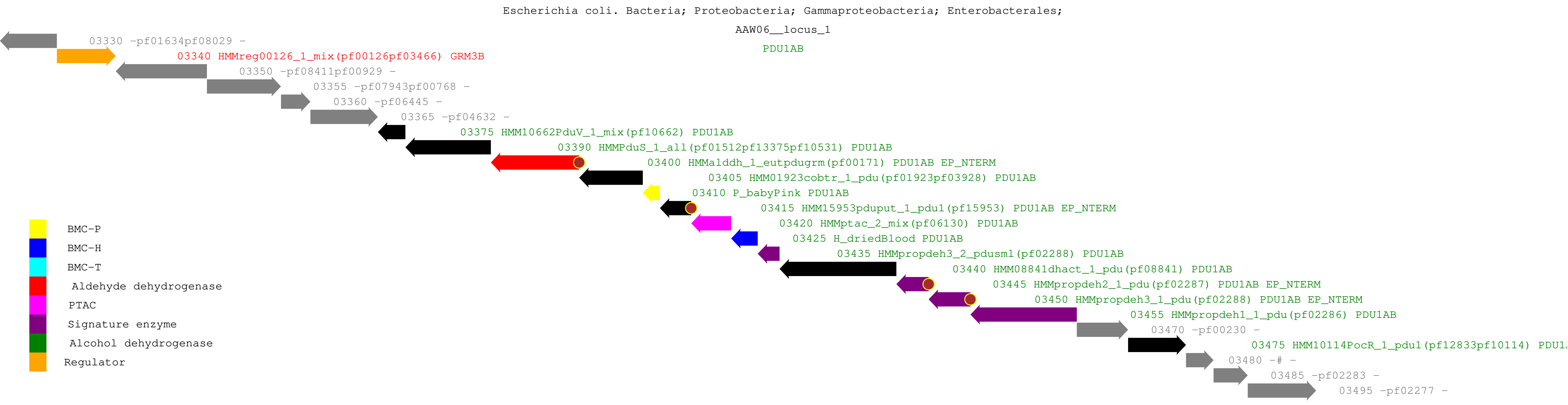

Escherichia coli. Bacteria; Proteobacteria; Gammaproteobacteria; Enterobacterales;
AAW06__locus_1
PDU1AB
03330 -pf01634pf08029 -
03340 HMMreg00126_1_mix(pf00126pf03466) GRM3B
03350 -pf08411pf00929 -
03355 -pf07943pf00768 -
03360 -pf06445 -
03365 -pf04632 -
03375 HMM10662PduV_1_mix(pf10662) PDU1AB
03390 HMMPduS_1_all(pf01512pf13375pf10531) PDU1AB
03400 HMMalddh_1_eutpdugrm(pf00171) PDU1AB EP_NTERM
03405 HMM01923cobtr_1_pdu(pf01923pf03928) PDU1AB
03410 P_babyPink PDU1AB
03415 HMM15953pduput_1_pdu1(pf15953) PDU1AB EP_NTERM
03420 HMMptac_2_mix(pf06130) PDU1AB
03425 H_driedBlood PDU1AB
03435 HMMpropdeh3_2_pdusml(pf02288) PDU1AB
03440 HMM08841dhact_1_pdu(pf08841) PDU1AB
03445 HMMpropdeh2_1_pdu(pf02287) PDU1AB EP_NTERM
03450 HMMpropdeh3_1_pdu(pf02288) PDU1AB EP_NTERM
03455 HMMpropdeh1_1_pdu(pf02286) PDU1AB
03470 -pf00230 -
03475 HMM10114PocR_1_pdu1(pf12833pf10114) PDU1
03480 -# -
03485 -pf02283 -
03495 -pf02277 -
BMC-P
BMC-H
BMC-T
Aldehyde dehydrogenase
PTAC
Signature enzyme
Alcohol dehydrogenase
Regulator

Vibrio parahaemolyticus. Bacteria; Proteobacteria; Gammaproteobacteria; Vibrionales;

AAY51__locus_1

PDU1AB

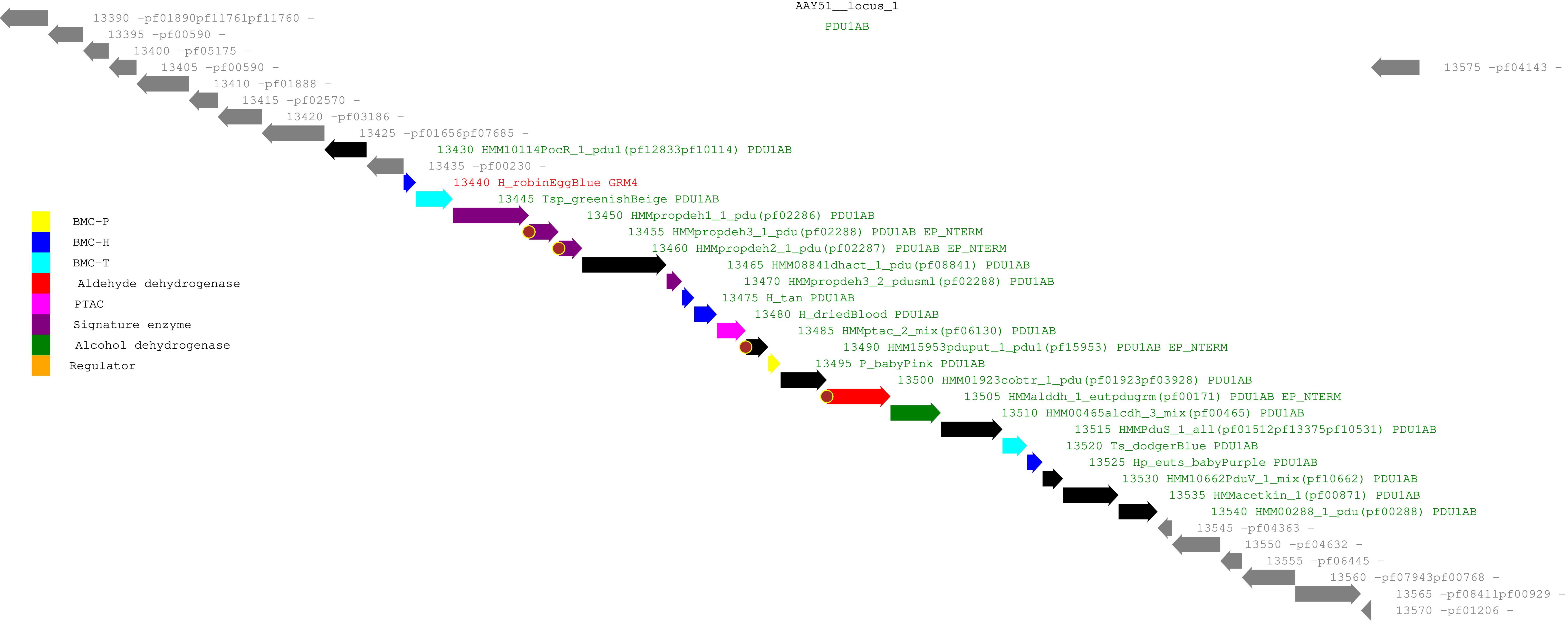

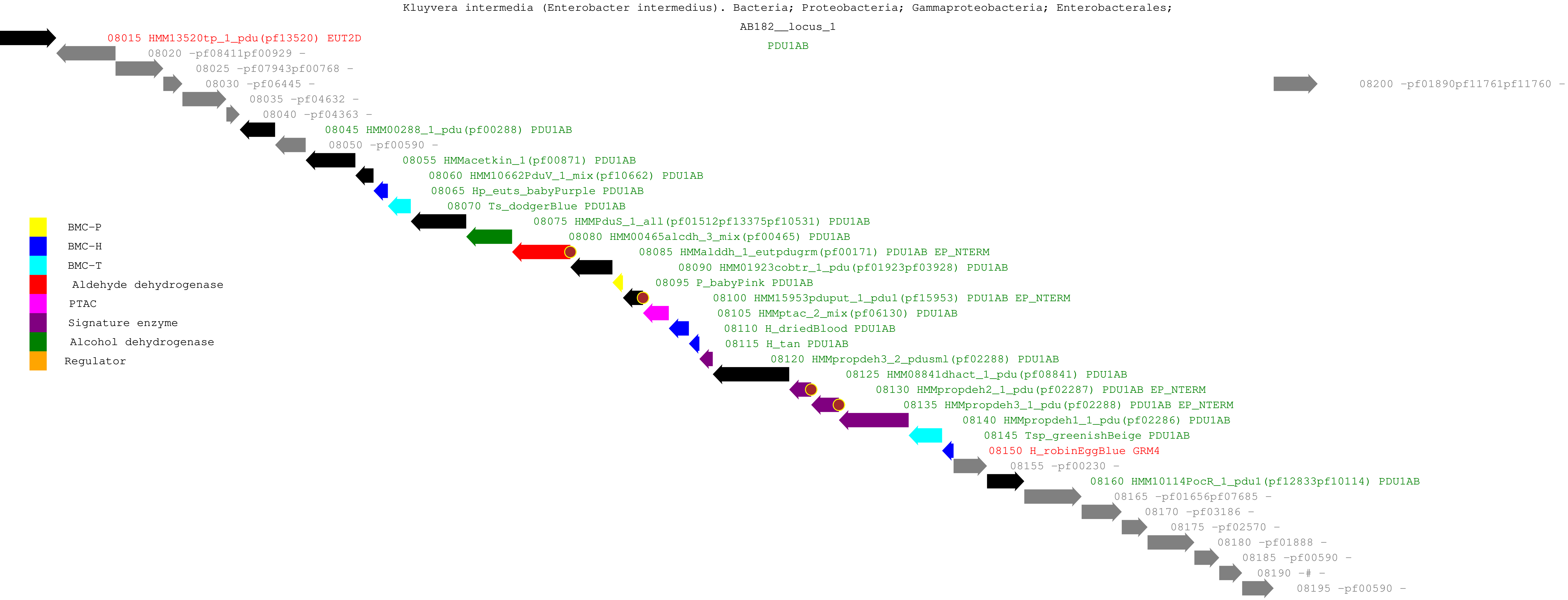
Kluyvera intermedia (Enterobacter intermedius). Bacteria; Proteobacteria; Gammaproteobacteria; Enterobacterales;
AB182__locus_1
PDU1AB
08015 HMM13520tp_1_pdu(pf13520) EUT2D
08020 -pf08411pf00929 -
08025 -pf07943pf00768 -
08030 -pf06445 -
08035 -pf04632 -
08040 -pf04363 -
08045 HMM00288_1_pdu(pf00288) PDU1AB
08050 -pf00590 -
08055 HMMacetkin_1(pf00871) PDU1AB
08060 HMM10662PduV_1_mix(pf10662) PDU1AB
08065 Hp_euts_babyPurple PDU1AB
08070 Ts_dodgerBlue PDU1AB
08075 HMMPduS_1_all(pf01512pf13375pf10531) PDU1AB
08080 HMM00465alcdh_3_mix(pf00465) PDU1AB
08085 HMMalddh_1_eutpdugrm(pf00171) PDU1AB EP_NTERM
08090 HMM01923cobtr_1_pdu(pf01923pf03928) PDU1AB
08095 P_babyPink PDU1AB
08100 HMM15953pduput_1_pdu1(pf15953) PDU1AB EP_NTERM
08105 HMMptac_2_mix(pf06130) PDU1AB
08110 H_driedBlood PDU1AB
08115 H_tan PDU1AB
08120 HMMpropdeh3_2_pdusml(pf02288) PDU1AB
08125 HMM08841dhact_1_pdu(pf08841) PDU1AB
08130 HMMpropdeh2_1_pdu(pf02287) PDU1AB EP_NTERM
08135 HMMpropdeh3_1_pdu(pf02288) PDU1AB EP_NTERM
08140 HMMpropdeh1_1_pdu(pf02286) PDU1AB
08145 Tsp_greenishBeige PDU1AB
08150 H_robinEggBlue GRM4
08155 -pf00230 -
08160 HMM10114PocR_1_pdu1(pf12833pf10114) PDU1AB
08165 -pf01656pf07685 -
08170 -pf03186 -
08175 -pf02570 -
08180 -pf01888 -
08185 -pf00590 -
08190 -# -
08195 -pf00590 -
08200 -pf01890pf11761pf11760 -
BMC-P
BMC-H
BMC-T
Aldehyde dehydrogenase
PTAC
Signature enzyme
Alcohol dehydrogenase
Regulator

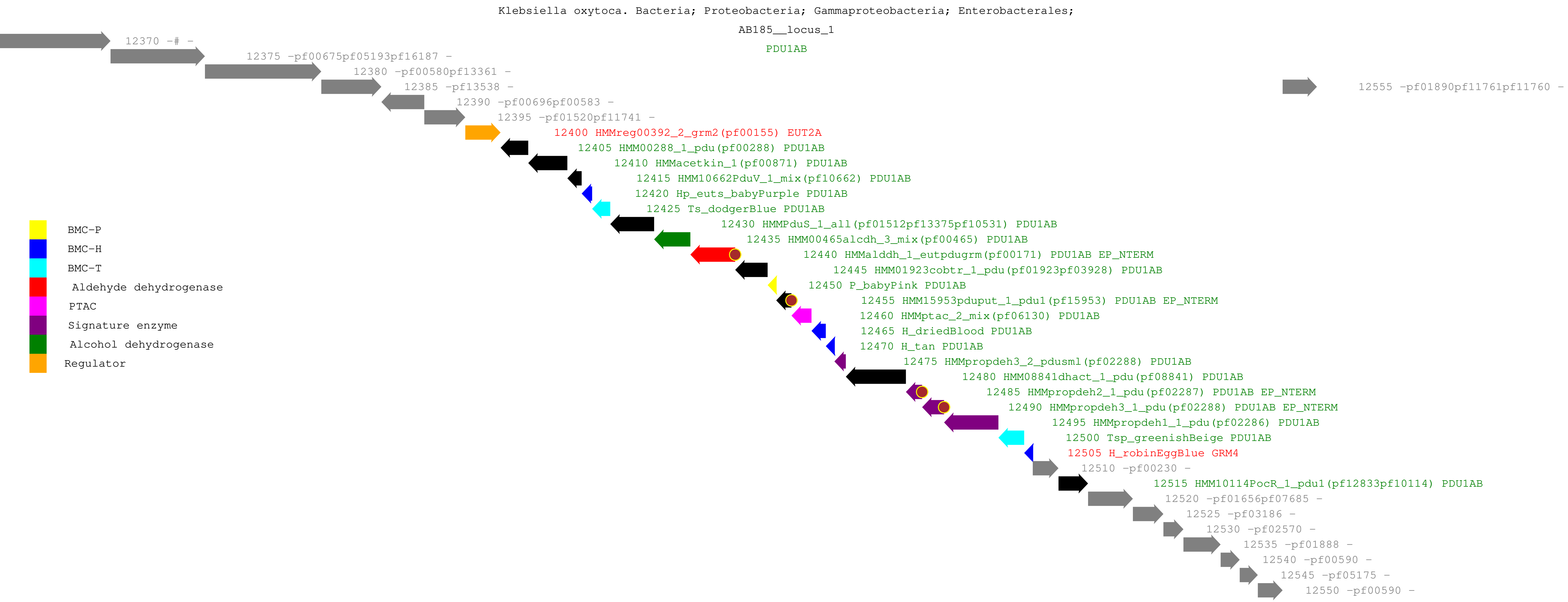

Klebsiella oxytoca. Bacteria; Proteobacteria; Gammaproteobacteria; Enterobacterales;
AB185__locus_1
PDU1AB
12370 -# -
12375 -pf00675pf05193pf16187 -
12380 -pf00580pf13361 -
12385 -pf13538 -
12555 -pf01890pf11761pf11760 -
12390 -pf00696pf00583 -
12395 -pf01520pf11741 -
12400 HMMreg00392_2_grm2(pf00155) EUT2A
12405 HMM00288_1_pdu(pf00288) PDU1AB
12410 HMMacetkin_1(pf00871) PDU1AB
12415 HMM10662PduV_1_mix(pf10662) PDU1AB
12420 Hp_euts_babyPurple PDU1AB
12425 Ts_dodgerBlue PDU1AB
12430 HMMPduS_1_all(pf01512pf13375pf10531) PDU1AB
12435 HMM00465alcdh_3_mix(pf00465) PDU1AB
12440 HMMaldh_1_eutpdugrm(pf00171) PDU1AB EP_NTERM
12445 HMM01923cobtr_1_pdu(pf01923pf03928) PDU1AB
12450 P_babyPink PDU1AB
12455 HMM15953pduput_1_pdu1(pf15953) PDU1AB EP_NTERM
12460 HMMptac_2_mix(pf06130) PDU1AB
12465 H_driedBlood PDU1AB
12470 H_tan PDU1AB
12475 HMMpropdeh3_2_pdusml(pf02288) PDU1AB
12480 HMM08841dhact_1_pdu(pf08841) PDU1AB
12485 HMMpropdeh2_1_pdu(pf02287) PDU1AB EP_NTERM
12490 HMMpropdeh3_1_pdu(pf02288) PDU1AB EP_NTERM
12495 HMMpropdeh1_1_pdu(pf02286) PDU1AB
12500 Tsp_greenishBeige PDU1AB
12505 H_robinEggBlue GRM4
12510 -pf00230 -
12515 HMM10114PocR_1_pdu1(pf12833pf10114) PDU1AB
12520 -pf01656pf07685 -
12525 -pf03186 -
12530 -pf02570 -
12535 -pf01888 -
12540 -pf00590 -
12545 -pf05175 -
12550 -pf00590 -
BMC-P
BMC-H
BMC-T
Aldehyde dehydrogenase
PTAC
Signature enzyme
Alcohol dehydrogenase
Regulator

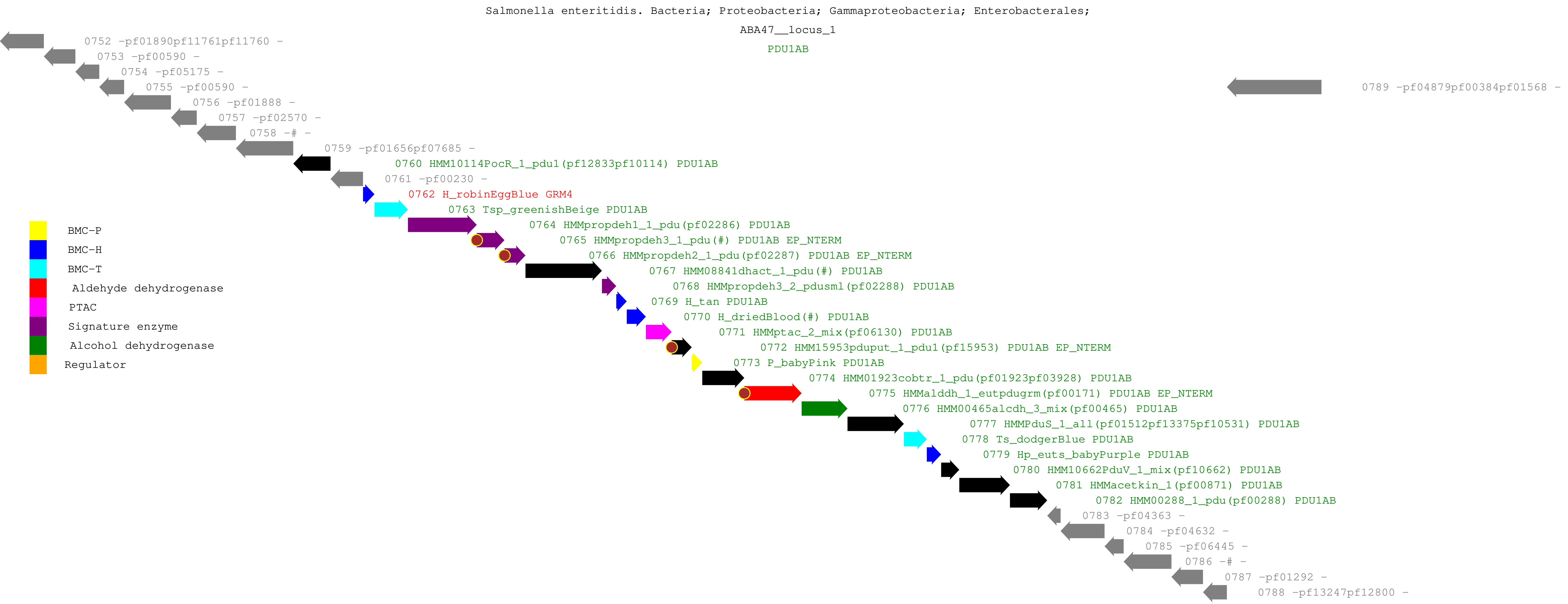

Salmonella enteritidis. Bacteria; Proteobacteria; Gammaproteobacteria; Enterobacterales;
ABA47__locus_1
PDU1AB
0752 -pf01890pf11761pf11760 -
0753 -pf00590 -
0754 -pf05175 -
0755 -pf00590 -
0756 -pf01888 -
0757 -pf02570 -
0758 -# -
0759 -pf01656pf07685 -
0760 HMM10114PocR_1_pdu1(pf12833pf10114) PDU1AB
0761 -pf00230 -
0762 H_robinEggBlue GRM4
0763 Tsp_greenishBeige PDU1AB
0764 HMMpropdeh1_1_pdu(pf02286) PDU1AB
0765 HMMpropdeh3_1_pdu(#) PDU1AB EP_NTERM
0766 HMMpropdeh2_1_pdu(pf02287) PDU1AB EP_NTERM
0767 HMM08841dhact_1_pdu(#) PDU1AB
0768 HMMpropdeh3_2_pdusml(pf02288) PDU1AB
0769 H_tan PDU1AB
0770 H_driedBlood(#) PDU1AB
0771 HMMptac_2_mix(pf06130) PDU1AB
0772 HMM15953pduput_1_pdu1(pf15953) PDU1AB EP_NTERM
0773 P_babyPink PDU1AB
0774 HMM01923cobtr_1_pdu(pf01923pf03928) PDU1AB
0775 HMMalddh_1_eutpdugrm(pf00171) PDU1AB EP_NTERM
0776 HMM00465alcdh_3_mix(pf00465) PDU1AB
0777 HMMPduS_1_all(pf01512pf13375pf10531) PDU1AB
0778 Ts_dodgerBlue PDU1AB
0779 Hp_euts_babyPurple PDU1AB
0780 HMM10662PduV_1_mix(pf10662) PDU1AB
0781 HMMacetkin_1(pf00871) PDU1AB
0782 HMM00288_1_pdu(pf00288) PDU1AB
0783 -pf04363 -
0784 -pf04632 -
0785 -pf06445 -
0786 -# -
0787 -pf01292 -
0788 -pf13247pf12800 -
0789 -pf04879pf00384pf01568 -
BMC-P
BMC-H
BMC-T
Aldehyde dehydrogenase
PTAC
Signature enzyme
Alcohol dehydrogenase
Regulator

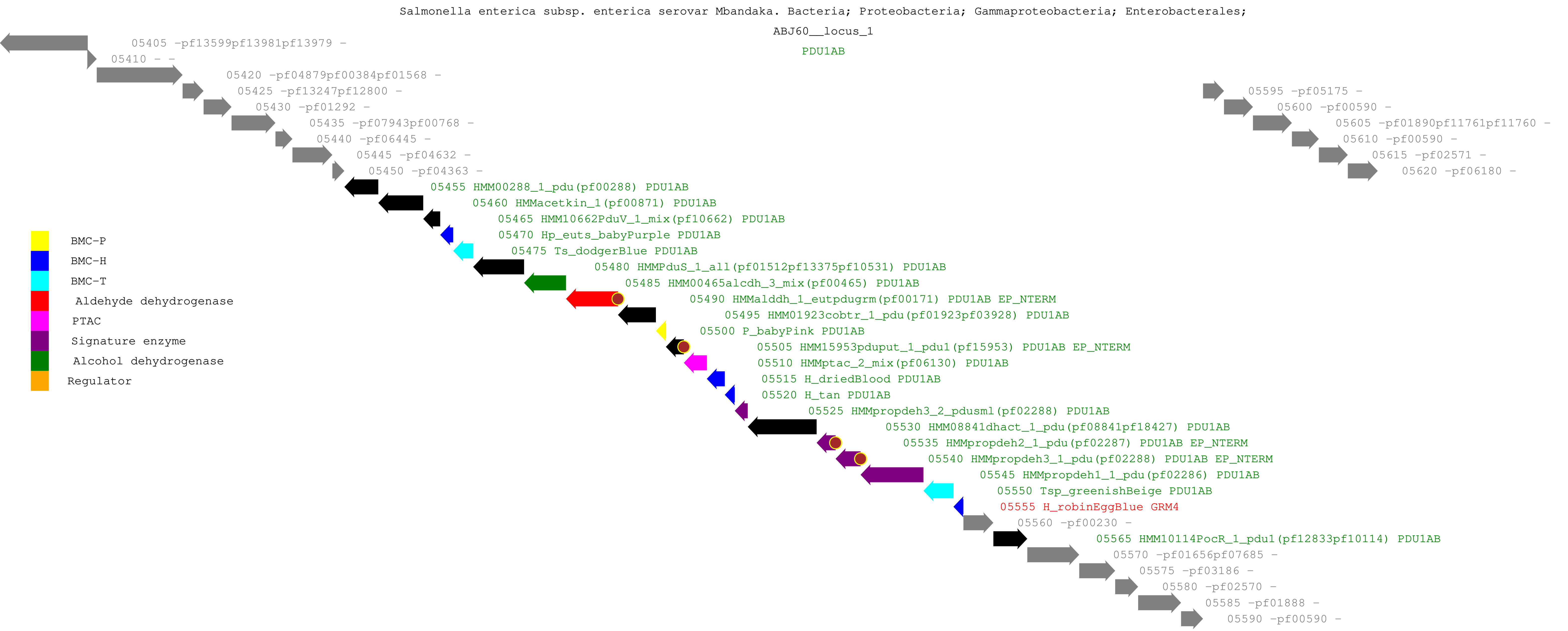
Salmonella enterica subsp. enterica serovar Mbandaka. Bacteria; Proteobacteria; Gammaproteobacteria; Enterobacterales;
ABJ60__locus_1
PDU1AB
05405 -pf13599pf13981pf13979 -
05410 - -
05420 -pf04879pf00384pf01568 -
05425 -pf13247pf12800 -
05430 -pf01292 -
05435 -pf07943pf00768 -
05440 -pf06445 -
05445 -pf04632 -
05450 -pf04363 -
05455 HMM00288_1_pdu(pf00288) PDU1AB
05460 HMMacetkin_1(pf00871) PDU1AB
05465 HMM10662PduV_1_mix(pf10662) PDU1AB
05470 Hp_euts_babyPurple PDU1AB
05475 Ts_dodgerBlue PDU1AB
05480 HMMPduS_1_all(pf01512pf13375pf10531) PDU1AB
05485 HMM00465alcdh_3_mix(pf00465) PDU1AB
05490 HMMalddh_1_eutpdugrm(pf00171) PDU1AB EP_NTERM
05495 HMM01923cobtr_1_pdu(pf01923pf03928) PDU1AB
05500 P_babyPink PDU1AB
05505 HMM15953pduput_1_pdu1(pf15953) PDU1AB EP_NTERM
05510 HMMptac_2_mix(pf06130) PDU1AB
05515 H_driedBlood PDU1AB
05520 H_tan PDU1AB
05525 HMMpropdeh3_2_pdusml(pf02288) PDU1AB
05530 HMM08841dhact_1_pdu(pf08841pf18427) PDU1AB
05535 HMMpropdeh2_1_pdu(pf02287) PDU1AB EP_NTERM
05540 HMMpropdeh3_1_pdu(pf02288) PDU1AB EP_NTERM
05545 HMMpropdeh1_1_pdu(pf02286) PDU1AB
05550 Tsp_greenishBeige PDU1AB
05555 H_robinEggBlue GRM4
05560 -pf00230 -
05565 HMM10114PocR_1_pdu1(pf12833pf10114) PDU1AB
05570 -pf01656pf07685 -
05575 -pf03186 -
05580 -pf02570 -
05585 -pf01888 -
05590 -pf00590 -
05595 -pf05175 -
05600 -pf00590 -
05605 -pf01890pf11761pf11760 -
05610 -pf00590 -
05615 -pf02571 -
05620 -pf06180 -
BMC-P
BMC-H
BMC-T
Aldehyde dehydrogenase
PTAC
Signature enzyme
Alcohol dehydrogenase
Regulator

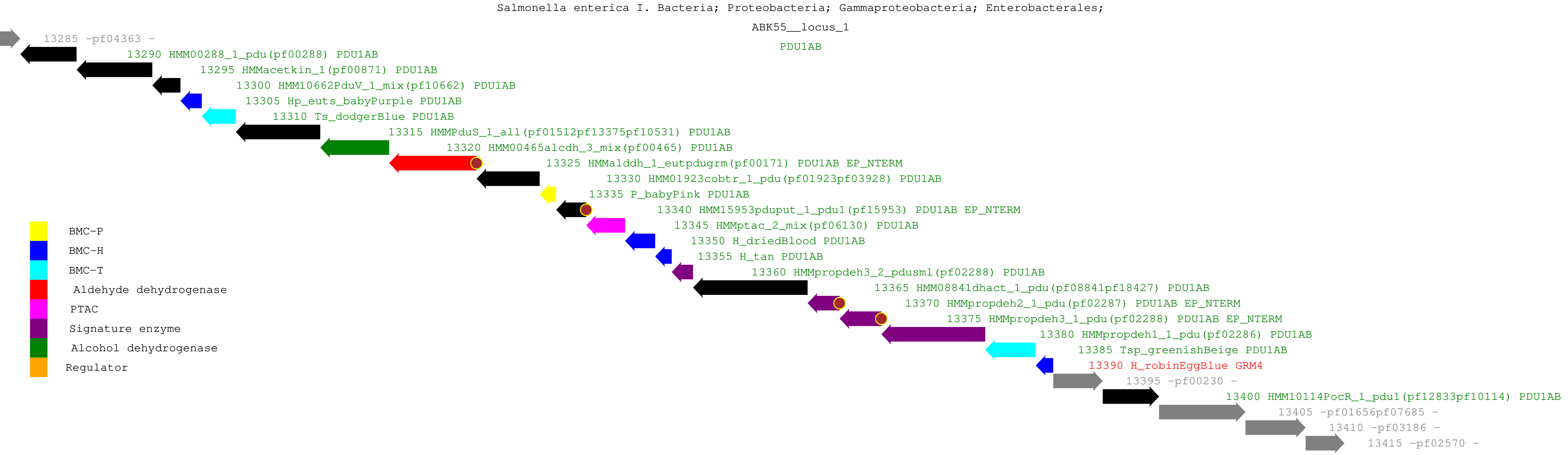

Salmonella enterica I. Bacteria; Proteobacteria; Gammaproteobacteria; Enterobacterales;
ABK55__locus_1
PDU1AB
13285 -pf04363 -
13290 HMM00288_1_pdu(pf00288) PDU1AB
13295 HMMacetkin_1(pf00871) PDU1AB
13300 HMM10662PduV_1_mix(pf10662) PDU1AB
13305 Hp_euts_babyPurple PDU1AB
13310 Ts_dodgerBlue PDU1AB
13315 HMMPduS_1_all(pf01512pf13375pf10531) PDU1AB
13320 HMM00465alcdh_3_mix(pf00465) PDU1AB
13325 HMMaldh_1_eutpdugrm(pf00171) PDU1AB EP_NTERM
13330 HMM01923cobtr_1_pdu(pf01923pf03928) PDU1AB
13335 P_babyPink PDU1AB
13340 HMM15953pduput_1_pdu1(pf15953) PDU1AB EP_NTERM
13345 HMMptac_2_mix(pf06130) PDU1AB
13350 H_driedBlood PDU1AB
13355 H_tan PDU1AB
13360 HMMpropdeh3_2_pdusml(pf02288) PDU1AB
13365 HMM08841dhact_1_pdu(pf08841pf18427) PDU1AB
13370 HMMpropdeh2_1_pdu(pf02287) PDU1AB EP_NTERM
13375 HMMpropdeh3_1_pdu(pf02288) PDU1AB EP_NTERM
13380 HMMpropdeh1_1_pdu(pf02286) PDU1AB
13385 Tsp_greenishBeige PDU1AB
13390 H_robinEggBlue GRM4
13395 -pf00230 -
13400 HMM10114PocR_1_pdu1(pf12833pf10114) PDU1AB
13405 -pf01656pf07685 -
13410 -pf03186 -
13415 -pf02570 -
BMC-P
BMC-H
BMC-T
Aldehyde dehydrogenase
PTAC
Signature enzyme
Alcohol dehydrogenase
Regulator

Salmonella senftenberg. Bacteria; Proteobacteria; Gammaproteobacteria; Enterobacterales;

ABM45__locus_1

PDU1AB

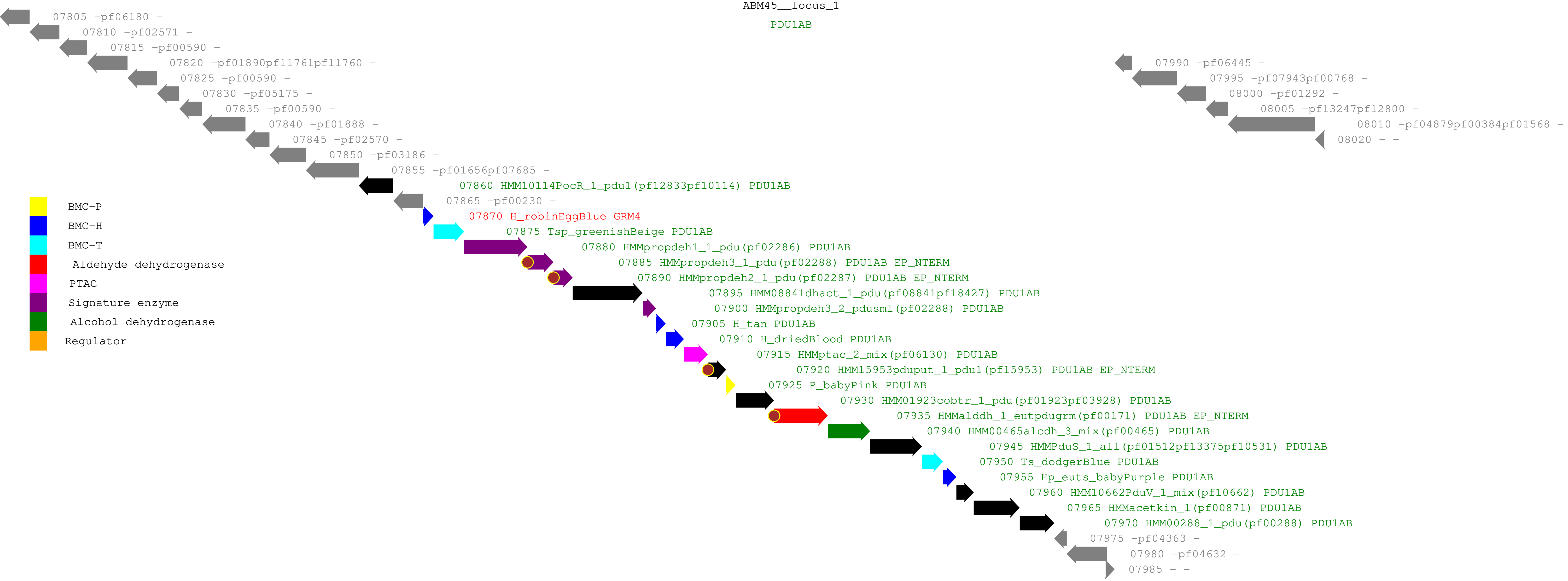

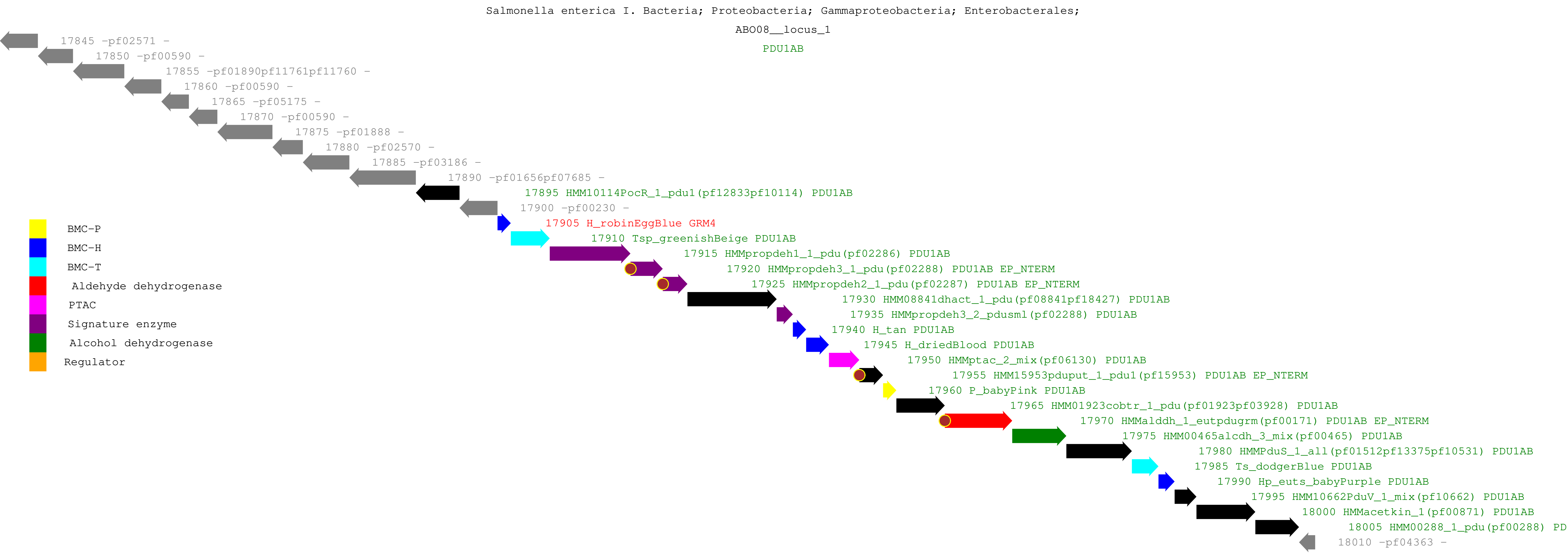
Salmonella enterica I. Bacteria; Proteobacteria; Gammaproteobacteria; Enterobacterales;
ABO08__locus_1
PDU1AB
17845 -pf02571 -
17850 -pf00590 -
17855 -pf01890pf11761pf11760 -
17860 -pf00590 -
17865 -pf05175 -
17870 -pf00590 -
17875 -pf01888 -
17880 -pf02570 -
17885 -pf03186 -
17890 -pf01656pf07685 -
17895 HMM10114PocR_1_pdu1(pf12833pf10114) PDU1AB
17900 -pf00230 -
17905 H_robinEggBlue GRM4
17910 Tsp_greenishBeige PDU1AB
17915 HMMpropdeh1_1_pdu(pf02286) PDU1AB
17920 HMMpropdeh3_1_pdu(pf02288) PDU1AB EP_NTERM
17925 HMMpropdeh2_1_pdu(pf02287) PDU1AB EP_NTERM
17930 HMM08841dhact_1_pdu(pf08841pf18427) PDU1AB
17935 HMMpropdeh3_2_pdusml(pf02288) PDU1AB
17940 H_tan PDU1AB
17945 H_driedBlood PDU1AB
17950 HMMptac_2_mix(pf06130) PDU1AB
17955 HMM15953pduput_1_pdu1(pf15953) PDU1AB EP_NTERM
17960 P_babyPink PDU1AB
17965 HMM01923cobtr_1_pdu(pf01923pf03928) PDU1AB
17970 HMMalddh_1_eutpdugrm(pf00171) PDU1AB EP_NTERM
17975 HMM00465alcdh_3_mix(pf00465) PDU1AB
17980 HMMPduS_1_all(pf01512pf13375pf10531) PDU1AB
17985 Ts_dodgerBlue PDU1AB
17990 Hp_euts_babyPurple PDU1AB
17995 HMM10662PduV_1_mix(pf10662) PDU1AB
18000 HMMacetkin_1(pf00871) PDU1AB
18005 HMM00288_1_pdu(pf00288) PD
18010 -pf04363 -
BMC-P
BMC-H
BMC-T
Aldehyde dehydrogenase
PTAC
Signature enzyme
Alcohol dehydrogenase
Regulator

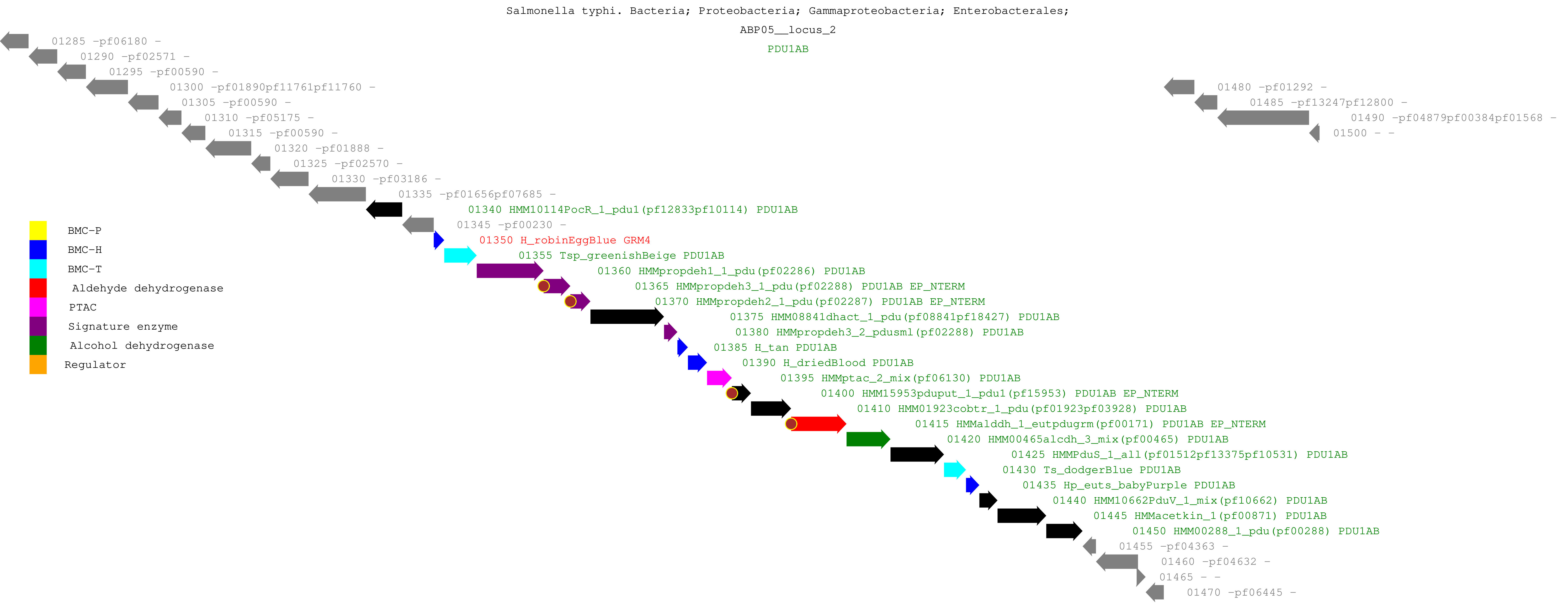
Salmonella typhi. Bacteria; Proteobacteria; Gammaproteobacteria; Enterobacterales;
ABP05__locus_2
PDU1AB
01285 -pf06180 -
01290 -pf02571 -
01295 -pf00590 -
01300 -pf01890pf11761pf11760 -
01305 -pf00590 -
01310 -pf05175 -
01315 -pf00590 -
01320 -pf01888 -
01325 -pf02570 -
01330 -pf03186 -
01335 -pf01656pf07685 -
01340 HMM10114PocR_1_pdu1(pf12833pf10114) PDU1AB
01345 -pf00230 -
01350 H_robinEggBlue GRM4
01355 Tsp_greenishBeige PDU1AB
01360 HMMpropdeh1_1_pdu(pf02286) PDU1AB
01365 HMMpropdeh3_1_pdu(pf02288) PDU1AB EP_NTERM
01370 HMMpropdeh2_1_pdu(pf02287) PDU1AB EP_NTERM
01375 HMM08841dhact_1_pdu(pf08841pf18427) PDU1AB
01380 HMMpropdeh3_2_pdusml(pf02288) PDU1AB
01385 H_tan PDU1AB
01390 H_driedBlood PDU1AB
01395 HMMptac_2_mix(pf06130) PDU1AB
01400 HMM15953pduput_1_pdu1(pf15953) PDU1AB EP_NTERM
01410 HMM01923cobtr_1_pdu(pf01923pf03928) PDU1AB
01415 HMMalddh_1_eutpdugrm(pf00171) PDU1AB EP_NTERM
01420 HMM00465alcdh_3_mix(pf00465) PDU1AB
01425 HMMPduS_1_all(pf01512pf13375pf10531) PDU1AB
01430 Ts_dodgerBlue PDU1AB
01435 Hp_euts_babyPurple PDU1AB
01440 HMM10662PduV_1_mix(pf10662) PDU1AB
01445 HMMacetkin_1(pf00871) PDU1AB
01450 HMM00288_1_pdu(pf00288) PDU1AB
01455 -pf04363 -
01460 -pf04632 -
01465 - -
01470 -pf06445 -
01480 -pf01292 -
01485 -pf13247pf12800 -
01490 -pf04879pf00384pf01568 -
01500 - -
BMC-P
BMC-H
BMC-T
Aldehyde dehydrogenase
PTAC
Signature enzyme
Alcohol dehydrogenase
Regulator

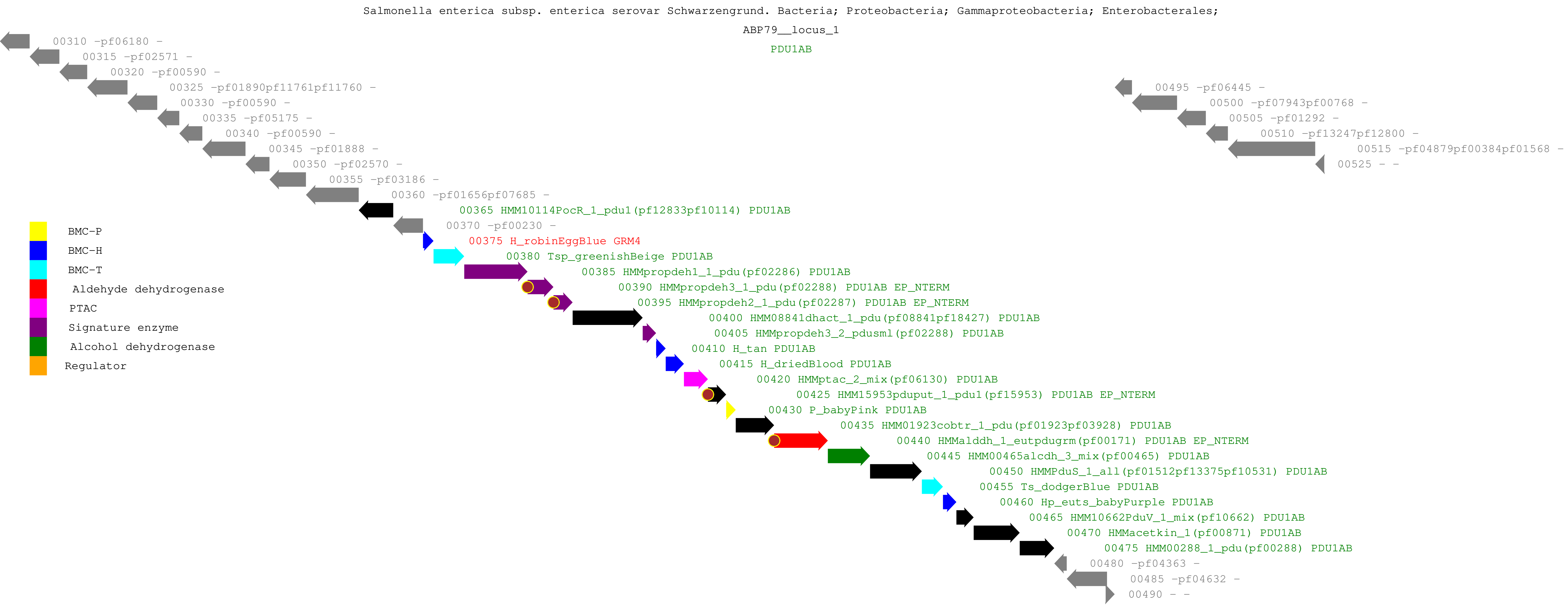
Salmonella enterica subsp. enterica serovar Schwarzengrund. Bacteria; Proteobacteria; Gammaproteobacteria; Enterobacterales;
ABP79__locus_1
PDU1AB
00310 -pf06180 -
00315 -pf02571 -
00320 -pf00590 -
00325 -pf01890pf11761pf11760 -
00330 -pf00590 -
00335 -pf05175 -
00340 -pf00590 -
00345 -pf01888 -
00350 -pf02570 -
00355 -pf03186 -
00360 -pf01656pf07685 -
00365 HMM10114PocR_1_pdu1(pf12833pf10114) PDU1AB
00370 -pf00230 -
00375 H_robinEggBlue GRM4
00380 Tsp_greenishBeige PDU1AB
00385 HMMpropdeh1_1_pdu(pf02286) PDU1AB
00390 HMMpropdeh3_1_pdu(pf02288) PDU1AB EP_NTERM
00395 HMMpropdeh2_1_pdu(pf02287) PDU1AB EP_NTERM
00400 HMM08841dhact_1_pdu(pf08841pf18427) PDU1AB
00405 HMMpropdeh3_2_pdusml(pf02288) PDU1AB
00410 H_tan PDU1AB
00415 H_driedBlood PDU1AB
00420 HMMptac_2_mix(pf06130) PDU1AB
00425 HMM15953pduput_1_pdu1(pf15953) PDU1AB EP_NTERM
00430 P_babyPink PDU1AB
00435 HMM01923cobtr_1_pdu(pf01923pf03928) PDU1AB
00440 HMMalddh_1_eutpdugrm(pf00171) PDU1AB EP_NTERM
00445 HMM00465alcdh_3_mix(pf00465) PDU1AB
00450 HMMPduS_1_all(pf01512pf13375pf10531) PDU1AB
00455 Ts_dodgerBlue PDU1AB
00460 Hp_euts_babyPurple PDU1AB
00465 HMM10662PduV_1_mix(pf10662) PDU1AB
00470 HMMacetkin_1(pf00871) PDU1AB
00475 HMM00288_1_pdu(pf00288) PDU1AB
00480 -pf04363 -
00485 -pf04632 -
00490 - -
00495 -pf06445 -
00500 -pf07943pf00768 -
00505 -pf01292 -
00510 -pf13247pf12800 -
00515 -pf04879pf00384pf01568 -
00525 - -
BMC-P
BMC-H
BMC-T
Aldehyde dehydrogenase
PTAC
Signature enzyme
Alcohol dehydrogenase
Regulator

Enterobacter sp. GN02600. Bacteria; Proteobacteria; Gammaproteobacteria; Enterobacterales;

ABR34__locus_1

PDU1AB

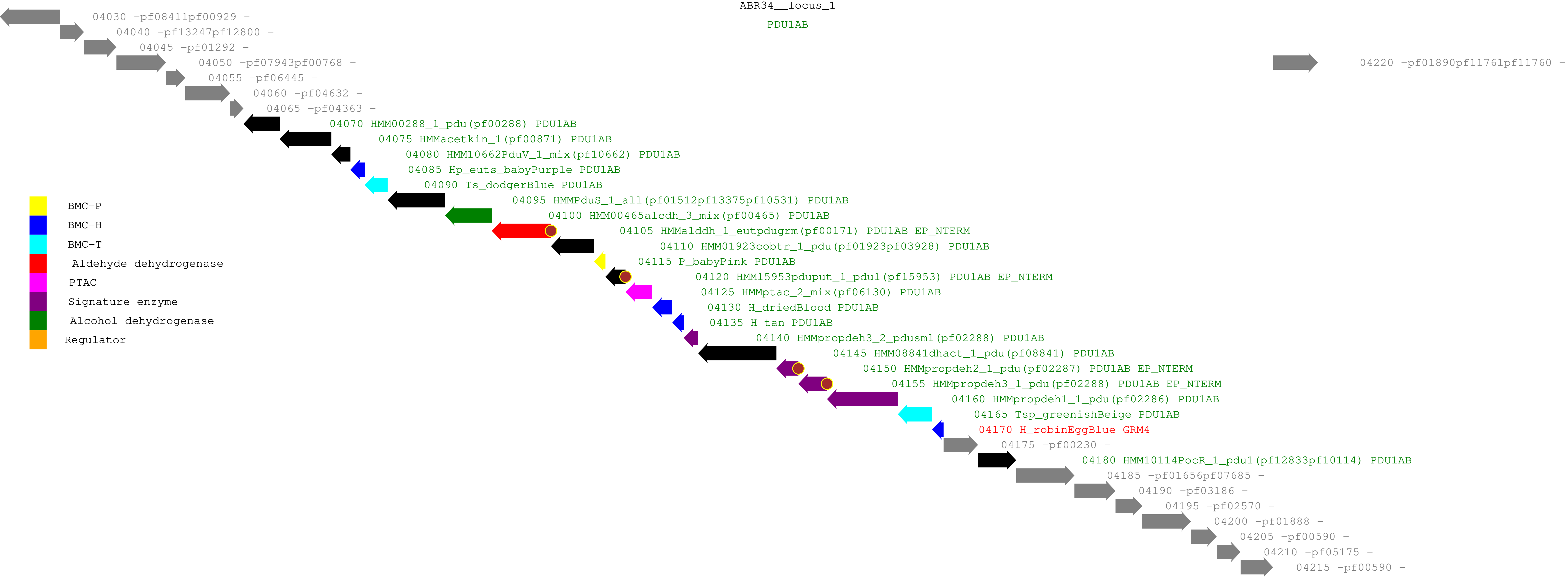

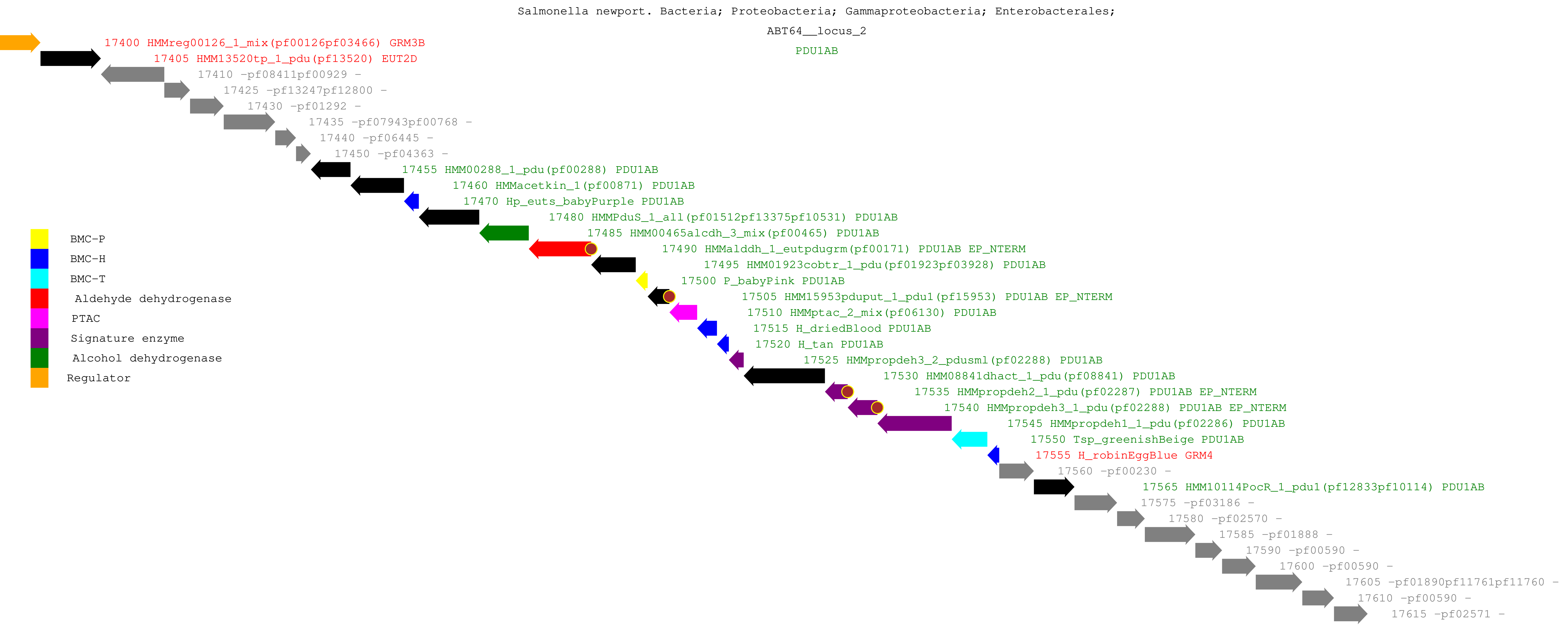
Salmonella newport. Bacteria; Proteobacteria; Gammaproteobacteria; Enterobacterales;
ABT64__locus_2
PDU1AB
17400 HMMreg00126_1_mix(pf00126pf03466) GRM3B
17405 HMM13520tp_1_pdu(pf13520) EUT2D
17410 -pf08411pf00929 -
17425 -pf13247pf12800 -
17430 -pf01292 -
17435 -pf07943pf00768 -
17440 -pf06445 -
17450 -pf04363 -
17455 HMM00288_1_pdu(pf00288) PDU1AB
17460 HMMacetkin_1(pf00871) PDU1AB
17470 Hp_euts_babyPurple PDU1AB
17480 HMMPduS_1_all(pf01512pf13375pf10531) PDU1AB
17485 HMM00465alcdh_3_mix(pf00465) PDU1AB
17490 HMMalddh_1_eutpdugrm(pf00171) PDU1AB EP_NTERM
17495 HMM01923cobtr_1_pdu(pf01923pf03928) PDU1AB
17500 P_babyPink PDU1AB
17505 HMM15953pduput_1_pdu1(pf15953) PDU1AB EP_NTERM
17510 HMMptac_2_mix(pf06130) PDU1AB
17515 H_driedBlood PDU1AB
17520 H_tan PDU1AB
17525 HMMpropdeh3_2_pdusml(pf02288) PDU1AB
17530 HMM08841dhact_1_pdu(pf08841) PDU1AB
17535 HMMpropdeh2_1_pdu(pf02287) PDU1AB EP_NTERM
17540 HMMpropdeh3_1_pdu(pf02288) PDU1AB EP_NTERM
17545 HMMpropdeh1_1_pdu(pf02286) PDU1AB
17550 Tsp_greenishBeige PDU1AB
17555 H_robinEggBlue GRM4
17560 -pf00230 -
17565 HMM10114PocR_1_pdu1(pf12833pf10114) PDU1AB
17575 -pf03186 -
17580 -pf02570 -
17585 -pf01888 -
17590 -pf00590 -
17600 -pf00590 -
17605 -pf01890pf11761pf11760 -
17610 -pf00590 -
17615 -pf02571 -
BMC-P
BMC-H
BMC-T
Aldehyde dehydrogenase
PTAC
Signature enzyme
Alcohol dehydrogenase
Regulator

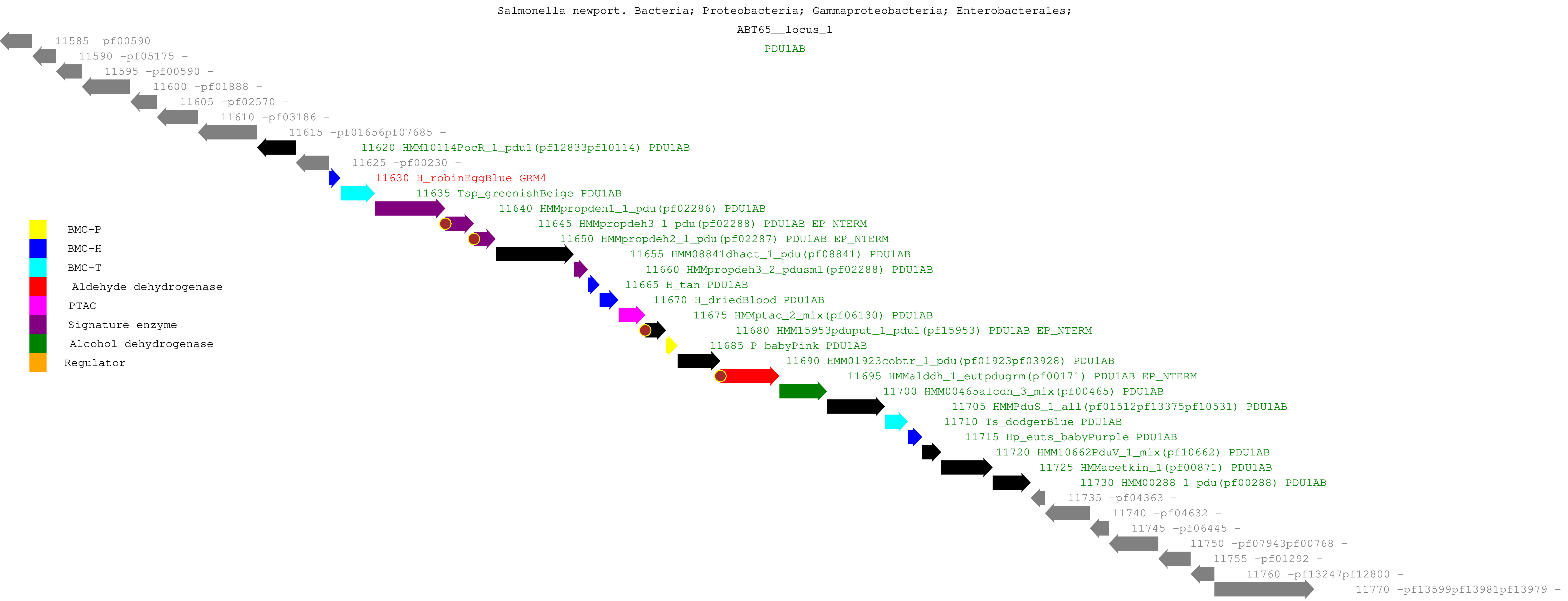

Salmonella newport. Bacteria; Proteobacteria; Gammaproteobacteria; Enterobacterales;
ABT65__locus_1
PDU1AB
11585 -pf00590 -
11590 -pf05175 -
11595 -pf00590 -
11600 -pf01888 -
11605 -pf02570 -
11610 -pf03186 -
11615 -pf01656pf07685 -
11620 HMM10114PocR_1_pdu1(pf12833pf10114) PDU1AB
11625 -pf00230 -
11630 H_robinEggBlue GRM4
11635 Tsp_greenishBeige PDU1AB
11640 HMMpropdeh1_1_pdu(pf02286) PDU1AB
11645 HMMpropdeh3_1_pdu(pf02288) PDU1AB EP_NTERM
11650 HMMpropdeh2_1_pdu(pf02287) PDU1AB EP_NTERM
11655 HMM08841dhact_1_pdu(pf08841) PDU1AB
11660 HMMpropdeh3_2_pdusml(pf02288) PDU1AB
11665 H_tan PDU1AB
11670 H_driedBlood PDU1AB
11675 HMMptac_2_mix(pf06130) PDU1AB
11680 HMM15953pduput_1_pdu1(pf15953) PDU1AB EP_NTERM
11685 P_babyPink PDU1AB
11690 HMM01923cobtr_1_pdu(pf01923pf03928) PDU1AB
11695 HMMalddh_1_eutpdugrm(pf00171) PDU1AB EP_NTERM
11700 HMM00465alcdh_3_mix(pf00465) PDU1AB
11705 HMMPduS_1_all(pf01512pf13375pf10531) PDU1AB
11710 Ts_dodgerBlue PDU1AB
11715 Hp_euts_babyPurple PDU1AB
11720 HMM10662PduV_1_mix(pf10662) PDU1AB
11725 HMMacetkin_1(pf00871) PDU1AB
11730 HMM00288_1_pdu(pf00288) PDU1AB
11735 -pf04363 -
11740 -pf04632 -
11745 -pf06445 -
11750 -pf07943pf00768 -
11755 -pf01292 -
11760 -pf13247pf12800 -
11770 -pf13599pf13981pf13979 -
BMC-P
BMC-H
BMC-T
Aldehyde dehydrogenase
PTAC
Signature enzyme
Alcohol dehydrogenase
Regulator

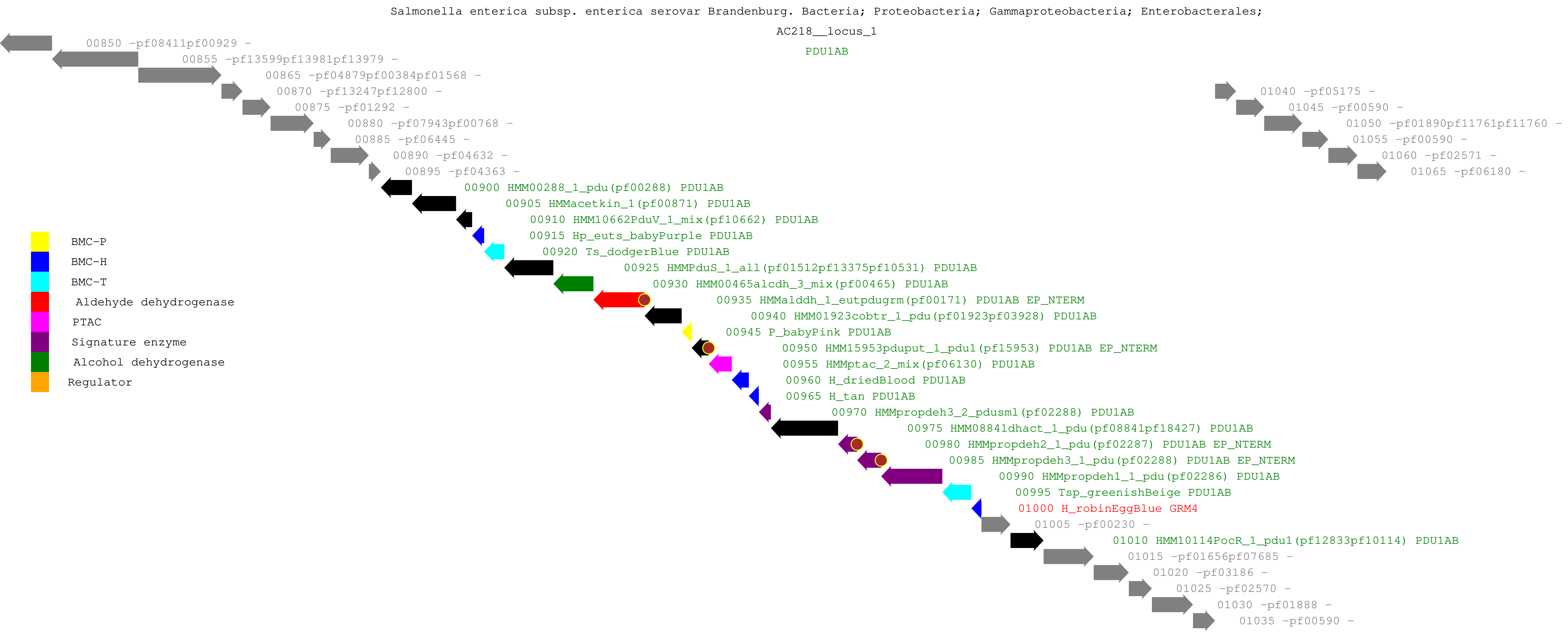

Salmonella enterica subsp. enterica serovar Brandenburg. Bacteria; Proteobacteria; Gammaproteobacteria; Enterobacterales;
AC218__locus_1
PDU1AB
00850 -pf08411pf00929 -
00855 -pf13599pf13981pf13979 -
00865 -pf04879pf00384pf01568 -
00870 -pf13247pf12800 -
00875 -pf01292 -
00880 -pf07943pf00768 -
00885 -pf06445 -
00890 -pf04632 -
00895 -pf04363 -
00900 HMM00288_1_pdu(pf00288) PDU1AB
00905 HMMacetkin_1(pf00871) PDU1AB
00910 HMM10662PduV_1_mix(pf10662) PDU1AB
00915 Hp_euts_babyPurple PDU1AB
00920 Ts_dodgerBlue PDU1AB
00925 HMMPduS_1_all(pf01512pf13375pf10531) PDU1AB
00930 HMM00465alcdh_3_mix(pf00465) PDU1AB
00935 HMMaldh_1_eutpdugrm(pf00171) PDU1AB EP_NTERM
00940 HMM01923cobtr_1_pdu(pf01923pf03928) PDU1AB
00945 P_babyPink PDU1AB
00950 HMM15953pduput_1_pdu1(pf15953) PDU1AB EP_NTERM
00955 HMMptac_2_mix(pf06130) PDU1AB
00960 H_driedBlood PDU1AB
00965 H_tan PDU1AB
00970 HMMpropdeh3_2_pdusml(pf02288) PDU1AB
00975 HMM08841dhact_1_pdu(pf08841pf18427) PDU1AB
00980 HMMpropdeh2_1_pdu(pf02287) PDU1AB EP_NTERM
00985 HMMpropdeh3_1_pdu(pf02288) PDU1AB EP_NTERM
00990 HMMpropdeh1_1_pdu(pf02286) PDU1AB
00995 Tsp_greenishBeige PDU1AB
01000 H_robinEggBlue GRM4
01005 -pf00230 -
01010 HMM10114PocR_1_pdu1(pf12833pf10114) PDU1AB
01015 -pf01656pf07685 -
01020 -pf03186 -
01025 -pf02570 -
01030 -pf01888 -
01035 -pf00590 -
01040 -pf05175 -
01045 -pf00590 -
01050 -pf01890pf11761pf11760 -
01055 -pf00590 -
01060 -pf02571 -
01065 -pf06180 -
BMC-P
BMC-H
BMC-T
Aldehyde dehydrogenase
PTAC
Signature enzyme
Alcohol dehydrogenase
Regulator

Escherichia coli 1-176-05_S3_C2. Bacteria; Proteobacteria; Gammaproteobacteria; Enterobacterales;

AC26__locus_1

PDU1AB

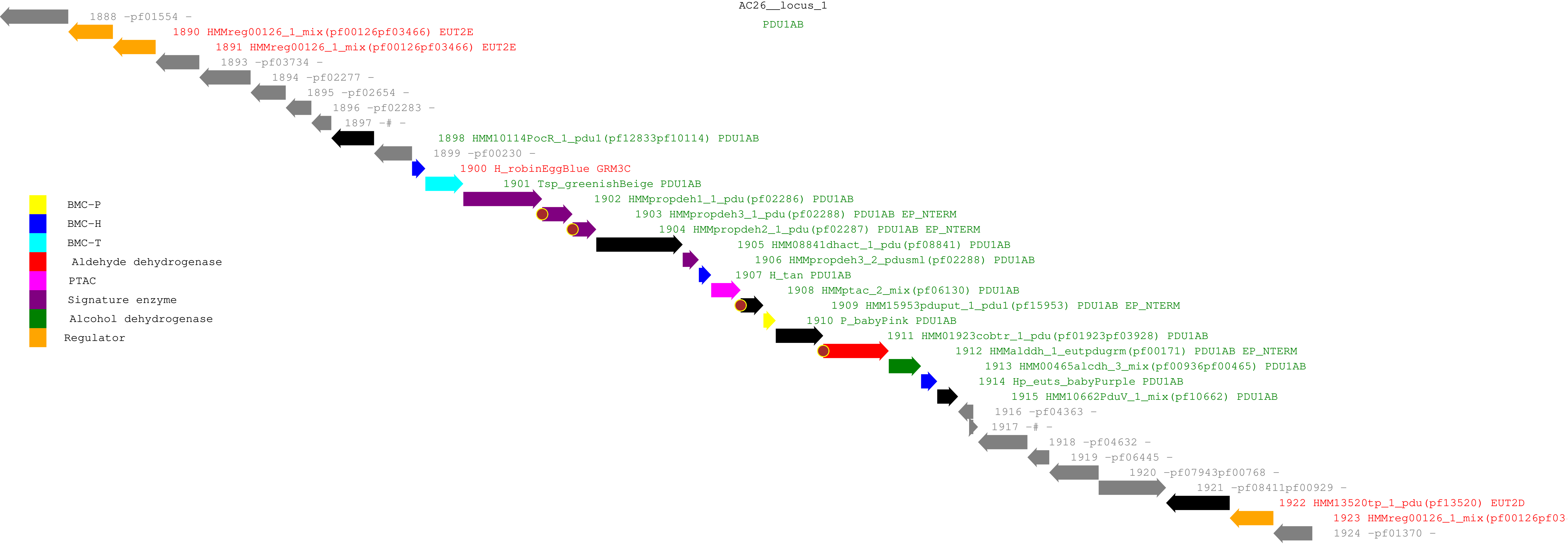

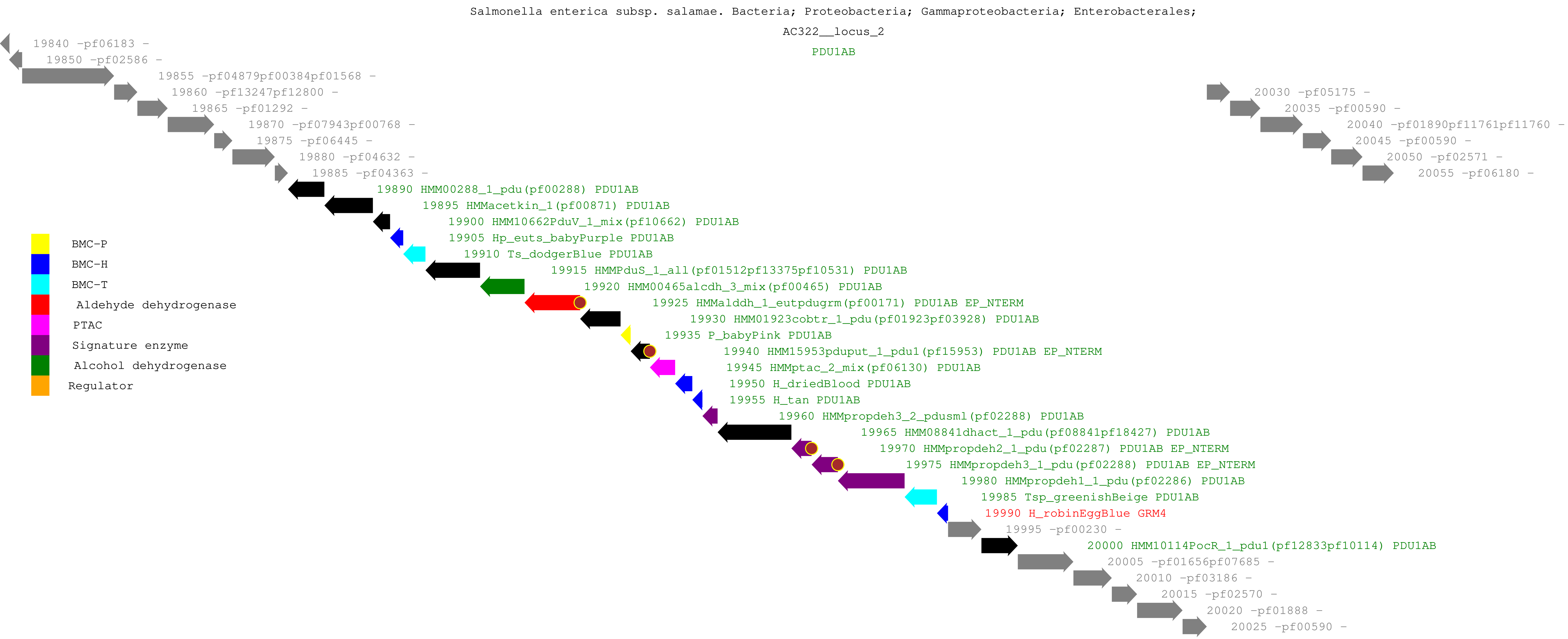
Salmonella enterica subsp. salamae. Bacteria; Proteobacteria; Gammaproteobacteria; Enterobacterales;
AC322__locus_2
PDU1AB
19840 -pf06183 -
19850 -pf02586 -
19855 -pf04879pf00384pf01568 -
19860 -pf13247pf12800 -
19865 -pf01292 -
19870 -pf07943pf00768 -
19875 -pf06445 -
19880 -pf04632 -
19885 -pf04363 -
19890 HMM00288_1_pdu(pf00288) PDU1AB
19895 HMMacetkin_1(pf00871) PDU1AB
19900 HMM10662PduV_1_mix(pf10662) PDU1AB
19905 Hp_euts_babyPurple PDU1AB
19910 Ts_dodgerBlue PDU1AB
19915 HMMPduS_1_all(pf01512pf13375pf10531) PDU1AB
19920 HMM00465alcdh_3_mix(pf00465) PDU1AB
19925 HMMalddh_1_eutpdugrm(pf00171) PDU1AB EP_NTERM
19930 HMM01923cobtr_1_pdu(pf01923pf03928) PDU1AB
19935 P_babyPink PDU1AB
19940 HMM15953pduput_1_pdu1(pf15953) PDU1AB EP_NTERM
19945 HMMptac_2_mix(pf06130) PDU1AB
19950 H_driedBlood PDU1AB
19955 H_tan PDU1AB
19960 HMMpropdeh3_2_pdusml(pf02288) PDU1AB
19965 HMM08841dhact_1_pdu(pf08841pf18427) PDU1AB
19970 HMMpropdeh2_1_pdu(pf02287) PDU1AB EP_NTERM
19975 HMMpropdeh3_1_pdu(pf02288) PDU1AB EP_NTERM
19980 HMMpropdeh1_1_pdu(pf02286) PDU1AB
19985 Tsp_greenishBeige PDU1AB
19990 H_robinEggBlue GRM4
19995 -pf00230 -
20000 HMM10114PocR_1_pdu1(pf12833pf10114) PDU1AB
20005 -pf01656pf07685 -
20010 -pf03186 -
20015 -pf02570 -
20020 -pf01888 -
20025 -pf00590 -
20030 -pf05175 -
20035 -pf00590 -
20040 -pf01890pf11761pf11760 -
20045 -pf00590 -
20050 -pf02571 -
20055 -pf06180 -
BMC-P
BMC-H
BMC-T
Aldehyde dehydrogenase
PTAC
Signature enzyme
Alcohol dehydrogenase
Regulator

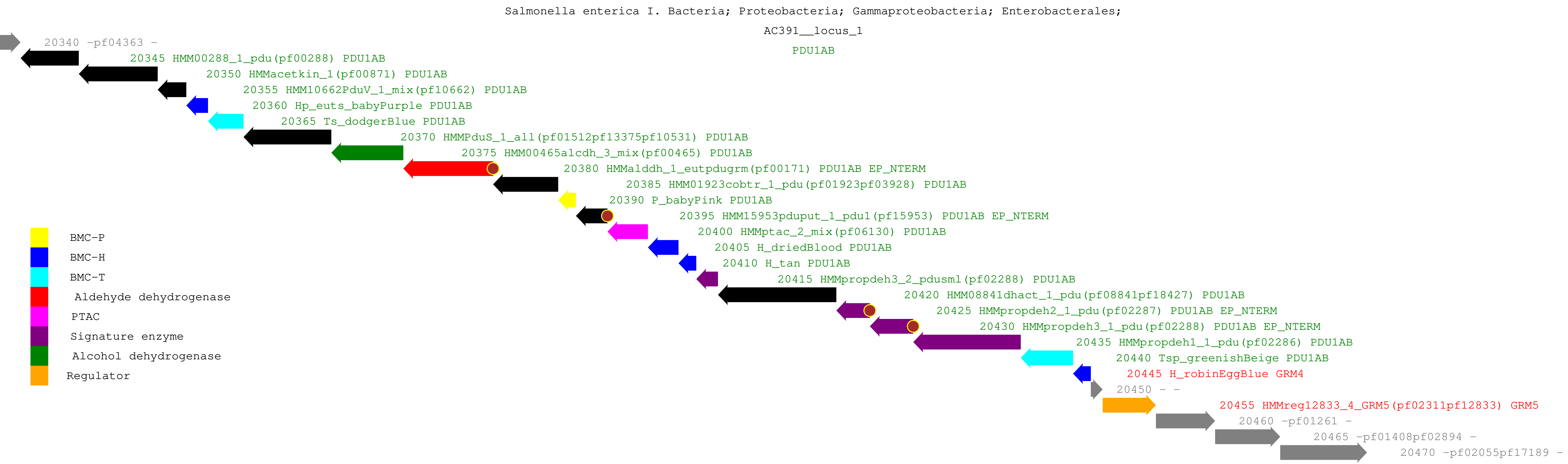

Salmonella enterica I. Bacteria; Proteobacteria; Gammaproteobacteria; Enterobacterales;
AC391__locus_1
PDU1AB
20340 -pf04363 -
20345 HMM00288_1_pdu(pf00288) PDU1AB
20350 HMMacetkin_1(pf00871) PDU1AB
20355 HMM10662PduV_1_mix(pf10662) PDU1AB
20360 Hp_euts_babyPurple PDU1AB
20365 Ts_dodgerBlue PDU1AB
20370 HMMPduS_1_all(pf01512pf13375pf10531) PDU1AB
20375 HMM00465alcdh_3_mix(pf00465) PDU1AB
20380 HMMalddh_1_eutpdugrm(pf00171) PDU1AB EP_NTERM
20385 HMM01923cobtr_1_pdu(pf01923pf03928) PDU1AB
20390 P_babyPink PDU1AB
20395 HMM15953pduput_1_pdu1(pf15953) PDU1AB EP_NTERM
20400 HMMptac_2_mix(pf06130) PDU1AB
20405 H_driedBlood PDU1AB
20410 H_tan PDU1AB
20415 HMMpropdeh3_2_pdusml(pf02288) PDU1AB
20420 HMM08841dhact_1_pdu(pf08841pf18427) PDU1AB
20425 HMMpropdeh2_1_pdu(pf02287) PDU1AB EP_NTERM
20430 HMMpropdeh3_1_pdu(pf02288) PDU1AB EP_NTERM
20435 HMMpropdeh1_1_pdu(pf02286) PDU1AB
20440 Tsp_greenishBeige PDU1AB
20445 H_robinEggBlue GRM4
20450 - -
20455 HMMreg12833_4_GRM5(pf02311pf12833) GRM5
20460 -pf01261 -
20465 -pf01408pf02894 -
20470 -pf02055pf17189 -
BMC-P
BMC-H
BMC-T
Aldehyde dehydrogenase
PTAC
Signature enzyme
Alcohol dehydrogenase
Regulator

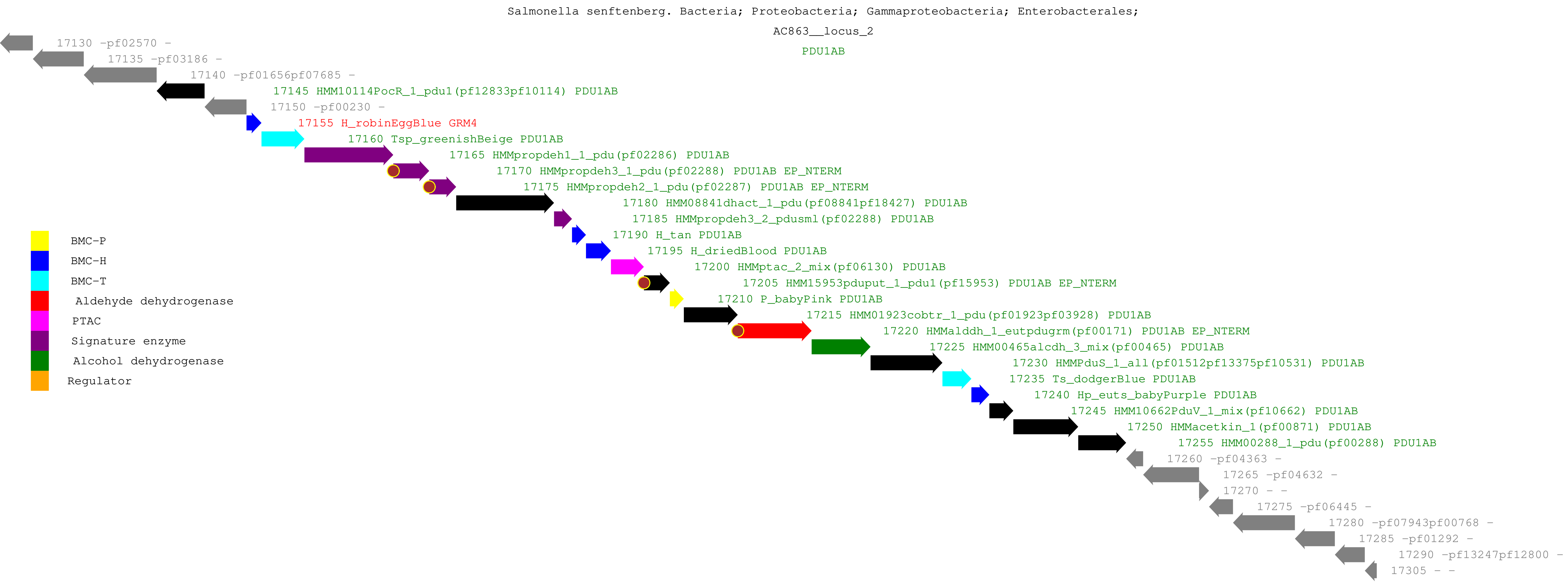
Salmonella senftenberg. Bacteria; Proteobacteria; Gammaproteobacteria; Enterobacterales;
AC863__locus_2
PDU1AB
17130 -pf02570 -
17135 -pf03186 -
17140 -pf01656pf07685 -
17145 HMM10114PocR_1_pdu1(pf12833pf10114) PDU1AB
17150 -pf00230 -
17155 H_robinEggBlue GRM4
17160 Tsp_greenishBeige PDU1AB
17165 HMMpropdeh1_1_pdu(pf02286) PDU1AB
17170 HMMpropdeh3_1_pdu(pf02288) PDU1AB EP_NTERM
17175 HMMpropdeh2_1_pdu(pf02287) PDU1AB EP_NTERM
17180 HMM08841dhact_1_pdu(pf08841pf18427) PDU1AB
17185 HMMpropdeh3_2_pdusml(pf02288) PDU1AB
17190 H_tan PDU1AB
17195 H_driedBlood PDU1AB
17200 HMMptac_2_mix(pf06130) PDU1AB
17205 HMM15953pduput_1_pdu1(pf15953) PDU1AB EP_NTERM
17210 P_babyPink PDU1AB
17215 HMM01923cobtr_1_pdu(pf01923pf03928) PDU1AB
17220 HMMalddh_1_eutpdugrm(pf00171) PDU1AB EP_NTERM
17225 HMM00465alcdh_3_mix(pf00465) PDU1AB
17230 HMMPduS_1_all(pf01512pf13375pf10531) PDU1AB
17235 Ts_dodgerBlue PDU1AB
17240 Hp_euts_babyPurple PDU1AB
17245 HMM10662PduV_1_mix(pf10662) PDU1AB
17250 HMMacetkin_1(pf00871) PDU1AB
17255 HMM00288_1_pdu(pf00288) PDU1AB
17260 -pf04363 -
17265 -pf04632 -
17270 - -
17275 -pf06445 -
17280 -pf07943pf00768 -
17285 -pf01292 -
17290 -pf13247pf12800 -
17305 - -
BMC-P
BMC-H
BMC-T
Aldehyde dehydrogenase
PTAC
Signature enzyme
Alcohol dehydrogenase
Regulator

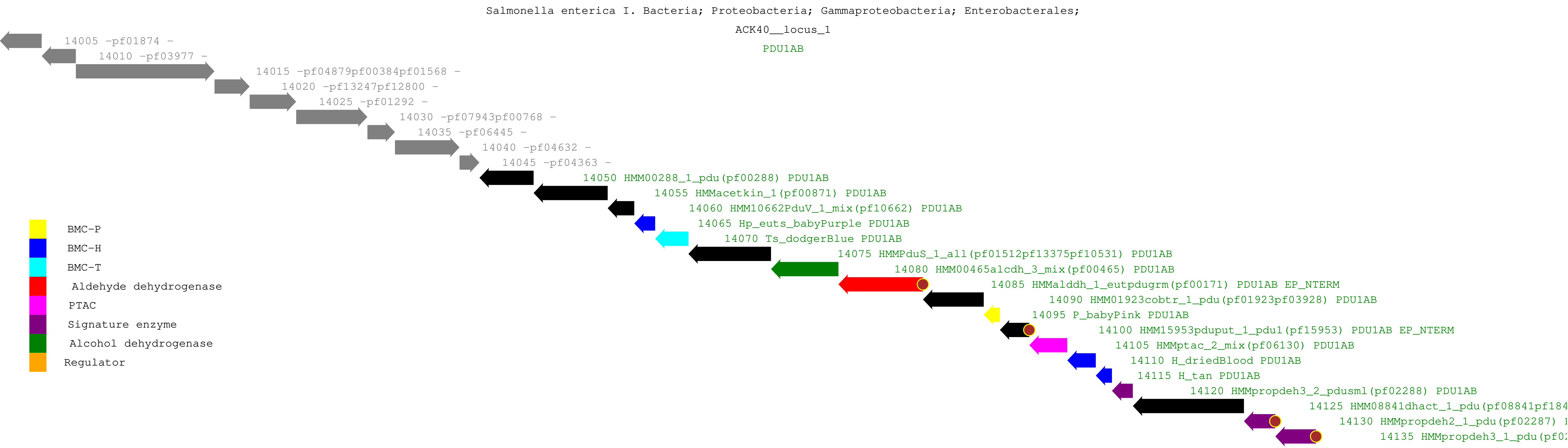

Salmonella enterica I. Bacteria; Proteobacteria; Gammaproteobacteria; Enterobacterales;
ACK40__locus_1
PDU1AB
14005 -pf01874 -
14010 -pf03977 -
14015 -pf04879pf00384pf01568 -
14020 -pf13247pf12800 -
14025 -pf01292 -
14030 -pf07943pf00768 -
14035 -pf06445 -
14040 -pf04632 -
14045 -pf04363 -
14050 HMM00288_1_pdu(pf00288) PDU1AB
14055 HMMacetkin_1(pf00871) PDU1AB
14060 HMM10662PduV_1_mix(pf10662) PDU1AB
14065 Hp_euts_babyPurple PDU1AB
14070 Ts_dodgerBlue PDU1AB
14075 HMMPduS_1_all(pf01512pf13375pf10531) PDU1AB
14080 HMM00465alcdh_3_mix(pf00465) PDU1AB
14085 HMMalddh_1_eutpdugrm(pf00171) PDU1AB EP_NTERM
14090 HMM01923cobtr_1_pdu(pf01923pf03928) PDU1AB
14095 P_babyPink PDU1AB
14100 HMM15953pduput_1_pdu1(pf15953) PDU1AB EP_NTERM
14105 HMMptac_2_mix(pf06130) PDU1AB
14110 H_driedBlood PDU1AB
14115 H_tan PDU1AB
14120 HMMpropdeh3_2_pdusml(pf02288) PDU1AB
14125 HMM08841dhact_1_pdu(pf08841pf184
14130 HMMpropdeh2_1_pdu(pf02287) I
14135 HMMpropdeh3_1_pdu(pf0
BMC-P
BMC-H
BMC-T
Aldehyde dehydrogenase
PTAC
Signature enzyme
Alcohol dehydrogenase
Regulator

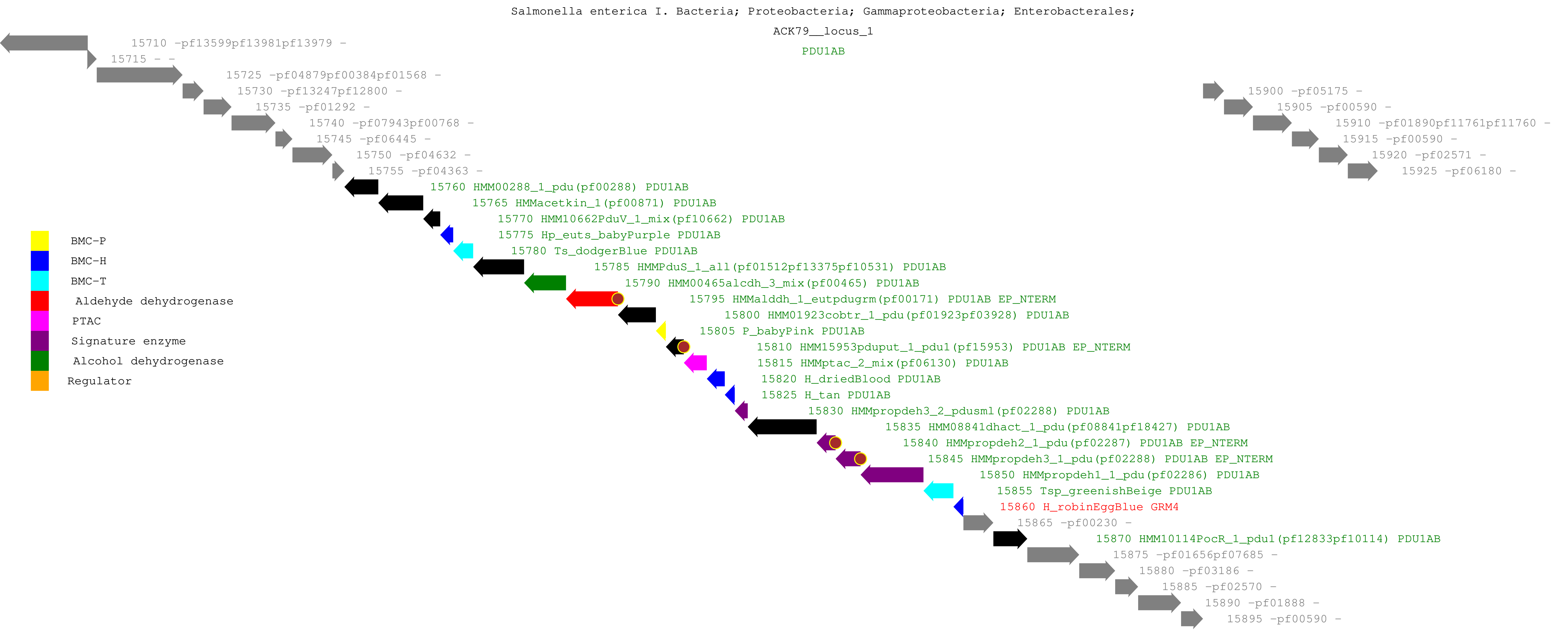
Salmonella enterica I. Bacteria; Proteobacteria; Gammaproteobacteria; Enterobacterales;
ACK79__locus_1
PDU1AB
15710 -pf13599pf13981pf13979 -
15715 - -
15725 -pf04879pf00384pf01568 -
15730 -pf13247pf12800 -
15735 -pf01292 -
15740 -pf07943pf00768 -
15745 -pf06445 -
15750 -pf04632 -
15755 -pf04363 -
15760 HMM00288_1_pdu(pf00288) PDU1AB
15765 HMMacetkin_1(pf00871) PDU1AB
15770 HMM10662PduV_1_mix(pf10662) PDU1AB
15775 Hp_euts_babyPurple PDU1AB
15780 Ts_dodgerBlue PDU1AB
15785 HMMPduS_1_all(pf01512pf13375pf10531) PDU1AB
15790 HMM00465alcdh_3_mix(pf00465) PDU1AB
15795 HMMalddh_1_eutpdugrm(pf00171) PDU1AB EP_NTERM
15800 HMM01923cobtr_1_pdu(pf01923pf03928) PDU1AB
15805 P_babyPink PDU1AB
15810 HMM15953pduput_1_pdu1(pf15953) PDU1AB EP_NTERM
15815 HMMptac_2_mix(pf06130) PDU1AB
15820 H_driedBlood PDU1AB
15825 H_tan PDU1AB
15830 HMMpropdeh3_2_pdusml(pf02288) PDU1AB
15835 HMM08841dhact_1_pdu(pf08841pf18427) PDU1AB
15840 HMMpropdeh2_1_pdu(pf02287) PDU1AB EP_NTERM
15845 HMMpropdeh3_1_pdu(pf02288) PDU1AB EP_NTERM
15850 HMMpropdeh1_1_pdu(pf02286) PDU1AB
15855 Tsp_greenishBeige PDU1AB
15860 H_robinEggBlue GRM4
15865 -pf00230 -
15870 HMM10114PocR_1_pdu1(pf12833pf10114) PDU1AB
15875 -pf01656pf07685 -
15880 -pf03186 -
15885 -pf02570 -
15890 -pf01888 -
15895 -pf00590 -
15900 -pf05175 -
15905 -pf00590 -
15910 -pf01890pf11761pf11760 -
15915 -pf00590 -
15920 -pf02571 -
15925 -pf06180 -
BMC-P
BMC-H
BMC-T
Aldehyde dehydrogenase
PTAC
Signature enzyme
Alcohol dehydrogenase
Regulator

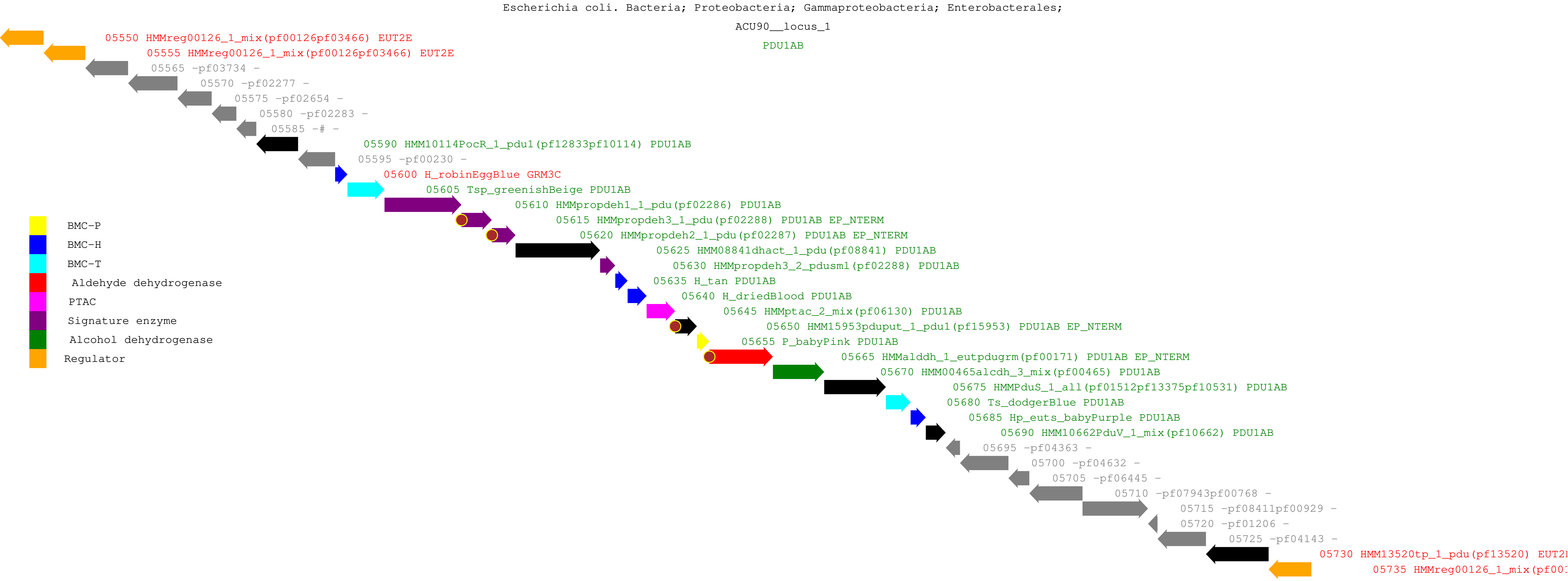
Escherichia coli. Bacteria; Proteobacteria; Gammaproteobacteria; Enterobacterales;
ACU90__locus_1
PDU1AB
05550 HMMreg00126_1_mix(pf00126pf03466) EUT2E
05555 HMMreg00126_1_mix(pf00126pf03466) EUT2E
05565 -pf03734 -
05570 -pf02277 -
05575 -pf02654 -
05580 -pf02283 -
05585 -# -
05590 HMM10114PocR_1_pdu1(pf12833pf10114) PDU1AB
05595 -pf00230 -
05600 H_robinEggBlue GRM3C
05605 Tsp_greenishBeige PDU1AB
05610 HMMpropdeh1_1_pdu(pf02286) PDU1AB
05615 HMMpropdeh3_1_pdu(pf02288) PDU1AB EP_NTERM
05620 HMMpropdeh2_1_pdu(pf02287) PDU1AB EP_NTERM
05625 HMM08841dhact_1_pdu(pf08841) PDU1AB
05630 HMMpropdeh3_2_pdusml(pf02288) PDU1AB
05635 H_tan PDU1AB
05640 H_driedBlood PDU1AB
05645 HMMptac_2_mix(pf06130) PDU1AB
05650 HMM15953pduput_1_pdu1(pf15953) PDU1AB EP_NTERM
05655 P_babyPink PDU1AB
05665 HMMalddh_1_eutpdugrm(pf00171) PDU1AB EP_NTERM
05670 HMM00465alcdh_3_mix(pf00465) PDU1AB
05675 HMMPduS_1_all(pf01512pf13375pf10531) PDU1AB
05680 Ts_dodgerBlue PDU1AB
05685 Hp_euts_babyPurple PDU1AB
05690 HMM10662PduV_1_mix(pf10662) PDU1AB
05695 -pf04363 -
05700 -pf04632 -
05705 -pf06445 -
05710 -pf07943pf00768 -
05715 -pf08411pf00929 -
05720 -pf01206 -
05725 -pf04143 -
05730 HMM13520tp_1_pdu(pf13520) EUT2
05735 HMMreg00126_1_mix(pf00
BMC-P
BMC-H
BMC-T
Aldehyde dehydrogenase
PTAC
Signature enzyme
Alcohol dehydrogenase
Regulator

Salmonella enterica I. Bacteria; Proteobacteria; Gammaproteobacteria; Enterobacterales;

ACW36__locus_2

PDU1AB

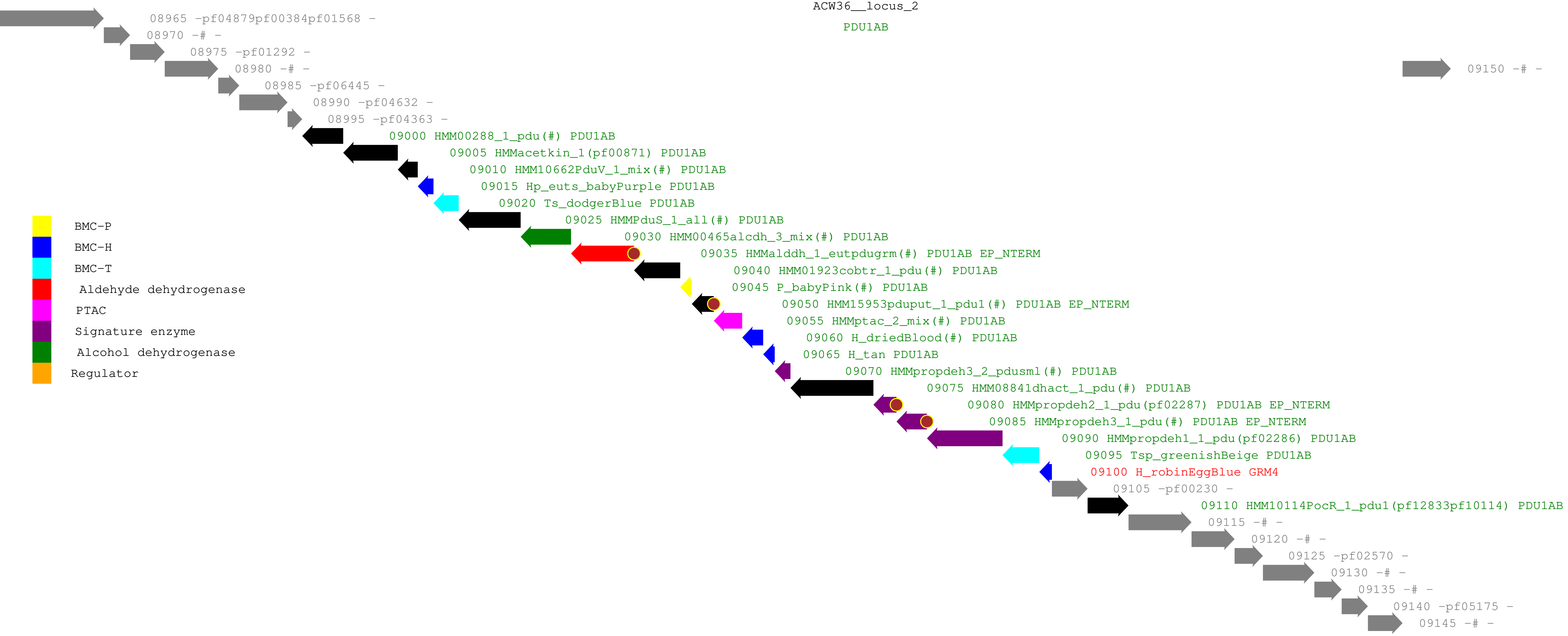

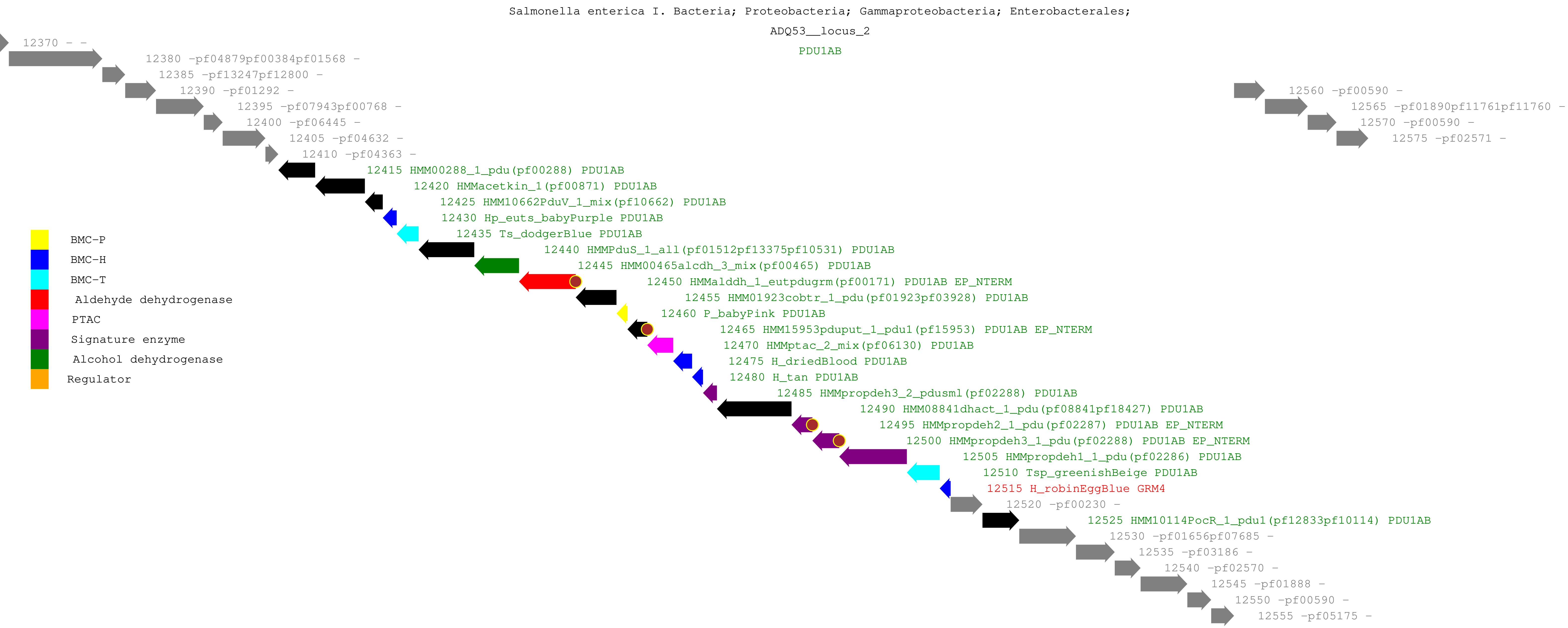
Salmonella enterica I. Bacteria; Proteobacteria; Gammaproteobacteria; Enterobacterales;
ADQ53__locus_2
PDU1AB
12370 - -
12380 -pf04879pf00384pf01568 -
12385 -pf13247pf12800 -
12390 -pf01292 -
12395 -pf07943pf00768 -
12400 -pf06445 -
12405 -pf04632 -
12410 -pf04363 -
12415 HMM00288_1_pdu(pf00288) PDU1AB
12420 HMMacetkin_1(pf00871) PDU1AB
12425 HMM10662PduV_1_mix(pf10662) PDU1AB
12430 Hp_euts_babyPurple PDU1AB
12435 Ts_dodgerBlue PDU1AB
12440 HMMPduS_1_all(pf01512pf13375pf10531) PDU1AB
12445 HMM00465alcdh_3_mix(pf00465) PDU1AB
12450 HMMaldh_1_eutpdugrm(pf00171) PDU1AB EP_NTERM
12455 HMM01923cobtr_1_pdu(pf01923pf03928) PDU1AB
12460 P_babyPink PDU1AB
12465 HMM15953pduput_1_pdu1(pf15953) PDU1AB EP_NTERM
12470 HMMptac_2_mix(pf06130) PDU1AB
12475 H_driedBlood PDU1AB
12480 H_tan PDU1AB
12485 HMMpropdeh3_2_pdusml(pf02288) PDU1AB
12490 HMM08841dhact_1_pdu(pf08841pf18427) PDU1AB
12495 HMMpropdeh2_1_pdu(pf02287) PDU1AB EP_NTERM
12500 HMMpropdeh3_1_pdu(pf02288) PDU1AB EP_NTERM
12505 HMMpropdeh1_1_pdu(pf02286) PDU1AB
12510 Tsp_greenishBeige PDU1AB
12515 H_robinEggBlue GRM4
12520 -pf00230 -
12525 HMM10114PocR_1_pdu1(pf12833pf10114) PDU1AB
12530 -pf01656pf07685 -
12535 -pf03186 -
12540 -pf02570 -
12545 -pf01888 -
12550 -pf00590 -
12555 -pf05175 -
12560 -pf00590 -
12565 -pf01890pf11761pf11760 -
12570 -pf00590 -
12575 -pf02571 -
BMC-P
BMC-H
BMC-T
Aldehyde dehydrogenase
PTAC
Signature enzyme
Alcohol dehydrogenase
Regulator

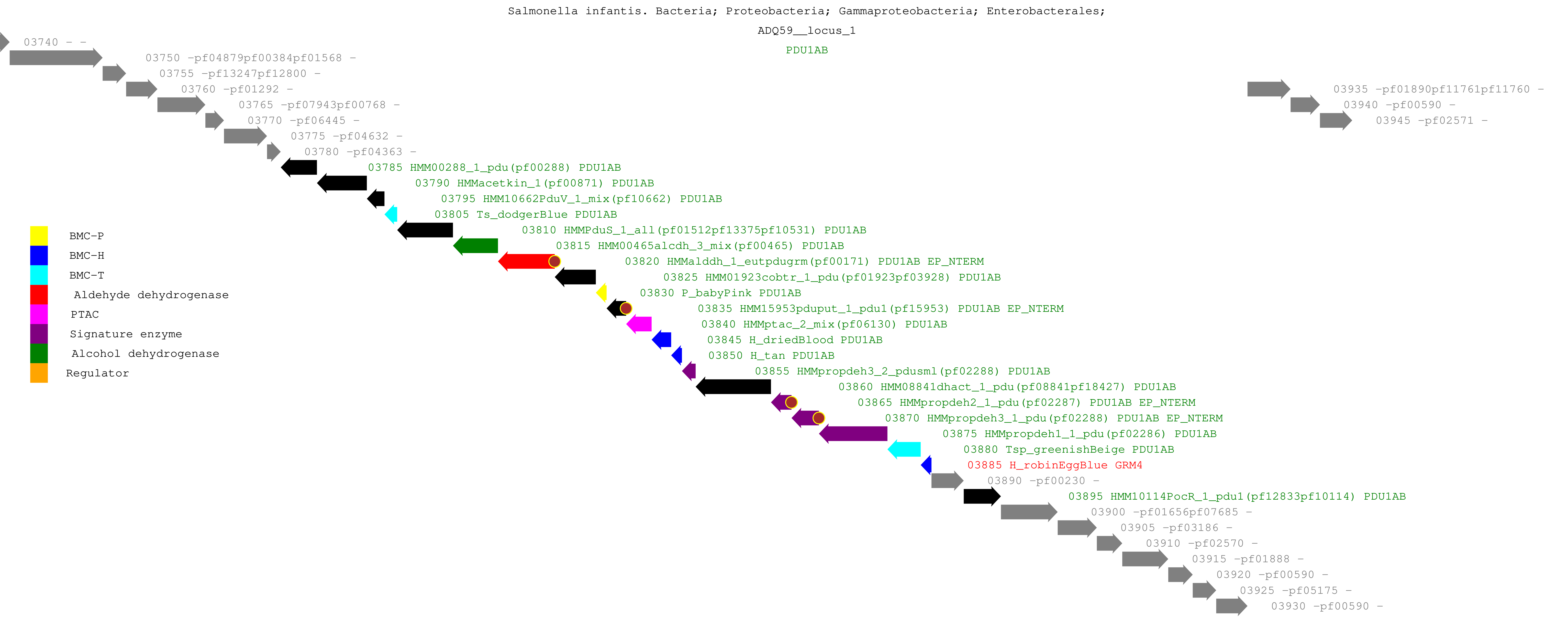
Salmonella infantis. Bacteria; Proteobacteria; Gammaproteobacteria; Enterobacterales;
ADQ59__locus_1
PDU1AB
03740 - -
03750 -pf04879pf00384pf01568 -
03755 -pf13247pf12800 -
03760 -pf01292 -
03765 -pf07943pf00768 -
03770 -pf06445 -
03775 -pf04632 -
03780 -pf04363 -
03785 HMM00288_1_pdu(pf00288) PDU1AB
03790 HMMacetkin_1(pf00871) PDU1AB
03795 HMM10662PduV_1_mix(pf10662) PDU1AB
03805 Ts_dodgerBlue PDU1AB
03810 HMMPduS_1_all(pf01512pf13375pf10531) PDU1AB
03815 HMM00465alcdh_3_mix(pf00465) PDU1AB
03820 HMMaldh_1_eutpdugrm(pf00171) PDU1AB EP_NTERM
03825 HMM01923cobtr_1_pdu(pf01923pf03928) PDU1AB
03830 P_babyPink PDU1AB
03835 HMM15953pduput_1_pdu1(pf15953) PDU1AB EP_NTERM
03840 HMMptac_2_mix(pf06130) PDU1AB
03845 H_driedBlood PDU1AB
03850 H_tan PDU1AB
03855 HMMpropdeh3_2_pdusml(pf02288) PDU1AB
03860 HMM08841dhact_1_pdu(pf08841pf18427) PDU1AB
03865 HMMpropdeh2_1_pdu(pf02287) PDU1AB EP_NTERM
03870 HMMpropdeh3_1_pdu(pf02288) PDU1AB EP_NTERM
03875 HMMpropdeh1_1_pdu(pf02286) PDU1AB
03880 Tsp_greenishBeige PDU1AB
03885 H_robinEggBlue GRM4
03890 -pf00230 -
03895 HMM10114PocR_1_pdu1(pf12833pf10114) PDU1AB
03900 -pf01656pf07685 -
03905 -pf03186 -
03910 -pf02570 -
03915 -pf01888 -
03920 -pf00590 -
03925 -pf05175 -
03930 -pf00590 -
03935 -pf01890pf11761pf11760 -
03940 -pf00590 -
03945 -pf02571 -
BMC-P
BMC-H
BMC-T
Aldehyde dehydrogenase
PTAC
Signature enzyme
Alcohol dehydrogenase
Regulator

Salmonella enterica (Salmonella choleraesuis). Bacteria; Proteobacteria; Gammaproteobacteria; Enterobacterales;

ADQ66__locus_1

PDU1AB

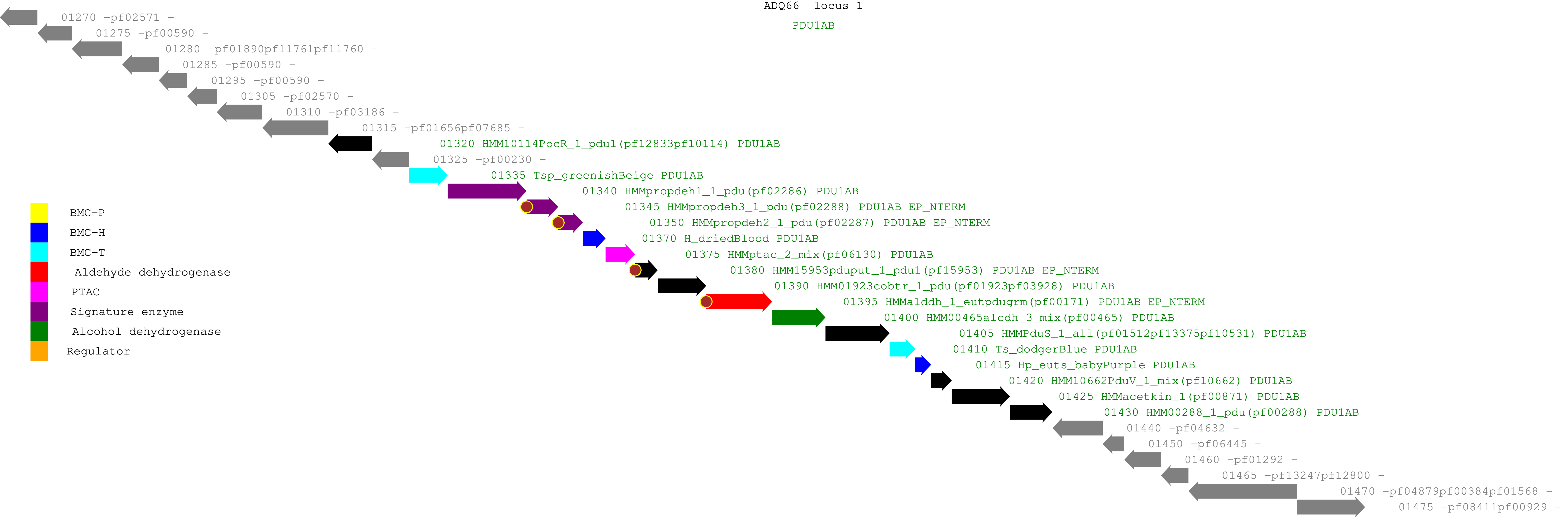

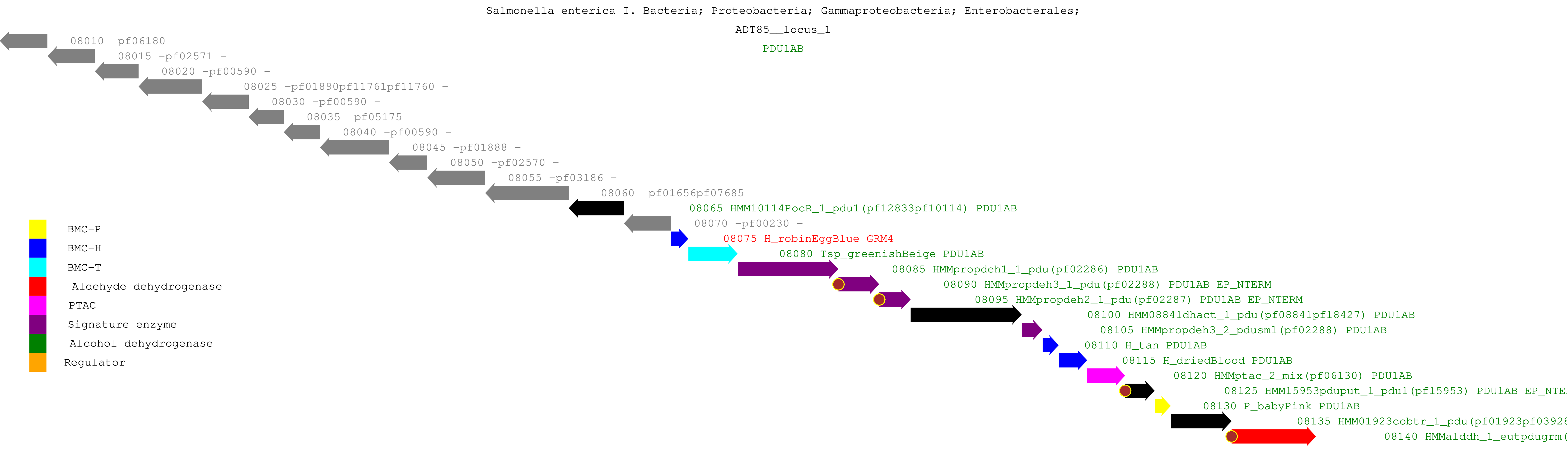
Salmonella enterica I. Bacteria; Proteobacteria; Gammaproteobacteria; Enterobacterales;
ADT85__locus_1
PDU1AB
08010 -pf06180 -
08015 -pf02571 -
08020 -pf00590 -
08025 -pf01890pf11761pf11760 -
08030 -pf00590 -
08035 -pf05175 -
08040 -pf00590 -
08045 -pf01888 -
08050 -pf02570 -
08055 -pf03186 -
08060 -pf01656pf07685 -
08065 HMM10114PocR_1_pdu1(pf12833pf10114) PDU1AB
08070 -pf00230 -
08075 H_robinEggBlue GRM4
08080 Tsp_greenishBeige PDU1AB
08085 HMMpropdeh1_1_pdu(pf02286) PDU1AB
08090 HMMpropdeh3_1_pdu(pf02288) PDU1AB EP_NTERM
08095 HMMpropdeh2_1_pdu(pf02287) PDU1AB EP_NTERM
08100 HMM08841dhact_1_pdu(pf08841pf18427) PDU1AB
08105 HMMpropdeh3_2_pdusml(pf02288) PDU1AB
08110 H_tan PDU1AB
08115 H_driedBlood PDU1AB
08120 HMMptac_2_mix(pf06130) PDU1AB
08125 HMM15953pduput_1_pdu1(pf15953) PDU1AB EP_NTE
08130 P_babyPink PDU1AB
08135 HMM01923cobtr_1_pdu(pf01923pf0392
08140 HMMalddh_1_eutpdugrm(
BMC-P
BMC-H
BMC-T
Aldehyde dehydrogenase
PTAC
Signature enzyme
Alcohol dehydrogenase
Regulator

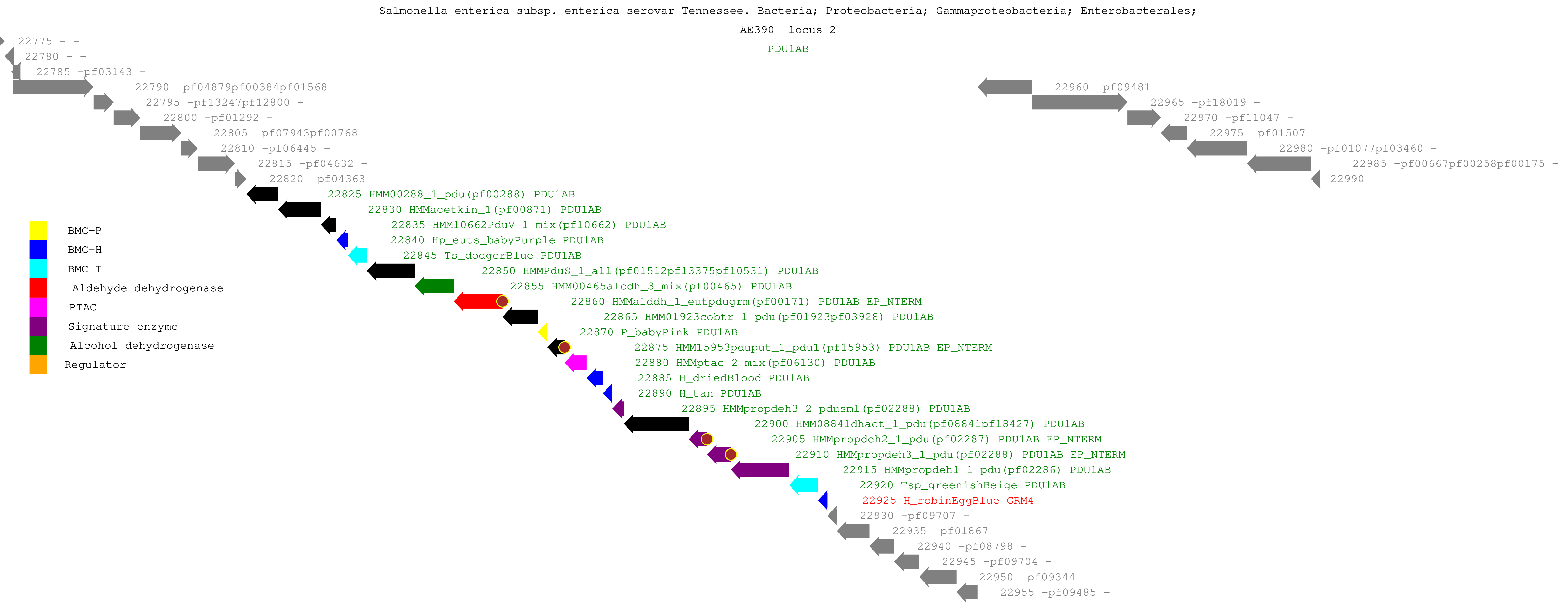
Salmonella enterica subsp. enterica serovar Tennessee. Bacteria; Proteobacteria; Gammaproteobacteria; Enterobacterales;
AE390__locus_2
PDU1AB
22775 - -
22780 - -
22785 -pf03143 -
22790 -pf04879pf00384pf01568 -
22795 -pf13247pf12800 -
22800 -pf01292 -
22805 -pf07943pf00768 -
22810 -pf06445 -
22815 -pf04632 -
22820 -pf04363 -
22825 HMM00288_1_pdu(pf00288) PDU1AB
22830 HMMacetkin_1(pf00871) PDU1AB
22835 HMM10662PduV_1_mix(pf10662) PDU1AB
22840 Hp_euts_babyPurple PDU1AB
22845 Ts_dodgerBlue PDU1AB
22850 HMMPduS_1_all(pf01512pf13375pf10531) PDU1AB
22855 HMM00465alcdh_3_mix(pf00465) PDU1AB
22860 HMMaldh_1_eutpdugrm(pf00171) PDU1AB EP_NTERM
22865 HMM01923cobtr_1_pdu(pf01923pf03928) PDU1AB
22870 P_babyPink PDU1AB
22875 HMM15953pduput_1_pdu1(pf15953) PDU1AB EP_NTERM
22880 HMMptac_2_mix(pf06130) PDU1AB
22885 H_driedBlood PDU1AB
22890 H_tan PDU1AB
22895 HMMpropdeh3_2_pdusml(pf02288) PDU1AB
22900 HMM08841dhact_1_pdu(pf08841pf18427) PDU1AB
22905 HMMpropdeh2_1_pdu(pf02287) PDU1AB EP_NTERM
22910 HMMpropdeh3_1_pdu(pf02288) PDU1AB EP_NTERM
22915 HMMpropdeh1_1_pdu(pf02286) PDU1AB
22920 Tsp_greenishBeige PDU1AB
22925 H_robinEggBlue GRM4
22930 -pf09707 -
22935 -pf01867 -
22940 -pf08798 -
22945 -pf09704 -
22950 -pf09344 -
22955 -pf09485 -
22960 -pf09481 -
22965 -pf18019 -
22970 -pf11047 -
22975 -pf01507 -
22980 -pf01077pf03460 -
22985 -pf00667pf00258pf00175 -
22990 - -
BMC-P
BMC-H
BMC-T
Aldehyde dehydrogenase
PTAC
Signature enzyme
Alcohol dehydrogenase
Regulator

Salmonella enterica subsp. enterica serovar Heidelberg. Bacteria; Proteobacteria; Gammaproteobacteria; Enterobacterales;

AEV26__locus_2

PDU1AB

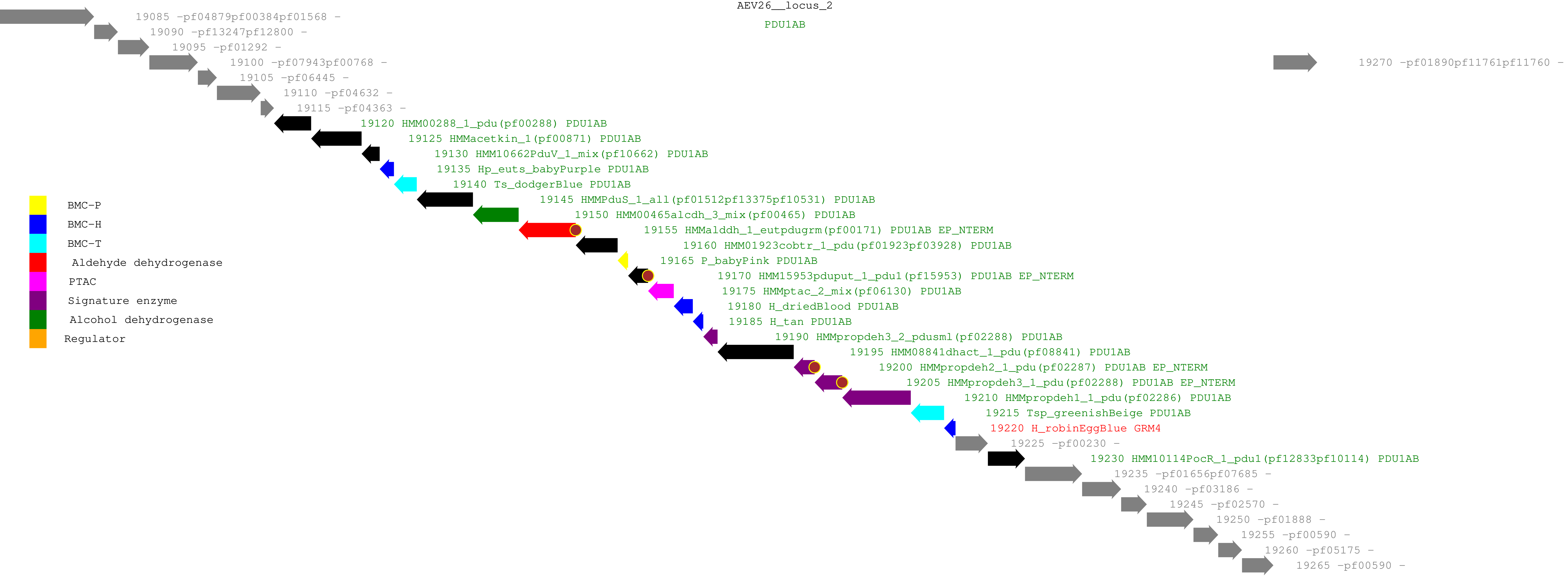

Salmonella enterica subsp. enterica serovar Johannesburg. Bacteria; Proteobacteria; Gammaproteobacteria; Enterobacterales;

AEX30__locus_2

PDU1AB

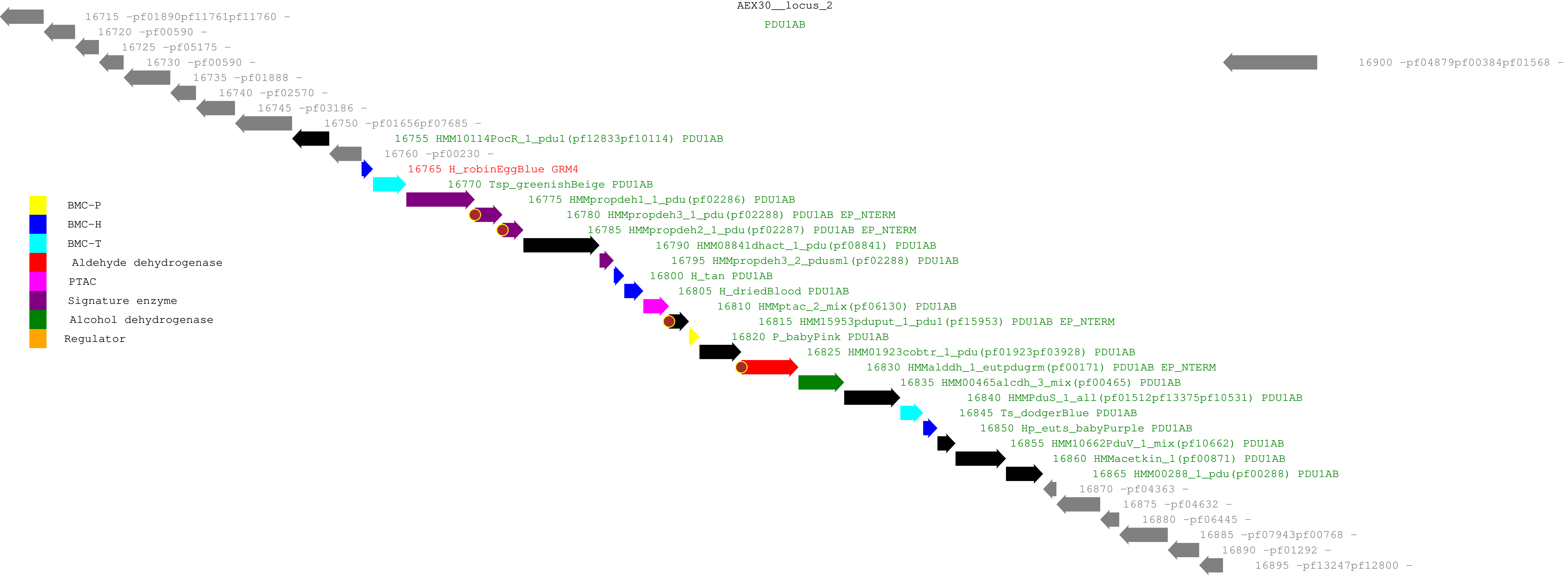

Citrobacter sp. MGH 55. Bacteria; Proteobacteria; Gammaproteobacteria; Enterobacterales;

AF41__locus_2

PDU1AB

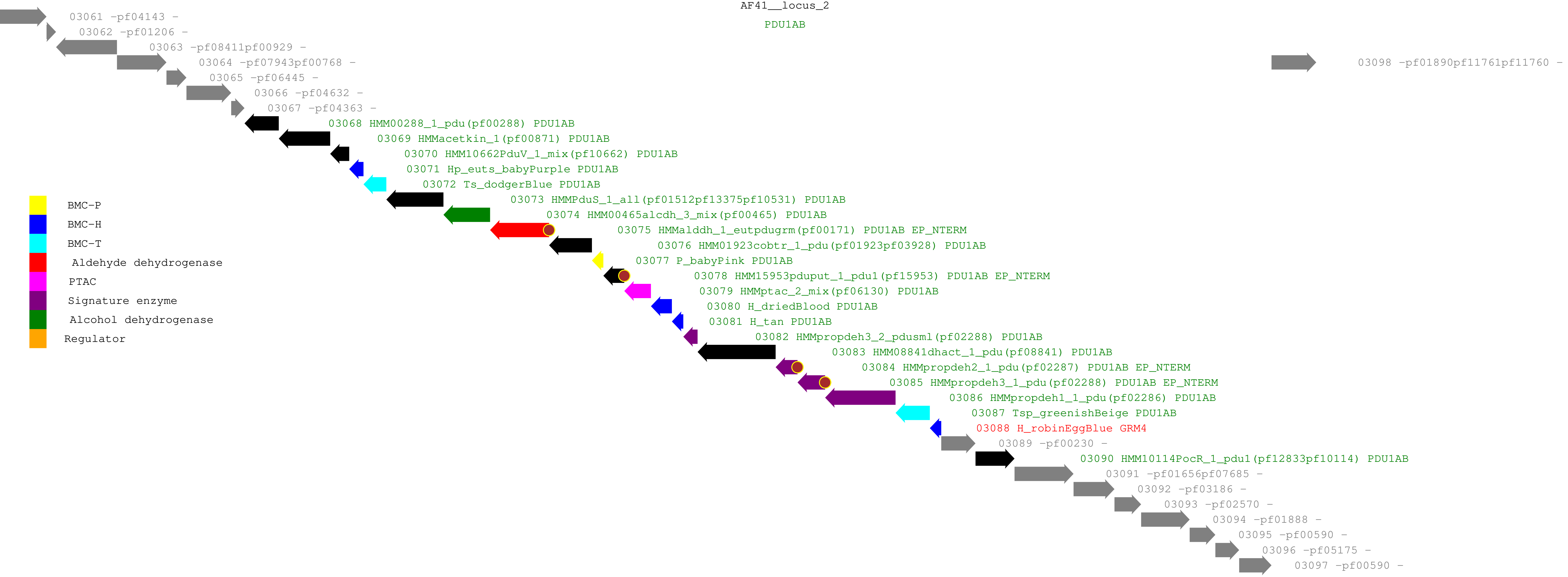

Citrobacter freundii MGH 56. Bacteria; Proteobacteria; Gammaproteobacteria; Enterobacterales;

AF42__locus_1

PDU1AB

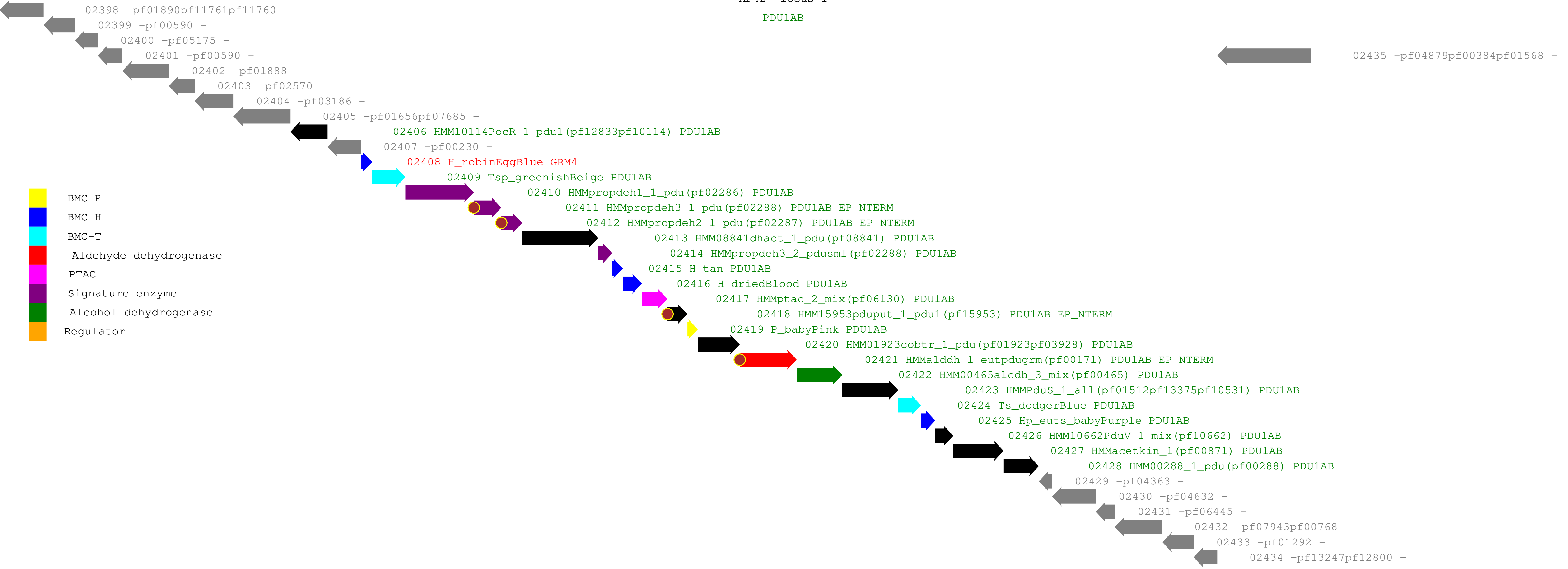

Salmonella typhimurium. Bacteria; Proteobacteria; Gammaproteobacteria; Enterobacterales;

AF480__locus_1

PDU1AB

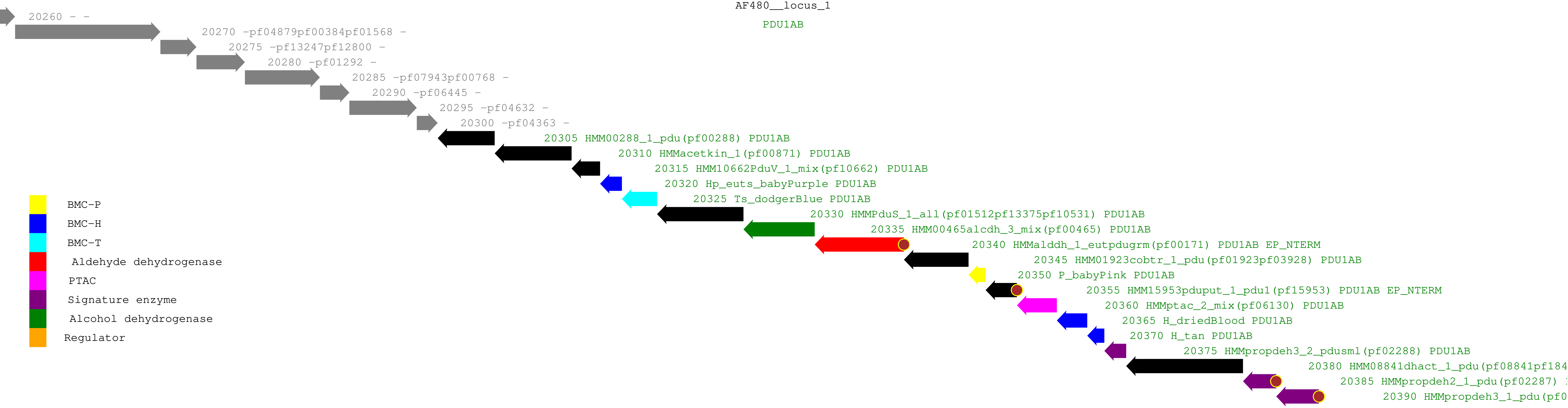

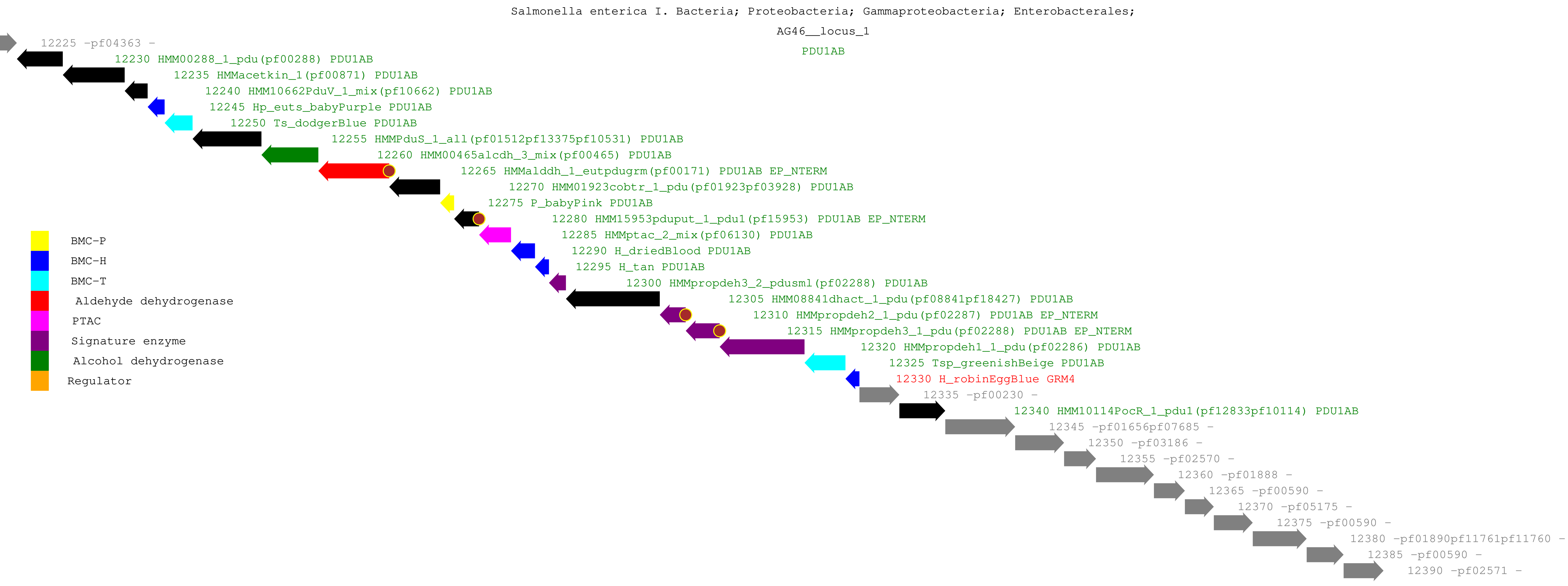

Salmonella enterica I. Bacteria; Proteobacteria; Gammaproteobacteria; Enterobacterales;
AG46__locus_1
PDU1AB
12225 -pf04363 -
12230 HMM00288_1_pdu(pf00288) PDU1AB
12235 HMMacetkin_1(pf00871) PDU1AB
12240 HMM10662PduV_1_mix(pf10662) PDU1AB
12245 Hp_euts_babyPurple PDU1AB
12250 Ts_dodgerBlue PDU1AB
12255 HMMPduS_1_all(pf01512pf13375pf10531) PDU1AB
12260 HMM00465alcdh_3_mix(pf00465) PDU1AB
12265 HMMalddh_1_eutpdugrm(pf00171) PDU1AB EP_NTERM
12270 HMM01923cobtr_1_pdu(pf01923pf03928) PDU1AB
12275 P_babyPink PDU1AB
12280 HMM15953pduput_1_pdu1(pf15953) PDU1AB EP_NTERM
12285 HMMptac_2_mix(pf06130) PDU1AB
12290 H_driedBlood PDU1AB
12295 H_tan PDU1AB
12300 HMMpropdeh3_2_pdusml(pf02288) PDU1AB
12305 HMM08841dhact_1_pdu(pf08841pf18427) PDU1AB
12310 HMMpropdeh2_1_pdu(pf02287) PDU1AB EP_NTERM
12315 HMMpropdeh3_1_pdu(pf02288) PDU1AB EP_NTERM
12320 HMMpropdeh1_1_pdu(pf02286) PDU1AB
12325 Tsp_greenishBeige PDU1AB
12330 H_robinEggBlue GRM4
12335 -pf00230 -
12340 HMM10114PocR_1_pdu1(pf12833pf10114) PDU1AB
12345 -pf01656pf07685 -
12350 -pf03186 -
12355 -pf02570 -
12360 -pf01888 -
12365 -pf00590 -
12370 -pf05175 -
12375 -pf00590 -
12380 -pf01890pf11761pf11760 -
12385 -pf00590 -
12390 -pf02571 -
BMC-P
BMC-H
BMC-T
Aldehyde dehydrogenase
PTAC
Signature enzyme
Alcohol dehydrogenase
Regulator

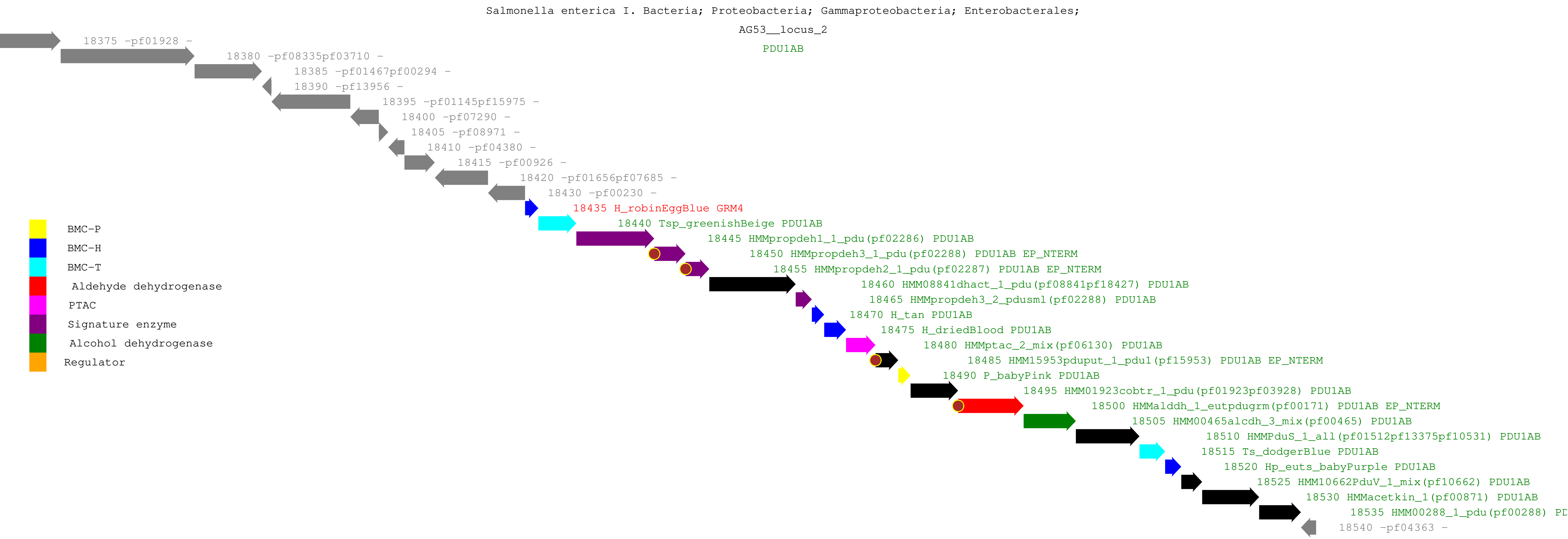
Salmonella enterica I. Bacteria; Proteobacteria; Gammaproteobacteria; Enterobacterales;
AG53__locus_2
PDU1AB
18375 -pf01928 -
18380 -pf08335pf03710 -
18385 -pf01467pf00294 -
18390 -pf13956 -
18395 -pf01145pf15975 -
18400 -pf07290 -
18405 -pf08971 -
18410 -pf04380 -
18415 -pf00926 -
18420 -pf01656pf07685 -
18430 -pf00230 -
18435 H_robinEggBlue GRM4
18440 Tsp_greenishBeige PDU1AB
18445 HMMpropdeh1_1_pdu(pf02286) PDU1AB
18450 HMMpropdeh3_1_pdu(pf02288) PDU1AB EP_NTERM
18455 HMMpropdeh2_1_pdu(pf02287) PDU1AB EP_NTERM
18460 HMM08841dhact_1_pdu(pf08841pf18427) PDU1AB
18465 HMMpropdeh3_2_pdusml(pf02288) PDU1AB
18470 H_tan PDU1AB
18475 H_driedBlood PDU1AB
18480 HMMptac_2_mix(pf06130) PDU1AB
18485 HMM15953pduput_1_pdu1(pf15953) PDU1AB EP_NTERM
18490 P_babyPink PDU1AB
18495 HMM01923cobtr_1_pdu(pf01923pf03928) PDU1AB
18500 HMMaldh_1_eutpdugrm(pf00171) PDU1AB EP_NTERM
18505 HMM00465alcdh_3_mix(pf00465) PDU1AB
18510 HMMPduS_1_all(pf01512pf13375pf10531) PDU1AB
18515 Ts_dodgerBlue PDU1AB
18520 Hp_euts_babyPurple PDU1AB
18525 HMM10662PduV_1_mix(pf10662) PDU1AB
18530 HMMacetkin_1(pf00871) PDU1AB
18535 HMM00288_1_pdu(pf00288) PD
18540 -pf04363 -
BMC-P
BMC-H
BMC-T
Aldehyde dehydrogenase
PTAC
Signature enzyme
Alcohol dehydrogenase
Regulator

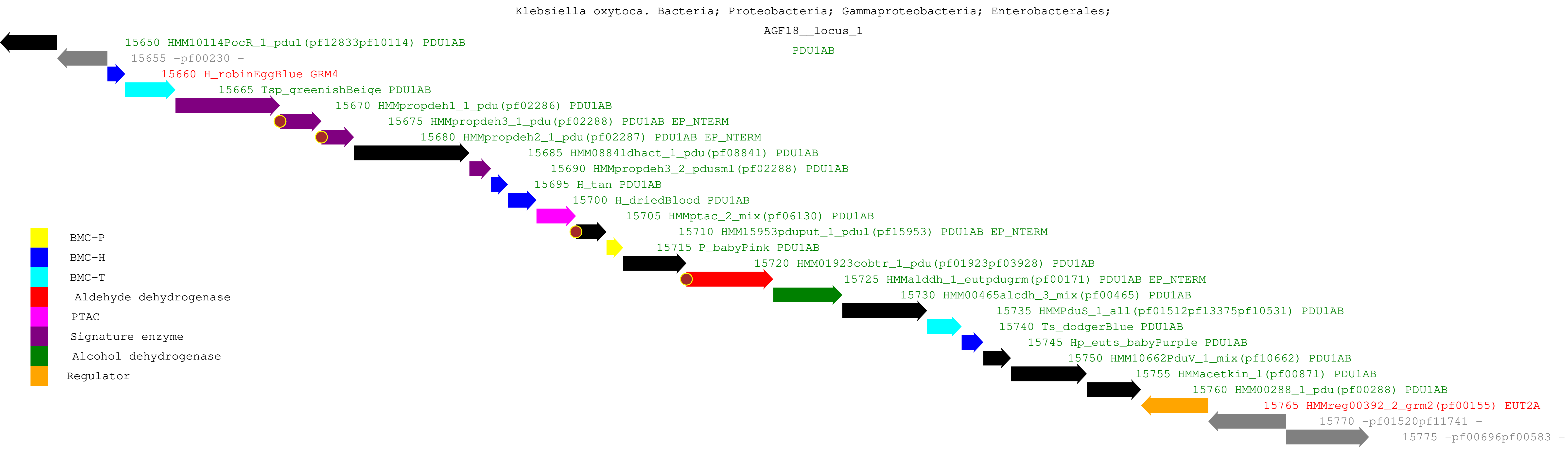
Klebsiella oxytoca. Bacteria; Proteobacteria; Gammaproteobacteria; Enterobacterales;
AGF18__locus_1
PDU1AB
15650 HMM10114PocR_1_pdu1(pf12833pf10114) PDU1AB
15655 -pf00230 -
15660 H_robinEggBlue GRM4
15665 Tsp_greenishBeige PDU1AB
15670 HMMpropdeh1_1_pdu(pf02286) PDU1AB
15675 HMMpropdeh3_1_pdu(pf02288) PDU1AB EP_NTERM
15680 HMMpropdeh2_1_pdu(pf02287) PDU1AB EP_NTERM
15685 HMM08841dhact_1_pdu(pf08841) PDU1AB
15690 HMMpropdeh3_2_pdusml(pf02288) PDU1AB
15695 H_tan PDU1AB
15700 H_driedBlood PDU1AB
15705 HMMptac_2_mix(pf06130) PDU1AB
15710 HMM15953pduput_1_pdu1(pf15953) PDU1AB EP_NTERM
15715 P_babyPink PDU1AB
15720 HMM01923cobtr_1_pdu(pf01923pf03928) PDU1AB
15725 HMMalddh_1_eutpdugrm(pf00171) PDU1AB EP_NTERM
15730 HMM00465alcdh_3_mix(pf00465) PDU1AB
15735 HMMPduS_1_all(pf01512pf13375pf10531) PDU1AB
15740 Ts_dodgerBlue PDU1AB
15745 Hp_euts_babyPurple PDU1AB
15750 HMM10662PduV_1_mix(pf10662) PDU1AB
15755 HMMacetkin_1(pf00871) PDU1AB
15760 HMM00288_1_pdu(pf00288) PDU1AB
15765 HMMreg00392_2_grm2(pf00155) EUT2A
15770 -pf01520pf11741 -
15775 -pf00696pf00583 -
BMC-P
BMC-H
BMC-T
Aldehyde dehydrogenase
PTAC
Signature enzyme
Alcohol dehydrogenase
Regulator

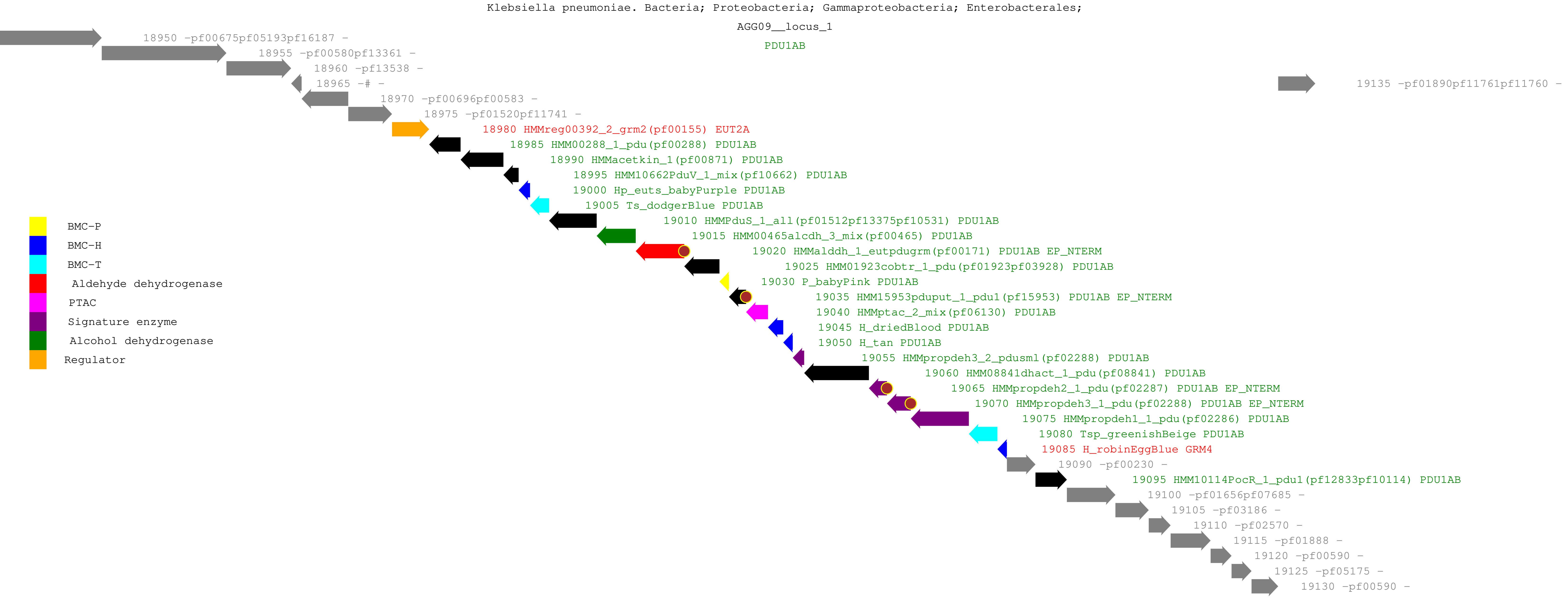
Klebsiella pneumoniae. Bacteria; Proteobacteria; Gammaproteobacteria; Enterobacterales;
AGG09__locus_1
PDU1AB
18950 -pf00675pf05193pf16187 -
18955 -pf00580pf13361 -
18960 -pf13538 -
18965 -# -
18970 -pf00696pf00583 -
18975 -pf01520pf11741 -
18980 HMMreg00392_2_grm2(pf00155) EUT2A
18985 HMM00288_1_pdu(pf00288) PDU1AB
18990 HMMacetkin_1(pf00871) PDU1AB
18995 HMM10662PduV_1_mix(pf10662) PDU1AB
19000 Hp_euts_babyPurple PDU1AB
19005 Ts_dodgerBlue PDU1AB
19010 HMMPduS_1_all(pf01512pf13375pf10531) PDU1AB
19015 HMM00465alcdh_3_mix(pf00465) PDU1AB
19020 HMMaldh_1_eutpdugrm(pf00171) PDU1AB EP_NTERM
19025 HMM01923cobtr_1_pdu(pf01923pf03928) PDU1AB
19030 P_babyPink PDU1AB
19035 HMM15953pduput_1_pdu1(pf15953) PDU1AB EP_NTERM
19040 HMMptac_2_mix(pf06130) PDU1AB
19045 H_driedBlood PDU1AB
19050 H_tan PDU1AB
19055 HMMpropdeh3_2_pdusml(pf02288) PDU1AB
19060 HMM08841dhact_1_pdu(pf08841) PDU1AB
19065 HMMpropdeh2_1_pdu(pf02287) PDU1AB EP_NTERM
19070 HMMpropdeh3_1_pdu(pf02288) PDU1AB EP_NTERM
19075 HMMpropdeh1_1_pdu(pf02286) PDU1AB
19080 Tsp_greenishBeige PDU1AB
19085 H_robinEggBlue GRM4
19090 -pf00230 -
19095 HMM10114PocR_1_pdu1(pf12833pf10114) PDU1AB
19100 -pf01656pf07685 -
19105 -pf03186 -
19110 -pf02570 -
19115 -pf01888 -
19120 -pf00590 -
19125 -pf05175 -
19130 -pf00590 -
19135 -pf01890pf11761pf11760 -
BMC-P
BMC-H
BMC-T
Aldehyde dehydrogenase
PTAC
Signature enzyme
Alcohol dehydrogenase
Regulator

Klebsiella michiganensis. Bacteria; Proteobacteria; Gammaproteobacteria; Enterobacterales;

AGG97__locus_3

PDU1AB

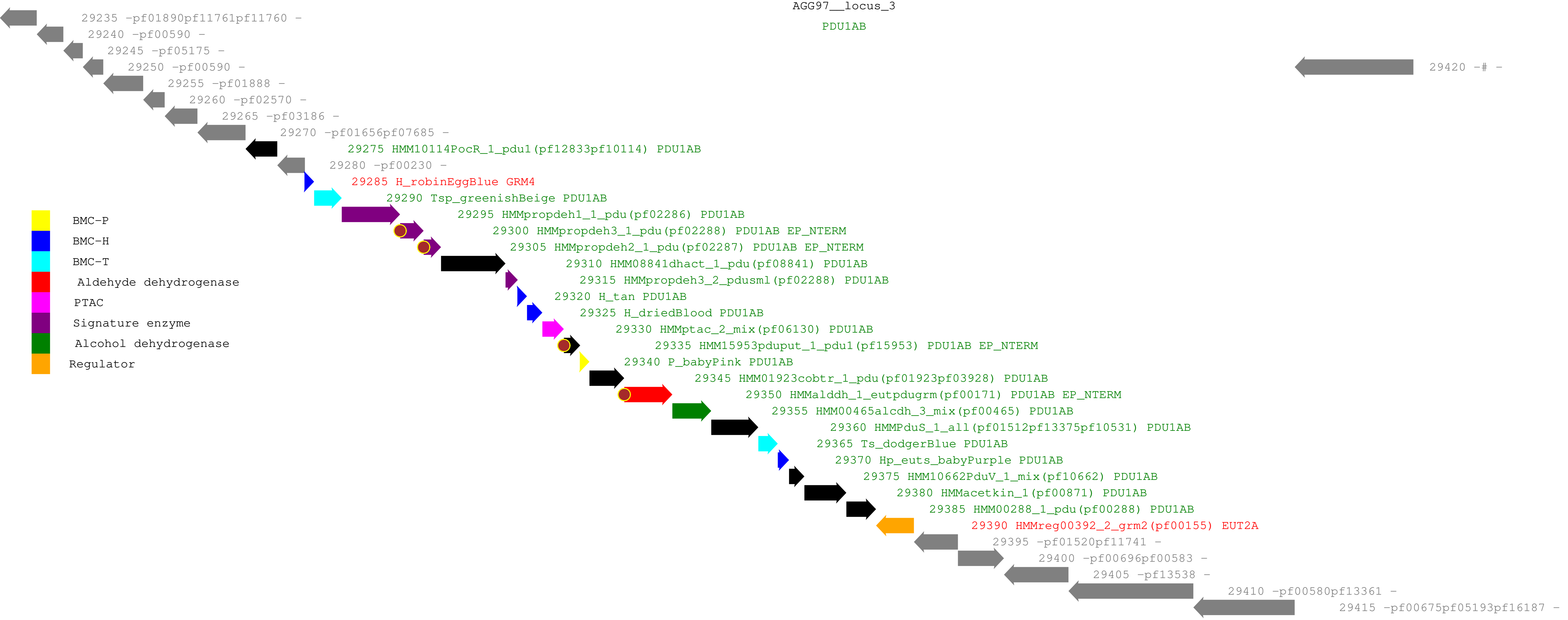

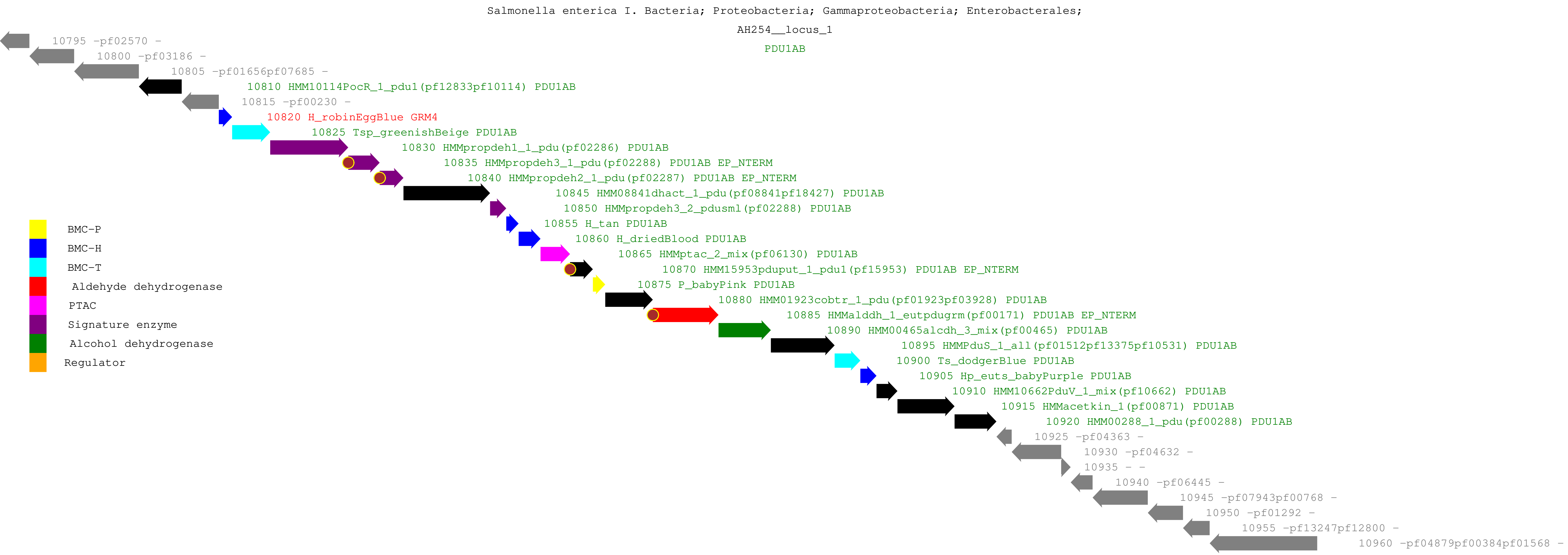
Salmonella enterica I. Bacteria; Proteobacteria; Gammaproteobacteria; Enterobacterales;
AH254__locus_1
PDU1AB
10795 -pf02570 -
10800 -pf03186 -
10805 -pf01656pf07685 -
10810 HMM10114PocR_1_pdu1(pf12833pf10114) PDU1AB
10815 -pf00230 -
10820 H_robinEggBlue GRM4
10825 Tsp_greenishBeige PDU1AB
10830 HMMpropdeh1_1_pdu(pf02286) PDU1AB
10835 HMMpropdeh3_1_pdu(pf02288) PDU1AB EP_NTERM
10840 HMMpropdeh2_1_pdu(pf02287) PDU1AB EP_NTERM
10845 HMM08841dhact_1_pdu(pf08841pf18427) PDU1AB
10850 HMMpropdeh3_2_pdusml(pf02288) PDU1AB
10855 H_tan PDU1AB
10860 H_driedBlood PDU1AB
10865 HMMptac_2_mix(pf06130) PDU1AB
10870 HMM15953pduput_1_pdu1(pf15953) PDU1AB EP_NTERM
10875 P_babyPink PDU1AB
10880 HMM01923cobtr_1_pdu(pf01923pf03928) PDU1AB
10885 HMMalddh_1_eutpdugrm(pf00171) PDU1AB EP_NTERM
10890 HMM00465alcdh_3_mix(pf00465) PDU1AB
10895 HMMPduS_1_all(pf01512pf13375pf10531) PDU1AB
10900 Ts_dodgerBlue PDU1AB
10905 Hp_euts_babyPurple PDU1AB
10910 HMM10662PduV_1_mix(pf10662) PDU1AB
10915 HMMacetkin_1(pf00871) PDU1AB
10920 HMM00288_1_pdu(pf00288) PDU1AB
10925 -pf04363 -
10930 -pf04632 -
10935 - -
10940 -pf06445 -
10945 -pf07943pf00768 -
10950 -pf01292 -
10955 -pf13247pf12800 -
10960 -pf04879pf00384pf01568 -
BMC-P
BMC-H
BMC-T
Aldehyde dehydrogenase
PTAC
Signature enzyme
Alcohol dehydrogenase
Regulator

Salmonella enterica subsp. enterica serovar Kentucky. Bacteria; Proteobacteria; Gammaproteobacteria; Enterobacterales;

AH329__locus_1

PDU1AB

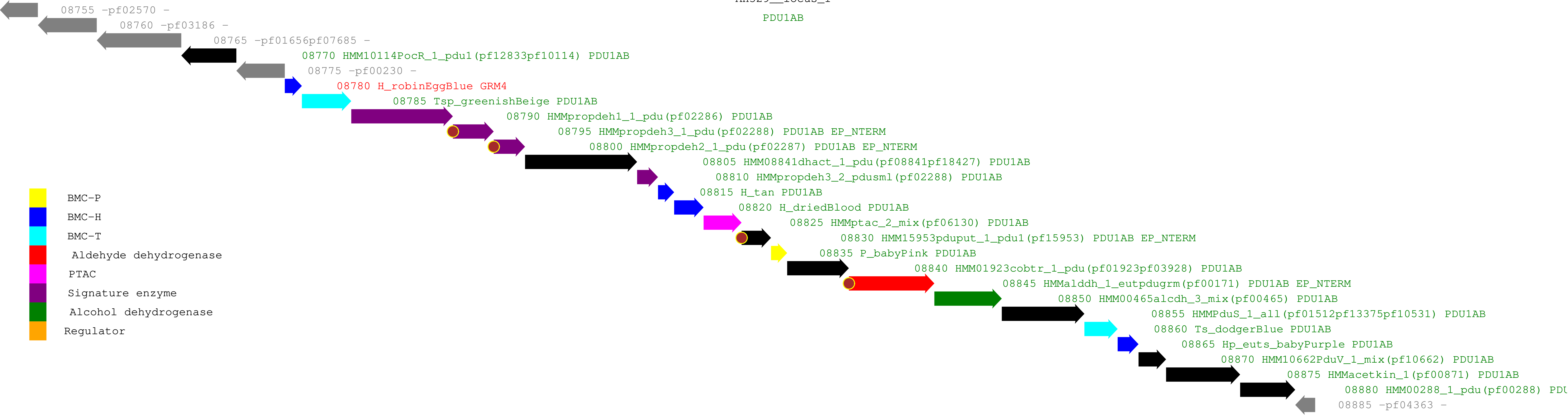

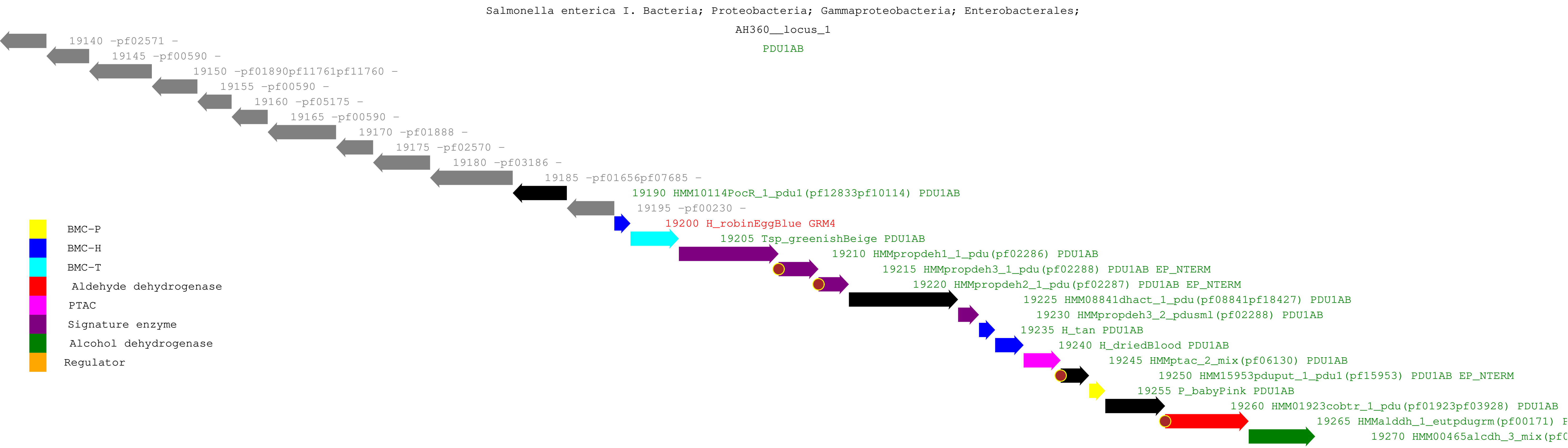
Salmonella enterica I. Bacteria; Proteobacteria; Gammaproteobacteria; Enterobacterales;
AH360__locus_1
PDU1AB
19140 -pf02571 -
19145 -pf00590 -
19150 -pf01890pf11761pf11760 -
19155 -pf00590 -
19160 -pf05175 -
19165 -pf00590 -
19170 -pf01888 -
19175 -pf02570 -
19180 -pf03186 -
19185 -pf01656pf07685 -
19190 HMM10114PocR_1_pdu1(pf12833pf10114) PDU1AB
19195 -pf00230 -
19200 H_robinEggBlue GRM4
19205 Tsp_greenishBeige PDU1AB
19210 HMMpropdeh1_1_pdu(pf02286) PDU1AB
19215 HMMpropdeh3_1_pdu(pf02288) PDU1AB EP_NTERM
19220 HMMpropdeh2_1_pdu(pf02287) PDU1AB EP_NTERM
19225 HMM08841dhact_1_pdu(pf08841pf18427) PDU1AB
19230 HMMpropdeh3_2_pdusml(pf02288) PDU1AB
19235 H_tan PDU1AB
19240 H_driedBlood PDU1AB
19245 HMMptac_2_mix(pf06130) PDU1AB
19250 HMM15953pduput_1_pdu1(pf15953) PDU1AB EP_NTERM
19255 P_babyPink PDU1AB
19260 HMM01923cobtr_1_pdu(pf01923pf03928) PDU1AB
19265 HMMalddh_1_eutpdugrm(pf00171)
19270 HMM00465alcdh_3_mix(pf0
BMC-P
BMC-H
BMC-T
Aldehyde dehydrogenase
PTAC
Signature enzyme
Alcohol dehydrogenase
Regulator

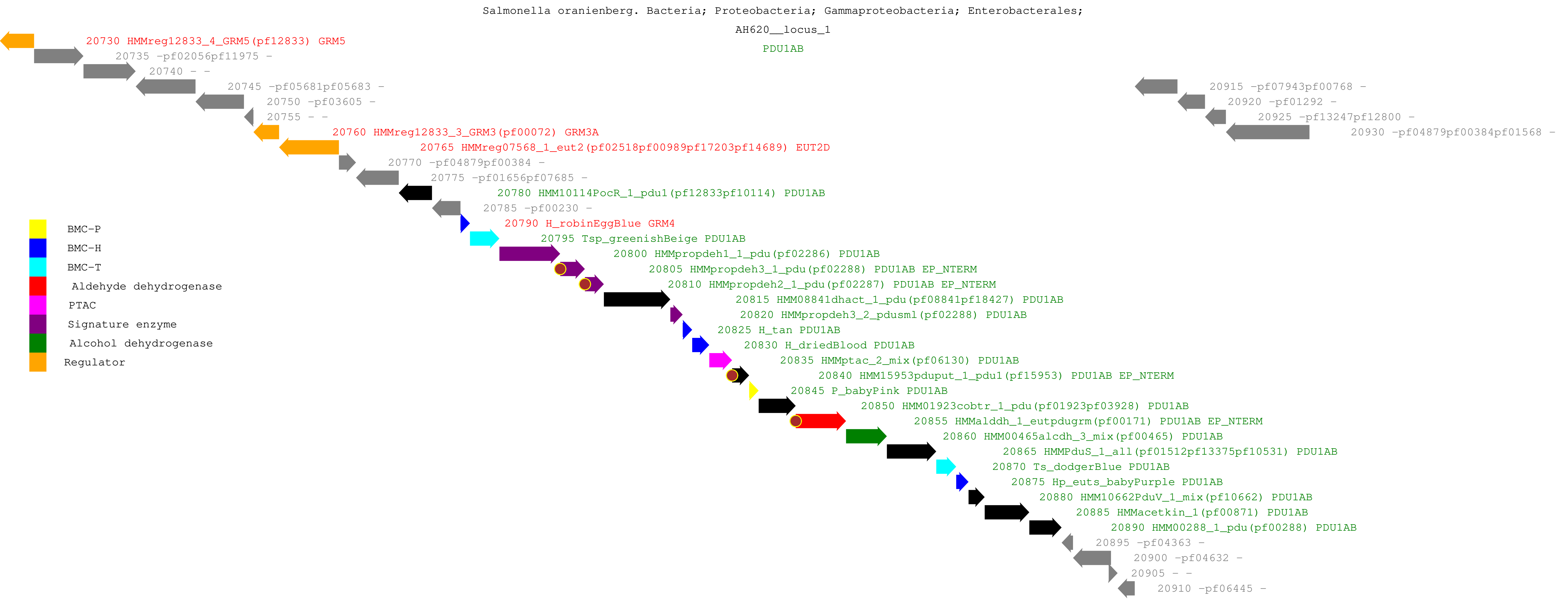
Salmonella oranienberg. Bacteria; Proteobacteria; Gammaproteobacteria; Enterobacterales;
AH620__locus_1
PDU1AB
20730 HMMreg12833_4_GRM5(pf12833) GRM5
20735 -pf02056pf11975 -
20740 - -
20745 -pf05681pf05683 -
20750 -pf03605 -
20755 - -
20760 HMMreg12833_3_GRM3(pf00072) GRM3A
20765 HMMreg07568_1_eut2(pf02518pf00989pf17203pf14689) EUT2D
20770 -pf04879pf00384 -
20775 -pf01656pf07685 -
20780 HMM10114PocR_1_pdu1(pf12833pf10114) PDU1AB
20785 -pf00230 -
20790 H_robinEggBlue GRM4
20795 Tsp_greenishBeige PDU1AB
20800 HMMpropdeh1_1_pdu(pf02286) PDU1AB
20805 HMMpropdeh3_1_pdu(pf02288) PDU1AB EP_NTERM
20810 HMMpropdeh2_1_pdu(pf02287) PDU1AB EP_NTERM
20815 HMM08841dhact_1_pdu(pf08841pf18427) PDU1AB
20820 HMMpropdeh3_2_pdusml(pf02288) PDU1AB
20825 H_tan PDU1AB
20830 H_driedBlood PDU1AB
20835 HMMptac_2_mix(pf06130) PDU1AB
20840 HMM15953pduput_1_pdu1(pf15953) PDU1AB EP_NTERM
20845 P_babyPink PDU1AB
20850 HMM01923cobtr_1_pdu(pf01923pf03928) PDU1AB
20855 HMMalddh_1_eutpdugrm(pf00171) PDU1AB EP_NTERM
20860 HMM00465alcdh_3_mix(pf00465) PDU1AB
20865 HMMPduS_1_all(pf01512pf13375pf10531) PDU1AB
20870 Ts_dodgerBlue PDU1AB
20875 Hp_euts_babyPurple PDU1AB
20880 HMM10662PduV_1_mix(pf10662) PDU1AB
20885 HMMacetkin_1(pf00871) PDU1AB
20890 HMM00288_1_pdu(pf00288) PDU1AB
20895 -pf04363 -
20900 -pf04632 -
20905 - -
20910 -pf06445 -
20915 -pf07943pf00768 -
20920 -pf01292 -
20925 -pf13247pf12800 -
20930 -pf04879pf00384pf01568 -
BMC-P
BMC-H
BMC-T
Aldehyde dehydrogenase
PTAC
Signature enzyme
Alcohol dehydrogenase
Regulator

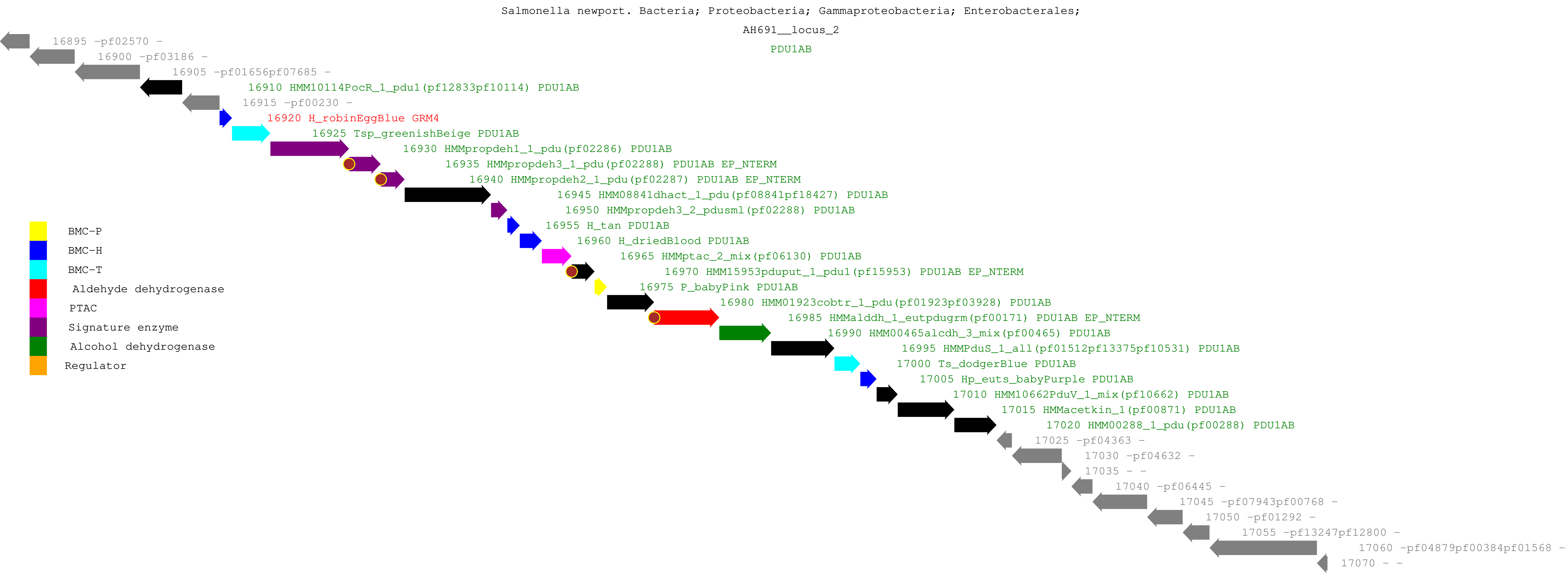

Salmonella newport. Bacteria; Proteobacteria; Gammaproteobacteria; Enterobacterales;
AH691__locus_2
PDU1AB
16895 -pf02570 -
16900 -pf03186 -
16905 -pf01656pf07685 -
16910 HMM10114PocR_1_pdu1(pf12833pf10114) PDU1AB
16915 -pf00230 -
16920 H_robinEggBlue GRM4
16925 Tsp_greenishBeige PDU1AB
16930 HMMpropdeh1_1_pdu(pf02286) PDU1AB
16935 HMMpropdeh3_1_pdu(pf02288) PDU1AB EP_NTERM
16940 HMMpropdeh2_1_pdu(pf02287) PDU1AB EP_NTERM
16945 HMM08841dhact_1_pdu(pf08841pf18427) PDU1AB
16950 HMMpropdeh3_2_pdusml(pf02288) PDU1AB
16955 H_tan PDU1AB
16960 H_driedBlood PDU1AB
16965 HMMptac_2_mix(pf06130) PDU1AB
16970 HMM15953pduput_1_pdu1(pf15953) PDU1AB EP_NTERM
16975 P_babyPink PDU1AB
16980 HMM01923cobtr_1_pdu(pf01923pf03928) PDU1AB
16985 HMMalddh_1_eutpdugrm(pf00171) PDU1AB EP_NTERM
16990 HMM00465alcdh_3_mix(pf00465) PDU1AB
16995 HMMPduS_1_all(pf01512pf13375pf10531) PDU1AB
17000 Ts_dodgerBlue PDU1AB
17005 Hp_euts_babyPurple PDU1AB
17010 HMM10662PduV_1_mix(pf10662) PDU1AB
17015 HMMacetkin_1(pf00871) PDU1AB
17020 HMM00288_1_pdu(pf00288) PDU1AB
17025 -pf04363 -
17030 -pf04632 -
17035 - -
17040 -pf06445 -
17045 -pf07943pf00768 -
17050 -pf01292 -
17055 -pf13247pf12800 -
17060 -pf04879pf00384pf01568 -
17070 - -
BMC-P
BMC-H
BMC-T
Aldehyde dehydrogenase
PTAC
Signature enzyme
Alcohol dehydrogenase
Regulator

Salmonella enterica I. Bacteria; Proteobacteria; Gammaproteobacteria; Enterobacterales;

AH874__locus_1

PDU1AB

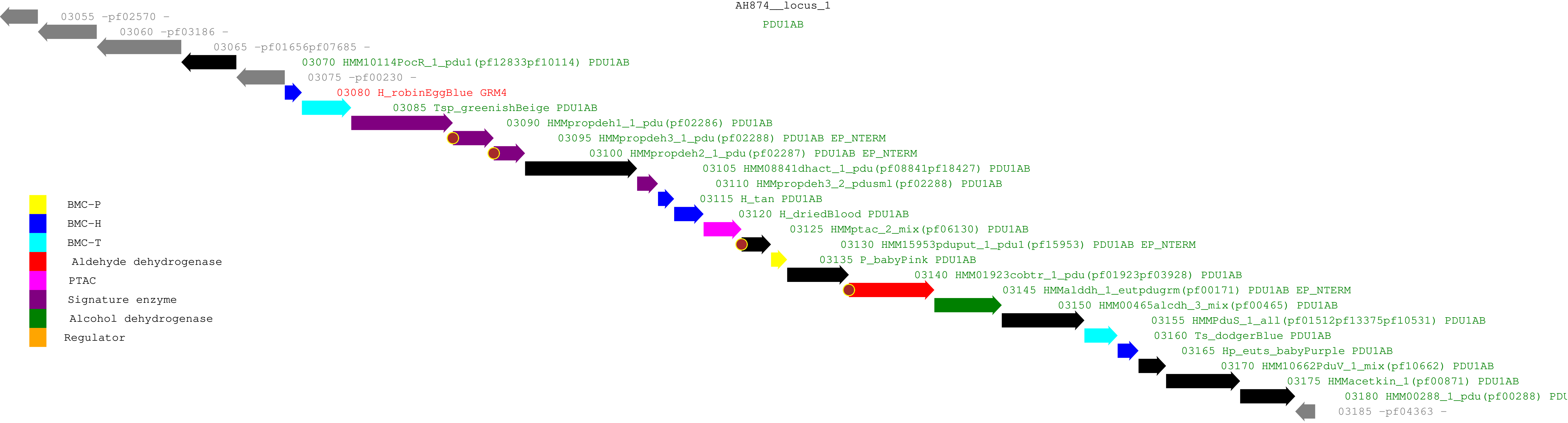

Salmonella enterica subsp. enterica serovar Havana. Bacteria; Proteobacteria; Gammaproteobacteria; Enterobacterales;

AH902__locus_2

PDU1AB

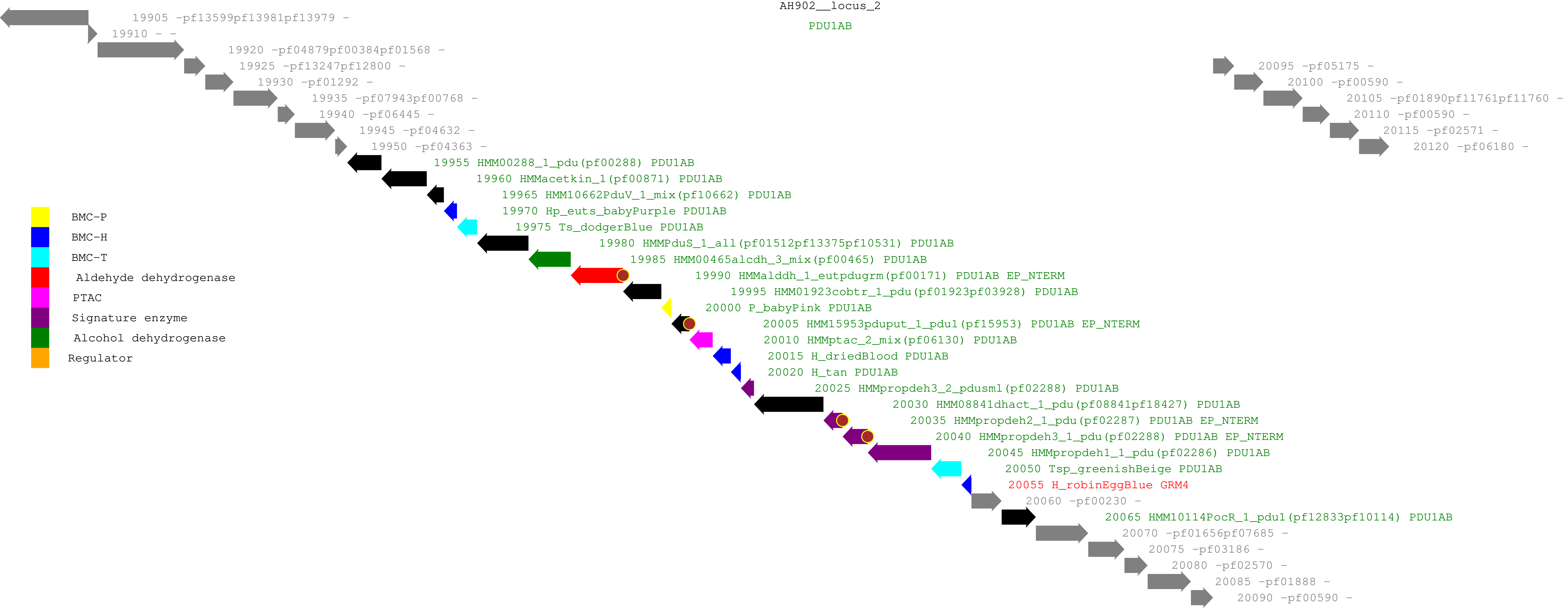

Salmonella enterica subsp. enterica serovar Paratyphi B. Bacteria; Proteobacteria; Gammaproteobacteria; Enterobacterales;

AHS25__locus_1

PDU1AB

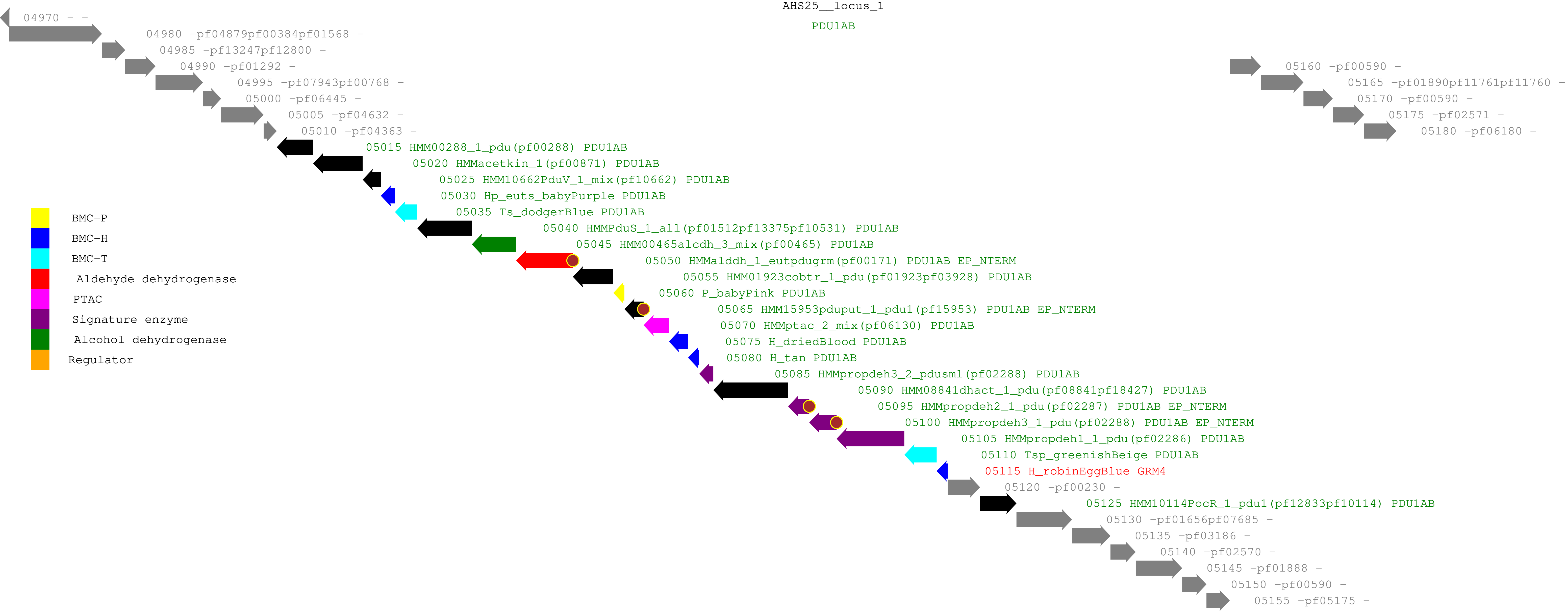

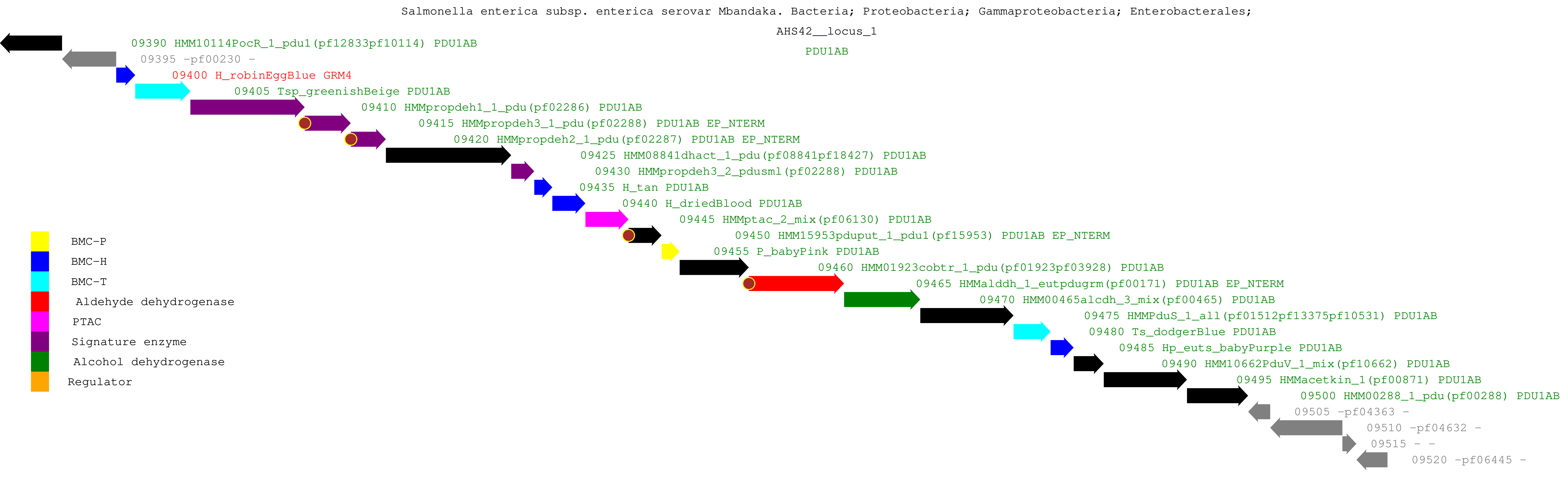
Salmonella enterica subsp. enterica serovar Mbandaka. Bacteria; Proteobacteria; Gammaproteobacteria; Enterobacterales;
AHS42__locus_1
PDU1AB
09390 HMM10114PocR_1_pdu1(pf12833pf10114) PDU1AB
09395 -pf00230 -
09400 H_robinEggBlue GRM4
09405 Tsp_greenishBeige PDU1AB
09410 HMMpropdeh1_1_pdu(pf02286) PDU1AB
09415 HMMpropdeh3_1_pdu(pf02288) PDU1AB EP_NTERM
09420 HMMpropdeh2_1_pdu(pf02287) PDU1AB EP_NTERM
09425 HMM08841dhact_1_pdu(pf08841pf18427) PDU1AB
09430 HMMpropdeh3_2_pdusml(pf02288) PDU1AB
09435 H_tan PDU1AB
09440 H_driedBlood PDU1AB
09445 HMMptac_2_mix(pf06130) PDU1AB
09450 HMM15953pduput_1_pdu1(pf15953) PDU1AB EP_NTERM
09455 P_babyPink PDU1AB
09460 HMM01923cobtr_1_pdu(pf01923pf03928) PDU1AB
09465 HMMalddh_1_eutpdugrm(pf00171) PDU1AB EP_NTERM
09470 HMM00465alcdh_3_mix(pf00465) PDU1AB
09475 HMMPduS_1_all(pf01512pf13375pf10531) PDU1AB
09480 Ts_dodgerBlue PDU1AB
09485 Hp_euts_babyPurple PDU1AB
09490 HMM10662PduV_1_mix(pf10662) PDU1AB
09495 HMMacetkin_1(pf00871) PDU1AB
09500 HMM00288_1_pdu(pf00288) PDU1AB
09505 -pf04363 -
09510 -pf04632 -
09515 - -
09520 -pf06445 -
BMC-P
BMC-H
BMC-T
Aldehyde dehydrogenase
PTAC
Signature enzyme
Alcohol dehydrogenase
Regulator

Salmonella enterica I. Bacteria; Proteobacteria; Gammaproteobacteria; Enterobacterales;

AHS94__locus_1

PDU1AB

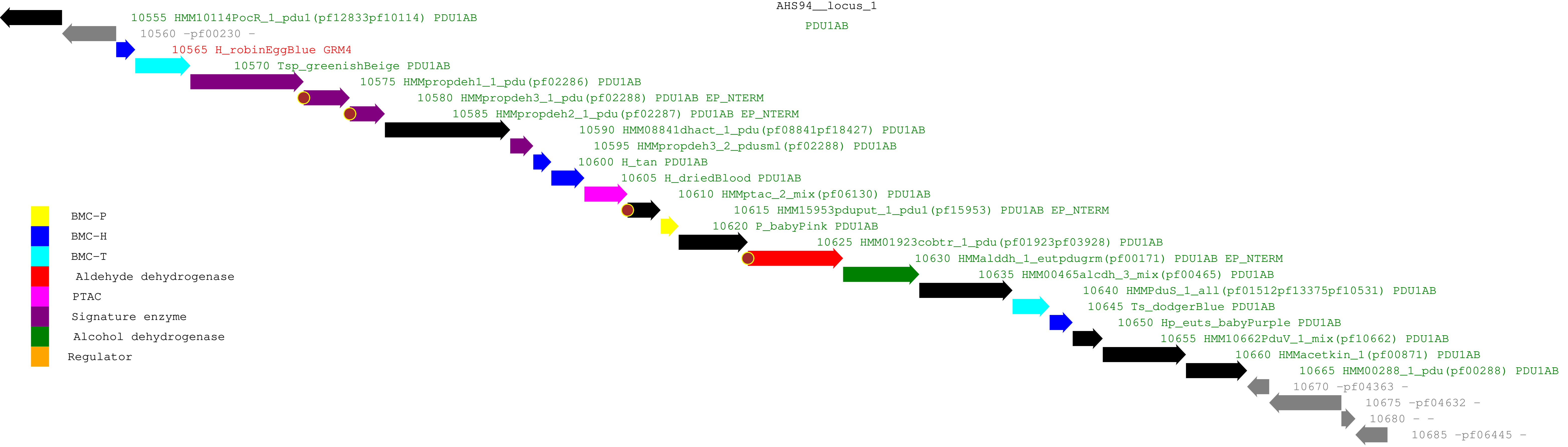

Salmonella oranienberg. Bacteria; Proteobacteria; Gammaproteobacteria; Enterobacterales;

AHW54__locus_1

PDU1AB

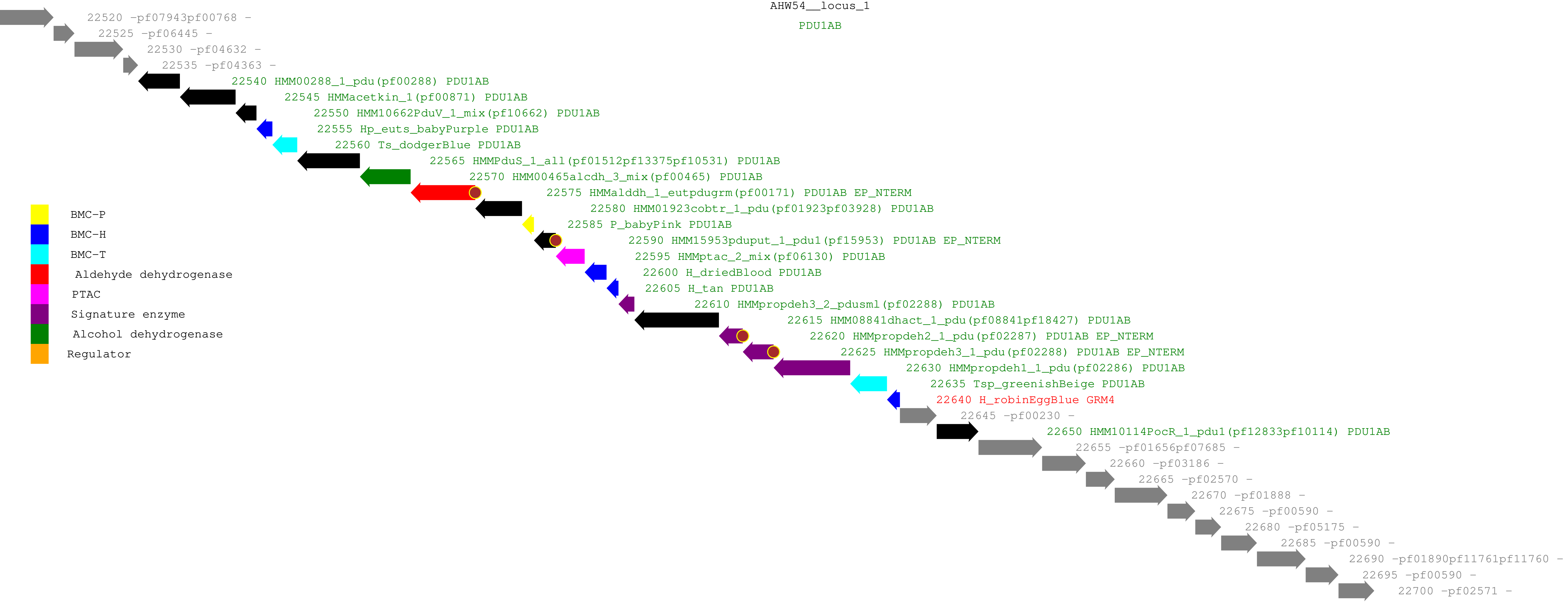

Salmonella enterica I. Bacteria; Proteobacteria; Gammaproteobacteria; Enterobacterales;

AHY46__locus_1

PDU1AB

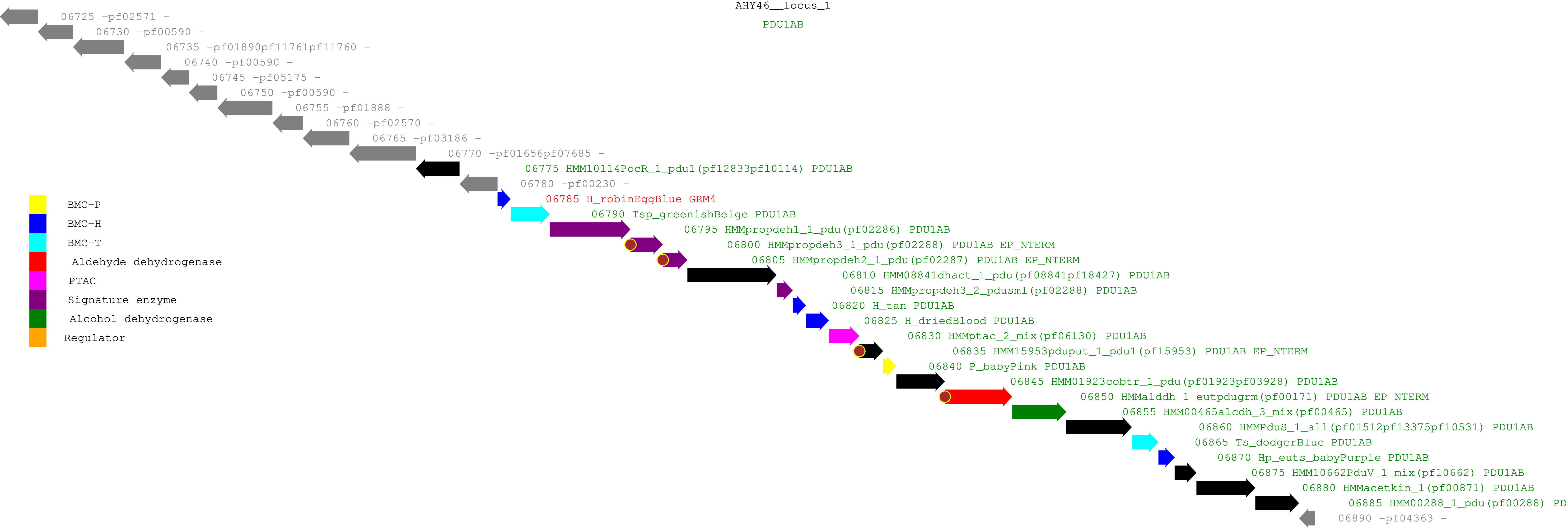

Salmonella enterica I. Bacteria; Proteobacteria; Gammaproteobacteria; Enterobacterales;

AIA22__locus_1

PDU1AB

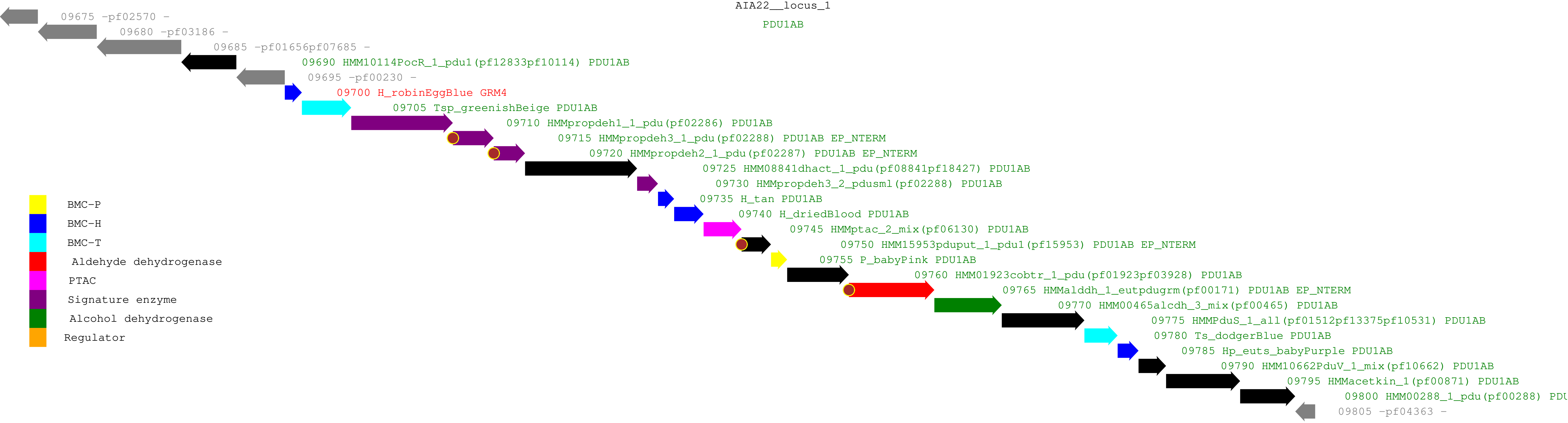

Salmonella enterica I. Bacteria; Proteobacteria; Gammaproteobacteria; Enterobacterales;

AIA94__locus_2

PDU1AB

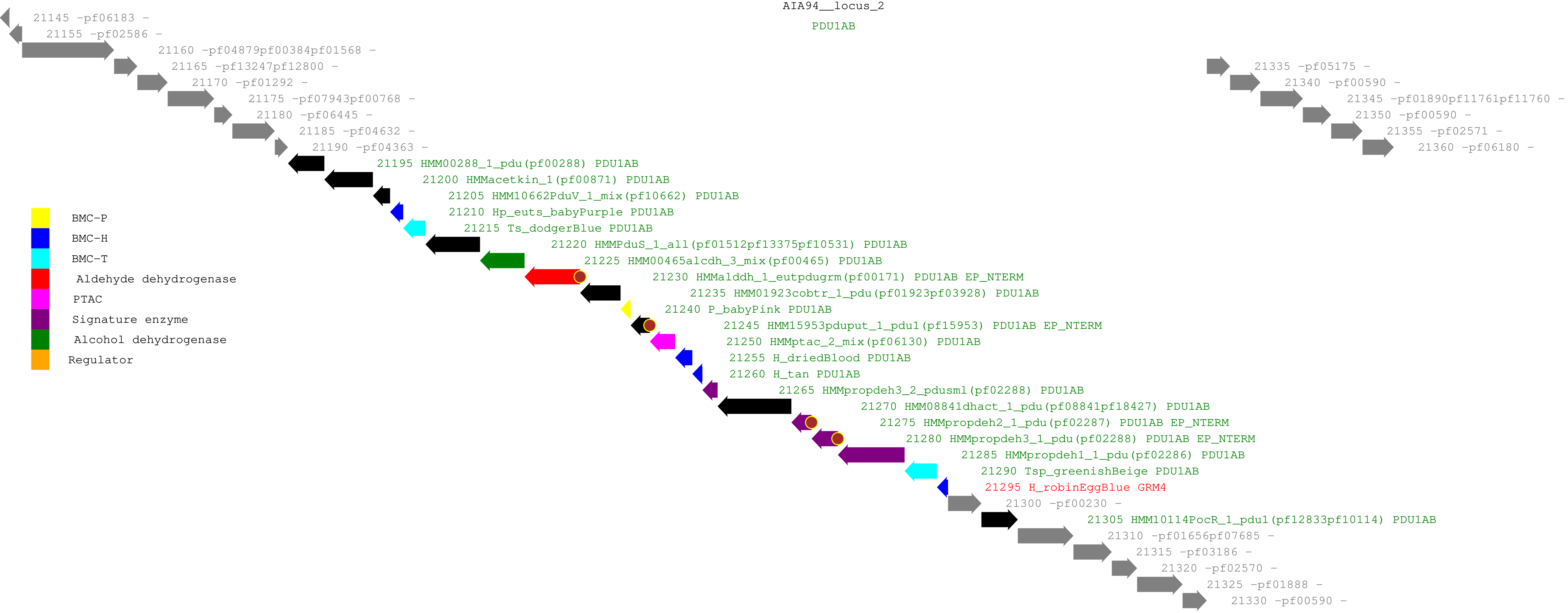

Salmonella enterica (Salmonella choleraesuis). Bacteria; Proteobacteria; Gammaproteobacteria; Enterobacterales;

AIF45__locus_1

PDU1AB

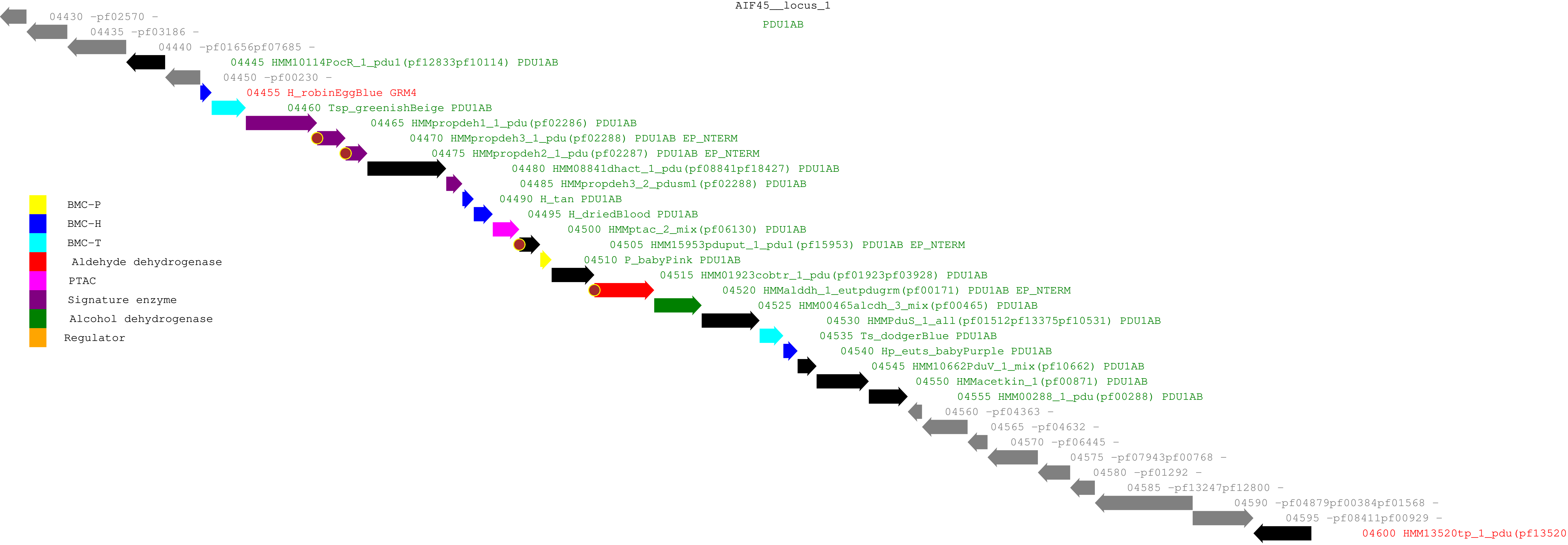

Salmonella thompson. Bacteria; Proteobacteria; Gammaproteobacteria; Enterobacterales;

AIG11__locus_2

PDU1AB

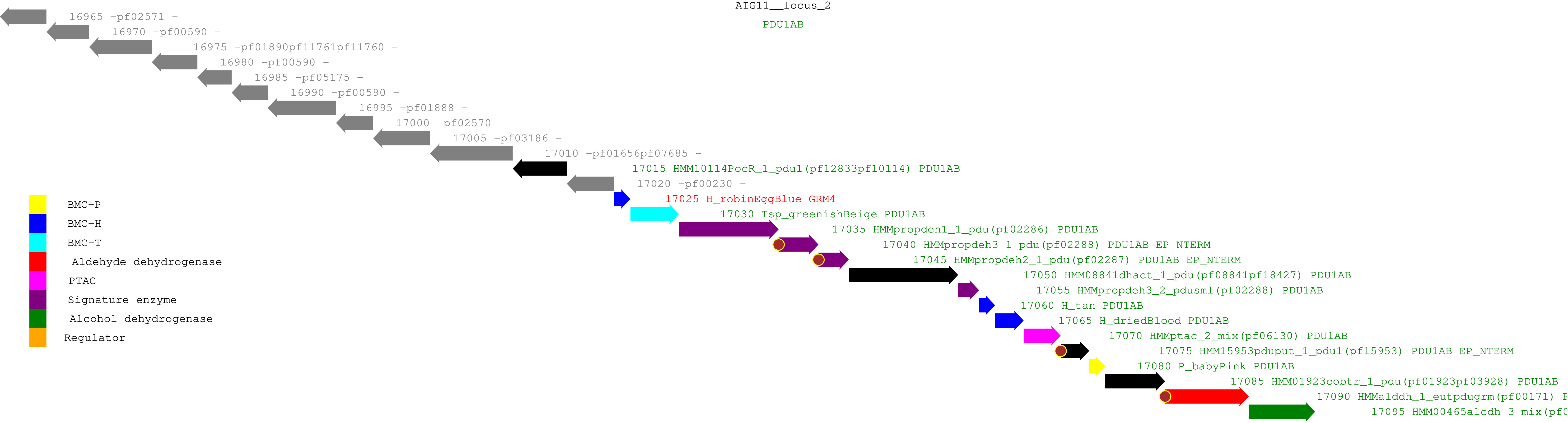

Salmonella enterica subsp. enterica serovar Pomona. Bacteria; Proteobacteria; Gammaproteobacteria; Enterobacterales;

AIG86__locus_2

PDU1AB

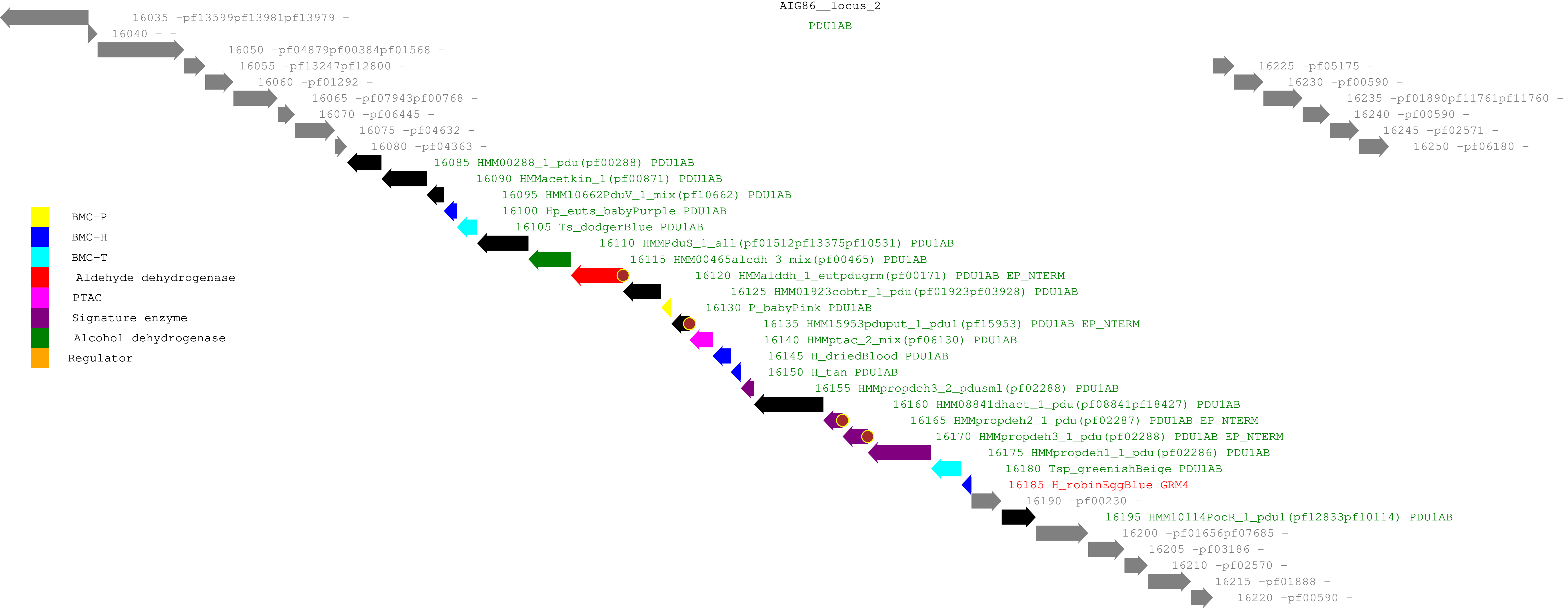

Salmonella enterica subsp. enterica serovar Havana. Bacteria; Proteobacteria; Gammaproteobacteria; Enterobacterales;

AIH56__locus_1

PDU1AB

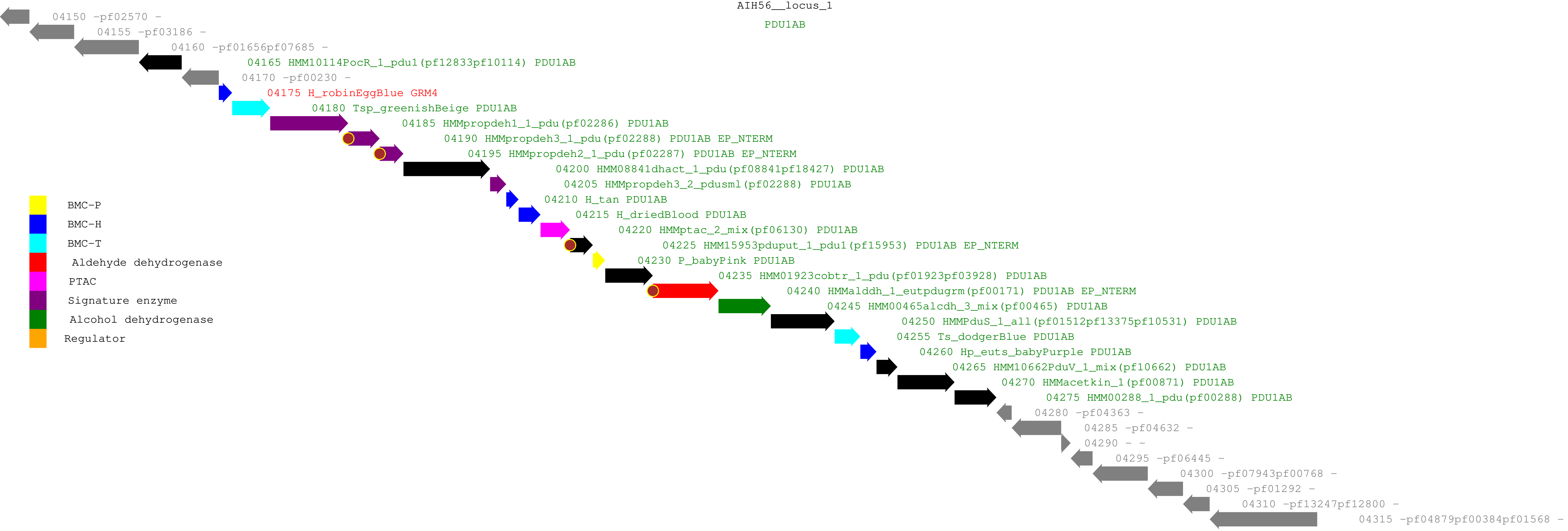

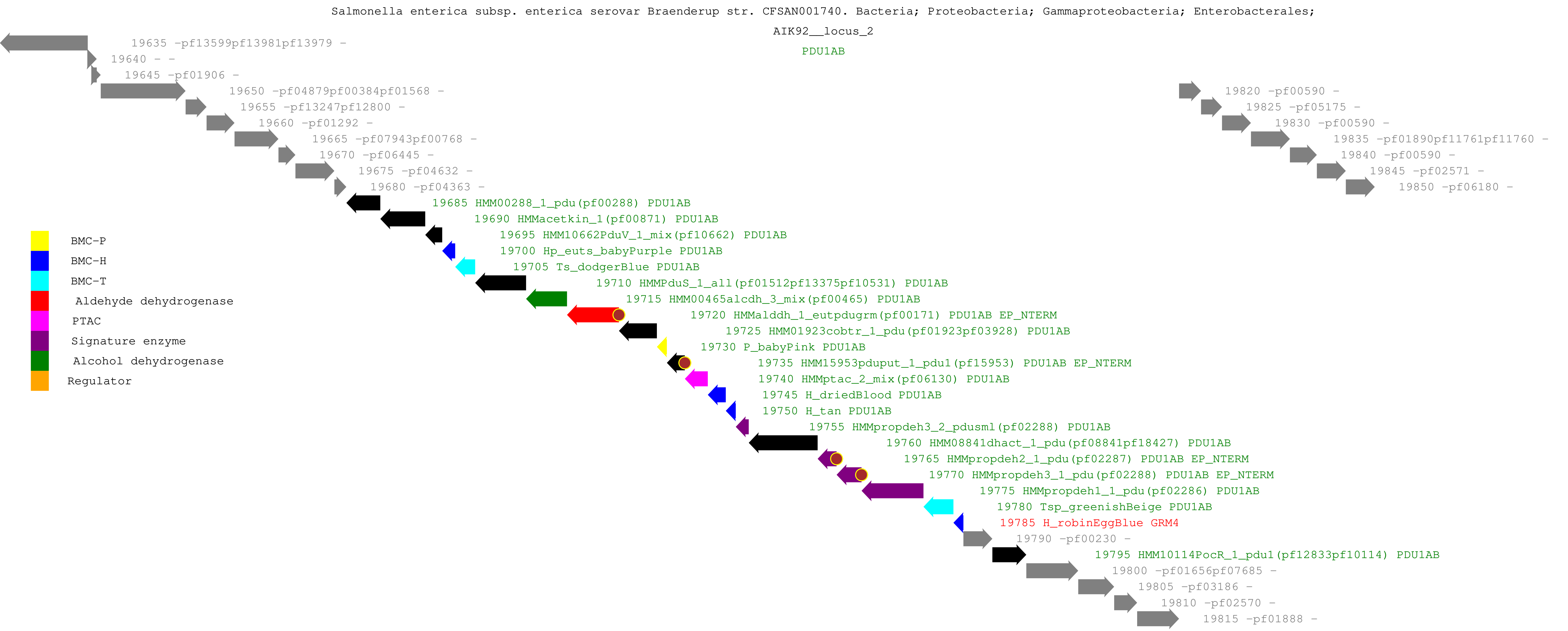
Salmonella enterica subsp. enterica serovar Braenderup str. CFSAN001740. Bacteria; Proteobacteria; Gammaproteobacteria; Enterobacterales;
AIK92__locus_2
PDU1AB
19635 -pf13599pf13981pf13979 -
19640 - -
19645 -pf01906 -
19650 -pf04879pf00384pf01568 -
19655 -pf13247pf12800 -
19660 -pf01292 -
19665 -pf07943pf00768 -
19670 -pf06445 -
19675 -pf04632 -
19680 -pf04363 -
19685 HMM00288_1_pdu(pf00288) PDU1AB
19690 HMMacetkin_1(pf00871) PDU1AB
19695 HMM10662PduV_1_mix(pf10662) PDU1AB
19700 Hp_euts_babyPurple PDU1AB
19705 Ts_dodgerBlue PDU1AB
19710 HMMPduS_1_all(pf01512pf13375pf10531) PDU1AB
19715 HMM00465alcdh_3_mix(pf00465) PDU1AB
19720 HMMalddh_1_eutpdugrm(pf00171) PDU1AB EP_NTERM
19725 HMM01923cobtr_1_pdu(pf01923pf03928) PDU1AB
19730 P_babyPink PDU1AB
19735 HMM15953pduput_1_pdu1(pf15953) PDU1AB EP_NTERM
19740 HMMptac_2_mix(pf06130) PDU1AB
19745 H_driedBlood PDU1AB
19750 H_tan PDU1AB
19755 HMMpropdeh3_2_pdusml(pf02288) PDU1AB
19760 HMM08841dhact_1_pdu(pf08841pf18427) PDU1AB
19765 HMMpropdeh2_1_pdu(pf02287) PDU1AB EP_NTERM
19770 HMMpropdeh3_1_pdu(pf02288) PDU1AB EP_NTERM
19775 HMMpropdeh1_1_pdu(pf02286) PDU1AB
19780 Tsp_greenishBeige PDU1AB
19785 H_robinEggBlue GRM4
19790 -pf00230 -
19795 HMM10114PocR_1_pdu1(pf12833pf10114) PDU1AB
19800 -pf01656pf07685 -
19805 -pf03186 -
19810 -pf02570 -
19815 -pf01888 -
19820 -pf00590 -
19825 -pf05175 -
19830 -pf00590 -
19835 -pf01890pf11761pf11760 -
19840 -pf00590 -
19845 -pf02571 -
19850 -pf06180 -
BMC-P
BMC-H
BMC-T
Aldehyde dehydrogenase
PTAC
Signature enzyme
Alcohol dehydrogenase
Regulator

Salmonella enterica subsp. arizonae serovar 41:z4,z23:- str. 01-0089. Bacteria; Proteobacteria; Gammaproteobacteria; Enterobacterales;

AIT14__locus_2

PDU1AB

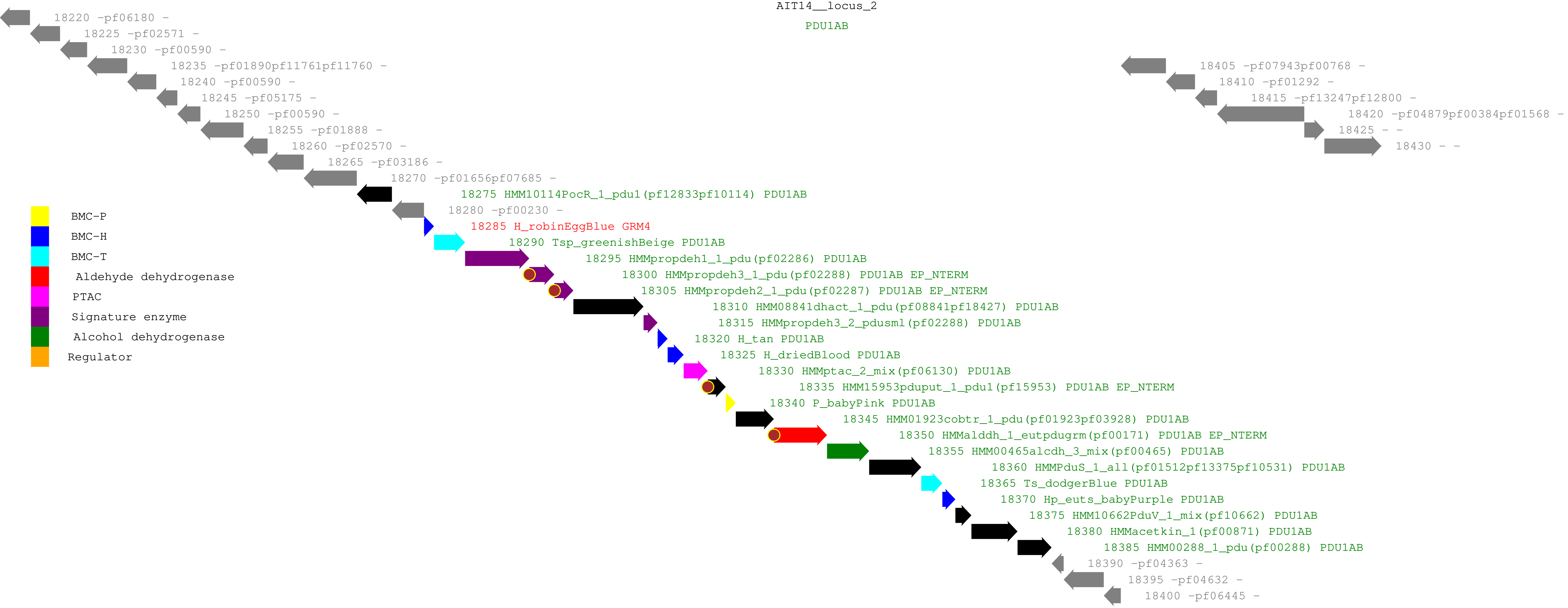

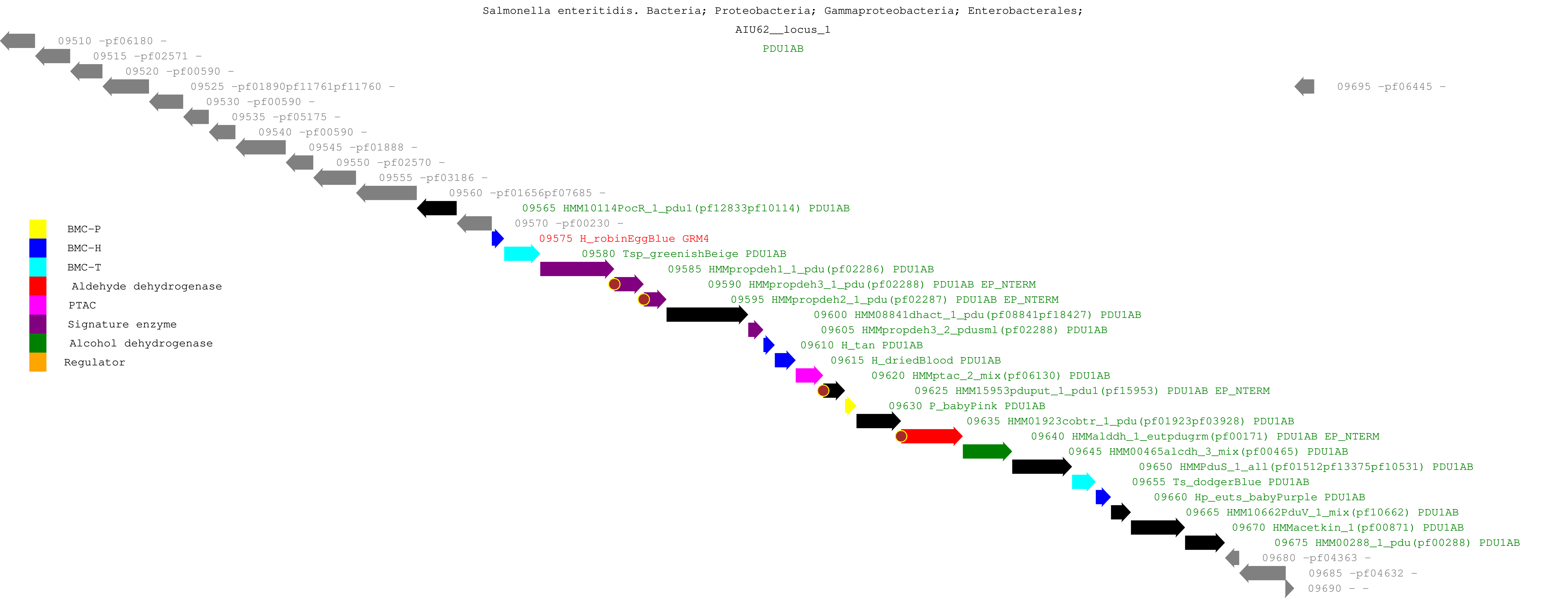
Salmonella enteritidis. Bacteria; Proteobacteria; Gammaproteobacteria; Enterobacterales;
AIU62__locus_1
PDU1AB
09510 -pf06180 -
09515 -pf02571 -
09520 -pf00590 -
09525 -pf01890pf11761pf11760 -
09530 -pf00590 -
09535 -pf05175 -
09540 -pf00590 -
09545 -pf01888 -
09550 -pf02570 -
09555 -pf03186 -
09560 -pf01656pf07685 -
09565 HMM10114PocR_1_pdu1(pf12833pf10114) PDU1AB
09570 -pf00230 -
09575 H_robinEggBlue GRM4
09580 Tsp_greenishBeige PDU1AB
09585 HMMpropdeh1_1_pdu(pf02286) PDU1AB
09590 HMMpropdeh3_1_pdu(pf02288) PDU1AB EP_NTERM
09595 HMMpropdeh2_1_pdu(pf02287) PDU1AB EP_NTERM
09600 HMM08841dhact_1_pdu(pf08841pf18427) PDU1AB
09605 HMMpropdeh3_2_pdusml(pf02288) PDU1AB
09610 H_tan PDU1AB
09615 H_driedBlood PDU1AB
09620 HMMptac_2_mix(pf06130) PDU1AB
09625 HMM15953pduput_1_pdu1(pf15953) PDU1AB EP_NTERM
09630 P_babyPink PDU1AB
09635 HMM01923cobtr_1_pdu(pf01923pf03928) PDU1AB
09640 HMMalddh_1_eutpdugrm(pf00171) PDU1AB EP_NTERM
09645 HMM00465alcdh_3_mix(pf00465) PDU1AB
09650 HMMPduS_1_all(pf01512pf13375pf10531) PDU1AB
09655 Ts_dodgerBlue PDU1AB
09660 Hp_euts_babyPurple PDU1AB
09665 HMM10662PduV_1_mix(pf10662) PDU1AB
09670 HMMacetkin_1(pf00871) PDU1AB
09675 HMM00288_1_pdu(pf00288) PDU1AB
09680 -pf04363 -
09685 -pf04632 -
09690 - -
09695 -pf06445 -
BMC-P
BMC-H
BMC-T
Aldehyde dehydrogenase
PTAC
Signature enzyme
Alcohol dehydrogenase
Regulator

Escherichia coli. Bacteria; Proteobacteria; Gammaproteobacteria; Enterobacterales;

AJ318__locus_2

PDU1AB

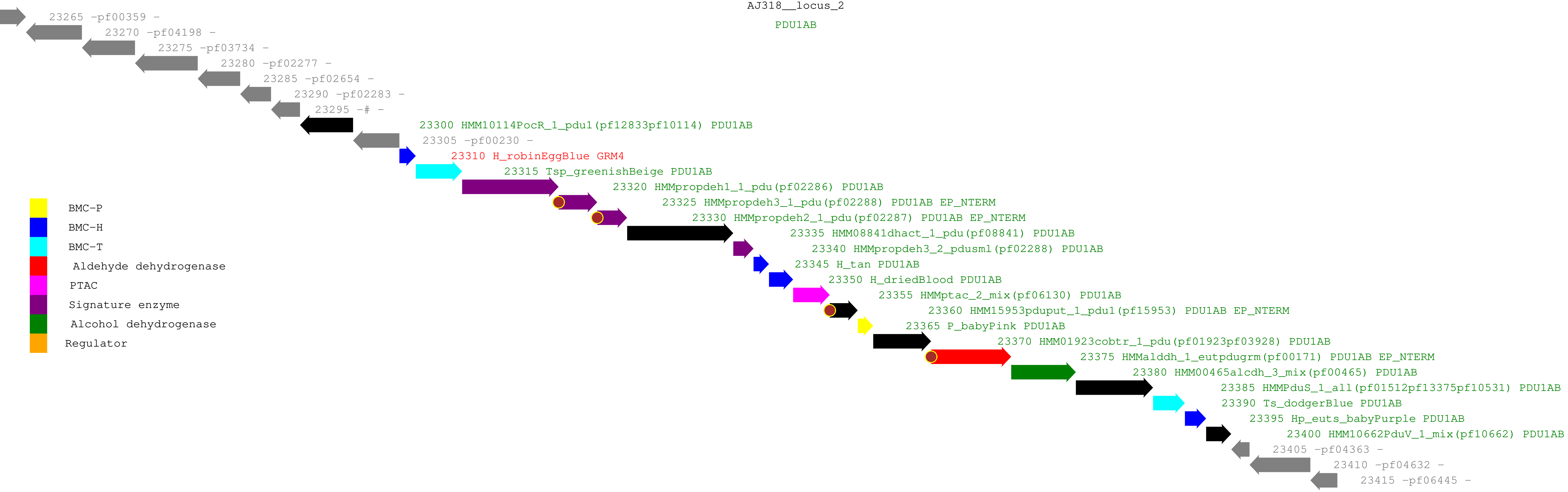

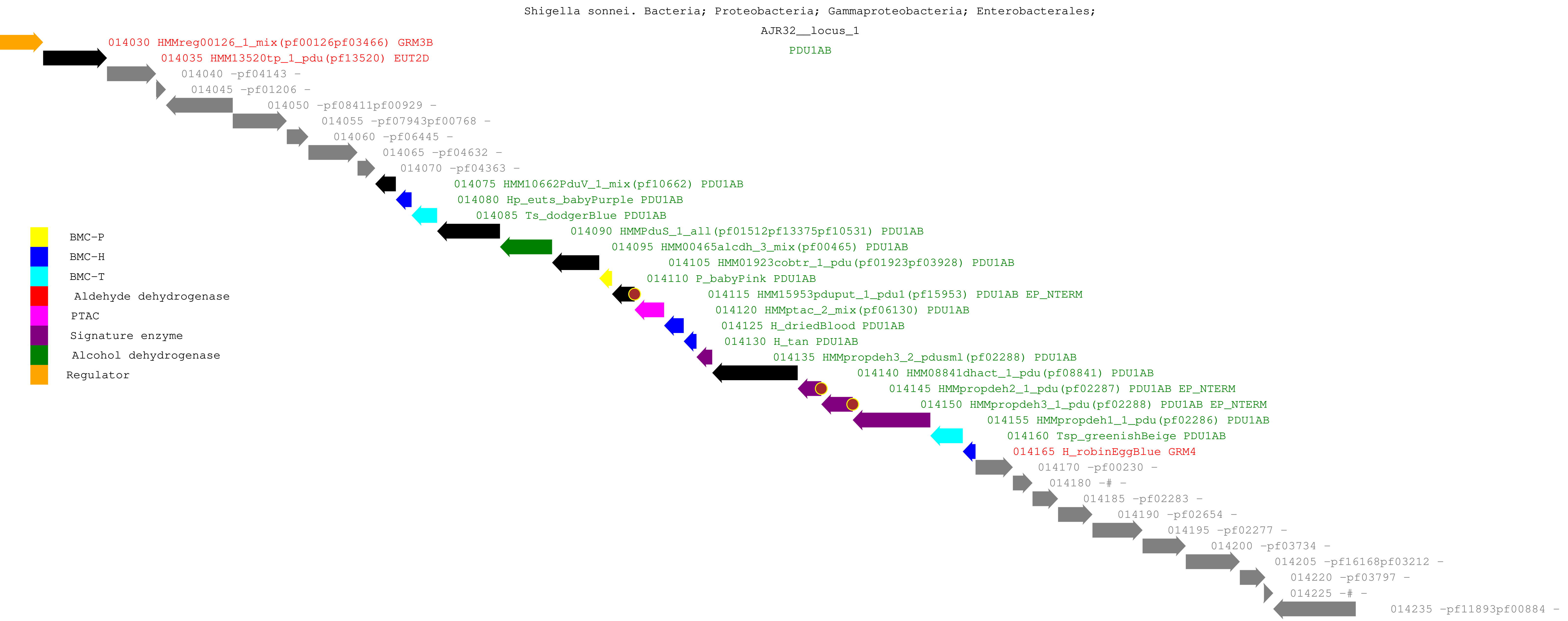
Shigella sonnei. Bacteria; Proteobacteria; Gammaproteobacteria; Enterobacterales;
AJR32__locus_1
PDU1AB
014030 HMMreg00126_1_mix(pf00126pf03466) GRM3B
014035 HMM13520tp_1_pdu(pf13520) EUT2D
014040 -pf04143 -
014045 -pf01206 -
014050 -pf08411pf00929 -
014055 -pf07943pf00768 -
014060 -pf06445 -
014065 -pf04632 -
014070 -pf04363 -
014075 HMM10662PduV_1_mix(pf10662) PDU1AB
014080 Hp_euts_babyPurple PDU1AB
014085 Ts_dodgerBlue PDU1AB
014090 HMMPduS_1_all(pf01512pf13375pf10531) PDU1AB
014095 HMM00465alcdh_3_mix(pf00465) PDU1AB
014105 HMM01923cobtr_1_pdu(pf01923pf03928) PDU1AB
014110 P_babyPink PDU1AB
014115 HMM15953pduput_1_pdu1(pf15953) PDU1AB EP_NTERM
014120 HMMptac_2_mix(pf06130) PDU1AB
014125 H_driedBlood PDU1AB
014130 H_tan PDU1AB
014135 HMMpropdeh3_2_pdusml(pf02288) PDU1AB
014140 HMM08841dhact_1_pdu(pf08841) PDU1AB
014145 HMMpropdeh2_1_pdu(pf02287) PDU1AB EP_NTERM
014150 HMMpropdeh3_1_pdu(pf02288) PDU1AB EP_NTERM
014155 HMMpropdeh1_1_pdu(pf02286) PDU1AB
014160 Tsp_greenishBeige PDU1AB
014165 H_robinEggBlue GRM4
014170 -pf00230 -
014180 -# -
014185 -pf02283 -
014190 -pf02654 -
014195 -pf02277 -
014200 -pf03734 -
014205 -pf16168pf03212 -
014220 -pf03797 -
014225 -# -
014235 -pf11893pf00884 -
BMC-P
BMC-H
BMC-T
Aldehyde dehydrogenase
PTAC
Signature enzyme
Alcohol dehydrogenase
Regulator

Citrobacter amalonaticus. Bacteria; Proteobacteria; Gammaproteobacteria; Enterobacterales;

AL012__locus_2

PDU1AB

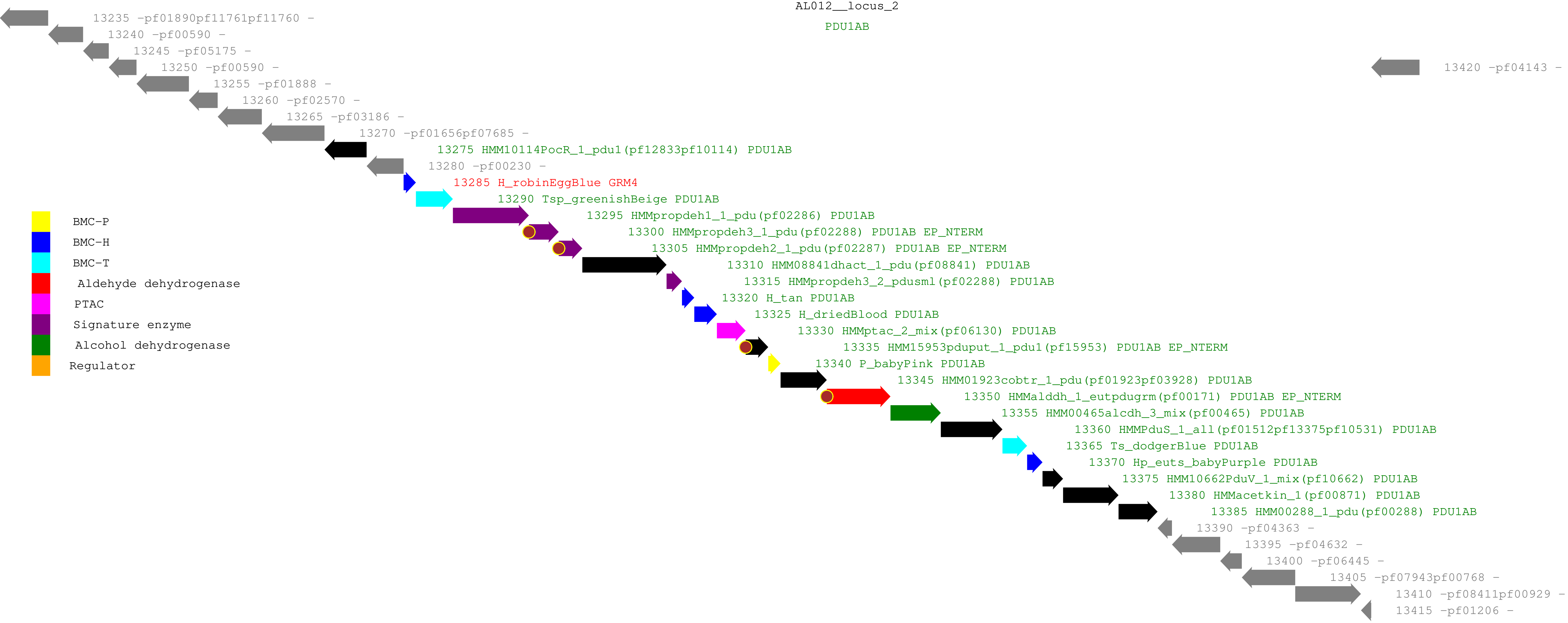

Salmonella choleraesuis. Bacteria; Proteobacteria; Gammaproteobacteria; Enterobacterales;

AL032__locus_1

PDU1AB

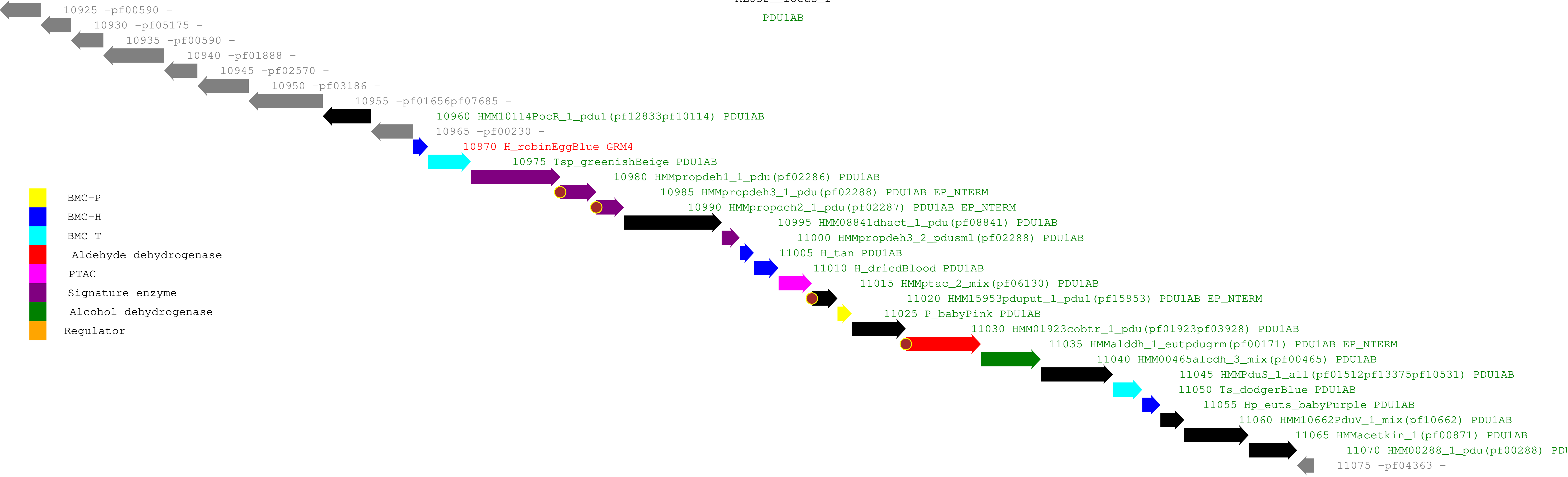

Salmonella enterica (Salmonella choleraesuis). Bacteria; Proteobacteria; Gammaproteobacteria; Enterobacterales;

AL463__locus_1

PDU1AB

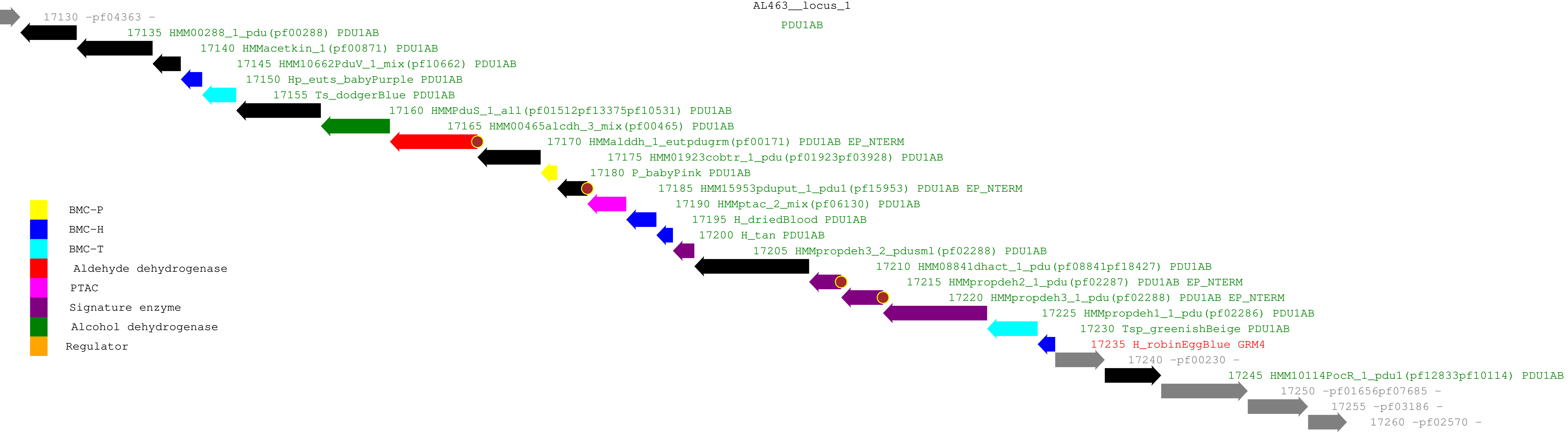

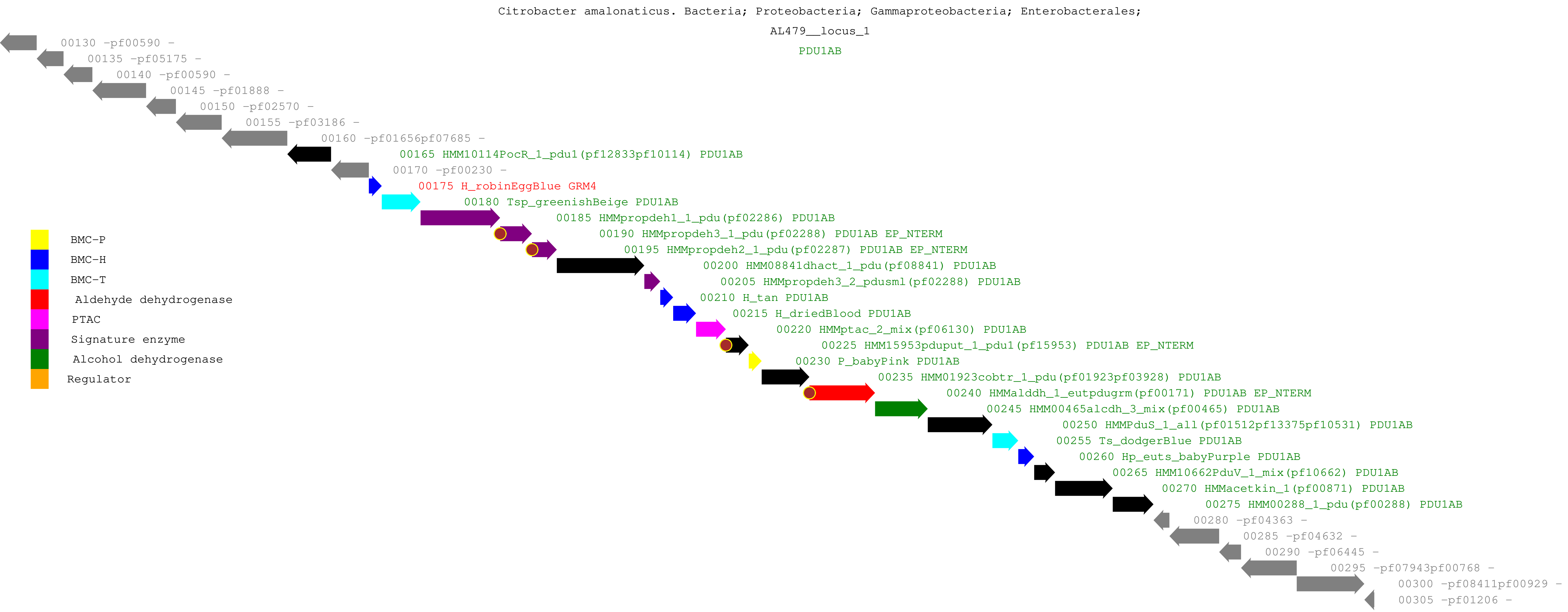

Citrobacter amalonaticus. Bacteria; Proteobacteria; Gammaproteobacteria; Enterobacterales;
AL479__locus_1
PDU1AB
00130 -pf00590 -
00135 -pf05175 -
00140 -pf00590 -
00145 -pf01888 -
00150 -pf02570 -
00155 -pf03186 -
00160 -pf01656pf07685 -
00165 HMM10114PocR_1_pdu1(pf12833pf10114) PDU1AB
00170 -pf00230 -
00175 H_robinEggBlue GRM4
00180 Tsp_greenishBeige PDU1AB
00185 HMMpropdeh1_1_pdu(pf02286) PDU1AB
00190 HMMpropdeh3_1_pdu(pf02288) PDU1AB EP_NTERM
00195 HMMpropdeh2_1_pdu(pf02287) PDU1AB EP_NTERM
00200 HMM08841dhact_1_pdu(pf08841) PDU1AB
00205 HMMpropdeh3_2_pdusml(pf02288) PDU1AB
00210 H_tan PDU1AB
00215 H_driedBlood PDU1AB
00220 HMMptac_2_mix(pf06130) PDU1AB
00225 HMM15953pduput_1_pdu1(pf15953) PDU1AB EP_NTERM
00230 P_babyPink PDU1AB
00235 HMM01923cobtr_1_pdu(pf01923pf03928) PDU1AB
00240 HMMalddh_1_eutpdugrm(pf00171) PDU1AB EP_NTERM
00245 HMM00465alcdh_3_mix(pf00465) PDU1AB
00250 HMMPduS_1_all(pf01512pf13375pf10531) PDU1AB
00255 Ts_dodgerBlue PDU1AB
00260 Hp_euts_babyPurple PDU1AB
00265 HMM10662PduV_1_mix(pf10662) PDU1AB
00270 HMMacetkin_1(pf00871) PDU1AB
00275 HMM00288_1_pdu(pf00288) PDU1AB
00280 -pf04363 -
00285 -pf04632 -
00290 -pf06445 -
00295 -pf07943pf00768 -
00300 -pf08411pf00929 -
00305 -pf01206 -
BMC-P
BMC-H
BMC-T
Aldehyde dehydrogenase
PTAC
Signature enzyme
Alcohol dehydrogenase
Regulator

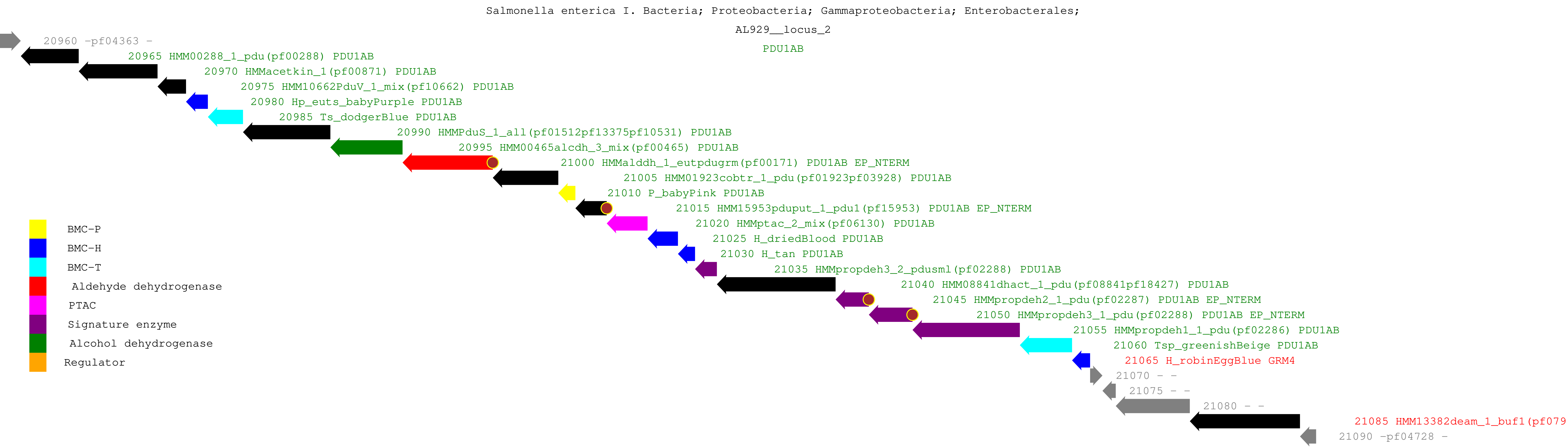

Salmonella enterica I. Bacteria; Proteobacteria; Gammaproteobacteria; Enterobacterales;
AL929__locus_2
PDU1AB
20960 -pf04363 -
20965 HMM00288_1_pdu(pf00288) PDU1AB
20970 HMMacetkin_1(pf00871) PDU1AB
20975 HMM10662PduV_1_mix(pf10662) PDU1AB
20980 Hp_euts_babyPurple PDU1AB
20985 Ts_dodgerBlue PDU1AB
20990 HMMPduS_1_all(pf01512pf13375pf10531) PDU1AB
20995 HMM00465alcdh_3_mix(pf00465) PDU1AB
21000 HMMalddh_1_eutpdugrm(pf00171) PDU1AB EP_NTERM
21005 HMM01923cobtr_1_pdu(pf01923pf03928) PDU1AB
21010 P_babyPink PDU1AB
21015 HMM15953pduput_1_pdu1(pf15953) PDU1AB EP_NTERM
21020 HMMptac_2_mix(pf06130) PDU1AB
21025 H_driedBlood PDU1AB
21030 H_tan PDU1AB
21035 HMMpropdeh3_2_pdusml(pf02288) PDU1AB
21040 HMM08841dhact_1_pdu(pf08841pf18427) PDU1AB
21045 HMMpropdeh2_1_pdu(pf02287) PDU1AB EP_NTERM
21050 HMMpropdeh3_1_pdu(pf02288) PDU1AB EP_NTERM
21055 HMMpropdeh1_1_pdu(pf02286) PDU1AB
21060 Tsp_greenishBeige PDU1AB
21065 H_robinEggBlue GRM4
21070 - -
21075 - -
21080 - -
21085 HMM13382deam_1_buf1(pf079
21090 -pf04728 -
BMC-P
BMC-H
BMC-T
Aldehyde dehydrogenase
PTAC
Signature enzyme
Alcohol dehydrogenase
Regulator

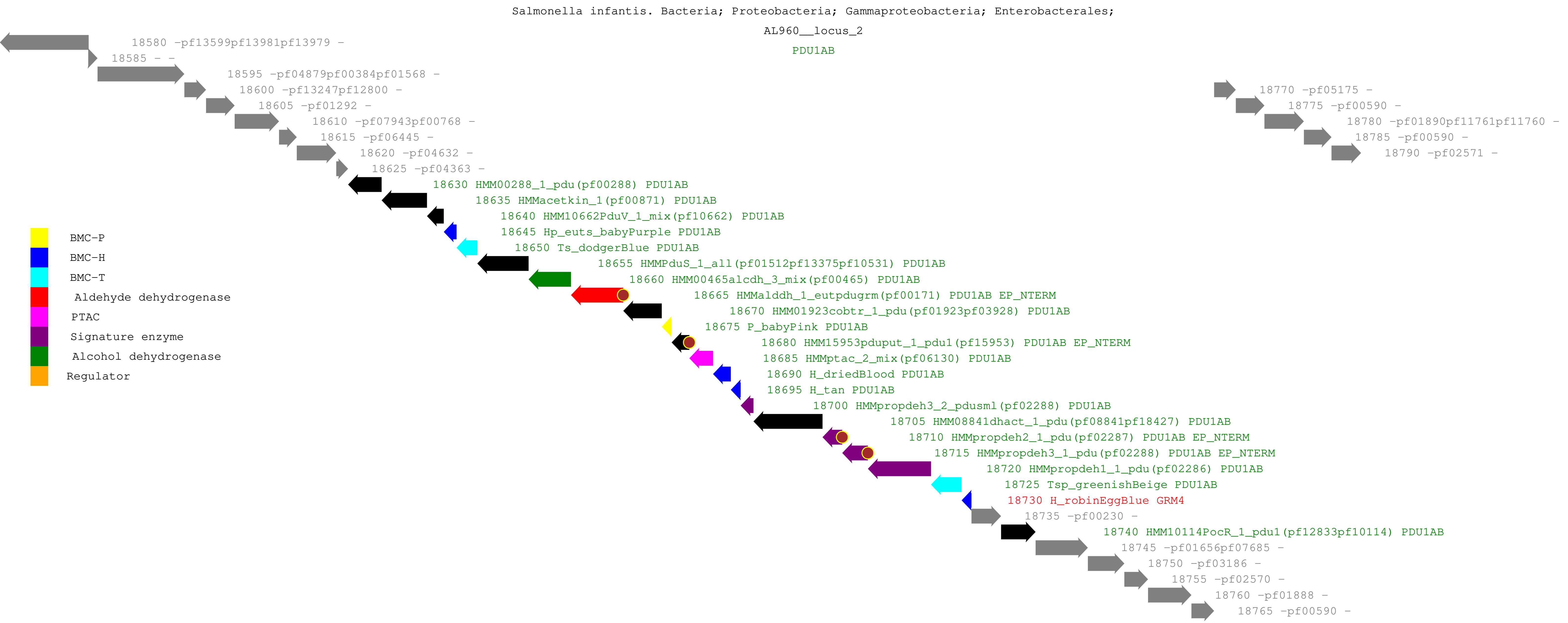
Salmonella infantis. Bacteria; Proteobacteria; Gammaproteobacteria; Enterobacterales;
AL960__locus_2
PDU1AB
18580 -pf13599pf13981pf13979 -
18585 - -
18595 -pf04879pf00384pf01568 -
18600 -pf13247pf12800 -
18605 -pf01292 -
18610 -pf07943pf00768 -
18615 -pf06445 -
18620 -pf04632 -
18625 -pf04363 -
18630 HMM00288_1_pdu(pf00288) PDU1AB
18635 HMMacetkin_1(pf00871) PDU1AB
18640 HMM10662PduV_1_mix(pf10662) PDU1AB
18645 Hp_euts_babyPurple PDU1AB
18650 Ts_dodgerBlue PDU1AB
18655 HMMPduS_1_all(pf01512pf13375pf10531) PDU1AB
18660 HMM00465alcdh_3_mix(pf00465) PDU1AB
18665 HMMalddh_1_eutpdugrm(pf00171) PDU1AB EP_NTERM
18670 HMM01923cobtr_1_pdu(pf01923pf03928) PDU1AB
18675 P_babyPink PDU1AB
18680 HMM15953pduput_1_pdu1(pf15953) PDU1AB EP_NTERM
18685 HMMptac_2_mix(pf06130) PDU1AB
18690 H_driedBlood PDU1AB
18695 H_tan PDU1AB
18700 HMMpropdeh3_2_pdusml(pf02288) PDU1AB
18705 HMM08841dhact_1_pdu(pf08841pf18427) PDU1AB
18710 HMMpropdeh2_1_pdu(pf02287) PDU1AB EP_NTERM
18715 HMMpropdeh3_1_pdu(pf02288) PDU1AB EP_NTERM
18720 HMMpropdeh1_1_pdu(pf02286) PDU1AB
18725 Tsp_greenishBeige PDU1AB
18730 H_robinEggBlue GRM4
18735 -pf00230 -
18740 HMM10114PocR_1_pdu1(pf12833pf10114) PDU1AB
18745 -pf01656pf07685 -
18750 -pf03186 -
18755 -pf02570 -
18760 -pf01888 -
18765 -pf00590 -
18770 -pf05175 -
18775 -pf00590 -
18780 -pf01890pf11761pf11760 -
18785 -pf00590 -
18790 -pf02571 -
BMC-P
BMC-H
BMC-T
Aldehyde dehydrogenase
PTAC
Signature enzyme
Alcohol dehydrogenase
Regulator

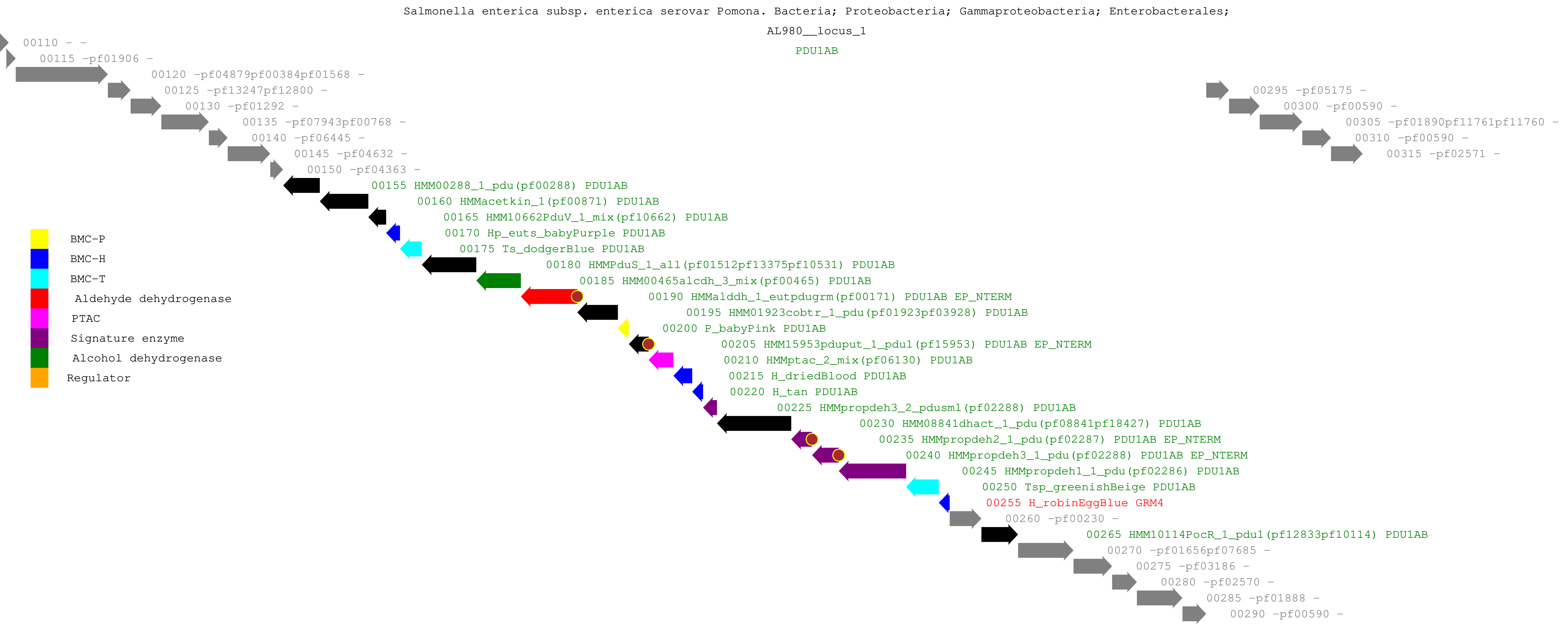
Salmonella enterica subsp. enterica serovar Pomona. Bacteria; Proteobacteria; Gammaproteobacteria; Enterobacterales;
AL980__locus_1
PDU1AB
00110 - -
00115 -pf01906 -
00120 -pf04879pf00384pf01568 -
00125 -pf13247pf12800 -
00130 -pf01292 -
00135 -pf07943pf00768 -
00140 -pf06445 -
00145 -pf04632 -
00150 -pf04363 -
00155 HMM00288_1_pdu(pf00288) PDU1AB
00160 HMMacetkin_1(pf00871) PDU1AB
00165 HMM10662PduV_1_mix(pf10662) PDU1AB
00170 Hp_euts_babyPurple PDU1AB
00175 Ts_dodgerBlue PDU1AB
00180 HMMPduS_1_all(pf01512pf13375pf10531) PDU1AB
00185 HMM00465alcdh_3_mix(pf00465) PDU1AB
00190 HMMalddh_1_eutpdugrm(pf00171) PDU1AB EP_NTERM
00195 HMM01923cobtr_1_pdu(pf01923pf03928) PDU1AB
00200 P_babyPink PDU1AB
00205 HMM15953pduput_1_pdu1(pf15953) PDU1AB EP_NTERM
00210 HMMptac_2_mix(pf06130) PDU1AB
00215 H_driedBlood PDU1AB
00220 H_tan PDU1AB
00225 HMMpropdeh3_2_pdusml(pf02288) PDU1AB
00230 HMM08841dhact_1_pdu(pf08841pf18427) PDU1AB
00235 HMMpropdeh2_1_pdu(pf02287) PDU1AB EP_NTERM
00240 HMMpropdeh3_1_pdu(pf02288) PDU1AB EP_NTERM
00245 HMMpropdeh1_1_pdu(pf02286) PDU1AB
00250 Tsp_greenishBeige PDU1AB
00255 H_robinEggBlue GRM4
00260 -pf00230 -
00265 HMM10114PocR_1_pdu1(pf12833pf10114) PDU1AB
00270 -pf01656pf07685 -
00275 -pf03186 -
00280 -pf02570 -
00285 -pf01888 -
00290 -pf00590 -
00295 -pf05175 -
00300 -pf00590 -
00305 -pf01890pf11761pf11760 -
00310 -pf00590 -
00315 -pf02571 -
BMC-P
BMC-H
BMC-T
Aldehyde dehydrogenase
PTAC
Signature enzyme
Alcohol dehydrogenase
Regulator

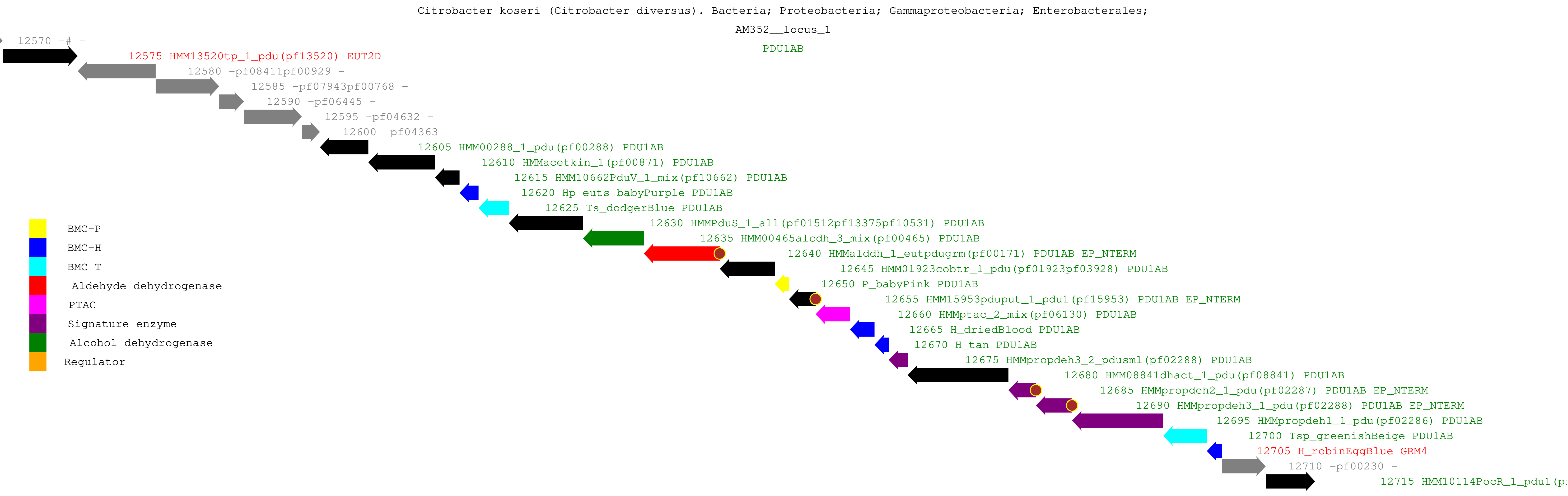
Citrobacter koseri (Citrobacter diversus). Bacteria; Proteobacteria; Gammaproteobacteria; Enterobacterales;
AM352__locus_1
PDU1AB
12570 -# -
12575 HMM13520tp_1_pdu(pf13520) EUT2D
12580 -pf08411pf00929 -
12585 -pf07943pf00768 -
12590 -pf06445 -
12595 -pf04632 -
12600 -pf04363 -
12605 HMM00288_1_pdu(pf00288) PDU1AB
12610 HMMacetkin_1(pf00871) PDU1AB
12615 HMM10662PduV_1_mix(pf10662) PDU1AB
12620 Hp_euts_babyPurple PDU1AB
12625 Ts_dodgerBlue PDU1AB
12630 HMMPduS_1_all(pf01512pf13375pf10531) PDU1AB
12635 HMM00465alcdh_3_mix(pf00465) PDU1AB
12640 HMMalddh_1_eutpdugrm(pf00171) PDU1AB EP_NTERM
12645 HMM01923cobtr_1_pdu(pf01923pf03928) PDU1AB
12650 P_babyPink PDU1AB
12655 HMM15953pduput_1_pdu1(pf15953) PDU1AB EP_NTERM
12660 HMMptac_2_mix(pf06130) PDU1AB
12665 H_driedBlood PDU1AB
12670 H_tan PDU1AB
12675 HMMpropdeh3_2_pdusml(pf02288) PDU1AB
12680 HMM08841dhact_1_pdu(pf08841) PDU1AB
12685 HMMpropdeh2_1_pdu(pf02287) PDU1AB EP_NTERM
12690 HMMpropdeh3_1_pdu(pf02288) PDU1AB EP_NTERM
12695 HMMpropdeh1_1_pdu(pf02286) PDU1AB
12700 Tsp_greenishBeige PDU1AB
12705 H_robinEggBlue GRM4
12710 -pf00230 -
12715 HMM10114PocR_1_pdu1(p
BMC-P
BMC-H
BMC-T
Aldehyde dehydrogenase
PTAC
Signature enzyme
Alcohol dehydrogenase
Regulator

Salmonella typhimurium. Bacteria; Proteobacteria; Gammaproteobacteria; Enterobacterales;

AM358__locus_2

PDU1AB

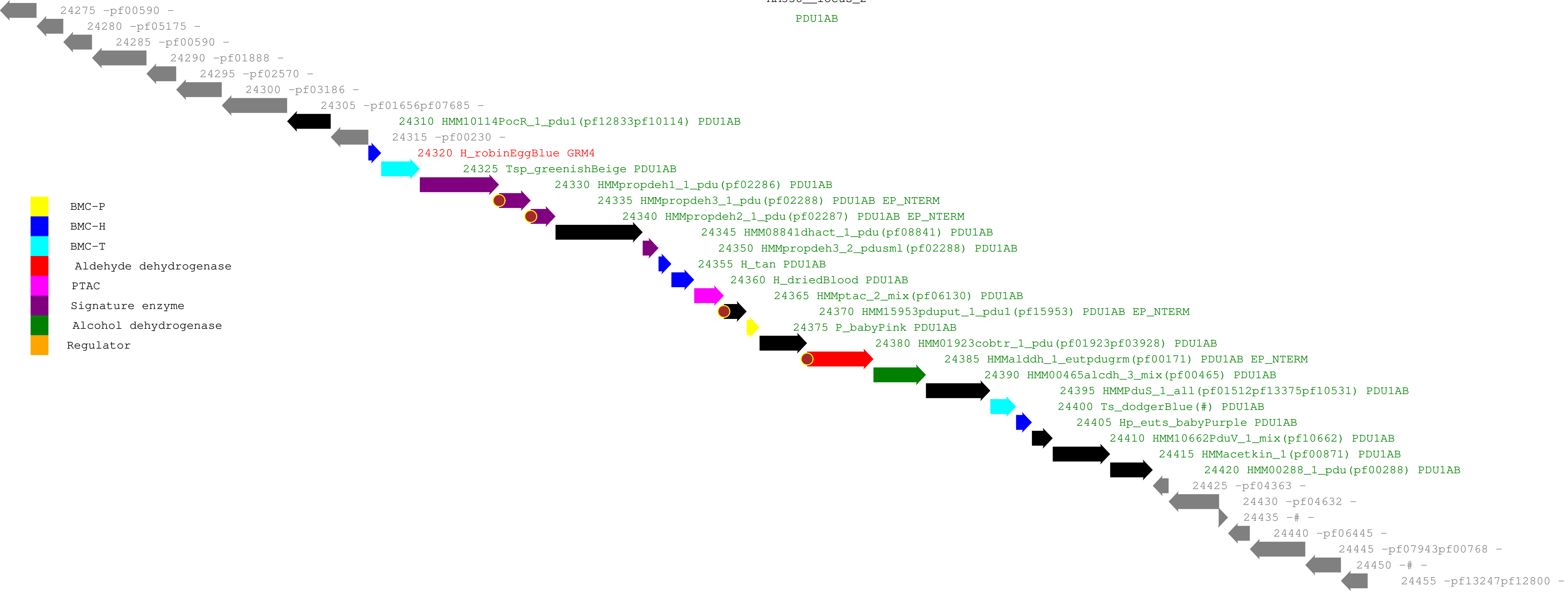

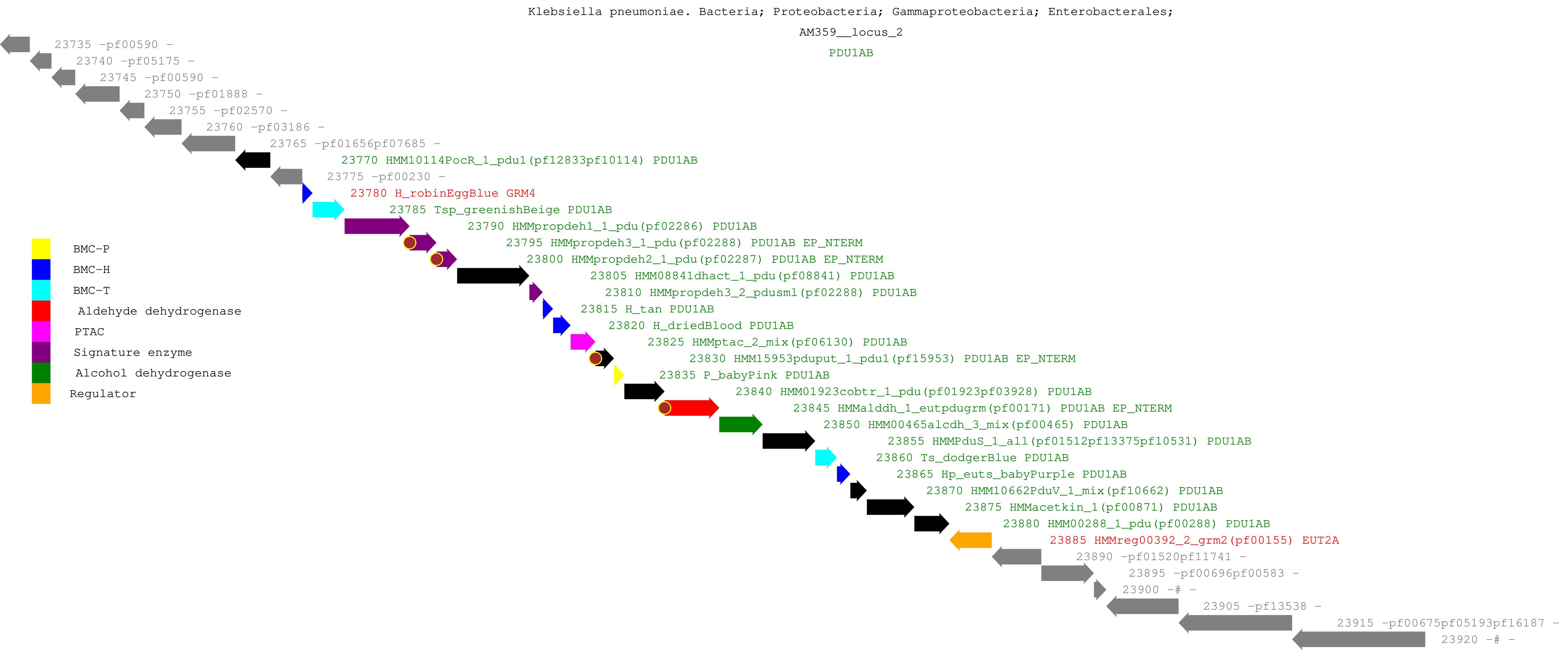
Klebsiella pneumoniae. Bacteria; Proteobacteria; Gammaproteobacteria; Enterobacterales;
AM359__locus_2
PDU1AB
23735 -pf00590 -
23740 -pf05175 -
23745 -pf00590 -
23750 -pf01888 -
23755 -pf02570 -
23760 -pf03186 -
23765 -pf01656pf07685 -
23770 HMM10114PocR_1_pdu1(pf12833pf10114) PDU1AB
23775 -pf00230 -
23780 H_robinEggBlue GRM4
23785 Tsp_greenishBeige PDU1AB
23790 HMMpropdeh1_1_pdu(pf02286) PDU1AB
23795 HMMpropdeh3_1_pdu(pf02288) PDU1AB EP_NTERM
23800 HMMpropdeh2_1_pdu(pf02287) PDU1AB EP_NTERM
23805 HMM08841dhact_1_pdu(pf08841) PDU1AB
23810 HMMpropdeh3_2_pdusml(pf02288) PDU1AB
23815 H_tan PDU1AB
23820 H_driedBlood PDU1AB
23825 HMMptac_2_mix(pf06130) PDU1AB
23830 HMM15953pduput_1_pdu1(pf15953) PDU1AB EP_NTERM
23835 P_babyPink PDU1AB
23840 HMM01923cobtr_1_pdu(pf01923pf03928) PDU1AB
23845 HMMalddh_1_eutpdugrm(pf00171) PDU1AB EP_NTERM
23850 HMM00465alcdh_3_mix(pf00465) PDU1AB
23855 HMMPduS_1_all(pf01512pf13375pf10531) PDU1AB
23860 Ts_dodgerBlue PDU1AB
23865 Hp_euts_babyPurple PDU1AB
23870 HMM10662PduV_1_mix(pf10662) PDU1AB
23875 HMMacetkin_1(pf00871) PDU1AB
23880 HMM00288_1_pdu(pf00288) PDU1AB
23885 HMMreg00392_2_grm2(pf00155) EUT2A
23890 -pf01520pf11741 -
23895 -pf00696pf00583 -
23900 -# -
23905 -pf13538 -
23915 -pf00675pf05193pf16187 -
23920 -# -
BMC-P
BMC-H
BMC-T
Aldehyde dehydrogenase
PTAC
Signature enzyme
Alcohol dehydrogenase
Regulator

Klebsiella pneumoniae. Bacteria; Proteobacteria; Gammaproteobacteria; Enterobacterales;

AM390__locus_2

PDU1AB

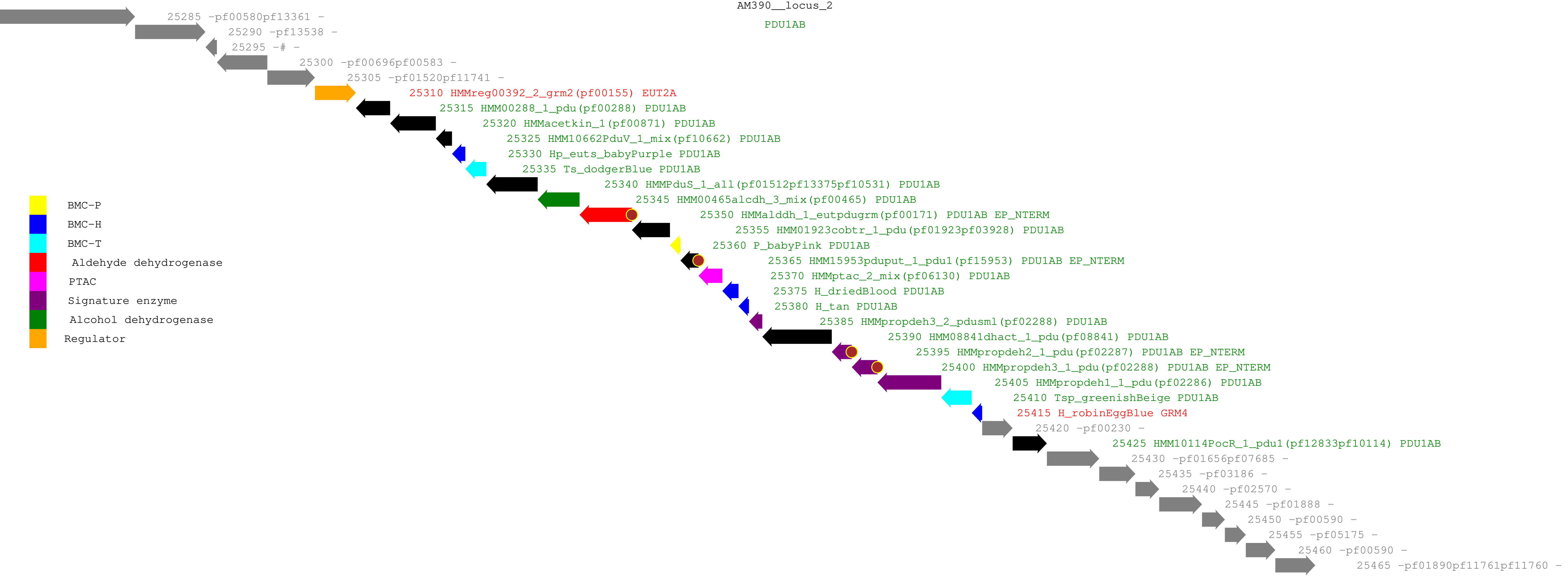

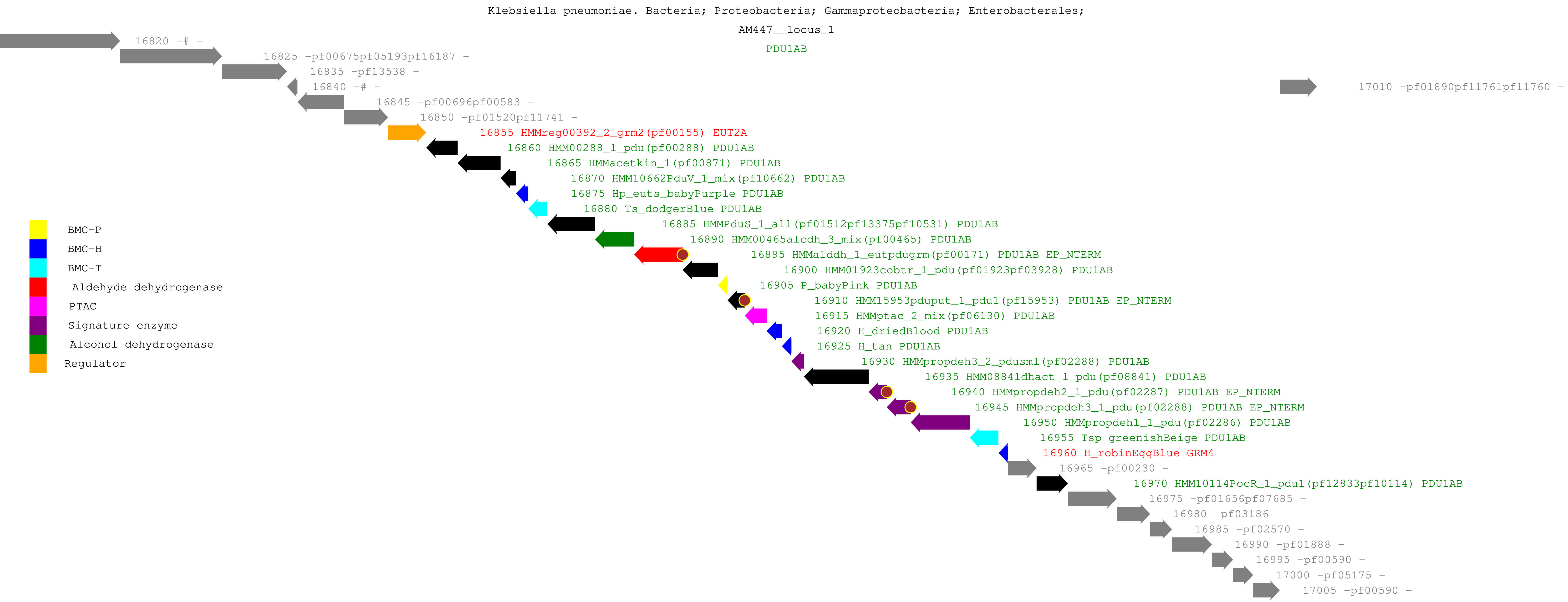

Klebsiella pneumoniae. Bacteria; Proteobacteria; Gammaproteobacteria; Enterobacterales;
AM447__locus_1
PDU1AB
16820 -# -
16825 -pf00675pf05193pf16187 -
16835 -pf13538 -
16840 -# -
16845 -pf00696pf00583 -
16850 -pf01520pf11741 -
16855 HMMreg00392_2_grm2(pf00155) EUT2A
16860 HMM00288_1_pdu(pf00288) PDU1AB
16865 HMMacetkin_1(pf00871) PDU1AB
16870 HMM10662PduV_1_mix(pf10662) PDU1AB
16875 Hp_euts_babyPurple PDU1AB
16880 Ts_dodgerBlue PDU1AB
16885 HMMPduS_1_all(pf01512pf13375pf10531) PDU1AB
16890 HMM00465alcdh_3_mix(pf00465) PDU1AB
16895 HMMalddh_1_eutpdugrm(pf00171) PDU1AB EP_NTERM
16900 HMM01923cobtr_1_pdu(pf01923pf03928) PDU1AB
16905 P_babyPink PDU1AB
16910 HMM15953pduput_1_pdu1(pf15953) PDU1AB EP_NTERM
16915 HMMptac_2_mix(pf06130) PDU1AB
16920 H_driedBlood PDU1AB
16925 H_tan PDU1AB
16930 HMMpropdeh3_2_pdusml(pf02288) PDU1AB
16935 HMM08841dhact_1_pdu(pf08841) PDU1AB
16940 HMMpropdeh2_1_pdu(pf02287) PDU1AB EP_NTERM
16945 HMMpropdeh3_1_pdu(pf02288) PDU1AB EP_NTERM
16950 HMMpropdeh1_1_pdu(pf02286) PDU1AB
16955 Tsp_greenishBeige PDU1AB
16960 H_robinEggBlue GRM4
16965 -pf00230 -
16970 HMM10114PocR_1_pdu1(pf12833pf10114) PDU1AB
16975 -pf01656pf07685 -
16980 -pf03186 -
16985 -pf02570 -
16990 -pf01888 -
16995 -pf00590 -
17000 -pf05175 -
17005 -pf00590 -
17010 -pf01890pf11761pf11760 -
BMC-P
BMC-H
BMC-T
Aldehyde dehydrogenase
PTAC
Signature enzyme
Alcohol dehydrogenase
Regulator

Salmonella enterica subsp. enterica serovar Brandenburg. Bacteria; Proteobacteria; Gammaproteobacteria; Enterobacterales;

AMA33__locus_2

PDU1AB

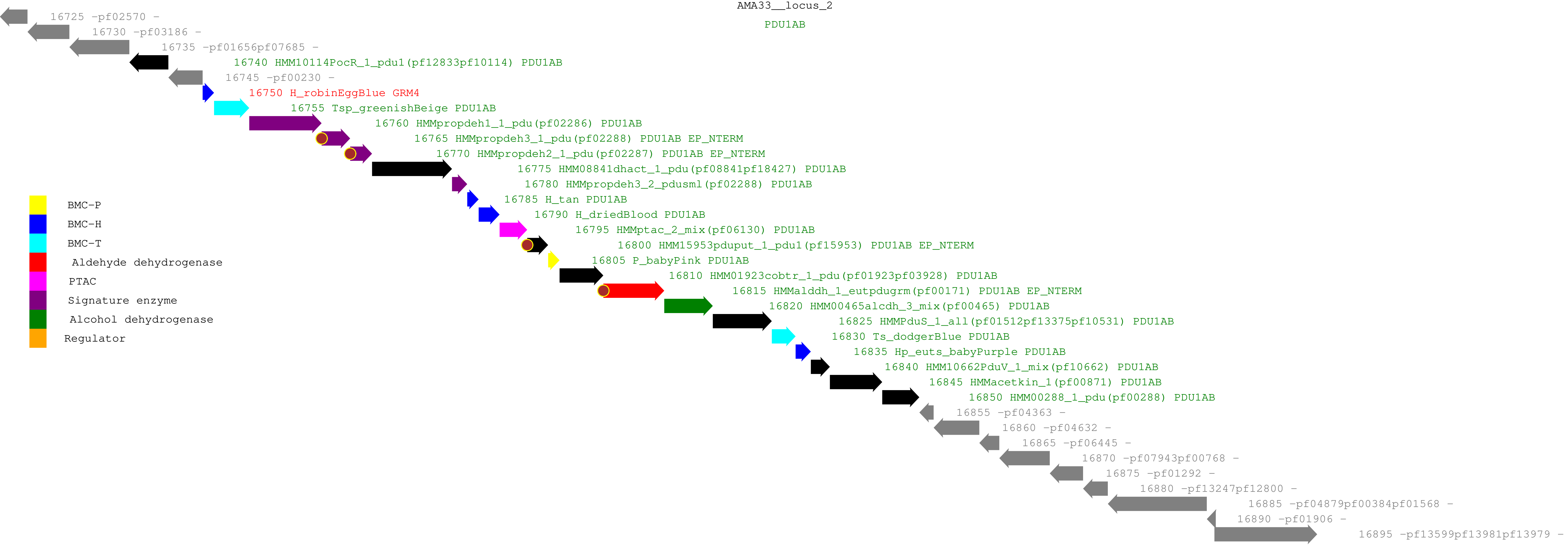

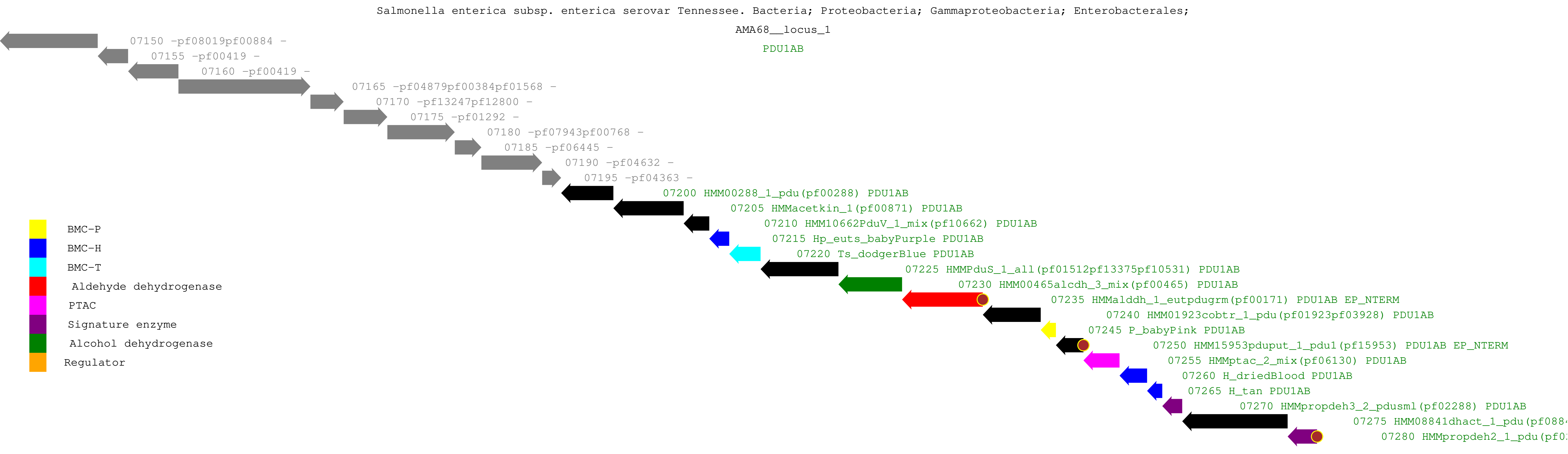
Salmonella enterica subsp. enterica serovar Tennessee. Bacteria; Proteobacteria; Gammaproteobacteria; Enterobacterales;
AMA68__locus_1
PDU1AB
07150 -pf08019pf00884 -
07155 -pf00419 -
07160 -pf00419 -
07165 -pf04879pf00384pf01568 -
07170 -pf13247pf12800 -
07175 -pf01292 -
07180 -pf07943pf00768 -
07185 -pf06445 -
07190 -pf04632 -
07195 -pf04363 -
07200 HMM00288_1_pdu(pf00288) PDU1AB
07205 HMMacetkin_1(pf00871) PDU1AB
07210 HMM10662PduV_1_mix(pf10662) PDU1AB
07215 Hp_euts_babyPurple PDU1AB
07220 Ts_dodgerBlue PDU1AB
07225 HMMPduS_1_all(pf01512pf13375pf10531) PDU1AB
07230 HMM00465alcdh_3_mix(pf00465) PDU1AB
07235 HMMalddh_1_eutpdugrm(pf00171) PDU1AB EP_NTERM
07240 HMM01923cobtr_1_pdu(pf01923pf03928) PDU1AB
07245 P_babyPink PDU1AB
07250 HMM15953pduput_1_pdu1(pf15953) PDU1AB EP_NTERM
07255 HMMptac_2_mix(pf06130) PDU1AB
07260 H_driedBlood PDU1AB
07265 H_tan PDU1AB
07270 HMMpropdeh3_2_pdusml(pf02288) PDU1AB
07275 HMM08841dhact_1_pdu(pf088
07280 HMMpropdeh2_1_pdu(pf0
BMC-P
BMC-H
BMC-T
Aldehyde dehydrogenase
PTAC
Signature enzyme
Alcohol dehydrogenase
Regulator

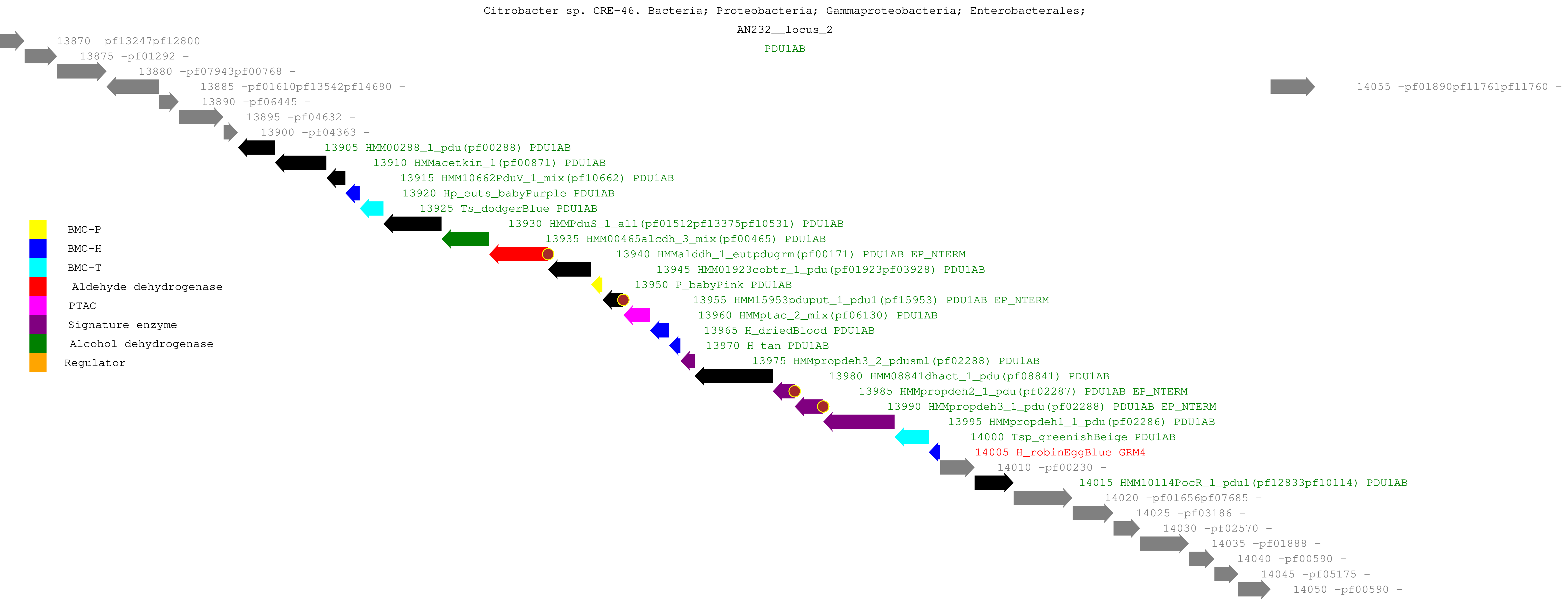

Citrobacter sp. CRE-46. Bacteria; Proteobacteria; Gammaproteobacteria; Enterobacterales;
AN232__locus_2
PDU1AB
13870 -pf13247pf12800 -
13875 -pf01292 -
13880 -pf07943pf00768 -
13885 -pf01610pf13542pf14690 -
13890 -pf06445 -
13895 -pf04632 -
13900 -pf04363 -
13905 HMM00288_1_pdu(pf00288) PDU1AB
13910 HMMacetkin_1(pf00871) PDU1AB
13915 HMM10662PduV_1_mix(pf10662) PDU1AB
13920 Hp_euts_babyPurple PDU1AB
13925 Ts_dodgerBlue PDU1AB
13930 HMMPduS_1_all(pf01512pf13375pf10531) PDU1AB
13935 HMM00465alcdh_3_mix(pf00465) PDU1AB
13940 HMMalddh_1_eutpdugrm(pf00171) PDU1AB EP_NTERM
13945 HMM01923cobtr_1_pdu(pf01923pf03928) PDU1AB
13950 P_babyPink PDU1AB
13955 HMM15953pduput_1_pdu1(pf15953) PDU1AB EP_NTERM
13960 HMMptac_2_mix(pf06130) PDU1AB
13965 H_driedBlood PDU1AB
13970 H_tan PDU1AB
13975 HMMpropdeh3_2_pdusml(pf02288) PDU1AB
13980 HMM08841dhact_1_pdu(pf08841) PDU1AB
13985 HMMpropdeh2_1_pdu(pf02287) PDU1AB EP_NTERM
13990 HMMpropdeh3_1_pdu(pf02288) PDU1AB EP_NTERM
13995 HMMpropdeh1_1_pdu(pf02286) PDU1AB
14000 Tsp_greenishBeige PDU1AB
14005 H_robinEggBlue GRM4
14010 -pf00230 -
14015 HMM10114PocR_1_pdu1(pf12833pf10114) PDU1AB
14020 -pf01656pf07685 -
14025 -pf03186 -
14030 -pf02570 -
14035 -pf01888 -
14040 -pf00590 -
14045 -pf05175 -
14050 -pf00590 -
14055 -pf01890pf11761pf11760 -
BMC-P
BMC-H
BMC-T
Aldehyde dehydrogenase
PTAC
Signature enzyme
Alcohol dehydrogenase
Regulator

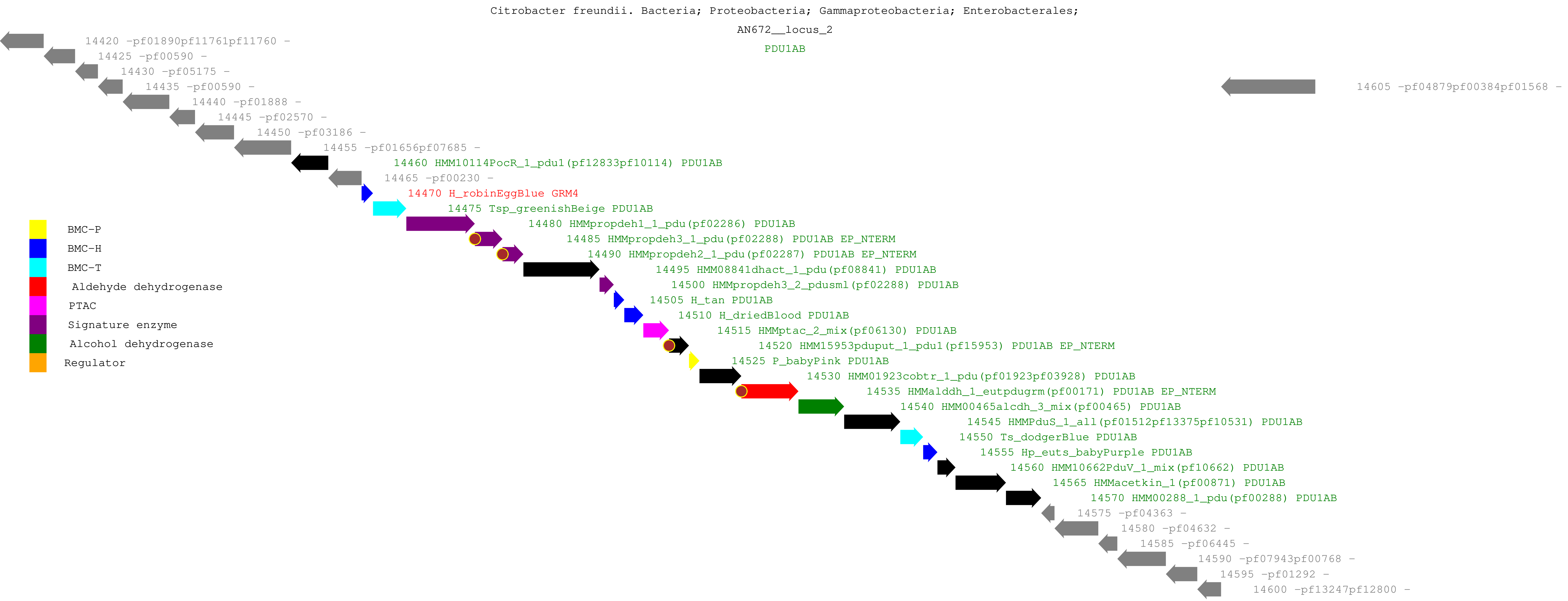
Citrobacter freundii. Bacteria; Proteobacteria; Gammaproteobacteria; Enterobacterales;
AN672__locus_2
PDU1AB
14420 -pf01890pf11761pf11760 -
14425 -pf00590 -
14430 -pf05175 -
14435 -pf00590 -
14440 -pf01888 -
14445 -pf02570 -
14450 -pf03186 -
14455 -pf01656pf07685 -
14460 HMM10114PocR_1_pdu1(pf12833pf10114) PDU1AB
14465 -pf00230 -
14470 H_robinEggBlue GRM4
14475 Tsp_greenishBeige PDU1AB
14480 HMMpropdeh1_1_pdu(pf02286) PDU1AB
14485 HMMpropdeh3_1_pdu(pf02288) PDU1AB EP_NTERM
14490 HMMpropdeh2_1_pdu(pf02287) PDU1AB EP_NTERM
14495 HMM08841dhact_1_pdu(pf08841) PDU1AB
14500 HMMpropdeh3_2_pdusml(pf02288) PDU1AB
14505 H_tan PDU1AB
14510 H_driedBlood PDU1AB
14515 HMMptac_2_mix(pf06130) PDU1AB
14520 HMM15953pduput_1_pdu1(pf15953) PDU1AB EP_NTERM
14525 P_babyPink PDU1AB
14530 HMM01923cobtr_1_pdu(pf01923pf03928) PDU1AB
14535 HMMalddh_1_eutpdugrm(pf00171) PDU1AB EP_NTERM
14540 HMM00465alcdh_3_mix(pf00465) PDU1AB
14545 HMMPduS_1_all(pf01512pf13375pf10531) PDU1AB
14550 Ts_dodgerBlue PDU1AB
14555 Hp_euts_babyPurple PDU1AB
14560 HMM10662PduV_1_mix(pf10662) PDU1AB
14565 HMMacetkin_1(pf00871) PDU1AB
14570 HMM00288_1_pdu(pf00288) PDU1AB
14575 -pf04363 -
14580 -pf04632 -
14585 -pf06445 -
14590 -pf07943pf00768 -
14595 -pf01292 -
14600 -pf13247pf12800 -
14605 -pf04879pf00384pf01568 -
BMC-P
BMC-H
BMC-T
Aldehyde dehydrogenase
PTAC
Signature enzyme
Alcohol dehydrogenase
Regulator

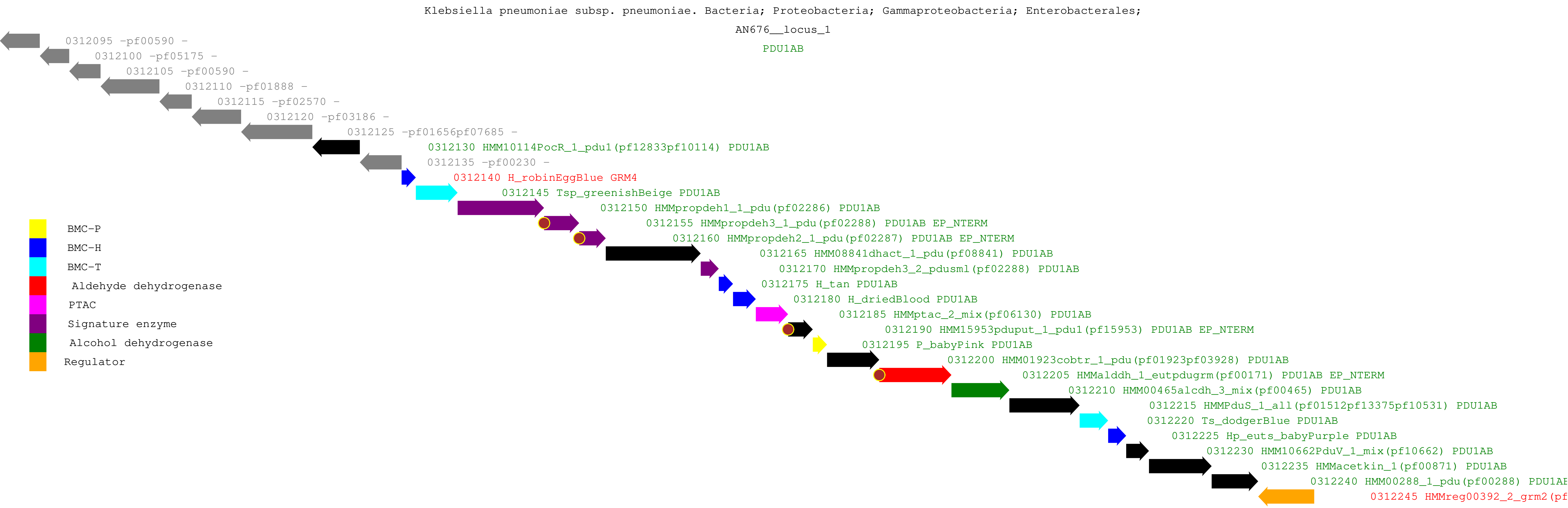
Klebsiella pneumoniae subsp. pneumoniae. Bacteria; Proteobacteria; Gammaproteobacteria; Enterobacterales;
AN676__locus_1
PDU1AB
0312095 -pf00590 -
0312100 -pf05175 -
0312105 -pf00590 -
0312110 -pf01888 -
0312115 -pf02570 -
0312120 -pf03186 -
0312125 -pf01656pf07685 -
0312130 HMM10114PocR_1_pdu1(pf12833pf10114) PDU1AB
0312135 -pf00230 -
0312140 H_robinEggBlue GRM4
0312145 Tsp_greenishBeige PDU1AB
0312150 HMMpropdeh1_1_pdu(pf02286) PDU1AB
0312155 HMMpropdeh3_1_pdu(pf02288) PDU1AB EP_NTERM
0312160 HMMpropdeh2_1_pdu(pf02287) PDU1AB EP_NTERM
0312165 HMM08841dhact_1_pdu(pf08841) PDU1AB
0312170 HMMpropdeh3_2_pdusml(pf02288) PDU1AB
0312175 H_tan PDU1AB
0312180 H_driedBlood PDU1AB
0312185 HMMptac_2_mix(pf06130) PDU1AB
0312190 HMM15953pduput_1_pdu1(pf15953) PDU1AB EP_NTERM
0312195 P_babyPink PDU1AB
0312200 HMM01923cobtr_1_pdu(pf01923pf03928) PDU1AB
0312205 HMMaldh_1_eutpdugrm(pf00171) PDU1AB EP_NTERM
0312210 HMM00465alcdh_3_mix(pf00465) PDU1AB
0312215 HMMPduS_1_all(pf01512pf13375pf10531) PDU1AB
0312220 Ts_dodgerBlue PDU1AB
0312225 Hp_euts_babyPurple PDU1AB
0312230 HMM10662PduV_1_mix(pf10662) PDU1AB
0312235 HMMacetkin_1(pf00871) PDU1AB
0312240 HMM00288_1_pdu(pf00288) PDU1A
0312245 HMMreg00392_2_grm2(pf
BMC-P
BMC-H
BMC-T
Aldehyde dehydrogenase
PTAC
Signature enzyme
Alcohol dehydrogenase
Regulator

Citrobacter freundii. Bacteria; Proteobacteria; Gammaproteobacteria; Enterobacterales;

AN688__locus_2

PDU1AB

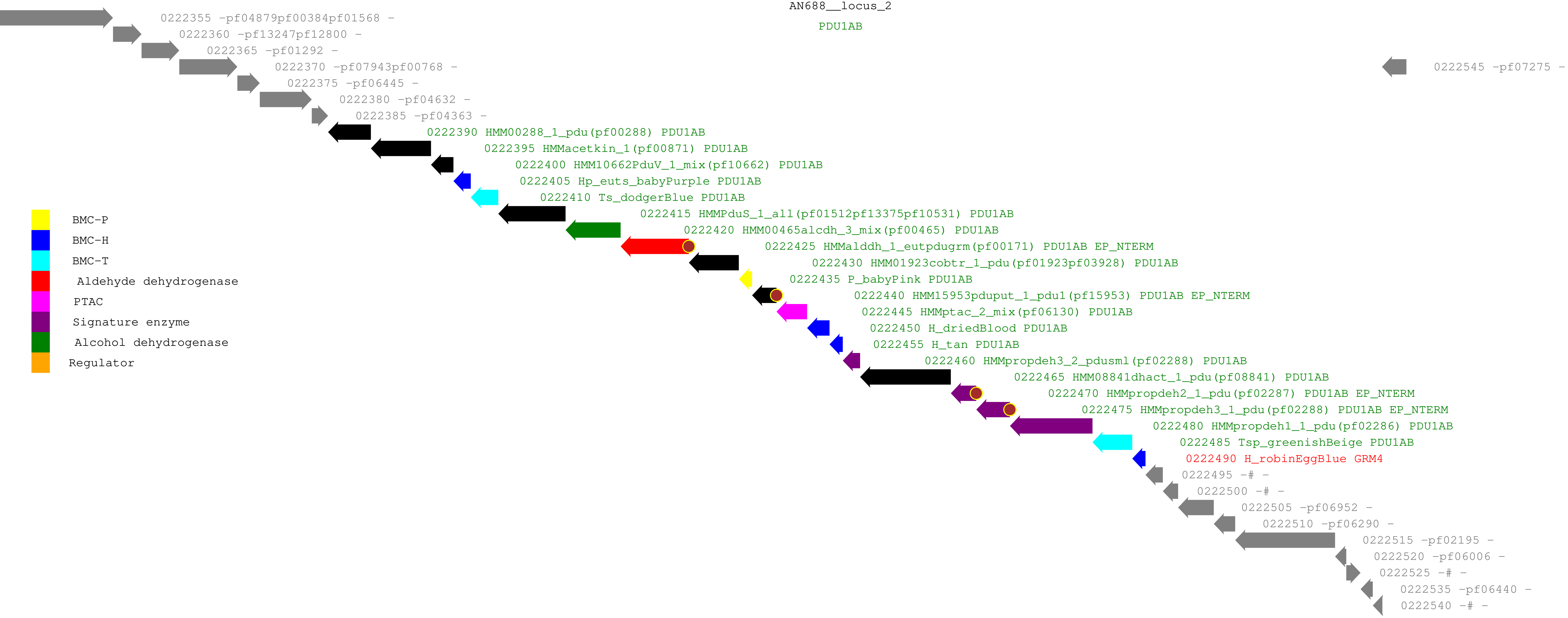

Salmonella enterica I. Bacteria; Proteobacteria; Gammaproteobacteria; Enterobacterales;

AO411__locus_2_main_sat

PDU1AB

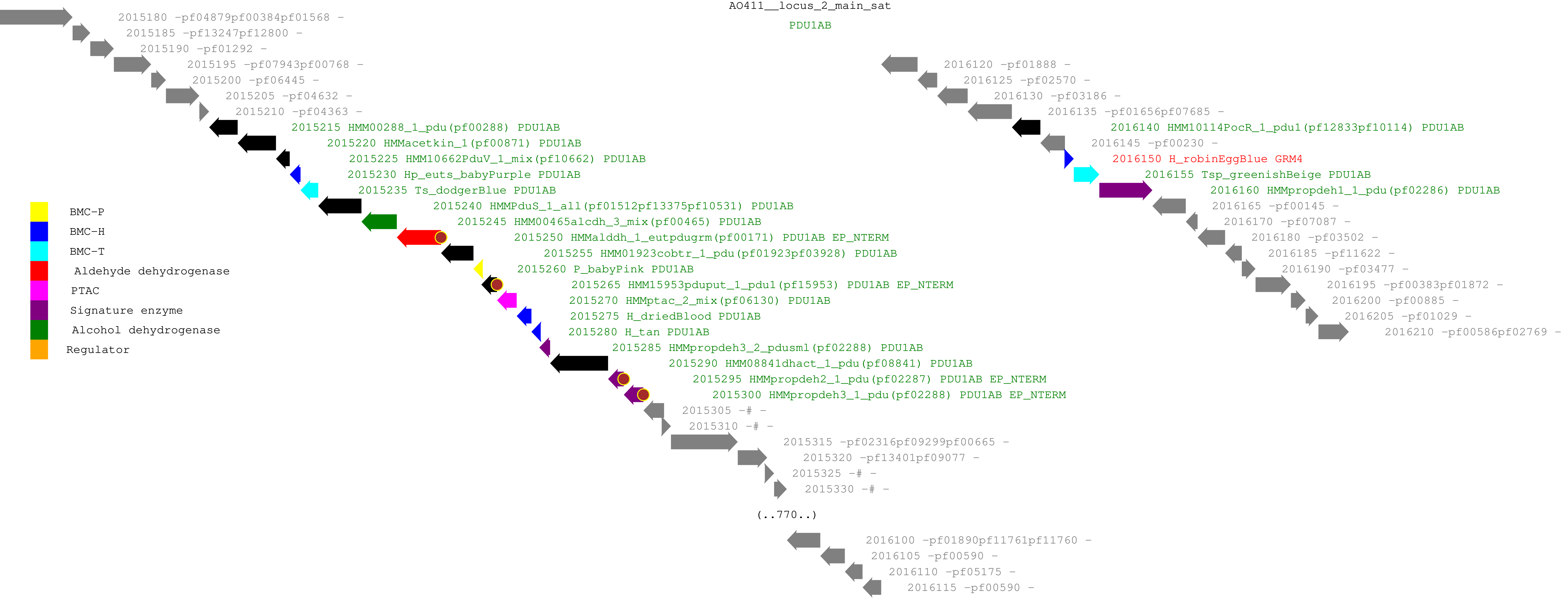

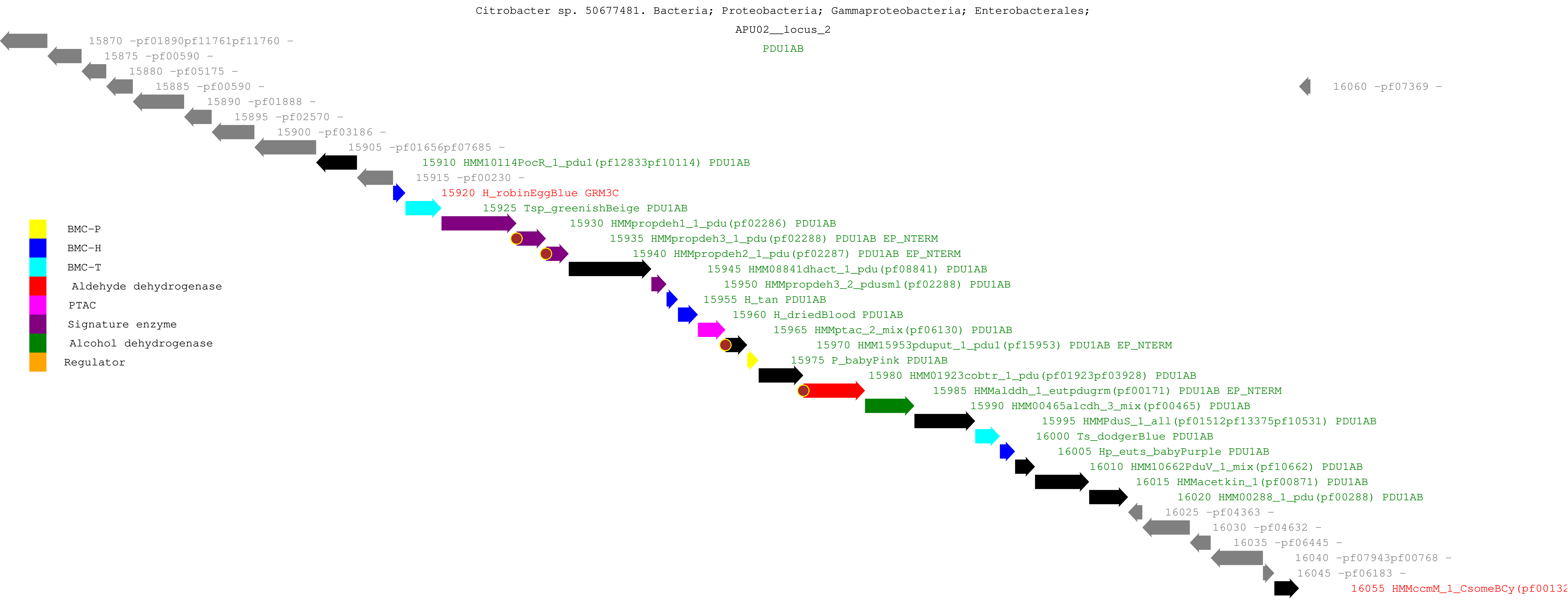
Citrobacter sp. 50677481. Bacteria; Proteobacteria; Gammaproteobacteria; Enterobacterales;
APU02__locus_2
PDU1AB
15870 -pf01890pf11761pf11760 -
15875 -pf00590 -
15880 -pf05175 -
15885 -pf00590 -
15890 -pf01888 -
15895 -pf02570 -
15900 -pf03186 -
15905 -pf01656pf07685 -
15910 HMM10114PocR_1_pdu1(pf12833pf10114) PDU1AB
15915 -pf00230 -
15920 H_robinEggBlue GRM3C
15925 Tsp_greenishBeige PDU1AB
15930 HMMpropdeh1_1_pdu(pf02286) PDU1AB
15935 HMMpropdeh3_1_pdu(pf02288) PDU1AB EP_NTERM
15940 HMMpropdeh2_1_pdu(pf02287) PDU1AB EP_NTERM
15945 HMM08841dhact_1_pdu(pf08841) PDU1AB
15950 HMMpropdeh3_2_pdusml(pf02288) PDU1AB
15955 H_tan PDU1AB
15960 H_driedBlood PDU1AB
15965 HMMptac_2_mix(pf06130) PDU1AB
15970 HMM15953pduput_1_pdu1(pf15953) PDU1AB EP_NTERM
15975 P_babyPink PDU1AB
15980 HMM01923cobtr_1_pdu(pf01923pf03928) PDU1AB
15985 HMMalddh_1_eutpdugrm(pf00171) PDU1AB EP_NTERM
15990 HMM00465alcdh_3_mix(pf00465) PDU1AB
15995 HMMPduS_1_all(pf01512pf13375pf10531) PDU1AB
16000 Ts_dodgerBlue PDU1AB
16005 Hp_euts_babyPurple PDU1AB
16010 HMM10662PduV_1_mix(pf10662) PDU1AB
16015 HMMacetkin_1(pf00871) PDU1AB
16020 HMM00288_1_pdu(pf00288) PDU1AB
16025 -pf04363 -
16030 -pf04632 -
16035 -pf06445 -
16040 -pf07943pf00768 -
16045 -pf06183 -
16055 HMMccmM_1_CsomeBCy(pf00132
16060 -pf07369 -
BMC-P
BMC-H
BMC-T
Aldehyde dehydrogenase
PTAC
Signature enzyme
Alcohol dehydrogenase
Regulator

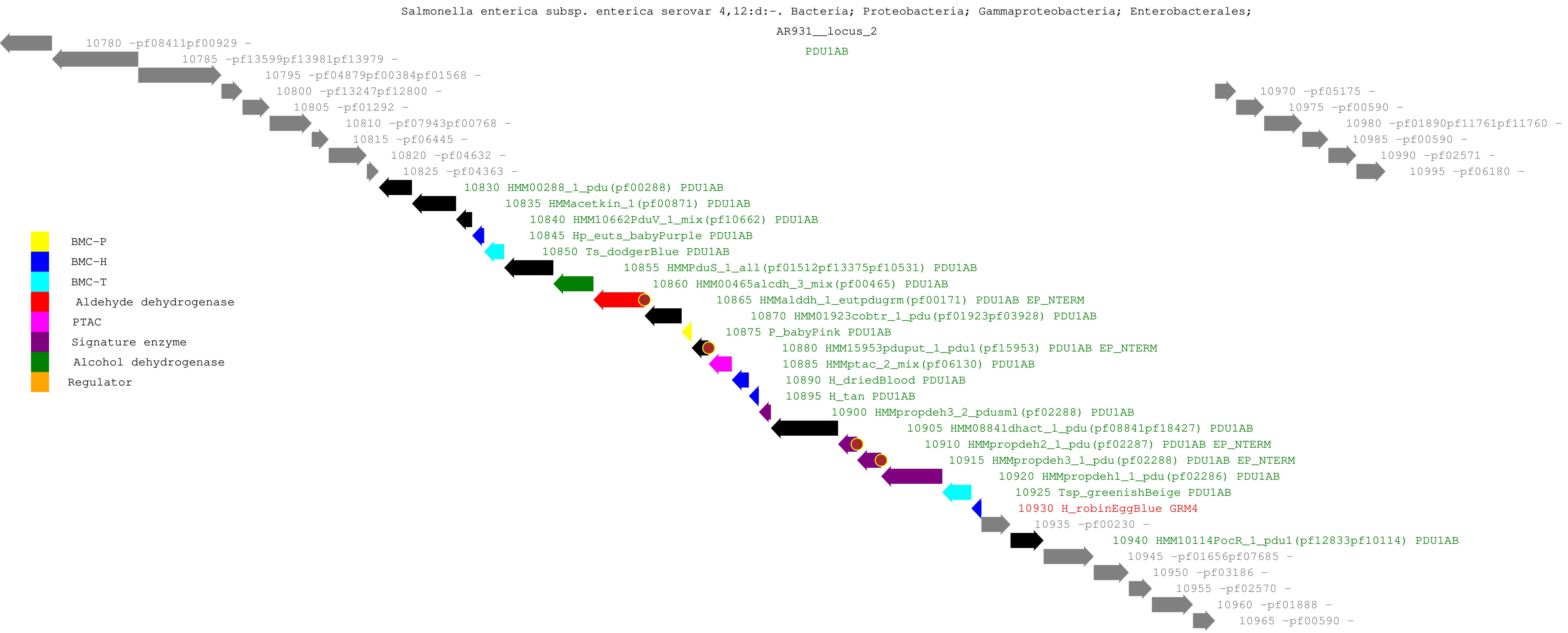

Salmonella enterica subsp. enterica serovar 4,12:d:-. Bacteria; Proteobacteria; Gammaproteobacteria; Enterobacterales;
AR931__locus_2
PDU1AB
10780 -pf08411pf00929 -
10785 -pf13599pf13981pf13979 -
10795 -pf04879pf00384pf01568 -
10800 -pf13247pf12800 -
10805 -pf01292 -
10810 -pf07943pf00768 -
10815 -pf06445 -
10820 -pf04632 -
10825 -pf04363 -
10830 HMM00288_1_pdu(pf00288) PDU1AB
10835 HMMacetkin_1(pf00871) PDU1AB
10840 HMM10662PduV_1_mix(pf10662) PDU1AB
10845 Hp_euts_babyPurple PDU1AB
10850 Ts_dodgerBlue PDU1AB
10855 HMMPduS_1_all(pf01512pf13375pf10531) PDU1AB
10860 HMM00465alcdh_3_mix(pf00465) PDU1AB
10865 HMMalddh_1_eutpdugrm(pf00171) PDU1AB EP_NTERM
10870 HMM01923cobtr_1_pdu(pf01923pf03928) PDU1AB
10875 P_babyPink PDU1AB
10880 HMM15953pduput_1_pdu1(pf15953) PDU1AB EP_NTERM
10885 HMMptac_2_mix(pf06130) PDU1AB
10890 H_driedBlood PDU1AB
10895 H_tan PDU1AB
10900 HMMpropdeh3_2_pdusml(pf02288) PDU1AB
10905 HMM08841dhact_1_pdu(pf08841pf18427) PDU1AB
10910 HMMpropdeh2_1_pdu(pf02287) PDU1AB EP_NTERM
10915 HMMpropdeh3_1_pdu(pf02288) PDU1AB EP_NTERM
10920 HMMpropdeh1_1_pdu(pf02286) PDU1AB
10925 Tsp_greenishBeige PDU1AB
10930 H_robinEggBlue GRM4
10935 -pf00230 -
10940 HMM10114PocR_1_pdu1(pf12833pf10114) PDU1AB
10945 -pf01656pf07685 -
10950 -pf03186 -
10955 -pf02570 -
10960 -pf01888 -
10965 -pf00590 -
10970 -pf05175 -
10975 -pf00590 -
10980 -pf01890pf11761pf11760 -
10985 -pf00590 -
10990 -pf02571 -
10995 -pf06180 -
BMC-P
BMC-H
BMC-T
Aldehyde dehydrogenase
PTAC
Signature enzyme
Alcohol dehydrogenase
Regulator

Salmonella enterica subsp. enterica serovar Heidelberg. Bacteria; Proteobacteria; Gammaproteobacteria; Enterobacterales;

ARF21__locus_1

PDU1AB

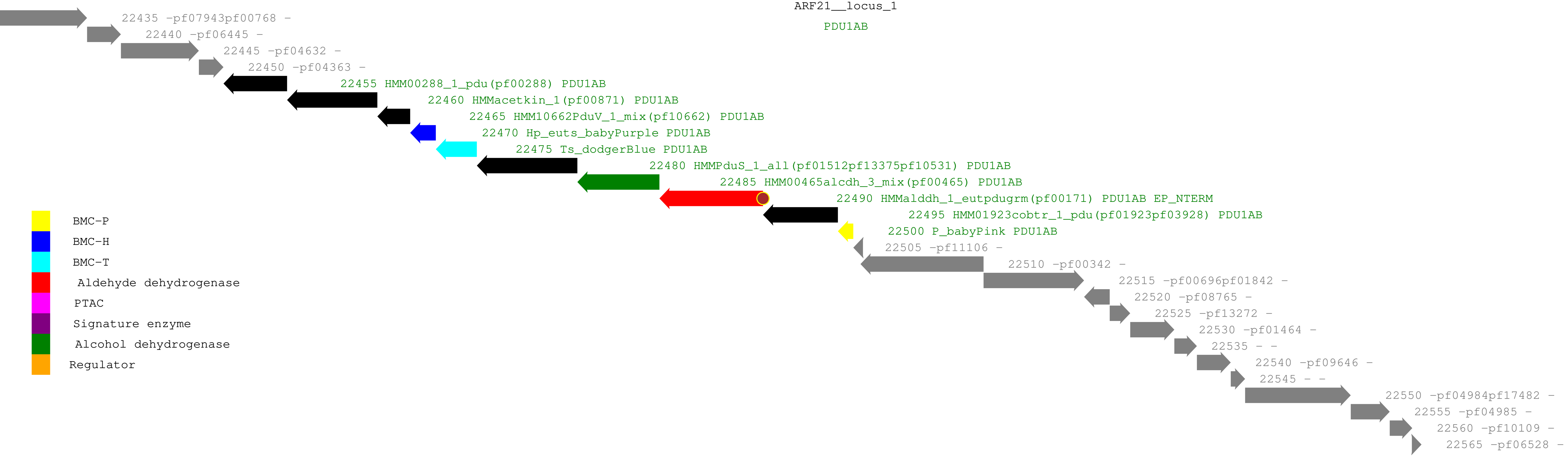

Salmonella enterica subsp. enterica serovar 4,5,12:r:-. Bacteria; Proteobacteria; Gammaproteobacteria; Enterobacterales;

ASI96__locus_1

PDU1AB

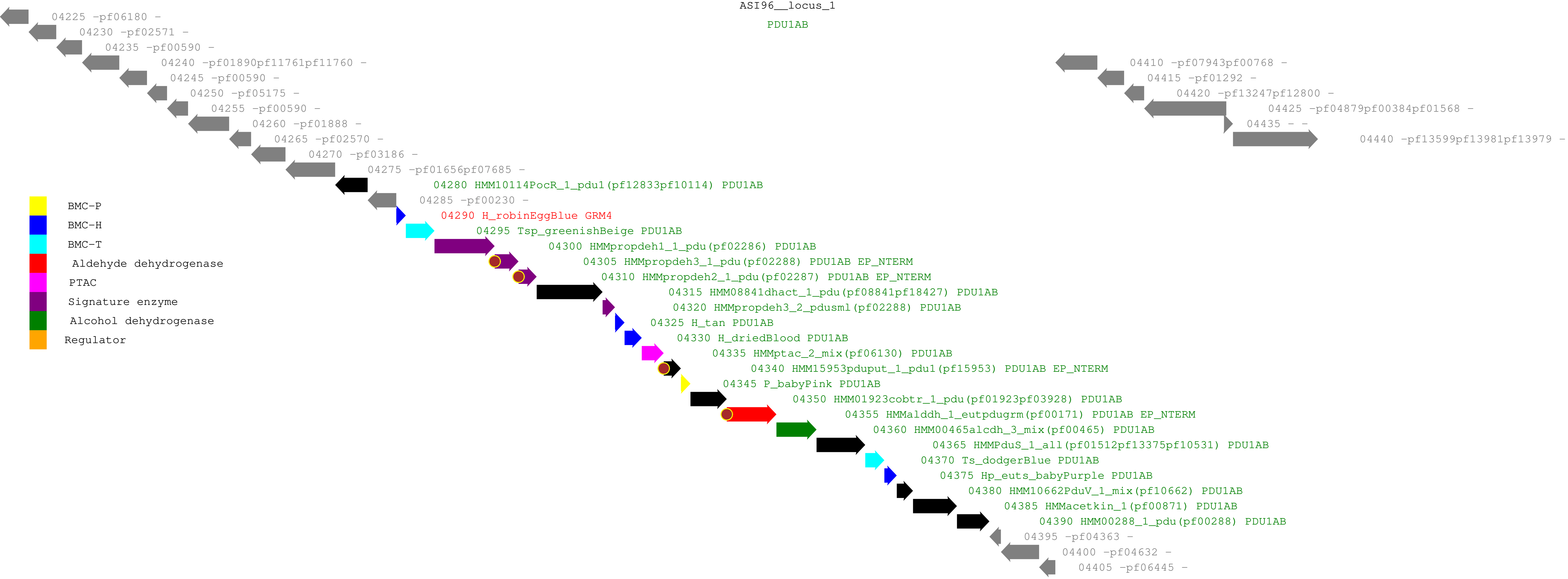

Salmonella enterica (Salmonella choleraesuis). Bacteria; Proteobacteria; Gammaproteobacteria; Enterobacterales;

ASQ14__locus_1

PDU1AB

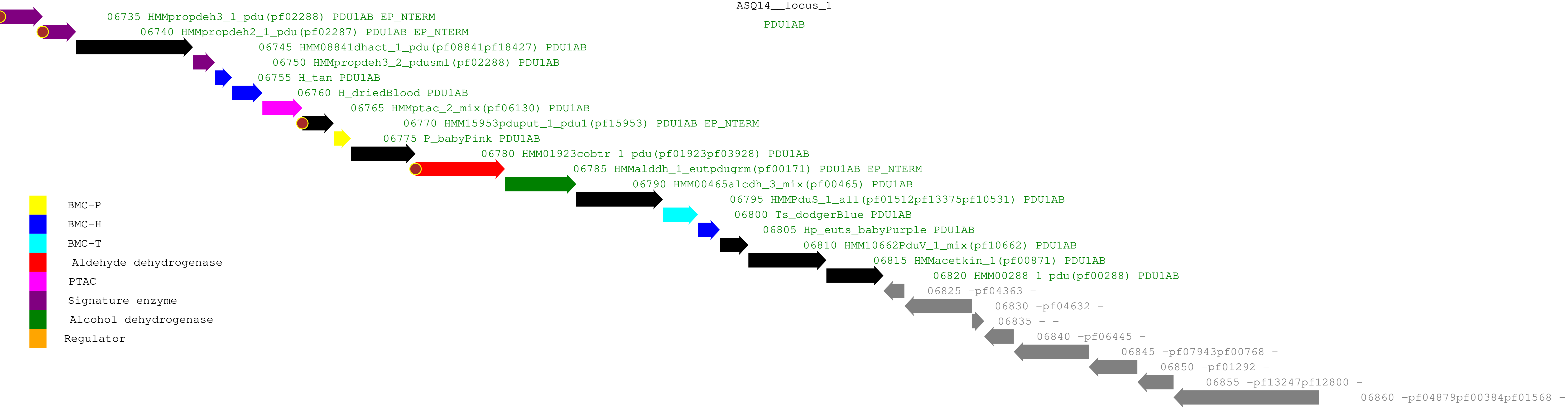

Salmonella enterica (Salmonella choleraesuis). Bacteria; Proteobacteria; Gammaproteobacteria; Enterobacterales;

ATP91__locus_2

PDU1AB

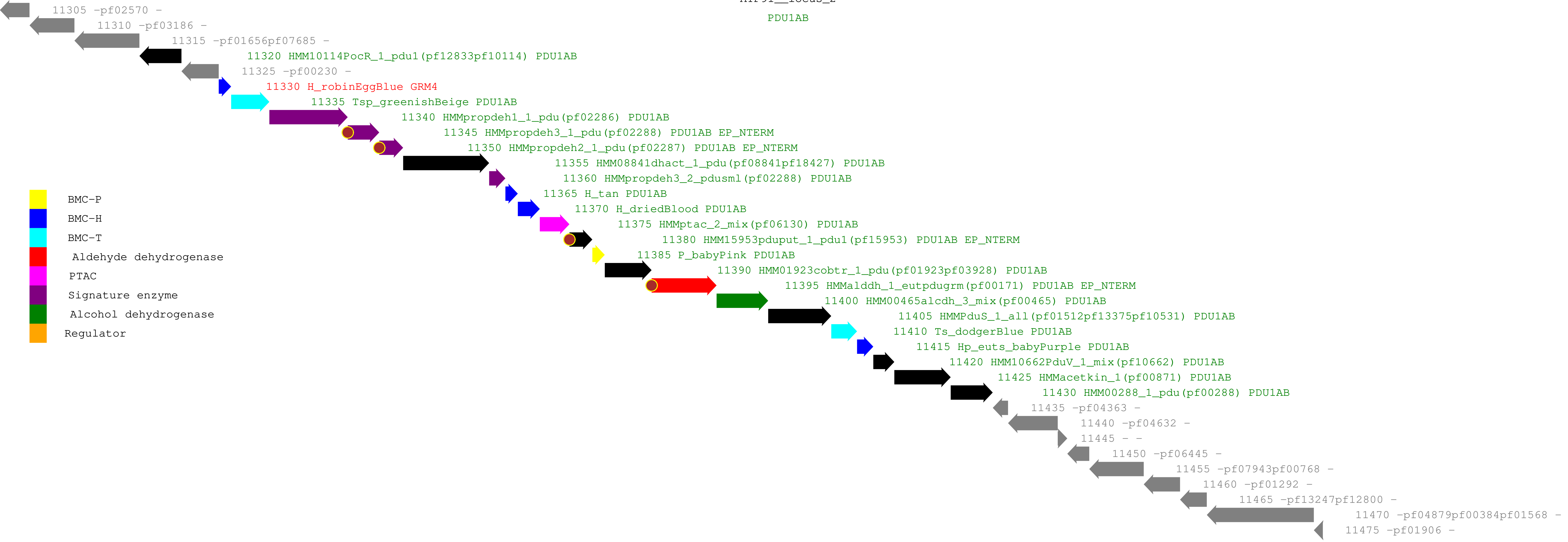

Salmonella enterica (Salmonella choleraesuis). Bacteria; Proteobacteria; Gammaproteobacteria; Enterobacterales;

ATQ15__locus_1

PDU1AB

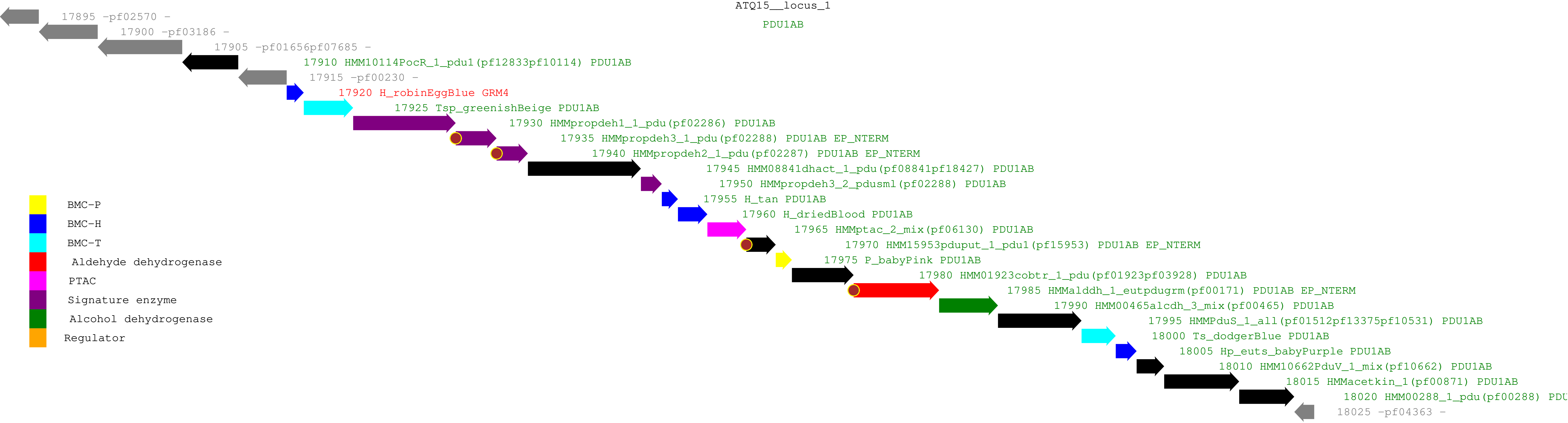

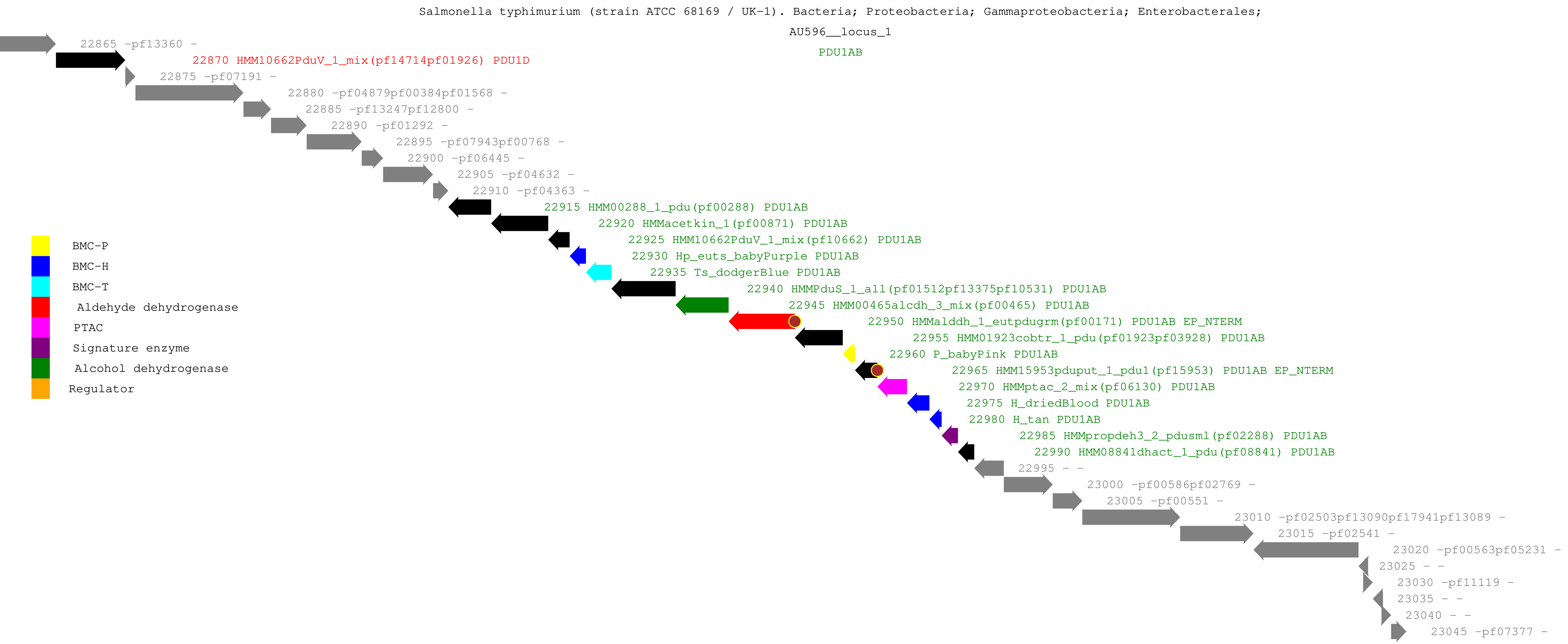
Salmonella typhimurium (strain ATCC 68169 / UK-1). Bacteria; Proteobacteria; Gammaproteobacteria; Enterobacterales;
AU596__locus_1
PDU1AB
22865 -pf13360 -
22870 HMM10662PduV_1_mix(pf14714pf01926) PDU1D
22875 -pf07191 -
22880 -pf04879pf00384pf01568 -
22885 -pf13247pf12800 -
22890 -pf01292 -
22895 -pf07943pf00768 -
22900 -pf06445 -
22905 -pf04632 -
22910 -pf04363 -
22915 HMM00288_1_pdu(pf00288) PDU1AB
22920 HMMacetkin_1(pf00871) PDU1AB
22925 HMM10662PduV_1_mix(pf10662) PDU1AB
22930 Hp_euts_babyPurple PDU1AB
22935 Ts_dodgerBlue PDU1AB
22940 HMMPduS_1_all(pf01512pf13375pf10531) PDU1AB
22945 HMM00465alcdh_3_mix(pf00465) PDU1AB
22950 HMMalddh_1_eutpdugrm(pf00171) PDU1AB EP_NTERM
22955 HMM01923cobtr_1_pdu(pf01923pf03928) PDU1AB
22960 P_babyPink PDU1AB
22965 HMM15953pduput_1_pdu1(pf15953) PDU1AB EP_NTERM
22970 HMMptac_2_mix(pf06130) PDU1AB
22975 H_driedBlood PDU1AB
22980 H_tan PDU1AB
22985 HMMpropdeh3_2_pdusml(pf02288) PDU1AB
22990 HMM08841dhact_1_pdu(pf08841) PDU1AB
22995 - -
23000 -pf00586pf02769 -
23005 -pf00551 -
23010 -pf02503pf13090pf17941pf13089 -
23015 -pf02541 -
23020 -pf00563pf05231 -
23025 - -
23030 -pf11119 -
23035 - -
23040 - -
23045 -pf07377 -
BMC-P
BMC-H
BMC-T
Aldehyde dehydrogenase
PTAC
Signature enzyme
Alcohol dehydrogenase
Regulator

Salmonella enterica subsp. enterica serovar Sekondi. Bacteria; Proteobacteria; Gammaproteobacteria; Enterobacterales;

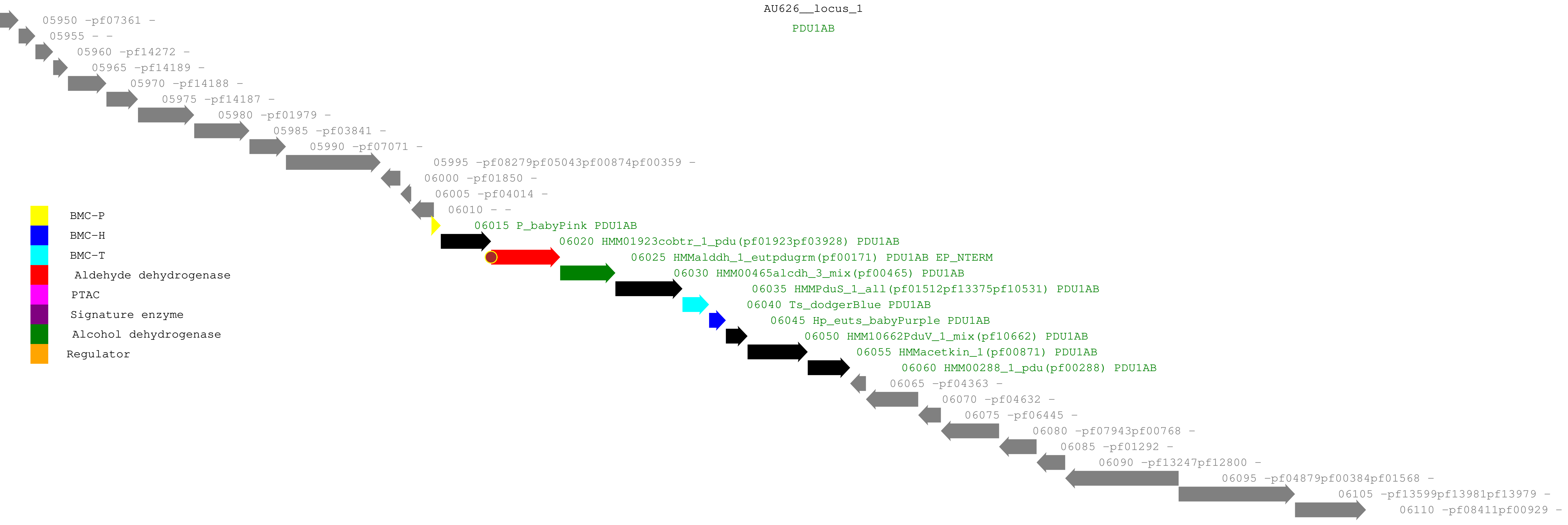

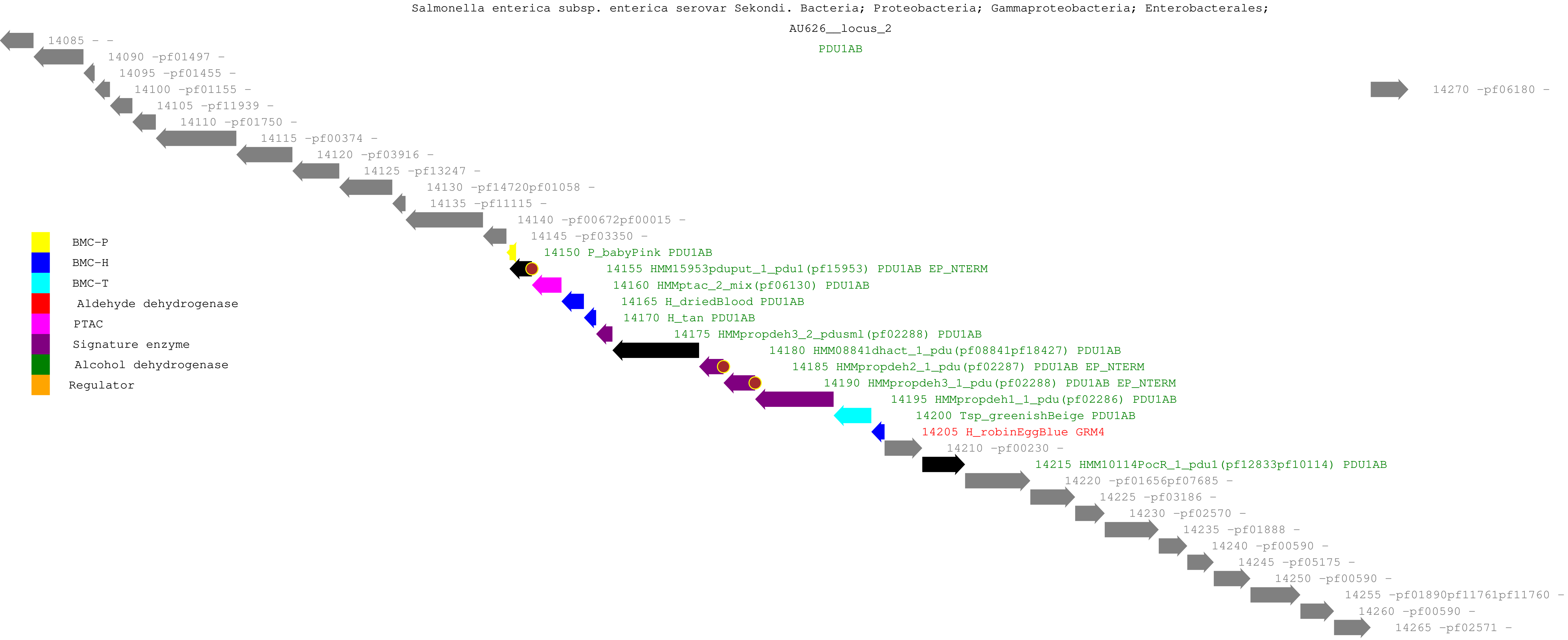
Salmonella enterica subsp. enterica serovar Sekondi. Bacteria; Proteobacteria; Gammaproteobacteria; Enterobacterales;
AU626__locus_2
PDU1AB
14085 - -
14090 -pf01497 -
14095 -pf01455 -
14100 -pf01155 -
14105 -pf11939 -
14110 -pf01750 -
14115 -pf00374 -
14120 -pf03916 -
14125 -pf13247 -
14130 -pf14720pf01058 -
14135 -pf11115 -
14140 -pf00672pf00015 -
14145 -pf03350 -
14150 P_babyPink PDU1AB
14155 HMM15953pduput_1_pdu1(pf15953) PDU1AB EP_NTERM
14160 HMMptac_2_mix(pf06130) PDU1AB
14165 H_driedBlood PDU1AB
14170 H_tan PDU1AB
14175 HMMpropdeh3_2_pdusml(pf02288) PDU1AB
14180 HMM08841dhact_1_pdu(pf08841pf18427) PDU1AB
14185 HMMpropdeh2_1_pdu(pf02287) PDU1AB EP_NTERM
14190 HMMpropdeh3_1_pdu(pf02288) PDU1AB EP_NTERM
14195 HMMpropdeh1_1_pdu(pf02286) PDU1AB
14200 Tsp_greenishBeige PDU1AB
14205 H_robinEggBlue GRM4
14210 -pf00230 -
14215 HMM10114PocR_1_pdu1(pf12833pf10114) PDU1AB
14220 -pf01656pf07685 -
14225 -pf03186 -
14230 -pf02570 -
14235 -pf01888 -
14240 -pf00590 -
14245 -pf05175 -
14250 -pf00590 -
14255 -pf01890pf11761pf11760 -
14260 -pf00590 -
14265 -pf02571 -
14270 -pf06180 -
BMC-P
BMC-H
BMC-T
Aldehyde dehydrogenase
PTAC
Signature enzyme
Alcohol dehydrogenase
Regulator

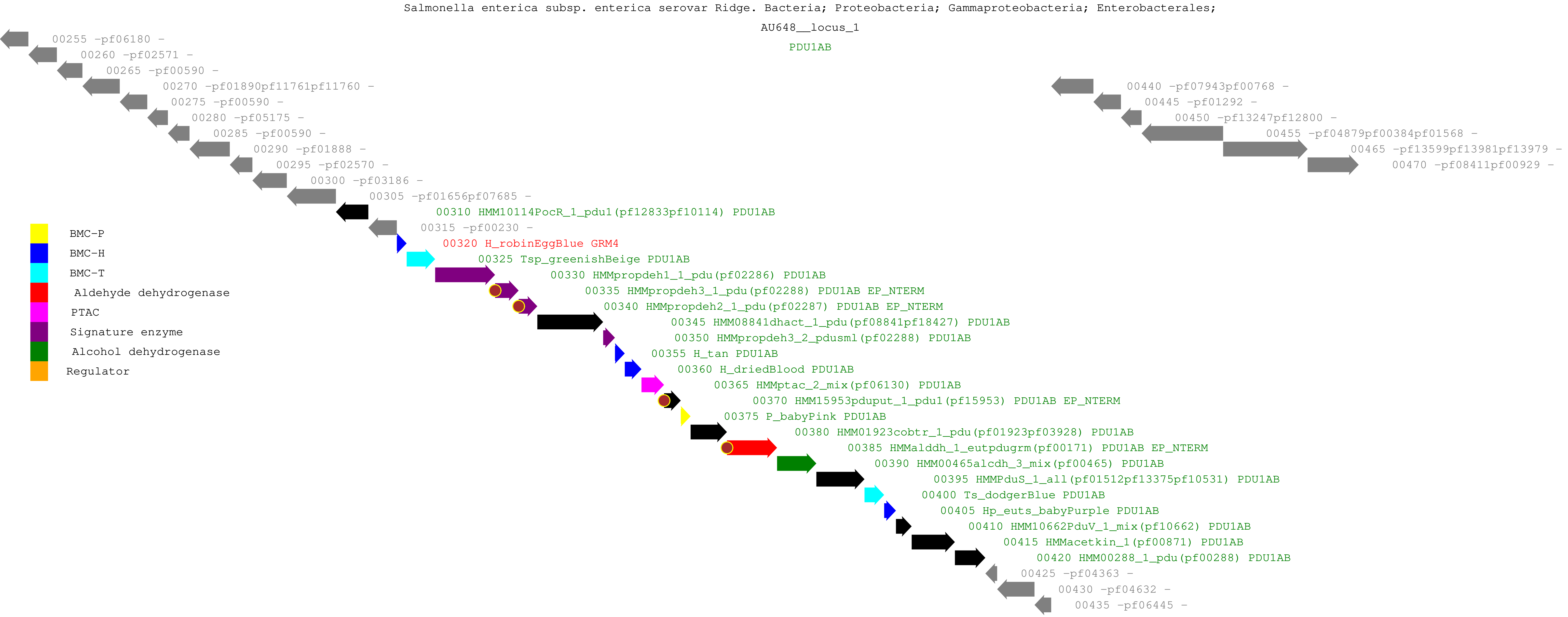
Salmonella enterica subsp. enterica serovar Ridge. Bacteria; Proteobacteria; Gammaproteobacteria; Enterobacterales;
AU648__locus_1
PDU1AB
00255 -pf06180 -
00260 -pf02571 -
00265 -pf00590 -
00270 -pf01890pf11761pf11760 -
00275 -pf00590 -
00280 -pf05175 -
00285 -pf00590 -
00290 -pf01888 -
00295 -pf02570 -
00300 -pf03186 -
00305 -pf01656pf07685 -
00310 HMM10114PocR_1_pdu1(pf12833pf10114) PDU1AB
00315 -pf00230 -
00320 H_robinEggBlue GRM4
00325 Tsp_greenishBeige PDU1AB
00330 HMMpropdeh1_1_pdu(pf02286) PDU1AB
00335 HMMpropdeh3_1_pdu(pf02288) PDU1AB EP_NTERM
00340 HMMpropdeh2_1_pdu(pf02287) PDU1AB EP_NTERM
00345 HMM08841dhact_1_pdu(pf08841pf18427) PDU1AB
00350 HMMpropdeh3_2_pdusml(pf02288) PDU1AB
00355 H_tan PDU1AB
00360 H_driedBlood PDU1AB
00365 HMMptac_2_mix(pf06130) PDU1AB
00370 HMM15953pduput_1_pdu1(pf15953) PDU1AB EP_NTERM
00375 P_babyPink PDU1AB
00380 HMM01923cobtr_1_pdu(pf01923pf03928) PDU1AB
00385 HMMaldh_1_eutpdugrm(pf00171) PDU1AB EP_NTERM
00390 HMM00465alcdh_3_mix(pf00465) PDU1AB
00395 HMMPduS_1_all(pf01512pf13375pf10531) PDU1AB
00400 Ts_dodgerBlue PDU1AB
00405 Hp_euts_babyPurple PDU1AB
00410 HMM10662PduV_1_mix(pf10662) PDU1AB
00415 HMMacetkin_1(pf00871) PDU1AB
00420 HMM00288_1_pdu(pf00288) PDU1AB
00425 -pf04363 -
00430 -pf04632 -
00435 -pf06445 -
00440 -pf07943pf00768 -
00445 -pf01292 -
00450 -pf13247pf12800 -
00455 -pf04879pf00384pf01568 -
00465 -pf13599pf13981pf13979 -
00470 -pf08411pf00929 -
BMC-P
BMC-H
BMC-T
Aldehyde dehydrogenase
PTAC
Signature enzyme
Alcohol dehydrogenase
Regulator

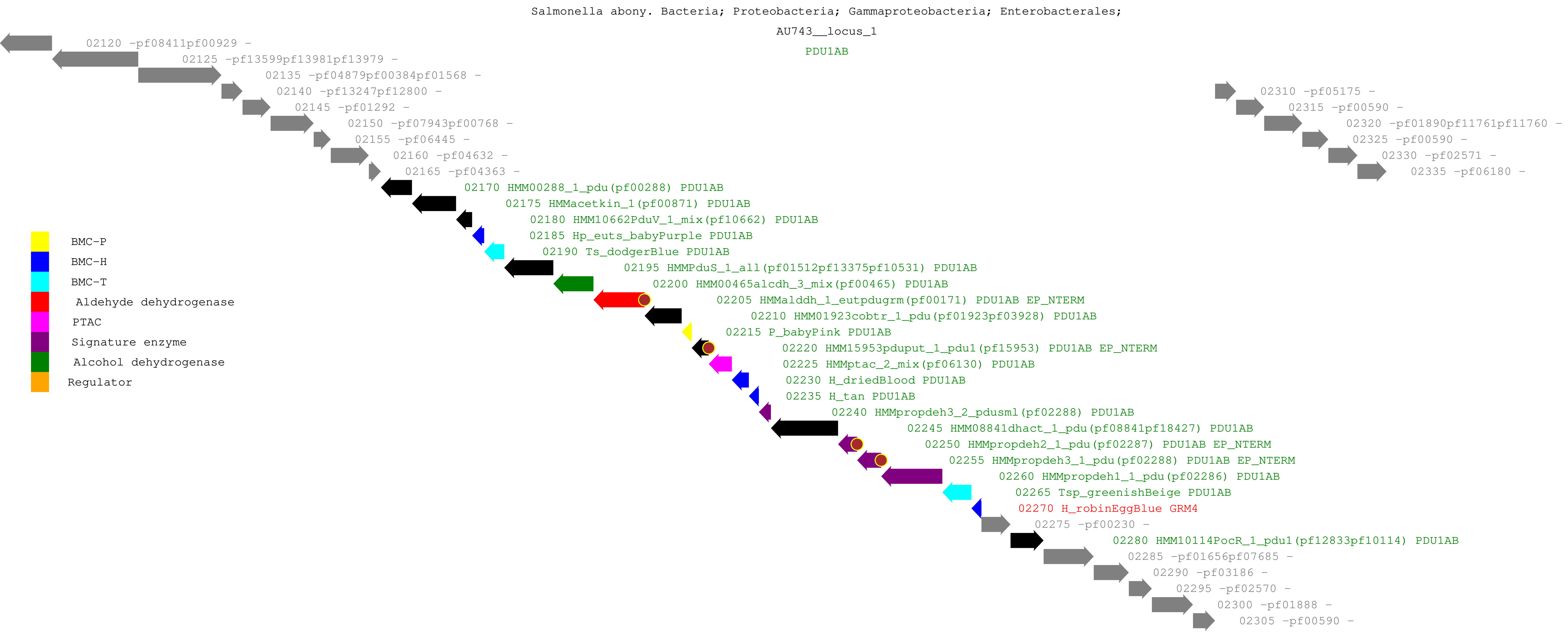

Salmonella abony. Bacteria; Proteobacteria; Gammaproteobacteria; Enterobacterales;
AU743__locus_1
PDU1AB
02120 -pf08411pf00929 -
02125 -pf13599pf13981pf13979 -
02135 -pf04879pf00384pf01568 -
02140 -pf13247pf12800 -
02145 -pf01292 -
02150 -pf07943pf00768 -
02155 -pf06445 -
02160 -pf04632 -
02165 -pf04363 -
02170 HMM00288_1_pdu(pf00288) PDU1AB
02175 HMMacetkin_1(pf00871) PDU1AB
02180 HMM10662PduV_1_mix(pf10662) PDU1AB
02185 Hp_euts_babyPurple PDU1AB
02190 Ts_dodgerBlue PDU1AB
02195 HMMPduS_1_all(pf01512pf13375pf10531) PDU1AB
02200 HMM00465alcdh_3_mix(pf00465) PDU1AB
02205 HMMalddh_1_eutpdugrm(pf00171) PDU1AB EP_NTERM
02210 HMM01923cobtr_1_pdu(pf01923pf03928) PDU1AB
02215 P_babyPink PDU1AB
02220 HMM15953pduput_1_pdu1(pf15953) PDU1AB EP_NTERM
02225 HMMptac_2_mix(pf06130) PDU1AB
02230 H_driedBlood PDU1AB
02235 H_tan PDU1AB
02240 HMMpropdeh3_2_pdusml(pf02288) PDU1AB
02245 HMM08841dhact_1_pdu(pf08841pf18427) PDU1AB
02250 HMMpropdeh2_1_pdu(pf02287) PDU1AB EP_NTERM
02255 HMMpropdeh3_1_pdu(pf02288) PDU1AB EP_NTERM
02260 HMMpropdeh1_1_pdu(pf02286) PDU1AB
02265 Tsp_greenishBeige PDU1AB
02270 H_robinEggBlue GRM4
02275 -pf00230 -
02280 HMM10114PocR_1_pdu1(pf12833pf10114) PDU1AB
02285 -pf01656pf07685 -
02290 -pf03186 -
02295 -pf02570 -
02300 -pf01888 -
02305 -pf00590 -
02310 -pf05175 -
02315 -pf00590 -
02320 -pf01890pf11761pf11760 -
02325 -pf00590 -
02330 -pf02571 -
02335 -pf06180 -
BMC-P
BMC-H
BMC-T
Aldehyde dehydrogenase
PTAC
Signature enzyme
Alcohol dehydrogenase
Regulator

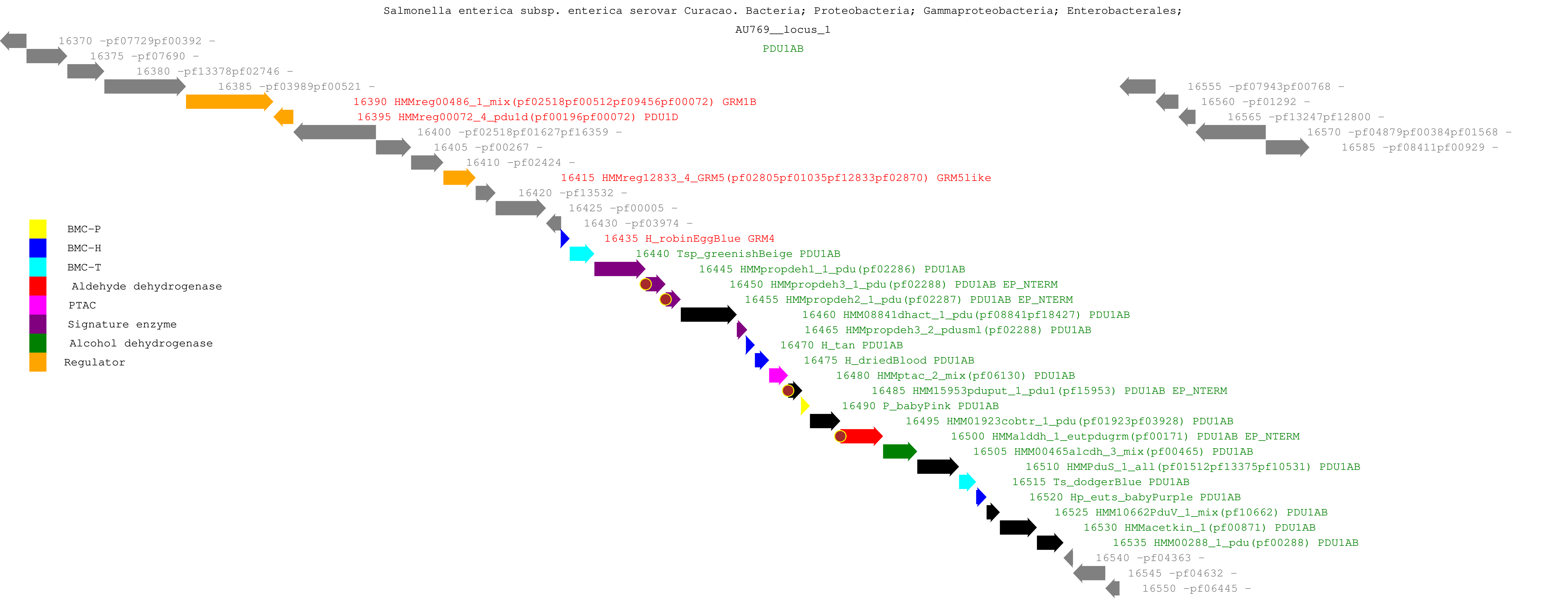

Salmonella enterica subsp. enterica serovar Curacao. Bacteria; Proteobacteria; Gammaproteobacteria; Enterobacterales;
AU769__locus_1
PDU1AB
16370 -pf07729pf00392 -
16375 -pf07690 -
16380 -pf13378pf02746 -
16385 -pf03989pf00521 -
16390 HMMreg00486_1_mix(pf02518pf00512pf09456pf00072) GRM1B
16395 HMMreg00072_4_pdu1d(pf00196pf00072) PDU1D
16400 -pf02518pf01627pf16359 -
16405 -pf00267 -
16410 -pf02424 -
16415 HMMreg12833_4_GRM5(pf02805pf01035pf12833pf02870) GRM5like
16420 -pf13532 -
16425 -pf00005 -
16430 -pf03974 -
16435 H_robinEggBlue GRM4
16440 Tsp_greenishBeige PDU1AB
16445 HMMpropdeh1_1_pdu(pf02286) PDU1AB
16450 HMMpropdeh3_1_pdu(pf02288) PDU1AB EP_NTERM
16455 HMMpropdeh2_1_pdu(pf02287) PDU1AB EP_NTERM
16460 HMM08841dhact_1_pdu(pf08841pf18427) PDU1AB
16465 HMMpropdeh3_2_pdusml(pf02288) PDU1AB
16470 H_tan PDU1AB
16475 H_driedBlood PDU1AB
16480 HMMptac_2_mix(pf06130) PDU1AB
16485 HMM15953pduput_1_pdu1(pf15953) PDU1AB EP_NTERM
16490 P_babyPink PDU1AB
16495 HMM01923cobtr_1_pdu(pf01923pf03928) PDU1AB
16500 HMMaldh_1_eutpdugrm(pf00171) PDU1AB EP_NTERM
16505 HMM00465alcdh_3_mix(pf00465) PDU1AB
16510 HMMPduS_1_all(pf01512pf13375pf10531) PDU1AB
16515 Ts_dodgerBlue PDU1AB
16520 Hp_euts_babyPurple PDU1AB
16525 HMM10662PduV_1_mix(pf10662) PDU1AB
16530 HMMacetkin_1(pf00871) PDU1AB
16535 HMM00288_1_pdu(pf00288) PDU1AB
16540 -pf04363 -
16545 -pf04632 -
16550 -pf06445 -
16555 -pf07943pf00768 -
16560 -pf01292 -
16565 -pf13247pf12800 -
16570 -pf04879pf00384pf01568 -
16585 -pf08411pf00929 -
BMC-P
BMC-H
BMC-T
Aldehyde dehydrogenase
PTAC
Signature enzyme
Alcohol dehydrogenase
Regulator

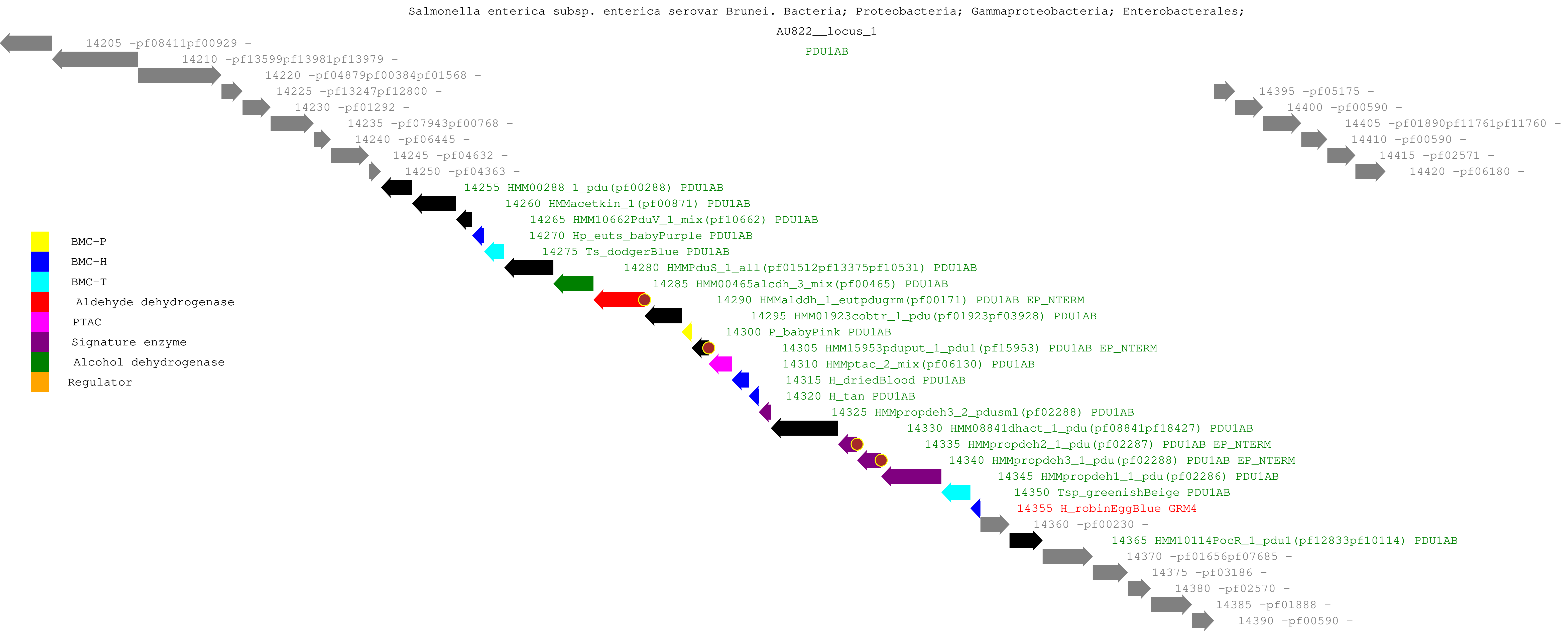

Salmonella enterica subsp. enterica serovar Brunei. Bacteria; Proteobacteria; Gammaproteobacteria; Enterobacterales;
AU822__locus_1
PDU1AB
14205 -pf08411pf00929 -
14210 -pf13599pf13981pf13979 -
14220 -pf04879pf00384pf01568 -
14225 -pf13247pf12800 -
14230 -pf01292 -
14235 -pf07943pf00768 -
14240 -pf06445 -
14245 -pf04632 -
14250 -pf04363 -
14255 HMM00288_1_pdu(pf00288) PDU1AB
14260 HMMacetkin_1(pf00871) PDU1AB
14265 HMM10662PduV_1_mix(pf10662) PDU1AB
14270 Hp_euts_babyPurple PDU1AB
14275 Ts_dodgerBlue PDU1AB
14280 HMMPduS_1_all(pf01512pf13375pf10531) PDU1AB
14285 HMM00465alcdh_3_mix(pf00465) PDU1AB
14290 HMMalddh_1_eutpdugrm(pf00171) PDU1AB EP_NTERM
14295 HMM01923cobtr_1_pdu(pf01923pf03928) PDU1AB
14300 P_babyPink PDU1AB
14305 HMM15953pduput_1_pdu1(pf15953) PDU1AB EP_NTERM
14310 HMMptac_2_mix(pf06130) PDU1AB
14315 H_driedBlood PDU1AB
14320 H_tan PDU1AB
14325 HMMpropdeh3_2_pdusml(pf02288) PDU1AB
14330 HMM08841dhact_1_pdu(pf08841pf18427) PDU1AB
14335 HMMpropdeh2_1_pdu(pf02287) PDU1AB EP_NTERM
14340 HMMpropdeh3_1_pdu(pf02288) PDU1AB EP_NTERM
14345 HMMpropdeh1_1_pdu(pf02286) PDU1AB
14350 Tsp_greenishBeige PDU1AB
14355 H_robinEggBlue GRM4
14360 -pf00230 -
14365 HMM10114PocR_1_pdu1(pf12833pf10114) PDU1AB
14370 -pf01656pf07685 -
14375 -pf03186 -
14380 -pf02570 -
14385 -pf01888 -
14390 -pf00590 -
14395 -pf05175 -
14400 -pf00590 -
14405 -pf01890pf11761pf11760 -
14410 -pf00590 -
14415 -pf02571 -
14420 -pf06180 -
BMC-P
BMC-H
BMC-T
Aldehyde dehydrogenase
PTAC
Signature enzyme
Alcohol dehydrogenase
Regulator

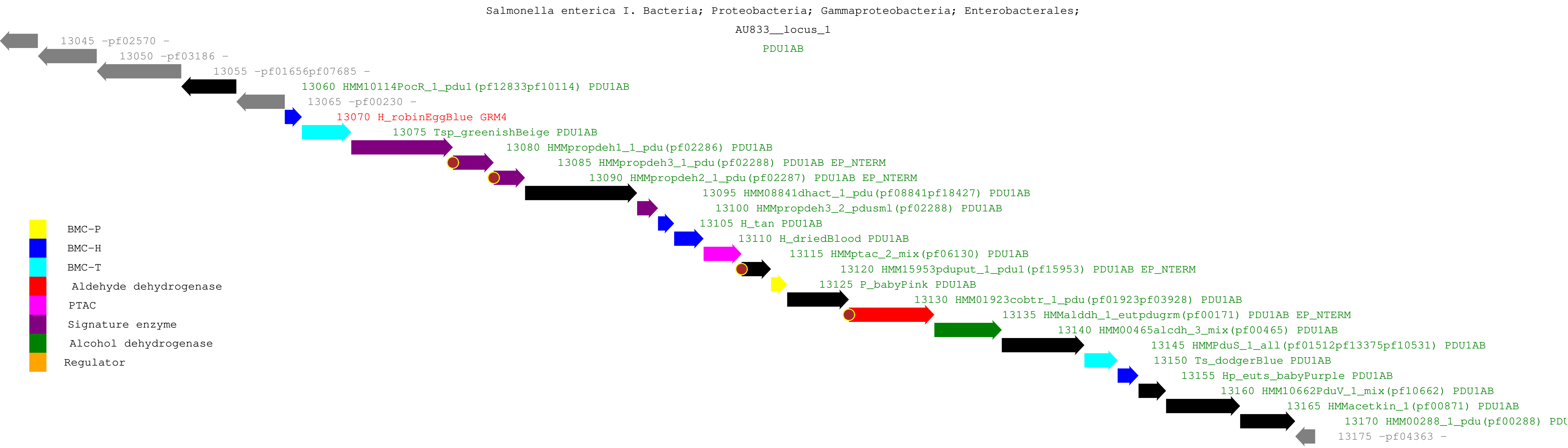

Salmonella enterica I. Bacteria; Proteobacteria; Gammaproteobacteria; Enterobacterales;
AU833__locus_1
PDU1AB
13045 -pf02570 -
13050 -pf03186 -
13055 -pf01656pf07685 -
13060 HMM10114PocR_1_pdu1(pf12833pf10114) PDU1AB
13065 -pf00230 -
13070 H_robinEggBlue GRM4
13075 Tsp_greenishBeige PDU1AB
13080 HMMpropdeh1_1_pdu(pf02286) PDU1AB
13085 HMMpropdeh3_1_pdu(pf02288) PDU1AB EP_NTERM
13090 HMMpropdeh2_1_pdu(pf02287) PDU1AB EP_NTERM
13095 HMM08841dhact_1_pdu(pf08841pf18427) PDU1AB
13100 HMMpropdeh3_2_pdusml(pf02288) PDU1AB
13105 H_tan PDU1AB
13110 H_driedBlood PDU1AB
13115 HMMptac_2_mix(pf06130) PDU1AB
13120 HMM15953pduput_1_pdu1(pf15953) PDU1AB EP_NTERM
13125 P_babyPink PDU1AB
13130 HMM01923cobtr_1_pdu(pf01923pf03928) PDU1AB
13135 HMMalddh_1_eutpdugrm(pf00171) PDU1AB EP_NTERM
13140 HMM00465alcdh_3_mix(pf00465) PDU1AB
13145 HMMPduS_1_all(pf01512pf13375pf10531) PDU1AB
13150 Ts_dodgerBlue PDU1AB
13155 Hp_euts_babyPurple PDU1AB
13160 HMM10662PduV_1_mix(pf10662) PDU1AB
13165 HMMacetkin_1(pf00871) PDU1AB
13170 HMM00288_1_pdu(pf00288) PDU
13175 -pf04363 -
BMC-P
BMC-H
BMC-T
Aldehyde dehydrogenase
PTAC
Signature enzyme
Alcohol dehydrogenase
Regulator

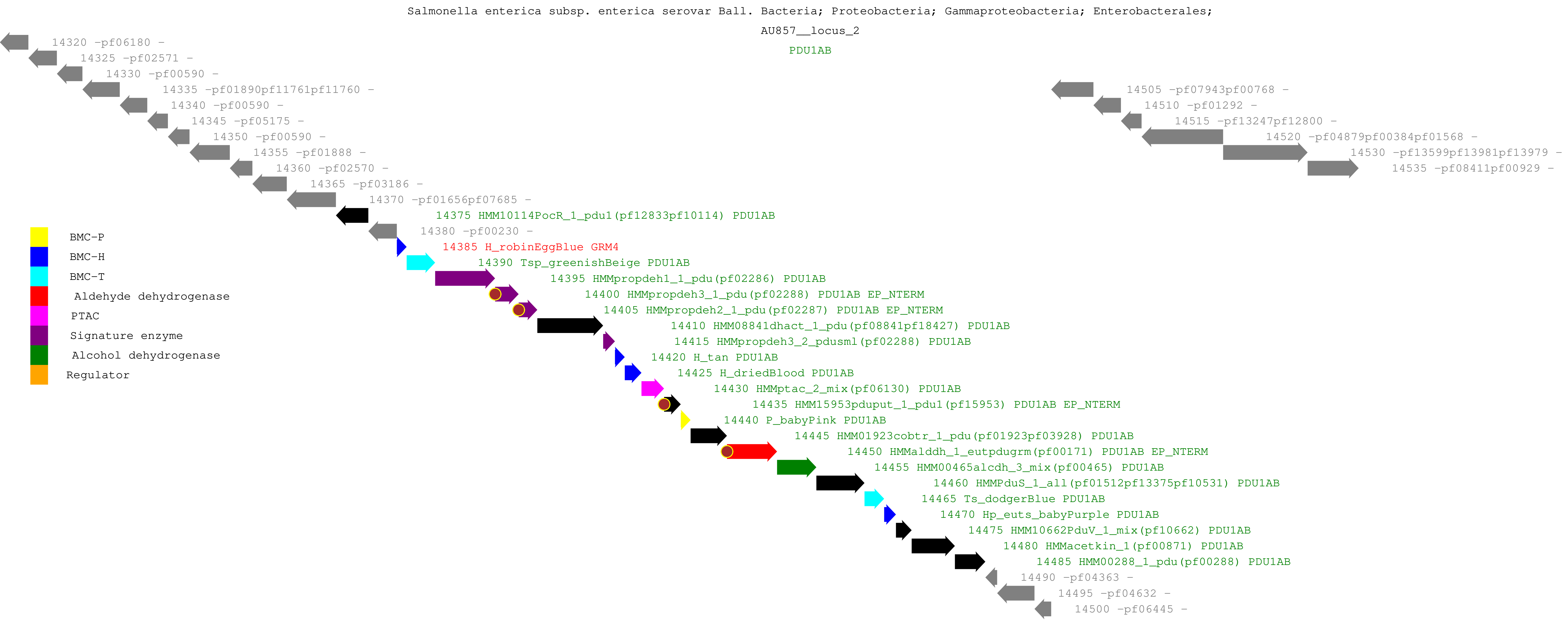

Salmonella enterica subsp. enterica serovar Ball. Bacteria; Proteobacteria; Gammaproteobacteria; Enterobacterales;
AU857__locus_2
PDU1AB
14320 -pf06180 -
14325 -pf02571 -
14330 -pf00590 -
14335 -pf01890pf11761pf11760 -
14340 -pf00590 -
14345 -pf05175 -
14350 -pf00590 -
14355 -pf01888 -
14360 -pf02570 -
14365 -pf03186 -
14370 -pf01656pf07685 -
14375 HMM10114PocR_1_pdu1(pf12833pf10114) PDU1AB
14380 -pf00230 -
14385 H_robinEggBlue GRM4
14390 Tsp_greenishBeige PDU1AB
14395 HMMpropdeh1_1_pdu(pf02286) PDU1AB
14400 HMMpropdeh3_1_pdu(pf02288) PDU1AB EP_NTERM
14405 HMMpropdeh2_1_pdu(pf02287) PDU1AB EP_NTERM
14410 HMM08841dhact_1_pdu(pf08841pf18427) PDU1AB
14415 HMMpropdeh3_2_pdusml(pf02288) PDU1AB
14420 H_tan PDU1AB
14425 H_driedBlood PDU1AB
14430 HMMptac_2_mix(pf06130) PDU1AB
14435 HMM15953pduput_1_pdu1(pf15953) PDU1AB EP_NTERM
14440 P_babyPink PDU1AB
14445 HMM01923cobtr_1_pdu(pf01923pf03928) PDU1AB
14450 HMMaldd_1_eutpdugrm(pf00171) PDU1AB EP_NTERM
14455 HMM00465alcdh_3_mix(pf00465) PDU1AB
14460 HMMPduS_1_all(pf01512pf13375pf10531) PDU1AB
14465 Ts_dodgerBlue PDU1AB
14470 Hp_euts_babyPurple PDU1AB
14475 HMM10662PduV_1_mix(pf10662) PDU1AB
14480 HMMacetkin_1(pf00871) PDU1AB
14485 HMM00288_1_pdu(pf00288) PDU1AB
14490 -pf04363 -
14495 -pf04632 -
14500 -pf06445 -
14505 -pf07943pf00768 -
14510 -pf01292 -
14515 -pf13247pf12800 -
14520 -pf04879pf00384pf01568 -
14530 -pf13599pf13981pf13979 -
14535 -pf08411pf00929 -
BMC-P
BMC-H
BMC-T
Aldehyde dehydrogenase
PTAC
Signature enzyme
Alcohol dehydrogenase
Regulator

Salmonella enterica subsp. enterica serovar Ibadan. Bacteria; Proteobacteria; Gammaproteobacteria; Enterobacterales;

AU867__locus_1

PDU1AB

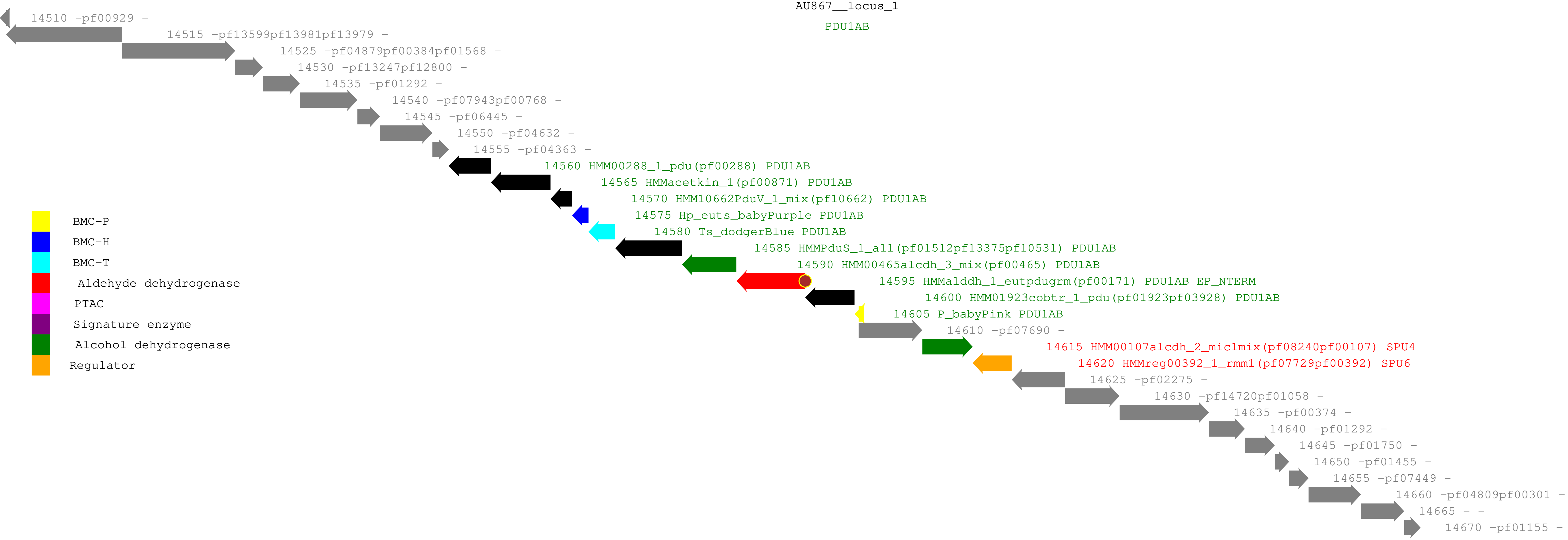

Salmonella enterica subsp. enterica serovar Kottbus. Bacteria; Proteobacteria; Gammaproteobacteria; Enterobacterales;

AU884__locus_1

PDU1AB

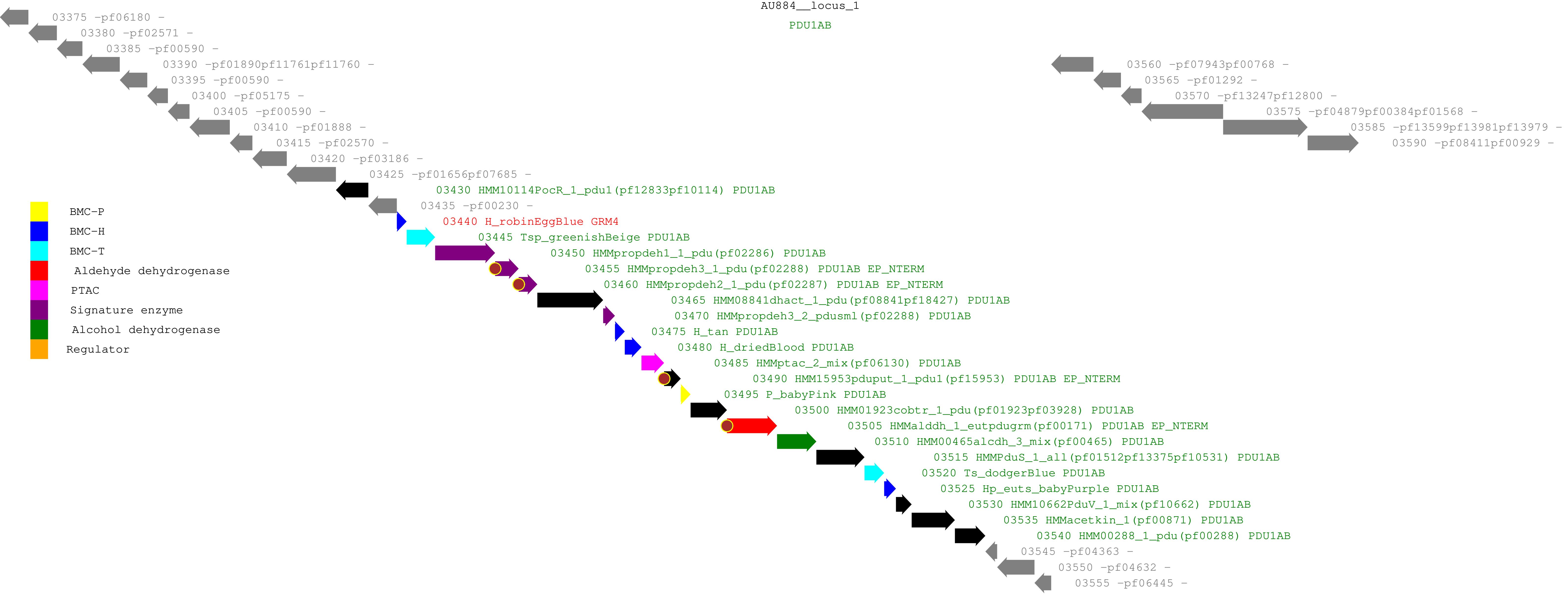

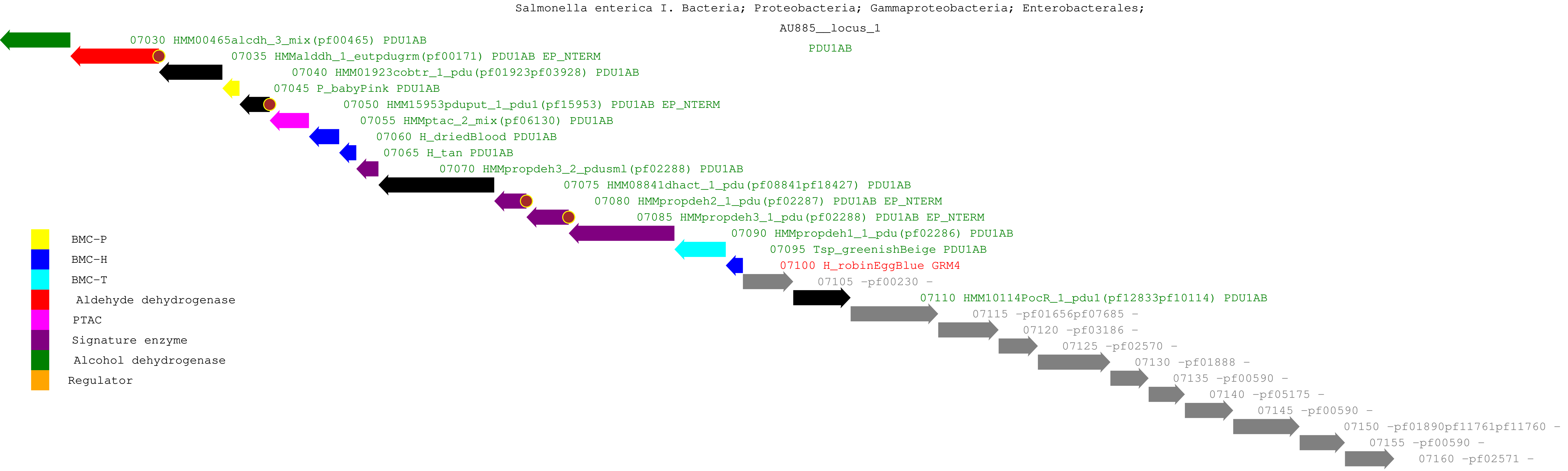
Salmonella enterica I. Bacteria; Proteobacteria; Gammaproteobacteria; Enterobacterales;
AU885__locus_1
PDU1AB
07030 HMM00465alcdh_3_mix(pf00465) PDU1AB
07035 HMMalddh_1_eutpdugrm(pf00171) PDU1AB EP_NTERM
07040 HMM01923cobtr_1_pdu(pf01923pf03928) PDU1AB
07045 P_babyPink PDU1AB
07050 HMM15953pduput_1_pdu1(pf15953) PDU1AB EP_NTERM
07055 HMMptac_2_mix(pf06130) PDU1AB
07060 H_driedBlood PDU1AB
07065 H_tan PDU1AB
07070 HMMpropdeh3_2_pdusml(pf02288) PDU1AB
07075 HMM08841dhact_1_pdu(pf08841pf18427) PDU1AB
07080 HMMpropdeh2_1_pdu(pf02287) PDU1AB EP_NTERM
07085 HMMpropdeh3_1_pdu(pf02288) PDU1AB EP_NTERM
07090 HMMpropdeh1_1_pdu(pf02286) PDU1AB
07095 Tsp_greenishBeige PDU1AB
07100 H_robinEggBlue GRM4
07105 -pf00230 -
07110 HMM10114PocR_1_pdu1(pf12833pf10114) PDU1AB
07115 -pf01656pf07685 -
07120 -pf03186 -
07125 -pf02570 -
07130 -pf01888 -
07135 -pf00590 -
07140 -pf05175 -
07145 -pf00590 -
07150 -pf01890pf11761pf11760 -
07155 -pf00590 -
07160 -pf02571 -
BMC-P
BMC-H
BMC-T
Aldehyde dehydrogenase
PTAC
Signature enzyme
Alcohol dehydrogenase
Regulator

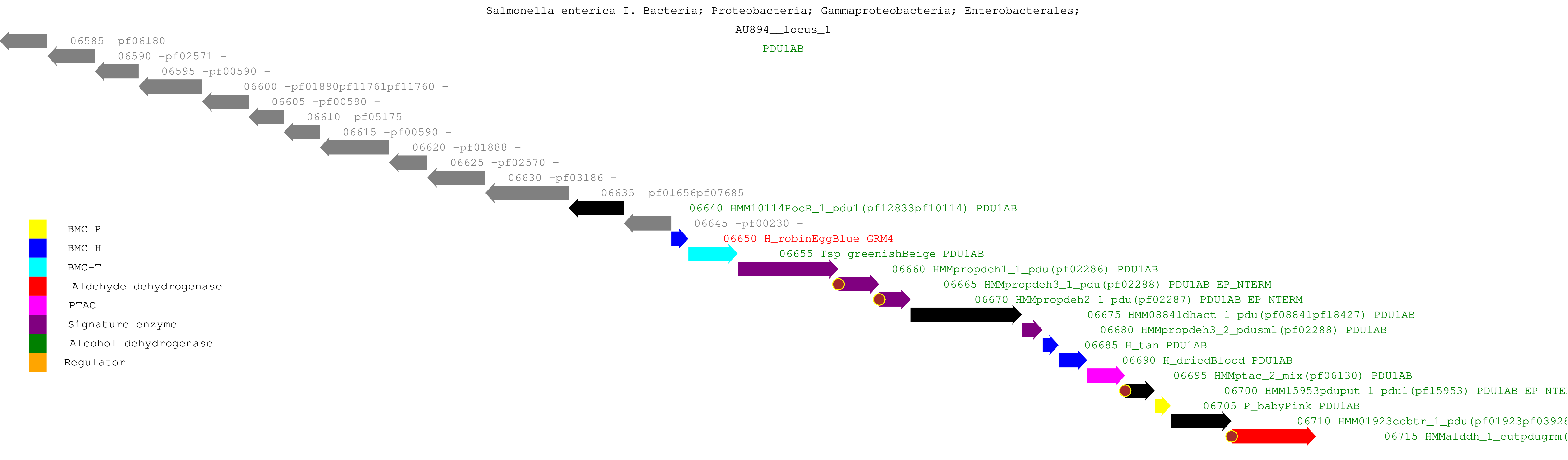

Salmonella enterica I. Bacteria; Proteobacteria; Gammaproteobacteria; Enterobacterales;
AU894__locus_1
PDU1AB
06585 -pf06180 -
06590 -pf02571 -
06595 -pf00590 -
06600 -pf01890pf11761pf11760 -
06605 -pf00590 -
06610 -pf05175 -
06615 -pf00590 -
06620 -pf01888 -
06625 -pf02570 -
06630 -pf03186 -
06635 -pf01656pf07685 -
06640 HMM10114PocR_1_pdu1(pf12833pf10114) PDU1AB
06645 -pf00230 -
06650 H_robinEggBlue GRM4
06655 Tsp_greenishBeige PDU1AB
06660 HMMpropdeh1_1_pdu(pf02286) PDU1AB
06665 HMMpropdeh3_1_pdu(pf02288) PDU1AB EP_NTERM
06670 HMMpropdeh2_1_pdu(pf02287) PDU1AB EP_NTERM
06675 HMM08841dhact_1_pdu(pf08841pf18427) PDU1AB
06680 HMMpropdeh3_2_pdusml(pf02288) PDU1AB
06685 H_tan PDU1AB
06690 H_driedBlood PDU1AB
06695 HMMptac_2_mix(pf06130) PDU1AB
06700 HMM15953pduput_1_pdu1(pf15953) PDU1AB EP_NTE
06705 P_babyPink PDU1AB
06710 HMM01923cobtr_1_pdu(pf01923pf0392
06715 HMMalddh_1_eutpdugrm(
BMC-P
BMC-H
BMC-T
Aldehyde dehydrogenase
PTAC
Signature enzyme
Alcohol dehydrogenase
Regulator

Salmonella enterica subsp. enterica serovar Ago. Bacteria; Proteobacteria; Gammaproteobacteria; Enterobacterales;

AU897__locus_1

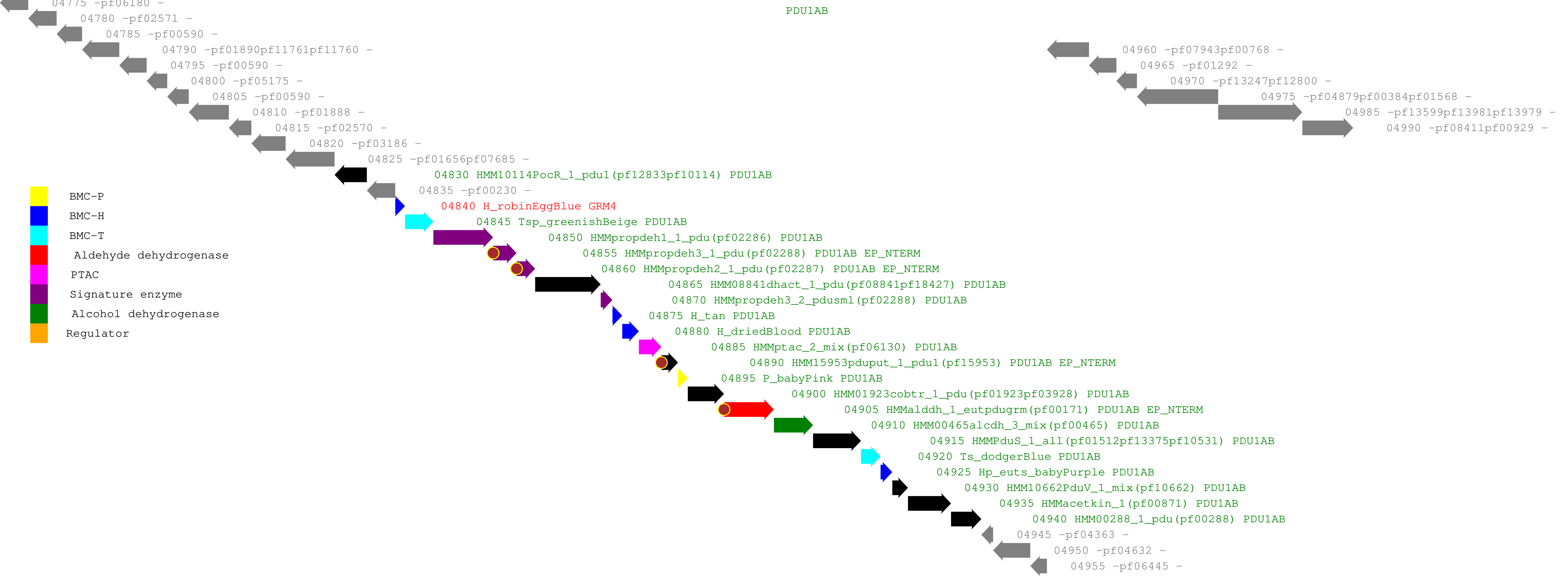

Salmonella enterica subsp. enterica serovar Remo. Bacteria; Proteobacteria; Gammaproteobacteria; Enterobacterales;

AU902__locus_2

PDU1AB

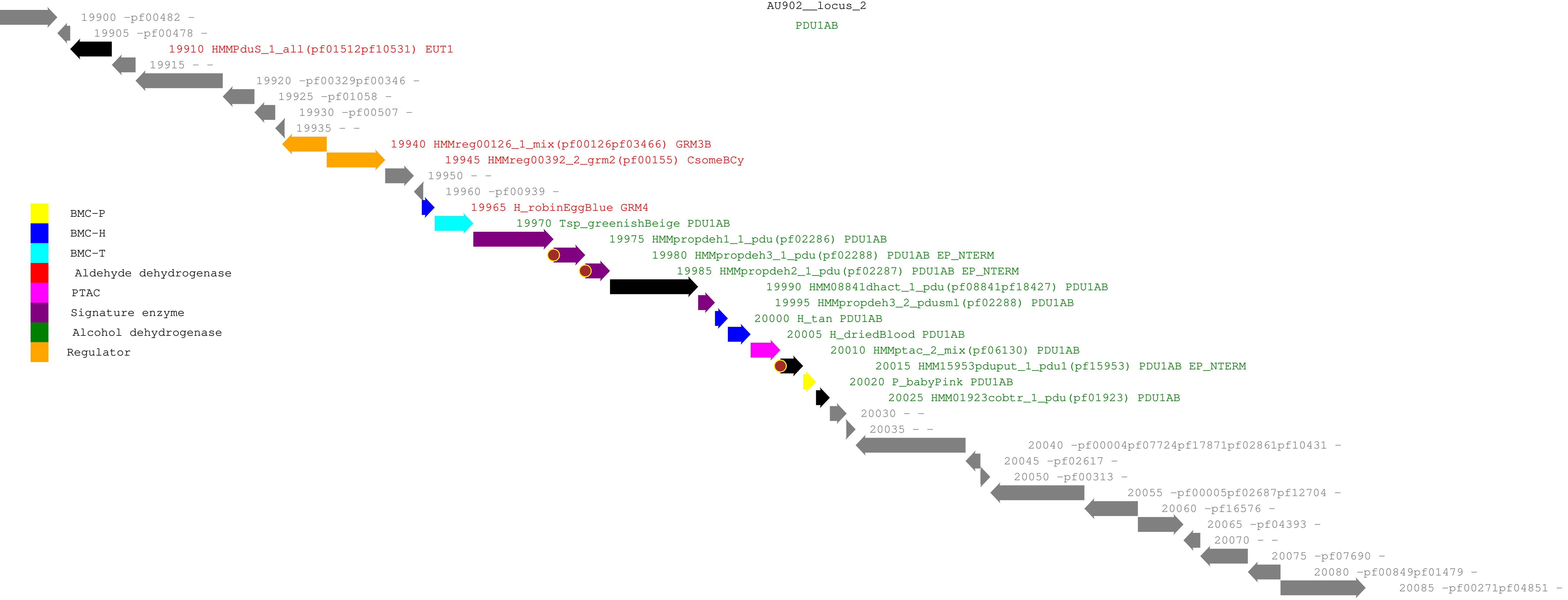

Salmonella enterica subsp. enterica serovar Eastbourne. Bacteria; Proteobacteria; Gammaproteobacteria; Enterobacterales;

AU911__locus_1

PDU1AB

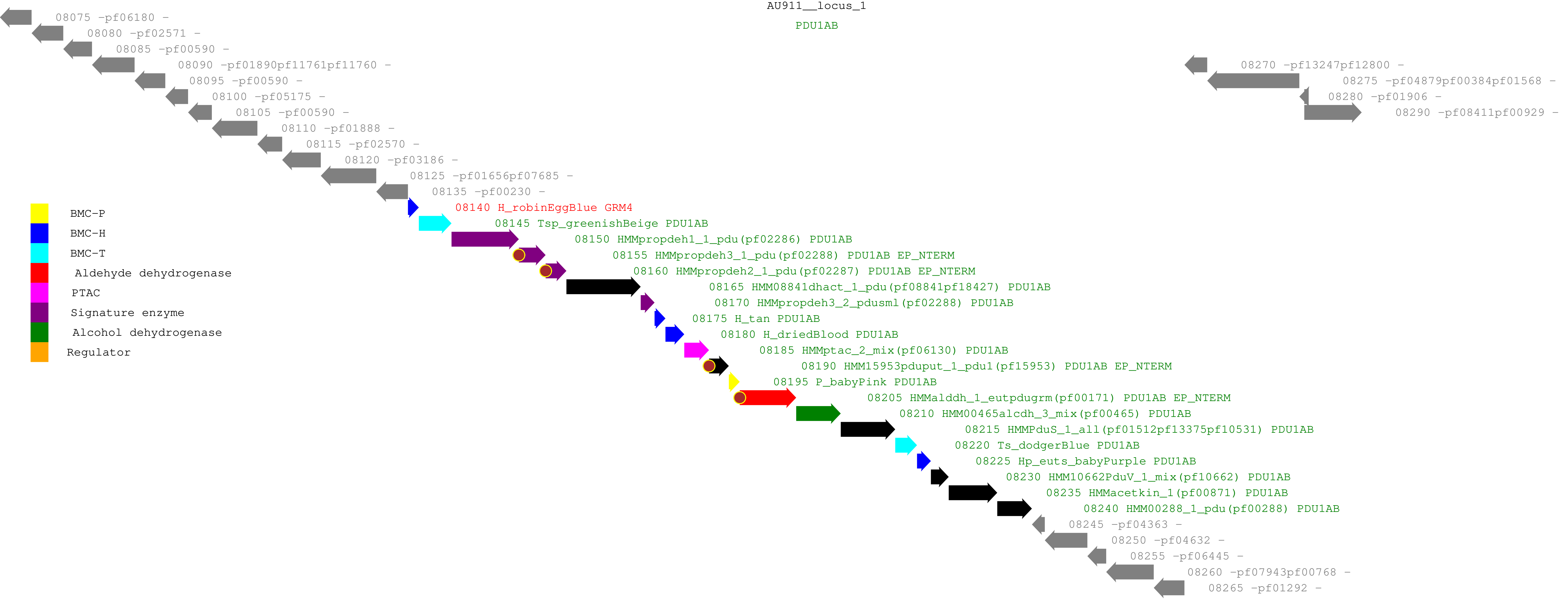

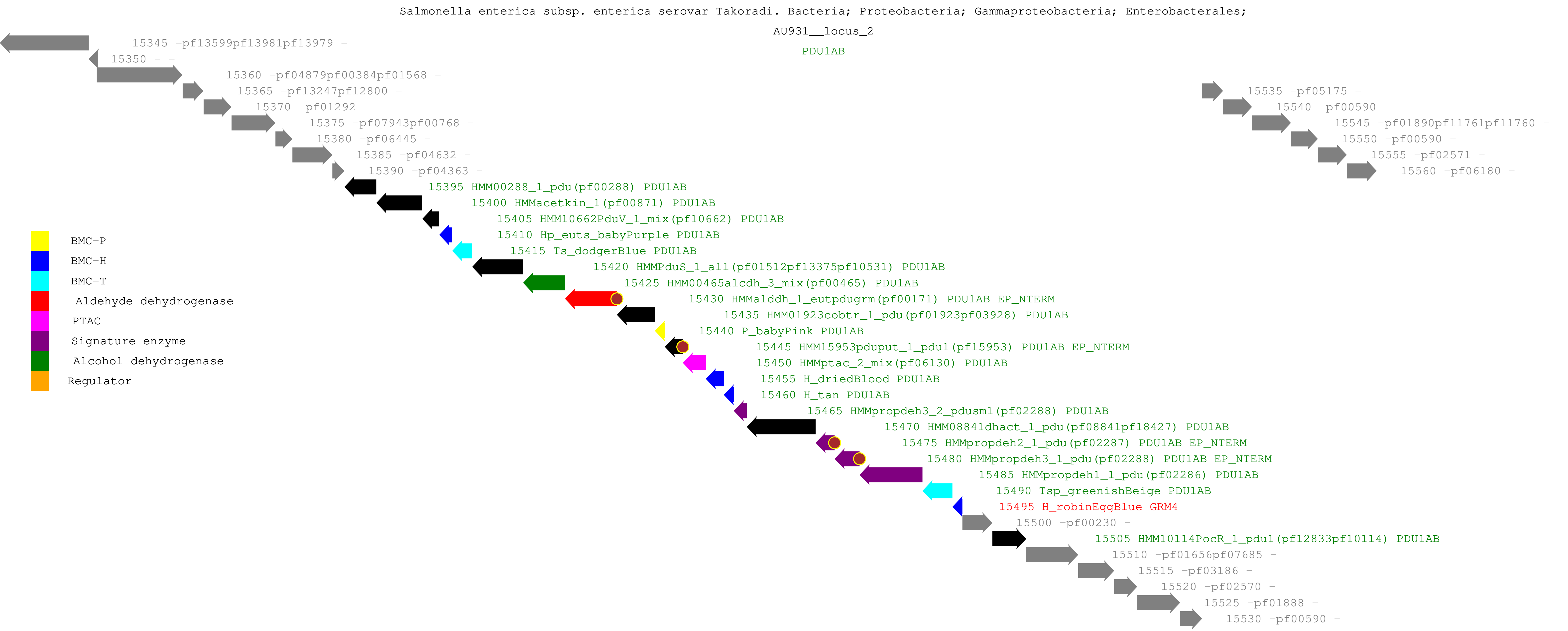
Salmonella enterica subsp. enterica serovar Takoradi. Bacteria; Proteobacteria; Gammaproteobacteria; Enterobacterales;
AU931__locus_2
PDU1AB
15345 -pf13599pf13981pf13979 -
15350 - -
15360 -pf04879pf00384pf01568 -
15365 -pf13247pf12800 -
15370 -pf01292 -
15375 -pf07943pf00768 -
15380 -pf06445 -
15385 -pf04632 -
15390 -pf04363 -
15395 HMM00288_1_pdu(pf00288) PDU1AB
15400 HMMacetkin_1(pf00871) PDU1AB
15405 HMM10662PduV_1_mix(pf10662) PDU1AB
15410 Hp_euts_babyPurple PDU1AB
15415 Ts_dodgerBlue PDU1AB
15420 HMMPduS_1_all(pf01512pf13375pf10531) PDU1AB
15425 HMM00465alcdh_3_mix(pf00465) PDU1AB
15430 HMMalddh_1_eutpdugrm(pf00171) PDU1AB EP_NTERM
15435 HMM01923cobtr_1_pdu(pf01923pf03928) PDU1AB
15440 P_babyPink PDU1AB
15445 HMM15953pduput_1_pdu1(pf15953) PDU1AB EP_NTERM
15450 HMMptac_2_mix(pf06130) PDU1AB
15455 H_driedBlood PDU1AB
15460 H_tan PDU1AB
15465 HMMpropdeh3_2_pdusml(pf02288) PDU1AB
15470 HMM08841dhact_1_pdu(pf08841pf18427) PDU1AB
15475 HMMpropdeh2_1_pdu(pf02287) PDU1AB EP_NTERM
15480 HMMpropdeh3_1_pdu(pf02288) PDU1AB EP_NTERM
15485 HMMpropdeh1_1_pdu(pf02286) PDU1AB
15490 Tsp_greenishBeige PDU1AB
15495 H_robinEggBlue GRM4
15500 -pf00230 -
15505 HMM10114PocR_1_pdu1(pf12833pf10114) PDU1AB
15510 -pf01656pf07685 -
15515 -pf03186 -
15520 -pf02570 -
15525 -pf01888 -
15530 -pf00590 -
15535 -pf05175 -
15540 -pf00590 -
15545 -pf01890pf11761pf11760 -
15550 -pf00590 -
15555 -pf02571 -
15560 -pf06180 -
BMC-P
BMC-H
BMC-T
Aldehyde dehydrogenase
PTAC
Signature enzyme
Alcohol dehydrogenase
Regulator

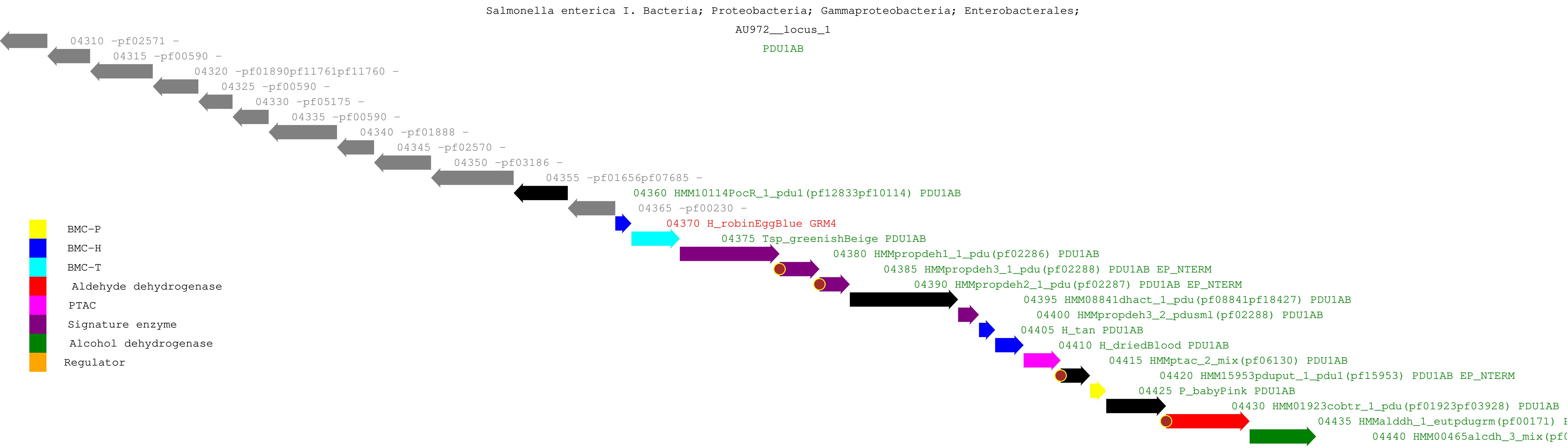
Salmonella enterica I. Bacteria; Proteobacteria; Gammaproteobacteria; Enterobacterales;
AU972__locus_1
PDU1AB
04310 -pf02571 -
04315 -pf00590 -
04320 -pf01890pf11761pf11760 -
04325 -pf00590 -
04330 -pf05175 -
04335 -pf00590 -
04340 -pf01888 -
04345 -pf02570 -
04350 -pf03186 -
04355 -pf01656pf07685 -
04360 HMM10114PocR_1_pdu1(pf12833pf10114) PDU1AB
04365 -pf00230 -
04370 H_robinEggBlue GRM4
04375 Tsp_greenishBeige PDU1AB
04380 HMMpropdeh1_1_pdu(pf02286) PDU1AB
04385 HMMpropdeh3_1_pdu(pf02288) PDU1AB EP_NTERM
04390 HMMpropdeh2_1_pdu(pf02287) PDU1AB EP_NTERM
04395 HMM08841dhact_1_pdu(pf08841pf18427) PDU1AB
04400 HMMpropdeh3_2_pdusml(pf02288) PDU1AB
04405 H_tan PDU1AB
04410 H_driedBlood PDU1AB
04415 HMMptac_2_mix(pf06130) PDU1AB
04420 HMM15953pduput_1_pdu1(pf15953) PDU1AB EP_NTERM
04425 P_babyPink PDU1AB
04430 HMM01923cobtr_1_pdu(pf01923pf03928) PDU1AB
04435 HMMalddh_1_eutpdugrm(pf00171)
04440 HMM00465alcdh_3_mix(pf0
BMC-P
BMC-H
BMC-T
Aldehyde dehydrogenase
PTAC
Signature enzyme
Alcohol dehydrogenase
Regulator

Salmonella enterica subsp. enterica serovar Telelkebir. Bacteria; Proteobacteria; Gammaproteobacteria; Enterobacterales;

AU975__locus_1

PDU1AB

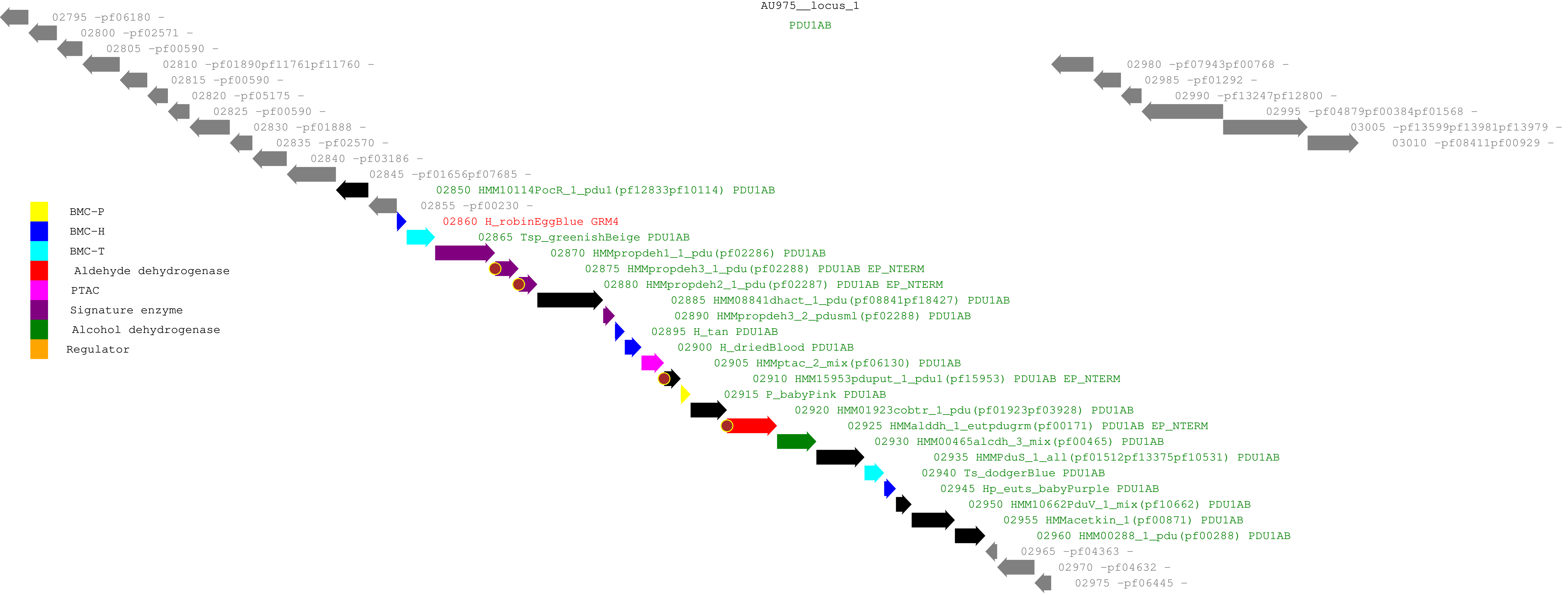

Salmonella paratyphi B (strain ATCC BAA-1250 / SPB7). Bacteria; Proteobacteria; Gammaproteobacteria; Enterobacterales;

AU980__locus_1

PDU1AB

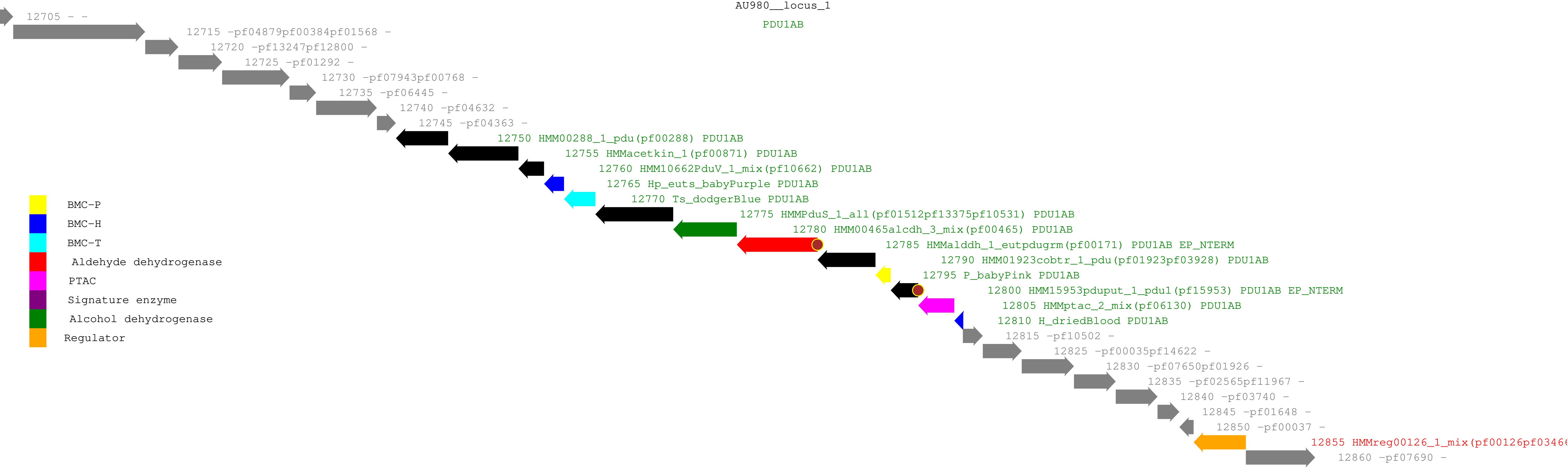

Salmonella enterica subsp. enterica serovar Senftenberg str. Bacteria; Proteobacteria; Gammaproteobacteria; Enterobacterales;

AV984__locus_2

PDU1AB

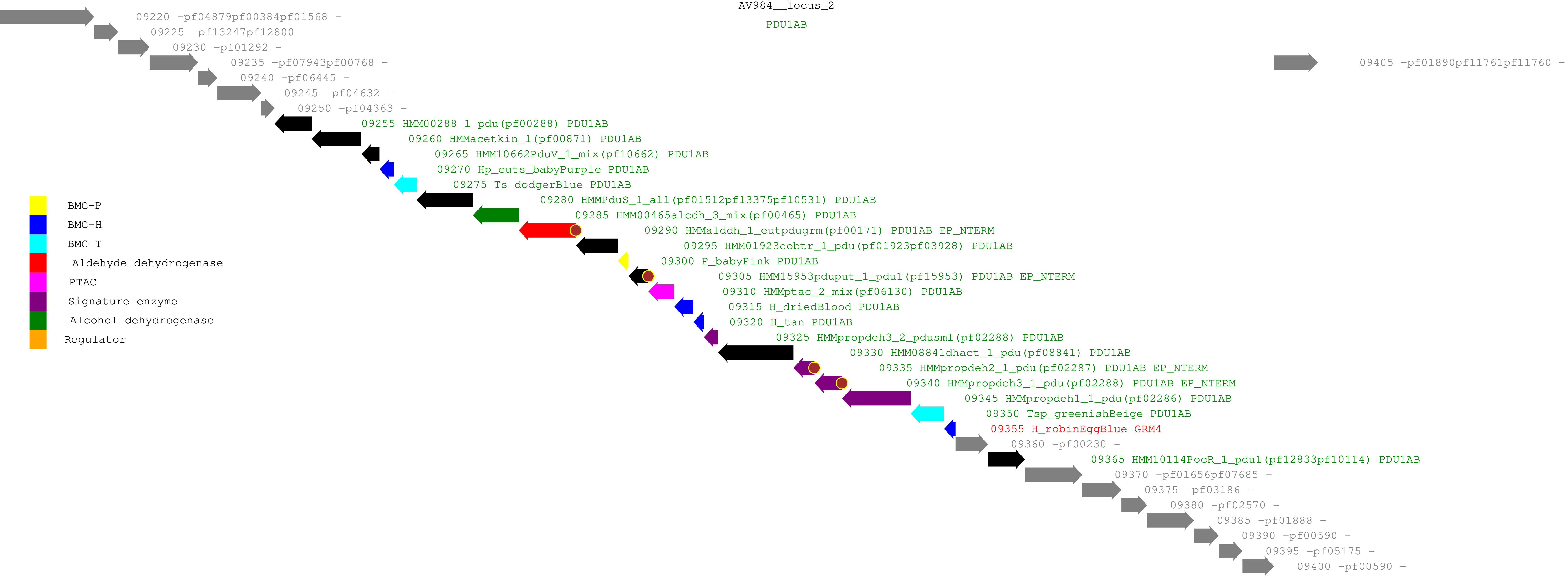

Salmonella enterica subsp. enterica serovar Litchfield. Bacteria; Proteobacteria; Gammaproteobacteria; Enterobacterales;

AVA37__locus_2

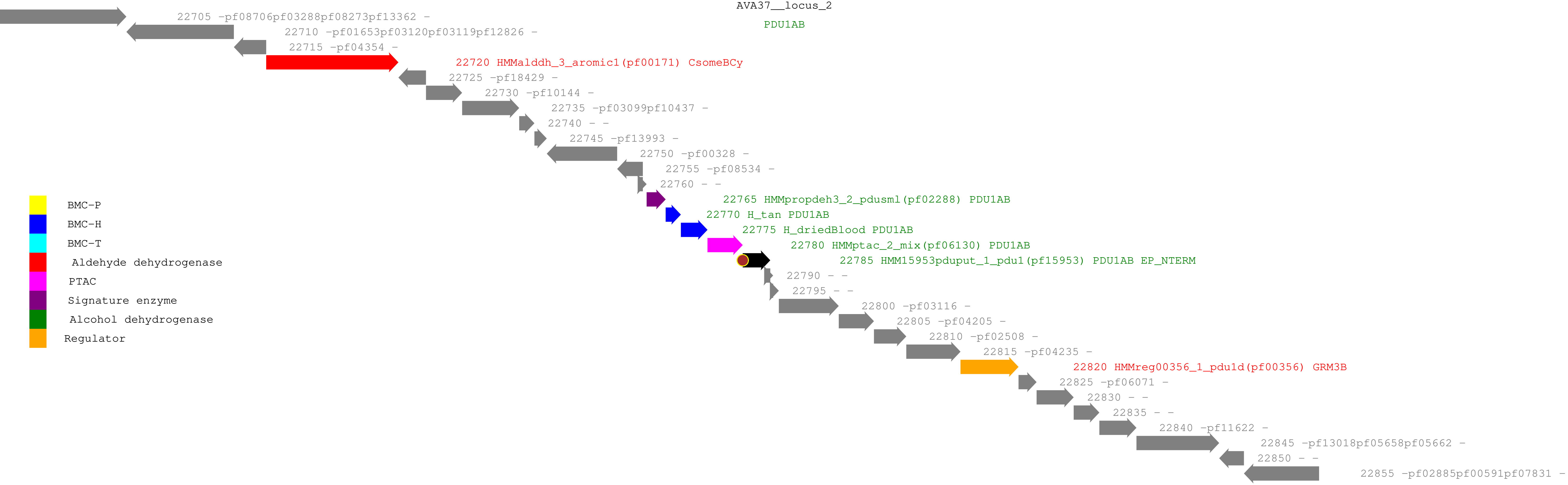

Salmonella enterica subsp. enterica serovar Bredeney. Bacteria; Proteobacteria; Gammaproteobacteria; Enterobacterales;

AVA51__locus_2

PDU1AB

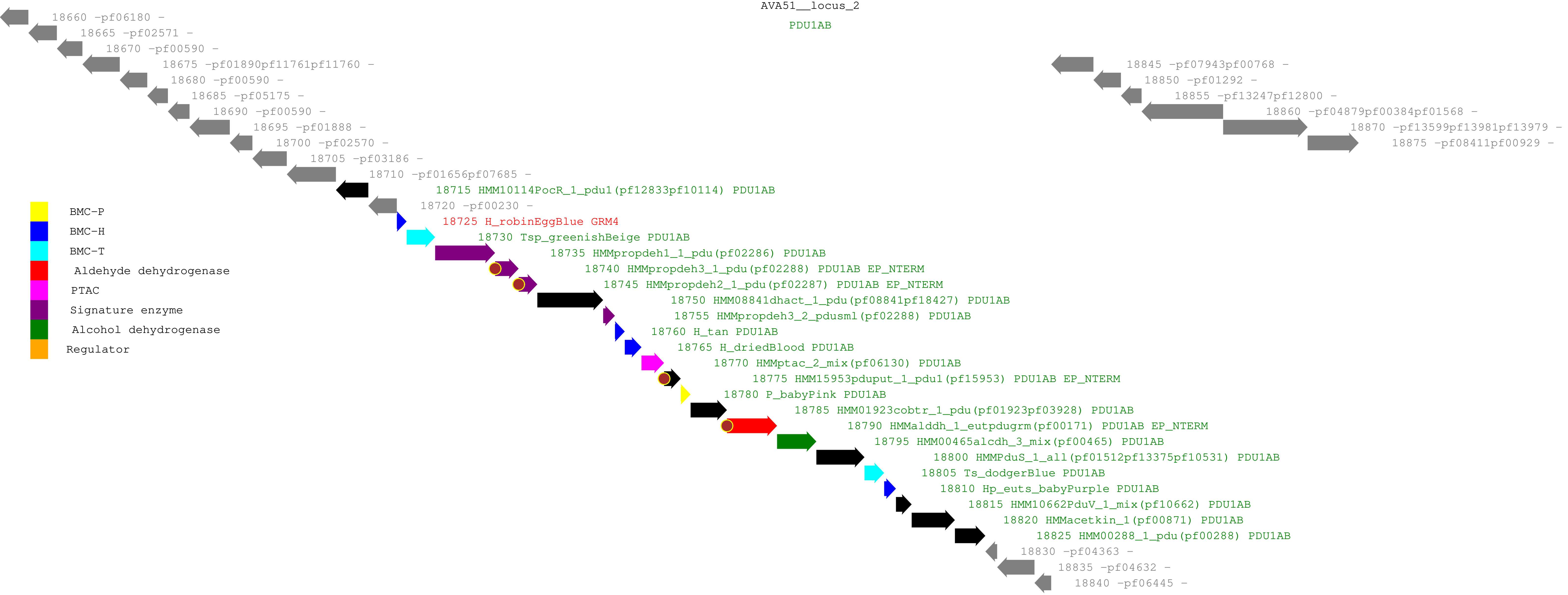

Salmonella enterica subsp. enterica serovar Altona. Bacteria; Proteobacteria; Gammaproteobacteria; Enterobacterales;

AVA57__locus_1

PDU1AB

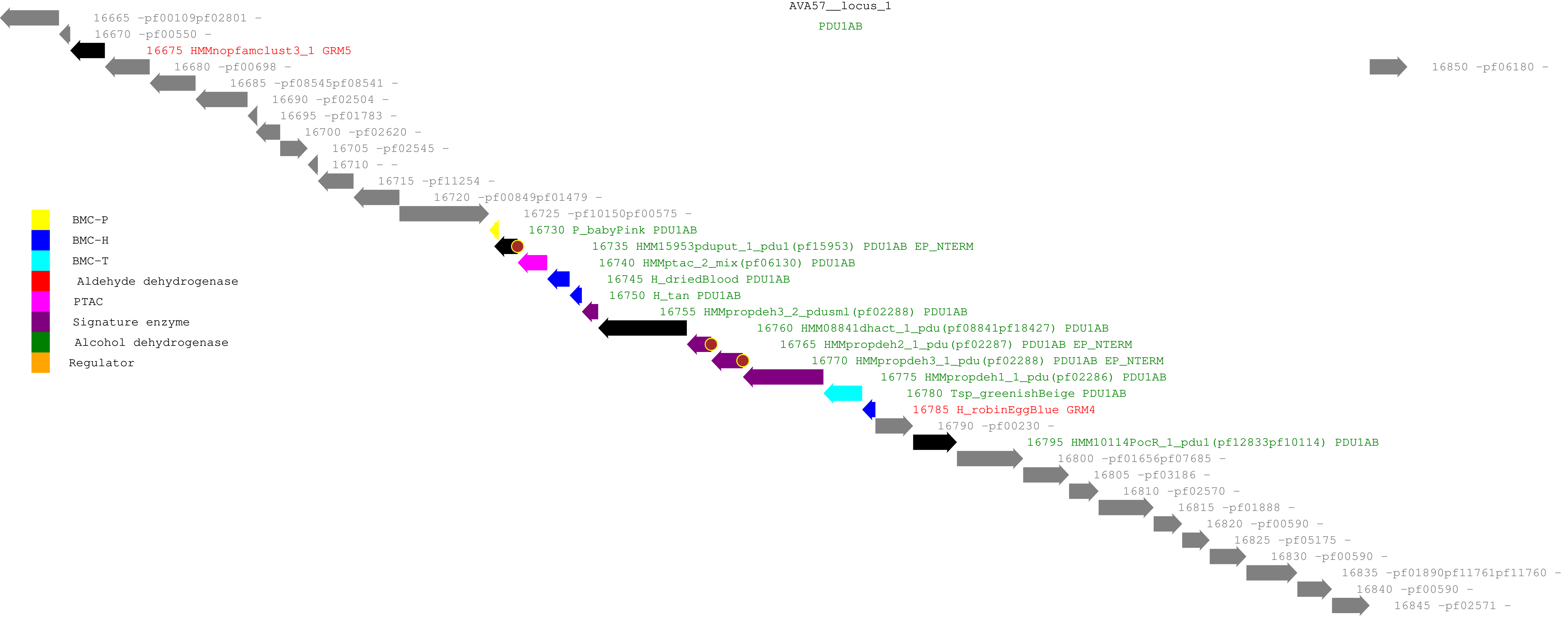

Salmonella enterica subsp. enterica serovar Altona. Bacteria; Proteobacteria; Gammaproteobacteria; Enterobacterales;

AVA57__locus_2

PDU1AB

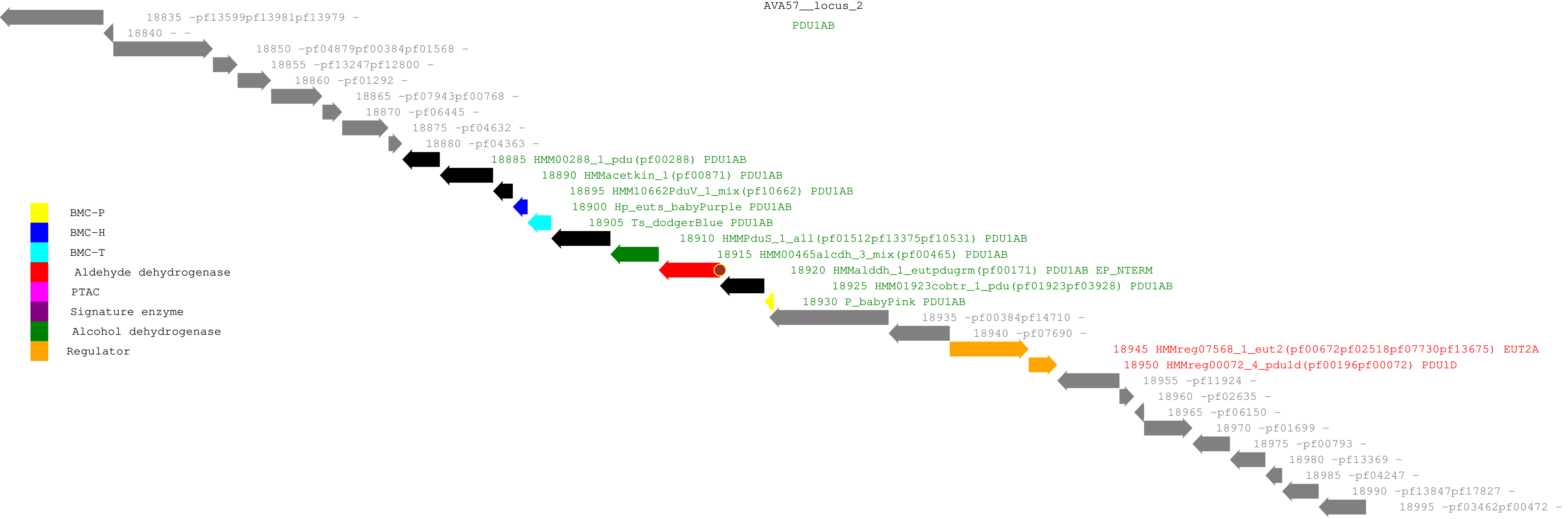

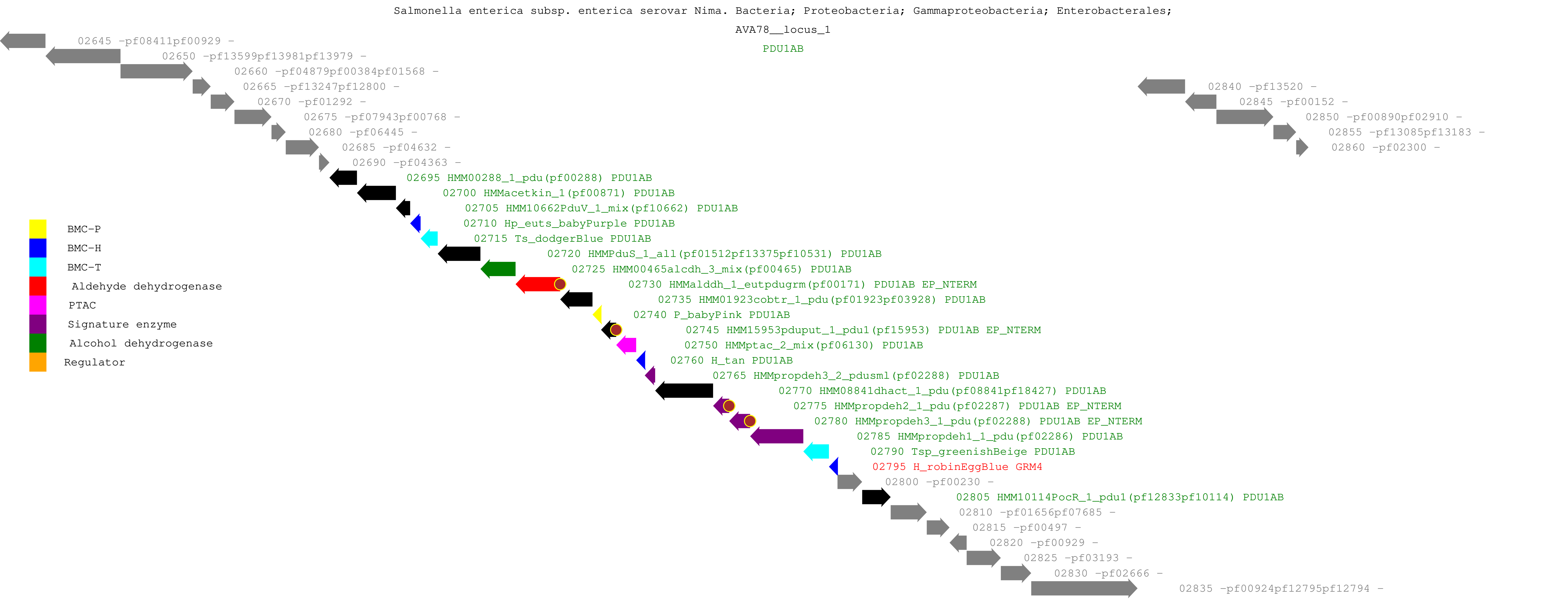
Salmonella enterica subsp. enterica serovar Nima. Bacteria; Proteobacteria; Gammaproteobacteria; Enterobacterales;
AVA78__locus_1
PDU1AB
02645 -pf08411pf00929 -
02650 -pf13599pf13981pf13979 -
02660 -pf04879pf00384pf01568 -
02665 -pf13247pf12800 -
02670 -pf01292 -
02675 -pf07943pf00768 -
02680 -pf06445 -
02685 -pf04632 -
02690 -pf04363 -
02695 HMM00288_1_pdu(pf00288) PDU1AB
02700 HMMacetkin_1(pf00871) PDU1AB
02705 HMM10662PduV_1_mix(pf10662) PDU1AB
02710 Hp_euts_babyPurple PDU1AB
02715 Ts_dodgerBlue PDU1AB
02720 HMMPduS_1_all(pf01512pf13375pf10531) PDU1AB
02725 HMM00465alcdh_3_mix(pf00465) PDU1AB
02730 HMMalddh_1_eutpdugrm(pf00171) PDU1AB EP_NTERM
02735 HMM01923cobtr_1_pdu(pf01923pf03928) PDU1AB
02740 P_babyPink PDU1AB
02745 HMM15953pduput_1_pdu1(pf15953) PDU1AB EP_NTERM
02750 HMMptac_2_mix(pf06130) PDU1AB
02760 H_tan PDU1AB
02765 HMMpropdeh3_2_pdusml(pf02288) PDU1AB
02770 HMM08841dhact_1_pdu(pf08841pf18427) PDU1AB
02775 HMMpropdeh2_1_pdu(pf02287) PDU1AB EP_NTERM
02780 HMMpropdeh3_1_pdu(pf02288) PDU1AB EP_NTERM
02785 HMMpropdeh1_1_pdu(pf02286) PDU1AB
02790 Tsp_greenishBeige PDU1AB
02795 H_robinEggBlue GRM4
02800 -pf00230 -
02805 HMM10114PocR_1_pdu1(pf12833pf10114) PDU1AB
02810 -pf01656pf07685 -
02815 -pf00497 -
02820 -pf00929 -
02825 -pf03193 -
02830 -pf02666 -
02835 -pf00924pf12795pf12794 -
02840 -pf13520 -
02845 -pf00152 -
02850 -pf00890pf02910 -
02855 -pf13085pf13183 -
02860 -pf02300 -
BMC-P
BMC-H
BMC-T
Aldehyde dehydrogenase
PTAC
Signature enzyme
Alcohol dehydrogenase
Regulator

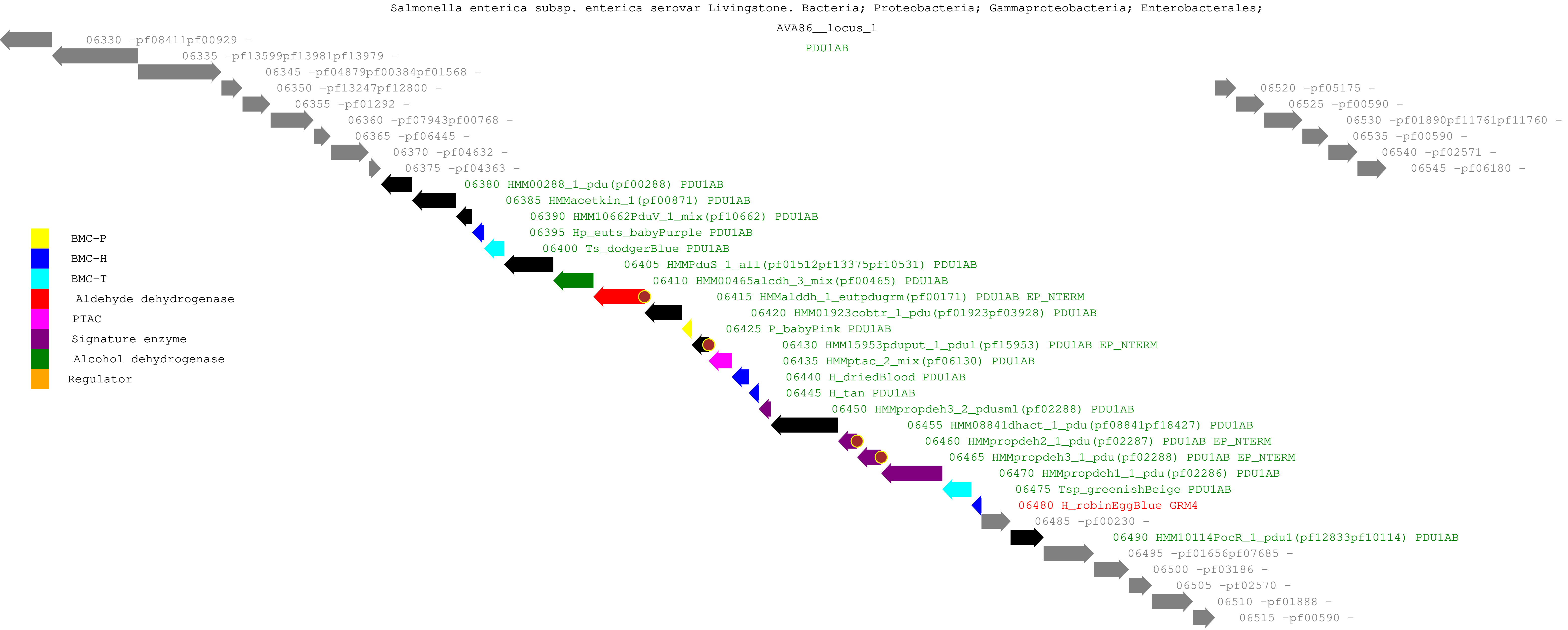
Salmonella enterica subsp. enterica serovar Livingstone. Bacteria; Proteobacteria; Gammaproteobacteria; Enterobacterales;
AVA86__locus_1
PDU1AB
06330 -pf08411pf00929 -
06335 -pf13599pf13981pf13979 -
06345 -pf04879pf00384pf01568 -
06350 -pf13247pf12800 -
06355 -pf01292 -
06360 -pf07943pf00768 -
06365 -pf06445 -
06370 -pf04632 -
06375 -pf04363 -
06380 HMM00288_1_pdu(pf00288) PDU1AB
06385 HMMacetkin_1(pf00871) PDU1AB
06390 HMM10662PduV_1_mix(pf10662) PDU1AB
06395 Hp_euts_babyPurple PDU1AB
06400 Ts_dodgerBlue PDU1AB
06405 HMMPduS_1_all(pf01512pf13375pf10531) PDU1AB
06410 HMM00465alcdh_3_mix(pf00465) PDU1AB
06415 HMMalddh_1_eutpdugrm(pf00171) PDU1AB EP_NTERM
06420 HMM01923cobtr_1_pdu(pf01923pf03928) PDU1AB
06425 P_babyPink PDU1AB
06430 HMM15953pduput_1_pdu1(pf15953) PDU1AB EP_NTERM
06435 HMMptac_2_mix(pf06130) PDU1AB
06440 H_driedBlood PDU1AB
06445 H_tan PDU1AB
06450 HMMpropdeh3_2_pdusml(pf02288) PDU1AB
06455 HMM08841dhact_1_pdu(pf08841pf18427) PDU1AB
06460 HMMpropdeh2_1_pdu(pf02287) PDU1AB EP_NTERM
06465 HMMpropdeh3_1_pdu(pf02288) PDU1AB EP_NTERM
06470 HMMpropdeh1_1_pdu(pf02286) PDU1AB
06475 Tsp_greenishBeige PDU1AB
06480 H_robinEggBlue GRM4
06485 -pf00230 -
06490 HMM10114PocR_1_pdu1(pf12833pf10114) PDU1AB
06495 -pf01656pf07685 -
06500 -pf03186 -
06505 -pf02570 -
06510 -pf01888 -
06515 -pf00590 -
06520 -pf05175 -
06525 -pf00590 -
06530 -pf01890pf11761pf11760 -
06535 -pf00590 -
06540 -pf02571 -
06545 -pf06180 -
BMC-P
BMC-H
BMC-T
Aldehyde dehydrogenase
PTAC
Signature enzyme
Alcohol dehydrogenase
Regulator

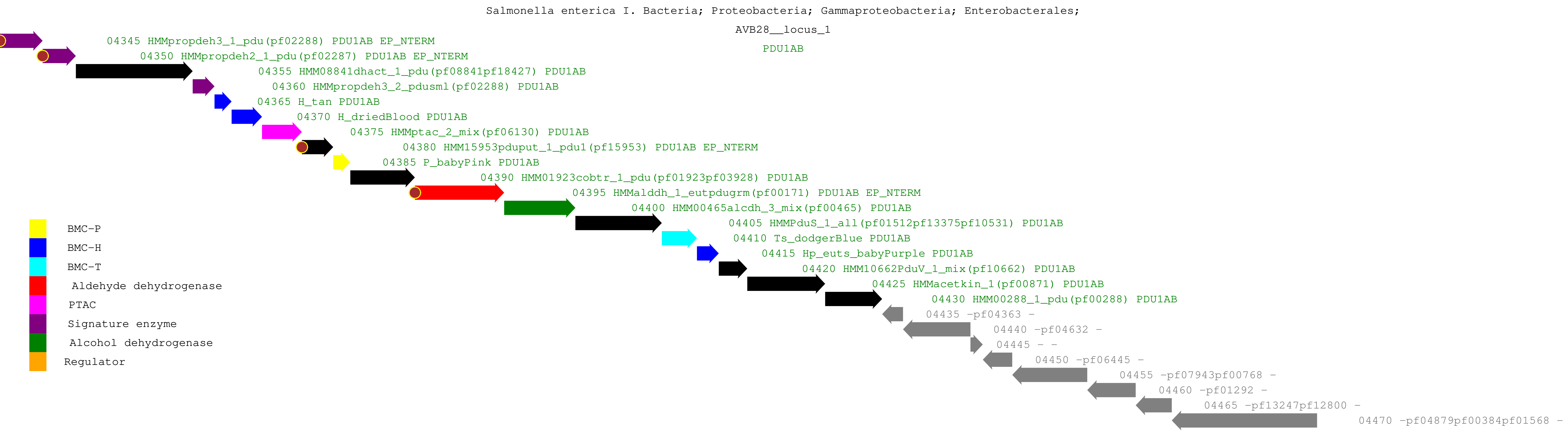
Salmonella enterica I. Bacteria; Proteobacteria; Gammaproteobacteria; Enterobacterales;
AVB28__locus_1
PDU1AB
04345 HMMpropdeh3_1_pdu(pf02288) PDU1AB EP_NTERM
04350 HMMpropdeh2_1_pdu(pf02287) PDU1AB EP_NTERM
04355 HMM08841dhact_1_pdu(pf08841pf18427) PDU1AB
04360 HMMpropdeh3_2_pdusml(pf02288) PDU1AB
04365 H_tan PDU1AB
04370 H_driedBlood PDU1AB
04375 HMMptac_2_mix(pf06130) PDU1AB
04380 HMM15953pduput_1_pdu1(pf15953) PDU1AB EP_NTERM
04385 P_babyPink PDU1AB
04390 HMM01923cobtr_1_pdu(pf01923pf03928) PDU1AB
04395 HMMalddh_1_eutpdugrm(pf00171) PDU1AB EP_NTERM
04400 HMM00465alcdh_3_mix(pf00465) PDU1AB
04405 HMMPduS_1_all(pf01512pf13375pf10531) PDU1AB
04410 Ts_dodgerBlue PDU1AB
04415 Hp_euts_babyPurple PDU1AB
04420 HMM10662PduV_1_mix(pf10662) PDU1AB
04425 HMMacetkin_1(pf00871) PDU1AB
04430 HMM00288_1_pdu(pf00288) PDU1AB
04435 -pf04363 -
04440 -pf04632 -
04445 - -
04450 -pf06445 -
04455 -pf07943pf00768 -
04460 -pf01292 -
04465 -pf13247pf12800 -
04470 -pf04879pf00384pf01568 -
BMC-P
BMC-H
BMC-T
Aldehyde dehydrogenase
PTAC
Signature enzyme
Alcohol dehydrogenase
Regulator

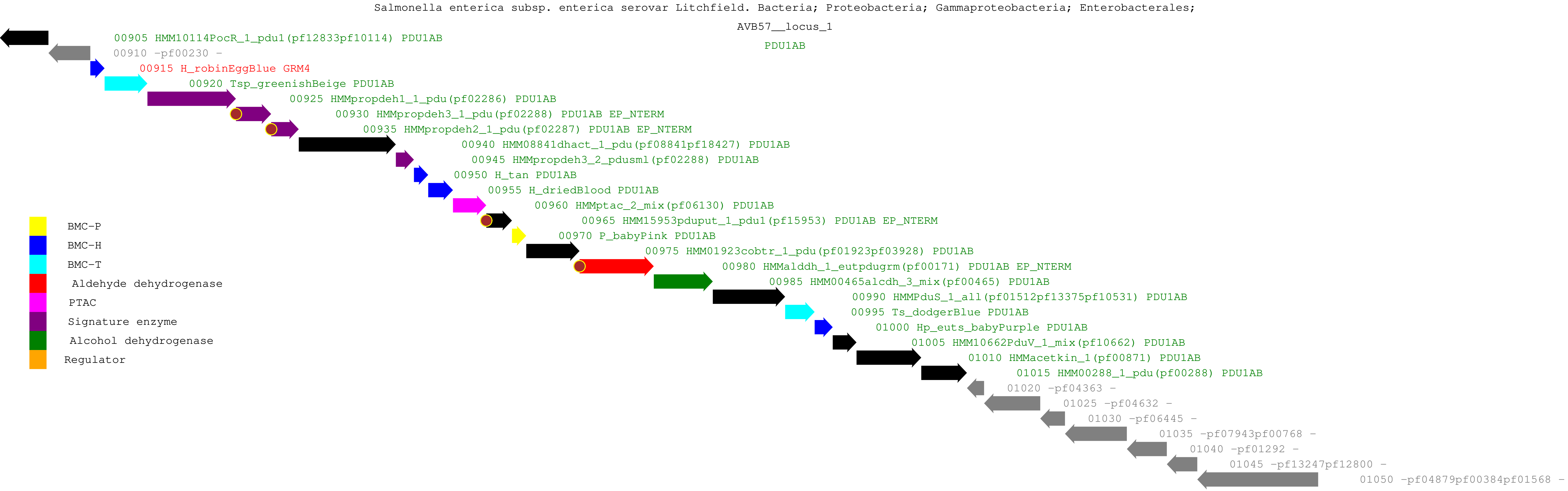
Salmonella enterica subsp. enterica serovar Litchfield. Bacteria; Proteobacteria; Gammaproteobacteria; Enterobacterales;
AVB57__locus_1
PDU1AB
00905 HMM10114PocR_1_pdu1(pf12833pf10114) PDU1AB
00910 -pf00230 -
00915 H_robinEggBlue GRM4
00920 Tsp_greenishBeige PDU1AB
00925 HMMpropdeh1_1_pdu(pf02286) PDU1AB
00930 HMMpropdeh3_1_pdu(pf02288) PDU1AB EP_NTERM
00935 HMMpropdeh2_1_pdu(pf02287) PDU1AB EP_NTERM
00940 HMM08841dhact_1_pdu(pf08841pf18427) PDU1AB
00945 HMMpropdeh3_2_pdusml(pf02288) PDU1AB
00950 H_tan PDU1AB
00955 H_driedBlood PDU1AB
00960 HMMptac_2_mix(pf06130) PDU1AB
00965 HMM15953pduput_1_pdu1(pf15953) PDU1AB EP_NTERM
00970 P_babyPink PDU1AB
00975 HMM01923cobtr_1_pdu(pf01923pf03928) PDU1AB
00980 HMMalddh_1_eutpdugrm(pf00171) PDU1AB EP_NTERM
00985 HMM00465alcdh_3_mix(pf00465) PDU1AB
00990 HMMPduS_1_all(pf01512pf13375pf10531) PDU1AB
00995 Ts_dodgerBlue PDU1AB
01000 Hp_euts_babyPurple PDU1AB
01005 HMM10662PduV_1_mix(pf10662) PDU1AB
01010 HMMacetkin_1(pf00871) PDU1AB
01015 HMM00288_1_pdu(pf00288) PDU1AB
01020 -pf04363 -
01025 -pf04632 -
01030 -pf06445 -
01035 -pf07943pf00768 -
01040 -pf01292 -
01045 -pf13247pf12800 -
01050 -pf04879pf00384pf01568 -
BMC-P
BMC-H
BMC-T
Aldehyde dehydrogenase
PTAC
Signature enzyme
Alcohol dehydrogenase
Regulator

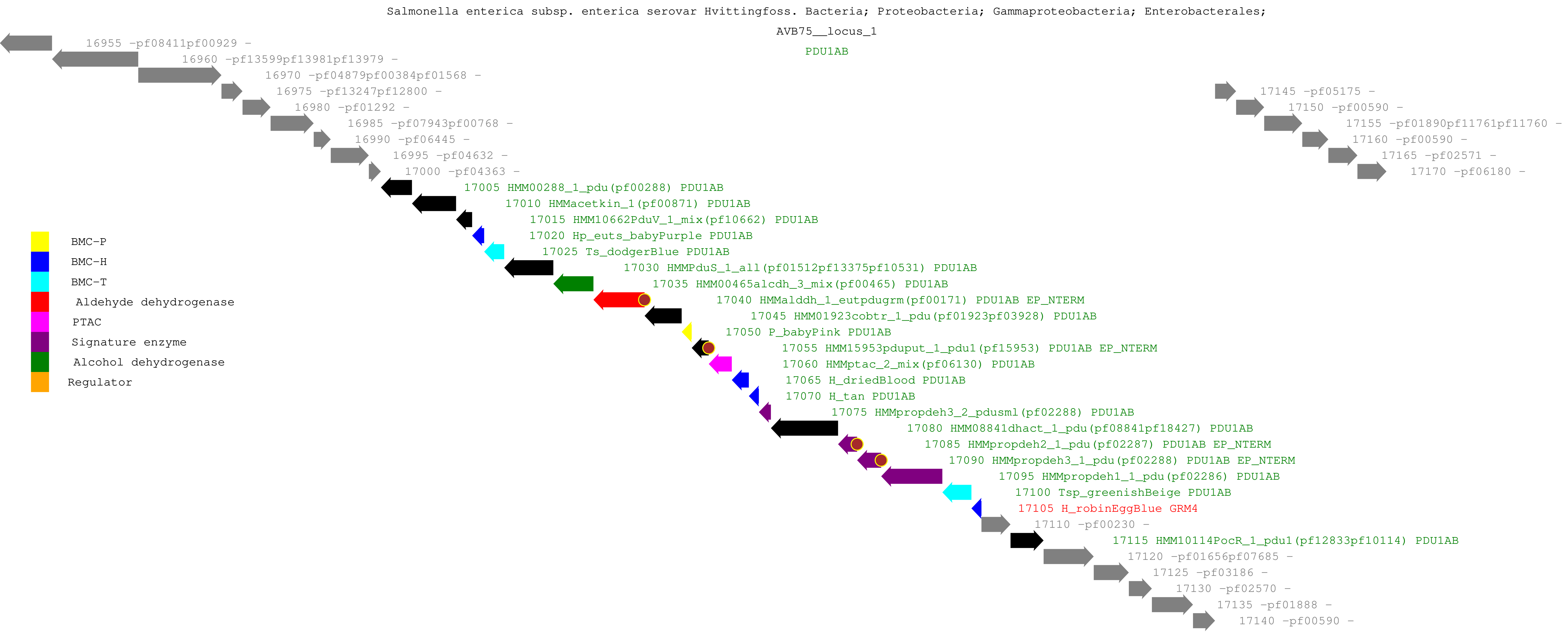
Salmonella enterica subsp. enterica serovar Hvittingfoss. Bacteria; Proteobacteria; Gammaproteobacteria; Enterobacterales;
AVB75__locus_1
PDU1AB
16955 -pf08411pf00929 -
16960 -pf13599pf13981pf13979 -
16970 -pf04879pf00384pf01568 -
16975 -pf13247pf12800 -
16980 -pf01292 -
16985 -pf07943pf00768 -
16990 -pf06445 -
16995 -pf04632 -
17000 -pf04363 -
17005 HMM00288_1_pdu(pf00288) PDU1AB
17010 HMMacetkin_1(pf00871) PDU1AB
17015 HMM10662PduV_1_mix(pf10662) PDU1AB
17020 Hp_euts_babyPurple PDU1AB
17025 Ts_dodgerBlue PDU1AB
17030 HMMPduS_1_all(pf01512pf13375pf10531) PDU1AB
17035 HMM00465alcdh_3_mix(pf00465) PDU1AB
17040 HMMalddh_1_eutpdugrm(pf00171) PDU1AB EP_NTERM
17045 HMM01923cobtr_1_pdu(pf01923pf03928) PDU1AB
17050 P_babyPink PDU1AB
17055 HMM15953pduput_1_pdu1(pf15953) PDU1AB EP_NTERM
17060 HMMptac_2_mix(pf06130) PDU1AB
17065 H_driedBlood PDU1AB
17070 H_tan PDU1AB
17075 HMMpropdeh3_2_pdusml(pf02288) PDU1AB
17080 HMM08841dhact_1_pdu(pf08841pf18427) PDU1AB
17085 HMMpropdeh2_1_pdu(pf02287) PDU1AB EP_NTERM
17090 HMMpropdeh3_1_pdu(pf02288) PDU1AB EP_NTERM
17095 HMMpropdeh1_1_pdu(pf02286) PDU1AB
17100 Tsp_greenishBeige PDU1AB
17105 H_robinEggBlue GRM4
17110 -pf00230 -
17115 HMM10114PocR_1_pdu1(pf12833pf10114) PDU1AB
17120 -pf01656pf07685 -
17125 -pf03186 -
17130 -pf02570 -
17135 -pf01888 -
17140 -pf00590 -
17145 -pf05175 -
17150 -pf00590 -
17155 -pf01890pf11761pf11760 -
17160 -pf00590 -
17165 -pf02571 -
17170 -pf06180 -
BMC-P
BMC-H
BMC-T
Aldehyde dehydrogenase
PTAC
Signature enzyme
Alcohol dehydrogenase
Regulator

Salmonella enterica subsp. enterica serovar Vitkin. Bacteria; Proteobacteria; Gammaproteobacteria; Enterobacterales;

AVB78__locus_2

PDU1AB

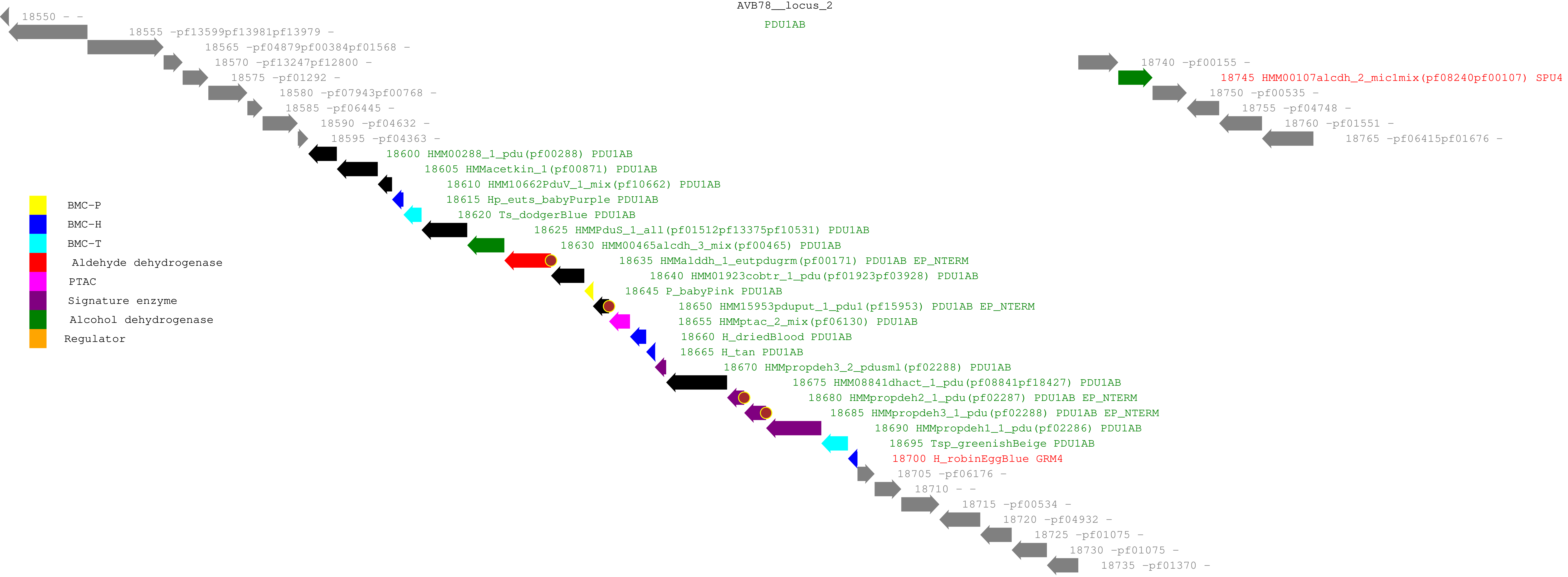

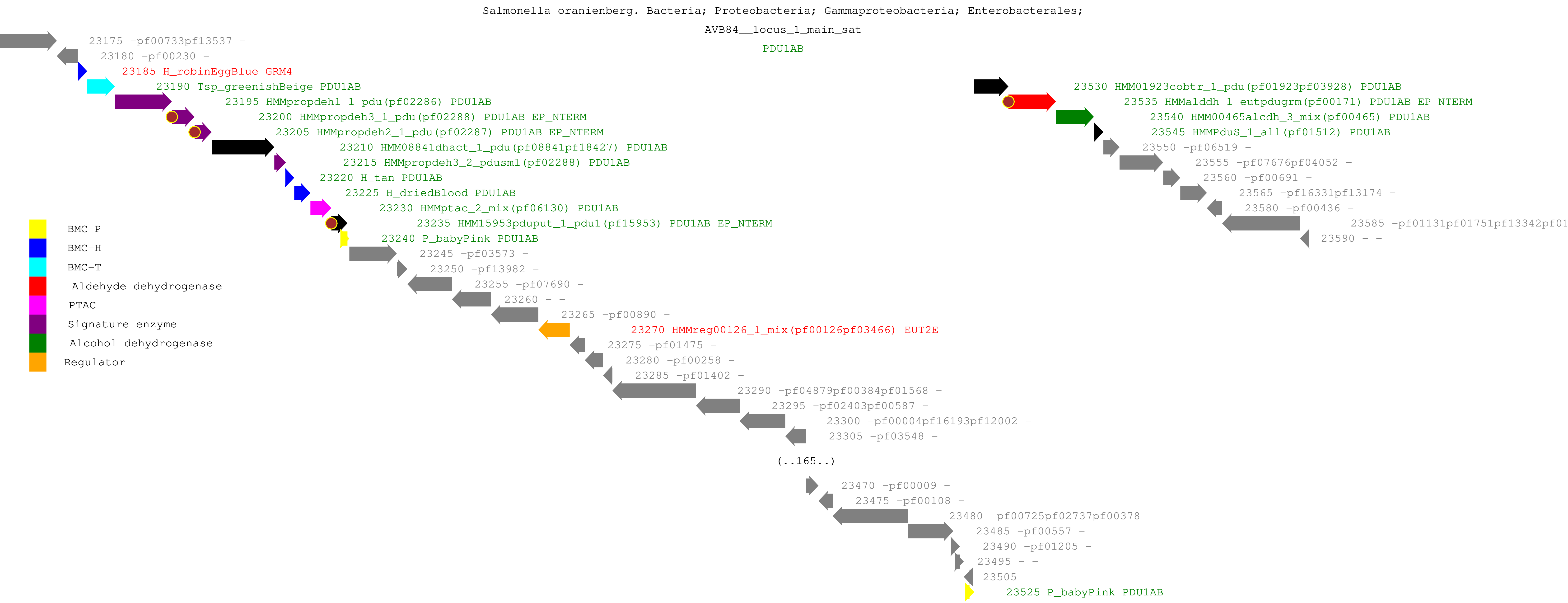
Salmonella oranienberg. Bacteria; Proteobacteria; Gammaproteobacteria; Enterobacterales;
AVB84__locus_1_main_sat
PDU1AB
23175 -pf00733pf13537 -
23180 -pf00230 -
23185 H_robinEggBlue GRM4
23190 Tsp_greenishBeige PDU1AB
23195 HMMpropdeh1_1_pdu(pf02286) PDU1AB
23200 HMMpropdeh3_1_pdu(pf02288) PDU1AB EP_NTERM
23205 HMMpropdeh2_1_pdu(pf02287) PDU1AB EP_NTERM
23210 HMM08841dhact_1_pdu(pf08841pf18427) PDU1AB
23215 HMMpropdeh3_2_pdusml(pf02288) PDU1AB
23220 H_tan PDU1AB
23225 H_driedBlood PDU1AB
23230 HMMptac_2_mix(pf06130) PDU1AB
23235 HMM15953pduput_1_pdu1(pf15953) PDU1AB EP_NTERM
23240 P_babyPink PDU1AB
23245 -pf03573 -
23250 -pf13982 -
23255 -pf07690 -
23260 - -
23265 -pf00890 -
23270 HMMreg00126_1_mix(pf00126pf03466) EUT2E
23275 -pf01475 -
23280 -pf00258 -
23285 -pf01402 -
23290 -pf04879pf00384pf01568 -
23295 -pf02403pf00587 -
23300 -pf00004pf16193pf12002 -
23305 -pf03548 -
(..165..)
23470 -pf00009 -
23475 -pf00108 -
23480 -pf00725pf02737pf00378 -
23485 -pf00557 -
23490 -pf01205 -
23495 - -
23505 - -
23525 P_babyPink PDU1AB
23530 HMM01923cobtr_1_pdu(pf01923pf03928) PDU1AB
23535 HMMalddh_1_eutpdugrm(pf00171) PDU1AB EP_NTERM
23540 HMM00465alcdh_3_mix(pf00465) PDU1AB
23545 HMMPduS_1_all(pf01512) PDU1AB
23550 -pf06519 -
23555 -pf07676pf04052 -
23560 -pf00691 -
23565 -pf16331pf13174 -
23580 -pf00436 -
23585 -pf01131pf01751pf13342pf01
23590 - -
BMC-P
BMC-H
BMC-T
Aldehyde dehydrogenase
PTAC
Signature enzyme
Alcohol dehydrogenase
Regulator

Salmonella enterica subsp. enterica serovar Vitkin. Bacteria; Proteobacteria; Gammaproteobacteria; Enterobacterales;

AVB85__locus_1

PDU1AB

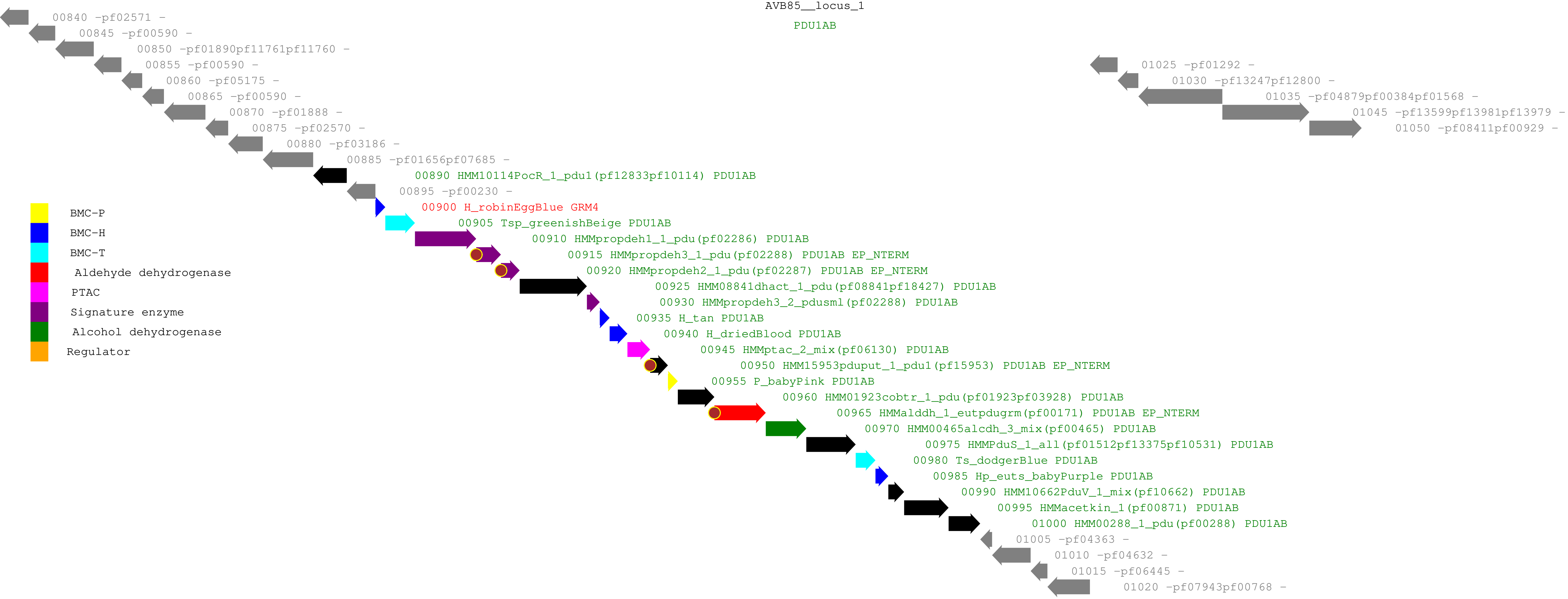

Salmonella enterica subsp. enterica serovar Ajiobo. Bacteria; Proteobacteria; Gammaproteobacteria; Enterobacterales;

AVB95__locus_1

PDU1AB

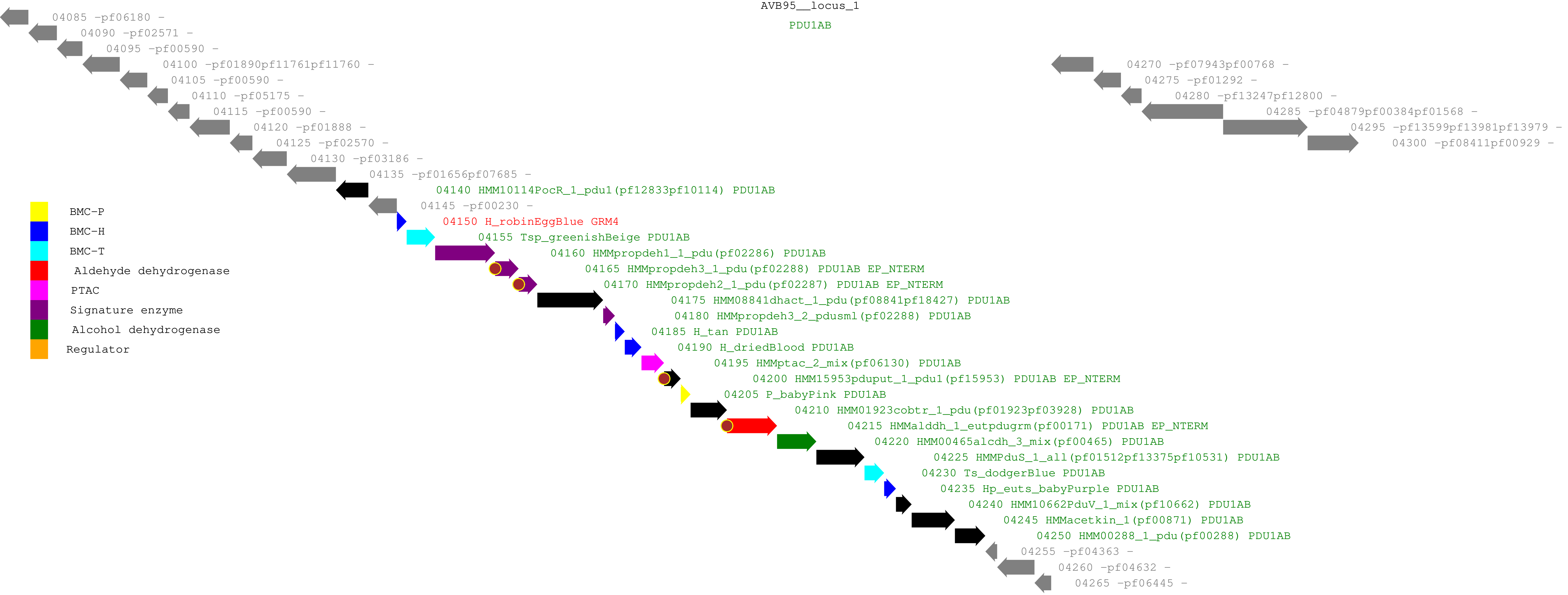

Salmonella enterica subsp. enterica serovar Kambole. Bacteria; Proteobacteria; Gammaproteobacteria; Enterobacterales;

AVC51__locus_2

PDU1AB

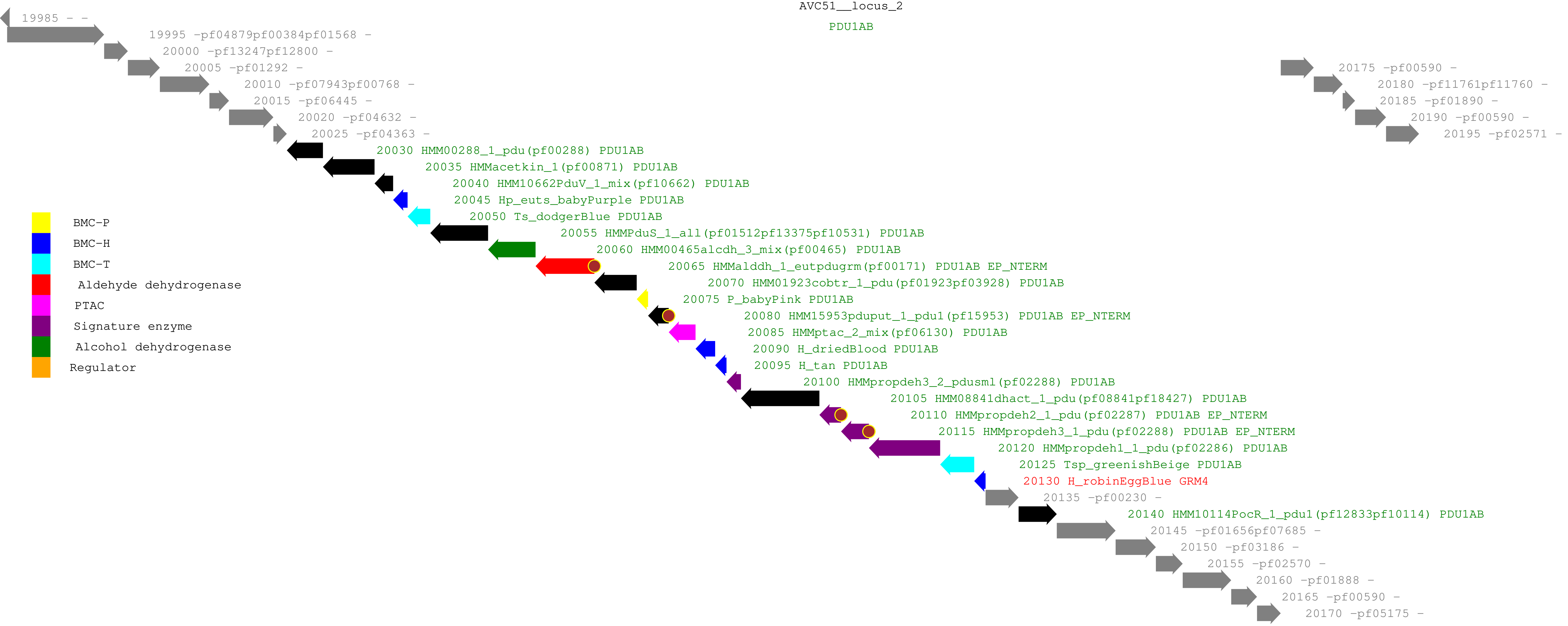

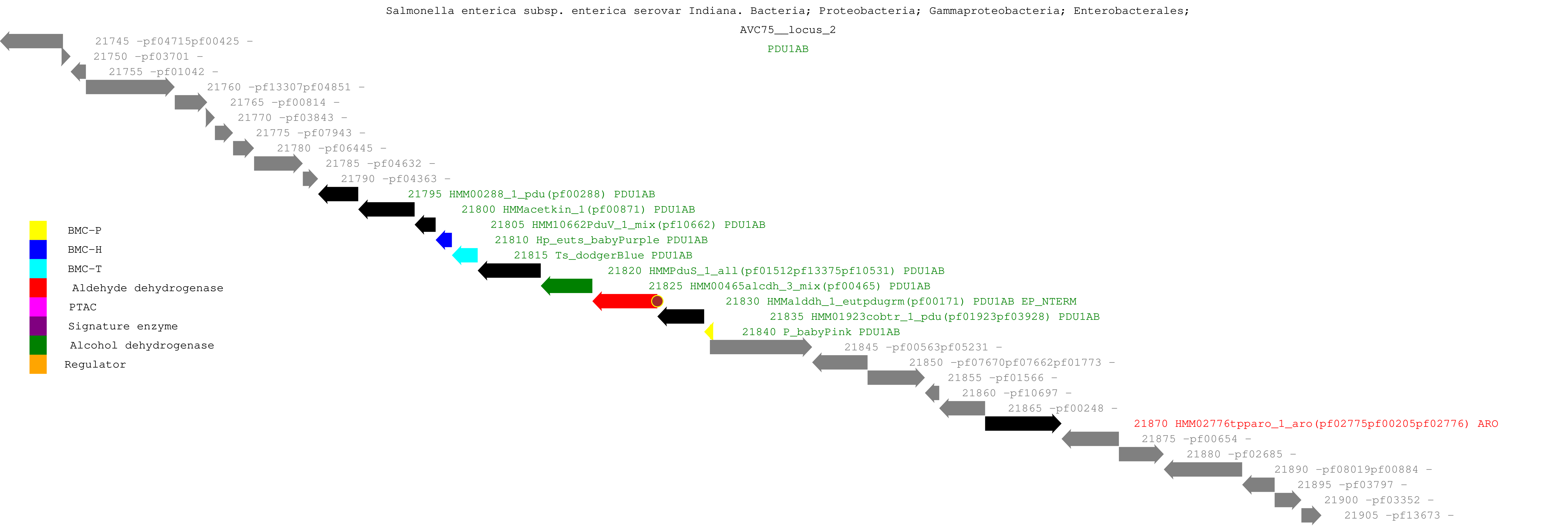
Salmonella enterica subsp. enterica serovar Indiana. Bacteria; Proteobacteria; Gammaproteobacteria; Enterobacterales;
AVC75__locus_2
PDU1AB
21745 -pf04715pf00425 -
21750 -pf03701 -
21755 -pf01042 -
21760 -pf13307pf04851 -
21765 -pf00814 -
21770 -pf03843 -
21775 -pf07943 -
21780 -pf06445 -
21785 -pf04632 -
21790 -pf04363 -
21795 HMM00288_1_pdu(pf00288) PDU1AB
21800 HMMacetkin_1(pf00871) PDU1AB
21805 HMM10662PduV_1_mix(pf10662) PDU1AB
21810 Hp_euts_babyPurple PDU1AB
21815 Ts_dodgerBlue PDU1AB
21820 HMMPduS_1_all(pf01512pf13375pf10531) PDU1AB
21825 HMM00465alcdh_3_mix(pf00465) PDU1AB
21830 HMMaldh_1_eutpdugrm(pf00171) PDU1AB EP_NTERM
21835 HMM01923cobtr_1_pdu(pf01923pf03928) PDU1AB
21840 P_babyPink PDU1AB
21845 -pf00563pf05231 -
21850 -pf07670pf07662pf01773 -
21855 -pf01566 -
21860 -pf10697 -
21865 -pf00248 -
21870 HMM02776tpparo_1_aro(pf02775pf00205pf02776) ARO
21875 -pf00654 -
21880 -pf02685 -
21890 -pf08019pf00884 -
21895 -pf03797 -
21900 -pf03352 -
21905 -pf13673 -
BMC-P
BMC-H
BMC-T
Aldehyde dehydrogenase
PTAC
Signature enzyme
Alcohol dehydrogenase
Regulator

Salmonella enterica subsp. enterica serovar Agama. Bacteria; Proteobacteria; Gammaproteobacteria; Enterobacterales;

AVC84__locus_1

PDU1AB

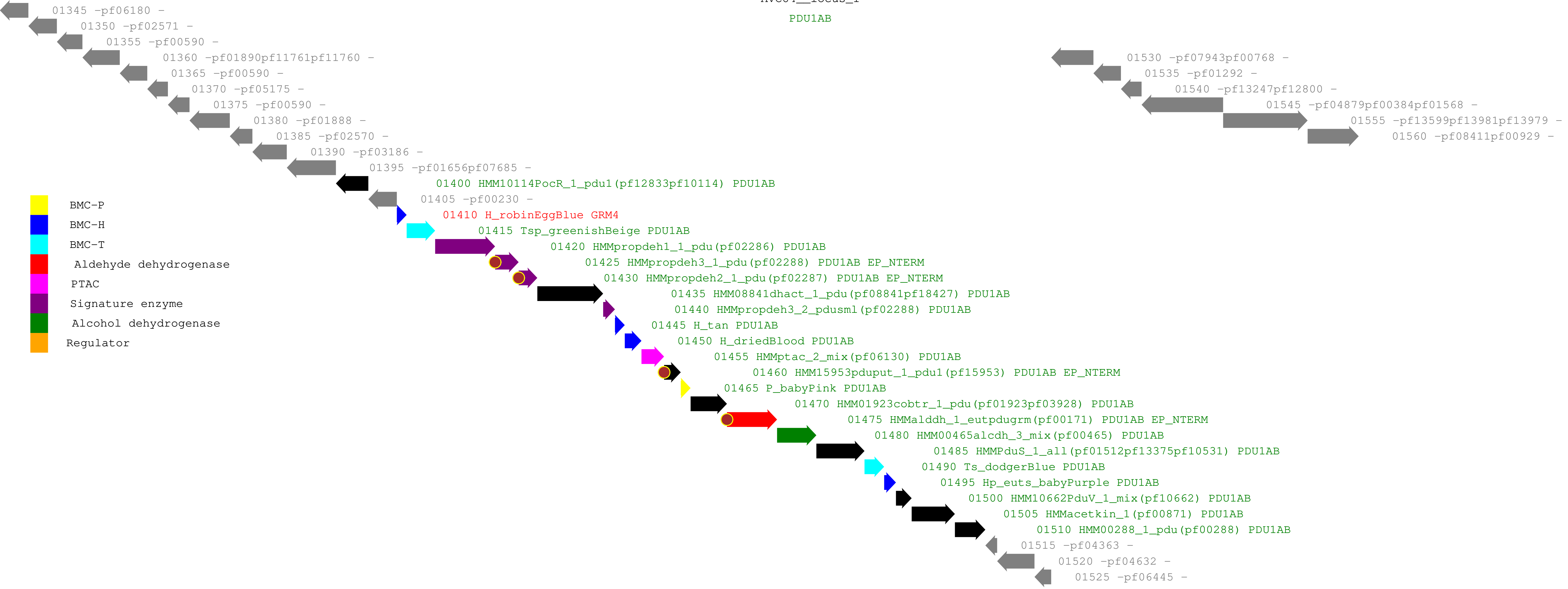

Salmonella enterica subsp. enterica serovar Javiana. Bacteria; Proteobacteria; Gammaproteobacteria; Enterobacterales;

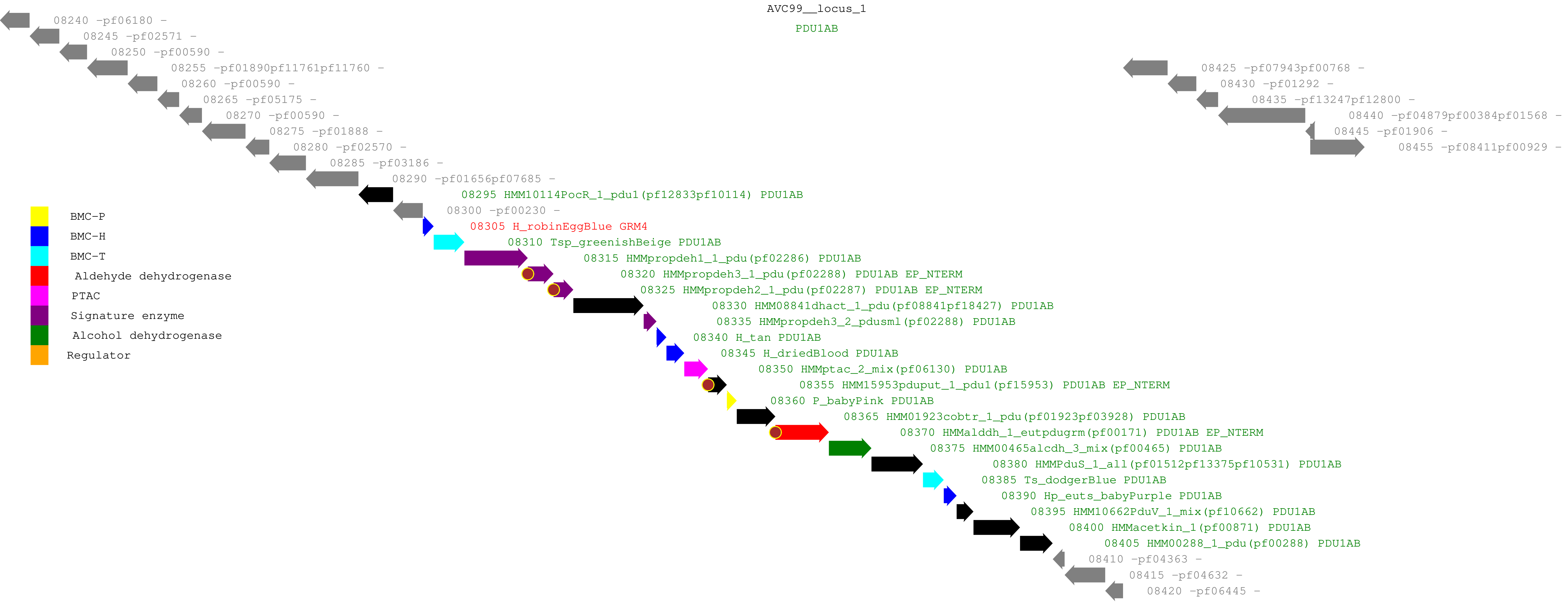

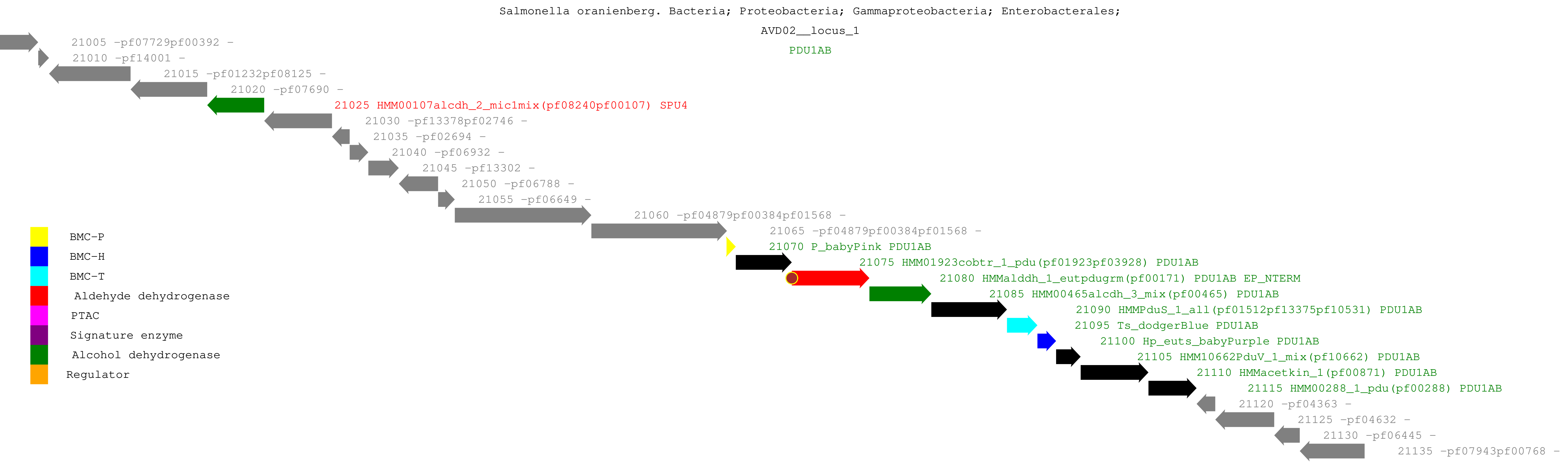

Salmonella oranienberg. Bacteria; Proteobacteria; Gammaproteobacteria; Enterobacterales;
AVD02__locus_1
PDU1AB
21005 -pf07729pf00392 -
21010 -pf14001 -
21015 -pf01232pf08125 -
21020 -pf07690 -
21025 HMM00107alcdh_2_mic1mix(pf08240pf00107) SPU4
21030 -pf13378pf02746 -
21035 -pf02694 -
21040 -pf06932 -
21045 -pf13302 -
21050 -pf06788 -
21055 -pf06649 -
21060 -pf04879pf00384pf01568 -
21065 -pf04879pf00384pf01568 -
21070 P_babyPink PDU1AB
21075 HMM01923cobtr_1_pdu(pf01923pf03928) PDU1AB
21080 HMMalddh_1_eutpdugrm(pf00171) PDU1AB EP_NTERM
21085 HMM00465alcdh_3_mix(pf00465) PDU1AB
21090 HMMPduS_1_all(pf01512pf13375pf10531) PDU1AB
21095 Ts_dodgerBlue PDU1AB
21100 Hp_euts_babyPurple PDU1AB
21105 HMM10662PduV_1_mix(pf10662) PDU1AB
21110 HMMacetkin_1(pf00871) PDU1AB
21115 HMM00288_1_pdu(pf00288) PDU1AB
21120 -pf04363 -
21125 -pf04632 -
21130 -pf06445 -
21135 -pf07943pf00768 -
BMC-P
BMC-H
BMC-T
Aldehyde dehydrogenase
PTAC
Signature enzyme
Alcohol dehydrogenase
Regulator

Salmonella oranienberg. Bacteria; Proteobacteria; Gammaproteobacteria; Enterobacterales;

AVD02__locus_2

PDU1AB

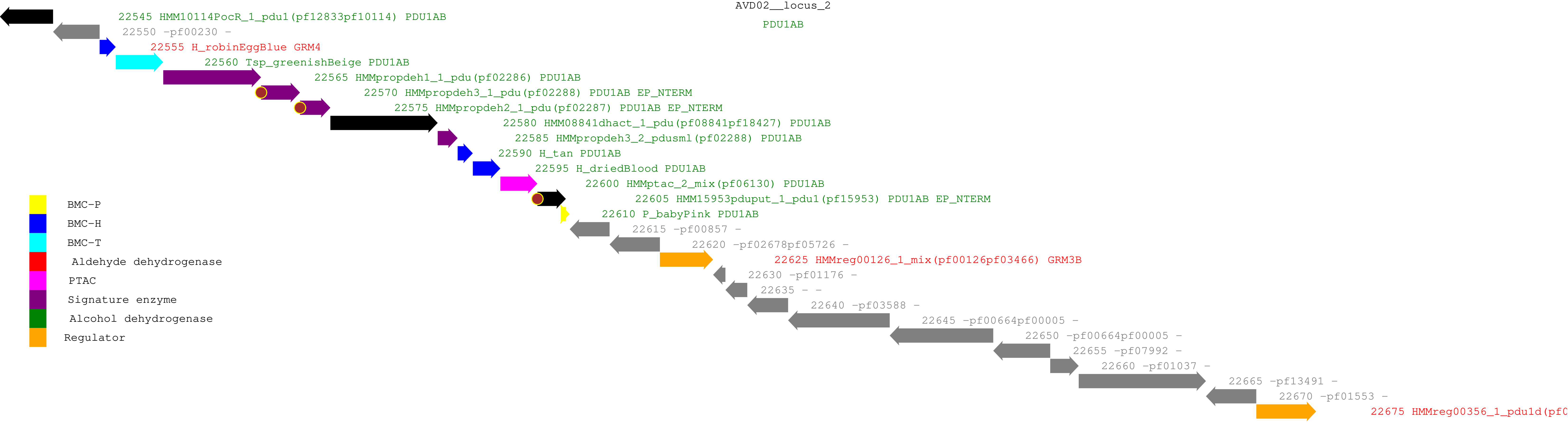

Salmonella enterica subsp. enterica serovar Chester. Bacteria; Proteobacteria; Gammaproteobacteria; Enterobacterales;

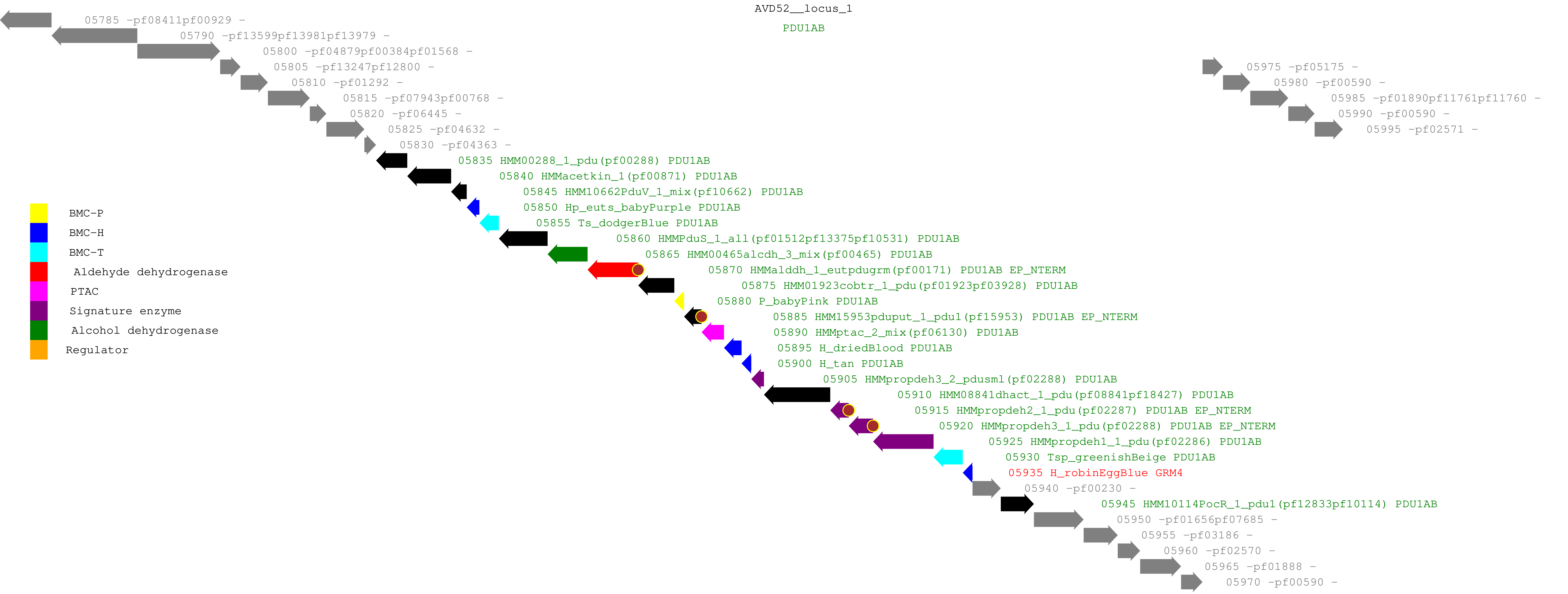

Salmonella enterica subsp. enterica serovar Alachua. Bacteria; Proteobacteria; Gammaproteobacteria; Enterobacterales;

AVD57__locus_1

PDU1AB

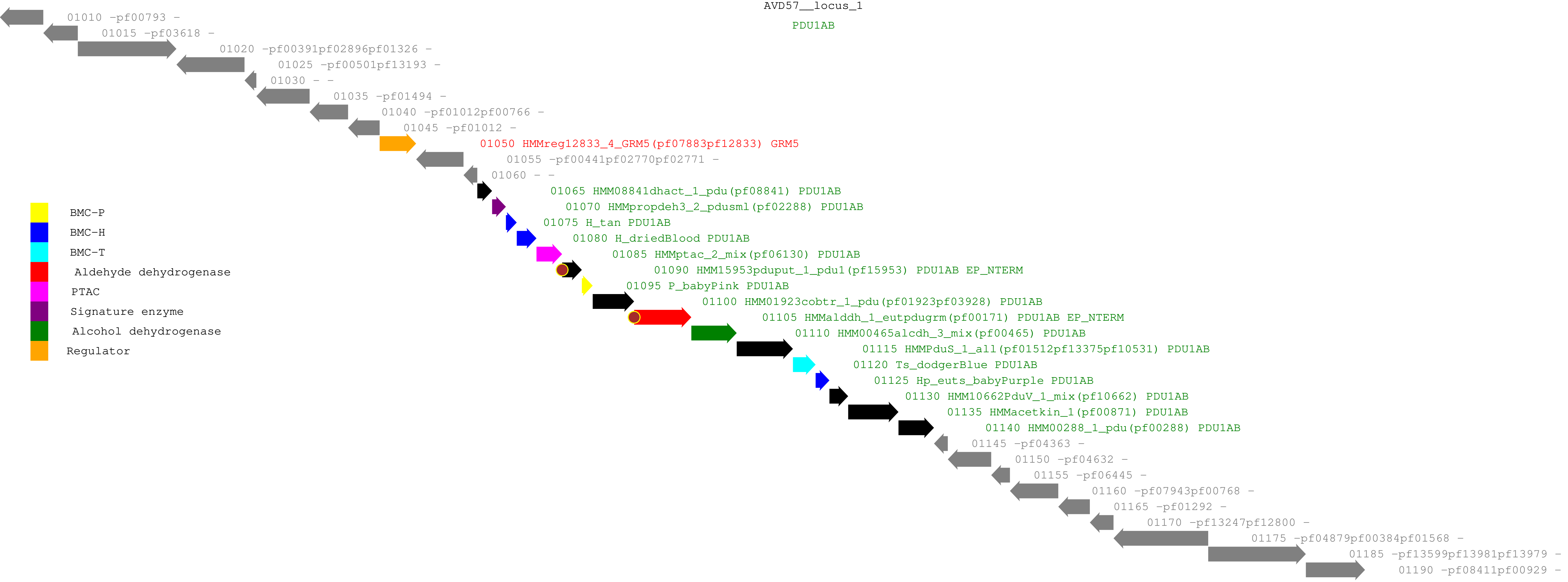

Salmonella enterica subsp. enterica serovar Indiana. Bacteria; Proteobacteria; Gammaproteobacteria; Enterobacterales;

AVE11__locus_1

PDU1AB

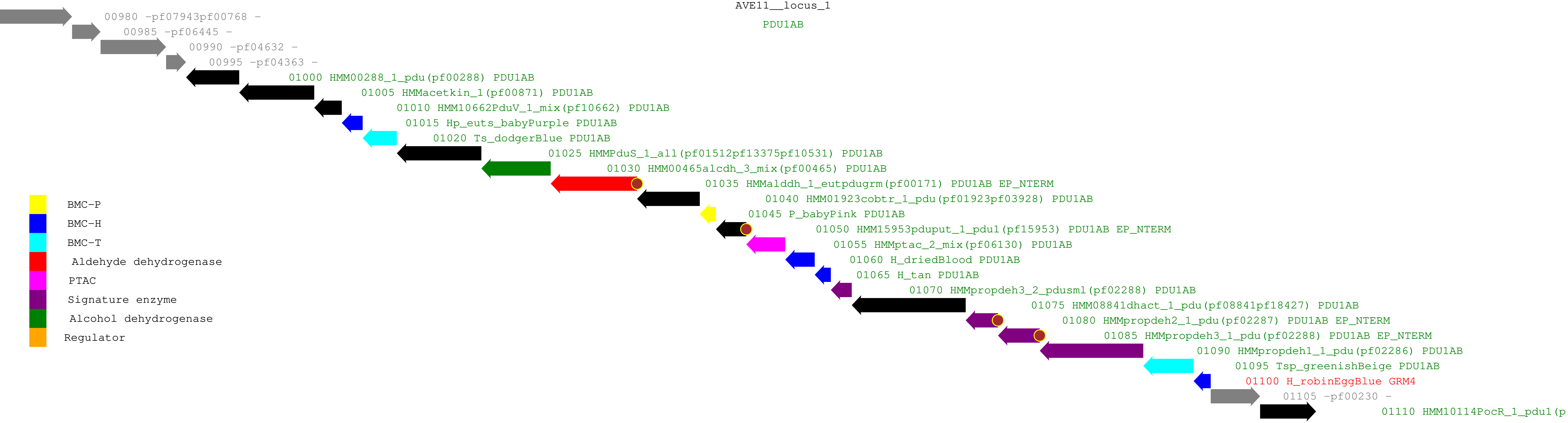

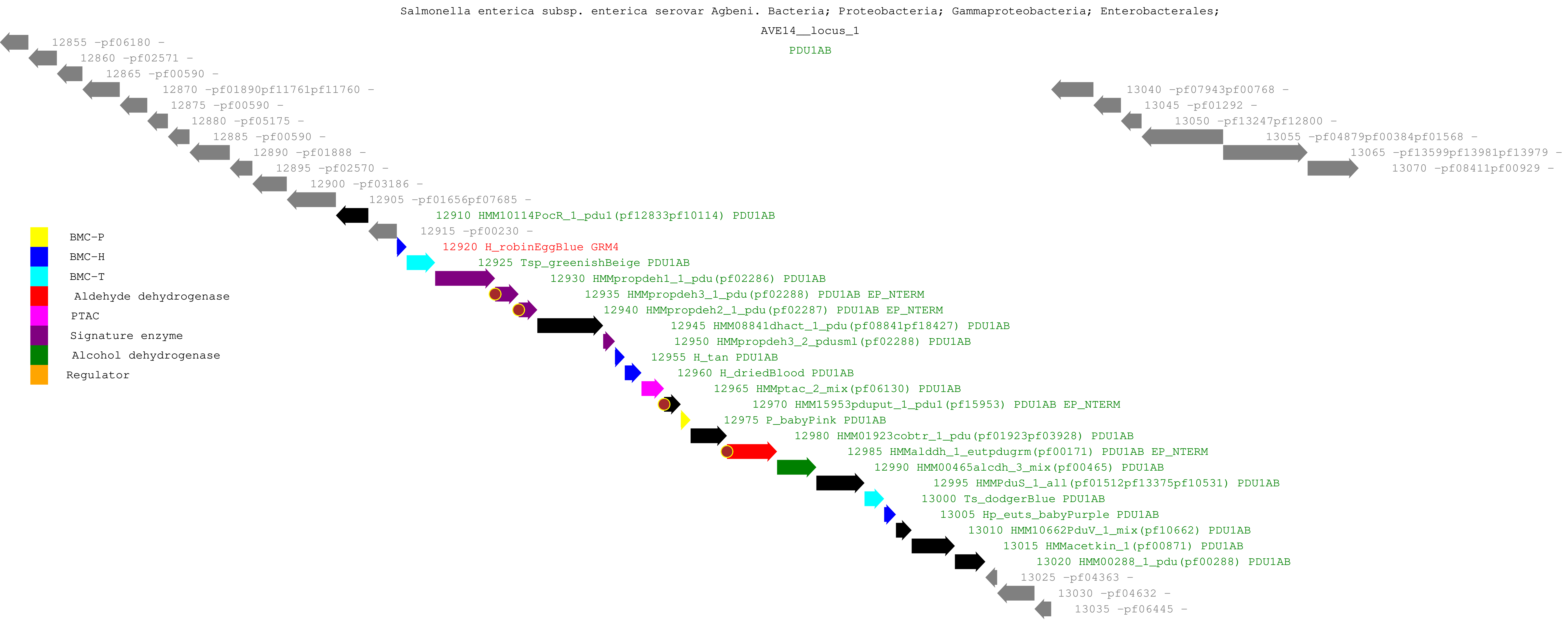
Salmonella enterica subsp. enterica serovar Agbeni. Bacteria; Proteobacteria; Gammaproteobacteria; Enterobacterales;
AVE14__locus_1
PDU1AB
12855 -pf06180 -
12860 -pf02571 -
12865 -pf00590 -
12870 -pf01890pf11761pf11760 -
12875 -pf00590 -
12880 -pf05175 -
12885 -pf00590 -
12890 -pf01888 -
12895 -pf02570 -
12900 -pf03186 -
12905 -pf01656pf07685 -
12910 HMM10114PocR_1_pdu1(pf12833pf10114) PDU1AB
12915 -pf00230 -
12920 H_robinEggBlue GRM4
12925 Tsp_greenishBeige PDU1AB
12930 HMMpropdeh1_1_pdu(pf02286) PDU1AB
12935 HMMpropdeh3_1_pdu(pf02288) PDU1AB EP_NTERM
12940 HMMpropdeh2_1_pdu(pf02287) PDU1AB EP_NTERM
12945 HMM08841dhact_1_pdu(pf08841pf18427) PDU1AB
12950 HMMpropdeh3_2_pdusml(pf02288) PDU1AB
12955 H_tan PDU1AB
12960 H_driedBlood PDU1AB
12965 HMMptac_2_mix(pf06130) PDU1AB
12970 HMM15953pduput_1_pdu1(pf15953) PDU1AB EP_NTERM
12975 P_babyPink PDU1AB
12980 HMM01923cobtr_1_pdu(pf01923pf03928) PDU1AB
12985 HMMaldd h_1_eutpdugrm(pf00171) PDU1AB EP_NTERM
12990 HMM00465alcdh_3_mix(pf00465) PDU1AB
12995 HMMPduS_1_all(pf01512pf13375pf10531) PDU1AB
13000 Ts_dodgerBlue PDU1AB
13005 Hp_euts_babyPurple PDU1AB
13010 HMM10662PduV_1_mix(pf10662) PDU1AB
13015 HMMacetkin_1(pf00871) PDU1AB
13020 HMM00288_1_pdu(pf00288) PDU1AB
13025 -pf04363 -
13030 -pf04632 -
13035 -pf06445 -
13040 -pf07943pf00768 -
13045 -pf01292 -
13050 -pf13247pf12800 -
13055 -pf04879pf00384pf01568 -
13065 -pf13599pf13981pf13979 -
13070 -pf08411pf00929 -
BMC-P
BMC-H
BMC-T
Aldehyde dehydrogenase
PTAC
Signature enzyme
Alcohol dehydrogenase
Regulator

Salmonella enterica subsp. enterica serovar Havana. Bacteria; Proteobacteria; Gammaproteobacteria; Enterobacterales;

AVF28__locus_1

PDU1AB

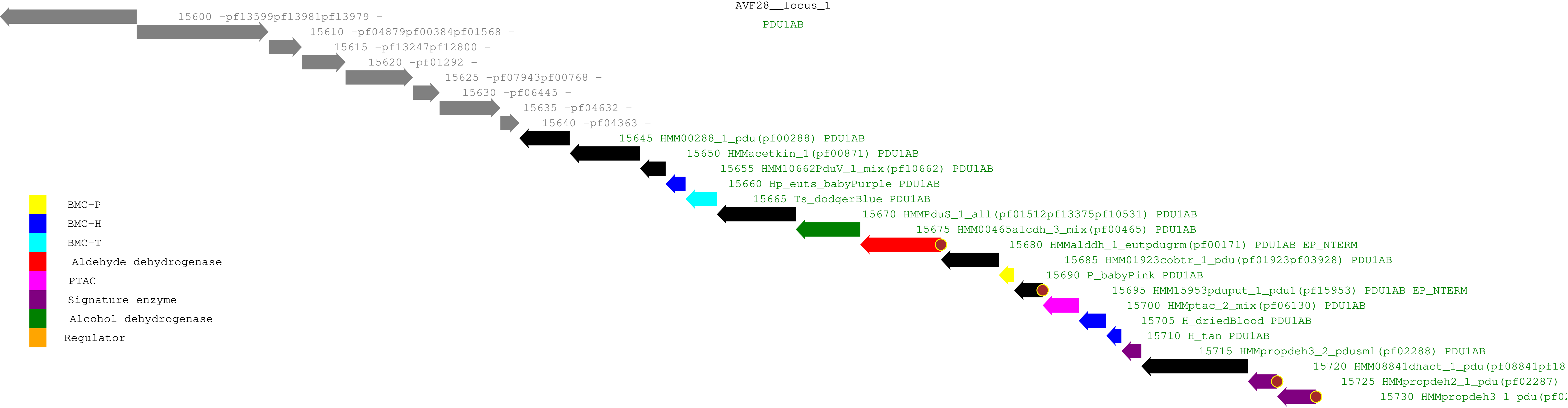

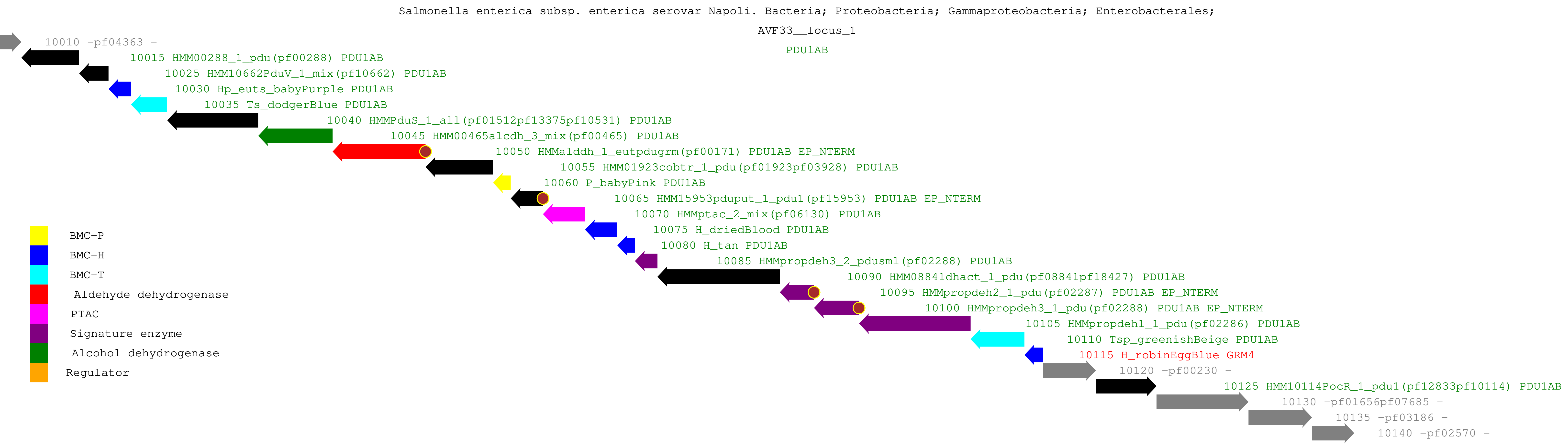

Salmonella enterica subsp. enterica serovar Napoli. Bacteria; Proteobacteria; Gammaproteobacteria; Enterobacterales;
AVF33__locus_1
PDU1AB
10010 -pf04363 -
10015 HMM00288_1_pdu(pf00288) PDU1AB
10025 HMM10662PduV_1_mix(pf10662) PDU1AB
10030 Hp_euts_babyPurple PDU1AB
10035 Ts_dodgerBlue PDU1AB
10040 HMMPduS_1_all(pf01512pf13375pf10531) PDU1AB
10045 HMM00465alcdh_3_mix(pf00465) PDU1AB
10050 HMMalddh_1_eutpdugrm(pf00171) PDU1AB EP_NTERM
10055 HMM01923cobtr_1_pdu(pf01923pf03928) PDU1AB
10060 P_babyPink PDU1AB
10065 HMM15953pduput_1_pdu1(pf15953) PDU1AB EP_NTERM
10070 HMMptac_2_mix(pf06130) PDU1AB
10075 H_driedBlood PDU1AB
10080 H_tan PDU1AB
10085 HMMpropdeh3_2_pdusml(pf02288) PDU1AB
10090 HMM08841dhact_1_pdu(pf08841pf18427) PDU1AB
10095 HMMpropdeh2_1_pdu(pf02287) PDU1AB EP_NTERM
10100 HMMpropdeh3_1_pdu(pf02288) PDU1AB EP_NTERM
10105 HMMpropdeh1_1_pdu(pf02286) PDU1AB
10110 Tsp_greenishBeige PDU1AB
10115 H_robinEggBlue GRM4
10120 -pf00230 -
10125 HMM10114PocR_1_pdu1(pf12833pf10114) PDU1AB
10130 -pf01656pf07685 -
10135 -pf03186 -
10140 -pf02570 -
BMC-P
BMC-H
BMC-T
Aldehyde dehydrogenase
PTAC
Signature enzyme
Alcohol dehydrogenase
Regulator

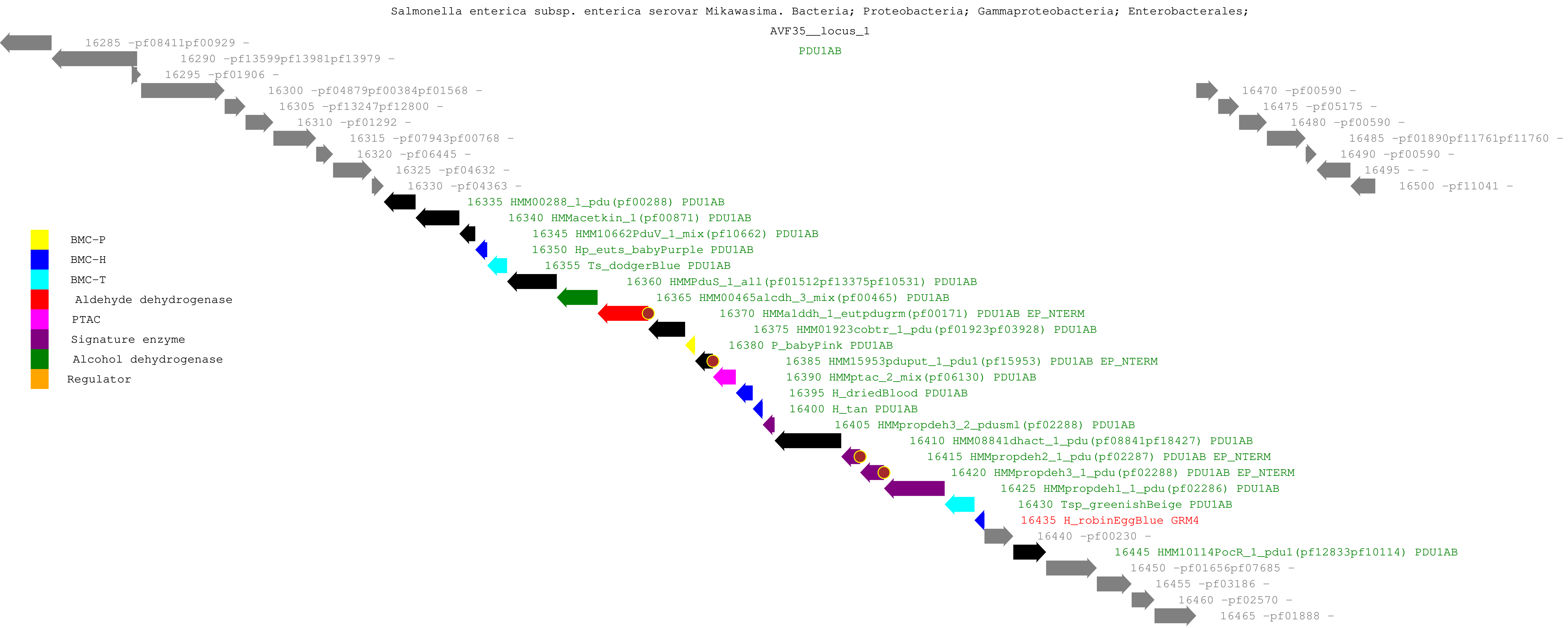
Salmonella enterica subsp. enterica serovar Mikawasima. Bacteria; Proteobacteria; Gammaproteobacteria; Enterobacterales;
AVF35__locus_1
PDU1AB
16285 -pf08411pf00929 -
16290 -pf13599pf13981pf13979 -
16295 -pf01906 -
16300 -pf04879pf00384pf01568 -
16305 -pf13247pf12800 -
16310 -pf01292 -
16315 -pf07943pf00768 -
16320 -pf06445 -
16325 -pf04632 -
16330 -pf04363 -
16335 HMM00288_1_pdu(pf00288) PDU1AB
16340 HMMacetkin_1(pf00871) PDU1AB
16345 HMM10662PduV_1_mix(pf10662) PDU1AB
16350 Hp_euts_babyPurple PDU1AB
16355 Ts_dodgerBlue PDU1AB
16360 HMMPduS_1_all(pf01512pf13375pf10531) PDU1AB
16365 HMM00465alcdh_3_mix(pf00465) PDU1AB
16370 HMMalddh_1_eutpdugrm(pf00171) PDU1AB EP_NTERM
16375 HMM01923cobtr_1_pdu(pf01923pf03928) PDU1AB
16380 P_babyPink PDU1AB
16385 HMM15953pduput_1_pdu1(pf15953) PDU1AB EP_NTERM
16390 HMMptac_2_mix(pf06130) PDU1AB
16395 H_driedBlood PDU1AB
16400 H_tan PDU1AB
16405 HMMpropdeh3_2_pdusml(pf02288) PDU1AB
16410 HMM08841dhact_1_pdu(pf08841pf18427) PDU1AB
16415 HMMpropdeh2_1_pdu(pf02287) PDU1AB EP_NTERM
16420 HMMpropdeh3_1_pdu(pf02288) PDU1AB EP_NTERM
16425 HMMpropdeh1_1_pdu(pf02286) PDU1AB
16430 Tsp_greenishBeige PDU1AB
16435 H_robinEggBlue GRM4
16440 -pf00230 -
16445 HMM10114PocR_1_pdu1(pf12833pf10114) PDU1AB
16450 -pf01656pf07685 -
16455 -pf03186 -
16460 -pf02570 -
16465 -pf01888 -
16470 -pf00590 -
16475 -pf05175 -
16480 -pf00590 -
16485 -pf01890pf11761pf11760 -
16490 -pf00590 -
16495 - -
16500 -pf11041 -
BMC-P
BMC-H
BMC-T
Aldehyde dehydrogenase
PTAC
Signature enzyme
Alcohol dehydrogenase
Regulator

Salmonella enterica subsp. enterica serovar Kottbus. Bacteria; Proteobacteria; Gammaproteobacteria; Enterobacterales;

AVF53__locus_1

PDU1AB

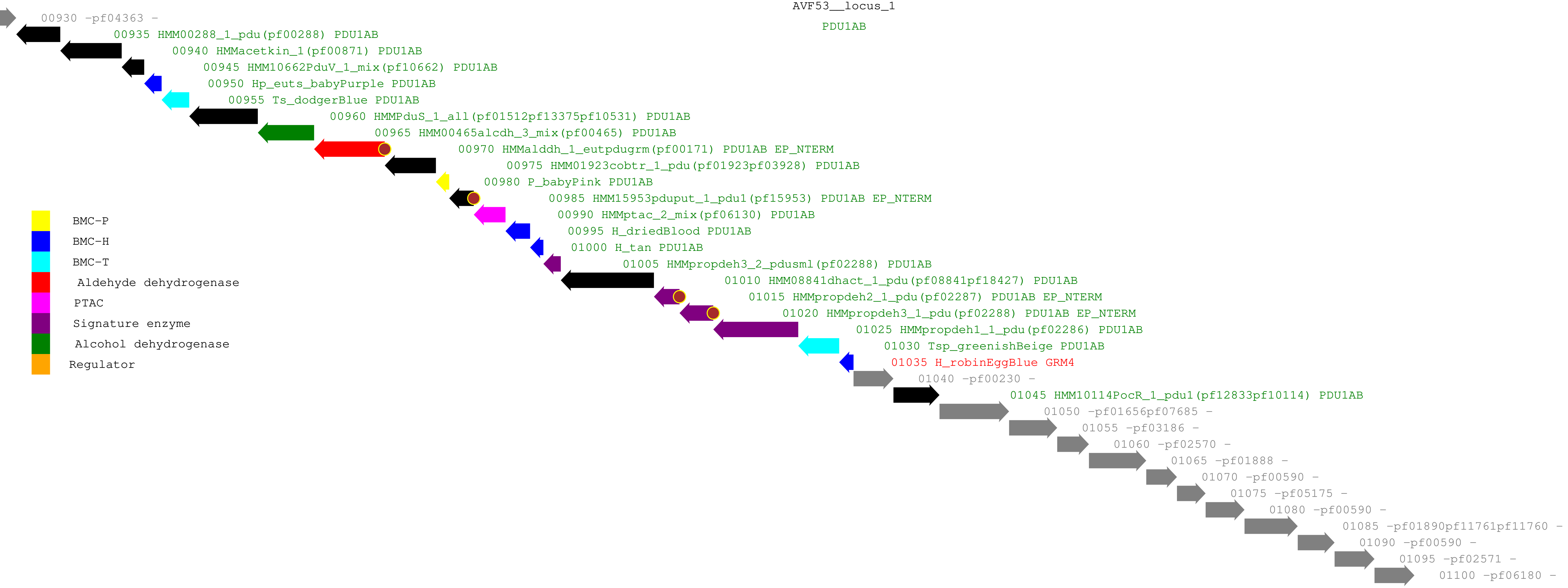

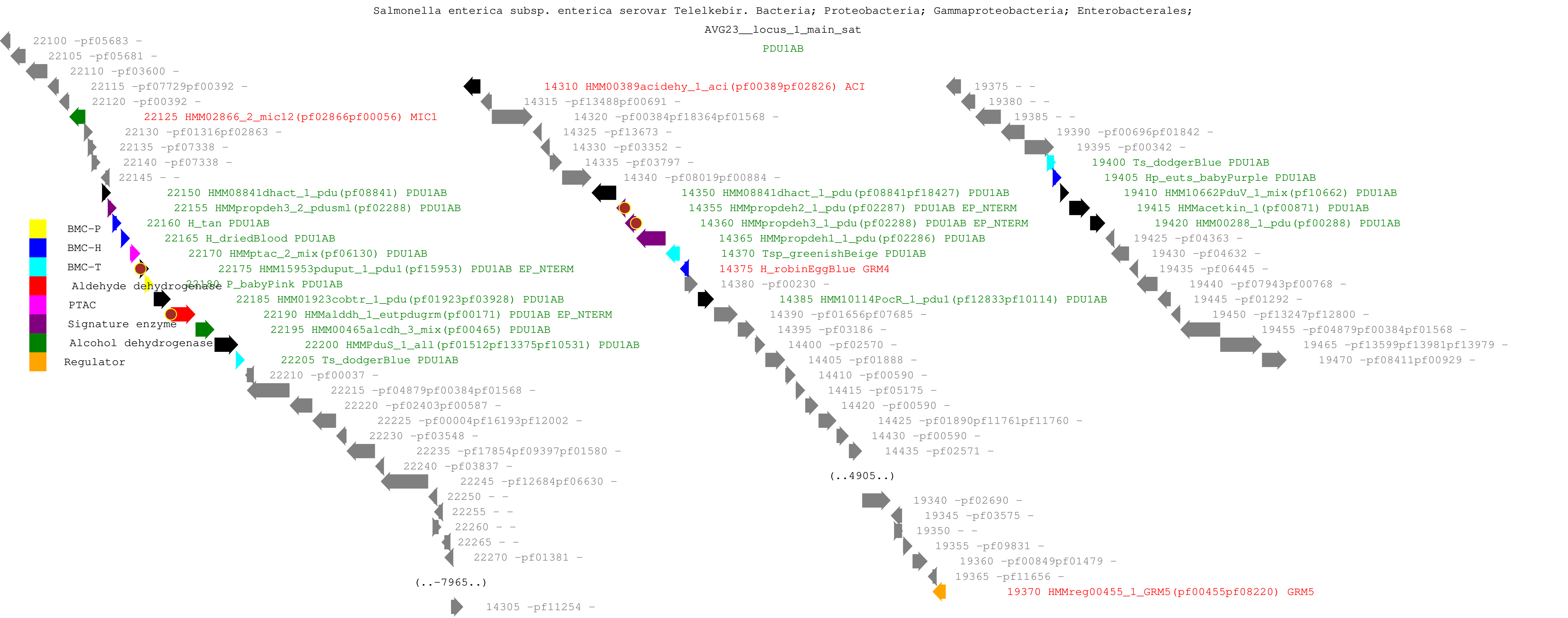
Salmonella enterica subsp. enterica serovar Telelkebir. Bacteria; Proteobacteria; Gammaproteobacteria; Enterobacterales;
AVG23__locus_1_main_sat
PDU1AB
22100 -pf05683 -
22105 -pf05681 -
22110 -pf03600 -
22115 -pf07729pf00392 -
22120 -pf00392 -
22125 HMM02866_2_mic12(pf02866pf00056) MIC1
22130 -pf01316pf02863 -
22135 -pf07338 -
22140 -pf07338 -
22145 - -
22150 HMM08841dhact_1_pdu(pf08841) PDU1AB
22155 HMMpropdeh3_2_pdusml(pf02288) PDU1AB
22160 H_tan PDU1AB
22165 H_driedBlood PDU1AB
22170 HMMptac_2_mix(pf06130) PDU1AB
22175 HMM15953pduput_1_pdu1(pf15953) PDU1AB EP_NTERM
22180 P_babyPink PDU1AB
22185 HMM01923cobtr_1_pdu(pf01923pf03928) PDU1AB
22190 HMMalddh_1_eutpdugrm(pf00171) PDU1AB EP_NTERM
22195 HMM00465alcdh_3_mix(pf00465) PDU1AB
22200 HMMPduS_1_all(pf01512pf13375pf10531) PDU1AB
22205 Ts_dodgerBlue PDU1AB
22210 -pf00037 -
22215 -pf04879pf00384pf01568 -
22220 -pf02403pf00587 -
22225 -pf00004pf16193pf12002 -
22230 -pf03548 -
22235 -pf17854pf09397pf01580 -
22240 -pf03837 -
22245 -pf12684pf06630 -
22250 - -
22255 - -
22260 - -
22265 - -
22270 -pf01381 -
(..-7965..)
14305 -pf11254 -
BMC-P
BMC-H
BMC-T
Aldehyde dehydrogenase
PTAC
Signature enzyme
Alcohol dehydrogenase
Regulator
14310 HMM00389acidehy_1_aci(pf00389pf02826) ACI
14315 -pf13488pf00691 -
14320 -pf00384pf18364pf01568 -
14325 -pf13673 -
14330 -pf03352 -
14335 -pf03797 -
14340 -pf08019pf00884 -
14350 HMM08841dhact_1_pdu(pf08841pf18427) PDU1AB
14355 HMMpropdeh2_1_pdu(pf02287) PDU1AB EP_NTERM
14360 HMMpropdeh3_1_pdu(pf02288) PDU1AB EP_NTERM
14365 HMMpropdeh1_1_pdu(pf02286) PDU1AB
14370 Tsp_greenishBeige PDU1AB
14375 H_robinEggBlue GRM4
14380 -pf00230 -
14385 HMM10114PocR_1_pdu1(pf12833pf10114) PDU1AB
14390 -pf01656pf07685 -
14395 -pf03186 -
14400 -pf02570 -
14405 -pf01888 -
14410 -pf00590 -
14415 -pf05175 -
14420 -pf00590 -
14425 -pf01890pf11761pf11760 -
14430 -pf00590 -
14435 -pf02571 -
(..4905..)
19340 -pf02690 -
19345 -pf03575 -
19350 - -
19355 -pf09831 -
19360 -pf00849pf01479 -
19365 -pf11656 -
19370 HMMreg00455_1_GRM5(pf00455pf08220) GRM5
19375 - -
19380 - -
19385 - -
19390 -pf00696pf01842 -
19395 -pf00342 -
19400 Ts_dodgerBlue PDU1AB
19405 Hp_euts_babyPurple PDU1AB
19410 HMM10662PduV_1_mix(pf10662) PDU1AB
19415 HMMacetkin_1(pf00871) PDU1AB
19420 HMM00288_1_pdu(pf00288) PDU1AB
19425 -pf04363 -
19430 -pf04632 -
19435 -pf06445 -
19440 -pf07943pf00768 -
19445 -pf01292 -
19450 -pf13247pf12800 -
19455 -pf04879pf00384pf01568 -
19465 -pf13599pf13981pf13979 -
19470 -pf08411pf00929 -

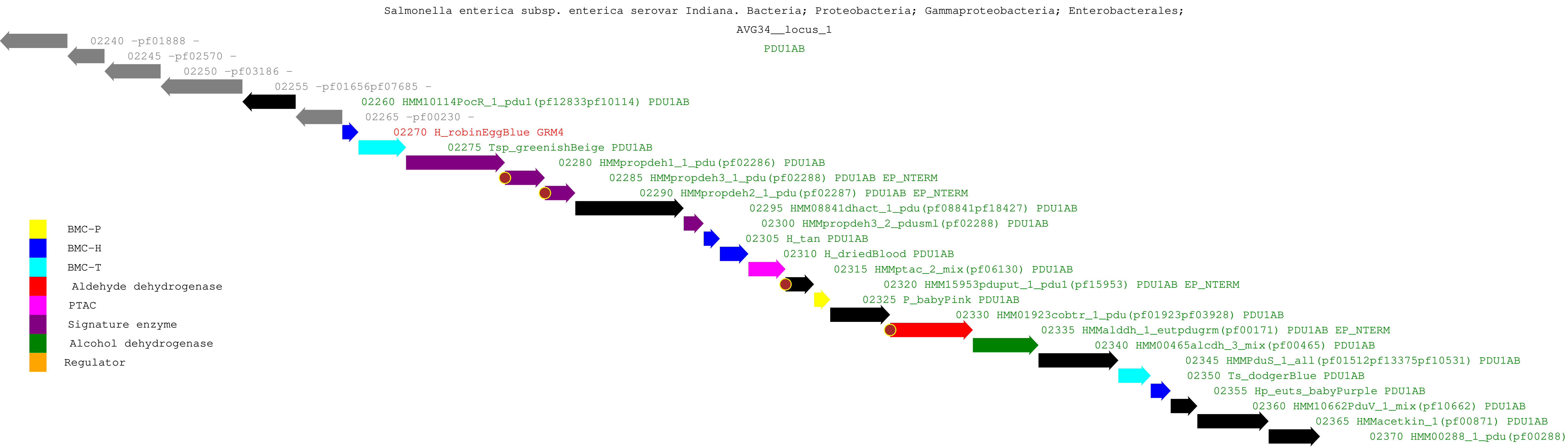
Salmonella enterica subsp. enterica serovar Indiana. Bacteria; Proteobacteria; Gammaproteobacteria; Enterobacterales;
AVG34__locus_1
PDU1AB
02240 -pf01888 -
02245 -pf02570 -
02250 -pf03186 -
02255 -pf01656pf07685 -
02260 HMM10114PocR_1_pdu1(pf12833pf10114) PDU1AB
02265 -pf00230 -
02270 H_robinEggBlue GRM4
02275 Tsp_greenishBeige PDU1AB
02280 HMMpropdeh1_1_pdu(pf02286) PDU1AB
02285 HMMpropdeh3_1_pdu(pf02288) PDU1AB EP_NTERM
02290 HMMpropdeh2_1_pdu(pf02287) PDU1AB EP_NTERM
02295 HMM08841dhact_1_pdu(pf08841pf18427) PDU1AB
02300 HMMpropdeh3_2_pdusml(pf02288) PDU1AB
02305 H_tan PDU1AB
02310 H_driedBlood PDU1AB
02315 HMMptac_2_mix(pf06130) PDU1AB
02320 HMM15953pduput_1_pdu1(pf15953) PDU1AB EP_NTERM
02325 P_babyPink PDU1AB
02330 HMM01923cobtr_1_pdu(pf01923pf03928) PDU1AB
02335 HMMalddh_1_eutpdugrm(pf00171) PDU1AB EP_NTERM
02340 HMM00465alcdh_3_mix(pf00465) PDU1AB
02345 HMMPduS_1_all(pf01512pf13375pf10531) PDU1AB
02350 Ts_dodgerBlue PDU1AB
02355 Hp_euts_babyPurple PDU1AB
02360 HMM10662PduV_1_mix(pf10662) PDU1AB
02365 HMMacetkin_1(pf00871) PDU1AB
02370 HMM00288_1_pdu(pf00288)
BMC-P
BMC-H
BMC-T
Aldehyde dehydrogenase
PTAC
Signature enzyme
Alcohol dehydrogenase
Regulator

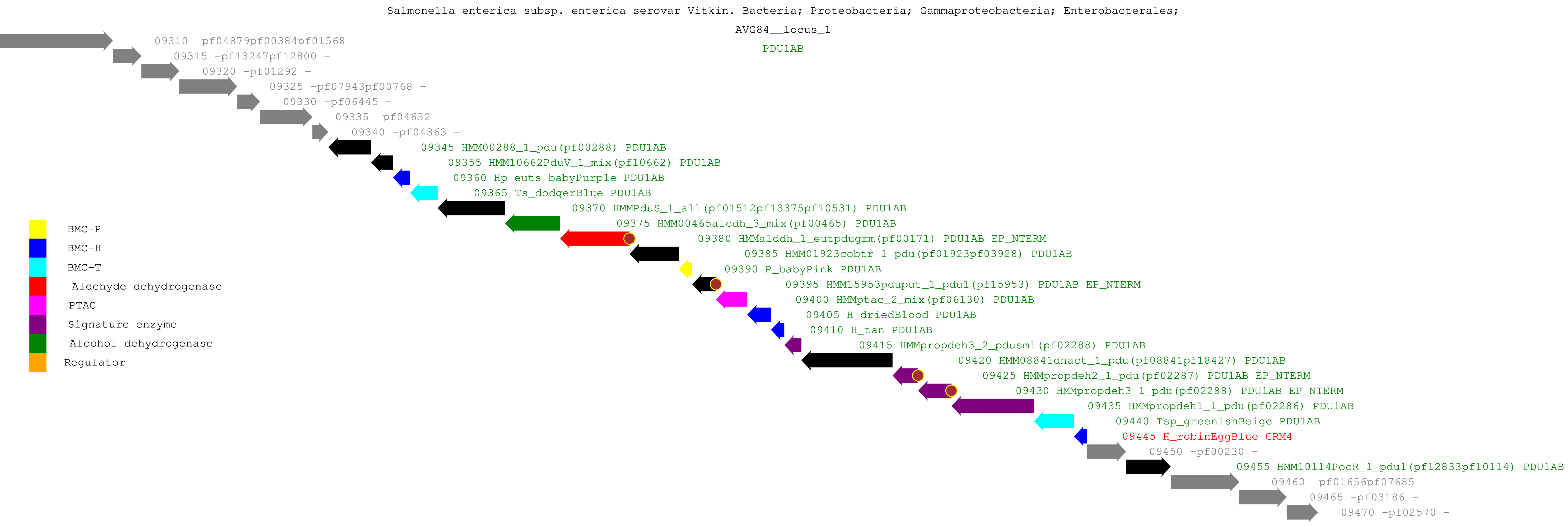
Salmonella enterica subsp. enterica serovar Vitkin. Bacteria; Proteobacteria; Gammaproteobacteria; Enterobacterales;
AVG84__locus_1
PDU1AB
09310 -pf04879pf00384pf01568 -
09315 -pf13247pf12800 -
09320 -pf01292 -
09325 -pf07943pf00768 -
09330 -pf06445 -
09335 -pf04632 -
09340 -pf04363 -
09345 HMM00288_1_pdu(pf00288) PDU1AB
09355 HMM10662PduV_1_mix(pf10662) PDU1AB
09360 Hp_euts_babyPurple PDU1AB
09365 Ts_dodgerBlue PDU1AB
09370 HMMPduS_1_all(pf01512pf13375pf10531) PDU1AB
09375 HMM00465alcdh_3_mix(pf00465) PDU1AB
09380 HMMalddh_1_eutpdugrm(pf00171) PDU1AB EP_NTERM
09385 HMM01923cobtr_1_pdu(pf01923pf03928) PDU1AB
09390 P_babyPink PDU1AB
09395 HMM15953pduput_1_pdu1(pf15953) PDU1AB EP_NTERM
09400 HMMptac_2_mix(pf06130) PDU1AB
09405 H_driedBlood PDU1AB
09410 H_tan PDU1AB
09415 HMMpropdeh3_2_pdusml(pf02288) PDU1AB
09420 HMM08841dhact_1_pdu(pf08841pf18427) PDU1AB
09425 HMMpropdeh2_1_pdu(pf02287) PDU1AB EP_NTERM
09430 HMMpropdeh3_1_pdu(pf02288) PDU1AB EP_NTERM
09435 HMMpropdeh1_1_pdu(pf02286) PDU1AB
09440 Tsp_greenishBeige PDU1AB
09445 H_robinEggBlue GRM4
09450 -pf00230 -
09455 HMM10114PocR_1_pdu1(pf12833pf10114) PDU1AB
09460 -pf01656pf07685 -
09465 -pf03186 -
09470 -pf02570 -
BMC-P
BMC-H
BMC-T
Aldehyde dehydrogenase
PTAC
Signature enzyme
Alcohol dehydrogenase
Regulator

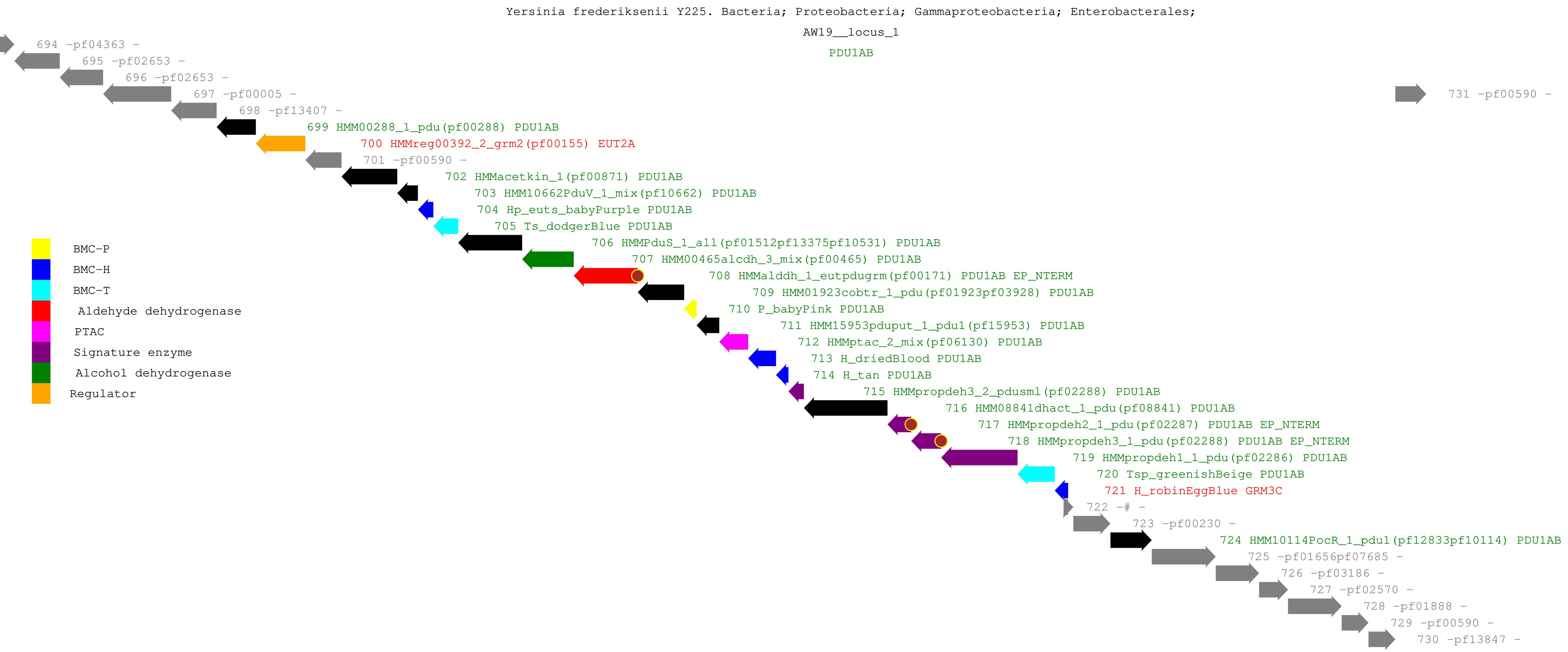

Yersinia frederiksenii Y225. Bacteria; Proteobacteria; Gammaproteobacteria; Enterobacterales;
AW19__locus_1
PDU1AB
694 -pf04363 -
695 -pf02653 -
696 -pf02653 -
697 -pf00005 -
698 -pf13407 -
699 HMM00288_1_pdu(pf00288) PDU1AB
700 HMMreg00392_2_grm2(pf00155) EUT2A
701 -pf00590 -
702 HMMacetkin_1(pf00871) PDU1AB
703 HMM10662PduV_1_mix(pf10662) PDU1AB
704 Hp_euts_babyPurple PDU1AB
705 Ts_dodgerBlue PDU1AB
706 HMMPduS_1_all(pf01512pf13375pf10531) PDU1AB
707 HMM00465alcdh_3_mix(pf00465) PDU1AB
708 HMMaldh_1_eutpdugrm(pf00171) PDU1AB EP_NTERM
709 HMM01923cobtr_1_pdu(pf01923pf03928) PDU1AB
710 P_babyPink PDU1AB
711 HMM15953pduput_1_pdu1(pf15953) PDU1AB
712 HMMptac_2_mix(pf06130) PDU1AB
713 H_driedBlood PDU1AB
714 H_tan PDU1AB
715 HMMpropdeh3_2_pdusml(pf02288) PDU1AB
716 HMM08841dhact_1_pdu(pf08841) PDU1AB
717 HMMpropdeh2_1_pdu(pf02287) PDU1AB EP_NTERM
718 HMMpropdeh3_1_pdu(pf02288) PDU1AB EP_NTERM
719 HMMpropdeh1_1_pdu(pf02286) PDU1AB
720 Tsp_greenishBeige PDU1AB
721 H_robinEggBlue GRM3C
722 -# -
723 -pf00230 -
724 HMM10114PocR_1_pdu1(pf12833pf10114) PDU1AB
725 -pf01656pf07685 -
726 -pf03186 -
727 -pf02570 -
728 -pf01888 -
729 -pf00590 -
730 -pf13847 -
731 -pf00590 -
BMC-P
BMC-H
BMC-T
Aldehyde dehydrogenase
PTAC
Signature enzyme
Alcohol dehydrogenase
Regulator

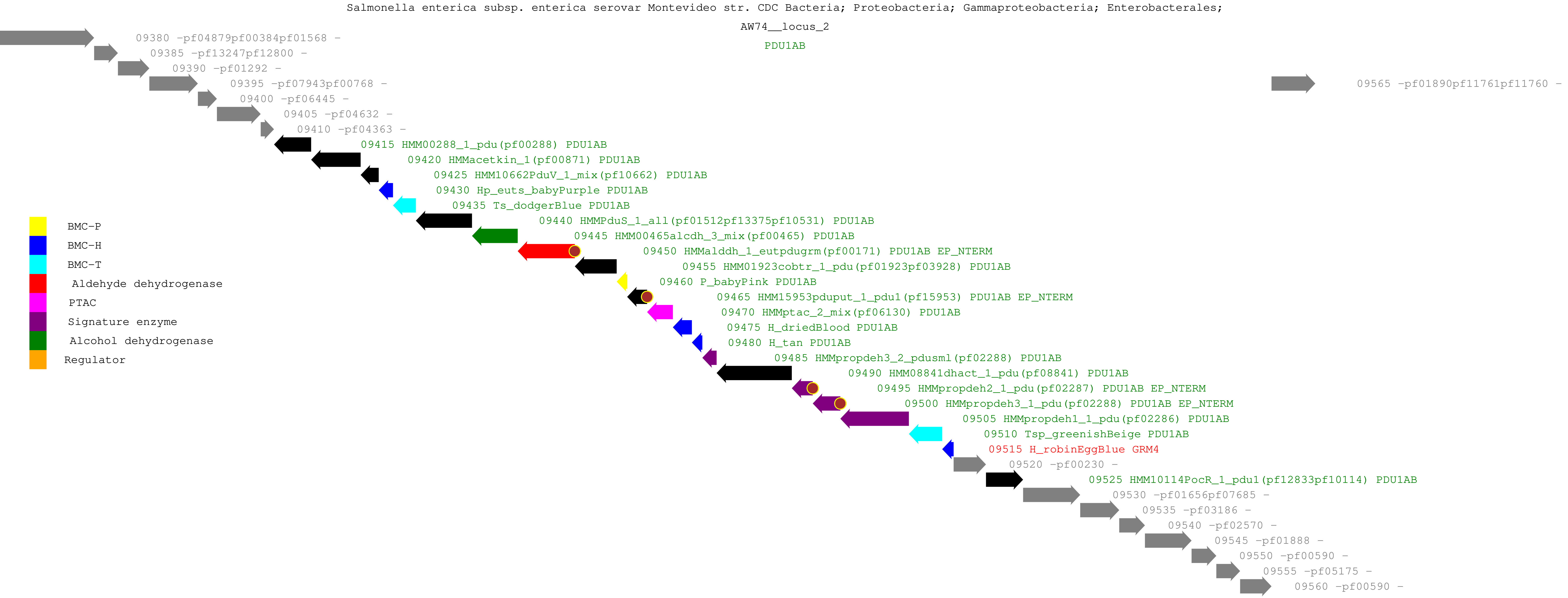
Salmonella enterica subsp. enterica serovar Montevideo str. CDC Bacteria; Proteobacteria; Gammaproteobacteria; Enterobacterales;
AW74__locus_2
PDU1AB
09380 -pf04879pf00384pf01568 -
09385 -pf13247pf12800 -
09390 -pf01292 -
09395 -pf07943pf00768 -
09400 -pf06445 -
09405 -pf04632 -
09410 -pf04363 -
09415 HMM00288_1_pdu(pf00288) PDU1AB
09420 HMMacetkin_1(pf00871) PDU1AB
09425 HMM10662PduV_1_mix(pf10662) PDU1AB
09430 Hp_euts_babyPurple PDU1AB
09435 Ts_dodgerBlue PDU1AB
09440 HMMPduS_1_all(pf01512pf13375pf10531) PDU1AB
09445 HMM00465alcdh_3_mix(pf00465) PDU1AB
09450 HMMalddh_1_eutpdugrm(pf00171) PDU1AB EP_NTERM
09455 HMM01923cobtr_1_pdu(pf01923pf03928) PDU1AB
09460 P_babyPink PDU1AB
09465 HMM15953pduput_1_pdu1(pf15953) PDU1AB EP_NTERM
09470 HMMptac_2_mix(pf06130) PDU1AB
09475 H_driedBlood PDU1AB
09480 H_tan PDU1AB
09485 HMMpropdeh3_2_pdusml(pf02288) PDU1AB
09490 HMM08841dhact_1_pdu(pf08841) PDU1AB
09495 HMMpropdeh2_1_pdu(pf02287) PDU1AB EP_NTERM
09500 HMMpropdeh3_1_pdu(pf02288) PDU1AB EP_NTERM
09505 HMMpropdeh1_1_pdu(pf02286) PDU1AB
09510 Tsp_greenishBeige PDU1AB
09515 H_robinEggBlue GRM4
09520 -pf00230 -
09525 HMM10114PocR_1_pdu1(pf12833pf10114) PDU1AB
09530 -pf01656pf07685 -
09535 -pf03186 -
09540 -pf02570 -
09545 -pf01888 -
09550 -pf00590 -
09555 -pf05175 -
09560 -pf00590 -
09565 -pf01890pf11761pf11760 -
BMC-P
BMC-H
BMC-T
Aldehyde dehydrogenase
PTAC
Signature enzyme
Alcohol dehydrogenase
Regulator

Escherichia coli. Bacteria; Proteobacteria; Gammaproteobacteria; Enterobacterales;

AWG90__locus_1

PDU1AB

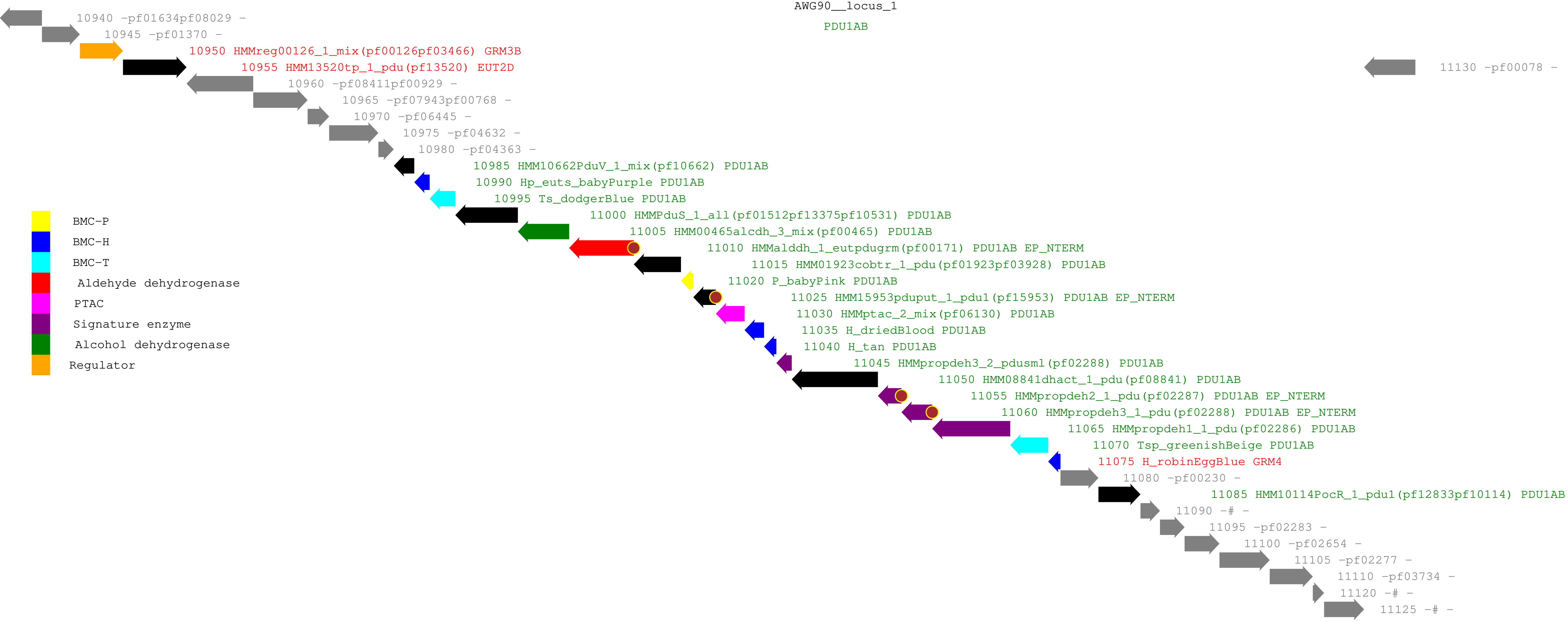

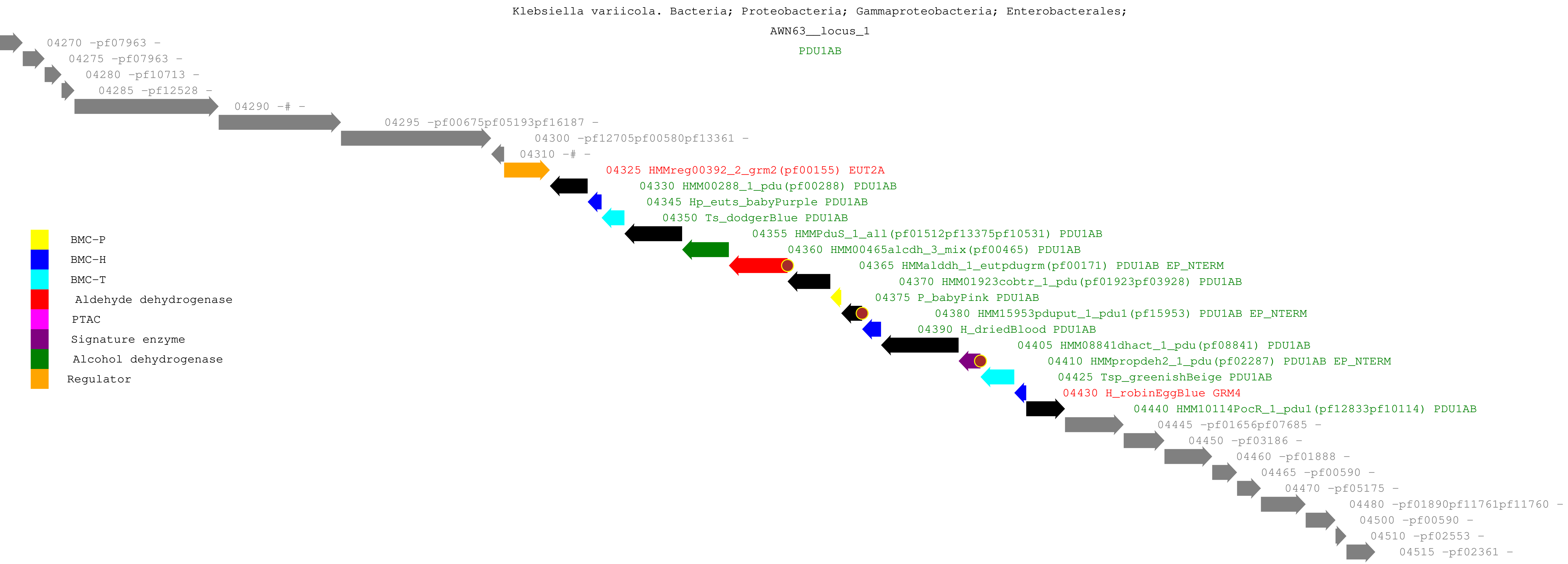
Klebsiella variicola. Bacteria; Proteobacteria; Gammaproteobacteria; Enterobacterales;
AWN63__locus_1
PDU1AB
04270 -pf07963 -
04275 -pf07963 -
04280 -pf10713 -
04285 -pf12528 -
04290 -# -
04295 -pf00675pf05193pf16187 -
04300 -pf12705pf00580pf13361 -
04310 -# -
04325 HMMreg00392_2_grm2(pf00155) EUT2A
04330 HMM00288_1_pdu(pf00288) PDU1AB
04345 Hp_euts_babyPurple PDU1AB
04350 Ts_dodgerBlue PDU1AB
04355 HMMPduS_1_all(pf01512pf13375pf10531) PDU1AB
04360 HMM00465alcdh_3_mix(pf00465) PDU1AB
04365 HMMalddh_1_eutpdugrm(pf00171) PDU1AB EP_NTERM
04370 HMM01923cobtr_1_pdu(pf01923pf03928) PDU1AB
04375 P_babyPink PDU1AB
04380 HMM15953pduput_1_pdu1(pf15953) PDU1AB EP_NTERM
04390 H_driedBlood PDU1AB
04405 HMM08841dhact_1_pdu(pf08841) PDU1AB
04410 HMMpropdeh2_1_pdu(pf02287) PDU1AB EP_NTERM
04425 Tsp_greenishBeige PDU1AB
04430 H_robinEggBlue GRM4
04440 HMM10114PocR_1_pdu1(pf12833pf10114) PDU1AB
04445 -pf01656pf07685 -
04450 -pf03186 -
04460 -pf01888 -
04465 -pf00590 -
04470 -pf05175 -
04480 -pf01890pf11761pf11760 -
04500 -pf00590 -
04510 -pf02553 -
04515 -pf02361 -
BMC-P
BMC-H
BMC-T
Aldehyde dehydrogenase
PTAC
Signature enzyme
Alcohol dehydrogenase
Regulator

Salmonella enterica I. Bacteria; Proteobacteria; Gammaproteobacteria; Enterobacterales;

AXM50__locus_1

PDU1AB

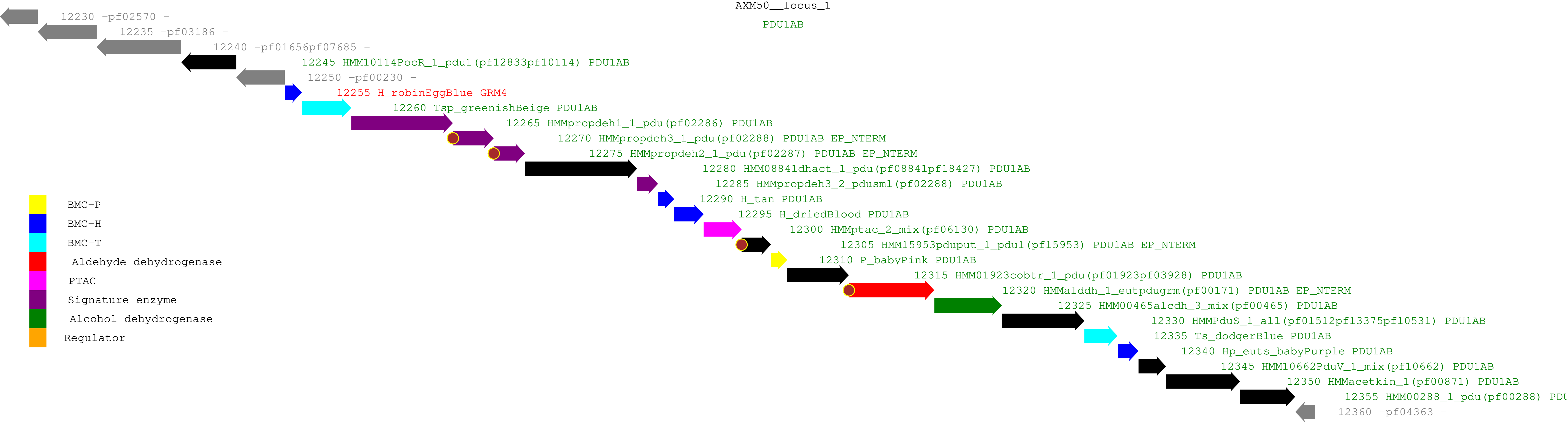

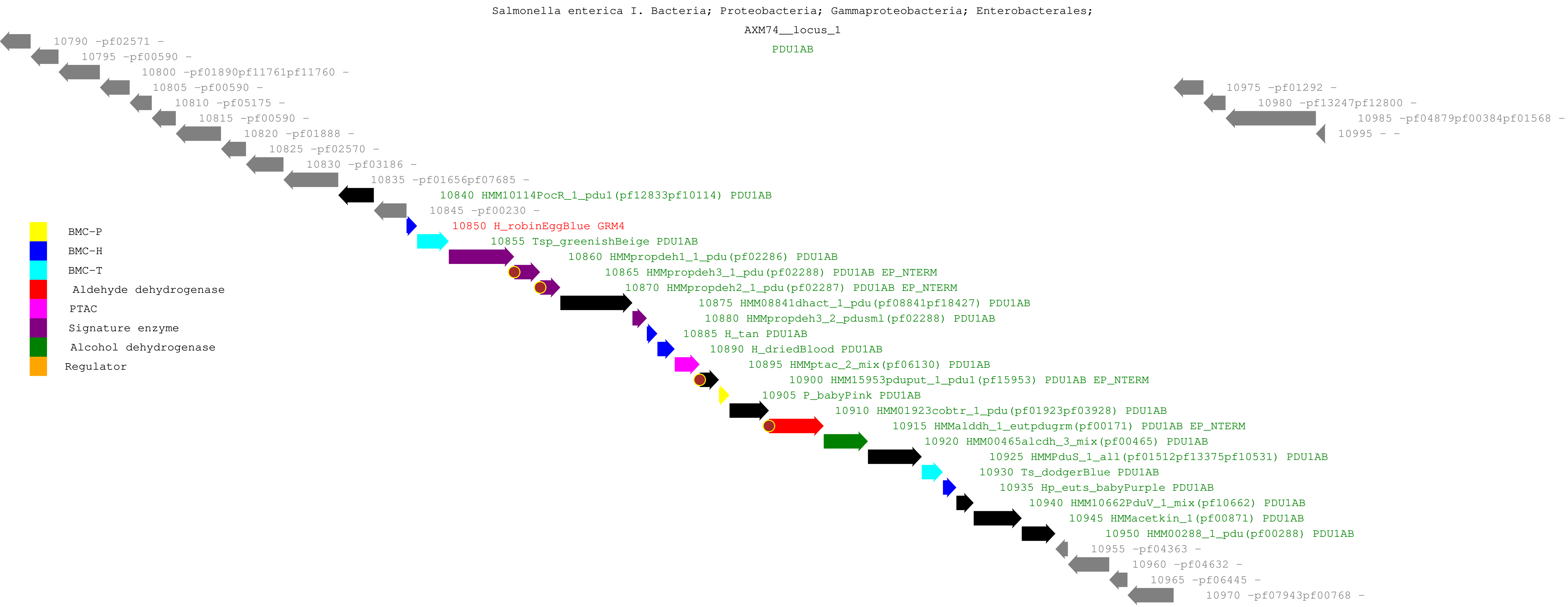
Salmonella enterica I. Bacteria; Proteobacteria; Gammaproteobacteria; Enterobacterales;
AXM74__locus_1
PDU1AB
10790 -pf02571 -
10795 -pf00590 -
10800 -pf01890pf11761pf11760 -
10805 -pf00590 -
10810 -pf05175 -
10815 -pf00590 -
10820 -pf01888 -
10825 -pf02570 -
10830 -pf03186 -
10835 -pf01656pf07685 -
10840 HMM10114PocR_1_pdu1(pf12833pf10114) PDU1AB
10845 -pf00230 -
10850 H_robinEggBlue GRM4
10855 Tsp_greenishBeige PDU1AB
10860 HMMpropdeh1_1_pdu(pf02286) PDU1AB
10865 HMMpropdeh3_1_pdu(pf02288) PDU1AB EP_NTERM
10870 HMMpropdeh2_1_pdu(pf02287) PDU1AB EP_NTERM
10875 HMM08841dhact_1_pdu(pf08841pf18427) PDU1AB
10880 HMMpropdeh3_2_pdusml(pf02288) PDU1AB
10885 H_tan PDU1AB
10890 H_driedBlood PDU1AB
10895 HMMptac_2_mix(pf06130) PDU1AB
10900 HMM15953pduput_1_pdu1(pf15953) PDU1AB EP_NTERM
10905 P_babyPink PDU1AB
10910 HMM01923cobtr_1_pdu(pf01923pf03928) PDU1AB
10915 HMMalddh_1_eutpdugrm(pf00171) PDU1AB EP_NTERM
10920 HMM00465alcdh_3_mix(pf00465) PDU1AB
10925 HMMPduS_1_all(pf01512pf13375pf10531) PDU1AB
10930 Ts_dodgerBlue PDU1AB
10935 Hp_euts_babyPurple PDU1AB
10940 HMM10662PduV_1_mix(pf10662) PDU1AB
10945 HMMacetkin_1(pf00871) PDU1AB
10950 HMM00288_1_pdu(pf00288) PDU1AB
10955 -pf04363 -
10960 -pf04632 -
10965 -pf06445 -
10970 -pf07943pf00768 -
10975 -pf01292 -
10980 -pf13247pf12800 -
10985 -pf04879pf00384pf01568 -
10995 - -
BMC-P
BMC-H
BMC-T
Aldehyde dehydrogenase
PTAC
Signature enzyme
Alcohol dehydrogenase
Regulator

Salmonella enterica I. Bacteria; Proteobacteria; Gammaproteobacteria; Enterobacterales;

AXN72__locus_1

PDU1AB

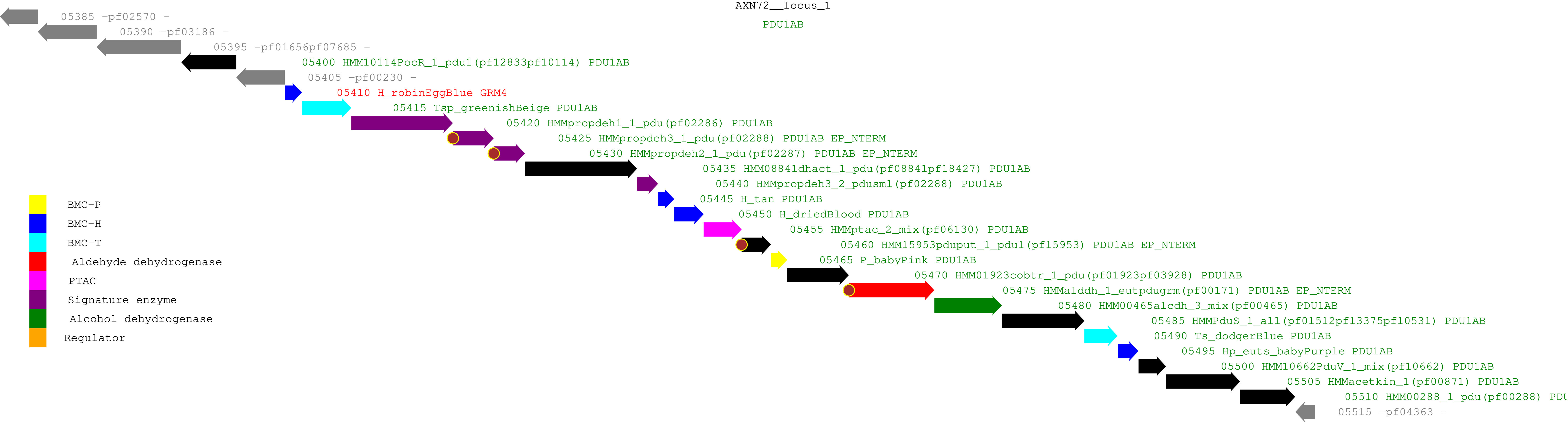

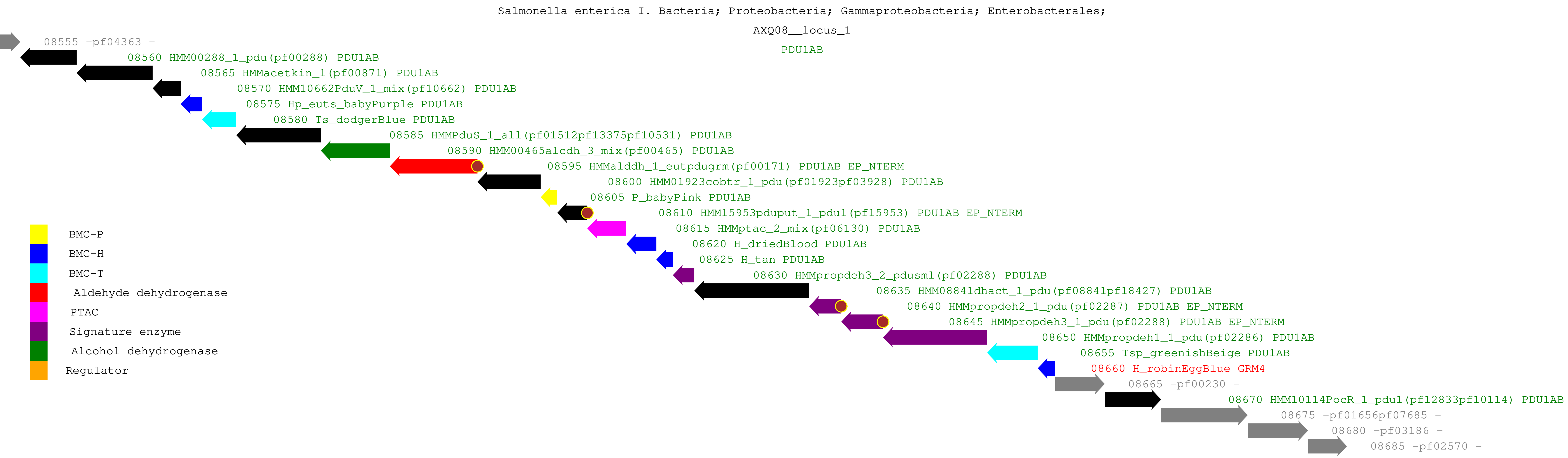
Salmonella enterica I. Bacteria; Proteobacteria; Gammaproteobacteria; Enterobacterales;
AXQ08__locus_1
PDU1AB
08555 -pf04363 -
08560 HMM00288_1_pdu(pf00288) PDU1AB
08565 HMMacetkin_1(pf00871) PDU1AB
08570 HMM10662PduV_1_mix(pf10662) PDU1AB
08575 Hp_euts_babyPurple PDU1AB
08580 Ts_dodgerBlue PDU1AB
08585 HMMPduS_1_all(pf01512pf13375pf10531) PDU1AB
08590 HMM00465alcdh_3_mix(pf00465) PDU1AB
08595 HMMalddh_1_eutpdugrm(pf00171) PDU1AB EP_NTERM
08600 HMM01923cobtr_1_pdu(pf01923pf03928) PDU1AB
08605 P_babyPink PDU1AB
08610 HMM15953pduput_1_pdu1(pf15953) PDU1AB EP_NTERM
08615 HMMptac_2_mix(pf06130) PDU1AB
08620 H_driedBlood PDU1AB
08625 H_tan PDU1AB
08630 HMMpropdeh3_2_pdusml(pf02288) PDU1AB
08635 HMM08841dhact_1_pdu(pf08841pf18427) PDU1AB
08640 HMMpropdeh2_1_pdu(pf02287) PDU1AB EP_NTERM
08645 HMMpropdeh3_1_pdu(pf02288) PDU1AB EP_NTERM
08650 HMMpropdeh1_1_pdu(pf02286) PDU1AB
08655 Tsp_greenishBeige PDU1AB
08660 H_robinEggBlue GRM4
08665 -pf00230 -
08670 HMM10114PocR_1_pdu1(pf12833pf10114) PDU1AB
08675 -pf01656pf07685 -
08680 -pf03186 -
08685 -pf02570 -
BMC-P
BMC-H
BMC-T
Aldehyde dehydrogenase
PTAC
Signature enzyme
Alcohol dehydrogenase
Regulator

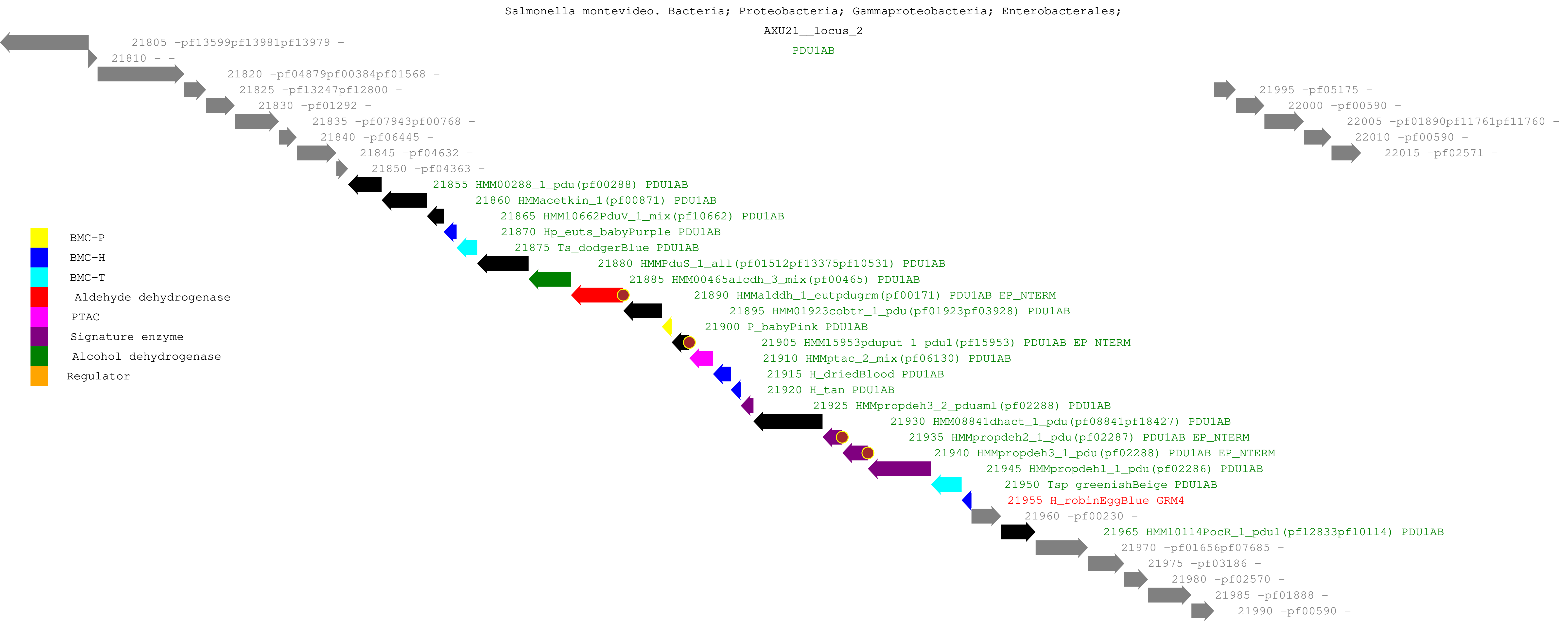
Salmonella montevideo. Bacteria; Proteobacteria; Gammaproteobacteria; Enterobacterales;
AXU21__locus_2
PDU1AB
21805 -pf13599pf13981pf13979 -
21810 - -
21820 -pf04879pf00384pf01568 -
21825 -pf13247pf12800 -
21830 -pf01292 -
21835 -pf07943pf00768 -
21840 -pf06445 -
21845 -pf04632 -
21850 -pf04363 -
21855 HMM00288_1_pdu(pf00288) PDU1AB
21860 HMMacetkin_1(pf00871) PDU1AB
21865 HMM10662PduV_1_mix(pf10662) PDU1AB
21870 Hp_euts_babyPurple PDU1AB
21875 Ts_dodgerBlue PDU1AB
21880 HMMPduS_1_all(pf01512pf13375pf10531) PDU1AB
21885 HMM00465alcdh_3_mix(pf00465) PDU1AB
21890 HMMalddh_1_eutpdugrm(pf00171) PDU1AB EP_NTERM
21895 HMM01923cobtr_1_pdu(pf01923pf03928) PDU1AB
21900 P_babyPink PDU1AB
21905 HMM15953pduput_1_pdu1(pf15953) PDU1AB EP_NTERM
21910 HMMptac_2_mix(pf06130) PDU1AB
21915 H_driedBlood PDU1AB
21920 H_tan PDU1AB
21925 HMMpropdeh3_2_pdusml(pf02288) PDU1AB
21930 HMM08841dhact_1_pdu(pf08841pf18427) PDU1AB
21935 HMMpropdeh2_1_pdu(pf02287) PDU1AB EP_NTERM
21940 HMMpropdeh3_1_pdu(pf02288) PDU1AB EP_NTERM
21945 HMMpropdeh1_1_pdu(pf02286) PDU1AB
21950 Tsp_greenishBeige PDU1AB
21955 H_robinEggBlue GRM4
21960 -pf00230 -
21965 HMM10114PocR_1_pdu1(pf12833pf10114) PDU1AB
21970 -pf01656pf07685 -
21975 -pf03186 -
21980 -pf02570 -
21985 -pf01888 -
21990 -pf00590 -
21995 -pf05175 -
22000 -pf00590 -
22005 -pf01890pf11761pf11760 -
22010 -pf00590 -
22015 -pf02571 -
BMC-P
BMC-H
BMC-T
Aldehyde dehydrogenase
PTAC
Signature enzyme
Alcohol dehydrogenase
Regulator

Citrobacter freundii. Bacteria; Proteobacteria; Gammaproteobacteria; Enterobacterales;

AZ002__locus_2

PDU1AB

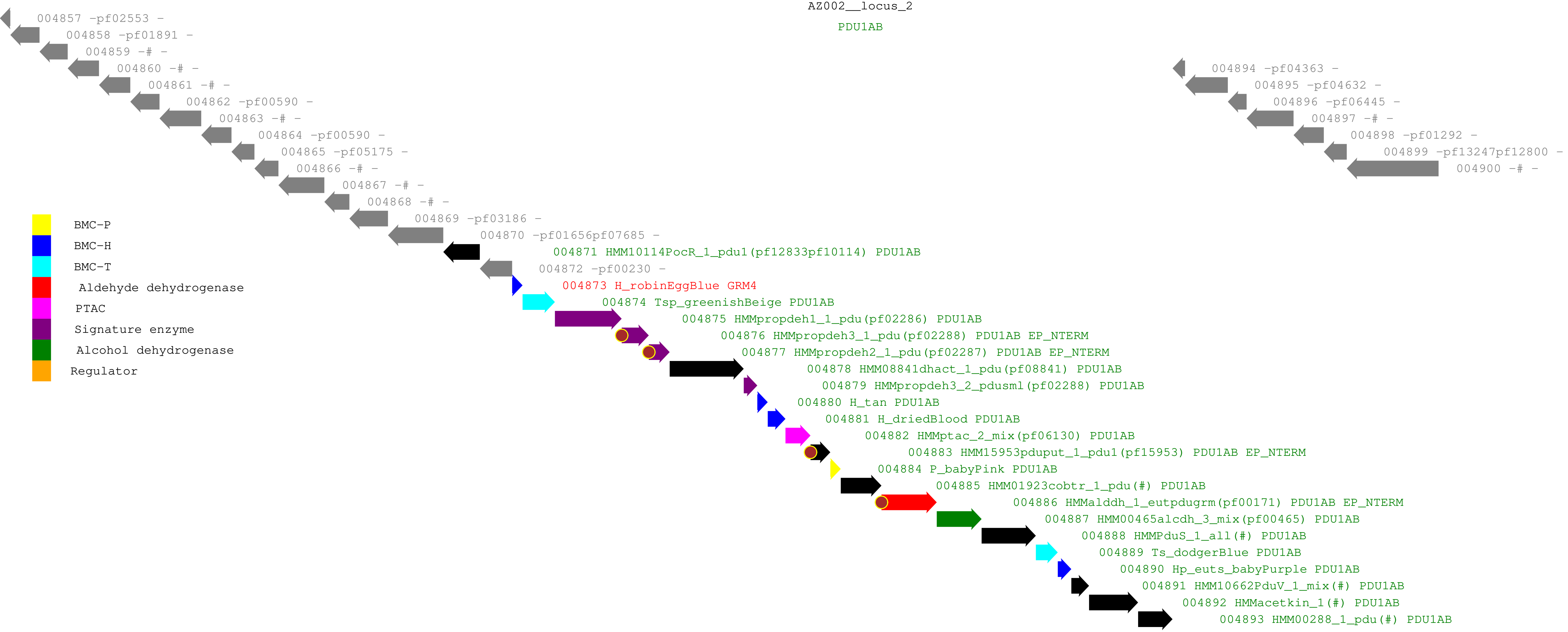

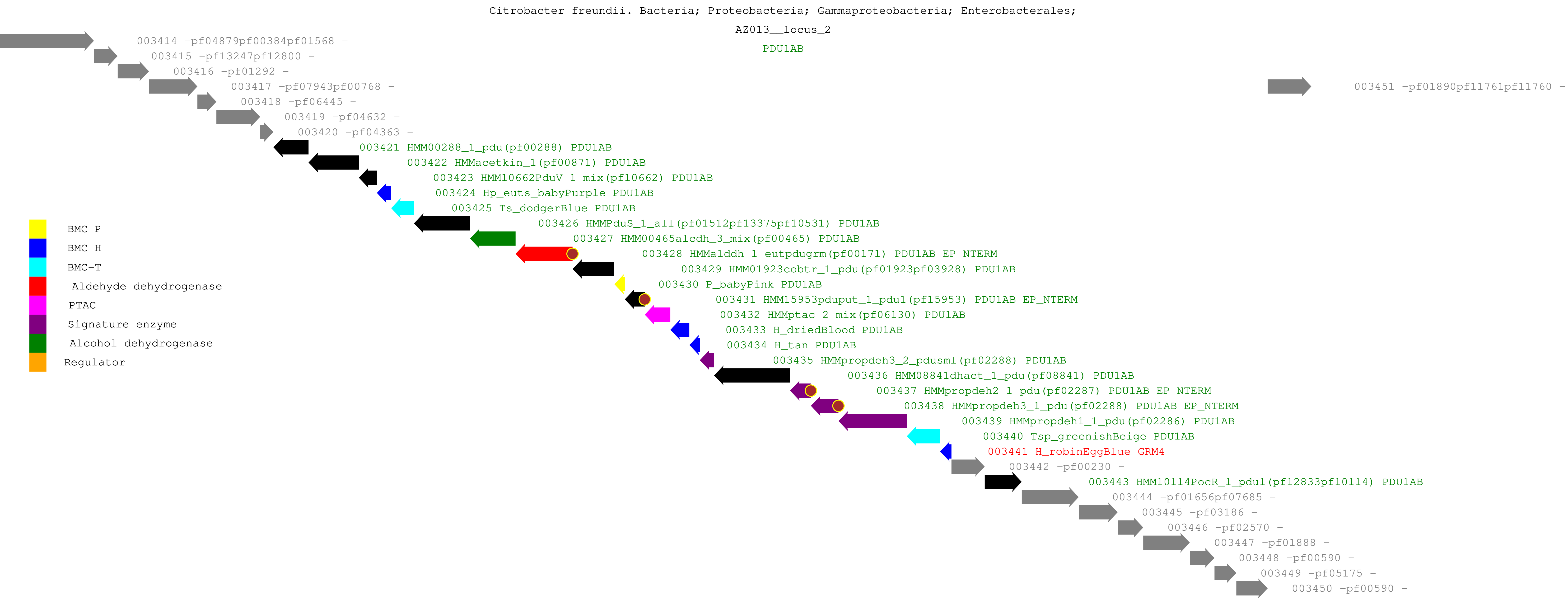
Citrobacter freundii. Bacteria; Proteobacteria; Gammaproteobacteria; Enterobacterales;
AZ013__locus_2
PDU1AB
003414 -pf04879pf00384pf01568 -
003415 -pf13247pf12800 -
003416 -pf01292 -
003417 -pf07943pf00768 -
003418 -pf06445 -
003419 -pf04632 -
003420 -pf04363 -
003421 HMM00288_1_pdu(pf00288) PDU1AB
003422 HMMacetkin_1(pf00871) PDU1AB
003423 HMM10662PduV_1_mix(pf10662) PDU1AB
003424 Hp_euts_babyPurple PDU1AB
003425 Ts_dodgerBlue PDU1AB
003426 HMMPduS_1_all(pf01512pf13375pf10531) PDU1AB
003427 HMM00465alcdh_3_mix(pf00465) PDU1AB
003428 HMMalddh_1_eutpdugrm(pf00171) PDU1AB EP_NTERM
003429 HMM01923cobtr_1_pdu(pf01923pf03928) PDU1AB
003430 P_babyPink PDU1AB
003431 HMM15953pduput_1_pdu1(pf15953) PDU1AB EP_NTERM
003432 HMMptac_2_mix(pf06130) PDU1AB
003433 H_driedBlood PDU1AB
003434 H_tan PDU1AB
003435 HMMpropdeh3_2_pdusml(pf02288) PDU1AB
003436 HMM08841dhact_1_pdu(pf08841) PDU1AB
003437 HMMpropdeh2_1_pdu(pf02287) PDU1AB EP_NTERM
003438 HMMpropdeh3_1_pdu(pf02288) PDU1AB EP_NTERM
003439 HMMpropdeh1_1_pdu(pf02286) PDU1AB
003440 Tsp_greenishBeige PDU1AB
003441 H_robinEggBlue GRM4
003442 -pf00230 -
003443 HMM10114PocR_1_pdu1(pf12833pf10114) PDU1AB
003444 -pf01656pf07685 -
003445 -pf03186 -
003446 -pf02570 -
003447 -pf01888 -
003448 -pf00590 -
003449 -pf05175 -
003450 -pf00590 -
003451 -pf01890pf11761pf11760 -
BMC-P
BMC-H
BMC-T
Aldehyde dehydrogenase
PTAC
Signature enzyme
Alcohol dehydrogenase
Regulator

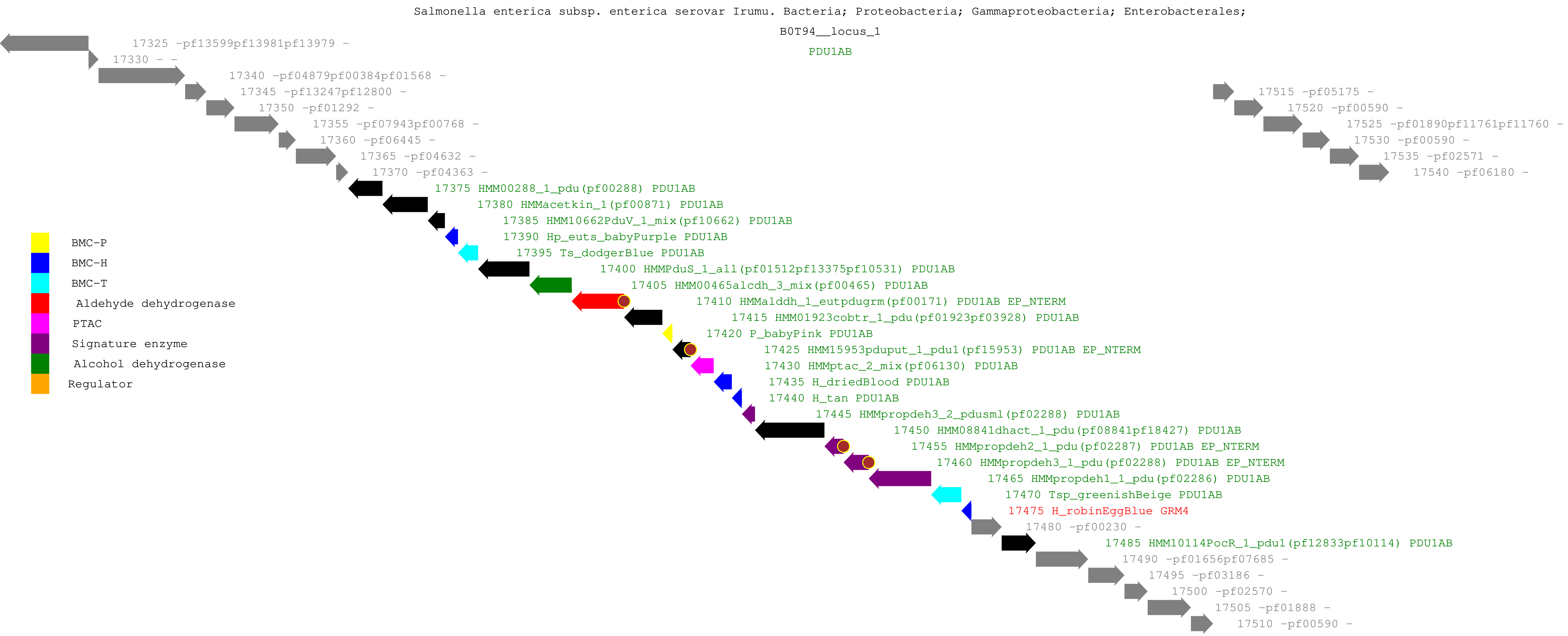
Salmonella enterica subsp. enterica serovar Irumu. Bacteria; Proteobacteria; Gammaproteobacteria; Enterobacterales;
B0T94__locus_1
PDU1AB
17325 -pf13599pf13981pf13979 -
17330 - -
17340 -pf04879pf00384pf01568 -
17345 -pf13247pf12800 -
17350 -pf01292 -
17355 -pf07943pf00768 -
17360 -pf06445 -
17365 -pf04632 -
17370 -pf04363 -
17375 HMM00288_1_pdu(pf00288) PDU1AB
17380 HMMacetkin_1(pf00871) PDU1AB
17385 HMM10662PduV_1_mix(pf10662) PDU1AB
17390 Hp_euts_babyPurple PDU1AB
17395 Ts_dodgerBlue PDU1AB
17400 HMMPduS_1_all(pf01512pf13375pf10531) PDU1AB
17405 HMM00465alcdh_3_mix(pf00465) PDU1AB
17410 HMMalddh_1_eutpdugrm(pf00171) PDU1AB EP_NTERM
17415 HMM01923cobtr_1_pdu(pf01923pf03928) PDU1AB
17420 P_babyPink PDU1AB
17425 HMM15953pduput_1_pdu1(pf15953) PDU1AB EP_NTERM
17430 HMMptac_2_mix(pf06130) PDU1AB
17435 H_driedBlood PDU1AB
17440 H_tan PDU1AB
17445 HMMpropdeh3_2_pdusml(pf02288) PDU1AB
17450 HMM08841dhact_1_pdu(pf08841pf18427) PDU1AB
17455 HMMpropdeh2_1_pdu(pf02287) PDU1AB EP_NTERM
17460 HMMpropdeh3_1_pdu(pf02288) PDU1AB EP_NTERM
17465 HMMpropdeh1_1_pdu(pf02286) PDU1AB
17470 Tsp_greenishBeige PDU1AB
17475 H_robinEggBlue GRM4
17480 -pf00230 -
17485 HMM10114PocR_1_pdu1(pf12833pf10114) PDU1AB
17490 -pf01656pf07685 -
17495 -pf03186 -
17500 -pf02570 -
17505 -pf01888 -
17510 -pf00590 -
17515 -pf05175 -
17520 -pf00590 -
17525 -pf01890pf11761pf11760 -
17530 -pf00590 -
17535 -pf02571 -
17540 -pf06180 -
BMC-P
BMC-H
BMC-T
Aldehyde dehydrogenase
PTAC
Signature enzyme
Alcohol dehydrogenase
Regulator

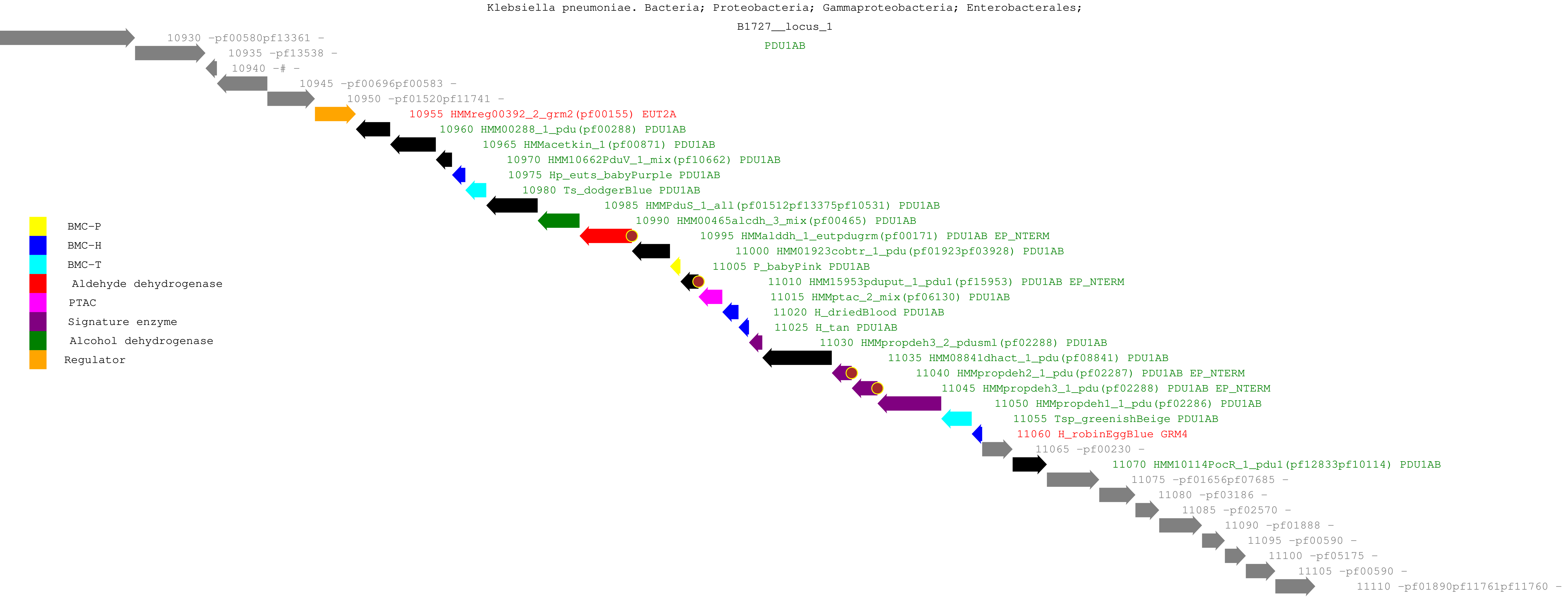
Klebsiella pneumoniae. Bacteria; Proteobacteria; Gammaproteobacteria; Enterobacterales;
B1727__locus_1
PDU1AB
10930 -pf00580pf13361 -
10935 -pf13538 -
10940 -# -
10945 -pf00696pf00583 -
10950 -pf01520pf11741 -
10955 HMMreg00392_2_grm2(pf00155) EUT2A
10960 HMM00288_1_pdu(pf00288) PDU1AB
10965 HMMacetkin_1(pf00871) PDU1AB
10970 HMM10662PduV_1_mix(pf10662) PDU1AB
10975 Hp_euts_babyPurple PDU1AB
10980 Ts_dodgerBlue PDU1AB
10985 HMMPduS_1_all(pf01512pf13375pf10531) PDU1AB
10990 HMM00465alcdh_3_mix(pf00465) PDU1AB
10995 HMMalddh_1_eutpdugrm(pf00171) PDU1AB EP_NTERM
11000 HMM01923cobtr_1_pdu(pf01923pf03928) PDU1AB
11005 P_babyPink PDU1AB
11010 HMM15953pduput_1_pdu1(pf15953) PDU1AB EP_NTERM
11015 HMMptac_2_mix(pf06130) PDU1AB
11020 H_driedBlood PDU1AB
11025 H_tan PDU1AB
11030 HMMpropdeh3_2_pdusml(pf02288) PDU1AB
11035 HMM08841dhact_1_pdu(pf08841) PDU1AB
11040 HMMpropdeh2_1_pdu(pf02287) PDU1AB EP_NTERM
11045 HMMpropdeh3_1_pdu(pf02288) PDU1AB EP_NTERM
11050 HMMpropdeh1_1_pdu(pf02286) PDU1AB
11055 Tsp_greenishBeige PDU1AB
11060 H_robinEggBlue GRM4
11065 -pf00230 -
11070 HMM10114PocR_1_pdu1(pf12833pf10114) PDU1AB
11075 -pf01656pf07685 -
11080 -pf03186 -
11085 -pf02570 -
11090 -pf01888 -
11095 -pf00590 -
11100 -pf05175 -
11105 -pf00590 -
11110 -pf01890pf11761pf11760 -
BMC-P
BMC-H
BMC-T
Aldehyde dehydrogenase
PTAC
Signature enzyme
Alcohol dehydrogenase
Regulator

Escherichia coli. Bacteria; Proteobacteria; Gammaproteobacteria; Enterobacterales;

B1K96__locus_1

PDU1AB

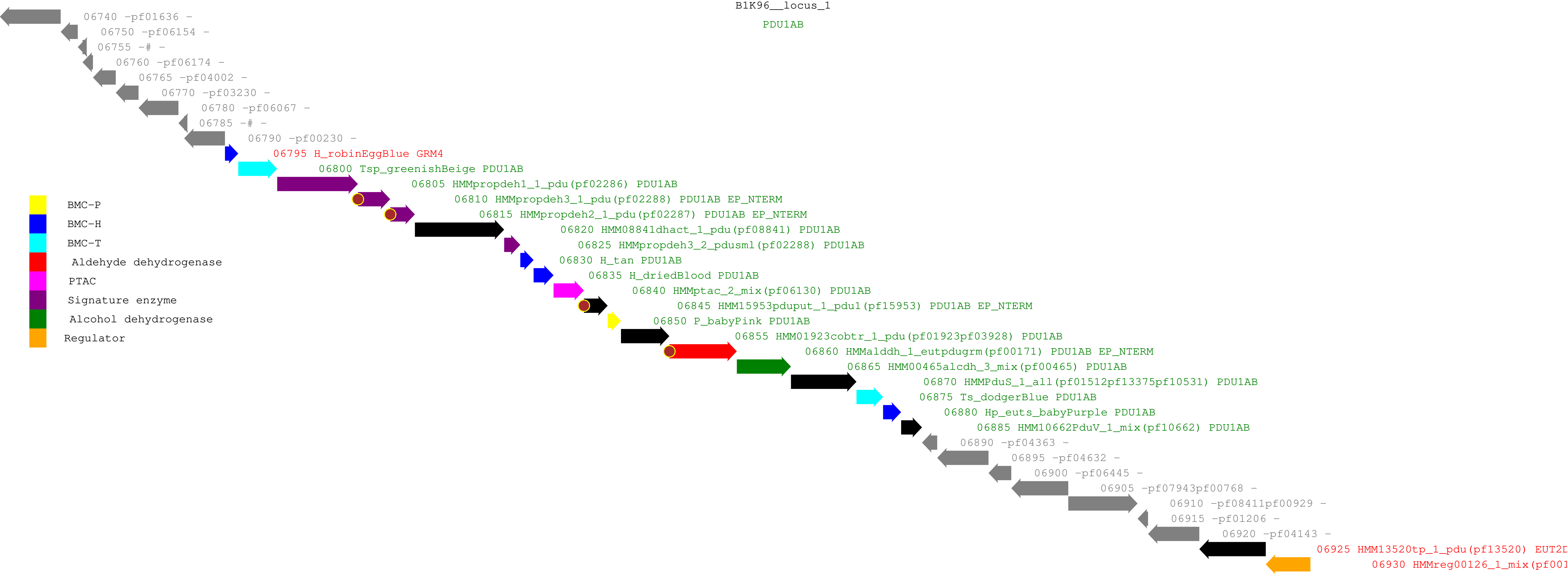

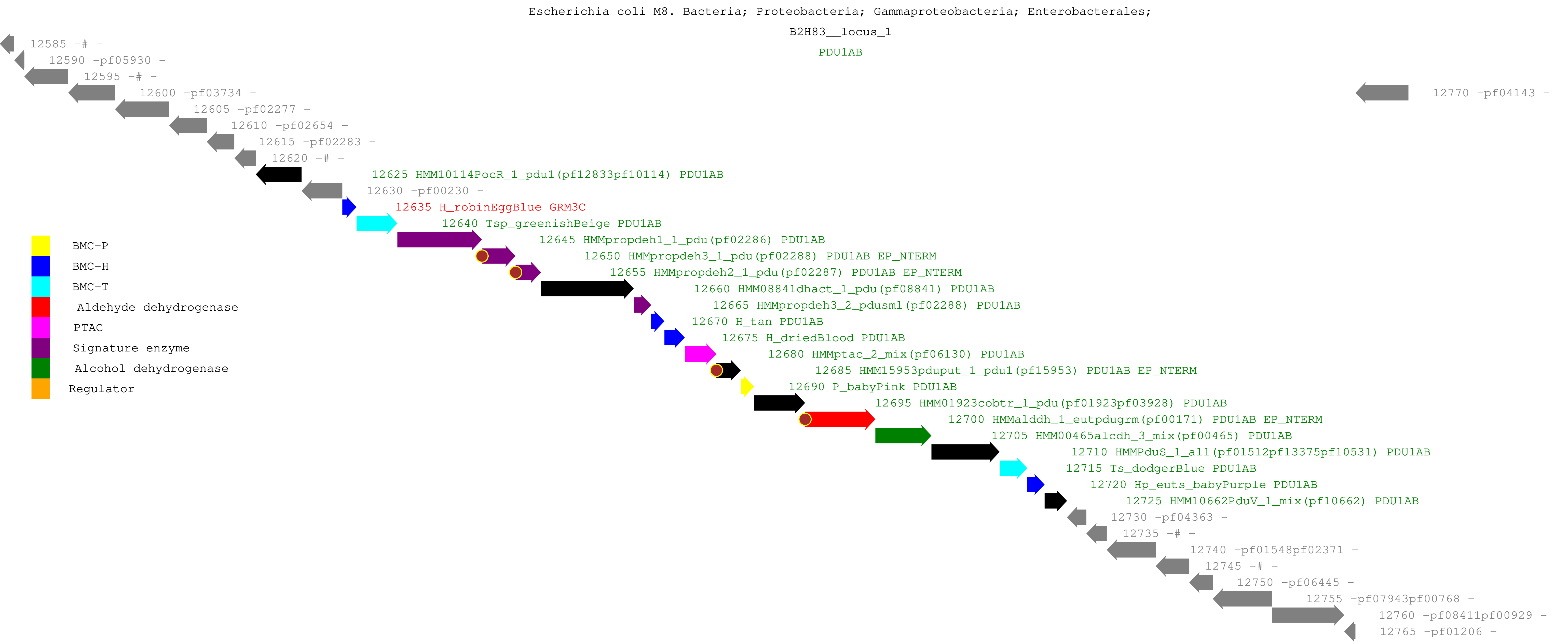
Escherichia coli M8. Bacteria; Proteobacteria; Gammaproteobacteria; Enterobacterales;
B2H83__locus_1
PDU1AB
12585 -# -
12590 -pf05930 -
12595 -# -
12600 -pf03734 -
12605 -pf02277 -
12610 -pf02654 -
12615 -pf02283 -
12620 -# -
12625 HMM10114PocR_1_pdu1(pf12833pf10114) PDU1AB
12630 -pf00230 -
12635 H_robinEggBlue GRM3C
12640 Tsp_greenishBeige PDU1AB
12645 HMMpropdeh1_1_pdu(pf02286) PDU1AB
12650 HMMpropdeh3_1_pdu(pf02288) PDU1AB EP_NTERM
12655 HMMpropdeh2_1_pdu(pf02287) PDU1AB EP_NTERM
12660 HMM08841dhact_1_pdu(pf08841) PDU1AB
12665 HMMpropdeh3_2_pdusml(pf02288) PDU1AB
12670 H_tan PDU1AB
12675 H_driedBlood PDU1AB
12680 HMMptac_2_mix(pf06130) PDU1AB
12685 HMM15953pduput_1_pdu1(pf15953) PDU1AB EP_NTERM
12690 P_babyPink PDU1AB
12695 HMM01923cobtr_1_pdu(pf01923pf03928) PDU1AB
12700 HMMalddh_1_eutpdugrm(pf00171) PDU1AB EP_NTERM
12705 HMM00465alcdh_3_mix(pf00465) PDU1AB
12710 HMMPduS_1_all(pf01512pf13375pf10531) PDU1AB
12715 Ts_dodgerBlue PDU1AB
12720 Hp_euts_babyPurple PDU1AB
12725 HMM10662PduV_1_mix(pf10662) PDU1AB
12730 -pf04363 -
12735 -# -
12740 -pf01548pf02371 -
12745 -# -
12750 -pf06445 -
12755 -pf07943pf00768 -
12760 -pf08411pf00929 -
12765 -pf01206 -
12770 -pf04143 -
BMC-P
BMC-H
BMC-T
Aldehyde dehydrogenase
PTAC
Signature enzyme
Alcohol dehydrogenase
Regulator

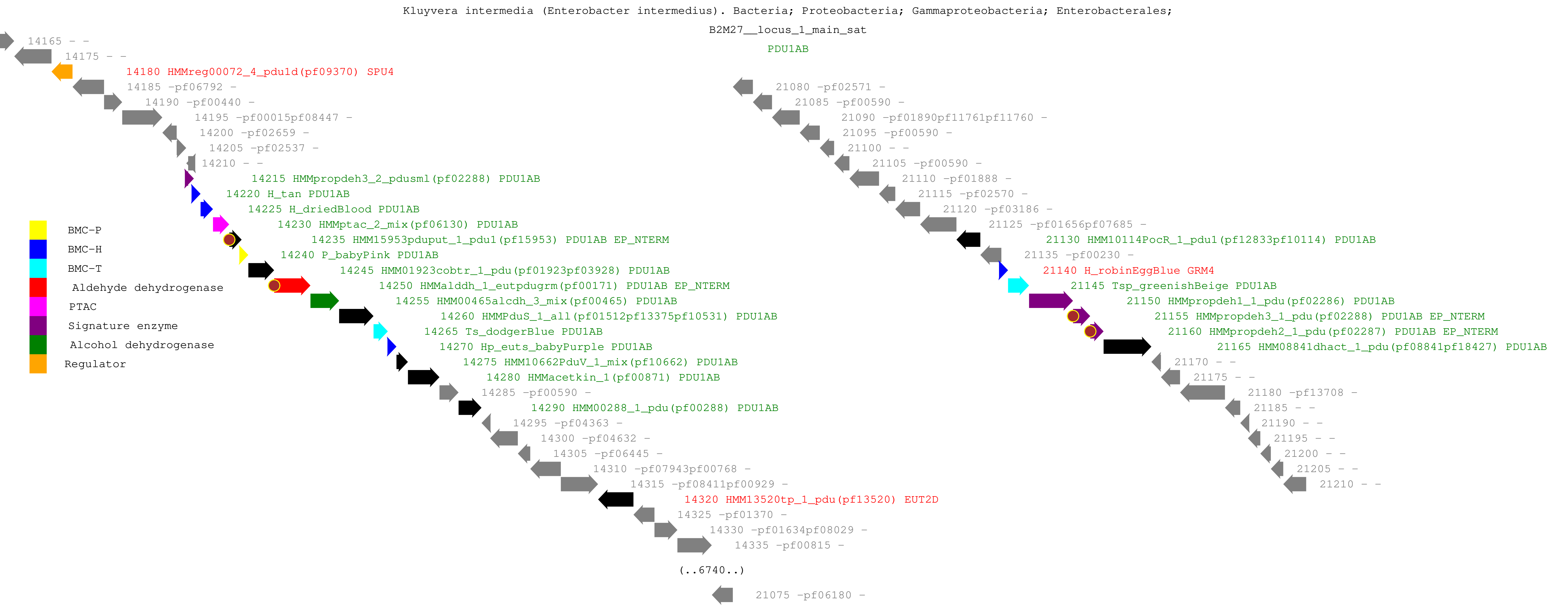
Kluyvera intermedia (Enterobacter intermedius). Bacteria; Proteobacteria; Gammaproteobacteria; Enterobacterales;
B2M27__locus_1_main_sat
PDU1AB
14165 - -
14175 - -
14180 HMMreg00072_4_pdu1d(pf09370) SPU4
14185 -pf06792 -
14190 -pf00440 -
14195 -pf00015pf08447 -
14200 -pf02659 -
14205 -pf02537 -
14210 - -
14215 HMMpropdeh3_2_pdusml(pf02288) PDU1AB
14220 H_tan PDU1AB
14225 H_driedBlood PDU1AB
14230 HMMptac_2_mix(pf06130) PDU1AB
14235 HMM15953pduput_1_pdu1(pf15953) PDU1AB EP_NTERM
14240 P_babyPink PDU1AB
14245 HMM01923cobtr_1_pdu(pf01923pf03928) PDU1AB
14250 HMMalddh_1_eutpdugrm(pf00171) PDU1AB EP_NTERM
14255 HMM00465alcdh_3_mix(pf00465) PDU1AB
14260 HMMPduS_1_all(pf01512pf13375pf10531) PDU1AB
14265 Ts_dodgerBlue PDU1AB
14270 Hp_euts_babyPurple PDU1AB
14275 HMM10662PduV_1_mix(pf10662) PDU1AB
14280 HMMacetkin_1(pf00871) PDU1AB
14285 -pf00590 -
14290 HMM00288_1_pdu(pf00288) PDU1AB
14295 -pf04363 -
14300 -pf04632 -
14305 -pf06445 -
14310 -pf07943pf00768 -
14315 -pf08411pf00929 -
14320 HMM13520tp_1_pdu(pf13520) EUT2D
14325 -pf01370 -
14330 -pf01634pf08029 -
14335 -pf00815 -
(..6740..)
21075 -pf06180 -
21080 -pf02571 -
21085 -pf00590 -
21090 -pf01890pf11761pf11760 -
21095 -pf00590 -
21100 - -
21105 -pf00590 -
21110 -pf01888 -
21115 -pf02570 -
21120 -pf03186 -
21125 -pf01656pf07685 -
21130 HMM10114PocR_1_pdu1(pf12833pf10114) PDU1AB
21135 -pf00230 -
21140 H_robinEggBlue GRM4
21145 Tsp_greenishBeige PDU1AB
21150 HMMpropdeh1_1_pdu(pf02286) PDU1AB
21155 HMMpropdeh3_1_pdu(pf02288) PDU1AB EP_NTERM
21160 HMMpropdeh2_1_pdu(pf02287) PDU1AB EP_NTERM
21165 HMM08841dhact_1_pdu(pf08841pf18427) PDU1AB
21170 - -
21175 - -
21180 -pf13708 -
21185 - -
21190 - -
21195 - -
21200 - -
21205 - -
21210 - -
BMC-P
BMC-H
BMC-T
Aldehyde dehydrogenase
PTAC
Signature enzyme
Alcohol dehydrogenase
Regulator

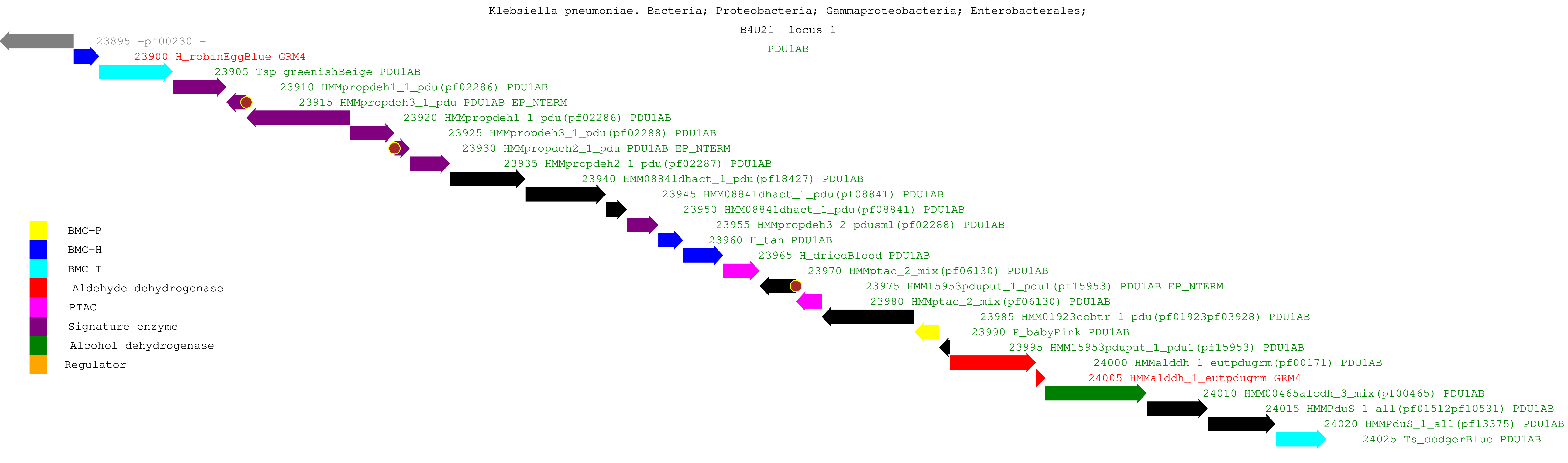
Klebsiella pneumoniae. Bacteria; Proteobacteria; Gammaproteobacteria; Enterobacterales;
B4U21__locus_1
PDU1AB
23895 -pf00230 -
23900 H_robinEggBlue GRM4
23905 Tsp_greenishBeige PDU1AB
23910 HMMpropdeh1_1_pdu(pf02286) PDU1AB
23915 HMMpropdeh3_1_pdu PDU1AB EP_NTERM
23920 HMMpropdeh1_1_pdu(pf02286) PDU1AB
23925 HMMpropdeh3_1_pdu(pf02288) PDU1AB
23930 HMMpropdeh2_1_pdu PDU1AB EP_NTERM
23935 HMMpropdeh2_1_pdu(pf02287) PDU1AB
23940 HMM08841dhact_1_pdu(pf18427) PDU1AB
23945 HMM08841dhact_1_pdu(pf08841) PDU1AB
23950 HMM08841dhact_1_pdu(pf08841) PDU1AB
23955 HMMpropdeh3_2_pdusml(pf02288) PDU1AB
23960 H_tan PDU1AB
23965 H_driedBlood PDU1AB
23970 HMMptac_2_mix(pf06130) PDU1AB
23975 HMM15953pduput_1_pdu1(pf15953) PDU1AB EP_NTERM
23980 HMMptac_2_mix(pf06130) PDU1AB
23985 HMM01923cobtr_1_pdu(pf01923pf03928) PDU1AB
23990 P_babyPink PDU1AB
23995 HMM15953pduput_1_pdu1(pf15953) PDU1AB
24000 HMMalddh_1_eutpdugrm(pf00171) PDU1AB
24005 HMMalddh_1_eutpdugrm GRM4
24010 HMM00465alcdh_3_mix(pf00465) PDU1AB
24015 HMMPduS_1_all(pf01512pf10531) PDU1AB
24020 HMMPduS_1_all(pf13375) PDU1AB
24025 Ts_dodgerBlue PDU1AB
BMC-P
BMC-H
BMC-T
Aldehyde dehydrogenase
PTAC
Signature enzyme
Alcohol dehydrogenase
Regulator

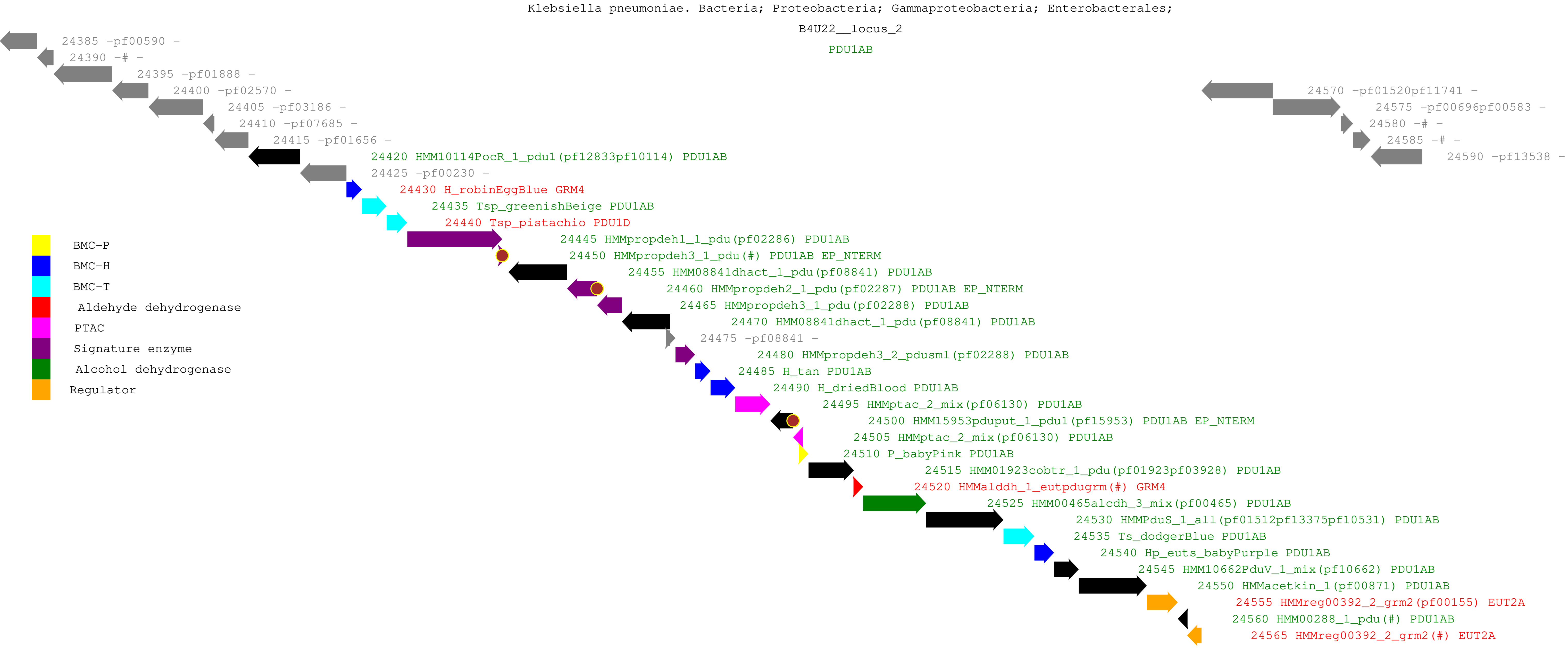
Klebsiella pneumoniae. Bacteria; Proteobacteria; Gammaproteobacteria; Enterobacterales;
B4U22__locus_2
PDU1AB
24385 -pf00590 -
24390 -# -
24395 -pf01888 -
24400 -pf02570 -
24405 -pf03186 -
24410 -pf07685 -
24415 -pf01656 -
24420 HMM10114PocR_1_pdu1(pf12833pf10114) PDU1AB
24425 -pf00230 -
24430 H_robinEggBlue GRM4
24435 Tsp_greenishBeige PDU1AB
24440 Tsp_pistachio PDU1D
24445 HMMpropdeh1_1_pdu(pf02286) PDU1AB
24450 HMMpropdeh3_1_pdu(#) PDU1AB EP_NTERM
24455 HMM08841dhact_1_pdu(pf08841) PDU1AB
24460 HMMpropdeh2_1_pdu(pf02287) PDU1AB EP_NTERM
24465 HMMpropdeh3_1_pdu(pf02288) PDU1AB
24470 HMM08841dhact_1_pdu(pf08841) PDU1AB
24475 -pf08841 -
24480 HMMpropdeh3_2_pdusml(pf02288) PDU1AB
24485 H_tan PDU1AB
24490 H_driedBlood PDU1AB
24495 HMMptac_2_mix(pf06130) PDU1AB
24500 HMM15953pduput_1_pdu1(pf15953) PDU1AB EP_NTERM
24505 HMMptac_2_mix(pf06130) PDU1AB
24510 P_babyPink PDU1AB
24515 HMM01923cobtr_1_pdu(pf01923pf03928) PDU1AB
24520 HMMalddh_1_eutpdugrm(#) GRM4
24525 HMM00465alcdh_3_mix(pf00465) PDU1AB
24530 HMMPduS_1_all(pf01512pf13375pf10531) PDU1AB
24535 Ts_dodgerBlue PDU1AB
24540 Hp_euts_babyPurple PDU1AB
24545 HMM10662PduV_1_mix(pf10662) PDU1AB
24550 HMMacetkin_1(pf00871) PDU1AB
24555 HMMreg00392_2_grm2(pf00155) EUT2A
24560 HMM00288_1_pdu(#) PDU1AB
24565 HMMreg00392_2_grm2(#) EUT2A
24570 -pf01520pf11741 -
24575 -pf00696pf00583 -
24580 -# -
24585 -# -
24590 -pf13538 -
BMC-P
BMC-H
BMC-T
Aldehyde dehydrogenase
PTAC
Signature enzyme
Alcohol dehydrogenase
Regulator

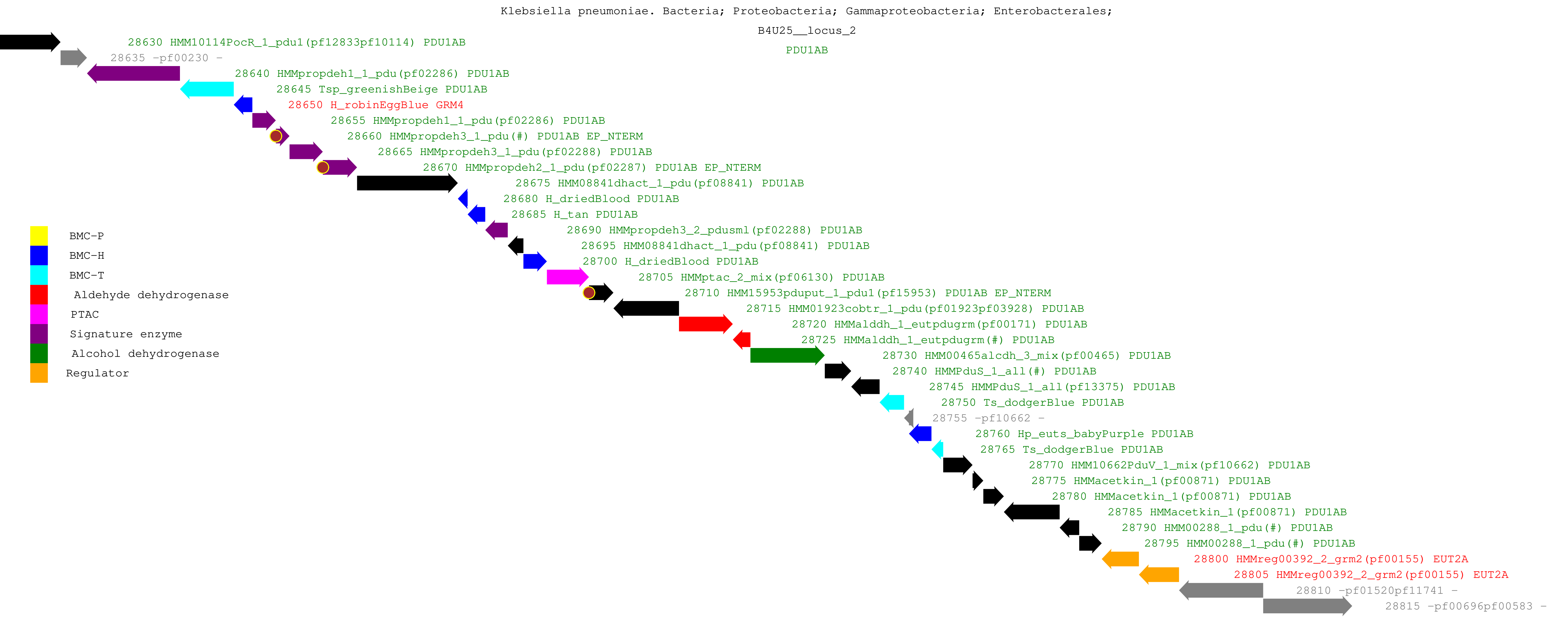
Klebsiella pneumoniae. Bacteria; Proteobacteria; Gammaproteobacteria; Enterobacterales;
B4U25__locus_2
PDU1AB
28630 HMM10114PocR_1_pdu1(pf12833pf10114) PDU1AB
28635 -pf00230 -
28640 HMMpropdeh1_1_pdu(pf02286) PDU1AB
28645 Tsp_greenishBeige PDU1AB
28650 H_robinEggBlue GRM4
28655 HMMpropdeh1_1_pdu(pf02286) PDU1AB
28660 HMMpropdeh3_1_pdu(#) PDU1AB EP_NTERM
28665 HMMpropdeh3_1_pdu(pf02288) PDU1AB
28670 HMMpropdeh2_1_pdu(pf02287) PDU1AB EP_NTERM
28675 HMM08841dhact_1_pdu(pf08841) PDU1AB
28680 H_driedBlood PDU1AB
28685 H_tan PDU1AB
28690 HMMpropdeh3_2_pdusml(pf02288) PDU1AB
28695 HMM08841dhact_1_pdu(pf08841) PDU1AB
28700 H_driedBlood PDU1AB
28705 HMMptac_2_mix(pf06130) PDU1AB
28710 HMM15953pduput_1_pdu1(pf15953) PDU1AB EP_NTERM
28715 HMM01923cobtr_1_pdu(pf01923pf03928) PDU1AB
28720 HMMalddh_1_eutpdugrm(pf00171) PDU1AB
28725 HMMalddh_1_eutpdugrm(#) PDU1AB
28730 HMM00465alcdh_3_mix(pf00465) PDU1AB
28740 HMMPduS_1_all(#) PDU1AB
28745 HMMPduS_1_all(pf13375) PDU1AB
28750 Ts_dodgerBlue PDU1AB
28755 -pf10662 -
28760 Hp_euts_babyPurple PDU1AB
28765 Ts_dodgerBlue PDU1AB
28770 HMM10662PduV_1_mix(pf10662) PDU1AB
28775 HMMacetkin_1(pf00871) PDU1AB
28780 HMMacetkin_1(pf00871) PDU1AB
28785 HMMacetkin_1(pf00871) PDU1AB
28790 HMM00288_1_pdu(#) PDU1AB
28795 HMM00288_1_pdu(#) PDU1AB
28800 HMMreg00392_2_grm2(pf00155) EUT2A
28805 HMMreg00392_2_grm2(pf00155) EUT2A
28810 -pf01520pf11741 -
28815 -pf00696pf00583 -
BMC-P
BMC-H
BMC-T
Aldehyde dehydrogenase
PTAC
Signature enzyme
Alcohol dehydrogenase
Regulator

Klebsiella pneumoniae. Bacteria; Proteobacteria; Gammaproteobacteria; Enterobacterales;

B4U27__locus_2

PDU1AB

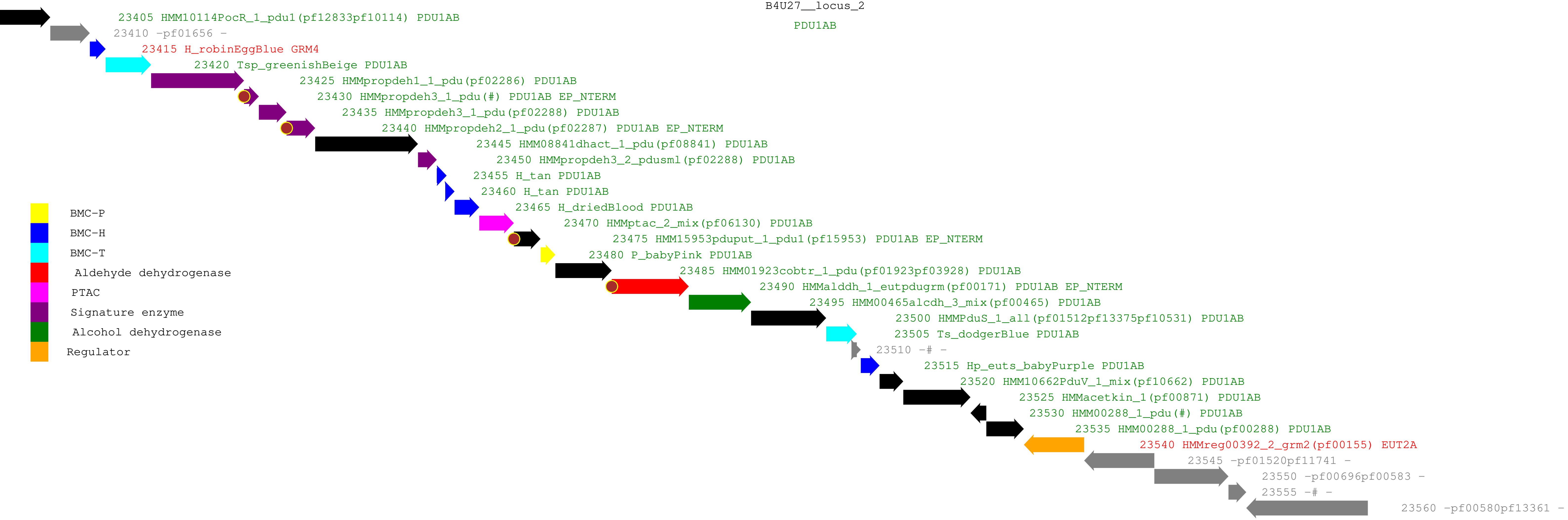

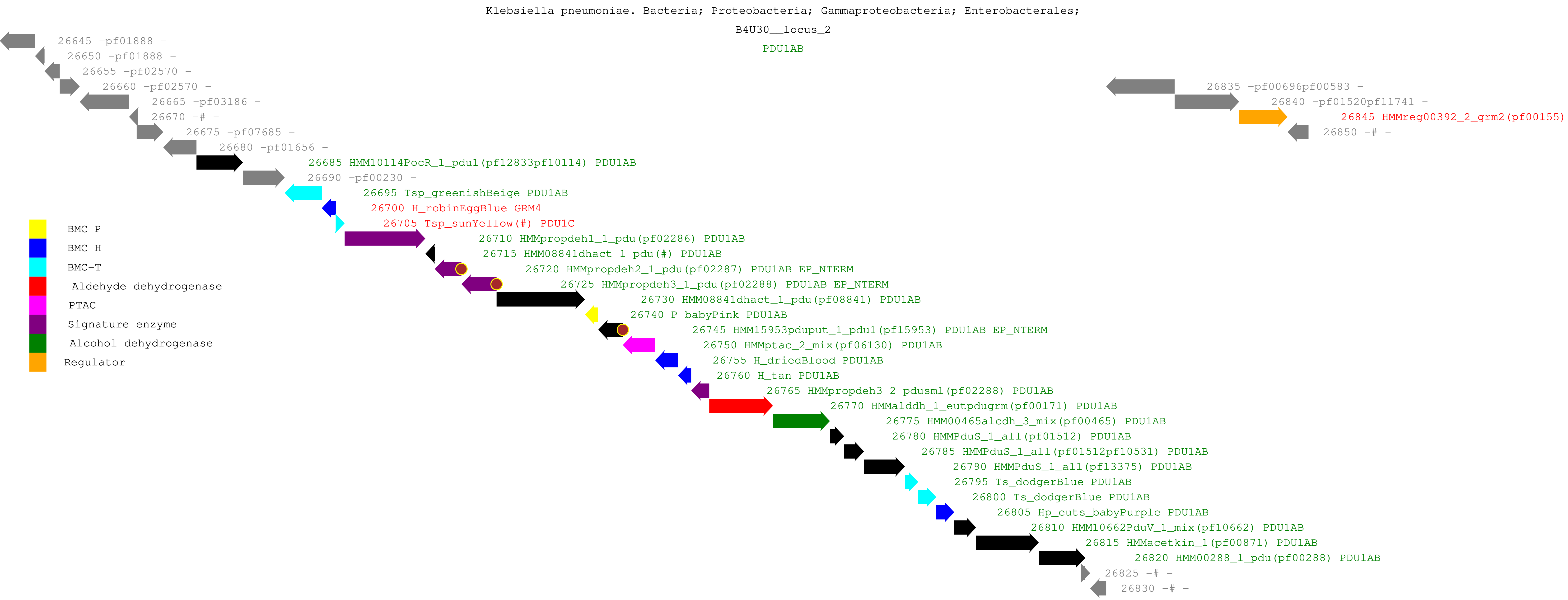
Klebsiella pneumoniae. Bacteria; Proteobacteria; Gammaproteobacteria; Enterobacterales;
B4U30__locus_2
PDU1AB
26645 -pf01888 -
26650 -pf01888 -
26655 -pf02570 -
26660 -pf02570 -
26665 -pf03186 -
26670 -# -
26675 -pf07685 -
26680 -pf01656 -
26685 HMM10114PocR_1_pdu1(pf12833pf10114) PDU1AB
26690 -pf00230 -
26695 Tsp_greenishBeige PDU1AB
26700 H_robinEggBlue GRM4
26705 Tsp_sunYellow(#) PDU1C
26710 HMMpropdeh1_1_pdu(pf02286) PDU1AB
26715 HMM08841dhact_1_pdu(#) PDU1AB
26720 HMMpropdeh2_1_pdu(pf02287) PDU1AB EP_NTERM
26725 HMMpropdeh3_1_pdu(pf02288) PDU1AB EP_NTERM
26730 HMM08841dhact_1_pdu(pf08841) PDU1AB
26740 P_babyPink PDU1AB
26745 HMM15953pduput_1_pdu1(pf15953) PDU1AB EP_NTERM
26750 HMMptac_2_mix(pf06130) PDU1AB
26755 H_driedBlood PDU1AB
26760 H_tan PDU1AB
26765 HMMpropdeh3_2_pdusml(pf02288) PDU1AB
26770 HMMalddh_1_eutpdugrm(pf00171) PDU1AB
26775 HMM00465alcdh_3_mix(pf00465) PDU1AB
26780 HMMPduS_1_all(pf01512) PDU1AB
26785 HMMPduS_1_all(pf01512pf10531) PDU1AB
26790 HMMPduS_1_all(pf13375) PDU1AB
26795 Ts_dodgerBlue PDU1AB
26800 Ts_dodgerBlue PDU1AB
26805 Hp_euts_babyPurple PDU1AB
26810 HMM10662PduV_1_mix(pf10662) PDU1AB
26815 HMMacetkin_1(pf00871) PDU1AB
26820 HMM00288_1_pdu(pf00288) PDU1AB
26825 -# -
26830 -# -
26835 -pf00696pf00583 -
26840 -pf01520pf11741 -
26845 HMMreg00392_2_grm2(pf00155)
26850 -# -
BMC-P
BMC-H
BMC-T
Aldehyde dehydrogenase
PTAC
Signature enzyme
Alcohol dehydrogenase
Regulator

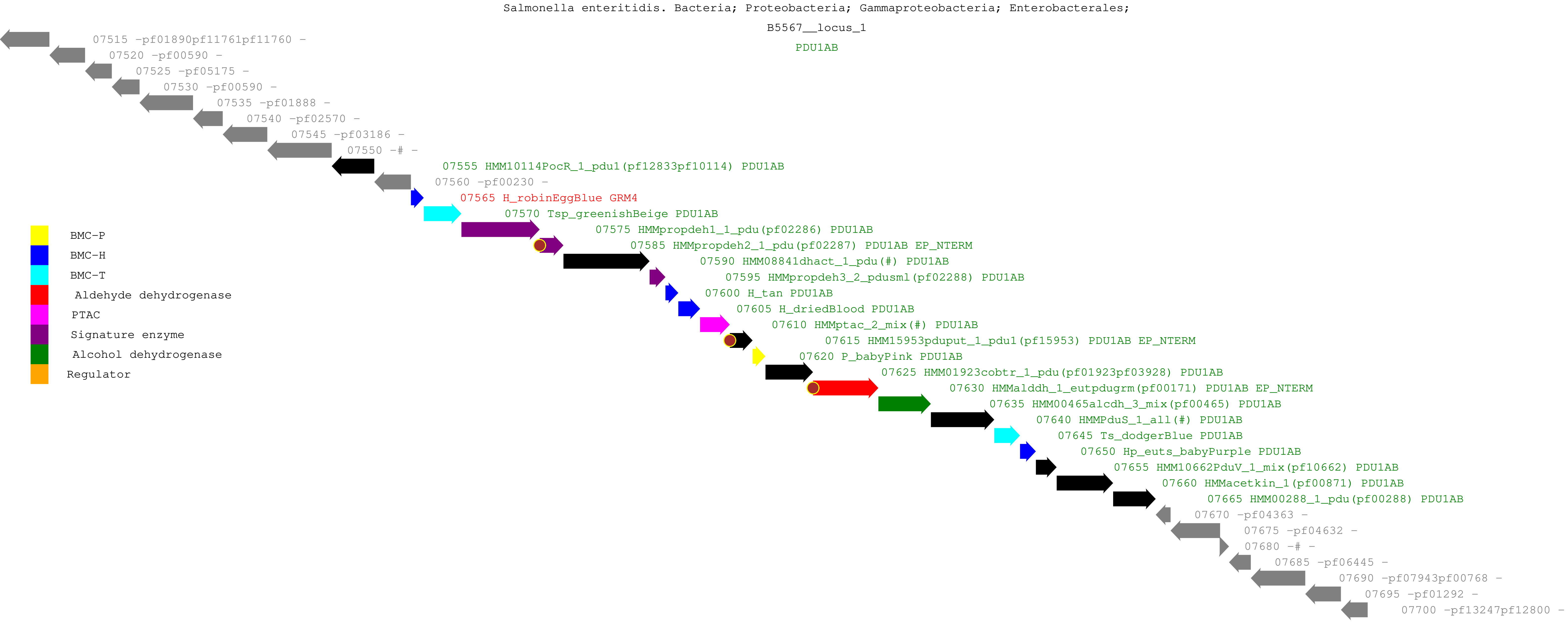
Salmonella enteritidis. Bacteria; Proteobacteria; Gammaproteobacteria; Enterobacterales;
B5567__locus_1
PDU1AB
07515 -pf01890pf11761pf11760 -
07520 -pf00590 -
07525 -pf05175 -
07530 -pf00590 -
07535 -pf01888 -
07540 -pf02570 -
07545 -pf03186 -
07550 -# -
07555 HMM10114PocR_1_pdu1(pf12833pf10114) PDU1AB
07560 -pf00230 -
07565 H_robinEggBlue GRM4
07570 Tsp_greenishBeige PDU1AB
07575 HMMpropdeh1_1_pdu(pf02286) PDU1AB
07585 HMMpropdeh2_1_pdu(pf02287) PDU1AB EP_NTERM
07590 HMM08841dhact_1_pdu(#) PDU1AB
07595 HMMpropdeh3_2_pdusml(pf02288) PDU1AB
07600 H_tan PDU1AB
07605 H_driedBlood PDU1AB
07610 HMMptac_2_mix(#) PDU1AB
07615 HMM15953pduput_1_pdu1(pf15953) PDU1AB EP_NTERM
07620 P_babyPink PDU1AB
07625 HMM01923cobtr_1_pdu(pf01923pf03928) PDU1AB
07630 HMMalddh_1_eutpdugrm(pf00171) PDU1AB EP_NTERM
07635 HMM00465alcdh_3_mix(pf00465) PDU1AB
07640 HMMPduS_1_all(#) PDU1AB
07645 Ts_dodgerBlue PDU1AB
07650 Hp_euts_babyPurple PDU1AB
07655 HMM10662PduV_1_mix(pf10662) PDU1AB
07660 HMMacetkin_1(pf00871) PDU1AB
07665 HMM00288_1_pdu(pf00288) PDU1AB
07670 -pf04363 -
07675 -pf04632 -
07680 -# -
07685 -pf06445 -
07690 -pf07943pf00768 -
07695 -pf01292 -
07700 -pf13247pf12800 -
BMC-P
BMC-H
BMC-T
Aldehyde dehydrogenase
PTAC
Signature enzyme
Alcohol dehydrogenase
Regulator

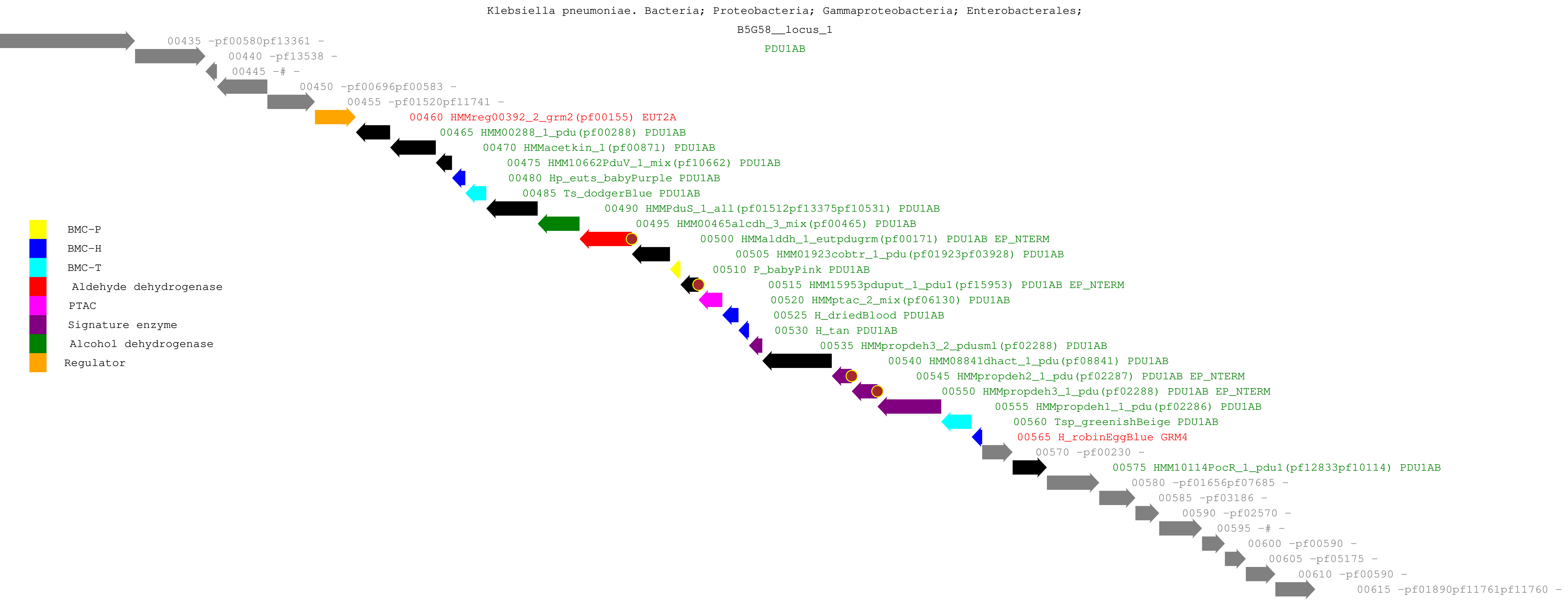
Klebsiella pneumoniae. Bacteria; Proteobacteria; Gammaproteobacteria; Enterobacterales;
B5G58__locus_1
PDU1AB
00435 -pf00580pf13361 -
00440 -pf13538 -
00445 -# -
00450 -pf00696pf00583 -
00455 -pf01520pf11741 -
00460 HMMreg00392_2_grm2(pf00155) EUT2A
00465 HMM00288_1_pdu(pf00288) PDU1AB
00470 HMMacetkin_1(pf00871) PDU1AB
00475 HMM10662PduV_1_mix(pf10662) PDU1AB
00480 Hp_euts_babyPurple PDU1AB
00485 Ts_dodgerBlue PDU1AB
00490 HMMPduS_1_all(pf01512pf13375pf10531) PDU1AB
00495 HMM00465alcdh_3_mix(pf00465) PDU1AB
00500 HMMalddh_1_eutpdugrm(pf00171) PDU1AB EP_NTERM
00505 HMM01923cobtr_1_pdu(pf01923pf03928) PDU1AB
00510 P_babyPink PDU1AB
00515 HMM15953pduput_1_pdu1(pf15953) PDU1AB EP_NTERM
00520 HMMptac_2_mix(pf06130) PDU1AB
00525 H_driedBlood PDU1AB
00530 H_tan PDU1AB
00535 HMMpropdeh3_2_pdusml(pf02288) PDU1AB
00540 HMM08841dhact_1_pdu(pf08841) PDU1AB
00545 HMMpropdeh2_1_pdu(pf02287) PDU1AB EP_NTERM
00550 HMMpropdeh3_1_pdu(pf02288) PDU1AB EP_NTERM
00555 HMMpropdeh1_1_pdu(pf02286) PDU1AB
00560 Tsp_greenishBeige PDU1AB
00565 H_robinEggBlue GRM4
00570 -pf00230 -
00575 HMM10114PocR_1_pdu1(pf12833pf10114) PDU1AB
00580 -pf01656pf07685 -
00585 -pf03186 -
00590 -pf02570 -
00595 -# -
00600 -pf00590 -
00605 -pf05175 -
00610 -pf00590 -
00615 -pf01890pf11761pf11760 -
BMC-P
BMC-H
BMC-T
Aldehyde dehydrogenase
PTAC
Signature enzyme
Alcohol dehydrogenase
Regulator

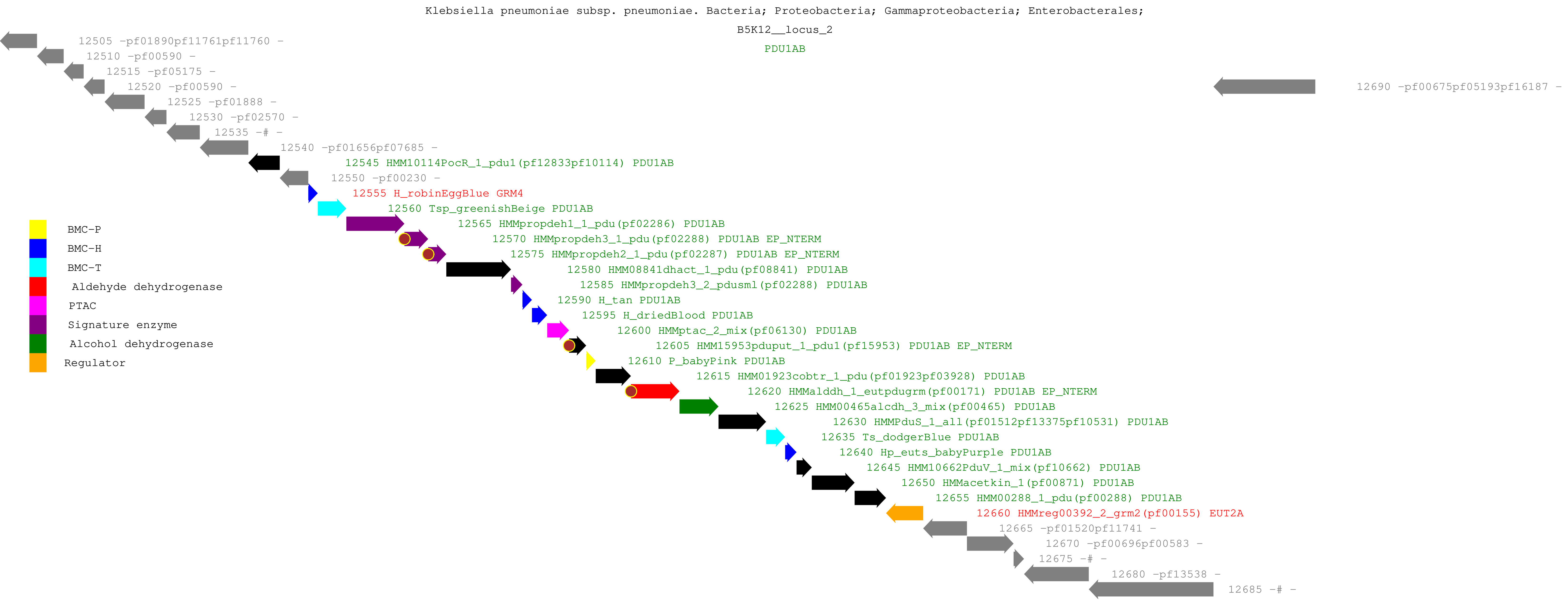

Klebsiella pneumoniae subsp. pneumoniae. Bacteria; Proteobacteria; Gammaproteobacteria; Enterobacterales;
B5K12__locus_2
PDU1AB
12505 -pf01890pf11761pf11760 -
12510 -pf00590 -
12515 -pf05175 -
12520 -pf00590 -
12525 -pf01888 -
12530 -pf02570 -
12535 -# -
12540 -pf01656pf07685 -
12545 HMM10114PocR_1_pdu1(pf12833pf10114) PDU1AB
12550 -pf00230 -
12555 H_robinEggBlue GRM4
12560 Tsp_greenishBeige PDU1AB
12565 HMMpropdeh1_1_pdu(pf02286) PDU1AB
12570 HMMpropdeh3_1_pdu(pf02288) PDU1AB EP_NTERM
12575 HMMpropdeh2_1_pdu(pf02287) PDU1AB EP_NTERM
12580 HMM08841dhact_1_pdu(pf08841) PDU1AB
12585 HMMpropdeh3_2_pdusml(pf02288) PDU1AB
12590 H_tan PDU1AB
12595 H_driedBlood PDU1AB
12600 HMMptac_2_mix(pf06130) PDU1AB
12605 HMM15953pduput_1_pdu1(pf15953) PDU1AB EP_NTERM
12610 P_babyPink PDU1AB
12615 HMM01923cobtr_1_pdu(pf01923pf03928) PDU1AB
12620 HMMaldh_1_eutpdugrm(pf00171) PDU1AB EP_NTERM
12625 HMM00465alcdh_3_mix(pf00465) PDU1AB
12630 HMMPduS_1_all(pf01512pf13375pf10531) PDU1AB
12635 Ts_dodgerBlue PDU1AB
12640 Hp_euts_babyPurple PDU1AB
12645 HMM10662PduV_1_mix(pf10662) PDU1AB
12650 HMMacetkin_1(pf00871) PDU1AB
12655 HMM00288_1_pdu(pf00288) PDU1AB
12660 HMMreg00392_2_grm2(pf00155) EUT2A
12665 -pf01520pf11741 -
12670 -pf00696pf00583 -
12675 -# -
12680 -pf13538 -
12685 -# -
12690 -pf00675pf05193pf16187 -
BMC-P
BMC-H
BMC-T
Aldehyde dehydrogenase
PTAC
Signature enzyme
Alcohol dehydrogenase
Regulator

Salmonella rubislaw. Bacteria; Proteobacteria; Gammaproteobacteria; Enterobacterales;

B6G08__locus_1

PDU1AB

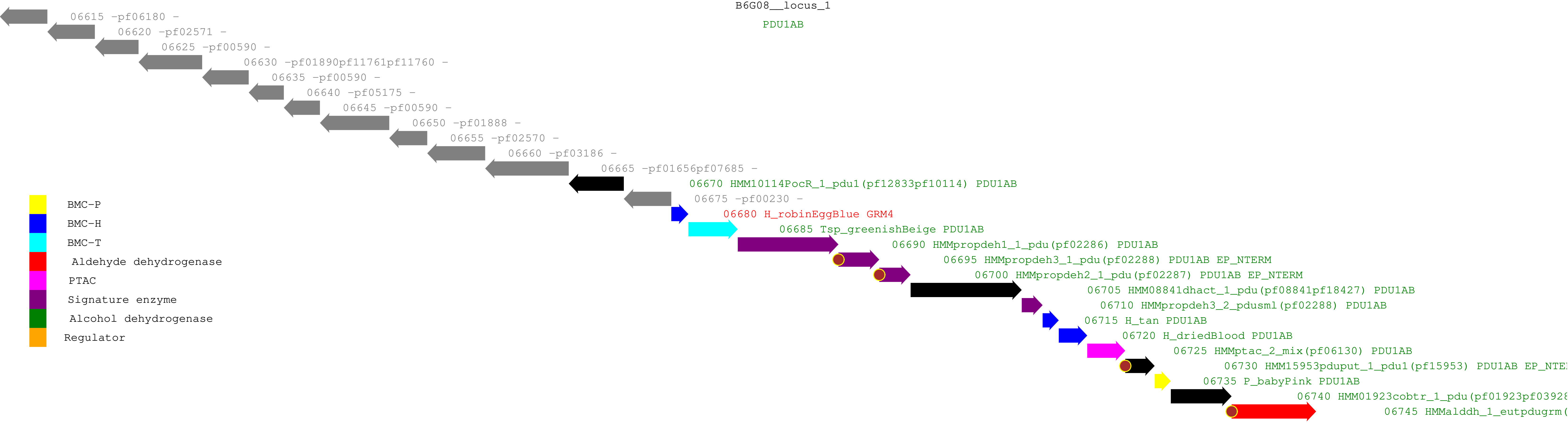

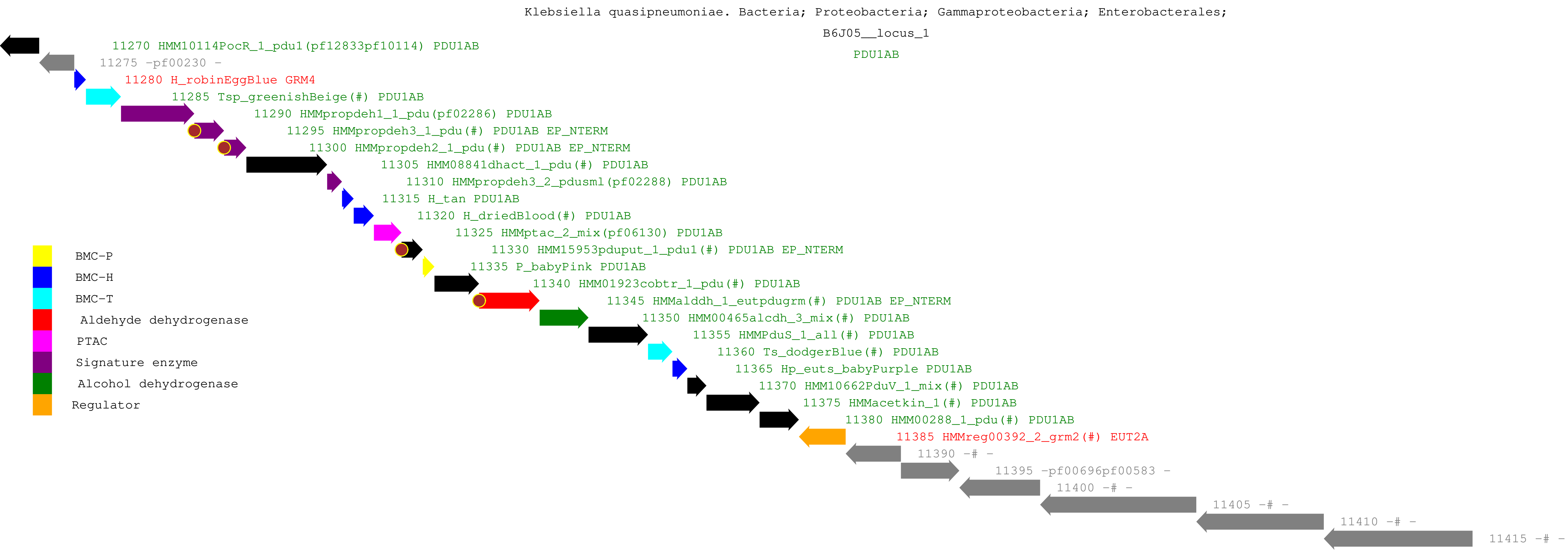

Klebsiella quasipneumoniae. Bacteria; Proteobacteria; Gammaproteobacteria; Enterobacterales;
B6J05__locus_1
PDU1AB
11270 HMM10114PocR_1_pdu1(pf12833pf10114) PDU1AB
11275 -pf00230 -
11280 H_robinEggBlue GRM4
11285 Tsp_greenishBeige(#) PDU1AB
11290 HMMpropdeh1_1_pdu(pf02286) PDU1AB
11295 HMMpropdeh3_1_pdu(#) PDU1AB EP_NTERM
11300 HMMpropdeh2_1_pdu(#) PDU1AB EP_NTERM
11305 HMM08841dhact_1_pdu(#) PDU1AB
11310 HMMpropdeh3_2_pdusml(pf02288) PDU1AB
11315 H_tan PDU1AB
11320 H_driedBlood(#) PDU1AB
11325 HMMptac_2_mix(pf06130) PDU1AB
11330 HMM15953pduput_1_pdu1(#) PDU1AB EP_NTERM
11335 P_babyPink PDU1AB
11340 HMM01923cobtr_1_pdu(#) PDU1AB
11345 HMMalddh_1_eutpdugrm(#) PDU1AB EP_NTERM
11350 HMM00465alcdh_3_mix(#) PDU1AB
11355 HMMPduS_1_all(#) PDU1AB
11360 Ts_dodgerBlue(#) PDU1AB
11365 Hp_euts_babyPurple PDU1AB
11370 HMM10662PduV_1_mix(#) PDU1AB
11375 HMMacetkin_1(#) PDU1AB
11380 HMM00288_1_pdu(#) PDU1AB
11385 HMMreg00392_2_grm2(#) EUT2A
11390 -# -
11395 -pf00696pf00583 -
11400 -# -
11405 -# -
11410 -# -
11415 -# -
BMC-P
BMC-H
BMC-T
Aldehyde dehydrogenase
PTAC
Signature enzyme
Alcohol dehydrogenase
Regulator

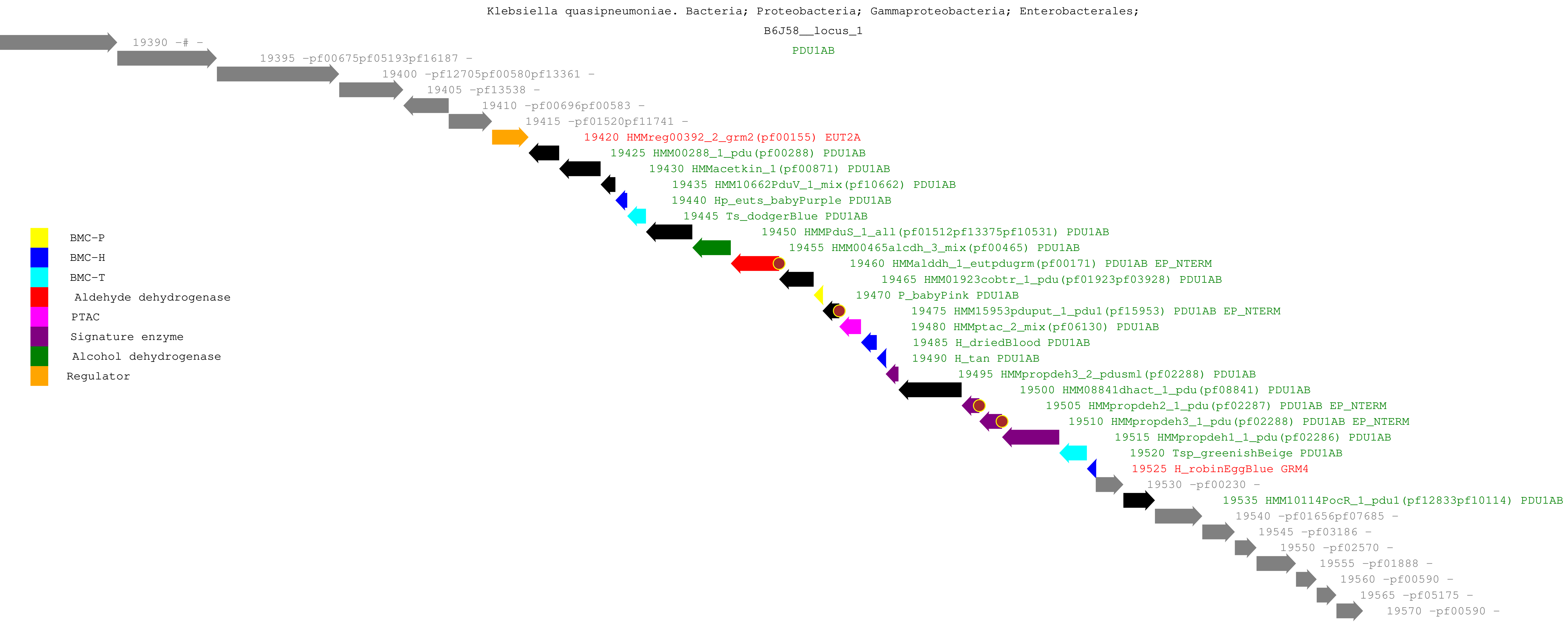
Klebsiella quasipneumoniae. Bacteria; Proteobacteria; Gammaproteobacteria; Enterobacterales;
B6J58__locus_1
PDU1AB
19390 -# -
19395 -pf00675pf05193pf16187 -
19400 -pf12705pf00580pf13361 -
19405 -pf13538 -
19410 -pf00696pf00583 -
19415 -pf01520pf11741 -
19420 HMMreg00392_2_grm2(pf00155) EUT2A
19425 HMM00288_1_pdu(pf00288) PDU1AB
19430 HMMacetkin_1(pf00871) PDU1AB
19435 HMM10662PduV_1_mix(pf10662) PDU1AB
19440 Hp_euts_babyPurple PDU1AB
19445 Ts_dodgerBlue PDU1AB
19450 HMMPduS_1_all(pf01512pf13375pf10531) PDU1AB
19455 HMM00465alcdh_3_mix(pf00465) PDU1AB
19460 HMMalddh_1_eutpdugrm(pf00171) PDU1AB EP_NTERM
19465 HMM01923cobtr_1_pdu(pf01923pf03928) PDU1AB
19470 P_babyPink PDU1AB
19475 HMM15953pduput_1_pdu1(pf15953) PDU1AB EP_NTERM
19480 HMMptac_2_mix(pf06130) PDU1AB
19485 H_driedBlood PDU1AB
19490 H_tan PDU1AB
19495 HMMpropdeh3_2_pdusml(pf02288) PDU1AB
19500 HMM08841dhact_1_pdu(pf08841) PDU1AB
19505 HMMpropdeh2_1_pdu(pf02287) PDU1AB EP_NTERM
19510 HMMpropdeh3_1_pdu(pf02288) PDU1AB EP_NTERM
19515 HMMpropdeh1_1_pdu(pf02286) PDU1AB
19520 Tsp_greenishBeige PDU1AB
19525 H_robinEggBlue GRM4
19530 -pf00230 -
19535 HMM10114PocR_1_pdu1(pf12833pf10114) PDU1AB
19540 -pf01656pf07685 -
19545 -pf03186 -
19550 -pf02570 -
19555 -pf01888 -
19560 -pf00590 -
19565 -pf05175 -
19570 -pf00590 -
BMC-P
BMC-H
BMC-T
Aldehyde dehydrogenase
PTAC
Signature enzyme
Alcohol dehydrogenase
Regulator

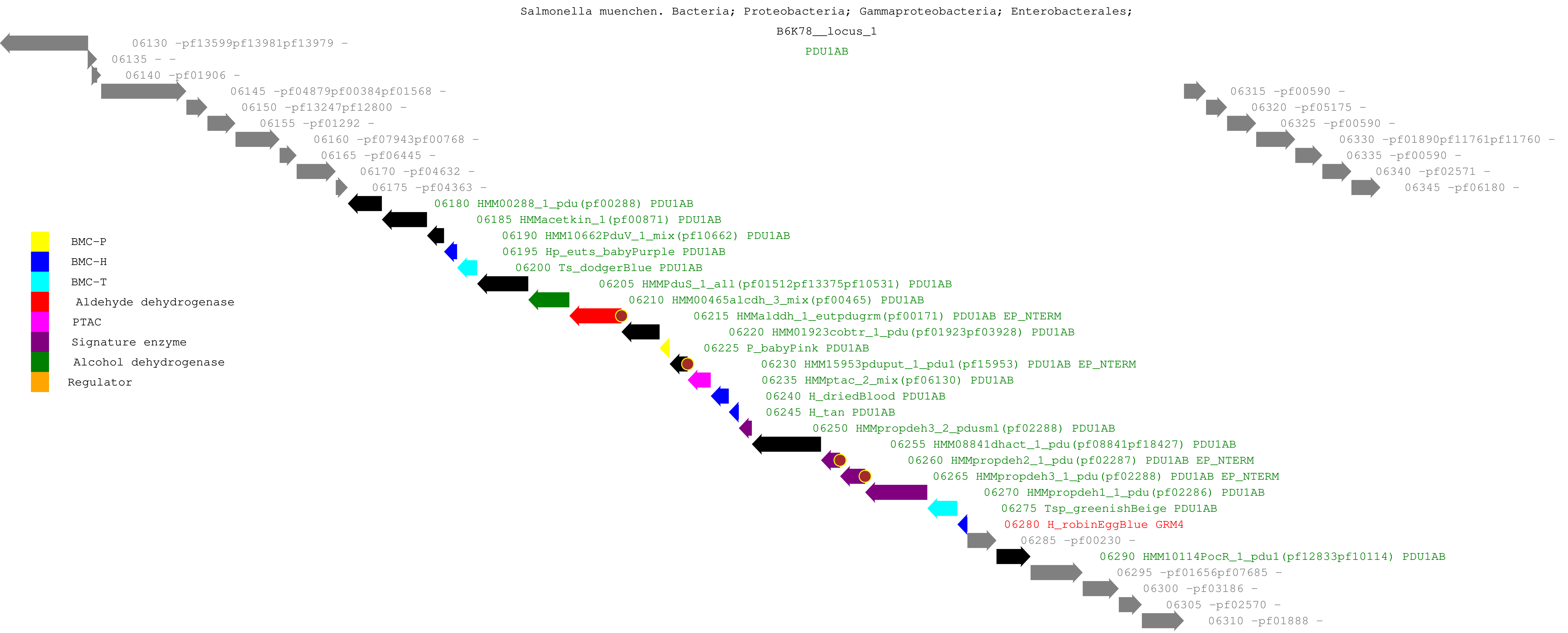
Salmonella muenchen. Bacteria; Proteobacteria; Gammaproteobacteria; Enterobacterales;
B6K78__locus_1
PDU1AB
06130 -pf13599pf13981pf13979 -
06135 - -
06140 -pf01906 -
06145 -pf04879pf00384pf01568 -
06150 -pf13247pf12800 -
06155 -pf01292 -
06160 -pf07943pf00768 -
06165 -pf06445 -
06170 -pf04632 -
06175 -pf04363 -
06180 HMM00288_1_pdu(pf00288) PDU1AB
06185 HMMacetkin_1(pf00871) PDU1AB
06190 HMM10662PduV_1_mix(pf10662) PDU1AB
06195 Hp_euts_babyPurple PDU1AB
06200 Ts_dodgerBlue PDU1AB
06205 HMMPduS_1_all(pf01512pf13375pf10531) PDU1AB
06210 HMM00465alcdh_3_mix(pf00465) PDU1AB
06215 HMMalddh_1_eutpdugrm(pf00171) PDU1AB EP_NTERM
06220 HMM01923cobtr_1_pdu(pf01923pf03928) PDU1AB
06225 P_babyPink PDU1AB
06230 HMM15953pduput_1_pdu1(pf15953) PDU1AB EP_NTERM
06235 HMMptac_2_mix(pf06130) PDU1AB
06240 H_driedBlood PDU1AB
06245 H_tan PDU1AB
06250 HMMpropdeh3_2_pdusml(pf02288) PDU1AB
06255 HMM08841dhact_1_pdu(pf08841pf18427) PDU1AB
06260 HMMpropdeh2_1_pdu(pf02287) PDU1AB EP_NTERM
06265 HMMpropdeh3_1_pdu(pf02288) PDU1AB EP_NTERM
06270 HMMpropdeh1_1_pdu(pf02286) PDU1AB
06275 Tsp_greenishBeige PDU1AB
06280 H_robinEggBlue GRM4
06285 -pf00230 -
06290 HMM10114PocR_1_pdu1(pf12833pf10114) PDU1AB
06295 -pf01656pf07685 -
06300 -pf03186 -
06305 -pf02570 -
06310 -pf01888 -
06315 -pf00590 -
06320 -pf05175 -
06325 -pf00590 -
06330 -pf01890pf11761pf11760 -
06335 -pf00590 -
06340 -pf02571 -
06345 -pf06180 -
BMC-P
BMC-H
BMC-T
Aldehyde dehydrogenase
PTAC
Signature enzyme
Alcohol dehydrogenase
Regulator

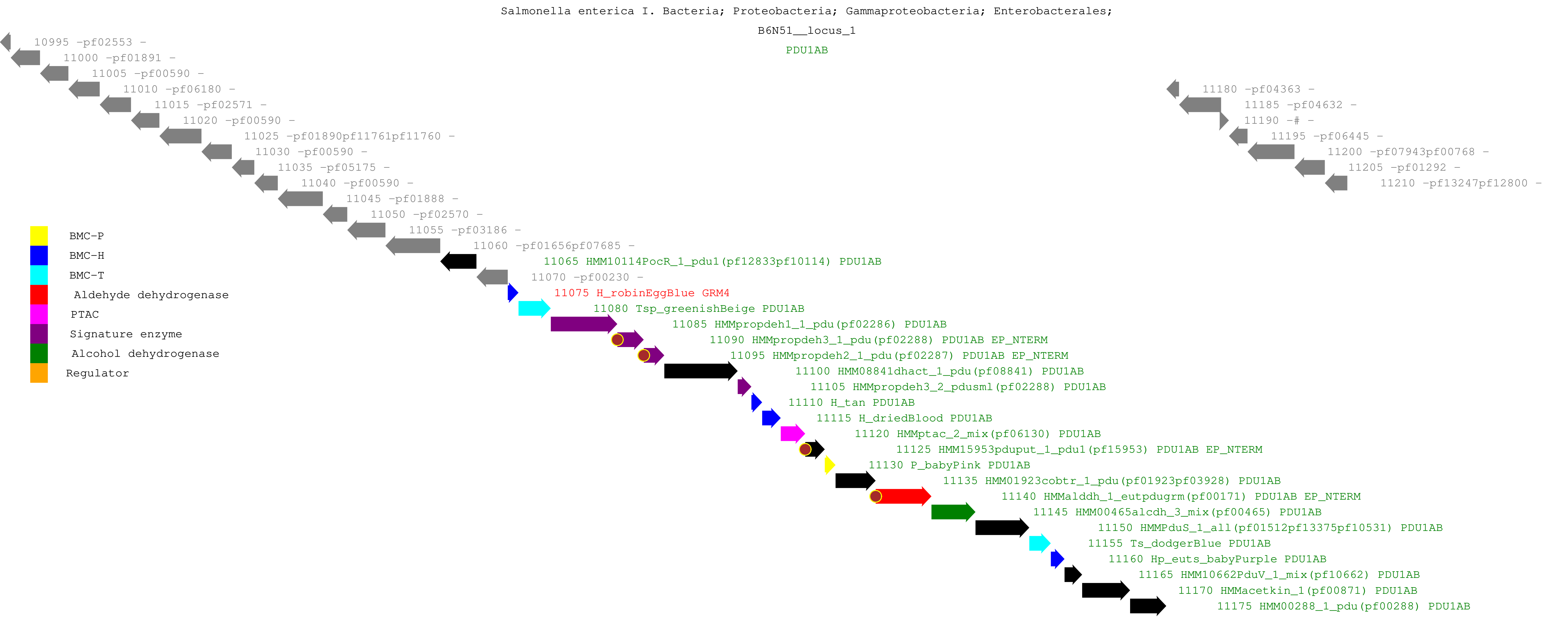
Salmonella enterica I. Bacteria; Proteobacteria; Gammaproteobacteria; Enterobacterales;
B6N51__locus_1
PDU1AB
10995 -pf02553 -
11000 -pf01891 -
11005 -pf00590 -
11010 -pf06180 -
11015 -pf02571 -
11020 -pf00590 -
11025 -pf01890pf11761pf11760 -
11030 -pf00590 -
11035 -pf05175 -
11040 -pf00590 -
11045 -pf01888 -
11050 -pf02570 -
11055 -pf03186 -
11060 -pf01656pf07685 -
11065 HMM10114PocR_1_pdu1(pf12833pf10114) PDU1AB
11070 -pf00230 -
11075 H_robinEggBlue GRM4
11080 Tsp_greenishBeige PDU1AB
11085 HMMpropdeh1_1_pdu(pf02286) PDU1AB
11090 HMMpropdeh3_1_pdu(pf02288) PDU1AB EP_NTERM
11095 HMMpropdeh2_1_pdu(pf02287) PDU1AB EP_NTERM
11100 HMM08841dhact_1_pdu(pf08841) PDU1AB
11105 HMMpropdeh3_2_pdusml(pf02288) PDU1AB
11110 H_tan PDU1AB
11115 H_driedBlood PDU1AB
11120 HMMptac_2_mix(pf06130) PDU1AB
11125 HMM15953pduput_1_pdu1(pf15953) PDU1AB EP_NTERM
11130 P_babyPink PDU1AB
11135 HMM01923cobtr_1_pdu(pf01923pf03928) PDU1AB
11140 HMMalddh_1_eutpdugrm(pf00171) PDU1AB EP_NTERM
11145 HMM00465alcdh_3_mix(pf00465) PDU1AB
11150 HMMPduS_1_all(pf01512pf13375pf10531) PDU1AB
11155 Ts_dodgerBlue PDU1AB
11160 Hp_euts_babyPurple PDU1AB
11165 HMM10662PduV_1_mix(pf10662) PDU1AB
11170 HMMacetkin_1(pf00871) PDU1AB
11175 HMM00288_1_pdu(pf00288) PDU1AB
11180 -pf04363 -
11185 -pf04632 -
11190 -# -
11195 -pf06445 -
11200 -pf07943pf00768 -
11205 -pf01292 -
11210 -pf13247pf12800 -
BMC-P
BMC-H
BMC-T
Aldehyde dehydrogenase
PTAC
Signature enzyme
Alcohol dehydrogenase
Regulator

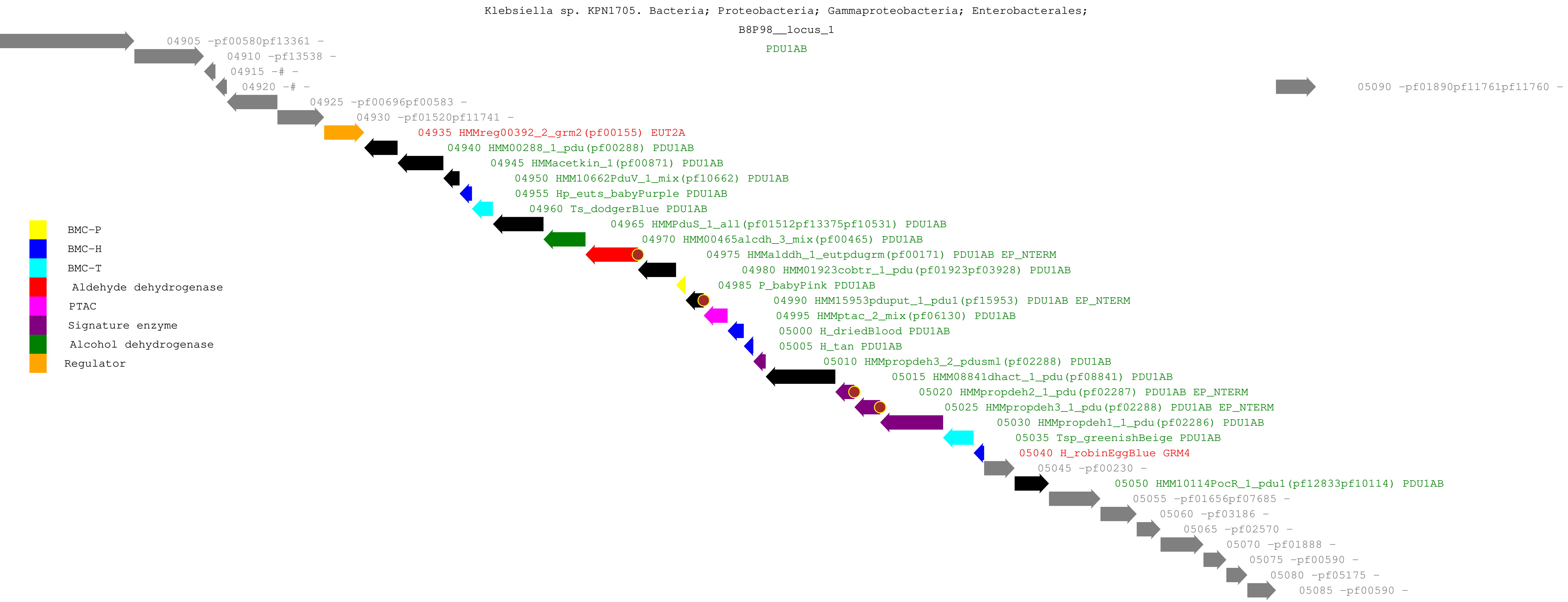
Klebsiella sp. KPN1705. Bacteria; Proteobacteria; Gammaproteobacteria; Enterobacterales;
B8P98__locus_1
PDU1AB
04905 -pf00580pf13361 -
04910 -pf13538 -
04915 -# -
04920 -# -
04925 -pf00696pf00583 -
04930 -pf01520pf11741 -
04935 HMMreg00392_2_grm2(pf00155) EUT2A
04940 HMM00288_1_pdu(pf00288) PDU1AB
04945 HMMacetkin_1(pf00871) PDU1AB
04950 HMM10662PduV_1_mix(pf10662) PDU1AB
04955 Hp_euts_babyPurple PDU1AB
04960 Ts_dodgerBlue PDU1AB
04965 HMMPduS_1_all(pf01512pf13375pf10531) PDU1AB
04970 HMM00465alcdh_3_mix(pf00465) PDU1AB
04975 HMMalddh_1_eutpdugrm(pf00171) PDU1AB EP_NTERM
04980 HMM01923cobtr_1_pdu(pf01923pf03928) PDU1AB
04985 P_babyPink PDU1AB
04990 HMM15953pduput_1_pdu1(pf15953) PDU1AB EP_NTERM
04995 HMMptac_2_mix(pf06130) PDU1AB
05000 H_driedBlood PDU1AB
05005 H_tan PDU1AB
05010 HMMpropdeh3_2_pdusml(pf02288) PDU1AB
05015 HMM08841dhact_1_pdu(pf08841) PDU1AB
05020 HMMpropdeh2_1_pdu(pf02287) PDU1AB EP_NTERM
05025 HMMpropdeh3_1_pdu(pf02288) PDU1AB EP_NTERM
05030 HMMpropdeh1_1_pdu(pf02286) PDU1AB
05035 Tsp_greenishBeige PDU1AB
05040 H_robinEggBlue GRM4
05045 -pf00230 -
05050 HMM10114PocR_1_pdu1(pf12833pf10114) PDU1AB
05055 -pf01656pf07685 -
05060 -pf03186 -
05065 -pf02570 -
05070 -pf01888 -
05075 -pf00590 -
05080 -pf05175 -
05085 -pf00590 -
05090 -pf01890pf11761pf11760 -
BMC-P
BMC-H
BMC-T
Aldehyde dehydrogenase
PTAC
Signature enzyme
Alcohol dehydrogenase
Regulator

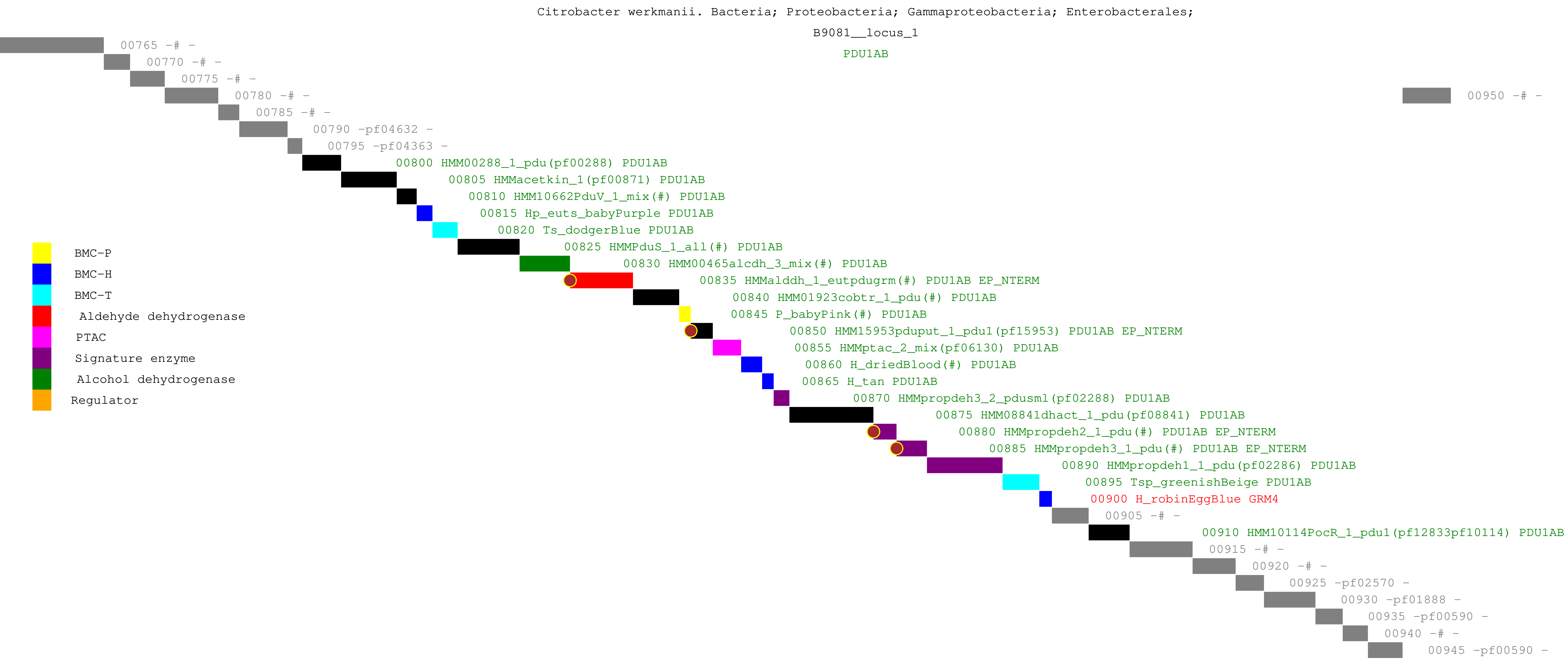
Citrobacter werkmanii. Bacteria; Proteobacteria; Gammaproteobacteria; Enterobacterales;
B9081__locus_1
PDU1AB
00765 -# -
00770 -# -
00775 -# -
00780 -# -
00785 -# -
00790 -pf04632 -
00795 -pf04363 -
00800 HMM00288_1_pdu(pf00288) PDU1AB
00805 HMMacetkin_1(pf00871) PDU1AB
00810 HMM10662PduV_1_mix(#) PDU1AB
00815 Hp_euts_babyPurple PDU1AB
00820 Ts_dodgerBlue PDU1AB
00825 HMMPduS_1_all(#) PDU1AB
00830 HMM00465alcdh_3_mix(#) PDU1AB
00835 HMMalddh_1_eutpdugrm(#) PDU1AB EP_NTERM
00840 HMM01923cobtr_1_pdu(#) PDU1AB
00845 P_babyPink(#) PDU1AB
00850 HMM15953pduput_1_pdu1(pf15953) PDU1AB EP_NTERM
00855 HMMptac_2_mix(pf06130) PDU1AB
00860 H_driedBlood(#) PDU1AB
00865 H_tan PDU1AB
00870 HMMpropdeh3_2_pdusml(pf02288) PDU1AB
00875 HMM08841dhact_1_pdu(pf08841) PDU1AB
00880 HMMpropdeh2_1_pdu(#) PDU1AB EP_NTERM
00885 HMMpropdeh3_1_pdu(#) PDU1AB EP_NTERM
00890 HMMpropdeh1_1_pdu(pf02286) PDU1AB
00895 Tsp_greenishBeige PDU1AB
00900 H_robinEggBlue GRM4
00905 -# -
00910 HMM10114PocR_1_pdu1(pf12833pf10114) PDU1AB
00915 -# -
00920 -# -
00925 -pf02570 -
00930 -pf01888 -
00935 -pf00590 -
00940 -# -
00945 -pf00590 -
00950 -# -
BMC-P
BMC-H
BMC-T
Aldehyde dehydrogenase
PTAC
Signature enzyme
Alcohol dehydrogenase
Regulator

Citrobacter sp. L55. Bacteria; Proteobacteria; Gammaproteobacteria; Enterobacterales;

B9P82__locus_1

PDU1AB

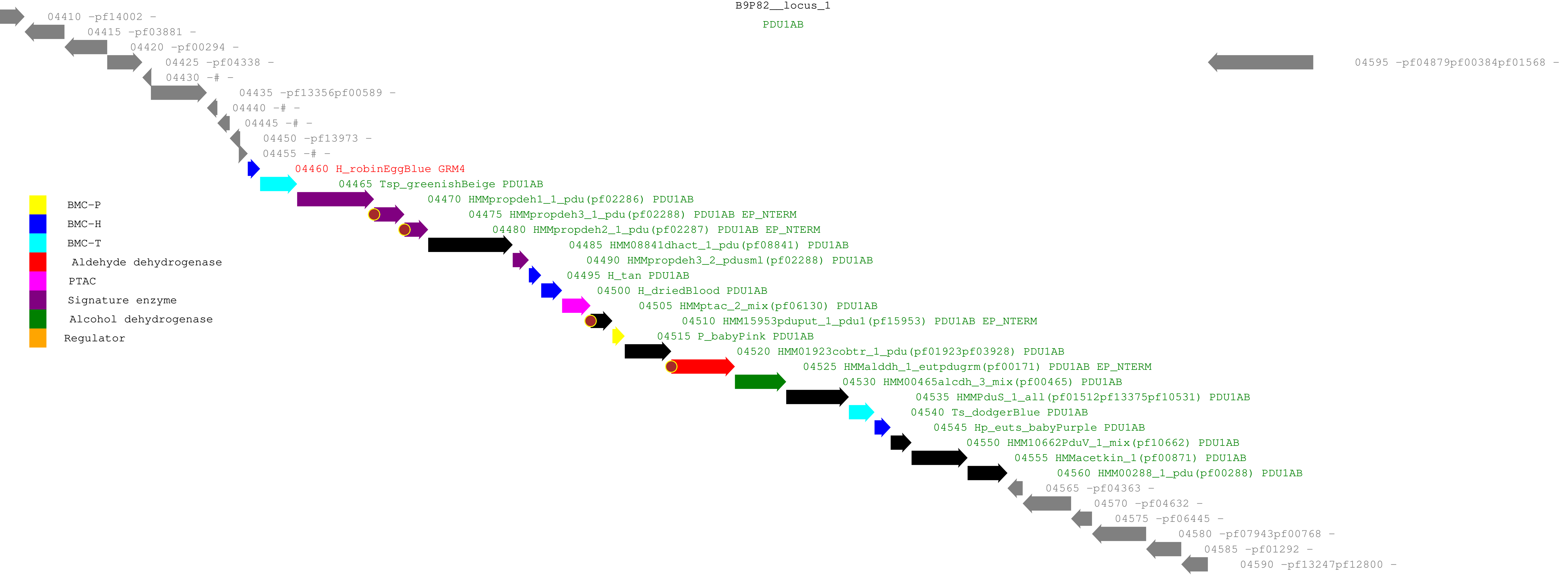

Citrobacter freundii. Bacteria; Proteobacteria; Gammaproteobacteria; Enterobacterales;

B9P90__locus_1

PDU1AB

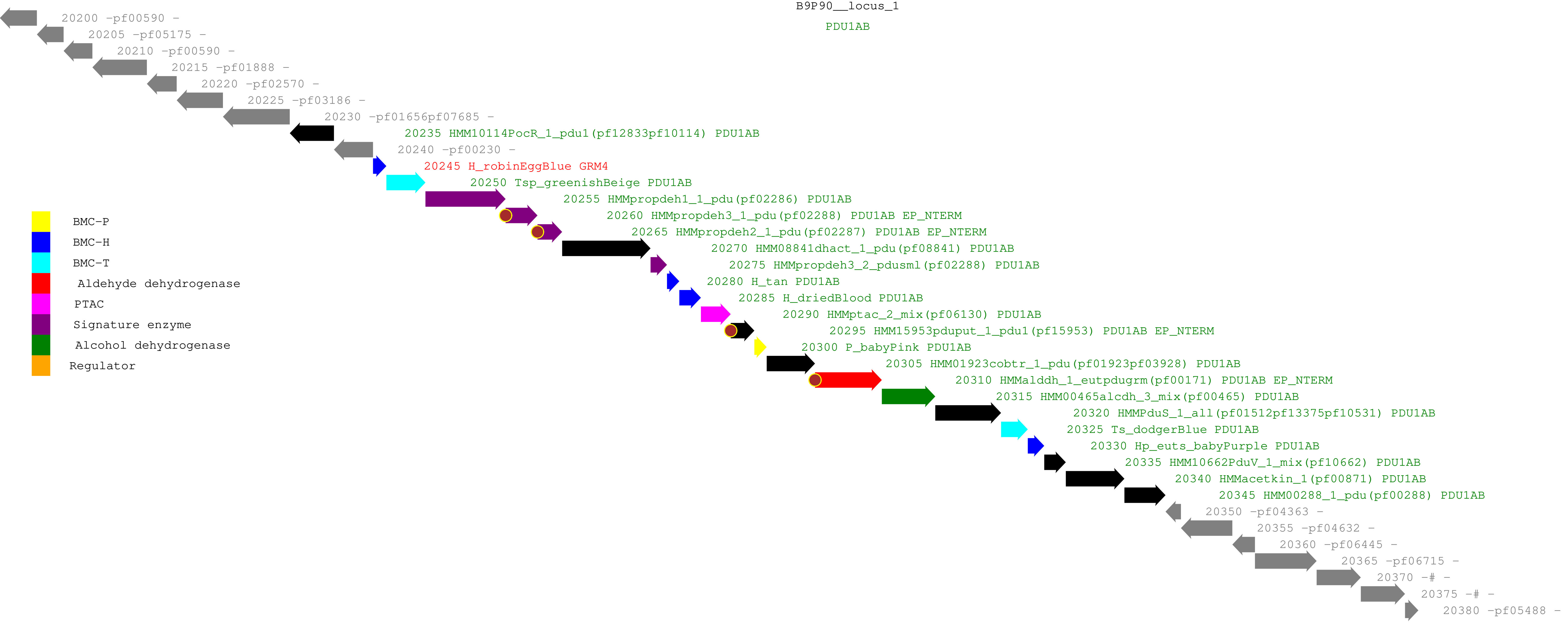

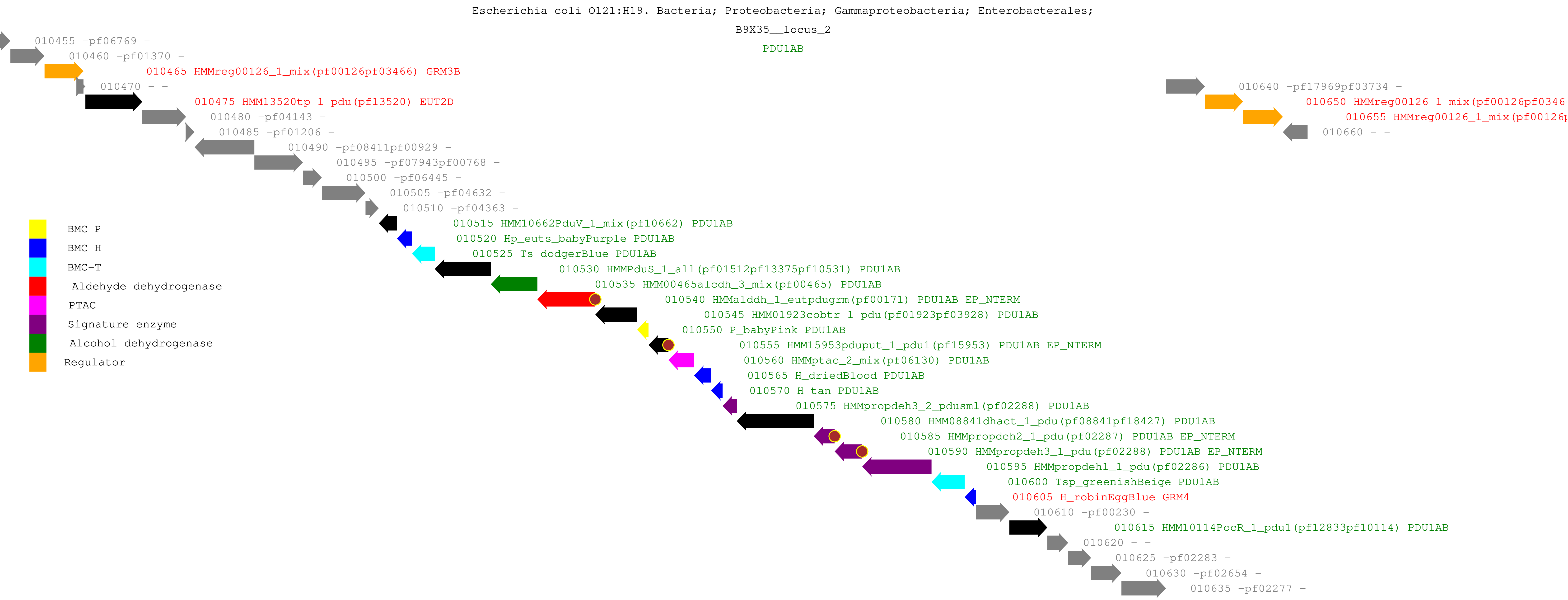
Escherichia coli O121:H19. Bacteria; Proteobacteria; Gammaproteobacteria; Enterobacterales;
B9X35__locus_2
PDU1AB
010455 -pf06769 -
010460 -pf01370 -
010465 HMMreg00126_1_mix(pf00126pf03466) GRM3B
010470 - -
010475 HMM13520tp_1_pdu(pf13520) EUT2D
010480 -pf04143 -
010485 -pf01206 -
010490 -pf08411pf00929 -
010495 -pf07943pf00768 -
010500 -pf06445 -
010505 -pf04632 -
010510 -pf04363 -
010515 HMM10662PduV_1_mix(pf10662) PDU1AB
010520 Hp_euts_babyPurple PDU1AB
010525 Ts_dodgerBlue PDU1AB
010530 HMMPduS_1_all(pf01512pf13375pf10531) PDU1AB
010535 HMM00465alcdh_3_mix(pf00465) PDU1AB
010540 HMMalddh_1_eutpdugrm(pf00171) PDU1AB EP_NTERM
010545 HMM01923cobtr_1_pdu(pf01923pf03928) PDU1AB
010550 P_babyPink PDU1AB
010555 HMM15953pduput_1_pdu1(pf15953) PDU1AB EP_NTERM
010560 HMMptac_2_mix(pf06130) PDU1AB
010565 H_driedBlood PDU1AB
010570 H_tan PDU1AB
010575 HMMpropdeh3_2_pdusml(pf02288) PDU1AB
010580 HMM08841dhact_1_pdu(pf08841pf18427) PDU1AB
010585 HMMpropdeh2_1_pdu(pf02287) PDU1AB EP_NTERM
010590 HMMpropdeh3_1_pdu(pf02288) PDU1AB EP_NTERM
010595 HMMpropdeh1_1_pdu(pf02286) PDU1AB
010600 Tsp_greenishBeige PDU1AB
010605 H_robinEggBlue GRM4
010610 -pf00230 -
010615 HMM10114PocR_1_pdu1(pf12833pf10114) PDU1AB
010620 - -
010625 -pf02283 -
010630 -pf02654 -
010635 -pf02277 -
010640 -pf17969pf03734 -
010650 HMMreg00126_1_mix(pf00126pf0346
010655 HMMreg00126_1_mix(pf00126p
010660 - -
BMC-P
BMC-H
BMC-T
Aldehyde dehydrogenase
PTAC
Signature enzyme
Alcohol dehydrogenase
Regulator

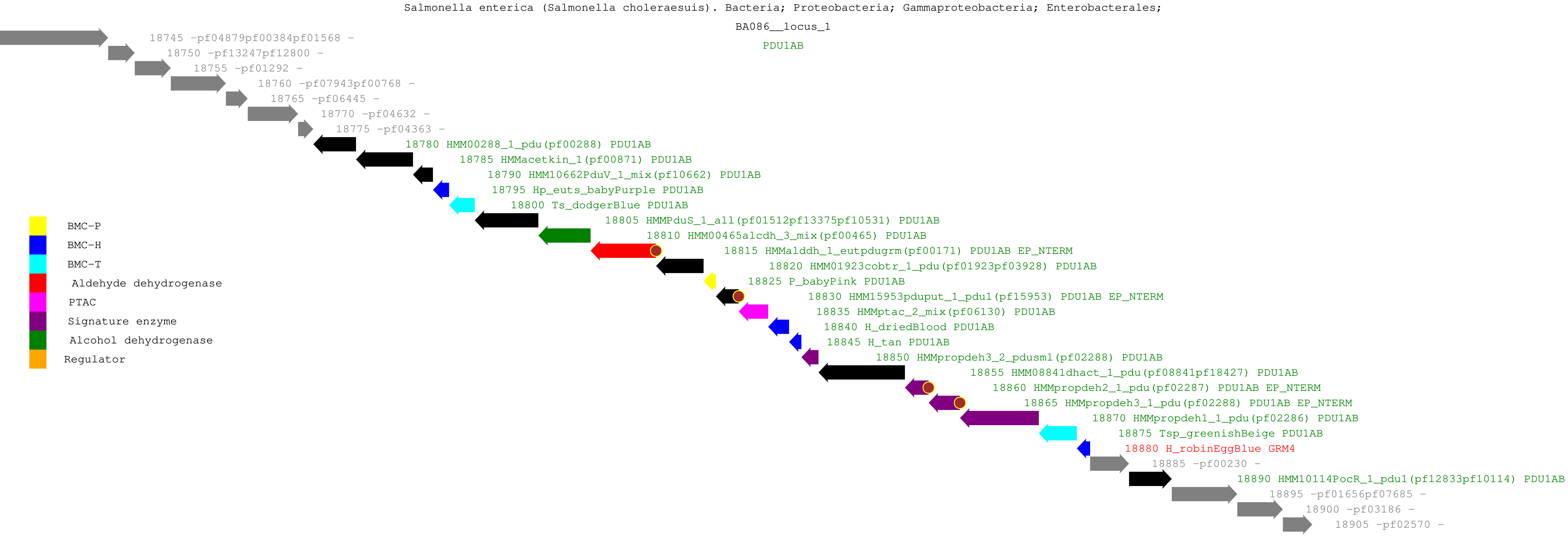
Salmonella enterica (Salmonella choleraesuis). Bacteria; Proteobacteria; Gammaproteobacteria; Enterobacterales;
BA086__locus_1
PDU1AB
18745 -pf04879pf00384pf01568 -
18750 -pf13247pf12800 -
18755 -pf01292 -
18760 -pf07943pf00768 -
18765 -pf06445 -
18770 -pf04632 -
18775 -pf04363 -
18780 HMM00288_1_pdu(pf00288) PDU1AB
18785 HMMacetkin_1(pf00871) PDU1AB
18790 HMM10662PduV_1_mix(pf10662) PDU1AB
18795 Hp_euts_babyPurple PDU1AB
18800 Ts_dodgerBlue PDU1AB
18805 HMMPduS_1_all(pf01512pf13375pf10531) PDU1AB
18810 HMM00465alcdh_3_mix(pf00465) PDU1AB
18815 HMMalddh_1_eutpdugrm(pf00171) PDU1AB EP_NTERM
18820 HMM01923cobtr_1_pdu(pf01923pf03928) PDU1AB
18825 P_babyPink PDU1AB
18830 HMM15953pduput_1_pdu1(pf15953) PDU1AB EP_NTERM
18835 HMMptac_2_mix(pf06130) PDU1AB
18840 H_driedBlood PDU1AB
18845 H_tan PDU1AB
18850 HMMpropdeh3_2_pdusml(pf02288) PDU1AB
18855 HMM08841dhact_1_pdu(pf08841pf18427) PDU1AB
18860 HMMpropdeh2_1_pdu(pf02287) PDU1AB EP_NTERM
18865 HMMpropdeh3_1_pdu(pf02288) PDU1AB EP_NTERM
18870 HMMpropdeh1_1_pdu(pf02286) PDU1AB
18875 Tsp_greenishBeige PDU1AB
18880 H_robinEggBlue GRM4
18885 -pf00230 -
18890 HMM10114PocR_1_pdu1(pf12833pf10114) PDU1AB
18895 -pf01656pf07685 -
18900 -pf03186 -
18905 -pf02570 -
BMC-P
BMC-H
BMC-T
Aldehyde dehydrogenase
PTAC
Signature enzyme
Alcohol dehydrogenase
Regulator

Salmonella choleraesuis. Bacteria; Proteobacteria; Gammaproteobacteria; Enterobacterales;

BA753__locus_1

PDU1AB

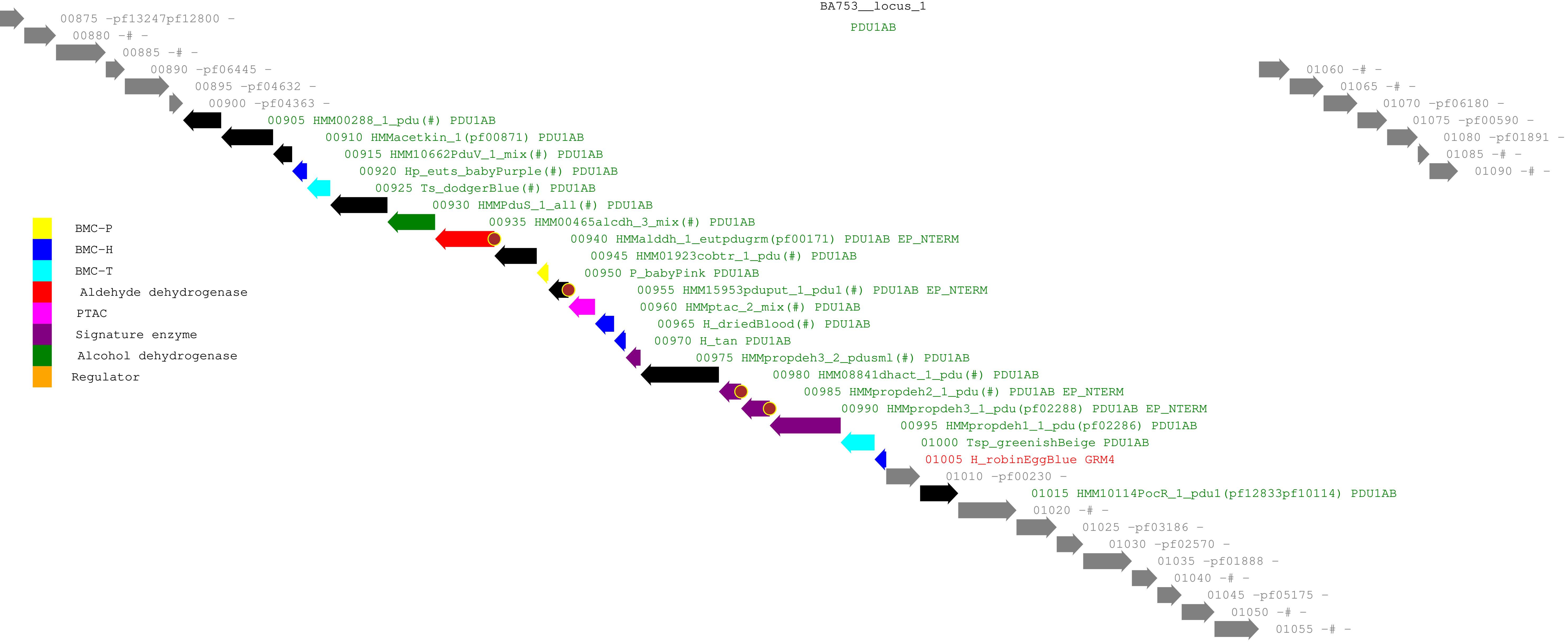

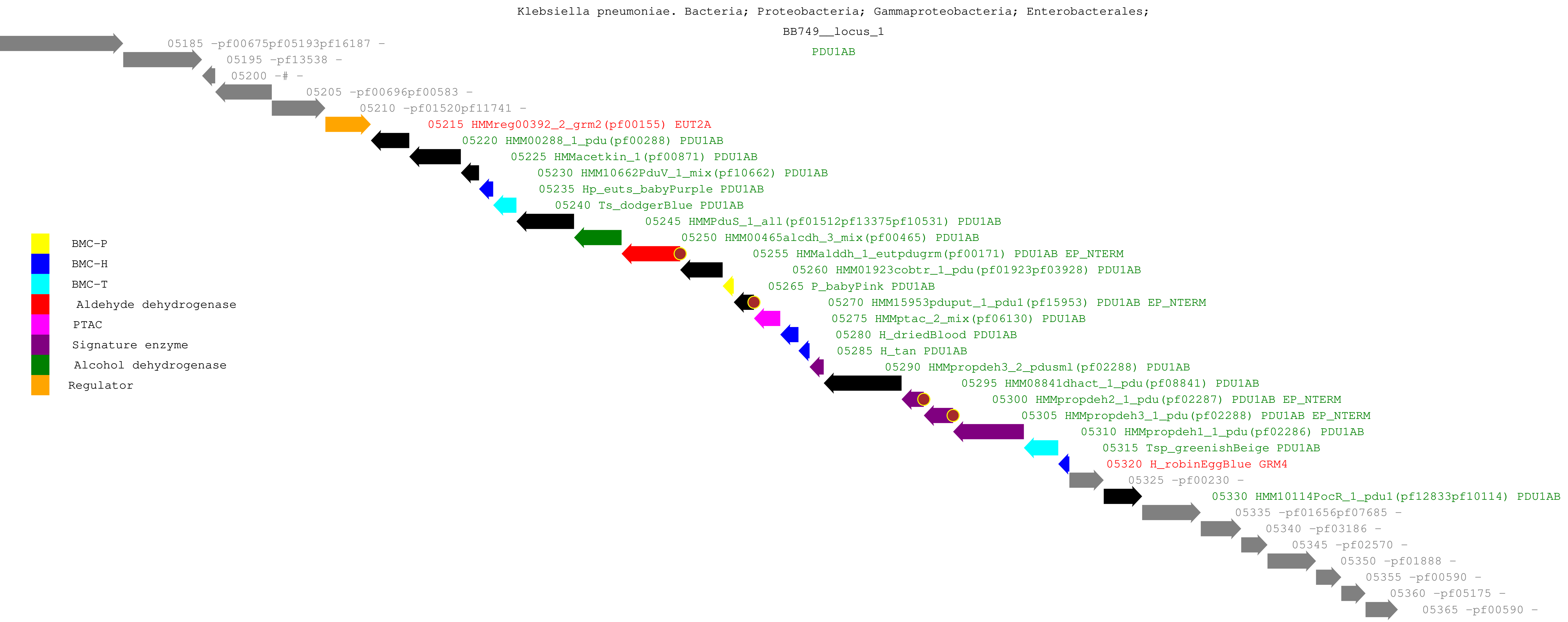
Klebsiella pneumoniae. Bacteria; Proteobacteria; Gammaproteobacteria; Enterobacterales;
BB749__locus_1
PDU1AB
05185 -pf00675pf05193pf16187 -
05195 -pf13538 -
05200 -# -
05205 -pf00696pf00583 -
05210 -pf01520pf11741 -
05215 HMMreg00392_2_grm2(pf00155) EUT2A
05220 HMM00288_1_pdu(pf00288) PDU1AB
05225 HMMacetkin_1(pf00871) PDU1AB
05230 HMM10662PduV_1_mix(pf10662) PDU1AB
05235 Hp_euts_babyPurple PDU1AB
05240 Ts_dodgerBlue PDU1AB
05245 HMMPduS_1_all(pf01512pf13375pf10531) PDU1AB
05250 HMM00465alcdh_3_mix(pf00465) PDU1AB
05255 HMMaldh_1_eutpdugrm(pf00171) PDU1AB EP_NTERM
05260 HMM01923cobtr_1_pdu(pf01923pf03928) PDU1AB
05265 P_babyPink PDU1AB
05270 HMM15953pduput_1_pdu1(pf15953) PDU1AB EP_NTERM
05275 HMMptac_2_mix(pf06130) PDU1AB
05280 H_driedBlood PDU1AB
05285 H_tan PDU1AB
05290 HMMpropdeh3_2_pdusml(pf02288) PDU1AB
05295 HMM08841dhact_1_pdu(pf08841) PDU1AB
05300 HMMpropdeh2_1_pdu(pf02287) PDU1AB EP_NTERM
05305 HMMpropdeh3_1_pdu(pf02288) PDU1AB EP_NTERM
05310 HMMpropdeh1_1_pdu(pf02286) PDU1AB
05315 Tsp_greenishBeige PDU1AB
05320 H_robinEggBlue GRM4
05325 -pf00230 -
05330 HMM10114PocR_1_pdu1(pf12833pf10114) PDU1AB
05335 -pf01656pf07685 -
05340 -pf03186 -
05345 -pf02570 -
05350 -pf01888 -
05355 -pf00590 -
05360 -pf05175 -
05365 -pf00590 -
BMC-P
BMC-H
BMC-T
Aldehyde dehydrogenase
PTAC
Signature enzyme
Alcohol dehydrogenase
Regulator

Klebsiella pneumoniae. Bacteria; Proteobacteria; Gammaproteobacteria; Enterobacterales;

BB785__locus_1

PDU1AB

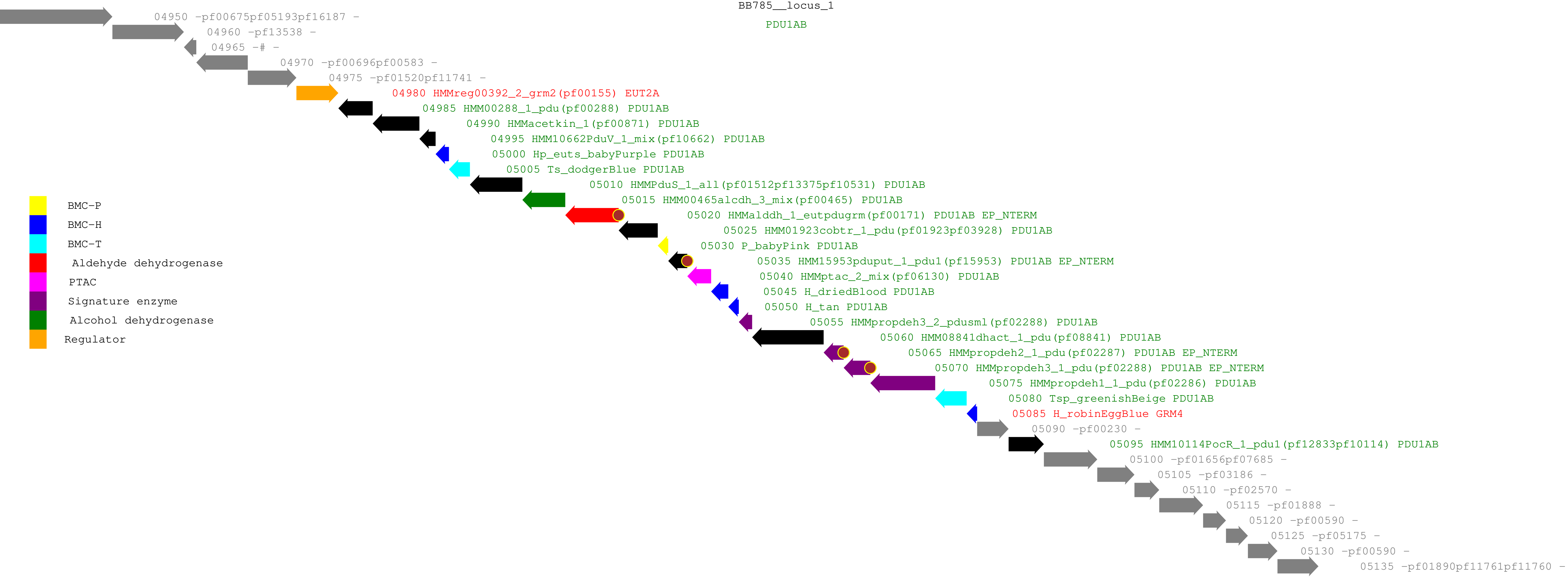

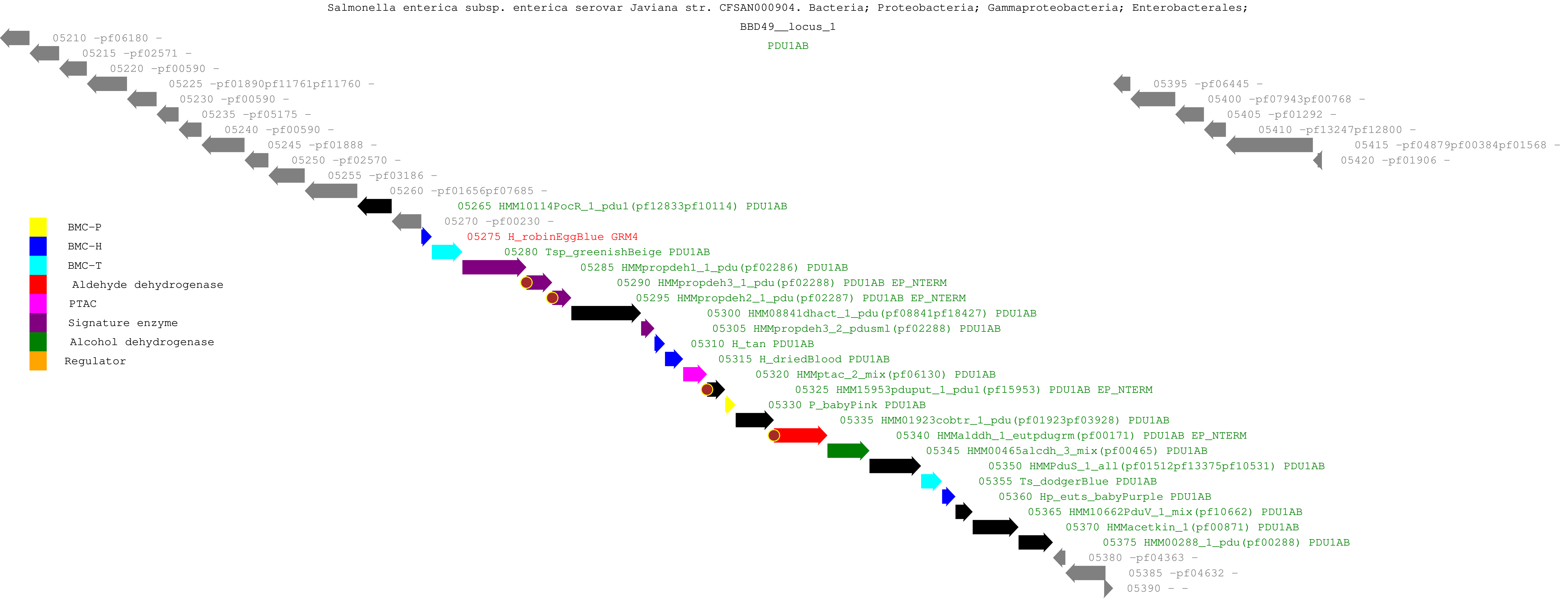
Salmonella enterica subsp. enterica serovar Javiana str. CFSAN000904. Bacteria; Proteobacteria; Gammaproteobacteria; Enterobacterales;
BBD49__locus_1
PDU1AB
05210 -pf06180 -
05215 -pf02571 -
05220 -pf00590 -
05225 -pf01890pf11761pf11760 -
05230 -pf00590 -
05235 -pf05175 -
05240 -pf00590 -
05245 -pf01888 -
05250 -pf02570 -
05255 -pf03186 -
05260 -pf01656pf07685 -
05265 HMM10114PocR_1_pdu1(pf12833pf10114) PDU1AB
05270 -pf00230 -
05275 H_robinEggBlue GRM4
05280 Tsp_greenishBeige PDU1AB
05285 HMMpropdeh1_1_pdu(pf02286) PDU1AB
05290 HMMpropdeh3_1_pdu(pf02288) PDU1AB EP_NTERM
05295 HMMpropdeh2_1_pdu(pf02287) PDU1AB EP_NTERM
05300 HMM08841dhact_1_pdu(pf08841pf18427) PDU1AB
05305 HMMpropdeh3_2_pdusml(pf02288) PDU1AB
05310 H_tan PDU1AB
05315 H_driedBlood PDU1AB
05320 HMMptac_2_mix(pf06130) PDU1AB
05325 HMM15953pduput_1_pdu1(pf15953) PDU1AB EP_NTERM
05330 P_babyPink PDU1AB
05335 HMM01923cobtr_1_pdu(pf01923pf03928) PDU1AB
05340 HMMalddh_1_eutpdugrm(pf00171) PDU1AB EP_NTERM
05345 HMM00465alcdh_3_mix(pf00465) PDU1AB
05350 HMMPduS_1_all(pf01512pf13375pf10531) PDU1AB
05355 Ts_dodgerBlue PDU1AB
05360 Hp_euts_babyPurple PDU1AB
05365 HMM10662PduV_1_mix(pf10662) PDU1AB
05370 HMMacetkin_1(pf00871) PDU1AB
05375 HMM00288_1_pdu(pf00288) PDU1AB
05380 -pf04363 -
05385 -pf04632 -
05390 - -
05395 -pf06445 -
05400 -pf07943pf00768 -
05405 -pf01292 -
05410 -pf13247pf12800 -
05415 -pf04879pf00384pf01568 -
05420 -pf01906 -
BMC-P
BMC-H
BMC-T
Aldehyde dehydrogenase
PTAC
Signature enzyme
Alcohol dehydrogenase
Regulator

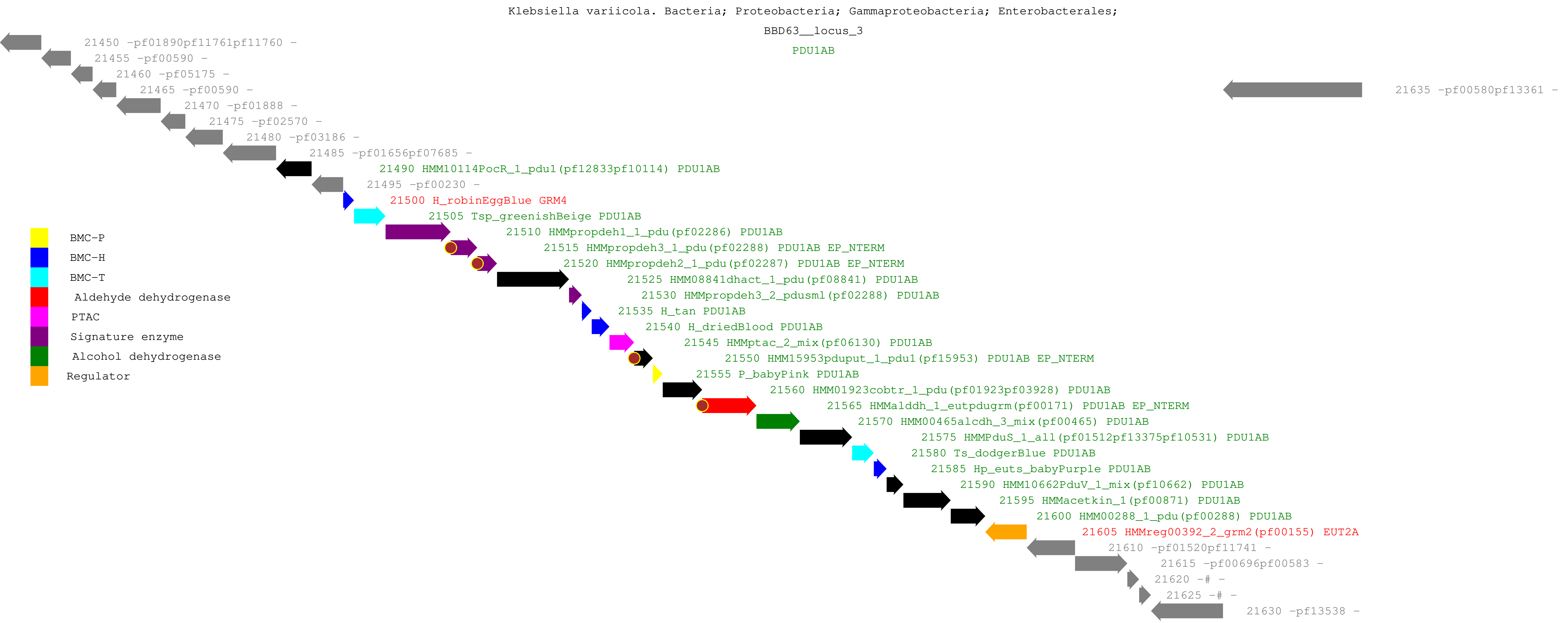

Klebsiella variicola. Bacteria; Proteobacteria; Gammaproteobacteria; Enterobacterales;
BBD63__locus_3
PDU1AB
21450 -pf01890pf11761pf11760 -
21455 -pf00590 -
21460 -pf05175 -
21465 -pf00590 -
21470 -pf01888 -
21475 -pf02570 -
21480 -pf03186 -
21485 -pf01656pf07685 -
21490 HMM10114PocR_1_pdu1(pf12833pf10114) PDU1AB
21495 -pf00230 -
21500 H_robinEggBlue GRM4
21505 Tsp_greenishBeige PDU1AB
21510 HMMpropdeh1_1_pdu(pf02286) PDU1AB
21515 HMMpropdeh3_1_pdu(pf02288) PDU1AB EP_NTERM
21520 HMMpropdeh2_1_pdu(pf02287) PDU1AB EP_NTERM
21525 HMM08841dhact_1_pdu(pf08841) PDU1AB
21530 HMMpropdeh3_2_pdusml(pf02288) PDU1AB
21535 H_tan PDU1AB
21540 H_driedBlood PDU1AB
21545 HMMptac_2_mix(pf06130) PDU1AB
21550 HMM15953pduput_1_pdu1(pf15953) PDU1AB EP_NTERM
21555 P_babyPink PDU1AB
21560 HMM01923cobtr_1_pdu(pf01923pf03928) PDU1AB
21565 HMMalddh_1_eutpdugrm(pf00171) PDU1AB EP_NTERM
21570 HMM00465alcdh_3_mix(pf00465) PDU1AB
21575 HMMPduS_1_all(pf01512pf13375pf10531) PDU1AB
21580 Ts_dodgerBlue PDU1AB
21585 Hp_euts_babyPurple PDU1AB
21590 HMM10662PduV_1_mix(pf10662) PDU1AB
21595 HMMacetkin_1(pf00871) PDU1AB
21600 HMM00288_1_pdu(pf00288) PDU1AB
21605 HMMreg00392_2_grm2(pf00155) EUT2A
21610 -pf01520pf11741 -
21615 -pf00696pf00583 -
21620 -# -
21625 -# -
21630 -pf13538 -
21635 -pf00580pf13361 -
BMC-P
BMC-H
BMC-T
Aldehyde dehydrogenase
PTAC
Signature enzyme
Alcohol dehydrogenase
Regulator

Salmonella enteritidis. Bacteria; Proteobacteria; Gammaproteobacteria; Enterobacterales;

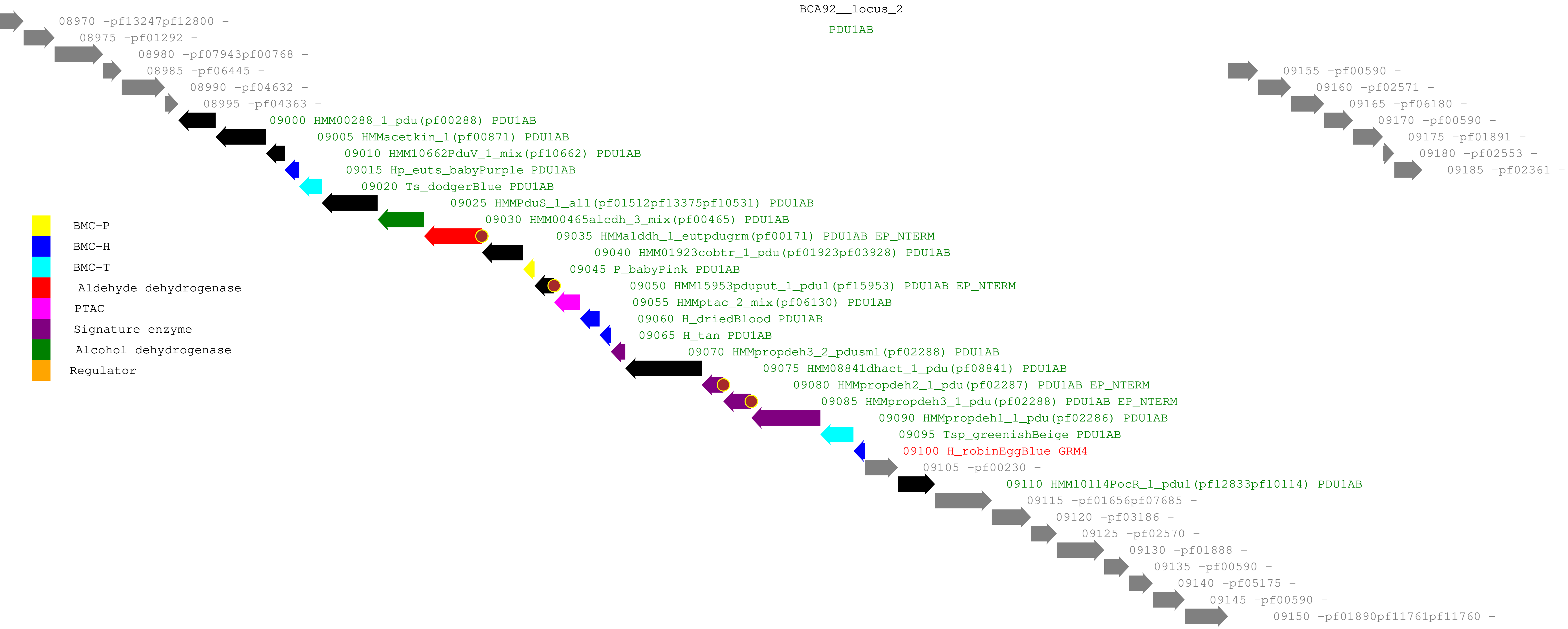

Klebsiella pneumoniae. Bacteria; Proteobacteria; Gammaproteobacteria; Enterobacterales;

BCB67__locus_1

PDU1AB

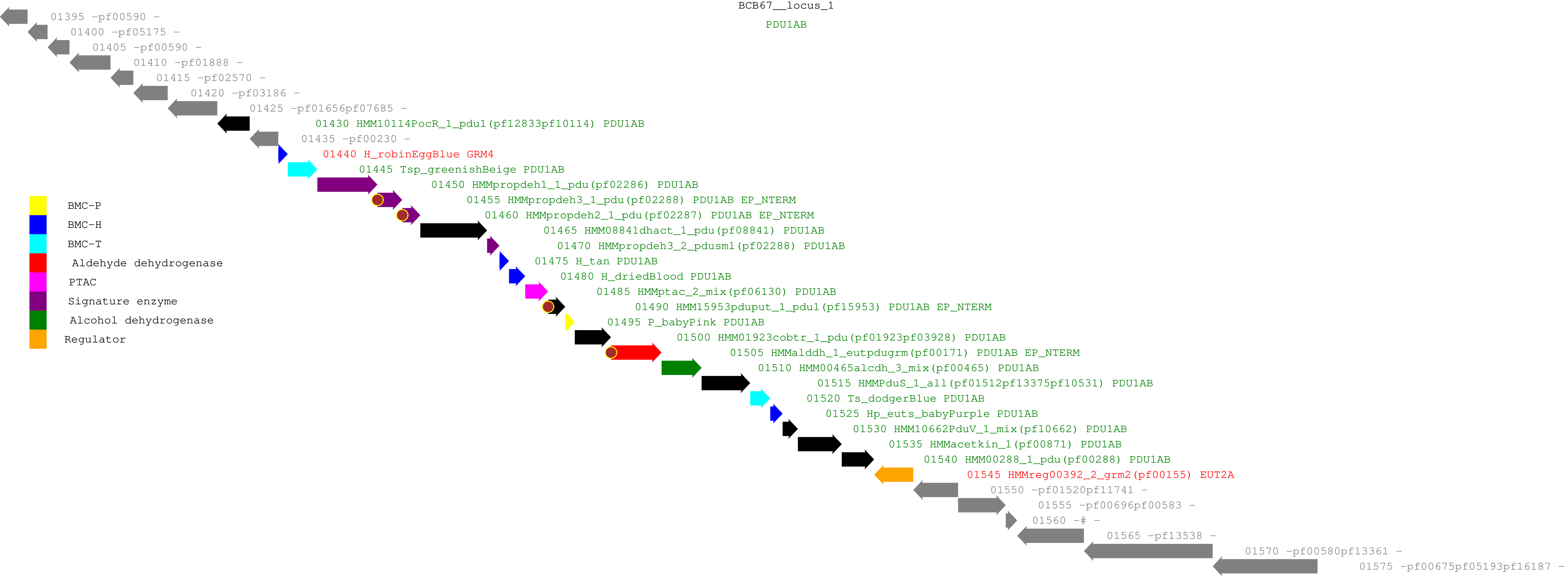

Salmonella enterica subsp. enterica serovar Altona. Bacteria; Proteobacteria; Gammaproteobacteria; Enterobacterales;

BCP30__locus_1

PDU1AB

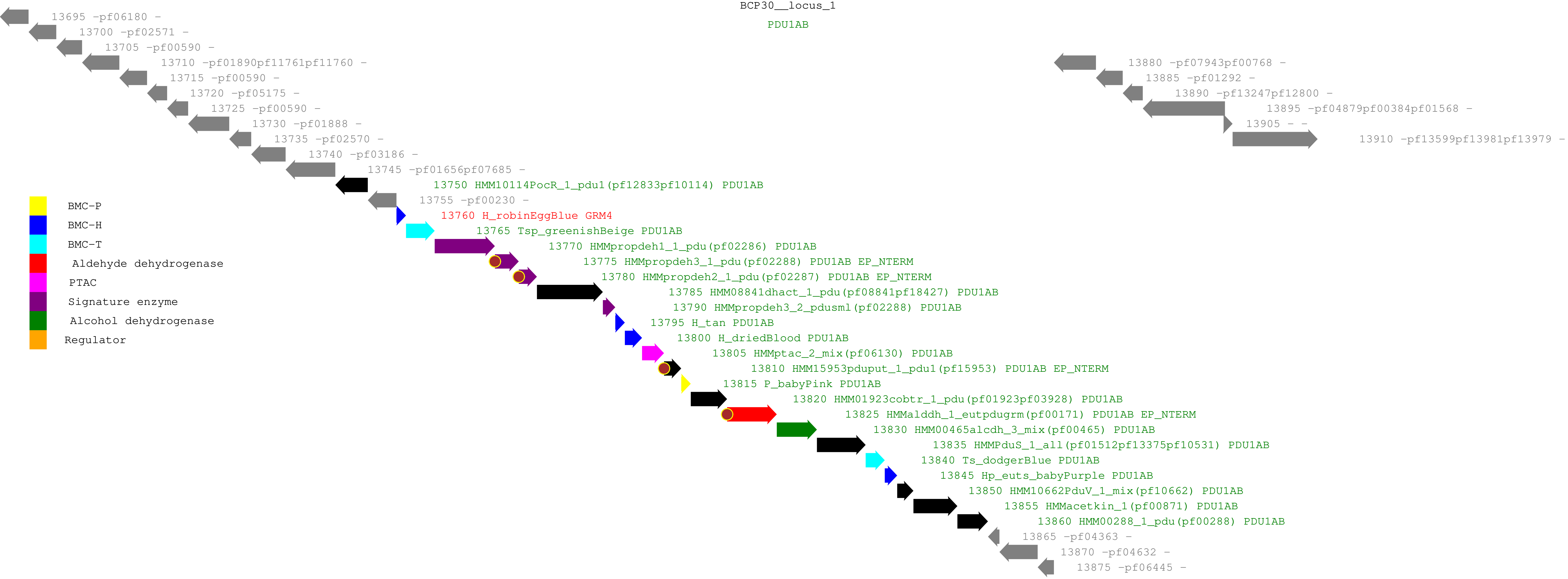

Klebsiella quasipneumoniae. Bacteria; Proteobacteria; Gammaproteobacteria; Enterobacterales;

BCY82__locus_2

PDU1AB

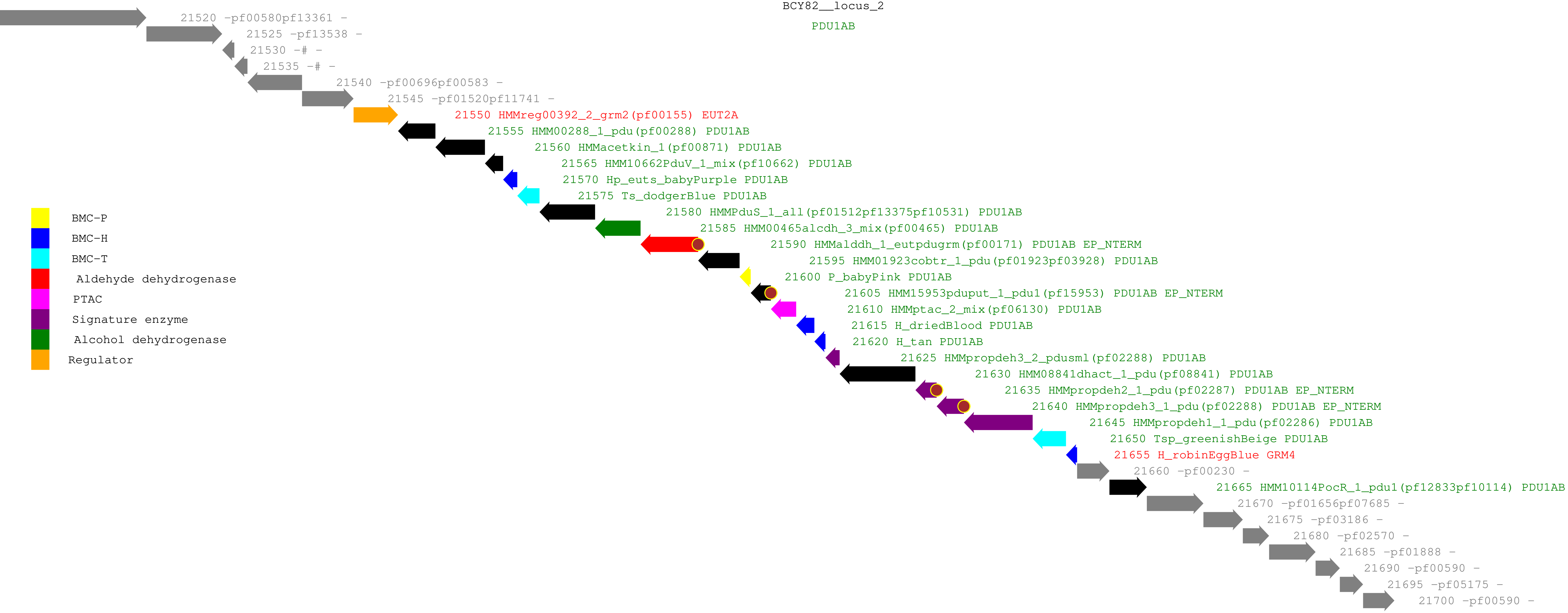

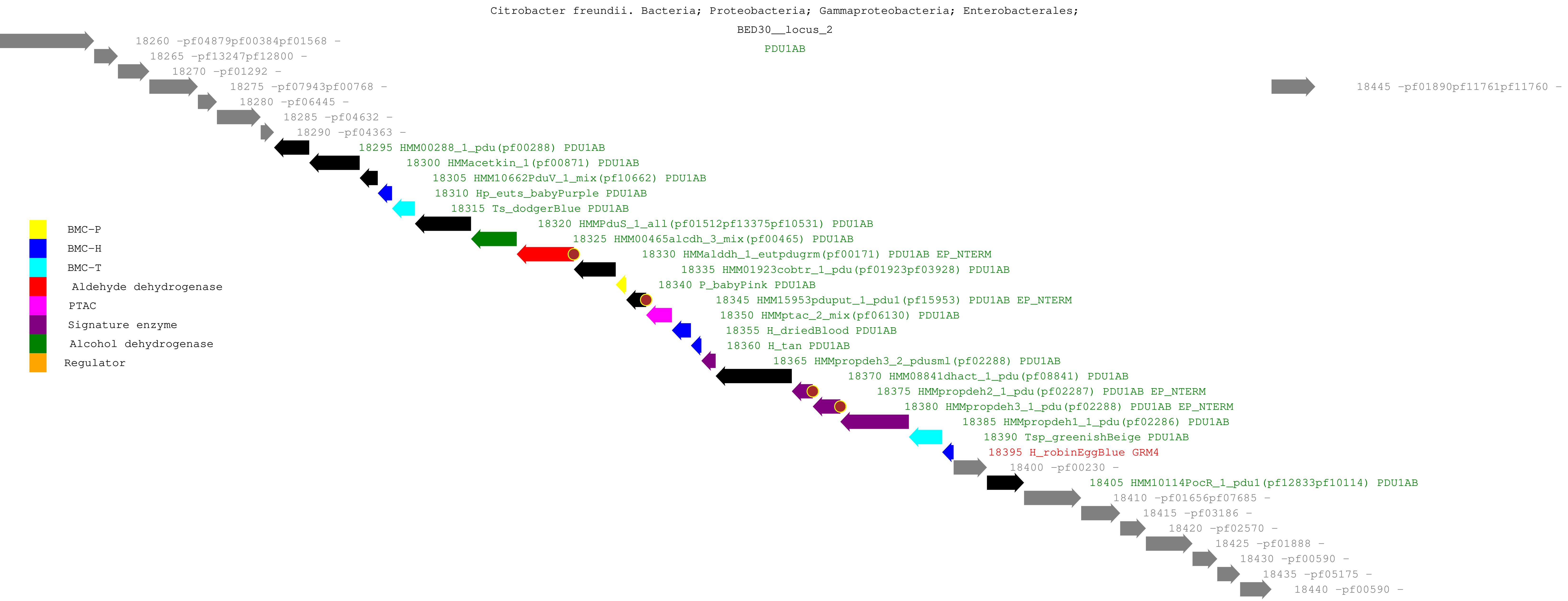
Citrobacter freundii. Bacteria; Proteobacteria; Gammaproteobacteria; Enterobacterales;
BED30__locus_2
PDU1AB
18260 -pf04879pf00384pf01568 -
18265 -pf13247pf12800 -
18270 -pf01292 -
18275 -pf07943pf00768 -
18280 -pf06445 -
18285 -pf04632 -
18290 -pf04363 -
18295 HMM00288_1_pdu(pf00288) PDU1AB
18300 HMMacetkin_1(pf00871) PDU1AB
18305 HMM10662PduV_1_mix(pf10662) PDU1AB
18310 Hp_euts_babyPurple PDU1AB
18315 Ts_dodgerBlue PDU1AB
18320 HMMPduS_1_all(pf01512pf13375pf10531) PDU1AB
18325 HMM00465alcdh_3_mix(pf00465) PDU1AB
18330 HMMalddh_1_eutpdugrm(pf00171) PDU1AB EP_NTERM
18335 HMM01923cobtr_1_pdu(pf01923pf03928) PDU1AB
18340 P_babyPink PDU1AB
18345 HMM15953pduput_1_pdu1(pf15953) PDU1AB EP_NTERM
18350 HMMptac_2_mix(pf06130) PDU1AB
18355 H_driedBlood PDU1AB
18360 H_tan PDU1AB
18365 HMMpropdeh3_2_pdusml(pf02288) PDU1AB
18370 HMM08841dhact_1_pdu(pf08841) PDU1AB
18375 HMMpropdeh2_1_pdu(pf02287) PDU1AB EP_NTERM
18380 HMMpropdeh3_1_pdu(pf02288) PDU1AB EP_NTERM
18385 HMMpropdeh1_1_pdu(pf02286) PDU1AB
18390 Tsp_greenishBeige PDU1AB
18395 H_robinEggBlue GRM4
18400 -pf00230 -
18405 HMM10114PocR_1_pdu1(pf12833pf10114) PDU1AB
18410 -pf01656pf07685 -
18415 -pf03186 -
18420 -pf02570 -
18425 -pf01888 -
18430 -pf00590 -
18435 -pf05175 -
18440 -pf00590 -
18445 -pf01890pf11761pf11760 -
BMC-P
BMC-H
BMC-T
Aldehyde dehydrogenase
PTAC
Signature enzyme
Alcohol dehydrogenase
Regulator

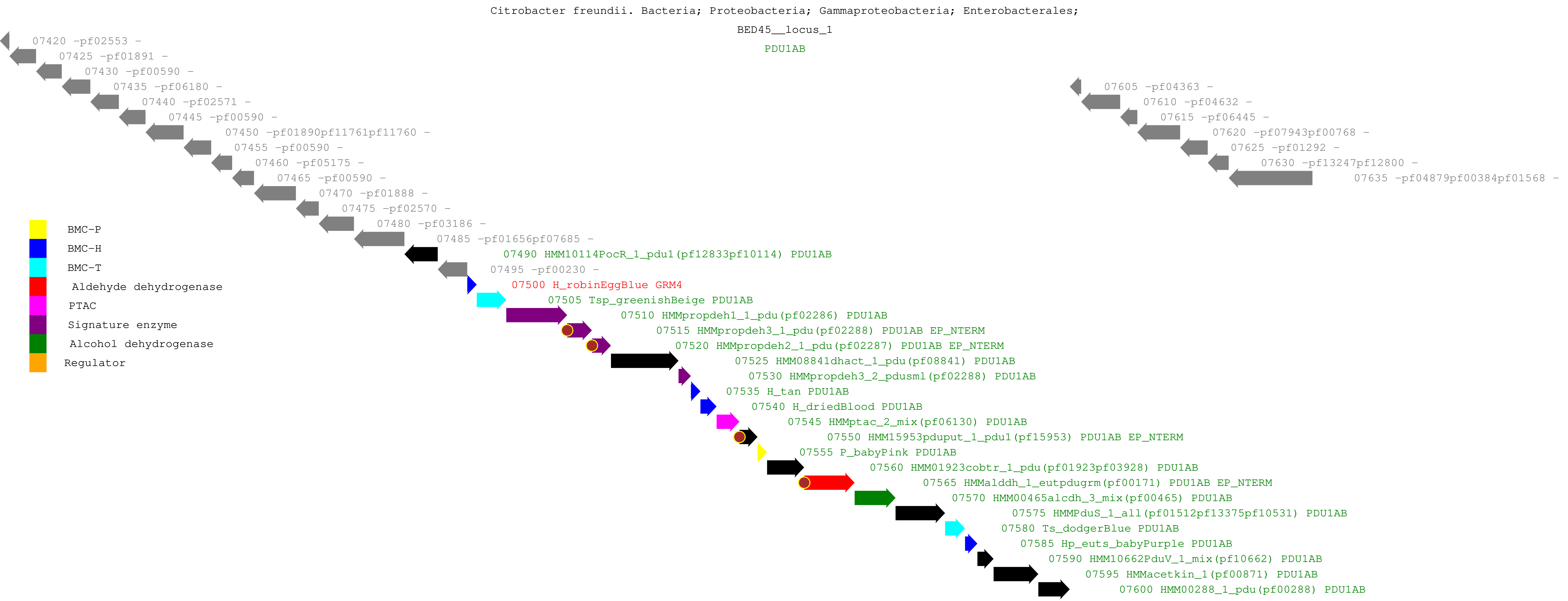
Citrobacter freundii. Bacteria; Proteobacteria; Gammaproteobacteria; Enterobacterales;
BED45__locus_1
PDU1AB
07420 -pf02553 -
07425 -pf01891 -
07430 -pf00590 -
07435 -pf06180 -
07440 -pf02571 -
07445 -pf00590 -
07450 -pf01890pf11761pf11760 -
07455 -pf00590 -
07460 -pf05175 -
07465 -pf00590 -
07470 -pf01888 -
07475 -pf02570 -
07480 -pf03186 -
07485 -pf01656pf07685 -
07490 HMM10114PocR_1_pdu1(pf12833pf10114) PDU1AB
07495 -pf00230 -
07500 H_robinEggBlue GRM4
07505 Tsp_greenishBeige PDU1AB
07510 HMMpropdeh1_1_pdu(pf02286) PDU1AB
07515 HMMpropdeh3_1_pdu(pf02288) PDU1AB EP_NTERM
07520 HMMpropdeh2_1_pdu(pf02287) PDU1AB EP_NTERM
07525 HMM08841dhact_1_pdu(pf08841) PDU1AB
07530 HMMpropdeh3_2_pdusml(pf02288) PDU1AB
07535 H_tan PDU1AB
07540 H_driedBlood PDU1AB
07545 HMMptac_2_mix(pf06130) PDU1AB
07550 HMM15953pduput_1_pdu1(pf15953) PDU1AB EP_NTERM
07555 P_babyPink PDU1AB
07560 HMM01923cobtr_1_pdu(pf01923pf03928) PDU1AB
07565 HMMaldh_1_eutpdugrm(pf00171) PDU1AB EP_NTERM
07570 HMM00465alcdh_3_mix(pf00465) PDU1AB
07575 HMMPduS_1_all(pf01512pf13375pf10531) PDU1AB
07580 Ts_dodgerBlue PDU1AB
07585 Hp_euts_babyPurple PDU1AB
07590 HMM10662PduV_1_mix(pf10662) PDU1AB
07595 HMMacetkin_1(pf00871) PDU1AB
07600 HMM00288_1_pdu(pf00288) PDU1AB
07605 -pf04363 -
07610 -pf04632 -
07615 -pf06445 -
07620 -pf07943pf00768 -
07625 -pf01292 -
07630 -pf13247pf12800 -
07635 -pf04879pf00384pf01568 -
BMC-P
BMC-H
BMC-T
Aldehyde dehydrogenase
PTAC
Signature enzyme
Alcohol dehydrogenase
Regulator

Citrobacter freundii. Bacteria; Proteobacteria; Gammaproteobacteria; Enterobacterales;

BEH70__locus_1

PDU1AB

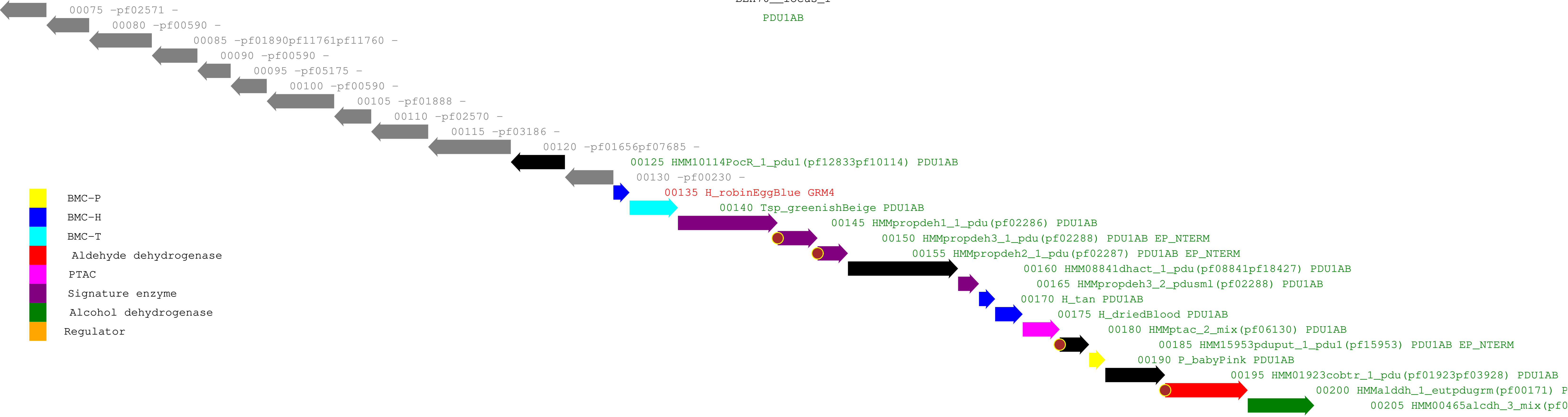

Citrobacter freundii. Bacteria; Proteobacteria; Gammaproteobacteria; Enterobacterales;

BEH73__locus_1

PDU1AB

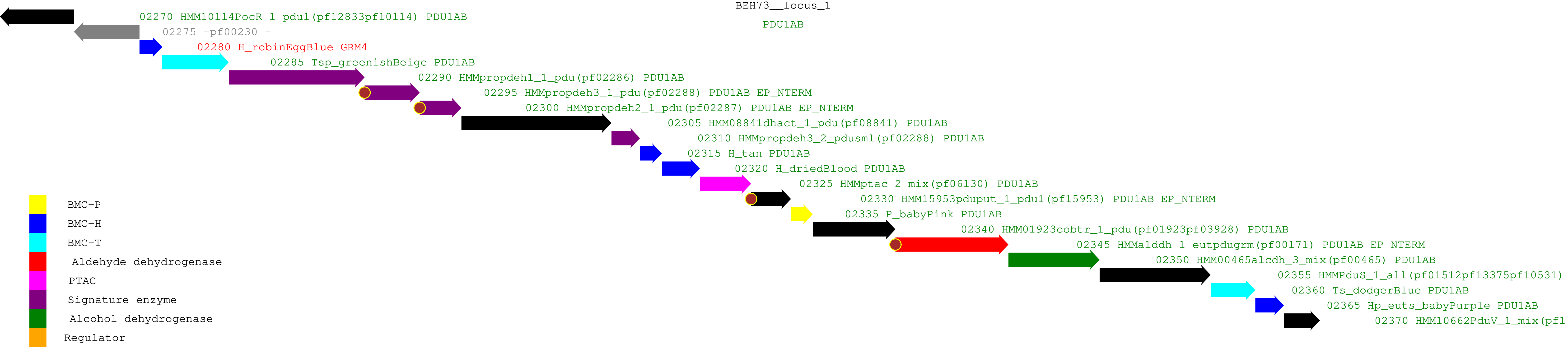

Salmonella enterica subsp. enterica serovar Grumpensis. Bacteria; Proteobacteria; Gammaproteobacteria; Enterobacterales;

BEL51__locus_2

PDU1AB

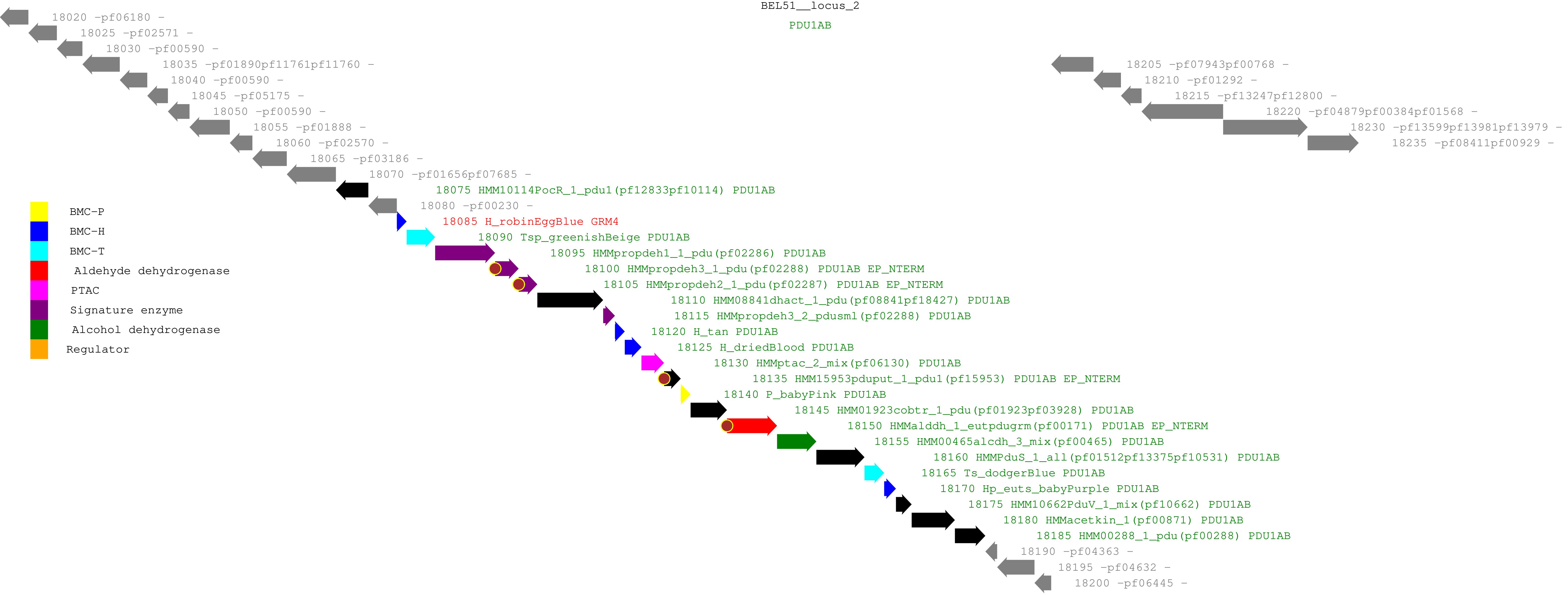

Escherichia sp. MOD1-EC5451. Bacteria; Proteobacteria; Gammaproteobacteria; Enterobacterales;

BEM40__locus_2

PDU1AB

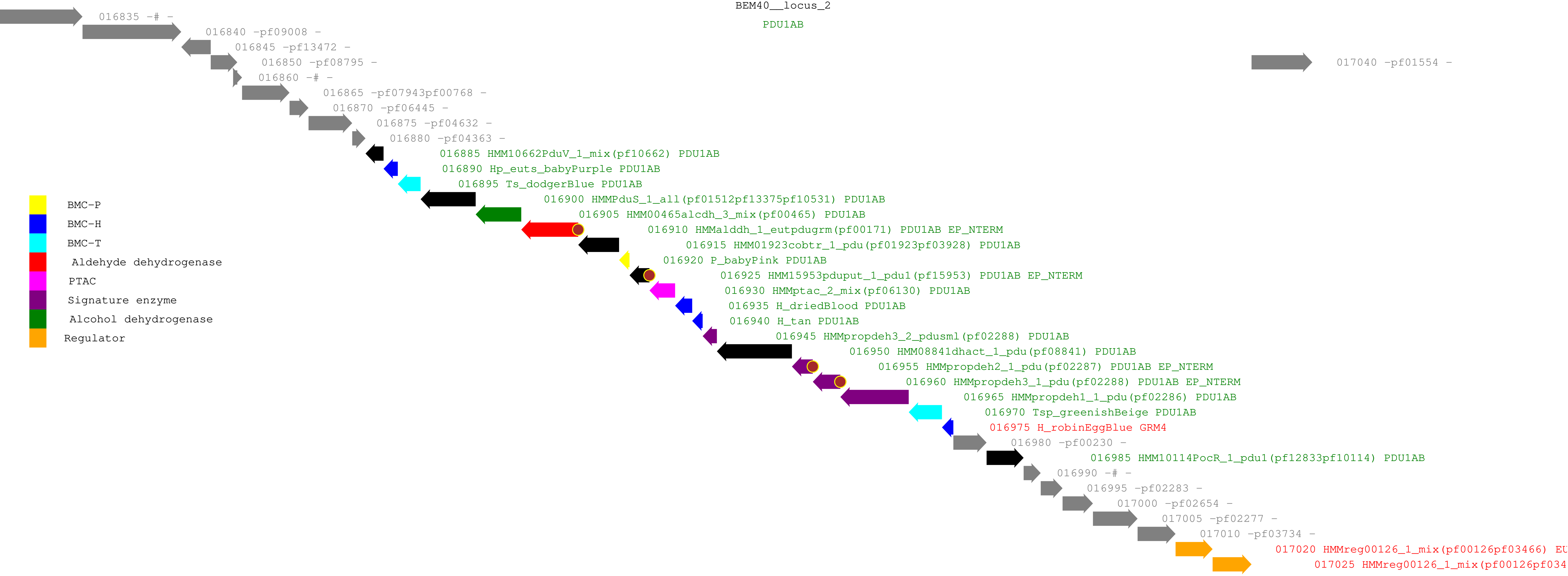

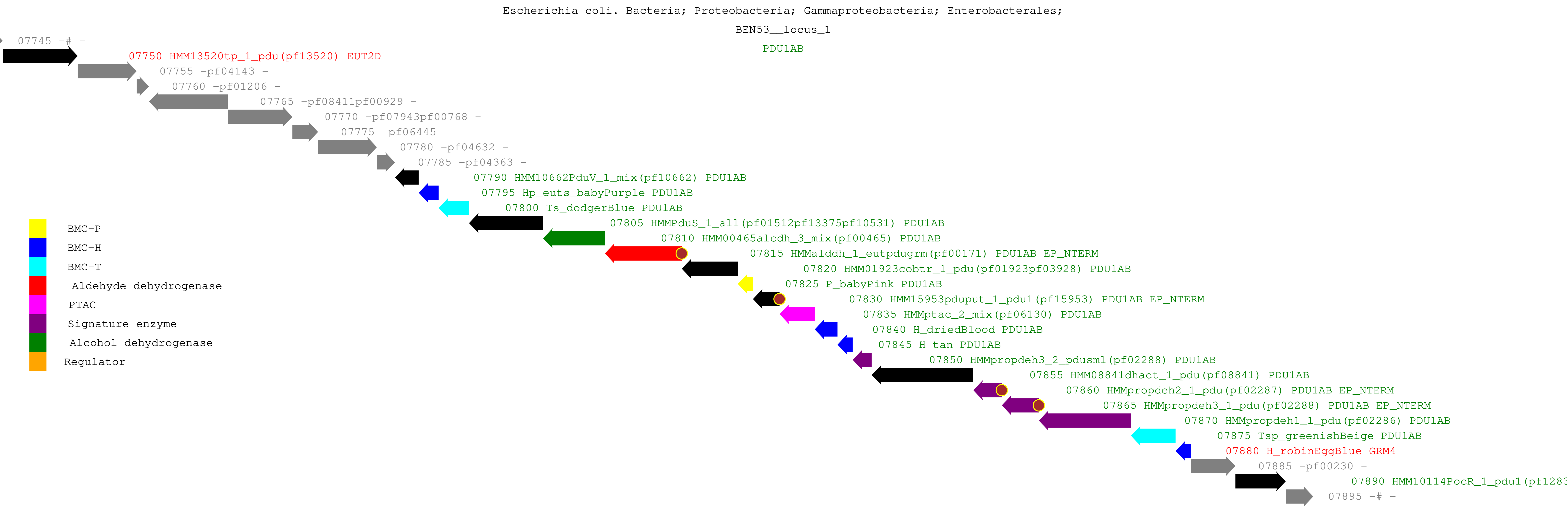

Escherichia coli. Bacteria; Proteobacteria; Gammaproteobacteria; Enterobacterales;
BEN53__locus_1
PDU1AB
07745 -# -
07750 HMM13520tp_1_pdu(pf13520) EUT2D
07755 -pf04143 -
07760 -pf01206 -
07765 -pf08411pf00929 -
07770 -pf07943pf00768 -
07775 -pf06445 -
07780 -pf04632 -
07785 -pf04363 -
07790 HMM10662PduV_1_mix(pf10662) PDU1AB
07795 Hp_euts_babyPurple PDU1AB
07800 Ts_dodgerBlue PDU1AB
07805 HMMPduS_1_all(pf01512pf13375pf10531) PDU1AB
07810 HMM00465alcdh_3_mix(pf00465) PDU1AB
07815 HMMalddh_1_eutpdugrm(pf00171) PDU1AB EP_NTERM
07820 HMM01923cobtr_1_pdu(pf01923pf03928) PDU1AB
07825 P_babyPink PDU1AB
07830 HMM15953pduput_1_pdu1(pf15953) PDU1AB EP_NTERM
07835 HMMptac_2_mix(pf06130) PDU1AB
07840 H_driedBlood PDU1AB
07845 H_tan PDU1AB
07850 HMMpropdeh3_2_pdusml(pf02288) PDU1AB
07855 HMM08841dhact_1_pdu(pf08841) PDU1AB
07860 HMMpropdeh2_1_pdu(pf02287) PDU1AB EP_NTERM
07865 HMMpropdeh3_1_pdu(pf02288) PDU1AB EP_NTERM
07870 HMMpropdeh1_1_pdu(pf02286) PDU1AB
07875 Tsp_greenishBeige PDU1AB
07880 H_robinEggBlue GRM4
07885 -pf00230 -
07890 HMM10114PocR_1_pdu1(pf1283
07895 -# -
BMC-P
BMC-H
BMC-T
Aldehyde dehydrogenase
PTAC
Signature enzyme
Alcohol dehydrogenase
Regulator

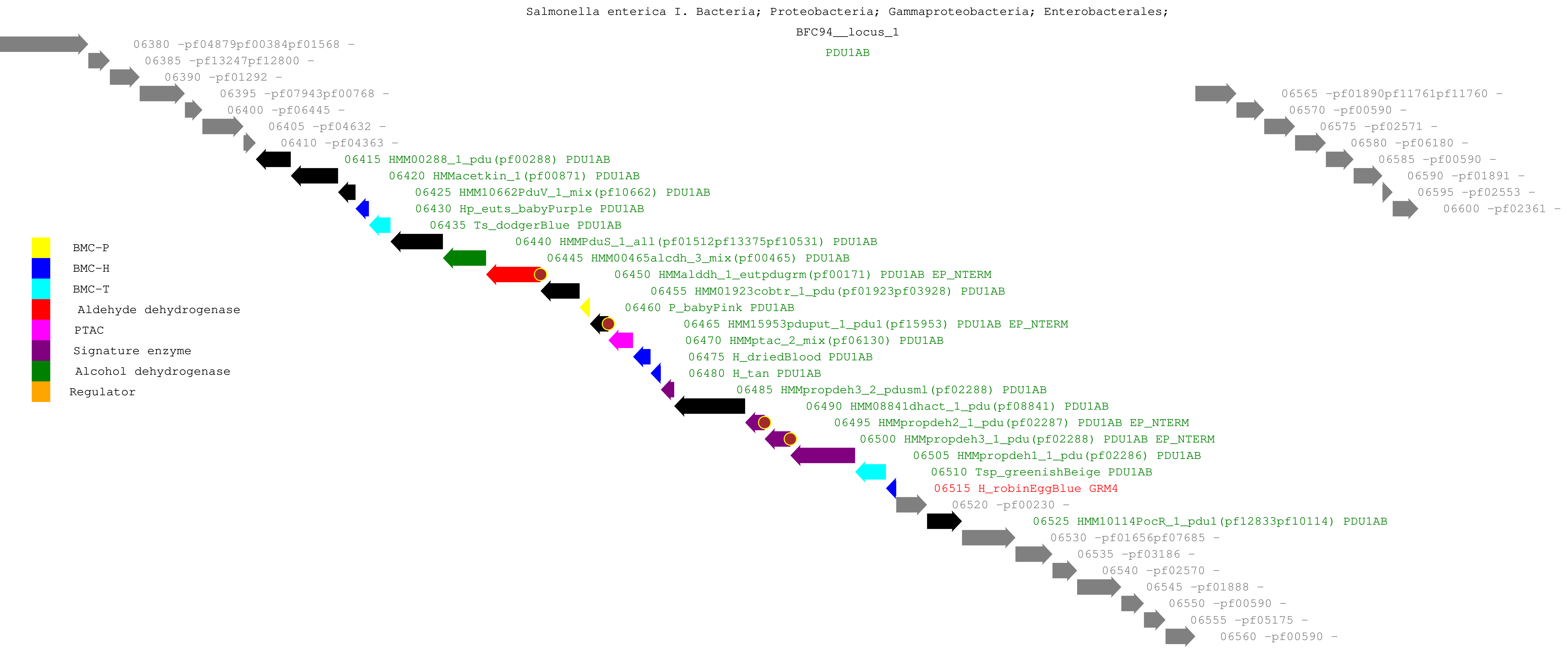
Salmonella enterica I. Bacteria; Proteobacteria; Gammaproteobacteria; Enterobacterales;
BFC94__locus_1
PDU1AB
06380 -pf04879pf00384pf01568 -
06385 -pf13247pf12800 -
06390 -pf01292 -
06395 -pf07943pf00768 -
06400 -pf06445 -
06405 -pf04632 -
06410 -pf04363 -
06415 HMM00288_1_pdu(pf00288) PDU1AB
06420 HMMacetkin_1(pf00871) PDU1AB
06425 HMM10662PduV_1_mix(pf10662) PDU1AB
06430 Hp_euts_babyPurple PDU1AB
06435 Ts_dodgerBlue PDU1AB
06440 HMMPduS_1_all(pf01512pf13375pf10531) PDU1AB
06445 HMM00465alcdh_3_mix(pf00465) PDU1AB
06450 HMMalddh_1_eutpdugrm(pf00171) PDU1AB EP_NTERM
06455 HMM01923cobtr_1_pdu(pf01923pf03928) PDU1AB
06460 P_babyPink PDU1AB
06465 HMM15953pduput_1_pdu1(pf15953) PDU1AB EP_NTERM
06470 HMMptac_2_mix(pf06130) PDU1AB
06475 H_driedBlood PDU1AB
06480 H_tan PDU1AB
06485 HMMpropdeh3_2_pdusml(pf02288) PDU1AB
06490 HMM08841dhact_1_pdu(pf08841) PDU1AB
06495 HMMpropdeh2_1_pdu(pf02287) PDU1AB EP_NTERM
06500 HMMpropdeh3_1_pdu(pf02288) PDU1AB EP_NTERM
06505 HMMpropdeh1_1_pdu(pf02286) PDU1AB
06510 Tsp_greenishBeige PDU1AB
06515 H_robinEggBlue GRM4
06520 -pf00230 -
06525 HMM10114PocR_1_pdu1(pf12833pf10114) PDU1AB
06530 -pf01656pf07685 -
06535 -pf03186 -
06540 -pf02570 -
06545 -pf01888 -
06550 -pf00590 -
06555 -pf05175 -
06560 -pf00590 -
06565 -pf01890pf11761pf11760 -
06570 -pf00590 -
06575 -pf02571 -
06580 -pf06180 -
06585 -pf00590 -
06590 -pf01891 -
06595 -pf02553 -
06600 -pf02361 -
BMC-P
BMC-H
BMC-T
Aldehyde dehydrogenase
PTAC
Signature enzyme
Alcohol dehydrogenase
Regulator

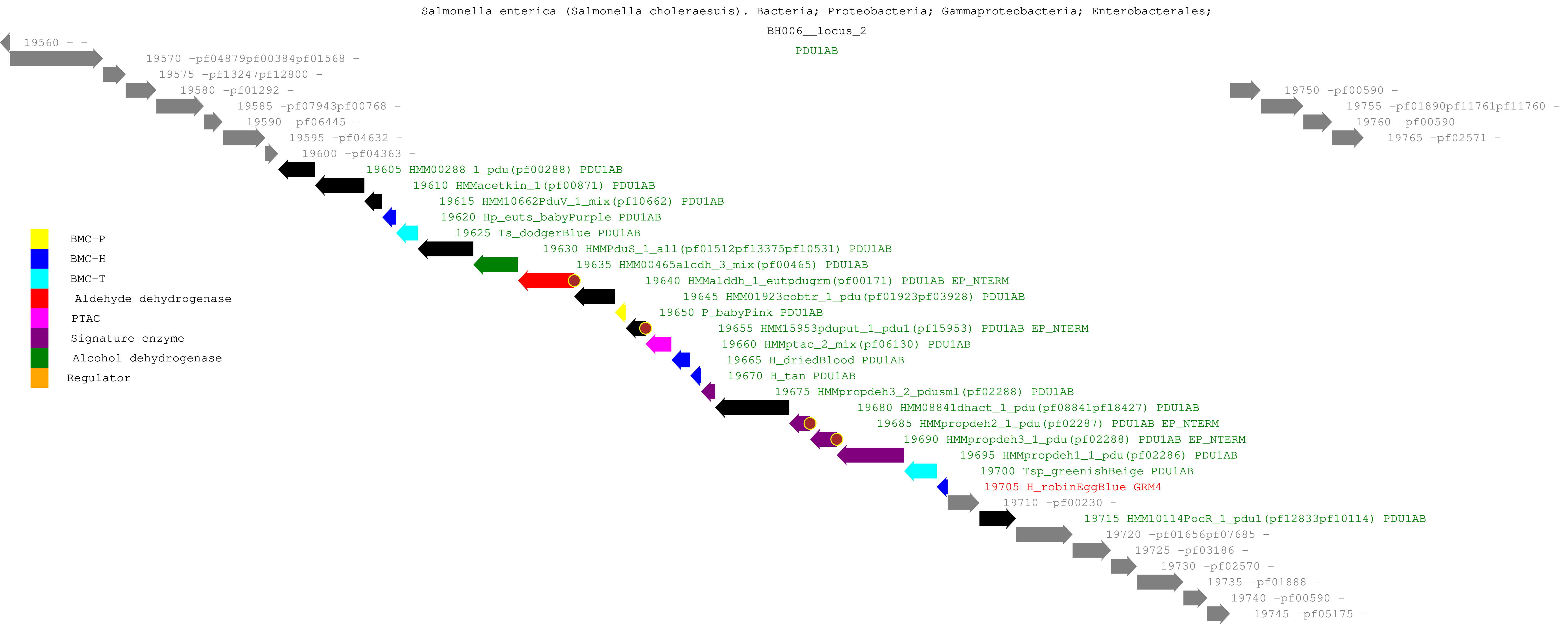
Salmonella enterica (Salmonella choleraesuis). Bacteria; Proteobacteria; Gammaproteobacteria; Enterobacterales;
BH006__locus_2
PDU1AB
19560 - -
19570 -pf04879pf00384pf01568 -
19575 -pf13247pf12800 -
19580 -pf01292 -
19585 -pf07943pf00768 -
19590 -pf06445 -
19595 -pf04632 -
19600 -pf04363 -
19605 HMM00288_1_pdu(pf00288) PDU1AB
19610 HMMacetkin_1(pf00871) PDU1AB
19615 HMM10662PduV_1_mix(pf10662) PDU1AB
19620 Hp_euts_babyPurple PDU1AB
19625 Ts_dodgerBlue PDU1AB
19630 HMMPduS_1_all(pf01512pf13375pf10531) PDU1AB
19635 HMM00465alcdh_3_mix(pf00465) PDU1AB
19640 HMMalddh_1_eutpdugrm(pf00171) PDU1AB EP_NTERM
19645 HMM01923cobtr_1_pdu(pf01923pf03928) PDU1AB
19650 P_babyPink PDU1AB
19655 HMM15953pduput_1_pdu1(pf15953) PDU1AB EP_NTERM
19660 HMMptac_2_mix(pf06130) PDU1AB
19665 H_driedBlood PDU1AB
19670 H_tan PDU1AB
19675 HMMpropdeh3_2_pdusml(pf02288) PDU1AB
19680 HMM08841dhact_1_pdu(pf08841pf18427) PDU1AB
19685 HMMpropdeh2_1_pdu(pf02287) PDU1AB EP_NTERM
19690 HMMpropdeh3_1_pdu(pf02288) PDU1AB EP_NTERM
19695 HMMpropdeh1_1_pdu(pf02286) PDU1AB
19700 Tsp_greenishBeige PDU1AB
19705 H_robinEggBlue GRM4
19710 -pf00230 -
19715 HMM10114PocR_1_pdu1(pf12833pf10114) PDU1AB
19720 -pf01656pf07685 -
19725 -pf03186 -
19730 -pf02570 -
19735 -pf01888 -
19740 -pf00590 -
19745 -pf05175 -
19750 -pf00590 -
19755 -pf01890pf11761pf11760 -
19760 -pf00590 -
19765 -pf02571 -
BMC-P
BMC-H
BMC-T
Aldehyde dehydrogenase
PTAC
Signature enzyme
Alcohol dehydrogenase
Regulator

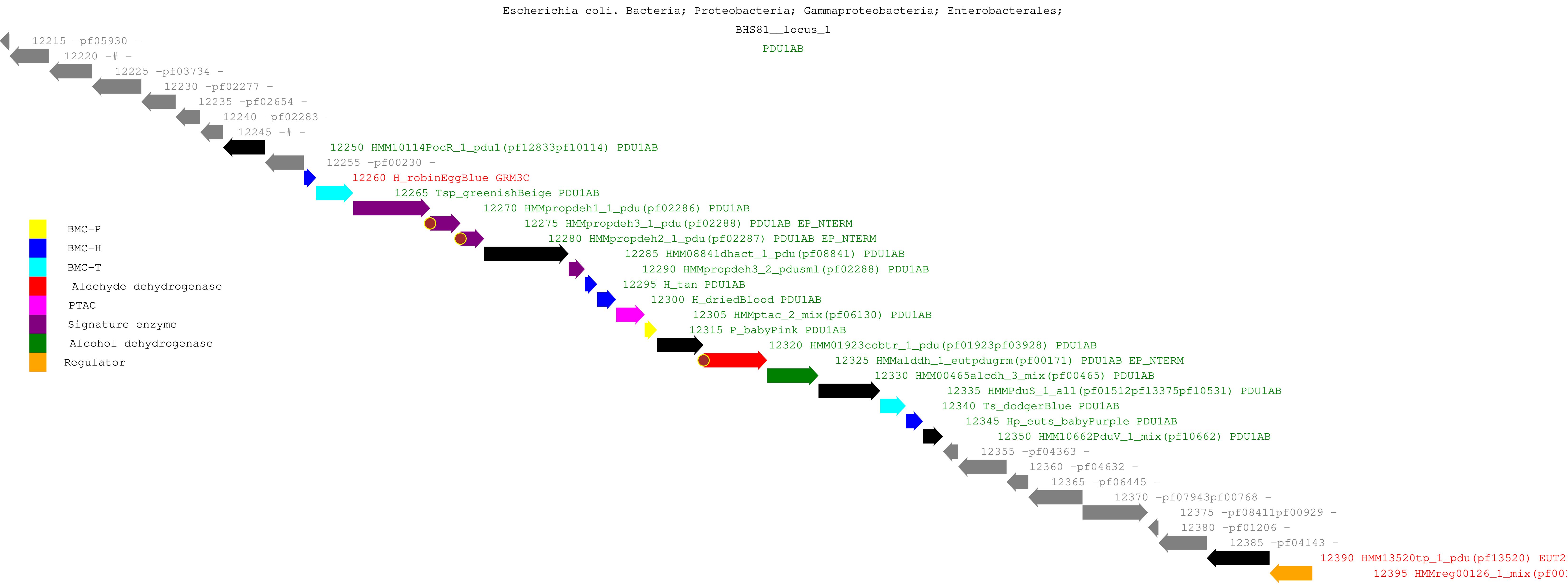
Escherichia coli. Bacteria; Proteobacteria; Gammaproteobacteria; Enterobacterales;
BHS81__locus_1
PDU1AB
12215 -pf05930 -
12220 -# -
12225 -pf03734 -
12230 -pf02277 -
12235 -pf02654 -
12240 -pf02283 -
12245 -# -
12250 HMM10114PocR_1_pdu1(pf12833pf10114) PDU1AB
12255 -pf00230 -
12260 H_robinEggBlue GRM3C
12265 Tsp_greenishBeige PDU1AB
12270 HMMpropdeh1_1_pdu(pf02286) PDU1AB
12275 HMMpropdeh3_1_pdu(pf02288) PDU1AB EP_NTERM
12280 HMMpropdeh2_1_pdu(pf02287) PDU1AB EP_NTERM
12285 HMM08841dhact_1_pdu(pf08841) PDU1AB
12290 HMMpropdeh3_2_pdusml(pf02288) PDU1AB
12295 H_tan PDU1AB
12300 H_driedBlood PDU1AB
12305 HMMptac_2_mix(pf06130) PDU1AB
12315 P_babyPink PDU1AB
12320 HMM01923cobtr_1_pdu(pf01923pf03928) PDU1AB
12325 HMMalddh_1_eutpdugrm(pf00171) PDU1AB EP_NTERM
12330 HMM00465alcdh_3_mix(pf00465) PDU1AB
12335 HMMPduS_1_all(pf01512pf13375pf10531) PDU1AB
12340 Ts_dodgerBlue PDU1AB
12345 Hp_euts_babyPurple PDU1AB
12350 HMM10662PduV_1_mix(pf10662) PDU1AB
12355 -pf04363 -
12360 -pf04632 -
12365 -pf06445 -
12370 -pf07943pf00768 -
12375 -pf08411pf00929 -
12380 -pf01206 -
12385 -pf04143 -
12390 HMM13520tp_1_pdu(pf13520) EUT2
12395 HMMreg00126_1_mix(pf00
BMC-P
BMC-H
BMC-T
Aldehyde dehydrogenase
PTAC
Signature enzyme
Alcohol dehydrogenase
Regulator

Kluyvera intestini. Bacteria; Proteobacteria; Gammaproteobacteria; Enterobacterales;

BHT19__locus_2

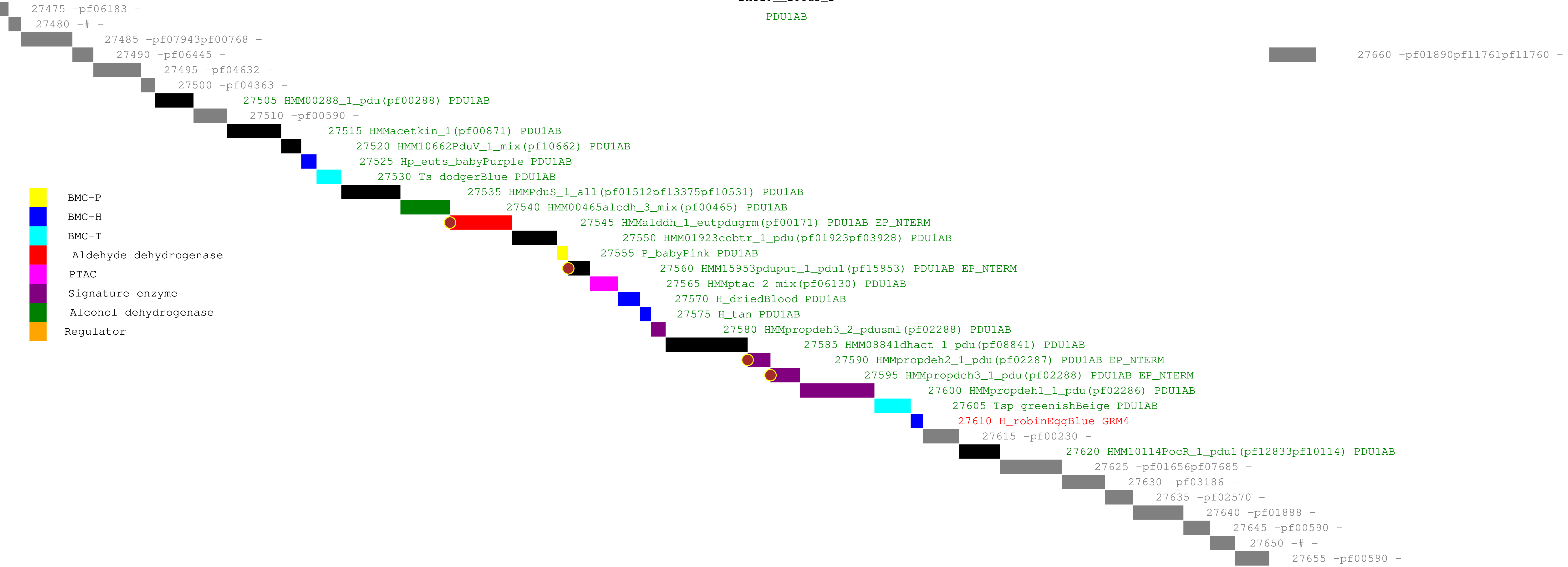

Escherichia coli. Bacteria; Proteobacteria; Gammaproteobacteria; Enterobacterales;

BIQ85__locus_1

PDU1AB

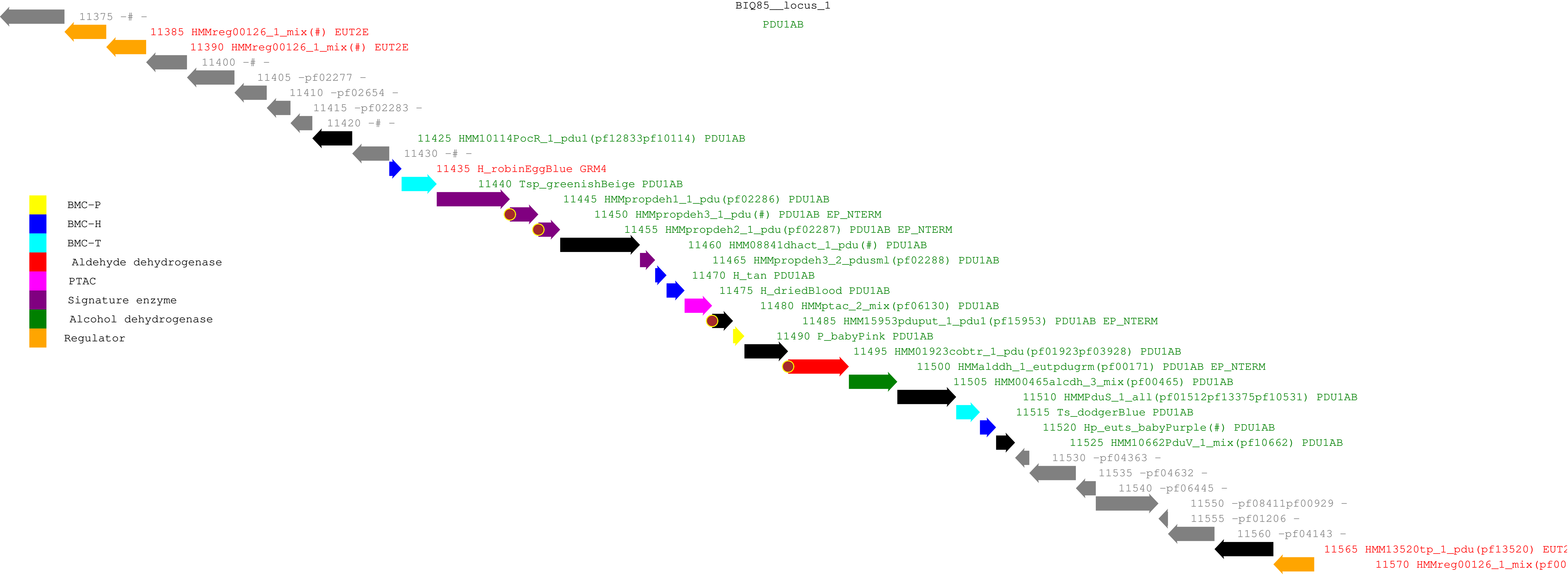

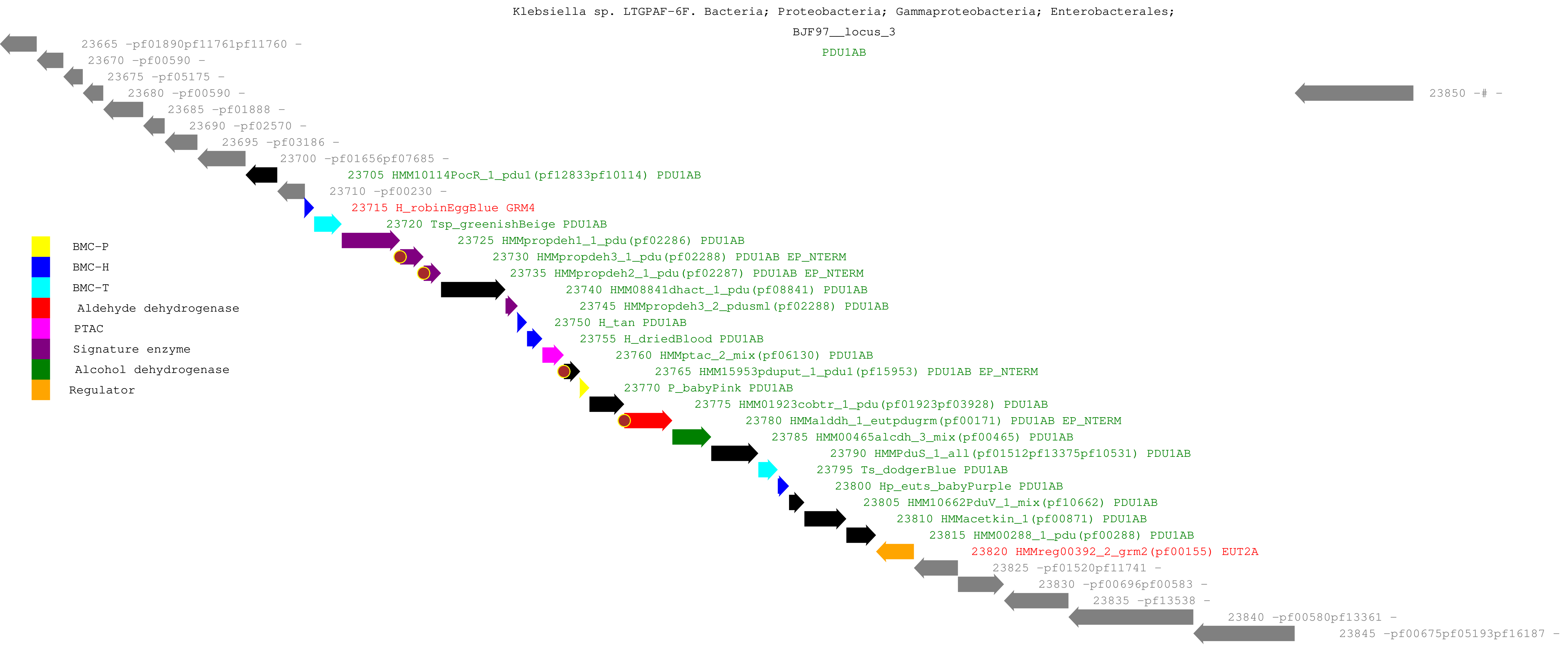
Klebsiella sp. LTGPAF-6F. Bacteria; Proteobacteria; Gammaproteobacteria; Enterobacterales;
BJF97__locus_3
PDU1AB
23665 -pf01890pf11761pf11760 -
23670 -pf00590 -
23675 -pf05175 -
23680 -pf00590 -
23685 -pf01888 -
23690 -pf02570 -
23695 -pf03186 -
23700 -pf01656pf07685 -
23705 HMM10114PocR_1_pdu1(pf12833pf10114) PDU1AB
23710 -pf00230 -
23715 H_robinEggBlue GRM4
23720 Tsp_greenishBeige PDU1AB
23725 HMMpropdeh1_1_pdu(pf02286) PDU1AB
23730 HMMpropdeh3_1_pdu(pf02288) PDU1AB EP_NTERM
23735 HMMpropdeh2_1_pdu(pf02287) PDU1AB EP_NTERM
23740 HMM08841dhact_1_pdu(pf08841) PDU1AB
23745 HMMpropdeh3_2_pdusml(pf02288) PDU1AB
23750 H_tan PDU1AB
23755 H_driedBlood PDU1AB
23760 HMMptac_2_mix(pf06130) PDU1AB
23765 HMM15953pduput_1_pdu1(pf15953) PDU1AB EP_NTERM
23770 P_babyPink PDU1AB
23775 HMM01923cobtr_1_pdu(pf01923pf03928) PDU1AB
23780 HMMalddh_1_eutpdugrm(pf00171) PDU1AB EP_NTERM
23785 HMM00465alcdh_3_mix(pf00465) PDU1AB
23790 HMMPduS_1_all(pf01512pf13375pf10531) PDU1AB
23795 Ts_dodgerBlue PDU1AB
23800 Hp_euts_babyPurple PDU1AB
23805 HMM10662PduV_1_mix(pf10662) PDU1AB
23810 HMMacetkin_1(pf00871) PDU1AB
23815 HMM00288_1_pdu(pf00288) PDU1AB
23820 HMMreg00392_2_grm2(pf00155) EUT2A
23825 -pf01520pf11741 -
23830 -pf00696pf00583 -
23835 -pf13538 -
23840 -pf00580pf13361 -
23845 -pf00675pf05193pf16187 -
23850 -# -
BMC-P
BMC-H
BMC-T
Aldehyde dehydrogenase
PTAC
Signature enzyme
Alcohol dehydrogenase
Regulator

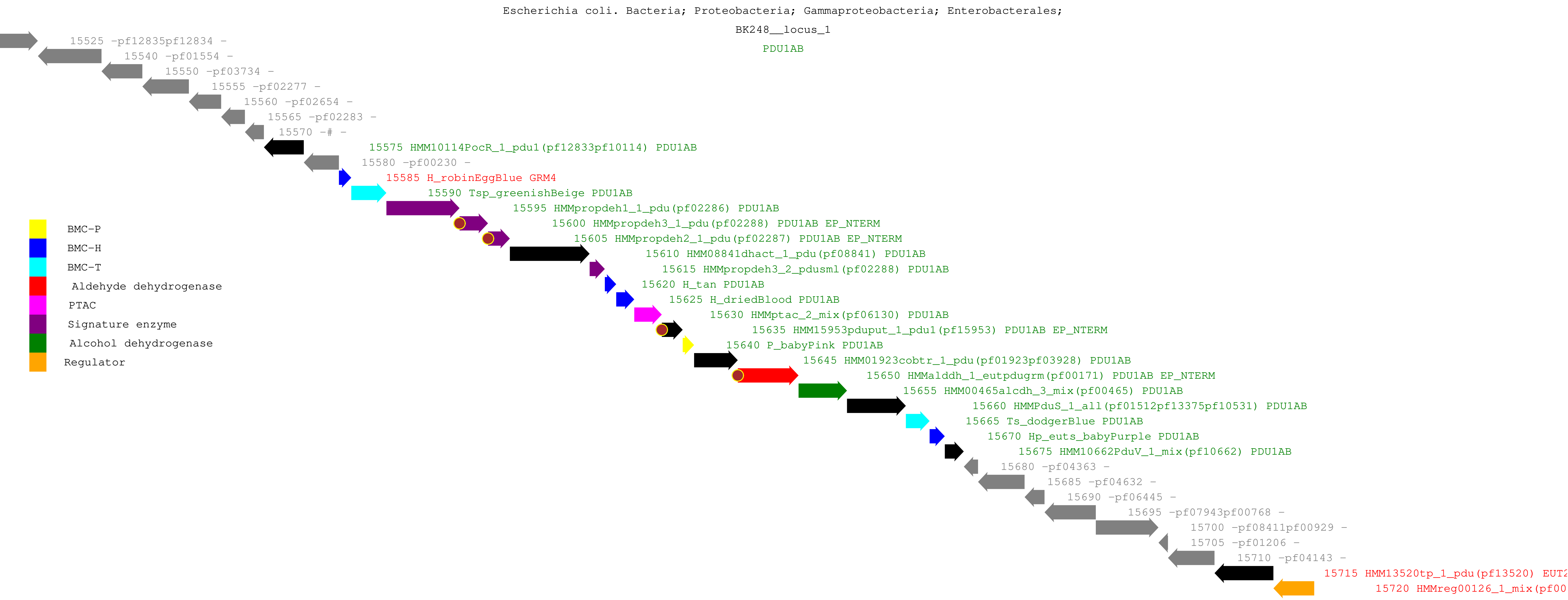

Escherichia coli. Bacteria; Proteobacteria; Gammaproteobacteria; Enterobacterales;
BK248__locus_1
PDU1AB
15525 -pf12835pf12834 -
15540 -pf01554 -
15550 -pf03734 -
15555 -pf02277 -
15560 -pf02654 -
15565 -pf02283 -
15570 -# -
15575 HMM10114PocR_1_pdu1(pf12833pf10114) PDU1AB
15580 -pf00230 -
15585 H_robinEggBlue GRM4
15590 Tsp_greenishBeige PDU1AB
15595 HMMpropdeh1_1_pdu(pf02286) PDU1AB
15600 HMMpropdeh3_1_pdu(pf02288) PDU1AB EP_NTERM
15605 HMMpropdeh2_1_pdu(pf02287) PDU1AB EP_NTERM
15610 HMM08841dhact_1_pdu(pf08841) PDU1AB
15615 HMMpropdeh3_2_pdusml(pf02288) PDU1AB
15620 H_tan PDU1AB
15625 H_driedBlood PDU1AB
15630 HMMptac_2_mix(pf06130) PDU1AB
15635 HMM15953pduput_1_pdu1(pf15953) PDU1AB EP_NTERM
15640 P_babyPink PDU1AB
15645 HMM01923cobtr_1_pdu(pf01923pf03928) PDU1AB
15650 HMMalddh_1_eutpdugrm(pf00171) PDU1AB EP_NTERM
15655 HMM00465alcdh_3_mix(pf00465) PDU1AB
15660 HMMPduS_1_all(pf01512pf13375pf10531) PDU1AB
15665 Ts_dodgerBlue PDU1AB
15670 Hp_euts_babyPurple PDU1AB
15675 HMM10662PduV_1_mix(pf10662) PDU1AB
15680 -pf04363 -
15685 -pf04632 -
15690 -pf06445 -
15695 -pf07943pf00768 -
15700 -pf08411pf00929 -
15705 -pf01206 -
15710 -pf04143 -
15715 HMM13520tp_1_pdu(pf13520) EUT2
15720 HMMreg00126_1_mix(pf00
BMC-P
BMC-H
BMC-T
Aldehyde dehydrogenase
PTAC
Signature enzyme
Alcohol dehydrogenase
Regulator

Escherichia coli. Bacteria; Proteobacteria; Gammaproteobacteria; Enterobacterales;

BK334__locus_1

PDU1AB

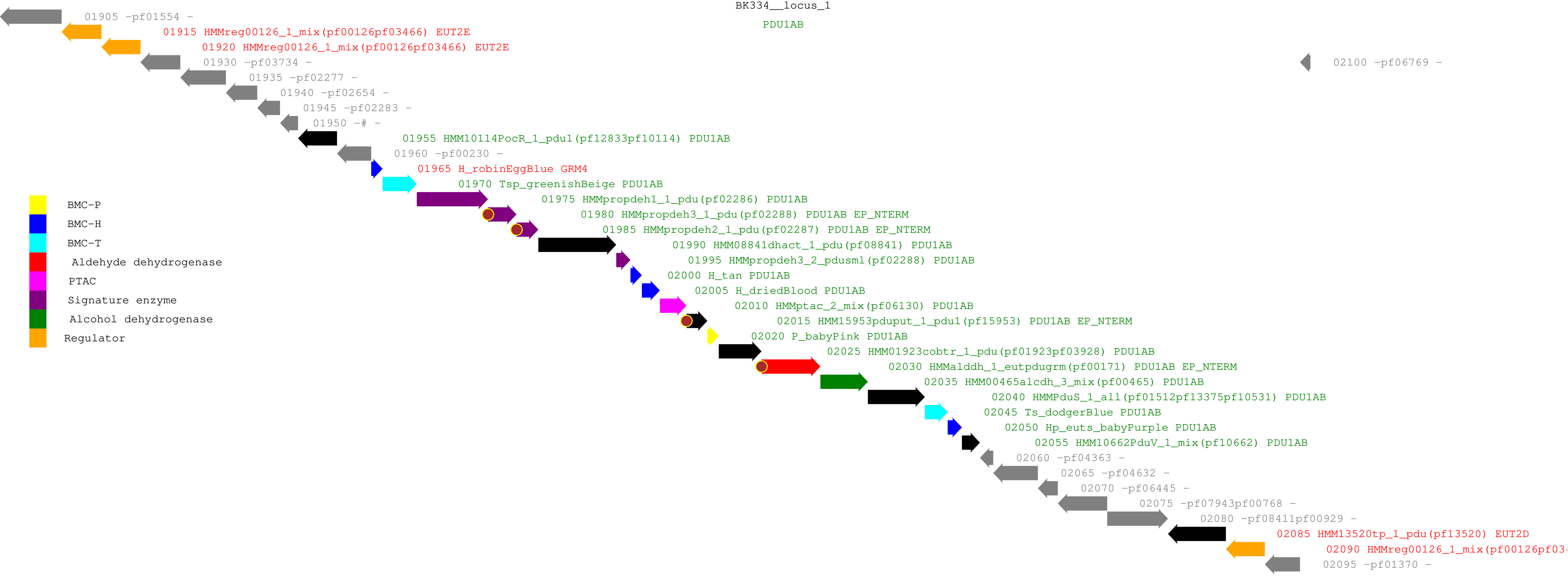

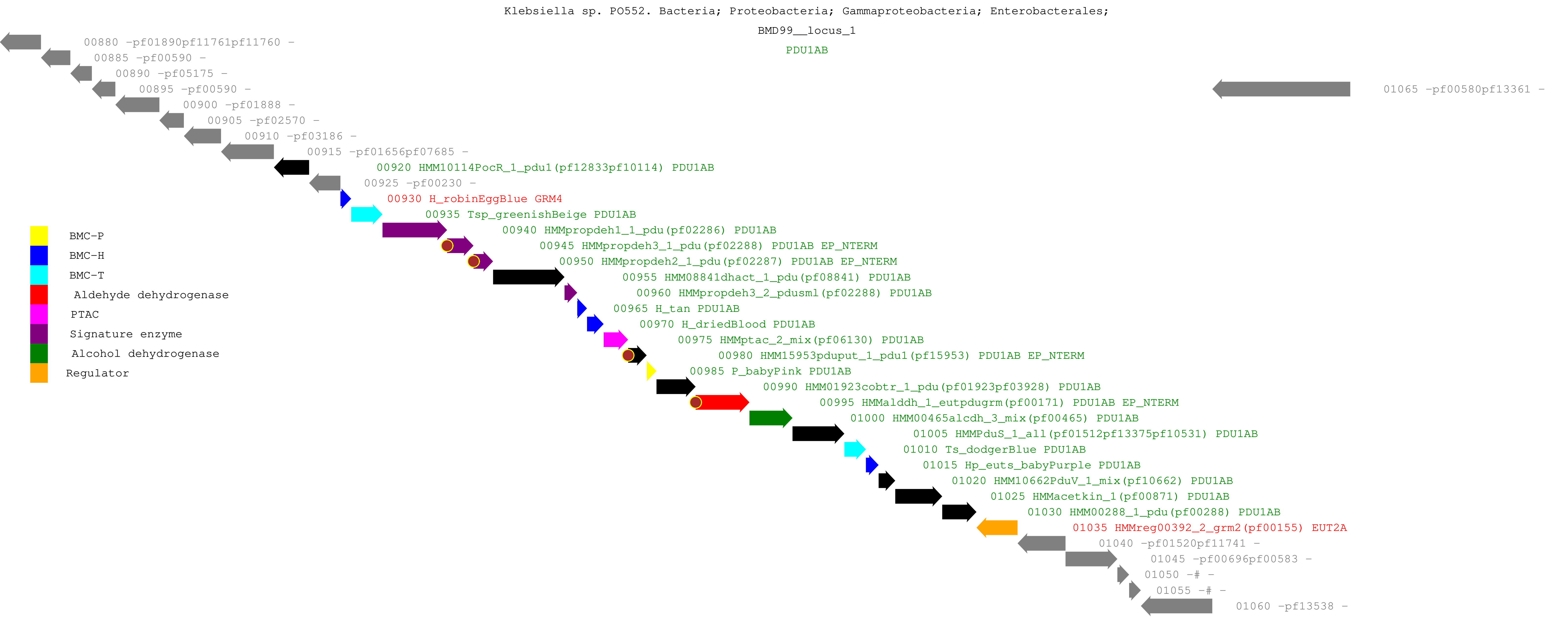
Klebsiella sp. PO552. Bacteria; Proteobacteria; Gammaproteobacteria; Enterobacterales;
BMD99__locus_1
PDU1AB
00880 -pf01890pf11761pf11760 -
00885 -pf00590 -
00890 -pf05175 -
00895 -pf00590 -
00900 -pf01888 -
00905 -pf02570 -
00910 -pf03186 -
00915 -pf01656pf07685 -
00920 HMM10114PocR_1_pdu1(pf12833pf10114) PDU1AB
00925 -pf00230 -
00930 H_robinEggBlue GRM4
00935 Tsp_greenishBeige PDU1AB
00940 HMMpropdeh1_1_pdu(pf02286) PDU1AB
00945 HMMpropdeh3_1_pdu(pf02288) PDU1AB EP_NTERM
00950 HMMpropdeh2_1_pdu(pf02287) PDU1AB EP_NTERM
00955 HMM08841dhact_1_pdu(pf08841) PDU1AB
00960 HMMpropdeh3_2_pdusml(pf02288) PDU1AB
00965 H_tan PDU1AB
00970 H_driedBlood PDU1AB
00975 HMMptac_2_mix(pf06130) PDU1AB
00980 HMM15953pduput_1_pdu1(pf15953) PDU1AB EP_NTERM
00985 P_babyPink PDU1AB
00990 HMM01923cobtr_1_pdu(pf01923pf03928) PDU1AB
00995 HMMalddh_1_eutpdugrm(pf00171) PDU1AB EP_NTERM
01000 HMM00465alcdh_3_mix(pf00465) PDU1AB
01005 HMMPduS_1_all(pf01512pf13375pf10531) PDU1AB
01010 Ts_dodgerBlue PDU1AB
01015 Hp_euts_babyPurple PDU1AB
01020 HMM10662PduV_1_mix(pf10662) PDU1AB
01025 HMMacetkin_1(pf00871) PDU1AB
01030 HMM00288_1_pdu(pf00288) PDU1AB
01035 HMMreg00392_2_grm2(pf00155) EUT2A
01040 -pf01520pf11741 -
01045 -pf00696pf00583 -
01050 -# -
01055 -# -
01060 -pf13538 -
01065 -pf00580pf13361 -
BMC-P
BMC-H
BMC-T
Aldehyde dehydrogenase
PTAC
Signature enzyme
Alcohol dehydrogenase
Regulator

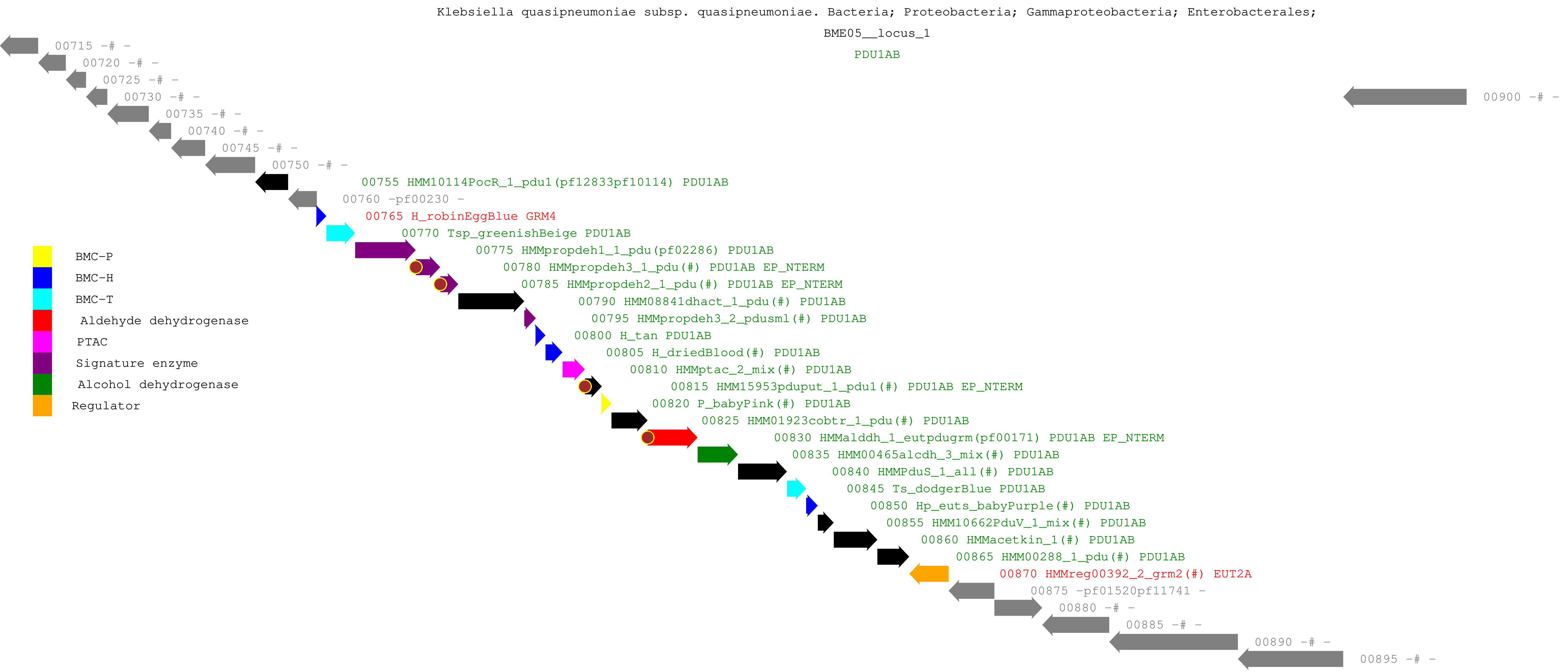
Klebsiella quasipneumoniae subsp. quasipneumoniae. Bacteria; Proteobacteria; Gammaproteobacteria; Enterobacterales;
BME05__locus_1
PDU1AB
00715 -# -
00720 -# -
00725 -# -
00730 -# -
00735 -# -
00740 -# -
00745 -# -
00750 -# -
00755 HMM10114PocR_1_pdu1(pf12833pf10114) PDU1AB
00760 -pf00230 -
00765 H_robinEggBlue GRM4
00770 Tsp_greenishBeige PDU1AB
00775 HMMpropdeh1_1_pdu(pf02286) PDU1AB
00780 HMMpropdeh3_1_pdu(#) PDU1AB EP_NTERM
00785 HMMpropdeh2_1_pdu(#) PDU1AB EP_NTERM
00790 HMM08841dhact_1_pdu(#) PDU1AB
00795 HMMpropdeh3_2_pdusml(#) PDU1AB
00800 H_tan PDU1AB
00805 H_driedBlood(#) PDU1AB
00810 HMMptac_2_mix(#) PDU1AB
00815 HMM15953pduput_1_pdu1(#) PDU1AB EP_NTERM
00820 P_babyPink(#) PDU1AB
00825 HMM01923cobtr_1_pdu(#) PDU1AB
00830 HMMalddh_1_eutpdugrm(pf00171) PDU1AB EP_NTERM
00835 HMM00465alcdh_3_mix(#) PDU1AB
00840 HMMPduS_1_all(#) PDU1AB
00845 Ts_dodgerBlue PDU1AB
00850 Hp_euts_babyPurple(#) PDU1AB
00855 HMM10662PduV_1_mix(#) PDU1AB
00860 HMMacetkin_1(#) PDU1AB
00865 HMM00288_1_pdu(#) PDU1AB
00870 HMMreg00392_2_grm2(#) EUT2A
00875 -pf01520pf11741 -
00880 -# -
00885 -# -
00890 -# -
00895 -# -
00900 -# -
BMC-P
BMC-H
BMC-T
Aldehyde dehydrogenase
PTAC
Signature enzyme
Alcohol dehydrogenase
Regulator

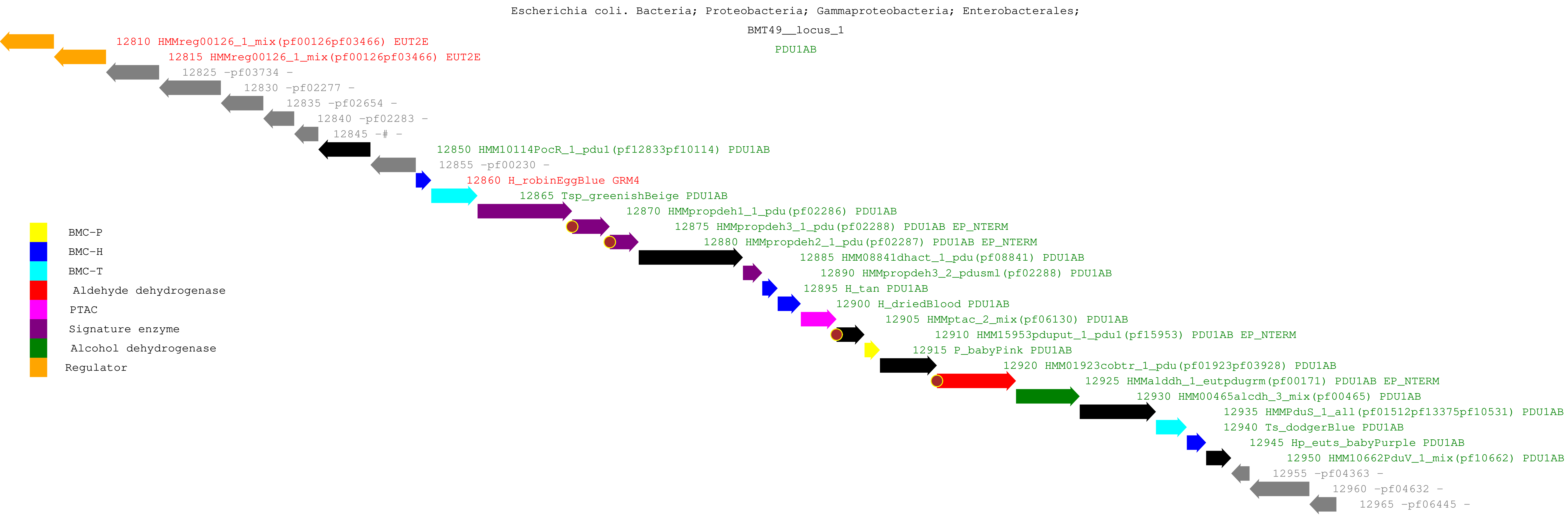

Escherichia coli. Bacteria; Proteobacteria; Gammaproteobacteria; Enterobacterales;
BMT49__locus_1
PDU1AB
12810 HMMreg00126_1_mix(pf00126pf03466) EUT2E
12815 HMMreg00126_1_mix(pf00126pf03466) EUT2E
12825 -pf03734 -
12830 -pf02277 -
12835 -pf02654 -
12840 -pf02283 -
12845 -# -
12850 HMM10114PocR_1_pdu1(pf12833pf10114) PDU1AB
12855 -pf00230 -
12860 H_robinEggBlue GRM4
12865 Tsp_greenishBeige PDU1AB
12870 HMMpropdeh1_1_pdu(pf02286) PDU1AB
12875 HMMpropdeh3_1_pdu(pf02288) PDU1AB EP_NTERM
12880 HMMpropdeh2_1_pdu(pf02287) PDU1AB EP_NTERM
12885 HMM08841dhact_1_pdu(pf08841) PDU1AB
12890 HMMpropdeh3_2_pdusml(pf02288) PDU1AB
12895 H_tan PDU1AB
12900 H_driedBlood PDU1AB
12905 HMMptac_2_mix(pf06130) PDU1AB
12910 HMM15953pduput_1_pdu1(pf15953) PDU1AB EP_NTERM
12915 P_babyPink PDU1AB
12920 HMM01923cobtr_1_pdu(pf01923pf03928) PDU1AB
12925 HMMalddh_1_eutpdugrm(pf00171) PDU1AB EP_NTERM
12930 HMM00465alcdh_3_mix(pf00465) PDU1AB
12935 HMMPduS_1_all(pf01512pf13375pf10531) PDU1AB
12940 Ts_dodgerBlue PDU1AB
12945 Hp_euts_babyPurple PDU1AB
12950 HMM10662PduV_1_mix(pf10662) PDU1AB
12955 -pf04363 -
12960 -pf04632 -
12965 -pf06445 -
BMC-P
BMC-H
BMC-T
Aldehyde dehydrogenase
PTAC
Signature enzyme
Alcohol dehydrogenase
Regulator

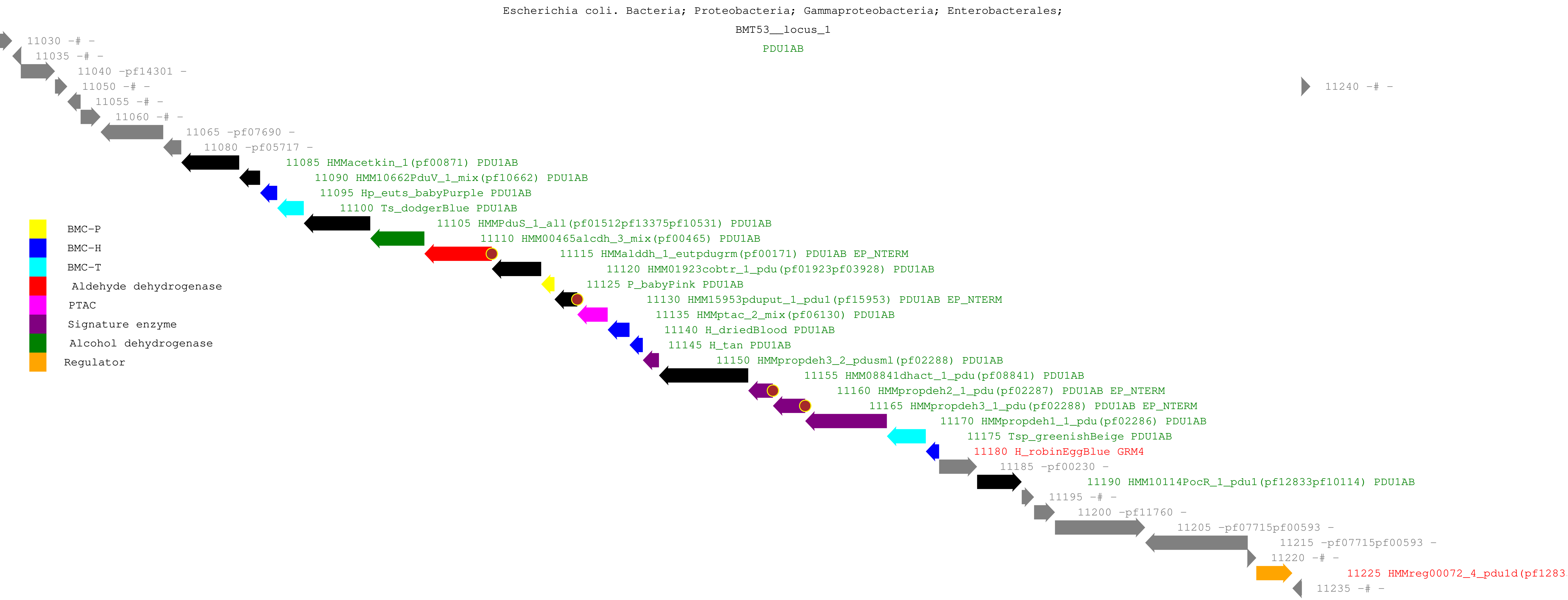
Escherichia coli. Bacteria; Proteobacteria; Gammaproteobacteria; Enterobacterales;
BMT53__locus_1
PDU1AB
11030 -# -
11035 -# -
11040 -pf14301 -
11050 -# -
11055 -# -
11060 -# -
11065 -pf07690 -
11080 -pf05717 -
11085 HMMacetkin_1(pf00871) PDU1AB
11090 HMM10662PduV_1_mix(pf10662) PDU1AB
11095 Hp_euts_babyPurple PDU1AB
11100 Ts_dodgerBlue PDU1AB
11105 HMMPduS_1_all(pf01512pf13375pf10531) PDU1AB
11110 HMM00465alcdh_3_mix(pf00465) PDU1AB
11115 HMMaldh_1_eutpdugrm(pf00171) PDU1AB EP_NTERM
11120 HMM01923cobtr_1_pdu(pf01923pf03928) PDU1AB
11125 P_babyPink PDU1AB
11130 HMM15953pduput_1_pdu1(pf15953) PDU1AB EP_NTERM
11135 HMMptac_2_mix(pf06130) PDU1AB
11140 H_driedBlood PDU1AB
11145 H_tan PDU1AB
11150 HMMpropdeh3_2_pdusml(pf02288) PDU1AB
11155 HMM08841dhact_1_pdu(pf08841) PDU1AB
11160 HMMpropdeh2_1_pdu(pf02287) PDU1AB EP_NTERM
11165 HMMpropdeh3_1_pdu(pf02288) PDU1AB EP_NTERM
11170 HMMpropdeh1_1_pdu(pf02286) PDU1AB
11175 Tsp_greenishBeige PDU1AB
11180 H_robinEggBlue GRM4
11185 -pf00230 -
11190 HMM10114PocR_1_pdu1(pf12833pf10114) PDU1AB
11195 -# -
11200 -pf11760 -
11205 -pf07715pf00593 -
11215 -pf07715pf00593 -
11220 -# -
11225 HMMreg00072_4_pdu1d(pf1283
11235 -# -
11240 -# -
BMC-P
BMC-H
BMC-T
Aldehyde dehydrogenase
PTAC
Signature enzyme
Alcohol dehydrogenase
Regulator

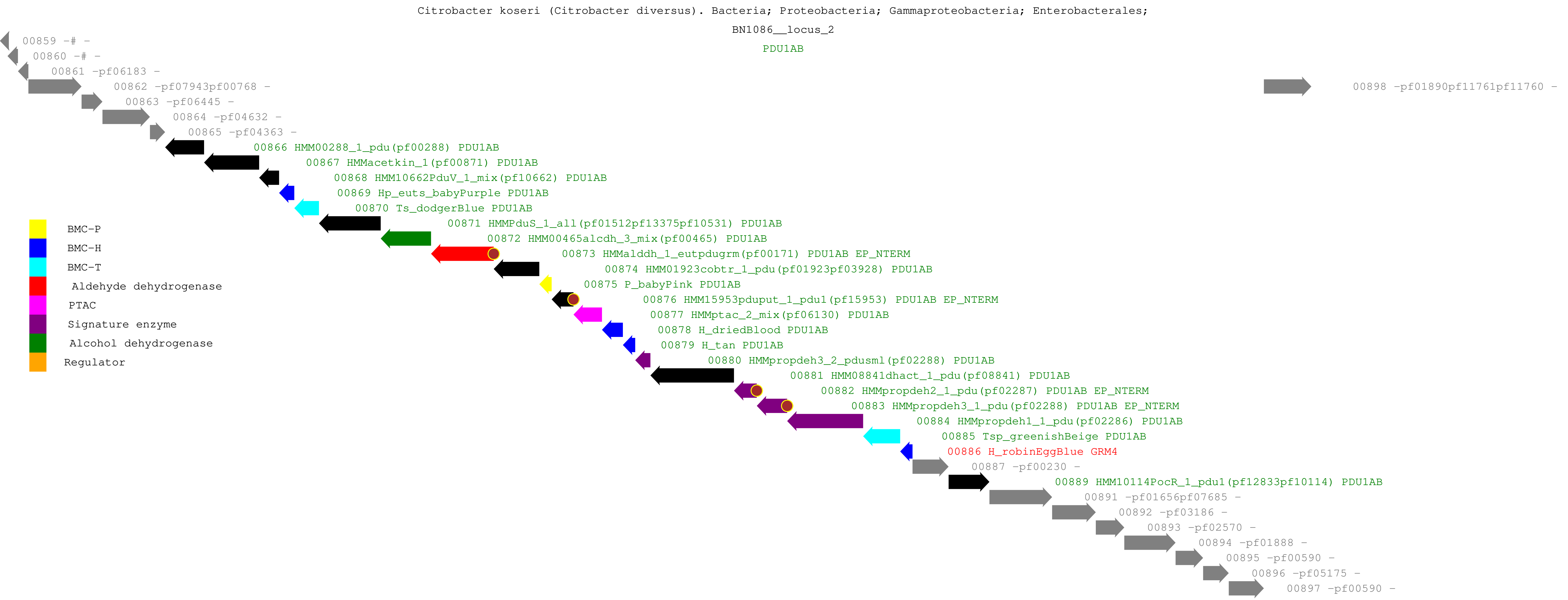
Citrobacter koseri (Citrobacter diversus). Bacteria; Proteobacteria; Gammaproteobacteria; Enterobacterales;
BN1086__locus_2
PDU1AB
00859 -# -
00860 -# -
00861 -pf06183 -
00862 -pf07943pf00768 -
00863 -pf06445 -
00864 -pf04632 -
00865 -pf04363 -
00866 HMM00288_1_pdu(pf00288) PDU1AB
00867 HMMacetkin_1(pf00871) PDU1AB
00868 HMM10662PduV_1_mix(pf10662) PDU1AB
00869 Hp_euts_babyPurple PDU1AB
00870 Ts_dodgerBlue PDU1AB
00871 HMMPduS_1_all(pf01512pf13375pf10531) PDU1AB
00872 HMM00465alcdh_3_mix(pf00465) PDU1AB
00873 HMMalddh_1_eutpdugrm(pf00171) PDU1AB EP_NTERM
00874 HMM01923cobtr_1_pdu(pf01923pf03928) PDU1AB
00875 P_babyPink PDU1AB
00876 HMM15953pduput_1_pdu1(pf15953) PDU1AB EP_NTERM
00877 HMMptac_2_mix(pf06130) PDU1AB
00878 H_driedBlood PDU1AB
00879 H_tan PDU1AB
00880 HMMpropdeh3_2_pdusml(pf02288) PDU1AB
00881 HMM08841dhact_1_pdu(pf08841) PDU1AB
00882 HMMpropdeh2_1_pdu(pf02287) PDU1AB EP_NTERM
00883 HMMpropdeh3_1_pdu(pf02288) PDU1AB EP_NTERM
00884 HMMpropdeh1_1_pdu(pf02286) PDU1AB
00885 Tsp_greenishBeige PDU1AB
00886 H_robinEggBlue GRM4
00887 -pf00230 -
00889 HMM10114PocR_1_pdu1(pf12833pf10114) PDU1AB
00891 -pf01656pf07685 -
00892 -pf03186 -
00893 -pf02570 -
00894 -pf01888 -
00895 -pf00590 -
00896 -pf05175 -
00897 -pf00590 -
00898 -pf01890pf11761pf11760 -
BMC-P
BMC-H
BMC-T
Aldehyde dehydrogenase
PTAC
Signature enzyme
Alcohol dehydrogenase
Regulator

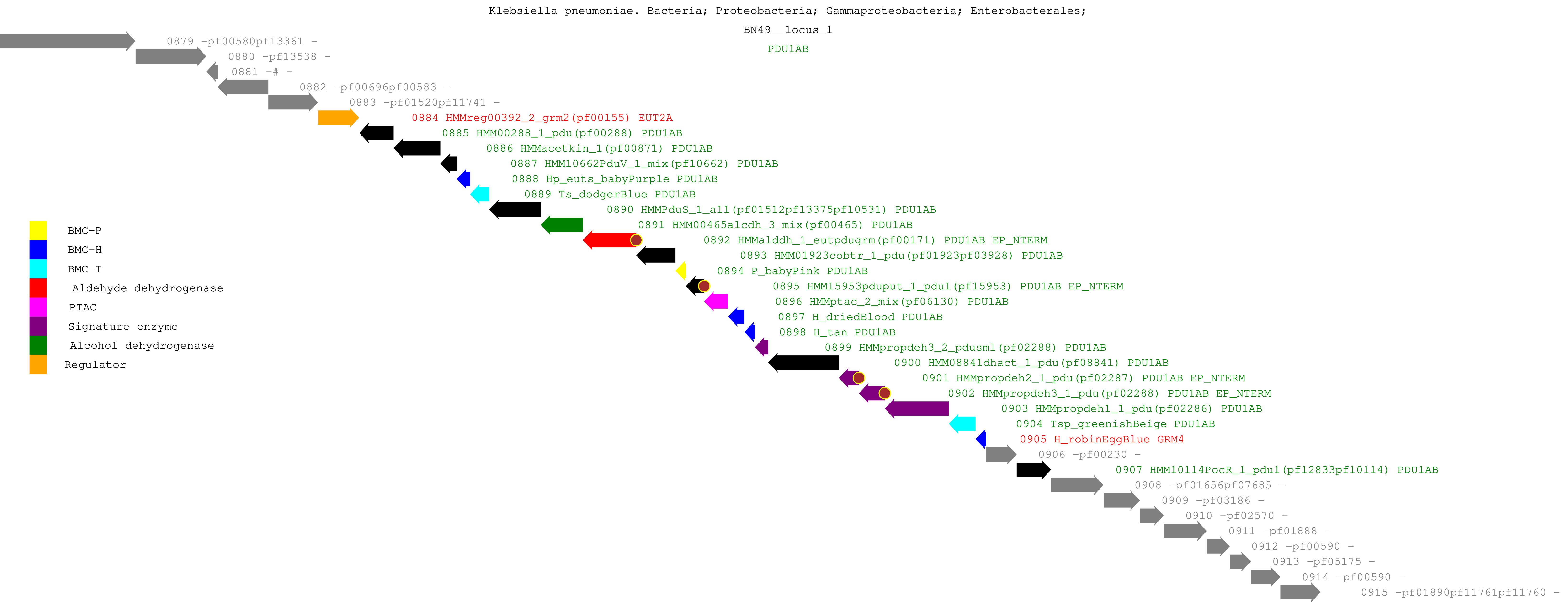

Klebsiella pneumoniae. Bacteria; Proteobacteria; Gammaproteobacteria; Enterobacterales;
BN49__locus_1
PDU1AB
0879 -pf00580pf13361 -
0880 -pf13538 -
0881 -# -
0882 -pf00696pf00583 -
0883 -pf01520pf11741 -
0884 HMMreg00392_2_grm2(pf00155) EUT2A
0885 HMM00288_1_pdu(pf00288) PDU1AB
0886 HMMacetkin_1(pf00871) PDU1AB
0887 HMM10662PduV_1_mix(pf10662) PDU1AB
0888 Hp_euts_babyPurple PDU1AB
0889 Ts_dodgerBlue PDU1AB
0890 HMMPduS_1_all(pf01512pf13375pf10531) PDU1AB
0891 HMM00465alcdh_3_mix(pf00465) PDU1AB
0892 HMMalddh_1_eutpdugrm(pf00171) PDU1AB EP_NTERM
0893 HMM01923cobtr_1_pdu(pf01923pf03928) PDU1AB
0894 P_babyPink PDU1AB
0895 HMM15953pduput_1_pdu1(pf15953) PDU1AB EP_NTERM
0896 HMMptac_2_mix(pf06130) PDU1AB
0897 H_driedBlood PDU1AB
0898 H_tan PDU1AB
0899 HMMpropdeh3_2_pdusml(pf02288) PDU1AB
0900 HMM08841dhact_1_pdu(pf08841) PDU1AB
0901 HMMpropdeh2_1_pdu(pf02287) PDU1AB EP_NTERM
0902 HMMpropdeh3_1_pdu(pf02288) PDU1AB EP_NTERM
0903 HMMpropdeh1_1_pdu(pf02286) PDU1AB
0904 Tsp_greenishBeige PDU1AB
0905 H_robinEggBlue GRM4
0906 -pf00230 -
0907 HMM10114PocR_1_pdu1(pf12833pf10114) PDU1AB
0908 -pf01656pf07685 -
0909 -pf03186 -
0910 -pf02570 -
0911 -pf01888 -
0912 -pf00590 -
0913 -pf05175 -
0914 -pf00590 -
0915 -pf01890pf11761pf11760 -
BMC-P
BMC-H
BMC-T
Aldehyde dehydrogenase
PTAC
Signature enzyme
Alcohol dehydrogenase
Regulator

Escherichia coli O127:H6. Bacteria; Proteobacteria; Gammaproteobacteria; Enterobacterales;

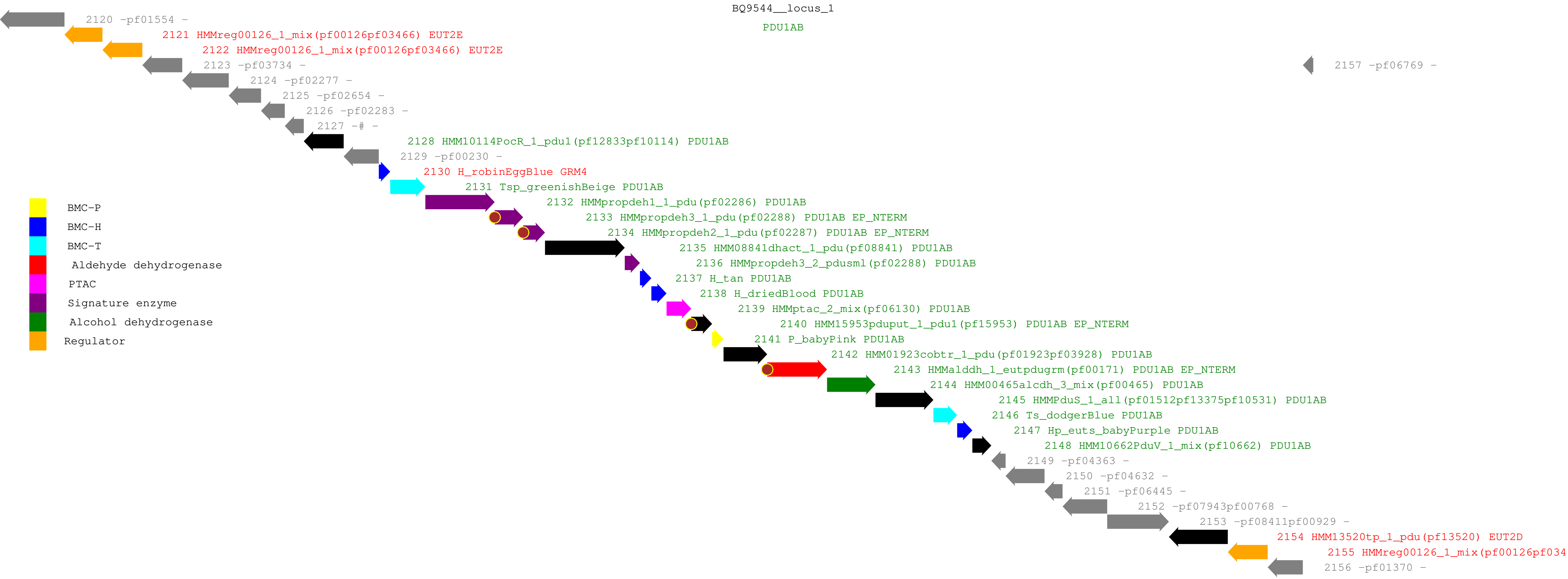

Salmonella enteritidis. Bacteria; Proteobacteria; Gammaproteobacteria; Enterobacterales;

BTN74__locus_1

PDU1AB

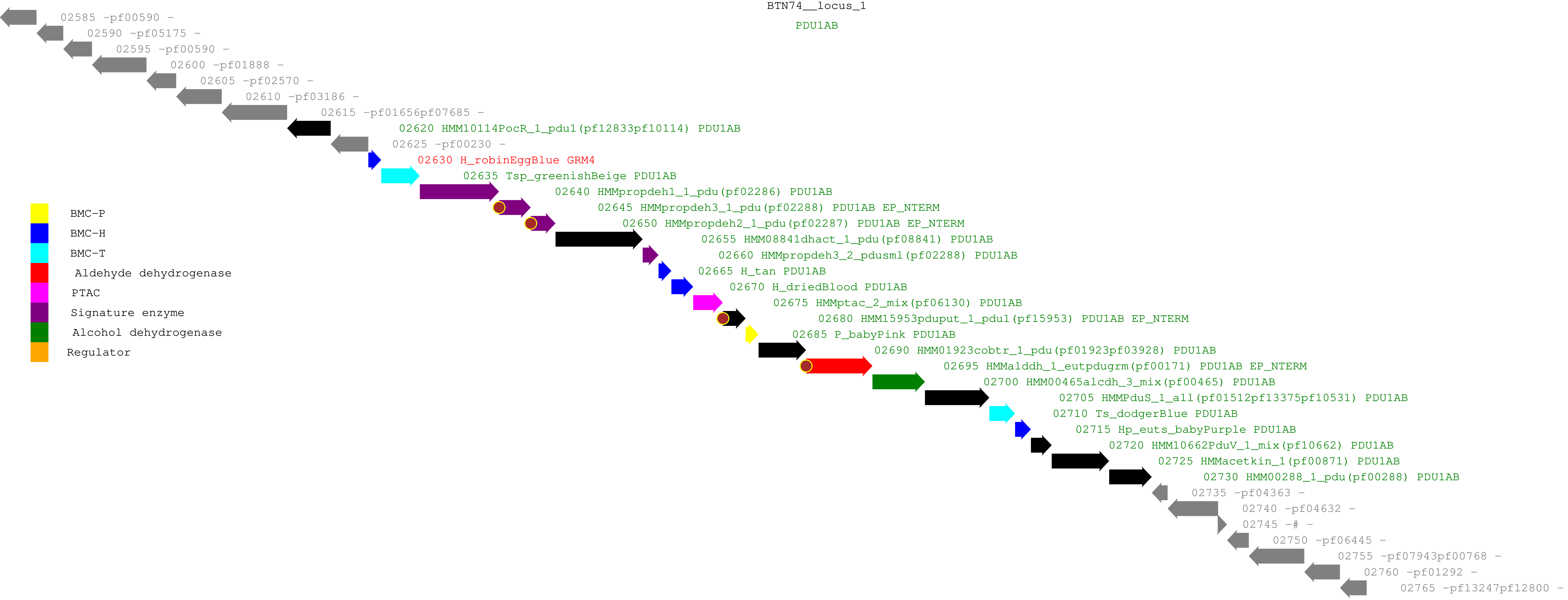

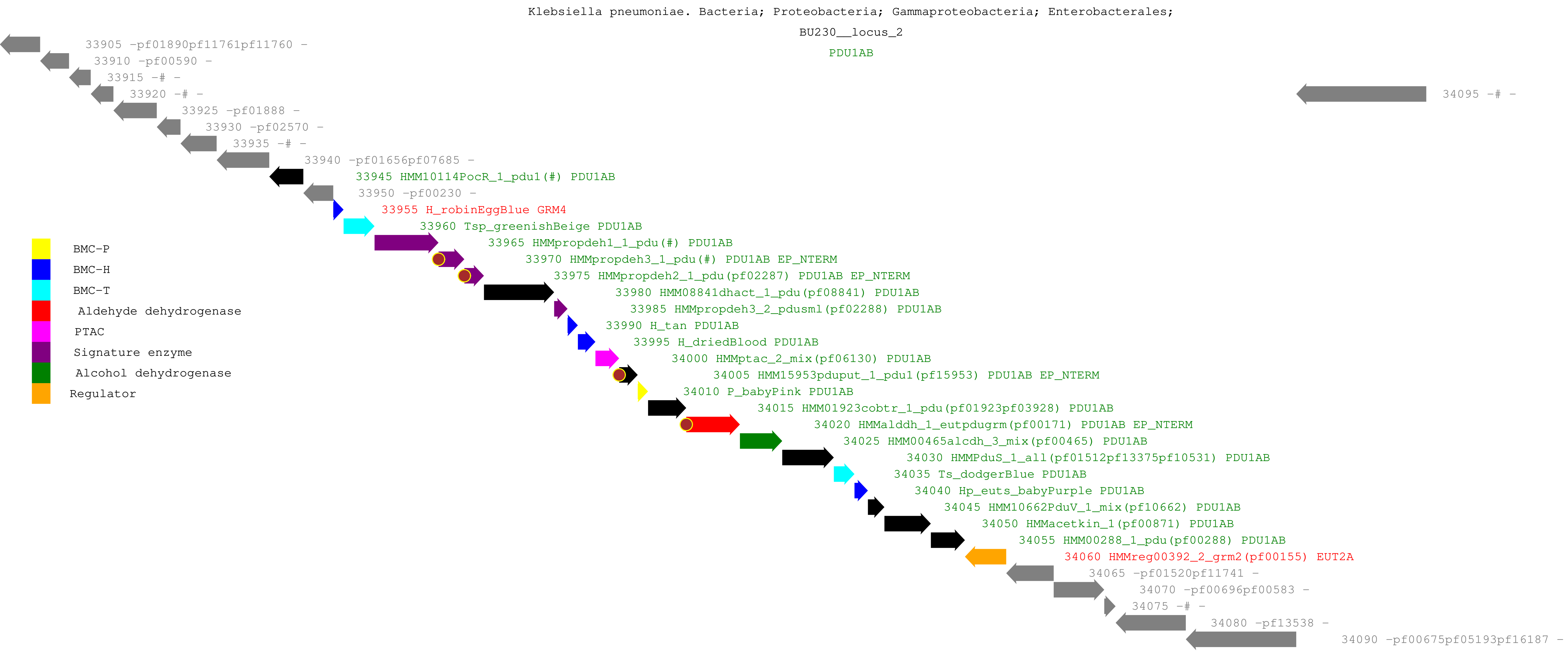
Klebsiella pneumoniae. Bacteria; Proteobacteria; Gammaproteobacteria; Enterobacterales;
BU230__locus_2
PDU1AB
33905 -pf01890pf11761pf11760 -
33910 -pf00590 -
33915 -# -
33920 -# -
33925 -pf01888 -
33930 -pf02570 -
33935 -# -
33940 -pf01656pf07685 -
33945 HMM10114PocR_1_pdu1(#) PDU1AB
33950 -pf00230 -
33955 H_robinEggBlue GRM4
33960 Tsp_greenishBeige PDU1AB
33965 HMMpropdeh1_1_pdu(#) PDU1AB
33970 HMMpropdeh3_1_pdu(#) PDU1AB EP_NTERM
33975 HMMpropdeh2_1_pdu(pf02287) PDU1AB EP_NTERM
33980 HMM08841dhact_1_pdu(pf08841) PDU1AB
33985 HMMpropdeh3_2_pdusml(pf02288) PDU1AB
33990 H_tan PDU1AB
33995 H_driedBlood PDU1AB
34000 HMMptac_2_mix(pf06130) PDU1AB
34005 HMM15953pduput_1_pdu1(pf15953) PDU1AB EP_NTERM
34010 P_babyPink PDU1AB
34015 HMM01923cobtr_1_pdu(pf01923pf03928) PDU1AB
34020 HMMalddh_1_eutpdugrm(pf00171) PDU1AB EP_NTERM
34025 HMM00465alcdh_3_mix(pf00465) PDU1AB
34030 HMMPduS_1_all(pf01512pf13375pf10531) PDU1AB
34035 Ts_dodgerBlue PDU1AB
34040 Hp_euts_babyPurple PDU1AB
34045 HMM10662PduV_1_mix(pf10662) PDU1AB
34050 HMMacetkin_1(pf00871) PDU1AB
34055 HMM00288_1_pdu(pf00288) PDU1AB
34060 HMMreg00392_2_grm2(pf00155) EUT2A
34065 -pf01520pf11741 -
34070 -pf00696pf00583 -
34075 -# -
34080 -pf13538 -
34090 -pf00675pf05193pf16187 -
34095 -# -
BMC-P
BMC-H
BMC-T
Aldehyde dehydrogenase
PTAC
Signature enzyme
Alcohol dehydrogenase
Regulator

Escherichia coli O128:H2 str. 2011C-3317. Bacteria; Proteobacteria; Gammaproteobacteria; Enterobacterales;

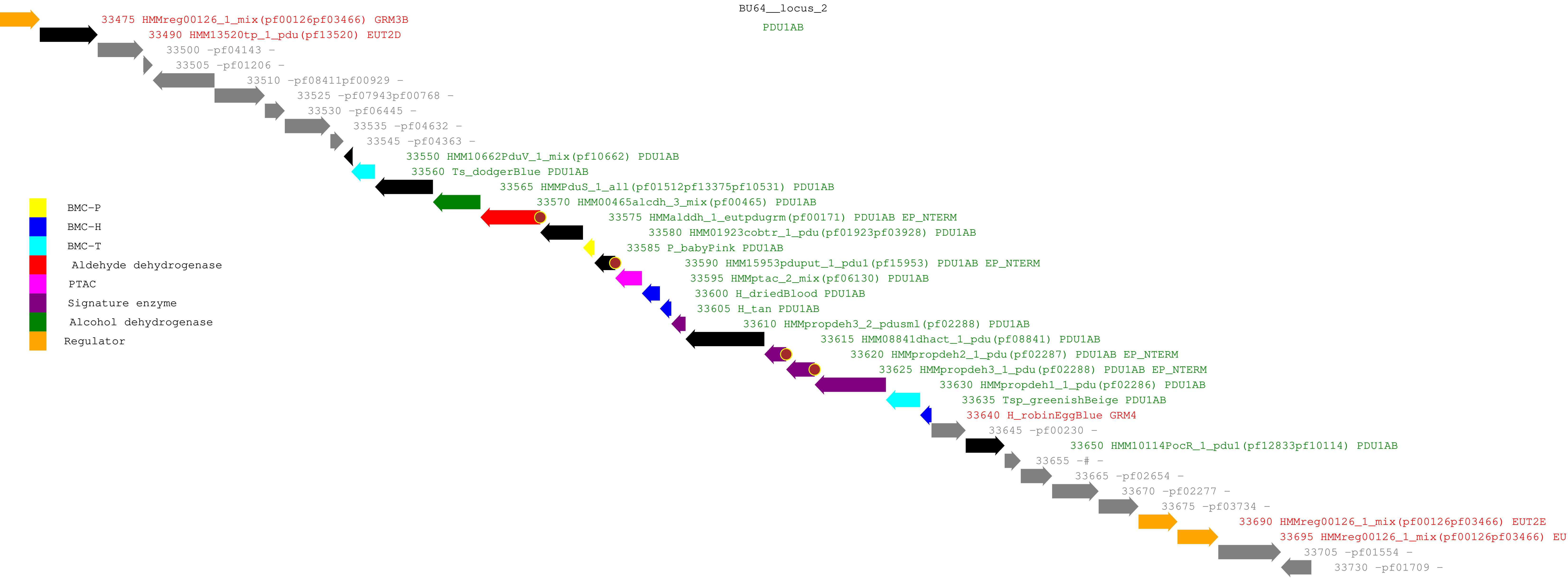

Klebsiella variicola. Bacteria; Proteobacteria; Gammaproteobacteria; Enterobacterales;

BUE65__locus_2

PDU1AB

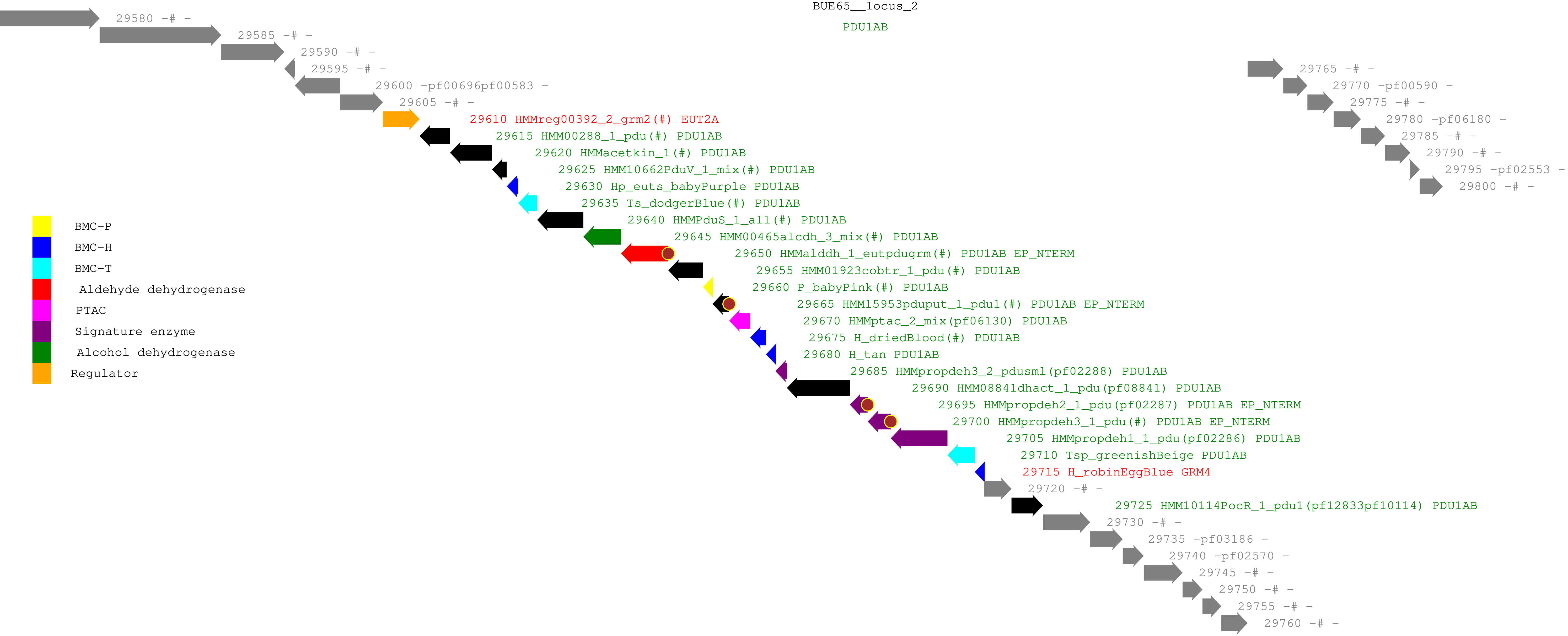

Salmonella enterica subsp. diarizonae serovar 60:r:e,n,x,z15. Bacteria; Proteobacteria; Gammaproteobacteria; Enterobacterales;

BUJ13__locus_2

PDU1AB

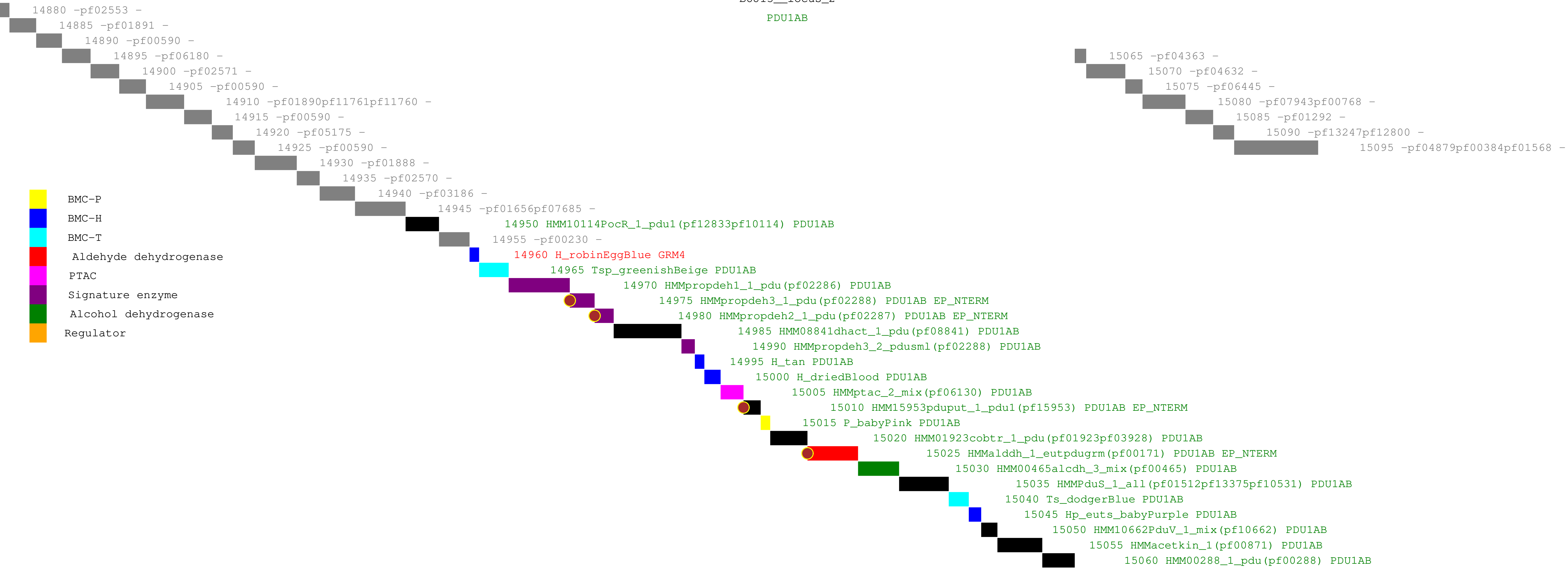

Salmonella rubislaw. Bacteria; Proteobacteria; Gammaproteobacteria; Enterobacterales;

BUJ14__locus_2

PDU1AB

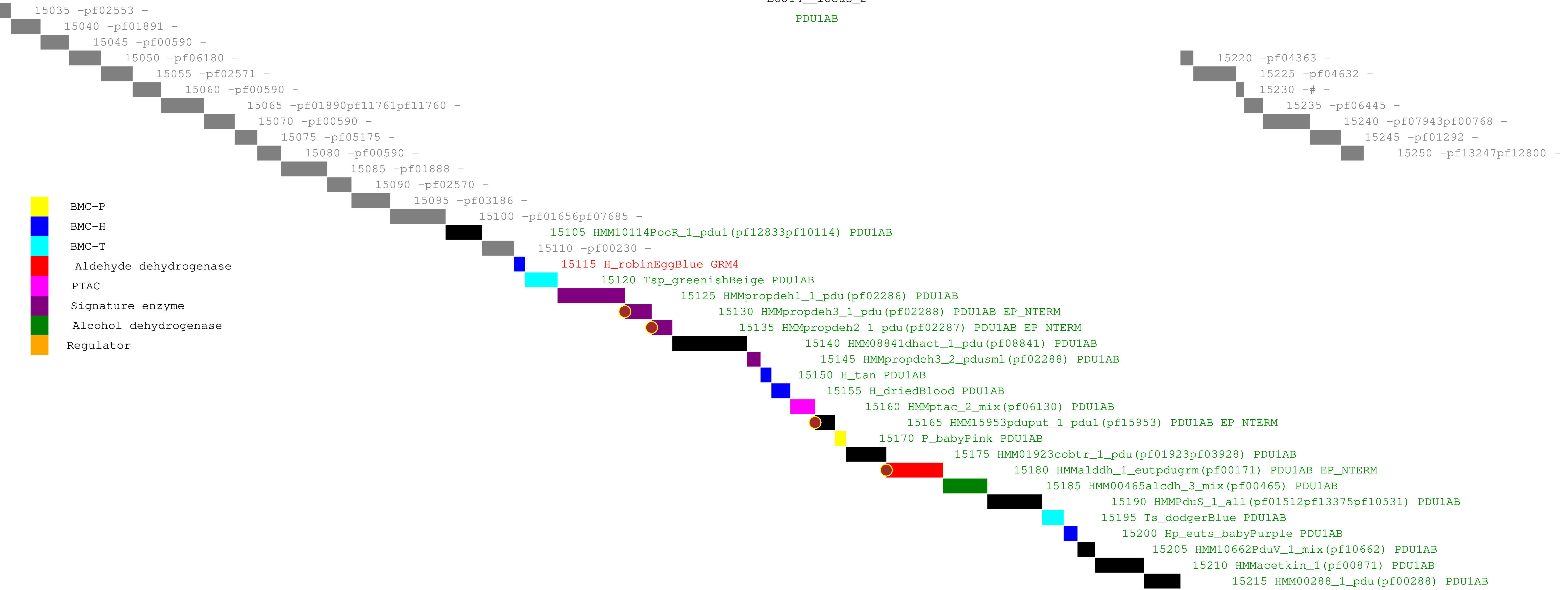

Salmonella newport. Bacteria; Proteobacteria; Gammaproteobacteria; Enterobacterales;

BUJ15__locus_2

PDU1AB

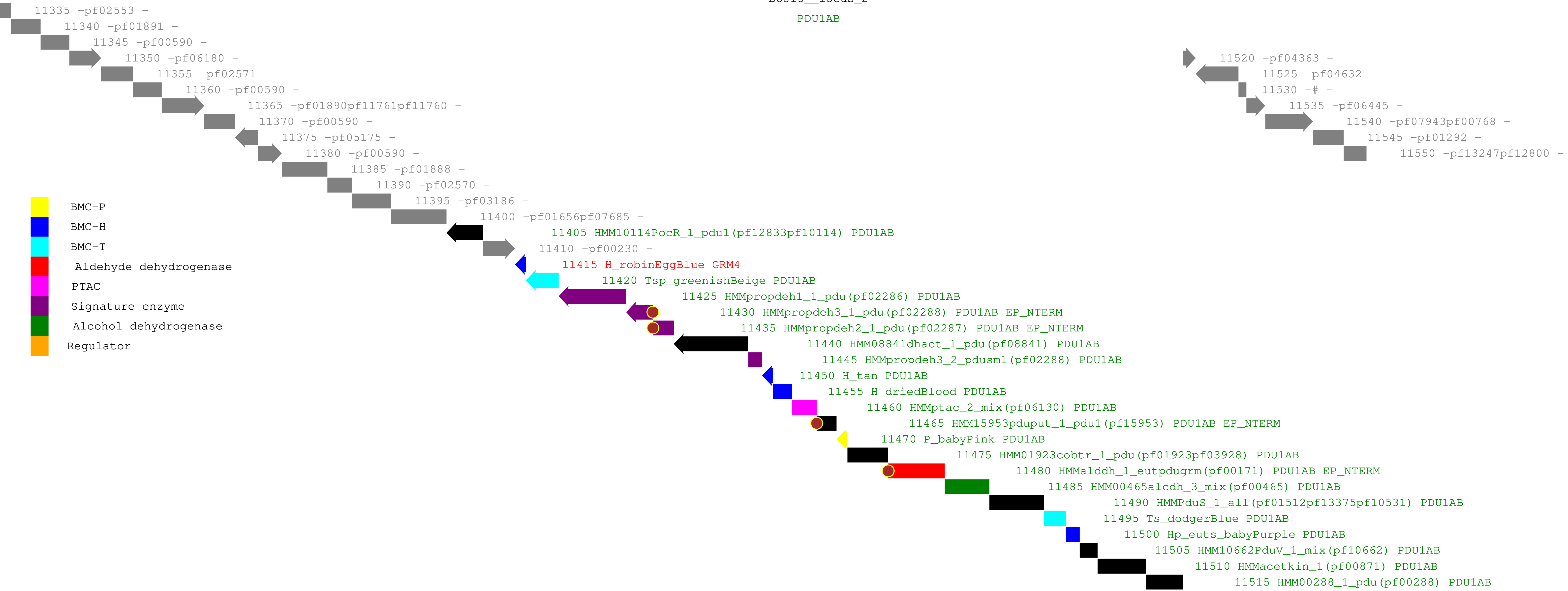

Salmonella choleraesuis. Bacteria; Proteobacteria; Gammaproteobacteria; Enterobacterales;

BUJ19__locus_2

PDU1AB

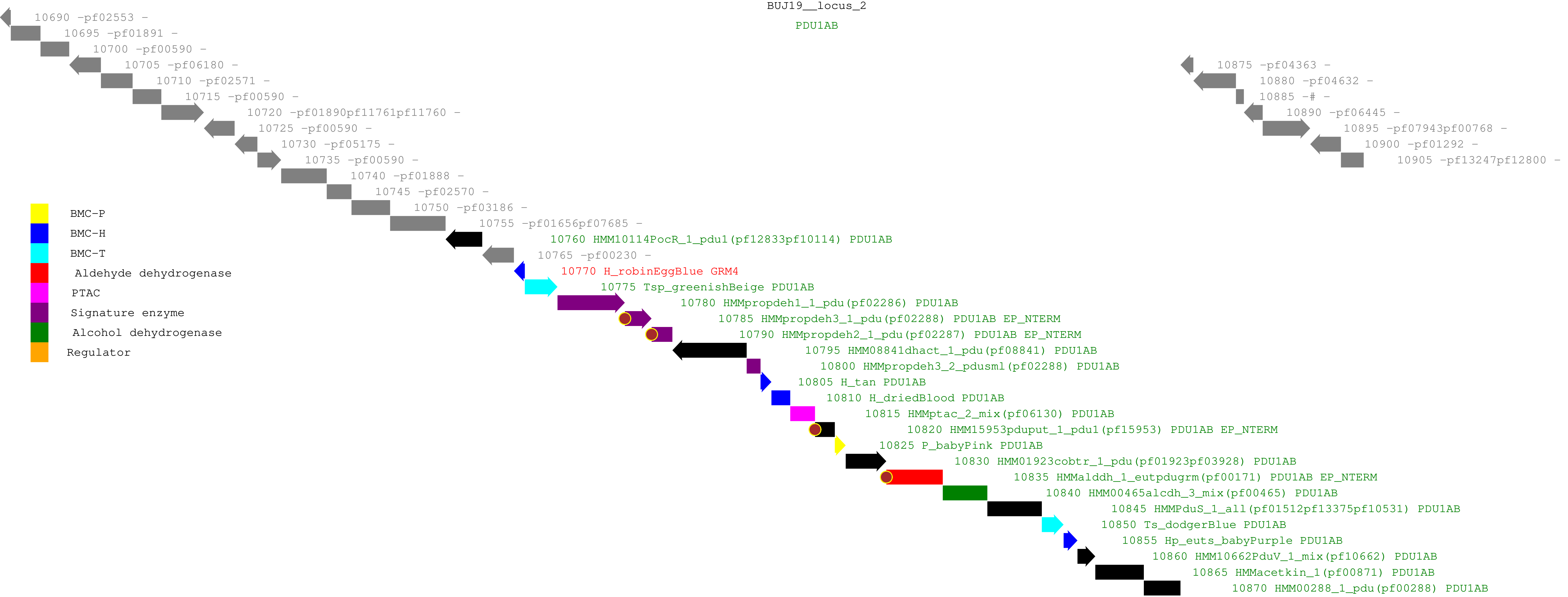

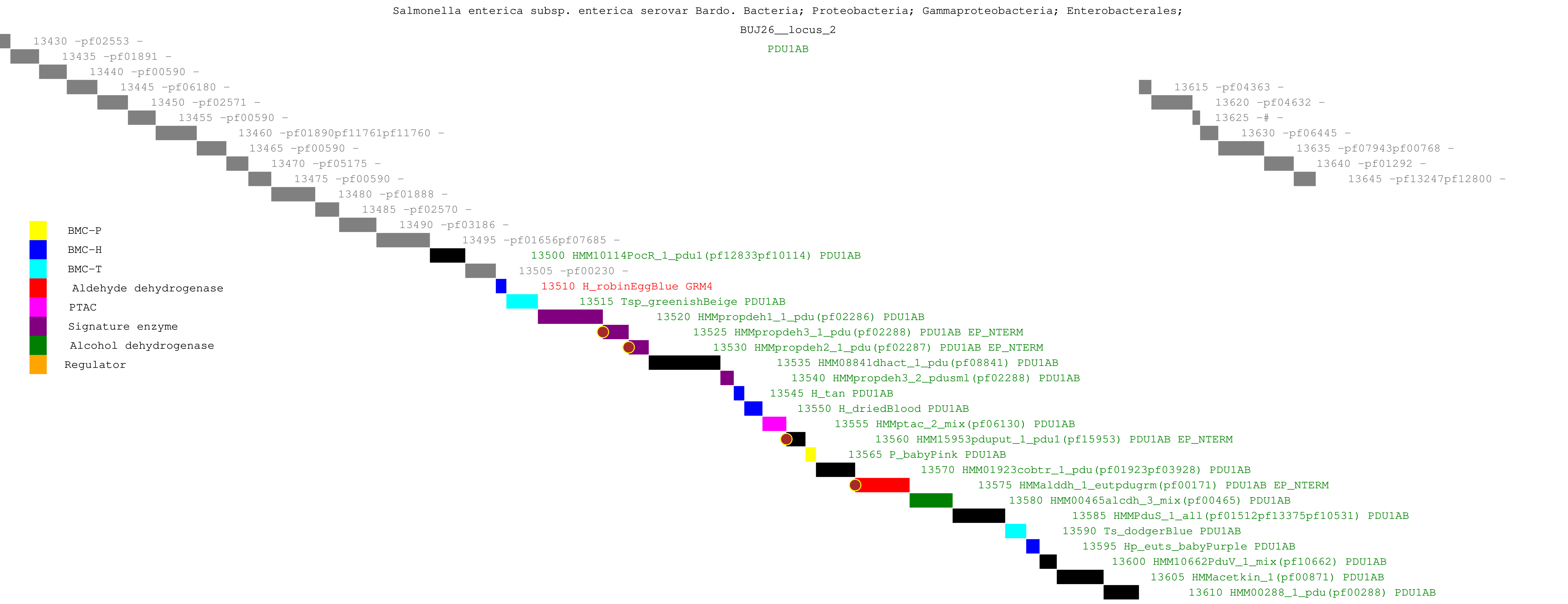
Salmonella enterica subsp. enterica serovar Bardo. Bacteria; Proteobacteria; Gammaproteobacteria; Enterobacterales;
BUJ26__locus_2
PDU1AB
13430 -pf02553 -
13435 -pf01891 -
13440 -pf00590 -
13445 -pf06180 -
13450 -pf02571 -
13455 -pf00590 -
13460 -pf01890pf11761pf11760 -
13465 -pf00590 -
13470 -pf05175 -
13475 -pf00590 -
13480 -pf01888 -
13485 -pf02570 -
13490 -pf03186 -
13495 -pf01656pf07685 -
13500 HMM10114PocR_1_pdu1(pf12833pf10114) PDU1AB
13505 -pf00230 -
13510 H_robinEggBlue GRM4
13515 Tsp_greenishBeige PDU1AB
13520 HMMpropdeh1_1_pdu(pf02286) PDU1AB
13525 HMMpropdeh3_1_pdu(pf02288) PDU1AB EP_NTERM
13530 HMMpropdeh2_1_pdu(pf02287) PDU1AB EP_NTERM
13535 HMM08841dhact_1_pdu(pf08841) PDU1AB
13540 HMMpropdeh3_2_pdusml(pf02288) PDU1AB
13545 H_tan PDU1AB
13550 H_driedBlood PDU1AB
13555 HMMptac_2_mix(pf06130) PDU1AB
13560 HMM15953pduput_1_pdu1(pf15953) PDU1AB EP_NTERM
13565 P_babyPink PDU1AB
13570 HMM01923cobtr_1_pdu(pf01923pf03928) PDU1AB
13575 HMMalddh_1_eutpdugrm(pf00171) PDU1AB EP_NTERM
13580 HMM00465alcdh_3_mix(pf00465) PDU1AB
13585 HMMPduS_1_all(pf01512pf13375pf10531) PDU1AB
13590 Ts_dodgerBlue PDU1AB
13595 Hp_euts_babyPurple PDU1AB
13600 HMM10662PduV_1_mix(pf10662) PDU1AB
13605 HMMacetkin_1(pf00871) PDU1AB
13610 HMM00288_1_pdu(pf00288) PDU1AB
13615 -pf04363 -
13620 -pf04632 -
13625 -# -
13630 -pf06445 -
13635 -pf07943pf00768 -
13640 -pf01292 -
13645 -pf13247pf12800 -
BMC-P
BMC-H
BMC-T
Aldehyde dehydrogenase
PTAC
Signature enzyme
Alcohol dehydrogenase
Regulator

Salmonella enterica subsp. enterica serovar Hartford. Bacteria; Proteobacteria; Gammaproteobacteria; Enterobacterales;

BUJ27__locus_2

PDU1AB

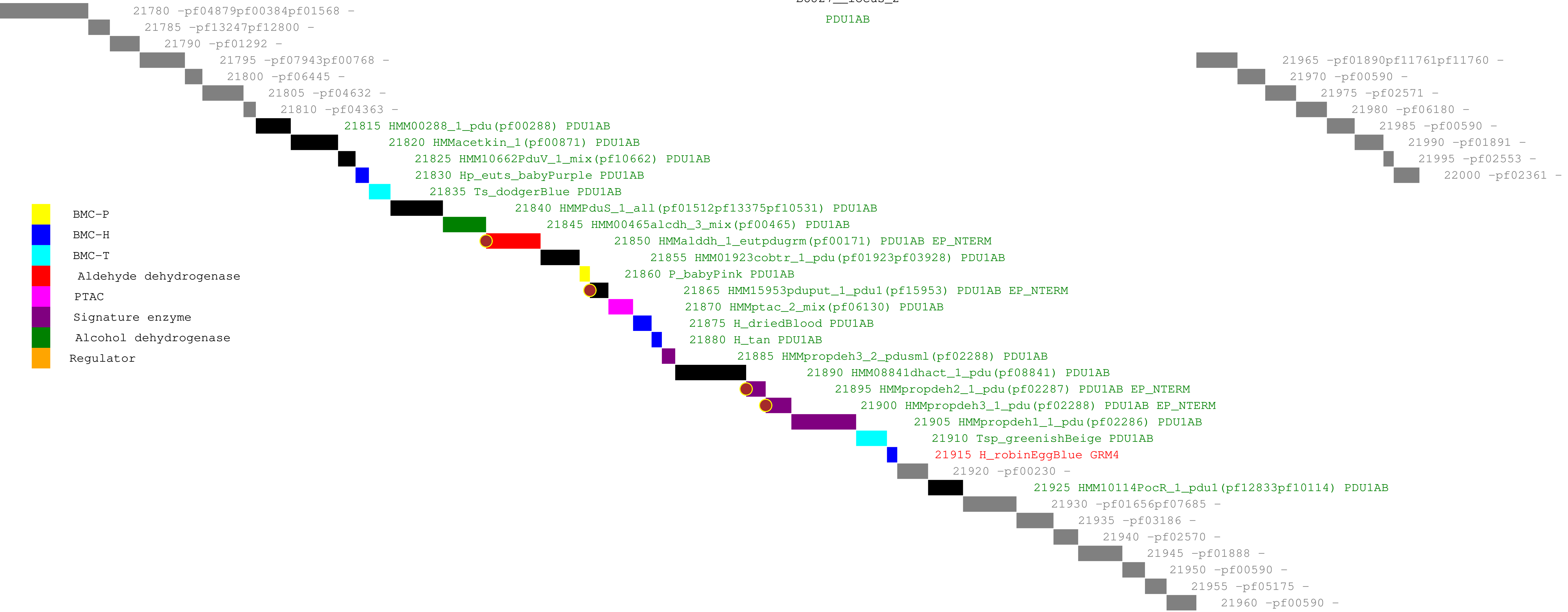

Salmonella muenchen. Bacteria; Proteobacteria; Gammaproteobacteria; Enterobacterales;

BUJ29__locus_1

PDU1AB

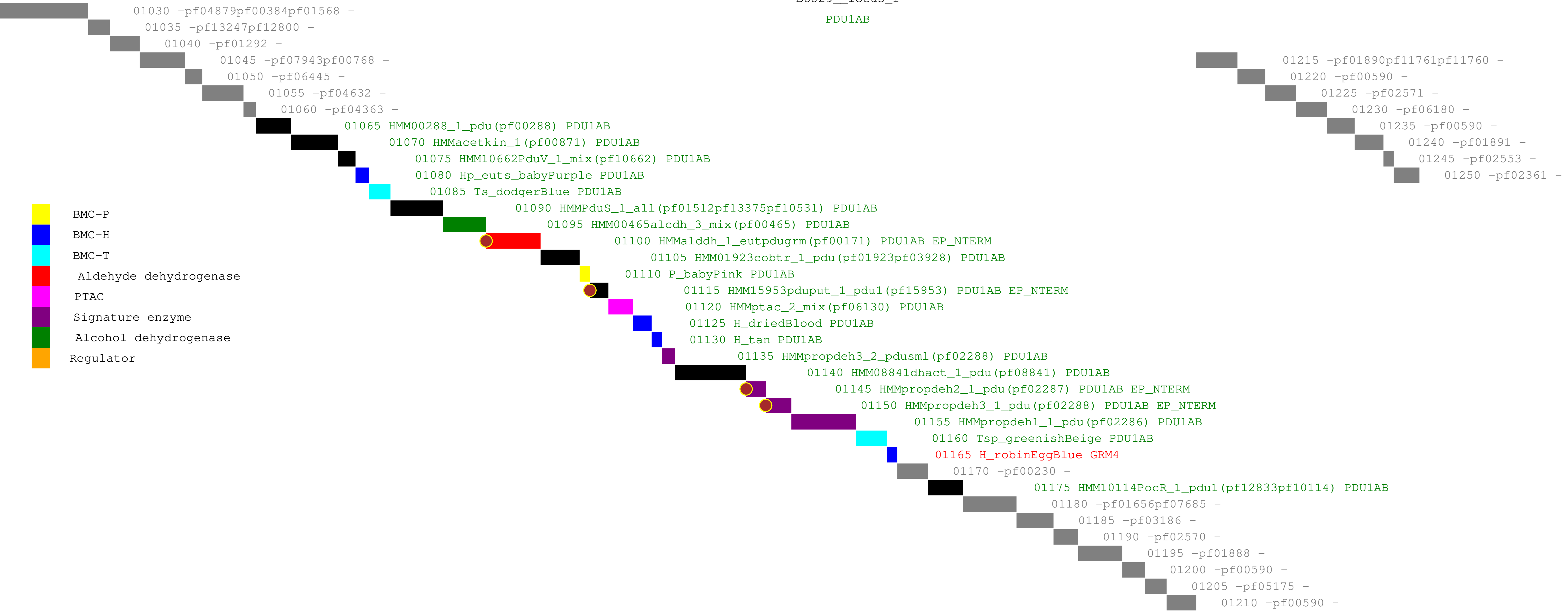

Escherichia coli. Bacteria; Proteobacteria; Gammaproteobacteria; Enterobacterales;

BvCmsKKP036__locus_1

PDU1AB

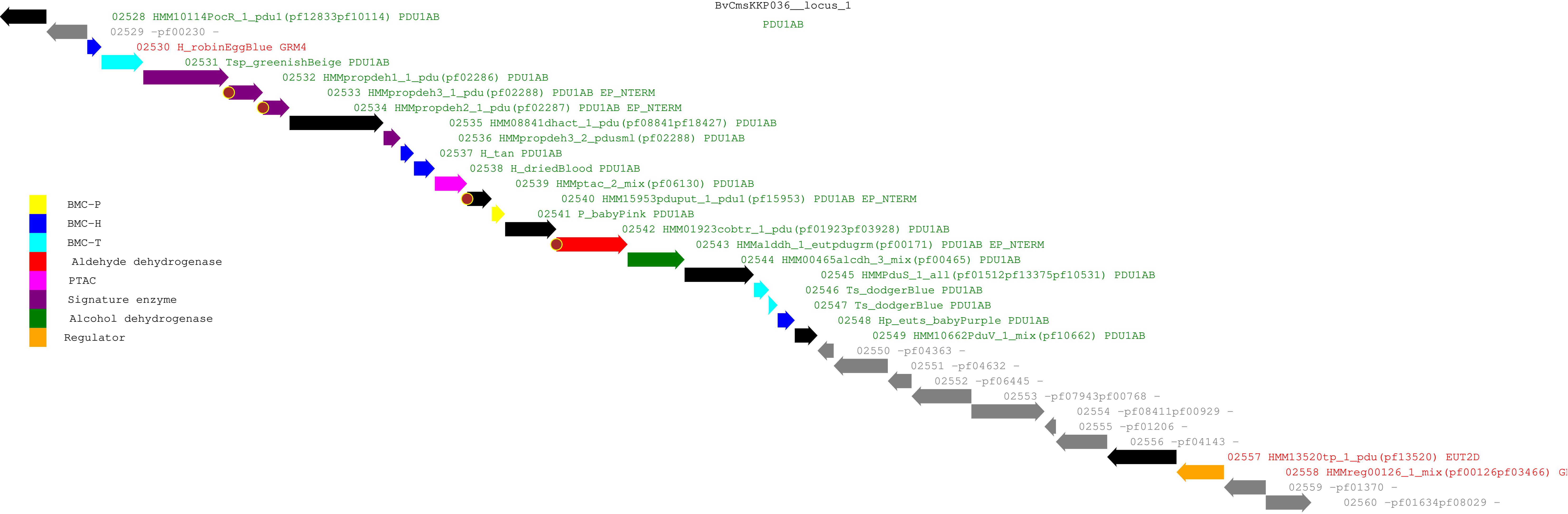

Escherichia coli. Bacteria; Proteobacteria; Gammaproteobacteria; Enterobacterales;

BvCmsOUP014__locus_1

PDU1AB

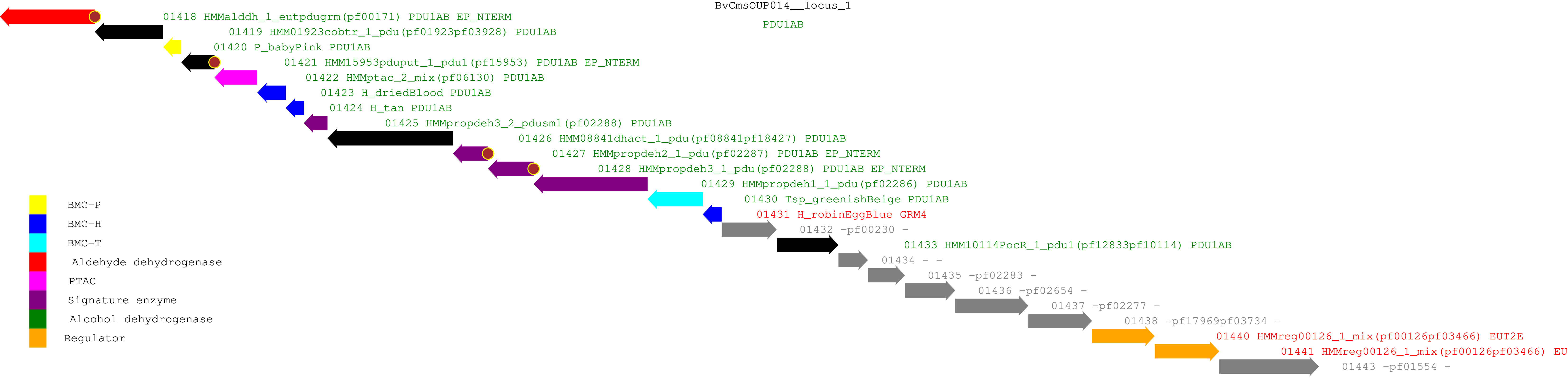

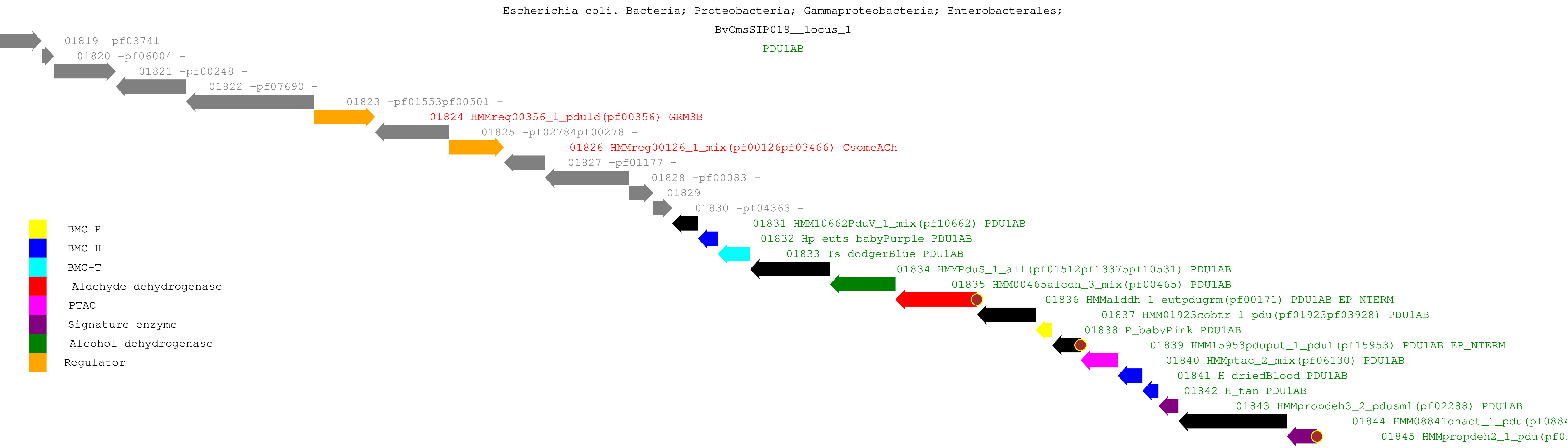
Escherichia coli. Bacteria; Proteobacteria; Gammaproteobacteria; Enterobacterales;
BvCmsSIP019__locus_1
PDU1AB
01819 -pf03741 -
01820 -pf06004 -
01821 -pf00248 -
01822 -pf07690 -
01823 -pf01553pf00501 -
01824 HMMreg00356_1_pdu1d(pf00356) GRM3B
01825 -pf02784pf00278 -
01826 HMMreg00126_1_mix(pf00126pf03466) CsomeACh
01827 -pf01177 -
01828 -pf00083 -
01829 - -
01830 -pf04363 -
01831 HMM10662PduV_1_mix(pf10662) PDU1AB
01832 Hp_euts_babyPurple PDU1AB
01833 Ts_dodgerBlue PDU1AB
01834 HMMPduS_1_all(pf01512pf13375pf10531) PDU1AB
01835 HMM00465alcdh_3_mix(pf00465) PDU1AB
01836 HMMalddh_1_eutpdugrm(pf00171) PDU1AB EP_NTERM
01837 HMM01923cobtr_1_pdu(pf01923pf03928) PDU1AB
01838 P_babyPink PDU1AB
01839 HMM15953pduput_1_pdu1(pf15953) PDU1AB EP_NTERM
01840 HMMptac_2_mix(pf06130) PDU1AB
01841 H_driedBlood PDU1AB
01842 H_tan PDU1AB
01843 HMMpropdeh3_2_pdusml(pf02288) PDU1AB
01844 HMM08841dhact_1_pdu(pf0884
01845 HMMpropdeh2_1_pdu(pf0
BMC-P
BMC-H
BMC-T
Aldehyde dehydrogenase
PTAC
Signature enzyme
Alcohol dehydrogenase
Regulator

Klebsiella pneumoniae. Bacteria; Proteobacteria; Gammaproteobacteria; Enterobacterales;

BVX91__locus_1

PDU1AB

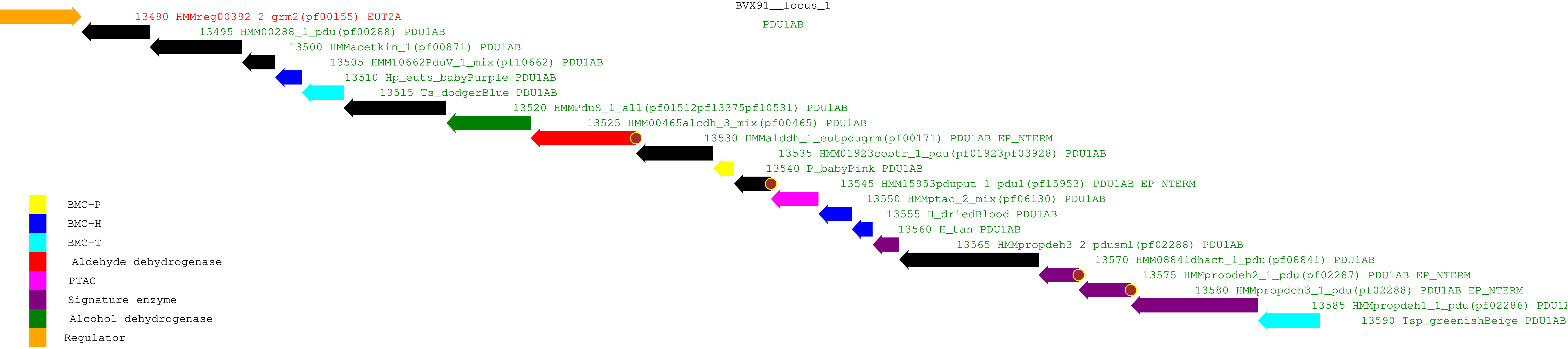

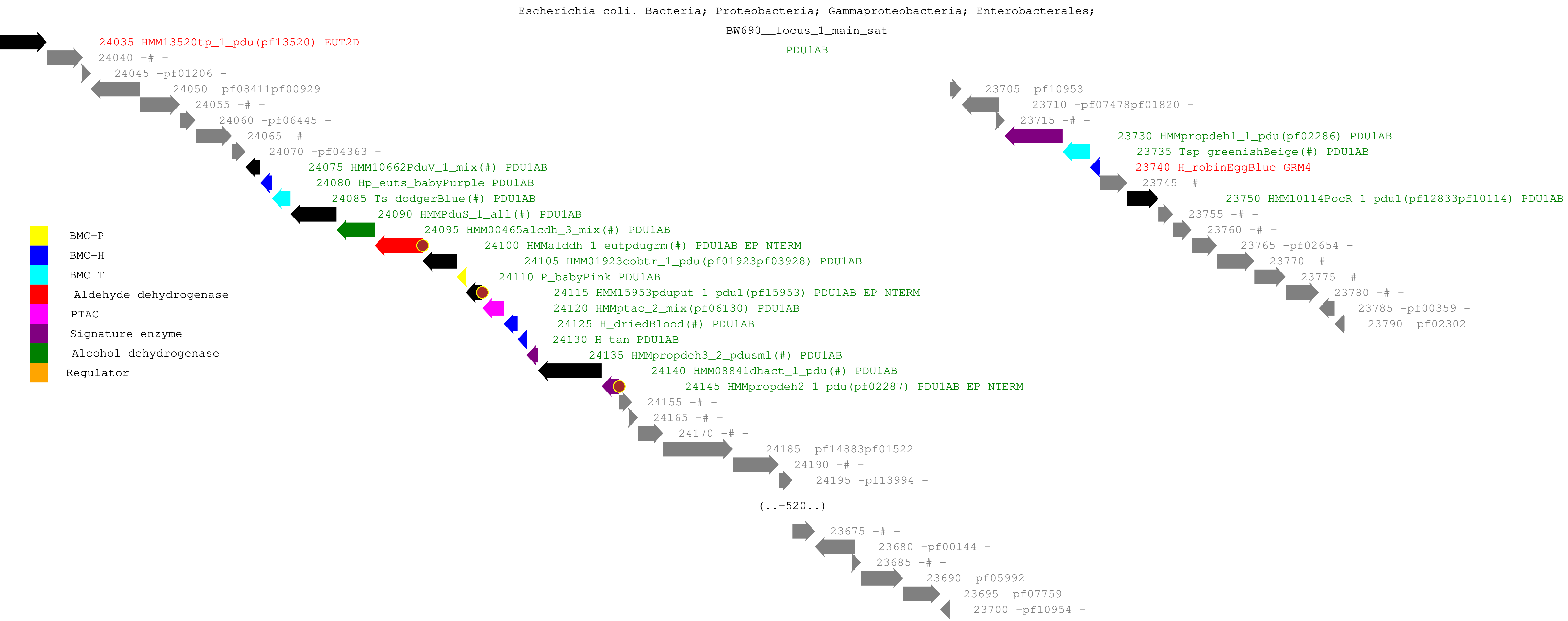
Escherichia coli. Bacteria; Proteobacteria; Gammaproteobacteria; Enterobacterales;
BW690__locus_1_main_sat
PDU1AB
24035 HMM13520tp_1_pdu(pf13520) EUT2D
24040 -# -
24045 -pf01206 -
24050 -pf08411pf00929 -
24055 -# -
24060 -pf06445 -
24065 -# -
24070 -pf04363 -
24075 HMM10662PduV_1_mix(#) PDU1AB
24080 Hp_euts_babyPurple PDU1AB
24085 Ts_dodgerBlue(#) PDU1AB
24090 HMMPduS_1_all(#) PDU1AB
24095 HMM00465alcdh_3_mix(#) PDU1AB
24100 HMMalddh_1_eutpdugrm(#) PDU1AB EP_NTERM
24105 HMM01923cobtr_1_pdu(pf01923pf03928) PDU1AB
24110 P_babyPink PDU1AB
24115 HMM15953pduput_1_pdu1(pf15953) PDU1AB EP_NTERM
24120 HMMptac_2_mix(pf06130) PDU1AB
24125 H_driedBlood(#) PDU1AB
24130 H_tan PDU1AB
24135 HMMpropdeh3_2_pdusml(#) PDU1AB
24140 HMM08841dhact_1_pdu(#) PDU1AB
24145 HMMpropdeh2_1_pdu(pf02287) PDU1AB EP_NTERM
24155 -# -
24165 -# -
24170 -# -
24185 -pf14883pf01522 -
24190 -# -
24195 -pf13994 -
(..-520..)
23675 -# -
23680 -pf00144 -
23685 -# -
23690 -pf05992 -
23695 -pf07759 -
23700 -pf10954 -
23705 -pf10953 -
23710 -pf07478pf01820 -
23715 -# -
23730 HMMpropdeh1_1_pdu(pf02286) PDU1AB
23735 Tsp_greenishBeige(#) PDU1AB
23740 H_robinEggBlue GRM4
23745 -# -
23750 HMM10114PocR_1_pdu1(pf12833pf10114) PDU1AB
23755 -# -
23760 -# -
23765 -pf02654 -
23770 -# -
23775 -# -
23780 -# -
23785 -pf00359 -
23790 -pf02302 -
BMC-P
BMC-H
BMC-T
Aldehyde dehydrogenase
PTAC
Signature enzyme
Alcohol dehydrogenase
Regulator

Citrobacter braakii. Bacteria; Proteobacteria; Gammaproteobacteria; Enterobacterales;

BWD41__locus_2

PDU1AB

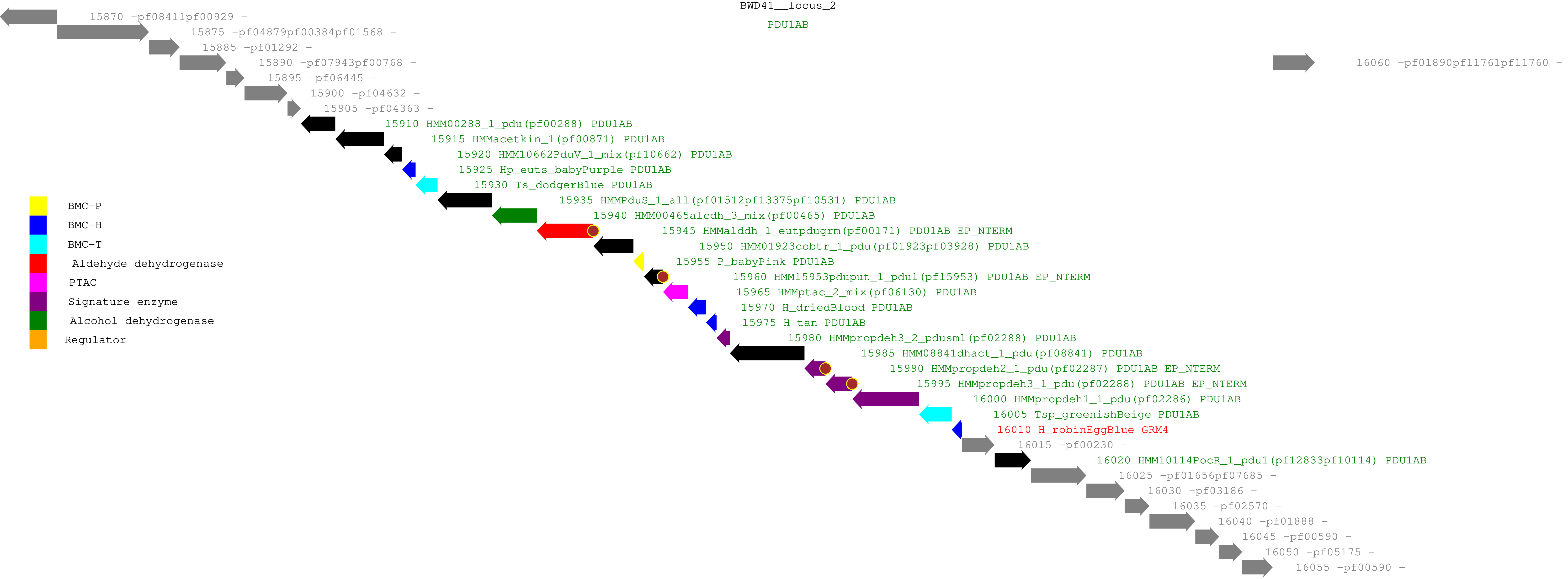

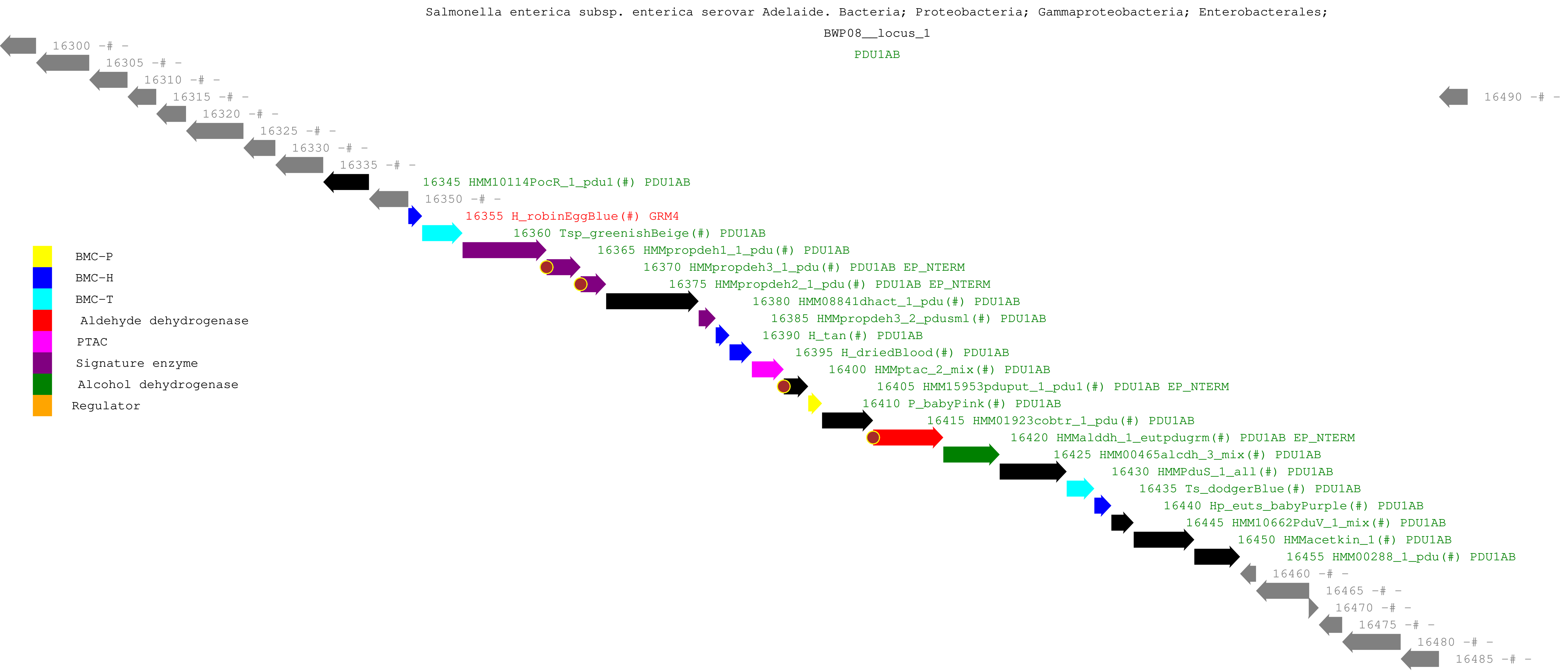

Salmonella enterica subsp. enterica serovar Adelaide. Bacteria; Proteobacteria; Gammaproteobacteria; Enterobacterales;
BWP08__locus_1
PDU1AB
16300 -# -
16305 -# -
16310 -# -
16315 -# -
16320 -# -
16325 -# -
16330 -# -
16335 -# -
16345 HMM10114PocR_1_pdu1(#) PDU1AB
16350 -# -
16355 H_robinEggBlue(#) GRM4
16360 Tsp_greenishBeige(#) PDU1AB
16365 HMMpropdeh1_1_pdu(#) PDU1AB
16370 HMMpropdeh3_1_pdu(#) PDU1AB EP_NTERM
16375 HMMpropdeh2_1_pdu(#) PDU1AB EP_NTERM
16380 HMM08841dhact_1_pdu(#) PDU1AB
16385 HMMpropdeh3_2_pdusml(#) PDU1AB
16390 H_tan(#) PDU1AB
16395 H_driedBlood(#) PDU1AB
16400 HMMptac_2_mix(#) PDU1AB
16405 HMM15953pduput_1_pdu1(#) PDU1AB EP_NTERM
16410 P_babyPink(#) PDU1AB
16415 HMM01923cobtr_1_pdu(#) PDU1AB
16420 HMMaldd h_1_eutpdugrm(#) PDU1AB EP_NTERM
16425 HMM00465alcdh_3_mix(#) PDU1AB
16430 HMMPduS_1_all(#) PDU1AB
16435 Ts_dodgerBlue(#) PDU1AB
16440 Hp_euts_babyPurple(#) PDU1AB
16445 HMM10662PduV_1_mix(#) PDU1AB
16450 HMMacetkin_1(#) PDU1AB
16455 HMM00288_1_pdu(#) PDU1AB
16460 -# -
16465 -# -
16470 -# -
16475 -# -
16480 -# -
16485 -# -
16490 -# -
BMC-P
BMC-H
BMC-T
Aldehyde dehydrogenase
PTAC
Signature enzyme
Alcohol dehydrogenase
Regulator

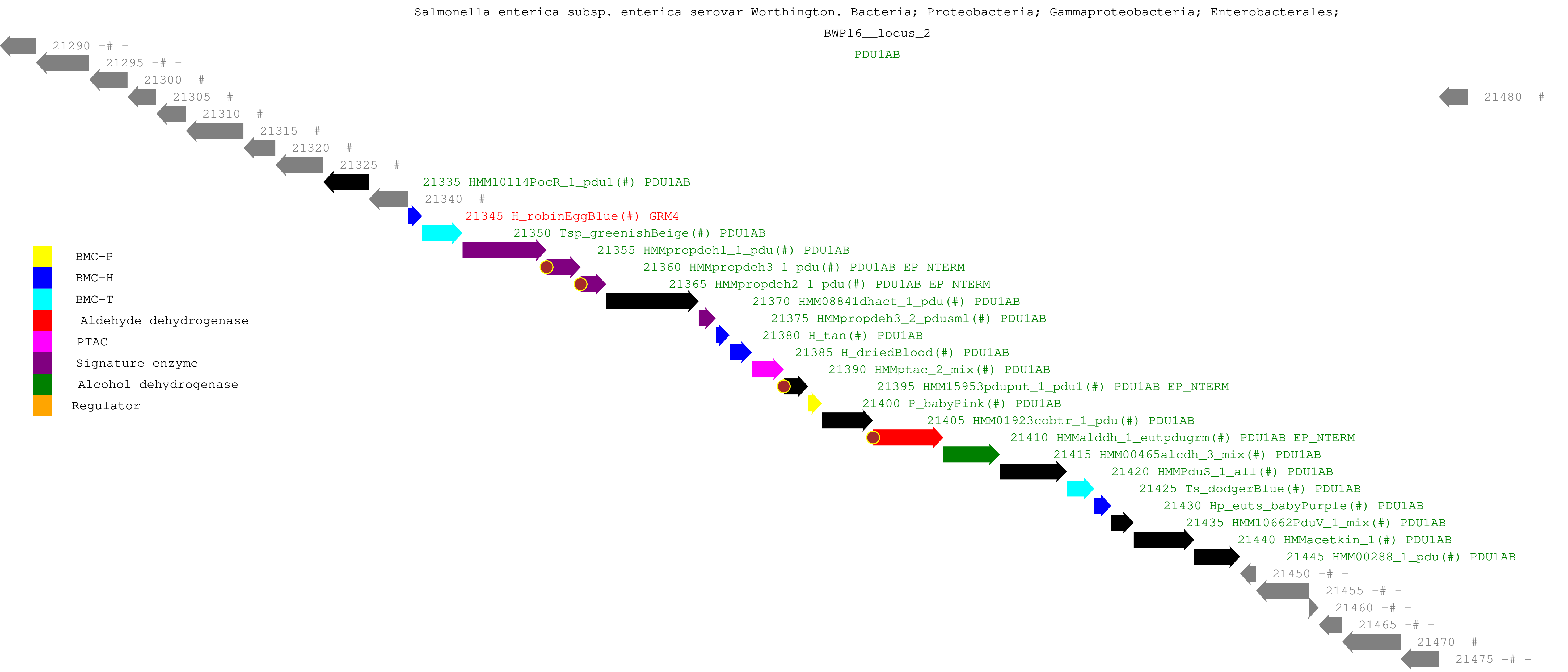
Salmonella enterica subsp. enterica serovar Worthington. Bacteria; Proteobacteria; Gammaproteobacteria; Enterobacterales;
BWP16__locus_2
PDU1AB
21290 -# -
21295 -# -
21300 -# -
21305 -# -
21310 -# -
21315 -# -
21320 -# -
21325 -# -
21335 HMM10114PocR_1_pdu1(#) PDU1AB
21340 -# -
21345 H_robinEggBlue(#) GRM4
21350 Tsp_greenishBeige(#) PDU1AB
21355 HMMpropdeh1_1_pdu(#) PDU1AB
21360 HMMpropdeh3_1_pdu(#) PDU1AB EP_NTERM
21365 HMMpropdeh2_1_pdu(#) PDU1AB EP_NTERM
21370 HMM08841dhact_1_pdu(#) PDU1AB
21375 HMMpropdeh3_2_pdusml(#) PDU1AB
21380 H_tan(#) PDU1AB
21385 H_driedBlood(#) PDU1AB
21390 HMMptac_2_mix(#) PDU1AB
21395 HMM15953pduput_1_pdu1(#) PDU1AB EP_NTERM
21400 P_babyPink(#) PDU1AB
21405 HMM01923cobtr_1_pdu(#) PDU1AB
21410 HMMalddh_1_eutpdugrm(#) PDU1AB EP_NTERM
21415 HMM00465alcdh_3_mix(#) PDU1AB
21420 HMMPduS_1_all(#) PDU1AB
21425 Ts_dodgerBlue(#) PDU1AB
21430 Hp_euts_babyPurple(#) PDU1AB
21435 HMM10662PduV_1_mix(#) PDU1AB
21440 HMMacetkin_1(#) PDU1AB
21445 HMM00288_1_pdu(#) PDU1AB
21450 -# -
21455 -# -
21460 -# -
21465 -# -
21470 -# -
21475 -# -
21480 -# -
BMC-P
BMC-H
BMC-T
Aldehyde dehydrogenase
PTAC
Signature enzyme
Alcohol dehydrogenase
Regulator

Escherichia coli. Bacteria; Proteobacteria; Gammaproteobacteria; Enterobacterales;

BWP18__locus_1

PDU1AB

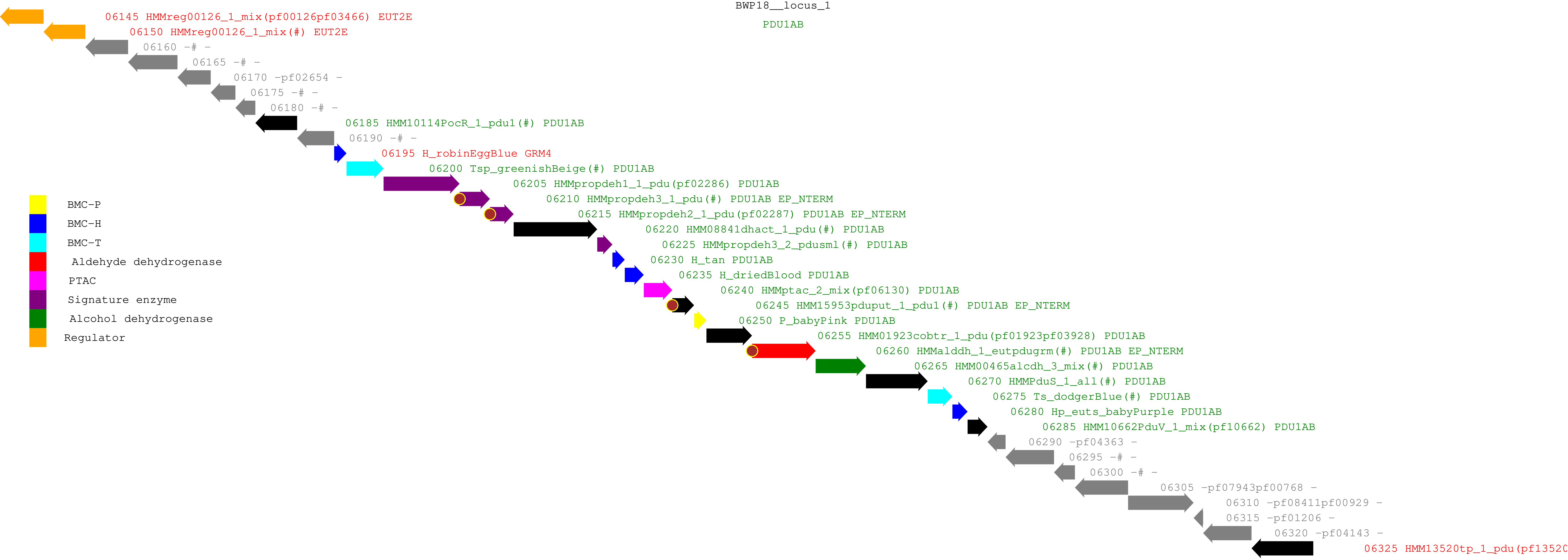

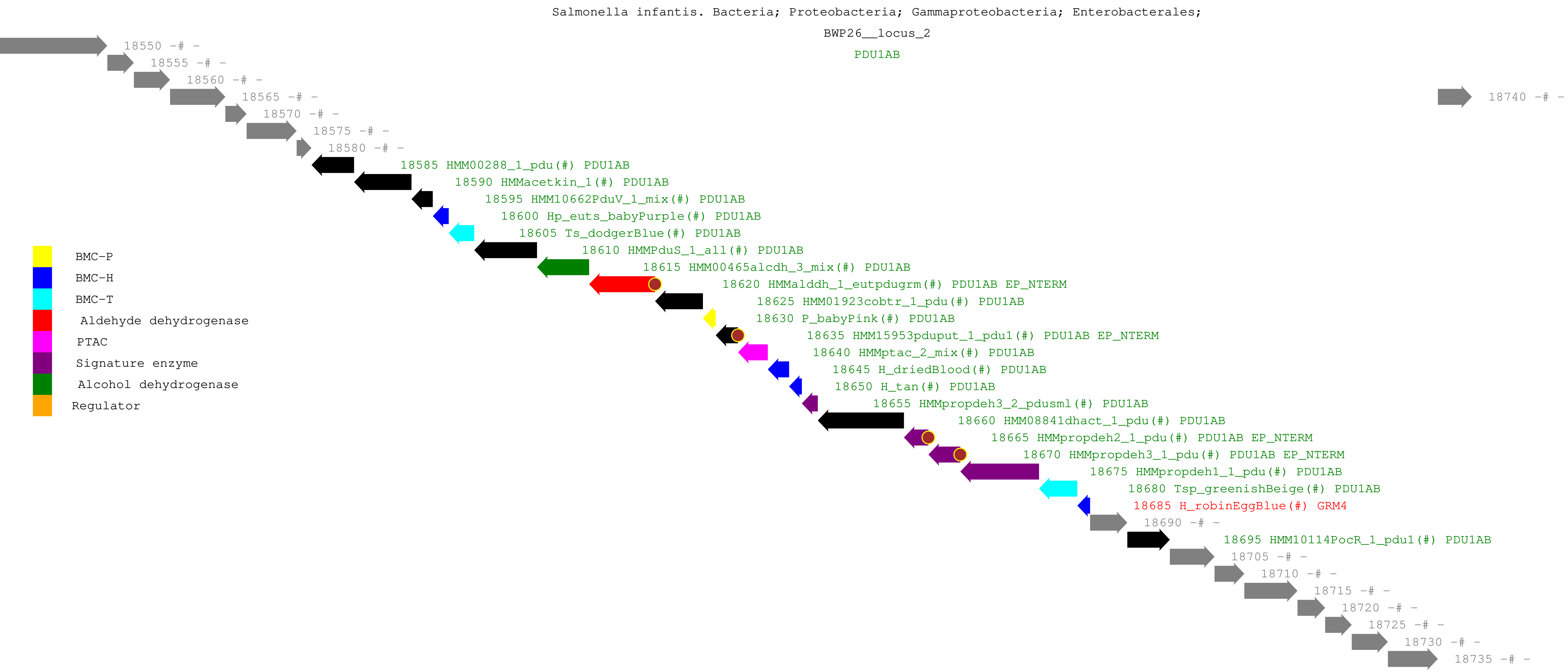

Salmonella infantis. Bacteria; Proteobacteria; Gammaproteobacteria; Enterobacterales;
BWP26__locus_2
PDU1AB
18550 -# -
18555 -# -
18560 -# -
18565 -# -
18570 -# -
18575 -# -
18580 -# -
18585 HMM00288_1_pdu(#) PDU1AB
18590 HMMacetkin_1(#) PDU1AB
18595 HMM10662PduV_1_mix(#) PDU1AB
18600 Hp_euts_babyPurple(#) PDU1AB
18605 Ts_dodgerBlue(#) PDU1AB
18610 HMMPduS_1_all(#) PDU1AB
18615 HMM00465alcdh_3_mix(#) PDU1AB
18620 HMMalddh_1_eutpdugrm(#) PDU1AB EP_NTERM
18625 HMM01923cobtr_1_pdu(#) PDU1AB
18630 P_babyPink(#) PDU1AB
18635 HMM15953pduput_1_pdu1(#) PDU1AB EP_NTERM
18640 HMMptac_2_mix(#) PDU1AB
18645 H_driedBlood(#) PDU1AB
18650 H_tan(#) PDU1AB
18655 HMMpropdeh3_2_pdusml(#) PDU1AB
18660 HMM08841dhact_1_pdu(#) PDU1AB
18665 HMMpropdeh2_1_pdu(#) PDU1AB EP_NTERM
18670 HMMpropdeh3_1_pdu(#) PDU1AB EP_NTERM
18675 HMMpropdeh1_1_pdu(#) PDU1AB
18680 Tsp_greenishBeige(#) PDU1AB
18685 H_robinEggBlue(#) GRM4
18690 -# -
18695 HMM10114PocR_1_pdu1(#) PDU1AB
18705 -# -
18710 -# -
18715 -# -
18720 -# -
18725 -# -
18730 -# -
18735 -# -
18740 -# -
BMC-P
BMC-H
BMC-T
Aldehyde dehydrogenase
PTAC
Signature enzyme
Alcohol dehydrogenase
Regulator

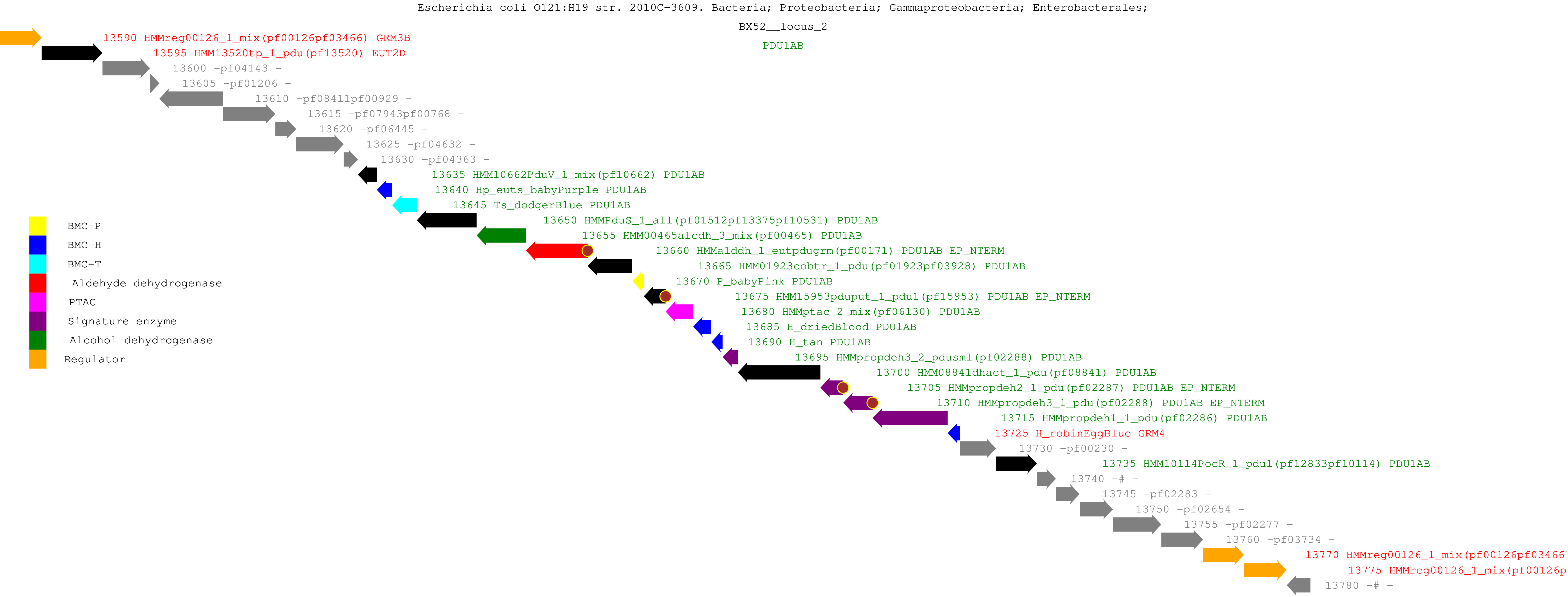
Escherichia coli O121:H19 str. 2010C-3609. Bacteria; Proteobacteria; Gammaproteobacteria; Enterobacterales;
BX52__locus_2
PDU1AB
13590 HMMreg00126_1_mix(pf00126pf03466) GRM3B
13595 HMM13520tp_1_pdu(pf13520) EUT2D
13600 -pf04143 -
13605 -pf01206 -
13610 -pf08411pf00929 -
13615 -pf07943pf00768 -
13620 -pf06445 -
13625 -pf04632 -
13630 -pf04363 -
13635 HMM10662PduV_1_mix(pf10662) PDU1AB
13640 Hp_euts_babyPurple PDU1AB
13645 Ts_dodgerBlue PDU1AB
13650 HMMPduS_1_all(pf01512pf13375pf10531) PDU1AB
13655 HMM00465alcdh_3_mix(pf00465) PDU1AB
13660 HMMaldddh_1_eutpdugrm(pf00171) PDU1AB EP_NTERM
13665 HMM01923cobtr_1_pdu(pf01923pf03928) PDU1AB
13670 P_babyPink PDU1AB
13675 HMM15953pduput_1_pdu1(pf15953) PDU1AB EP_NTERM
13680 HMMptac_2_mix(pf06130) PDU1AB
13685 H_driedBlood PDU1AB
13690 H_tan PDU1AB
13695 HMMpropdeh3_2_pdusml(pf02288) PDU1AB
13700 HMM08841dhact_1_pdu(pf08841) PDU1AB
13705 HMMpropdeh2_1_pdu(pf02287) PDU1AB EP_NTERM
13710 HMMpropdeh3_1_pdu(pf02288) PDU1AB EP_NTERM
13715 HMMpropdeh1_1_pdu(pf02286) PDU1AB
13725 H_robinEggBlue GRM4
13730 -pf00230 -
13735 HMM10114PocR_1_pdu1(pf12833pf10114) PDU1AB
13740 -# -
13745 -pf02283 -
13750 -pf02654 -
13755 -pf02277 -
13760 -pf03734 -
13770 HMMreg00126_1_mix(pf00126pf03466)
13775 HMMreg00126_1_mix(pf00126p
13780 -# -
BMC-P
BMC-H
BMC-T
Aldehyde dehydrogenase
PTAC
Signature enzyme
Alcohol dehydrogenase
Regulator

Salmonella enterica subsp. enterica serovar Java. Bacteria; Proteobacteria; Gammaproteobacteria; Enterobacterales;

BXN20__locus_1

PDU1AB

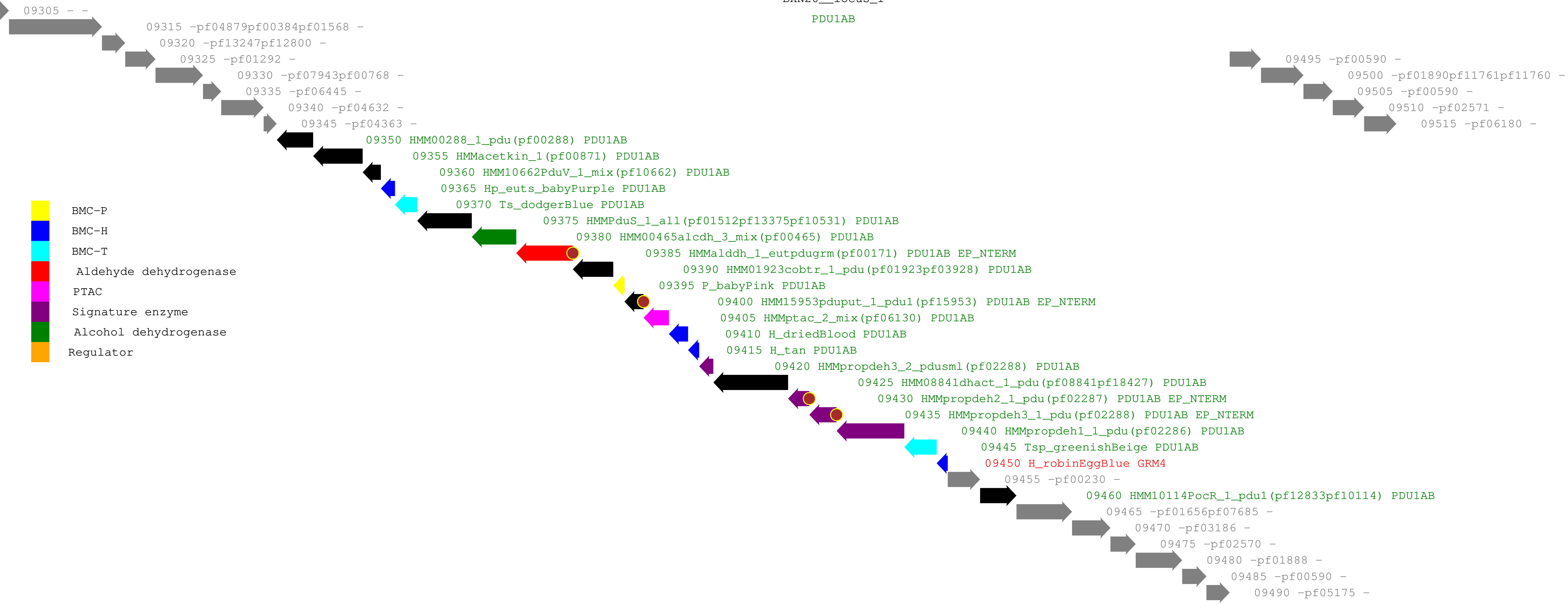

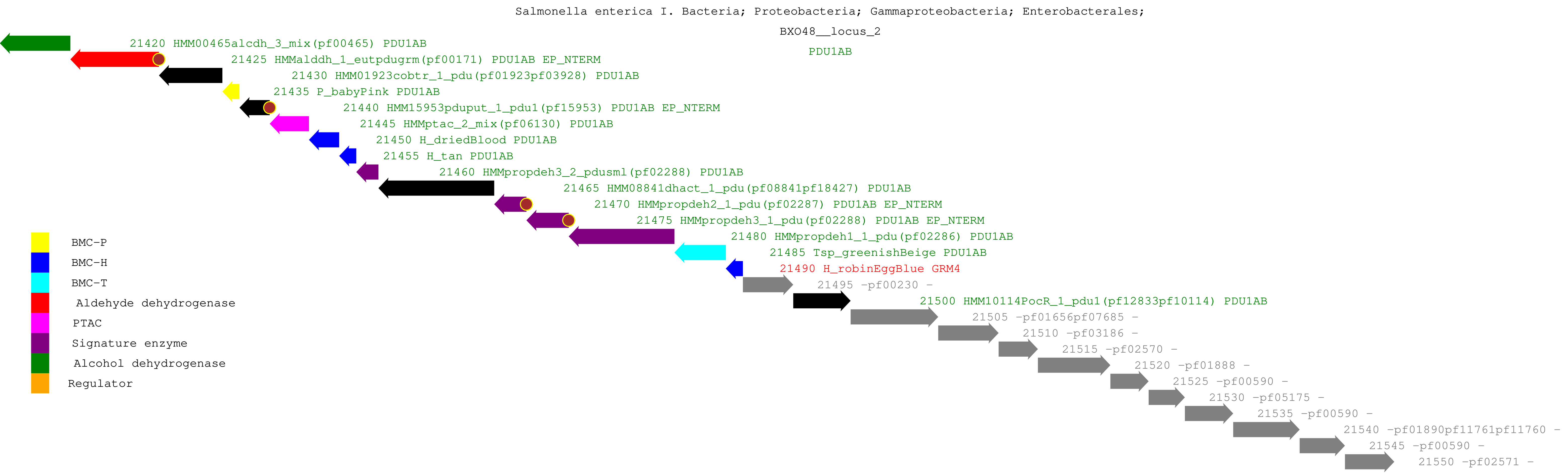
Salmonella enterica I. Bacteria; Proteobacteria; Gammaproteobacteria; Enterobacterales;
BX048__locus_2
PDU1AB
21420 HMM00465alcdh_3_mix(pf00465) PDU1AB
21425 HMMalddh_1_eutpdugrm(pf00171) PDU1AB EP_NTERM
21430 HMM01923cobtr_1_pdu(pf01923pf03928) PDU1AB
21435 P_babyPink PDU1AB
21440 HMM15953pduput_1_pdu1(pf15953) PDU1AB EP_NTERM
21445 HMMptac_2_mix(pf06130) PDU1AB
21450 H_driedBlood PDU1AB
21455 H_tan PDU1AB
21460 HMMpropdeh3_2_pdusml(pf02288) PDU1AB
21465 HMM08841dhact_1_pdu(pf08841pf18427) PDU1AB
21470 HMMpropdeh2_1_pdu(pf02287) PDU1AB EP_NTERM
21475 HMMpropdeh3_1_pdu(pf02288) PDU1AB EP_NTERM
21480 HMMpropdeh1_1_pdu(pf02286) PDU1AB
21485 Tsp_greenishBeige PDU1AB
21490 H_robinEggBlue GRM4
21495 -pf00230 -
21500 HMM10114PocR_1_pdu1(pf12833pf10114) PDU1AB
21505 -pf01656pf07685 -
21510 -pf03186 -
21515 -pf02570 -
21520 -pf01888 -
21525 -pf00590 -
21530 -pf05175 -
21535 -pf00590 -
21540 -pf01890pf11761pf11760 -
21545 -pf00590 -
21550 -pf02571 -
BMC-P
BMC-H
BMC-T
Aldehyde dehydrogenase
PTAC
Signature enzyme
Alcohol dehydrogenase
Regulator

Escherichia coli. Bacteria; Proteobacteria; Gammaproteobacteria; Enterobacterales;

BXT93__locus_1

PDU1AB

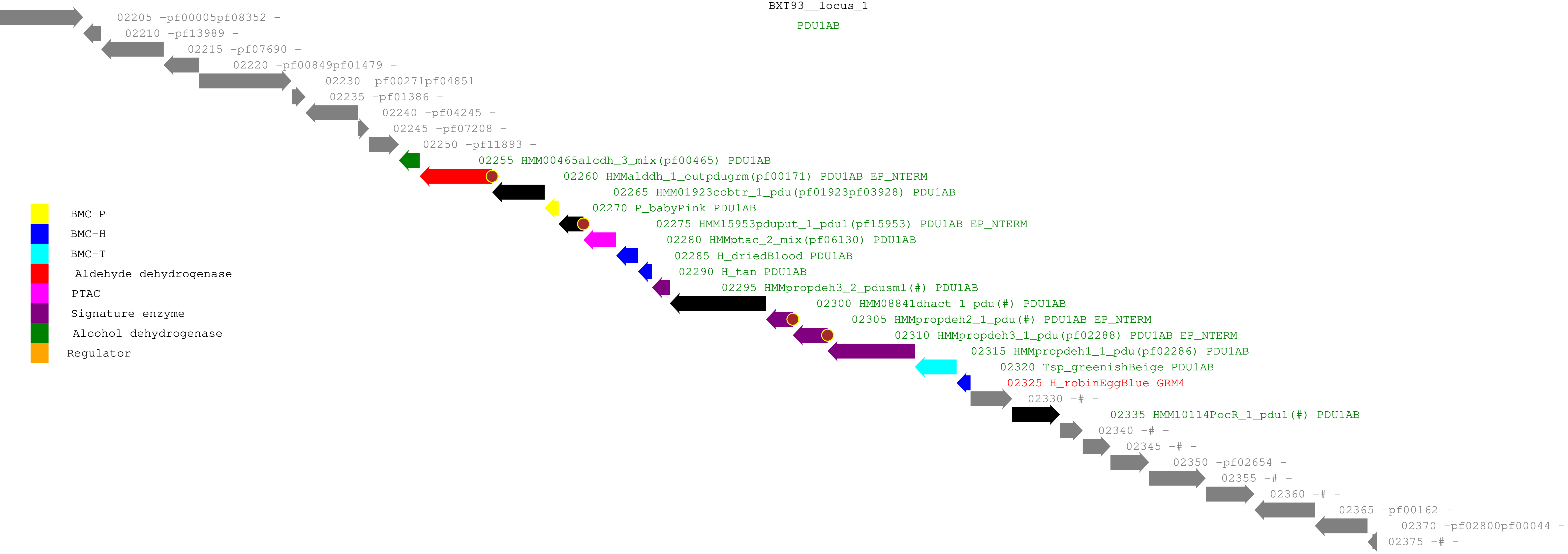

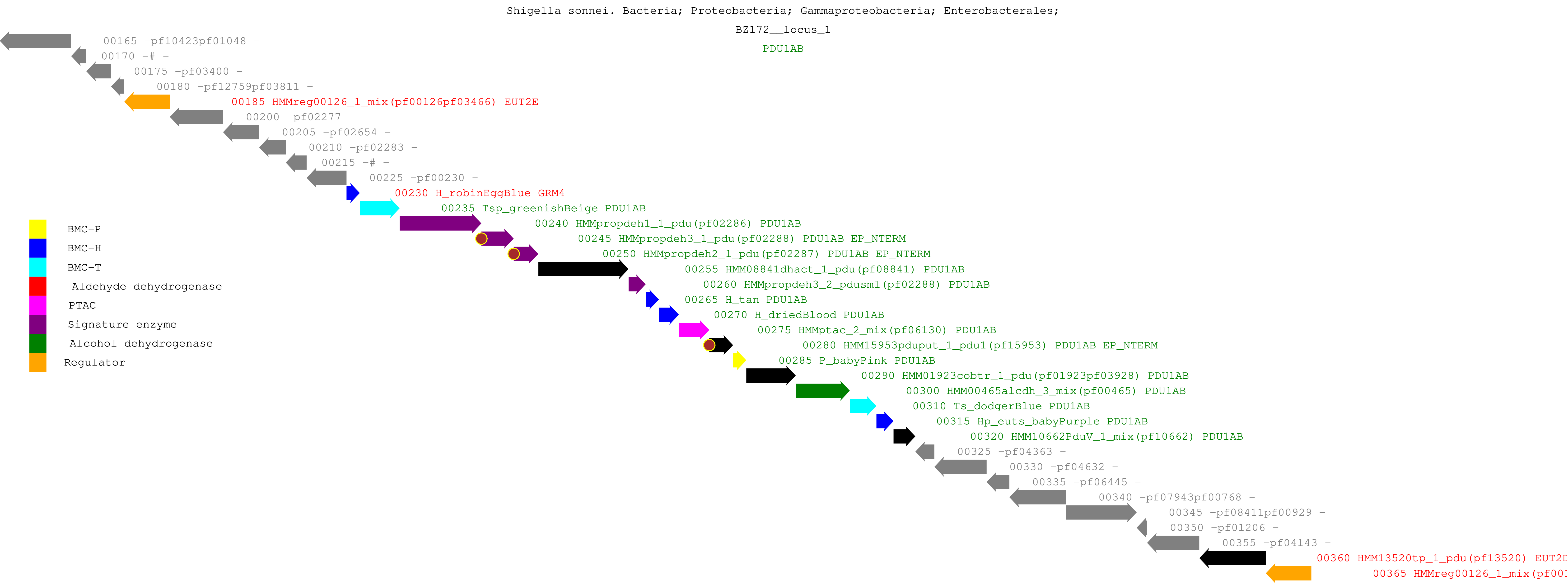
Shigella sonnei. Bacteria; Proteobacteria; Gammaproteobacteria; Enterobacterales;
BZ172__locus_1
PDU1AB
00165 -pf10423pf01048 -
00170 -# -
00175 -pf03400 -
00180 -pf12759pf03811 -
00185 HMMreg00126_1_mix(pf00126pf03466) EUT2E
00200 -pf02277 -
00205 -pf02654 -
00210 -pf02283 -
00215 -# -
00225 -pf00230 -
00230 H_robinEggBlue GRM4
00235 Tsp_greenishBeige PDU1AB
00240 HMMpropdeh1_1_pdu(pf02286) PDU1AB
00245 HMMpropdeh3_1_pdu(pf02288) PDU1AB EP_NTERM
00250 HMMpropdeh2_1_pdu(pf02287) PDU1AB EP_NTERM
00255 HMM08841dhact_1_pdu(pf08841) PDU1AB
00260 HMMpropdeh3_2_pdusml(pf02288) PDU1AB
00265 H_tan PDU1AB
00270 H_driedBlood PDU1AB
00275 HMMptac_2_mix(pf06130) PDU1AB
00280 HMM15953pduput_1_pdu1(pf15953) PDU1AB EP_NTERM
00285 P_babyPink PDU1AB
00290 HMM01923cobtr_1_pdu(pf01923pf03928) PDU1AB
00300 HMM00465alcdh_3_mix(pf00465) PDU1AB
00310 Ts_dodgerBlue PDU1AB
00315 Hp_euts_babyPurple PDU1AB
00320 HMM10662PduV_1_mix(pf10662) PDU1AB
00325 -pf04363 -
00330 -pf04632 -
00335 -pf06445 -
00340 -pf07943pf00768 -
00345 -pf08411pf00929 -
00350 -pf01206 -
00355 -pf04143 -
00360 HMM13520tp_1_pdu(pf13520) EUT2D
00365 HMMreg00126_1_mix(pf00
BMC-P
BMC-H
BMC-T
Aldehyde dehydrogenase
PTAC
Signature enzyme
Alcohol dehydrogenase
Regulator

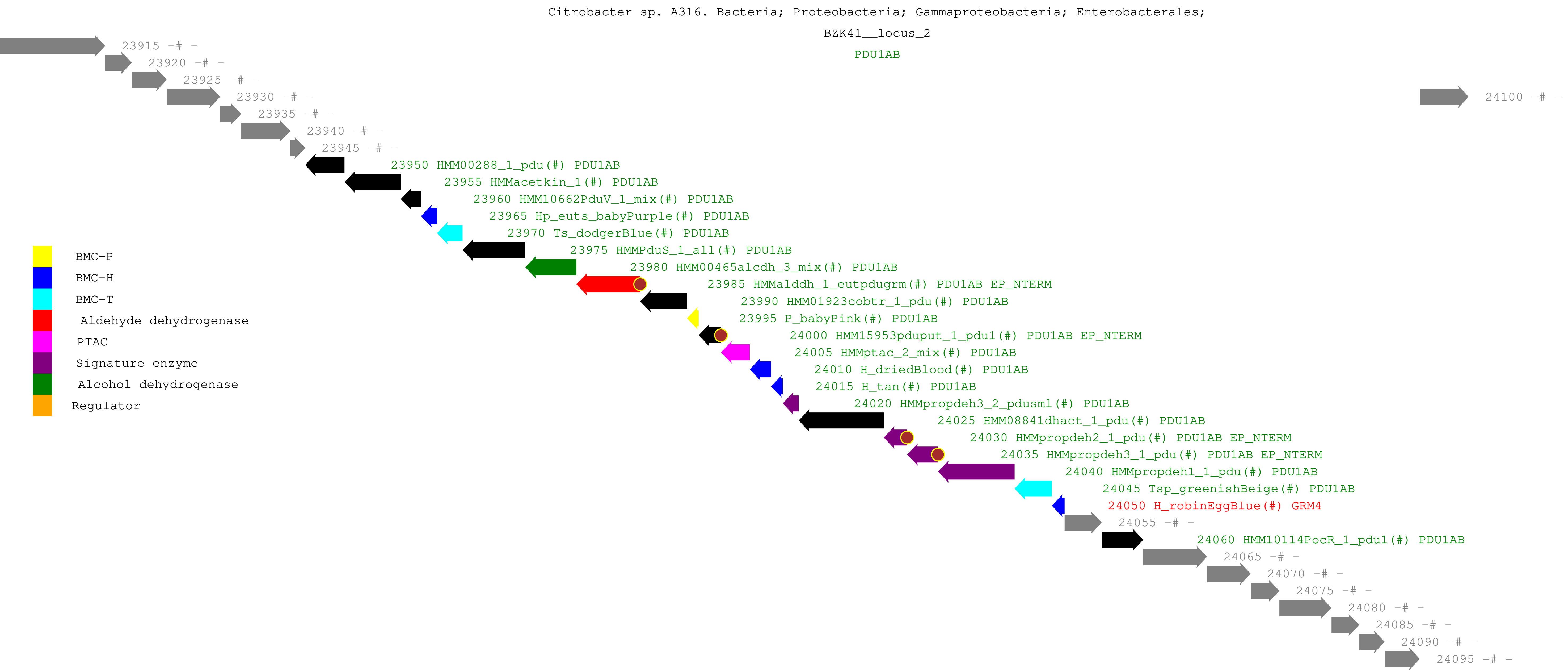

Citrobacter sp. A316. Bacteria; Proteobacteria; Gammaproteobacteria; Enterobacterales;
BZK41__locus_2
PDU1AB
23915 -# -
23920 -# -
23925 -# -
23930 -# -
23935 -# -
23940 -# -
23945 -# -
23950 HMM00288_1_pdu(#) PDU1AB
23955 HMMacetkin_1(#) PDU1AB
23960 HMM10662PduV_1_mix(#) PDU1AB
23965 Hp_euts_babyPurple(#) PDU1AB
23970 Ts_dodgerBlue(#) PDU1AB
23975 HMMPduS_1_all(#) PDU1AB
23980 HMM00465alcdh_3_mix(#) PDU1AB
23985 HMMalddh_1_eutpdugrm(#) PDU1AB EP_NTERM
23990 HMM01923cobtr_1_pdu(#) PDU1AB
23995 P_babyPink(#) PDU1AB
24000 HMM15953pduput_1_pdu1(#) PDU1AB EP_NTERM
24005 HMMptac_2_mix(#) PDU1AB
24010 H_driedBlood(#) PDU1AB
24015 H_tan(#) PDU1AB
24020 HMMpropdeh3_2_pdusml(#) PDU1AB
24025 HMM08841dhact_1_pdu(#) PDU1AB
24030 HMMpropdeh2_1_pdu(#) PDU1AB EP_NTERM
24035 HMMpropdeh3_1_pdu(#) PDU1AB EP_NTERM
24040 HMMpropdeh1_1_pdu(#) PDU1AB
24045 Tsp_greenishBeige(#) PDU1AB
24050 H_robinEggBlue(#) GRM4
24055 -# -
24060 HMM10114PocR_1_pdu1(#) PDU1AB
24065 -# -
24070 -# -
24075 -# -
24080 -# -
24085 -# -
24090 -# -
24095 -# -
24100 -# -
BMC-P
BMC-H
BMC-T
Aldehyde dehydrogenase
PTAC
Signature enzyme
Alcohol dehydrogenase
Regulator

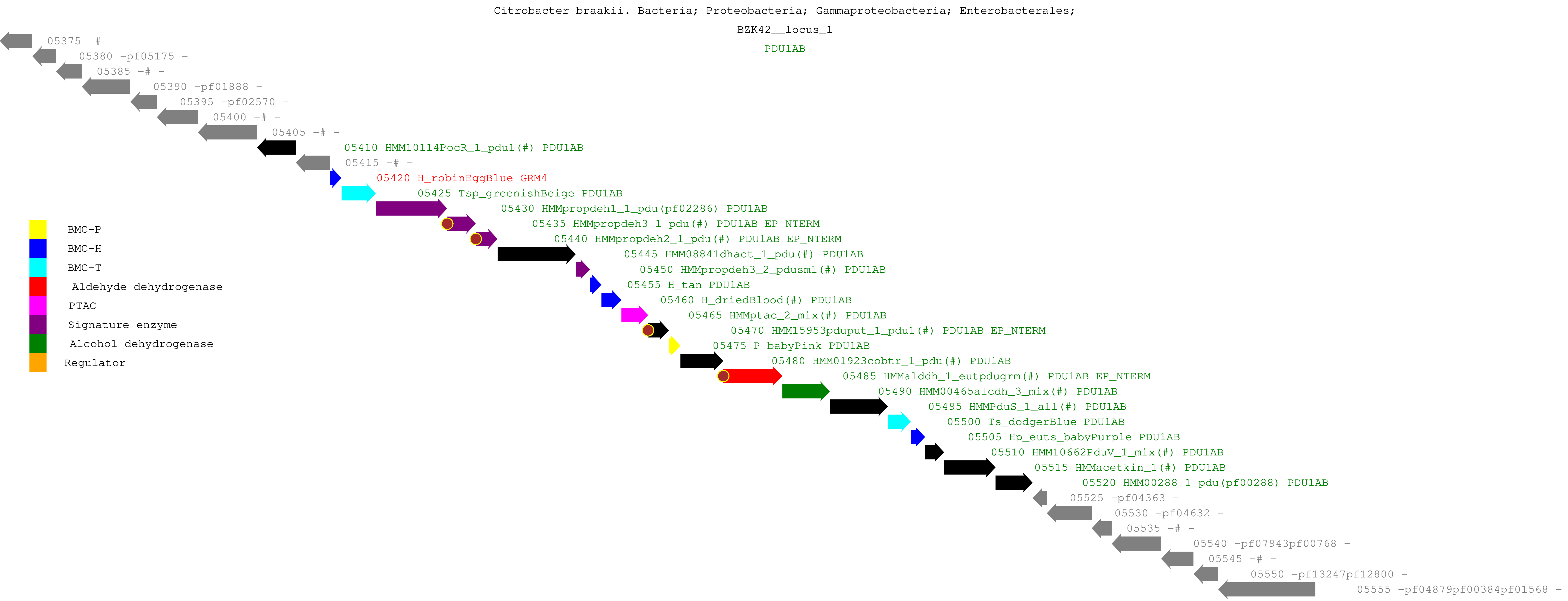

Citrobacter braakii. Bacteria; Proteobacteria; Gammaproteobacteria; Enterobacterales;
BZK42__locus_1
PDU1AB
05375 -# -
05380 -pf05175 -
05385 -# -
05390 -pf01888 -
05395 -pf02570 -
05400 -# -
05405 -# -
05410 HMM10114PocR_1_pdu1(#) PDU1AB
05415 -# -
05420 H_robinEggBlue GRM4
05425 Tsp_greenishBeige PDU1AB
05430 HMMpropdeh1_1_pdu(pf02286) PDU1AB
05435 HMMpropdeh3_1_pdu(#) PDU1AB EP_NTERM
05440 HMMpropdeh2_1_pdu(#) PDU1AB EP_NTERM
05445 HMM08841dhact_1_pdu(#) PDU1AB
05450 HMMpropdeh3_2_pdusml(#) PDU1AB
05455 H_tan PDU1AB
05460 H_driedBlood(#) PDU1AB
05465 HMMptac_2_mix(#) PDU1AB
05470 HMM15953pduput_1_pdu1(#) PDU1AB EP_NTERM
05475 P_babyPink PDU1AB
05480 HMM01923cobtr_1_pdu(#) PDU1AB
05485 HMMalddh_1_eutpdugrm(#) PDU1AB EP_NTERM
05490 HMM00465alcdh_3_mix(#) PDU1AB
05495 HMMPduS_1_all(#) PDU1AB
05500 Ts_dodgerBlue PDU1AB
05505 Hp_euts_babyPurple PDU1AB
05510 HMM10662PduV_1_mix(#) PDU1AB
05515 HMMacetkin_1(#) PDU1AB
05520 HMM00288_1_pdu(pf00288) PDU1AB
05525 -pf04363 -
05530 -pf04632 -
05535 -# -
05540 -pf07943pf00768 -
05545 -# -
05550 -pf13247pf12800 -
05555 -pf04879pf00384pf01568 -
BMC-P
BMC-H
BMC-T
Aldehyde dehydrogenase
PTAC
Signature enzyme
Alcohol dehydrogenase
Regulator

Escherichia coli. Bacteria; Proteobacteria; Gammaproteobacteria; Enterobacterales;

BZL31__locus_1

PDU1AB

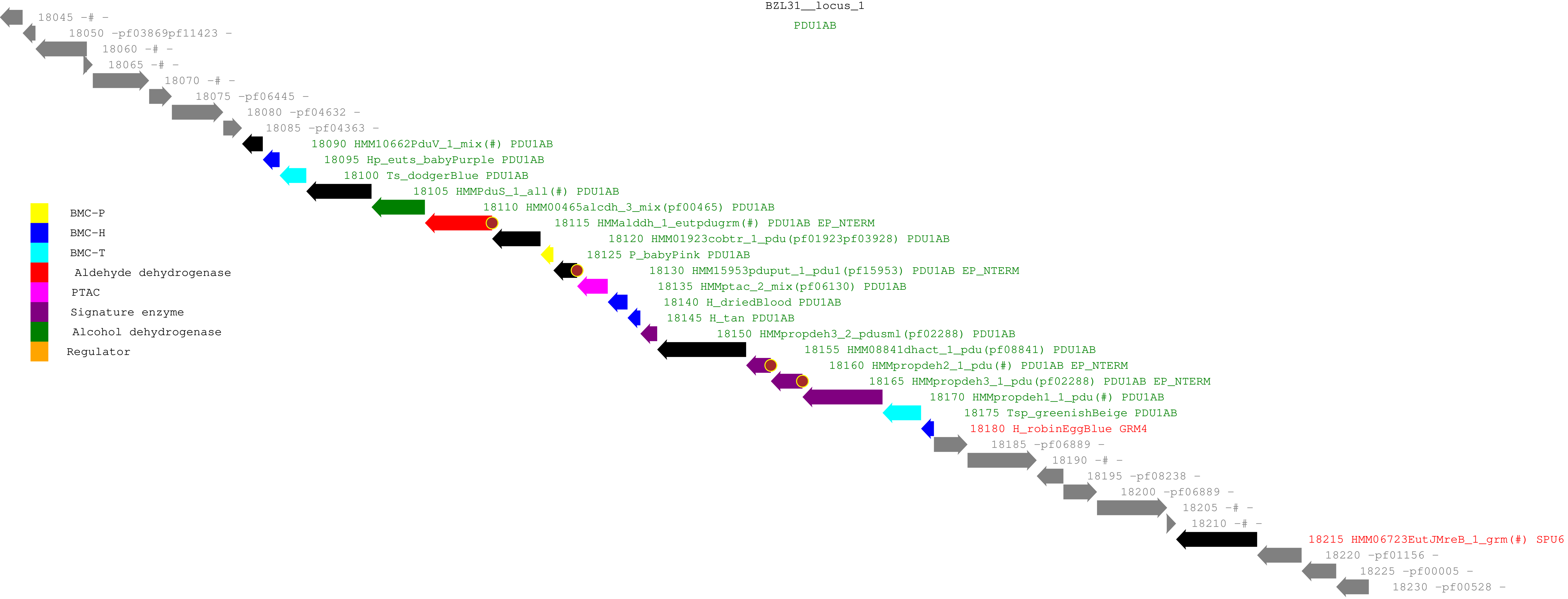

Salmonella enterica I. Bacteria; Proteobacteria; Gammaproteobacteria; Enterobacterales;

BZW54__locus_1

PDU1AB

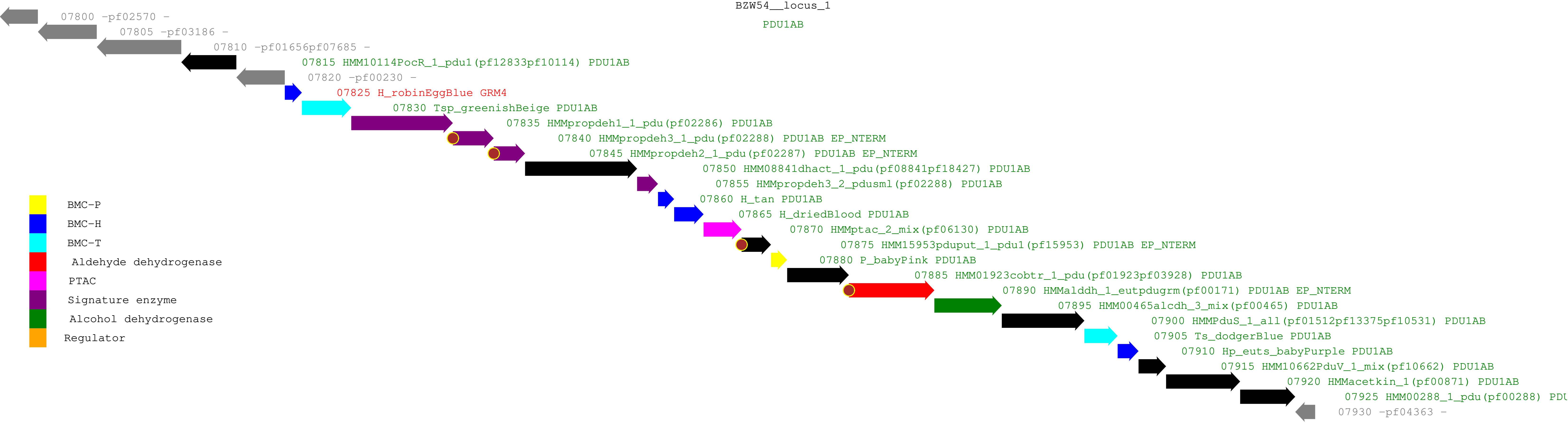

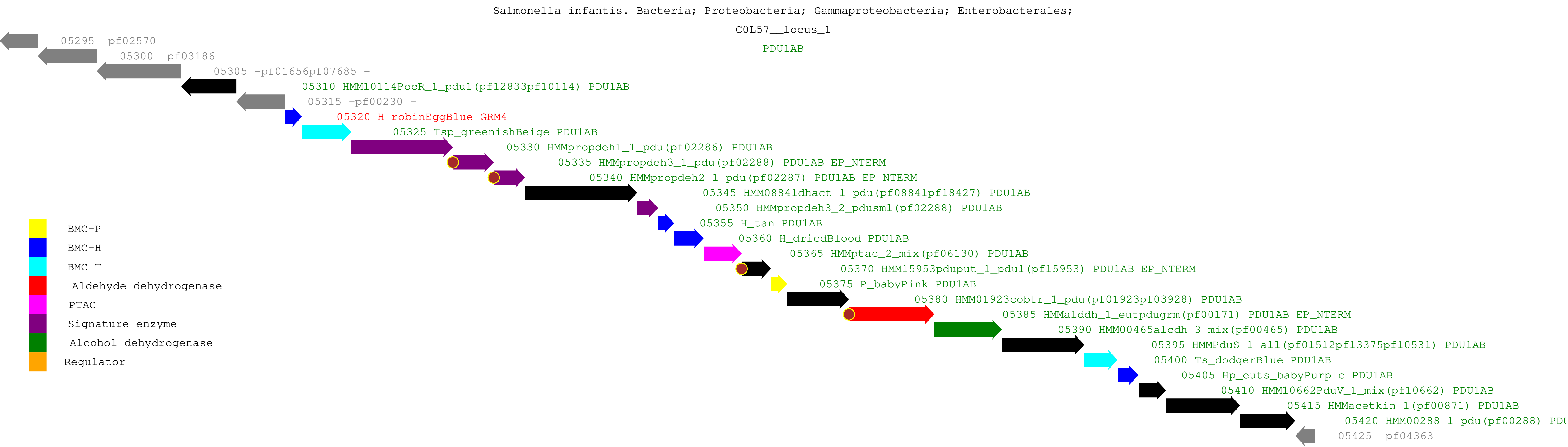
Salmonella infantis. Bacteria; Proteobacteria; Gammaproteobacteria; Enterobacterales;
C0L57__locus_1
PDU1AB
05295 -pf02570 -
05300 -pf03186 -
05305 -pf01656pf07685 -
05310 HMM10114PocR_1_pdu1(pf12833pf10114) PDU1AB
05315 -pf00230 -
05320 H_robinEggBlue GRM4
05325 Tsp_greenishBeige PDU1AB
05330 HMMpropdeh1_1_pdu(pf02286) PDU1AB
05335 HMMpropdeh3_1_pdu(pf02288) PDU1AB EP_NTERM
05340 HMMpropdeh2_1_pdu(pf02287) PDU1AB EP_NTERM
05345 HMM08841dhact_1_pdu(pf08841pf18427) PDU1AB
05350 HMMpropdeh3_2_pdusml(pf02288) PDU1AB
05355 H_tan PDU1AB
05360 H_driedBlood PDU1AB
05365 HMMptac_2_mix(pf06130) PDU1AB
05370 HMM15953pduput_1_pdu1(pf15953) PDU1AB EP_NTERM
05375 P_babyPink PDU1AB
05380 HMM01923cobtr_1_pdu(pf01923pf03928) PDU1AB
05385 HMMalddh_1_eutpdugrm(pf00171) PDU1AB EP_NTERM
05390 HMM00465alcdh_3_mix(pf00465) PDU1AB
05395 HMMPduS_1_all(pf01512pf13375pf10531) PDU1AB
05400 Ts_dodgerBlue PDU1AB
05405 Hp_euts_babyPurple PDU1AB
05410 HMM10662PduV_1_mix(pf10662) PDU1AB
05415 HMMacetkin_1(pf00871) PDU1AB
05420 HMM00288_1_pdu(pf00288) PDU
05425 -pf04363 -
BMC-P
BMC-H
BMC-T
Aldehyde dehydrogenase
PTAC
Signature enzyme
Alcohol dehydrogenase
Regulator

Escherichia marmotae. Bacteria; Proteobacteria; Gammaproteobacteria; Enterobacterales;

C1192__locus_1

PDU1AB

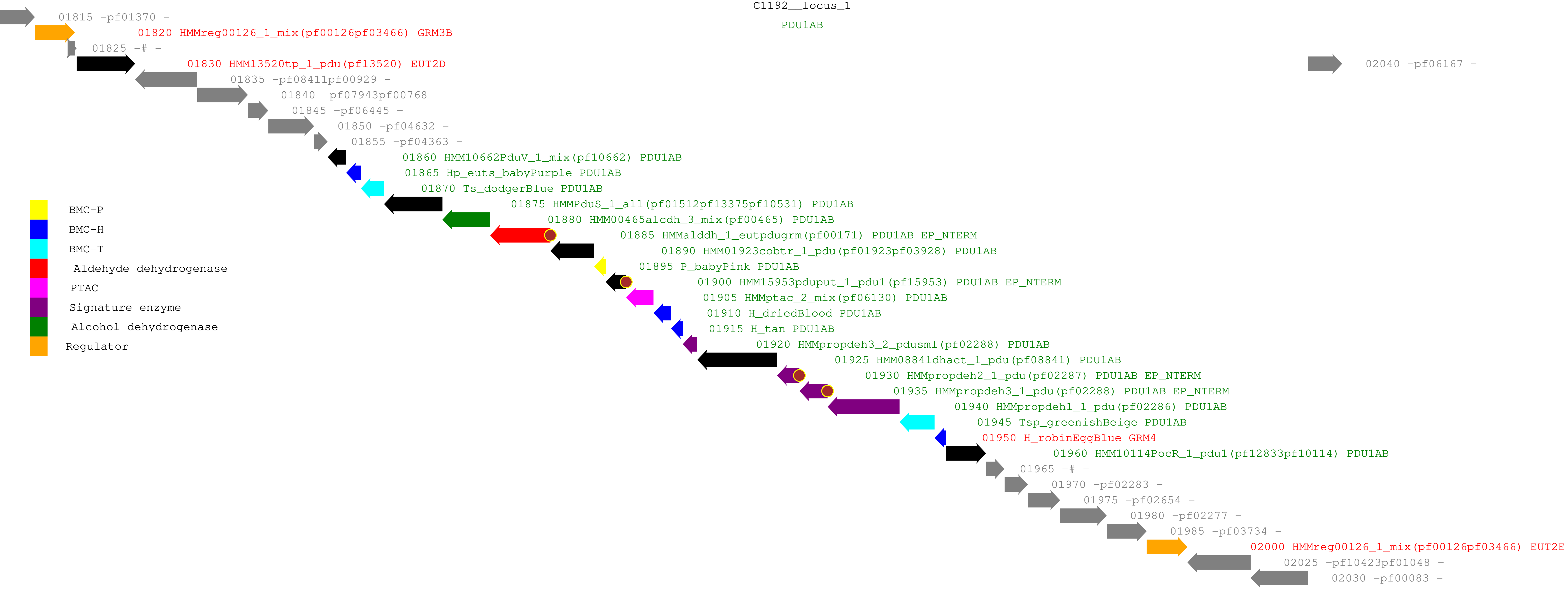

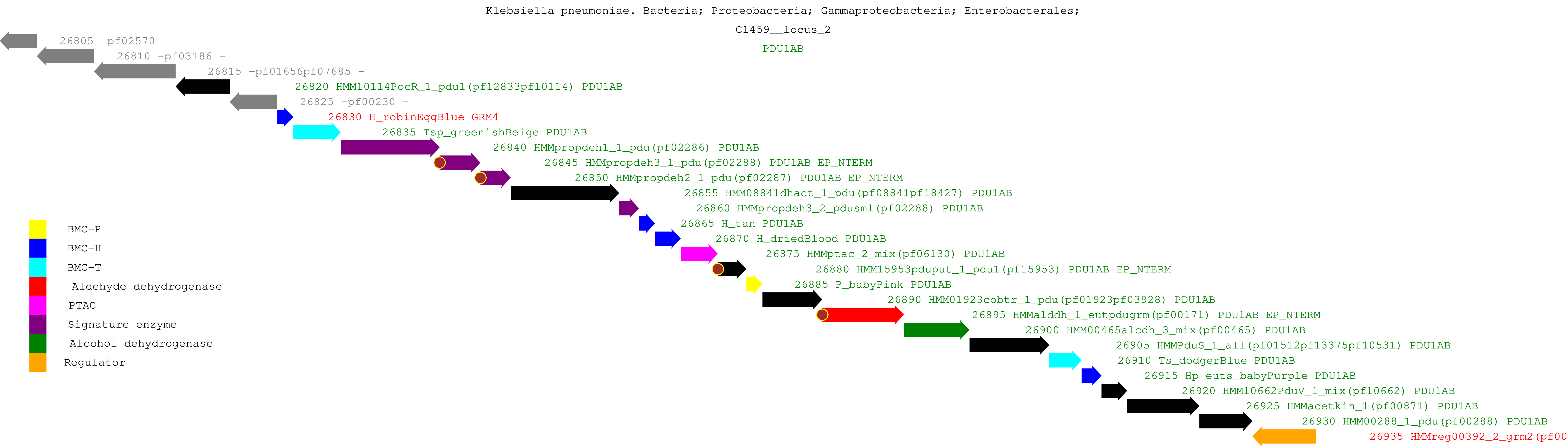
Klebsiella pneumoniae. Bacteria; Proteobacteria; Gammaproteobacteria; Enterobacterales;
C1459__locus_2
PDU1AB
26805 -pf02570 -
26810 -pf03186 -
26815 -pf01656pf07685 -
26820 HMM10114PocR_1_pdu1(pf12833pf10114) PDU1AB
26825 -pf00230 -
26830 H_robinEggBlue GRM4
26835 Tsp_greenishBeige PDU1AB
26840 HMMpropdeh1_1_pdu(pf02286) PDU1AB
26845 HMMpropdeh3_1_pdu(pf02288) PDU1AB EP_NTERM
26850 HMMpropdeh2_1_pdu(pf02287) PDU1AB EP_NTERM
26855 HMM08841dhact_1_pdu(pf08841pf18427) PDU1AB
26860 HMMpropdeh3_2_pdusml(pf02288) PDU1AB
26865 H_tan PDU1AB
26870 H_driedBlood PDU1AB
26875 HMMptac_2_mix(pf06130) PDU1AB
26880 HMM15953pduput_1_pdu1(pf15953) PDU1AB EP_NTERM
26885 P_babyPink PDU1AB
26890 HMM01923cobtr_1_pdu(pf01923pf03928) PDU1AB
26895 HMMalddh_1_eutpdugrm(pf00171) PDU1AB EP_NTERM
26900 HMM00465alcdh_3_mix(pf00465) PDU1AB
26905 HMMPduS_1_all(pf01512pf13375pf10531) PDU1AB
26910 Ts_dodgerBlue PDU1AB
26915 Hp_euts_babyPurple PDU1AB
26920 HMM10662PduV_1_mix(pf10662) PDU1AB
26925 HMMacetkin_1(pf00871) PDU1AB
26930 HMM00288_1_pdu(pf00288) PDU1AB
26935 HMMreg00392_2_grm2(pf00
BMC-P
BMC-H
BMC-T
Aldehyde dehydrogenase
PTAC
Signature enzyme
Alcohol dehydrogenase
Regulator

Escherichia coli. Bacteria; Proteobacteria; Gammaproteobacteria; Enterobacterales;

C1I39__locus_1

PDU1AB

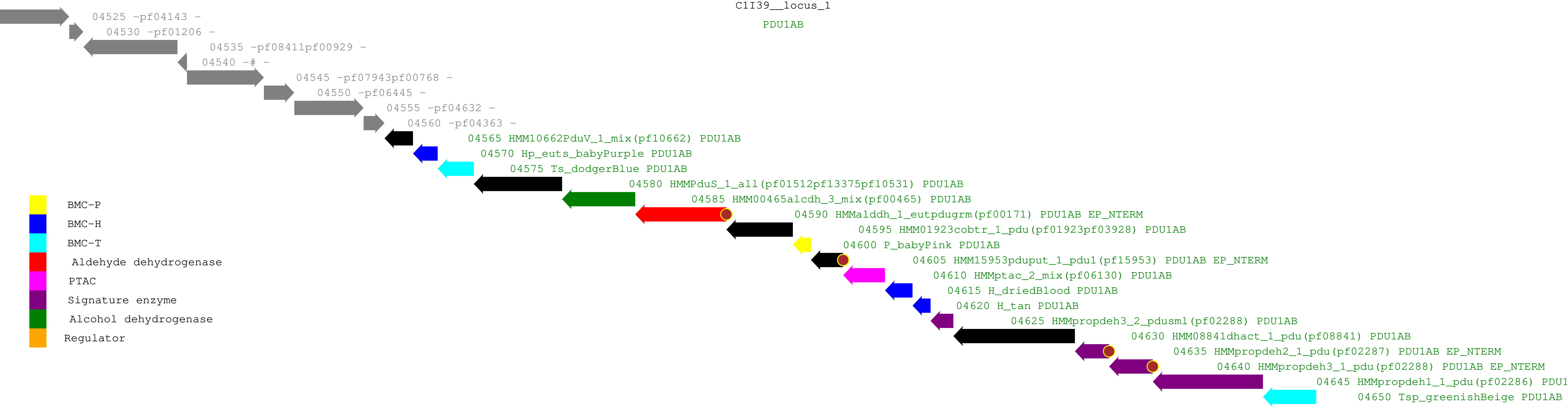

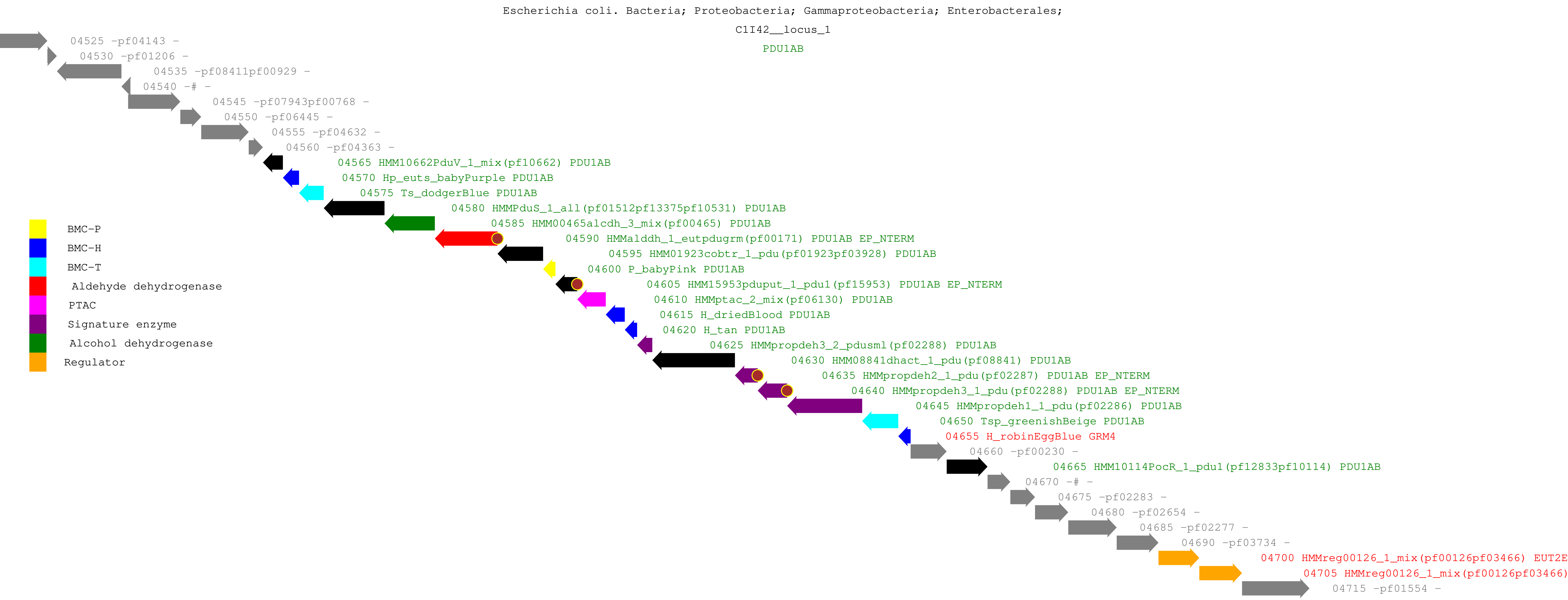
Escherichia coli. Bacteria; Proteobacteria; Gammaproteobacteria; Enterobacterales;
C1I42__locus_1
PDU1AB
04525 -pf04143 -
04530 -pf01206 -
04535 -pf08411pf00929 -
04540 -# -
04545 -pf07943pf00768 -
04550 -pf06445 -
04555 -pf04632 -
04560 -pf04363 -
04565 HMM10662PduV_1_mix(pf10662) PDU1AB
04570 Hp_euts_babyPurple PDU1AB
04575 Ts_dodgerBlue PDU1AB
04580 HMMPduS_1_all(pf01512pf13375pf10531) PDU1AB
04585 HMM00465alcdh_3_mix(pf00465) PDU1AB
04590 HMMaldh_1_eutpdugrm(pf00171) PDU1AB EP_NTERM
04595 HMM01923cobtr_1_pdu(pf01923pf03928) PDU1AB
04600 P_babyPink PDU1AB
04605 HMM15953pduput_1_pdu1(pf15953) PDU1AB EP_NTERM
04610 HMMptac_2_mix(pf06130) PDU1AB
04615 H_driedBlood PDU1AB
04620 H_tan PDU1AB
04625 HMMpropdeh3_2_pdusml(pf02288) PDU1AB
04630 HMM08841dhact_1_pdu(pf08841) PDU1AB
04635 HMMpropdeh2_1_pdu(pf02287) PDU1AB EP_NTERM
04640 HMMpropdeh3_1_pdu(pf02288) PDU1AB EP_NTERM
04645 HMMpropdeh1_1_pdu(pf02286) PDU1AB
04650 Tsp_greenishBeige PDU1AB
04655 H_robinEggBlue GRM4
04660 -pf00230 -
04665 HMM10114PocR_1_pdu1(pf12833pf10114) PDU1AB
04670 -# -
04675 -pf02283 -
04680 -pf02654 -
04685 -pf02277 -
04690 -pf03734 -
04700 HMMreg00126_1_mix(pf00126pf03466) EUT2E
04705 HMMreg00126_1_mix(pf00126pf03466)
04715 -pf01554 -
BMC-P
BMC-H
BMC-T
Aldehyde dehydrogenase
PTAC
Signature enzyme
Alcohol dehydrogenase
Regulator

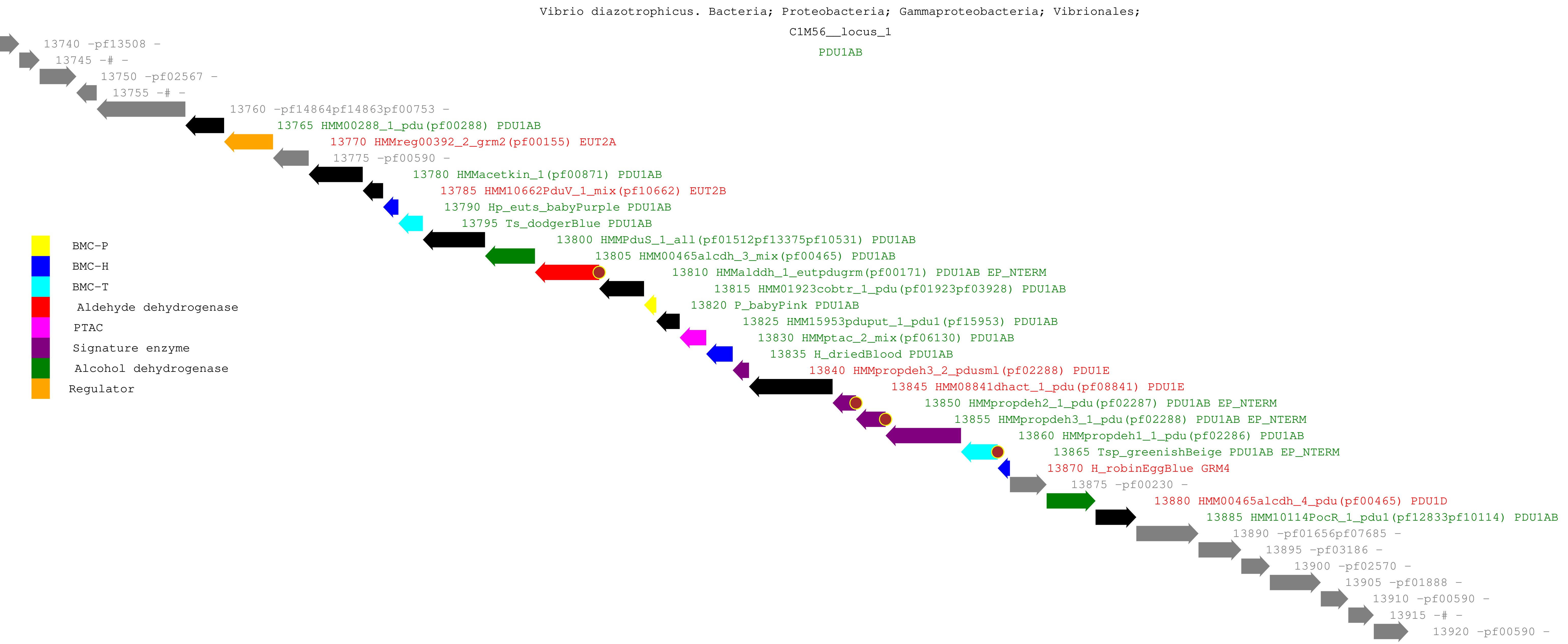
Vibrio diazotrophicus. Bacteria; Proteobacteria; Gammaproteobacteria; Vibrionales;
C1M56__locus_1
PDU1AB
13740 -pf13508 -
13745 -# -
13750 -pf02567 -
13755 -# -
13760 -pf14864pf14863pf00753 -
13765 HMM00288_1_pdu(pf00288) PDU1AB
13770 HMMreg00392_2_grm2(pf00155) EUT2A
13775 -pf00590 -
13780 HMMacetkin_1(pf00871) PDU1AB
13785 HMM10662PduV_1_mix(pf10662) EUT2B
13790 Hp_euts_babyPurple PDU1AB
13795 Ts_dodgerBlue PDU1AB
13800 HMMPduS_1_all(pf01512pf13375pf10531) PDU1AB
13805 HMM00465alcdh_3_mix(pf00465) PDU1AB
13810 HMMalddh_1_eutpdugrm(pf00171) PDU1AB EP_NTERM
13815 HMM01923cobtr_1_pdu(pf01923pf03928) PDU1AB
13820 P_babyPink PDU1AB
13825 HMM15953pduput_1_pdu1(pf15953) PDU1AB
13830 HMMptac_2_mix(pf06130) PDU1AB
13835 H_driedBlood PDU1AB
13840 HMMpropdeh3_2_pdusml(pf02288) PDU1E
13845 HMM08841dhact_1_pdu(pf08841) PDU1E
13850 HMMpropdeh2_1_pdu(pf02287) PDU1AB EP_NTERM
13855 HMMpropdeh3_1_pdu(pf02288) PDU1AB EP_NTERM
13860 HMMpropdeh1_1_pdu(pf02286) PDU1AB
13865 Tsp_greenishBeige PDU1AB EP_NTERM
13870 H_robinEggBlue GRM4
13875 -pf00230 -
13880 HMM00465alcdh_4_pdu(pf00465) PDU1D
13885 HMM10114PocR_1_pdu1(pf12833pf10114) PDU1AB
13890 -pf01656pf07685 -
13895 -pf03186 -
13900 -pf02570 -
13905 -pf01888 -
13910 -pf00590 -
13915 -# -
13920 -pf00590 -
BMC-P
BMC-H
BMC-T
Aldehyde dehydrogenase
PTAC
Signature enzyme
Alcohol dehydrogenase
Regulator

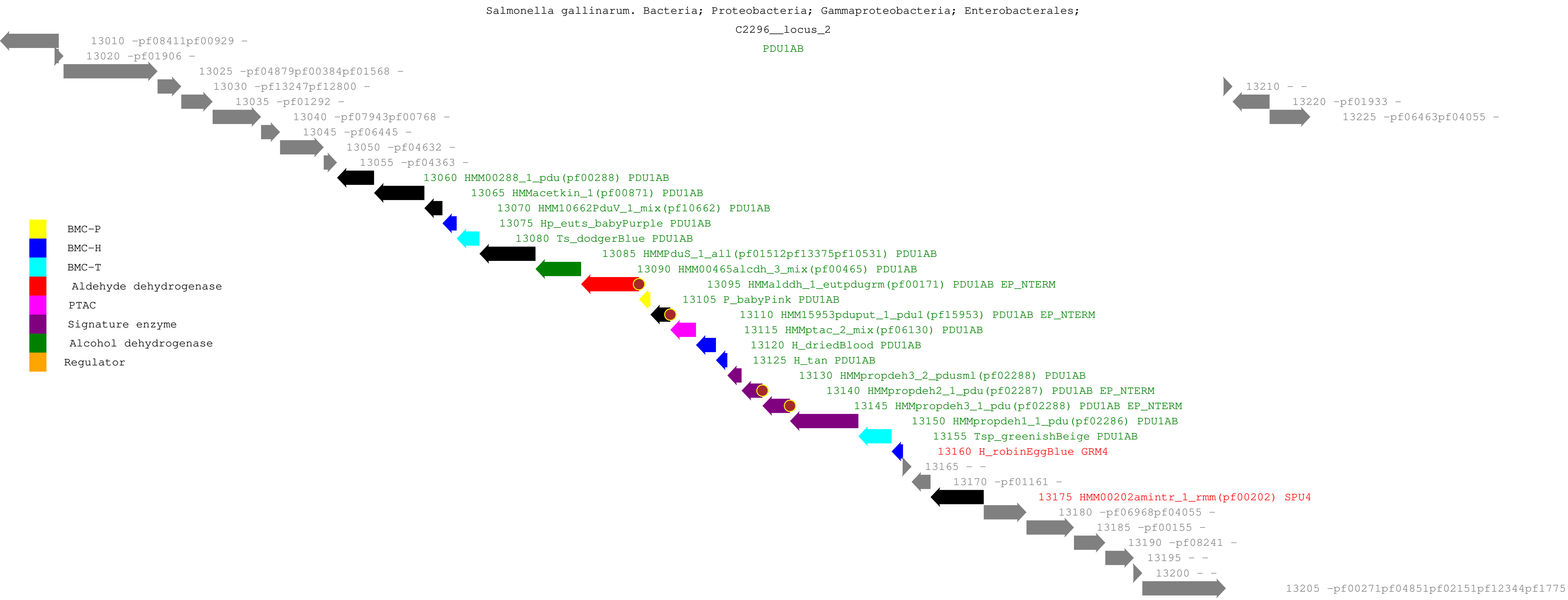
Salmonella gallinarum. Bacteria; Proteobacteria; Gammaproteobacteria; Enterobacterales;
C2296__locus_2
PDU1AB
13010 -pf08411pf00929 -
13020 -pf01906 -
13025 -pf04879pf00384pf01568 -
13030 -pf13247pf12800 -
13035 -pf01292 -
13040 -pf07943pf00768 -
13045 -pf06445 -
13050 -pf04632 -
13055 -pf04363 -
13060 HMM00288_1_pdu(pf00288) PDU1AB
13065 HMMacetkin_1(pf00871) PDU1AB
13070 HMM10662PduV_1_mix(pf10662) PDU1AB
13075 Hp_euts_babyPurple PDU1AB
13080 Ts_dodgerBlue PDU1AB
13085 HMMPduS_1_all(pf01512pf13375pf10531) PDU1AB
13090 HMM00465alcdh_3_mix(pf00465) PDU1AB
13095 HMMalddh_1_eutpdugrm(pf00171) PDU1AB EP_NTERM
13105 P_babyPink PDU1AB
13110 HMM15953pduput_1_pdu1(pf15953) PDU1AB EP_NTERM
13115 HMMptac_2_mix(pf06130) PDU1AB
13120 H_driedBlood PDU1AB
13125 H_tan PDU1AB
13130 HMMpropdeh3_2_pdusml(pf02288) PDU1AB
13140 HMMpropdeh2_1_pdu(pf02287) PDU1AB EP_NTERM
13145 HMMpropdeh3_1_pdu(pf02288) PDU1AB EP_NTERM
13150 HMMpropdeh1_1_pdu(pf02286) PDU1AB
13155 Tsp_greenishBeige PDU1AB
13160 H_robinEggBlue GRM4
13165 - -
13170 -pf01161 -
13175 HMM00202amintr_1_rmm(pf00202) SPU4
13180 -pf06968pf04055 -
13185 -pf00155 -
13190 -pf08241 -
13195 - -
13200 - -
13205 -pf00271pf04851pf02151pf12344pf1775
13210 - -
13220 -pf01933 -
13225 -pf06463pf04055 -
BMC-P
BMC-H
BMC-T
Aldehyde dehydrogenase
PTAC
Signature enzyme
Alcohol dehydrogenase
Regulator

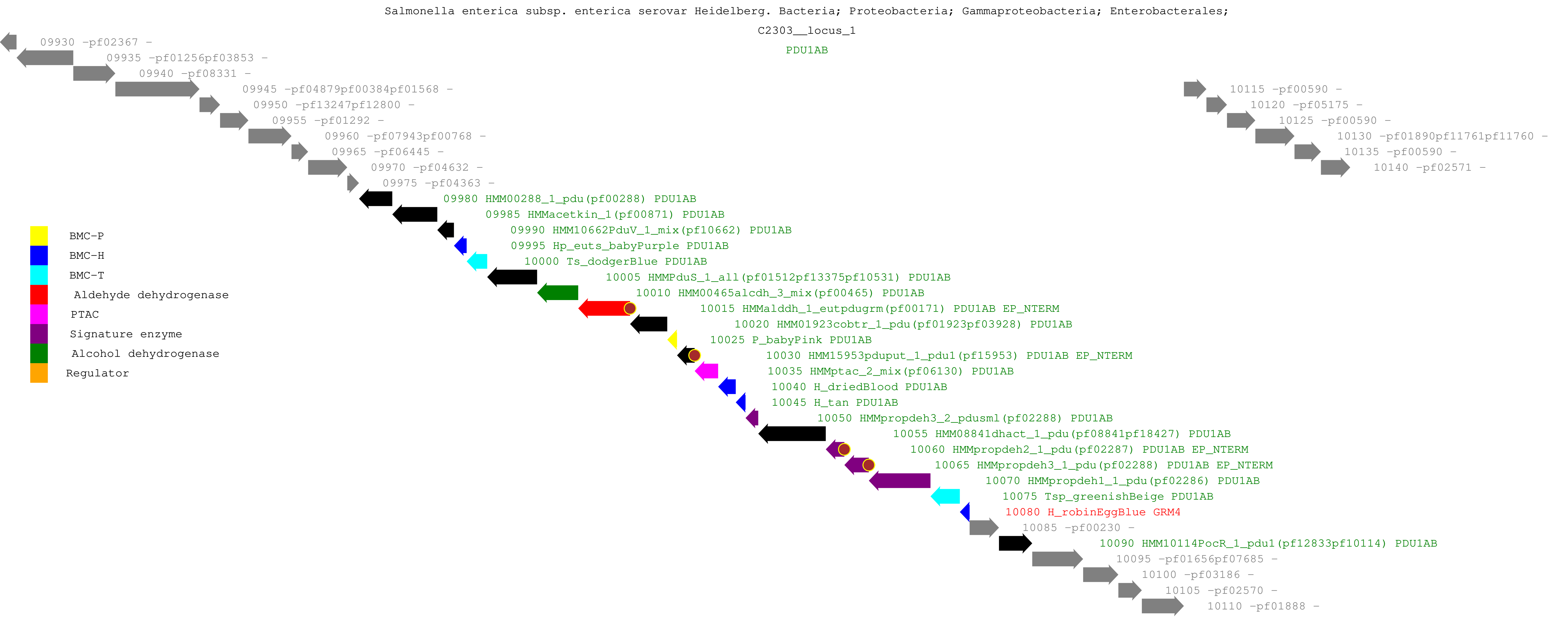
Salmonella enterica subsp. enterica serovar Heidelberg. Bacteria; Proteobacteria; Gammaproteobacteria; Enterobacterales;
C2303__locus_1
PDU1AB
09930 -pf02367 -
09935 -pf01256pf03853 -
09940 -pf08331 -
09945 -pf04879pf00384pf01568 -
09950 -pf13247pf12800 -
09955 -pf01292 -
09960 -pf07943pf00768 -
09965 -pf06445 -
09970 -pf04632 -
09975 -pf04363 -
09980 HMM00288_1_pdu(pf00288) PDU1AB
09985 HMMacetkin_1(pf00871) PDU1AB
09990 HMM10662PduV_1_mix(pf10662) PDU1AB
09995 Hp_euts_babyPurple PDU1AB
10000 Ts_dodgerBlue PDU1AB
10005 HMMPduS_1_all(pf01512pf13375pf10531) PDU1AB
10010 HMM00465alcdh_3_mix(pf00465) PDU1AB
10015 HMMalddh_1_eutpdugrm(pf00171) PDU1AB EP_NTERM
10020 HMM01923cobtr_1_pdu(pf01923pf03928) PDU1AB
10025 P_babyPink PDU1AB
10030 HMM15953pduput_1_pdu1(pf15953) PDU1AB EP_NTERM
10035 HMMptac_2_mix(pf06130) PDU1AB
10040 H_driedBlood PDU1AB
10045 H_tan PDU1AB
10050 HMMpropdeh3_2_pdusml(pf02288) PDU1AB
10055 HMM08841dhact_1_pdu(pf08841pf18427) PDU1AB
10060 HMMpropdeh2_1_pdu(pf02287) PDU1AB EP_NTERM
10065 HMMpropdeh3_1_pdu(pf02288) PDU1AB EP_NTERM
10070 HMMpropdeh1_1_pdu(pf02286) PDU1AB
10075 Tsp_greenishBeige PDU1AB
10080 H_robinEggBlue GRM4
10085 -pf00230 -
10090 HMM10114PocR_1_pdu1(pf12833pf10114) PDU1AB
10095 -pf01656pf07685 -
10100 -pf03186 -
10105 -pf02570 -
10110 -pf01888 -
10115 -pf00590 -
10120 -pf05175 -
10125 -pf00590 -
10130 -pf01890pf11761pf11760 -
10135 -pf00590 -
10140 -pf02571 -
BMC-P
BMC-H
BMC-T
Aldehyde dehydrogenase
PTAC
Signature enzyme
Alcohol dehydrogenase
Regulator

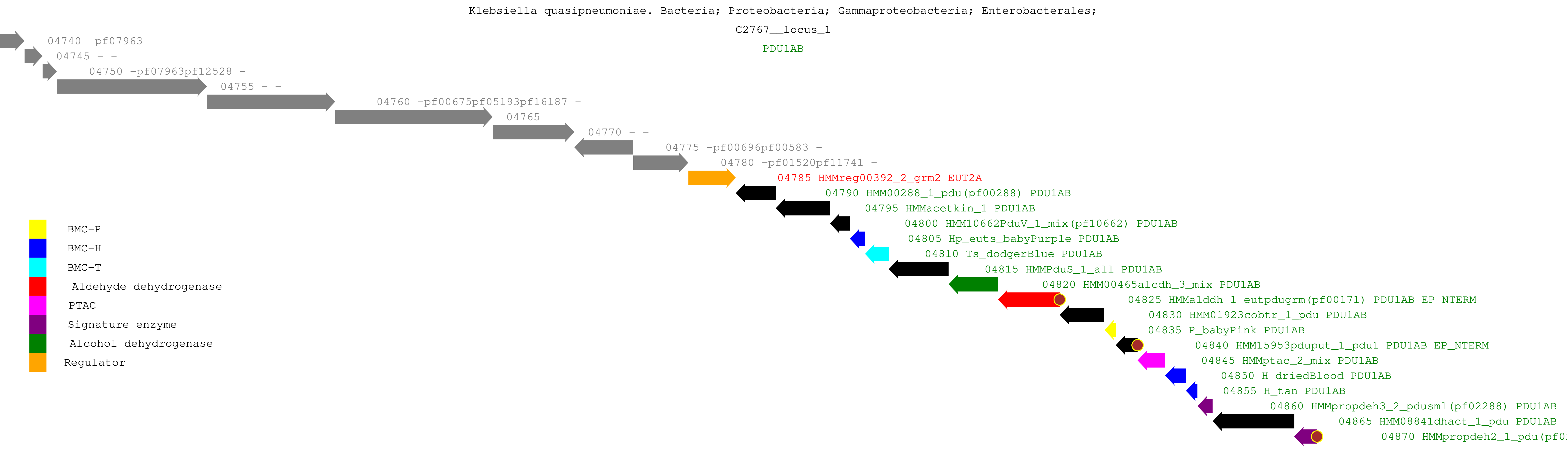
Klebsiella quasipneumoniae. Bacteria; Proteobacteria; Gammaproteobacteria; Enterobacterales;
C2767__locus_1
PDU1AB
04740 -pf07963 -
04745 - -
04750 -pf07963pf12528 -
04755 - -
04760 -pf00675pf05193pf16187 -
04765 - -
04770 - -
04775 -pf00696pf00583 -
04780 -pf01520pf11741 -
04785 HMMreg00392_2_grm2 EUT2A
04790 HMM00288_1_pdu(pf00288) PDU1AB
04795 HMMacetkin_1 PDU1AB
04800 HMM10662PduV_1_mix(pf10662) PDU1AB
04805 Hp_euts_babyPurple PDU1AB
04810 Ts_dodgerBlue PDU1AB
04815 HMMPduS_1_all PDU1AB
04820 HMM00465alcdh_3_mix PDU1AB
04825 HMMaldh_1_eutpdugrm(pf00171) PDU1AB EP_NTERM
04830 HMM01923cobtr_1_pdu PDU1AB
04835 P_babyPink PDU1AB
04840 HMM15953pduput_1_pdu1 PDU1AB EP_NTERM
04845 HMMptac_2_mix PDU1AB
04850 H_driedBlood PDU1AB
04855 H_tan PDU1AB
04860 HMMpropdeh3_2_pdusml(pf02288) PDU1AB
04865 HMM08841dhact_1_pdu PDU1AB
04870 HMMpropdeh2_1_pdu(pf0
BMC-P
BMC-H
BMC-T
Aldehyde dehydrogenase
PTAC
Signature enzyme
Alcohol dehydrogenase
Regulator

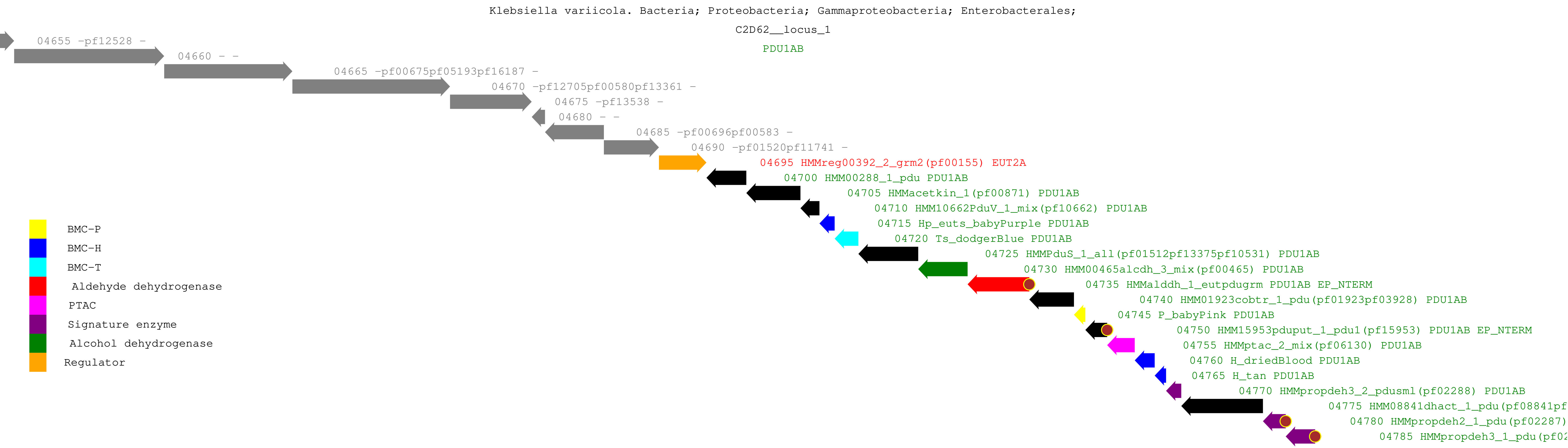
Klebsiella variicola. Bacteria; Proteobacteria; Gammaproteobacteria; Enterobacterales;
C2D62__locus_1
PDU1AB
04655 -pf12528 -
04660 - -
04665 -pf00675pf05193pf16187 -
04670 -pf12705pf00580pf13361 -
04675 -pf13538 -
04680 - -
04685 -pf00696pf00583 -
04690 -pf01520pf11741 -
04695 HMMreg00392_2_grm2(pf00155) EUT2A
04700 HMM00288_1_pdu PDU1AB
04705 HMMacetkin_1(pf00871) PDU1AB
04710 HMM10662PduV_1_mix(pf10662) PDU1AB
04715 Hp_euts_babyPurple PDU1AB
04720 Ts_dodgerBlue PDU1AB
04725 HMMPduS_1_all(pf01512pf13375pf10531) PDU1AB
04730 HMM00465alcdh_3_mix(pf00465) PDU1AB
04735 HMMalddh_1_eutpdugrm PDU1AB EP_NTERM
04740 HMM01923cobtr_1_pdu(pf01923pf03928) PDU1AB
04745 P_babyPink PDU1AB
04750 HMM15953pduput_1_pdu1(pf15953) PDU1AB EP_NTERM
04755 HMMptac_2_mix(pf06130) PDU1AB
04760 H_driedBlood PDU1AB
04765 H_tan PDU1AB
04770 HMMpropdeh3_2_pdusml(pf02288) PDU1AB
04775 HMM08841dhact_1_pdu(pf08841pf
04780 HMMpropdeh2_1_pdu(pf02287)
04785 HMMpropdeh3_1_pdu(pf02
BMC-P
BMC-H
BMC-T
Aldehyde dehydrogenase
PTAC
Signature enzyme
Alcohol dehydrogenase
Regulator

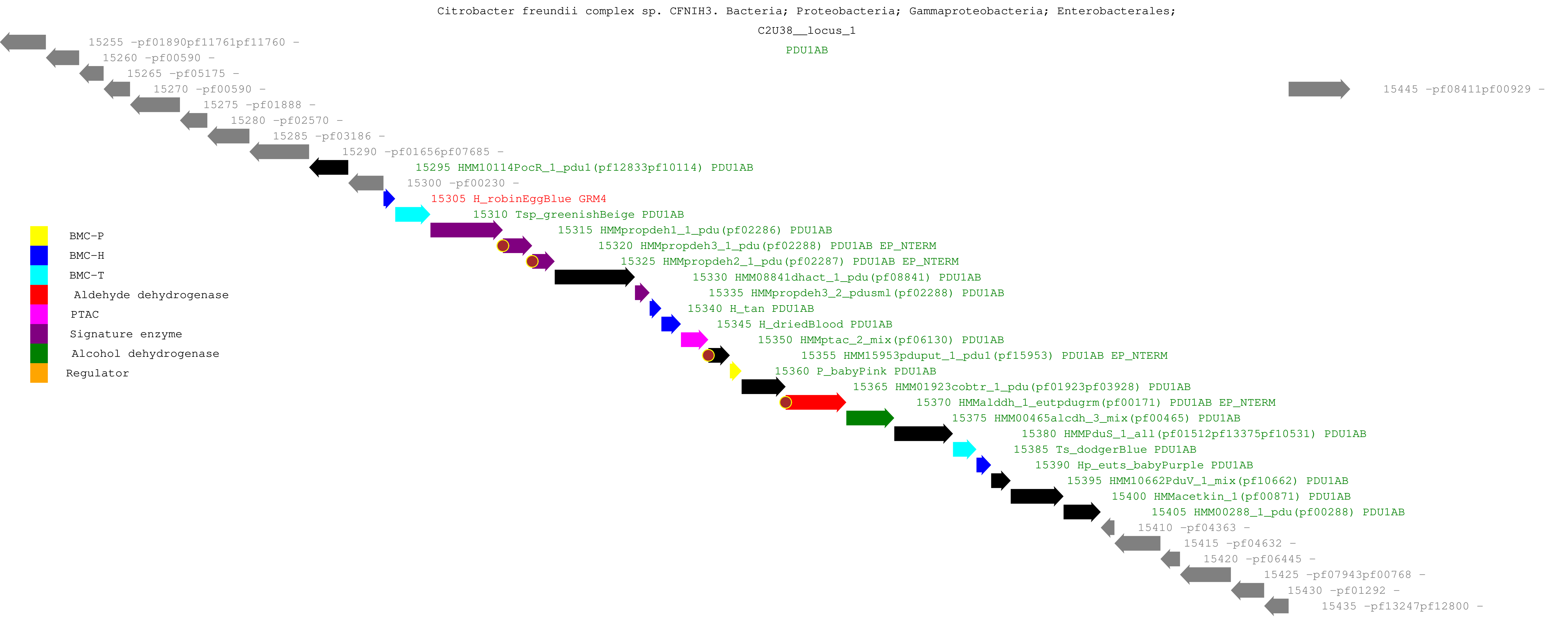
Citrobacter freundii complex sp. CFNIH3. Bacteria; Proteobacteria; Gammaproteobacteria; Enterobacterales;
C2U38__locus_1
PDU1AB
15255 -pf01890pf11761pf11760 -
15260 -pf00590 -
15265 -pf05175 -
15270 -pf00590 -
15275 -pf01888 -
15280 -pf02570 -
15285 -pf03186 -
15290 -pf01656pf07685 -
15295 HMM10114PocR_1_pdu1(pf12833pf10114) PDU1AB
15300 -pf00230 -
15305 H_robinEggBlue GRM4
15310 Tsp_greenishBeige PDU1AB
15315 HMMpropdeh1_1_pdu(pf02286) PDU1AB
15320 HMMpropdeh3_1_pdu(pf02288) PDU1AB EP_NTERM
15325 HMMpropdeh2_1_pdu(pf02287) PDU1AB EP_NTERM
15330 HMM08841dhact_1_pdu(pf08841) PDU1AB
15335 HMMpropdeh3_2_pdusml(pf02288) PDU1AB
15340 H_tan PDU1AB
15345 H_driedBlood PDU1AB
15350 HMMptac_2_mix(pf06130) PDU1AB
15355 HMM15953pduput_1_pdu1(pf15953) PDU1AB EP_NTERM
15360 P_babyPink PDU1AB
15365 HMM01923cobtr_1_pdu(pf01923pf03928) PDU1AB
15370 HMMalddh_1_eutpdugrm(pf00171) PDU1AB EP_NTERM
15375 HMM00465alcdh_3_mix(pf00465) PDU1AB
15380 HMMPduS_1_all(pf01512pf13375pf10531) PDU1AB
15385 Ts_dodgerBlue PDU1AB
15390 Hp_euts_babyPurple PDU1AB
15395 HMM10662PduV_1_mix(pf10662) PDU1AB
15400 HMMacetkin_1(pf00871) PDU1AB
15405 HMM00288_1_pdu(pf00288) PDU1AB
15410 -pf04363 -
15415 -pf04632 -
15420 -pf06445 -
15425 -pf07943pf00768 -
15430 -pf01292 -
15435 -pf13247pf12800 -
15445 -pf08411pf00929 -
BMC-P
BMC-H
BMC-T
Aldehyde dehydrogenase
PTAC
Signature enzyme
Alcohol dehydrogenase
Regulator

Citrobacter freundii complex sp. CFNIH4. Bacteria; Proteobacteria; Gammaproteobacteria; Enterobacterales;

C2U41__locus_1

PDU1AB

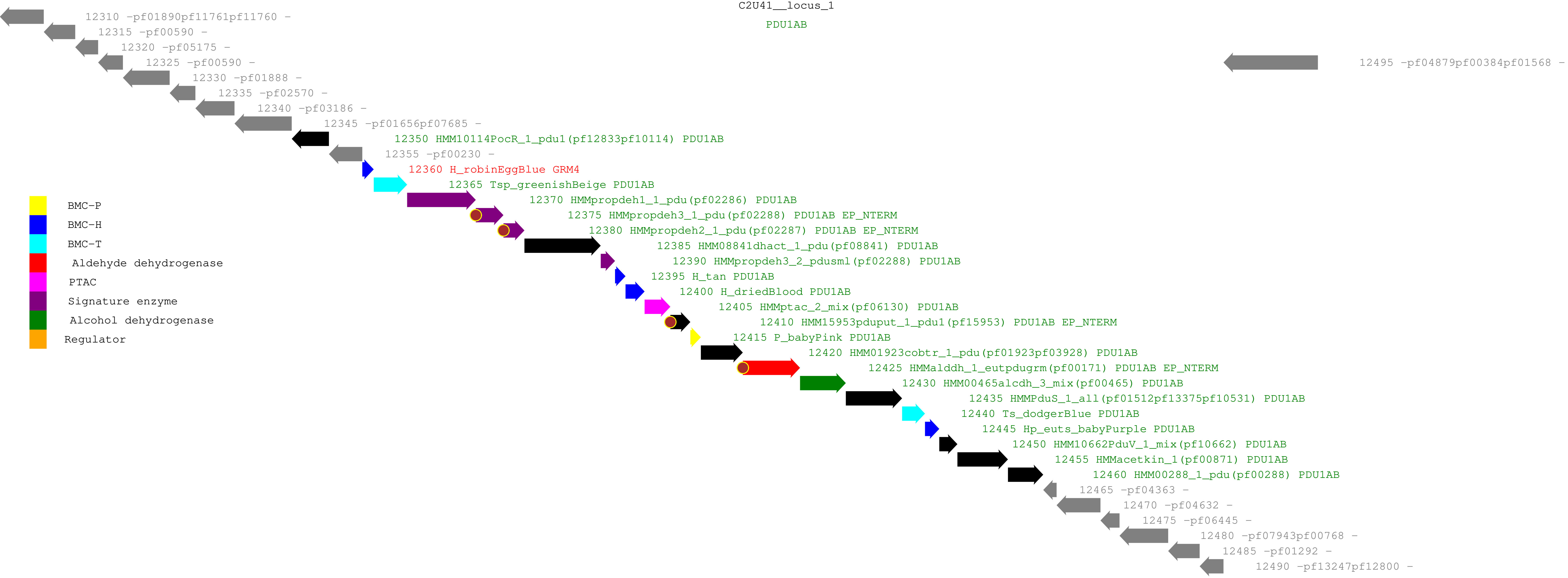

Klebsiella oxytoca. Bacteria; Proteobacteria; Gammaproteobacteria; Enterobacterales;

C2U42__locus_1

PDU1AB

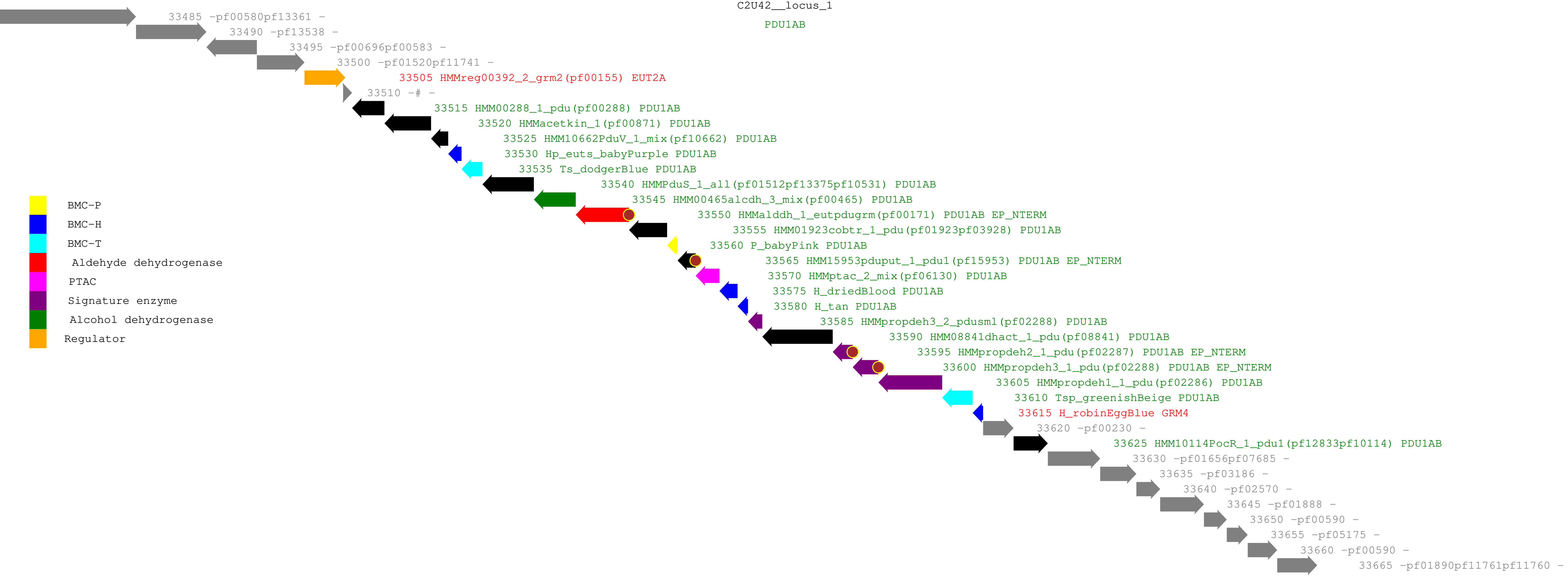

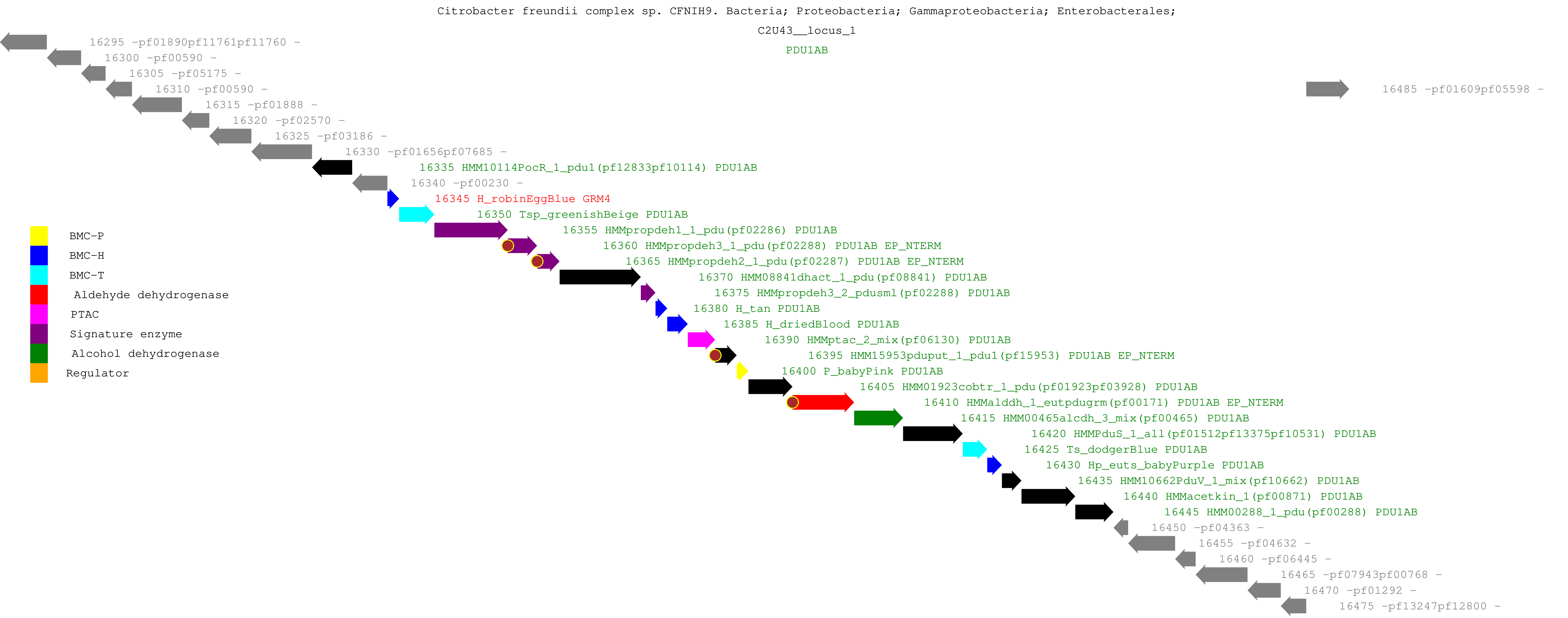

Citrobacter freundii complex sp. CFNIH9. Bacteria; Proteobacteria; Gammaproteobacteria; Enterobacterales;
C2U43__locus_1
PDU1AB
16295 -pf01890pf11761pf11760 -
16300 -pf00590 -
16305 -pf05175 -
16310 -pf00590 -
16315 -pf01888 -
16320 -pf02570 -
16325 -pf03186 -
16330 -pf01656pf07685 -
16335 HMM10114PocR_1_pdu1(pf12833pf10114) PDU1AB
16340 -pf00230 -
16345 H_robinEggBlue GRM4
16350 Tsp_greenishBeige PDU1AB
16355 HMMpropdeh1_1_pdu(pf02286) PDU1AB
16360 HMMpropdeh3_1_pdu(pf02288) PDU1AB EP_NTERM
16365 HMMpropdeh2_1_pdu(pf02287) PDU1AB EP_NTERM
16370 HMM08841dhact_1_pdu(pf08841) PDU1AB
16375 HMMpropdeh3_2_pdusml(pf02288) PDU1AB
16380 H_tan PDU1AB
16385 H_driedBlood PDU1AB
16390 HMMptac_2_mix(pf06130) PDU1AB
16395 HMM15953pduput_1_pdu1(pf15953) PDU1AB EP_NTERM
16400 P_babyPink PDU1AB
16405 HMM01923cobtr_1_pdu(pf01923pf03928) PDU1AB
16410 HMMalddh_1_eutpdugrm(pf00171) PDU1AB EP_NTERM
16415 HMM00465alcdh_3_mix(pf00465) PDU1AB
16420 HMMPduS_1_all(pf01512pf13375pf10531) PDU1AB
16425 Ts_dodgerBlue PDU1AB
16430 Hp_euts_babyPurple PDU1AB
16435 HMM10662PduV_1_mix(pf10662) PDU1AB
16440 HMMacetkin_1(pf00871) PDU1AB
16445 HMM00288_1_pdu(pf00288) PDU1AB
16450 -pf04363 -
16455 -pf04632 -
16460 -pf06445 -
16465 -pf07943pf00768 -
16470 -pf01292 -
16475 -pf13247pf12800 -
16485 -pf01609pf05598 -
BMC-P
BMC-H
BMC-T
Aldehyde dehydrogenase
PTAC
Signature enzyme
Alcohol dehydrogenase
Regulator

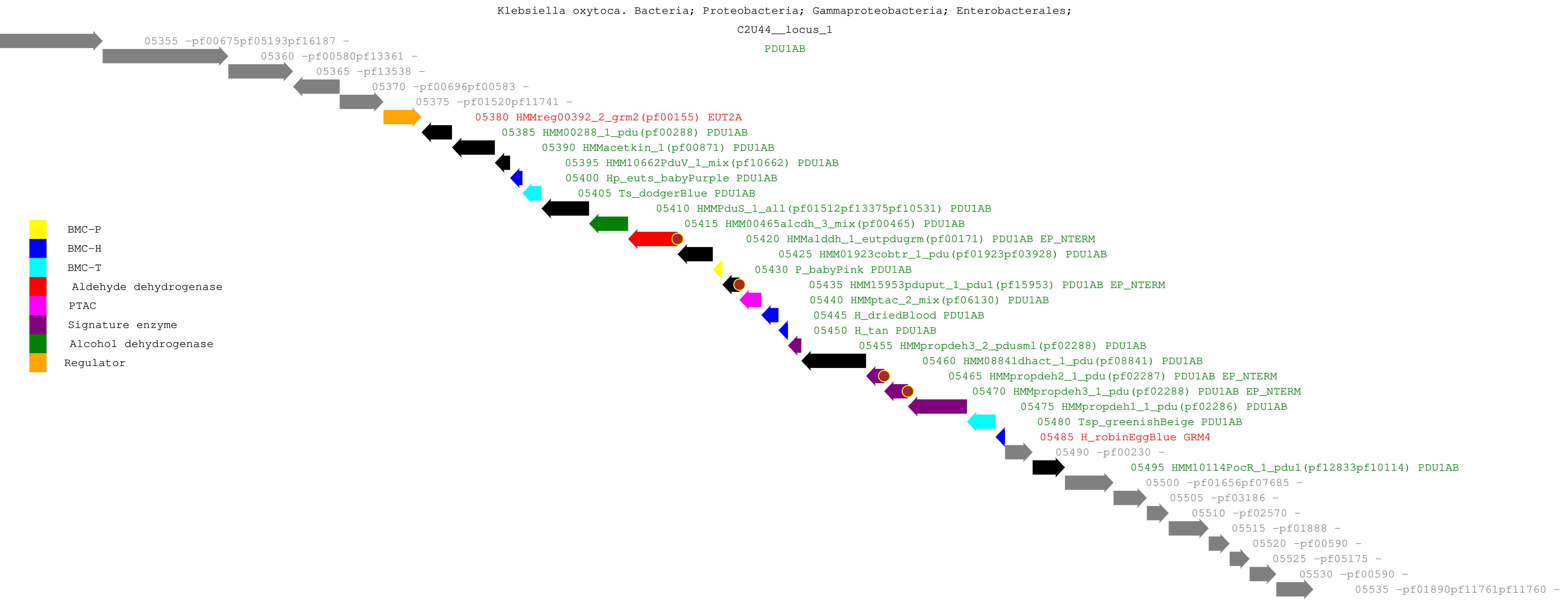

Klebsiella oxytoca. Bacteria; Proteobacteria; Gammaproteobacteria; Enterobacterales;
C2U44__locus_1
PDU1AB
05355 -pf00675pf05193pf16187 -
05360 -pf00580pf13361 -
05365 -pf13538 -
05370 -pf00696pf00583 -
05375 -pf01520pf11741 -
05380 HMMreg00392_2_grm2(pf00155) EUT2A
05385 HMM00288_1_pdu(pf00288) PDU1AB
05390 HMMacetkin_1(pf00871) PDU1AB
05395 HMM10662PduV_1_mix(pf10662) PDU1AB
05400 Hp_euts_babyPurple PDU1AB
05405 Ts_dodgerBlue PDU1AB
05410 HMMPduS_1_all(pf01512pf13375pf10531) PDU1AB
05415 HMM00465alcdh_3_mix(pf00465) PDU1AB
05420 HMMalddh_1_eutpdugrm(pf00171) PDU1AB EP_NTERM
05425 HMM01923cobtr_1_pdu(pf01923pf03928) PDU1AB
05430 P_babyPink PDU1AB
05435 HMM15953pduput_1_pdu1(pf15953) PDU1AB EP_NTERM
05440 HMMptac_2_mix(pf06130) PDU1AB
05445 H_driedBlood PDU1AB
05450 H_tan PDU1AB
05455 HMMpropdeh3_2_pdusml(pf02288) PDU1AB
05460 HMM08841dhact_1_pdu(pf08841) PDU1AB
05465 HMMpropdeh2_1_pdu(pf02287) PDU1AB EP_NTERM
05470 HMMpropdeh3_1_pdu(pf02288) PDU1AB EP_NTERM
05475 HMMpropdeh1_1_pdu(pf02286) PDU1AB
05480 Tsp_greenishBeige PDU1AB
05485 H_robinEggBlue GRM4
05490 -pf00230 -
05495 HMM10114PocR_1_pdu1(pf12833pf10114) PDU1AB
05500 -pf01656pf07685 -
05505 -pf03186 -
05510 -pf02570 -
05515 -pf01888 -
05520 -pf00590 -
05525 -pf05175 -
05530 -pf00590 -
05535 -pf01890pf11761pf11760 -
BMC-P
BMC-H
BMC-T
Aldehyde dehydrogenase
PTAC
Signature enzyme
Alcohol dehydrogenase
Regulator

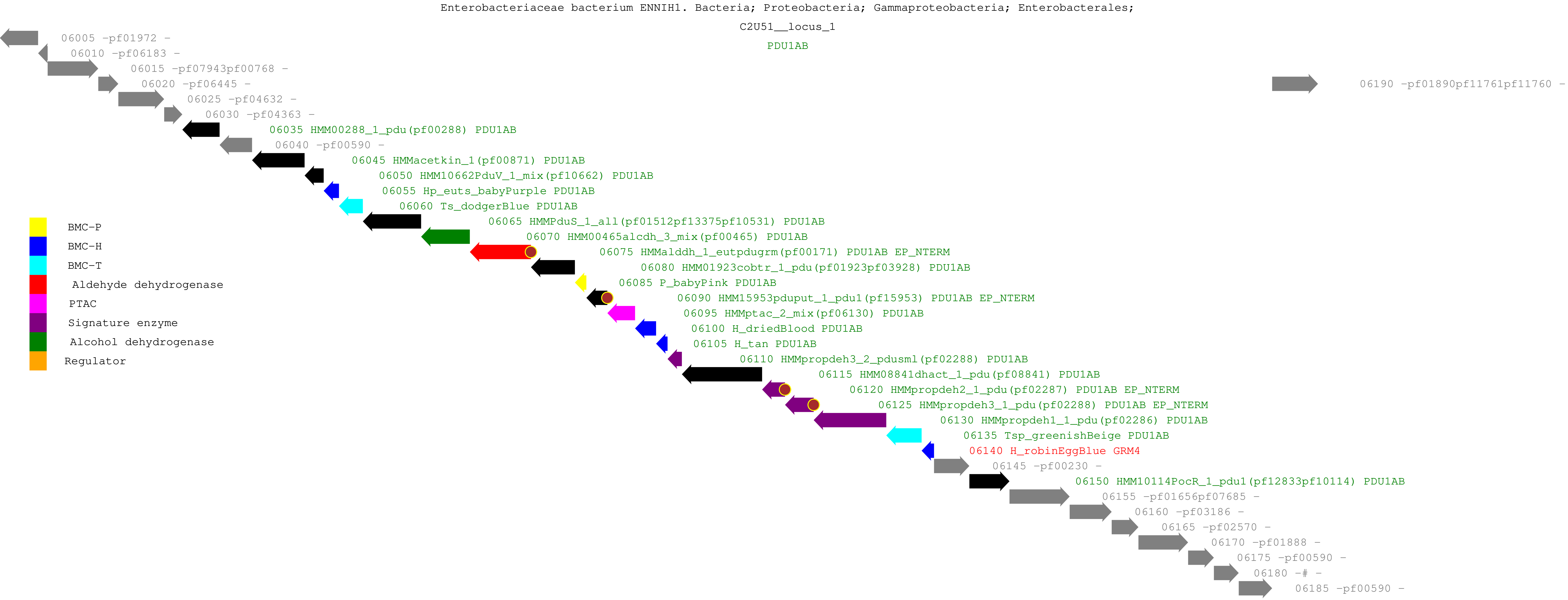
Enterobacteriaceae bacterium ENNIH1. Bacteria; Proteobacteria; Gammaproteobacteria; Enterobacterales;
C2U51__locus_1
PDU1AB
06005 -pf01972 -
06010 -pf06183 -
06015 -pf07943pf00768 -
06020 -pf06445 -
06190 -pf01890pf11761pf11760 -
06025 -pf04632 -
06030 -pf04363 -
06035 HMM00288_1_pdu(pf00288) PDU1AB
06040 -pf00590 -
06045 HMMacetkin_1(pf00871) PDU1AB
06050 HMM10662PduV_1_mix(pf10662) PDU1AB
06055 Hp_euts_babyPurple PDU1AB
06060 Ts_dodgerBlue PDU1AB
06065 HMMPduS_1_all(pf01512pf13375pf10531) PDU1AB
06070 HMM00465alcdh_3_mix(pf00465) PDU1AB
06075 HMMalddh_1_eutpdugrm(pf00171) PDU1AB EP_NTERM
06080 HMM01923cobtr_1_pdu(pf01923pf03928) PDU1AB
06085 P_babyPink PDU1AB
06090 HMM15953pduput_1_pdu1(pf15953) PDU1AB EP_NTERM
06095 HMMptac_2_mix(pf06130) PDU1AB
06100 H_driedBlood PDU1AB
06105 H_tan PDU1AB
06110 HMMpropdeh3_2_pdusml(pf02288) PDU1AB
06115 HMM08841dhact_1_pdu(pf08841) PDU1AB
06120 HMMpropdeh2_1_pdu(pf02287) PDU1AB EP_NTERM
06125 HMMpropdeh3_1_pdu(pf02288) PDU1AB EP_NTERM
06130 HMMpropdeh1_1_pdu(pf02286) PDU1AB
06135 Tsp_greenishBeige PDU1AB
06140 H_robinEggBlue GRM4
06145 -pf00230 -
06150 HMM10114PocR_1_pdu1(pf12833pf10114) PDU1AB
06155 -pf01656pf07685 -
06160 -pf03186 -
06165 -pf02570 -
06170 -pf01888 -
06175 -pf00590 -
06180 -# -
06185 -pf00590 -
BMC-P
BMC-H
BMC-T
Aldehyde dehydrogenase
PTAC
Signature enzyme
Alcohol dehydrogenase
Regulator

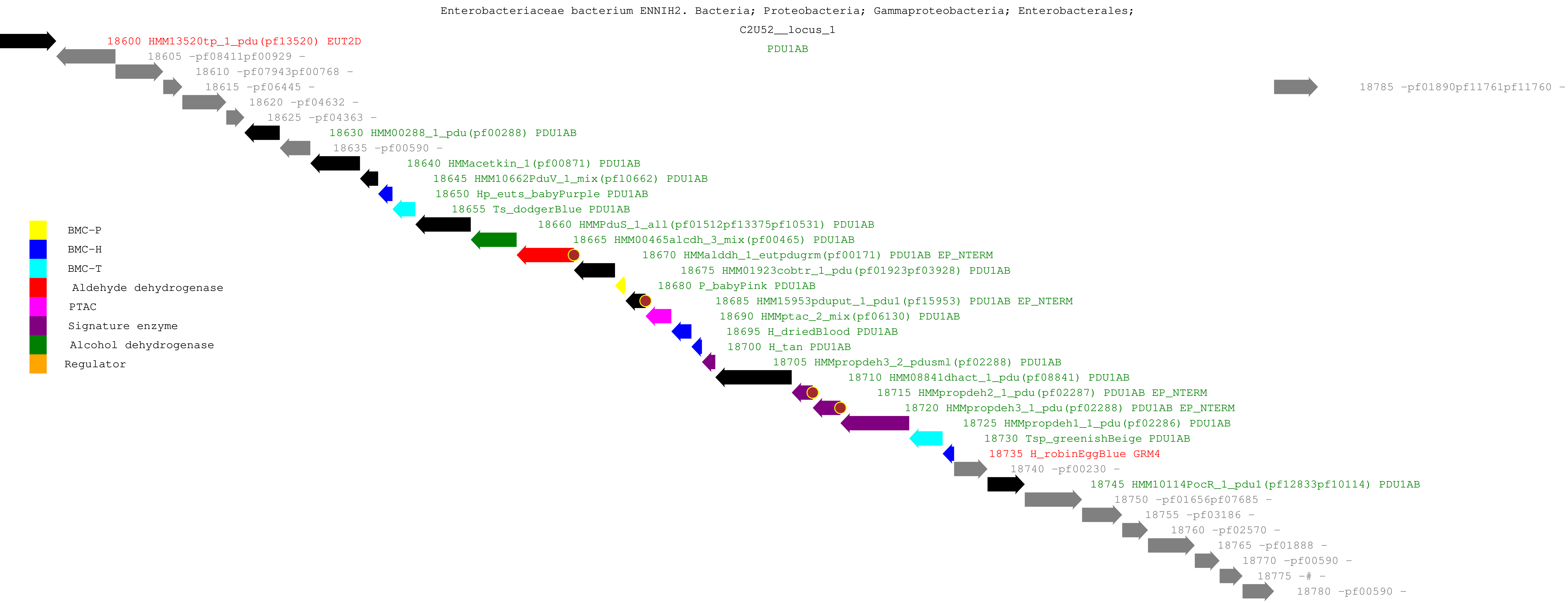
Enterobacteriaceae bacterium ENNIH2. Bacteria; Proteobacteria; Gammaproteobacteria; Enterobacterales;
C2U52__locus_1
PDU1AB
18600 HMM13520tp_1_pdu(pf13520) EUT2D
18605 -pf08411pf00929 -
18610 -pf07943pf00768 -
18615 -pf06445 -
18620 -pf04632 -
18625 -pf04363 -
18630 HMM00288_1_pdu(pf00288) PDU1AB
18635 -pf00590 -
18640 HMMacetkin_1(pf00871) PDU1AB
18645 HMM10662PduV_1_mix(pf10662) PDU1AB
18650 Hp_euts_babyPurple PDU1AB
18655 Ts_dodgerBlue PDU1AB
18660 HMMPduS_1_all(pf01512pf13375pf10531) PDU1AB
18665 HMM00465alcdh_3_mix(pf00465) PDU1AB
18670 HMMalddh_1_eutpdugrm(pf00171) PDU1AB EP_NTERM
18675 HMM01923cobtr_1_pdu(pf01923pf03928) PDU1AB
18680 P_babyPink PDU1AB
18685 HMM15953pduput_1_pdu1(pf15953) PDU1AB EP_NTERM
18690 HMMptac_2_mix(pf06130) PDU1AB
18695 H_driedBlood PDU1AB
18700 H_tan PDU1AB
18705 HMMpropdeh3_2_pdusml(pf02288) PDU1AB
18710 HMM08841dhact_1_pdu(pf08841) PDU1AB
18715 HMMpropdeh2_1_pdu(pf02287) PDU1AB EP_NTERM
18720 HMMpropdeh3_1_pdu(pf02288) PDU1AB EP_NTERM
18725 HMMpropdeh1_1_pdu(pf02286) PDU1AB
18730 Tsp_greenishBeige PDU1AB
18735 H_robinEggBlue GRM4
18740 -pf00230 -
18745 HMM10114PocR_1_pdu1(pf12833pf10114) PDU1AB
18750 -pf01656pf07685 -
18755 -pf03186 -
18760 -pf02570 -
18765 -pf01888 -
18770 -pf00590 -
18775 -# -
18780 -pf00590 -
18785 -pf01890pf11761pf11760 -
BMC-P
BMC-H
BMC-T
Aldehyde dehydrogenase
PTAC
Signature enzyme
Alcohol dehydrogenase
Regulator

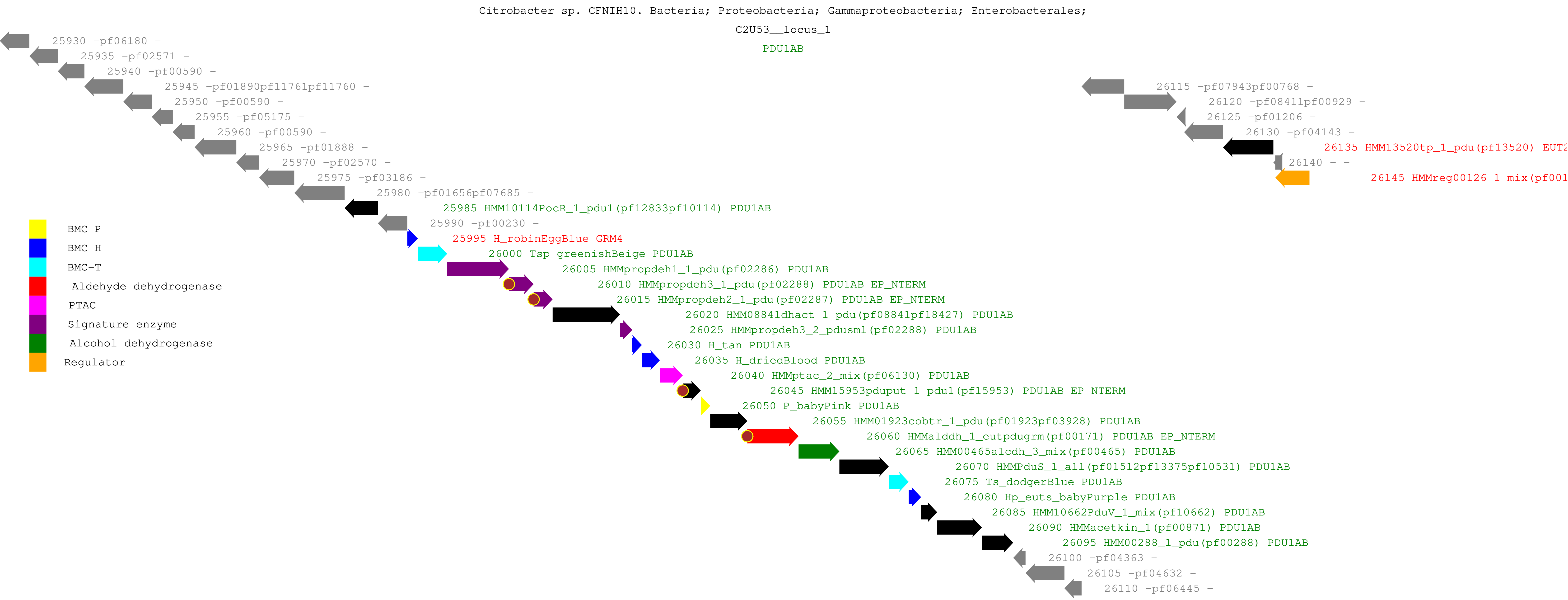
Citrobacter sp. CFNIH10. Bacteria; Proteobacteria; Gammaproteobacteria; Enterobacterales;
C2U53__locus_1
PDU1AB
25930 -pf06180 -
25935 -pf02571 -
25940 -pf00590 -
25945 -pf01890pf11761pf11760 -
25950 -pf00590 -
25955 -pf05175 -
25960 -pf00590 -
25965 -pf01888 -
25970 -pf02570 -
25975 -pf03186 -
25980 -pf01656pf07685 -
25985 HMM10114PocR_1_pdu1(pf12833pf10114) PDU1AB
25990 -pf00230 -
25995 H_robinEggBlue GRM4
26000 Tsp_greenishBeige PDU1AB
26005 HMMpropdeh1_1_pdu(pf02286) PDU1AB
26010 HMMpropdeh3_1_pdu(pf02288) PDU1AB EP_NTERM
26015 HMMpropdeh2_1_pdu(pf02287) PDU1AB EP_NTERM
26020 HMM08841dhact_1_pdu(pf08841pf18427) PDU1AB
26025 HMMpropdeh3_2_pdusml(pf02288) PDU1AB
26030 H_tan PDU1AB
26035 H_driedBlood PDU1AB
26040 HMMptac_2_mix(pf06130) PDU1AB
26045 HMM15953pduput_1_pdu1(pf15953) PDU1AB EP_NTERM
26050 P_babyPink PDU1AB
26055 HMM01923cobtr_1_pdu(pf01923pf03928) PDU1AB
26060 HMMalddh_1_eutpdugrm(pf00171) PDU1AB EP_NTERM
26065 HMM00465alcdh_3_mix(pf00465) PDU1AB
26070 HMMPduS_1_all(pf01512pf13375pf10531) PDU1AB
26075 Ts_dodgerBlue PDU1AB
26080 Hp_euts_babyPurple PDU1AB
26085 HMM10662PduV_1_mix(pf10662) PDU1AB
26090 HMMacetkin_1(pf00871) PDU1AB
26095 HMM00288_1_pdu(pf00288) PDU1AB
26100 -pf04363 -
26105 -pf04632 -
26110 -pf06445 -
26115 -pf07943pf00768 -
26120 -pf08411pf00929 -
26125 -pf01206 -
26130 -pf04143 -
26135 HMM13520tp_1_pdu(pf13520) EUT2
26140 - -
26145 HMMreg00126_1_mix(pf001
BMC-P
BMC-H
BMC-T
Aldehyde dehydrogenase
PTAC
Signature enzyme
Alcohol dehydrogenase
Regulator

Citrobacter amalonaticus. Bacteria; Proteobacteria; Gammaproteobacteria; Enterobacterales;

C3430__locus_1

PDU1AB

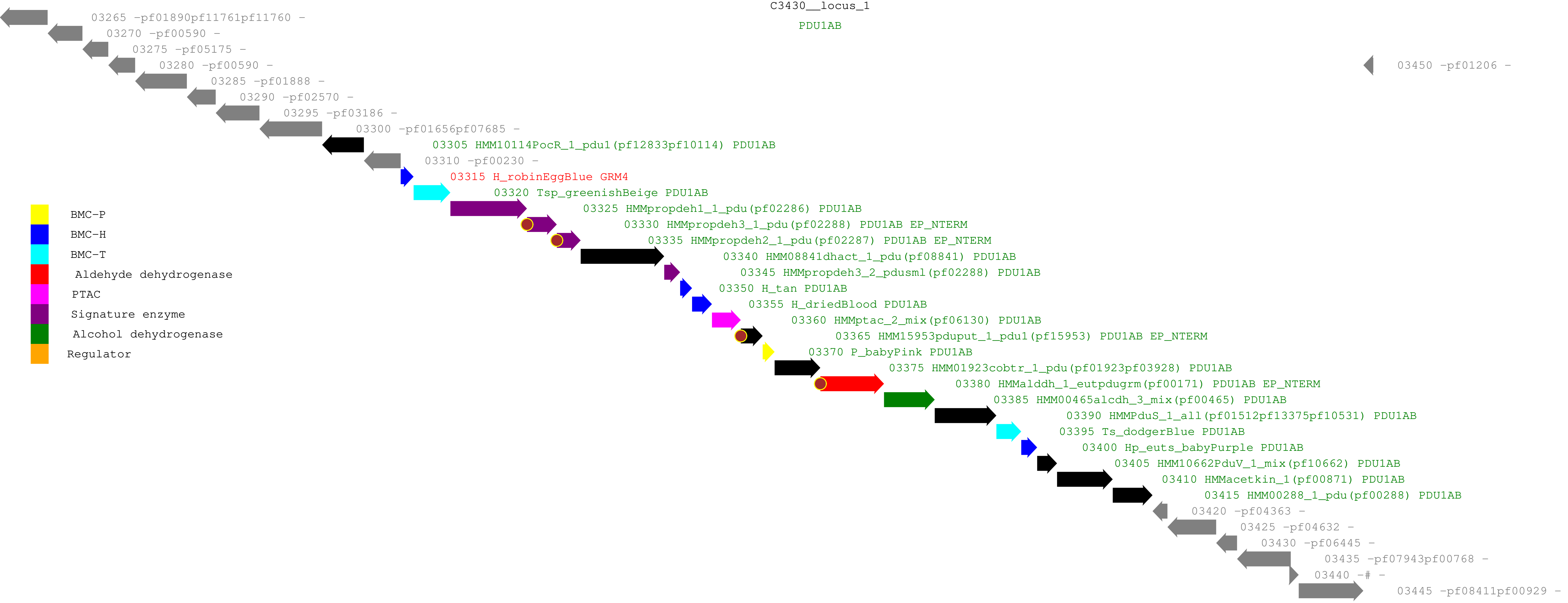

Citrobacter freundii. Bacteria; Proteobacteria; Gammaproteobacteria; Enterobacterales;

C3433__locus_1

PDU1AB

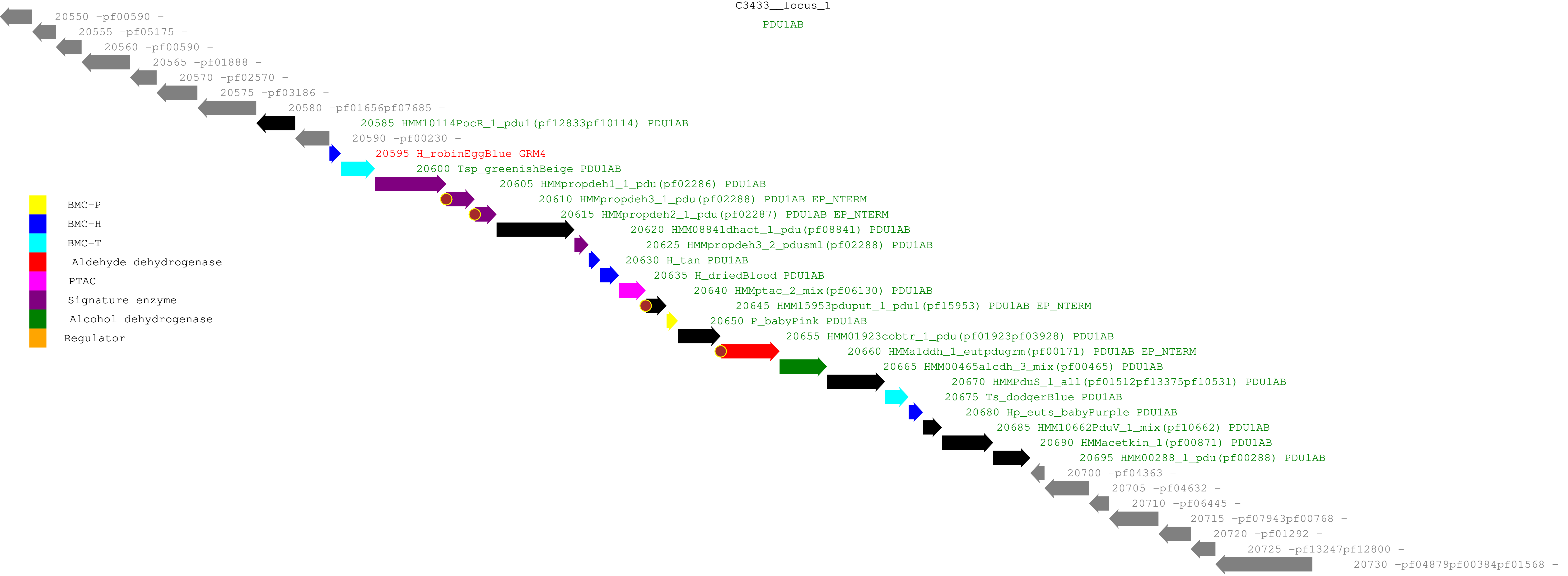

Salmonella enterica I. Bacteria; Proteobacteria; Gammaproteobacteria; Enterobacterales;

C3010__locus_1

PDU1AB

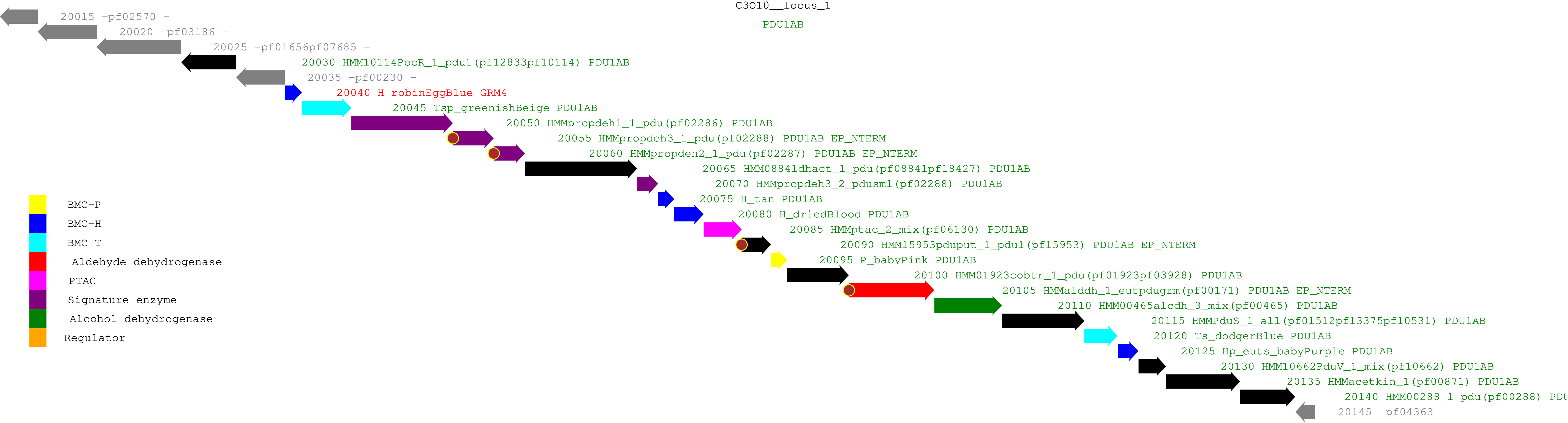

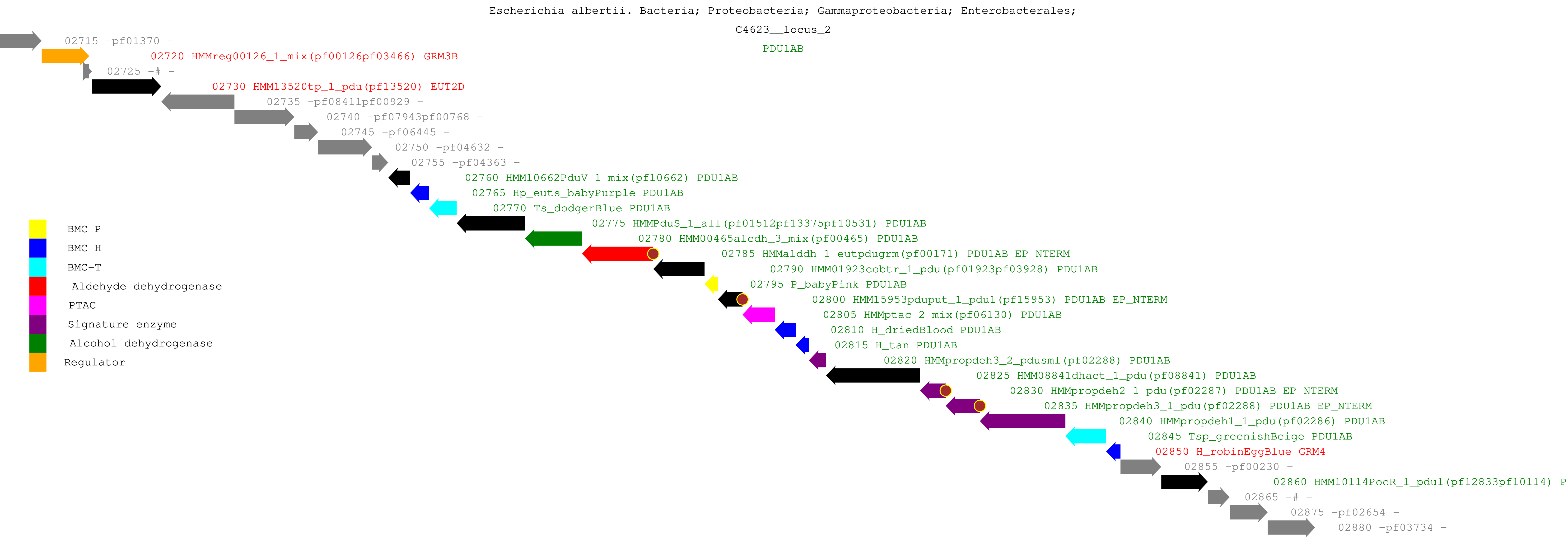
Escherichia albertii. Bacteria; Proteobacteria; Gammaproteobacteria; Enterobacterales;
C4623__locus_2
PDU1AB
02715 -pf01370 -
02720 HMMreg00126_1_mix(pf00126pf03466) GRM3B
02725 -# -
02730 HMM13520tp_1_pdu(pf13520) EUT2D
02735 -pf08411pf00929 -
02740 -pf07943pf00768 -
02745 -pf06445 -
02750 -pf04632 -
02755 -pf04363 -
02760 HMM10662PduV_1_mix(pf10662) PDU1AB
02765 Hp_euts_babyPurple PDU1AB
02770 Ts_dodgerBlue PDU1AB
02775 HMMPduS_1_all(pf01512pf13375pf10531) PDU1AB
02780 HMM00465alcdh_3_mix(pf00465) PDU1AB
02785 HMMaldh_1_eutpdugrm(pf00171) PDU1AB EP_NTERM
02790 HMM01923cobtr_1_pdu(pf01923pf03928) PDU1AB
02795 P_babyPink PDU1AB
02800 HMM15953pduput_1_pdu1(pf15953) PDU1AB EP_NTERM
02805 HMMptac_2_mix(pf06130) PDU1AB
02810 H_driedBlood PDU1AB
02815 H_tan PDU1AB
02820 HMMpropdeh3_2_pdusml(pf02288) PDU1AB
02825 HMM08841dhact_1_pdu(pf08841) PDU1AB
02830 HMMpropdeh2_1_pdu(pf02287) PDU1AB EP_NTERM
02835 HMMpropdeh3_1_pdu(pf02288) PDU1AB EP_NTERM
02840 HMMpropdeh1_1_pdu(pf02286) PDU1AB
02845 Tsp_greenishBeige PDU1AB
02850 H_robinEggBlue GRM4
02855 -pf00230 -
02860 HMM10114PocR_1_pdu1(pf12833pf10114) P
02865 -# -
02875 -pf02654 -
02880 -pf03734 -
BMC-P
BMC-H
BMC-T
Aldehyde dehydrogenase
PTAC
Signature enzyme
Alcohol dehydrogenase
Regulator

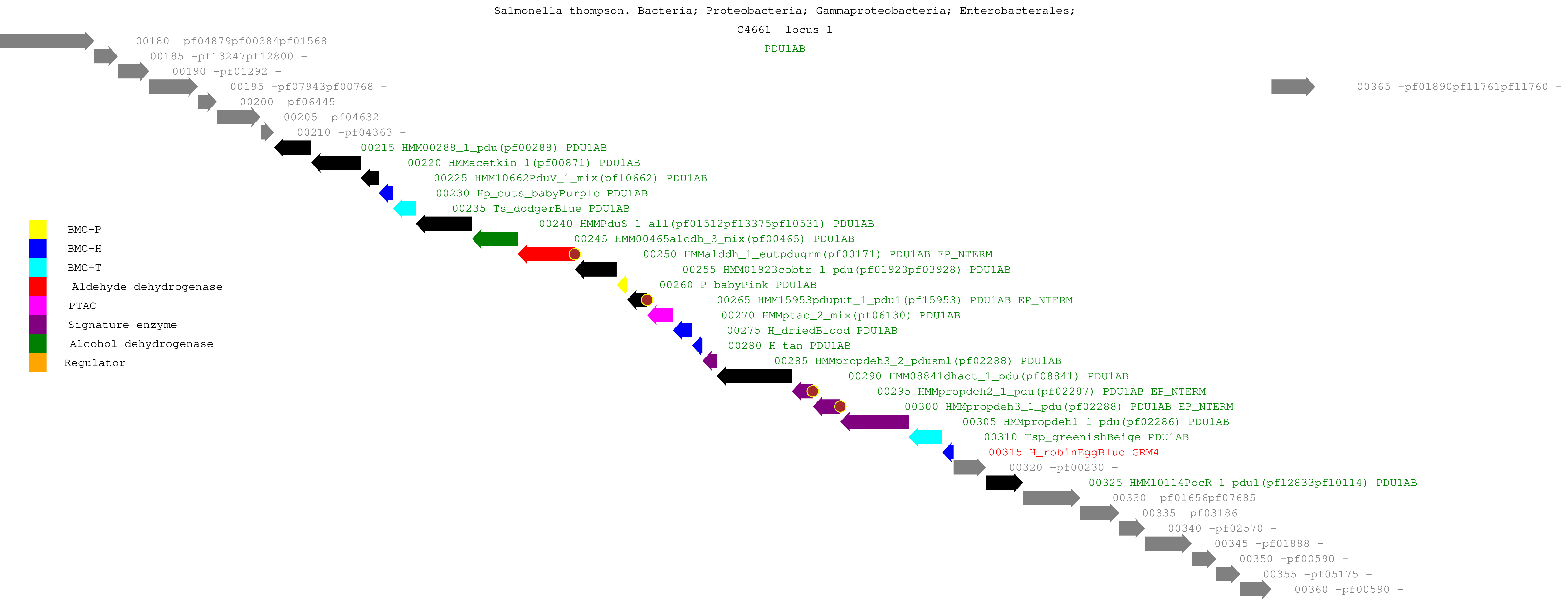

Salmonella thompson. Bacteria; Proteobacteria; Gammaproteobacteria; Enterobacterales;
C4661__locus_1
PDU1AB
00180 -pf04879pf00384pf01568 -
00185 -pf13247pf12800 -
00190 -pf01292 -
00195 -pf07943pf00768 -
00200 -pf06445 -
00205 -pf04632 -
00210 -pf04363 -
00215 HMM00288_1_pdu(pf00288) PDU1AB
00220 HMMacetkin_1(pf00871) PDU1AB
00225 HMM10662PduV_1_mix(pf10662) PDU1AB
00230 Hp_euts_babyPurple PDU1AB
00235 Ts_dodgerBlue PDU1AB
00240 HMMPduS_1_all(pf01512pf13375pf10531) PDU1AB
00245 HMM00465alcdh_3_mix(pf00465) PDU1AB
00250 HMMalddh_1_eutpdugrm(pf00171) PDU1AB EP_NTERM
00255 HMM01923cobtr_1_pdu(pf01923pf03928) PDU1AB
00260 P_babyPink PDU1AB
00265 HMM15953pduput_1_pdu1(pf15953) PDU1AB EP_NTERM
00270 HMMptac_2_mix(pf06130) PDU1AB
00275 H_driedBlood PDU1AB
00280 H_tan PDU1AB
00285 HMMpropdeh3_2_pdusml(pf02288) PDU1AB
00290 HMM08841dhact_1_pdu(pf08841) PDU1AB
00295 HMMpropdeh2_1_pdu(pf02287) PDU1AB EP_NTERM
00300 HMMpropdeh3_1_pdu(pf02288) PDU1AB EP_NTERM
00305 HMMpropdeh1_1_pdu(pf02286) PDU1AB
00310 Tsp_greenishBeige PDU1AB
00315 H_robinEggBlue GRM4
00320 -pf00230 -
00325 HMM10114PocR_1_pdu1(pf12833pf10114) PDU1AB
00330 -pf01656pf07685 -
00335 -pf03186 -
00340 -pf02570 -
00345 -pf01888 -
00350 -pf00590 -
00355 -pf05175 -
00360 -pf00590 -
00365 -pf01890pf11761pf11760 -
BMC-P
BMC-H
BMC-T
Aldehyde dehydrogenase
PTAC
Signature enzyme
Alcohol dehydrogenase
Regulator

Salmonella enterica subsp. enterica serovar Bareilly. Bacteria; Proteobacteria; Gammaproteobacteria; Enterobacterales;

C4680__locus_2

PDU1AB

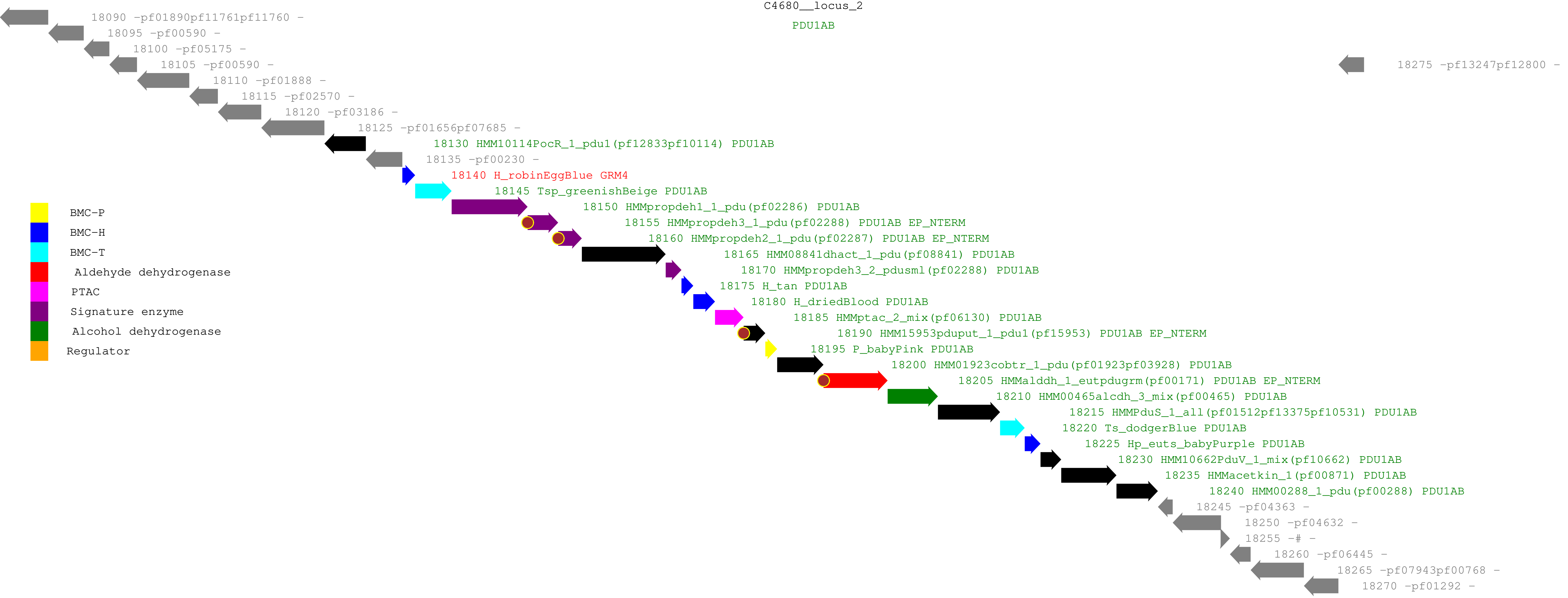

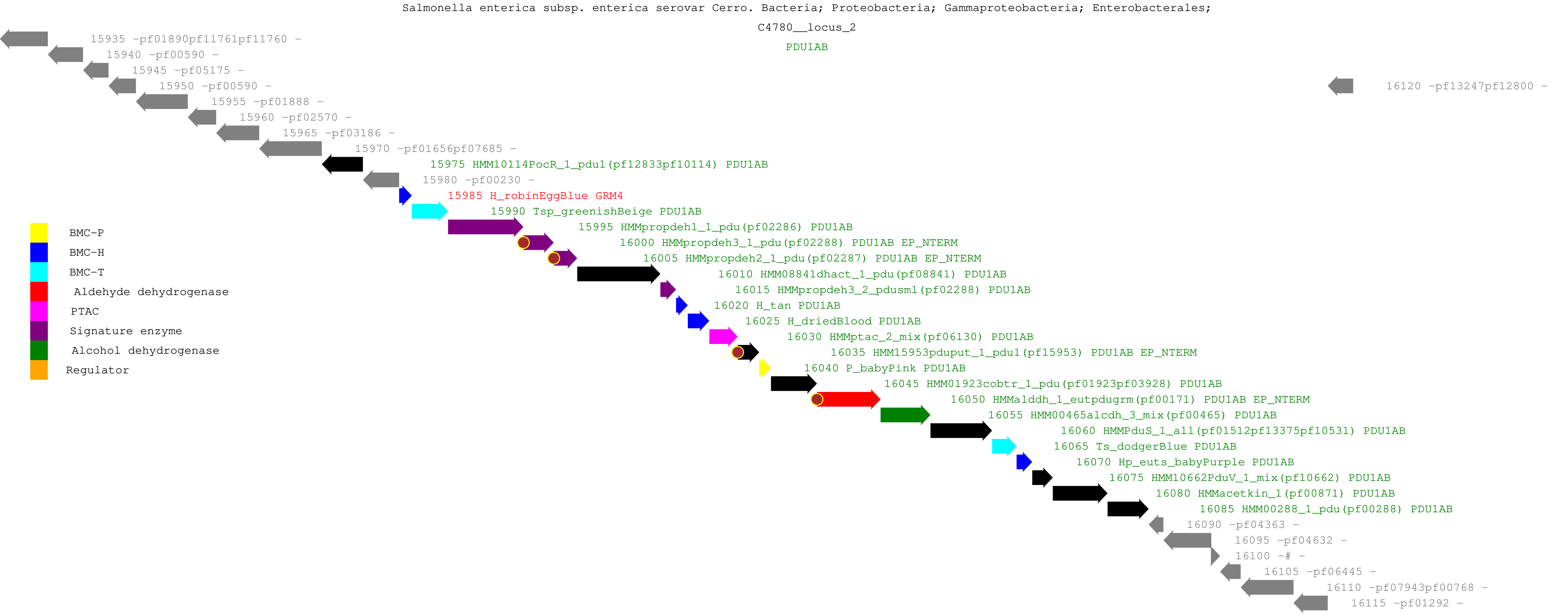

Salmonella enterica subsp. enterica serovar Cerro. Bacteria; Proteobacteria; Gammaproteobacteria; Enterobacterales;
C4780__locus_2
PDU1AB
15935 -pf01890pf11761pf11760 -
15940 -pf00590 -
15945 -pf05175 -
15950 -pf00590 -
15955 -pf01888 -
15960 -pf02570 -
15965 -pf03186 -
15970 -pf01656pf07685 -
15975 HMM10114PocR_1_pdu1(pf12833pf10114) PDU1AB
15980 -pf00230 -
15985 H_robinEggBlue GRM4
15990 Tsp_greenishBeige PDU1AB
15995 HMMpropdeh1_1_pdu(pf02286) PDU1AB
16000 HMMpropdeh3_1_pdu(pf02288) PDU1AB EP_NTERM
16005 HMMpropdeh2_1_pdu(pf02287) PDU1AB EP_NTERM
16010 HMM08841dhact_1_pdu(pf08841) PDU1AB
16015 HMMpropdeh3_2_pdusml(pf02288) PDU1AB
16020 H_tan PDU1AB
16025 H_driedBlood PDU1AB
16030 HMMptac_2_mix(pf06130) PDU1AB
16035 HMM15953pduput_1_pdu1(pf15953) PDU1AB EP_NTERM
16040 P_babyPink PDU1AB
16045 HMM01923cobtr_1_pdu(pf01923pf03928) PDU1AB
16050 HMMaldddh_1_eutpdugrm(pf00171) PDU1AB EP_NTERM
16055 HMM00465alcdh_3_mix(pf00465) PDU1AB
16060 HMMPduS_1_all(pf01512pf13375pf10531) PDU1AB
16065 Ts_dodgerBlue PDU1AB
16070 Hp_euts_babyPurple PDU1AB
16075 HMM10662PduV_1_mix(pf10662) PDU1AB
16080 HMMacetkin_1(pf00871) PDU1AB
16085 HMM00288_1_pdu(pf00288) PDU1AB
16090 -pf04363 -
16095 -pf04632 -
16100 -# -
16105 -pf06445 -
16110 -pf07943pf00768 -
16115 -pf01292 -
16120 -pf13247pf12800 -
BMC-P
BMC-H
BMC-T
Aldehyde dehydrogenase
PTAC
Signature enzyme
Alcohol dehydrogenase
Regulator

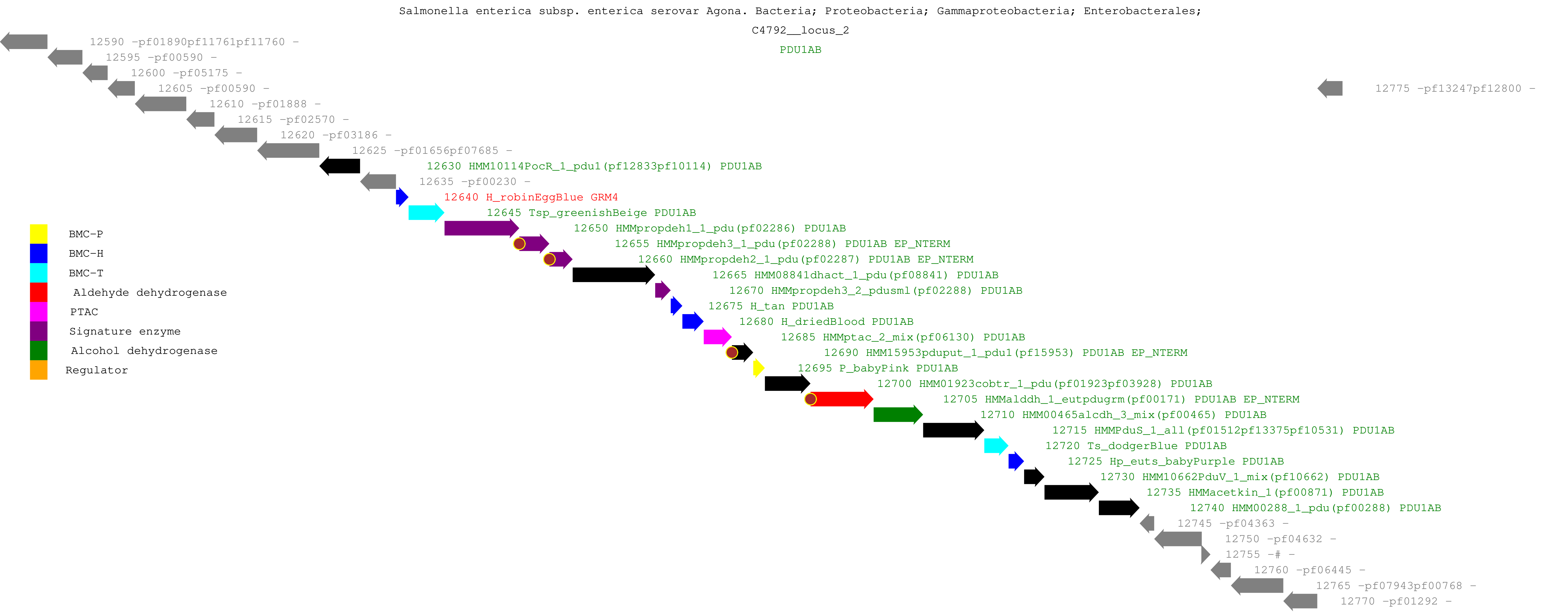
Salmonella enterica subsp. enterica serovar Agona. Bacteria; Proteobacteria; Gammaproteobacteria; Enterobacterales;
C4792__locus_2
PDU1AB
12590 -pf01890pf11761pf11760 -
12595 -pf00590 -
12600 -pf05175 -
12605 -pf00590 -
12610 -pf01888 -
12615 -pf02570 -
12620 -pf03186 -
12625 -pf01656pf07685 -
12630 HMM10114PocR_1_pdu1(pf12833pf10114) PDU1AB
12635 -pf00230 -
12640 H_robinEggBlue GRM4
12645 Tsp_greenishBeige PDU1AB
12650 HMMpropdeh1_1_pdu(pf02286) PDU1AB
12655 HMMpropdeh3_1_pdu(pf02288) PDU1AB EP_NTERM
12660 HMMpropdeh2_1_pdu(pf02287) PDU1AB EP_NTERM
12665 HMM08841dhact_1_pdu(pf08841) PDU1AB
12670 HMMpropdeh3_2_pdusml(pf02288) PDU1AB
12675 H_tan PDU1AB
12680 H_driedBlood PDU1AB
12685 HMMptac_2_mix(pf06130) PDU1AB
12690 HMM15953pduput_1_pdu1(pf15953) PDU1AB EP_NTERM
12695 P_babyPink PDU1AB
12700 HMM01923cobtr_1_pdu(pf01923pf03928) PDU1AB
12705 HMMalddh_1_eutpdugrm(pf00171) PDU1AB EP_NTERM
12710 HMM00465alcdh_3_mix(pf00465) PDU1AB
12715 HMMPduS_1_all(pf01512pf13375pf10531) PDU1AB
12720 Ts_dodgerBlue PDU1AB
12725 Hp_euts_babyPurple PDU1AB
12730 HMM10662PduV_1_mix(pf10662) PDU1AB
12735 HMMacetkin_1(pf00871) PDU1AB
12740 HMM00288_1_pdu(pf00288) PDU1AB
12745 -pf04363 -
12750 -pf04632 -
12755 -# -
12760 -pf06445 -
12765 -pf07943pf00768 -
12770 -pf01292 -
12775 -pf13247pf12800 -
BMC-P
BMC-H
BMC-T
Aldehyde dehydrogenase
PTAC
Signature enzyme
Alcohol dehydrogenase
Regulator

Salmonella choleraesuis. Bacteria; Proteobacteria; Gammaproteobacteria; Enterobacterales;

C4801__locus_2

PDU1AB

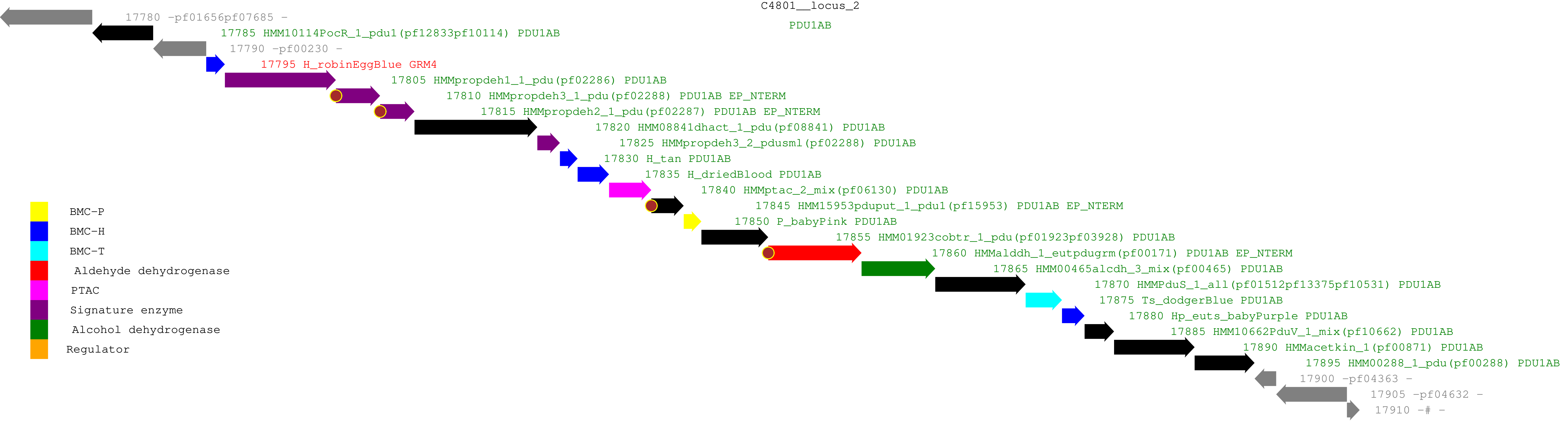

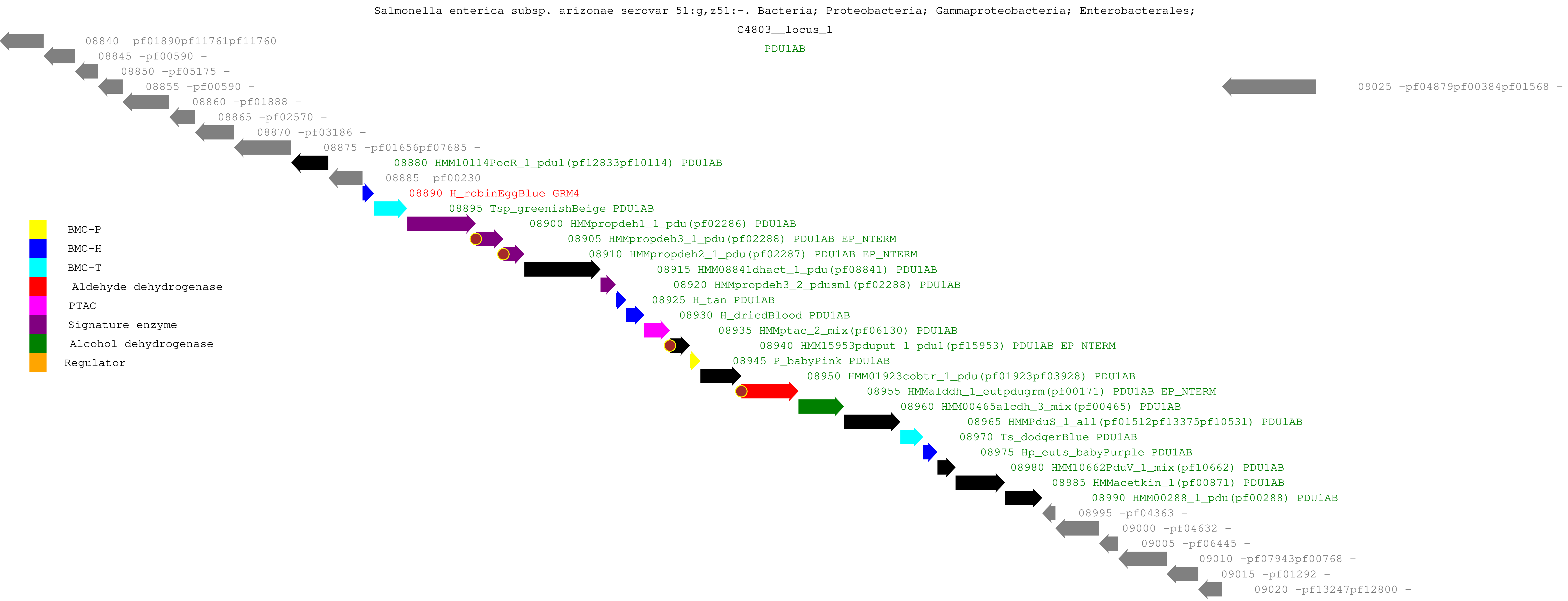
Salmonella enterica subsp. arizonae serovar 51:g,z51:-. Bacteria; Proteobacteria; Gammaproteobacteria; Enterobacterales;
C4803__locus_1
PDU1AB
08840 -pf01890pf11761pf11760 -
08845 -pf00590 -
08850 -pf05175 -
08855 -pf00590 -
08860 -pf01888 -
08865 -pf02570 -
08870 -pf03186 -
08875 -pf01656pf07685 -
08880 HMM10114PocR_1_pdu1(pf12833pf10114) PDU1AB
08885 -pf00230 -
08890 H_robinEggBlue GRM4
08895 Tsp_greenishBeige PDU1AB
08900 HMMpropdeh1_1_pdu(pf02286) PDU1AB
08905 HMMpropdeh3_1_pdu(pf02288) PDU1AB EP_NTERM
08910 HMMpropdeh2_1_pdu(pf02287) PDU1AB EP_NTERM
08915 HMM08841dhact_1_pdu(pf08841) PDU1AB
08920 HMMpropdeh3_2_pdusml(pf02288) PDU1AB
08925 H_tan PDU1AB
08930 H_driedBlood PDU1AB
08935 HMMptac_2_mix(pf06130) PDU1AB
08940 HMM15953pduput_1_pdu1(pf15953) PDU1AB EP_NTERM
08945 P_babyPink PDU1AB
08950 HMM01923cobtr_1_pdu(pf01923pf03928) PDU1AB
08955 HMMalddh_1_eutpdugrm(pf00171) PDU1AB EP_NTERM
08960 HMM00465alcdh_3_mix(pf00465) PDU1AB
08965 HMMPduS_1_all(pf01512pf13375pf10531) PDU1AB
08970 Ts_dodgerBlue PDU1AB
08975 Hp_euts_babyPurple PDU1AB
08980 HMM10662PduV_1_mix(pf10662) PDU1AB
08985 HMMacetkin_1(pf00871) PDU1AB
08990 HMM00288_1_pdu(pf00288) PDU1AB
08995 -pf04363 -
09000 -pf04632 -
09005 -pf06445 -
09010 -pf07943pf00768 -
09015 -pf01292 -
09020 -pf13247pf12800 -
09025 -pf04879pf00384pf01568 -
BMC-P
BMC-H
BMC-T
Aldehyde dehydrogenase
PTAC
Signature enzyme
Alcohol dehydrogenase
Regulator

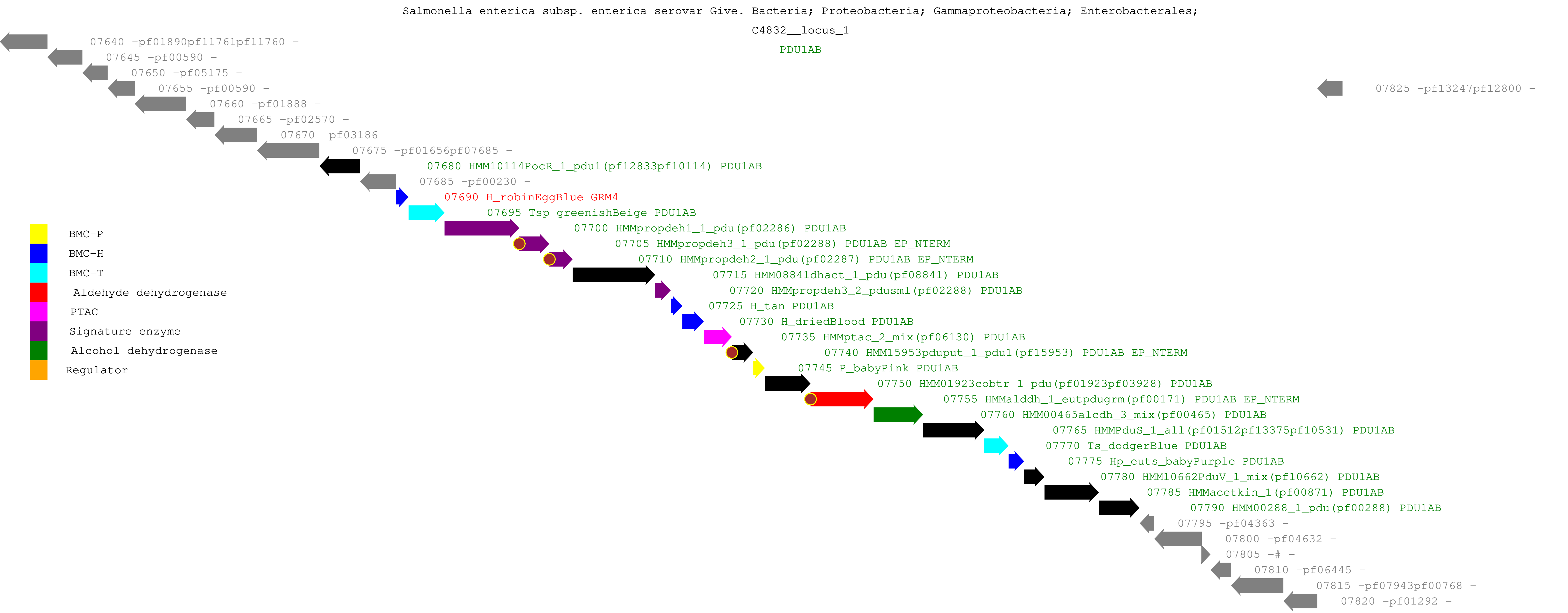
Salmonella enterica subsp. enterica serovar Give. Bacteria; Proteobacteria; Gammaproteobacteria; Enterobacterales;
C4832__locus_1
PDU1AB
07640 -pf01890pf11761pf11760 -
07645 -pf00590 -
07650 -pf05175 -
07655 -pf00590 -
07660 -pf01888 -
07665 -pf02570 -
07670 -pf03186 -
07675 -pf01656pf07685 -
07680 HMM10114PocR_1_pdu1(pf12833pf10114) PDU1AB
07685 -pf00230 -
07690 H_robinEggBlue GRM4
07695 Tsp_greenishBeige PDU1AB
07700 HMMpropdeh1_1_pdu(pf02286) PDU1AB
07705 HMMpropdeh3_1_pdu(pf02288) PDU1AB EP_NTERM
07710 HMMpropdeh2_1_pdu(pf02287) PDU1AB EP_NTERM
07715 HMM08841dhact_1_pdu(pf08841) PDU1AB
07720 HMMpropdeh3_2_pdusml(pf02288) PDU1AB
07725 H_tan PDU1AB
07730 H_driedBlood PDU1AB
07735 HMMptac_2_mix(pf06130) PDU1AB
07740 HMM15953pduput_1_pdu1(pf15953) PDU1AB EP_NTERM
07745 P_babyPink PDU1AB
07750 HMM01923cobtr_1_pdu(pf01923pf03928) PDU1AB
07755 HMMalddh_1_eutpdugrm(pf00171) PDU1AB EP_NTERM
07760 HMM00465alcdh_3_mix(pf00465) PDU1AB
07765 HMMPduS_1_all(pf01512pf13375pf10531) PDU1AB
07770 Ts_dodgerBlue PDU1AB
07775 Hp_euts_babyPurple PDU1AB
07780 HMM10662PduV_1_mix(pf10662) PDU1AB
07785 HMMacetkin_1(pf00871) PDU1AB
07790 HMM00288_1_pdu(pf00288) PDU1AB
07795 -pf04363 -
07800 -pf04632 -
07805 -# -
07810 -pf06445 -
07815 -pf07943pf00768 -
07820 -pf01292 -
07825 -pf13247pf12800 -
BMC-P
BMC-H
BMC-T
Aldehyde dehydrogenase
PTAC
Signature enzyme
Alcohol dehydrogenase
Regulator

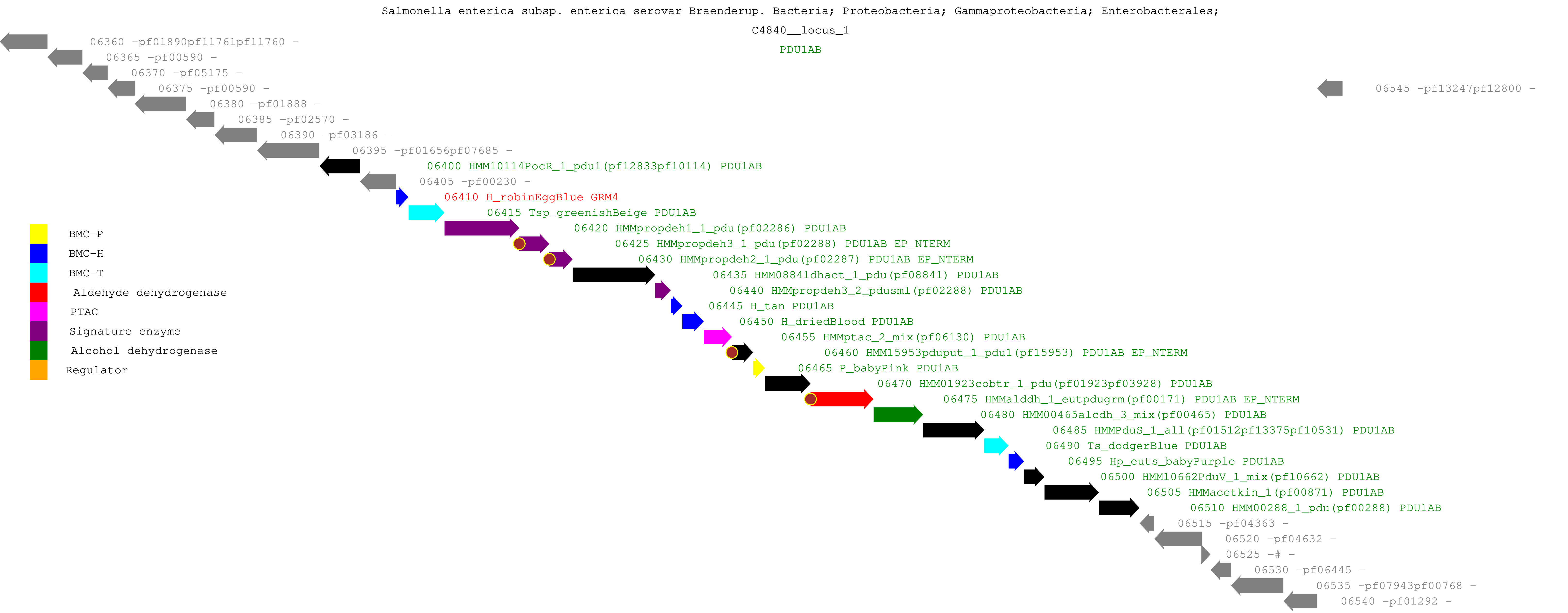
Salmonella enterica subsp. enterica serovar Braenderup. Bacteria; Proteobacteria; Gammaproteobacteria; Enterobacterales;
C4840__locus_1
PDU1AB
06360 -pf01890pf11761pf11760 -
06365 -pf00590 -
06370 -pf05175 -
06375 -pf00590 -
06380 -pf01888 -
06385 -pf02570 -
06390 -pf03186 -
06395 -pf01656pf07685 -
06400 HMM10114PocR_1_pdu1(pf12833pf10114) PDU1AB
06405 -pf00230 -
06410 H_robinEggBlue GRM4
06415 Tsp_greenishBeige PDU1AB
06420 HMMpropdeh1_1_pdu(pf02286) PDU1AB
06425 HMMpropdeh3_1_pdu(pf02288) PDU1AB EP_NTERM
06430 HMMpropdeh2_1_pdu(pf02287) PDU1AB EP_NTERM
06435 HMM08841dhact_1_pdu(pf08841) PDU1AB
06440 HMMpropdeh3_2_pdusml(pf02288) PDU1AB
06445 H_tan PDU1AB
06450 H_driedBlood PDU1AB
06455 HMMptac_2_mix(pf06130) PDU1AB
06460 HMM15953pduput_1_pdu1(pf15953) PDU1AB EP_NTERM
06465 P_babyPink PDU1AB
06470 HMM01923cobtr_1_pdu(pf01923pf03928) PDU1AB
06475 HMMalddh_1_eutpdugrm(pf00171) PDU1AB EP_NTERM
06480 HMM00465alcdh_3_mix(pf00465) PDU1AB
06485 HMMPduS_1_all(pf01512pf13375pf10531) PDU1AB
06490 Ts_dodgerBlue PDU1AB
06495 Hp_euts_babyPurple PDU1AB
06500 HMM10662PduV_1_mix(pf10662) PDU1AB
06505 HMMacetkin_1(pf00871) PDU1AB
06510 HMM00288_1_pdu(pf00288) PDU1AB
06515 -pf04363 -
06520 -pf04632 -
06525 -# -
06530 -pf06445 -
06535 -pf07943pf00768 -
06540 -pf01292 -
06545 -pf13247pf12800 -
BMC-P
BMC-H
BMC-T
Aldehyde dehydrogenase
PTAC
Signature enzyme
Alcohol dehydrogenase
Regulator

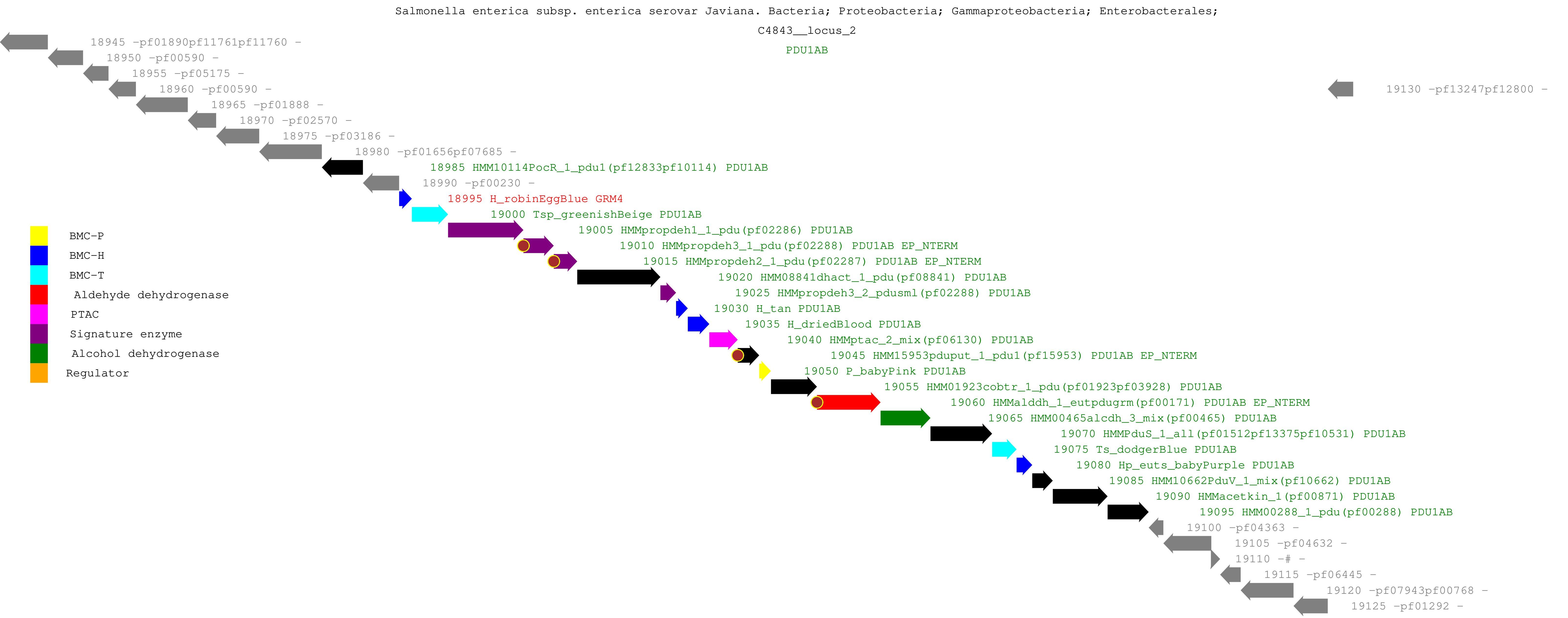
Salmonella enterica subsp. enterica serovar Javiana. Bacteria; Proteobacteria; Gammaproteobacteria; Enterobacterales;
C4843__locus_2
PDU1AB
18945 -pf01890pf11761pf11760 -
18950 -pf00590 -
18955 -pf05175 -
18960 -pf00590 -
18965 -pf01888 -
18970 -pf02570 -
18975 -pf03186 -
18980 -pf01656pf07685 -
18985 HMM10114PocR_1_pdu1(pf12833pf10114) PDU1AB
18990 -pf00230 -
18995 H_robinEggBlue GRM4
19000 Tsp_greenishBeige PDU1AB
19005 HMMpropdeh1_1_pdu(pf02286) PDU1AB
19010 HMMpropdeh3_1_pdu(pf02288) PDU1AB EP_NTERM
19015 HMMpropdeh2_1_pdu(pf02287) PDU1AB EP_NTERM
19020 HMM08841dhact_1_pdu(pf08841) PDU1AB
19025 HMMpropdeh3_2_pdusml(pf02288) PDU1AB
19030 H_tan PDU1AB
19035 H_driedBlood PDU1AB
19040 HMMptac_2_mix(pf06130) PDU1AB
19045 HMM15953pduput_1_pdu1(pf15953) PDU1AB EP_NTERM
19050 P_babyPink PDU1AB
19055 HMM01923cobtr_1_pdu(pf01923pf03928) PDU1AB
19060 HMMaldddh_1_eutpdugrm(pf00171) PDU1AB EP_NTERM
19065 HMM00465alcdh_3_mix(pf00465) PDU1AB
19070 HMMPduS_1_all(pf01512pf13375pf10531) PDU1AB
19075 Ts_dodgerBlue PDU1AB
19080 Hp_euts_babyPurple PDU1AB
19085 HMM10662PduV_1_mix(pf10662) PDU1AB
19090 HMMacetkin_1(pf00871) PDU1AB
19095 HMM00288_1_pdu(pf00288) PDU1AB
19100 -pf04363 -
19105 -pf04632 -
19110 -# -
19115 -pf06445 -
19120 -pf07943pf00768 -
19125 -pf01292 -
19130 -pf13247pf12800 -
BMC-P
BMC-H
BMC-T
Aldehyde dehydrogenase
PTAC
Signature enzyme
Alcohol dehydrogenase
Regulator

Salmonella infantis. Bacteria; Proteobacteria; Gammaproteobacteria; Enterobacterales;

C4846__locus_2

PDU1AB

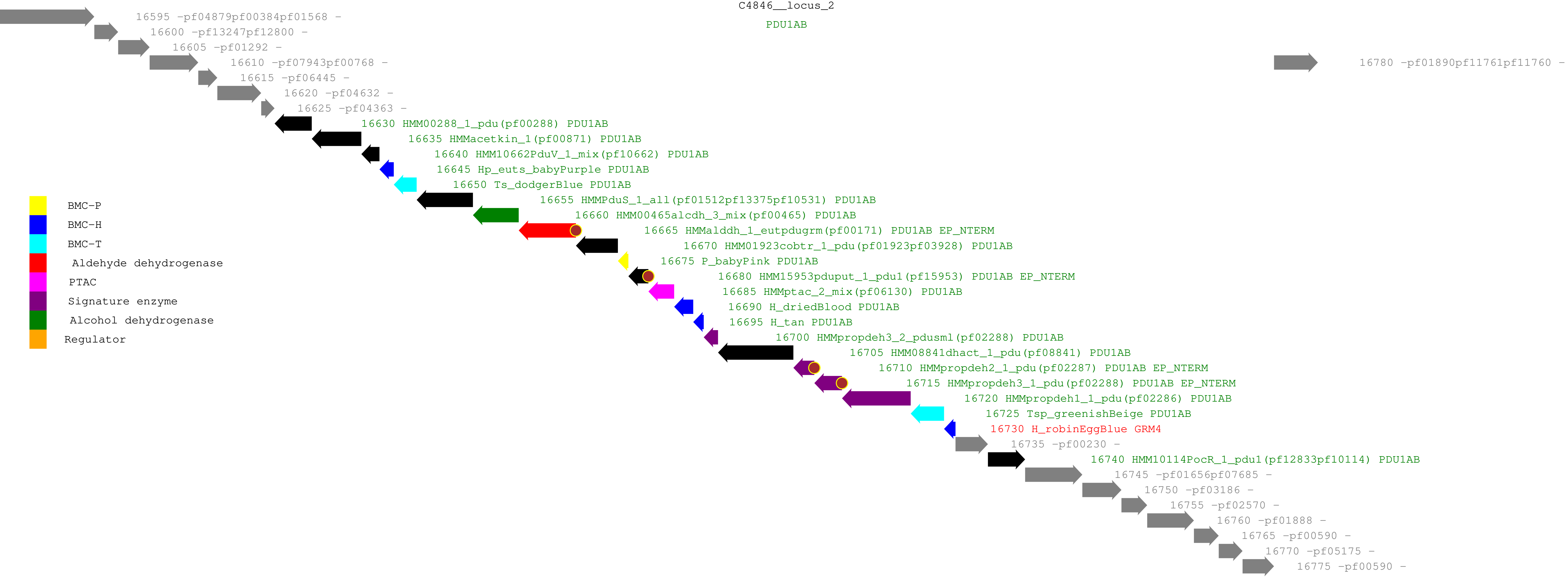

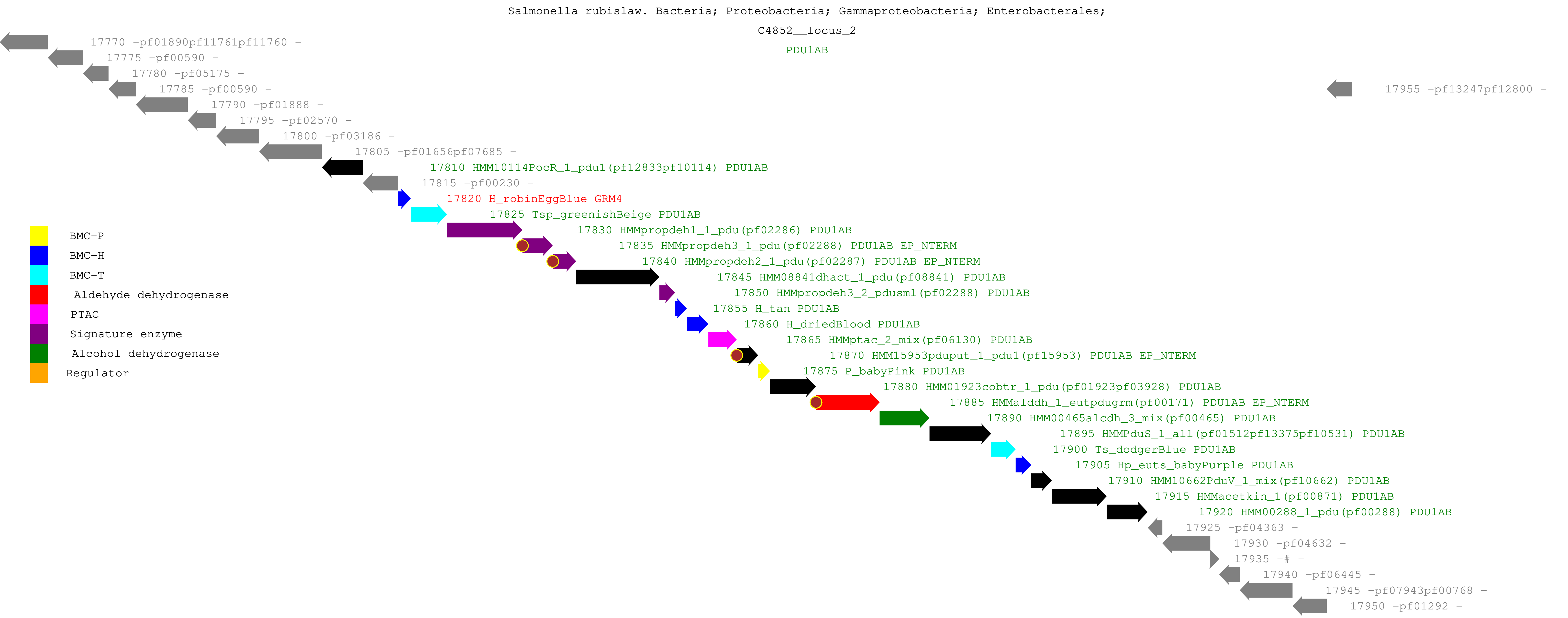

Salmonella rubislaw. Bacteria; Proteobacteria; Gammaproteobacteria; Enterobacterales;
C4852__locus_2
PDU1AB
17770 -pf01890pf11761pf11760 -
17775 -pf00590 -
17780 -pf05175 -
17785 -pf00590 -
17790 -pf01888 -
17795 -pf02570 -
17800 -pf03186 -
17805 -pf01656pf07685 -
17810 HMM10114PocR_1_pdu1(pf12833pf10114) PDU1AB
17815 -pf00230 -
17820 H_robinEggBlue GRM4
17825 Tsp_greenishBeige PDU1AB
17830 HMMpropdeh1_1_pdu(pf02286) PDU1AB
17835 HMMpropdeh3_1_pdu(pf02288) PDU1AB EP_NTERM
17840 HMMpropdeh2_1_pdu(pf02287) PDU1AB EP_NTERM
17845 HMM08841dhact_1_pdu(pf08841) PDU1AB
17850 HMMpropdeh3_2_pdusml(pf02288) PDU1AB
17855 H_tan PDU1AB
17860 H_driedBlood PDU1AB
17865 HMMptac_2_mix(pf06130) PDU1AB
17870 HMM15953pduput_1_pdu1(pf15953) PDU1AB EP_NTERM
17875 P_babyPink PDU1AB
17880 HMM01923cobtr_1_pdu(pf01923pf03928) PDU1AB
17885 HMMalddh_1_eutpdugrm(pf00171) PDU1AB EP_NTERM
17890 HMM00465alcdh_3_mix(pf00465) PDU1AB
17895 HMMPduS_1_all(pf01512pf13375pf10531) PDU1AB
17900 Ts_dodgerBlue PDU1AB
17905 Hp_euts_babyPurple PDU1AB
17910 HMM10662PduV_1_mix(pf10662) PDU1AB
17915 HMMacetkin_1(pf00871) PDU1AB
17920 HMM00288_1_pdu(pf00288) PDU1AB
17925 -pf04363 -
17930 -pf04632 -
17935 -# -
17940 -pf06445 -
17945 -pf07943pf00768 -
17950 -pf01292 -
17955 -pf13247pf12800 -
BMC-P
BMC-H
BMC-T
Aldehyde dehydrogenase
PTAC
Signature enzyme
Alcohol dehydrogenase
Regulator

Salmonella enterica subsp. enterica serovar Gaminara. Bacteria; Proteobacteria; Gammaproteobacteria; Enterobacterales;

C4855__locus_2

PDU1AB

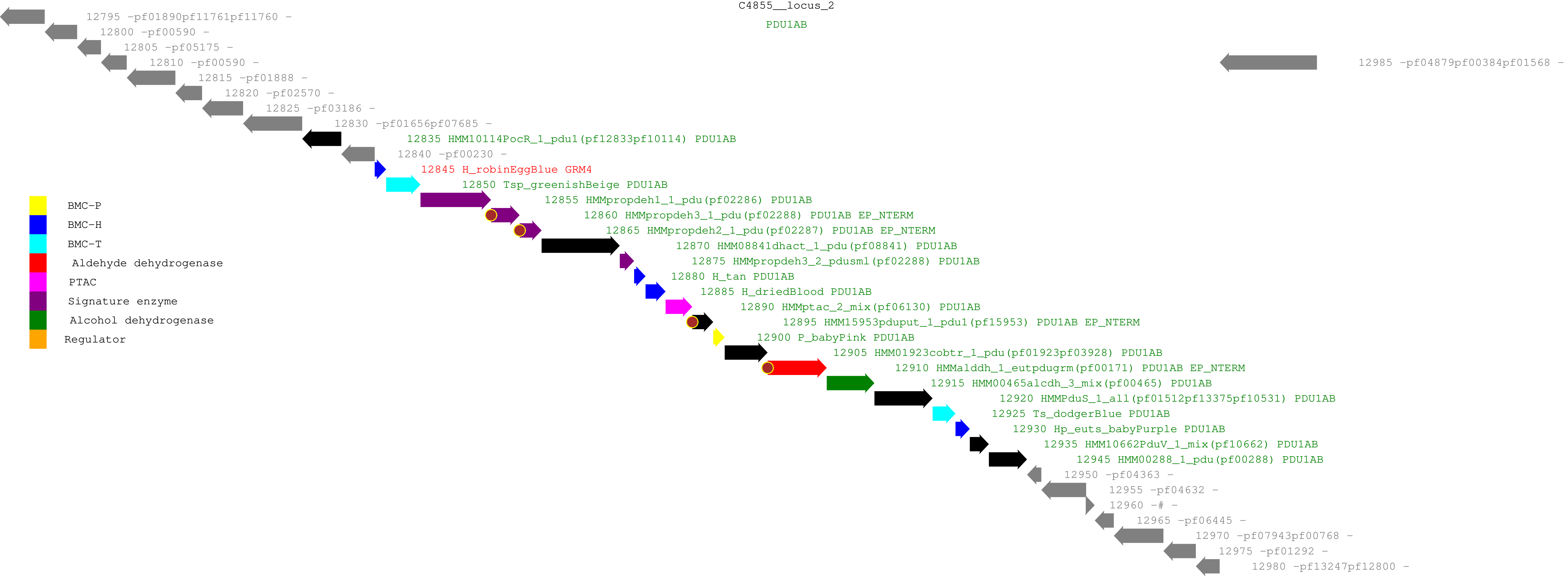

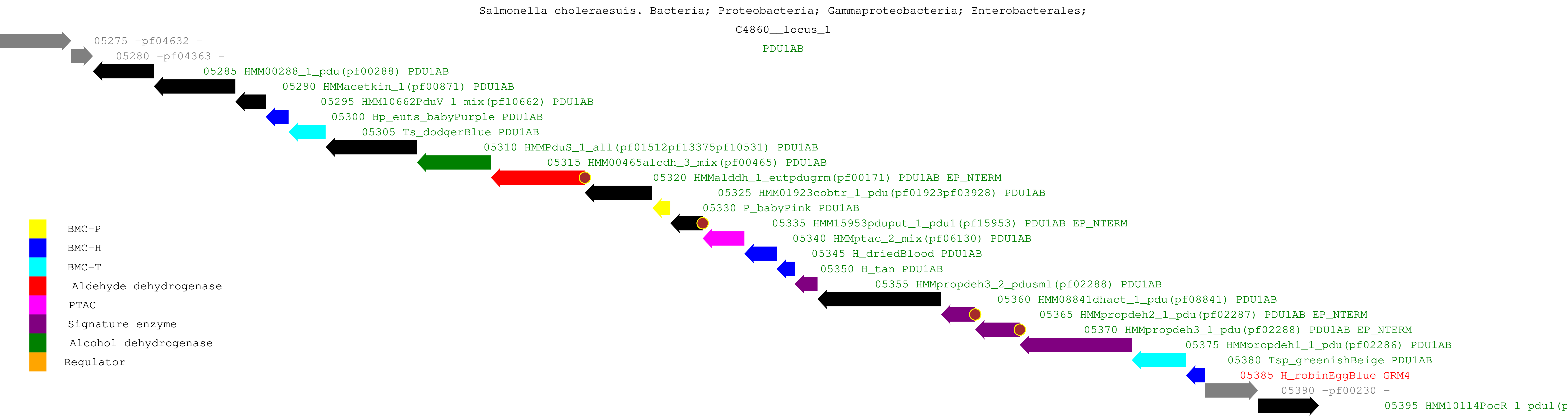

Salmonella choleraesuis. Bacteria; Proteobacteria; Gammaproteobacteria; Enterobacterales;
C4860__locus_1
PDU1AB
05275 -pf04632 -
05280 -pf04363 -
05285 HMM00288_1_pdu(pf00288) PDU1AB
05290 HMMacetkin_1(pf00871) PDU1AB
05295 HMM10662PduV_1_mix(pf10662) PDU1AB
05300 Hp_euts_babyPurple PDU1AB
05305 Ts_dodgerBlue PDU1AB
05310 HMMPduS_1_all(pf01512pf13375pf10531) PDU1AB
05315 HMM00465alcdh_3_mix(pf00465) PDU1AB
05320 HMMalddh_1_eutpdugrm(pf00171) PDU1AB EP_NTERM
05325 HMM01923cobtr_1_pdu(pf01923pf03928) PDU1AB
05330 P_babyPink PDU1AB
05335 HMM15953pduput_1_pdu1(pf15953) PDU1AB EP_NTERM
05340 HMMptac_2_mix(pf06130) PDU1AB
05345 H_driedBlood PDU1AB
05350 H_tan PDU1AB
05355 HMMpropdeh3_2_pdusml(pf02288) PDU1AB
05360 HMM08841dhact_1_pdu(pf08841) PDU1AB
05365 HMMpropdeh2_1_pdu(pf02287) PDU1AB EP_NTERM
05370 HMMpropdeh3_1_pdu(pf02288) PDU1AB EP_NTERM
05375 HMMpropdeh1_1_pdu(pf02286) PDU1AB
05380 Tsp_greenishBeige PDU1AB
05385 H_robinEggBlue GRM4
05390 -pf00230 -
05395 HMM10114PocR_1_pdu1(p
BMC-P
BMC-H
BMC-T
Aldehyde dehydrogenase
PTAC
Signature enzyme
Alcohol dehydrogenase
Regulator

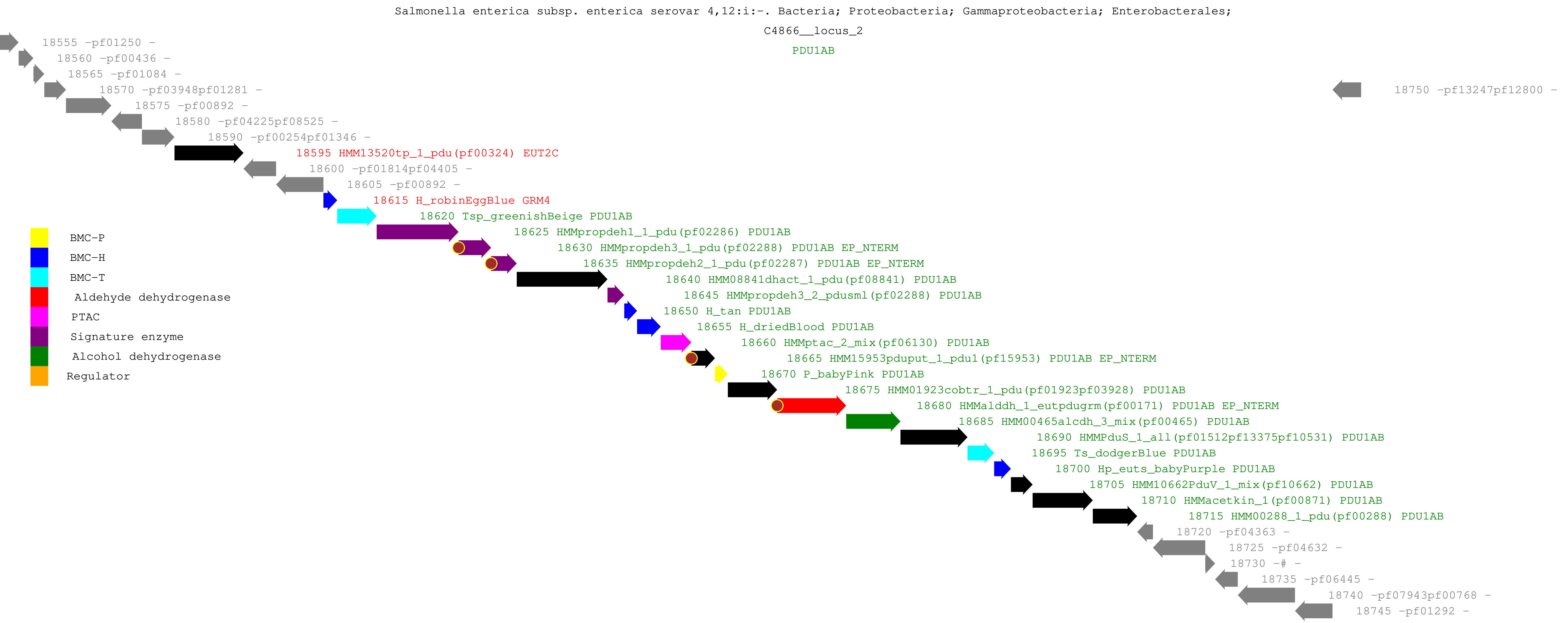

Salmonella enterica subsp. enterica serovar 4,12:i:-. Bacteria; Proteobacteria; Gammaproteobacteria; Enterobacterales;
C4866__locus_2
PDU1AB
18555 -pf01250 -
18560 -pf00436 -
18565 -pf01084 -
18570 -pf03948pf01281 -
18575 -pf00892 -
18580 -pf04225pf08525 -
18590 -pf00254pf01346 -
18595 HMM13520tp_1_pdu(pf00324) EUT2C
18600 -pf01814pf04405 -
18605 -pf00892 -
18615 H_robinEggBlue GRM4
18620 Tsp_greenishBeige PDU1AB
18625 HMMpropdeh1_1_pdu(pf02286) PDU1AB
18630 HMMpropdeh3_1_pdu(pf02288) PDU1AB EP_NTERM
18635 HMMpropdeh2_1_pdu(pf02287) PDU1AB EP_NTERM
18640 HMM08841dhact_1_pdu(pf08841) PDU1AB
18645 HMMpropdeh3_2_pdusml(pf02288) PDU1AB
18650 H_tan PDU1AB
18655 H_driedBlood PDU1AB
18660 HMMptac_2_mix(pf06130) PDU1AB
18665 HMM15953pduput_1_pdul(pf15953) PDU1AB EP_NTERM
18670 P_babyPink PDU1AB
18675 HMM01923cobtr_1_pdu(pf01923pf03928) PDU1AB
18680 HMMalddh_1_eutpdugrm(pf00171) PDU1AB EP_NTERM
18685 HMM00465alcdh_3_mix(pf00465) PDU1AB
18690 HMMPduS_1_all(pf01512pf13375pf10531) PDU1AB
18695 Ts_dodgerBlue PDU1AB
18700 Hp_euts_babyPurple PDU1AB
18705 HMM10662PduV_1_mix(pf10662) PDU1AB
18710 HMMacetkin_1(pf00871) PDU1AB
18715 HMM00288_1_pdu(pf00288) PDU1AB
18720 -pf04363 -
18725 -pf04632 -
18730 -# -
18735 -pf06445 -
18740 -pf07943pf00768 -
18745 -pf01292 -
18750 -pf13247pf12800 -
BMC-P
BMC-H
BMC-T
Aldehyde dehydrogenase
PTAC
Signature enzyme
Alcohol dehydrogenase
Regulator

Salmonella anatum. Bacteria; Proteobacteria; Gammaproteobacteria; Enterobacterales;

C4876__locus_2

PDU1AB

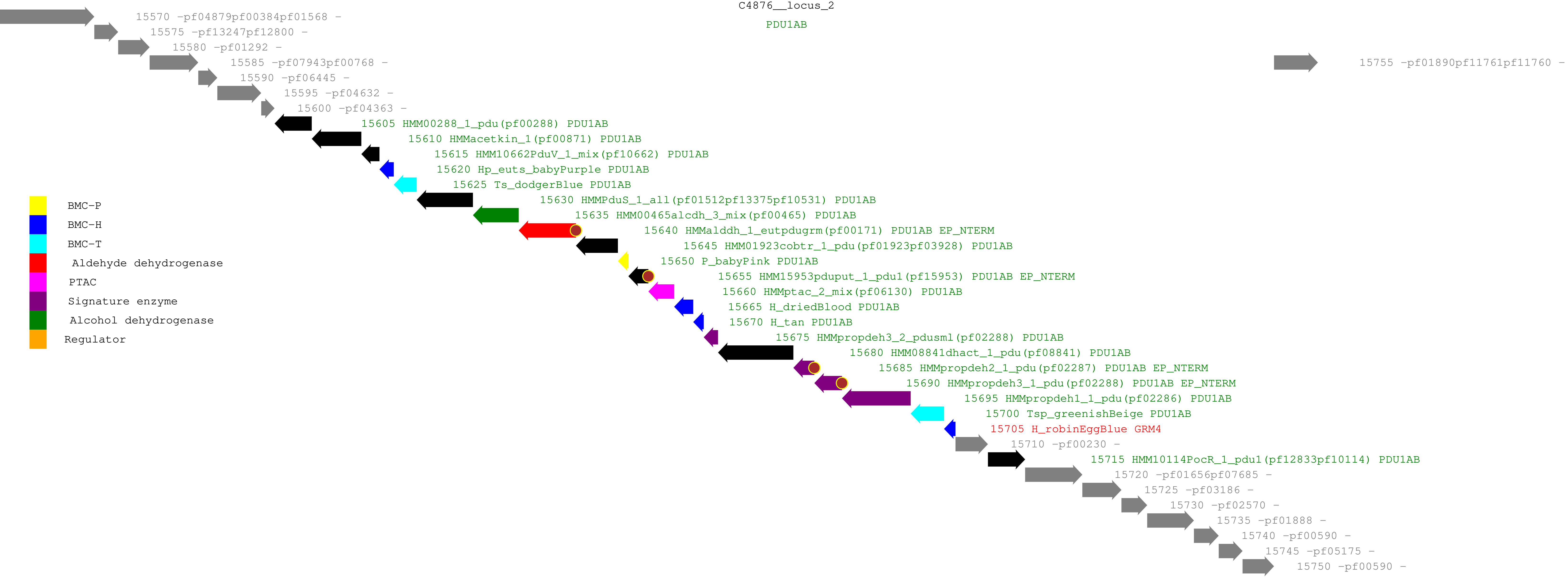

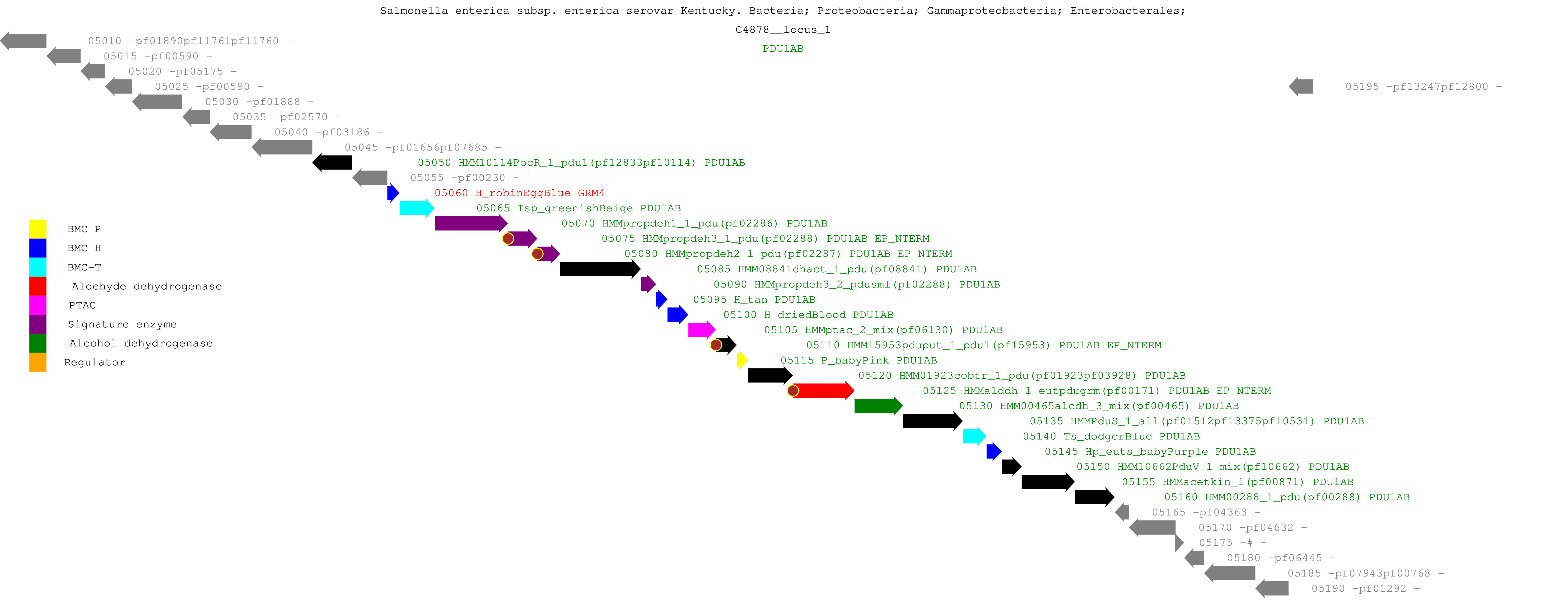
Salmonella enterica subsp. enterica serovar Kentucky. Bacteria; Proteobacteria; Gammaproteobacteria; Enterobacterales;
C4878__locus_1
PDU1AB
05010 -pf01890pf11761pf11760 -
05015 -pf00590 -
05020 -pf05175 -
05025 -pf00590 -
05030 -pf01888 -
05035 -pf02570 -
05040 -pf03186 -
05045 -pf01656pf07685 -
05050 HMM10114PocR_1_pdu1(pf12833pf10114) PDU1AB
05055 -pf00230 -
05060 H_robinEggBlue GRM4
05065 Tsp_greenishBeige PDU1AB
05070 HMMpropdeh1_1_pdu(pf02286) PDU1AB
05075 HMMpropdeh3_1_pdu(pf02288) PDU1AB EP_NTERM
05080 HMMpropdeh2_1_pdu(pf02287) PDU1AB EP_NTERM
05085 HMM08841dhact_1_pdu(pf08841) PDU1AB
05090 HMMpropdeh3_2_pdusml(pf02288) PDU1AB
05095 H_tan PDU1AB
05100 H_driedBlood PDU1AB
05105 HMMptac_2_mix(pf06130) PDU1AB
05110 HMM15953pduput_1_pdu1(pf15953) PDU1AB EP_NTERM
05115 P_babyPink PDU1AB
05120 HMM01923cobtr_1_pdu(pf01923pf03928) PDU1AB
05125 HMMaldh_1_eutpdugrm(pf00171) PDU1AB EP_NTERM
05130 HMM00465alcdh_3_mix(pf00465) PDU1AB
05135 HMMPduS_1_all(pf01512pf13375pf10531) PDU1AB
05140 Ts_dodgerBlue PDU1AB
05145 Hp_euts_babyPurple PDU1AB
05150 HMM10662PduV_1_mix(pf10662) PDU1AB
05155 HMMacetkin_1(pf00871) PDU1AB
05160 HMM00288_1_pdu(pf00288) PDU1AB
05165 -pf04363 -
05170 -pf04632 -
05175 -# -
05180 -pf06445 -
05185 -pf07943pf00768 -
05190 -pf01292 -
05195 -pf13247pf12800 -
BMC-P
BMC-H
BMC-T
Aldehyde dehydrogenase
PTAC
Signature enzyme
Alcohol dehydrogenase
Regulator

Salmonella enterica subsp. enterica serovar Teko. Bacteria; Proteobacteria; Gammaproteobacteria; Enterobacterales;

C4V46__locus_1

PDU1AB

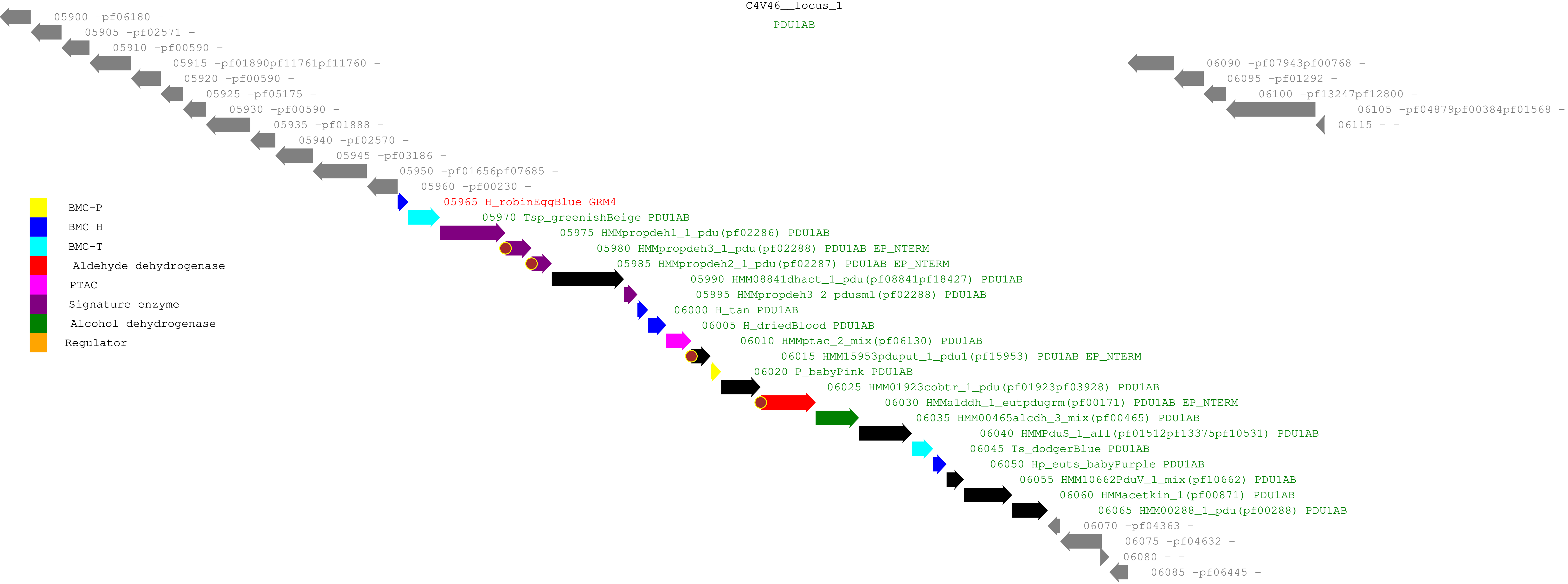

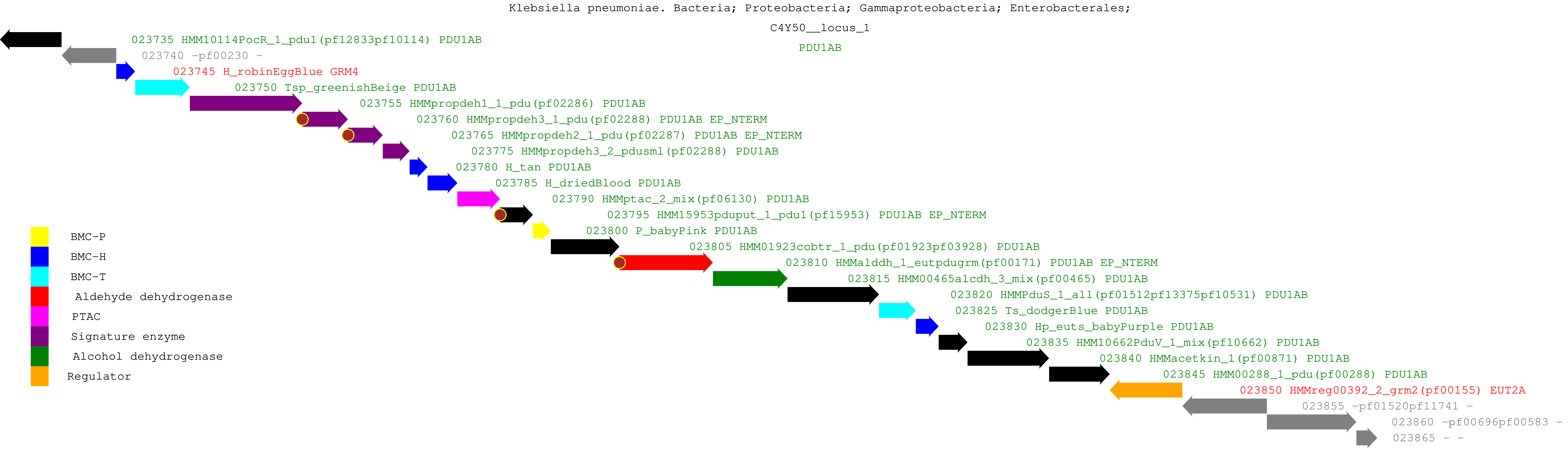

Klebsiella pneumoniae. Bacteria; Proteobacteria; Gammaproteobacteria; Enterobacterales;
C4Y50__locus_1
PDU1AB
023735 HMM10114PocR_1_pdu1(pf12833pf10114) PDU1AB
023740 -pf00230 -
023745 H_robinEggBlue GRM4
023750 Tsp_greenishBeige PDU1AB
023755 HMMpropdeh1_1_pdu(pf02286) PDU1AB
023760 HMMpropdeh3_1_pdu(pf02288) PDU1AB EP_NTERM
023765 HMMpropdeh2_1_pdu(pf02287) PDU1AB EP_NTERM
023775 HMMpropdeh3_2_pdusml(pf02288) PDU1AB
023780 H_tan PDU1AB
023785 H_driedBlood PDU1AB
023790 HMMptac_2_mix(pf06130) PDU1AB
023795 HMM15953pduput_1_pdu1(pf15953) PDU1AB EP_NTERM
023800 P_babyPink PDU1AB
023805 HMM01923cobtr_1_pdu(pf01923pf03928) PDU1AB
023810 HMMalddh_1_eutpdugrm(pf00171) PDU1AB EP_NTERM
023815 HMM00465alcdh_3_mix(pf00465) PDU1AB
023820 HMMPduS_1_all(pf01512pf13375pf10531) PDU1AB
023825 Ts_dodgerBlue PDU1AB
023830 Hp_euts_babyPurple PDU1AB
023835 HMM10662PduV_1_mix(pf10662) PDU1AB
023840 HMMacetkin_1(pf00871) PDU1AB
023845 HMM00288_1_pdu(pf00288) PDU1AB
023850 HMMreg00392_2_grm2(pf00155) EUT2A
023855 -pf01520pf11741 -
023860 -pf00696pf00583 -
023865 - -
BMC-P
BMC-H
BMC-T
Aldehyde dehydrogenase
PTAC
Signature enzyme
Alcohol dehydrogenase
Regulator

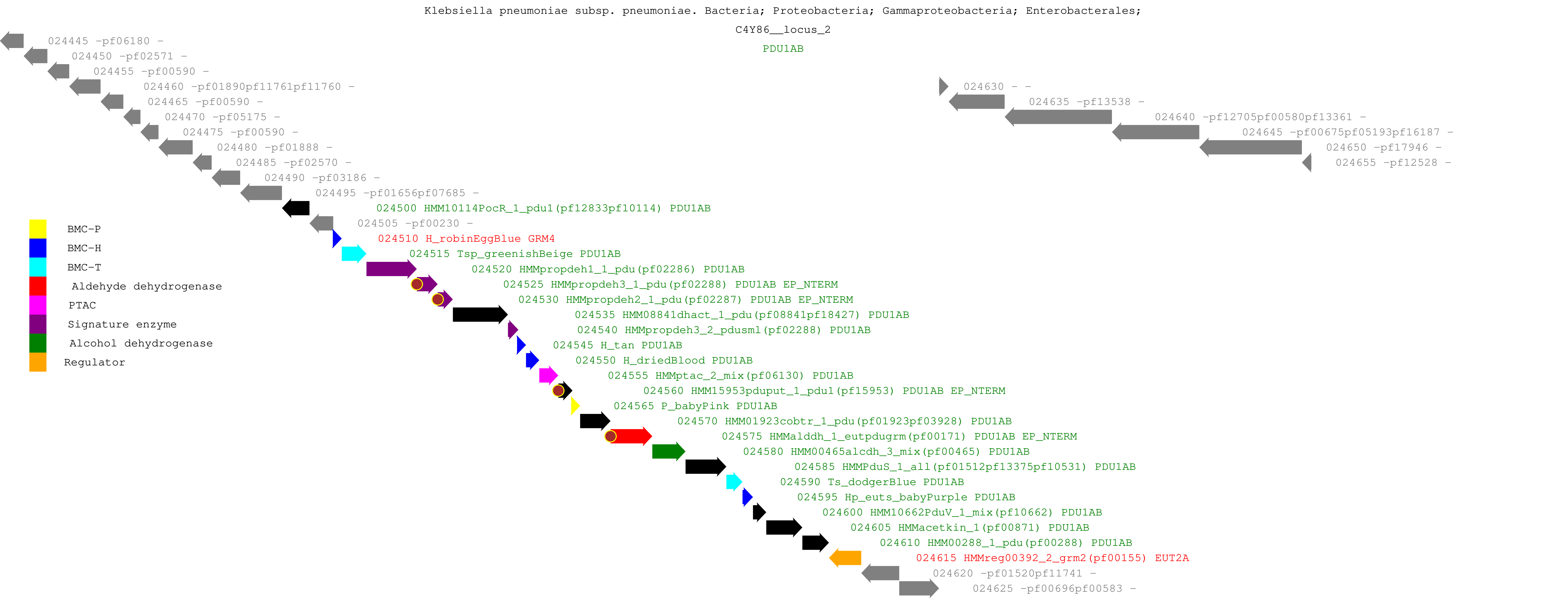
Klebsiella pneumoniae subsp. pneumoniae. Bacteria; Proteobacteria; Gammaproteobacteria; Enterobacterales;
C4Y86__locus_2
PDU1AB
024445 -pf06180 -
024450 -pf02571 -
024455 -pf00590 -
024460 -pf01890pf11761pf11760 -
024465 -pf00590 -
024470 -pf05175 -
024475 -pf00590 -
024480 -pf01888 -
024485 -pf02570 -
024490 -pf03186 -
024495 -pf01656pf07685 -
024500 HMM10114PocR_1_pdu1(pf12833pf10114) PDU1AB
024505 -pf00230 -
024510 H_robinEggBlue GRM4
024515 Tsp_greenishBeige PDU1AB
024520 HMMpropdeh1_1_pdu(pf02286) PDU1AB
024525 HMMpropdeh3_1_pdu(pf02288) PDU1AB EP_NTERM
024530 HMMpropdeh2_1_pdu(pf02287) PDU1AB EP_NTERM
024535 HMM08841dhact_1_pdu(pf08841pf18427) PDU1AB
024540 HMMpropdeh3_2_pdusml(pf02288) PDU1AB
024545 H_tan PDU1AB
024550 H_driedBlood PDU1AB
024555 HMMptac_2_mix(pf06130) PDU1AB
024560 HMM15953pduput_1_pdu1(pf15953) PDU1AB EP_NTERM
024565 P_babyPink PDU1AB
024570 HMM01923cobtr_1_pdu(pf01923pf03928) PDU1AB
024575 HMMalddh_1_eutpdugrm(pf00171) PDU1AB EP_NTERM
024580 HMM00465alcdh_3_mix(pf00465) PDU1AB
024585 HMMPduS_1_all(pf01512pf13375pf10531) PDU1AB
024590 Ts_dodgerBlue PDU1AB
024595 Hp_euts_babyPurple PDU1AB
024600 HMM10662PduV_1_mix(pf10662) PDU1AB
024605 HMMacetkin_1(pf00871) PDU1AB
024610 HMM00288_1_pdu(pf00288) PDU1AB
024615 HMMreg00392_2_grm2(pf00155) EUT2A
024620 -pf01520pf11741 -
024625 -pf00696pf00583 -
024630 - -
024635 -pf13538 -
024640 -pf12705pf00580pf13361 -
024645 -pf00675pf05193pf16187 -
024650 -pf17946 -
024655 -pf12528 -
BMC-P
BMC-H
BMC-T
Aldehyde dehydrogenase
PTAC
Signature enzyme
Alcohol dehydrogenase
Regulator

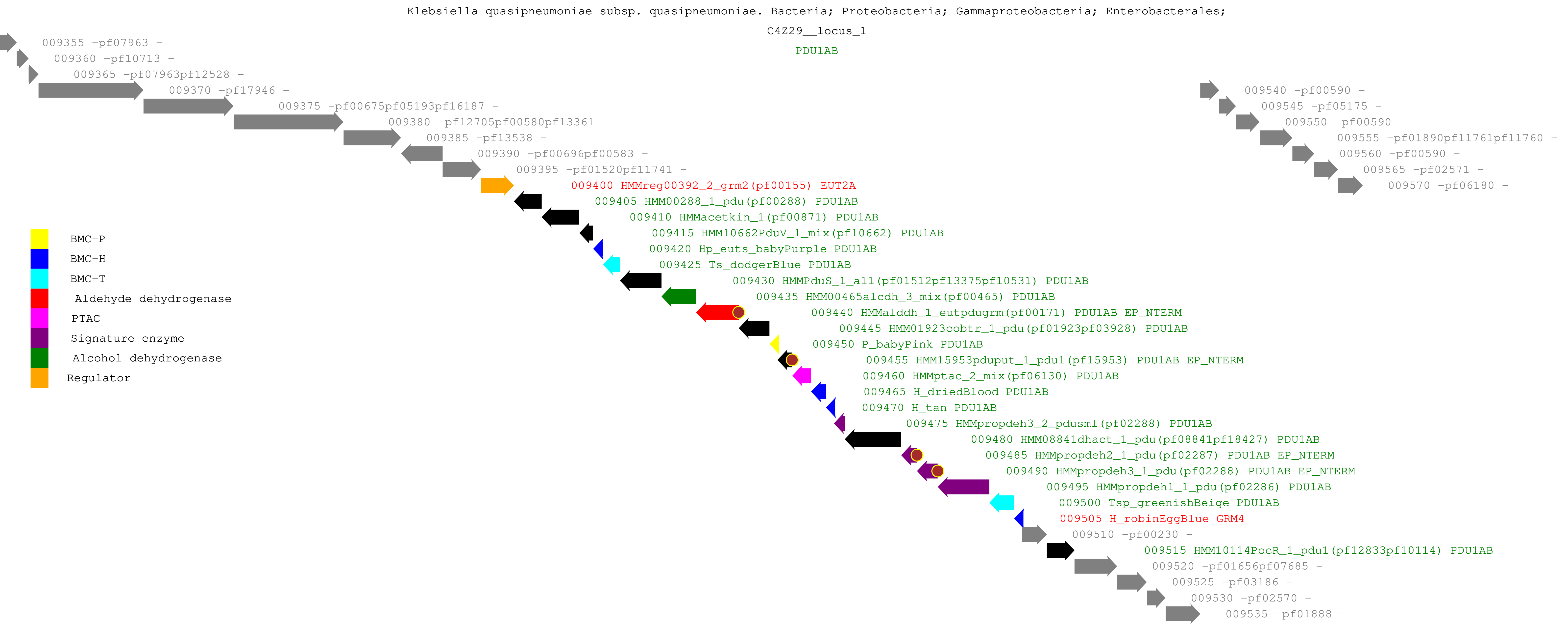
Klebsiella quasipneumoniae subsp. quasipneumoniae. Bacteria; Proteobacteria; Gammaproteobacteria; Enterobacterales;
C4Z29__locus_1
PDU1AB
009355 -pf07963 -
009360 -pf10713 -
009365 -pf07963pf12528 -
009370 -pf17946 -
009375 -pf00675pf05193pf16187 -
009380 -pf12705pf00580pf13361 -
009385 -pf13538 -
009390 -pf00696pf00583 -
009395 -pf01520pf11741 -
009400 HMMreg00392_2_grm2(pf00155) EUT2A
009405 HMM00288_1_pdu(pf00288) PDU1AB
009410 HMMacetkin_1(pf00871) PDU1AB
009415 HMM10662PduV_1_mix(pf10662) PDU1AB
009420 Hp_euts_babyPurple PDU1AB
009425 Ts_dodgerBlue PDU1AB
009430 HMMPduS_1_all(pf01512pf13375pf10531) PDU1AB
009435 HMM00465alcdh_3_mix(pf00465) PDU1AB
009440 HMMalddh_1_eutpdugrm(pf00171) PDU1AB EP_NTERM
009445 HMM01923cobtr_1_pdu(pf01923pf03928) PDU1AB
009450 P_babyPink PDU1AB
009455 HMM15953pduput_1_pdu1(pf15953) PDU1AB EP_NTERM
009460 HMMptac_2_mix(pf06130) PDU1AB
009465 H_driedBlood PDU1AB
009470 H_tan PDU1AB
009475 HMMpropdeh3_2_pdusml(pf02288) PDU1AB
009480 HMM08841dhact_1_pdu(pf08841pf18427) PDU1AB
009485 HMMpropdeh2_1_pdu(pf02287) PDU1AB EP_NTERM
009490 HMMpropdeh3_1_pdu(pf02288) PDU1AB EP_NTERM
009495 HMMpropdeh1_1_pdu(pf02286) PDU1AB
009500 Tsp_greenishBeige PDU1AB
009505 H_robinEggBlue GRM4
009510 -pf00230 -
009515 HMM10114PocR_1_pdu1(pf12833pf10114) PDU1AB
009520 -pf01656pf07685 -
009525 -pf03186 -
009530 -pf02570 -
009535 -pf01888 -
009540 -pf00590 -
009545 -pf05175 -
009550 -pf00590 -
009555 -pf01890pf11761pf11760 -
009560 -pf00590 -
009565 -pf02571 -
009570 -pf06180 -
BMC-P
BMC-H
BMC-T
Aldehyde dehydrogenase
PTAC
Signature enzyme
Alcohol dehydrogenase
Regulator

Klebsiella pneumoniae subsp. pneumoniae. Bacteria; Proteobacteria; Gammaproteobacteria; Enterobacterales;

C4Z38__locus_1

PDU1AB

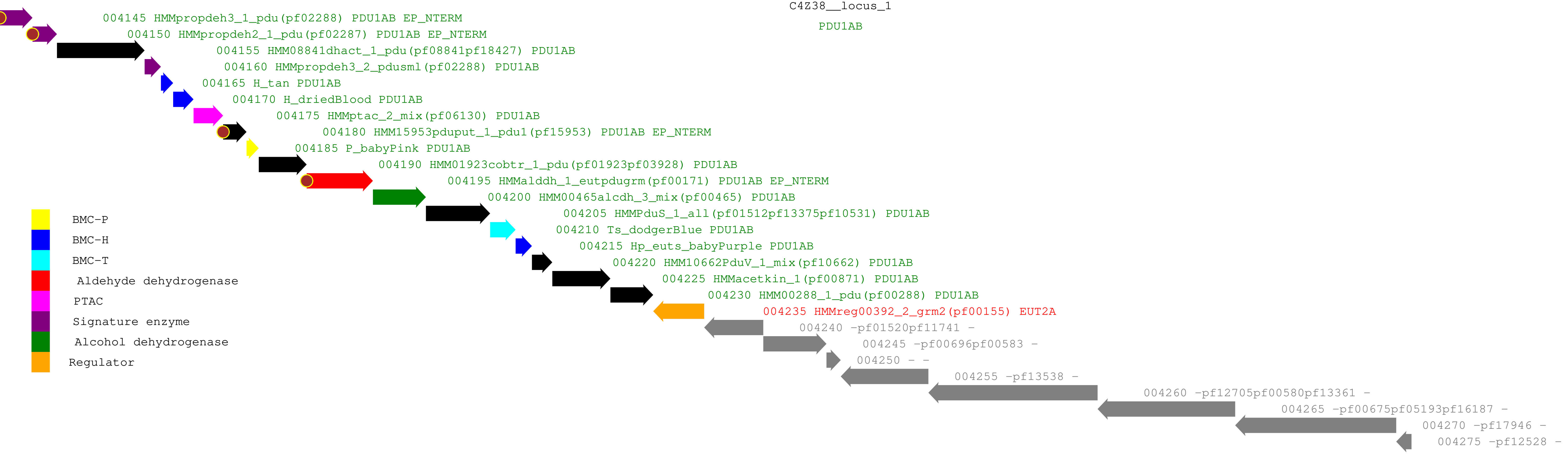

Salmonella enterica subsp. enterica serovar Concord. Bacteria; Proteobacteria; Gammaproteobacteria; Enterobacterales;

C6651__locus_2

PDU1AB

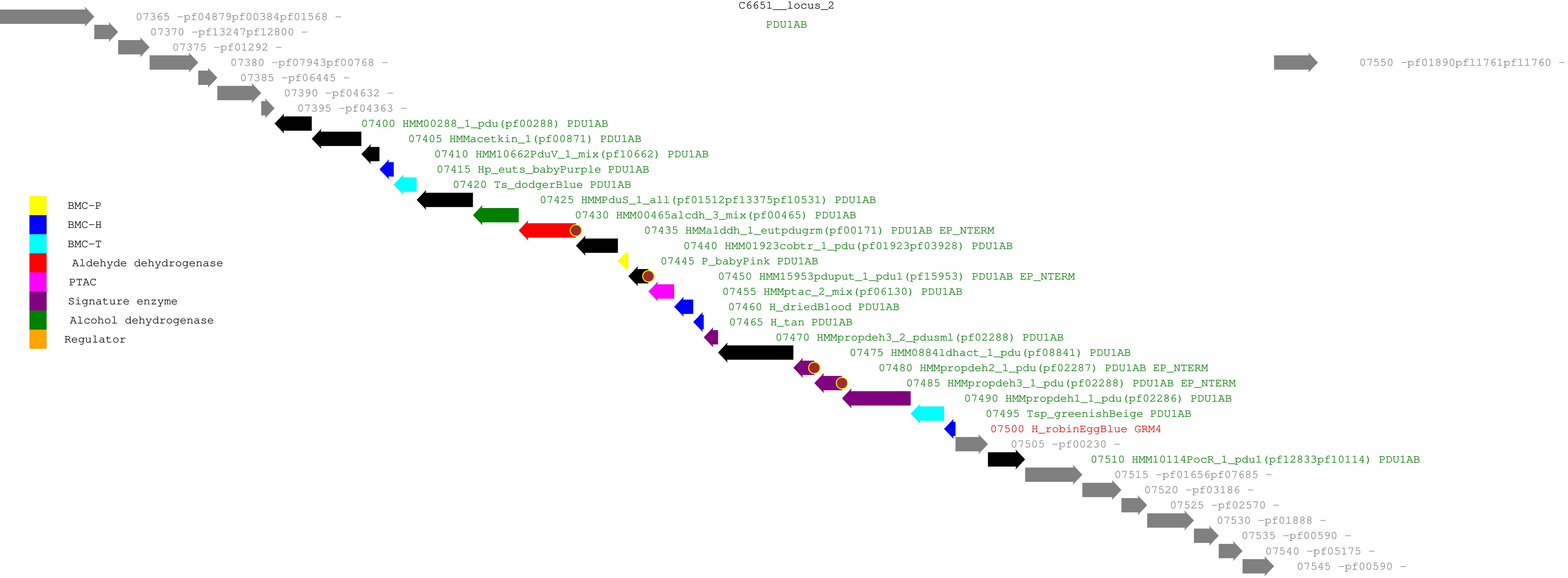

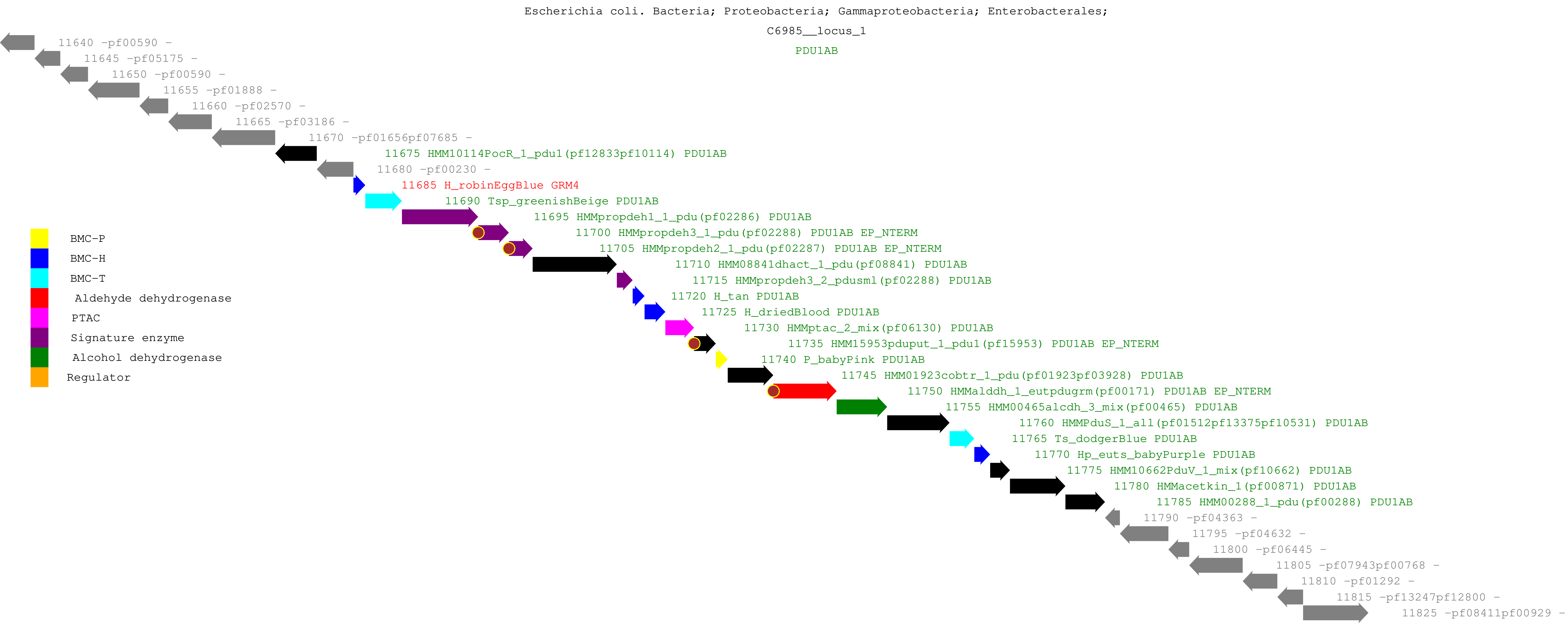
Escherichia coli. Bacteria; Proteobacteria; Gammaproteobacteria; Enterobacterales;
C6985__locus_1
PDU1AB
11640 -pf00590 -
11645 -pf05175 -
11650 -pf00590 -
11655 -pf01888 -
11660 -pf02570 -
11665 -pf03186 -
11670 -pf01656pf07685 -
11675 HMM10114PocR_1_pdu1(pf12833pf10114) PDU1AB
11680 -pf00230 -
11685 H_robinEggBlue GRM4
11690 Tsp_greenishBeige PDU1AB
11695 HMMpropdeh1_1_pdu(pf02286) PDU1AB
11700 HMMpropdeh3_1_pdu(pf02288) PDU1AB EP_NTERM
11705 HMMpropdeh2_1_pdu(pf02287) PDU1AB EP_NTERM
11710 HMM08841dhact_1_pdu(pf08841) PDU1AB
11715 HMMpropdeh3_2_pdusml(pf02288) PDU1AB
11720 H_tan PDU1AB
11725 H_driedBlood PDU1AB
11730 HMMptac_2_mix(pf06130) PDU1AB
11735 HMM15953pduput_1_pdu1(pf15953) PDU1AB EP_NTERM
11740 P_babyPink PDU1AB
11745 HMM01923cobtr_1_pdu(pf01923pf03928) PDU1AB
11750 HMMalddh_1_eutpdugrm(pf00171) PDU1AB EP_NTERM
11755 HMM00465alcdh_3_mix(pf00465) PDU1AB
11760 HMMPduS_1_all(pf01512pf13375pf10531) PDU1AB
11765 Ts_dodgerBlue PDU1AB
11770 Hp_euts_babyPurple PDU1AB
11775 HMM10662PduV_1_mix(pf10662) PDU1AB
11780 HMMacetkin_1(pf00871) PDU1AB
11785 HMM00288_1_pdu(pf00288) PDU1AB
11790 -pf04363 -
11795 -pf04632 -
11800 -pf06445 -
11805 -pf07943pf00768 -
11810 -pf01292 -
11815 -pf13247pf12800 -
11825 -pf08411pf00929 -
BMC-P
BMC-H
BMC-T
Aldehyde dehydrogenase
PTAC
Signature enzyme
Alcohol dehydrogenase
Regulator

Escherichia sp. 4726-5. Bacteria; Proteobacteria; Gammaproteobacteria; Enterobacterales;

C7B04__locus_1

PDU1AB

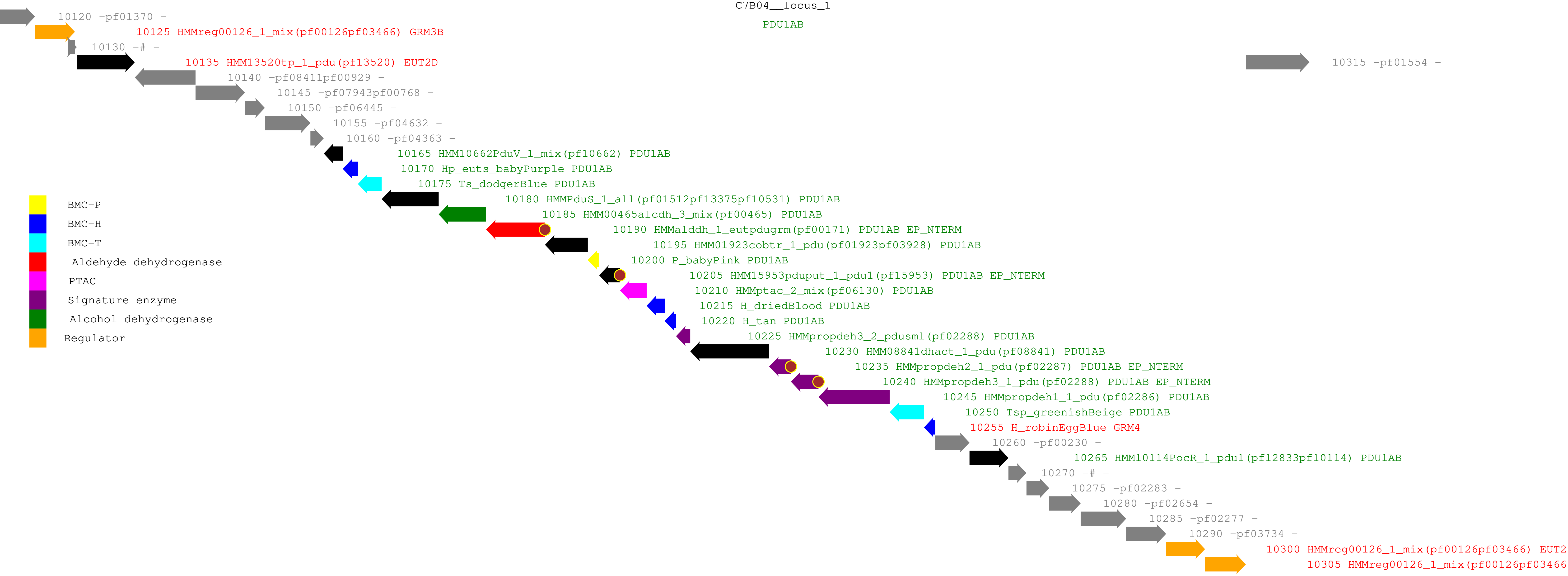

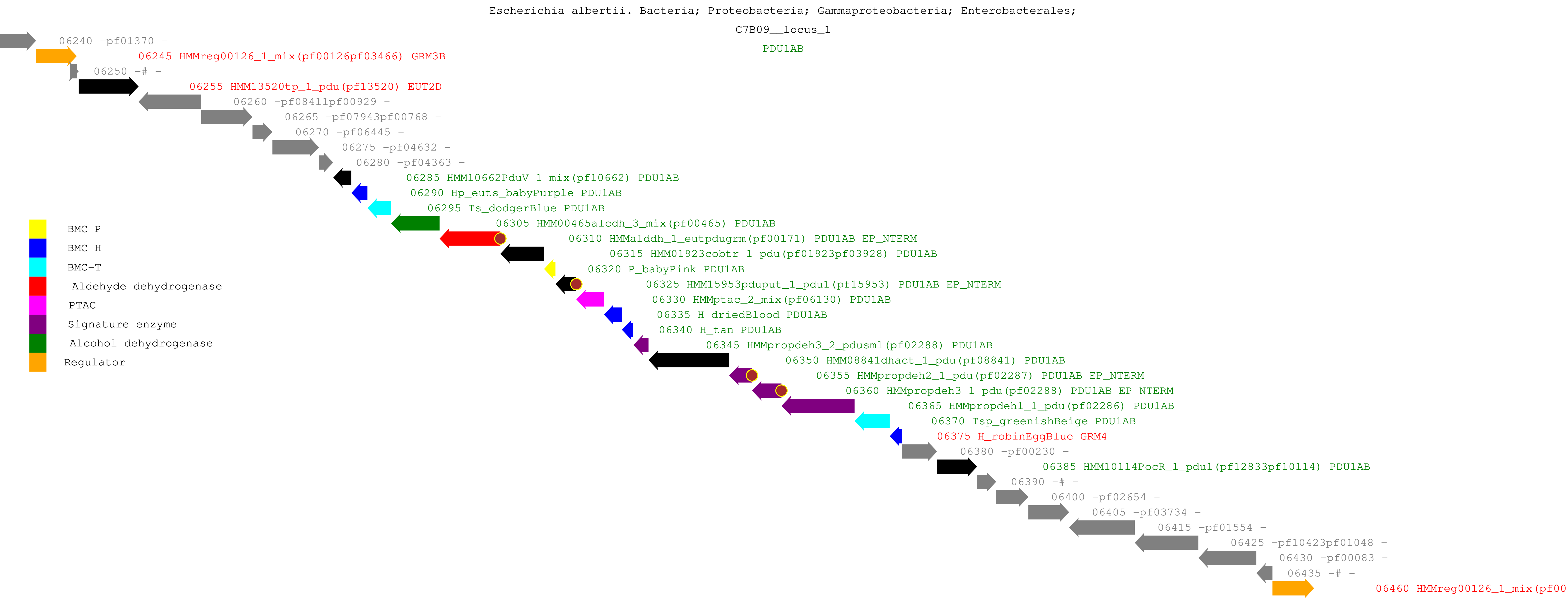
Escherichia albertii. Bacteria; Proteobacteria; Gammaproteobacteria; Enterobacterales;
C7B09__locus_1
PDU1AB
06240 -pf01370 -
06245 HMMreg00126_1_mix(pf00126pf03466) GRM3B
06250 -# -
06255 HMM13520tp_1_pdu(pf13520) EUT2D
06260 -pf08411pf00929 -
06265 -pf07943pf00768 -
06270 -pf06445 -
06275 -pf04632 -
06280 -pf04363 -
06285 HMM10662PduV_1_mix(pf10662) PDU1AB
06290 Hp_euts_babyPurple PDU1AB
06295 Ts_dodgerBlue PDU1AB
06305 HMM00465alcdh_3_mix(pf00465) PDU1AB
06310 HMMalddh_1_eutpdugrm(pf00171) PDU1AB EP_NTERM
06315 HMM01923cobtr_1_pdu(pf01923pf03928) PDU1AB
06320 P_babyPink PDU1AB
06325 HMM15953pduput_1_pdu1(pf15953) PDU1AB EP_NTERM
06330 HMMptac_2_mix(pf06130) PDU1AB
06335 H_driedBlood PDU1AB
06340 H_tan PDU1AB
06345 HMMpropdeh3_2_pdusml(pf02288) PDU1AB
06350 HMM08841dhact_1_pdu(pf08841) PDU1AB
06355 HMMpropdeh2_1_pdu(pf02287) PDU1AB EP_NTERM
06360 HMMpropdeh3_1_pdu(pf02288) PDU1AB EP_NTERM
06365 HMMpropdeh1_1_pdu(pf02286) PDU1AB
06370 Tsp_greenishBeige PDU1AB
06375 H_robinEggBlue GRM4
06380 -pf00230 -
06385 HMM10114PocR_1_pdu1(pf12833pf10114) PDU1AB
06390 -# -
06400 -pf02654 -
06405 -pf03734 -
06415 -pf01554 -
06425 -pf10423pf01048 -
06430 -pf00083 -
06435 -# -
06460 HMMreg00126_1_mix(pf00
BMC-P
BMC-H
BMC-T
Aldehyde dehydrogenase
PTAC
Signature enzyme
Alcohol dehydrogenase
Regulator

Salmonella enterica subsp. enterica serovar 4,5,12:b:-. Bacteria; Proteobacteria; Gammaproteobacteria; Enterobacterales;

C7S56__locus_1

PDU1AB

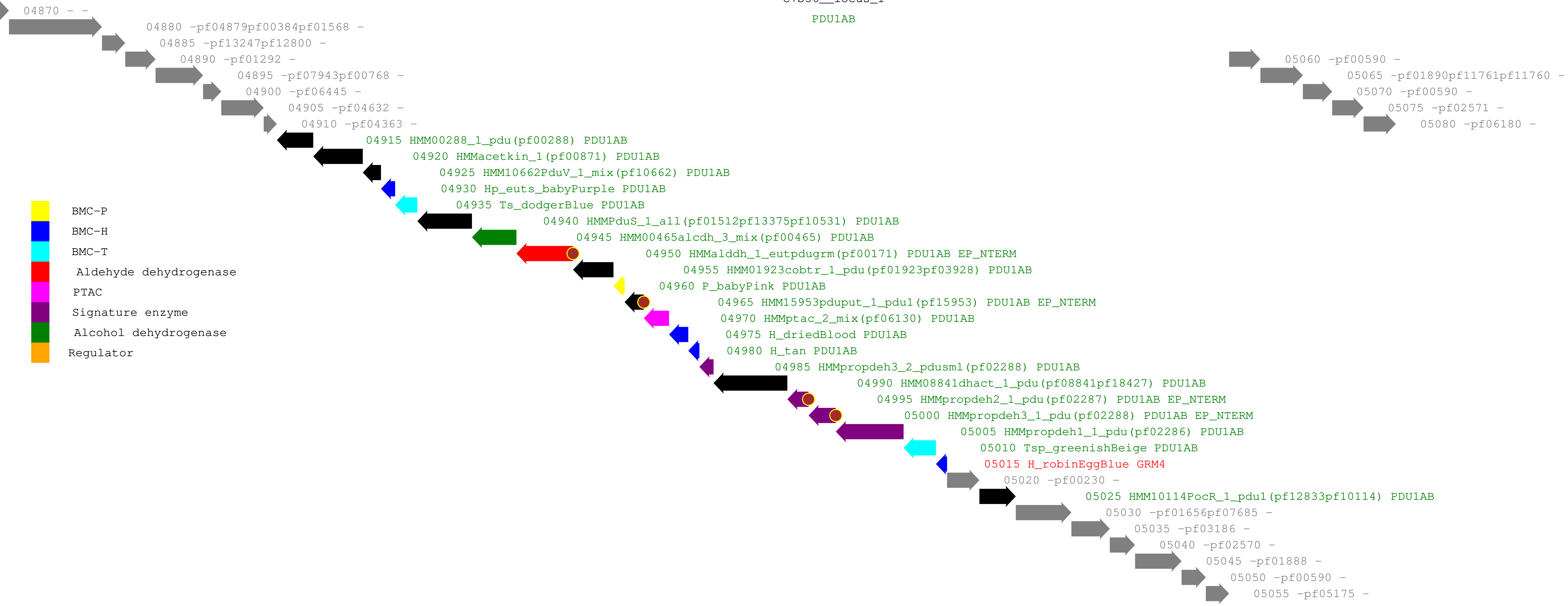

Klebsiella pneumoniae. Bacteria; Proteobacteria; Gammaproteobacteria; Enterobacterales;

C8P71__locus_1

PDU1AB

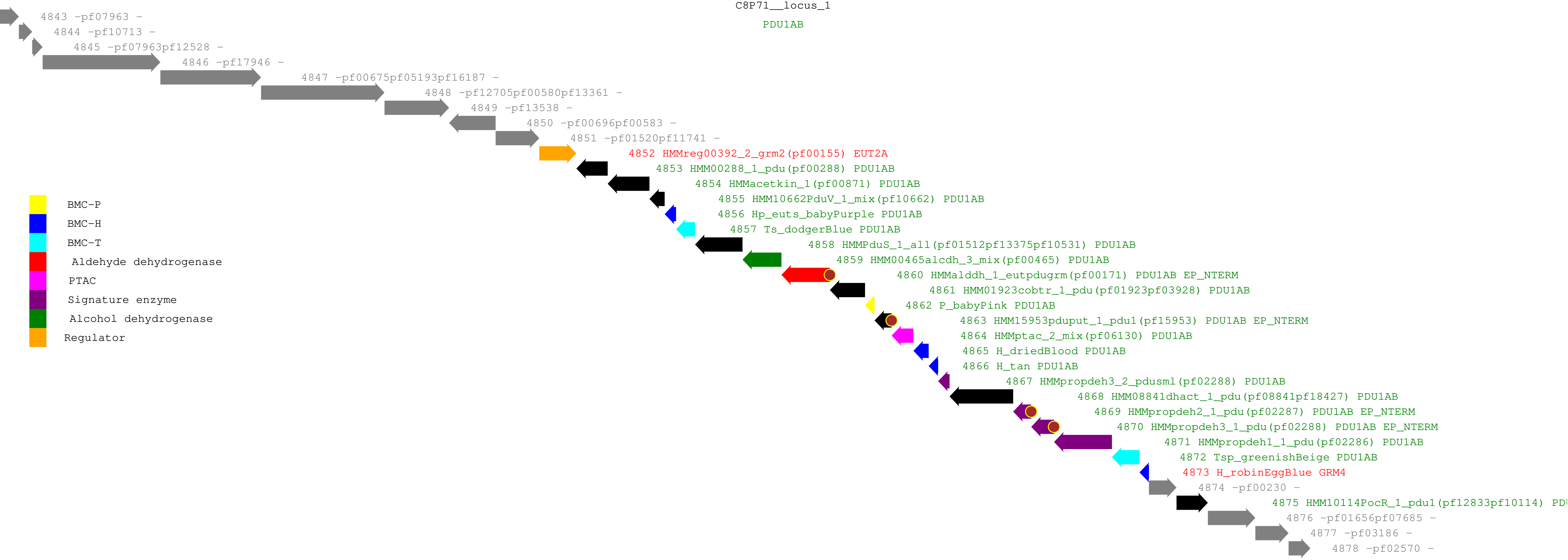

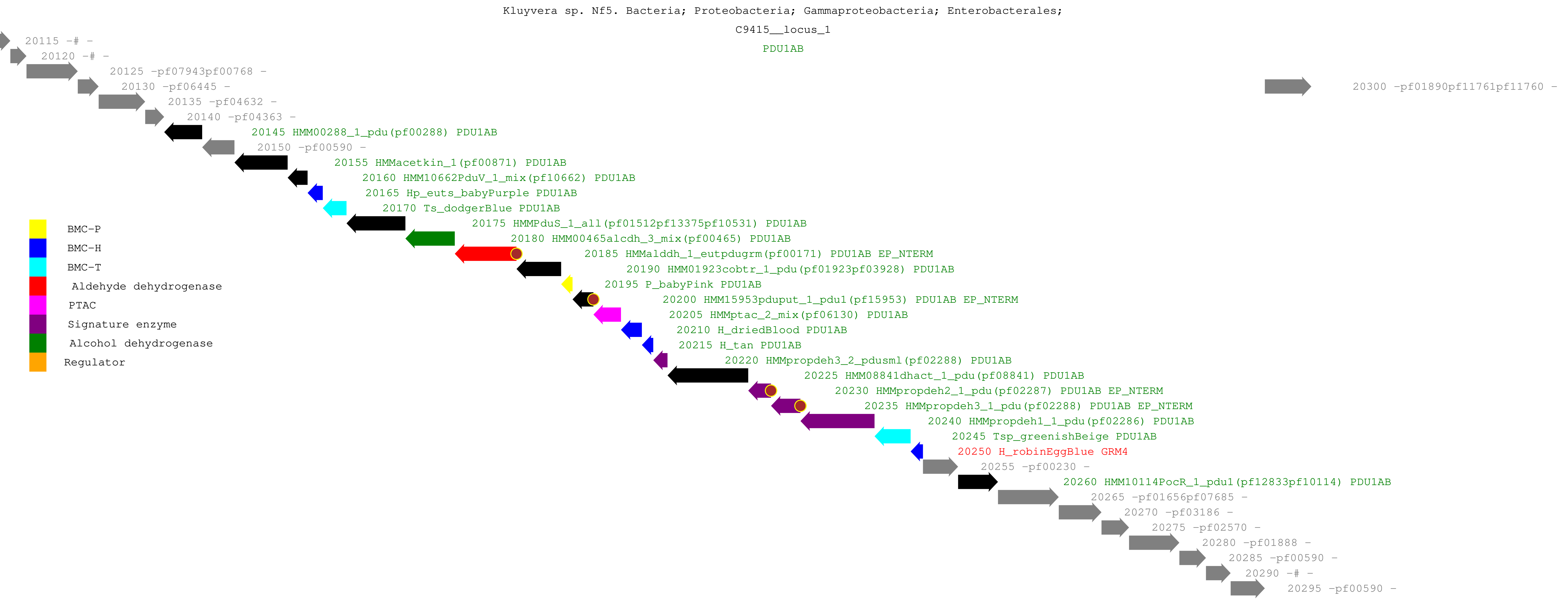

Kluyvera sp. Nf5. Bacteria; Proteobacteria; Gammaproteobacteria; Enterobacterales;
C9415__locus_1
PDU1AB
20115 -# -
20120 -# -
20125 -pf07943pf00768 -
20130 -pf06445 -
20135 -pf04632 -
20140 -pf04363 -
20145 HMM00288_1_pdu(pf00288) PDU1AB
20150 -pf00590 -
20155 HMMacetkin_1(pf00871) PDU1AB
20160 HMM10662PduV_1_mix(pf10662) PDU1AB
20165 Hp_euts_babyPurple PDU1AB
20170 Ts_dodgerBlue PDU1AB
20175 HMMPduS_1_all(pf01512pf13375pf10531) PDU1AB
20180 HMM00465alcdh_3_mix(pf00465) PDU1AB
20185 HMMalddh_1_eutpdugrm(pf00171) PDU1AB EP_NTERM
20190 HMM01923cobtr_1_pdu(pf01923pf03928) PDU1AB
20195 P_babyPink PDU1AB
20200 HMM15953pduput_1_pdu1(pf15953) PDU1AB EP_NTERM
20205 HMMptac_2_mix(pf06130) PDU1AB
20210 H_driedBlood PDU1AB
20215 H_tan PDU1AB
20220 HMMpropdeh3_2_pdusml(pf02288) PDU1AB
20225 HMM08841dhact_1_pdu(pf08841) PDU1AB
20230 HMMpropdeh2_1_pdu(pf02287) PDU1AB EP_NTERM
20235 HMMpropdeh3_1_pdu(pf02288) PDU1AB EP_NTERM
20240 HMMpropdeh1_1_pdu(pf02286) PDU1AB
20245 Tsp_greenishBeige PDU1AB
20250 H_robinEggBlue GRM4
20255 -pf00230 -
20260 HMM10114PocR_1_pdu1(pf12833pf10114) PDU1AB
20265 -pf01656pf07685 -
20270 -pf03186 -
20275 -pf02570 -
20280 -pf01888 -
20285 -pf00590 -
20290 -# -
20295 -pf00590 -
20300 -pf01890pf11761pf11760 -
BMC-P
BMC-H
BMC-T
Aldehyde dehydrogenase
PTAC
Signature enzyme
Alcohol dehydrogenase
Regulator

Salmonella enterica subsp. enterica serovar Poona. Bacteria; Proteobacteria; Gammaproteobacteria; Enterobacterales;

C9F07__locus_1

PDU1AB

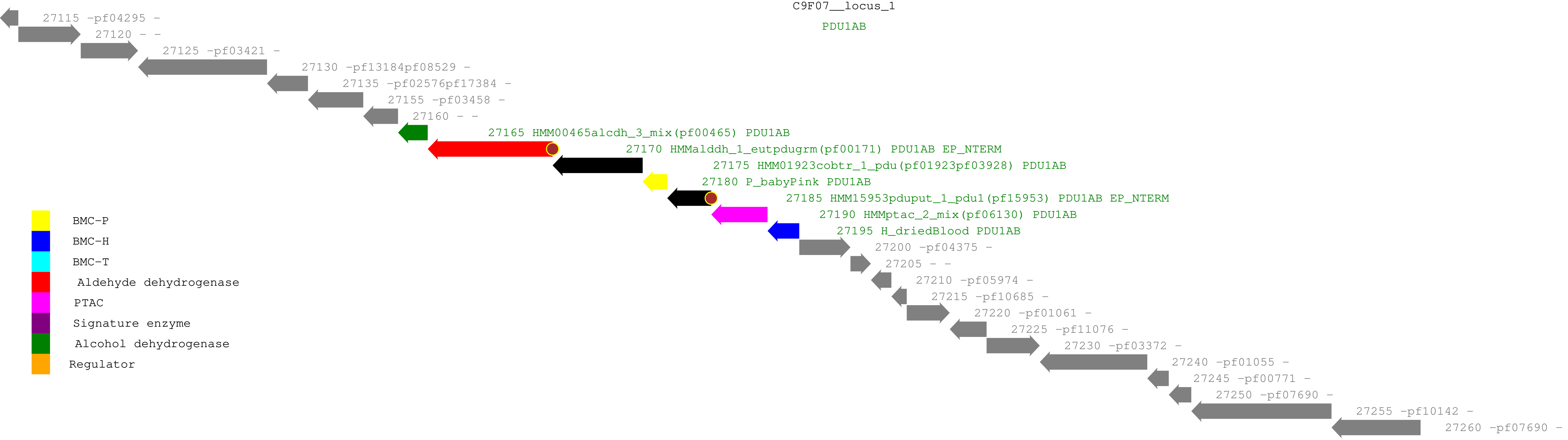

Salmonella enteritidis. Bacteria; Proteobacteria; Gammaproteobacteria; Enterobacterales;

CA285__locus_1

PDU1AB

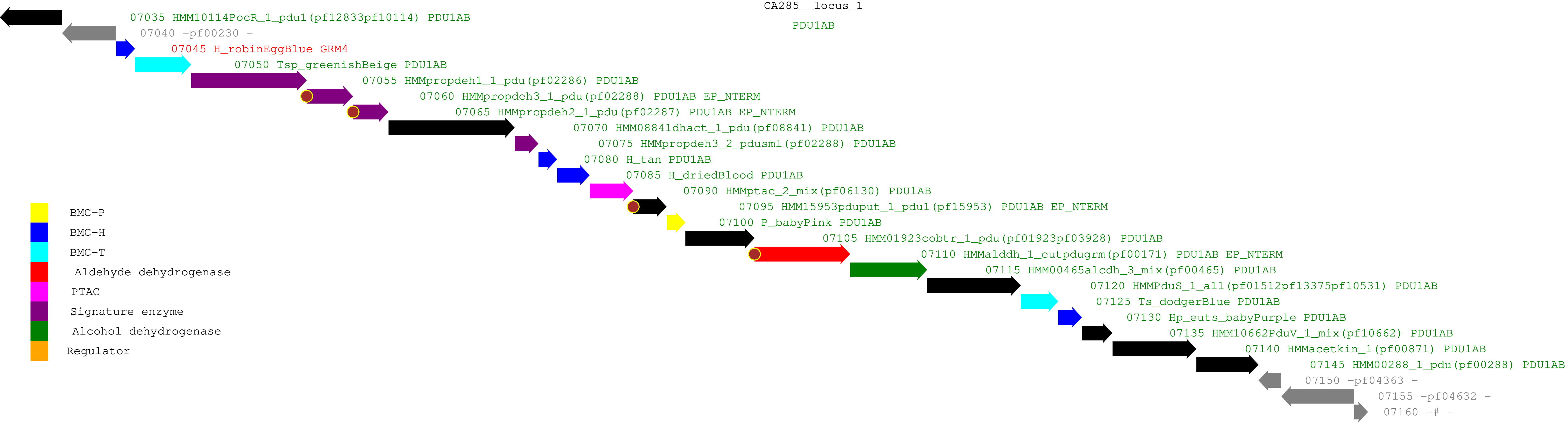

Salmonella enterica subsp. enterica serovar Paratyphi B str. SARA61. Bacteria; Proteobacteria; Gammaproteobacteria; Enterobacterales;

CAH10__locus_2

PDU1AB

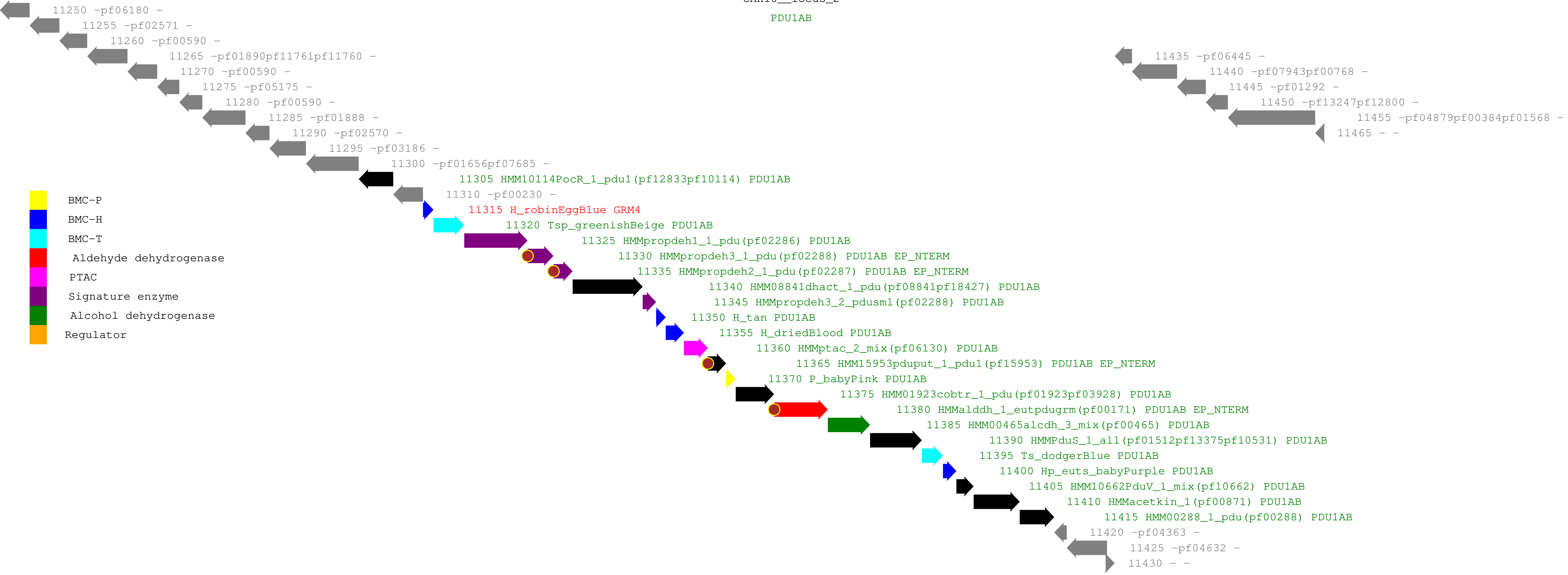

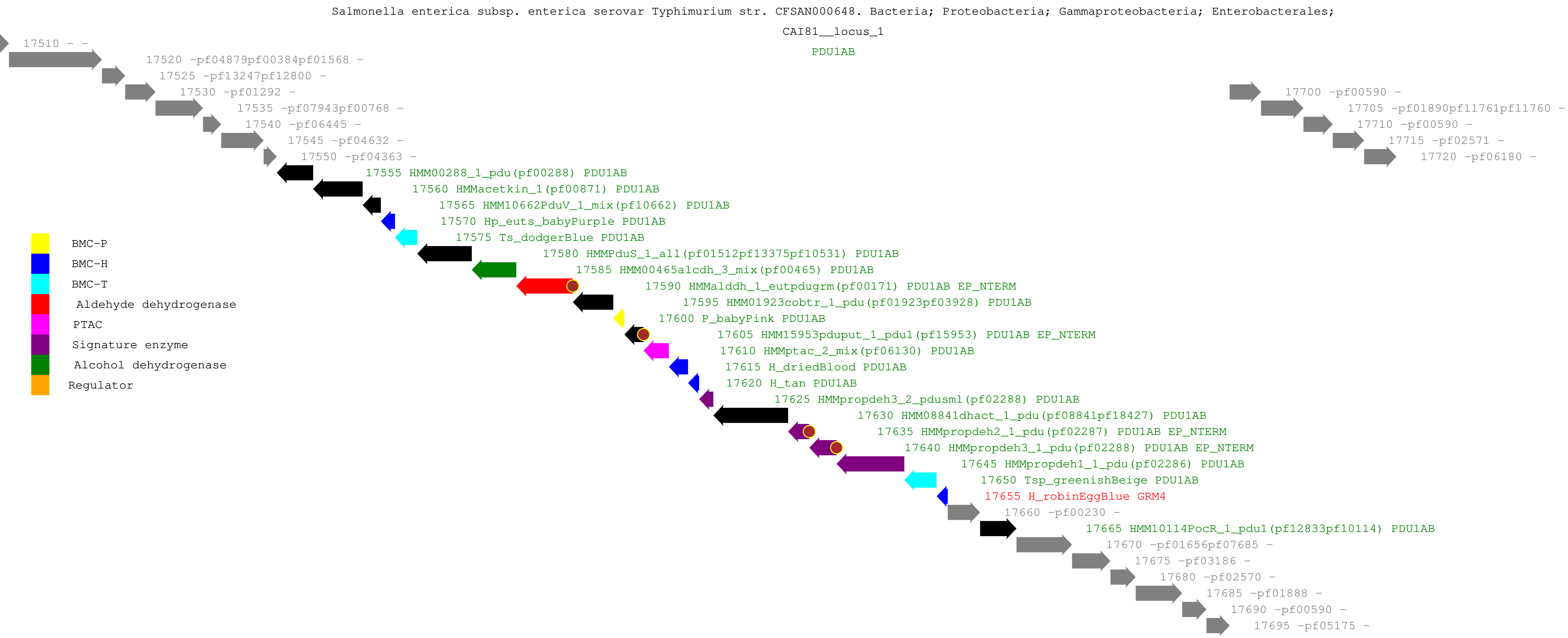

Salmonella enterica subsp. enterica serovar Typhimurium str. CFSAN000648. Bacteria; Proteobacteria; Gammaproteobacteria; Enterobacterales;
CAI81__locus_1
PDU1AB
17510 - -
17520 -pf04879pf00384pf01568 -
17525 -pf13247pf12800 -
17530 -pf01292 -
17535 -pf07943pf00768 -
17540 -pf06445 -
17545 -pf04632 -
17550 -pf04363 -
17555 HMM00288_1_pdu(pf00288) PDU1AB
17560 HMMacetkin_1(pf00871) PDU1AB
17565 HMM10662PduV_1_mix(pf10662) PDU1AB
17570 Hp_euts_babyPurple PDU1AB
17575 Ts_dodgerBlue PDU1AB
17580 HMMPduS_1_all(pf01512pf13375pf10531) PDU1AB
17585 HMM00465alcdh_3_mix(pf00465) PDU1AB
17590 HMMalddh_1_eutpdugrm(pf00171) PDU1AB EP_NTERM
17595 HMM01923cobtr_1_pdu(pf01923pf03928) PDU1AB
17600 P_babyPink PDU1AB
17605 HMM15953pduput_1_pdu1(pf15953) PDU1AB EP_NTERM
17610 HMMptac_2_mix(pf06130) PDU1AB
17615 H_driedBlood PDU1AB
17620 H_tan PDU1AB
17625 HMMpropdeh3_2_pdusml(pf02288) PDU1AB
17630 HMM08841dhact_1_pdu(pf08841pf18427) PDU1AB
17635 HMMpropdeh2_1_pdu(pf02287) PDU1AB EP_NTERM
17640 HMMpropdeh3_1_pdu(pf02288) PDU1AB EP_NTERM
17645 HMMpropdeh1_1_pdu(pf02286) PDU1AB
17650 Tsp_greenishBeige PDU1AB
17655 H_robinEggBlue GRM4
17660 -pf00230 -
17665 HMM10114PocR_1_pdu1(pf12833pf10114) PDU1AB
17670 -pf01656pf07685 -
17675 -pf03186 -
17680 -pf02570 -
17685 -pf01888 -
17690 -pf00590 -
17695 -pf05175 -
17700 -pf00590 -
17705 -pf01890pf11761pf11760 -
17710 -pf00590 -
17715 -pf02571 -
17720 -pf06180 -
BMC-P
BMC-H
BMC-T
Aldehyde dehydrogenase
PTAC
Signature enzyme
Alcohol dehydrogenase
Regulator

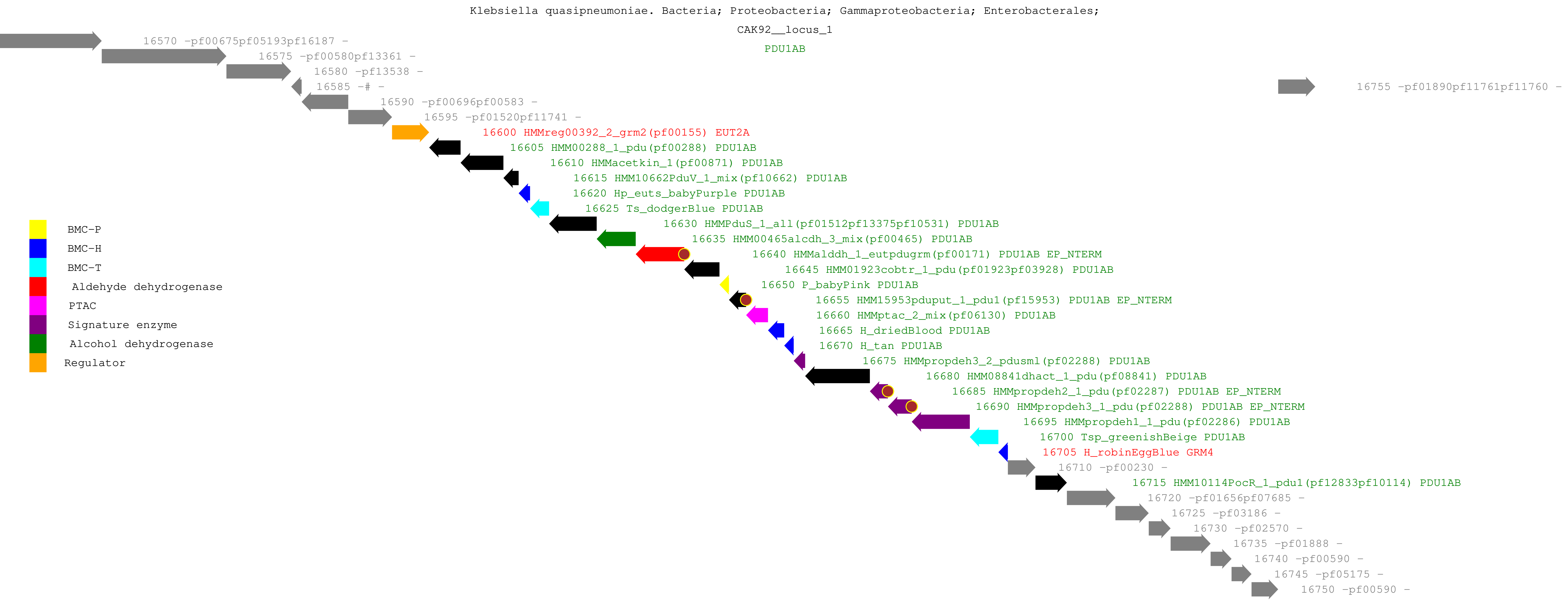
Klebsiella quasipneumoniae. Bacteria; Proteobacteria; Gammaproteobacteria; Enterobacterales;
CAK92__locus_1
PDU1AB
16570 -pf00675pf05193pf16187 -
16575 -pf00580pf13361 -
16580 -pf13538 -
16585 -# -
16590 -pf00696pf00583 -
16595 -pf01520pf11741 -
16600 HMMreg00392_2_grm2(pf00155) EUT2A
16605 HMM00288_1_pdu(pf00288) PDU1AB
16610 HMMacetkin_1(pf00871) PDU1AB
16615 HMM10662PduV_1_mix(pf10662) PDU1AB
16620 Hp_euts_babyPurple PDU1AB
16625 Ts_dodgerBlue PDU1AB
16630 HMMPduS_1_all(pf01512pf13375pf10531) PDU1AB
16635 HMM00465alcdh_3_mix(pf00465) PDU1AB
16640 HMMaldh_1_eutpdugrm(pf00171) PDU1AB EP_NTERM
16645 HMM01923cobtr_1_pdu(pf01923pf03928) PDU1AB
16650 P_babyPink PDU1AB
16655 HMM15953pduput_1_pdu1(pf15953) PDU1AB EP_NTERM
16660 HMMptac_2_mix(pf06130) PDU1AB
16665 H_driedBlood PDU1AB
16670 H_tan PDU1AB
16675 HMMpropdeh3_2_pdusml(pf02288) PDU1AB
16680 HMM08841dhact_1_pdu(pf08841) PDU1AB
16685 HMMpropdeh2_1_pdu(pf02287) PDU1AB EP_NTERM
16690 HMMpropdeh3_1_pdu(pf02288) PDU1AB EP_NTERM
16695 HMMpropdeh1_1_pdu(pf02286) PDU1AB
16700 Tsp_greenishBeige PDU1AB
16705 H_robinEggBlue GRM4
16710 -pf00230 -
16715 HMM10114PocR_1_pdu1(pf12833pf10114) PDU1AB
16720 -pf01656pf07685 -
16725 -pf03186 -
16730 -pf02570 -
16735 -pf01888 -
16740 -pf00590 -
16745 -pf05175 -
16750 -pf00590 -
16755 -pf01890pf11761pf11760 -
BMC-P
BMC-H
BMC-T
Aldehyde dehydrogenase
PTAC
Signature enzyme
Alcohol dehydrogenase
Regulator

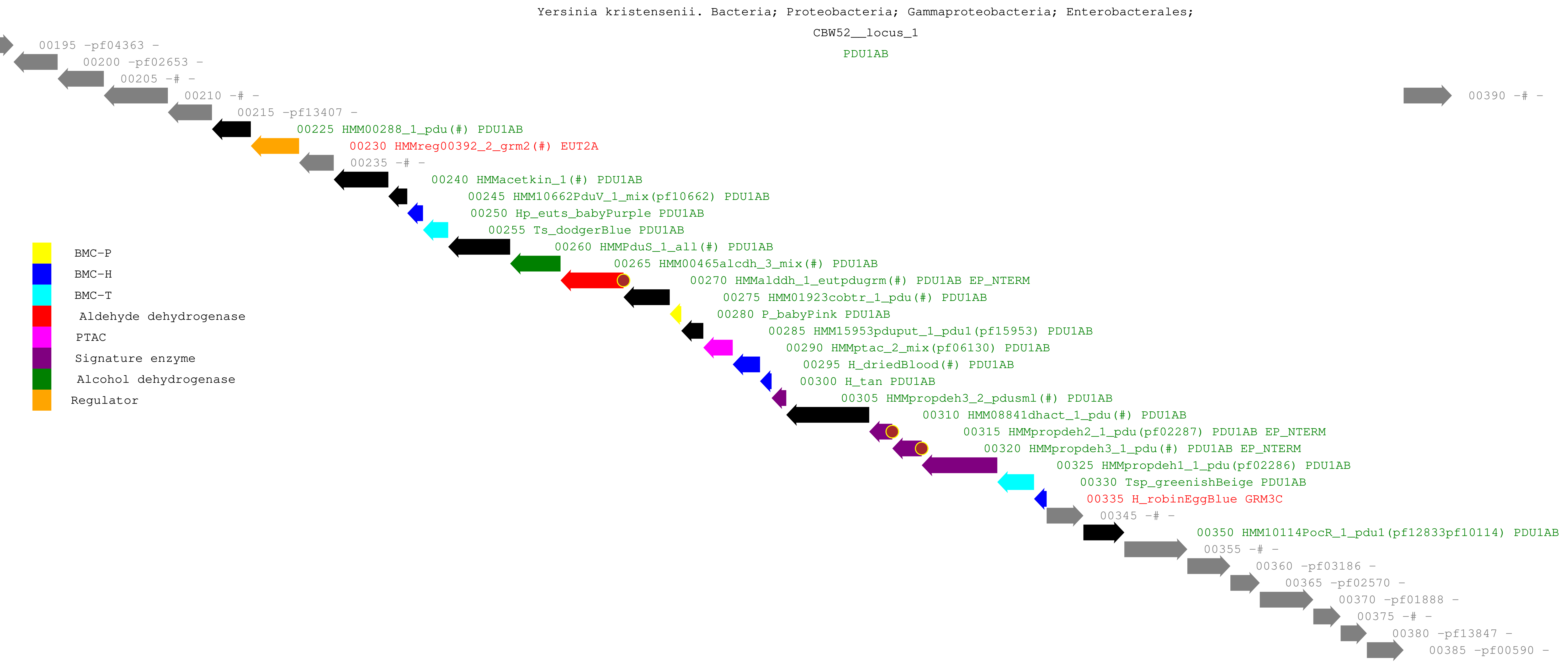
Yersinia kristensenii. Bacteria; Proteobacteria; Gammaproteobacteria; Enterobacterales;
CBW52__locus_1
PDU1AB
00195 -pf04363 -
00200 -pf02653 -
00205 -# -
00210 -# -
00215 -pf13407 -
00225 HMM00288_1_pdu(#) PDU1AB
00230 HMMreg00392_2_grm2(#) EUT2A
00235 -# -
00240 HMMacetkin_1(#) PDU1AB
00245 HMM10662PduV_1_mix(pf10662) PDU1AB
00250 Hp_euts_babyPurple PDU1AB
00255 Ts_dodgerBlue PDU1AB
00260 HMMPduS_1_all(#) PDU1AB
00265 HMM00465alcdh_3_mix(#) PDU1AB
00270 HMMalddh_1_eutpdugrm(#) PDU1AB EP_NTERM
00275 HMM01923cobtr_1_pdu(#) PDU1AB
00280 P_babyPink PDU1AB
00285 HMM15953pduput_1_pdu1(pf15953) PDU1AB
00290 HMMptac_2_mix(pf06130) PDU1AB
00295 H_driedBlood(#) PDU1AB
00300 H_tan PDU1AB
00305 HMMpropdeh3_2_pdusml(#) PDU1AB
00310 HMM08841dhact_1_pdu(#) PDU1AB
00315 HMMpropdeh2_1_pdu(pf02287) PDU1AB EP_NTERM
00320 HMMpropdeh3_1_pdu(#) PDU1AB EP_NTERM
00325 HMMpropdeh1_1_pdu(pf02286) PDU1AB
00330 Tsp_greenishBeige PDU1AB
00335 H_robinEggBlue GRM3C
00345 -# -
00350 HMM10114PocR_1_pdu1(pf12833pf10114) PDU1AB
00355 -# -
00360 -pf03186 -
00365 -pf02570 -
00370 -pf01888 -
00375 -# -
00380 -pf13847 -
00385 -pf00590 -
00390 -# -
BMC-P
BMC-H
BMC-T
Aldehyde dehydrogenase
PTAC
Signature enzyme
Alcohol dehydrogenase
Regulator

Yersinia frederiksenii. Bacteria; Proteobacteria; Gammaproteobacteria; Enterobacterales;
CBW53__locus_1
PDU1AB

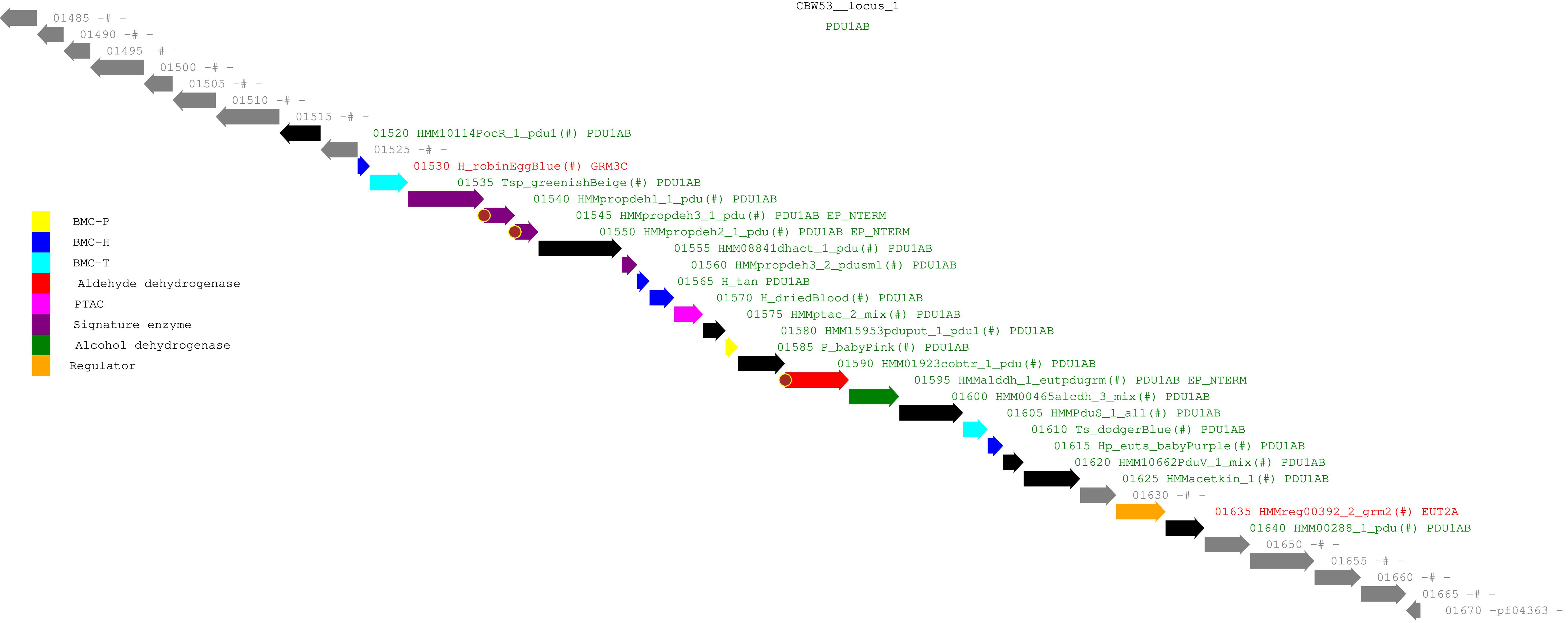

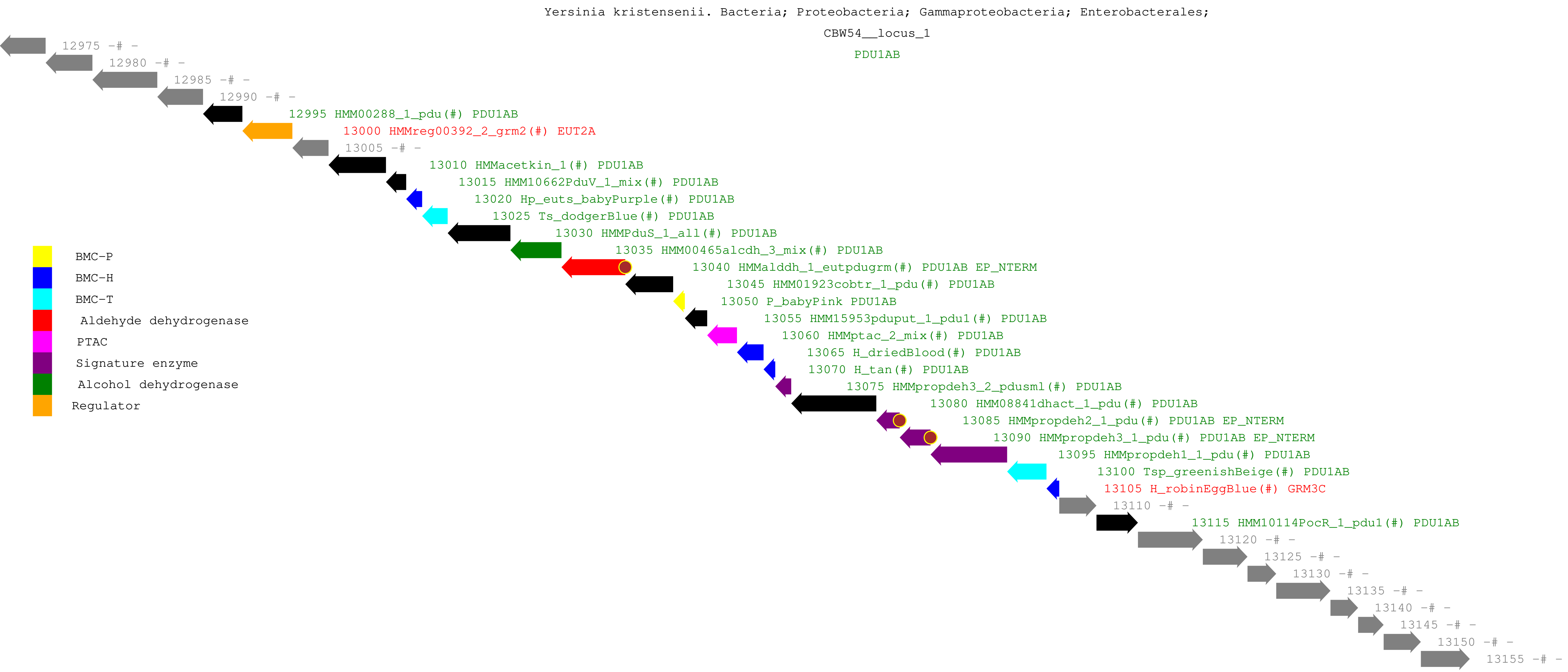
Yersinia kristensenii. Bacteria; Proteobacteria; Gammaproteobacteria; Enterobacterales;
CBW54__locus_1
PDU1AB
12975 -# -
12980 -# -
12985 -# -
12990 -# -
12995 HMM00288_1_pdu(#) PDU1AB
13000 HMMreg00392_2_grm2(#) EUT2A
13005 -# -
13010 HMMacetkin_1(#) PDU1AB
13015 HMM10662PduV_1_mix(#) PDU1AB
13020 Hp_euts_babyPurple(#) PDU1AB
13025 Ts_dodgerBlue(#) PDU1AB
13030 HMMPduS_1_all(#) PDU1AB
13035 HMM00465alcdh_3_mix(#) PDU1AB
13040 HMMalddh_1_eutpdugrm(#) PDU1AB EP_NTERM
13045 HMM01923cobtr_1_pdu(#) PDU1AB
13050 P_babyPink PDU1AB
13055 HMM15953pduput_1_pdu1(#) PDU1AB
13060 HMMptac_2_mix(#) PDU1AB
13065 H_driedBlood(#) PDU1AB
13070 H_tan(#) PDU1AB
13075 HMMpropdeh3_2_pdusml(#) PDU1AB
13080 HMM08841dhact_1_pdu(#) PDU1AB
13085 HMMpropdeh2_1_pdu(#) PDU1AB EP_NTERM
13090 HMMpropdeh3_1_pdu(#) PDU1AB EP_NTERM
13095 HMMpropdeh1_1_pdu(#) PDU1AB
13100 Tsp_greenishBeige(#) PDU1AB
13105 H_robinEggBlue(#) GRM3C
13110 -# -
13115 HMM10114PocR_1_pdu1(#) PDU1AB
13120 -# -
13125 -# -
13130 -# -
13135 -# -
13140 -# -
13145 -# -
13150 -# -
13155 -# -
BMC-P
BMC-H
BMC-T
Aldehyde dehydrogenase
PTAC
Signature enzyme
Alcohol dehydrogenase
Regulator

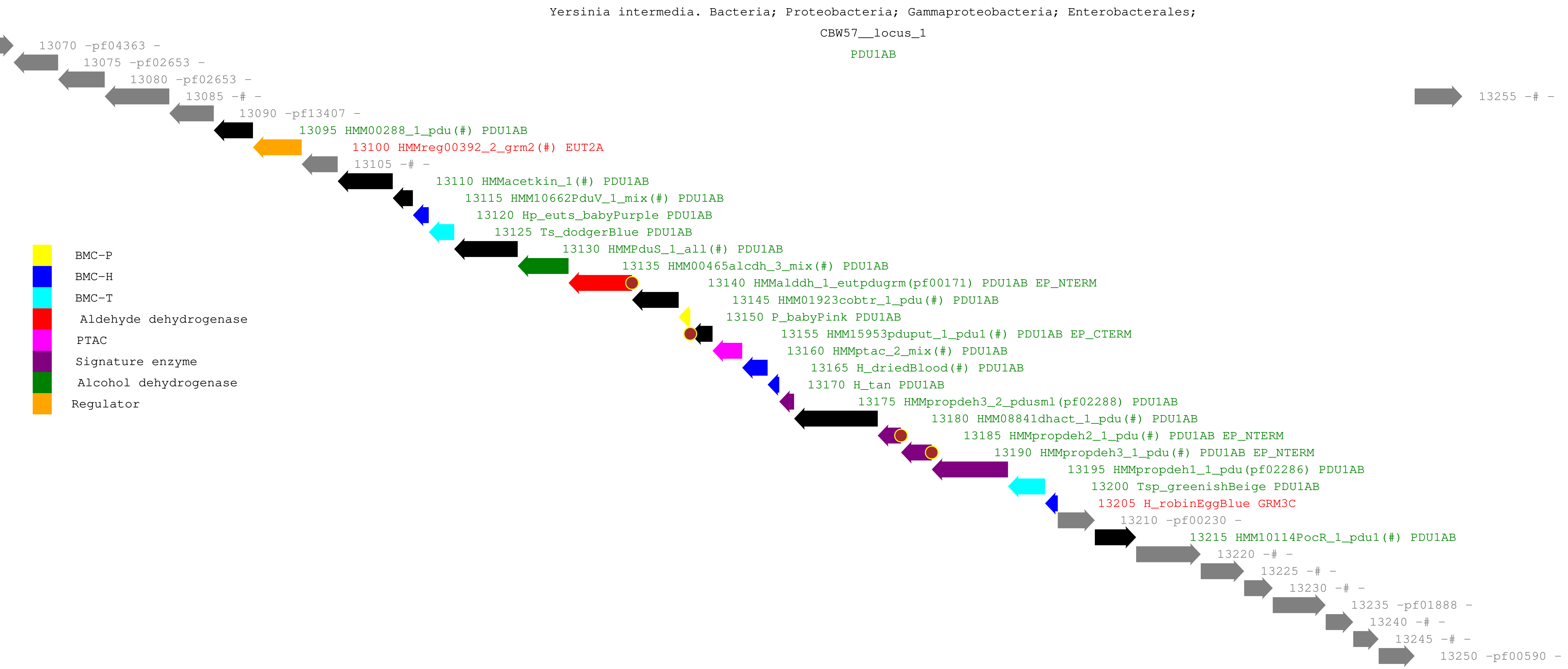
Yersinia intermedia. Bacteria; Proteobacteria; Gammaproteobacteria; Enterobacterales;
CBW57__locus_1
PDU1AB
13070 -pf04363 -
13075 -pf02653 -
13080 -pf02653 -
13085 -# -
13090 -pf13407 -
13095 HMM00288_1_pdu(#) PDU1AB
13100 HMMreg00392_2_grm2(#) EUT2A
13105 -# -
13110 HMMacetkin_1(#) PDU1AB
13115 HMM10662PduV_1_mix(#) PDU1AB
13120 Hp_euts_babyPurple PDU1AB
13125 Ts_dodgerBlue PDU1AB
13130 HMMPduS_1_all(#) PDU1AB
13135 HMM00465alcdh_3_mix(#) PDU1AB
13140 HMMalddh_1_eutpdugrm(pf00171) PDU1AB EP_NTERM
13145 HMM01923cobtr_1_pdu(#) PDU1AB
13150 P_babyPink PDU1AB
13155 HMM15953pduput_1_pdu1(#) PDU1AB EP_CTERM
13160 HMMptac_2_mix(#) PDU1AB
13165 H_driedBlood(#) PDU1AB
13170 H_tan PDU1AB
13175 HMMpropdeh3_2_pdusml(pf02288) PDU1AB
13180 HMM08841dhact_1_pdu(#) PDU1AB
13185 HMMpropdeh2_1_pdu(#) PDU1AB EP_NTERM
13190 HMMpropdeh3_1_pdu(#) PDU1AB EP_NTERM
13195 HMMpropdeh1_1_pdu(pf02286) PDU1AB
13200 Tsp_greenishBeige PDU1AB
13205 H_robinEggBlue GRM3C
13210 -pf00230 -
13215 HMM10114PocR_1_pdu1(#) PDU1AB
13220 -# -
13225 -# -
13230 -# -
13235 -pf01888 -
13240 -# -
13245 -# -
13250 -pf00590 -
13255 -# -
BMC-P
BMC-H
BMC-T
Aldehyde dehydrogenase
PTAC
Signature enzyme
Alcohol dehydrogenase
Regulator

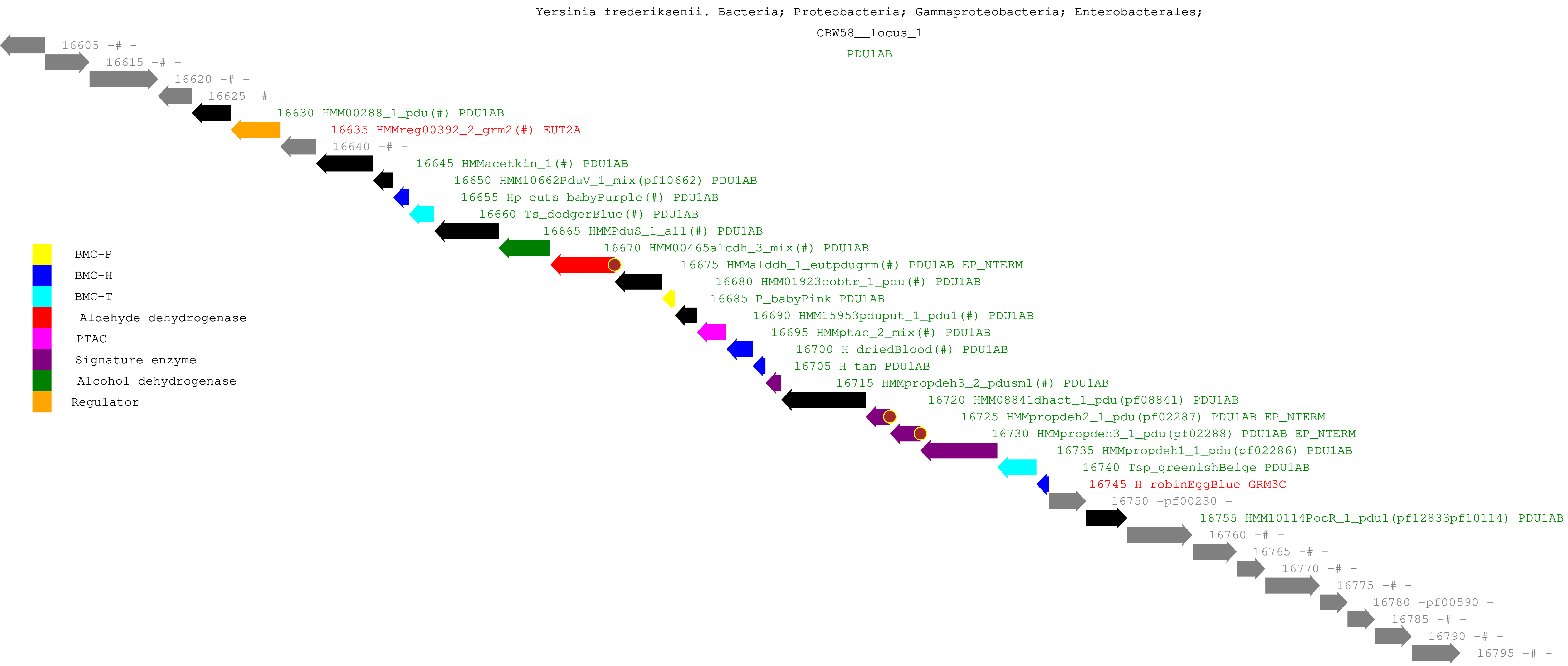
Yersinia frederiksenii. Bacteria; Proteobacteria; Gammaproteobacteria; Enterobacterales;
CBW58__locus_1
PDU1AB
16605 -# -
16615 -# -
16620 -# -
16625 -# -
16630 HMM00288_1_pdu(#) PDU1AB
16635 HMMreg00392_2_grm2(#) EUT2A
16640 -# -
16645 HMMacetkin_1(#) PDU1AB
16650 HMM10662PduV_1_mix(pf10662) PDU1AB
16655 Hp_euts_babyPurple(#) PDU1AB
16660 Ts_dodgerBlue(#) PDU1AB
16665 HMMPduS_1_all(#) PDU1AB
16670 HMM00465alcdh_3_mix(#) PDU1AB
16675 HMMalddh_1_eutpdugrm(#) PDU1AB EP_NTERM
16680 HMM01923cobtr_1_pdu(#) PDU1AB
16685 P_babyPink PDU1AB
16690 HMM15953pduput_1_pdu1(#) PDU1AB
16695 HMMptac_2_mix(#) PDU1AB
16700 H_driedBlood(#) PDU1AB
16705 H_tan PDU1AB
16715 HMMpropdeh3_2_pdusml(#) PDU1AB
16720 HMM08841dhact_1_pdu(pf08841) PDU1AB
16725 HMMpropdeh2_1_pdu(pf02287) PDU1AB EP_NTERM
16730 HMMpropdeh3_1_pdu(pf02288) PDU1AB EP_NTERM
16735 HMMpropdeh1_1_pdu(pf02286) PDU1AB
16740 Tsp_greenishBeige PDU1AB
16745 H_robinEggBlue GRM3C
16750 -pf00230 -
16755 HMM10114PocR_1_pdu1(pf12833pf10114) PDU1AB
16760 -# -
16765 -# -
16770 -# -
16775 -# -
16780 -pf00590 -
16785 -# -
16790 -# -
16795 -# -
BMC-P
BMC-H
BMC-T
Aldehyde dehydrogenase
PTAC
Signature enzyme
Alcohol dehydrogenase
Regulator

Salmonella enterica I. Bacteria; Proteobacteria; Gammaproteobacteria; Enterobacterales;

CC530__locus_1

PDU1AB

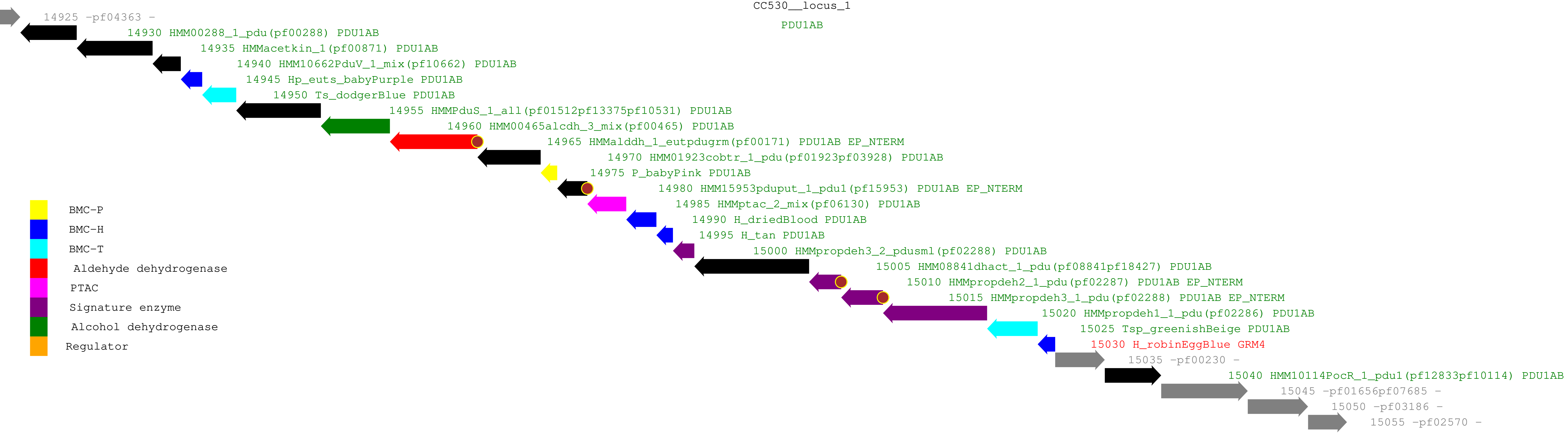

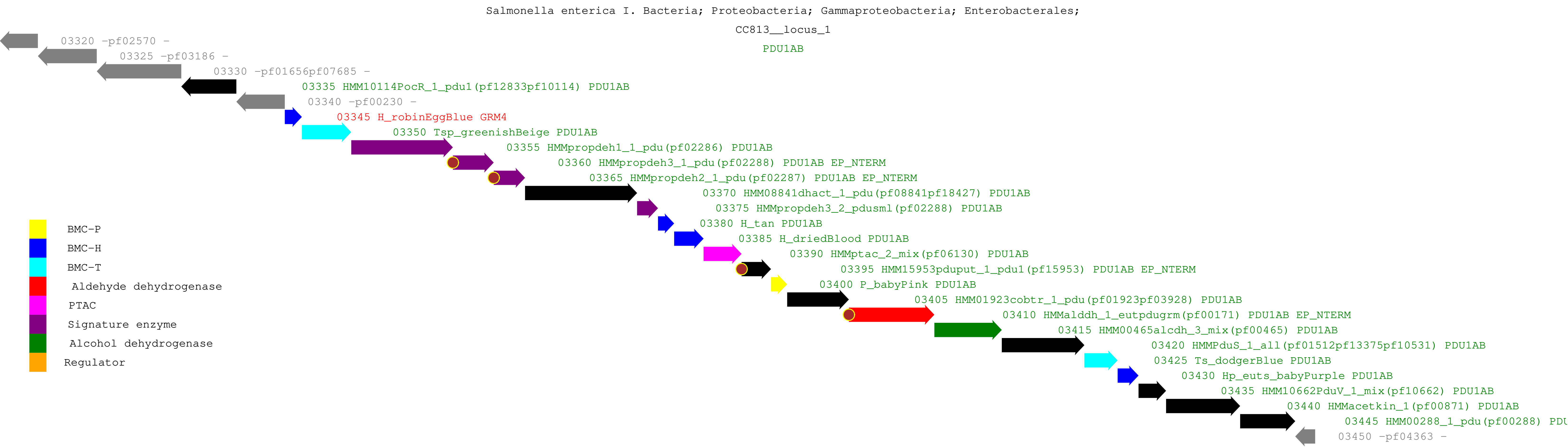
Salmonella enterica I. Bacteria; Proteobacteria; Gammaproteobacteria; Enterobacterales;
CC813__locus_1
PDU1AB
03320 -pf02570 -
03325 -pf03186 -
03330 -pf01656pf07685 -
03335 HMM10114PocR_1_pdu1(pf12833pf10114) PDU1AB
03340 -pf00230 -
03345 H_robinEggBlue GRM4
03350 Tsp_greenishBeige PDU1AB
03355 HMMpropdeh1_1_pdu(pf02286) PDU1AB
03360 HMMpropdeh3_1_pdu(pf02288) PDU1AB EP_NTERM
03365 HMMpropdeh2_1_pdu(pf02287) PDU1AB EP_NTERM
03370 HMM08841dhact_1_pdu(pf08841pf18427) PDU1AB
03375 HMMpropdeh3_2_pdusml(pf02288) PDU1AB
03380 H_tan PDU1AB
03385 H_driedBlood PDU1AB
03390 HMMptac_2_mix(pf06130) PDU1AB
03395 HMM15953pduput_1_pdu1(pf15953) PDU1AB EP_NTERM
03400 P_babyPink PDU1AB
03405 HMM01923cobtr_1_pdu(pf01923pf03928) PDU1AB
03410 HMMalddh_1_eutpdugrm(pf00171) PDU1AB EP_NTERM
03415 HMM00465alcdh_3_mix(pf00465) PDU1AB
03420 HMMPduS_1_all(pf01512pf13375pf10531) PDU1AB
03425 Ts_dodgerBlue PDU1AB
03430 Hp_euts_babyPurple PDU1AB
03435 HMM10662PduV_1_mix(pf10662) PDU1AB
03440 HMMacetkin_1(pf00871) PDU1AB
03445 HMM00288_1_pdu(pf00288) PDU
03450 -pf04363 -
BMC-P
BMC-H
BMC-T
Aldehyde dehydrogenase
PTAC
Signature enzyme
Alcohol dehydrogenase
Regulator

Enterobacter sp. 10-1. Bacteria; Proteobacteria; Gammaproteobacteria; Enterobacterales;

CD006__locus_1

PDU1AB

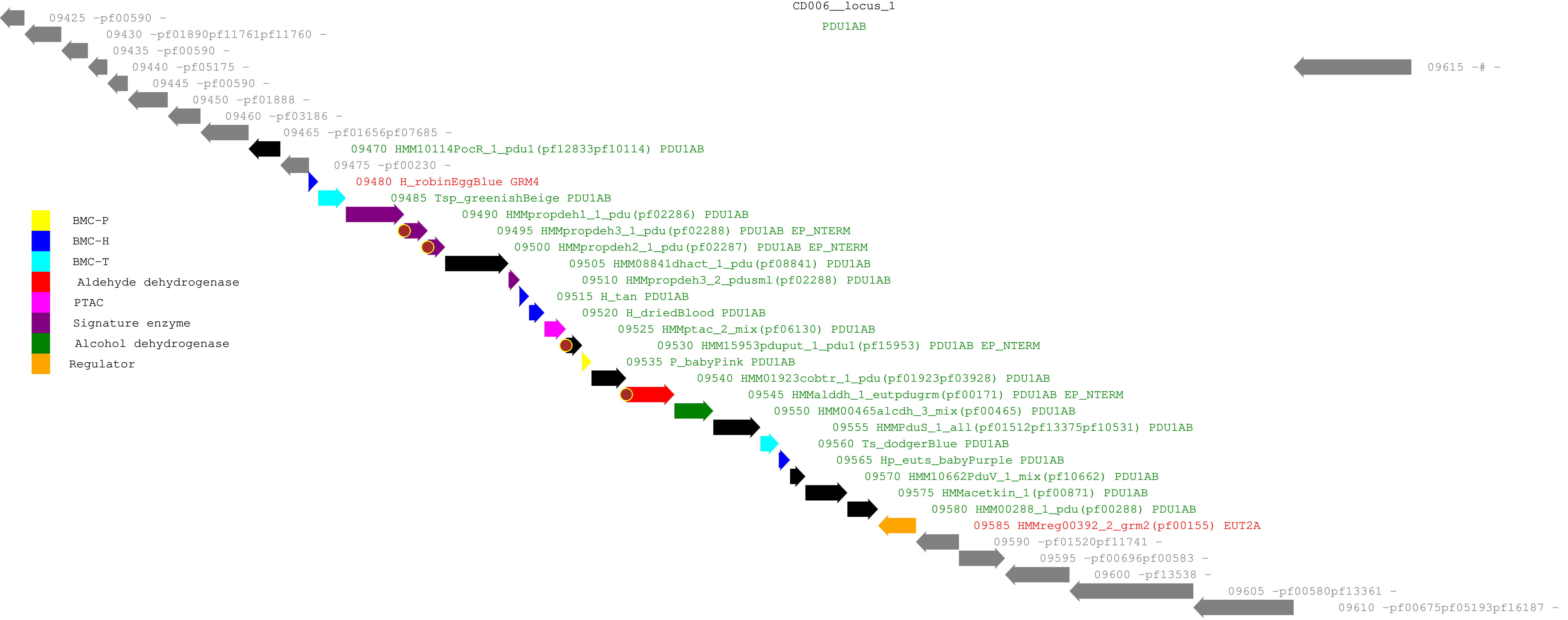

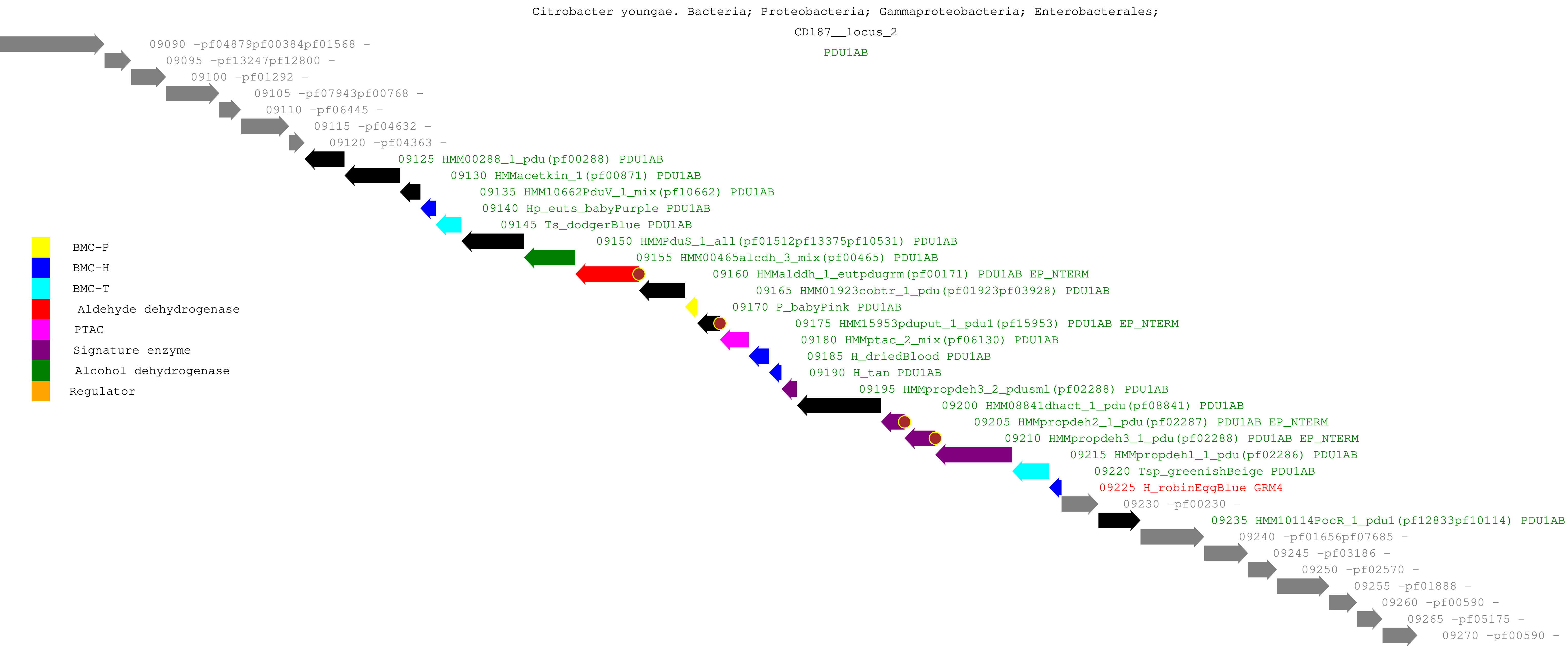

Citrobacter youngae. Bacteria; Proteobacteria; Gammaproteobacteria; Enterobacterales;
CD187__locus_2
PDU1AB
09090 -pf04879pf00384pf01568 -
09095 -pf13247pf12800 -
09100 -pf01292 -
09105 -pf07943pf00768 -
09110 -pf06445 -
09115 -pf04632 -
09120 -pf04363 -
09125 HMM00288_1_pdu(pf00288) PDU1AB
09130 HMMacetkin_1(pf00871) PDU1AB
09135 HMM10662PduV_1_mix(pf10662) PDU1AB
09140 Hp_euts_babyPurple PDU1AB
09145 Ts_dodgerBlue PDU1AB
09150 HMMPduS_1_all(pf01512pf13375pf10531) PDU1AB
09155 HMM00465alcdh_3_mix(pf00465) PDU1AB
09160 HMMalddh_1_eutpdugrm(pf00171) PDU1AB EP_NTERM
09165 HMM01923cobtr_1_pdu(pf01923pf03928) PDU1AB
09170 P_babyPink PDU1AB
09175 HMM15953pduput_1_pdu1(pf15953) PDU1AB EP_NTERM
09180 HMMptac_2_mix(pf06130) PDU1AB
09185 H_driedBlood PDU1AB
09190 H_tan PDU1AB
09195 HMMpropdeh3_2_pdusml(pf02288) PDU1AB
09200 HMM08841dhact_1_pdu(pf08841) PDU1AB
09205 HMMpropdeh2_1_pdu(pf02287) PDU1AB EP_NTERM
09210 HMMpropdeh3_1_pdu(pf02288) PDU1AB EP_NTERM
09215 HMMpropdeh1_1_pdu(pf02286) PDU1AB
09220 Tsp_greenishBeige PDU1AB
09225 H_robinEggBlue GRM4
09230 -pf00230 -
09235 HMM10114PocR_1_pdu1(pf12833pf10114) PDU1AB
09240 -pf01656pf07685 -
09245 -pf03186 -
09250 -pf02570 -
09255 -pf01888 -
09260 -pf00590 -
09265 -pf05175 -
09270 -pf00590 -
BMC-P
BMC-H
BMC-T
Aldehyde dehydrogenase
PTAC
Signature enzyme
Alcohol dehydrogenase
Regulator

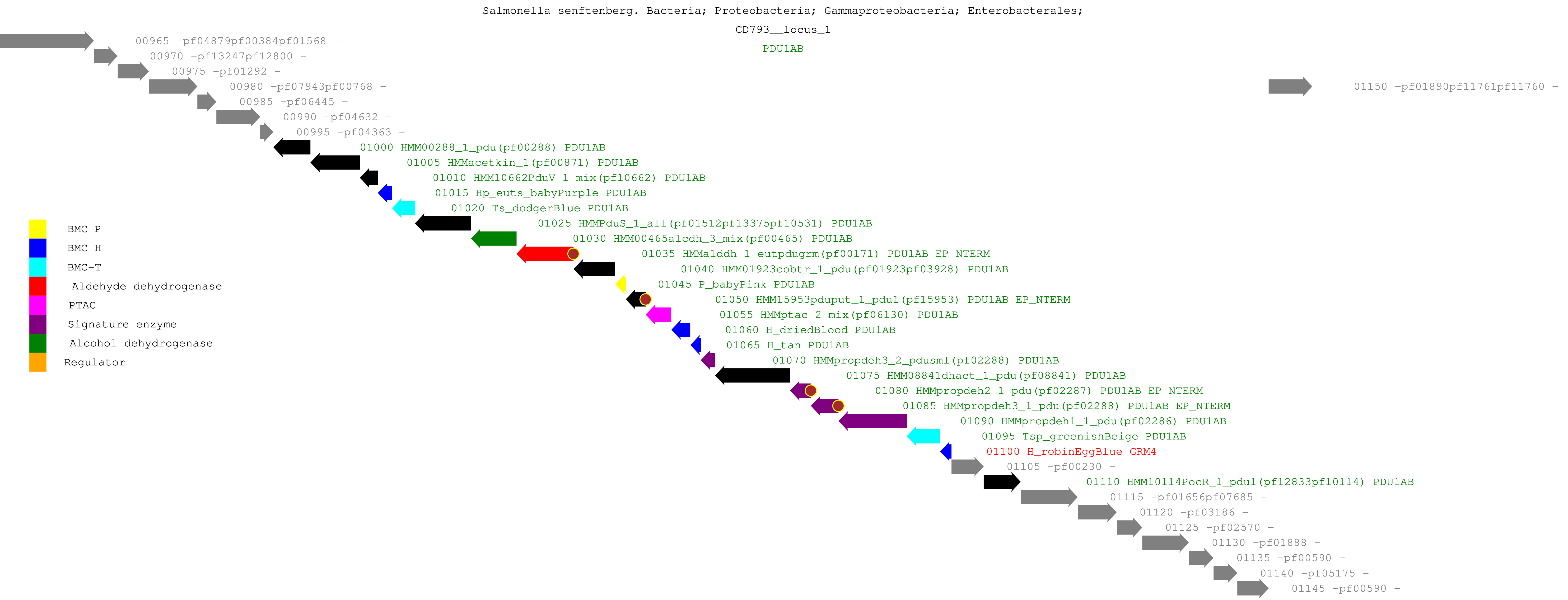
Salmonella senftenberg. Bacteria; Proteobacteria; Gammaproteobacteria; Enterobacterales;
CD793__locus_1
PDU1AB
00965 -pf04879pf00384pf01568 -
00970 -pf13247pf12800 -
00975 -pf01292 -
00980 -pf07943pf00768 -
00985 -pf06445 -
00990 -pf04632 -
00995 -pf04363 -
01000 HMM00288_1_pdu(pf00288) PDU1AB
01005 HMMacetkin_1(pf00871) PDU1AB
01010 HMM10662PduV_1_mix(pf10662) PDU1AB
01015 Hp_euts_babyPurple PDU1AB
01020 Ts_dodgerBlue PDU1AB
01025 HMMPduS_1_all(pf01512pf13375pf10531) PDU1AB
01030 HMM00465alcdh_3_mix(pf00465) PDU1AB
01035 HMMalddh_1_eutpdugrm(pf00171) PDU1AB EP_NTERM
01040 HMM01923cobtr_1_pdu(pf01923pf03928) PDU1AB
01045 P_babyPink PDU1AB
01050 HMM15953pduput_1_pdu1(pf15953) PDU1AB EP_NTERM
01055 HMMptac_2_mix(pf06130) PDU1AB
01060 H_driedBlood PDU1AB
01065 H_tan PDU1AB
01070 HMMpropdeh3_2_pdusml(pf02288) PDU1AB
01075 HMM08841dhact_1_pdu(pf08841) PDU1AB
01080 HMMpropdeh2_1_pdu(pf02287) PDU1AB EP_NTERM
01085 HMMpropdeh3_1_pdu(pf02288) PDU1AB EP_NTERM
01090 HMMpropdeh1_1_pdu(pf02286) PDU1AB
01095 Tsp_greenishBeige PDU1AB
01100 H_robinEggBlue GRM4
01105 -pf00230 -
01110 HMM10114PocR_1_pdu1(pf12833pf10114) PDU1AB
01115 -pf01656pf07685 -
01120 -pf03186 -
01125 -pf02570 -
01130 -pf01888 -
01135 -pf00590 -
01140 -pf05175 -
01145 -pf00590 -
01150 -pf01890pf11761pf11760 -
BMC-P
BMC-H
BMC-T
Aldehyde dehydrogenase
PTAC
Signature enzyme
Alcohol dehydrogenase
Regulator

Salmonella enterica subsp. enterica serovar Schwarzengrund. Bacteria; Proteobacteria; Gammaproteobacteria; Enterobacterales;

CDO01__locus_2

PDU1AB

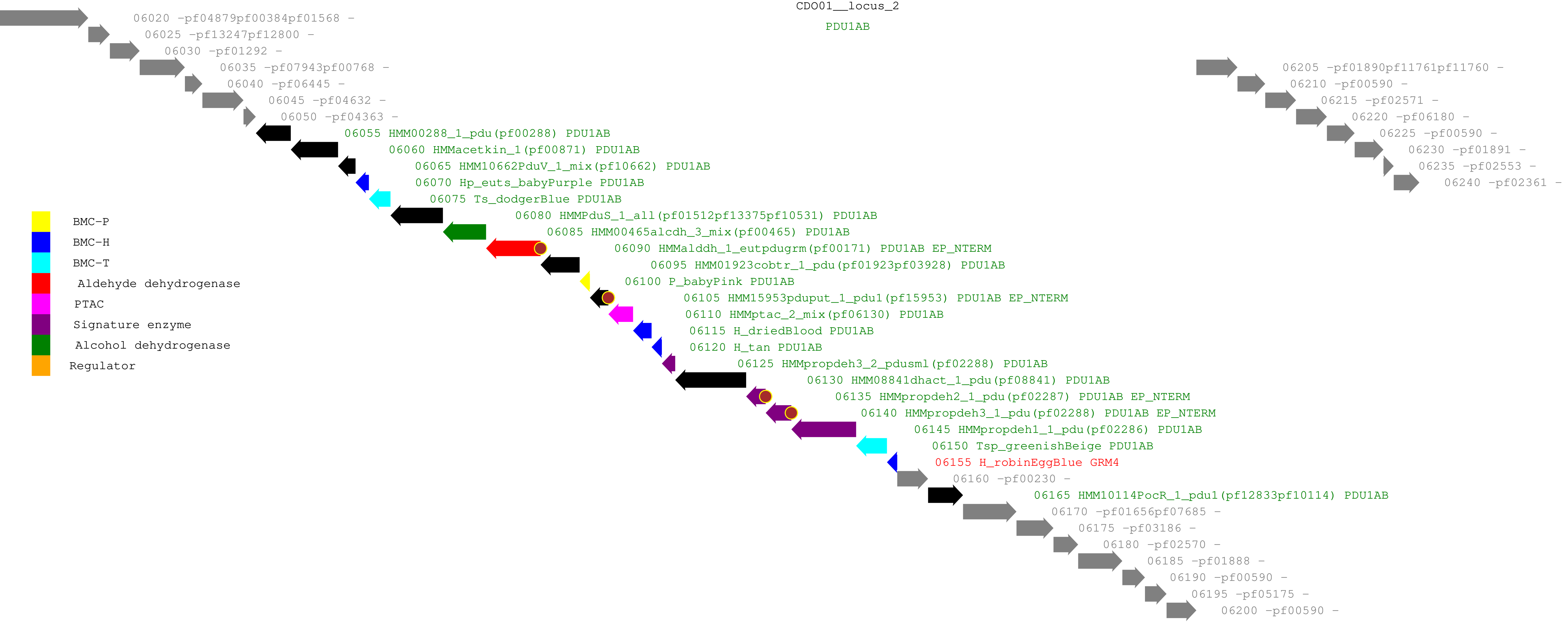

Salmonella enterica subsp. enterica serovar Enteritidis str. CFSAN000052. Bacteria; Proteobacteria; Gammaproteobacteria; Enterobacterales;

CDR59__locus_1

PDU1AB

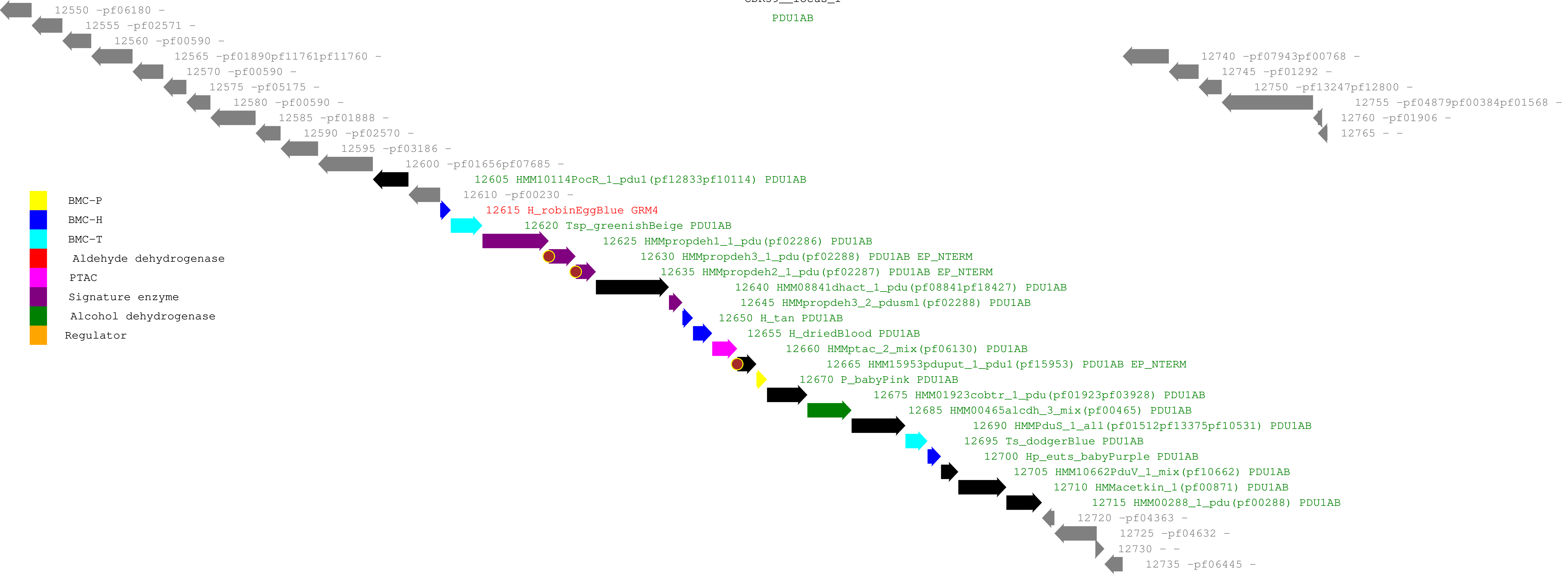

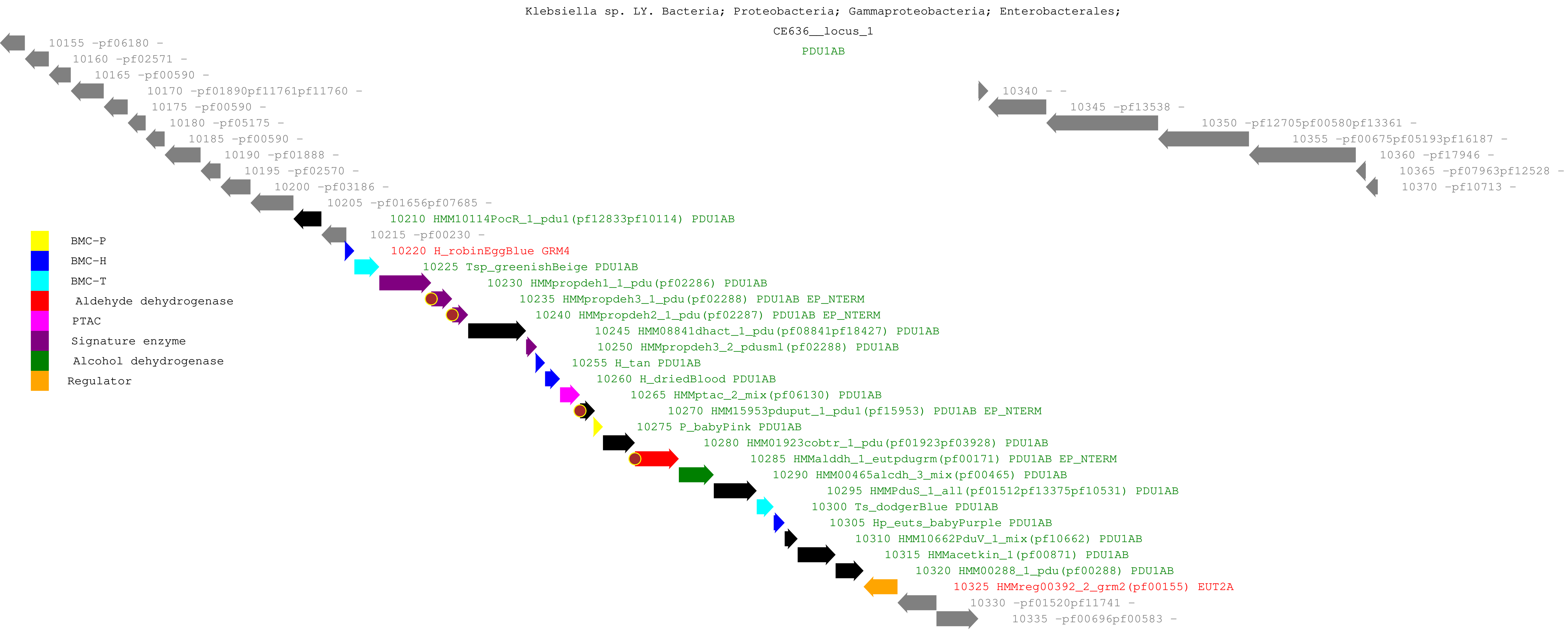

Klebsiella sp. LY. Bacteria; Proteobacteria; Gammaproteobacteria; Enterobacterales;
CE636__locus_1
PDU1AB
10155 -pf06180 -
10160 -pf02571 -
10165 -pf00590 -
10170 -pf01890pf11761pf11760 -
10175 -pf00590 -
10180 -pf05175 -
10185 -pf00590 -
10190 -pf01888 -
10195 -pf02570 -
10200 -pf03186 -
10205 -pf01656pf07685 -
10210 HMM10114PocR_1_pdu1(pf12833pf10114) PDU1AB
10215 -pf00230 -
10220 H_robinEggBlue GRM4
10225 Tsp_greenishBeige PDU1AB
10230 HMMpropdeh1_1_pdu(pf02286) PDU1AB
10235 HMMpropdeh3_1_pdu(pf02288) PDU1AB EP_NTERM
10240 HMMpropdeh2_1_pdu(pf02287) PDU1AB EP_NTERM
10245 HMM08841dhact_1_pdu(pf08841pf18427) PDU1AB
10250 HMMpropdeh3_2_pdusml(pf02288) PDU1AB
10255 H_tan PDU1AB
10260 H_driedBlood PDU1AB
10265 HMMptac_2_mix(pf06130) PDU1AB
10270 HMM15953pduput_1_pdu1(pf15953) PDU1AB EP_NTERM
10275 P_babyPink PDU1AB
10280 HMM01923cobtr_1_pdu(pf01923pf03928) PDU1AB
10285 HMMalddh_1_eutpdugrm(pf00171) PDU1AB EP_NTERM
10290 HMM00465alcdh_3_mix(pf00465) PDU1AB
10295 HMMPduS_1_all(pf01512pf13375pf10531) PDU1AB
10300 Ts_dodgerBlue PDU1AB
10305 Hp_euts_babyPurple PDU1AB
10310 HMM10662PduV_1_mix(pf10662) PDU1AB
10315 HMMacetkin_1(pf00871) PDU1AB
10320 HMM00288_1_pdu(pf00288) PDU1AB
10325 HMMreg00392_2_grm2(pf00155) EUT2A
10330 -pf01520pf11741 -
10335 -pf00696pf00583 -
10340 - -
10345 -pf13538 -
10350 -pf12705pf00580pf13361 -
10355 -pf00675pf05193pf16187 -
10360 -pf17946 -
10365 -pf07963pf12528 -
10370 -pf10713 -
BMC-P
BMC-H
BMC-T
Aldehyde dehydrogenase
PTAC
Signature enzyme
Alcohol dehydrogenase
Regulator

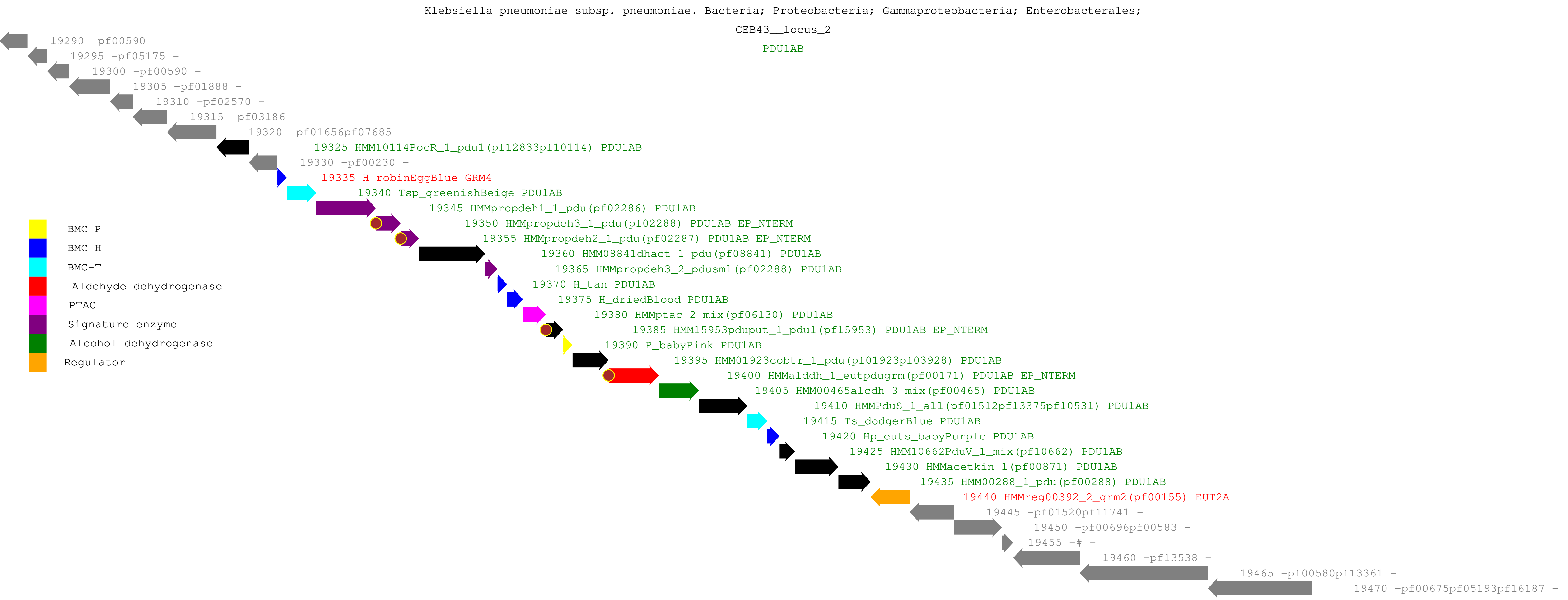
Klebsiella pneumoniae subsp. pneumoniae. Bacteria; Proteobacteria; Gammaproteobacteria; Enterobacterales;
CEB43__locus_2
PDU1AB
19290 -pf00590 -
19295 -pf05175 -
19300 -pf00590 -
19305 -pf01888 -
19310 -pf02570 -
19315 -pf03186 -
19320 -pf01656pf07685 -
19325 HMM10114PocR_1_pdu1(pf12833pf10114) PDU1AB
19330 -pf00230 -
19335 H_robinEggBlue GRM4
19340 Tsp_greenishBeige PDU1AB
19345 HMMpropdeh1_1_pdu(pf02286) PDU1AB
19350 HMMpropdeh3_1_pdu(pf02288) PDU1AB EP_NTERM
19355 HMMpropdeh2_1_pdu(pf02287) PDU1AB EP_NTERM
19360 HMM08841dhact_1_pdu(pf08841) PDU1AB
19365 HMMpropdeh3_2_pdusml(pf02288) PDU1AB
19370 H_tan PDU1AB
19375 H_driedBlood PDU1AB
19380 HMMptac_2_mix(pf06130) PDU1AB
19385 HMM15953pduput_1_pdu1(pf15953) PDU1AB EP_NTERM
19390 P_babyPink PDU1AB
19395 HMM01923cobtr_1_pdu(pf01923pf03928) PDU1AB
19400 HMMalddh_1_eutpdugrm(pf00171) PDU1AB EP_NTERM
19405 HMM00465alcdh_3_mix(pf00465) PDU1AB
19410 HMMPduS_1_all(pf01512pf13375pf10531) PDU1AB
19415 Ts_dodgerBlue PDU1AB
19420 Hp_euts_babyPurple PDU1AB
19425 HMM10662PduV_1_mix(pf10662) PDU1AB
19430 HMMacetkin_1(pf00871) PDU1AB
19435 HMM00288_1_pdu(pf00288) PDU1AB
19440 HMMreg00392_2_grm2(pf00155) EUT2A
19445 -pf01520pf11741 -
19450 -pf00696pf00583 -
19455 -# -
19460 -pf13538 -
19465 -pf00580pf13361 -
19470 -pf00675pf05193pf16187 -
BMC-P
BMC-H
BMC-T
Aldehyde dehydrogenase
PTAC
Signature enzyme
Alcohol dehydrogenase
Regulator

Klebsiella pneumoniae subsp. pneumoniae. Bacteria; Proteobacteria; Gammaproteobacteria; Enterobacterales;

CEB45__locus_1

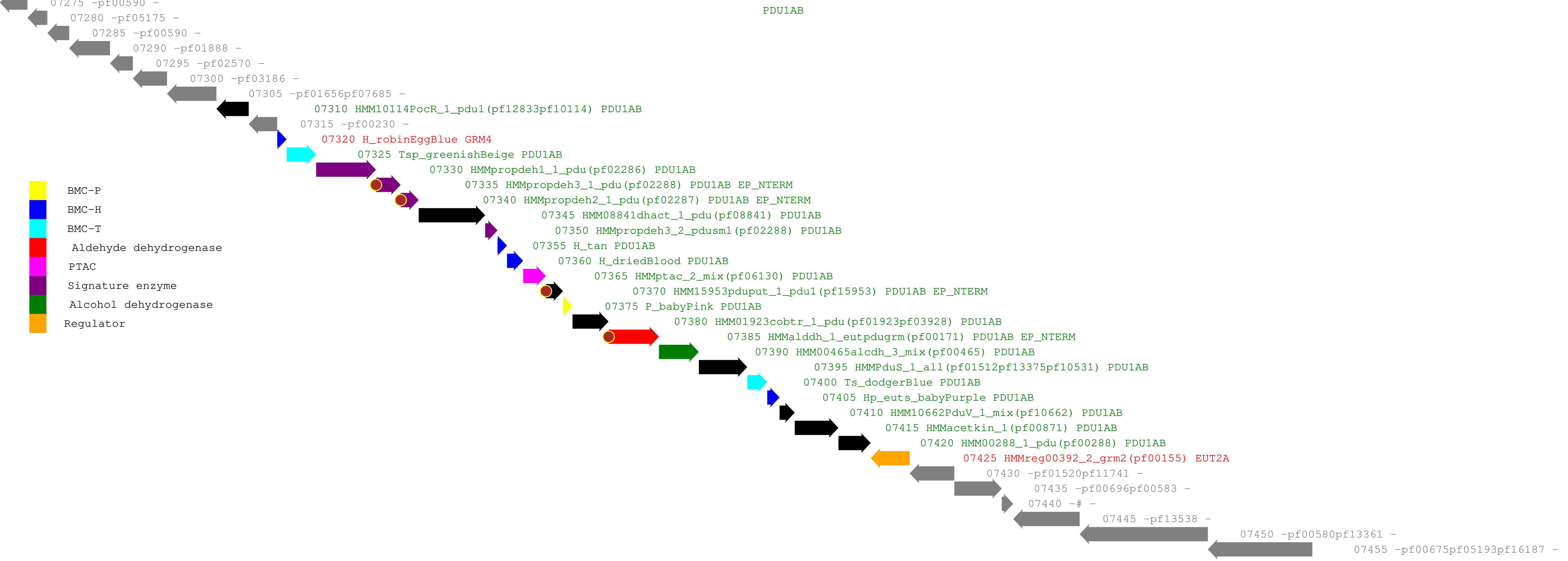

Klebsiella pneumoniae subsp. pneumoniae. Bacteria; Proteobacteria; Gammaproteobacteria; Enterobacterales;

CEB46__locus_1

PDU1AB

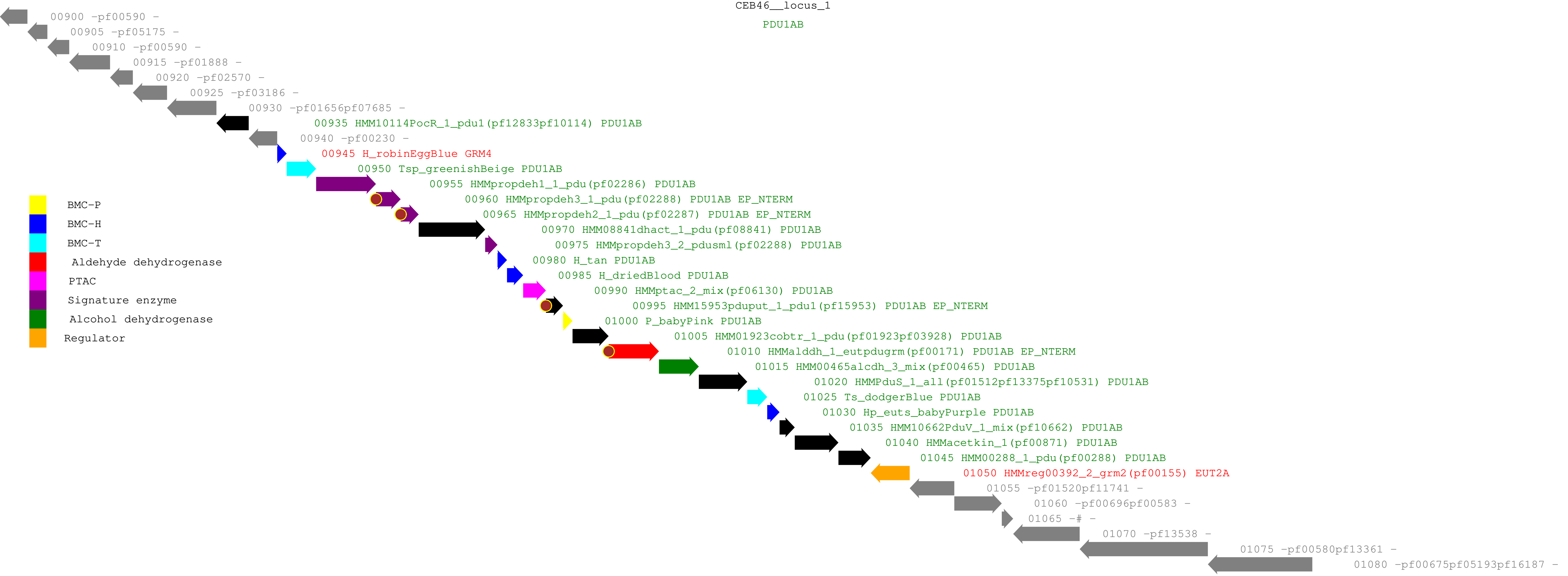

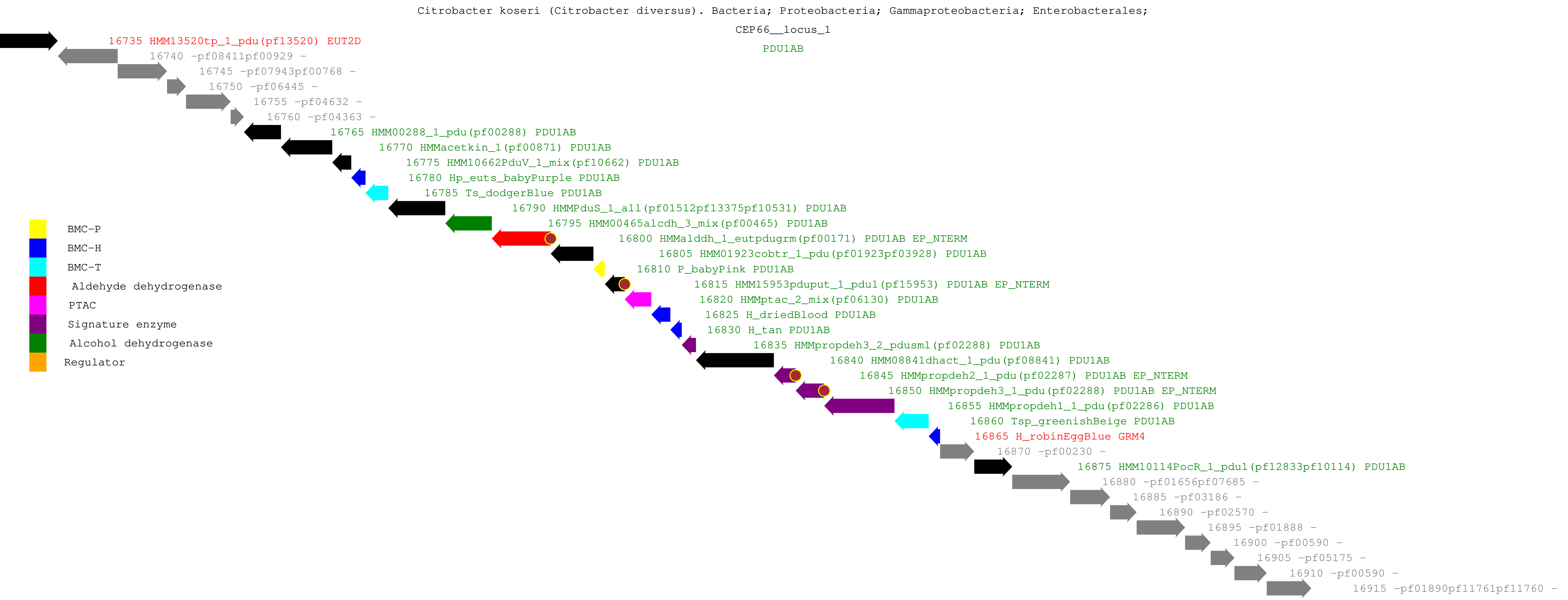
Citrobacter koseri (Citrobacter diversus). Bacteria; Proteobacteria; Gammaproteobacteria; Enterobacterales;
CEP66__locus_1
PDU1AB
16735 HMM13520tp_1_pdu(pf13520) EUT2D
16740 -pf08411pf00929 -
16745 -pf07943pf00768 -
16750 -pf06445 -
16755 -pf04632 -
16760 -pf04363 -
16765 HMM00288_1_pdu(pf00288) PDU1AB
16770 HMMacetkin_1(pf00871) PDU1AB
16775 HMM10662PduV_1_mix(pf10662) PDU1AB
16780 Hp_euts_babyPurple PDU1AB
16785 Ts_dodgerBlue PDU1AB
16790 HMMPduS_1_all(pf01512pf13375pf10531) PDU1AB
16795 HMM00465alcdh_3_mix(pf00465) PDU1AB
16800 HMMaldh_1_eutpdugrm(pf00171) PDU1AB EP_NTERM
16805 HMM01923cobtr_1_pdu(pf01923pf03928) PDU1AB
16810 P_babyPink PDU1AB
16815 HMM15953pduput_1_pdu1(pf15953) PDU1AB EP_NTERM
16820 HMMptac_2_mix(pf06130) PDU1AB
16825 H_driedBlood PDU1AB
16830 H_tan PDU1AB
16835 HMMpropdeh3_2_pdusml(pf02288) PDU1AB
16840 HMM08841dhact_1_pdu(pf08841) PDU1AB
16845 HMMpropdeh2_1_pdu(pf02287) PDU1AB EP_NTERM
16850 HMMpropdeh3_1_pdu(pf02288) PDU1AB EP_NTERM
16855 HMMpropdeh1_1_pdu(pf02286) PDU1AB
16860 Tsp_greenishBeige PDU1AB
16865 H_robinEggBlue GRM4
16870 -pf00230 -
16875 HMM10114PocR_1_pdu1(pf12833pf10114) PDU1AB
16880 -pf01656pf07685 -
16885 -pf03186 -
16890 -pf02570 -
16895 -pf01888 -
16900 -pf00590 -
16905 -pf05175 -
16910 -pf00590 -
16915 -pf01890pf11761pf11760 -
BMC-P
BMC-H
BMC-T
Aldehyde dehydrogenase
PTAC
Signature enzyme
Alcohol dehydrogenase
Regulator

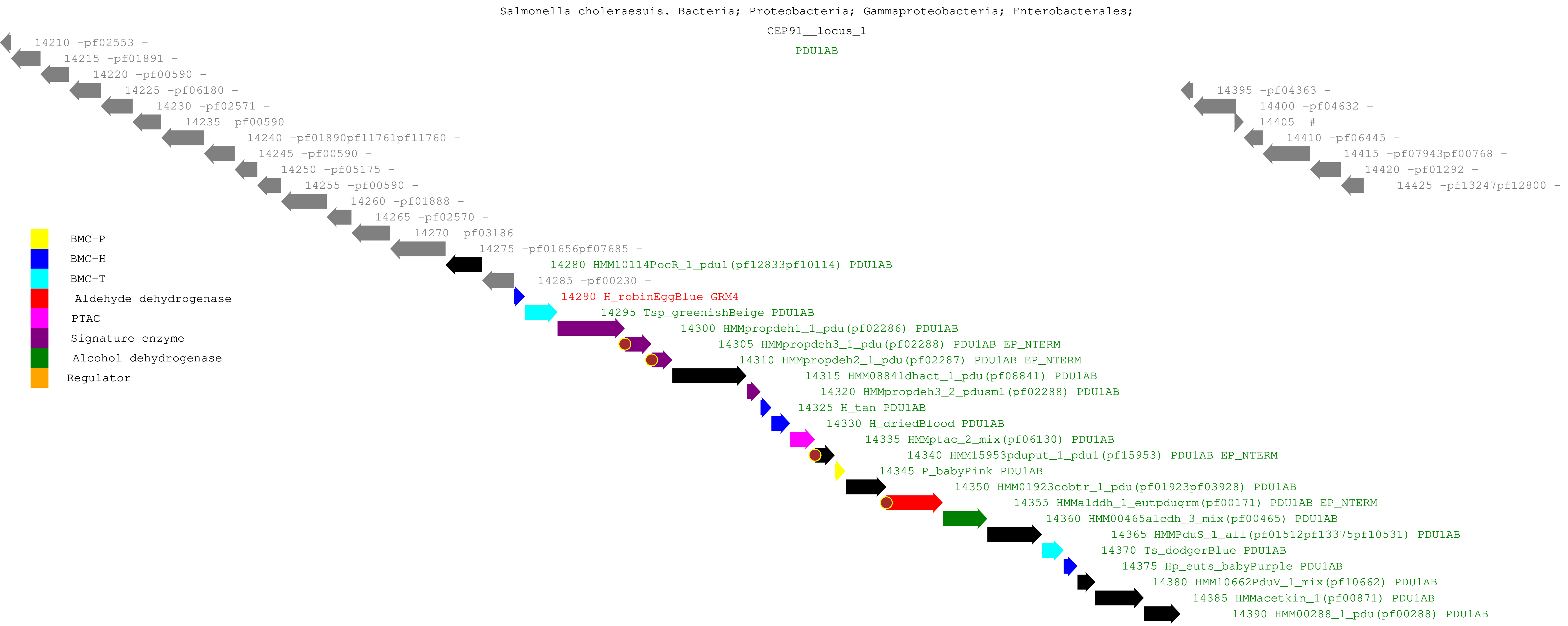

Salmonella choleraesuis. Bacteria; Proteobacteria; Gammaproteobacteria; Enterobacterales;
CEP91__locus_1
PDU1AB
14210 -pf02553 -
14215 -pf01891 -
14220 -pf00590 -
14225 -pf06180 -
14230 -pf02571 -
14235 -pf00590 -
14240 -pf01890pf11761pf11760 -
14245 -pf00590 -
14250 -pf05175 -
14255 -pf00590 -
14260 -pf01888 -
14265 -pf02570 -
14270 -pf03186 -
14275 -pf01656pf07685 -
14280 HMM10114PocR_1_pdu1(pf12833pf10114) PDU1AB
14285 -pf00230 -
14290 H_robinEggBlue GRM4
14295 Tsp_greenishBeige PDU1AB
14300 HMMpropdeh1_1_pdu(pf02286) PDU1AB
14305 HMMpropdeh3_1_pdu(pf02288) PDU1AB EP_NTERM
14310 HMMpropdeh2_1_pdu(pf02287) PDU1AB EP_NTERM
14315 HMM08841dhact_1_pdu(pf08841) PDU1AB
14320 HMMpropdeh3_2_pdusml(pf02288) PDU1AB
14325 H_tan PDU1AB
14330 H_driedBlood PDU1AB
14335 HMMptac_2_mix(pf06130) PDU1AB
14340 HMM15953pduput_1_pdu1(pf15953) PDU1AB EP_NTERM
14345 P_babyPink PDU1AB
14350 HMM01923cobtr_1_pdu(pf01923pf03928) PDU1AB
14355 HMMalddh_1_eutpdugrm(pf00171) PDU1AB EP_NTERM
14360 HMM00465alcdh_3_mix(pf00465) PDU1AB
14365 HMMPduS_1_all(pf01512pf13375pf10531) PDU1AB
14370 Ts_dodgerBlue PDU1AB
14375 Hp_euts_babyPurple PDU1AB
14380 HMM10662PduV_1_mix(pf10662) PDU1AB
14385 HMMacetkin_1(pf00871) PDU1AB
14390 HMM00288_1_pdu(pf00288) PDU1AB
14395 -pf04363 -
14400 -pf04632 -
14405 -# -
14410 -pf06445 -
14415 -pf07943pf00768 -
14420 -pf01292 -
14425 -pf13247pf12800 -
BMC-P
BMC-H
BMC-T
Aldehyde dehydrogenase
PTAC
Signature enzyme
Alcohol dehydrogenase
Regulator

Salmonella typhimurium. Bacteria; Proteobacteria; Gammaproteobacteria; Enterobacterales;

CET98__locus_1

PDU1AB

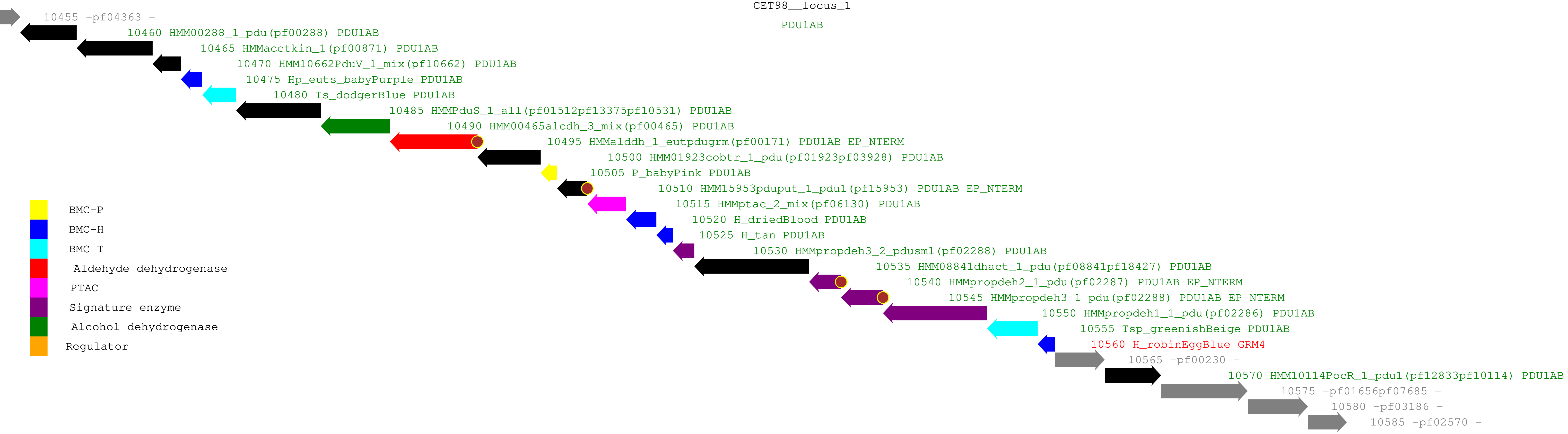

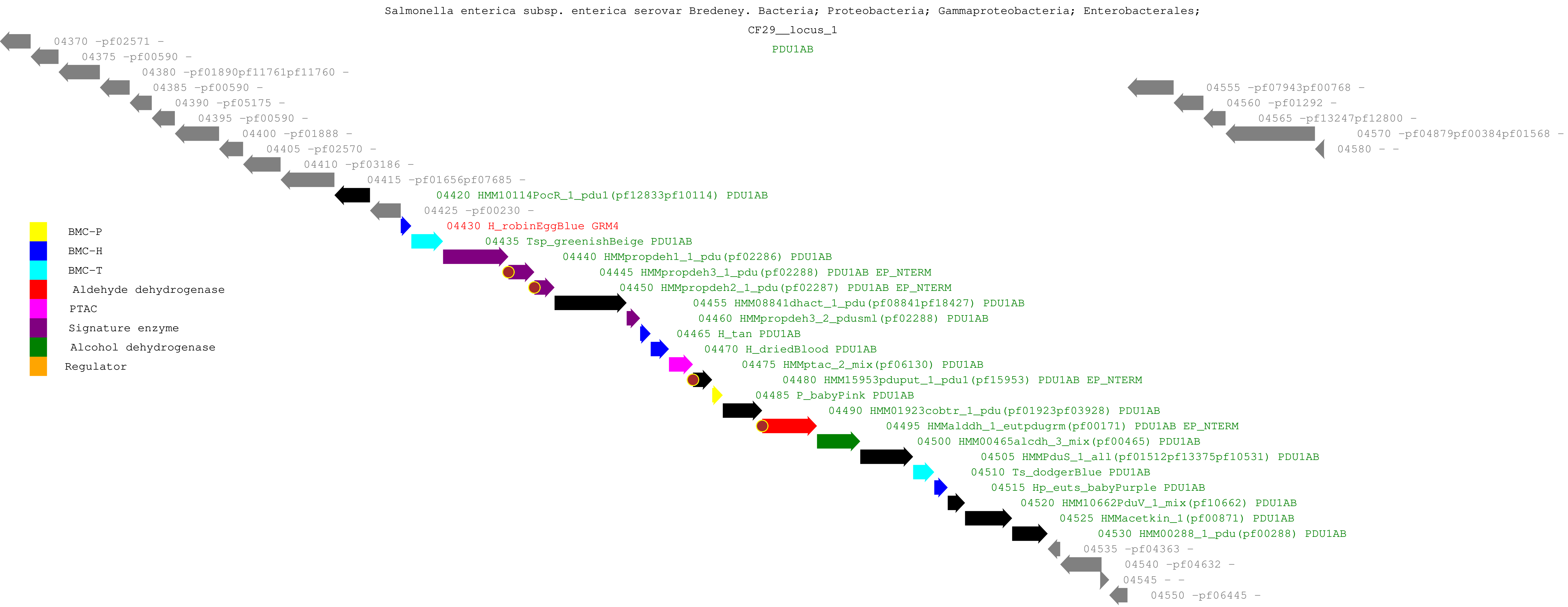
Salmonella enterica subsp. enterica serovar Bredeney. Bacteria; Proteobacteria; Gammaproteobacteria; Enterobacterales;
CF29__locus_1
PDU1AB
04370 -pf02571 -
04375 -pf00590 -
04380 -pf01890pf11761pf11760 -
04385 -pf00590 -
04390 -pf05175 -
04395 -pf00590 -
04400 -pf01888 -
04405 -pf02570 -
04410 -pf03186 -
04415 -pf01656pf07685 -
04420 HMM10114PocR_1_pdu1(pf12833pf10114) PDU1AB
04425 -pf00230 -
04430 H_robinEggBlue GRM4
04435 Tsp_greenishBeige PDU1AB
04440 HMMpropdeh1_1_pdu(pf02286) PDU1AB
04445 HMMpropdeh3_1_pdu(pf02288) PDU1AB EP_NTERM
04450 HMMpropdeh2_1_pdu(pf02287) PDU1AB EP_NTERM
04455 HMM08841dhact_1_pdu(pf08841pf18427) PDU1AB
04460 HMMpropdeh3_2_pdusml(pf02288) PDU1AB
04465 H_tan PDU1AB
04470 H_driedBlood PDU1AB
04475 HMMptac_2_mix(pf06130) PDU1AB
04480 HMM15953pduput_1_pdu1(pf15953) PDU1AB EP_NTERM
04485 P_babyPink PDU1AB
04490 HMM01923cobtr_1_pdu(pf01923pf03928) PDU1AB
04495 HMMalddh_1_eutpdugrm(pf00171) PDU1AB EP_NTERM
04500 HMM00465alcdh_3_mix(pf00465) PDU1AB
04505 HMMPduS_1_all(pf01512pf13375pf10531) PDU1AB
04510 Ts_dodgerBlue PDU1AB
04515 Hp_euts_babyPurple PDU1AB
04520 HMM10662PduV_1_mix(pf10662) PDU1AB
04525 HMMacetkin_1(pf00871) PDU1AB
04530 HMM00288_1_pdu(pf00288) PDU1AB
04535 -pf04363 -
04540 -pf04632 -
04545 - -
04550 -pf06445 -
04555 -pf07943pf00768 -
04560 -pf01292 -
04565 -pf13247pf12800 -
04570 -pf04879pf00384pf01568 -
04580 - -
BMC-P
BMC-H
BMC-T
Aldehyde dehydrogenase
PTAC
Signature enzyme
Alcohol dehydrogenase
Regulator

Citrobacter freundii. Bacteria; Proteobacteria; Gammaproteobacteria; Enterobacterales;

CFA70__locus_1

PDU1AB

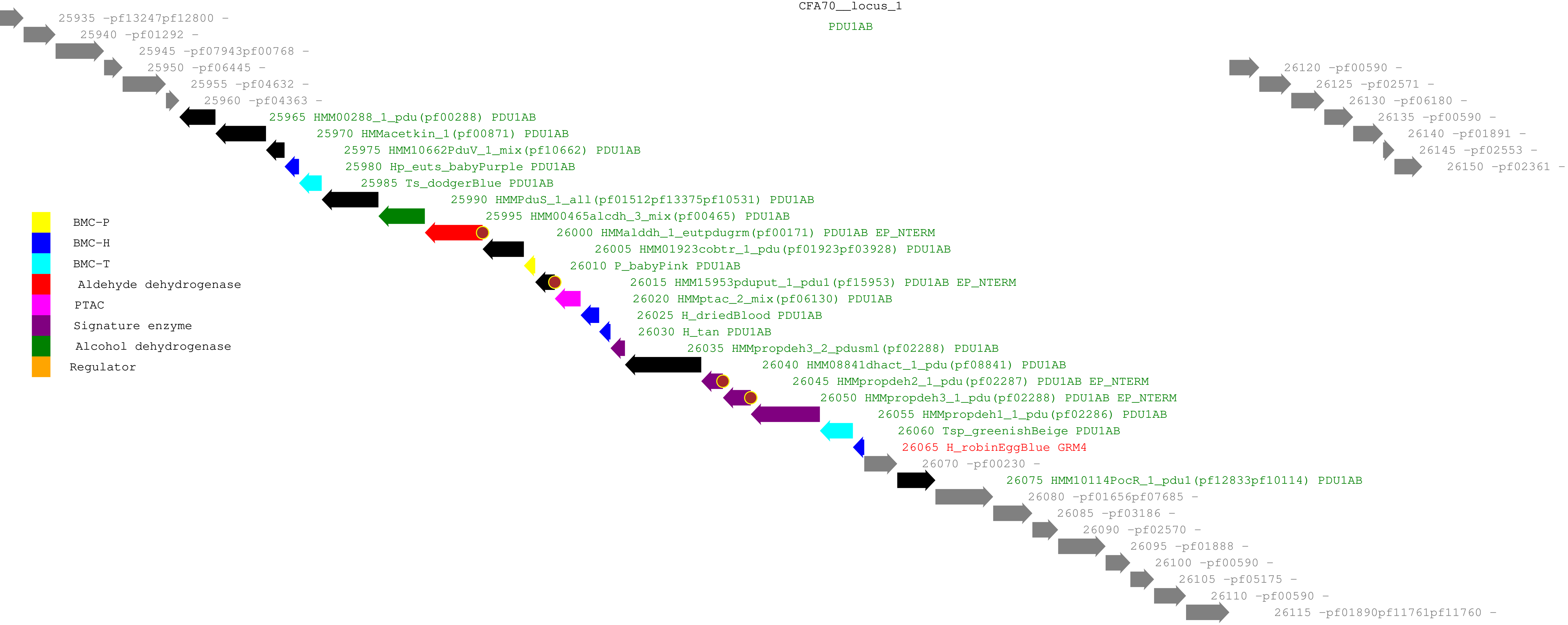

Citrobacter freundii. Bacteria; Proteobacteria; Gammaproteobacteria; Enterobacterales;

CfB38__locus_1

PDU1AB

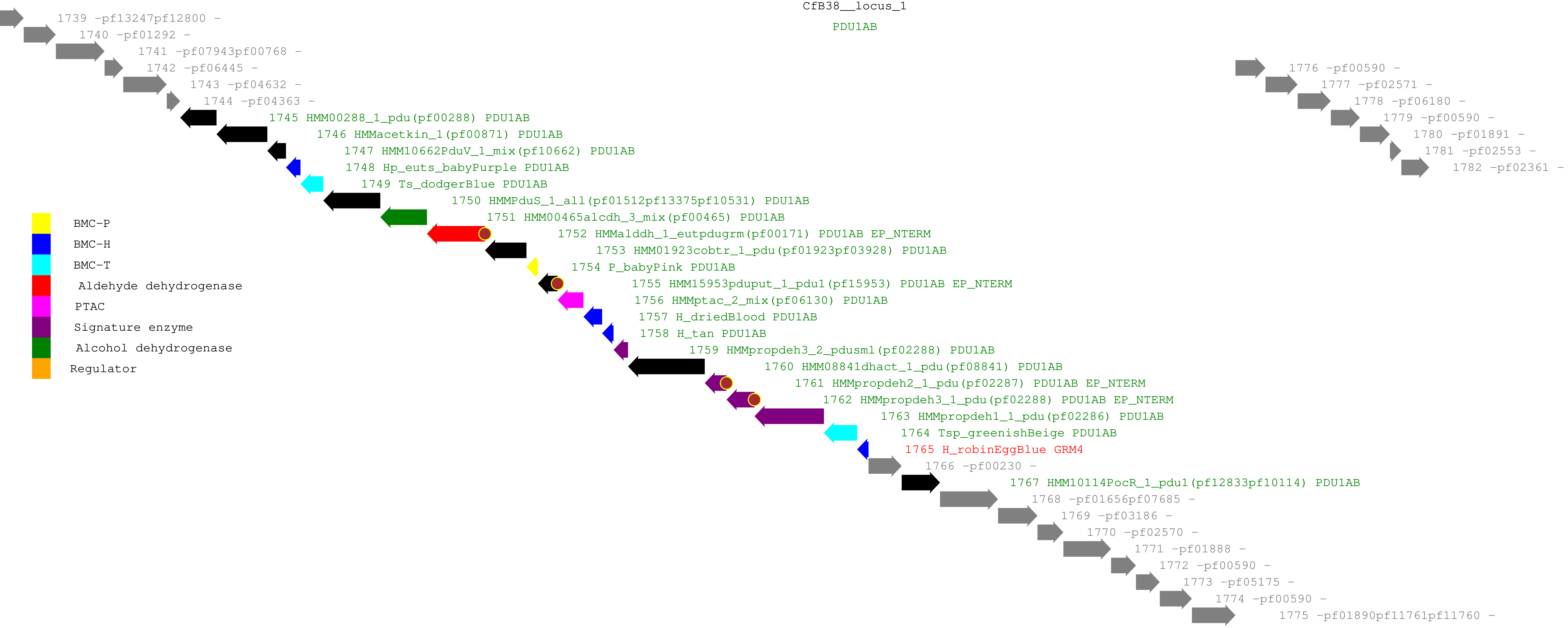

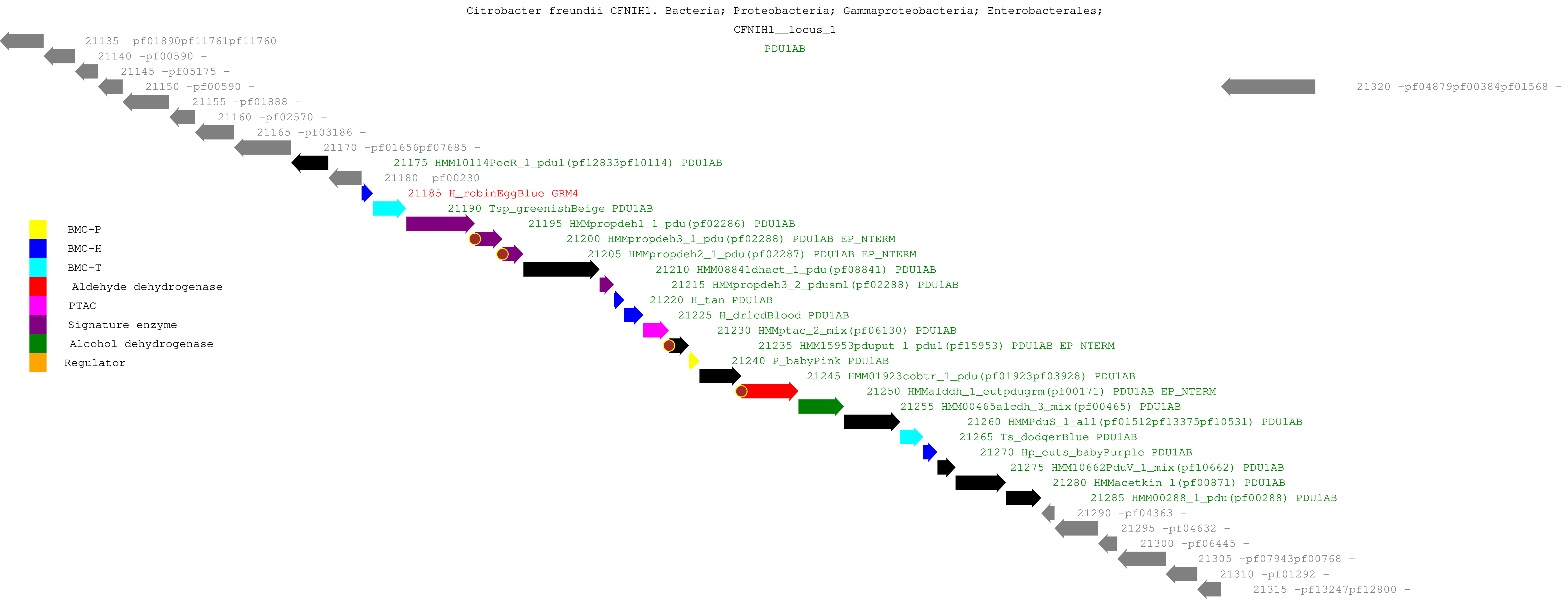
Citrobacter freundii CFNIH1. Bacteria; Proteobacteria; Gammaproteobacteria; Enterobacterales;
CFNIH1__locus_1
PDU1AB
21135 -pf01890pf11761pf11760 -
21140 -pf00590 -
21145 -pf05175 -
21150 -pf00590 -
21155 -pf01888 -
21160 -pf02570 -
21165 -pf03186 -
21170 -pf01656pf07685 -
21175 HMM10114PocR_1_pdu1(pf12833pf10114) PDU1AB
21180 -pf00230 -
21185 H_robinEggBlue GRM4
21190 Tsp_greenishBeige PDU1AB
21195 HMMpropdeh1_1_pdu(pf02286) PDU1AB
21200 HMMpropdeh3_1_pdu(pf02288) PDU1AB EP_NTERM
21205 HMMpropdeh2_1_pdu(pf02287) PDU1AB EP_NTERM
21210 HMM08841dhact_1_pdu(pf08841) PDU1AB
21215 HMMpropdeh3_2_pdusml(pf02288) PDU1AB
21220 H_tan PDU1AB
21225 H_driedBlood PDU1AB
21230 HMMptac_2_mix(pf06130) PDU1AB
21235 HMM15953pduput_1_pdu1(pf15953) PDU1AB EP_NTERM
21240 P_babyPink PDU1AB
21245 HMM01923cobtr_1_pdu(pf01923pf03928) PDU1AB
21250 HMMalddh_1_eutpdugrm(pf00171) PDU1AB EP_NTERM
21255 HMM00465alcdh_3_mix(pf00465) PDU1AB
21260 HMMPduS_1_all(pf01512pf13375pf10531) PDU1AB
21265 Ts_dodgerBlue PDU1AB
21270 Hp_euts_babyPurple PDU1AB
21275 HMM10662PduV_1_mix(pf10662) PDU1AB
21280 HMMacetkin_1(pf00871) PDU1AB
21285 HMM00288_1_pdu(pf00288) PDU1AB
21290 -pf04363 -
21295 -pf04632 -
21300 -pf06445 -
21305 -pf07943pf00768 -
21310 -pf01292 -
21315 -pf13247pf12800 -
21320 -pf04879pf00384pf01568 -
BMC-P
BMC-H
BMC-T
Aldehyde dehydrogenase
PTAC
Signature enzyme
Alcohol dehydrogenase
Regulator

Salmonella enterica subsp. enterica serovar Choleraesuis str. CFSAN000514. Bacteria; Proteobacteria; Gammaproteobacteria; Enterobacterales;

CFSAN000514__locus_2

PDU1AB

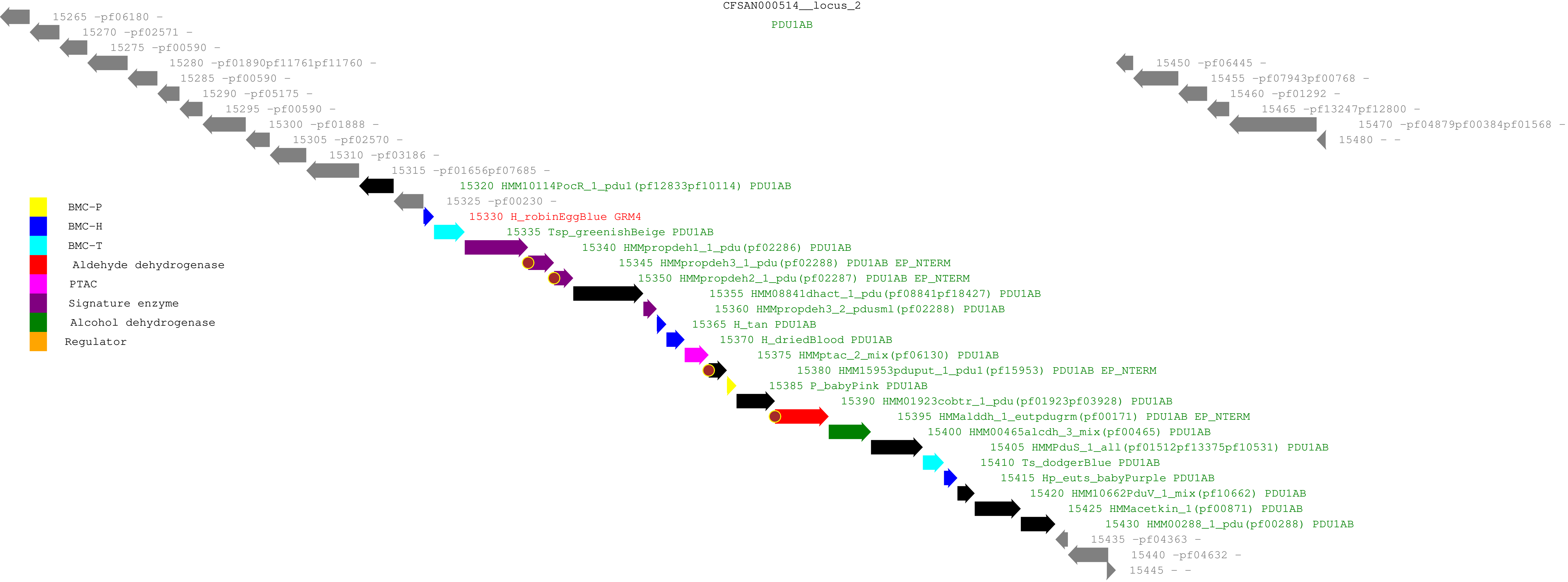

Salmonella enterica subsp. salamae str. CFSAN000559. Bacteria; Proteobacteria; Gammaproteobacteria; Enterobacterales;

CFSAN000559__locus_2

PDU1AB

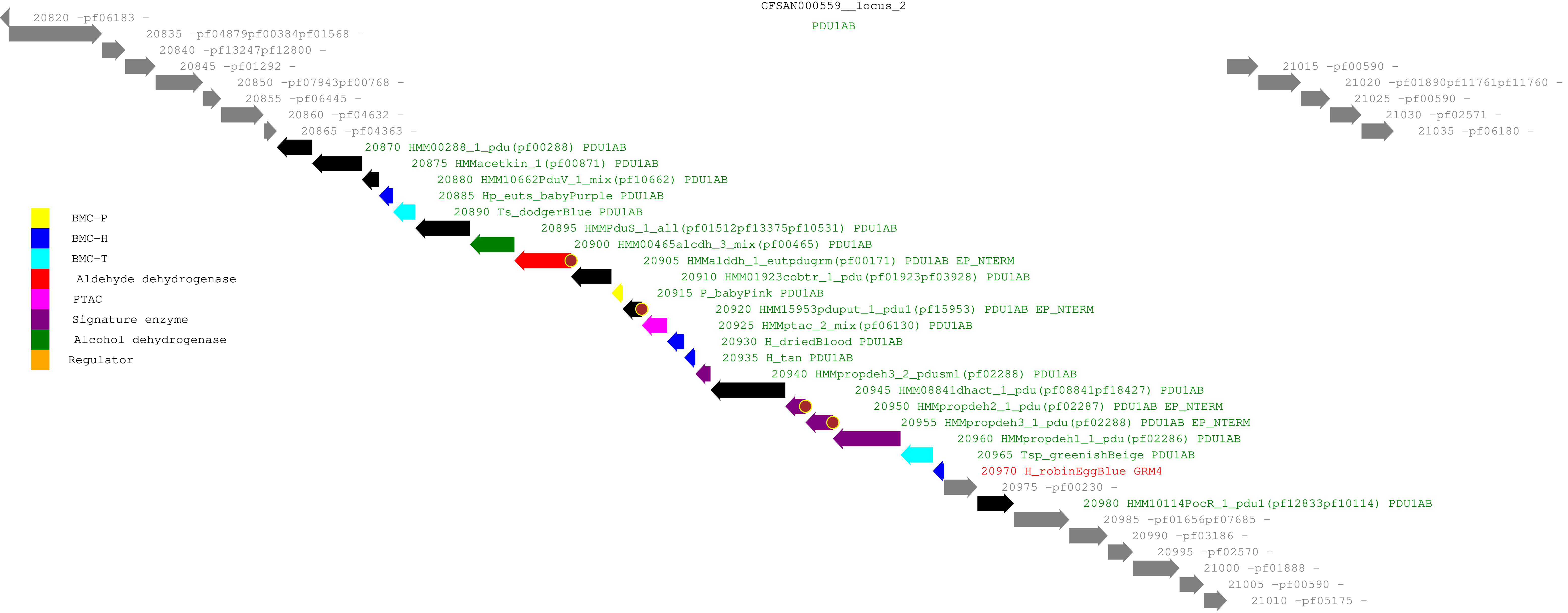

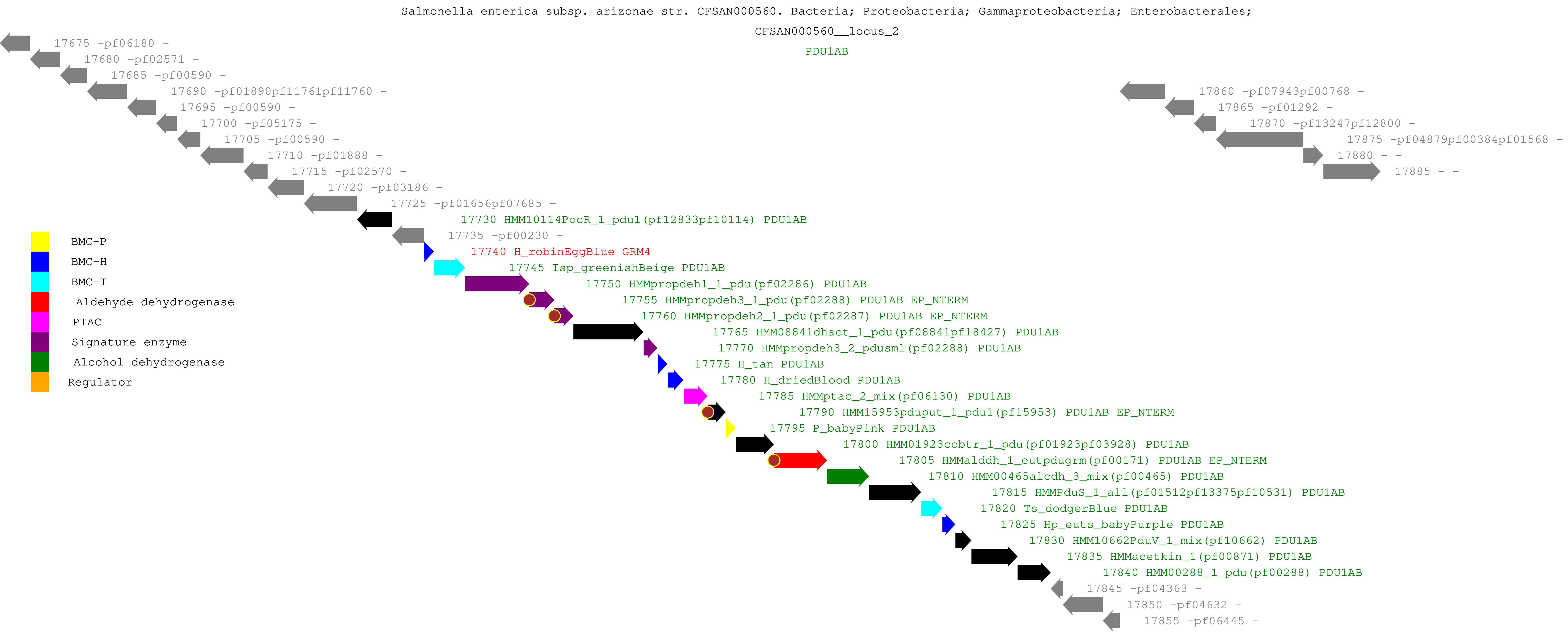
Salmonella enterica subsp. arizonae str. CFSAN000560. Bacteria; Proteobacteria; Gammaproteobacteria; Enterobacterales;
CFSAN000560__locus_2
PDU1AB
17675 -pf06180 -
17680 -pf02571 -
17685 -pf00590 -
17690 -pf01890pf11761pf11760 -
17695 -pf00590 -
17700 -pf05175 -
17705 -pf00590 -
17710 -pf01888 -
17715 -pf02570 -
17720 -pf03186 -
17725 -pf01656pf07685 -
17730 HMM10114PocR_1_pdu1(pf12833pf10114) PDU1AB
17735 -pf00230 -
17740 H_robinEggBlue GRM4
17745 Tsp_greenishBeige PDU1AB
17750 HMMpropdeh1_1_pdu(pf02286) PDU1AB
17755 HMMpropdeh3_1_pdu(pf02288) PDU1AB EP_NTERM
17760 HMMpropdeh2_1_pdu(pf02287) PDU1AB EP_NTERM
17765 HMM08841dhact_1_pdu(pf08841pf18427) PDU1AB
17770 HMMpropdeh3_2_pdusml(pf02288) PDU1AB
17775 H_tan PDU1AB
17780 H_driedBlood PDU1AB
17785 HMMptac_2_mix(pf06130) PDU1AB
17790 HMM15953pduput_1_pdu1(pf15953) PDU1AB EP_NTERM
17795 P_babyPink PDU1AB
17800 HMM01923cobtr_1_pdu(pf01923pf03928) PDU1AB
17805 HMMalddh_1_eutpdugrm(pf00171) PDU1AB EP_NTERM
17810 HMM00465alcdh_3_mix(pf00465) PDU1AB
17815 HMMPduS_1_all(pf01512pf13375pf10531) PDU1AB
17820 Ts_dodgerBlue PDU1AB
17825 Hp_euts_babyPurple PDU1AB
17830 HMM10662PduV_1_mix(pf10662) PDU1AB
17835 HMMacetkin_1(pf00871) PDU1AB
17840 HMM00288_1_pdu(pf00288) PDU1AB
17845 -pf04363 -
17850 -pf04632 -
17855 -pf06445 -
17860 -pf07943pf00768 -
17865 -pf01292 -
17870 -pf13247pf12800 -
17875 -pf04879pf00384pf01568 -
17880 - -
17885 - -
BMC-P
BMC-H
BMC-T
Aldehyde dehydrogenase
PTAC
Signature enzyme
Alcohol dehydrogenase
Regulator

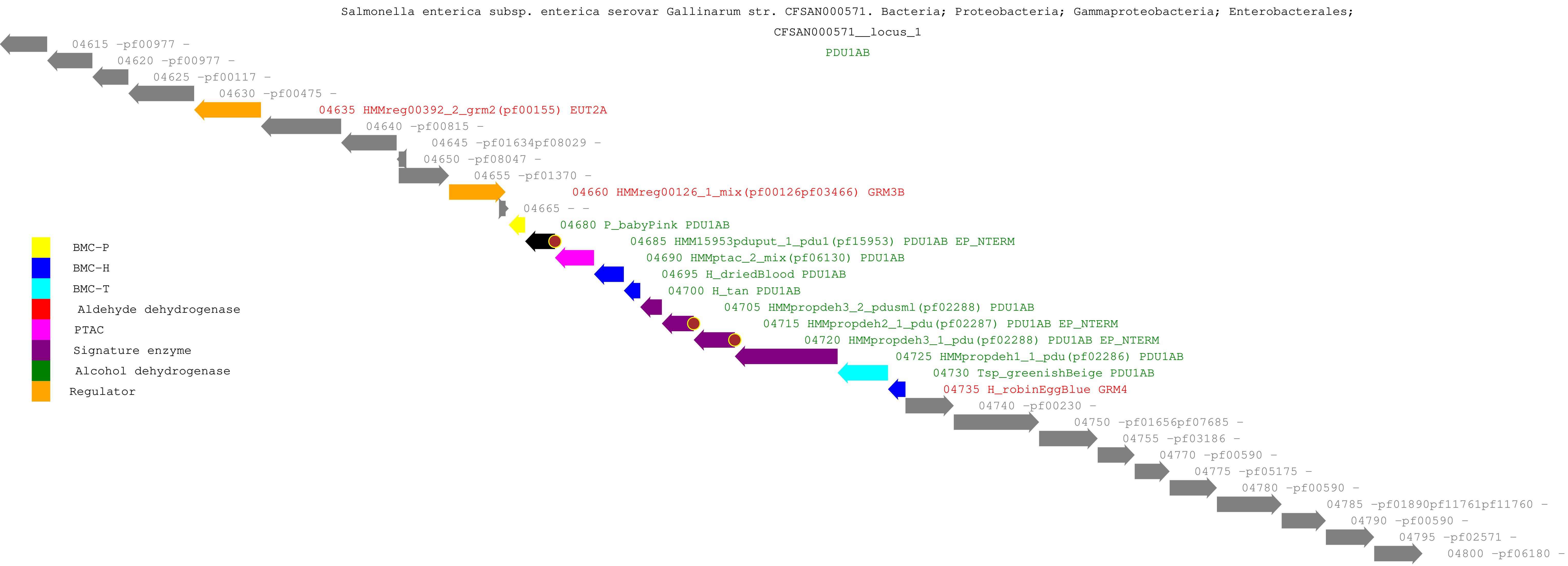

Salmonella enterica subsp. enterica serovar Gallinarum str. CFSAN000571. Bacteria; Proteobacteria; Gammaproteobacteria; Enterobacterales;
CFSAN000571__locus_1
PDU1AB
04615 -pf00977 -
04620 -pf00977 -
04625 -pf00117 -
04630 -pf00475 -
04635 HMMreg00392_2_grm2(pf00155) EUT2A
04640 -pf00815 -
04645 -pf01634pf08029 -
04650 -pf08047 -
04655 -pf01370 -
04660 HMMreg00126_1_mix(pf00126pf03466) GRM3B
04665 - -
04680 P_babyPink PDU1AB
04685 HMM15953pduput_1_pdu1(pf15953) PDU1AB EP_NTERM
04690 HMMptac_2_mix(pf06130) PDU1AB
04695 H_driedBlood PDU1AB
04700 H_tan PDU1AB
04705 HMMpropdeh3_2_pdusml(pf02288) PDU1AB
04715 HMMpropdeh2_1_pdu(pf02287) PDU1AB EP_NTERM
04720 HMMpropdeh3_1_pdu(pf02288) PDU1AB EP_NTERM
04725 HMMpropdeh1_1_pdu(pf02286) PDU1AB
04730 Tsp_greenishBeige PDU1AB
04735 H_robinEggBlue GRM4
04740 -pf00230 -
04750 -pf01656pf07685 -
04755 -pf03186 -
04770 -pf00590 -
04775 -pf05175 -
04780 -pf00590 -
04785 -pf01890pf11761pf11760 -
04790 -pf00590 -
04795 -pf02571 -
04800 -pf06180 -
BMC-P
BMC-H
BMC-T
Aldehyde dehydrogenase
PTAC
Signature enzyme
Alcohol dehydrogenase
Regulator

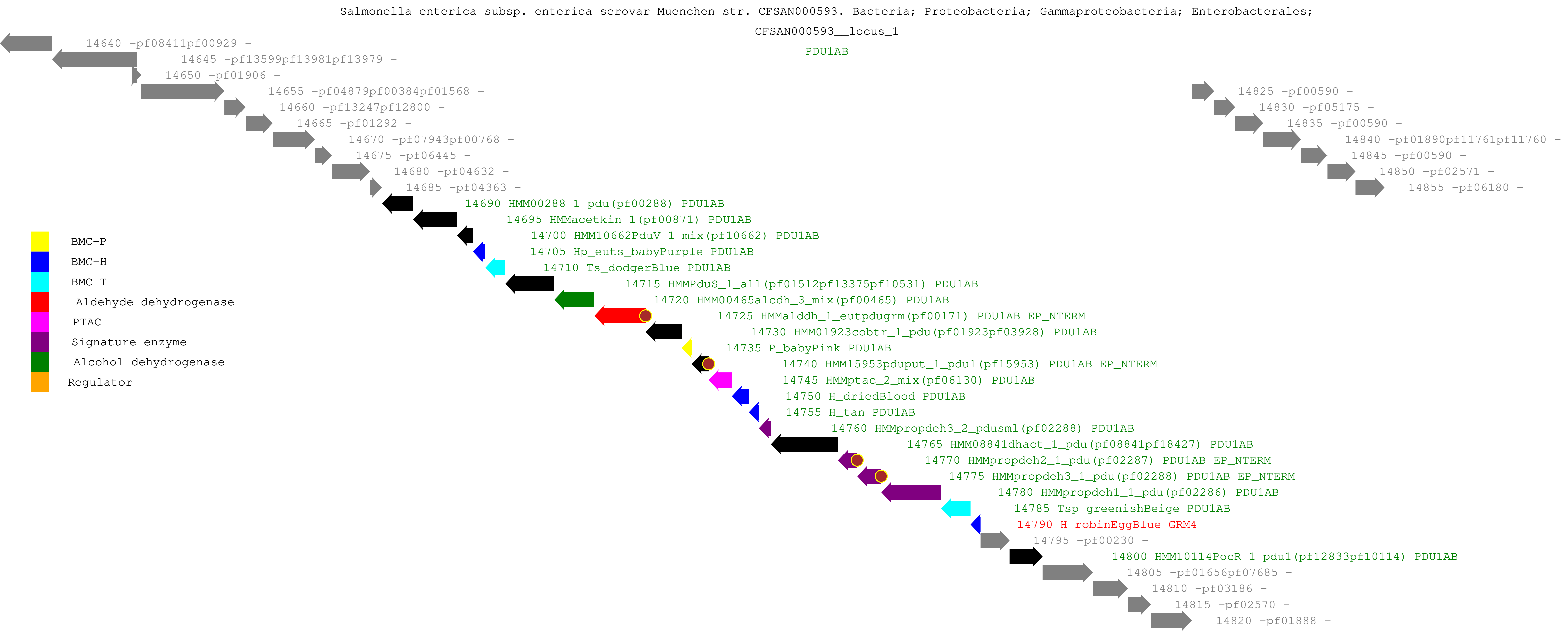
Salmonella enterica subsp. enterica serovar Muenchen str. CFSAN000593. Bacteria; Proteobacteria; Gammaproteobacteria; Enterobacterales;
CFSAN000593__locus_1
PDU1AB
14640 -pf08411pf00929 -
14645 -pf13599pf13981pf13979 -
14650 -pf01906 -
14655 -pf04879pf00384pf01568 -
14660 -pf13247pf12800 -
14665 -pf01292 -
14670 -pf07943pf00768 -
14675 -pf06445 -
14680 -pf04632 -
14685 -pf04363 -
14690 HMM00288_1_pdu(pf00288) PDU1AB
14695 HMMacetkin_1(pf00871) PDU1AB
14700 HMM10662PduV_1_mix(pf10662) PDU1AB
14705 Hp_euts_babyPurple PDU1AB
14710 Ts_dodgerBlue PDU1AB
14715 HMMPduS_1_all(pf01512pf13375pf10531) PDU1AB
14720 HMM00465alcdh_3_mix(pf00465) PDU1AB
14725 HMMalddh_1_eutpdugrm(pf00171) PDU1AB EP_NTERM
14730 HMM01923cobtr_1_pdu(pf01923pf03928) PDU1AB
14735 P_babyPink PDU1AB
14740 HMM15953pduput_1_pdu1(pf15953) PDU1AB EP_NTERM
14745 HMMptac_2_mix(pf06130) PDU1AB
14750 H_driedBlood PDU1AB
14755 H_tan PDU1AB
14760 HMMpropdeh3_2_pdusml(pf02288) PDU1AB
14765 HMM08841dhact_1_pdu(pf08841pf18427) PDU1AB
14770 HMMpropdeh2_1_pdu(pf02287) PDU1AB EP_NTERM
14775 HMMpropdeh3_1_pdu(pf02288) PDU1AB EP_NTERM
14780 HMMpropdeh1_1_pdu(pf02286) PDU1AB
14785 Tsp_greenishBeige PDU1AB
14790 H_robinEggBlue GRM4
14795 -pf00230 -
14800 HMM10114PocR_1_pdu1(pf12833pf10114) PDU1AB
14805 -pf01656pf07685 -
14810 -pf03186 -
14815 -pf02570 -
14820 -pf01888 -
14825 -pf00590 -
14830 -pf05175 -
14835 -pf00590 -
14840 -pf01890pf11761pf11760 -
14845 -pf00590 -
14850 -pf02571 -
14855 -pf06180 -
BMC-P
BMC-H
BMC-T
Aldehyde dehydrogenase
PTAC
Signature enzyme
Alcohol dehydrogenase
Regulator

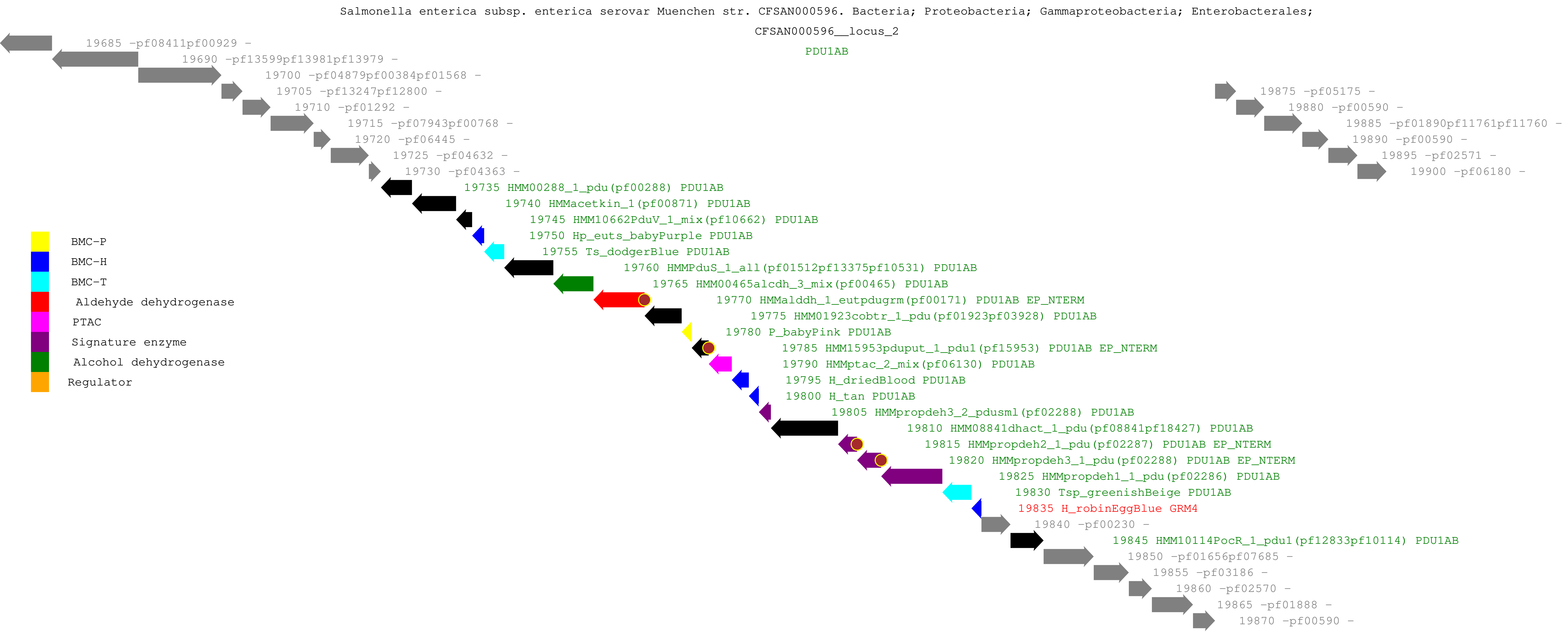
Salmonella enterica subsp. enterica serovar Muenchen str. CFSAN000596. Bacteria; Proteobacteria; Gammaproteobacteria; Enterobacterales;
CFSAN000596__locus_2
PDU1AB
19685 -pf08411pf00929 -
19690 -pf13599pf13981pf13979 -
19700 -pf04879pf00384pf01568 -
19705 -pf13247pf12800 -
19710 -pf01292 -
19715 -pf07943pf00768 -
19720 -pf06445 -
19725 -pf04632 -
19730 -pf04363 -
19735 HMM00288_1_pdu(pf00288) PDU1AB
19740 HMMacetkin_1(pf00871) PDU1AB
19745 HMM10662PduV_1_mix(pf10662) PDU1AB
19750 Hp_euts_babyPurple PDU1AB
19755 Ts_dodgerBlue PDU1AB
19760 HMMPduS_1_all(pf01512pf13375pf10531) PDU1AB
19765 HMM00465alcdh_3_mix(pf00465) PDU1AB
19770 HMMalddh_1_eutpdugrm(pf00171) PDU1AB EP_NTERM
19775 HMM01923cobtr_1_pdu(pf01923pf03928) PDU1AB
19780 P_babyPink PDU1AB
19785 HMM15953pduput_1_pdu1(pf15953) PDU1AB EP_NTERM
19790 HMMptac_2_mix(pf06130) PDU1AB
19795 H_driedBlood PDU1AB
19800 H_tan PDU1AB
19805 HMMpropdeh3_2_pdusml(pf02288) PDU1AB
19810 HMM08841dhact_1_pdu(pf08841pf18427) PDU1AB
19815 HMMpropdeh2_1_pdu(pf02287) PDU1AB EP_NTERM
19820 HMMpropdeh3_1_pdu(pf02288) PDU1AB EP_NTERM
19825 HMMpropdeh1_1_pdu(pf02286) PDU1AB
19830 Tsp_greenishBeige PDU1AB
19835 H_robinEggBlue GRM4
19840 -pf00230 -
19845 HMM10114PocR_1_pdu1(pf12833pf10114) PDU1AB
19850 -pf01656pf07685 -
19855 -pf03186 -
19860 -pf02570 -
19865 -pf01888 -
19870 -pf00590 -
19875 -pf05175 -
19880 -pf00590 -
19885 -pf01890pf11761pf11760 -
19890 -pf00590 -
19895 -pf02571 -
19900 -pf06180 -
BMC-P
BMC-H
BMC-T
Aldehyde dehydrogenase
PTAC
Signature enzyme
Alcohol dehydrogenase
Regulator

Salmonella enterica subsp. enterica serovar Panama str. CFSAN000600. Bacteria; Proteobacteria; Gammaproteobacteria; Enterobacterales;

CFSAN000600__locus_1

PDU1AB

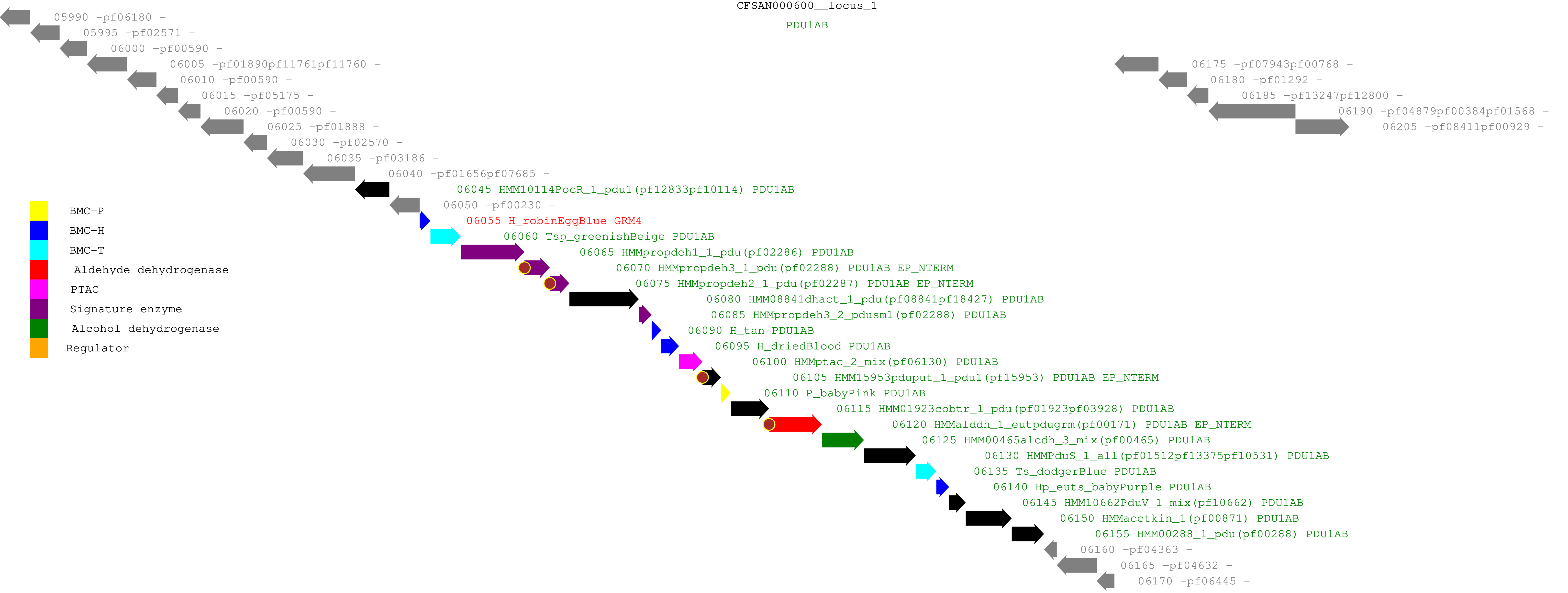

Salmonella enterica subsp. enterica serovar Saintpaul str. CFSAN000618. Bacteria; Proteobacteria; Gammaproteobacteria; Enterobacterales;

CFSAN000618__locus_2

PDU1AB

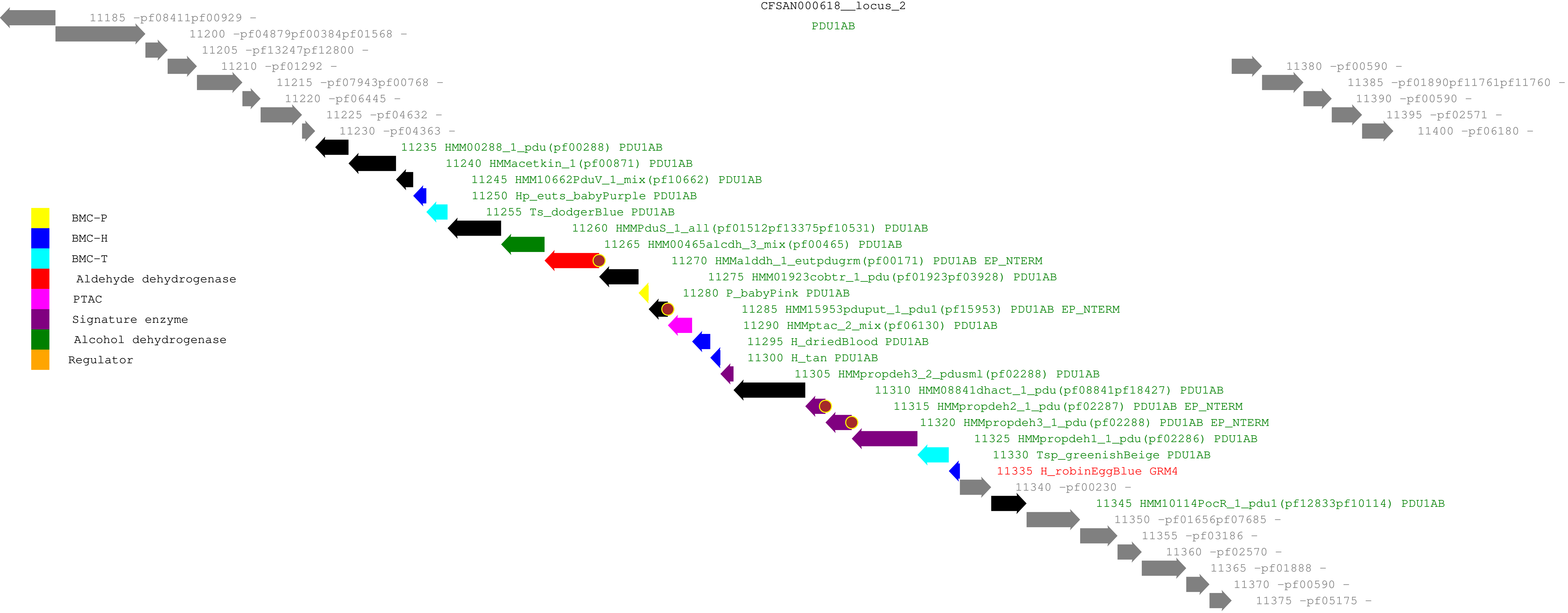

Salmonella enterica subsp. enterica serovar Saintpaul str. CFSAN000619. Bacteria; Proteobacteria; Gammaproteobacteria; Enterobacterales;

CFSAN000619__locus_1

PDU1AB

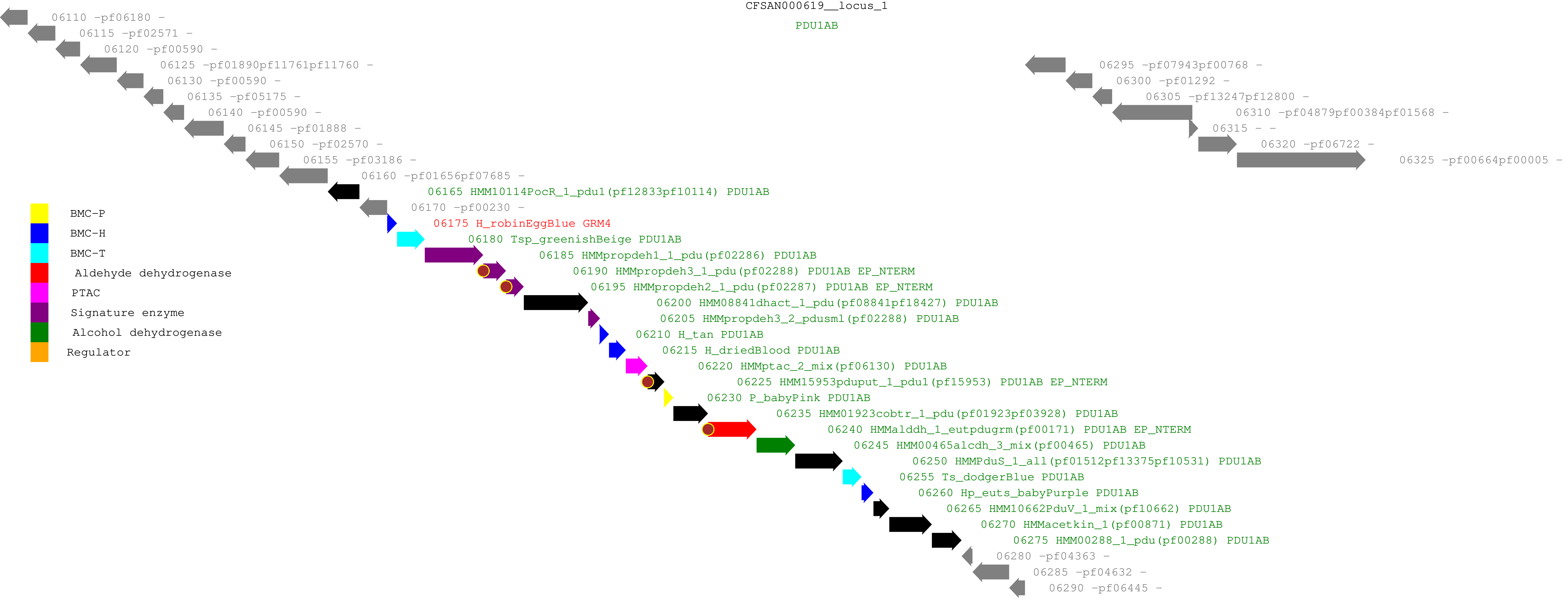

Salmonella enterica subsp. enterica serovar Stanley str. CFSAN000623. Bacteria; Proteobacteria; Gammaproteobacteria; Enterobacterales;

CFSAN000623__locus_1

PDU1AB

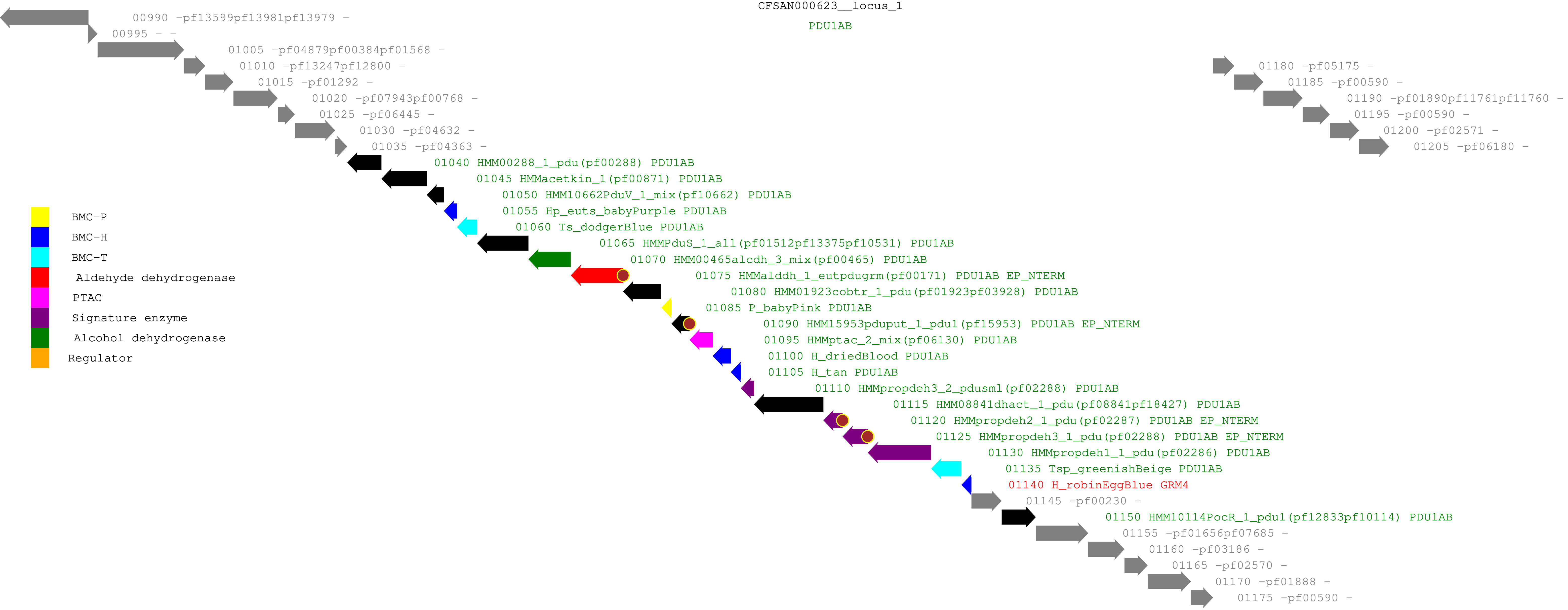

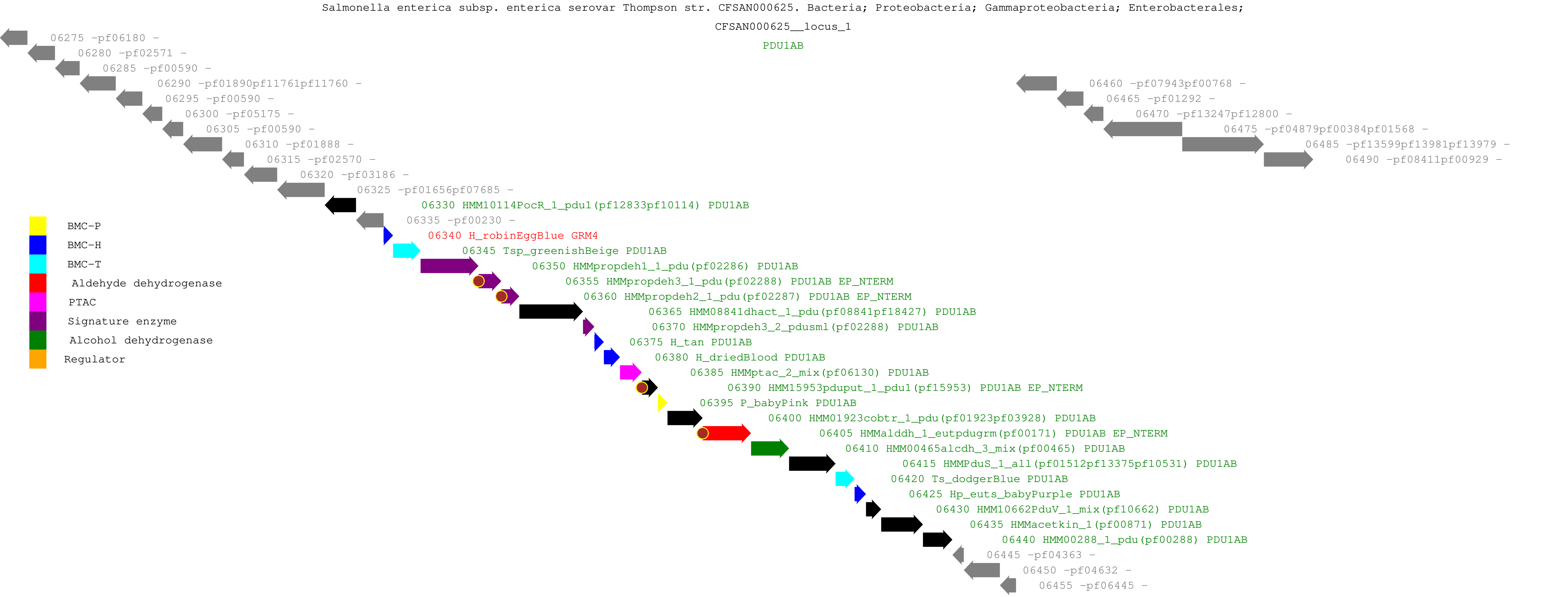

Salmonella enterica subsp. enterica serovar Thompson str. CFSAN000625. Bacteria; Proteobacteria; Gammaproteobacteria; Enterobacterales;
CFSAN000625__locus_1
PDU1AB
06275 -pf06180 -
06280 -pf02571 -
06285 -pf00590 -
06290 -pf01890pf11761pf11760 -
06295 -pf00590 -
06300 -pf05175 -
06305 -pf00590 -
06310 -pf01888 -
06315 -pf02570 -
06320 -pf03186 -
06325 -pf01656pf07685 -
06330 HMM10114PocR_1_pdu1(pf12833pf10114) PDU1AB
06335 -pf00230 -
06340 H_robinEggBlue GRM4
06345 Tsp_greenishBeige PDU1AB
06350 HMMpropdeh1_1_pdu(pf02286) PDU1AB
06355 HMMpropdeh3_1_pdu(pf02288) PDU1AB EP_NTERM
06360 HMMpropdeh2_1_pdu(pf02287) PDU1AB EP_NTERM
06365 HMM08841dhact_1_pdu(pf08841pf18427) PDU1AB
06370 HMMpropdeh3_2_pdusml(pf02288) PDU1AB
06375 H_tan PDU1AB
06380 H_driedBlood PDU1AB
06385 HMMptac_2_mix(pf06130) PDU1AB
06390 HMM15953pduput_1_pdu1(pf15953) PDU1AB EP_NTERM
06395 P_babyPink PDU1AB
06400 HMM01923cobtr_1_pdu(pf01923pf03928) PDU1AB
06405 HMMalddh_1_eutpdugrm(pf00171) PDU1AB EP_NTERM
06410 HMM00465alcdh_3_mix(pf00465) PDU1AB
06415 HMMPduS_1_all(pf01512pf13375pf10531) PDU1AB
06420 Ts_dodgerBlue PDU1AB
06425 Hp_euts_babyPurple PDU1AB
06430 HMM10662PduV_1_mix(pf10662) PDU1AB
06435 HMMacetkin_1(pf00871) PDU1AB
06440 HMM00288_1_pdu(pf00288) PDU1AB
06445 -pf04363 -
06450 -pf04632 -
06455 -pf06445 -
06460 -pf07943pf00768 -
06465 -pf01292 -
06470 -pf13247pf12800 -
06475 -pf04879pf00384pf01568 -
06485 -pf13599pf13981pf13979 -
06490 -pf08411pf00929 -
BMC-P
BMC-H
BMC-T
Aldehyde dehydrogenase
PTAC
Signature enzyme
Alcohol dehydrogenase
Regulator

Salmonella enterica subsp. enterica serovar Cerro str. CFSAN001590. Bacteria; Proteobacteria; Gammaproteobacteria; Enterobacterales;

CFSAN001590__locus_1

PDU1AB

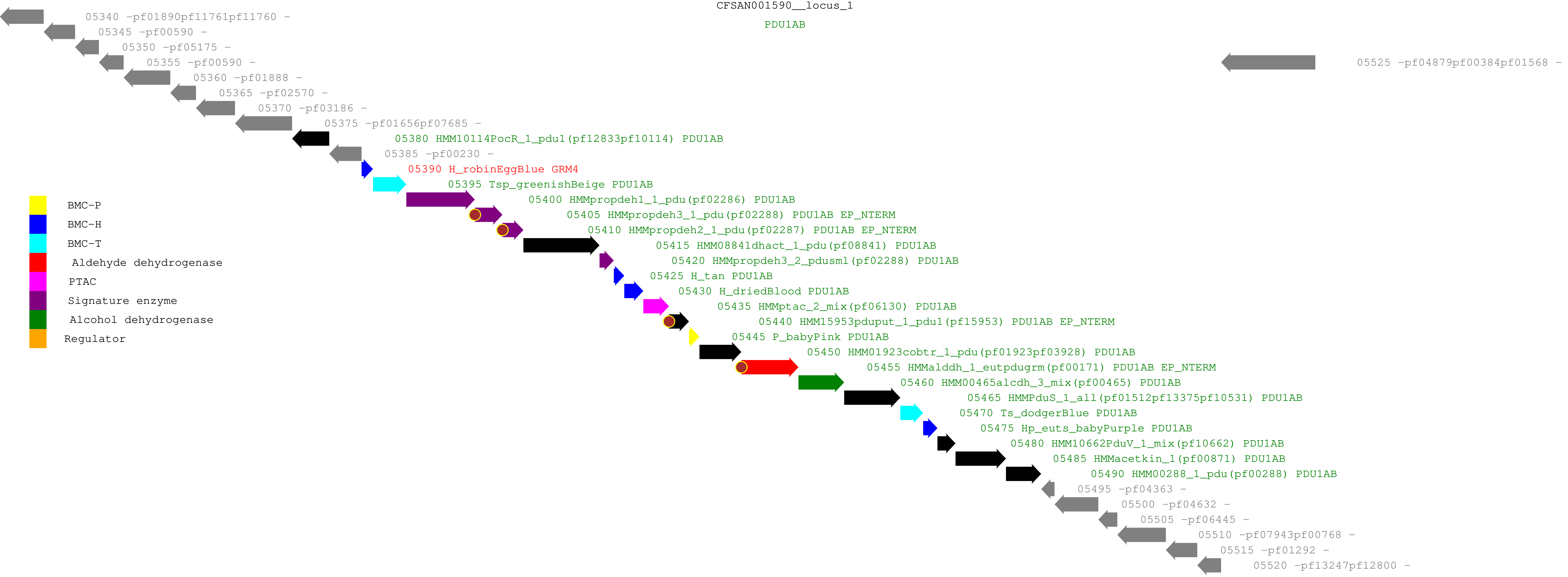

Salmonella enterica subsp. enterica serovar Cubana str. CFSAN002050. Bacteria; Proteobacteria; Gammaproteobacteria; Enterobacterales;

CFSAN002050__locus_1

PDU1AB

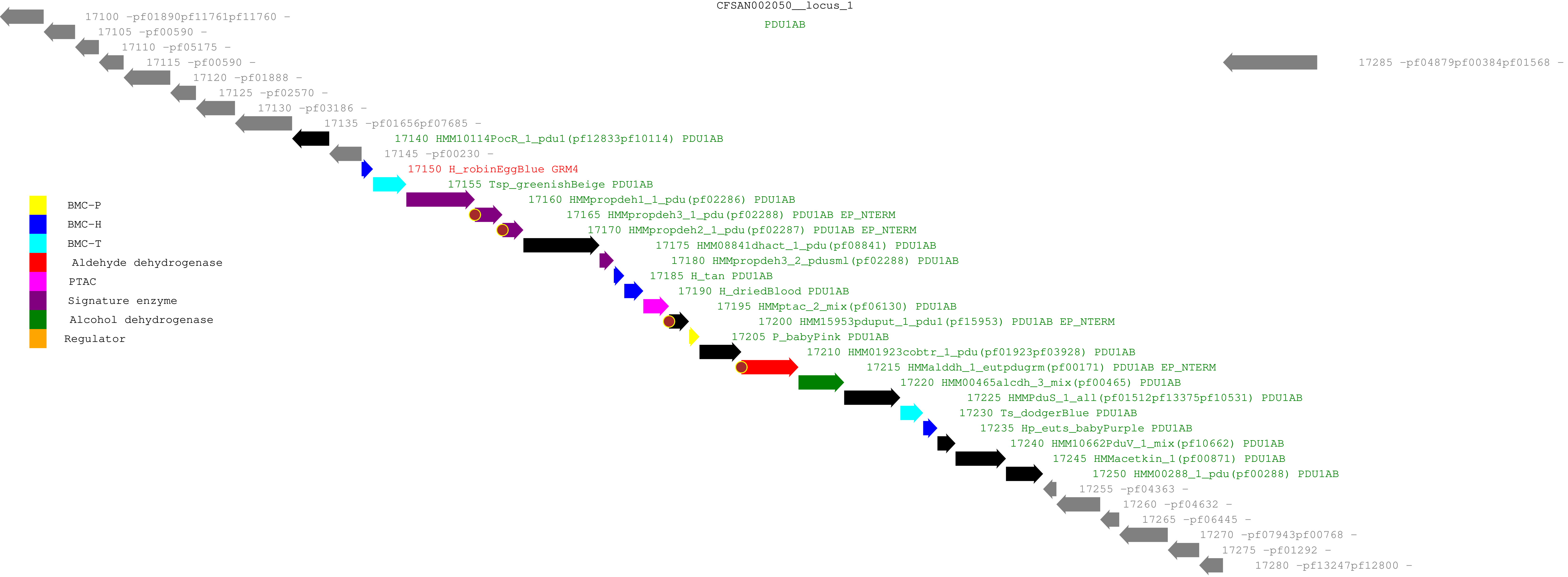

Salmonella enterica subsp. enterica serovar Indiana. Bacteria; Proteobacteria; Gammaproteobacteria; Enterobacterales;

CGL53__locus_2

PDU1AB

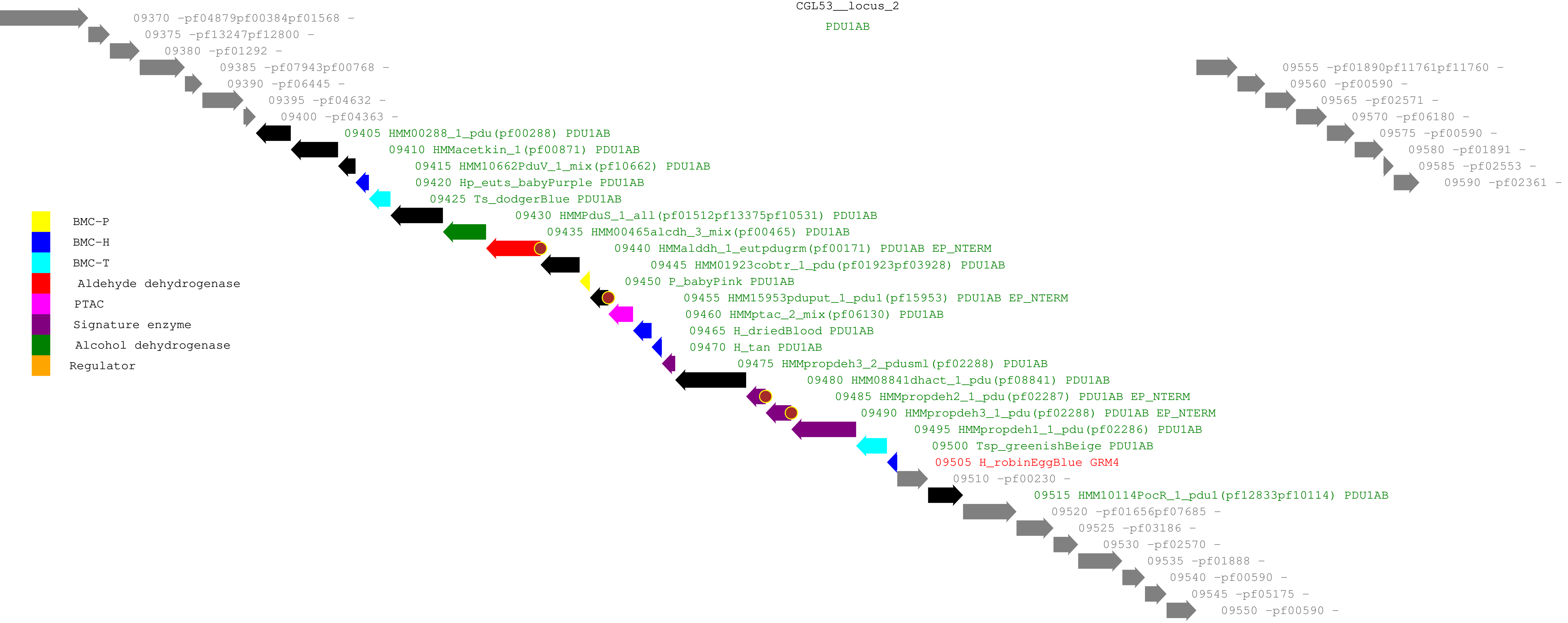

Salmonella enterica subsp. enterica serovar Heidelberg. Bacteria; Proteobacteria; Gammaproteobacteria; Enterobacterales;

CH228__locus_2

PDU1AB

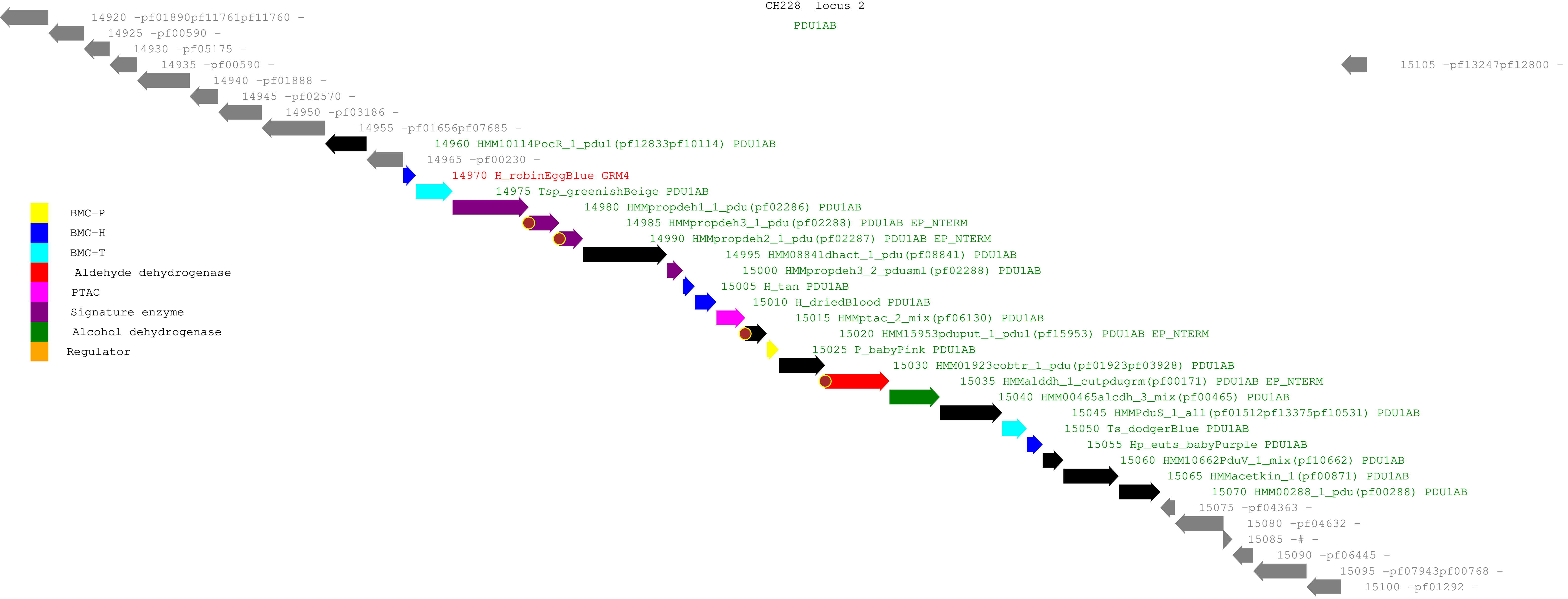

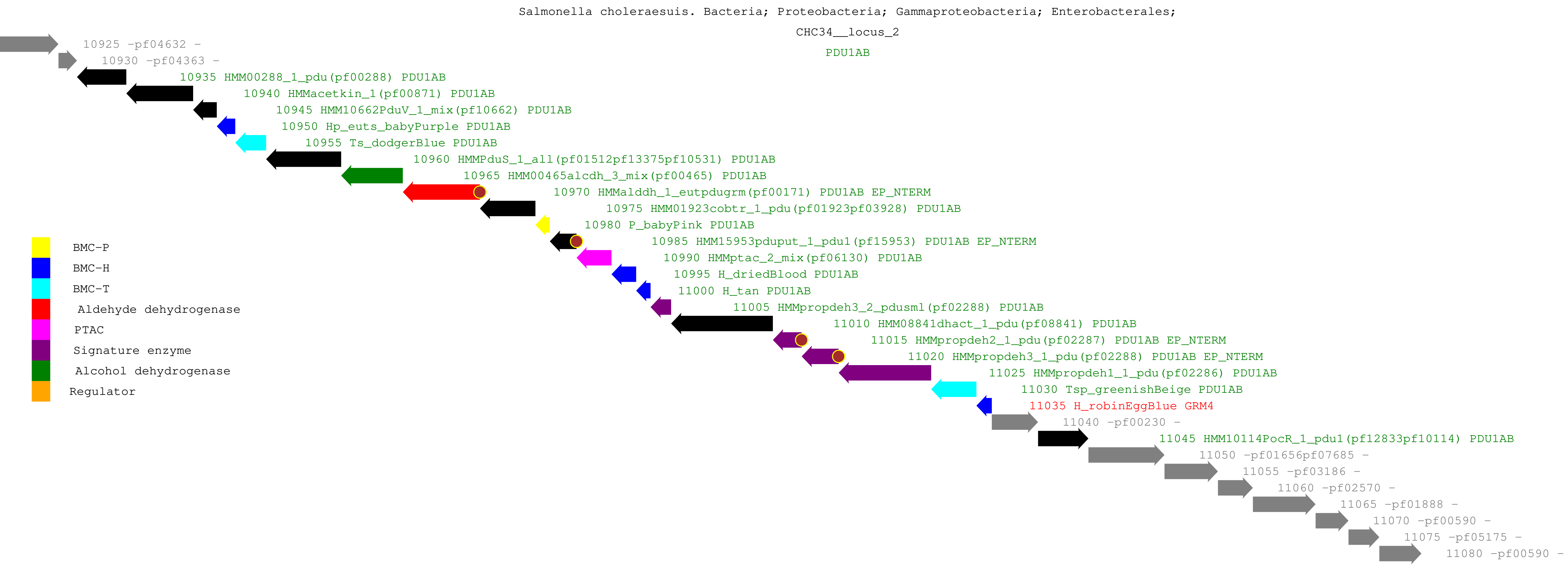
Salmonella choleraesuis. Bacteria; Proteobacteria; Gammaproteobacteria; Enterobacterales;
CHC34__locus_2
PDU1AB
10925 -pf04632 -
10930 -pf04363 -
10935 HMM00288_1_pdu(pf00288) PDU1AB
10940 HMMacetkin_1(pf00871) PDU1AB
10945 HMM10662PduV_1_mix(pf10662) PDU1AB
10950 Hp_euts_babyPurple PDU1AB
10955 Ts_dodgerBlue PDU1AB
10960 HMMPduS_1_all(pf01512pf13375pf10531) PDU1AB
10965 HMM00465alcdh_3_mix(pf00465) PDU1AB
10970 HMMalddh_1_eutpdugrm(pf00171) PDU1AB EP_NTERM
10975 HMM01923cobtr_1_pdu(pf01923pf03928) PDU1AB
10980 P_babyPink PDU1AB
10985 HMM15953pduput_1_pdu1(pf15953) PDU1AB EP_NTERM
10990 HMMptac_2_mix(pf06130) PDU1AB
10995 H_driedBlood PDU1AB
11000 H_tan PDU1AB
11005 HMMpropdeh3_2_pdusml(pf02288) PDU1AB
11010 HMM08841dhact_1_pdu(pf08841) PDU1AB
11015 HMMpropdeh2_1_pdu(pf02287) PDU1AB EP_NTERM
11020 HMMpropdeh3_1_pdu(pf02288) PDU1AB EP_NTERM
11025 HMMpropdeh1_1_pdu(pf02286) PDU1AB
11030 Tsp_greenishBeige PDU1AB
11035 H_robinEggBlue GRM4
11040 -pf00230 -
11045 HMM10114PocR_1_pdu1(pf12833pf10114) PDU1AB
11050 -pf01656pf07685 -
11055 -pf03186 -
11060 -pf02570 -
11065 -pf01888 -
11070 -pf00590 -
11075 -pf05175 -
11080 -pf00590 -
BMC-P
BMC-H
BMC-T
Aldehyde dehydrogenase
PTAC
Signature enzyme
Alcohol dehydrogenase
Regulator

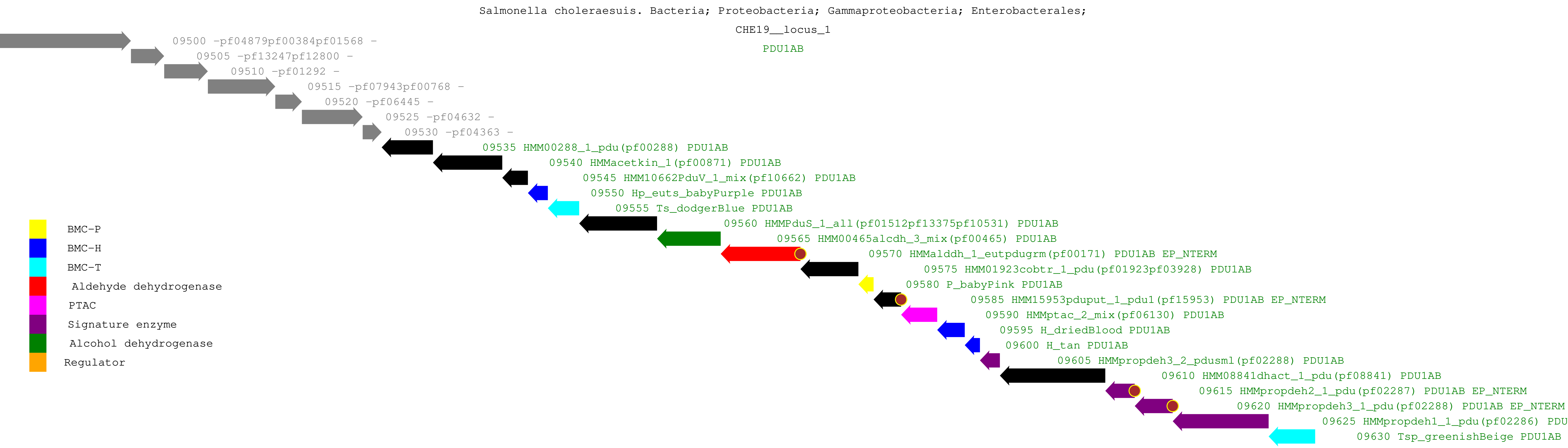
Salmonella choleraesuis. Bacteria; Proteobacteria; Gammaproteobacteria; Enterobacterales;
CHE19__locus_1
PDU1AB
09500 -pf04879pf00384pf01568 -
09505 -pf13247pf12800 -
09510 -pf01292 -
09515 -pf07943pf00768 -
09520 -pf06445 -
09525 -pf04632 -
09530 -pf04363 -
09535 HMM00288_1_pdu(pf00288) PDU1AB
09540 HMMacetkin_1(pf00871) PDU1AB
09545 HMM10662PduV_1_mix(pf10662) PDU1AB
09550 Hp_euts_babyPurple PDU1AB
09555 Ts_dodgerBlue PDU1AB
09560 HMMPduS_1_all(pf01512pf13375pf10531) PDU1AB
09565 HMM00465alcdh_3_mix(pf00465) PDU1AB
09570 HMMalddh_1_eutpdugrm(pf00171) PDU1AB EP_NTERM
09575 HMM01923cobtr_1_pdu(pf01923pf03928) PDU1AB
09580 P_babyPink PDU1AB
09585 HMM15953pduput_1_pdu1(pf15953) PDU1AB EP_NTERM
09590 HMMptac_2_mix(pf06130) PDU1AB
09595 H_driedBlood PDU1AB
09600 H_tan PDU1AB
09605 HMMpropdeh3_2_pdusml(pf02288) PDU1AB
09610 HMM08841dhact_1_pdu(pf08841) PDU1AB
09615 HMMpropdeh2_1_pdu(pf02287) PDU1AB EP_NTERM
09620 HMMpropdeh3_1_pdu(pf02288) PDU1AB EP_NTERM
09625 HMMpropdeh1_1_pdu(pf02286) PDU
09630 Tsp_greenishBeige PDU1AB
BMC-P
BMC-H
BMC-T
Aldehyde dehydrogenase
PTAC
Signature enzyme
Alcohol dehydrogenase
Regulator

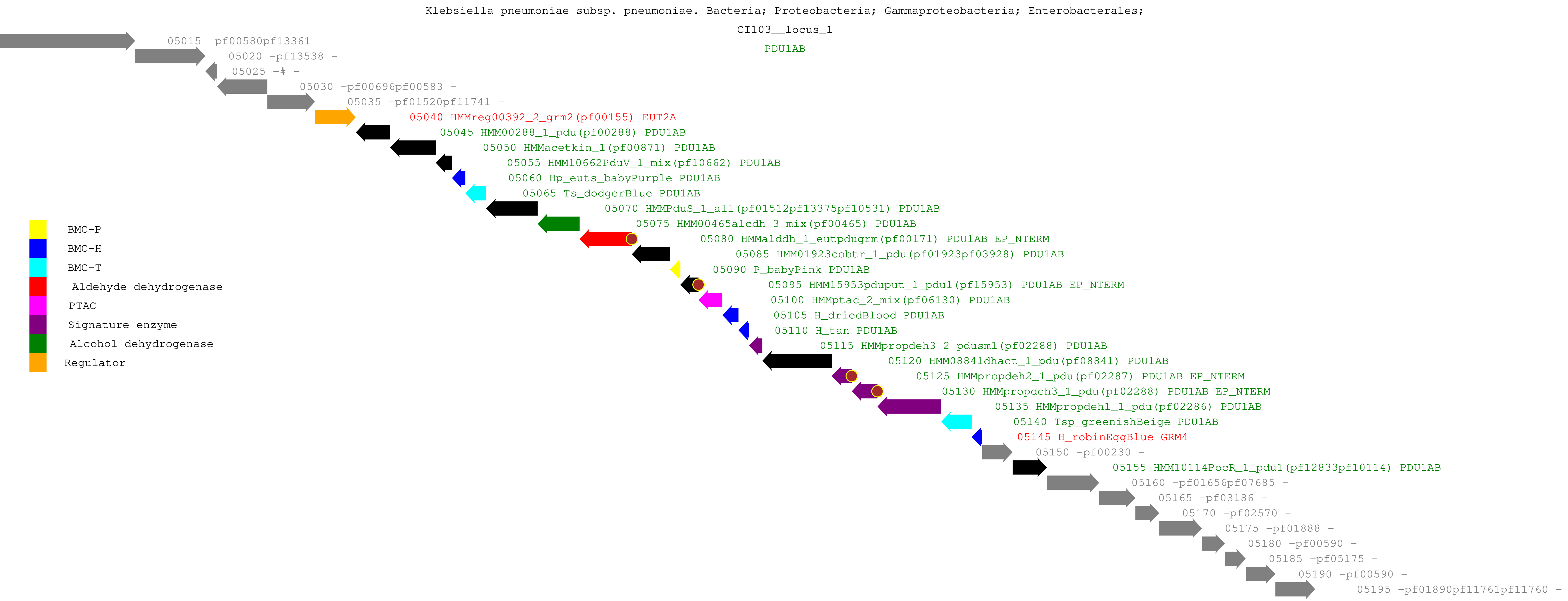

Klebsiella pneumoniae subsp. pneumoniae. Bacteria; Proteobacteria; Gammaproteobacteria; Enterobacterales;
CI103__locus_1
PDU1AB
05015 -pf00580pf13361 -
05020 -pf13538 -
05025 -# -
05030 -pf00696pf00583 -
05035 -pf01520pf11741 -
05040 HMMreg00392_2_grm2(pf00155) EUT2A
05045 HMM00288_1_pdu(pf00288) PDU1AB
05050 HMMacetkin_1(pf00871) PDU1AB
05055 HMM10662PduV_1_mix(pf10662) PDU1AB
05060 Hp_euts_babyPurple PDU1AB
05065 Ts_dodgerBlue PDU1AB
05070 HMMPduS_1_all(pf01512pf13375pf10531) PDU1AB
05075 HMM00465alcdh_3_mix(pf00465) PDU1AB
05080 HMMalddh_1_eutpdugrm(pf00171) PDU1AB EP_NTERM
05085 HMM01923cobtr_1_pdu(pf01923pf03928) PDU1AB
05090 P_babyPink PDU1AB
05095 HMM15953pduput_1_pdu1(pf15953) PDU1AB EP_NTERM
05100 HMMptac_2_mix(pf06130) PDU1AB
05105 H_driedBlood PDU1AB
05110 H_tan PDU1AB
05115 HMMpropdeh3_2_pdusml(pf02288) PDU1AB
05120 HMM08841dhact_1_pdu(pf08841) PDU1AB
05125 HMMpropdeh2_1_pdu(pf02287) PDU1AB EP_NTERM
05130 HMMpropdeh3_1_pdu(pf02288) PDU1AB EP_NTERM
05135 HMMpropdeh1_1_pdu(pf02286) PDU1AB
05140 Tsp_greenishBeige PDU1AB
05145 H_robinEggBlue GRM4
05150 -pf00230 -
05155 HMM10114PocR_1_pdu1(pf12833pf10114) PDU1AB
05160 -pf01656pf07685 -
05165 -pf03186 -
05170 -pf02570 -
05175 -pf01888 -
05180 -pf00590 -
05185 -pf05175 -
05190 -pf00590 -
05195 -pf01890pf11761pf11760 -
BMC-P
BMC-H
BMC-T
Aldehyde dehydrogenase
PTAC
Signature enzyme
Alcohol dehydrogenase
Regulator

Citrobacter farmeri. Bacteria; Proteobacteria; Gammaproteobacteria; Enterobacterales;

CI104__locus_1

PDU1AB

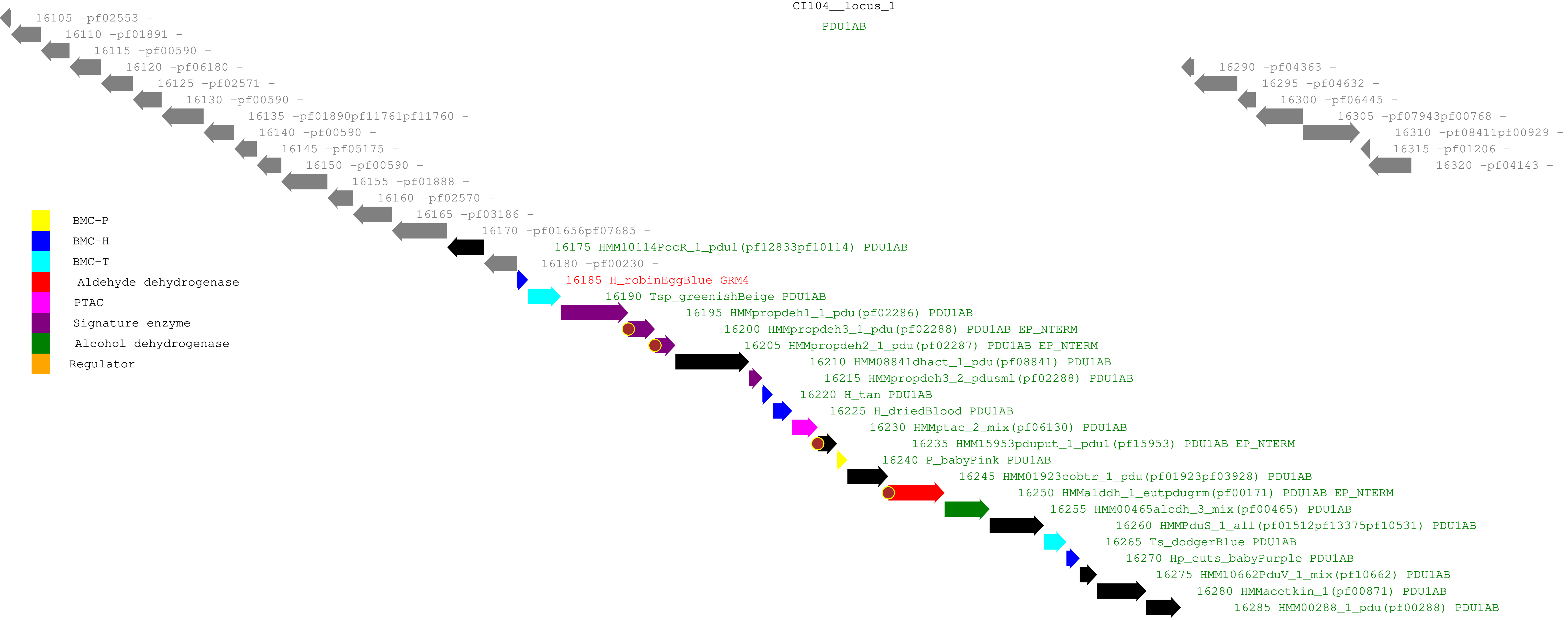

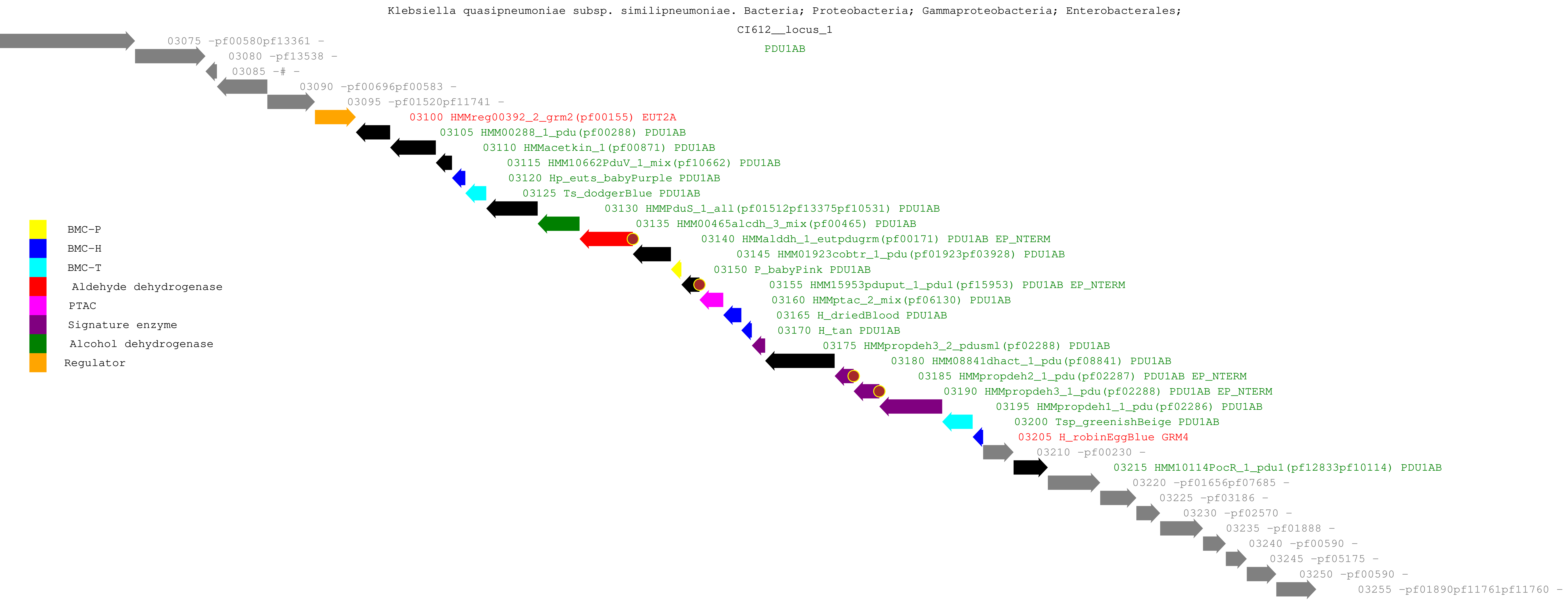
Klebsiella quasipneumoniae subsp. similipneumoniae. Bacteria; Proteobacteria; Gammaproteobacteria; Enterobacterales;
CI612__locus_1
PDU1AB
03075 -pf00580pf13361 -
03080 -pf13538 -
03085 -# -
03090 -pf00696pf00583 -
03095 -pf01520pf11741 -
03100 HMMreg00392_2_grm2(pf00155) EUT2A
03105 HMM00288_1_pdu(pf00288) PDU1AB
03110 HMMacetkin_1(pf00871) PDU1AB
03115 HMM10662PduV_1_mix(pf10662) PDU1AB
03120 Hp_euts_babyPurple PDU1AB
03125 Ts_dodgerBlue PDU1AB
03130 HMMPduS_1_all(pf01512pf13375pf10531) PDU1AB
03135 HMM00465alcdh_3_mix(pf00465) PDU1AB
03140 HMMalddh_1_eutpdugrm(pf00171) PDU1AB EP_NTERM
03145 HMM01923cobtr_1_pdu(pf01923pf03928) PDU1AB
03150 P_babyPink PDU1AB
03155 HMM15953pduput_1_pdu1(pf15953) PDU1AB EP_NTERM
03160 HMMptac_2_mix(pf06130) PDU1AB
03165 H_driedBlood PDU1AB
03170 H_tan PDU1AB
03175 HMMpropdeh3_2_pdusml(pf02288) PDU1AB
03180 HMM08841dhact_1_pdu(pf08841) PDU1AB
03185 HMMpropdeh2_1_pdu(pf02287) PDU1AB EP_NTERM
03190 HMMpropdeh3_1_pdu(pf02288) PDU1AB EP_NTERM
03195 HMMpropdeh1_1_pdu(pf02286) PDU1AB
03200 Tsp_greenishBeige PDU1AB
03205 H_robinEggBlue GRM4
03210 -pf00230 -
03215 HMM10114PocR_1_pdu1(pf12833pf10114) PDU1AB
03220 -pf01656pf07685 -
03225 -pf03186 -
03230 -pf02570 -
03235 -pf01888 -
03240 -pf00590 -
03245 -pf05175 -
03250 -pf00590 -
03255 -pf01890pf11761pf11760 -
BMC-P
BMC-H
BMC-T
Aldehyde dehydrogenase
PTAC
Signature enzyme
Alcohol dehydrogenase
Regulator

Shigella sonnei. Bacteria; Proteobacteria; Gammaproteobacteria; Enterobacterales;

CI641__locus_1

PDU1AB

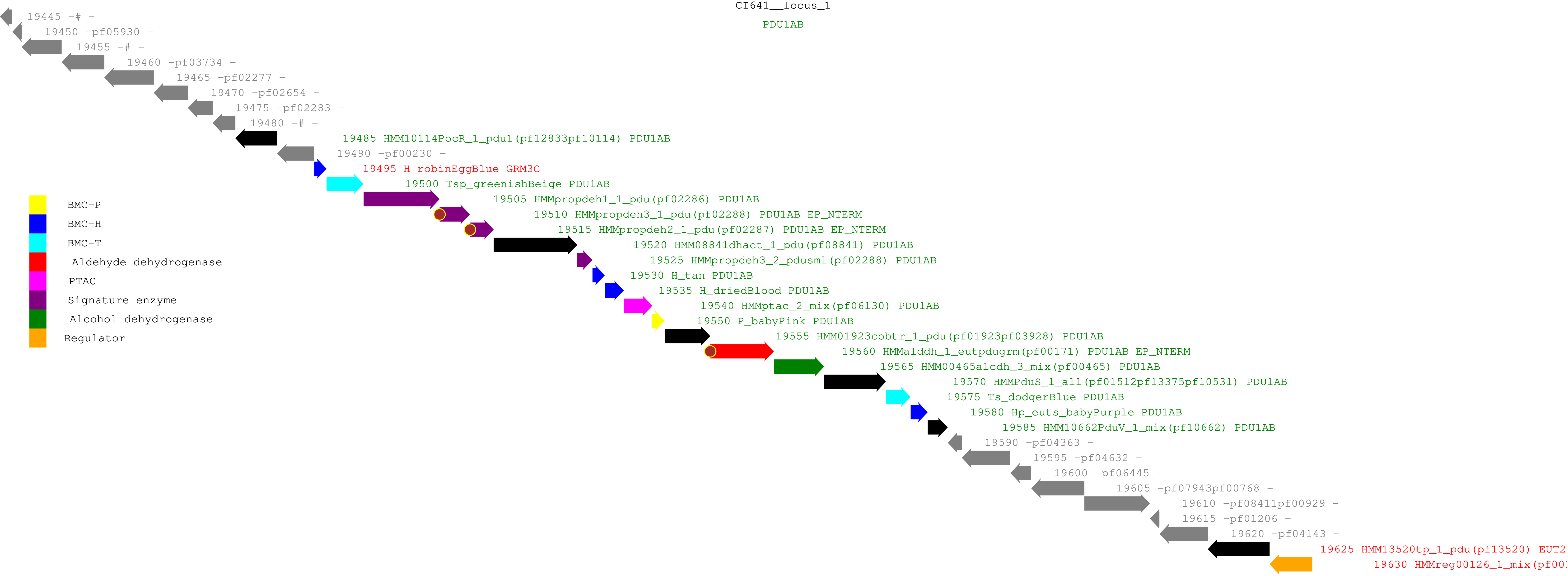

Shigella sonnei. Bacteria; Proteobacteria; Gammaproteobacteria; Enterobacterales;

CI642__locus_1

PDU1AB

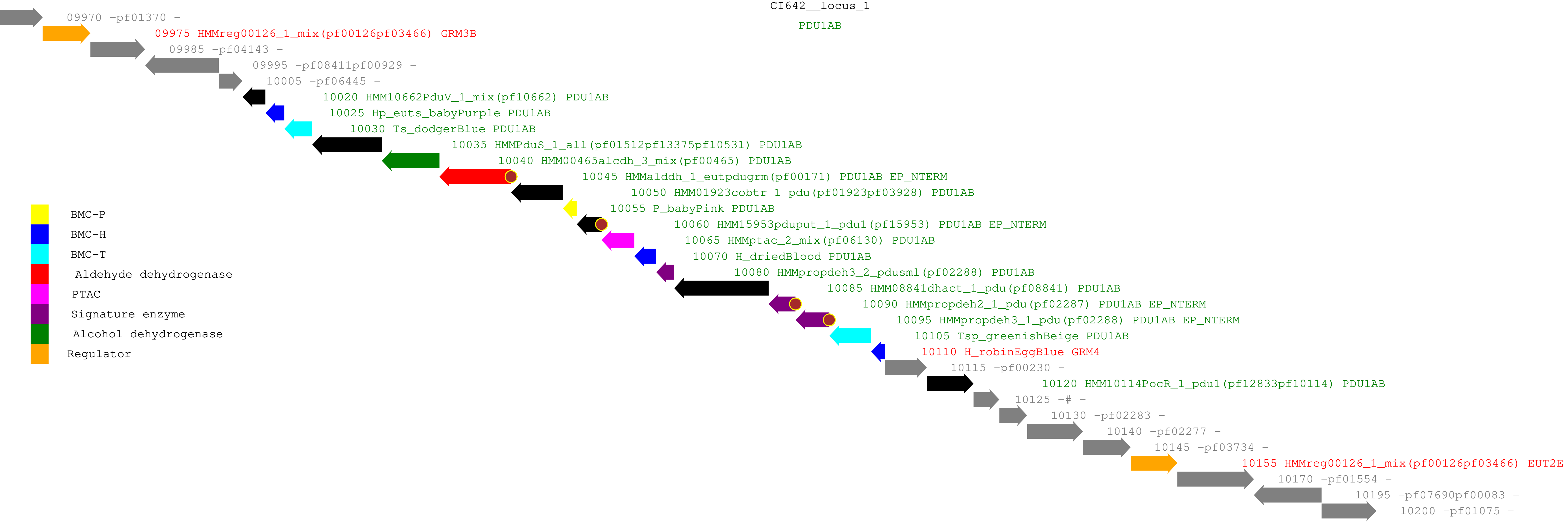

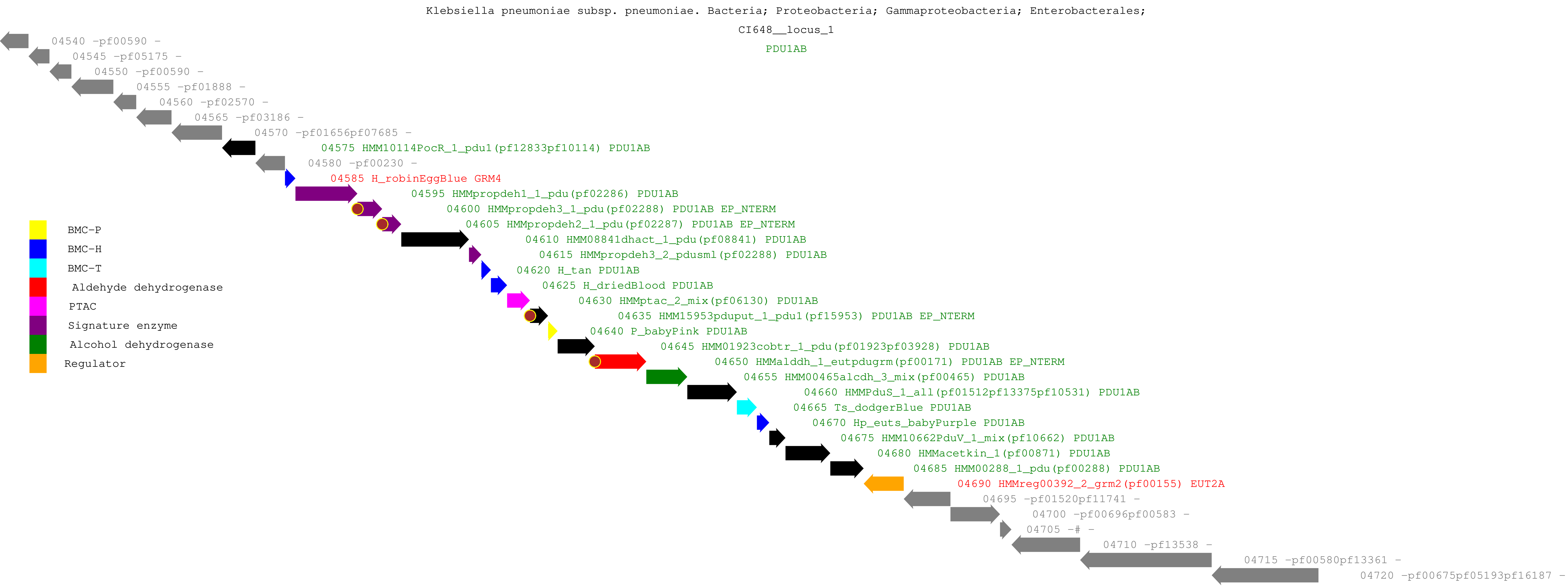

Klebsiella pneumoniae subsp. pneumoniae. Bacteria; Proteobacteria; Gammaproteobacteria; Enterobacterales;
CI648__locus_1
PDU1AB
04540 -pf00590 -
04545 -pf05175 -
04550 -pf00590 -
04555 -pf01888 -
04560 -pf02570 -
04565 -pf03186 -
04570 -pf01656pf07685 -
04575 HMM10114PocR_1_pdu1(pf12833pf10114) PDU1AB
04580 -pf00230 -
04585 H_robinEggBlue GRM4
04595 HMMpropdeh1_1_pdu(pf02286) PDU1AB
04600 HMMpropdeh3_1_pdu(pf02288) PDU1AB EP_NTERM
04605 HMMpropdeh2_1_pdu(pf02287) PDU1AB EP_NTERM
04610 HMM08841dhact_1_pdu(pf08841) PDU1AB
04615 HMMpropdeh3_2_pdusml(pf02288) PDU1AB
04620 H_tan PDU1AB
04625 H_driedBlood PDU1AB
04630 HMMptac_2_mix(pf06130) PDU1AB
04635 HMM15953pduput_1_pdu1(pf15953) PDU1AB EP_NTERM
04640 P_babyPink PDU1AB
04645 HMM01923cobtr_1_pdu(pf01923pf03928) PDU1AB
04650 HMMalddh_1_eutpdugrm(pf00171) PDU1AB EP_NTERM
04655 HMM00465alcdh_3_mix(pf00465) PDU1AB
04660 HMMPduS_1_all(pf01512pf13375pf10531) PDU1AB
04665 Ts_dodgerBlue PDU1AB
04670 Hp_euts_babyPurple PDU1AB
04675 HMM10662PduV_1_mix(pf10662) PDU1AB
04680 HMMacetkin_1(pf00871) PDU1AB
04685 HMM00288_1_pdu(pf00288) PDU1AB
04690 HMMreg00392_2_grm2(pf00155) EUT2A
04695 -pf01520pf11741 -
04700 -pf00696pf00583 -
04705 -# -
04710 -pf13538 -
04715 -pf00580pf13361 -
04720 -pf00675pf05193pf16187 -
BMC-P
BMC-H
BMC-T
Aldehyde dehydrogenase
PTAC
Signature enzyme
Alcohol dehydrogenase
Regulator

Klebsiella quasipneumoniae subsp. similipneumoniae. Bacteria; Proteobacteria; Gammaproteobacteria; Enterobacterales;

CI664__locus_2

PDU1AB

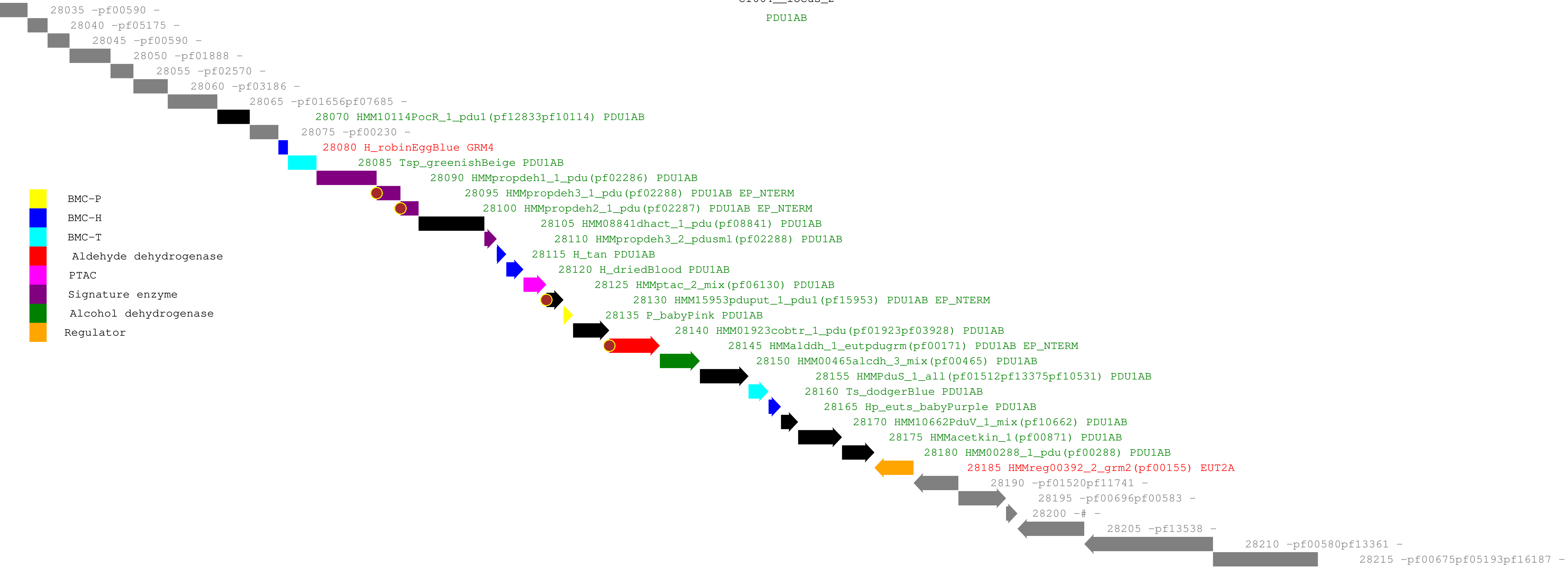

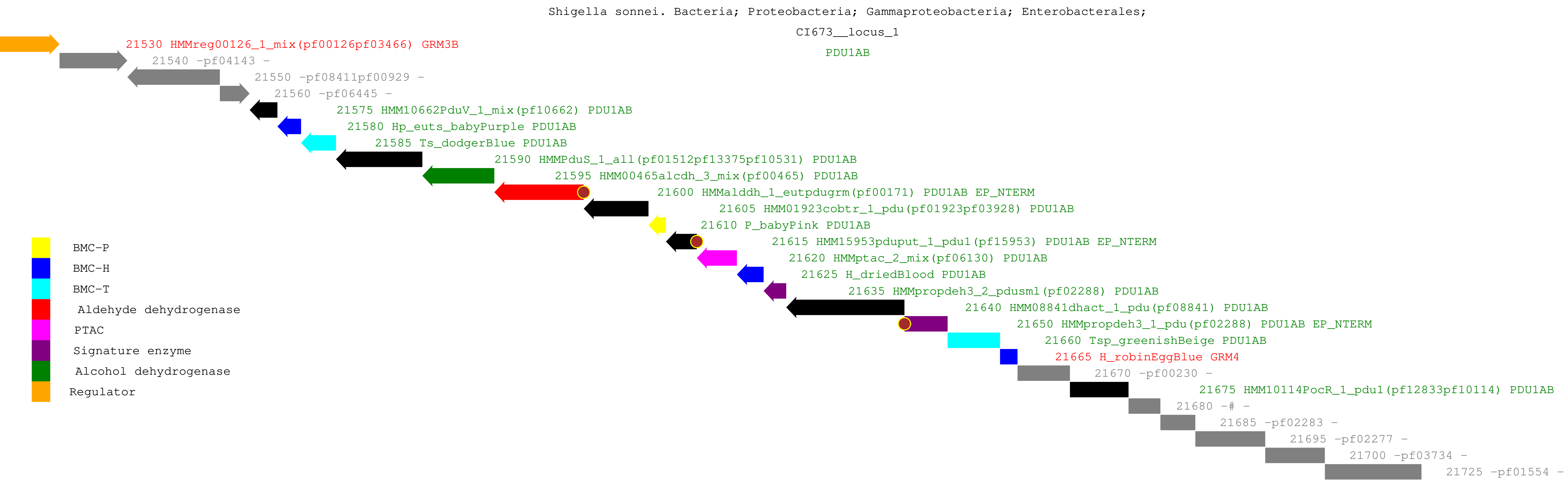
Shigella sonnei. Bacteria; Proteobacteria; Gammaproteobacteria; Enterobacterales;
CI673__locus_1
PDU1AB
21530 HMMreg00126_1_mix(pf00126pf03466) GRM3B
21540 -pf04143 -
21550 -pf08411pf00929 -
21560 -pf06445 -
21575 HMM10662PduV_1_mix(pf10662) PDU1AB
21580 Hp_euts_babyPurple PDU1AB
21585 Ts_dodgerBlue PDU1AB
21590 HMMPduS_1_all(pf01512pf13375pf10531) PDU1AB
21595 HMM00465alcdh_3_mix(pf00465) PDU1AB
21600 HMMalddh_1_eutpdugrm(pf00171) PDU1AB EP_NTERM
21605 HMM01923cobtr_1_pdu(pf01923pf03928) PDU1AB
21610 P_babyPink PDU1AB
21615 HMM15953pduput_1_pdu1(pf15953) PDU1AB EP_NTERM
21620 HMMptac_2_mix(pf06130) PDU1AB
21625 H_driedBlood PDU1AB
21635 HMMpropdeh3_2_pdusml(pf02288) PDU1AB
21640 HMM08841dhact_1_pdu(pf08841) PDU1AB
21650 HMMpropdeh3_1_pdu(pf02288) PDU1AB EP_NTERM
21660 Tsp_greenishBeige PDU1AB
21665 H_robinEggBlue GRM4
21670 -pf00230 -
21675 HMM10114PocR_1_pdu1(pf12833pf10114) PDU1AB
21680 -# -
21685 -pf02283 -
21695 -pf02277 -
21700 -pf03734 -
21725 -pf01554 -
BMC-P
BMC-H
BMC-T
Aldehyde dehydrogenase
PTAC
Signature enzyme
Alcohol dehydrogenase
Regulator

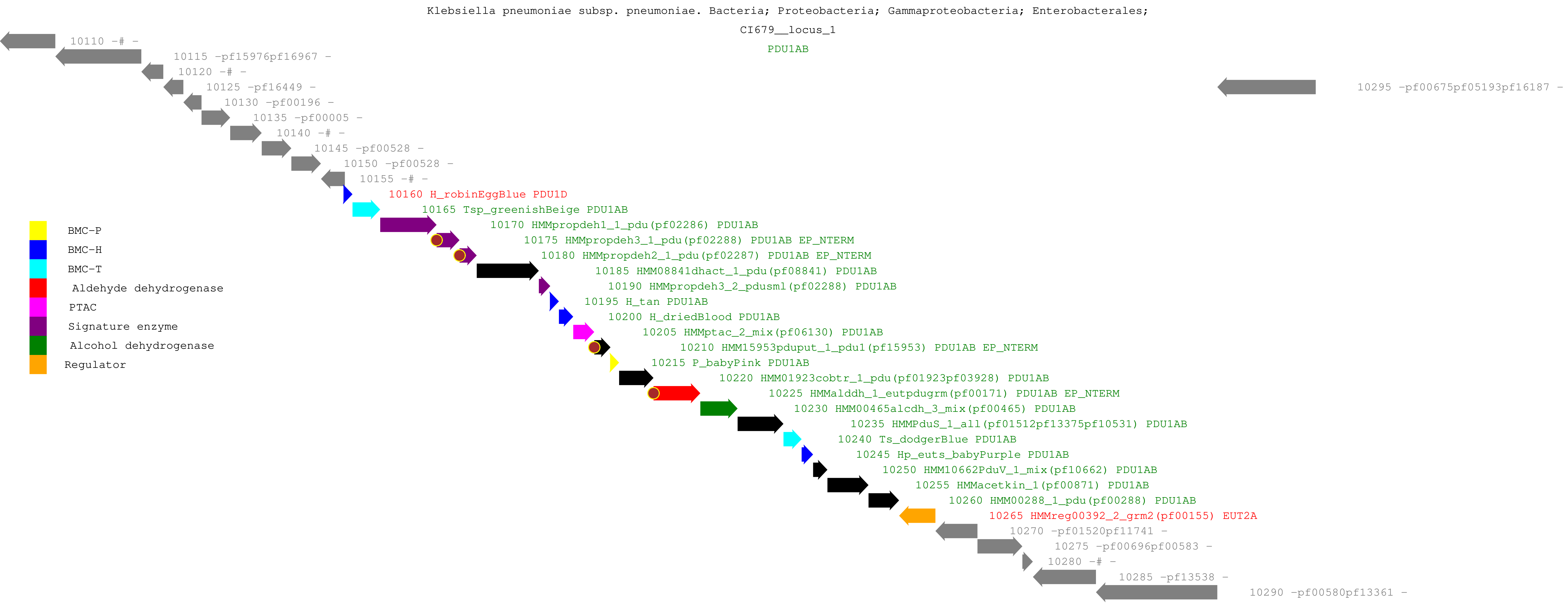

Klebsiella pneumoniae subsp. pneumoniae. Bacteria; Proteobacteria; Gammaproteobacteria; Enterobacterales;
CI679__locus_1
PDU1AB
10110 -# -
10115 -pf15976pf16967 -
10120 -# -
10125 -pf16449 -
10130 -pf00196 -
10135 -pf00005 -
10140 -# -
10145 -pf00528 -
10150 -pf00528 -
10155 -# -
10160 H_robinEggBlue PDU1D
10165 Tsp_greenishBeige PDU1AB
10170 HMMpropdeh1_1_pdu(pf02286) PDU1AB
10175 HMMpropdeh3_1_pdu(pf02288) PDU1AB EP_NTERM
10180 HMMpropdeh2_1_pdu(pf02287) PDU1AB EP_NTERM
10185 HMM08841dhact_1_pdu(pf08841) PDU1AB
10190 HMMpropdeh3_2_pdusml(pf02288) PDU1AB
10195 H_tan PDU1AB
10200 H_driedBlood PDU1AB
10205 HMMptac_2_mix(pf06130) PDU1AB
10210 HMM15953pduput_1_pdu1(pf15953) PDU1AB EP_NTERM
10215 P_babyPink PDU1AB
10220 HMM01923cobtr_1_pdu(pf01923pf03928) PDU1AB
10225 HMMalddh_1_eutpdugrm(pf00171) PDU1AB EP_NTERM
10230 HMM00465alcdh_3_mix(pf00465) PDU1AB
10235 HMMPduS_1_all(pf01512pf13375pf10531) PDU1AB
10240 Ts_dodgerBlue PDU1AB
10245 Hp_euts_babyPurple PDU1AB
10250 HMM10662PduV_1_mix(pf10662) PDU1AB
10255 HMMacetkin_1(pf00871) PDU1AB
10260 HMM00288_1_pdu(pf00288) PDU1AB
10265 HMMreg00392_2_grm2(pf00155) EUT2A
10270 -pf01520pf11741 -
10275 -pf00696pf00583 -
10280 -# -
10285 -pf13538 -
10290 -pf00580pf13361 -
10295 -pf00675pf05193pf16187 -
BMC-P
BMC-H
BMC-T
Aldehyde dehydrogenase
PTAC
Signature enzyme
Alcohol dehydrogenase
Regulator

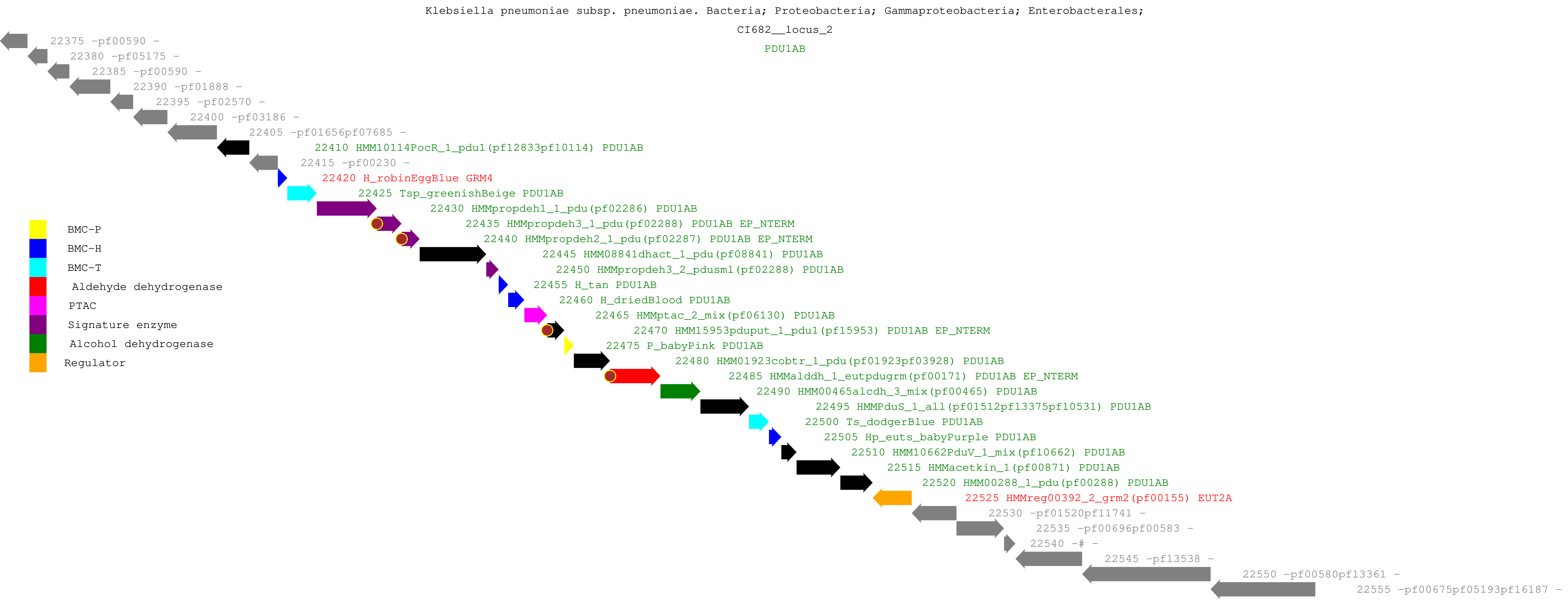
Klebsiella pneumoniae subsp. pneumoniae. Bacteria; Proteobacteria; Gammaproteobacteria; Enterobacterales;
CI682__locus_2
PDU1AB
22375 -pf00590 -
22380 -pf05175 -
22385 -pf00590 -
22390 -pf01888 -
22395 -pf02570 -
22400 -pf03186 -
22405 -pf01656pf07685 -
22410 HMM10114PocR_1_pdu1(pf12833pf10114) PDU1AB
22415 -pf00230 -
22420 H_robinEggBlue GRM4
22425 Tsp_greenishBeige PDU1AB
22430 HMMpropdeh1_1_pdu(pf02286) PDU1AB
22435 HMMpropdeh3_1_pdu(pf02288) PDU1AB EP_NTERM
22440 HMMpropdeh2_1_pdu(pf02287) PDU1AB EP_NTERM
22445 HMM08841dhact_1_pdu(pf08841) PDU1AB
22450 HMMpropdeh3_2_pdusml(pf02288) PDU1AB
22455 H_tan PDU1AB
22460 H_driedBlood PDU1AB
22465 HMMptac_2_mix(pf06130) PDU1AB
22470 HMM15953pduput_1_pdu1(pf15953) PDU1AB EP_NTERM
22475 P_babyPink PDU1AB
22480 HMM01923cobtr_1_pdu(pf01923pf03928) PDU1AB
22485 HMMalddh_1_eutpdugrm(pf00171) PDU1AB EP_NTERM
22490 HMM00465alcdh_3_mix(pf00465) PDU1AB
22495 HMMPduS_1_all(pf01512pf13375pf10531) PDU1AB
22500 Ts_dodgerBlue PDU1AB
22505 Hp_euts_babyPurple PDU1AB
22510 HMM10662PduV_1_mix(pf10662) PDU1AB
22515 HMMacetkin_1(pf00871) PDU1AB
22520 HMM00288_1_pdu(pf00288) PDU1AB
22525 HMMreg00392_2_grm2(pf00155) EUT2A
22530 -pf01520pf11741 -
22535 -pf00696pf00583 -
22540 -# -
22545 -pf13538 -
22550 -pf00580pf13361 -
22555 -pf00675pf05193pf16187 -
BMC-P
BMC-H
BMC-T
Aldehyde dehydrogenase
PTAC
Signature enzyme
Alcohol dehydrogenase
Regulator

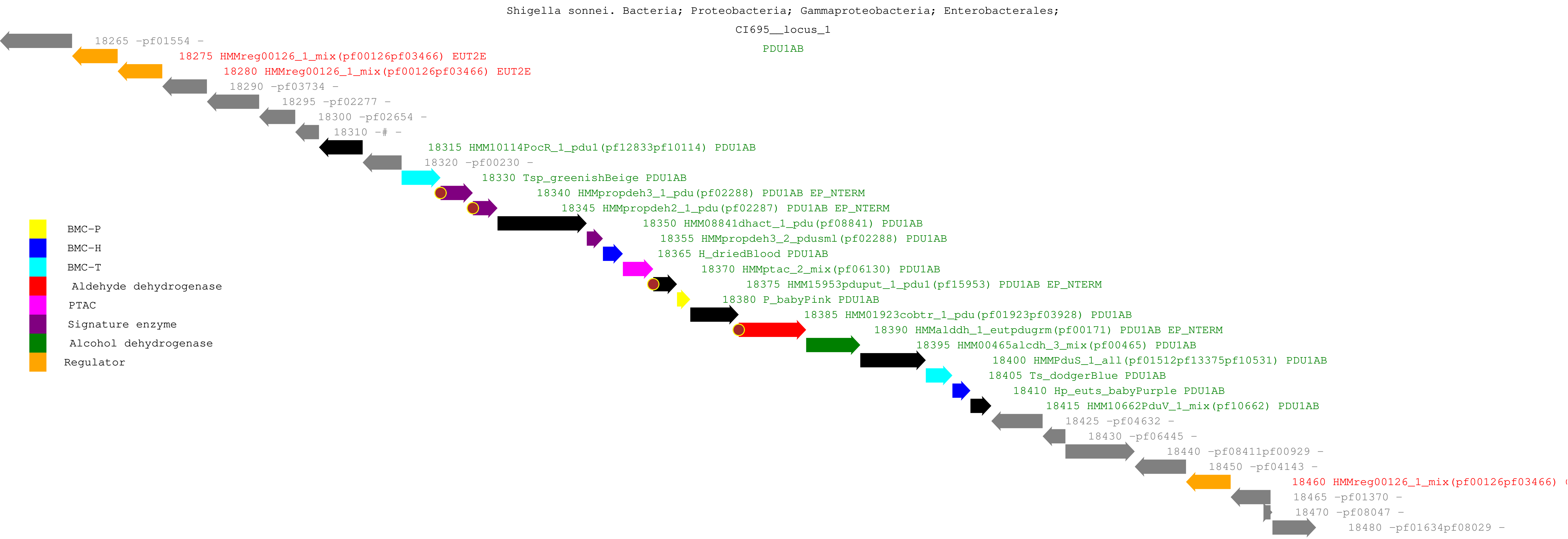
Shigella sonnei. Bacteria; Proteobacteria; Gammaproteobacteria; Enterobacterales;
CI695__locus_1
PDU1AB
18265 -pf01554 -
18275 HMMreg00126_1_mix(pf00126pf03466) EUT2E
18280 HMMreg00126_1_mix(pf00126pf03466) EUT2E
18290 -pf03734 -
18295 -pf02277 -
18300 -pf02654 -
18310 -# -
18315 HMM10114PocR_1_pdu1(pf12833pf10114) PDU1AB
18320 -pf00230 -
18330 Tsp_greenishBeige PDU1AB
18340 HMMpropdeh3_1_pdu(pf02288) PDU1AB EP_NTERM
18345 HMMpropdeh2_1_pdu(pf02287) PDU1AB EP_NTERM
18350 HMM08841dhact_1_pdu(pf08841) PDU1AB
18355 HMMpropdeh3_2_pdusml(pf02288) PDU1AB
18365 H_driedBlood PDU1AB
18370 HMMptac_2_mix(pf06130) PDU1AB
18375 HMM15953pduput_1_pdu1(pf15953) PDU1AB EP_NTERM
18380 P_babyPink PDU1AB
18385 HMM01923cobtr_1_pdu(pf01923pf03928) PDU1AB
18390 HMMaldh_1_eutpdugrm(pf00171) PDU1AB EP_NTERM
18395 HMM00465alcdh_3_mix(pf00465) PDU1AB
18400 HMMPduS_1_all(pf01512pf13375pf10531) PDU1AB
18405 Ts_dodgerBlue PDU1AB
18410 Hp_euts_babyPurple PDU1AB
18415 HMM10662PduV_1_mix(pf10662) PDU1AB
18425 -pf04632 -
18430 -pf06445 -
18440 -pf08411pf00929 -
18450 -pf04143 -
18460 HMMreg00126_1_mix(pf00126pf03466)
18465 -pf01370 -
18470 -pf08047 -
18480 -pf01634pf08029 -
BMC-P
BMC-H
BMC-T
Aldehyde dehydrogenase
PTAC
Signature enzyme
Alcohol dehydrogenase
Regulator

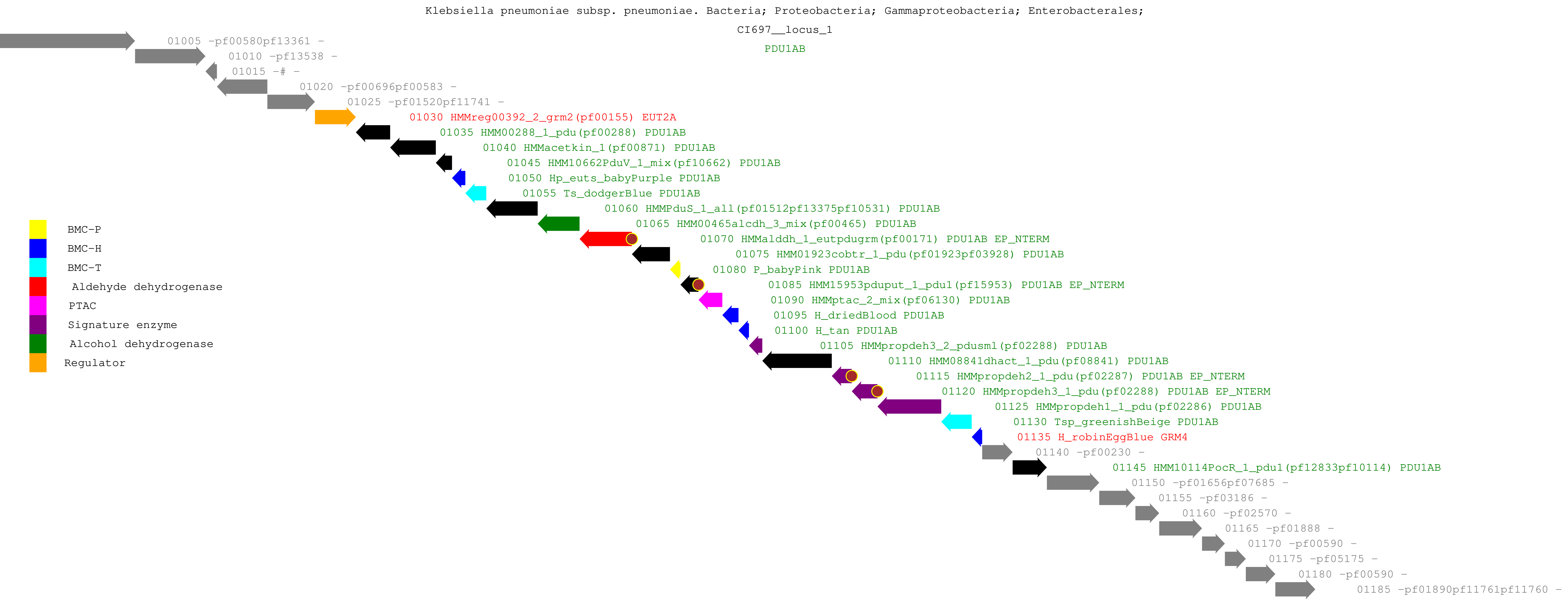
Klebsiella pneumoniae subsp. pneumoniae. Bacteria; Proteobacteria; Gammaproteobacteria; Enterobacterales;
CI697__locus_1
PDU1AB
01005 -pf00580pf13361 -
01010 -pf13538 -
01015 -# -
01020 -pf00696pf00583 -
01025 -pf01520pf11741 -
01030 HMMreg00392_2_grm2(pf00155) EUT2A
01035 HMM00288_1_pdu(pf00288) PDU1AB
01040 HMMacetkin_1(pf00871) PDU1AB
01045 HMM10662PduV_1_mix(pf10662) PDU1AB
01050 Hp_euts_babyPurple PDU1AB
01055 Ts_dodgerBlue PDU1AB
01060 HMMPduS_1_all(pf01512pf13375pf10531) PDU1AB
01065 HMM00465alcdh_3_mix(pf00465) PDU1AB
01070 HMMalddh_1_eutpdugrm(pf00171) PDU1AB EP_NTERM
01075 HMM01923cobtr_1_pdu(pf01923pf03928) PDU1AB
01080 P_babyPink PDU1AB
01085 HMM15953pduput_1_pdu1(pf15953) PDU1AB EP_NTERM
01090 HMMptac_2_mix(pf06130) PDU1AB
01095 H_driedBlood PDU1AB
01100 H_tan PDU1AB
01105 HMMpropdeh3_2_pdusml(pf02288) PDU1AB
01110 HMM08841dhact_1_pdu(pf08841) PDU1AB
01115 HMMpropdeh2_1_pdu(pf02287) PDU1AB EP_NTERM
01120 HMMpropdeh3_1_pdu(pf02288) PDU1AB EP_NTERM
01125 HMMpropdeh1_1_pdu(pf02286) PDU1AB
01130 Tsp_greenishBeige PDU1AB
01135 H_robinEggBlue GRM4
01140 -pf00230 -
01145 HMM10114PocR_1_pdu1(pf12833pf10114) PDU1AB
01150 -pf01656pf07685 -
01155 -pf03186 -
01160 -pf02570 -
01165 -pf01888 -
01170 -pf00590 -
01175 -pf05175 -
01180 -pf00590 -
01185 -pf01890pf11761pf11760 -
BMC-P
BMC-H
BMC-T
Aldehyde dehydrogenase
PTAC
Signature enzyme
Alcohol dehydrogenase
Regulator

Klebsiella pneumoniae subsp. pneumoniae. Bacteria; Proteobacteria; Gammaproteobacteria; Enterobacterales;

CI739__locus_1

PDU1AB

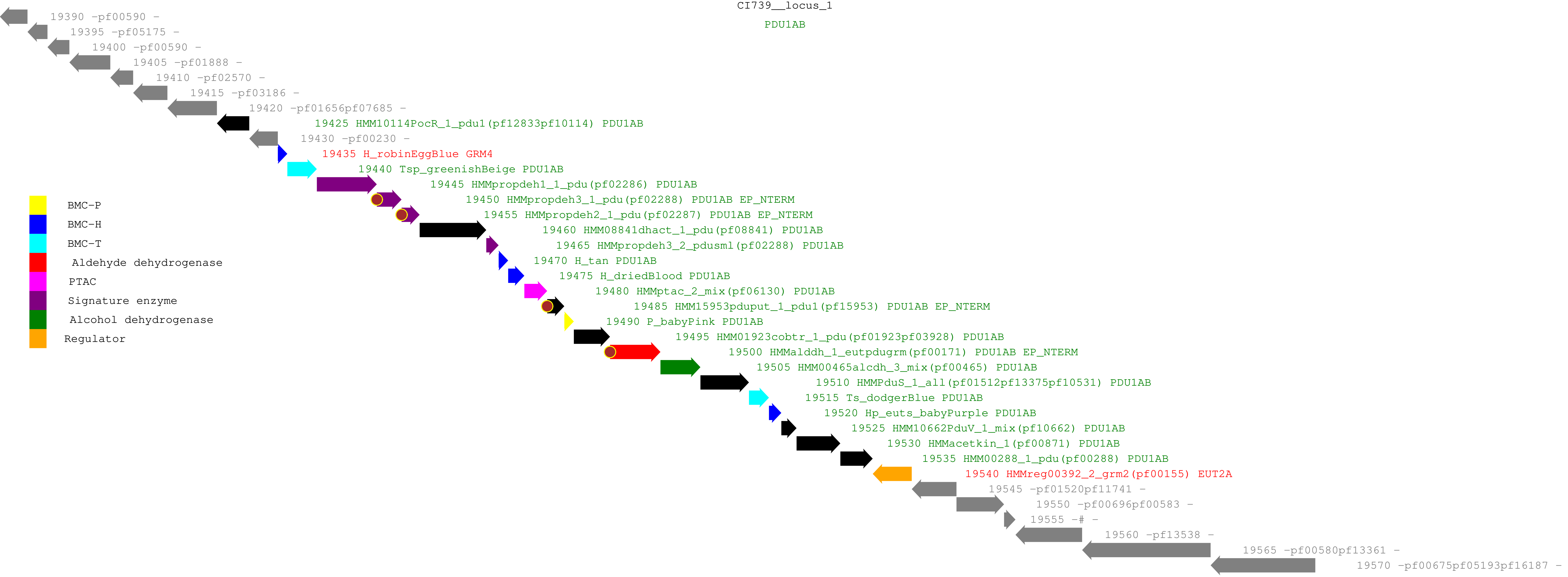

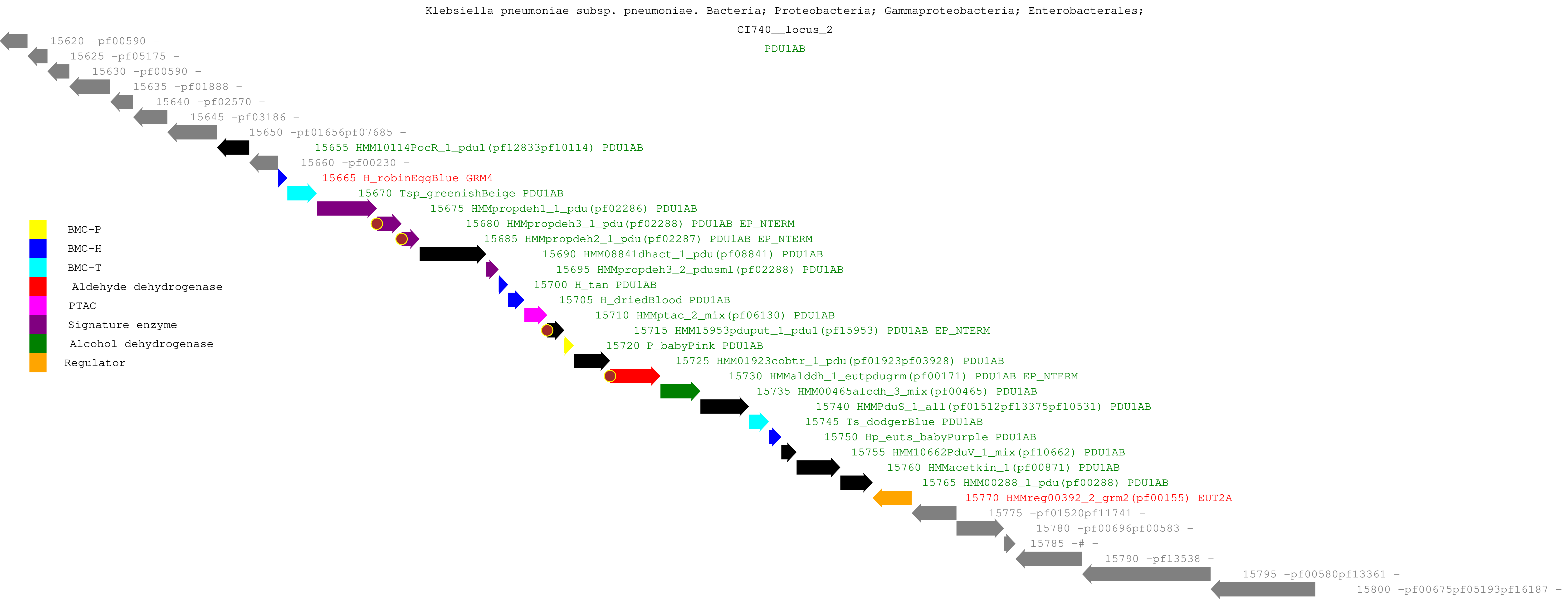

Klebsiella pneumoniae subsp. pneumoniae. Bacteria; Proteobacteria; Gammaproteobacteria; Enterobacterales;
CI740__locus_2
PDU1AB
15620 -pf00590 -
15625 -pf05175 -
15630 -pf00590 -
15635 -pf01888 -
15640 -pf02570 -
15645 -pf03186 -
15650 -pf01656pf07685 -
15655 HMM10114PocR_1_pdu1(pf12833pf10114) PDU1AB
15660 -pf00230 -
15665 H_robinEggBlue GRM4
15670 Tsp_greenishBeige PDU1AB
15675 HMMpropdeh1_1_pdu(pf02286) PDU1AB
15680 HMMpropdeh3_1_pdu(pf02288) PDU1AB EP_NTERM
15685 HMMpropdeh2_1_pdu(pf02287) PDU1AB EP_NTERM
15690 HMM08841dhact_1_pdu(pf08841) PDU1AB
15695 HMMpropdeh3_2_pdusml(pf02288) PDU1AB
15700 H_tan PDU1AB
15705 H_driedBlood PDU1AB
15710 HMMptac_2_mix(pf06130) PDU1AB
15715 HMM15953pduput_1_pdu1(pf15953) PDU1AB EP_NTERM
15720 P_babyPink PDU1AB
15725 HMM01923cobtr_1_pdu(pf01923pf03928) PDU1AB
15730 HMMalddh_1_eutpdugrm(pf00171) PDU1AB EP_NTERM
15735 HMM00465alcdh_3_mix(pf00465) PDU1AB
15740 HMMPduS_1_all(pf01512pf13375pf10531) PDU1AB
15745 Ts_dodgerBlue PDU1AB
15750 Hp_euts_babyPurple PDU1AB
15755 HMM10662PduV_1_mix(pf10662) PDU1AB
15760 HMMacetkin_1(pf00871) PDU1AB
15765 HMM00288_1_pdu(pf00288) PDU1AB
15770 HMMreg00392_2_grm2(pf00155) EUT2A
15775 -pf01520pf11741 -
15780 -pf00696pf00583 -
15785 -# -
15790 -pf13538 -
15795 -pf00580pf13361 -
15800 -pf00675pf05193pf16187 -
BMC-P
BMC-H
BMC-T
Aldehyde dehydrogenase
PTAC
Signature enzyme
Alcohol dehydrogenase
Regulator

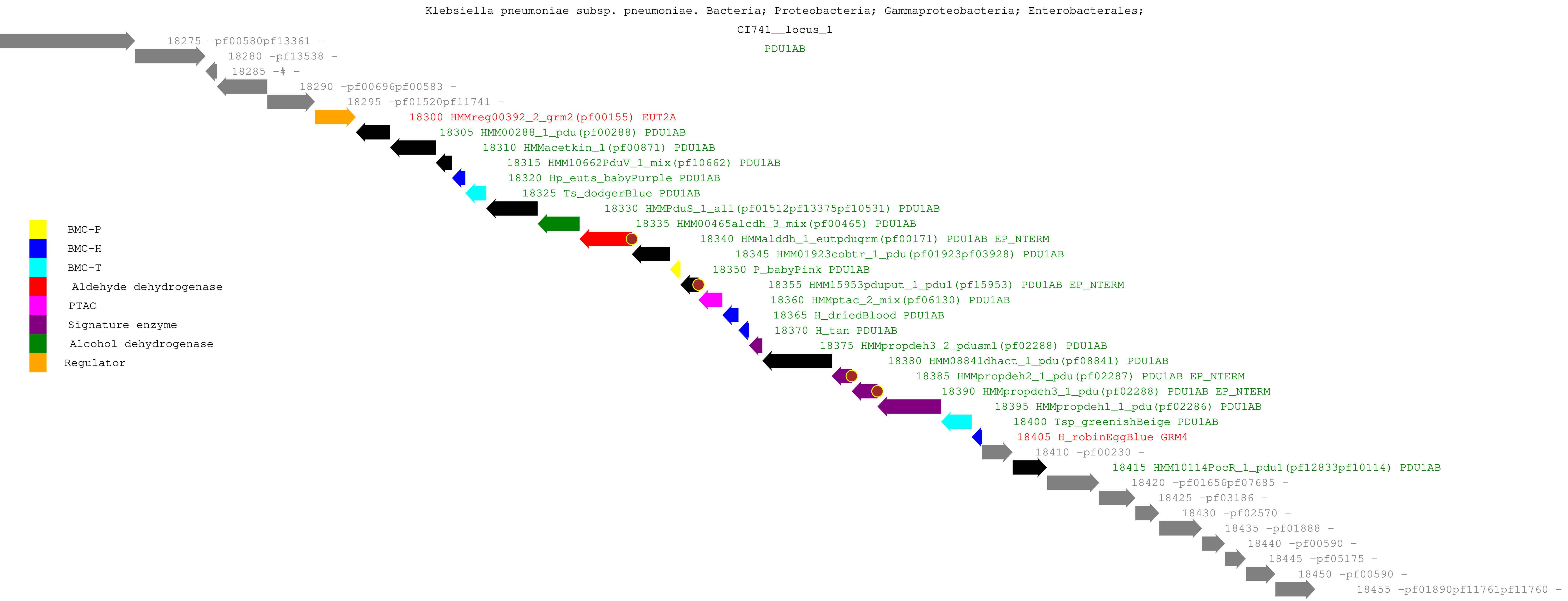

Klebsiella pneumoniae subsp. pneumoniae. Bacteria; Proteobacteria; Gammaproteobacteria; Enterobacterales;
CI741__locus_1
PDU1AB
18275 -pf00580pf13361 -
18280 -pf13538 -
18285 -# -
18290 -pf00696pf00583 -
18295 -pf01520pf11741 -
18300 HMMreg00392_2_grm2(pf00155) EUT2A
18305 HMM00288_1_pdu(pf00288) PDU1AB
18310 HMMacetkin_1(pf00871) PDU1AB
18315 HMM10662PduV_1_mix(pf10662) PDU1AB
18320 Hp_euts_babyPurple PDU1AB
18325 Ts_dodgerBlue PDU1AB
18330 HMMPduS_1_all(pf01512pf13375pf10531) PDU1AB
18335 HMM00465alcdh_3_mix(pf00465) PDU1AB
18340 HMMalddh_1_eutpdugrm(pf00171) PDU1AB EP_NTERM
18345 HMM01923cobtr_1_pdu(pf01923pf03928) PDU1AB
18350 P_babyPink PDU1AB
18355 HMM15953pduput_1_pdu1(pf15953) PDU1AB EP_NTERM
18360 HMMptac_2_mix(pf06130) PDU1AB
18365 H_driedBlood PDU1AB
18370 H_tan PDU1AB
18375 HMMpropdeh3_2_pdusml(pf02288) PDU1AB
18380 HMM08841dhact_1_pdu(pf08841) PDU1AB
18385 HMMpropdeh2_1_pdu(pf02287) PDU1AB EP_NTERM
18390 HMMpropdeh3_1_pdu(pf02288) PDU1AB EP_NTERM
18395 HMMpropdeh1_1_pdu(pf02286) PDU1AB
18400 Tsp_greenishBeige PDU1AB
18405 H_robinEggBlue GRM4
18410 -pf00230 -
18415 HMM10114PocR_1_pdu1(pf12833pf10114) PDU1AB
18420 -pf01656pf07685 -
18425 -pf03186 -
18430 -pf02570 -
18435 -pf01888 -
18440 -pf00590 -
18445 -pf05175 -
18450 -pf00590 -
18455 -pf01890pf11761pf11760 -
BMC-P
BMC-H
BMC-T
Aldehyde dehydrogenase
PTAC
Signature enzyme
Alcohol dehydrogenase
Regulator

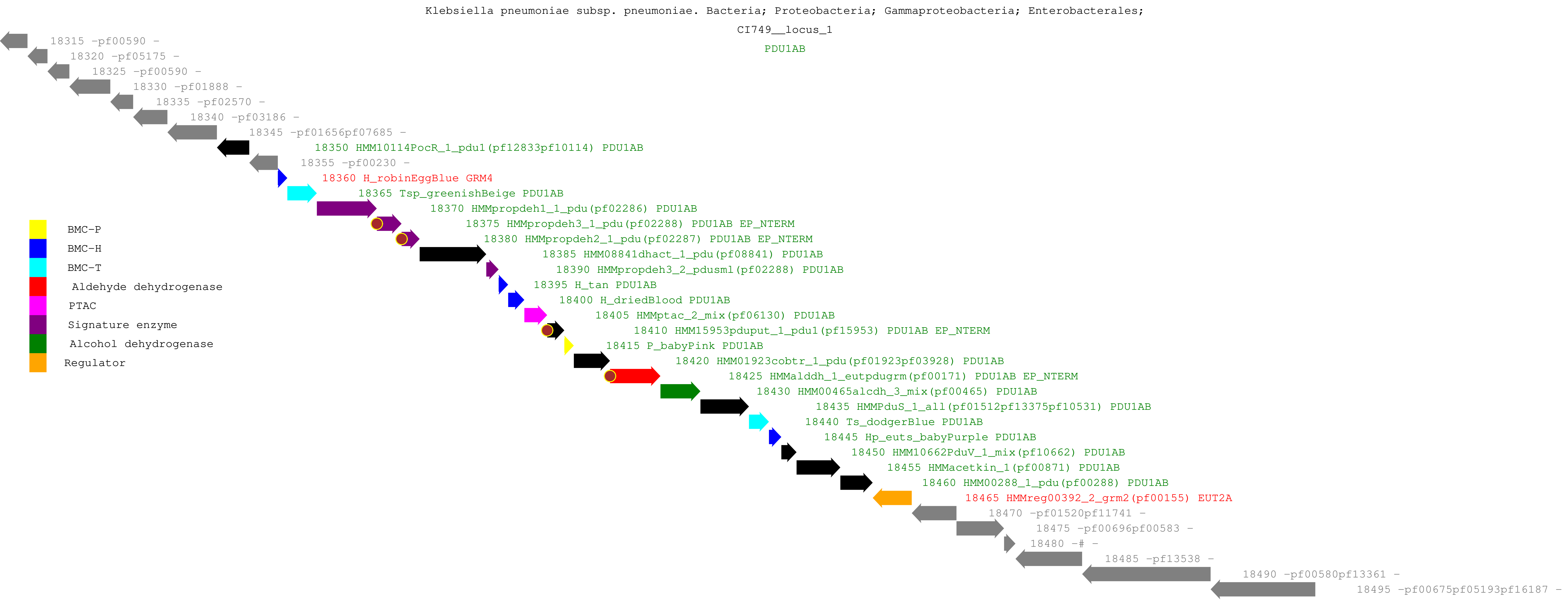
Klebsiella pneumoniae subsp. pneumoniae. Bacteria; Proteobacteria; Gammaproteobacteria; Enterobacterales;
CI749__locus_1
PDU1AB
18315 -pf00590 -
18320 -pf05175 -
18325 -pf00590 -
18330 -pf01888 -
18335 -pf02570 -
18340 -pf03186 -
18345 -pf01656pf07685 -
18350 HMM10114PocR_1_pdu1(pf12833pf10114) PDU1AB
18355 -pf00230 -
18360 H_robinEggBlue GRM4
18365 Tsp_greenishBeige PDU1AB
18370 HMMpropdeh1_1_pdu(pf02286) PDU1AB
18375 HMMpropdeh3_1_pdu(pf02288) PDU1AB EP_NTERM
18380 HMMpropdeh2_1_pdu(pf02287) PDU1AB EP_NTERM
18385 HMM08841dhact_1_pdu(pf08841) PDU1AB
18390 HMMpropdeh3_2_pdusml(pf02288) PDU1AB
18395 H_tan PDU1AB
18400 H_driedBlood PDU1AB
18405 HMMptac_2_mix(pf06130) PDU1AB
18410 HMM15953pduput_1_pdu1(pf15953) PDU1AB EP_NTERM
18415 P_babyPink PDU1AB
18420 HMM01923cobtr_1_pdu(pf01923pf03928) PDU1AB
18425 HMMaldh_1_eutpdugrm(pf00171) PDU1AB EP_NTERM
18430 HMM00465alcdh_3_mix(pf00465) PDU1AB
18435 HMMPduS_1_all(pf01512pf13375pf10531) PDU1AB
18440 Ts_dodgerBlue PDU1AB
18445 Hp_euts_babyPurple PDU1AB
18450 HMM10662PduV_1_mix(pf10662) PDU1AB
18455 HMMacetkin_1(pf00871) PDU1AB
18460 HMM00288_1_pdu(pf00288) PDU1AB
18465 HMMreg00392_2_grm2(pf00155) EUT2A
18470 -pf01520pf11741 -
18475 -pf00696pf00583 -
18480 -# -
18485 -pf13538 -
18490 -pf00580pf13361 -
18495 -pf00675pf05193pf16187 -
BMC-P
BMC-H
BMC-T
Aldehyde dehydrogenase
PTAC
Signature enzyme
Alcohol dehydrogenase
Regulator

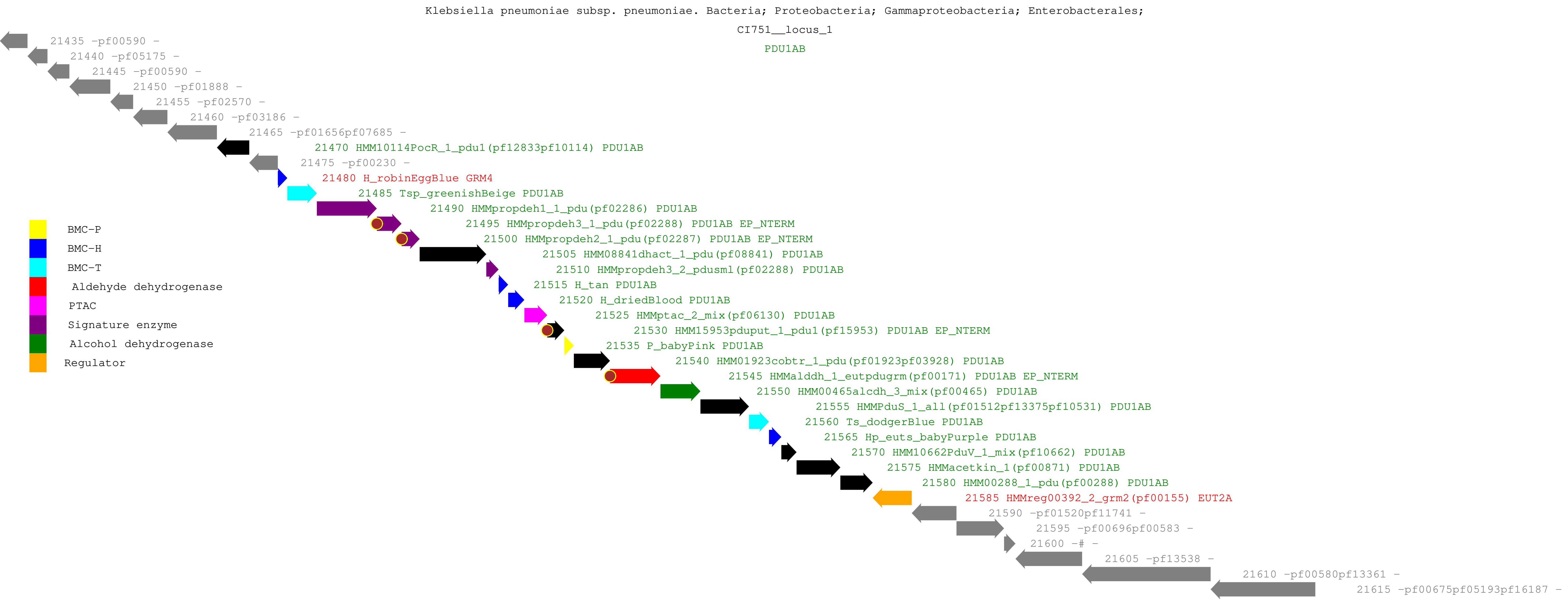
Klebsiella pneumoniae subsp. pneumoniae. Bacteria; Proteobacteria; Gammaproteobacteria; Enterobacterales;
CI751__locus_1
PDU1AB
21435 -pf00590 -
21440 -pf05175 -
21445 -pf00590 -
21450 -pf01888 -
21455 -pf02570 -
21460 -pf03186 -
21465 -pf01656pf07685 -
21470 HMM10114PocR_1_pdu1(pf12833pf10114) PDU1AB
21475 -pf00230 -
21480 H_robinEggBlue GRM4
21485 Tsp_greenishBeige PDU1AB
21490 HMMpropdeh1_1_pdu(pf02286) PDU1AB
21495 HMMpropdeh3_1_pdu(pf02288) PDU1AB EP_NTERM
21500 HMMpropdeh2_1_pdu(pf02287) PDU1AB EP_NTERM
21505 HMM08841dhact_1_pdu(pf08841) PDU1AB
21510 HMMpropdeh3_2_pdusml(pf02288) PDU1AB
21515 H_tan PDU1AB
21520 H_driedBlood PDU1AB
21525 HMMptac_2_mix(pf06130) PDU1AB
21530 HMM15953pduput_1_pdu1(pf15953) PDU1AB EP_NTERM
21535 P_babyPink PDU1AB
21540 HMM01923cobtr_1_pdu(pf01923pf03928) PDU1AB
21545 HMMalddh_1_eutpdugrm(pf00171) PDU1AB EP_NTERM
21550 HMM00465alcdh_3_mix(pf00465) PDU1AB
21555 HMMPduS_1_all(pf01512pf13375pf10531) PDU1AB
21560 Ts_dodgerBlue PDU1AB
21565 Hp_euts_babyPurple PDU1AB
21570 HMM10662PduV_1_mix(pf10662) PDU1AB
21575 HMMacetkin_1(pf00871) PDU1AB
21580 HMM00288_1_pdu(pf00288) PDU1AB
21585 HMMreg00392_2_grm2(pf00155) EUT2A
21590 -pf01520pf11741 -
21595 -pf00696pf00583 -
21600 -# -
21605 -pf13538 -
21610 -pf00580pf13361 -
21615 -pf00675pf05193pf16187 -
BMC-P
BMC-H
BMC-T
Aldehyde dehydrogenase
PTAC
Signature enzyme
Alcohol dehydrogenase
Regulator

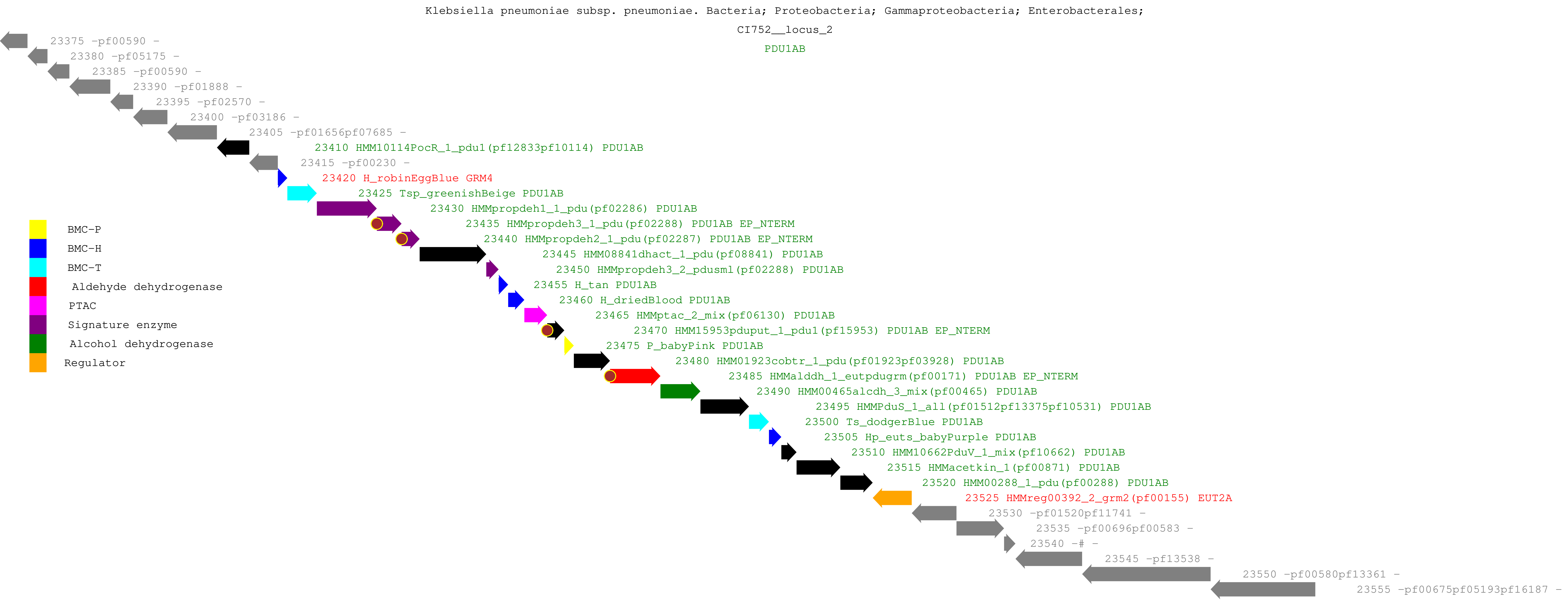
Klebsiella pneumoniae subsp. pneumoniae. Bacteria; Proteobacteria; Gammaproteobacteria; Enterobacterales;
CI752__locus_2
PDU1AB
23375 -pf00590 -
23380 -pf05175 -
23385 -pf00590 -
23390 -pf01888 -
23395 -pf02570 -
23400 -pf03186 -
23405 -pf01656pf07685 -
23410 HMM10114PocR_1_pdu1(pf12833pf10114) PDU1AB
23415 -pf00230 -
23420 H_robinEggBlue GRM4
23425 Tsp_greenishBeige PDU1AB
23430 HMMpropdeh1_1_pdu(pf02286) PDU1AB
23435 HMMpropdeh3_1_pdu(pf02288) PDU1AB EP_NTERM
23440 HMMpropdeh2_1_pdu(pf02287) PDU1AB EP_NTERM
23445 HMM08841dhact_1_pdu(pf08841) PDU1AB
23450 HMMpropdeh3_2_pdusml(pf02288) PDU1AB
23455 H_tan PDU1AB
23460 H_driedBlood PDU1AB
23465 HMMptac_2_mix(pf06130) PDU1AB
23470 HMM15953pduput_1_pdu1(pf15953) PDU1AB EP_NTERM
23475 P_babyPink PDU1AB
23480 HMM01923cobtr_1_pdu(pf01923pf03928) PDU1AB
23485 HMMalddh_1_eutpdugrm(pf00171) PDU1AB EP_NTERM
23490 HMM00465alcdh_3_mix(pf00465) PDU1AB
23495 HMMPduS_1_all(pf01512pf13375pf10531) PDU1AB
23500 Ts_dodgerBlue PDU1AB
23505 Hp_euts_babyPurple PDU1AB
23510 HMM10662PduV_1_mix(pf10662) PDU1AB
23515 HMMacetkin_1(pf00871) PDU1AB
23520 HMM00288_1_pdu(pf00288) PDU1AB
23525 HMMreg00392_2_grm2(pf00155) EUT2A
23530 -pf01520pf11741 -
23535 -pf00696pf00583 -
23540 -# -
23545 -pf13538 -
23550 -pf00580pf13361 -
23555 -pf00675pf05193pf16187 -
BMC-P
BMC-H
BMC-T
Aldehyde dehydrogenase
PTAC
Signature enzyme
Alcohol dehydrogenase
Regulator

Salmonella choleraesuis. Bacteria; Proteobacteria; Gammaproteobacteria; Enterobacterales;

CIC26__locus_1

PDU1AB

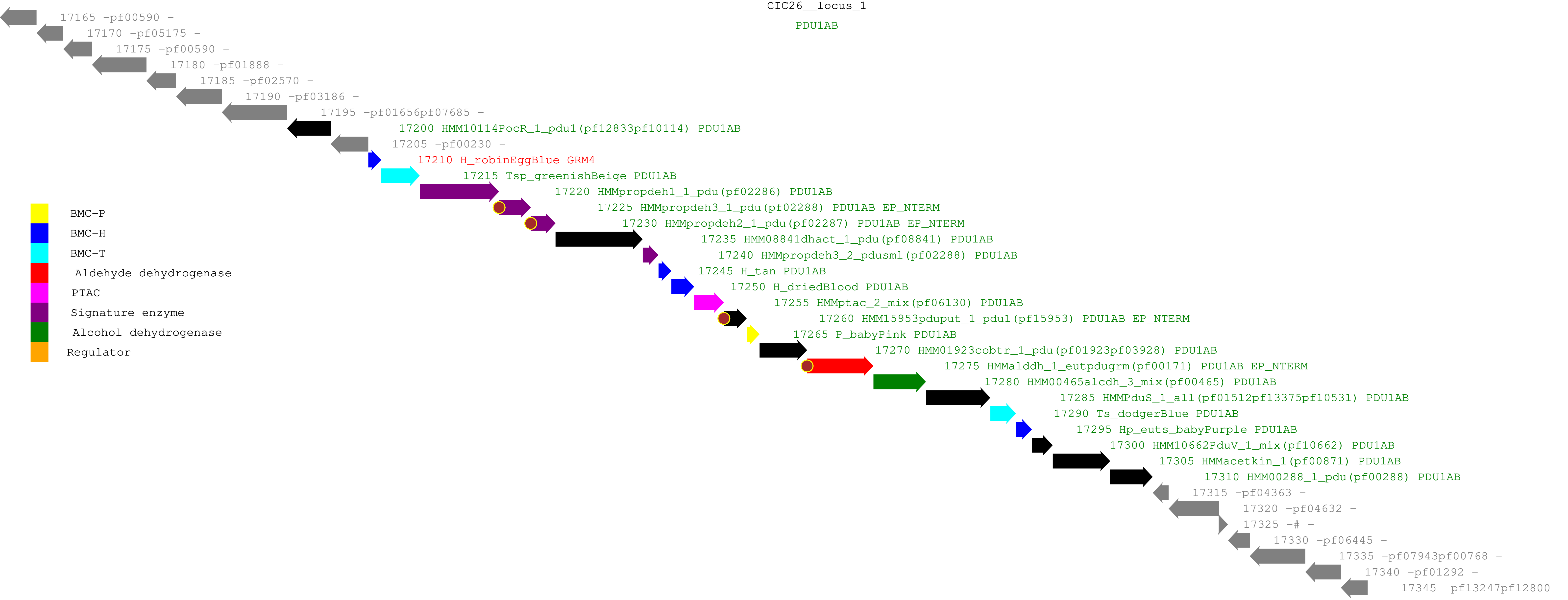

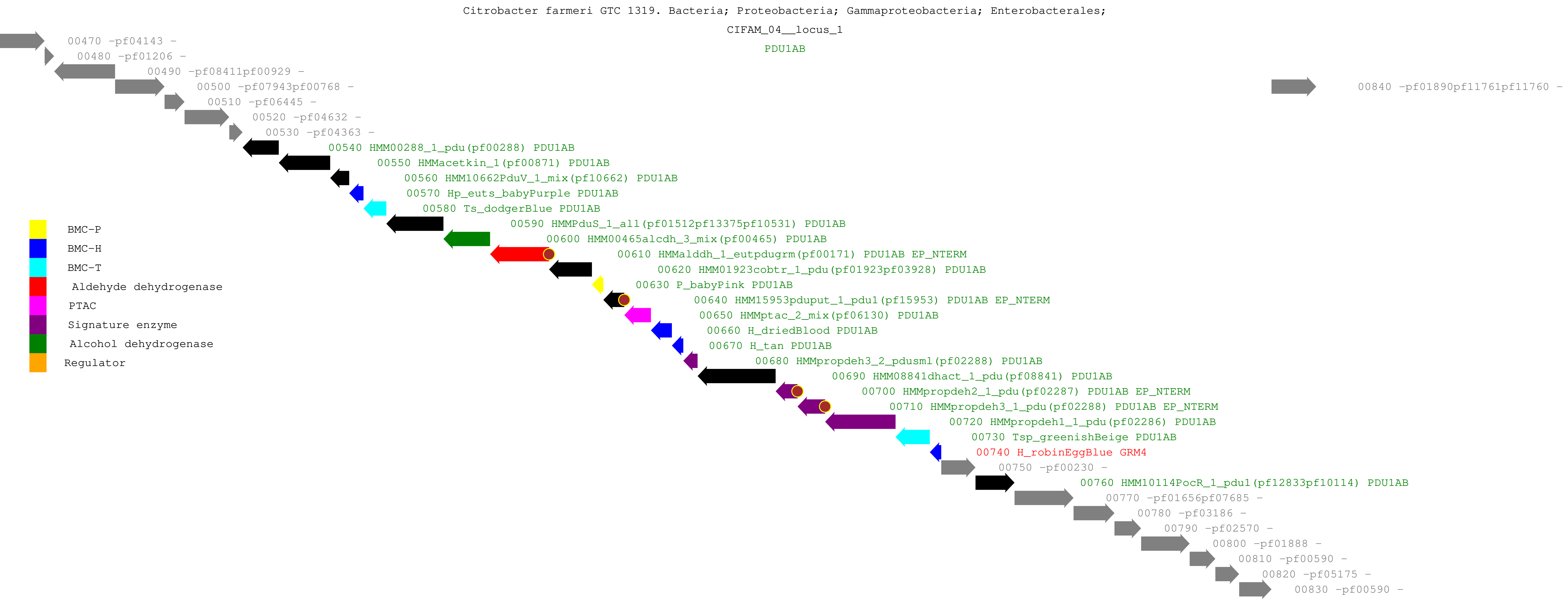
Citrobacter farmeri GTC 1319. Bacteria; Proteobacteria; Gammaproteobacteria; Enterobacterales;
CIFAM_04__locus_1
PDU1AB
00470 -pf04143 -
00480 -pf01206 -
00490 -pf08411pf00929 -
00500 -pf07943pf00768 -
00510 -pf06445 -
00520 -pf04632 -
00530 -pf04363 -
00540 HMM00288_1_pdu(pf00288) PDU1AB
00550 HMMacetkin_1(pf00871) PDU1AB
00560 HMM10662PduV_1_mix(pf10662) PDU1AB
00570 Hp_euts_babyPurple PDU1AB
00580 Ts_dodgerBlue PDU1AB
00590 HMMPduS_1_all(pf01512pf13375pf10531) PDU1AB
00600 HMM00465alcdh_3_mix(pf00465) PDU1AB
00610 HMMaldddh_1_eutpdugrm(pf00171) PDU1AB EP_NTERM
00620 HMM01923cobtr_1_pdu(pf01923pf03928) PDU1AB
00630 P_babyPink PDU1AB
00640 HMM15953pduput_1_pdu1(pf15953) PDU1AB EP_NTERM
00650 HMMptac_2_mix(pf06130) PDU1AB
00660 H_driedBlood PDU1AB
00670 H_tan PDU1AB
00680 HMMpropdeh3_2_pdusml(pf02288) PDU1AB
00690 HMM08841dhact_1_pdu(pf08841) PDU1AB
00700 HMMpropdeh2_1_pdu(pf02287) PDU1AB EP_NTERM
00710 HMMpropdeh3_1_pdu(pf02288) PDU1AB EP_NTERM
00720 HMMpropdeh1_1_pdu(pf02286) PDU1AB
00730 Tsp_greenishBeige PDU1AB
00740 H_robinEggBlue GRM4
00750 -pf00230 -
00760 HMM10114PocR_1_pdu1(pf12833pf10114) PDU1AB
00770 -pf01656pf07685 -
00780 -pf03186 -
00790 -pf02570 -
00800 -pf01888 -
00810 -pf00590 -
00820 -pf05175 -
00830 -pf00590 -
00840 -pf01890pf11761pf11760 -
BMC-P
BMC-H
BMC-T
Aldehyde dehydrogenase
PTAC
Signature enzyme
Alcohol dehydrogenase
Regulator

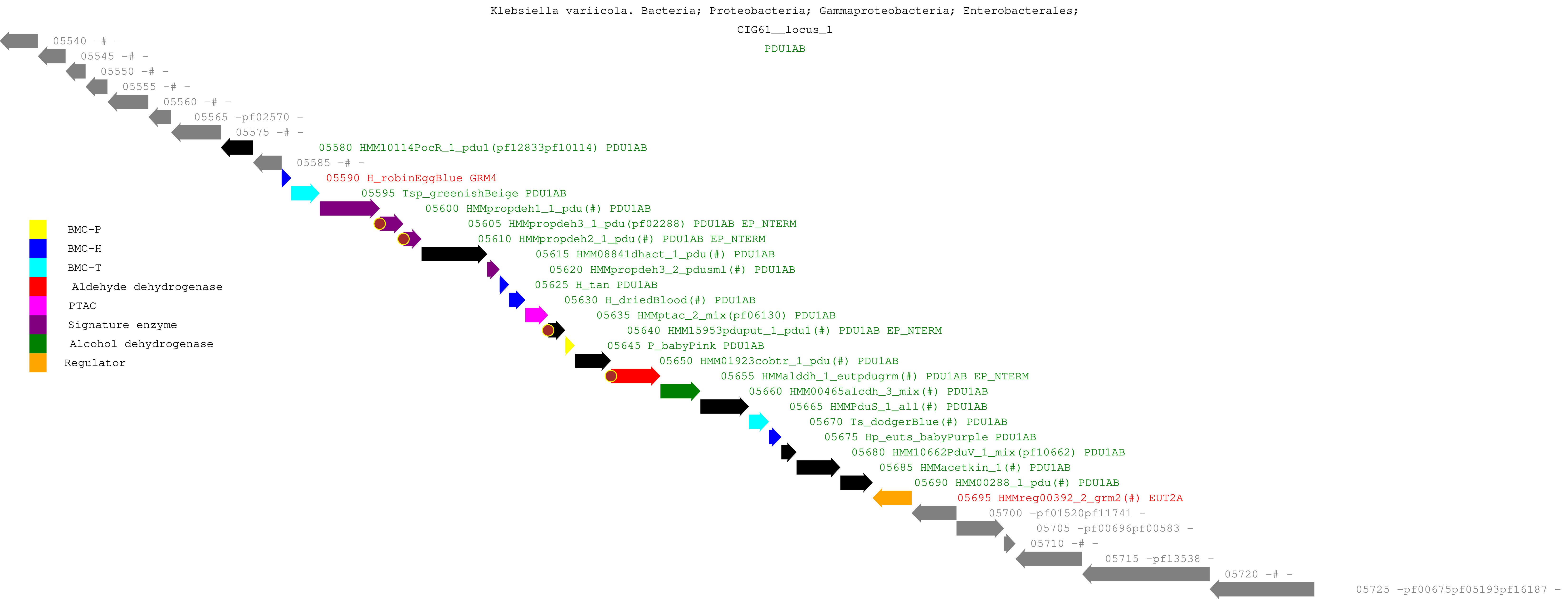

Klebsiella variicola. Bacteria; Proteobacteria; Gammaproteobacteria; Enterobacterales;
CIG61__locus_1
PDU1AB
05540 -# -
05545 -# -
05550 -# -
05555 -# -
05560 -# -
05565 -pf02570 -
05575 -# -
05580 HMM10114PocR_1_pdu1(pf12833pf10114) PDU1AB
05585 -# -
05590 H_robinEggBlue GRM4
05595 Tsp_greenishBeige PDU1AB
05600 HMMpropdeh1_1_pdu(#) PDU1AB
05605 HMMpropdeh3_1_pdu(pf02288) PDU1AB EP_NTERM
05610 HMMpropdeh2_1_pdu(#) PDU1AB EP_NTERM
05615 HMM08841dhact_1_pdu(#) PDU1AB
05620 HMMpropdeh3_2_pdusml(#) PDU1AB
05625 H_tan PDU1AB
05630 H_driedBlood(#) PDU1AB
05635 HMMptac_2_mix(pf06130) PDU1AB
05640 HMM15953pduput_1_pdu1(#) PDU1AB EP_NTERM
05645 P_babyPink PDU1AB
05650 HMM01923cobtr_1_pdu(#) PDU1AB
05655 HMMalddh_1_eutpdugrm(#) PDU1AB EP_NTERM
05660 HMM00465alcdh_3_mix(#) PDU1AB
05665 HMMPduS_1_all(#) PDU1AB
05670 Ts_dodgerBlue(#) PDU1AB
05675 Hp_euts_babyPurple PDU1AB
05680 HMM10662PduV_1_mix(pf10662) PDU1AB
05685 HMMacetkin_1(#) PDU1AB
05690 HMM00288_1_pdu(#) PDU1AB
05695 HMMreg00392_2_grm2(#) EUT2A
05700 -pf01520pf11741 -
05705 -pf00696pf00583 -
05710 -# -
05715 -pf13538 -
05720 -# -
05725 -pf00675pf05193pf16187 -
BMC-P
BMC-H
BMC-T
Aldehyde dehydrogenase
PTAC
Signature enzyme
Alcohol dehydrogenase
Regulator

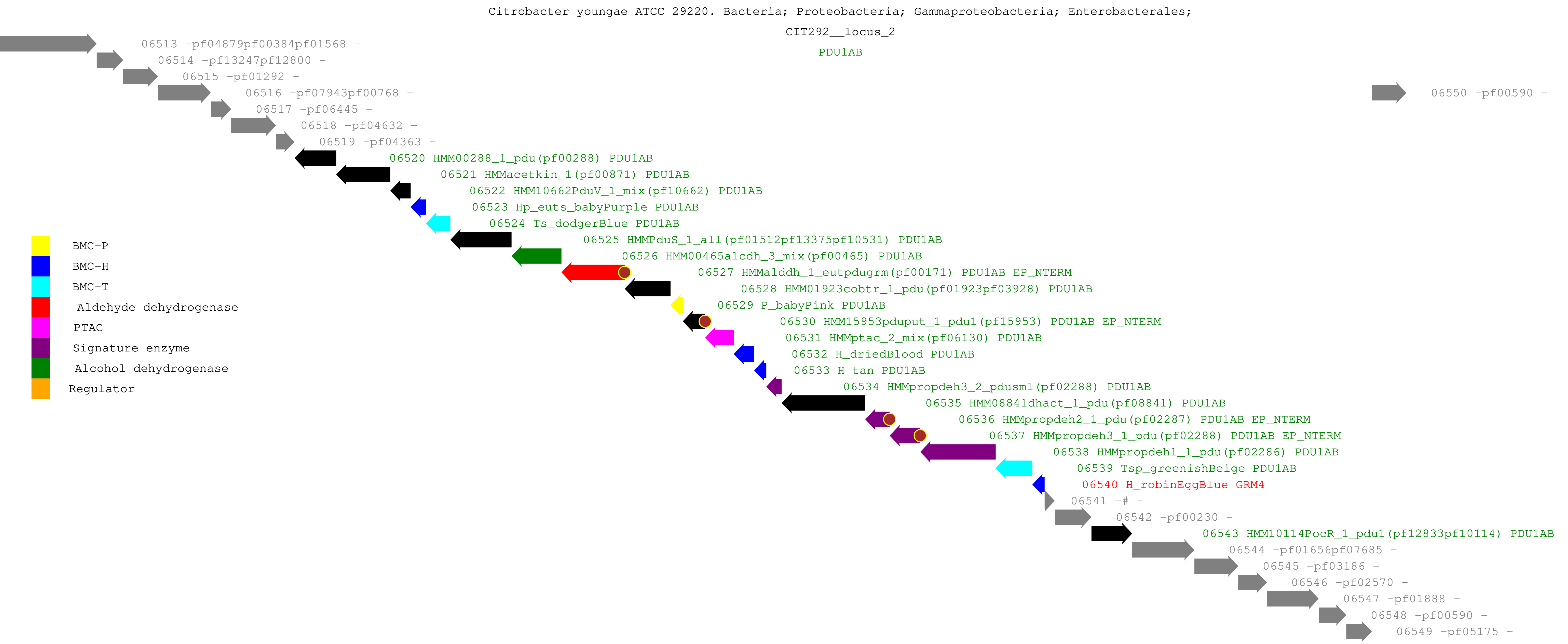

Citrobacter youngae ATCC 29220. Bacteria; Proteobacteria; Gammaproteobacteria; Enterobacterales;
CIT292__locus_2
PDU1AB
06513 -pf04879pf00384pf01568 -
06514 -pf13247pf12800 -
06515 -pf01292 -
06516 -pf07943pf00768 -
06517 -pf06445 -
06518 -pf04632 -
06519 -pf04363 -
06520 HMM00288_1_pdu(pf00288) PDU1AB
06521 HMMacetkin_1(pf00871) PDU1AB
06522 HMM10662PduV_1_mix(pf10662) PDU1AB
06523 Hp_euts_babyPurple PDU1AB
06524 Ts_dodgerBlue PDU1AB
06525 HMMPduS_1_all(pf01512pf13375pf10531) PDU1AB
06526 HMM00465alcdh_3_mix(pf00465) PDU1AB
06527 HMMalddh_1_eutpdugrm(pf00171) PDU1AB EP_NTERM
06528 HMM01923cobtr_1_pdu(pf01923pf03928) PDU1AB
06529 P_babyPink PDU1AB
06530 HMM15953pduput_1_pdu1(pf15953) PDU1AB EP_NTERM
06531 HMMptac_2_mix(pf06130) PDU1AB
06532 H_driedBlood PDU1AB
06533 H_tan PDU1AB
06534 HMMpropdeh3_2_pdusml(pf02288) PDU1AB
06535 HMM08841dhact_1_pdu(pf08841) PDU1AB
06536 HMMpropdeh2_1_pdu(pf02287) PDU1AB EP_NTERM
06537 HMMpropdeh3_1_pdu(pf02288) PDU1AB EP_NTERM
06538 HMMpropdeh1_1_pdu(pf02286) PDU1AB
06539 Tsp_greenishBeige PDU1AB
06540 H_robinEggBlue GRM4
06541 -# -
06542 -pf00230 -
06543 HMM10114PocR_1_pdu1(pf12833pf10114) PDU1AB
06544 -pf01656pf07685 -
06545 -pf03186 -
06546 -pf02570 -
06547 -pf01888 -
06548 -pf00590 -
06549 -pf05175 -
06550 -pf00590 -
BMC-P
BMC-H
BMC-T
Aldehyde dehydrogenase
PTAC
Signature enzyme
Alcohol dehydrogenase
Regulator

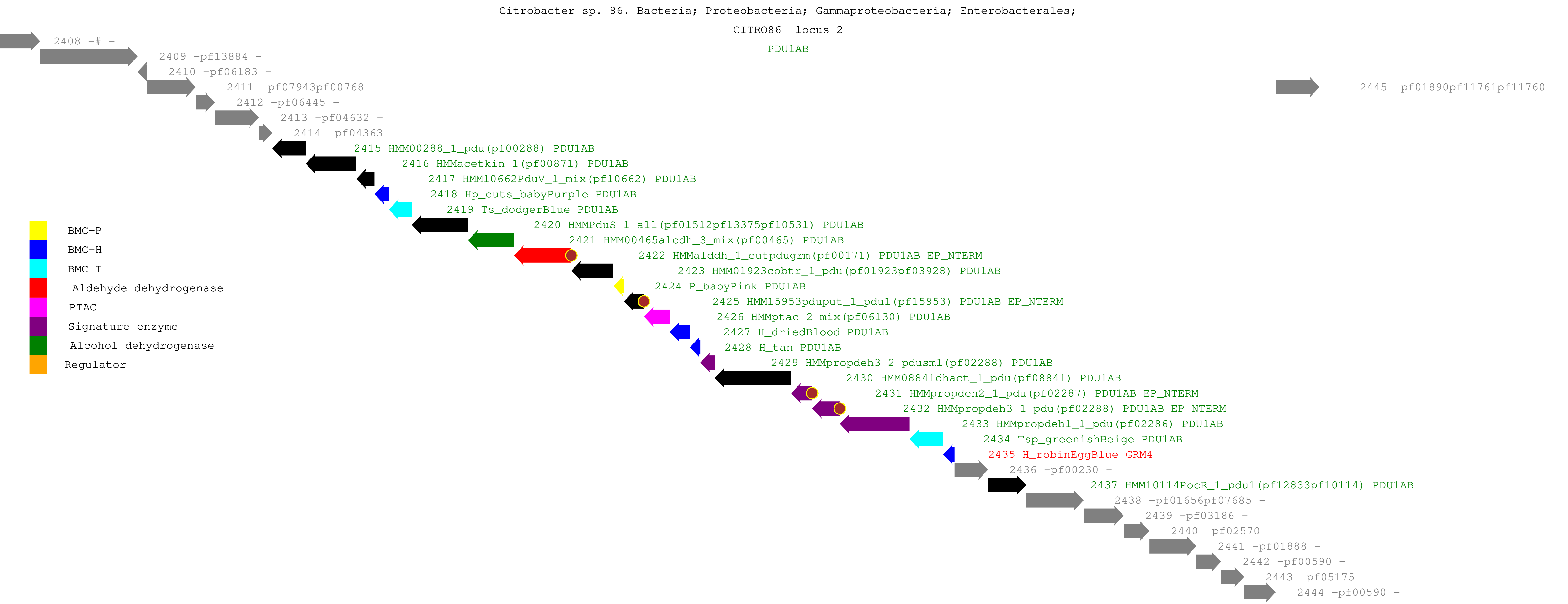
Citrobacter sp. 86. Bacteria; Proteobacteria; Gammaproteobacteria; Enterobacterales;
CITRO86__locus_2
PDU1AB
2408 -# -
2409 -pf13884 -
2410 -pf06183 -
2411 -pf07943pf00768 -
2412 -pf06445 -
2413 -pf04632 -
2414 -pf04363 -
2415 HMM00288_1_pdu(pf00288) PDU1AB
2416 HMMacetkin_1(pf00871) PDU1AB
2417 HMM10662PduV_1_mix(pf10662) PDU1AB
2418 Hp_euts_babyPurple PDU1AB
2419 Ts_dodgerBlue PDU1AB
2420 HMMPduS_1_all(pf01512pf13375pf10531) PDU1AB
2421 HMM00465alcdh_3_mix(pf00465) PDU1AB
2422 HMMalddh_1_eutpdugrm(pf00171) PDU1AB EP_NTERM
2423 HMM01923cobtr_1_pdu(pf01923pf03928) PDU1AB
2424 P_babyPink PDU1AB
2425 HMM15953pduput_1_pdu1(pf15953) PDU1AB EP_NTERM
2426 HMMptac_2_mix(pf06130) PDU1AB
2427 H_driedBlood PDU1AB
2428 H_tan PDU1AB
2429 HMMpropdeh3_2_pdusml(pf02288) PDU1AB
2430 HMM08841dhact_1_pdu(pf08841) PDU1AB
2431 HMMpropdeh2_1_pdu(pf02287) PDU1AB EP_NTERM
2432 HMMpropdeh3_1_pdu(pf02288) PDU1AB EP_NTERM
2433 HMMpropdeh1_1_pdu(pf02286) PDU1AB
2434 Tsp_greenishBeige PDU1AB
2435 H_robinEggBlue GRM4
2436 -pf00230 -
2437 HMM10114PocR_1_pdu1(pf12833pf10114) PDU1AB
2438 -pf01656pf07685 -
2439 -pf03186 -
2440 -pf02570 -
2441 -pf01888 -
2442 -pf00590 -
2443 -pf05175 -
2444 -pf00590 -
2445 -pf01890pf11761pf11760 -
BMC-P
BMC-H
BMC-T
Aldehyde dehydrogenase
PTAC
Signature enzyme
Alcohol dehydrogenase
Regulator

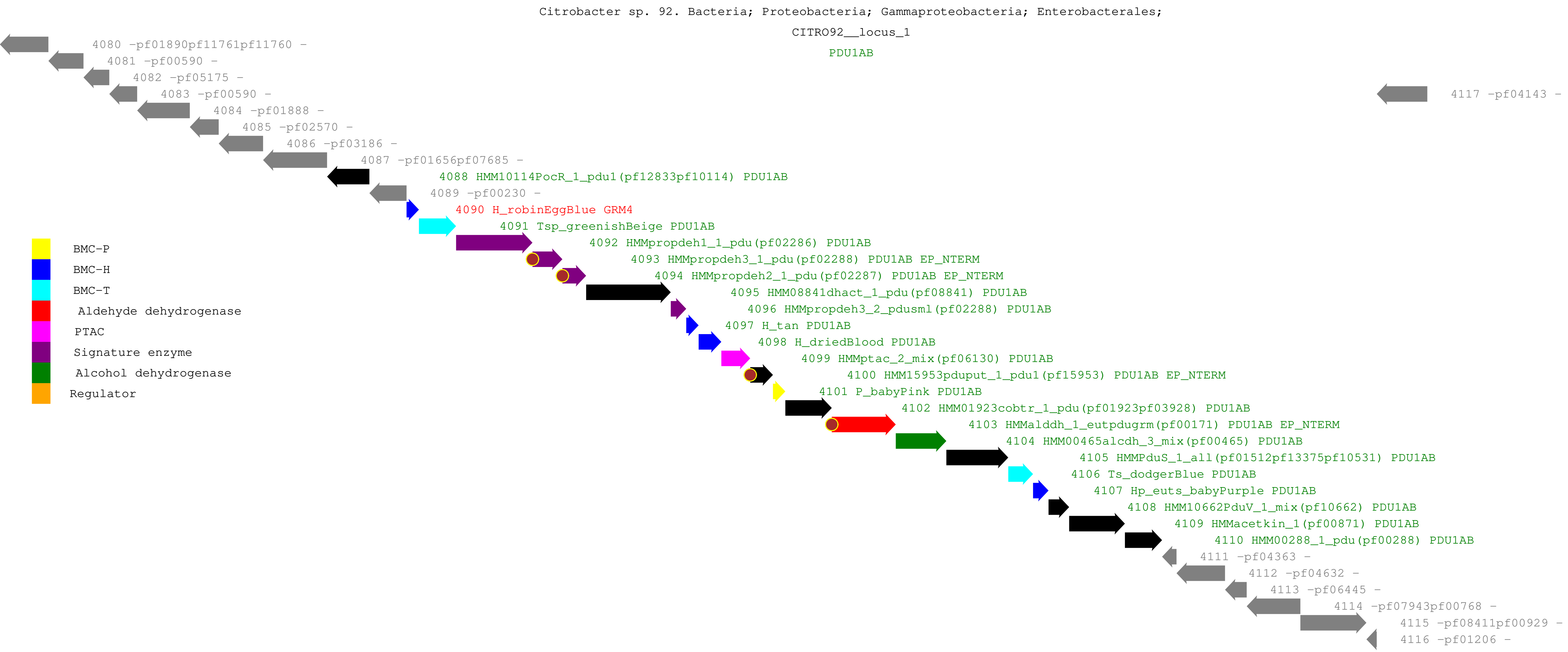
Citrobacter sp. 92. Bacteria; Proteobacteria; Gammaproteobacteria; Enterobacterales;
CITRO92__locus_1
PDU1AB
4080 -pf01890pf11761pf11760 -
4081 -pf00590 -
4082 -pf05175 -
4083 -pf00590 -
4084 -pf01888 -
4085 -pf02570 -
4086 -pf03186 -
4087 -pf01656pf07685 -
4088 HMM10114PocR_1_pdu1(pf12833pf10114) PDU1AB
4089 -pf00230 -
4090 H_robinEggBlue GRM4
4091 Tsp_greenishBeige PDU1AB
4092 HMMpropdeh1_1_pdu(pf02286) PDU1AB
4093 HMMpropdeh3_1_pdu(pf02288) PDU1AB EP_NTERM
4094 HMMpropdeh2_1_pdu(pf02287) PDU1AB EP_NTERM
4095 HMM08841dhact_1_pdu(pf08841) PDU1AB
4096 HMMpropdeh3_2_pdusml(pf02288) PDU1AB
4097 H_tan PDU1AB
4098 H_driedBlood PDU1AB
4099 HMMptac_2_mix(pf06130) PDU1AB
4100 HMM15953pduput_1_pdu1(pf15953) PDU1AB EP_NTERM
4101 P_babyPink PDU1AB
4102 HMM01923cobtr_1_pdu(pf01923pf03928) PDU1AB
4103 HMMaldh_1_eutpdugrm(pf00171) PDU1AB EP_NTERM
4104 HMM00465alcdh_3_mix(pf00465) PDU1AB
4105 HMMPduS_1_all(pf01512pf13375pf10531) PDU1AB
4106 Ts_dodgerBlue PDU1AB
4107 Hp_euts_babyPurple PDU1AB
4108 HMM10662PduV_1_mix(pf10662) PDU1AB
4109 HMMacetkin_1(pf00871) PDU1AB
4110 HMM00288_1_pdu(pf00288) PDU1AB
4111 -pf04363 -
4112 -pf04632 -
4113 -pf06445 -
4114 -pf07943pf00768 -
4115 -pf08411pf00929 -
4116 -pf01206 -
4117 -pf04143 -
BMC-P
BMC-H
BMC-T
Aldehyde dehydrogenase
PTAC
Signature enzyme
Alcohol dehydrogenase
Regulator

Citrobacter werkmanii NBRC 105721. Bacteria; Proteobacteria; Gammaproteobacteria; Enterobacterales;

CIWKM_03__locus_1

PDU1AB

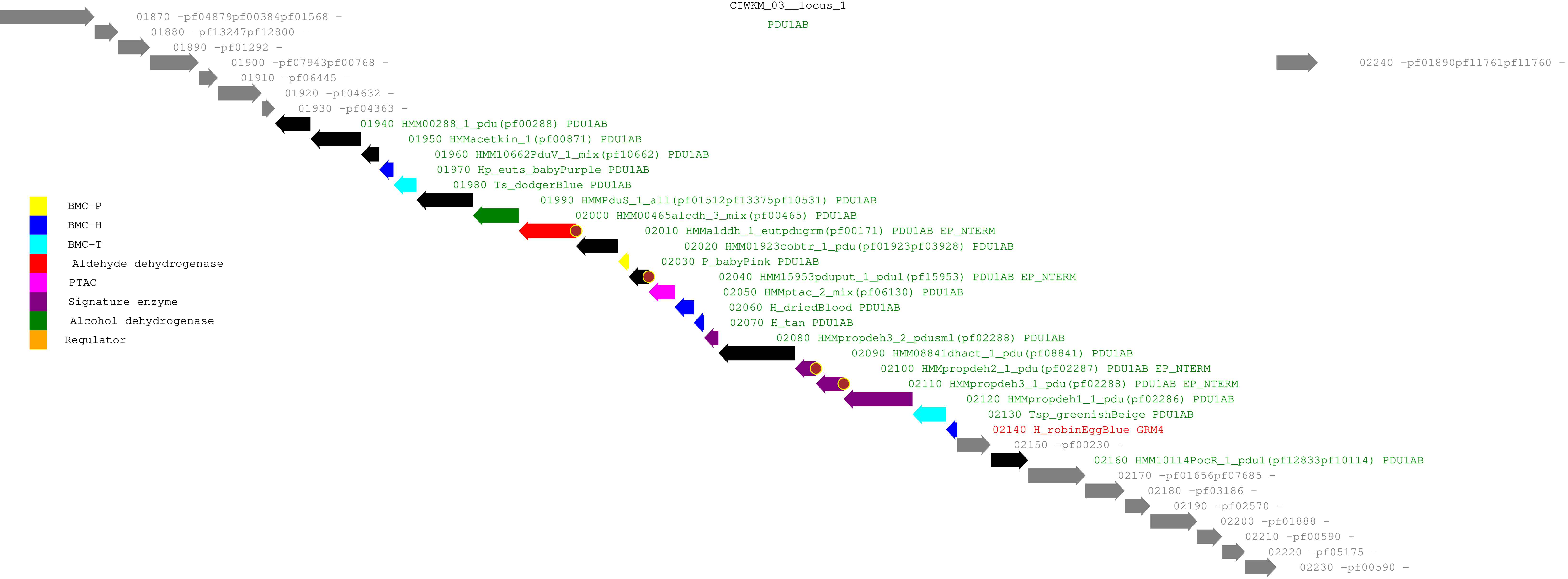

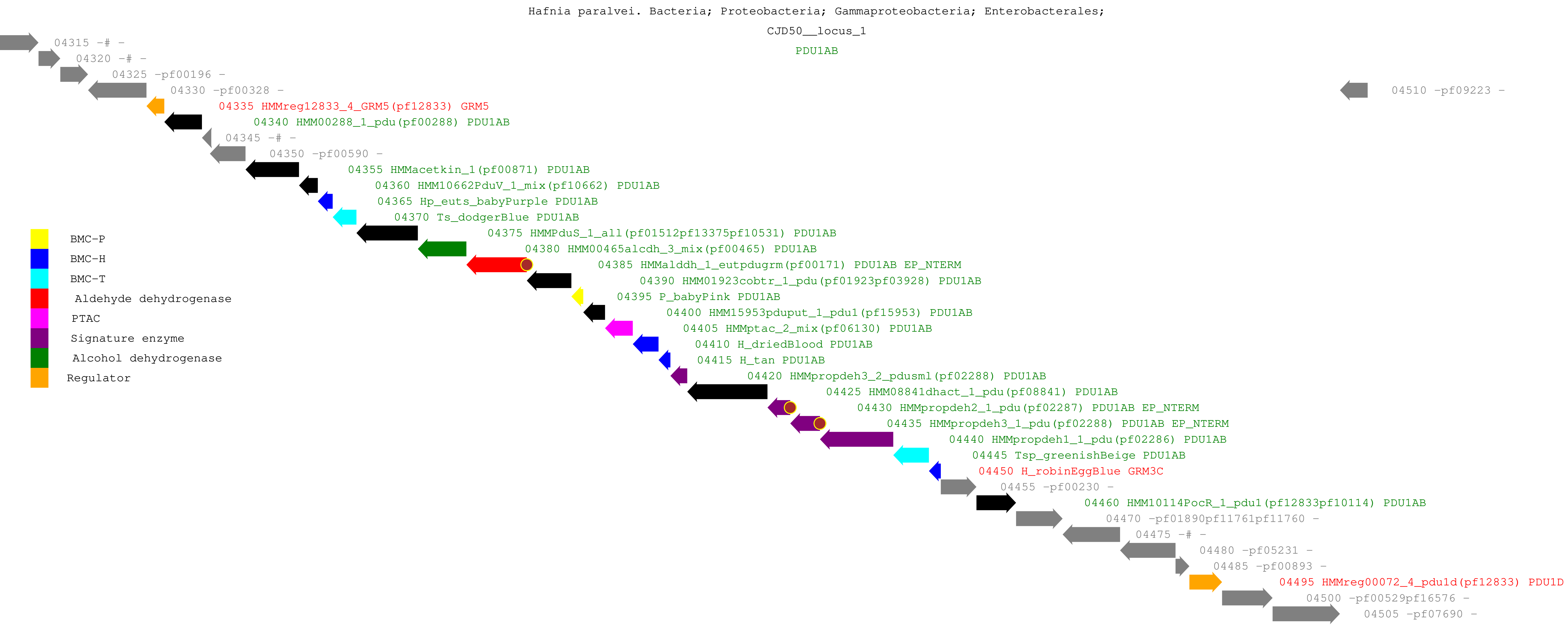
Hafnia paralvei. Bacteria; Proteobacteria; Gammaproteobacteria; Enterobacterales;
CJD50__locus_1
PDU1AB
04315 -# -
04320 -# -
04325 -pf00196 -
04330 -pf00328 -
04335 HMMreg12833_4_GRM5(pf12833) GRM5
04340 HMM00288_1_pdu(pf00288) PDU1AB
04345 -# -
04350 -pf00590 -
04355 HMMacetkin_1(pf00871) PDU1AB
04360 HMM10662PduV_1_mix(pf10662) PDU1AB
04365 Hp_euts_babyPurple PDU1AB
04370 Ts_dodgerBlue PDU1AB
04375 HMMPduS_1_all(pf01512pf13375pf10531) PDU1AB
04380 HMM00465alcdh_3_mix(pf00465) PDU1AB
04385 HMMalddh_1_eutpdugrm(pf00171) PDU1AB EP_NTERM
04390 HMM01923cobtr_1_pdu(pf01923pf03928) PDU1AB
04395 P_babyPink PDU1AB
04400 HMM15953pduput_1_pdu1(pf15953) PDU1AB
04405 HMMptac_2_mix(pf06130) PDU1AB
04410 H_driedBlood PDU1AB
04415 H_tan PDU1AB
04420 HMMpropdeh3_2_pdusml(pf02288) PDU1AB
04425 HMM08841dhact_1_pdu(pf08841) PDU1AB
04430 HMMpropdeh2_1_pdu(pf02287) PDU1AB EP_NTERM
04435 HMMpropdeh3_1_pdu(pf02288) PDU1AB EP_NTERM
04440 HMMpropdeh1_1_pdu(pf02286) PDU1AB
04445 Tsp_greenishBeige PDU1AB
04450 H_robinEggBlue GRM3C
04455 -pf00230 -
04460 HMM10114PocR_1_pdu1(pf12833pf10114) PDU1AB
04470 -pf01890pf11761pf11760 -
04475 -# -
04480 -pf05231 -
04485 -pf00893 -
04495 HMMreg00072_4_pdu1d(pf12833) PDU1D
04500 -pf00529pf16576 -
04505 -pf07690 -
04510 -pf09223 -
BMC-P
BMC-H
BMC-T
Aldehyde dehydrogenase
PTAC
Signature enzyme
Alcohol dehydrogenase
Regulator

Salmonella enterica subsp. enterica serovar Schwarzengrund. Bacteria; Proteobacteria; Gammaproteobacteria; Enterobacterales;

CJS39__locus_1

PDU1AB

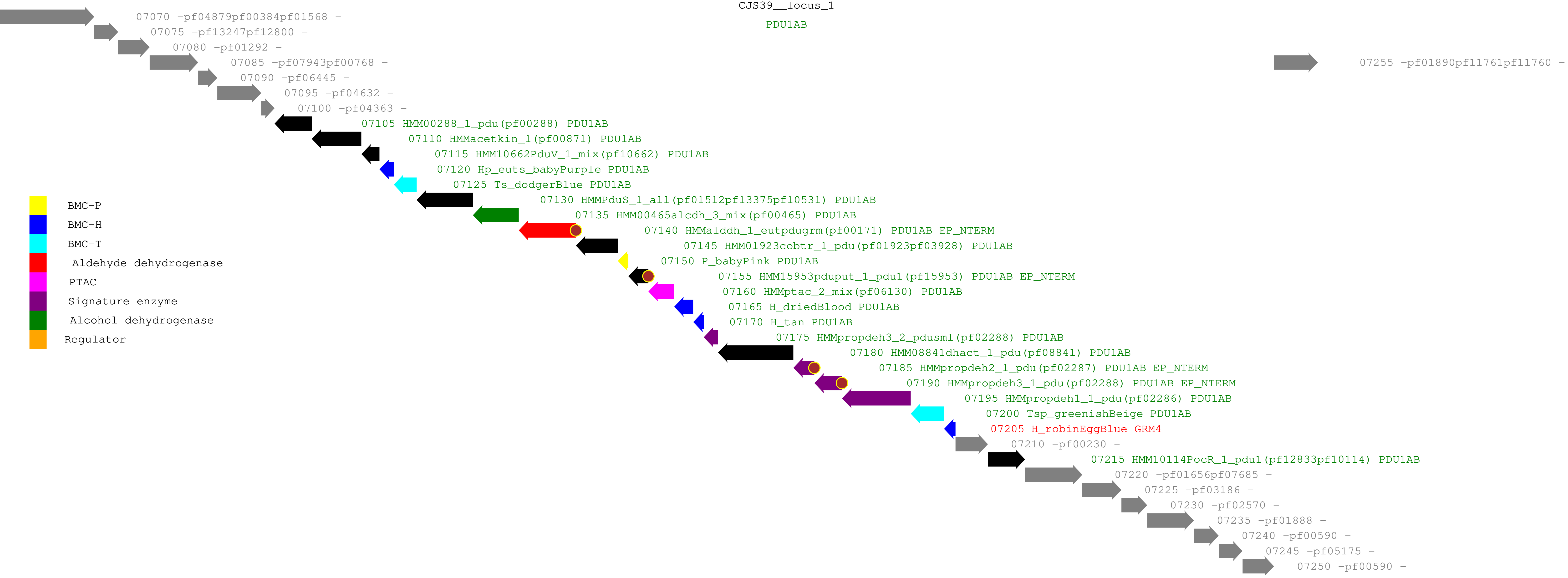

Salmonella enterica subsp. enterica serovar Kentucky. Bacteria; Proteobacteria; Gammaproteobacteria; Enterobacterales;

CJS41__locus_1

PDU1AB

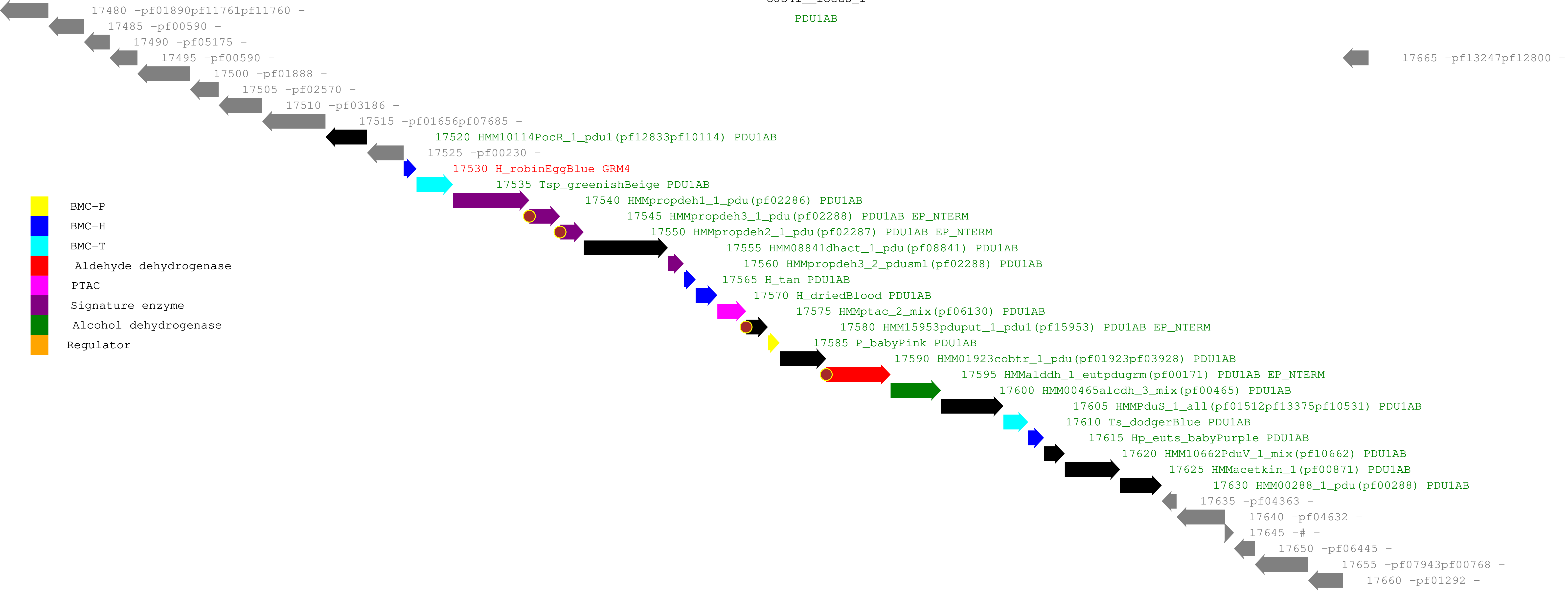

Salmonella enterica subsp. enterica serovar Saintpaul. Bacteria; Proteobacteria; Gammaproteobacteria; Enterobacterales;

CK947__locus_2

PDU1AB

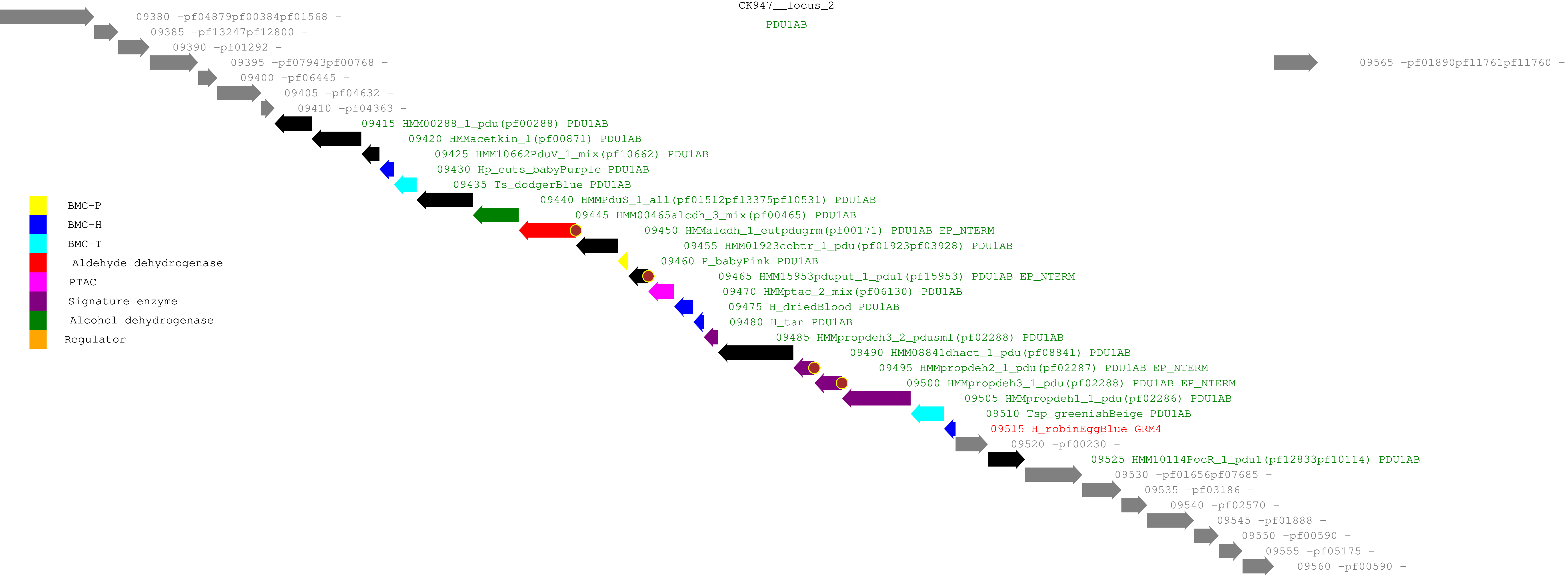

Citrobacter koseri (strain ATCC BAA-895 / CDC 4225-83 / SGSC4696). Bacteria; Proteobacteria; Gammaproteobacteria; Enterobacterales;

CKO__locus_2

PDU1AB

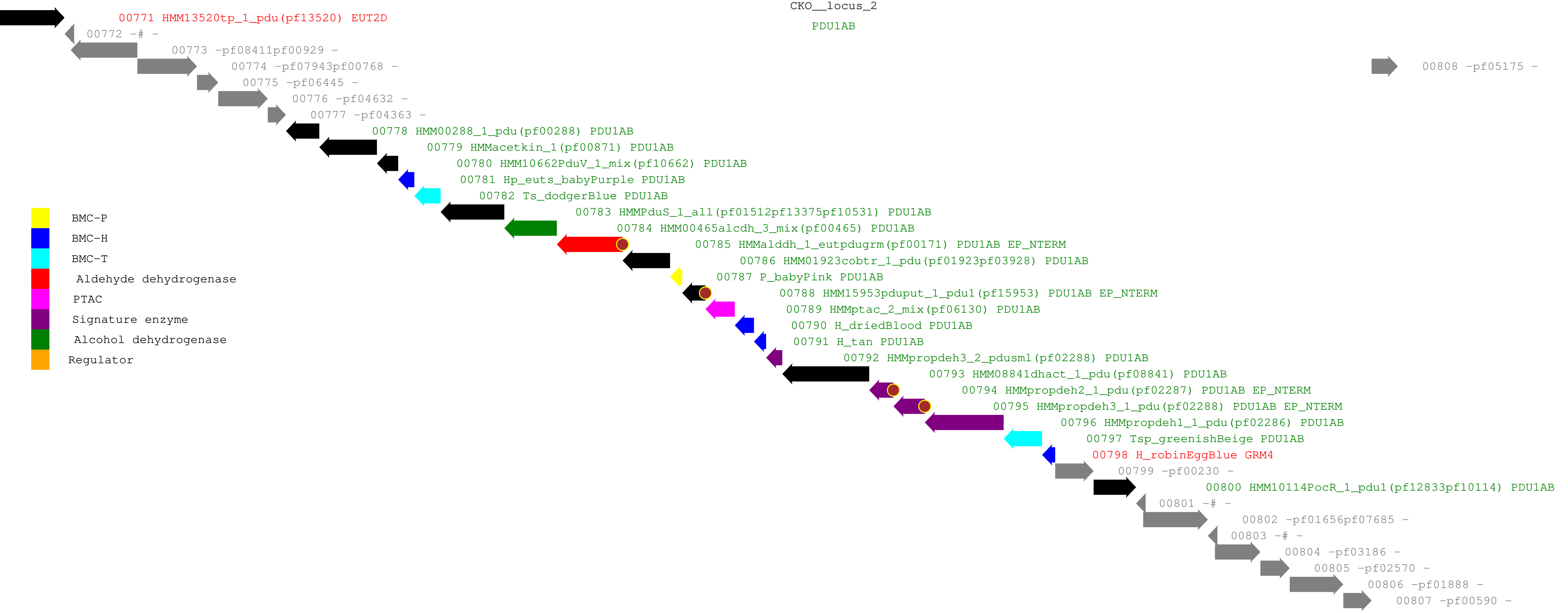

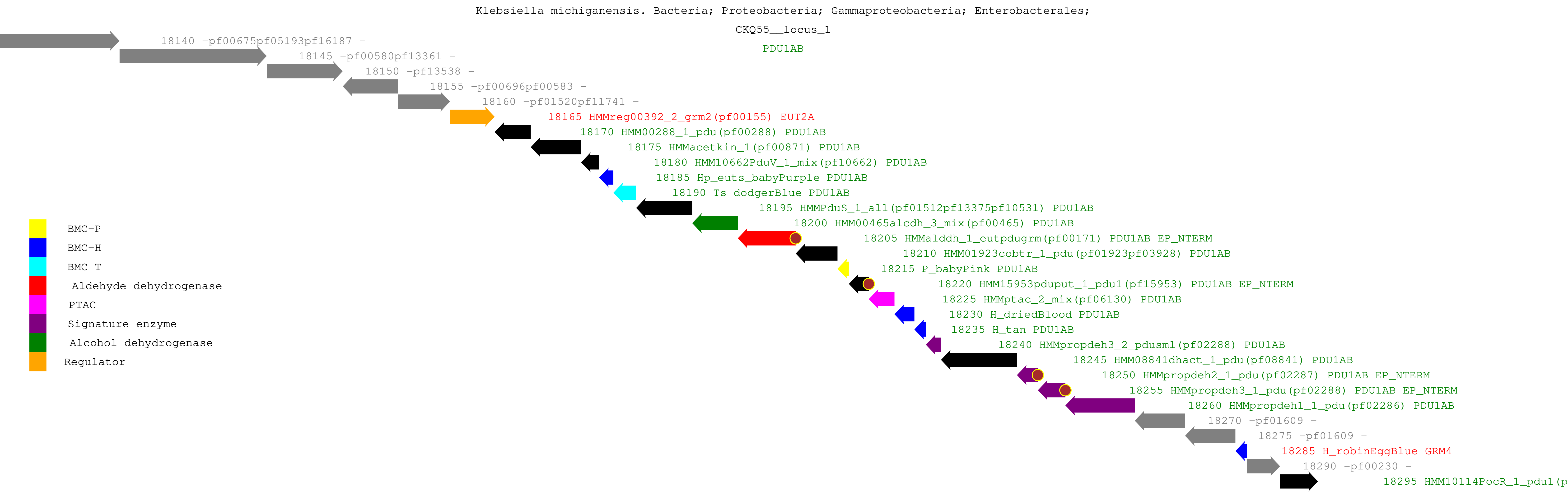
Klebsiella michiganensis. Bacteria; Proteobacteria; Gammaproteobacteria; Enterobacterales;
CKQ55__locus_1
PDU1AB
18140 -pf00675pf05193pf16187 -
18145 -pf00580pf13361 -
18150 -pf13538 -
18155 -pf00696pf00583 -
18160 -pf01520pf11741 -
18165 HMMreg00392_2_grm2(pf00155) EUT2A
18170 HMM00288_1_pdu(pf00288) PDU1AB
18175 HMMacetkin_1(pf00871) PDU1AB
18180 HMM10662PduV_1_mix(pf10662) PDU1AB
18185 Hp_euts_babyPurple PDU1AB
18190 Ts_dodgerBlue PDU1AB
18195 HMMPduS_1_all(pf01512pf13375pf10531) PDU1AB
18200 HMM00465alcdh_3_mix(pf00465) PDU1AB
18205 HMMalddh_1_eutpdugrm(pf00171) PDU1AB EP_NTERM
18210 HMM01923cobtr_1_pdu(pf01923pf03928) PDU1AB
18215 P_babyPink PDU1AB
18220 HMM15953pduput_1_pdu1(pf15953) PDU1AB EP_NTERM
18225 HMMptac_2_mix(pf06130) PDU1AB
18230 H_driedBlood PDU1AB
18235 H_tan PDU1AB
18240 HMMpropdeh3_2_pdusml(pf02288) PDU1AB
18245 HMM08841dhact_1_pdu(pf08841) PDU1AB
18250 HMMpropdeh2_1_pdu(pf02287) PDU1AB EP_NTERM
18255 HMMpropdeh3_1_pdu(pf02288) PDU1AB EP_NTERM
18260 HMMpropdeh1_1_pdu(pf02286) PDU1AB
18270 -pf01609 -
18275 -pf01609 -
18285 H_robinEggBlue GRM4
18290 -pf00230 -
18295 HMM10114PocR_1_pdu1(p
BMC-P
BMC-H
BMC-T
Aldehyde dehydrogenase
PTAC
Signature enzyme
Alcohol dehydrogenase
Regulator

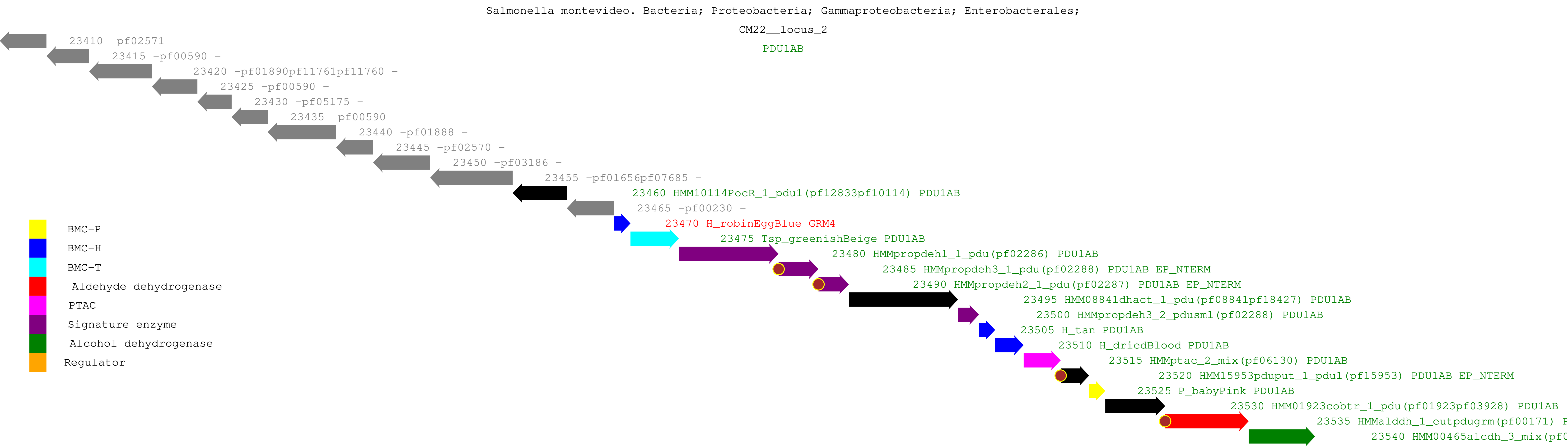

Salmonella montevideo. Bacteria; Proteobacteria; Gammaproteobacteria; Enterobacterales;
CM22__locus_2
PDU1AB
23410 -pf02571 -
23415 -pf00590 -
23420 -pf01890pf11761pf11760 -
23425 -pf00590 -
23430 -pf05175 -
23435 -pf00590 -
23440 -pf01888 -
23445 -pf02570 -
23450 -pf03186 -
23455 -pf01656pf07685 -
23460 HMM10114PocR_1_pdu1(pf12833pf10114) PDU1AB
23465 -pf00230 -
23470 H_robinEggBlue GRM4
23475 Tsp_greenishBeige PDU1AB
23480 HMMpropdeh1_1_pdu(pf02286) PDU1AB
23485 HMMpropdeh3_1_pdu(pf02288) PDU1AB EP_NTERM
23490 HMMpropdeh2_1_pdu(pf02287) PDU1AB EP_NTERM
23495 HMM08841dhact_1_pdu(pf08841pf18427) PDU1AB
23500 HMMpropdeh3_2_pdusml(pf02288) PDU1AB
23505 H_tan PDU1AB
23510 H_driedBlood PDU1AB
23515 HMMptac_2_mix(pf06130) PDU1AB
23520 HMM15953pduput_1_pdu1(pf15953) PDU1AB EP_NTERM
23525 P_babyPink PDU1AB
23530 HMM01923cobtr_1_pdu(pf01923pf03928) PDU1AB
23535 HMMalddh_1_eutpdugrm(pf00171) P
23540 HMM00465alcdh_3_mix(pf0
BMC-P
BMC-H
BMC-T
Aldehyde dehydrogenase
PTAC
Signature enzyme
Alcohol dehydrogenase
Regulator

Salmonella diarizonae. Bacteria; Proteobacteria; Gammaproteobacteria; Enterobacterales;

CNQ75__locus_2

PDU1AB

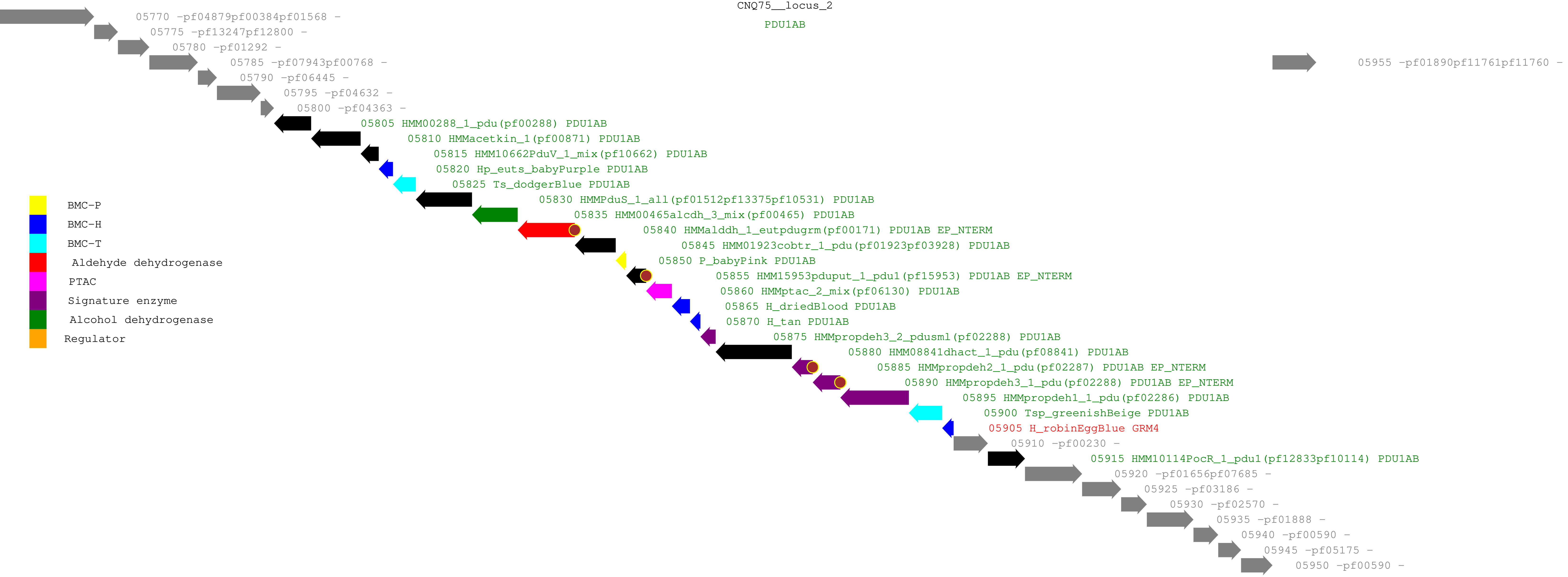

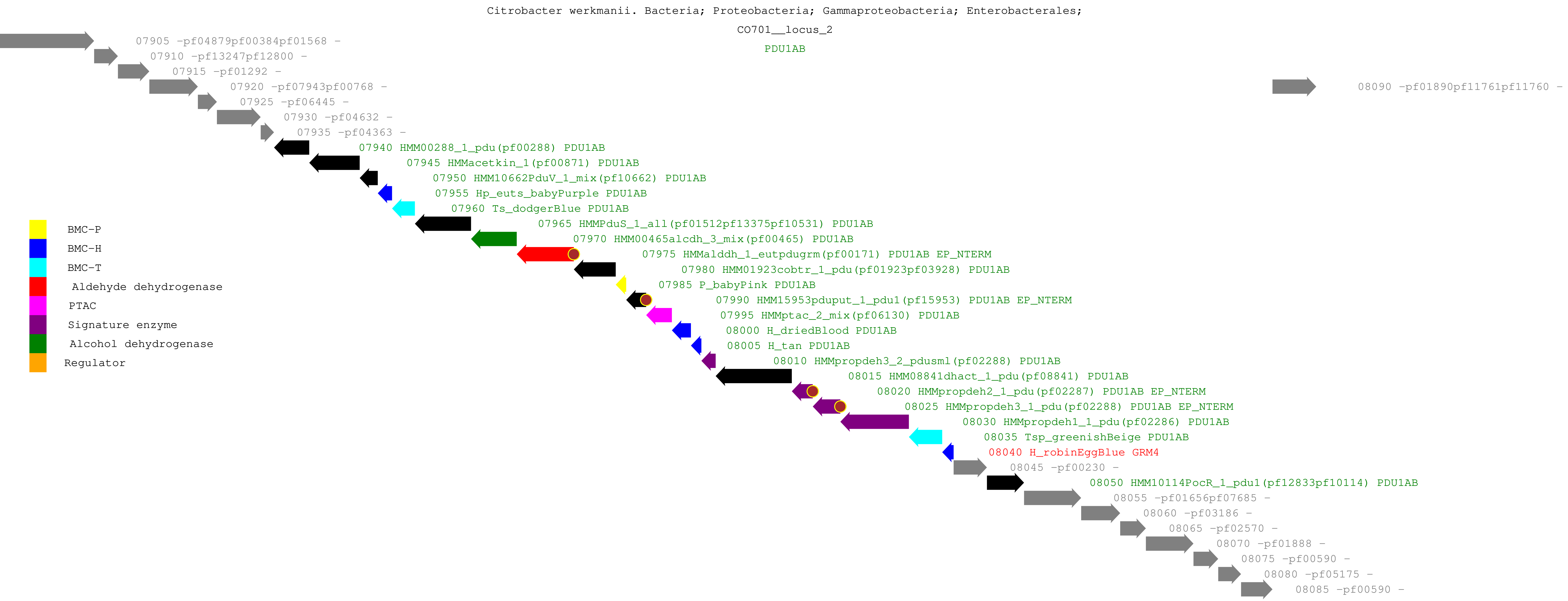
Citrobacter werkmanii. Bacteria; Proteobacteria; Gammaproteobacteria; Enterobacterales;
CO701__locus_2
PDU1AB
07905 -pf04879pf00384pf01568 -
07910 -pf13247pf12800 -
07915 -pf01292 -
07920 -pf07943pf00768 -
07925 -pf06445 -
07930 -pf04632 -
07935 -pf04363 -
07940 HMM00288_1_pdu(pf00288) PDU1AB
07945 HMMacetkin_1(pf00871) PDU1AB
07950 HMM10662PduV_1_mix(pf10662) PDU1AB
07955 Hp_euts_babyPurple PDU1AB
07960 Ts_dodgerBlue PDU1AB
07965 HMMPduS_1_all(pf01512pf13375pf10531) PDU1AB
07970 HMM00465alcdh_3_mix(pf00465) PDU1AB
07975 HMMalddh_1_eutpdugrm(pf00171) PDU1AB EP_NTERM
07980 HMM01923cobtr_1_pdu(pf01923pf03928) PDU1AB
07985 P_babyPink PDU1AB
07990 HMM15953pduput_1_pdu1(pf15953) PDU1AB EP_NTERM
07995 HMMptac_2_mix(pf06130) PDU1AB
08000 H_driedBlood PDU1AB
08005 H_tan PDU1AB
08010 HMMpropdeh3_2_pdusml(pf02288) PDU1AB
08015 HMM08841dhact_1_pdu(pf08841) PDU1AB
08020 HMMpropdeh2_1_pdu(pf02287) PDU1AB EP_NTERM
08025 HMMpropdeh3_1_pdu(pf02288) PDU1AB EP_NTERM
08030 HMMpropdeh1_1_pdu(pf02286) PDU1AB
08035 Tsp_greenishBeige PDU1AB
08040 H_robinEggBlue GRM4
08045 -pf00230 -
08050 HMM10114PocR_1_pdu1(pf12833pf10114) PDU1AB
08055 -pf01656pf07685 -
08060 -pf03186 -
08065 -pf02570 -
08070 -pf01888 -
08075 -pf00590 -
08080 -pf05175 -
08085 -pf00590 -
08090 -pf01890pf11761pf11760 -
BMC-P
BMC-H
BMC-T
Aldehyde dehydrogenase
PTAC
Signature enzyme
Alcohol dehydrogenase
Regulator

Klebsiella quasipneumoniae. Bacteria; Proteobacteria; Gammaproteobacteria; Enterobacterales;

CP911__locus_1

PDU1AB

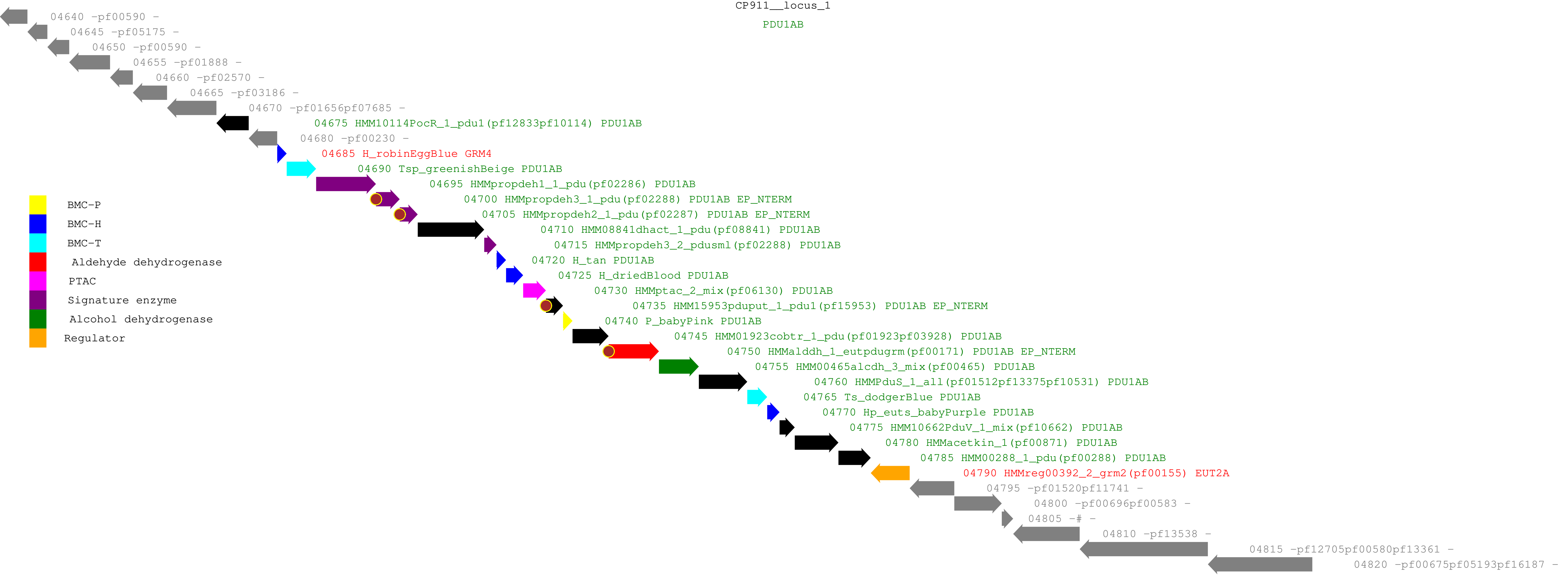

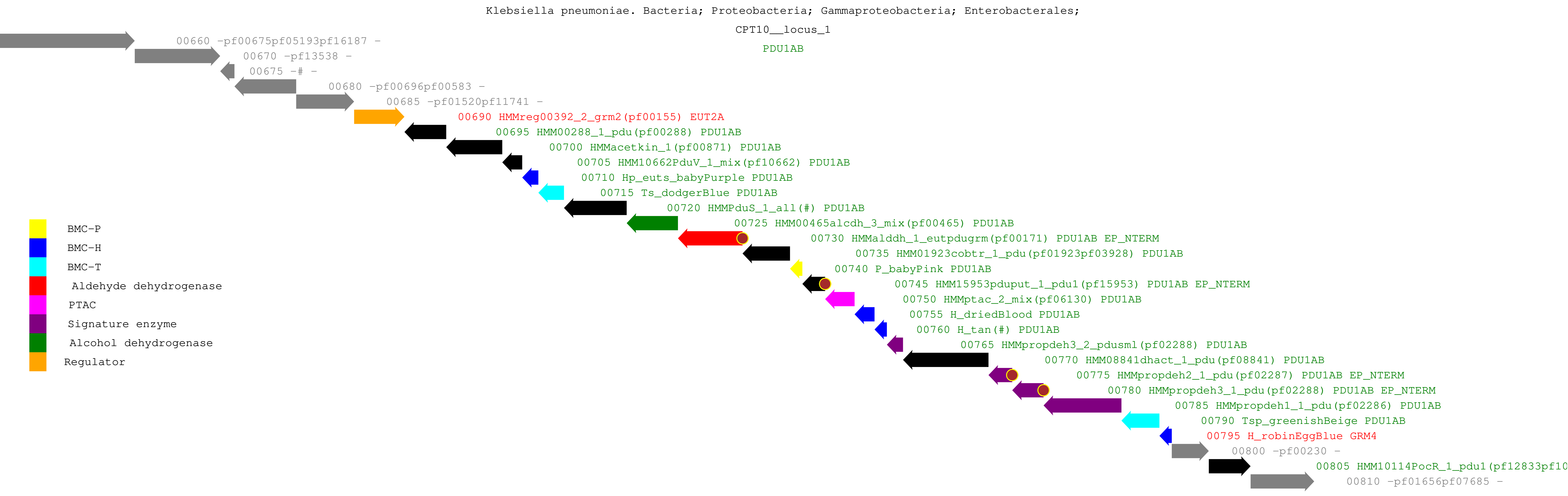
Klebsiella pneumoniae. Bacteria; Proteobacteria; Gammaproteobacteria; Enterobacterales;
CPT10__locus_1
PDU1AB
00660 -pf00675pf05193pf16187 -
00670 -pf13538 -
00675 -# -
00680 -pf00696pf00583 -
00685 -pf01520pf11741 -
00690 HMMreg00392_2_grm2(pf00155) EUT2A
00695 HMM00288_1_pdu(pf00288) PDU1AB
00700 HMMacetkin_1(pf00871) PDU1AB
00705 HMM10662PduV_1_mix(pf10662) PDU1AB
00710 Hp_euts_babyPurple PDU1AB
00715 Ts_dodgerBlue PDU1AB
00720 HMMPduS_1_all(#) PDU1AB
00725 HMM00465alcdh_3_mix(pf00465) PDU1AB
00730 HMMalddh_1_eutpdugrm(pf00171) PDU1AB EP_NTERM
00735 HMM01923cobtr_1_pdu(pf01923pf03928) PDU1AB
00740 P_babyPink PDU1AB
00745 HMM15953pduput_1_pdu1(pf15953) PDU1AB EP_NTERM
00750 HMMptac_2_mix(pf06130) PDU1AB
00755 H_driedBlood PDU1AB
00760 H_tan(#) PDU1AB
00765 HMMpropdeh3_2_pdusml(pf02288) PDU1AB
00770 HMM08841dhact_1_pdu(pf08841) PDU1AB
00775 HMMpropdeh2_1_pdu(pf02287) PDU1AB EP_NTERM
00780 HMMpropdeh3_1_pdu(pf02288) PDU1AB EP_NTERM
00785 HMMpropdeh1_1_pdu(pf02286) PDU1AB
00790 Tsp_greenishBeige PDU1AB
00795 H_robinEggBlue GRM4
00800 -pf00230 -
00805 HMM10114PocR_1_pdu1(pf12833pf10
00810 -pf01656pf07685 -
BMC-P
BMC-H
BMC-T
Aldehyde dehydrogenase
PTAC
Signature enzyme
Alcohol dehydrogenase
Regulator

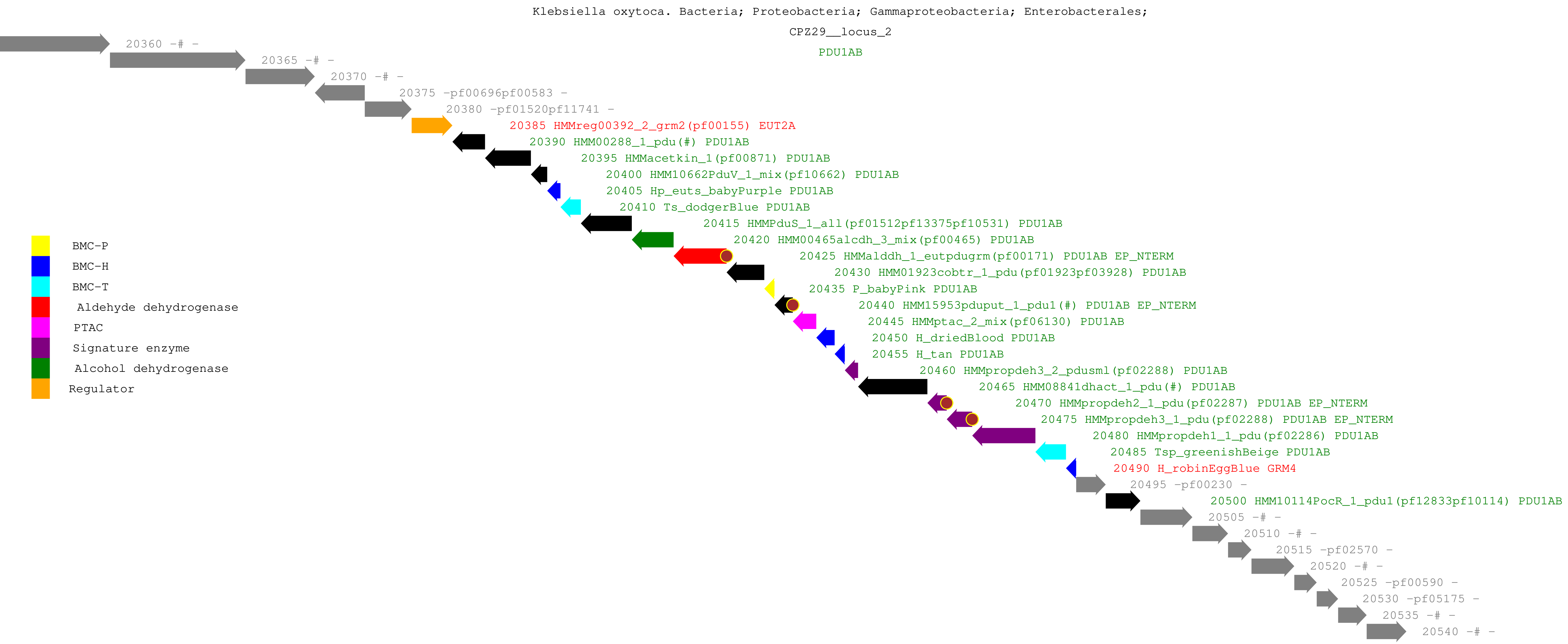
Klebsiella oxytoca. Bacteria; Proteobacteria; Gammaproteobacteria; Enterobacterales;
CPZ29__locus_2
PDU1AB
20360 -# -
20365 -# -
20370 -# -
20375 -pf00696pf00583 -
20380 -pf01520pf11741 -
20385 HMMreg00392_2_grm2(pf00155) EUT2A
20390 HMM00288_1_pdu(#) PDU1AB
20395 HMMacetkin_1(pf00871) PDU1AB
20400 HMM10662PduV_1_mix(pf10662) PDU1AB
20405 Hp_euts_babyPurple PDU1AB
20410 Ts_dodgerBlue PDU1AB
20415 HMMPduS_1_all(pf01512pf13375pf10531) PDU1AB
20420 HMM00465alcdh_3_mix(pf00465) PDU1AB
20425 HMMalddh_1_eutpdugrm(pf00171) PDU1AB EP_NTERM
20430 HMM01923cobtr_1_pdu(pf01923pf03928) PDU1AB
20435 P_babyPink PDU1AB
20440 HMM15953pduput_1_pdu1(#) PDU1AB EP_NTERM
20445 HMMptac_2_mix(pf06130) PDU1AB
20450 H_driedBlood PDU1AB
20455 H_tan PDU1AB
20460 HMMpropdeh3_2_pdusml(pf02288) PDU1AB
20465 HMM08841dhact_1_pdu(#) PDU1AB
20470 HMMpropdeh2_1_pdu(pf02287) PDU1AB EP_NTERM
20475 HMMpropdeh3_1_pdu(pf02288) PDU1AB EP_NTERM
20480 HMMpropdeh1_1_pdu(pf02286) PDU1AB
20485 Tsp_greenishBeige PDU1AB
20490 H_robinEggBlue GRM4
20495 -pf00230 -
20500 HMM10114PocR_1_pdu1(pf12833pf10114) PDU1AB
20505 -# -
20510 -# -
20515 -pf02570 -
20520 -# -
20525 -pf00590 -
20530 -pf05175 -
20535 -# -
20540 -# -
BMC-P
BMC-H
BMC-T
Aldehyde dehydrogenase
PTAC
Signature enzyme
Alcohol dehydrogenase
Regulator

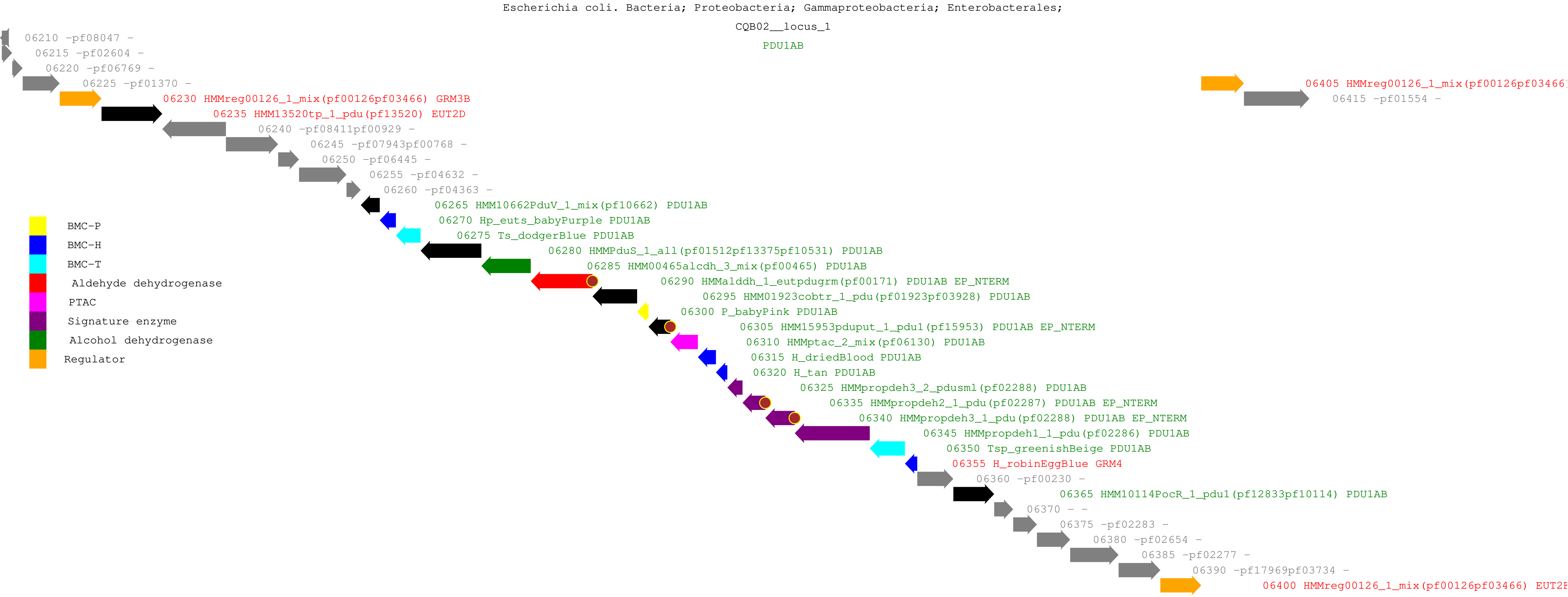
Escherichia coli. Bacteria; Proteobacteria; Gammaproteobacteria; Enterobacterales;
CQB02__locus_1
PDU1AB
06210 -pf08047 -
06215 -pf02604 -
06220 -pf06769 -
06225 -pf01370 -
06230 HMMreg00126_1_mix(pf00126pf03466) GRM3B
06235 HMM13520tp_1_pdu(pf13520) EUT2D
06240 -pf08411pf00929 -
06245 -pf07943pf00768 -
06250 -pf06445 -
06255 -pf04632 -
06260 -pf04363 -
06265 HMM10662PduV_1_mix(pf10662) PDU1AB
06270 Hp_euts_babyPurple PDU1AB
06275 Ts_dodgerBlue PDU1AB
06280 HMMPduS_1_all(pf01512pf13375pf10531) PDU1AB
06285 HMM00465alcdh_3_mix(pf00465) PDU1AB
06290 HMMalddh_1_eutpdugrm(pf00171) PDU1AB EP_NTERM
06295 HMM01923cobtr_1_pdu(pf01923pf03928) PDU1AB
06300 P_babyPink PDU1AB
06305 HMM15953pduput_1_pdu1(pf15953) PDU1AB EP_NTERM
06310 HMMptac_2_mix(pf06130) PDU1AB
06315 H_driedBlood PDU1AB
06320 H_tan PDU1AB
06325 HMMpropdeh3_2_pdusml(pf02288) PDU1AB
06335 HMMpropdeh2_1_pdu(pf02287) PDU1AB EP_NTERM
06340 HMMpropdeh3_1_pdu(pf02288) PDU1AB EP_NTERM
06345 HMMpropdeh1_1_pdu(pf02286) PDU1AB
06350 Tsp_greenishBeige PDU1AB
06355 H_robinEggBlue GRM4
06360 -pf00230 -
06365 HMM10114PocR_1_pdu1(pf12833pf10114) PDU1AB
06370 - -
06375 -pf02283 -
06380 -pf02654 -
06385 -pf02277 -
06390 -pf17969pf03734 -
06400 HMMreg00126_1_mix(pf00126pf03466) EUT2I
06405 HMMreg00126_1_mix(pf00126pf03466)
06415 -pf01554 -
BMC-P
BMC-H
BMC-T
Aldehyde dehydrogenase
PTAC
Signature enzyme
Alcohol dehydrogenase
Regulator

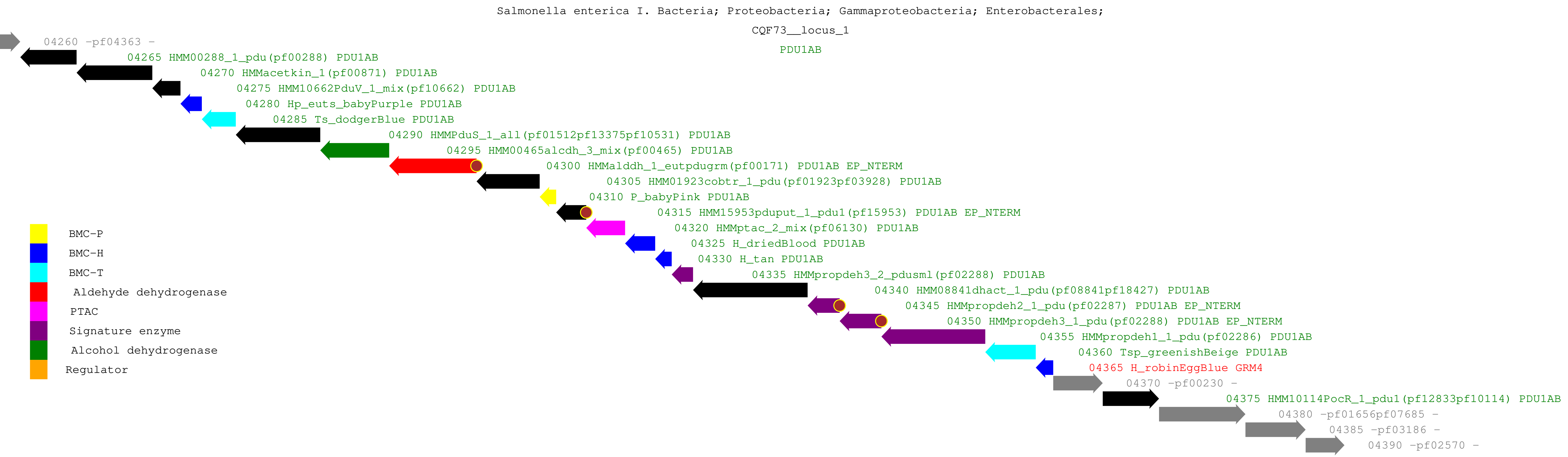
Salmonella enterica I. Bacteria; Proteobacteria; Gammaproteobacteria; Enterobacterales;
CQF73__locus_1
PDU1AB
04260 -pf04363 -
04265 HMM00288_1_pdu(pf00288) PDU1AB
04270 HMMacetkin_1(pf00871) PDU1AB
04275 HMM10662PduV_1_mix(pf10662) PDU1AB
04280 Hp_euts_babyPurple PDU1AB
04285 Ts_dodgerBlue PDU1AB
04290 HMMPduS_1_all(pf01512pf13375pf10531) PDU1AB
04295 HMM00465alcdh_3_mix(pf00465) PDU1AB
04300 HMMalddh_1_eutpdugrm(pf00171) PDU1AB EP_NTERM
04305 HMM01923cobtr_1_pdu(pf01923pf03928) PDU1AB
04310 P_babyPink PDU1AB
04315 HMM15953pduput_1_pdu1(pf15953) PDU1AB EP_NTERM
04320 HMMptac_2_mix(pf06130) PDU1AB
04325 H_driedBlood PDU1AB
04330 H_tan PDU1AB
04335 HMMpropdeh3_2_pdusml(pf02288) PDU1AB
04340 HMM08841dhact_1_pdu(pf08841pf18427) PDU1AB
04345 HMMpropdeh2_1_pdu(pf02287) PDU1AB EP_NTERM
04350 HMMpropdeh3_1_pdu(pf02288) PDU1AB EP_NTERM
04355 HMMpropdeh1_1_pdu(pf02286) PDU1AB
04360 Tsp_greenishBeige PDU1AB
04365 H_robinEggBlue GRM4
04370 -pf00230 -
04375 HMM10114PocR_1_pdu1(pf12833pf10114) PDU1AB
04380 -pf01656pf07685 -
04385 -pf03186 -
04390 -pf02570 -
BMC-P
BMC-H
BMC-T
Aldehyde dehydrogenase
PTAC
Signature enzyme
Alcohol dehydrogenase
Regulator

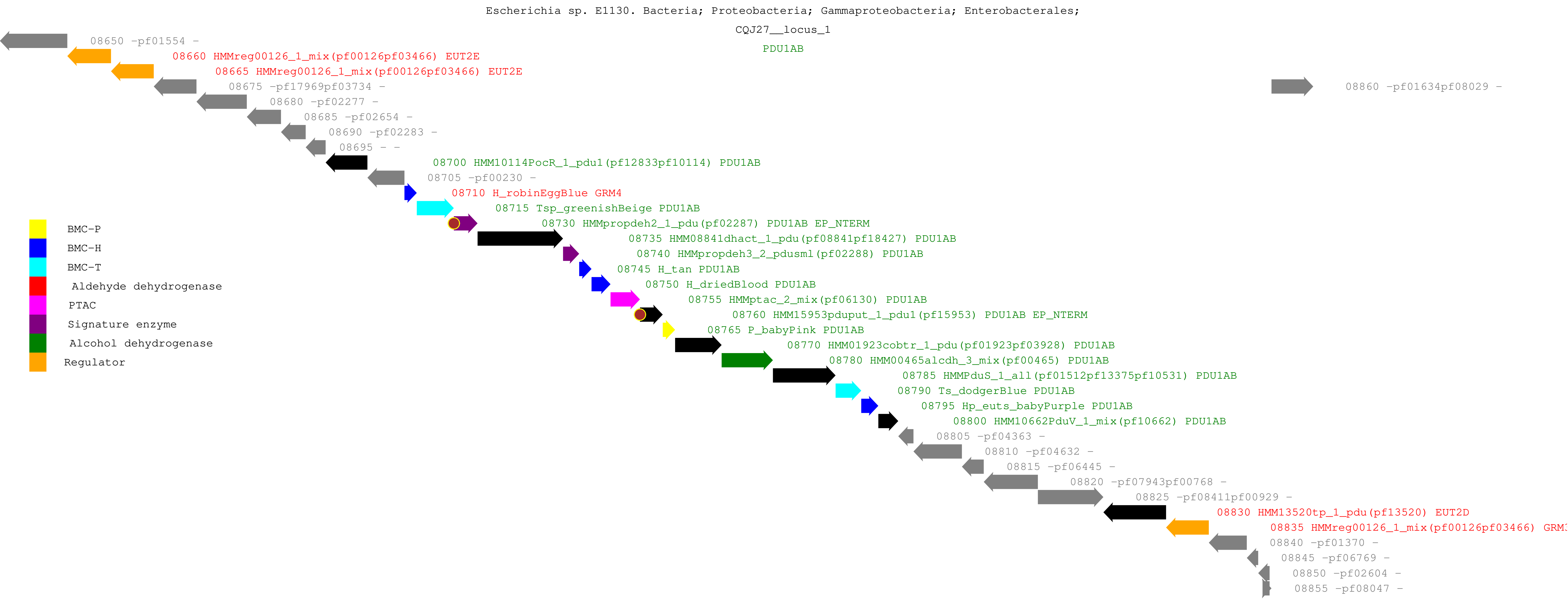
Escherichia sp. E1130. Bacteria; Proteobacteria; Gammaproteobacteria; Enterobacterales;
CQJ27__locus_1
PDU1AB
08650 -pf01554 -
08660 HMMreg00126_1_mix(pf00126pf03466) EUT2E
08665 HMMreg00126_1_mix(pf00126pf03466) EUT2E
08675 -pf17969pf03734 -
08860 -pf01634pf08029 -
08680 -pf02277 -
08685 -pf02654 -
08690 -pf02283 -
08695 - -
08700 HMM10114PocR_1_pdu1(pf12833pf10114) PDU1AB
08705 -pf00230 -
08710 H_robinEggBlue GRM4
08715 Tsp_greenishBeige PDU1AB
08730 HMMpropdeh2_1_pdu(pf02287) PDU1AB EP_NTERM
08735 HMM08841dhact_1_pdu(pf08841pf18427) PDU1AB
08740 HMMpropdeh3_2_pdusml(pf02288) PDU1AB
08745 H_tan PDU1AB
08750 H_driedBlood PDU1AB
08755 HMMptac_2_mix(pf06130) PDU1AB
08760 HMM15953pduput_1_pdu1(pf15953) PDU1AB EP_NTERM
08765 P_babyPink PDU1AB
08770 HMM01923cobtr_1_pdu(pf01923pf03928) PDU1AB
08780 HMM00465alcdh_3_mix(pf00465) PDU1AB
08785 HMMPduS_1_all(pf01512pf13375pf10531) PDU1AB
08790 Ts_dodgerBlue PDU1AB
08795 Hp_euts_babyPurple PDU1AB
08800 HMM10662PduV_1_mix(pf10662) PDU1AB
08805 -pf04363 -
08810 -pf04632 -
08815 -pf06445 -
08820 -pf07943pf00768 -
08825 -pf08411pf00929 -
08830 HMM13520tp_1_pdu(pf13520) EUT2D
08835 HMMreg00126_1_mix(pf00126pf03466) GRM
08840 -pf01370 -
08845 -pf06769 -
08850 -pf02604 -
08855 -pf08047 -
BMC-P
BMC-H
BMC-T
Aldehyde dehydrogenase
PTAC
Signature enzyme
Alcohol dehydrogenase
Regulator

Escherichia sp. E2562. Bacteria; Proteobacteria; Gammaproteobacteria; Enterobacterales;

CQJ28__locus_1

PDU1AB

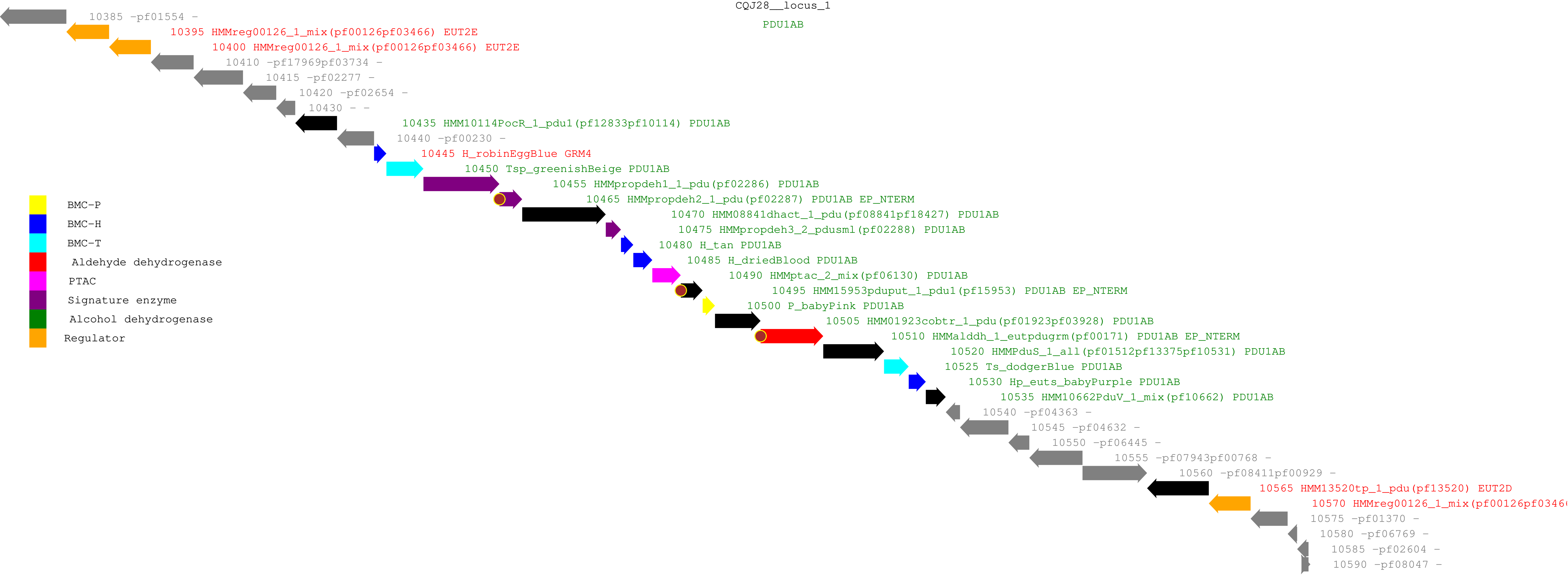

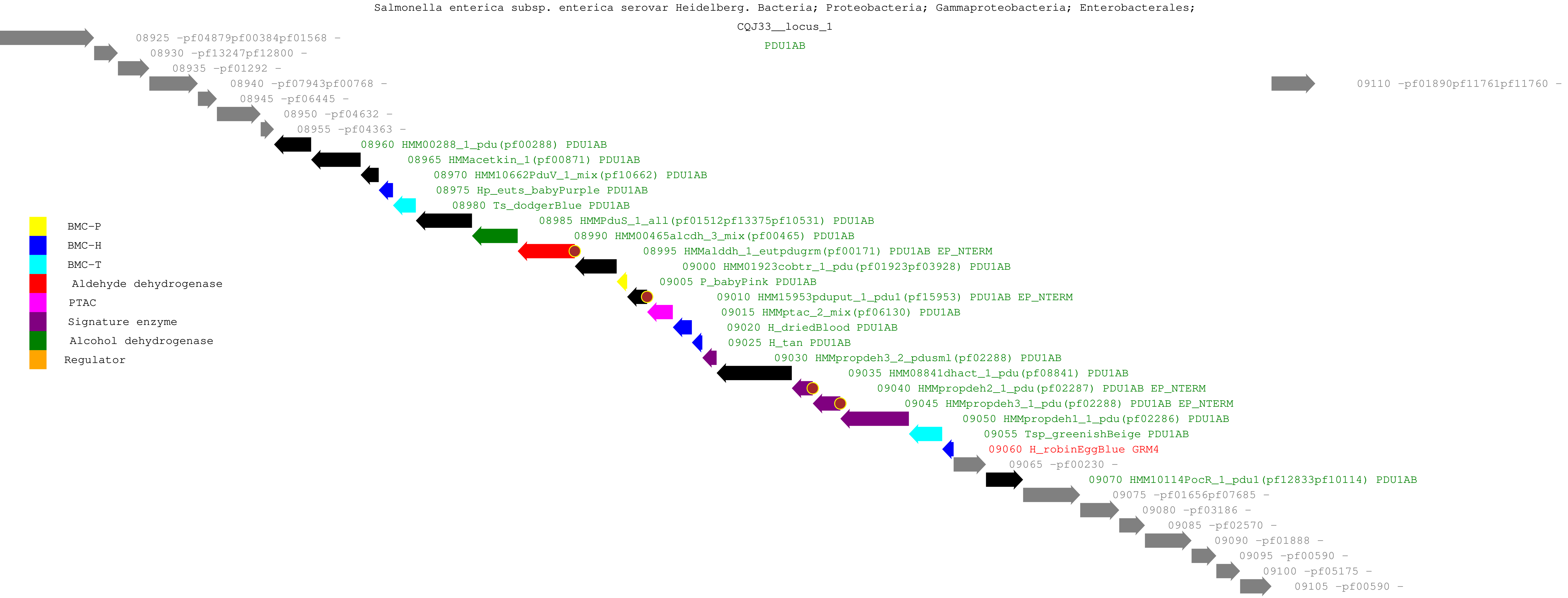
Salmonella enterica subsp. enterica serovar Heidelberg. Bacteria; Proteobacteria; Gammaproteobacteria; Enterobacterales;
CQJ33__locus_1
PDU1AB
08925 -pf04879pf00384pf01568 -
08930 -pf13247pf12800 -
08935 -pf01292 -
08940 -pf07943pf00768 -
08945 -pf06445 -
08950 -pf04632 -
08955 -pf04363 -
08960 HMM00288_1_pdu(pf00288) PDU1AB
08965 HMMacetkin_1(pf00871) PDU1AB
08970 HMM10662PduV_1_mix(pf10662) PDU1AB
08975 Hp_euts_babyPurple PDU1AB
08980 Ts_dodgerBlue PDU1AB
08985 HMMPduS_1_all(pf01512pf13375pf10531) PDU1AB
08990 HMM00465alcdh_3_mix(pf00465) PDU1AB
08995 HMMalddh_1_eutpdugrm(pf00171) PDU1AB EP_NTERM
09000 HMM01923cobtr_1_pdu(pf01923pf03928) PDU1AB
09005 P_babyPink PDU1AB
09010 HMM15953pduput_1_pdu1(pf15953) PDU1AB EP_NTERM
09015 HMMptac_2_mix(pf06130) PDU1AB
09020 H_driedBlood PDU1AB
09025 H_tan PDU1AB
09030 HMMpropdeh3_2_pdusml(pf02288) PDU1AB
09035 HMM08841dhact_1_pdu(pf08841) PDU1AB
09040 HMMpropdeh2_1_pdu(pf02287) PDU1AB EP_NTERM
09045 HMMpropdeh3_1_pdu(pf02288) PDU1AB EP_NTERM
09050 HMMpropdeh1_1_pdu(pf02286) PDU1AB
09055 Tsp_greenishBeige PDU1AB
09060 H_robinEggBlue GRM4
09065 -pf00230 -
09070 HMM10114PocR_1_pdu1(pf12833pf10114) PDU1AB
09075 -pf01656pf07685 -
09080 -pf03186 -
09085 -pf02570 -
09090 -pf01888 -
09095 -pf00590 -
09100 -pf05175 -
09105 -pf00590 -
09110 -pf01890pf11761pf11760 -
BMC-P
BMC-H
BMC-T
Aldehyde dehydrogenase
PTAC
Signature enzyme
Alcohol dehydrogenase
Regulator

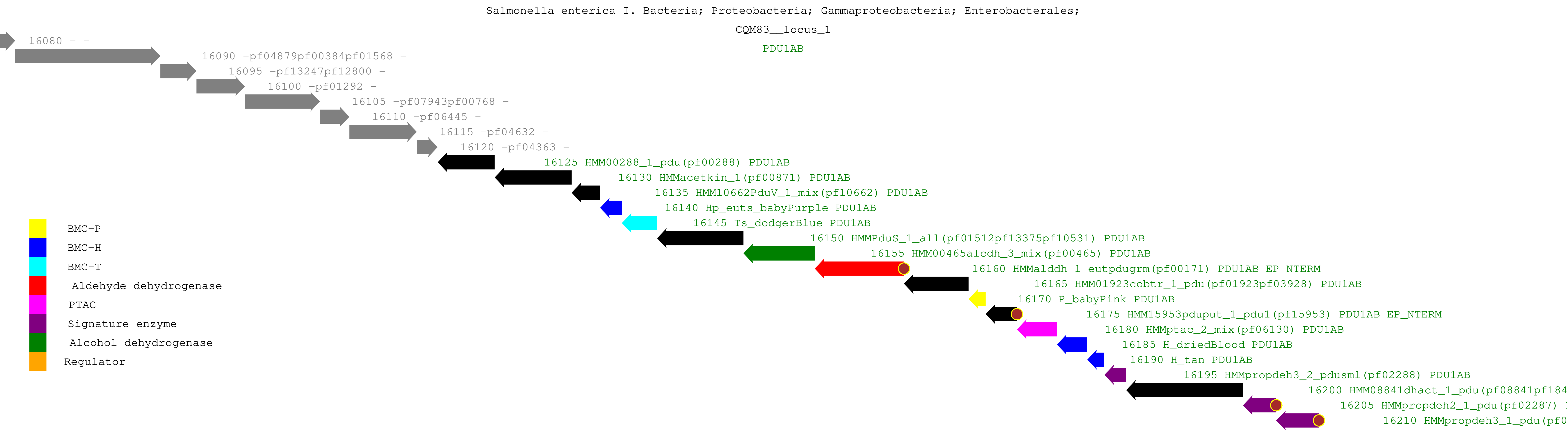
Salmonella enterica I. Bacteria; Proteobacteria; Gammaproteobacteria; Enterobacterales;
CQM83__locus_1
PDU1AB
16080 - -
16090 -pf04879pf00384pf01568 -
16095 -pf13247pf12800 -
16100 -pf01292 -
16105 -pf07943pf00768 -
16110 -pf06445 -
16115 -pf04632 -
16120 -pf04363 -
16125 HMM00288_1_pdu(pf00288) PDU1AB
16130 HMMacetkin_1(pf00871) PDU1AB
16135 HMM10662PduV_1_mix(pf10662) PDU1AB
16140 Hp_euts_babyPurple PDU1AB
16145 Ts_dodgerBlue PDU1AB
16150 HMMPduS_1_all(pf01512pf13375pf10531) PDU1AB
16155 HMM00465alcdh_3_mix(pf00465) PDU1AB
16160 HMMaldh_1_eutpdugrm(pf00171) PDU1AB EP_NTERM
16165 HMM01923cobtr_1_pdu(pf01923pf03928) PDU1AB
16170 P_babyPink PDU1AB
16175 HMM15953pduput_1_pdu1(pf15953) PDU1AB EP_NTERM
16180 HMMptac_2_mix(pf06130) PDU1AB
16185 H_driedBlood PDU1AB
16190 H_tan PDU1AB
16195 HMMpropdeh3_2_pdusml(pf02288) PDU1AB
16200 HMM08841dhact_1_pdu(pf08841pf184
16205 HMMpropdeh2_1_pdu(pf02287)
16210 HMMpropdeh3_1_pdu(pf0
BMC-P
BMC-H
BMC-T
Aldehyde dehydrogenase
PTAC
Signature enzyme
Alcohol dehydrogenase
Regulator

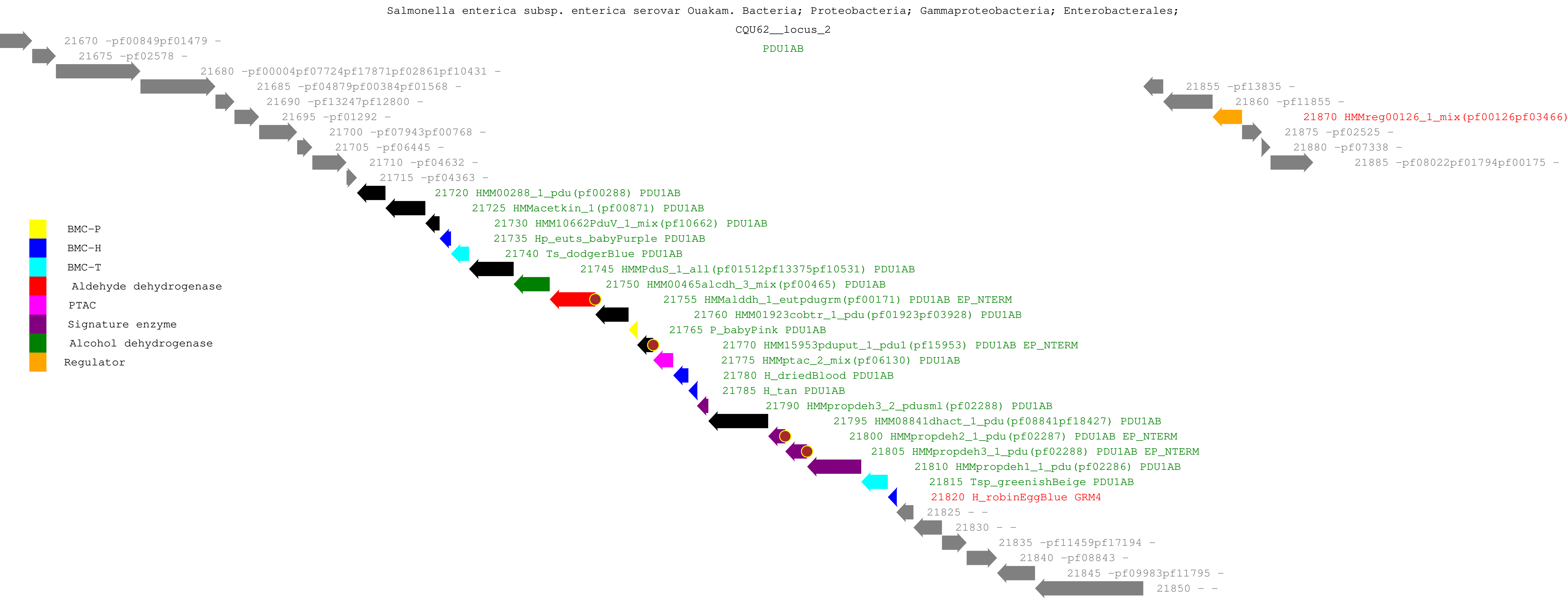
Salmonella enterica subsp. enterica serovar Ouakam. Bacteria; Proteobacteria; Gammaproteobacteria; Enterobacterales;
CQU62__locus_2
PDU1AB
21670 -pf00849pf01479 -
21675 -pf02578 -
21680 -pf00004pf07724pf17871pf02861pf10431 -
21685 -pf04879pf00384pf01568 -
21690 -pf13247pf12800 -
21695 -pf01292 -
21700 -pf07943pf00768 -
21705 -pf06445 -
21710 -pf04632 -
21715 -pf04363 -
21720 HMM00288_1_pdu(pf00288) PDU1AB
21725 HMMacetkin_1(pf00871) PDU1AB
21730 HMM10662PduV_1_mix(pf10662) PDU1AB
21735 Hp_euts_babyPurple PDU1AB
21740 Ts_dodgerBlue PDU1AB
21745 HMMPduS_1_all(pf01512pf13375pf10531) PDU1AB
21750 HMM00465alcdh_3_mix(pf00465) PDU1AB
21755 HMMalddh_1_eutpdugrm(pf00171) PDU1AB EP_NTERM
21760 HMM01923cobtr_1_pdu(pf01923pf03928) PDU1AB
21765 P_babyPink PDU1AB
21770 HMM15953pduput_1_pdu1(pf15953) PDU1AB EP_NTERM
21775 HMMptac_2_mix(pf06130) PDU1AB
21780 H_driedBlood PDU1AB
21785 H_tan PDU1AB
21790 HMMpropdeh3_2_pdusml(pf02288) PDU1AB
21795 HMM08841dhact_1_pdu(pf08841pf18427) PDU1AB
21800 HMMpropdeh2_1_pdu(pf02287) PDU1AB EP_NTERM
21805 HMMpropdeh3_1_pdu(pf02288) PDU1AB EP_NTERM
21810 HMMpropdeh1_1_pdu(pf02286) PDU1AB
21815 Tsp_greenishBeige PDU1AB
21820 H_robinEggBlue GRM4
21825 - -
21830 - -
21835 -pf11459pf17194 -
21840 -pf08843 -
21845 -pf09983pf11795 -
21850 - -
21855 -pf13835 -
21860 -pf11855 -
21870 HMMreg00126_1_mix(pf00126pf03466)
21875 -pf02525 -
21880 -pf07338 -
21885 -pf08022pf01794pf00175 -
BMC-P
BMC-H
BMC-T
Aldehyde dehydrogenase
PTAC
Signature enzyme
Alcohol dehydrogenase
Regulator

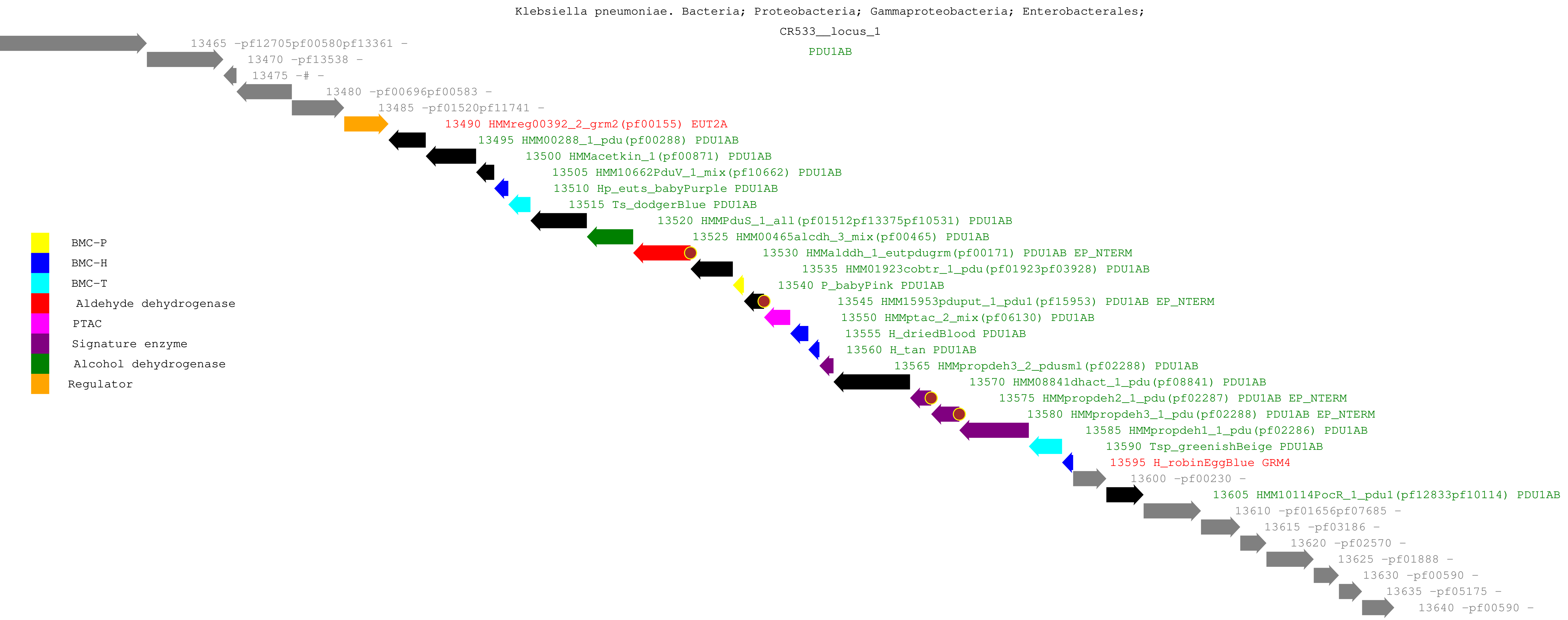
Klebsiella pneumoniae. Bacteria; Proteobacteria; Gammaproteobacteria; Enterobacterales;
CR533__locus_1
PDU1AB
13465 -pf12705pf00580pf13361 -
13470 -pf13538 -
13475 -# -
13480 -pf00696pf00583 -
13485 -pf01520pf11741 -
13490 HMMreg00392_2_grm2(pf00155) EUT2A
13495 HMM00288_1_pdu(pf00288) PDU1AB
13500 HMMacetkin_1(pf00871) PDU1AB
13505 HMM10662PduV_1_mix(pf10662) PDU1AB
13510 Hp_euts_babyPurple PDU1AB
13515 Ts_dodgerBlue PDU1AB
13520 HMMPduS_1_all(pf01512pf13375pf10531) PDU1AB
13525 HMM00465alcdh_3_mix(pf00465) PDU1AB
13530 HMMalddh_1_eutpdugrm(pf00171) PDU1AB EP_NTERM
13535 HMM01923cobtr_1_pdu(pf01923pf03928) PDU1AB
13540 P_babyPink PDU1AB
13545 HMM15953pduput_1_pdu1(pf15953) PDU1AB EP_NTERM
13550 HMMptac_2_mix(pf06130) PDU1AB
13555 H_driedBlood PDU1AB
13560 H_tan PDU1AB
13565 HMMpropdeh3_2_pdusml(pf02288) PDU1AB
13570 HMM08841dhact_1_pdu(pf08841) PDU1AB
13575 HMMpropdeh2_1_pdu(pf02287) PDU1AB EP_NTERM
13580 HMMpropdeh3_1_pdu(pf02288) PDU1AB EP_NTERM
13585 HMMpropdeh1_1_pdu(pf02286) PDU1AB
13590 Tsp_greenishBeige PDU1AB
13595 H_robinEggBlue GRM4
13600 -pf00230 -
13605 HMM10114PocR_1_pdu1(pf12833pf10114) PDU1AB
13610 -pf01656pf07685 -
13615 -pf03186 -
13620 -pf02570 -
13625 -pf01888 -
13630 -pf00590 -
13635 -pf05175 -
13640 -pf00590 -
BMC-P
BMC-H
BMC-T
Aldehyde dehydrogenase
PTAC
Signature enzyme
Alcohol dehydrogenase
Regulator

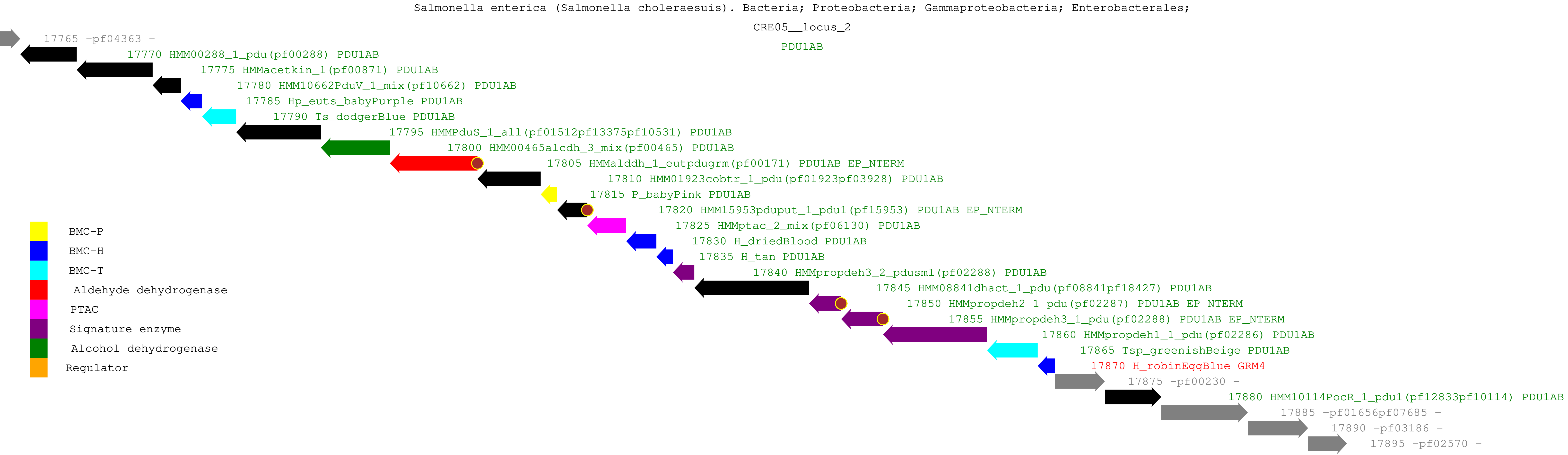
Salmonella enterica (Salmonella choleraesuis). Bacteria; Proteobacteria; Gammaproteobacteria; Enterobacterales;
CRE05__locus_2
PDU1AB
17765 -pf04363 -
17770 HMM00288_1_pdu(pf00288) PDU1AB
17775 HMMacetkin_1(pf00871) PDU1AB
17780 HMM10662PduV_1_mix(pf10662) PDU1AB
17785 Hp_euts_babyPurple PDU1AB
17790 Ts_dodgerBlue PDU1AB
17795 HMMPduS_1_all(pf01512pf13375pf10531) PDU1AB
17800 HMM00465alcdh_3_mix(pf00465) PDU1AB
17805 HMMalddh_1_eutpdugrm(pf00171) PDU1AB EP_NTERM
17810 HMM01923cobtr_1_pdu(pf01923pf03928) PDU1AB
17815 P_babyPink PDU1AB
17820 HMM15953pduput_1_pdu1(pf15953) PDU1AB EP_NTERM
17825 HMMptac_2_mix(pf06130) PDU1AB
17830 H_driedBlood PDU1AB
17835 H_tan PDU1AB
17840 HMMpropdeh3_2_pdusml(pf02288) PDU1AB
17845 HMM08841dhact_1_pdu(pf08841pf18427) PDU1AB
17850 HMMpropdeh2_1_pdu(pf02287) PDU1AB EP_NTERM
17855 HMMpropdeh3_1_pdu(pf02288) PDU1AB EP_NTERM
17860 HMMpropdeh1_1_pdu(pf02286) PDU1AB
17865 Tsp_greenishBeige PDU1AB
17870 H_robinEggBlue GRM4
17875 -pf00230 -
17880 HMM10114PocR_1_pdu1(pf12833pf10114) PDU1AB
17885 -pf01656pf07685 -
17890 -pf03186 -
17895 -pf02570 -
BMC-P
BMC-H
BMC-T
Aldehyde dehydrogenase
PTAC
Signature enzyme
Alcohol dehydrogenase
Regulator

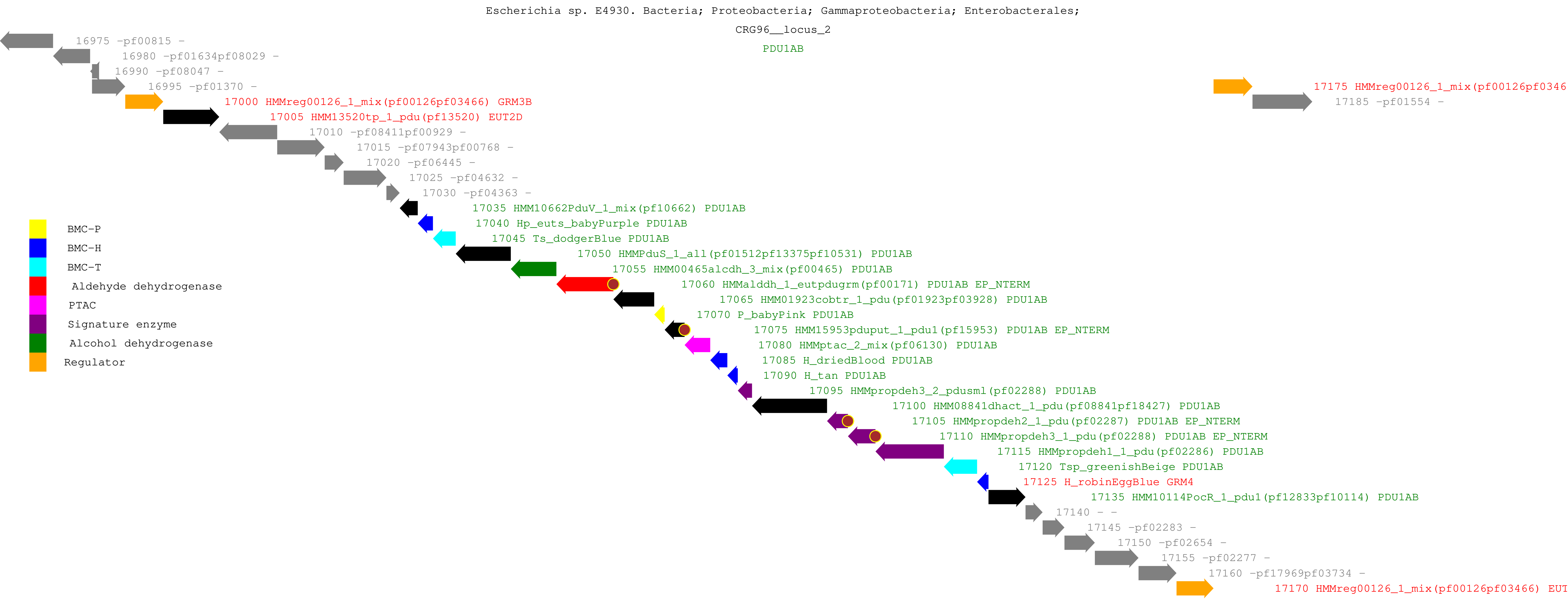
Escherichia sp. E4930. Bacteria; Proteobacteria; Gammaproteobacteria; Enterobacterales;
CRG96__locus_2
PDU1AB
16975 -pf00815 -
16980 -pf01634pf08029 -
16990 -pf08047 -
16995 -pf01370 -
17000 HMMreg00126_1_mix(pf00126pf03466) GRM3B
17005 HMM13520tp_1_pdu(pf13520) EUT2D
17010 -pf08411pf00929 -
17015 -pf07943pf00768 -
17020 -pf06445 -
17025 -pf04632 -
17030 -pf04363 -
17035 HMM10662PduV_1_mix(pf10662) PDU1AB
17040 Hp_euts_babyPurple PDU1AB
17045 Ts_dodgerBlue PDU1AB
17050 HMMPduS_1_all(pf01512pf13375pf10531) PDU1AB
17055 HMM00465alcdh_3_mix(pf00465) PDU1AB
17060 HMMaldh_1_eutpdugrm(pf00171) PDU1AB EP_NTERM
17065 HMM01923cobtr_1_pdu(pf01923pf03928) PDU1AB
17070 P_babyPink PDU1AB
17075 HMM15953pduput_1_pdu1(pf15953) PDU1AB EP_NTERM
17080 HMMptac_2_mix(pf06130) PDU1AB
17085 H_driedBlood PDU1AB
17090 H_tan PDU1AB
17095 HMMpropdeh3_2_pdusml(pf02288) PDU1AB
17100 HMM08841dhact_1_pdu(pf08841pf18427) PDU1AB
17105 HMMpropdeh2_1_pdu(pf02287) PDU1AB EP_NTERM
17110 HMMpropdeh3_1_pdu(pf02288) PDU1AB EP_NTERM
17115 HMMpropdeh1_1_pdu(pf02286) PDU1AB
17120 Tsp_greenishBeige PDU1AB
17125 H_robinEggBlue GRM4
17135 HMM10114PocR_1_pdu1(pf12833pf10114) PDU1AB
17140 - -
17145 -pf02283 -
17150 -pf02654 -
17155 -pf02277 -
17160 -pf17969pf03734 -
17170 HMMreg00126_1_mix(pf00126pf03466) EUT
17175 HMMreg00126_1_mix(pf00126pf0346
17185 -pf01554 -
BMC-P
BMC-H
BMC-T
Aldehyde dehydrogenase
PTAC
Signature enzyme
Alcohol dehydrogenase
Regulator

Escherichia sp. E3659. Bacteria; Proteobacteria; Gammaproteobacteria; Enterobacterales;

CRI65__locus_2

PDU1AB

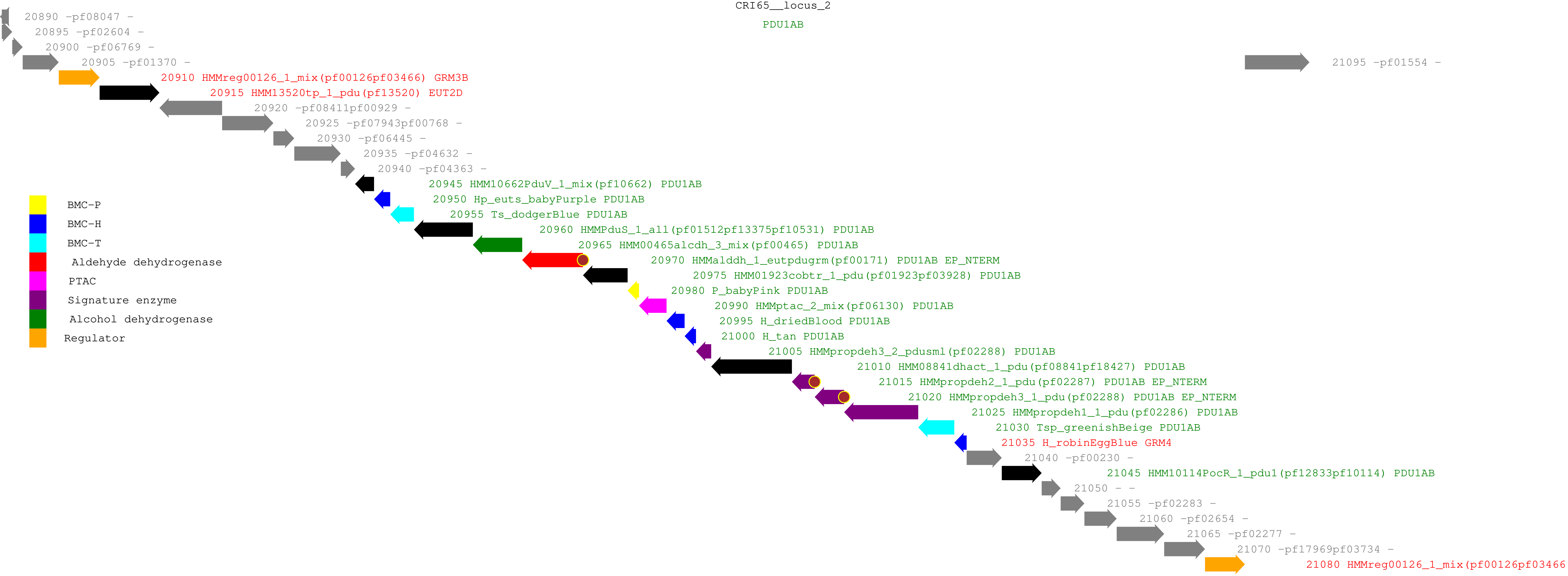

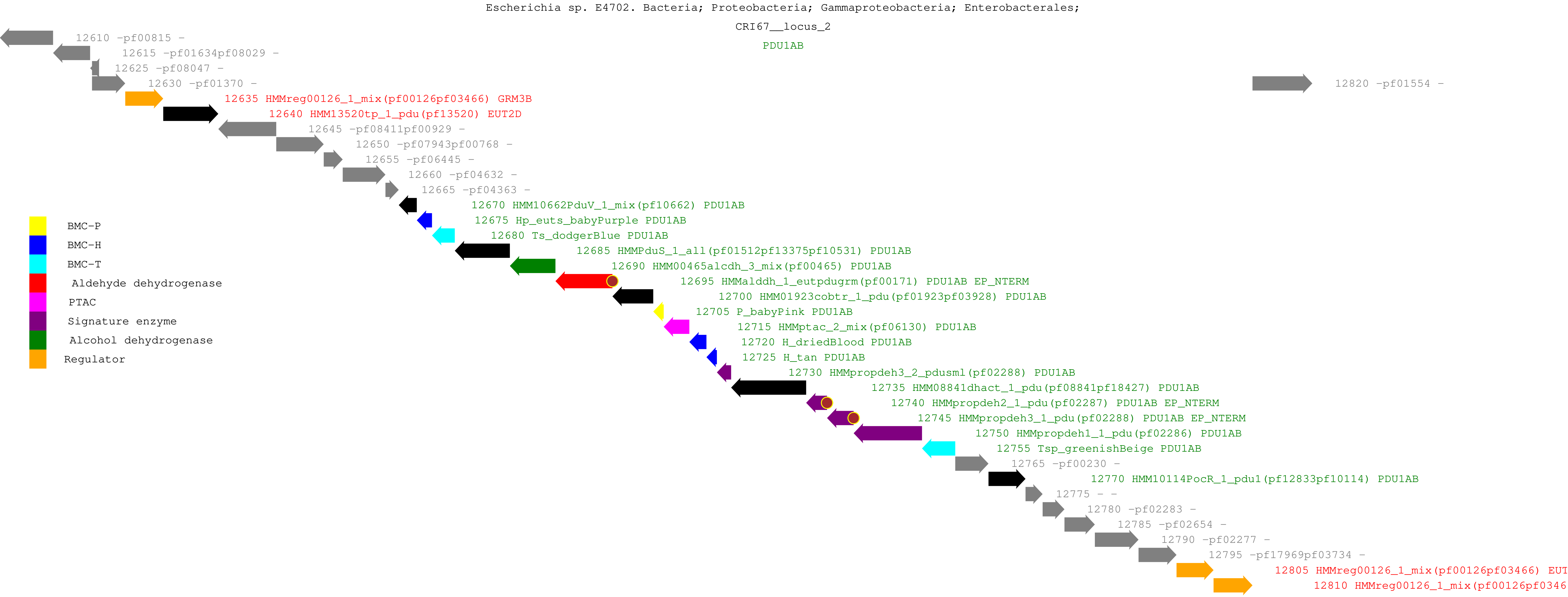

Escherichia sp. E4702. Bacteria; Proteobacteria; Gammaproteobacteria; Enterobacterales;
CRI67__locus_2
PDU1AB
12610 -pf00815 -
12615 -pf01634pf08029 -
12625 -pf08047 -
12630 -pf01370 -
12820 -pf01554 -
12635 HMMreg00126_1_mix(pf00126pf03466) GRM3B
12640 HMM13520tp_1_pdu(pf13520) EUT2D
12645 -pf08411pf00929 -
12650 -pf07943pf00768 -
12655 -pf06445 -
12660 -pf04632 -
12665 -pf04363 -
12670 HMM10662PduV_1_mix(pf10662) PDU1AB
12675 Hp_euts_babyPurple PDU1AB
12680 Ts_dodgerBlue PDU1AB
12685 HMMPduS_1_all(pf01512pf13375pf10531) PDU1AB
12690 HMM00465alcdh_3_mix(pf00465) PDU1AB
12695 HMMalddh_1_eutpdugrm(pf00171) PDU1AB EP_NTERM
12700 HMM01923cobtr_1_pdu(pf01923pf03928) PDU1AB
12705 P_babyPink PDU1AB
12715 HMMptac_2_mix(pf06130) PDU1AB
12720 H_driedBlood PDU1AB
12725 H_tan PDU1AB
12730 HMMpropdeh3_2_pdusml(pf02288) PDU1AB
12735 HMM08841dhact_1_pdu(pf08841pf18427) PDU1AB
12740 HMMpropdeh2_1_pdu(pf02287) PDU1AB EP_NTERM
12745 HMMpropdeh3_1_pdu(pf02288) PDU1AB EP_NTERM
12750 HMMpropdeh1_1_pdu(pf02286) PDU1AB
12755 Tsp_greenishBeige PDU1AB
12765 -pf00230 -
12770 HMM10114PocR_1_pdu1(pf12833pf10114) PDU1AB
12775 - -
12780 -pf02283 -
12785 -pf02654 -
12790 -pf02277 -
12795 -pf17969pf03734 -
12805 HMMreg00126_1_mix(pf00126pf03466) EUT
12810 HMMreg00126_1_mix(pf00126pf0346
BMC-P
BMC-H
BMC-T
Aldehyde dehydrogenase
PTAC
Signature enzyme
Alcohol dehydrogenase
Regulator

Escherichia sp. E4742. Bacteria; Proteobacteria; Gammaproteobacteria; Enterobacterales;

CRI69__locus_2

PDU1AB

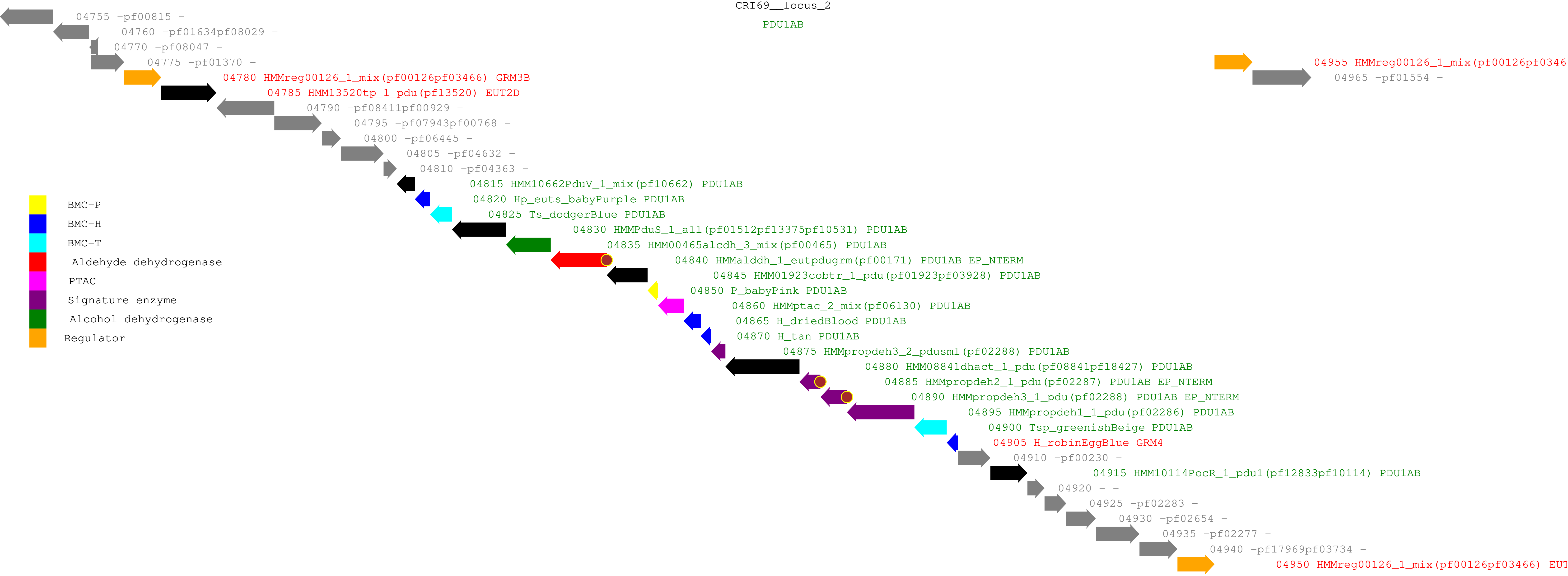

Yersinia kristensenii. Bacteria; Proteobacteria; Gammaproteobacteria; Enterobacterales;

CRM81__locus_1

PDU1AB

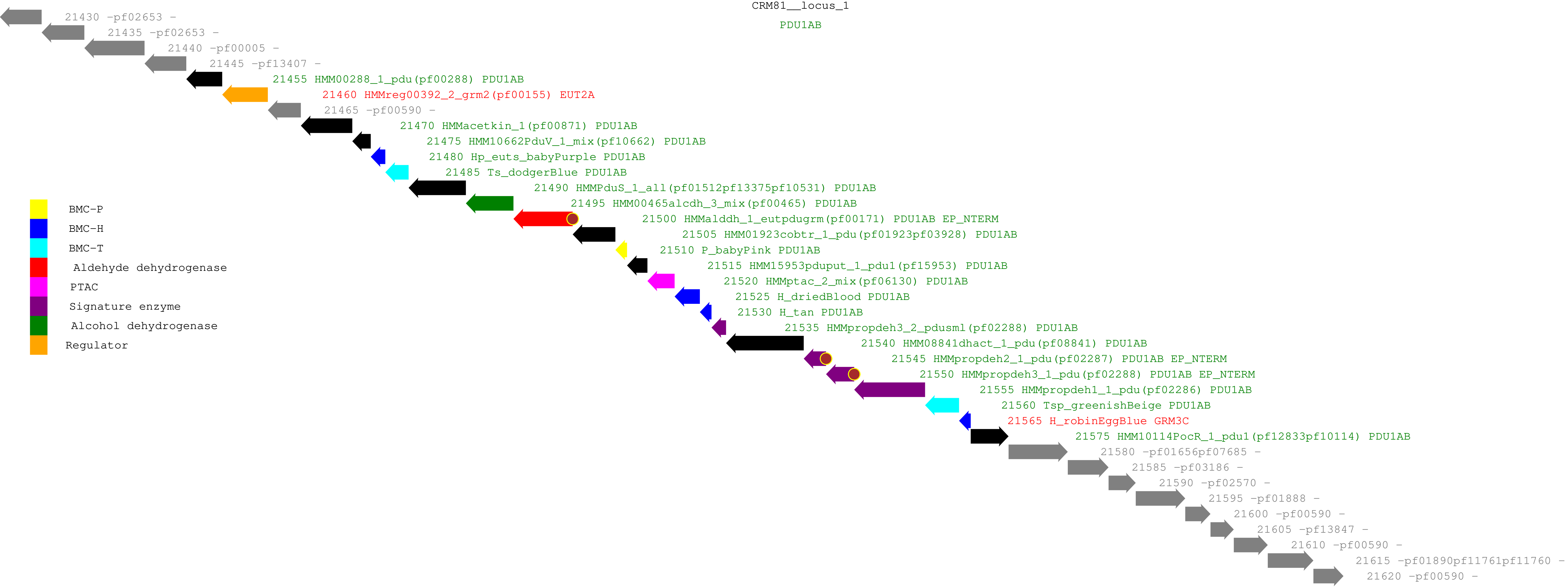

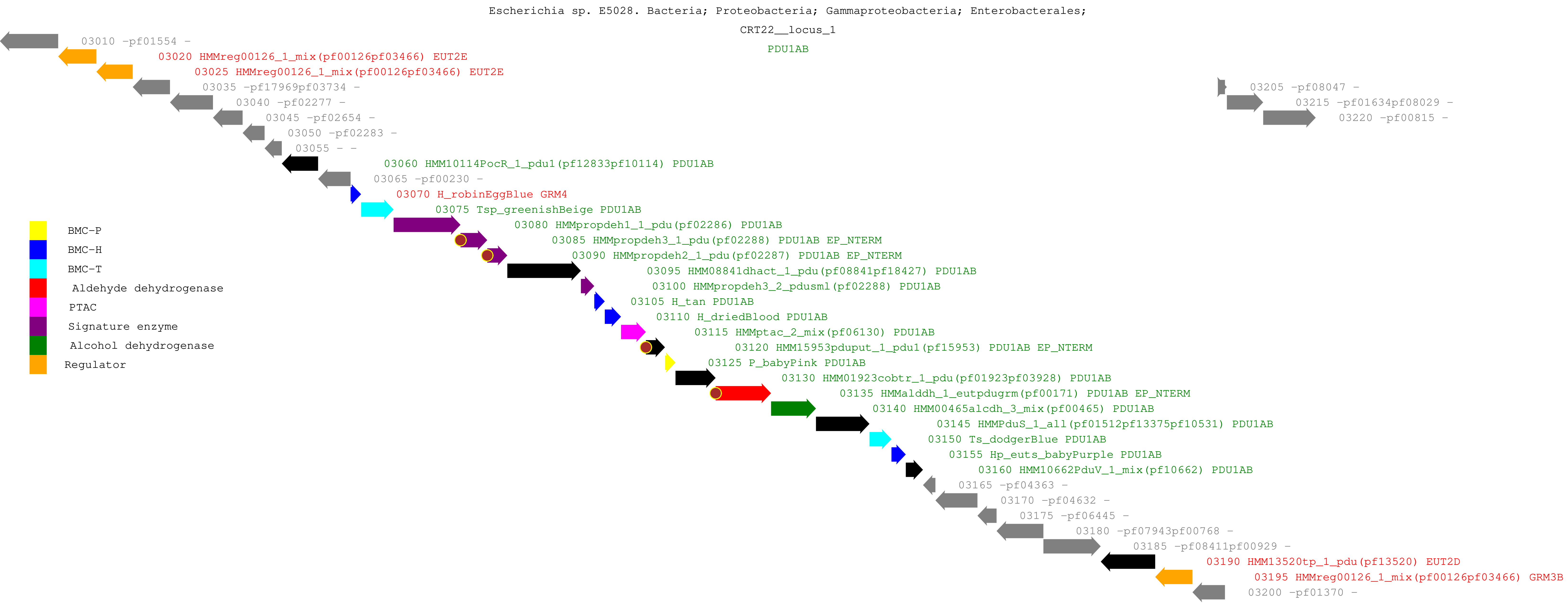
Escherichia sp. E5028. Bacteria; Proteobacteria; Gammaproteobacteria; Enterobacterales;
CRT22__locus_1
PDU1AB
03010 -pf01554 -
03020 HMMreg00126_1_mix(pf00126pf03466) EUT2E
03025 HMMreg00126_1_mix(pf00126pf03466) EUT2E
03035 -pf17969pf03734 -
03040 -pf02277 -
03045 -pf02654 -
03050 -pf02283 -
03055 - -
03060 HMM10114PocR_1_pdu1(pf12833pf10114) PDU1AB
03065 -pf00230 -
03070 H_robinEggBlue GRM4
03075 Tsp_greenishBeige PDU1AB
03080 HMMpropdeh1_1_pdu(pf02286) PDU1AB
03085 HMMpropdeh3_1_pdu(pf02288) PDU1AB EP_NTERM
03090 HMMpropdeh2_1_pdu(pf02287) PDU1AB EP_NTERM
03095 HMM08841dhact_1_pdu(pf08841pf18427) PDU1AB
03100 HMMpropdeh3_2_pdusml(pf02288) PDU1AB
03105 H_tan PDU1AB
03110 H_driedBlood PDU1AB
03115 HMMptac_2_mix(pf06130) PDU1AB
03120 HMM15953pduput_1_pdu1(pf15953) PDU1AB EP_NTERM
03125 P_babyPink PDU1AB
03130 HMM01923cobtr_1_pdu(pf01923pf03928) PDU1AB
03135 HMMalddh_1_eutpdugrm(pf00171) PDU1AB EP_NTERM
03140 HMM00465alcdh_3_mix(pf00465) PDU1AB
03145 HMMPduS_1_all(pf01512pf13375pf10531) PDU1AB
03150 Ts_dodgerBlue PDU1AB
03155 Hp_euts_babyPurple PDU1AB
03160 HMM10662PduV_1_mix(pf10662) PDU1AB
03165 -pf04363 -
03170 -pf04632 -
03175 -pf06445 -
03180 -pf07943pf00768 -
03185 -pf08411pf00929 -
03190 HMM13520tp_1_pdu(pf13520) EUT2D
03195 HMMreg00126_1_mix(pf00126pf03466) GRM3B
03200 -pf01370 -
03205 -pf08047 -
03215 -pf01634pf08029 -
03220 -pf00815 -
BMC-P
BMC-H
BMC-T
Aldehyde dehydrogenase
PTAC
Signature enzyme
Alcohol dehydrogenase
Regulator

Escherichia sp. E4385. Bacteria; Proteobacteria; Gammaproteobacteria; Enterobacterales;

CRU79__locus_1

PDU1AB

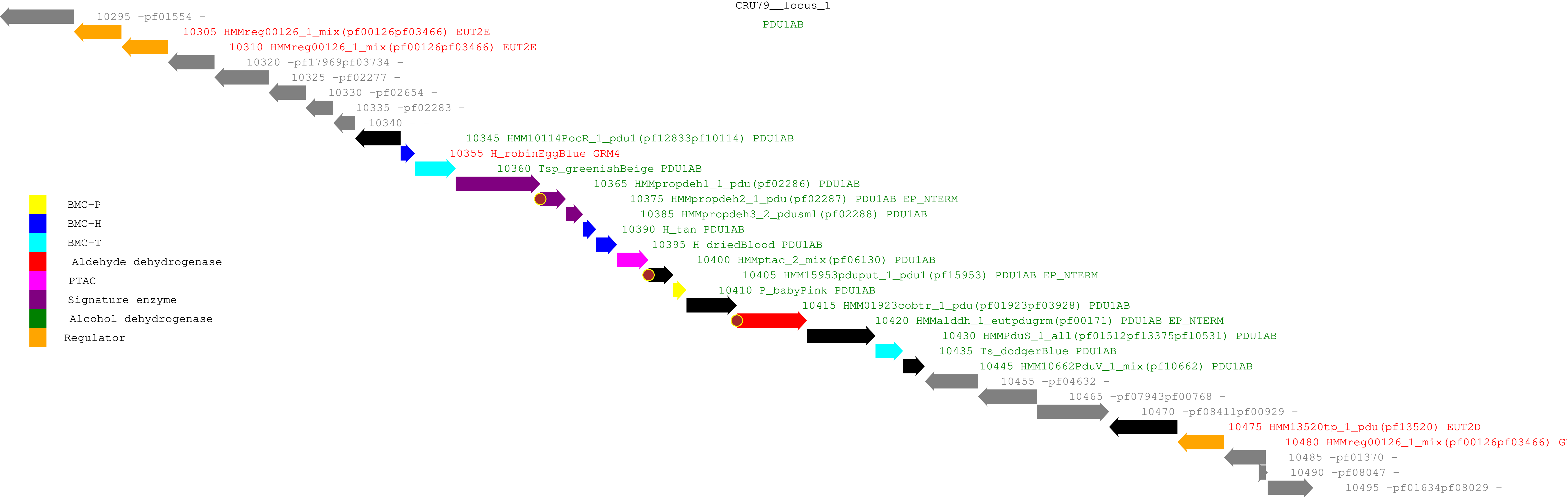

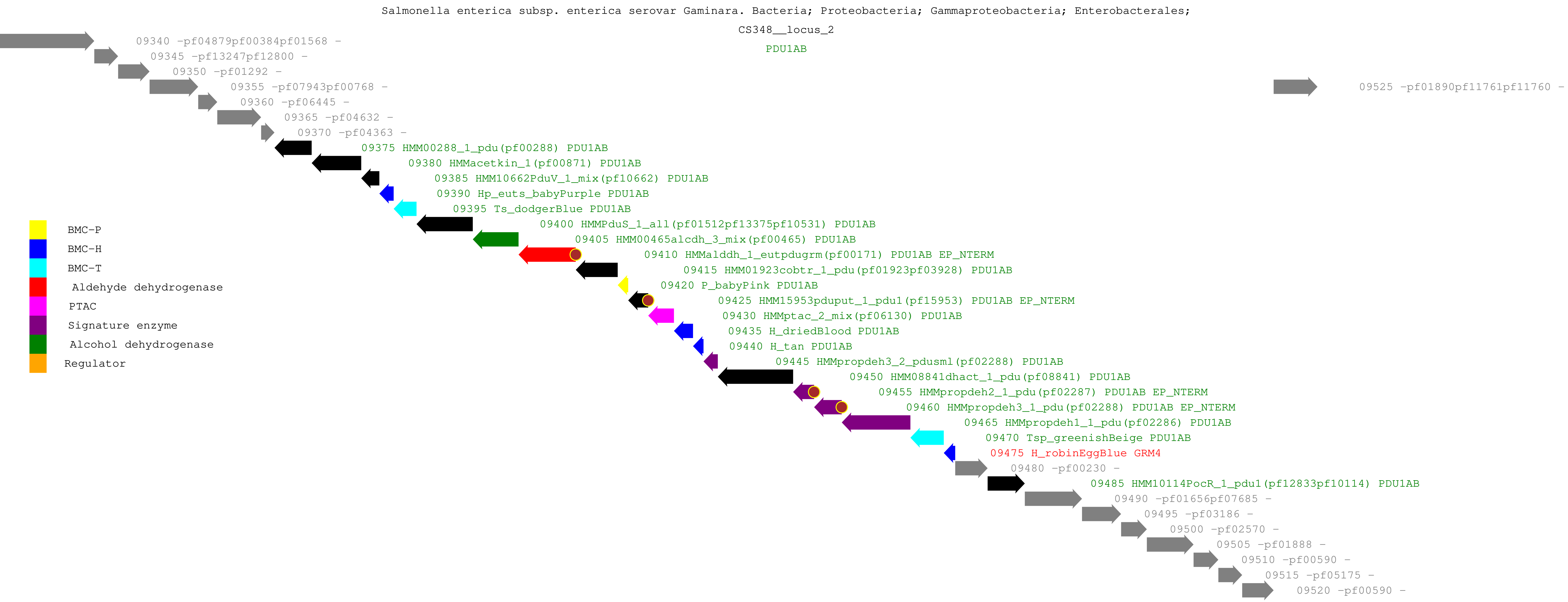
Salmonella enterica subsp. enterica serovar Gaminara. Bacteria; Proteobacteria; Gammaproteobacteria; Enterobacterales;
CS348__locus_2
PDU1AB
09340 -pf04879pf00384pf01568 -
09345 -pf13247pf12800 -
09350 -pf01292 -
09355 -pf07943pf00768 -
09360 -pf06445 -
09365 -pf04632 -
09370 -pf04363 -
09375 HMM00288_1_pdu(pf00288) PDU1AB
09380 HMMacetkin_1(pf00871) PDU1AB
09385 HMM10662PduV_1_mix(pf10662) PDU1AB
09390 Hp_euts_babyPurple PDU1AB
09395 Ts_dodgerBlue PDU1AB
09400 HMMPduS_1_all(pf01512pf13375pf10531) PDU1AB
09405 HMM00465alcdh_3_mix(pf00465) PDU1AB
09410 HMMalddh_1_eutpdugrm(pf00171) PDU1AB EP_NTERM
09415 HMM01923cobtr_1_pdu(pf01923pf03928) PDU1AB
09420 P_babyPink PDU1AB
09425 HMM15953pduput_1_pdu1(pf15953) PDU1AB EP_NTERM
09430 HMMptac_2_mix(pf06130) PDU1AB
09435 H_driedBlood PDU1AB
09440 H_tan PDU1AB
09445 HMMpropdeh3_2_pdusml(pf02288) PDU1AB
09450 HMM08841dhact_1_pdu(pf08841) PDU1AB
09455 HMMpropdeh2_1_pdu(pf02287) PDU1AB EP_NTERM
09460 HMMpropdeh3_1_pdu(pf02288) PDU1AB EP_NTERM
09465 HMMpropdeh1_1_pdu(pf02286) PDU1AB
09470 Tsp_greenishBeige PDU1AB
09475 H_robinEggBlue GRM4
09480 -pf00230 -
09485 HMM10114PocR_1_pdu1(pf12833pf10114) PDU1AB
09490 -pf01656pf07685 -
09495 -pf03186 -
09500 -pf02570 -
09505 -pf01888 -
09510 -pf00590 -
09515 -pf05175 -
09520 -pf00590 -
09525 -pf01890pf11761pf11760 -
BMC-P
BMC-H
BMC-T
Aldehyde dehydrogenase
PTAC
Signature enzyme
Alcohol dehydrogenase
Regulator

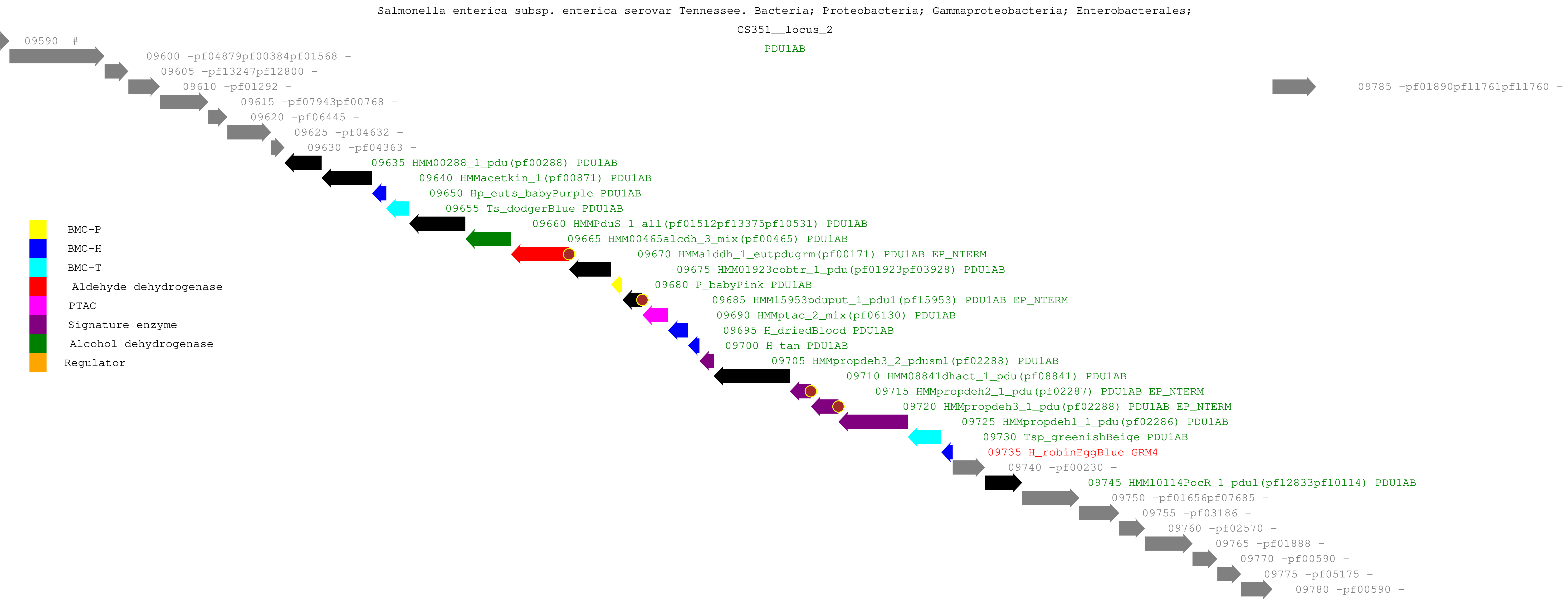
Salmonella enterica subsp. enterica serovar Tennessee. Bacteria; Proteobacteria; Gammaproteobacteria; Enterobacterales;
CS351__locus_2
PDU1AB
09590 -# -
09600 -pf04879pf00384pf01568 -
09605 -pf13247pf12800 -
09610 -pf01292 -
09615 -pf07943pf00768 -
09620 -pf06445 -
09625 -pf04632 -
09630 -pf04363 -
09635 HMM00288_1_pdu(pf00288) PDU1AB
09640 HMMacetkin_1(pf00871) PDU1AB
09650 Hp_euts_babyPurple PDU1AB
09655 Ts_dodgerBlue PDU1AB
09660 HMMPduS_1_all(pf01512pf13375pf10531) PDU1AB
09665 HMM00465alcdh_3_mix(pf00465) PDU1AB
09670 HMMalddh_1_eutpdugrm(pf00171) PDU1AB EP_NTERM
09675 HMM01923cobtr_1_pdu(pf01923pf03928) PDU1AB
09680 P_babyPink PDU1AB
09685 HMM15953pduput_1_pdu1(pf15953) PDU1AB EP_NTERM
09690 HMMptac_2_mix(pf06130) PDU1AB
09695 H_driedBlood PDU1AB
09700 H_tan PDU1AB
09705 HMMpropdeh3_2_pdusml(pf02288) PDU1AB
09710 HMM08841dhact_1_pdu(pf08841) PDU1AB
09715 HMMpropdeh2_1_pdu(pf02287) PDU1AB EP_NTERM
09720 HMMpropdeh3_1_pdu(pf02288) PDU1AB EP_NTERM
09725 HMMpropdeh1_1_pdu(pf02286) PDU1AB
09730 Tsp_greenishBeige PDU1AB
09735 H_robinEggBlue GRM4
09740 -pf00230 -
09745 HMM10114PocR_1_pdu1(pf12833pf10114) PDU1AB
09750 -pf01656pf07685 -
09755 -pf03186 -
09760 -pf02570 -
09765 -pf01888 -
09770 -pf00590 -
09775 -pf05175 -
09780 -pf00590 -
09785 -pf01890pf11761pf11760 -
BMC-P
BMC-H
BMC-T
Aldehyde dehydrogenase
PTAC
Signature enzyme
Alcohol dehydrogenase
Regulator

Yersinia bercovieri. Bacteria; Proteobacteria; Gammaproteobacteria; Enterobacterales;

CS533__locus_1

PDU1AB

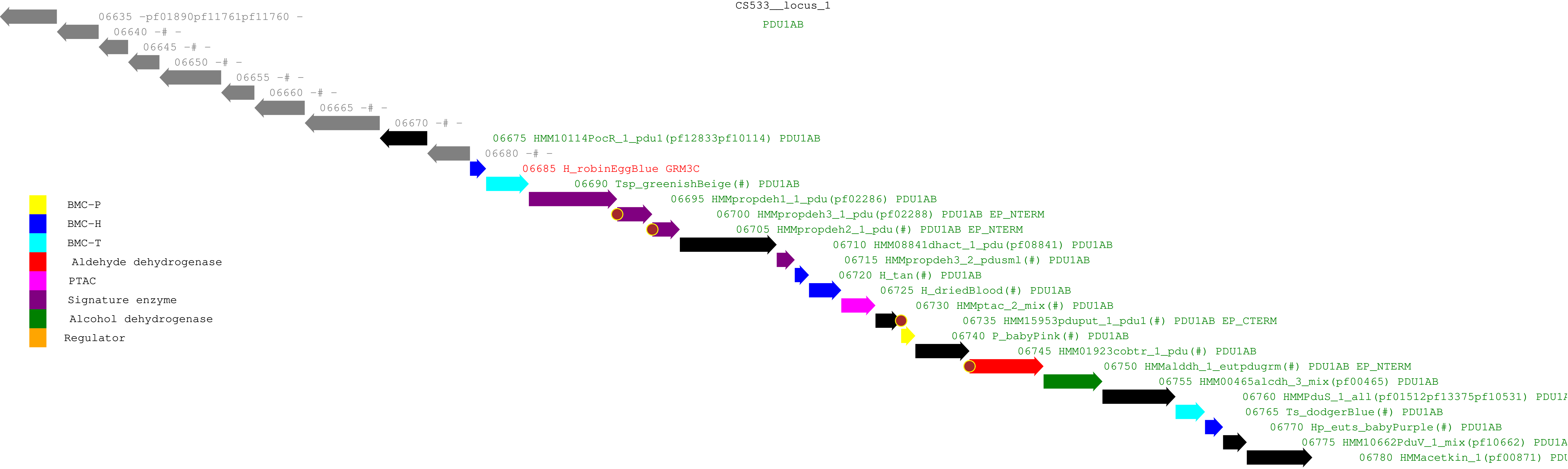

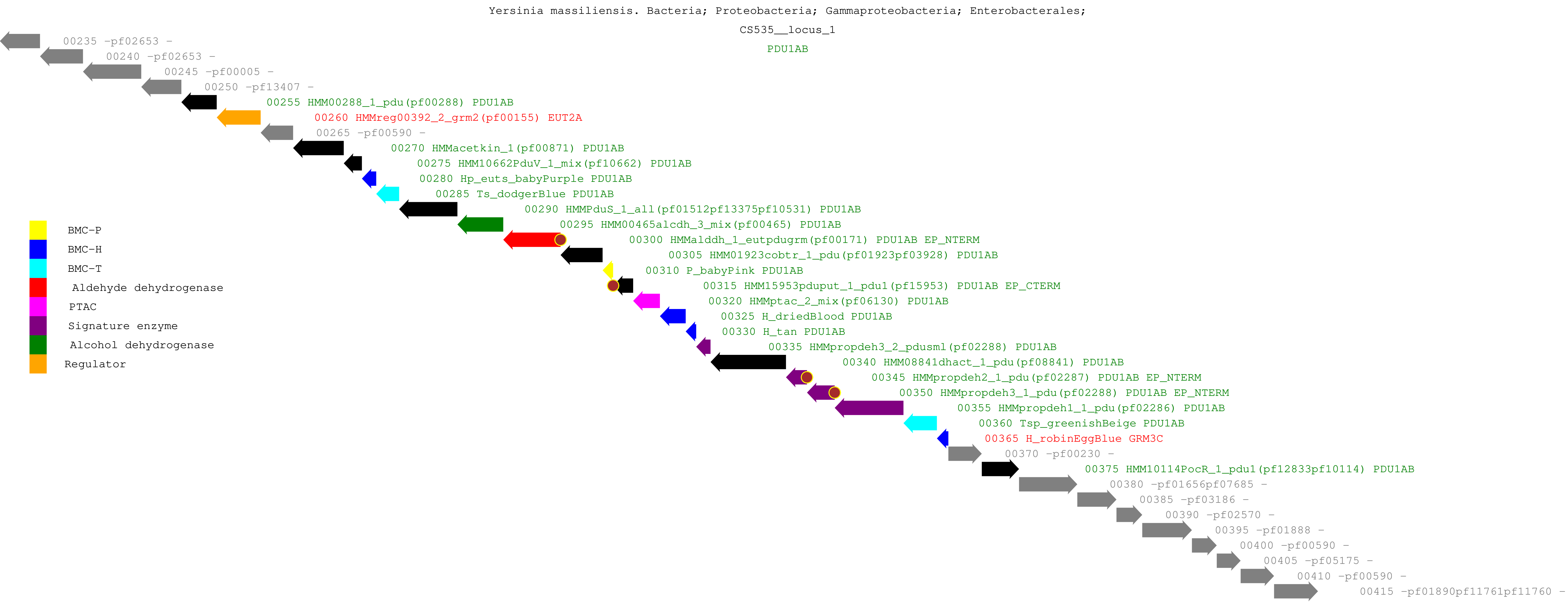
Yersinia massiliensis. Bacteria; Proteobacteria; Gammaproteobacteria; Enterobacterales;
CS535__locus_1
PDU1AB
00235 -pf02653 -
00240 -pf02653 -
00245 -pf00005 -
00250 -pf13407 -
00255 HMM00288_1_pdu(pf00288) PDU1AB
00260 HMMreg00392_2_grm2(pf00155) EUT2A
00265 -pf00590 -
00270 HMMacetkin_1(pf00871) PDU1AB
00275 HMM10662PduV_1_mix(pf10662) PDU1AB
00280 Hp_euts_babyPurple PDU1AB
00285 Ts_dodgerBlue PDU1AB
00290 HMMPduS_1_all(pf01512pf13375pf10531) PDU1AB
00295 HMM00465alcdh_3_mix(pf00465) PDU1AB
00300 HMMalddh_1_eutpdugrm(pf00171) PDU1AB EP_NTERM
00305 HMM01923cobtr_1_pdu(pf01923pf03928) PDU1AB
00310 P_babyPink PDU1AB
00315 HMM15953pduput_1_pdu1(pf15953) PDU1AB EP_CTERM
00320 HMMptac_2_mix(pf06130) PDU1AB
00325 H_driedBlood PDU1AB
00330 H_tan PDU1AB
00335 HMMpropdeh3_2_pdusml(pf02288) PDU1AB
00340 HMM08841dhact_1_pdu(pf08841) PDU1AB
00345 HMMpropdeh2_1_pdu(pf02287) PDU1AB EP_NTERM
00350 HMMpropdeh3_1_pdu(pf02288) PDU1AB EP_NTERM
00355 HMMpropdeh1_1_pdu(pf02286) PDU1AB
00360 Tsp_greenishBeige PDU1AB
00365 H_robinEggBlue GRM3C
00370 -pf00230 -
00375 HMM10114PocR_1_pdu1(pf12833pf10114) PDU1AB
00380 -pf01656pf07685 -
00385 -pf03186 -
00390 -pf02570 -
00395 -pf01888 -
00400 -pf00590 -
00405 -pf05175 -
00410 -pf00590 -
00415 -pf01890pf11761pf11760 -
BMC-P
BMC-H
BMC-T
Aldehyde dehydrogenase
PTAC
Signature enzyme
Alcohol dehydrogenase
Regulator

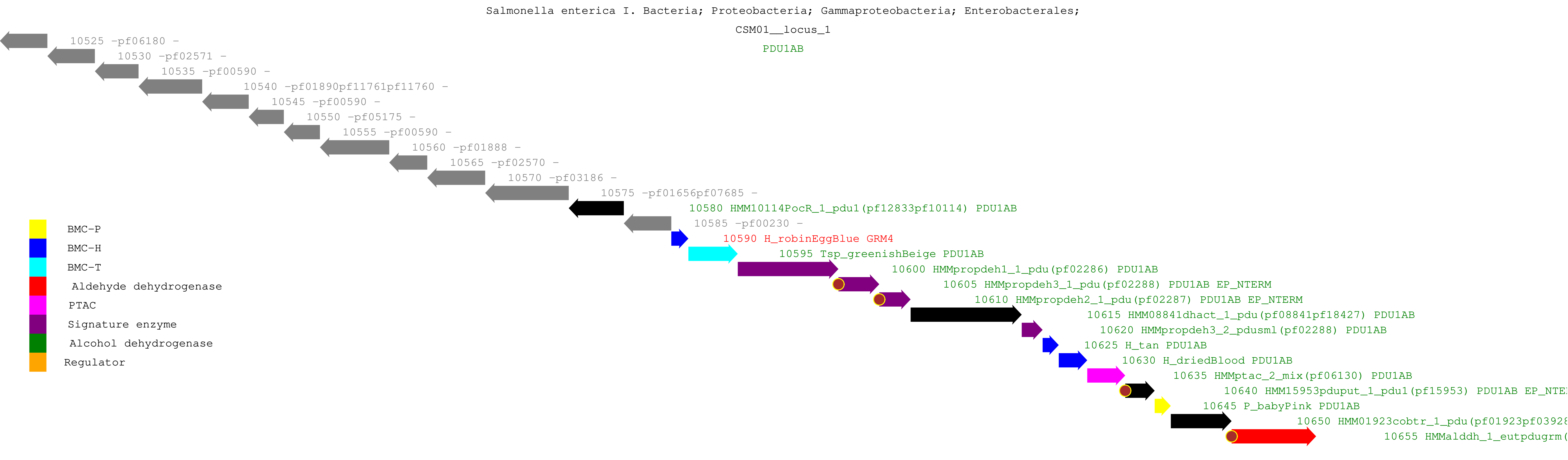
Salmonella enterica I. Bacteria; Proteobacteria; Gammaproteobacteria; Enterobacterales;
CSM01__locus_1
PDU1AB
10525 -pf06180 -
10530 -pf02571 -
10535 -pf00590 -
10540 -pf01890pf11761pf11760 -
10545 -pf00590 -
10550 -pf05175 -
10555 -pf00590 -
10560 -pf01888 -
10565 -pf02570 -
10570 -pf03186 -
10575 -pf01656pf07685 -
10580 HMM10114PocR_1_pdu1(pf12833pf10114) PDU1AB
10585 -pf00230 -
10590 H_robinEggBlue GRM4
10595 Tsp_greenishBeige PDU1AB
10600 HMMpropdeh1_1_pdu(pf02286) PDU1AB
10605 HMMpropdeh3_1_pdu(pf02288) PDU1AB EP_NTERM
10610 HMMpropdeh2_1_pdu(pf02287) PDU1AB EP_NTERM
10615 HMM08841dhact_1_pdu(pf08841pf18427) PDU1AB
10620 HMMpropdeh3_2_pdusml(pf02288) PDU1AB
10625 H_tan PDU1AB
10630 H_driedBlood PDU1AB
10635 HMMptac_2_mix(pf06130) PDU1AB
10640 HMM15953pduput_1_pdu1(pf15953) PDU1AB EP_NTE
10645 P_babyPink PDU1AB
10650 HMM01923cobtr_1_pdu(pf01923pf0392
10655 HMMalddh_1_eutpdugrm(
BMC-P
BMC-H
BMC-T
Aldehyde dehydrogenase
PTAC
Signature enzyme
Alcohol dehydrogenase
Regulator

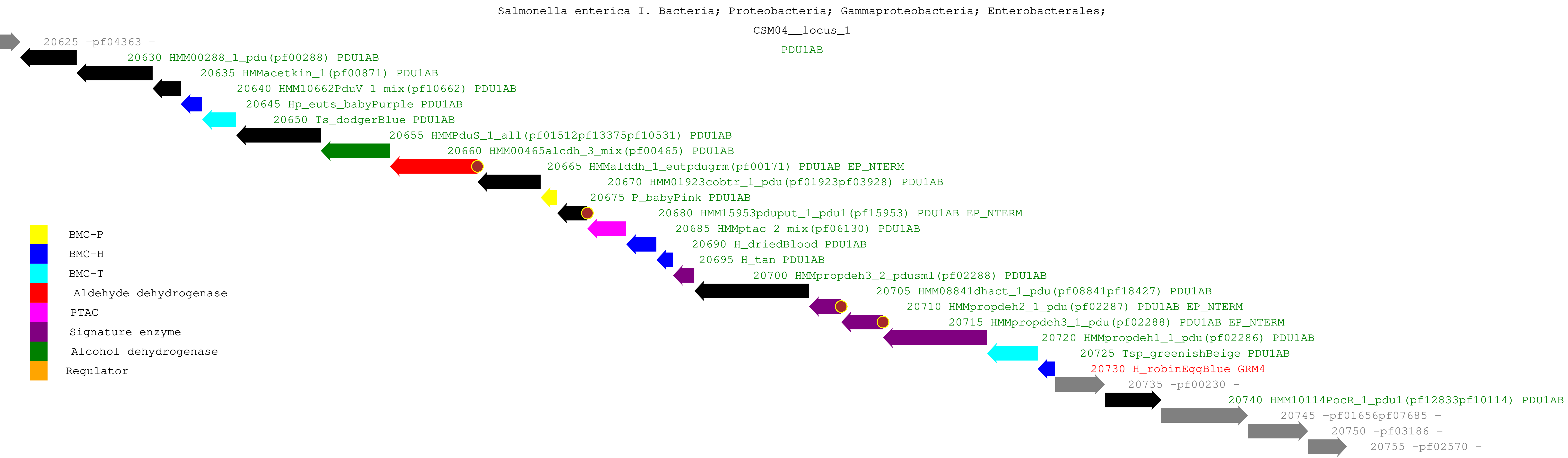
Salmonella enterica I. Bacteria; Proteobacteria; Gammaproteobacteria; Enterobacterales;
CSM04__locus_1
PDU1AB
20625 -pf04363 -
20630 HMM00288_1_pdu(pf00288) PDU1AB
20635 HMMacetkin_1(pf00871) PDU1AB
20640 HMM10662PduV_1_mix(pf10662) PDU1AB
20645 Hp_euts_babyPurple PDU1AB
20650 Ts_dodgerBlue PDU1AB
20655 HMMPduS_1_all(pf01512pf13375pf10531) PDU1AB
20660 HMM00465alcdh_3_mix(pf00465) PDU1AB
20665 HMMalddh_1_eutpdugrm(pf00171) PDU1AB EP_NTERM
20670 HMM01923cobtr_1_pdu(pf01923pf03928) PDU1AB
20675 P_babyPink PDU1AB
20680 HMM15953pduput_1_pdu1(pf15953) PDU1AB EP_NTERM
20685 HMMptac_2_mix(pf06130) PDU1AB
20690 H_driedBlood PDU1AB
20695 H_tan PDU1AB
20700 HMMpropdeh3_2_pdusml(pf02288) PDU1AB
20705 HMM08841dhact_1_pdu(pf08841pf18427) PDU1AB
20710 HMMpropdeh2_1_pdu(pf02287) PDU1AB EP_NTERM
20715 HMMpropdeh3_1_pdu(pf02288) PDU1AB EP_NTERM
20720 HMMpropdeh1_1_pdu(pf02286) PDU1AB
20725 Tsp_greenishBeige PDU1AB
20730 H_robinEggBlue GRM4
20735 -pf00230 -
20740 HMM10114PocR_1_pdu1(pf12833pf10114) PDU1AB
20745 -pf01656pf07685 -
20750 -pf03186 -
20755 -pf02570 -
BMC-P
BMC-H
BMC-T
Aldehyde dehydrogenase
PTAC
Signature enzyme
Alcohol dehydrogenase
Regulator

Salmonella enterica subsp. enterica serovar Weltevreden. Bacteria; Proteobacteria; Gammaproteobacteria; Enterobacterales;

CSR62__locus_2

PDU1AB

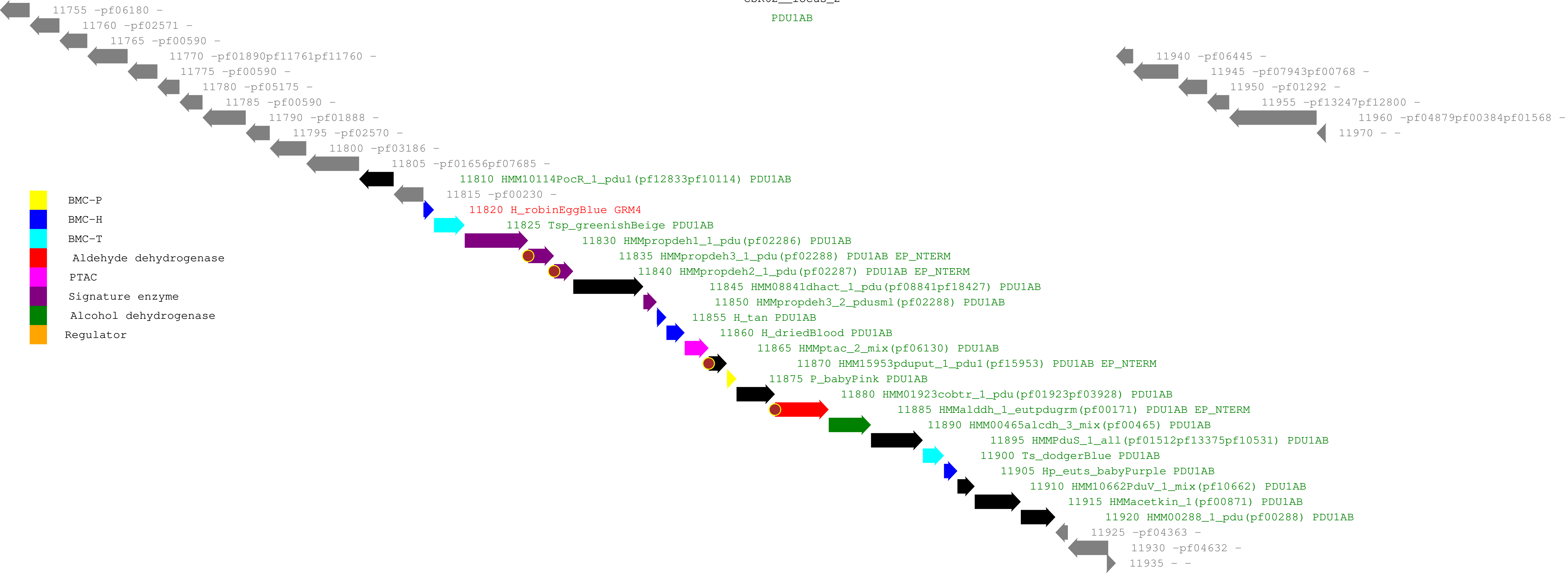

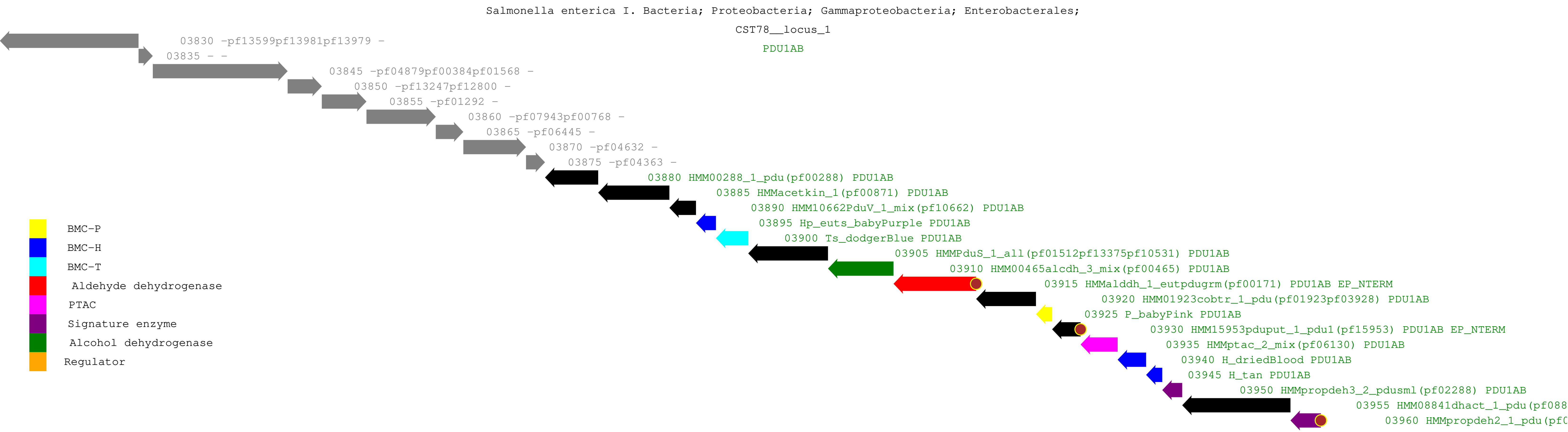
Salmonella enterica I. Bacteria; Proteobacteria; Gammaproteobacteria; Enterobacterales;
CST78__locus_1
PDU1AB
03830 -pf13599pf13981pf13979 -
03835 - -
03845 -pf04879pf00384pf01568 -
03850 -pf13247pf12800 -
03855 -pf01292 -
03860 -pf07943pf00768 -
03865 -pf06445 -
03870 -pf04632 -
03875 -pf04363 -
03880 HMM00288_1_pdu(pf00288) PDU1AB
03885 HMMacetkin_1(pf00871) PDU1AB
03890 HMM10662PduV_1_mix(pf10662) PDU1AB
03895 Hp_euts_babyPurple PDU1AB
03900 Ts_dodgerBlue PDU1AB
03905 HMMPduS_1_all(pf01512pf13375pf10531) PDU1AB
03910 HMM00465alcdh_3_mix(pf00465) PDU1AB
03915 HMMalddh_1_eutpdugrm(pf00171) PDU1AB EP_NTERM
03920 HMM01923cobtr_1_pdu(pf01923pf03928) PDU1AB
03925 P_babyPink PDU1AB
03930 HMM15953pduput_1_pdu1(pf15953) PDU1AB EP_NTERM
03935 HMMptac_2_mix(pf06130) PDU1AB
03940 H_driedBlood PDU1AB
03945 H_tan PDU1AB
03950 HMMpropdeh3_2_pdusml(pf02288) PDU1AB
03955 HMM08841dhact_1_pdu(pf088
03960 HMMpropdeh2_1_pdu(pf0
BMC-P
BMC-H
BMC-T
Aldehyde dehydrogenase
PTAC
Signature enzyme
Alcohol dehydrogenase
Regulator

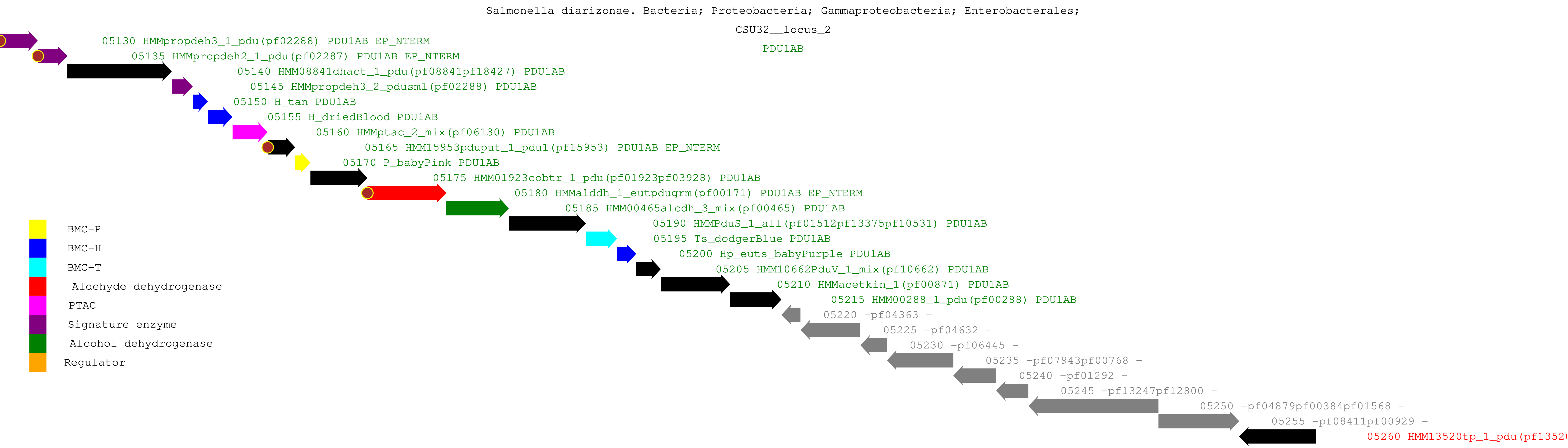
Salmonella diarizonae. Bacteria; Proteobacteria; Gammaproteobacteria; Enterobacterales;
CSU32__locus_2
PDU1AB
05130 HMMpropdeh3_1_pdu(pf02288) PDU1AB EP_NTERM
05135 HMMpropdeh2_1_pdu(pf02287) PDU1AB EP_NTERM
05140 HMM08841dhact_1_pdu(pf08841pf18427) PDU1AB
05145 HMMpropdeh3_2_pdusml(pf02288) PDU1AB
05150 H_tan PDU1AB
05155 H_driedBlood PDU1AB
05160 HMMptac_2_mix(pf06130) PDU1AB
05165 HMM15953pduput_1_pdu1(pf15953) PDU1AB EP_NTERM
05170 P_babyPink PDU1AB
05175 HMM01923cobtr_1_pdu(pf01923pf03928) PDU1AB
05180 HMMalddh_1_eutpdugrm(pf00171) PDU1AB EP_NTERM
05185 HMM00465alcdh_3_mix(pf00465) PDU1AB
05190 HMMPduS_1_all(pf01512pf13375pf10531) PDU1AB
05195 Ts_dodgerBlue PDU1AB
05200 Hp_euts_babyPurple PDU1AB
05205 HMM10662PduV_1_mix(pf10662) PDU1AB
05210 HMMacetkin_1(pf00871) PDU1AB
05215 HMM00288_1_pdu(pf00288) PDU1AB
05220 -pf04363 -
05225 -pf04632 -
05230 -pf06445 -
05235 -pf07943pf00768 -
05240 -pf01292 -
05245 -pf13247pf12800 -
05250 -pf04879pf00384pf01568 -
05255 -pf08411pf00929 -
05260 HMM13520tp_1_pdu(pf1352
BMC-P
BMC-H
BMC-T
Aldehyde dehydrogenase
PTAC
Signature enzyme
Alcohol dehydrogenase
Regulator

Salmonella enterica subsp. enterica serovar Javiana. Bacteria; Proteobacteria; Gammaproteobacteria; Enterobacterales;

CT060__locus_1

PDU1AB

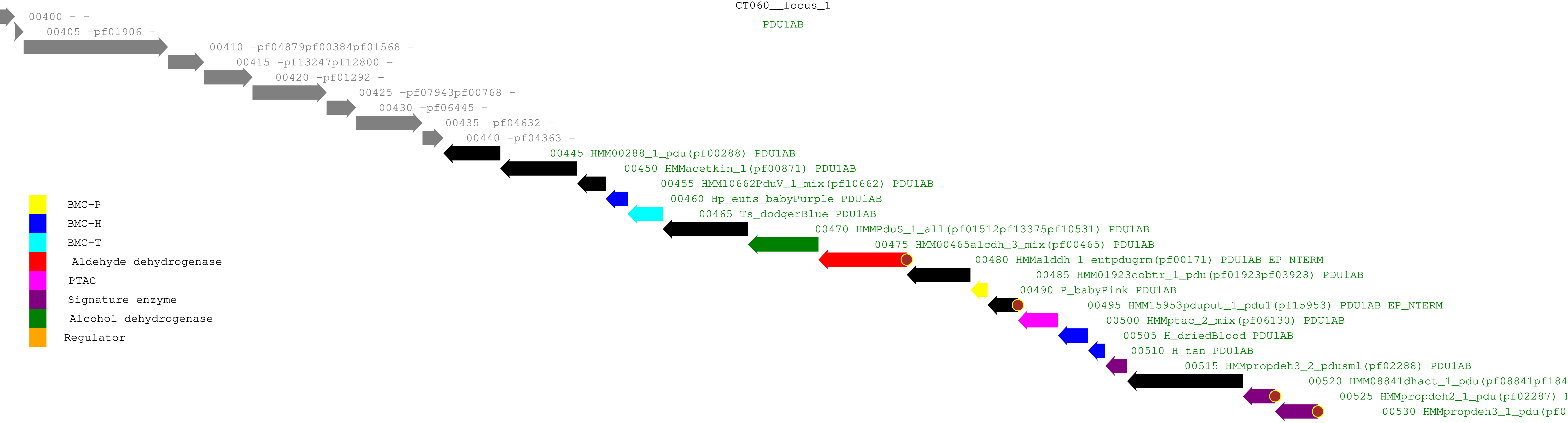

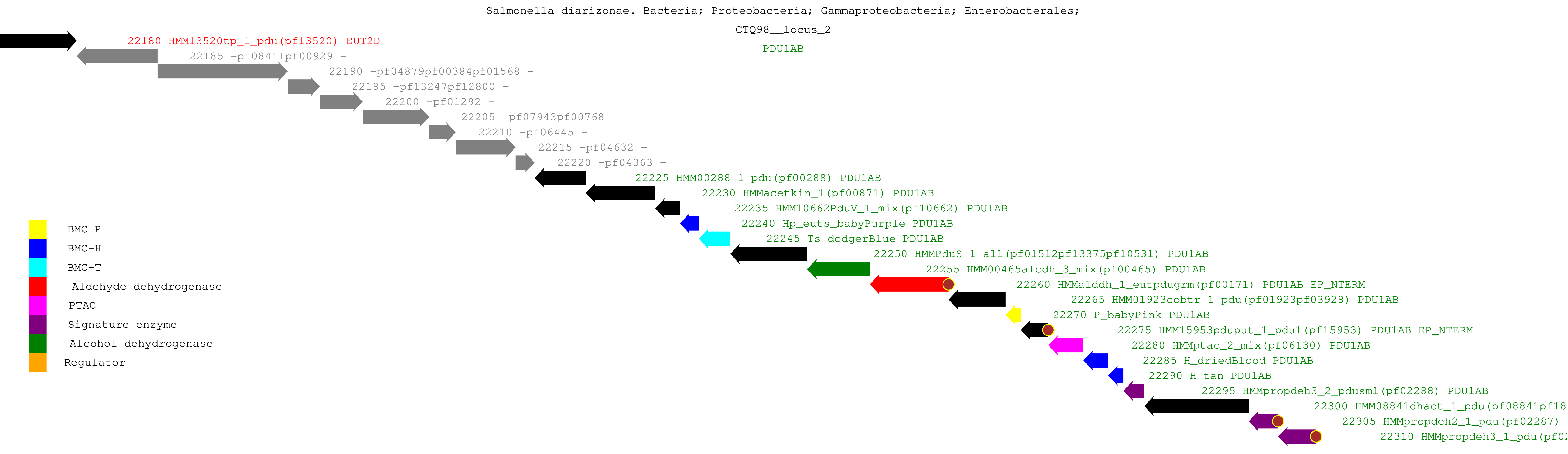

Salmonella diarizonae. Bacteria; Proteobacteria; Gammaproteobacteria; Enterobacterales;
CTQ98__locus_2
PDU1AB
22180 HMM13520tp_1_pdu(pf13520) EUT2D
22185 -pf08411pf00929 -
22190 -pf04879pf00384pf01568 -
22195 -pf13247pf12800 -
22200 -pf01292 -
22205 -pf07943pf00768 -
22210 -pf06445 -
22215 -pf04632 -
22220 -pf04363 -
22225 HMM00288_1_pdu(pf00288) PDU1AB
22230 HMMacetkin_1(pf00871) PDU1AB
22235 HMM10662PduV_1_mix(pf10662) PDU1AB
22240 Hp_euts_babyPurple PDU1AB
22245 Ts_dodgerBlue PDU1AB
22250 HMMPduS_1_all(pf01512pf13375pf10531) PDU1AB
22255 HMM00465alcdh_3_mix(pf00465) PDU1AB
22260 HMMalddh_1_eutpdugrm(pf00171) PDU1AB EP_NTERM
22265 HMM01923cobtr_1_pdu(pf01923pf03928) PDU1AB
22270 P_babyPink PDU1AB
22275 HMM15953pduput_1_pdu1(pf15953) PDU1AB EP_NTERM
22280 HMMptac_2_mix(pf06130) PDU1AB
22285 H_driedBlood PDU1AB
22290 H_tan PDU1AB
22295 HMMpropdeh3_2_pdusml(pf02288) PDU1AB
22300 HMM08841dhact_1_pdu(pf08841pf18
22305 HMMpropdeh2_1_pdu(pf02287)
22310 HMMpropdeh3_1_pdu(pf0
BMC-P
BMC-H
BMC-T
Aldehyde dehydrogenase
PTAC
Signature enzyme
Alcohol dehydrogenase
Regulator

Enterobacteriaceae bacterium S05. Bacteria; Proteobacteria; Gammaproteobacteria; Enterobacterales;

CUC76__locus_2

PDU1AB

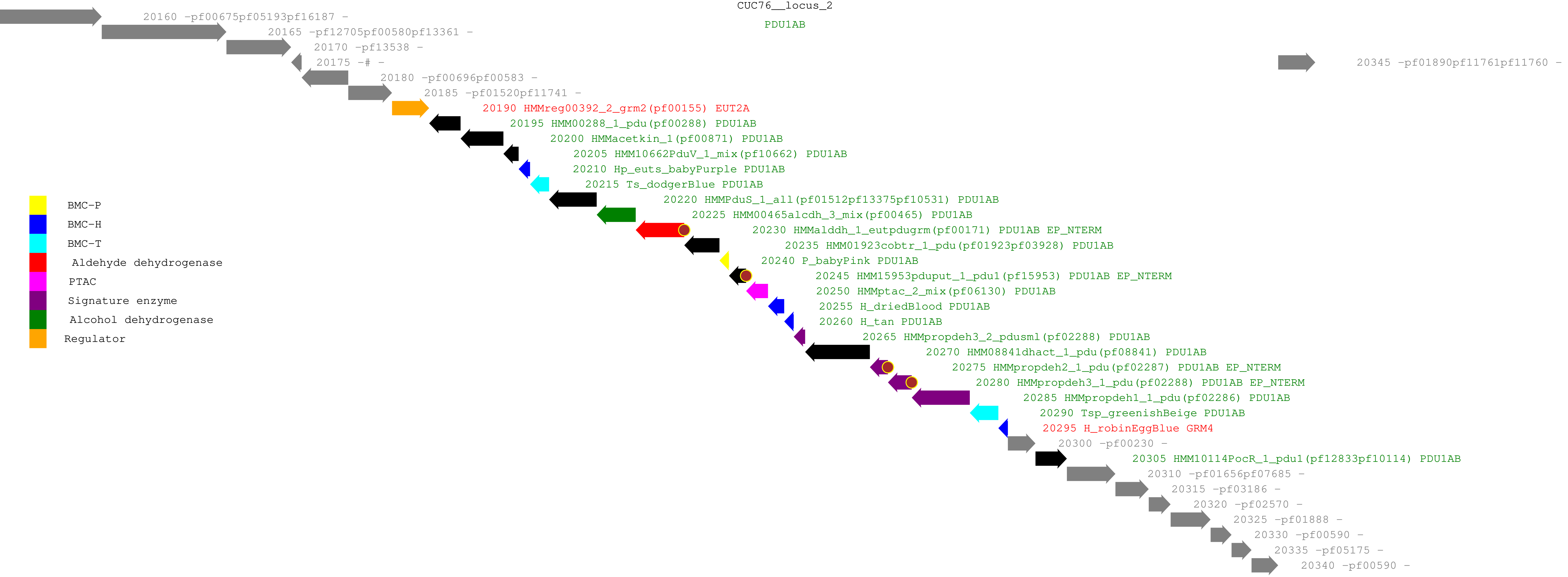

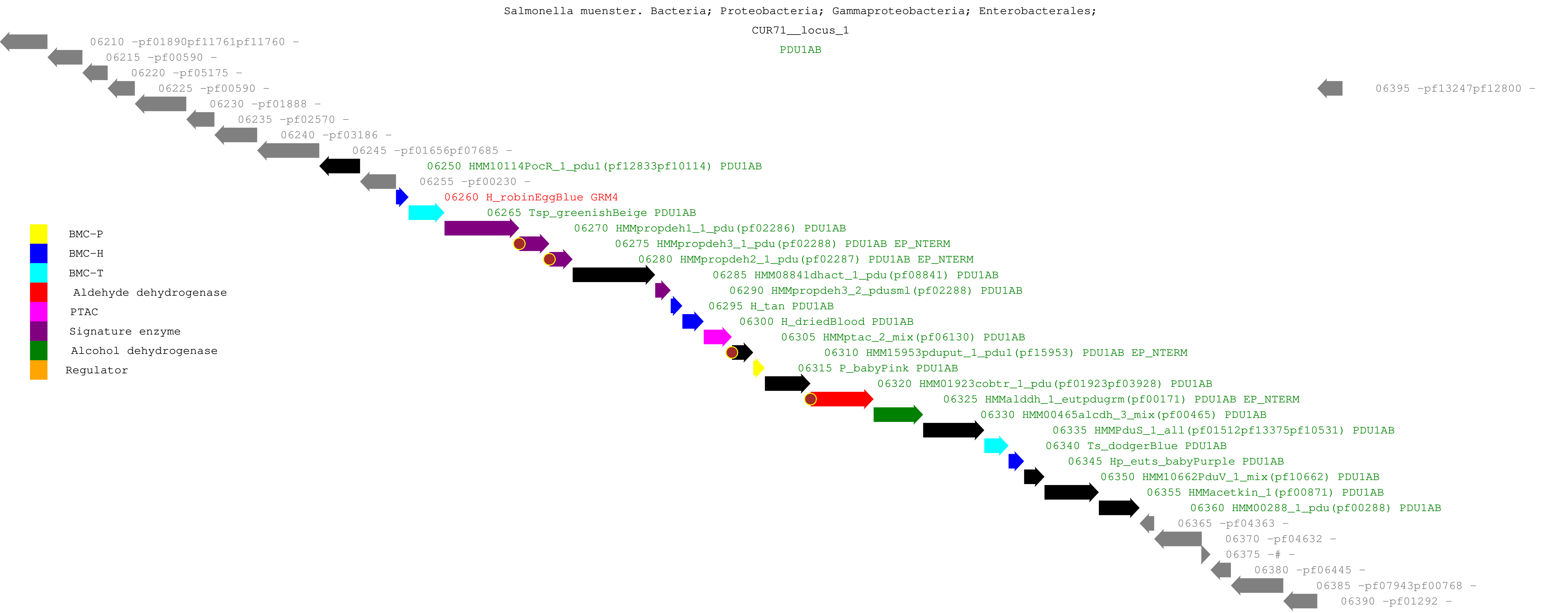
Salmonella muenster. Bacteria; Proteobacteria; Gammaproteobacteria; Enterobacterales;
CUR71__locus_1
PDU1AB
06210 -pf01890pf11761pf11760 -
06215 -pf00590 -
06220 -pf05175 -
06225 -pf00590 -
06230 -pf01888 -
06235 -pf02570 -
06240 -pf03186 -
06245 -pf01656pf07685 -
06250 HMM10114PocR_1_pdu1(pf12833pf10114) PDU1AB
06255 -pf00230 -
06260 H_robinEggBlue GRM4
06265 Tsp_greenishBeige PDU1AB
06270 HMMpropdeh1_1_pdu(pf02286) PDU1AB
06275 HMMpropdeh3_1_pdu(pf02288) PDU1AB EP_NTERM
06280 HMMpropdeh2_1_pdu(pf02287) PDU1AB EP_NTERM
06285 HMM08841dhact_1_pdu(pf08841) PDU1AB
06290 HMMpropdeh3_2_pdusml(pf02288) PDU1AB
06295 H_tan PDU1AB
06300 H_driedBlood PDU1AB
06305 HMMptac_2_mix(pf06130) PDU1AB
06310 HMM15953pduput_1_pdu1(pf15953) PDU1AB EP_NTERM
06315 P_babyPink PDU1AB
06320 HMM01923cobtr_1_pdu(pf01923pf03928) PDU1AB
06325 HMMalddh_1_eutpdugrm(pf00171) PDU1AB EP_NTERM
06330 HMM00465alcdh_3_mix(pf00465) PDU1AB
06335 HMMPduS_1_all(pf01512pf13375pf10531) PDU1AB
06340 Ts_dodgerBlue PDU1AB
06345 Hp_euts_babyPurple PDU1AB
06350 HMM10662PduV_1_mix(pf10662) PDU1AB
06355 HMMacetkin_1(pf00871) PDU1AB
06360 HMM00288_1_pdu(pf00288) PDU1AB
06365 -pf04363 -
06370 -pf04632 -
06375 -# -
06380 -pf06445 -
06385 -pf07943pf00768 -
06390 -pf01292 -
06395 -pf13247pf12800 -
BMC-P
BMC-H
BMC-T
Aldehyde dehydrogenase
PTAC
Signature enzyme
Alcohol dehydrogenase
Regulator

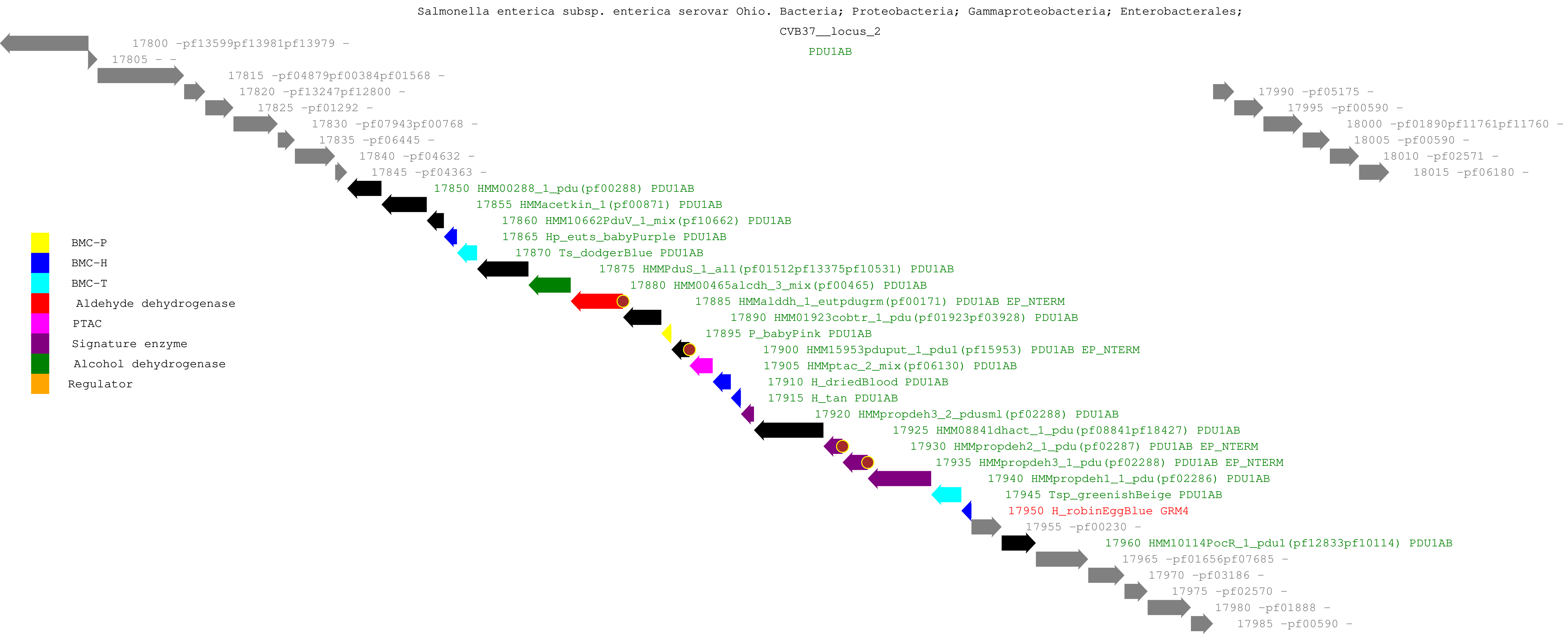

Salmonella enterica subsp. enterica serovar Ohio. Bacteria; Proteobacteria; Gammaproteobacteria; Enterobacterales;
CVB37__locus_2
PDU1AB
17800 -pf13599pf13981pf13979 -
17805 - -
17815 -pf04879pf00384pf01568 -
17820 -pf13247pf12800 -
17825 -pf01292 -
17830 -pf07943pf00768 -
17835 -pf06445 -
17840 -pf04632 -
17845 -pf04363 -
17850 HMM00288_1_pdu(pf00288) PDU1AB
17855 HMMacetkin_1(pf00871) PDU1AB
17860 HMM10662PduV_1_mix(pf10662) PDU1AB
17865 Hp_euts_babyPurple PDU1AB
17870 Ts_dodgerBlue PDU1AB
17875 HMMPduS_1_all(pf01512pf13375pf10531) PDU1AB
17880 HMM00465alcdh_3_mix(pf00465) PDU1AB
17885 HMMalddh_1_eutpdugrm(pf00171) PDU1AB EP_NTERM
17890 HMM01923cobtr_1_pdu(pf01923pf03928) PDU1AB
17895 P_babyPink PDU1AB
17900 HMM15953pduput_1_pdu1(pf15953) PDU1AB EP_NTERM
17905 HMMptac_2_mix(pf06130) PDU1AB
17910 H_driedBlood PDU1AB
17915 H_tan PDU1AB
17920 HMMpropdeh3_2_pdusml(pf02288) PDU1AB
17925 HMM08841dhact_1_pdu(pf08841pf18427) PDU1AB
17930 HMMpropdeh2_1_pdu(pf02287) PDU1AB EP_NTERM
17935 HMMpropdeh3_1_pdu(pf02288) PDU1AB EP_NTERM
17940 HMMpropdeh1_1_pdu(pf02286) PDU1AB
17945 Tsp_greenishBeige PDU1AB
17950 H_robinEggBlue GRM4
17955 -pf00230 -
17960 HMM10114PocR_1_pdu1(pf12833pf10114) PDU1AB
17965 -pf01656pf07685 -
17970 -pf03186 -
17975 -pf02570 -
17980 -pf01888 -
17985 -pf00590 -
17990 -pf05175 -
17995 -pf00590 -
18000 -pf01890pf11761pf11760 -
18005 -pf00590 -
18010 -pf02571 -
18015 -pf06180 -
BMC-P
BMC-H
BMC-T
Aldehyde dehydrogenase
PTAC
Signature enzyme
Alcohol dehydrogenase
Regulator

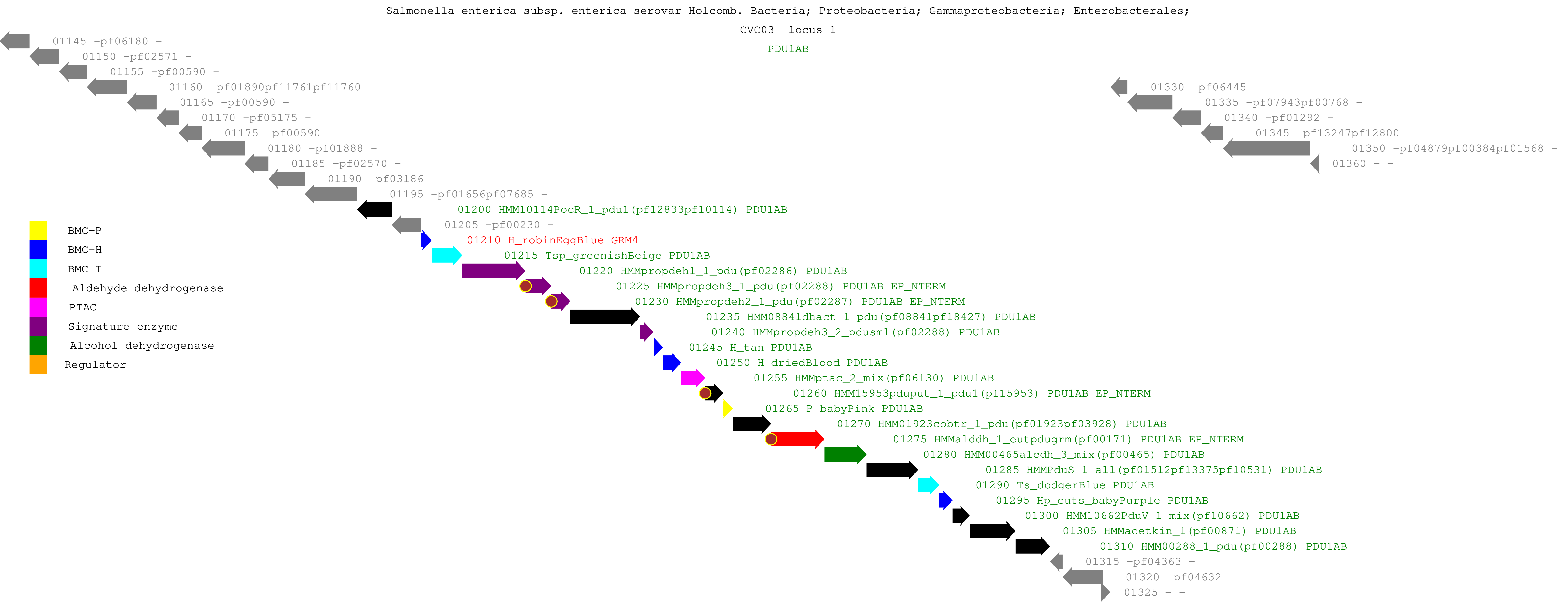

Salmonella enterica subsp. enterica serovar Holcomb. Bacteria; Proteobacteria; Gammaproteobacteria; Enterobacterales;
CVC03__locus_1
PDU1AB
01145 -pf06180 -
01150 -pf02571 -
01155 -pf00590 -
01160 -pf01890pf11761pf11760 -
01165 -pf00590 -
01170 -pf05175 -
01175 -pf00590 -
01180 -pf01888 -
01185 -pf02570 -
01190 -pf03186 -
01195 -pf01656pf07685 -
01200 HMM10114PocR_1_pdu1(pf12833pf10114) PDU1AB
01205 -pf00230 -
01210 H_robinEggBlue GRM4
01215 Tsp_greenishBeige PDU1AB
01220 HMMpropdeh1_1_pdu(pf02286) PDU1AB
01225 HMMpropdeh3_1_pdu(pf02288) PDU1AB EP_NTERM
01230 HMMpropdeh2_1_pdu(pf02287) PDU1AB EP_NTERM
01235 HMM08841dhact_1_pdu(pf08841pf18427) PDU1AB
01240 HMMpropdeh3_2_pdusml(pf02288) PDU1AB
01245 H_tan PDU1AB
01250 H_driedBlood PDU1AB
01255 HMMptac_2_mix(pf06130) PDU1AB
01260 HMM15953pduput_1_pdu1(pf15953) PDU1AB EP_NTERM
01265 P_babyPink PDU1AB
01270 HMM01923cobtr_1_pdu(pf01923pf03928) PDU1AB
01275 HMMalddh_1_eutpdugrm(pf00171) PDU1AB EP_NTERM
01280 HMM00465alcdh_3_mix(pf00465) PDU1AB
01285 HMMPduS_1_all(pf01512pf13375pf10531) PDU1AB
01290 Ts_dodgerBlue PDU1AB
01295 Hp_euts_babyPurple PDU1AB
01300 HMM10662PduV_1_mix(pf10662) PDU1AB
01305 HMMacetkin_1(pf00871) PDU1AB
01310 HMM00288_1_pdu(pf00288) PDU1AB
01315 -pf04363 -
01320 -pf04632 -
01325 - -
01330 -pf06445 -
01335 -pf07943pf00768 -
01340 -pf01292 -
01345 -pf13247pf12800 -
01350 -pf04879pf00384pf01568 -
01360 - -
BMC-P
BMC-H
BMC-T
Aldehyde dehydrogenase
PTAC
Signature enzyme
Alcohol dehydrogenase
Regulator

Salmonella enterica I. Bacteria; Proteobacteria; Gammaproteobacteria; Enterobacterales;

CVR97__locus_1

PDU1AB

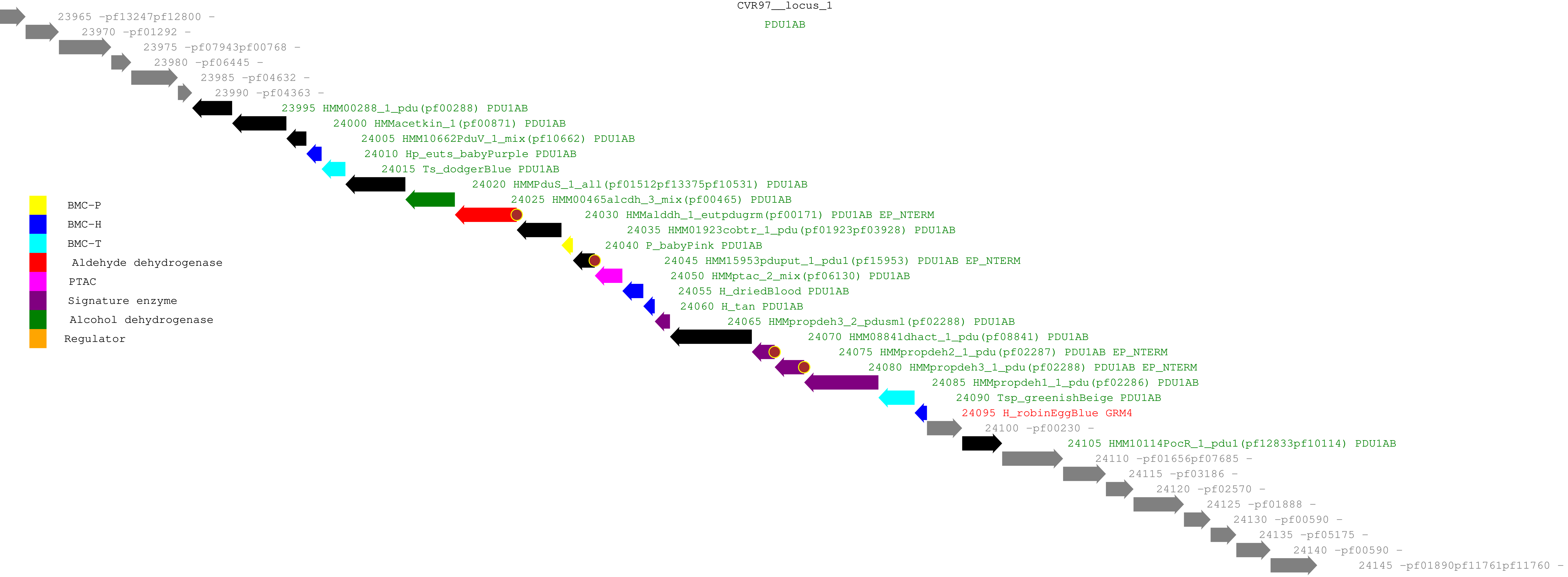

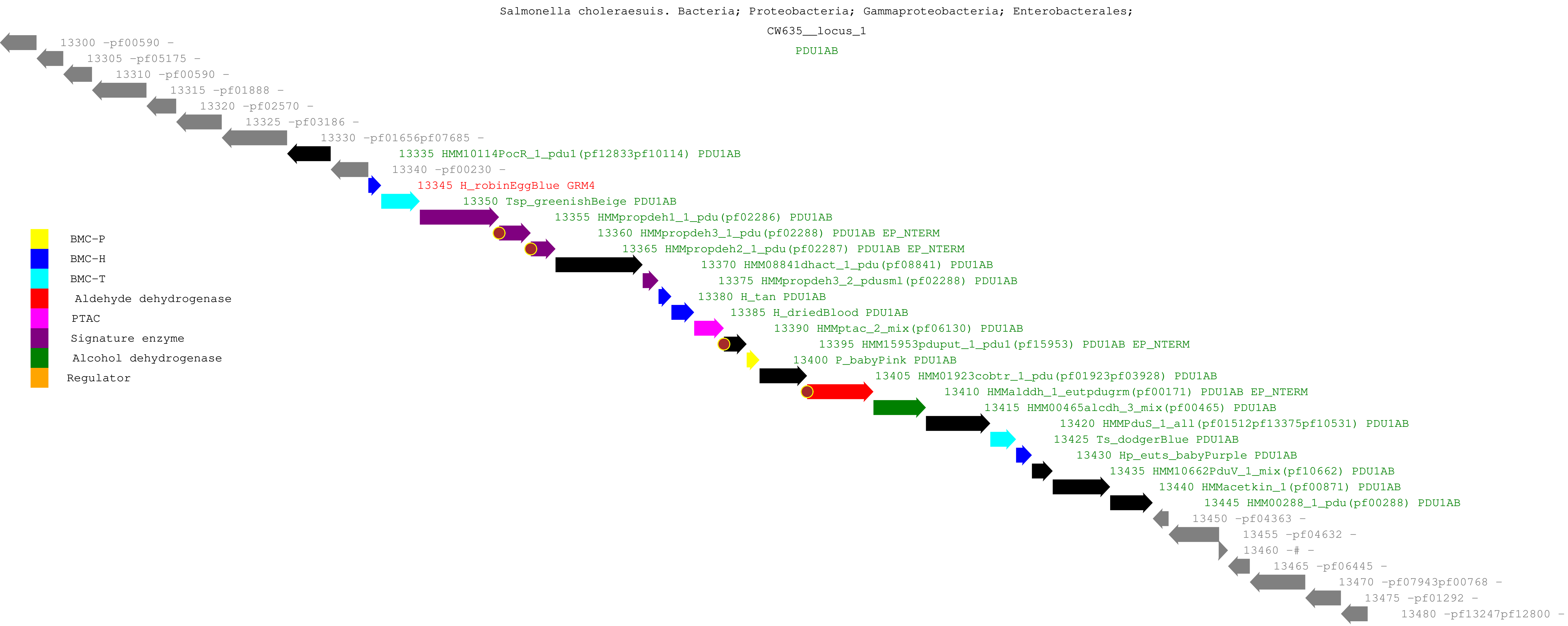

Salmonella choleraesuis. Bacteria; Proteobacteria; Gammaproteobacteria; Enterobacterales;
CW635__locus_1
PDU1AB
13300 -pf00590 -
13305 -pf05175 -
13310 -pf00590 -
13315 -pf01888 -
13320 -pf02570 -
13325 -pf03186 -
13330 -pf01656pf07685 -
13335 HMM10114PocR_1_pdu1(pf12833pf10114) PDU1AB
13340 -pf00230 -
13345 H_robinEggBlue GRM4
13350 Tsp_greenishBeige PDU1AB
13355 HMMpropdeh1_1_pdu(pf02286) PDU1AB
13360 HMMpropdeh3_1_pdu(pf02288) PDU1AB EP_NTERM
13365 HMMpropdeh2_1_pdu(pf02287) PDU1AB EP_NTERM
13370 HMM08841dhact_1_pdu(pf08841) PDU1AB
13375 HMMpropdeh3_2_pdusml(pf02288) PDU1AB
13380 H_tan PDU1AB
13385 H_driedBlood PDU1AB
13390 HMMptac_2_mix(pf06130) PDU1AB
13395 HMM15953pduput_1_pdu1(pf15953) PDU1AB EP_NTERM
13400 P_babyPink PDU1AB
13405 HMM01923cobtr_1_pdu(pf01923pf03928) PDU1AB
13410 HMMaldh_1_eutpdugrm(pf00171) PDU1AB EP_NTERM
13415 HMM00465alcdh_3_mix(pf00465) PDU1AB
13420 HMMPduS_1_all(pf01512pf13375pf10531) PDU1AB
13425 Ts_dodgerBlue PDU1AB
13430 Hp_euts_babyPurple PDU1AB
13435 HMM10662PduV_1_mix(pf10662) PDU1AB
13440 HMMacetkin_1(pf00871) PDU1AB
13445 HMM00288_1_pdu(pf00288) PDU1AB
13450 -pf04363 -
13455 -pf04632 -
13460 -# -
13465 -pf06445 -
13470 -pf07943pf00768 -
13475 -pf01292 -
13480 -pf13247pf12800 -
BMC-P
BMC-H
BMC-T
Aldehyde dehydrogenase
PTAC
Signature enzyme
Alcohol dehydrogenase
Regulator

Salmonella enterica I. Bacteria; Proteobacteria; Gammaproteobacteria; Enterobacterales;

CWG53__locus_1

PDU1AB

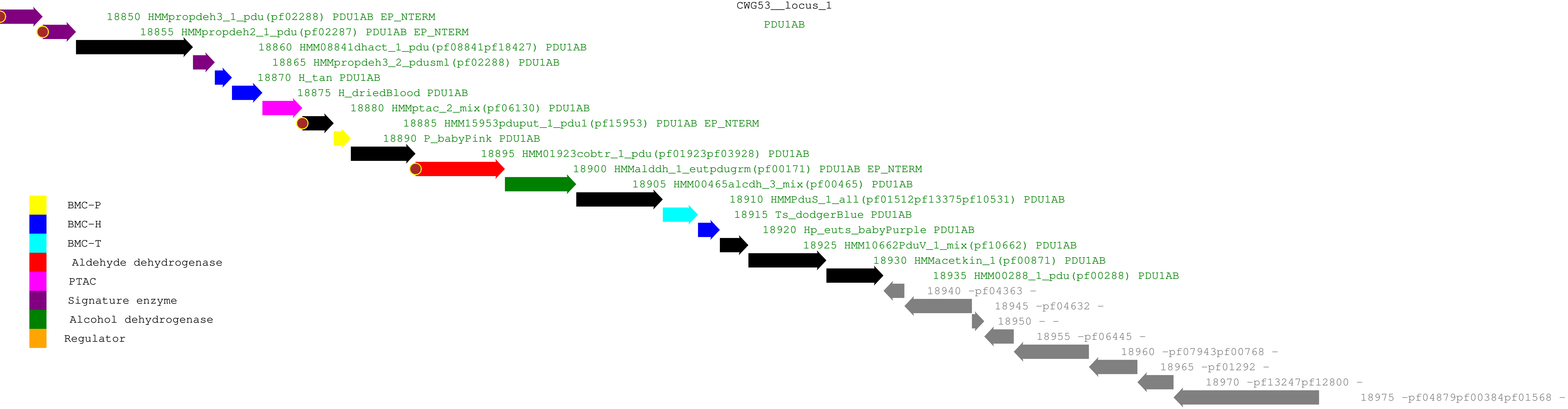

Salmonella enterica I. Bacteria; Proteobacteria; Gammaproteobacteria; Enterobacterales;

CWG97__locus_1

PDU1AB

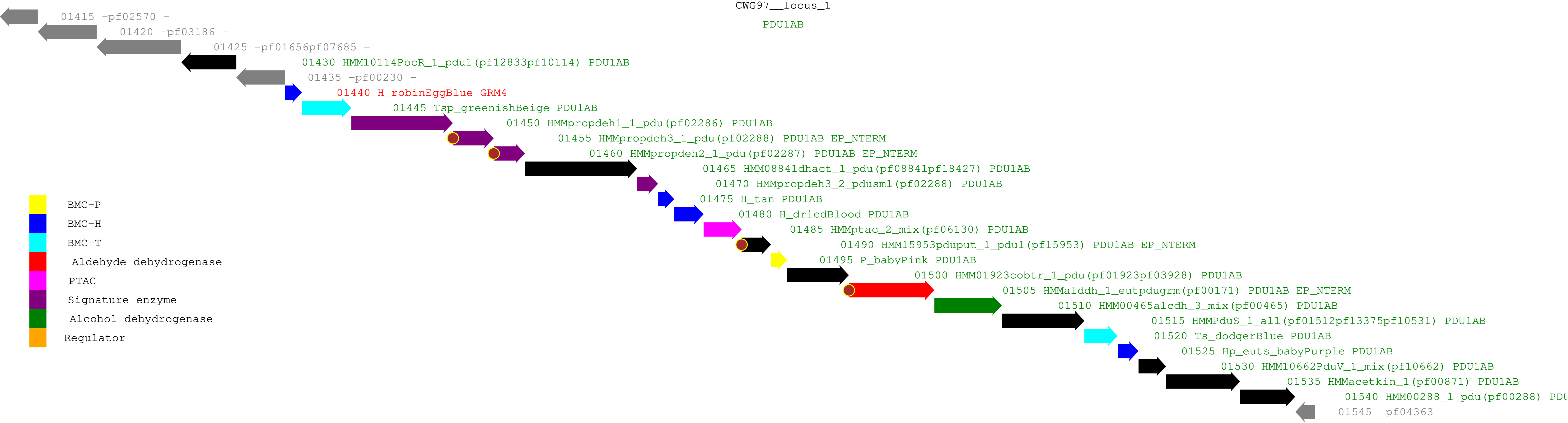

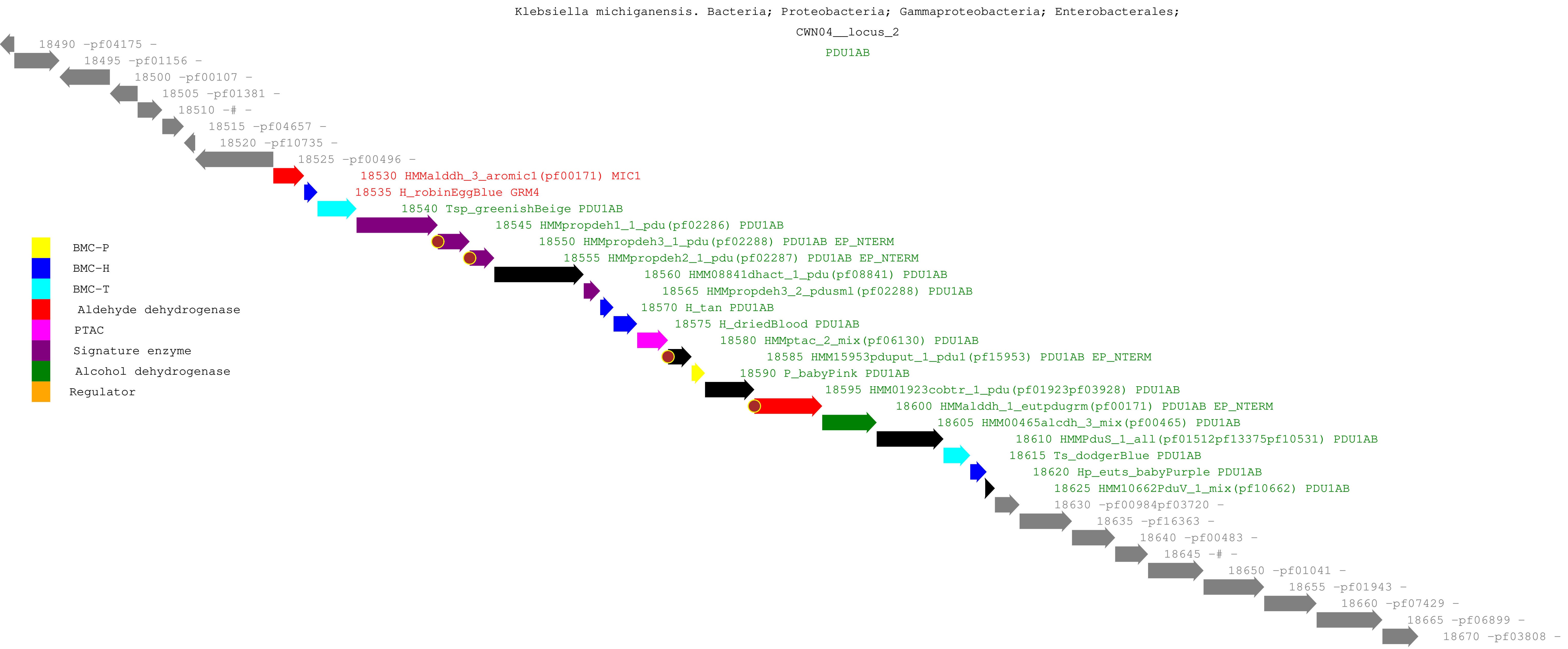
Klebsiella michiganensis. Bacteria; Proteobacteria; Gammaproteobacteria; Enterobacterales;
CWN04__locus_2
PDU1AB
18490 -pf04175 -
18495 -pf01156 -
18500 -pf00107 -
18505 -pf01381 -
18510 -# -
18515 -pf04657 -
18520 -pf10735 -
18525 -pf00496 -
18530 HMMalddh_3_aromic1(pf00171) MIC1
18535 H_robinEggBlue GRM4
18540 Tsp_greenishBeige PDU1AB
18545 HMMpropdeh1_1_pdu(pf02286) PDU1AB
18550 HMMpropdeh3_1_pdu(pf02288) PDU1AB EP_NTERM
18555 HMMpropdeh2_1_pdu(pf02287) PDU1AB EP_NTERM
18560 HMM08841dhact_1_pdu(pf08841) PDU1AB
18565 HMMpropdeh3_2_pdusml(pf02288) PDU1AB
18570 H_tan PDU1AB
18575 H_driedBlood PDU1AB
18580 HMMptac_2_mix(pf06130) PDU1AB
18585 HMM15953pduput_1_pdu1(pf15953) PDU1AB EP_NTERM
18590 P_babyPink PDU1AB
18595 HMM01923cobtr_1_pdu(pf01923pf03928) PDU1AB
18600 HMMalddh_1_eutpdugrm(pf00171) PDU1AB EP_NTERM
18605 HMM00465alcdh_3_mix(pf00465) PDU1AB
18610 HMMPduS_1_all(pf01512pf13375pf10531) PDU1AB
18615 Ts_dodgerBlue PDU1AB
18620 Hp_euts_babyPurple PDU1AB
18625 HMM10662PduV_1_mix(pf10662) PDU1AB
18630 -pf00984pf03720 -
18635 -pf16363 -
18640 -pf00483 -
18645 -# -
18650 -pf01041 -
18655 -pf01943 -
18660 -pf07429 -
18665 -pf06899 -
18670 -pf03808 -
BMC-P
BMC-H
BMC-T
Aldehyde dehydrogenase
PTAC
Signature enzyme
Alcohol dehydrogenase
Regulator

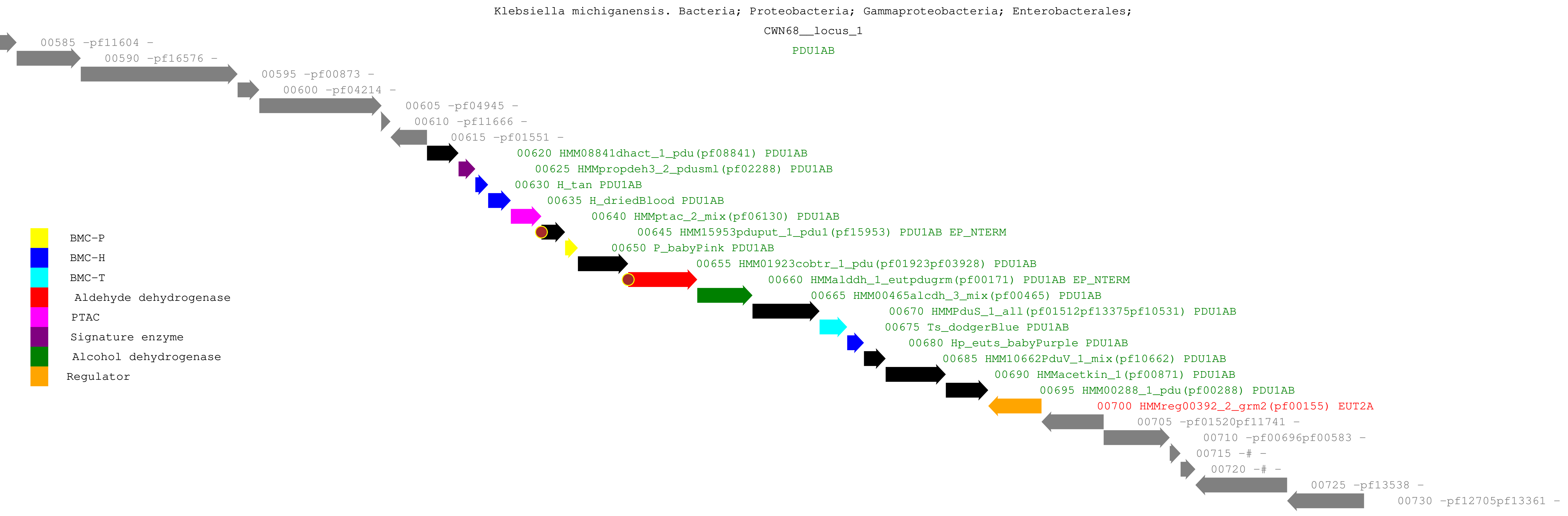

Klebsiella michiganensis. Bacteria; Proteobacteria; Gammaproteobacteria; Enterobacterales;
CWN68__locus_1
PDU1AB
00585 -pf11604 -
00590 -pf16576 -
00595 -pf00873 -
00600 -pf04214 -
00605 -pf04945 -
00610 -pf11666 -
00615 -pf01551 -
00620 HMM08841dhact_1_pdu(pf08841) PDU1AB
00625 HMMpropdeh3_2_pdusml(pf02288) PDU1AB
00630 H_tan PDU1AB
00635 H_driedBlood PDU1AB
00640 HMMptac_2_mix(pf06130) PDU1AB
00645 HMM15953pduput_1_pdu1(pf15953) PDU1AB EP_NTERM
00650 P_babyPink PDU1AB
00655 HMM01923cobtr_1_pdu(pf01923pf03928) PDU1AB
00660 HMMalddh_1_eutpdugrm(pf00171) PDU1AB EP_NTERM
00665 HMM00465alcdh_3_mix(pf00465) PDU1AB
00670 HMMPduS_1_all(pf01512pf13375pf10531) PDU1AB
00675 Ts_dodgerBlue PDU1AB
00680 Hp_euts_babyPurple PDU1AB
00685 HMM10662PduV_1_mix(pf10662) PDU1AB
00690 HMMacetkin_1(pf00871) PDU1AB
00695 HMM00288_1_pdu(pf00288) PDU1AB
00700 HMMreg00392_2_grm2(pf00155) EUT2A
00705 -pf01520pf11741 -
00710 -pf00696pf00583 -
00715 -# -
00720 -# -
00725 -pf13538 -
00730 -pf12705pf13361 -
BMC-P
BMC-H
BMC-T
Aldehyde dehydrogenase
PTAC
Signature enzyme
Alcohol dehydrogenase
Regulator

Salmonella enterica subsp. enterica serovar Heidelberg. Bacteria; Proteobacteria; Gammaproteobacteria; Enterobacterales;

CWX81__locus_fragmented

PDU1AB

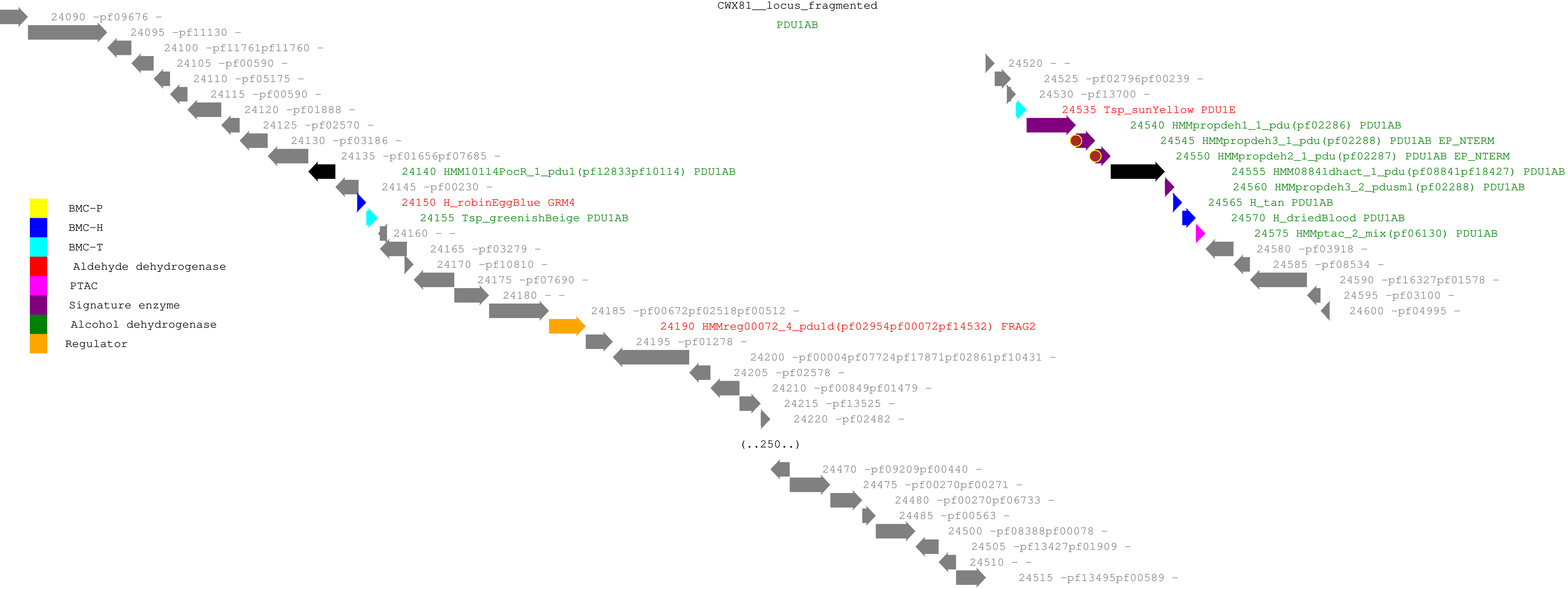

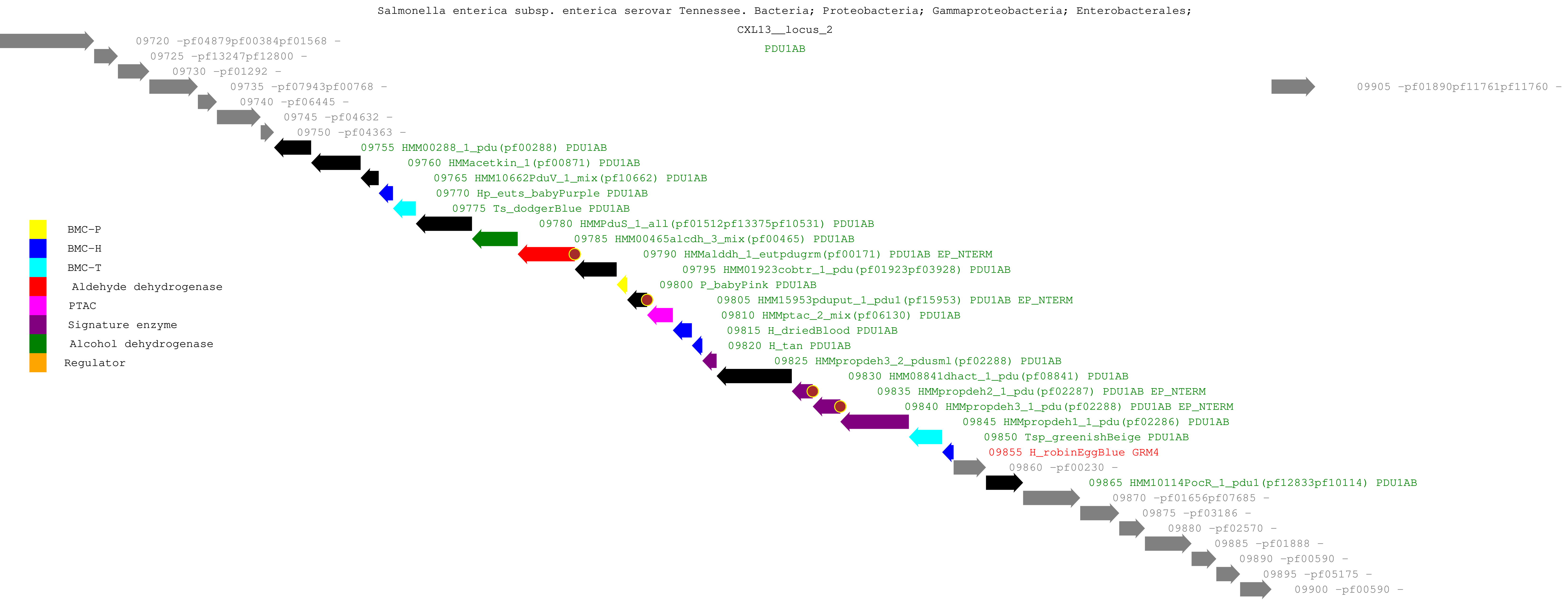
Salmonella enterica subsp. enterica serovar Tennessee. Bacteria; Proteobacteria; Gammaproteobacteria; Enterobacterales;
CXL13__locus_2
PDU1AB
09720 -pf04879pf00384pf01568 -
09725 -pf13247pf12800 -
09730 -pf01292 -
09735 -pf07943pf00768 -
09740 -pf06445 -
09745 -pf04632 -
09750 -pf04363 -
09755 HMM00288_1_pdu(pf00288) PDU1AB
09760 HMMacetkin_1(pf00871) PDU1AB
09765 HMM10662PduV_1_mix(pf10662) PDU1AB
09770 Hp_euts_babyPurple PDU1AB
09775 Ts_dodgerBlue PDU1AB
09780 HMMPduS_1_all(pf01512pf13375pf10531) PDU1AB
09785 HMM00465alcdh_3_mix(pf00465) PDU1AB
09790 HMMalddh_1_eutpdugrm(pf00171) PDU1AB EP_NTERM
09795 HMM01923cobtr_1_pdu(pf01923pf03928) PDU1AB
09800 P_babyPink PDU1AB
09805 HMM15953pduput_1_pdu1(pf15953) PDU1AB EP_NTERM
09810 HMMptac_2_mix(pf06130) PDU1AB
09815 H_driedBlood PDU1AB
09820 H_tan PDU1AB
09825 HMMpropdeh3_2_pdusml(pf02288) PDU1AB
09830 HMM08841dhact_1_pdu(pf08841) PDU1AB
09835 HMMpropdeh2_1_pdu(pf02287) PDU1AB EP_NTERM
09840 HMMpropdeh3_1_pdu(pf02288) PDU1AB EP_NTERM
09845 HMMpropdeh1_1_pdu(pf02286) PDU1AB
09850 Tsp_greenishBeige PDU1AB
09855 H_robinEggBlue GRM4
09860 -pf00230 -
09865 HMM10114PocR_1_pdu1(pf12833pf10114) PDU1AB
09870 -pf01656pf07685 -
09875 -pf03186 -
09880 -pf02570 -
09885 -pf01888 -
09890 -pf00590 -
09895 -pf05175 -
09900 -pf00590 -
09905 -pf01890pf11761pf11760 -
BMC-P
BMC-H
BMC-T
Aldehyde dehydrogenase
PTAC
Signature enzyme
Alcohol dehydrogenase
Regulator

Escherichia albertii. Bacteria; Proteobacteria; Gammaproteobacteria; Enterobacterales;

CXP54__locus_1

PDU1AB

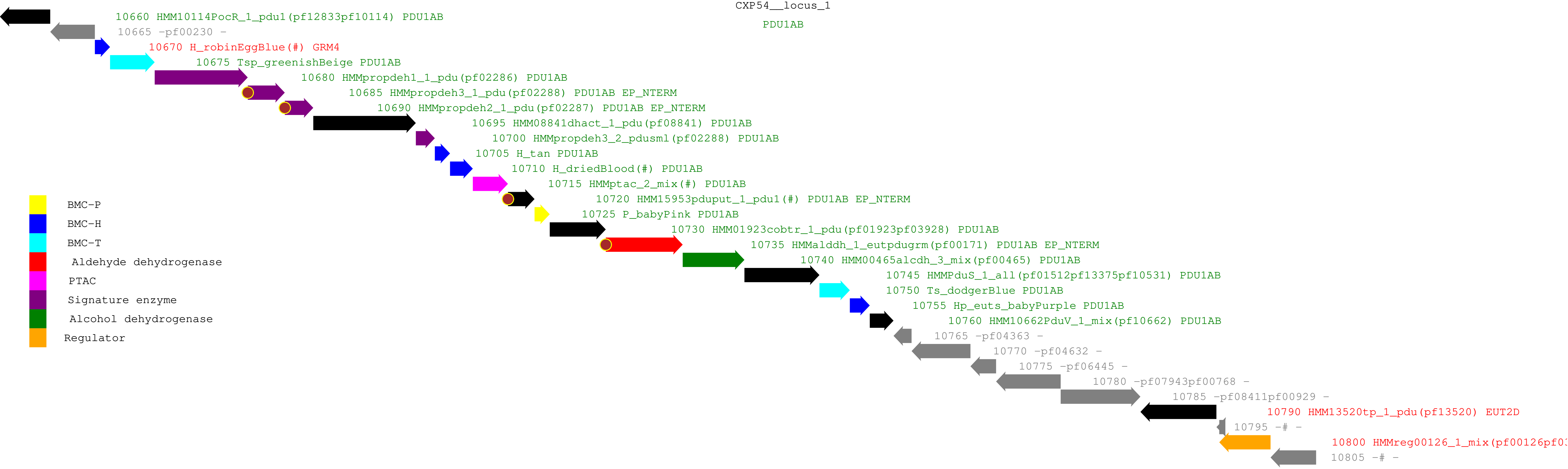

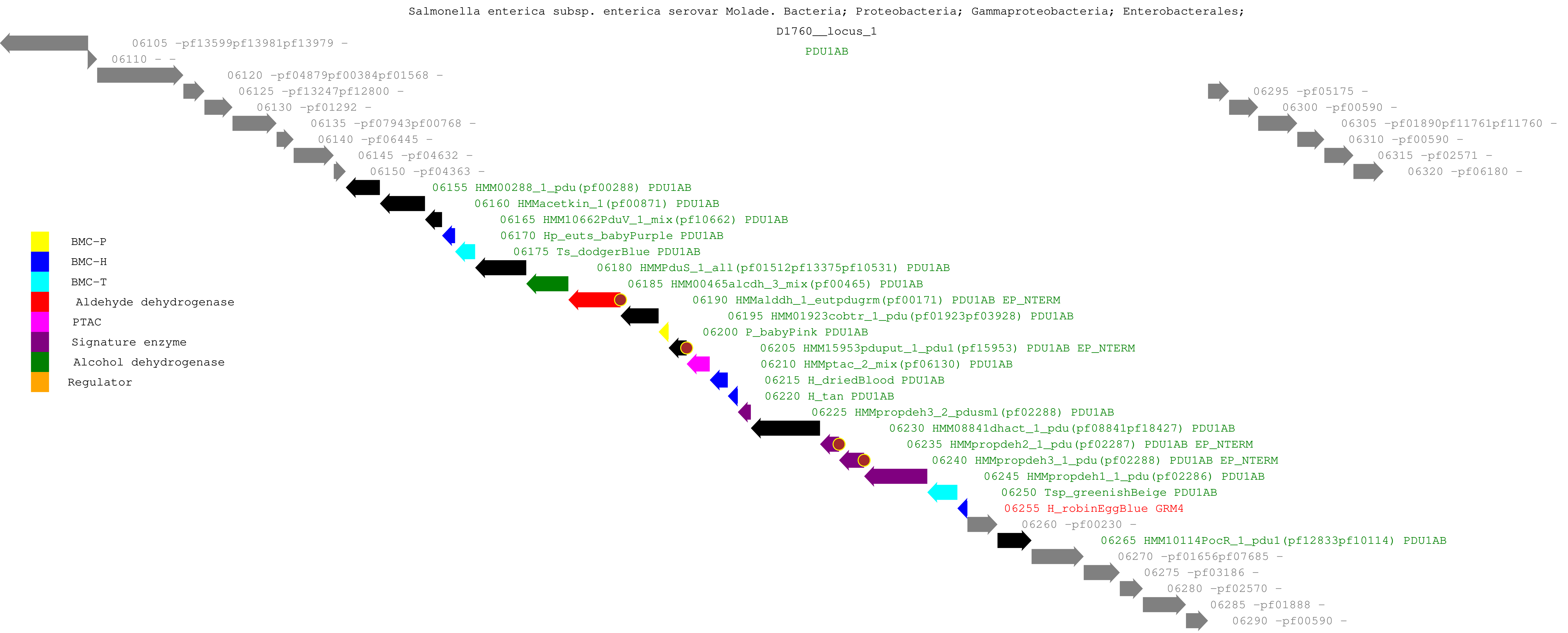

Salmonella enterica subsp. enterica serovar Molade. Bacteria; Proteobacteria; Gammaproteobacteria; Enterobacterales;
D1760__locus_1
PDU1AB
06105 -pf13599pf13981pf13979 -
06110 - -
06120 -pf04879pf00384pf01568 -
06125 -pf13247pf12800 -
06130 -pf01292 -
06135 -pf07943pf00768 -
06140 -pf06445 -
06145 -pf04632 -
06150 -pf04363 -
06155 HMM00288_1_pdu(pf00288) PDU1AB
06160 HMMacetkin_1(pf00871) PDU1AB
06165 HMM10662PduV_1_mix(pf10662) PDU1AB
06170 Hp_euts_babyPurple PDU1AB
06175 Ts_dodgerBlue PDU1AB
06180 HMMPduS_1_all(pf01512pf13375pf10531) PDU1AB
06185 HMM00465alcdh_3_mix(pf00465) PDU1AB
06190 HMMalddh_1_eutpdugrm(pf00171) PDU1AB EP_NTERM
06195 HMM01923cobtr_1_pdu(pf01923pf03928) PDU1AB
06200 P_babyPink PDU1AB
06205 HMM15953pduput_1_pdu1(pf15953) PDU1AB EP_NTERM
06210 HMMptac_2_mix(pf06130) PDU1AB
06215 H_driedBlood PDU1AB
06220 H_tan PDU1AB
06225 HMMpropdeh3_2_pdusml(pf02288) PDU1AB
06230 HMM08841dhact_1_pdu(pf08841pf18427) PDU1AB
06235 HMMpropdeh2_1_pdu(pf02287) PDU1AB EP_NTERM
06240 HMMpropdeh3_1_pdu(pf02288) PDU1AB EP_NTERM
06245 HMMpropdeh1_1_pdu(pf02286) PDU1AB
06250 Tsp_greenishBeige PDU1AB
06255 H_robinEggBlue GRM4
06260 -pf00230 -
06265 HMM10114PocR_1_pdu1(pf12833pf10114) PDU1AB
06270 -pf01656pf07685 -
06275 -pf03186 -
06280 -pf02570 -
06285 -pf01888 -
06290 -pf00590 -
06295 -pf05175 -
06300 -pf00590 -
06305 -pf01890pf11761pf11760 -
06310 -pf00590 -
06315 -pf02571 -
06320 -pf06180 -
BMC-P
BMC-H
BMC-T
Aldehyde dehydrogenase
PTAC
Signature enzyme
Alcohol dehydrogenase
Regulator

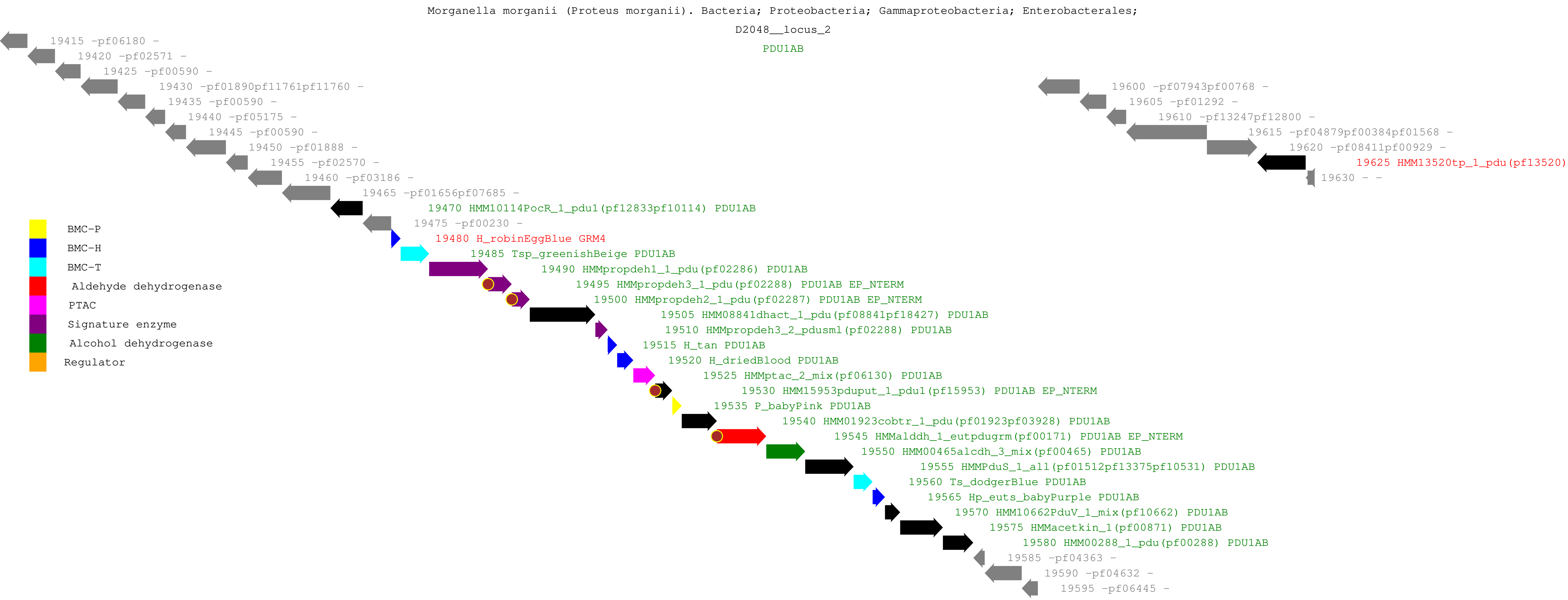
Morganella morganii (Proteus morganii). Bacteria; Proteobacteria; Gammaproteobacteria; Enterobacterales;
D2048__locus_2
PDU1AB
19415 -pf06180 -
19420 -pf02571 -
19425 -pf00590 -
19430 -pf01890pf11761pf11760 -
19435 -pf00590 -
19440 -pf05175 -
19445 -pf00590 -
19450 -pf01888 -
19455 -pf02570 -
19460 -pf03186 -
19465 -pf01656pf07685 -
19470 HMM10114PocR_1_pdu1(pf12833pf10114) PDU1AB
19475 -pf00230 -
19480 H_robinEggBlue GRM4
19485 Tsp_greenishBeige PDU1AB
19490 HMMpropdeh1_1_pdu(pf02286) PDU1AB
19495 HMMpropdeh3_1_pdu(pf02288) PDU1AB EP_NTERM
19500 HMMpropdeh2_1_pdu(pf02287) PDU1AB EP_NTERM
19505 HMM08841dhact_1_pdu(pf08841pf18427) PDU1AB
19510 HMMpropdeh3_2_pdusml(pf02288) PDU1AB
19515 H_tan PDU1AB
19520 H_driedBlood PDU1AB
19525 HMMptac_2_mix(pf06130) PDU1AB
19530 HMM15953pduput_1_pdu1(pf15953) PDU1AB EP_NTERM
19535 P_babyPink PDU1AB
19540 HMM01923cobtr_1_pdu(pf01923pf03928) PDU1AB
19545 HMMalddh_1_eutpdugrm(pf00171) PDU1AB EP_NTERM
19550 HMM00465alcdh_3_mix(pf00465) PDU1AB
19555 HMMPduS_1_all(pf01512pf13375pf10531) PDU1AB
19560 Ts_dodgerBlue PDU1AB
19565 Hp_euts_babyPurple PDU1AB
19570 HMM10662PduV_1_mix(pf10662) PDU1AB
19575 HMMacetkin_1(pf00871) PDU1AB
19580 HMM00288_1_pdu(pf00288) PDU1AB
19585 -pf04363 -
19590 -pf04632 -
19595 -pf06445 -
19600 -pf07943pf00768 -
19605 -pf01292 -
19610 -pf13247pf12800 -
19615 -pf04879pf00384pf01568 -
19620 -pf08411pf00929 -
19625 HMM13520tp_1_pdu(pf13520)
19630 - -
BMC-P
BMC-H
BMC-T
Aldehyde dehydrogenase
PTAC
Signature enzyme
Alcohol dehydrogenase
Regulator

Escherichia coli. Bacteria; Proteobacteria; Gammaproteobacteria; Enterobacterales;

D3822__locus_1_main_sat

PDU1AB

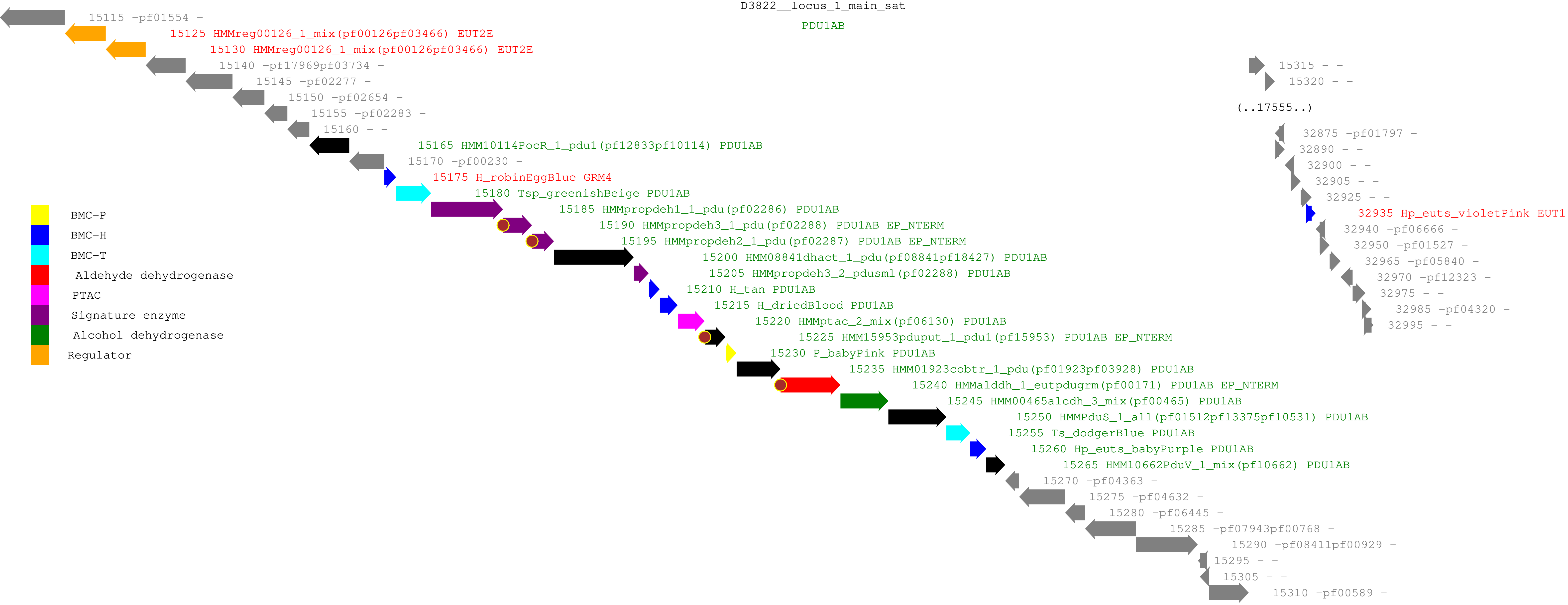

Citrobacter portucalensis. Bacteria; Proteobacteria; Gammaproteobacteria; Enterobacterales;

D3H66__locus_1

PDU1AB

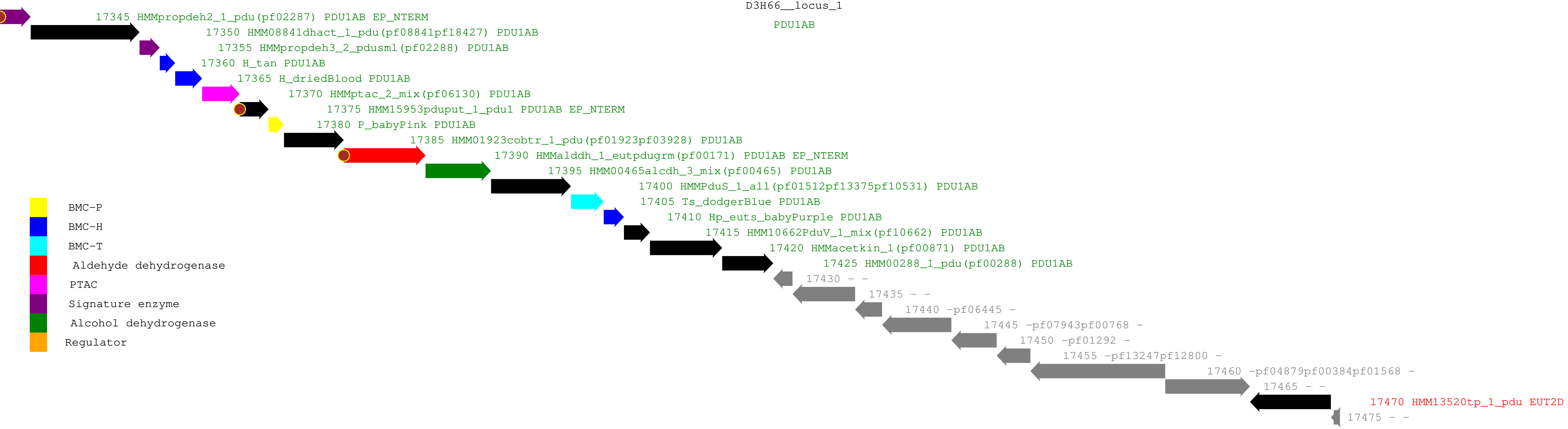

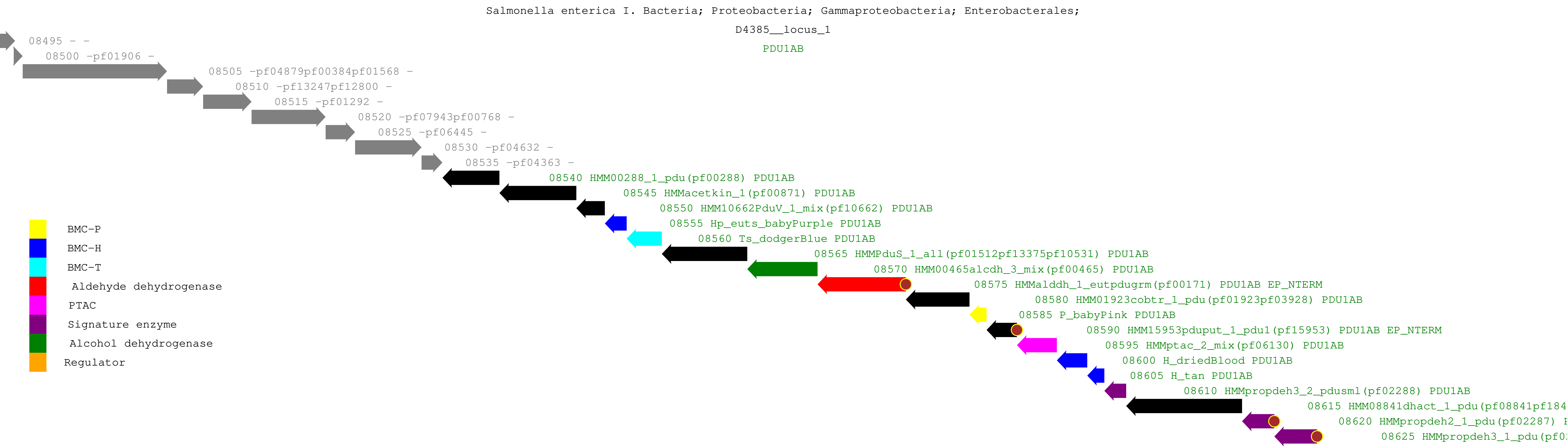

Salmonella enterica I. Bacteria; Proteobacteria; Gammaproteobacteria; Enterobacterales;
D4385__locus_1
PDU1AB
08495 - -
08500 -pf01906 -
08505 -pf04879pf00384pf01568 -
08510 -pf13247pf12800 -
08515 -pf01292 -
08520 -pf07943pf00768 -
08525 -pf06445 -
08530 -pf04632 -
08535 -pf04363 -
08540 HMM00288_1_pdu(pf00288) PDU1AB
08545 HMMacetkin_1(pf00871) PDU1AB
08550 HMM10662PduV_1_mix(pf10662) PDU1AB
08555 Hp_euts_babyPurple PDU1AB
08560 Ts_dodgerBlue PDU1AB
08565 HMMPduS_1_all(pf01512pf13375pf10531) PDU1AB
08570 HMM00465alcdh_3_mix(pf00465) PDU1AB
08575 HMMaldh_1_eutpdugrm(pf00171) PDU1AB EP_NTERM
08580 HMM01923cobtr_1_pdu(pf01923pf03928) PDU1AB
08585 P_babyPink PDU1AB
08590 HMM15953pduput_1_pdu1(pf15953) PDU1AB EP_NTERM
08595 HMMptac_2_mix(pf06130) PDU1AB
08600 H_driedBlood PDU1AB
08605 H_tan PDU1AB
08610 HMMpropdeh3_2_pdusml(pf02288) PDU1AB
08615 HMM08841dhact_1_pdu(pf08841pf184
08620 HMMpropdeh2_1_pdu(pf02287) P
08625 HMMpropdeh3_1_pdu(pf0
BMC-P
BMC-H
BMC-T
Aldehyde dehydrogenase
PTAC
Signature enzyme
Alcohol dehydrogenase
Regulator

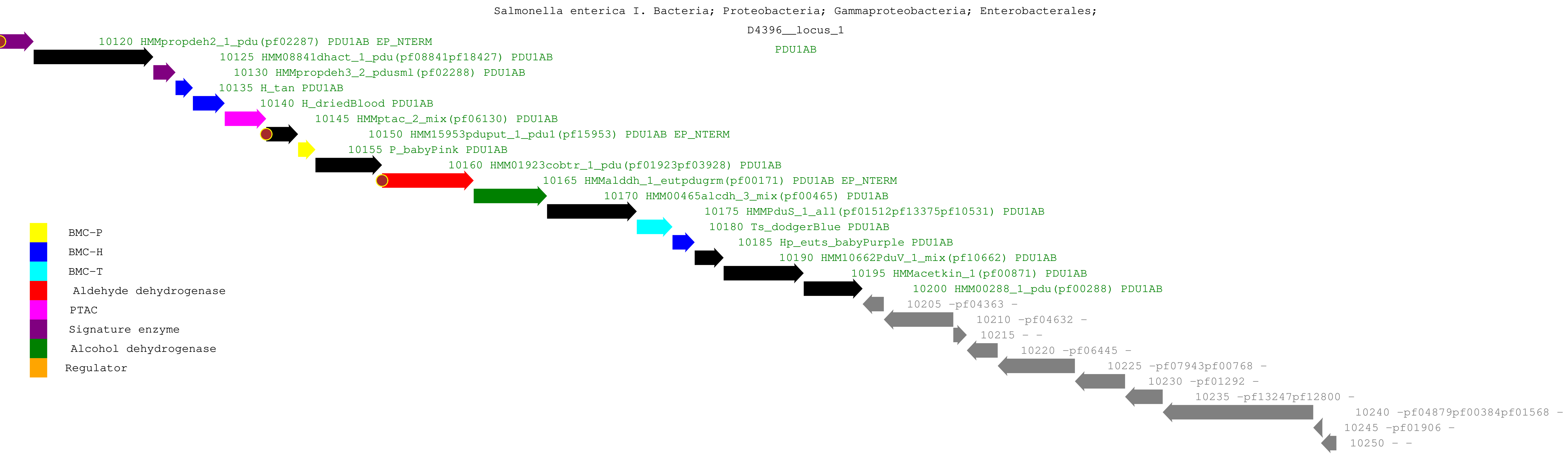

Salmonella enterica I. Bacteria; Proteobacteria; Gammaproteobacteria; Enterobacterales;
D4396__locus_1
PDU1AB
10120 HMMpropdeh2_1_pdu(pf02287) PDU1AB EP_NTERM
10125 HMM08841dhact_1_pdu(pf08841pf18427) PDU1AB
10130 HMMpropdeh3_2_pdusml(pf02288) PDU1AB
10135 H_tan PDU1AB
10140 H_driedBlood PDU1AB
10145 HMMptac_2_mix(pf06130) PDU1AB
10150 HMM15953pduput_1_pdu1(pf15953) PDU1AB EP_NTERM
10155 P_babyPink PDU1AB
10160 HMM01923cobtr_1_pdu(pf01923pf03928) PDU1AB
10165 HMMaldh_1_eutpdugrm(pf00171) PDU1AB EP_NTERM
10170 HMM00465alcdh_3_mix(pf00465) PDU1AB
10175 HMMPduS_1_all(pf01512pf13375pf10531) PDU1AB
10180 Ts_dodgerBlue PDU1AB
10185 Hp_euts_babyPurple PDU1AB
10190 HMM10662PduV_1_mix(pf10662) PDU1AB
10195 HMMacetkin_1(pf00871) PDU1AB
10200 HMM00288_1_pdu(pf00288) PDU1AB
10205 -pf04363 -
10210 -pf04632 -
10215 - -
10220 -pf06445 -
10225 -pf07943pf00768 -
10230 -pf01292 -
10235 -pf13247pf12800 -
10240 -pf04879pf00384pf01568 -
10245 -pf01906 -
10250 - -
BMC-P
BMC-H
BMC-T
Aldehyde dehydrogenase
PTAC
Signature enzyme
Alcohol dehydrogenase
Regulator

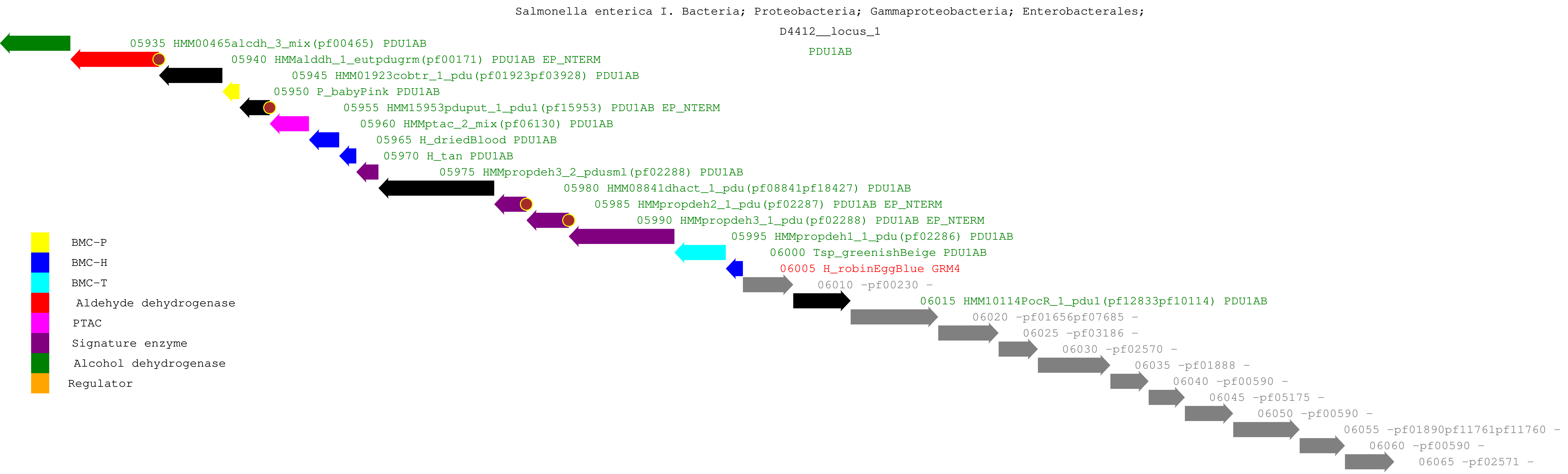
Salmonella enterica I. Bacteria; Proteobacteria; Gammaproteobacteria; Enterobacterales;
D4412__locus_1
PDU1AB
05935 HMM00465alcdh_3_mix(pf00465) PDU1AB
05940 HMMalddh_1_eutpdugrm(pf00171) PDU1AB EP_NTERM
05945 HMM01923cobtr_1_pdu(pf01923pf03928) PDU1AB
05950 P_babyPink PDU1AB
05955 HMM15953pduput_1_pdu1(pf15953) PDU1AB EP_NTERM
05960 HMMptac_2_mix(pf06130) PDU1AB
05965 H_driedBlood PDU1AB
05970 H_tan PDU1AB
05975 HMMpropdeh3_2_pdusml(pf02288) PDU1AB
05980 HMM08841dhact_1_pdu(pf08841pf18427) PDU1AB
05985 HMMpropdeh2_1_pdu(pf02287) PDU1AB EP_NTERM
05990 HMMpropdeh3_1_pdu(pf02288) PDU1AB EP_NTERM
05995 HMMpropdeh1_1_pdu(pf02286) PDU1AB
06000 Tsp_greenishBeige PDU1AB
06005 H_robinEggBlue GRM4
06010 -pf00230 -
06015 HMM10114PocR_1_pdu1(pf12833pf10114) PDU1AB
06020 -pf01656pf07685 -
06025 -pf03186 -
06030 -pf02570 -
06035 -pf01888 -
06040 -pf00590 -
06045 -pf05175 -
06050 -pf00590 -
06055 -pf01890pf11761pf11760 -
06060 -pf00590 -
06065 -pf02571 -
BMC-P
BMC-H
BMC-T
Aldehyde dehydrogenase
PTAC
Signature enzyme
Alcohol dehydrogenase
Regulator

Salmonella enterica I. Bacteria; Proteobacteria; Gammaproteobacteria; Enterobacterales;

D4428__locus_1

PDU1AB

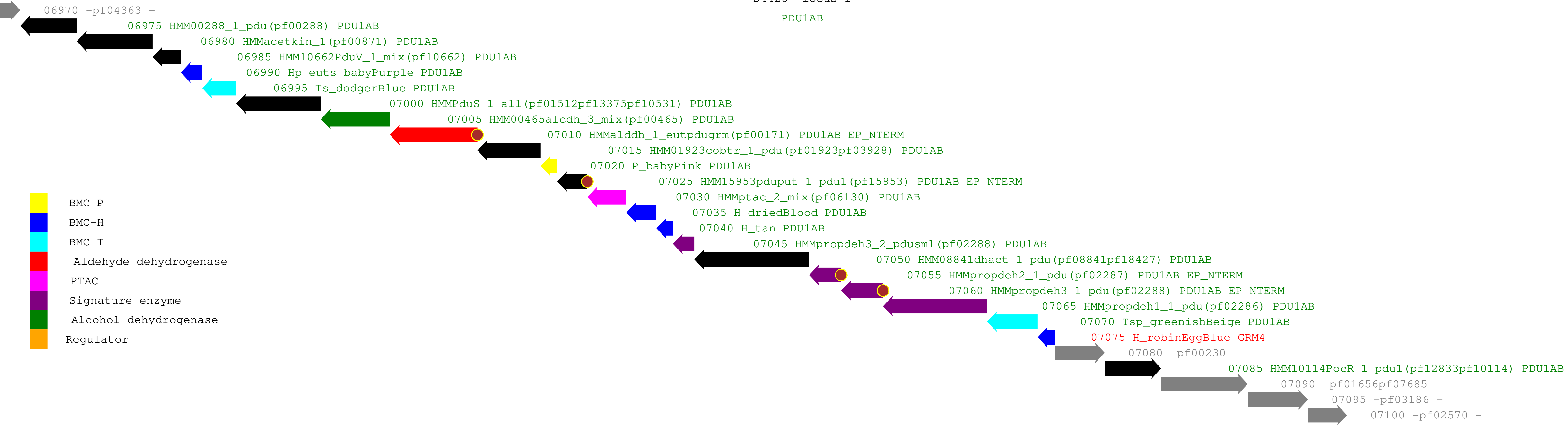

Salmonella enterica I. Bacteria; Proteobacteria; Gammaproteobacteria; Enterobacterales;

D4445__locus_1

PDU1AB

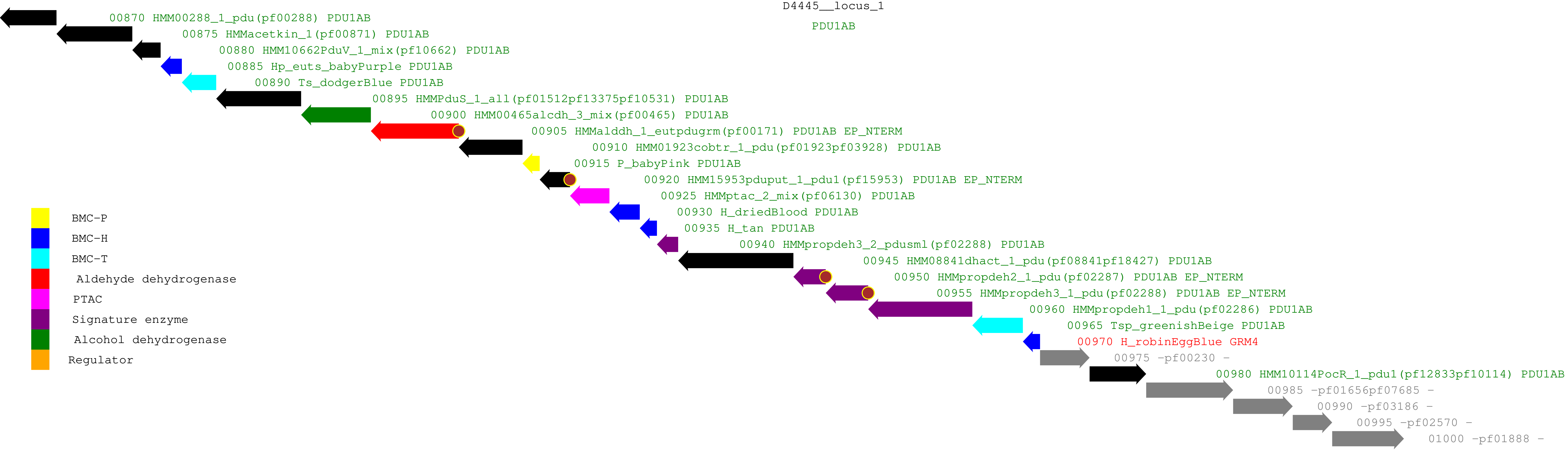

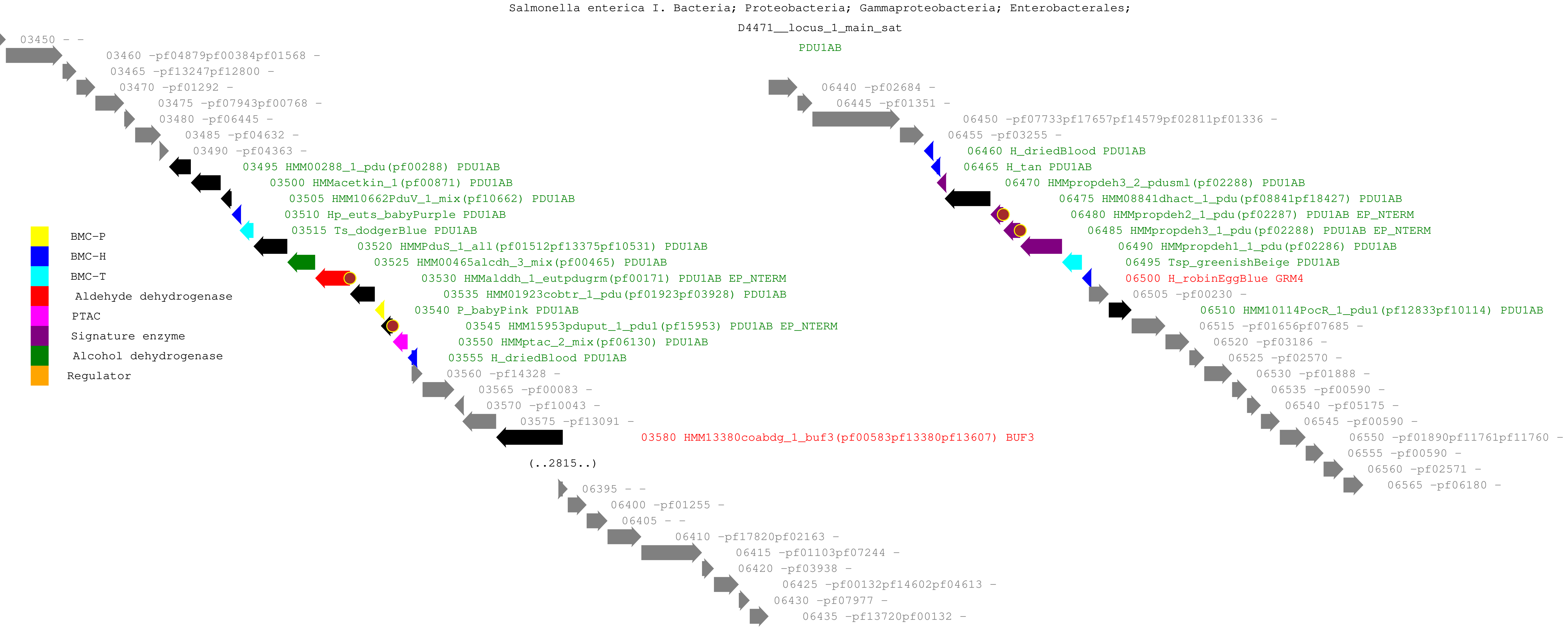
Salmonella enterica I. Bacteria; Proteobacteria; Gammaproteobacteria; Enterobacterales;
D4471__locus_1_main_sat
PDU1AB
03450 - -
03460 -pf04879pf00384pf01568 -
03465 -pf13247pf12800 -
03470 -pf01292 -
03475 -pf07943pf00768 -
03480 -pf06445 -
03485 -pf04632 -
03490 -pf04363 -
03495 HMM00288_1_pdu(pf00288) PDU1AB
03500 HMMacetkin_1(pf00871) PDU1AB
03505 HMM10662PduV_1_mix(pf10662) PDU1AB
03510 Hp_euts_babyPurple PDU1AB
03515 Ts_dodgerBlue PDU1AB
03520 HMMPduS_1_all(pf01512pf13375pf10531) PDU1AB
03525 HMM00465alcdh_3_mix(pf00465) PDU1AB
03530 HMMaldh_1_eutpdugrm(pf00171) PDU1AB EP_NTERM
03535 HMM01923cobtr_1_pdu(pf01923pf03928) PDU1AB
03540 P_babyPink PDU1AB
03545 HMM15953pduput_1_pdu1(pf15953) PDU1AB EP_NTERM
03550 HMMptac_2_mix(pf06130) PDU1AB
03555 H_driedBlood PDU1AB
03560 -pf14328 -
03565 -pf00083 -
03570 -pf10043 -
03575 -pf13091 -
03580 HMM13380coabdg_1_buf3(pf00583pf13380pf13607) BUF3
(..2815..)
06395 - -
06400 -pf01255 -
06405 - -
06410 -pf17820pf02163 -
06415 -pf01103pf07244 -
06420 -pf03938 -
06425 -pf00132pf14602pf04613 -
06430 -pf07977 -
06435 -pf13720pf00132 -
06440 -pf02684 -
06445 -pf01351 -
06450 -pf07733pf17657pf14579pf02811pf01336 -
06455 -pf03255 -
06460 H_driedBlood PDU1AB
06465 H_tan PDU1AB
06470 HMMpropdeh3_2_pdusml(pf02288) PDU1AB
06475 HMM08841dhact_1_pdu(pf08841pf18427) PDU1AB
06480 HMMpropdeh2_1_pdu(pf02287) PDU1AB EP_NTERM
06485 HMMpropdeh3_1_pdu(pf02288) PDU1AB EP_NTERM
06490 HMMpropdeh1_1_pdu(pf02286) PDU1AB
06495 Tsp_greenishBeige PDU1AB
06500 H_robinEggBlue GRM4
06505 -pf00230 -
06510 HMM10114PocR_1_pdu1(pf12833pf10114) PDU1AB
06515 -pf01656pf07685 -
06520 -pf03186 -
06525 -pf02570 -
06530 -pf01888 -
06535 -pf00590 -
06540 -pf05175 -
06545 -pf00590 -
06550 -pf01890pf11761pf11760 -
06555 -pf00590 -
06560 -pf02571 -
06565 -pf06180 -
BMC-P
BMC-H
BMC-T
Aldehyde dehydrogenase
PTAC
Signature enzyme
Alcohol dehydrogenase
Regulator

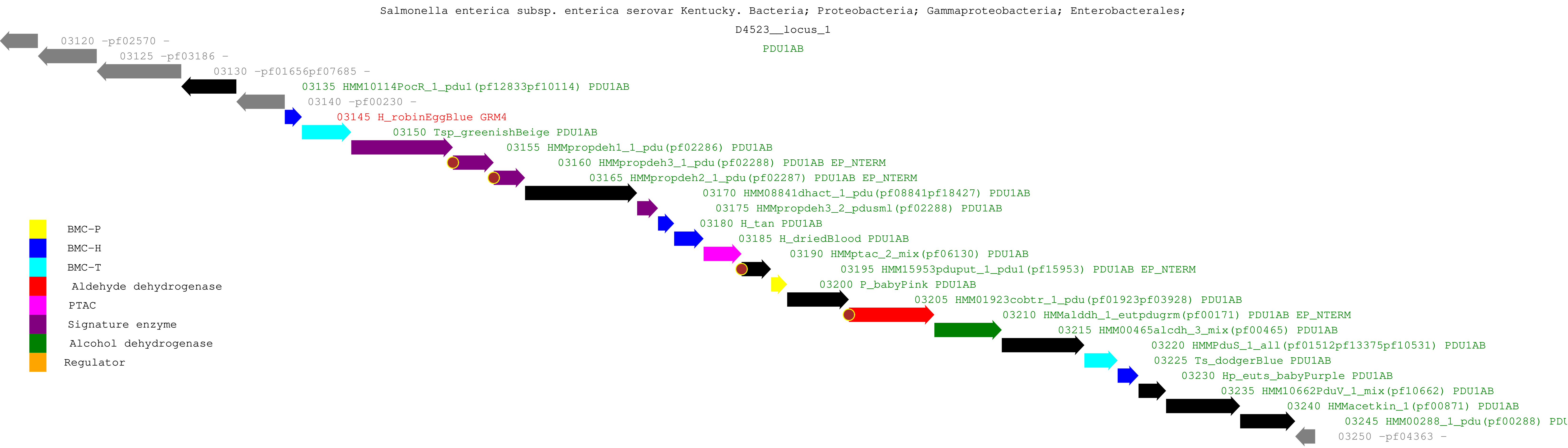

Salmonella enterica subsp. enterica serovar Kentucky. Bacteria; Proteobacteria; Gammaproteobacteria; Enterobacterales;
D4523__locus_1
PDU1AB
03120 -pf02570 -
03125 -pf03186 -
03130 -pf01656pf07685 -
03135 HMM10114PocR_1_pdu1(pf12833pf10114) PDU1AB
03140 -pf00230 -
03145 H_robinEggBlue GRM4
03150 Tsp_greenishBeige PDU1AB
03155 HMMpropdeh1_1_pdu(pf02286) PDU1AB
03160 HMMpropdeh3_1_pdu(pf02288) PDU1AB EP_NTERM
03165 HMMpropdeh2_1_pdu(pf02287) PDU1AB EP_NTERM
03170 HMM08841dhact_1_pdu(pf08841pf18427) PDU1AB
03175 HMMpropdeh3_2_pdusml(pf02288) PDU1AB
03180 H_tan PDU1AB
03185 H_driedBlood PDU1AB
03190 HMMptac_2_mix(pf06130) PDU1AB
03195 HMM15953pduput_1_pdu1(pf15953) PDU1AB EP_NTERM
03200 P_babyPink PDU1AB
03205 HMM01923cobtr_1_pdu(pf01923pf03928) PDU1AB
03210 HMMalddh_1_eutpdugrm(pf00171) PDU1AB EP_NTERM
03215 HMM00465alcdh_3_mix(pf00465) PDU1AB
03220 HMMPduS_1_all(pf01512pf13375pf10531) PDU1AB
03225 Ts_dodgerBlue PDU1AB
03230 Hp_euts_babyPurple PDU1AB
03235 HMM10662PduV_1_mix(pf10662) PDU1AB
03240 HMMacetkin_1(pf00871) PDU1AB
03245 HMM00288_1_pdu(pf00288) PDU
03250 -pf04363 -
BMC-P
BMC-H
BMC-T
Aldehyde dehydrogenase
PTAC
Signature enzyme
Alcohol dehydrogenase
Regulator

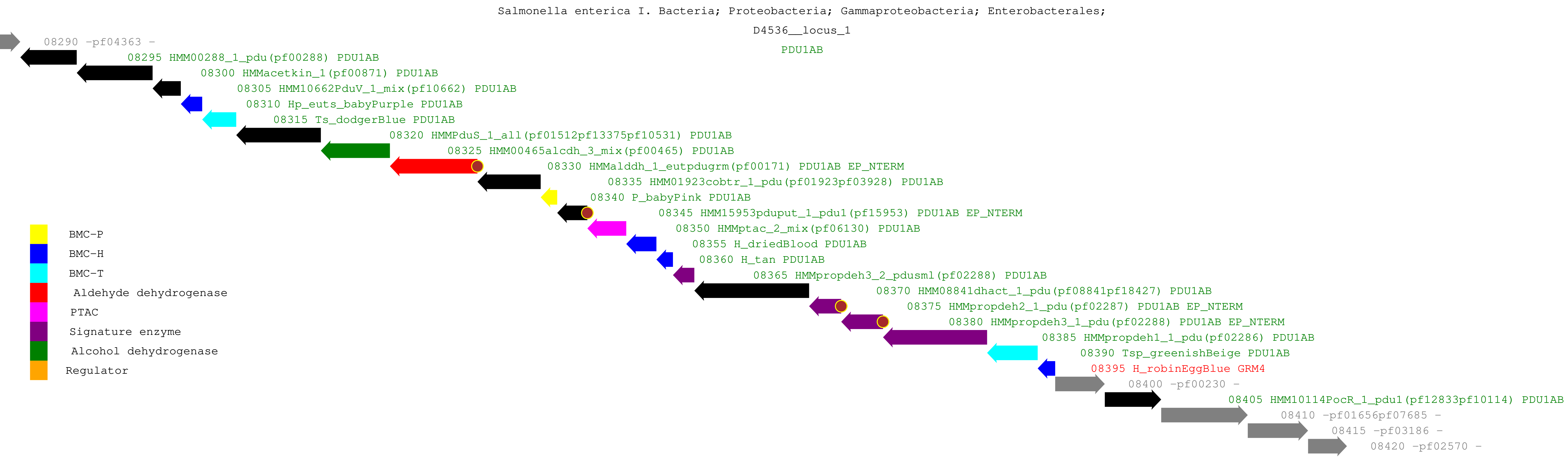
Salmonella enterica I. Bacteria; Proteobacteria; Gammaproteobacteria; Enterobacterales;
D4536__locus_1
PDU1AB
08290 -pf04363 -
08295 HMM00288_1_pdu(pf00288) PDU1AB
08300 HMMacetkin_1(pf00871) PDU1AB
08305 HMM10662PduV_1_mix(pf10662) PDU1AB
08310 Hp_euts_babyPurple PDU1AB
08315 Ts_dodgerBlue PDU1AB
08320 HMMPduS_1_all(pf01512pf13375pf10531) PDU1AB
08325 HMM00465alcdh_3_mix(pf00465) PDU1AB
08330 HMMalddh_1_eutpdugrm(pf00171) PDU1AB EP_NTERM
08335 HMM01923cobtr_1_pdu(pf01923pf03928) PDU1AB
08340 P_babyPink PDU1AB
08345 HMM15953pduput_1_pdu1(pf15953) PDU1AB EP_NTERM
08350 HMMptac_2_mix(pf06130) PDU1AB
08355 H_driedBlood PDU1AB
08360 H_tan PDU1AB
08365 HMMpropdeh3_2_pdusml(pf02288) PDU1AB
08370 HMM08841dhact_1_pdu(pf08841pf18427) PDU1AB
08375 HMMpropdeh2_1_pdu(pf02287) PDU1AB EP_NTERM
08380 HMMpropdeh3_1_pdu(pf02288) PDU1AB EP_NTERM
08385 HMMpropdeh1_1_pdu(pf02286) PDU1AB
08390 Tsp_greenishBeige PDU1AB
08395 H_robinEggBlue GRM4
08400 -pf00230 -
08405 HMM10114PocR_1_pdu1(pf12833pf10114) PDU1AB
08410 -pf01656pf07685 -
08415 -pf03186 -
08420 -pf02570 -
BMC-P
BMC-H
BMC-T
Aldehyde dehydrogenase
PTAC
Signature enzyme
Alcohol dehydrogenase
Regulator

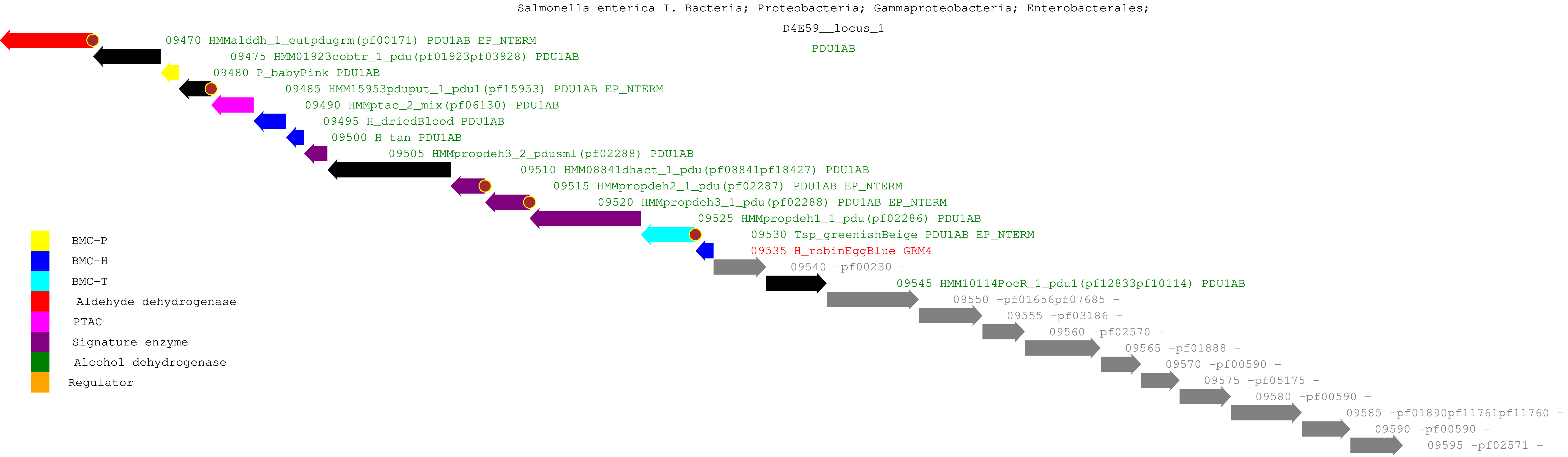
Salmonella enterica I. Bacteria; Proteobacteria; Gammaproteobacteria; Enterobacterales;
D4E59__locus_1
PDU1AB
09470 HMMalddh_1_eutpdugrm(pf00171) PDU1AB EP_NTERM
09475 HMM01923cobtr_1_pdu(pf01923pf03928) PDU1AB
09480 P_babyPink PDU1AB
09485 HMM15953pduput_1_pdu1(pf15953) PDU1AB EP_NTERM
09490 HMMptac_2_mix(pf06130) PDU1AB
09495 H_driedBlood PDU1AB
09500 H_tan PDU1AB
09505 HMMpropdeh3_2_pdusml(pf02288) PDU1AB
09510 HMM08841dhact_1_pdu(pf08841pf18427) PDU1AB
09515 HMMpropdeh2_1_pdu(pf02287) PDU1AB EP_NTERM
09520 HMMpropdeh3_1_pdu(pf02288) PDU1AB EP_NTERM
09525 HMMpropdeh1_1_pdu(pf02286) PDU1AB
09530 Tsp_greenishBeige PDU1AB EP_NTERM
09535 H_robinEggBlue GRM4
09540 -pf00230 -
09545 HMM10114PocR_1_pdu1(pf12833pf10114) PDU1AB
09550 -pf01656pf07685 -
09555 -pf03186 -
09560 -pf02570 -
09565 -pf01888 -
09570 -pf00590 -
09575 -pf05175 -
09580 -pf00590 -
09585 -pf01890pf11761pf11760 -
09590 -pf00590 -
09595 -pf02571 -
BMC-P
BMC-H
BMC-T
Aldehyde dehydrogenase
PTAC
Signature enzyme
Alcohol dehydrogenase
Regulator

Salmonella enterica I. Bacteria; Proteobacteria; Gammaproteobacteria; Enterobacterales;

D4F19__locus_1

PDU1AB

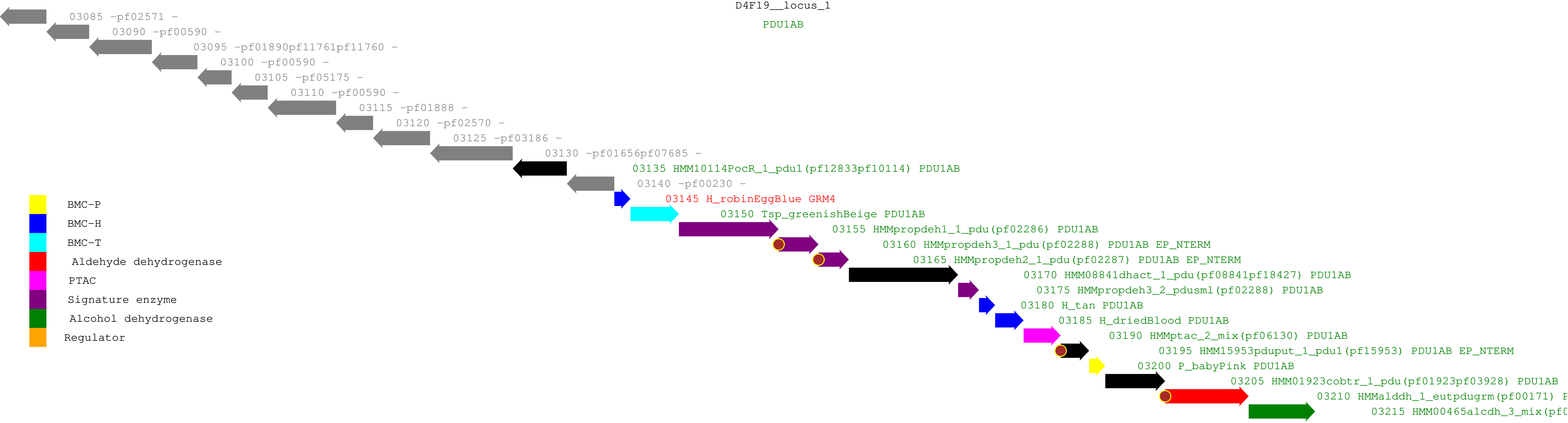

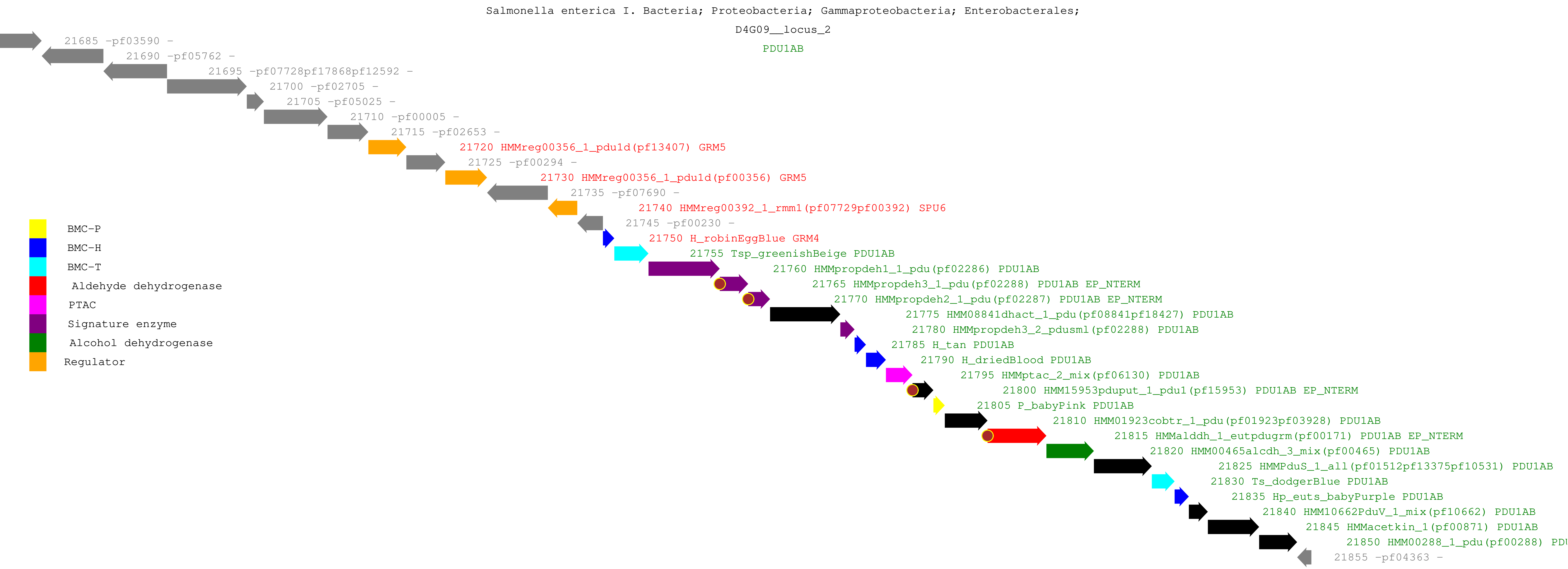
Salmonella enterica I. Bacteria; Proteobacteria; Gammaproteobacteria; Enterobacterales;
D4G09__locus_2
PDU1AB
21685 -pf03590 -
21690 -pf05762 -
21695 -pf07728pf17868pf12592 -
21700 -pf02705 -
21705 -pf05025 -
21710 -pf00005 -
21715 -pf02653 -
21720 HMMreg00356_1_pdu1d(pf13407) GRM5
21725 -pf00294 -
21730 HMMreg00356_1_pdu1d(pf00356) GRM5
21735 -pf07690 -
21740 HMMreg00392_1_rmm1(pf07729pf00392) SPU6
21745 -pf00230 -
21750 H_robinEggBlue GRM4
21755 Tsp_greenishBeige PDU1AB
21760 HMMpropdeh1_1_pdu(pf02286) PDU1AB
21765 HMMpropdeh3_1_pdu(pf02288) PDU1AB EP_NTERM
21770 HMMpropdeh2_1_pdu(pf02287) PDU1AB EP_NTERM
21775 HMM08841dhact_1_pdu(pf08841pf18427) PDU1AB
21780 HMMpropdeh3_2_pdusml(pf02288) PDU1AB
21785 H_tan PDU1AB
21790 H_driedBlood PDU1AB
21795 HMMptac_2_mix(pf06130) PDU1AB
21800 HMM15953pduput_1_pdul(pf15953) PDU1AB EP_NTERM
21805 P_babyPink PDU1AB
21810 HMM01923cobtr_1_pdu(pf01923pf03928) PDU1AB
21815 HMMaldh_1_eutpdugrm(pf00171) PDU1AB EP_NTERM
21820 HMM00465alcdh_3_mix(pf00465) PDU1AB
21825 HMMPduS_1_all(pf01512pf13375pf10531) PDU1AB
21830 Ts_dodgerBlue PDU1AB
21835 Hp_euts_babyPurple PDU1AB
21840 HMM10662PduV_1_mix(pf10662) PDU1AB
21845 HMMacetkin_1(pf00871) PDU1AB
21850 HMM00288_1_pdu(pf00288) PDU
21855 -pf04363 -
BMC-P
BMC-H
BMC-T
Aldehyde dehydrogenase
PTAC
Signature enzyme
Alcohol dehydrogenase
Regulator

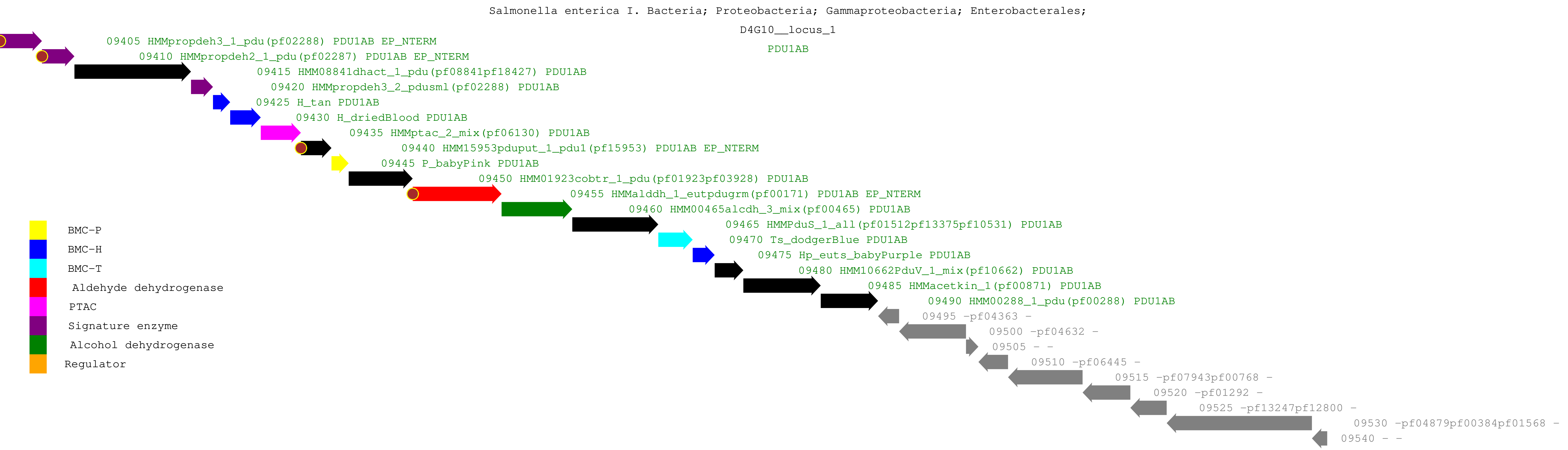
Salmonella enterica I. Bacteria; Proteobacteria; Gammaproteobacteria; Enterobacterales;
D4G10__locus_1
PDU1AB
09405 HMMpropdeh3_1_pdu(pf02288) PDU1AB EP_NTERM
09410 HMMpropdeh2_1_pdu(pf02287) PDU1AB EP_NTERM
09415 HMM08841dhact_1_pdu(pf08841pf18427) PDU1AB
09420 HMMpropdeh3_2_pdusml(pf02288) PDU1AB
09425 H_tan PDU1AB
09430 H_driedBlood PDU1AB
09435 HMMptac_2_mix(pf06130) PDU1AB
09440 HMM15953pduput_1_pdu1(pf15953) PDU1AB EP_NTERM
09445 P_babyPink PDU1AB
09450 HMM01923cobtr_1_pdu(pf01923pf03928) PDU1AB
09455 HMMalddh_1_eutpdugrm(pf00171) PDU1AB EP_NTERM
09460 HMM00465alcdh_3_mix(pf00465) PDU1AB
09465 HMMPduS_1_all(pf01512pf13375pf10531) PDU1AB
09470 Ts_dodgerBlue PDU1AB
09475 Hp_euts_babyPurple PDU1AB
09480 HMM10662PduV_1_mix(pf10662) PDU1AB
09485 HMMacetkin_1(pf00871) PDU1AB
09490 HMM00288_1_pdu(pf00288) PDU1AB
09495 -pf04363 -
09500 -pf04632 -
09505 - -
09510 -pf06445 -
09515 -pf07943pf00768 -
09520 -pf01292 -
09525 -pf13247pf12800 -
09530 -pf04879pf00384pf01568 -
09540 - -
BMC-P
BMC-H
BMC-T
Aldehyde dehydrogenase
PTAC
Signature enzyme
Alcohol dehydrogenase
Regulator

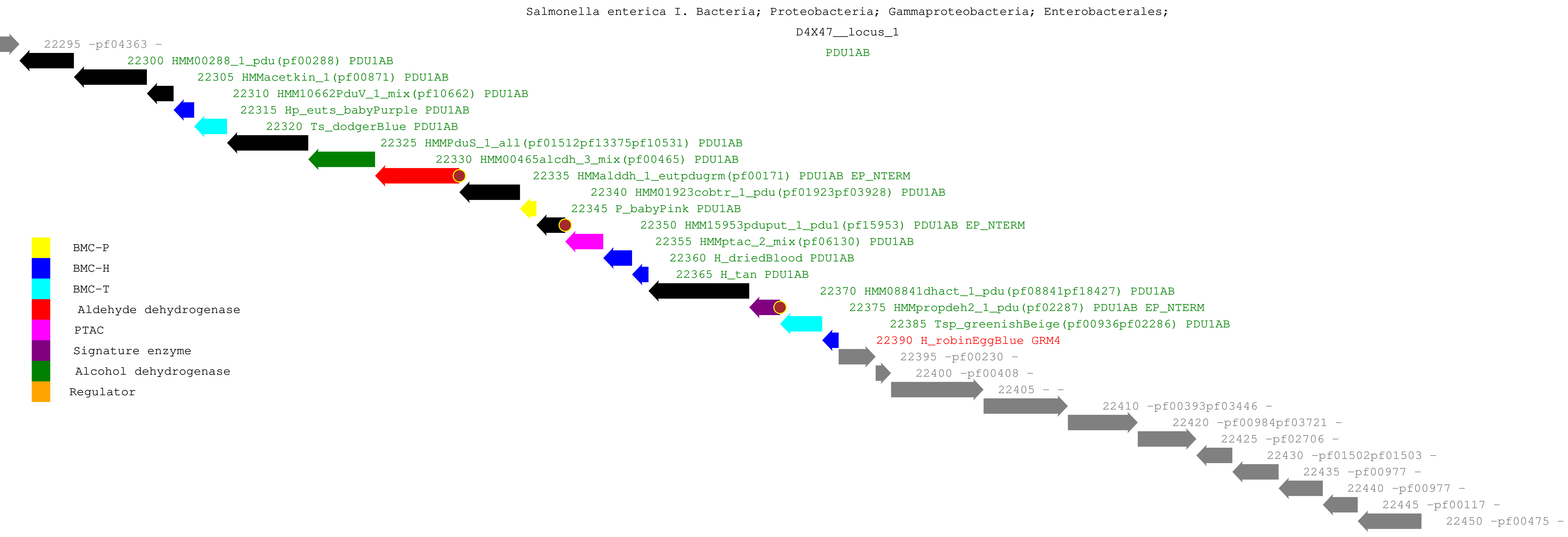
Salmonella enterica I. Bacteria; Proteobacteria; Gammaproteobacteria; Enterobacterales;
D4X47__locus_1
PDU1AB
22295 -pf04363 -
22300 HMM00288_1_pdu(pf00288) PDU1AB
22305 HMMacetkin_1(pf00871) PDU1AB
22310 HMM10662PduV_1_mix(pf10662) PDU1AB
22315 Hp_euts_babyPurple PDU1AB
22320 Ts_dodgerBlue PDU1AB
22325 HMMPduS_1_all(pf01512pf13375pf10531) PDU1AB
22330 HMM00465alcdh_3_mix(pf00465) PDU1AB
22335 HMMalddh_1_eutpdugrm(pf00171) PDU1AB EP_NTERM
22340 HMM01923cobtr_1_pdu(pf01923pf03928) PDU1AB
22345 P_babyPink PDU1AB
22350 HMM15953pduput_1_pdu1(pf15953) PDU1AB EP_NTERM
22355 HMMptac_2_mix(pf06130) PDU1AB
22360 H_driedBlood PDU1AB
22365 H_tan PDU1AB
22370 HMM08841dhact_1_pdu(pf08841pf18427) PDU1AB
22375 HMMpropdeh2_1_pdu(pf02287) PDU1AB EP_NTERM
22385 Tsp_greenishBeige(pf00936pf02286) PDU1AB
22390 H_robinEggBlue GRM4
22395 -pf00230 -
22400 -pf00408 -
22405 - -
22410 -pf00393pf03446 -
22420 -pf00984pf03721 -
22425 -pf02706 -
22430 -pf01502pf01503 -
22435 -pf00977 -
22440 -pf00977 -
22445 -pf00117 -
22450 -pf00475 -
BMC-P
BMC-H
BMC-T
Aldehyde dehydrogenase
PTAC
Signature enzyme
Alcohol dehydrogenase
Regulator

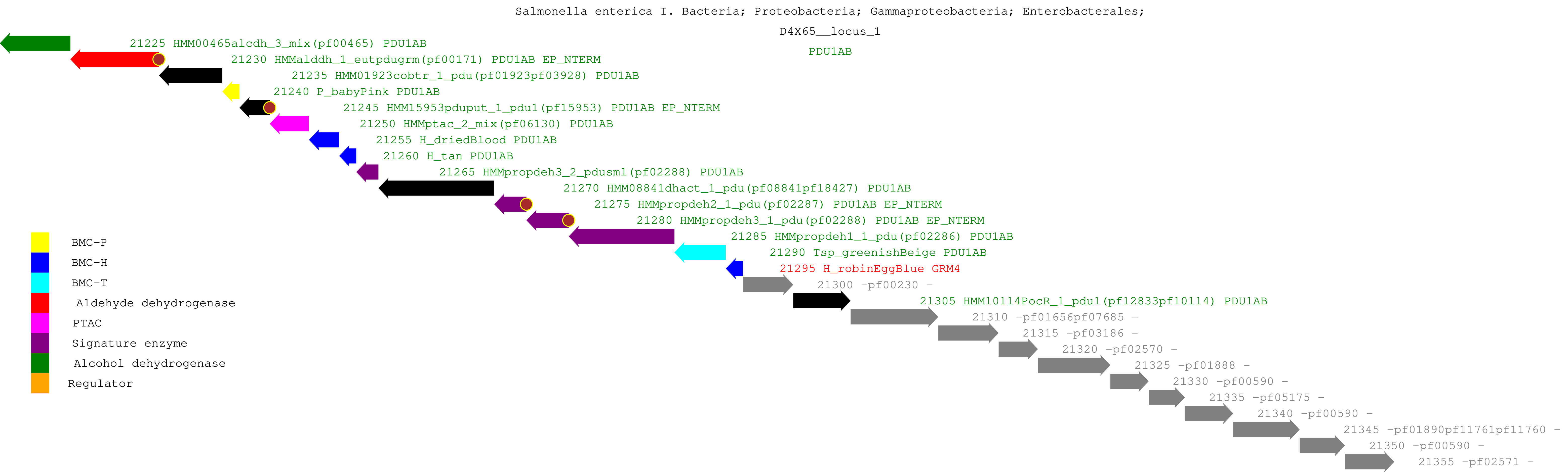

Salmonella enterica I. Bacteria; Proteobacteria; Gammaproteobacteria; Enterobacterales;
D4X65__locus_1
PDU1AB
21225 HMM00465alcdh_3_mix(pf00465) PDU1AB
21230 HMMalddh_1_eutpdugrm(pf00171) PDU1AB EP_NTERM
21235 HMM01923cobtr_1_pdu(pf01923pf03928) PDU1AB
21240 P_babyPink PDU1AB
21245 HMM15953pduput_1_pdu1(pf15953) PDU1AB EP_NTERM
21250 HMMptac_2_mix(pf06130) PDU1AB
21255 H_driedBlood PDU1AB
21260 H_tan PDU1AB
21265 HMMpropdeh3_2_pdusml(pf02288) PDU1AB
21270 HMM08841dhact_1_pdu(pf08841pf18427) PDU1AB
21275 HMMpropdeh2_1_pdu(pf02287) PDU1AB EP_NTERM
21280 HMMpropdeh3_1_pdu(pf02288) PDU1AB EP_NTERM
21285 HMMpropdeh1_1_pdu(pf02286) PDU1AB
21290 Tsp_greenishBeige PDU1AB
21295 H_robinEggBlue GRM4
21300 -pf00230 -
21305 HMM10114PocR_1_pdu1(pf12833pf10114) PDU1AB
21310 -pf01656pf07685 -
21315 -pf03186 -
21320 -pf02570 -
21325 -pf01888 -
21330 -pf00590 -
21335 -pf05175 -
21340 -pf00590 -
21345 -pf01890pf11761pf11760 -
21350 -pf00590 -
21355 -pf02571 -
BMC-P
BMC-H
BMC-T
Aldehyde dehydrogenase
PTAC
Signature enzyme
Alcohol dehydrogenase
Regulator

Salmonella enterica I. Bacteria; Proteobacteria; Gammaproteobacteria; Enterobacterales;

D4Y00__locus_1

PDU1AB

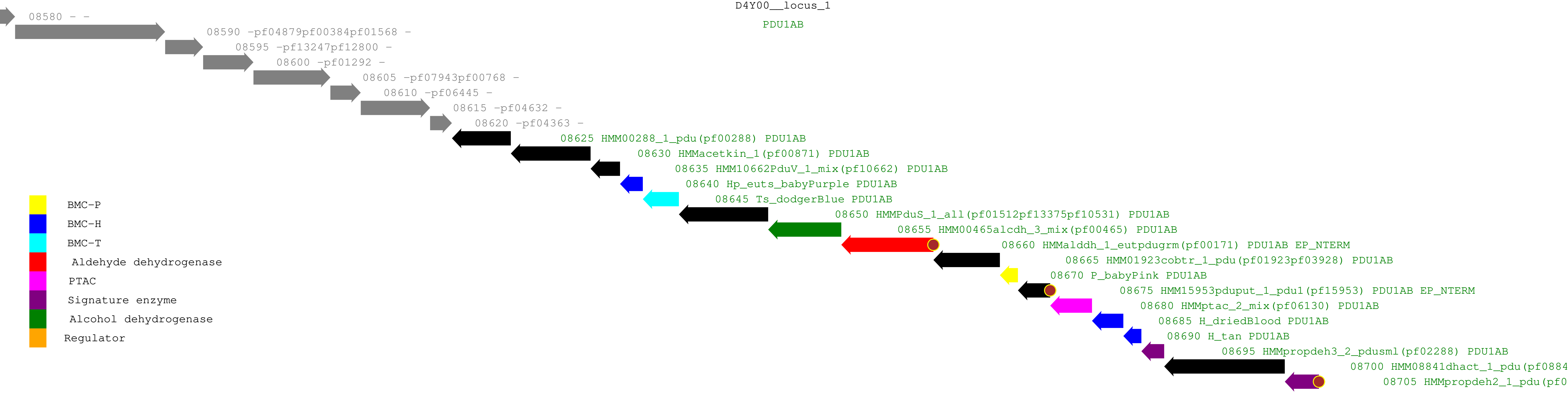

Salmonella enterica I. Bacteria; Proteobacteria; Gammaproteobacteria; Enterobacterales;

D4Y56__locus_1

PDU1AB

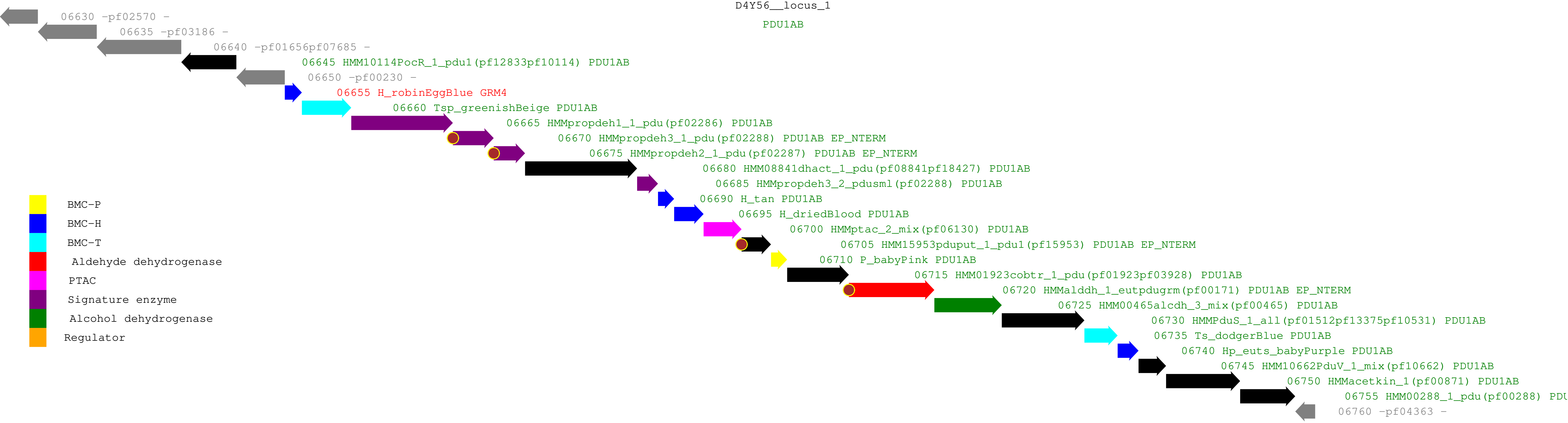

Salmonella enterica I. Bacteria; Proteobacteria; Gammaproteobacteria; Enterobacterales;

D4Y61__locus_1

PDU1AB

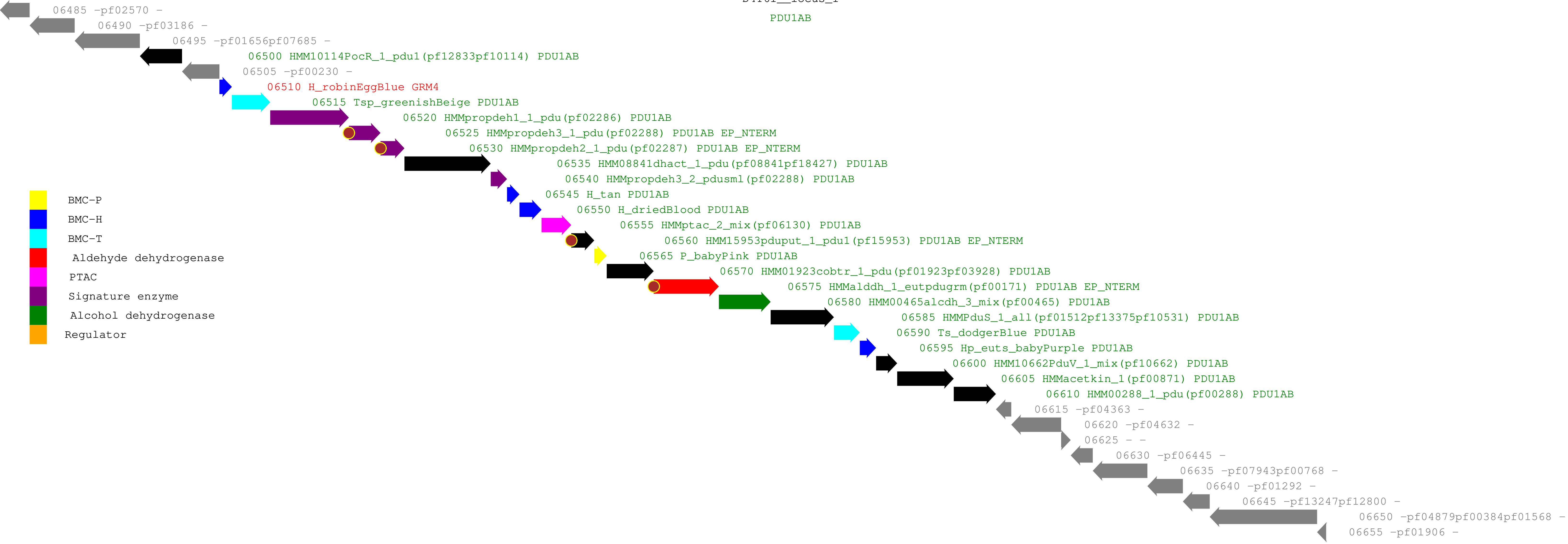

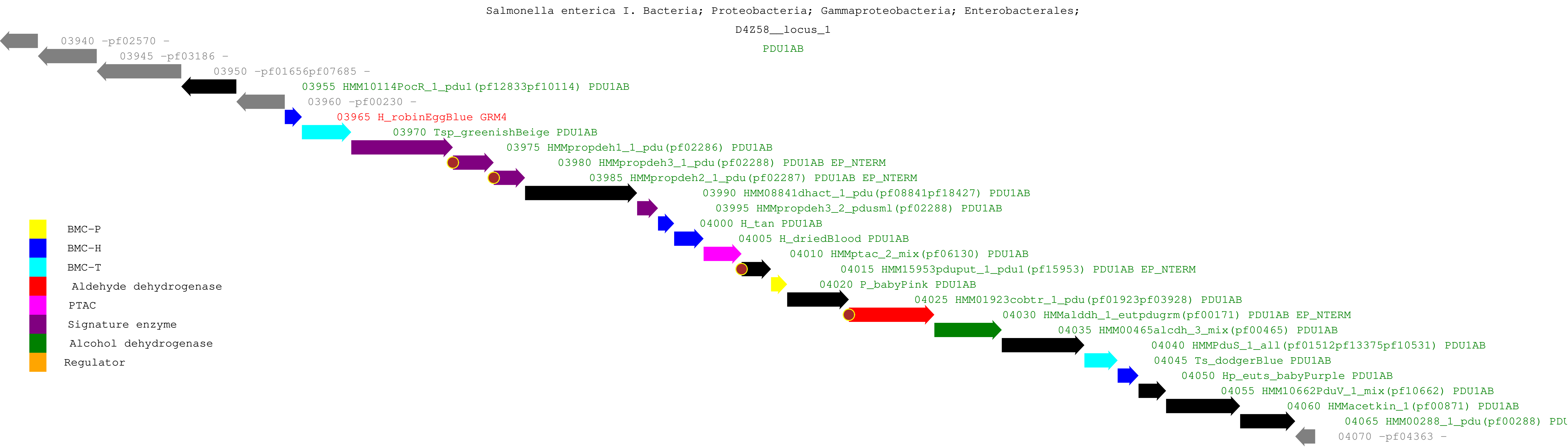
Salmonella enterica I. Bacteria; Proteobacteria; Gammaproteobacteria; Enterobacterales;
D4Z58__locus_1
PDU1AB
03940 -pf02570 -
03945 -pf03186 -
03950 -pf01656pf07685 -
03955 HMM10114PocR_1_pdu1(pf12833pf10114) PDU1AB
03960 -pf00230 -
03965 H_robinEggBlue GRM4
03970 Tsp_greenishBeige PDU1AB
03975 HMMpropdeh1_1_pdu(pf02286) PDU1AB
03980 HMMpropdeh3_1_pdu(pf02288) PDU1AB EP_NTERM
03985 HMMpropdeh2_1_pdu(pf02287) PDU1AB EP_NTERM
03990 HMM08841dhact_1_pdu(pf08841pf18427) PDU1AB
03995 HMMpropdeh3_2_pdusml(pf02288) PDU1AB
04000 H_tan PDU1AB
04005 H_driedBlood PDU1AB
04010 HMMptac_2_mix(pf06130) PDU1AB
04015 HMM15953pduput_1_pdu1(pf15953) PDU1AB EP_NTERM
04020 P_babyPink PDU1AB
04025 HMM01923cobtr_1_pdu(pf01923pf03928) PDU1AB
04030 HMMalddh_1_eutpdugrm(pf00171) PDU1AB EP_NTERM
04035 HMM00465alcdh_3_mix(pf00465) PDU1AB
04040 HMMPduS_1_all(pf01512pf13375pf10531) PDU1AB
04045 Ts_dodgerBlue PDU1AB
04050 Hp_euts_babyPurple PDU1AB
04055 HMM10662PduV_1_mix(pf10662) PDU1AB
04060 HMMacetkin_1(pf00871) PDU1AB
04065 HMM00288_1_pdu(pf00288) PDU
04070 -pf04363 -
BMC-P
BMC-H
BMC-T
Aldehyde dehydrogenase
PTAC
Signature enzyme
Alcohol dehydrogenase
Regulator

Salmonella muenchen. Bacteria; Proteobacteria; Gammaproteobacteria; Enterobacterales;

D4Z66__locus_1

PDU1AB

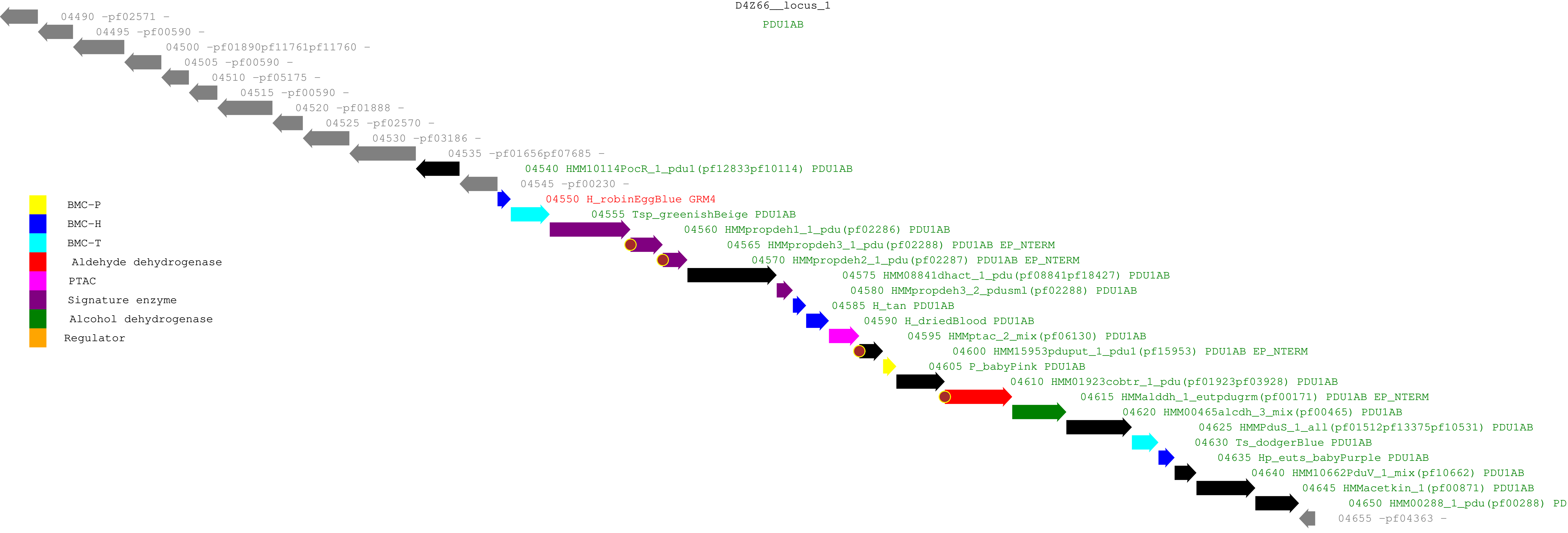

Salmonella enterica I. Bacteria; Proteobacteria; Gammaproteobacteria; Enterobacterales;

D5A36__locus_1

PDU1AB

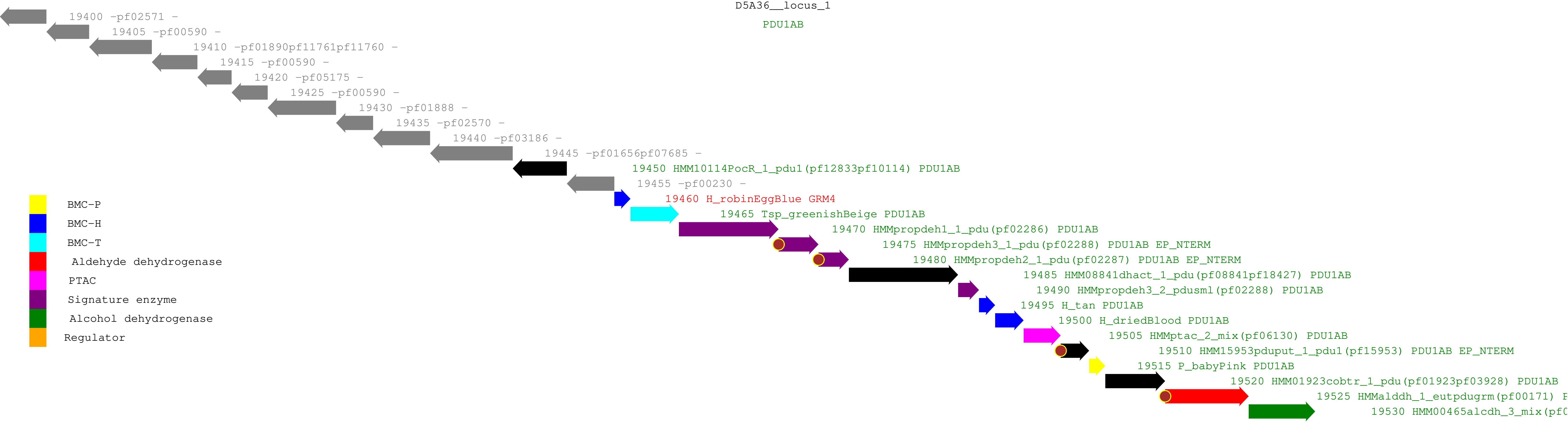

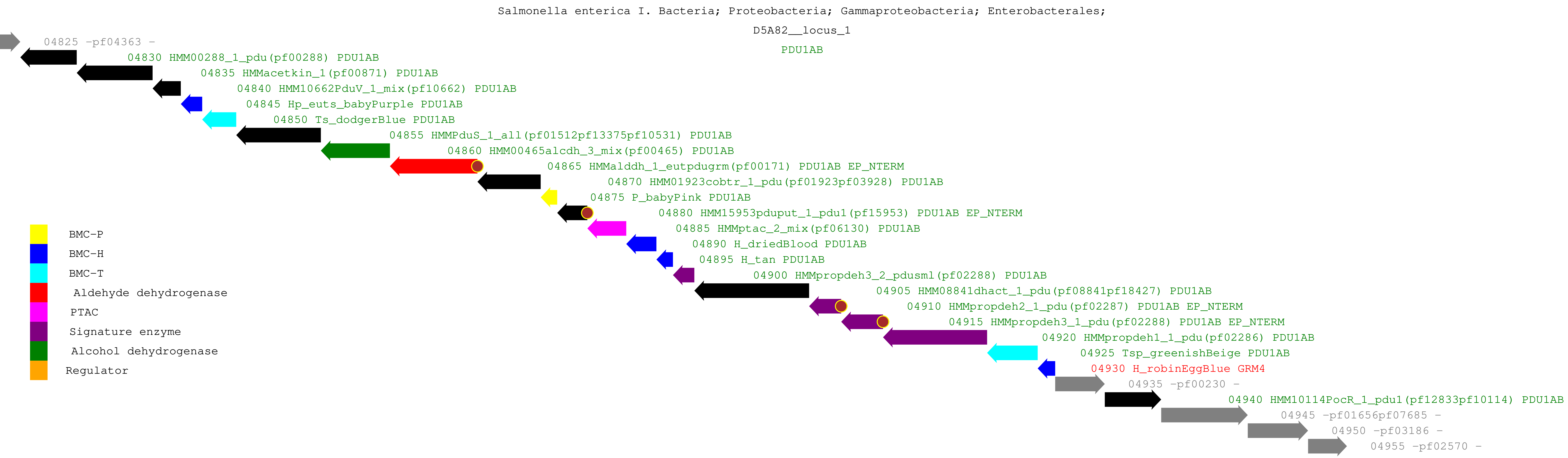
Salmonella enterica I. Bacteria; Proteobacteria; Gammaproteobacteria; Enterobacterales;
D5A82__locus_1
PDU1AB
04825 -pf04363 -
04830 HMM00288_1_pdu(pf00288) PDU1AB
04835 HMMacetkin_1(pf00871) PDU1AB
04840 HMM10662PduV_1_mix(pf10662) PDU1AB
04845 Hp_euts_babyPurple PDU1AB
04850 Ts_dodgerBlue PDU1AB
04855 HMMPduS_1_all(pf01512pf13375pf10531) PDU1AB
04860 HMM00465alcdh_3_mix(pf00465) PDU1AB
04865 HMMalddh_1_eutpdugrm(pf00171) PDU1AB EP_NTERM
04870 HMM01923cobtr_1_pdu(pf01923pf03928) PDU1AB
04875 P_babyPink PDU1AB
04880 HMM15953pduput_1_pdu1(pf15953) PDU1AB EP_NTERM
04885 HMMptac_2_mix(pf06130) PDU1AB
04890 H_driedBlood PDU1AB
04895 H_tan PDU1AB
04900 HMMpropdeh3_2_pdusml(pf02288) PDU1AB
04905 HMM08841dhact_1_pdu(pf08841pf18427) PDU1AB
04910 HMMpropdeh2_1_pdu(pf02287) PDU1AB EP_NTERM
04915 HMMpropdeh3_1_pdu(pf02288) PDU1AB EP_NTERM
04920 HMMpropdeh1_1_pdu(pf02286) PDU1AB
04925 Tsp_greenishBeige PDU1AB
04930 H_robinEggBlue GRM4
04935 -pf00230 -
04940 HMM10114PocR_1_pdu1(pf12833pf10114) PDU1AB
04945 -pf01656pf07685 -
04950 -pf03186 -
04955 -pf02570 -
BMC-P
BMC-H
BMC-T
Aldehyde dehydrogenase
PTAC
Signature enzyme
Alcohol dehydrogenase
Regulator

Salmonella enterica I. Bacteria; Proteobacteria; Gammaproteobacteria; Enterobacterales;

D5B67__locus_1

PDU1AB

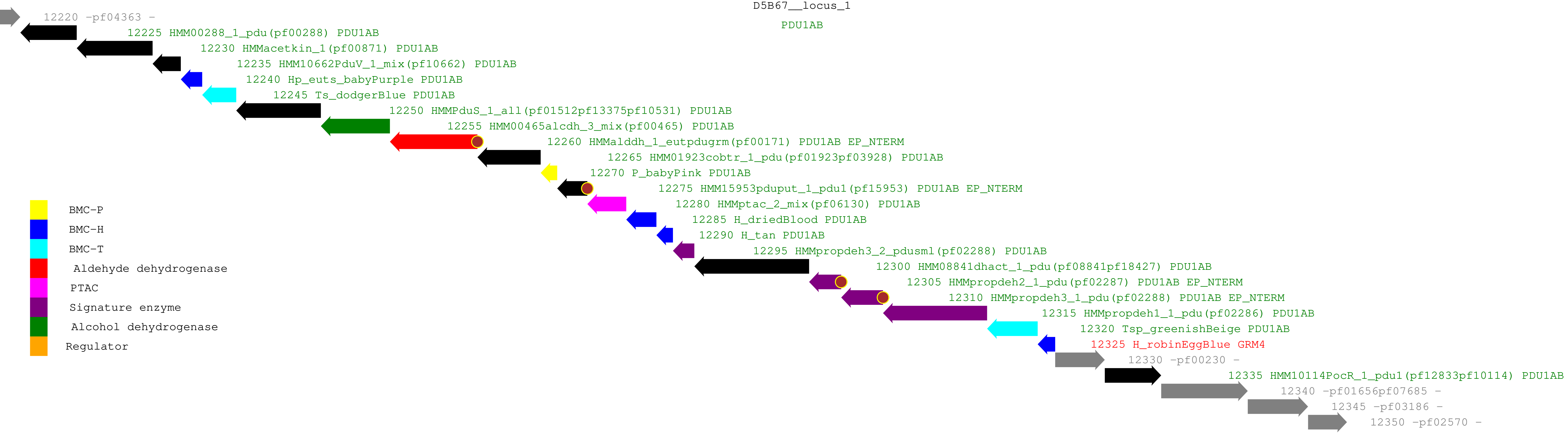

Yersinia hibernica. Bacteria; Proteobacteria; Gammaproteobacteria; Enterobacterales;

D5F51__locus_1

PDU1AB

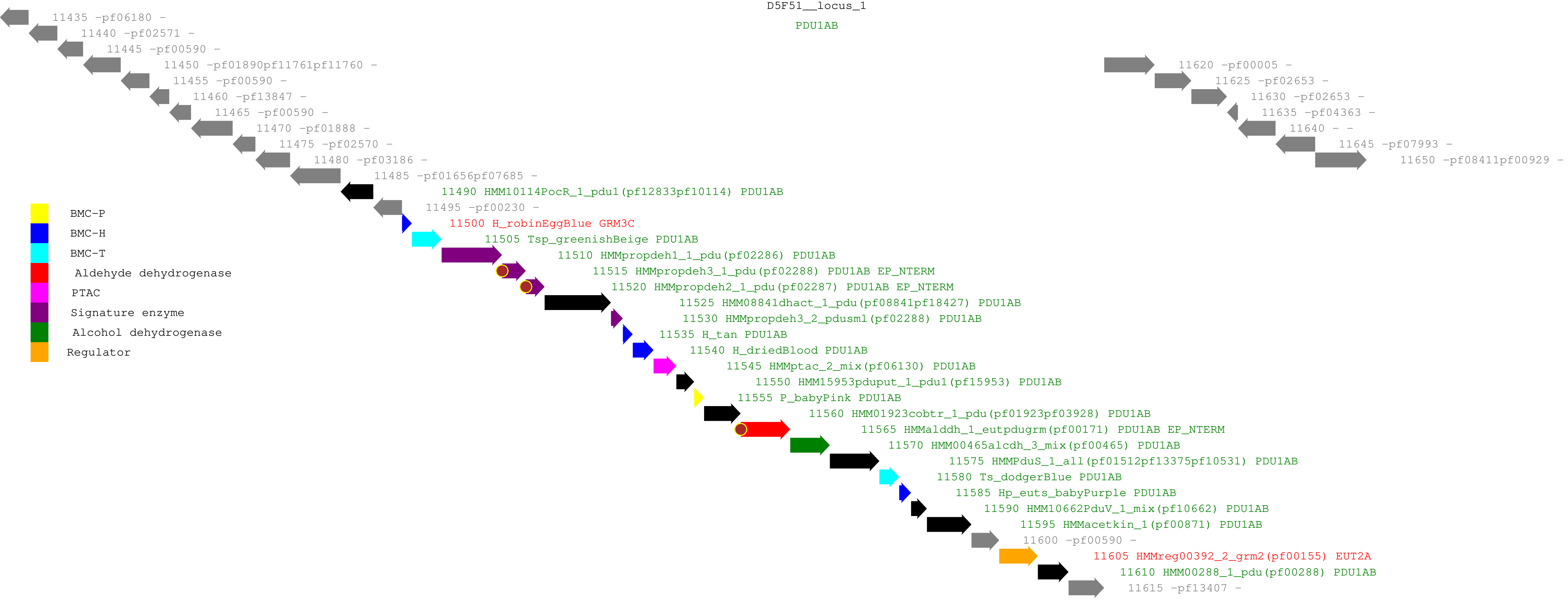

Salmonella enterica I. Bacteria; Proteobacteria; Gammaproteobacteria; Enterobacterales;

D5080__locus_1

PDU1AB

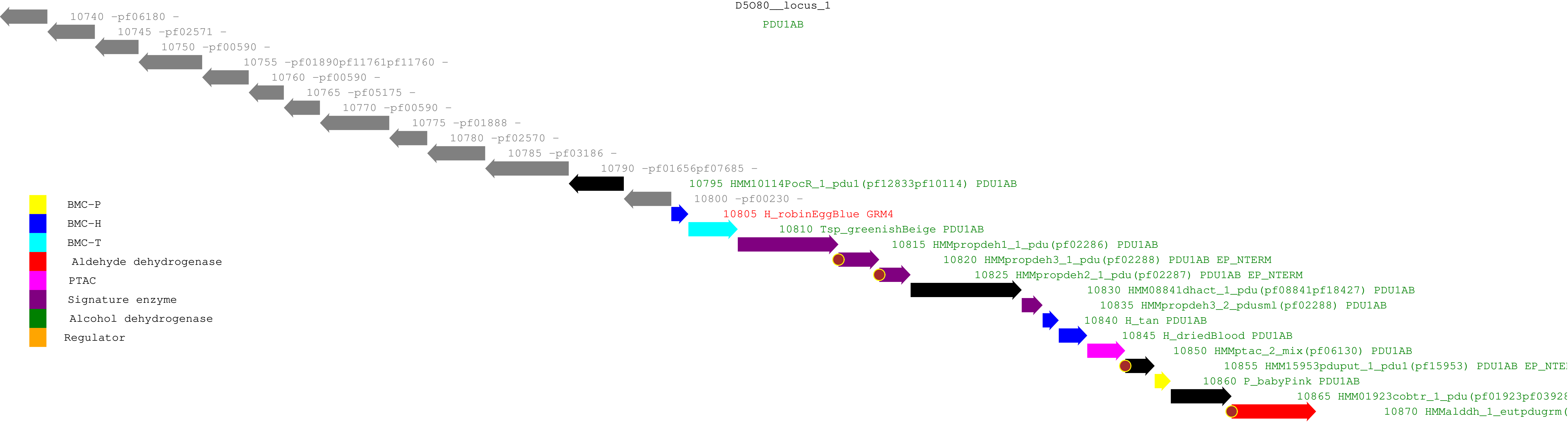

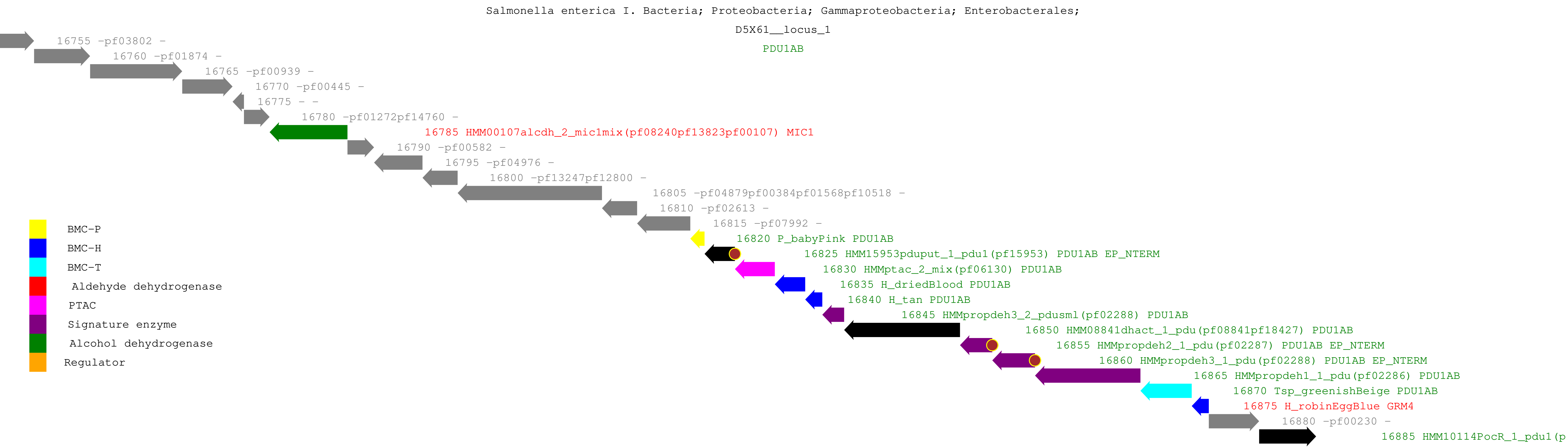
Salmonella enterica I. Bacteria; Proteobacteria; Gammaproteobacteria; Enterobacterales;
D5X61__locus_1
PDU1AB
16755 -pf03802 -
16760 -pf01874 -
16765 -pf00939 -
16770 -pf00445 -
16775 - -
16780 -pf01272pf14760 -
16785 HMM00107alcdh_2_mic1mix(pf08240pf13823pf00107) MIC1
16790 -pf00582 -
16795 -pf04976 -
16800 -pf13247pf12800 -
16805 -pf04879pf00384pf01568pf10518 -
16810 -pf02613 -
16815 -pf07992 -
16820 P_babyPink PDU1AB
16825 HMM15953pduput_1_pdu1(pf15953) PDU1AB EP_NTERM
16830 HMMptac_2_mix(pf06130) PDU1AB
16835 H_driedBlood PDU1AB
16840 H_tan PDU1AB
16845 HMMpropdeh3_2_pdusml(pf02288) PDU1AB
16850 HMM08841dhact_1_pdu(pf08841pf18427) PDU1AB
16855 HMMpropdeh2_1_pdu(pf02287) PDU1AB EP_NTERM
16860 HMMpropdeh3_1_pdu(pf02288) PDU1AB EP_NTERM
16865 HMMpropdeh1_1_pdu(pf02286) PDU1AB
16870 Tsp_greenishBeige PDU1AB
16875 H_robinEggBlue GRM4
16880 -pf00230 -
16885 HMM10114PocR_1_pdu1(p
BMC-P
BMC-H
BMC-T
Aldehyde dehydrogenase
PTAC
Signature enzyme
Alcohol dehydrogenase
Regulator

Salmonella enterica subsp. enterica serovar Stanley. Bacteria; Proteobacteria; Gammaproteobacteria; Enterobacterales;

D6410__locus_1

PDU1AB

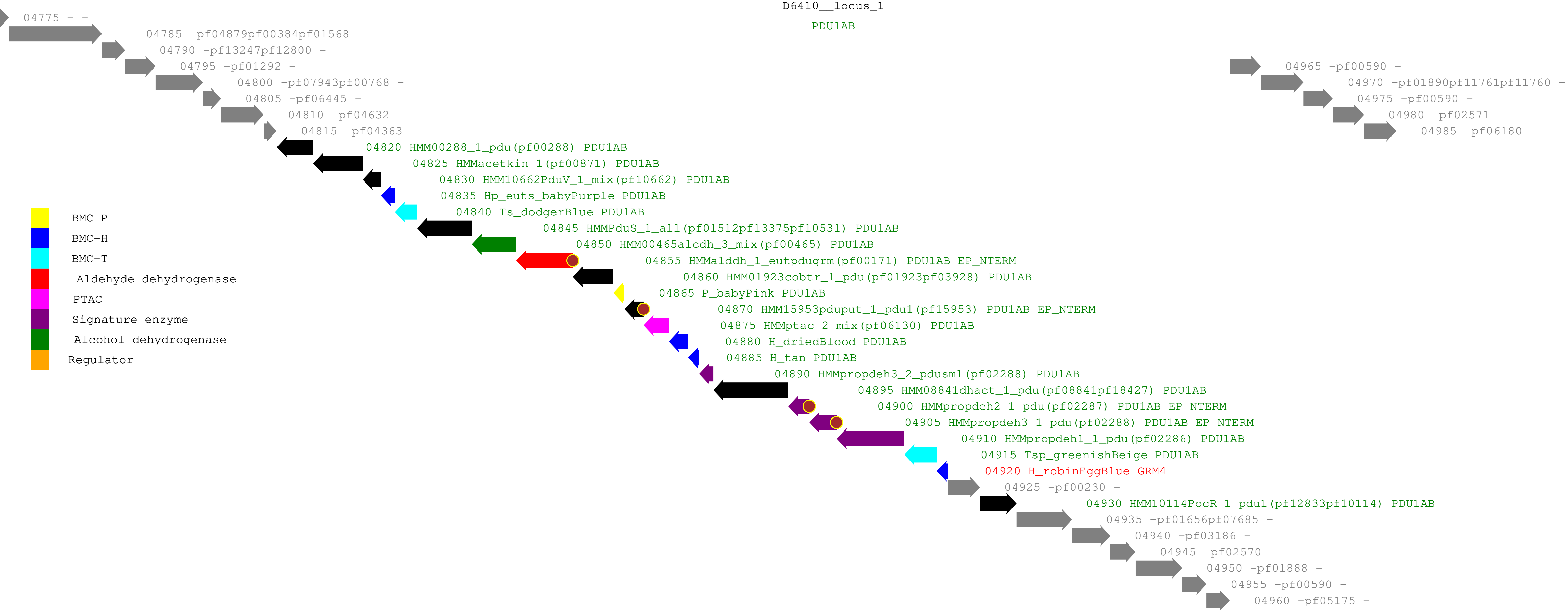

Salmonella enterica subsp. enterica serovar Schwarzengrund. Bacteria; Proteobacteria; Gammaproteobacteria; Enterobacterales;

D6A15__locus_2

PDU1AB

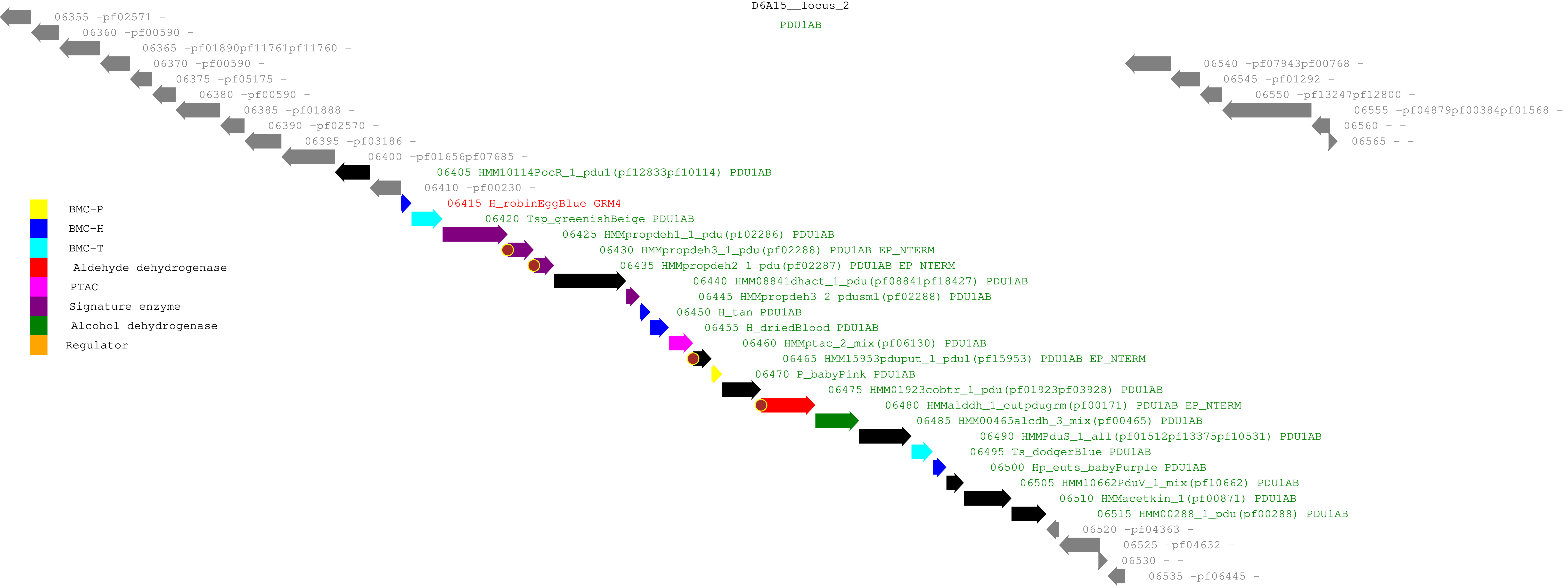

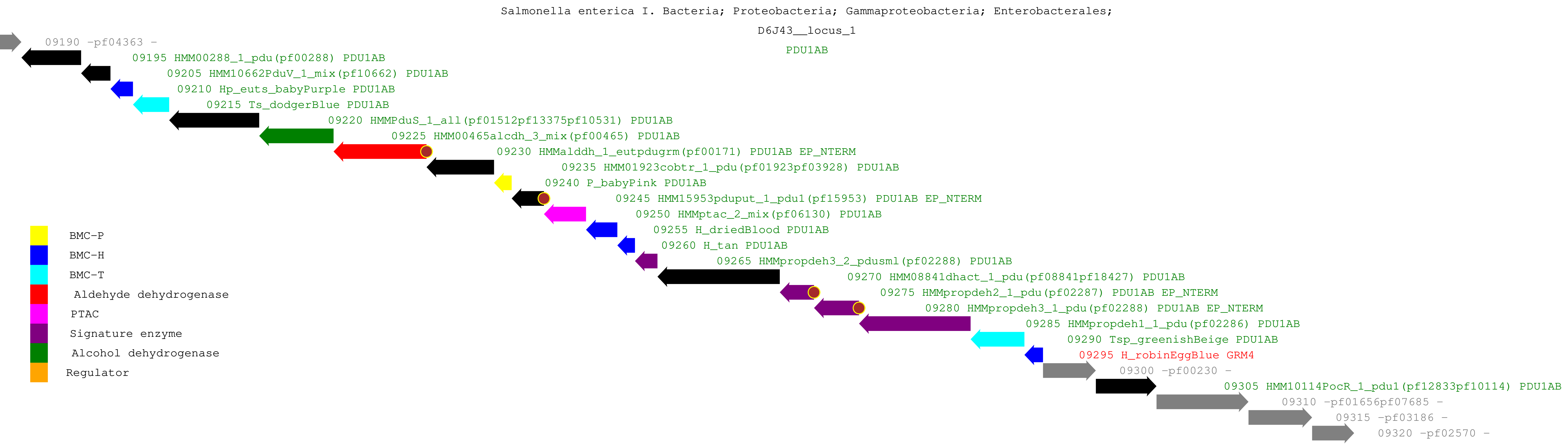

Salmonella enterica I. Bacteria; Proteobacteria; Gammaproteobacteria; Enterobacterales;
D6J43__locus_1
PDU1AB
09190 -pf04363 -
09195 HMM00288_1_pdu(pf00288) PDU1AB
09205 HMM10662PduV_1_mix(pf10662) PDU1AB
09210 Hp_euts_babyPurple PDU1AB
09215 Ts_dodgerBlue PDU1AB
09220 HMMPduS_1_all(pf01512pf13375pf10531) PDU1AB
09225 HMM00465alcdh_3_mix(pf00465) PDU1AB
09230 HMMalddh_1_eutpdugrm(pf00171) PDU1AB EP_NTERM
09235 HMM01923cobtr_1_pdu(pf01923pf03928) PDU1AB
09240 P_babyPink PDU1AB
09245 HMM15953pduput_1_pdu1(pf15953) PDU1AB EP_NTERM
09250 HMMptac_2_mix(pf06130) PDU1AB
09255 H_driedBlood PDU1AB
09260 H_tan PDU1AB
09265 HMMpropdeh3_2_pdusml(pf02288) PDU1AB
09270 HMM08841dhact_1_pdu(pf08841pf18427) PDU1AB
09275 HMMpropdeh2_1_pdu(pf02287) PDU1AB EP_NTERM
09280 HMMpropdeh3_1_pdu(pf02288) PDU1AB EP_NTERM
09285 HMMpropdeh1_1_pdu(pf02286) PDU1AB
09290 Tsp_greenishBeige PDU1AB
09295 H_robinEggBlue GRM4
09300 -pf00230 -
09305 HMM10114PocR_1_pdu1(pf12833pf10114) PDU1AB
09310 -pf01656pf07685 -
09315 -pf03186 -
09320 -pf02570 -
BMC-P
BMC-H
BMC-T
Aldehyde dehydrogenase
PTAC
Signature enzyme
Alcohol dehydrogenase
Regulator

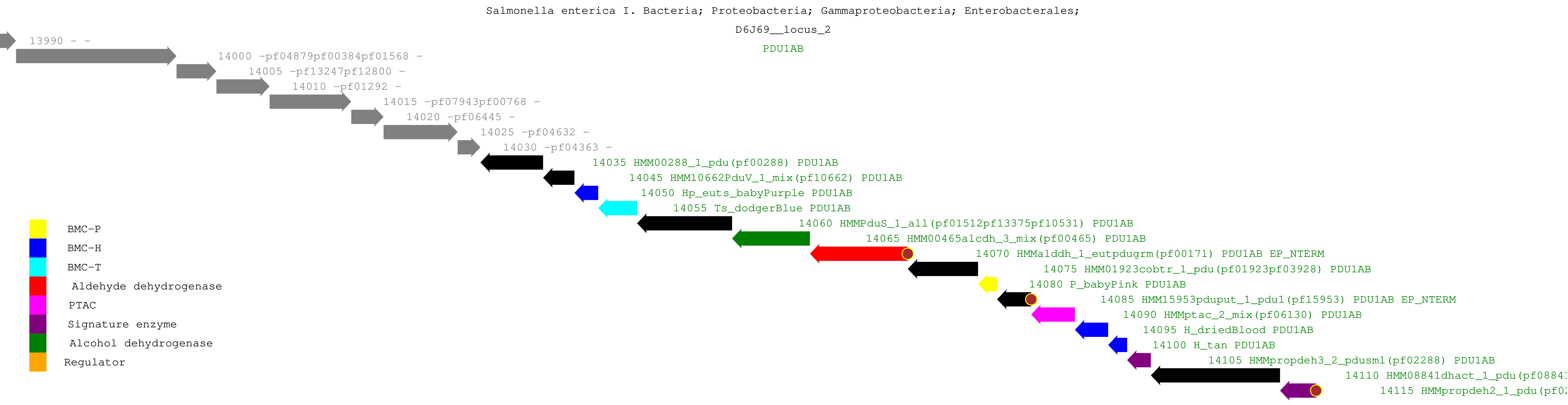
Salmonella enterica I. Bacteria; Proteobacteria; Gammaproteobacteria; Enterobacterales;
D6J69__locus_2
PDU1AB
13990 - -
14000 -pf04879pf00384pf01568 -
14005 -pf13247pf12800 -
14010 -pf01292 -
14015 -pf07943pf00768 -
14020 -pf06445 -
14025 -pf04632 -
14030 -pf04363 -
14035 HMM00288_1_pdu(pf00288) PDU1AB
14045 HMM10662PduV_1_mix(pf10662) PDU1AB
14050 Hp_euts_babyPurple PDU1AB
14055 Ts_dodgerBlue PDU1AB
14060 HMMPduS_1_all(pf01512pf13375pf10531) PDU1AB
14065 HMM00465alcdh_3_mix(pf00465) PDU1AB
14070 HMMalddh_1_eutpdugrm(pf00171) PDU1AB EP_NTERM
14075 HMM01923cobtr_1_pdu(pf01923pf03928) PDU1AB
14080 P_babyPink PDU1AB
14085 HMM15953pduput_1_pdu1(pf15953) PDU1AB EP_NTERM
14090 HMMptac_2_mix(pf06130) PDU1AB
14095 H_driedBlood PDU1AB
14100 H_tan PDU1AB
14105 HMMpropdeh3_2_pdusml(pf02288) PDU1AB
BMC-P
BMC-H
BMC-T
Aldehyde dehydrogenase
PTAC
Signature enzyme
Alcohol dehydrogenase
Regulator

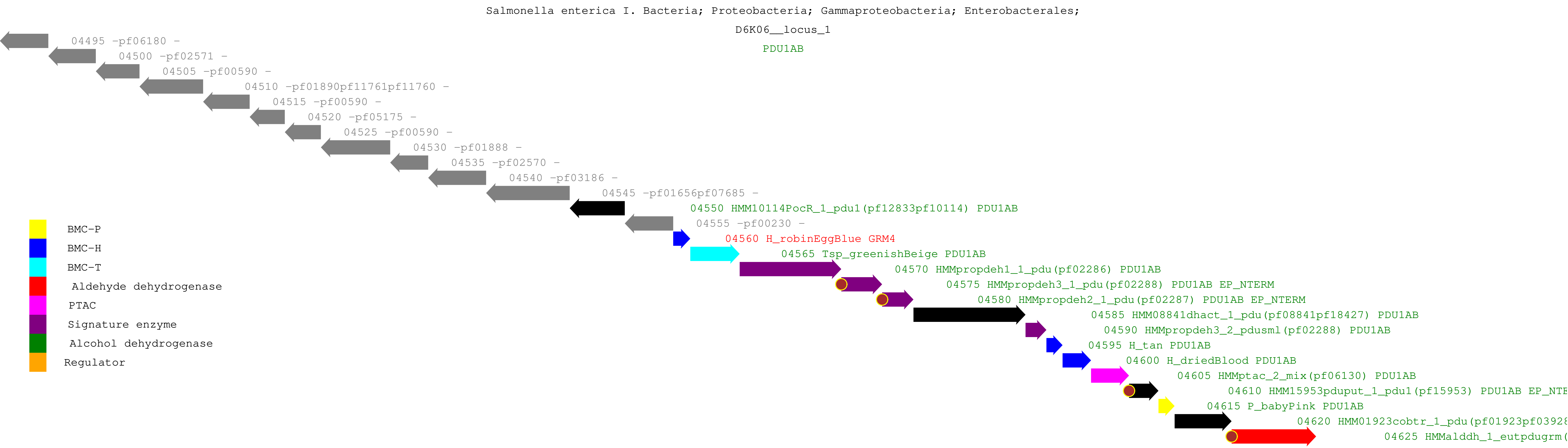
Salmonella enterica I. Bacteria; Proteobacteria; Gammaproteobacteria; Enterobacterales;
D6K06__locus_1
PDU1AB
04495 -pf06180 -
04500 -pf02571 -
04505 -pf00590 -
04510 -pf01890pf11761pf11760 -
04515 -pf00590 -
04520 -pf05175 -
04525 -pf00590 -
04530 -pf01888 -
04535 -pf02570 -
04540 -pf03186 -
04545 -pf01656pf07685 -
04550 HMM10114PocR_1_pdu1(pf12833pf10114) PDU1AB
04555 -pf00230 -
04560 H_robinEggBlue GRM4
04565 Tsp_greenishBeige PDU1AB
04570 HMMpropdeh1_1_pdu(pf02286) PDU1AB
04575 HMMpropdeh3_1_pdu(pf02288) PDU1AB EP_NTERM
04580 HMMpropdeh2_1_pdu(pf02287) PDU1AB EP_NTERM
04585 HMM08841dhact_1_pdu(pf08841pf18427) PDU1AB
04590 HMMpropdeh3_2_pdusml(pf02288) PDU1AB
04595 H_tan PDU1AB
04600 H_driedBlood PDU1AB
04605 HMMptac_2_mix(pf06130) PDU1AB
04610 HMM15953pduput_1_pdu1(pf15953) PDU1AB EP_NTE
04615 P_babyPink PDU1AB
04620 HMM01923cobtr_1_pdu(pf01923pf03928
04625 HMMalddh_1_eutpdugrm(
BMC-P
BMC-H
BMC-T
Aldehyde dehydrogenase
PTAC
Signature enzyme
Alcohol dehydrogenase
Regulator

Salmonella enterica subsp. enterica serovar Java. Bacteria; Proteobacteria; Gammaproteobacteria; Enterobacterales;

D6K54__locus_2

PDU1AB

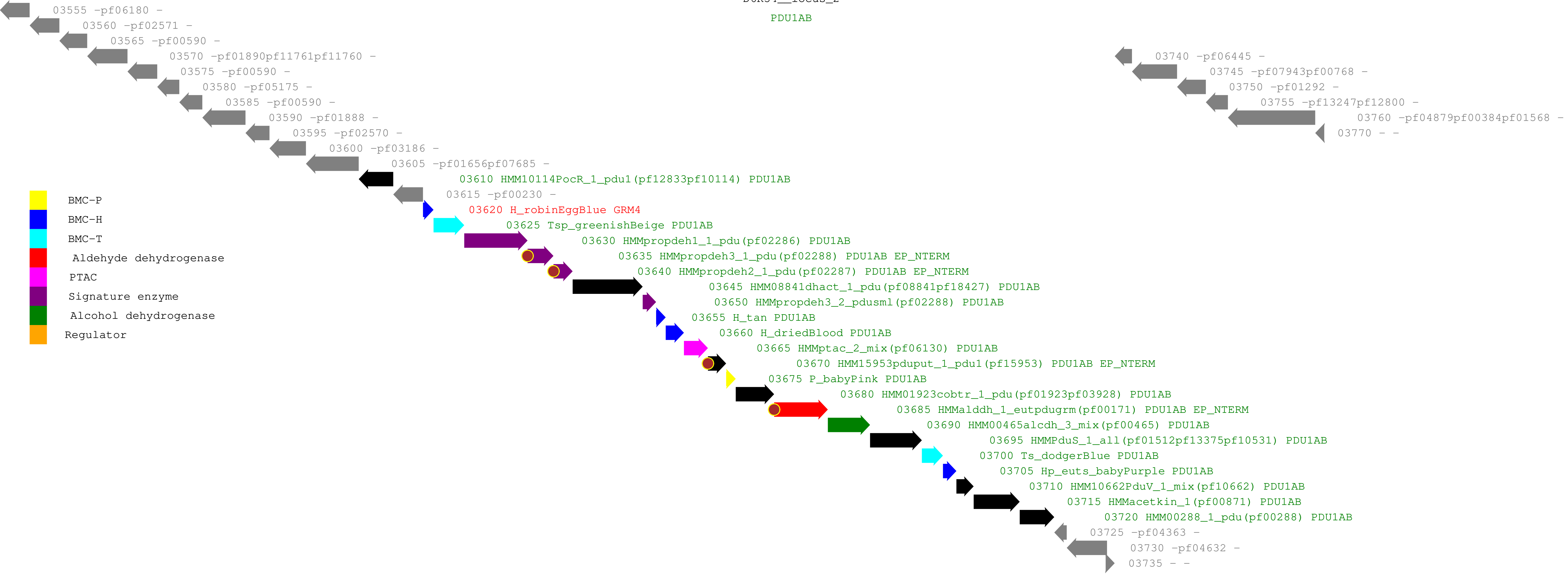

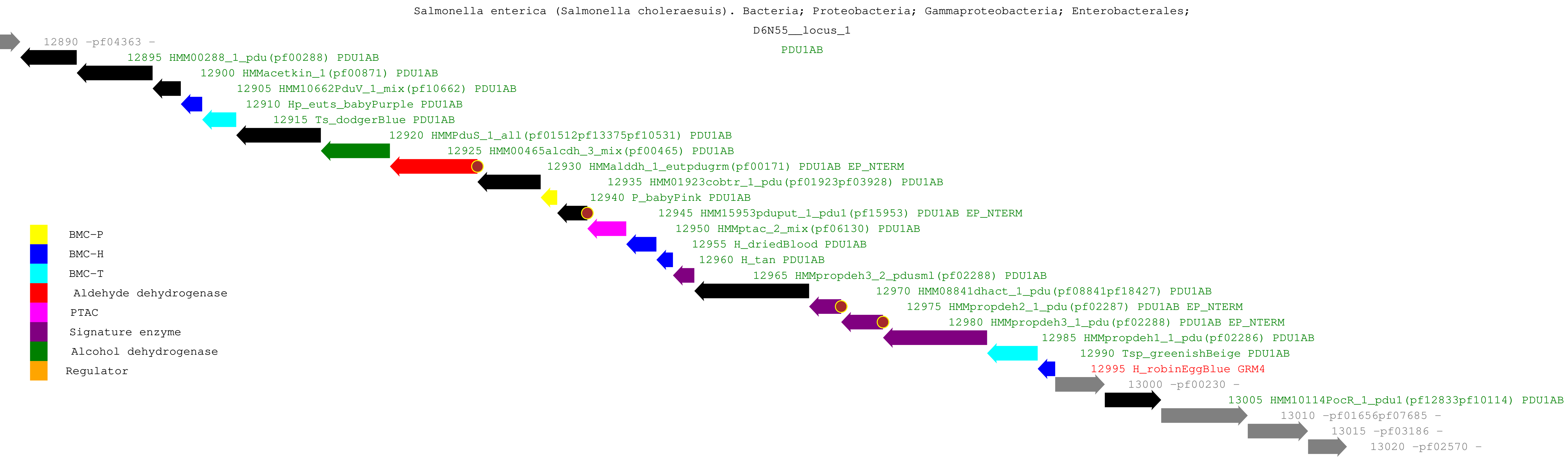
Salmonella enterica (Salmonella choleraesuis). Bacteria; Proteobacteria; Gammaproteobacteria; Enterobacterales;
D6N55__locus_1
PDU1AB
12890 -pf04363 -
12895 HMM00288_1_pdu(pf00288) PDU1AB
12900 HMMacetkin_1(pf00871) PDU1AB
12905 HMM10662PduV_1_mix(pf10662) PDU1AB
12910 Hp_euts_babyPurple PDU1AB
12915 Ts_dodgerBlue PDU1AB
12920 HMMPduS_1_all(pf01512pf13375pf10531) PDU1AB
12925 HMM00465alcdh_3_mix(pf00465) PDU1AB
12930 HMMalddh_1_eutpdugrm(pf00171) PDU1AB EP_NTERM
12935 HMM01923cobtr_1_pdu(pf01923pf03928) PDU1AB
12940 P_babyPink PDU1AB
12945 HMM15953pduput_1_pdu1(pf15953) PDU1AB EP_NTERM
12950 HMMptac_2_mix(pf06130) PDU1AB
12955 H_driedBlood PDU1AB
12960 H_tan PDU1AB
12965 HMMpropdeh3_2_pdusml(pf02288) PDU1AB
12970 HMM08841dhact_1_pdu(pf08841pf18427) PDU1AB
12975 HMMpropdeh2_1_pdu(pf02287) PDU1AB EP_NTERM
12980 HMMpropdeh3_1_pdu(pf02288) PDU1AB EP_NTERM
12985 HMMpropdeh1_1_pdu(pf02286) PDU1AB
12990 Tsp_greenishBeige PDU1AB
12995 H_robinEggBlue GRM4
13000 -pf00230 -
13005 HMM10114PocR_1_pdu1(pf12833pf10114) PDU1AB
13010 -pf01656pf07685 -
13015 -pf03186 -
13020 -pf02570 -
BMC-P
BMC-H
BMC-T
Aldehyde dehydrogenase
PTAC
Signature enzyme
Alcohol dehydrogenase
Regulator

Salmonella enterica I. Bacteria; Proteobacteria; Gammaproteobacteria; Enterobacterales;

D6S13__locus_1

PDU1AB

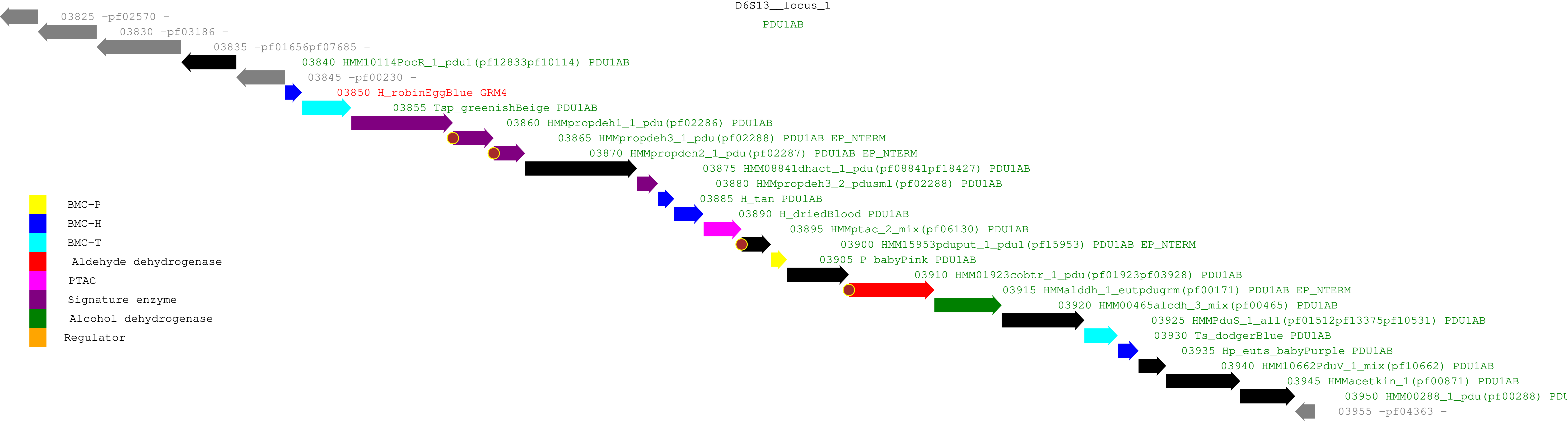

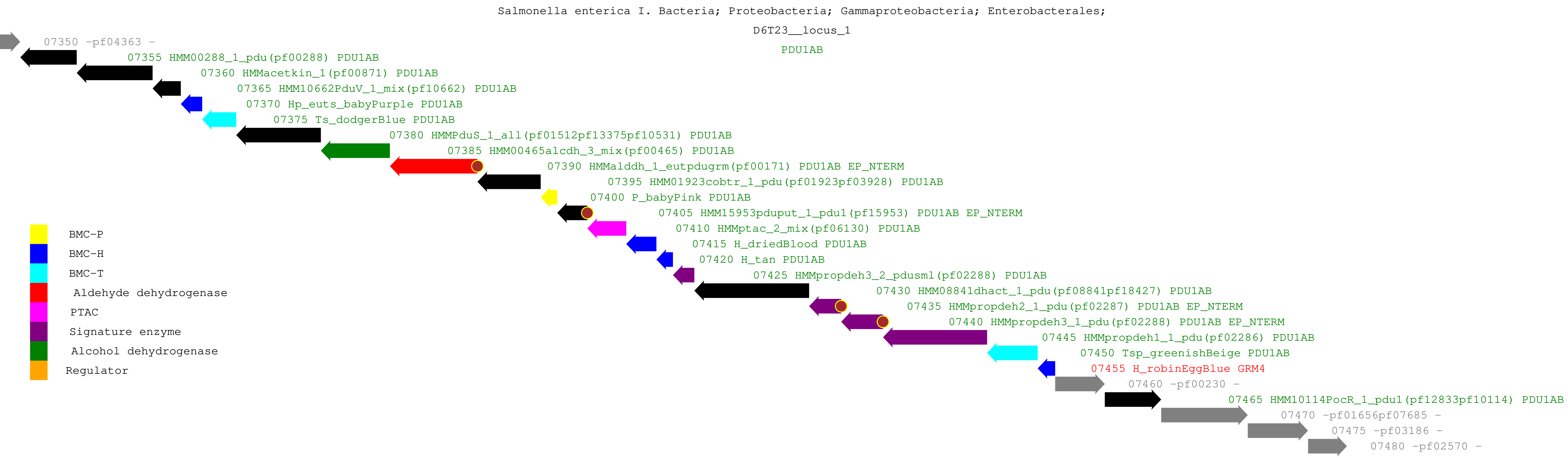
Salmonella enterica I. Bacteria; Proteobacteria; Gammaproteobacteria; Enterobacterales;
D6T23__locus_1
PDU1AB
07350 -pf04363 -
07355 HMM00288_1_pdu(pf00288) PDU1AB
07360 HMMacetkin_1(pf00871) PDU1AB
07365 HMM10662PduV_1_mix(pf10662) PDU1AB
07370 Hp_euts_babyPurple PDU1AB
07375 Ts_dodgerBlue PDU1AB
07380 HMMPduS_1_all(pf01512pf13375pf10531) PDU1AB
07385 HMM00465alcdh_3_mix(pf00465) PDU1AB
07390 HMMalddh_1_eutpdugrm(pf00171) PDU1AB EP_NTERM
07395 HMM01923cobtr_1_pdu(pf01923pf03928) PDU1AB
07400 P_babyPink PDU1AB
07405 HMM15953pduput_1_pdu1(pf15953) PDU1AB EP_NTERM
07410 HMMptac_2_mix(pf06130) PDU1AB
07415 H_driedBlood PDU1AB
07420 H_tan PDU1AB
07425 HMMpropdeh3_2_pdusml(pf02288) PDU1AB
07430 HMM08841dhact_1_pdu(pf08841pf18427) PDU1AB
07435 HMMpropdeh2_1_pdu(pf02287) PDU1AB EP_NTERM
07440 HMMpropdeh3_1_pdu(pf02288) PDU1AB EP_NTERM
07445 HMMpropdeh1_1_pdu(pf02286) PDU1AB
07450 Tsp_greenishBeige PDU1AB
07455 H_robinEggBlue GRM4
07460 -pf00230 -
07465 HMM10114PocR_1_pdu1(pf12833pf10114) PDU1AB
07470 -pf01656pf07685 -
07475 -pf03186 -
07480 -pf02570 -
BMC-P
BMC-H
BMC-T
Aldehyde dehydrogenase
PTAC
Signature enzyme
Alcohol dehydrogenase
Regulator

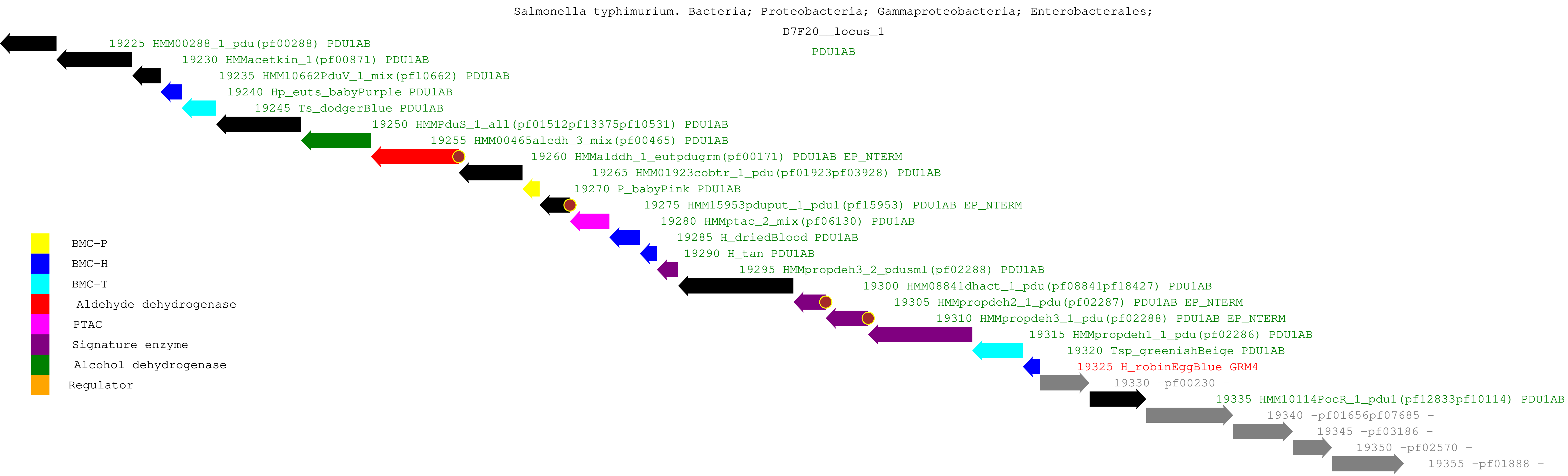

Salmonella typhimurium. Bacteria; Proteobacteria; Gammaproteobacteria; Enterobacterales;
D7F20__locus_1
PDU1AB
19225 HMM00288_1_pdu(pf00288) PDU1AB
19230 HMMacetkin_1(pf00871) PDU1AB
19235 HMM10662PduV_1_mix(pf10662) PDU1AB
19240 Hp_euts_babyPurple PDU1AB
19245 Ts_dodgerBlue PDU1AB
19250 HMMPduS_1_all(pf01512pf13375pf10531) PDU1AB
19255 HMM00465alcdh_3_mix(pf00465) PDU1AB
19260 HMMalddh_1_eutpdugrm(pf00171) PDU1AB EP_NTERM
19265 HMM01923cobtr_1_pdu(pf01923pf03928) PDU1AB
19270 P_babyPink PDU1AB
19275 HMM15953pduput_1_pdu1(pf15953) PDU1AB EP_NTERM
19280 HMMptac_2_mix(pf06130) PDU1AB
19285 H_driedBlood PDU1AB
19290 H_tan PDU1AB
19295 HMMpropdeh3_2_pdusml(pf02288) PDU1AB
19300 HMM08841dhact_1_pdu(pf08841pf18427) PDU1AB
19305 HMMpropdeh2_1_pdu(pf02287) PDU1AB EP_NTERM
19310 HMMpropdeh3_1_pdu(pf02288) PDU1AB EP_NTERM
19315 HMMpropdeh1_1_pdu(pf02286) PDU1AB
19320 Tsp_greenishBeige PDU1AB
19325 H_robinEggBlue GRM4
19330 -pf00230 -
19335 HMM10114PocR_1_pdu1(pf12833pf10114) PDU1AB
19340 -pf01656pf07685 -
19345 -pf03186 -
19350 -pf02570 -
19355 -pf01888 -
BMC-P
BMC-H
BMC-T
Aldehyde dehydrogenase
PTAC
Signature enzyme
Alcohol dehydrogenase
Regulator

Salmonella typhimurium. Bacteria; Proteobacteria; Gammaproteobacteria; Enterobacterales;

D7H43__locus_1

PDU1AB

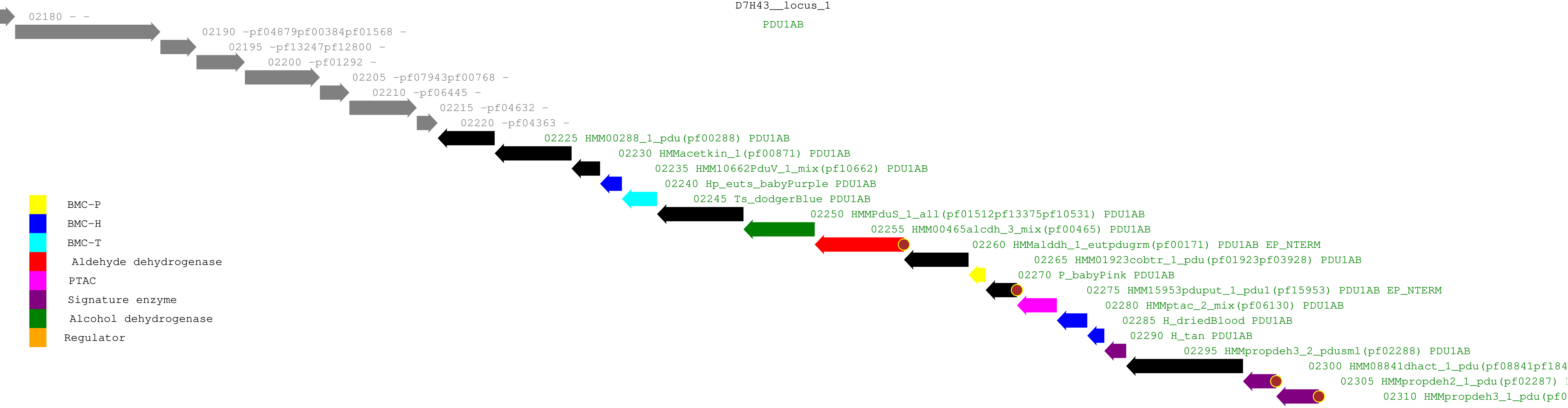

Citrobacter sp. MH181794. Bacteria; Proteobacteria; Gammaproteobacteria; Enterobacterales;

D7I40__locus_1

PDU1AB

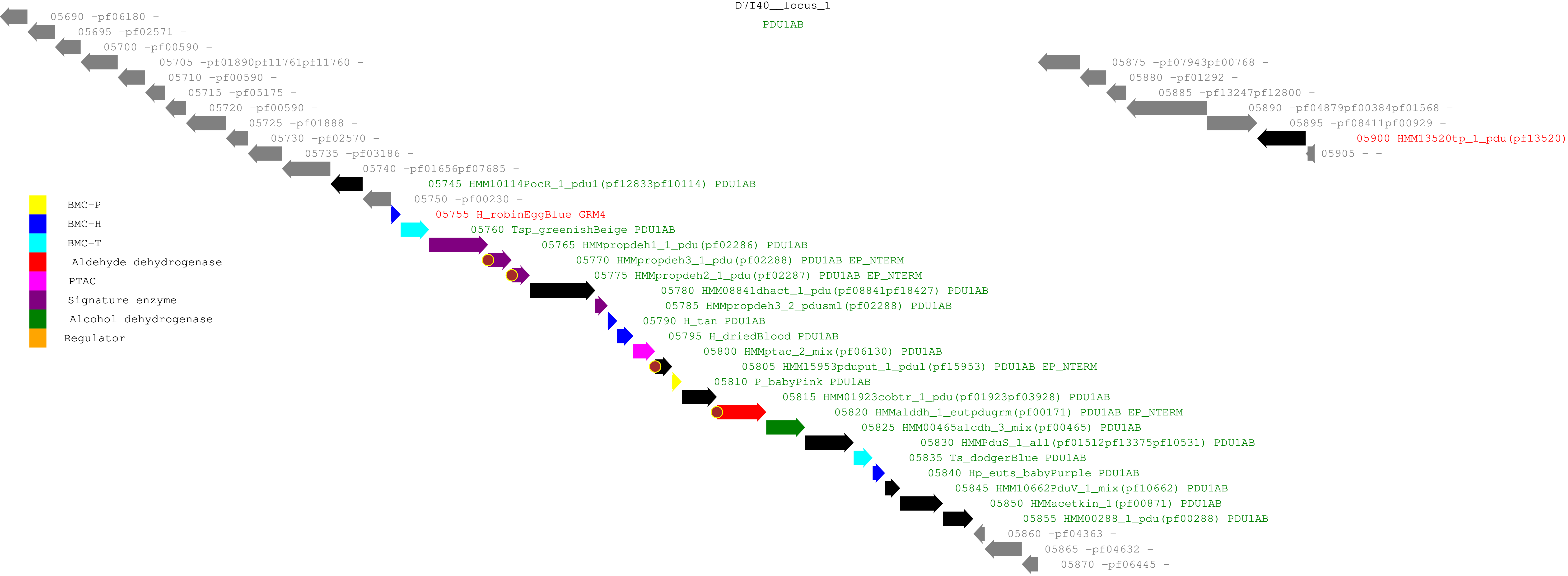

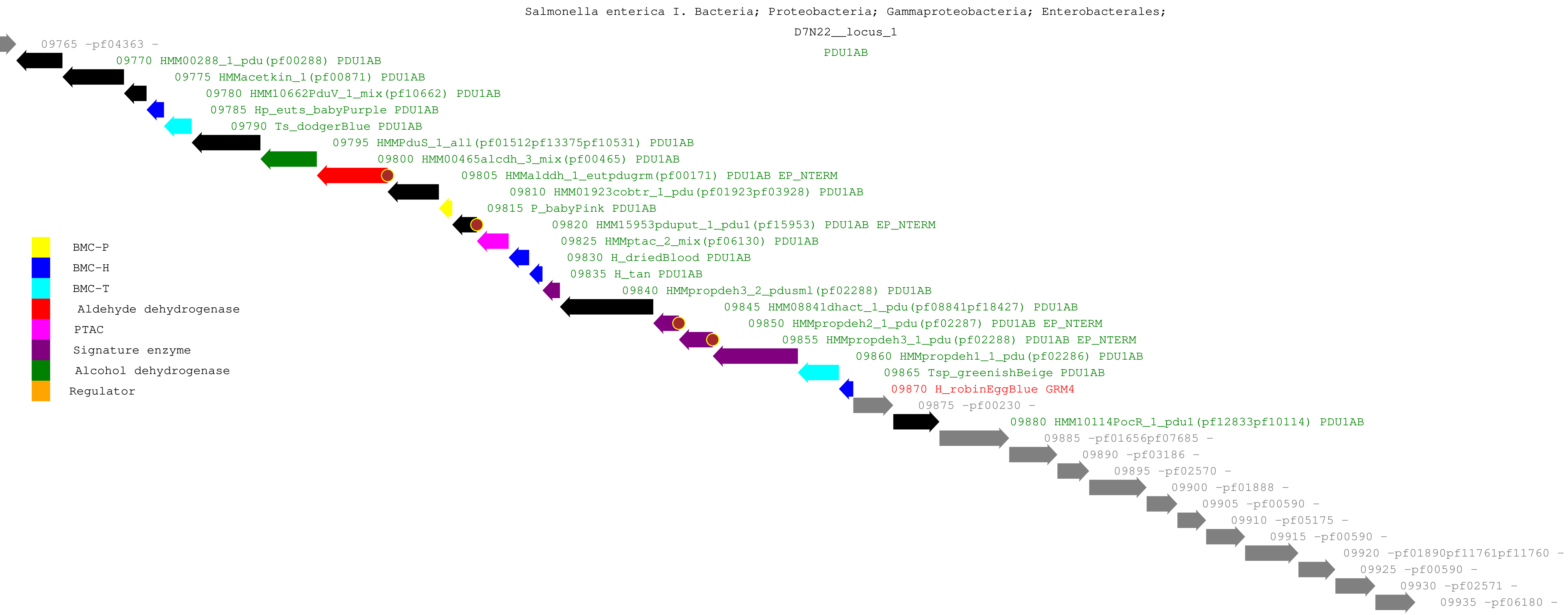
Salmonella enterica I. Bacteria; Proteobacteria; Gammaproteobacteria; Enterobacterales;
D7N22__locus_1
PDU1AB
09765 -pf04363 -
09770 HMM00288_1_pdu(pf00288) PDU1AB
09775 HMMacetkin_1(pf00871) PDU1AB
09780 HMM10662PduV_1_mix(pf10662) PDU1AB
09785 Hp_euts_babyPurple PDU1AB
09790 Ts_dodgerBlue PDU1AB
09795 HMMPduS_1_all(pf01512pf13375pf10531) PDU1AB
09800 HMM00465alcdh_3_mix(pf00465) PDU1AB
09805 HMMalddh_1_eutpdugrm(pf00171) PDU1AB EP_NTERM
09810 HMM01923cobtr_1_pdu(pf01923pf03928) PDU1AB
09815 P_babyPink PDU1AB
09820 HMM15953pduput_1_pdu1(pf15953) PDU1AB EP_NTERM
09825 HMMptac_2_mix(pf06130) PDU1AB
09830 H_driedBlood PDU1AB
09835 H_tan PDU1AB
09840 HMMpropdeh3_2_pdusml(pf02288) PDU1AB
09845 HMM08841dhact_1_pdu(pf08841pf18427) PDU1AB
09850 HMMpropdeh2_1_pdu(pf02287) PDU1AB EP_NTERM
09855 HMMpropdeh3_1_pdu(pf02288) PDU1AB EP_NTERM
09860 HMMpropdeh1_1_pdu(pf02286) PDU1AB
09865 Tsp_greenishBeige PDU1AB
09870 H_robinEggBlue GRM4
09875 -pf00230 -
09880 HMM10114PocR_1_pdu1(pf12833pf10114) PDU1AB
09885 -pf01656pf07685 -
09890 -pf03186 -
09895 -pf02570 -
09900 -pf01888 -
09905 -pf00590 -
09910 -pf05175 -
09915 -pf00590 -
09920 -pf01890pf11761pf11760 -
09925 -pf00590 -
09930 -pf02571 -
09935 -pf06180 -
BMC-P
BMC-H
BMC-T
Aldehyde dehydrogenase
PTAC
Signature enzyme
Alcohol dehydrogenase
Regulator

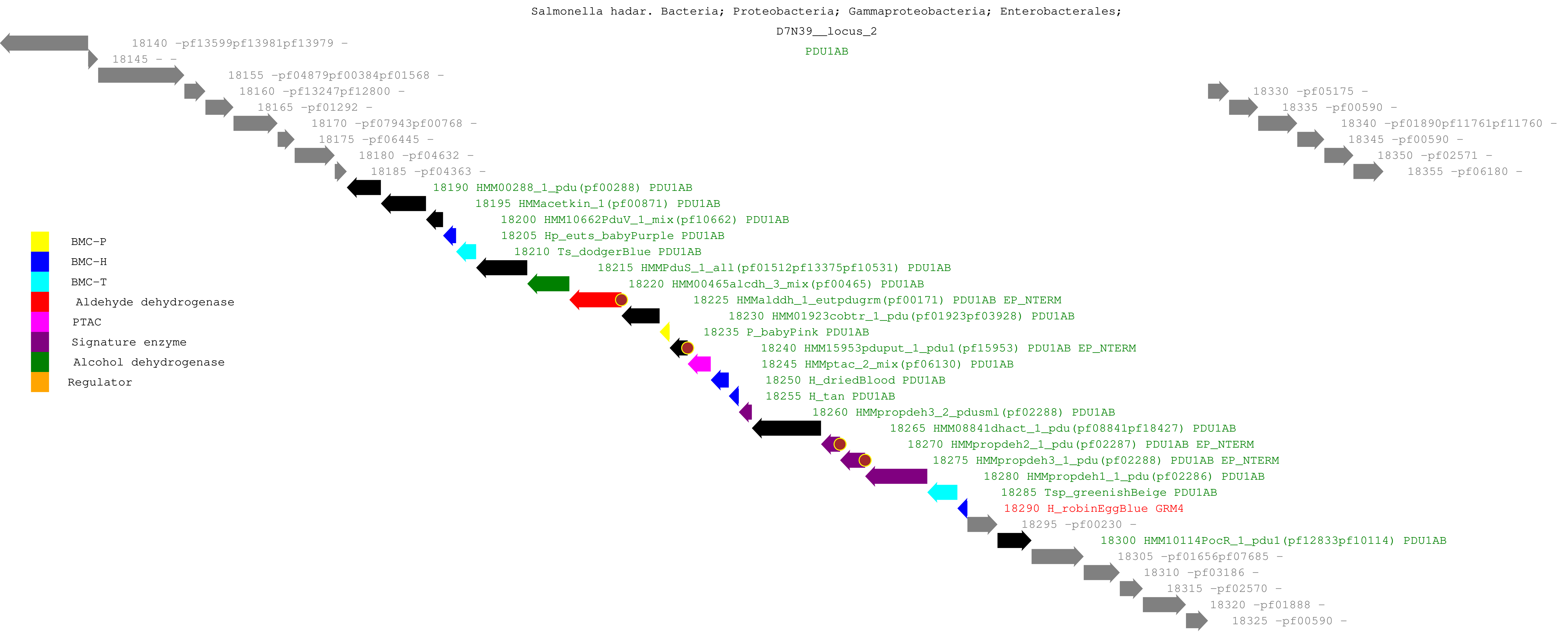

Salmonella hadar. Bacteria; Proteobacteria; Gammaproteobacteria; Enterobacterales;
D7N39__locus_2
PDU1AB
18140 -pf13599pf13981pf13979 -
18145 - -
18155 -pf04879pf00384pf01568 -
18160 -pf13247pf12800 -
18165 -pf01292 -
18170 -pf07943pf00768 -
18175 -pf06445 -
18180 -pf04632 -
18185 -pf04363 -
18190 HMM00288_1_pdu(pf00288) PDU1AB
18195 HMMacetkin_1(pf00871) PDU1AB
18200 HMM10662PduV_1_mix(pf10662) PDU1AB
18205 Hp_euts_babyPurple PDU1AB
18210 Ts_dodgerBlue PDU1AB
18215 HMMPduS_1_all(pf01512pf13375pf10531) PDU1AB
18220 HMM00465alcdh_3_mix(pf00465) PDU1AB
18225 HMMalddh_1_eutpdugrm(pf00171) PDU1AB EP_NTERM
18230 HMM01923cobtr_1_pdu(pf01923pf03928) PDU1AB
18235 P_babyPink PDU1AB
18240 HMM15953pduput_1_pdu1(pf15953) PDU1AB EP_NTERM
18245 HMMptac_2_mix(pf06130) PDU1AB
18250 H_driedBlood PDU1AB
18255 H_tan PDU1AB
18260 HMMpropdeh3_2_pdusml(pf02288) PDU1AB
18265 HMM08841dhact_1_pdu(pf08841pf18427) PDU1AB
18270 HMMpropdeh2_1_pdu(pf02287) PDU1AB EP_NTERM
18275 HMMpropdeh3_1_pdu(pf02288) PDU1AB EP_NTERM
18280 HMMpropdeh1_1_pdu(pf02286) PDU1AB
18285 Tsp_greenishBeige PDU1AB
18290 H_robinEggBlue GRM4
18295 -pf00230 -
18300 HMM10114PocR_1_pdu1(pf12833pf10114) PDU1AB
18305 -pf01656pf07685 -
18310 -pf03186 -
18315 -pf02570 -
18320 -pf01888 -
18325 -pf00590 -
18330 -pf05175 -
18335 -pf00590 -
18340 -pf01890pf11761pf11760 -
18345 -pf00590 -
18350 -pf02571 -
18355 -pf06180 -
BMC-P
BMC-H
BMC-T
Aldehyde dehydrogenase
PTAC
Signature enzyme
Alcohol dehydrogenase
Regulator

Salmonella enterica I. Bacteria; Proteobacteria; Gammaproteobacteria; Enterobacterales;

D7N47__locus_1

PDU1AB

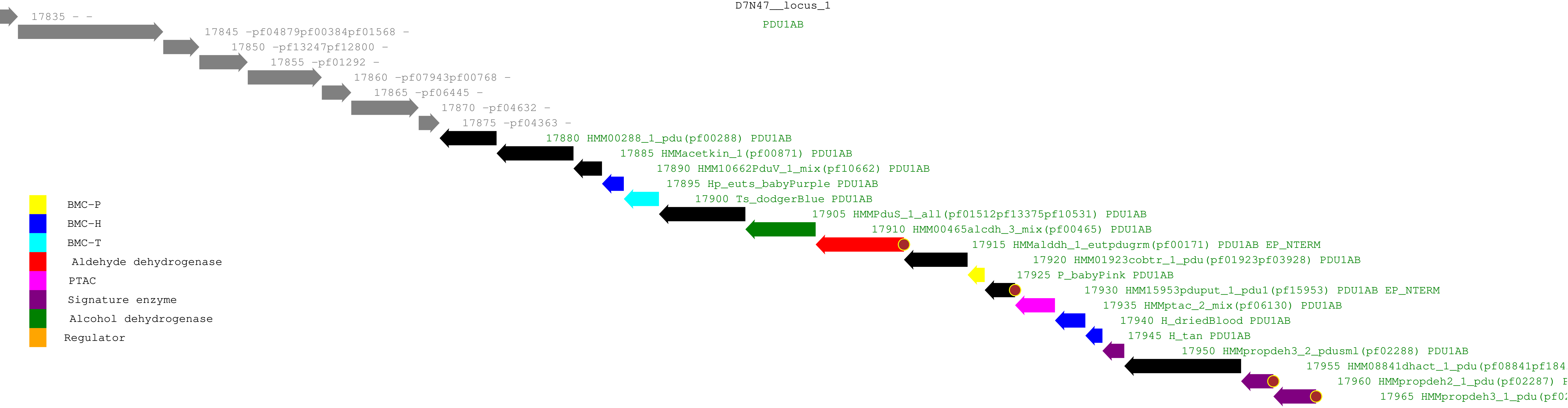

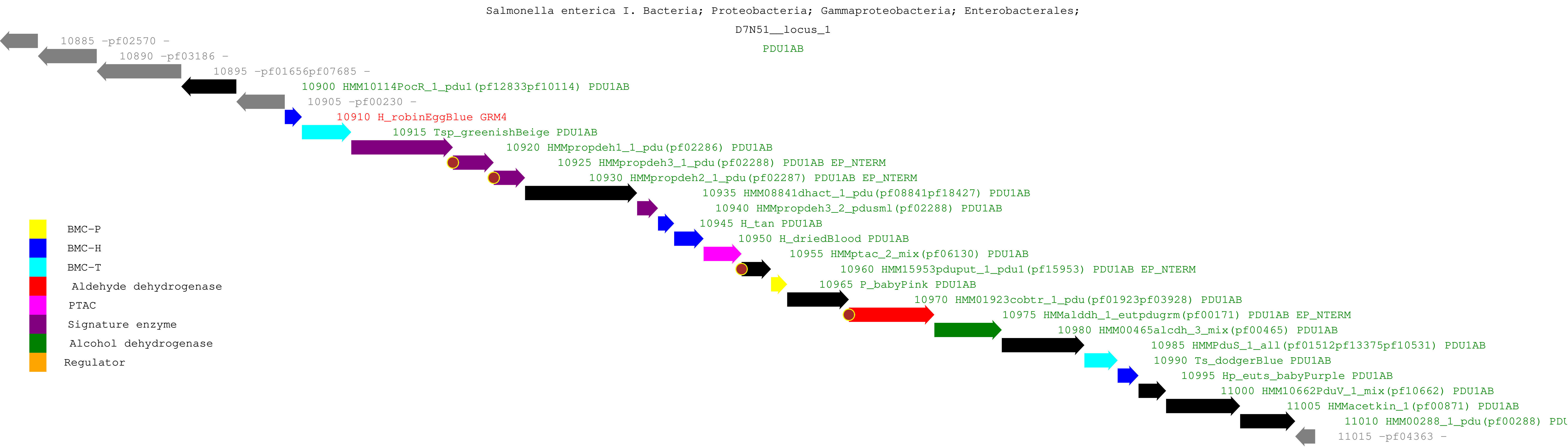

Salmonella enterica I. Bacteria; Proteobacteria; Gammaproteobacteria; Enterobacterales;
D7N51__locus_1
PDU1AB
10885 -pf02570 -
10890 -pf03186 -
10895 -pf01656pf07685 -
10900 HMM10114PocR_1_pdu1(pf12833pf10114) PDU1AB
10905 -pf00230 -
10910 H_robinEggBlue GRM4
10915 Tsp_greenishBeige PDU1AB
10920 HMMpropdeh1_1_pdu(pf02286) PDU1AB
10925 HMMpropdeh3_1_pdu(pf02288) PDU1AB EP_NTERM
10930 HMMpropdeh2_1_pdu(pf02287) PDU1AB EP_NTERM
10935 HMM08841dhact_1_pdu(pf08841pf18427) PDU1AB
10940 HMMpropdeh3_2_pdusml(pf02288) PDU1AB
10945 H_tan PDU1AB
10950 H_driedBlood PDU1AB
10955 HMMptac_2_mix(pf06130) PDU1AB
10960 HMM15953pduput_1_pdu1(pf15953) PDU1AB EP_NTERM
10965 P_babyPink PDU1AB
10970 HMM01923cobtr_1_pdu(pf01923pf03928) PDU1AB
10975 HMMalddh_1_eutpdugrm(pf00171) PDU1AB EP_NTERM
10980 HMM00465alcdh_3_mix(pf00465) PDU1AB
10985 HMMPduS_1_all(pf01512pf13375pf10531) PDU1AB
10990 Ts_dodgerBlue PDU1AB
10995 Hp_euts_babyPurple PDU1AB
11000 HMM10662PduV_1_mix(pf10662) PDU1AB
11005 HMMacetkin_1(pf00871) PDU1AB
11010 HMM00288_1_pdu(pf00288) PDU
11015 -pf04363 -
BMC-P
BMC-H
BMC-T
Aldehyde dehydrogenase
PTAC
Signature enzyme
Alcohol dehydrogenase
Regulator

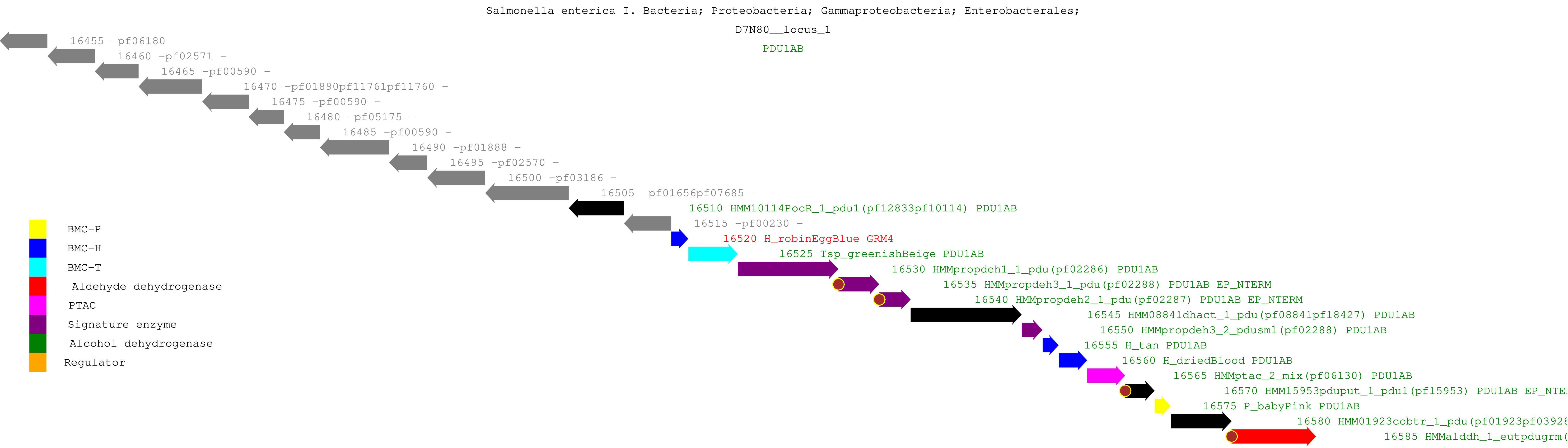
Salmonella enterica I. Bacteria; Proteobacteria; Gammaproteobacteria; Enterobacterales;
D7N80__locus_1
PDU1AB
16455 -pf06180 -
16460 -pf02571 -
16465 -pf00590 -
16470 -pf01890pf11761pf11760 -
16475 -pf00590 -
16480 -pf05175 -
16485 -pf00590 -
16490 -pf01888 -
16495 -pf02570 -
16500 -pf03186 -
16505 -pf01656pf07685 -
16510 HMM10114PocR_1_pdu1(pf12833pf10114) PDU1AB
16515 -pf00230 -
16520 H_robinEggBlue GRM4
16525 Tsp_greenishBeige PDU1AB
16530 HMMpropdeh1_1_pdu(pf02286) PDU1AB
16535 HMMpropdeh3_1_pdu(pf02288) PDU1AB EP_NTERM
16540 HMMpropdeh2_1_pdu(pf02287) PDU1AB EP_NTERM
16545 HMM08841dhact_1_pdu(pf08841pf18427) PDU1AB
16550 HMMpropdeh3_2_pdusml(pf02288) PDU1AB
16555 H_tan PDU1AB
16560 H_driedBlood PDU1AB
16565 HMMptac_2_mix(pf06130) PDU1AB
16570 HMM15953pduput_1_pdu1(pf15953) PDU1AB EP_NTE
16575 P_babyPink PDU1AB
16580 HMM01923cobtr_1_pdu(pf01923pf0392
16585 HMMalddh_1_eutpdugrm(
BMC-P
BMC-H
BMC-T
Aldehyde dehydrogenase
PTAC
Signature enzyme
Alcohol dehydrogenase
Regulator

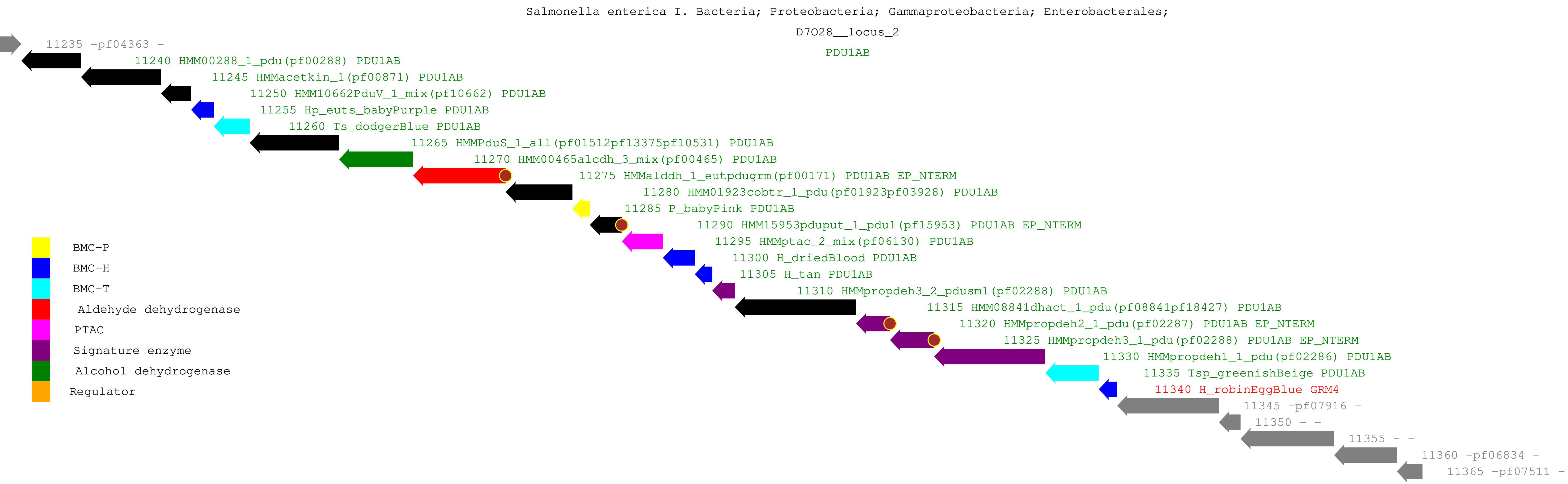
Salmonella enterica I. Bacteria; Proteobacteria; Gammaproteobacteria; Enterobacterales;
D7028__locus_2
PDU1AB
11235 -pf04363 -
11240 HMM00288_1_pdu(pf00288) PDU1AB
11245 HMMacetkin_1(pf00871) PDU1AB
11250 HMM10662PduV_1_mix(pf10662) PDU1AB
11255 Hp_euts_babyPurple PDU1AB
11260 Ts_dodgerBlue PDU1AB
11265 HMMPduS_1_all(pf01512pf13375pf10531) PDU1AB
11270 HMM00465alcdh_3_mix(pf00465) PDU1AB
11275 HMMalddh_1_eutpdugrm(pf00171) PDU1AB EP_NTERM
11280 HMM01923cobtr_1_pdu(pf01923pf03928) PDU1AB
11285 P_babyPink PDU1AB
11290 HMM15953pduput_1_pdu1(pf15953) PDU1AB EP_NTERM
11295 HMMptac_2_mix(pf06130) PDU1AB
11300 H_driedBlood PDU1AB
11305 H_tan PDU1AB
11310 HMMpropdeh3_2_pdusml(pf02288) PDU1AB
11315 HMM08841dhact_1_pdu(pf08841pf18427) PDU1AB
11320 HMMpropdeh2_1_pdu(pf02287) PDU1AB EP_NTERM
11325 HMMpropdeh3_1_pdu(pf02288) PDU1AB EP_NTERM
11330 HMMpropdeh1_1_pdu(pf02286) PDU1AB
11335 Tsp_greenishBeige PDU1AB
11340 H_robinEggBlue GRM4
11345 -pf07916 -
11350 - -
11355 - -
11360 -pf06834 -
11365 -pf07511 -
BMC-P
BMC-H
BMC-T
Aldehyde dehydrogenase
PTAC
Signature enzyme
Alcohol dehydrogenase
Regulator

Salmonella enterica subsp. enterica serovar Saintpaul. Bacteria; Proteobacteria; Gammaproteobacteria; Enterobacterales;

D7055__locus_2

PDU1AB

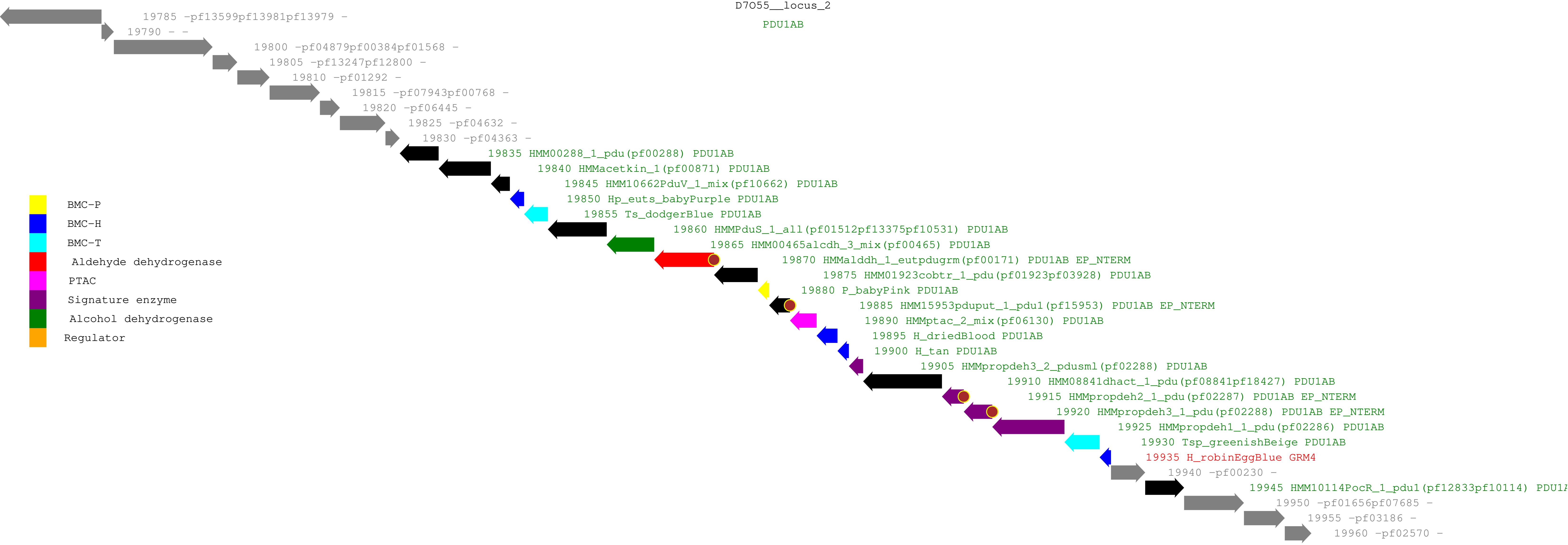

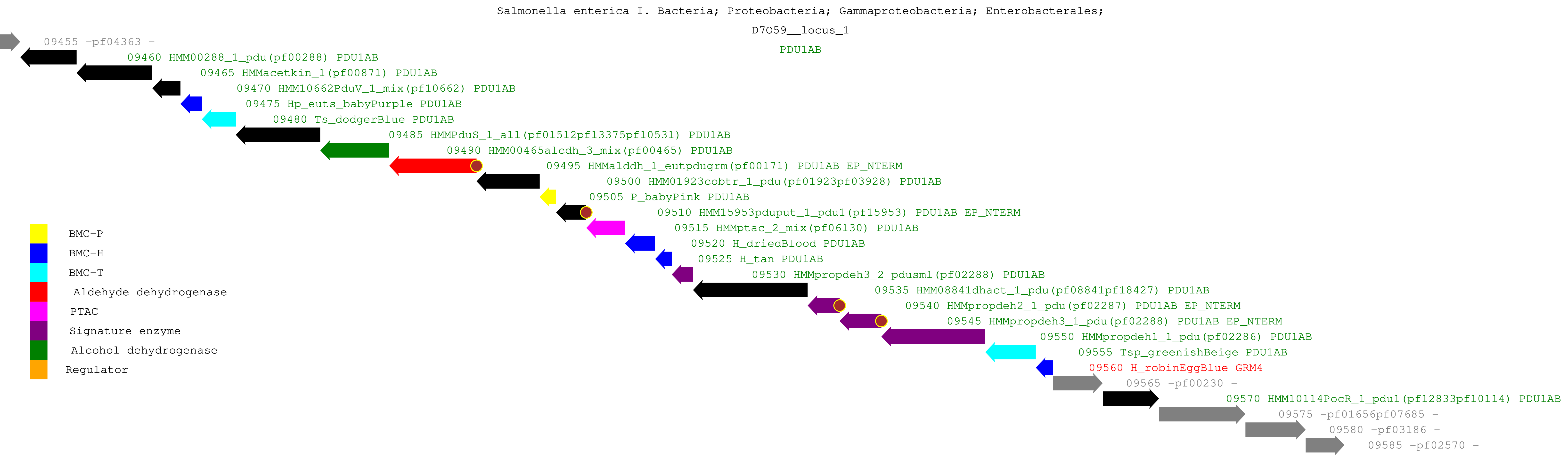

Salmonella enterica I. Bacteria; Proteobacteria; Gammaproteobacteria; Enterobacterales;
D7059__locus_1
PDU1AB
09455 -pf04363 -
09460 HMM00288_1_pdu(pf00288) PDU1AB
09465 HMMacetkin_1(pf00871) PDU1AB
09470 HMM10662PduV_1_mix(pf10662) PDU1AB
09475 Hp_euts_babyPurple PDU1AB
09480 Ts_dodgerBlue PDU1AB
09485 HMMPduS_1_all(pf01512pf13375pf10531) PDU1AB
09490 HMM00465alcdh_3_mix(pf00465) PDU1AB
09495 HMMalddh_1_eutpdugrm(pf00171) PDU1AB EP_NTERM
09500 HMM01923cobtr_1_pdu(pf01923pf03928) PDU1AB
09505 P_babyPink PDU1AB
09510 HMM15953pduput_1_pdu1(pf15953) PDU1AB EP_NTERM
09515 HMMptac_2_mix(pf06130) PDU1AB
09520 H_driedBlood PDU1AB
09525 H_tan PDU1AB
09530 HMMpropdeh3_2_pdusml(pf02288) PDU1AB
09535 HMM08841dhact_1_pdu(pf08841pf18427) PDU1AB
09540 HMMpropdeh2_1_pdu(pf02287) PDU1AB EP_NTERM
09545 HMMpropdeh3_1_pdu(pf02288) PDU1AB EP_NTERM
09550 HMMpropdeh1_1_pdu(pf02286) PDU1AB
09555 Tsp_greenishBeige PDU1AB
09560 H_robinEggBlue GRM4
09565 -pf00230 -
09570 HMM10114PocR_1_pdu1(pf12833pf10114) PDU1AB
09575 -pf01656pf07685 -
09580 -pf03186 -
09585 -pf02570 -
BMC-P
BMC-H
BMC-T
Aldehyde dehydrogenase
PTAC
Signature enzyme
Alcohol dehydrogenase
Regulator

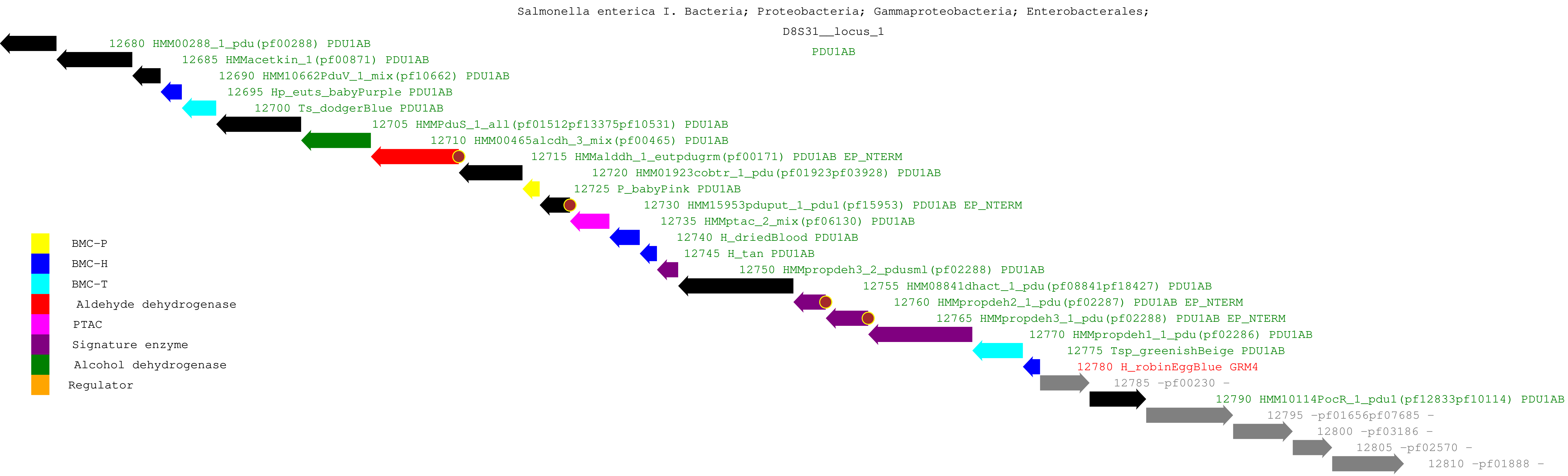
Salmonella enterica I. Bacteria; Proteobacteria; Gammaproteobacteria; Enterobacterales;
D8S31__locus_1
PDU1AB
12680 HMM00288_1_pdu(pf00288) PDU1AB
12685 HMMacetkin_1(pf00871) PDU1AB
12690 HMM10662PduV_1_mix(pf10662) PDU1AB
12695 Hp_euts_babyPurple PDU1AB
12700 Ts_dodgerBlue PDU1AB
12705 HMMPduS_1_all(pf01512pf13375pf10531) PDU1AB
12710 HMM00465alcdh_3_mix(pf00465) PDU1AB
12715 HMMalddh_1_eutpdugrm(pf00171) PDU1AB EP_NTERM
12720 HMM01923cobtr_1_pdu(pf01923pf03928) PDU1AB
12725 P_babyPink PDU1AB
12730 HMM15953pduput_1_pdu1(pf15953) PDU1AB EP_NTERM
12735 HMMptac_2_mix(pf06130) PDU1AB
12740 H_driedBlood PDU1AB
12745 H_tan PDU1AB
12750 HMMpropdeh3_2_pdusml(pf02288) PDU1AB
12755 HMM08841dhact_1_pdu(pf08841pf18427) PDU1AB
12760 HMMpropdeh2_1_pdu(pf02287) PDU1AB EP_NTERM
12765 HMMpropdeh3_1_pdu(pf02288) PDU1AB EP_NTERM
12770 HMMpropdeh1_1_pdu(pf02286) PDU1AB
12775 Tsp_greenishBeige PDU1AB
12780 H_robinEggBlue GRM4
12785 -pf00230 -
12790 HMM10114PocR_1_pdu1(pf12833pf10114) PDU1AB
12795 -pf01656pf07685 -
12800 -pf03186 -
12805 -pf02570 -
12810 -pf01888 -
BMC-P
BMC-H
BMC-T
Aldehyde dehydrogenase
PTAC
Signature enzyme
Alcohol dehydrogenase
Regulator

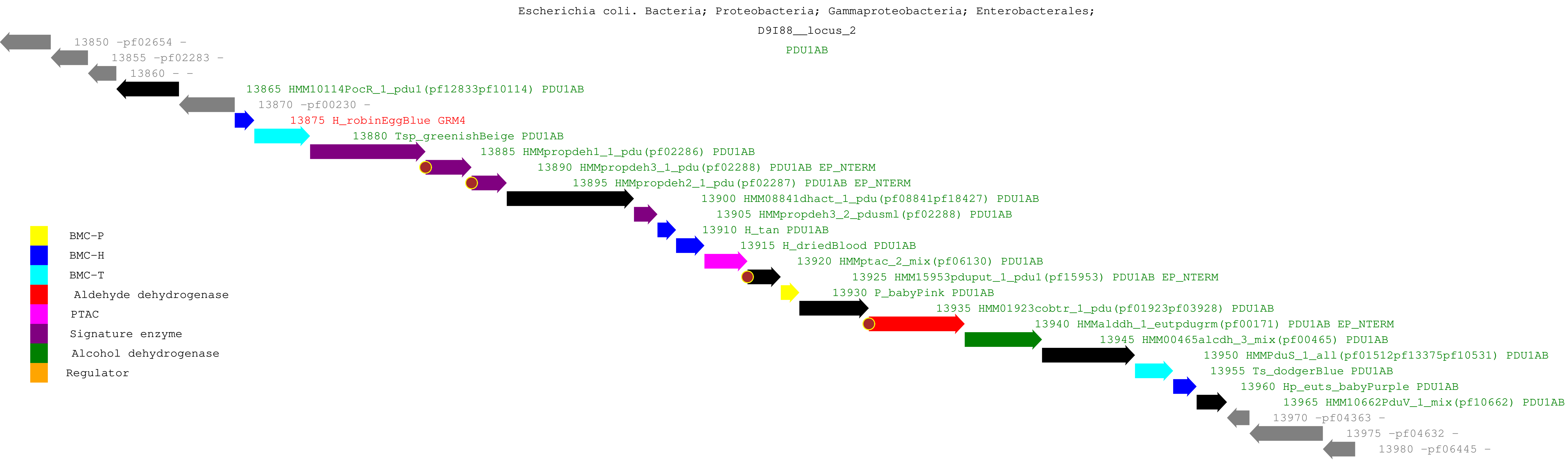
Escherichia coli. Bacteria; Proteobacteria; Gammaproteobacteria; Enterobacterales;
D9I88__locus_2
PDU1AB
13850 -pf02654 -
13855 -pf02283 -
13860 - -
13865 HMM10114PocR_1_pdu1(pf12833pf10114) PDU1AB
13870 -pf00230 -
13875 H_robinEggBlue GRM4
13880 Tsp_greenishBeige PDU1AB
13885 HMMpropdeh1_1_pdu(pf02286) PDU1AB
13890 HMMpropdeh3_1_pdu(pf02288) PDU1AB EP_NTERM
13895 HMMpropdeh2_1_pdu(pf02287) PDU1AB EP_NTERM
13900 HMM08841dhact_1_pdu(pf08841pf18427) PDU1AB
13905 HMMpropdeh3_2_pdusml(pf02288) PDU1AB
13910 H_tan PDU1AB
13915 H_driedBlood PDU1AB
13920 HMMptac_2_mix(pf06130) PDU1AB
13925 HMM15953pduput_1_pdu1(pf15953) PDU1AB EP_NTERM
13930 P_babyPink PDU1AB
13935 HMM01923cobtr_1_pdu(pf01923pf03928) PDU1AB
13940 HMMalddh_1_eutpdugrm(pf00171) PDU1AB EP_NTERM
13945 HMM00465alcdh_3_mix(pf00465) PDU1AB
13950 HMMPduS_1_all(pf01512pf13375pf10531) PDU1AB
13955 Ts_dodgerBlue PDU1AB
13960 Hp_euts_babyPurple PDU1AB
13965 HMM10662PduV_1_mix(pf10662) PDU1AB
13970 -pf04363 -
13975 -pf04632 -
13980 -pf06445 -
BMC-P
BMC-H
BMC-T
Aldehyde dehydrogenase
PTAC
Signature enzyme
Alcohol dehydrogenase
Regulator

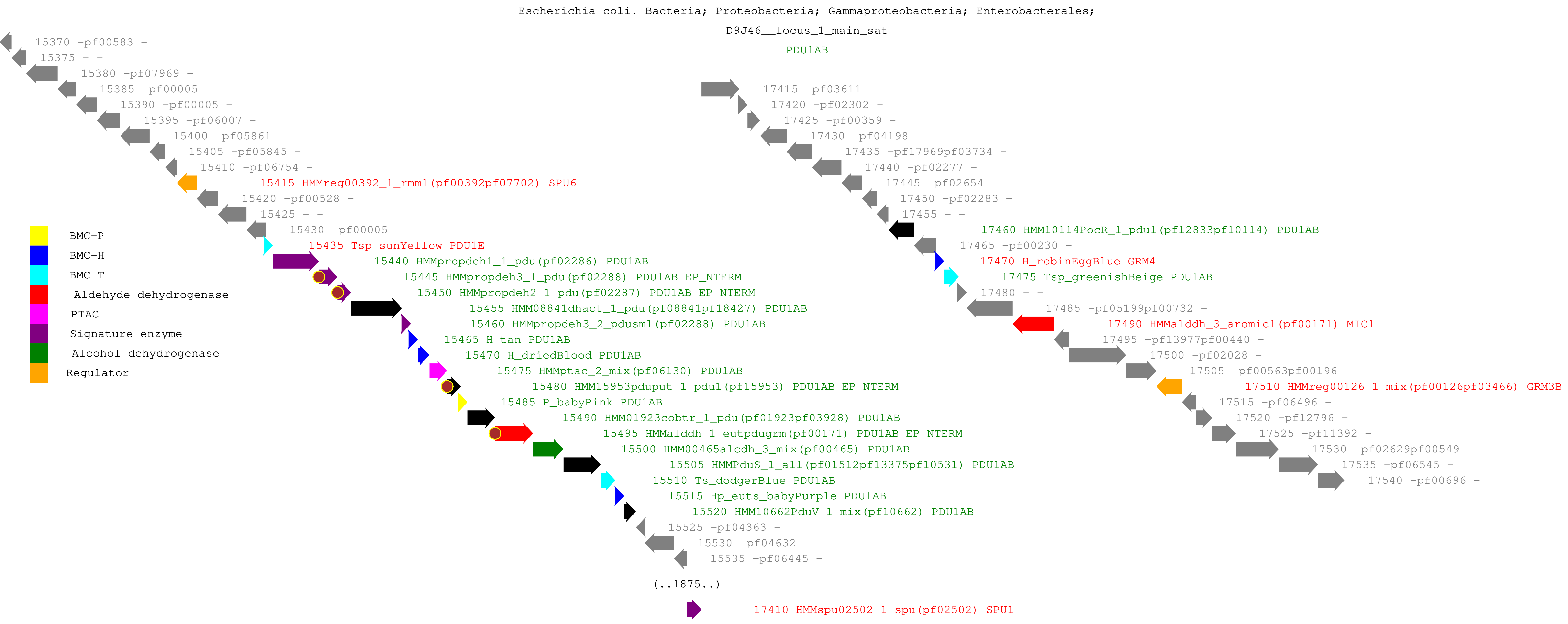
Escherichia coli. Bacteria; Proteobacteria; Gammaproteobacteria; Enterobacterales;
D9J46__locus_1_main_sat
PDU1AB
15370 -pf00583 -
15375 - -
15380 -pf07969 -
15385 -pf00005 -
15390 -pf00005 -
15395 -pf06007 -
15400 -pf05861 -
15405 -pf05845 -
15410 -pf06754 -
15415 HMMreg00392_1_rmm1(pf00392pf07702) SPU6
15420 -pf00528 -
15425 - -
15430 -pf00005 -
15435 Tsp_sunYellow PDU1E
15440 HMMpropdeh1_1_pdu(pf02286) PDU1AB
15445 HMMpropdeh3_1_pdu(pf02288) PDU1AB EP_NTERM
15450 HMMpropdeh2_1_pdu(pf02287) PDU1AB EP_NTERM
15455 HMM08841dhact_1_pdu(pf08841pf18427) PDU1AB
15460 HMMpropdeh3_2_pdusml(pf02288) PDU1AB
15465 H_tan PDU1AB
15470 H_driedBlood PDU1AB
15475 HMMptac_2_mix(pf06130) PDU1AB
15480 HMM15953pduput_1_pdu1(pf15953) PDU1AB EP_NTERM
15485 P_babyPink PDU1AB
15490 HMM01923cobtr_1_pdu(pf01923pf03928) PDU1AB
15495 HMMalddh_1_eutpdugrm(pf00171) PDU1AB EP_NTERM
15500 HMM00465alcdh_3_mix(pf00465) PDU1AB
15505 HMMPduS_1_all(pf01512pf13375pf10531) PDU1AB
15510 Ts_dodgerBlue PDU1AB
15515 Hp_euts_babyPurple PDU1AB
15520 HMM10662PduV_1_mix(pf10662) PDU1AB
15525 -pf04363 -
15530 -pf04632 -
15535 -pf06445 -
(..1875..)
17410 HMMspu02502_1_spu(pf02502) SPU1
BMC-P
BMC-H
BMC-T
Aldehyde dehydrogenase
PTAC
Signature enzyme
Alcohol dehydrogenase
Regulator
17415 -pf03611 -
17420 -pf02302 -
17425 -pf00359 -
17430 -pf04198 -
17435 -pf17969pf03734 -
17440 -pf02277 -
17445 -pf02654 -
17450 -pf02283 -
17455 - -
17460 HMM10114PocR_1_pdu1(pf12833pf10114) PDU1AB
17465 -pf00230 -
17470 H_robinEggBlue GRM4
17475 Tsp_greenishBeige PDU1AB
17480 - -
17485 -pf05199pf00732 -
17490 HMMaldh_3_aromic1(pf00171) MIC1
17495 -pf13977pf00440 -
17500 -pf02028 -
17505 -pf00563pf00196 -
17510 HMMreg00126_1_mix(pf00126pf03466) GRM3B
17515 -pf06496 -
17520 -pf12796 -
17525 -pf11392 -
17530 -pf02629pf00549 -
17535 -pf06545 -
17540 -pf00696 -

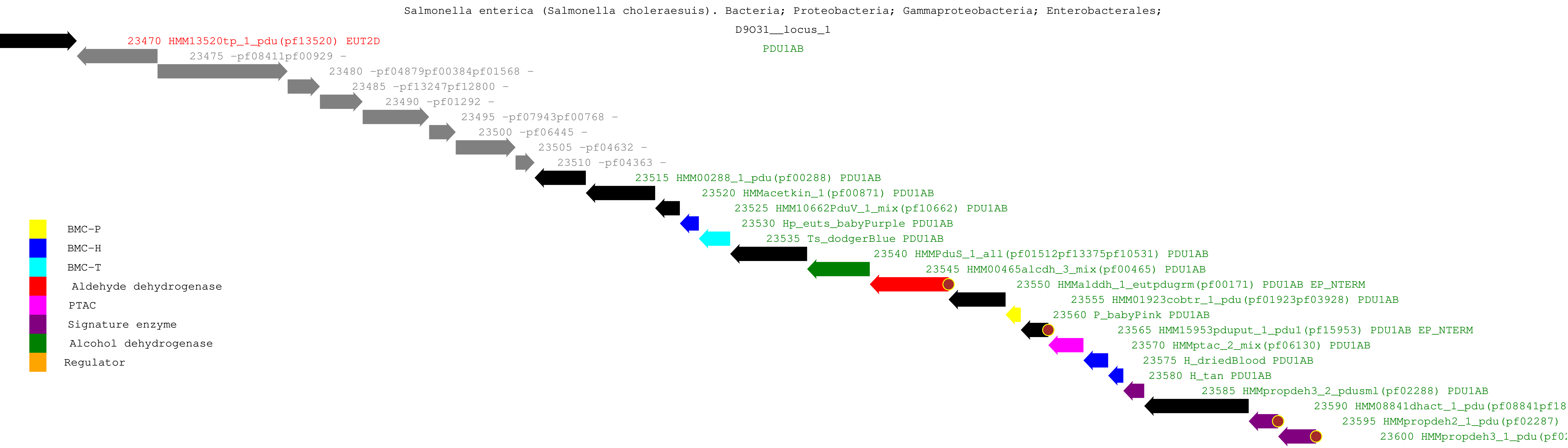
Salmonella enterica (Salmonella choleraesuis). Bacteria; Proteobacteria; Gammaproteobacteria; Enterobacterales;
D9031__locus_1
PDU1AB
23470 HMM13520tp_1_pdu(pf13520) EUT2D
23475 -pf08411pf00929 -
23480 -pf04879pf00384pf01568 -
23485 -pf13247pf12800 -
23490 -pf01292 -
23495 -pf07943pf00768 -
23500 -pf06445 -
23505 -pf04632 -
23510 -pf04363 -
23515 HMM00288_1_pdu(pf00288) PDU1AB
23520 HMMacetkin_1(pf00871) PDU1AB
23525 HMM10662PduV_1_mix(pf10662) PDU1AB
23530 Hp_euts_babyPurple PDU1AB
23535 Ts_dodgerBlue PDU1AB
23540 HMMPduS_1_all(pf01512pf13375pf10531) PDU1AB
23545 HMM00465alcdh_3_mix(pf00465) PDU1AB
23550 HMMalddh_1_eutpdugrm(pf00171) PDU1AB EP_NTERM
23555 HMM01923cobtr_1_pdu(pf01923pf03928) PDU1AB
23560 P_babyPink PDU1AB
23565 HMM15953pduput_1_pdu1(pf15953) PDU1AB EP_NTERM
23570 HMMptac_2_mix(pf06130) PDU1AB
23575 H_driedBlood PDU1AB
23580 H_tan PDU1AB
23585 HMMpropdeh3_2_pdusml(pf02288) PDU1AB
23590 HMM08841dhact_1_pdu(pf08841pf18
23595 HMMpropdeh2_1_pdu(pf02287)
23600 HMMpropdeh3_1_pdu(pf0
BMC-P
BMC-H
BMC-T
Aldehyde dehydrogenase
PTAC
Signature enzyme
Alcohol dehydrogenase
Regulator

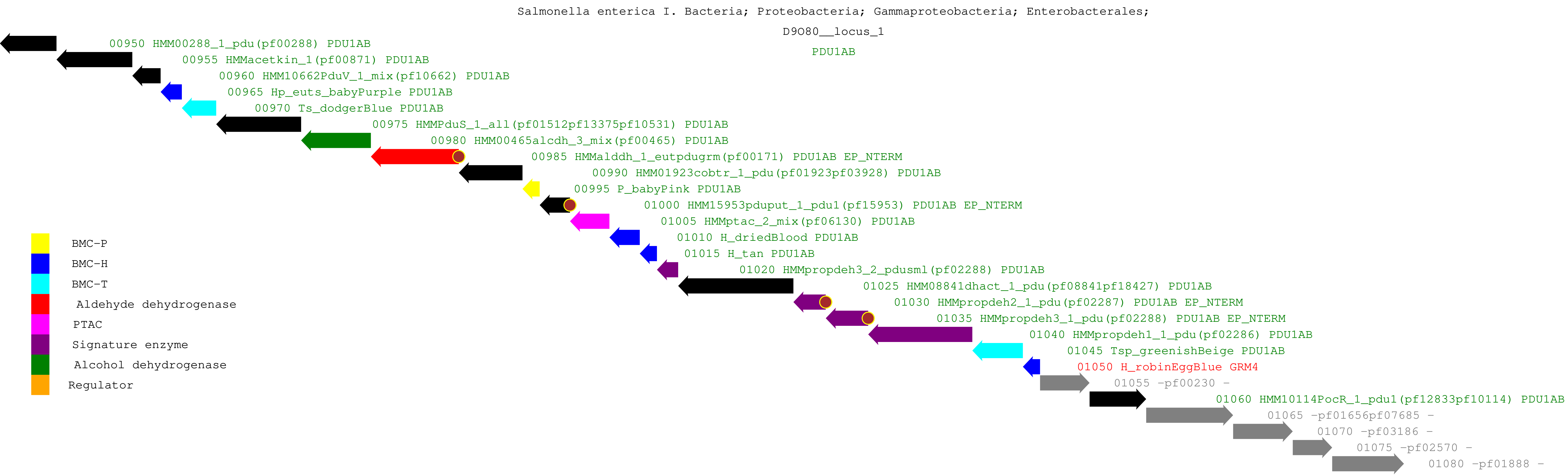

Salmonella enterica I. Bacteria; Proteobacteria; Gammaproteobacteria; Enterobacterales;
D9080__locus_1
PDU1AB
00950 HMM00288_1_pdu(pf00288) PDU1AB
00955 HMMacetkin_1(pf00871) PDU1AB
00960 HMM10662PduV_1_mix(pf10662) PDU1AB
00965 Hp_euts_babyPurple PDU1AB
00970 Ts_dodgerBlue PDU1AB
00975 HMMPduS_1_all(pf01512pf13375pf10531) PDU1AB
00980 HMM00465alcdh_3_mix(pf00465) PDU1AB
00985 HMMalddh_1_eutpdugrm(pf00171) PDU1AB EP_NTERM
00990 HMM01923cobtr_1_pdu(pf01923pf03928) PDU1AB
00995 P_babyPink PDU1AB
01000 HMM15953pduput_1_pdu1(pf15953) PDU1AB EP_NTERM
01005 HMMptac_2_mix(pf06130) PDU1AB
01010 H_driedBlood PDU1AB
01015 H_tan PDU1AB
01020 HMMpropdeh3_2_pdusml(pf02288) PDU1AB
01025 HMM08841dhact_1_pdu(pf08841pf18427) PDU1AB
01030 HMMpropdeh2_1_pdu(pf02287) PDU1AB EP_NTERM
01035 HMMpropdeh3_1_pdu(pf02288) PDU1AB EP_NTERM
01040 HMMpropdeh1_1_pdu(pf02286) PDU1AB
01045 Tsp_greenishBeige PDU1AB
01050 H_robinEggBlue GRM4
01055 -pf00230 -
01060 HMM10114PocR_1_pdu1(pf12833pf10114) PDU1AB
01065 -pf01656pf07685 -
01070 -pf03186 -
01075 -pf02570 -
01080 -pf01888 -
BMC-P
BMC-H
BMC-T
Aldehyde dehydrogenase
PTAC
Signature enzyme
Alcohol dehydrogenase
Regulator

Salmonella enterica I. Bacteria; Proteobacteria; Gammaproteobacteria; Enterobacterales;

D9086__locus_1

PDU1AB

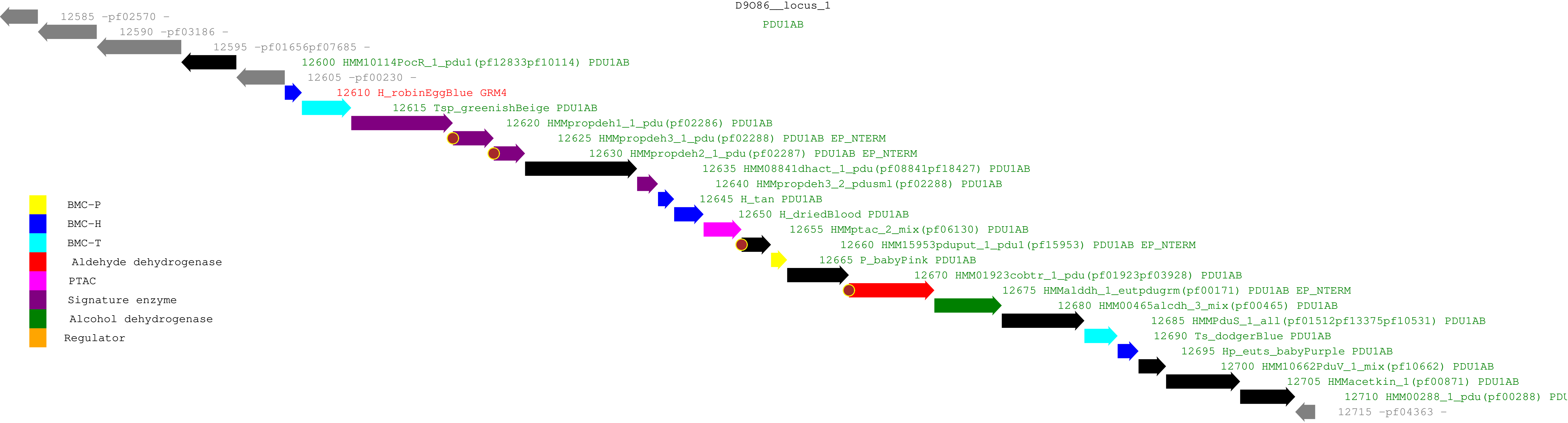

Salmonella enterica I. Bacteria; Proteobacteria; Gammaproteobacteria; Enterobacterales;

D9095__locus_1

PDU1AB

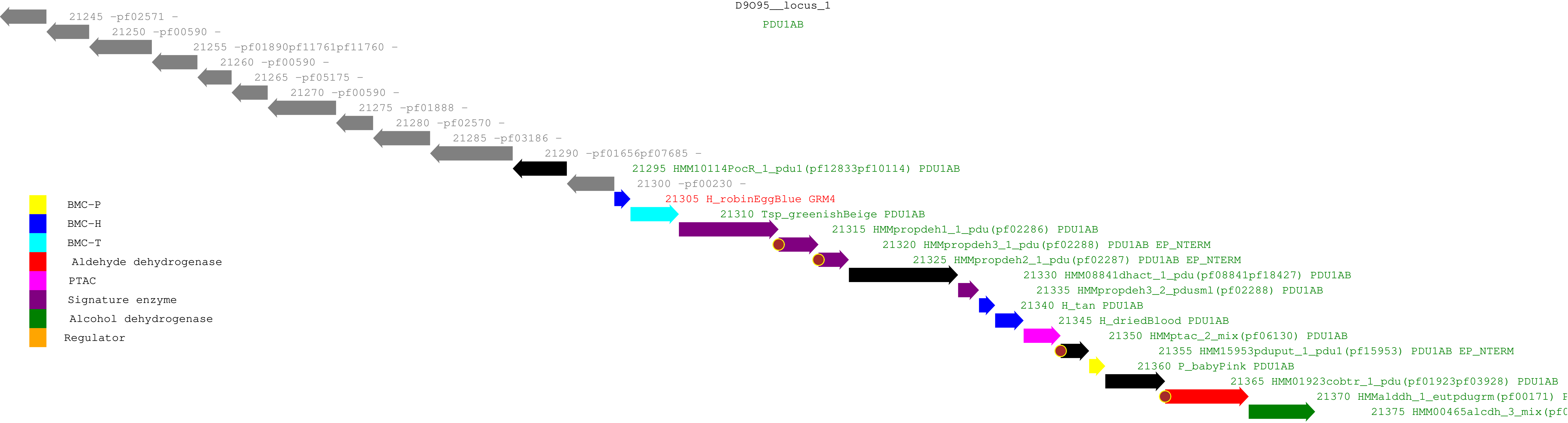

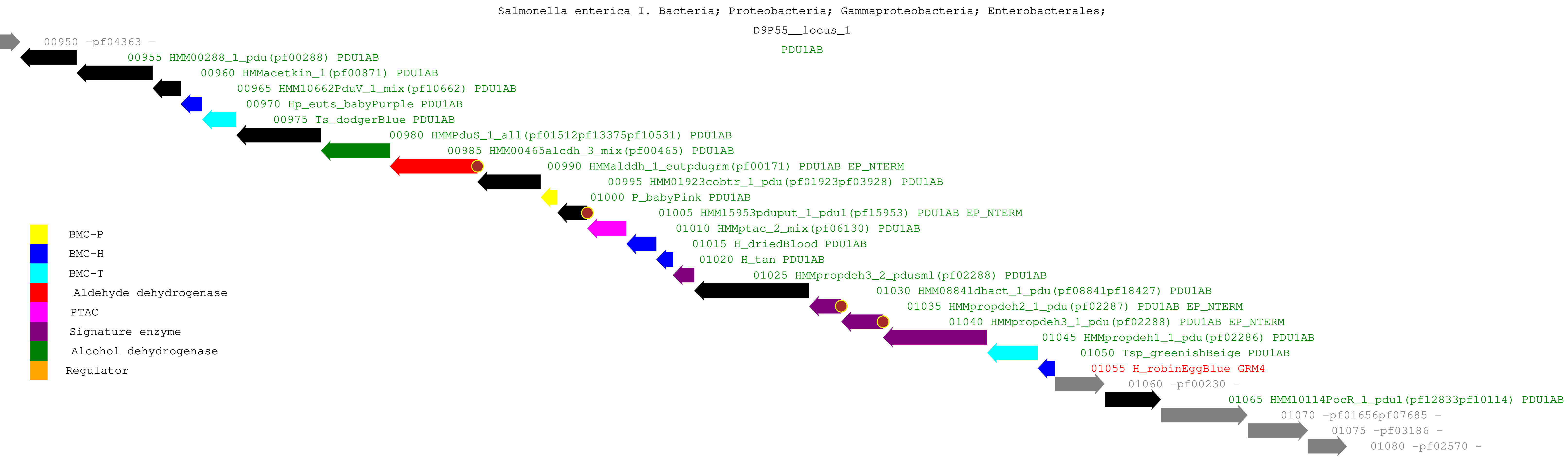
Salmonella enterica I. Bacteria; Proteobacteria; Gammaproteobacteria; Enterobacterales;
D9P55__locus_1
PDU1AB
00950 -pf04363 -
00955 HMM00288_1_pdu(pf00288) PDU1AB
00960 HMMacetkin_1(pf00871) PDU1AB
00965 HMM10662PduV_1_mix(pf10662) PDU1AB
00970 Hp_euts_babyPurple PDU1AB
00975 Ts_dodgerBlue PDU1AB
00980 HMMPduS_1_all(pf01512pf13375pf10531) PDU1AB
00985 HMM00465alcdh_3_mix(pf00465) PDU1AB
00990 HMMalddh_1_eutpdugrm(pf00171) PDU1AB EP_NTERM
00995 HMM01923cobtr_1_pdu(pf01923pf03928) PDU1AB
01000 P_babyPink PDU1AB
01005 HMM15953pduput_1_pdu1(pf15953) PDU1AB EP_NTERM
01010 HMMptac_2_mix(pf06130) PDU1AB
01015 H_driedBlood PDU1AB
01020 H_tan PDU1AB
01025 HMMpropdeh3_2_pdusml(pf02288) PDU1AB
01030 HMM08841dhact_1_pdu(pf08841pf18427) PDU1AB
01035 HMMpropdeh2_1_pdu(pf02287) PDU1AB EP_NTERM
01040 HMMpropdeh3_1_pdu(pf02288) PDU1AB EP_NTERM
01045 HMMpropdeh1_1_pdu(pf02286) PDU1AB
01050 Tsp_greenishBeige PDU1AB
01055 H_robinEggBlue GRM4
01060 -pf00230 -
01065 HMM10114PocR_1_pdu1(pf12833pf10114) PDU1AB
01070 -pf01656pf07685 -
01075 -pf03186 -
01080 -pf02570 -
BMC-P
BMC-H
BMC-T
Aldehyde dehydrogenase
PTAC
Signature enzyme
Alcohol dehydrogenase
Regulator

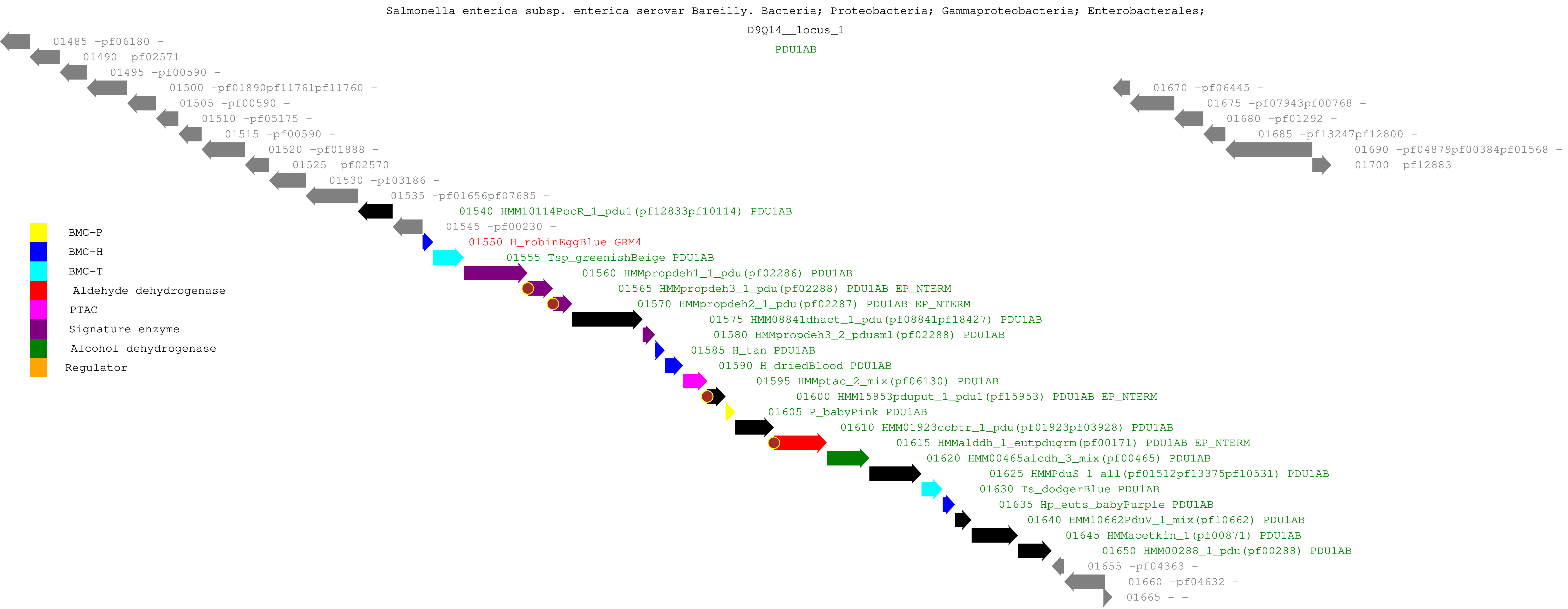

Salmonella enterica subsp. enterica serovar Bareilly. Bacteria; Proteobacteria; Gammaproteobacteria; Enterobacterales;
D9Q14__locus_1
PDU1AB
01485 -pf06180 -
01490 -pf02571 -
01495 -pf00590 -
01500 -pf01890pf11761pf11760 -
01505 -pf00590 -
01510 -pf05175 -
01515 -pf00590 -
01520 -pf01888 -
01525 -pf02570 -
01530 -pf03186 -
01535 -pf01656pf07685 -
01540 HMM10114PocR_1_pdu1(pf12833pf10114) PDU1AB
01545 -pf00230 -
01550 H_robinEggBlue GRM4
01555 Tsp_greenishBeige PDU1AB
01560 HMMpropdeh1_1_pdu(pf02286) PDU1AB
01565 HMMpropdeh3_1_pdu(pf02288) PDU1AB EP_NTERM
01570 HMMpropdeh2_1_pdu(pf02287) PDU1AB EP_NTERM
01575 HMM08841dhact_1_pdu(pf08841pf18427) PDU1AB
01580 HMMpropdeh3_2_pdusml(pf02288) PDU1AB
01585 H_tan PDU1AB
01590 H_driedBlood PDU1AB
01595 HMMptac_2_mix(pf06130) PDU1AB
01600 HMM15953pduput_1_pdu1(pf15953) PDU1AB EP_NTERM
01605 P_babyPink PDU1AB
01610 HMM01923cobtr_1_pdu(pf01923pf03928) PDU1AB
01615 HMMalddh_1_eutpdugrm(pf00171) PDU1AB EP_NTERM
01620 HMM00465alcdh_3_mix(pf00465) PDU1AB
01625 HMMPduS_1_all(pf01512pf13375pf10531) PDU1AB
01630 Ts_dodgerBlue PDU1AB
01635 Hp_euts_babyPurple PDU1AB
01640 HMM10662PduV_1_mix(pf10662) PDU1AB
01645 HMMacetkin_1(pf00871) PDU1AB
01650 HMM00288_1_pdu(pf00288) PDU1AB
01655 -pf04363 -
01660 -pf04632 -
01665 - -
01670 -pf06445 -
01675 -pf07943pf00768 -
01680 -pf01292 -
01685 -pf13247pf12800 -
01690 -pf04879pf00384pf01568 -
01700 -pf12883 -
BMC-P
BMC-H
BMC-T
Aldehyde dehydrogenase
PTAC
Signature enzyme
Alcohol dehydrogenase
Regulator

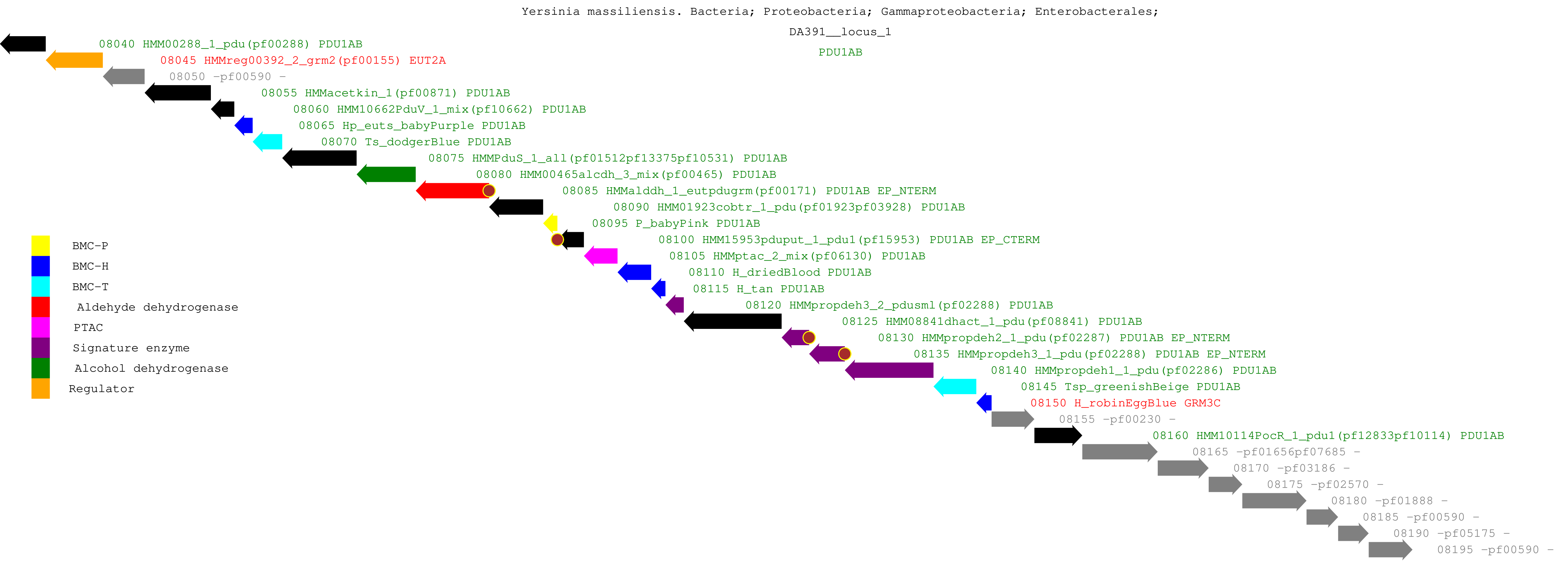
Yersinia massiliensis. Bacteria; Proteobacteria; Gammaproteobacteria; Enterobacterales;
DA391__locus_1
PDU1AB
08040 HMM00288_1_pdu(pf00288) PDU1AB
08045 HMMreg00392_2_grm2(pf00155) EUT2A
08050 -pf00590 -
08055 HMMacetkin_1(pf00871) PDU1AB
08060 HMM10662PduV_1_mix(pf10662) PDU1AB
08065 Hp_euts_babyPurple PDU1AB
08070 Ts_dodgerBlue PDU1AB
08075 HMMPduS_1_all(pf01512pf13375pf10531) PDU1AB
08080 HMM00465alcdh_3_mix(pf00465) PDU1AB
08085 HMMalddh_1_eutpdugrm(pf00171) PDU1AB EP_NTERM
08090 HMM01923cobtr_1_pdu(pf01923pf03928) PDU1AB
08095 P_babyPink PDU1AB
08100 HMM15953pduput_1_pdu1(pf15953) PDU1AB EP_CTERM
08105 HMMptac_2_mix(pf06130) PDU1AB
08110 H_driedBlood PDU1AB
08115 H_tan PDU1AB
08120 HMMpropdeh3_2_pdusml(pf02288) PDU1AB
08125 HMM08841dhact_1_pdu(pf08841) PDU1AB
08130 HMMpropdeh2_1_pdu(pf02287) PDU1AB EP_NTERM
08135 HMMpropdeh3_1_pdu(pf02288) PDU1AB EP_NTERM
08140 HMMpropdeh1_1_pdu(pf02286) PDU1AB
08145 Tsp_greenishBeige PDU1AB
08150 H_robinEggBlue GRM3C
08155 -pf00230 -
08160 HMM10114PocR_1_pdu1(pf12833pf10114) PDU1AB
08165 -pf01656pf07685 -
08170 -pf03186 -
08175 -pf02570 -
08180 -pf01888 -
08185 -pf00590 -
08190 -pf05175 -
08195 -pf00590 -
BMC-P
BMC-H
BMC-T
Aldehyde dehydrogenase
PTAC
Signature enzyme
Alcohol dehydrogenase
Regulator

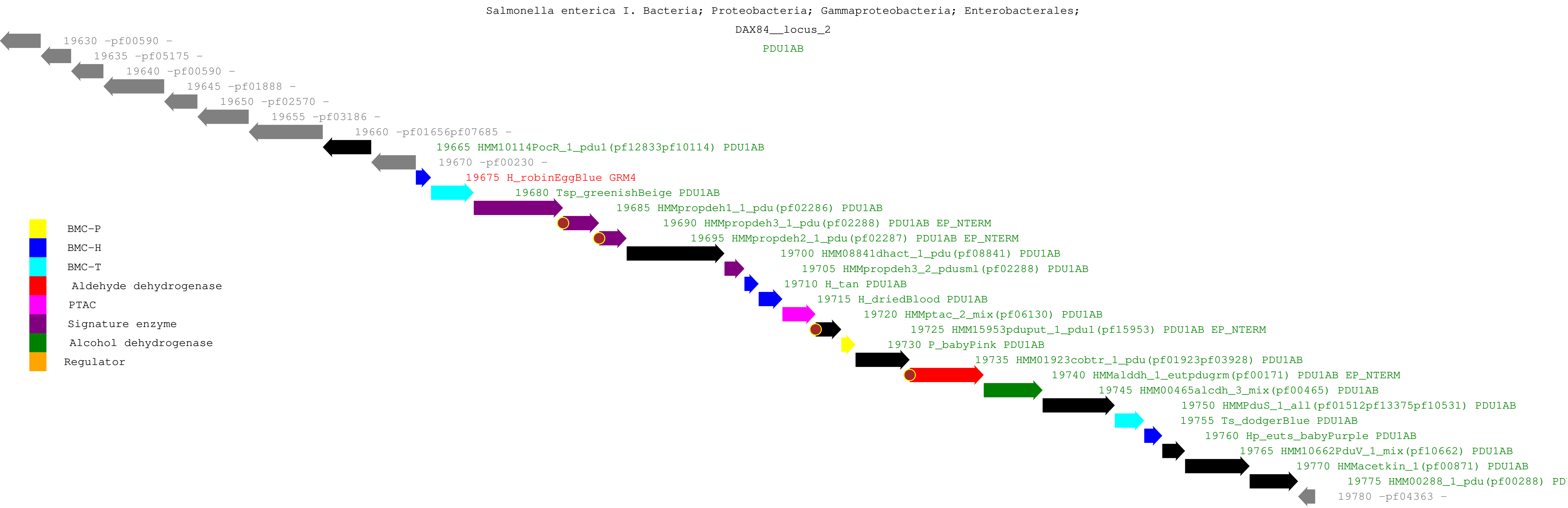

Salmonella enterica I. Bacteria; Proteobacteria; Gammaproteobacteria; Enterobacterales;
DAX84__locus_2
PDU1AB
19630 -pf00590 -
19635 -pf05175 -
19640 -pf00590 -
19645 -pf01888 -
19650 -pf02570 -
19655 -pf03186 -
19660 -pf01656pf07685 -
19665 HMM10114PocR_1_pdu1(pf12833pf10114) PDU1AB
19670 -pf00230 -
19675 H_robinEggBlue GRM4
19680 Tsp_greenishBeige PDU1AB
19685 HMMpropdeh1_1_pdu(pf02286) PDU1AB
19690 HMMpropdeh3_1_pdu(pf02288) PDU1AB EP_NTERM
19695 HMMpropdeh2_1_pdu(pf02287) PDU1AB EP_NTERM
19700 HMM08841dhact_1_pdu(pf08841) PDU1AB
19705 HMMpropdeh3_2_pdusml(pf02288) PDU1AB
19710 H_tan PDU1AB
19715 H_driedBlood PDU1AB
19720 HMMptac_2_mix(pf06130) PDU1AB
19725 HMM15953pduput_1_pdu1(pf15953) PDU1AB EP_NTERM
19730 P_babyPink PDU1AB
19735 HMM01923cobtr_1_pdu(pf01923pf03928) PDU1AB
19740 HMMaldh_1_eutpdugrm(pf00171) PDU1AB EP_NTERM
19745 HMM00465alcdh_3_mix(pf00465) PDU1AB
19750 HMMPduS_1_all(pf01512pf13375pf10531) PDU1AB
19755 Ts_dodgerBlue PDU1AB
19760 Hp_euts_babyPurple PDU1AB
19765 HMM10662PduV_1_mix(pf10662) PDU1AB
19770 HMMacetkin_1(pf00871) PDU1AB
19775 HMM00288_1_pdu(pf00288) PD
19780 -pf04363 -
BMC-P
BMC-H
BMC-T
Aldehyde dehydrogenase
PTAC
Signature enzyme
Alcohol dehydrogenase
Regulator

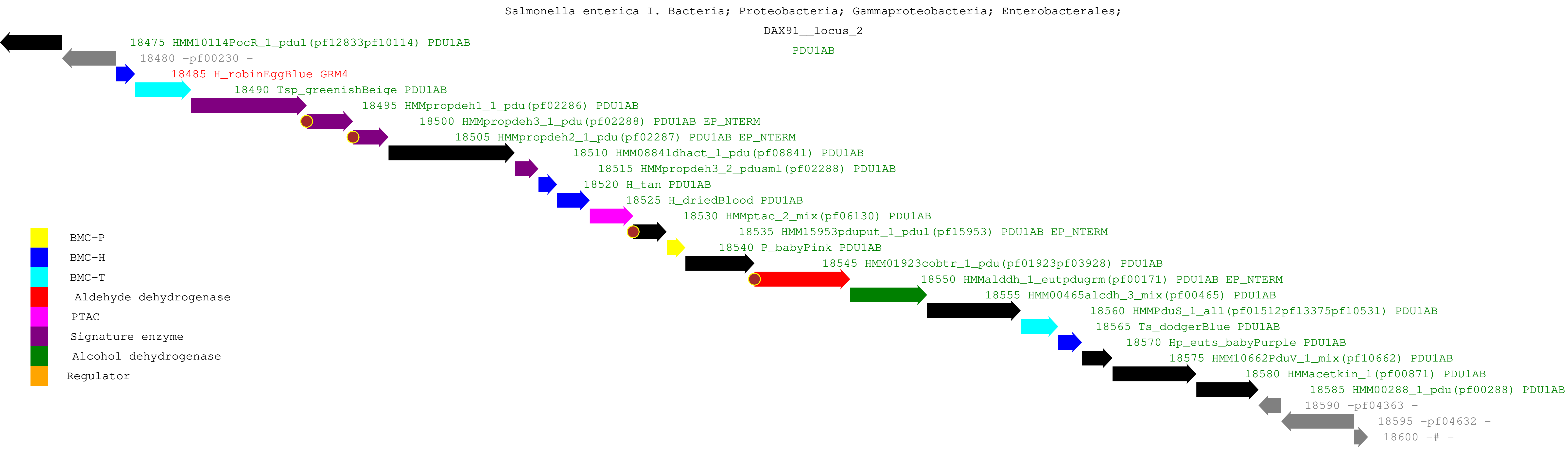
Salmonella enterica I. Bacteria; Proteobacteria; Gammaproteobacteria; Enterobacterales;
DAX91__locus_2
PDU1AB
18475 HMM10114PocR_1_pdu1(pf12833pf10114) PDU1AB
18480 -pf00230 -
18485 H_robinEggBlue GRM4
18490 Tsp_greenishBeige PDU1AB
18495 HMMpropdeh1_1_pdu(pf02286) PDU1AB
18500 HMMpropdeh3_1_pdu(pf02288) PDU1AB EP_NTERM
18505 HMMpropdeh2_1_pdu(pf02287) PDU1AB EP_NTERM
18510 HMM08841dhact_1_pdu(pf08841) PDU1AB
18515 HMMpropdeh3_2_pdusml(pf02288) PDU1AB
18520 H_tan PDU1AB
18525 H_driedBlood PDU1AB
18530 HMMptac_2_mix(pf06130) PDU1AB
18535 HMM15953pduput_1_pdu1(pf15953) PDU1AB EP_NTERM
18540 P_babyPink PDU1AB
18545 HMM01923cobtr_1_pdu(pf01923pf03928) PDU1AB
18550 HMMalddh_1_eutpdugrm(pf00171) PDU1AB EP_NTERM
18555 HMM00465alcdh_3_mix(pf00465) PDU1AB
18560 HMMPduS_1_all(pf01512pf13375pf10531) PDU1AB
18565 Ts_dodgerBlue PDU1AB
18570 Hp_euts_babyPurple PDU1AB
18575 HMM10662PduV_1_mix(pf10662) PDU1AB
18580 HMMacetkin_1(pf00871) PDU1AB
18585 HMM00288_1_pdu(pf00288) PDU1AB
18590 -pf04363 -
18595 -pf04632 -
18600 -# -
BMC-P
BMC-H
BMC-T
Aldehyde dehydrogenase
PTAC
Signature enzyme
Alcohol dehydrogenase
Regulator

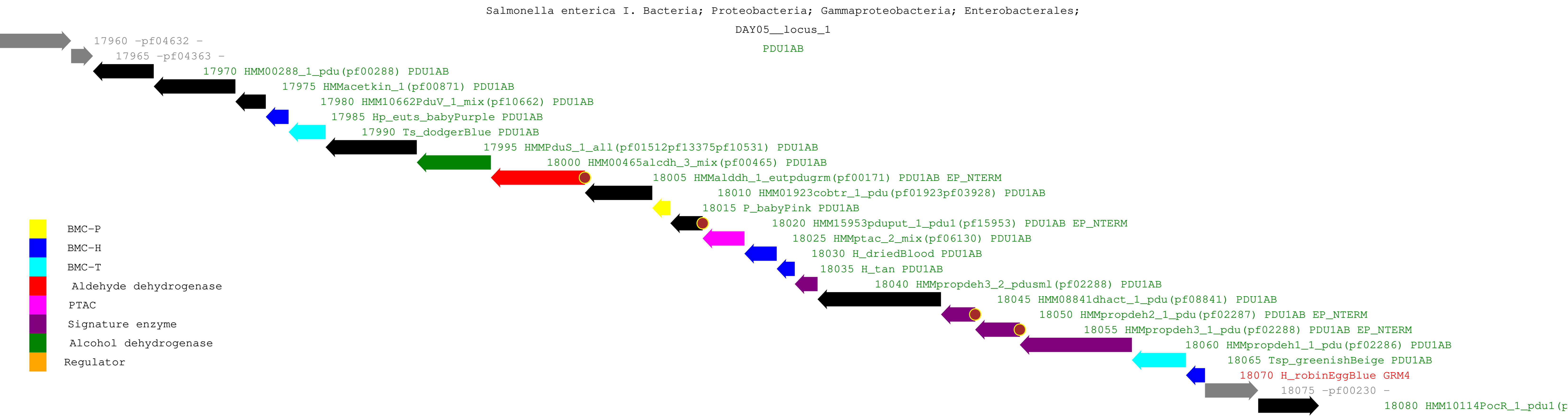
Salmonella enterica I. Bacteria; Proteobacteria; Gammaproteobacteria; Enterobacterales;
DAY05__locus_1
PDU1AB
17960 -pf04632 -
17965 -pf04363 -
17970 HMM00288_1_pdu(pf00288) PDU1AB
17975 HMMacetkin_1(pf00871) PDU1AB
17980 HMM10662PduV_1_mix(pf10662) PDU1AB
17985 Hp_euts_babyPurple PDU1AB
17990 Ts_dodgerBlue PDU1AB
17995 HMMPduS_1_all(pf01512pf13375pf10531) PDU1AB
18000 HMM00465alcdh_3_mix(pf00465) PDU1AB
18005 HMMalddh_1_eutpdugrm(pf00171) PDU1AB EP_NTERM
18010 HMM01923cobtr_1_pdu(pf01923pf03928) PDU1AB
18015 P_babyPink PDU1AB
18020 HMM15953pduput_1_pdu1(pf15953) PDU1AB EP_NTERM
18025 HMMptac_2_mix(pf06130) PDU1AB
18030 H_driedBlood PDU1AB
18035 H_tan PDU1AB
18040 HMMpropdeh3_2_pdusml(pf02288) PDU1AB
18045 HMM08841dhact_1_pdu(pf08841) PDU1AB
18050 HMMpropdeh2_1_pdu(pf02287) PDU1AB EP_NTERM
18055 HMMpropdeh3_1_pdu(pf02288) PDU1AB EP_NTERM
18060 HMMpropdeh1_1_pdu(pf02286) PDU1AB
18065 Tsp_greenishBeige PDU1AB
18070 H_robinEggBlue GRM4
18075 -pf00230 -
18080 HMM10114PocR_1_pdu1(p
BMC-P
BMC-H
BMC-T
Aldehyde dehydrogenase
PTAC
Signature enzyme
Alcohol dehydrogenase
Regulator

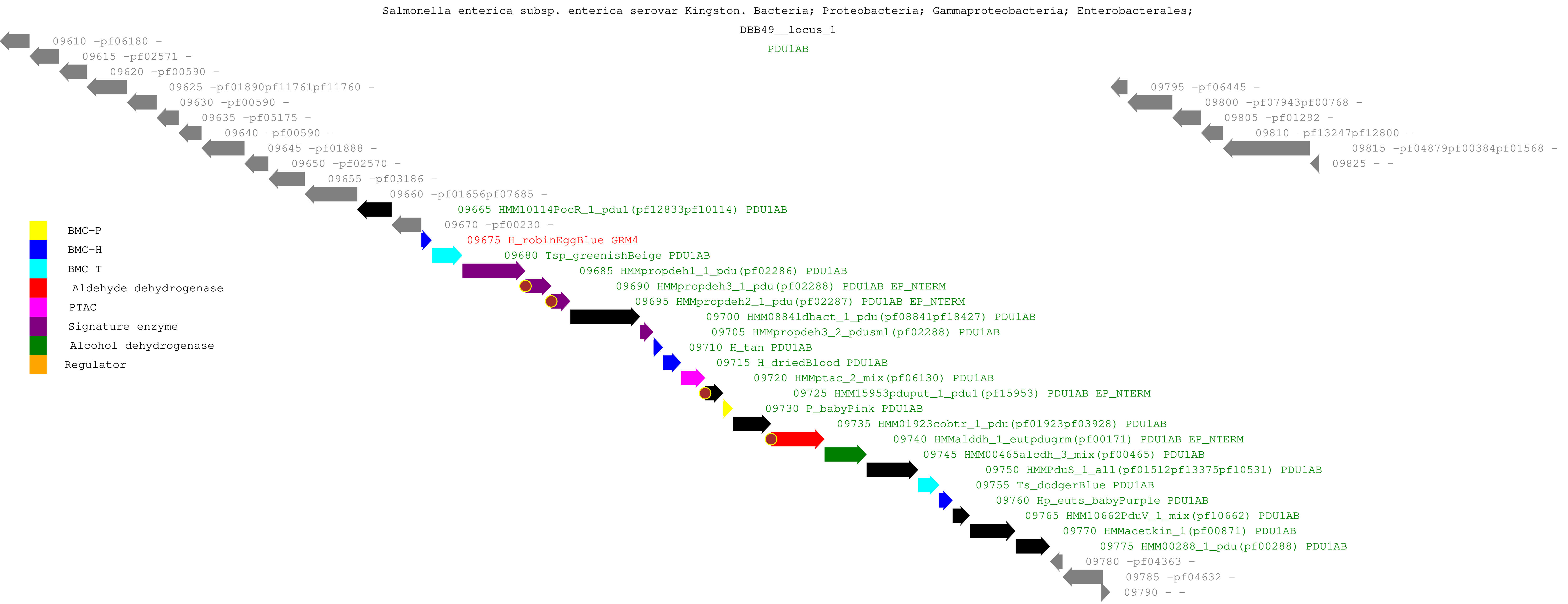

Salmonella enterica subsp. enterica serovar Kingston. Bacteria; Proteobacteria; Gammaproteobacteria; Enterobacterales;
DBB49__locus_1
PDU1AB
09610 -pf06180 -
09615 -pf02571 -
09620 -pf00590 -
09625 -pf01890pf11761pf11760 -
09630 -pf00590 -
09635 -pf05175 -
09640 -pf00590 -
09645 -pf01888 -
09650 -pf02570 -
09655 -pf03186 -
09660 -pf01656pf07685 -
09665 HMM10114PocR_1_pdu1(pf12833pf10114) PDU1AB
09670 -pf00230 -
09675 H_robinEggBlue GRM4
09680 Tsp_greenishBeige PDU1AB
09685 HMMpropdeh1_1_pdu(pf02286) PDU1AB
09690 HMMpropdeh3_1_pdu(pf02288) PDU1AB EP_NTERM
09695 HMMpropdeh2_1_pdu(pf02287) PDU1AB EP_NTERM
09700 HMM08841dhact_1_pdu(pf08841pf18427) PDU1AB
09705 HMMpropdeh3_2_pdusml(pf02288) PDU1AB
09710 H_tan PDU1AB
09715 H_driedBlood PDU1AB
09720 HMMptac_2_mix(pf06130) PDU1AB
09725 HMM15953pduput_1_pdu1(pf15953) PDU1AB EP_NTERM
09730 P_babyPink PDU1AB
09735 HMM01923cobtr_1_pdu(pf01923pf03928) PDU1AB
09740 HMMalddh_1_eutpdugrm(pf00171) PDU1AB EP_NTERM
09745 HMM00465alcdh_3_mix(pf00465) PDU1AB
09750 HMMPduS_1_all(pf01512pf13375pf10531) PDU1AB
09755 Ts_dodgerBlue PDU1AB
09760 Hp_euts_babyPurple PDU1AB
09765 HMM10662PduV_1_mix(pf10662) PDU1AB
09770 HMMacetkin_1(pf00871) PDU1AB
09775 HMM00288_1_pdu(pf00288) PDU1AB
09780 -pf04363 -
09785 -pf04632 -
09790 - -
09795 -pf06445 -
09800 -pf07943pf00768 -
09805 -pf01292 -
09810 -pf13247pf12800 -
09815 -pf04879pf00384pf01568 -
09825 - -
BMC-P
BMC-H
BMC-T
Aldehyde dehydrogenase
PTAC
Signature enzyme
Alcohol dehydrogenase
Regulator

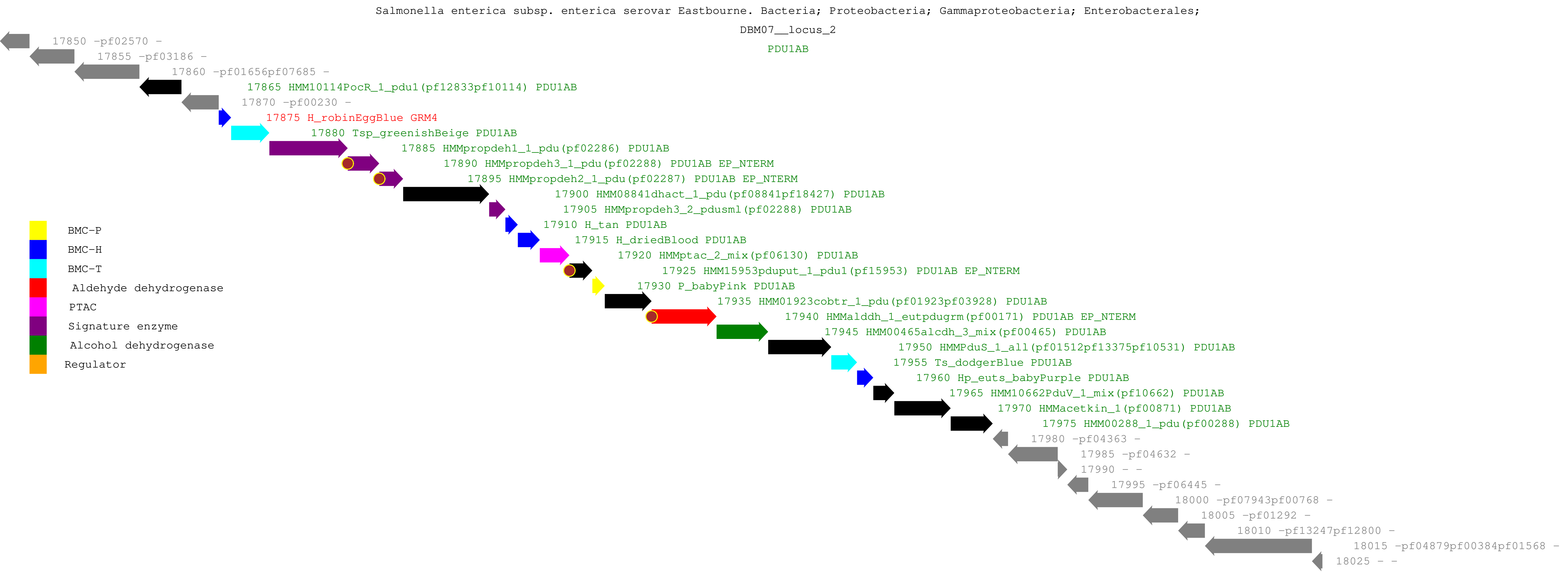

Salmonella enterica subsp. enterica serovar Eastbourne. Bacteria; Proteobacteria; Gammaproteobacteria; Enterobacterales;
DBM07__locus_2
PDU1AB
17850 -pf02570 -
17855 -pf03186 -
17860 -pf01656pf07685 -
17865 HMM10114PocR_1_pdu1(pf12833pf10114) PDU1AB
17870 -pf00230 -
17875 H_robinEggBlue GRM4
17880 Tsp_greenishBeige PDU1AB
17885 HMMpropdeh1_1_pdu(pf02286) PDU1AB
17890 HMMpropdeh3_1_pdu(pf02288) PDU1AB EP_NTERM
17895 HMMpropdeh2_1_pdu(pf02287) PDU1AB EP_NTERM
17900 HMM08841dhact_1_pdu(pf08841pf18427) PDU1AB
17905 HMMpropdeh3_2_pdusml(pf02288) PDU1AB
17910 H_tan PDU1AB
17915 H_driedBlood PDU1AB
17920 HMMptac_2_mix(pf06130) PDU1AB
17925 HMM15953pduput_1_pdu1(pf15953) PDU1AB EP_NTERM
17930 P_babyPink PDU1AB
17935 HMM01923cobtr_1_pdu(pf01923pf03928) PDU1AB
17940 HMMalddh_1_eutpdugrm(pf00171) PDU1AB EP_NTERM
17945 HMM00465alcdh_3_mix(pf00465) PDU1AB
17950 HMMPduS_1_all(pf01512pf13375pf10531) PDU1AB
17955 Ts_dodgerBlue PDU1AB
17960 Hp_euts_babyPurple PDU1AB
17965 HMM10662PduV_1_mix(pf10662) PDU1AB
17970 HMMacetkin_1(pf00871) PDU1AB
17975 HMM00288_1_pdu(pf00288) PDU1AB
17980 -pf04363 -
17985 -pf04632 -
17990 - -
17995 -pf06445 -
18000 -pf07943pf00768 -
18005 -pf01292 -
18010 -pf13247pf12800 -
18015 -pf04879pf00384pf01568 -
18025 - -
BMC-P
BMC-H
BMC-T
Aldehyde dehydrogenase
PTAC
Signature enzyme
Alcohol dehydrogenase
Regulator

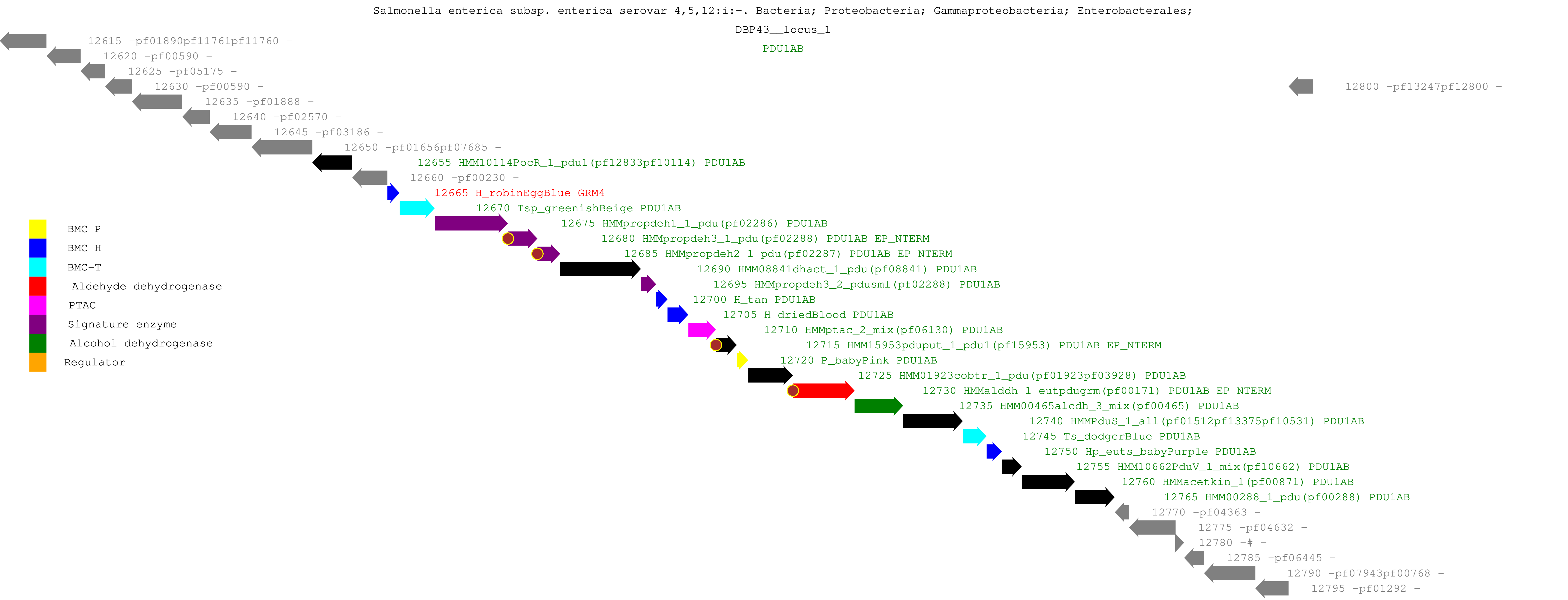

Salmonella enterica subsp. enterica serovar 4,5,12:i:-. Bacteria; Proteobacteria; Gammaproteobacteria; Enterobacterales;
DBP43__locus_1
PDU1AB
12615 -pf01890pf11761pf11760 -
12620 -pf00590 -
12625 -pf05175 -
12630 -pf00590 -
12635 -pf01888 -
12640 -pf02570 -
12645 -pf03186 -
12650 -pf01656pf07685 -
12655 HMM10114PocR_1_pdu1(pf12833pf10114) PDU1AB
12660 -pf00230 -
12665 H_robinEggBlue GRM4
12670 Tsp_greenishBeige PDU1AB
12675 HMMpropdeh1_1_pdu(pf02286) PDU1AB
12680 HMMpropdeh3_1_pdu(pf02288) PDU1AB EP_NTERM
12685 HMMpropdeh2_1_pdu(pf02287) PDU1AB EP_NTERM
12690 HMM08841dhact_1_pdu(pf08841) PDU1AB
12695 HMMpropdeh3_2_pdusml(pf02288) PDU1AB
12700 H_tan PDU1AB
12705 H_driedBlood PDU1AB
12710 HMMptac_2_mix(pf06130) PDU1AB
12715 HMM15953pduput_1_pdu1(pf15953) PDU1AB EP_NTERM
12720 P_babyPink PDU1AB
12725 HMM01923cobtr_1_pdu(pf01923pf03928) PDU1AB
12730 HMMalddh_1_eutpdugrm(pf00171) PDU1AB EP_NTERM
12735 HMM00465alcdh_3_mix(pf00465) PDU1AB
12740 HMMPduS_1_all(pf01512pf13375pf10531) PDU1AB
12745 Ts_dodgerBlue PDU1AB
12750 Hp_euts_babyPurple PDU1AB
12755 HMM10662PduV_1_mix(pf10662) PDU1AB
12760 HMMacetkin_1(pf00871) PDU1AB
12765 HMM00288_1_pdu(pf00288) PDU1AB
12770 -pf04363 -
12775 -pf04632 -
12780 -# -
12785 -pf06445 -
12790 -pf07943pf00768 -
12795 -pf01292 -
12800 -pf13247pf12800 -
BMC-P
BMC-H
BMC-T
Aldehyde dehydrogenase
PTAC
Signature enzyme
Alcohol dehydrogenase
Regulator

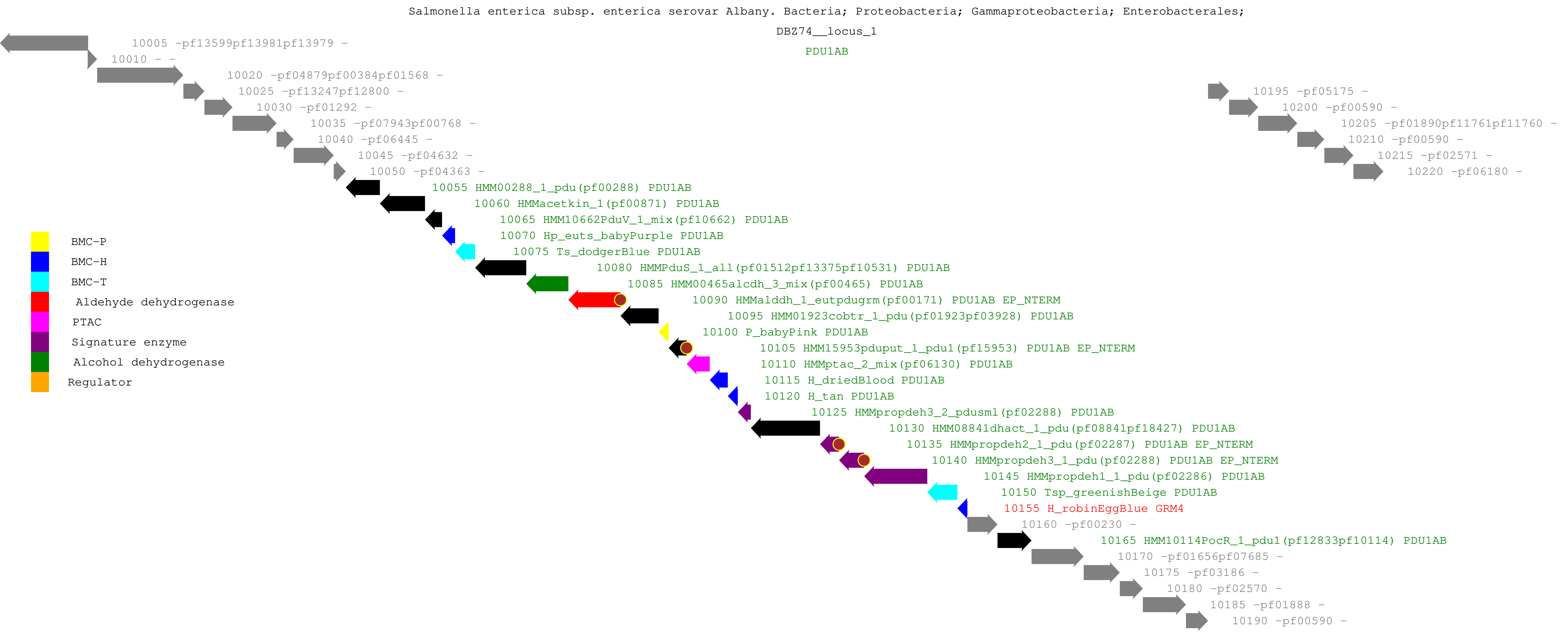
Salmonella enterica subsp. enterica serovar Albany. Bacteria; Proteobacteria; Gammaproteobacteria; Enterobacterales;
DBZ74__locus_1
PDU1AB
10005 -pf13599pf13981pf13979 -
10010 - -
10020 -pf04879pf00384pf01568 -
10025 -pf13247pf12800 -
10030 -pf01292 -
10035 -pf07943pf00768 -
10040 -pf06445 -
10045 -pf04632 -
10050 -pf04363 -
10055 HMM00288_1_pdu(pf00288) PDU1AB
10060 HMMacetkin_1(pf00871) PDU1AB
10065 HMM10662PduV_1_mix(pf10662) PDU1AB
10070 Hp_euts_babyPurple PDU1AB
10075 Ts_dodgerBlue PDU1AB
10080 HMMPduS_1_all(pf01512pf13375pf10531) PDU1AB
10085 HMM00465alcdh_3_mix(pf00465) PDU1AB
10090 HMMalddh_1_eutpdugrm(pf00171) PDU1AB EP_NTERM
10095 HMM01923cobtr_1_pdu(pf01923pf03928) PDU1AB
10100 P_babyPink PDU1AB
10105 HMM15953pduput_1_pdu1(pf15953) PDU1AB EP_NTERM
10110 HMMptac_2_mix(pf06130) PDU1AB
10115 H_driedBlood PDU1AB
10120 H_tan PDU1AB
10125 HMMpropdeh3_2_pdusml(pf02288) PDU1AB
10130 HMM08841dhact_1_pdu(pf08841pf18427) PDU1AB
10135 HMMpropdeh2_1_pdu(pf02287) PDU1AB EP_NTERM
10140 HMMpropdeh3_1_pdu(pf02288) PDU1AB EP_NTERM
10145 HMMpropdeh1_1_pdu(pf02286) PDU1AB
10150 Tsp_greenishBeige PDU1AB
10155 H_robinEggBlue GRM4
10160 -pf00230 -
10165 HMM10114PocR_1_pdu1(pf12833pf10114) PDU1AB
10170 -pf01656pf07685 -
10175 -pf03186 -
10180 -pf02570 -
10185 -pf01888 -
10190 -pf00590 -
10195 -pf05175 -
10200 -pf00590 -
10205 -pf01890pf11761pf11760 -
10210 -pf00590 -
10215 -pf02571 -
10220 -pf06180 -
BMC-P
BMC-H
BMC-T
Aldehyde dehydrogenase
PTAC
Signature enzyme
Alcohol dehydrogenase
Regulator

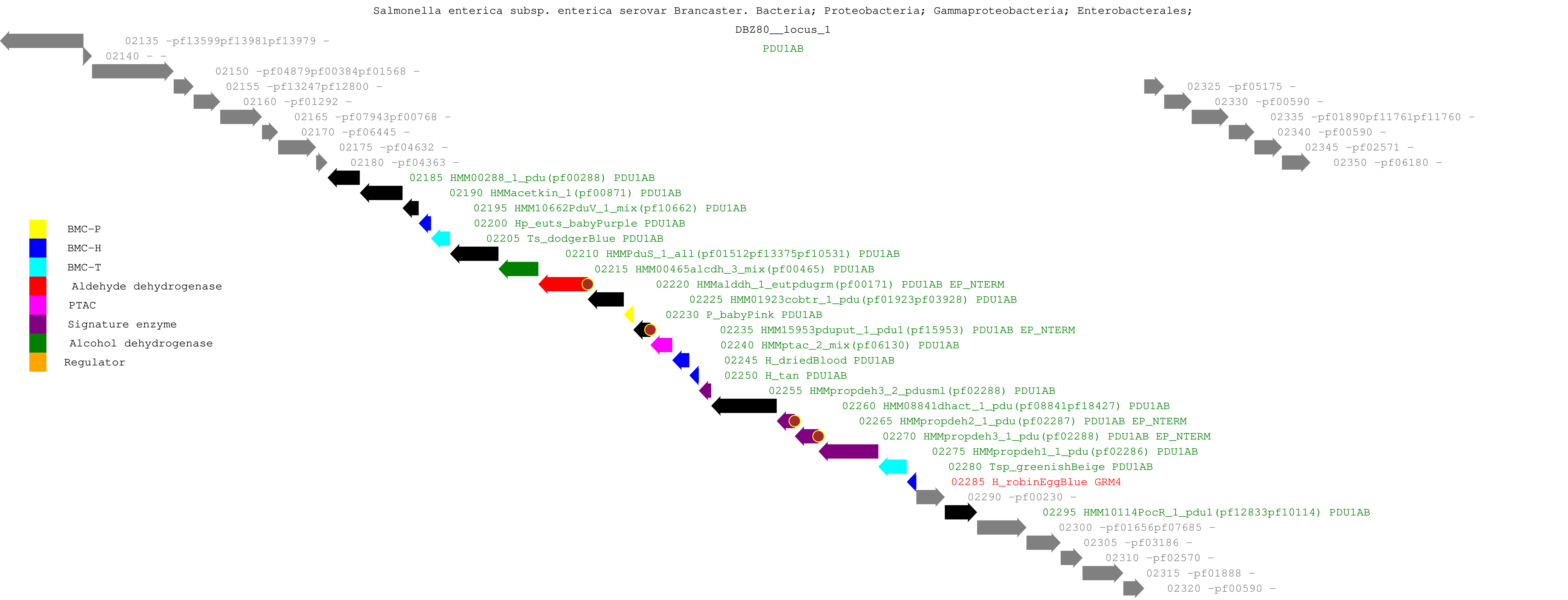

Salmonella enterica subsp. enterica serovar Brancaster. Bacteria; Proteobacteria; Gammaproteobacteria; Enterobacterales;
DBZ80__locus_1
PDU1AB
02135 -pf13599pf13981pf13979 -
02140 - -
02150 -pf04879pf00384pf01568 -
02155 -pf13247pf12800 -
02160 -pf01292 -
02165 -pf07943pf00768 -
02170 -pf06445 -
02175 -pf04632 -
02180 -pf04363 -
02185 HMM00288_1_pdu(pf00288) PDU1AB
02190 HMMacetkin_1(pf00871) PDU1AB
02195 HMM10662PduV_1_mix(pf10662) PDU1AB
02200 Hp_euts_babyPurple PDU1AB
02205 Ts_dodgerBlue PDU1AB
02210 HMMPduS_1_all(pf01512pf13375pf10531) PDU1AB
02215 HMM00465alcdh_3_mix(pf00465) PDU1AB
02220 HMMalddh_1_eutpdugrm(pf00171) PDU1AB EP_NTERM
02225 HMM01923cobtr_1_pdu(pf01923pf03928) PDU1AB
02230 P_babyPink PDU1AB
02235 HMM15953pduput_1_pdu1(pf15953) PDU1AB EP_NTERM
02240 HMMptac_2_mix(pf06130) PDU1AB
02245 H_driedBlood PDU1AB
02250 H_tan PDU1AB
02255 HMMpropdeh3_2_pdusml(pf02288) PDU1AB
02260 HMM08841dhact_1_pdu(pf08841pf18427) PDU1AB
02265 HMMpropdeh2_1_pdu(pf02287) PDU1AB EP_NTERM
02270 HMMpropdeh3_1_pdu(pf02288) PDU1AB EP_NTERM
02275 HMMpropdeh1_1_pdu(pf02286) PDU1AB
02280 Tsp_greenishBeige PDU1AB
02285 H_robinEggBlue GRM4
02290 -pf00230 -
02295 HMM10114PocR_1_pdu1(pf12833pf10114) PDU1AB
02300 -pf01656pf07685 -
02305 -pf03186 -
02310 -pf02570 -
02315 -pf01888 -
02320 -pf00590 -
02325 -pf05175 -
02330 -pf00590 -
02335 -pf01890pf11761pf11760 -
02340 -pf00590 -
02345 -pf02571 -
02350 -pf06180 -
BMC-P
BMC-H
BMC-T
Aldehyde dehydrogenase
PTAC
Signature enzyme
Alcohol dehydrogenase
Regulator

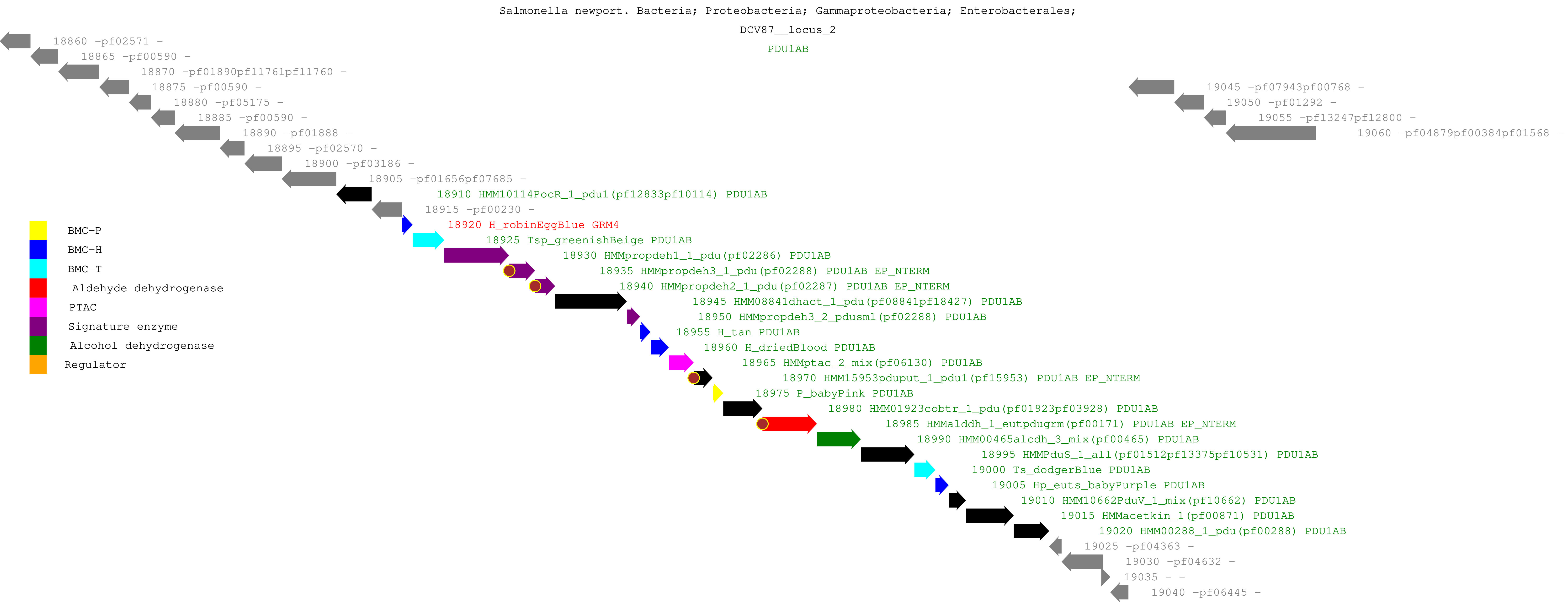

Salmonella newport. Bacteria; Proteobacteria; Gammaproteobacteria; Enterobacterales;
DCV87__locus_2
PDU1AB
18860 -pf02571 -
18865 -pf00590 -
18870 -pf01890pf11761pf11760 -
18875 -pf00590 -
18880 -pf05175 -
18885 -pf00590 -
18890 -pf01888 -
18895 -pf02570 -
18900 -pf03186 -
18905 -pf01656pf07685 -
18910 HMM10114PocR_1_pdu1(pf12833pf10114) PDU1AB
18915 -pf00230 -
18920 H_robinEggBlue GRM4
18925 Tsp_greenishBeige PDU1AB
18930 HMMpropdeh1_1_pdu(pf02286) PDU1AB
18935 HMMpropdeh3_1_pdu(pf02288) PDU1AB EP_NTERM
18940 HMMpropdeh2_1_pdu(pf02287) PDU1AB EP_NTERM
18945 HMM08841dhact_1_pdu(pf08841pf18427) PDU1AB
18950 HMMpropdeh3_2_pdusml(pf02288) PDU1AB
18955 H_tan PDU1AB
18960 H_driedBlood PDU1AB
18965 HMMptac_2_mix(pf06130) PDU1AB
18970 HMM15953pduput_1_pdu1(pf15953) PDU1AB EP_NTERM
18975 P_babyPink PDU1AB
18980 HMM01923cobtr_1_pdu(pf01923pf03928) PDU1AB
18985 HMMalddh_1_eutpdugrm(pf00171) PDU1AB EP_NTERM
18990 HMM00465alcdh_3_mix(pf00465) PDU1AB
18995 HMMPduS_1_all(pf01512pf13375pf10531) PDU1AB
19000 Ts_dodgerBlue PDU1AB
19005 Hp_euts_babyPurple PDU1AB
19010 HMM10662PduV_1_mix(pf10662) PDU1AB
19015 HMMacetkin_1(pf00871) PDU1AB
19020 HMM00288_1_pdu(pf00288) PDU1AB
19025 -pf04363 -
19030 -pf04632 -
19035 - -
19040 -pf06445 -
19045 -pf07943pf00768 -
19050 -pf01292 -
19055 -pf13247pf12800 -
19060 -pf04879pf00384pf01568 -
BMC-P
BMC-H
BMC-T
Aldehyde dehydrogenase
PTAC
Signature enzyme
Alcohol dehydrogenase
Regulator

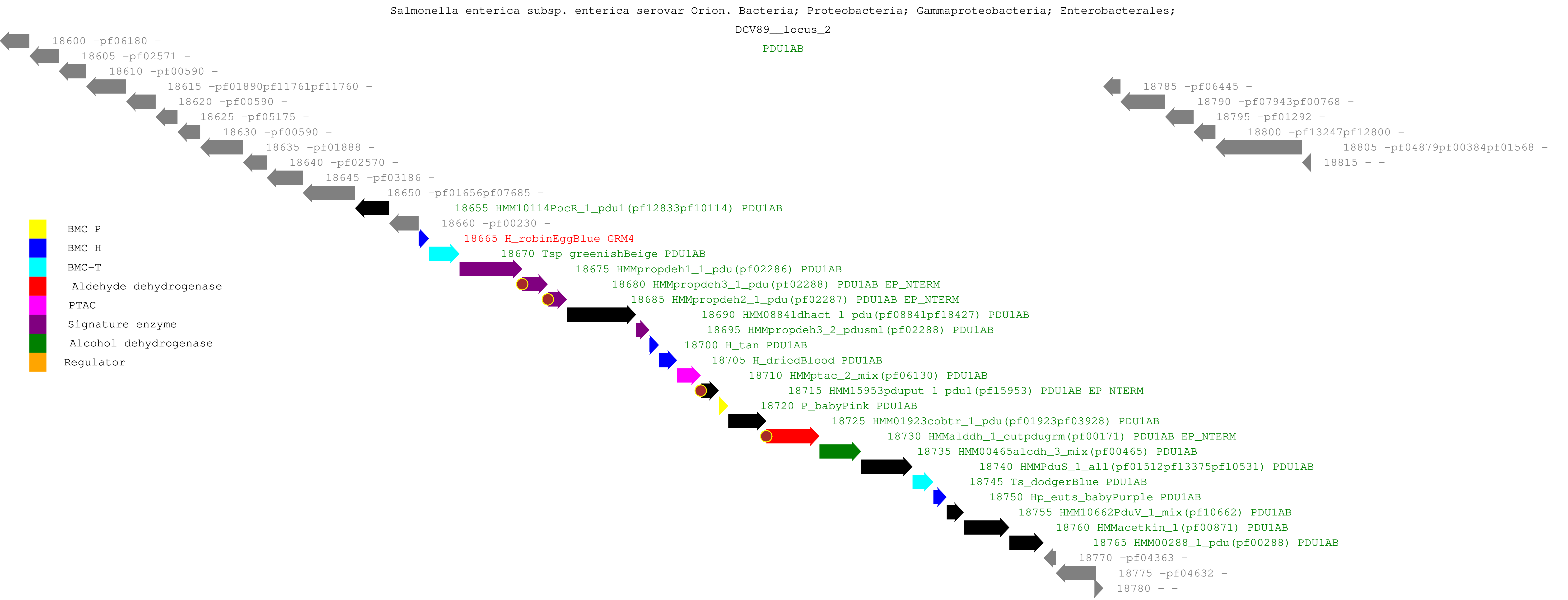
Salmonella enterica subsp. enterica serovar Orion. Bacteria; Proteobacteria; Gammaproteobacteria; Enterobacterales;
DCV89__locus_2
PDU1AB
18600 -pf06180 -
18605 -pf02571 -
18610 -pf00590 -
18615 -pf01890pf11761pf11760 -
18620 -pf00590 -
18625 -pf05175 -
18630 -pf00590 -
18635 -pf01888 -
18640 -pf02570 -
18645 -pf03186 -
18650 -pf01656pf07685 -
18655 HMM10114PocR_1_pdu1(pf12833pf10114) PDU1AB
18660 -pf00230 -
18665 H_robinEggBlue GRM4
18670 Tsp_greenishBeige PDU1AB
18675 HMMpropdeh1_1_pdu(pf02286) PDU1AB
18680 HMMpropdeh3_1_pdu(pf02288) PDU1AB EP_NTERM
18685 HMMpropdeh2_1_pdu(pf02287) PDU1AB EP_NTERM
18690 HMM08841dhact_1_pdu(pf08841pf18427) PDU1AB
18695 HMMpropdeh3_2_pdusml(pf02288) PDU1AB
18700 H_tan PDU1AB
18705 H_driedBlood PDU1AB
18710 HMMptac_2_mix(pf06130) PDU1AB
18715 HMM15953pduput_1_pdu1(pf15953) PDU1AB EP_NTERM
18720 P_babyPink PDU1AB
18725 HMM01923cobtr_1_pdu(pf01923pf03928) PDU1AB
18730 HMMalddh_1_eutpdugrm(pf00171) PDU1AB EP_NTERM
18735 HMM00465alcdh_3_mix(pf00465) PDU1AB
18740 HMMPduS_1_all(pf01512pf13375pf10531) PDU1AB
18745 Ts_dodgerBlue PDU1AB
18750 Hp_euts_babyPurple PDU1AB
18755 HMM10662PduV_1_mix(pf10662) PDU1AB
18760 HMMacetkin_1(pf00871) PDU1AB
18765 HMM00288_1_pdu(pf00288) PDU1AB
18770 -pf04363 -
18775 -pf04632 -
18780 - -
18785 -pf06445 -
18790 -pf07943pf00768 -
18795 -pf01292 -
18800 -pf13247pf12800 -
18805 -pf04879pf00384pf01568 -
18815 - -
BMC-P
BMC-H
BMC-T
Aldehyde dehydrogenase
PTAC
Signature enzyme
Alcohol dehydrogenase
Regulator

Escherichia coli. Bacteria; Proteobacteria; Gammaproteobacteria; Enterobacterales;

DD647__locus_1

PDU1AB

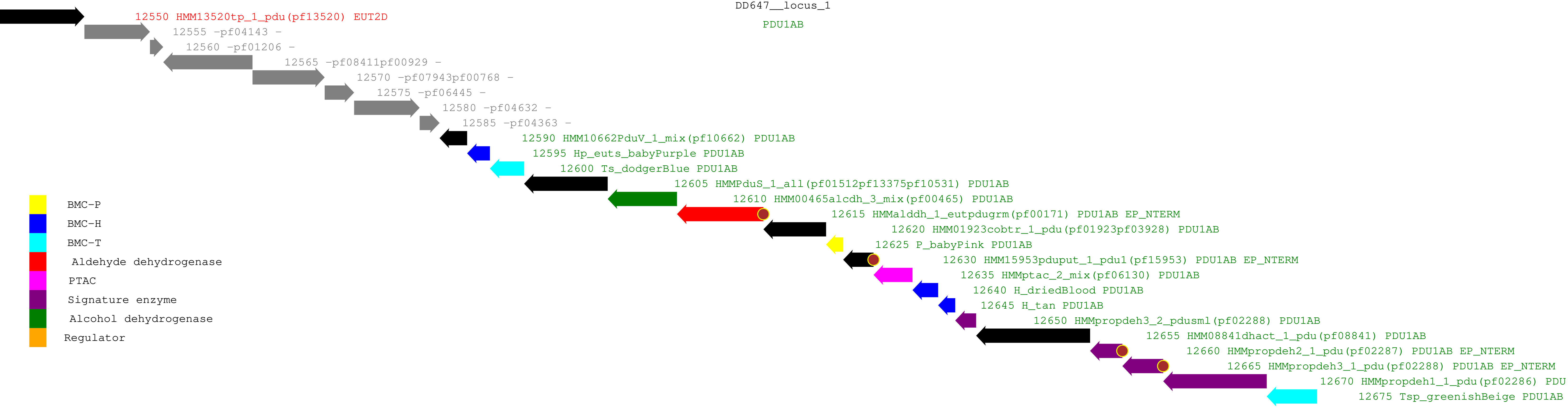

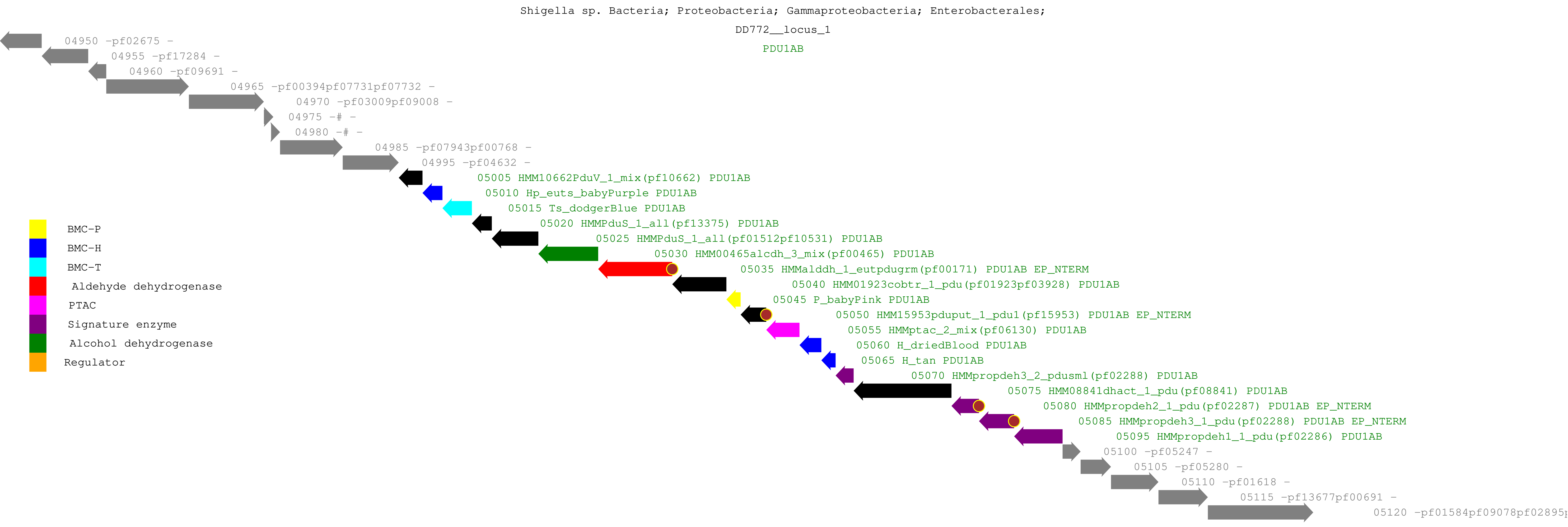
Shigella sp. Bacteria; Proteobacteria; Gammaproteobacteria; Enterobacterales;
DD772__locus_1
PDU1AB
04950 -pf02675 -
04955 -pf17284 -
04960 -pf09691 -
04965 -pf00394pf07731pf07732 -
04970 -pf03009pf09008 -
04975 -# -
04980 -# -
04985 -pf07943pf00768 -
04995 -pf04632 -
05005 HMM10662PduV_1_mix(pf10662) PDU1AB
05010 Hp_euts_babyPurple PDU1AB
05015 Ts_dodgerBlue PDU1AB
05020 HMMPduS_1_all(pf13375) PDU1AB
05025 HMMPduS_1_all(pf01512pf10531) PDU1AB
05030 HMM00465alcdh_3_mix(pf00465) PDU1AB
05035 HMMalddh_1_eutpdugrm(pf00171) PDU1AB EP_NTERM
05040 HMM01923cobtr_1_pdu(pf01923pf03928) PDU1AB
05045 P_babyPink PDU1AB
05050 HMM15953pduput_1_pdu1(pf15953) PDU1AB EP_NTERM
05055 HMMptac_2_mix(pf06130) PDU1AB
05060 H_driedBlood PDU1AB
05065 H_tan PDU1AB
05070 HMMpropdeh3_2_pdusml(pf02288) PDU1AB
05075 HMM08841dhact_1_pdu(pf08841) PDU1AB
05080 HMMpropdeh2_1_pdu(pf02287) PDU1AB EP_NTERM
05085 HMMpropdeh3_1_pdu(pf02288) PDU1AB EP_NTERM
05095 HMMpropdeh1_1_pdu(pf02286) PDU1AB
05100 -pf05247 -
05105 -pf05280 -
05110 -pf01618 -
05115 -pf13677pf00691 -
05120 -pf01584pf09078pf02895
BMC-P
BMC-H
BMC-T
Aldehyde dehydrogenase
PTAC
Signature enzyme
Alcohol dehydrogenase
Regulator

Salmonella typhimurium. Bacteria; Proteobacteria; Gammaproteobacteria; Enterobacterales;

DD95__locus_2

PDU1AB

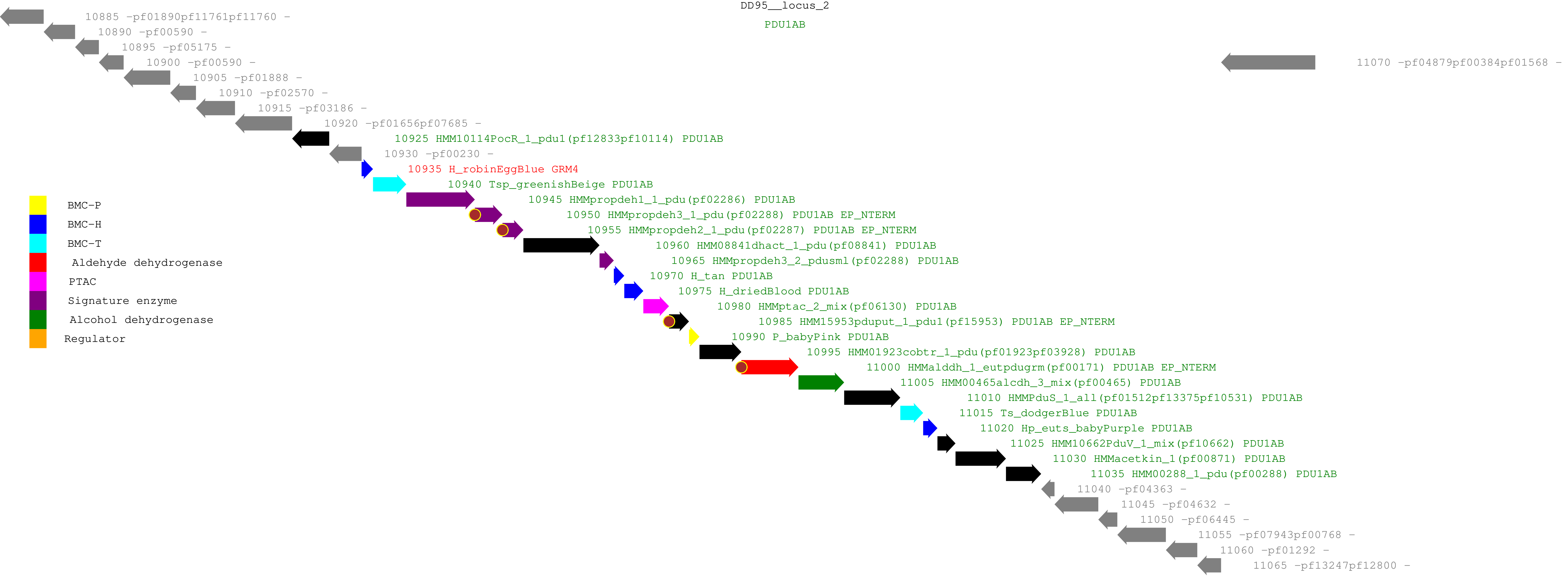

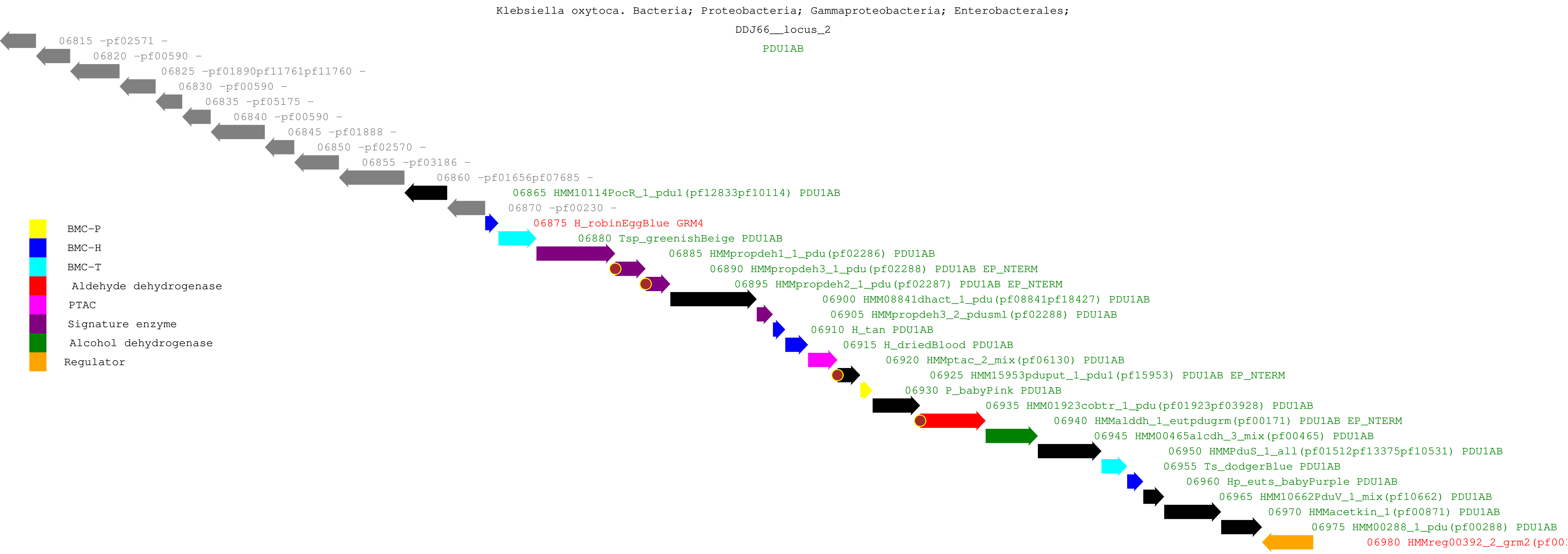
Klebsiella oxytoca. Bacteria; Proteobacteria; Gammaproteobacteria; Enterobacterales;
DDJ66__locus_2
PDU1AB
06815 -pf02571 -
06820 -pf00590 -
06825 -pf01890pf11761pf11760 -
06830 -pf00590 -
06835 -pf05175 -
06840 -pf00590 -
06845 -pf01888 -
06850 -pf02570 -
06855 -pf03186 -
06860 -pf01656pf07685 -
06865 HMM10114PocR_1_pdu1(pf12833pf10114) PDU1AB
06870 -pf00230 -
06875 H_robinEggBlue GRM4
06880 Tsp_greenishBeige PDU1AB
06885 HMMpropdeh1_1_pdu(pf02286) PDU1AB
06890 HMMpropdeh3_1_pdu(pf02288) PDU1AB EP_NTERM
06895 HMMpropdeh2_1_pdu(pf02287) PDU1AB EP_NTERM
06900 HMM08841dhact_1_pdu(pf08841pf18427) PDU1AB
06905 HMMpropdeh3_2_pdusml(pf02288) PDU1AB
06910 H_tan PDU1AB
06915 H_driedBlood PDU1AB
06920 HMMptac_2_mix(pf06130) PDU1AB
06925 HMM15953pduput_1_pdu1(pf15953) PDU1AB EP_NTERM
06930 P_babyPink PDU1AB
06935 HMM01923cobtr_1_pdu(pf01923pf03928) PDU1AB
06940 HMMalddh_1_eutpdugrm(pf00171) PDU1AB EP_NTERM
06945 HMM00465alcdh_3_mix(pf00465) PDU1AB
06950 HMMPduS_1_all(pf01512pf13375pf10531) PDU1AB
06955 Ts_dodgerBlue PDU1AB
06960 Hp_euts_babyPurple PDU1AB
06965 HMM10662PduV_1_mix(pf10662) PDU1AB
06970 HMMacetkin_1(pf00871) PDU1AB
06975 HMM00288_1_pdu(pf00288) PDU1AB
06980 HMMreg00392_2_grm2(pf00
BMC-P
BMC-H
BMC-T
Aldehyde dehydrogenase
PTAC
Signature enzyme
Alcohol dehydrogenase
Regulator

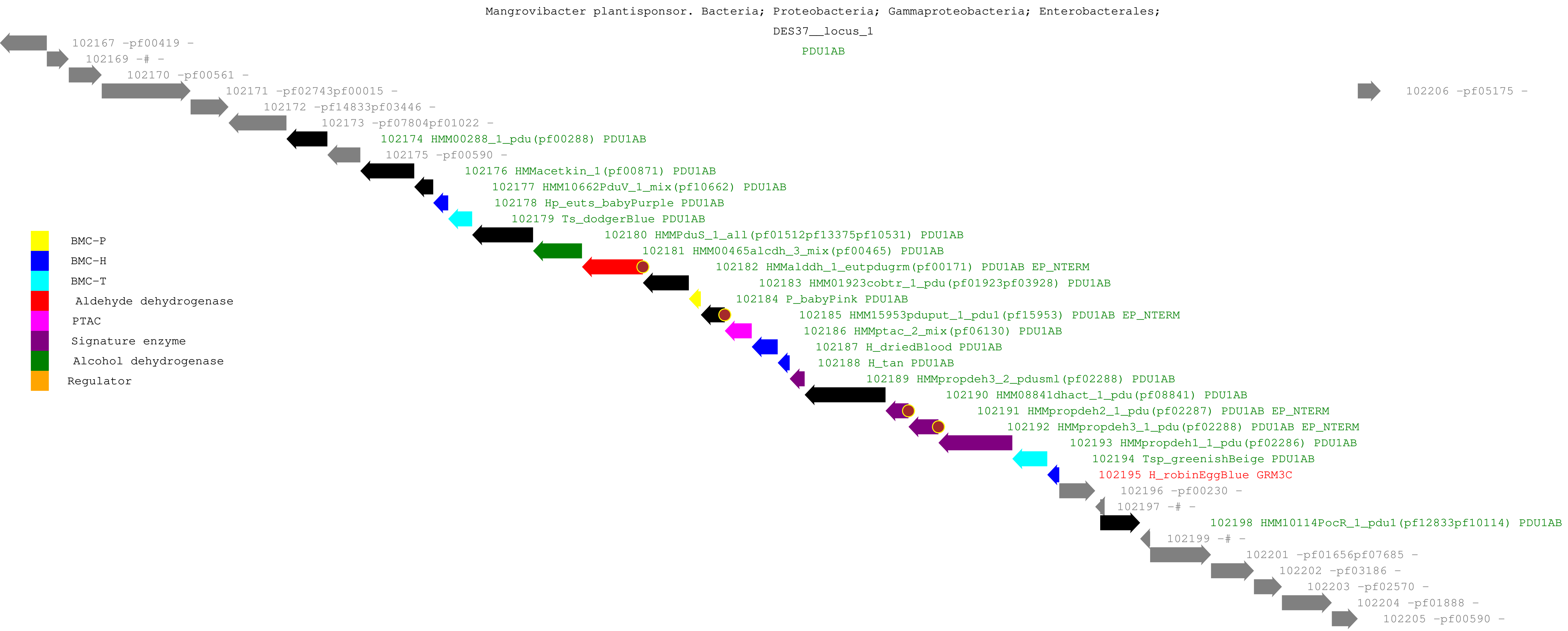

Mangrovibacter plantisponsor. Bacteria; Proteobacteria; Gammaproteobacteria; Enterobacterales;
DES37__locus_1
PDU1AB
102167 -pf00419 -
102169 -# -
102170 -pf00561 -
102171 -pf02743pf00015 -
102172 -pf14833pf03446 -
102173 -pf07804pf01022 -
102174 HMM00288_1_pdu(pf00288) PDU1AB
102175 -pf00590 -
102176 HMMacetkin_1(pf00871) PDU1AB
102177 HMM10662PduV_1_mix(pf10662) PDU1AB
102178 Hp_euts_babyPurple PDU1AB
102179 Ts_dodgerBlue PDU1AB
102180 HMMPduS_1_all(pf01512pf13375pf10531) PDU1AB
102181 HMM00465alcdh_3_mix(pf00465) PDU1AB
102182 HMMalddh_1_eutpdugrm(pf00171) PDU1AB EP_NTERM
102183 HMM01923cobtr_1_pdu(pf01923pf03928) PDU1AB
102184 P_babyPink PDU1AB
102185 HMM15953pduput_1_pdu1(pf15953) PDU1AB EP_NTERM
102186 HMMptac_2_mix(pf06130) PDU1AB
102187 H_driedBlood PDU1AB
102188 H_tan PDU1AB
102189 HMMpropdeh3_2_pdusml(pf02288) PDU1AB
102190 HMM08841dhact_1_pdu(pf08841) PDU1AB
102191 HMMpropdeh2_1_pdu(pf02287) PDU1AB EP_NTERM
102192 HMMpropdeh3_1_pdu(pf02288) PDU1AB EP_NTERM
102193 HMMpropdeh1_1_pdu(pf02286) PDU1AB
102194 Tsp_greenishBeige PDU1AB
102195 H_robinEggBlue GRM3C
102196 -pf00230 -
102197 -# -
102198 HMM10114PocR_1_pdu1(pf12833pf10114) PDU1AB
102199 -# -
102201 -pf01656pf07685 -
102202 -pf03186 -
102203 -pf02570 -
102204 -pf01888 -
102205 -pf00590 -
102206 -pf05175 -
BMC-P
BMC-H
BMC-T
Aldehyde dehydrogenase
PTAC
Signature enzyme
Alcohol dehydrogenase
Regulator

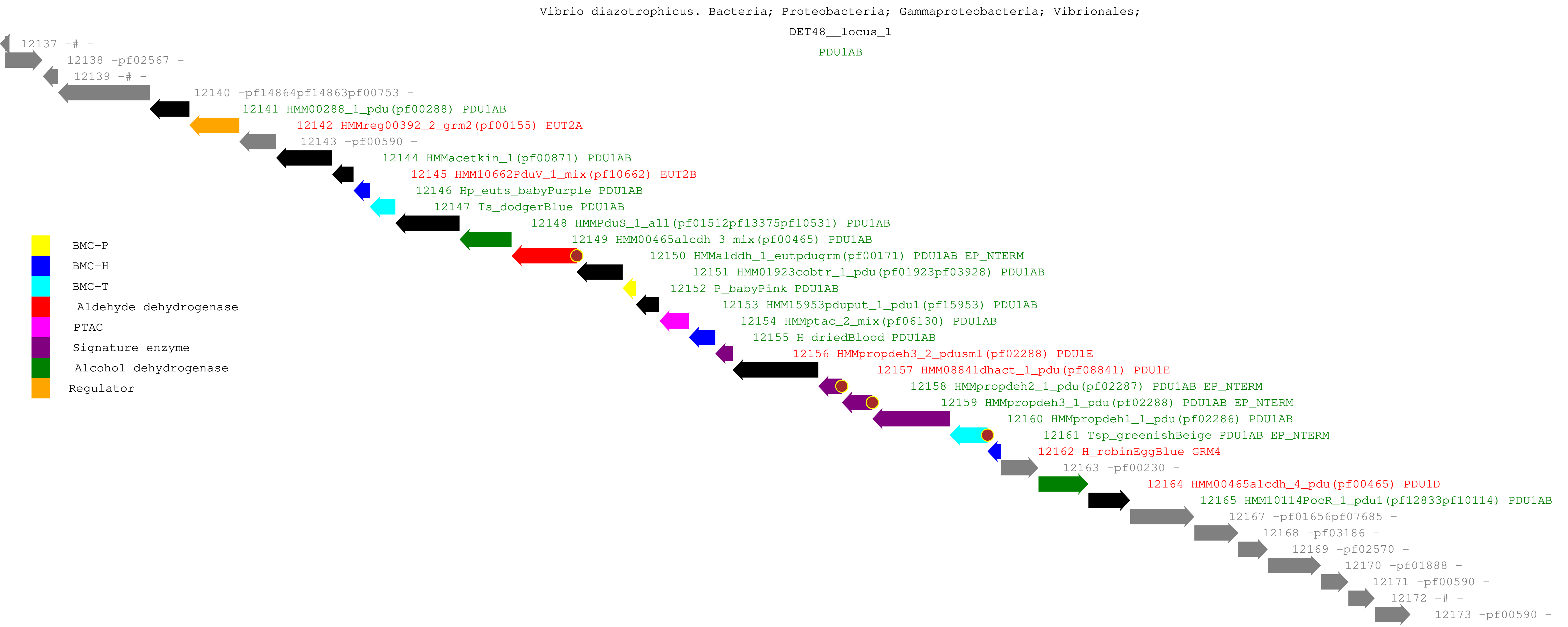
Vibrio diazotrophicus. Bacteria; Proteobacteria; Gammaproteobacteria; Vibrionales;
DET48__locus_1
PDU1AB
12137 -# -
12138 -pf02567 -
12139 -# -
12140 -pf14864pf14863pf00753 -
12141 HMM00288_1_pdu(pf00288) PDU1AB
12142 HMMreg00392_2_grm2(pf00155) EUT2A
12143 -pf00590 -
12144 HMMacetkin_1(pf00871) PDU1AB
12145 HMM10662PduV_1_mix(pf10662) EUT2B
12146 Hp_euts_babyPurple PDU1AB
12147 Ts_dodgerBlue PDU1AB
12148 HMMPduS_1_all(pf01512pf13375pf10531) PDU1AB
12149 HMM00465alcdh_3_mix(pf00465) PDU1AB
12150 HMMalddh_1_eutpdugrm(pf00171) PDU1AB EP_NTERM
12151 HMM01923cobtr_1_pdu(pf01923pf03928) PDU1AB
12152 P_babyPink PDU1AB
12153 HMM15953pduput_1_pdu1(pf15953) PDU1AB
12154 HMMptac_2_mix(pf06130) PDU1AB
12155 H_driedBlood PDU1AB
12156 HMMpropdeh3_2_pdusml(pf02288) PDU1E
12157 HMM08841dhact_1_pdu(pf08841) PDU1E
12158 HMMpropdeh2_1_pdu(pf02287) PDU1AB EP_NTERM
12159 HMMpropdeh3_1_pdu(pf02288) PDU1AB EP_NTERM
12160 HMMpropdeh1_1_pdu(pf02286) PDU1AB
12161 Tsp_greenishBeige PDU1AB EP_NTERM
12162 H_robinEggBlue GRM4
12163 -pf00230 -
12164 HMM00465alcdh_4_pdu(pf00465) PDU1D
12165 HMM10114PocR_1_pdu1(pf12833pf10114) PDU1AB
12167 -pf01656pf07685 -
12168 -pf03186 -
12169 -pf02570 -
12170 -pf01888 -
12171 -pf00590 -
12172 -# -
12173 -pf00590 -
BMC-P
BMC-H
BMC-T
Aldehyde dehydrogenase
PTAC
Signature enzyme
Alcohol dehydrogenase
Regulator

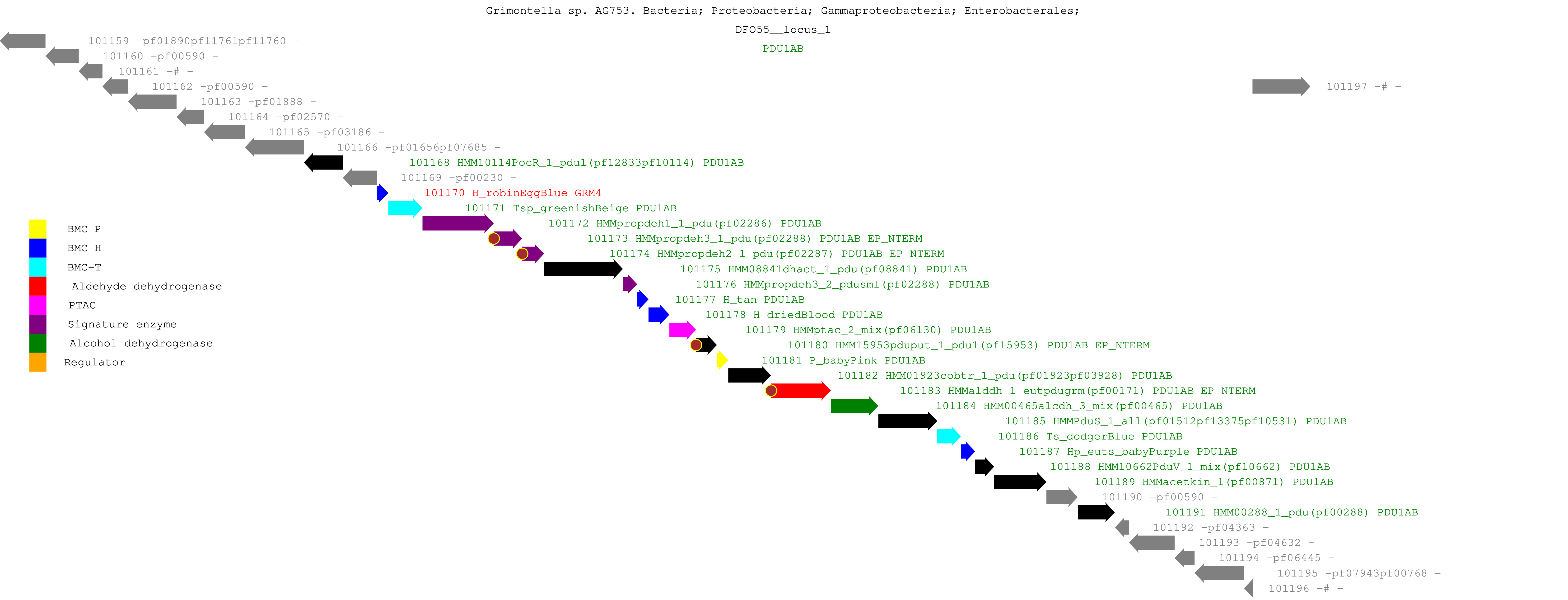

Grimontella sp. AG753. Bacteria; Proteobacteria; Gammaproteobacteria; Enterobacterales;
DFO55__locus_1
PDU1AB
101159 -pf01890pf11761pf11760 -
101160 -pf00590 -
101161 -# -
101162 -pf00590 -
101163 -pf01888 -
101164 -pf02570 -
101165 -pf03186 -
101166 -pf01656pf07685 -
101168 HMM10114PocR_1_pdu1(pf12833pf10114) PDU1AB
101169 -pf00230 -
101170 H_robinEggBlue GRM4
101171 Tsp_greenishBeige PDU1AB
101172 HMMpropdeh1_1_pdu(pf02286) PDU1AB
101173 HMMpropdeh3_1_pdu(pf02288) PDU1AB EP_NTERM
101174 HMMpropdeh2_1_pdu(pf02287) PDU1AB EP_NTERM
101175 HMM08841dhact_1_pdu(pf08841) PDU1AB
101176 HMMpropdeh3_2_pdusml(pf02288) PDU1AB
101177 H_tan PDU1AB
101178 H_driedBlood PDU1AB
101179 HMMptac_2_mix(pf06130) PDU1AB
101180 HMM15953pduput_1_pdu1(pf15953) PDU1AB EP_NTERM
101181 P_babyPink PDU1AB
101182 HMM01923cobtr_1_pdu(pf01923pf03928) PDU1AB
101183 HMMaldh_1_eutpdugrm(pf00171) PDU1AB EP_NTERM
101184 HMM00465alcdh_3_mix(pf00465) PDU1AB
101185 HMMPduS_1_all(pf01512pf13375pf10531) PDU1AB
101186 Ts_dodgerBlue PDU1AB
101187 Hp_euts_babyPurple PDU1AB
101188 HMM10662PduV_1_mix(pf10662) PDU1AB
101189 HMMacetkin_1(pf00871) PDU1AB
101190 -pf00590 -
101191 HMM00288_1_pdu(pf00288) PDU1AB
101192 -pf04363 -
101193 -pf04632 -
101194 -pf06445 -
101195 -pf07943pf00768 -
101196 -# -
101197 -# -
BMC-P
BMC-H
BMC-T
Aldehyde dehydrogenase
PTAC
Signature enzyme
Alcohol dehydrogenase
Regulator

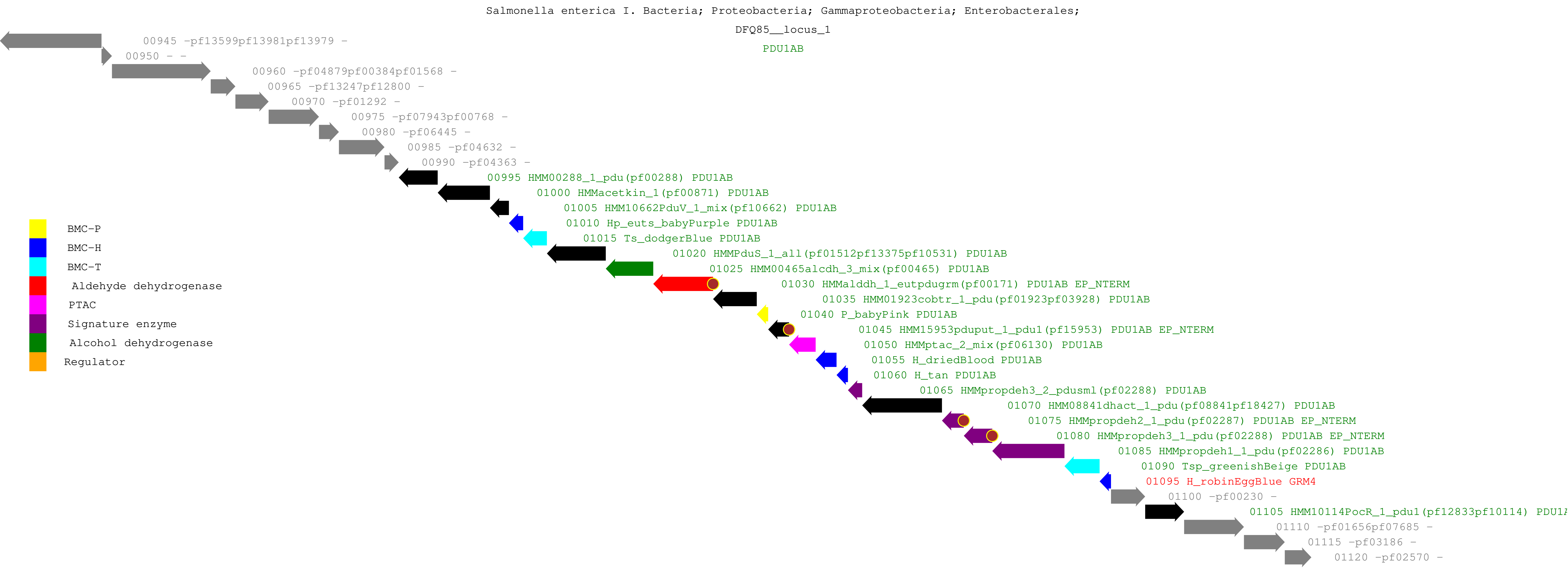
Salmonella enterica I. Bacteria; Proteobacteria; Gammaproteobacteria; Enterobacterales;
DFQ85__locus_1
PDU1AB
00945 -pf13599pf13981pf13979 -
00950 - -
00960 -pf04879pf00384pf01568 -
00965 -pf13247pf12800 -
00970 -pf01292 -
00975 -pf07943pf00768 -
00980 -pf06445 -
00985 -pf04632 -
00990 -pf04363 -
00995 HMM00288_1_pdu(pf00288) PDU1AB
01000 HMMacetkin_1(pf00871) PDU1AB
01005 HMM10662PduV_1_mix(pf10662) PDU1AB
01010 Hp_euts_babyPurple PDU1AB
01015 Ts_dodgerBlue PDU1AB
01020 HMMPduS_1_all(pf01512pf13375pf10531) PDU1AB
01025 HMM00465alcdh_3_mix(pf00465) PDU1AB
01030 HMMalddh_1_eutpdugrm(pf00171) PDU1AB EP_NTERM
01035 HMM01923cobtr_1_pdu(pf01923pf03928) PDU1AB
01040 P_babyPink PDU1AB
01045 HMM15953pduput_1_pdu1(pf15953) PDU1AB EP_NTERM
01050 HMMptac_2_mix(pf06130) PDU1AB
01055 H_driedBlood PDU1AB
01060 H_tan PDU1AB
01065 HMMpropdeh3_2_pdusml(pf02288) PDU1AB
01070 HMM08841dhact_1_pdu(pf08841pf18427) PDU1AB
01075 HMMpropdeh2_1_pdu(pf02287) PDU1AB EP_NTERM
01080 HMMpropdeh3_1_pdu(pf02288) PDU1AB EP_NTERM
01085 HMMpropdeh1_1_pdu(pf02286) PDU1AB
01090 Tsp_greenishBeige PDU1AB
01095 H_robinEggBlue GRM4
01100 -pf00230 -
01105 HMM10114PocR_1_pdu1(pf12833pf10114) PDU1A
01110 -pf01656pf07685 -
01115 -pf03186 -
01120 -pf02570 -
BMC-P
BMC-H
BMC-T
Aldehyde dehydrogenase
PTAC
Signature enzyme
Alcohol dehydrogenase
Regulator

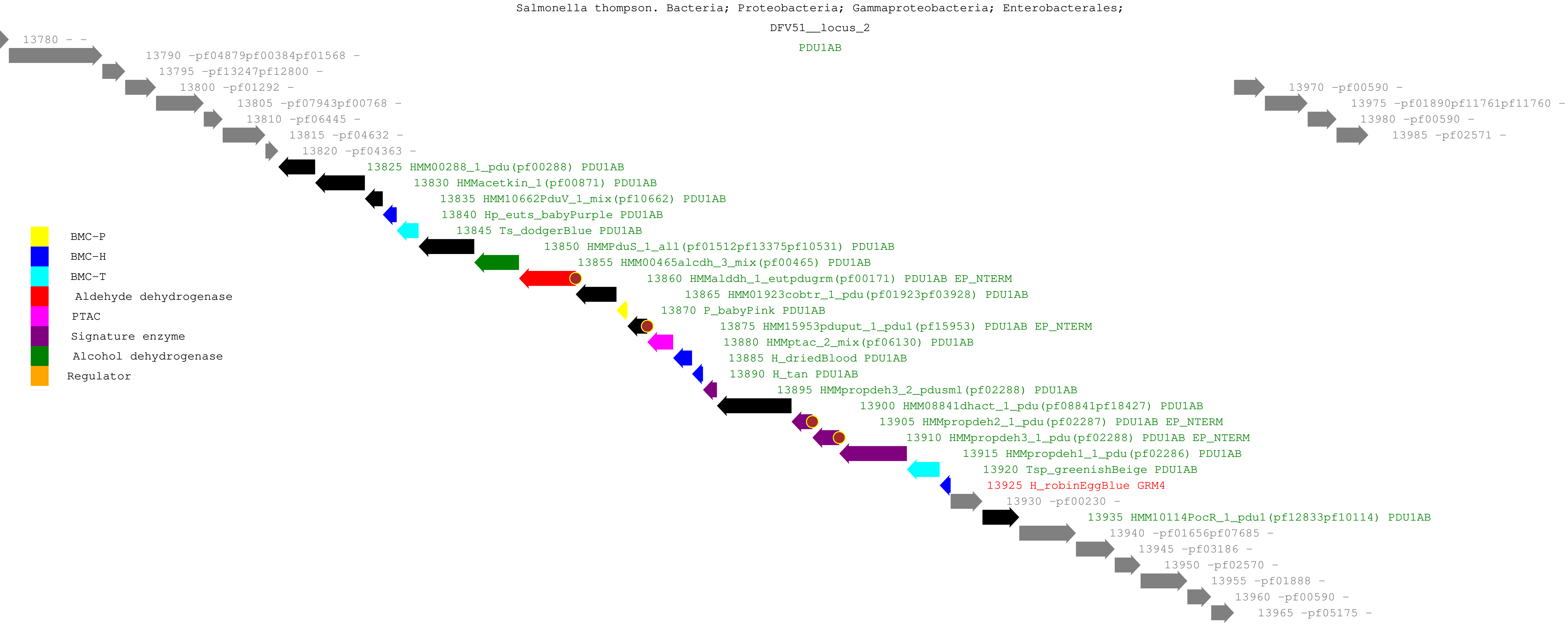

Salmonella thompson. Bacteria; Proteobacteria; Gammaproteobacteria; Enterobacterales;
DFV51__locus_2
PDU1AB
13780 - -
13790 -pf04879pf00384pf01568 -
13795 -pf13247pf12800 -
13800 -pf01292 -
13805 -pf07943pf00768 -
13810 -pf06445 -
13815 -pf04632 -
13820 -pf04363 -
13825 HMM00288_1_pdu(pf00288) PDU1AB
13830 HMMacetkin_1(pf00871) PDU1AB
13835 HMM10662PduV_1_mix(pf10662) PDU1AB
13840 Hp_euts_babyPurple PDU1AB
13845 Ts_dodgerBlue PDU1AB
13850 HMMPduS_1_all(pf01512pf13375pf10531) PDU1AB
13855 HMM00465alcdh_3_mix(pf00465) PDU1AB
13860 HMMaldh_1_eutpdugrm(pf00171) PDU1AB EP_NTERM
13865 HMM01923cobtr_1_pdu(pf01923pf03928) PDU1AB
13870 P_babyPink PDU1AB
13875 HMM15953pduput_1_pdu1(pf15953) PDU1AB EP_NTERM
13880 HMMptac_2_mix(pf06130) PDU1AB
13885 H_driedBlood PDU1AB
13890 H_tan PDU1AB
13895 HMMpropdeh3_2_pdusml(pf02288) PDU1AB
13900 HMM08841dhact_1_pdu(pf08841pf18427) PDU1AB
13905 HMMpropdeh2_1_pdu(pf02287) PDU1AB EP_NTERM
13910 HMMpropdeh3_1_pdu(pf02288) PDU1AB EP_NTERM
13915 HMMpropdeh1_1_pdu(pf02286) PDU1AB
13920 Tsp_greenishBeige PDU1AB
13925 H_robinEggBlue GRM4
13930 -pf00230 -
13935 HMM10114PocR_1_pdu1(pf12833pf10114) PDU1AB
13940 -pf01656pf07685 -
13945 -pf03186 -
13950 -pf02570 -
13955 -pf01888 -
13960 -pf00590 -
13965 -pf05175 -
13970 -pf00590 -
13975 -pf01890pf11761pf11760 -
13980 -pf00590 -
13985 -pf02571 -
BMC-P
BMC-H
BMC-T
Aldehyde dehydrogenase
PTAC
Signature enzyme
Alcohol dehydrogenase
Regulator

Salmonella enterica I. Bacteria; Proteobacteria; Gammaproteobacteria; Enterobacterales;

DJ273__locus_1

PDU1AB

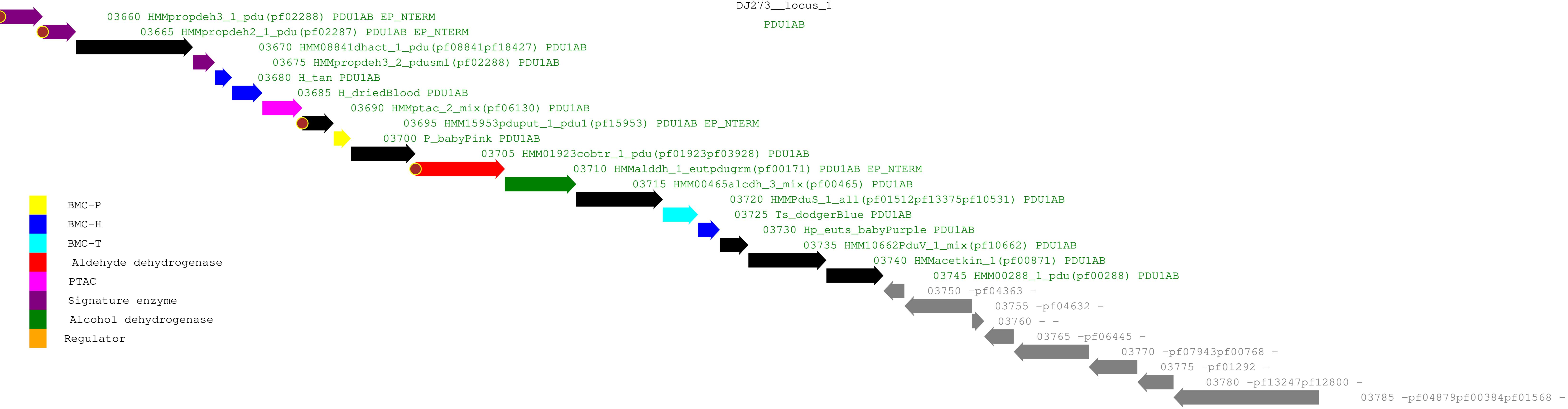

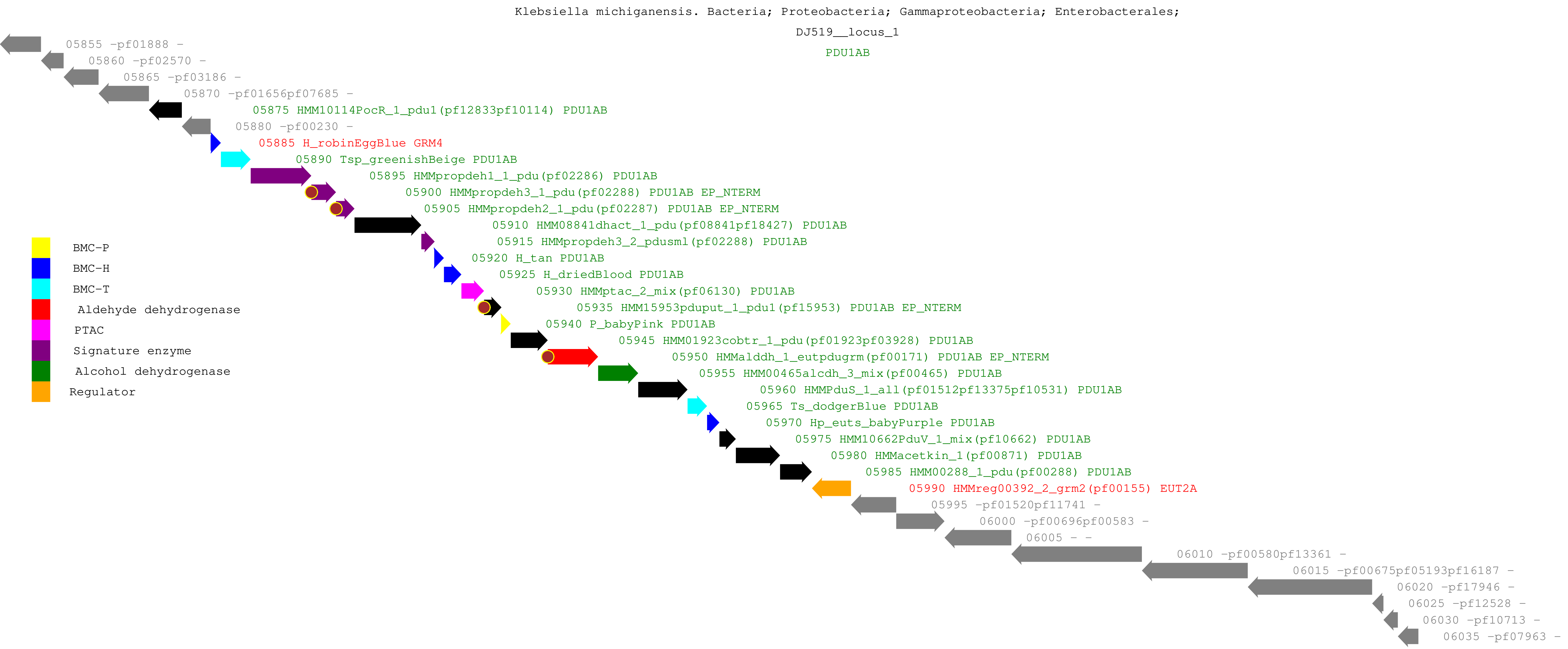
Klebsiella michiganensis. Bacteria; Proteobacteria; Gammaproteobacteria; Enterobacterales;
DJ519__locus_1
PDU1AB
05855 -pf01888 -
05860 -pf02570 -
05865 -pf03186 -
05870 -pf01656pf07685 -
05875 HMM10114PocR_1_pdu1(pf12833pf10114) PDU1AB
05880 -pf00230 -
05885 H_robinEggBlue GRM4
05890 Tsp_greenishBeige PDU1AB
05895 HMMpropdeh1_1_pdu(pf02286) PDU1AB
05900 HMMpropdeh3_1_pdu(pf02288) PDU1AB EP_NTERM
05905 HMMpropdeh2_1_pdu(pf02287) PDU1AB EP_NTERM
05910 HMM08841dhact_1_pdu(pf08841pf18427) PDU1AB
05915 HMMpropdeh3_2_pdusml(pf02288) PDU1AB
05920 H_tan PDU1AB
05925 H_driedBlood PDU1AB
05930 HMMptac_2_mix(pf06130) PDU1AB
05935 HMM15953pduput_1_pdu1(pf15953) PDU1AB EP_NTERM
05940 P_babyPink PDU1AB
05945 HMM01923cobtr_1_pdu(pf01923pf03928) PDU1AB
05950 HMMalddh_1_eutpdugrm(pf00171) PDU1AB EP_NTERM
05955 HMM00465alcdh_3_mix(pf00465) PDU1AB
05960 HMMPduS_1_all(pf01512pf13375pf10531) PDU1AB
05965 Ts_dodgerBlue PDU1AB
05970 Hp_euts_babyPurple PDU1AB
05975 HMM10662PduV_1_mix(pf10662) PDU1AB
05980 HMMacetkin_1(pf00871) PDU1AB
05985 HMM00288_1_pdu(pf00288) PDU1AB
05990 HMMreg00392_2_grm2(pf00155) EUT2A
05995 -pf01520pf11741 -
06000 -pf00696pf00583 -
06005 - -
06010 -pf00580pf13361 -
06015 -pf00675pf05193pf16187 -
06020 -pf17946 -
06025 -pf12528 -
06030 -pf10713 -
06035 -pf07963 -
BMC-P
BMC-H
BMC-T
Aldehyde dehydrogenase
PTAC
Signature enzyme
Alcohol dehydrogenase
Regulator

Klebsiella grimontii. Bacteria; Proteobacteria; Gammaproteobacteria; Enterobacterales;

DJ542__locus_3

PDU1AB

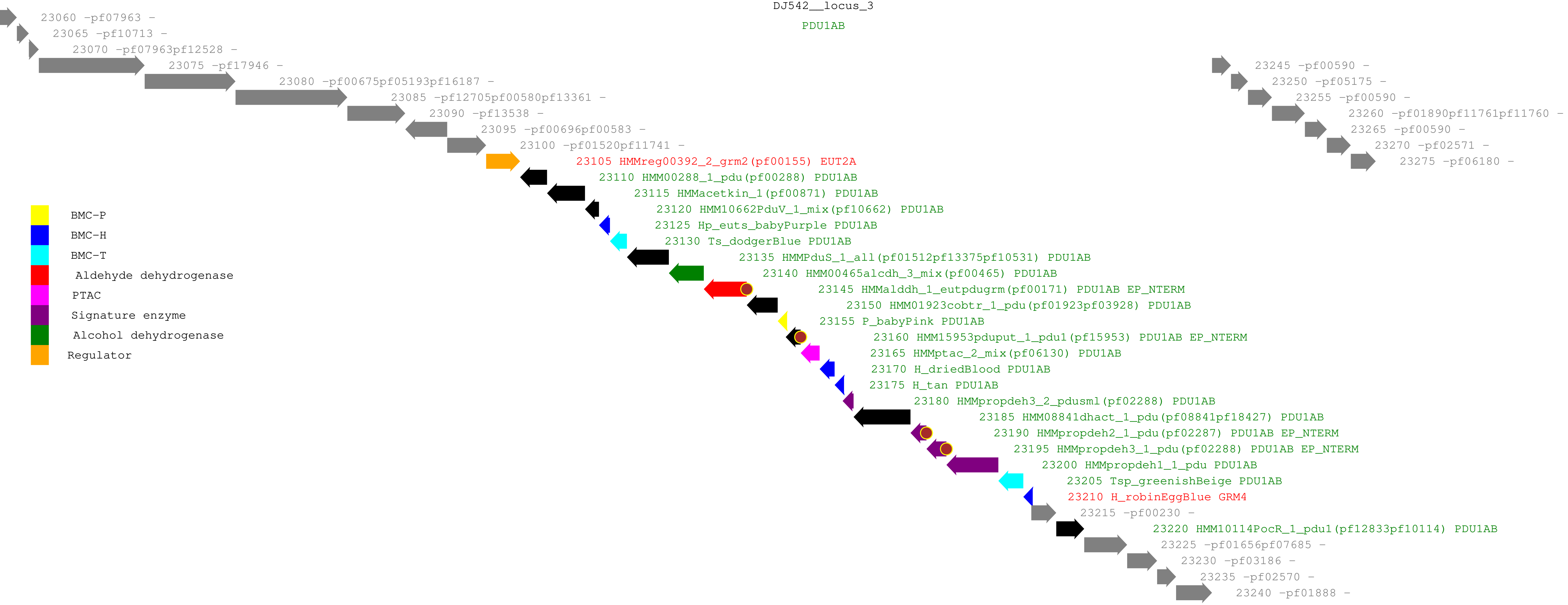

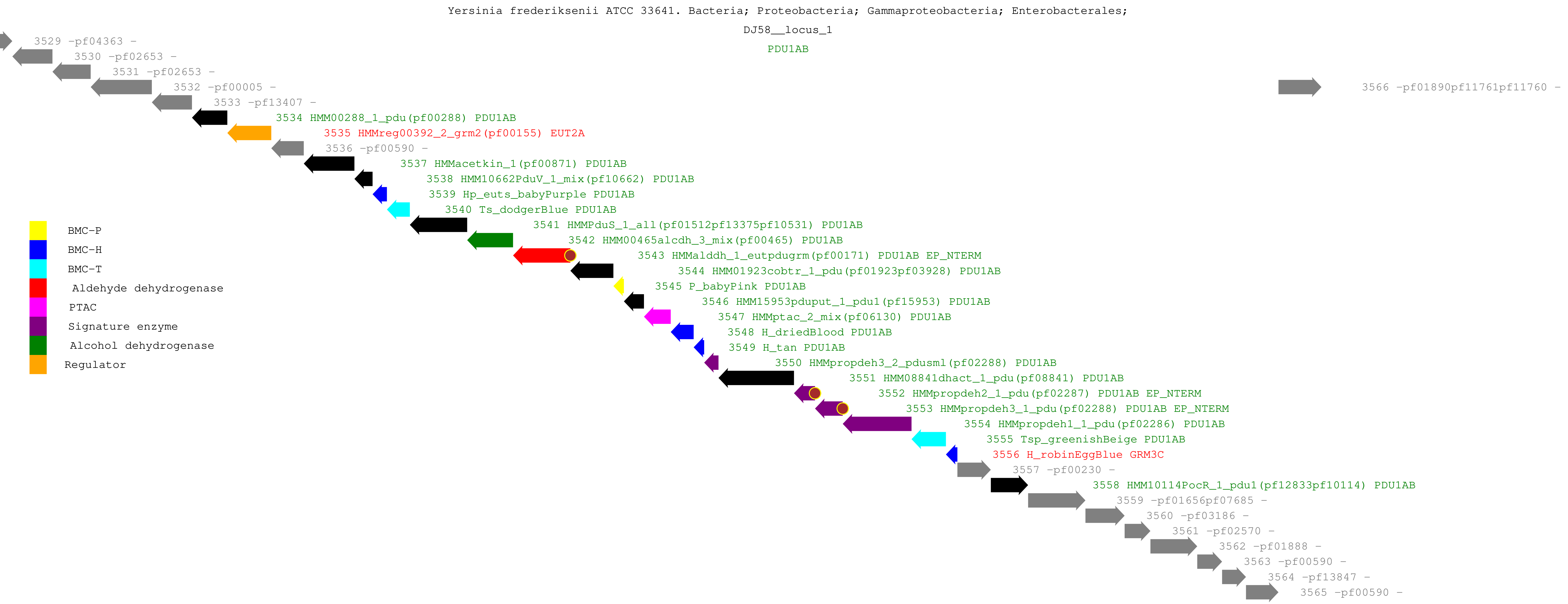

Yersinia frederiksenii ATCC 33641. Bacteria; Proteobacteria; Gammaproteobacteria; Enterobacterales;
DJ58__locus_1
PDU1AB
3529 -pf04363 -
3530 -pf02653 -
3531 -pf02653 -
3532 -pf00005 -
3566 -pf01890pf11761pf11760 -
3533 -pf13407 -
3534 HMM00288_1_pdu(pf00288) PDU1AB
3535 HMMreg00392_2_grm2(pf00155) EUT2A
3536 -pf00590 -
3537 HMMacetkin_1(pf00871) PDU1AB
3538 HMM10662PduV_1_mix(pf10662) PDU1AB
3539 Hp_euts_babyPurple PDU1AB
3540 Ts_dodgerBlue PDU1AB
3541 HMMPduS_1_all(pf01512pf13375pf10531) PDU1AB
3542 HMM00465alcdh_3_mix(pf00465) PDU1AB
3543 HMMalddh_1_eutpdugrm(pf00171) PDU1AB EP_NTERM
3544 HMM01923cobtr_1_pdu(pf01923pf03928) PDU1AB
3545 P_babyPink PDU1AB
3546 HMM15953pduput_1_pdu1(pf15953) PDU1AB
3547 HMMptac_2_mix(pf06130) PDU1AB
3548 H_driedBlood PDU1AB
3549 H_tan PDU1AB
3550 HMMpropdeh3_2_pdusml(pf02288) PDU1AB
3551 HMM08841dhact_1_pdu(pf08841) PDU1AB
3552 HMMpropdeh2_1_pdu(pf02287) PDU1AB EP_NTERM
3553 HMMpropdeh3_1_pdu(pf02288) PDU1AB EP_NTERM
3554 HMMpropdeh1_1_pdu(pf02286) PDU1AB
3555 Tsp_greenishBeige PDU1AB
3556 H_robinEggBlue GRM3C
3557 -pf00230 -
3558 HMM10114PocR_1_pdu1(pf12833pf10114) PDU1AB
3559 -pf01656pf07685 -
3560 -pf03186 -
3561 -pf02570 -
3562 -pf01888 -
3563 -pf00590 -
3564 -pf13847 -
3565 -pf00590 -
BMC-P
BMC-H
BMC-T
Aldehyde dehydrogenase
PTAC
Signature enzyme
Alcohol dehydrogenase
Regulator

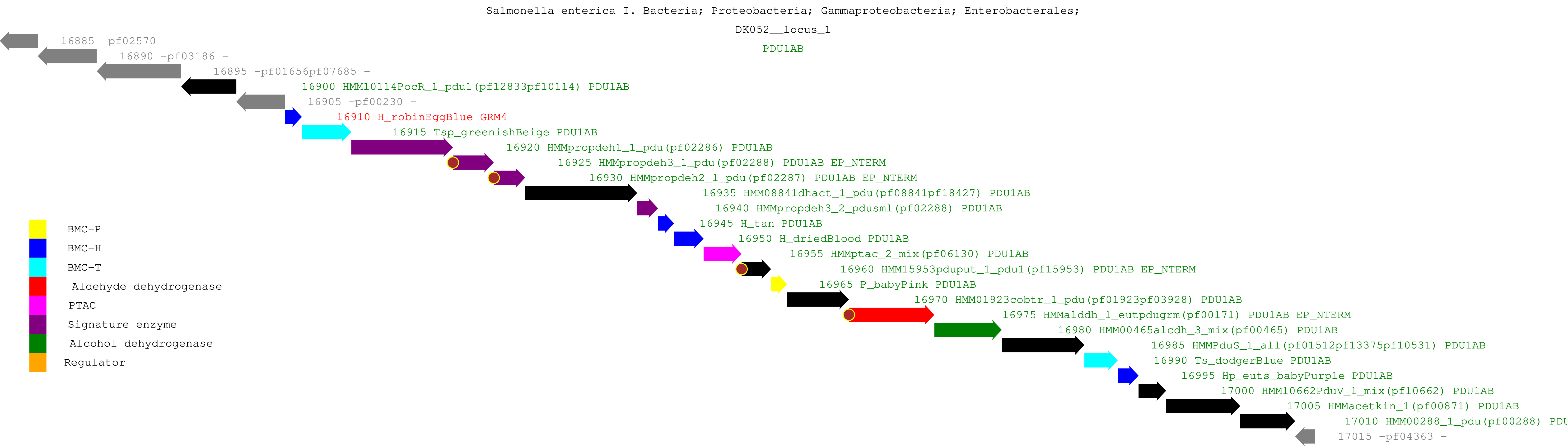
Salmonella enterica I. Bacteria; Proteobacteria; Gammaproteobacteria; Enterobacterales;
DK052__locus_1
PDU1AB
16885 -pf02570 -
16890 -pf03186 -
16895 -pf01656pf07685 -
16900 HMM10114PocR_1_pdu1(pf12833pf10114) PDU1AB
16905 -pf00230 -
16910 H_robinEggBlue GRM4
16915 Tsp_greenishBeige PDU1AB
16920 HMMpropdeh1_1_pdu(pf02286) PDU1AB
16925 HMMpropdeh3_1_pdu(pf02288) PDU1AB EP_NTERM
16930 HMMpropdeh2_1_pdu(pf02287) PDU1AB EP_NTERM
16935 HMM08841dhact_1_pdu(pf08841pf18427) PDU1AB
16940 HMMpropdeh3_2_pdusml(pf02288) PDU1AB
16945 H_tan PDU1AB
16950 H_driedBlood PDU1AB
16955 HMMptac_2_mix(pf06130) PDU1AB
16960 HMM15953pduput_1_pdu1(pf15953) PDU1AB EP_NTERM
16965 P_babyPink PDU1AB
16970 HMM01923cobtr_1_pdu(pf01923pf03928) PDU1AB
16975 HMMalddh_1_eutpdugrm(pf00171) PDU1AB EP_NTERM
16980 HMM00465alcdh_3_mix(pf00465) PDU1AB
16985 HMMPduS_1_all(pf01512pf13375pf10531) PDU1AB
16990 Ts_dodgerBlue PDU1AB
16995 Hp_euts_babyPurple PDU1AB
17000 HMM10662PduV_1_mix(pf10662) PDU1AB
17005 HMMacetkin_1(pf00871) PDU1AB
17010 HMM00288_1_pdu(pf00288) PDU[illegible]
17015 -pf04363 -
BMC-P
BMC-H
BMC-T
Aldehyde dehydrogenase
PTAC
Signature enzyme
Alcohol dehydrogenase
Regulator

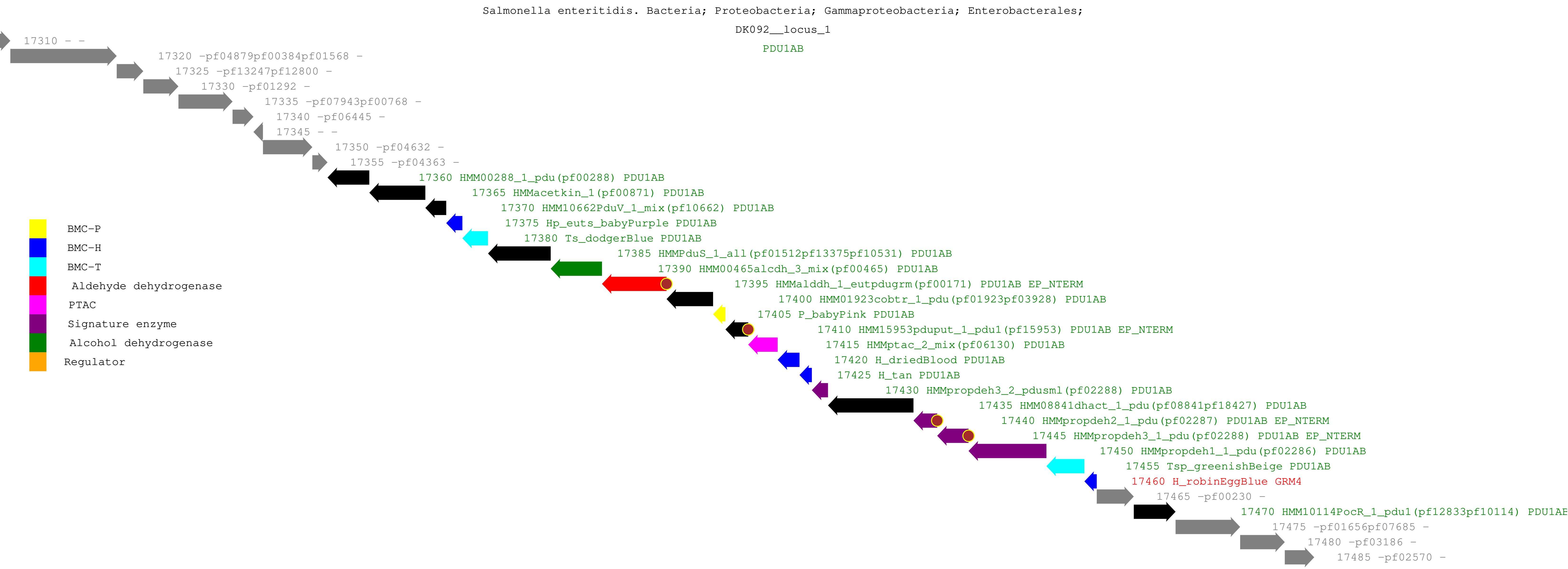
Salmonella enteritidis. Bacteria; Proteobacteria; Gammaproteobacteria; Enterobacterales;
DK092__locus_1
PDU1AB
17310 - -
17320 -pf04879pf00384pf01568 -
17325 -pf13247pf12800 -
17330 -pf01292 -
17335 -pf07943pf00768 -
17340 -pf06445 -
17345 - -
17350 -pf04632 -
17355 -pf04363 -
17360 HMM00288_1_pdu(pf00288) PDU1AB
17365 HMMacetkin_1(pf00871) PDU1AB
17370 HMM10662PduV_1_mix(pf10662) PDU1AB
17375 Hp_euts_babyPurple PDU1AB
17380 Ts_dodgerBlue PDU1AB
17385 HMMPduS_1_all(pf01512pf13375pf10531) PDU1AB
17390 HMM00465alcdh_3_mix(pf00465) PDU1AB
17395 HMMalddh_1_eutpdugrm(pf00171) PDU1AB EP_NTERM
17400 HMM01923cobtr_1_pdu(pf01923pf03928) PDU1AB
17405 P_babyPink PDU1AB
17410 HMM15953pduput_1_pdu1(pf15953) PDU1AB EP_NTERM
17415 HMMptac_2_mix(pf06130) PDU1AB
17420 H_driedBlood PDU1AB
17425 H_tan PDU1AB
17430 HMMpropdeh3_2_pdusml(pf02288) PDU1AB
17435 HMM08841dhact_1_pdu(pf08841pf18427) PDU1AB
17440 HMMpropdeh2_1_pdu(pf02287) PDU1AB EP_NTERM
17445 HMMpropdeh3_1_pdu(pf02288) PDU1AB EP_NTERM
17450 HMMpropdeh1_1_pdu(pf02286) PDU1AB
17455 Tsp_greenishBeige PDU1AB
17460 H_robinEggBlue GRM4
17465 -pf00230 -
17470 HMM10114PocR_1_pdu1(pf12833pf10114) PDU1AB
17475 -pf01656pf07685 -
17480 -pf03186 -
17485 -pf02570 -
BMC-P
BMC-H
BMC-T
Aldehyde dehydrogenase
PTAC
Signature enzyme
Alcohol dehydrogenase
Regulator

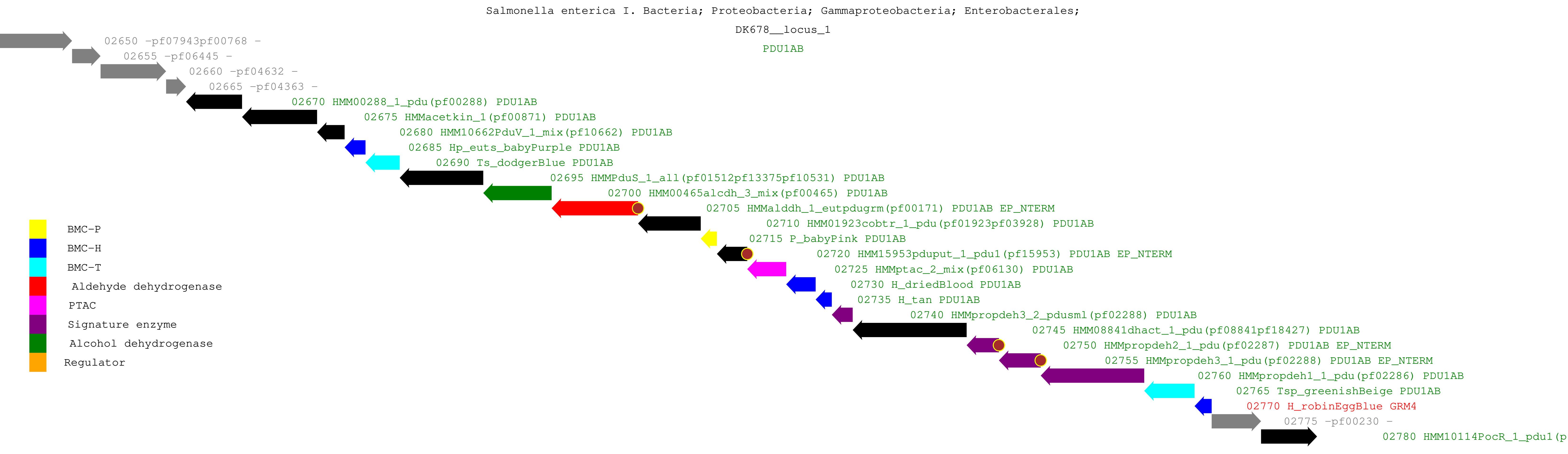
Salmonella enterica I. Bacteria; Proteobacteria; Gammaproteobacteria; Enterobacterales;
DK678__locus_1
PDU1AB
02650 -pf07943pf00768 -
02655 -pf06445 -
02660 -pf04632 -
02665 -pf04363 -
02670 HMM00288_1_pdu(pf00288) PDU1AB
02675 HMMacetkin_1(pf00871) PDU1AB
02680 HMM10662PduV_1_mix(pf10662) PDU1AB
02685 Hp_euts_babyPurple PDU1AB
02690 Ts_dodgerBlue PDU1AB
02695 HMMPduS_1_all(pf01512pf13375pf10531) PDU1AB
02700 HMM00465alcdh_3_mix(pf00465) PDU1AB
02705 HMMalddh_1_eutpdugrm(pf00171) PDU1AB EP_NTERM
02710 HMM01923cobtr_1_pdu(pf01923pf03928) PDU1AB
02715 P_babyPink PDU1AB
02720 HMM15953pduput_1_pdu1(pf15953) PDU1AB EP_NTERM
02725 HMMptac_2_mix(pf06130) PDU1AB
02730 H_driedBlood PDU1AB
02735 H_tan PDU1AB
02740 HMMpropdeh3_2_pdusml(pf02288) PDU1AB
02745 HMM08841dhact_1_pdu(pf08841pf18427) PDU1AB
02750 HMMpropdeh2_1_pdu(pf02287) PDU1AB EP_NTERM
02755 HMMpropdeh3_1_pdu(pf02288) PDU1AB EP_NTERM
02760 HMMpropdeh1_1_pdu(pf02286) PDU1AB
02765 Tsp_greenishBeige PDU1AB
02770 H_robinEggBlue GRM4
02775 -pf00230 -
02780 HMM10114PocR_1_pdu1(p
BMC-P
BMC-H
BMC-T
Aldehyde dehydrogenase
PTAC
Signature enzyme
Alcohol dehydrogenase
Regulator

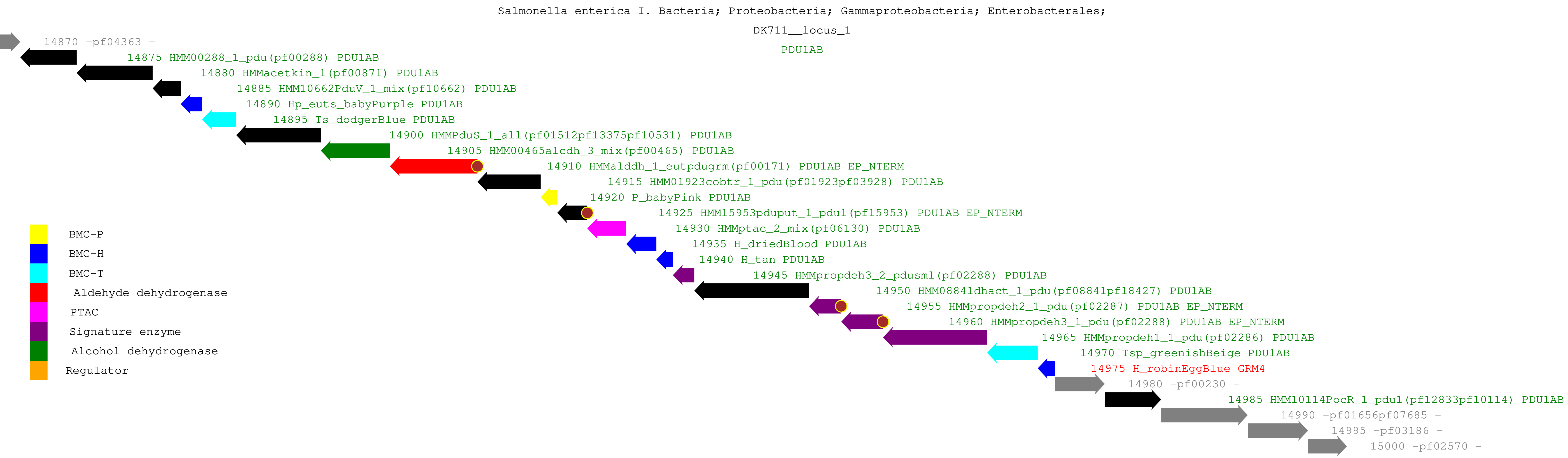
Salmonella enterica I. Bacteria; Proteobacteria; Gammaproteobacteria; Enterobacterales;
DK711__locus_1
PDU1AB
14870 -pf04363 -
14875 HMM00288_1_pdu(pf00288) PDU1AB
14880 HMMacetkin_1(pf00871) PDU1AB
14885 HMM10662PduV_1_mix(pf10662) PDU1AB
14890 Hp_euts_babyPurple PDU1AB
14895 Ts_dodgerBlue PDU1AB
14900 HMMPduS_1_all(pf01512pf13375pf10531) PDU1AB
14905 HMM00465alcdh_3_mix(pf00465) PDU1AB
14910 HMMalddh_1_eutpdugrm(pf00171) PDU1AB EP_NTERM
14915 HMM01923cobtr_1_pdu(pf01923pf03928) PDU1AB
14920 P_babyPink PDU1AB
14925 HMM15953pduput_1_pdu1(pf15953) PDU1AB EP_NTERM
14930 HMMptac_2_mix(pf06130) PDU1AB
14935 H_driedBlood PDU1AB
14940 H_tan PDU1AB
14945 HMMpropdeh3_2_pdusml(pf02288) PDU1AB
14950 HMM08841dhact_1_pdu(pf08841pf18427) PDU1AB
14955 HMMpropdeh2_1_pdu(pf02287) PDU1AB EP_NTERM
14960 HMMpropdeh3_1_pdu(pf02288) PDU1AB EP_NTERM
14965 HMMpropdeh1_1_pdu(pf02286) PDU1AB
14970 Tsp_greenishBeige PDU1AB
14975 H_robinEggBlue GRM4
14980 -pf00230 -
14985 HMM10114PocR_1_pdu1(pf12833pf10114) PDU1AB
14990 -pf01656pf07685 -
14995 -pf03186 -
15000 -pf02570 -
BMC-P
BMC-H
BMC-T
Aldehyde dehydrogenase
PTAC
Signature enzyme
Alcohol dehydrogenase
Regulator

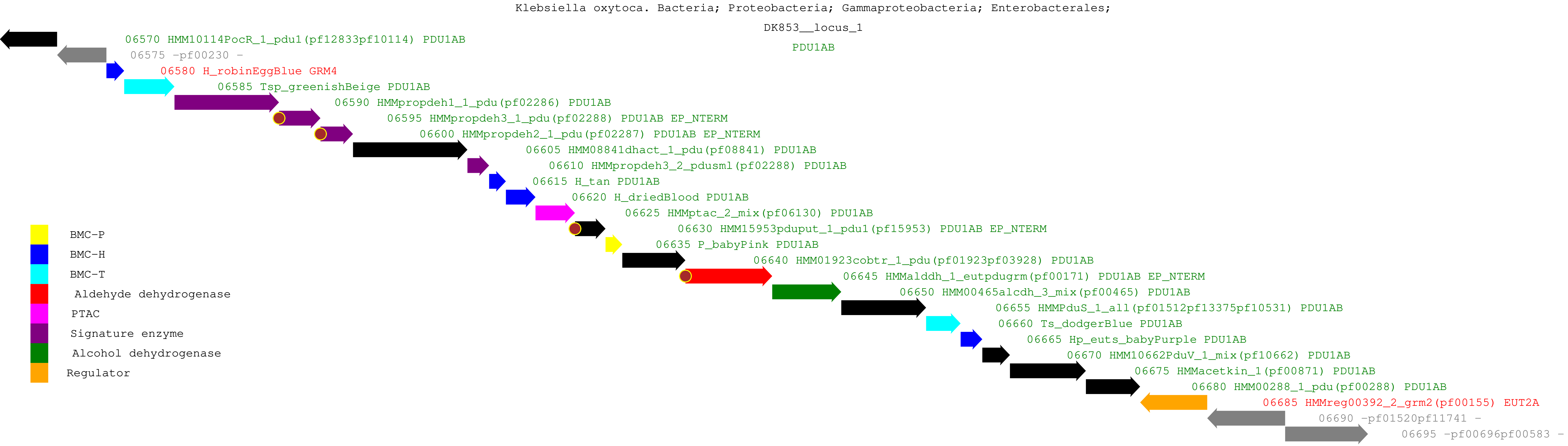

Klebsiella oxytoca. Bacteria; Proteobacteria; Gammaproteobacteria; Enterobacterales;
DK853__locus_1
PDU1AB
06570 HMM10114PocR_1_pdu1(pf12833pf10114) PDU1AB
06575 -pf00230 -
06580 H_robinEggBlue GRM4
06585 Tsp_greenishBeige PDU1AB
06590 HMMpropdeh1_1_pdu(pf02286) PDU1AB
06595 HMMpropdeh3_1_pdu(pf02288) PDU1AB EP_NTERM
06600 HMMpropdeh2_1_pdu(pf02287) PDU1AB EP_NTERM
06605 HMM08841dhact_1_pdu(pf08841) PDU1AB
06610 HMMpropdeh3_2_pdusml(pf02288) PDU1AB
06615 H_tan PDU1AB
06620 H_driedBlood PDU1AB
06625 HMMptac_2_mix(pf06130) PDU1AB
06630 HMM15953pduput_1_pdu1(pf15953) PDU1AB EP_NTERM
06635 P_babyPink PDU1AB
06640 HMM01923cobtr_1_pdu(pf01923pf03928) PDU1AB
06645 HMMalddh_1_eutpdugrm(pf00171) PDU1AB EP_NTERM
06650 HMM00465alcdh_3_mix(pf00465) PDU1AB
06655 HMMPduS_1_all(pf01512pf13375pf10531) PDU1AB
06660 Ts_dodgerBlue PDU1AB
06665 Hp_euts_babyPurple PDU1AB
06670 HMM10662PduV_1_mix(pf10662) PDU1AB
06675 HMMacetkin_1(pf00871) PDU1AB
06680 HMM00288_1_pdu(pf00288) PDU1AB
06685 HMMreg00392_2_grm2(pf00155) EUT2A
06690 -pf01520pf11741 -
06695 -pf00696pf00583 -
BMC-P
BMC-H
BMC-T
Aldehyde dehydrogenase
PTAC
Signature enzyme
Alcohol dehydrogenase
Regulator

Salmonella enterica subsp. enterica serovar Mishmarhaemek. Bacteria; Proteobacteria; Gammaproteobacteria; Enterobacterales;

DKA99__locus_2

PDU1AB

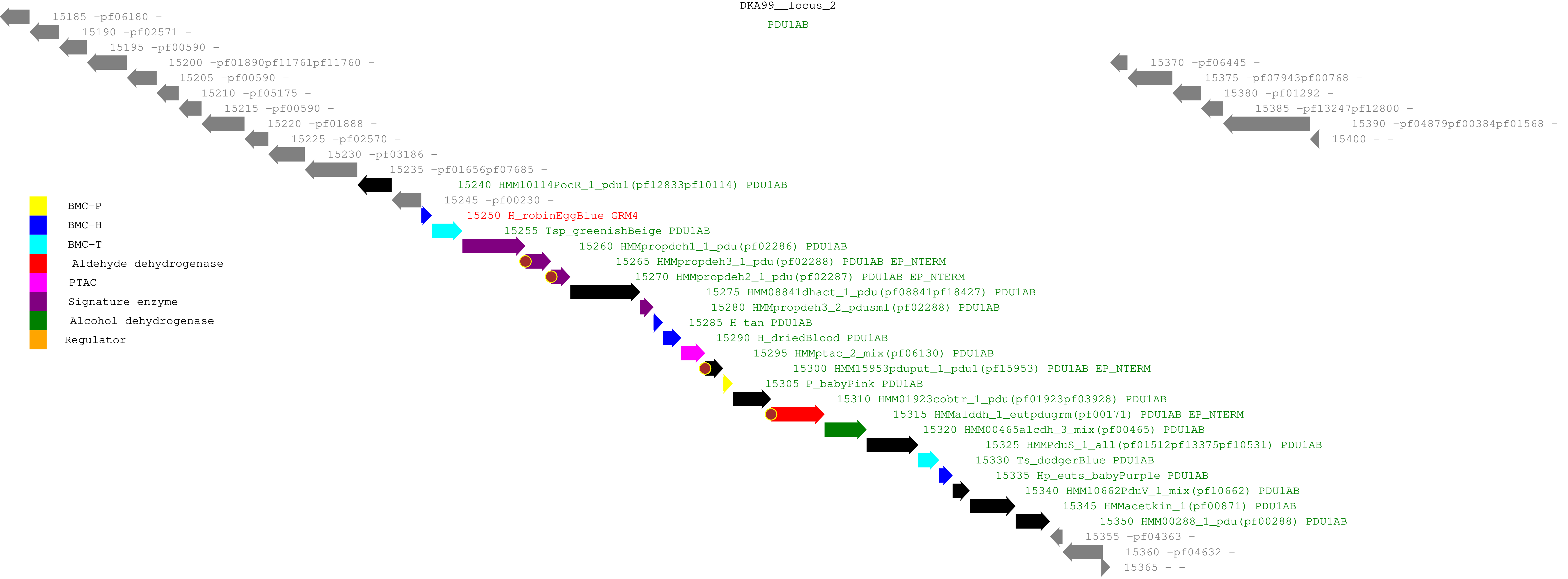

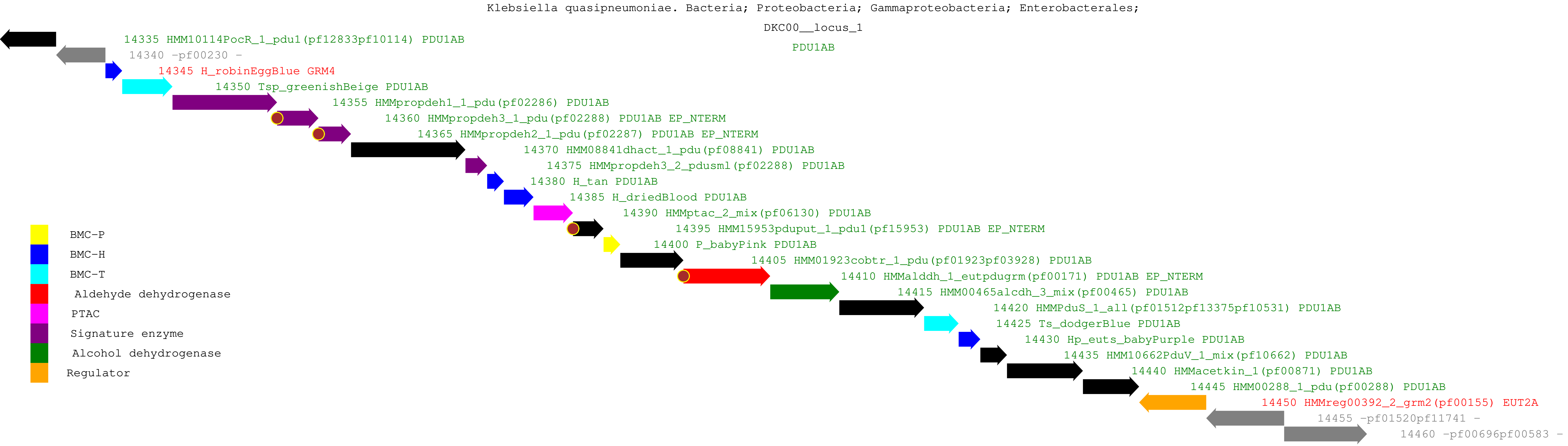
Klebsiella quasipneumoniae. Bacteria; Proteobacteria; Gammaproteobacteria; Enterobacterales;
DKC00__locus_1
PDU1AB
14335 HMM10114PocR_1_pdu1(pf12833pf10114) PDU1AB
14340 -pf00230 -
14345 H_robinEggBlue GRM4
14350 Tsp_greenishBeige PDU1AB
14355 HMMpropdeh1_1_pdu(pf02286) PDU1AB
14360 HMMpropdeh3_1_pdu(pf02288) PDU1AB EP_NTERM
14365 HMMpropdeh2_1_pdu(pf02287) PDU1AB EP_NTERM
14370 HMM08841dhact_1_pdu(pf08841) PDU1AB
14375 HMMpropdeh3_2_pdusml(pf02288) PDU1AB
14380 H_tan PDU1AB
14385 H_driedBlood PDU1AB
14390 HMMptac_2_mix(pf06130) PDU1AB
14395 HMM15953pduput_1_pdu1(pf15953) PDU1AB EP_NTERM
14400 P_babyPink PDU1AB
14405 HMM01923cobtr_1_pdu(pf01923pf03928) PDU1AB
14410 HMMalddh_1_eutpdugrm(pf00171) PDU1AB EP_NTERM
14415 HMM00465alcdh_3_mix(pf00465) PDU1AB
14420 HMMPduS_1_all(pf01512pf13375pf10531) PDU1AB
14425 Ts_dodgerBlue PDU1AB
14430 Hp_euts_babyPurple PDU1AB
14435 HMM10662PduV_1_mix(pf10662) PDU1AB
14440 HMMacetkin_1(pf00871) PDU1AB
14445 HMM00288_1_pdu(pf00288) PDU1AB
14450 HMMreg00392_2_grm2(pf00155) EUT2A
14455 -pf01520pf11741 -
14460 -pf00696pf00583 -
BMC-P
BMC-H
BMC-T
Aldehyde dehydrogenase
PTAC
Signature enzyme
Alcohol dehydrogenase
Regulator

Salmonella enterica subsp. enterica serovar Lexington. Bacteria; Proteobacteria; Gammaproteobacteria; Enterobacterales;

DKG82__locus_1

PDU1AB

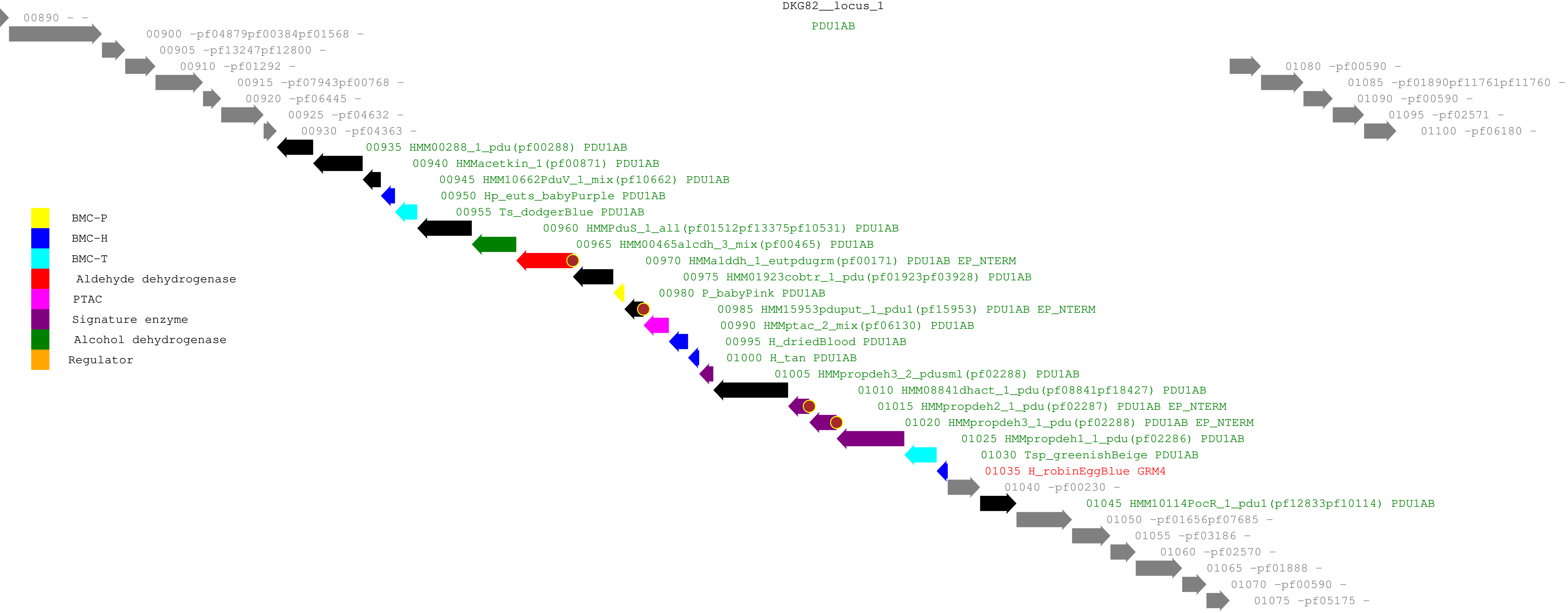

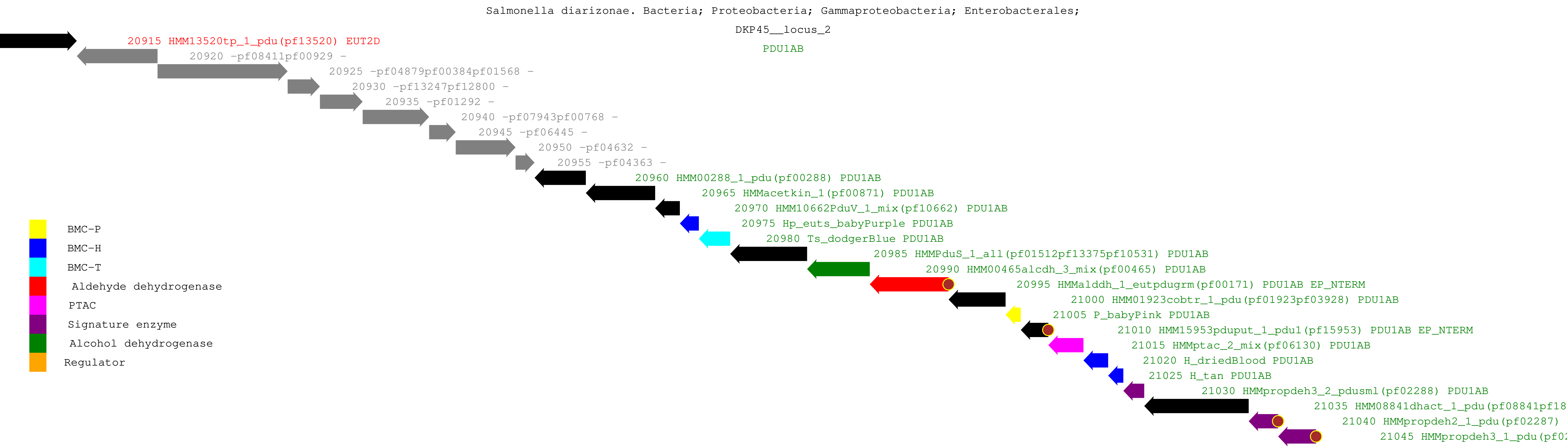
Salmonella diarizonae. Bacteria; Proteobacteria; Gammaproteobacteria; Enterobacterales;
DKP45__locus_2
PDU1AB
20915 HMM13520tp_1_pdu(pf13520) EUT2D
20920 -pf08411pf00929 -
20925 -pf04879pf00384pf01568 -
20930 -pf13247pf12800 -
20935 -pf01292 -
20940 -pf07943pf00768 -
20945 -pf06445 -
20950 -pf04632 -
20955 -pf04363 -
20960 HMM00288_1_pdu(pf00288) PDU1AB
20965 HMMacetkin_1(pf00871) PDU1AB
20970 HMM10662PduV_1_mix(pf10662) PDU1AB
20975 Hp_euts_babyPurple PDU1AB
20980 Ts_dodgerBlue PDU1AB
20985 HMMPduS_1_all(pf01512pf13375pf10531) PDU1AB
20990 HMM00465alcdh_3_mix(pf00465) PDU1AB
20995 HMMalddh_1_eutpdugrm(pf00171) PDU1AB EP_NTERM
21000 HMM01923cobtr_1_pdu(pf01923pf03928) PDU1AB
21005 P_babyPink PDU1AB
21010 HMM15953pduput_1_pdu1(pf15953) PDU1AB EP_NTERM
21015 HMMptac_2_mix(pf06130) PDU1AB
21020 H_driedBlood PDU1AB
21025 H_tan PDU1AB
21030 HMMpropdeh3_2_pdusml(pf02288) PDU1AB
21035 HMM08841dhact_1_pdu(pf08841pf18
21040 HMMpropdeh2_1_pdu(pf02287)
21045 HMMpropdeh3_1_pdu(pf0
BMC-P
BMC-H
BMC-T
Aldehyde dehydrogenase
PTAC
Signature enzyme
Alcohol dehydrogenase
Regulator

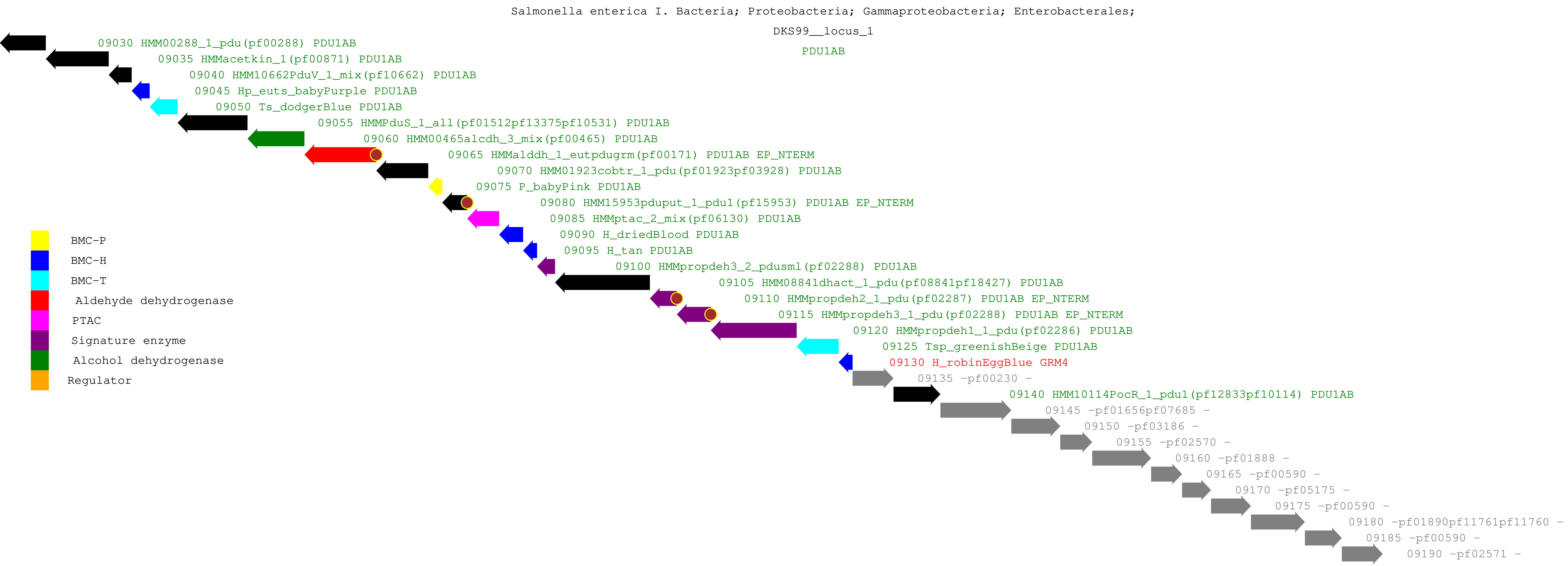
Salmonella enterica I. Bacteria; Proteobacteria; Gammaproteobacteria; Enterobacterales;
DKS99__locus_1
PDU1AB
09030 HMM00288_1_pdu(pf00288) PDU1AB
09035 HMMacetkin_1(pf00871) PDU1AB
09040 HMM10662PduV_1_mix(pf10662) PDU1AB
09045 Hp_euts_babyPurple PDU1AB
09050 Ts_dodgerBlue PDU1AB
09055 HMMPduS_1_all(pf01512pf13375pf10531) PDU1AB
09060 HMM00465alcdh_3_mix(pf00465) PDU1AB
09065 HMMalddh_1_eutpdugrm(pf00171) PDU1AB EP_NTERM
09070 HMM01923cobtr_1_pdu(pf01923pf03928) PDU1AB
09075 P_babyPink PDU1AB
09080 HMM15953pduput_1_pdu1(pf15953) PDU1AB EP_NTERM
09085 HMMptac_2_mix(pf06130) PDU1AB
09090 H_driedBlood PDU1AB
09095 H_tan PDU1AB
09100 HMMpropdeh3_2_pdusml(pf02288) PDU1AB
09105 HMM08841dhact_1_pdu(pf08841pf18427) PDU1AB
09110 HMMpropdeh2_1_pdu(pf02287) PDU1AB EP_NTERM
09115 HMMpropdeh3_1_pdu(pf02288) PDU1AB EP_NTERM
09120 HMMpropdeh1_1_pdu(pf02286) PDU1AB
09125 Tsp_greenishBeige PDU1AB
09130 H_robinEggBlue GRM4
09135 -pf00230 -
09140 HMM10114PocR_1_pdu1(pf12833pf10114) PDU1AB
09145 -pf01656pf07685 -
09150 -pf03186 -
09155 -pf02570 -
09160 -pf01888 -
09165 -pf00590 -
09170 -pf05175 -
09175 -pf00590 -
09180 -pf01890pf11761pf11760 -
09185 -pf00590 -
09190 -pf02571 -
BMC-P
BMC-H
BMC-T
Aldehyde dehydrogenase
PTAC
Signature enzyme
Alcohol dehydrogenase
Regulator

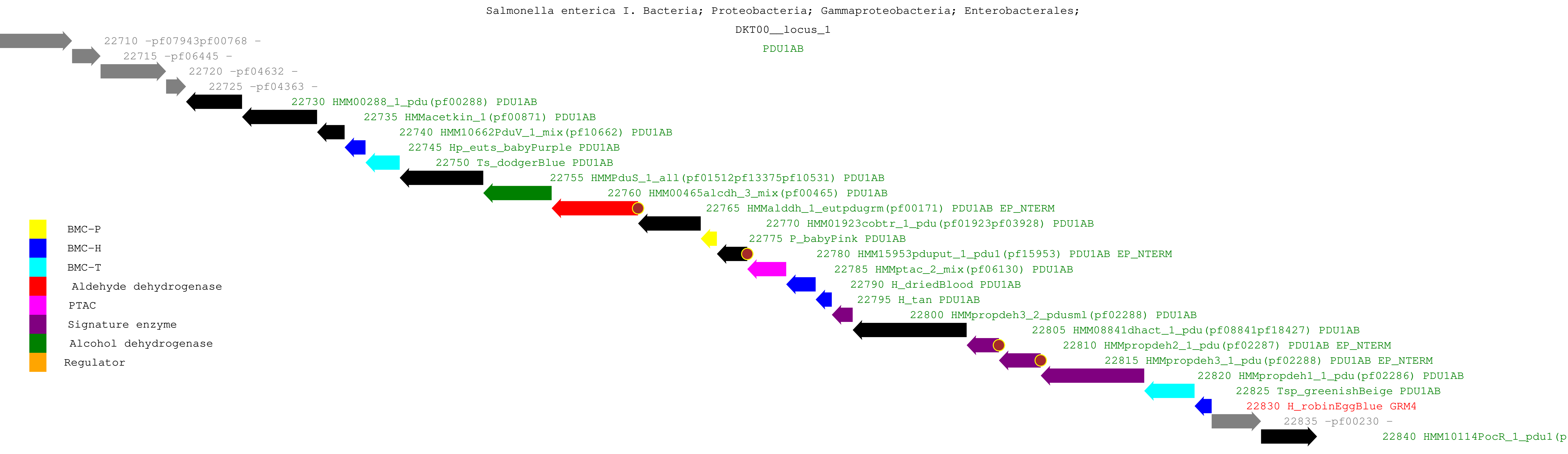
Salmonella enterica I. Bacteria; Proteobacteria; Gammaproteobacteria; Enterobacterales;
DKT00__locus_1
PDU1AB
22710 -pf07943pf00768 -
22715 -pf06445 -
22720 -pf04632 -
22725 -pf04363 -
22730 HMM00288_1_pdu(pf00288) PDU1AB
22735 HMMacetkin_1(pf00871) PDU1AB
22740 HMM10662PduV_1_mix(pf10662) PDU1AB
22745 Hp_euts_babyPurple PDU1AB
22750 Ts_dodgerBlue PDU1AB
22755 HMMPduS_1_all(pf01512pf13375pf10531) PDU1AB
22760 HMM00465alcdh_3_mix(pf00465) PDU1AB
22765 HMMaldh_1_eutpdugrm(pf00171) PDU1AB EP_NTERM
22770 HMM01923cobtr_1_pdu(pf01923pf03928) PDU1AB
22775 P_babyPink PDU1AB
22780 HMM15953pduput_1_pdu1(pf15953) PDU1AB EP_NTERM
22785 HMMptac_2_mix(pf06130) PDU1AB
22790 H_driedBlood PDU1AB
22795 H_tan PDU1AB
22800 HMMpropdeh3_2_pdusml(pf02288) PDU1AB
22805 HMM08841dhact_1_pdu(pf08841pf18427) PDU1AB
22810 HMMpropdeh2_1_pdu(pf02287) PDU1AB EP_NTERM
22815 HMMpropdeh3_1_pdu(pf02288) PDU1AB EP_NTERM
22820 HMMpropdeh1_1_pdu(pf02286) PDU1AB
22825 Tsp_greenishBeige PDU1AB
22830 H_robinEggBlue GRM4
22835 -pf00230 -
22840 HMM10114PocR_1_pdu1(p
BMC-P
BMC-H
BMC-T
Aldehyde dehydrogenase
PTAC
Signature enzyme
Alcohol dehydrogenase
Regulator

Salmonella enterica subsp. enterica serovar Mountpleasant. Bacteria; Proteobacteria; Gammaproteobacteria; Enterobacterales;

DKT33__locus_1

PDU1AB

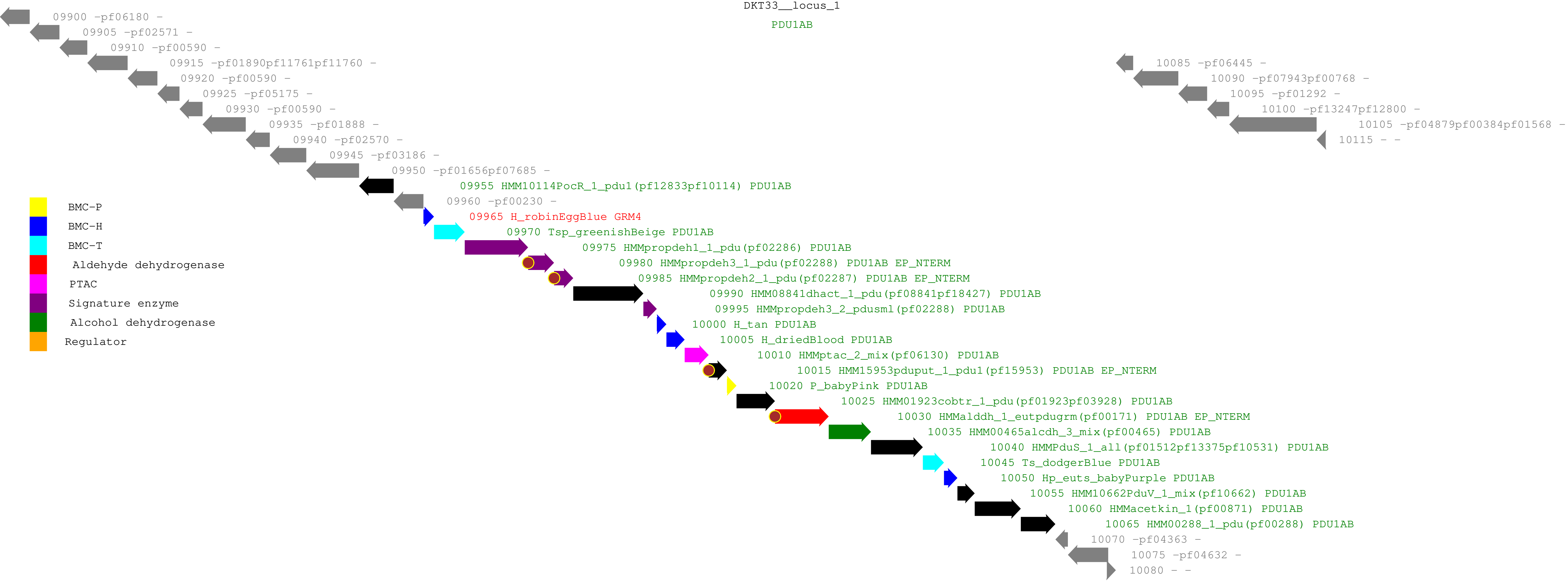

Salmonella typhimurium. Bacteria; Proteobacteria; Gammaproteobacteria; Enterobacterales;

DKU57__locus_1

PDU1AB

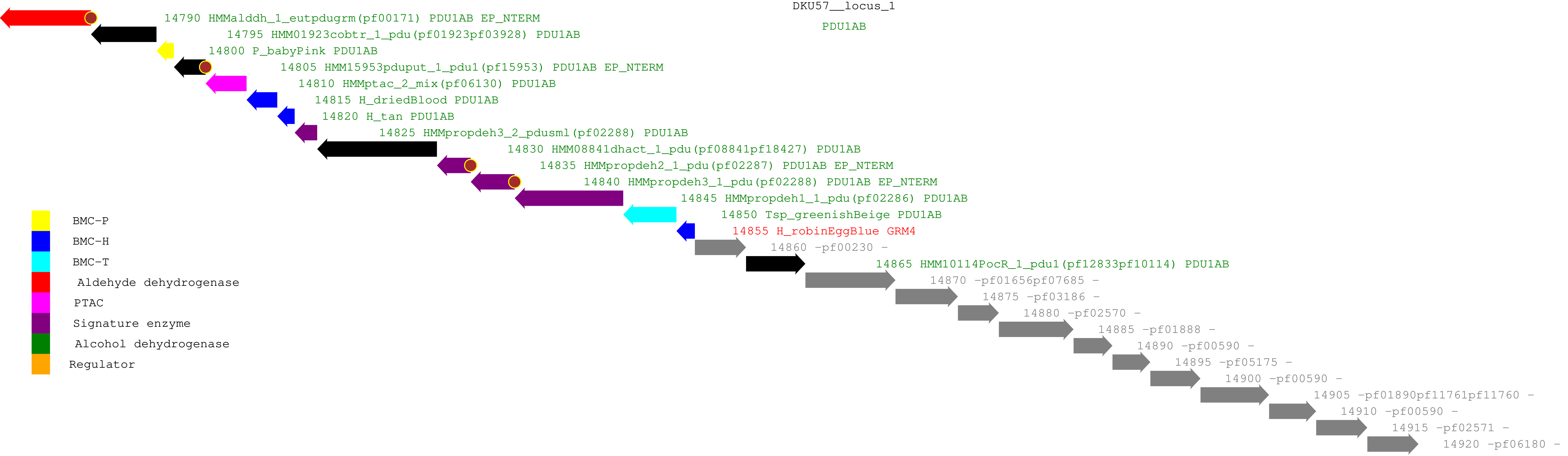

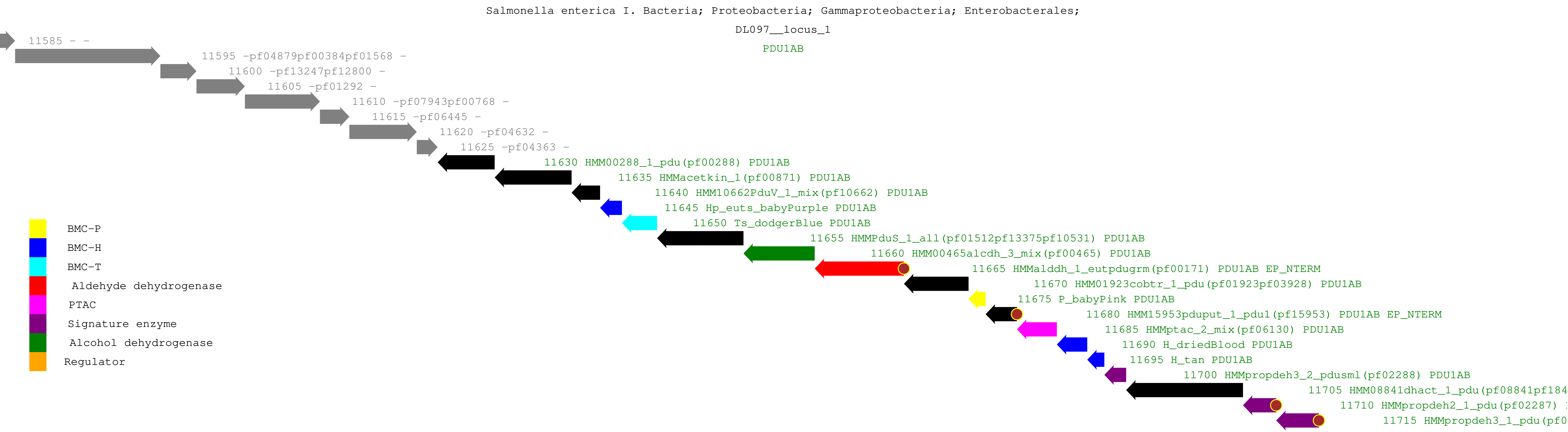
Salmonella enterica I. Bacteria; Proteobacteria; Gammaproteobacteria; Enterobacterales;
DL097__locus_1
PDU1AB
11585 - -
11595 -pf04879pf00384pf01568 -
11600 -pf13247pf12800 -
11605 -pf01292 -
11610 -pf07943pf00768 -
11615 -pf06445 -
11620 -pf04632 -
11625 -pf04363 -
11630 HMM00288_1_pdu(pf00288) PDU1AB
11635 HMMacetkin_1(pf00871) PDU1AB
11640 HMM10662PduV_1_mix(pf10662) PDU1AB
11645 Hp_euts_babyPurple PDU1AB
11650 Ts_dodgerBlue PDU1AB
11655 HMMPduS_1_all(pf01512pf13375pf10531) PDU1AB
11660 HMM00465alcdh_3_mix(pf00465) PDU1AB
11665 HMMaldh_1_eutpdugrm(pf00171) PDU1AB EP_NTERM
11670 HMM01923cobtr_1_pdu(pf01923pf03928) PDU1AB
11675 P_babyPink PDU1AB
11680 HMM15953pduput_1_pdu1(pf15953) PDU1AB EP_NTERM
11685 HMMptac_2_mix(pf06130) PDU1AB
11690 H_driedBlood PDU1AB
11695 H_tan PDU1AB
11700 HMMpropdeh3_2_pdusml(pf02288) PDU1AB
11705 HMM08841dhact_1_pdu(pf08841pf184
11710 HMMpropdeh2_1_pdu(pf02287)
11715 HMMpropdeh3_1_pdu(pf0
BMC-P
BMC-H
BMC-T
Aldehyde dehydrogenase
PTAC
Signature enzyme
Alcohol dehydrogenase
Regulator

Salmonella enterica I. Bacteria; Proteobacteria; Gammaproteobacteria; Enterobacterales;

DL111__locus_1

PDU1AB

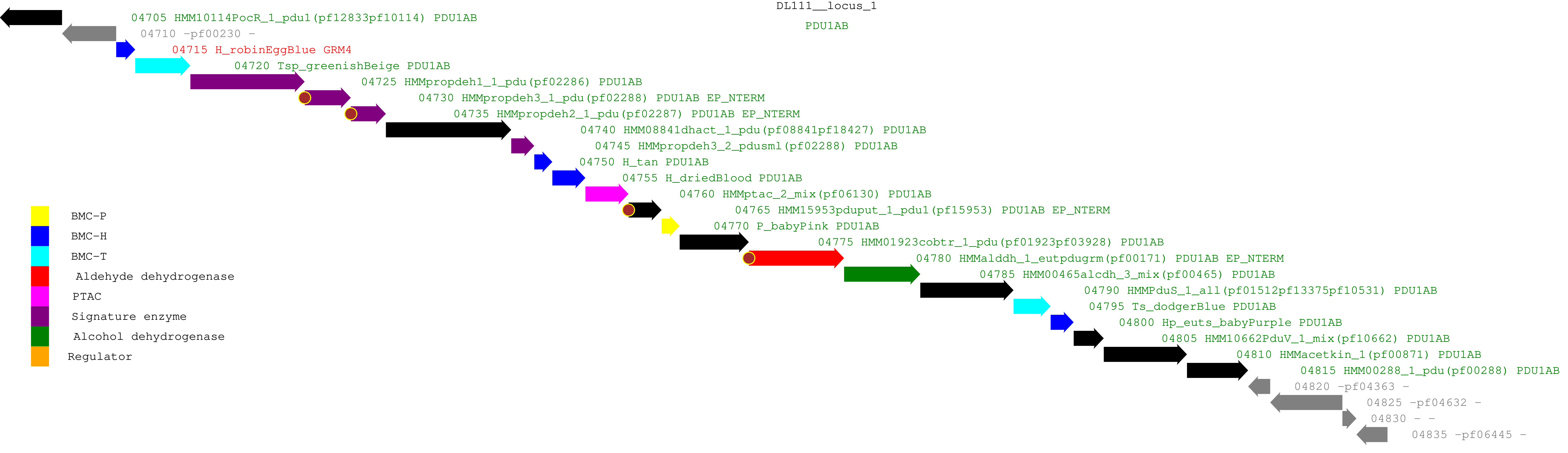

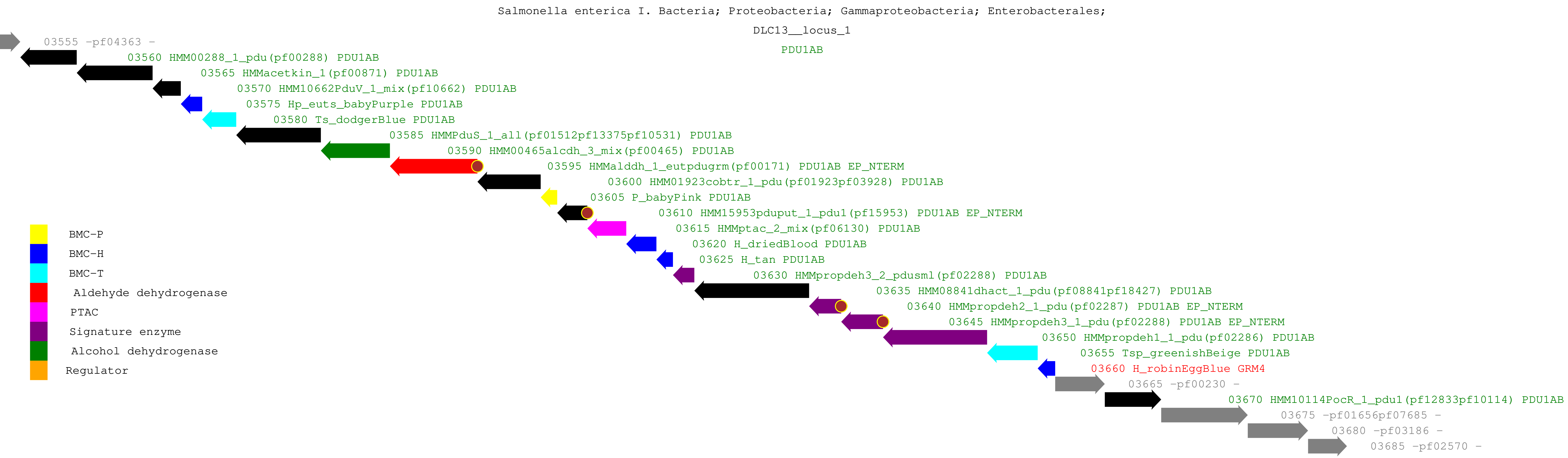
Salmonella enterica I. Bacteria; Proteobacteria; Gammaproteobacteria; Enterobacterales;
DLC13__locus_1
PDU1AB
03555 -pf04363 -
03560 HMM00288_1_pdu(pf00288) PDU1AB
03565 HMMacetkin_1(pf00871) PDU1AB
03570 HMM10662PduV_1_mix(pf10662) PDU1AB
03575 Hp_euts_babyPurple PDU1AB
03580 Ts_dodgerBlue PDU1AB
03585 HMMPduS_1_all(pf01512pf13375pf10531) PDU1AB
03590 HMM00465alcdh_3_mix(pf00465) PDU1AB
03595 HMMalddh_1_eutpdugrm(pf00171) PDU1AB EP_NTERM
03600 HMM01923cobtr_1_pdu(pf01923pf03928) PDU1AB
03605 P_babyPink PDU1AB
03610 HMM15953pduput_1_pdu1(pf15953) PDU1AB EP_NTERM
03615 HMMptac_2_mix(pf06130) PDU1AB
03620 H_driedBlood PDU1AB
03625 H_tan PDU1AB
03630 HMMpropdeh3_2_pdusml(pf02288) PDU1AB
03635 HMM08841dhact_1_pdu(pf08841pf18427) PDU1AB
03640 HMMpropdeh2_1_pdu(pf02287) PDU1AB EP_NTERM
03645 HMMpropdeh3_1_pdu(pf02288) PDU1AB EP_NTERM
03650 HMMpropdeh1_1_pdu(pf02286) PDU1AB
03655 Tsp_greenishBeige PDU1AB
03660 H_robinEggBlue GRM4
03665 -pf00230 -
03670 HMM10114PocR_1_pdu1(pf12833pf10114) PDU1AB
03675 -pf01656pf07685 -
03680 -pf03186 -
03685 -pf02570 -
BMC-P
BMC-H
BMC-T
Aldehyde dehydrogenase
PTAC
Signature enzyme
Alcohol dehydrogenase
Regulator

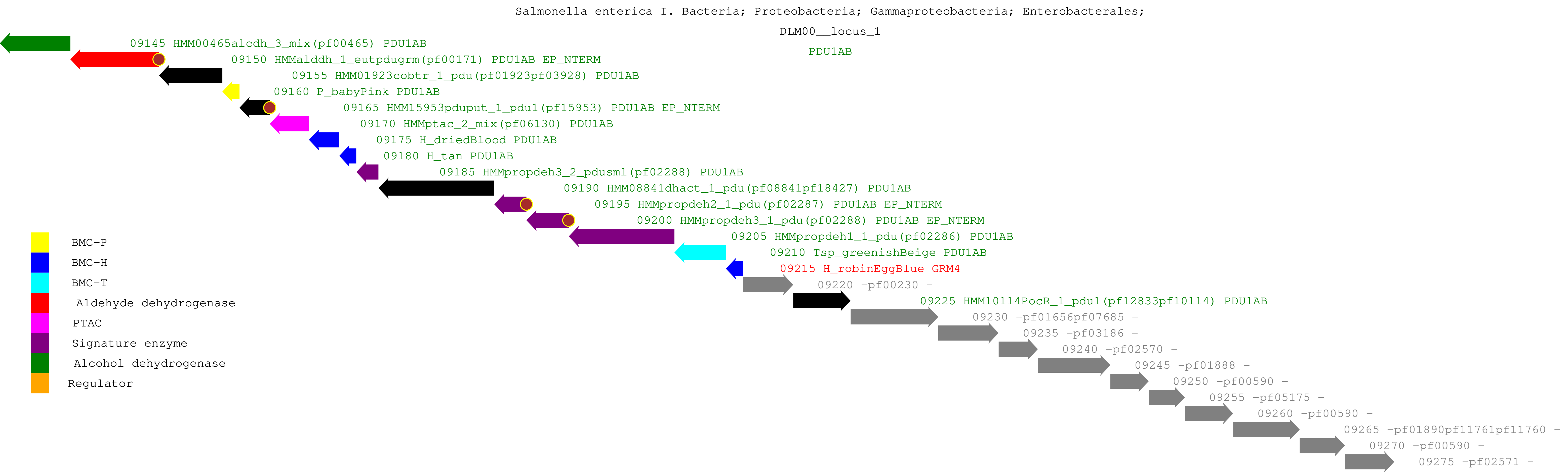
Salmonella enterica I. Bacteria; Proteobacteria; Gammaproteobacteria; Enterobacterales;
DLM00__locus_1
PDU1AB
09145 HMM00465alcdh_3_mix(pf00465) PDU1AB
09150 HMMalddh_1_eutpdugrm(pf00171) PDU1AB EP_NTERM
09155 HMM01923cobtr_1_pdu(pf01923pf03928) PDU1AB
09160 P_babyPink PDU1AB
09165 HMM15953pduput_1_pdu1(pf15953) PDU1AB EP_NTERM
09170 HMMptac_2_mix(pf06130) PDU1AB
09175 H_driedBlood PDU1AB
09180 H_tan PDU1AB
09185 HMMpropdeh3_2_pdusml(pf02288) PDU1AB
09190 HMM08841dhact_1_pdu(pf08841pf18427) PDU1AB
09195 HMMpropdeh2_1_pdu(pf02287) PDU1AB EP_NTERM
09200 HMMpropdeh3_1_pdu(pf02288) PDU1AB EP_NTERM
09205 HMMpropdeh1_1_pdu(pf02286) PDU1AB
09210 Tsp_greenishBeige PDU1AB
09215 H_robinEggBlue GRM4
09220 -pf00230 -
09225 HMM10114PocR_1_pdu1(pf12833pf10114) PDU1AB
09230 -pf01656pf07685 -
09235 -pf03186 -
09240 -pf02570 -
09245 -pf01888 -
09250 -pf00590 -
09255 -pf05175 -
09260 -pf00590 -
09265 -pf01890pf11761pf11760 -
09270 -pf00590 -
09275 -pf02571 -
BMC-P
BMC-H
BMC-T
Aldehyde dehydrogenase
PTAC
Signature enzyme
Alcohol dehydrogenase
Regulator

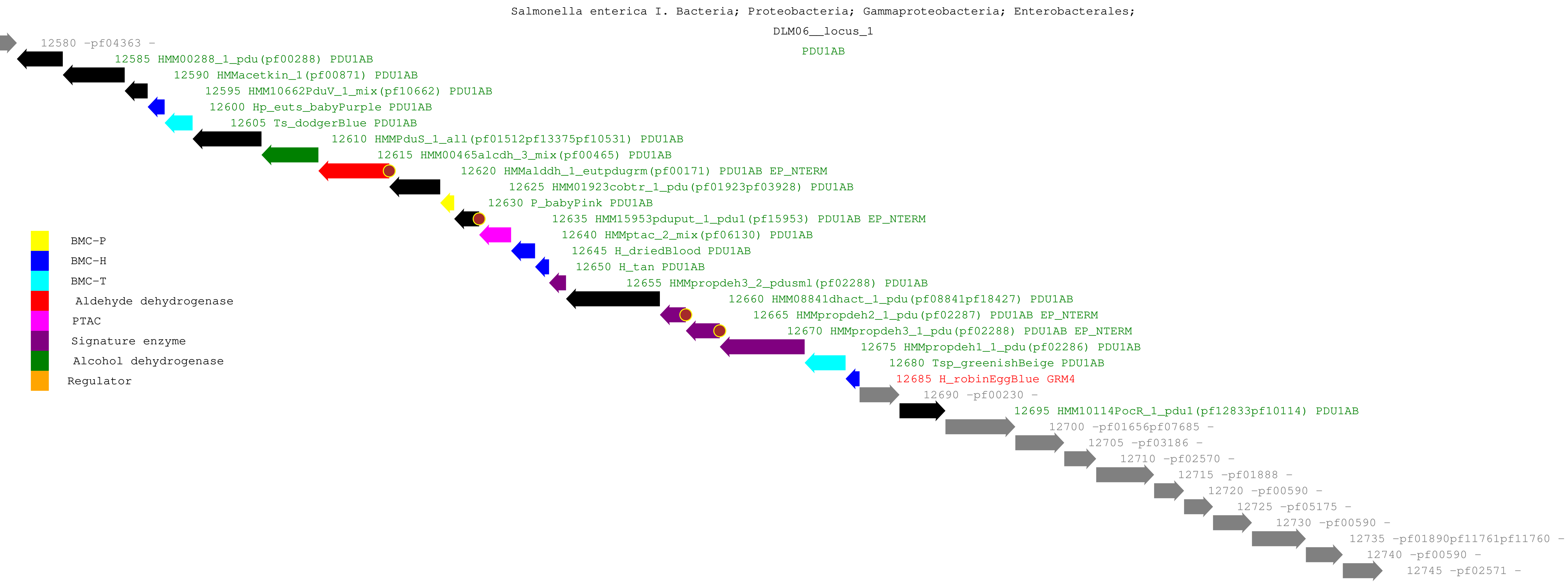
Salmonella enterica I. Bacteria; Proteobacteria; Gammaproteobacteria; Enterobacterales;
DLM06__locus_1
PDU1AB
12580 -pf04363 -
12585 HMM00288_1_pdu(pf00288) PDU1AB
12590 HMMacetkin_1(pf00871) PDU1AB
12595 HMM10662PduV_1_mix(pf10662) PDU1AB
12600 Hp_euts_babyPurple PDU1AB
12605 Ts_dodgerBlue PDU1AB
12610 HMMPduS_1_all(pf01512pf13375pf10531) PDU1AB
12615 HMM00465alcdh_3_mix(pf00465) PDU1AB
12620 HMMalddh_1_eutpdugrm(pf00171) PDU1AB EP_NTERM
12625 HMM01923cobtr_1_pdu(pf01923pf03928) PDU1AB
12630 P_babyPink PDU1AB
12635 HMM15953pduput_1_pdu1(pf15953) PDU1AB EP_NTERM
12640 HMMptac_2_mix(pf06130) PDU1AB
12645 H_driedBlood PDU1AB
12650 H_tan PDU1AB
12655 HMMpropdeh3_2_pdusml(pf02288) PDU1AB
12660 HMM08841dhact_1_pdu(pf08841pf18427) PDU1AB
12665 HMMpropdeh2_1_pdu(pf02287) PDU1AB EP_NTERM
12670 HMMpropdeh3_1_pdu(pf02288) PDU1AB EP_NTERM
12675 HMMpropdeh1_1_pdu(pf02286) PDU1AB
12680 Tsp_greenishBeige PDU1AB
12685 H_robinEggBlue GRM4
12690 -pf00230 -
12695 HMM10114PocR_1_pdu1(pf12833pf10114) PDU1AB
12700 -pf01656pf07685 -
12705 -pf03186 -
12710 -pf02570 -
12715 -pf01888 -
12720 -pf00590 -
12725 -pf05175 -
12730 -pf00590 -
12735 -pf01890pf11761pf11760 -
12740 -pf00590 -
12745 -pf02571 -
BMC-P
BMC-H
BMC-T
Aldehyde dehydrogenase
PTAC
Signature enzyme
Alcohol dehydrogenase
Regulator

Klebsiella pneumoniae. Bacteria; Proteobacteria; Gammaproteobacteria; Enterobacterales;

DM059__locus_2

PDU1AB

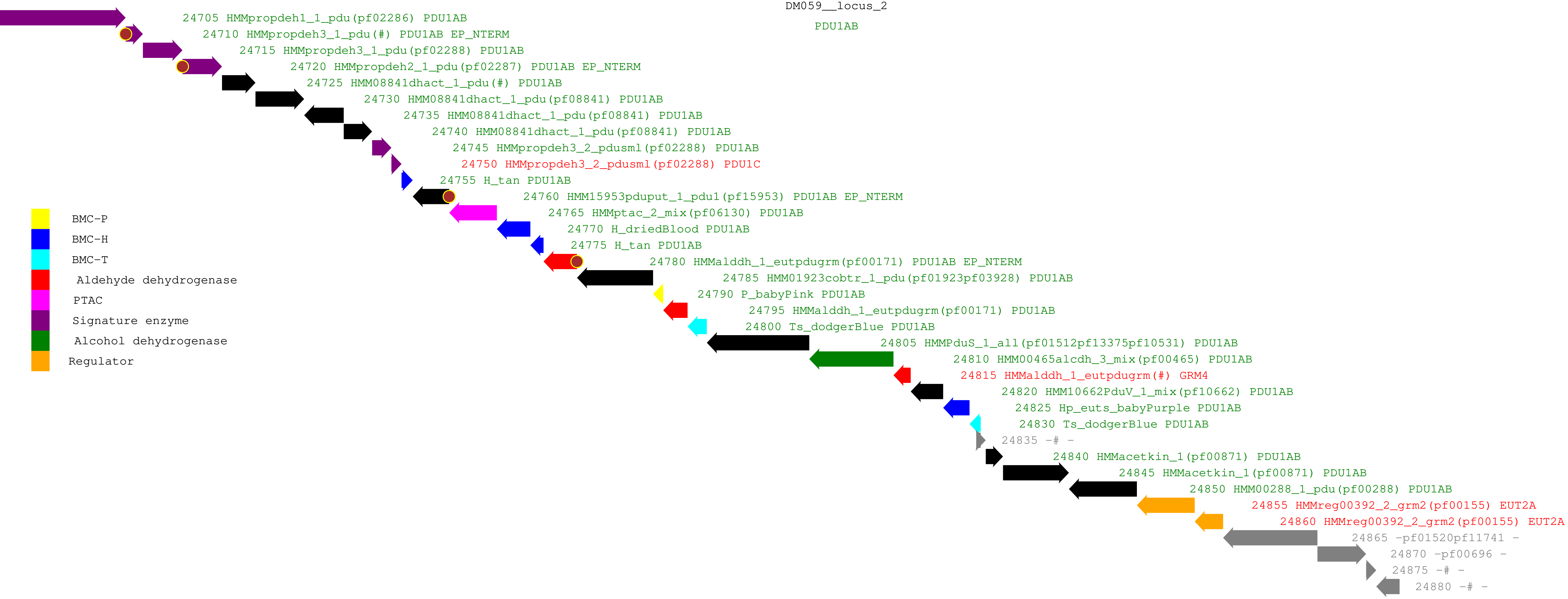

Klebsiella pneumoniae. Bacteria; Proteobacteria; Gammaproteobacteria; Enterobacterales;

DM062__locus_2

PDU1AB

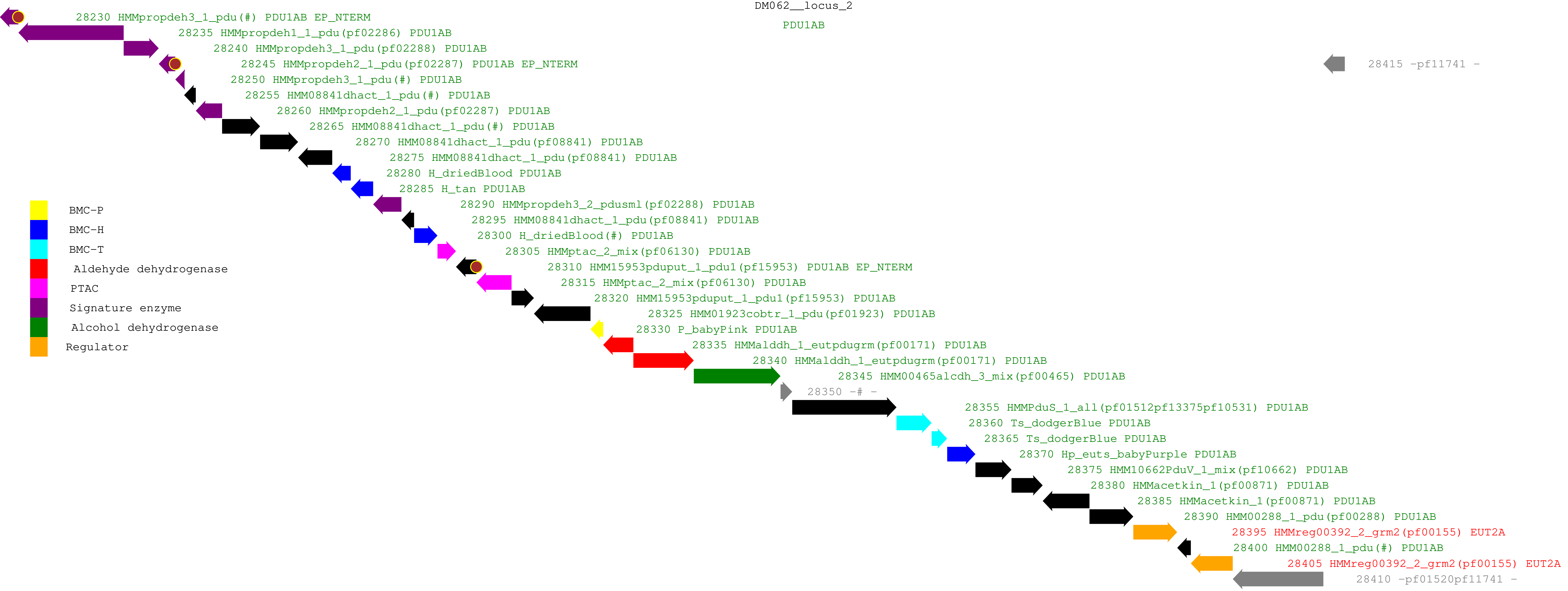

Escherichia coli. Bacteria; Proteobacteria; Gammaproteobacteria; Enterobacterales;

DM463__locus_1_main_sat

PDU1AB

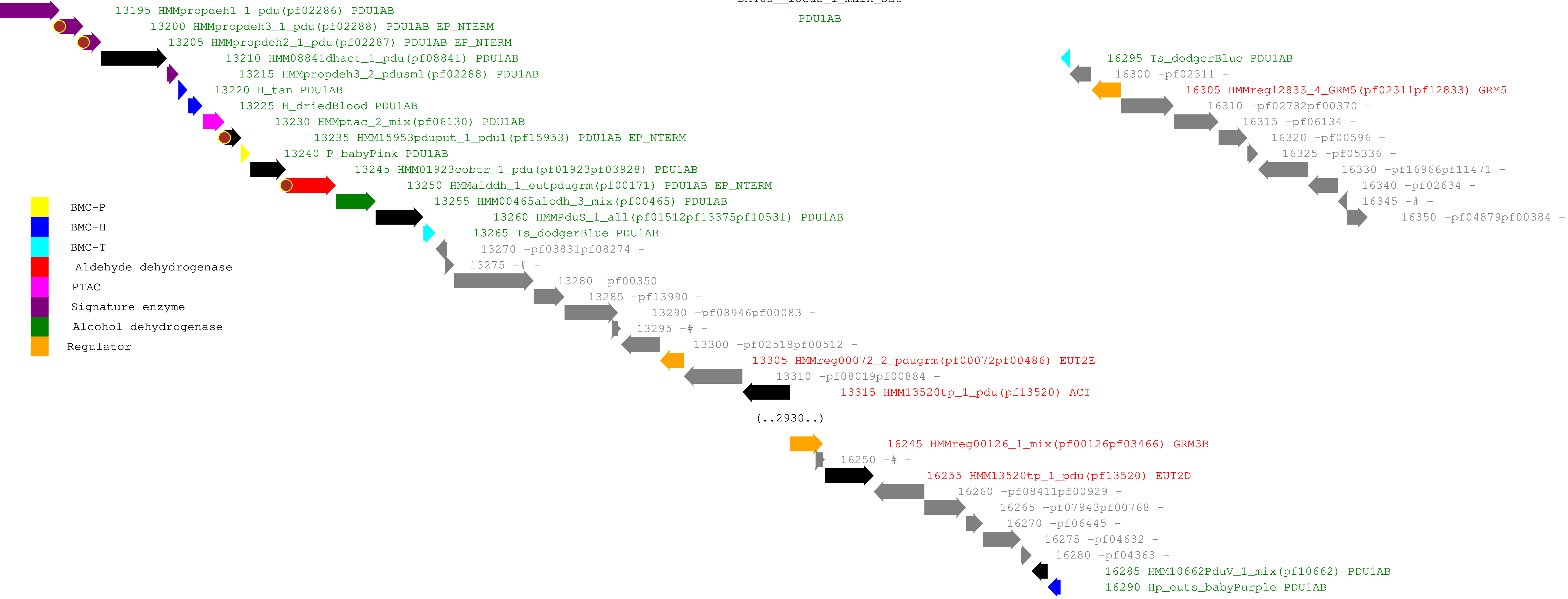

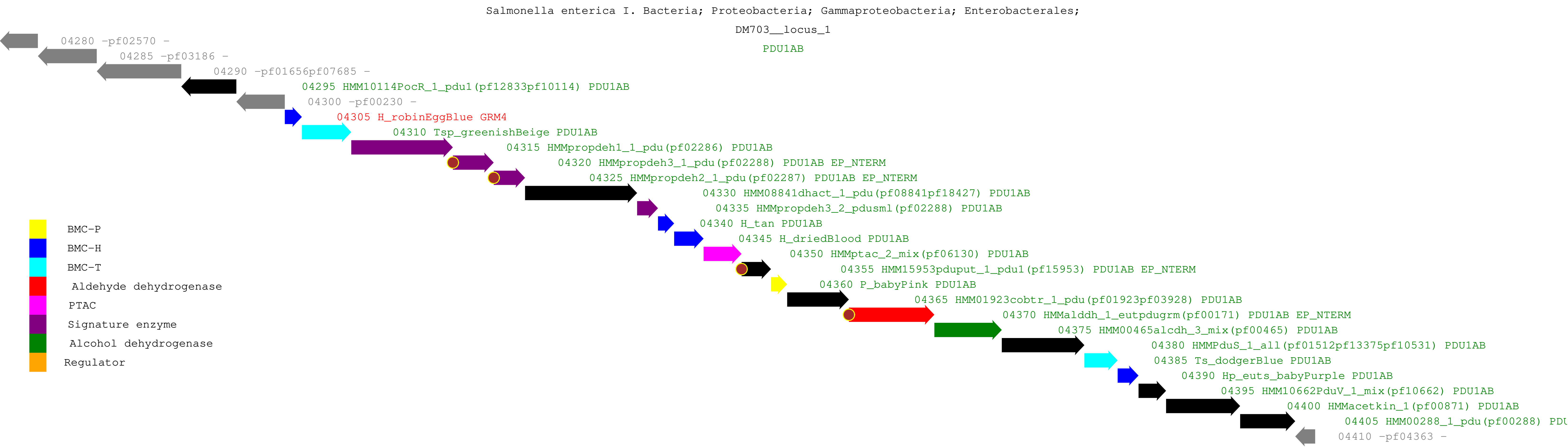
Salmonella enterica I. Bacteria; Proteobacteria; Gammaproteobacteria; Enterobacterales;
DM703__locus_1
PDU1AB
04280 -pf02570 -
04285 -pf03186 -
04290 -pf01656pf07685 -
04295 HMM10114PocR_1_pdu1(pf12833pf10114) PDU1AB
04300 -pf00230 -
04305 H_robinEggBlue GRM4
04310 Tsp_greenishBeige PDU1AB
04315 HMMpropdeh1_1_pdu(pf02286) PDU1AB
04320 HMMpropdeh3_1_pdu(pf02288) PDU1AB EP_NTERM
04325 HMMpropdeh2_1_pdu(pf02287) PDU1AB EP_NTERM
04330 HMM08841dhact_1_pdu(pf08841pf18427) PDU1AB
04335 HMMpropdeh3_2_pdusml(pf02288) PDU1AB
04340 H_tan PDU1AB
04345 H_driedBlood PDU1AB
04350 HMMptac_2_mix(pf06130) PDU1AB
04355 HMM15953pduput_1_pdu1(pf15953) PDU1AB EP_NTERM
04360 P_babyPink PDU1AB
04365 HMM01923cobtr_1_pdu(pf01923pf03928) PDU1AB
04370 HMMalddh_1_eutpdugrm(pf00171) PDU1AB EP_NTERM
04375 HMM00465alcdh_3_mix(pf00465) PDU1AB
04380 HMMPduS_1_all(pf01512pf13375pf10531) PDU1AB
04385 Ts_dodgerBlue PDU1AB
04390 Hp_euts_babyPurple PDU1AB
04395 HMM10662PduV_1_mix(pf10662) PDU1AB
04400 HMMacetkin_1(pf00871) PDU1AB
04405 HMM00288_1_pdu(pf00288) PDU
04410 -pf04363 -
BMC-P
BMC-H
BMC-T
Aldehyde dehydrogenase
PTAC
Signature enzyme
Alcohol dehydrogenase
Regulator

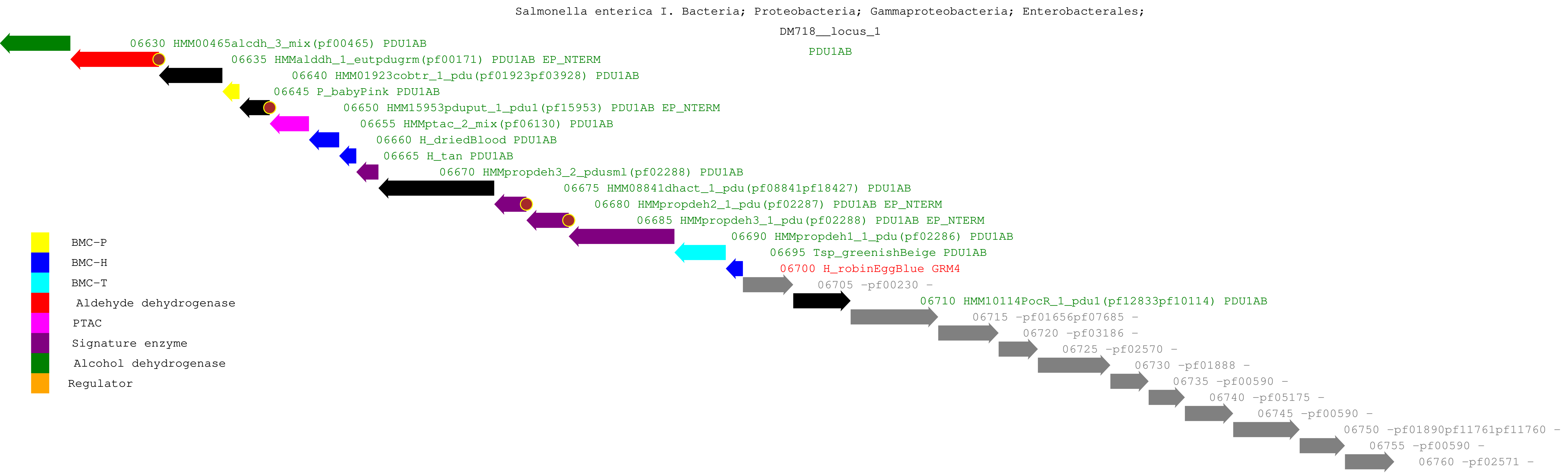
Salmonella enterica I. Bacteria; Proteobacteria; Gammaproteobacteria; Enterobacterales;
DM718__locus_1
PDU1AB
06630 HMM00465alcdh_3_mix(pf00465) PDU1AB
06635 HMMalddh_1_eutpdugrm(pf00171) PDU1AB EP_NTERM
06640 HMM01923cobtr_1_pdu(pf01923pf03928) PDU1AB
06645 P_babyPink PDU1AB
06650 HMM15953pduput_1_pdu1(pf15953) PDU1AB EP_NTERM
06655 HMMptac_2_mix(pf06130) PDU1AB
06660 H_driedBlood PDU1AB
06665 H_tan PDU1AB
06670 HMMpropdeh3_2_pdusml(pf02288) PDU1AB
06675 HMM08841dhact_1_pdu(pf08841pf18427) PDU1AB
06680 HMMpropdeh2_1_pdu(pf02287) PDU1AB EP_NTERM
06685 HMMpropdeh3_1_pdu(pf02288) PDU1AB EP_NTERM
06690 HMMpropdeh1_1_pdu(pf02286) PDU1AB
06695 Tsp_greenishBeige PDU1AB
06700 H_robinEggBlue GRM4
06705 -pf00230 -
06710 HMM10114PocR_1_pdu1(pf12833pf10114) PDU1AB
06715 -pf01656pf07685 -
06720 -pf03186 -
06725 -pf02570 -
06730 -pf01888 -
06735 -pf00590 -
06740 -pf05175 -
06745 -pf00590 -
06750 -pf01890pf11761pf11760 -
06755 -pf00590 -
06760 -pf02571 -
BMC-P
BMC-H
BMC-T
Aldehyde dehydrogenase
PTAC
Signature enzyme
Alcohol dehydrogenase
Regulator

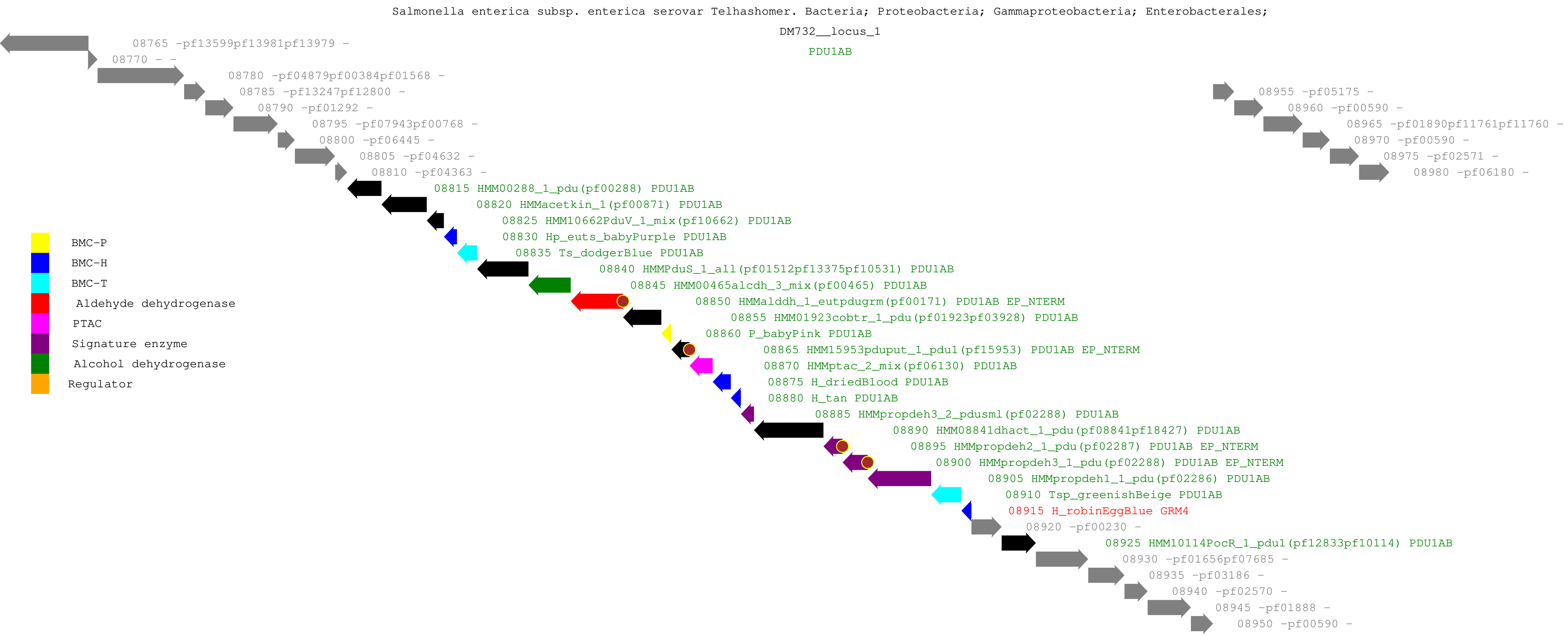
Salmonella enterica subsp. enterica serovar Telhashomer. Bacteria; Proteobacteria; Gammaproteobacteria; Enterobacterales;
DM732__locus_1
PDU1AB
08765 -pf13599pf13981pf13979 -
08770 - -
08780 -pf04879pf00384pf01568 -
08785 -pf13247pf12800 -
08790 -pf01292 -
08795 -pf07943pf00768 -
08800 -pf06445 -
08805 -pf04632 -
08810 -pf04363 -
08815 HMM00288_1_pdu(pf00288) PDU1AB
08820 HMMacetkin_1(pf00871) PDU1AB
08825 HMM10662PduV_1_mix(pf10662) PDU1AB
08830 Hp_euts_babyPurple PDU1AB
08835 Ts_dodgerBlue PDU1AB
08840 HMMPduS_1_all(pf01512pf13375pf10531) PDU1AB
08845 HMM00465alcdh_3_mix(pf00465) PDU1AB
08850 HMMalddh_1_eutpdugrm(pf00171) PDU1AB EP_NTERM
08855 HMM01923cobtr_1_pdu(pf01923pf03928) PDU1AB
08860 P_babyPink PDU1AB
08865 HMM15953pduput_1_pdu1(pf15953) PDU1AB EP_NTERM
08870 HMMptac_2_mix(pf06130) PDU1AB
08875 H_driedBlood PDU1AB
08880 H_tan PDU1AB
08885 HMMpropdeh3_2_pdusml(pf02288) PDU1AB
08890 HMM08841dhact_1_pdu(pf08841pf18427) PDU1AB
08895 HMMpropdeh2_1_pdu(pf02287) PDU1AB EP_NTERM
08900 HMMpropdeh3_1_pdu(pf02288) PDU1AB EP_NTERM
08905 HMMpropdeh1_1_pdu(pf02286) PDU1AB
08910 Tsp_greenishBeige PDU1AB
08915 H_robinEggBlue GRM4
08920 -pf00230 -
08925 HMM10114PocR_1_pdu1(pf12833pf10114) PDU1AB
08930 -pf01656pf07685 -
08935 -pf03186 -
08940 -pf02570 -
08945 -pf01888 -
08950 -pf00590 -
08955 -pf05175 -
08960 -pf00590 -
08965 -pf01890pf11761pf11760 -
08970 -pf00590 -
08975 -pf02571 -
08980 -pf06180 -
BMC-P
BMC-H
BMC-T
Aldehyde dehydrogenase
PTAC
Signature enzyme
Alcohol dehydrogenase
Regulator

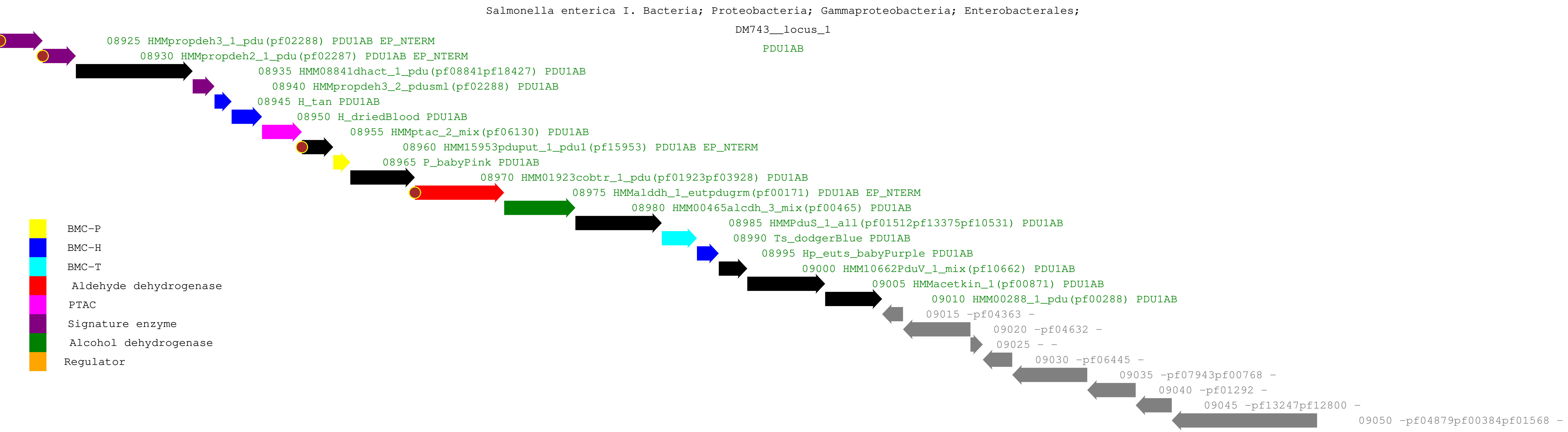

Salmonella enterica I. Bacteria; Proteobacteria; Gammaproteobacteria; Enterobacterales;
DM743__locus_1
PDU1AB
08925 HMMpropdeh3_1_pdu(pf02288) PDU1AB EP_NTERM
08930 HMMpropdeh2_1_pdu(pf02287) PDU1AB EP_NTERM
08935 HMM08841dhact_1_pdu(pf08841pf18427) PDU1AB
08940 HMMpropdeh3_2_pdusml(pf02288) PDU1AB
08945 H_tan PDU1AB
08950 H_driedBlood PDU1AB
08955 HMMptac_2_mix(pf06130) PDU1AB
08960 HMM15953pduput_1_pdu1(pf15953) PDU1AB EP_NTERM
08965 P_babyPink PDU1AB
08970 HMM01923cobtr_1_pdu(pf01923pf03928) PDU1AB
08975 HMMalddh_1_eutpdugrm(pf00171) PDU1AB EP_NTERM
08980 HMM00465alcdh_3_mix(pf00465) PDU1AB
08985 HMMPduS_1_all(pf01512pf13375pf10531) PDU1AB
08990 Ts_dodgerBlue PDU1AB
08995 Hp_euts_babyPurple PDU1AB
09000 HMM10662PduV_1_mix(pf10662) PDU1AB
09005 HMMacetkin_1(pf00871) PDU1AB
09010 HMM00288_1_pdu(pf00288) PDU1AB
09015 -pf04363 -
09020 -pf04632 -
09025 - -
09030 -pf06445 -
09035 -pf07943pf00768 -
09040 -pf01292 -
09045 -pf13247pf12800 -
09050 -pf04879pf00384pf01568 -
BMC-P
BMC-H
BMC-T
Aldehyde dehydrogenase
PTAC
Signature enzyme
Alcohol dehydrogenase
Regulator

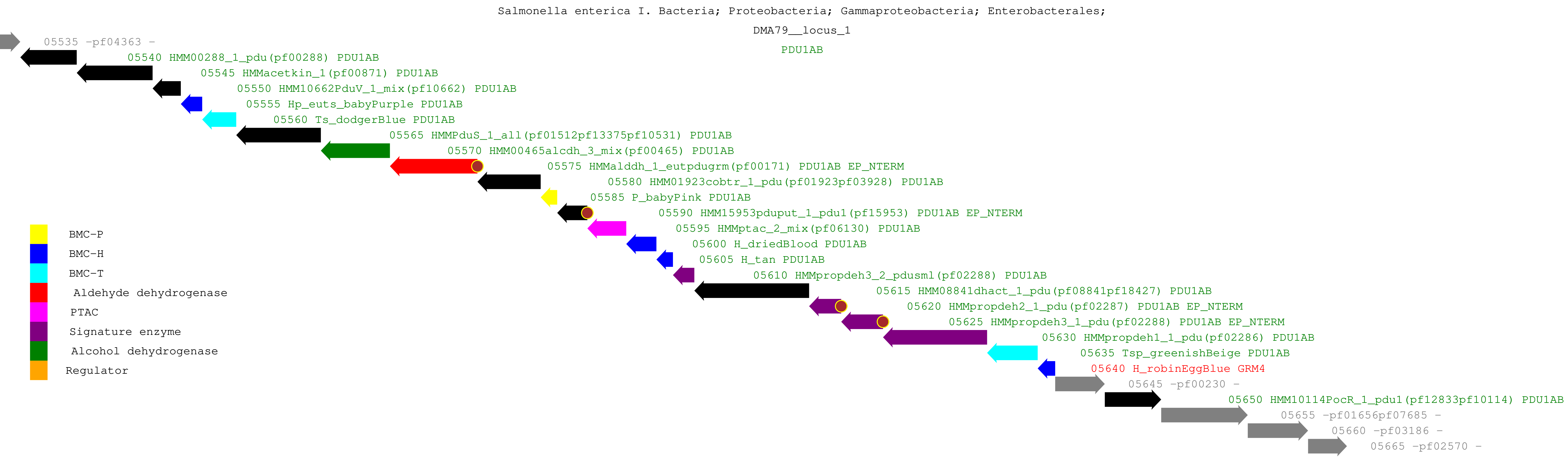
Salmonella enterica I. Bacteria; Proteobacteria; Gammaproteobacteria; Enterobacterales;
DMA79__locus_1
PDU1AB
05535 -pf04363 -
05540 HMM00288_1_pdu(pf00288) PDU1AB
05545 HMMacetkin_1(pf00871) PDU1AB
05550 HMM10662PduV_1_mix(pf10662) PDU1AB
05555 Hp_euts_babyPurple PDU1AB
05560 Ts_dodgerBlue PDU1AB
05565 HMMPduS_1_all(pf01512pf13375pf10531) PDU1AB
05570 HMM00465alcdh_3_mix(pf00465) PDU1AB
05575 HMMalddh_1_eutpdugrm(pf00171) PDU1AB EP_NTERM
05580 HMM01923cobtr_1_pdu(pf01923pf03928) PDU1AB
05585 P_babyPink PDU1AB
05590 HMM15953pduput_1_pdu1(pf15953) PDU1AB EP_NTERM
05595 HMMptac_2_mix(pf06130) PDU1AB
05600 H_driedBlood PDU1AB
05605 H_tan PDU1AB
05610 HMMpropdeh3_2_pdusml(pf02288) PDU1AB
05615 HMM08841dhact_1_pdu(pf08841pf18427) PDU1AB
05620 HMMpropdeh2_1_pdu(pf02287) PDU1AB EP_NTERM
05625 HMMpropdeh3_1_pdu(pf02288) PDU1AB EP_NTERM
05630 HMMpropdeh1_1_pdu(pf02286) PDU1AB
05635 Tsp_greenishBeige PDU1AB
05640 H_robinEggBlue GRM4
05645 -pf00230 -
05650 HMM10114PocR_1_pdu1(pf12833pf10114) PDU1AB
05655 -pf01656pf07685 -
05660 -pf03186 -
05665 -pf02570 -
BMC-P
BMC-H
BMC-T
Aldehyde dehydrogenase
PTAC
Signature enzyme
Alcohol dehydrogenase
Regulator

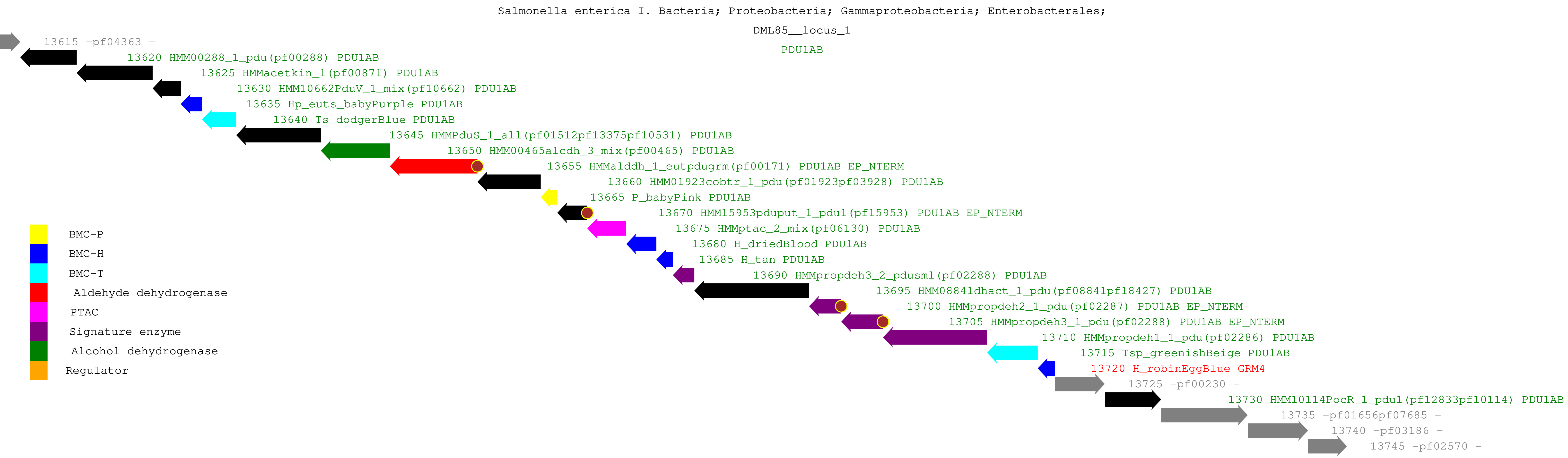
Salmonella enterica I. Bacteria; Proteobacteria; Gammaproteobacteria; Enterobacterales;
DML85__locus_1
PDU1AB
13615 -pf04363 -
13620 HMM00288_1_pdu(pf00288) PDU1AB
13625 HMMacetkin_1(pf00871) PDU1AB
13630 HMM10662PduV_1_mix(pf10662) PDU1AB
13635 Hp_euts_babyPurple PDU1AB
13640 Ts_dodgerBlue PDU1AB
13645 HMMPduS_1_all(pf01512pf13375pf10531) PDU1AB
13650 HMM00465alcdh_3_mix(pf00465) PDU1AB
13655 HMMalddh_1_eutpdugrm(pf00171) PDU1AB EP_NTERM
13660 HMM01923cobtr_1_pdu(pf01923pf03928) PDU1AB
13665 P_babyPink PDU1AB
13670 HMM15953pduput_1_pdu1(pf15953) PDU1AB EP_NTERM
13675 HMMptac_2_mix(pf06130) PDU1AB
13680 H_driedBlood PDU1AB
13685 H_tan PDU1AB
13690 HMMpropdeh3_2_pdusml(pf02288) PDU1AB
13695 HMM08841dhact_1_pdu(pf08841pf18427) PDU1AB
13700 HMMpropdeh2_1_pdu(pf02287) PDU1AB EP_NTERM
13705 HMMpropdeh3_1_pdu(pf02288) PDU1AB EP_NTERM
13710 HMMpropdeh1_1_pdu(pf02286) PDU1AB
13715 Tsp_greenishBeige PDU1AB
13720 H_robinEggBlue GRM4
13725 -pf00230 -
13730 HMM10114PocR_1_pdu1(pf12833pf10114) PDU1AB
13735 -pf01656pf07685 -
13740 -pf03186 -
13745 -pf02570 -
BMC-P
BMC-H
BMC-T
Aldehyde dehydrogenase
PTAC
Signature enzyme
Alcohol dehydrogenase
Regulator

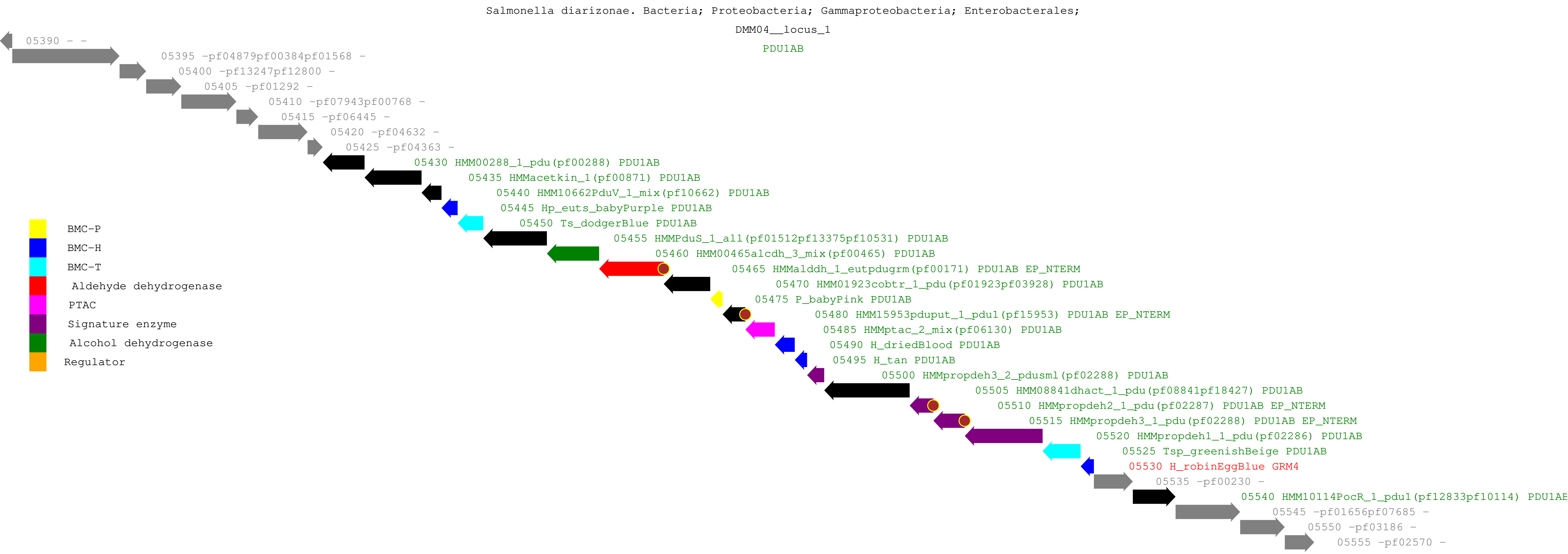
Salmonella diarizonae. Bacteria; Proteobacteria; Gammaproteobacteria; Enterobacterales;
DMM04__locus_1
PDU1AB
05390 - -
05395 -pf04879pf00384pf01568 -
05400 -pf13247pf12800 -
05405 -pf01292 -
05410 -pf07943pf00768 -
05415 -pf06445 -
05420 -pf04632 -
05425 -pf04363 -
05430 HMM00288_1_pdu(pf00288) PDU1AB
05435 HMMacetkin_1(pf00871) PDU1AB
05440 HMM10662PduV_1_mix(pf10662) PDU1AB
05445 Hp_euts_babyPurple PDU1AB
05450 Ts_dodgerBlue PDU1AB
05455 HMMPduS_1_all(pf01512pf13375pf10531) PDU1AB
05460 HMM00465alcdh_3_mix(pf00465) PDU1AB
05465 HMMaldh_1_eutpdugrm(pf00171) PDU1AB EP_NTERM
05470 HMM01923cobtr_1_pdu(pf01923pf03928) PDU1AB
05475 P_babyPink PDU1AB
05480 HMM15953pduput_1_pdu1(pf15953) PDU1AB EP_NTERM
05485 HMMptac_2_mix(pf06130) PDU1AB
05490 H_driedBlood PDU1AB
05495 H_tan PDU1AB
05500 HMMpropdeh3_2_pdusml(pf02288) PDU1AB
05505 HMM08841dhact_1_pdu(pf08841pf18427) PDU1AB
05510 HMMpropdeh2_1_pdu(pf02287) PDU1AB EP_NTERM
05515 HMMpropdeh3_1_pdu(pf02288) PDU1AB EP_NTERM
05520 HMMpropdeh1_1_pdu(pf02286) PDU1AB
05525 Tsp_greenishBeige PDU1AB
05530 H_robinEggBlue GRM4
05535 -pf00230 -
05540 HMM10114PocR_1_pdu1(pf12833pf10114) PDU1A
05545 -pf01656pf07685 -
05550 -pf03186 -
05555 -pf02570 -
BMC-P
BMC-H
BMC-T
Aldehyde dehydrogenase
PTAC
Signature enzyme
Alcohol dehydrogenase
Regulator

Salmonella enterica I. Bacteria; Proteobacteria; Gammaproteobacteria; Enterobacterales;

DMO79__locus_1

PDU1AB

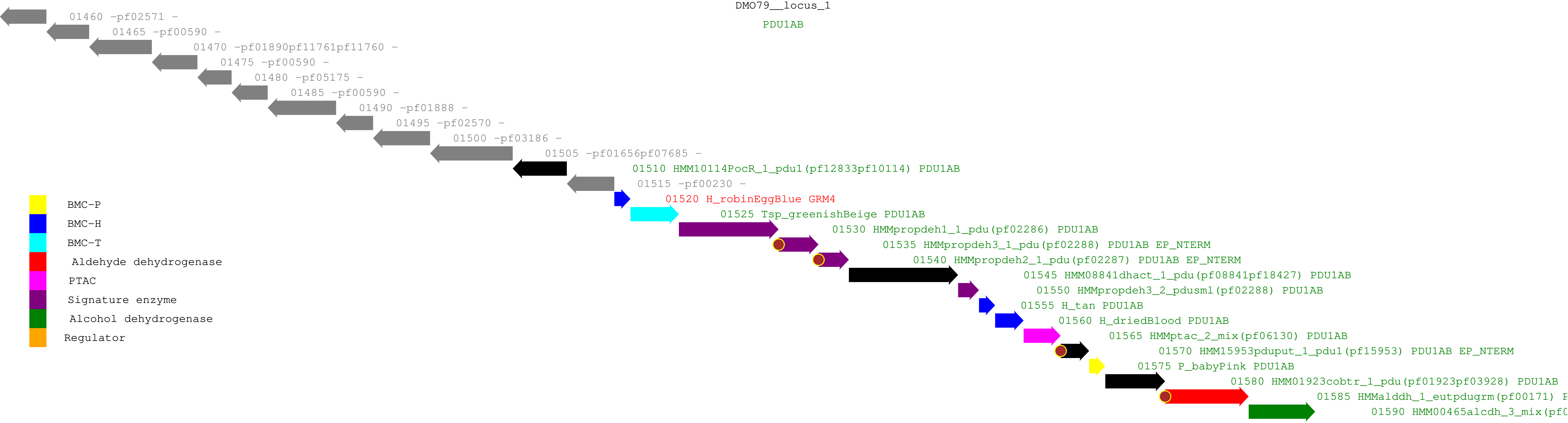

Klebsiella variicola. Bacteria; Proteobacteria; Gammaproteobacteria; Enterobacterales;

DMS35__locus_2

PDU1AB

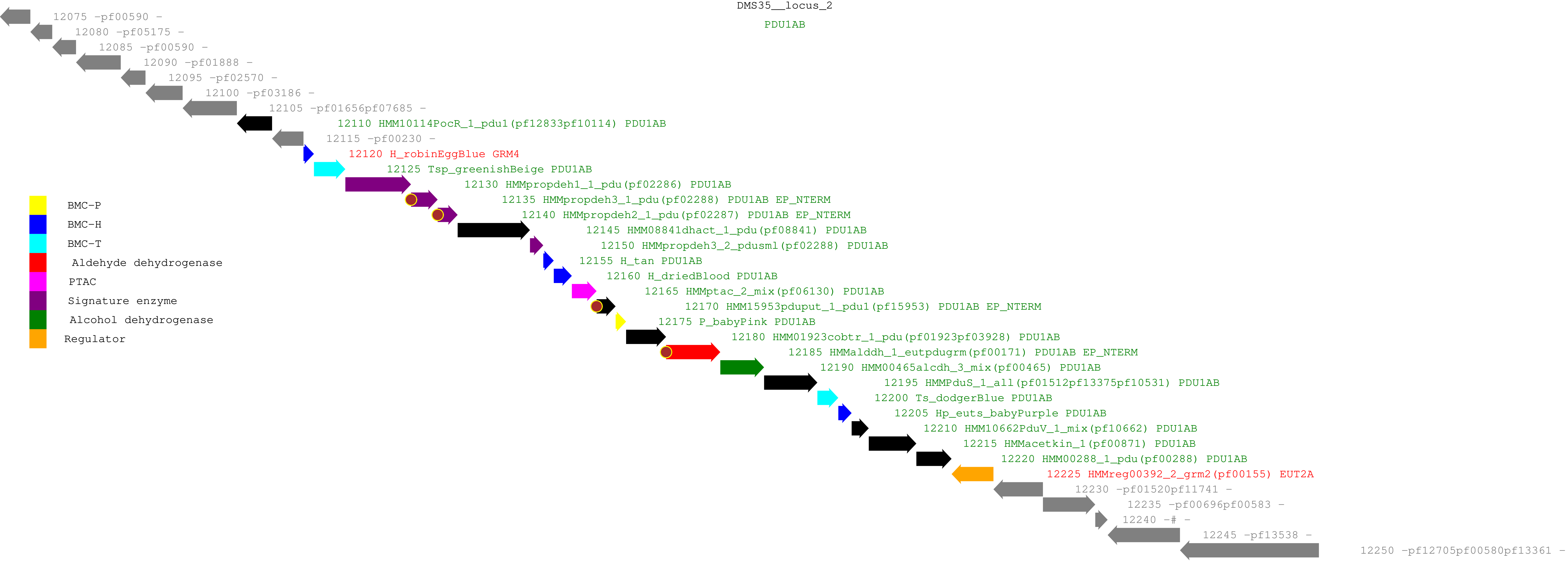

Salmonella enterica I. Bacteria; Proteobacteria; Gammaproteobacteria; Enterobacterales;

DMU73__locus_1

PDU1AB

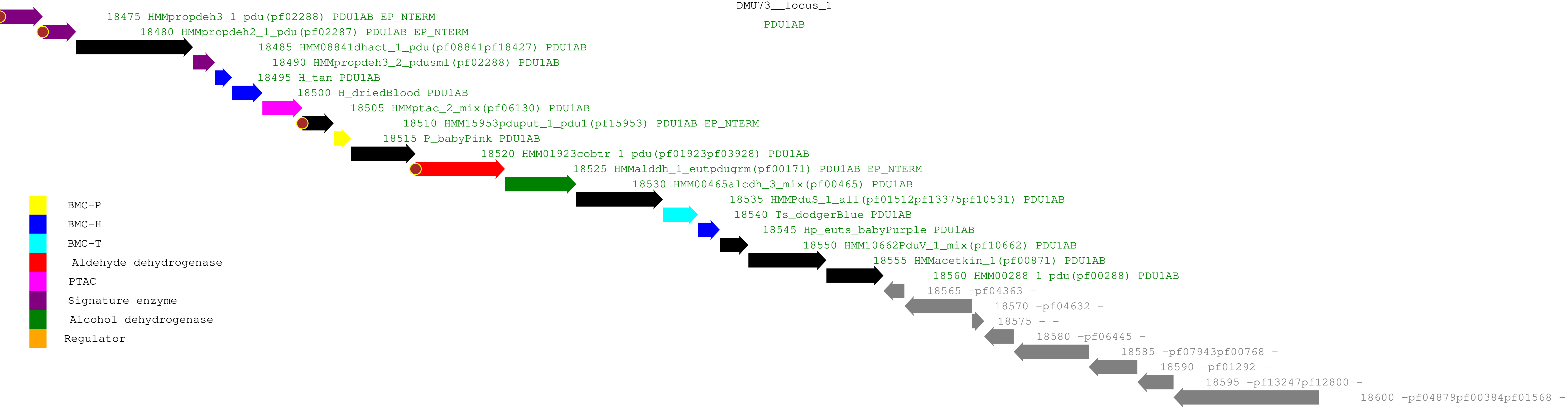

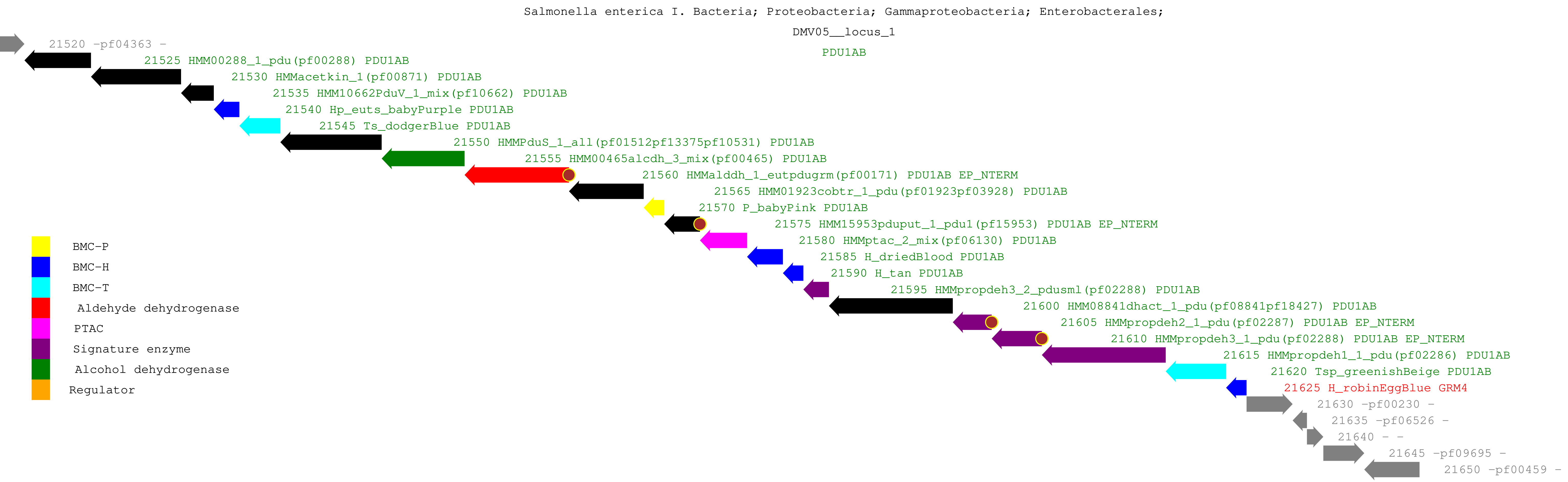

Salmonella enterica I. Bacteria; Proteobacteria; Gammaproteobacteria; Enterobacterales;
DMV05__locus_1
PDU1AB
21520 -pf04363 -
21525 HMM00288_1_pdu(pf00288) PDU1AB
21530 HMMacetkin_1(pf00871) PDU1AB
21535 HMM10662PduV_1_mix(pf10662) PDU1AB
21540 Hp_euts_babyPurple PDU1AB
21545 Ts_dodgerBlue PDU1AB
21550 HMMPduS_1_all(pf01512pf13375pf10531) PDU1AB
21555 HMM00465alcdh_3_mix(pf00465) PDU1AB
21560 HMMalddh_1_eutpdugrm(pf00171) PDU1AB EP_NTERM
21565 HMM01923cobtr_1_pdu(pf01923pf03928) PDU1AB
21570 P_babyPink PDU1AB
21575 HMM15953pduput_1_pdu1(pf15953) PDU1AB EP_NTERM
21580 HMMptac_2_mix(pf06130) PDU1AB
21585 H_driedBlood PDU1AB
21590 H_tan PDU1AB
21595 HMMpropdeh3_2_pdusml(pf02288) PDU1AB
21600 HMM08841dhact_1_pdu(pf08841pf18427) PDU1AB
21605 HMMpropdeh2_1_pdu(pf02287) PDU1AB EP_NTERM
21610 HMMpropdeh3_1_pdu(pf02288) PDU1AB EP_NTERM
21615 HMMpropdeh1_1_pdu(pf02286) PDU1AB
21620 Tsp_greenishBeige PDU1AB
21625 H_robinEggBlue GRM4
21630 -pf00230 -
21635 -pf06526 -
21640 - -
21645 -pf09695 -
21650 -pf00459 -
BMC-P
BMC-H
BMC-T
Aldehyde dehydrogenase
PTAC
Signature enzyme
Alcohol dehydrogenase
Regulator

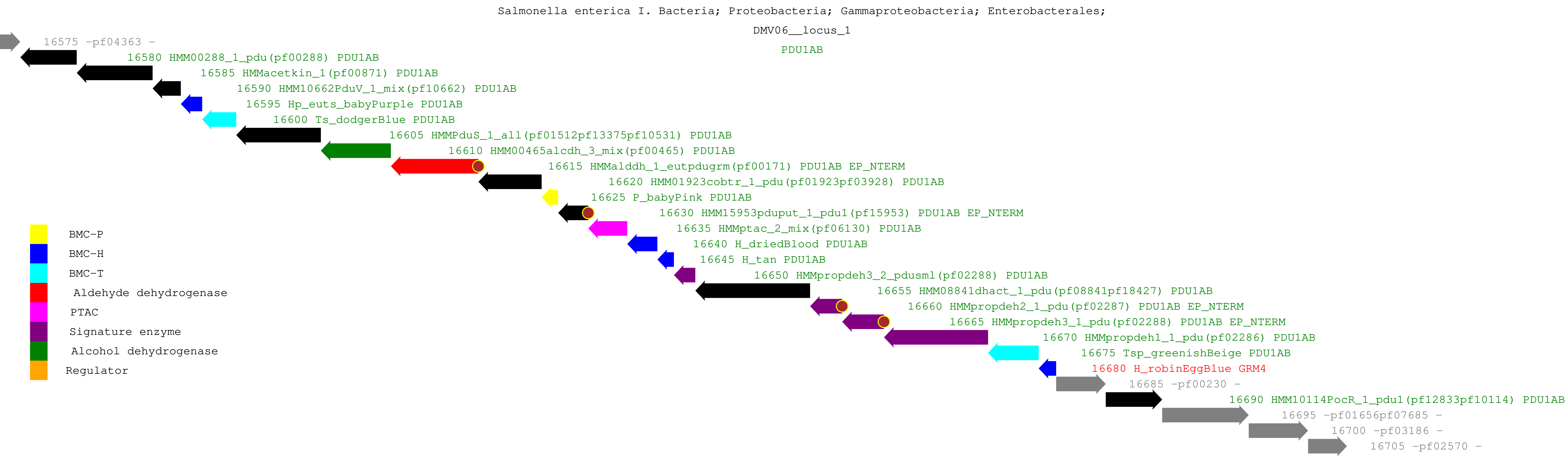

Salmonella enterica I. Bacteria; Proteobacteria; Gammaproteobacteria; Enterobacterales;
DMV06__locus_1
PDU1AB
16575 -pf04363 -
16580 HMM00288_1_pdu(pf00288) PDU1AB
16585 HMMacetkin_1(pf00871) PDU1AB
16590 HMM10662PduV_1_mix(pf10662) PDU1AB
16595 Hp_euts_babyPurple PDU1AB
16600 Ts_dodgerBlue PDU1AB
16605 HMMPduS_1_all(pf01512pf13375pf10531) PDU1AB
16610 HMM00465alcdh_3_mix(pf00465) PDU1AB
16615 HMMalddh_1_eutpdugrm(pf00171) PDU1AB EP_NTERM
16620 HMM01923cobtr_1_pdu(pf01923pf03928) PDU1AB
16625 P_babyPink PDU1AB
16630 HMM15953pduput_1_pdu1(pf15953) PDU1AB EP_NTERM
16635 HMMptac_2_mix(pf06130) PDU1AB
16640 H_driedBlood PDU1AB
16645 H_tan PDU1AB
16650 HMMpropdeh3_2_pdusml(pf02288) PDU1AB
16655 HMM08841dhact_1_pdu(pf08841pf18427) PDU1AB
16660 HMMpropdeh2_1_pdu(pf02287) PDU1AB EP_NTERM
16665 HMMpropdeh3_1_pdu(pf02288) PDU1AB EP_NTERM
16670 HMMpropdeh1_1_pdu(pf02286) PDU1AB
16675 Tsp_greenishBeige PDU1AB
16680 H_robinEggBlue GRM4
16685 -pf00230 -
16690 HMM10114PocR_1_pdu1(pf12833pf10114) PDU1AB
16695 -pf01656pf07685 -
16700 -pf03186 -
16705 -pf02570 -
BMC-P
BMC-H
BMC-T
Aldehyde dehydrogenase
PTAC
Signature enzyme
Alcohol dehydrogenase
Regulator

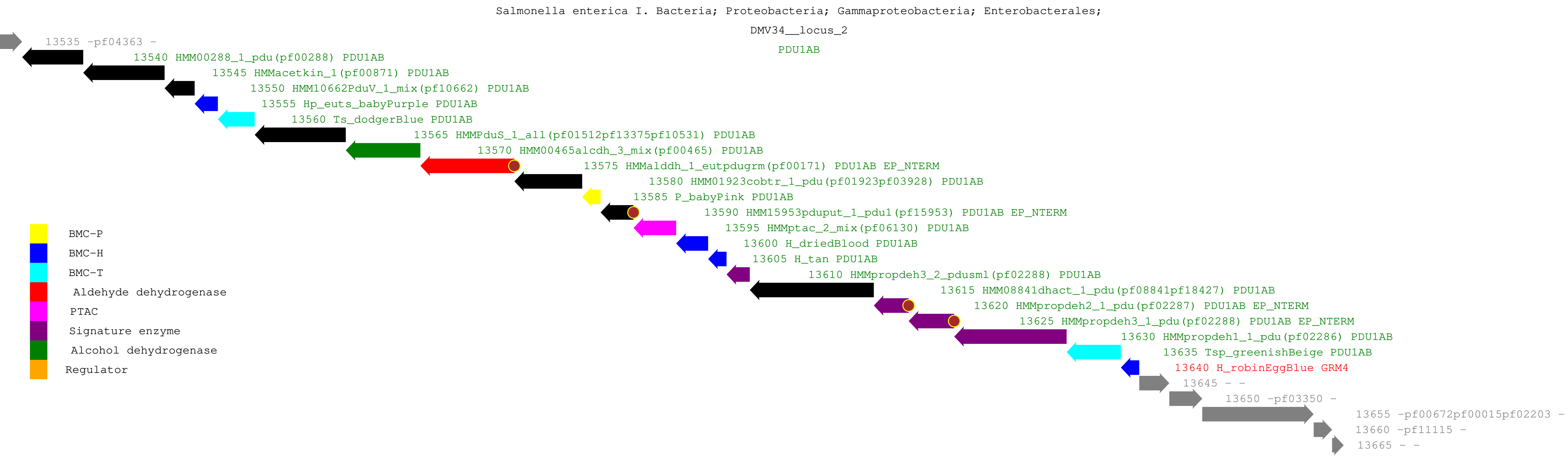
Salmonella enterica I. Bacteria; Proteobacteria; Gammaproteobacteria; Enterobacterales;
DMV34__locus_2
PDU1AB
13535 -pf04363 -
13540 HMM00288_1_pdu(pf00288) PDU1AB
13545 HMMacetkin_1(pf00871) PDU1AB
13550 HMM10662PduV_1_mix(pf10662) PDU1AB
13555 Hp_euts_babyPurple PDU1AB
13560 Ts_dodgerBlue PDU1AB
13565 HMMPduS_1_all(pf01512pf13375pf10531) PDU1AB
13570 HMM00465alcdh_3_mix(pf00465) PDU1AB
13575 HMMalddh_1_eutpdugrm(pf00171) PDU1AB EP_NTERM
13580 HMM01923cobtr_1_pdu(pf01923pf03928) PDU1AB
13585 P_babyPink PDU1AB
13590 HMM15953pduput_1_pdu1(pf15953) PDU1AB EP_NTERM
13595 HMMptac_2_mix(pf06130) PDU1AB
13600 H_driedBlood PDU1AB
13605 H_tan PDU1AB
13610 HMMpropdeh3_2_pdusml(pf02288) PDU1AB
13615 HMM08841dhact_1_pdu(pf08841pf18427) PDU1AB
13620 HMMpropdeh2_1_pdu(pf02287) PDU1AB EP_NTERM
13625 HMMpropdeh3_1_pdu(pf02288) PDU1AB EP_NTERM
13630 HMMpropdeh1_1_pdu(pf02286) PDU1AB
13635 Tsp_greenishBeige PDU1AB
13640 H_robinEggBlue GRM4
13645 - -
13650 -pf03350 -
13655 -pf00672pf00015pf02203 -
13660 -pf11115 -
13665 - -
BMC-P
BMC-H
BMC-T
Aldehyde dehydrogenase
PTAC
Signature enzyme
Alcohol dehydrogenase
Regulator

Klebsiella pneumoniae. Bacteria; Proteobacteria; Gammaproteobacteria; Enterobacterales;

DN589__locus_2

PDU1AB

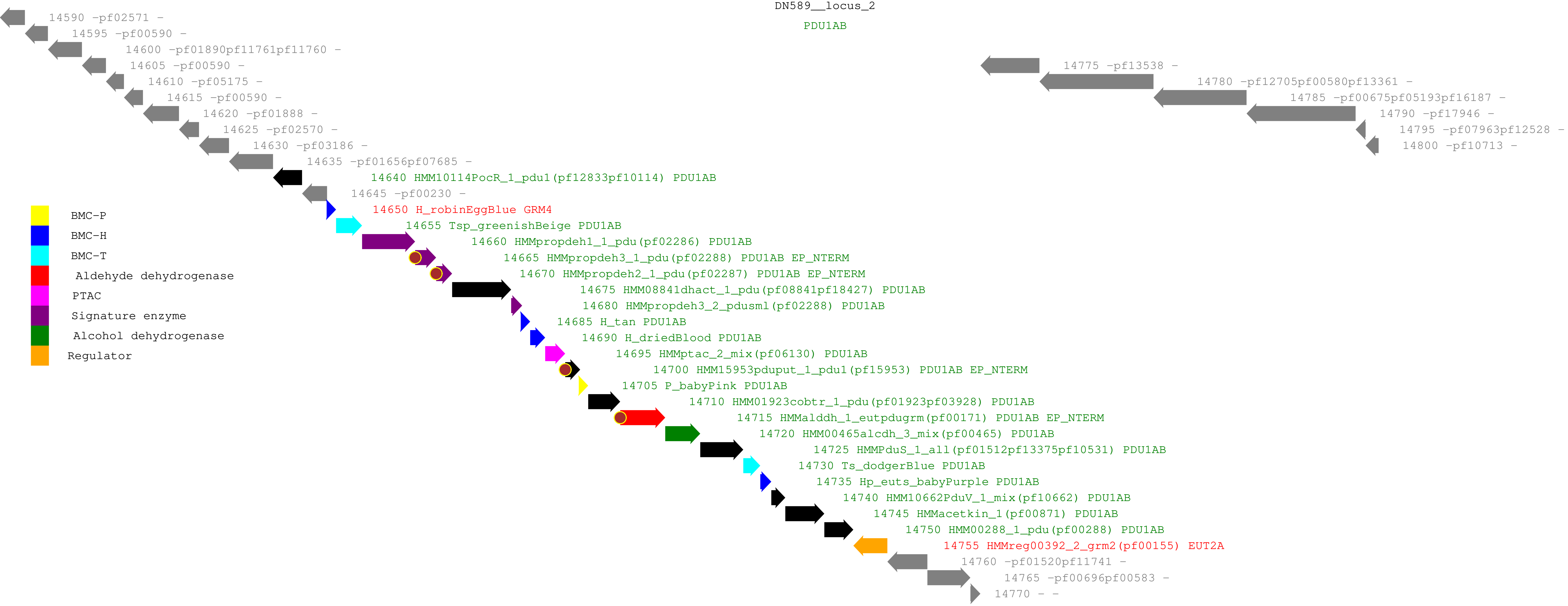

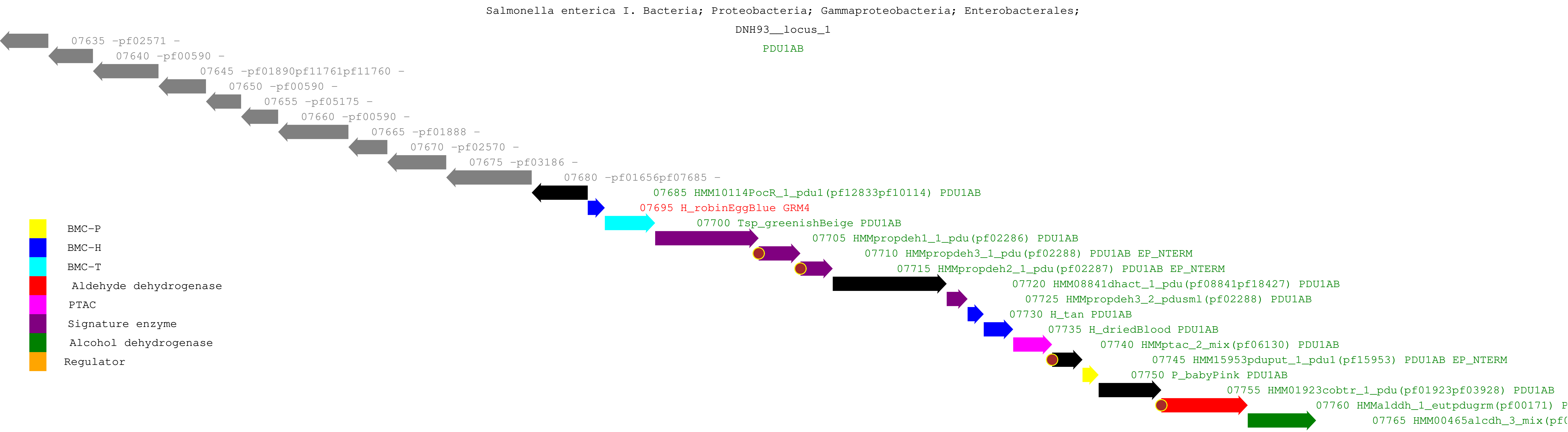
Salmonella enterica I. Bacteria; Proteobacteria; Gammaproteobacteria; Enterobacterales;
DNH93__locus_1
PDU1AB
07635 -pf02571 -
07640 -pf00590 -
07645 -pf01890pf11761pf11760 -
07650 -pf00590 -
07655 -pf05175 -
07660 -pf00590 -
07665 -pf01888 -
07670 -pf02570 -
07675 -pf03186 -
07680 -pf01656pf07685 -
07685 HMM10114PocR_1_pdu1(pf12833pf10114) PDU1AB
07695 H_robinEggBlue GRM4
07700 Tsp_greenishBeige PDU1AB
07705 HMMpropdeh1_1_pdu(pf02286) PDU1AB
07710 HMMpropdeh3_1_pdu(pf02288) PDU1AB EP_NTERM
07715 HMMpropdeh2_1_pdu(pf02287) PDU1AB EP_NTERM
07720 HMM08841dhact_1_pdu(pf08841pf18427) PDU1AB
07725 HMMpropdeh3_2_pdusml(pf02288) PDU1AB
07730 H_tan PDU1AB
07735 H_driedBlood PDU1AB
07740 HMMptac_2_mix(pf06130) PDU1AB
07745 HMM15953pduput_1_pdu1(pf15953) PDU1AB EP_NTERM
07750 P_babyPink PDU1AB
07755 HMM01923cobtr_1_pdu(pf01923pf03928) PDU1AB
07760 HMMalddh_1_eutpdugrm(pf00171) P
07765 HMM00465alcdh_3_mix(pf(
BMC-P
BMC-H
BMC-T
Aldehyde dehydrogenase
PTAC
Signature enzyme
Alcohol dehydrogenase
Regulator

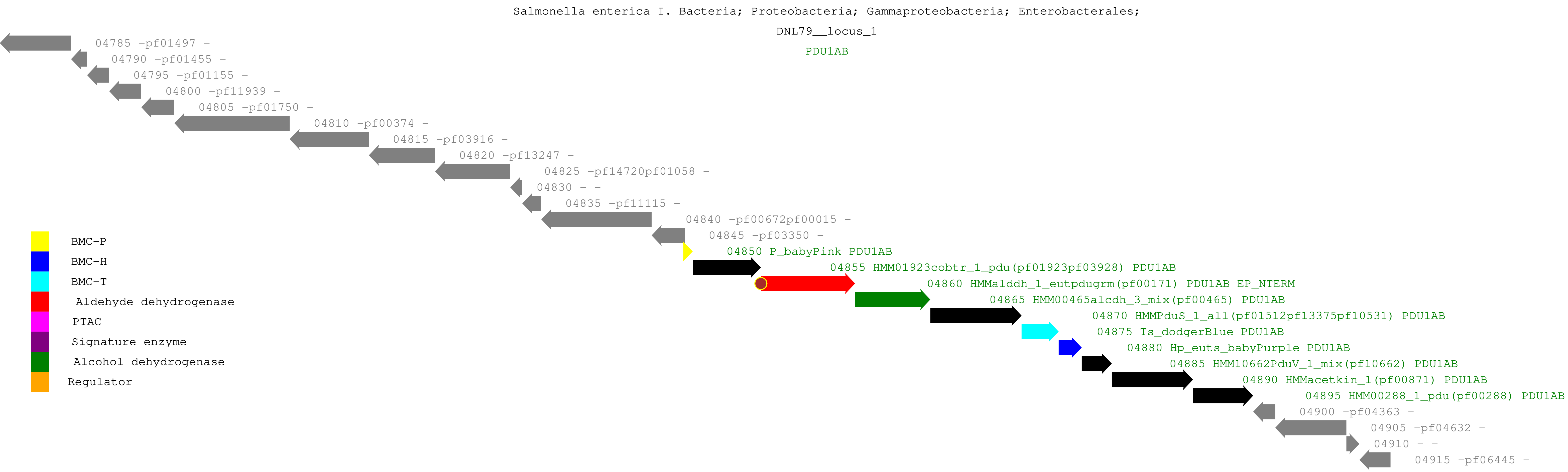
Salmonella enterica I. Bacteria; Proteobacteria; Gammaproteobacteria; Enterobacterales;
DNL79__locus_1
PDU1AB
04785 -pf01497 -
04790 -pf01455 -
04795 -pf01155 -
04800 -pf11939 -
04805 -pf01750 -
04810 -pf00374 -
04815 -pf03916 -
04820 -pf13247 -
04825 -pf14720pf01058 -
04830 - -
04835 -pf11115 -
04840 -pf00672pf00015 -
04845 -pf03350 -
04850 P_babyPink PDU1AB
04855 HMM01923cobtr_1_pdu(pf01923pf03928) PDU1AB
04860 HMMalddh_1_eutpdugrm(pf00171) PDU1AB EP_NTERM
04865 HMM00465alcdh_3_mix(pf00465) PDU1AB
04870 HMMPduS_1_all(pf01512pf13375pf10531) PDU1AB
04875 Ts_dodgerBlue PDU1AB
04880 Hp_euts_babyPurple PDU1AB
04885 HMM10662PduV_1_mix(pf10662) PDU1AB
04890 HMMacetkin_1(pf00871) PDU1AB
04895 HMM00288_1_pdu(pf00288) PDU1AB
04900 -pf04363 -
04905 -pf04632 -
04910 - -
04915 -pf06445 -
BMC-P
BMC-H
BMC-T
Aldehyde dehydrogenase
PTAC
Signature enzyme
Alcohol dehydrogenase
Regulator

Salmonella enterica I. Bacteria; Proteobacteria; Gammaproteobacteria; Enterobacterales;

DNL79__locus_2

PDU1AB

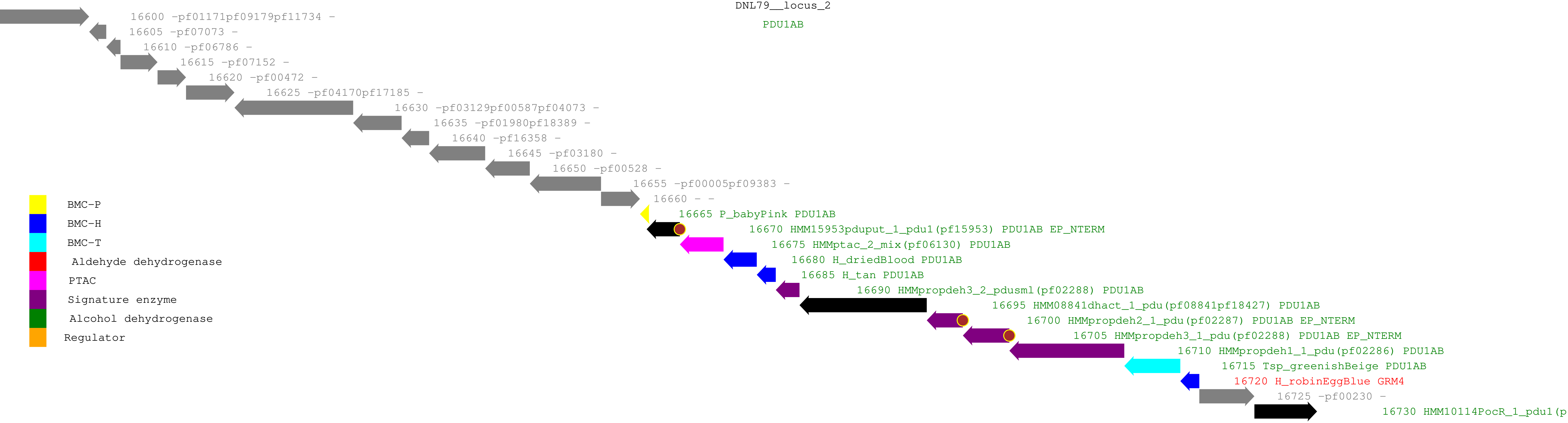

Salmonella enterica I. Bacteria; Proteobacteria; Gammaproteobacteria; Enterobacterales;

DNM39__locus_1

PDU1AB

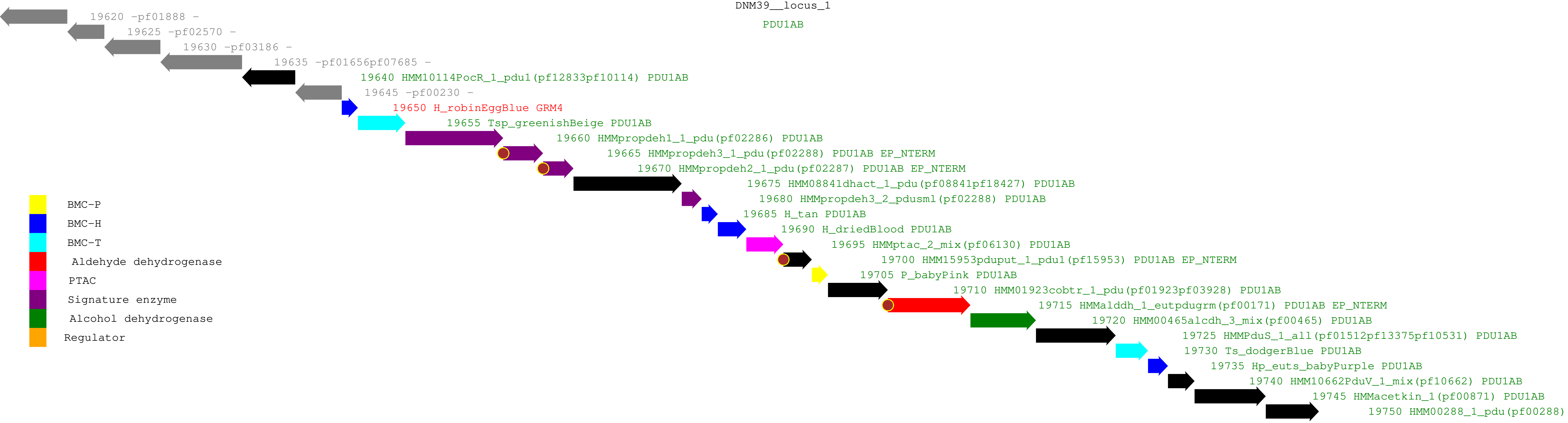

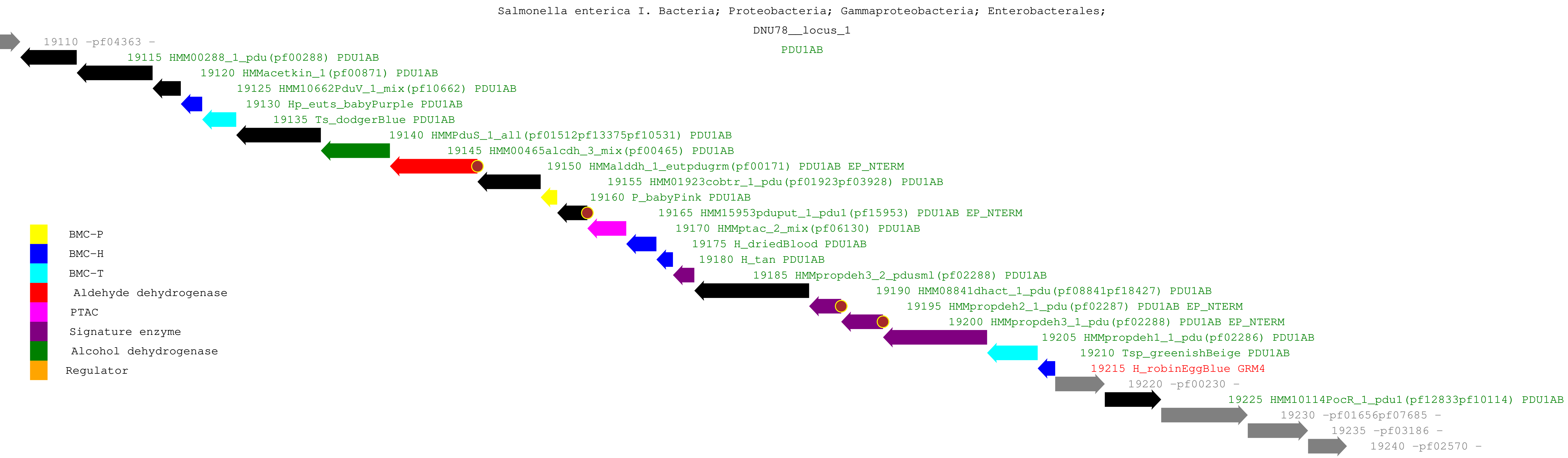
Salmonella enterica I. Bacteria; Proteobacteria; Gammaproteobacteria; Enterobacterales;
DNU78__locus_1
PDU1AB
19110 -pf04363 -
19115 HMM00288_1_pdu(pf00288) PDU1AB
19120 HMMacetkin_1(pf00871) PDU1AB
19125 HMM10662PduV_1_mix(pf10662) PDU1AB
19130 Hp_euts_babyPurple PDU1AB
19135 Ts_dodgerBlue PDU1AB
19140 HMMPduS_1_all(pf01512pf13375pf10531) PDU1AB
19145 HMM00465alcdh_3_mix(pf00465) PDU1AB
19150 HMMalddh_1_eutpdugrm(pf00171) PDU1AB EP_NTERM
19155 HMM01923cobtr_1_pdu(pf01923pf03928) PDU1AB
19160 P_babyPink PDU1AB
19165 HMM15953pduput_1_pdu1(pf15953) PDU1AB EP_NTERM
19170 HMMptac_2_mix(pf06130) PDU1AB
19175 H_driedBlood PDU1AB
19180 H_tan PDU1AB
19185 HMMpropdeh3_2_pdusml(pf02288) PDU1AB
19190 HMM08841dhact_1_pdu(pf08841pf18427) PDU1AB
19195 HMMpropdeh2_1_pdu(pf02287) PDU1AB EP_NTERM
19200 HMMpropdeh3_1_pdu(pf02288) PDU1AB EP_NTERM
19205 HMMpropdeh1_1_pdu(pf02286) PDU1AB
19210 Tsp_greenishBeige PDU1AB
19215 H_robinEggBlue GRM4
19220 -pf00230 -
19225 HMM10114PocR_1_pdu1(pf12833pf10114) PDU1AB
19230 -pf01656pf07685 -
19235 -pf03186 -
19240 -pf02570 -
BMC-P
BMC-H
BMC-T
Aldehyde dehydrogenase
PTAC
Signature enzyme
Alcohol dehydrogenase
Regulator

Salmonella enterica I. Bacteria; Proteobacteria; Gammaproteobacteria; Enterobacterales;

DNV24__locus_2

PDU1AB

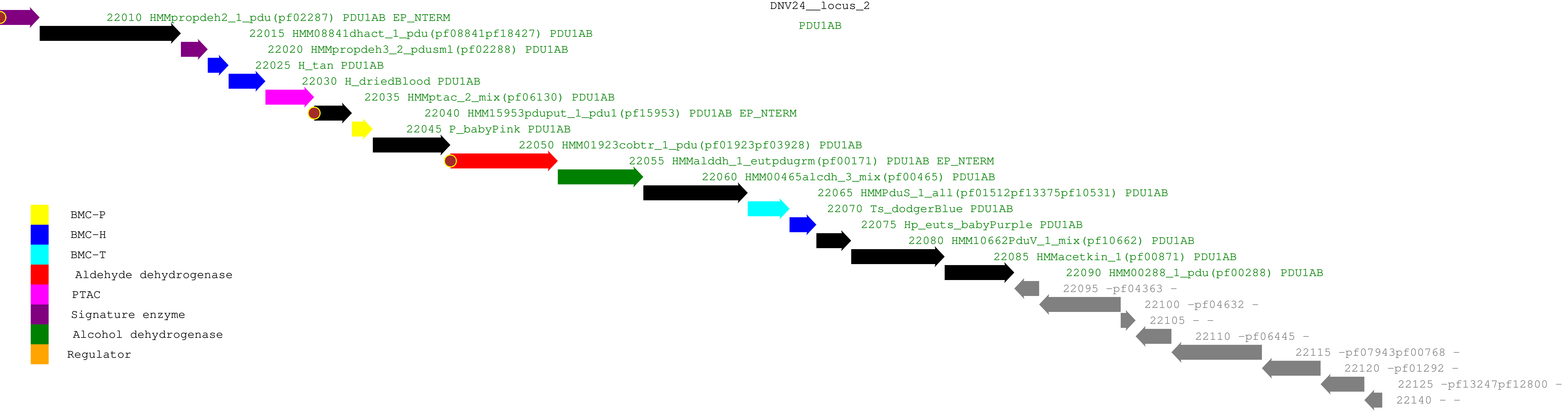

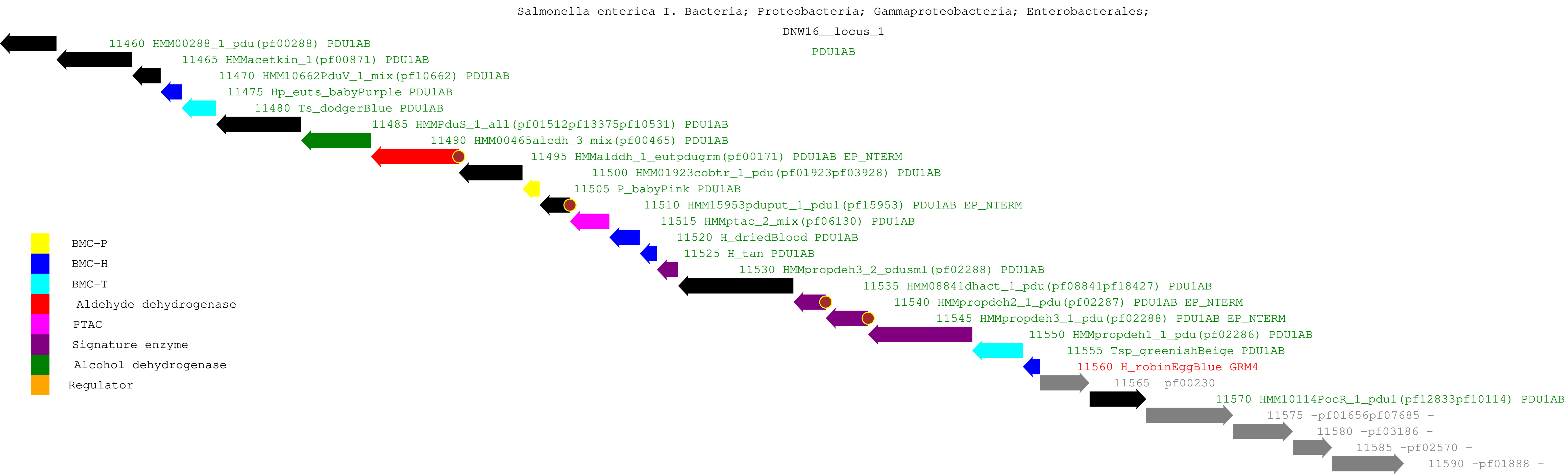
Salmonella enterica I. Bacteria; Proteobacteria; Gammaproteobacteria; Enterobacterales;
DNW16__locus_1
PDU1AB
11460 HMM00288_1_pdu(pf00288) PDU1AB
11465 HMMacetkin_1(pf00871) PDU1AB
11470 HMM10662PduV_1_mix(pf10662) PDU1AB
11475 Hp_euts_babyPurple PDU1AB
11480 Ts_dodgerBlue PDU1AB
11485 HMMPduS_1_all(pf01512pf13375pf10531) PDU1AB
11490 HMM00465alcdh_3_mix(pf00465) PDU1AB
11495 HMMalddh_1_eutpdugrm(pf00171) PDU1AB EP_NTERM
11500 HMM01923cobtr_1_pdu(pf01923pf03928) PDU1AB
11505 P_babyPink PDU1AB
11510 HMM15953pduput_1_pdu1(pf15953) PDU1AB EP_NTERM
11515 HMMptac_2_mix(pf06130) PDU1AB
11520 H_driedBlood PDU1AB
11525 H_tan PDU1AB
11530 HMMpropdeh3_2_pdusml(pf02288) PDU1AB
11535 HMM08841dhact_1_pdu(pf08841pf18427) PDU1AB
11540 HMMpropdeh2_1_pdu(pf02287) PDU1AB EP_NTERM
11545 HMMpropdeh3_1_pdu(pf02288) PDU1AB EP_NTERM
11550 HMMpropdeh1_1_pdu(pf02286) PDU1AB
11555 Tsp_greenishBeige PDU1AB
11560 H_robinEggBlue GRM4
11565 -pf00230 -
11570 HMM10114PocR_1_pdu1(pf12833pf10114) PDU1AB
11575 -pf01656pf07685 -
11580 -pf03186 -
11585 -pf02570 -
11590 -pf01888 -
BMC-P
BMC-H
BMC-T
Aldehyde dehydrogenase
PTAC
Signature enzyme
Alcohol dehydrogenase
Regulator

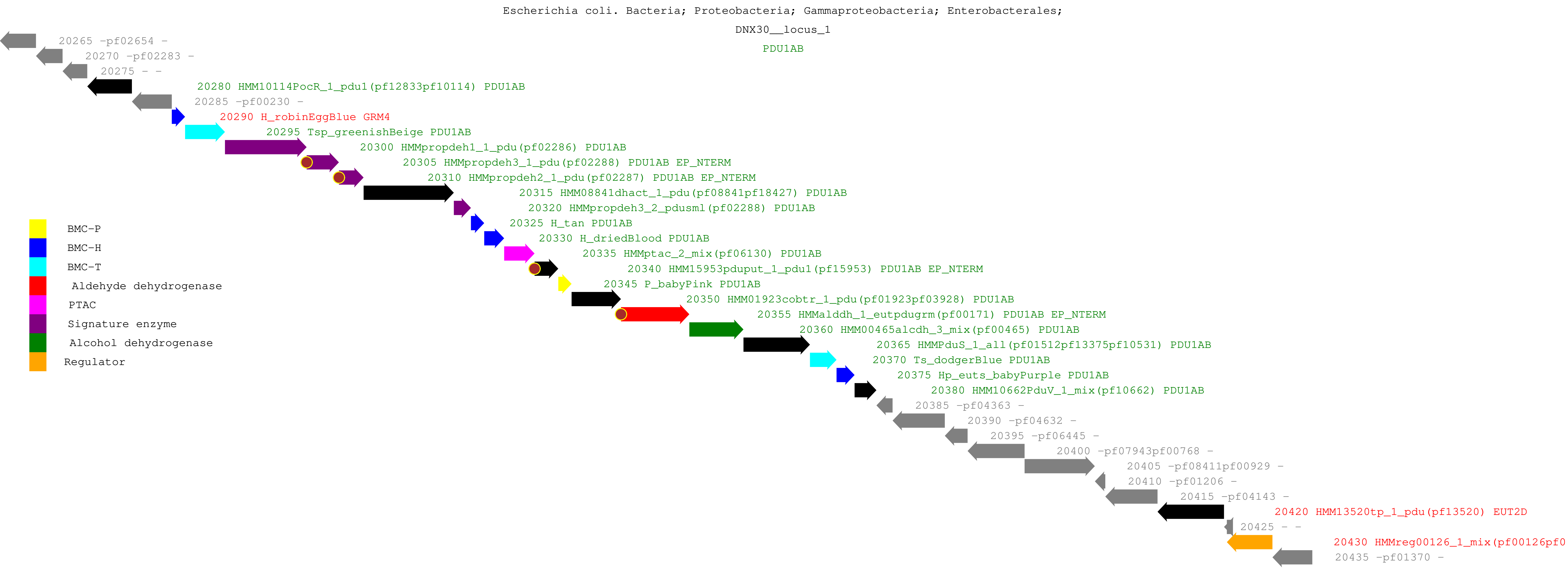

Escherichia coli. Bacteria; Proteobacteria; Gammaproteobacteria; Enterobacterales;
DNX30__locus_1
PDU1AB
20265 -pf02654 -
20270 -pf02283 -
20275 - -
20280 HMM10114PocR_1_pdu1(pf12833pf10114) PDU1AB
20285 -pf00230 -
20290 H_robinEggBlue GRM4
20295 Tsp_greenishBeige PDU1AB
20300 HMMpropdeh1_1_pdu(pf02286) PDU1AB
20305 HMMpropdeh3_1_pdu(pf02288) PDU1AB EP_NTERM
20310 HMMpropdeh2_1_pdu(pf02287) PDU1AB EP_NTERM
20315 HMM08841dhact_1_pdu(pf08841pf18427) PDU1AB
20320 HMMpropdeh3_2_pdusml(pf02288) PDU1AB
20325 H_tan PDU1AB
20330 H_driedBlood PDU1AB
20335 HMMptac_2_mix(pf06130) PDU1AB
20340 HMM15953pduput_1_pdu1(pf15953) PDU1AB EP_NTERM
20345 P_babyPink PDU1AB
20350 HMM01923cobtr_1_pdu(pf01923pf03928) PDU1AB
20355 HMMalddh_1_eutpdugrm(pf00171) PDU1AB EP_NTERM
20360 HMM00465alcdh_3_mix(pf00465) PDU1AB
20365 HMMPduS_1_all(pf01512pf13375pf10531) PDU1AB
20370 Ts_dodgerBlue PDU1AB
20375 Hp_euts_babyPurple PDU1AB
20380 HMM10662PduV_1_mix(pf10662) PDU1AB
20385 -pf04363 -
20390 -pf04632 -
20395 -pf06445 -
20400 -pf07943pf00768 -
20405 -pf08411pf00929 -
20410 -pf01206 -
20415 -pf04143 -
20420 HMM13520tp_1_pdu(pf13520) EUT2D
20425 - -
20430 HMMreg00126_1_mix(pf00126pf0
20435 -pf01370 -
BMC-P
BMC-H
BMC-T
Aldehyde dehydrogenase
PTAC
Signature enzyme
Alcohol dehydrogenase
Regulator

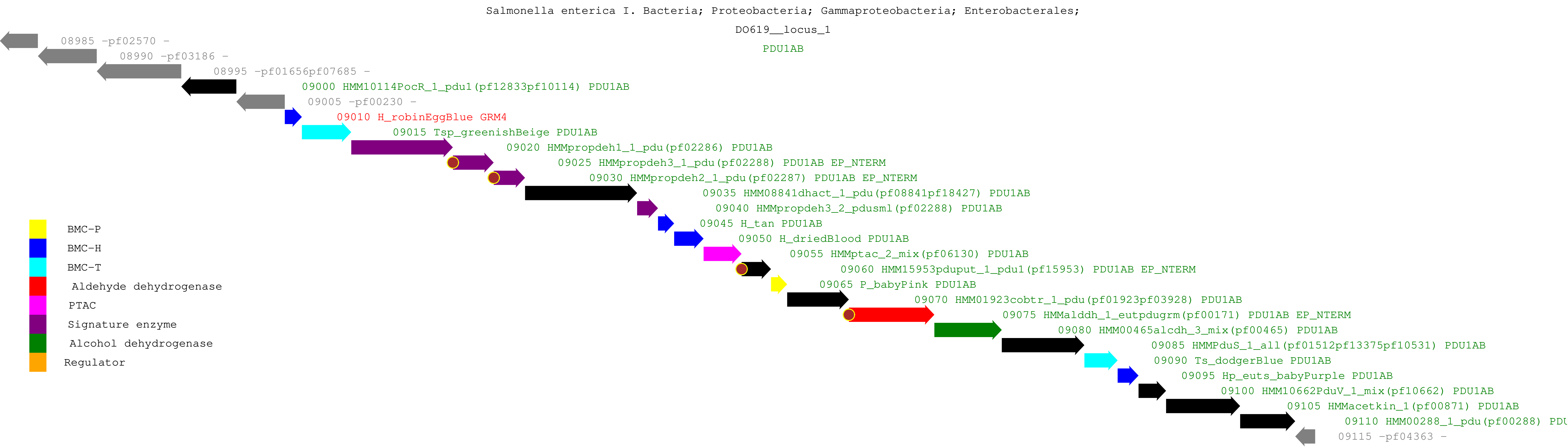
Salmonella enterica I. Bacteria; Proteobacteria; Gammaproteobacteria; Enterobacterales;
DO619__locus_1
PDU1AB
08985 -pf02570 -
08990 -pf03186 -
08995 -pf01656pf07685 -
09000 HMM10114PocR_1_pdu1(pf12833pf10114) PDU1AB
09005 -pf00230 -
09010 H_robinEggBlue GRM4
09015 Tsp_greenishBeige PDU1AB
09020 HMMpropdeh1_1_pdu(pf02286) PDU1AB
09025 HMMpropdeh3_1_pdu(pf02288) PDU1AB EP_NTERM
09030 HMMpropdeh2_1_pdu(pf02287) PDU1AB EP_NTERM
09035 HMM08841dhact_1_pdu(pf08841pf18427) PDU1AB
09040 HMMpropdeh3_2_pdusml(pf02288) PDU1AB
09045 H_tan PDU1AB
09050 H_driedBlood PDU1AB
09055 HMMptac_2_mix(pf06130) PDU1AB
09060 HMM15953pduput_1_pdu1(pf15953) PDU1AB EP_NTERM
09065 P_babyPink PDU1AB
09070 HMM01923cobtr_1_pdu(pf01923pf03928) PDU1AB
09075 HMMaldh_1_eutpdugrm(pf00171) PDU1AB EP_NTERM
09080 HMM00465alcdh_3_mix(pf00465) PDU1AB
09085 HMMPduS_1_all(pf01512pf13375pf10531) PDU1AB
09090 Ts_dodgerBlue PDU1AB
09095 Hp_euts_babyPurple PDU1AB
09100 HMM10662PduV_1_mix(pf10662) PDU1AB
09105 HMMacetkin_1(pf00871) PDU1AB
09110 HMM00288_1_pdu(pf00288) PDU
09115 -pf04363 -
BMC-P
BMC-H
BMC-T
Aldehyde dehydrogenase
PTAC
Signature enzyme
Alcohol dehydrogenase
Regulator

Salmonella enterica subsp. salamae. Bacteria; Proteobacteria; Gammaproteobacteria; Enterobacterales;

DO628__locus_2

PDU1AB

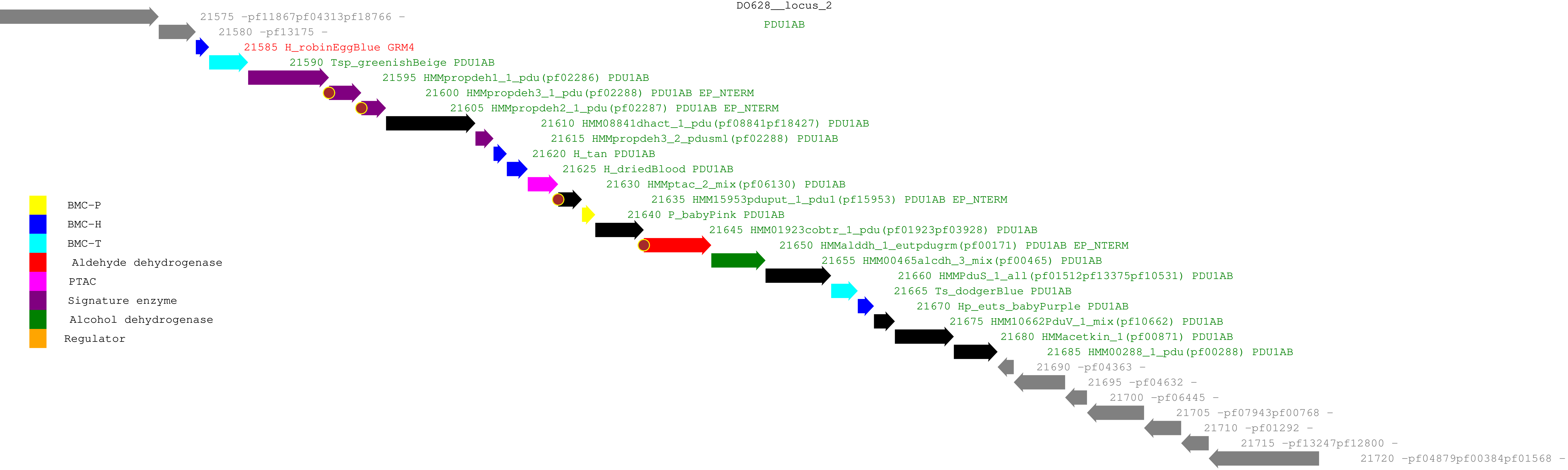

Salmonella enterica subsp. enterica serovar Weltevreden. Bacteria; Proteobacteria; Gammaproteobacteria; Enterobacterales;

DO955__locus_1

PDU1AB

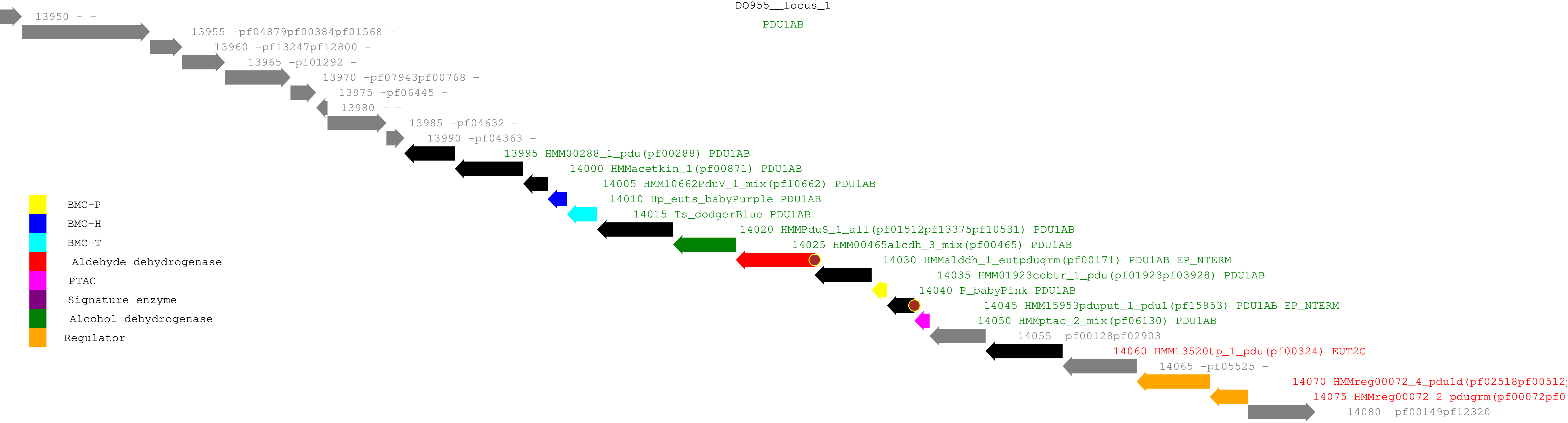

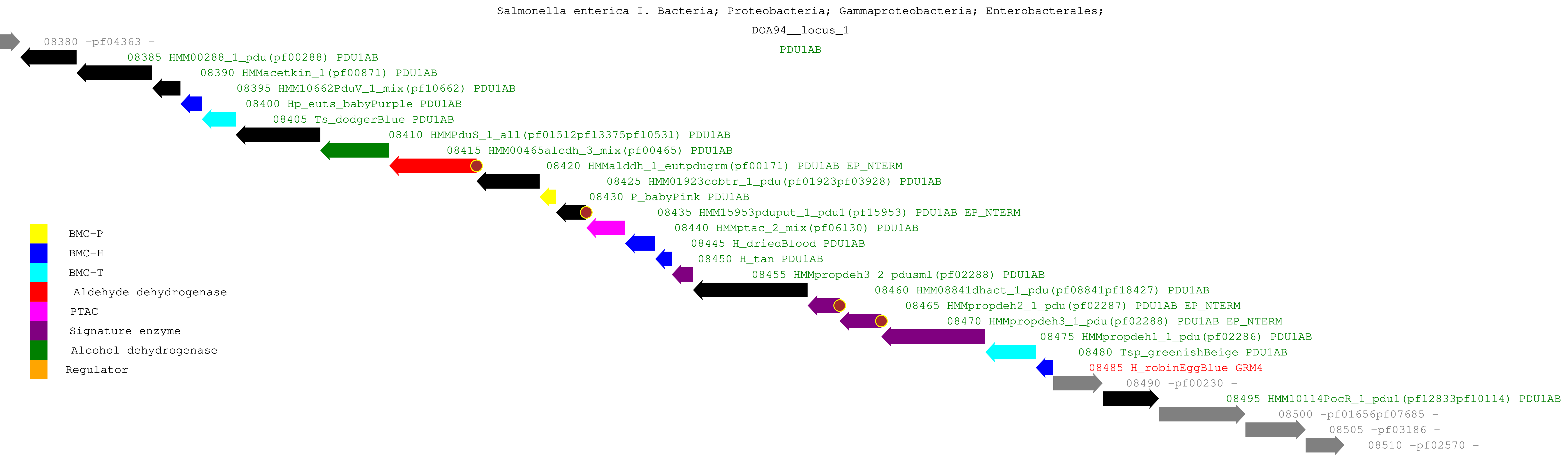
Salmonella enterica I. Bacteria; Proteobacteria; Gammaproteobacteria; Enterobacterales;
DOA94__locus_1
PDU1AB
08380 -pf04363 -
08385 HMM00288_1_pdu(pf00288) PDU1AB
08390 HMMacetkin_1(pf00871) PDU1AB
08395 HMM10662PduV_1_mix(pf10662) PDU1AB
08400 Hp_euts_babyPurple PDU1AB
08405 Ts_dodgerBlue PDU1AB
08410 HMMPduS_1_all(pf01512pf13375pf10531) PDU1AB
08415 HMM00465alcdh_3_mix(pf00465) PDU1AB
08420 HMMalddh_1_eutpdugrm(pf00171) PDU1AB EP_NTERM
08425 HMM01923cobtr_1_pdu(pf01923pf03928) PDU1AB
08430 P_babyPink PDU1AB
08435 HMM15953pduput_1_pdu1(pf15953) PDU1AB EP_NTERM
08440 HMMptac_2_mix(pf06130) PDU1AB
08445 H_driedBlood PDU1AB
08450 H_tan PDU1AB
08455 HMMpropdeh3_2_pdusml(pf02288) PDU1AB
08460 HMM08841dhact_1_pdu(pf08841pf18427) PDU1AB
08465 HMMpropdeh2_1_pdu(pf02287) PDU1AB EP_NTERM
08470 HMMpropdeh3_1_pdu(pf02288) PDU1AB EP_NTERM
08475 HMMpropdeh1_1_pdu(pf02286) PDU1AB
08480 Tsp_greenishBeige PDU1AB
08485 H_robinEggBlue GRM4
08490 -pf00230 -
08495 HMM10114PocR_1_pdu1(pf12833pf10114) PDU1AB
08500 -pf01656pf07685 -
08505 -pf03186 -
08510 -pf02570 -
BMC-P
BMC-H
BMC-T
Aldehyde dehydrogenase
PTAC
Signature enzyme
Alcohol dehydrogenase
Regulator

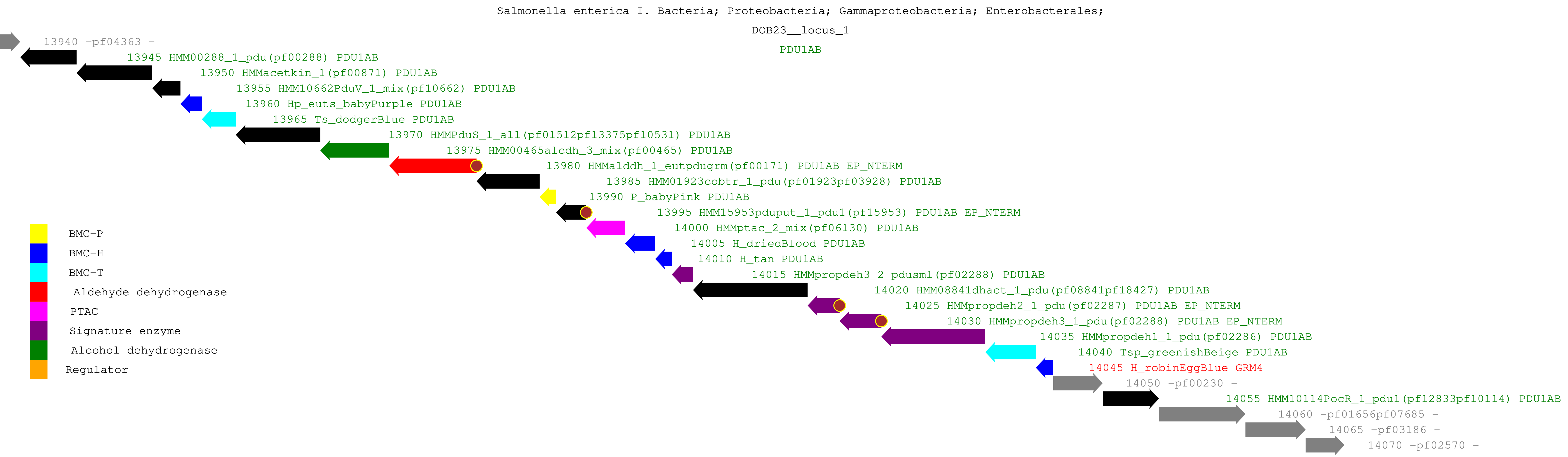
Salmonella enterica I. Bacteria; Proteobacteria; Gammaproteobacteria; Enterobacterales;
DOB23__locus_1
PDU1AB
13940 -pf04363 -
13945 HMM00288_1_pdu(pf00288) PDU1AB
13950 HMMacetkin_1(pf00871) PDU1AB
13955 HMM10662PduV_1_mix(pf10662) PDU1AB
13960 Hp_euts_babyPurple PDU1AB
13965 Ts_dodgerBlue PDU1AB
13970 HMMPduS_1_all(pf01512pf13375pf10531) PDU1AB
13975 HMM00465alcdh_3_mix(pf00465) PDU1AB
13980 HMMalddh_1_eutpdugrm(pf00171) PDU1AB EP_NTERM
13985 HMM01923cobtr_1_pdu(pf01923pf03928) PDU1AB
13990 P_babyPink PDU1AB
13995 HMM15953pduput_1_pdu1(pf15953) PDU1AB EP_NTERM
14000 HMMptac_2_mix(pf06130) PDU1AB
14005 H_driedBlood PDU1AB
14010 H_tan PDU1AB
14015 HMMpropdeh3_2_pdusml(pf02288) PDU1AB
14020 HMM08841dhact_1_pdu(pf08841pf18427) PDU1AB
14025 HMMpropdeh2_1_pdu(pf02287) PDU1AB EP_NTERM
14030 HMMpropdeh3_1_pdu(pf02288) PDU1AB EP_NTERM
14035 HMMpropdeh1_1_pdu(pf02286) PDU1AB
14040 Tsp_greenishBeige PDU1AB
14045 H_robinEggBlue GRM4
14050 -pf00230 -
14055 HMM10114PocR_1_pdu1(pf12833pf10114) PDU1AB
14060 -pf01656pf07685 -
14065 -pf03186 -
14070 -pf02570 -
BMC-P
BMC-H
BMC-T
Aldehyde dehydrogenase
PTAC
Signature enzyme
Alcohol dehydrogenase
Regulator

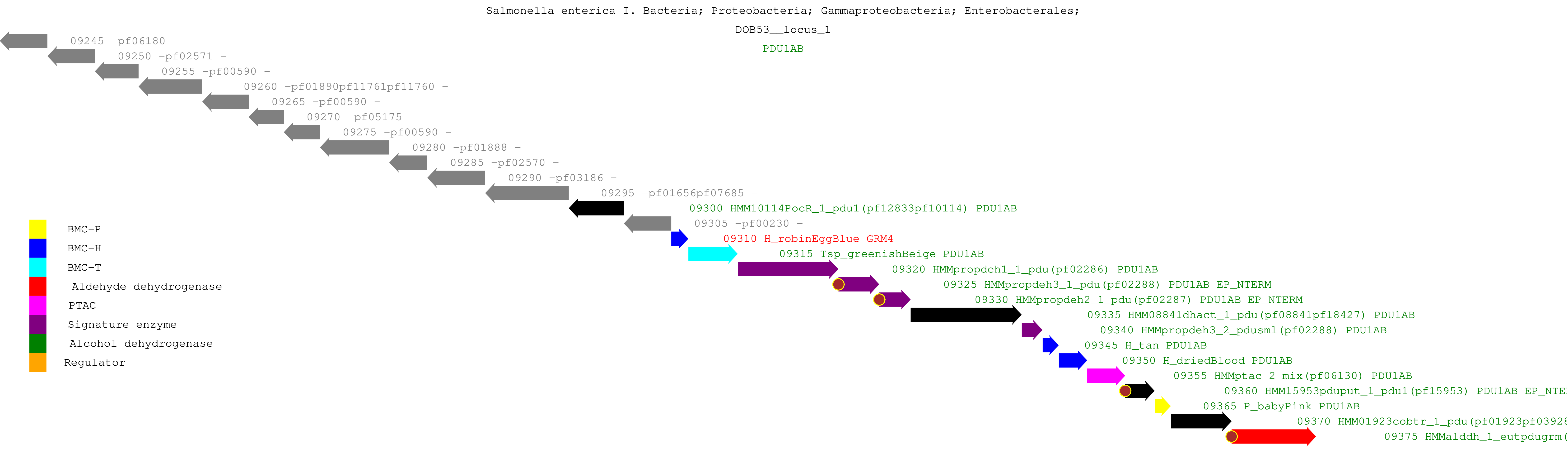
Salmonella enterica I. Bacteria; Proteobacteria; Gammaproteobacteria; Enterobacterales;
DOB53__locus_1
PDU1AB
09245 -pf06180 -
09250 -pf02571 -
09255 -pf00590 -
09260 -pf01890pf11761pf11760 -
09265 -pf00590 -
09270 -pf05175 -
09275 -pf00590 -
09280 -pf01888 -
09285 -pf02570 -
09290 -pf03186 -
09295 -pf01656pf07685 -
09300 HMM10114PocR_1_pdu1(pf12833pf10114) PDU1AB
09305 -pf00230 -
09310 H_robinEggBlue GRM4
09315 Tsp_greenishBeige PDU1AB
09320 HMMpropdeh1_1_pdu(pf02286) PDU1AB
09325 HMMpropdeh3_1_pdu(pf02288) PDU1AB EP_NTERM
09330 HMMpropdeh2_1_pdu(pf02287) PDU1AB EP_NTERM
09335 HMM08841dhact_1_pdu(pf08841pf18427) PDU1AB
09340 HMMpropdeh3_2_pdusml(pf02288) PDU1AB
09345 H_tan PDU1AB
09350 H_driedBlood PDU1AB
09355 HMMptac_2_mix(pf06130) PDU1AB
09360 HMM15953pduput_1_pdu1(pf15953) PDU1AB EP_NTE
09365 P_babyPink PDU1AB
09370 HMM01923cobtr_1_pdu(pf01923pf0392
09375 HMMalddh_1_eutpdugrm(
BMC-P
BMC-H
BMC-T
Aldehyde dehydrogenase
PTAC
Signature enzyme
Alcohol dehydrogenase
Regulator

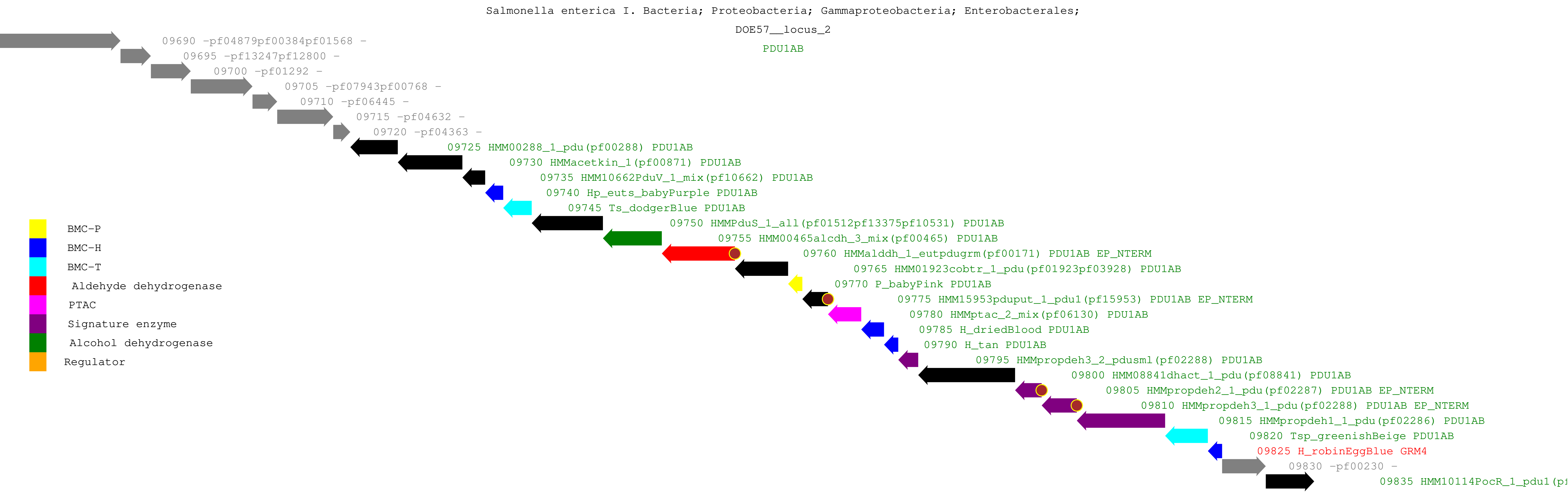
Salmonella enterica I. Bacteria; Proteobacteria; Gammaproteobacteria; Enterobacterales;
DOE57__locus_2
PDU1AB
09690 -pf04879pf00384pf01568 -
09695 -pf13247pf12800 -
09700 -pf01292 -
09705 -pf07943pf00768 -
09710 -pf06445 -
09715 -pf04632 -
09720 -pf04363 -
09725 HMM00288_1_pdu(pf00288) PDU1AB
09730 HMMacetkin_1(pf00871) PDU1AB
09735 HMM10662PduV_1_mix(pf10662) PDU1AB
09740 Hp_euts_babyPurple PDU1AB
09745 Ts_dodgerBlue PDU1AB
09750 HMMPduS_1_all(pf01512pf13375pf10531) PDU1AB
09755 HMM00465alcdh_3_mix(pf00465) PDU1AB
09760 HMMaldh_1_eutpdugrm(pf00171) PDU1AB EP_NTERM
09765 HMM01923cobtr_1_pdu(pf01923pf03928) PDU1AB
09770 P_babyPink PDU1AB
09775 HMM15953pduput_1_pdu1(pf15953) PDU1AB EP_NTERM
09780 HMMptac_2_mix(pf06130) PDU1AB
09785 H_driedBlood PDU1AB
09790 H_tan PDU1AB
09795 HMMpropdeh3_2_pdusml(pf02288) PDU1AB
09800 HMM08841dhact_1_pdu(pf08841) PDU1AB
09805 HMMpropdeh2_1_pdu(pf02287) PDU1AB EP_NTERM
09810 HMMpropdeh3_1_pdu(pf02288) PDU1AB EP_NTERM
09815 HMMpropdeh1_1_pdu(pf02286) PDU1AB
09820 Tsp_greenishBeige PDU1AB
09825 H_robinEggBlue GRM4
09830 -pf00230 -
09835 HMM10114PocR_1_pdu1(p
BMC-P
BMC-H
BMC-T
Aldehyde dehydrogenase
PTAC
Signature enzyme
Alcohol dehydrogenase
Regulator

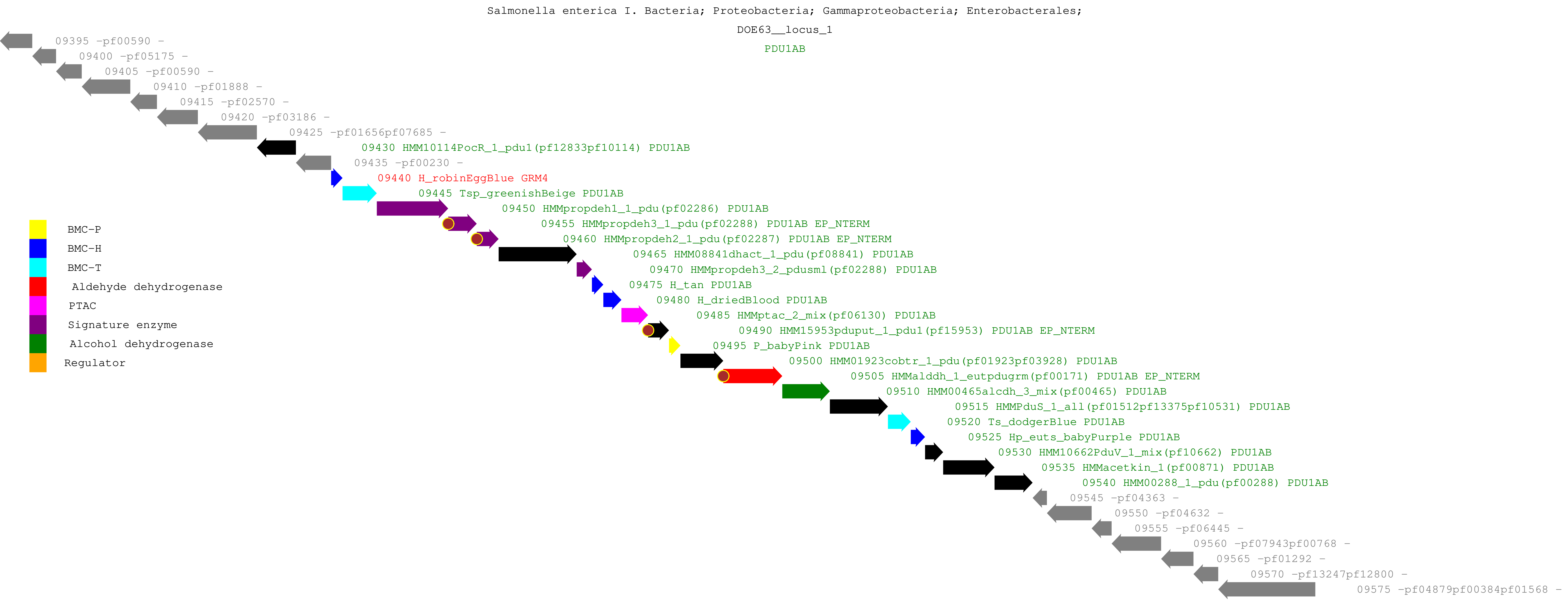
Salmonella enterica I. Bacteria; Proteobacteria; Gammaproteobacteria; Enterobacterales;
DOE63__locus_1
PDU1AB
09395 -pf00590 -
09400 -pf05175 -
09405 -pf00590 -
09410 -pf01888 -
09415 -pf02570 -
09420 -pf03186 -
09425 -pf01656pf07685 -
09430 HMM10114PocR_1_pdu1(pf12833pf10114) PDU1AB
09435 -pf00230 -
09440 H_robinEggBlue GRM4
09445 Tsp_greenishBeige PDU1AB
09450 HMMpropdeh1_1_pdu(pf02286) PDU1AB
09455 HMMpropdeh3_1_pdu(pf02288) PDU1AB EP_NTERM
09460 HMMpropdeh2_1_pdu(pf02287) PDU1AB EP_NTERM
09465 HMM08841dhact_1_pdu(pf08841) PDU1AB
09470 HMMpropdeh3_2_pdusml(pf02288) PDU1AB
09475 H_tan PDU1AB
09480 H_driedBlood PDU1AB
09485 HMMptac_2_mix(pf06130) PDU1AB
09490 HMM15953pduput_1_pdu1(pf15953) PDU1AB EP_NTERM
09495 P_babyPink PDU1AB
09500 HMM01923cobtr_1_pdu(pf01923pf03928) PDU1AB
09505 HMMalddh_1_eutpdugrm(pf00171) PDU1AB EP_NTERM
09510 HMM00465alcdh_3_mix(pf00465) PDU1AB
09515 HMMPduS_1_all(pf01512pf13375pf10531) PDU1AB
09520 Ts_dodgerBlue PDU1AB
09525 Hp_euts_babyPurple PDU1AB
09530 HMM10662PduV_1_mix(pf10662) PDU1AB
09535 HMMacetkin_1(pf00871) PDU1AB
09540 HMM00288_1_pdu(pf00288) PDU1AB
09545 -pf04363 -
09550 -pf04632 -
09555 -pf06445 -
09560 -pf07943pf00768 -
09565 -pf01292 -
09570 -pf13247pf12800 -
09575 -pf04879pf00384pf01568 -
BMC-P
BMC-H
BMC-T
Aldehyde dehydrogenase
PTAC
Signature enzyme
Alcohol dehydrogenase
Regulator

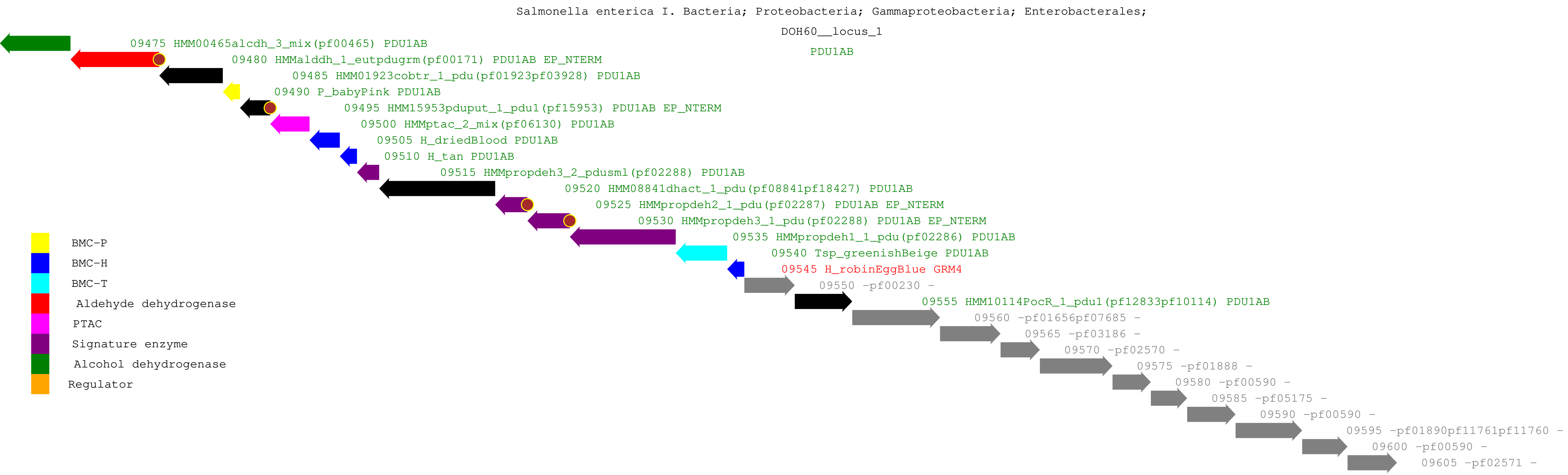

Salmonella enterica I. Bacteria; Proteobacteria; Gammaproteobacteria; Enterobacterales;
DOH60__locus_1
PDU1AB
09475 HMM00465alcdh_3_mix(pf00465) PDU1AB
09480 HMMalddh_1_eutpdugrm(pf00171) PDU1AB EP_NTERM
09485 HMM01923cobtr_1_pdu(pf01923pf03928) PDU1AB
09490 P_babyPink PDU1AB
09495 HMM15953pduput_1_pdu1(pf15953) PDU1AB EP_NTERM
09500 HMMptac_2_mix(pf06130) PDU1AB
09505 H_driedBlood PDU1AB
09510 H_tan PDU1AB
09515 HMMpropdeh3_2_pdusml(pf02288) PDU1AB
09520 HMM08841dhact_1_pdu(pf08841pf18427) PDU1AB
09525 HMMpropdeh2_1_pdu(pf02287) PDU1AB EP_NTERM
09530 HMMpropdeh3_1_pdu(pf02288) PDU1AB EP_NTERM
09535 HMMpropdeh1_1_pdu(pf02286) PDU1AB
09540 Tsp_greenishBeige PDU1AB
09545 H_robinEggBlue GRM4
09550 -pf00230 -
09555 HMM10114PocR_1_pdu1(pf12833pf10114) PDU1AB
09560 -pf01656pf07685 -
09565 -pf03186 -
09570 -pf02570 -
09575 -pf01888 -
09580 -pf00590 -
09585 -pf05175 -
09590 -pf00590 -
09595 -pf01890pf11761pf11760 -
09600 -pf00590 -
09605 -pf02571 -
BMC-P
BMC-H
BMC-T
Aldehyde dehydrogenase
PTAC
Signature enzyme
Alcohol dehydrogenase
Regulator

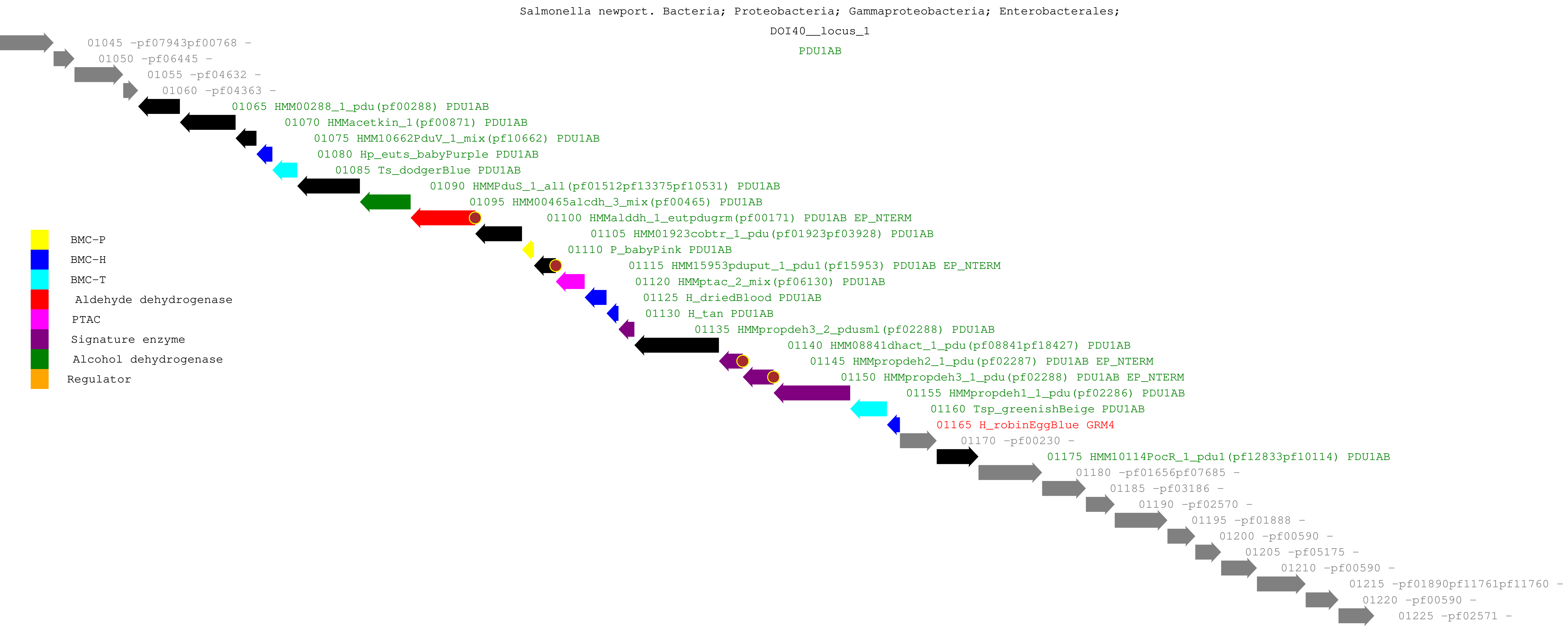
Salmonella newport. Bacteria; Proteobacteria; Gammaproteobacteria; Enterobacterales;
DOI40__locus_1
PDU1AB
01045 -pf07943pf00768 -
01050 -pf06445 -
01055 -pf04632 -
01060 -pf04363 -
01065 HMM00288_1_pdu(pf00288) PDU1AB
01070 HMMacetkin_1(pf00871) PDU1AB
01075 HMM10662PduV_1_mix(pf10662) PDU1AB
01080 Hp_euts_babyPurple PDU1AB
01085 Ts_dodgerBlue PDU1AB
01090 HMMPduS_1_all(pf01512pf13375pf10531) PDU1AB
01095 HMM00465alcdh_3_mix(pf00465) PDU1AB
01100 HMMaldh_1_eutpdugrm(pf00171) PDU1AB EP_NTERM
01105 HMM01923cobtr_1_pdu(pf01923pf03928) PDU1AB
01110 P_babyPink PDU1AB
01115 HMM15953pduput_1_pdu1(pf15953) PDU1AB EP_NTERM
01120 HMMptac_2_mix(pf06130) PDU1AB
01125 H_driedBlood PDU1AB
01130 H_tan PDU1AB
01135 HMMpropdeh3_2_pdusml(pf02288) PDU1AB
01140 HMM08841dhact_1_pdu(pf08841pf18427) PDU1AB
01145 HMMpropdeh2_1_pdu(pf02287) PDU1AB EP_NTERM
01150 HMMpropdeh3_1_pdu(pf02288) PDU1AB EP_NTERM
01155 HMMpropdeh1_1_pdu(pf02286) PDU1AB
01160 Tsp_greenishBeige PDU1AB
01165 H_robinEggBlue GRM4
01170 -pf00230 -
01175 HMM10114PocR_1_pdu1(pf12833pf10114) PDU1AB
01180 -pf01656pf07685 -
01185 -pf03186 -
01190 -pf02570 -
01195 -pf01888 -
01200 -pf00590 -
01205 -pf05175 -
01210 -pf00590 -
01215 -pf01890pf11761pf11760 -
01220 -pf00590 -
01225 -pf02571 -
BMC-P
BMC-H
BMC-T
Aldehyde dehydrogenase
PTAC
Signature enzyme
Alcohol dehydrogenase
Regulator

Salmonella enterica I. Bacteria; Proteobacteria; Gammaproteobacteria; Enterobacterales;

DOW79__locus_1

PDU1AB

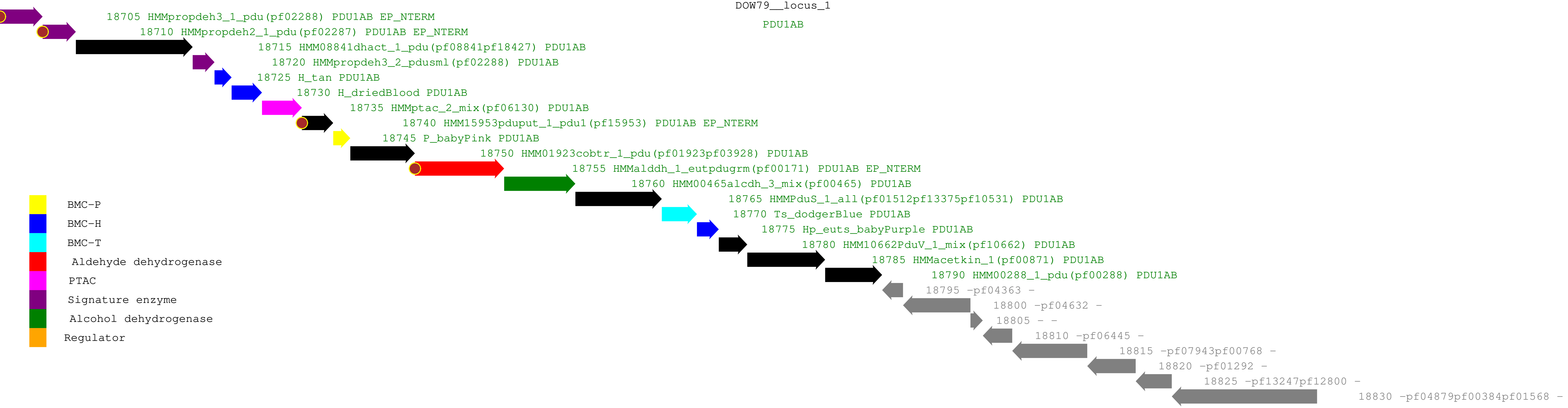

Salmonella enterica subsp. enterica serovar Give. Bacteria; Proteobacteria; Gammaproteobacteria; Enterobacterales;

DOW88__locus_1

PDU1AB

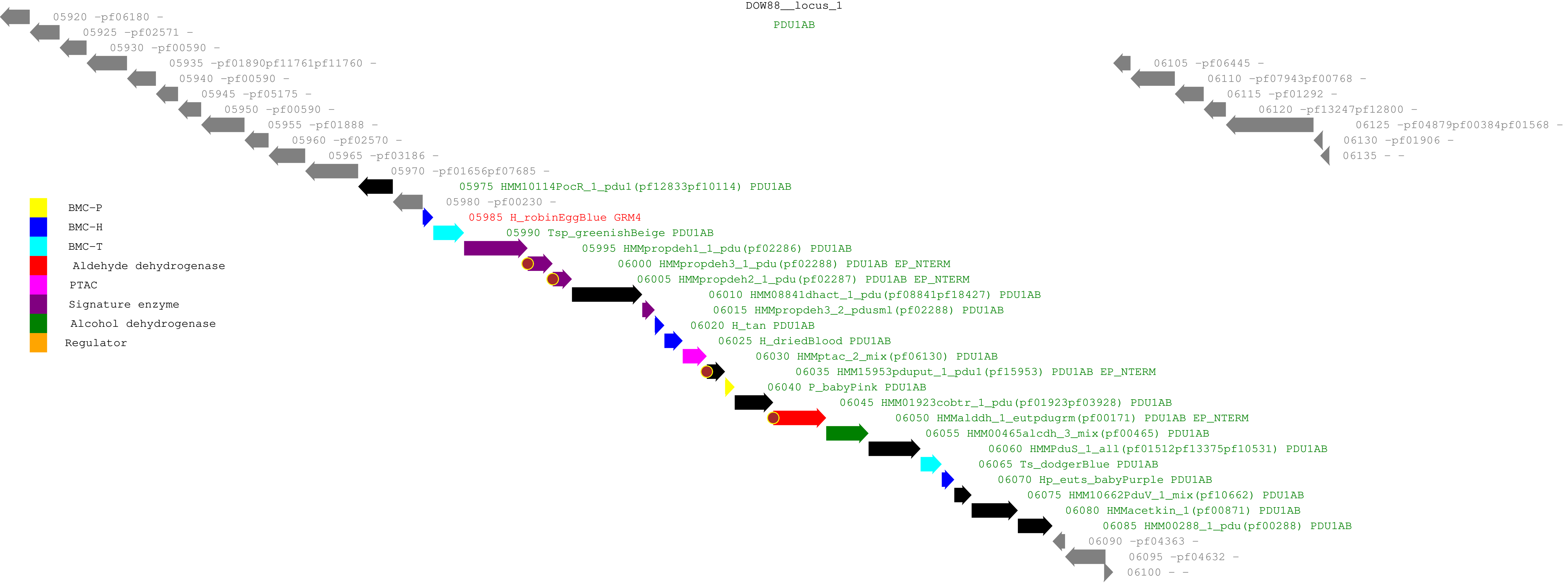

Salmonella diarizonae. Bacteria; Proteobacteria; Gammaproteobacteria; Enterobacterales;

DP705__locus_2

PDU1AB

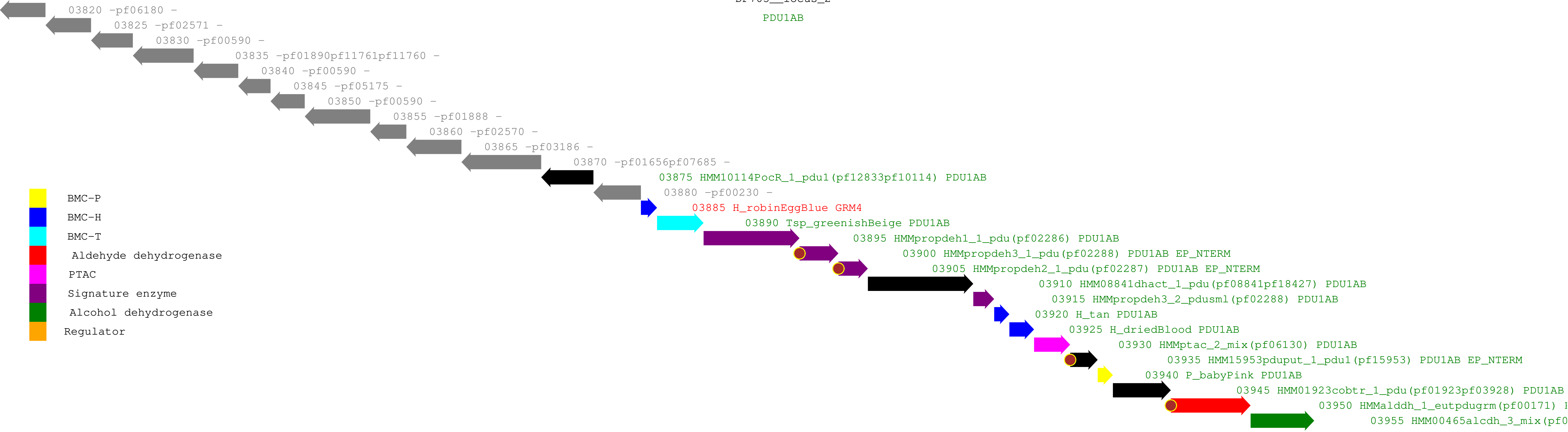

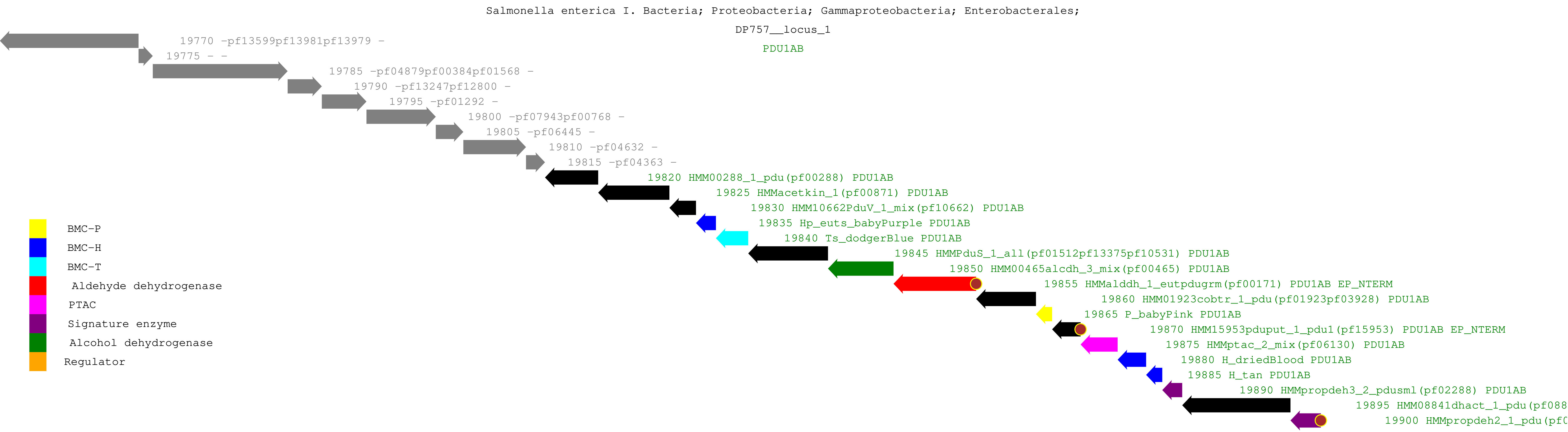
Salmonella enterica I. Bacteria; Proteobacteria; Gammaproteobacteria; Enterobacterales;
DP757__locus_1
PDU1AB
19770 -pf13599pf13981pf13979 -
19775 - -
19785 -pf04879pf00384pf01568 -
19790 -pf13247pf12800 -
19795 -pf01292 -
19800 -pf07943pf00768 -
19805 -pf06445 -
19810 -pf04632 -
19815 -pf04363 -
19820 HMM00288_1_pdu(pf00288) PDU1AB
19825 HMMacetkin_1(pf00871) PDU1AB
19830 HMM10662PduV_1_mix(pf10662) PDU1AB
19835 Hp_euts_babyPurple PDU1AB
19840 Ts_dodgerBlue PDU1AB
19845 HMMPduS_1_all(pf01512pf13375pf10531) PDU1AB
19850 HMM00465alcdh_3_mix(pf00465) PDU1AB
19855 HMMalddh_1_eutpdugrm(pf00171) PDU1AB EP_NTERM
19860 HMM01923cobtr_1_pdu(pf01923pf03928) PDU1AB
19865 P_babyPink PDU1AB
19870 HMM15953pduput_1_pdu1(pf15953) PDU1AB EP_NTERM
19875 HMMptac_2_mix(pf06130) PDU1AB
19880 H_driedBlood PDU1AB
19885 H_tan PDU1AB
19890 HMMpropdeh3_2_pdusml(pf02288) PDU1AB
19895 HMM08841dhact_1_pdu(pf088
19900 HMMpropdeh2_1_pdu(pf0
BMC-P
BMC-H
BMC-T
Aldehyde dehydrogenase
PTAC
Signature enzyme
Alcohol dehydrogenase
Regulator

Salmonella enterica subsp. arizonae. Bacteria; Proteobacteria; Gammaproteobacteria; Enterobacterales;

DP773__locus_2

PDU1AB

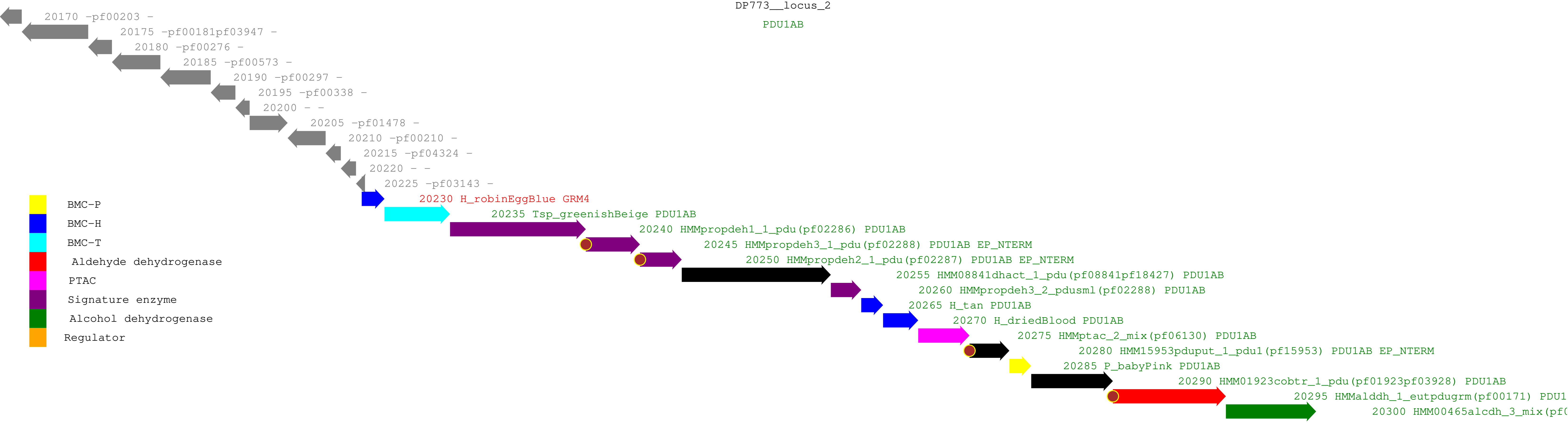

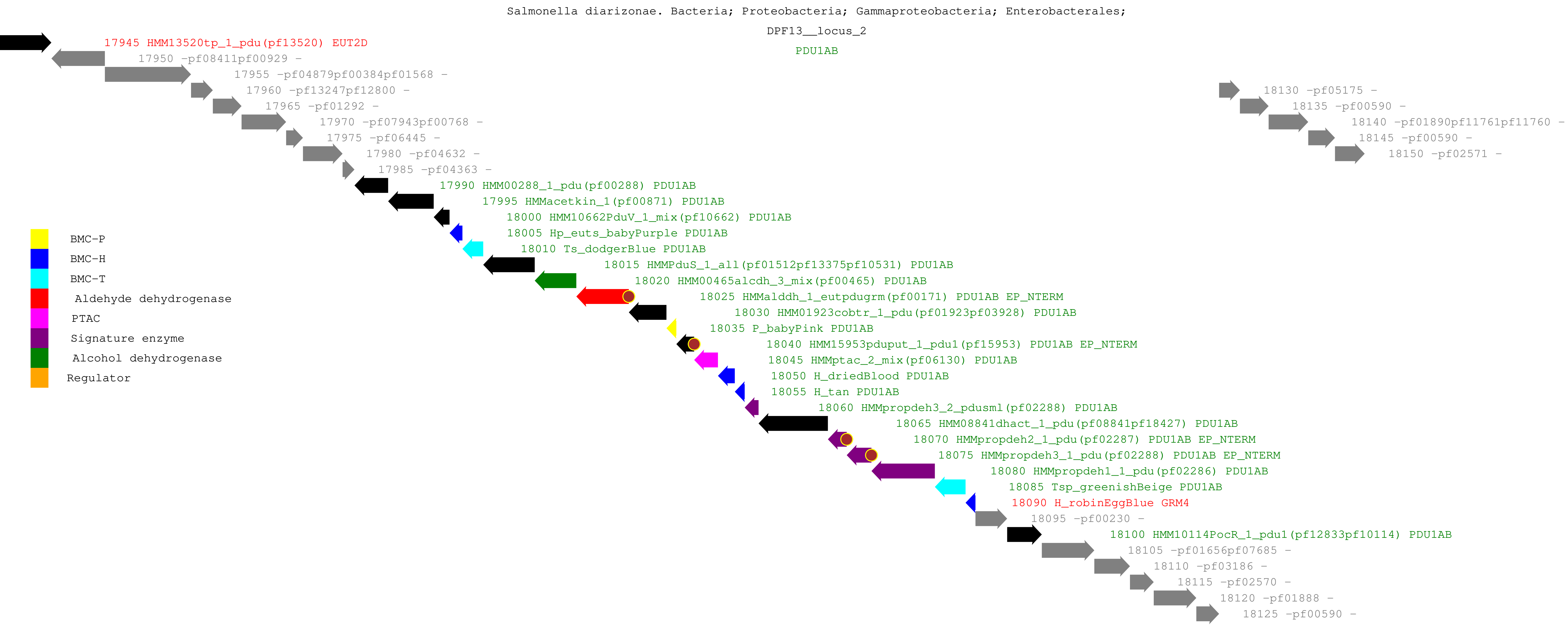
Salmonella diarizonae. Bacteria; Proteobacteria; Gammaproteobacteria; Enterobacterales;
DPF13__locus_2
PDU1AB
17945 HMM13520tp_1_pdu(pf13520) EUT2D
17950 -pf08411pf00929 -
17955 -pf04879pf00384pf01568 -
17960 -pf13247pf12800 -
17965 -pf01292 -
17970 -pf07943pf00768 -
17975 -pf06445 -
17980 -pf04632 -
17985 -pf04363 -
17990 HMM00288_1_pdu(pf00288) PDU1AB
17995 HMMacetkin_1(pf00871) PDU1AB
18000 HMM10662PduV_1_mix(pf10662) PDU1AB
18005 Hp_euts_babyPurple PDU1AB
18010 Ts_dodgerBlue PDU1AB
18015 HMMPduS_1_all(pf01512pf13375pf10531) PDU1AB
18020 HMM00465alcdh_3_mix(pf00465) PDU1AB
18025 HMMalddh_1_eutpdugrm(pf00171) PDU1AB EP_NTERM
18030 HMM01923cobtr_1_pdu(pf01923pf03928) PDU1AB
18035 P_babyPink PDU1AB
18040 HMM15953pduput_1_pdu1(pf15953) PDU1AB EP_NTERM
18045 HMMptac_2_mix(pf06130) PDU1AB
18050 H_driedBlood PDU1AB
18055 H_tan PDU1AB
18060 HMMpropdeh3_2_pdusml(pf02288) PDU1AB
18065 HMM08841dhact_1_pdu(pf08841pf18427) PDU1AB
18070 HMMpropdeh2_1_pdu(pf02287) PDU1AB EP_NTERM
18075 HMMpropdeh3_1_pdu(pf02288) PDU1AB EP_NTERM
18080 HMMpropdeh1_1_pdu(pf02286) PDU1AB
18085 Tsp_greenishBeige PDU1AB
18090 H_robinEggBlue GRM4
18095 -pf00230 -
18100 HMM10114PocR_1_pdu1(pf12833pf10114) PDU1AB
18105 -pf01656pf07685 -
18110 -pf03186 -
18115 -pf02570 -
18120 -pf01888 -
18125 -pf00590 -
18130 -pf05175 -
18135 -pf00590 -
18140 -pf01890pf11761pf11760 -
18145 -pf00590 -
18150 -pf02571 -
BMC-P
BMC-H
BMC-T
Aldehyde dehydrogenase
PTAC
Signature enzyme
Alcohol dehydrogenase
Regulator

Salmonella enterica I. Bacteria; Proteobacteria; Gammaproteobacteria; Enterobacterales;

DPK19__locus_1

PDU1AB

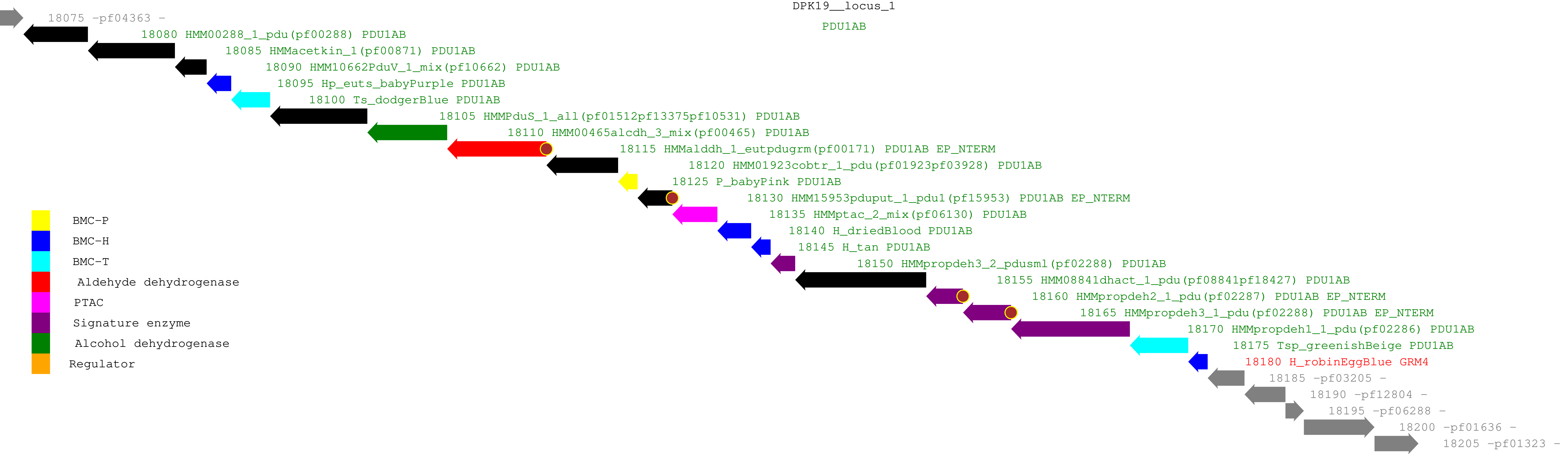

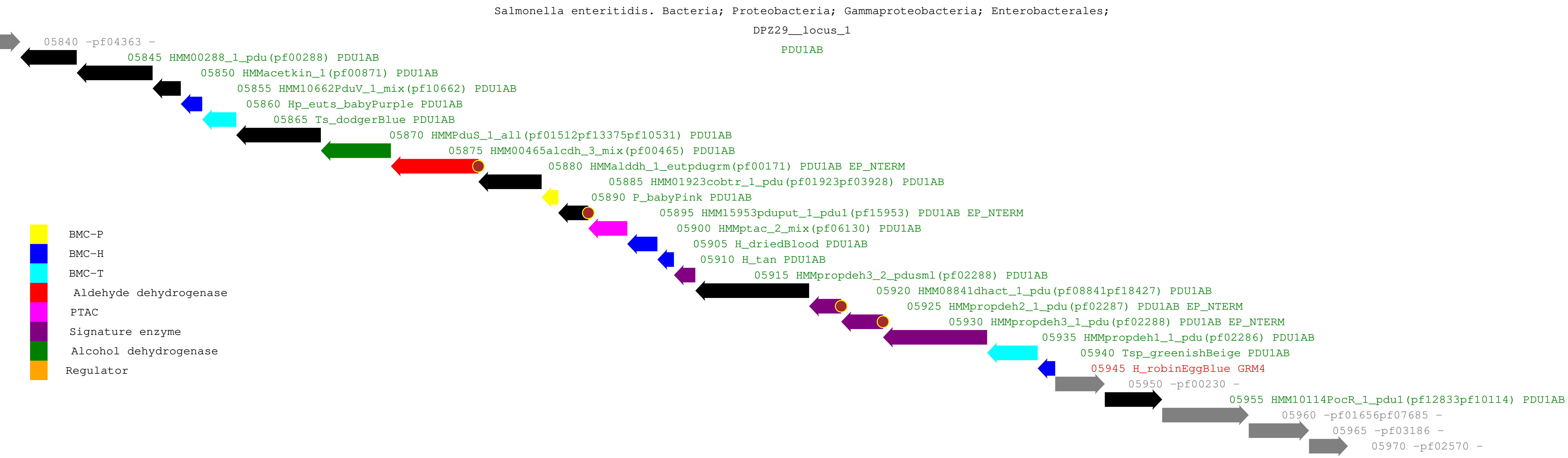
Salmonella enteritidis. Bacteria; Proteobacteria; Gammaproteobacteria; Enterobacterales;
DPZ29__locus_1
PDU1AB
05840 -pf04363 -
05845 HMM00288_1_pdu(pf00288) PDU1AB
05850 HMMacetkin_1(pf00871) PDU1AB
05855 HMM10662PduV_1_mix(pf10662) PDU1AB
05860 Hp_euts_babyPurple PDU1AB
05865 Ts_dodgerBlue PDU1AB
05870 HMMPduS_1_all(pf01512pf13375pf10531) PDU1AB
05875 HMM00465alcdh_3_mix(pf00465) PDU1AB
05880 HMMalddh_1_eutpdugrm(pf00171) PDU1AB EP_NTERM
05885 HMM01923cobtr_1_pdu(pf01923pf03928) PDU1AB
05890 P_babyPink PDU1AB
05895 HMM15953pduput_1_pdu1(pf15953) PDU1AB EP_NTERM
05900 HMMptac_2_mix(pf06130) PDU1AB
05905 H_driedBlood PDU1AB
05910 H_tan PDU1AB
05915 HMMpropdeh3_2_pdusml(pf02288) PDU1AB
05920 HMM08841dhact_1_pdu(pf08841pf18427) PDU1AB
05925 HMMpropdeh2_1_pdu(pf02287) PDU1AB EP_NTERM
05930 HMMpropdeh3_1_pdu(pf02288) PDU1AB EP_NTERM
05935 HMMpropdeh1_1_pdu(pf02286) PDU1AB
05940 Tsp_greenishBeige PDU1AB
05945 H_robinEggBlue GRM4
05950 -pf00230 -
05955 HMM10114PocR_1_pdu1(pf12833pf10114) PDU1AB
05960 -pf01656pf07685 -
05965 -pf03186 -
05970 -pf02570 -
BMC-P
BMC-H
BMC-T
Aldehyde dehydrogenase
PTAC
Signature enzyme
Alcohol dehydrogenase
Regulator

Salmonella enterica subsp. salamae. Bacteria; Proteobacteria; Gammaproteobacteria; Enterobacterales;

DQ004__locus_2

PDU1AB

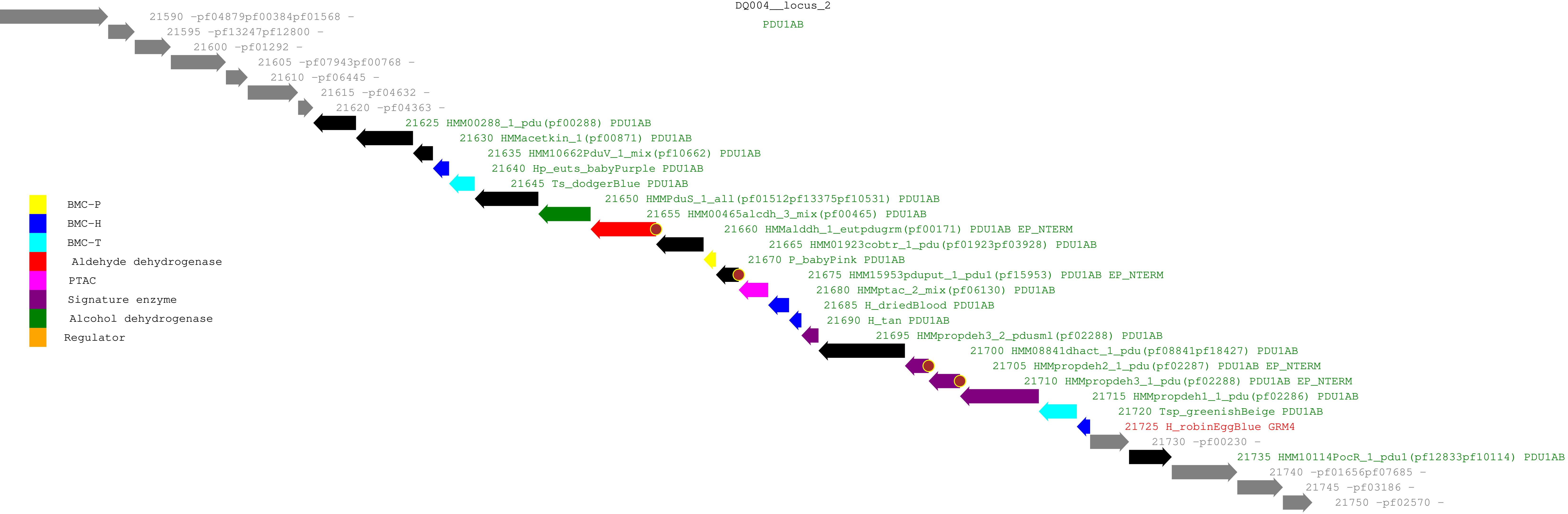

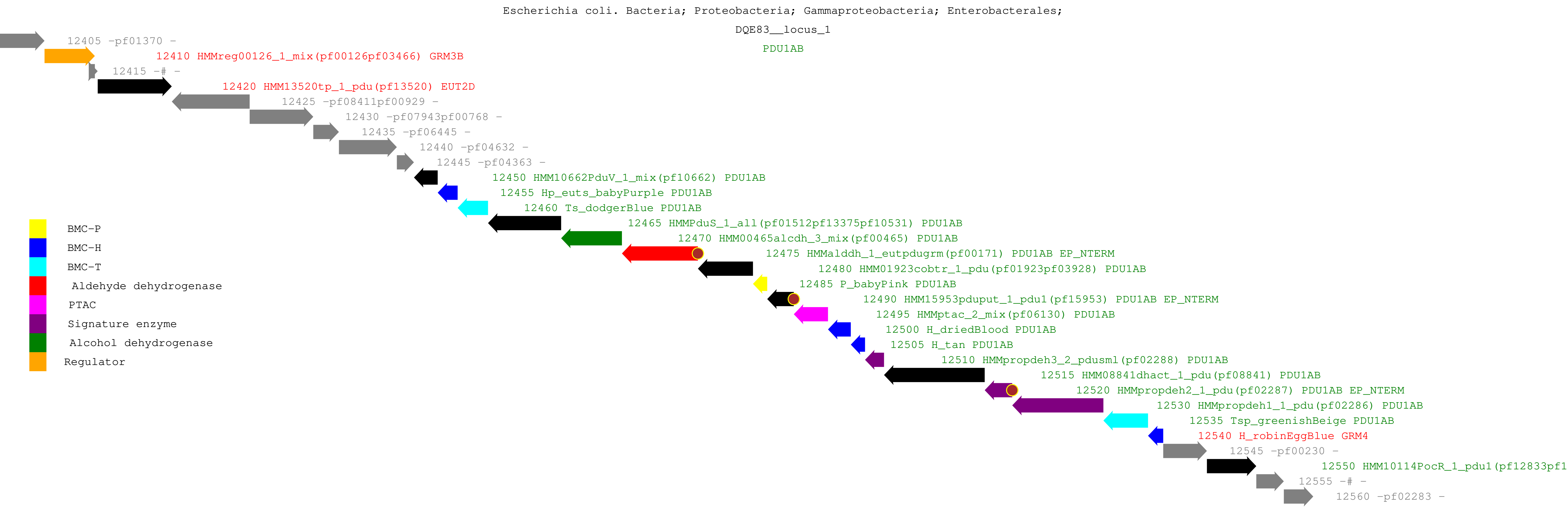

Escherichia coli. Bacteria; Proteobacteria; Gammaproteobacteria; Enterobacterales;
DQE83__locus_1
PDU1AB
12405 -pf01370 -
12410 HMMreg00126_1_mix(pf00126pf03466) GRM3B
12415 -# -
12420 HMM13520tp_1_pdu(pf13520) EUT2D
12425 -pf08411pf00929 -
12430 -pf07943pf00768 -
12435 -pf06445 -
12440 -pf04632 -
12445 -pf04363 -
12450 HMM10662PduV_1_mix(pf10662) PDU1AB
12455 Hp_euts_babyPurple PDU1AB
12460 Ts_dodgerBlue PDU1AB
12465 HMMPduS_1_all(pf01512pf13375pf10531) PDU1AB
12470 HMM00465alcdh_3_mix(pf00465) PDU1AB
12475 HMMaldh_1_eutpdugrm(pf00171) PDU1AB EP_NTERM
12480 HMM01923cobtr_1_pdu(pf01923pf03928) PDU1AB
12485 P_babyPink PDU1AB
12490 HMM15953pduput_1_pdu1(pf15953) PDU1AB EP_NTERM
12495 HMMptac_2_mix(pf06130) PDU1AB
12500 H_driedBlood PDU1AB
12505 H_tan PDU1AB
12510 HMMpropdeh3_2_pdusml(pf02288) PDU1AB
12515 HMM08841dhact_1_pdu(pf08841) PDU1AB
12520 HMMpropdeh2_1_pdu(pf02287) PDU1AB EP_NTERM
12530 HMMpropdeh1_1_pdu(pf02286) PDU1AB
12535 Tsp_greenishBeige PDU1AB
12540 H_robinEggBlue GRM4
12545 -pf00230 -
12550 HMM10114PocR_1_pdu1(pf12833pf1
12555 -# -
12560 -pf02283 -
BMC-P
BMC-H
BMC-T
Aldehyde dehydrogenase
PTAC
Signature enzyme
Alcohol dehydrogenase
Regulator

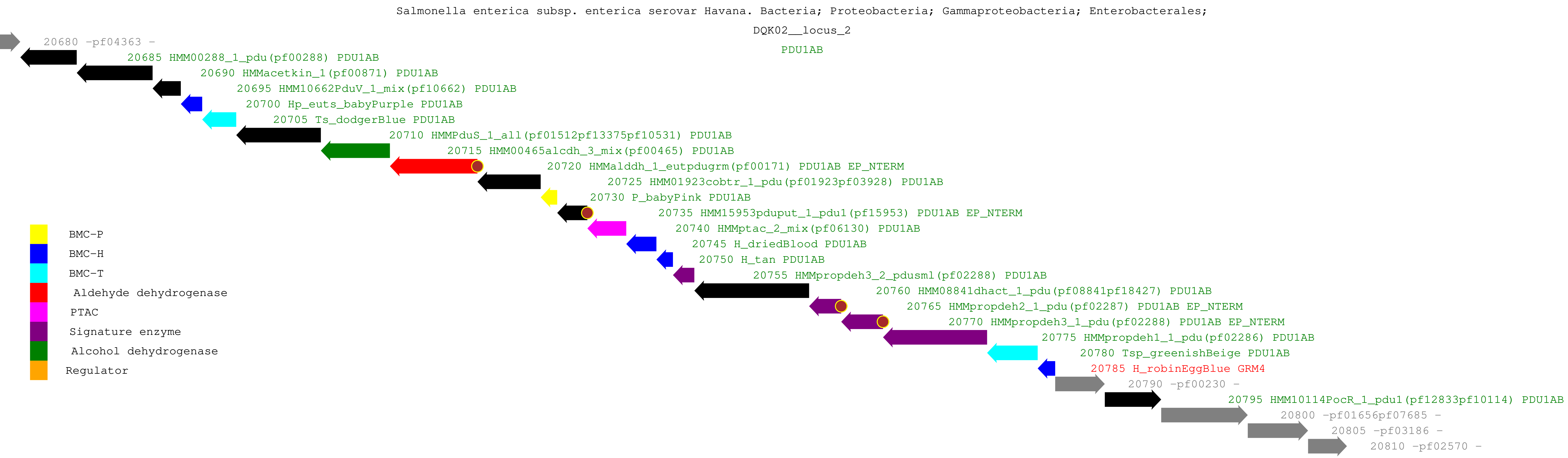
Salmonella enterica subsp. enterica serovar Havana. Bacteria; Proteobacteria; Gammaproteobacteria; Enterobacterales;
DQK02__locus_2
PDU1AB
20680 -pf04363 -
20685 HMM00288_1_pdu(pf00288) PDU1AB
20690 HMMacetkin_1(pf00871) PDU1AB
20695 HMM10662PduV_1_mix(pf10662) PDU1AB
20700 Hp_euts_babyPurple PDU1AB
20705 Ts_dodgerBlue PDU1AB
20710 HMMPduS_1_all(pf01512pf13375pf10531) PDU1AB
20715 HMM00465alcdh_3_mix(pf00465) PDU1AB
20720 HMMalddh_1_eutpdugrm(pf00171) PDU1AB EP_NTERM
20725 HMM01923cobtr_1_pdu(pf01923pf03928) PDU1AB
20730 P_babyPink PDU1AB
20735 HMM15953pduput_1_pdu1(pf15953) PDU1AB EP_NTERM
20740 HMMptac_2_mix(pf06130) PDU1AB
20745 H_driedBlood PDU1AB
20750 H_tan PDU1AB
20755 HMMpropdeh3_2_pdusml(pf02288) PDU1AB
20760 HMM08841dhact_1_pdu(pf08841pf18427) PDU1AB
20765 HMMpropdeh2_1_pdu(pf02287) PDU1AB EP_NTERM
20770 HMMpropdeh3_1_pdu(pf02288) PDU1AB EP_NTERM
20775 HMMpropdeh1_1_pdu(pf02286) PDU1AB
20780 Tsp_greenishBeige PDU1AB
20785 H_robinEggBlue GRM4
20790 -pf00230 -
20795 HMM10114PocR_1_pdu1(pf12833pf10114) PDU1AB
20800 -pf01656pf07685 -
20805 -pf03186 -
20810 -pf02570 -
BMC-P
BMC-H
BMC-T
Aldehyde dehydrogenase
PTAC
Signature enzyme
Alcohol dehydrogenase
Regulator

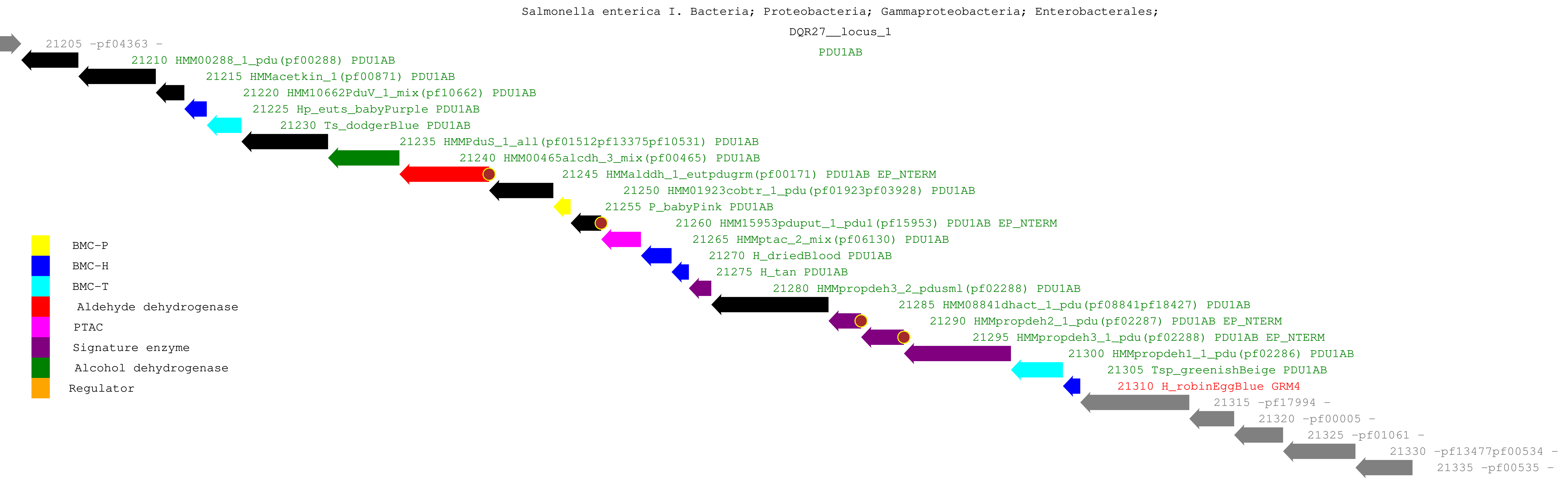
Salmonella enterica I. Bacteria; Proteobacteria; Gammaproteobacteria; Enterobacterales;
DQR27__locus_1
PDU1AB
21205 -pf04363 -
21210 HMM00288_1_pdu(pf00288) PDU1AB
21215 HMMacetkin_1(pf00871) PDU1AB
21220 HMM10662PduV_1_mix(pf10662) PDU1AB
21225 Hp_euts_babyPurple PDU1AB
21230 Ts_dodgerBlue PDU1AB
21235 HMMPduS_1_all(pf01512pf13375pf10531) PDU1AB
21240 HMM00465alcdh_3_mix(pf00465) PDU1AB
21245 HMMalddh_1_eutpdugrm(pf00171) PDU1AB EP_NTERM
21250 HMM01923cobtr_1_pdu(pf01923pf03928) PDU1AB
21255 P_babyPink PDU1AB
21260 HMM15953pduput_1_pdu1(pf15953) PDU1AB EP_NTERM
21265 HMMptac_2_mix(pf06130) PDU1AB
21270 H_driedBlood PDU1AB
21275 H_tan PDU1AB
21280 HMMpropdeh3_2_pdusml(pf02288) PDU1AB
21285 HMM08841dhact_1_pdu(pf08841pf18427) PDU1AB
21290 HMMpropdeh2_1_pdu(pf02287) PDU1AB EP_NTERM
21295 HMMpropdeh3_1_pdu(pf02288) PDU1AB EP_NTERM
21300 HMMpropdeh1_1_pdu(pf02286) PDU1AB
21305 Tsp_greenishBeige PDU1AB
21310 H_robinEggBlue GRM4
21315 -pf17994 -
21320 -pf00005 -
21325 -pf01061 -
21330 -pf13477pf00534 -
21335 -pf00535 -
BMC-P
BMC-H
BMC-T
Aldehyde dehydrogenase
PTAC
Signature enzyme
Alcohol dehydrogenase
Regulator

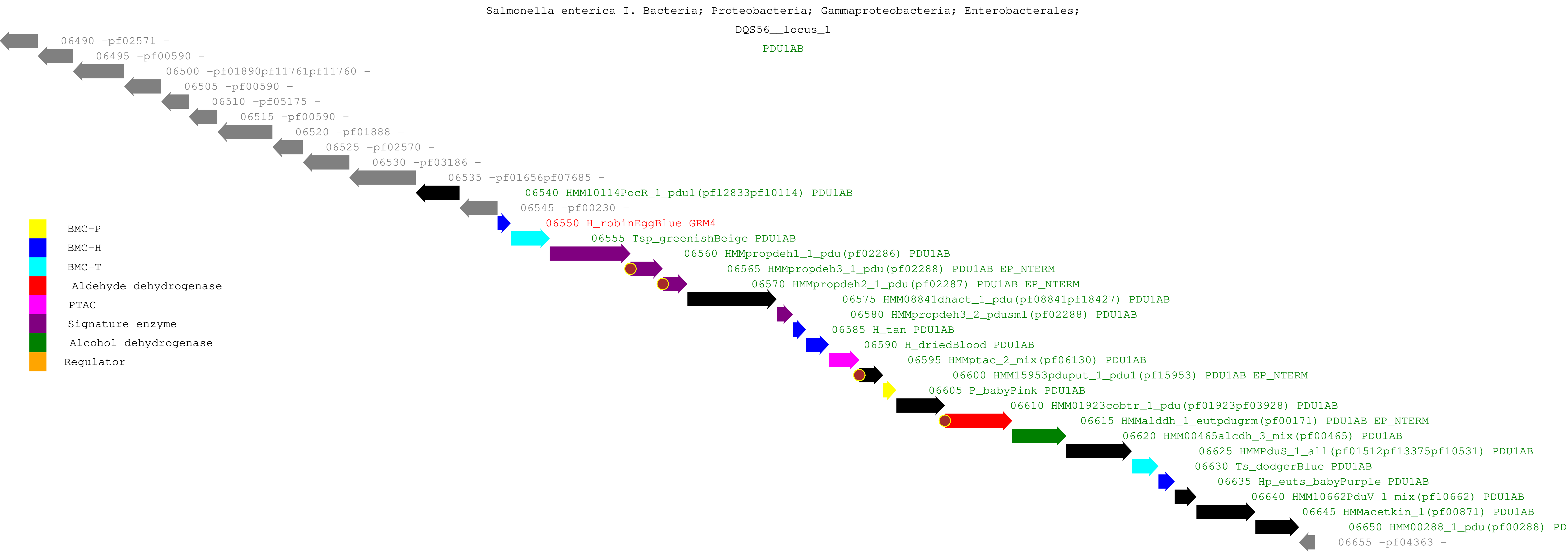
Salmonella enterica I. Bacteria; Proteobacteria; Gammaproteobacteria; Enterobacterales;
DQS56__locus_1
PDU1AB
06490 -pf02571 -
06495 -pf00590 -
06500 -pf01890pf11761pf11760 -
06505 -pf00590 -
06510 -pf05175 -
06515 -pf00590 -
06520 -pf01888 -
06525 -pf02570 -
06530 -pf03186 -
06535 -pf01656pf07685 -
06540 HMM10114PocR_1_pdu1(pf12833pf10114) PDU1AB
06545 -pf00230 -
06550 H_robinEggBlue GRM4
06555 Tsp_greenishBeige PDU1AB
06560 HMMpropdeh1_1_pdu(pf02286) PDU1AB
06565 HMMpropdeh3_1_pdu(pf02288) PDU1AB EP_NTERM
06570 HMMpropdeh2_1_pdu(pf02287) PDU1AB EP_NTERM
06575 HMM08841dhact_1_pdu(pf08841pf18427) PDU1AB
06580 HMMpropdeh3_2_pdusml(pf02288) PDU1AB
06585 H_tan PDU1AB
06590 H_driedBlood PDU1AB
06595 HMMptac_2_mix(pf06130) PDU1AB
06600 HMM15953pduput_1_pdu1(pf15953) PDU1AB EP_NTERM
06605 P_babyPink PDU1AB
06610 HMM01923cobtr_1_pdu(pf01923pf03928) PDU1AB
06615 HMMalddh_1_eutpdugrm(pf00171) PDU1AB EP_NTERM
06620 HMM00465alcdh_3_mix(pf00465) PDU1AB
06625 HMMPduS_1_all(pf01512pf13375pf10531) PDU1AB
06630 Ts_dodgerBlue PDU1AB
06635 Hp_euts_babyPurple PDU1AB
06640 HMM10662PduV_1_mix(pf10662) PDU1AB
06645 HMMacetkin_1(pf00871) PDU1AB
06650 HMM00288_1_pdu(pf00288) PD
06655 -pf04363 -
BMC-P
BMC-H
BMC-T
Aldehyde dehydrogenase
PTAC
Signature enzyme
Alcohol dehydrogenase
Regulator

Salmonella enterica I. Bacteria; Proteobacteria; Gammaproteobacteria; Enterobacterales;

DQS71__locus_1

PDU1AB

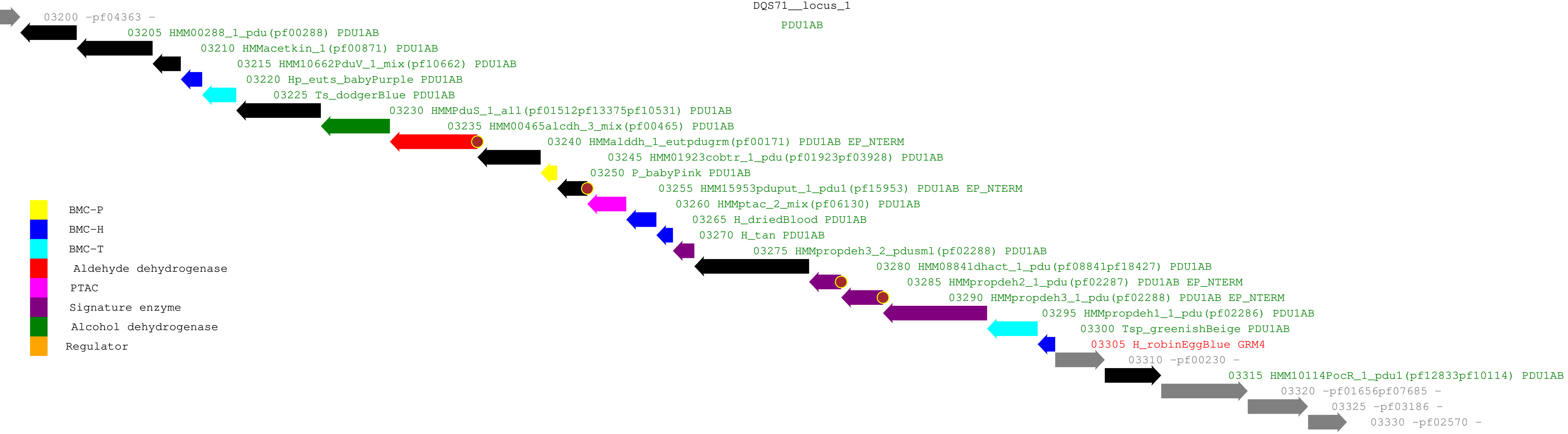

Salmonella enterica I. Bacteria; Proteobacteria; Gammaproteobacteria; Enterobacterales;

DQZ68__locus_1

PDU1AB

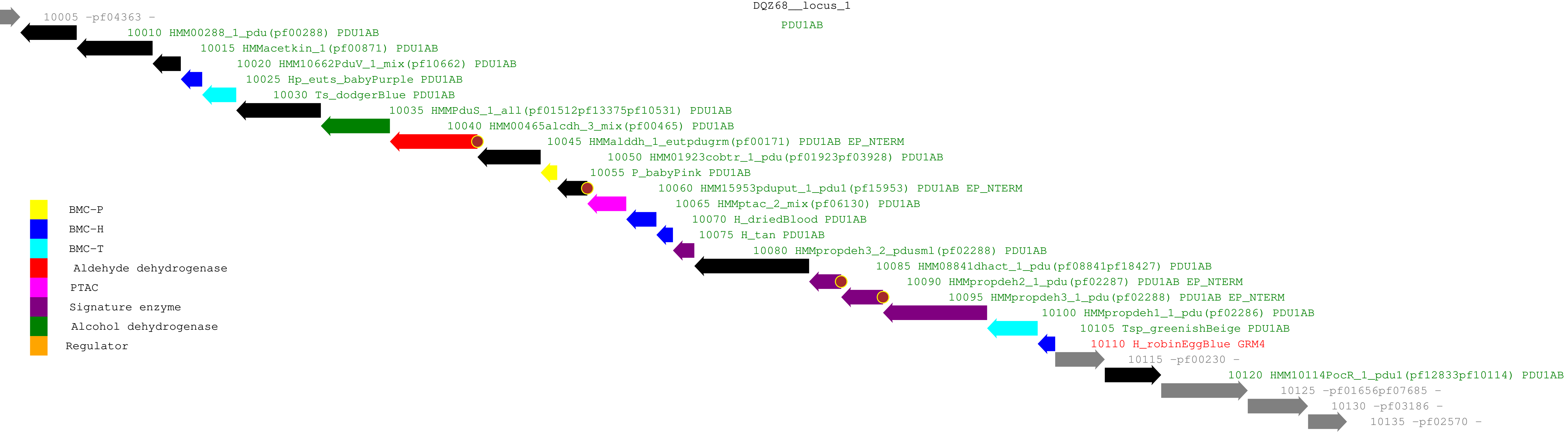

Salmonella enterica I. Bacteria; Proteobacteria; Gammaproteobacteria; Enterobacterales;

DRE09__locus_1

PDU1AB

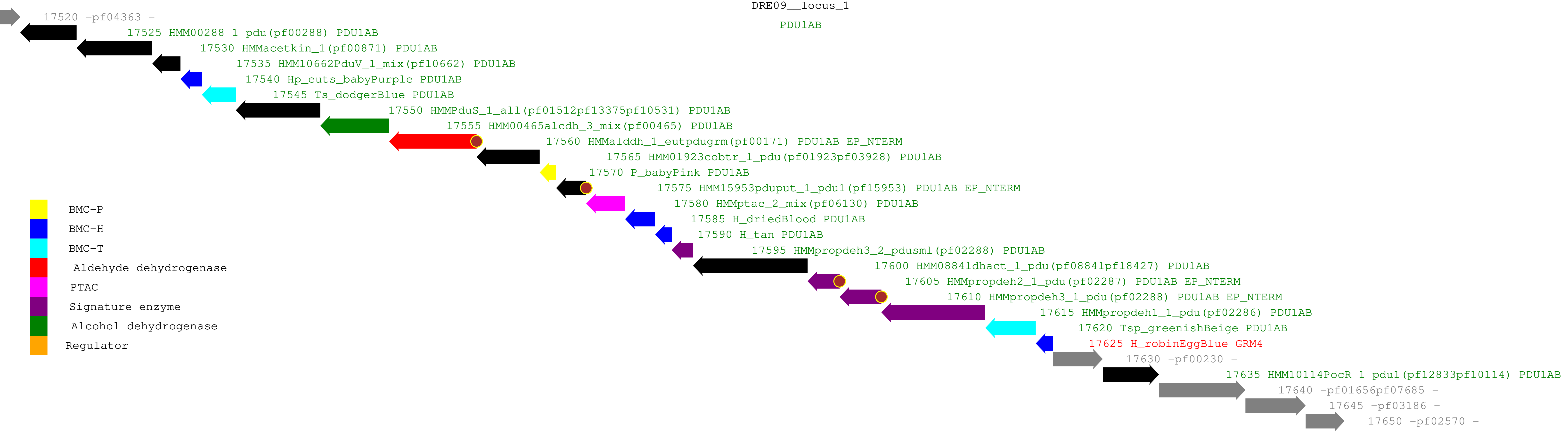

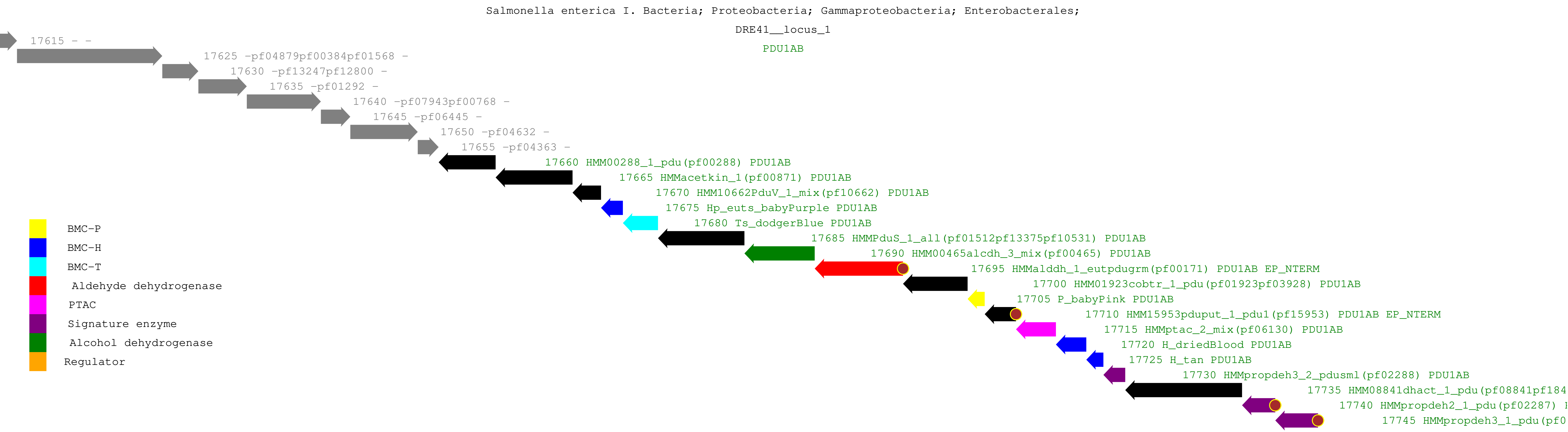
Salmonella enterica I. Bacteria; Proteobacteria; Gammaproteobacteria; Enterobacterales;
DRE41__locus_1
PDU1AB
17615 - -
17625 -pf04879pf00384pf01568 -
17630 -pf13247pf12800 -
17635 -pf01292 -
17640 -pf07943pf00768 -
17645 -pf06445 -
17650 -pf04632 -
17655 -pf04363 -
17660 HMM00288_1_pdu(pf00288) PDU1AB
17665 HMMacetkin_1(pf00871) PDU1AB
17670 HMM10662PduV_1_mix(pf10662) PDU1AB
17675 Hp_euts_babyPurple PDU1AB
17680 Ts_dodgerBlue PDU1AB
17685 HMMPduS_1_all(pf01512pf13375pf10531) PDU1AB
17690 HMM00465alcdh_3_mix(pf00465) PDU1AB
17695 HMMalddh_1_eutpdugrm(pf00171) PDU1AB EP_NTERM
17700 HMM01923cobtr_1_pdu(pf01923pf03928) PDU1AB
17705 P_babyPink PDU1AB
17710 HMM15953pduput_1_pdu1(pf15953) PDU1AB EP_NTERM
17715 HMMptac_2_mix(pf06130) PDU1AB
17720 H_driedBlood PDU1AB
17725 H_tan PDU1AB
17730 HMMpropdeh3_2_pdusml(pf02288) PDU1AB
17735 HMM08841dhact_1_pdu(pf08841pf184
17740 HMMpropdeh2_1_pdu(pf02287) I
17745 HMMpropdeh3_1_pdu(pf0
BMC-P
BMC-H
BMC-T
Aldehyde dehydrogenase
PTAC
Signature enzyme
Alcohol dehydrogenase
Regulator

Salmonella enterica I. Bacteria; Proteobacteria; Gammaproteobacteria; Enterobacterales;

DRK91__locus_1

PDU1AB

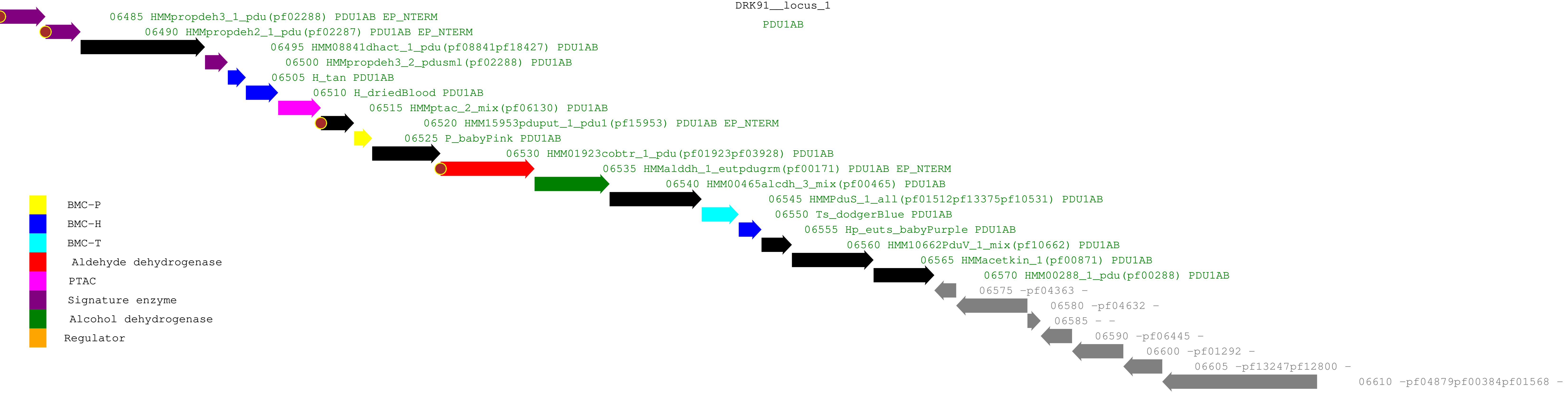

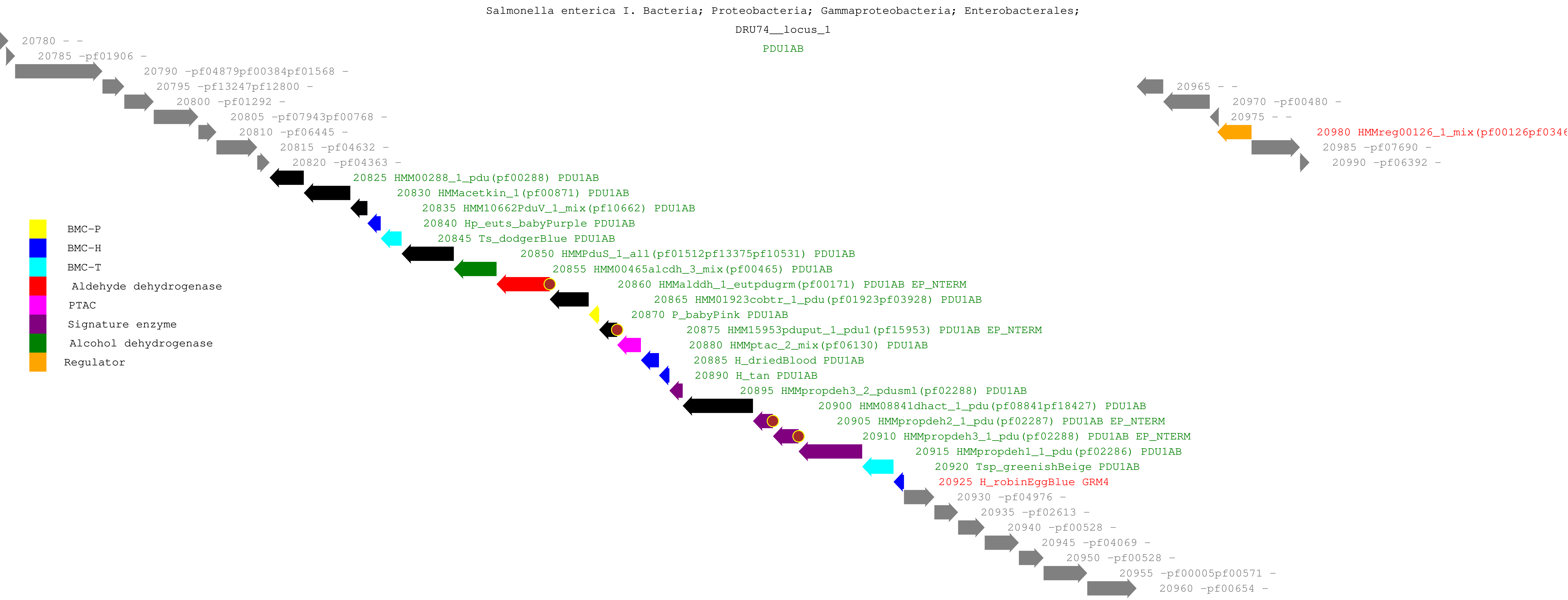

Salmonella enterica I. Bacteria; Proteobacteria; Gammaproteobacteria; Enterobacterales;
DRU74__locus_1
PDU1AB
20780 - -
20785 -pf01906 -
20790 -pf04879pf00384pf01568 -
20795 -pf13247pf12800 -
20800 -pf01292 -
20805 -pf07943pf00768 -
20810 -pf06445 -
20815 -pf04632 -
20820 -pf04363 -
20825 HMM00288_1_pdu(pf00288) PDU1AB
20830 HMMacetkin_1(pf00871) PDU1AB
20835 HMM10662PduV_1_mix(pf10662) PDU1AB
20840 Hp_euts_babyPurple PDU1AB
20845 Ts_dodgerBlue PDU1AB
20850 HMMPduS_1_all(pf01512pf13375pf10531) PDU1AB
20855 HMM00465alcdh_3_mix(pf00465) PDU1AB
20860 HMMalddh_1_eutpdugrm(pf00171) PDU1AB EP_NTERM
20865 HMM01923cobtr_1_pdu(pf01923pf03928) PDU1AB
20870 P_babyPink PDU1AB
20875 HMM15953pduput_1_pdu1(pf15953) PDU1AB EP_NTERM
20880 HMMptac_2_mix(pf06130) PDU1AB
20885 H_driedBlood PDU1AB
20890 H_tan PDU1AB
20895 HMMpropdeh3_2_pdusml(pf02288) PDU1AB
20900 HMM08841dhact_1_pdu(pf08841pf18427) PDU1AB
20905 HMMpropdeh2_1_pdu(pf02287) PDU1AB EP_NTERM
20910 HMMpropdeh3_1_pdu(pf02288) PDU1AB EP_NTERM
20915 HMMpropdeh1_1_pdu(pf02286) PDU1AB
20920 Tsp_greenishBeige PDU1AB
20925 H_robinEggBlue GRM4
20930 -pf04976 -
20935 -pf02613 -
20940 -pf00528 -
20945 -pf04069 -
20950 -pf00528 -
20955 -pf00005pf00571 -
20960 -pf00654 -
20965 - -
20970 -pf00480 -
20975 - -
20980 HMMreg00126_1_mix(pf00126pf0346
20985 -pf07690 -
20990 -pf06392 -
BMC-P
BMC-H
BMC-T
Aldehyde dehydrogenase
PTAC
Signature enzyme
Alcohol dehydrogenase
Regulator

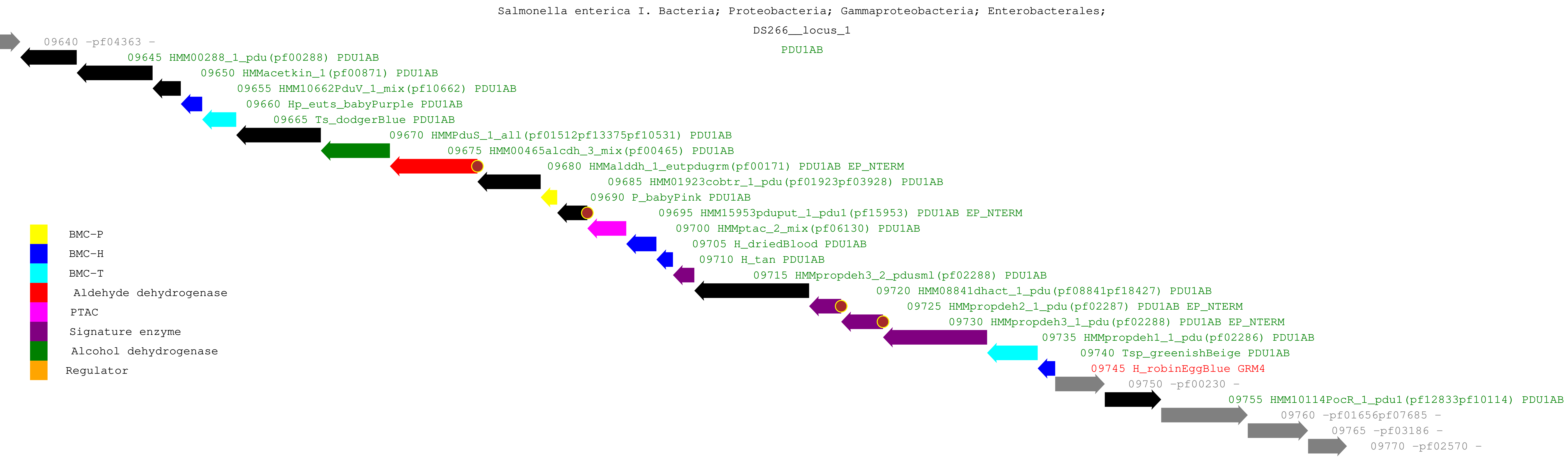
Salmonella enterica I. Bacteria; Proteobacteria; Gammaproteobacteria; Enterobacterales;
DS266__locus_1
PDU1AB
09640 -pf04363 -
09645 HMM00288_1_pdu(pf00288) PDU1AB
09650 HMMacetkin_1(pf00871) PDU1AB
09655 HMM10662PduV_1_mix(pf10662) PDU1AB
09660 Hp_euts_babyPurple PDU1AB
09665 Ts_dodgerBlue PDU1AB
09670 HMMPduS_1_all(pf01512pf13375pf10531) PDU1AB
09675 HMM00465alcdh_3_mix(pf00465) PDU1AB
09680 HMMalddh_1_eutpdugrm(pf00171) PDU1AB EP_NTERM
09685 HMM01923cobtr_1_pdu(pf01923pf03928) PDU1AB
09690 P_babyPink PDU1AB
09695 HMM15953pduput_1_pdu1(pf15953) PDU1AB EP_NTERM
09700 HMMptac_2_mix(pf06130) PDU1AB
09705 H_driedBlood PDU1AB
09710 H_tan PDU1AB
09715 HMMpropdeh3_2_pdusml(pf02288) PDU1AB
09720 HMM08841dhact_1_pdu(pf08841pf18427) PDU1AB
09725 HMMpropdeh2_1_pdu(pf02287) PDU1AB EP_NTERM
09730 HMMpropdeh3_1_pdu(pf02288) PDU1AB EP_NTERM
09735 HMMpropdeh1_1_pdu(pf02286) PDU1AB
09740 Tsp_greenishBeige PDU1AB
09745 H_robinEggBlue GRM4
09750 -pf00230 -
09755 HMM10114PocR_1_pdu1(pf12833pf10114) PDU1AB
09760 -pf01656pf07685 -
09765 -pf03186 -
09770 -pf02570 -
BMC-P
BMC-H
BMC-T
Aldehyde dehydrogenase
PTAC
Signature enzyme
Alcohol dehydrogenase
Regulator

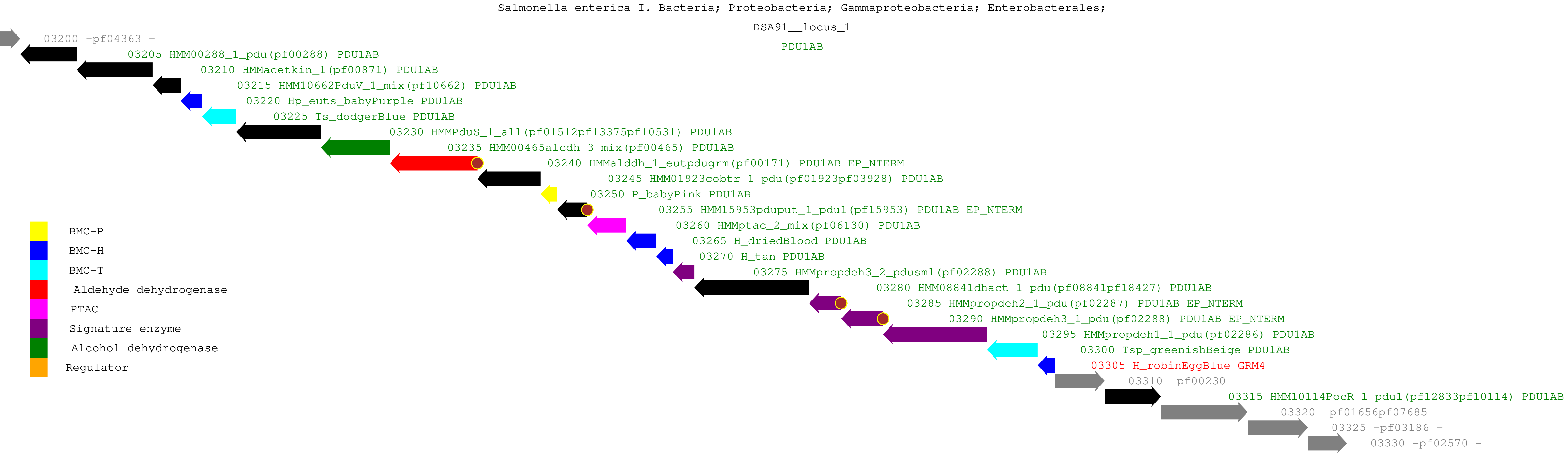
Salmonella enterica I. Bacteria; Proteobacteria; Gammaproteobacteria; Enterobacterales;
DSA91__locus_1
PDU1AB
03200 -pf04363 -
03205 HMM00288_1_pdu(pf00288) PDU1AB
03210 HMMacetkin_1(pf00871) PDU1AB
03215 HMM10662PduV_1_mix(pf10662) PDU1AB
03220 Hp_euts_babyPurple PDU1AB
03225 Ts_dodgerBlue PDU1AB
03230 HMMPduS_1_all(pf01512pf13375pf10531) PDU1AB
03235 HMM00465alcdh_3_mix(pf00465) PDU1AB
03240 HMMalddh_1_eutpdugrm(pf00171) PDU1AB EP_NTERM
03245 HMM01923cobtr_1_pdu(pf01923pf03928) PDU1AB
03250 P_babyPink PDU1AB
03255 HMM15953pduput_1_pdu1(pf15953) PDU1AB EP_NTERM
03260 HMMptac_2_mix(pf06130) PDU1AB
03265 H_driedBlood PDU1AB
03270 H_tan PDU1AB
03275 HMMpropdeh3_2_pdusml(pf02288) PDU1AB
03280 HMM08841dhact_1_pdu(pf08841pf18427) PDU1AB
03285 HMMpropdeh2_1_pdu(pf02287) PDU1AB EP_NTERM
03290 HMMpropdeh3_1_pdu(pf02288) PDU1AB EP_NTERM
03295 HMMpropdeh1_1_pdu(pf02286) PDU1AB
03300 Tsp_greenishBeige PDU1AB
03305 H_robinEggBlue GRM4
03310 -pf00230 -
03315 HMM10114PocR_1_pdu1(pf12833pf10114) PDU1AB
03320 -pf01656pf07685 -
03325 -pf03186 -
03330 -pf02570 -
BMC-P
BMC-H
BMC-T
Aldehyde dehydrogenase
PTAC
Signature enzyme
Alcohol dehydrogenase
Regulator

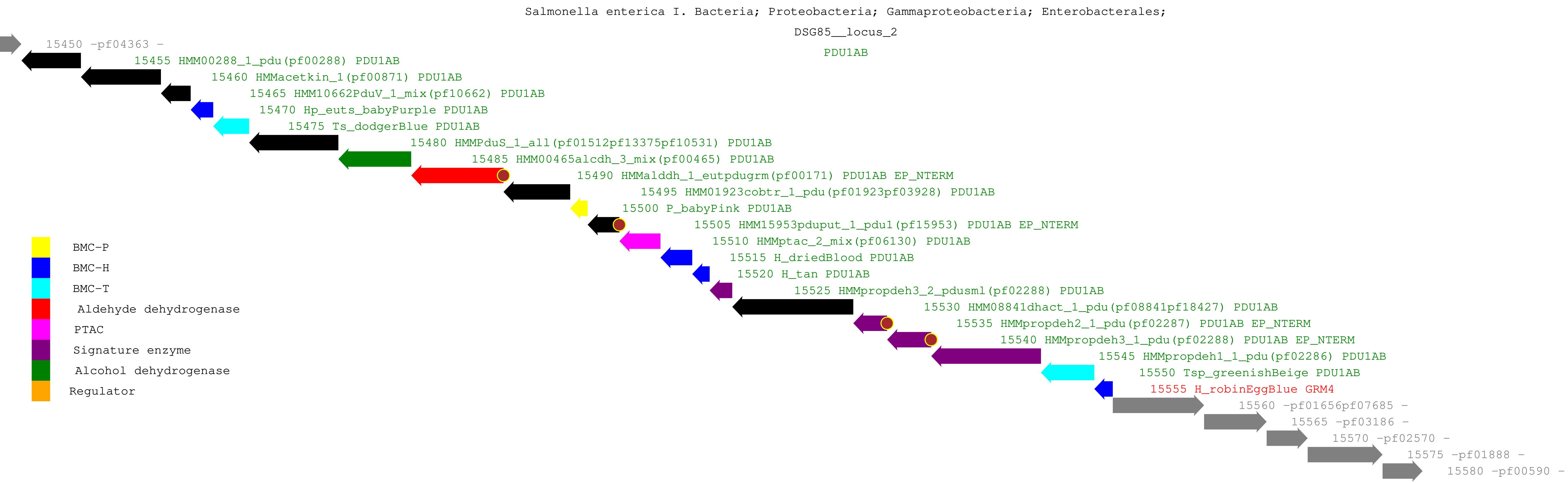
Salmonella enterica I. Bacteria; Proteobacteria; Gammaproteobacteria; Enterobacterales;
DSG85__locus_2
PDU1AB
15450 -pf04363 -
15455 HMM00288_1_pdu(pf00288) PDU1AB
15460 HMMacetkin_1(pf00871) PDU1AB
15465 HMM10662PduV_1_mix(pf10662) PDU1AB
15470 Hp_euts_babyPurple PDU1AB
15475 Ts_dodgerBlue PDU1AB
15480 HMMPduS_1_all(pf01512pf13375pf10531) PDU1AB
15485 HMM00465alcdh_3_mix(pf00465) PDU1AB
15490 HMMalddh_1_eutpdugrm(pf00171) PDU1AB EP_NTERM
15495 HMM01923cobtr_1_pdu(pf01923pf03928) PDU1AB
15500 P_babyPink PDU1AB
15505 HMM15953pduput_1_pdu1(pf15953) PDU1AB EP_NTERM
15510 HMMptac_2_mix(pf06130) PDU1AB
15515 H_driedBlood PDU1AB
15520 H_tan PDU1AB
15525 HMMpropdeh3_2_pdusml(pf02288) PDU1AB
15530 HMM08841dhact_1_pdu(pf08841pf18427) PDU1AB
15535 HMMpropdeh2_1_pdu(pf02287) PDU1AB EP_NTERM
15540 HMMpropdeh3_1_pdu(pf02288) PDU1AB EP_NTERM
15545 HMMpropdeh1_1_pdu(pf02286) PDU1AB
15550 Tsp_greenishBeige PDU1AB
15555 H_robinEggBlue GRM4
15560 -pf01656pf07685 -
15565 -pf03186 -
15570 -pf02570 -
15575 -pf01888 -
15580 -pf00590 -
BMC-P
BMC-H
BMC-T
Aldehyde dehydrogenase
PTAC
Signature enzyme
Alcohol dehydrogenase
Regulator

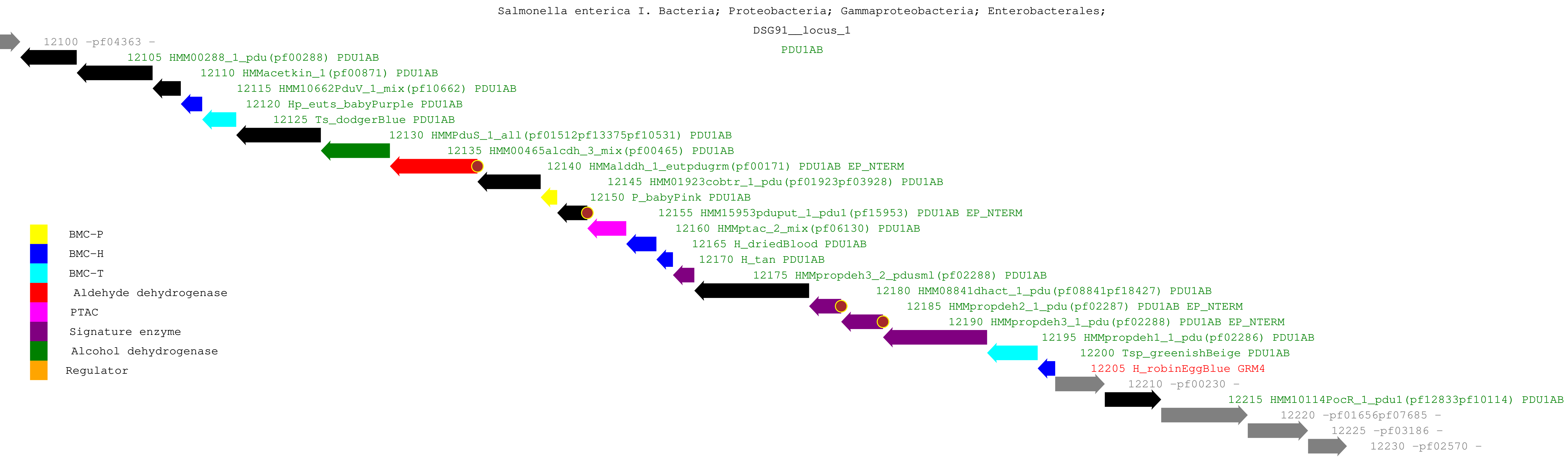

Salmonella enterica I. Bacteria; Proteobacteria; Gammaproteobacteria; Enterobacterales;
DSG91__locus_1
PDU1AB
12100 -pf04363 -
12105 HMM00288_1_pdu(pf00288) PDU1AB
12110 HMMacetkin_1(pf00871) PDU1AB
12115 HMM10662PduV_1_mix(pf10662) PDU1AB
12120 Hp_euts_babyPurple PDU1AB
12125 Ts_dodgerBlue PDU1AB
12130 HMMPduS_1_all(pf01512pf13375pf10531) PDU1AB
12135 HMM00465alcdh_3_mix(pf00465) PDU1AB
12140 HMMalddh_1_eutpdugrm(pf00171) PDU1AB EP_NTERM
12145 HMM01923cobtr_1_pdu(pf01923pf03928) PDU1AB
12150 P_babyPink PDU1AB
12155 HMM15953pduput_1_pdu1(pf15953) PDU1AB EP_NTERM
12160 HMMptac_2_mix(pf06130) PDU1AB
12165 H_driedBlood PDU1AB
12170 H_tan PDU1AB
12175 HMMpropdeh3_2_pdusml(pf02288) PDU1AB
12180 HMM08841dhact_1_pdu(pf08841pf18427) PDU1AB
12185 HMMpropdeh2_1_pdu(pf02287) PDU1AB EP_NTERM
12190 HMMpropdeh3_1_pdu(pf02288) PDU1AB EP_NTERM
12195 HMMpropdeh1_1_pdu(pf02286) PDU1AB
12200 Tsp_greenishBeige PDU1AB
12205 H_robinEggBlue GRM4
12210 -pf00230 -
12215 HMM10114PocR_1_pdu1(pf12833pf10114) PDU1AB
12220 -pf01656pf07685 -
12225 -pf03186 -
12230 -pf02570 -
BMC-P
BMC-H
BMC-T
Aldehyde dehydrogenase
PTAC
Signature enzyme
Alcohol dehydrogenase
Regulator

Salmonella enterica I. Bacteria; Proteobacteria; Gammaproteobacteria; Enterobacterales;

DSM93__locus_1

PDU1AB

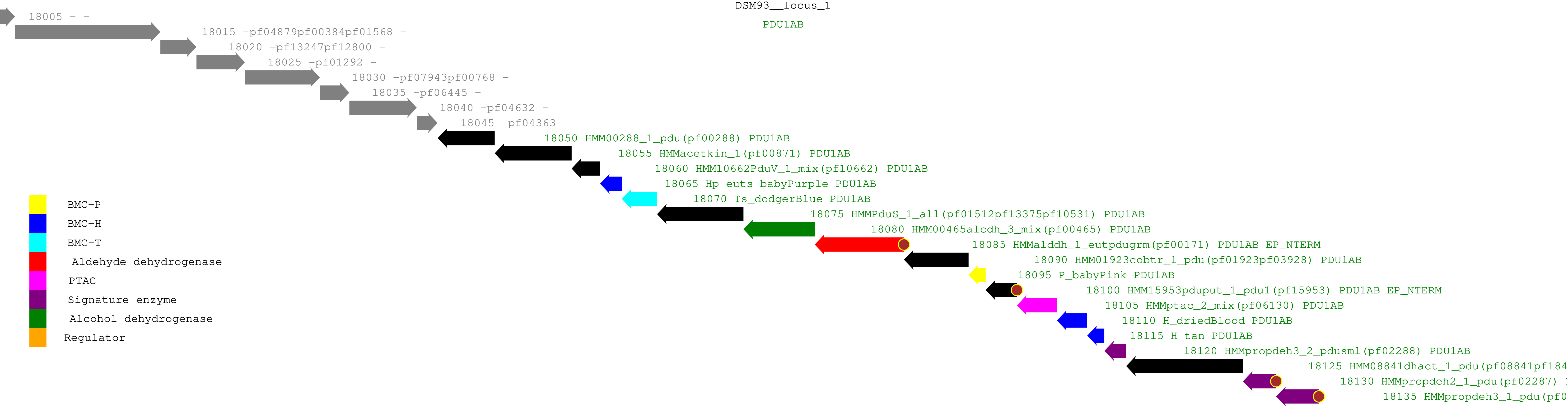

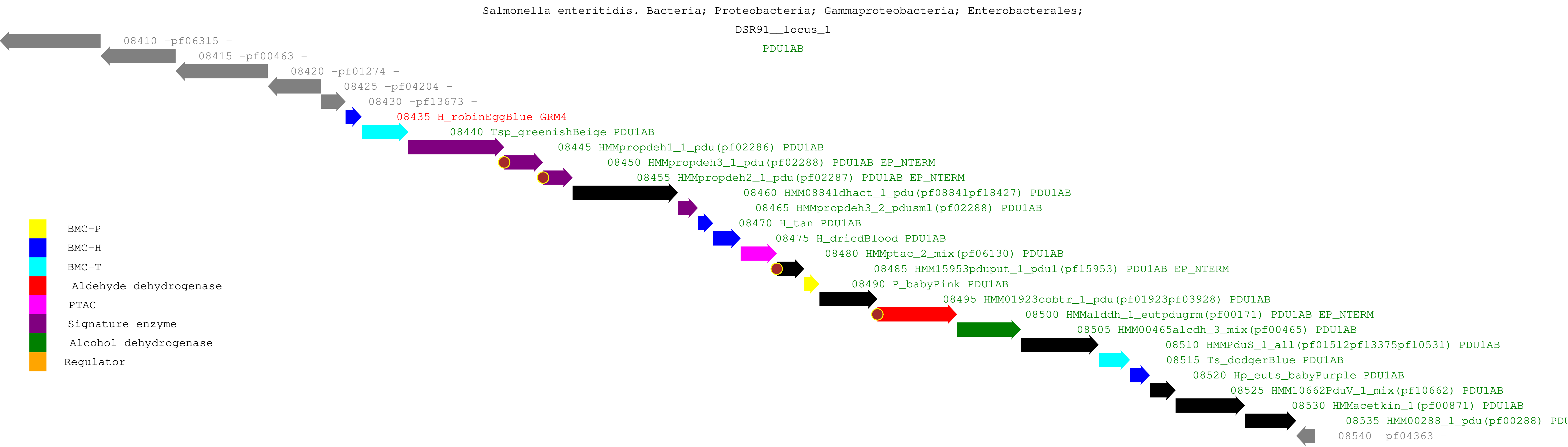
Salmonella enteritidis. Bacteria; Proteobacteria; Gammaproteobacteria; Enterobacterales;
DSR91__locus_1
PDU1AB
08410 -pf06315 -
08415 -pf00463 -
08420 -pf01274 -
08425 -pf04204 -
08430 -pf13673 -
08435 H_robinEggBlue GRM4
08440 Tsp_greenishBeige PDU1AB
08445 HMMpropdeh1_1_pdu(pf02286) PDU1AB
08450 HMMpropdeh3_1_pdu(pf02288) PDU1AB EP_NTERM
08455 HMMpropdeh2_1_pdu(pf02287) PDU1AB EP_NTERM
08460 HMM08841dhact_1_pdu(pf08841pf18427) PDU1AB
08465 HMMpropdeh3_2_pdusml(pf02288) PDU1AB
08470 H_tan PDU1AB
08475 H_driedBlood PDU1AB
08480 HMMptac_2_mix(pf06130) PDU1AB
08485 HMM15953pduput_1_pdu1(pf15953) PDU1AB EP_NTERM
08490 P_babyPink PDU1AB
08495 HMM01923cobtr_1_pdu(pf01923pf03928) PDU1AB
08500 HMMalddh_1_eutpdugrm(pf00171) PDU1AB EP_NTERM
08505 HMM00465alcdh_3_mix(pf00465) PDU1AB
08510 HMMPduS_1_all(pf01512pf13375pf10531) PDU1AB
08515 Ts_dodgerBlue PDU1AB
08520 Hp_euts_babyPurple PDU1AB
08525 HMM10662PduV_1_mix(pf10662) PDU1AB
08530 HMMacetkin_1(pf00871) PDU1AB
08535 HMM00288_1_pdu(pf00288) PDU
08540 -pf04363 -
BMC-P
BMC-H
BMC-T
Aldehyde dehydrogenase
PTAC
Signature enzyme
Alcohol dehydrogenase
Regulator

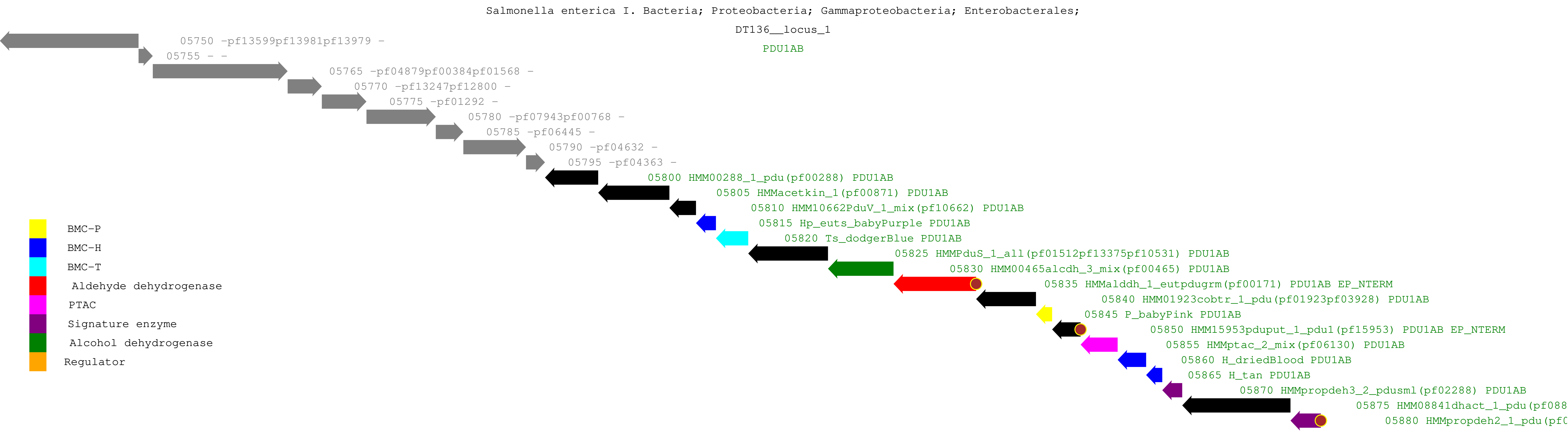
Salmonella enterica I. Bacteria; Proteobacteria; Gammaproteobacteria; Enterobacterales;
DT136__locus_1
PDU1AB
05750 -pf13599pf13981pf13979 -
05755 - -
05765 -pf04879pf00384pf01568 -
05770 -pf13247pf12800 -
05775 -pf01292 -
05780 -pf07943pf00768 -
05785 -pf06445 -
05790 -pf04632 -
05795 -pf04363 -
05800 HMM00288_1_pdu(pf00288) PDU1AB
05805 HMMacetkin_1(pf00871) PDU1AB
05810 HMM10662PduV_1_mix(pf10662) PDU1AB
05815 Hp_euts_babyPurple PDU1AB
05820 Ts_dodgerBlue PDU1AB
05825 HMMPduS_1_all(pf01512pf13375pf10531) PDU1AB
05830 HMM00465alcdh_3_mix(pf00465) PDU1AB
05835 HMMalddh_1_eutpdugrm(pf00171) PDU1AB EP_NTERM
05840 HMM01923cobtr_1_pdu(pf01923pf03928) PDU1AB
05845 P_babyPink PDU1AB
05850 HMM15953pduput_1_pdu1(pf15953) PDU1AB EP_NTERM
05855 HMMptac_2_mix(pf06130) PDU1AB
05860 H_driedBlood PDU1AB
05865 H_tan PDU1AB
05870 HMMpropdeh3_2_pdusml(pf02288) PDU1AB
05875 HMM08841dhact_1_pdu(pf088
05880 HMMpropdeh2_1_pdu(pf0
BMC-P
BMC-H
BMC-T
Aldehyde dehydrogenase
PTAC
Signature enzyme
Alcohol dehydrogenase
Regulator

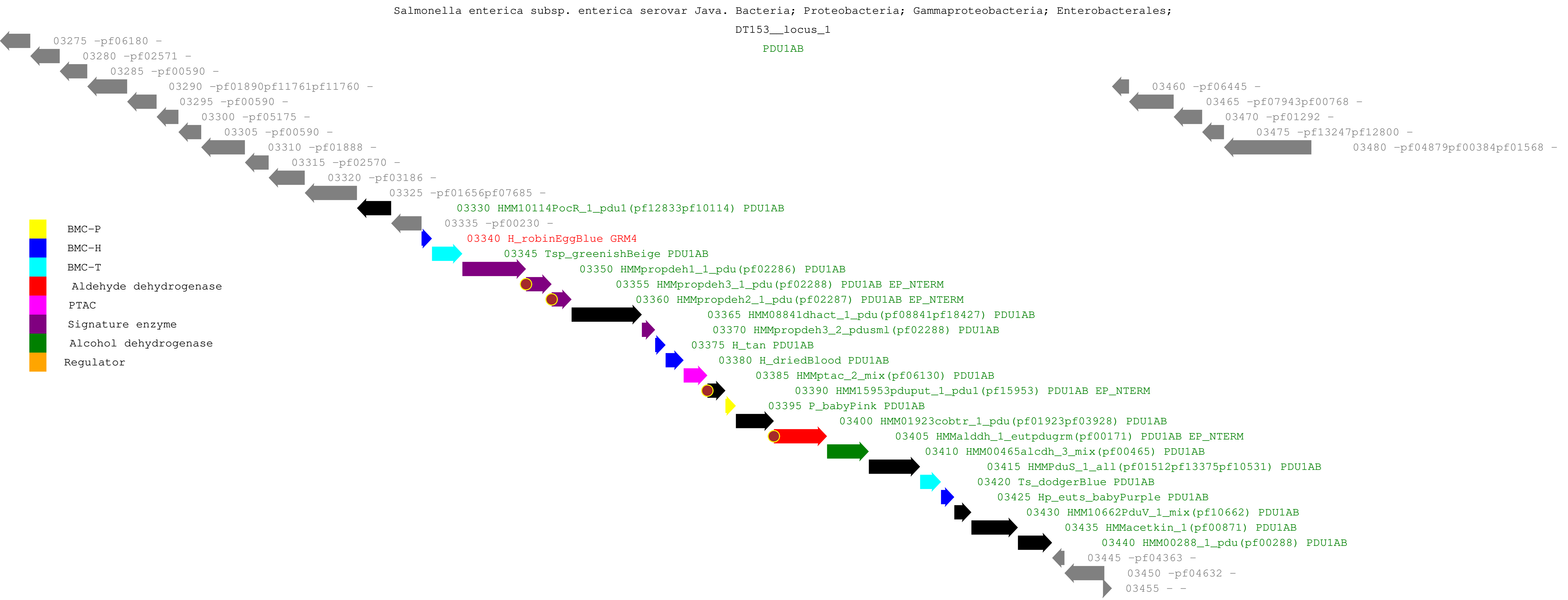

Salmonella enterica subsp. enterica serovar Java. Bacteria; Proteobacteria; Gammaproteobacteria; Enterobacterales;
DT153__locus_1
PDU1AB
03275 -pf06180 -
03280 -pf02571 -
03285 -pf00590 -
03290 -pf01890pf11761pf11760 -
03295 -pf00590 -
03300 -pf05175 -
03305 -pf00590 -
03310 -pf01888 -
03315 -pf02570 -
03320 -pf03186 -
03325 -pf01656pf07685 -
03330 HMM10114PocR_1_pdu1(pf12833pf10114) PDU1AB
03335 -pf00230 -
03340 H_robinEggBlue GRM4
03345 Tsp_greenishBeige PDU1AB
03350 HMMpropdeh1_1_pdu(pf02286) PDU1AB
03355 HMMpropdeh3_1_pdu(pf02288) PDU1AB EP_NTERM
03360 HMMpropdeh2_1_pdu(pf02287) PDU1AB EP_NTERM
03365 HMM08841dhact_1_pdu(pf08841pf18427) PDU1AB
03370 HMMpropdeh3_2_pdusml(pf02288) PDU1AB
03375 H_tan PDU1AB
03380 H_driedBlood PDU1AB
03385 HMMptac_2_mix(pf06130) PDU1AB
03390 HMM15953pduput_1_pdu1(pf15953) PDU1AB EP_NTERM
03395 P_babyPink PDU1AB
03400 HMM01923cobtr_1_pdu(pf01923pf03928) PDU1AB
03405 HMMalddh_1_eutpdugrm(pf00171) PDU1AB EP_NTERM
03410 HMM00465alcdh_3_mix(pf00465) PDU1AB
03415 HMMPduS_1_all(pf01512pf13375pf10531) PDU1AB
03420 Ts_dodgerBlue PDU1AB
03425 Hp_euts_babyPurple PDU1AB
03430 HMM10662PduV_1_mix(pf10662) PDU1AB
03435 HMMacetkin_1(pf00871) PDU1AB
03440 HMM00288_1_pdu(pf00288) PDU1AB
03445 -pf04363 -
03450 -pf04632 -
03455 - -
03460 -pf06445 -
03465 -pf07943pf00768 -
03470 -pf01292 -
03475 -pf13247pf12800 -
03480 -pf04879pf00384pf01568 -
BMC-P
BMC-H
BMC-T
Aldehyde dehydrogenase
PTAC
Signature enzyme
Alcohol dehydrogenase
Regulator

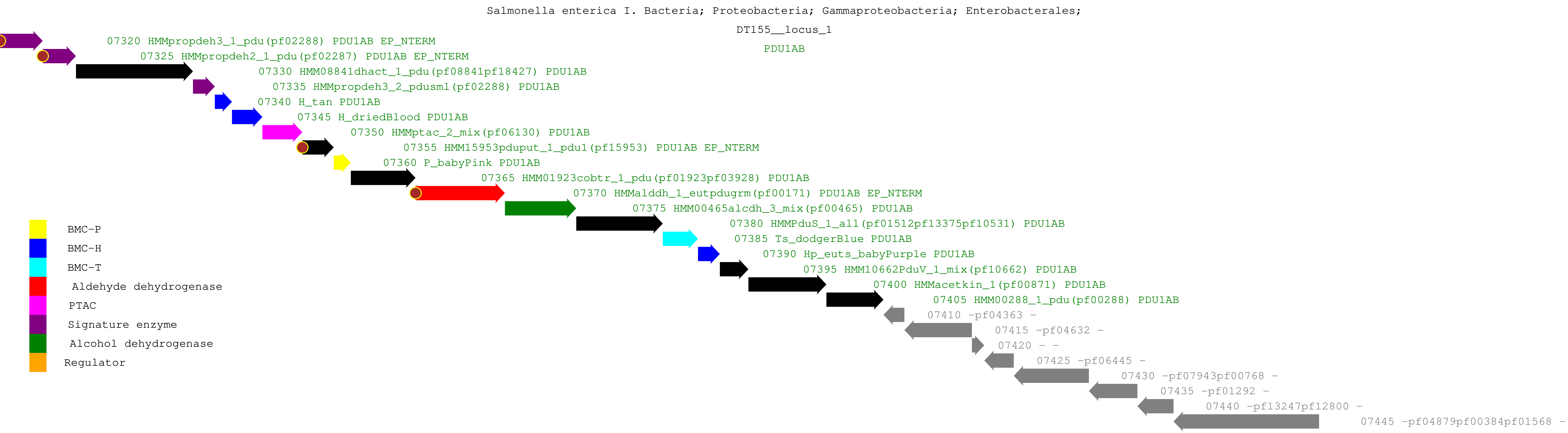
Salmonella enterica I. Bacteria; Proteobacteria; Gammaproteobacteria; Enterobacterales;
DT155__locus_1
PDU1AB
07320 HMMpropdeh3_1_pdu(pf02288) PDU1AB EP_NTERM
07325 HMMpropdeh2_1_pdu(pf02287) PDU1AB EP_NTERM
07330 HMM08841dhact_1_pdu(pf08841pf18427) PDU1AB
07335 HMMpropdeh3_2_pdusml(pf02288) PDU1AB
07340 H_tan PDU1AB
07345 H_driedBlood PDU1AB
07350 HMMptac_2_mix(pf06130) PDU1AB
07355 HMM15953pduput_1_pdu1(pf15953) PDU1AB EP_NTERM
07360 P_babyPink PDU1AB
07365 HMM01923cobtr_1_pdu(pf01923pf03928) PDU1AB
07370 HMMaldh_1_eutpdugrm(pf00171) PDU1AB EP_NTERM
07375 HMM00465alcdh_3_mix(pf00465) PDU1AB
07380 HMMPduS_1_all(pf01512pf13375pf10531) PDU1AB
07385 Ts_dodgerBlue PDU1AB
07390 Hp_euts_babyPurple PDU1AB
07395 HMM10662PduV_1_mix(pf10662) PDU1AB
07400 HMMacetkin_1(pf00871) PDU1AB
07405 HMM00288_1_pdu(pf00288) PDU1AB
07410 -pf04363 -
07415 -pf04632 -
07420 - -
07425 -pf06445 -
07430 -pf07943pf00768 -
07435 -pf01292 -
07440 -pf13247pf12800 -
07445 -pf04879pf00384pf01568 -
BMC-P
BMC-H
BMC-T
Aldehyde dehydrogenase
PTAC
Signature enzyme
Alcohol dehydrogenase
Regulator

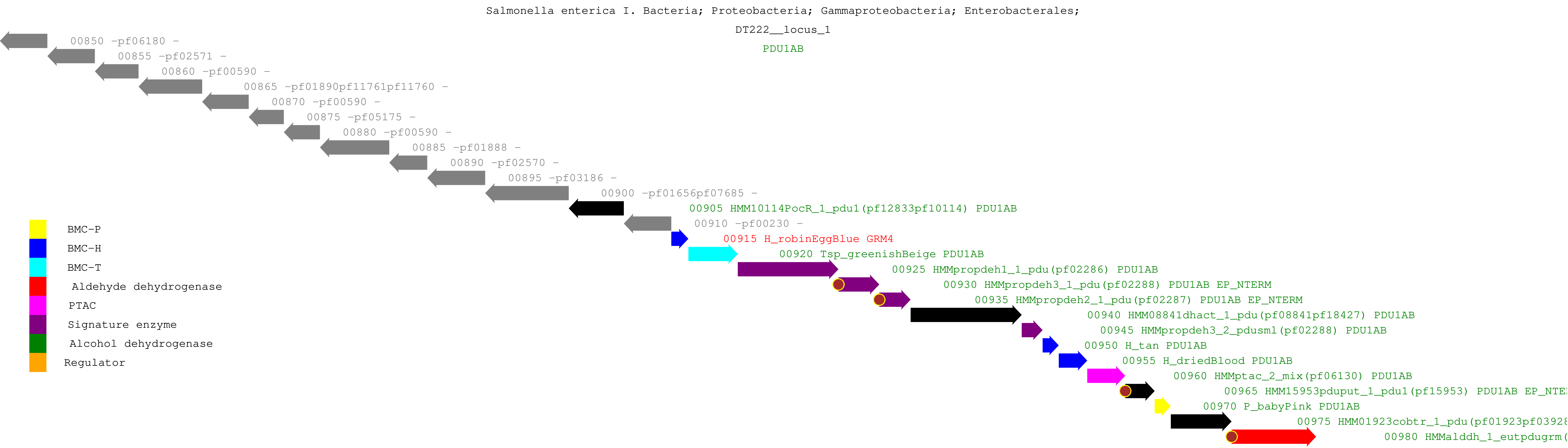
Salmonella enterica I. Bacteria; Proteobacteria; Gammaproteobacteria; Enterobacterales;
DT222__locus_1
PDU1AB
00850 -pf06180 -
00855 -pf02571 -
00860 -pf00590 -
00865 -pf01890pf11761pf11760 -
00870 -pf00590 -
00875 -pf05175 -
00880 -pf00590 -
00885 -pf01888 -
00890 -pf02570 -
00895 -pf03186 -
00900 -pf01656pf07685 -
00905 HMM10114PocR_1_pdu1(pf12833pf10114) PDU1AB
00910 -pf00230 -
00915 H_robinEggBlue GRM4
00920 Tsp_greenishBeige PDU1AB
00925 HMMpropdeh1_1_pdu(pf02286) PDU1AB
00930 HMMpropdeh3_1_pdu(pf02288) PDU1AB EP_NTERM
00935 HMMpropdeh2_1_pdu(pf02287) PDU1AB EP_NTERM
00940 HMM08841dhact_1_pdu(pf08841pf18427) PDU1AB
00945 HMMpropdeh3_2_pdusml(pf02288) PDU1AB
00950 H_tan PDU1AB
00955 H_driedBlood PDU1AB
00960 HMMptac_2_mix(pf06130) PDU1AB
00965 HMM15953pduput_1_pdu1(pf15953) PDU1AB EP_NTE
00970 P_babyPink PDU1AB
00975 HMM01923cobtr_1_pdu(pf01923pf0392
00980 HMMalddh_1_eutpdugrm(
BMC-P
BMC-H
BMC-T
Aldehyde dehydrogenase
PTAC
Signature enzyme
Alcohol dehydrogenase
Regulator

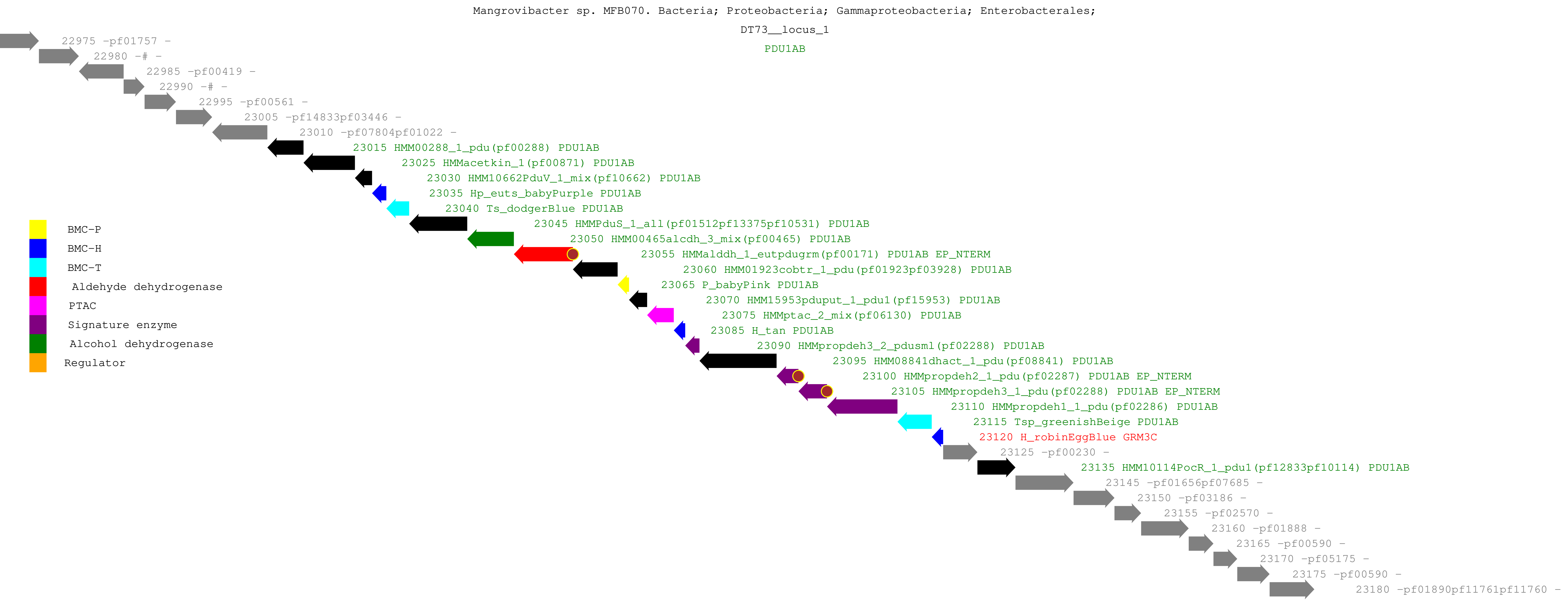
Mangrovibacter sp. MFB070. Bacteria; Proteobacteria; Gammaproteobacteria; Enterobacterales;
DT73__locus_1
PDU1AB
22975 -pf01757 -
22980 -# -
22985 -pf00419 -
22990 -# -
22995 -pf00561 -
23005 -pf14833pf03446 -
23010 -pf07804pf01022 -
23015 HMM00288_1_pdu(pf00288) PDU1AB
23025 HMMacetkin_1(pf00871) PDU1AB
23030 HMM10662PduV_1_mix(pf10662) PDU1AB
23035 Hp_euts_babyPurple PDU1AB
23040 Ts_dodgerBlue PDU1AB
23045 HMMPduS_1_all(pf01512pf13375pf10531) PDU1AB
23050 HMM00465alcdh_3_mix(pf00465) PDU1AB
23055 HMMalddh_1_eutpdugrm(pf00171) PDU1AB EP_NTERM
23060 HMM01923cobtr_1_pdu(pf01923pf03928) PDU1AB
23065 P_babyPink PDU1AB
23070 HMM15953pduput_1_pdu1(pf15953) PDU1AB
23075 HMMptac_2_mix(pf06130) PDU1AB
23085 H_tan PDU1AB
23090 HMMpropdeh3_2_pdusml(pf02288) PDU1AB
23095 HMM08841dhact_1_pdu(pf08841) PDU1AB
23100 HMMpropdeh2_1_pdu(pf02287) PDU1AB EP_NTERM
23105 HMMpropdeh3_1_pdu(pf02288) PDU1AB EP_NTERM
23110 HMMpropdeh1_1_pdu(pf02286) PDU1AB
23115 Tsp_greenishBeige PDU1AB
23120 H_robinEggBlue GRM3C
23125 -pf00230 -
23135 HMM10114PocR_1_pdu1(pf12833pf10114) PDU1AB
23145 -pf01656pf07685 -
23150 -pf03186 -
23155 -pf02570 -
23160 -pf01888 -
23165 -pf00590 -
23170 -pf05175 -
23175 -pf00590 -
23180 -pf01890pf11761pf11760 -
BMC-P
BMC-H
BMC-T
Aldehyde dehydrogenase
PTAC
Signature enzyme
Alcohol dehydrogenase
Regulator

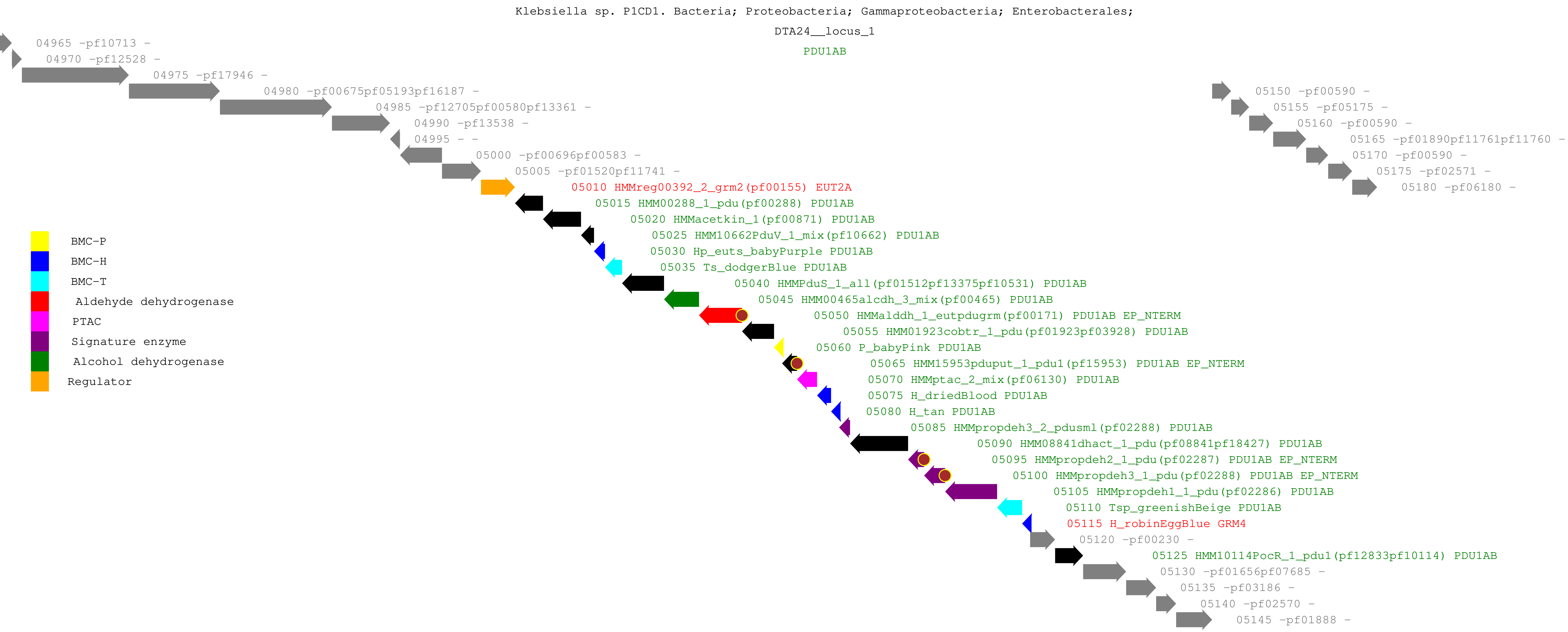
Klebsiella sp. P1CD1. Bacteria; Proteobacteria; Gammaproteobacteria; Enterobacterales;
DTA24__locus_1
PDU1AB
04965 -pf10713 -
04970 -pf12528 -
04975 -pf17946 -
04980 -pf00675pf05193pf16187 -
04985 -pf12705pf00580pf13361 -
04990 -pf13538 -
04995 - -
05000 -pf00696pf00583 -
05005 -pf01520pf11741 -
05010 HMMreg00392_2_grm2(pf00155) EUT2A
05015 HMM00288_1_pdu(pf00288) PDU1AB
05020 HMMacetkin_1(pf00871) PDU1AB
05025 HMM10662PduV_1_mix(pf10662) PDU1AB
05030 Hp_euts_babyPurple PDU1AB
05035 Ts_dodgerBlue PDU1AB
05040 HMMPduS_1_all(pf01512pf13375pf10531) PDU1AB
05045 HMM00465alcdh_3_mix(pf00465) PDU1AB
05050 HMMalddh_1_eutpdugrm(pf00171) PDU1AB EP_NTERM
05055 HMM01923cobtr_1_pdu(pf01923pf03928) PDU1AB
05060 P_babyPink PDU1AB
05065 HMM15953pduput_1_pdu1(pf15953) PDU1AB EP_NTERM
05070 HMMptac_2_mix(pf06130) PDU1AB
05075 H_driedBlood PDU1AB
05080 H_tan PDU1AB
05085 HMMpropdeh3_2_pdusml(pf02288) PDU1AB
05090 HMM08841dhact_1_pdu(pf08841pf18427) PDU1AB
05095 HMMpropdeh2_1_pdu(pf02287) PDU1AB EP_NTERM
05100 HMMpropdeh3_1_pdu(pf02288) PDU1AB EP_NTERM
05105 HMMpropdeh1_1_pdu(pf02286) PDU1AB
05110 Tsp_greenishBeige PDU1AB
05115 H_robinEggBlue GRM4
05120 -pf00230 -
05125 HMM10114PocR_1_pdu1(pf12833pf10114) PDU1AB
05130 -pf01656pf07685 -
05135 -pf03186 -
05140 -pf02570 -
05145 -pf01888 -
05150 -pf00590 -
05155 -pf05175 -
05160 -pf00590 -
05165 -pf01890pf11761pf11760 -
05170 -pf00590 -
05175 -pf02571 -
05180 -pf06180 -
BMC-P
BMC-H
BMC-T
Aldehyde dehydrogenase
PTAC
Signature enzyme
Alcohol dehydrogenase
Regulator

Salmonella enterica I. Bacteria; Proteobacteria; Gammaproteobacteria; Enterobacterales;

DTF39__locus_1

PDU1AB

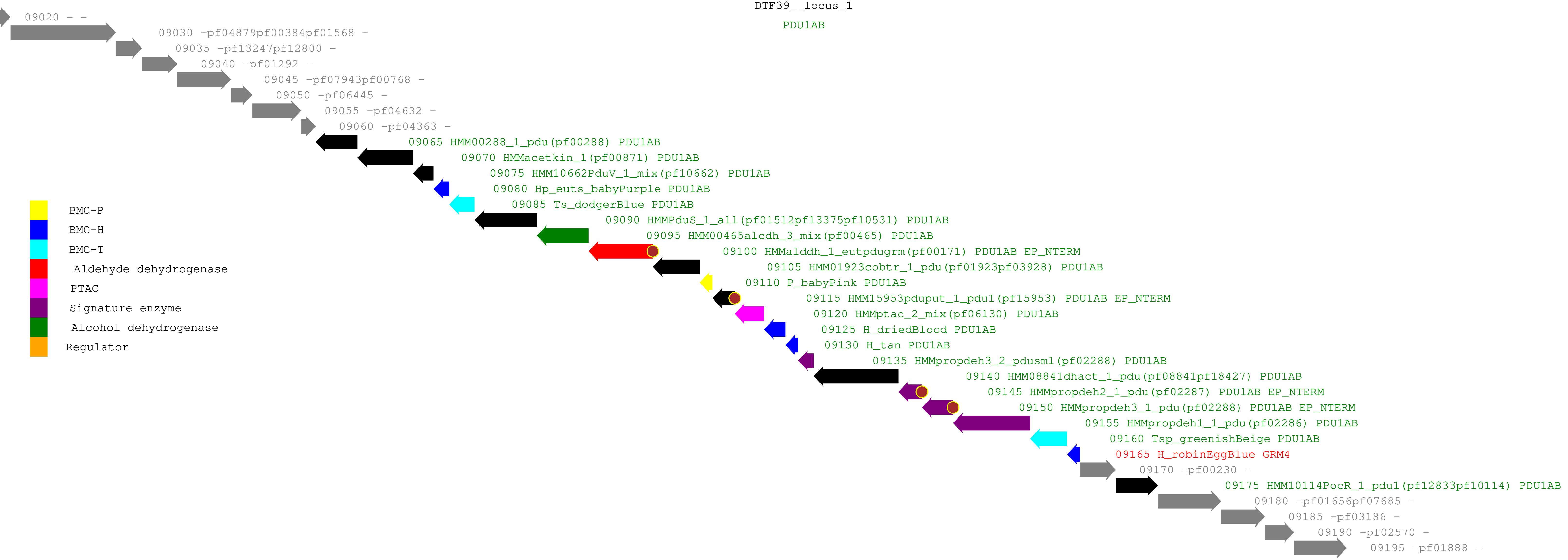

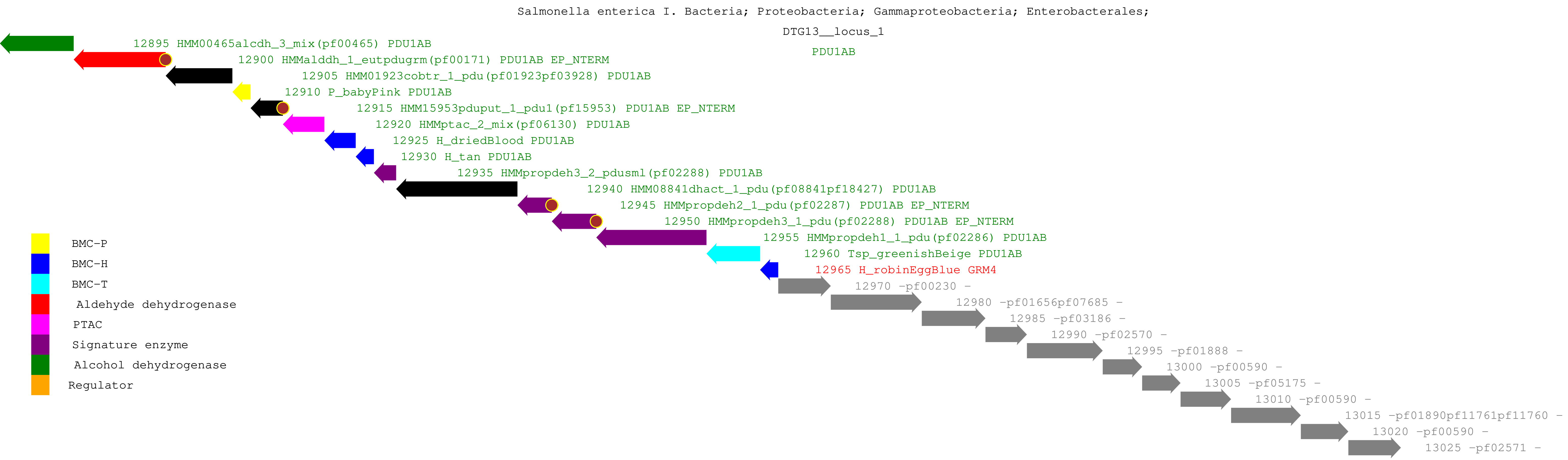
Salmonella enterica I. Bacteria; Proteobacteria; Gammaproteobacteria; Enterobacterales;
DTG13__locus_1
PDU1AB
12895 HMM00465alcdh_3_mix(pf00465) PDU1AB
12900 HMMalddh_1_eutpdugrm(pf00171) PDU1AB EP_NTERM
12905 HMM01923cobtr_1_pdu(pf01923pf03928) PDU1AB
12910 P_babyPink PDU1AB
12915 HMM15953pduput_1_pdu1(pf15953) PDU1AB EP_NTERM
12920 HMMptac_2_mix(pf06130) PDU1AB
12925 H_driedBlood PDU1AB
12930 H_tan PDU1AB
12935 HMMpropdeh3_2_pdusml(pf02288) PDU1AB
12940 HMM08841dhact_1_pdu(pf08841pf18427) PDU1AB
12945 HMMpropdeh2_1_pdu(pf02287) PDU1AB EP_NTERM
12950 HMMpropdeh3_1_pdu(pf02288) PDU1AB EP_NTERM
12955 HMMpropdeh1_1_pdu(pf02286) PDU1AB
12960 Tsp_greenishBeige PDU1AB
12965 H_robinEggBlue GRM4
12970 -pf00230 -
12980 -pf01656pf07685 -
12985 -pf03186 -
12990 -pf02570 -
12995 -pf01888 -
13000 -pf00590 -
13005 -pf05175 -
13010 -pf00590 -
13015 -pf01890pf11761pf11760 -
13020 -pf00590 -
13025 -pf02571 -
BMC-P
BMC-H
BMC-T
Aldehyde dehydrogenase
PTAC
Signature enzyme
Alcohol dehydrogenase
Regulator

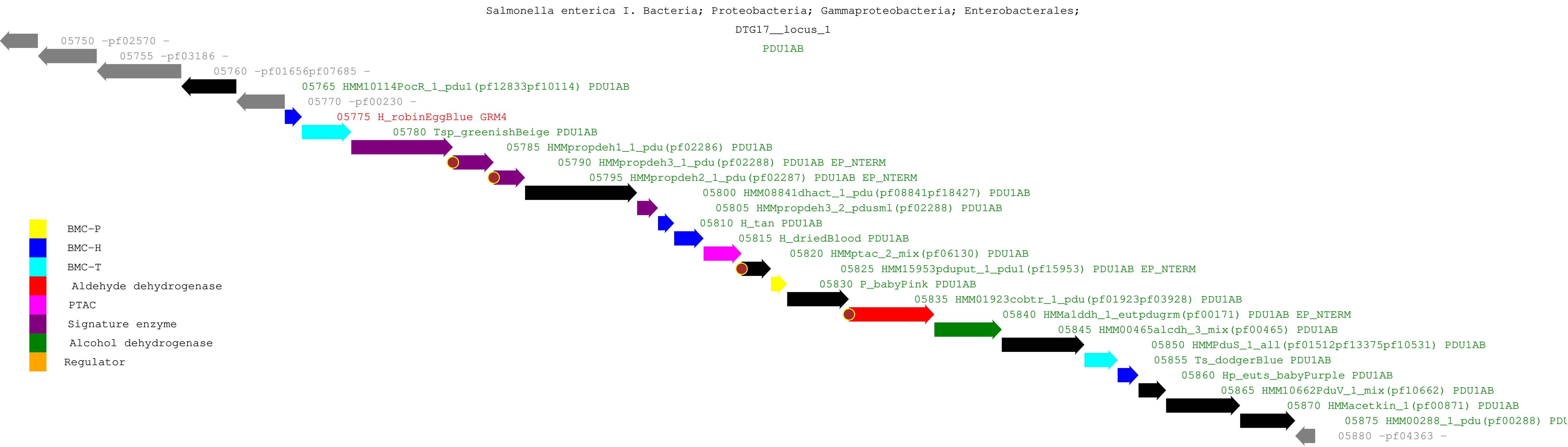
Salmonella enterica I. Bacteria; Proteobacteria; Gammaproteobacteria; Enterobacterales;
DTG17__locus_1
PDU1AB
05750 -pf02570 -
05755 -pf03186 -
05760 -pf01656pf07685 -
05765 HMM10114PocR_1_pdu1(pf12833pf10114) PDU1AB
05770 -pf00230 -
05775 H_robinEggBlue GRM4
05780 Tsp_greenishBeige PDU1AB
05785 HMMpropdeh1_1_pdu(pf02286) PDU1AB
05790 HMMpropdeh3_1_pdu(pf02288) PDU1AB EP_NTERM
05795 HMMpropdeh2_1_pdu(pf02287) PDU1AB EP_NTERM
05800 HMM08841dhact_1_pdu(pf08841pf18427) PDU1AB
05805 HMMpropdeh3_2_pdusml(pf02288) PDU1AB
05810 H_tan PDU1AB
05815 H_driedBlood PDU1AB
05820 HMMptac_2_mix(pf06130) PDU1AB
05825 HMM15953pduput_1_pdu1(pf15953) PDU1AB EP_NTERM
05830 P_babyPink PDU1AB
05835 HMM01923cobtr_1_pdu(pf01923pf03928) PDU1AB
05840 HMMalddh_1_eutpdugrm(pf00171) PDU1AB EP_NTERM
05845 HMM00465alcdh_3_mix(pf00465) PDU1AB
05850 HMMPduS_1_all(pf01512pf13375pf10531) PDU1AB
05855 Ts_dodgerBlue PDU1AB
05860 Hp_euts_babyPurple PDU1AB
05865 HMM10662PduV_1_mix(pf10662) PDU1AB
05870 HMMacetkin_1(pf00871) PDU1AB
05875 HMM00288_1_pdu(pf00288) PDU
05880 -pf04363 -
BMC-P
BMC-H
BMC-T
Aldehyde dehydrogenase
PTAC
Signature enzyme
Alcohol dehydrogenase
Regulator

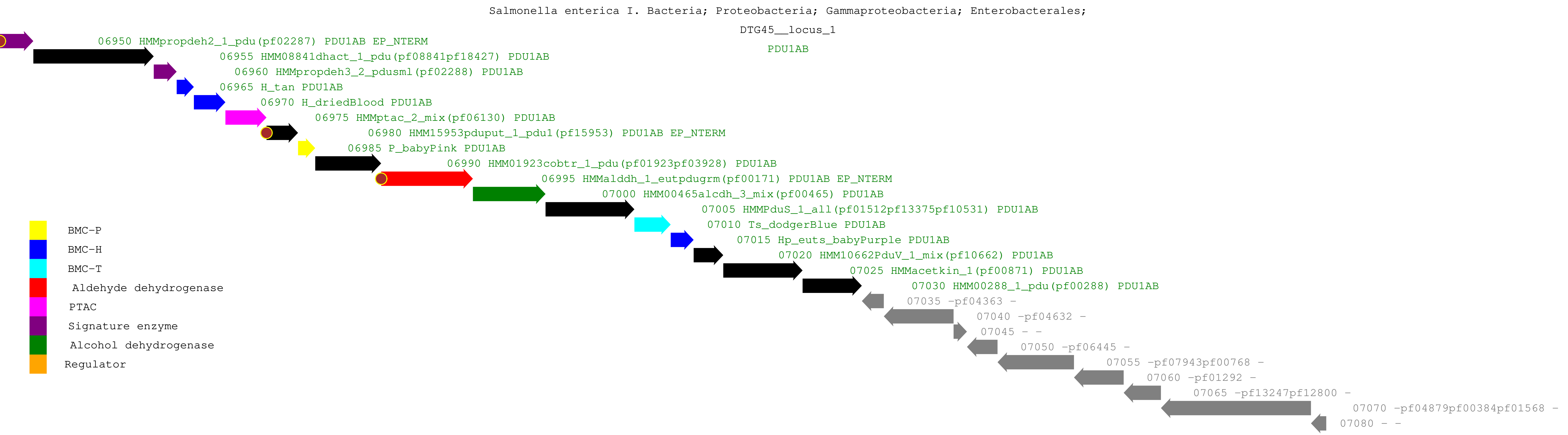
Salmonella enterica I. Bacteria; Proteobacteria; Gammaproteobacteria; Enterobacterales;
DTG45__locus_1
PDU1AB
06950 HMMpropdeh2_1_pdu(pf02287) PDU1AB EP_NTERM
06955 HMM08841dhact_1_pdu(pf08841pf18427) PDU1AB
06960 HMMpropdeh3_2_pdusml(pf02288) PDU1AB
06965 H_tan PDU1AB
06970 H_driedBlood PDU1AB
06975 HMMptac_2_mix(pf06130) PDU1AB
06980 HMM15953pduput_1_pdu1(pf15953) PDU1AB EP_NTERM
06985 P_babyPink PDU1AB
06990 HMM01923cobtr_1_pdu(pf01923pf03928) PDU1AB
06995 HMMalddh_1_eutpdugrm(pf00171) PDU1AB EP_NTERM
07000 HMM00465alcdh_3_mix(pf00465) PDU1AB
07005 HMMPduS_1_all(pf01512pf13375pf10531) PDU1AB
07010 Ts_dodgerBlue PDU1AB
07015 Hp_euts_babyPurple PDU1AB
07020 HMM10662PduV_1_mix(pf10662) PDU1AB
07025 HMMacetkin_1(pf00871) PDU1AB
07030 HMM00288_1_pdu(pf00288) PDU1AB
07035 -pf04363 -
07040 -pf04632 -
07045 - -
07050 -pf06445 -
07055 -pf07943pf00768 -
07060 -pf01292 -
07065 -pf13247pf12800 -
07070 -pf04879pf00384pf01568 -
07080 - -
BMC-P
BMC-H
BMC-T
Aldehyde dehydrogenase
PTAC
Signature enzyme
Alcohol dehydrogenase
Regulator

Salmonella enterica subsp. enterica serovar Heidelberg. Bacteria; Proteobacteria; Gammaproteobacteria; Enterobacterales;

DTG66__locus_1

PDU1AB

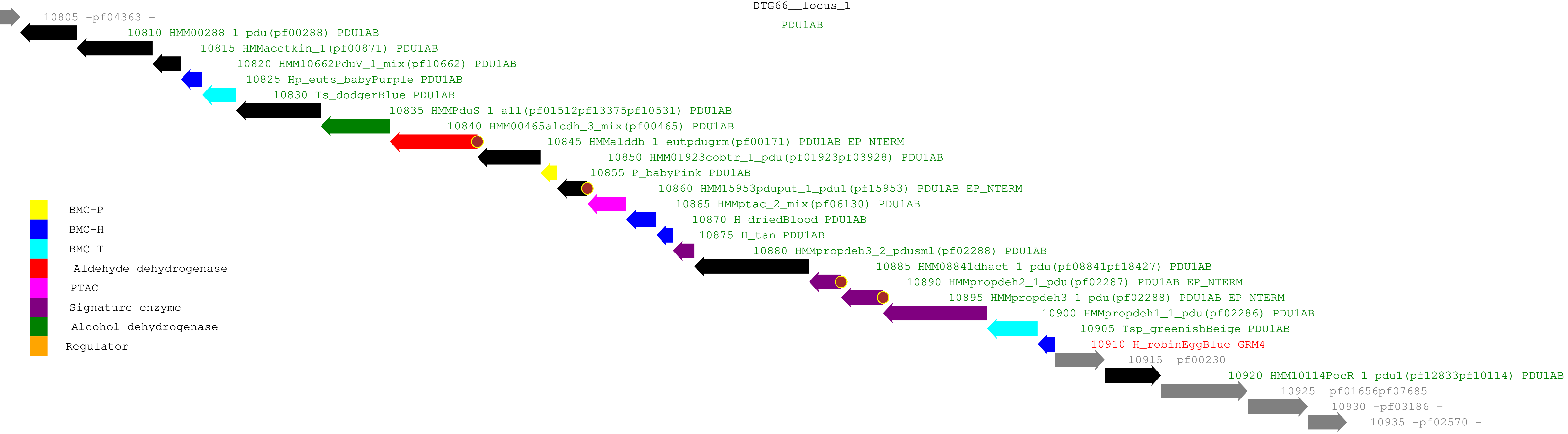

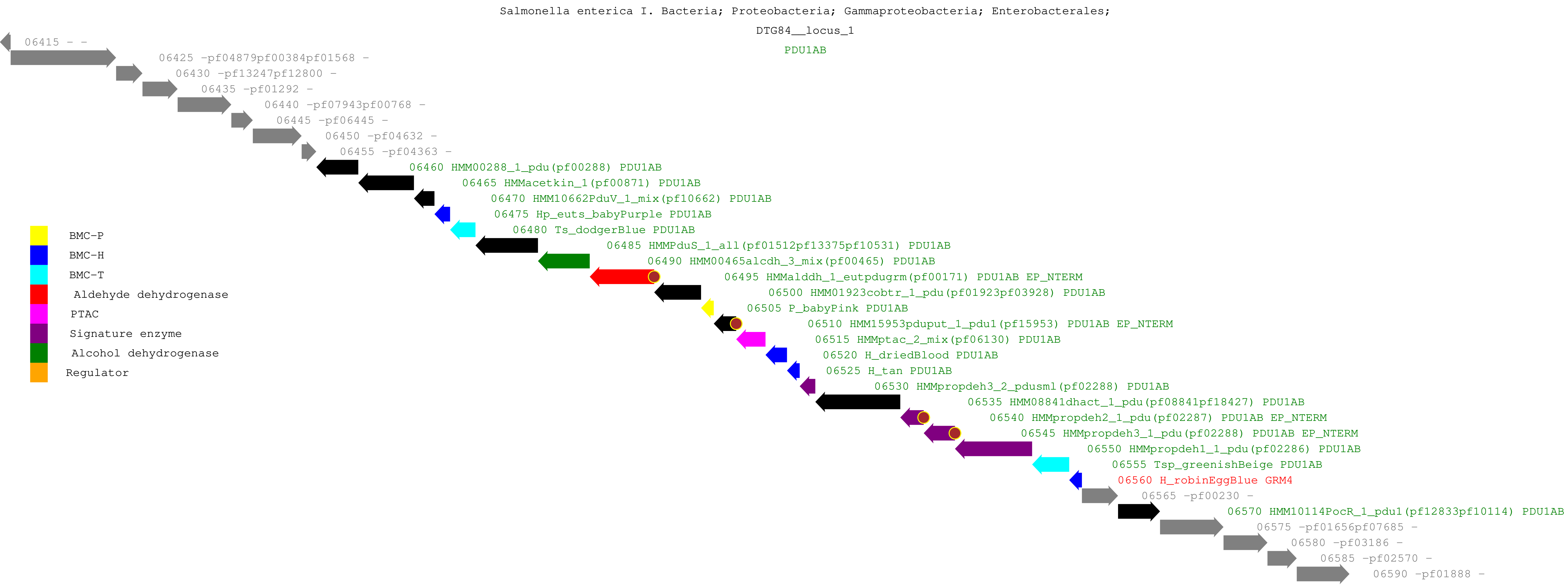
Salmonella enterica I. Bacteria; Proteobacteria; Gammaproteobacteria; Enterobacterales;
DTG84__locus_1
PDU1AB
06415 - -
06425 -pf04879pf00384pf01568 -
06430 -pf13247pf12800 -
06435 -pf01292 -
06440 -pf07943pf00768 -
06445 -pf06445 -
06450 -pf04632 -
06455 -pf04363 -
06460 HMM00288_1_pdu(pf00288) PDU1AB
06465 HMMacetkin_1(pf00871) PDU1AB
06470 HMM10662PduV_1_mix(pf10662) PDU1AB
06475 Hp_euts_babyPurple PDU1AB
06480 Ts_dodgerBlue PDU1AB
06485 HMMPduS_1_all(pf01512pf13375pf10531) PDU1AB
06490 HMM00465alcdh_3_mix(pf00465) PDU1AB
06495 HMMalddh_1_eutpdugrm(pf00171) PDU1AB EP_NTERM
06500 HMM01923cobtr_1_pdu(pf01923pf03928) PDU1AB
06505 P_babyPink PDU1AB
06510 HMM15953pduput_1_pdu1(pf15953) PDU1AB EP_NTERM
06515 HMMptac_2_mix(pf06130) PDU1AB
06520 H_driedBlood PDU1AB
06525 H_tan PDU1AB
06530 HMMpropdeh3_2_pdusml(pf02288) PDU1AB
06535 HMM08841dhact_1_pdu(pf08841pf18427) PDU1AB
06540 HMMpropdeh2_1_pdu(pf02287) PDU1AB EP_NTERM
06545 HMMpropdeh3_1_pdu(pf02288) PDU1AB EP_NTERM
06550 HMMpropdeh1_1_pdu(pf02286) PDU1AB
06555 Tsp_greenishBeige PDU1AB
06560 H_robinEggBlue GRM4
06565 -pf00230 -
06570 HMM10114PocR_1_pdu1(pf12833pf10114) PDU1AB
06575 -pf01656pf07685 -
06580 -pf03186 -
06585 -pf02570 -
06590 -pf01888 -
BMC-P
BMC-H
BMC-T
Aldehyde dehydrogenase
PTAC
Signature enzyme
Alcohol dehydrogenase
Regulator

Salmonella enteritidis. Bacteria; Proteobacteria; Gammaproteobacteria; Enterobacterales;

DTI44__locus_2

PDU1AB

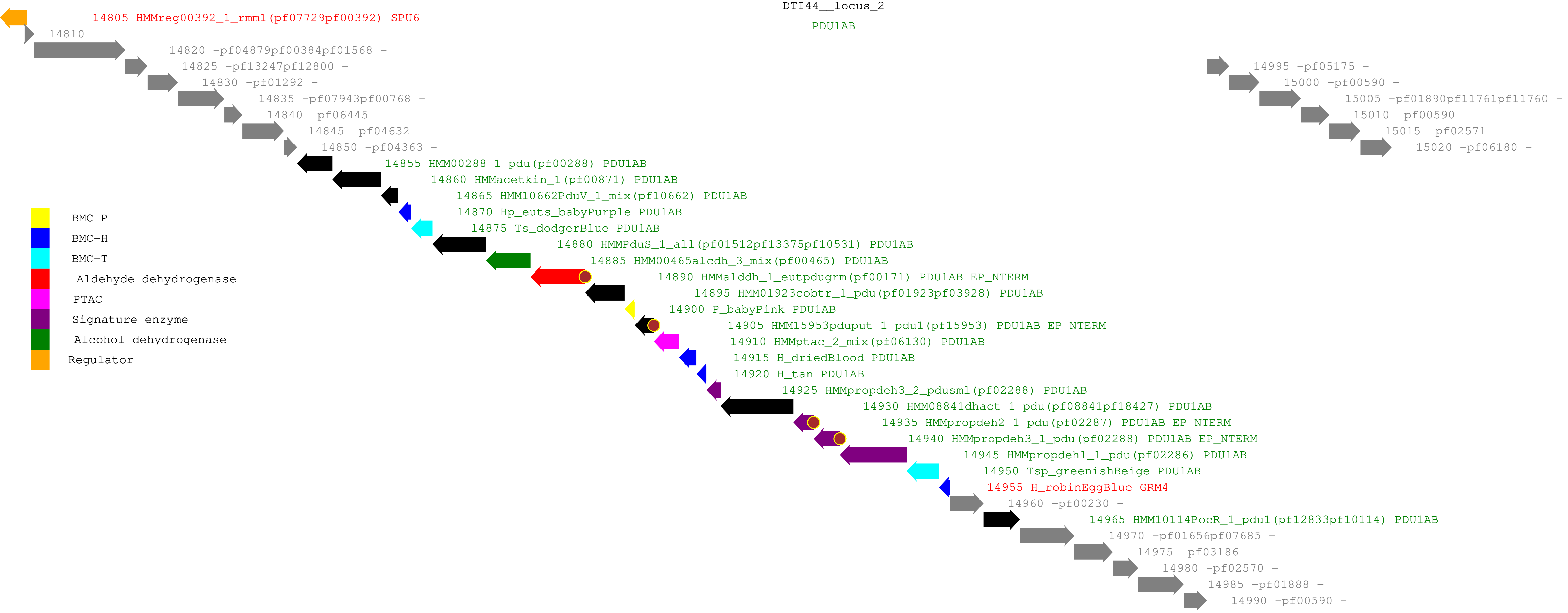

Escherichia coli. Bacteria; Proteobacteria; Gammaproteobacteria; Enterobacterales;

DTM45__locus_1

PDU1AB

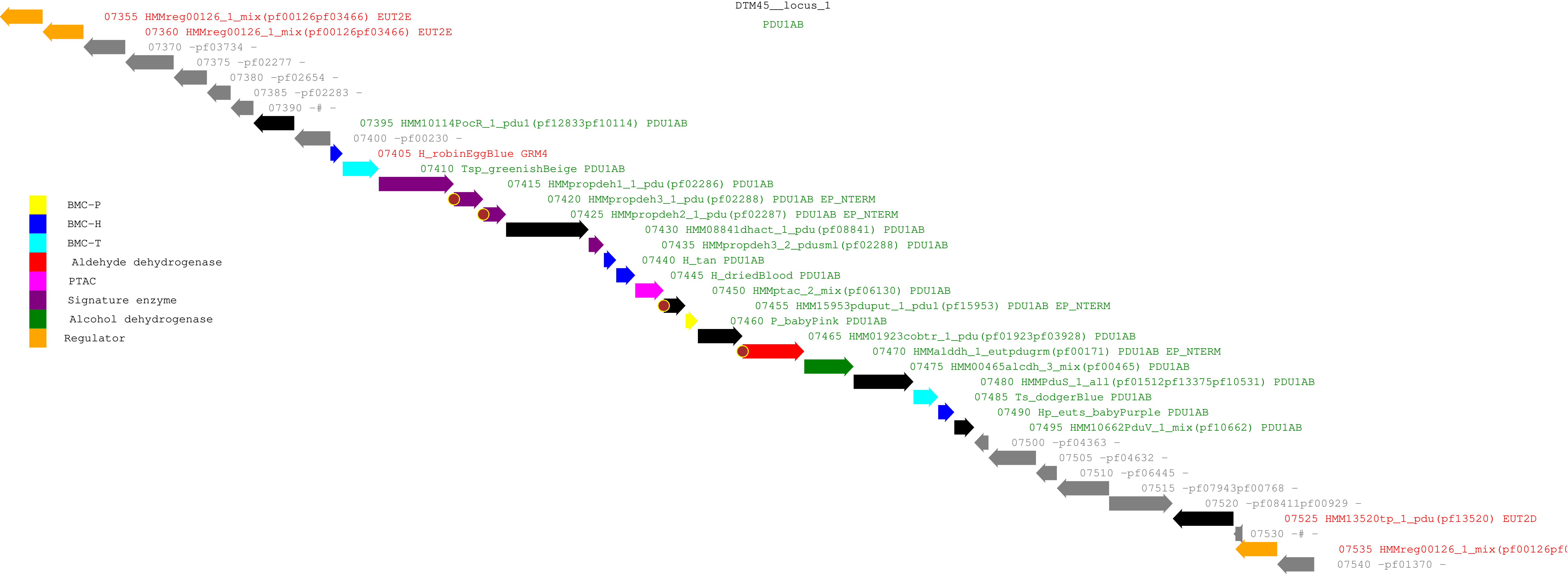

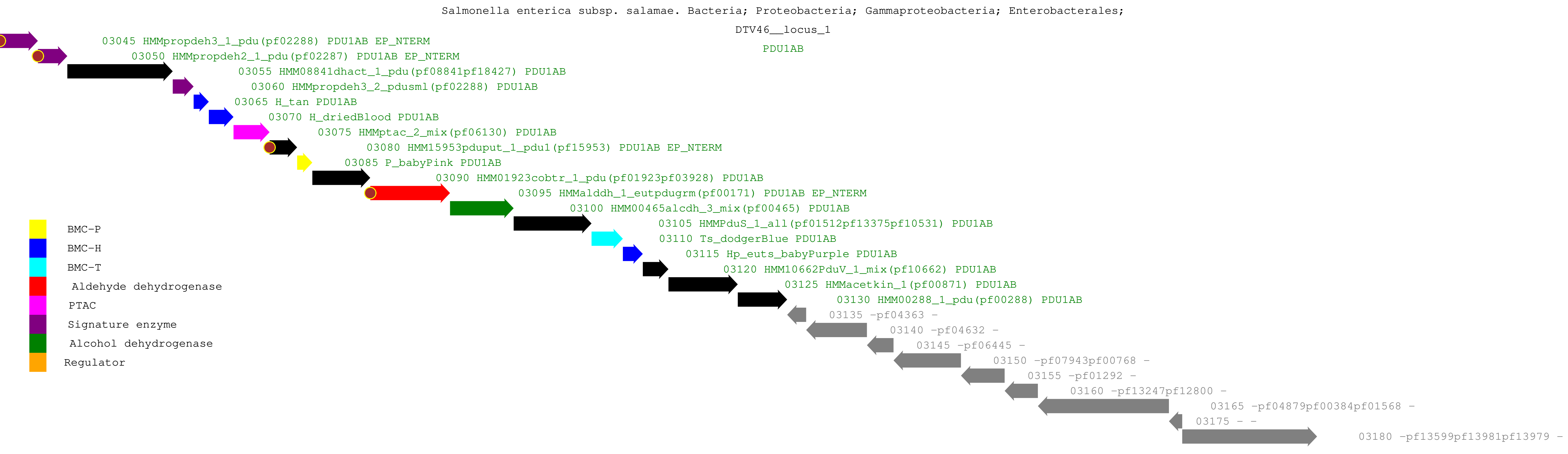
Salmonella enterica subsp. salamae. Bacteria; Proteobacteria; Gammaproteobacteria; Enterobacterales;
DTV46__locus_1
PDU1AB
03045 HMMpropdeh3_1_pdu(pf02288) PDU1AB EP_NTERM
03050 HMMpropdeh2_1_pdu(pf02287) PDU1AB EP_NTERM
03055 HMM08841dhact_1_pdu(pf08841pf18427) PDU1AB
03060 HMMpropdeh3_2_pdusml(pf02288) PDU1AB
03065 H_tan PDU1AB
03070 H_driedBlood PDU1AB
03075 HMMptac_2_mix(pf06130) PDU1AB
03080 HMM15953pduput_1_pdu1(pf15953) PDU1AB EP_NTERM
03085 P_babyPink PDU1AB
03090 HMM01923cobtr_1_pdu(pf01923pf03928) PDU1AB
03095 HMMalddh_1_eutpdugrm(pf00171) PDU1AB EP_NTERM
03100 HMM00465alcdh_3_mix(pf00465) PDU1AB
03105 HMMPduS_1_all(pf01512pf13375pf10531) PDU1AB
03110 Ts_dodgerBlue PDU1AB
03115 Hp_euts_babyPurple PDU1AB
03120 HMM10662PduV_1_mix(pf10662) PDU1AB
03125 HMMacetkin_1(pf00871) PDU1AB
03130 HMM00288_1_pdu(pf00288) PDU1AB
03135 -pf04363 -
03140 -pf04632 -
03145 -pf06445 -
03150 -pf07943pf00768 -
03155 -pf01292 -
03160 -pf13247pf12800 -
03165 -pf04879pf00384pf01568 -
03175 - -
03180 -pf13599pf13981pf13979 -
BMC-P
BMC-H
BMC-T
Aldehyde dehydrogenase
PTAC
Signature enzyme
Alcohol dehydrogenase
Regulator

Salmonella enterica I. Bacteria; Proteobacteria; Gammaproteobacteria; Enterobacterales;

DU161__locus_2

PDU1AB

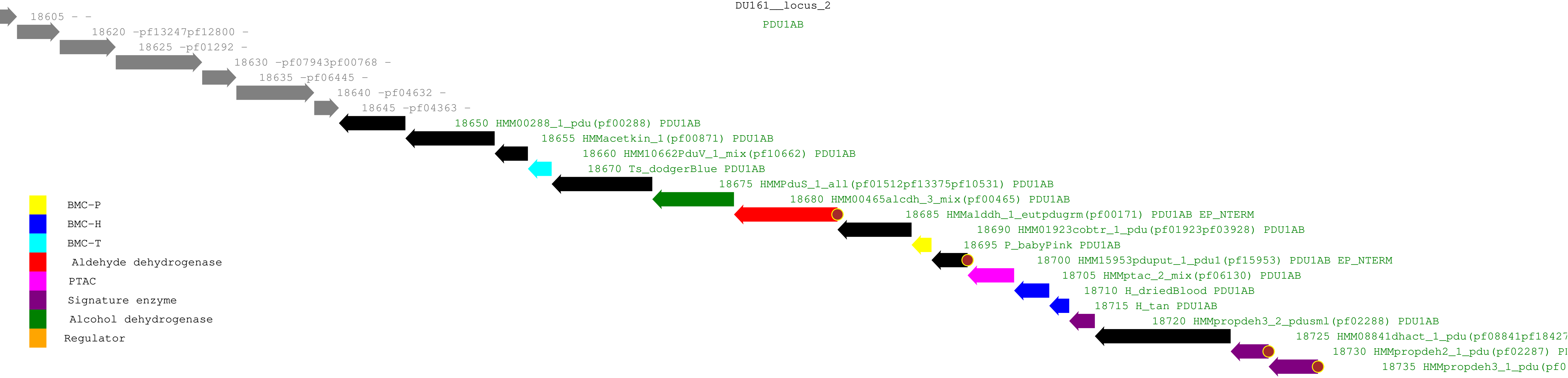

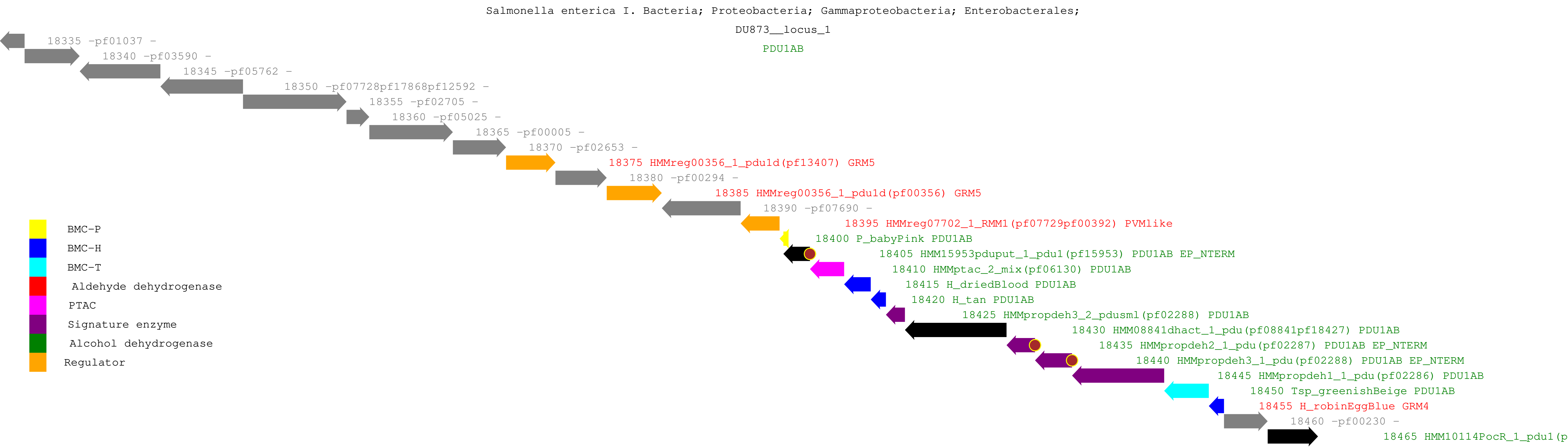

Salmonella enterica I. Bacteria; Proteobacteria; Gammaproteobacteria; Enterobacterales;
DU873__locus_1
PDU1AB
18335 -pf01037 -
18340 -pf03590 -
18345 -pf05762 -
18350 -pf07728pf17868pf12592 -
18355 -pf02705 -
18360 -pf05025 -
18365 -pf00005 -
18370 -pf02653 -
18375 HMMreg00356_1_pdu1d(pf13407) GRM5
18380 -pf00294 -
18385 HMMreg00356_1_pdu1d(pf00356) GRM5
18390 -pf07690 -
18395 HMMreg07702_1_RMM1(pf07729pf00392) PVMlike
18400 P_babyPink PDU1AB
18405 HMM15953pduput_1_pdu1(pf15953) PDU1AB EP_NTERM
18410 HMMptac_2_mix(pf06130) PDU1AB
18415 H_driedBlood PDU1AB
18420 H_tan PDU1AB
18425 HMMpropdeh3_2_pdusml(pf02288) PDU1AB
18430 HMM08841dhact_1_pdu(pf08841pf18427) PDU1AB
18435 HMMpropdeh2_1_pdu(pf02287) PDU1AB EP_NTERM
18440 HMMpropdeh3_1_pdu(pf02288) PDU1AB EP_NTERM
18445 HMMpropdeh1_1_pdu(pf02286) PDU1AB
18450 Tsp_greenishBeige PDU1AB
18455 H_robinEggBlue GRM4
18460 -pf00230 -
18465 HMM10114PocR_1_pdu1(p
BMC-P
BMC-H
BMC-T
Aldehyde dehydrogenase
PTAC
Signature enzyme
Alcohol dehydrogenase
Regulator

Salmonella enterica I. Bacteria; Proteobacteria; Gammaproteobacteria; Enterobacterales;

DU873__locus_2

PDU1AB

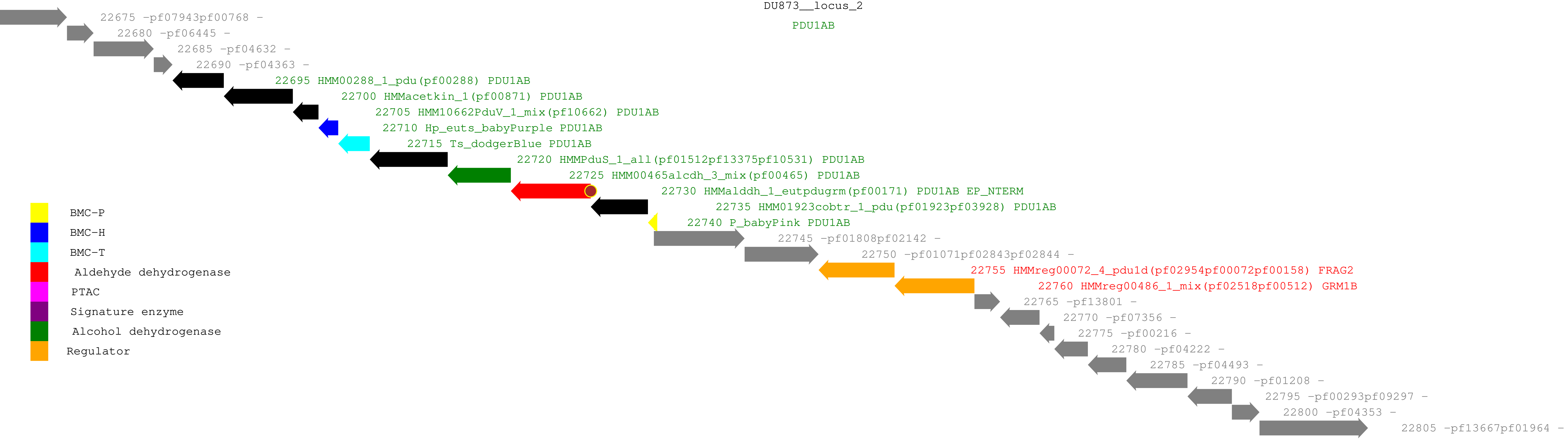

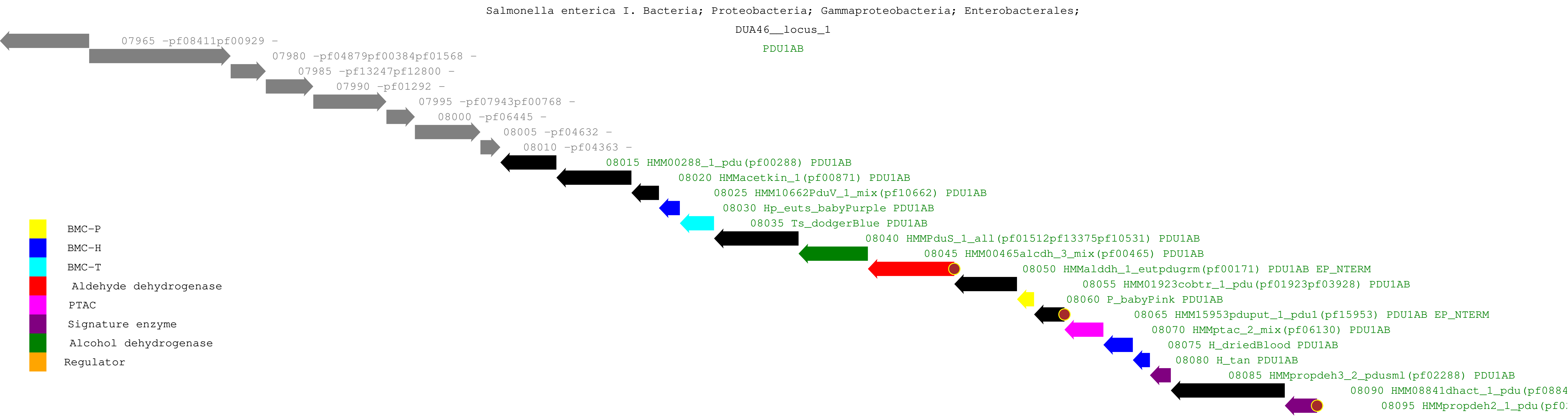

Salmonella enterica I. Bacteria; Proteobacteria; Gammaproteobacteria; Enterobacterales;
DUA46__locus_1
PDU1AB
07965 -pf08411pf00929 -
07980 -pf04879pf00384pf01568 -
07985 -pf13247pf12800 -
07990 -pf01292 -
07995 -pf07943pf00768 -
08000 -pf06445 -
08005 -pf04632 -
08010 -pf04363 -
08015 HMM00288_1_pdu(pf00288) PDU1AB
08020 HMMacetkin_1(pf00871) PDU1AB
08025 HMM10662PduV_1_mix(pf10662) PDU1AB
08030 Hp_euts_babyPurple PDU1AB
08035 Ts_dodgerBlue PDU1AB
08040 HMMPduS_1_all(pf01512pf13375pf10531) PDU1AB
08045 HMM00465alcdh_3_mix(pf00465) PDU1AB
08050 HMMalddh_1_eutpdugrm(pf00171) PDU1AB EP_NTERM
08055 HMM01923cobtr_1_pdu(pf01923pf03928) PDU1AB
08060 P_babyPink PDU1AB
08065 HMM15953pduput_1_pdu1(pf15953) PDU1AB EP_NTERM
08070 HMMptac_2_mix(pf06130) PDU1AB
08075 H_driedBlood PDU1AB
08080 H_tan PDU1AB
08085 HMMpropdeh3_2_pdusml(pf02288) PDU1AB
08090 HMM08841dhact_1_pdu(pf0884
08095 HMMpropdeh2_1_pdu(pf0
BMC-P
BMC-H
BMC-T
Aldehyde dehydrogenase
PTAC
Signature enzyme
Alcohol dehydrogenase
Regulator

Salmonella enterica I. Bacteria; Proteobacteria; Gammaproteobacteria; Enterobacterales;

DUC08__locus_1

PDU1AB

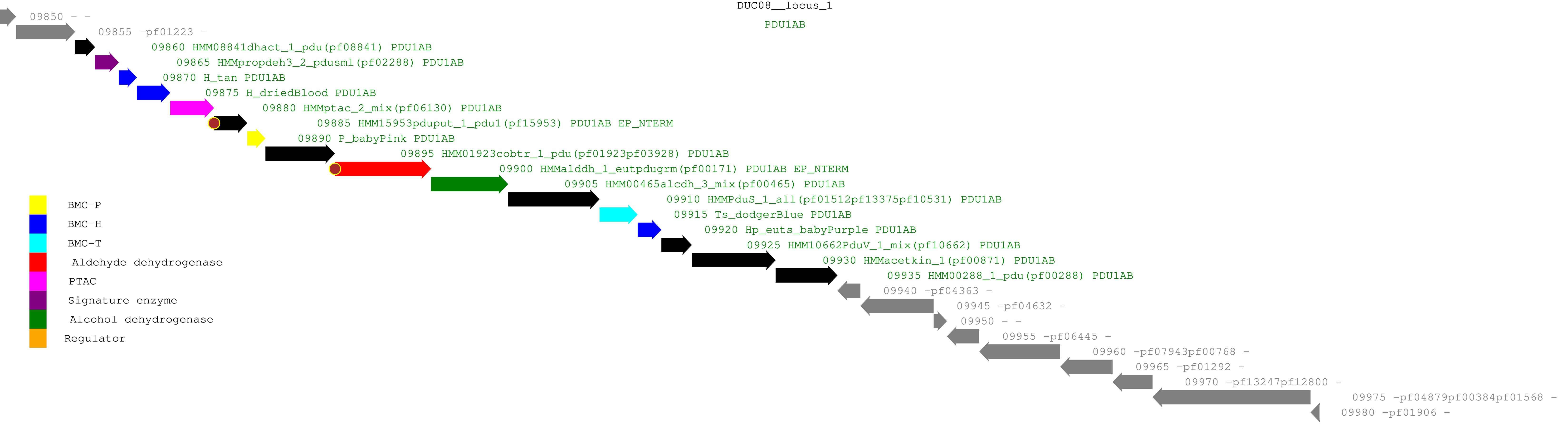

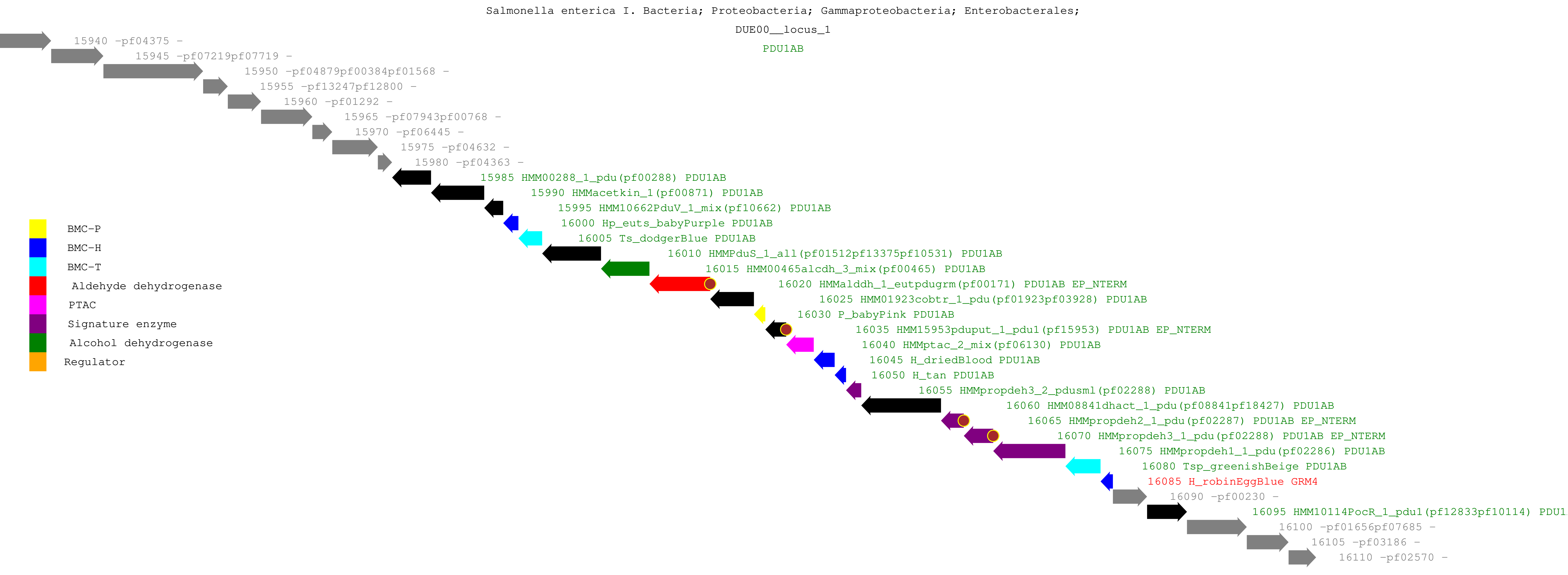
Salmonella enterica I. Bacteria; Proteobacteria; Gammaproteobacteria; Enterobacterales;
DUE00__locus_1
PDU1AB
15940 -pf04375 -
15945 -pf07219pf07719 -
15950 -pf04879pf00384pf01568 -
15955 -pf13247pf12800 -
15960 -pf01292 -
15965 -pf07943pf00768 -
15970 -pf06445 -
15975 -pf04632 -
15980 -pf04363 -
15985 HMM00288_1_pdu(pf00288) PDU1AB
15990 HMMacetkin_1(pf00871) PDU1AB
15995 HMM10662PduV_1_mix(pf10662) PDU1AB
16000 Hp_euts_babyPurple PDU1AB
16005 Ts_dodgerBlue PDU1AB
16010 HMMPduS_1_all(pf01512pf13375pf10531) PDU1AB
16015 HMM00465alcdh_3_mix(pf00465) PDU1AB
16020 HMMalddh_1_eutpdugrm(pf00171) PDU1AB EP_NTERM
16025 HMM01923cobtr_1_pdu(pf01923pf03928) PDU1AB
16030 P_babyPink PDU1AB
16035 HMM15953pduput_1_pdu1(pf15953) PDU1AB EP_NTERM
16040 HMMptac_2_mix(pf06130) PDU1AB
16045 H_driedBlood PDU1AB
16050 H_tan PDU1AB
16055 HMMpropdeh3_2_pdusml(pf02288) PDU1AB
16060 HMM08841dhact_1_pdu(pf08841pf18427) PDU1AB
16065 HMMpropdeh2_1_pdu(pf02287) PDU1AB EP_NTERM
16070 HMMpropdeh3_1_pdu(pf02288) PDU1AB EP_NTERM
16075 HMMpropdeh1_1_pdu(pf02286) PDU1AB
16080 Tsp_greenishBeige PDU1AB
16085 H_robinEggBlue GRM4
16090 -pf00230 -
16095 HMM10114PocR_1_pdu1(pf12833pf10114) PDU1
16100 -pf01656pf07685 -
16105 -pf03186 -
16110 -pf02570 -
BMC-P
BMC-H
BMC-T
Aldehyde dehydrogenase
PTAC
Signature enzyme
Alcohol dehydrogenase
Regulator

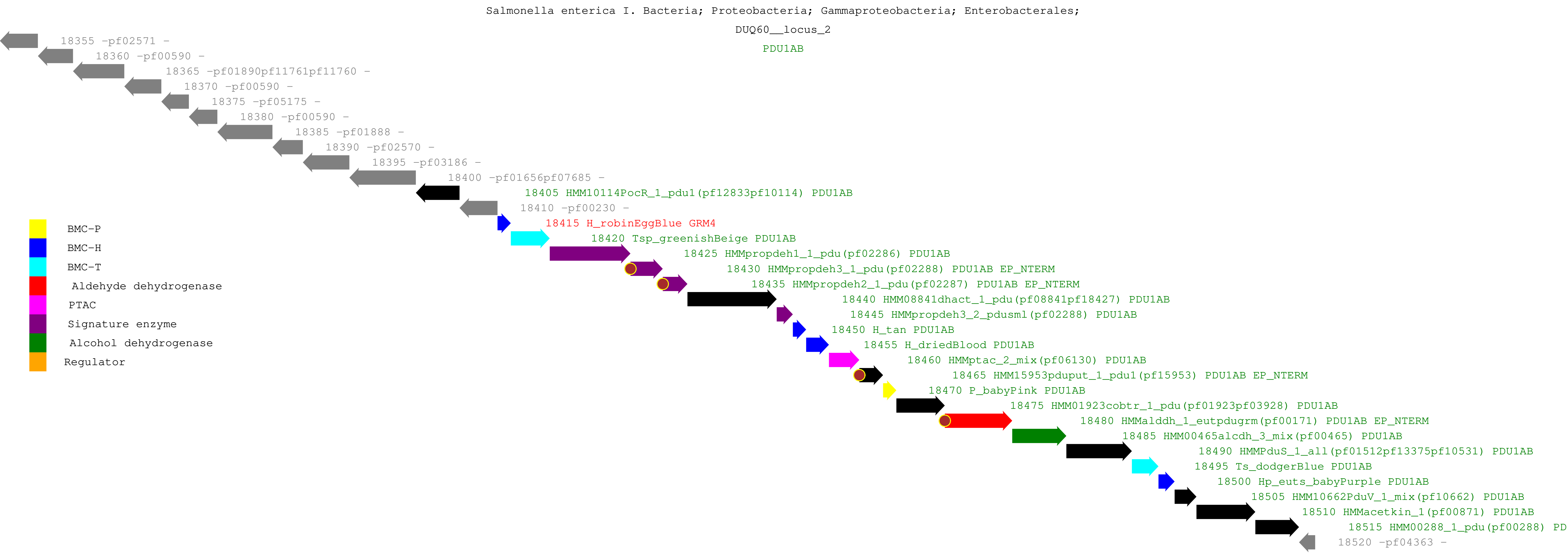

Salmonella enterica I. Bacteria; Proteobacteria; Gammaproteobacteria; Enterobacterales;
DUQ60__locus_2
PDU1AB
18355 -pf02571 -
18360 -pf00590 -
18365 -pf01890pf11761pf11760 -
18370 -pf00590 -
18375 -pf05175 -
18380 -pf00590 -
18385 -pf01888 -
18390 -pf02570 -
18395 -pf03186 -
18400 -pf01656pf07685 -
18405 HMM10114PocR_1_pdu1(pf12833pf10114) PDU1AB
18410 -pf00230 -
18415 H_robinEggBlue GRM4
18420 Tsp_greenishBeige PDU1AB
18425 HMMpropdeh1_1_pdu(pf02286) PDU1AB
18430 HMMpropdeh3_1_pdu(pf02288) PDU1AB EP_NTERM
18435 HMMpropdeh2_1_pdu(pf02287) PDU1AB EP_NTERM
18440 HMM08841dhact_1_pdu(pf08841pf18427) PDU1AB
18445 HMMpropdeh3_2_pdusml(pf02288) PDU1AB
18450 H_tan PDU1AB
18455 H_driedBlood PDU1AB
18460 HMMptac_2_mix(pf06130) PDU1AB
18465 HMM15953pduput_1_pdu1(pf15953) PDU1AB EP_NTERM
18470 P_babyPink PDU1AB
18475 HMM01923cobtr_1_pdu(pf01923pf03928) PDU1AB
18480 HMMalddh_1_eutpdugrm(pf00171) PDU1AB EP_NTERM
18485 HMM00465alcdh_3_mix(pf00465) PDU1AB
18490 HMMPduS_1_all(pf01512pf13375pf10531) PDU1AB
18495 Ts_dodgerBlue PDU1AB
18500 Hp_euts_babyPurple PDU1AB
18505 HMM10662PduV_1_mix(pf10662) PDU1AB
18510 HMMacetkin_1(pf00871) PDU1AB
18515 HMM00288_1_pdu(pf00288) PD
18520 -pf04363 -
BMC-P
BMC-H
BMC-T
Aldehyde dehydrogenase
PTAC
Signature enzyme
Alcohol dehydrogenase
Regulator

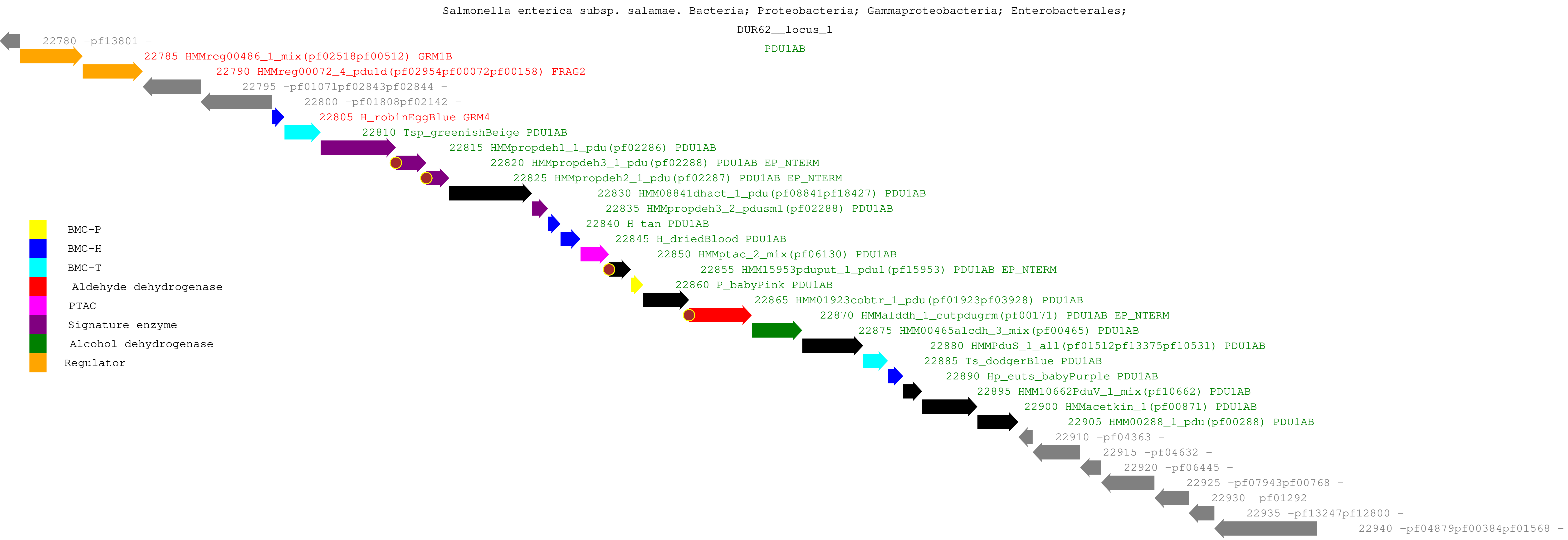

Salmonella enterica subsp. salamae. Bacteria; Proteobacteria; Gammaproteobacteria; Enterobacterales;
DUR62__locus_1
PDU1AB
22780 -pf13801 -
22785 HMMreg00486_1_mix(pf02518pf00512) GRM1B
22790 HMMreg00072_4_pdu1d(pf02954pf00072pf00158) FRAG2
22795 -pf01071pf02843pf02844 -
22800 -pf01808pf02142 -
22805 H_robinEggBlue GRM4
22810 Tsp_greenishBeige PDU1AB
22815 HMMpropdeh1_1_pdu(pf02286) PDU1AB
22820 HMMpropdeh3_1_pdu(pf02288) PDU1AB EP_NTERM
22825 HMMpropdeh2_1_pdu(pf02287) PDU1AB EP_NTERM
22830 HMM08841dhact_1_pdu(pf08841pf18427) PDU1AB
22835 HMMpropdeh3_2_pdusml(pf02288) PDU1AB
22840 H_tan PDU1AB
22845 H_driedBlood PDU1AB
22850 HMMptac_2_mix(pf06130) PDU1AB
22855 HMM15953pduput_1_pdu1(pf15953) PDU1AB EP_NTERM
22860 P_babyPink PDU1AB
22865 HMM01923cobtr_1_pdu(pf01923pf03928) PDU1AB
22870 HMMalddh_1_eutpdugrm(pf00171) PDU1AB EP_NTERM
22875 HMM00465alcdh_3_mix(pf00465) PDU1AB
22880 HMMPduS_1_all(pf01512pf13375pf10531) PDU1AB
22885 Ts_dodgerBlue PDU1AB
22890 Hp_euts_babyPurple PDU1AB
22895 HMM10662PduV_1_mix(pf10662) PDU1AB
22900 HMMacetkin_1(pf00871) PDU1AB
22905 HMM00288_1_pdu(pf00288) PDU1AB
22910 -pf04363 -
22915 -pf04632 -
22920 -pf06445 -
22925 -pf07943pf00768 -
22930 -pf01292 -
22935 -pf13247pf12800 -
22940 -pf04879pf00384pf01568 -
BMC-P
BMC-H
BMC-T
Aldehyde dehydrogenase
PTAC
Signature enzyme
Alcohol dehydrogenase
Regulator

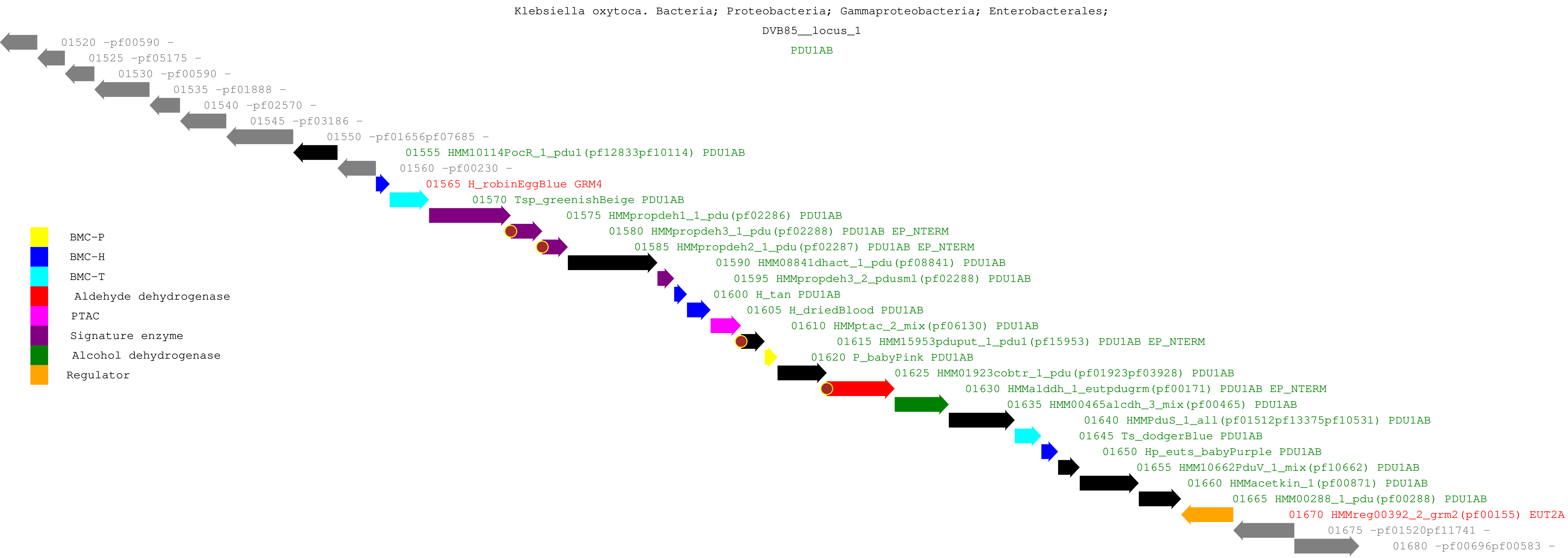
Klebsiella oxytoca. Bacteria; Proteobacteria; Gammaproteobacteria; Enterobacterales;
DVB85__locus_1
PDU1AB
01520 -pf00590 -
01525 -pf05175 -
01530 -pf00590 -
01535 -pf01888 -
01540 -pf02570 -
01545 -pf03186 -
01550 -pf01656pf07685 -
01555 HMM10114PocR_1_pdu1(pf12833pf10114) PDU1AB
01560 -pf00230 -
01565 H_robinEggBlue GRM4
01570 Tsp_greenishBeige PDU1AB
01575 HMMpropdeh1_1_pdu(pf02286) PDU1AB
01580 HMMpropdeh3_1_pdu(pf02288) PDU1AB EP_NTERM
01585 HMMpropdeh2_1_pdu(pf02287) PDU1AB EP_NTERM
01590 HMM08841dhact_1_pdu(pf08841) PDU1AB
01595 HMMpropdeh3_2_pdusml(pf02288) PDU1AB
01600 H_tan PDU1AB
01605 H_driedBlood PDU1AB
01610 HMMptac_2_mix(pf06130) PDU1AB
01615 HMM15953pduput_1_pdu1(pf15953) PDU1AB EP_NTERM
01620 P_babyPink PDU1AB
01625 HMM01923cobtr_1_pdu(pf01923pf03928) PDU1AB
01630 HMMalddh_1_eutpdugrm(pf00171) PDU1AB EP_NTERM
01635 HMM00465alcdh_3_mix(pf00465) PDU1AB
01640 HMMPduS_1_all(pf01512pf13375pf10531) PDU1AB
01645 Ts_dodgerBlue PDU1AB
01650 Hp_euts_babyPurple PDU1AB
01655 HMM10662PduV_1_mix(pf10662) PDU1AB
01660 HMMacetkin_1(pf00871) PDU1AB
01665 HMM00288_1_pdu(pf00288) PDU1AB
01670 HMMreg00392_2_grm2(pf00155) EUT2A
01675 -pf01520pf11741 -
01680 -pf00696pf00583 -
BMC-P
BMC-H
BMC-T
Aldehyde dehydrogenase
PTAC
Signature enzyme
Alcohol dehydrogenase
Regulator

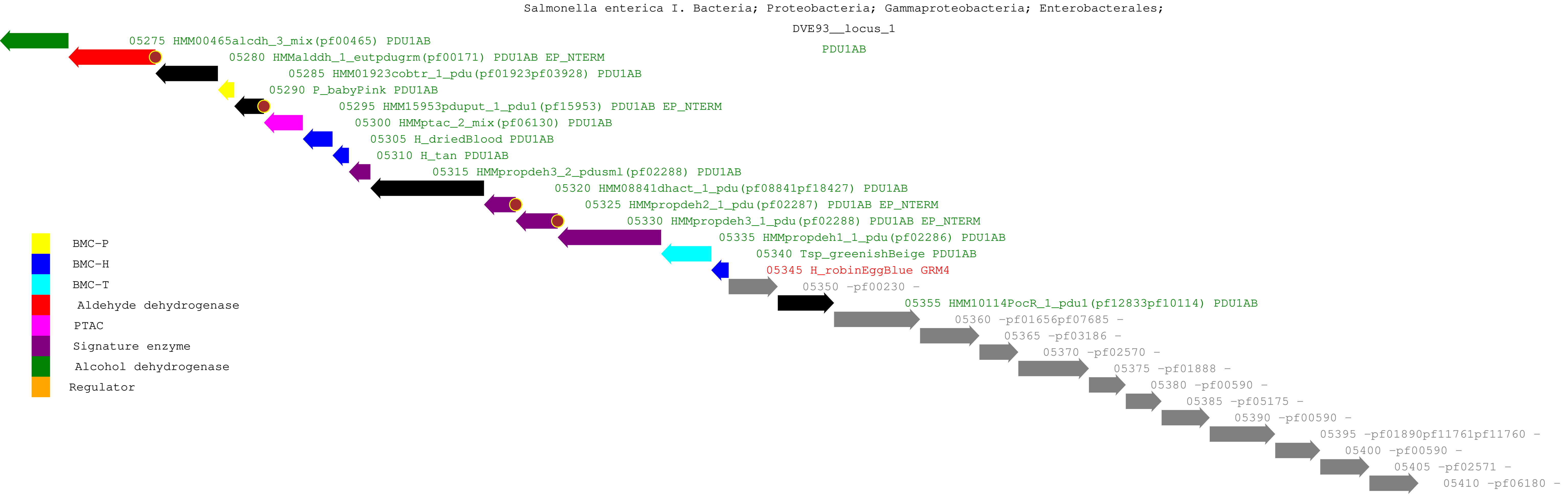
Salmonella enterica I. Bacteria; Proteobacteria; Gammaproteobacteria; Enterobacterales;
DVE93__locus_1
PDU1AB
05275 HMM00465alcdh_3_mix(pf00465) PDU1AB
05280 HMMaldh_1_eutpdugrm(pf00171) PDU1AB EP_NTERM
05285 HMM01923cobtr_1_pdu(pf01923pf03928) PDU1AB
05290 P_babyPink PDU1AB
05295 HMM15953pduput_1_pdu1(pf15953) PDU1AB EP_NTERM
05300 HMMptac_2_mix(pf06130) PDU1AB
05305 H_driedBlood PDU1AB
05310 H_tan PDU1AB
05315 HMMpropdeh3_2_pdusml(pf02288) PDU1AB
05320 HMM08841dhact_1_pdu(pf08841pf18427) PDU1AB
05325 HMMpropdeh2_1_pdu(pf02287) PDU1AB EP_NTERM
05330 HMMpropdeh3_1_pdu(pf02288) PDU1AB EP_NTERM
05335 HMMpropdeh1_1_pdu(pf02286) PDU1AB
05340 Tsp_greenishBeige PDU1AB
05345 H_robinEggBlue GRM4
05350 -pf00230 -
05355 HMM10114PocR_1_pdu1(pf12833pf10114) PDU1AB
05360 -pf01656pf07685 -
05365 -pf03186 -
05370 -pf02570 -
05375 -pf01888 -
05380 -pf00590 -
05385 -pf05175 -
05390 -pf00590 -
05395 -pf01890pf11761pf11760 -
05400 -pf00590 -
05405 -pf02571 -
05410 -pf06180 -
BMC-P
BMC-H
BMC-T
Aldehyde dehydrogenase
PTAC
Signature enzyme
Alcohol dehydrogenase
Regulator

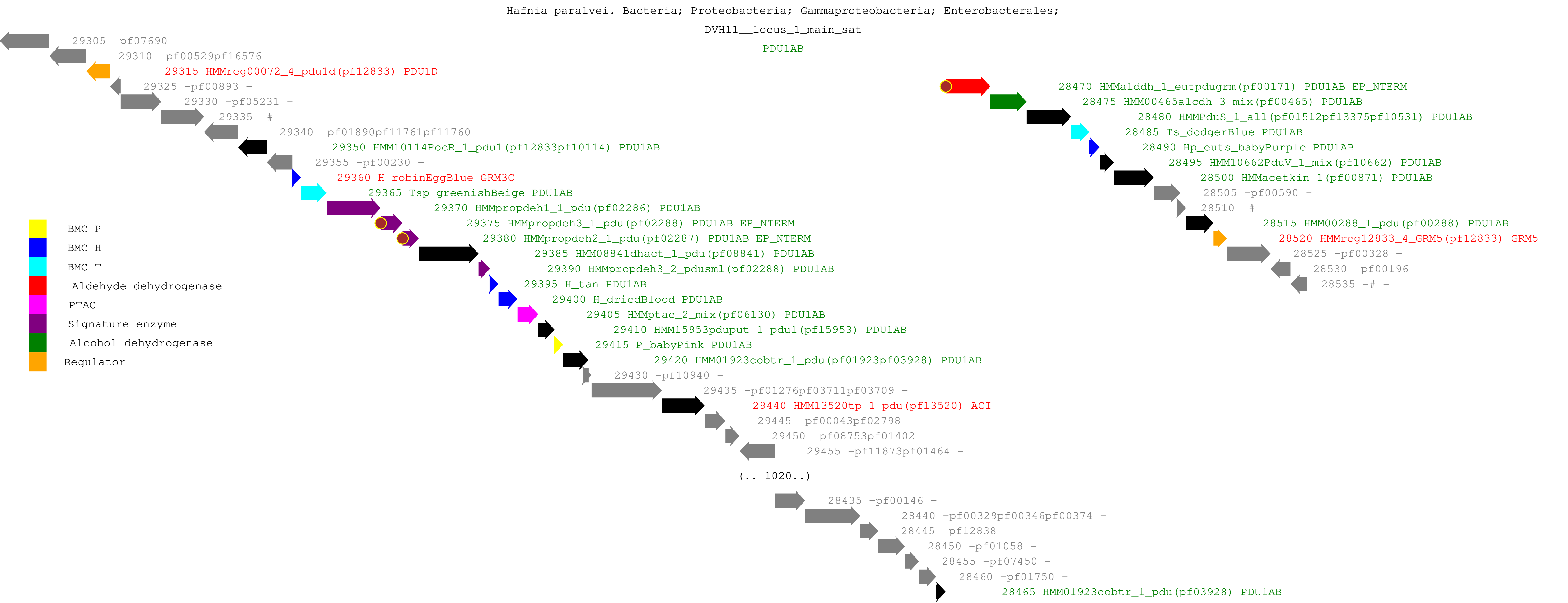
Hafnia paralvei. Bacteria; Proteobacteria; Gammaproteobacteria; Enterobacterales;
DVH11__locus_1_main_sat
PDU1AB
29305 -pf07690 -
29310 -pf00529pf16576 -
29315 HMMreg00072_4_pdu1d(pf12833) PDU1D
29325 -pf00893 -
29330 -pf05231 -
29335 -# -
29340 -pf01890pf11761pf11760 -
29350 HMM10114PocR_1_pdu1(pf12833pf10114) PDU1AB
29355 -pf00230 -
29360 H_robinEggBlue GRM3C
29365 Tsp_greenishBeige PDU1AB
29370 HMMpropdeh1_1_pdu(pf02286) PDU1AB
29375 HMMpropdeh3_1_pdu(pf02288) PDU1AB EP_NTERM
29380 HMMpropdeh2_1_pdu(pf02287) PDU1AB EP_NTERM
29385 HMM08841dhact_1_pdu(pf08841) PDU1AB
29390 HMMpropdeh3_2_pdusml(pf02288) PDU1AB
29395 H_tan PDU1AB
29400 H_driedBlood PDU1AB
29405 HMMptac_2_mix(pf06130) PDU1AB
29410 HMM15953pduput_1_pdu1(pf15953) PDU1AB
29415 P_babyPink PDU1AB
29420 HMM01923cobtr_1_pdu(pf01923pf03928) PDU1AB
29430 -pf10940 -
29435 -pf01276pf03711pf03709 -
29440 HMM13520tp_1_pdu(pf13520) ACI
29445 -pf00043pf02798 -
29450 -pf08753pf01402 -
29455 -pf11873pf01464 -
(..-1020..)
28435 -pf00146 -
28440 -pf00329pf00346pf00374 -
28445 -pf12838 -
28450 -pf01058 -
28455 -pf07450 -
28460 -pf01750 -
28465 HMM01923cobtr_1_pdu(pf03928) PDU1AB
28470 HMMalddh_1_eutpdugrm(pf00171) PDU1AB EP_NTERM
28475 HMM00465alcdh_3_mix(pf00465) PDU1AB
28480 HMMPduS_1_all(pf01512pf13375pf10531) PDU1AB
28485 Ts_dodgerBlue PDU1AB
28490 Hp_euts_babyPurple PDU1AB
28495 HMM10662PduV_1_mix(pf10662) PDU1AB
28500 HMMacetkin_1(pf00871) PDU1AB
28505 -pf00590 -
28510 -# -
28515 HMM00288_1_pdu(pf00288) PDU1AB
28520 HMMreg12833_4_GRM5(pf12833) GRM5
28525 -pf00328 -
28530 -pf00196 -
28535 -# -
BMC-P
BMC-H
BMC-T
Aldehyde dehydrogenase
PTAC
Signature enzyme
Alcohol dehydrogenase
Regulator

Escherichia coli. Bacteria; Proteobacteria; Gammaproteobacteria; Enterobacterales;

DXF93__locus_1

PDU1AB

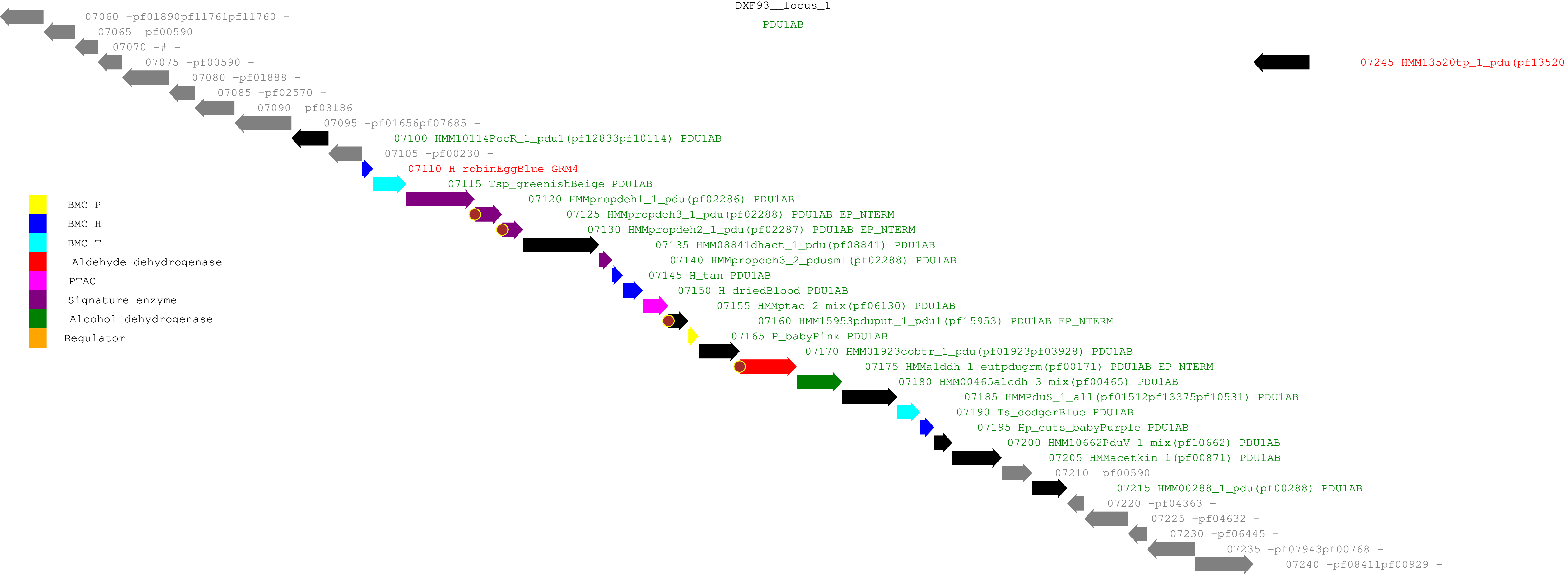

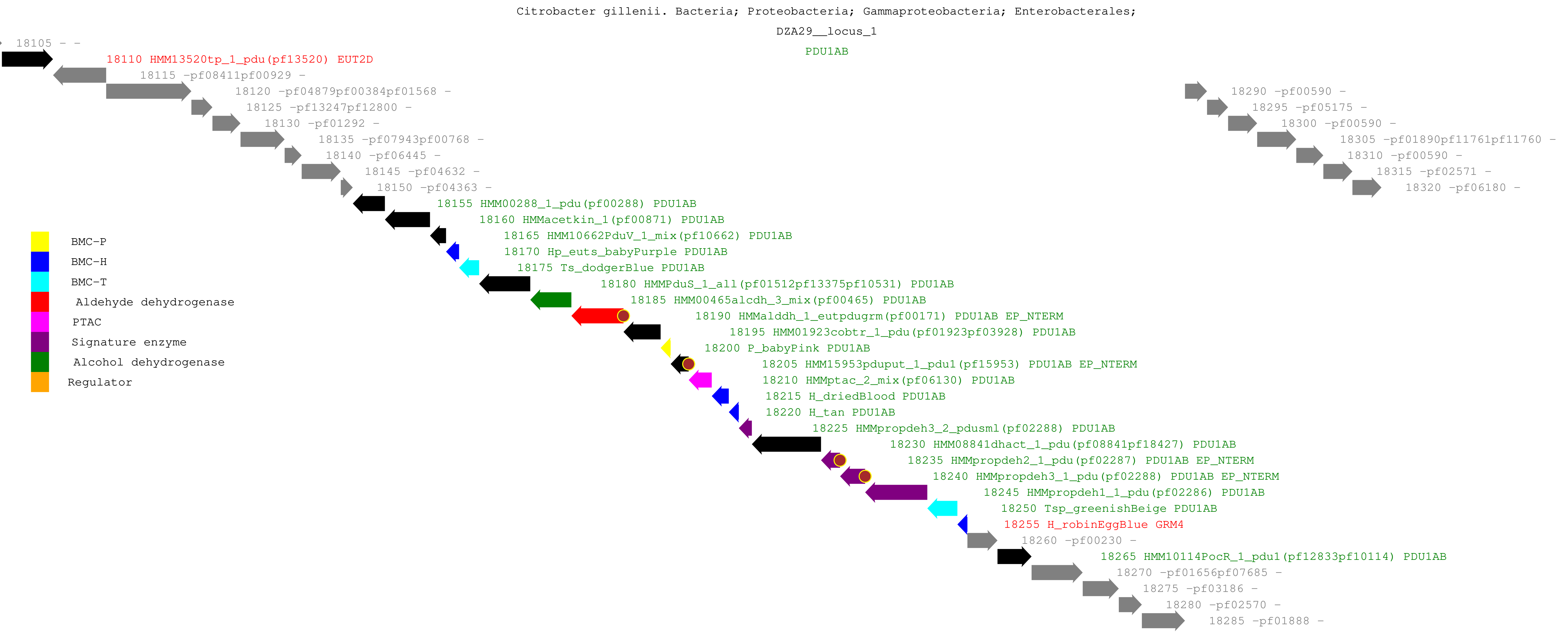

Citrobacter gillenii. Bacteria; Proteobacteria; Gammaproteobacteria; Enterobacterales;
DZA29__locus_1
PDU1AB
18105 - -
18110 HMM13520tp_1_pdu(pf13520) EUT2D
18115 -pf08411pf00929 -
18120 -pf04879pf00384pf01568 -
18125 -pf13247pf12800 -
18130 -pf01292 -
18135 -pf07943pf00768 -
18140 -pf06445 -
18145 -pf04632 -
18150 -pf04363 -
18155 HMM00288_1_pdu(pf00288) PDU1AB
18160 HMMacetkin_1(pf00871) PDU1AB
18165 HMM10662PduV_1_mix(pf10662) PDU1AB
18170 Hp_euts_babyPurple PDU1AB
18175 Ts_dodgerBlue PDU1AB
18180 HMMPduS_1_all(pf01512pf13375pf10531) PDU1AB
18185 HMM00465alcdh_3_mix(pf00465) PDU1AB
18190 HMMalddh_1_eutpdugrm(pf00171) PDU1AB EP_NTERM
18195 HMM01923cobtr_1_pdu(pf01923pf03928) PDU1AB
18200 P_babyPink PDU1AB
18205 HMM15953pduput_1_pdu1(pf15953) PDU1AB EP_NTERM
18210 HMMptac_2_mix(pf06130) PDU1AB
18215 H_driedBlood PDU1AB
18220 H_tan PDU1AB
18225 HMMpropdeh3_2_pdusml(pf02288) PDU1AB
18230 HMM08841dhact_1_pdu(pf08841pf18427) PDU1AB
18235 HMMpropdeh2_1_pdu(pf02287) PDU1AB EP_NTERM
18240 HMMpropdeh3_1_pdu(pf02288) PDU1AB EP_NTERM
18245 HMMpropdeh1_1_pdu(pf02286) PDU1AB
18250 Tsp_greenishBeige PDU1AB
18255 H_robinEggBlue GRM4
18260 -pf00230 -
18265 HMM10114PocR_1_pdu1(pf12833pf10114) PDU1AB
18270 -pf01656pf07685 -
18275 -pf03186 -
18280 -pf02570 -
18285 -pf01888 -
18290 -pf00590 -
18295 -pf05175 -
18300 -pf00590 -
18305 -pf01890pf11761pf11760 -
18310 -pf00590 -
18315 -pf02571 -
18320 -pf06180 -
BMC-P
BMC-H
BMC-T
Aldehyde dehydrogenase
PTAC
Signature enzyme
Alcohol dehydrogenase
Regulator

Salmonella enterica subsp. enterica serovar Tilene. Bacteria; Proteobacteria; Gammaproteobacteria; Enterobacterales;

E0539__locus_2

PDU1AB

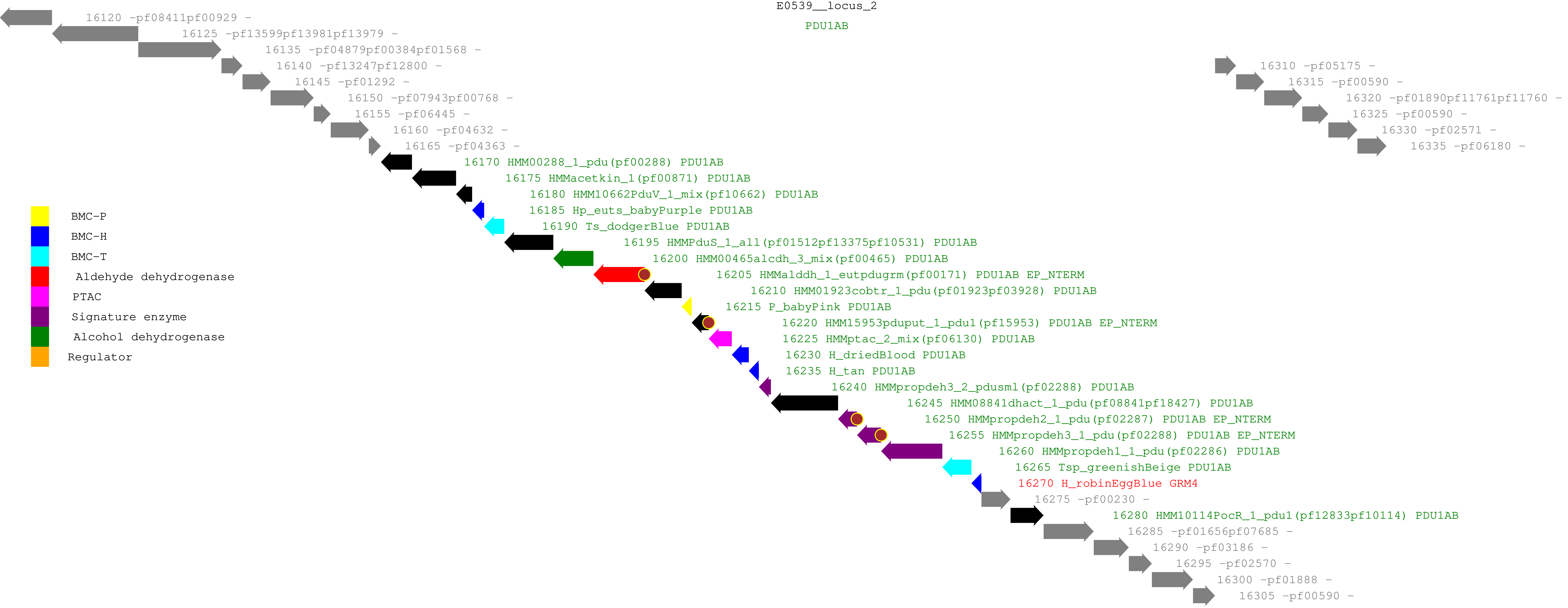

Salmonella enterica subsp. enterica serovar London. Bacteria; Proteobacteria; Gammaproteobacteria; Enterobacterales;

E0541__locus_1

PDU1AB

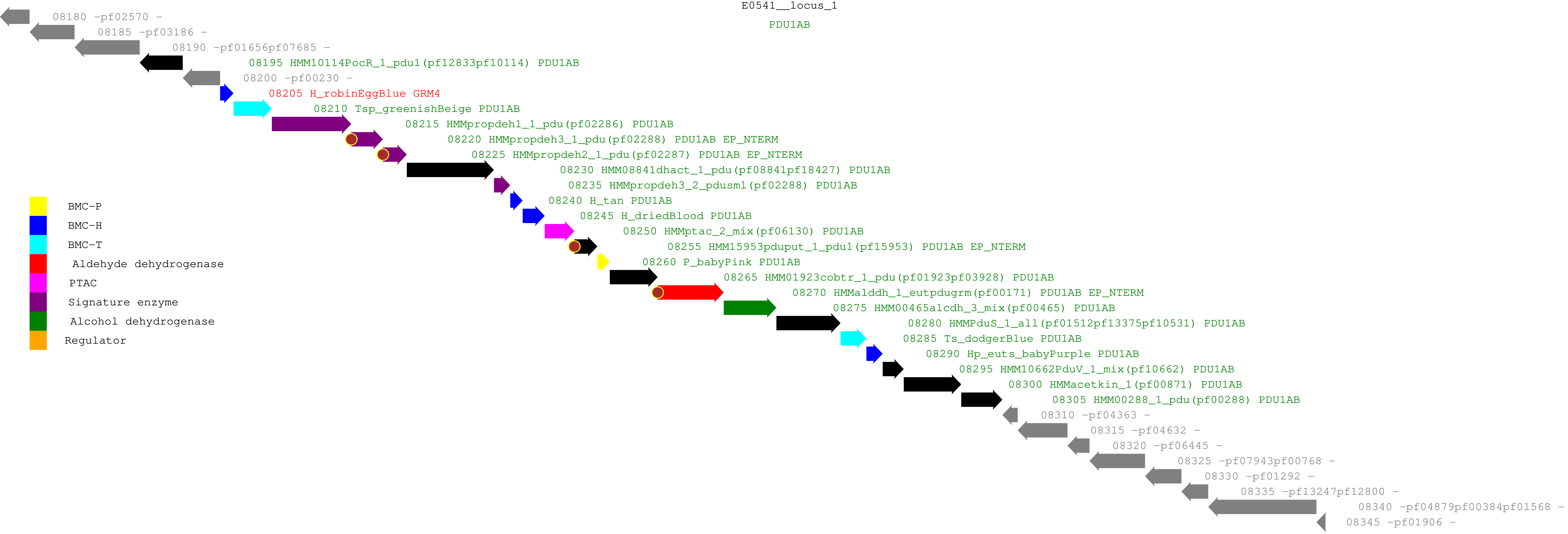

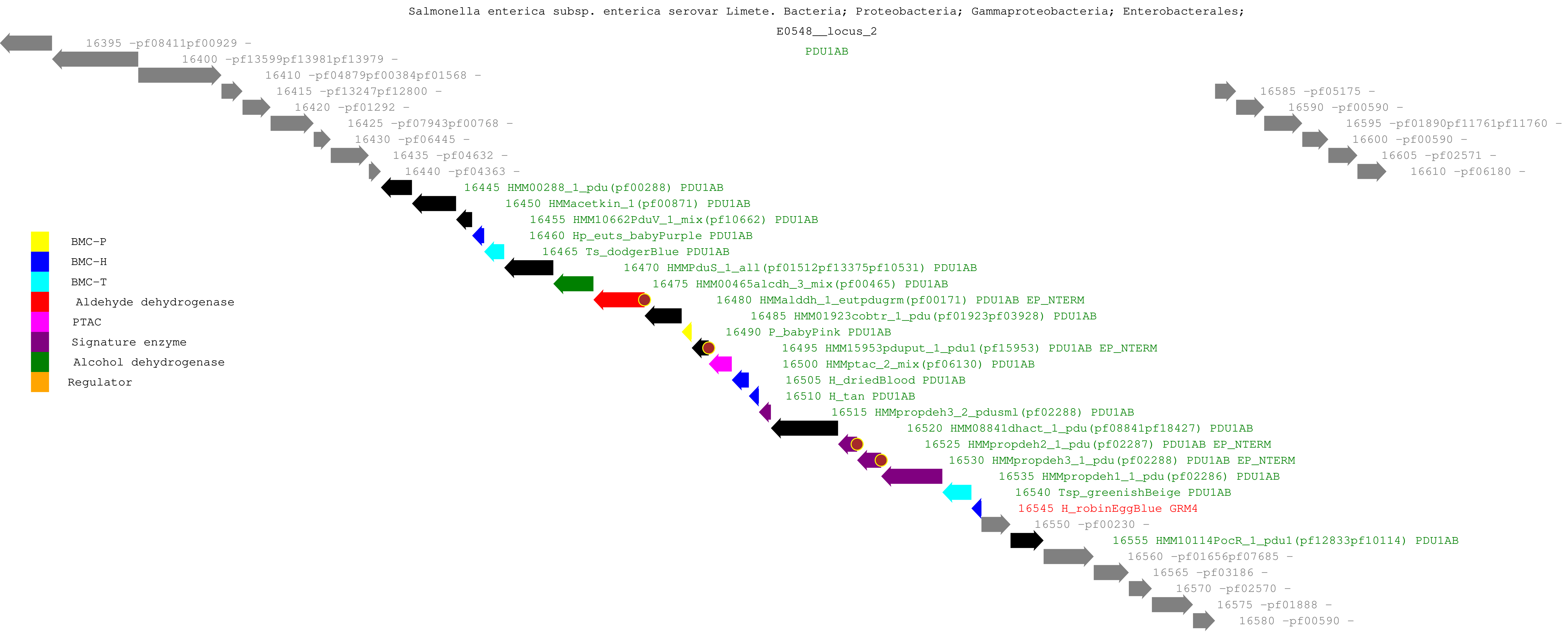
Salmonella enterica subsp. enterica serovar Limete. Bacteria; Proteobacteria; Gammaproteobacteria; Enterobacterales;
E0548__locus_2
PDU1AB
16395 -pf08411pf00929 -
16400 -pf13599pf13981pf13979 -
16410 -pf04879pf00384pf01568 -
16415 -pf13247pf12800 -
16420 -pf01292 -
16425 -pf07943pf00768 -
16430 -pf06445 -
16435 -pf04632 -
16440 -pf04363 -
16445 HMM00288_1_pdu(pf00288) PDU1AB
16450 HMMacetkin_1(pf00871) PDU1AB
16455 HMM10662PduV_1_mix(pf10662) PDU1AB
16460 Hp_euts_babyPurple PDU1AB
16465 Ts_dodgerBlue PDU1AB
16470 HMMPduS_1_all(pf01512pf13375pf10531) PDU1AB
16475 HMM00465alcdh_3_mix(pf00465) PDU1AB
16480 HMMalddh_1_eutpdugrm(pf00171) PDU1AB EP_NTERM
16485 HMM01923cobtr_1_pdu(pf01923pf03928) PDU1AB
16490 P_babyPink PDU1AB
16495 HMM15953pduput_1_pdu1(pf15953) PDU1AB EP_NTERM
16500 HMMptac_2_mix(pf06130) PDU1AB
16505 H_driedBlood PDU1AB
16510 H_tan PDU1AB
16515 HMMpropdeh3_2_pdusml(pf02288) PDU1AB
16520 HMM08841dhact_1_pdu(pf08841pf18427) PDU1AB
16525 HMMpropdeh2_1_pdu(pf02287) PDU1AB EP_NTERM
16530 HMMpropdeh3_1_pdu(pf02288) PDU1AB EP_NTERM
16535 HMMpropdeh1_1_pdu(pf02286) PDU1AB
16540 Tsp_greenishBeige PDU1AB
16545 H_robinEggBlue GRM4
16550 -pf00230 -
16555 HMM10114PocR_1_pdu1(pf12833pf10114) PDU1AB
16560 -pf01656pf07685 -
16565 -pf03186 -
16570 -pf02570 -
16575 -pf01888 -
16580 -pf00590 -
16585 -pf05175 -
16590 -pf00590 -
16595 -pf01890pf11761pf11760 -
16600 -pf00590 -
16605 -pf02571 -
16610 -pf06180 -
BMC-P
BMC-H
BMC-T
Aldehyde dehydrogenase
PTAC
Signature enzyme
Alcohol dehydrogenase
Regulator

Salmonella enterica subsp. enterica serovar Richmond. Bacteria; Proteobacteria; Gammaproteobacteria; Enterobacterales;

E0577__locus_2

PDU1AB

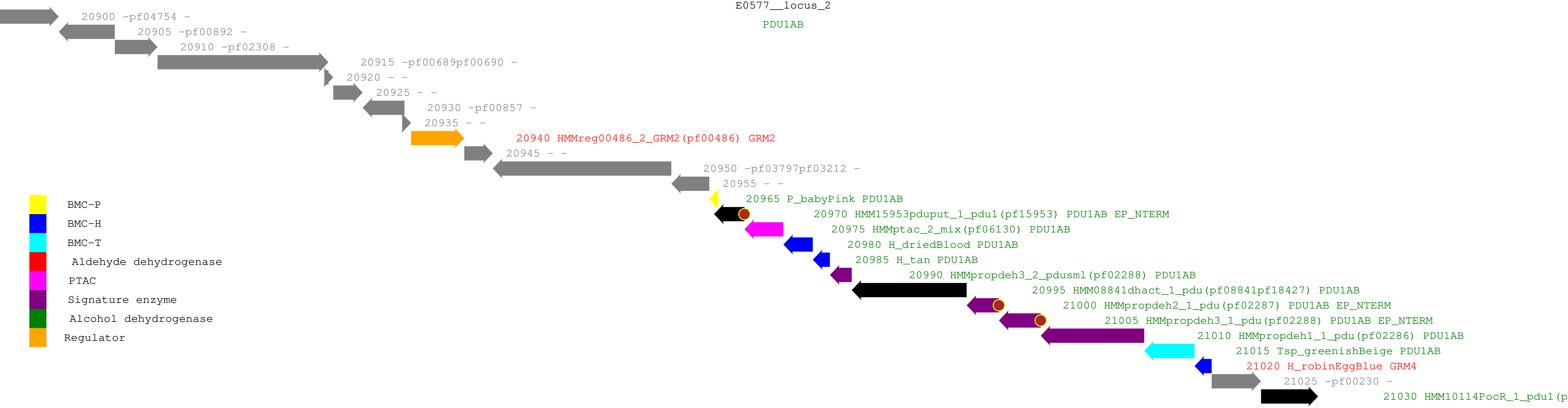

Salmonella enterica subsp. enterica serovar Richmond. Bacteria; Proteobacteria; Gammaproteobacteria; Enterobacterales;

E0577__locus_3

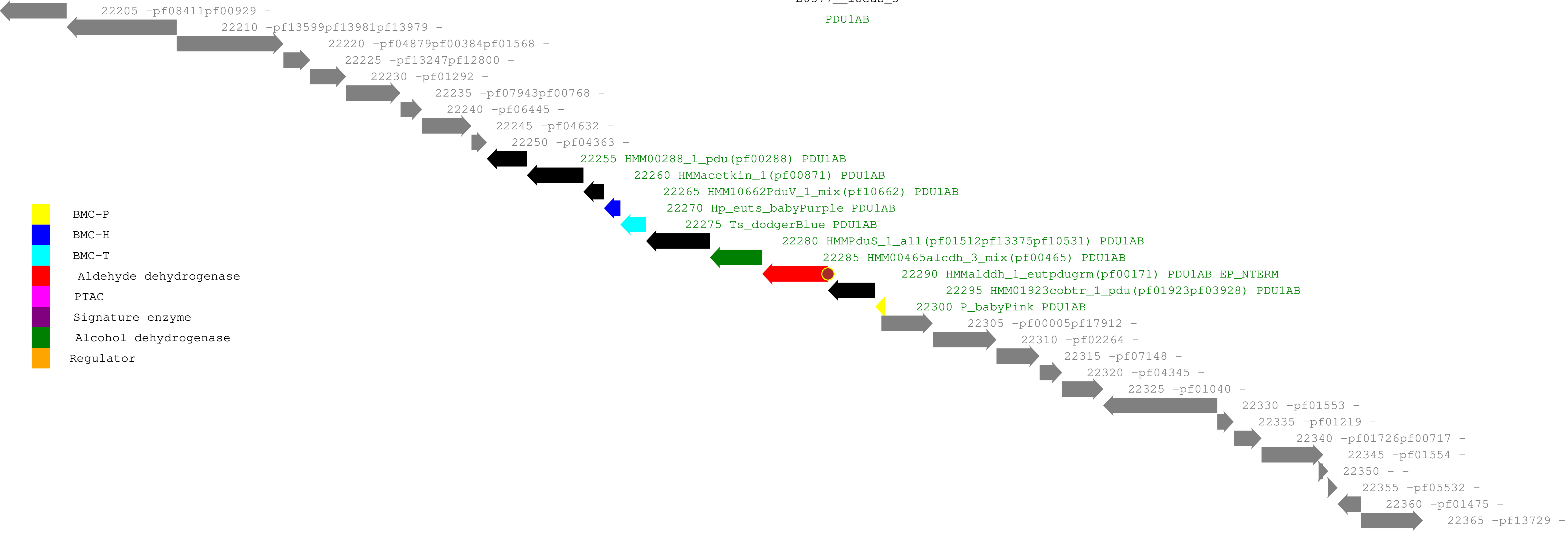

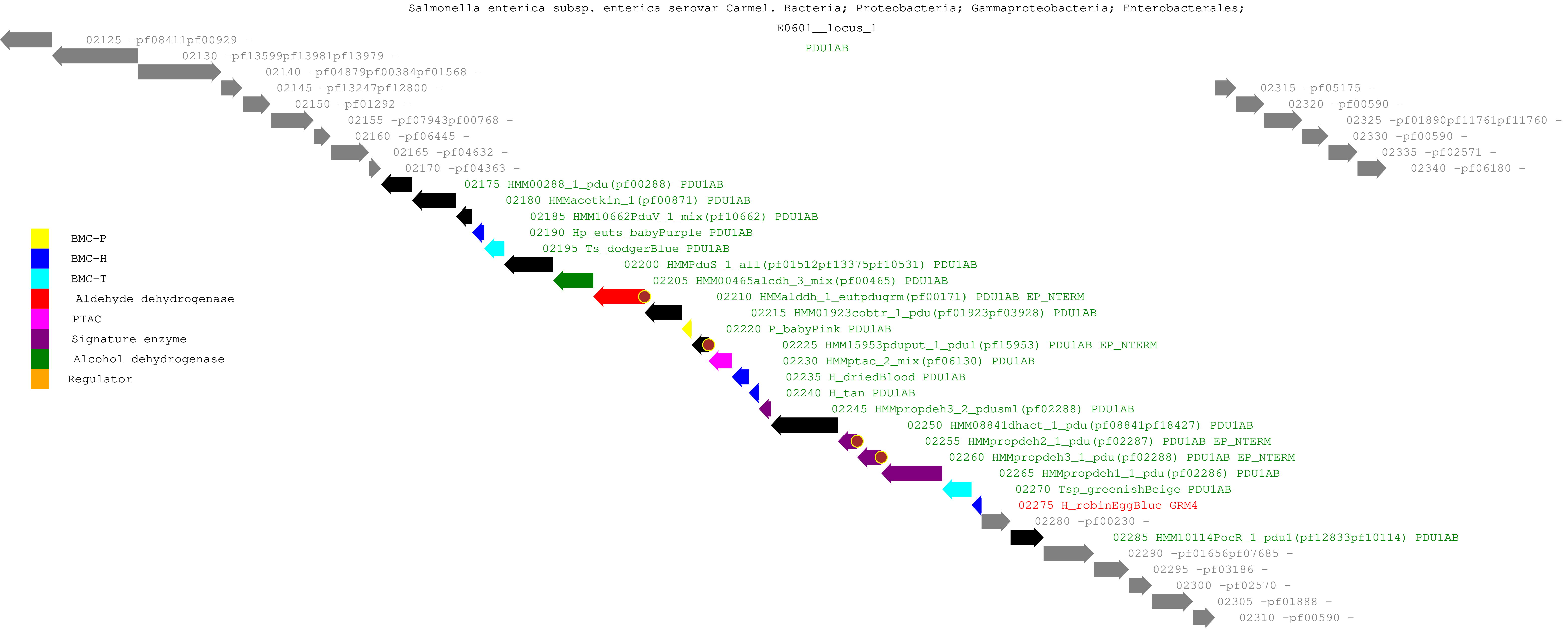
Salmonella enterica subsp. enterica serovar Carmel. Bacteria; Proteobacteria; Gammaproteobacteria; Enterobacterales;
E0601__locus_1
PDU1AB
02125 -pf08411pf00929 -
02130 -pf13599pf13981pf13979 -
02140 -pf04879pf00384pf01568 -
02145 -pf13247pf12800 -
02150 -pf01292 -
02155 -pf07943pf00768 -
02160 -pf06445 -
02165 -pf04632 -
02170 -pf04363 -
02175 HMM00288_1_pdu(pf00288) PDU1AB
02180 HMMacetkin_1(pf00871) PDU1AB
02185 HMM10662PduV_1_mix(pf10662) PDU1AB
02190 Hp_euts_babyPurple PDU1AB
02195 Ts_dodgerBlue PDU1AB
02200 HMMPduS_1_all(pf01512pf13375pf10531) PDU1AB
02205 HMM00465alcdh_3_mix(pf00465) PDU1AB
02210 HMMalddh_1_eutpdugrm(pf00171) PDU1AB EP_NTERM
02215 HMM01923cobtr_1_pdu(pf01923pf03928) PDU1AB
02220 P_babyPink PDU1AB
02225 HMM15953pduput_1_pdu1(pf15953) PDU1AB EP_NTERM
02230 HMMptac_2_mix(pf06130) PDU1AB
02235 H_driedBlood PDU1AB
02240 H_tan PDU1AB
02245 HMMpropdeh3_2_pdusml(pf02288) PDU1AB
02250 HMM08841dhact_1_pdu(pf08841pf18427) PDU1AB
02255 HMMpropdeh2_1_pdu(pf02287) PDU1AB EP_NTERM
02260 HMMpropdeh3_1_pdu(pf02288) PDU1AB EP_NTERM
02265 HMMpropdeh1_1_pdu(pf02286) PDU1AB
02270 Tsp_greenishBeige PDU1AB
02275 H_robinEggBlue GRM4
02280 -pf00230 -
02285 HMM10114PocR_1_pdu1(pf12833pf10114) PDU1AB
02290 -pf01656pf07685 -
02295 -pf03186 -
02300 -pf02570 -
02305 -pf01888 -
02310 -pf00590 -
02315 -pf05175 -
02320 -pf00590 -
02325 -pf01890pf11761pf11760 -
02330 -pf00590 -
02335 -pf02571 -
02340 -pf06180 -
BMC-P
BMC-H
BMC-T
Aldehyde dehydrogenase
PTAC
Signature enzyme
Alcohol dehydrogenase
Regulator

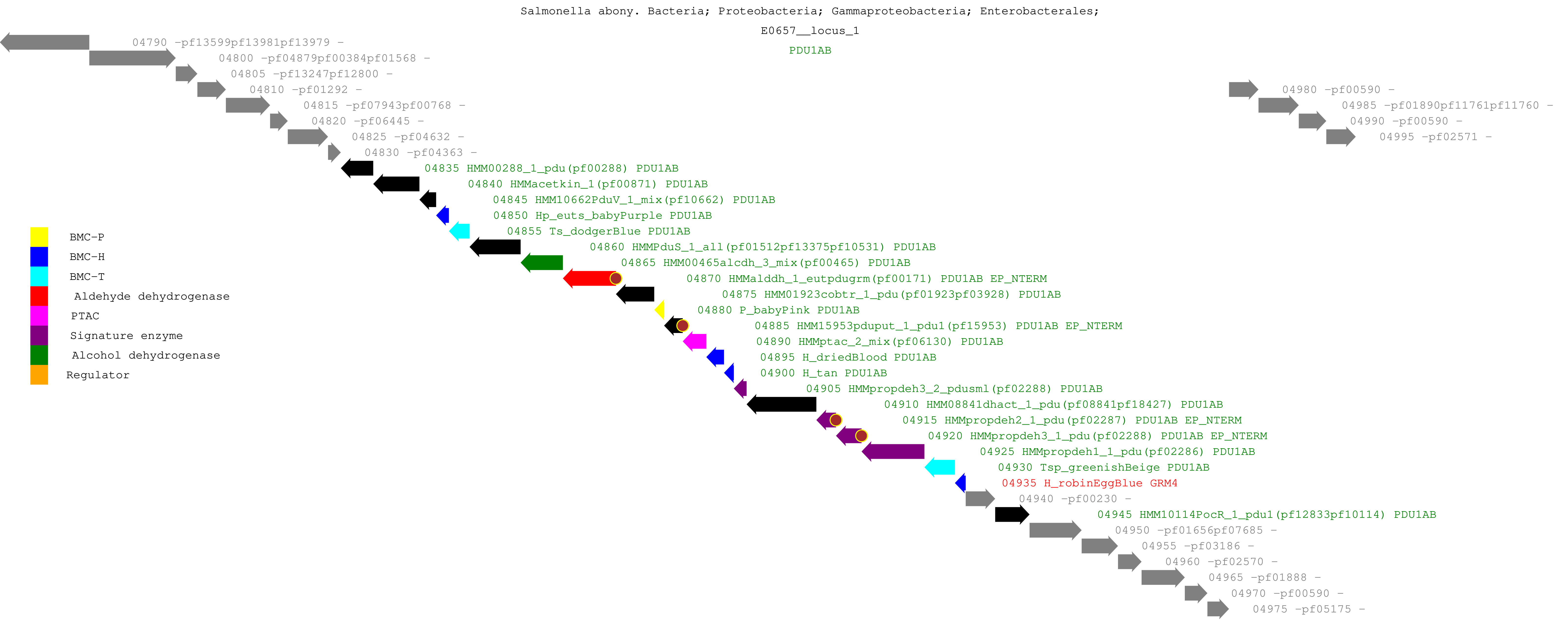
Salmonella abony. Bacteria; Proteobacteria; Gammaproteobacteria; Enterobacterales;
E0657__locus_1
PDU1AB
04790 -pf13599pf13981pf13979 -
04800 -pf04879pf00384pf01568 -
04805 -pf13247pf12800 -
04810 -pf01292 -
04815 -pf07943pf00768 -
04820 -pf06445 -
04825 -pf04632 -
04830 -pf04363 -
04835 HMM00288_1_pdu(pf00288) PDU1AB
04840 HMMacetkin_1(pf00871) PDU1AB
04845 HMM10662PduV_1_mix(pf10662) PDU1AB
04850 Hp_euts_babyPurple PDU1AB
04855 Ts_dodgerBlue PDU1AB
04860 HMMPduS_1_all(pf01512pf13375pf10531) PDU1AB
04865 HMM00465alcdh_3_mix(pf00465) PDU1AB
04870 HMMaldh_1_eutpdugrm(pf00171) PDU1AB EP_NTERM
04875 HMM01923cobtr_1_pdu(pf01923pf03928) PDU1AB
04880 P_babyPink PDU1AB
04885 HMM15953pduput_1_pdu1(pf15953) PDU1AB EP_NTERM
04890 HMMptac_2_mix(pf06130) PDU1AB
04895 H_driedBlood PDU1AB
04900 H_tan PDU1AB
04905 HMMpropdeh3_2_pdusml(pf02288) PDU1AB
04910 HMM08841dhact_1_pdu(pf08841pf18427) PDU1AB
04915 HMMpropdeh2_1_pdu(pf02287) PDU1AB EP_NTERM
04920 HMMpropdeh3_1_pdu(pf02288) PDU1AB EP_NTERM
04925 HMMpropdeh1_1_pdu(pf02286) PDU1AB
04930 Tsp_greenishBeige PDU1AB
04935 H_robinEggBlue GRM4
04940 -pf00230 -
04945 HMM10114PocR_1_pdu1(pf12833pf10114) PDU1AB
04950 -pf01656pf07685 -
04955 -pf03186 -
04960 -pf02570 -
04965 -pf01888 -
04970 -pf00590 -
04975 -pf05175 -
04980 -pf00590 -
04985 -pf01890pf11761pf11760 -
04990 -pf00590 -
04995 -pf02571 -
BMC-P
BMC-H
BMC-T
Aldehyde dehydrogenase
PTAC
Signature enzyme
Alcohol dehydrogenase
Regulator

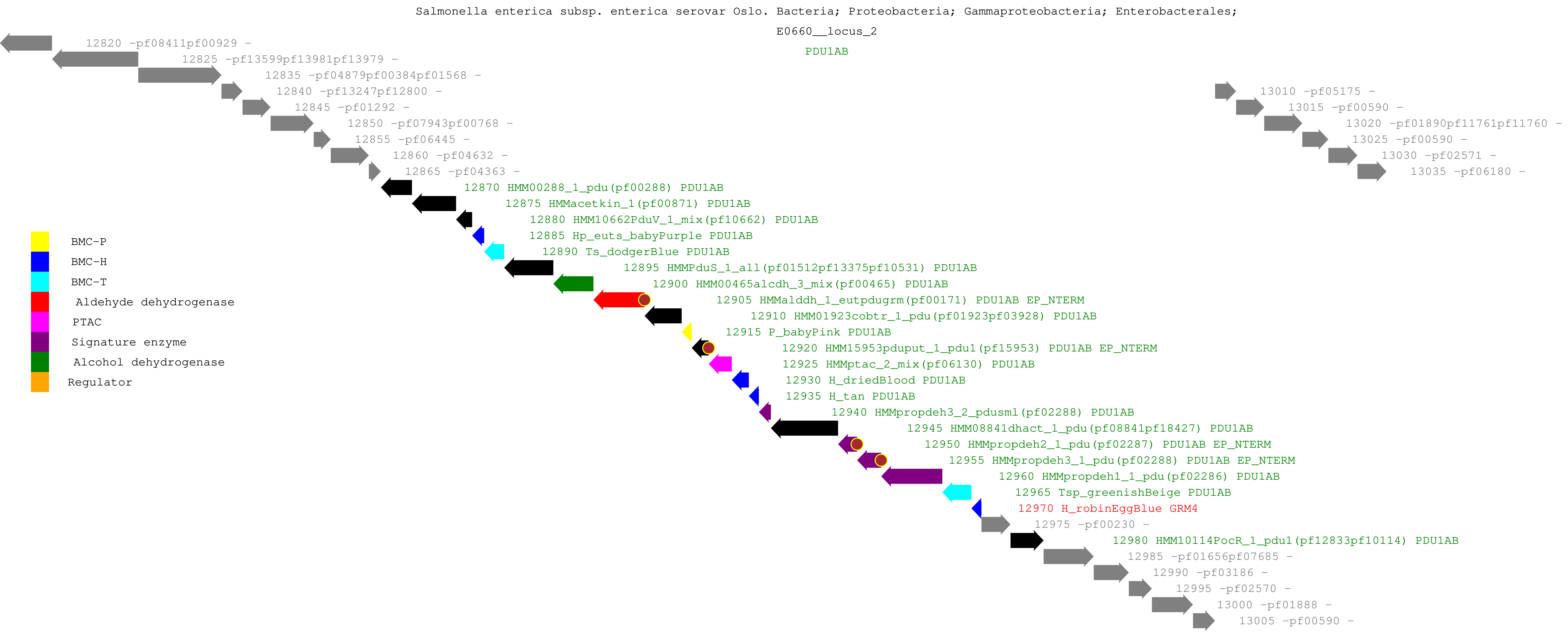
Salmonella enterica subsp. enterica serovar Oslo. Bacteria; Proteobacteria; Gammaproteobacteria; Enterobacterales;
E0660__locus_2
PDU1AB
12820 -pf08411pf00929 -
12825 -pf13599pf13981pf13979 -
12835 -pf04879pf00384pf01568 -
12840 -pf13247pf12800 -
12845 -pf01292 -
12850 -pf07943pf00768 -
12855 -pf06445 -
12860 -pf04632 -
12865 -pf04363 -
12870 HMM00288_1_pdu(pf00288) PDU1AB
12875 HMMacetkin_1(pf00871) PDU1AB
12880 HMM10662PduV_1_mix(pf10662) PDU1AB
12885 Hp_euts_babyPurple PDU1AB
12890 Ts_dodgerBlue PDU1AB
12895 HMMPduS_1_all(pf01512pf13375pf10531) PDU1AB
12900 HMM00465alcdh_3_mix(pf00465) PDU1AB
12905 HMMalddh_1_eutpdugrm(pf00171) PDU1AB EP_NTERM
12910 HMM01923cobtr_1_pdu(pf01923pf03928) PDU1AB
12915 P_babyPink PDU1AB
12920 HMM15953pduput_1_pdu1(pf15953) PDU1AB EP_NTERM
12925 HMMptac_2_mix(pf06130) PDU1AB
12930 H_driedBlood PDU1AB
12935 H_tan PDU1AB
12940 HMMpropdeh3_2_pdusml(pf02288) PDU1AB
12945 HMM08841dhact_1_pdu(pf08841pf18427) PDU1AB
12950 HMMpropdeh2_1_pdu(pf02287) PDU1AB EP_NTERM
12955 HMMpropdeh3_1_pdu(pf02288) PDU1AB EP_NTERM
12960 HMMpropdeh1_1_pdu(pf02286) PDU1AB
12965 Tsp_greenishBeige PDU1AB
12970 H_robinEggBlue GRM4
12975 -pf00230 -
12980 HMM10114PocR_1_pdu1(pf12833pf10114) PDU1AB
12985 -pf01656pf07685 -
12990 -pf03186 -
12995 -pf02570 -
13000 -pf01888 -
13005 -pf00590 -
13010 -pf05175 -
13015 -pf00590 -
13020 -pf01890pf11761pf11760 -
13025 -pf00590 -
13030 -pf02571 -
13035 -pf06180 -
BMC-P
BMC-H
BMC-T
Aldehyde dehydrogenase
PTAC
Signature enzyme
Alcohol dehydrogenase
Regulator

Salmonella berta. Bacteria; Proteobacteria; Gammaproteobacteria; Enterobacterales;

E0917__locus_1

PDU1AB

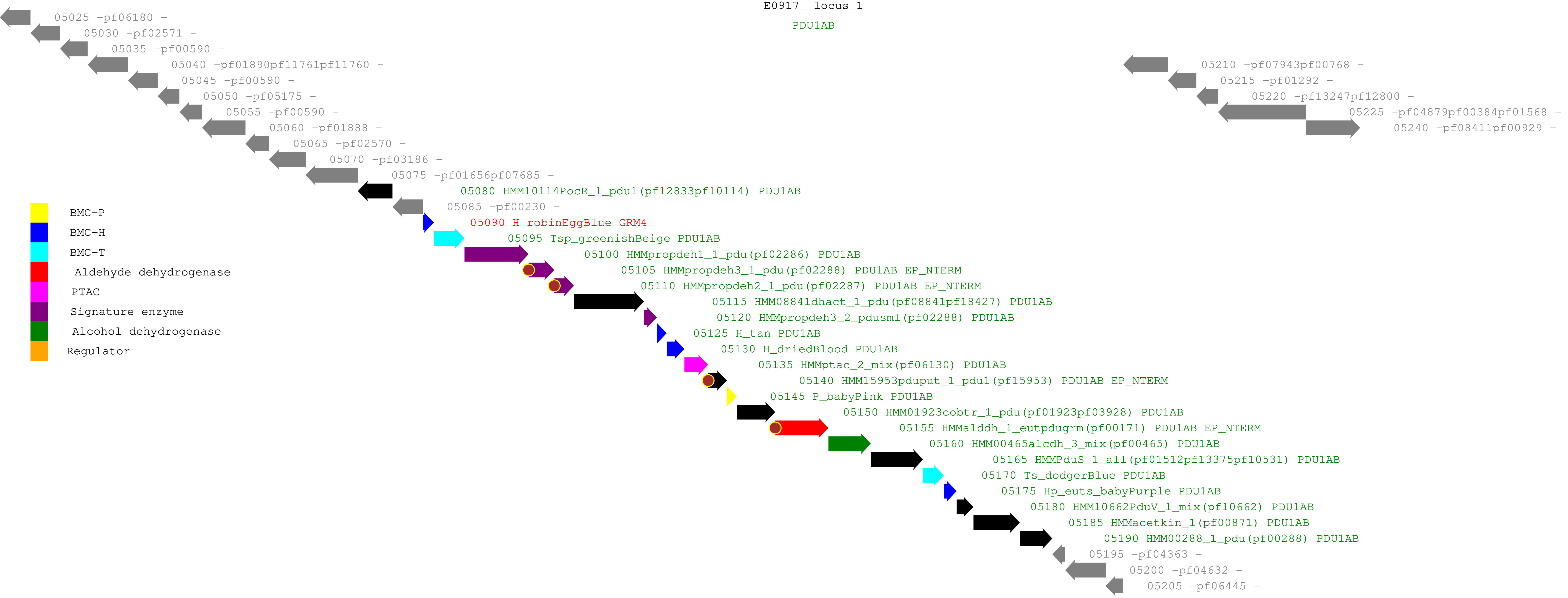

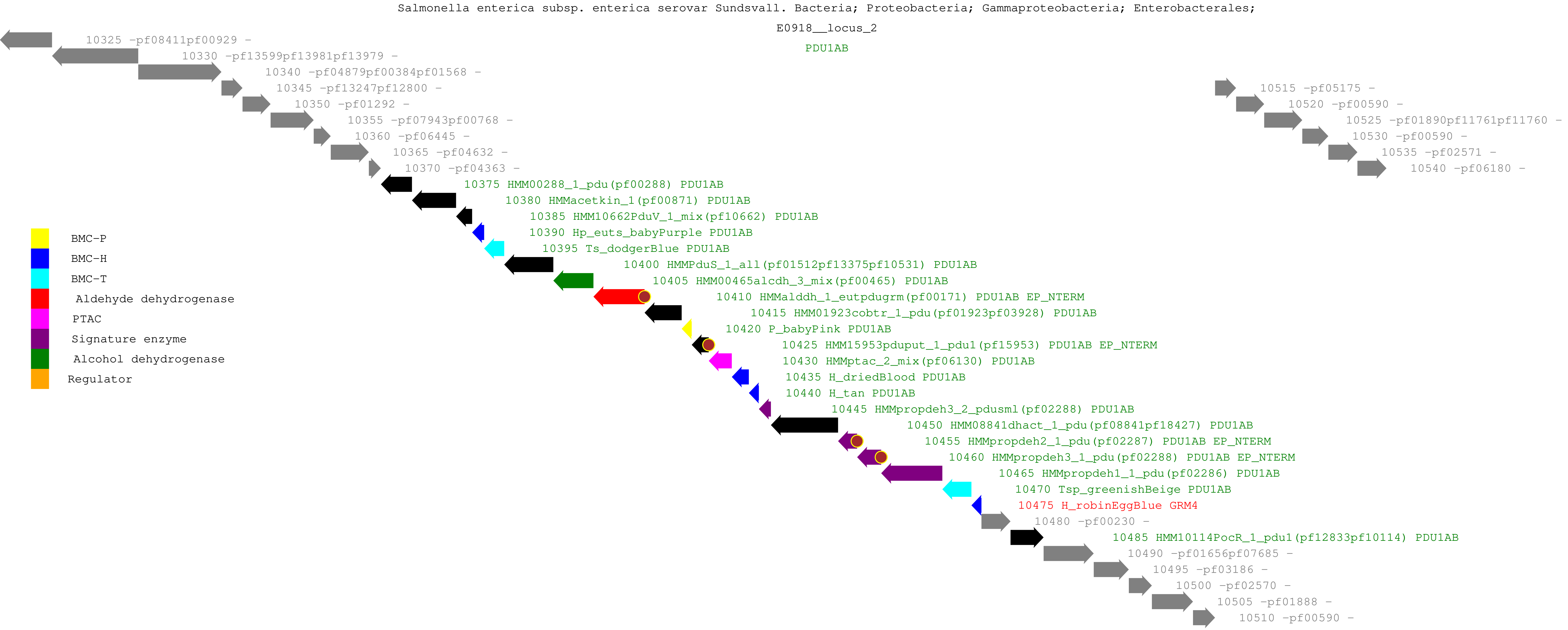
Salmonella enterica subsp. enterica serovar Sundsvall. Bacteria; Proteobacteria; Gammaproteobacteria; Enterobacterales;
E0918__locus_2
PDU1AB
10325 -pf08411pf00929 -
10330 -pf13599pf13981pf13979 -
10340 -pf04879pf00384pf01568 -
10345 -pf13247pf12800 -
10350 -pf01292 -
10355 -pf07943pf00768 -
10360 -pf06445 -
10365 -pf04632 -
10370 -pf04363 -
10375 HMM00288_1_pdu(pf00288) PDU1AB
10380 HMMacetkin_1(pf00871) PDU1AB
10385 HMM10662PduV_1_mix(pf10662) PDU1AB
10390 Hp_euts_babyPurple PDU1AB
10395 Ts_dodgerBlue PDU1AB
10400 HMMPduS_1_all(pf01512pf13375pf10531) PDU1AB
10405 HMM00465alcdh_3_mix(pf00465) PDU1AB
10410 HMMalddh_1_eutpdugrm(pf00171) PDU1AB EP_NTERM
10415 HMM01923cobtr_1_pdu(pf01923pf03928) PDU1AB
10420 P_babyPink PDU1AB
10425 HMM15953pduput_1_pdu1(pf15953) PDU1AB EP_NTERM
10430 HMMptac_2_mix(pf06130) PDU1AB
10435 H_driedBlood PDU1AB
10440 H_tan PDU1AB
10445 HMMpropdeh3_2_pdusml(pf02288) PDU1AB
10450 HMM08841dhact_1_pdu(pf08841pf18427) PDU1AB
10455 HMMpropdeh2_1_pdu(pf02287) PDU1AB EP_NTERM
10460 HMMpropdeh3_1_pdu(pf02288) PDU1AB EP_NTERM
10465 HMMpropdeh1_1_pdu(pf02286) PDU1AB
10470 Tsp_greenishBeige PDU1AB
10475 H_robinEggBlue GRM4
10480 -pf00230 -
10485 HMM10114PocR_1_pdu1(pf12833pf10114) PDU1AB
10490 -pf01656pf07685 -
10495 -pf03186 -
10500 -pf02570 -
10505 -pf01888 -
10510 -pf00590 -
10515 -pf05175 -
10520 -pf00590 -
10525 -pf01890pf11761pf11760 -
10530 -pf00590 -
10535 -pf02571 -
10540 -pf06180 -
BMC-P
BMC-H
BMC-T
Aldehyde dehydrogenase
PTAC
Signature enzyme
Alcohol dehydrogenase
Regulator

Salmonella virchow. Bacteria; Proteobacteria; Gammaproteobacteria; Enterobacterales;

E0928__locus_1

PDU1AB

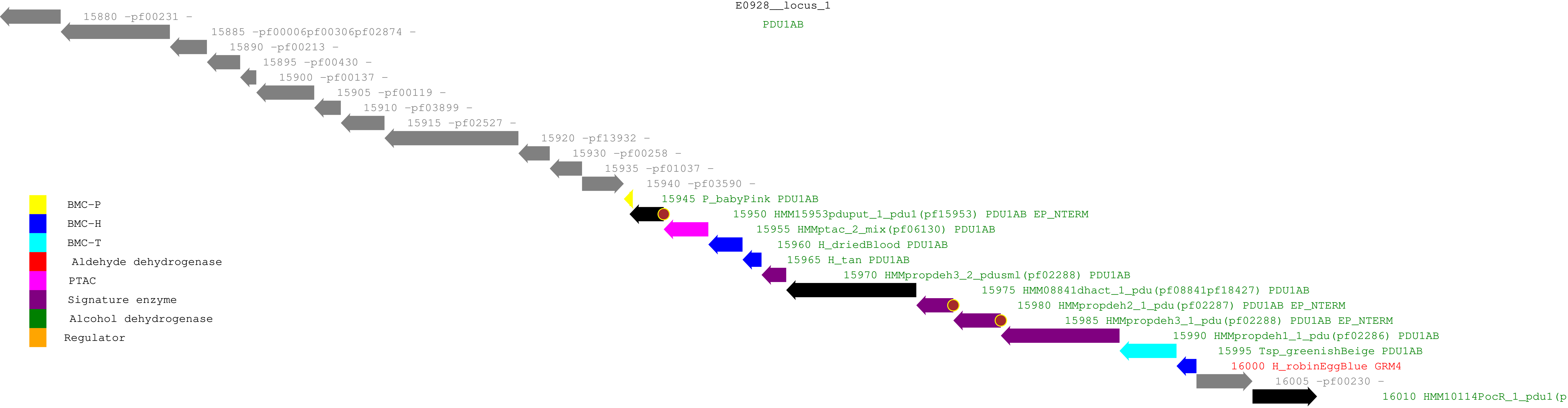

Salmonella enterica subsp. enterica serovar Szentes. Bacteria; Proteobacteria; Gammaproteobacteria; Enterobacterales;

E0931__locus_1

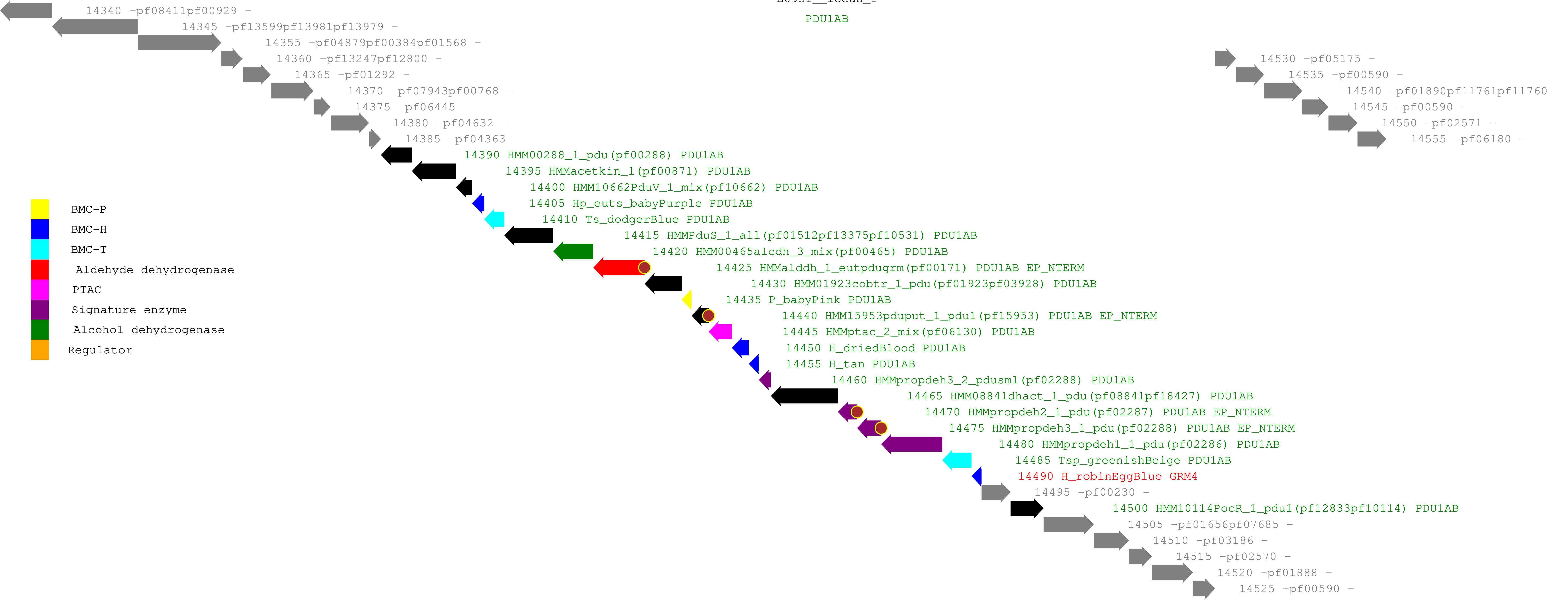

Salmonella enterica subsp. enterica serovar Mapo. Bacteria; Proteobacteria; Gammaproteobacteria; Enterobacterales;

E0940__locus_1

PDU1AB

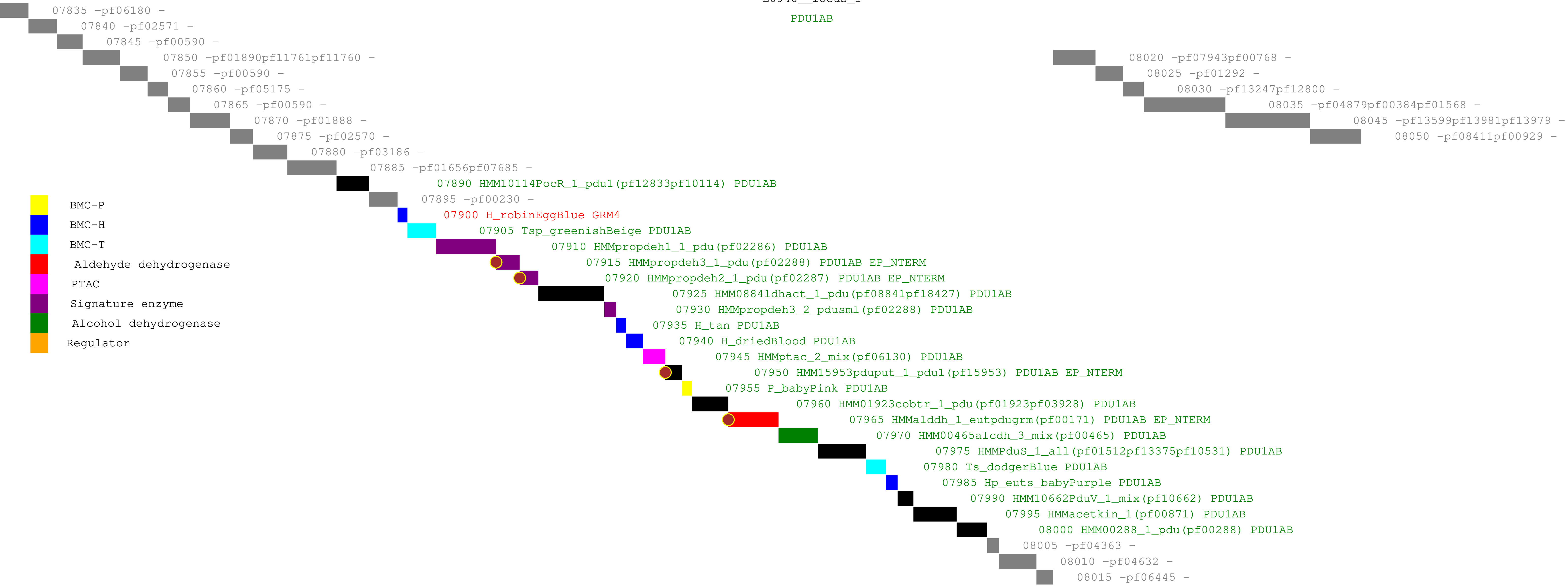

Salmonella enterica subsp. enterica serovar Manhattan. Bacteria; Proteobacteria; Gammaproteobacteria; Enterobacterales;

E0960__locus_1

PDU1AB

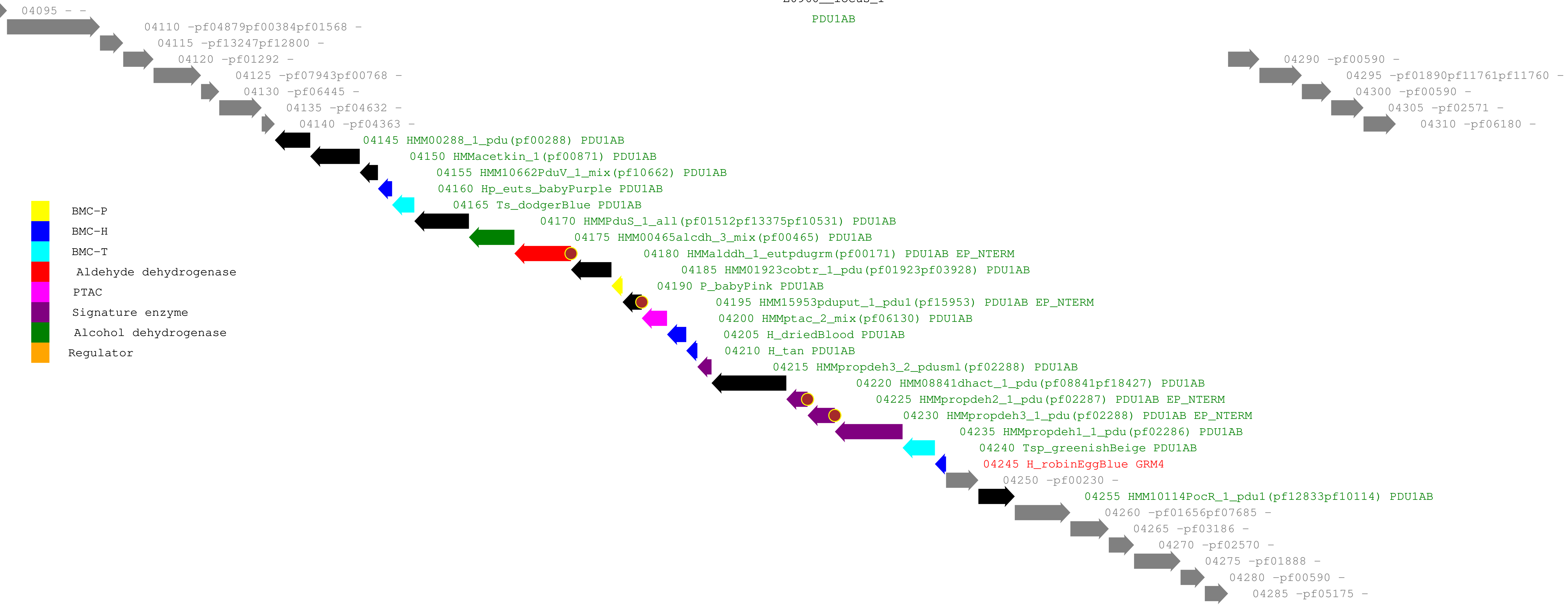

Salmonella enterica subsp. enterica serovar Livingstone. Bacteria; Proteobacteria; Gammaproteobacteria; Enterobacterales;

E0961__locus_1

PDU1AB

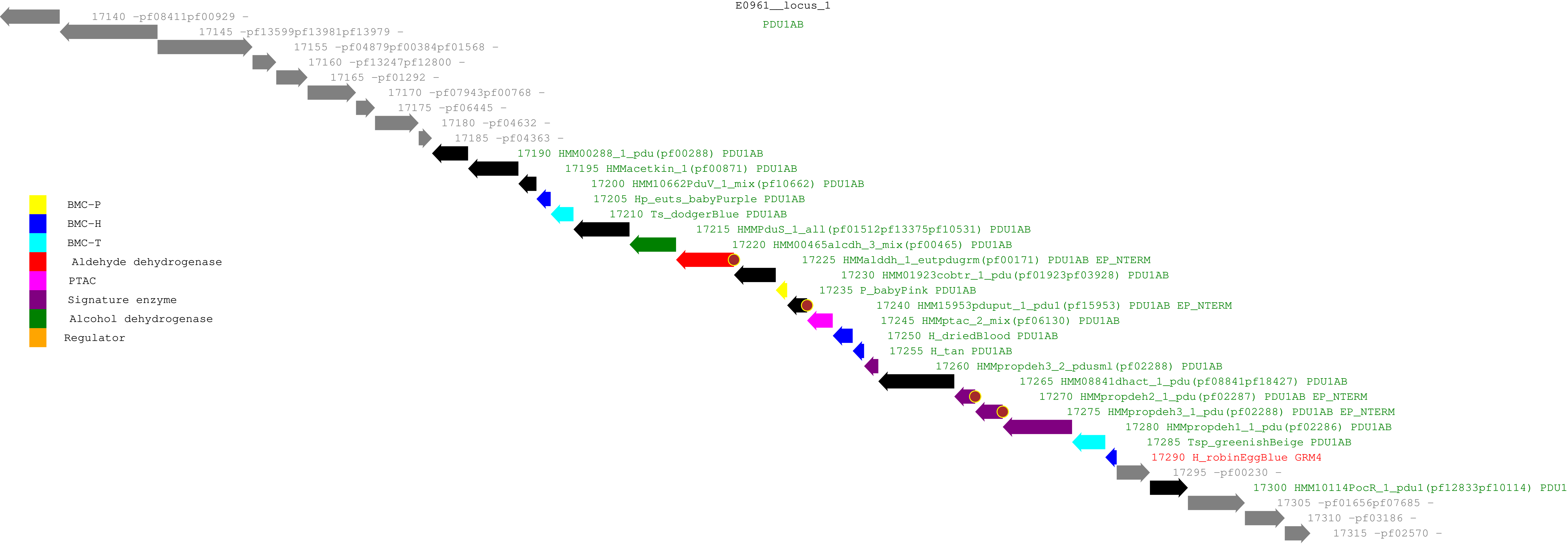

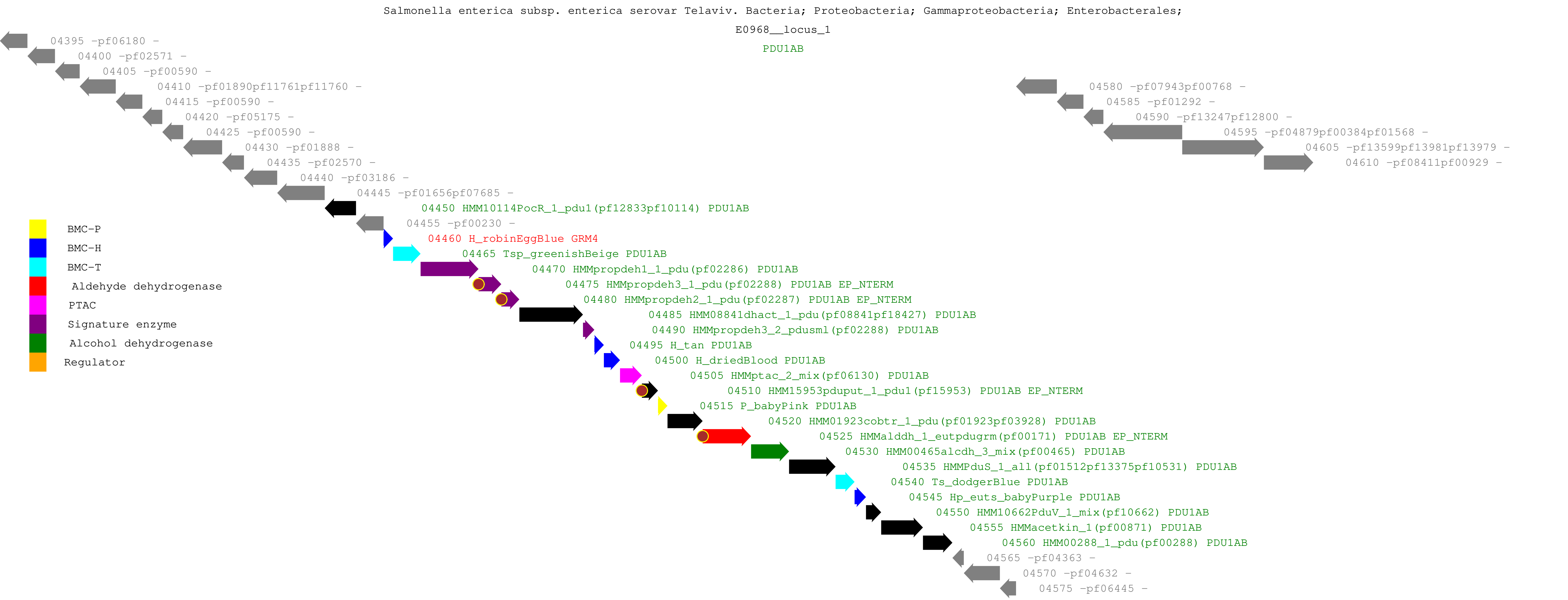
Salmonella enterica subsp. enterica serovar Telaviv. Bacteria; Proteobacteria; Gammaproteobacteria; Enterobacterales;
E0968__locus_1
PDU1AB
04395 -pf06180 -
04400 -pf02571 -
04405 -pf00590 -
04410 -pf01890pf11761pf11760 -
04415 -pf00590 -
04420 -pf05175 -
04425 -pf00590 -
04430 -pf01888 -
04435 -pf02570 -
04440 -pf03186 -
04445 -pf01656pf07685 -
04450 HMM10114PocR_1_pdu1(pf12833pf10114) PDU1AB
04455 -pf00230 -
04460 H_robinEggBlue GRM4
04465 Tsp_greenishBeige PDU1AB
04470 HMMpropdeh1_1_pdu(pf02286) PDU1AB
04475 HMMpropdeh3_1_pdu(pf02288) PDU1AB EP_NTERM
04480 HMMpropdeh2_1_pdu(pf02287) PDU1AB EP_NTERM
04485 HMM08841dhact_1_pdu(pf08841pf18427) PDU1AB
04490 HMMpropdeh3_2_pdusml(pf02288) PDU1AB
04495 H_tan PDU1AB
04500 H_driedBlood PDU1AB
04505 HMMptac_2_mix(pf06130) PDU1AB
04510 HMM15953pduput_1_pdu1(pf15953) PDU1AB EP_NTERM
04515 P_babyPink PDU1AB
04520 HMM01923cobtr_1_pdu(pf01923pf03928) PDU1AB
04525 HMMalddh_1_eutpdugrm(pf00171) PDU1AB EP_NTERM
04530 HMM00465alcdh_3_mix(pf00465) PDU1AB
04535 HMMPduS_1_all(pf01512pf13375pf10531) PDU1AB
04540 Ts_dodgerBlue PDU1AB
04545 Hp_euts_babyPurple PDU1AB
04550 HMM10662PduV_1_mix(pf10662) PDU1AB
04555 HMMacetkin_1(pf00871) PDU1AB
04560 HMM00288_1_pdu(pf00288) PDU1AB
04565 -pf04363 -
04570 -pf04632 -
04575 -pf06445 -
04580 -pf07943pf00768 -
04585 -pf01292 -
04590 -pf13247pf12800 -
04595 -pf04879pf00384pf01568 -
04605 -pf13599pf13981pf13979 -
04610 -pf08411pf00929 -
BMC-P
BMC-H
BMC-T
Aldehyde dehydrogenase
PTAC
Signature enzyme
Alcohol dehydrogenase
Regulator

Salmonella rubislaw. Bacteria; Proteobacteria; Gammaproteobacteria; Enterobacterales;

E0976__locus_1

PDU1AB

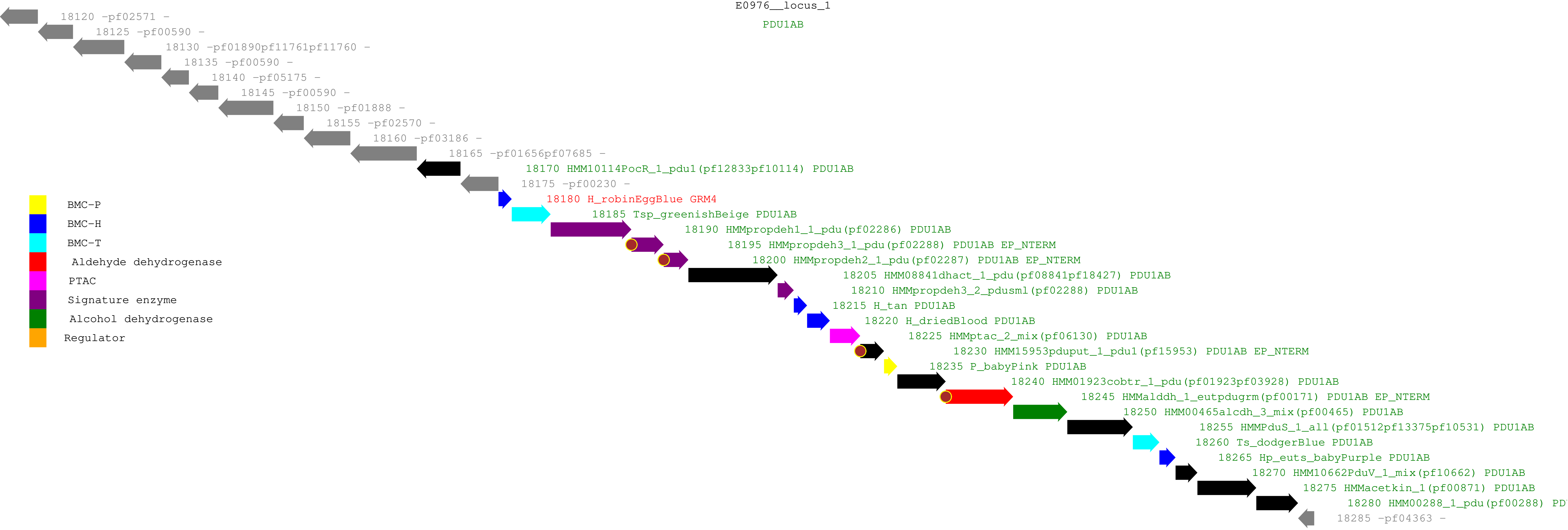

Salmonella enterica subsp. enterica serovar Waycross. Bacteria; Proteobacteria; Gammaproteobacteria; Enterobacterales;
E0980__locus_1
PDU1AB

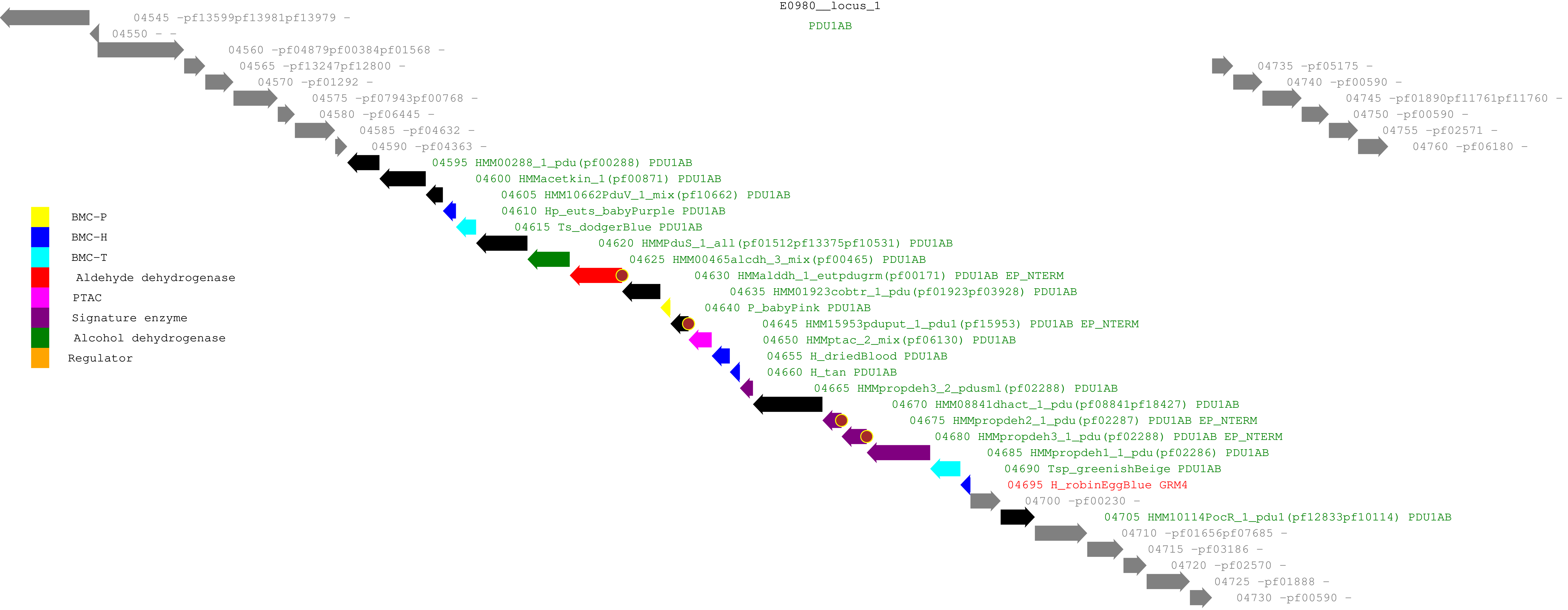

Salmonella enterica subsp. enterica serovar Oslo. Bacteria; Proteobacteria; Gammaproteobacteria; Enterobacterales;

E0985__locus_1

PDU1AB

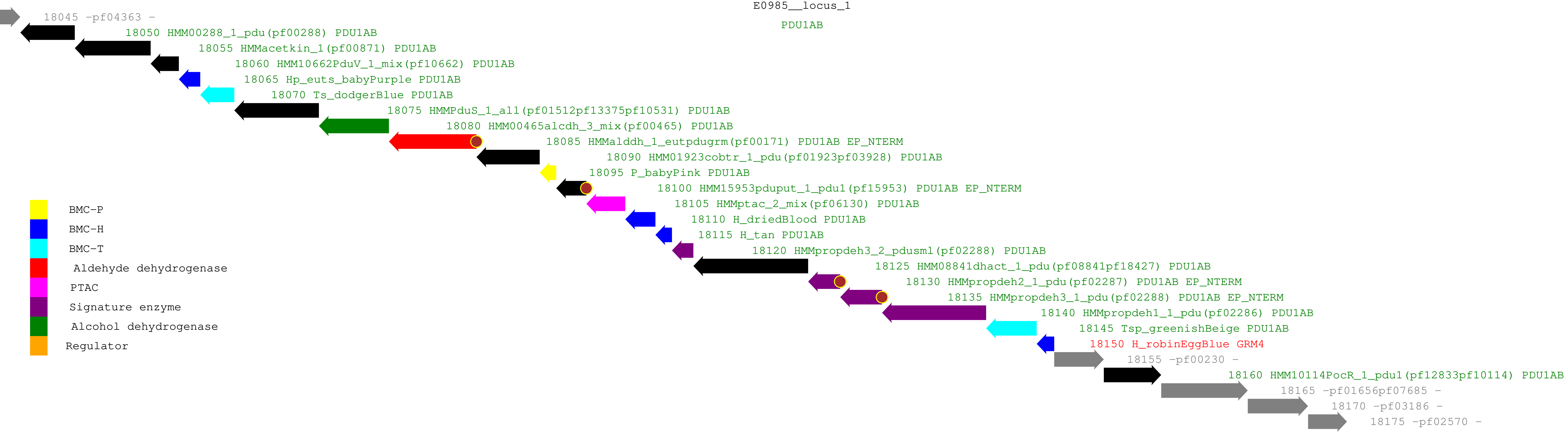

Salmonella enterica subsp. enterica serovar Newington. Bacteria; Proteobacteria; Gammaproteobacteria; Enterobacterales;

E0991__locus_1

PDU1AB

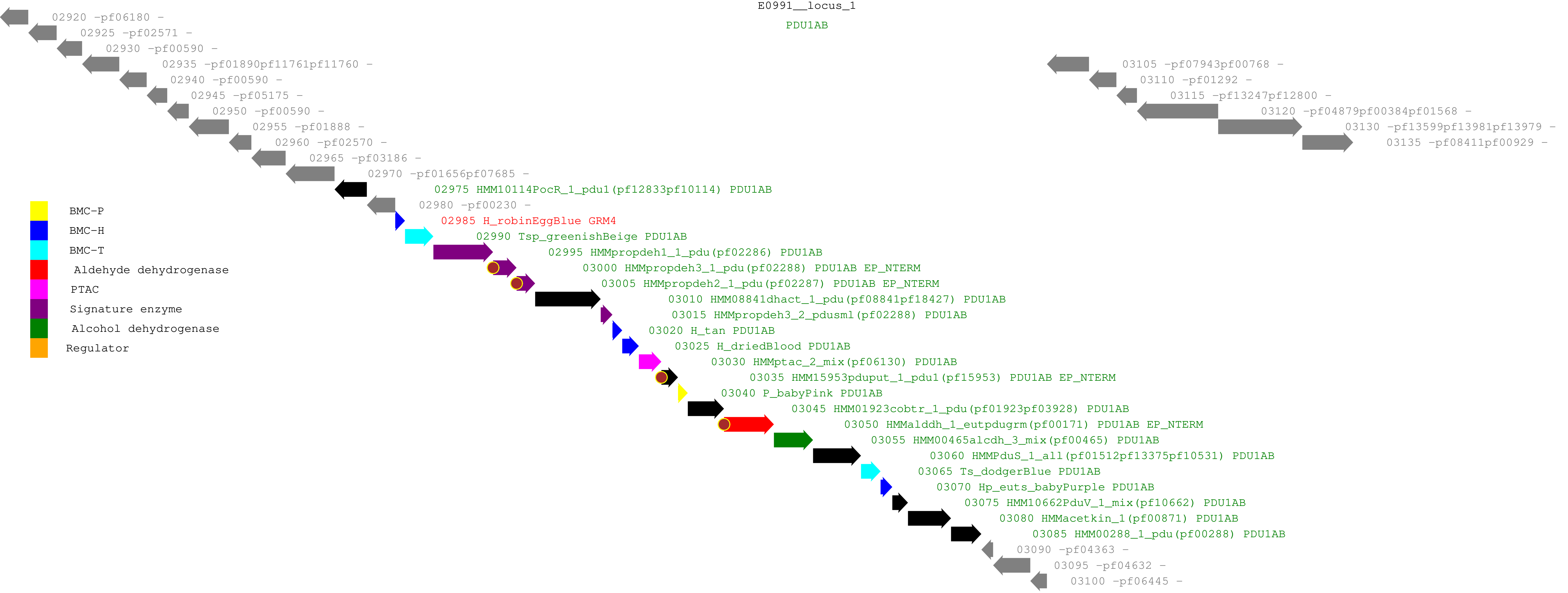

Salmonella enterica subsp. enterica serovar Urbana. Bacteria; Proteobacteria; Gammaproteobacteria; Enterobacterales;

E0999__locus_2

PDU1AB

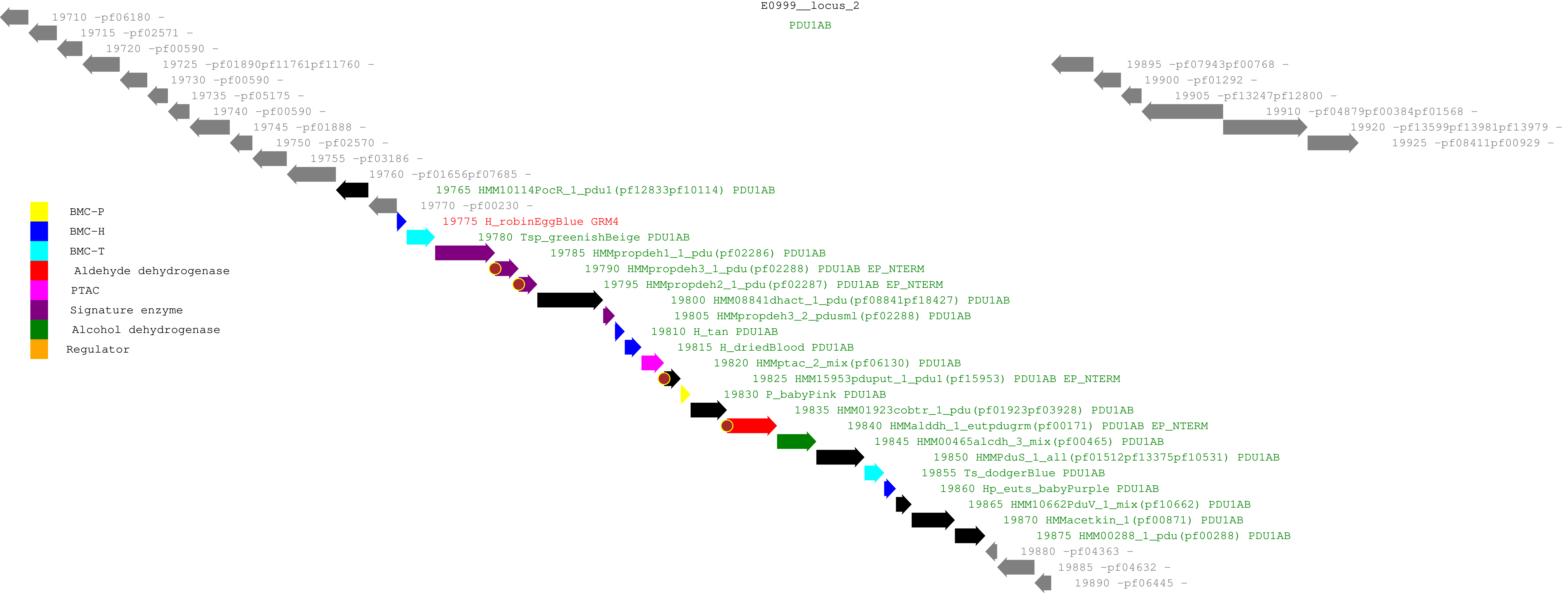

Salmonella enterica subsp. enterica serovar Kisangani. Bacteria; Proteobacteria; Gammaproteobacteria; Enterobacterales;

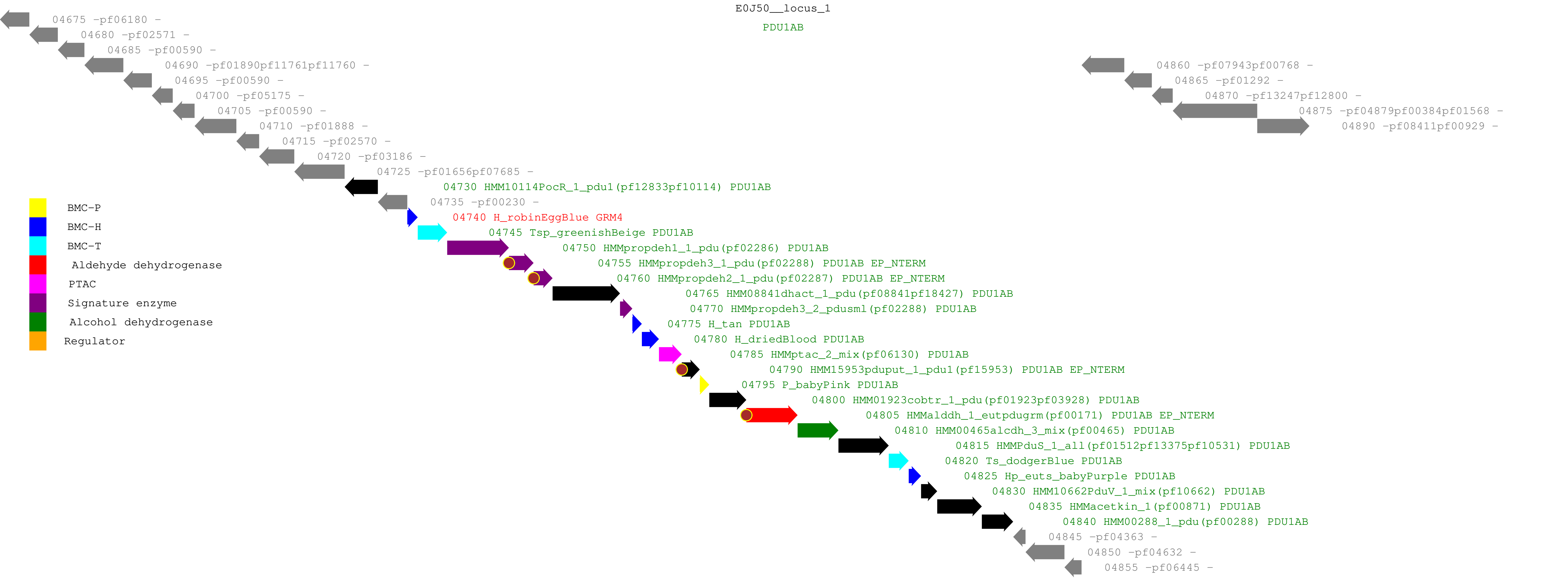

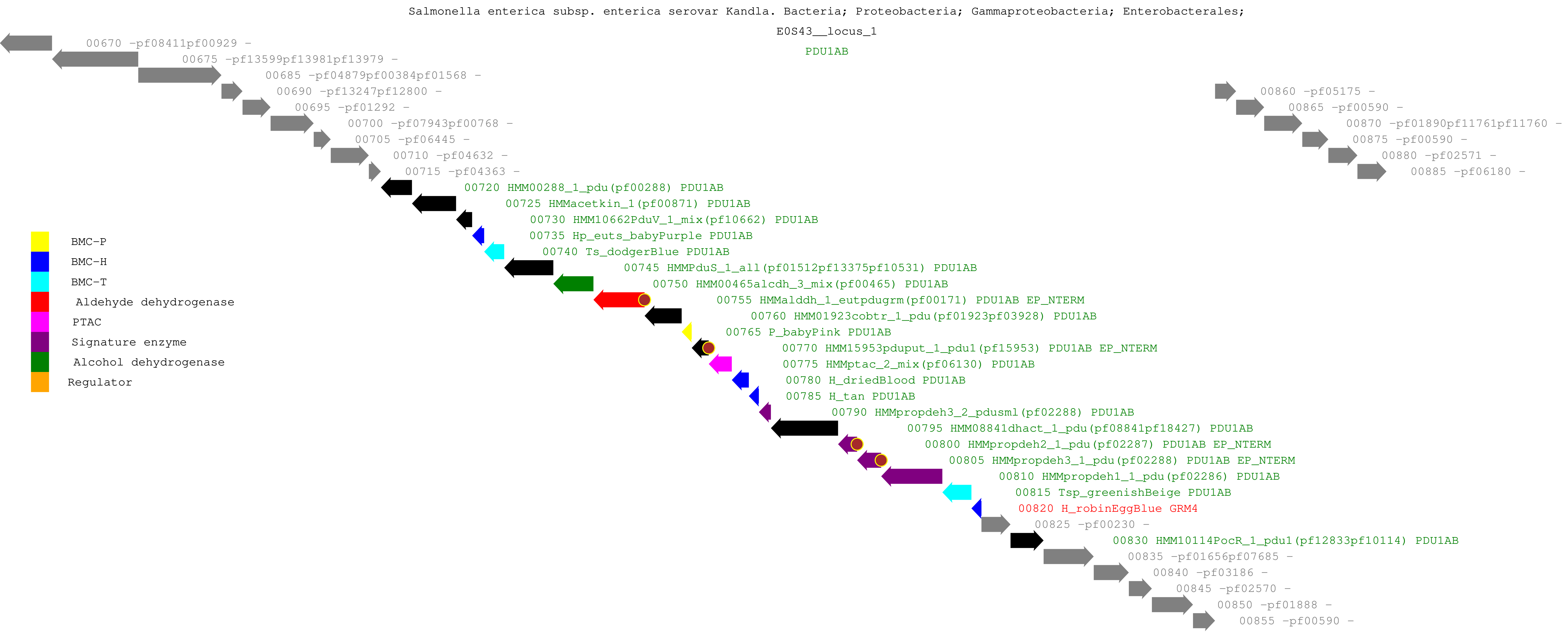

Salmonella enterica subsp. enterica serovar Kandla. Bacteria; Proteobacteria; Gammaproteobacteria; Enterobacterales;
E0S43__locus_1
PDU1AB
00670 -pf08411pf00929 -
00675 -pf13599pf13981pf13979 -
00685 -pf04879pf00384pf01568 -
00690 -pf13247pf12800 -
00695 -pf01292 -
00700 -pf07943pf00768 -
00705 -pf06445 -
00710 -pf04632 -
00715 -pf04363 -
00720 HMM00288_1_pdu(pf00288) PDU1AB
00725 HMMacetkin_1(pf00871) PDU1AB
00730 HMM10662PduV_1_mix(pf10662) PDU1AB
00735 Hp_euts_babyPurple PDU1AB
00740 Ts_dodgerBlue PDU1AB
00745 HMMPduS_1_all(pf01512pf13375pf10531) PDU1AB
00750 HMM00465alcdh_3_mix(pf00465) PDU1AB
00755 HMMalddh_1_eutpdugrm(pf00171) PDU1AB EP_NTERM
00760 HMM01923cobtr_1_pdu(pf01923pf03928) PDU1AB
00765 P_babyPink PDU1AB
00770 HMM15953pduput_1_pdu1(pf15953) PDU1AB EP_NTERM
00775 HMMptac_2_mix(pf06130) PDU1AB
00780 H_driedBlood PDU1AB
00785 H_tan PDU1AB
00790 HMMpropdeh3_2_pdusml(pf02288) PDU1AB
00795 HMM08841dhact_1_pdu(pf08841pf18427) PDU1AB
00800 HMMpropdeh2_1_pdu(pf02287) PDU1AB EP_NTERM
00805 HMMpropdeh3_1_pdu(pf02288) PDU1AB EP_NTERM
00810 HMMpropdeh1_1_pdu(pf02286) PDU1AB
00815 Tsp_greenishBeige PDU1AB
00820 H_robinEggBlue GRM4
00825 -pf00230 -
00830 HMM10114PocR_1_pdu1(pf12833pf10114) PDU1AB
00835 -pf01656pf07685 -
00840 -pf03186 -
00845 -pf02570 -
00850 -pf01888 -
00855 -pf00590 -
00860 -pf05175 -
00865 -pf00590 -
00870 -pf01890pf11761pf11760 -
00875 -pf00590 -
00880 -pf02571 -
00885 -pf06180 -
BMC-P
BMC-H
BMC-T
Aldehyde dehydrogenase
PTAC
Signature enzyme
Alcohol dehydrogenase
Regulator

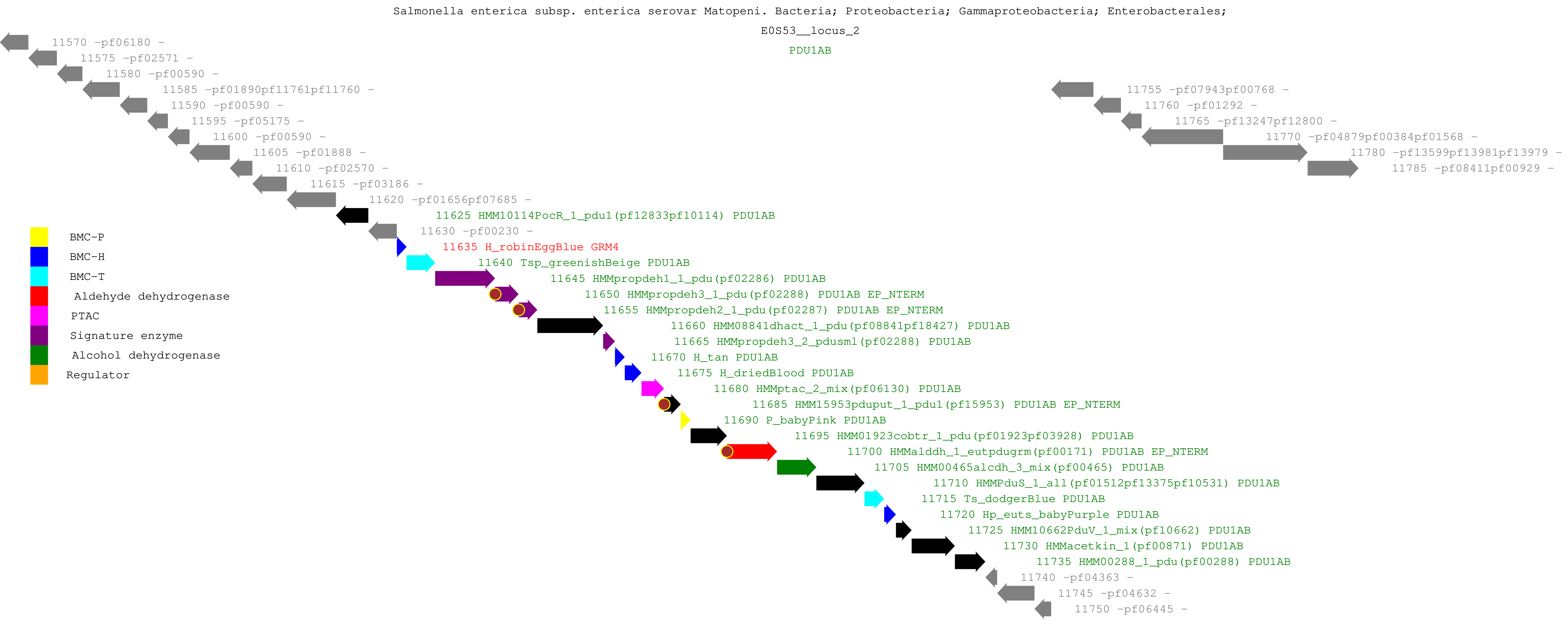
Salmonella enterica subsp. enterica serovar Matopeni. Bacteria; Proteobacteria; Gammaproteobacteria; Enterobacterales;
E0S53__locus_2
PDU1AB
11570 -pf06180 -
11575 -pf02571 -
11580 -pf00590 -
11585 -pf01890pf11761pf11760 -
11590 -pf00590 -
11595 -pf05175 -
11600 -pf00590 -
11605 -pf01888 -
11610 -pf02570 -
11615 -pf03186 -
11620 -pf01656pf07685 -
11625 HMM10114PocR_1_pdu1(pf12833pf10114) PDU1AB
11630 -pf00230 -
11635 H_robinEggBlue GRM4
11640 Tsp_greenishBeige PDU1AB
11645 HMMpropdeh1_1_pdu(pf02286) PDU1AB
11650 HMMpropdeh3_1_pdu(pf02288) PDU1AB EP_NTERM
11655 HMMpropdeh2_1_pdu(pf02287) PDU1AB EP_NTERM
11660 HMM08841dhact_1_pdu(pf08841pf18427) PDU1AB
11665 HMMpropdeh3_2_pdusml(pf02288) PDU1AB
11670 H_tan PDU1AB
11675 H_driedBlood PDU1AB
11680 HMMptac_2_mix(pf06130) PDU1AB
11685 HMM15953pduput_1_pdu1(pf15953) PDU1AB EP_NTERM
11690 P_babyPink PDU1AB
11695 HMM01923cobtr_1_pdu(pf01923pf03928) PDU1AB
11700 HMMaldddh_1_eutpdugrm(pf00171) PDU1AB EP_NTERM
11705 HMM00465alcdh_3_mix(pf00465) PDU1AB
11710 HMMPduS_1_all(pf01512pf13375pf10531) PDU1AB
11715 Ts_dodgerBlue PDU1AB
11720 Hp_euts_babyPurple PDU1AB
11725 HMM10662PduV_1_mix(pf10662) PDU1AB
11730 HMMacetkin_1(pf00871) PDU1AB
11735 HMM00288_1_pdu(pf00288) PDU1AB
11740 -pf04363 -
11745 -pf04632 -
11750 -pf06445 -
11755 -pf07943pf00768 -
11760 -pf01292 -
11765 -pf13247pf12800 -
11770 -pf04879pf00384pf01568 -
11780 -pf13599pf13981pf13979 -
11785 -pf08411pf00929 -
BMC-P
BMC-H
BMC-T
Aldehyde dehydrogenase
PTAC
Signature enzyme
Alcohol dehydrogenase
Regulator

Salmonella enterica subsp. enterica serovar Butantan. Bacteria; Proteobacteria; Gammaproteobacteria; Enterobacterales;

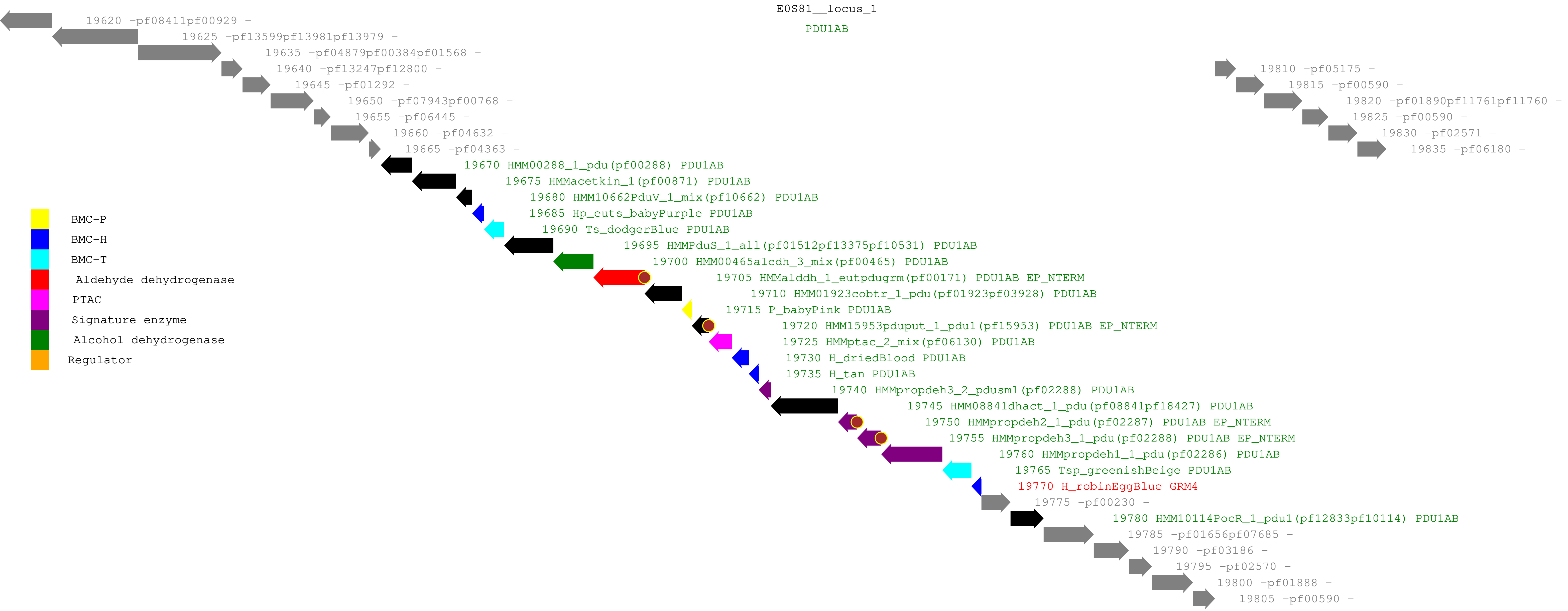

Salmonella enterica subsp. enterica serovar Manhattan. Bacteria; Proteobacteria; Gammaproteobacteria; Enterobacterales;

E0T26__locus_1

PDU1AB

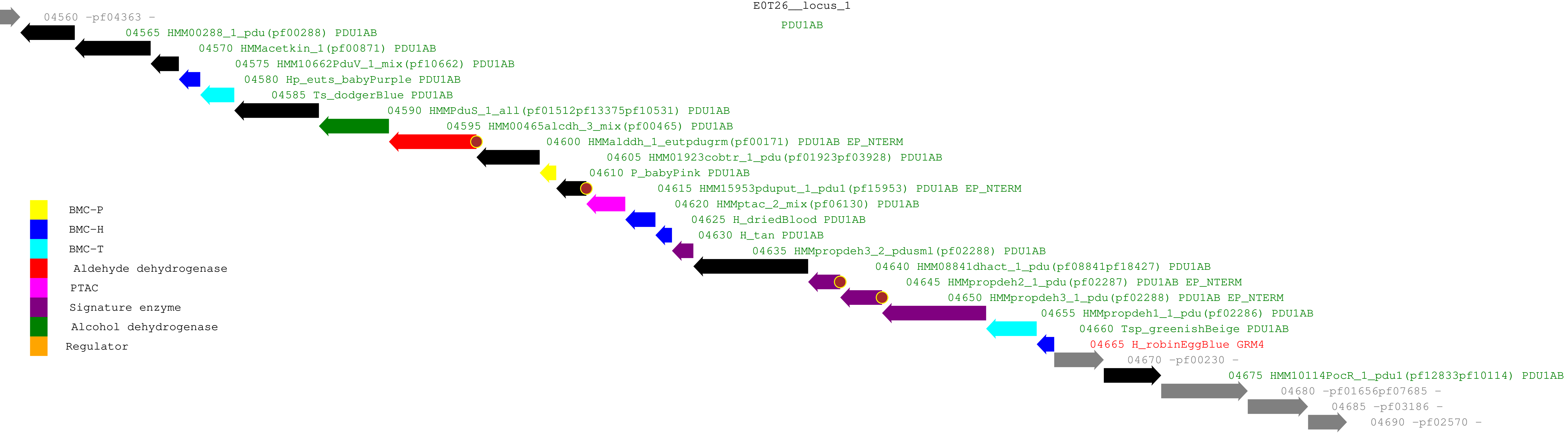

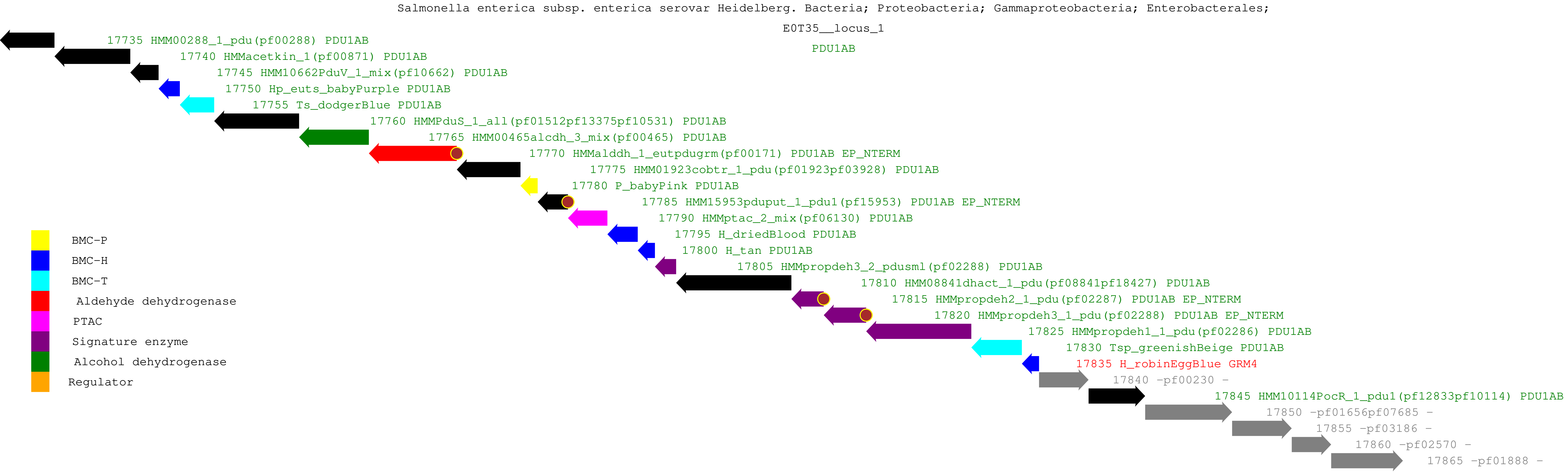
Salmonella enterica subsp. enterica serovar Heidelberg. Bacteria; Proteobacteria; Gammaproteobacteria; Enterobacterales;
E0T35__locus_1
PDU1AB
17735 HMM00288_1_pdu(pf00288) PDU1AB
17740 HMMacetkin_1(pf00871) PDU1AB
17745 HMM10662PduV_1_mix(pf10662) PDU1AB
17750 Hp_euts_babyPurple PDU1AB
17755 Ts_dodgerBlue PDU1AB
17760 HMMPduS_1_all(pf01512pf13375pf10531) PDU1AB
17765 HMM00465alcdh_3_mix(pf00465) PDU1AB
17770 HMMaldh_1_eutpdugrm(pf00171) PDU1AB EP_NTERM
17775 HMM01923cobtr_1_pdu(pf01923pf03928) PDU1AB
17780 P_babyPink PDU1AB
17785 HMM15953pduput_1_pdu1(pf15953) PDU1AB EP_NTERM
17790 HMMptac_2_mix(pf06130) PDU1AB
17795 H_driedBlood PDU1AB
17800 H_tan PDU1AB
17805 HMMpropdeh3_2_pdusml(pf02288) PDU1AB
17810 HMM08841dhact_1_pdu(pf08841pf18427) PDU1AB
17815 HMMpropdeh2_1_pdu(pf02287) PDU1AB EP_NTERM
17820 HMMpropdeh3_1_pdu(pf02288) PDU1AB EP_NTERM
17825 HMMpropdeh1_1_pdu(pf02286) PDU1AB
17830 Tsp_greenishBeige PDU1AB
17835 H_robinEggBlue GRM4
17840 -pf00230 -
17845 HMM10114PocR_1_pdu1(pf12833pf10114) PDU1AB
17850 -pf01656pf07685 -
17855 -pf03186 -
17860 -pf02570 -
17865 -pf01888 -
BMC-P
BMC-H
BMC-T
Aldehyde dehydrogenase
PTAC
Signature enzyme
Alcohol dehydrogenase
Regulator

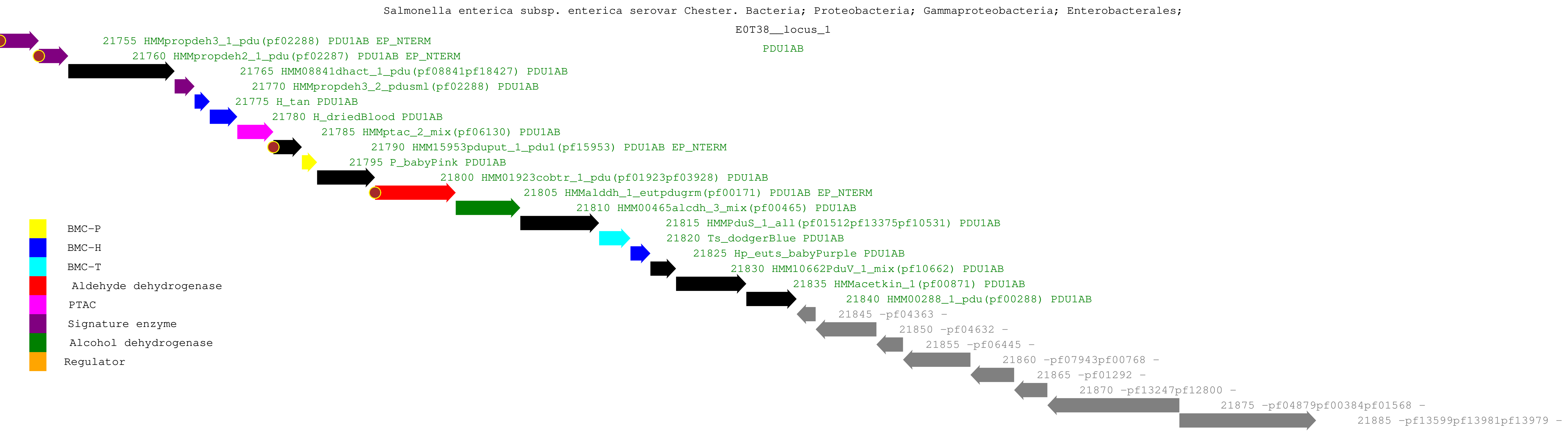

Salmonella enterica subsp. enterica serovar Chester. Bacteria; Proteobacteria; Gammaproteobacteria; Enterobacterales;
E0T38__locus_1
PDU1AB
21755 HMMpropdeh3_1_pdu(pf02288) PDU1AB EP_NTERM
21760 HMMpropdeh2_1_pdu(pf02287) PDU1AB EP_NTERM
21765 HMM08841dhact_1_pdu(pf08841pf18427) PDU1AB
21770 HMMpropdeh3_2_pdusml(pf02288) PDU1AB
21775 H_tan PDU1AB
21780 H_driedBlood PDU1AB
21785 HMMptac_2_mix(pf06130) PDU1AB
21790 HMM15953pduput_1_pdu1(pf15953) PDU1AB EP_NTERM
21795 P_babyPink PDU1AB
21800 HMM01923cobtr_1_pdu(pf01923pf03928) PDU1AB
21805 HMMalddh_1_eutpdugrm(pf00171) PDU1AB EP_NTERM
21810 HMM00465alcdh_3_mix(pf00465) PDU1AB
21815 HMMPduS_1_all(pf01512pf13375pf10531) PDU1AB
21820 Ts_dodgerBlue PDU1AB
21825 Hp_euts_babyPurple PDU1AB
21830 HMM10662PduV_1_mix(pf10662) PDU1AB
21835 HMMacetkin_1(pf00871) PDU1AB
21840 HMM00288_1_pdu(pf00288) PDU1AB
21845 -pf04363 -
21850 -pf04632 -
21855 -pf06445 -
21860 -pf07943pf00768 -
21865 -pf01292 -
21870 -pf13247pf12800 -
21875 -pf04879pf00384pf01568 -
21885 -pf13599pf13981pf13979 -
BMC-P
BMC-H
BMC-T
Aldehyde dehydrogenase
PTAC
Signature enzyme
Alcohol dehydrogenase
Regulator

Salmonella enterica subsp. enterica serovar Galiema. Bacteria; Proteobacteria; Gammaproteobacteria; Enterobacterales;

E0U28__locus_1

PDU1AB

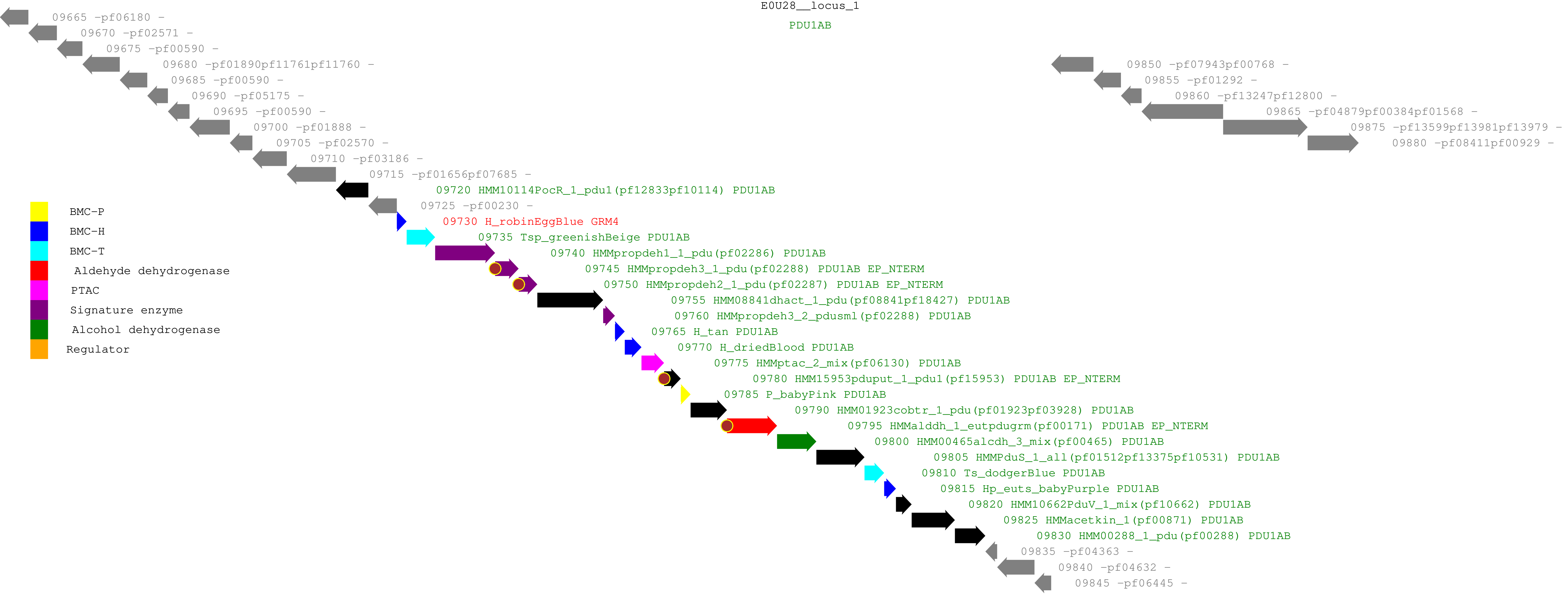

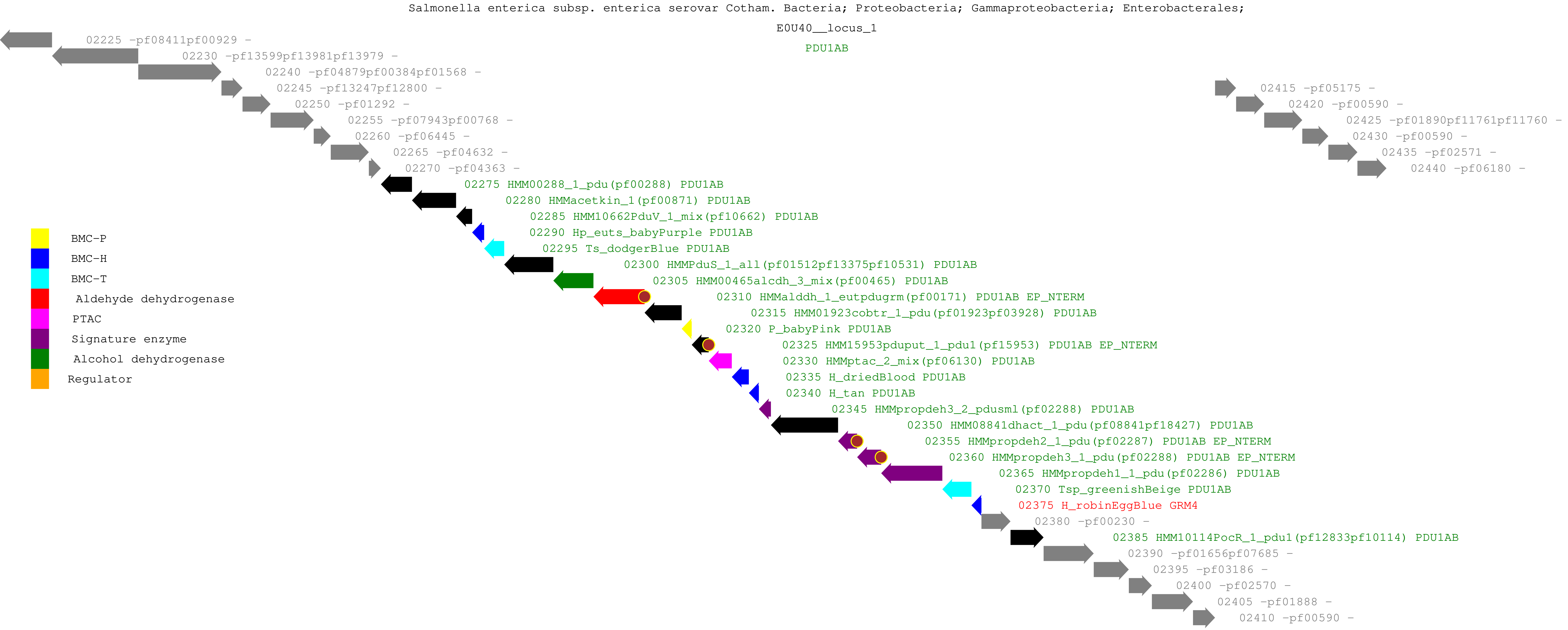
Salmonella enterica subsp. enterica serovar Cotham. Bacteria; Proteobacteria; Gammaproteobacteria; Enterobacterales;
E0U40__locus_1
PDU1AB
02225 -pf08411pf00929 -
02230 -pf13599pf13981pf13979 -
02240 -pf04879pf00384pf01568 -
02245 -pf13247pf12800 -
02250 -pf01292 -
02255 -pf07943pf00768 -
02260 -pf06445 -
02265 -pf04632 -
02270 -pf04363 -
02275 HMM00288_1_pdu(pf00288) PDU1AB
02280 HMMacetkin_1(pf00871) PDU1AB
02285 HMM10662PduV_1_mix(pf10662) PDU1AB
02290 Hp_euts_babyPurple PDU1AB
02295 Ts_dodgerBlue PDU1AB
02300 HMMPduS_1_all(pf01512pf13375pf10531) PDU1AB
02305 HMM00465alcdh_3_mix(pf00465) PDU1AB
02310 HMMalddh_1_eutpdugrm(pf00171) PDU1AB EP_NTERM
02315 HMM01923cobtr_1_pdu(pf01923pf03928) PDU1AB
02320 P_babyPink PDU1AB
02325 HMM15953pduput_1_pdu1(pf15953) PDU1AB EP_NTERM
02330 HMMptac_2_mix(pf06130) PDU1AB
02335 H_driedBlood PDU1AB
02340 H_tan PDU1AB
02345 HMMpropdeh3_2_pdusml(pf02288) PDU1AB
02350 HMM08841dhact_1_pdu(pf08841pf18427) PDU1AB
02355 HMMpropdeh2_1_pdu(pf02287) PDU1AB EP_NTERM
02360 HMMpropdeh3_1_pdu(pf02288) PDU1AB EP_NTERM
02365 HMMpropdeh1_1_pdu(pf02286) PDU1AB
02370 Tsp_greenishBeige PDU1AB
02375 H_robinEggBlue GRM4
02380 -pf00230 -
02385 HMM10114PocR_1_pdu1(pf12833pf10114) PDU1AB
02390 -pf01656pf07685 -
02395 -pf03186 -
02400 -pf02570 -
02405 -pf01888 -
02410 -pf00590 -
02415 -pf05175 -
02420 -pf00590 -
02425 -pf01890pf11761pf11760 -
02430 -pf00590 -
02435 -pf02571 -
02440 -pf06180 -
BMC-P
BMC-H
BMC-T
Aldehyde dehydrogenase
PTAC
Signature enzyme
Alcohol dehydrogenase
Regulator

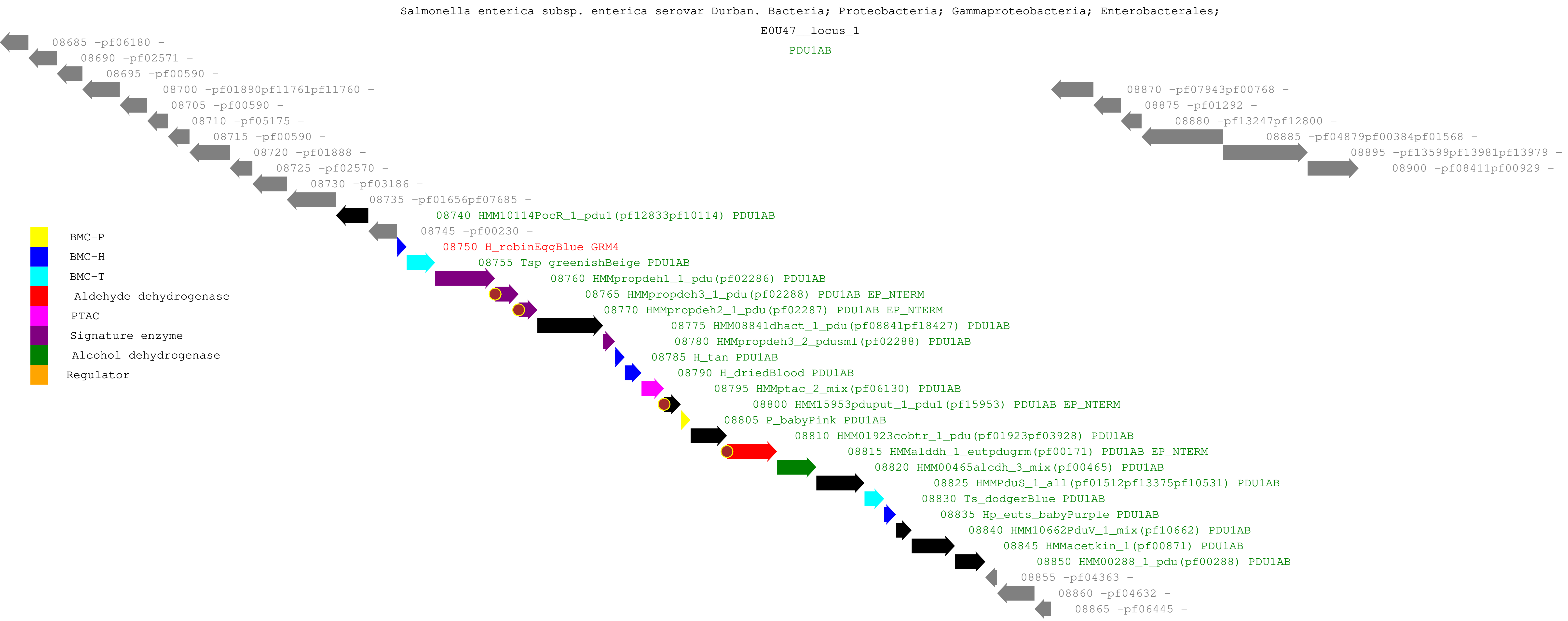
Salmonella enterica subsp. enterica serovar Durban. Bacteria; Proteobacteria; Gammaproteobacteria; Enterobacterales;
E0U47__locus_1
PDU1AB
08685 -pf06180 -
08690 -pf02571 -
08695 -pf00590 -
08700 -pf01890pf11761pf11760 -
08705 -pf00590 -
08710 -pf05175 -
08715 -pf00590 -
08720 -pf01888 -
08725 -pf02570 -
08730 -pf03186 -
08735 -pf01656pf07685 -
08740 HMM10114PocR_1_pdu1(pf12833pf10114) PDU1AB
08745 -pf00230 -
08750 H_robinEggBlue GRM4
08755 Tsp_greenishBeige PDU1AB
08760 HMMpropdeh1_1_pdu(pf02286) PDU1AB
08765 HMMpropdeh3_1_pdu(pf02288) PDU1AB EP_NTERM
08770 HMMpropdeh2_1_pdu(pf02287) PDU1AB EP_NTERM
08775 HMM08841dhact_1_pdu(pf08841pf18427) PDU1AB
08780 HMMpropdeh3_2_pdusml(pf02288) PDU1AB
08785 H_tan PDU1AB
08790 H_driedBlood PDU1AB
08795 HMMptac_2_mix(pf06130) PDU1AB
08800 HMM15953pduput_1_pdu1(pf15953) PDU1AB EP_NTERM
08805 P_babyPink PDU1AB
08810 HMM01923cobtr_1_pdu(pf01923pf03928) PDU1AB
08815 HMMalddh_1_eutpdugrm(pf00171) PDU1AB EP_NTERM
08820 HMM00465alcdh_3_mix(pf00465) PDU1AB
08825 HMMPduS_1_all(pf01512pf13375pf10531) PDU1AB
08830 Ts_dodgerBlue PDU1AB
08835 Hp_euts_babyPurple PDU1AB
08840 HMM10662PduV_1_mix(pf10662) PDU1AB
08845 HMMacetkin_1(pf00871) PDU1AB
08850 HMM00288_1_pdu(pf00288) PDU1AB
08855 -pf04363 -
08860 -pf04632 -
08865 -pf06445 -
08870 -pf07943pf00768 -
08875 -pf01292 -
08880 -pf13247pf12800 -
08885 -pf04879pf00384pf01568 -
08895 -pf13599pf13981pf13979 -
08900 -pf08411pf00929 -
BMC-P
BMC-H
BMC-T
Aldehyde dehydrogenase
PTAC
Signature enzyme
Alcohol dehydrogenase
Regulator

Salmonella enterica subsp. enterica serovar Haga. Bacteria; Proteobacteria; Gammaproteobacteria; Enterobacterales;

E0U59__locus_1

PDU1AB

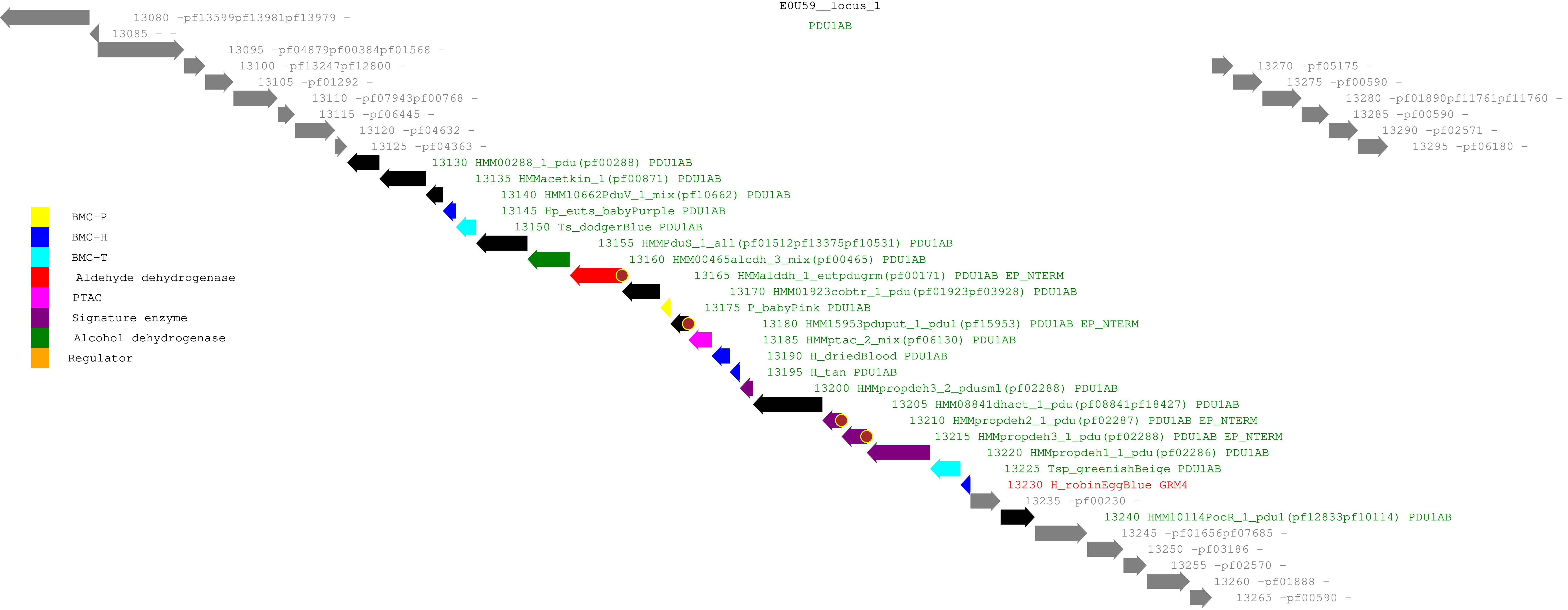

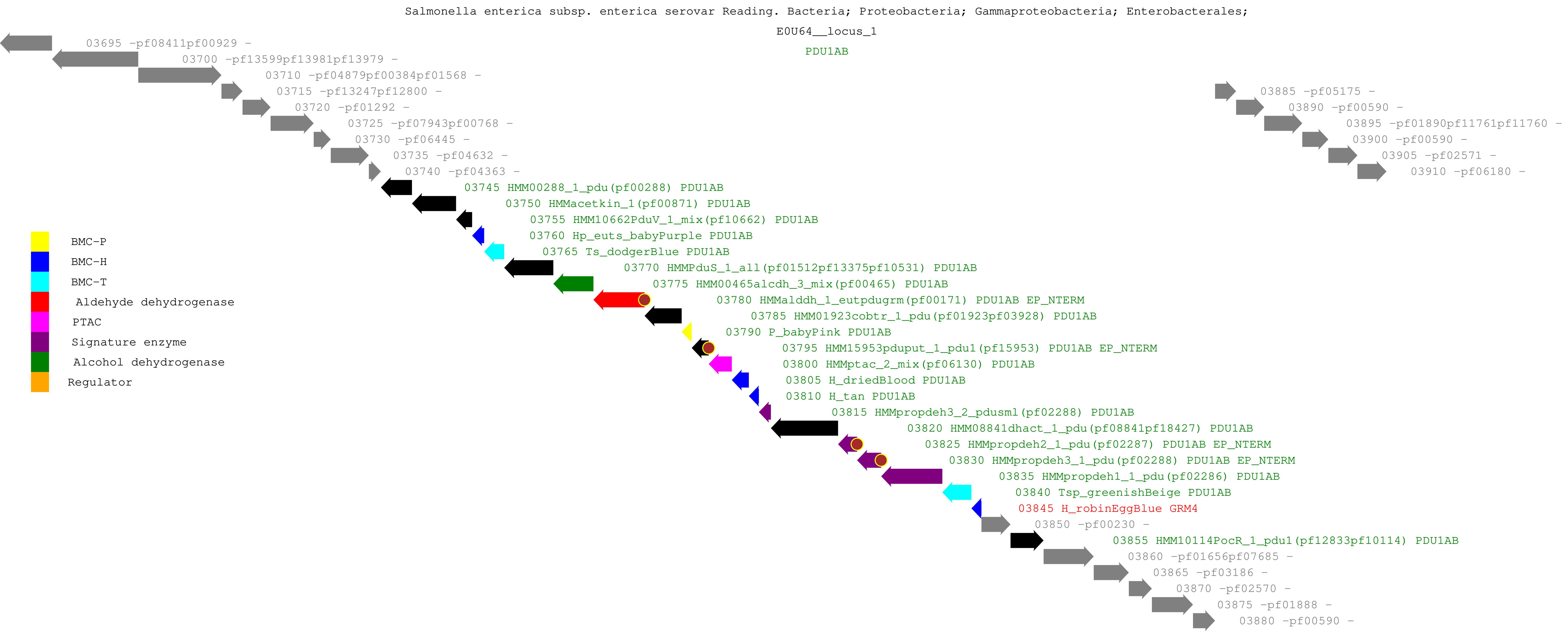
Salmonella enterica subsp. enterica serovar Reading. Bacteria; Proteobacteria; Gammaproteobacteria; Enterobacterales;
E0U64__locus_1
PDU1AB
03695 -pf08411pf00929 -
03700 -pf13599pf13981pf13979 -
03710 -pf04879pf00384pf01568 -
03715 -pf13247pf12800 -
03720 -pf01292 -
03725 -pf07943pf00768 -
03730 -pf06445 -
03735 -pf04632 -
03740 -pf04363 -
03745 HMM00288_1_pdu(pf00288) PDU1AB
03750 HMMacetkin_1(pf00871) PDU1AB
03755 HMM10662PduV_1_mix(pf10662) PDU1AB
03760 Hp_euts_babyPurple PDU1AB
03765 Ts_dodgerBlue PDU1AB
03770 HMMPduS_1_all(pf01512pf13375pf10531) PDU1AB
03775 HMM00465alcdh_3_mix(pf00465) PDU1AB
03780 HMMalddh_1_eutpdugrm(pf00171) PDU1AB EP_NTERM
03785 HMM01923cobtr_1_pdu(pf01923pf03928) PDU1AB
03790 P_babyPink PDU1AB
03795 HMM15953pduput_1_pdu1(pf15953) PDU1AB EP_NTERM
03800 HMMptac_2_mix(pf06130) PDU1AB
03805 H_driedBlood PDU1AB
03810 H_tan PDU1AB
03815 HMMpropdeh3_2_pdusml(pf02288) PDU1AB
03820 HMM08841dhact_1_pdu(pf08841pf18427) PDU1AB
03825 HMMpropdeh2_1_pdu(pf02287) PDU1AB EP_NTERM
03830 HMMpropdeh3_1_pdu(pf02288) PDU1AB EP_NTERM
03835 HMMpropdeh1_1_pdu(pf02286) PDU1AB
03840 Tsp_greenishBeige PDU1AB
03845 H_robinEggBlue GRM4
03850 -pf00230 -
03855 HMM10114PocR_1_pdu1(pf12833pf10114) PDU1AB
03860 -pf01656pf07685 -
03865 -pf03186 -
03870 -pf02570 -
03875 -pf01888 -
03880 -pf00590 -
03885 -pf05175 -
03890 -pf00590 -
03895 -pf01890pf11761pf11760 -
03900 -pf00590 -
03905 -pf02571 -
03910 -pf06180 -
BMC-P
BMC-H
BMC-T
Aldehyde dehydrogenase
PTAC
Signature enzyme
Alcohol dehydrogenase
Regulator

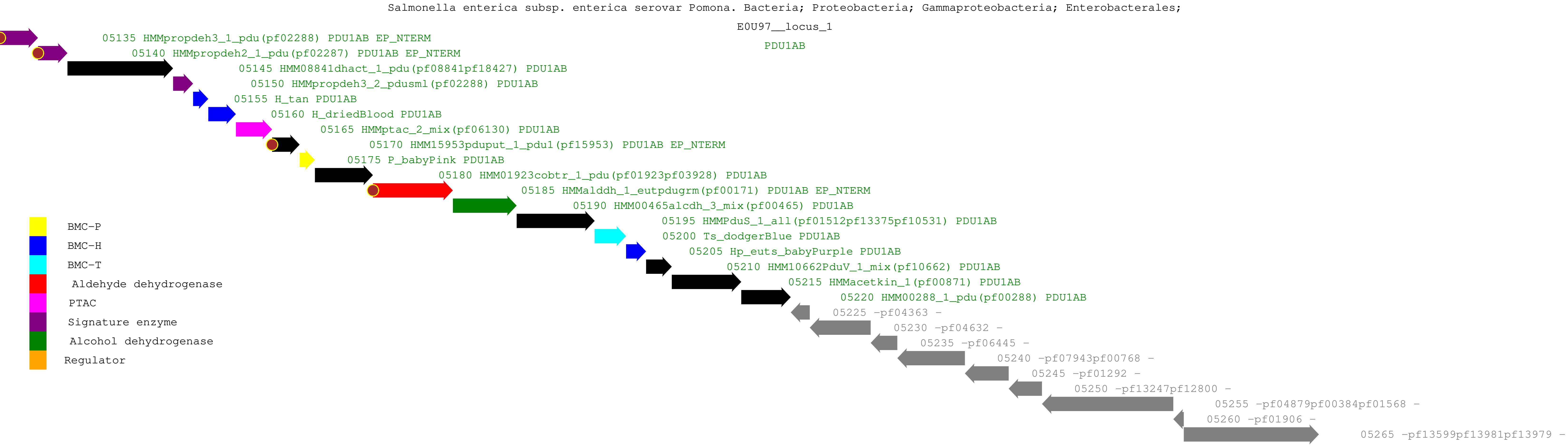

Salmonella enterica subsp. enterica serovar Pomona. Bacteria; Proteobacteria; Gammaproteobacteria; Enterobacterales;
E0U97__locus_1
PDU1AB
05135 HMMpropdeh3_1_pdu(pf02288) PDU1AB EP_NTERM
05140 HMMpropdeh2_1_pdu(pf02287) PDU1AB EP_NTERM
05145 HMM08841dhact_1_pdu(pf08841pf18427) PDU1AB
05150 HMMpropdeh3_2_pdusml(pf02288) PDU1AB
05155 H_tan PDU1AB
05160 H_driedBlood PDU1AB
05165 HMMptac_2_mix(pf06130) PDU1AB
05170 HMM15953pduput_1_pdu1(pf15953) PDU1AB EP_NTERM
05175 P_babyPink PDU1AB
05180 HMM01923cobtr_1_pdu(pf01923pf03928) PDU1AB
05185 HMMaldh_1_eutpdugrm(pf00171) PDU1AB EP_NTERM
05190 HMM00465alcdh_3_mix(pf00465) PDU1AB
05195 HMMPduS_1_all(pf01512pf13375pf10531) PDU1AB
05200 Ts_dodgerBlue PDU1AB
05205 Hp_euts_babyPurple PDU1AB
05210 HMM10662PduV_1_mix(pf10662) PDU1AB
05215 HMMacetkin_1(pf00871) PDU1AB
05220 HMM00288_1_pdu(pf00288) PDU1AB
05225 -pf04363 -
05230 -pf04632 -
05235 -pf06445 -
05240 -pf07943pf00768 -
05245 -pf01292 -
05250 -pf13247pf12800 -
05255 -pf04879pf00384pf01568 -
05260 -pf01906 -
05265 -pf13599pf13981pf13979 -
BMC-P
BMC-H
BMC-T
Aldehyde dehydrogenase
PTAC
Signature enzyme
Alcohol dehydrogenase
Regulator

Salmonella enterica subsp. enterica serovar Hvittingfoss. Bacteria; Proteobacteria; Gammaproteobacteria; Enterobacterales;

E0V05__locus_1

PDU1AB

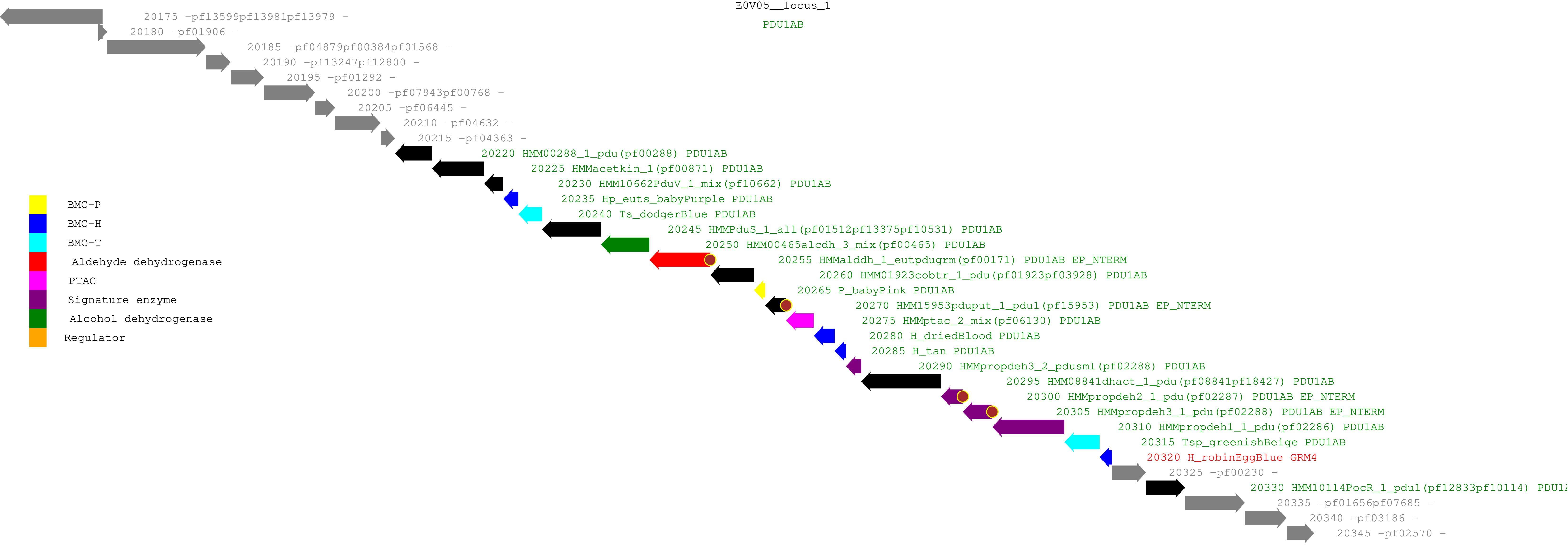

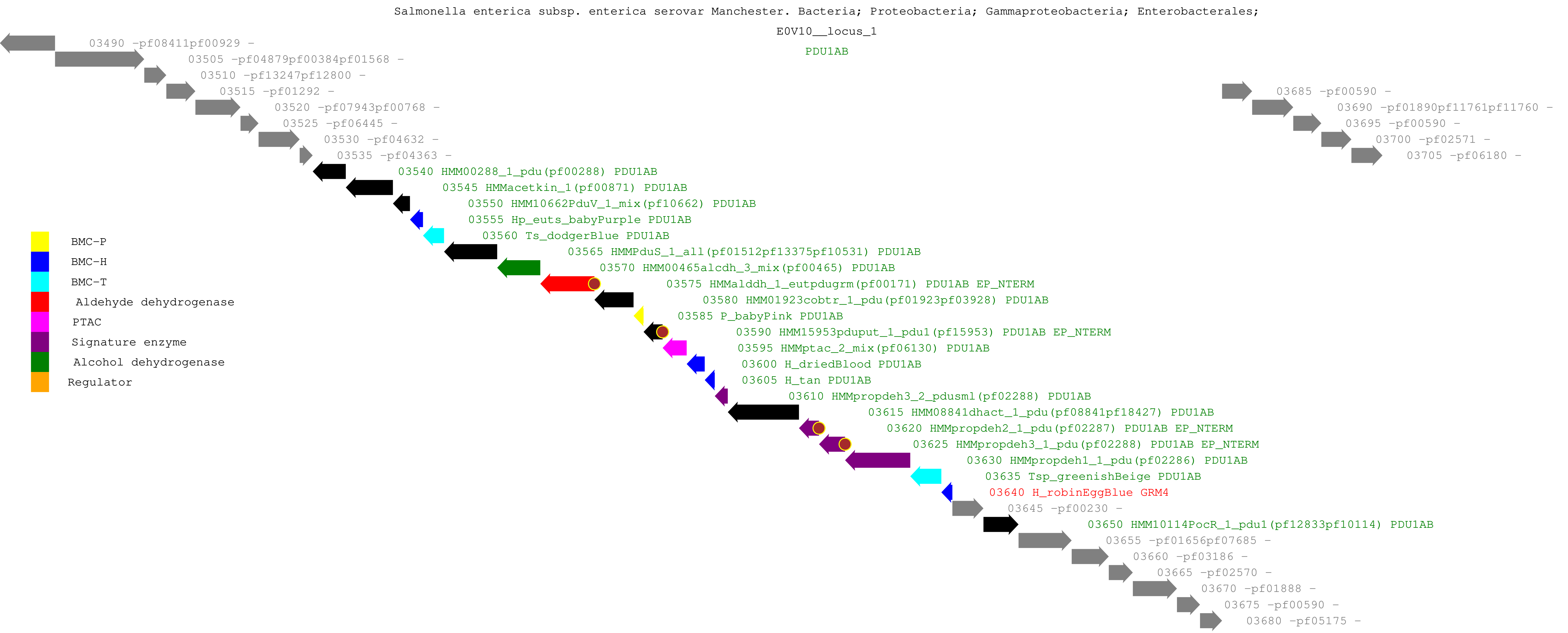

Salmonella enterica subsp. enterica serovar Manchester. Bacteria; Proteobacteria; Gammaproteobacteria; Enterobacterales;
E0V10__locus_1
PDU1AB
03490 -pf08411pf00929 -
03505 -pf04879pf00384pf01568 -
03510 -pf13247pf12800 -
03515 -pf01292 -
03520 -pf07943pf00768 -
03525 -pf06445 -
03530 -pf04632 -
03535 -pf04363 -
03540 HMM00288_1_pdu(pf00288) PDU1AB
03545 HMMacetkin_1(pf00871) PDU1AB
03550 HMM10662PduV_1_mix(pf10662) PDU1AB
03555 Hp_euts_babyPurple PDU1AB
03560 Ts_dodgerBlue PDU1AB
03565 HMMPduS_1_all(pf01512pf13375pf10531) PDU1AB
03570 HMM00465alcdh_3_mix(pf00465) PDU1AB
03575 HMMaldh_1_eutpdugrm(pf00171) PDU1AB EP_NTERM
03580 HMM01923cobtr_1_pdu(pf01923pf03928) PDU1AB
03585 P_babyPink PDU1AB
03590 HMM15953pduput_1_pdu1(pf15953) PDU1AB EP_NTERM
03595 HMMptac_2_mix(pf06130) PDU1AB
03600 H_driedBlood PDU1AB
03605 H_tan PDU1AB
03610 HMMpropdeh3_2_pdusml(pf02288) PDU1AB
03615 HMM08841dhact_1_pdu(pf08841pf18427) PDU1AB
03620 HMMpropdeh2_1_pdu(pf02287) PDU1AB EP_NTERM
03625 HMMpropdeh3_1_pdu(pf02288) PDU1AB EP_NTERM
03630 HMMpropdeh1_1_pdu(pf02286) PDU1AB
03635 Tsp_greenishBeige PDU1AB
03640 H_robinEggBlue GRM4
03645 -pf00230 -
03650 HMM10114PocR_1_pdu1(pf12833pf10114) PDU1AB
03655 -pf01656pf07685 -
03660 -pf03186 -
03665 -pf02570 -
03670 -pf01888 -
03675 -pf00590 -
03680 -pf05175 -
03685 -pf00590 -
03690 -pf01890pf11761pf11760 -
03695 -pf00590 -
03700 -pf02571 -
03705 -pf06180 -
BMC-P
BMC-H
BMC-T
Aldehyde dehydrogenase
PTAC
Signature enzyme
Alcohol dehydrogenase
Regulator

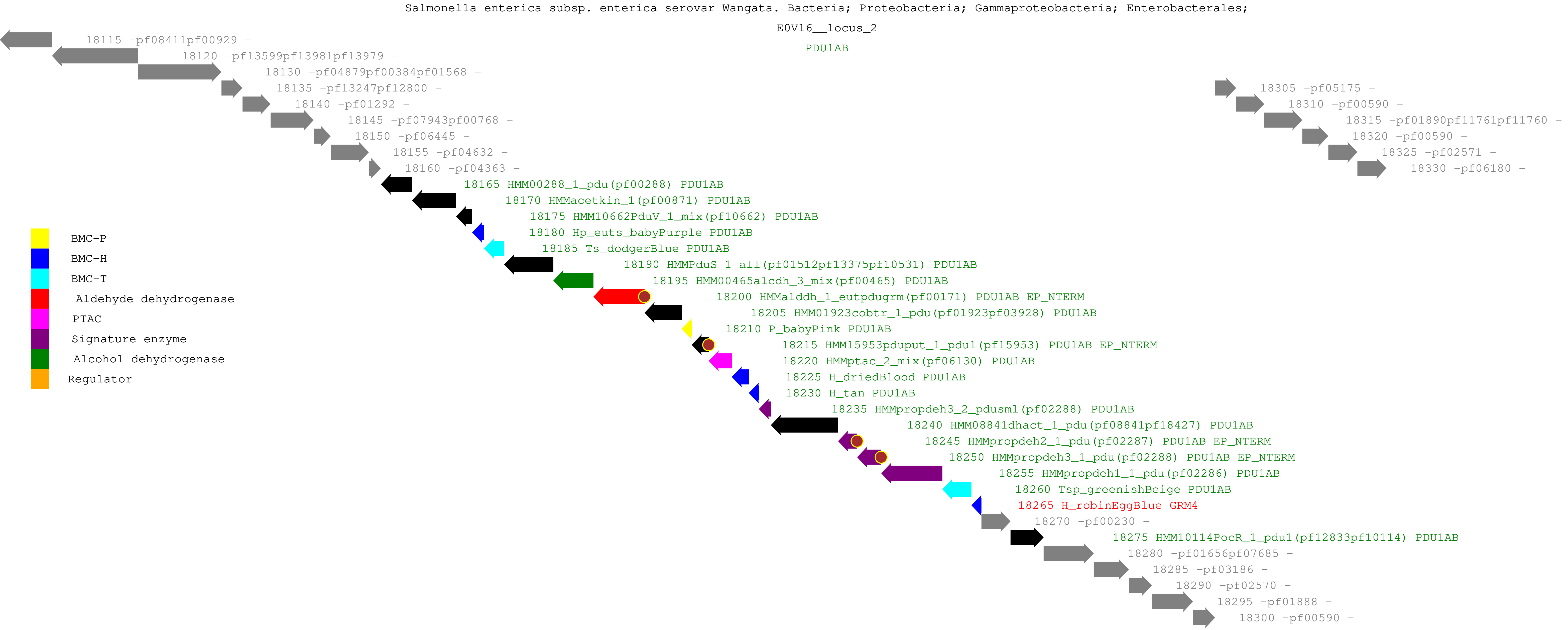
Salmonella enterica subsp. enterica serovar Wangata. Bacteria; Proteobacteria; Gammaproteobacteria; Enterobacterales;
E0V16__locus_2
PDU1AB
18115 -pf08411pf00929 -
18120 -pf13599pf13981pf13979 -
18130 -pf04879pf00384pf01568 -
18135 -pf13247pf12800 -
18140 -pf01292 -
18145 -pf07943pf00768 -
18150 -pf06445 -
18155 -pf04632 -
18160 -pf04363 -
18165 HMM00288_1_pdu(pf00288) PDU1AB
18170 HMMacetkin_1(pf00871) PDU1AB
18175 HMM10662PduV_1_mix(pf10662) PDU1AB
18180 Hp_euts_babyPurple PDU1AB
18185 Ts_dodgerBlue PDU1AB
18190 HMMPduS_1_all(pf01512pf13375pf10531) PDU1AB
18195 HMM00465alcdh_3_mix(pf00465) PDU1AB
18200 HMMalddh_1_eutpdugrm(pf00171) PDU1AB EP_NTERM
18205 HMM01923cobtr_1_pdu(pf01923pf03928) PDU1AB
18210 P_babyPink PDU1AB
18215 HMM15953pduput_1_pdu1(pf15953) PDU1AB EP_NTERM
18220 HMMptac_2_mix(pf06130) PDU1AB
18225 H_driedBlood PDU1AB
18230 H_tan PDU1AB
18235 HMMpropdeh3_2_pdusml(pf02288) PDU1AB
18240 HMM08841dhact_1_pdu(pf08841pf18427) PDU1AB
18245 HMMpropdeh2_1_pdu(pf02287) PDU1AB EP_NTERM
18250 HMMpropdeh3_1_pdu(pf02288) PDU1AB EP_NTERM
18255 HMMpropdeh1_1_pdu(pf02286) PDU1AB
18260 Tsp_greenishBeige PDU1AB
18265 H_robinEggBlue GRM4
18270 -pf00230 -
18275 HMM10114PocR_1_pdu1(pf12833pf10114) PDU1AB
18280 -pf01656pf07685 -
18285 -pf03186 -
18290 -pf02570 -
18295 -pf01888 -
18300 -pf00590 -
18305 -pf05175 -
18310 -pf00590 -
18315 -pf01890pf11761pf11760 -
18320 -pf00590 -
18325 -pf02571 -
18330 -pf06180 -
BMC-P
BMC-H
BMC-T
Aldehyde dehydrogenase
PTAC
Signature enzyme
Alcohol dehydrogenase
Regulator

Salmonella enterica subsp. enterica serovar Nottingham. Bacteria; Proteobacteria; Gammaproteobacteria; Enterobacterales;

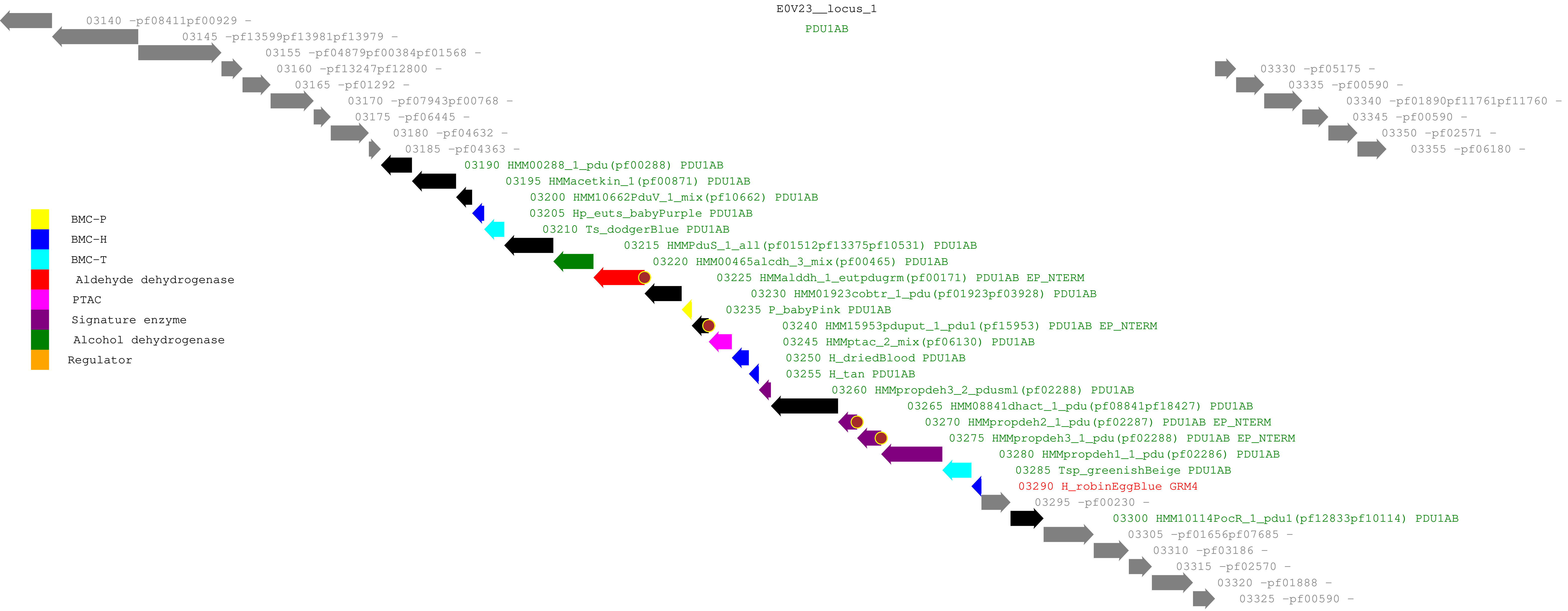

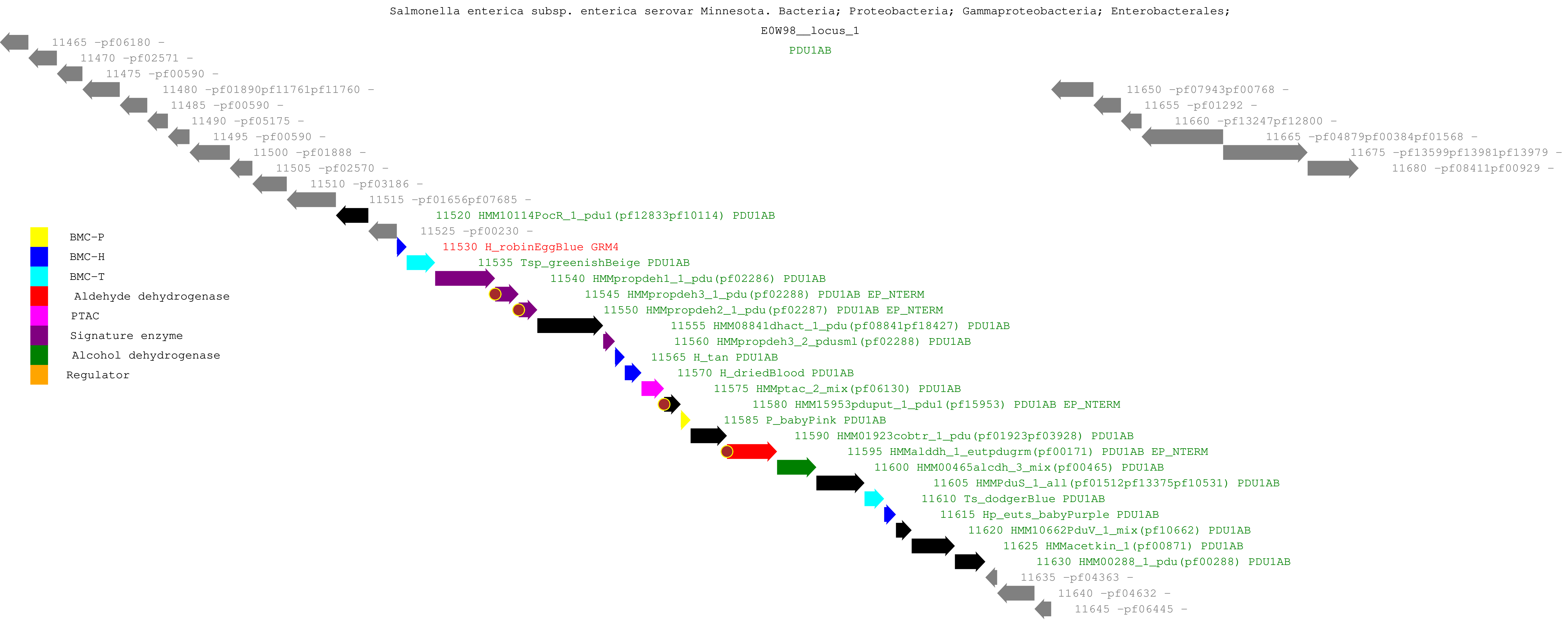
Salmonella enterica subsp. enterica serovar Minnesota. Bacteria; Proteobacteria; Gammaproteobacteria; Enterobacterales;
E0W98__locus_1
PDU1AB
11465 -pf06180 -
11470 -pf02571 -
11475 -pf00590 -
11480 -pf01890pf11761pf11760 -
11485 -pf00590 -
11490 -pf05175 -
11495 -pf00590 -
11500 -pf01888 -
11505 -pf02570 -
11510 -pf03186 -
11515 -pf01656pf07685 -
11520 HMM10114PocR_1_pdu1(pf12833pf10114) PDU1AB
11525 -pf00230 -
11530 H_robinEggBlue GRM4
11535 Tsp_greenishBeige PDU1AB
11540 HMMpropdeh1_1_pdu(pf02286) PDU1AB
11545 HMMpropdeh3_1_pdu(pf02288) PDU1AB EP_NTERM
11550 HMMpropdeh2_1_pdu(pf02287) PDU1AB EP_NTERM
11555 HMM08841dhact_1_pdu(pf08841pf18427) PDU1AB
11560 HMMpropdeh3_2_pdusml(pf02288) PDU1AB
11565 H_tan PDU1AB
11570 H_driedBlood PDU1AB
11575 HMMptac_2_mix(pf06130) PDU1AB
11580 HMM15953pduput_1_pdu1(pf15953) PDU1AB EP_NTERM
11585 P_babyPink PDU1AB
11590 HMM01923cobtr_1_pdu(pf01923pf03928) PDU1AB
11595 HMMalddh_1_eutpdugrm(pf00171) PDU1AB EP_NTERM
11600 HMM00465alcdh_3_mix(pf00465) PDU1AB
11605 HMMPduS_1_all(pf01512pf13375pf10531) PDU1AB
11610 Ts_dodgerBlue PDU1AB
11615 Hp_euts_babyPurple PDU1AB
11620 HMM10662PduV_1_mix(pf10662) PDU1AB
11625 HMMacetkin_1(pf00871) PDU1AB
11630 HMM00288_1_pdu(pf00288) PDU1AB
11635 -pf04363 -
11640 -pf04632 -
11645 -pf06445 -
11650 -pf07943pf00768 -
11655 -pf01292 -
11660 -pf13247pf12800 -
11665 -pf04879pf00384pf01568 -
11675 -pf13599pf13981pf13979 -
11680 -pf08411pf00929 -
BMC-P
BMC-H
BMC-T
Aldehyde dehydrogenase
PTAC
Signature enzyme
Alcohol dehydrogenase
Regulator

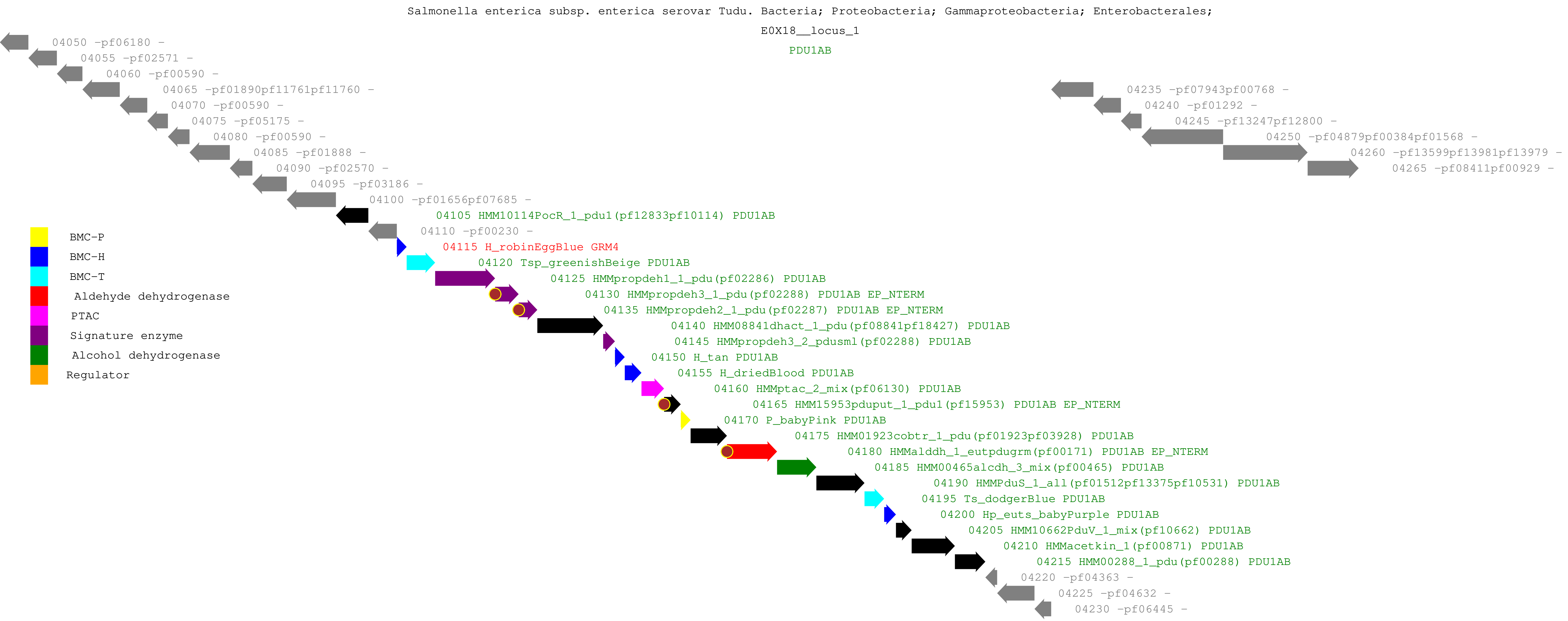
Salmonella enterica subsp. enterica serovar Tudu. Bacteria; Proteobacteria; Gammaproteobacteria; Enterobacterales;
E0X18__locus_1
PDU1AB
04050 -pf06180 -
04055 -pf02571 -
04060 -pf00590 -
04065 -pf01890pf11761pf11760 -
04070 -pf00590 -
04075 -pf05175 -
04080 -pf00590 -
04085 -pf01888 -
04090 -pf02570 -
04095 -pf03186 -
04100 -pf01656pf07685 -
04105 HMM10114PocR_1_pdu1(pf12833pf10114) PDU1AB
04110 -pf00230 -
04115 H_robinEggBlue GRM4
04120 Tsp_greenishBeige PDU1AB
04125 HMMpropdeh1_1_pdu(pf02286) PDU1AB
04130 HMMpropdeh3_1_pdu(pf02288) PDU1AB EP_NTERM
04135 HMMpropdeh2_1_pdu(pf02287) PDU1AB EP_NTERM
04140 HMM08841dhact_1_pdu(pf08841pf18427) PDU1AB
04145 HMMpropdeh3_2_pdusml(pf02288) PDU1AB
04150 H_tan PDU1AB
04155 H_driedBlood PDU1AB
04160 HMMptac_2_mix(pf06130) PDU1AB
04165 HMM15953pduput_1_pdu1(pf15953) PDU1AB EP_NTERM
04170 P_babyPink PDU1AB
04175 HMM01923cobtr_1_pdu(pf01923pf03928) PDU1AB
04180 HMMalddh_1_eutpdugrm(pf00171) PDU1AB EP_NTERM
04185 HMM00465alcdh_3_mix(pf00465) PDU1AB
04190 HMMPduS_1_all(pf01512pf13375pf10531) PDU1AB
04195 Ts_dodgerBlue PDU1AB
04200 Hp_euts_babyPurple PDU1AB
04205 HMM10662PduV_1_mix(pf10662) PDU1AB
04210 HMMacetkin_1(pf00871) PDU1AB
04215 HMM00288_1_pdu(pf00288) PDU1AB
04220 -pf04363 -
04225 -pf04632 -
04230 -pf06445 -
04235 -pf07943pf00768 -
04240 -pf01292 -
04245 -pf13247pf12800 -
04250 -pf04879pf00384pf01568 -
04260 -pf13599pf13981pf13979 -
04265 -pf08411pf00929 -
BMC-P
BMC-H
BMC-T
Aldehyde dehydrogenase
PTAC
Signature enzyme
Alcohol dehydrogenase
Regulator

Salmonella enterica subsp. enterica serovar Agama. Bacteria; Proteobacteria; Gammaproteobacteria; Enterobacterales;

E0X19__locus_1

PDU1AB

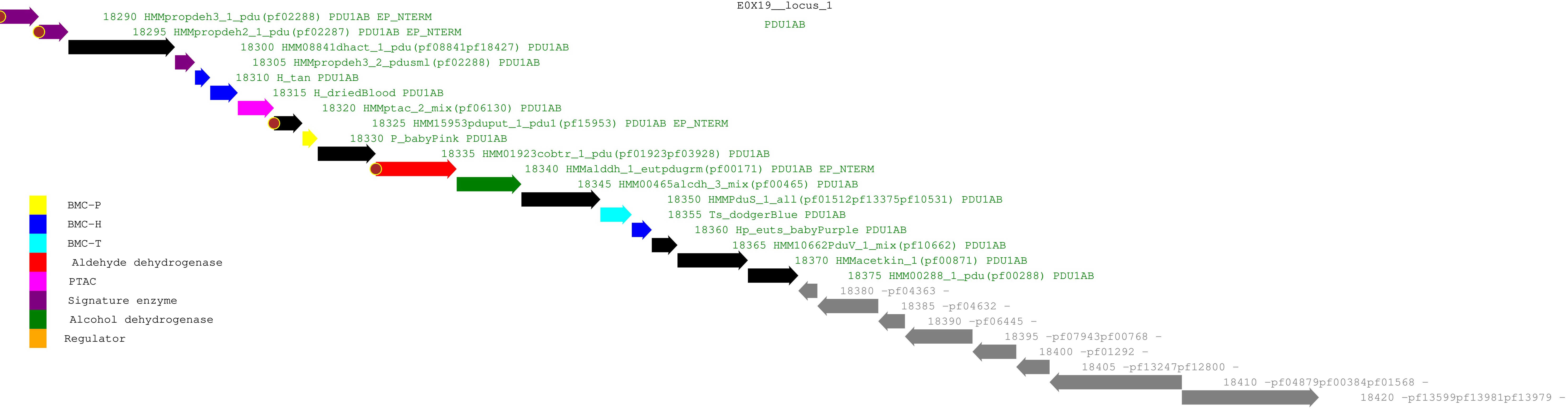

Salmonella potsdam. Bacteria; Proteobacteria; Gammaproteobacteria; Enterobacterales;

E0X61__locus_1

PDU1AB

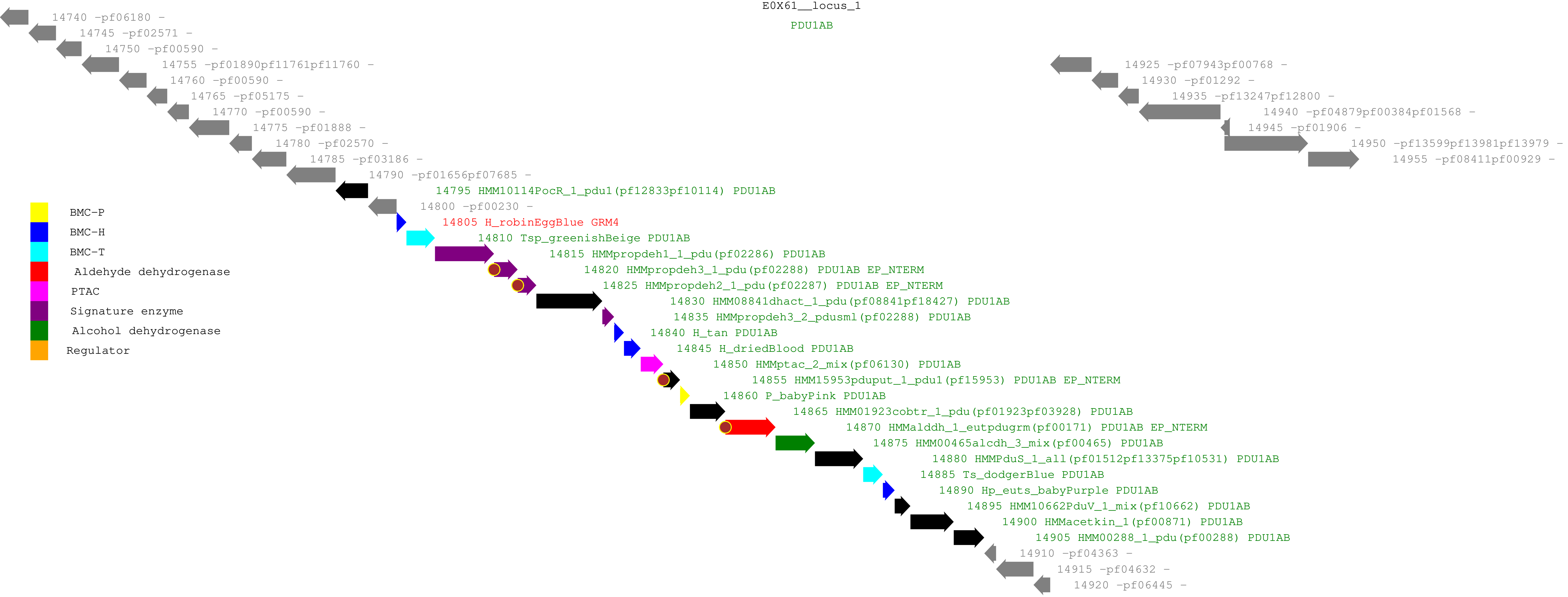

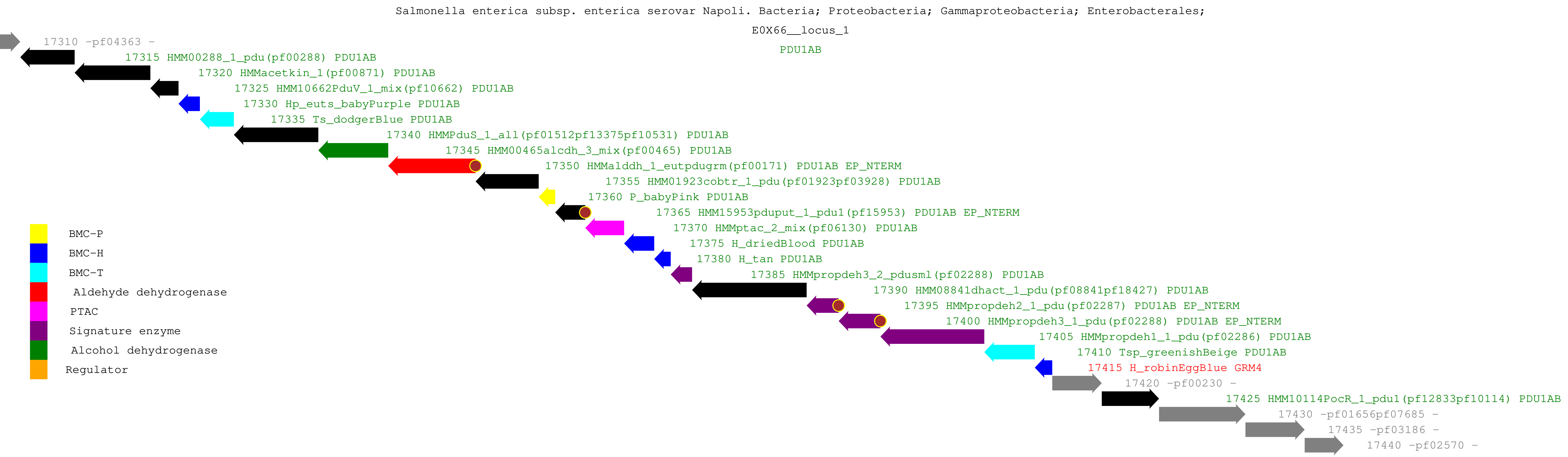
Salmonella enterica subsp. enterica serovar Napoli. Bacteria; Proteobacteria; Gammaproteobacteria; Enterobacterales;
E0X66__locus_1
PDU1AB
17310 -pf04363 -
17315 HMM00288_1_pdu(pf00288) PDU1AB
17320 HMMacetkin_1(pf00871) PDU1AB
17325 HMM10662PduV_1_mix(pf10662) PDU1AB
17330 Hp_euts_babyPurple PDU1AB
17335 Ts_dodgerBlue PDU1AB
17340 HMMPduS_1_all(pf01512pf13375pf10531) PDU1AB
17345 HMM00465alcdh_3_mix(pf00465) PDU1AB
17350 HMMaldddh_1_eutpdugrm(pf00171) PDU1AB EP_NTERM
17355 HMM01923cobtr_1_pdu(pf01923pf03928) PDU1AB
17360 P_babyPink PDU1AB
17365 HMM15953pduput_1_pdu1(pf15953) PDU1AB EP_NTERM
17370 HMMptac_2_mix(pf06130) PDU1AB
17375 H_driedBlood PDU1AB
17380 H_tan PDU1AB
17385 HMMpropdeh3_2_pdusml(pf02288) PDU1AB
17390 HMM08841dhact_1_pdu(pf08841pf18427) PDU1AB
17395 HMMpropdeh2_1_pdu(pf02287) PDU1AB EP_NTERM
17400 HMMpropdeh3_1_pdu(pf02288) PDU1AB EP_NTERM
17405 HMMpropdeh1_1_pdu(pf02286) PDU1AB
17410 Tsp_greenishBeige PDU1AB
17415 H_robinEggBlue GRM4
17420 -pf00230 -
17425 HMM10114PocR_1_pdu1(pf12833pf10114) PDU1AB
17430 -pf01656pf07685 -
17435 -pf03186 -
17440 -pf02570 -
BMC-P
BMC-H
BMC-T
Aldehyde dehydrogenase
PTAC
Signature enzyme
Alcohol dehydrogenase
Regulator

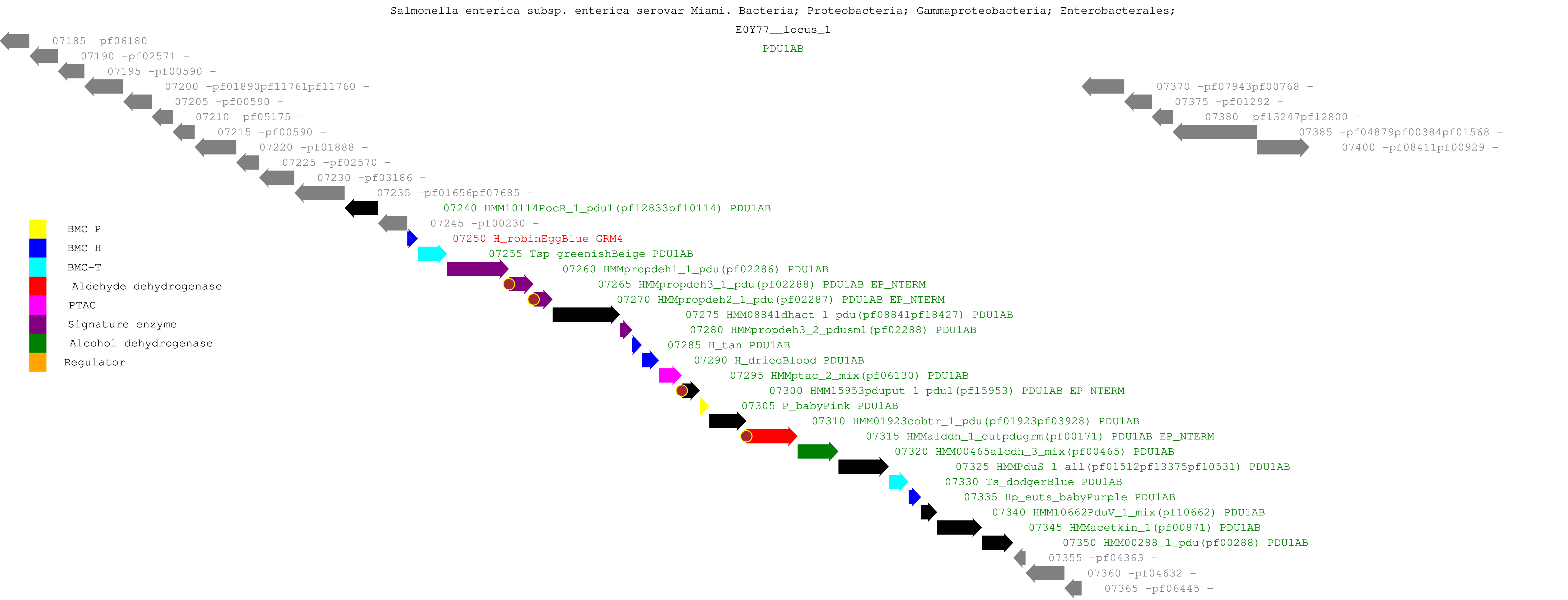
Salmonella enterica subsp. enterica serovar Miami. Bacteria; Proteobacteria; Gammaproteobacteria; Enterobacterales;
E0Y77__locus_1
PDU1AB
07185 -pf06180 -
07190 -pf02571 -
07195 -pf00590 -
07200 -pf01890pf11761pf11760 -
07205 -pf00590 -
07210 -pf05175 -
07215 -pf00590 -
07220 -pf01888 -
07225 -pf02570 -
07230 -pf03186 -
07235 -pf01656pf07685 -
07240 HMM10114PocR_1_pdu1(pf12833pf10114) PDU1AB
07245 -pf00230 -
07250 H_robinEggBlue GRM4
07255 Tsp_greenishBeige PDU1AB
07260 HMMpropdeh1_1_pdu(pf02286) PDU1AB
07265 HMMpropdeh3_1_pdu(pf02288) PDU1AB EP_NTERM
07270 HMMpropdeh2_1_pdu(pf02287) PDU1AB EP_NTERM
07275 HMM08841dhact_1_pdu(pf08841pf18427) PDU1AB
07280 HMMpropdeh3_2_pdusml(pf02288) PDU1AB
07285 H_tan PDU1AB
07290 H_driedBlood PDU1AB
07295 HMMptac_2_mix(pf06130) PDU1AB
07300 HMM15953pduput_1_pdu1(pf15953) PDU1AB EP_NTERM
07305 P_babyPink PDU1AB
07310 HMM01923cobtr_1_pdu(pf01923pf03928) PDU1AB
07315 HMMalddh_1_eutpdugrm(pf00171) PDU1AB EP_NTERM
07320 HMM00465alcdh_3_mix(pf00465) PDU1AB
07325 HMMPduS_1_all(pf01512pf13375pf10531) PDU1AB
07330 Ts_dodgerBlue PDU1AB
07335 Hp_euts_babyPurple PDU1AB
07340 HMM10662PduV_1_mix(pf10662) PDU1AB
07345 HMMacetkin_1(pf00871) PDU1AB
07350 HMM00288_1_pdu(pf00288) PDU1AB
07355 -pf04363 -
07360 -pf04632 -
07365 -pf06445 -
07370 -pf07943pf00768 -
07375 -pf01292 -
07380 -pf13247pf12800 -
07385 -pf04879pf00384pf01568 -
07400 -pf08411pf00929 -
BMC-P
BMC-H
BMC-T
Aldehyde dehydrogenase
PTAC
Signature enzyme
Alcohol dehydrogenase
Regulator

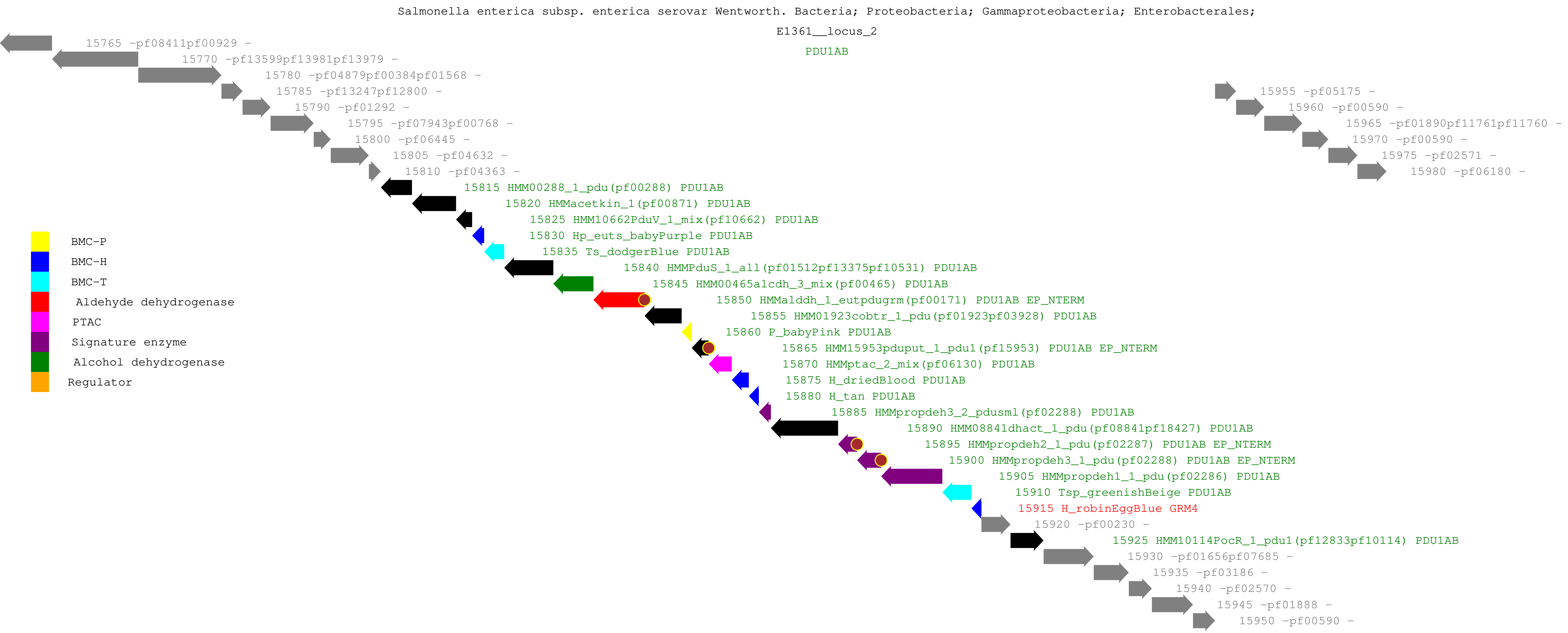

Salmonella enterica subsp. enterica serovar Wentworth. Bacteria; Proteobacteria; Gammaproteobacteria; Enterobacterales;
E1361__locus_2
PDU1AB
15765 -pf08411pf00929 -
15770 -pf13599pf13981pf13979 -
15780 -pf04879pf00384pf01568 -
15785 -pf13247pf12800 -
15790 -pf01292 -
15795 -pf07943pf00768 -
15800 -pf06445 -
15805 -pf04632 -
15810 -pf04363 -
15815 HMM00288_1_pdu(pf00288) PDU1AB
15820 HMMacetkin_1(pf00871) PDU1AB
15825 HMM10662PduV_1_mix(pf10662) PDU1AB
15830 Hp_euts_babyPurple PDU1AB
15835 Ts_dodgerBlue PDU1AB
15840 HMMPduS_1_all(pf01512pf13375pf10531) PDU1AB
15845 HMM00465alcdh_3_mix(pf00465) PDU1AB
15850 HMMalddh_1_eutpdugrm(pf00171) PDU1AB EP_NTERM
15855 HMM01923cobtr_1_pdu(pf01923pf03928) PDU1AB
15860 P_babyPink PDU1AB
15865 HMM15953pduput_1_pdu1(pf15953) PDU1AB EP_NTERM
15870 HMMptac_2_mix(pf06130) PDU1AB
15875 H_driedBlood PDU1AB
15880 H_tan PDU1AB
15885 HMMpropdeh3_2_pdusml(pf02288) PDU1AB
15890 HMM08841dhact_1_pdu(pf08841pf18427) PDU1AB
15895 HMMpropdeh2_1_pdu(pf02287) PDU1AB EP_NTERM
15900 HMMpropdeh3_1_pdu(pf02288) PDU1AB EP_NTERM
15905 HMMpropdeh1_1_pdu(pf02286) PDU1AB
15910 Tsp_greenishBeige PDU1AB
15915 H_robinEggBlue GRM4
15920 -pf00230 -
15925 HMM10114PocR_1_pdu1(pf12833pf10114) PDU1AB
15930 -pf01656pf07685 -
15935 -pf03186 -
15940 -pf02570 -
15945 -pf01888 -
15950 -pf00590 -
15955 -pf05175 -
15960 -pf00590 -
15965 -pf01890pf11761pf11760 -
15970 -pf00590 -
15975 -pf02571 -
15980 -pf06180 -
BMC-P
BMC-H
BMC-T
Aldehyde dehydrogenase
PTAC
Signature enzyme
Alcohol dehydrogenase
Regulator

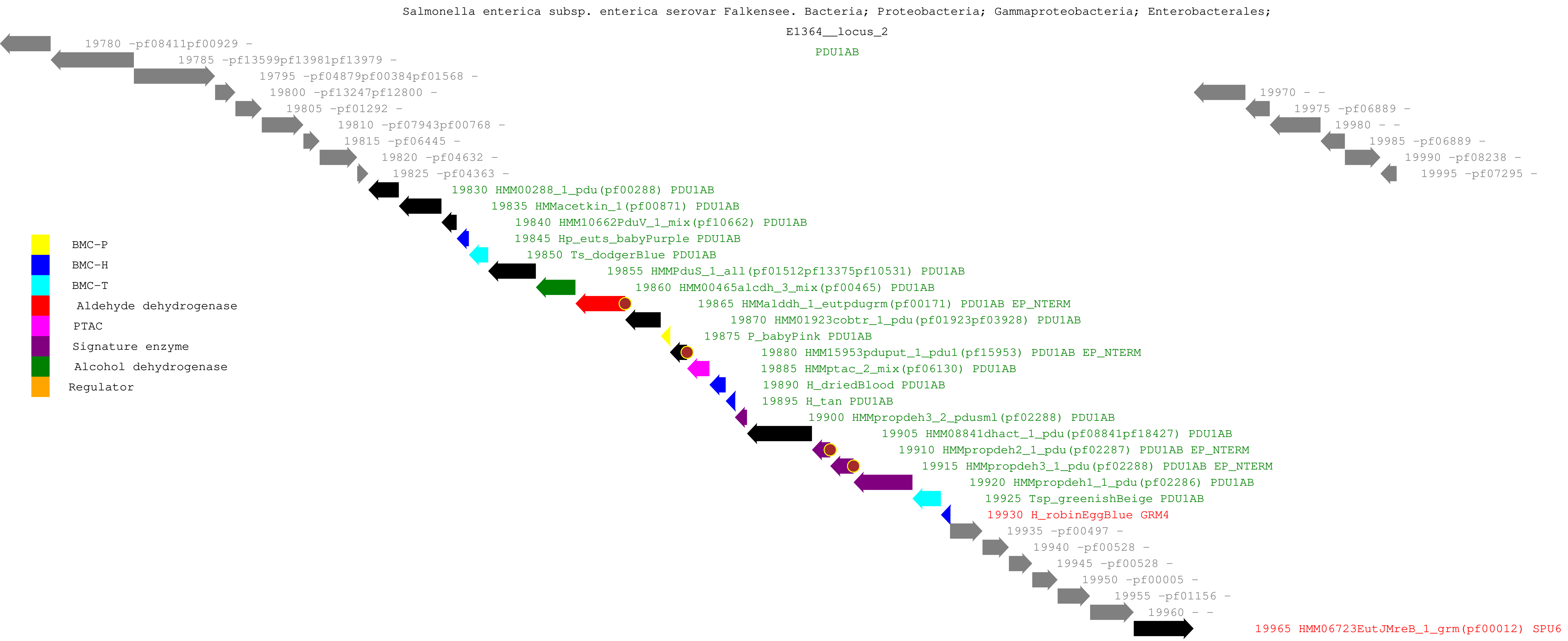
Salmonella enterica subsp. enterica serovar Falkensee. Bacteria; Proteobacteria; Gammaproteobacteria; Enterobacterales;
E1364__locus_2
PDU1AB
19780 -pf08411pf00929 -
19785 -pf13599pf13981pf13979 -
19795 -pf04879pf00384pf01568 -
19800 -pf13247pf12800 -
19805 -pf01292 -
19810 -pf07943pf00768 -
19815 -pf06445 -
19820 -pf04632 -
19825 -pf04363 -
19830 HMM00288_1_pdu(pf00288) PDU1AB
19835 HMMacetkin_1(pf00871) PDU1AB
19840 HMM10662PduV_1_mix(pf10662) PDU1AB
19845 Hp_euts_babyPurple PDU1AB
19850 Ts_dodgerBlue PDU1AB
19855 HMMPduS_1_all(pf01512pf13375pf10531) PDU1AB
19860 HMM00465alcdh_3_mix(pf00465) PDU1AB
19865 HMMalddh_1_eutpdugrm(pf00171) PDU1AB EP_NTERM
19870 HMM01923cobtr_1_pdu(pf01923pf03928) PDU1AB
19875 P_babyPink PDU1AB
19880 HMM15953pduput_1_pdu1(pf15953) PDU1AB EP_NTERM
19885 HMMptac_2_mix(pf06130) PDU1AB
19890 H_driedBlood PDU1AB
19895 H_tan PDU1AB
19900 HMMpropdeh3_2_pdusml(pf02288) PDU1AB
19905 HMM08841dhact_1_pdu(pf08841pf18427) PDU1AB
19910 HMMpropdeh2_1_pdu(pf02287) PDU1AB EP_NTERM
19915 HMMpropdeh3_1_pdu(pf02288) PDU1AB EP_NTERM
19920 HMMpropdeh1_1_pdu(pf02286) PDU1AB
19925 Tsp_greenishBeige PDU1AB
19930 H_robinEggBlue GRM4
19935 -pf00497 -
19940 -pf00528 -
19945 -pf00528 -
19950 -pf00005 -
19955 -pf01156 -
19960 - -
19965 HMM06723EutJMreB_1_grm(pf00012) SPU6
19970 - -
19975 -pf06889 -
19980 - -
19985 -pf06889 -
19990 -pf08238 -
19995 -pf07295 -
BMC-P
BMC-H
BMC-T
Aldehyde dehydrogenase
PTAC
Signature enzyme
Alcohol dehydrogenase
Regulator

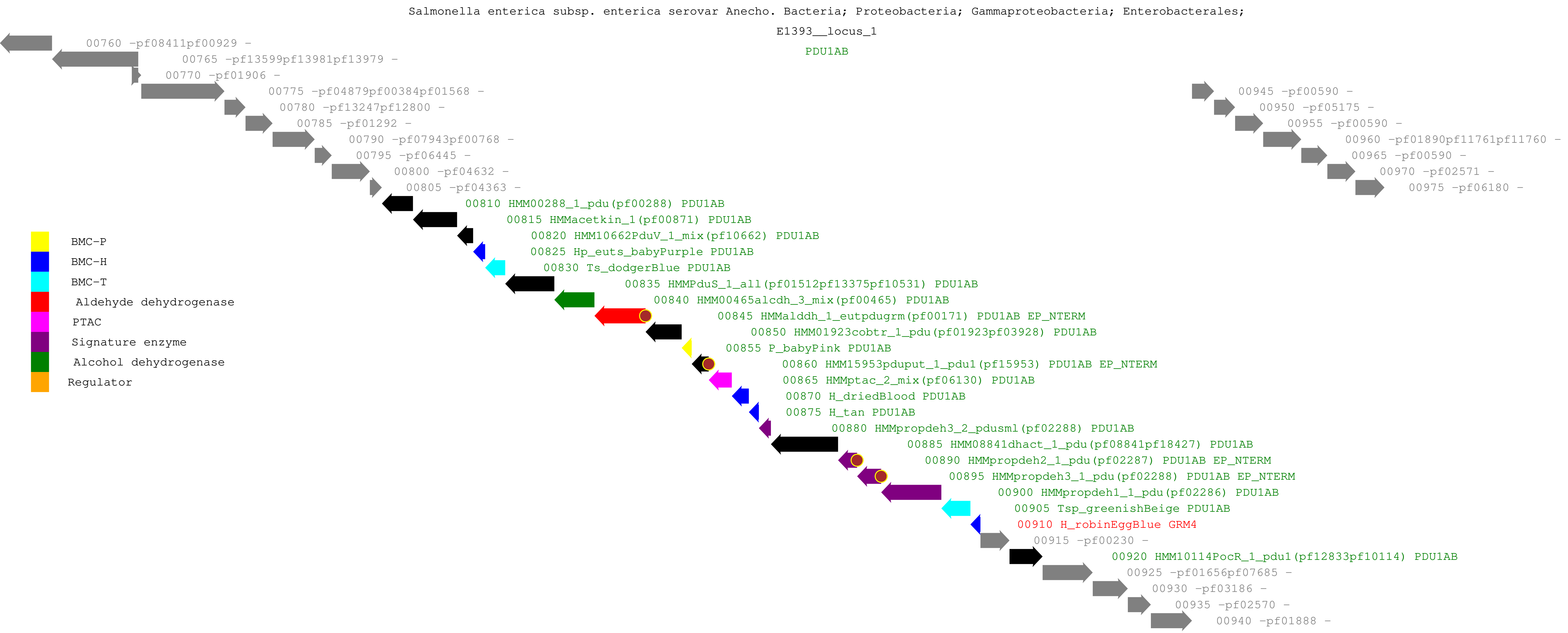

Salmonella enterica subsp. enterica serovar Anecho. Bacteria; Proteobacteria; Gammaproteobacteria; Enterobacterales;
E1393__locus_1
PDU1AB
00760 -pf08411pf00929 -
00765 -pf13599pf13981pf13979 -
00770 -pf01906 -
00775 -pf04879pf00384pf01568 -
00780 -pf13247pf12800 -
00785 -pf01292 -
00790 -pf07943pf00768 -
00795 -pf06445 -
00800 -pf04632 -
00805 -pf04363 -
00810 HMM00288_1_pdu(pf00288) PDU1AB
00815 HMMacetkin_1(pf00871) PDU1AB
00820 HMM10662PduV_1_mix(pf10662) PDU1AB
00825 Hp_euts_babyPurple PDU1AB
00830 Ts_dodgerBlue PDU1AB
00835 HMMPduS_1_all(pf01512pf13375pf10531) PDU1AB
00840 HMM00465alcdh_3_mix(pf00465) PDU1AB
00845 HMMalddh_1_eutpdugrm(pf00171) PDU1AB EP_NTERM
00850 HMM01923cobtr_1_pdu(pf01923pf03928) PDU1AB
00855 P_babyPink PDU1AB
00860 HMM15953pduput_1_pdu1(pf15953) PDU1AB EP_NTERM
00865 HMMptac_2_mix(pf06130) PDU1AB
00870 H_driedBlood PDU1AB
00875 H_tan PDU1AB
00880 HMMpropdeh3_2_pdusml(pf02288) PDU1AB
00885 HMM08841dhact_1_pdu(pf08841pf18427) PDU1AB
00890 HMMpropdeh2_1_pdu(pf02287) PDU1AB EP_NTERM
00895 HMMpropdeh3_1_pdu(pf02288) PDU1AB EP_NTERM
00900 HMMpropdeh1_1_pdu(pf02286) PDU1AB
00905 Tsp_greenishBeige PDU1AB
00910 H_robinEggBlue GRM4
00915 -pf00230 -
00920 HMM10114PocR_1_pdu1(pf12833pf10114) PDU1AB
00925 -pf01656pf07685 -
00930 -pf03186 -
00935 -pf02570 -
00940 -pf01888 -
00945 -pf00590 -
00950 -pf05175 -
00955 -pf00590 -
00960 -pf01890pf11761pf11760 -
00965 -pf00590 -
00970 -pf02571 -
00975 -pf06180 -
BMC-P
BMC-H
BMC-T
Aldehyde dehydrogenase
PTAC
Signature enzyme
Alcohol dehydrogenase
Regulator

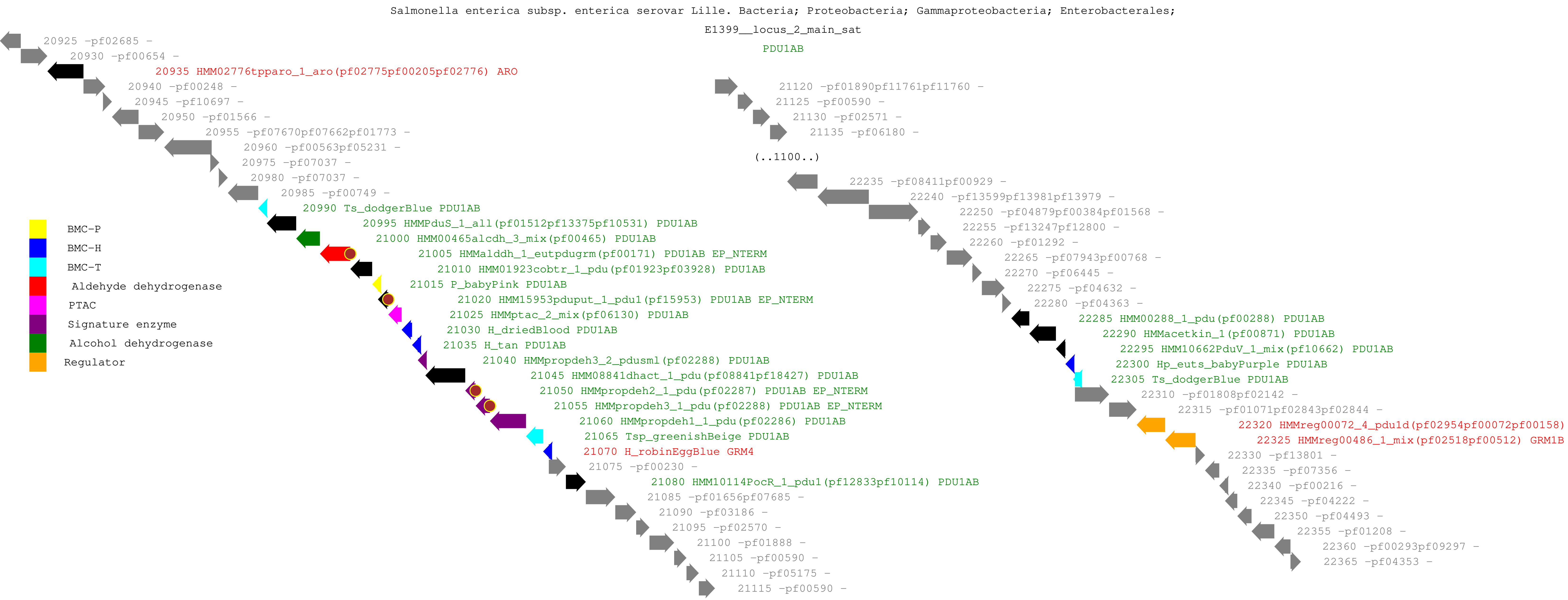
Salmonella enterica subsp. enterica serovar Lille. Bacteria; Proteobacteria; Gammaproteobacteria; Enterobacterales;
E1399__locus_2_main_sat
PDU1AB
20925 -pf02685 -
20930 -pf00654 -
20935 HMM02776tpparo_1_aro(pf02775pf00205pf02776) ARO
20940 -pf00248 -
20945 -pf10697 -
20950 -pf01566 -
20955 -pf07670pf07662pf01773 -
20960 -pf00563pf05231 -
20975 -pf07037 -
20980 -pf07037 -
20985 -pf00749 -
20990 Ts_dodgerBlue PDU1AB
20995 HMMPduS_1_all(pf01512pf13375pf10531) PDU1AB
21000 HMM00465alcdh_3_mix(pf00465) PDU1AB
21005 HMMalddh_1_eutpdugrm(pf00171) PDU1AB EP_NTERM
21010 HMM01923cobtr_1_pdu(pf01923pf03928) PDU1AB
21015 P_babyPink PDU1AB
21020 HMM15953pduput_1_pdu1(pf15953) PDU1AB EP_NTERM
21025 HMMptac_2_mix(pf06130) PDU1AB
21030 H_driedBlood PDU1AB
21035 H_tan PDU1AB
21040 HMMpropdeh3_2_pdusml(pf02288) PDU1AB
21045 HMM08841dhact_1_pdu(pf08841pf18427) PDU1AB
21050 HMMpropdeh2_1_pdu(pf02287) PDU1AB EP_NTERM
21055 HMMpropdeh3_1_pdu(pf02288) PDU1AB EP_NTERM
21060 HMMpropdeh1_1_pdu(pf02286) PDU1AB
21065 Tsp_greenishBeige PDU1AB
21070 H_robinEggBlue GRM4
21075 -pf00230 -
21080 HMM10114PocR_1_pdu1(pf12833pf10114) PDU1AB
21085 -pf01656pf07685 -
21090 -pf03186 -
21095 -pf02570 -
21100 -pf01888 -
21105 -pf00590 -
21110 -pf05175 -
21115 -pf00590 -
21120 -pf01890pf11761pf11760 -
21125 -pf00590 -
21130 -pf02571 -
21135 -pf06180 -
(..1100..)
22235 -pf08411pf00929 -
22240 -pf13599pf13981pf13979 -
22250 -pf04879pf00384pf01568 -
22255 -pf13247pf12800 -
22260 -pf01292 -
22265 -pf07943pf00768 -
22270 -pf06445 -
22275 -pf04632 -
22280 -pf04363 -
22285 HMM00288_1_pdu(pf00288) PDU1AB
22290 HMMacetkin_1(pf00871) PDU1AB
22295 HMM10662PduV_1_mix(pf10662) PDU1AB
22300 Hp_euts_babyPurple PDU1AB
22305 Ts_dodgerBlue PDU1AB
22310 -pf01808pf02142 -
22315 -pf01071pf02843pf02844 -
22320 HMMreg00072_4_pdu1d(pf02954pf00072pf00158)
22325 HMMreg00486_1_mix(pf02518pf00512) GRM1B
22330 -pf13801 -
22335 -pf07356 -
22340 -pf00216 -
22345 -pf04222 -
22350 -pf04493 -
22355 -pf01208 -
22360 -pf00293pf09297 -
22365 -pf04353 -
BMC-P
BMC-H
BMC-T
Aldehyde dehydrogenase
PTAC
Signature enzyme
Alcohol dehydrogenase
Regulator

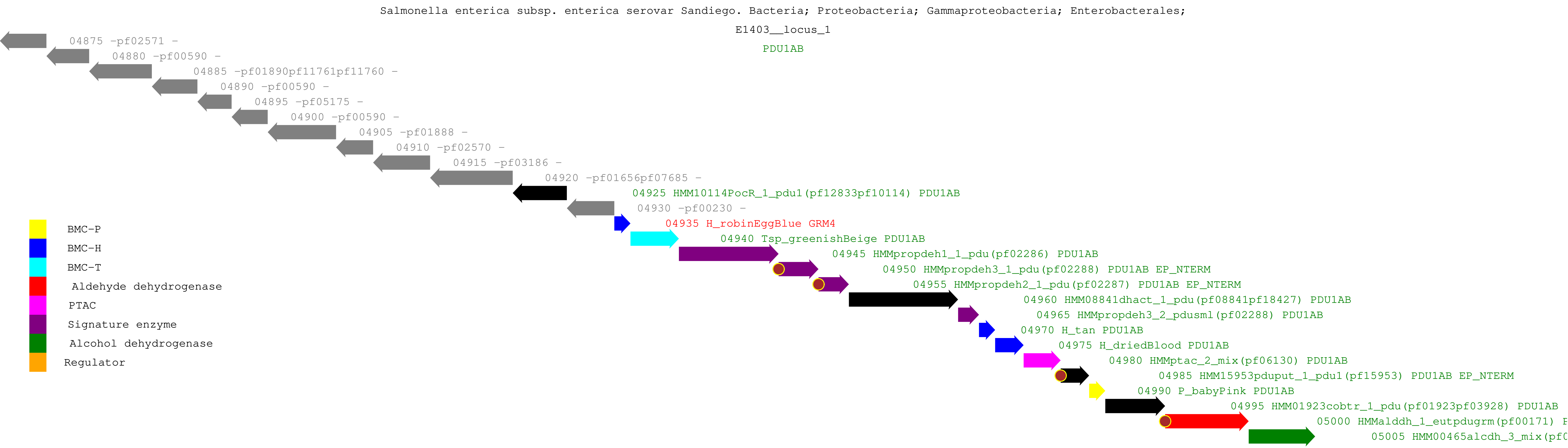
Salmonella enterica subsp. enterica serovar Sandiego. Bacteria; Proteobacteria; Gammaproteobacteria; Enterobacterales;
E1403__locus_1
PDU1AB
04875 -pf02571 -
04880 -pf00590 -
04885 -pf01890pf11761pf11760 -
04890 -pf00590 -
04895 -pf05175 -
04900 -pf00590 -
04905 -pf01888 -
04910 -pf02570 -
04915 -pf03186 -
04920 -pf01656pf07685 -
04925 HMM10114PocR_1_pdu1(pf12833pf10114) PDU1AB
04930 -pf00230 -
04935 H_robinEggBlue GRM4
04940 Tsp_greenishBeige PDU1AB
04945 HMMpropdeh1_1_pdu(pf02286) PDU1AB
04950 HMMpropdeh3_1_pdu(pf02288) PDU1AB EP_NTERM
04955 HMMpropdeh2_1_pdu(pf02287) PDU1AB EP_NTERM
04960 HMM08841dhact_1_pdu(pf08841pf18427) PDU1AB
04965 HMMpropdeh3_2_pdusml(pf02288) PDU1AB
04970 H_tan PDU1AB
04975 H_driedBlood PDU1AB
04980 HMMptac_2_mix(pf06130) PDU1AB
04985 HMM15953pduput_1_pdu1(pf15953) PDU1AB EP_NTERM
04990 P_babyPink PDU1AB
04995 HMM01923cobtr_1_pdu(pf01923pf03928) PDU1AB
05000 HMMalddh_1_eutpdugrm(pf00171) P
05005 HMM00465alcdh_3_mix(pf0
BMC-P
BMC-H
BMC-T
Aldehyde dehydrogenase
PTAC
Signature enzyme
Alcohol dehydrogenase
Regulator

Salmonella enterica subsp. enterica serovar Augustenborg. Bacteria; Proteobacteria; Gammaproteobacteria; Enterobacterales;

E1429__locus_1

PDU1AB

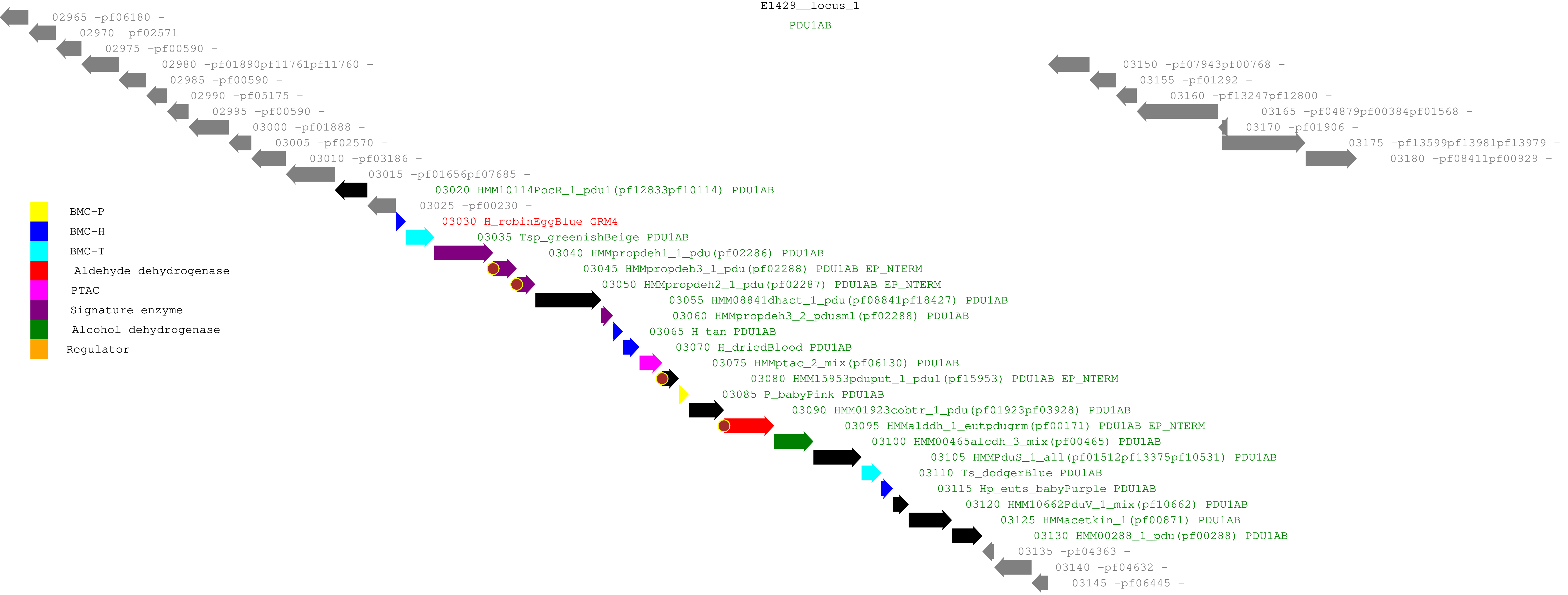

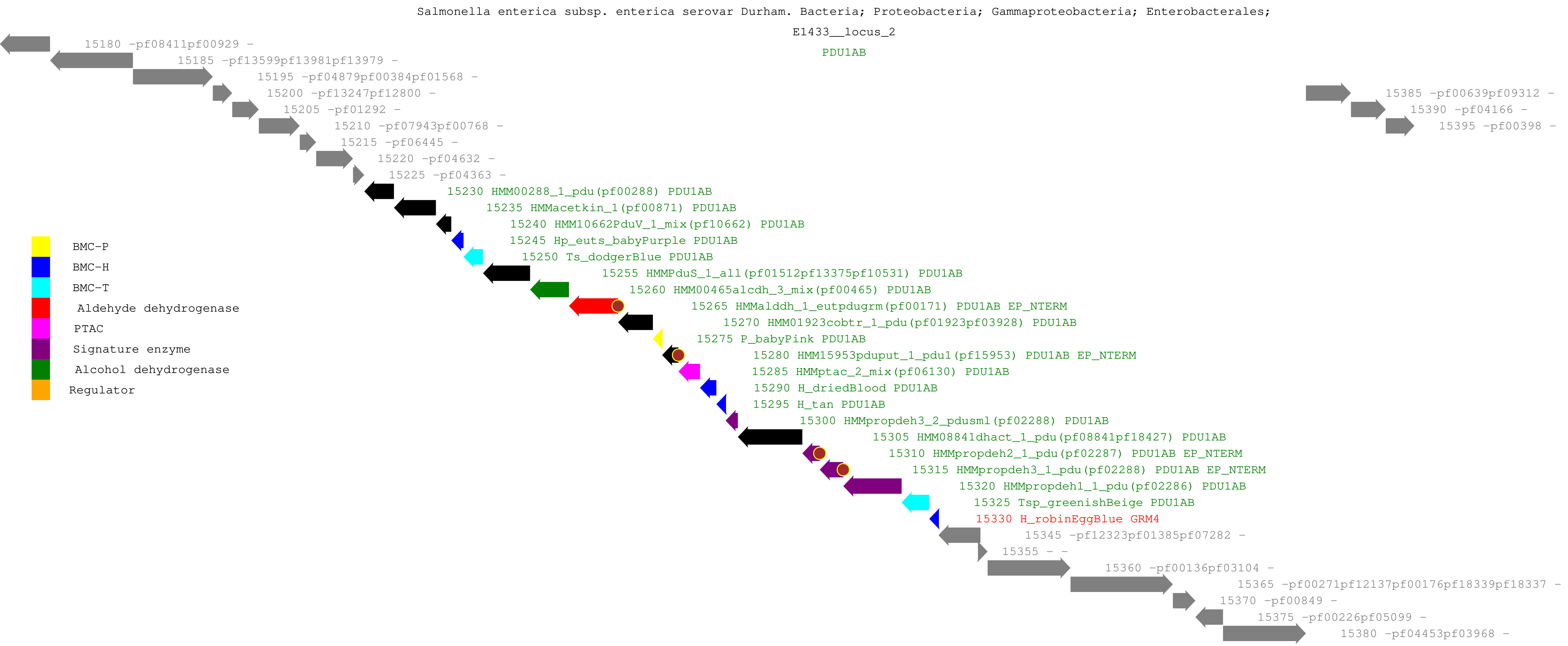

Salmonella enterica subsp. enterica serovar Durham. Bacteria; Proteobacteria; Gammaproteobacteria; Enterobacterales;
E1433__locus_2
PDU1AB
15180 -pf08411pf00929 -
15185 -pf13599pf13981pf13979 -
15195 -pf04879pf00384pf01568 -
15200 -pf13247pf12800 -
15205 -pf01292 -
15210 -pf07943pf00768 -
15215 -pf06445 -
15220 -pf04632 -
15225 -pf04363 -
15230 HMM00288_1_pdu(pf00288) PDU1AB
15235 HMMacetkin_1(pf00871) PDU1AB
15240 HMM10662PduV_1_mix(pf10662) PDU1AB
15245 Hp_euts_babyPurple PDU1AB
15250 Ts_dodgerBlue PDU1AB
15255 HMMPduS_1_all(pf01512pf13375pf10531) PDU1AB
15260 HMM00465alcdh_3_mix(pf00465) PDU1AB
15265 HMMalddh_1_eutpdugrm(pf00171) PDU1AB EP_NTERM
15270 HMM01923cobtr_1_pdu(pf01923pf03928) PDU1AB
15275 P_babyPink PDU1AB
15280 HMM15953pduput_1_pdu1(pf15953) PDU1AB EP_NTERM
15285 HMMptac_2_mix(pf06130) PDU1AB
15290 H_driedBlood PDU1AB
15295 H_tan PDU1AB
15300 HMMpropdeh3_2_pdusml(pf02288) PDU1AB
15305 HMM08841dhact_1_pdu(pf08841pf18427) PDU1AB
15310 HMMpropdeh2_1_pdu(pf02287) PDU1AB EP_NTERM
15315 HMMpropdeh3_1_pdu(pf02288) PDU1AB EP_NTERM
15320 HMMpropdeh1_1_pdu(pf02286) PDU1AB
15325 Tsp_greenishBeige PDU1AB
15330 H_robinEggBlue GRM4
15345 -pf12323pf01385pf07282 -
15355 - -
15360 -pf00136pf03104 -
15365 -pf00271pf12137pf00176pf18339pf18337 -
15370 -pf00849 -
15375 -pf00226pf05099 -
15380 -pf04453pf03968 -
15385 -pf00639pf09312 -
15390 -pf04166 -
15395 -pf00398 -
BMC-P
BMC-H
BMC-T
Aldehyde dehydrogenase
PTAC
Signature enzyme
Alcohol dehydrogenase
Regulator

Salmonella enterica subsp. enterica serovar Kimuenza. Bacteria; Proteobacteria; Gammaproteobacteria; Enterobacterales;

E1444__locus_2

PDU1AB

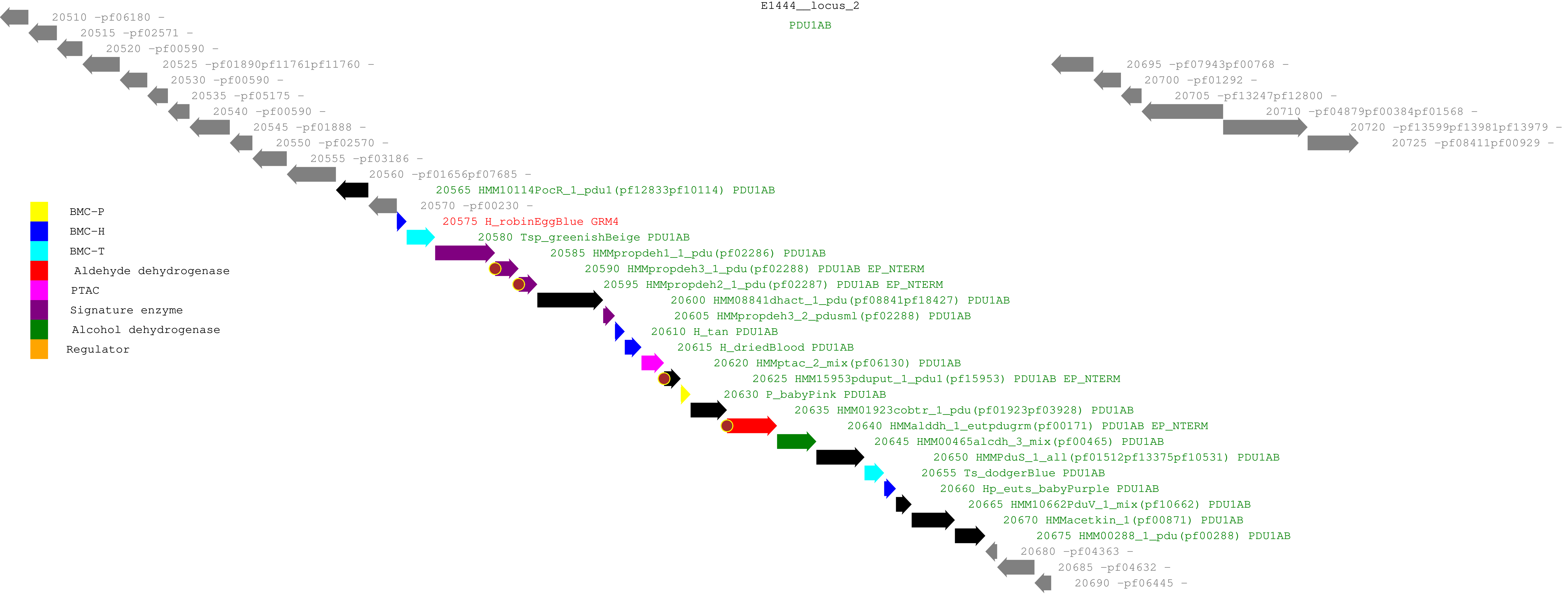

Salmonella enterica subsp. enterica serovar Caracas. Bacteria; Proteobacteria; Gammaproteobacteria; Enterobacterales;

E1460__locus_2

PDU1AB

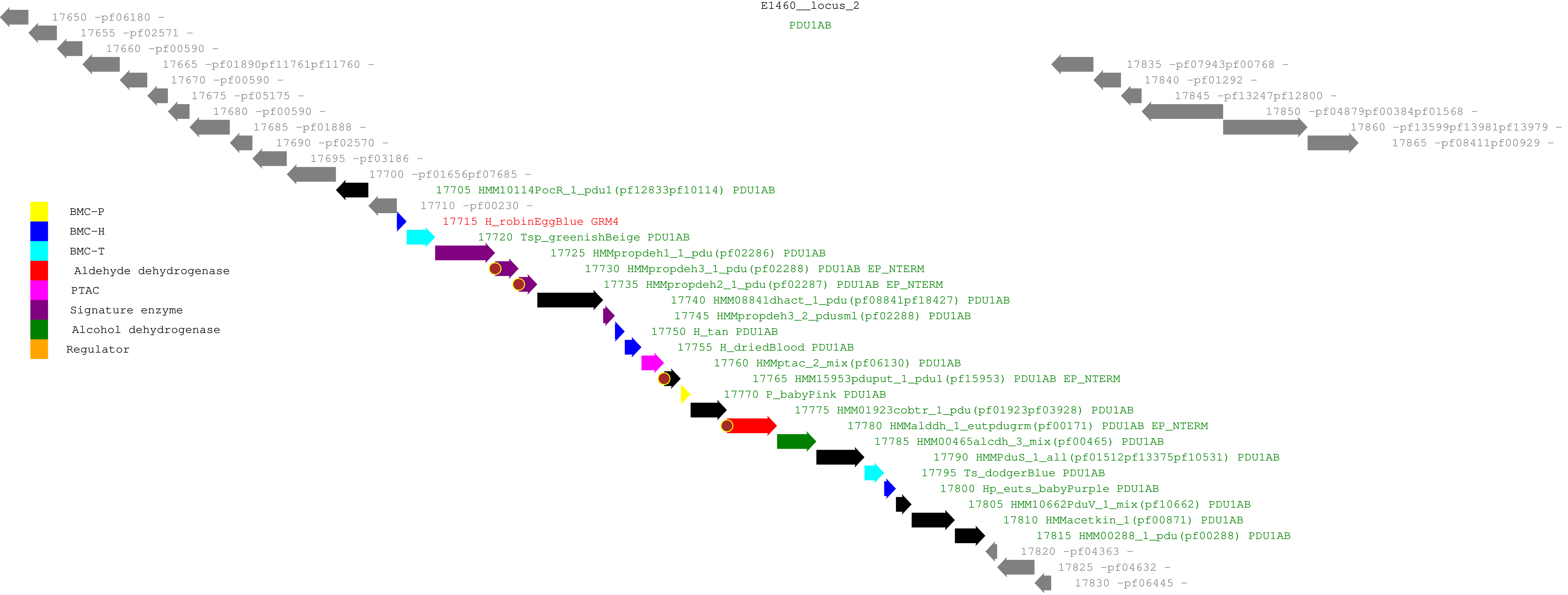

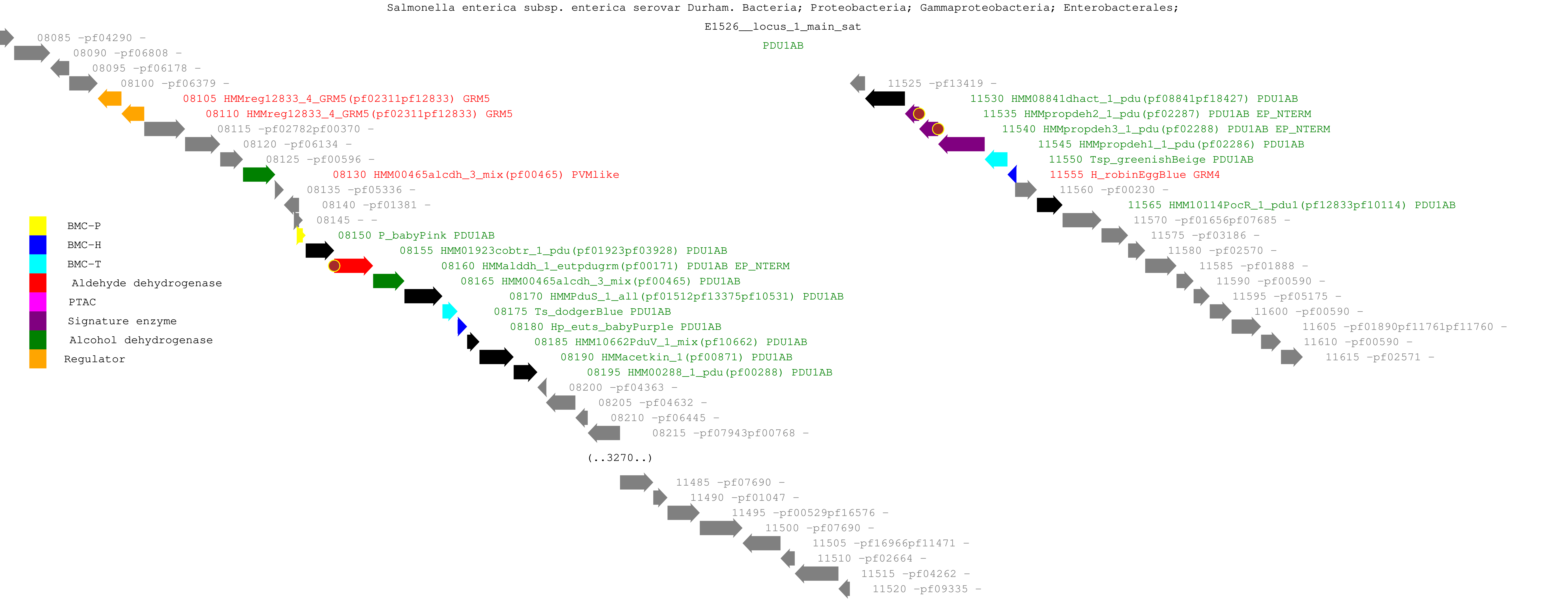
Salmonella enterica subsp. enterica serovar Durham. Bacteria; Proteobacteria; Gammaproteobacteria; Enterobacterales;
E1526__locus_1_main_sat
PDU1AB
08085 -pf04290 -
08090 -pf06808 -
08095 -pf06178 -
08100 -pf06379 -
08105 HMMreg12833_4_GRM5(pf02311pf12833) GRM5
08110 HMMreg12833_4_GRM5(pf02311pf12833) GRM5
08115 -pf02782pf00370 -
08120 -pf06134 -
08125 -pf00596 -
08130 HMM00465alcdh_3_mix(pf00465) PVMlike
08135 -pf05336 -
08140 -pf01381 -
08145 - -
08150 P_babyPink PDU1AB
08155 HMM01923cobtr_1_pdu(pf01923pf03928) PDU1AB
08160 HMMaldh_1_eutpdugrm(pf00171) PDU1AB EP_NTERM
08165 HMM00465alcdh_3_mix(pf00465) PDU1AB
08170 HMMPduS_1_all(pf01512pf13375pf10531) PDU1AB
08175 Ts_dodgerBlue PDU1AB
08180 Hp_euts_babyPurple PDU1AB
08185 HMM10662PduV_1_mix(pf10662) PDU1AB
08190 HMMacetkin_1(pf00871) PDU1AB
08195 HMM00288_1_pdu(pf00288) PDU1AB
08200 -pf04363 -
08205 -pf04632 -
08210 -pf06445 -
08215 -pf07943pf00768 -
(..3270..)
11485 -pf07690 -
11490 -pf01047 -
11495 -pf00529pf16576 -
11500 -pf07690 -
11505 -pf16966pf11471 -
11510 -pf02664 -
11515 -pf04262 -
11520 -pf09335 -
11525 -pf13419 -
11530 HMM08841dhact_1_pdu(pf08841pf18427) PDU1AB
11535 HMMpropdeh2_1_pdu(pf02287) PDU1AB EP_NTERM
11540 HMMpropdeh3_1_pdu(pf02288) PDU1AB EP_NTERM
11545 HMMpropdeh1_1_pdu(pf02286) PDU1AB
11550 Tsp_greenishBeige PDU1AB
11555 H_robinEggBlue GRM4
11560 -pf00230 -
11565 HMM10114PocR_1_pdu1(pf12833pf10114) PDU1AB
11570 -pf01656pf07685 -
11575 -pf03186 -
11580 -pf02570 -
11585 -pf01888 -
11590 -pf00590 -
11595 -pf05175 -
11600 -pf00590 -
11605 -pf01890pf11761pf11760 -
11610 -pf00590 -
11615 -pf02571 -
BMC-P
BMC-H
BMC-T
Aldehyde dehydrogenase
PTAC
Signature enzyme
Alcohol dehydrogenase
Regulator

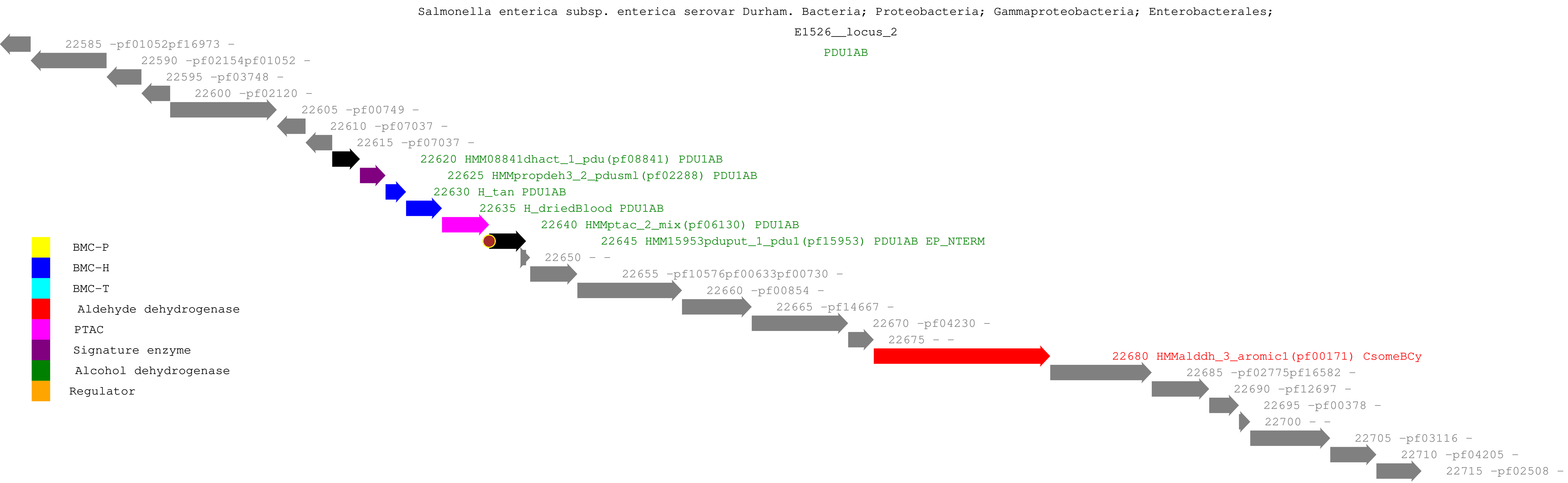

Salmonella enterica subsp. enterica serovar Durham. Bacteria; Proteobacteria; Gammaproteobacteria; Enterobacterales;
E1526__locus_2
PDU1AB
22585 -pf01052pf16973 -
22590 -pf02154pf01052 -
22595 -pf03748 -
22600 -pf02120 -
22605 -pf00749 -
22610 -pf07037 -
22615 -pf07037 -
22620 HMM08841dhact_1_pdu(pf08841) PDU1AB
22625 HMMpropdeh3_2_pdusml(pf02288) PDU1AB
22630 H_tan PDU1AB
22635 H_driedBlood PDU1AB
22640 HMMptac_2_mix(pf06130) PDU1AB
22645 HMM15953pduput_1_pdu1(pf15953) PDU1AB EP_NTERM
22650 - -
22655 -pf10576pf00633pf00730 -
22660 -pf00854 -
22665 -pf14667 -
22670 -pf04230 -
22675 - -
22680 HMMaldh_3_aromic1(pf00171) CsomeBCy
22685 -pf02775pf16582 -
22690 -pf12697 -
22695 -pf00378 -
22700 - -
22705 -pf03116 -
22710 -pf04205 -
22715 -pf02508 -
BMC-P
BMC-H
BMC-T
Aldehyde dehydrogenase
PTAC
Signature enzyme
Alcohol dehydrogenase
Regulator

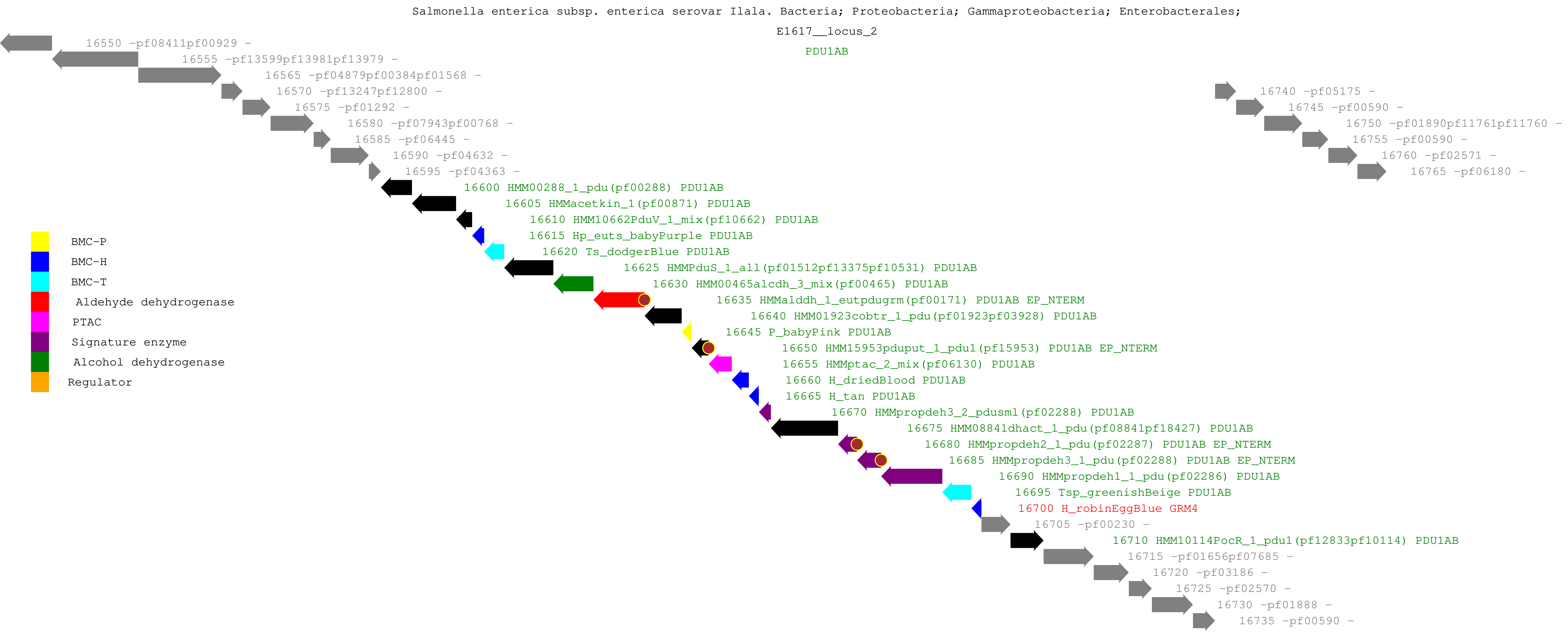
Salmonella enterica subsp. enterica serovar Ilala. Bacteria; Proteobacteria; Gammaproteobacteria; Enterobacterales;
E1617__locus_2
PDU1AB
16550 -pf08411pf00929 -
16555 -pf13599pf13981pf13979 -
16565 -pf04879pf00384pf01568 -
16570 -pf13247pf12800 -
16575 -pf01292 -
16580 -pf07943pf00768 -
16585 -pf06445 -
16590 -pf04632 -
16595 -pf04363 -
16600 HMM00288_1_pdu(pf00288) PDU1AB
16605 HMMacetkin_1(pf00871) PDU1AB
16610 HMM10662PduV_1_mix(pf10662) PDU1AB
16615 Hp_euts_babyPurple PDU1AB
16620 Ts_dodgerBlue PDU1AB
16625 HMMPduS_1_all(pf01512pf13375pf10531) PDU1AB
16630 HMM00465alcdh_3_mix(pf00465) PDU1AB
16635 HMMalddh_1_eutpdugrm(pf00171) PDU1AB EP_NTERM
16640 HMM01923cobtr_1_pdu(pf01923pf03928) PDU1AB
16645 P_babyPink PDU1AB
16650 HMM15953pduput_1_pdu1(pf15953) PDU1AB EP_NTERM
16655 HMMptac_2_mix(pf06130) PDU1AB
16660 H_driedBlood PDU1AB
16665 H_tan PDU1AB
16670 HMMpropdeh3_2_pdusml(pf02288) PDU1AB
16675 HMM08841dhact_1_pdu(pf08841pf18427) PDU1AB
16680 HMMpropdeh2_1_pdu(pf02287) PDU1AB EP_NTERM
16685 HMMpropdeh3_1_pdu(pf02288) PDU1AB EP_NTERM
16690 HMMpropdeh1_1_pdu(pf02286) PDU1AB
16695 Tsp_greenishBeige PDU1AB
16700 H_robinEggBlue GRM4
16705 -pf00230 -
16710 HMM10114PocR_1_pdu1(pf12833pf10114) PDU1AB
16715 -pf01656pf07685 -
16720 -pf03186 -
16725 -pf02570 -
16730 -pf01888 -
16735 -pf00590 -
16740 -pf05175 -
16745 -pf00590 -
16750 -pf01890pf11761pf11760 -
16755 -pf00590 -
16760 -pf02571 -
16765 -pf06180 -
BMC-P
BMC-H
BMC-T
Aldehyde dehydrogenase
PTAC
Signature enzyme
Alcohol dehydrogenase
Regulator

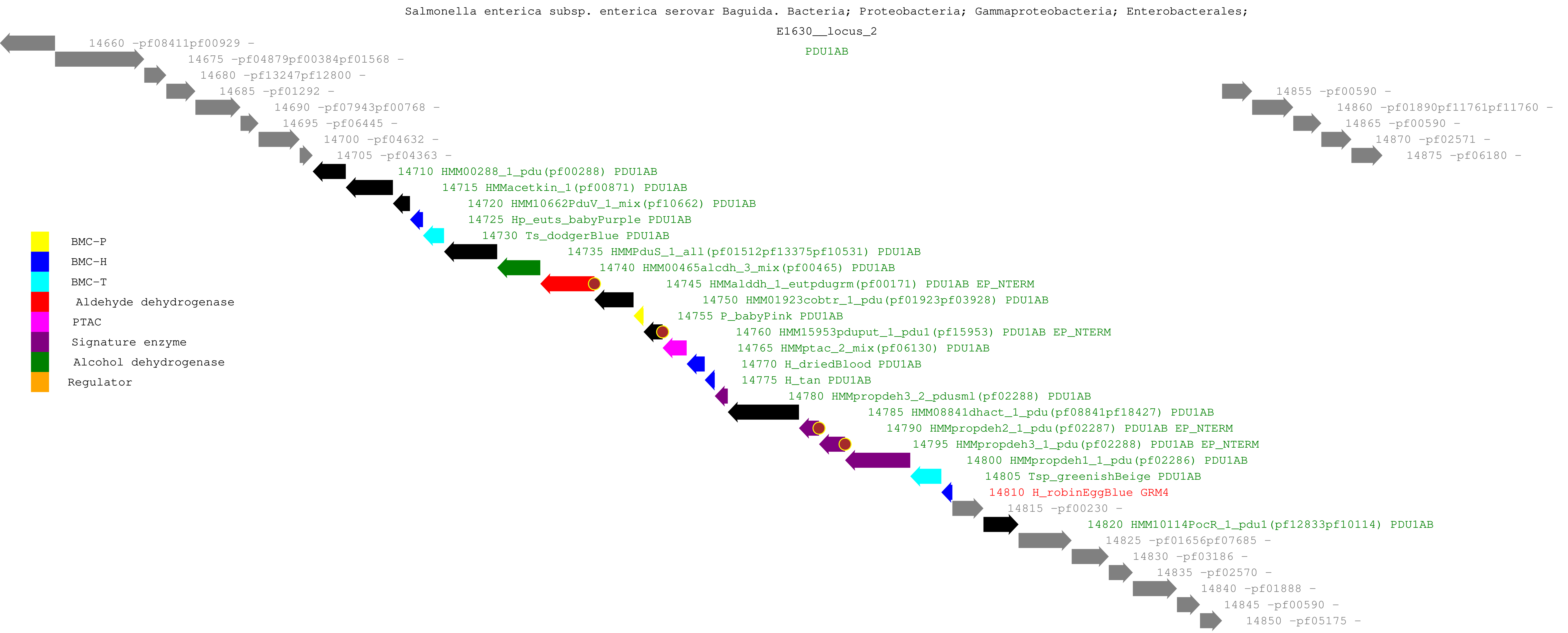
Salmonella enterica subsp. enterica serovar Baguida. Bacteria; Proteobacteria; Gammaproteobacteria; Enterobacterales;
E1630__locus_2
PDU1AB
14660 -pf08411pf00929 -
14675 -pf04879pf00384pf01568 -
14680 -pf13247pf12800 -
14685 -pf01292 -
14690 -pf07943pf00768 -
14695 -pf06445 -
14700 -pf04632 -
14705 -pf04363 -
14710 HMM00288_1_pdu(pf00288) PDU1AB
14715 HMMacetkin_1(pf00871) PDU1AB
14720 HMM10662PduV_1_mix(pf10662) PDU1AB
14725 Hp_euts_babyPurple PDU1AB
14730 Ts_dodgerBlue PDU1AB
14735 HMMPduS_1_all(pf01512pf13375pf10531) PDU1AB
14740 HMM00465alcdh_3_mix(pf00465) PDU1AB
14745 HMMaldh_1_eutpdugrm(pf00171) PDU1AB EP_NTERM
14750 HMM01923cobtr_1_pdu(pf01923pf03928) PDU1AB
14755 P_babyPink PDU1AB
14760 HMM15953pduput_1_pdu1(pf15953) PDU1AB EP_NTERM
14765 HMMptac_2_mix(pf06130) PDU1AB
14770 H_driedBlood PDU1AB
14775 H_tan PDU1AB
14780 HMMpropdeh3_2_pdusml(pf02288) PDU1AB
14785 HMM08841dhact_1_pdu(pf08841pf18427) PDU1AB
14790 HMMpropdeh2_1_pdu(pf02287) PDU1AB EP_NTERM
14795 HMMpropdeh3_1_pdu(pf02288) PDU1AB EP_NTERM
14800 HMMpropdeh1_1_pdu(pf02286) PDU1AB
14805 Tsp_greenishBeige PDU1AB
14810 H_robinEggBlue GRM4
14815 -pf00230 -
14820 HMM10114PocR_1_pdu1(pf12833pf10114) PDU1AB
14825 -pf01656pf07685 -
14830 -pf03186 -
14835 -pf02570 -
14840 -pf01888 -
14845 -pf00590 -
14850 -pf05175 -
14855 -pf00590 -
14860 -pf01890pf11761pf11760 -
14865 -pf00590 -
14870 -pf02571 -
14875 -pf06180 -
BMC-P
BMC-H
BMC-T
Aldehyde dehydrogenase
PTAC
Signature enzyme
Alcohol dehydrogenase
Regulator

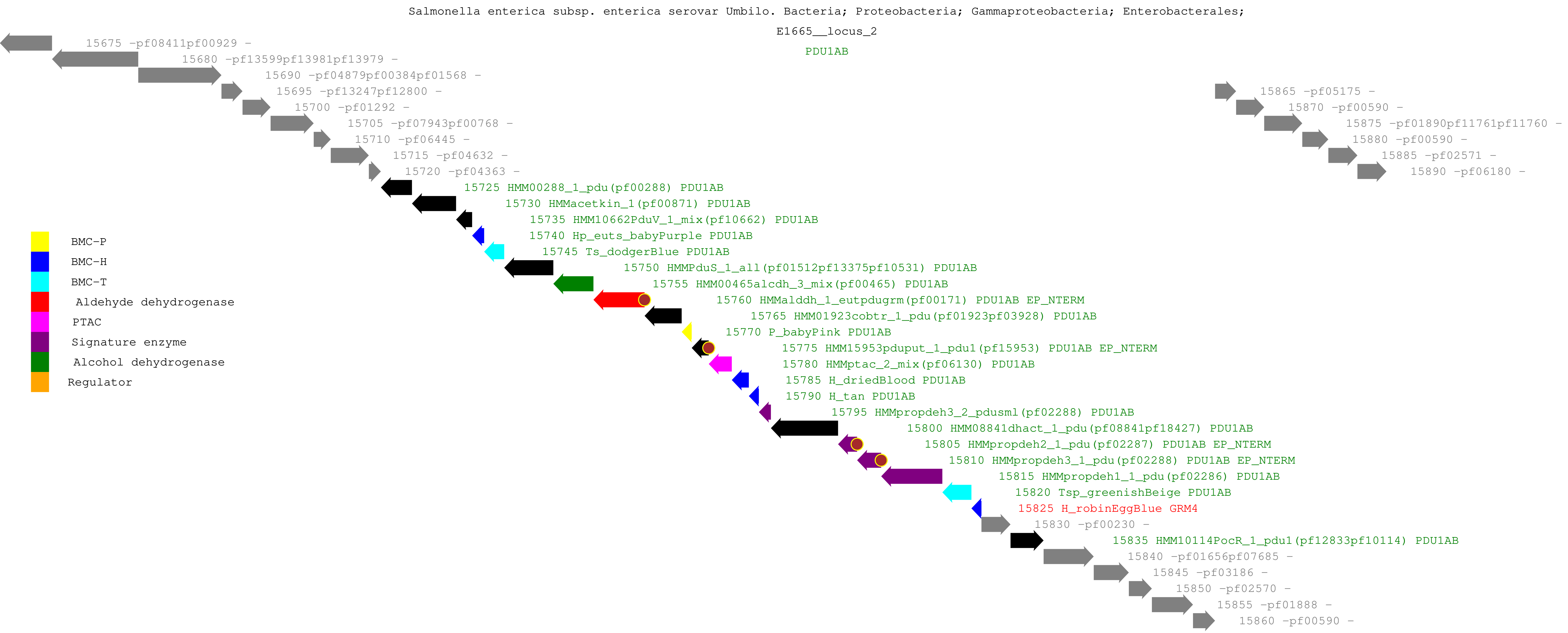

Salmonella enterica subsp. enterica serovar Umbilo. Bacteria; Proteobacteria; Gammaproteobacteria; Enterobacterales;
E1665__locus_2
PDU1AB
15675 -pf08411pf00929 -
15680 -pf13599pf13981pf13979 -
15690 -pf04879pf00384pf01568 -
15695 -pf13247pf12800 -
15700 -pf01292 -
15705 -pf07943pf00768 -
15710 -pf06445 -
15715 -pf04632 -
15720 -pf04363 -
15725 HMM00288_1_pdu(pf00288) PDU1AB
15730 HMMacetkin_1(pf00871) PDU1AB
15735 HMM10662PduV_1_mix(pf10662) PDU1AB
15740 Hp_euts_babyPurple PDU1AB
15745 Ts_dodgerBlue PDU1AB
15750 HMMPduS_1_all(pf01512pf13375pf10531) PDU1AB
15755 HMM00465alcdh_3_mix(pf00465) PDU1AB
15760 HMMalddh_1_eutpdugrm(pf00171) PDU1AB EP_NTERM
15765 HMM01923cobtr_1_pdu(pf01923pf03928) PDU1AB
15770 P_babyPink PDU1AB
15775 HMM15953pduput_1_pdu1(pf15953) PDU1AB EP_NTERM
15780 HMMptac_2_mix(pf06130) PDU1AB
15785 H_driedBlood PDU1AB
15790 H_tan PDU1AB
15795 HMMpropdeh3_2_pdusml(pf02288) PDU1AB
15800 HMM08841dhact_1_pdu(pf08841pf18427) PDU1AB
15805 HMMpropdeh2_1_pdu(pf02287) PDU1AB EP_NTERM
15810 HMMpropdeh3_1_pdu(pf02288) PDU1AB EP_NTERM
15815 HMMpropdeh1_1_pdu(pf02286) PDU1AB
15820 Tsp_greenishBeige PDU1AB
15825 H_robinEggBlue GRM4
15830 -pf00230 -
15835 HMM10114PocR_1_pdu1(pf12833pf10114) PDU1AB
15840 -pf01656pf07685 -
15845 -pf03186 -
15850 -pf02570 -
15855 -pf01888 -
15860 -pf00590 -
15865 -pf05175 -
15870 -pf00590 -
15875 -pf01890pf11761pf11760 -
15880 -pf00590 -
15885 -pf02571 -
15890 -pf06180 -
BMC-P
BMC-H
BMC-T
Aldehyde dehydrogenase
PTAC
Signature enzyme
Alcohol dehydrogenase
Regulator

Salmonella blockley. Bacteria; Proteobacteria; Gammaproteobacteria; Enterobacterales;

E1672__locus_1

PDU1AB

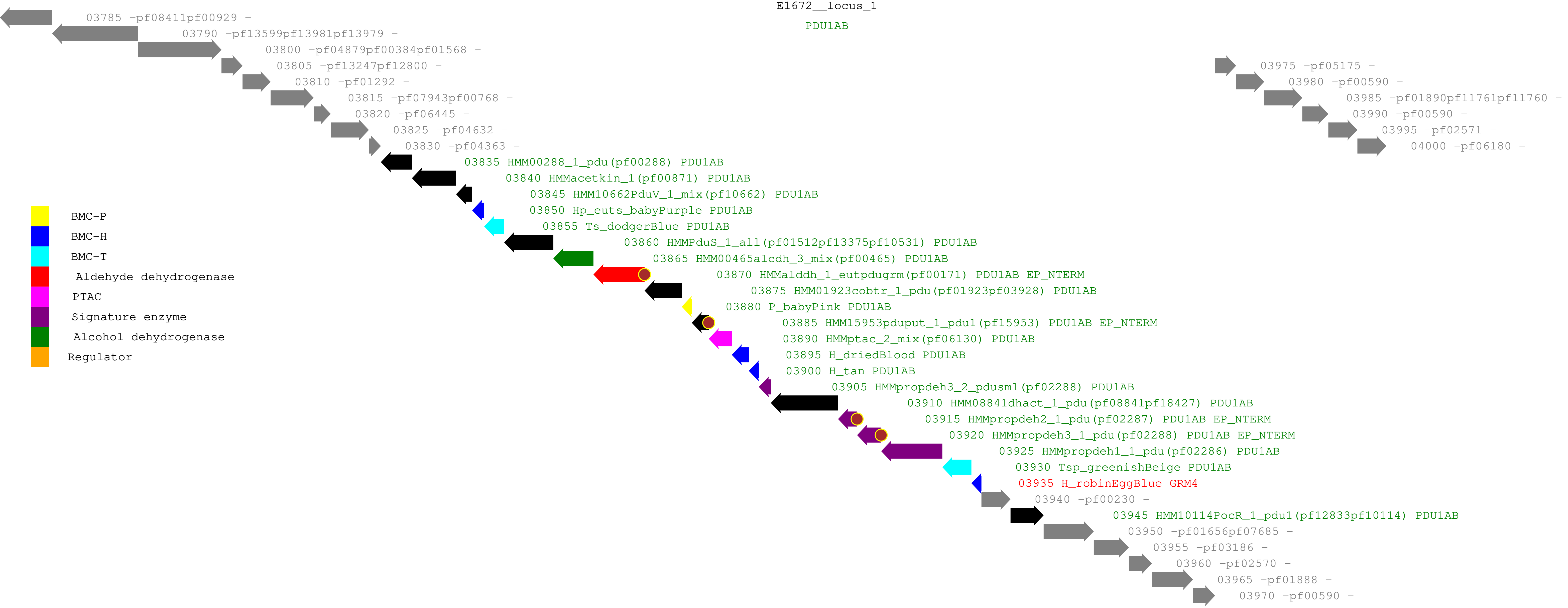

Salmonella enterica subsp. enterica serovar Duesseldorf. Bacteria; Proteobacteria; Gammaproteobacteria; Enterobacterales;

E1759__locus_2

PDU1AB

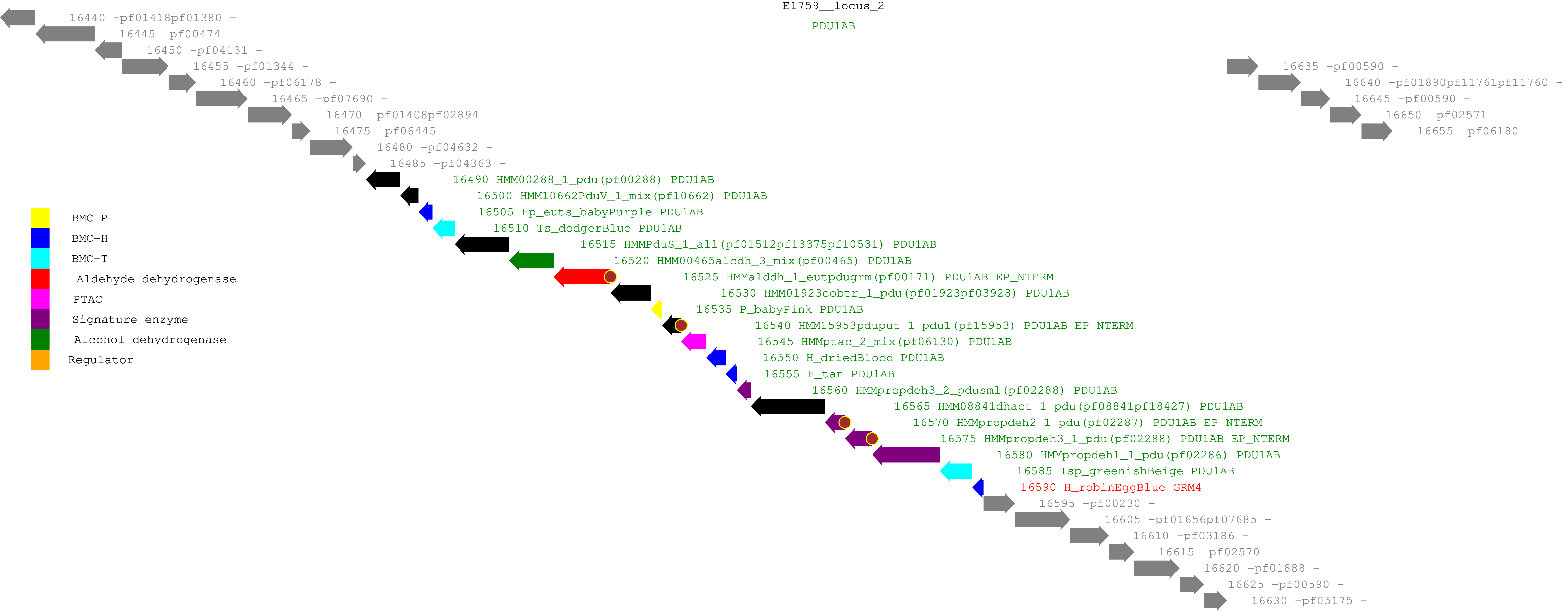

Salmonella enterica subsp. enterica serovar Nagoya. Bacteria; Proteobacteria; Gammaproteobacteria; Enterobacterales;

E1811__locus_2

PDU1AB

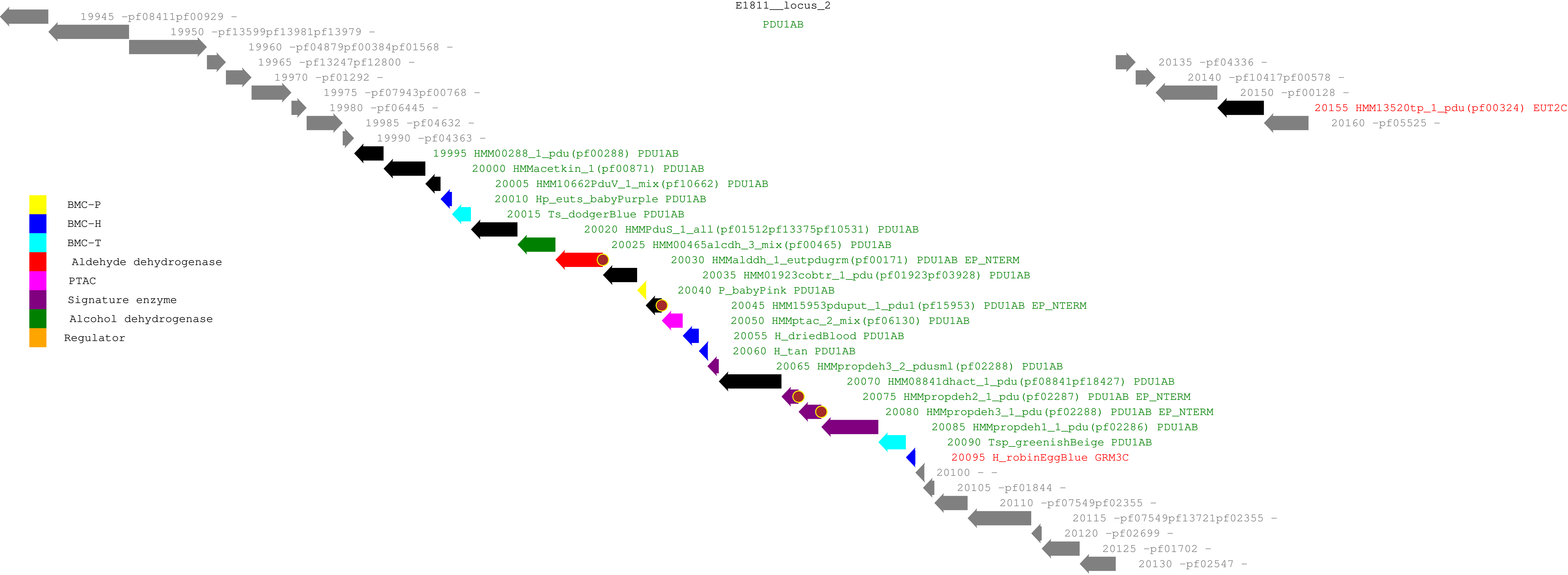

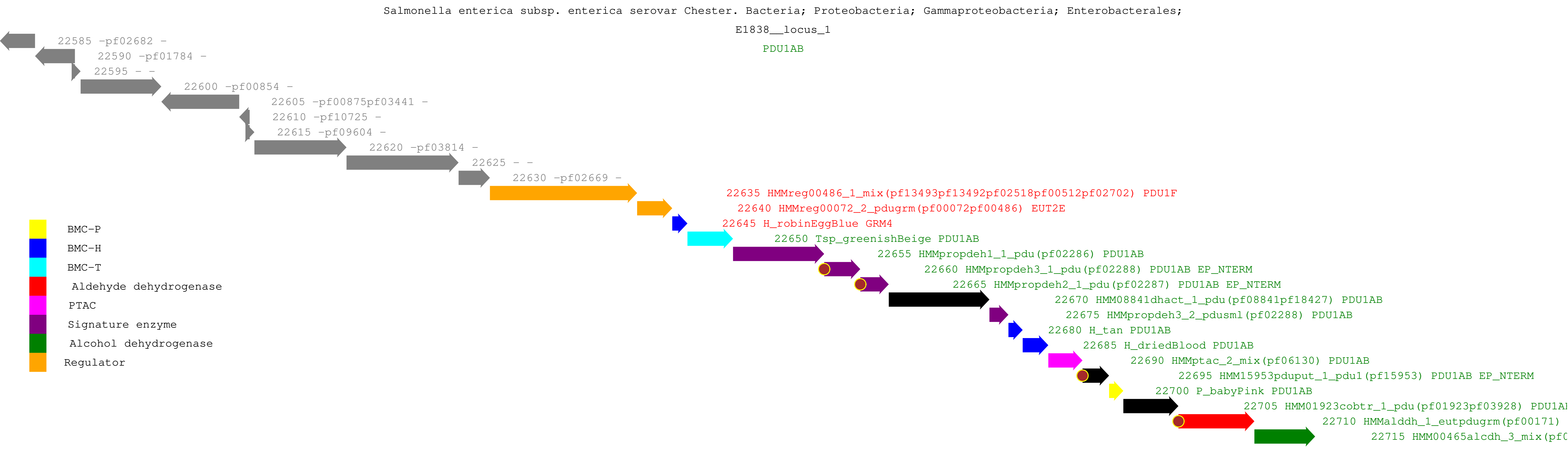

Salmonella enterica subsp. enterica serovar Chester. Bacteria; Proteobacteria; Gammaproteobacteria; Enterobacterales;
E1838__locus_1
PDU1AB
22585 -pf02682 -
22590 -pf01784 -
22595 - -
22600 -pf00854 -
22605 -pf00875pf03441 -
22610 -pf10725 -
22615 -pf09604 -
22620 -pf03814 -
22625 - -
22630 -pf02669 -
22635 HMMreg00486_1_mix(pf13493pf13492pf02518pf00512pf02702) PDU1F
22640 HMMreg00072_2_pdugrm(pf00072pf00486) EUT2E
22645 H_robinEggBlue GRM4
22650 Tsp_greenishBeige PDU1AB
22655 HMMpropdeh1_1_pdu(pf02286) PDU1AB
22660 HMMpropdeh3_1_pdu(pf02288) PDU1AB EP_NTERM
22665 HMMpropdeh2_1_pdu(pf02287) PDU1AB EP_NTERM
22670 HMM08841dhact_1_pdu(pf08841pf18427) PDU1AB
22675 HMMpropdeh3_2_pdusml(pf02288) PDU1AB
22680 H_tan PDU1AB
22685 H_driedBlood PDU1AB
22690 HMMptac_2_mix(pf06130) PDU1AB
22695 HMM15953pduput_1_pdu1(pf15953) PDU1AB EP_NTERM
22700 P_babyPink PDU1AB
22705 HMM01923cobtr_1_pdu(pf01923pf03928) PDU1AB
22710 HMMalddh_1_eutpdugrm(pf00171)
22715 HMM00465alcdh_3_mix(pf0
BMC-P
BMC-H
BMC-T
Aldehyde dehydrogenase
PTAC
Signature enzyme
Alcohol dehydrogenase
Regulator

Salmonella enterica subsp. enterica serovar Glostrup. Bacteria; Proteobacteria; Gammaproteobacteria; Enterobacterales;

E1853__locus_1

PDU1AB

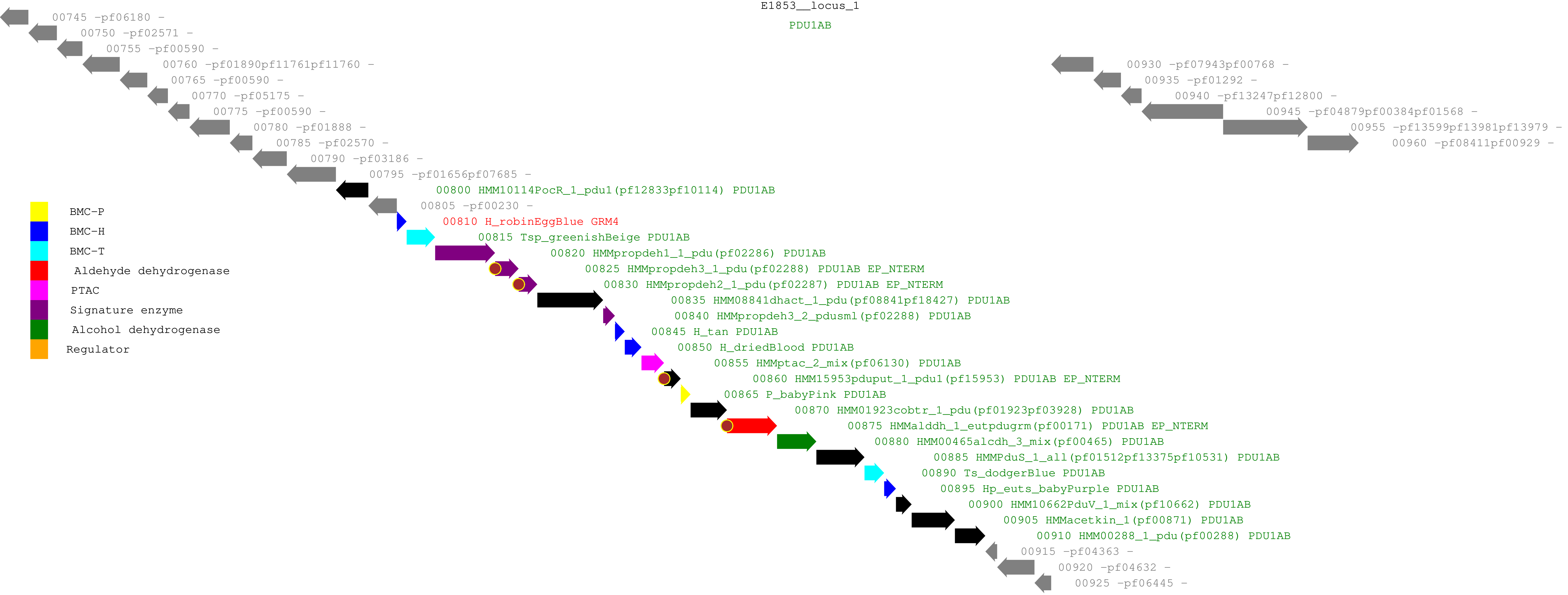

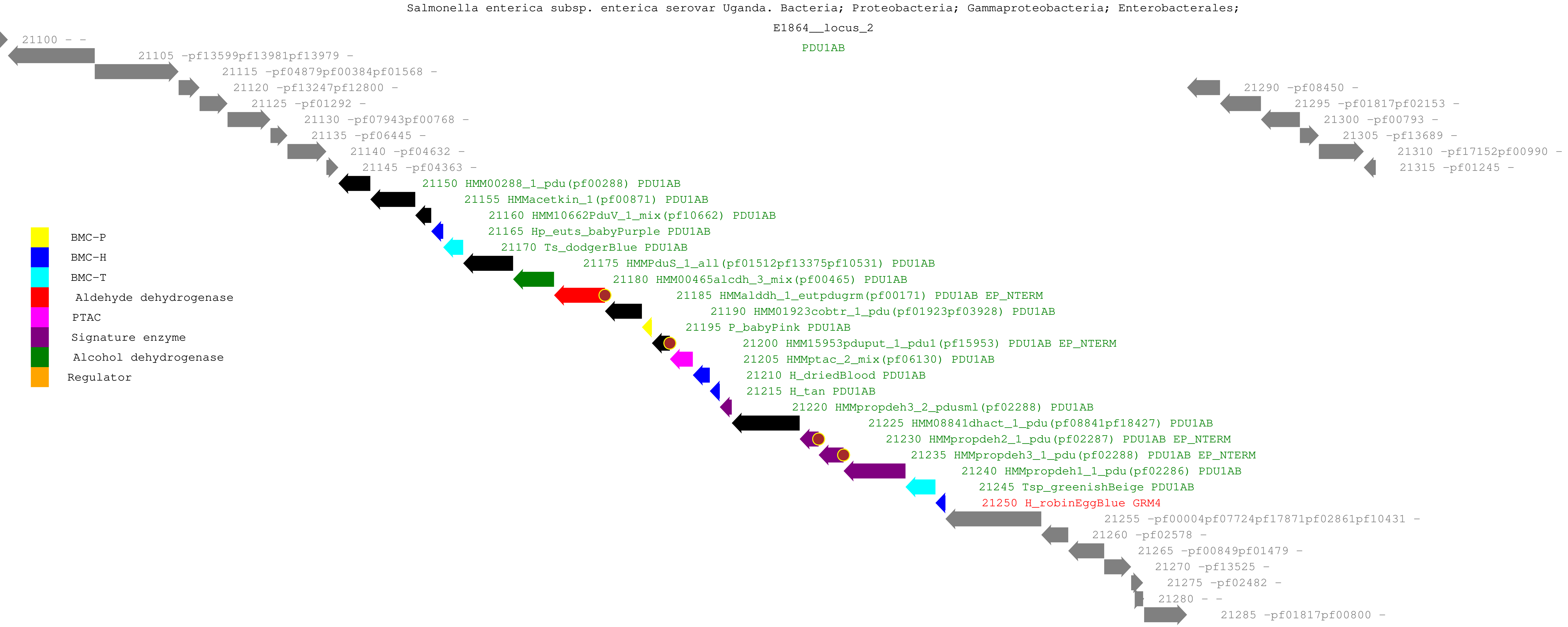

Salmonella enterica subsp. enterica serovar Uganda. Bacteria; Proteobacteria; Gammaproteobacteria; Enterobacterales;
E1864__locus_2
PDU1AB
21100 - -
21105 -pf13599pf13981pf13979 -
21115 -pf04879pf00384pf01568 -
21120 -pf13247pf12800 -
21125 -pf01292 -
21130 -pf07943pf00768 -
21135 -pf06445 -
21140 -pf04632 -
21145 -pf04363 -
21150 HMM00288_1_pdu(pf00288) PDU1AB
21155 HMMacetkin_1(pf00871) PDU1AB
21160 HMM10662PduV_1_mix(pf10662) PDU1AB
21165 Hp_euts_babyPurple PDU1AB
21170 Ts_dodgerBlue PDU1AB
21175 HMMPduS_1_all(pf01512pf13375pf10531) PDU1AB
21180 HMM00465alcdh_3_mix(pf00465) PDU1AB
21185 HMMalddh_1_eutpdugrm(pf00171) PDU1AB EP_NTERM
21190 HMM01923cobtr_1_pdu(pf01923pf03928) PDU1AB
21195 P_babyPink PDU1AB
21200 HMM15953pduput_1_pdu1(pf15953) PDU1AB EP_NTERM
21205 HMMptac_2_mix(pf06130) PDU1AB
21210 H_driedBlood PDU1AB
21215 H_tan PDU1AB
21220 HMMpropdeh3_2_pdusml(pf02288) PDU1AB
21225 HMM08841dhact_1_pdu(pf08841pf18427) PDU1AB
21230 HMMpropdeh2_1_pdu(pf02287) PDU1AB EP_NTERM
21235 HMMpropdeh3_1_pdu(pf02288) PDU1AB EP_NTERM
21240 HMMpropdeh1_1_pdu(pf02286) PDU1AB
21245 Tsp_greenishBeige PDU1AB
21250 H_robinEggBlue GRM4
21255 -pf00004pf07724pf17871pf02861pf10431 -
21260 -pf02578 -
21265 -pf00849pf01479 -
21270 -pf13525 -
21275 -pf02482 -
21280 - -
21285 -pf01817pf00800 -
21290 -pf08450 -
21295 -pf01817pf02153 -
21300 -pf00793 -
21305 -pf13689 -
21310 -pf17152pf00990 -
21315 -pf01245 -
BMC-P
BMC-H
BMC-T
Aldehyde dehydrogenase
PTAC
Signature enzyme
Alcohol dehydrogenase
Regulator

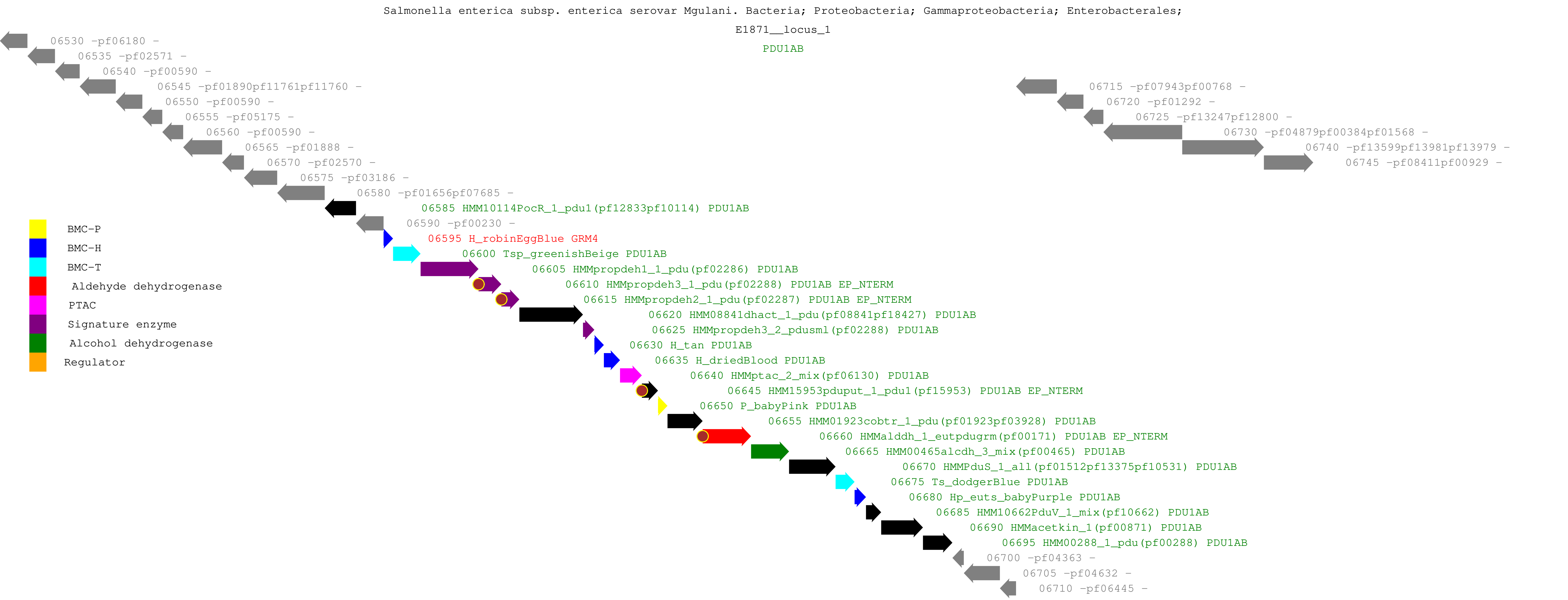
Salmonella enterica subsp. enterica serovar Mgulani. Bacteria; Proteobacteria; Gammaproteobacteria; Enterobacterales;
E1871__locus_1
PDU1AB
06530 -pf06180 -
06535 -pf02571 -
06540 -pf00590 -
06545 -pf01890pf11761pf11760 -
06550 -pf00590 -
06555 -pf05175 -
06560 -pf00590 -
06565 -pf01888 -
06570 -pf02570 -
06575 -pf03186 -
06580 -pf01656pf07685 -
06585 HMM10114PocR_1_pdu1(pf12833pf10114) PDU1AB
06590 -pf00230 -
06595 H_robinEggBlue GRM4
06600 Tsp_greenishBeige PDU1AB
06605 HMMpropdeh1_1_pdu(pf02286) PDU1AB
06610 HMMpropdeh3_1_pdu(pf02288) PDU1AB EP_NTERM
06615 HMMpropdeh2_1_pdu(pf02287) PDU1AB EP_NTERM
06620 HMM08841dhact_1_pdu(pf08841pf18427) PDU1AB
06625 HMMpropdeh3_2_pdusml(pf02288) PDU1AB
06630 H_tan PDU1AB
06635 H_driedBlood PDU1AB
06640 HMMptac_2_mix(pf06130) PDU1AB
06645 HMM15953pduput_1_pdu1(pf15953) PDU1AB EP_NTERM
06650 P_babyPink PDU1AB
06655 HMM01923cobtr_1_pdu(pf01923pf03928) PDU1AB
06660 HMMalddh_1_eutpdugrm(pf00171) PDU1AB EP_NTERM
06665 HMM00465alcdh_3_mix(pf00465) PDU1AB
06670 HMMPduS_1_all(pf01512pf13375pf10531) PDU1AB
06675 Ts_dodgerBlue PDU1AB
06680 Hp_euts_babyPurple PDU1AB
06685 HMM10662PduV_1_mix(pf10662) PDU1AB
06690 HMMacetkin_1(pf00871) PDU1AB
06695 HMM00288_1_pdu(pf00288) PDU1AB
06700 -pf04363 -
06705 -pf04632 -
06710 -pf06445 -
06715 -pf07943pf00768 -
06720 -pf01292 -
06725 -pf13247pf12800 -
06730 -pf04879pf00384pf01568 -
06740 -pf13599pf13981pf13979 -
06745 -pf08411pf00929 -
BMC-P
BMC-H
BMC-T
Aldehyde dehydrogenase
PTAC
Signature enzyme
Alcohol dehydrogenase
Regulator

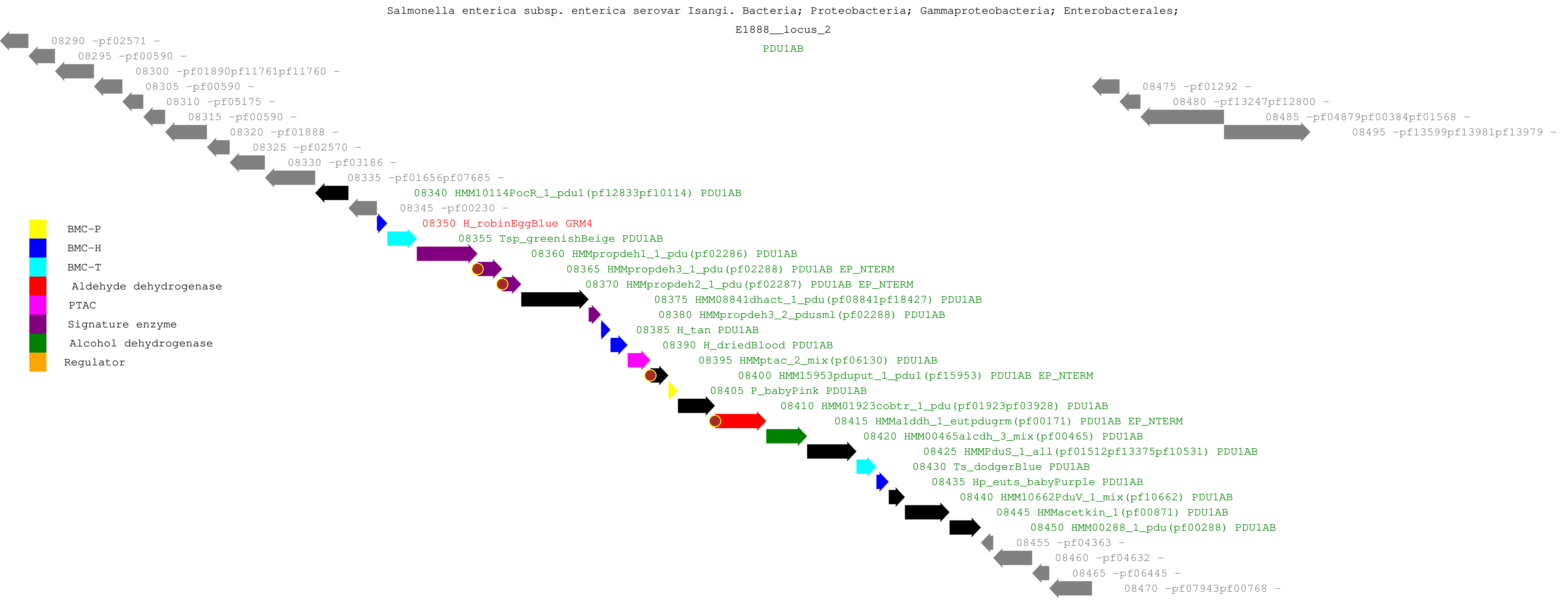
Salmonella enterica subsp. enterica serovar Isangi. Bacteria; Proteobacteria; Gammaproteobacteria; Enterobacterales;
E1888__locus_2
PDU1AB
08290 -pf02571 -
08295 -pf00590 -
08300 -pf01890pf11761pf11760 -
08305 -pf00590 -
08310 -pf05175 -
08315 -pf00590 -
08320 -pf01888 -
08325 -pf02570 -
08330 -pf03186 -
08335 -pf01656pf07685 -
08340 HMM10114PocR_1_pdu1(pf12833pf10114) PDU1AB
08345 -pf00230 -
08350 H_robinEggBlue GRM4
08355 Tsp_greenishBeige PDU1AB
08360 HMMpropdeh1_1_pdu(pf02286) PDU1AB
08365 HMMpropdeh3_1_pdu(pf02288) PDU1AB EP_NTERM
08370 HMMpropdeh2_1_pdu(pf02287) PDU1AB EP_NTERM
08375 HMM08841dhact_1_pdu(pf08841pf18427) PDU1AB
08380 HMMpropdeh3_2_pdusml(pf02288) PDU1AB
08385 H_tan PDU1AB
08390 H_driedBlood PDU1AB
08395 HMMptac_2_mix(pf06130) PDU1AB
08400 HMM15953pduput_1_pdu1(pf15953) PDU1AB EP_NTERM
08405 P_babyPink PDU1AB
08410 HMM01923cobtr_1_pdu(pf01923pf03928) PDU1AB
08415 HMMaldh_1_eutpdugrm(pf00171) PDU1AB EP_NTERM
08420 HMM00465alcdh_3_mix(pf00465) PDU1AB
08425 HMMPduS_1_all(pf01512pf13375pf10531) PDU1AB
08430 Ts_dodgerBlue PDU1AB
08435 Hp_euts_babyPurple PDU1AB
08440 HMM10662PduV_1_mix(pf10662) PDU1AB
08445 HMMacetkin_1(pf00871) PDU1AB
08450 HMM00288_1_pdu(pf00288) PDU1AB
08455 -pf04363 -
08460 -pf04632 -
08465 -pf06445 -
08470 -pf07943pf00768 -
08475 -pf01292 -
08480 -pf13247pf12800 -
08485 -pf04879pf00384pf01568 -
08495 -pf13599pf13981pf13979 -
BMC-P
BMC-H
BMC-T
Aldehyde dehydrogenase
PTAC
Signature enzyme
Alcohol dehydrogenase
Regulator

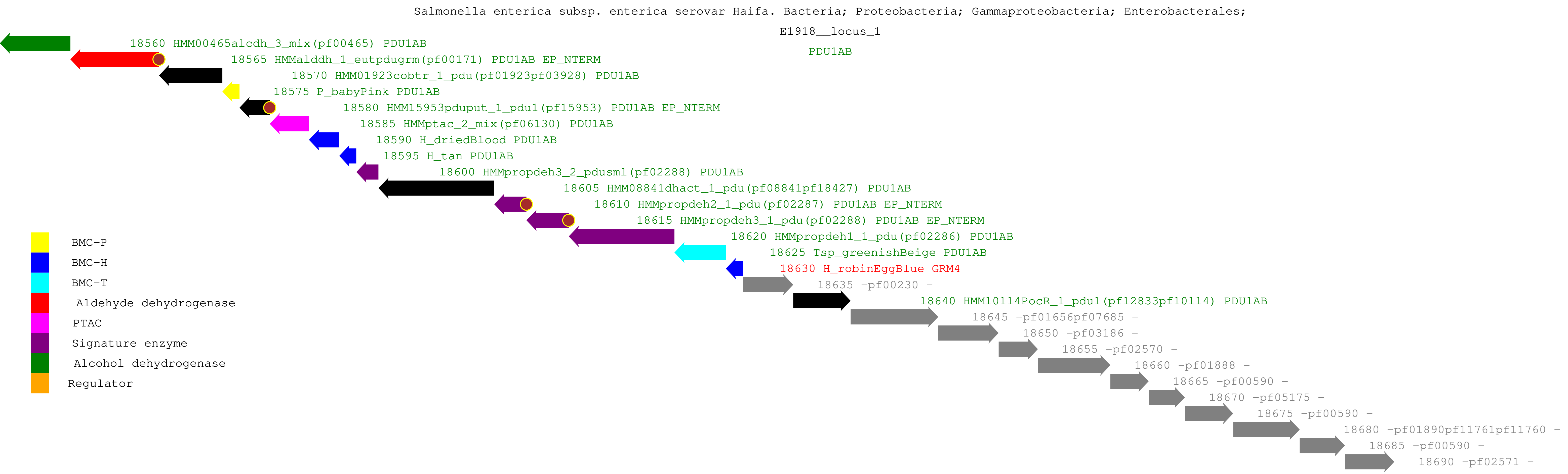
Salmonella enterica subsp. enterica serovar Haifa. Bacteria; Proteobacteria; Gammaproteobacteria; Enterobacterales;
E1918__locus_1
PDU1AB
18560 HMM00465alcdh_3_mix(pf00465) PDU1AB
18565 HMMalddh_1_eutpdugrm(pf00171) PDU1AB EP_NTERM
18570 HMM01923cobtr_1_pdu(pf01923pf03928) PDU1AB
18575 P_babyPink PDU1AB
18580 HMM15953pduput_1_pdu1(pf15953) PDU1AB EP_NTERM
18585 HMMptac_2_mix(pf06130) PDU1AB
18590 H_driedBlood PDU1AB
18595 H_tan PDU1AB
18600 HMMpropdeh3_2_pdusml(pf02288) PDU1AB
18605 HMM08841dhact_1_pdu(pf08841pf18427) PDU1AB
18610 HMMpropdeh2_1_pdu(pf02287) PDU1AB EP_NTERM
18615 HMMpropdeh3_1_pdu(pf02288) PDU1AB EP_NTERM
18620 HMMpropdeh1_1_pdu(pf02286) PDU1AB
18625 Tsp_greenishBeige PDU1AB
18630 H_robinEggBlue GRM4
18635 -pf00230 -
18640 HMM10114PocR_1_pdu1(pf12833pf10114) PDU1AB
18645 -pf01656pf07685 -
18650 -pf03186 -
18655 -pf02570 -
18660 -pf01888 -
18665 -pf00590 -
18670 -pf05175 -
18675 -pf00590 -
18680 -pf01890pf11761pf11760 -
18685 -pf00590 -
18690 -pf02571 -
BMC-P
BMC-H
BMC-T
Aldehyde dehydrogenase
PTAC
Signature enzyme
Alcohol dehydrogenase
Regulator

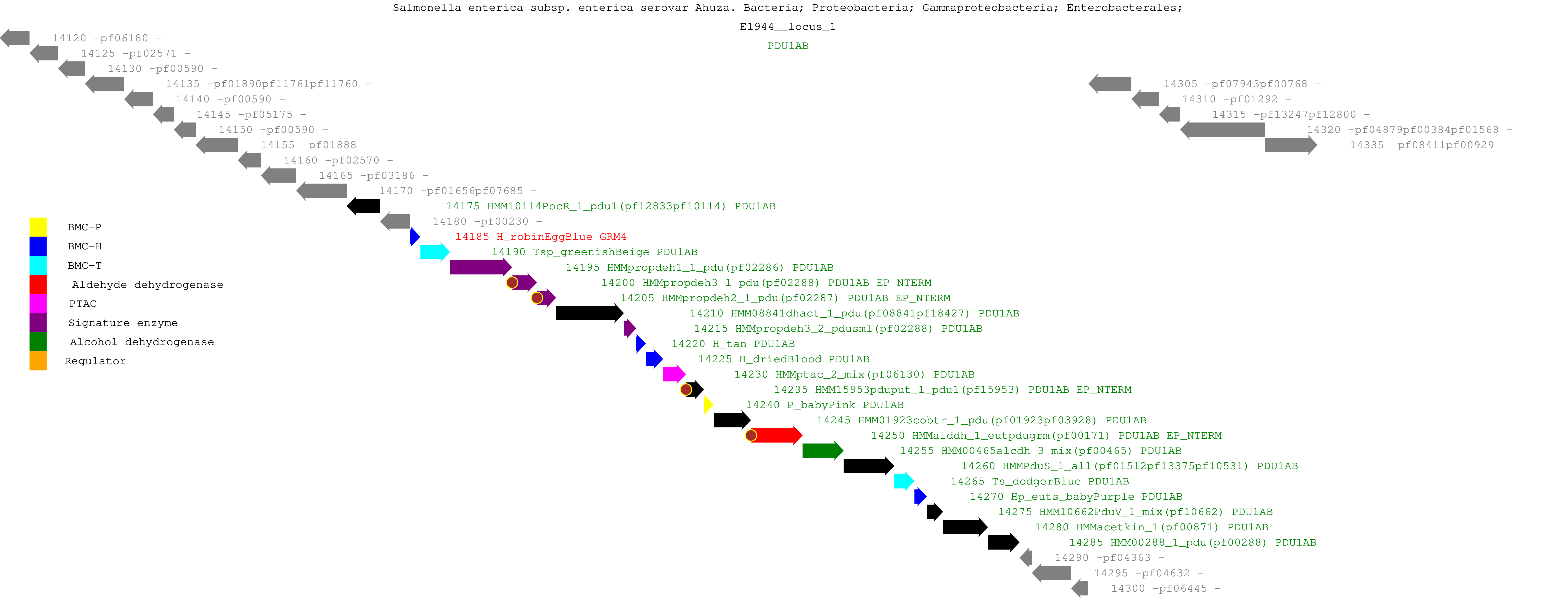

Salmonella enterica subsp. enterica serovar Ahuza. Bacteria; Proteobacteria; Gammaproteobacteria; Enterobacterales;
E1944__locus_1
PDU1AB
14120 -pf06180 -
14125 -pf02571 -
14130 -pf00590 -
14135 -pf01890pf11761pf11760 -
14140 -pf00590 -
14145 -pf05175 -
14150 -pf00590 -
14155 -pf01888 -
14160 -pf02570 -
14165 -pf03186 -
14170 -pf01656pf07685 -
14175 HMM10114PocR_1_pdu1(pf12833pf10114) PDU1AB
14180 -pf00230 -
14185 H_robinEggBlue GRM4
14190 Tsp_greenishBeige PDU1AB
14195 HMMpropdeh1_1_pdu(pf02286) PDU1AB
14200 HMMpropdeh3_1_pdu(pf02288) PDU1AB EP_NTERM
14205 HMMpropdeh2_1_pdu(pf02287) PDU1AB EP_NTERM
14210 HMM08841dhact_1_pdu(pf08841pf18427) PDU1AB
14215 HMMpropdeh3_2_pdusml(pf02288) PDU1AB
14220 H_tan PDU1AB
14225 H_driedBlood PDU1AB
14230 HMMptac_2_mix(pf06130) PDU1AB
14235 HMM15953pduput_1_pdu1(pf15953) PDU1AB EP_NTERM
14240 P_babyPink PDU1AB
14245 HMM01923cobtr_1_pdu(pf01923pf03928) PDU1AB
14250 HMMalddh_1_eutpdugrm(pf00171) PDU1AB EP_NTERM
14255 HMM00465alcdh_3_mix(pf00465) PDU1AB
14260 HMMPduS_1_all(pf01512pf13375pf10531) PDU1AB
14265 Ts_dodgerBlue PDU1AB
14270 Hp_euts_babyPurple PDU1AB
14275 HMM10662PduV_1_mix(pf10662) PDU1AB
14280 HMMacetkin_1(pf00871) PDU1AB
14285 HMM00288_1_pdu(pf00288) PDU1AB
14290 -pf04363 -
14295 -pf04632 -
14300 -pf06445 -
14305 -pf07943pf00768 -
14310 -pf01292 -
14315 -pf13247pf12800 -
14320 -pf04879pf00384pf01568 -
14335 -pf08411pf00929 -
BMC-P
BMC-H
BMC-T
Aldehyde dehydrogenase
PTAC
Signature enzyme
Alcohol dehydrogenase
Regulator

Salmonella enterica subsp. enterica serovar Cubana. Bacteria; Proteobacteria; Gammaproteobacteria; Enterobacterales;

E1A24__locus_1

PDU1AB

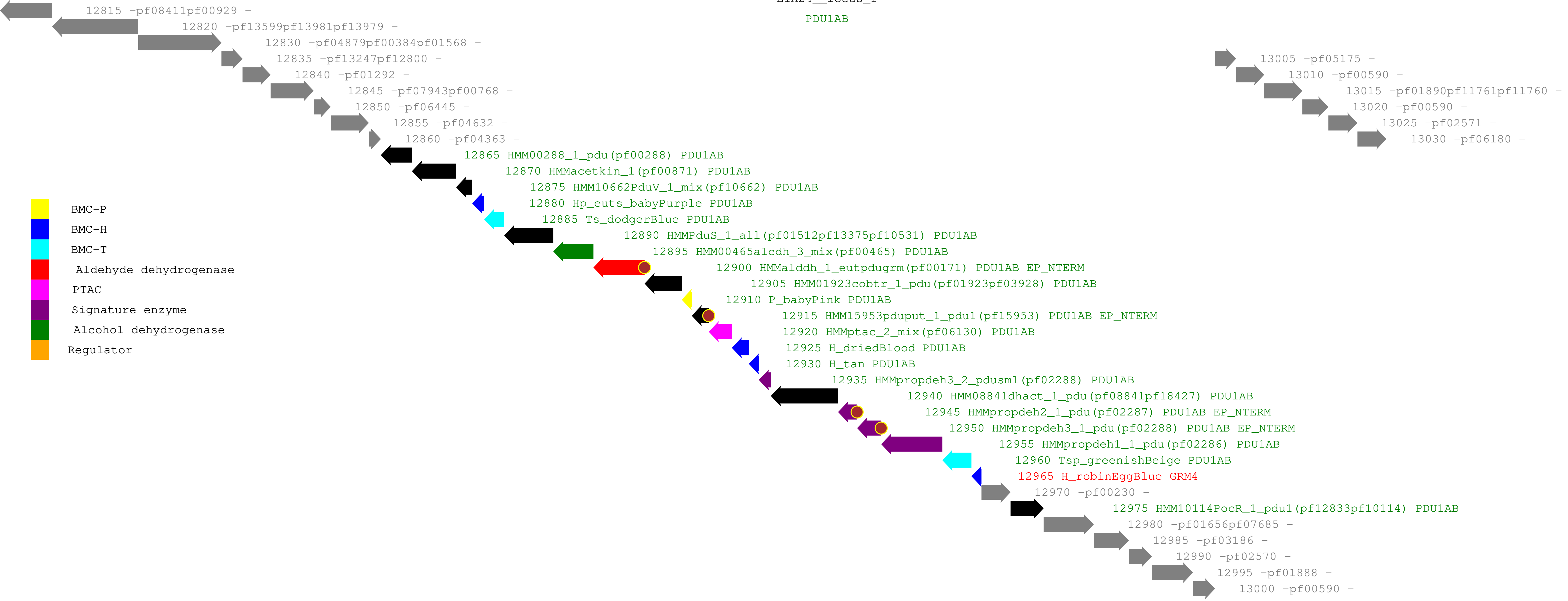

Citrobacter sp. LY-1. Bacteria; Proteobacteria; Gammaproteobacteria; Enterobacterales;

E1B03__locus_1

PDU1AB

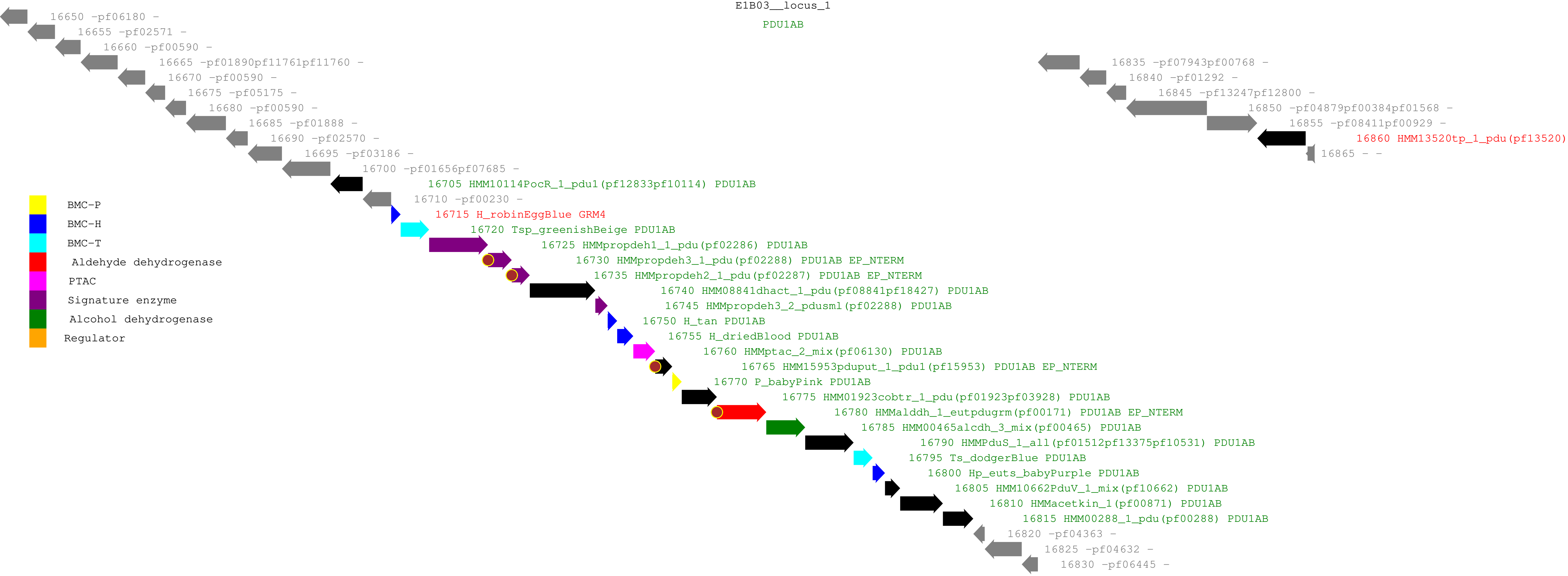

Salmonella moscow. Bacteria; Proteobacteria; Gammaproteobacteria; Enterobacterales;

E1B69__locus_2

PDU1AB

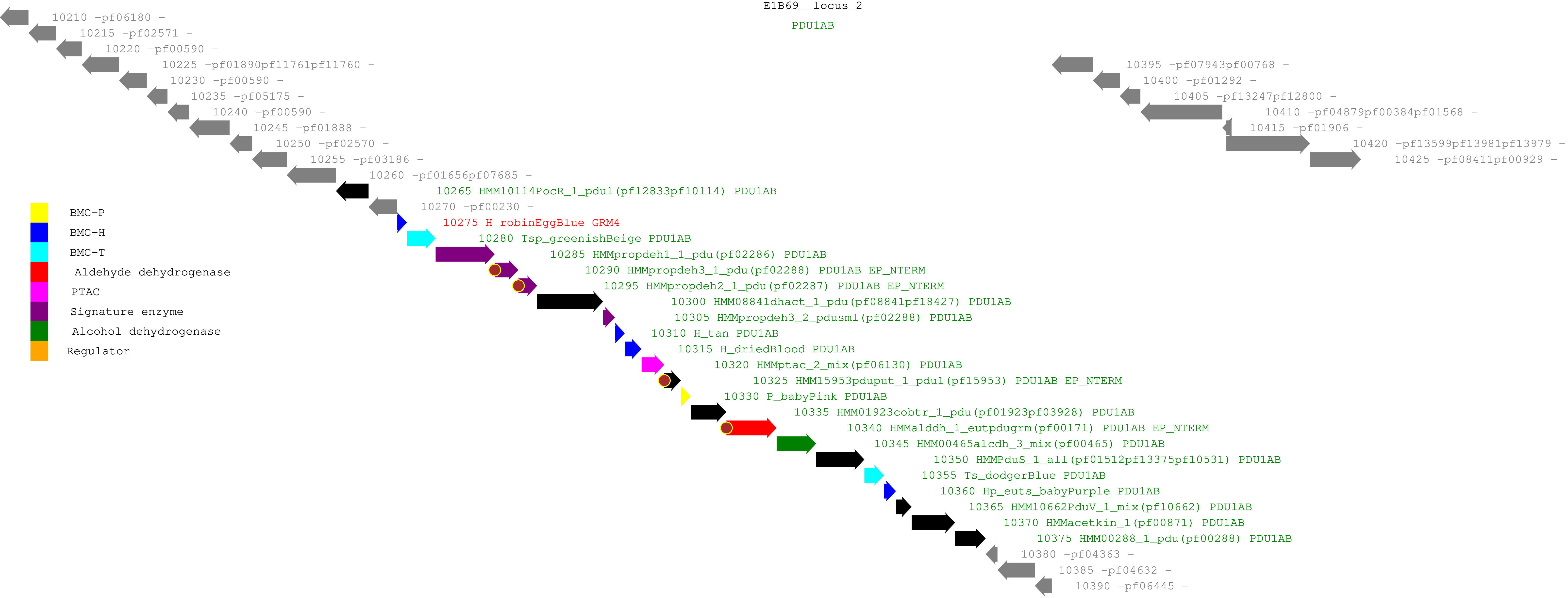

Salmonella enterica subsp. enterica serovar Bonariensis. Bacteria; Proteobacteria; Gammaproteobacteria; Enterobacterales;

E1C95__locus_1

PDU1AB

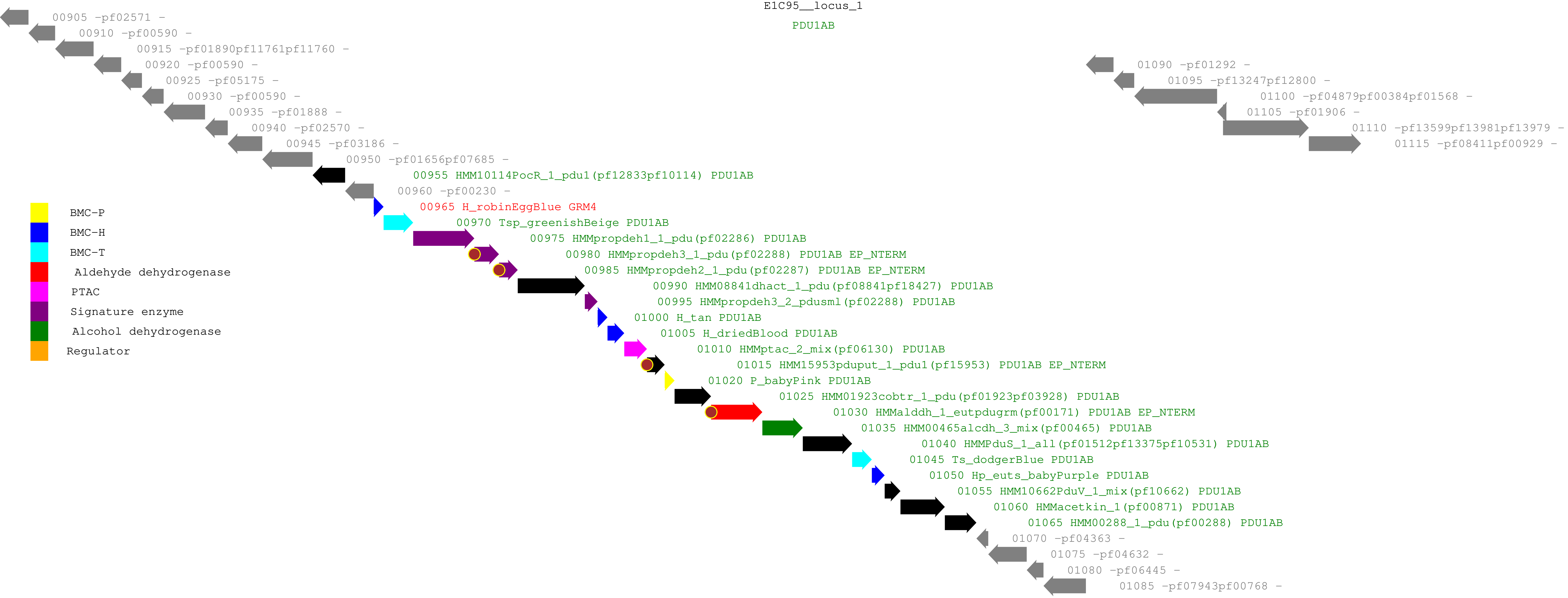

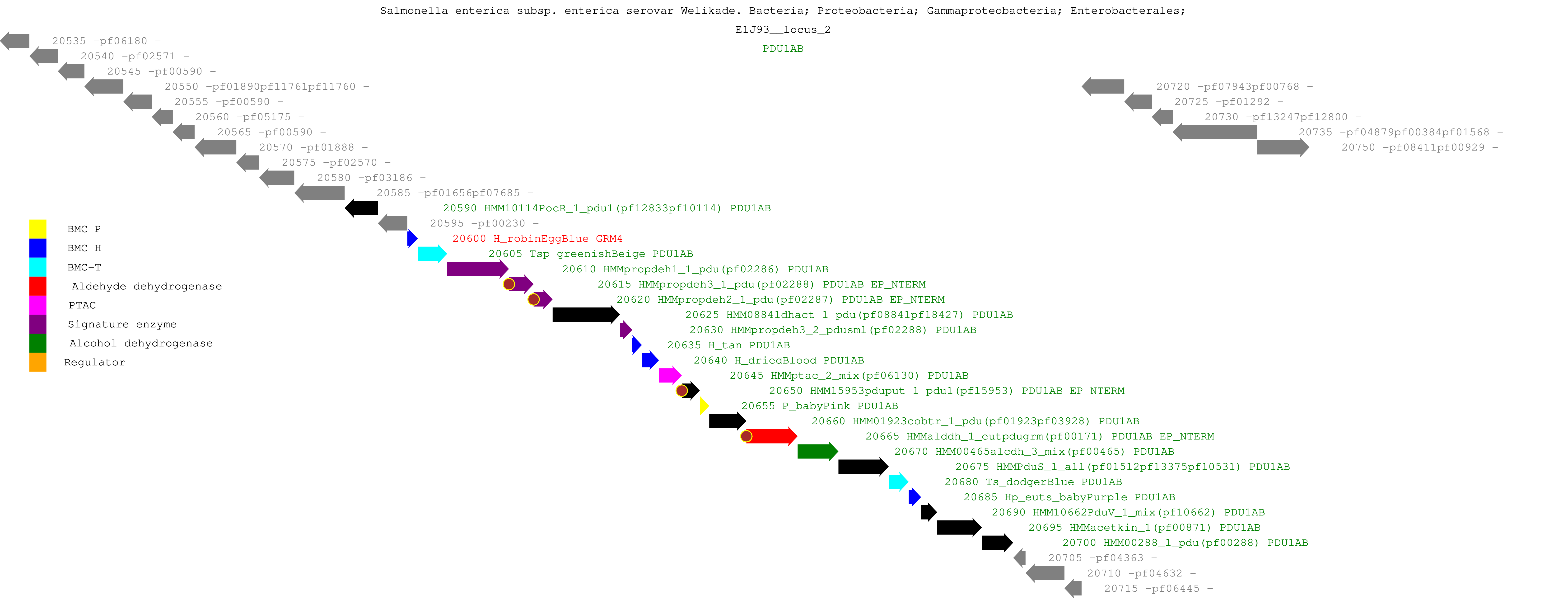

Salmonella enterica subsp. enterica serovar Welikade. Bacteria; Proteobacteria; Gammaproteobacteria; Enterobacterales;
E1J93__locus_2
PDU1AB
20535 -pf06180 -
20540 -pf02571 -
20545 -pf00590 -
20550 -pf01890pf11761pf11760 -
20555 -pf00590 -
20560 -pf05175 -
20565 -pf00590 -
20570 -pf01888 -
20575 -pf02570 -
20580 -pf03186 -
20585 -pf01656pf07685 -
20590 HMM10114PocR_1_pdu1(pf12833pf10114) PDU1AB
20595 -pf00230 -
20600 H_robinEggBlue GRM4
20605 Tsp_greenishBeige PDU1AB
20610 HMMpropdeh1_1_pdu(pf02286) PDU1AB
20615 HMMpropdeh3_1_pdu(pf02288) PDU1AB EP_NTERM
20620 HMMpropdeh2_1_pdu(pf02287) PDU1AB EP_NTERM
20625 HMM08841dhact_1_pdu(pf08841pf18427) PDU1AB
20630 HMMpropdeh3_2_pdusml(pf02288) PDU1AB
20635 H_tan PDU1AB
20640 H_driedBlood PDU1AB
20645 HMMptac_2_mix(pf06130) PDU1AB
20650 HMM15953pduput_1_pdu1(pf15953) PDU1AB EP_NTERM
20655 P_babyPink PDU1AB
20660 HMM01923cobtr_1_pdu(pf01923pf03928) PDU1AB
20665 HMMalddh_1_eutpdugrm(pf00171) PDU1AB EP_NTERM
20670 HMM00465alcdh_3_mix(pf00465) PDU1AB
20675 HMMPduS_1_all(pf01512pf13375pf10531) PDU1AB
20680 Ts_dodgerBlue PDU1AB
20685 Hp_euts_babyPurple PDU1AB
20690 HMM10662PduV_1_mix(pf10662) PDU1AB
20695 HMMacetkin_1(pf00871) PDU1AB
20700 HMM00288_1_pdu(pf00288) PDU1AB
20705 -pf04363 -
20710 -pf04632 -
20715 -pf06445 -
20720 -pf07943pf00768 -
20725 -pf01292 -
20730 -pf13247pf12800 -
20735 -pf04879pf00384pf01568 -
20750 -pf08411pf00929 -
BMC-P
BMC-H
BMC-T
Aldehyde dehydrogenase
PTAC
Signature enzyme
Alcohol dehydrogenase
Regulator

Salmonella enterica subsp. enterica serovar Amager. Bacteria; Proteobacteria; Gammaproteobacteria; Enterobacterales;

E1K22__locus_1

PDU1AB

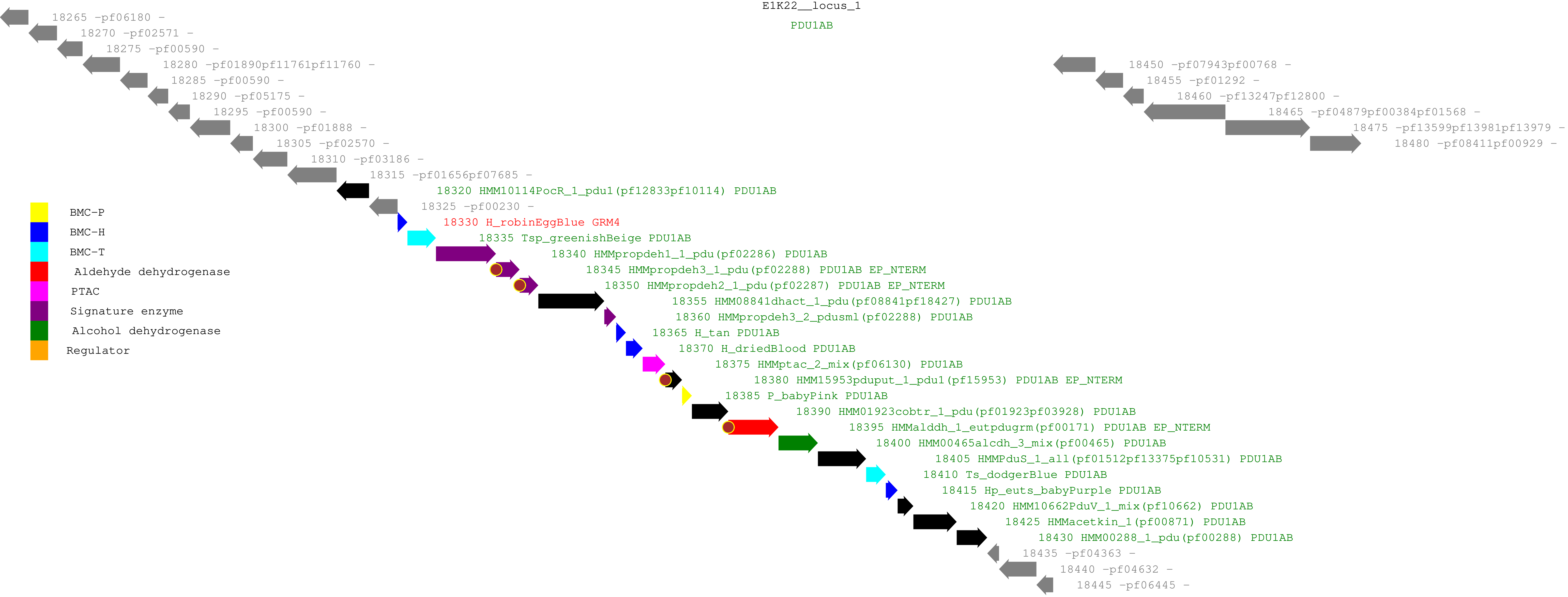

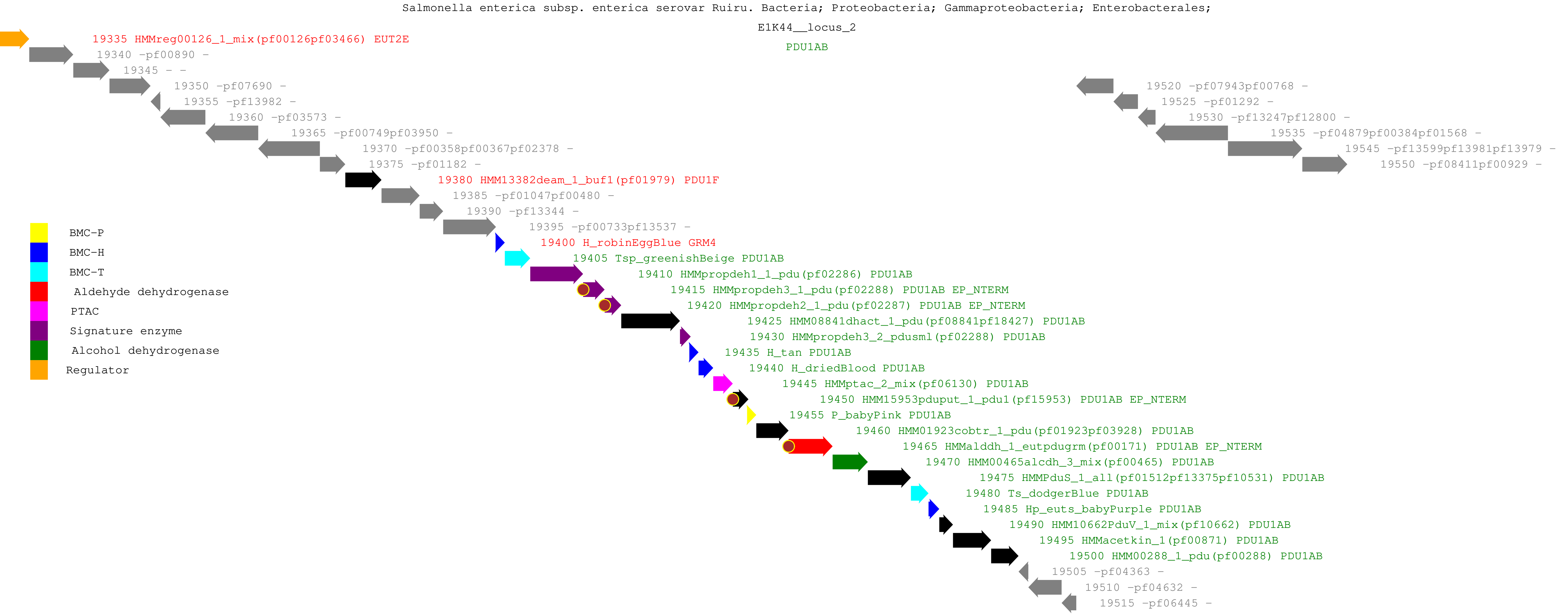

Salmonella enterica subsp. enterica serovar Ruiru. Bacteria; Proteobacteria; Gammaproteobacteria; Enterobacterales;
E1K44__locus_2
PDU1AB
19335 HMMreg00126_1_mix(pf00126pf03466) EUT2E
19340 -pf00890 -
19345 - -
19350 -pf07690 -
19355 -pf13982 -
19360 -pf03573 -
19365 -pf00749pf03950 -
19370 -pf00358pf00367pf02378 -
19375 -pf01182 -
19380 HMM13382deam_1_buf1(pf01979) PDU1F
19385 -pf01047pf00480 -
19390 -pf13344 -
19395 -pf00733pf13537 -
19400 H_robinEggBlue GRM4
19405 Tsp_greenishBeige PDU1AB
19410 HMMpropdeh1_1_pdu(pf02286) PDU1AB
19415 HMMpropdeh3_1_pdu(pf02288) PDU1AB EP_NTERM
19420 HMMpropdeh2_1_pdu(pf02287) PDU1AB EP_NTERM
19425 HMM08841dhact_1_pdu(pf08841pf18427) PDU1AB
19430 HMMpropdeh3_2_pdusml(pf02288) PDU1AB
19435 H_tan PDU1AB
19440 H_driedBlood PDU1AB
19445 HMMptac_2_mix(pf06130) PDU1AB
19450 HMM15953pduput_1_pdu1(pf15953) PDU1AB EP_NTERM
19455 P_babyPink PDU1AB
19460 HMM01923cobtr_1_pdu(pf01923pf03928) PDU1AB
19465 HMMalddh_1_eutpdugrm(pf00171) PDU1AB EP_NTERM
19470 HMM00465alcdh_3_mix(pf00465) PDU1AB
19475 HMMPduS_1_all(pf01512pf13375pf10531) PDU1AB
19480 Ts_dodgerBlue PDU1AB
19485 Hp_euts_babyPurple PDU1AB
19490 HMM10662PduV_1_mix(pf10662) PDU1AB
19495 HMMacetkin_1(pf00871) PDU1AB
19500 HMM00288_1_pdu(pf00288) PDU1AB
19505 -pf04363 -
19510 -pf04632 -
19515 -pf06445 -
19520 -pf07943pf00768 -
19525 -pf01292 -
19530 -pf13247pf12800 -
19535 -pf04879pf00384pf01568 -
19545 -pf13599pf13981pf13979 -
19550 -pf08411pf00929 -
BMC-P
BMC-H
BMC-T
Aldehyde dehydrogenase
PTAC
Signature enzyme
Alcohol dehydrogenase
Regulator

Salmonella enterica subsp. enterica serovar Coleypark. Bacteria; Proteobacteria; Gammaproteobacteria; Enterobacterales;

E1K87__locus_2

PDU1AB

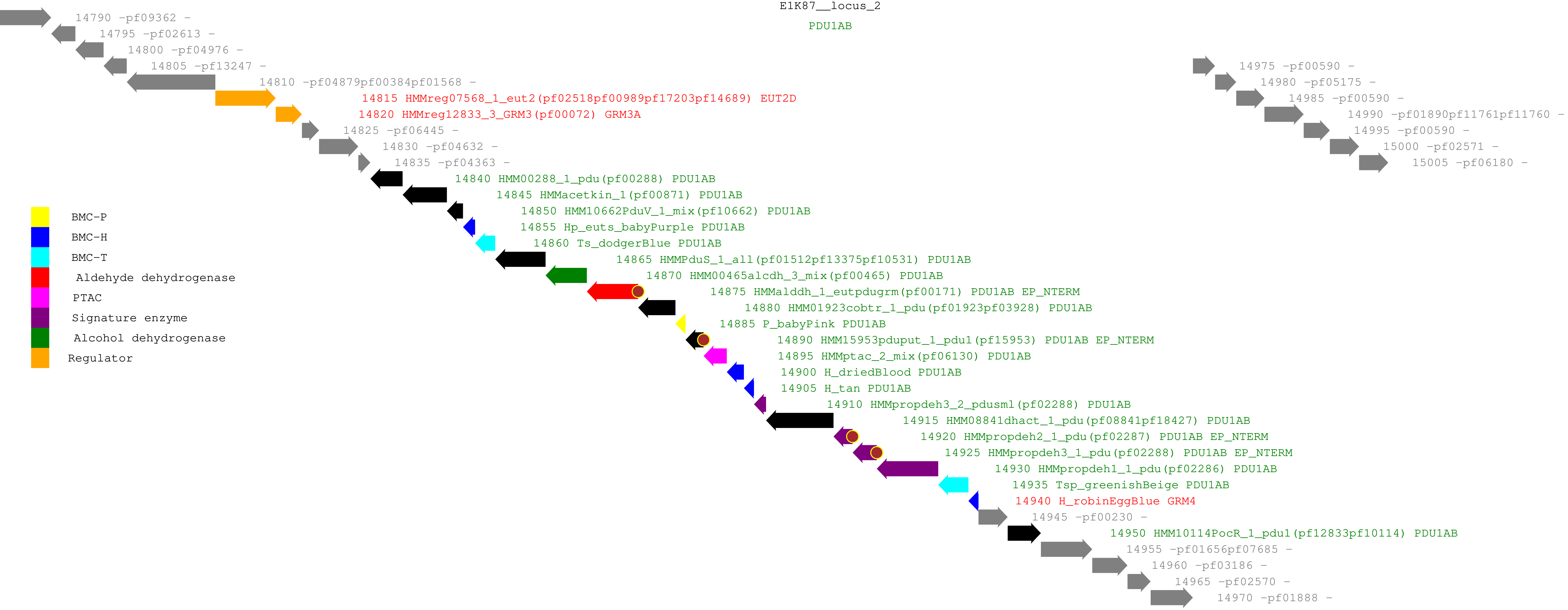

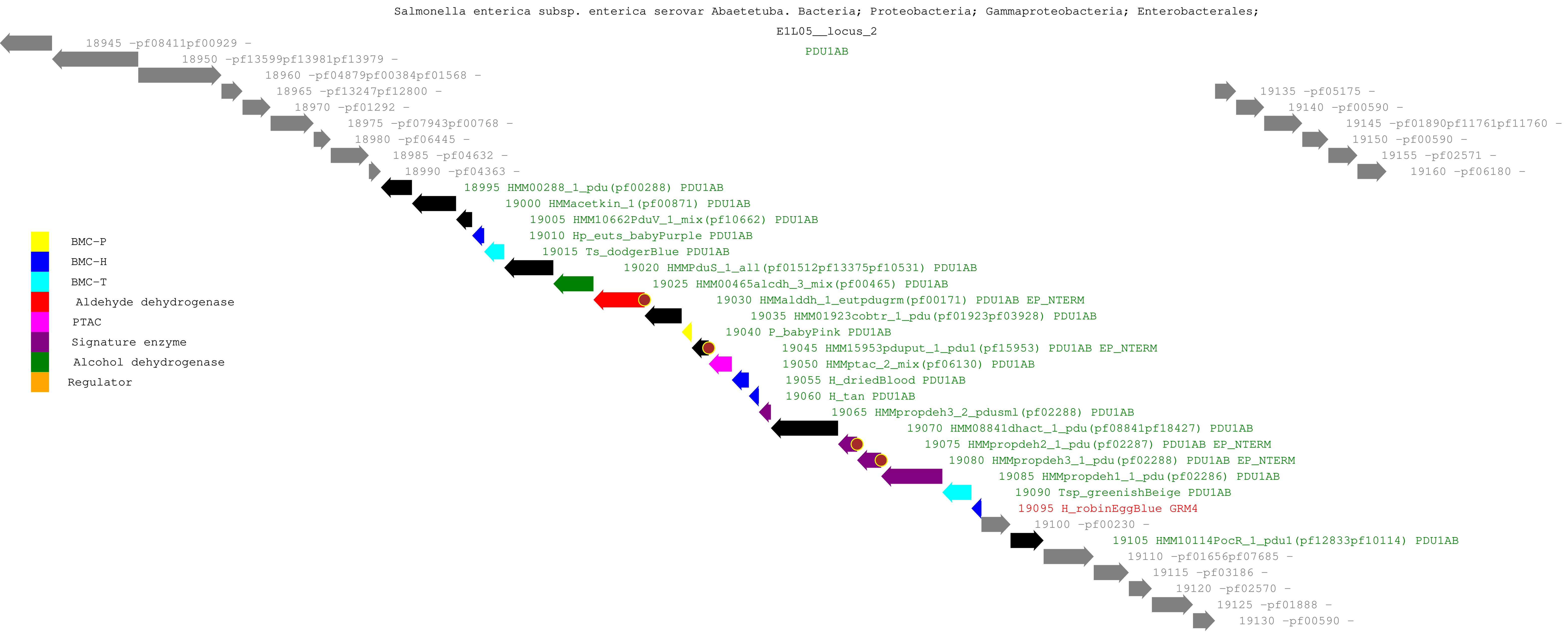

Salmonella enterica subsp. enterica serovar Abaetetuba. Bacteria; Proteobacteria; Gammaproteobacteria; Enterobacterales;
E1L05__locus_2
PDU1AB
18945 -pf08411pf00929 -
18950 -pf13599pf13981pf13979 -
18960 -pf04879pf00384pf01568 -
18965 -pf13247pf12800 -
18970 -pf01292 -
18975 -pf07943pf00768 -
18980 -pf06445 -
18985 -pf04632 -
18990 -pf04363 -
18995 HMM00288_1_pdu(pf00288) PDU1AB
19000 HMMacetkin_1(pf00871) PDU1AB
19005 HMM10662PduV_1_mix(pf10662) PDU1AB
19010 Hp_euts_babyPurple PDU1AB
19015 Ts_dodgerBlue PDU1AB
19020 HMMPduS_1_all(pf01512pf13375pf10531) PDU1AB
19025 HMM00465alcdh_3_mix(pf00465) PDU1AB
19030 HMMalddh_1_eutpdugrm(pf00171) PDU1AB EP_NTERM
19035 HMM01923cobtr_1_pdu(pf01923pf03928) PDU1AB
19040 P_babyPink PDU1AB
19045 HMM15953pduput_1_pdu1(pf15953) PDU1AB EP_NTERM
19050 HMMptac_2_mix(pf06130) PDU1AB
19055 H_driedBlood PDU1AB
19060 H_tan PDU1AB
19065 HMMpropdeh3_2_pdusml(pf02288) PDU1AB
19070 HMM08841dhact_1_pdu(pf08841pf18427) PDU1AB
19075 HMMpropdeh2_1_pdu(pf02287) PDU1AB EP_NTERM
19080 HMMpropdeh3_1_pdu(pf02288) PDU1AB EP_NTERM
19085 HMMpropdeh1_1_pdu(pf02286) PDU1AB
19090 Tsp_greenishBeige PDU1AB
19095 H_robinEggBlue GRM4
19100 -pf00230 -
19105 HMM10114PocR_1_pdu1(pf12833pf10114) PDU1AB
19110 -pf01656pf07685 -
19115 -pf03186 -
19120 -pf02570 -
19125 -pf01888 -
19130 -pf00590 -
19135 -pf05175 -
19140 -pf00590 -
19145 -pf01890pf11761pf11760 -
19150 -pf00590 -
19155 -pf02571 -
19160 -pf06180 -
BMC-P
BMC-H
BMC-T
Aldehyde dehydrogenase
PTAC
Signature enzyme
Alcohol dehydrogenase
Regulator

Salmonella enterica subsp. enterica serovar Everleigh. Bacteria; Proteobacteria; Gammaproteobacteria; Enterobacterales;

E1L13__locus_1

PDU1AB

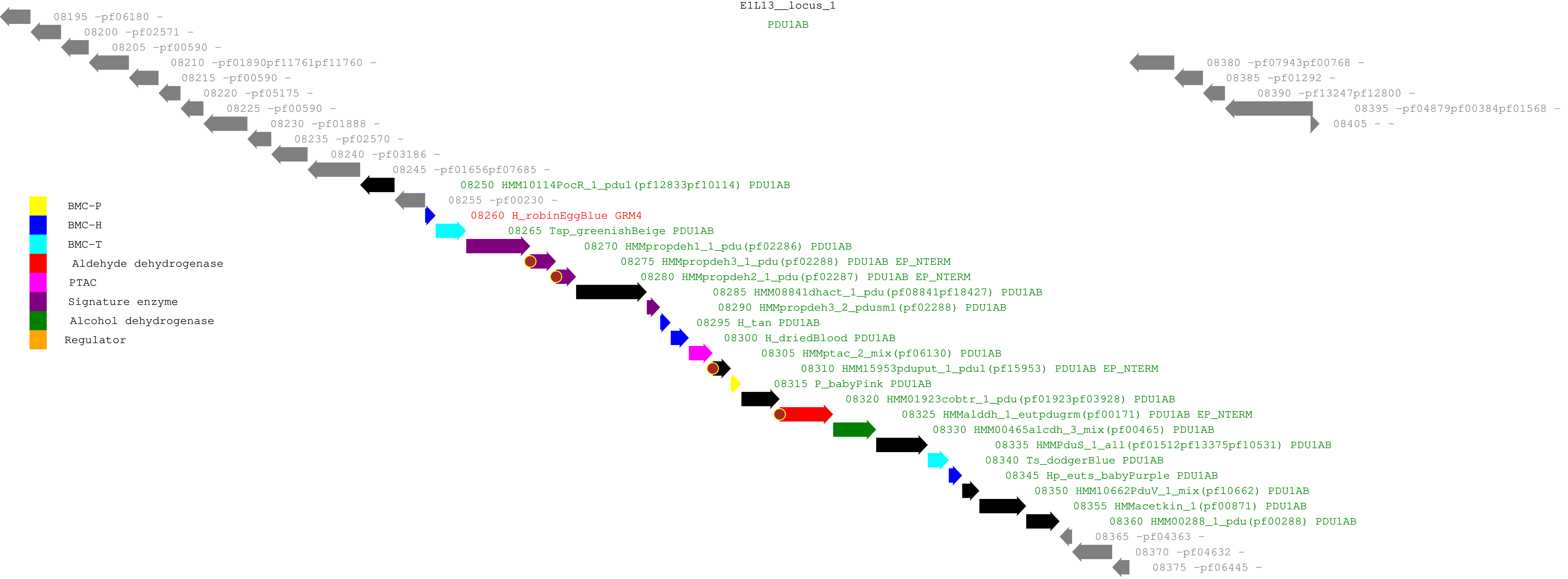

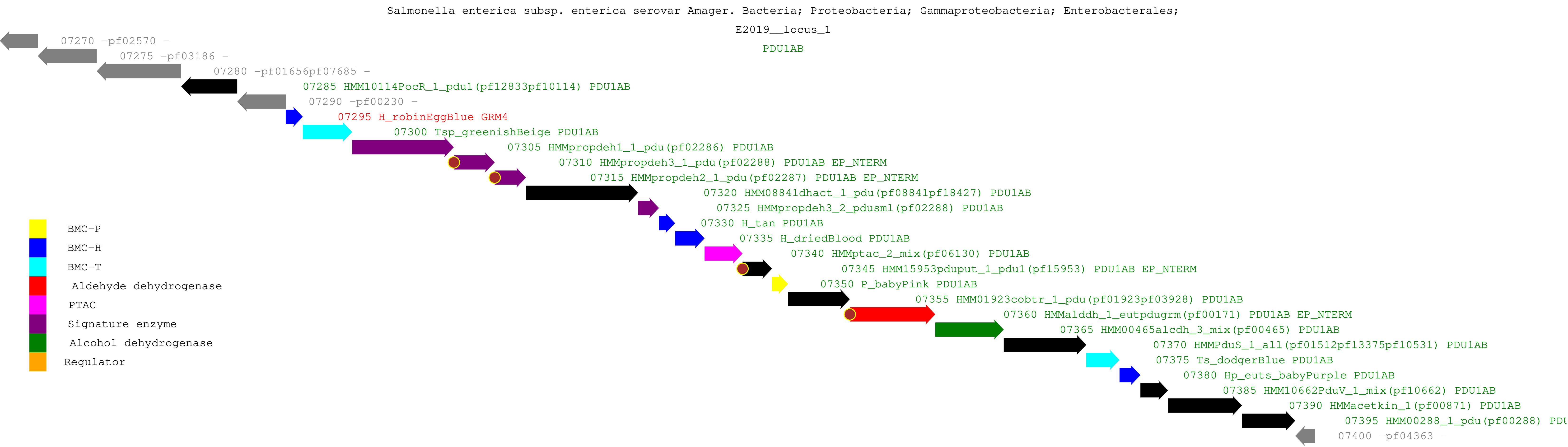

Salmonella enterica subsp. enterica serovar Amager. Bacteria; Proteobacteria; Gammaproteobacteria; Enterobacterales;
E2019__locus_1
PDU1AB
07270 -pf02570 -
07275 -pf03186 -
07280 -pf01656pf07685 -
07285 HMM10114PocR_1_pdu1(pf12833pf10114) PDU1AB
07290 -pf00230 -
07295 H_robinEggBlue GRM4
07300 Tsp_greenishBeige PDU1AB
07305 HMMpropdeh1_1_pdu(pf02286) PDU1AB
07310 HMMpropdeh3_1_pdu(pf02288) PDU1AB EP_NTERM
07315 HMMpropdeh2_1_pdu(pf02287) PDU1AB EP_NTERM
07320 HMM08841dhact_1_pdu(pf08841pf18427) PDU1AB
07325 HMMpropdeh3_2_pdusml(pf02288) PDU1AB
07330 H_tan PDU1AB
07335 H_driedBlood PDU1AB
07340 HMMptac_2_mix(pf06130) PDU1AB
07345 HMM15953pduput_1_pdu1(pf15953) PDU1AB EP_NTERM
07350 P_babyPink PDU1AB
07355 HMM01923cobtr_1_pdu(pf01923pf03928) PDU1AB
07360 HMMalddh_1_eutpdugrm(pf00171) PDU1AB EP_NTERM
07365 HMM00465alcdh_3_mix(pf00465) PDU1AB
07370 HMMPduS_1_all(pf01512pf13375pf10531) PDU1AB
07375 Ts_dodgerBlue PDU1AB
07380 Hp_euts_babyPurple PDU1AB
07385 HMM10662PduV_1_mix(pf10662) PDU1AB
07390 HMMacetkin_1(pf00871) PDU1AB
07395 HMM00288_1_pdu(pf00288) PDU
07400 -pf04363 -
BMC-P
BMC-H
BMC-T
Aldehyde dehydrogenase
PTAC
Signature enzyme
Alcohol dehydrogenase
Regulator

Salmonella oranienberg. Bacteria; Proteobacteria; Gammaproteobacteria; Enterobacterales;

E2040__locus_1

PDU1AB

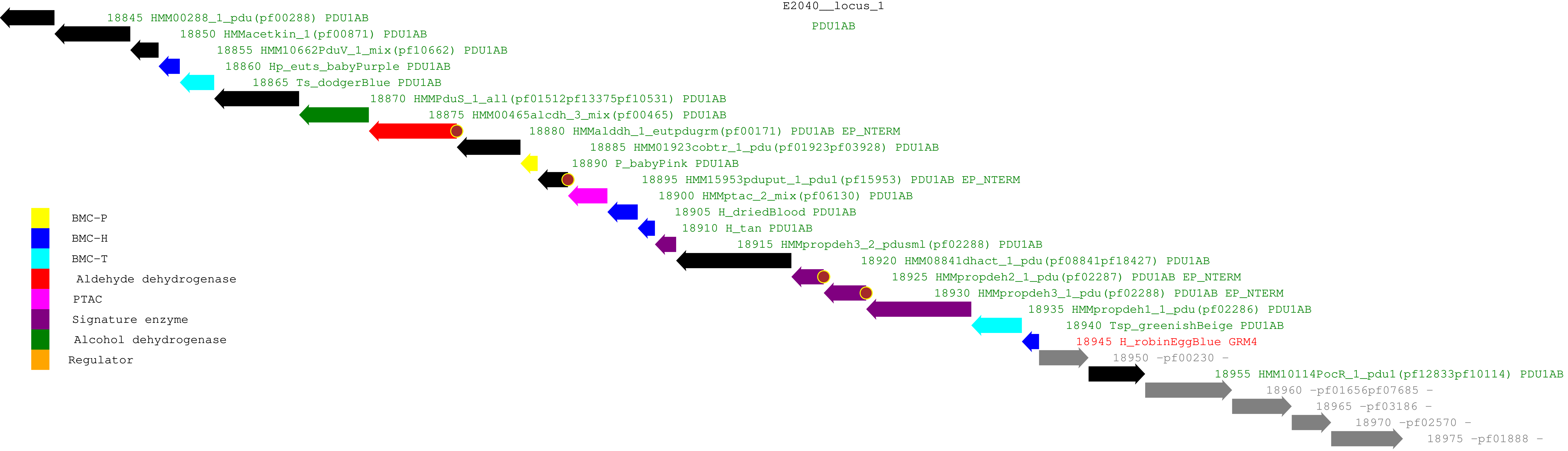

Salmonella enterica subsp. enterica serovar Wien. Bacteria; Proteobacteria; Gammaproteobacteria; Enterobacterales;

E2332__locus_1

PDU1AB

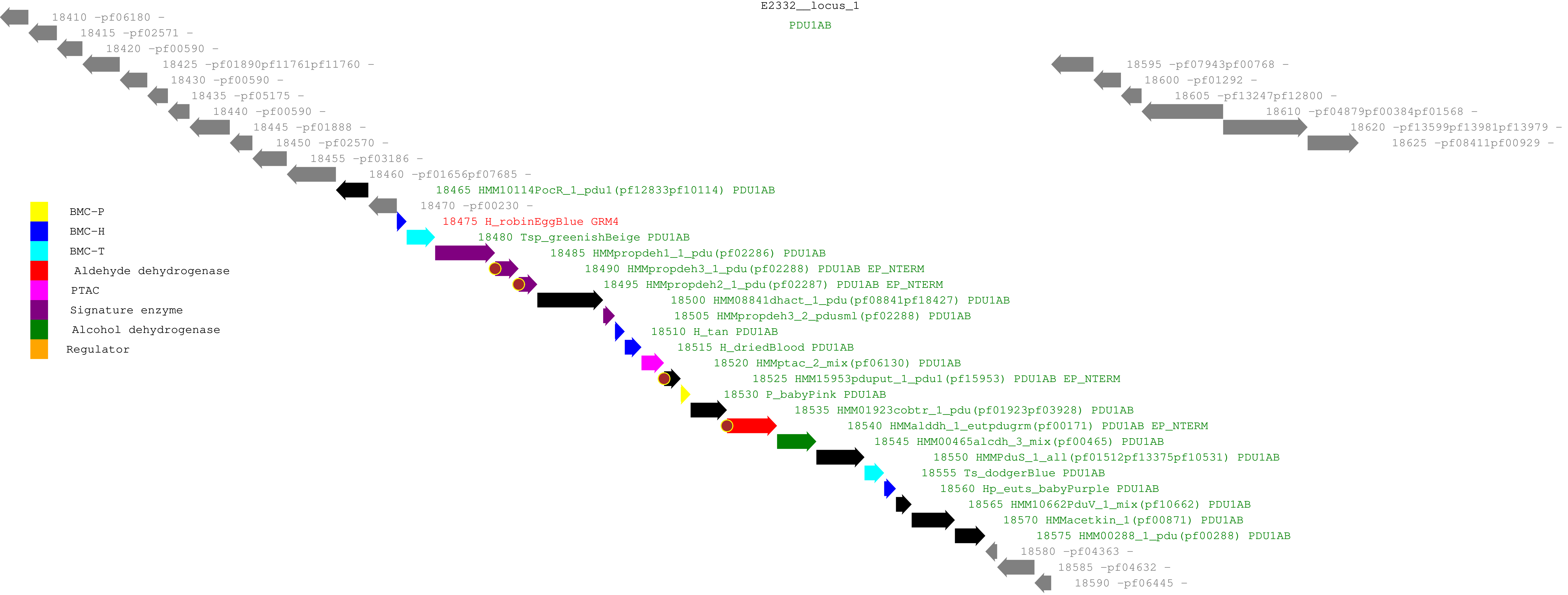

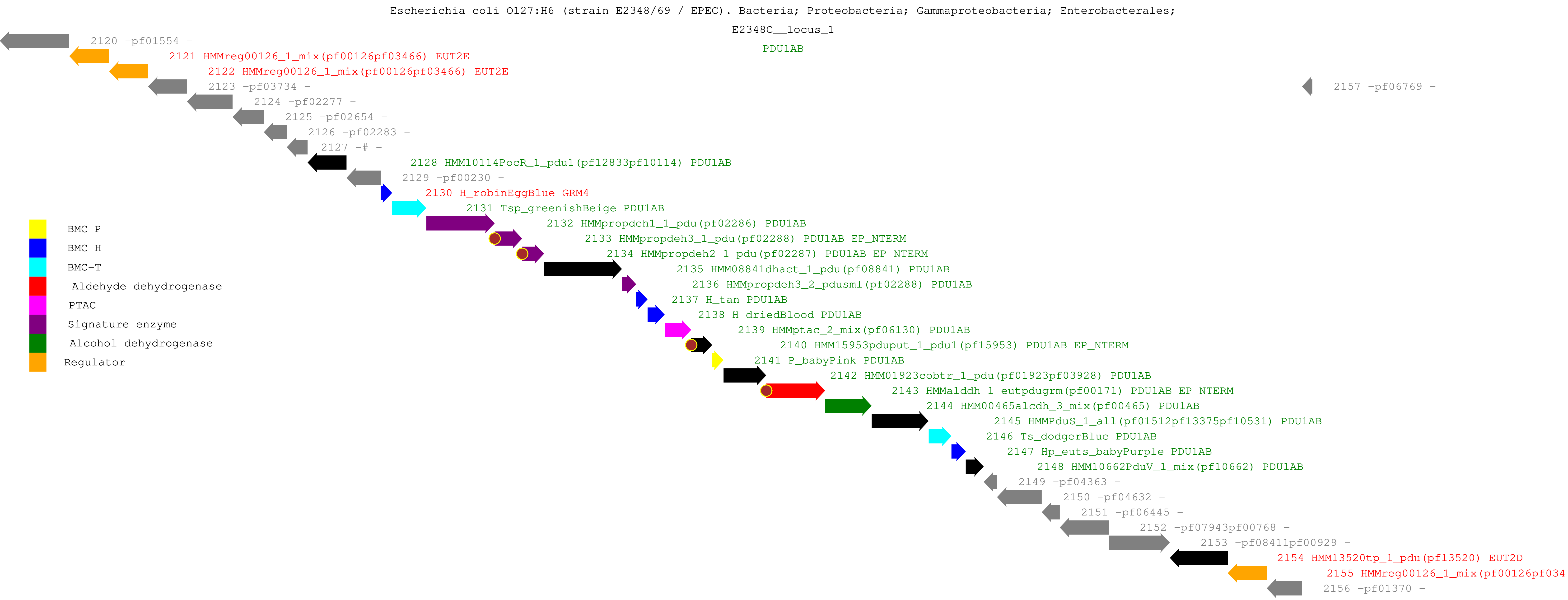
Escherichia coli O127:H6 (strain E2348/69 / EPEC). Bacteria; Proteobacteria; Gammaproteobacteria; Enterobacterales;
E2348C__locus_1
PDU1AB
2120 -pf01554 -
2121 HMMreg00126_1_mix(pf00126pf03466) EUT2E
2122 HMMreg00126_1_mix(pf00126pf03466) EUT2E
2123 -pf03734 -
2124 -pf02277 -
2125 -pf02654 -
2126 -pf02283 -
2127 -# -
2128 HMM10114PocR_1_pdu1(pf12833pf10114) PDU1AB
2129 -pf00230 -
2130 H_robinEggBlue GRM4
2131 Tsp_greenishBeige PDU1AB
2132 HMMpropdeh1_1_pdu(pf02286) PDU1AB
2133 HMMpropdeh3_1_pdu(pf02288) PDU1AB EP_NTERM
2134 HMMpropdeh2_1_pdu(pf02287) PDU1AB EP_NTERM
2135 HMM08841dhact_1_pdu(pf08841) PDU1AB
2136 HMMpropdeh3_2_pdusml(pf02288) PDU1AB
2137 H_tan PDU1AB
2138 H_driedBlood PDU1AB
2139 HMMptac_2_mix(pf06130) PDU1AB
2140 HMM15953pduput_1_pdu1(pf15953) PDU1AB EP_NTERM
2141 P_babyPink PDU1AB
2142 HMM01923cobtr_1_pdu(pf01923pf03928) PDU1AB
2143 HMMalddh_1_eutpdugrm(pf00171) PDU1AB EP_NTERM
2144 HMM00465alcdh_3_mix(pf00465) PDU1AB
2145 HMMPduS_1_all(pf01512pf13375pf10531) PDU1AB
2146 Ts_dodgerBlue PDU1AB
2147 Hp_euts_babyPurple PDU1AB
2148 HMM10662PduV_1_mix(pf10662) PDU1AB
2149 -pf04363 -
2150 -pf04632 -
2151 -pf06445 -
2152 -pf07943pf00768 -
2153 -pf08411pf00929 -
2154 HMM13520tp_1_pdu(pf13520) EUT2D
2155 HMMreg00126_1_mix(pf00126pf034
2156 -pf01370 -
2157 -pf06769 -
BMC-P
BMC-H
BMC-T
Aldehyde dehydrogenase
PTAC
Signature enzyme
Alcohol dehydrogenase
Regulator

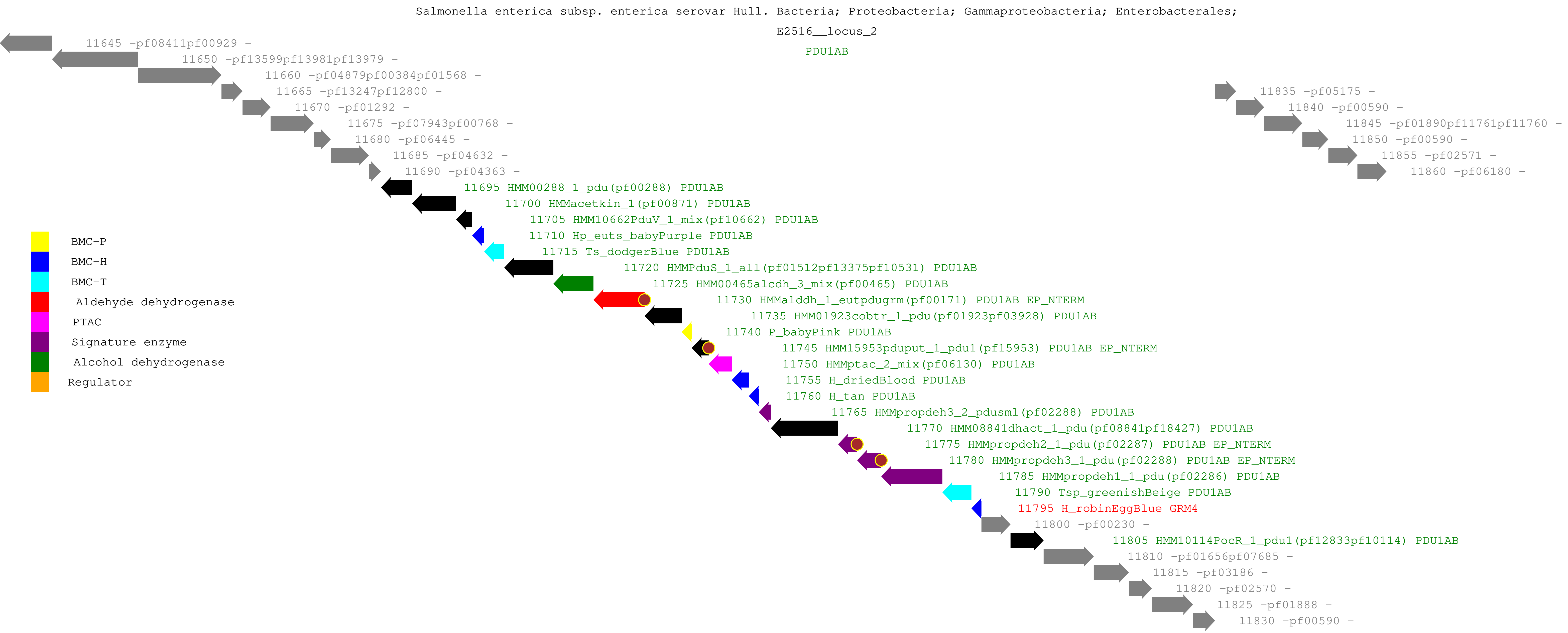
Salmonella enterica subsp. enterica serovar Hull. Bacteria; Proteobacteria; Gammaproteobacteria; Enterobacterales;
E2516__locus_2
PDU1AB
11645 -pf08411pf00929 -
11650 -pf13599pf13981pf13979 -
11660 -pf04879pf00384pf01568 -
11665 -pf13247pf12800 -
11670 -pf01292 -
11675 -pf07943pf00768 -
11680 -pf06445 -
11685 -pf04632 -
11690 -pf04363 -
11695 HMM00288_1_pdu(pf00288) PDU1AB
11700 HMMacetkin_1(pf00871) PDU1AB
11705 HMM10662PduV_1_mix(pf10662) PDU1AB
11710 Hp_euts_babyPurple PDU1AB
11715 Ts_dodgerBlue PDU1AB
11720 HMMPduS_1_all(pf01512pf13375pf10531) PDU1AB
11725 HMM00465alcdh_3_mix(pf00465) PDU1AB
11730 HMMalddh_1_eutpdugrm(pf00171) PDU1AB EP_NTERM
11735 HMM01923cobtr_1_pdu(pf01923pf03928) PDU1AB
11740 P_babyPink PDU1AB
11745 HMM15953pduput_1_pdu1(pf15953) PDU1AB EP_NTERM
11750 HMMptac_2_mix(pf06130) PDU1AB
11755 H_driedBlood PDU1AB
11760 H_tan PDU1AB
11765 HMMpropdeh3_2_pdusml(pf02288) PDU1AB
11770 HMM08841dhact_1_pdu(pf08841pf18427) PDU1AB
11775 HMMpropdeh2_1_pdu(pf02287) PDU1AB EP_NTERM
11780 HMMpropdeh3_1_pdu(pf02288) PDU1AB EP_NTERM
11785 HMMpropdeh1_1_pdu(pf02286) PDU1AB
11790 Tsp_greenishBeige PDU1AB
11795 H_robinEggBlue GRM4
11800 -pf00230 -
11805 HMM10114PocR_1_pdu1(pf12833pf10114) PDU1AB
11810 -pf01656pf07685 -
11815 -pf03186 -
11820 -pf02570 -
11825 -pf01888 -
11830 -pf00590 -
11835 -pf05175 -
11840 -pf00590 -
11845 -pf01890pf11761pf11760 -
11850 -pf00590 -
11855 -pf02571 -
11860 -pf06180 -
BMC-P
BMC-H
BMC-T
Aldehyde dehydrogenase
PTAC
Signature enzyme
Alcohol dehydrogenase
Regulator

Salmonella enterica subsp. enterica serovar Miami. Bacteria; Proteobacteria; Gammaproteobacteria; Enterobacterales;

E2D53__locus_2

PDU1AB

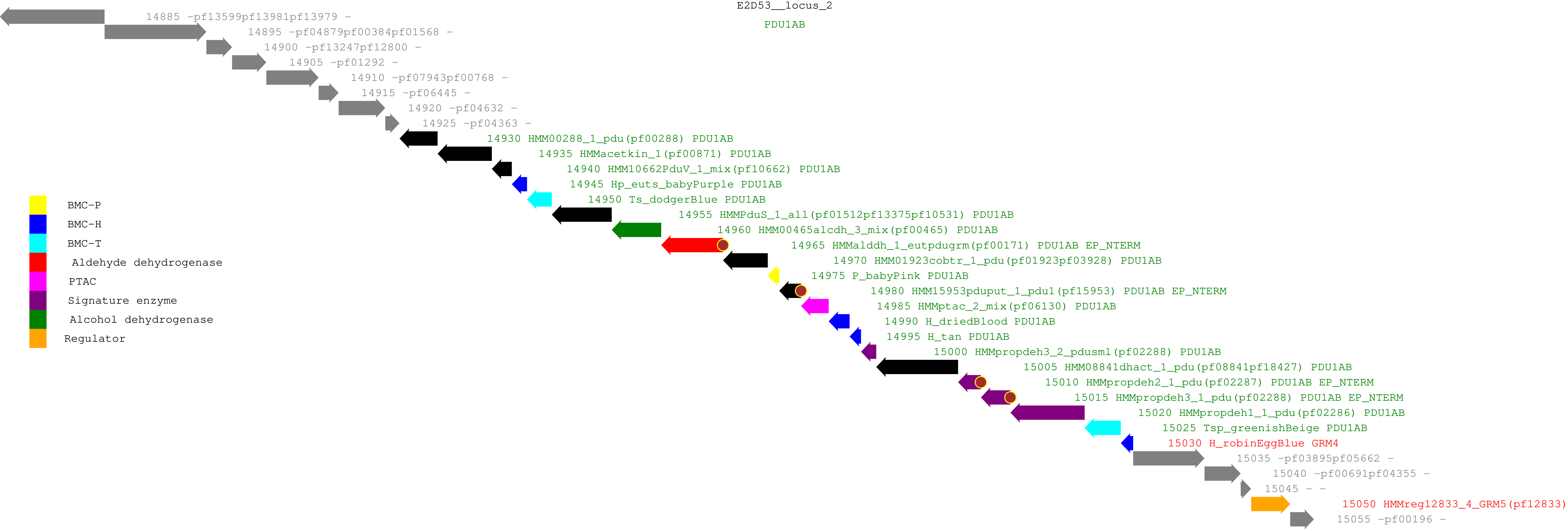

Salmonella enterica subsp. enterica serovar Kokomlemle. Bacteria; Proteobacteria; Gammaproteobacteria; Enterobacterales;

E2D59__locus_2

PDU1AB

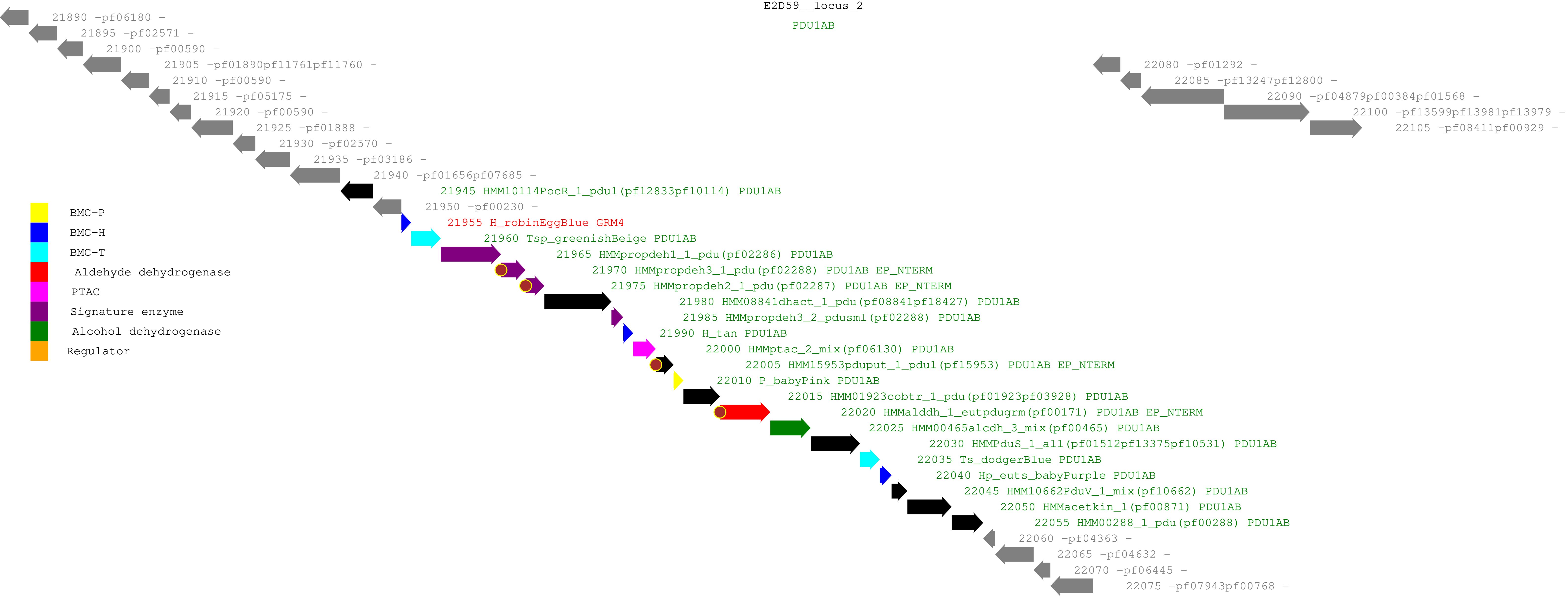

Salmonella enterica subsp. enterica serovar Reading. Bacteria; Proteobacteria; Gammaproteobacteria; Enterobacterales;

E2G14__locus_1

PDU1AB

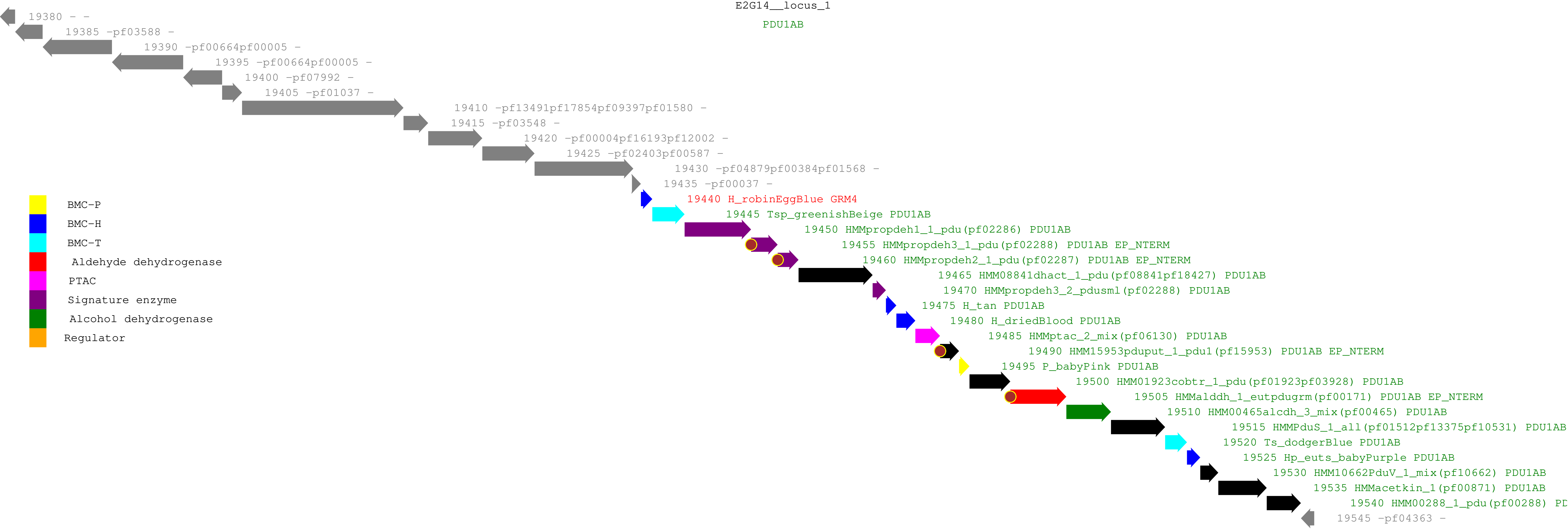

Salmonella enterica subsp. enterica serovar Chester. Bacteria; Proteobacteria; Gammaproteobacteria; Enterobacterales;

E2G30__locus_1

PDU1AB

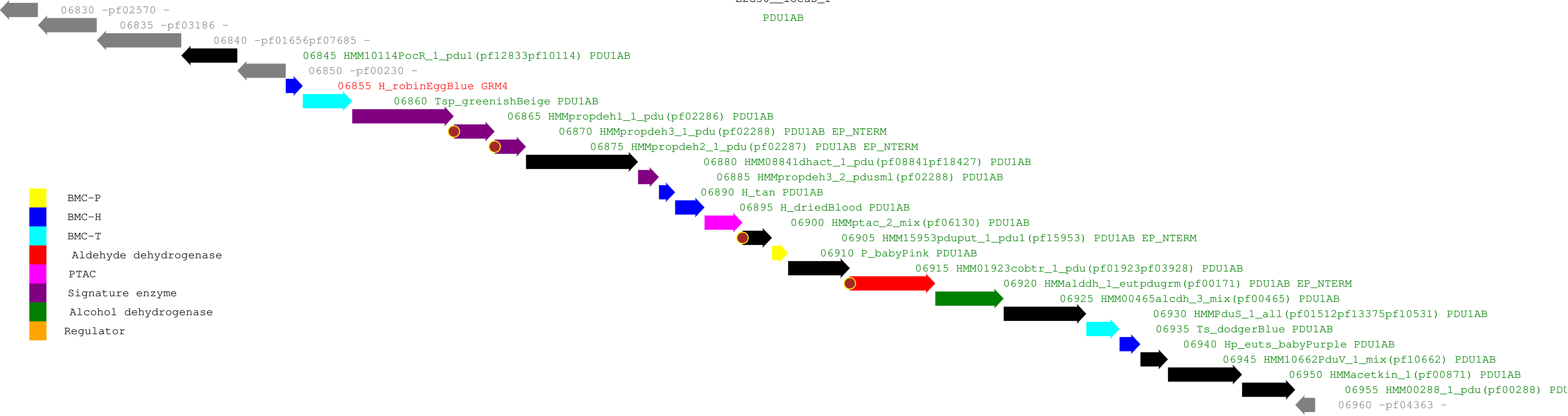

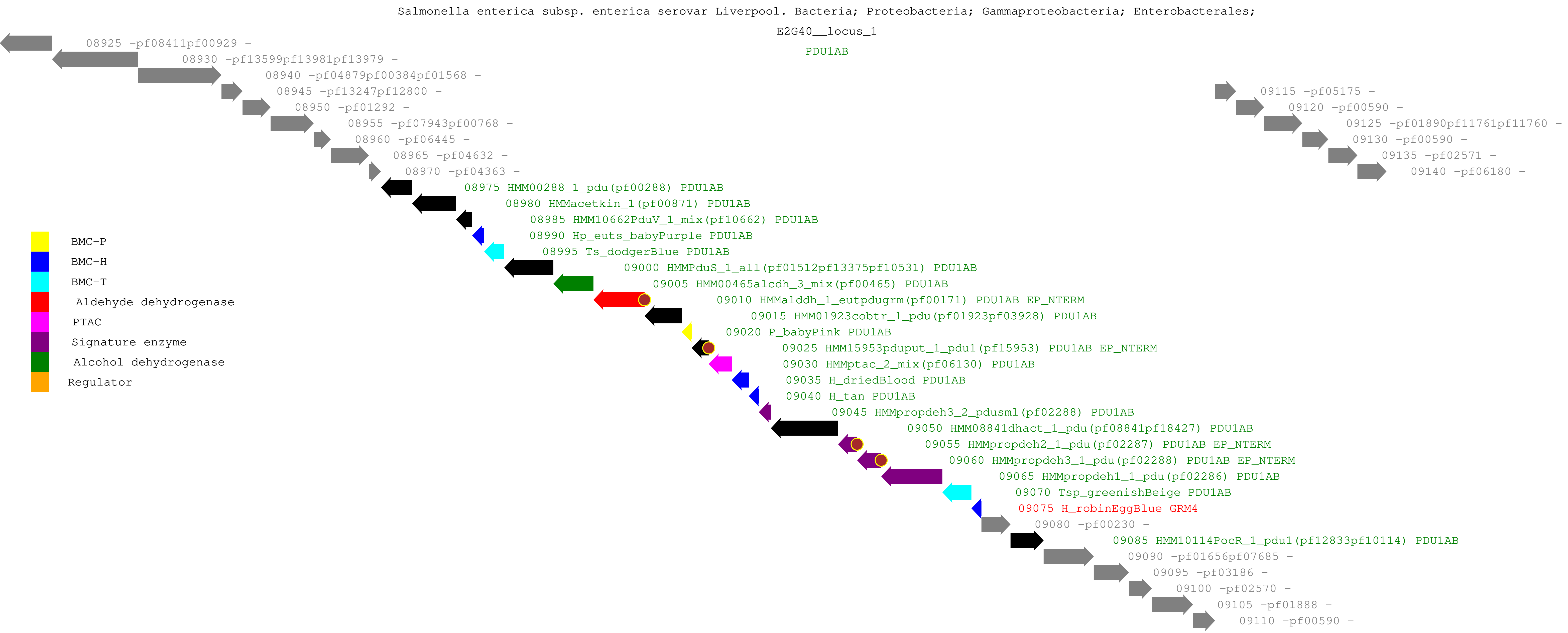

Salmonella enterica subsp. enterica serovar Liverpool. Bacteria; Proteobacteria; Gammaproteobacteria; Enterobacterales;
E2G40__locus_1
PDU1AB
08925 -pf08411pf00929 -
08930 -pf13599pf13981pf13979 -
08940 -pf04879pf00384pf01568 -
08945 -pf13247pf12800 -
08950 -pf01292 -
08955 -pf07943pf00768 -
08960 -pf06445 -
08965 -pf04632 -
08970 -pf04363 -
08975 HMM00288_1_pdu(pf00288) PDU1AB
08980 HMMacetkin_1(pf00871) PDU1AB
08985 HMM10662PduV_1_mix(pf10662) PDU1AB
08990 Hp_euts_babyPurple PDU1AB
08995 Ts_dodgerBlue PDU1AB
09000 HMMPduS_1_all(pf01512pf13375pf10531) PDU1AB
09005 HMM00465alcdh_3_mix(pf00465) PDU1AB
09010 HMMalddh_1_eutpdugrm(pf00171) PDU1AB EP_NTERM
09015 HMM01923cobtr_1_pdu(pf01923pf03928) PDU1AB
09020 P_babyPink PDU1AB
09025 HMM15953pduput_1_pdu1(pf15953) PDU1AB EP_NTERM
09030 HMMptac_2_mix(pf06130) PDU1AB
09035 H_driedBlood PDU1AB
09040 H_tan PDU1AB
09045 HMMpropdeh3_2_pdusml(pf02288) PDU1AB
09050 HMM08841dhact_1_pdu(pf08841pf18427) PDU1AB
09055 HMMpropdeh2_1_pdu(pf02287) PDU1AB EP_NTERM
09060 HMMpropdeh3_1_pdu(pf02288) PDU1AB EP_NTERM
09065 HMMpropdeh1_1_pdu(pf02286) PDU1AB
09070 Tsp_greenishBeige PDU1AB
09075 H_robinEggBlue GRM4
09080 -pf00230 -
09085 HMM10114PocR_1_pdu1(pf12833pf10114) PDU1AB
09090 -pf01656pf07685 -
09095 -pf03186 -
09100 -pf02570 -
09105 -pf01888 -
09110 -pf00590 -
09115 -pf05175 -
09120 -pf00590 -
09125 -pf01890pf11761pf11760 -
09130 -pf00590 -
09135 -pf02571 -
09140 -pf06180 -
BMC-P
BMC-H
BMC-T
Aldehyde dehydrogenase
PTAC
Signature enzyme
Alcohol dehydrogenase
Regulator

Salmonella enterica subsp. enterica serovar Stanleyville. Bacteria; Proteobacteria; Gammaproteobacteria; Enterobacterales;

E2G65__locus_2

PDU1AB

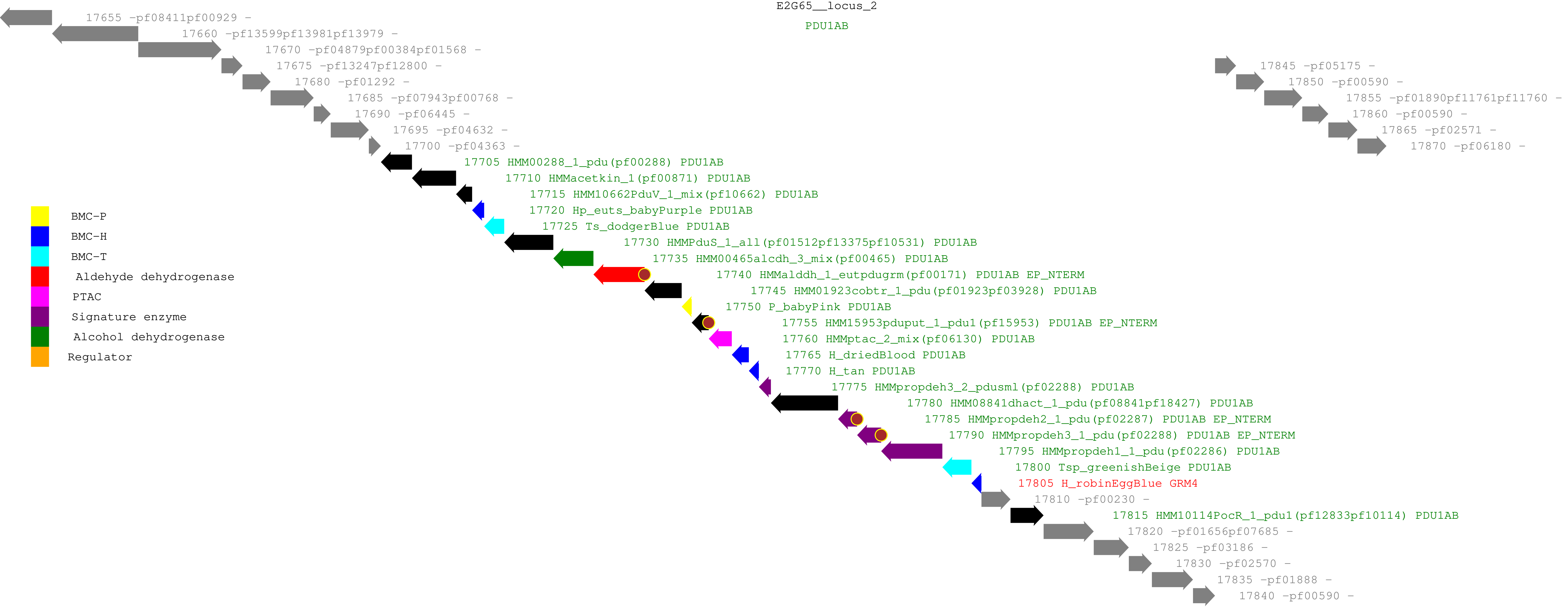

Salmonella enterica subsp. enterica serovar Wien. Bacteria; Proteobacteria; Gammaproteobacteria; Enterobacterales;

E2G78__locus_2

PDU1AB

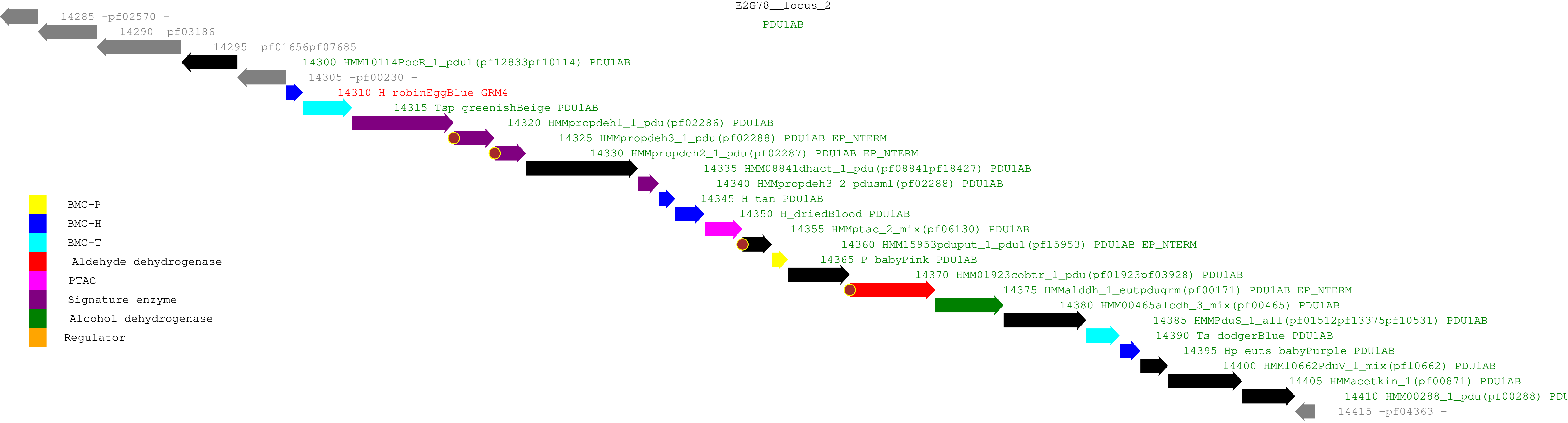

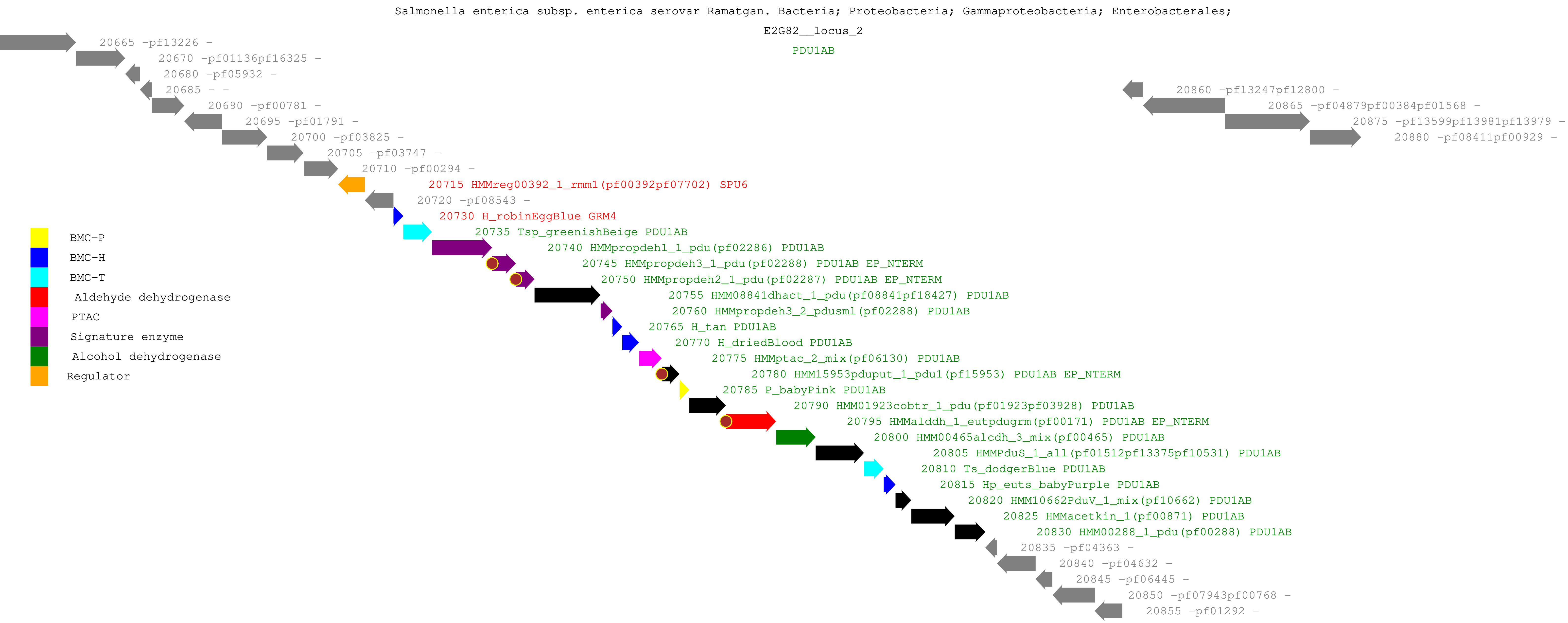
Salmonella enterica subsp. enterica serovar Ramatgan. Bacteria; Proteobacteria; Gammaproteobacteria; Enterobacterales;
E2G82__locus_2
PDU1AB
20665 -pf13226 -
20670 -pf01136pf16325 -
20680 -pf05932 -
20685 - -
20690 -pf00781 -
20695 -pf01791 -
20700 -pf03825 -
20705 -pf03747 -
20710 -pf00294 -
20715 HMMreg00392_1_rmm1(pf00392pf07702) SPU6
20720 -pf08543 -
20730 H_robinEggBlue GRM4
20735 Tsp_greenishBeige PDU1AB
20740 HMMpropdeh1_1_pdu(pf02286) PDU1AB
20745 HMMpropdeh3_1_pdu(pf02288) PDU1AB EP_NTERM
20750 HMMpropdeh2_1_pdu(pf02287) PDU1AB EP_NTERM
20755 HMM08841dhact_1_pdu(pf08841pf18427) PDU1AB
20760 HMMpropdeh3_2_pdusml(pf02288) PDU1AB
20765 H_tan PDU1AB
20770 H_driedBlood PDU1AB
20775 HMMptac_2_mix(pf06130) PDU1AB
20780 HMM15953pduput_1_pdu1(pf15953) PDU1AB EP_NTERM
20785 P_babyPink PDU1AB
20790 HMM01923cobtr_1_pdu(pf01923pf03928) PDU1AB
20795 HMMalddh_1_eutpdugrm(pf00171) PDU1AB EP_NTERM
20800 HMM00465alcdh_3_mix(pf00465) PDU1AB
20805 HMMPduS_1_all(pf01512pf13375pf10531) PDU1AB
20810 Ts_dodgerBlue PDU1AB
20815 Hp_euts_babyPurple PDU1AB
20820 HMM10662PduV_1_mix(pf10662) PDU1AB
20825 HMMacetkin_1(pf00871) PDU1AB
20830 HMM00288_1_pdu(pf00288) PDU1AB
20835 -pf04363 -
20840 -pf04632 -
20845 -pf06445 -
20850 -pf07943pf00768 -
20855 -pf01292 -
20860 -pf13247pf12800 -
20865 -pf04879pf00384pf01568 -
20875 -pf13599pf13981pf13979 -
20880 -pf08411pf00929 -
BMC-P
BMC-H
BMC-T
Aldehyde dehydrogenase
PTAC
Signature enzyme
Alcohol dehydrogenase
Regulator

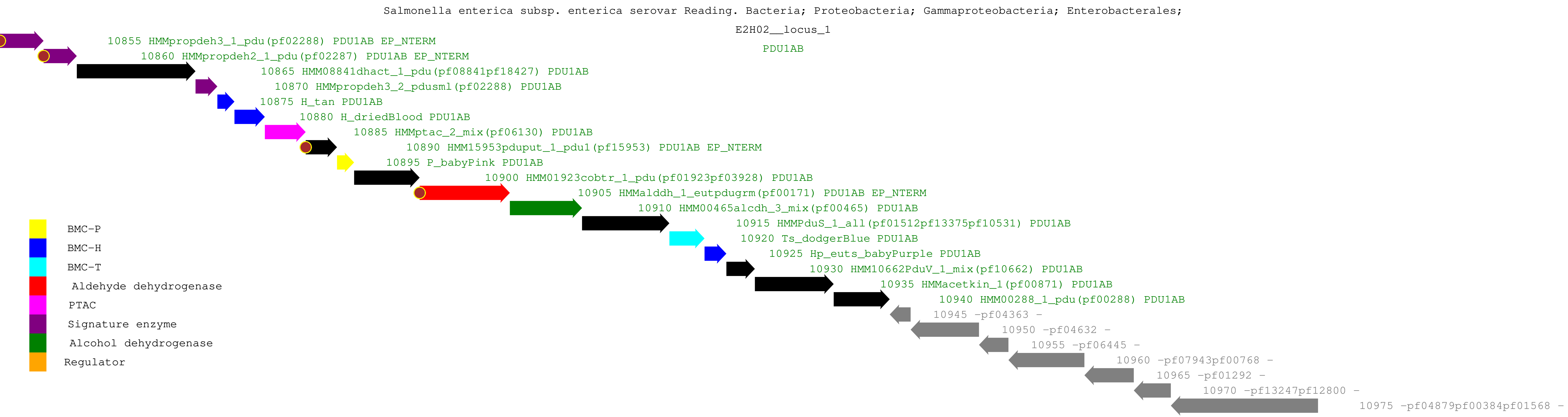
Salmonella enterica subsp. enterica serovar Reading. Bacteria; Proteobacteria; Gammaproteobacteria; Enterobacterales;
E2H02__locus_1
PDU1AB
10855 HMMpropdeh3_1_pdu(pf02288) PDU1AB EP_NTERM
10860 HMMpropdeh2_1_pdu(pf02287) PDU1AB EP_NTERM
10865 HMM08841dhact_1_pdu(pf08841pf18427) PDU1AB
10870 HMMpropdeh3_2_pdusml(pf02288) PDU1AB
10875 H_tan PDU1AB
10880 H_driedBlood PDU1AB
10885 HMMptac_2_mix(pf06130) PDU1AB
10890 HMM15953pduput_1_pdu1(pf15953) PDU1AB EP_NTERM
10895 P_babyPink PDU1AB
10900 HMM01923cobtr_1_pdu(pf01923pf03928) PDU1AB
10905 HMMalddh_1_eutpdugrm(pf00171) PDU1AB EP_NTERM
10910 HMM00465alcdh_3_mix(pf00465) PDU1AB
10915 HMMPduS_1_all(pf01512pf13375pf10531) PDU1AB
10920 Ts_dodgerBlue PDU1AB
10925 Hp_euts_babyPurple PDU1AB
10930 HMM10662PduV_1_mix(pf10662) PDU1AB
10935 HMMacetkin_1(pf00871) PDU1AB
10940 HMM00288_1_pdu(pf00288) PDU1AB
10945 -pf04363 -
10950 -pf04632 -
10955 -pf06445 -
10960 -pf07943pf00768 -
10965 -pf01292 -
10970 -pf13247pf12800 -
10975 -pf04879pf00384pf01568 -
BMC-P
BMC-H
BMC-T
Aldehyde dehydrogenase
PTAC
Signature enzyme
Alcohol dehydrogenase
Regulator

Salmonella enterica subsp. enterica serovar Volkmarsdorf. Bacteria; Proteobacteria; Gammaproteobacteria; Enterobacterales;

E2H20__locus_1

PDU1AB

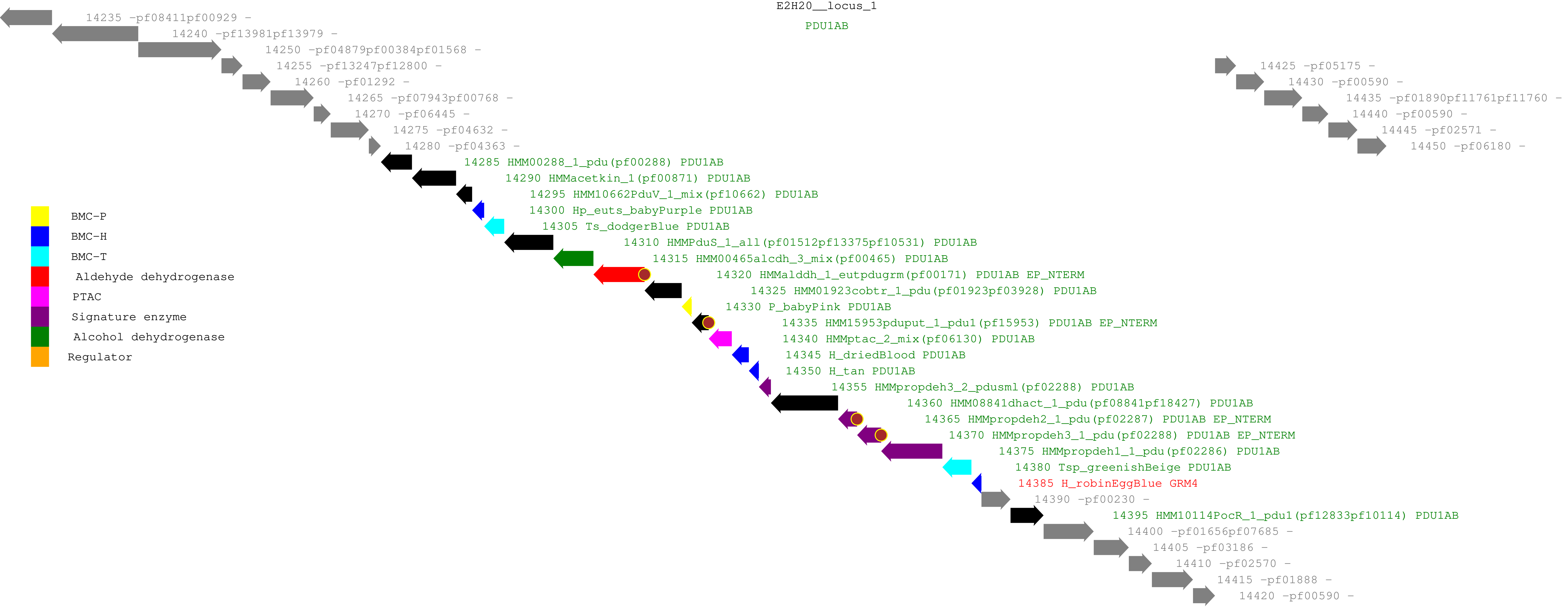

Salmonella enterica subsp. enterica serovar Alachua. Bacteria; Proteobacteria; Gammaproteobacteria; Enterobacterales;

E2H46__locus_1

PDU1AB

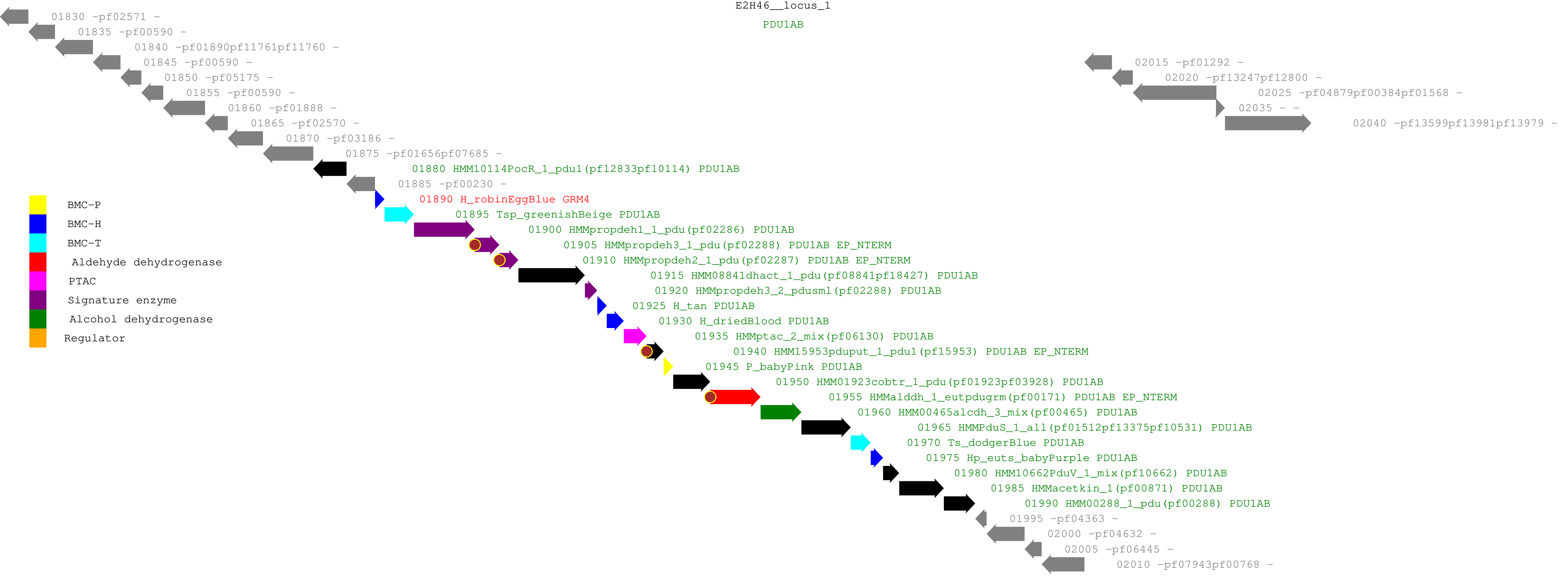

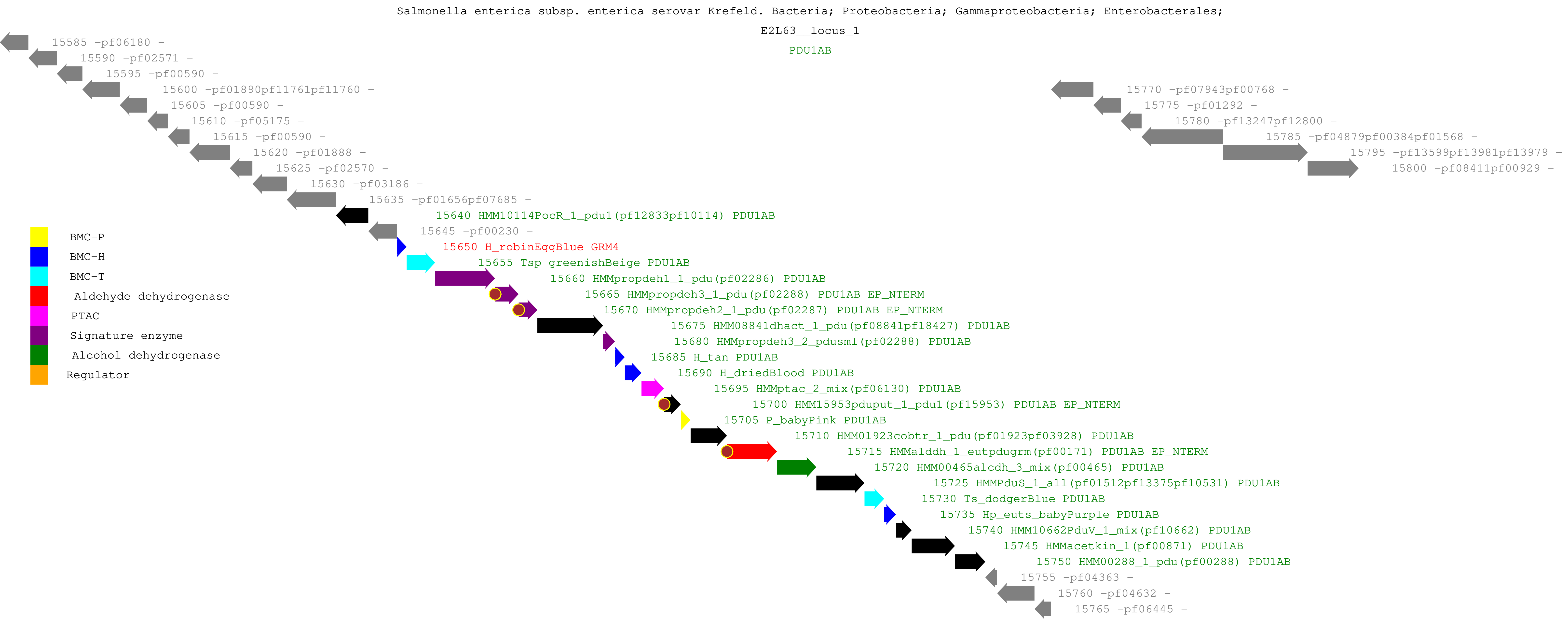

Salmonella enterica subsp. enterica serovar Krefeld. Bacteria; Proteobacteria; Gammaproteobacteria; Enterobacterales;
E2L63__locus_1
PDU1AB
15585 -pf06180 -
15590 -pf02571 -
15595 -pf00590 -
15600 -pf01890pf11761pf11760 -
15605 -pf00590 -
15610 -pf05175 -
15615 -pf00590 -
15620 -pf01888 -
15625 -pf02570 -
15630 -pf03186 -
15635 -pf01656pf07685 -
15640 HMM10114PocR_1_pdu1(pf12833pf10114) PDU1AB
15645 -pf00230 -
15650 H_robinEggBlue GRM4
15655 Tsp_greenishBeige PDU1AB
15660 HMMpropdeh1_1_pdu(pf02286) PDU1AB
15665 HMMpropdeh3_1_pdu(pf02288) PDU1AB EP_NTERM
15670 HMMpropdeh2_1_pdu(pf02287) PDU1AB EP_NTERM
15675 HMM08841dhact_1_pdu(pf08841pf18427) PDU1AB
15680 HMMpropdeh3_2_pdusml(pf02288) PDU1AB
15685 H_tan PDU1AB
15690 H_driedBlood PDU1AB
15695 HMMptac_2_mix(pf06130) PDU1AB
15700 HMM15953pduput_1_pdu1(pf15953) PDU1AB EP_NTERM
15705 P_babyPink PDU1AB
15710 HMM01923cobtr_1_pdu(pf01923pf03928) PDU1AB
15715 HMMalddh_1_eutpdugrm(pf00171) PDU1AB EP_NTERM
15720 HMM00465alcdh_3_mix(pf00465) PDU1AB
15725 HMMPduS_1_all(pf01512pf13375pf10531) PDU1AB
15730 Ts_dodgerBlue PDU1AB
15735 Hp_euts_babyPurple PDU1AB
15740 HMM10662PduV_1_mix(pf10662) PDU1AB
15745 HMMacetkin_1(pf00871) PDU1AB
15750 HMM00288_1_pdu(pf00288) PDU1AB
15755 -pf04363 -
15760 -pf04632 -
15765 -pf06445 -
15770 -pf07943pf00768 -
15775 -pf01292 -
15780 -pf13247pf12800 -
15785 -pf04879pf00384pf01568 -
15795 -pf13599pf13981pf13979 -
15800 -pf08411pf00929 -
BMC-P
BMC-H
BMC-T
Aldehyde dehydrogenase
PTAC
Signature enzyme
Alcohol dehydrogenase
Regulator

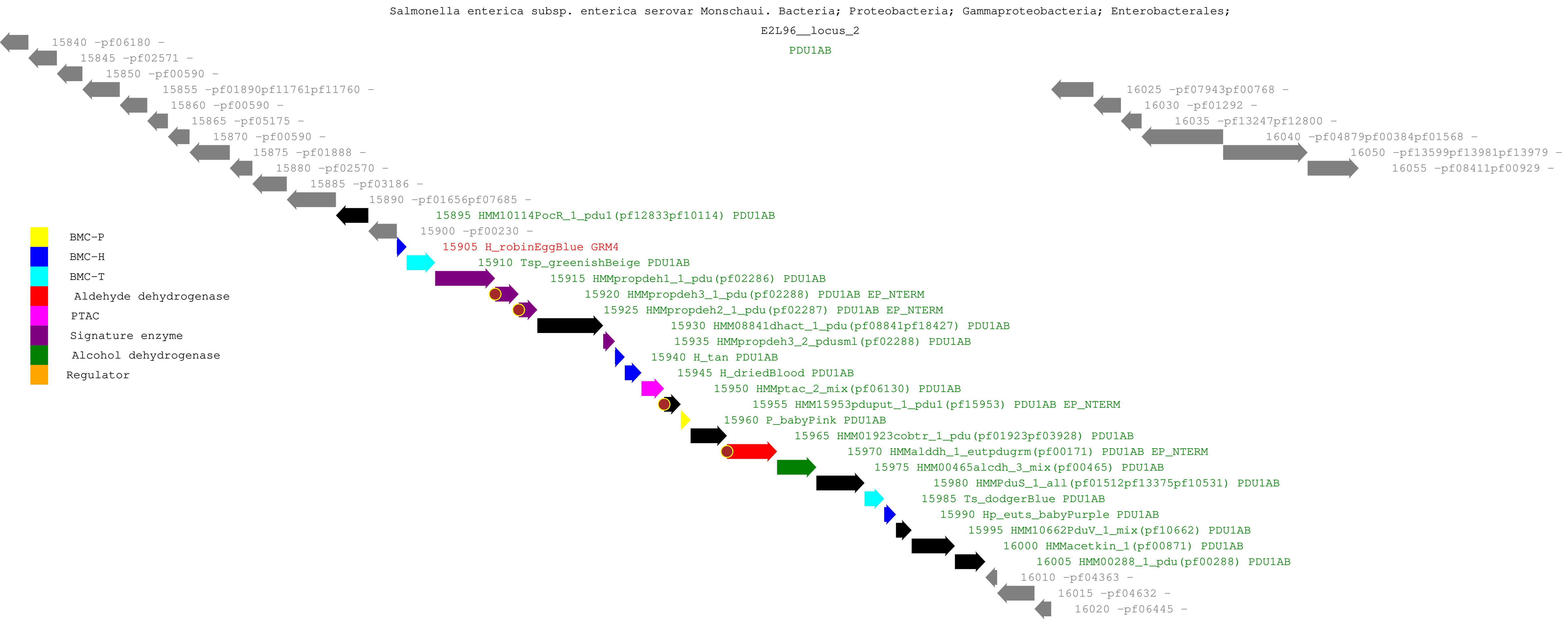
Salmonella enterica subsp. enterica serovar Monschaui. Bacteria; Proteobacteria; Gammaproteobacteria; Enterobacterales;
E2L96__locus_2
PDU1AB
15840 -pf06180 -
15845 -pf02571 -
15850 -pf00590 -
15855 -pf01890pf11761pf11760 -
15860 -pf00590 -
15865 -pf05175 -
15870 -pf00590 -
15875 -pf01888 -
15880 -pf02570 -
15885 -pf03186 -
15890 -pf01656pf07685 -
15895 HMM10114PocR_1_pdu1(pf12833pf10114) PDU1AB
15900 -pf00230 -
15905 H_robinEggBlue GRM4
15910 Tsp_greenishBeige PDU1AB
15915 HMMpropdeh1_1_pdu(pf02286) PDU1AB
15920 HMMpropdeh3_1_pdu(pf02288) PDU1AB EP_NTERM
15925 HMMpropdeh2_1_pdu(pf02287) PDU1AB EP_NTERM
15930 HMM08841dhact_1_pdu(pf08841pf18427) PDU1AB
15935 HMMpropdeh3_2_pdusml(pf02288) PDU1AB
15940 H_tan PDU1AB
15945 H_driedBlood PDU1AB
15950 HMMptac_2_mix(pf06130) PDU1AB
15955 HMM15953pduput_1_pdu1(pf15953) PDU1AB EP_NTERM
15960 P_babyPink PDU1AB
15965 HMM01923cobtr_1_pdu(pf01923pf03928) PDU1AB
15970 HMMalddh_1_eutpdugrm(pf00171) PDU1AB EP_NTERM
15975 HMM00465alcdh_3_mix(pf00465) PDU1AB
15980 HMMPduS_1_all(pf01512pf13375pf10531) PDU1AB
15985 Ts_dodgerBlue PDU1AB
15990 Hp_euts_babyPurple PDU1AB
15995 HMM10662PduV_1_mix(pf10662) PDU1AB
16000 HMMacetkin_1(pf00871) PDU1AB
16005 HMM00288_1_pdu(pf00288) PDU1AB
16010 -pf04363 -
16015 -pf04632 -
16020 -pf06445 -
16025 -pf07943pf00768 -
16030 -pf01292 -
16035 -pf13247pf12800 -
16040 -pf04879pf00384pf01568 -
16050 -pf13599pf13981pf13979 -
16055 -pf08411pf00929 -
BMC-P
BMC-H
BMC-T
Aldehyde dehydrogenase
PTAC
Signature enzyme
Alcohol dehydrogenase
Regulator

Salmonella enterica subsp. enterica serovar Arechavaleta. Bacteria; Proteobacteria; Gammaproteobacteria; Enterobacterales;

E2M22__locus_2

PDU1AB

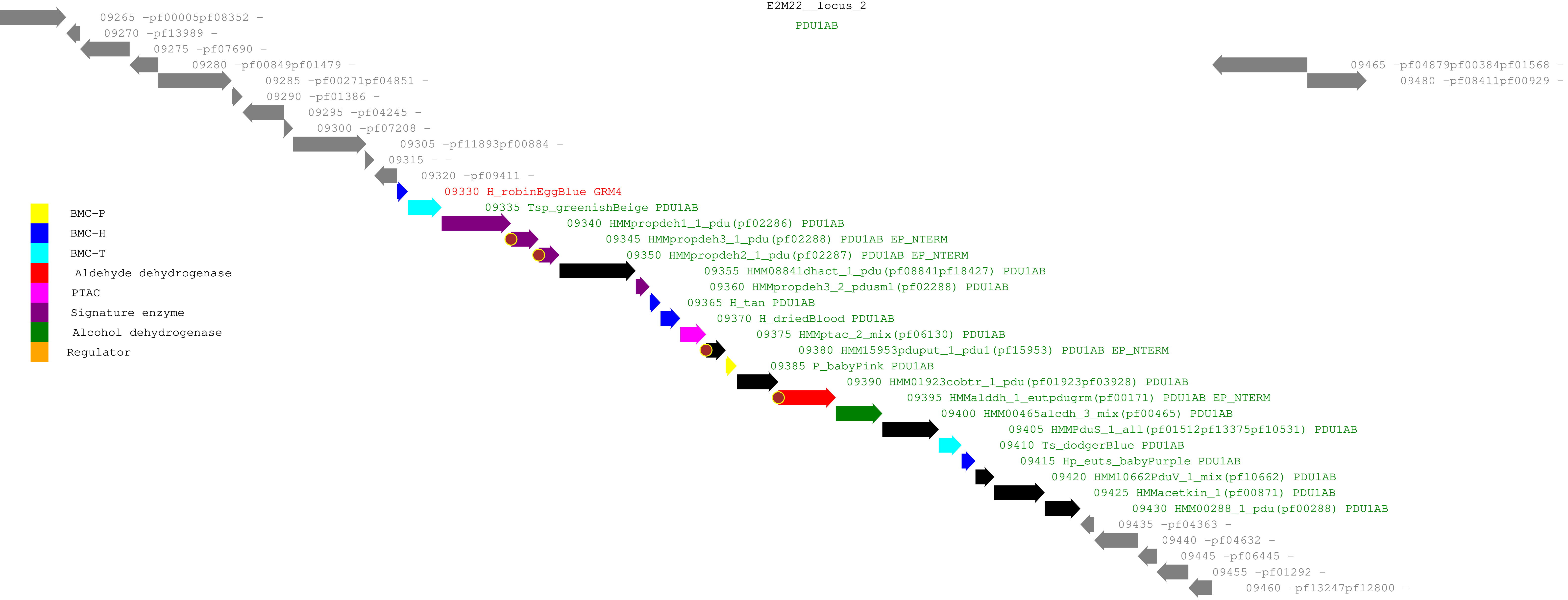

Salmonella enterica subsp. enterica serovar Nima. Bacteria; Proteobacteria; Gammaproteobacteria; Enterobacterales;

E2M36__locus_1

PDU1AB

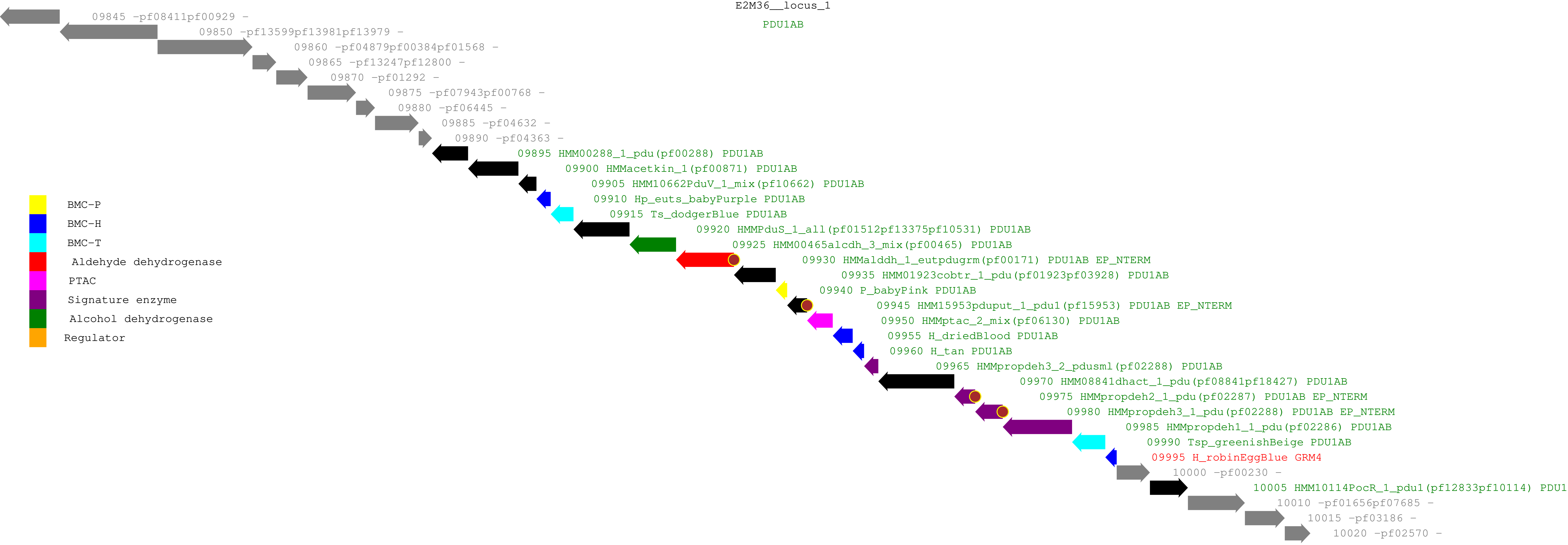

Salmonella enterica subsp. enterica serovar Othmarschen. Bacteria; Proteobacteria; Gammaproteobacteria; Enterobacterales;

E2M46__locus_1

PDU1AB

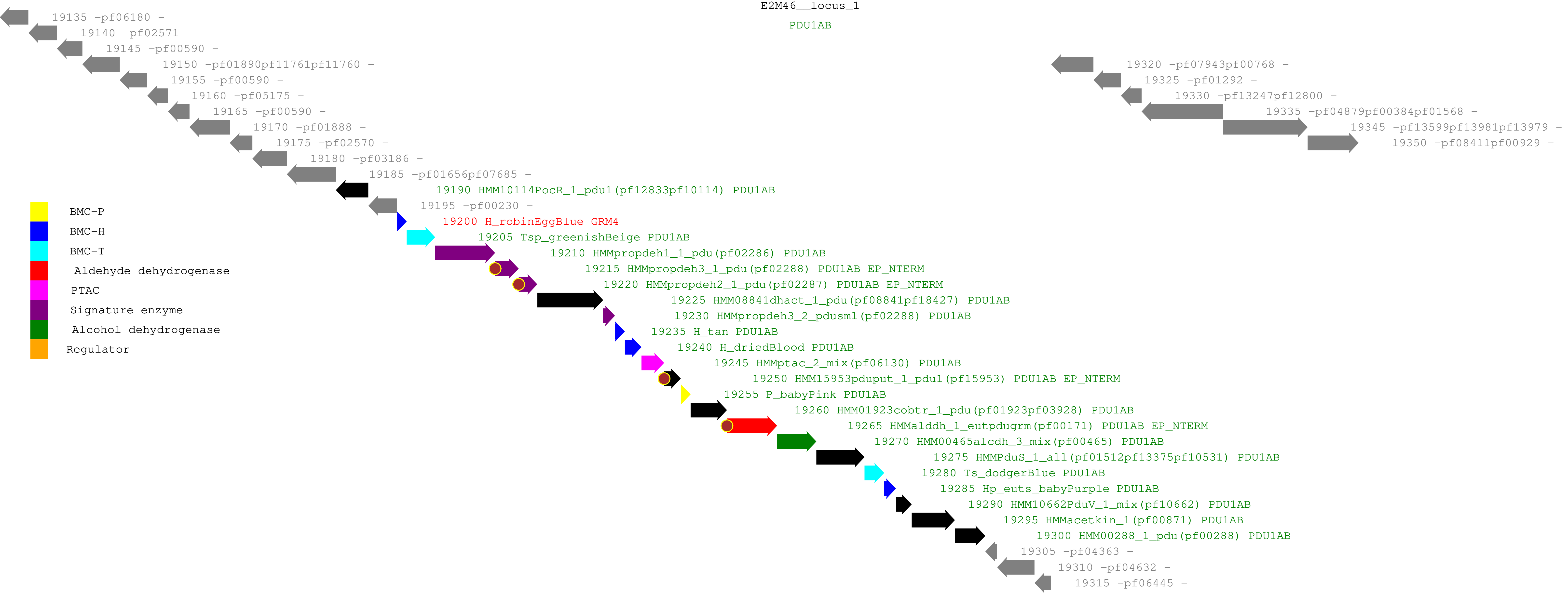

Salmonella enterica subsp. enterica serovar Amager. Bacteria; Proteobacteria; Gammaproteobacteria; Enterobacterales;

E2Q29__locus_1

PDU1AB

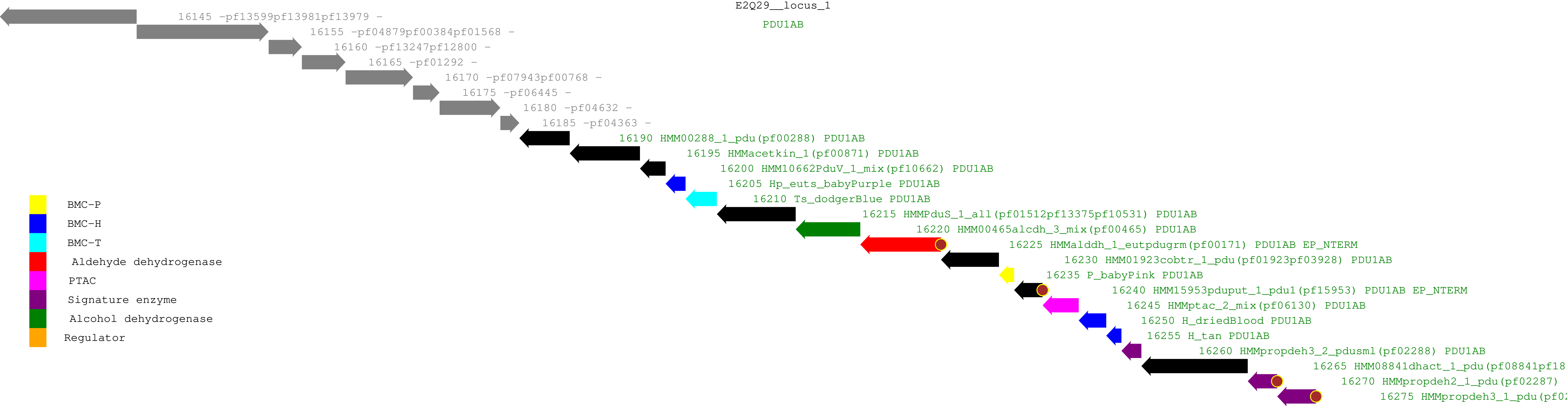

Salmonella enterica subsp. enterica serovar Durham. Bacteria; Proteobacteria; Gammaproteobacteria; Enterobacterales;

E2Q46__locus_1

PDU1AB

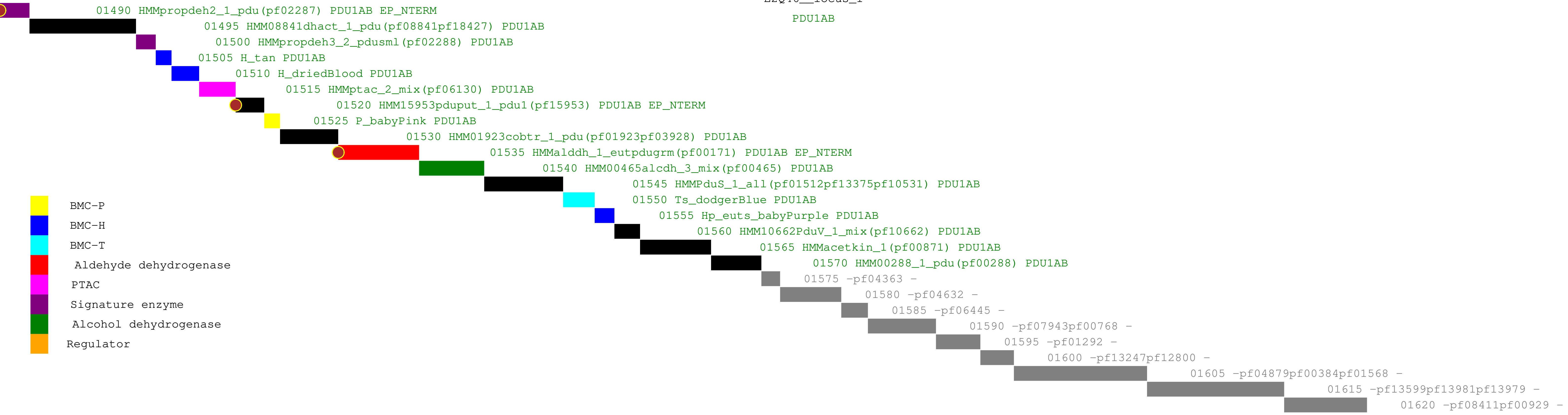

Citrobacter rodentium. Bacteria; Proteobacteria; Gammaproteobacteria; Enterobacterales;

E2R62__locus_1

PDU1AB

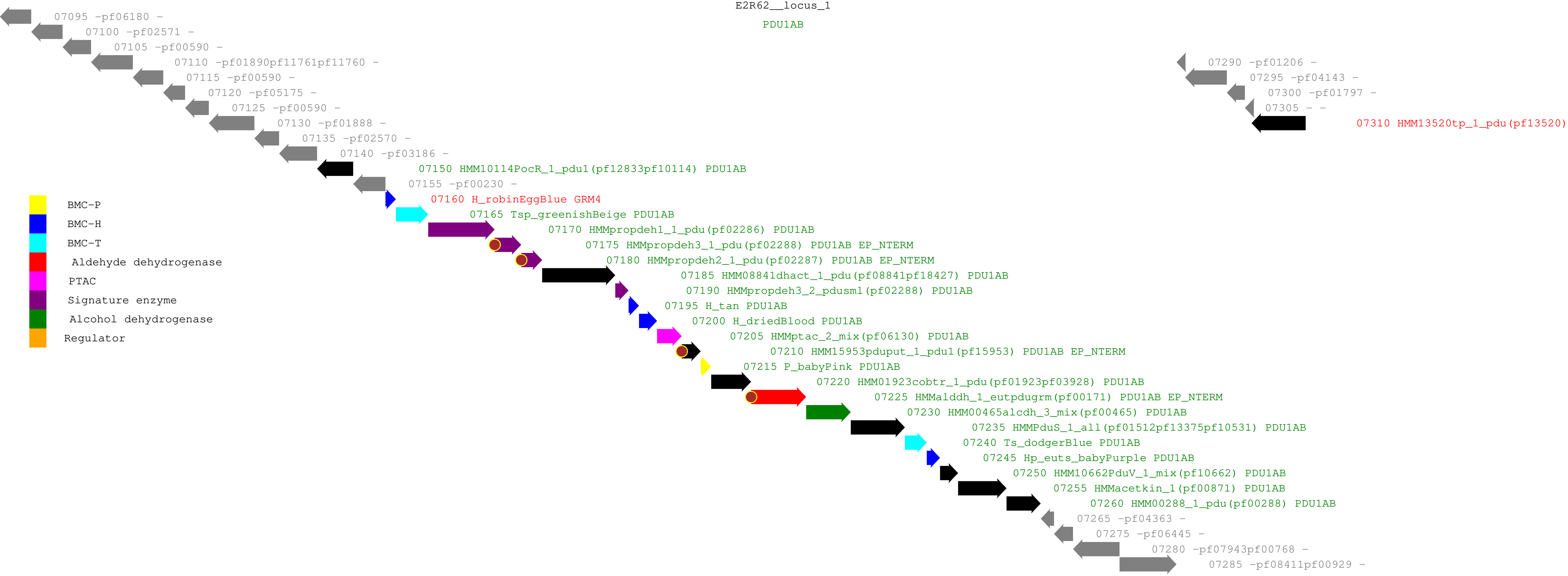

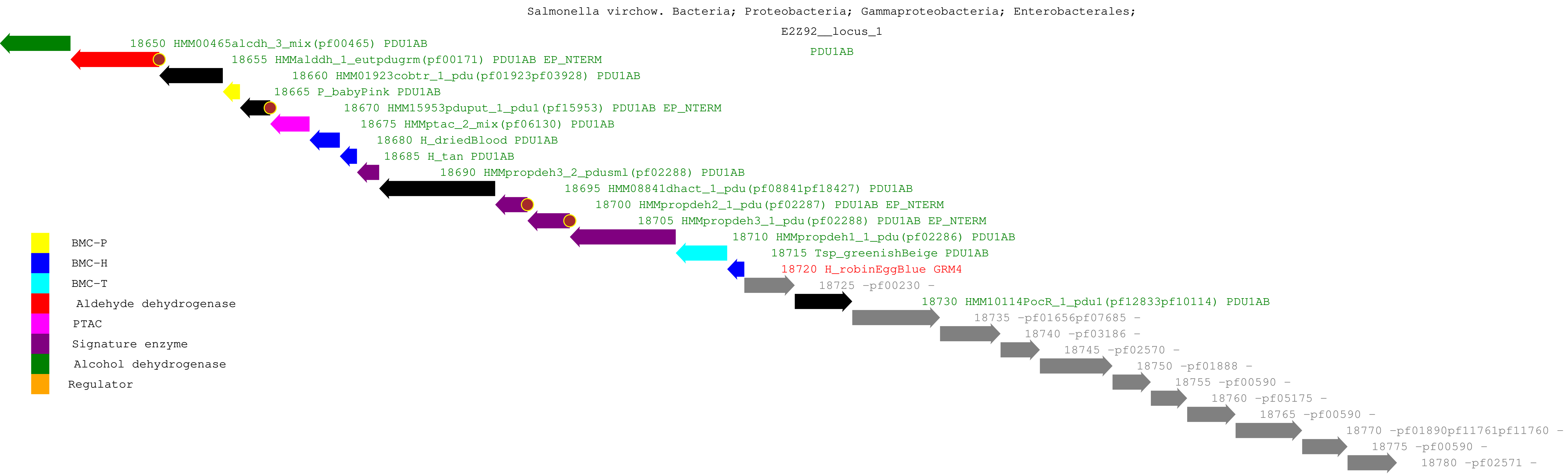
Salmonella virchow. Bacteria; Proteobacteria; Gammaproteobacteria; Enterobacterales;
E2Z92__locus_1
PDU1AB
18650 HMM00465alcdh_3_mix(pf00465) PDU1AB
18655 HMMalddh_1_eutpdugrm(pf00171) PDU1AB EP_NTERM
18660 HMM01923cobtr_1_pdu(pf01923pf03928) PDU1AB
18665 P_babyPink PDU1AB
18670 HMM15953pduput_1_pdu1(pf15953) PDU1AB EP_NTERM
18675 HMMptac_2_mix(pf06130) PDU1AB
18680 H_driedBlood PDU1AB
18685 H_tan PDU1AB
18690 HMMpropdeh3_2_pdusml(pf02288) PDU1AB
18695 HMM08841dhact_1_pdu(pf08841pf18427) PDU1AB
18700 HMMpropdeh2_1_pdu(pf02287) PDU1AB EP_NTERM
18705 HMMpropdeh3_1_pdu(pf02288) PDU1AB EP_NTERM
18710 HMMpropdeh1_1_pdu(pf02286) PDU1AB
18715 Tsp_greenishBeige PDU1AB
18720 H_robinEggBlue GRM4
18725 -pf00230 -
18730 HMM10114PocR_1_pdu1(pf12833pf10114) PDU1AB
18735 -pf01656pf07685 -
18740 -pf03186 -
18745 -pf02570 -
18750 -pf01888 -
18755 -pf00590 -
18760 -pf05175 -
18765 -pf00590 -
18770 -pf01890pf11761pf11760 -
18775 -pf00590 -
18780 -pf02571 -
BMC-P
BMC-H
BMC-T
Aldehyde dehydrogenase
PTAC
Signature enzyme
Alcohol dehydrogenase
Regulator

Salmonella enterica subsp. enterica serovar Chailey. Bacteria; Proteobacteria; Gammaproteobacteria; Enterobacterales;

E3314__locus_1

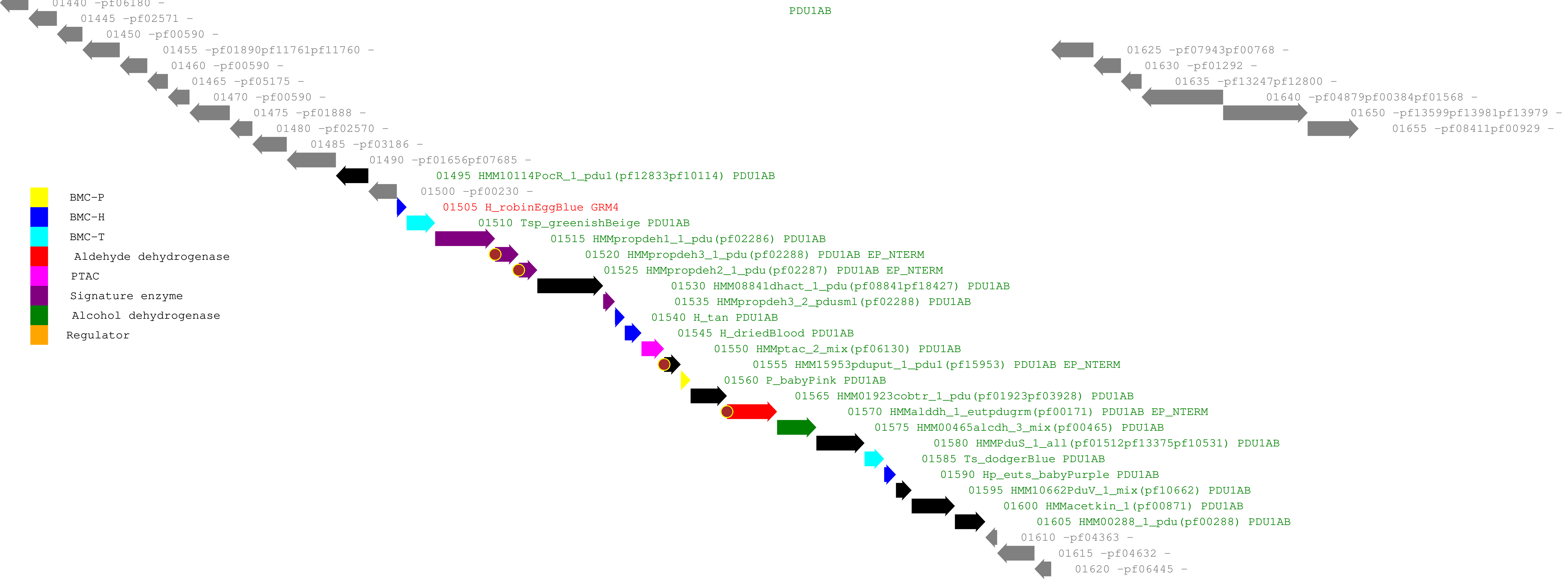

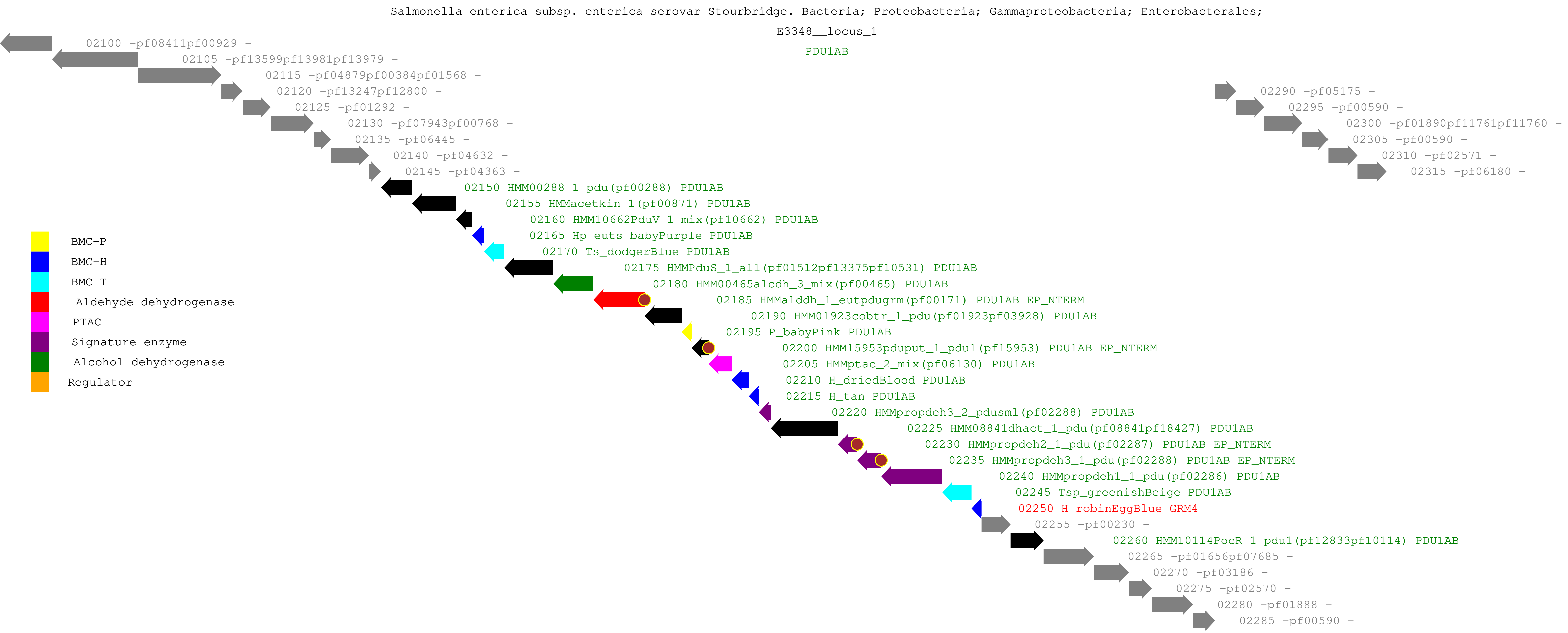

Salmonella enterica subsp. enterica serovar Stourbridge. Bacteria; Proteobacteria; Gammaproteobacteria; Enterobacterales;
E3348__locus_1
PDU1AB
02100 -pf08411pf00929 -
02105 -pf13599pf13981pf13979 -
02115 -pf04879pf00384pf01568 -
02120 -pf13247pf12800 -
02125 -pf01292 -
02130 -pf07943pf00768 -
02135 -pf06445 -
02140 -pf04632 -
02145 -pf04363 -
02150 HMM00288_1_pdu(pf00288) PDU1AB
02155 HMMacetkin_1(pf00871) PDU1AB
02160 HMM10662PduV_1_mix(pf10662) PDU1AB
02165 Hp_euts_babyPurple PDU1AB
02170 Ts_dodgerBlue PDU1AB
02175 HMMPduS_1_all(pf01512pf13375pf10531) PDU1AB
02180 HMM00465alcdh_3_mix(pf00465) PDU1AB
02185 HMMalddh_1_eutpdugrm(pf00171) PDU1AB EP_NTERM
02190 HMM01923cobtr_1_pdu(pf01923pf03928) PDU1AB
02195 P_babyPink PDU1AB
02200 HMM15953pduput_1_pdu1(pf15953) PDU1AB EP_NTERM
02205 HMMptac_2_mix(pf06130) PDU1AB
02210 H_driedBlood PDU1AB
02215 H_tan PDU1AB
02220 HMMpropdeh3_2_pdusml(pf02288) PDU1AB
02225 HMM08841dhact_1_pdu(pf08841pf18427) PDU1AB
02230 HMMpropdeh2_1_pdu(pf02287) PDU1AB EP_NTERM
02235 HMMpropdeh3_1_pdu(pf02288) PDU1AB EP_NTERM
02240 HMMpropdeh1_1_pdu(pf02286) PDU1AB
02245 Tsp_greenishBeige PDU1AB
02250 H_robinEggBlue GRM4
02255 -pf00230 -
02260 HMM10114PocR_1_pdu1(pf12833pf10114) PDU1AB
02265 -pf01656pf07685 -
02270 -pf03186 -
02275 -pf02570 -
02280 -pf01888 -
02285 -pf00590 -
02290 -pf05175 -
02295 -pf00590 -
02300 -pf01890pf11761pf11760 -
02305 -pf00590 -
02310 -pf02571 -
02315 -pf06180 -
BMC-P
BMC-H
BMC-T
Aldehyde dehydrogenase
PTAC
Signature enzyme
Alcohol dehydrogenase
Regulator

Salmonella enterica subsp. enterica serovar Brandenburg. Bacteria; Proteobacteria; Gammaproteobacteria; Enterobacterales;

E3354__locus_1

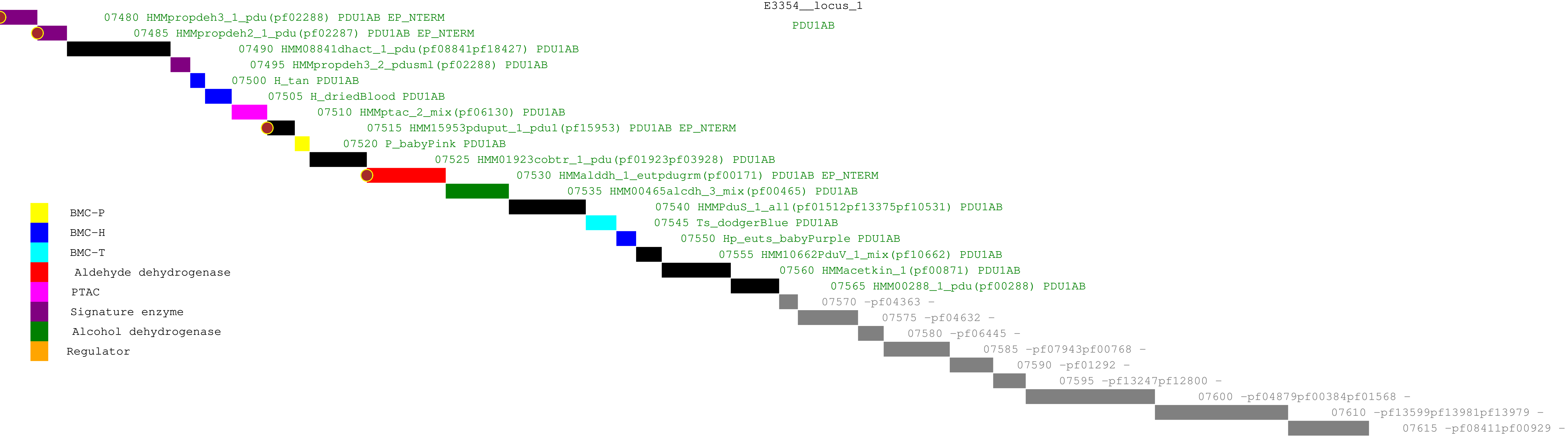

Salmonella enterica subsp. enterica serovar Waycross. Bacteria; Proteobacteria; Gammaproteobacteria; Enterobacterales;

E4339__locus_1

PDU1AB

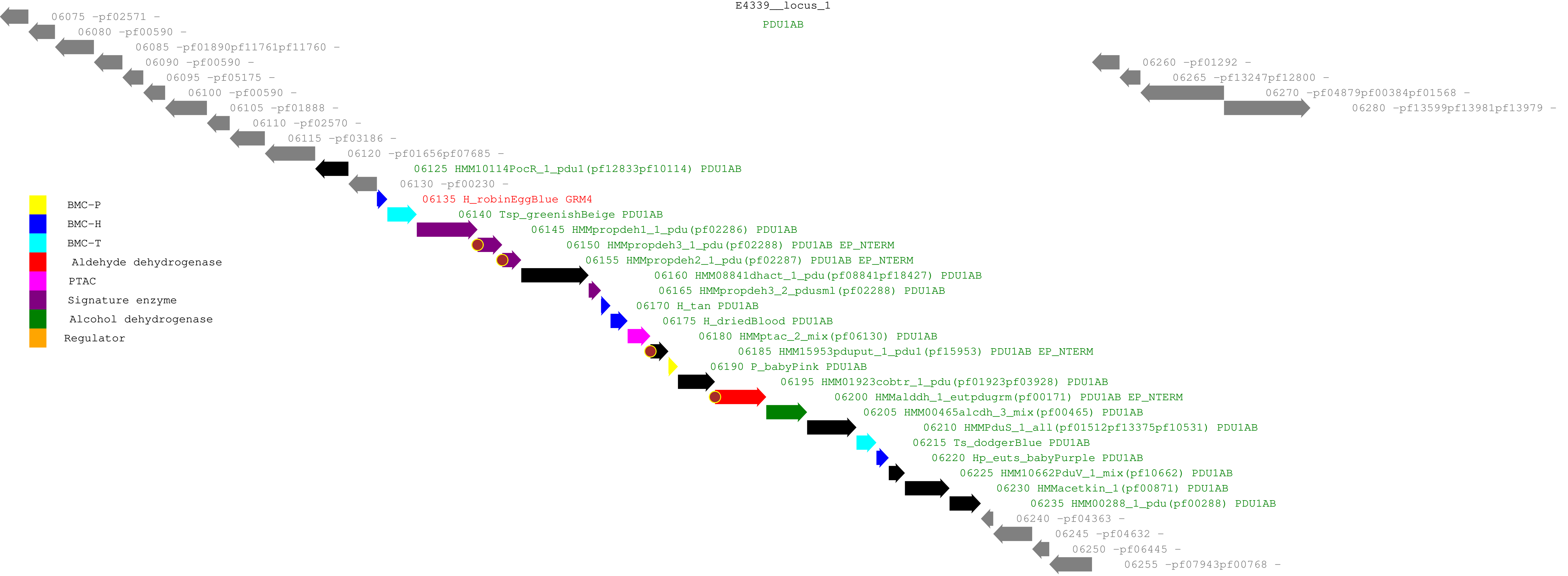

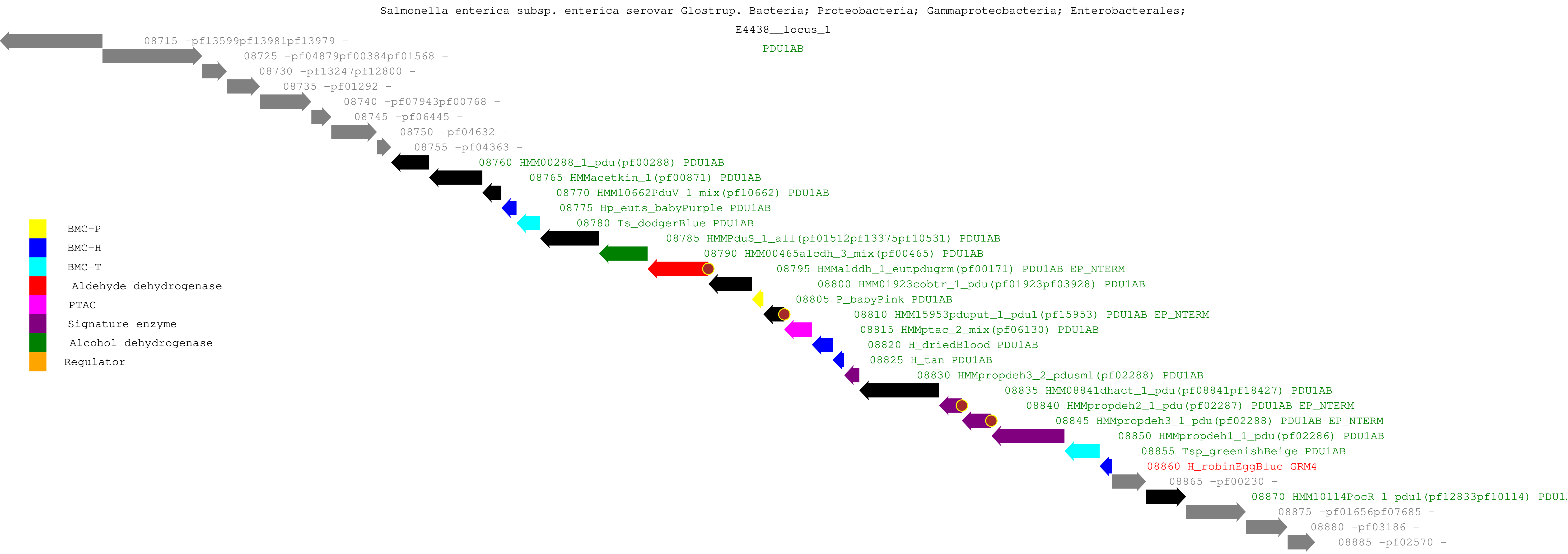

Salmonella enterica subsp. enterica serovar Glostrup. Bacteria; Proteobacteria; Gammaproteobacteria; Enterobacterales;
E4438__locus_1
PDU1AB
08715 -pf13599pf13981pf13979 -
08725 -pf04879pf00384pf01568 -
08730 -pf13247pf12800 -
08735 -pf01292 -
08740 -pf07943pf00768 -
08745 -pf06445 -
08750 -pf04632 -
08755 -pf04363 -
08760 HMM00288_1_pdu(pf00288) PDU1AB
08765 HMMacetkin_1(pf00871) PDU1AB
08770 HMM10662PduV_1_mix(pf10662) PDU1AB
08775 Hp_euts_babyPurple PDU1AB
08780 Ts_dodgerBlue PDU1AB
08785 HMMPduS_1_all(pf01512pf13375pf10531) PDU1AB
08790 HMM00465alcdh_3_mix(pf00465) PDU1AB
08795 HMMalddh_1_eutpdugrm(pf00171) PDU1AB EP_NTERM
08800 HMM01923cobtr_1_pdu(pf01923pf03928) PDU1AB
08805 P_babyPink PDU1AB
08810 HMM15953pduput_1_pdu1(pf15953) PDU1AB EP_NTERM
08815 HMMptac_2_mix(pf06130) PDU1AB
08820 H_driedBlood PDU1AB
08825 H_tan PDU1AB
08830 HMMpropdeh3_2_pdusml(pf02288) PDU1AB
08835 HMM08841dhact_1_pdu(pf08841pf18427) PDU1AB
08840 HMMpropdeh2_1_pdu(pf02287) PDU1AB EP_NTERM
08845 HMMpropdeh3_1_pdu(pf02288) PDU1AB EP_NTERM
08850 HMMpropdeh1_1_pdu(pf02286) PDU1AB
08855 Tsp_greenishBeige PDU1AB
08860 H_robinEggBlue GRM4
08865 -pf00230 -
08870 HMM10114PocR_1_pdu1(pf12833pf10114) PDU1
08875 -pf01656pf07685 -
08880 -pf03186 -
08885 -pf02570 -
BMC-P
BMC-H
BMC-T
Aldehyde dehydrogenase
PTAC
Signature enzyme
Alcohol dehydrogenase
Regulator

Salmonella enterica subsp. enterica serovar Apeyeme. Bacteria; Proteobacteria; Gammaproteobacteria; Enterobacterales;

E4443__locus_1

PDU1AB

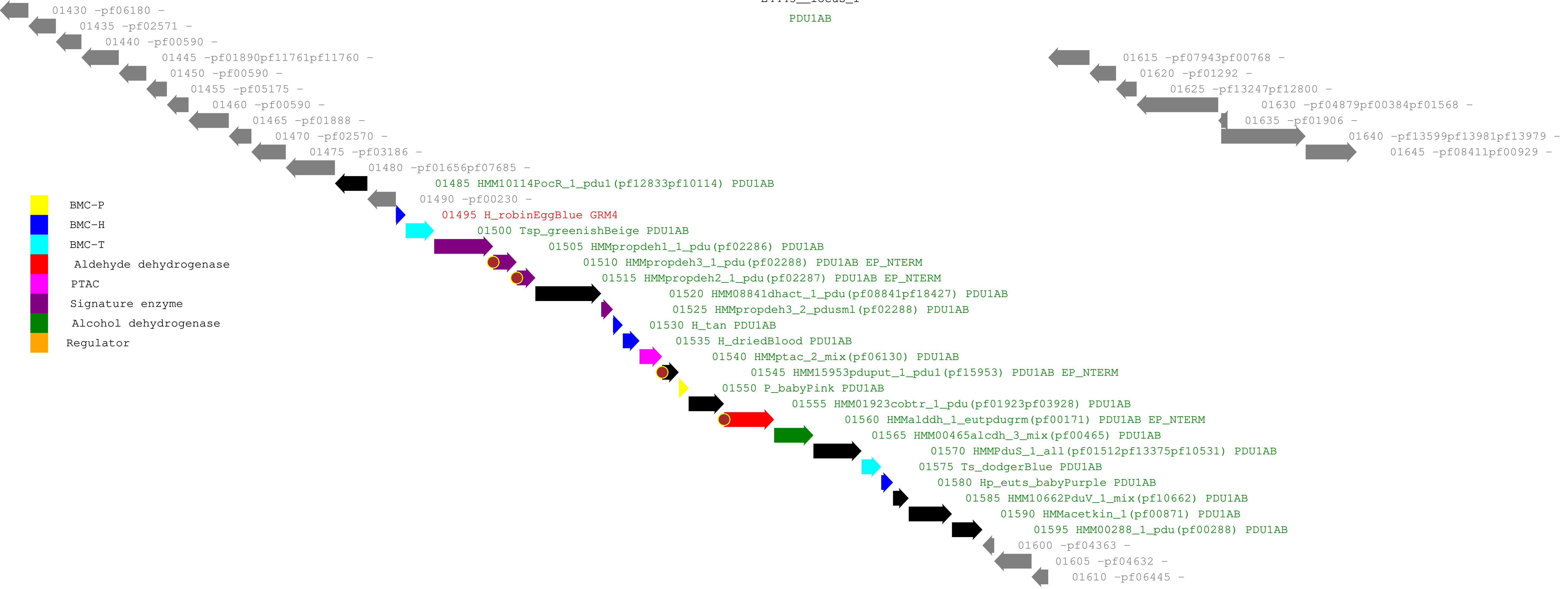

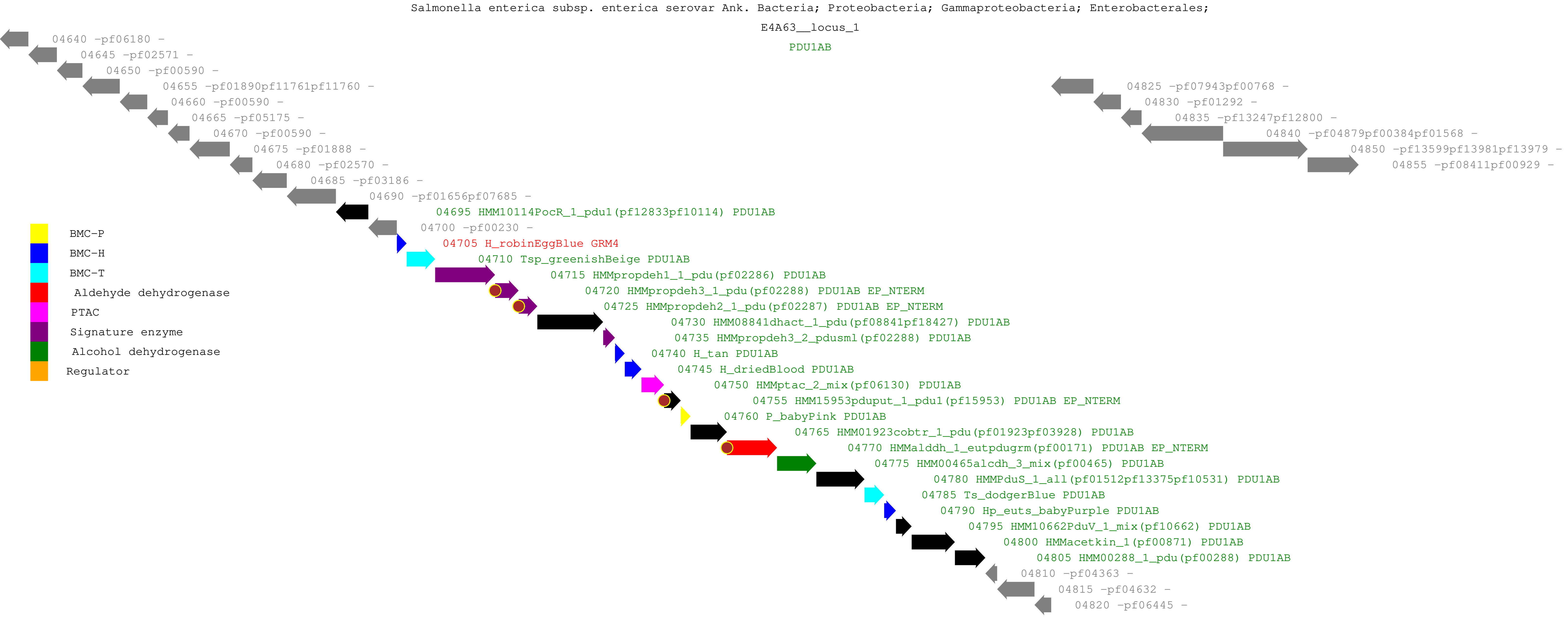

Salmonella enterica subsp. enterica serovar Ank. Bacteria; Proteobacteria; Gammaproteobacteria; Enterobacterales;
E4A63__locus_1
PDU1AB
04640 -pf06180 -
04645 -pf02571 -
04650 -pf00590 -
04655 -pf01890pf11761pf11760 -
04660 -pf00590 -
04665 -pf05175 -
04670 -pf00590 -
04675 -pf01888 -
04680 -pf02570 -
04685 -pf03186 -
04690 -pf01656pf07685 -
04695 HMM10114PocR_1_pdu1(pf12833pf10114) PDU1AB
04700 -pf00230 -
04705 H_robinEggBlue GRM4
04710 Tsp_greenishBeige PDU1AB
04715 HMMpropdeh1_1_pdu(pf02286) PDU1AB
04720 HMMpropdeh3_1_pdu(pf02288) PDU1AB EP_NTERM
04725 HMMpropdeh2_1_pdu(pf02287) PDU1AB EP_NTERM
04730 HMM08841dhact_1_pdu(pf08841pf18427) PDU1AB
04735 HMMpropdeh3_2_pdusml(pf02288) PDU1AB
04740 H_tan PDU1AB
04745 H_driedBlood PDU1AB
04750 HMMptac_2_mix(pf06130) PDU1AB
04755 HMM15953pduput_1_pdu1(pf15953) PDU1AB EP_NTERM
04760 P_babyPink PDU1AB
04765 HMM01923cobtr_1_pdu(pf01923pf03928) PDU1AB
04770 HMMalddh_1_eutpdugrm(pf00171) PDU1AB EP_NTERM
04775 HMM00465alcdh_3_mix(pf00465) PDU1AB
04780 HMMPduS_1_all(pf01512pf13375pf10531) PDU1AB
04785 Ts_dodgerBlue PDU1AB
04790 Hp_euts_babyPurple PDU1AB
04795 HMM10662PduV_1_mix(pf10662) PDU1AB
04800 HMMacetkin_1(pf00871) PDU1AB
04805 HMM00288_1_pdu(pf00288) PDU1AB
04810 -pf04363 -
04815 -pf04632 -
04820 -pf06445 -
04825 -pf07943pf00768 -
04830 -pf01292 -
04835 -pf13247pf12800 -
04840 -pf04879pf00384pf01568 -
04850 -pf13599pf13981pf13979 -
04855 -pf08411pf00929 -
BMC-P
BMC-H
BMC-T
Aldehyde dehydrogenase
PTAC
Signature enzyme
Alcohol dehydrogenase
Regulator

Salmonella enterica subsp. enterica serovar Adelaide. Bacteria; Proteobacteria; Gammaproteobacteria; Enterobacterales;

E4N46__locus_2

PDU1AB

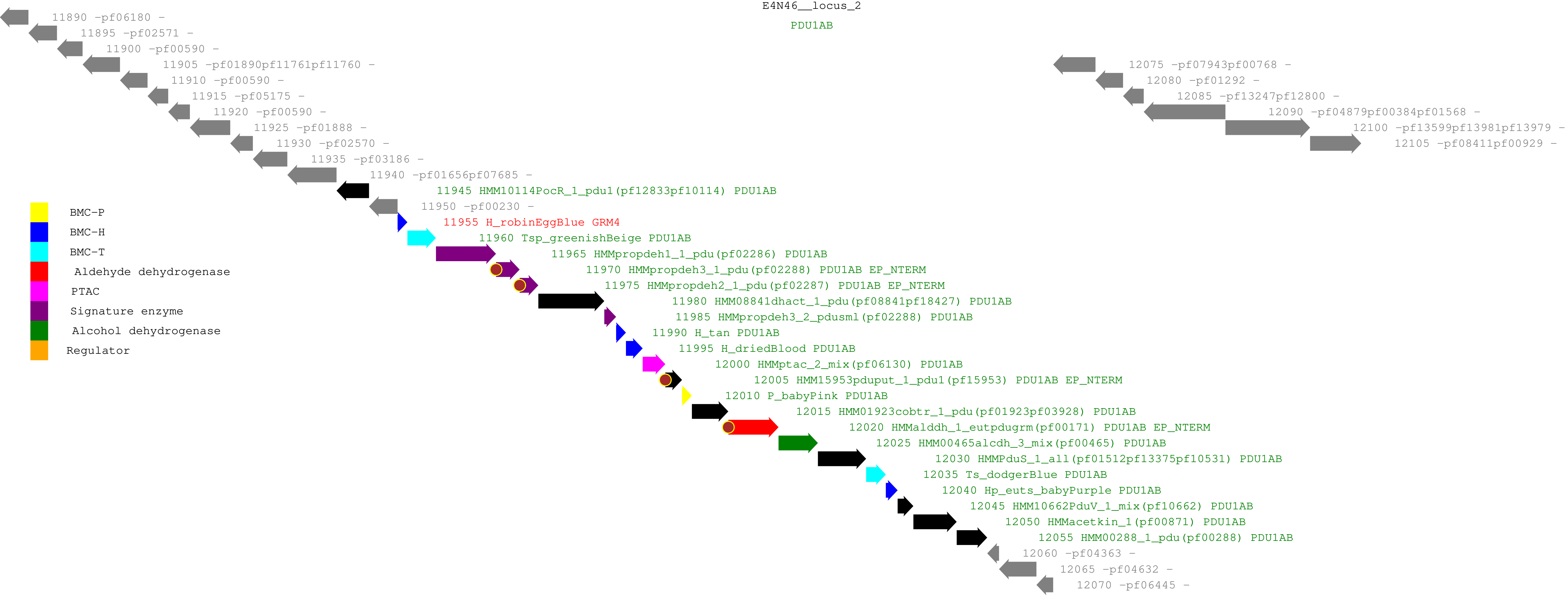

Salmonella potsdam. Bacteria; Proteobacteria; Gammaproteobacteria; Enterobacterales;

E4N53__locus_1

PDU1AB

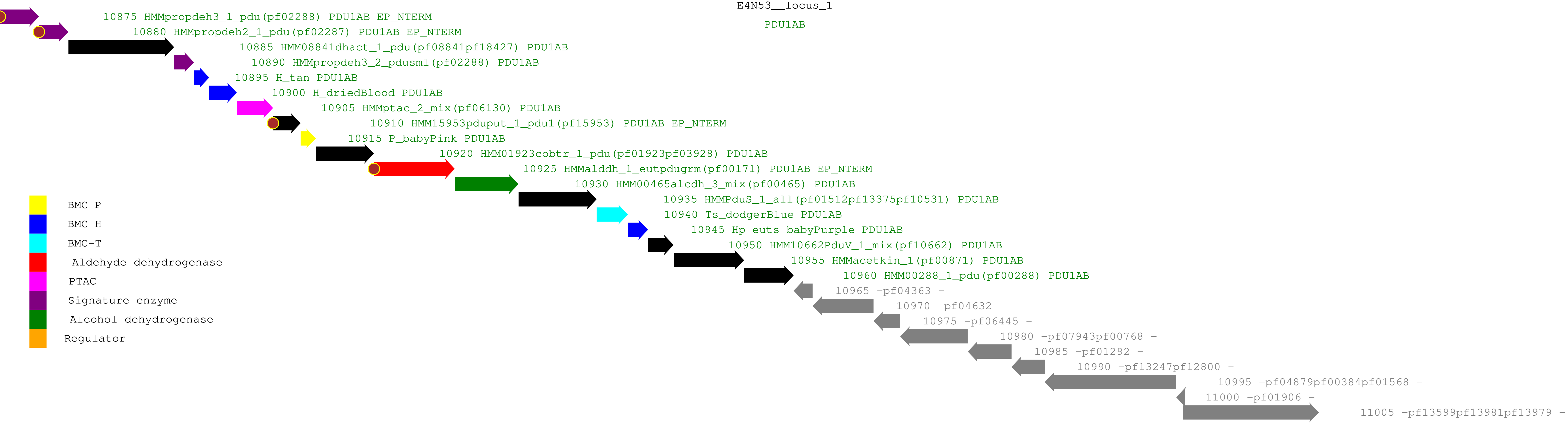

Citrobacter koseri (Citrobacter diversus). Bacteria; Proteobacteria; Gammaproteobacteria; Enterobacterales;

E4T98__locus_1

PDU1AB

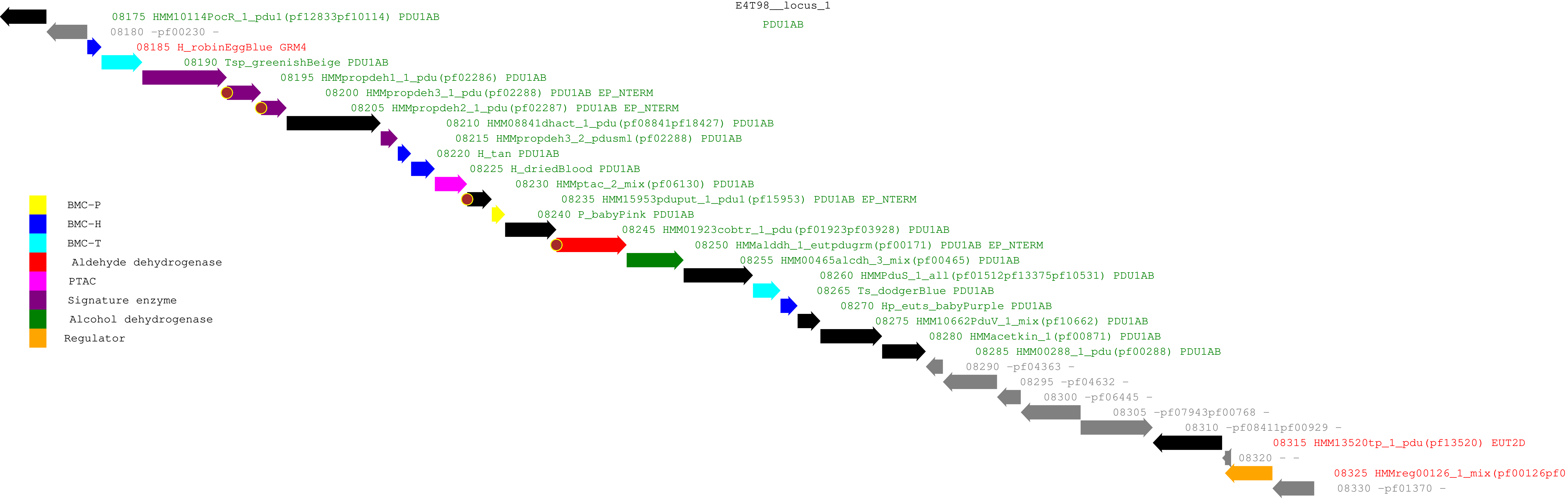

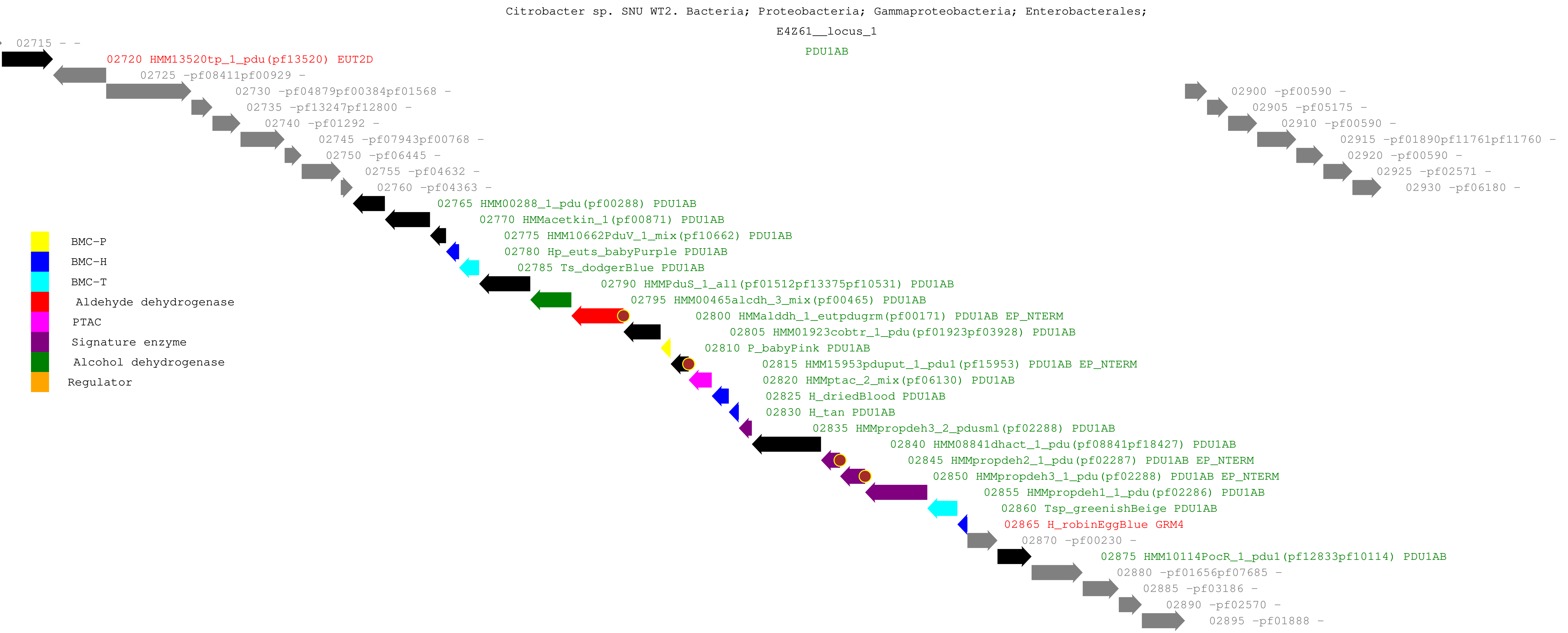
Citrobacter sp. SNU WT2. Bacteria; Proteobacteria; Gammaproteobacteria; Enterobacterales;
E4Z61__locus_1
PDU1AB
02715 - -
02720 HMM13520tp_1_pdu(pf13520) EUT2D
02725 -pf08411pf00929 -
02730 -pf04879pf00384pf01568 -
02735 -pf13247pf12800 -
02740 -pf01292 -
02745 -pf07943pf00768 -
02750 -pf06445 -
02755 -pf04632 -
02760 -pf04363 -
02765 HMM00288_1_pdu(pf00288) PDU1AB
02770 HMMacetkin_1(pf00871) PDU1AB
02775 HMM10662PduV_1_mix(pf10662) PDU1AB
02780 Hp_euts_babyPurple PDU1AB
02785 Ts_dodgerBlue PDU1AB
02790 HMMPduS_1_all(pf01512pf13375pf10531) PDU1AB
02795 HMM00465alcdh_3_mix(pf00465) PDU1AB
02800 HMMaldh_1_eutpdugrm(pf00171) PDU1AB EP_NTERM
02805 HMM01923cobtr_1_pdu(pf01923pf03928) PDU1AB
02810 P_babyPink PDU1AB
02815 HMM15953pduput_1_pdu1(pf15953) PDU1AB EP_NTERM
02820 HMMptac_2_mix(pf06130) PDU1AB
02825 H_driedBlood PDU1AB
02830 H_tan PDU1AB
02835 HMMpropdeh3_2_pdusml(pf02288) PDU1AB
02840 HMM08841dhact_1_pdu(pf08841pf18427) PDU1AB
02845 HMMpropdeh2_1_pdu(pf02287) PDU1AB EP_NTERM
02850 HMMpropdeh3_1_pdu(pf02288) PDU1AB EP_NTERM
02855 HMMpropdeh1_1_pdu(pf02286) PDU1AB
02860 Tsp_greenishBeige PDU1AB
02865 H_robinEggBlue GRM4
02870 -pf00230 -
02875 HMM10114PocR_1_pdu1(pf12833pf10114) PDU1AB
02880 -pf01656pf07685 -
02885 -pf03186 -
02890 -pf02570 -
02895 -pf01888 -
02900 -pf00590 -
02905 -pf05175 -
02910 -pf00590 -
02915 -pf01890pf11761pf11760 -
02920 -pf00590 -
02925 -pf02571 -
02930 -pf06180 -
BMC-P
BMC-H
BMC-T
Aldehyde dehydrogenase
PTAC
Signature enzyme
Alcohol dehydrogenase
Regulator

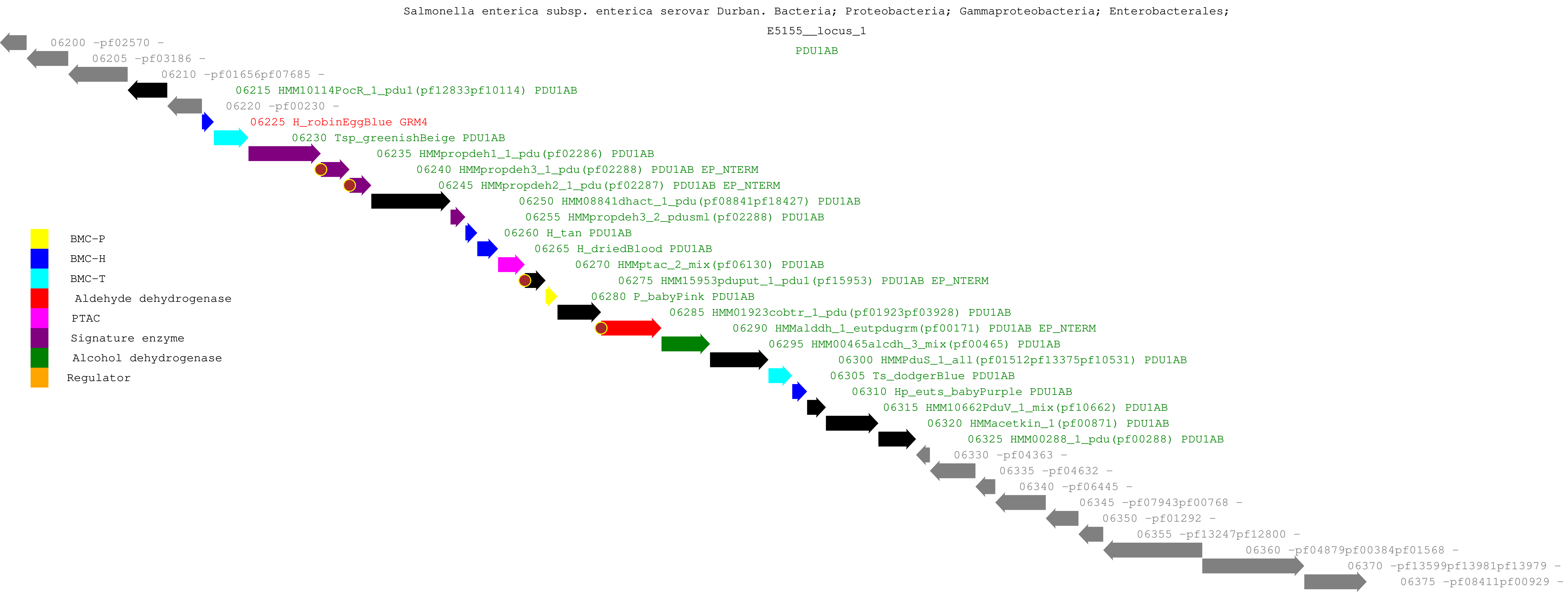
Salmonella enterica subsp. enterica serovar Durban. Bacteria; Proteobacteria; Gammaproteobacteria; Enterobacterales;
E5155__locus_1
PDU1AB
06200 -pf02570 -
06205 -pf03186 -
06210 -pf01656pf07685 -
06215 HMM10114PocR_1_pdu1(pf12833pf10114) PDU1AB
06220 -pf00230 -
06225 H_robinEggBlue GRM4
06230 Tsp_greenishBeige PDU1AB
06235 HMMpropdeh1_1_pdu(pf02286) PDU1AB
06240 HMMpropdeh3_1_pdu(pf02288) PDU1AB EP_NTERM
06245 HMMpropdeh2_1_pdu(pf02287) PDU1AB EP_NTERM
06250 HMM08841dhact_1_pdu(pf08841pf18427) PDU1AB
06255 HMMpropdeh3_2_pdusml(pf02288) PDU1AB
06260 H_tan PDU1AB
06265 H_driedBlood PDU1AB
06270 HMMptac_2_mix(pf06130) PDU1AB
06275 HMM15953pduput_1_pdu1(pf15953) PDU1AB EP_NTERM
06280 P_babyPink PDU1AB
06285 HMM01923cobtr_1_pdu(pf01923pf03928) PDU1AB
06290 HMMalddh_1_eutpdugrm(pf00171) PDU1AB EP_NTERM
06295 HMM00465alcdh_3_mix(pf00465) PDU1AB
06300 HMMPduS_1_all(pf01512pf13375pf10531) PDU1AB
06305 Ts_dodgerBlue PDU1AB
06310 Hp_euts_babyPurple PDU1AB
06315 HMM10662PduV_1_mix(pf10662) PDU1AB
06320 HMMacetkin_1(pf00871) PDU1AB
06325 HMM00288_1_pdu(pf00288) PDU1AB
06330 -pf04363 -
06335 -pf04632 -
06340 -pf06445 -
06345 -pf07943pf00768 -
06350 -pf01292 -
06355 -pf13247pf12800 -
06360 -pf04879pf00384pf01568 -
06370 -pf13599pf13981pf13979 -
06375 -pf08411pf00929 -
BMC-P
BMC-H
BMC-T
Aldehyde dehydrogenase
PTAC
Signature enzyme
Alcohol dehydrogenase
Regulator

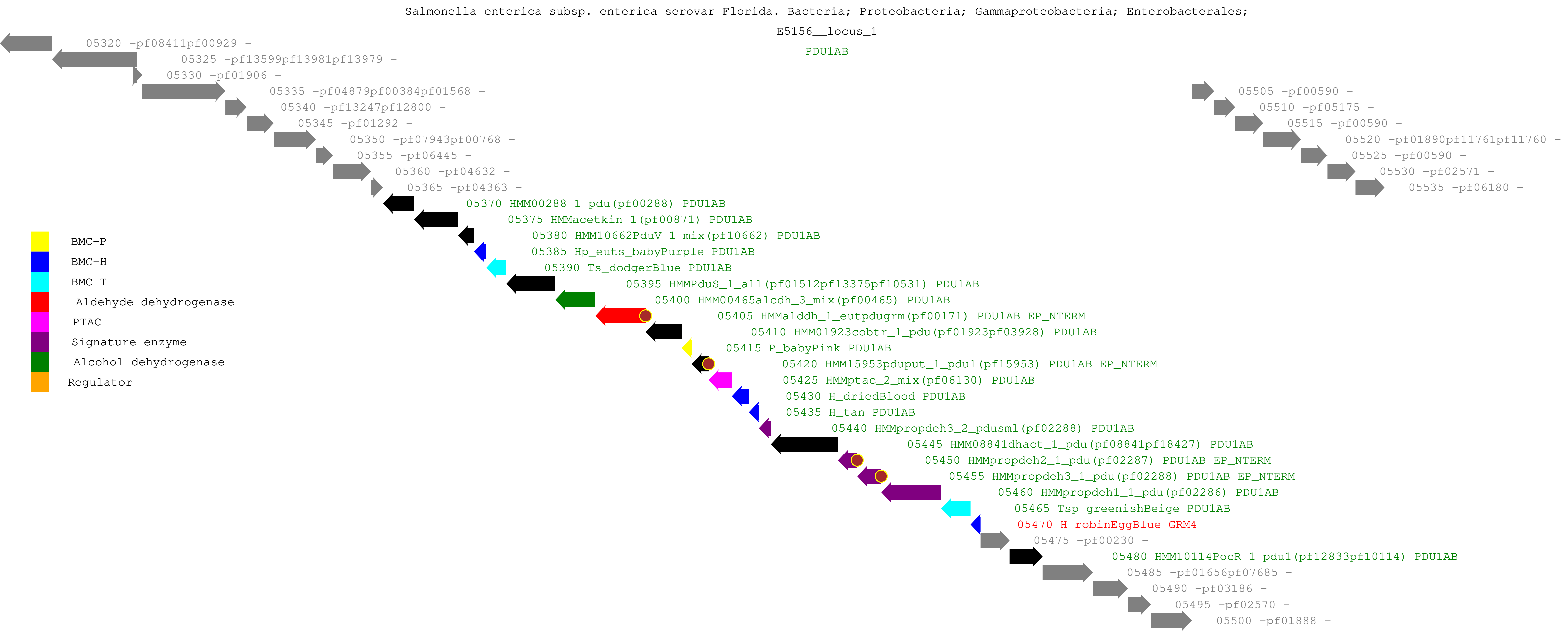

Salmonella enterica subsp. enterica serovar Florida. Bacteria; Proteobacteria; Gammaproteobacteria; Enterobacterales;
E5156__locus_1
PDU1AB
05320 -pf08411pf00929 -
05325 -pf13599pf13981pf13979 -
05330 -pf01906 -
05335 -pf04879pf00384pf01568 -
05340 -pf13247pf12800 -
05345 -pf01292 -
05350 -pf07943pf00768 -
05355 -pf06445 -
05360 -pf04632 -
05365 -pf04363 -
05370 HMM00288_1_pdu(pf00288) PDU1AB
05375 HMMacetkin_1(pf00871) PDU1AB
05380 HMM10662PduV_1_mix(pf10662) PDU1AB
05385 Hp_euts_babyPurple PDU1AB
05390 Ts_dodgerBlue PDU1AB
05395 HMMPduS_1_all(pf01512pf13375pf10531) PDU1AB
05400 HMM00465alcdh_3_mix(pf00465) PDU1AB
05405 HMMalddh_1_eutpdugrm(pf00171) PDU1AB EP_NTERM
05410 HMM01923cobtr_1_pdu(pf01923pf03928) PDU1AB
05415 P_babyPink PDU1AB
05420 HMM15953pduput_1_pdu1(pf15953) PDU1AB EP_NTERM
05425 HMMptac_2_mix(pf06130) PDU1AB
05430 H_driedBlood PDU1AB
05435 H_tan PDU1AB
05440 HMMpropdeh3_2_pdusml(pf02288) PDU1AB
05445 HMM08841dhact_1_pdu(pf08841pf18427) PDU1AB
05450 HMMpropdeh2_1_pdu(pf02287) PDU1AB EP_NTERM
05455 HMMpropdeh3_1_pdu(pf02288) PDU1AB EP_NTERM
05460 HMMpropdeh1_1_pdu(pf02286) PDU1AB
05465 Tsp_greenishBeige PDU1AB
05470 H_robinEggBlue GRM4
05475 -pf00230 -
05480 HMM10114PocR_1_pdu1(pf12833pf10114) PDU1AB
05485 -pf01656pf07685 -
05490 -pf03186 -
05495 -pf02570 -
05500 -pf01888 -
05505 -pf00590 -
05510 -pf05175 -
05515 -pf00590 -
05520 -pf01890pf11761pf11760 -
05525 -pf00590 -
05530 -pf02571 -
05535 -pf06180 -
BMC-P
BMC-H
BMC-T
Aldehyde dehydrogenase
PTAC
Signature enzyme
Alcohol dehydrogenase
Regulator

Salmonella enterica subsp. enterica serovar Westhampton. Bacteria; Proteobacteria; Gammaproteobacteria; Enterobacterales;

E5213__locus_2

PDU1AB

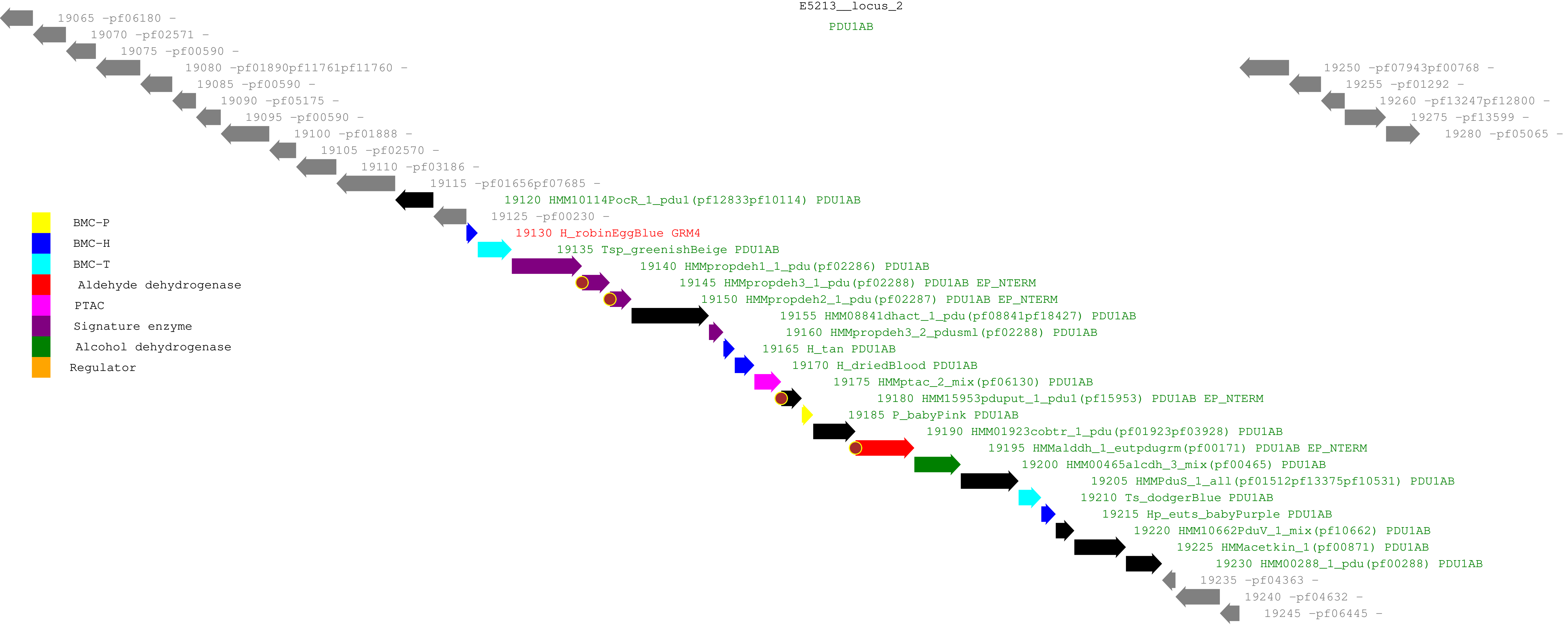

Citrobacter freundii. Bacteria; Proteobacteria; Gammaproteobacteria; Enterobacterales;

E5284__locus_2

PDU1AB

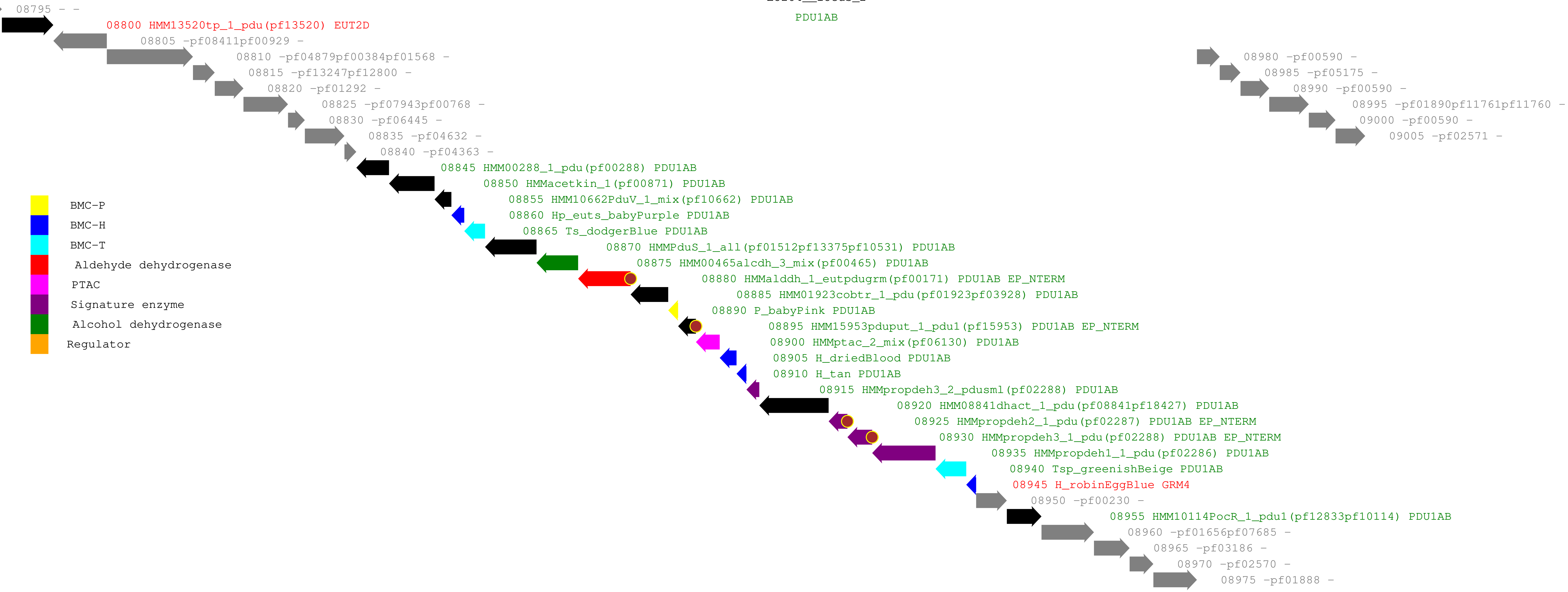

Salmonella enterica subsp. enterica serovar Lagos. Bacteria; Proteobacteria; Gammaproteobacteria; Enterobacterales;

E5J91__locus_1

PDU1AB

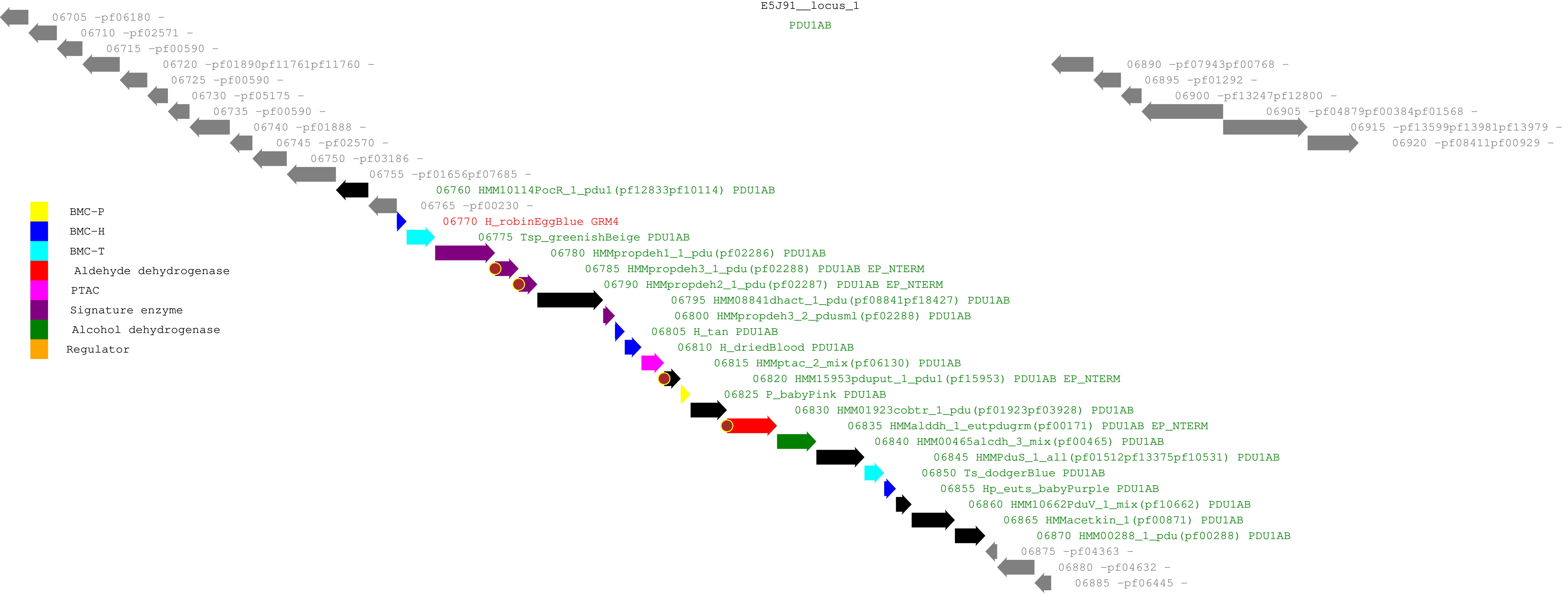

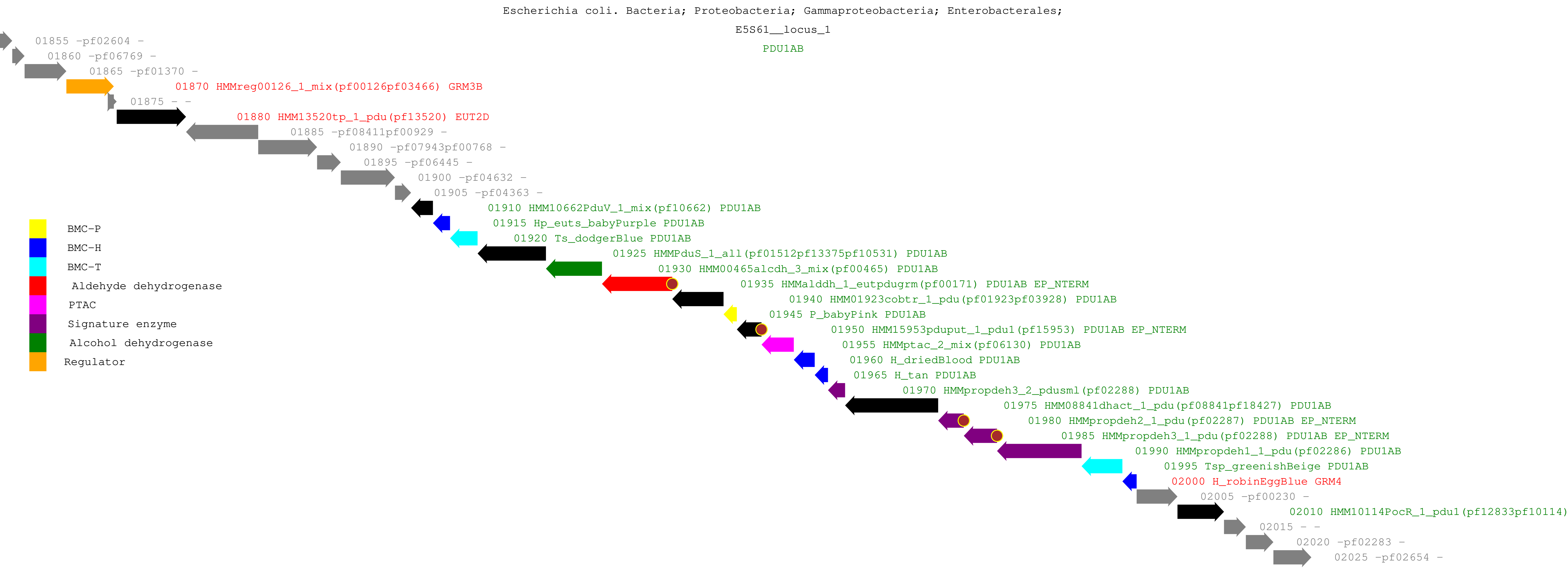
Escherichia coli. Bacteria; Proteobacteria; Gammaproteobacteria; Enterobacterales;
E5S61__locus_1
PDU1AB
01855 -pf02604 -
01860 -pf06769 -
01865 -pf01370 -
01870 HMMreg00126_1_mix(pf00126pf03466) GRM3B
01875 - -
01880 HMM13520tp_1_pdu(pf13520) EUT2D
01885 -pf08411pf00929 -
01890 -pf07943pf00768 -
01895 -pf06445 -
01900 -pf04632 -
01905 -pf04363 -
01910 HMM10662PduV_1_mix(pf10662) PDU1AB
01915 Hp_euts_babyPurple PDU1AB
01920 Ts_dodgerBlue PDU1AB
01925 HMMPduS_1_all(pf01512pf13375pf10531) PDU1AB
01930 HMM00465alcdh_3_mix(pf00465) PDU1AB
01935 HMMalddh_1_eutpdugrm(pf00171) PDU1AB EP_NTERM
01940 HMM01923cobtr_1_pdu(pf01923pf03928) PDU1AB
01945 P_babyPink PDU1AB
01950 HMM15953pduput_1_pdu1(pf15953) PDU1AB EP_NTERM
01955 HMMptac_2_mix(pf06130) PDU1AB
01960 H_driedBlood PDU1AB
01965 H_tan PDU1AB
01970 HMMpropdeh3_2_pdusml(pf02288) PDU1AB
01975 HMM08841dhact_1_pdu(pf08841pf18427) PDU1AB
01980 HMMpropdeh2_1_pdu(pf02287) PDU1AB EP_NTERM
01985 HMMpropdeh3_1_pdu(pf02288) PDU1AB EP_NTERM
01990 HMMpropdeh1_1_pdu(pf02286) PDU1AB
01995 Tsp_greenishBeige PDU1AB
02000 H_robinEggBlue GRM4
02005 -pf00230 -
02010 HMM10114PocR_1_pdu1(pf12833pf10114)
02015 - -
02020 -pf02283 -
02025 -pf02654 -
BMC-P
BMC-H
BMC-T
Aldehyde dehydrogenase
PTAC
Signature enzyme
Alcohol dehydrogenase
Regulator

Salmonella enterica subsp. enterica serovar Stanleyville. Bacteria; Proteobacteria; Gammaproteobacteria; Enterobacterales;

E5U21__locus_1

PDU1AB

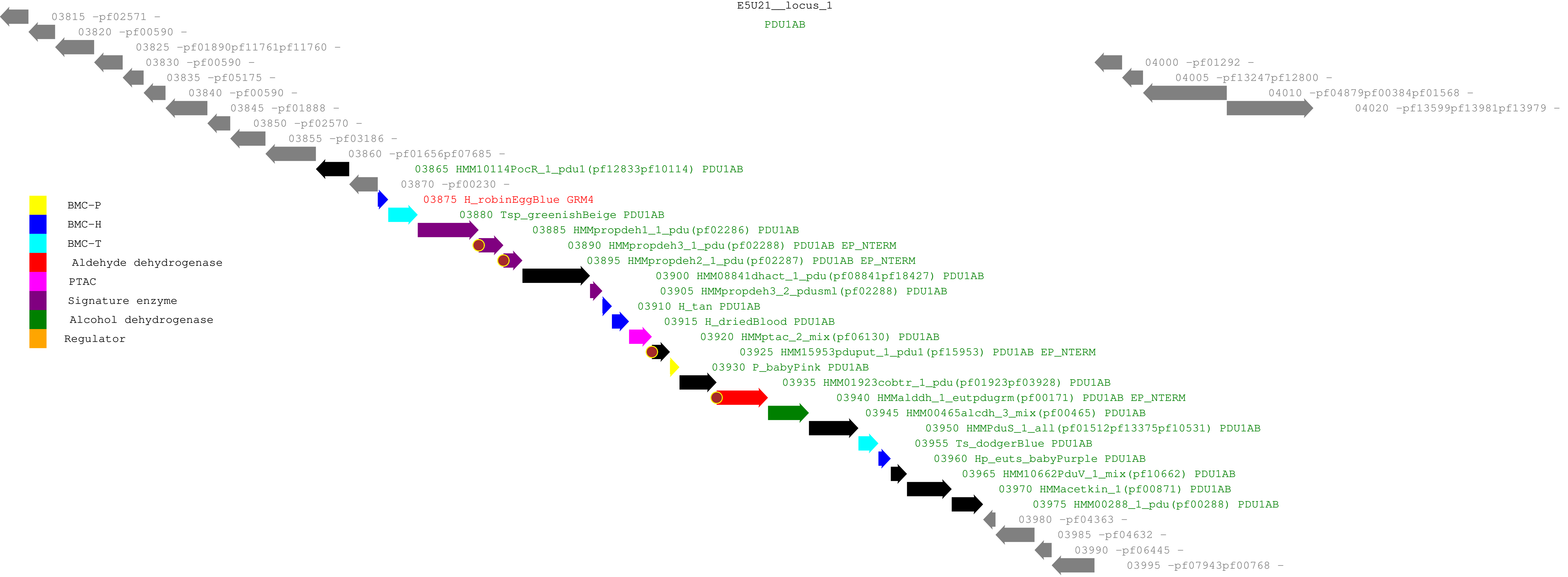

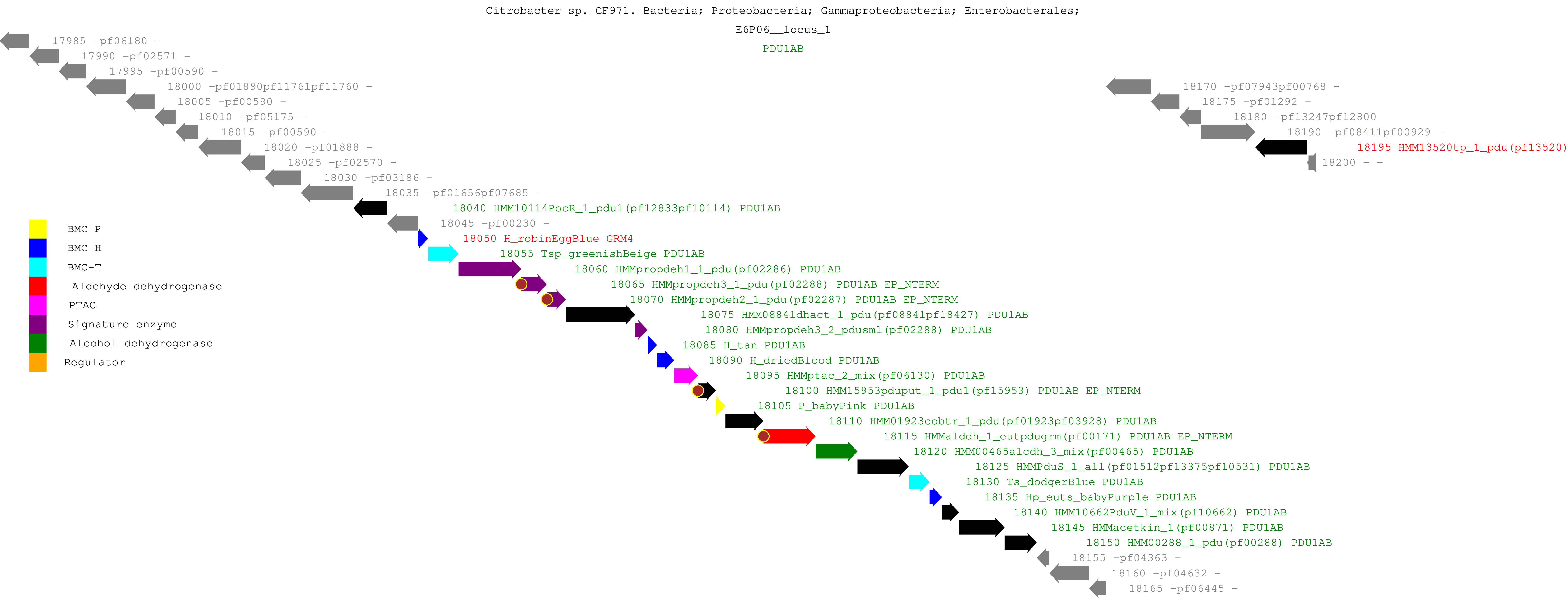
Citrobacter sp. CF971. Bacteria; Proteobacteria; Gammaproteobacteria; Enterobacterales;
E6P06__locus_1
PDU1AB
17985 -pf06180 -
17990 -pf02571 -
17995 -pf00590 -
18000 -pf01890pf11761pf11760 -
18005 -pf00590 -
18010 -pf05175 -
18015 -pf00590 -
18020 -pf01888 -
18025 -pf02570 -
18030 -pf03186 -
18035 -pf01656pf07685 -
18040 HMM10114PocR_1_pdu1(pf12833pf10114) PDU1AB
18045 -pf00230 -
18050 H_robinEggBlue GRM4
18055 Tsp_greenishBeige PDU1AB
18060 HMMpropdeh1_1_pdu(pf02286) PDU1AB
18065 HMMpropdeh3_1_pdu(pf02288) PDU1AB EP_NTERM
18070 HMMpropdeh2_1_pdu(pf02287) PDU1AB EP_NTERM
18075 HMM08841dhact_1_pdu(pf08841pf18427) PDU1AB
18080 HMMpropdeh3_2_pdusml(pf02288) PDU1AB
18085 H_tan PDU1AB
18090 H_driedBlood PDU1AB
18095 HMMptac_2_mix(pf06130) PDU1AB
18100 HMM15953pduput_1_pdu1(pf15953) PDU1AB EP_NTERM
18105 P_babyPink PDU1AB
18110 HMM01923cobtr_1_pdu(pf01923pf03928) PDU1AB
18115 HMMalddh_1_eutpdugrm(pf00171) PDU1AB EP_NTERM
18120 HMM00465alcdh_3_mix(pf00465) PDU1AB
18125 HMMPduS_1_all(pf01512pf13375pf10531) PDU1AB
18130 Ts_dodgerBlue PDU1AB
18135 Hp_euts_babyPurple PDU1AB
18140 HMM10662PduV_1_mix(pf10662) PDU1AB
18145 HMMacetkin_1(pf00871) PDU1AB
18150 HMM00288_1_pdu(pf00288) PDU1AB
18155 -pf04363 -
18160 -pf04632 -
18165 -pf06445 -
18170 -pf07943pf00768 -
18175 -pf01292 -
18180 -pf13247pf12800 -
18190 -pf08411pf00929 -
18195 HMM13520tp_1_pdu(pf13520)
18200 - -
BMC-P
BMC-H
BMC-T
Aldehyde dehydrogenase
PTAC
Signature enzyme
Alcohol dehydrogenase
Regulator

Salmonella enterica subsp. enterica serovar Berkeley. Bacteria; Proteobacteria; Gammaproteobacteria; Enterobacterales;

E6W86__locus_2

PDU1AB

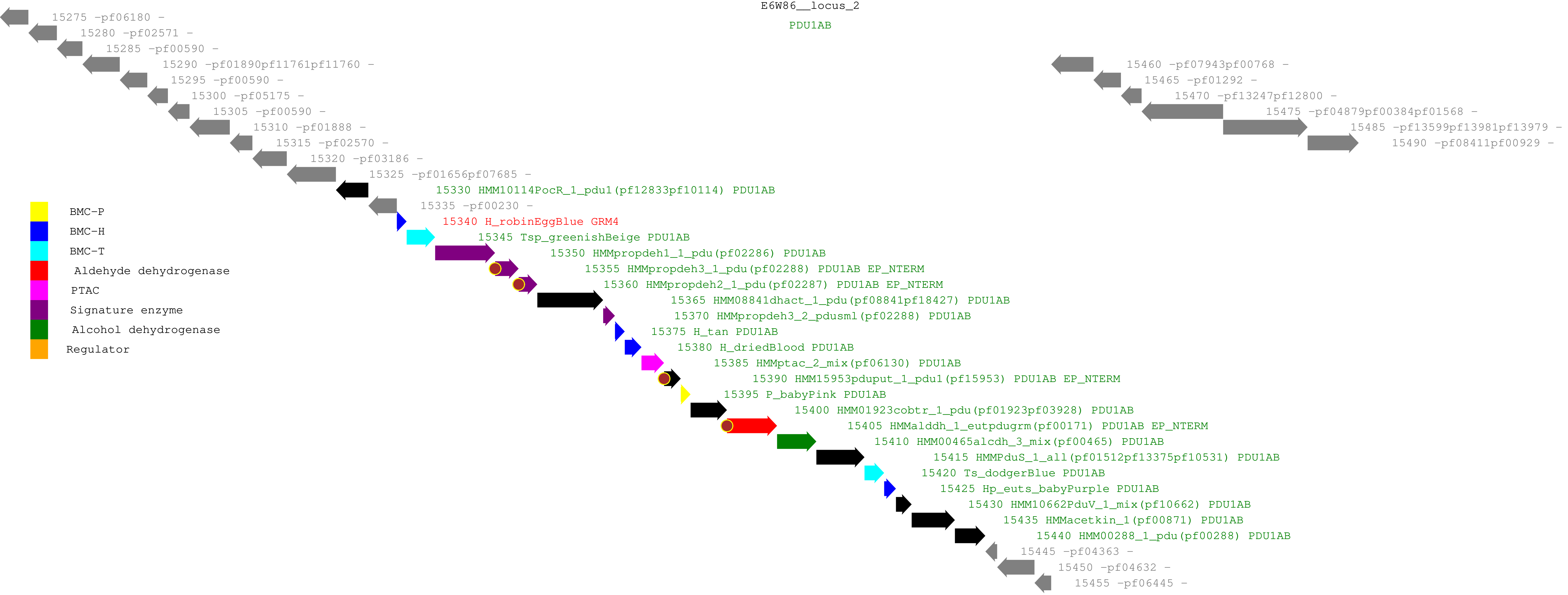

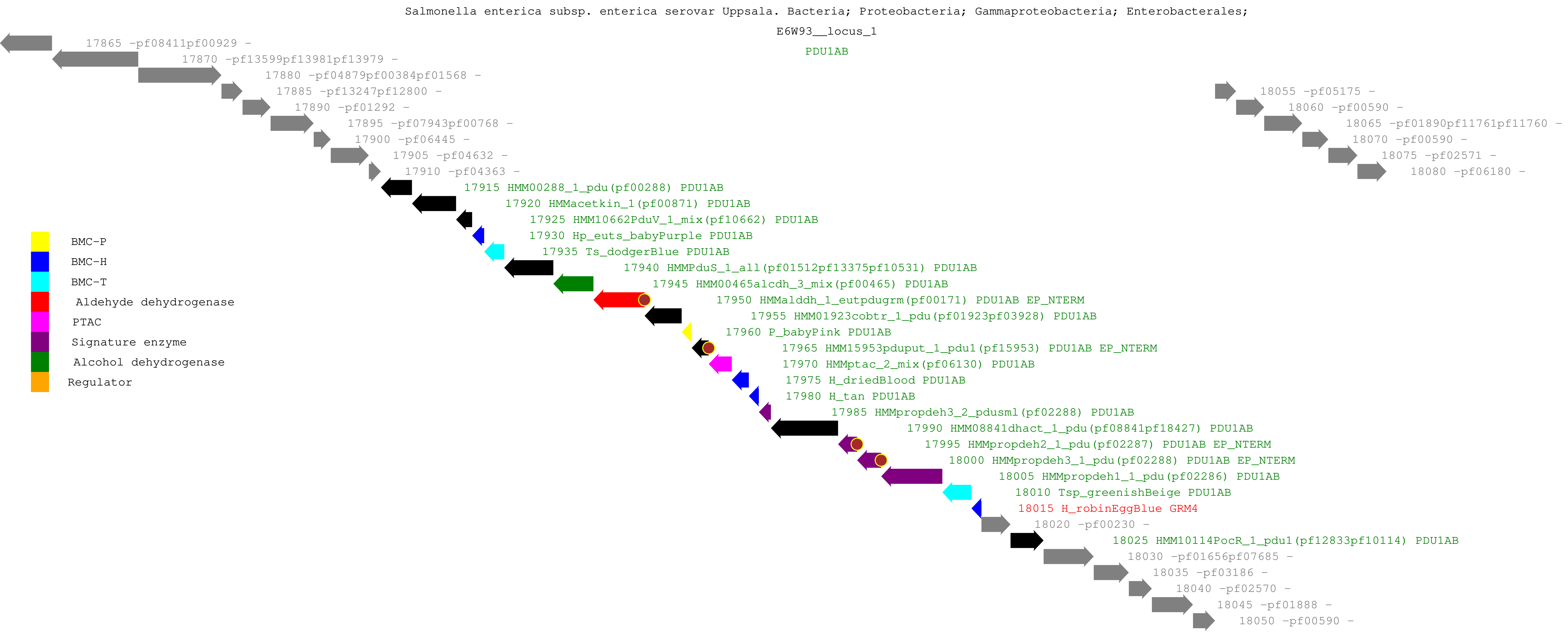

Salmonella enterica subsp. enterica serovar Uppsala. Bacteria; Proteobacteria; Gammaproteobacteria; Enterobacterales;
E6W93__locus_1
PDU1AB
17865 -pf08411pf00929 -
17870 -pf13599pf13981pf13979 -
17880 -pf04879pf00384pf01568 -
17885 -pf13247pf12800 -
17890 -pf01292 -
17895 -pf07943pf00768 -
17900 -pf06445 -
17905 -pf04632 -
17910 -pf04363 -
17915 HMM00288_1_pdu(pf00288) PDU1AB
17920 HMMacetkin_1(pf00871) PDU1AB
17925 HMM10662PduV_1_mix(pf10662) PDU1AB
17930 Hp_euts_babyPurple PDU1AB
17935 Ts_dodgerBlue PDU1AB
17940 HMMPduS_1_all(pf01512pf13375pf10531) PDU1AB
17945 HMM00465alcdh_3_mix(pf00465) PDU1AB
17950 HMMalddh_1_eutpdugrm(pf00171) PDU1AB EP_NTERM
17955 HMM01923cobtr_1_pdu(pf01923pf03928) PDU1AB
17960 P_babyPink PDU1AB
17965 HMM15953pduput_1_pdu1(pf15953) PDU1AB EP_NTERM
17970 HMMptac_2_mix(pf06130) PDU1AB
17975 H_driedBlood PDU1AB
17980 H_tan PDU1AB
17985 HMMpropdeh3_2_pdusml(pf02288) PDU1AB
17990 HMM08841dhact_1_pdu(pf08841pf18427) PDU1AB
17995 HMMpropdeh2_1_pdu(pf02287) PDU1AB EP_NTERM
18000 HMMpropdeh3_1_pdu(pf02288) PDU1AB EP_NTERM
18005 HMMpropdeh1_1_pdu(pf02286) PDU1AB
18010 Tsp_greenishBeige PDU1AB
18015 H_robinEggBlue GRM4
18020 -pf00230 -
18025 HMM10114PocR_1_pdu1(pf12833pf10114) PDU1AB
18030 -pf01656pf07685 -
18035 -pf03186 -
18040 -pf02570 -
18045 -pf01888 -
18050 -pf00590 -
18055 -pf05175 -
18060 -pf00590 -
18065 -pf01890pf11761pf11760 -
18070 -pf00590 -
18075 -pf02571 -
18080 -pf06180 -
BMC-P
BMC-H
BMC-T
Aldehyde dehydrogenase
PTAC
Signature enzyme
Alcohol dehydrogenase
Regulator

Salmonella enterica subsp. enterica serovar Poano. Bacteria; Proteobacteria; Gammaproteobacteria; Enterobacterales;

E6W94__locus_2

PDU1AB

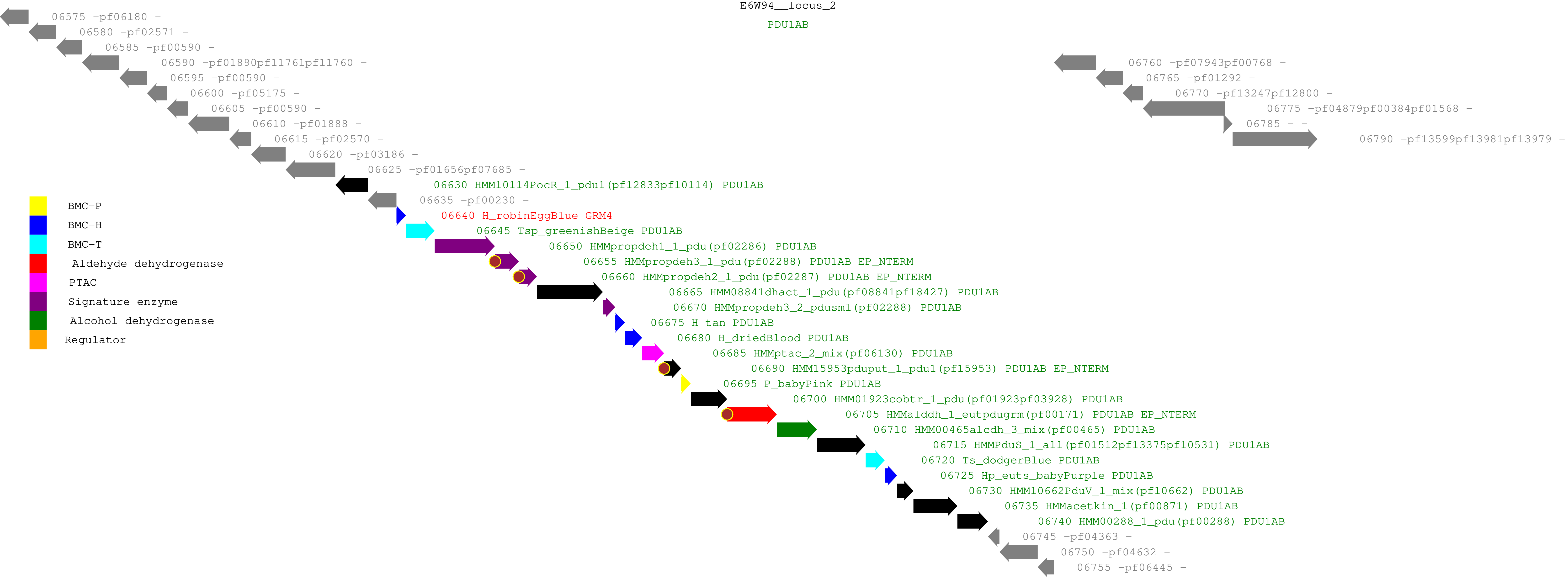

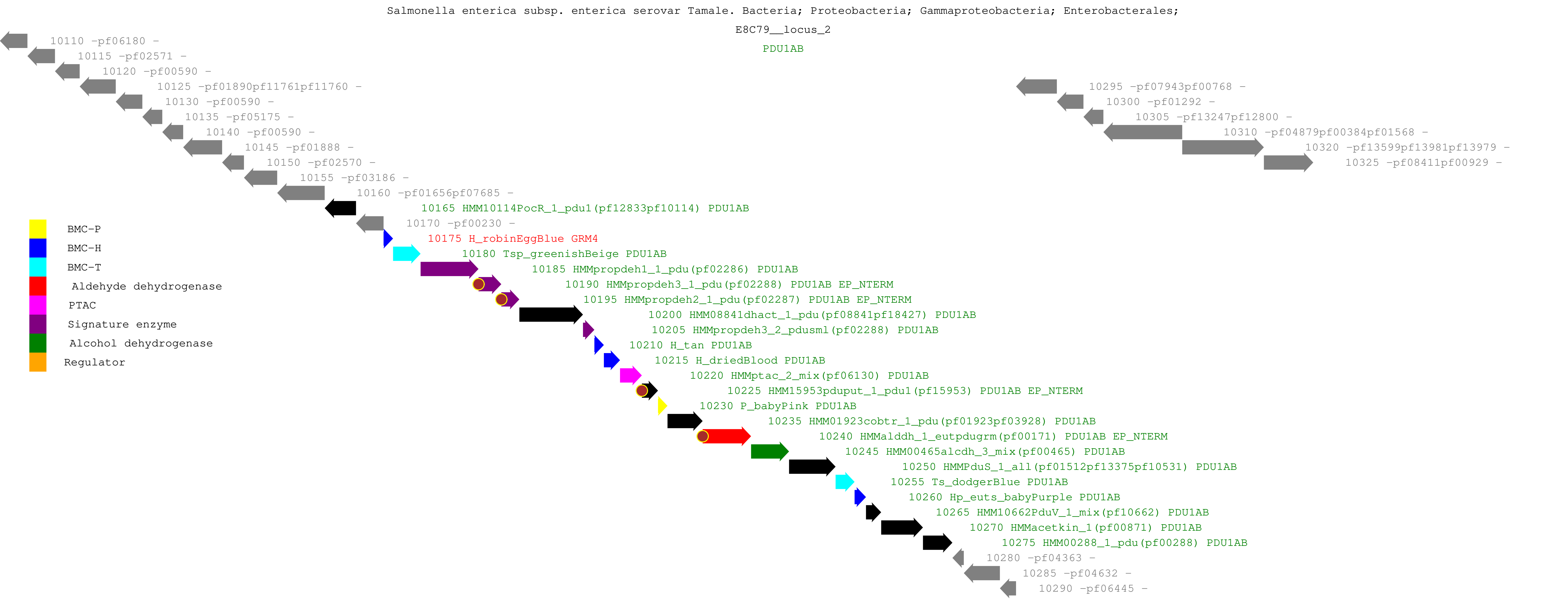
Salmonella enterica subsp. enterica serovar Tamale. Bacteria; Proteobacteria; Gammaproteobacteria; Enterobacterales;
E8C79__locus_2
PDU1AB
10110 -pf06180 -
10115 -pf02571 -
10120 -pf00590 -
10125 -pf01890pf11761pf11760 -
10130 -pf00590 -
10135 -pf05175 -
10140 -pf00590 -
10145 -pf01888 -
10150 -pf02570 -
10155 -pf03186 -
10160 -pf01656pf07685 -
10165 HMM10114PocR_1_pdu1(pf12833pf10114) PDU1AB
10170 -pf00230 -
10175 H_robinEggBlue GRM4
10180 Tsp_greenishBeige PDU1AB
10185 HMMpropdeh1_1_pdu(pf02286) PDU1AB
10190 HMMpropdeh3_1_pdu(pf02288) PDU1AB EP_NTERM
10195 HMMpropdeh2_1_pdu(pf02287) PDU1AB EP_NTERM
10200 HMM08841dhact_1_pdu(pf08841pf18427) PDU1AB
10205 HMMpropdeh3_2_pdusml(pf02288) PDU1AB
10210 H_tan PDU1AB
10215 H_driedBlood PDU1AB
10220 HMMptac_2_mix(pf06130) PDU1AB
10225 HMM15953pduput_1_pdu1(pf15953) PDU1AB EP_NTERM
10230 P_babyPink PDU1AB
10235 HMM01923cobtr_1_pdu(pf01923pf03928) PDU1AB
10240 HMMaldd h_1_eutpdugrm(pf00171) PDU1AB EP_NTERM
10245 HMM00465alcdh_3_mix(pf00465) PDU1AB
10250 HMMPduS_1_all(pf01512pf13375pf10531) PDU1AB
10255 Ts_dodgerBlue PDU1AB
10260 Hp_euts_babyPurple PDU1AB
10265 HMM10662PduV_1_mix(pf10662) PDU1AB
10270 HMMacetkin_1(pf00871) PDU1AB
10275 HMM00288_1_pdu(pf00288) PDU1AB
10280 -pf04363 -
10285 -pf04632 -
10290 -pf06445 -
10295 -pf07943pf00768 -
10300 -pf01292 -
10305 -pf13247pf12800 -
10310 -pf04879pf00384pf01568 -
10320 -pf13599pf13981pf13979 -
10325 -pf08411pf00929 -
BMC-P
BMC-H
BMC-T
Aldehyde dehydrogenase
PTAC
Signature enzyme
Alcohol dehydrogenase
Regulator

Salmonella enterica subsp. enterica serovar Amsterdam. Bacteria; Proteobacteria; Gammaproteobacteria; Enterobacterales;

E8Q13__locus_1

PDU1AB

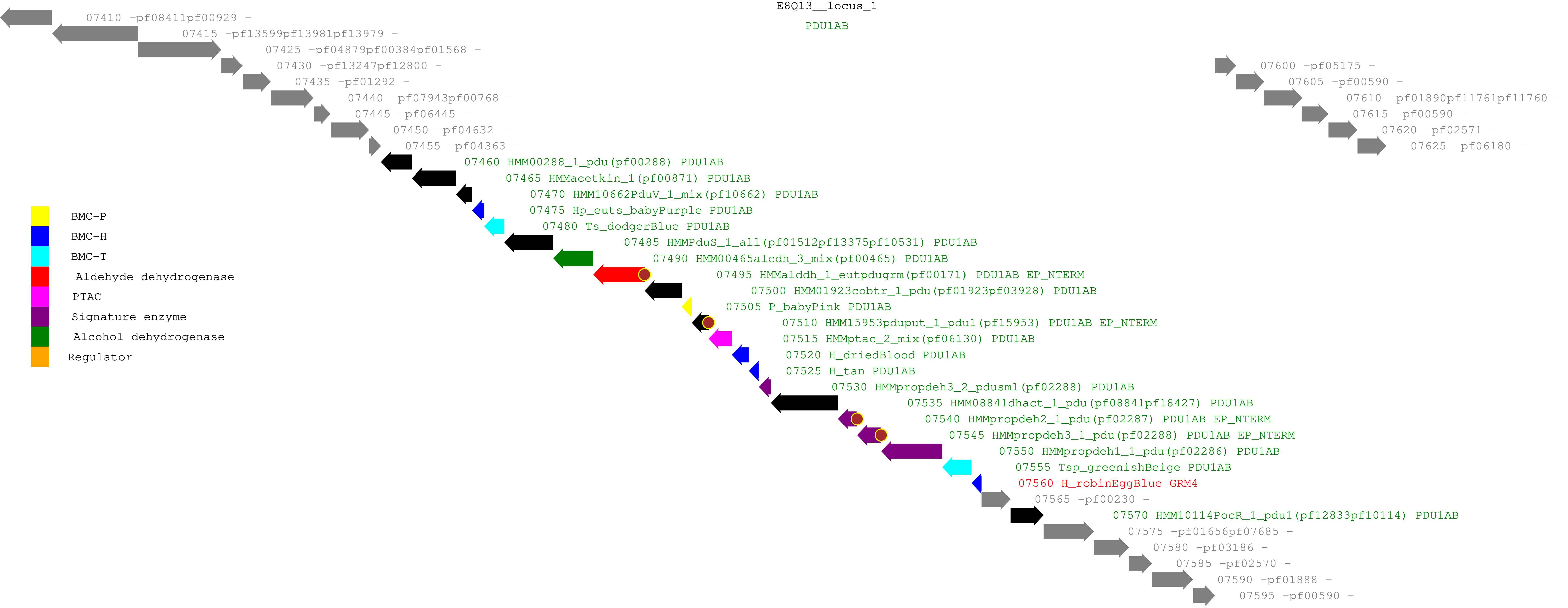

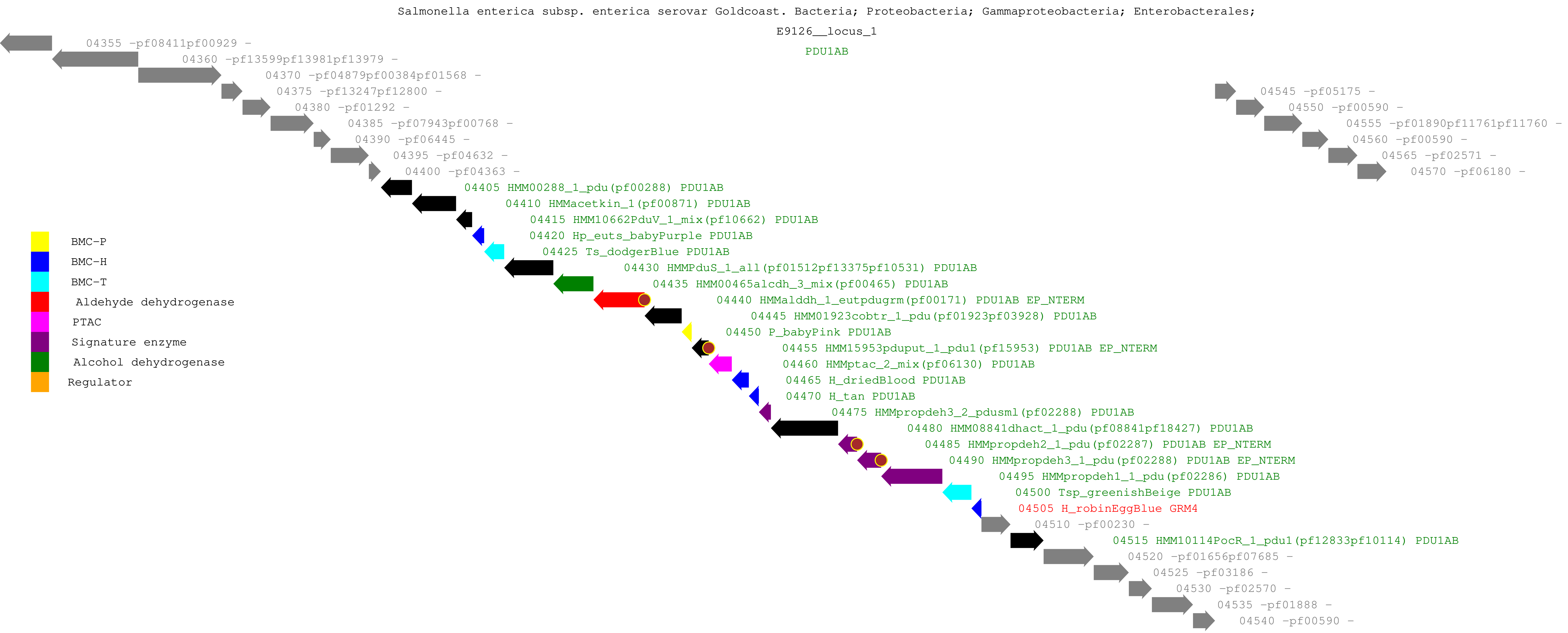

Salmonella enterica subsp. enterica serovar Goldcoast. Bacteria; Proteobacteria; Gammaproteobacteria; Enterobacterales;
E9126__locus_1
PDU1AB
04355 -pf08411pf00929 -
04360 -pf13599pf13981pf13979 -
04370 -pf04879pf00384pf01568 -
04375 -pf13247pf12800 -
04380 -pf01292 -
04385 -pf07943pf00768 -
04390 -pf06445 -
04395 -pf04632 -
04400 -pf04363 -
04405 HMM00288_1_pdu(pf00288) PDU1AB
04410 HMMacetkin_1(pf00871) PDU1AB
04415 HMM10662PduV_1_mix(pf10662) PDU1AB
04420 Hp_euts_babyPurple PDU1AB
04425 Ts_dodgerBlue PDU1AB
04430 HMMPduS_1_all(pf01512pf13375pf10531) PDU1AB
04435 HMM00465alcdh_3_mix(pf00465) PDU1AB
04440 HMMalddh_1_eutpdugrm(pf00171) PDU1AB EP_NTERM
04445 HMM01923cobtr_1_pdu(pf01923pf03928) PDU1AB
04450 P_babyPink PDU1AB
04455 HMM15953pduput_1_pdu1(pf15953) PDU1AB EP_NTERM
04460 HMMptac_2_mix(pf06130) PDU1AB
04465 H_driedBlood PDU1AB
04470 H_tan PDU1AB
04475 HMMpropdeh3_2_pdusml(pf02288) PDU1AB
04480 HMM08841dhact_1_pdu(pf08841pf18427) PDU1AB
04485 HMMpropdeh2_1_pdu(pf02287) PDU1AB EP_NTERM
04490 HMMpropdeh3_1_pdu(pf02288) PDU1AB EP_NTERM
04495 HMMpropdeh1_1_pdu(pf02286) PDU1AB
04500 Tsp_greenishBeige PDU1AB
04505 H_robinEggBlue GRM4
04510 -pf00230 -
04515 HMM10114PocR_1_pdu1(pf12833pf10114) PDU1AB
04520 -pf01656pf07685 -
04525 -pf03186 -
04530 -pf02570 -
04535 -pf01888 -
04540 -pf00590 -
04545 -pf05175 -
04550 -pf00590 -
04555 -pf01890pf11761pf11760 -
04560 -pf00590 -
04565 -pf02571 -
04570 -pf06180 -
BMC-P
BMC-H
BMC-T
Aldehyde dehydrogenase
PTAC
Signature enzyme
Alcohol dehydrogenase
Regulator

Klebsiella pneumoniae subsp. pneumoniae. Bacteria; Proteobacteria; Gammaproteobacteria; Enterobacterales;

E9154__locus_1

PDU1AB

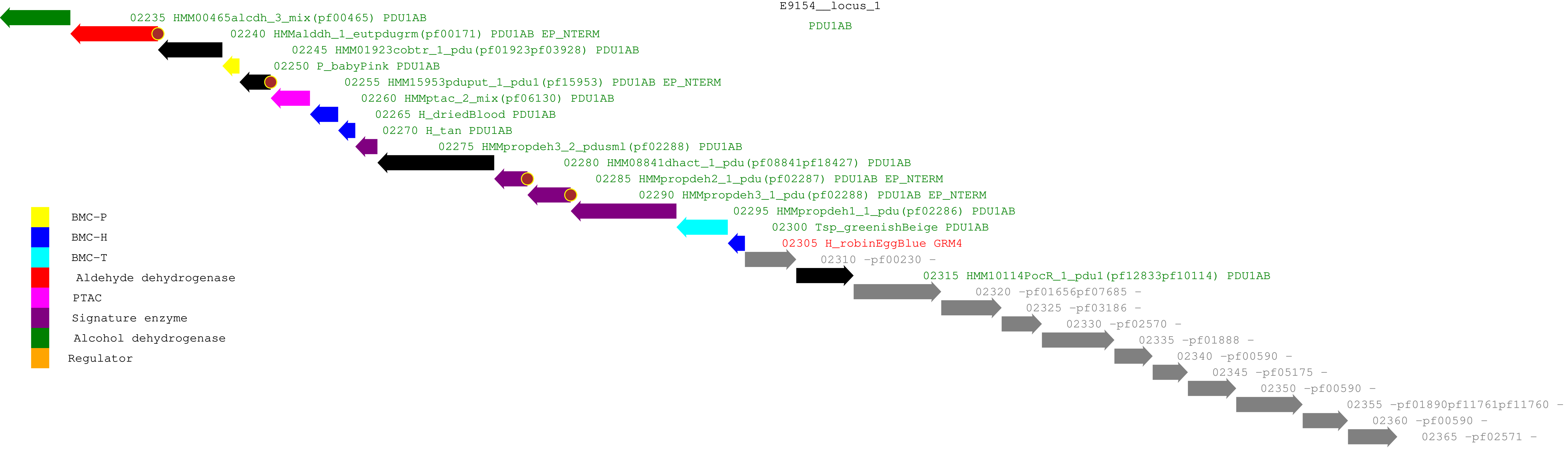

Klebsiella pneumoniae subsp. pneumoniae. Bacteria; Proteobacteria; Gammaproteobacteria; Enterobacterales;

E9161__locus_1

PDU1AB

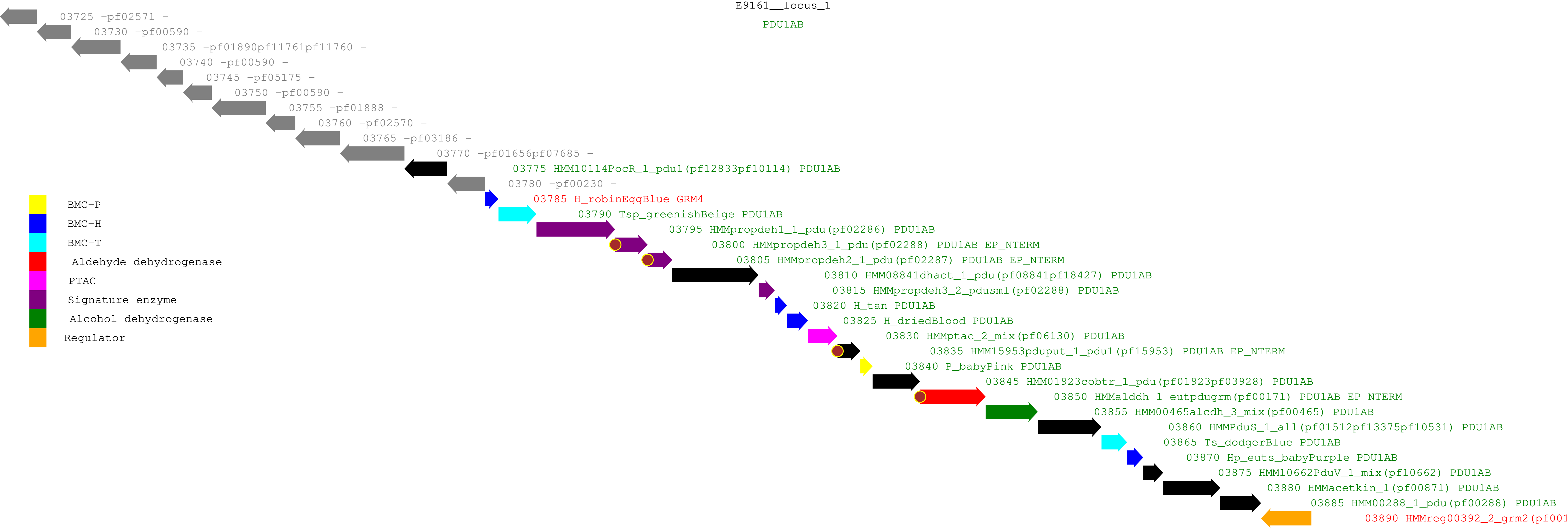

Escherichia albertii. Bacteria; Proteobacteria; Gammaproteobacteria; Enterobacterales;

EAB3F__locus_1

PDU1AB

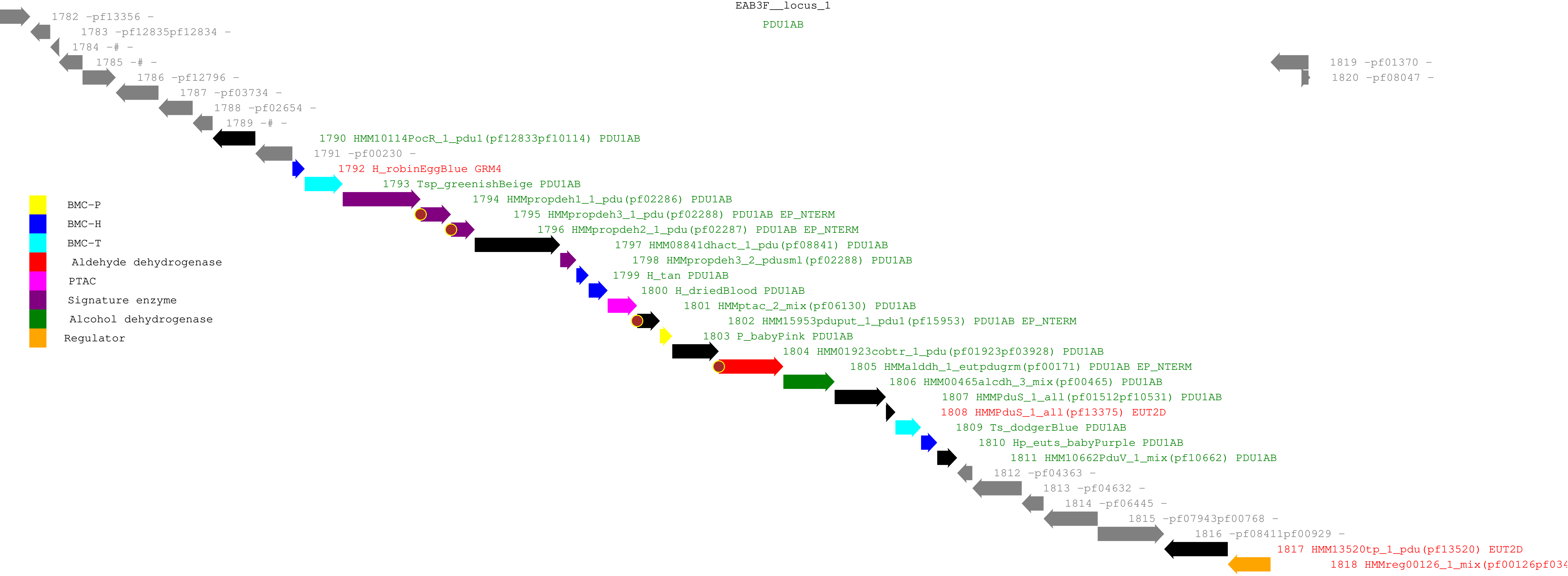

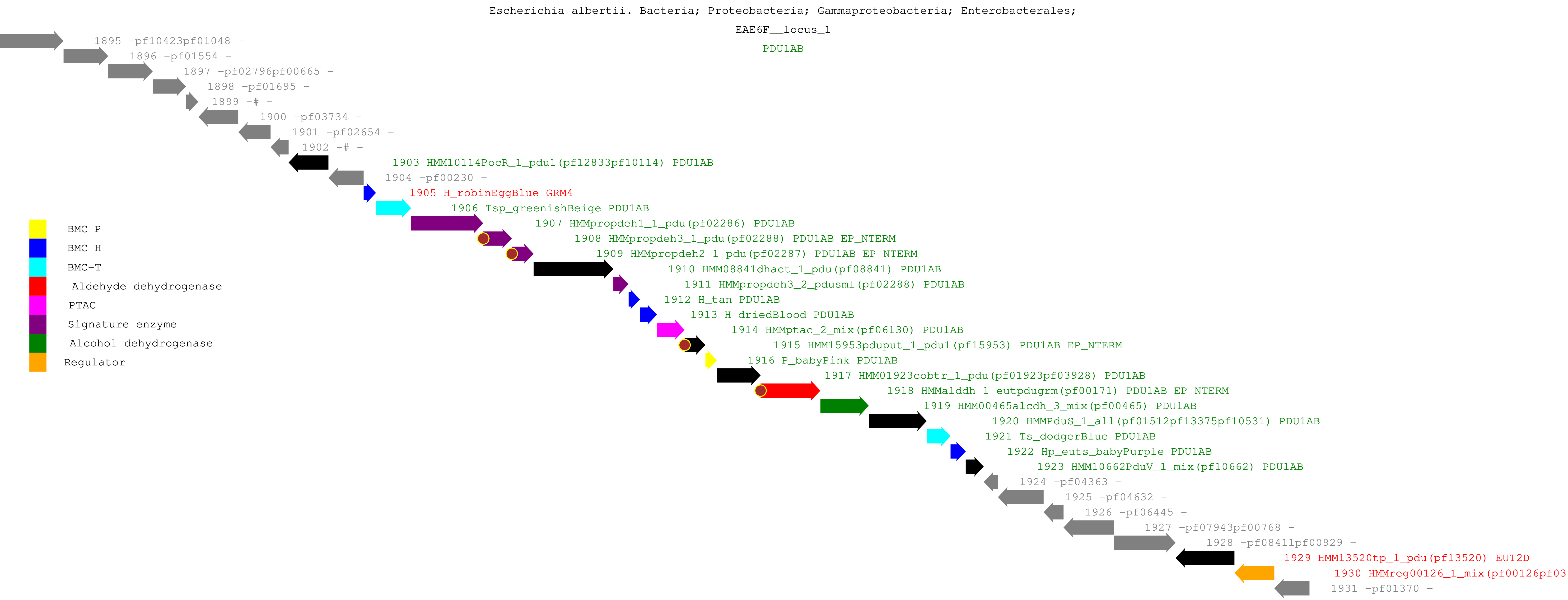
Escherichia albertii. Bacteria; Proteobacteria; Gammaproteobacteria; Enterobacterales;
EAE6F__locus_1
PDU1AB
1895 -pf10423pf01048 -
1896 -pf01554 -
1897 -pf02796pf00665 -
1898 -pf01695 -
1899 -# -
1900 -pf03734 -
1901 -pf02654 -
1902 -# -
1903 HMM10114PocR_1_pdu1(pf12833pf10114) PDU1AB
1904 -pf00230 -
1905 H_robinEggBlue GRM4
1906 Tsp_greenishBeige PDU1AB
1907 HMMpropdeh1_1_pdu(pf02286) PDU1AB
1908 HMMpropdeh3_1_pdu(pf02288) PDU1AB EP_NTERM
1909 HMMpropdeh2_1_pdu(pf02287) PDU1AB EP_NTERM
1910 HMM08841dhact_1_pdu(pf08841) PDU1AB
1911 HMMpropdeh3_2_pdusml(pf02288) PDU1AB
1912 H_tan PDU1AB
1913 H_driedBlood PDU1AB
1914 HMMptac_2_mix(pf06130) PDU1AB
1915 HMM15953pduput_1_pdu1(pf15953) PDU1AB EP_NTERM
1916 P_babyPink PDU1AB
1917 HMM01923cobtr_1_pdu(pf01923pf03928) PDU1AB
1918 HMMalddh_1_eutpdugrm(pf00171) PDU1AB EP_NTERM
1919 HMM00465alcdh_3_mix(pf00465) PDU1AB
1920 HMMPduS_1_all(pf01512pf13375pf10531) PDU1AB
1921 Ts_dodgerBlue PDU1AB
1922 Hp_euts_babyPurple PDU1AB
1923 HMM10662PduV_1_mix(pf10662) PDU1AB
1924 -pf04363 -
1925 -pf04632 -
1926 -pf06445 -
1927 -pf07943pf00768 -
1928 -pf08411pf00929 -
1929 HMM13520tp_1_pdu(pf13520) EUT2D
1930 HMMreg00126_1_mix(pf00126pf03
1931 -pf01370 -
BMC-P
BMC-H
BMC-T
Aldehyde dehydrogenase
PTAC
Signature enzyme
Alcohol dehydrogenase
Regulator

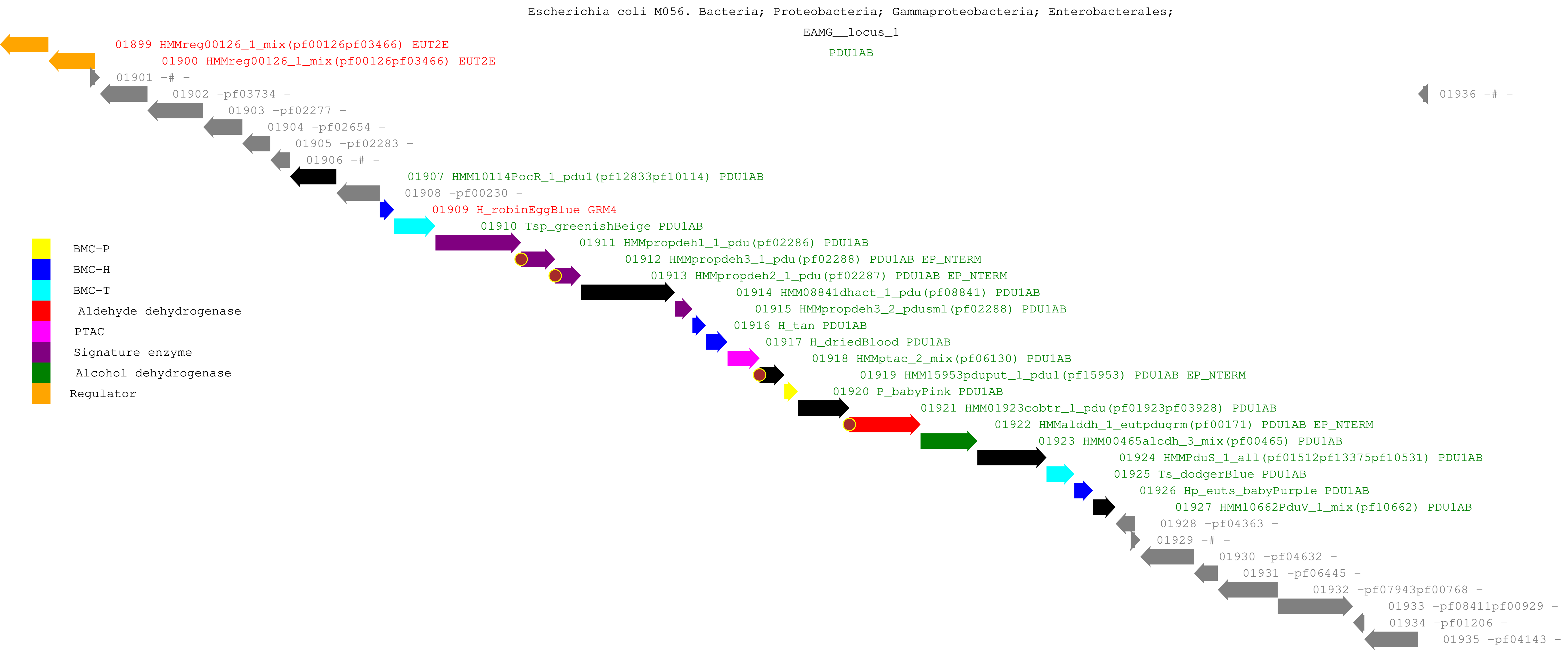
Escherichia coli M056. Bacteria; Proteobacteria; Gammaproteobacteria; Enterobacterales;
EAMG__locus_1
PDU1AB
01899 HMMreg00126_1_mix(pf00126pf03466) EUT2E
01900 HMMreg00126_1_mix(pf00126pf03466) EUT2E
01901 -# -
01902 -pf03734 -
01903 -pf02277 -
01904 -pf02654 -
01905 -pf02283 -
01906 -# -
01907 HMM10114PocR_1_pdu1(pf12833pf10114) PDU1AB
01908 -pf00230 -
01909 H_robinEggBlue GRM4
01910 Tsp_greenishBeige PDU1AB
01911 HMMpropdeh1_1_pdu(pf02286) PDU1AB
01912 HMMpropdeh3_1_pdu(pf02288) PDU1AB EP_NTERM
01913 HMMpropdeh2_1_pdu(pf02287) PDU1AB EP_NTERM
01914 HMM08841dhact_1_pdu(pf08841) PDU1AB
01915 HMMpropdeh3_2_pdusml(pf02288) PDU1AB
01916 H_tan PDU1AB
01917 H_driedBlood PDU1AB
01918 HMMptac_2_mix(pf06130) PDU1AB
01919 HMM15953pduput_1_pdu1(pf15953) PDU1AB EP_NTERM
01920 P_babyPink PDU1AB
01921 HMM01923cobtr_1_pdu(pf01923pf03928) PDU1AB
01922 HMMaldh_1_eutpdugrm(pf00171) PDU1AB EP_NTERM
01923 HMM00465alcdh_3_mix(pf00465) PDU1AB
01924 HMMPduS_1_all(pf01512pf13375pf10531) PDU1AB
01925 Ts_dodgerBlue PDU1AB
01926 Hp_euts_babyPurple PDU1AB
01927 HMM10662PduV_1_mix(pf10662) PDU1AB
01928 -pf04363 -
01929 -# -
01930 -pf04632 -
01931 -pf06445 -
01932 -pf07943pf00768 -
01933 -pf08411pf00929 -
01934 -pf01206 -
01935 -pf04143 -
01936 -# -
BMC-P
BMC-H
BMC-T
Aldehyde dehydrogenase
PTAC
Signature enzyme
Alcohol dehydrogenase
Regulator

Salmonella enterica (Salmonella choleraesuis). Bacteria; Proteobacteria; Gammaproteobacteria; Enterobacterales;

EAN84__locus_1

PDU1AB

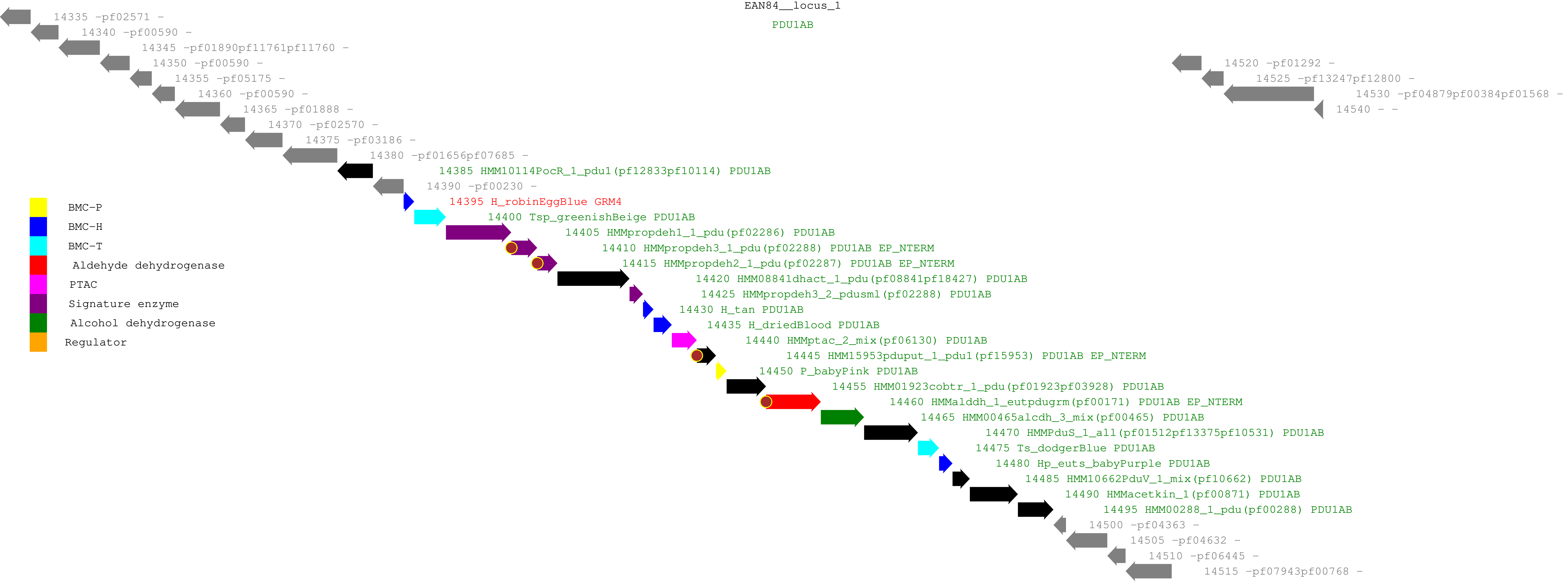

Escherichia albertii B156. Bacteria; Proteobacteria; Gammaproteobacteria; Enterobacterales;

EAPG__locus_1

PDU1AB

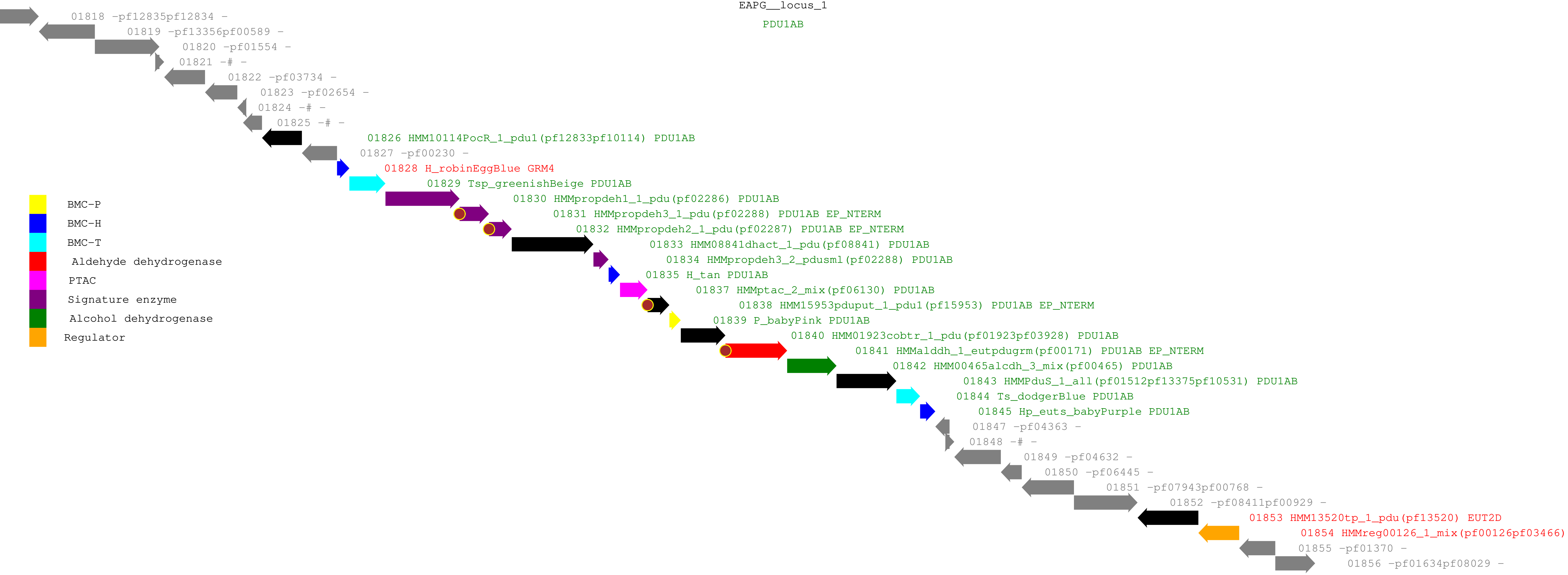

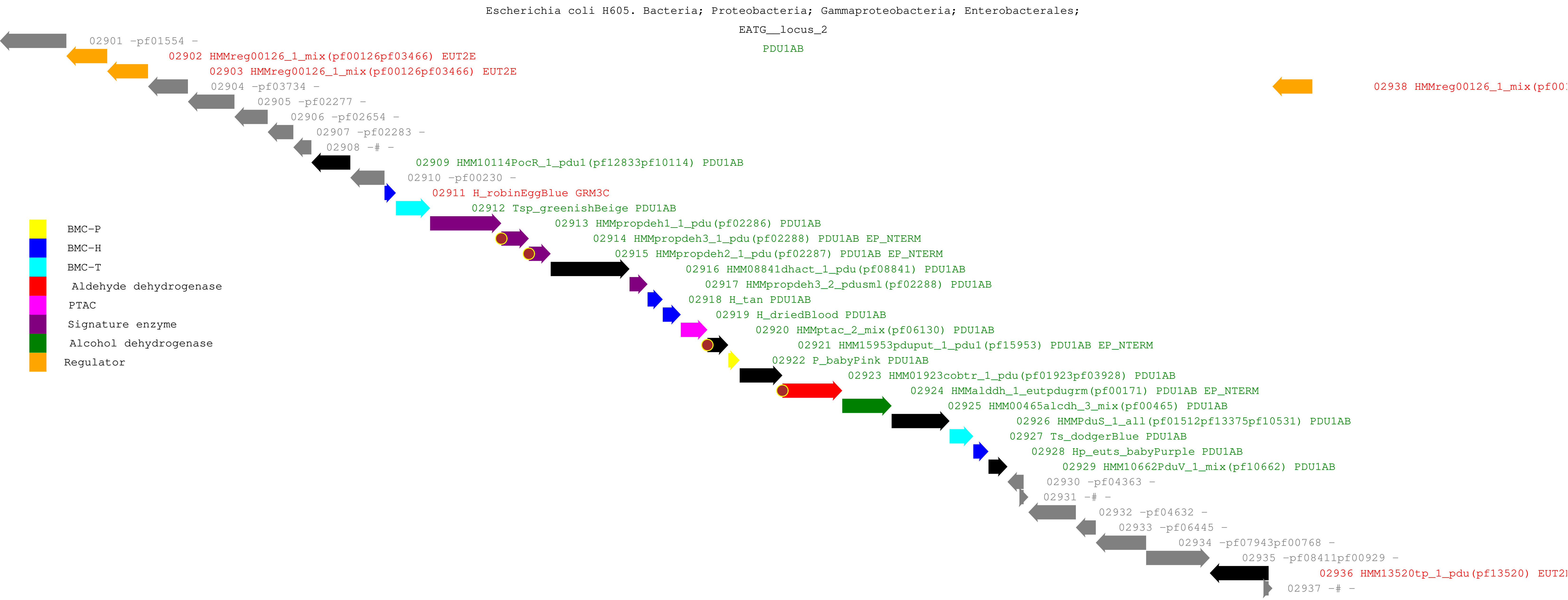

Escherichia coli H605. Bacteria; Proteobacteria; Gammaproteobacteria; Enterobacterales;
EATG__locus_2
PDU1AB
02901 -pf01554 -
02902 HMMreg00126_1_mix(pf00126pf03466) EUT2E
02903 HMMreg00126_1_mix(pf00126pf03466) EUT2E
02904 -pf03734 -
02905 -pf02277 -
02906 -pf02654 -
02907 -pf02283 -
02908 -# -
02909 HMM10114PocR_1_pdu1(pf12833pf10114) PDU1AB
02910 -pf00230 -
02911 H_robinEggBlue GRM3C
02912 Tsp_greenishBeige PDU1AB
02913 HMMpropdeh1_1_pdu(pf02286) PDU1AB
02914 HMMpropdeh3_1_pdu(pf02288) PDU1AB EP_NTERM
02915 HMMpropdeh2_1_pdu(pf02287) PDU1AB EP_NTERM
02916 HMM08841dhact_1_pdu(pf08841) PDU1AB
02917 HMMpropdeh3_2_pdusml(pf02288) PDU1AB
02918 H_tan PDU1AB
02919 H_driedBlood PDU1AB
02920 HMMptac_2_mix(pf06130) PDU1AB
02921 HMM15953pduput_1_pdu1(pf15953) PDU1AB EP_NTERM
02922 P_babyPink PDU1AB
02923 HMM01923cobtr_1_pdu(pf01923pf03928) PDU1AB
02924 HMMalddh_1_eutpdugrm(pf00171) PDU1AB EP_NTERM
02925 HMM00465alcdh_3_mix(pf00465) PDU1AB
02926 HMMPduS_1_all(pf01512pf13375pf10531) PDU1AB
02927 Ts_dodgerBlue PDU1AB
02928 Hp_euts_babyPurple PDU1AB
02929 HMM10662PduV_1_mix(pf10662) PDU1AB
02930 -pf04363 -
02931 -# -
02932 -pf04632 -
02933 -pf06445 -
02934 -pf07943pf00768 -
02935 -pf08411pf00929 -
02936 HMM13520tp_1_pdu(pf13520) EUT2
02937 -# -
02938 HMMreg00126_1_mix(pf00
BMC-P
BMC-H
BMC-T
Aldehyde dehydrogenase
PTAC
Signature enzyme
Alcohol dehydrogenase
Regulator

Escherichia coli TA249. Bacteria; Proteobacteria; Gammaproteobacteria; Enterobacterales;

EAZG__locus_1

PDU1AB

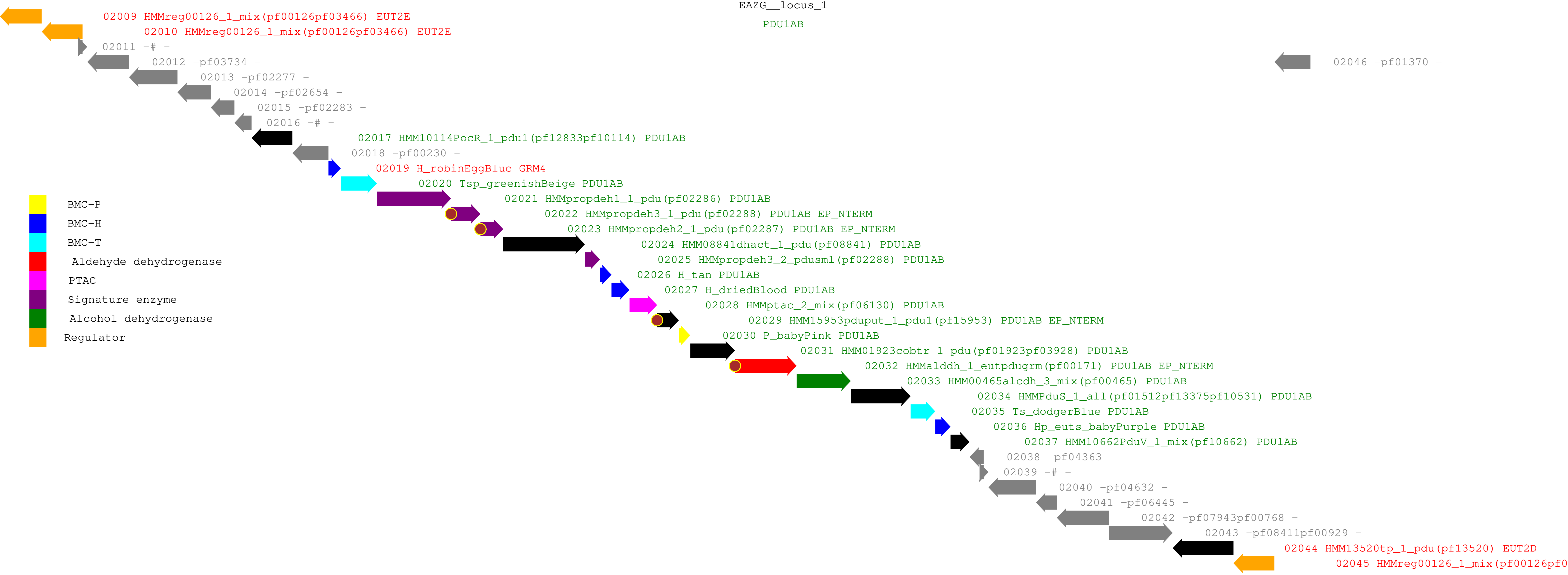

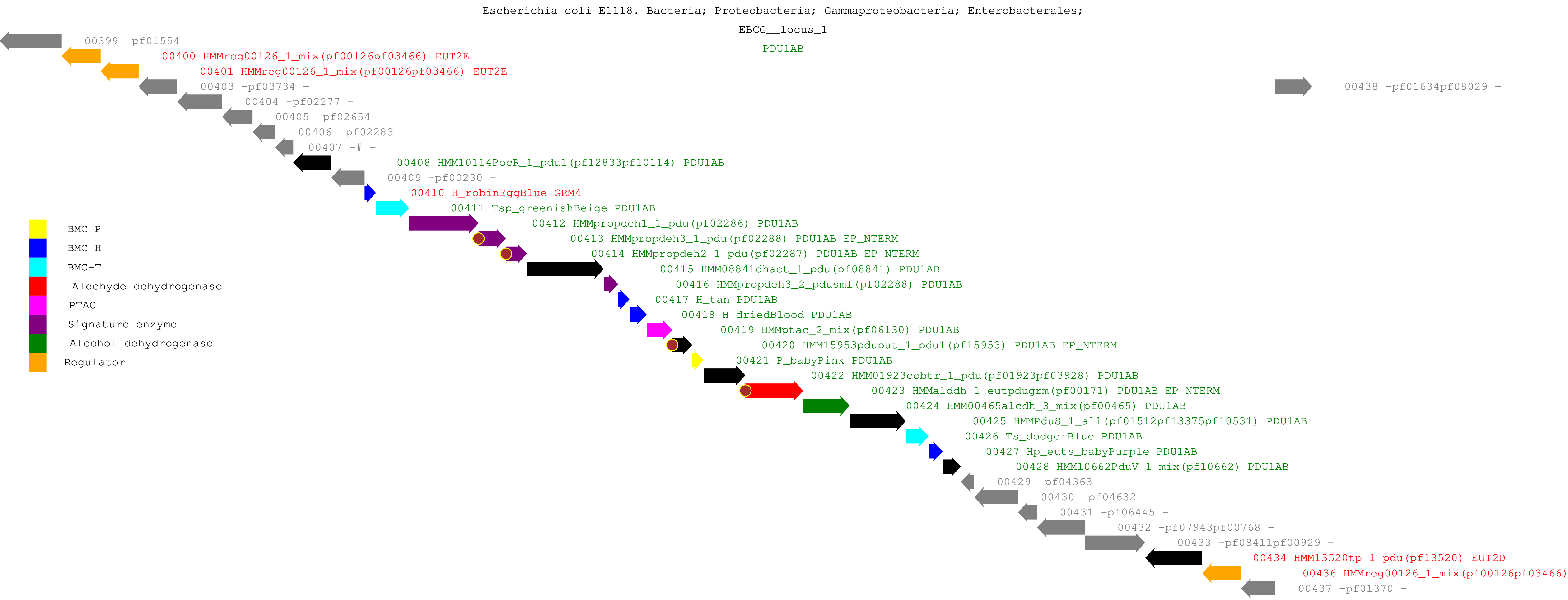
Escherichia coli E1118. Bacteria; Proteobacteria; Gammaproteobacteria; Enterobacterales;
EBCG__locus_1
PDU1AB
00399 -pf01554 -
00400 HMMreg00126_1_mix(pf00126pf03466) EUT2E
00401 HMMreg00126_1_mix(pf00126pf03466) EUT2E
00403 -pf03734 -
00404 -pf02277 -
00405 -pf02654 -
00406 -pf02283 -
00407 -# -
00408 HMM10114PocR_1_pdu1(pf12833pf10114) PDU1AB
00409 -pf00230 -
00410 H_robinEggBlue GRM4
00411 Tsp_greenishBeige PDU1AB
00412 HMMpropdeh1_1_pdu(pf02286) PDU1AB
00413 HMMpropdeh3_1_pdu(pf02288) PDU1AB EP_NTERM
00414 HMMpropdeh2_1_pdu(pf02287) PDU1AB EP_NTERM
00415 HMM08841dhact_1_pdu(pf08841) PDU1AB
00416 HMMpropdeh3_2_pdusml(pf02288) PDU1AB
00417 H_tan PDU1AB
00418 H_driedBlood PDU1AB
00419 HMMptac_2_mix(pf06130) PDU1AB
00420 HMM15953pduput_1_pdu1(pf15953) PDU1AB EP_NTERM
00421 P_babyPink PDU1AB
00422 HMM01923cobtr_1_pdu(pf01923pf03928) PDU1AB
00423 HMMalddh_1_eutpdugrm(pf00171) PDU1AB EP_NTERM
00424 HMM00465alcdh_3_mix(pf00465) PDU1AB
00425 HMMPduS_1_all(pf01512pf13375pf10531) PDU1AB
00426 Ts_dodgerBlue PDU1AB
00427 Hp_euts_babyPurple PDU1AB
00428 HMM10662PduV_1_mix(pf10662) PDU1AB
00429 -pf04363 -
00430 -pf04632 -
00431 -pf06445 -
00432 -pf07943pf00768 -
00433 -pf08411pf00929 -
00434 HMM13520tp_1_pdu(pf13520) EUT2D
00436 HMMreg00126_1_mix(pf00126pf03466)
00437 -pf01370 -
00438 -pf01634pf08029 -
BMC-P
BMC-H
BMC-T
Aldehyde dehydrogenase
PTAC
Signature enzyme
Alcohol dehydrogenase
Regulator

Salmonella enterica I. Bacteria; Proteobacteria; Gammaproteobacteria; Enterobacterales;

EBD05__locus_2

PDU1AB

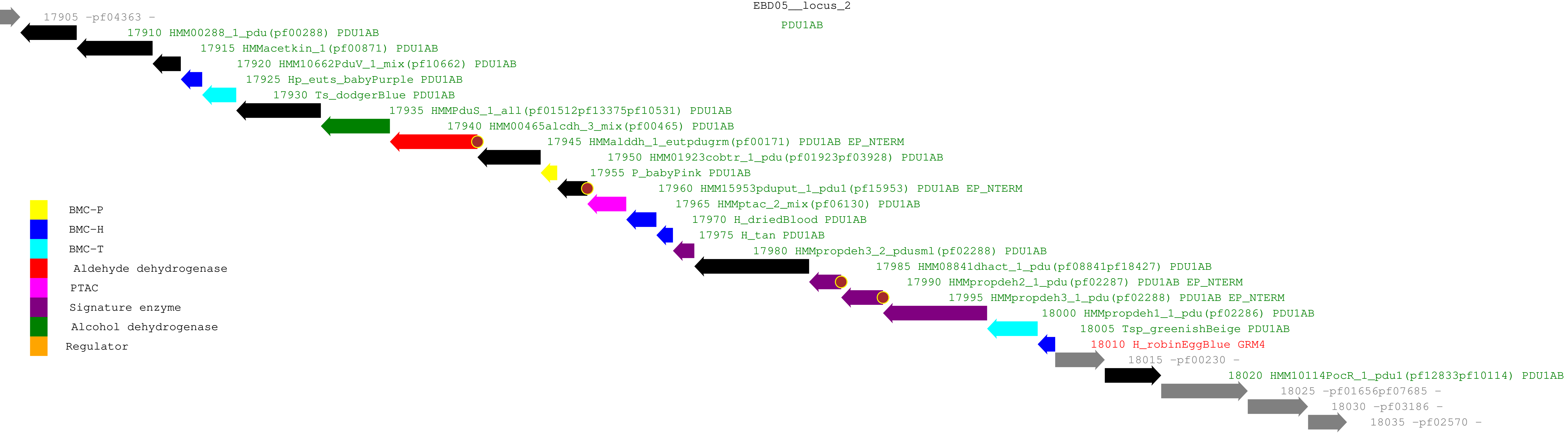

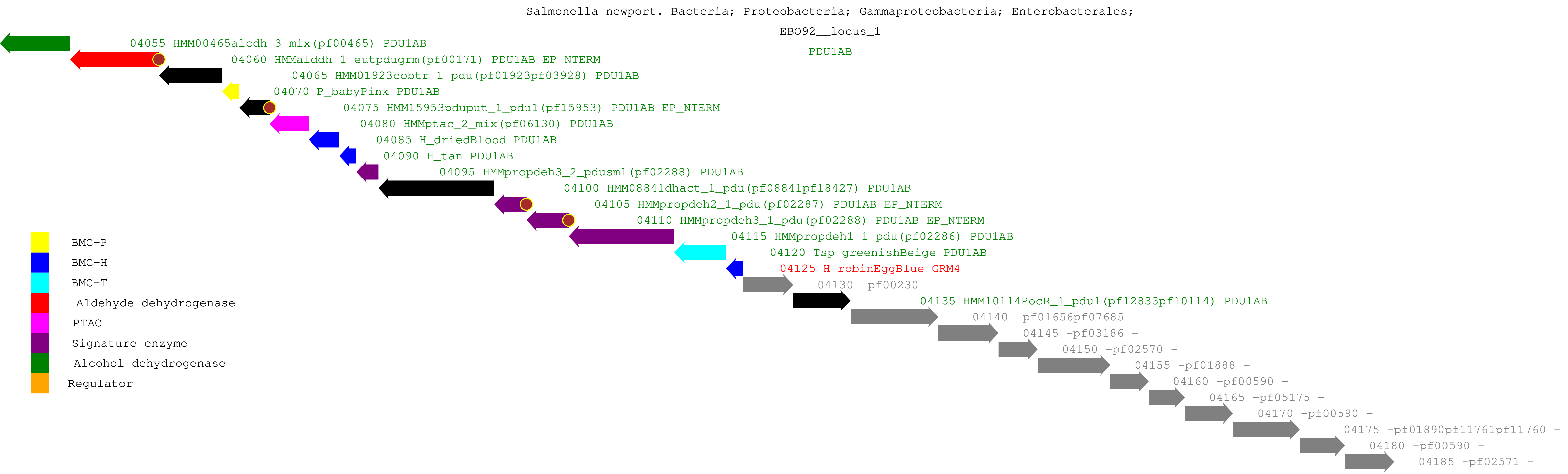

Salmonella newport. Bacteria; Proteobacteria; Gammaproteobacteria; Enterobacterales;
EBO92__locus_1
PDU1AB
04055 HMM00465alcdh_3_mix(pf00465) PDU1AB
04060 HMMalddh_1_eutpdugrm(pf00171) PDU1AB EP_NTERM
04065 HMM01923cobtr_1_pdu(pf01923pf03928) PDU1AB
04070 P_babyPink PDU1AB
04075 HMM15953pduput_1_pdu1(pf15953) PDU1AB EP_NTERM
04080 HMMptac_2_mix(pf06130) PDU1AB
04085 H_driedBlood PDU1AB
04090 H_tan PDU1AB
04095 HMMpropdeh3_2_pdusml(pf02288) PDU1AB
04100 HMM08841dhact_1_pdu(pf08841pf18427) PDU1AB
04105 HMMpropdeh2_1_pdu(pf02287) PDU1AB EP_NTERM
04110 HMMpropdeh3_1_pdu(pf02288) PDU1AB EP_NTERM
04115 HMMpropdeh1_1_pdu(pf02286) PDU1AB
04120 Tsp_greenishBeige PDU1AB
04125 H_robinEggBlue GRM4
04130 -pf00230 -
04135 HMM10114PocR_1_pdu1(pf12833pf10114) PDU1AB
04140 -pf01656pf07685 -
04145 -pf03186 -
04150 -pf02570 -
04155 -pf01888 -
04160 -pf00590 -
04165 -pf05175 -
04170 -pf00590 -
04175 -pf01890pf11761pf11760 -
04180 -pf00590 -
04185 -pf02571 -
BMC-P
BMC-H
BMC-T
Aldehyde dehydrogenase
PTAC
Signature enzyme
Alcohol dehydrogenase
Regulator

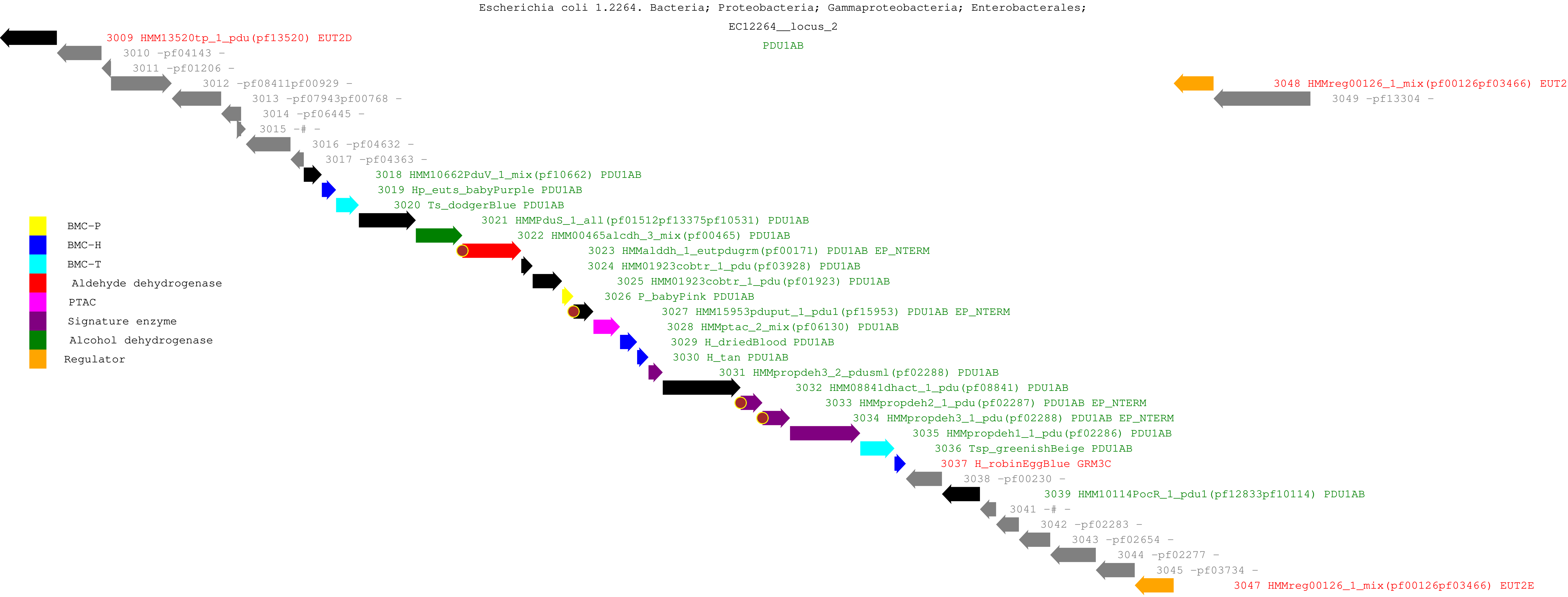
Escherichia coli 1.2264. Bacteria; Proteobacteria; Gammaproteobacteria; Enterobacterales;
EC12264__locus_2
PDU1AB
3009 HMM13520tp_1_pdu(pf13520) EUT2D
3010 -pf04143 -
3011 -pf01206 -
3012 -pf08411pf00929 -
3013 -pf07943pf00768 -
3014 -pf06445 -
3015 -# -
3016 -pf04632 -
3017 -pf04363 -
3018 HMM10662PduV_1_mix(pf10662) PDU1AB
3019 Hp_euts_babyPurple PDU1AB
3020 Ts_dodgerBlue PDU1AB
3021 HMMPduS_1_all(pf01512pf13375pf10531) PDU1AB
3022 HMM00465alcdh_3_mix(pf00465) PDU1AB
3023 HMMalddh_1_eutpdugrm(pf00171) PDU1AB EP_NTERM
3024 HMM01923cobtr_1_pdu(pf03928) PDU1AB
3025 HMM01923cobtr_1_pdu(pf01923) PDU1AB
3026 P_babyPink PDU1AB
3027 HMM15953pduput_1_pdu1(pf15953) PDU1AB EP_NTERM
3028 HMMptac_2_mix(pf06130) PDU1AB
3029 H_driedBlood PDU1AB
3030 H_tan PDU1AB
3031 HMMpropdeh3_2_pdusml(pf02288) PDU1AB
3032 HMM08841dhact_1_pdu(pf08841) PDU1AB
3033 HMMpropdeh2_1_pdu(pf02287) PDU1AB EP_NTERM
3034 HMMpropdeh3_1_pdu(pf02288) PDU1AB EP_NTERM
3035 HMMpropdeh1_1_pdu(pf02286) PDU1AB
3036 Tsp_greenishBeige PDU1AB
3037 H_robinEggBlue GRM3C
3038 -pf00230 -
3039 HMM10114PocR_1_pdu1(pf12833pf10114) PDU1AB
3041 -# -
3042 -pf02283 -
3043 -pf02654 -
3044 -pf02277 -
3045 -pf03734 -
3047 HMMreg00126_1_mix(pf00126pf03466) EUT2E
3048 HMMreg00126_1_mix(pf00126pf03466) EUT2
3049 -pf13304 -
BMC-P
BMC-H
BMC-T
Aldehyde dehydrogenase
PTAC
Signature enzyme
Alcohol dehydrogenase
Regulator

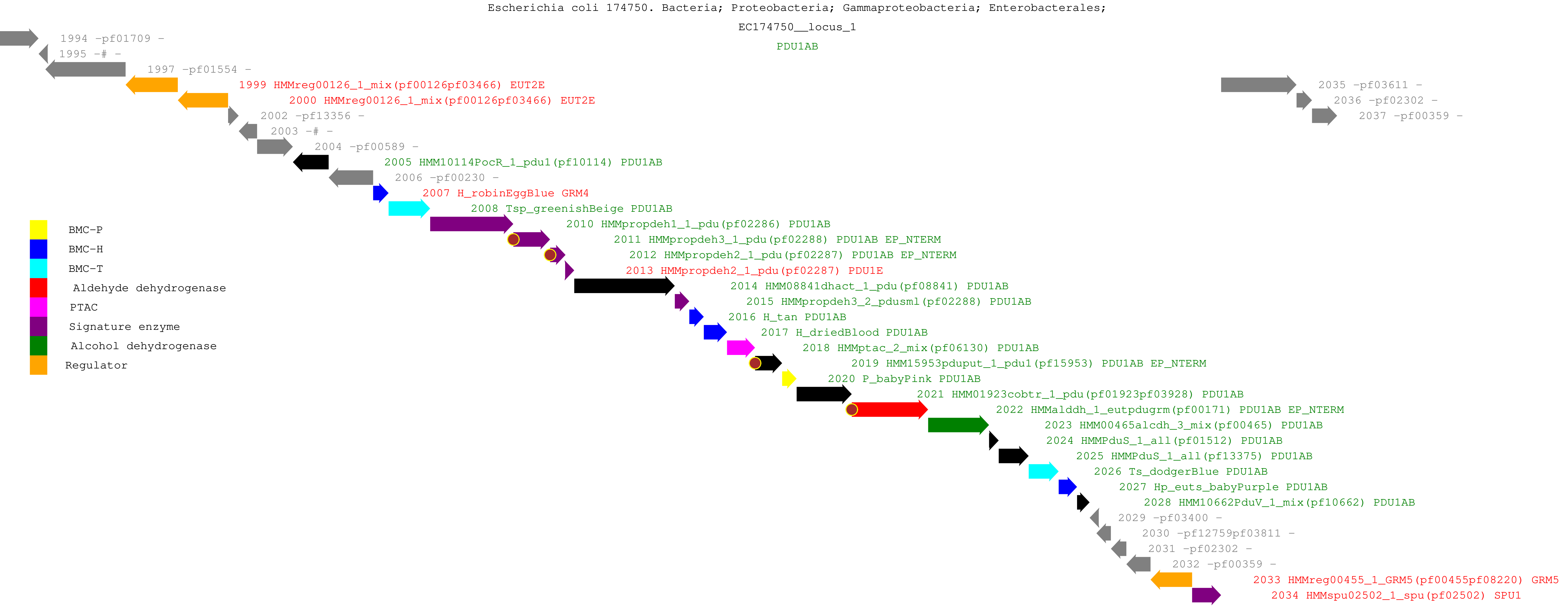

Escherichia coli 174750. Bacteria; Proteobacteria; Gammaproteobacteria; Enterobacterales;
EC174750__locus_1
PDU1AB
1994 -pf01709 -
1995 -# -
1997 -pf01554 -
1999 HMMreg00126_1_mix(pf00126pf03466) EUT2E
2000 HMMreg00126_1_mix(pf00126pf03466) EUT2E
2002 -pf13356 -
2003 -# -
2004 -pf00589 -
2005 HMM10114PocR_1_pdu1(pf10114) PDU1AB
2006 -pf00230 -
2007 H_robinEggBlue GRM4
2008 Tsp_greenishBeige PDU1AB
2010 HMMpropdeh1_1_pdu(pf02286) PDU1AB
2011 HMMpropdeh3_1_pdu(pf02288) PDU1AB EP_NTERM
2012 HMMpropdeh2_1_pdu(pf02287) PDU1AB EP_NTERM
2013 HMMpropdeh2_1_pdu(pf02287) PDU1E
2014 HMM08841dhact_1_pdu(pf08841) PDU1AB
2015 HMMpropdeh3_2_pdusml(pf02288) PDU1AB
2016 H_tan PDU1AB
2017 H_driedBlood PDU1AB
2018 HMMptac_2_mix(pf06130) PDU1AB
2019 HMM15953pduput_1_pdu1(pf15953) PDU1AB EP_NTERM
2020 P_babyPink PDU1AB
2021 HMM01923cobtr_1_pdu(pf01923pf03928) PDU1AB
2022 HMMaldh_1_eutpdugrm(pf00171) PDU1AB EP_NTERM
2023 HMM00465alcdh_3_mix(pf00465) PDU1AB
2024 HMMPduS_1_all(pf01512) PDU1AB
2025 HMMPduS_1_all(pf13375) PDU1AB
2026 Ts_dodgerBlue PDU1AB
2027 Hp_euts_babyPurple PDU1AB
2028 HMM10662PduV_1_mix(pf10662) PDU1AB
2029 -pf03400 -
2030 -pf12759pf03811 -
2031 -pf02302 -
2032 -pf00359 -
2033 HMMreg00455_1_GRM5(pf00455pf08220) GRM5
2034 HMMspu02502_1_spu(pf02502) SPU1
2035 -pf03611 -
2036 -pf02302 -
2037 -pf00359 -
BMC-P
BMC-H
BMC-T
Aldehyde dehydrogenase
PTAC
Signature enzyme
Alcohol dehydrogenase
Regulator

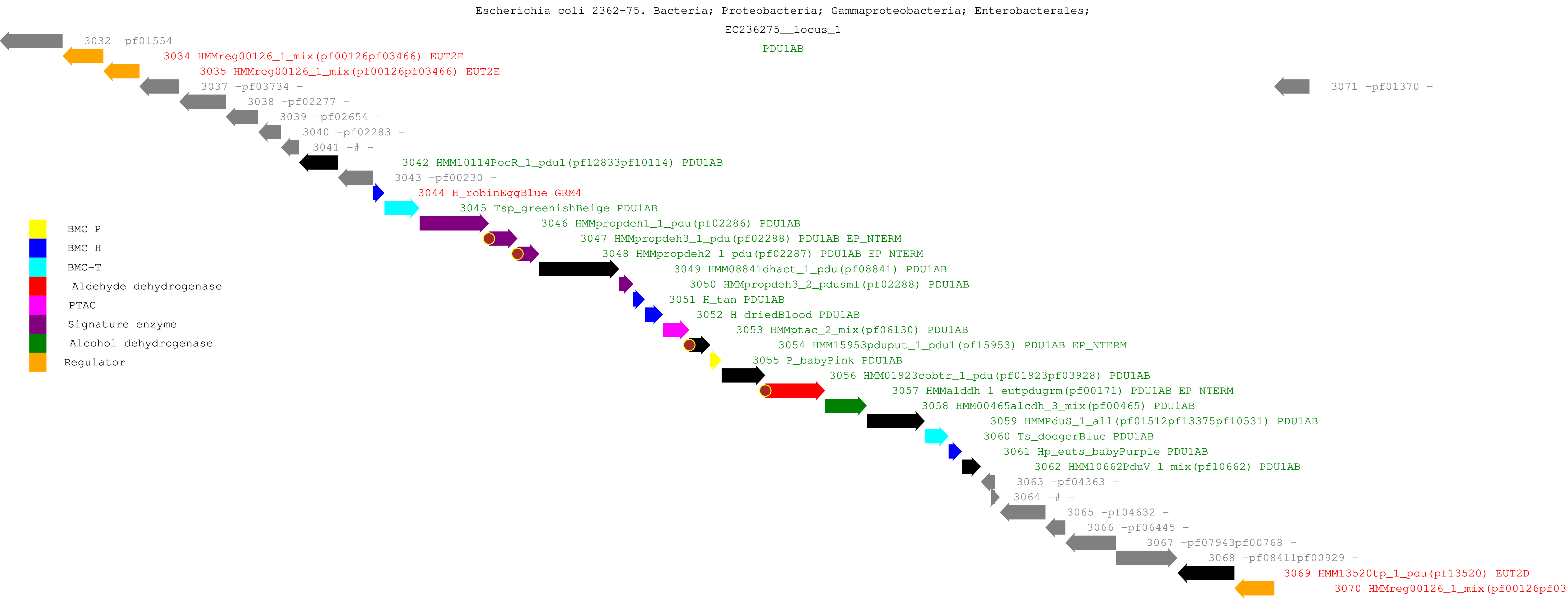
Escherichia coli 2362-75. Bacteria; Proteobacteria; Gammaproteobacteria; Enterobacterales;
EC236275__locus_1
PDU1AB
3032 -pf01554 -
3034 HMMreg00126_1_mix(pf00126pf03466) EUT2E
3035 HMMreg00126_1_mix(pf00126pf03466) EUT2E
3037 -pf03734 -
3038 -pf02277 -
3039 -pf02654 -
3040 -pf02283 -
3041 -# -
3042 HMM10114PocR_1_pdu1(pf12833pf10114) PDU1AB
3043 -pf00230 -
3044 H_robinEggBlue GRM4
3045 Tsp_greenishBeige PDU1AB
3046 HMMpropdeh1_1_pdu(pf02286) PDU1AB
3047 HMMpropdeh3_1_pdu(pf02288) PDU1AB EP_NTERM
3048 HMMpropdeh2_1_pdu(pf02287) PDU1AB EP_NTERM
3049 HMM08841dhact_1_pdu(pf08841) PDU1AB
3050 HMMpropdeh3_2_pdusml(pf02288) PDU1AB
3051 H_tan PDU1AB
3052 H_driedBlood PDU1AB
3053 HMMptac_2_mix(pf06130) PDU1AB
3054 HMM15953pduput_1_pdu1(pf15953) PDU1AB EP_NTERM
3055 P_babyPink PDU1AB
3056 HMM01923cobtr_1_pdu(pf01923pf03928) PDU1AB
3057 HMMalddh_1_eutpdugrm(pf00171) PDU1AB EP_NTERM
3058 HMM00465alcdh_3_mix(pf00465) PDU1AB
3059 HMMPduS_1_all(pf01512pf13375pf10531) PDU1AB
3060 Ts_dodgerBlue PDU1AB
3061 Hp_euts_babyPurple PDU1AB
3062 HMM10662PduV_1_mix(pf10662) PDU1AB
3063 -pf04363 -
3064 -# -
3065 -pf04632 -
3066 -pf06445 -
3067 -pf07943pf00768 -
3068 -pf08411pf00929 -
3069 HMM13520tp_1_pdu(pf13520) EUT2D
3070 HMMreg00126_1_mix(pf00126pf03
3071 -pf01370 -
BMC-P
BMC-H
BMC-T
Aldehyde dehydrogenase
PTAC
Signature enzyme
Alcohol dehydrogenase
Regulator

Escherichia coli 9.0111. Bacteria; Proteobacteria; Gammaproteobacteria; Enterobacterales;

EC90111__locus_2_main_sat

PDU1AB

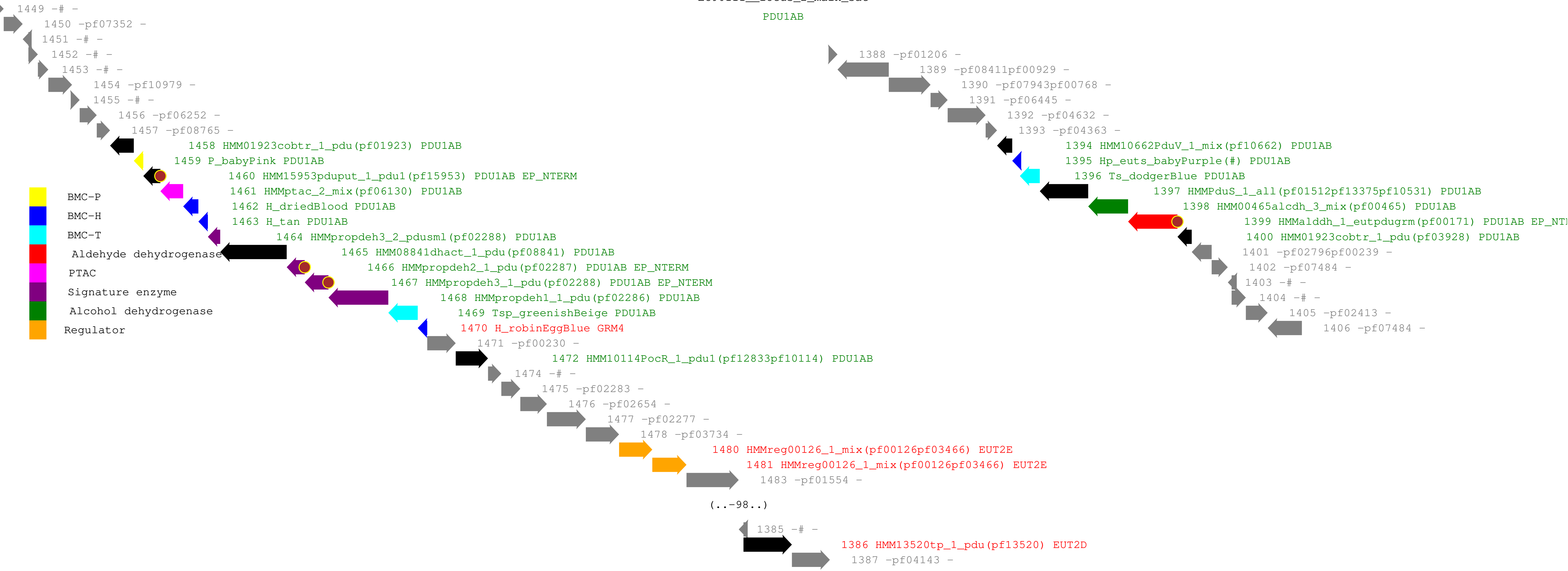

Escherichia fergusonii ECD227. Bacteria; Proteobacteria; Gammaproteobacteria; Enterobacterales;

ECD227__locus_2

PDU1AB

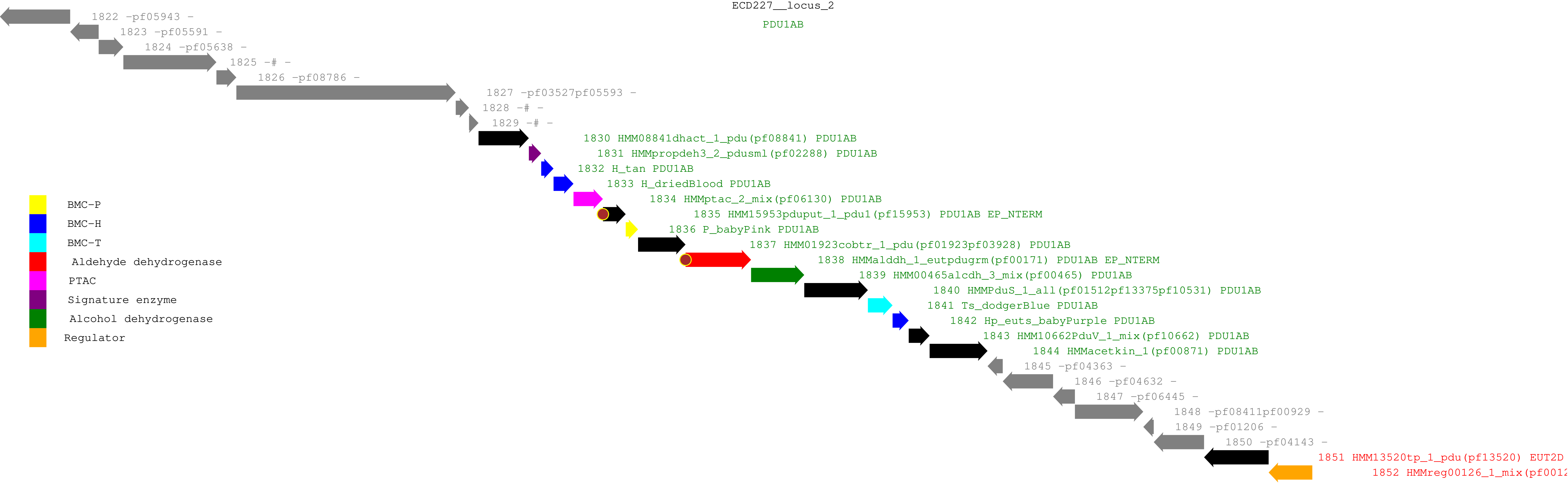

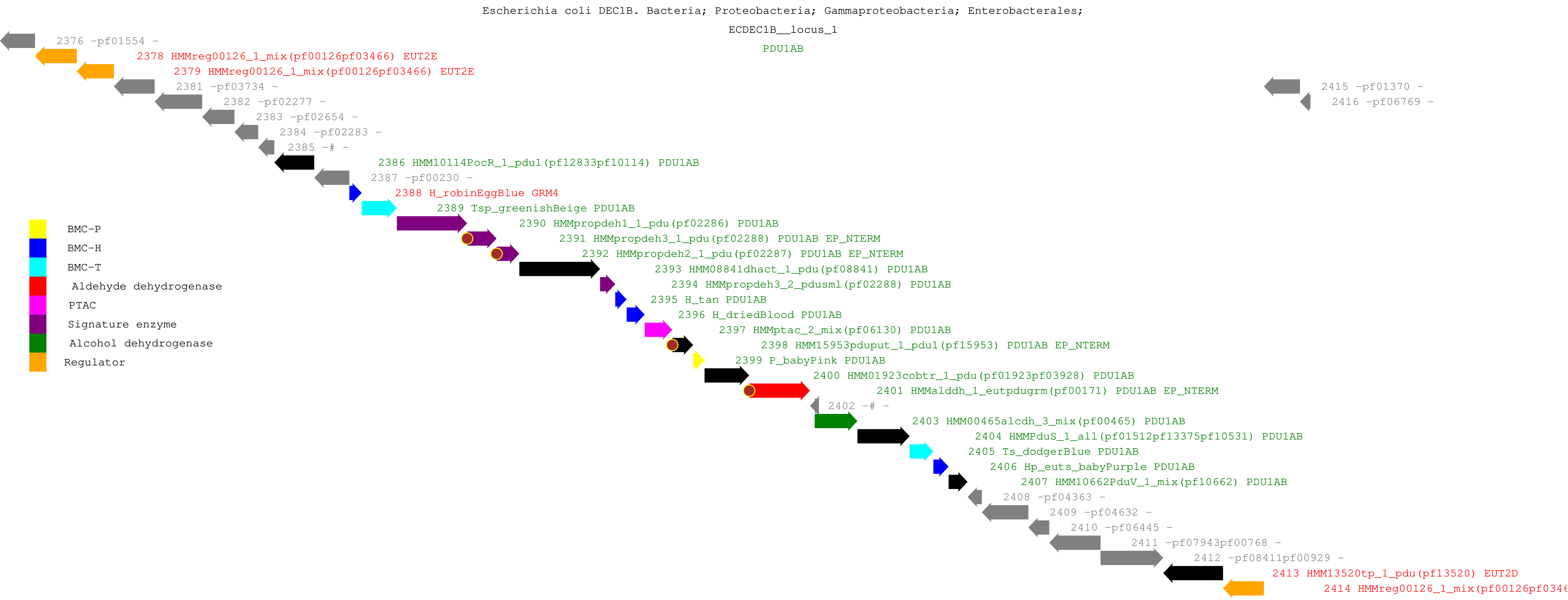
Escherichia coli DEC1B. Bacteria; Proteobacteria; Gammaproteobacteria; Enterobacterales;
ECDEC1B__locus_1
PDU1AB
2376 -pf01554 -
2378 HMMreg00126_1_mix(pf00126pf03466) EUT2E
2379 HMMreg00126_1_mix(pf00126pf03466) EUT2E
2381 -pf03734 -
2382 -pf02277 -
2383 -pf02654 -
2384 -pf02283 -
2385 -# -
2386 HMM10114PocR_1_pdu1(pf12833pf10114) PDU1AB
2387 -pf00230 -
2388 H_robinEggBlue GRM4
2389 Tsp_greenishBeige PDU1AB
2390 HMMpropdeh1_1_pdu(pf02286) PDU1AB
2391 HMMpropdeh3_1_pdu(pf02288) PDU1AB EP_NTERM
2392 HMMpropdeh2_1_pdu(pf02287) PDU1AB EP_NTERM
2393 HMM08841dhact_1_pdu(pf08841) PDU1AB
2394 HMMpropdeh3_2_pdusml(pf02288) PDU1AB
2395 H_tan PDU1AB
2396 H_driedBlood PDU1AB
2397 HMMptac_2_mix(pf06130) PDU1AB
2398 HMM15953pduput_1_pdu1(pf15953) PDU1AB EP_NTERM
2399 P_babyPink PDU1AB
2400 HMM01923cobtr_1_pdu(pf01923pf03928) PDU1AB
2401 HMMalddh_1_eutpdugrm(pf00171) PDU1AB EP_NTERM
2402 -# -
2403 HMM00465alcdh_3_mix(pf00465) PDU1AB
2404 HMMPduS_1_all(pf01512pf13375pf10531) PDU1AB
2405 Ts_dodgerBlue PDU1AB
2406 Hp_euts_babyPurple PDU1AB
2407 HMM10662PduV_1_mix(pf10662) PDU1AB
2408 -pf04363 -
2409 -pf04632 -
2410 -pf06445 -
2411 -pf07943pf00768 -
2412 -pf08411pf00929 -
2413 HMM13520tp_1_pdu(pf13520) EUT2D
2414 HMMreg00126_1_mix(pf00126pf034
2415 -pf01370 -
2416 -pf06769 -
BMC-P
BMC-H
BMC-T
Aldehyde dehydrogenase
PTAC
Signature enzyme
Alcohol dehydrogenase
Regulator

Escherichia coli DEC1C. Bacteria; Proteobacteria; Gammaproteobacteria; Enterobacterales;

ECDEC1C__locus_1

PDU1AB

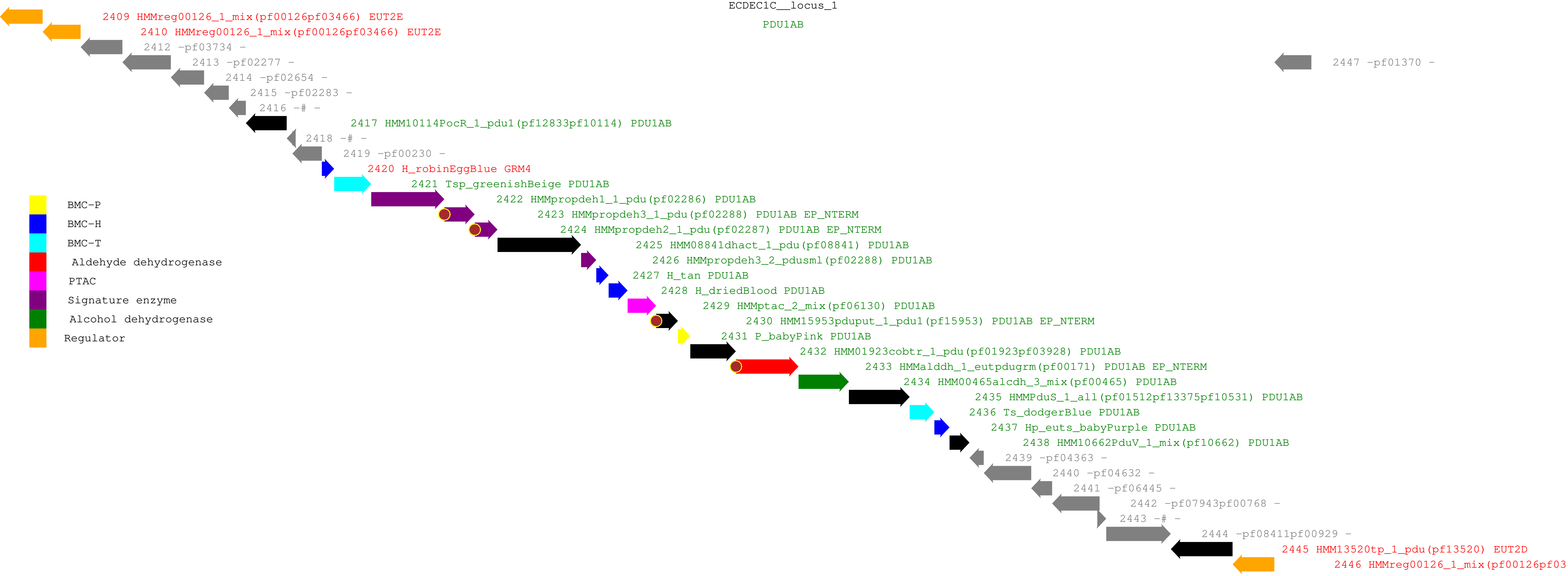

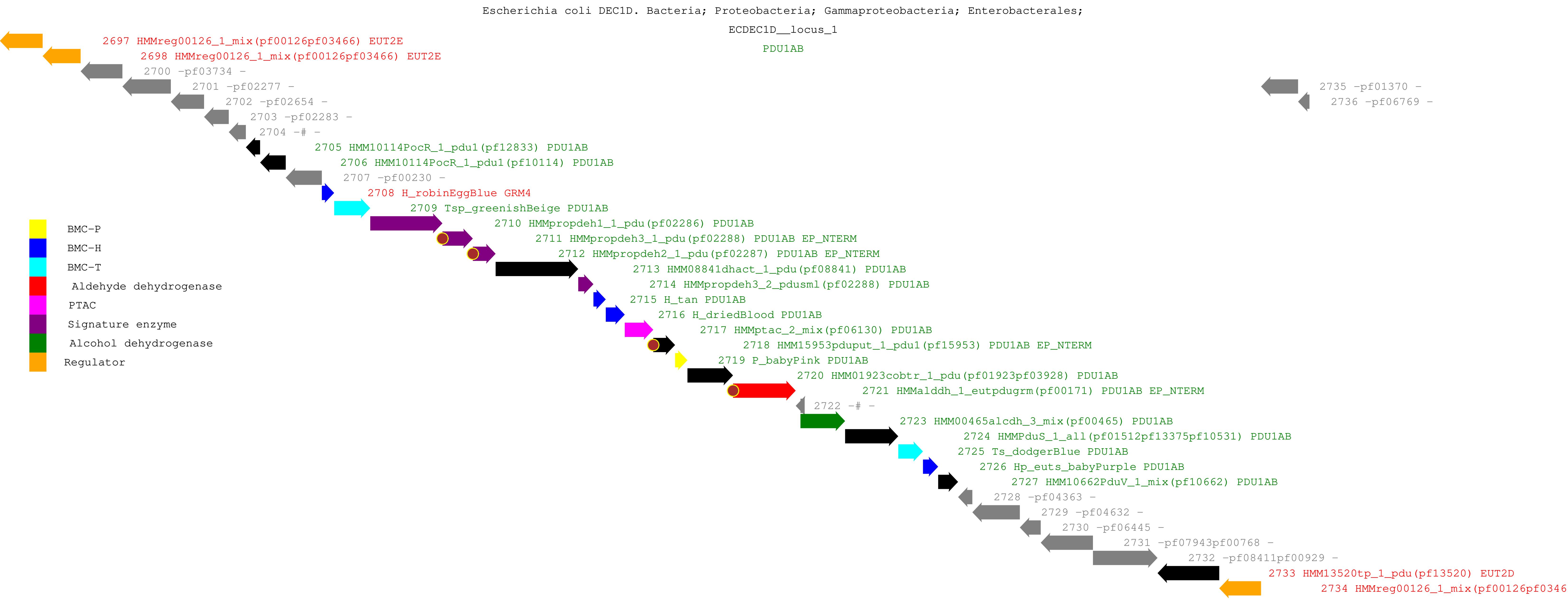

Escherichia coli DEC1D. Bacteria; Proteobacteria; Gammaproteobacteria; Enterobacterales;
ECDEC1D__locus_1
PDU1AB
2697 HMMreg00126_1_mix(pf00126pf03466) EUT2E
2698 HMMreg00126_1_mix(pf00126pf03466) EUT2E
2700 -pf03734 -
2701 -pf02277 -
2702 -pf02654 -
2703 -pf02283 -
2704 -# -
2705 HMM10114PocR_1_pdu1(pf12833) PDU1AB
2706 HMM10114PocR_1_pdu1(pf10114) PDU1AB
2707 -pf00230 -
2708 H_robinEggBlue GRM4
2709 Tsp_greenishBeige PDU1AB
2710 HMMpropdeh1_1_pdu(pf02286) PDU1AB
2711 HMMpropdeh3_1_pdu(pf02288) PDU1AB EP_NTERM
2712 HMMpropdeh2_1_pdu(pf02287) PDU1AB EP_NTERM
2713 HMM08841dhact_1_pdu(pf08841) PDU1AB
2714 HMMpropdeh3_2_pdusml(pf02288) PDU1AB
2715 H_tan PDU1AB
2716 H_driedBlood PDU1AB
2717 HMMptac_2_mix(pf06130) PDU1AB
2718 HMM15953pduput_1_pdu1(pf15953) PDU1AB EP_NTERM
2719 P_babyPink PDU1AB
2720 HMM01923cobtr_1_pdu(pf01923pf03928) PDU1AB
2721 HMMalddh_1_eutpdugrm(pf00171) PDU1AB EP_NTERM
2722 -# -
2723 HMM00465alcdh_3_mix(pf00465) PDU1AB
2724 HMMPduS_1_all(pf01512pf13375pf10531) PDU1AB
2725 Ts_dodgerBlue PDU1AB
2726 Hp_euts_babyPurple PDU1AB
2727 HMM10662PduV_1_mix(pf10662) PDU1AB
2728 -pf04363 -
2729 -pf04632 -
2730 -pf06445 -
2731 -pf07943pf00768 -
2732 -pf08411pf00929 -
2733 HMM13520tp_1_pdu(pf13520) EUT2D
2734 HMMreg00126_1_mix(pf00126pf0346
2735 -pf01370 -
2736 -pf06769 -
BMC-P
BMC-H
BMC-T
Aldehyde dehydrogenase
PTAC
Signature enzyme
Alcohol dehydrogenase
Regulator

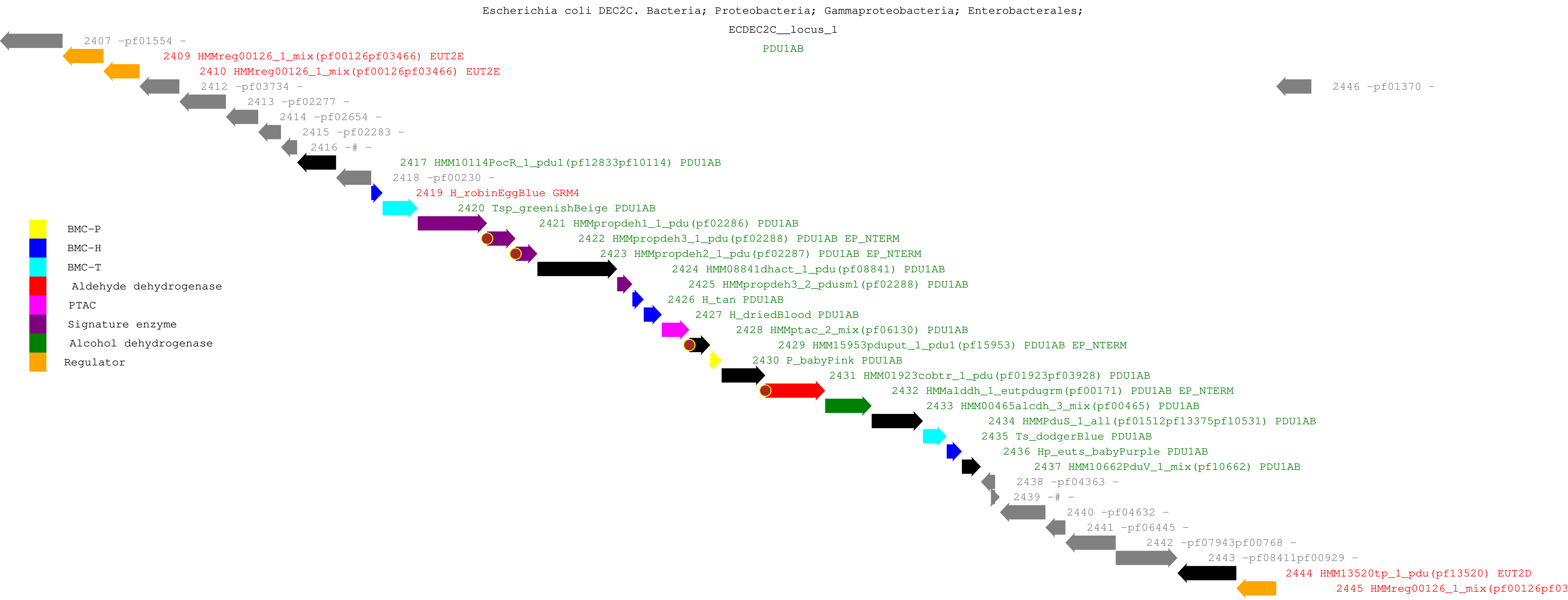
Escherichia coli DEC2C. Bacteria; Proteobacteria; Gammaproteobacteria; Enterobacterales;
ECDEC2C__locus_1
PDU1AB
2407 -pf01554 -
2409 HMMreg00126_1_mix(pf00126pf03466) EUT2E
2410 HMMreg00126_1_mix(pf00126pf03466) EUT2E
2412 -pf03734 -
2413 -pf02277 -
2414 -pf02654 -
2415 -pf02283 -
2416 -# -
2417 HMM10114PocR_1_pdu1(pf12833pf10114) PDU1AB
2418 -pf00230 -
2419 H_robinEggBlue GRM4
2420 Tsp_greenishBeige PDU1AB
2421 HMMpropdeh1_1_pdu(pf02286) PDU1AB
2422 HMMpropdeh3_1_pdu(pf02288) PDU1AB EP_NTERM
2423 HMMpropdeh2_1_pdu(pf02287) PDU1AB EP_NTERM
2424 HMM08841dhact_1_pdu(pf08841) PDU1AB
2425 HMMpropdeh3_2_pdusml(pf02288) PDU1AB
2426 H_tan PDU1AB
2427 H_driedBlood PDU1AB
2428 HMMptac_2_mix(pf06130) PDU1AB
2429 HMM15953pduput_1_pdu1(pf15953) PDU1AB EP_NTERM
2430 P_babyPink PDU1AB
2431 HMM01923cobtr_1_pdu(pf01923pf03928) PDU1AB
2432 HMMalddh_1_eutpdugrm(pf00171) PDU1AB EP_NTERM
2433 HMM00465alcdh_3_mix(pf00465) PDU1AB
2434 HMMPduS_1_all(pf01512pf13375pf10531) PDU1AB
2435 Ts_dodgerBlue PDU1AB
2436 Hp_euts_babyPurple PDU1AB
2437 HMM10662PduV_1_mix(pf10662) PDU1AB
2438 -pf04363 -
2439 -# -
2440 -pf04632 -
2441 -pf06445 -
2442 -pf07943pf00768 -
2443 -pf08411pf00929 -
2444 HMM13520tp_1_pdu(pf13520) EUT2D
2445 HMMreg00126_1_mix(pf00126pf03
2446 -pf01370 -
BMC-P
BMC-H
BMC-T
Aldehyde dehydrogenase
PTAC
Signature enzyme
Alcohol dehydrogenase
Regulator

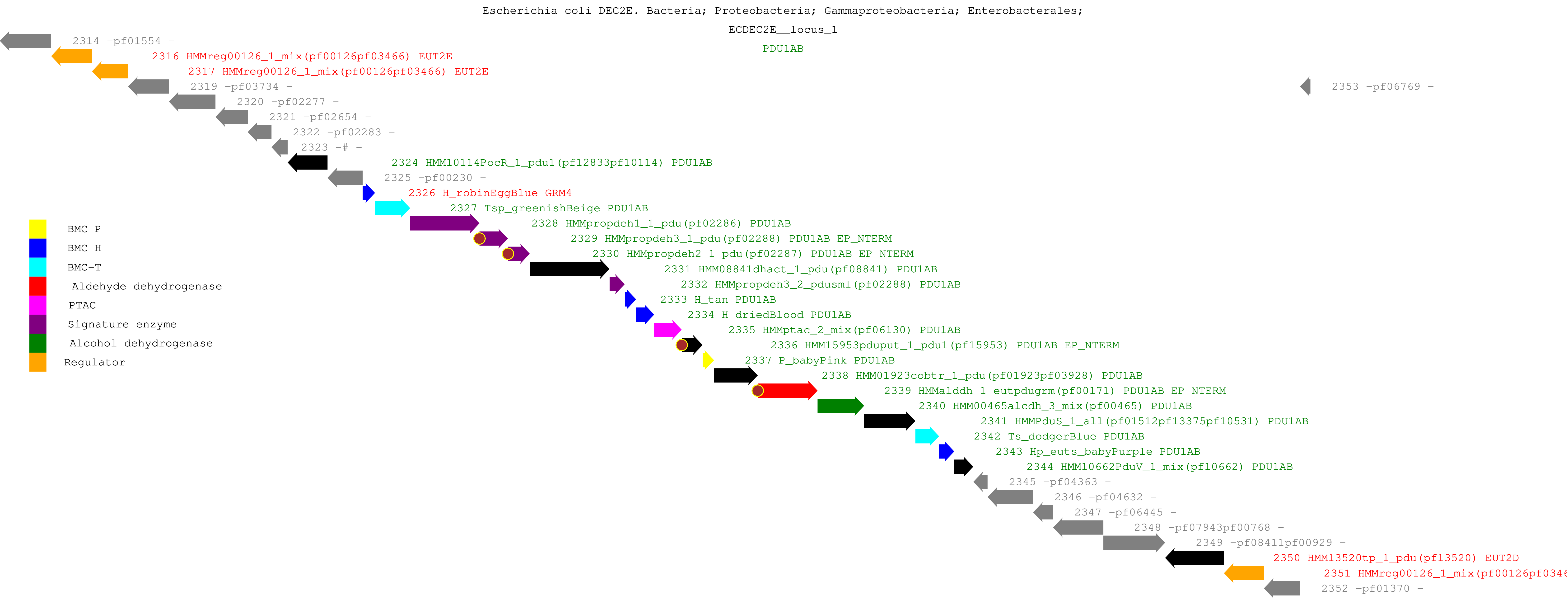
Escherichia coli DEC2E. Bacteria; Proteobacteria; Gammaproteobacteria; Enterobacterales;
ECDEC2E__locus_1
PDU1AB
2314 -pf01554 -
2316 HMMreg00126_1_mix(pf00126pf03466) EUT2E
2317 HMMreg00126_1_mix(pf00126pf03466) EUT2E
2319 -pf03734 -
2320 -pf02277 -
2321 -pf02654 -
2322 -pf02283 -
2323 -# -
2324 HMM10114PocR_1_pdu1(pf12833pf10114) PDU1AB
2325 -pf00230 -
2326 H_robinEggBlue GRM4
2327 Tsp_greenishBeige PDU1AB
2328 HMMpropdeh1_1_pdu(pf02286) PDU1AB
2329 HMMpropdeh3_1_pdu(pf02288) PDU1AB EP_NTERM
2330 HMMpropdeh2_1_pdu(pf02287) PDU1AB EP_NTERM
2331 HMM08841dhact_1_pdu(pf08841) PDU1AB
2332 HMMpropdeh3_2_pdusml(pf02288) PDU1AB
2333 H_tan PDU1AB
2334 H_driedBlood PDU1AB
2335 HMMptac_2_mix(pf06130) PDU1AB
2336 HMM15953pduput_1_pdu1(pf15953) PDU1AB EP_NTERM
2337 P_babyPink PDU1AB
2338 HMM01923cobtr_1_pdu(pf01923pf03928) PDU1AB
2339 HMMalddh_1_eutpdugrm(pf00171) PDU1AB EP_NTERM
2340 HMM00465alcdh_3_mix(pf00465) PDU1AB
2341 HMMPduS_1_all(pf01512pf13375pf10531) PDU1AB
2342 Ts_dodgerBlue PDU1AB
2343 Hp_euts_babyPurple PDU1AB
2344 HMM10662PduV_1_mix(pf10662) PDU1AB
2345 -pf04363 -
2346 -pf04632 -
2347 -pf06445 -
2348 -pf07943pf00768 -
2349 -pf08411pf00929 -
2350 HMM13520tp_1_pdu(pf13520) EUT2D
2351 HMMreg00126_1_mix(pf00126pf0346
2352 -pf01370 -
2353 -pf06769 -
BMC-P
BMC-H
BMC-T
Aldehyde dehydrogenase
PTAC
Signature enzyme
Alcohol dehydrogenase
Regulator

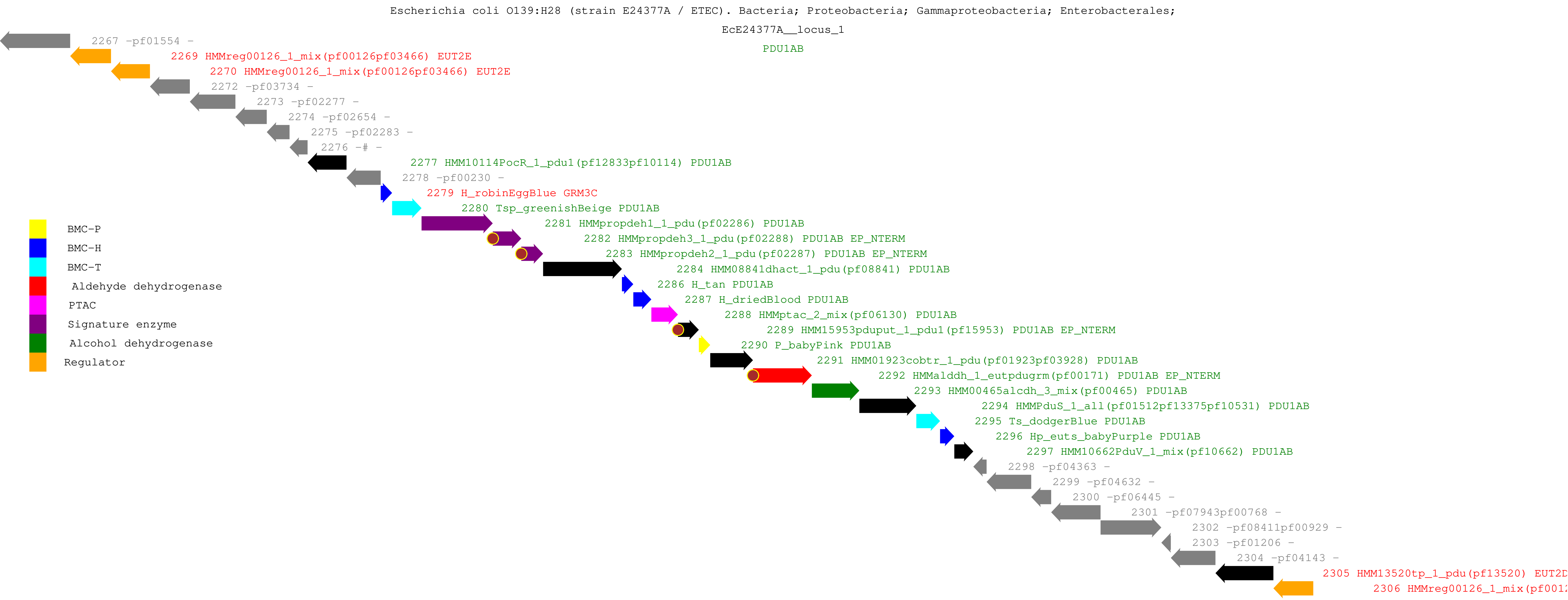
Escherichia coli O139:H28 (strain E24377A / ETEC). Bacteria; Proteobacteria; Gammaproteobacteria; Enterobacterales;
EcE24377A__locus_1
PDU1AB
2267 -pf01554 -
2269 HMMreg00126_1_mix(pf00126pf03466) EUT2E
2270 HMMreg00126_1_mix(pf00126pf03466) EUT2E
2272 -pf03734 -
2273 -pf02277 -
2274 -pf02654 -
2275 -pf02283 -
2276 -# -
2277 HMM10114PocR_1_pdu1(pf12833pf10114) PDU1AB
2278 -pf00230 -
2279 H_robinEggBlue GRM3C
2280 Tsp_greenishBeige PDU1AB
2281 HMMpropdeh1_1_pdu(pf02286) PDU1AB
2282 HMMpropdeh3_1_pdu(pf02288) PDU1AB EP_NTERM
2283 HMMpropdeh2_1_pdu(pf02287) PDU1AB EP_NTERM
2284 HMM08841dhact_1_pdu(pf08841) PDU1AB
2286 H_tan PDU1AB
2287 H_driedBlood PDU1AB
2288 HMMptac_2_mix(pf06130) PDU1AB
2289 HMM15953pduput_1_pdu1(pf15953) PDU1AB EP_NTERM
2290 P_babyPink PDU1AB
2291 HMM01923cobtr_1_pdu(pf01923pf03928) PDU1AB
2292 HMMalddh_1_eutpdugrm(pf00171) PDU1AB EP_NTERM
2293 HMM00465alcdh_3_mix(pf00465) PDU1AB
2294 HMMPduS_1_all(pf01512pf13375pf10531) PDU1AB
2295 Ts_dodgerBlue PDU1AB
2296 Hp_euts_babyPurple PDU1AB
2297 HMM10662PduV_1_mix(pf10662) PDU1AB
2298 -pf04363 -
2299 -pf04632 -
2300 -pf06445 -
2301 -pf07943pf00768 -
2302 -pf08411pf00929 -
2303 -pf01206 -
2304 -pf04143 -
2305 HMM13520tp_1_pdu(pf13520) EUT2D
2306 HMMreg00126_1_mix(pf001
BMC-P
BMC-H
BMC-T
Aldehyde dehydrogenase
PTAC
Signature enzyme
Alcohol dehydrogenase
Regulator

Escherichia coli B354. Bacteria; Proteobacteria; Gammaproteobacteria; Enterobacterales;

ECEG__locus_1

PDU1AB

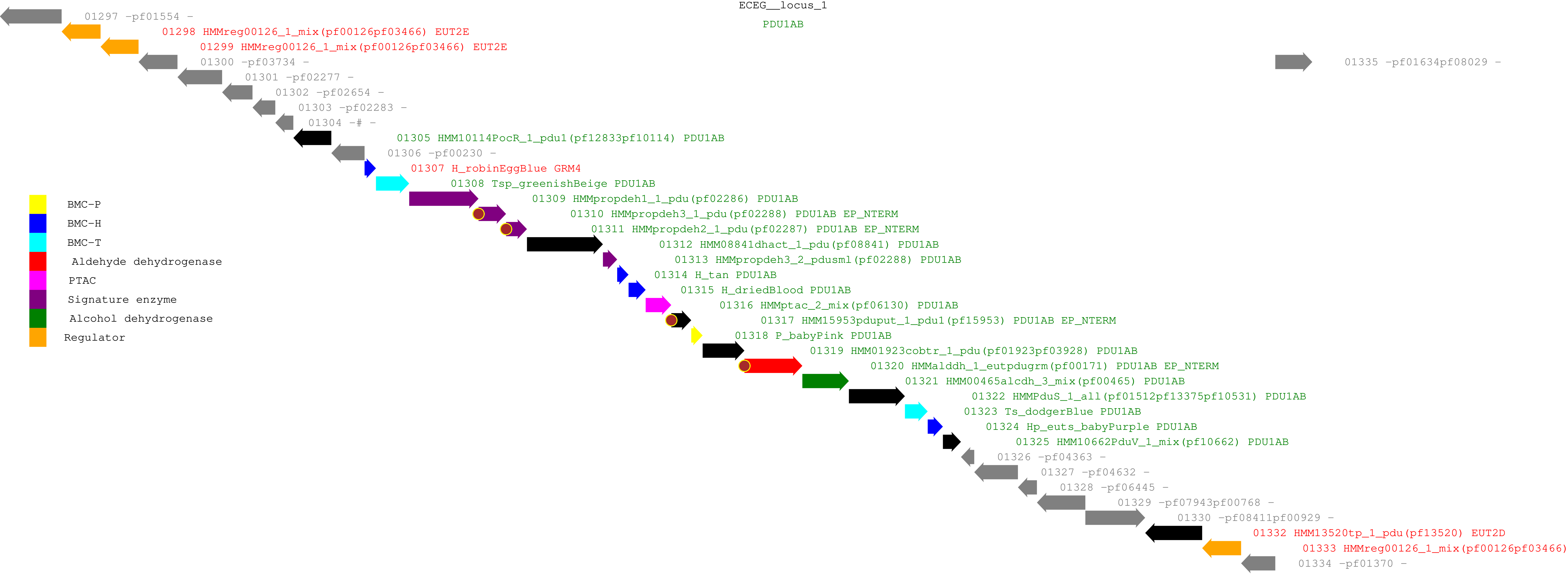

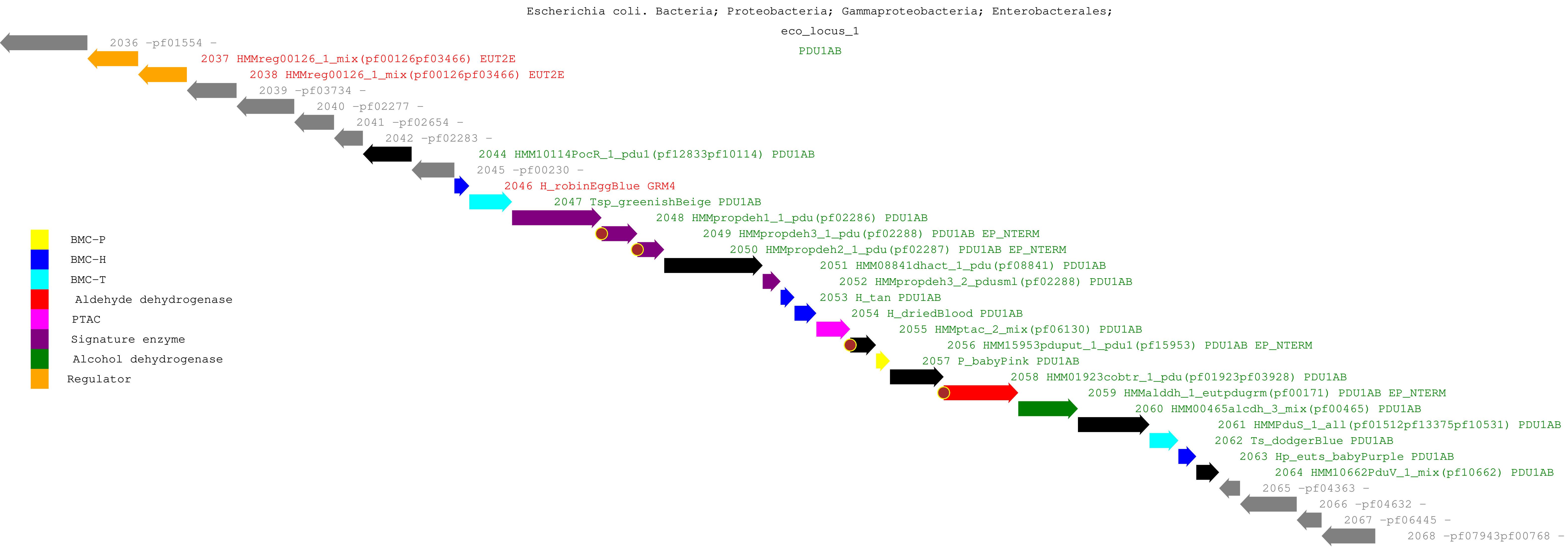
Escherichia coli. Bacteria; Proteobacteria; Gammaproteobacteria; Enterobacterales;
eco_locus_1
PDU1AB
2036 -pf01554 -
2037 HMMreg00126_1_mix(pf00126pf03466) EUT2E
2038 HMMreg00126_1_mix(pf00126pf03466) EUT2E
2039 -pf03734 -
2040 -pf02277 -
2041 -pf02654 -
2042 -pf02283 -
2044 HMM10114PocR_1_pdu1(pf12833pf10114) PDU1AB
2045 -pf00230 -
2046 H_robinEggBlue GRM4
2047 Tsp_greenishBeige PDU1AB
2048 HMMpropdeh1_1_pdu(pf02286) PDU1AB
2049 HMMpropdeh3_1_pdu(pf02288) PDU1AB EP_NTERM
2050 HMMpropdeh2_1_pdu(pf02287) PDU1AB EP_NTERM
2051 HMM08841dhact_1_pdu(pf08841) PDU1AB
2052 HMMpropdeh3_2_pdusml(pf02288) PDU1AB
2053 H_tan PDU1AB
2054 H_driedBlood PDU1AB
2055 HMMptac_2_mix(pf06130) PDU1AB
2056 HMM15953pduput_1_pdu1(pf15953) PDU1AB EP_NTERM
2057 P_babyPink PDU1AB
2058 HMM01923cobtr_1_pdu(pf01923pf03928) PDU1AB
2059 HMMalddh_1_eutpdugrm(pf00171) PDU1AB EP_NTERM
2060 HMM00465alcdh_3_mix(pf00465) PDU1AB
2061 HMMPduS_1_all(pf01512pf13375pf10531) PDU1AB
2062 Ts_dodgerBlue PDU1AB
2063 Hp_euts_babyPurple PDU1AB
2064 HMM10662PduV_1_mix(pf10662) PDU1AB
2065 -pf04363 -
2066 -pf04632 -
2067 -pf06445 -
2068 -pf07943pf00768 -
BMC-P
BMC-H
BMC-T
Aldehyde dehydrogenase
PTAC
Signature enzyme
Alcohol dehydrogenase
Regulator

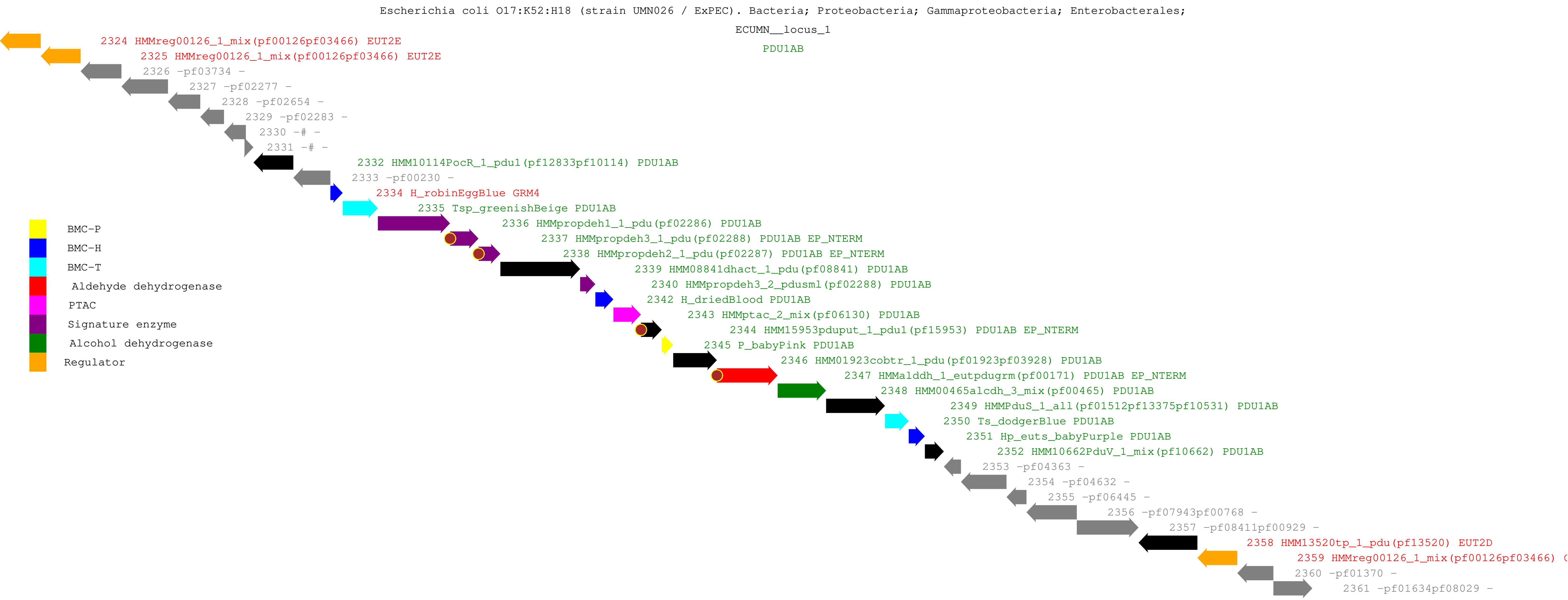
Escherichia coli O17:K52:H18 (strain UMN026 / ExPEC). Bacteria; Proteobacteria; Gammaproteobacteria; Enterobacterales;
ECUMN__locus_1
PDU1AB
2324 HMMreg00126_1_mix(pf00126pf03466) EUT2E
2325 HMMreg00126_1_mix(pf00126pf03466) EUT2E
2326 -pf03734 -
2327 -pf02277 -
2328 -pf02654 -
2329 -pf02283 -
2330 -# -
2331 -# -
2332 HMM10114PocR_1_pdu1(pf12833pf10114) PDU1AB
2333 -pf00230 -
2334 H_robinEggBlue GRM4
2335 Tsp_greenishBeige PDU1AB
2336 HMMpropdeh1_1_pdu(pf02286) PDU1AB
2337 HMMpropdeh3_1_pdu(pf02288) PDU1AB EP_NTERM
2338 HMMpropdeh2_1_pdu(pf02287) PDU1AB EP_NTERM
2339 HMM08841dhact_1_pdu(pf08841) PDU1AB
2340 HMMpropdeh3_2_pdusml(pf02288) PDU1AB
2342 H_driedBlood PDU1AB
2343 HMMptac_2_mix(pf06130) PDU1AB
2344 HMM15953pduput_1_pdu1(pf15953) PDU1AB EP_NTERM
2345 P_babyPink PDU1AB
2346 HMM01923cobtr_1_pdu(pf01923pf03928) PDU1AB
2347 HMMaldh_1_eutpdugrm(pf00171) PDU1AB EP_NTERM
2348 HMM00465alcdh_3_mix(pf00465) PDU1AB
2349 HMMPduS_1_all(pf01512pf13375pf10531) PDU1AB
2350 Ts_dodgerBlue PDU1AB
2351 Hp_euts_babyPurple PDU1AB
2352 HMM10662PduV_1_mix(pf10662) PDU1AB
2353 -pf04363 -
2354 -pf04632 -
2355 -pf06445 -
2356 -pf07943pf00768 -
2357 -pf08411pf00929 -
2358 HMM13520tp_1_pdu(pf13520) EUT2D
2359 HMMreg00126_1_mix(pf00126pf03466)
2360 -pf01370 -
2361 -pf01634pf08029 -
BMC-P
BMC-H
BMC-T
Aldehyde dehydrogenase
PTAC
Signature enzyme
Alcohol dehydrogenase
Regulator

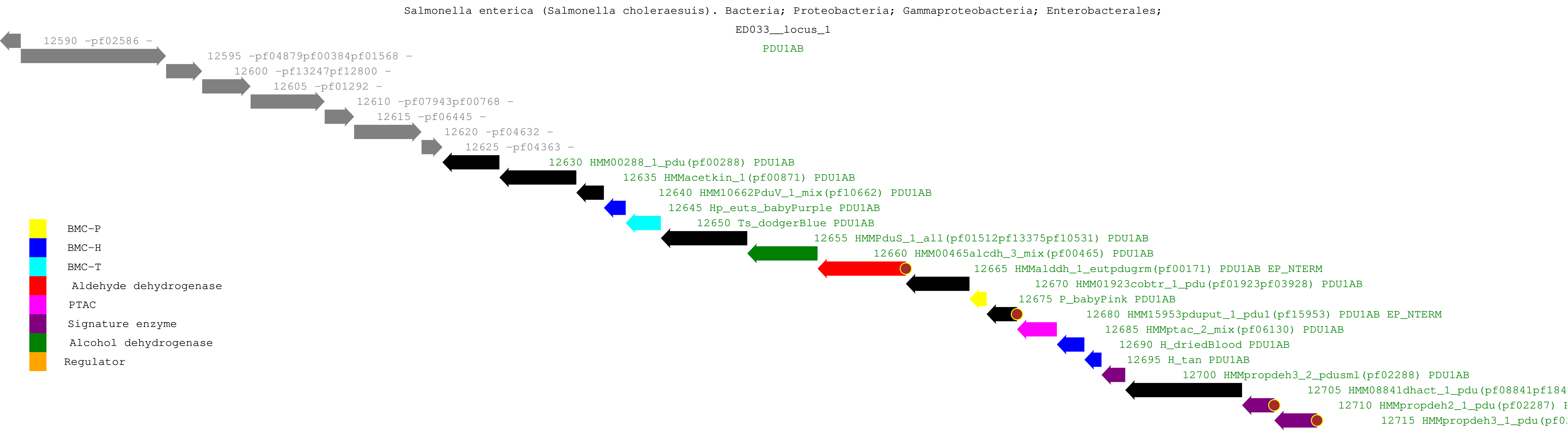
Salmonella enterica (Salmonella choleraesuis). Bacteria; Proteobacteria; Gammaproteobacteria; Enterobacterales;
ED033__locus_1
PDU1AB
12590 -pf02586 -
12595 -pf04879pf00384pf01568 -
12600 -pf13247pf12800 -
12605 -pf01292 -
12610 -pf07943pf00768 -
12615 -pf06445 -
12620 -pf04632 -
12625 -pf04363 -
12630 HMM00288_1_pdu(pf00288) PDU1AB
12635 HMMacetkin_1(pf00871) PDU1AB
12640 HMM10662PduV_1_mix(pf10662) PDU1AB
12645 Hp_euts_babyPurple PDU1AB
12650 Ts_dodgerBlue PDU1AB
12655 HMMPduS_1_all(pf01512pf13375pf10531) PDU1AB
12660 HMM00465alcdh_3_mix(pf00465) PDU1AB
12665 HMMaldh_1_eutpdugrm(pf00171) PDU1AB EP_NTERM
12670 HMM01923cobtr_1_pdu(pf01923pf03928) PDU1AB
12675 P_babyPink PDU1AB
12680 HMM15953pduput_1_pdu1(pf15953) PDU1AB EP_NTERM
12685 HMMptac_2_mix(pf06130) PDU1AB
12690 H_driedBlood PDU1AB
12695 H_tan PDU1AB
12700 HMMpropdeh3_2_pdusml(pf02288) PDU1AB
12705 HMM08841dhact_1_pdu(pf08841pf184
12710 HMMpropdeh2_1_pdu(pf02287)
12715 HMMpropdeh3_1_pdu(pf0
BMC-P
BMC-H
BMC-T
Aldehyde dehydrogenase
PTAC
Signature enzyme
Alcohol dehydrogenase
Regulator

Phytobacter diazotrophicus. Bacteria; Proteobacteria; Gammaproteobacteria; Enterobacterales;

EDC53__locus_1

PDU1AB

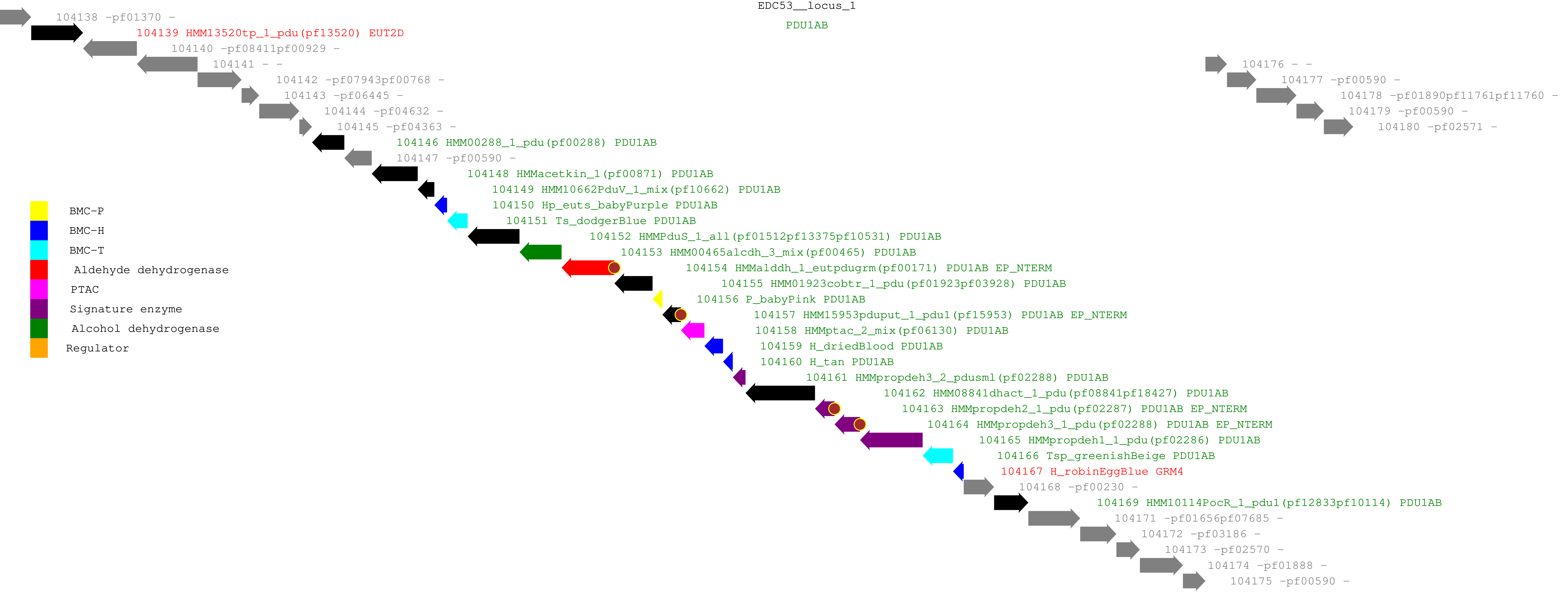

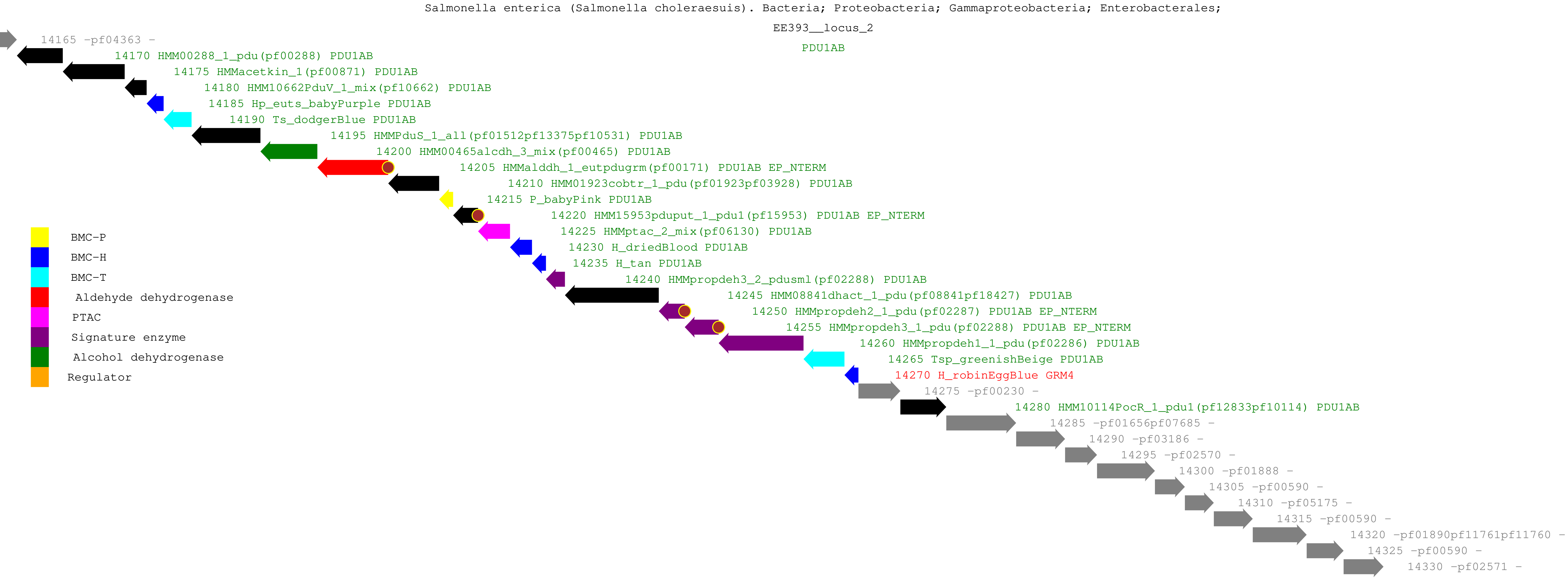

Salmonella enterica (Salmonella choleraesuis). Bacteria; Proteobacteria; Gammaproteobacteria; Enterobacterales;
EE393__locus_2
PDU1AB
14165 -pf04363 -
14170 HMM00288_1_pdu(pf00288) PDU1AB
14175 HMMacetkin_1(pf00871) PDU1AB
14180 HMM10662PduV_1_mix(pf10662) PDU1AB
14185 Hp_euts_babyPurple PDU1AB
14190 Ts_dodgerBlue PDU1AB
14195 HMMPduS_1_all(pf01512pf13375pf10531) PDU1AB
14200 HMM00465alcdh_3_mix(pf00465) PDU1AB
14205 HMMalddh_1_eutpdugrm(pf00171) PDU1AB EP_NTERM
14210 HMM01923cobtr_1_pdu(pf01923pf03928) PDU1AB
14215 P_babyPink PDU1AB
14220 HMM15953pduput_1_pdu1(pf15953) PDU1AB EP_NTERM
14225 HMMptac_2_mix(pf06130) PDU1AB
14230 H_driedBlood PDU1AB
14235 H_tan PDU1AB
14240 HMMpropdeh3_2_pdusml(pf02288) PDU1AB
14245 HMM08841dhact_1_pdu(pf08841pf18427) PDU1AB
14250 HMMpropdeh2_1_pdu(pf02287) PDU1AB EP_NTERM
14255 HMMpropdeh3_1_pdu(pf02288) PDU1AB EP_NTERM
14260 HMMpropdeh1_1_pdu(pf02286) PDU1AB
14265 Tsp_greenishBeige PDU1AB
14270 H_robinEggBlue GRM4
14275 -pf00230 -
14280 HMM10114PocR_1_pdu1(pf12833pf10114) PDU1AB
14285 -pf01656pf07685 -
14290 -pf03186 -
14295 -pf02570 -
14300 -pf01888 -
14305 -pf00590 -
14310 -pf05175 -
14315 -pf00590 -
14320 -pf01890pf11761pf11760 -
14325 -pf00590 -
14330 -pf02571 -
BMC-P
BMC-H
BMC-T
Aldehyde dehydrogenase
PTAC
Signature enzyme
Alcohol dehydrogenase
Regulator

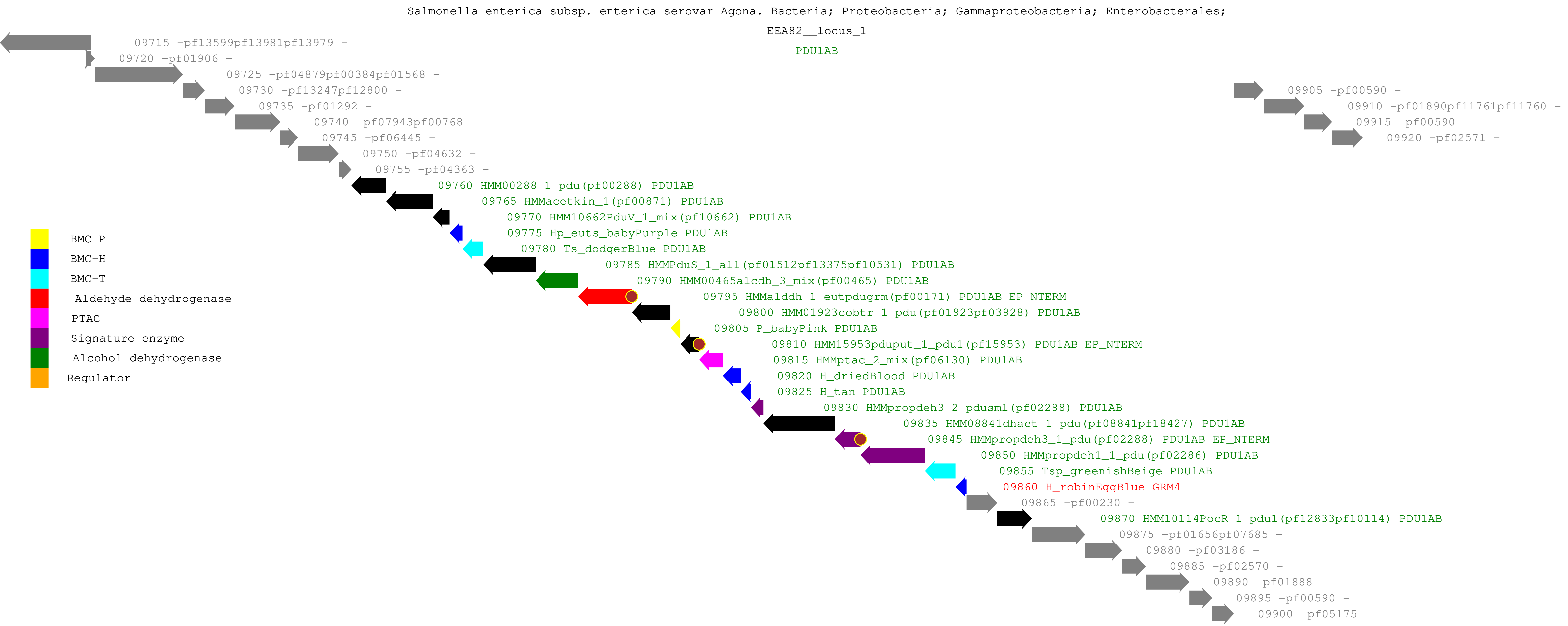

Salmonella enterica subsp. enterica serovar Agona. Bacteria; Proteobacteria; Gammaproteobacteria; Enterobacterales;
EEA82__locus_1
PDU1AB
09715 -pf13599pf13981pf13979 -
09720 -pf01906 -
09725 -pf04879pf00384pf01568 -
09730 -pf13247pf12800 -
09735 -pf01292 -
09740 -pf07943pf00768 -
09745 -pf06445 -
09750 -pf04632 -
09755 -pf04363 -
09760 HMM00288_1_pdu(pf00288) PDU1AB
09765 HMMacetkin_1(pf00871) PDU1AB
09770 HMM10662PduV_1_mix(pf10662) PDU1AB
09775 Hp_euts_babyPurple PDU1AB
09780 Ts_dodgerBlue PDU1AB
09785 HMMPduS_1_all(pf01512pf13375pf10531) PDU1AB
09790 HMM00465alcdh_3_mix(pf00465) PDU1AB
09795 HMMalddh_1_eutpdugrm(pf00171) PDU1AB EP_NTERM
09800 HMM01923cobtr_1_pdu(pf01923pf03928) PDU1AB
09805 P_babyPink PDU1AB
09810 HMM15953pduput_1_pdu1(pf15953) PDU1AB EP_NTERM
09815 HMMptac_2_mix(pf06130) PDU1AB
09820 H_driedBlood PDU1AB
09825 H_tan PDU1AB
09830 HMMpropdeh3_2_pdusml(pf02288) PDU1AB
09835 HMM08841dhact_1_pdu(pf08841pf18427) PDU1AB
09845 HMMpropdeh3_1_pdu(pf02288) PDU1AB EP_NTERM
09850 HMMpropdeh1_1_pdu(pf02286) PDU1AB
09855 Tsp_greenishBeige PDU1AB
09860 H_robinEggBlue GRM4
09865 -pf00230 -
09870 HMM10114PocR_1_pdu1(pf12833pf10114) PDU1AB
09875 -pf01656pf07685 -
09880 -pf03186 -
09885 -pf02570 -
09890 -pf01888 -
09895 -pf00590 -
09900 -pf05175 -
09905 -pf00590 -
09910 -pf01890pf11761pf11760 -
09915 -pf00590 -
09920 -pf02571 -
BMC-P
BMC-H
BMC-T
Aldehyde dehydrogenase
PTAC
Signature enzyme
Alcohol dehydrogenase
Regulator

Salmonella enterica (Salmonella choleraesuis). Bacteria; Proteobacteria; Gammaproteobacteria; Enterobacterales;

EEM01__locus_2

PDU1AB

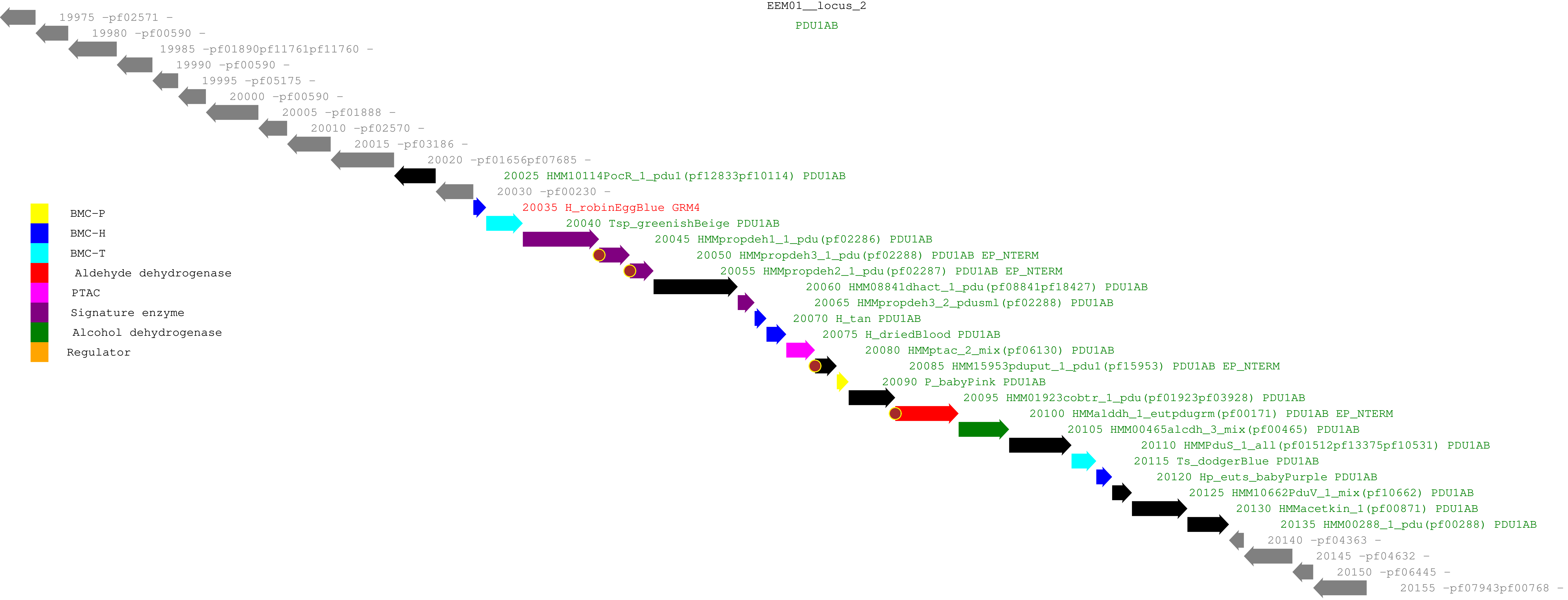

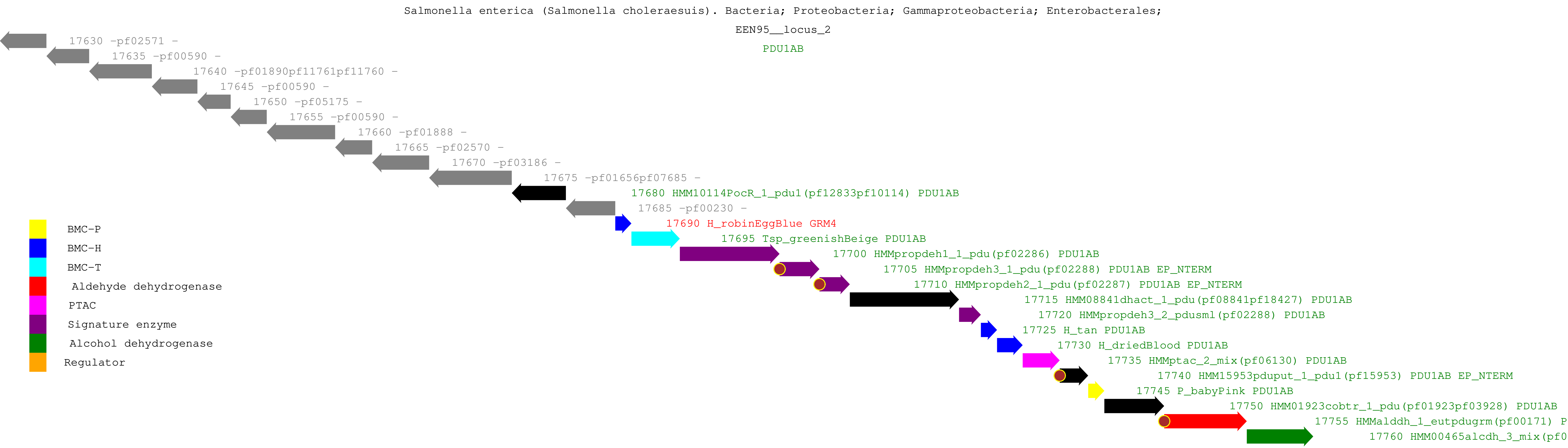

Salmonella enterica (Salmonella choleraesuis). Bacteria; Proteobacteria; Gammaproteobacteria; Enterobacterales;
EEN95__locus_2
PDU1AB
17630 -pf02571 -
17635 -pf00590 -
17640 -pf01890pf11761pf11760 -
17645 -pf00590 -
17650 -pf05175 -
17655 -pf00590 -
17660 -pf01888 -
17665 -pf02570 -
17670 -pf03186 -
17675 -pf01656pf07685 -
17680 HMM10114PocR_1_pdu1(pf12833pf10114) PDU1AB
17685 -pf00230 -
17690 H_robinEggBlue GRM4
17695 Tsp_greenishBeige PDU1AB
17700 HMMpropdeh1_1_pdu(pf02286) PDU1AB
17705 HMMpropdeh3_1_pdu(pf02288) PDU1AB EP_NTERM
17710 HMMpropdeh2_1_pdu(pf02287) PDU1AB EP_NTERM
17715 HMM08841dhact_1_pdu(pf08841pf18427) PDU1AB
17720 HMMpropdeh3_2_pdusml(pf02288) PDU1AB
17725 H_tan PDU1AB
17730 H_driedBlood PDU1AB
17735 HMMptac_2_mix(pf06130) PDU1AB
17740 HMM15953pduput_1_pdu1(pf15953) PDU1AB EP_NTERM
17745 P_babyPink PDU1AB
17750 HMM01923cobtr_1_pdu(pf01923pf03928) PDU1AB
17755 HMMalddh_1_eutpdugrm(pf00171) P
17760 HMM00465alcdh_3_mix(pf0
BMC-P
BMC-H
BMC-T
Aldehyde dehydrogenase
PTAC
Signature enzyme
Alcohol dehydrogenase
Regulator

Salmonella enterica subsp. enterica serovar Adelaide. Bacteria; Proteobacteria; Gammaproteobacteria; Enterobacterales;

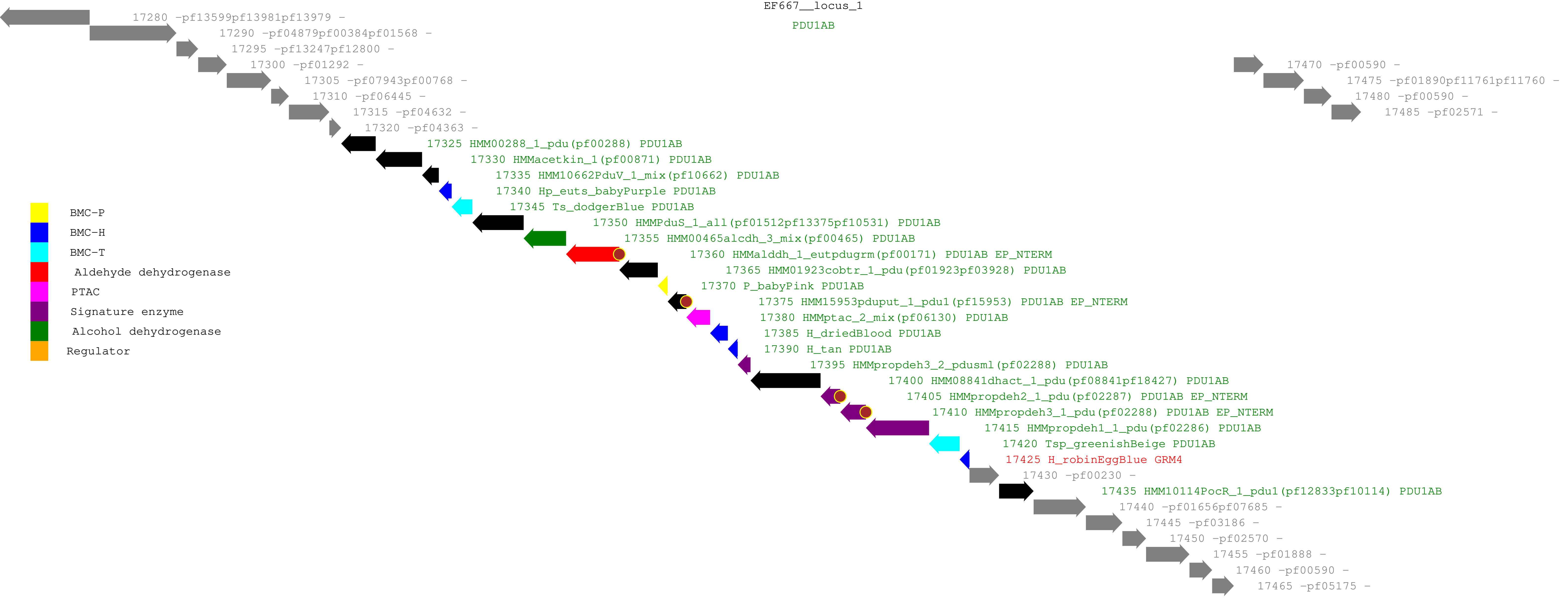

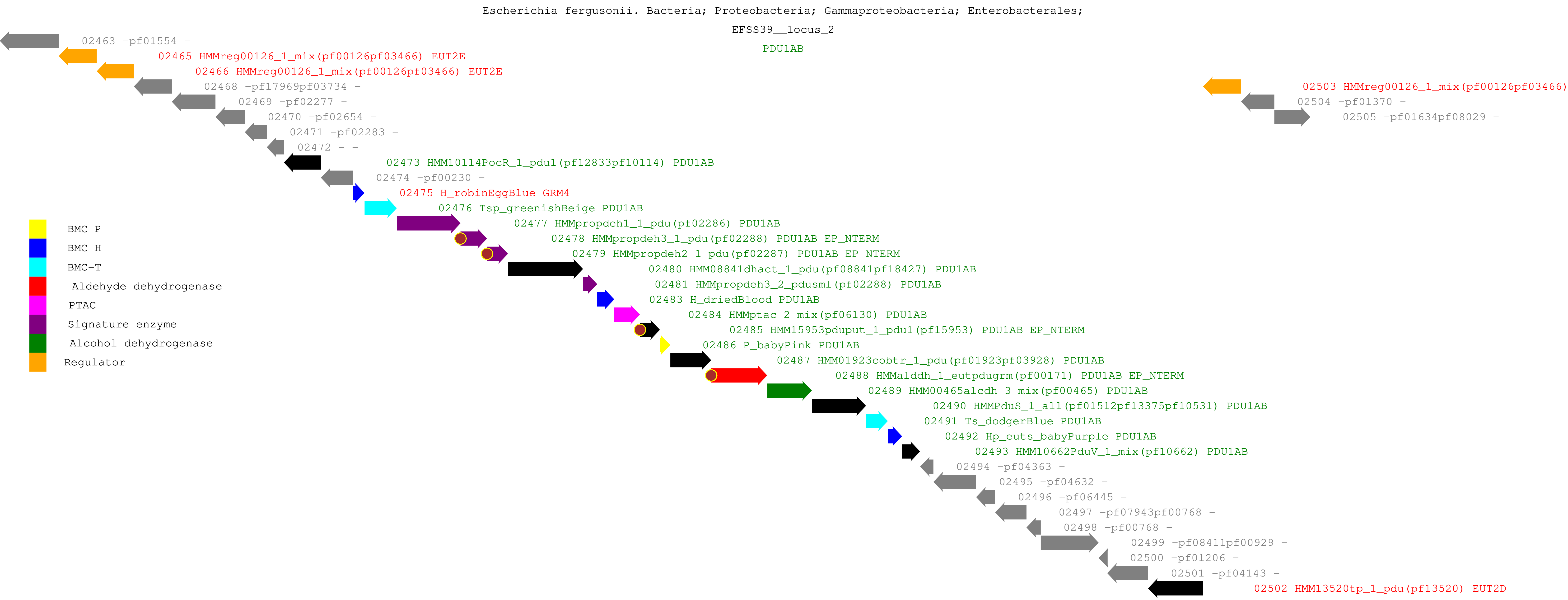

Escherichia fergusonii. Bacteria; Proteobacteria; Gammaproteobacteria; Enterobacterales;
EFSS39__locus_2
PDU1AB
02463 -pf01554 -
02465 HMMreg00126_1_mix(pf00126pf03466) EUT2E
02466 HMMreg00126_1_mix(pf00126pf03466) EUT2E
02468 -pf17969pf03734 -
02469 -pf02277 -
02470 -pf02654 -
02471 -pf02283 -
02472 - -
02473 HMM10114PocR_1_pdu1(pf12833pf10114) PDU1AB
02474 -pf00230 -
02475 H_robinEggBlue GRM4
02476 Tsp_greenishBeige PDU1AB
02477 HMMpropdeh1_1_pdu(pf02286) PDU1AB
02478 HMMpropdeh3_1_pdu(pf02288) PDU1AB EP_NTERM
02479 HMMpropdeh2_1_pdu(pf02287) PDU1AB EP_NTERM
02480 HMM08841dhact_1_pdu(pf08841pf18427) PDU1AB
02481 HMMpropdeh3_2_pdusml(pf02288) PDU1AB
02483 H_driedBlood PDU1AB
02484 HMMptac_2_mix(pf06130) PDU1AB
02485 HMM15953pduput_1_pdu1(pf15953) PDU1AB EP_NTERM
02486 P_babyPink PDU1AB
02487 HMM01923cobtr_1_pdu(pf01923pf03928) PDU1AB
02488 HMMaldd h_1_eutpdugrm(pf00171) PDU1AB EP_NTERM
02489 HMM00465alcdh_3_mix(pf00465) PDU1AB
02490 HMMPduS_1_all(pf01512pf13375pf10531) PDU1AB
02491 Ts_dodgerBlue PDU1AB
02492 Hp_euts_babyPurple PDU1AB
02493 HMM10662PduV_1_mix(pf10662) PDU1AB
02494 -pf04363 -
02495 -pf04632 -
02496 -pf06445 -
02497 -pf07943pf00768 -
02498 -pf00768 -
02499 -pf08411pf00929 -
02500 -pf01206 -
02501 -pf04143 -
02502 HMM13520tp_1_pdu(pf13520) EUT2D
02503 HMMreg00126_1_mix(pf00126pf03466)
02504 -pf01370 -
02505 -pf01634pf08029 -
BMC-P
BMC-H
BMC-T
Aldehyde dehydrogenase
PTAC
Signature enzyme
Alcohol dehydrogenase
Regulator

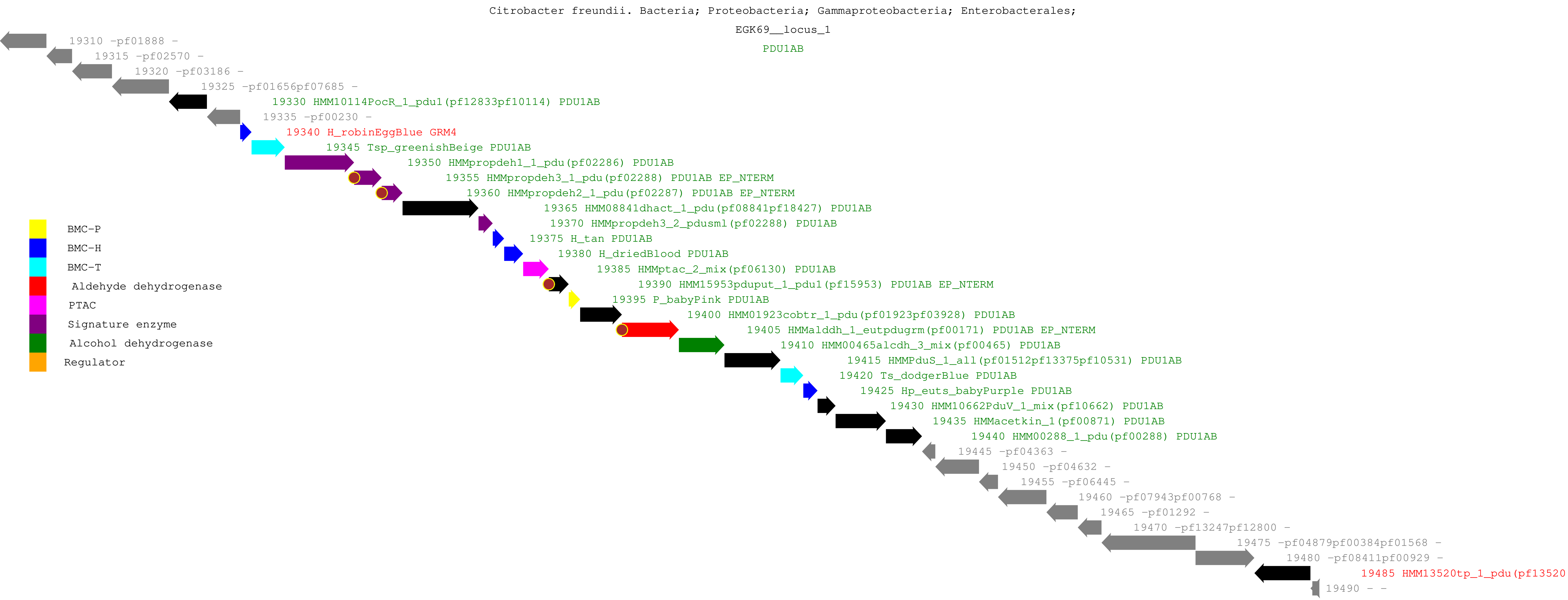
Citrobacter freundii. Bacteria; Proteobacteria; Gammaproteobacteria; Enterobacterales;
EGK69__locus_1
PDU1AB
19310 -pf01888 -
19315 -pf02570 -
19320 -pf03186 -
19325 -pf01656pf07685 -
19330 HMM10114PocR_1_pdu1(pf12833pf10114) PDU1AB
19335 -pf00230 -
19340 H_robinEggBlue GRM4
19345 Tsp_greenishBeige PDU1AB
19350 HMMpropdeh1_1_pdu(pf02286) PDU1AB
19355 HMMpropdeh3_1_pdu(pf02288) PDU1AB EP_NTERM
19360 HMMpropdeh2_1_pdu(pf02287) PDU1AB EP_NTERM
19365 HMM08841dhact_1_pdu(pf08841pf18427) PDU1AB
19370 HMMpropdeh3_2_pdusml(pf02288) PDU1AB
19375 H_tan PDU1AB
19380 H_driedBlood PDU1AB
19385 HMMptac_2_mix(pf06130) PDU1AB
19390 HMM15953pduput_1_pdu1(pf15953) PDU1AB EP_NTERM
19395 P_babyPink PDU1AB
19400 HMM01923cobtr_1_pdu(pf01923pf03928) PDU1AB
19405 HMMaldh_1_eutpdugrm(pf00171) PDU1AB EP_NTERM
19410 HMM00465alcdh_3_mix(pf00465) PDU1AB
19415 HMMPduS_1_all(pf01512pf13375pf10531) PDU1AB
19420 Ts_dodgerBlue PDU1AB
19425 Hp_euts_babyPurple PDU1AB
19430 HMM10662PduV_1_mix(pf10662) PDU1AB
19435 HMMacetkin_1(pf00871) PDU1AB
19440 HMM00288_1_pdu(pf00288) PDU1AB
19445 -pf04363 -
19450 -pf04632 -
19455 -pf06445 -
19460 -pf07943pf00768 -
19465 -pf01292 -
19470 -pf13247pf12800 -
19475 -pf04879pf00384pf01568 -
19480 -pf08411pf00929 -
19485 HMM13520tp_1_pdu(pf13520
19490 - -
BMC-P
BMC-H
BMC-T
Aldehyde dehydrogenase
PTAC
Signature enzyme
Alcohol dehydrogenase
Regulator

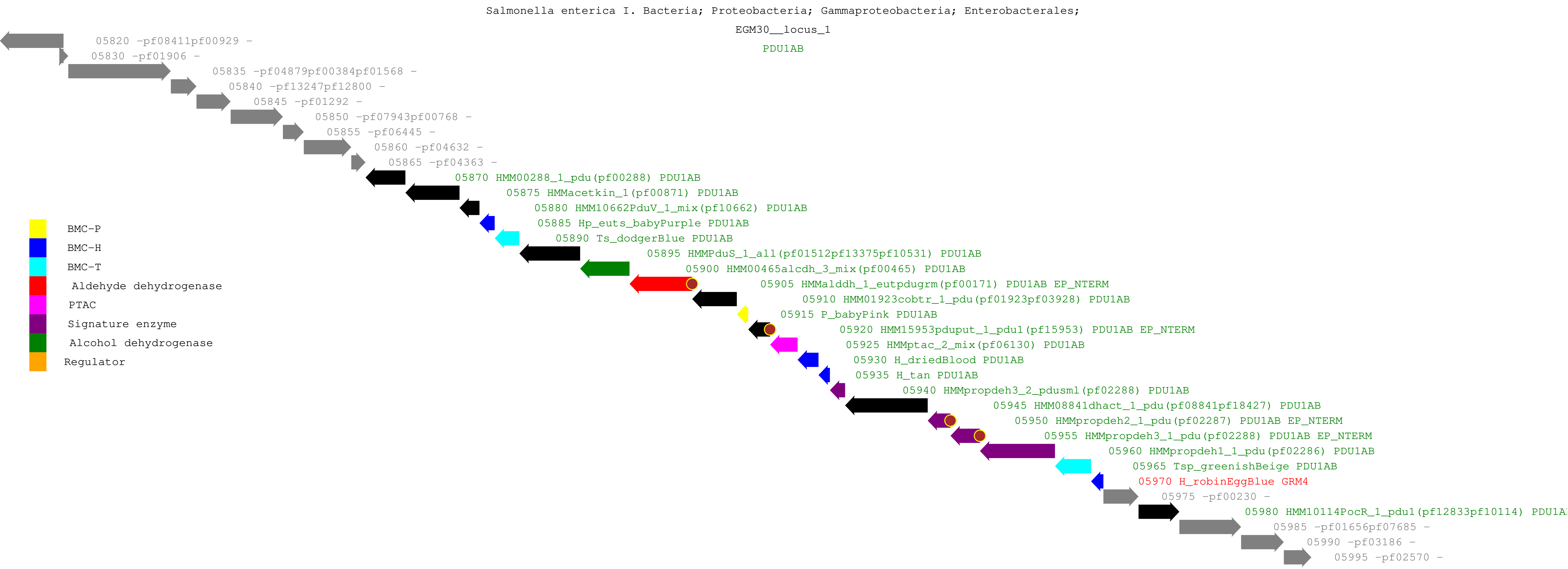

Salmonella enterica I. Bacteria; Proteobacteria; Gammaproteobacteria; Enterobacterales;
EGM30__locus_1
PDU1AB
05820 -pf08411pf00929 -
05830 -pf01906 -
05835 -pf04879pf00384pf01568 -
05840 -pf13247pf12800 -
05845 -pf01292 -
05850 -pf07943pf00768 -
05855 -pf06445 -
05860 -pf04632 -
05865 -pf04363 -
05870 HMM00288_1_pdu(pf00288) PDU1AB
05875 HMMacetkin_1(pf00871) PDU1AB
05880 HMM10662PduV_1_mix(pf10662) PDU1AB
05885 Hp_euts_babyPurple PDU1AB
05890 Ts_dodgerBlue PDU1AB
05895 HMMPduS_1_all(pf01512pf13375pf10531) PDU1AB
05900 HMM00465alcdh_3_mix(pf00465) PDU1AB
05905 HMMalddh_1_eutpdugrm(pf00171) PDU1AB EP_NTERM
05910 HMM01923cobtr_1_pdu(pf01923pf03928) PDU1AB
05915 P_babyPink PDU1AB
05920 HMM15953pduput_1_pdu1(pf15953) PDU1AB EP_NTERM
05925 HMMptac_2_mix(pf06130) PDU1AB
05930 H_driedBlood PDU1AB
05935 H_tan PDU1AB
05940 HMMpropdeh3_2_pdusml(pf02288) PDU1AB
05945 HMM08841dhact_1_pdu(pf08841pf18427) PDU1AB
05950 HMMpropdeh2_1_pdu(pf02287) PDU1AB EP_NTERM
05955 HMMpropdeh3_1_pdu(pf02288) PDU1AB EP_NTERM
05960 HMMpropdeh1_1_pdu(pf02286) PDU1AB
05965 Tsp_greenishBeige PDU1AB
05970 H_robinEggBlue GRM4
05975 -pf00230 -
05980 HMM10114PocR_1_pdu1(pf12833pf10114) PDU1A
05985 -pf01656pf07685 -
05990 -pf03186 -
05995 -pf02570 -
BMC-P
BMC-H
BMC-T
Aldehyde dehydrogenase
PTAC
Signature enzyme
Alcohol dehydrogenase
Regulator

Salmonella paratyphi A. Bacteria; Proteobacteria; Gammaproteobacteria; Enterobacterales;

EJI24__locus_2

PDU1AB

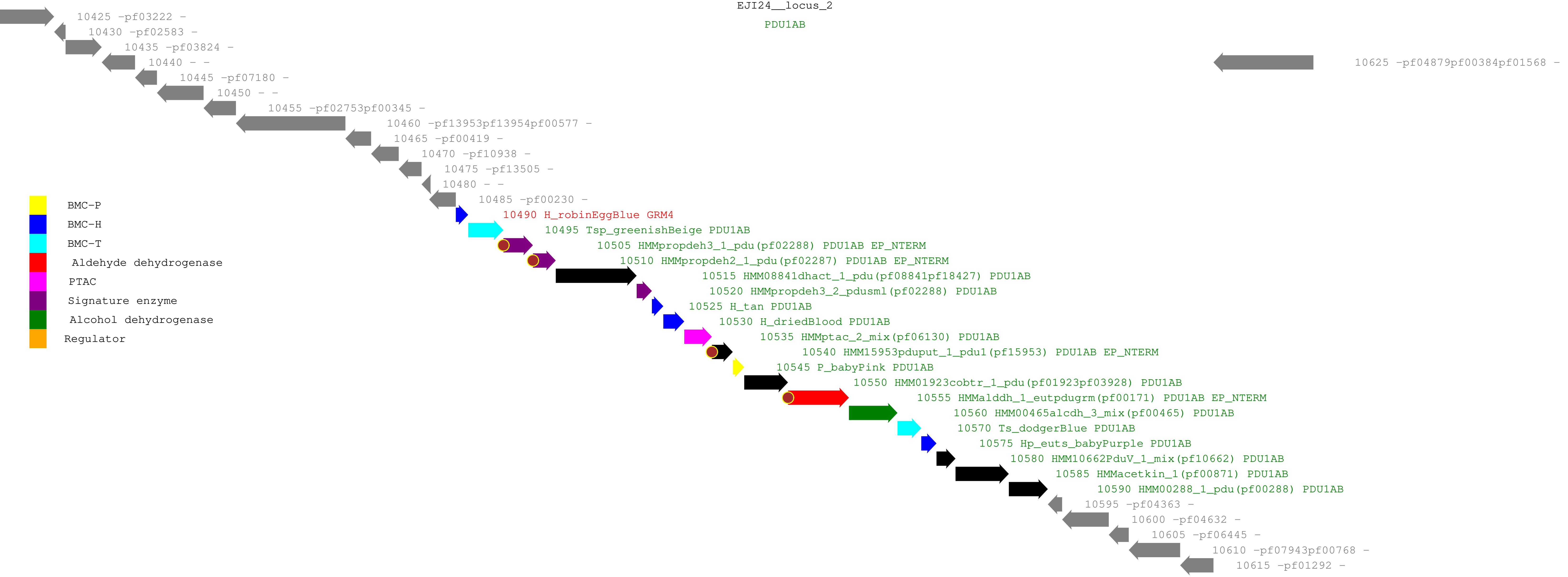

Salmonella enterica subsp. enterica serovar Karamoja. Bacteria; Proteobacteria; Gammaproteobacteria; Enterobacterales;

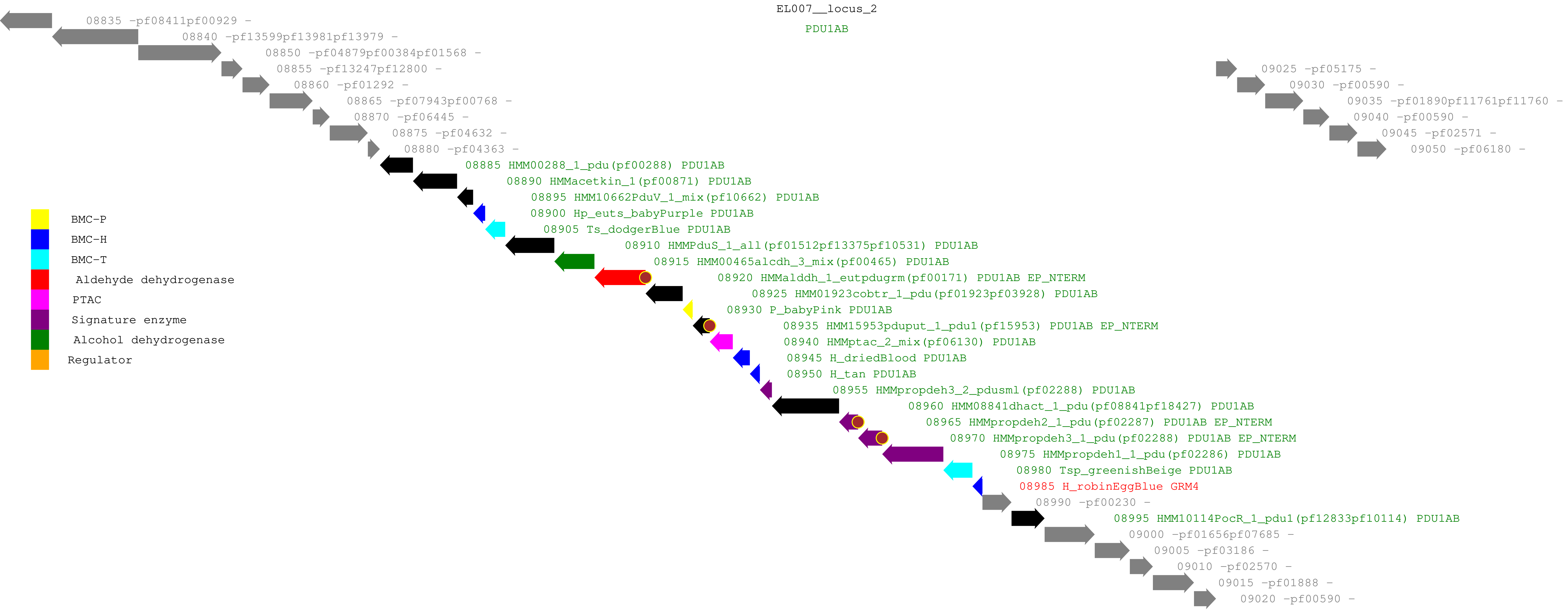

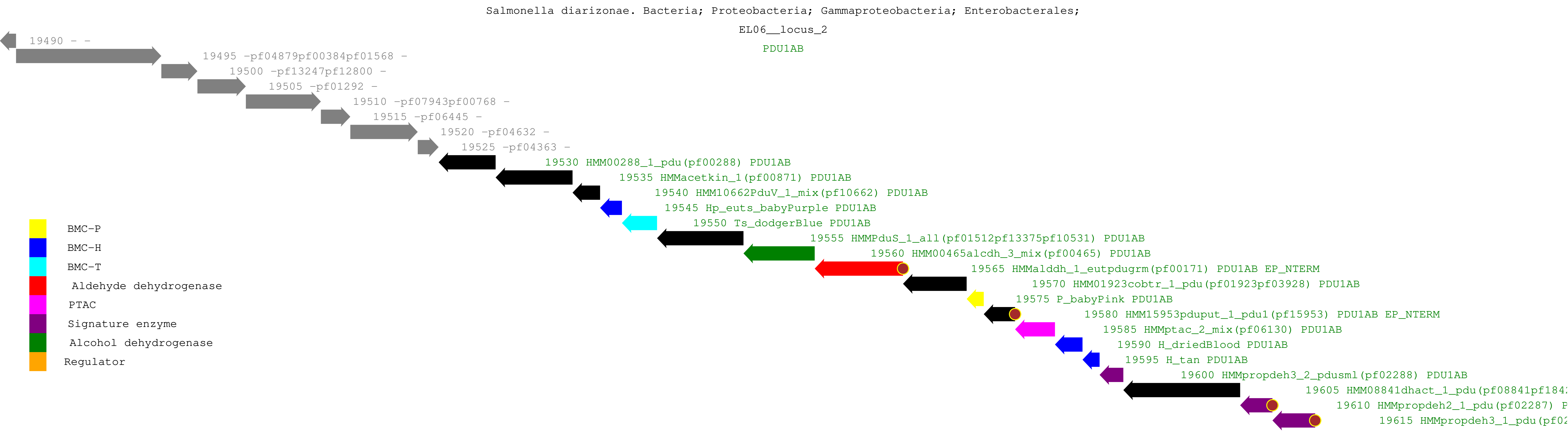
Salmonella diarizonae. Bacteria; Proteobacteria; Gammaproteobacteria; Enterobacterales;
EL06__locus_2
PDU1AB
19490 - -
19495 -pf04879pf00384pf01568 -
19500 -pf13247pf12800 -
19505 -pf01292 -
19510 -pf07943pf00768 -
19515 -pf06445 -
19520 -pf04632 -
19525 -pf04363 -
19530 HMM00288_1_pdu(pf00288) PDU1AB
19535 HMMacetkin_1(pf00871) PDU1AB
19540 HMM10662PduV_1_mix(pf10662) PDU1AB
19545 Hp_euts_babyPurple PDU1AB
19550 Ts_dodgerBlue PDU1AB
19555 HMMPduS_1_all(pf01512pf13375pf10531) PDU1AB
19560 HMM00465alcdh_3_mix(pf00465) PDU1AB
19565 HMMalddh_1_eutpdugrm(pf00171) PDU1AB EP_NTERM
19570 HMM01923cobtr_1_pdu(pf01923pf03928) PDU1AB
19575 P_babyPink PDU1AB
19580 HMM15953pduput_1_pdu1(pf15953) PDU1AB EP_NTERM
19585 HMMptac_2_mix(pf06130) PDU1AB
19590 H_driedBlood PDU1AB
19595 H_tan PDU1AB
19600 HMMpropdeh3_2_pdusml(pf02288) PDU1AB
19605 HMM08841dhact_1_pdu(pf08841pf184
19610 HMMpropdeh2_1_pdu(pf02287) P
19615 HMMpropdeh3_1_pdu(pf02
BMC-P
BMC-H
BMC-T
Aldehyde dehydrogenase
PTAC
Signature enzyme
Alcohol dehydrogenase
Regulator

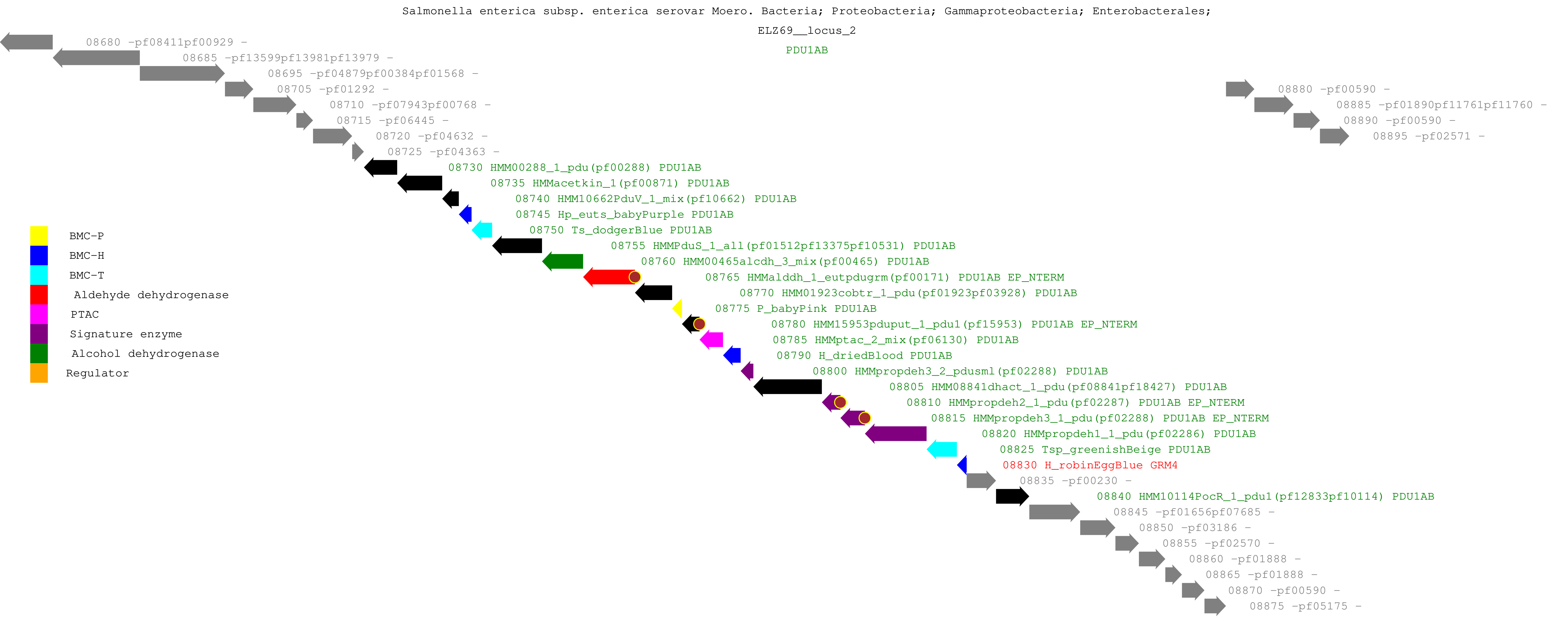
Salmonella enterica subsp. enterica serovar Moero. Bacteria; Proteobacteria; Gammaproteobacteria; Enterobacterales;
ELZ69__locus_2
PDU1AB
08680 -pf08411pf00929 -
08685 -pf13599pf13981pf13979 -
08695 -pf04879pf00384pf01568 -
08705 -pf01292 -
08710 -pf07943pf00768 -
08715 -pf06445 -
08720 -pf04632 -
08725 -pf04363 -
08730 HMM00288_1_pdu(pf00288) PDU1AB
08735 HMMacetkin_1(pf00871) PDU1AB
08740 HMM10662PduV_1_mix(pf10662) PDU1AB
08745 Hp_euts_babyPurple PDU1AB
08750 Ts_dodgerBlue PDU1AB
08755 HMMPduS_1_all(pf01512pf13375pf10531) PDU1AB
08760 HMM00465alcdh_3_mix(pf00465) PDU1AB
08765 HMMalddh_1_eutpdugrm(pf00171) PDU1AB EP_NTERM
08770 HMM01923cobtr_1_pdu(pf01923pf03928) PDU1AB
08775 P_babyPink PDU1AB
08780 HMM15953pduput_1_pdu1(pf15953) PDU1AB EP_NTERM
08785 HMMptac_2_mix(pf06130) PDU1AB
08790 H_driedBlood PDU1AB
08800 HMMpropdeh3_2_pdusml(pf02288) PDU1AB
08805 HMM08841dhact_1_pdu(pf08841pf18427) PDU1AB
08810 HMMpropdeh2_1_pdu(pf02287) PDU1AB EP_NTERM
08815 HMMpropdeh3_1_pdu(pf02288) PDU1AB EP_NTERM
08820 HMMpropdeh1_1_pdu(pf02286) PDU1AB
08825 Tsp_greenishBeige PDU1AB
08830 H_robinEggBlue GRM4
08835 -pf00230 -
08840 HMM10114PocR_1_pdu1(pf12833pf10114) PDU1AB
08845 -pf01656pf07685 -
08850 -pf03186 -
08855 -pf02570 -
08860 -pf01888 -
08865 -pf01888 -
08870 -pf00590 -
08875 -pf05175 -
08880 -pf00590 -
08885 -pf01890pf11761pf11760 -
08890 -pf00590 -
08895 -pf02571 -
BMC-P
BMC-H
BMC-T
Aldehyde dehydrogenase
PTAC
Signature enzyme
Alcohol dehydrogenase
Regulator

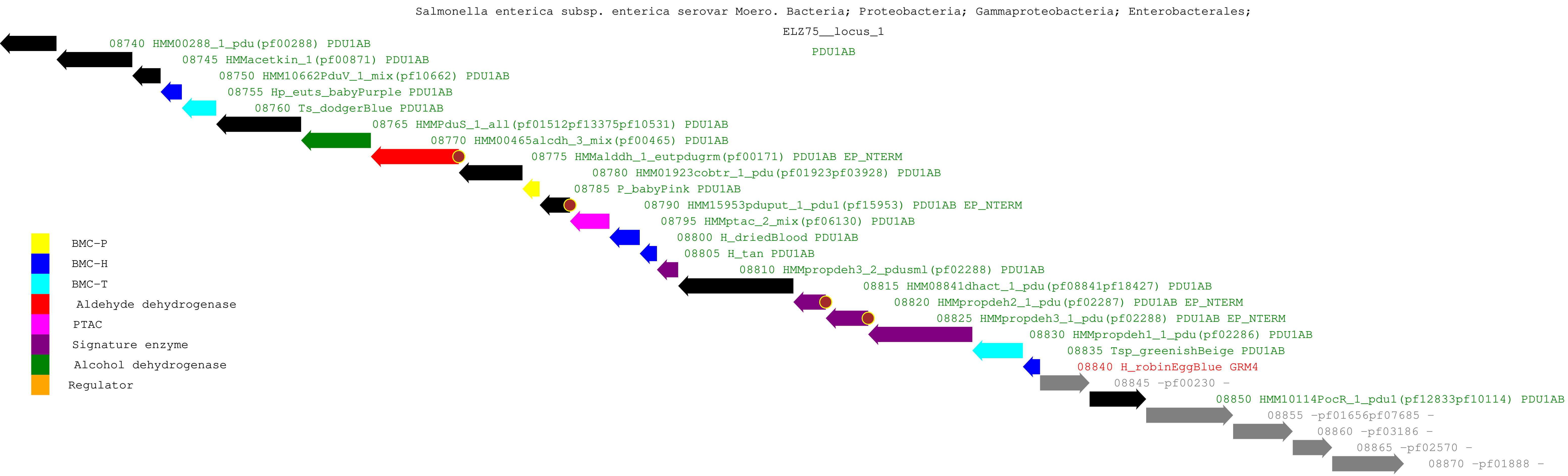
Salmonella enterica subsp. enterica serovar Moero. Bacteria; Proteobacteria; Gammaproteobacteria; Enterobacterales;
ELZ75__locus_1
PDU1AB
08740 HMM00288_1_pdu(pf00288) PDU1AB
08745 HMMacetkin_1(pf00871) PDU1AB
08750 HMM10662PduV_1_mix(pf10662) PDU1AB
08755 Hp_euts_babyPurple PDU1AB
08760 Ts_dodgerBlue PDU1AB
08765 HMMPduS_1_all(pf01512pf13375pf10531) PDU1AB
08770 HMM00465alcdh_3_mix(pf00465) PDU1AB
08775 HMMalddh_1_eutpdugrm(pf00171) PDU1AB EP_NTERM
08780 HMM01923cobtr_1_pdu(pf01923pf03928) PDU1AB
08785 P_babyPink PDU1AB
08790 HMM15953pduput_1_pdu1(pf15953) PDU1AB EP_NTERM
08795 HMMptac_2_mix(pf06130) PDU1AB
08800 H_driedBlood PDU1AB
08805 H_tan PDU1AB
08810 HMMpropdeh3_2_pdusml(pf02288) PDU1AB
08815 HMM08841dhact_1_pdu(pf08841pf18427) PDU1AB
08820 HMMpropdeh2_1_pdu(pf02287) PDU1AB EP_NTERM
08825 HMMpropdeh3_1_pdu(pf02288) PDU1AB EP_NTERM
08830 HMMpropdeh1_1_pdu(pf02286) PDU1AB
08835 Tsp_greenishBeige PDU1AB
08840 H_robinEggBlue GRM4
08845 -pf00230 -
08850 HMM10114PocR_1_pdu1(pf12833pf10114) PDU1AB
08855 -pf01656pf07685 -
08860 -pf03186 -
08865 -pf02570 -
08870 -pf01888 -
BMC-P
BMC-H
BMC-T
Aldehyde dehydrogenase
PTAC
Signature enzyme
Alcohol dehydrogenase
Regulator

Salmonella enterica subsp. enterica serovar 43:a:1,7. Bacteria; Proteobacteria; Gammaproteobacteria; Enterobacterales;

ELZ85__locus_2

PDU1AB

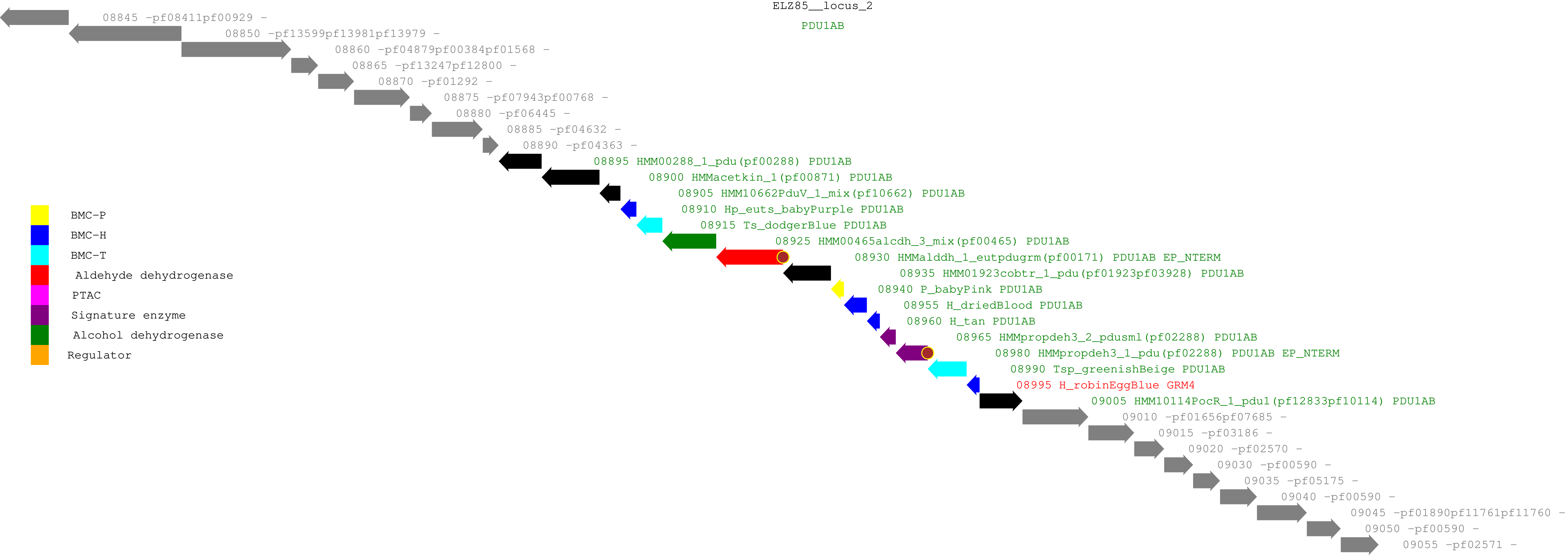

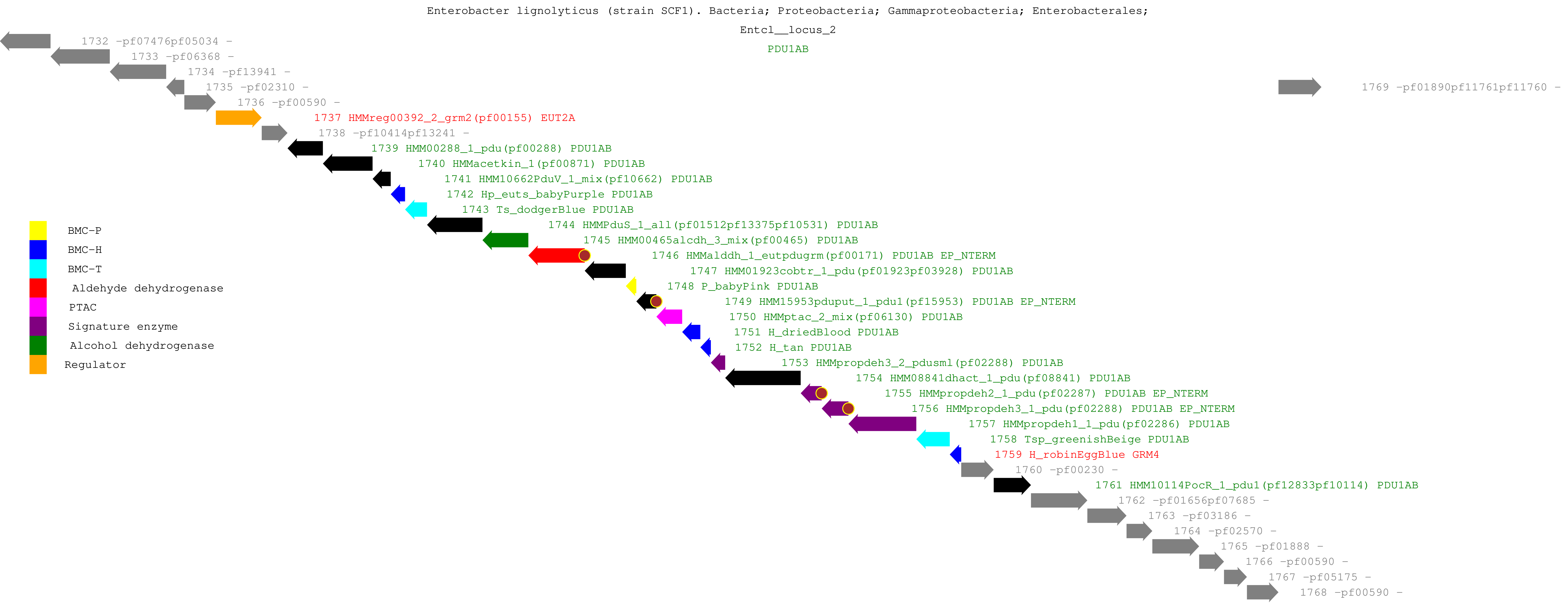
Enterobacter lignolyticus (strain SCF1). Bacteria; Proteobacteria; Gammaproteobacteria; Enterobacterales;
Entcl__locus_2
PDU1AB
1732 -pf07476pf05034 -
1733 -pf06368 -
1734 -pf13941 -
1735 -pf02310 -
1736 -pf00590 -
1737 HMMreg00392_2_grm2(pf00155) EUT2A
1738 -pf10414pf13241 -
1739 HMM00288_1_pdu(pf00288) PDU1AB
1740 HMMacetkin_1(pf00871) PDU1AB
1741 HMM10662PduV_1_mix(pf10662) PDU1AB
1742 Hp_euts_babyPurple PDU1AB
1743 Ts_dodgerBlue PDU1AB
1744 HMMPduS_1_all(pf01512pf13375pf10531) PDU1AB
1745 HMM00465alcdh_3_mix(pf00465) PDU1AB
1746 HMMalddh_1_eutpdugrm(pf00171) PDU1AB EP_NTERM
1747 HMM01923cobtr_1_pdu(pf01923pf03928) PDU1AB
1748 P_babyPink PDU1AB
1749 HMM15953pduput_1_pdu1(pf15953) PDU1AB EP_NTERM
1750 HMMptac_2_mix(pf06130) PDU1AB
1751 H_driedBlood PDU1AB
1752 H_tan PDU1AB
1753 HMMpropdeh3_2_pdusml(pf02288) PDU1AB
1754 HMM08841dhact_1_pdu(pf08841) PDU1AB
1755 HMMpropdeh2_1_pdu(pf02287) PDU1AB EP_NTERM
1756 HMMpropdeh3_1_pdu(pf02288) PDU1AB EP_NTERM
1757 HMMpropdeh1_1_pdu(pf02286) PDU1AB
1758 Tsp_greenishBeige PDU1AB
1759 H_robinEggBlue GRM4
1760 -pf00230 -
1761 HMM10114PocR_1_pdu1(pf12833pf10114) PDU1AB
1762 -pf01656pf07685 -
1763 -pf03186 -
1764 -pf02570 -
1765 -pf01888 -
1766 -pf00590 -
1767 -pf05175 -
1768 -pf00590 -
1769 -pf01890pf11761pf11760 -
BMC-P
BMC-H
BMC-T
Aldehyde dehydrogenase
PTAC
Signature enzyme
Alcohol dehydrogenase
Regulator

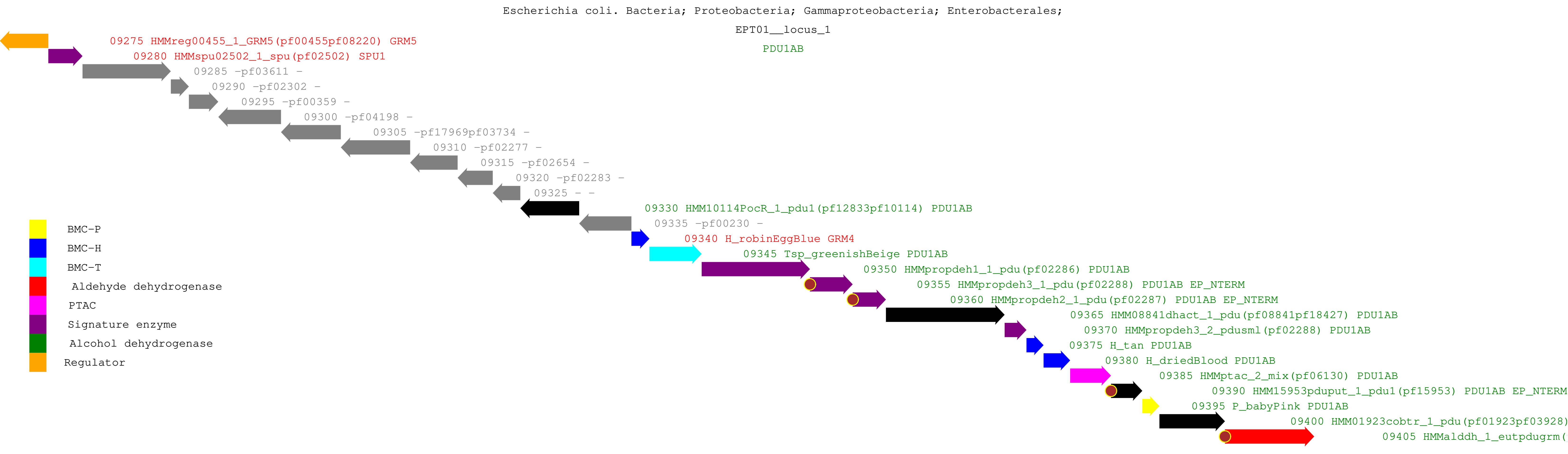
Escherichia coli. Bacteria; Proteobacteria; Gammaproteobacteria; Enterobacterales;
EPT01__locus_1
PDU1AB
09275 HMMreg00455_1_GRM5(pf00455pf08220) GRM5
09280 HMMspu02502_1_spu(pf02502) SPU1
09285 -pf03611 -
09290 -pf02302 -
09295 -pf00359 -
09300 -pf04198 -
09305 -pf17969pf03734 -
09310 -pf02277 -
09315 -pf02654 -
09320 -pf02283 -
09325 - -
09330 HMM10114PocR_1_pdu1(pf12833pf10114) PDU1AB
09335 -pf00230 -
09340 H_robinEggBlue GRM4
09345 Tsp_greenishBeige PDU1AB
09350 HMMpropdeh1_1_pdu(pf02286) PDU1AB
09355 HMMpropdeh3_1_pdu(pf02288) PDU1AB EP_NTERM
09360 HMMpropdeh2_1_pdu(pf02287) PDU1AB EP_NTERM
09365 HMM08841dhact_1_pdu(pf08841pf18427) PDU1AB
09370 HMMpropdeh3_2_pdusml(pf02288) PDU1AB
09375 H_tan PDU1AB
09380 H_driedBlood PDU1AB
09385 HMMptac_2_mix(pf06130) PDU1AB
09390 HMM15953pduput_1_pdu1(pf15953) PDU1AB EP_NTERM
09395 P_babyPink PDU1AB
09400 HMM01923cobtr_1_pdu(pf01923pf03928)
09405 HMMalddh_1_eutpdugrm(
BMC-P
BMC-H
BMC-T
Aldehyde dehydrogenase
PTAC
Signature enzyme
Alcohol dehydrogenase
Regulator

Klebsiella pneumoniae. Bacteria; Proteobacteria; Gammaproteobacteria; Enterobacterales;

EQH49__locus_1

PDU1AB

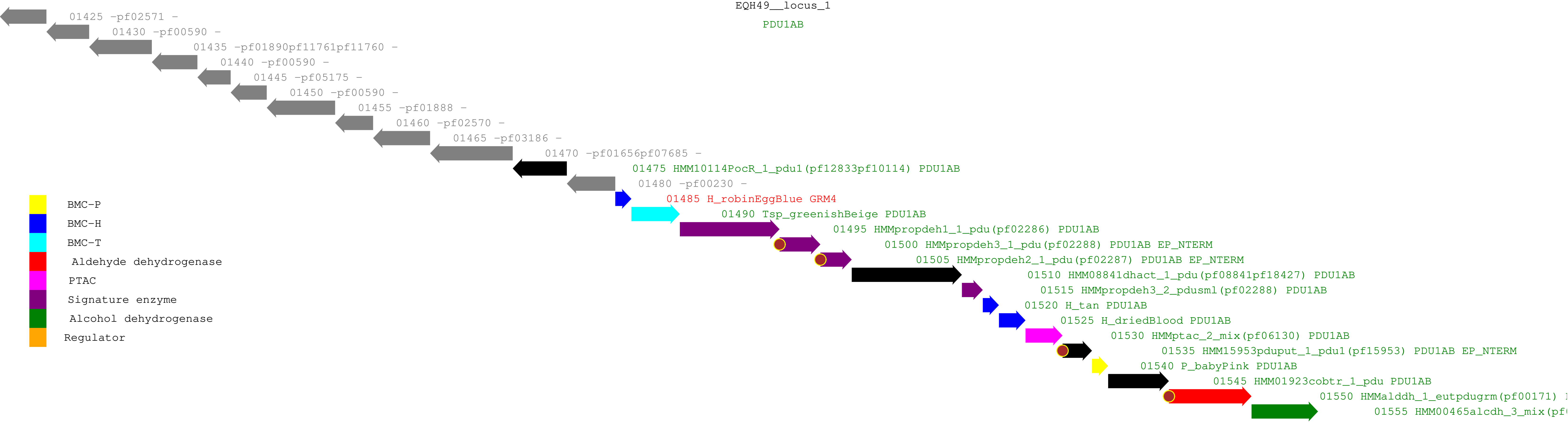

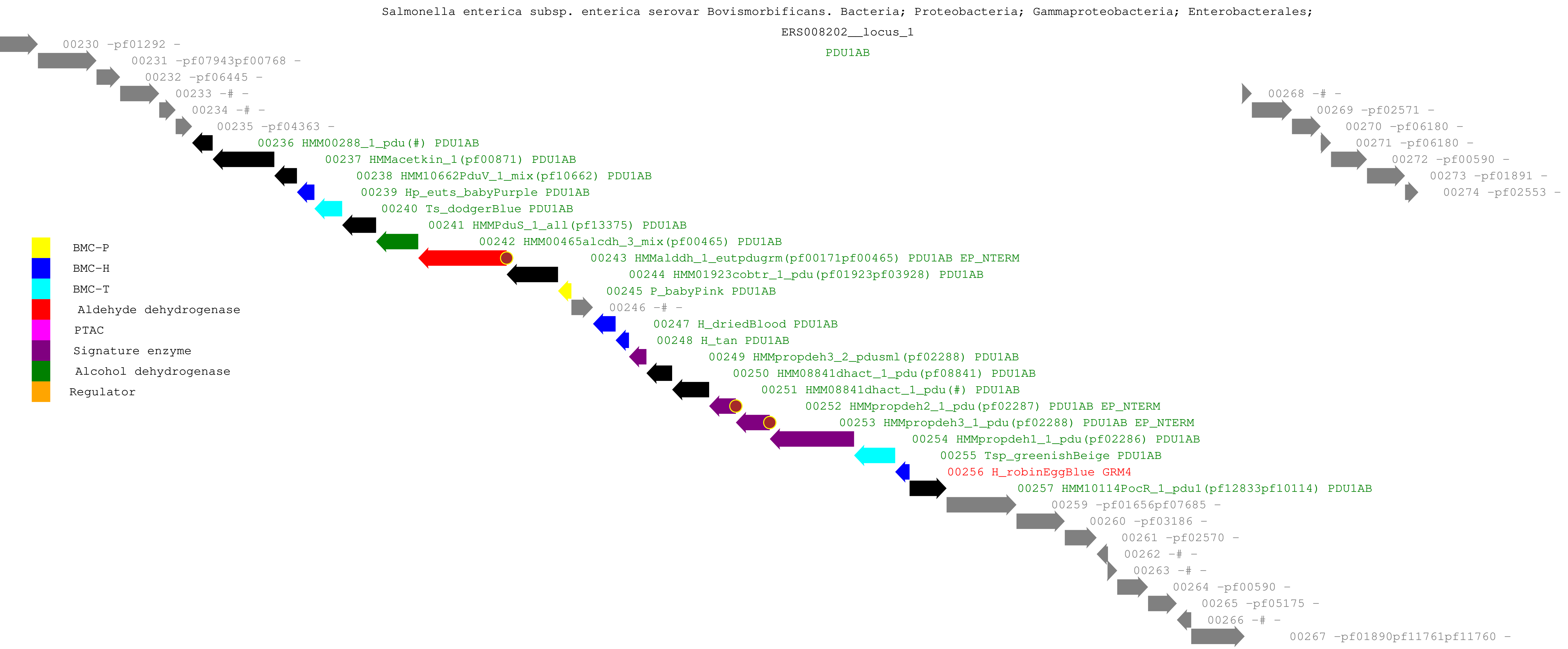
Salmonella enterica subsp. enterica serovar Bovismorbificans. Bacteria; Proteobacteria; Gammaproteobacteria; Enterobacterales;
ERS008202__locus_1
PDU1AB
00230 -pf01292 -
00231 -pf07943pf00768 -
00232 -pf06445 -
00233 -# -
00234 -# -
00235 -pf04363 -
00236 HMM00288_1_pdu(#) PDU1AB
00237 HMMacetkin_1(pf00871) PDU1AB
00238 HMM10662PduV_1_mix(pf10662) PDU1AB
00239 Hp_euts_babyPurple PDU1AB
00240 Ts_dodgerBlue PDU1AB
00241 HMMPduS_1_all(pf13375) PDU1AB
00242 HMM00465alcdh_3_mix(pf00465) PDU1AB
00243 HMMalddh_1_eutpdugrm(pf00171pf00465) PDU1AB EP_NTERM
00244 HMM01923cobtr_1_pdu(pf01923pf03928) PDU1AB
00245 P_babyPink PDU1AB
00246 -# -
00247 H_driedBlood PDU1AB
00248 H_tan PDU1AB
00249 HMMpropdeh3_2_pdusml(pf02288) PDU1AB
00250 HMM08841dhact_1_pdu(pf08841) PDU1AB
00251 HMM08841dhact_1_pdu(#) PDU1AB
00252 HMMpropdeh2_1_pdu(pf02287) PDU1AB EP_NTERM
00253 HMMpropdeh3_1_pdu(pf02288) PDU1AB EP_NTERM
00254 HMMpropdeh1_1_pdu(pf02286) PDU1AB
00255 Tsp_greenishBeige PDU1AB
00256 H_robinEggBlue GRM4
00257 HMM10114PocR_1_pdu1(pf12833pf10114) PDU1AB
00259 -pf01656pf07685 -
00260 -pf03186 -
00261 -pf02570 -
00262 -# -
00263 -# -
00264 -pf00590 -
00265 -pf05175 -
00266 -# -
00267 -pf01890pf11761pf11760 -
00268 -# -
00269 -pf02571 -
00270 -pf06180 -
00271 -pf06180 -
00272 -pf00590 -
00273 -pf01891 -
00274 -pf02553 -
BMC-P
BMC-H
BMC-T
Aldehyde dehydrogenase
PTAC
Signature enzyme
Alcohol dehydrogenase
Regulator

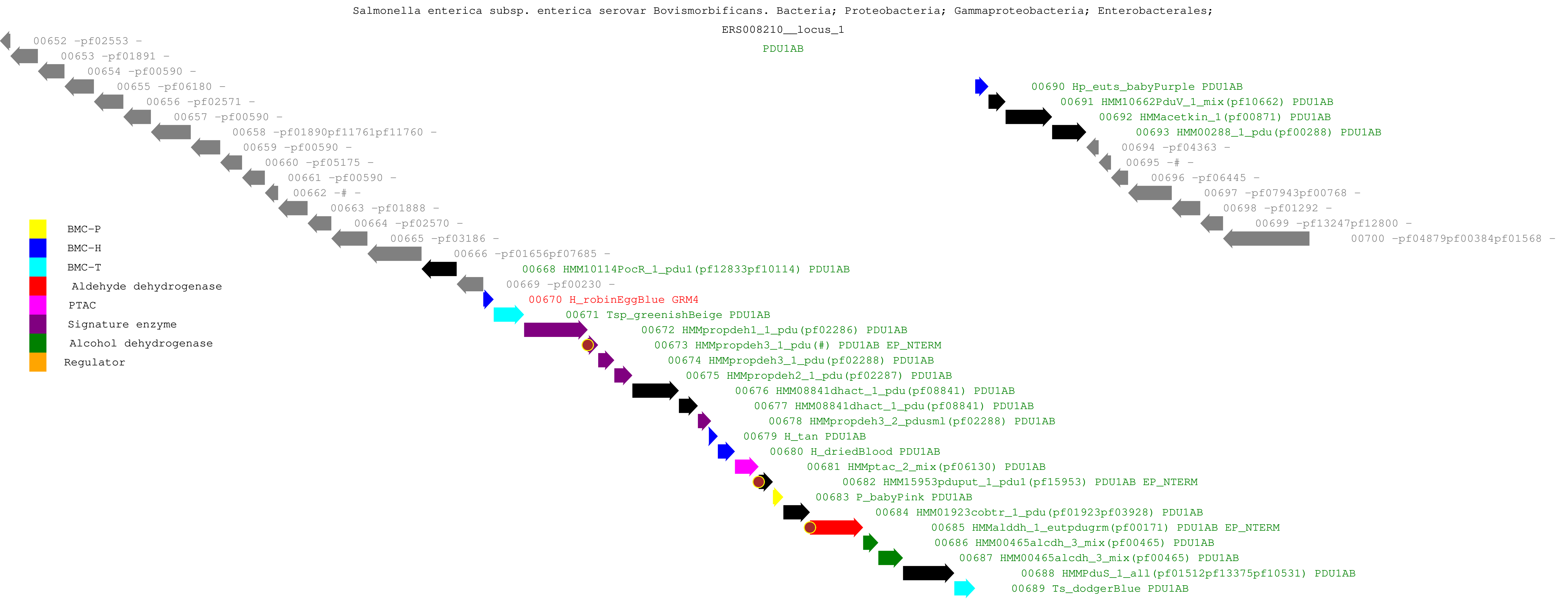
Salmonella enterica subsp. enterica serovar Bovismorbificans. Bacteria; Proteobacteria; Gammaproteobacteria; Enterobacterales;
ERS008210__locus_1
PDU1AB
00652 -pf02553 -
00653 -pf01891 -
00654 -pf00590 -
00655 -pf06180 -
00656 -pf02571 -
00657 -pf00590 -
00658 -pf01890pf11761pf11760 -
00659 -pf00590 -
00660 -pf05175 -
00661 -pf00590 -
00662 -# -
00663 -pf01888 -
00664 -pf02570 -
00665 -pf03186 -
00666 -pf01656pf07685 -
00668 HMM10114PocR_1_pdu1(pf12833pf10114) PDU1AB
00669 -pf00230 -
00670 H_robinEggBlue GRM4
00671 Tsp_greenishBeige PDU1AB
00672 HMMpropdeh1_1_pdu(pf02286) PDU1AB
00673 HMMpropdeh3_1_pdu(#) PDU1AB EP_NTERM
00674 HMMpropdeh3_1_pdu(pf02288) PDU1AB
00675 HMMpropdeh2_1_pdu(pf02287) PDU1AB
00676 HMM08841dhact_1_pdu(pf08841) PDU1AB
00677 HMM08841dhact_1_pdu(pf08841) PDU1AB
00678 HMMpropdeh3_2_pdusml(pf02288) PDU1AB
00679 H_tan PDU1AB
00680 H_driedBlood PDU1AB
00681 HMMptac_2_mix(pf06130) PDU1AB
00682 HMM15953pduput_1_pdu1(pf15953) PDU1AB EP_NTERM
00683 P_babyPink PDU1AB
00684 HMM01923cobtr_1_pdu(pf01923pf03928) PDU1AB
00685 HMMaldh_1_eutpdugrm(pf00171) PDU1AB EP_NTERM
00686 HMM00465alcdh_3_mix(pf00465) PDU1AB
00687 HMM00465alcdh_3_mix(pf00465) PDU1AB
00688 HMMPduS_1_all(pf01512pf13375pf10531) PDU1AB
00689 Ts_dodgerBlue PDU1AB
00690 Hp_euts_babyPurple PDU1AB
00691 HMM10662PduV_1_mix(pf10662) PDU1AB
00692 HMMacetkin_1(pf00871) PDU1AB
00693 HMM00288_1_pdu(pf00288) PDU1AB
00694 -pf04363 -
00695 -# -
00696 -pf06445 -
00697 -pf07943pf00768 -
00698 -pf01292 -
00699 -pf13247pf12800 -
00700 -pf04879pf00384pf01568 -
BMC-P
BMC-H
BMC-T
Aldehyde dehydrogenase
PTAC
Signature enzyme
Alcohol dehydrogenase
Regulator

Yersinia mollaretii. Bacteria; Proteobacteria; Gammaproteobacteria; Enterobacterales;

ERS008437__locus_1

PDU1AB

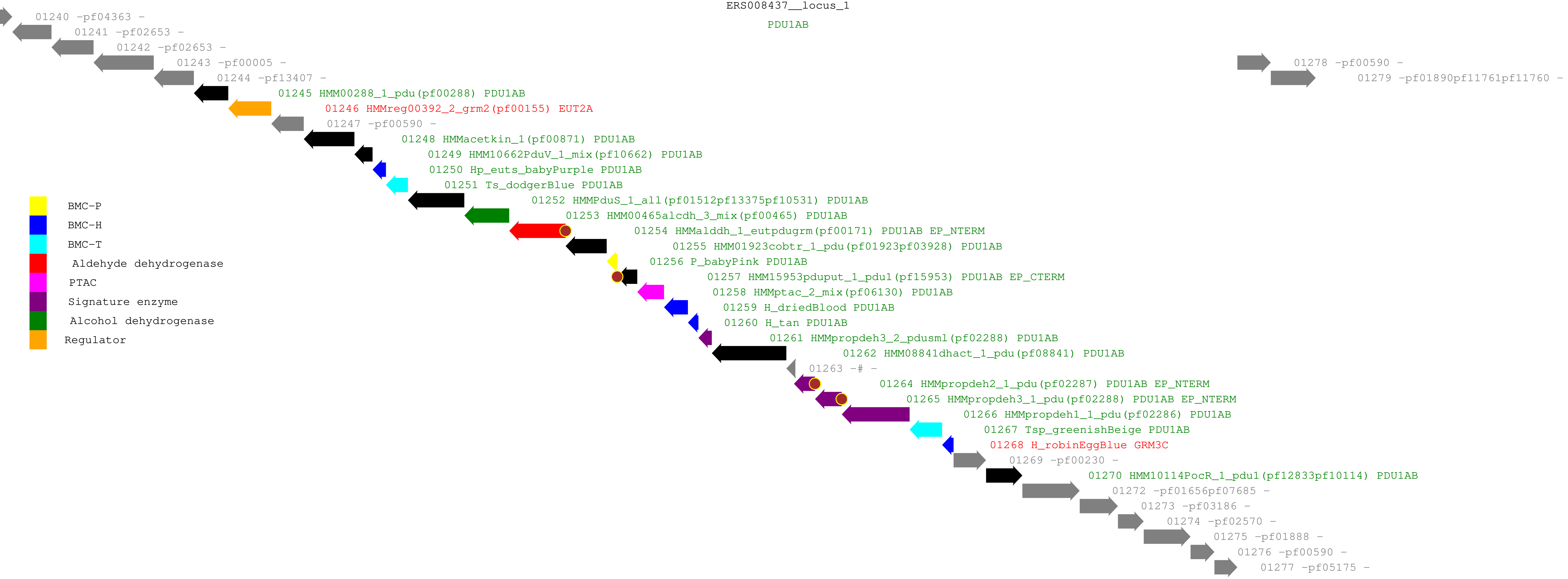

Yersinia kristensenii. Bacteria; Proteobacteria; Gammaproteobacteria; Enterobacterales;

ERS008460__locus_1

PDU1AB

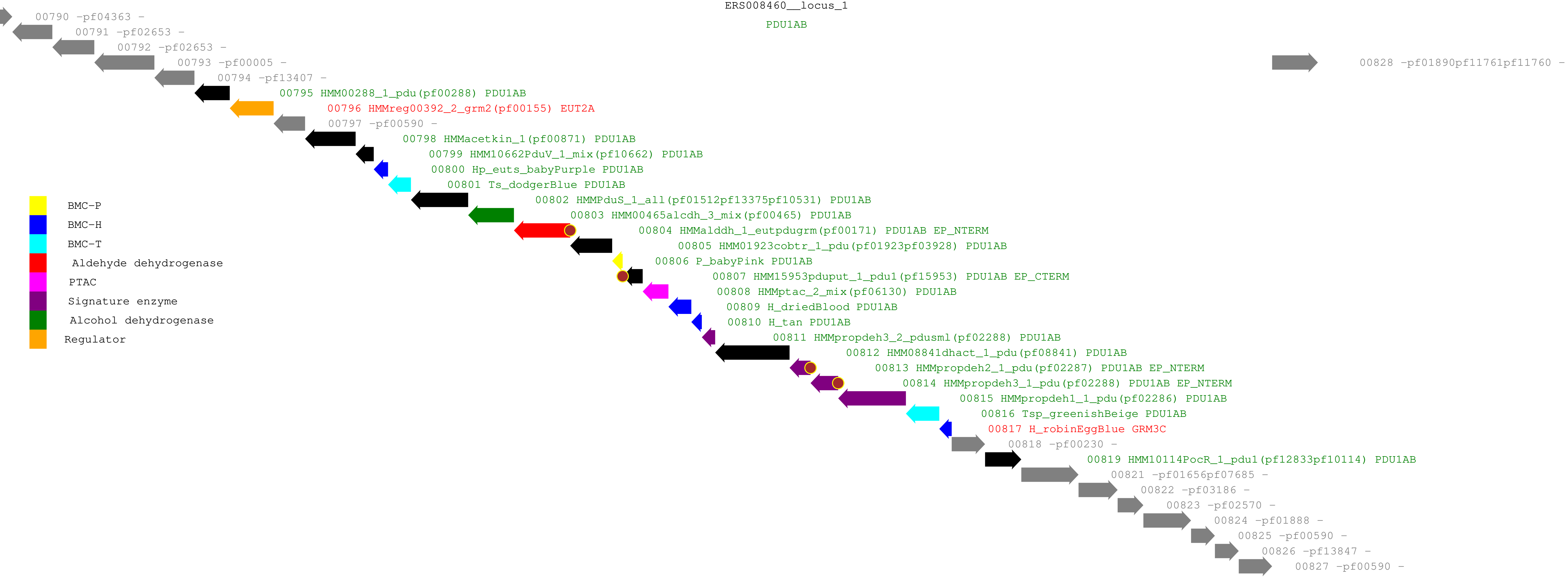

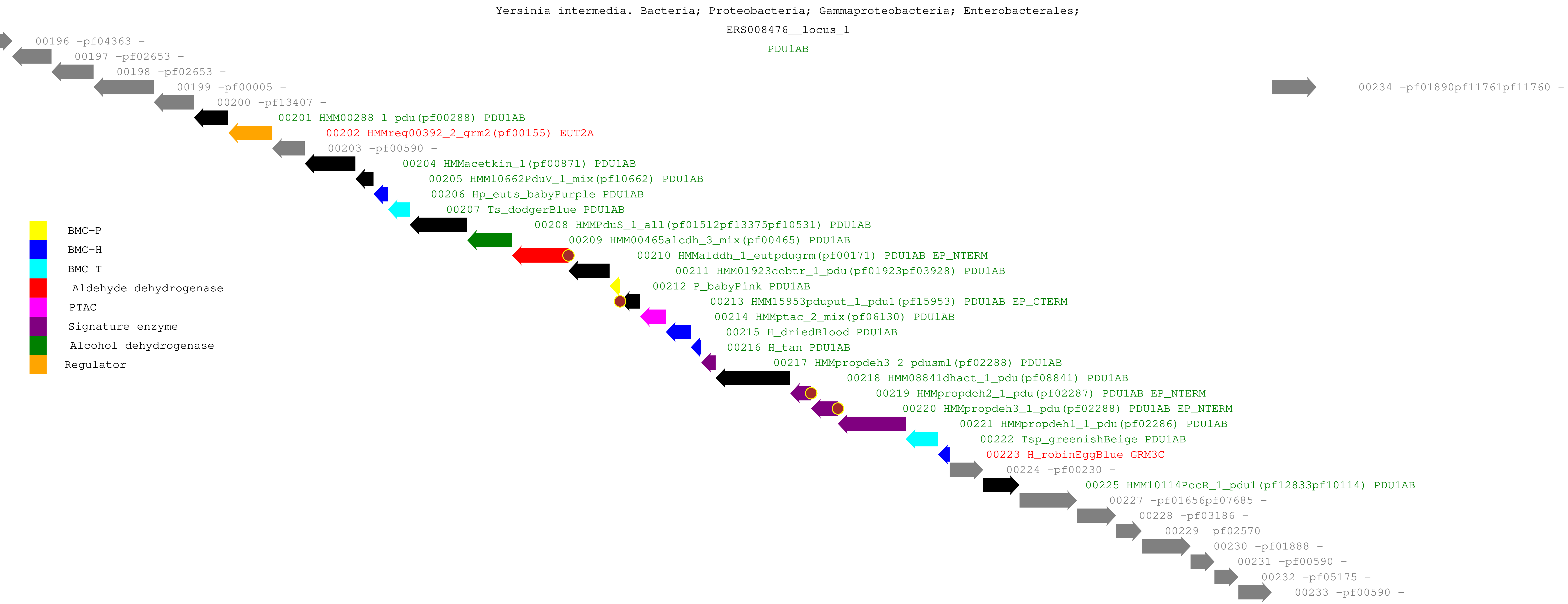
Yersinia intermedia. Bacteria; Proteobacteria; Gammaproteobacteria; Enterobacterales;
ERS008476__locus_1
PDU1AB
00196 -pf04363 -
00197 -pf02653 -
00198 -pf02653 -
00199 -pf00005 -
00200 -pf13407 -
00201 HMM00288_1_pdu(pf00288) PDU1AB
00202 HMMreg00392_2_grm2(pf00155) EUT2A
00203 -pf00590 -
00204 HMMacetkin_1(pf00871) PDU1AB
00205 HMM10662PduV_1_mix(pf10662) PDU1AB
00206 Hp_euts_babyPurple PDU1AB
00207 Ts_dodgerBlue PDU1AB
00208 HMMPduS_1_all(pf01512pf13375pf10531) PDU1AB
00209 HMM00465alcdh_3_mix(pf00465) PDU1AB
00210 HMMalddh_1_eutpdugrm(pf00171) PDU1AB EP_NTERM
00211 HMM01923cobtr_1_pdu(pf01923pf03928) PDU1AB
00212 P_babyPink PDU1AB
00213 HMM15953pduput_1_pdu1(pf15953) PDU1AB EP_CTERM
00214 HMMptac_2_mix(pf06130) PDU1AB
00215 H_driedBlood PDU1AB
00216 H_tan PDU1AB
00217 HMMpropdeh3_2_pdusml(pf02288) PDU1AB
00218 HMM08841dhact_1_pdu(pf08841) PDU1AB
00219 HMMpropdeh2_1_pdu(pf02287) PDU1AB EP_NTERM
00220 HMMpropdeh3_1_pdu(pf02288) PDU1AB EP_NTERM
00221 HMMpropdeh1_1_pdu(pf02286) PDU1AB
00222 Tsp_greenishBeige PDU1AB
00223 H_robinEggBlue GRM3C
00224 -pf00230 -
00225 HMM10114PocR_1_pdu1(pf12833pf10114) PDU1AB
00227 -pf01656pf07685 -
00228 -pf03186 -
00229 -pf02570 -
00230 -pf01888 -
00231 -pf00590 -
00232 -pf05175 -
00233 -pf00590 -
00234 -pf01890pf11761pf11760 -
BMC-P
BMC-H
BMC-T
Aldehyde dehydrogenase
PTAC
Signature enzyme
Alcohol dehydrogenase
Regulator

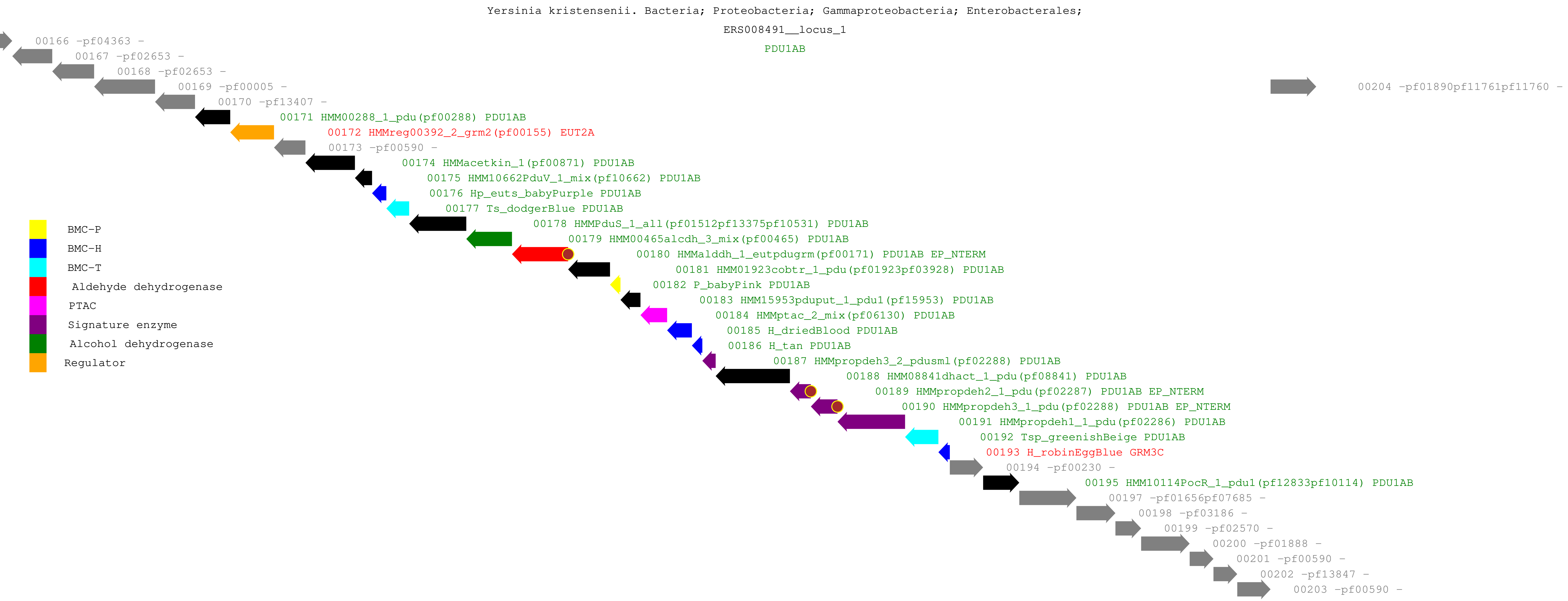

Yersinia kristensenii. Bacteria; Proteobacteria; Gammaproteobacteria; Enterobacterales;
ERS008491__locus_1
PDU1AB
00166 -pf04363 -
00167 -pf02653 -
00168 -pf02653 -
00169 -pf00005 -
00204 -pf01890pf11761pf11760 -
00170 -pf13407 -
00171 HMM00288_1_pdu(pf00288) PDU1AB
00172 HMMreg00392_2_grm2(pf00155) EUT2A
00173 -pf00590 -
00174 HMMacetkin_1(pf00871) PDU1AB
00175 HMM10662PduV_1_mix(pf10662) PDU1AB
00176 Hp_euts_babyPurple PDU1AB
00177 Ts_dodgerBlue PDU1AB
00178 HMMPduS_1_all(pf01512pf13375pf10531) PDU1AB
00179 HMM00465alcdh_3_mix(pf00465) PDU1AB
00180 HMMalddh_1_eutpdugrm(pf00171) PDU1AB EP_NTERM
00181 HMM01923cobtr_1_pdu(pf01923pf03928) PDU1AB
00182 P_babyPink PDU1AB
00183 HMM15953pduput_1_pdu1(pf15953) PDU1AB
00184 HMMptac_2_mix(pf06130) PDU1AB
00185 H_driedBlood PDU1AB
00186 H_tan PDU1AB
00187 HMMpropdeh3_2_pdusml(pf02288) PDU1AB
00188 HMM08841dhact_1_pdu(pf08841) PDU1AB
00189 HMMpropdeh2_1_pdu(pf02287) PDU1AB EP_NTERM
00190 HMMpropdeh3_1_pdu(pf02288) PDU1AB EP_NTERM
00191 HMMpropdeh1_1_pdu(pf02286) PDU1AB
00192 Tsp_greenishBeige PDU1AB
00193 H_robinEggBlue GRM3C
00194 -pf00230 -
00195 HMM10114PocR_1_pdu1(pf12833pf10114) PDU1AB
00197 -pf01656pf07685 -
00198 -pf03186 -
00199 -pf02570 -
00200 -pf01888 -
00201 -pf00590 -
00202 -pf13847 -
00203 -pf00590 -
BMC-P
BMC-H
BMC-T
Aldehyde dehydrogenase
PTAC
Signature enzyme
Alcohol dehydrogenase
Regulator

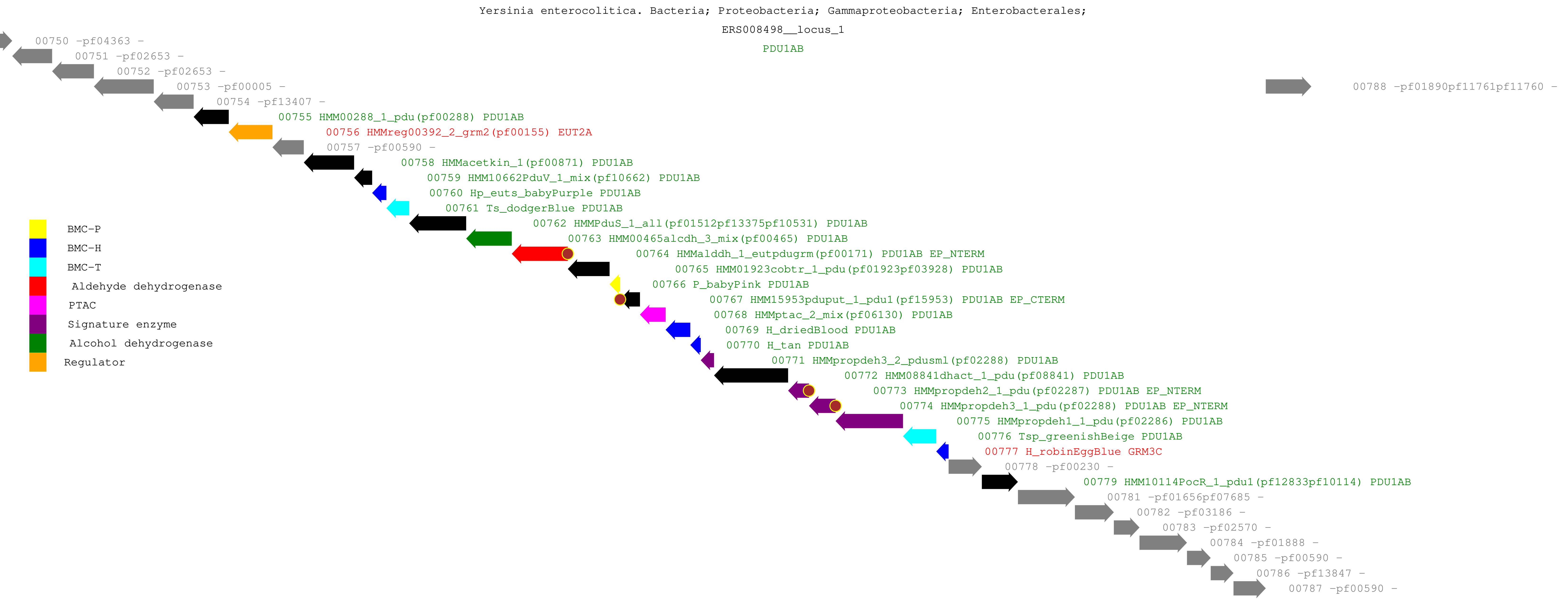
Yersinia enterocolitica. Bacteria; Proteobacteria; Gammaproteobacteria; Enterobacterales;
ERS008498__locus_1
PDU1AB
00750 -pf04363 -
00751 -pf02653 -
00752 -pf02653 -
00753 -pf00005 -
00754 -pf13407 -
00755 HMM00288_1_pdu(pf00288) PDU1AB
00756 HMMreg00392_2_grm2(pf00155) EUT2A
00757 -pf00590 -
00758 HMMacetkin_1(pf00871) PDU1AB
00759 HMM10662PduV_1_mix(pf10662) PDU1AB
00760 Hp_euts_babyPurple PDU1AB
00761 Ts_dodgerBlue PDU1AB
00762 HMMPduS_1_all(pf01512pf13375pf10531) PDU1AB
00763 HMM00465alcdh_3_mix(pf00465) PDU1AB
00764 HMMalddh_1_eutpdugrm(pf00171) PDU1AB EP_NTERM
00765 HMM01923cobtr_1_pdu(pf01923pf03928) PDU1AB
00766 P_babyPink PDU1AB
00767 HMM15953pduput_1_pdu1(pf15953) PDU1AB EP_CTERM
00768 HMMptac_2_mix(pf06130) PDU1AB
00769 H_driedBlood PDU1AB
00770 H_tan PDU1AB
00771 HMMpropdeh3_2_pdusml(pf02288) PDU1AB
00772 HMM08841dhact_1_pdu(pf08841) PDU1AB
00773 HMMpropdeh2_1_pdu(pf02287) PDU1AB EP_NTERM
00774 HMMpropdeh3_1_pdu(pf02288) PDU1AB EP_NTERM
00775 HMMpropdeh1_1_pdu(pf02286) PDU1AB
00776 Tsp_greenishBeige PDU1AB
00777 H_robinEggBlue GRM3C
00778 -pf00230 -
00779 HMM10114PocR_1_pdu1(pf12833pf10114) PDU1AB
00781 -pf01656pf07685 -
00782 -pf03186 -
00783 -pf02570 -
00784 -pf01888 -
00785 -pf00590 -
00786 -pf13847 -
00787 -pf00590 -
00788 -pf01890pf11761pf11760 -
BMC-P
BMC-H
BMC-T
Aldehyde dehydrogenase
PTAC
Signature enzyme
Alcohol dehydrogenase
Regulator

Yersinia kristensenii. Bacteria; Proteobacteria; Gammaproteobacteria; Enterobacterales;

ERS008500__locus_1

PDU1AB

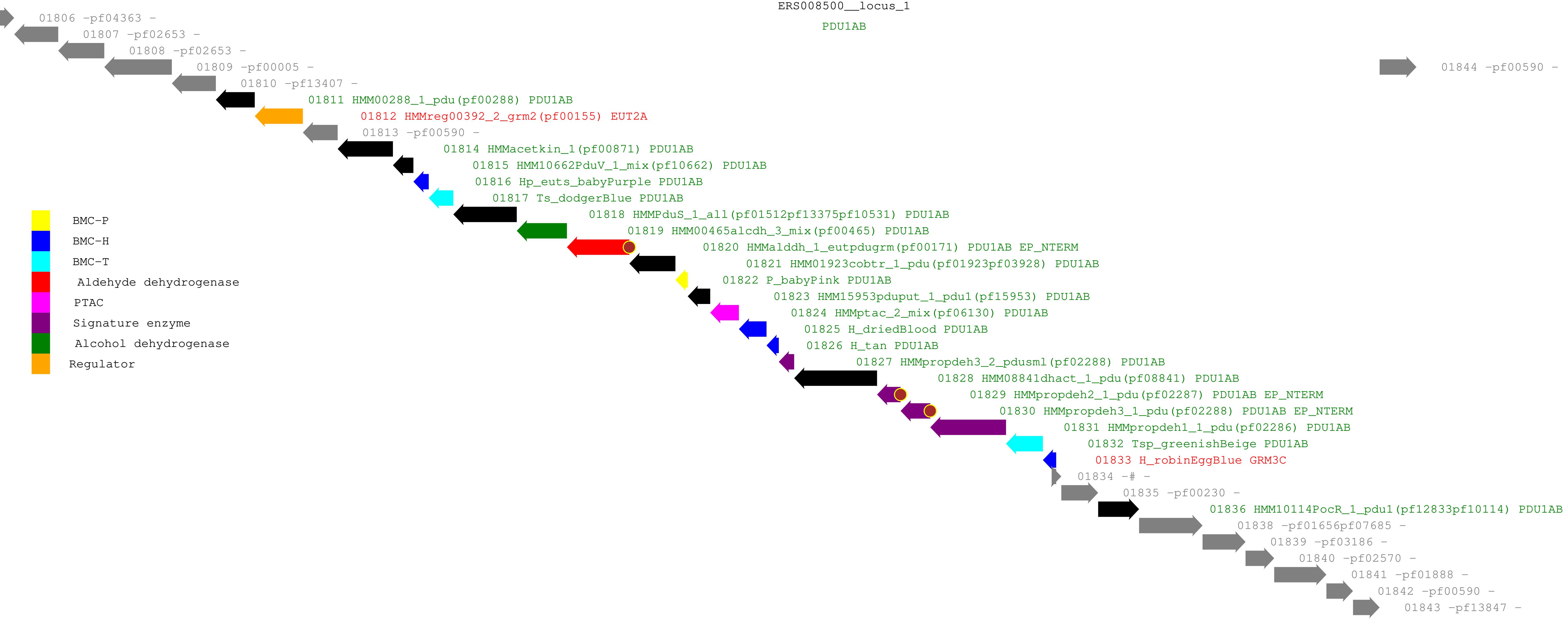

Yersinia frederiksenii. Bacteria; Proteobacteria; Gammaproteobacteria; Enterobacterales;

ERS008501__locus_1

PDU1AB

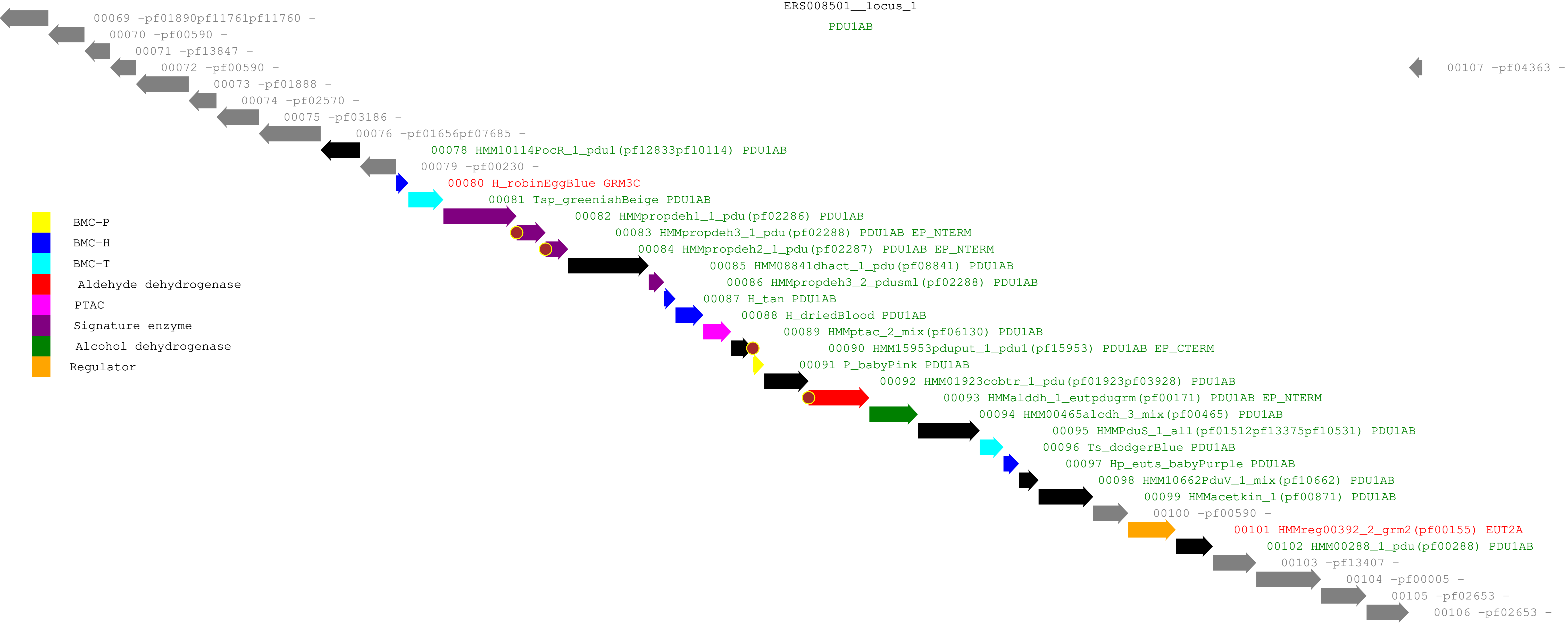

Yersinia bercovieri. Bacteria; Proteobacteria; Gammaproteobacteria; Enterobacterales;

ERS008506__locus_1

PDU1AB

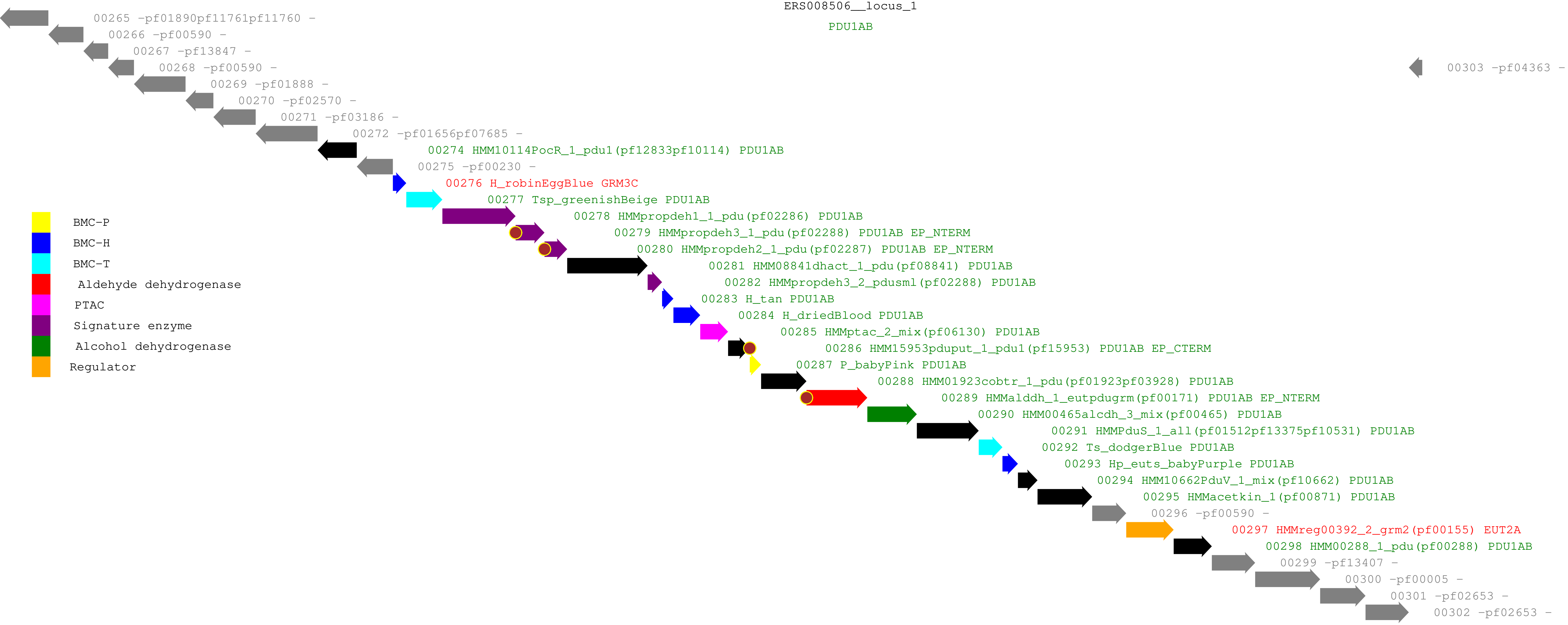

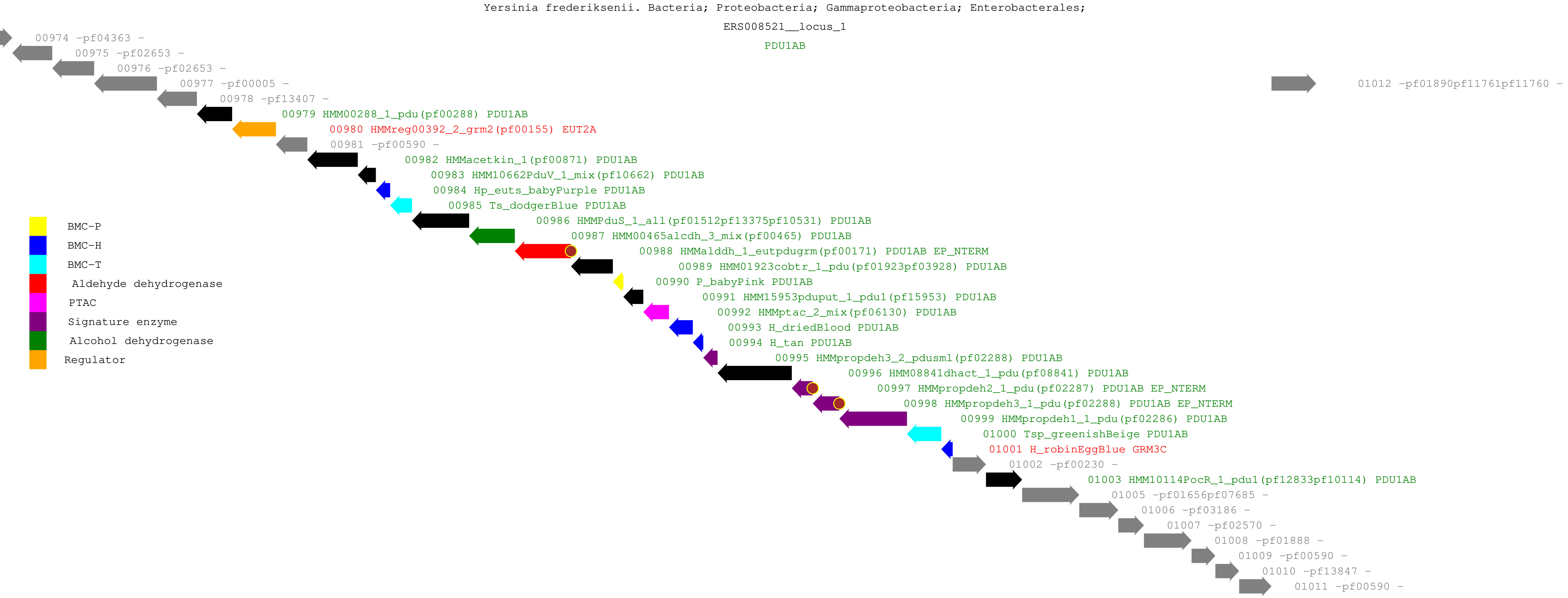
Yersinia frederiksenii. Bacteria; Proteobacteria; Gammaproteobacteria; Enterobacterales;
ERS008521__locus_1
PDU1AB
00974 -pf04363 -
00975 -pf02653 -
00976 -pf02653 -
00977 -pf00005 -
01012 -pf01890pf11761pf11760 -
00978 -pf13407 -
00979 HMM00288_1_pdu(pf00288) PDU1AB
00980 HMMreg00392_2_grm2(pf00155) EUT2A
00981 -pf00590 -
00982 HMMacetkin_1(pf00871) PDU1AB
00983 HMM10662PduV_1_mix(pf10662) PDU1AB
00984 Hp_euts_babyPurple PDU1AB
00985 Ts_dodgerBlue PDU1AB
00986 HMMPduS_1_all(pf01512pf13375pf10531) PDU1AB
00987 HMM00465alcdh_3_mix(pf00465) PDU1AB
00988 HMMaldh_1_eutpdugrm(pf00171) PDU1AB EP_NTERM
00989 HMM01923cobtr_1_pdu(pf01923pf03928) PDU1AB
00990 P_babyPink PDU1AB
00991 HMM15953pduput_1_pdu1(pf15953) PDU1AB
00992 HMMptac_2_mix(pf06130) PDU1AB
00993 H_driedBlood PDU1AB
00994 H_tan PDU1AB
00995 HMMpropdeh3_2_pdusml(pf02288) PDU1AB
00996 HMM08841dhact_1_pdu(pf08841) PDU1AB
00997 HMMpropdeh2_1_pdu(pf02287) PDU1AB EP_NTERM
00998 HMMpropdeh3_1_pdu(pf02288) PDU1AB EP_NTERM
00999 HMMpropdeh1_1_pdu(pf02286) PDU1AB
01000 Tsp_greenishBeige PDU1AB
01001 H_robinEggBlue GRM3C
01002 -pf00230 -
01003 HMM10114PocR_1_pdu1(pf12833pf10114) PDU1AB
01005 -pf01656pf07685 -
01006 -pf03186 -
01007 -pf02570 -
01008 -pf01888 -
01009 -pf00590 -
01010 -pf13847 -
01011 -pf00590 -
BMC-P
BMC-H
BMC-T
Aldehyde dehydrogenase
PTAC
Signature enzyme
Alcohol dehydrogenase
Regulator

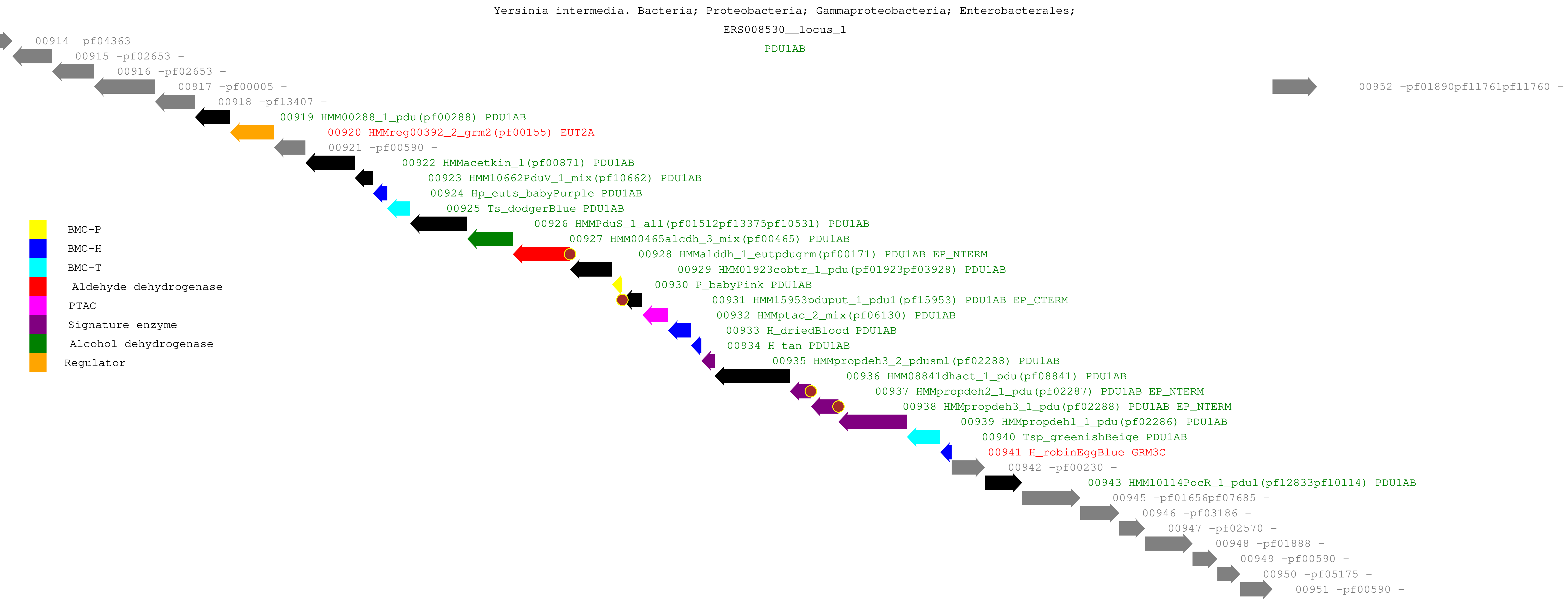
Yersinia intermedia. Bacteria; Proteobacteria; Gammaproteobacteria; Enterobacterales;
ERS008530__locus_1
PDU1AB
00914 -pf04363 -
00915 -pf02653 -
00916 -pf02653 -
00917 -pf00005 -
00918 -pf13407 -
00919 HMM00288_1_pdu(pf00288) PDU1AB
00920 HMMreg00392_2_grm2(pf00155) EUT2A
00921 -pf00590 -
00922 HMMacetkin_1(pf00871) PDU1AB
00923 HMM10662PduV_1_mix(pf10662) PDU1AB
00924 Hp_euts_babyPurple PDU1AB
00925 Ts_dodgerBlue PDU1AB
00926 HMMPduS_1_all(pf01512pf13375pf10531) PDU1AB
00927 HMM00465alcdh_3_mix(pf00465) PDU1AB
00928 HMMaldh_1_eutpdugrm(pf00171) PDU1AB EP_NTERM
00929 HMM01923cobtr_1_pdu(pf01923pf03928) PDU1AB
00930 P_babyPink PDU1AB
00931 HMM15953pduput_1_pdu1(pf15953) PDU1AB EP_CTERM
00932 HMMptac_2_mix(pf06130) PDU1AB
00933 H_driedBlood PDU1AB
00934 H_tan PDU1AB
00935 HMMpropdeh3_2_pdusml(pf02288) PDU1AB
00936 HMM08841dhact_1_pdu(pf08841) PDU1AB
00937 HMMpropdeh2_1_pdu(pf02287) PDU1AB EP_NTERM
00938 HMMpropdeh3_1_pdu(pf02288) PDU1AB EP_NTERM
00939 HMMpropdeh1_1_pdu(pf02286) PDU1AB
00940 Tsp_greenishBeige PDU1AB
00941 H_robinEggBlue GRM3C
00942 -pf00230 -
00943 HMM10114PocR_1_pdu1(pf12833pf10114) PDU1AB
00945 -pf01656pf07685 -
00946 -pf03186 -
00947 -pf02570 -
00948 -pf01888 -
00949 -pf00590 -
00950 -pf05175 -
00951 -pf00590 -
00952 -pf01890pf11761pf11760 -
BMC-P
BMC-H
BMC-T
Aldehyde dehydrogenase
PTAC
Signature enzyme
Alcohol dehydrogenase
Regulator

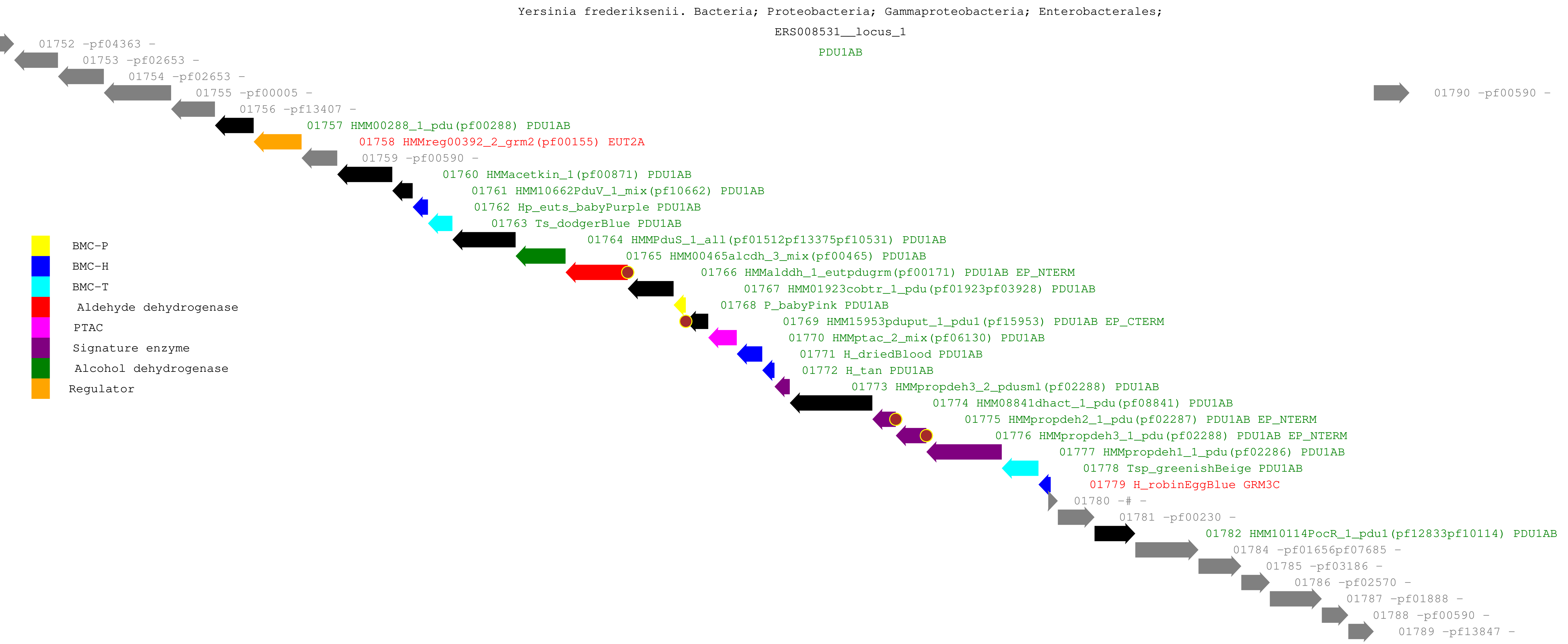
Yersinia frederiksenii. Bacteria; Proteobacteria; Gammaproteobacteria; Enterobacterales;
ERS008531__locus_1
PDU1AB
01752 -pf04363 -
01753 -pf02653 -
01754 -pf02653 -
01755 -pf00005 -
01790 -pf00590 -
01756 -pf13407 -
01757 HMM00288_1_pdu(pf00288) PDU1AB
01758 HMMreg00392_2_grm2(pf00155) EUT2A
01759 -pf00590 -
01760 HMMacetkin_1(pf00871) PDU1AB
01761 HMM10662PduV_1_mix(pf10662) PDU1AB
01762 Hp_euts_babyPurple PDU1AB
01763 Ts_dodgerBlue PDU1AB
01764 HMMPduS_1_all(pf01512pf13375pf10531) PDU1AB
01765 HMM00465alcdh_3_mix(pf00465) PDU1AB
01766 HMMalddh_1_eutpdugrm(pf00171) PDU1AB EP_NTERM
01767 HMM01923cobtr_1_pdu(pf01923pf03928) PDU1AB
01768 P_babyPink PDU1AB
01769 HMM15953pduput_1_pdu1(pf15953) PDU1AB EP_CTERM
01770 HMMptac_2_mix(pf06130) PDU1AB
01771 H_driedBlood PDU1AB
01772 H_tan PDU1AB
01773 HMMpropdeh3_2_pdusml(pf02288) PDU1AB
01774 HMM08841dhact_1_pdu(pf08841) PDU1AB
01775 HMMpropdeh2_1_pdu(pf02287) PDU1AB EP_NTERM
01776 HMMpropdeh3_1_pdu(pf02288) PDU1AB EP_NTERM
01777 HMMpropdeh1_1_pdu(pf02286) PDU1AB
01778 Tsp_greenishBeige PDU1AB
01779 H_robinEggBlue GRM3C
01780 -# -
01781 -pf00230 -
01782 HMM10114PocR_1_pdu1(pf12833pf10114) PDU1AB
01784 -pf01656pf07685 -
01785 -pf03186 -
01786 -pf02570 -
01787 -pf01888 -
01788 -pf00590 -
01789 -pf13847 -
BMC-P
BMC-H
BMC-T
Aldehyde dehydrogenase
PTAC
Signature enzyme
Alcohol dehydrogenase
Regulator

Yersinia frederiksenii. Bacteria; Proteobacteria; Gammaproteobacteria; Enterobacterales;

ERS008545__locus_1

PDU1AB

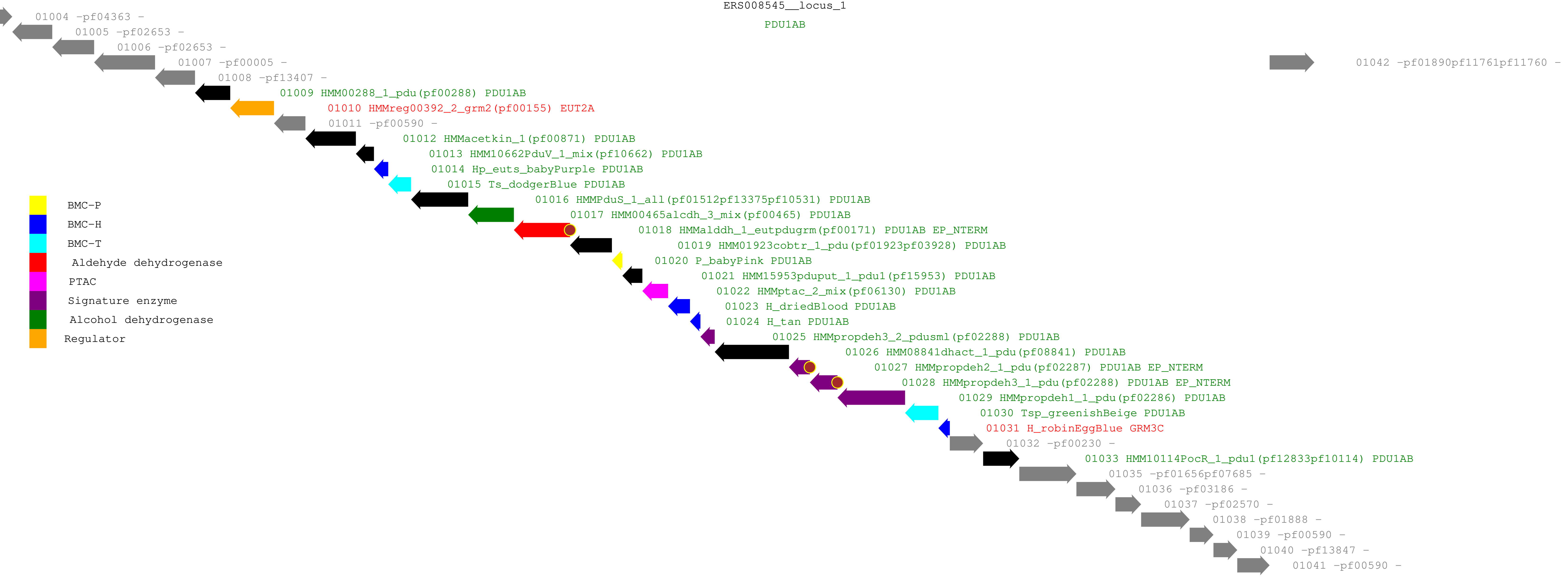

Yersinia intermedia. Bacteria; Proteobacteria; Gammaproteobacteria; Enterobacterales;

ERS008547__locus_1

PDU1AB

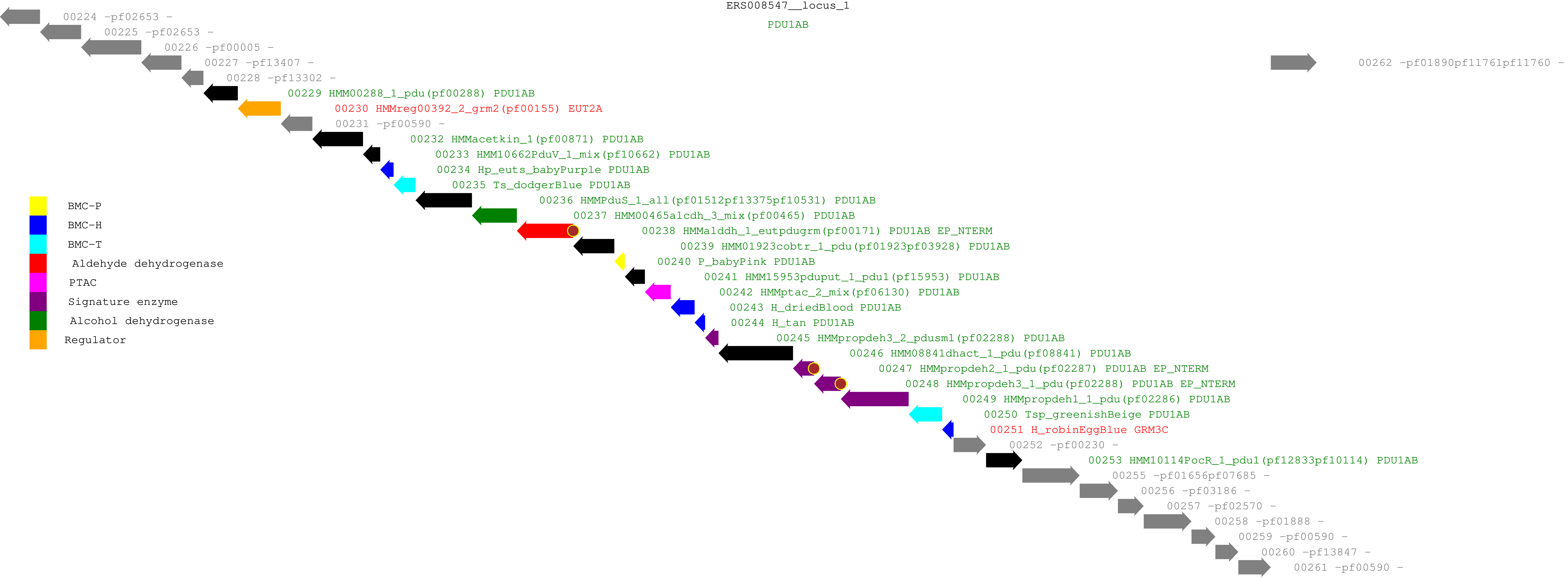

Yersinia mollaretii. Bacteria; Proteobacteria; Gammaproteobacteria; Enterobacterales;

ERS008557__locus_1

PDU1AB

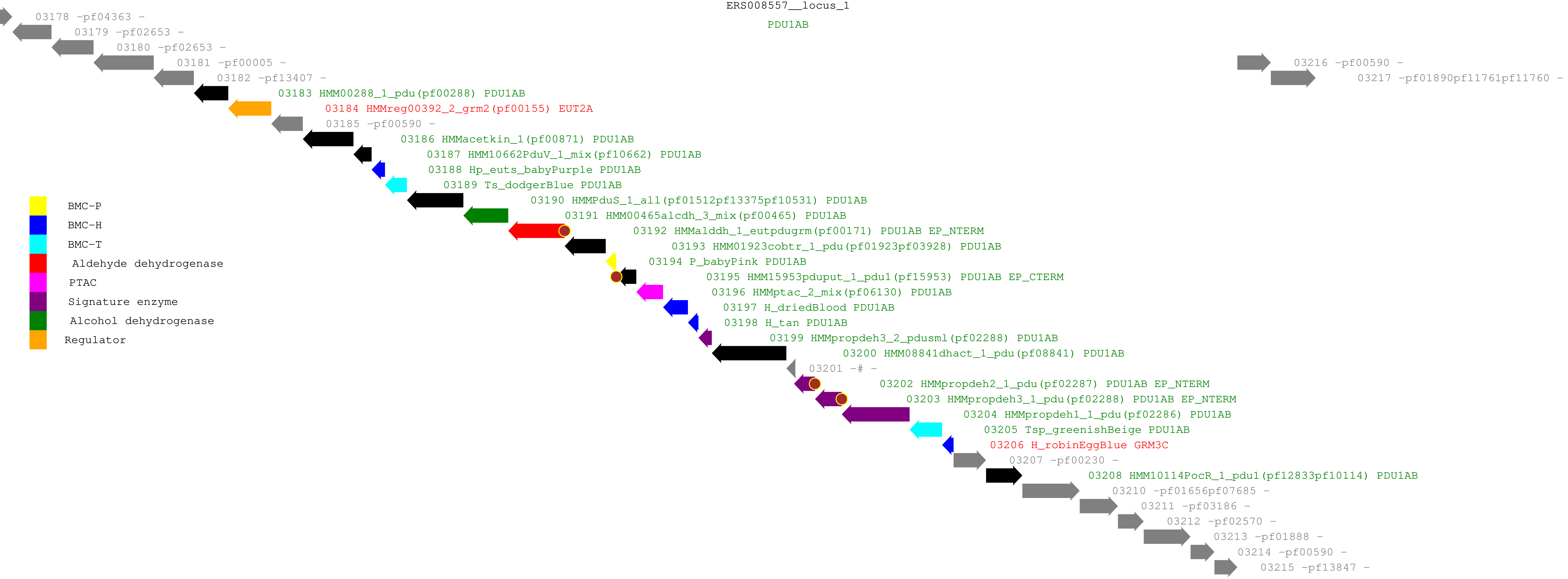

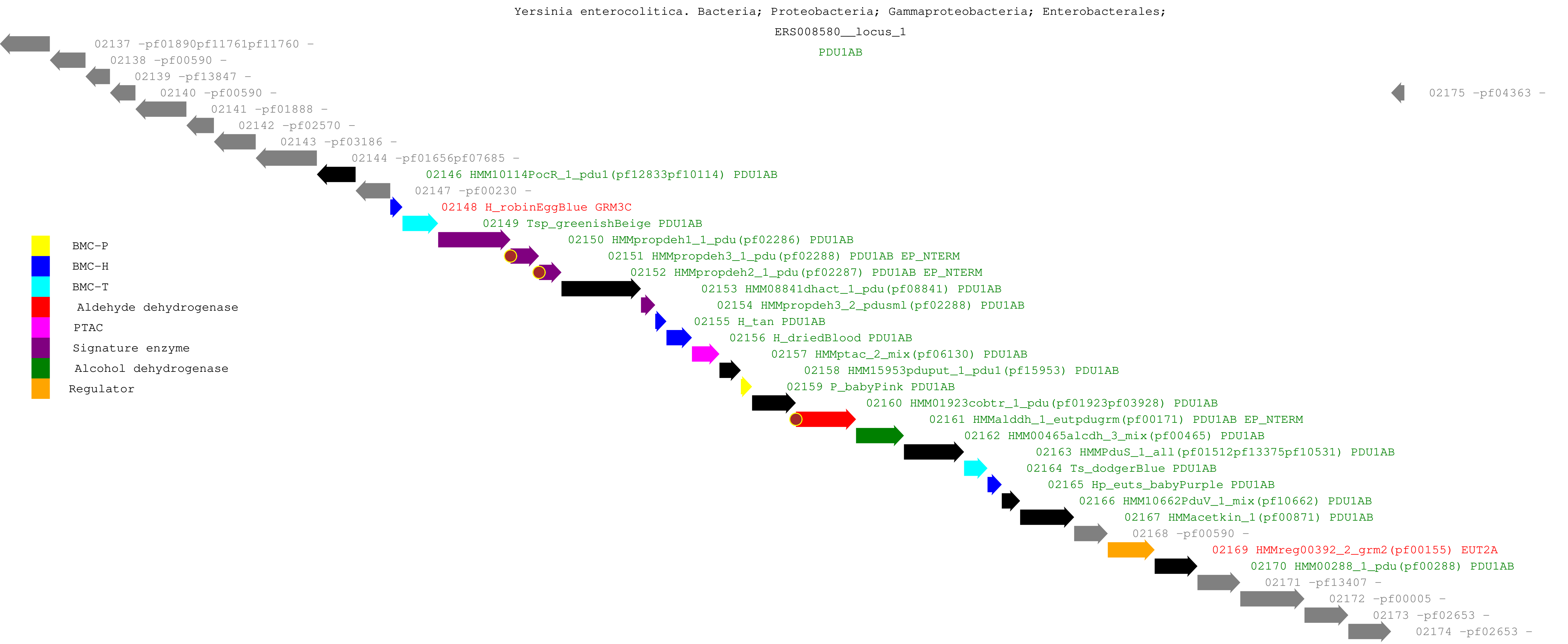
Yersinia enterocolitica. Bacteria; Proteobacteria; Gammaproteobacteria; Enterobacterales;
ERS008580__locus_1
PDU1AB
02137 -pf01890pf11761pf11760 -
02138 -pf00590 -
02139 -pf13847 -
02140 -pf00590 -
02141 -pf01888 -
02142 -pf02570 -
02143 -pf03186 -
02144 -pf01656pf07685 -
02146 HMM10114PocR_1_pdu1(pf12833pf10114) PDU1AB
02147 -pf00230 -
02148 H_robinEggBlue GRM3C
02149 Tsp_greenishBeige PDU1AB
02150 HMMpropdeh1_1_pdu(pf02286) PDU1AB
02151 HMMpropdeh3_1_pdu(pf02288) PDU1AB EP_NTERM
02152 HMMpropdeh2_1_pdu(pf02287) PDU1AB EP_NTERM
02153 HMM08841dhact_1_pdu(pf08841) PDU1AB
02154 HMMpropdeh3_2_pdusml(pf02288) PDU1AB
02155 H_tan PDU1AB
02156 H_driedBlood PDU1AB
02157 HMMptac_2_mix(pf06130) PDU1AB
02158 HMM15953pduput_1_pdu1(pf15953) PDU1AB
02159 P_babyPink PDU1AB
02160 HMM01923cobtr_1_pdu(pf01923pf03928) PDU1AB
02161 HMMalddh_1_eutpdugrm(pf00171) PDU1AB EP_NTERM
02162 HMM00465alcdh_3_mix(pf00465) PDU1AB
02163 HMMPduS_1_all(pf01512pf13375pf10531) PDU1AB
02164 Ts_dodgerBlue PDU1AB
02165 Hp_euts_babyPurple PDU1AB
02166 HMM10662PduV_1_mix(pf10662) PDU1AB
02167 HMMacetkin_1(pf00871) PDU1AB
02168 -pf00590 -
02169 HMMreg00392_2_grm2(pf00155) EUT2A
02170 HMM00288_1_pdu(pf00288) PDU1AB
02171 -pf13407 -
02172 -pf00005 -
02173 -pf02653 -
02174 -pf02653 -
02175 -pf04363 -
BMC-P
BMC-H
BMC-T
Aldehyde dehydrogenase
PTAC
Signature enzyme
Alcohol dehydrogenase
Regulator

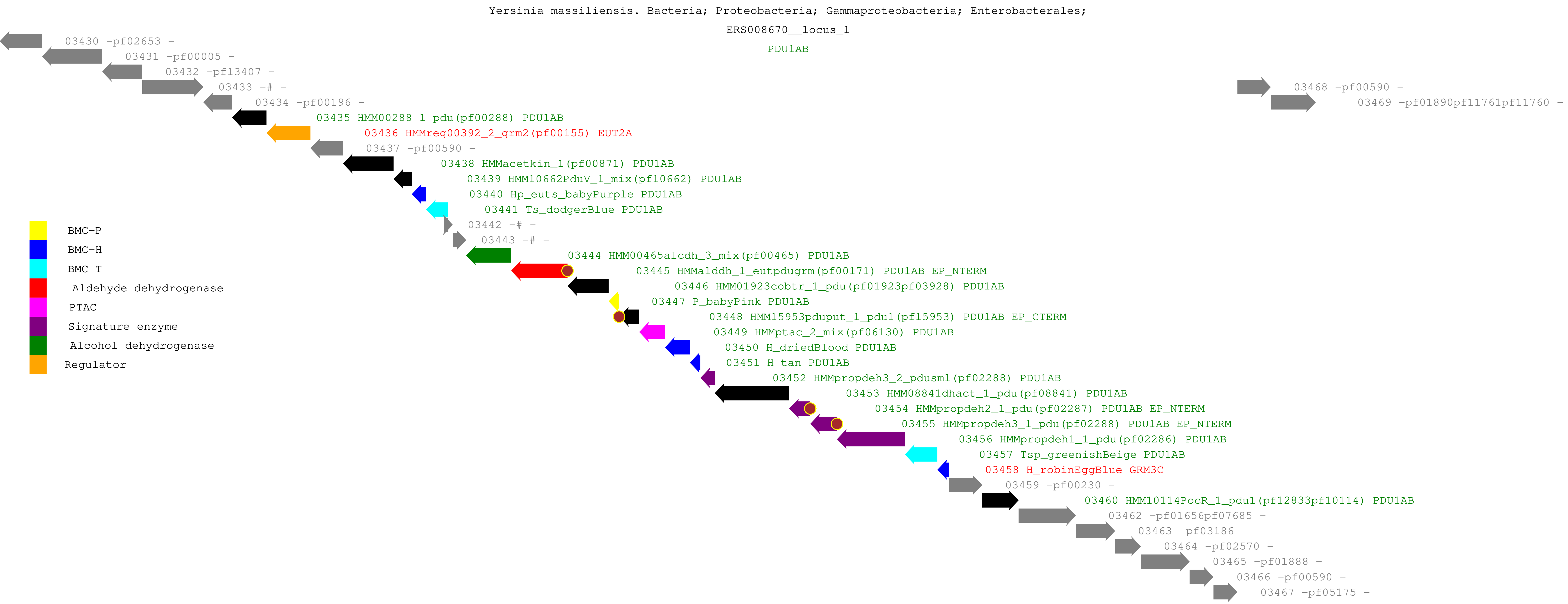
Yersinia massiliensis. Bacteria; Proteobacteria; Gammaproteobacteria; Enterobacterales;
ERS008670__locus_1
PDU1AB
03430 -pf02653 -
03431 -pf00005 -
03432 -pf13407 -
03433 -# -
03434 -pf00196 -
03435 HMM00288_1_pdu(pf00288) PDU1AB
03436 HMMreg00392_2_grm2(pf00155) EUT2A
03437 -pf00590 -
03438 HMMacetkin_1(pf00871) PDU1AB
03439 HMM10662PduV_1_mix(pf10662) PDU1AB
03440 Hp_euts_babyPurple PDU1AB
03441 Ts_dodgerBlue PDU1AB
03442 -# -
03443 -# -
03444 HMM00465alcdh_3_mix(pf00465) PDU1AB
03445 HMMalddh_1_eutpdugrm(pf00171) PDU1AB EP_NTERM
03446 HMM01923cobtr_1_pdu(pf01923pf03928) PDU1AB
03447 P_babyPink PDU1AB
03448 HMM15953pduput_1_pdu1(pf15953) PDU1AB EP_CTERM
03449 HMMptac_2_mix(pf06130) PDU1AB
03450 H_driedBlood PDU1AB
03451 H_tan PDU1AB
03452 HMMpropdeh3_2_pdusml(pf02288) PDU1AB
03453 HMM08841dhact_1_pdu(pf08841) PDU1AB
03454 HMMpropdeh2_1_pdu(pf02287) PDU1AB EP_NTERM
03455 HMMpropdeh3_1_pdu(pf02288) PDU1AB EP_NTERM
03456 HMMpropdeh1_1_pdu(pf02286) PDU1AB
03457 Tsp_greenishBeige PDU1AB
03458 H_robinEggBlue GRM3C
03459 -pf00230 -
03460 HMM10114PocR_1_pdu1(pf12833pf10114) PDU1AB
03462 -pf01656pf07685 -
03463 -pf03186 -
03464 -pf02570 -
03465 -pf01888 -
03466 -pf00590 -
03467 -pf05175 -
03468 -pf00590 -
03469 -pf01890pf11761pf11760 -
BMC-P
BMC-H
BMC-T
Aldehyde dehydrogenase
PTAC
Signature enzyme
Alcohol dehydrogenase
Regulator

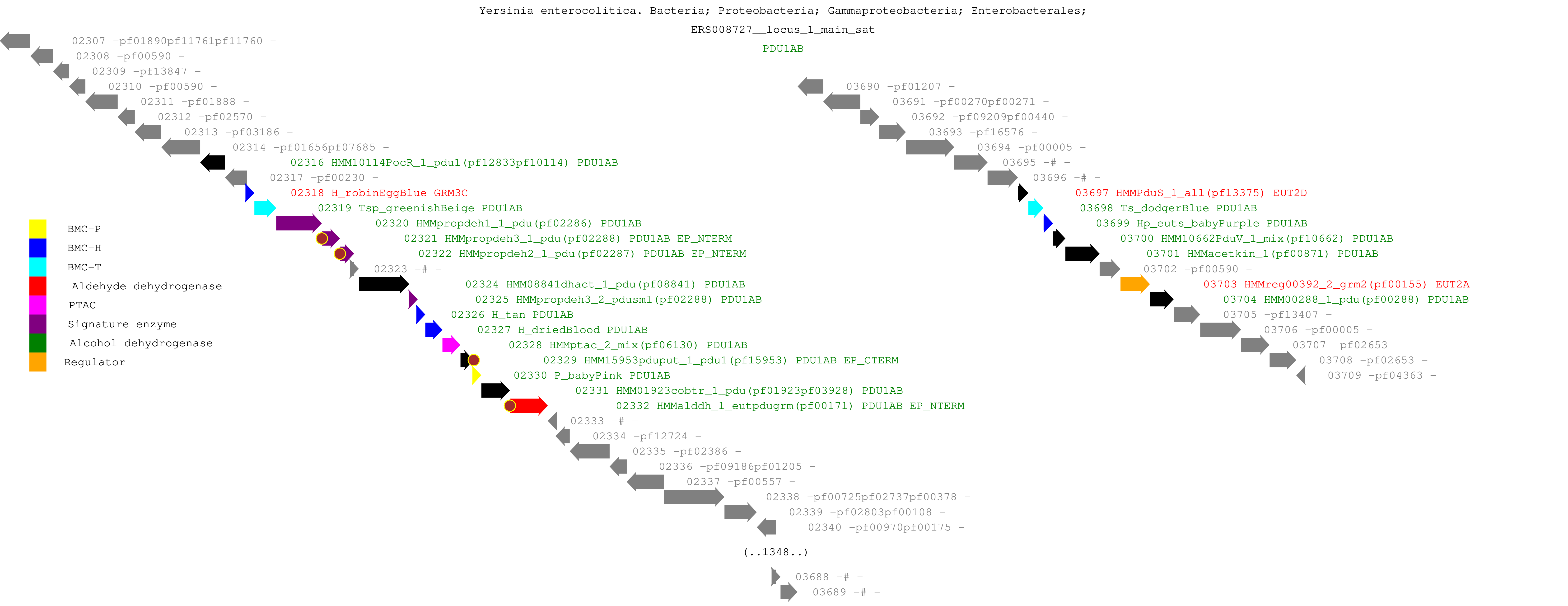
Yersinia enterocolitica. Bacteria; Proteobacteria; Gammaproteobacteria; Enterobacterales;
ERS008727__locus_1_main_sat
PDU1AB
02307 -pf01890pf11761pf11760 -
02308 -pf00590 -
02309 -pf13847 -
02310 -pf00590 -
02311 -pf01888 -
02312 -pf02570 -
02313 -pf03186 -
02314 -pf01656pf07685 -
02316 HMM10114PocR_1_pdu1(pf12833pf10114) PDU1AB
02317 -pf00230 -
02318 H_robinEggBlue GRM3C
02319 Tsp_greenishBeige PDU1AB
02320 HMMpropdeh1_1_pdu(pf02286) PDU1AB
02321 HMMpropdeh3_1_pdu(pf02288) PDU1AB EP_NTERM
02322 HMMpropdeh2_1_pdu(pf02287) PDU1AB EP_NTERM
02323 -# -
02324 HMM08841dhact_1_pdu(pf08841) PDU1AB
02325 HMMpropdeh3_2_pdusml(pf02288) PDU1AB
02326 H_tan PDU1AB
02327 H_driedBlood PDU1AB
02328 HMMptac_2_mix(pf06130) PDU1AB
02329 HMM15953pduput_1_pdu1(pf15953) PDU1AB EP_CTERM
02330 P_babyPink PDU1AB
02331 HMM01923cobtr_1_pdu(pf01923pf03928) PDU1AB
02332 HMMalddh_1_eutpdugrm(pf00171) PDU1AB EP_NTERM
02333 -# -
02334 -pf12724 -
02335 -pf02386 -
02336 -pf09186pf01205 -
02337 -pf00557 -
02338 -pf00725pf02737pf00378 -
02339 -pf02803pf00108 -
02340 -pf00970pf00175 -
(..1348..)
03688 -# -
03689 -# -
03690 -pf01207 -
03691 -pf00270pf00271 -
03692 -pf09209pf00440 -
03693 -pf16576 -
03694 -pf00005 -
03695 -# -
03696 -# -
03697 HMMPduS_1_all(pf13375) EUT2D
03698 Ts_dodgerBlue PDU1AB
03699 Hp_euts_babyPurple PDU1AB
03700 HMM10662PduV_1_mix(pf10662) PDU1AB
03701 HMMacetkin_1(pf00871) PDU1AB
03702 -pf00590 -
03703 HMMreg00392_2_grm2(pf00155) EUT2A
03704 HMM00288_1_pdu(pf00288) PDU1AB
03705 -pf13407 -
03706 -pf00005 -
03707 -pf02653 -
03708 -pf02653 -
03709 -pf04363 -
BMC-P
BMC-H
BMC-T
Aldehyde dehydrogenase
PTAC
Signature enzyme
Alcohol dehydrogenase
Regulator

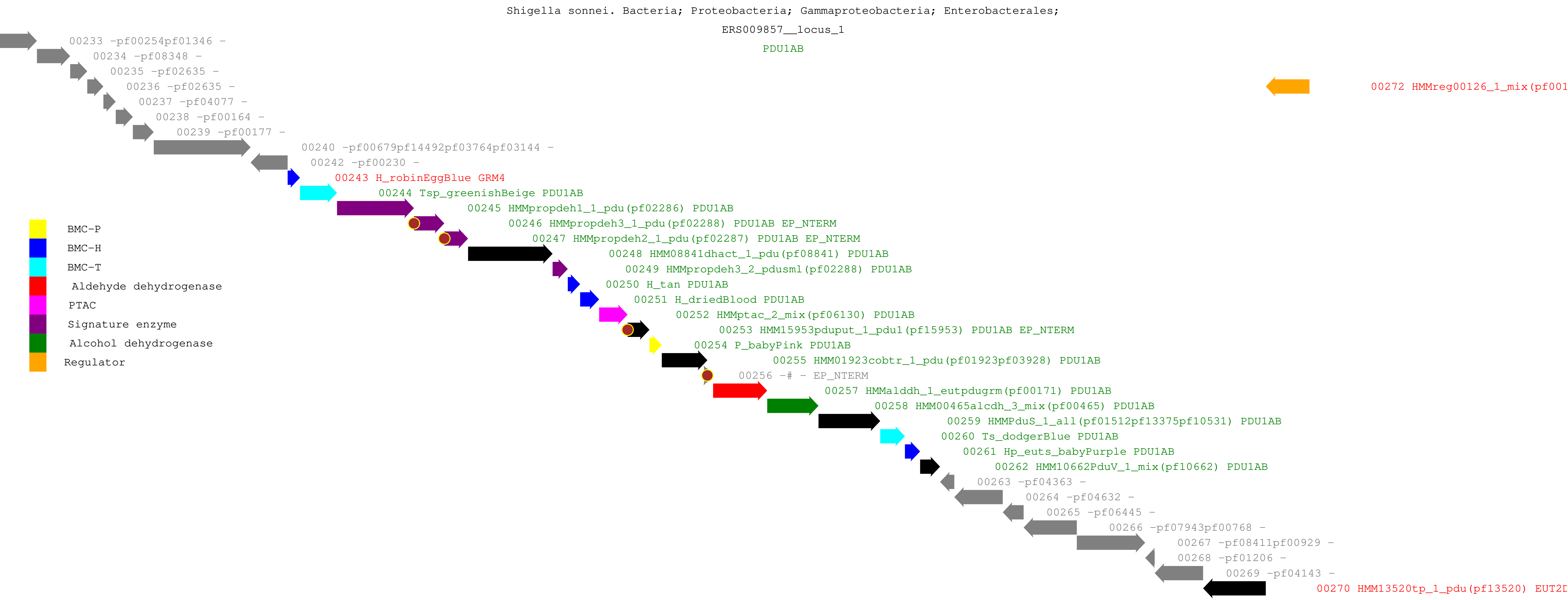
Shigella sonnei. Bacteria; Proteobacteria; Gammaproteobacteria; Enterobacterales;
ERS009857__locus_1
PDU1AB
00233 -pf00254pf01346 -
00234 -pf08348 -
00235 -pf02635 -
00236 -pf02635 -
00237 -pf04077 -
00238 -pf00164 -
00239 -pf00177 -
00240 -pf00679pf14492pf03764pf03144 -
00242 -pf00230 -
00243 H_robinEggBlue GRM4
00244 Tsp_greenishBeige PDU1AB
00245 HMMpropdeh1_1_pdu(pf02286) PDU1AB
00246 HMMpropdeh3_1_pdu(pf02288) PDU1AB EP_NTERM
00247 HMMpropdeh2_1_pdu(pf02287) PDU1AB EP_NTERM
00248 HMM08841dhact_1_pdu(pf08841) PDU1AB
00249 HMMpropdeh3_2_pdusml(pf02288) PDU1AB
00250 H_tan PDU1AB
00251 H_driedBlood PDU1AB
00252 HMMptac_2_mix(pf06130) PDU1AB
00253 HMM15953pduput_1_pdu1(pf15953) PDU1AB EP_NTERM
00254 P_babyPink PDU1AB
00255 HMM01923cobtr_1_pdu(pf01923pf03928) PDU1AB
00256 -# - EP_NTERM
00257 HMMalddh_1_eutpdugrm(pf00171) PDU1AB
00258 HMM00465alcdh_3_mix(pf00465) PDU1AB
00259 HMMPduS_1_all(pf01512pf13375pf10531) PDU1AB
00260 Ts_dodgerBlue PDU1AB
00261 Hp_euts_babyPurple PDU1AB
00262 HMM10662PduV_1_mix(pf10662) PDU1AB
00263 -pf04363 -
00264 -pf04632 -
00265 -pf06445 -
00266 -pf07943pf00768 -
00267 -pf08411pf00929 -
00268 -pf01206 -
00269 -pf04143 -
00270 HMM13520tp_1_pdu(pf13520) EUT2D
00272 HMMreg00126_1_mix(pf001
BMC-P
BMC-H
BMC-T
Aldehyde dehydrogenase
PTAC
Signature enzyme
Alcohol dehydrogenase
Regulator

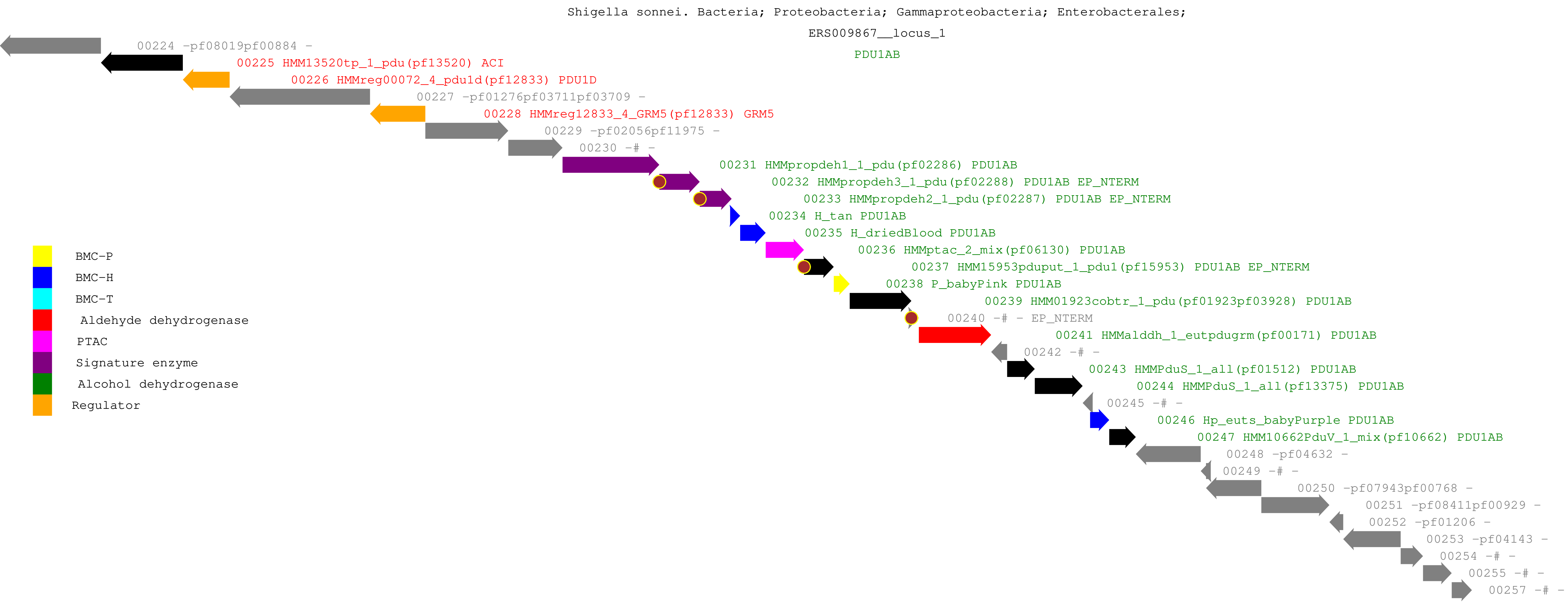

Shigella sonnei. Bacteria; Proteobacteria; Gammaproteobacteria; Enterobacterales;
ERS009867__locus_1
PDU1AB
00224 -pf08019pf00884 -
00225 HMM13520tp_1_pdu(pf13520) ACI
00226 HMMreg00072_4_pdu1d(pf12833) PDU1D
00227 -pf01276pf03711pf03709 -
00228 HMMreg12833_4_GRM5(pf12833) GRM5
00229 -pf02056pf11975 -
00230 -# -
00231 HMMpropdeh1_1_pdu(pf02286) PDU1AB
00232 HMMpropdeh3_1_pdu(pf02288) PDU1AB EP_NTERM
00233 HMMpropdeh2_1_pdu(pf02287) PDU1AB EP_NTERM
00234 H_tan PDU1AB
00235 H_driedBlood PDU1AB
00236 HMMptac_2_mix(pf06130) PDU1AB
00237 HMM15953pduput_1_pdu1(pf15953) PDU1AB EP_NTERM
00238 P_babyPink PDU1AB
00239 HMM01923cobtr_1_pdu(pf01923pf03928) PDU1AB
00240 -# - EP_NTERM
00241 HMMalddh_1_eutpdugrm(pf00171) PDU1AB
00242 -# -
00243 HMMPduS_1_all(pf01512) PDU1AB
00244 HMMPduS_1_all(pf13375) PDU1AB
00245 -# -
00246 Hp_euts_babyPurple PDU1AB
00247 HMM10662PduV_1_mix(pf10662) PDU1AB
00248 -pf04632 -
00249 -# -
00250 -pf07943pf00768 -
00251 -pf08411pf00929 -
00252 -pf01206 -
00253 -pf04143 -
00254 -# -
00255 -# -
00257 -# -
BMC-P
BMC-H
BMC-T
Aldehyde dehydrogenase
PTAC
Signature enzyme
Alcohol dehydrogenase
Regulator

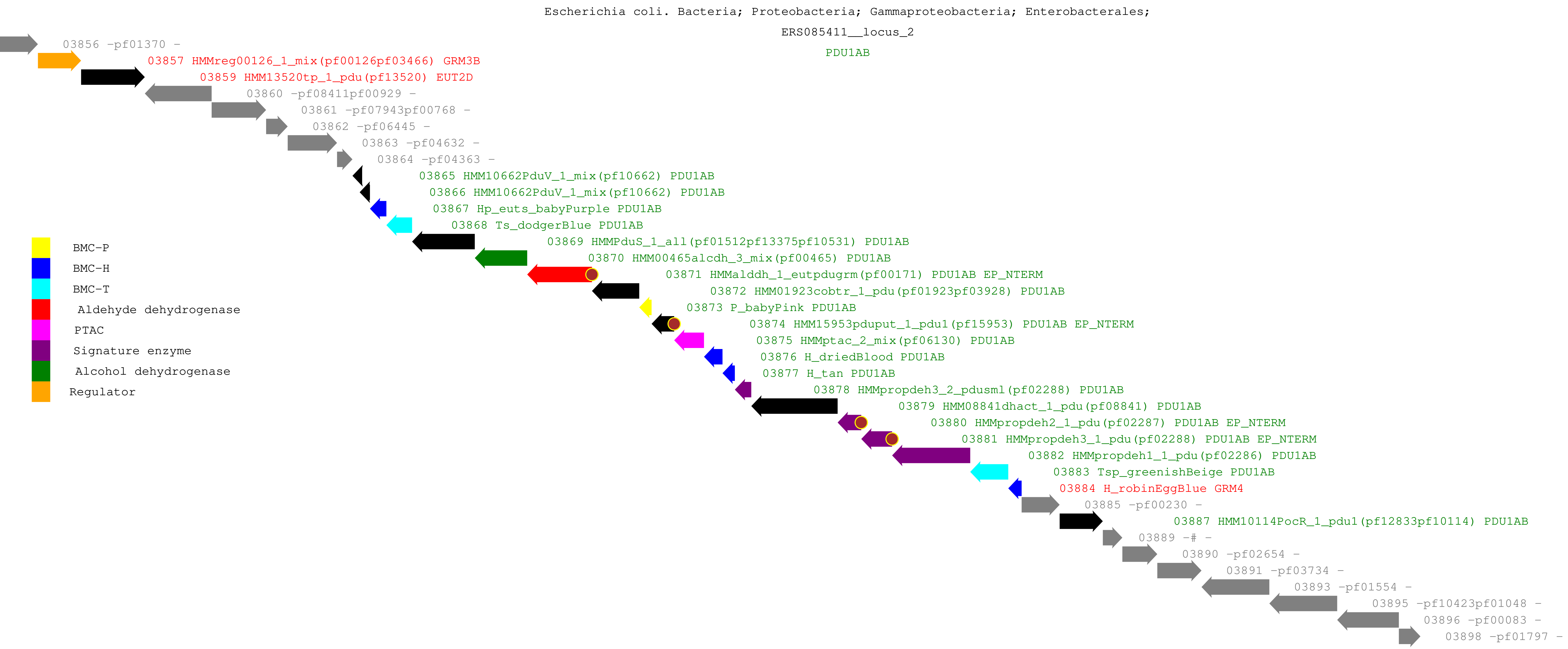
Escherichia coli. Bacteria; Proteobacteria; Gammaproteobacteria; Enterobacterales;
ERS085411__locus_2
PDU1AB
03856 -pf01370 -
03857 HMMreg00126_1_mix(pf00126pf03466) GRM3B
03859 HMM13520tp_1_pdu(pf13520) EUT2D
03860 -pf08411pf00929 -
03861 -pf07943pf00768 -
03862 -pf06445 -
03863 -pf04632 -
03864 -pf04363 -
03865 HMM10662PduV_1_mix(pf10662) PDU1AB
03866 HMM10662PduV_1_mix(pf10662) PDU1AB
03867 Hp_euts_babyPurple PDU1AB
03868 Ts_dodgerBlue PDU1AB
03869 HMMPduS_1_all(pf01512pf13375pf10531) PDU1AB
03870 HMM00465alcdh_3_mix(pf00465) PDU1AB
03871 HMMalddh_1_eutpdugrm(pf00171) PDU1AB EP_NTERM
03872 HMM01923cobtr_1_pdu(pf01923pf03928) PDU1AB
03873 P_babyPink PDU1AB
03874 HMM15953pduput_1_pdu1(pf15953) PDU1AB EP_NTERM
03875 HMMptac_2_mix(pf06130) PDU1AB
03876 H_driedBlood PDU1AB
03877 H_tan PDU1AB
03878 HMMpropdeh3_2_pdusml(pf02288) PDU1AB
03879 HMM08841dhact_1_pdu(pf08841) PDU1AB
03880 HMMpropdeh2_1_pdu(pf02287) PDU1AB EP_NTERM
03881 HMMpropdeh3_1_pdu(pf02288) PDU1AB EP_NTERM
03882 HMMpropdeh1_1_pdu(pf02286) PDU1AB
03883 Tsp_greenishBeige PDU1AB
03884 H_robinEggBlue GRM4
03885 -pf00230 -
03887 HMM10114PocR_1_pdu1(pf12833pf10114) PDU1AB
03889 -# -
03890 -pf02654 -
03891 -pf03734 -
03893 -pf01554 -
03895 -pf10423pf01048 -
03896 -pf00083 -
03898 -pf01797 -
BMC-P
BMC-H
BMC-T
Aldehyde dehydrogenase
PTAC
Signature enzyme
Alcohol dehydrogenase
Regulator

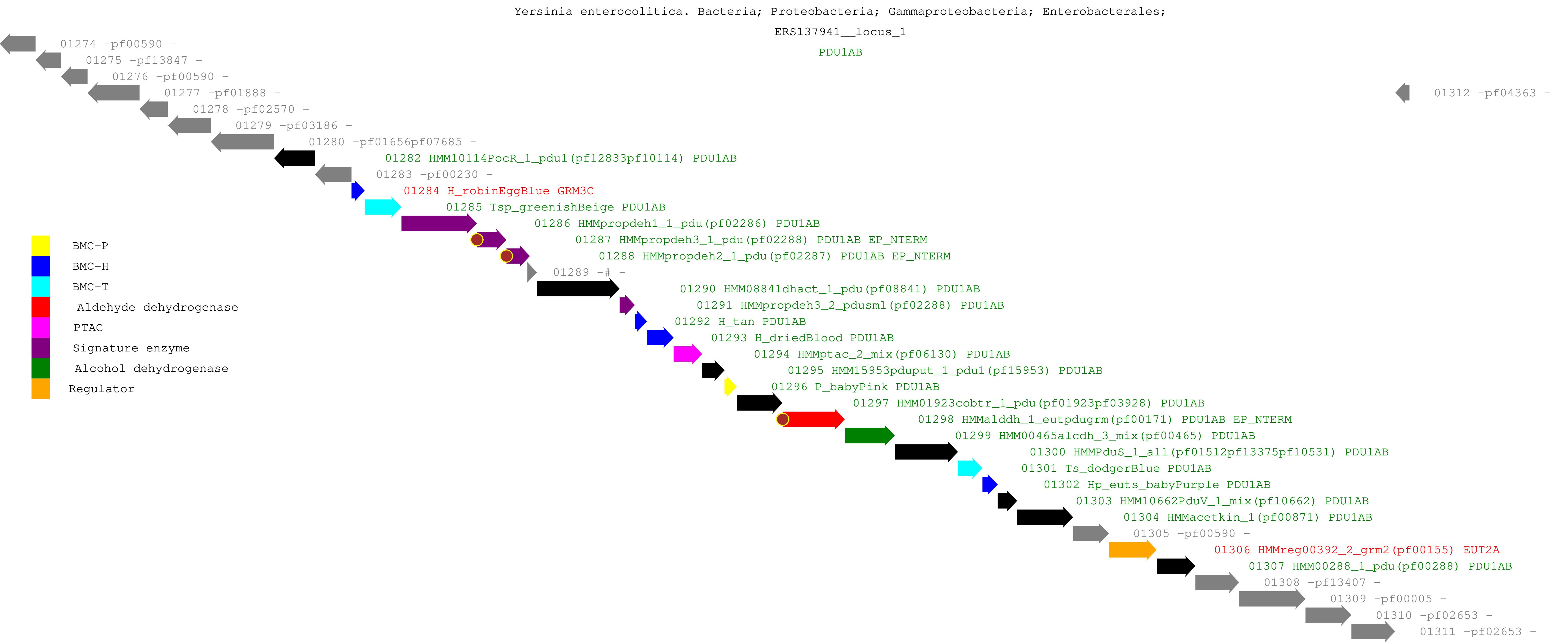
Yersinia enterocolitica. Bacteria; Proteobacteria; Gammaproteobacteria; Enterobacterales;
ERS137941__locus_1
PDU1AB
01274 -pf00590 -
01275 -pf13847 -
01276 -pf00590 -
01277 -pf01888 -
01278 -pf02570 -
01279 -pf03186 -
01280 -pf01656pf07685 -
01282 HMM10114PocR_1_pdu1(pf12833pf10114) PDU1AB
01283 -pf00230 -
01284 H_robinEggBlue GRM3C
01285 Tsp_greenishBeige PDU1AB
01286 HMMpropdeh1_1_pdu(pf02286) PDU1AB
01287 HMMpropdeh3_1_pdu(pf02288) PDU1AB EP_NTERM
01288 HMMpropdeh2_1_pdu(pf02287) PDU1AB EP_NTERM
01289 -# -
01290 HMM08841dhact_1_pdu(pf08841) PDU1AB
01291 HMMpropdeh3_2_pdusml(pf02288) PDU1AB
01292 H_tan PDU1AB
01293 H_driedBlood PDU1AB
01294 HMMptac_2_mix(pf06130) PDU1AB
01295 HMM15953pduput_1_pdu1(pf15953) PDU1AB
01296 P_babyPink PDU1AB
01297 HMM01923cobtr_1_pdu(pf01923pf03928) PDU1AB
01298 HMMalddh_1_eutpdugrm(pf00171) PDU1AB EP_NTERM
01299 HMM00465alcdh_3_mix(pf00465) PDU1AB
01300 HMMPduS_1_all(pf01512pf13375pf10531) PDU1AB
01301 Ts_dodgerBlue PDU1AB
01302 Hp_euts_babyPurple PDU1AB
01303 HMM10662PduV_1_mix(pf10662) PDU1AB
01304 HMMacetkin_1(pf00871) PDU1AB
01305 -pf00590 -
01306 HMMreg00392_2_grm2(pf00155) EUT2A
01307 HMM00288_1_pdu(pf00288) PDU1AB
01308 -pf13407 -
01309 -pf00005 -
01310 -pf02653 -
01311 -pf02653 -
01312 -pf04363 -
BMC-P
BMC-H
BMC-T
Aldehyde dehydrogenase
PTAC
Signature enzyme
Alcohol dehydrogenase
Regulator

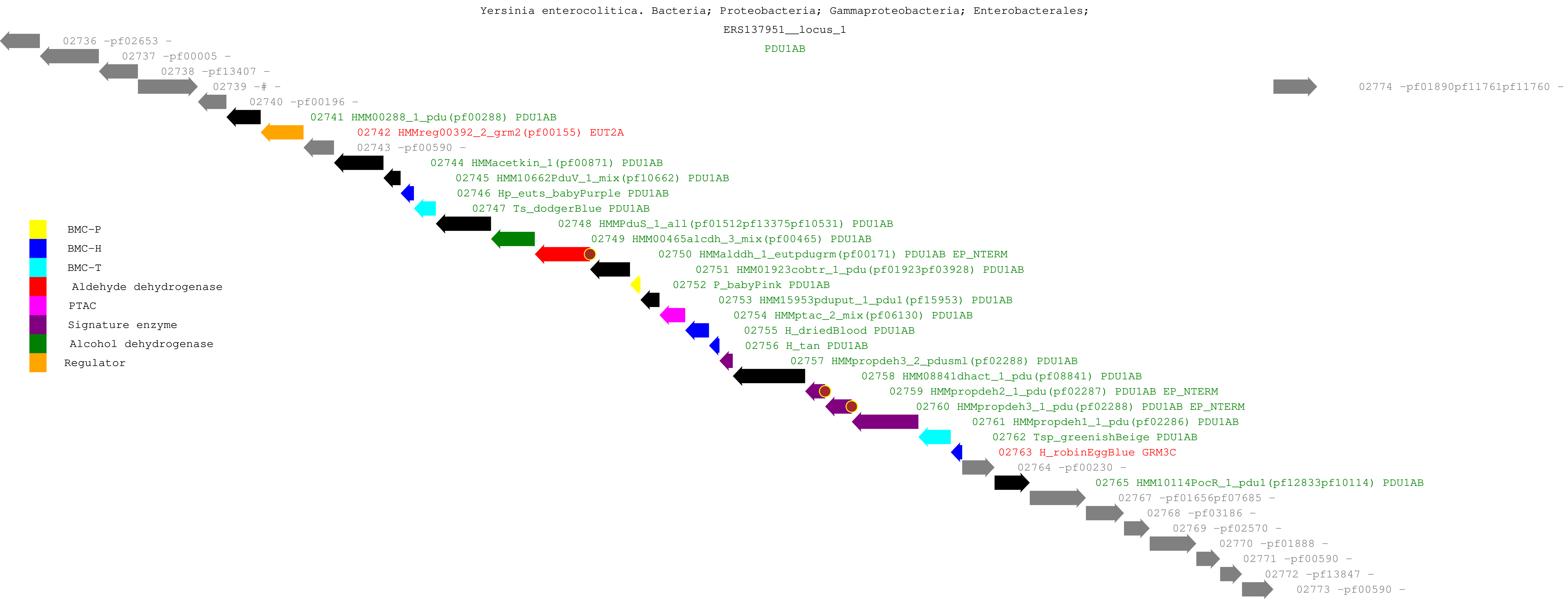
Yersinia enterocolitica. Bacteria; Proteobacteria; Gammaproteobacteria; Enterobacterales;
ERS137951__locus_1
PDU1AB
02736 -pf02653 -
02737 -pf00005 -
02738 -pf13407 -
02739 -# -
02740 -pf00196 -
02741 HMM00288_1_pdu(pf00288) PDU1AB
02742 HMMreg00392_2_grm2(pf00155) EUT2A
02743 -pf00590 -
02744 HMMacetkin_1(pf00871) PDU1AB
02745 HMM10662PduV_1_mix(pf10662) PDU1AB
02746 Hp_euts_babyPurple PDU1AB
02747 Ts_dodgerBlue PDU1AB
02748 HMMPduS_1_all(pf01512pf13375pf10531) PDU1AB
02749 HMM00465alcdh_3_mix(pf00465) PDU1AB
02750 HMMalddh_1_eutpdugrm(pf00171) PDU1AB EP_NTERM
02751 HMM01923cobtr_1_pdu(pf01923pf03928) PDU1AB
02752 P_babyPink PDU1AB
02753 HMM15953pduput_1_pdu1(pf15953) PDU1AB
02754 HMMptac_2_mix(pf06130) PDU1AB
02755 H_driedBlood PDU1AB
02756 H_tan PDU1AB
02757 HMMpropdeh3_2_pdusml(pf02288) PDU1AB
02758 HMM08841dhact_1_pdu(pf08841) PDU1AB
02759 HMMpropdeh2_1_pdu(pf02287) PDU1AB EP_NTERM
02760 HMMpropdeh3_1_pdu(pf02288) PDU1AB EP_NTERM
02761 HMMpropdeh1_1_pdu(pf02286) PDU1AB
02762 Tsp_greenishBeige PDU1AB
02763 H_robinEggBlue GRM3C
02764 -pf00230 -
02765 HMM10114PocR_1_pdu1(pf12833pf10114) PDU1AB
02767 -pf01656pf07685 -
02768 -pf03186 -
02769 -pf02570 -
02770 -pf01888 -
02771 -pf00590 -
02772 -pf13847 -
02773 -pf00590 -
02774 -pf01890pf11761pf11760 -
BMC-P
BMC-H
BMC-T
Aldehyde dehydrogenase
PTAC
Signature enzyme
Alcohol dehydrogenase
Regulator

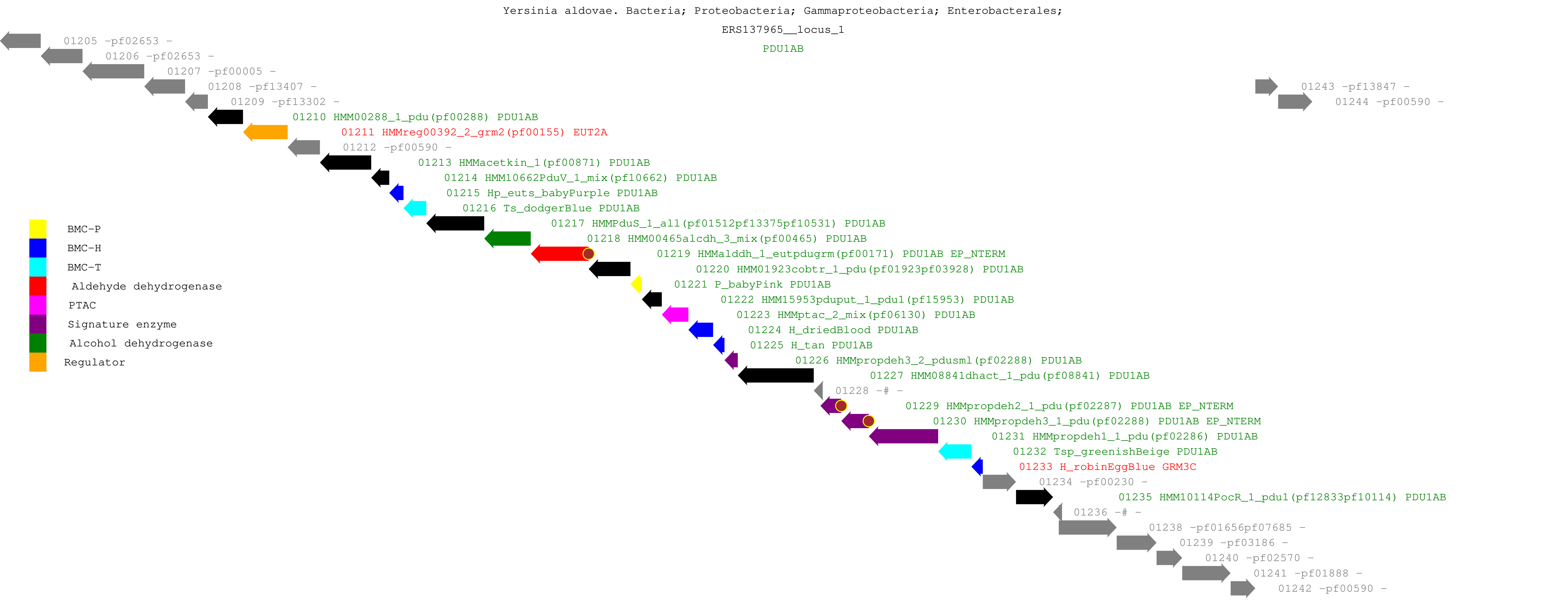

Yersinia aldovae. Bacteria; Proteobacteria; Gammaproteobacteria; Enterobacterales;
ERS137965__locus_1
PDU1AB
01205 -pf02653 -
01206 -pf02653 -
01207 -pf00005 -
01208 -pf13407 -
01209 -pf13302 -
01210 HMM00288_1_pdu(pf00288) PDU1AB
01211 HMMreg00392_2_grm2(pf00155) EUT2A
01212 -pf00590 -
01213 HMMacetkin_1(pf00871) PDU1AB
01214 HMM10662PduV_1_mix(pf10662) PDU1AB
01215 Hp_euts_babyPurple PDU1AB
01216 Ts_dodgerBlue PDU1AB
01217 HMMPduS_1_all(pf01512pf13375pf10531) PDU1AB
01218 HMM00465alcdh_3_mix(pf00465) PDU1AB
01219 HMMaldd h_1_eutpdugrm(pf00171) PDU1AB EP_NTERM
01220 HMM01923cobtr_1_pdu(pf01923pf03928) PDU1AB
01221 P_babyPink PDU1AB
01222 HMM15953pduput_1_pdu1(pf15953) PDU1AB
01223 HMMptac_2_mix(pf06130) PDU1AB
01224 H_driedBlood PDU1AB
01225 H_tan PDU1AB
01226 HMMpropdeh3_2_pdusml(pf02288) PDU1AB
01227 HMM08841dhact_1_pdu(pf08841) PDU1AB
01228 -# -
01229 HMMpropdeh2_1_pdu(pf02287) PDU1AB EP_NTERM
01230 HMMpropdeh3_1_pdu(pf02288) PDU1AB EP_NTERM
01231 HMMpropdeh1_1_pdu(pf02286) PDU1AB
01232 Tsp_greenishBeige PDU1AB
01233 H_robinEggBlue GRM3C
01234 -pf00230 -
01235 HMM10114PocR_1_pdu1(pf12833pf10114) PDU1AB
01236 -# -
01238 -pf01656pf07685 -
01239 -pf03186 -
01240 -pf02570 -
01241 -pf01888 -
01242 -pf00590 -
01243 -pf13847 -
01244 -pf00590 -
BMC-P
BMC-H
BMC-T
Aldehyde dehydrogenase
PTAC
Signature enzyme
Alcohol dehydrogenase
Regulator

Salmonella enterica subsp. enterica serovar Typhimurium str. DT104. Bacteria; Proteobacteria; Gammaproteobacteria; Enterobacterales;

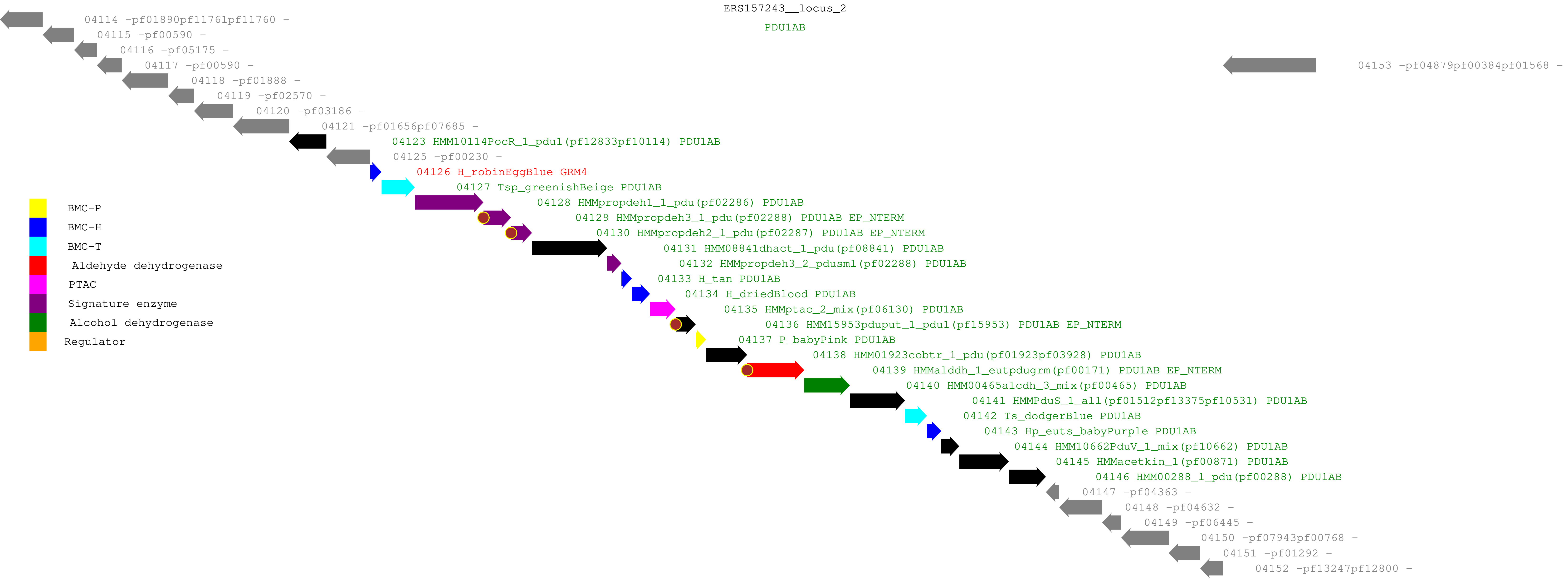

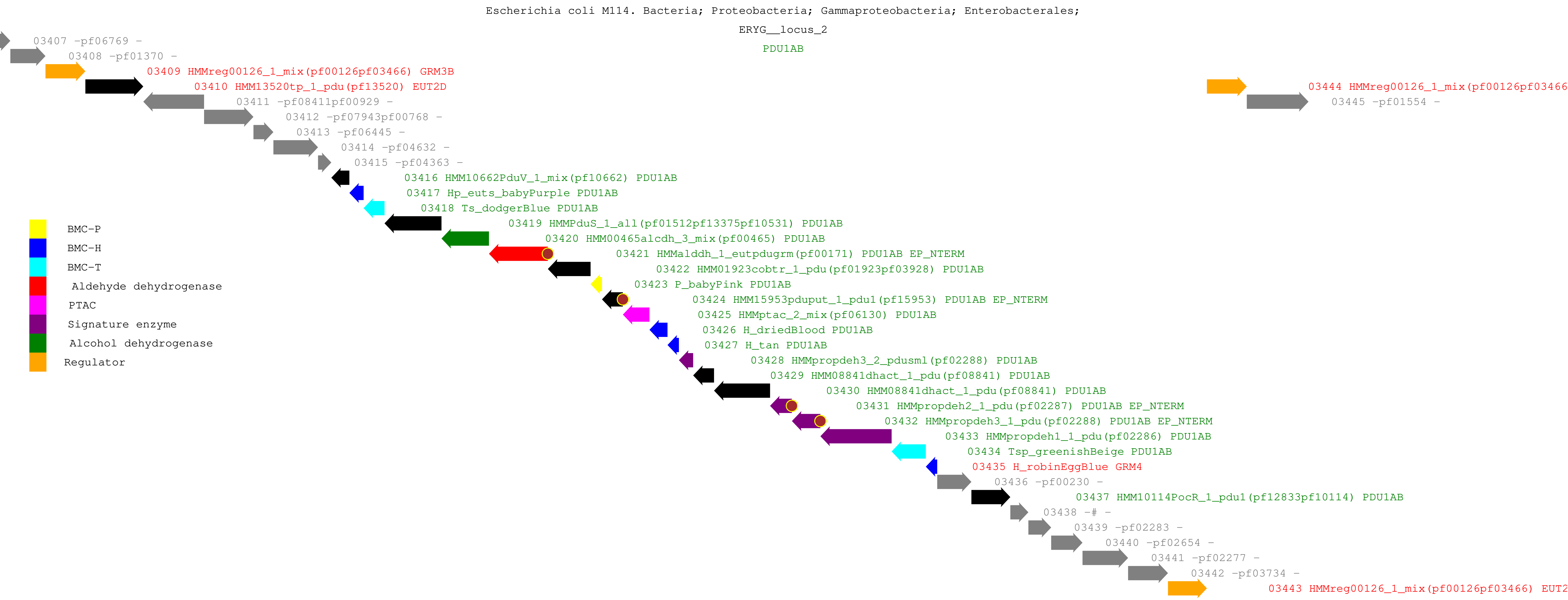

Escherichia coli M114. Bacteria; Proteobacteria; Gammaproteobacteria; Enterobacterales;
ERYG__locus_2
PDU1AB
03407 -pf06769 -
03408 -pf01370 -
03409 HMMreg00126_1_mix(pf00126pf03466) GRM3B
03410 HMM13520tp_1_pdu(pf13520) EUT2D
03411 -pf08411pf00929 -
03412 -pf07943pf00768 -
03413 -pf06445 -
03414 -pf04632 -
03415 -pf04363 -
03416 HMM10662PduV_1_mix(pf10662) PDU1AB
03417 Hp_euts_babyPurple PDU1AB
03418 Ts_dodgerBlue PDU1AB
03419 HMMPduS_1_all(pf01512pf13375pf10531) PDU1AB
03420 HMM00465alcdh_3_mix(pf00465) PDU1AB
03421 HMMaldh_1_eutpdugrm(pf00171) PDU1AB EP_NTERM
03422 HMM01923cobtr_1_pdu(pf01923pf03928) PDU1AB
03423 P_babyPink PDU1AB
03424 HMM15953pduput_1_pdu1(pf15953) PDU1AB EP_NTERM
03425 HMMptac_2_mix(pf06130) PDU1AB
03426 H_driedBlood PDU1AB
03427 H_tan PDU1AB
03428 HMMpropdeh3_2_pdusml(pf02288) PDU1AB
03429 HMM08841dhact_1_pdu(pf08841) PDU1AB
03430 HMM08841dhact_1_pdu(pf08841) PDU1AB
03431 HMMpropdeh2_1_pdu(pf02287) PDU1AB EP_NTERM
03432 HMMpropdeh3_1_pdu(pf02288) PDU1AB EP_NTERM
03433 HMMpropdeh1_1_pdu(pf02286) PDU1AB
03434 Tsp_greenishBeige PDU1AB
03435 H_robinEggBlue GRM4
03436 -pf00230 -
03437 HMM10114PocR_1_pdu1(pf12833pf10114) PDU1AB
03438 -# -
03439 -pf02283 -
03440 -pf02654 -
03441 -pf02277 -
03442 -pf03734 -
03443 HMMreg00126_1_mix(pf00126pf03466) EUT2
03444 HMMreg00126_1_mix(pf00126pf03466
03445 -pf01554 -
BMC-P
BMC-H
BMC-T
Aldehyde dehydrogenase
PTAC
Signature enzyme
Alcohol dehydrogenase
Regulator

Escherichia albertii (strain TW07627). Bacteria; Proteobacteria; Gammaproteobacteria; Enterobacterales;

ESCAB7627__locus_1

PDU1AB

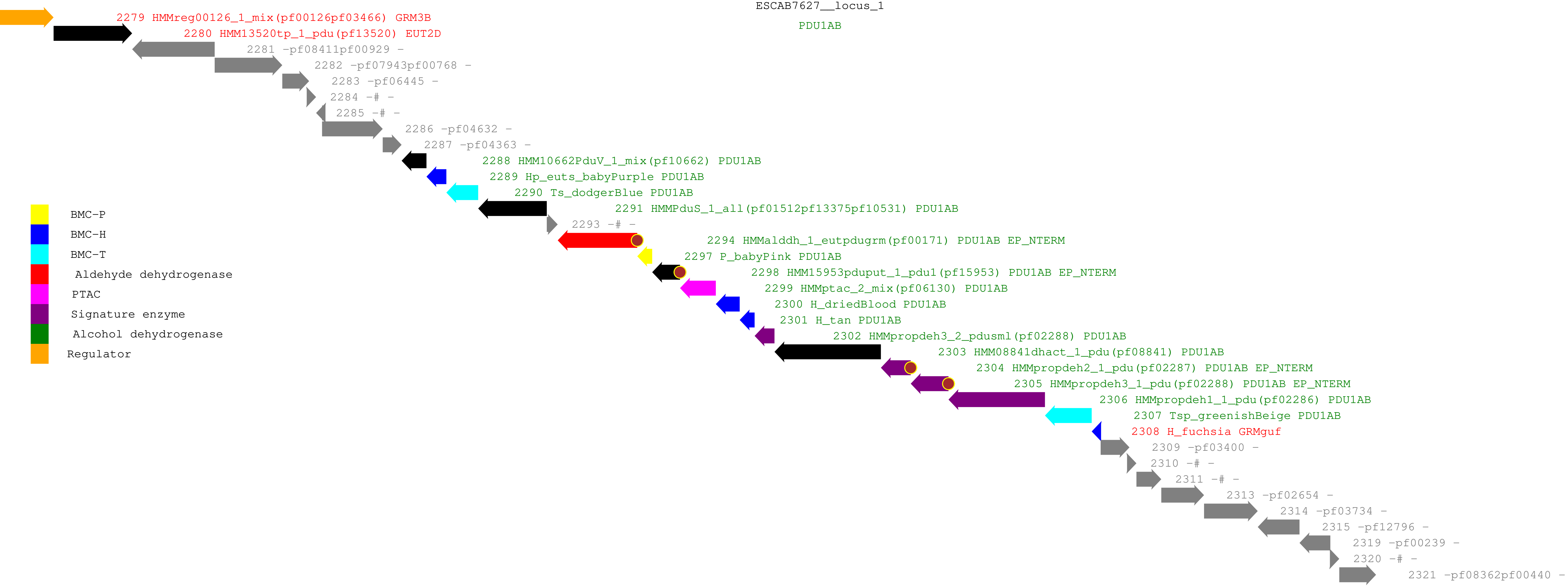

Salmonella enterica subsp. enterica serovar Monschaui. Bacteria; Proteobacteria; Gammaproteobacteria; Enterobacterales;

EU345__locus_1

PDU1AB

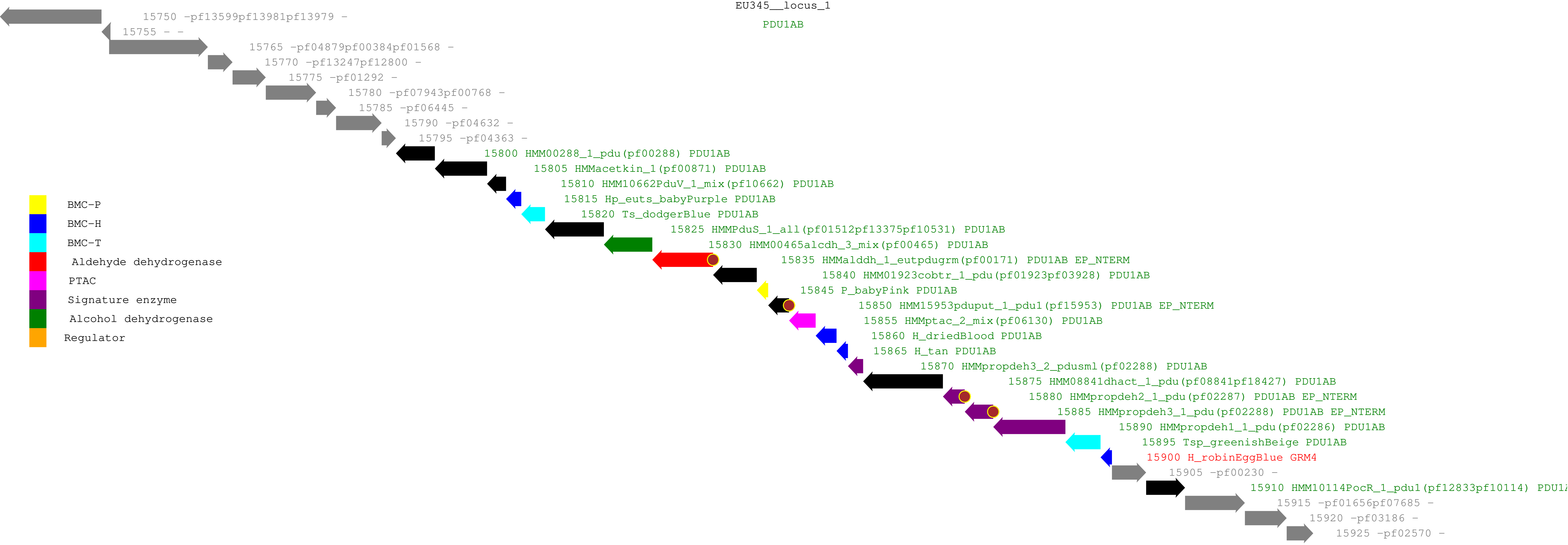

Salmonella enterica subsp. enterica serovar Bonn. Bacteria; Proteobacteria; Gammaproteobacteria; Enterobacterales;

EU363__locus_1

PDU1AB

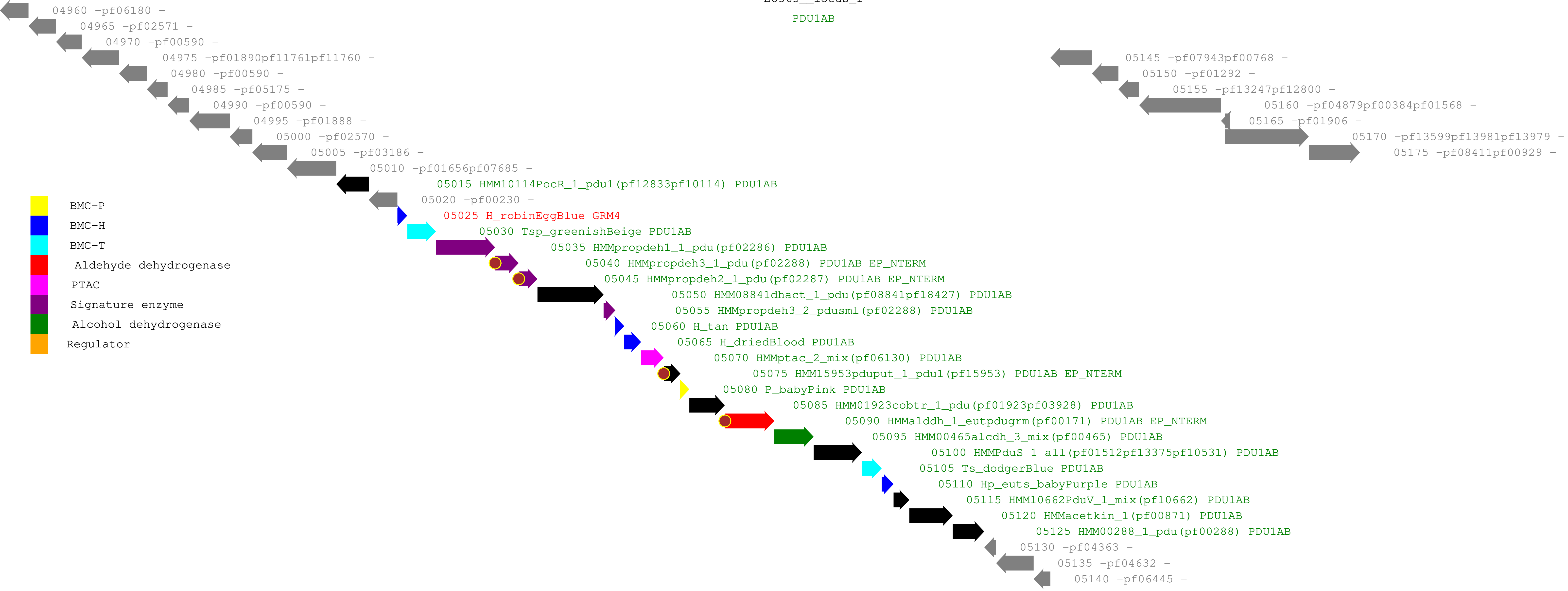

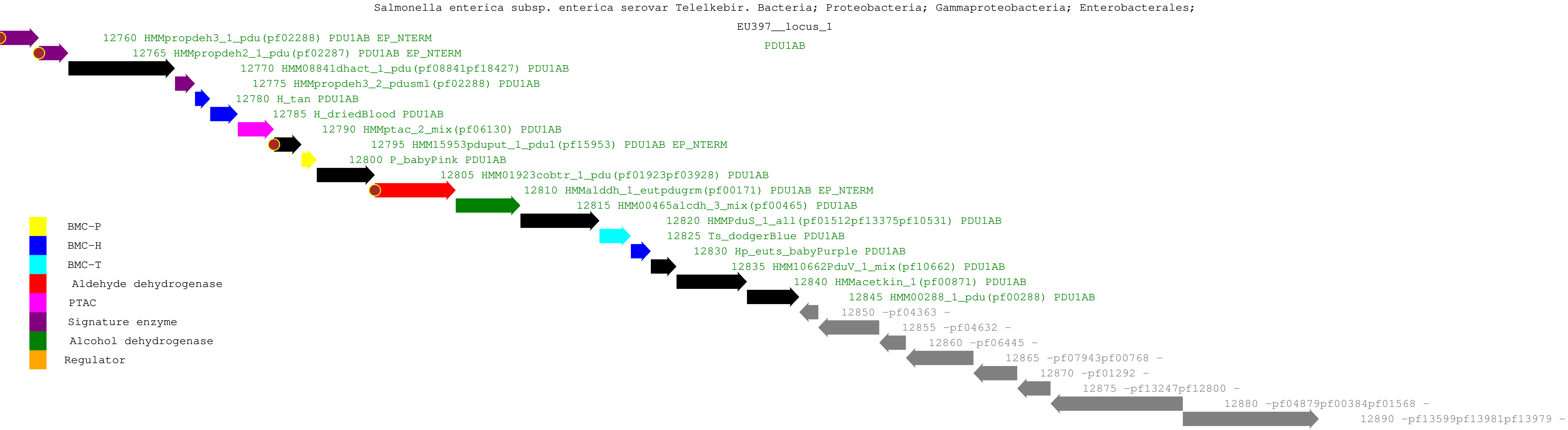

Salmonella enterica subsp. enterica serovar Telelkebir. Bacteria; Proteobacteria; Gammaproteobacteria; Enterobacterales;
EU397__locus_1
PDU1AB
12760 HMMpropdeh3_1_pdu(pf02288) PDU1AB EP_NTERM
12765 HMMpropdeh2_1_pdu(pf02287) PDU1AB EP_NTERM
12770 HMM08841dhact_1_pdu(pf08841pf18427) PDU1AB
12775 HMMpropdeh3_2_pdusml(pf02288) PDU1AB
12780 H_tan PDU1AB
12785 H_driedBlood PDU1AB
12790 HMMptac_2_mix(pf06130) PDU1AB
12795 HMM15953pduput_1_pdu1(pf15953) PDU1AB EP_NTERM
12800 P_babyPink PDU1AB
12805 HMM01923cobtr_1_pdu(pf01923pf03928) PDU1AB
12810 HMMaldh_1_eutpdugrm(pf00171) PDU1AB EP_NTERM
12815 HMM00465alcdh_3_mix(pf00465) PDU1AB
12820 HMMPduS_1_all(pf01512pf13375pf10531) PDU1AB
12825 Ts_dodgerBlue PDU1AB
12830 Hp_euts_babyPurple PDU1AB
12835 HMM10662PduV_1_mix(pf10662) PDU1AB
12840 HMMacetkin_1(pf00871) PDU1AB
12845 HMM00288_1_pdu(pf00288) PDU1AB
12850 -pf04363 -
12855 -pf04632 -
12860 -pf06445 -
12865 -pf07943pf00768 -
12870 -pf01292 -
12875 -pf13247pf12800 -
12880 -pf04879pf00384pf01568 -
12890 -pf13599pf13981pf13979 -
BMC-P
BMC-H
BMC-T
Aldehyde dehydrogenase
PTAC
Signature enzyme
Alcohol dehydrogenase
Regulator

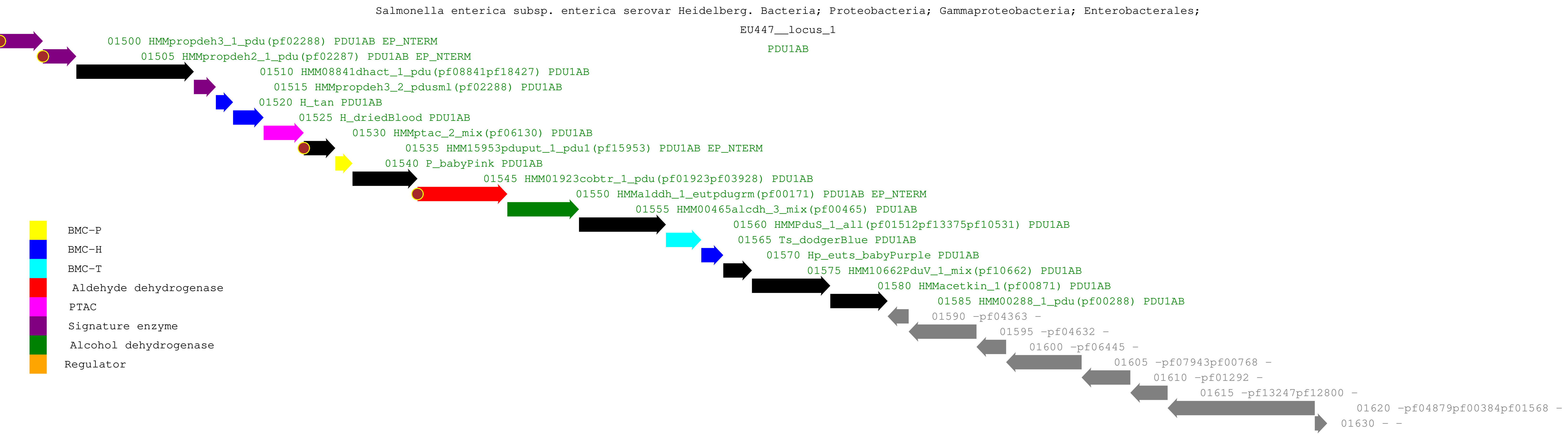
Salmonella enterica subsp. enterica serovar Heidelberg. Bacteria; Proteobacteria; Gammaproteobacteria; Enterobacterales;
EU447__locus_1
PDU1AB
01500 HMMpropdeh3_1_pdu(pf02288) PDU1AB EP_NTERM
01505 HMMpropdeh2_1_pdu(pf02287) PDU1AB EP_NTERM
01510 HMM08841dhact_1_pdu(pf08841pf18427) PDU1AB
01515 HMMpropdeh3_2_pdusml(pf02288) PDU1AB
01520 H_tan PDU1AB
01525 H_driedBlood PDU1AB
01530 HMMptac_2_mix(pf06130) PDU1AB
01535 HMM15953pduput_1_pdu1(pf15953) PDU1AB EP_NTERM
01540 P_babyPink PDU1AB
01545 HMM01923cobtr_1_pdu(pf01923pf03928) PDU1AB
01550 HMMalddh_1_eutpdugrm(pf00171) PDU1AB EP_NTERM
01555 HMM00465alcdh_3_mix(pf00465) PDU1AB
01560 HMMPduS_1_all(pf01512pf13375pf10531) PDU1AB
01565 Ts_dodgerBlue PDU1AB
01570 Hp_euts_babyPurple PDU1AB
01575 HMM10662PduV_1_mix(pf10662) PDU1AB
01580 HMMacetkin_1(pf00871) PDU1AB
01585 HMM00288_1_pdu(pf00288) PDU1AB
01590 -pf04363 -
01595 -pf04632 -
01600 -pf06445 -
01605 -pf07943pf00768 -
01610 -pf01292 -
01615 -pf13247pf12800 -
01620 -pf04879pf00384pf01568 -
01630 - -
BMC-P
BMC-H
BMC-T
Aldehyde dehydrogenase
PTAC
Signature enzyme
Alcohol dehydrogenase
Regulator

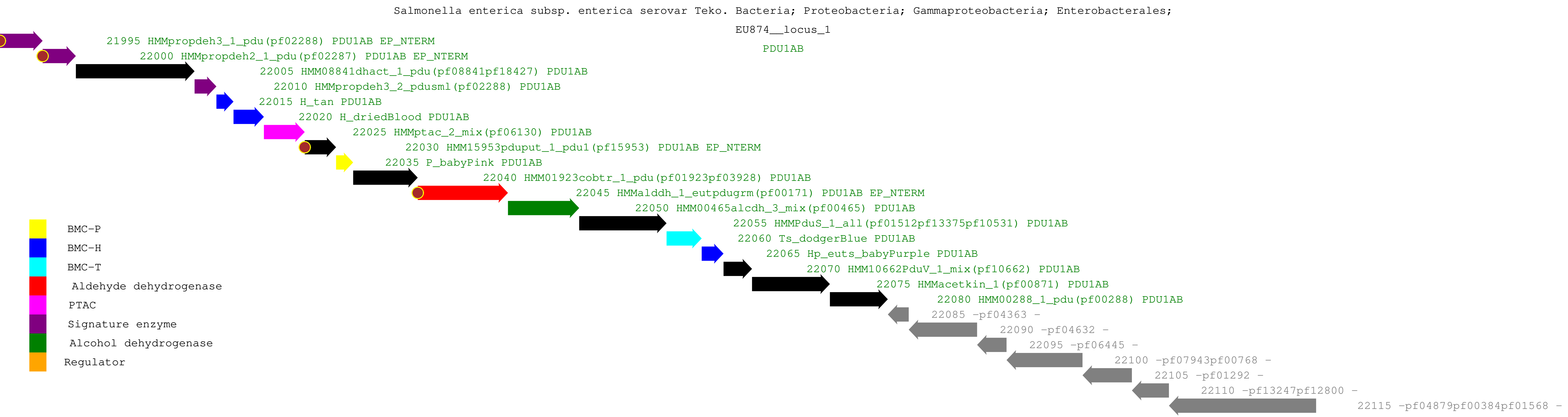

Salmonella enterica subsp. enterica serovar Teko. Bacteria; Proteobacteria; Gammaproteobacteria; Enterobacterales;
EU874__locus_1
PDU1AB
21995 HMMpropdeh3_1_pdu(pf02288) PDU1AB EP_NTERM
22000 HMMpropdeh2_1_pdu(pf02287) PDU1AB EP_NTERM
22005 HMM08841dhact_1_pdu(pf08841pf18427) PDU1AB
22010 HMMpropdeh3_2_pdusml(pf02288) PDU1AB
22015 H_tan PDU1AB
22020 H_driedBlood PDU1AB
22025 HMMptac_2_mix(pf06130) PDU1AB
22030 HMM15953pduput_1_pdu1(pf15953) PDU1AB EP_NTERM
22035 P_babyPink PDU1AB
22040 HMM01923cobtr_1_pdu(pf01923pf03928) PDU1AB
22045 HMMalddh_1_eutpdugrm(pf00171) PDU1AB EP_NTERM
22050 HMM00465alcdh_3_mix(pf00465) PDU1AB
22055 HMMPduS_1_all(pf01512pf13375pf10531) PDU1AB
22060 Ts_dodgerBlue PDU1AB
22065 Hp_euts_babyPurple PDU1AB
22070 HMM10662PduV_1_mix(pf10662) PDU1AB
22075 HMMacetkin_1(pf00871) PDU1AB
22080 HMM00288_1_pdu(pf00288) PDU1AB
22085 -pf04363 -
22090 -pf04632 -
22095 -pf06445 -
22100 -pf07943pf00768 -
22105 -pf01292 -
22110 -pf13247pf12800 -
22115 -pf04879pf00384pf01568 -
BMC-P
BMC-H
BMC-T
Aldehyde dehydrogenase
PTAC
Signature enzyme
Alcohol dehydrogenase
Regulator

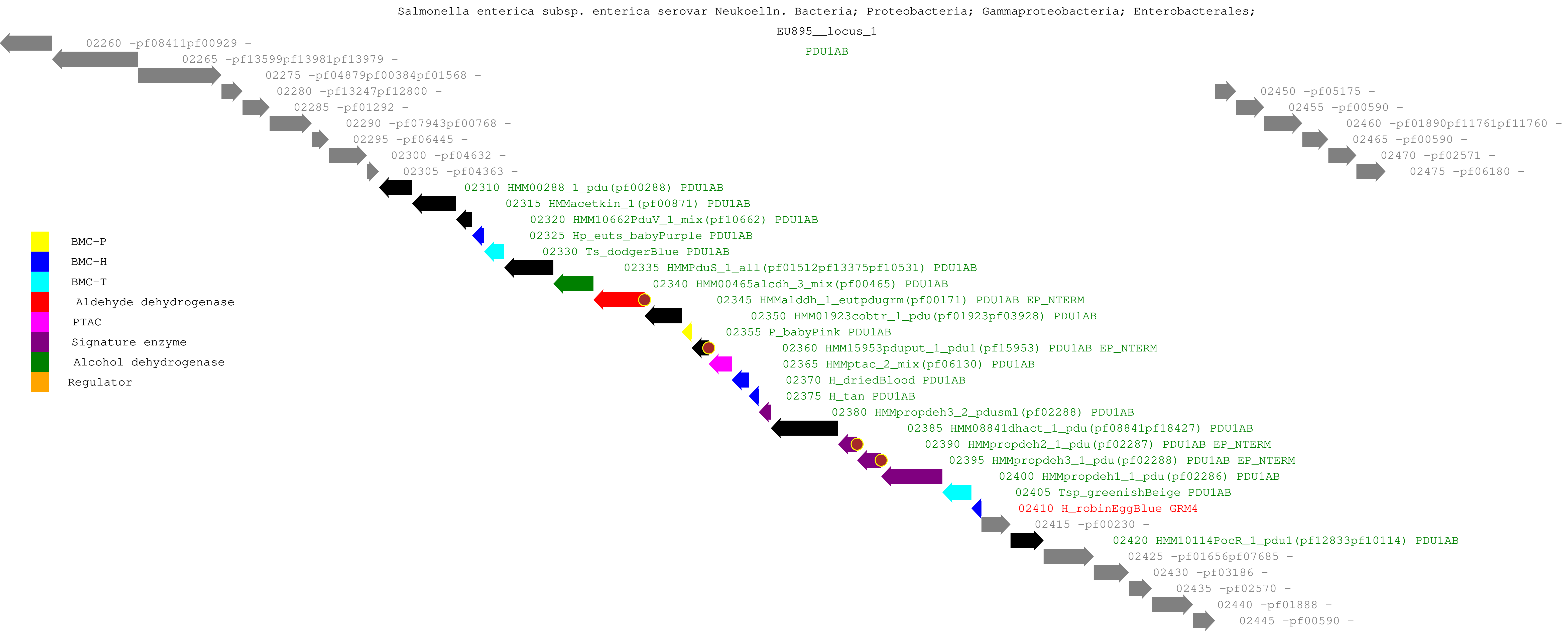

Salmonella enterica subsp. enterica serovar Neukoelln. Bacteria; Proteobacteria; Gammaproteobacteria; Enterobacterales;
EU895__locus_1
PDU1AB
02260 -pf08411pf00929 -
02265 -pf13599pf13981pf13979 -
02275 -pf04879pf00384pf01568 -
02280 -pf13247pf12800 -
02285 -pf01292 -
02290 -pf07943pf00768 -
02295 -pf06445 -
02300 -pf04632 -
02305 -pf04363 -
02310 HMM00288_1_pdu(pf00288) PDU1AB
02315 HMMacetkin_1(pf00871) PDU1AB
02320 HMM10662PduV_1_mix(pf10662) PDU1AB
02325 Hp_euts_babyPurple PDU1AB
02330 Ts_dodgerBlue PDU1AB
02335 HMMPduS_1_all(pf01512pf13375pf10531) PDU1AB
02340 HMM00465alcdh_3_mix(pf00465) PDU1AB
02345 HMMalddh_1_eutpdugrm(pf00171) PDU1AB EP_NTERM
02350 HMM01923cobtr_1_pdu(pf01923pf03928) PDU1AB
02355 P_babyPink PDU1AB
02360 HMM15953pduput_1_pdu1(pf15953) PDU1AB EP_NTERM
02365 HMMptac_2_mix(pf06130) PDU1AB
02370 H_driedBlood PDU1AB
02375 H_tan PDU1AB
02380 HMMpropdeh3_2_pdusml(pf02288) PDU1AB
02385 HMM08841dhact_1_pdu(pf08841pf18427) PDU1AB
02390 HMMpropdeh2_1_pdu(pf02287) PDU1AB EP_NTERM
02395 HMMpropdeh3_1_pdu(pf02288) PDU1AB EP_NTERM
02400 HMMpropdeh1_1_pdu(pf02286) PDU1AB
02405 Tsp_greenishBeige PDU1AB
02410 H_robinEggBlue GRM4
02415 -pf00230 -
02420 HMM10114PocR_1_pdu1(pf12833pf10114) PDU1AB
02425 -pf01656pf07685 -
02430 -pf03186 -
02435 -pf02570 -
02440 -pf01888 -
02445 -pf00590 -
02450 -pf05175 -
02455 -pf00590 -
02460 -pf01890pf11761pf11760 -
02465 -pf00590 -
02470 -pf02571 -
02475 -pf06180 -
BMC-P
BMC-H
BMC-T
Aldehyde dehydrogenase
PTAC
Signature enzyme
Alcohol dehydrogenase
Regulator

Salmonella enterica subsp. enterica serovar Stanleyville. Bacteria; Proteobacteria; Gammaproteobacteria; Enterobacterales;

EU930__locus_2

PDU1AB

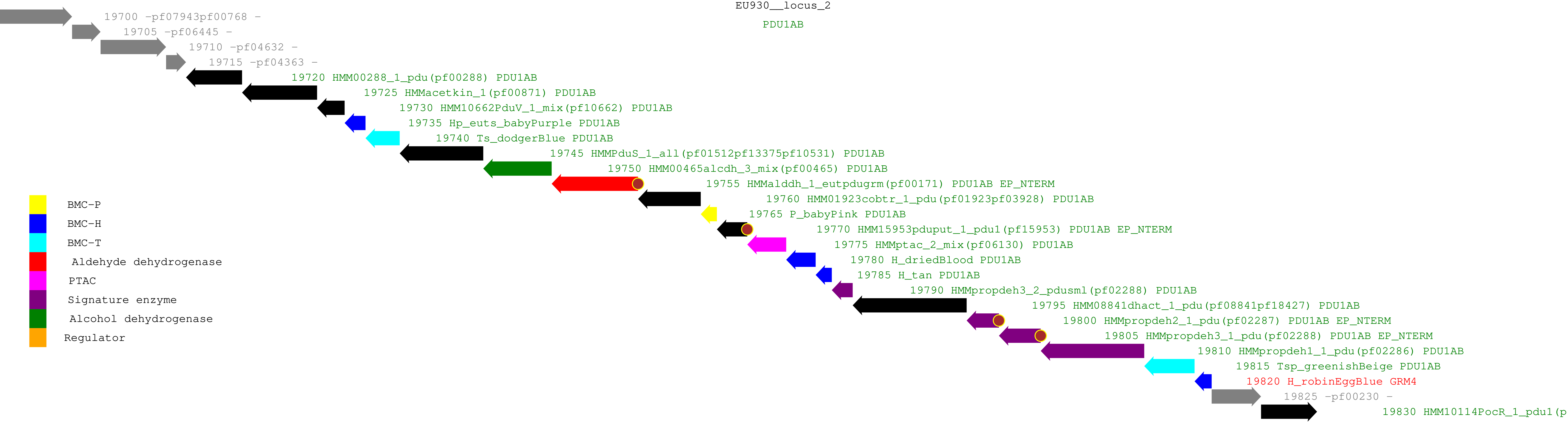

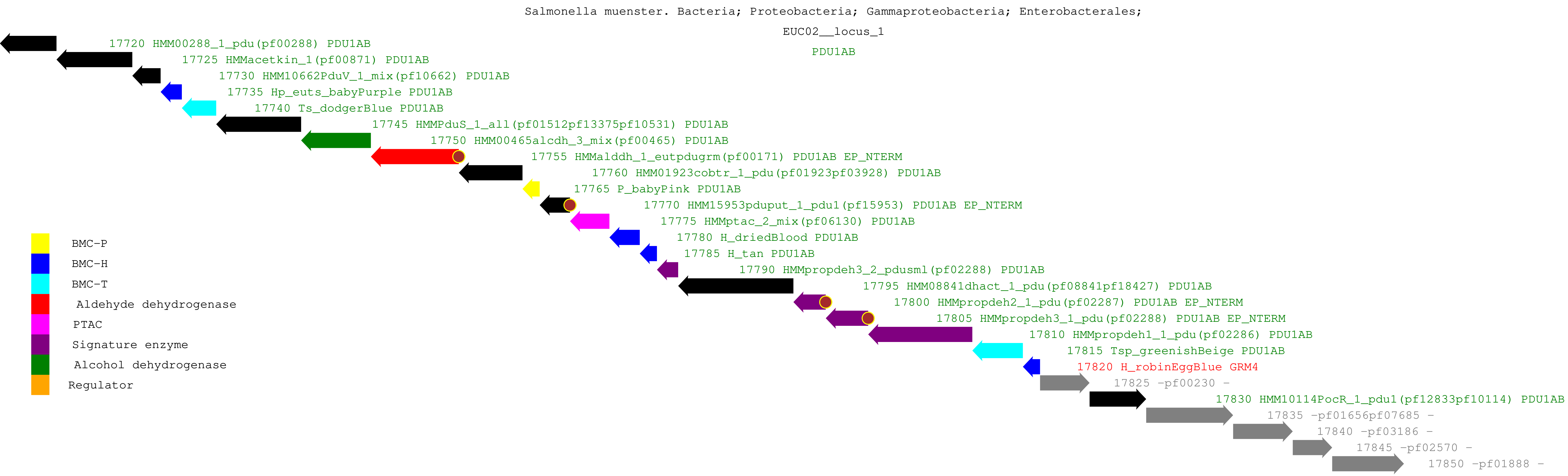
Salmonella muenster. Bacteria; Proteobacteria; Gammaproteobacteria; Enterobacterales;
EUC02__locus_1
PDU1AB
17720 HMM00288_1_pdu(pf00288) PDU1AB
17725 HMMacetkin_1(pf00871) PDU1AB
17730 HMM10662PduV_1_mix(pf10662) PDU1AB
17735 Hp_euts_babyPurple PDU1AB
17740 Ts_dodgerBlue PDU1AB
17745 HMMPduS_1_all(pf01512pf13375pf10531) PDU1AB
17750 HMM00465alcdh_3_mix(pf00465) PDU1AB
17755 HMMalddh_1_eutpdugrm(pf00171) PDU1AB EP_NTERM
17760 HMM01923cobtr_1_pdu(pf01923pf03928) PDU1AB
17765 P_babyPink PDU1AB
17770 HMM15953pduput_1_pdu1(pf15953) PDU1AB EP_NTERM
17775 HMMptac_2_mix(pf06130) PDU1AB
17780 H_driedBlood PDU1AB
17785 H_tan PDU1AB
17790 HMMpropdeh3_2_pdusml(pf02288) PDU1AB
17795 HMM08841dhact_1_pdu(pf08841pf18427) PDU1AB
17800 HMMpropdeh2_1_pdu(pf02287) PDU1AB EP_NTERM
17805 HMMpropdeh3_1_pdu(pf02288) PDU1AB EP_NTERM
17810 HMMpropdeh1_1_pdu(pf02286) PDU1AB
17815 Tsp_greenishBeige PDU1AB
17820 H_robinEggBlue GRM4
17825 -pf00230 -
17830 HMM10114PocR_1_pdu1(pf12833pf10114) PDU1AB
17835 -pf01656pf07685 -
17840 -pf03186 -
17845 -pf02570 -
17850 -pf01888 -
BMC-P
BMC-H
BMC-T
Aldehyde dehydrogenase
PTAC
Signature enzyme
Alcohol dehydrogenase
Regulator

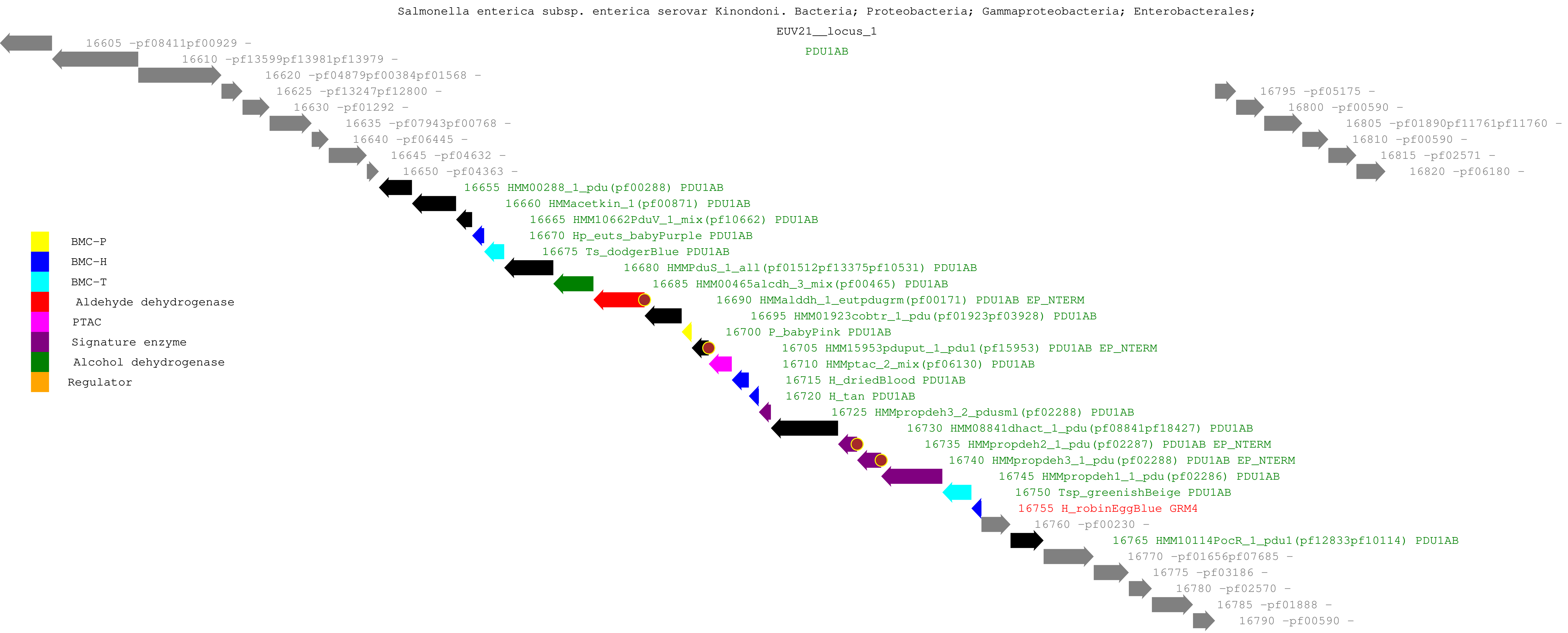

Salmonella enterica subsp. enterica serovar Kinondoni. Bacteria; Proteobacteria; Gammaproteobacteria; Enterobacterales;
EUV21__locus_1
PDU1AB
16605 -pf08411pf00929 -
16610 -pf13599pf13981pf13979 -
16620 -pf04879pf00384pf01568 -
16625 -pf13247pf12800 -
16630 -pf01292 -
16635 -pf07943pf00768 -
16640 -pf06445 -
16645 -pf04632 -
16650 -pf04363 -
16655 HMM00288_1_pdu(pf00288) PDU1AB
16660 HMMacetkin_1(pf00871) PDU1AB
16665 HMM10662PduV_1_mix(pf10662) PDU1AB
16670 Hp_euts_babyPurple PDU1AB
16675 Ts_dodgerBlue PDU1AB
16680 HMMPduS_1_all(pf01512pf13375pf10531) PDU1AB
16685 HMM00465alcdh_3_mix(pf00465) PDU1AB
16690 HMMalddh_1_eutpdugrm(pf00171) PDU1AB EP_NTERM
16695 HMM01923cobtr_1_pdu(pf01923pf03928) PDU1AB
16700 P_babyPink PDU1AB
16705 HMM15953pduput_1_pdu1(pf15953) PDU1AB EP_NTERM
16710 HMMptac_2_mix(pf06130) PDU1AB
16715 H_driedBlood PDU1AB
16720 H_tan PDU1AB
16725 HMMpropdeh3_2_pdusml(pf02288) PDU1AB
16730 HMM08841dhact_1_pdu(pf08841pf18427) PDU1AB
16735 HMMpropdeh2_1_pdu(pf02287) PDU1AB EP_NTERM
16740 HMMpropdeh3_1_pdu(pf02288) PDU1AB EP_NTERM
16745 HMMpropdeh1_1_pdu(pf02286) PDU1AB
16750 Tsp_greenishBeige PDU1AB
16755 H_robinEggBlue GRM4
16760 -pf00230 -
16765 HMM10114PocR_1_pdu1(pf12833pf10114) PDU1AB
16770 -pf01656pf07685 -
16775 -pf03186 -
16780 -pf02570 -
16785 -pf01888 -
16790 -pf00590 -
16795 -pf05175 -
16800 -pf00590 -
16805 -pf01890pf11761pf11760 -
16810 -pf00590 -
16815 -pf02571 -
16820 -pf06180 -
BMC-P
BMC-H
BMC-T
Aldehyde dehydrogenase
PTAC
Signature enzyme
Alcohol dehydrogenase
Regulator

Salmonella enterica subsp. enterica serovar Indiana. Bacteria; Proteobacteria; Gammaproteobacteria; Enterobacterales;

EUV68__locus_fragmented

PDU1AB

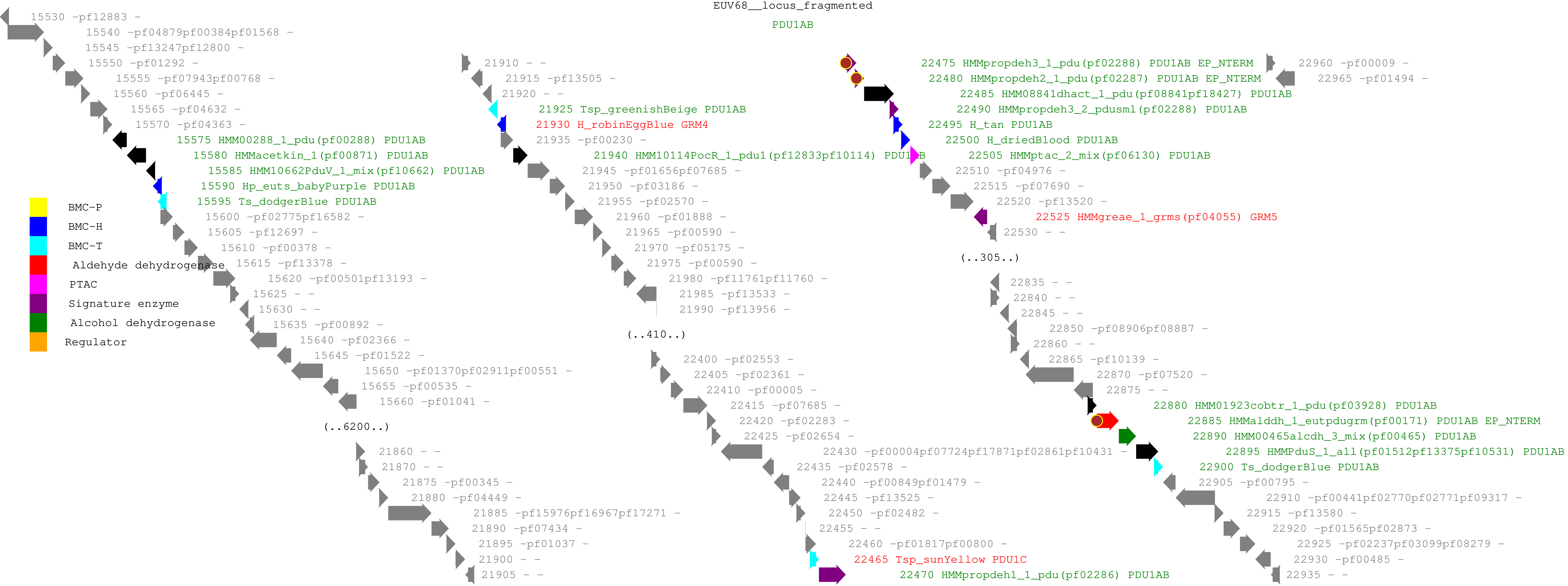

Salmonella enterica subsp. enterica serovar Carmel. Bacteria; Proteobacteria; Gammaproteobacteria; Enterobacterales;

EUW37__locus_1

PDU1AB

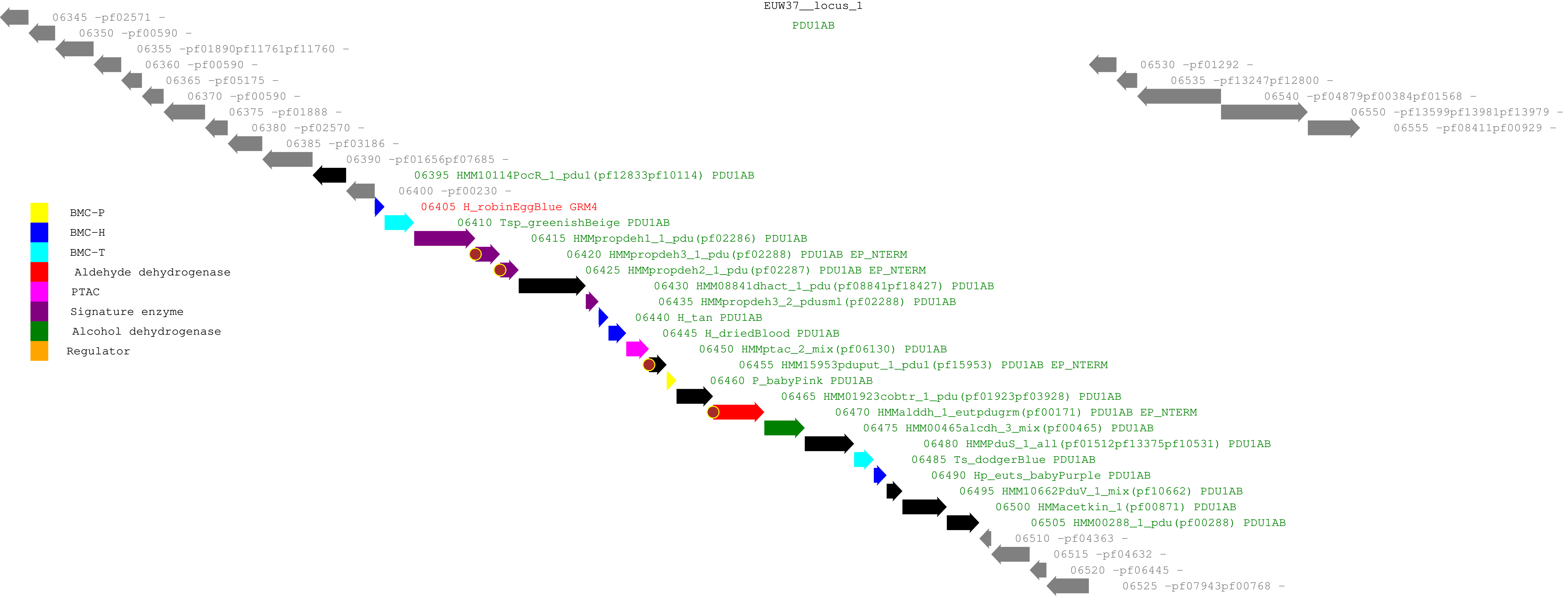

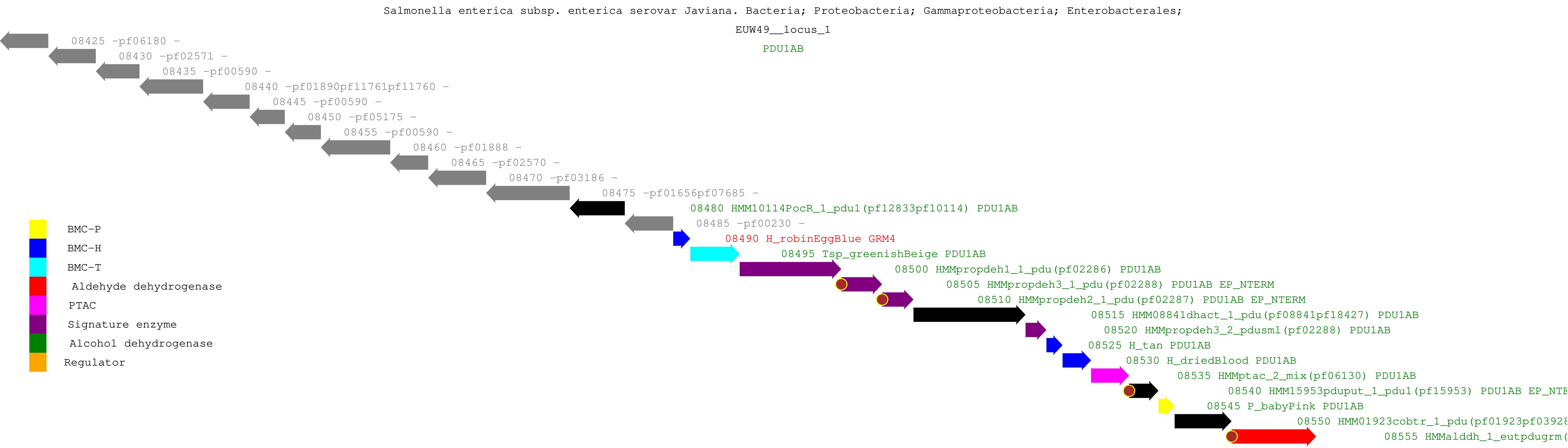
Salmonella enterica subsp. enterica serovar Javiana. Bacteria; Proteobacteria; Gammaproteobacteria; Enterobacterales;
EUW49__locus_1
PDU1AB
08425 -pf06180 -
08430 -pf02571 -
08435 -pf00590 -
08440 -pf01890pf11761pf11760 -
08445 -pf00590 -
08450 -pf05175 -
08455 -pf00590 -
08460 -pf01888 -
08465 -pf02570 -
08470 -pf03186 -
08475 -pf01656pf07685 -
08480 HMM10114PocR_1_pdu1(pf12833pf10114) PDU1AB
08485 -pf00230 -
08490 H_robinEggBlue GRM4
08495 Tsp_greenishBeige PDU1AB
08500 HMMpropdeh1_1_pdu(pf02286) PDU1AB
08505 HMMpropdeh3_1_pdu(pf02288) PDU1AB EP_NTERM
08510 HMMpropdeh2_1_pdu(pf02287) PDU1AB EP_NTERM
08515 HMM08841dhact_1_pdu(pf08841pf18427) PDU1AB
08520 HMMpropdeh3_2_pdusml(pf02288) PDU1AB
08525 H_tan PDU1AB
08530 H_driedBlood PDU1AB
08535 HMMptac_2_mix(pf06130) PDU1AB
08540 HMM15953pduput_1_pdu1(pf15953) PDU1AB EP_NTE
08545 P_babyPink PDU1AB
08550 HMM01923cobtr_1_pdu(pf01923pf0392
08555 HMMalddh_1_eutpdugrm(
BMC-P
BMC-H
BMC-T
Aldehyde dehydrogenase
PTAC
Signature enzyme
Alcohol dehydrogenase
Regulator

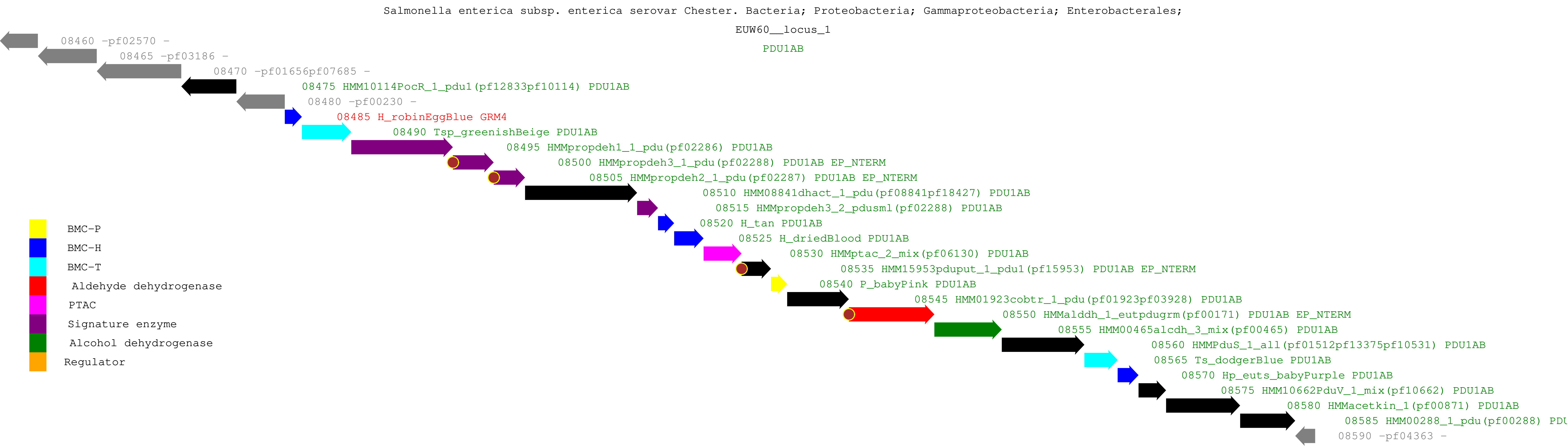
Salmonella enterica subsp. enterica serovar Chester. Bacteria; Proteobacteria; Gammaproteobacteria; Enterobacterales;
EUW60__locus_1
PDU1AB
08460 -pf02570 -
08465 -pf03186 -
08470 -pf01656pf07685 -
08475 HMM10114PocR_1_pdu1(pf12833pf10114) PDU1AB
08480 -pf00230 -
08485 H_robinEggBlue GRM4
08490 Tsp_greenishBeige PDU1AB
08495 HMMpropdeh1_1_pdu(pf02286) PDU1AB
08500 HMMpropdeh3_1_pdu(pf02288) PDU1AB EP_NTERM
08505 HMMpropdeh2_1_pdu(pf02287) PDU1AB EP_NTERM
08510 HMM08841dhact_1_pdu(pf08841pf18427) PDU1AB
08515 HMMpropdeh3_2_pdusml(pf02288) PDU1AB
08520 H_tan PDU1AB
08525 H_driedBlood PDU1AB
08530 HMMptac_2_mix(pf06130) PDU1AB
08535 HMM15953pduput_1_pdu1(pf15953) PDU1AB EP_NTERM
08540 P_babyPink PDU1AB
08545 HMM01923cobtr_1_pdu(pf01923pf03928) PDU1AB
08550 HMMaldh_1_eutpdugrm(pf00171) PDU1AB EP_NTERM
08555 HMM00465alcdh_3_mix(pf00465) PDU1AB
08560 HMMPduS_1_all(pf01512pf13375pf10531) PDU1AB
08565 Ts_dodgerBlue PDU1AB
08570 Hp_euts_babyPurple PDU1AB
08575 HMM10662PduV_1_mix(pf10662) PDU1AB
08580 HMMacetkin_1(pf00871) PDU1AB
08585 HMM00288_1_pdu(pf00288) PDU
08590 -pf04363 -
BMC-P
BMC-H
BMC-T
Aldehyde dehydrogenase
PTAC
Signature enzyme
Alcohol dehydrogenase
Regulator

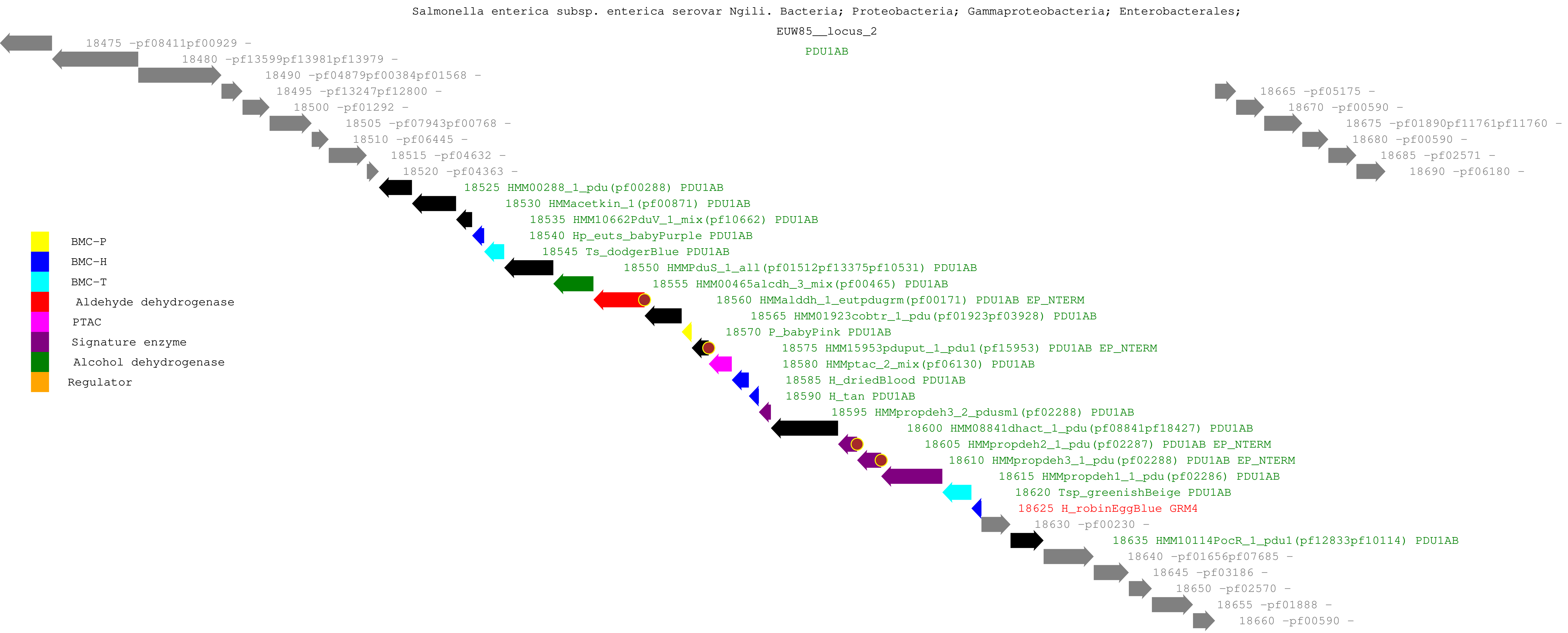
Salmonella enterica subsp. enterica serovar Ngili. Bacteria; Proteobacteria; Gammaproteobacteria; Enterobacterales;
EUW85__locus_2
PDU1AB
18475 -pf08411pf00929 -
18480 -pf13599pf13981pf13979 -
18490 -pf04879pf00384pf01568 -
18495 -pf13247pf12800 -
18500 -pf01292 -
18505 -pf07943pf00768 -
18510 -pf06445 -
18515 -pf04632 -
18520 -pf04363 -
18525 HMM00288_1_pdu(pf00288) PDU1AB
18530 HMMacetkin_1(pf00871) PDU1AB
18535 HMM10662PduV_1_mix(pf10662) PDU1AB
18540 Hp_euts_babyPurple PDU1AB
18545 Ts_dodgerBlue PDU1AB
18550 HMMPduS_1_all(pf01512pf13375pf10531) PDU1AB
18555 HMM00465alcdh_3_mix(pf00465) PDU1AB
18560 HMMalddh_1_eutpdugrm(pf00171) PDU1AB EP_NTERM
18565 HMM01923cobtr_1_pdu(pf01923pf03928) PDU1AB
18570 P_babyPink PDU1AB
18575 HMM15953pduput_1_pdu1(pf15953) PDU1AB EP_NTERM
18580 HMMptac_2_mix(pf06130) PDU1AB
18585 H_driedBlood PDU1AB
18590 H_tan PDU1AB
18595 HMMpropdeh3_2_pdusml(pf02288) PDU1AB
18600 HMM08841dhact_1_pdu(pf08841pf18427) PDU1AB
18605 HMMpropdeh2_1_pdu(pf02287) PDU1AB EP_NTERM
18610 HMMpropdeh3_1_pdu(pf02288) PDU1AB EP_NTERM
18615 HMMpropdeh1_1_pdu(pf02286) PDU1AB
18620 Tsp_greenishBeige PDU1AB
18625 H_robinEggBlue GRM4
18630 -pf00230 -
18635 HMM10114PocR_1_pdu1(pf12833pf10114) PDU1AB
18640 -pf01656pf07685 -
18645 -pf03186 -
18650 -pf02570 -
18655 -pf01888 -
18660 -pf00590 -
18665 -pf05175 -
18670 -pf00590 -
18675 -pf01890pf11761pf11760 -
18680 -pf00590 -
18685 -pf02571 -
18690 -pf06180 -
BMC-P
BMC-H
BMC-T
Aldehyde dehydrogenase
PTAC
Signature enzyme
Alcohol dehydrogenase
Regulator

Salmonella enterica subsp. enterica serovar Reading. Bacteria; Proteobacteria; Gammaproteobacteria; Enterobacterales;

EUX01__locus_1

PDU1AB

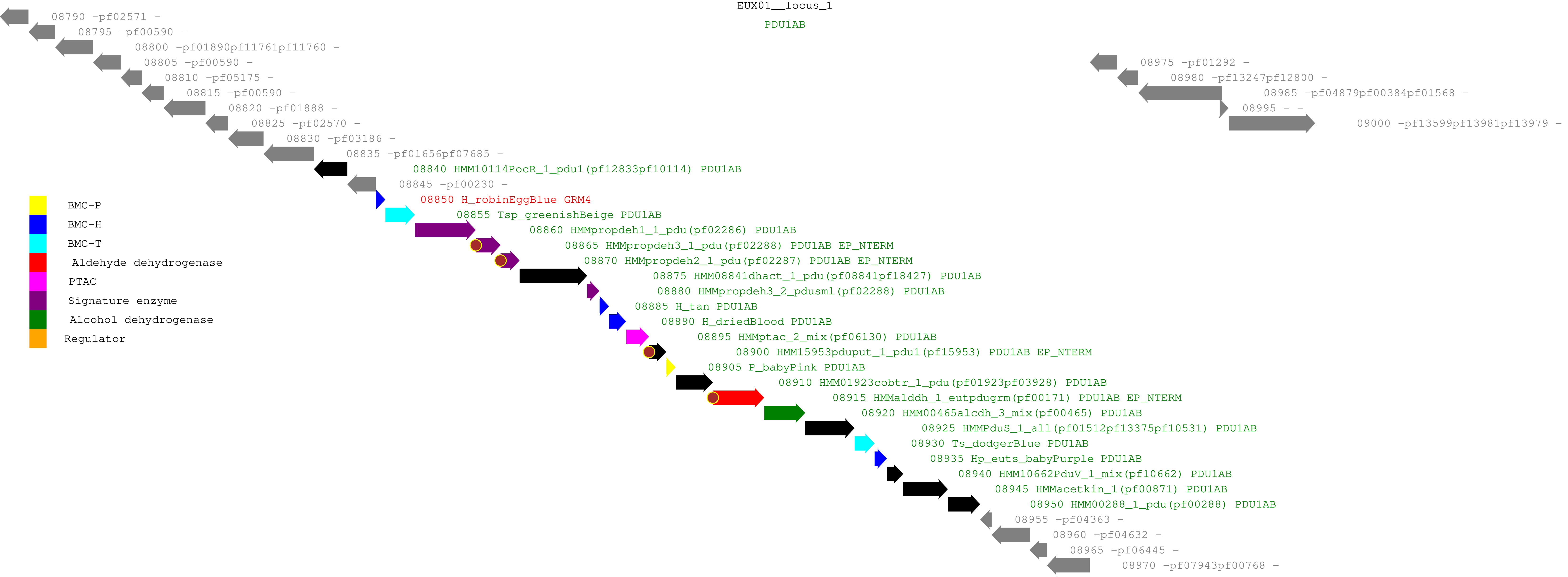

Salmonella enterica subsp. enterica serovar Ituri. Bacteria; Proteobacteria; Gammaproteobacteria; Enterobacterales;

EVG97__locus_2

PDU1AB

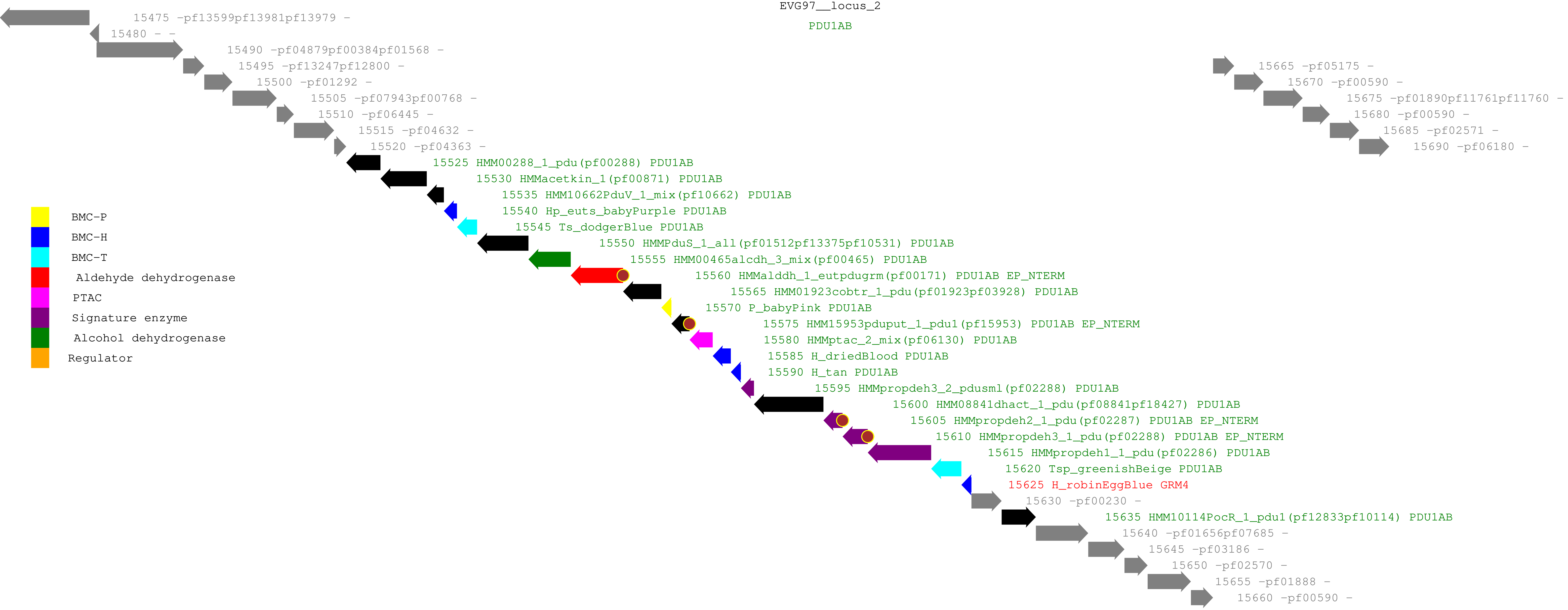

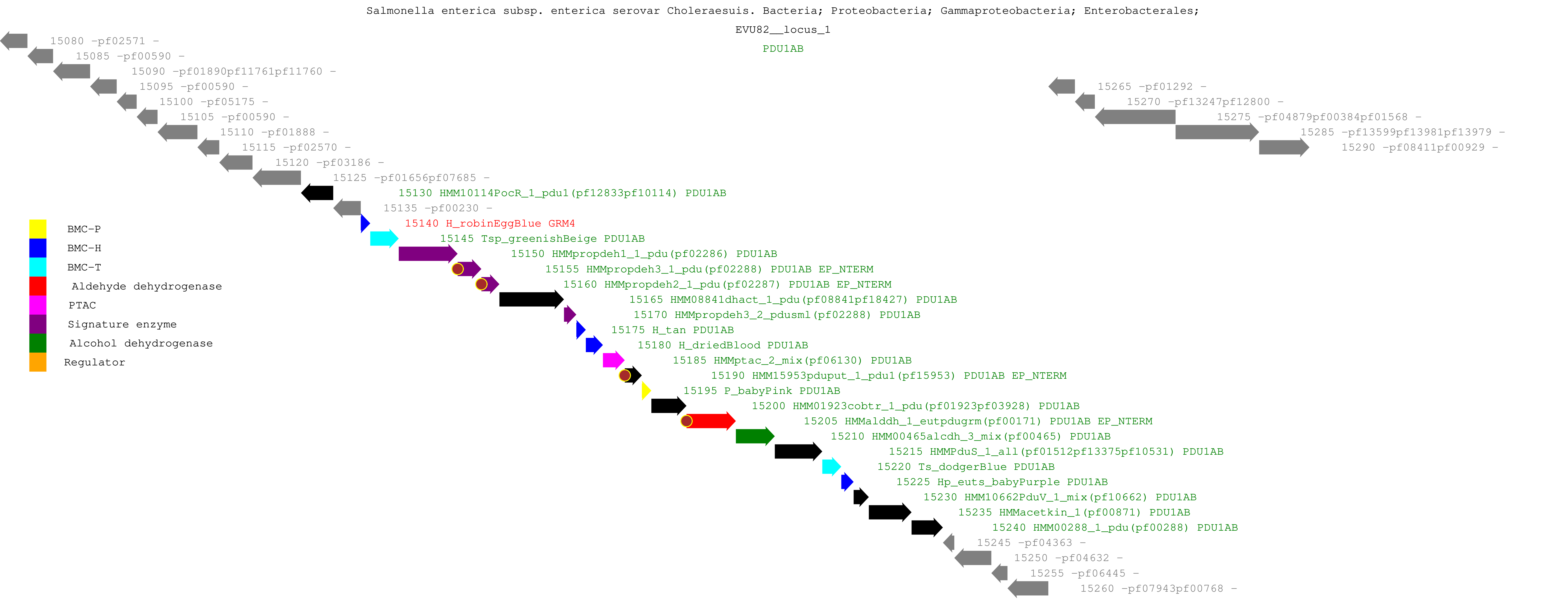
Salmonella enterica subsp. enterica serovar Choleraesuis. Bacteria; Proteobacteria; Gammaproteobacteria; Enterobacterales;
EVU82__locus_1
PDU1AB
15080 -pf02571 -
15085 -pf00590 -
15090 -pf01890pf11761pf11760 -
15095 -pf00590 -
15100 -pf05175 -
15105 -pf00590 -
15110 -pf01888 -
15115 -pf02570 -
15120 -pf03186 -
15125 -pf01656pf07685 -
15130 HMM10114PocR_1_pdu1(pf12833pf10114) PDU1AB
15135 -pf00230 -
15140 H_robinEggBlue GRM4
15145 Tsp_greenishBeige PDU1AB
15150 HMMpropdeh1_1_pdu(pf02286) PDU1AB
15155 HMMpropdeh3_1_pdu(pf02288) PDU1AB EP_NTERM
15160 HMMpropdeh2_1_pdu(pf02287) PDU1AB EP_NTERM
15165 HMM08841dhact_1_pdu(pf08841pf18427) PDU1AB
15170 HMMpropdeh3_2_pdusml(pf02288) PDU1AB
15175 H_tan PDU1AB
15180 H_driedBlood PDU1AB
15185 HMMptac_2_mix(pf06130) PDU1AB
15190 HMM15953pduput_1_pdu1(pf15953) PDU1AB EP_NTERM
15195 P_babyPink PDU1AB
15200 HMM01923cobtr_1_pdu(pf01923pf03928) PDU1AB
15205 HMMalddh_1_eutpdugrm(pf00171) PDU1AB EP_NTERM
15210 HMM00465alcdh_3_mix(pf00465) PDU1AB
15215 HMMPduS_1_all(pf01512pf13375pf10531) PDU1AB
15220 Ts_dodgerBlue PDU1AB
15225 Hp_euts_babyPurple PDU1AB
15230 HMM10662PduV_1_mix(pf10662) PDU1AB
15235 HMMacetkin_1(pf00871) PDU1AB
15240 HMM00288_1_pdu(pf00288) PDU1AB
15245 -pf04363 -
15250 -pf04632 -
15255 -pf06445 -
15260 -pf07943pf00768 -
15265 -pf01292 -
15270 -pf13247pf12800 -
15275 -pf04879pf00384pf01568 -
15285 -pf13599pf13981pf13979 -
15290 -pf08411pf00929 -
BMC-P
BMC-H
BMC-T
Aldehyde dehydrogenase
PTAC
Signature enzyme
Alcohol dehydrogenase
Regulator

Salmonella enterica subsp. enterica serovar Leoben. Bacteria; Proteobacteria; Gammaproteobacteria; Enterobacterales;

EW399__locus_1_main_sat

PDU1AB

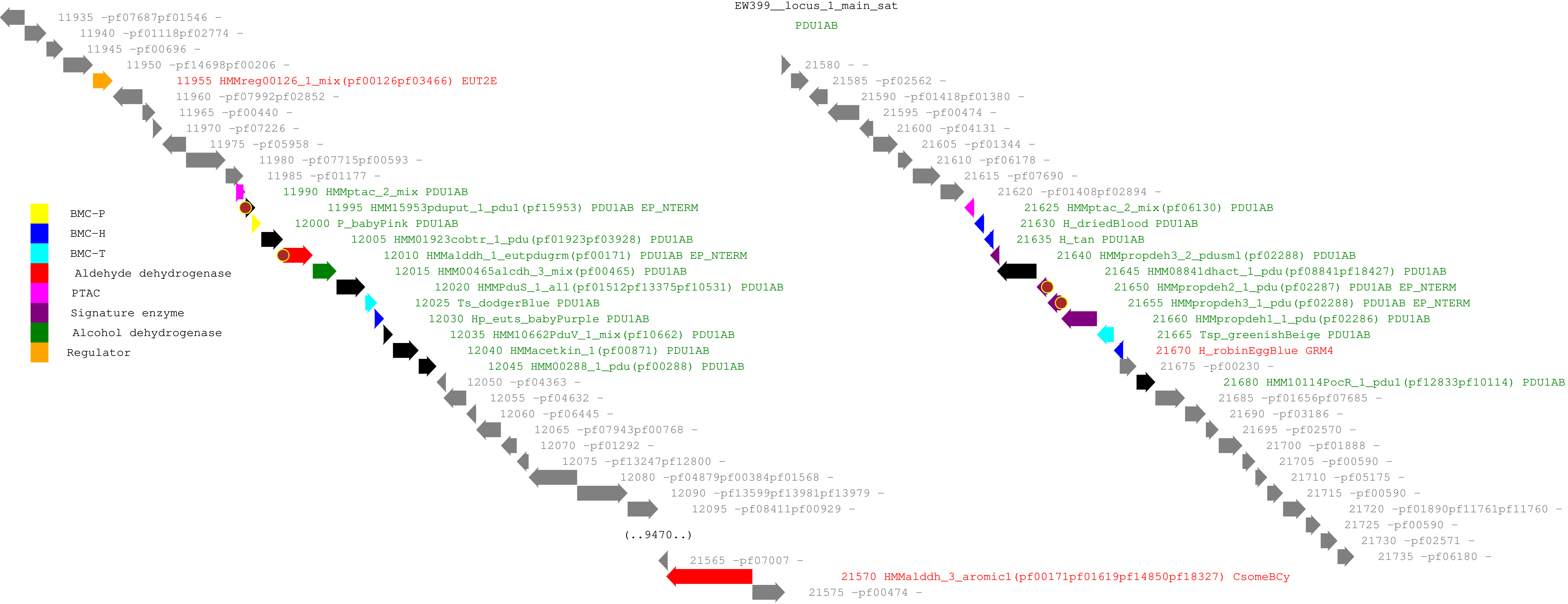

Salmonella enterica subsp. enterica serovar Carno. Bacteria; Proteobacteria; Gammaproteobacteria; Enterobacterales;

EW411__locus_1

PDU1AB

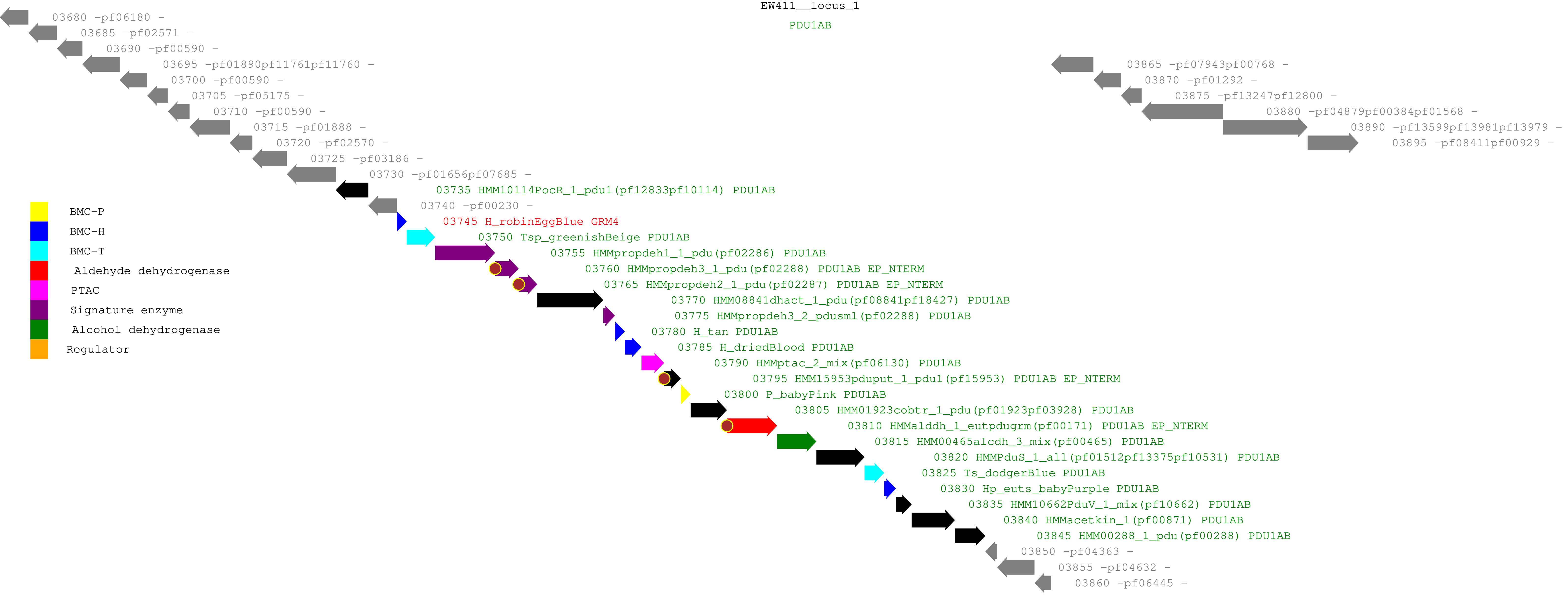

Salmonella enterica subsp. enterica serovar Kingabwa. Bacteria; Proteobacteria; Gammaproteobacteria; Enterobacterales;

EW439__locus_1_main_sat

PDU1AB

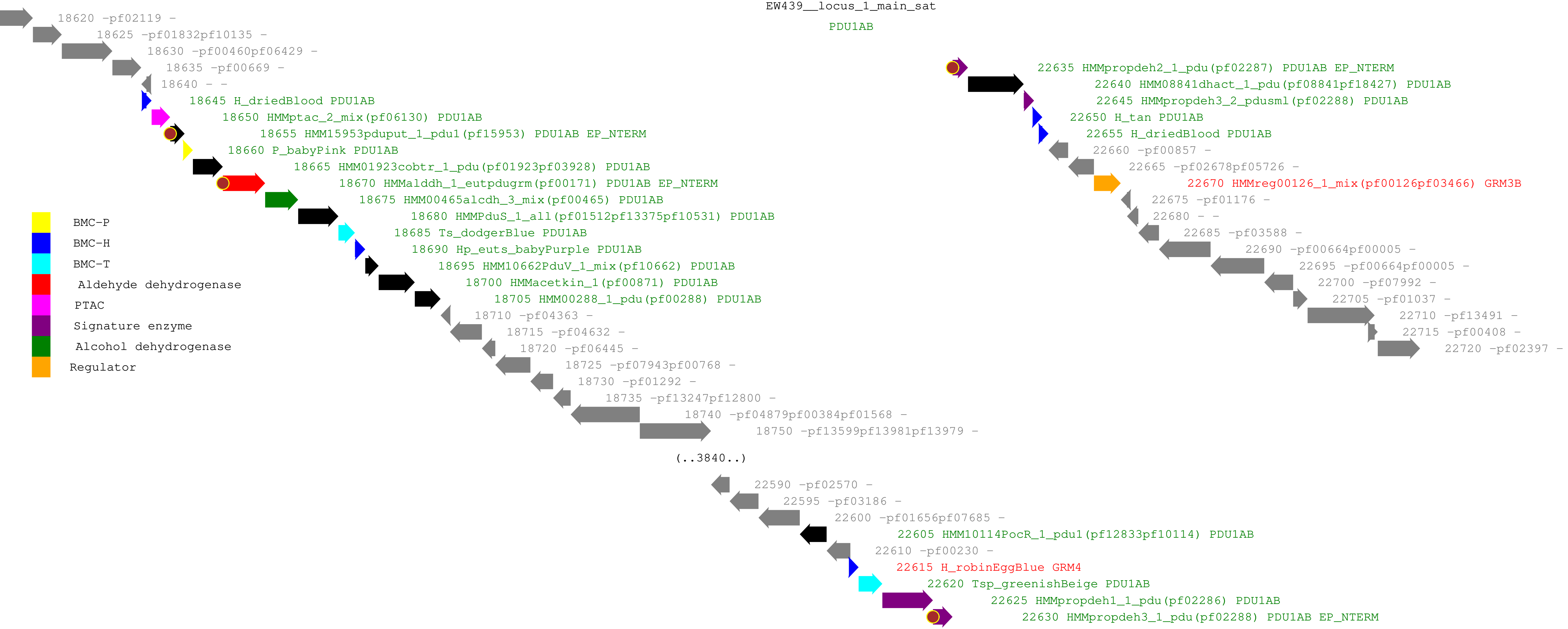

Salmonella enterica subsp. enterica serovar Singapore. Bacteria; Proteobacteria; Gammaproteobacteria; Enterobacterales;

EW464__locus_1

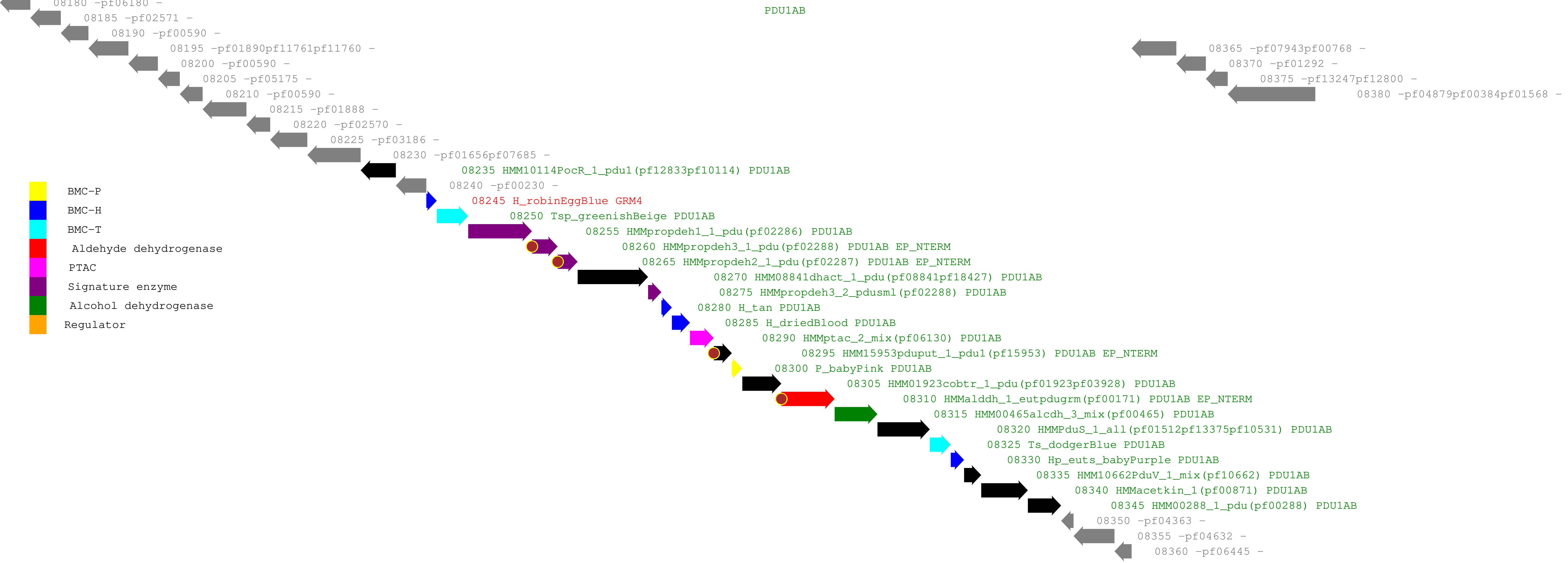

Salmonella enterica subsp. enterica serovar Stockholm. Bacteria; Proteobacteria; Gammaproteobacteria; Enterobacterales;

EW938__locus_1

PDU1AB

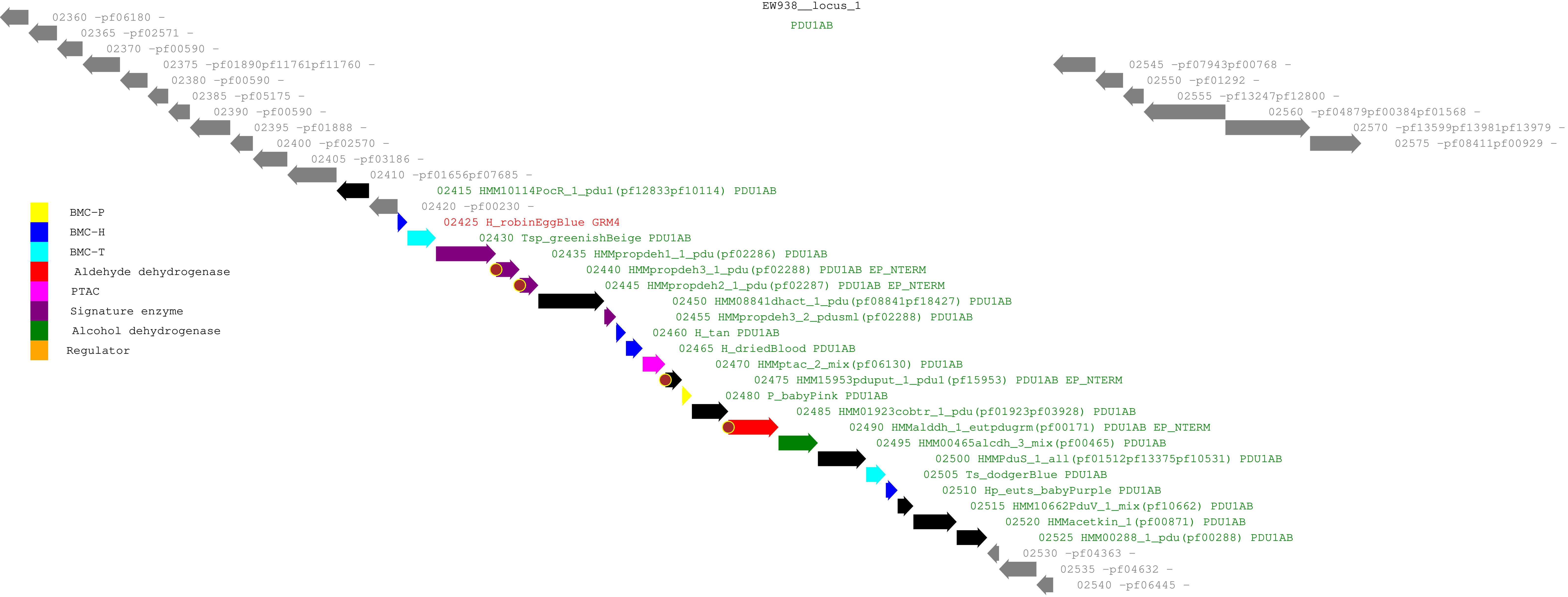

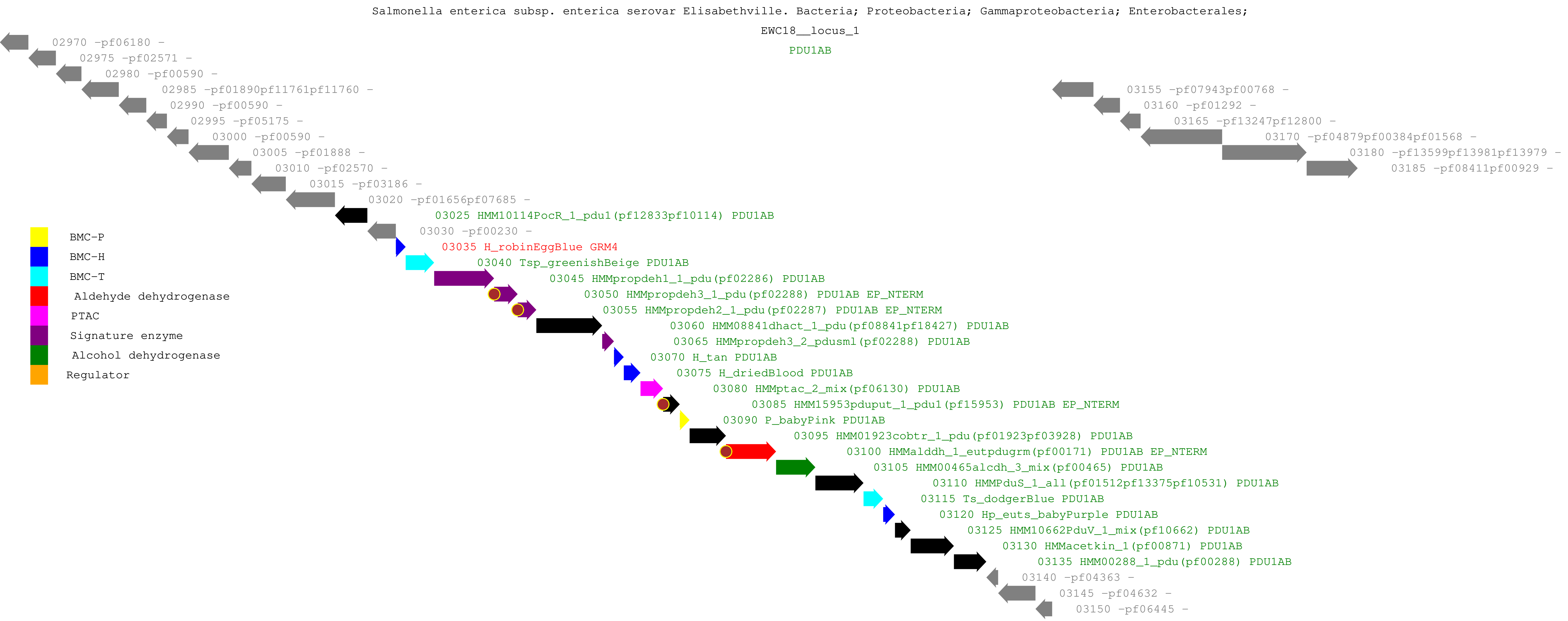

Salmonella enterica subsp. enterica serovar Elisabethville. Bacteria; Proteobacteria; Gammaproteobacteria; Enterobacterales;
EWC18__locus_1
PDU1AB
02970 -pf06180 -
02975 -pf02571 -
02980 -pf00590 -
02985 -pf01890pf11761pf11760 -
02990 -pf00590 -
02995 -pf05175 -
03000 -pf00590 -
03005 -pf01888 -
03010 -pf02570 -
03015 -pf03186 -
03020 -pf01656pf07685 -
03025 HMM10114PocR_1_pdu1(pf12833pf10114) PDU1AB
03030 -pf00230 -
03035 H_robinEggBlue GRM4
03040 Tsp_greenishBeige PDU1AB
03045 HMMpropdeh1_1_pdu(pf02286) PDU1AB
03050 HMMpropdeh3_1_pdu(pf02288) PDU1AB EP_NTERM
03055 HMMpropdeh2_1_pdu(pf02287) PDU1AB EP_NTERM
03060 HMM08841dhact_1_pdu(pf08841pf18427) PDU1AB
03065 HMMpropdeh3_2_pdusml(pf02288) PDU1AB
03070 H_tan PDU1AB
03075 H_driedBlood PDU1AB
03080 HMMptac_2_mix(pf06130) PDU1AB
03085 HMM15953pduput_1_pdu1(pf15953) PDU1AB EP_NTERM
03090 P_babyPink PDU1AB
03095 HMM01923cobtr_1_pdu(pf01923pf03928) PDU1AB
03100 HMMalddh_1_eutpdugrm(pf00171) PDU1AB EP_NTERM
03105 HMM00465alcdh_3_mix(pf00465) PDU1AB
03110 HMMPduS_1_all(pf01512pf13375pf10531) PDU1AB
03115 Ts_dodgerBlue PDU1AB
03120 Hp_euts_babyPurple PDU1AB
03125 HMM10662PduV_1_mix(pf10662) PDU1AB
03130 HMMacetkin_1(pf00871) PDU1AB
03135 HMM00288_1_pdu(pf00288) PDU1AB
03140 -pf04363 -
03145 -pf04632 -
03150 -pf06445 -
03155 -pf07943pf00768 -
03160 -pf01292 -
03165 -pf13247pf12800 -
03170 -pf04879pf00384pf01568 -
03180 -pf13599pf13981pf13979 -
03185 -pf08411pf00929 -
BMC-P
BMC-H
BMC-T
Aldehyde dehydrogenase
PTAC
Signature enzyme
Alcohol dehydrogenase
Regulator

Salmonella enterica subsp. enterica serovar Nottingham. Bacteria; Proteobacteria; Gammaproteobacteria; Enterobacterales;

EWC41__locus_2

PDU1AB

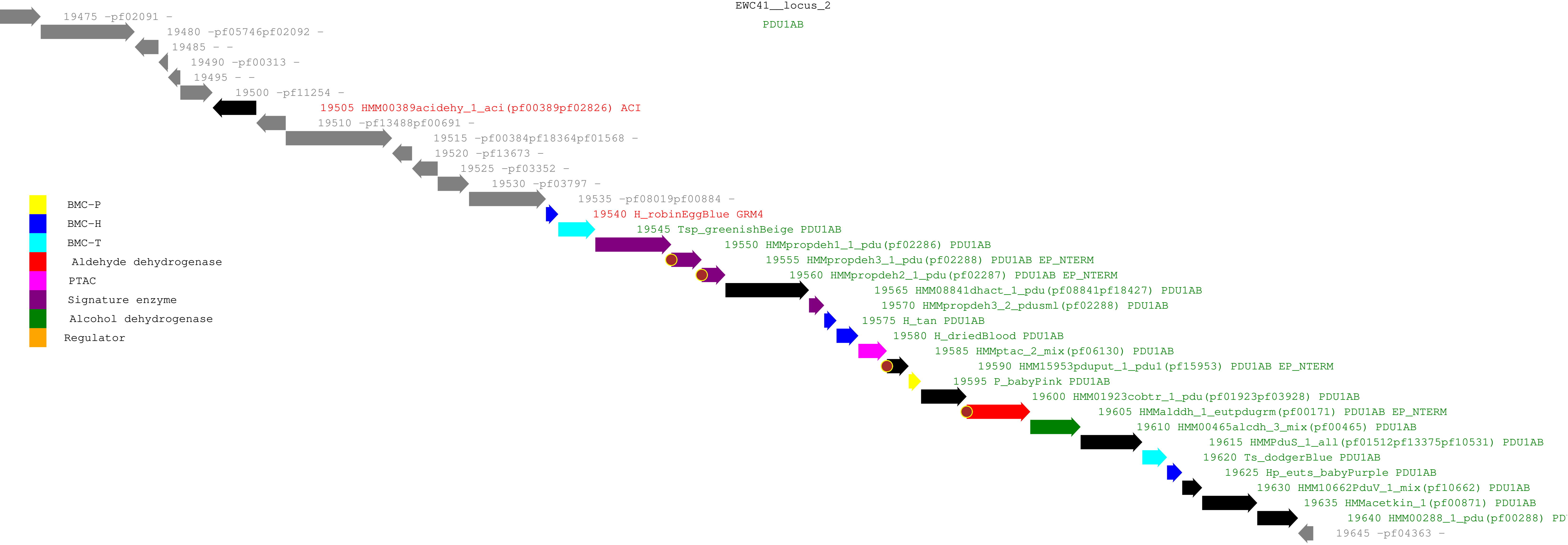

Salmonella enterica subsp. enterica serovar Adelaide. Bacteria; Proteobacteria; Gammaproteobacteria; Enterobacterales;

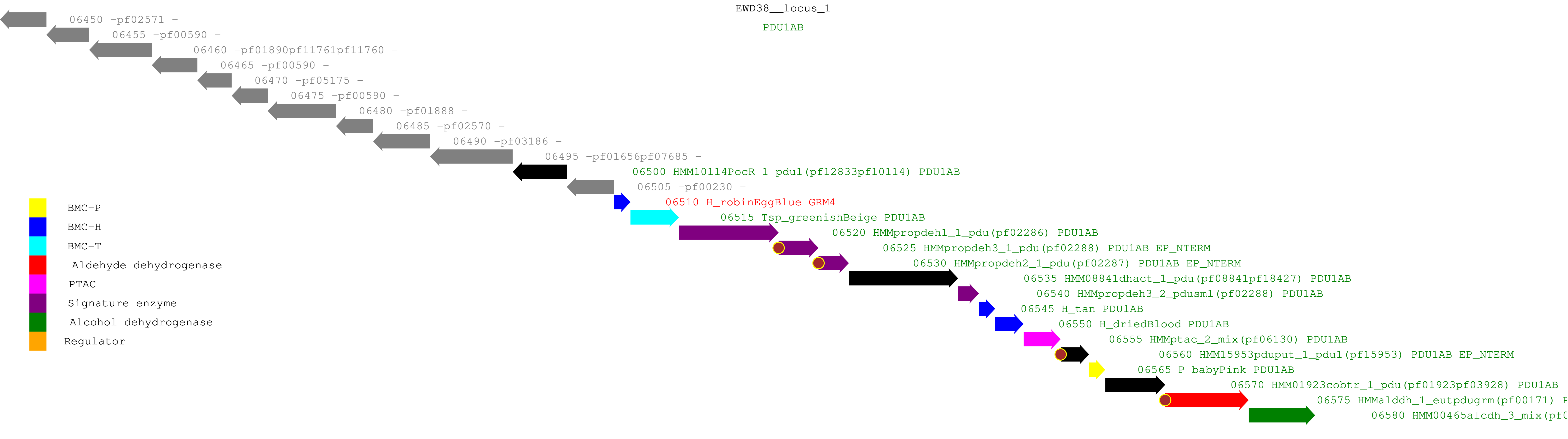

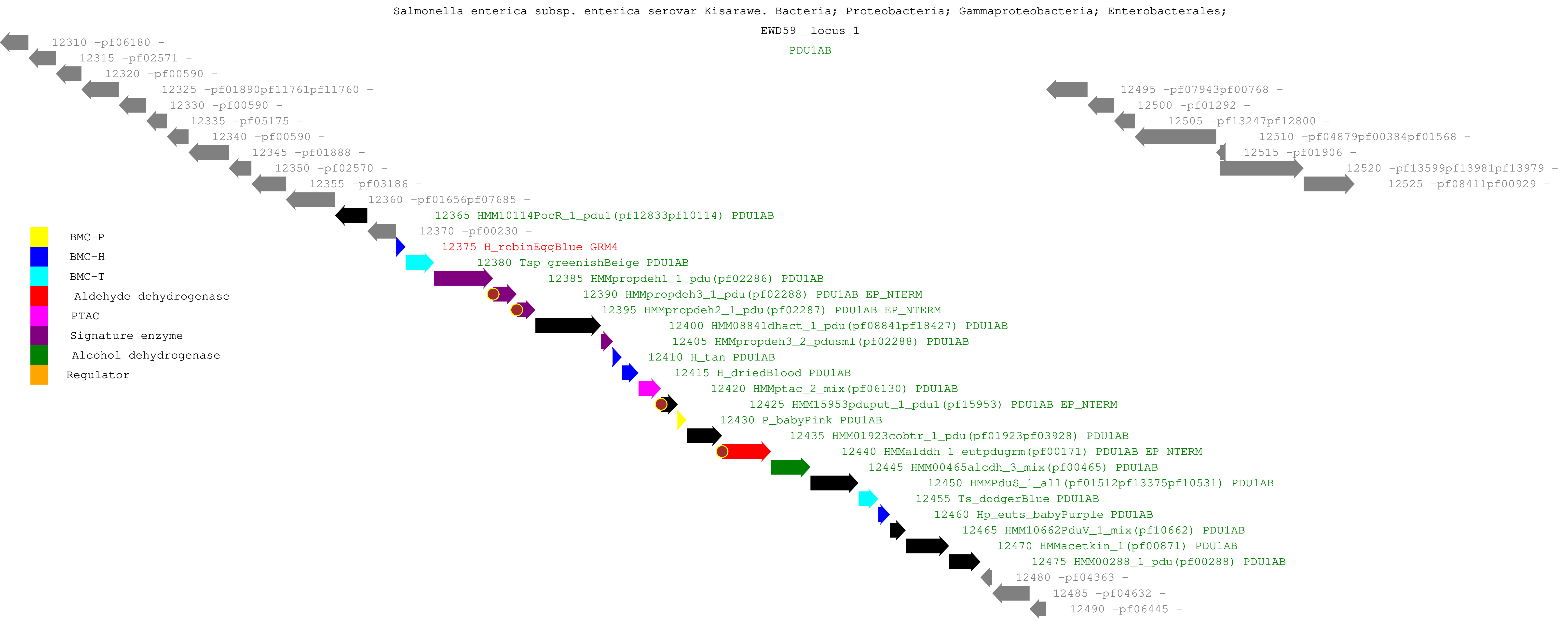
Salmonella enterica subsp. enterica serovar Kisarawe. Bacteria; Proteobacteria; Gammaproteobacteria; Enterobacterales;
EWD59__locus_1
PDU1AB
12310 -pf06180 -
12315 -pf02571 -
12320 -pf00590 -
12325 -pf01890pf11761pf11760 -
12330 -pf00590 -
12335 -pf05175 -
12340 -pf00590 -
12345 -pf01888 -
12350 -pf02570 -
12355 -pf03186 -
12360 -pf01656pf07685 -
12365 HMM10114PocR_1_pdu1(pf12833pf10114) PDU1AB
12370 -pf00230 -
12375 H_robinEggBlue GRM4
12380 Tsp_greenishBeige PDU1AB
12385 HMMpropdeh1_1_pdu(pf02286) PDU1AB
12390 HMMpropdeh3_1_pdu(pf02288) PDU1AB EP_NTERM
12395 HMMpropdeh2_1_pdu(pf02287) PDU1AB EP_NTERM
12400 HMM08841dhact_1_pdu(pf08841pf18427) PDU1AB
12405 HMMpropdeh3_2_pdusml(pf02288) PDU1AB
12410 H_tan PDU1AB
12415 H_driedBlood PDU1AB
12420 HMMptac_2_mix(pf06130) PDU1AB
12425 HMM15953pduput_1_pdu1(pf15953) PDU1AB EP_NTERM
12430 P_babyPink PDU1AB
12435 HMM01923cobtr_1_pdu(pf01923pf03928) PDU1AB
12440 HMMalddh_1_eutpdugrm(pf00171) PDU1AB EP_NTERM
12445 HMM00465alcdh_3_mix(pf00465) PDU1AB
12450 HMMPduS_1_all(pf01512pf13375pf10531) PDU1AB
12455 Ts_dodgerBlue PDU1AB
12460 Hp_euts_babyPurple PDU1AB
12465 HMM10662PduV_1_mix(pf10662) PDU1AB
12470 HMMacetkin_1(pf00871) PDU1AB
12475 HMM00288_1_pdu(pf00288) PDU1AB
12480 -pf04363 -
12485 -pf04632 -
12490 -pf06445 -
12495 -pf07943pf00768 -
12500 -pf01292 -
12505 -pf13247pf12800 -
12510 -pf04879pf00384pf01568 -
12515 -pf01906 -
12520 -pf13599pf13981pf13979 -
12525 -pf08411pf00929 -
BMC-P
BMC-H
BMC-T
Aldehyde dehydrogenase
PTAC
Signature enzyme
Alcohol dehydrogenase
Regulator

Salmonella enterica subsp. enterica serovar Praha. Bacteria; Proteobacteria; Gammaproteobacteria; Enterobacterales;

EWD62__locus_1

PDU1AB

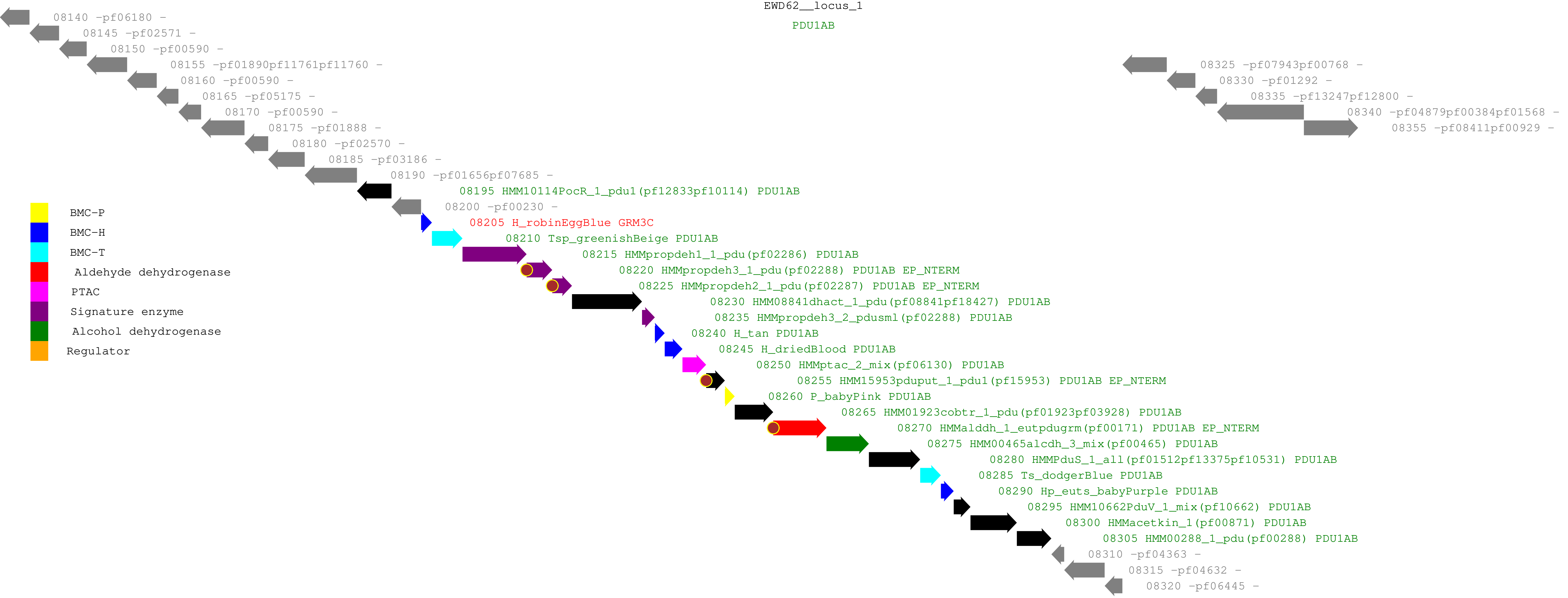

Salmonella enterica subsp. enterica serovar Vinohrady. Bacteria; Proteobacteria; Gammaproteobacteria; Enterobacterales;

EWF27__locus_1

PDU1AB

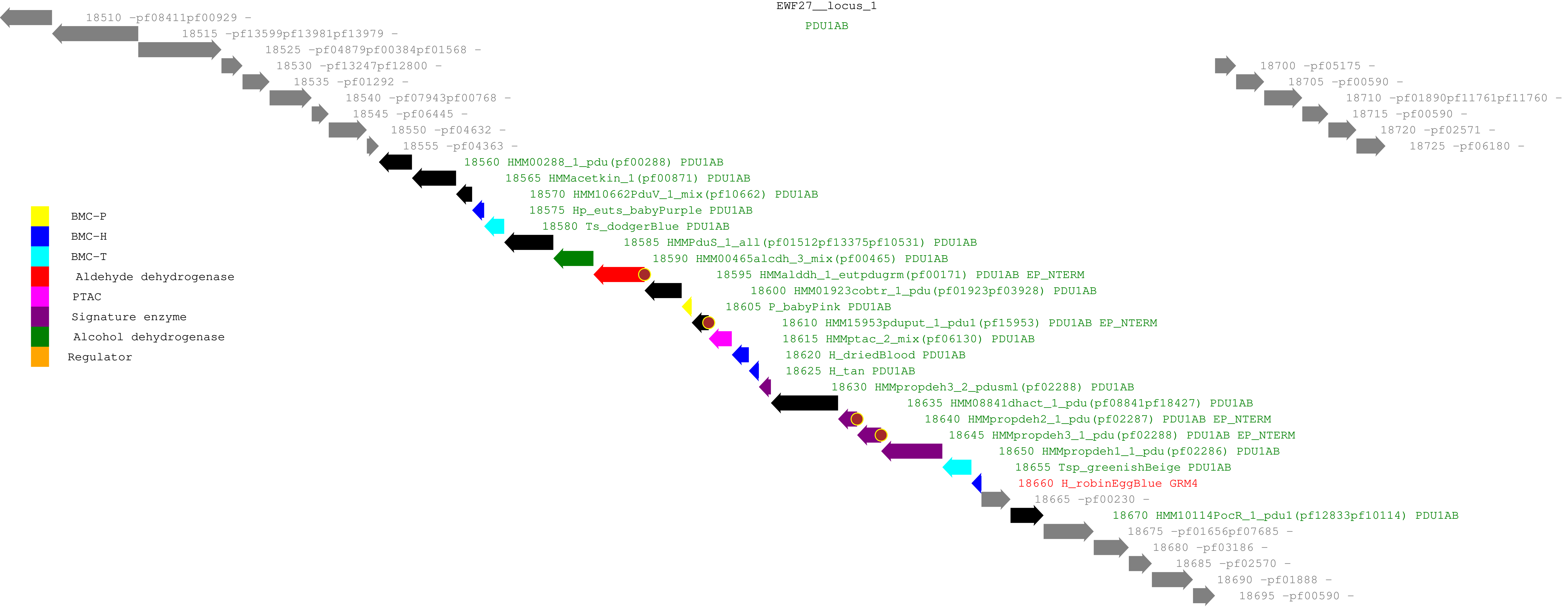

Salmonella enterica subsp. enterica serovar Herston. Bacteria; Proteobacteria; Gammaproteobacteria; Enterobacterales;

EWF40__locus_1

PDU1AB

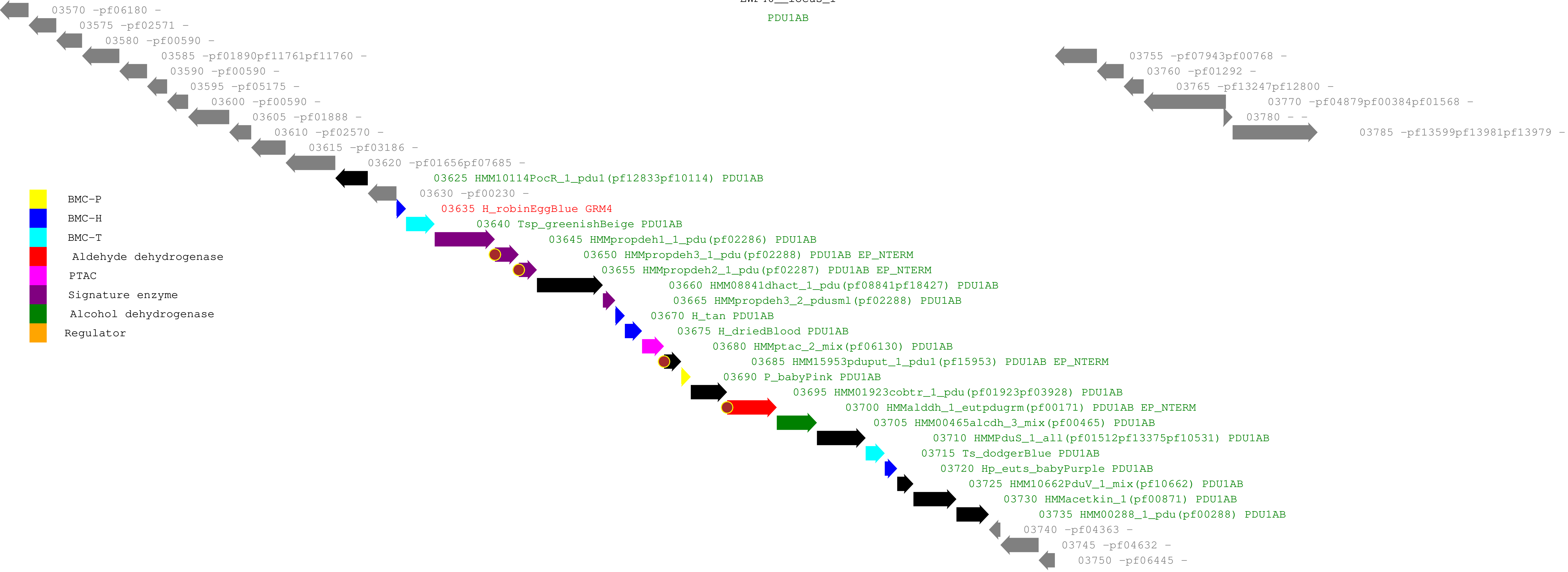

Salmonella enterica subsp. enterica serovar Choleraesuis. Bacteria; Proteobacteria; Gammaproteobacteria; Enterobacterales;

EWG00__locus_1

PDU1AB

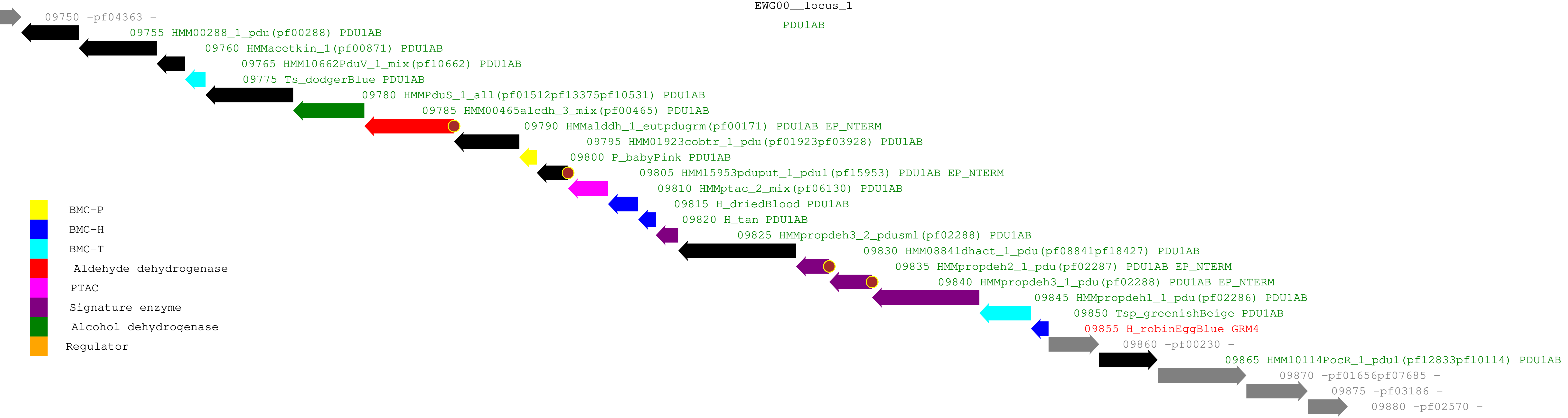

Salmonella enterica subsp. enterica serovar Zanzibar. Bacteria; Proteobacteria; Gammaproteobacteria; Enterobacterales;

EWG38__locus_2

PDU1AB

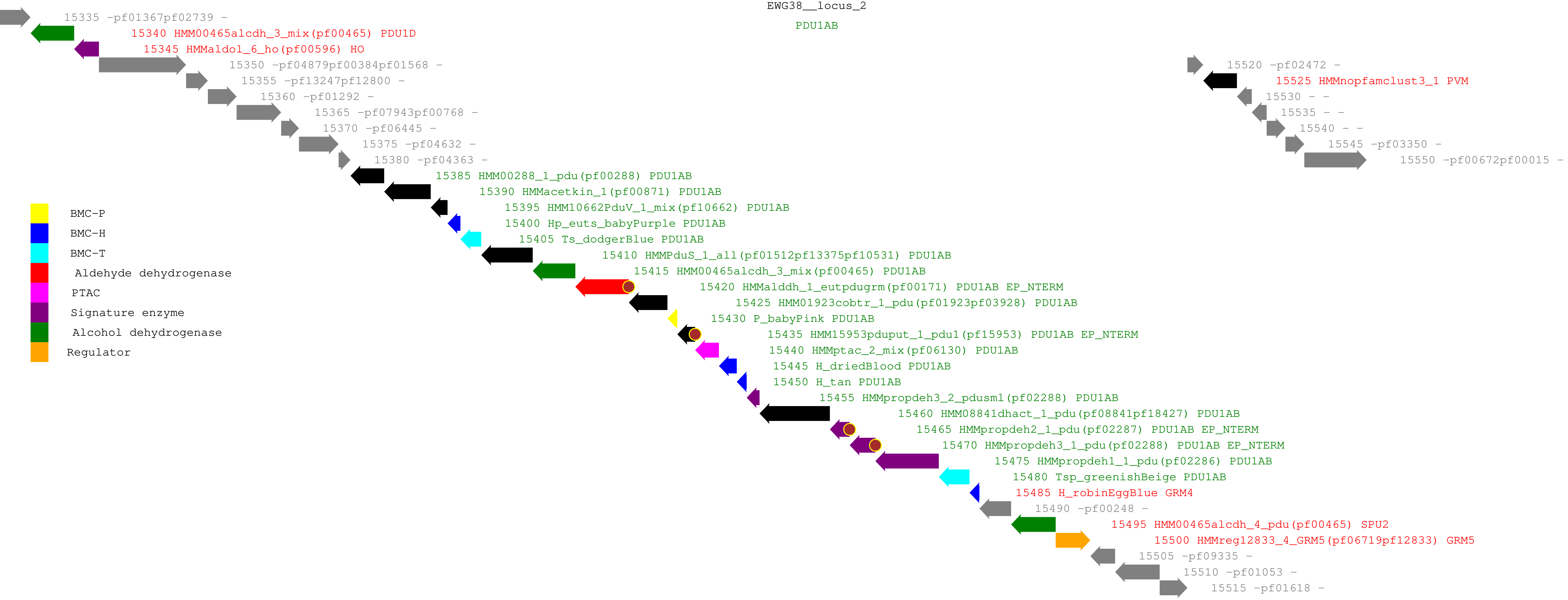

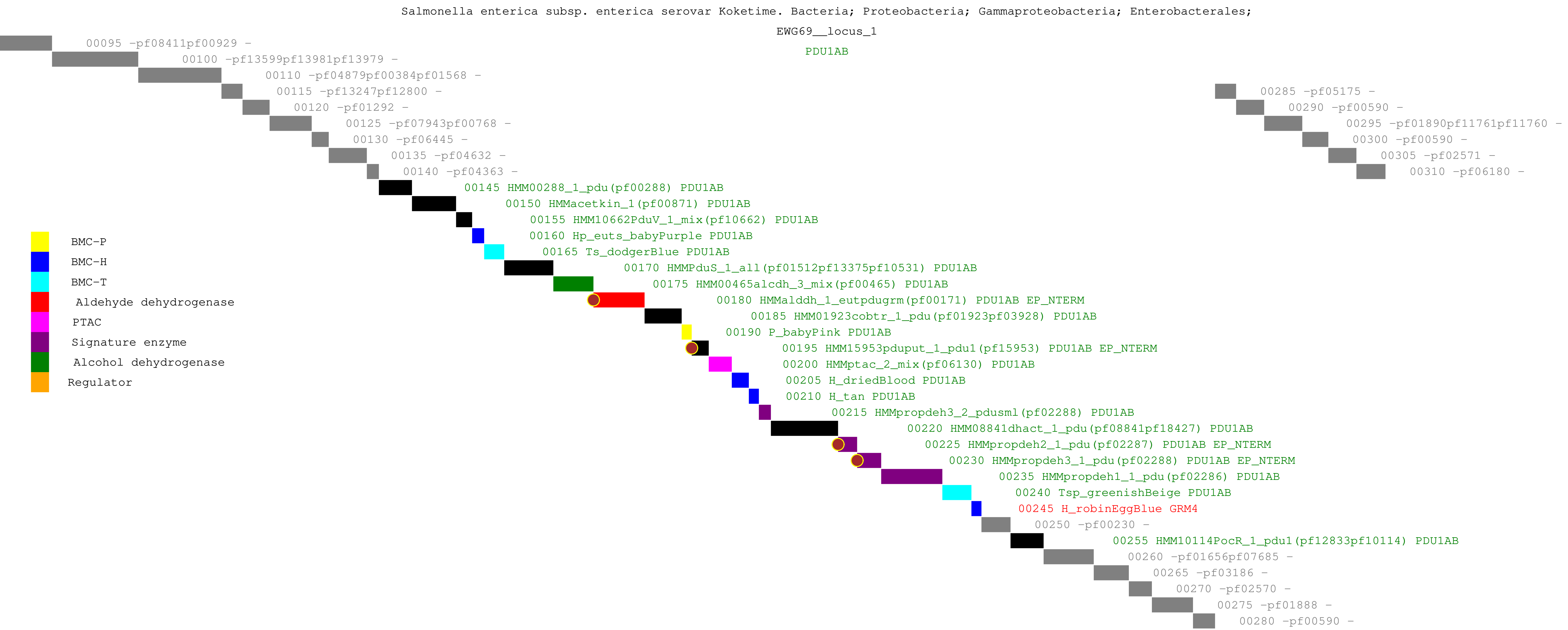
Salmonella enterica subsp. enterica serovar Koketime. Bacteria; Proteobacteria; Gammaproteobacteria; Enterobacterales;
EWG69__locus_1
PDU1AB
00095 -pf08411pf00929 -
00100 -pf13599pf13981pf13979 -
00110 -pf04879pf00384pf01568 -
00115 -pf13247pf12800 -
00120 -pf01292 -
00125 -pf07943pf00768 -
00130 -pf06445 -
00135 -pf04632 -
00140 -pf04363 -
00145 HMM00288_1_pdu(pf00288) PDU1AB
00150 HMMacetkin_1(pf00871) PDU1AB
00155 HMM10662PduV_1_mix(pf10662) PDU1AB
00160 Hp_euts_babyPurple PDU1AB
00165 Ts_dodgerBlue PDU1AB
00170 HMMPduS_1_all(pf01512pf13375pf10531) PDU1AB
00175 HMM00465alcdh_3_mix(pf00465) PDU1AB
00180 HMMalddh_1_eutpdugrm(pf00171) PDU1AB EP_NTERM
00185 HMM01923cobtr_1_pdu(pf01923pf03928) PDU1AB
00190 P_babyPink PDU1AB
00195 HMM15953pduput_1_pdu1(pf15953) PDU1AB EP_NTERM
00200 HMMptac_2_mix(pf06130) PDU1AB
00205 H_driedBlood PDU1AB
00210 H_tan PDU1AB
00215 HMMpropdeh3_2_pdusml(pf02288) PDU1AB
00220 HMM08841dhact_1_pdu(pf08841pf18427) PDU1AB
00225 HMMpropdeh2_1_pdu(pf02287) PDU1AB EP_NTERM
00230 HMMpropdeh3_1_pdu(pf02288) PDU1AB EP_NTERM
00235 HMMpropdeh1_1_pdu(pf02286) PDU1AB
00240 Tsp_greenishBeige PDU1AB
00245 H_robinEggBlue GRM4
00250 -pf00230 -
00255 HMM10114PocR_1_pdu1(pf12833pf10114) PDU1AB
00260 -pf01656pf07685 -
00265 -pf03186 -
00270 -pf02570 -
00275 -pf01888 -
00280 -pf00590 -
00285 -pf05175 -
00290 -pf00590 -
00295 -pf01890pf11761pf11760 -
00300 -pf00590 -
00305 -pf02571 -
00310 -pf06180 -
BMC-P
BMC-H
BMC-T
Aldehyde dehydrogenase
PTAC
Signature enzyme
Alcohol dehydrogenase
Regulator

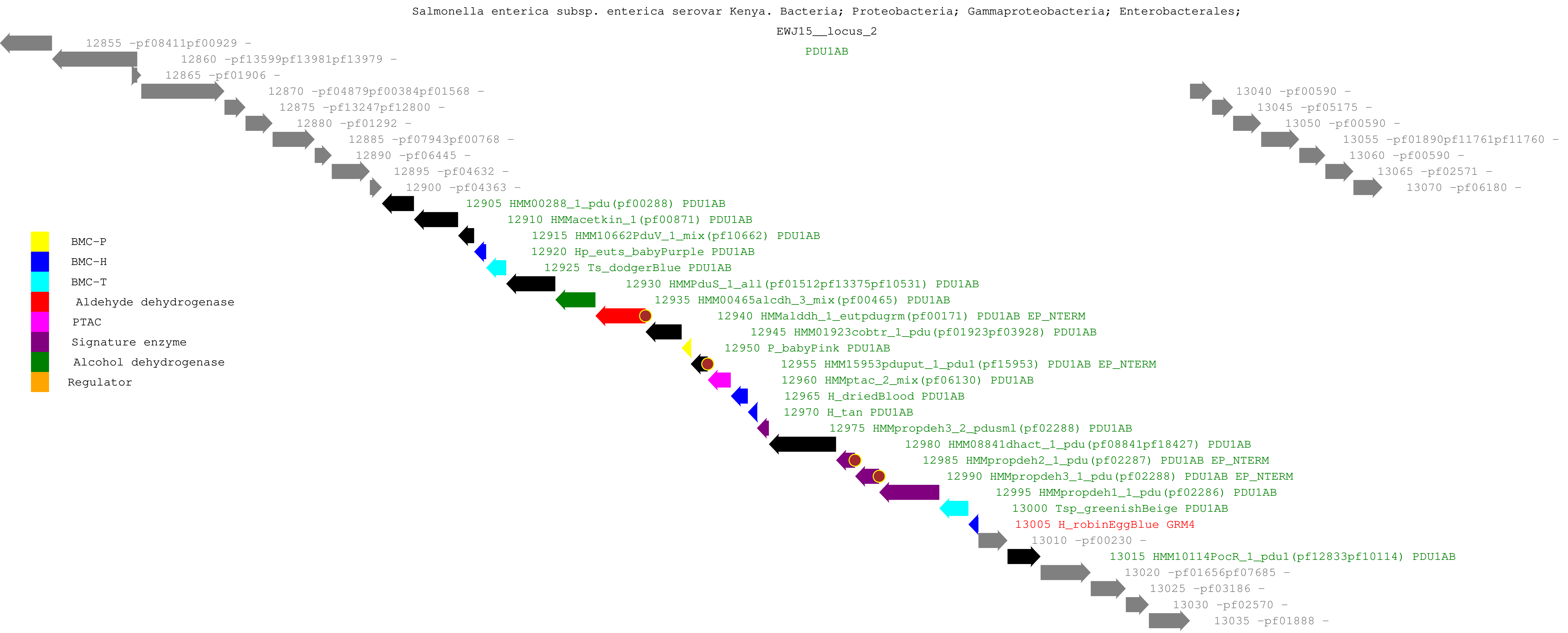
Salmonella enterica subsp. enterica serovar Kenya. Bacteria; Proteobacteria; Gammaproteobacteria; Enterobacterales;
EWJ15__locus_2
PDU1AB
12855 -pf08411pf00929 -
12860 -pf13599pf13981pf13979 -
12865 -pf01906 -
12870 -pf04879pf00384pf01568 -
12875 -pf13247pf12800 -
12880 -pf01292 -
12885 -pf07943pf00768 -
12890 -pf06445 -
12895 -pf04632 -
12900 -pf04363 -
12905 HMM00288_1_pdu(pf00288) PDU1AB
12910 HMMacetkin_1(pf00871) PDU1AB
12915 HMM10662PduV_1_mix(pf10662) PDU1AB
12920 Hp_euts_babyPurple PDU1AB
12925 Ts_dodgerBlue PDU1AB
12930 HMMPduS_1_all(pf01512pf13375pf10531) PDU1AB
12935 HMM00465alcdh_3_mix(pf00465) PDU1AB
12940 HMMalddh_1_eutpdugrm(pf00171) PDU1AB EP_NTERM
12945 HMM01923cobtr_1_pdu(pf01923pf03928) PDU1AB
12950 P_babyPink PDU1AB
12955 HMM15953pduput_1_pdu1(pf15953) PDU1AB EP_NTERM
12960 HMMptac_2_mix(pf06130) PDU1AB
12965 H_driedBlood PDU1AB
12970 H_tan PDU1AB
12975 HMMpropdeh3_2_pdusml(pf02288) PDU1AB
12980 HMM08841dhact_1_pdu(pf08841pf18427) PDU1AB
12985 HMMpropdeh2_1_pdu(pf02287) PDU1AB EP_NTERM
12990 HMMpropdeh3_1_pdu(pf02288) PDU1AB EP_NTERM
12995 HMMpropdeh1_1_pdu(pf02286) PDU1AB
13000 Tsp_greenishBeige PDU1AB
13005 H_robinEggBlue GRM4
13010 -pf00230 -
13015 HMM10114PocR_1_pdu1(pf12833pf10114) PDU1AB
13020 -pf01656pf07685 -
13025 -pf03186 -
13030 -pf02570 -
13035 -pf01888 -
13040 -pf00590 -
13045 -pf05175 -
13050 -pf00590 -
13055 -pf01890pf11761pf11760 -
13060 -pf00590 -
13065 -pf02571 -
13070 -pf06180 -
BMC-P
BMC-H
BMC-T
Aldehyde dehydrogenase
PTAC
Signature enzyme
Alcohol dehydrogenase
Regulator

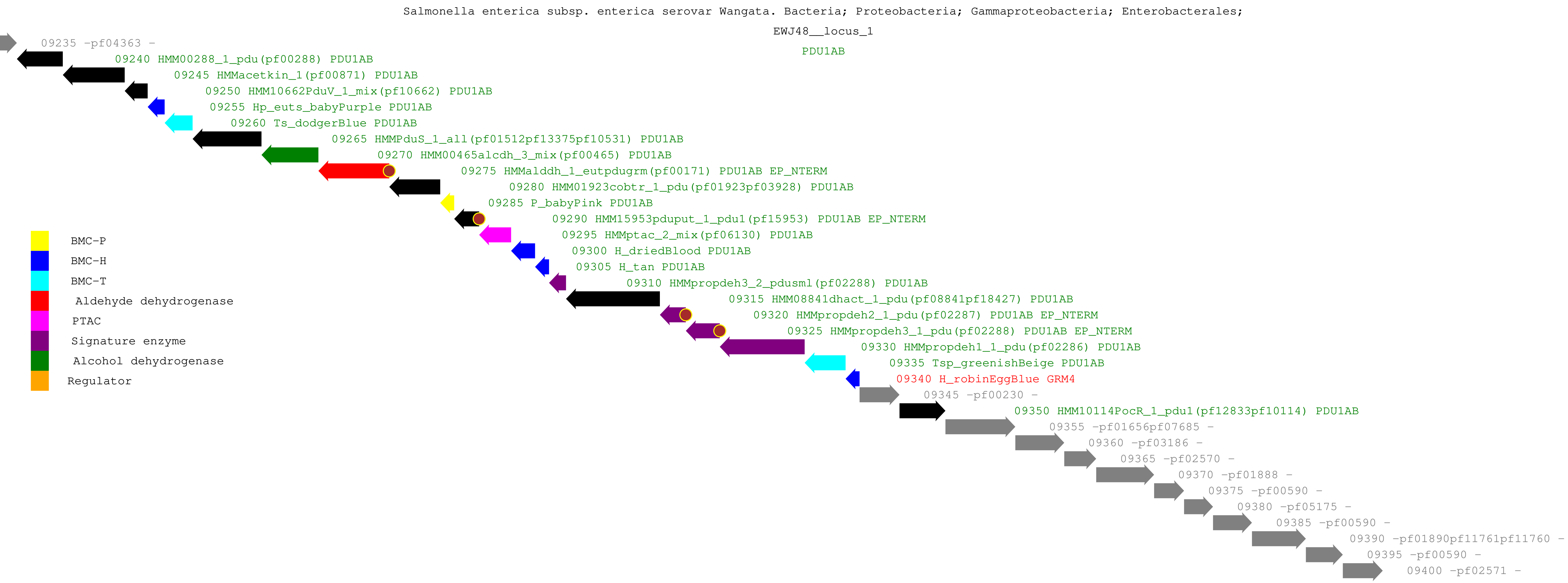
Salmonella enterica subsp. enterica serovar Wangata. Bacteria; Proteobacteria; Gammaproteobacteria; Enterobacterales;
EWJ48__locus_1
PDU1AB
09235 -pf04363 -
09240 HMM00288_1_pdu(pf00288) PDU1AB
09245 HMMacetkin_1(pf00871) PDU1AB
09250 HMM10662PduV_1_mix(pf10662) PDU1AB
09255 Hp_euts_babyPurple PDU1AB
09260 Ts_dodgerBlue PDU1AB
09265 HMMPduS_1_all(pf01512pf13375pf10531) PDU1AB
09270 HMM00465alcdh_3_mix(pf00465) PDU1AB
09275 HMMalddh_1_eutpdugrm(pf00171) PDU1AB EP_NTERM
09280 HMM01923cobtr_1_pdu(pf01923pf03928) PDU1AB
09285 P_babyPink PDU1AB
09290 HMM15953pduput_1_pdu1(pf15953) PDU1AB EP_NTERM
09295 HMMptac_2_mix(pf06130) PDU1AB
09300 H_driedBlood PDU1AB
09305 H_tan PDU1AB
09310 HMMpropdeh3_2_pdusml(pf02288) PDU1AB
09315 HMM08841dhact_1_pdu(pf08841pf18427) PDU1AB
09320 HMMpropdeh2_1_pdu(pf02287) PDU1AB EP_NTERM
09325 HMMpropdeh3_1_pdu(pf02288) PDU1AB EP_NTERM
09330 HMMpropdeh1_1_pdu(pf02286) PDU1AB
09335 Tsp_greenishBeige PDU1AB
09340 H_robinEggBlue GRM4
09345 -pf00230 -
09350 HMM10114PocR_1_pdu1(pf12833pf10114) PDU1AB
09355 -pf01656pf07685 -
09360 -pf03186 -
09365 -pf02570 -
09370 -pf01888 -
09375 -pf00590 -
09380 -pf05175 -
09385 -pf00590 -
09390 -pf01890pf11761pf11760 -
09395 -pf00590 -
09400 -pf02571 -
BMC-P
BMC-H
BMC-T
Aldehyde dehydrogenase
PTAC
Signature enzyme
Alcohol dehydrogenase
Regulator

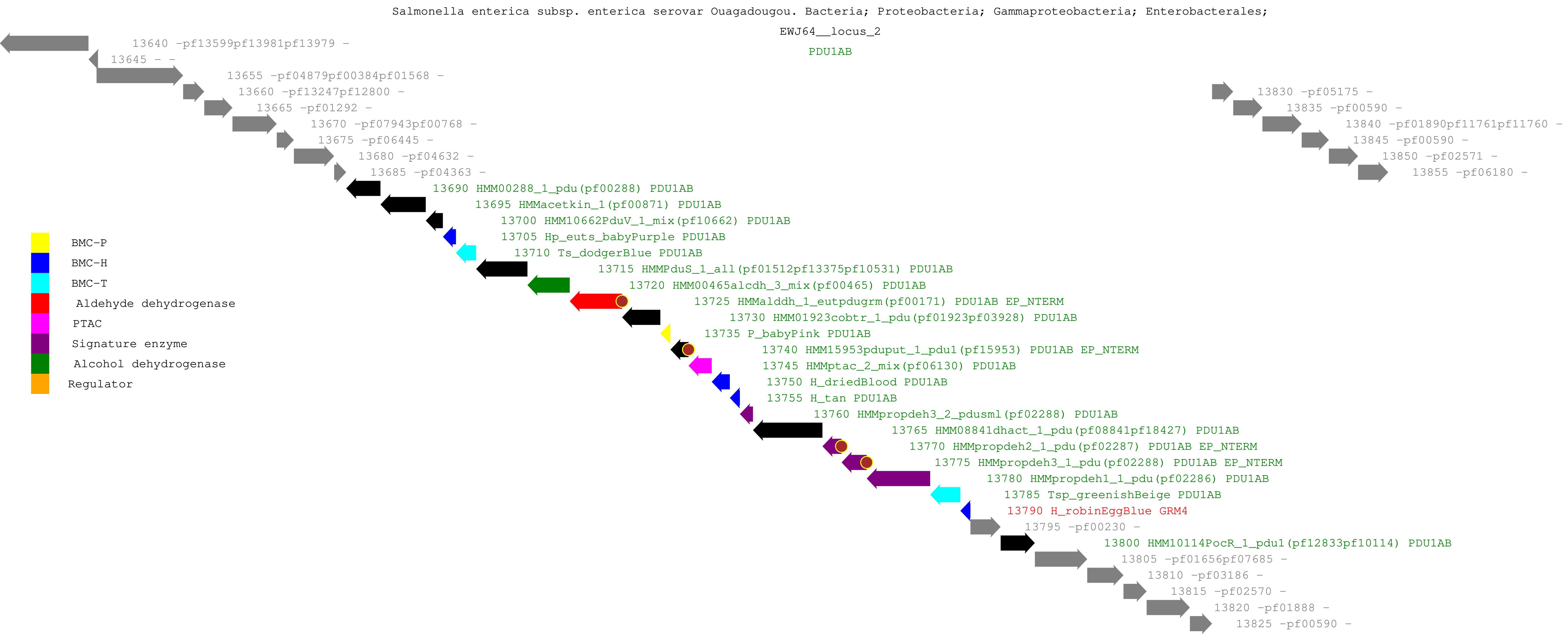
Salmonella enterica subsp. enterica serovar Ouagadougou. Bacteria; Proteobacteria; Gammaproteobacteria; Enterobacterales;
EWJ64__locus_2
PDU1AB
13640 -pf13599pf13981pf13979 -
13645 - -
13655 -pf04879pf00384pf01568 -
13660 -pf13247pf12800 -
13665 -pf01292 -
13670 -pf07943pf00768 -
13675 -pf06445 -
13680 -pf04632 -
13685 -pf04363 -
13690 HMM00288_1_pdu(pf00288) PDU1AB
13695 HMMacetkin_1(pf00871) PDU1AB
13700 HMM10662PduV_1_mix(pf10662) PDU1AB
13705 Hp_euts_babyPurple PDU1AB
13710 Ts_dodgerBlue PDU1AB
13715 HMMPduS_1_all(pf01512pf13375pf10531) PDU1AB
13720 HMM00465alcdh_3_mix(pf00465) PDU1AB
13725 HMMalddh_1_eutpdugrm(pf00171) PDU1AB EP_NTERM
13730 HMM01923cobtr_1_pdu(pf01923pf03928) PDU1AB
13735 P_babyPink PDU1AB
13740 HMM15953pduput_1_pdu1(pf15953) PDU1AB EP_NTERM
13745 HMMptac_2_mix(pf06130) PDU1AB
13750 H_driedBlood PDU1AB
13755 H_tan PDU1AB
13760 HMMpropdeh3_2_pdusml(pf02288) PDU1AB
13765 HMM08841dhact_1_pdu(pf08841pf18427) PDU1AB
13770 HMMpropdeh2_1_pdu(pf02287) PDU1AB EP_NTERM
13775 HMMpropdeh3_1_pdu(pf02288) PDU1AB EP_NTERM
13780 HMMpropdeh1_1_pdu(pf02286) PDU1AB
13785 Tsp_greenishBeige PDU1AB
13790 H_robinEggBlue GRM4
13795 -pf00230 -
13800 HMM10114PocR_1_pdu1(pf12833pf10114) PDU1AB
13805 -pf01656pf07685 -
13810 -pf03186 -
13815 -pf02570 -
13820 -pf01888 -
13825 -pf00590 -
13830 -pf05175 -
13835 -pf00590 -
13840 -pf01890pf11761pf11760 -
13845 -pf00590 -
13850 -pf02571 -
13855 -pf06180 -
BMC-P
BMC-H
BMC-T
Aldehyde dehydrogenase
PTAC
Signature enzyme
Alcohol dehydrogenase
Regulator

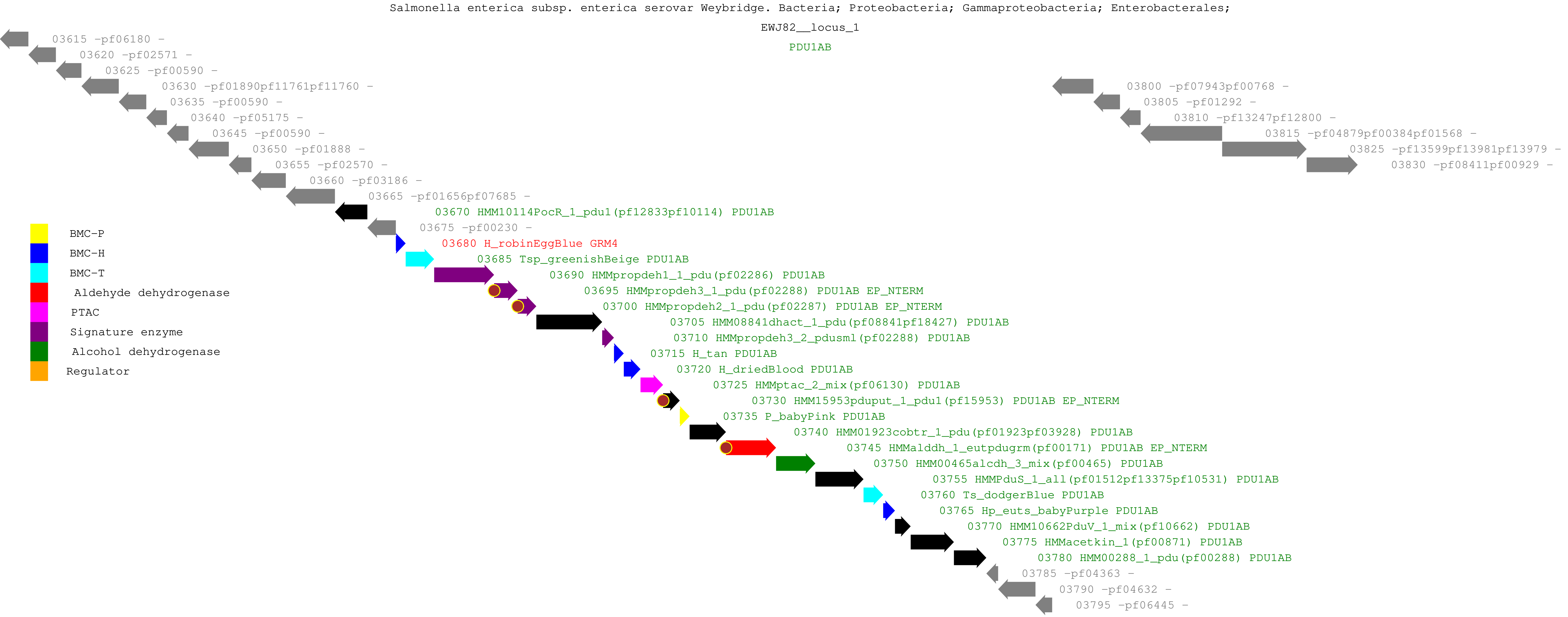
Salmonella enterica subsp. enterica serovar Weybridge. Bacteria; Proteobacteria; Gammaproteobacteria; Enterobacterales;
EWJ82__locus_1
PDU1AB
03615 -pf06180 -
03620 -pf02571 -
03625 -pf00590 -
03630 -pf01890pf11761pf11760 -
03635 -pf00590 -
03640 -pf05175 -
03645 -pf00590 -
03650 -pf01888 -
03655 -pf02570 -
03660 -pf03186 -
03665 -pf01656pf07685 -
03670 HMM10114PocR_1_pdu1(pf12833pf10114) PDU1AB
03675 -pf00230 -
03680 H_robinEggBlue GRM4
03685 Tsp_greenishBeige PDU1AB
03690 HMMpropdeh1_1_pdu(pf02286) PDU1AB
03695 HMMpropdeh3_1_pdu(pf02288) PDU1AB EP_NTERM
03700 HMMpropdeh2_1_pdu(pf02287) PDU1AB EP_NTERM
03705 HMM08841dhact_1_pdu(pf08841pf18427) PDU1AB
03710 HMMpropdeh3_2_pdusml(pf02288) PDU1AB
03715 H_tan PDU1AB
03720 H_driedBlood PDU1AB
03725 HMMptac_2_mix(pf06130) PDU1AB
03730 HMM15953pduput_1_pdu1(pf15953) PDU1AB EP_NTERM
03735 P_babyPink PDU1AB
03740 HMM01923cobtr_1_pdu(pf01923pf03928) PDU1AB
03745 HMMaldd h_1_eutpdugrm(pf00171) PDU1AB EP_NTERM
03750 HMM00465alcdh_3_mix(pf00465) PDU1AB
03755 HMMPduS_1_all(pf01512pf13375pf10531) PDU1AB
03760 Ts_dodgerBlue PDU1AB
03765 Hp_euts_babyPurple PDU1AB
03770 HMM10662PduV_1_mix(pf10662) PDU1AB
03775 HMMacetkin_1(pf00871) PDU1AB
03780 HMM00288_1_pdu(pf00288) PDU1AB
03785 -pf04363 -
03790 -pf04632 -
03795 -pf06445 -
03800 -pf07943pf00768 -
03805 -pf01292 -
03810 -pf13247pf12800 -
03815 -pf04879pf00384pf01568 -
03825 -pf13599pf13981pf13979 -
03830 -pf08411pf00929 -
BMC-P
BMC-H
BMC-T
Aldehyde dehydrogenase
PTAC
Signature enzyme
Alcohol dehydrogenase
Regulator

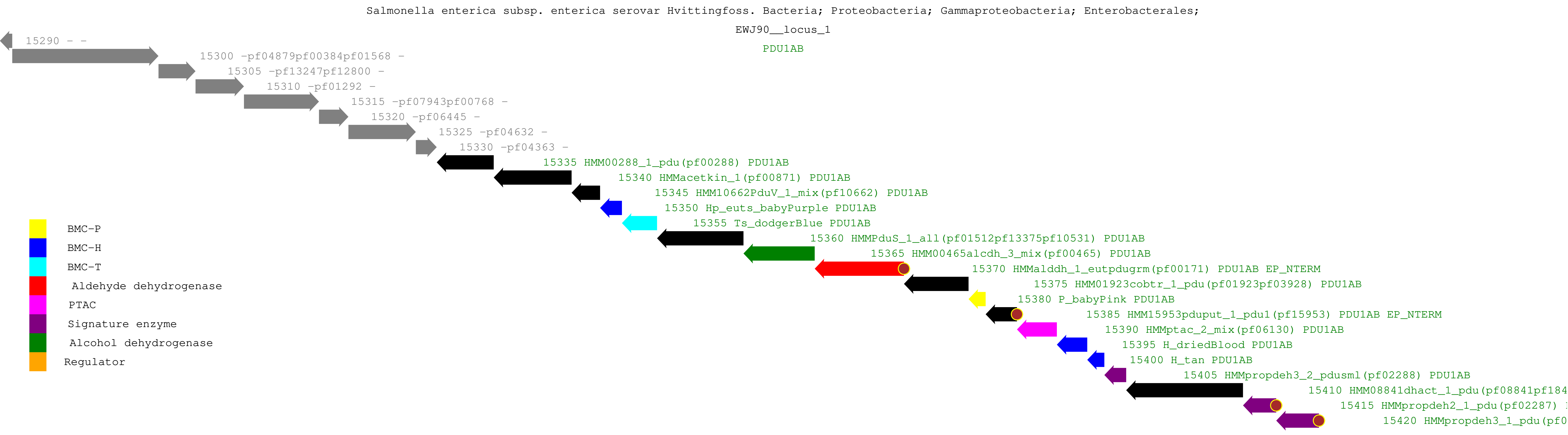
Salmonella enterica subsp. enterica serovar Hvittingfoss. Bacteria; Proteobacteria; Gammaproteobacteria; Enterobacterales;
EWJ90__locus_1
PDU1AB
15290 - -
15300 -pf04879pf00384pf01568 -
15305 -pf13247pf12800 -
15310 -pf01292 -
15315 -pf07943pf00768 -
15320 -pf06445 -
15325 -pf04632 -
15330 -pf04363 -
15335 HMM00288_1_pdu(pf00288) PDU1AB
15340 HMMacetkin_1(pf00871) PDU1AB
15345 HMM10662PduV_1_mix(pf10662) PDU1AB
15350 Hp_euts_babyPurple PDU1AB
15355 Ts_dodgerBlue PDU1AB
15360 HMMPduS_1_all(pf01512pf13375pf10531) PDU1AB
15365 HMM00465alcdh_3_mix(pf00465) PDU1AB
15370 HMMalddh_1_eutpdugrm(pf00171) PDU1AB EP_NTERM
15375 HMM01923cobtr_1_pdu(pf01923pf03928) PDU1AB
15380 P_babyPink PDU1AB
15385 HMM15953pduput_1_pdu1(pf15953) PDU1AB EP_NTERM
15390 HMMptac_2_mix(pf06130) PDU1AB
15395 H_driedBlood PDU1AB
15400 H_tan PDU1AB
15405 HMMpropdeh3_2_pdusml(pf02288) PDU1AB
15410 HMM08841dhact_1_pdu(pf08841pf184
15415 HMMpropdeh2_1_pdu(pf02287)
15420 HMMpropdeh3_1_pdu(pf0
BMC-P
BMC-H
BMC-T
Aldehyde dehydrogenase
PTAC
Signature enzyme
Alcohol dehydrogenase
Regulator

Salmonella virchow. Bacteria; Proteobacteria; Gammaproteobacteria; Enterobacterales;

EWK03__locus_1

PDU1AB

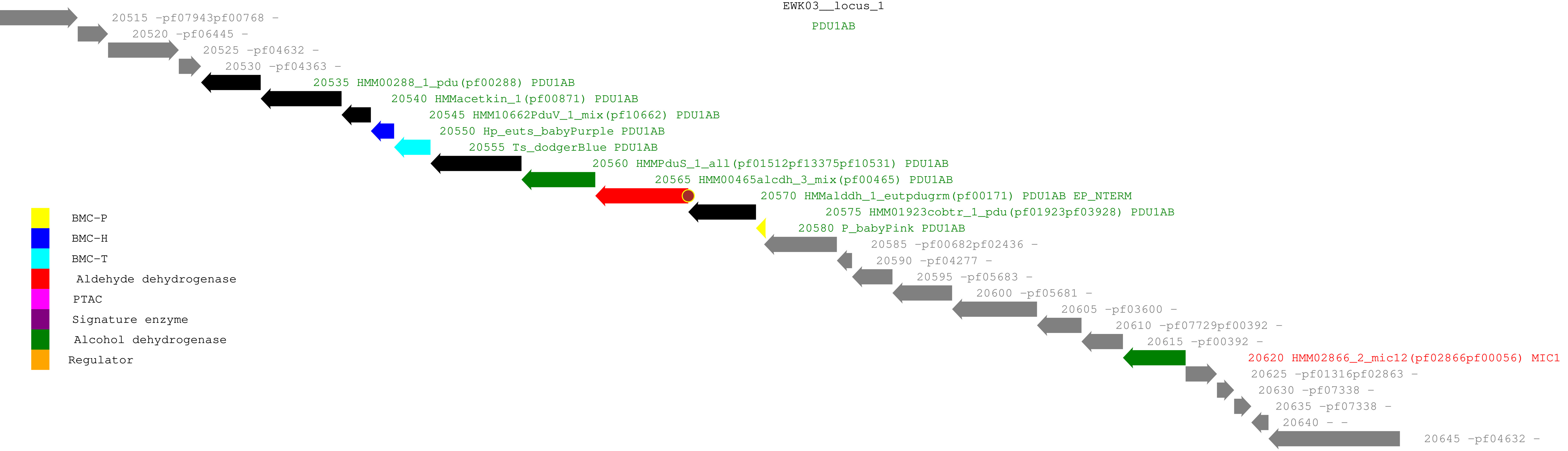

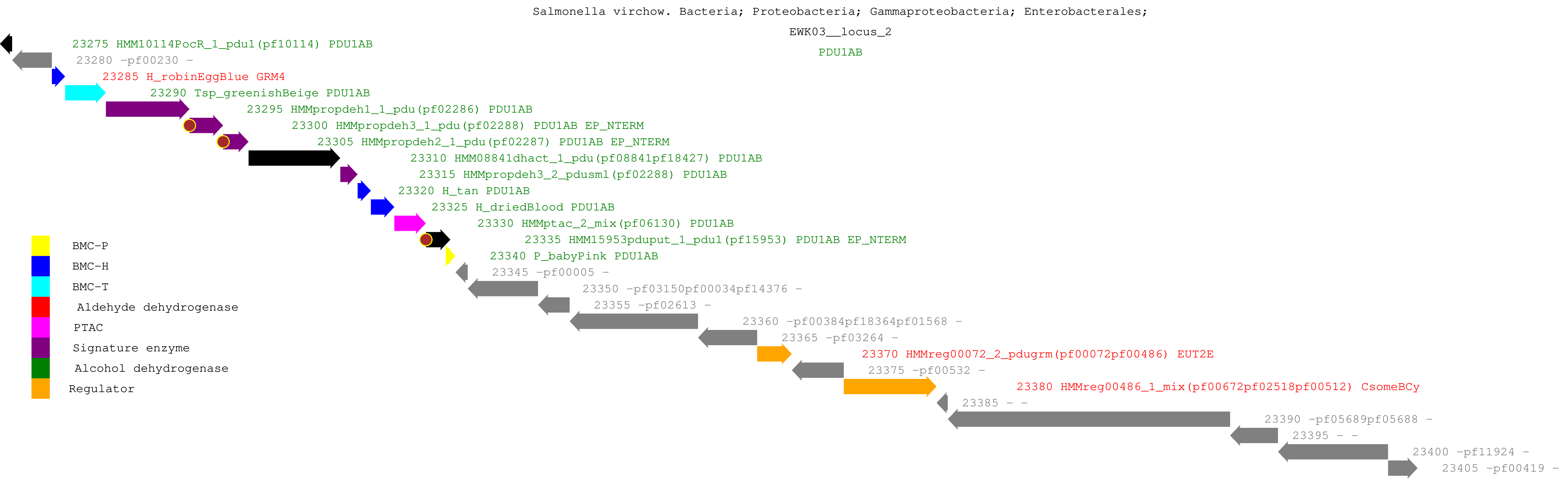
Salmonella virchow. Bacteria; Proteobacteria; Gammaproteobacteria; Enterobacterales;
EWK03__locus_2
PDU1AB
23275 HMM10114PocR_1_pdu1(pf10114) PDU1AB
23280 -pf00230 -
23285 H_robinEggBlue GRM4
23290 Tsp_greenishBeige PDU1AB
23295 HMMpropdeh1_1_pdu(pf02286) PDU1AB
23300 HMMpropdeh3_1_pdu(pf02288) PDU1AB EP_NTERM
23305 HMMpropdeh2_1_pdu(pf02287) PDU1AB EP_NTERM
23310 HMM08841dhact_1_pdu(pf08841pf18427) PDU1AB
23315 HMMpropdeh3_2_pdusml(pf02288) PDU1AB
23320 H_tan PDU1AB
23325 H_driedBlood PDU1AB
23330 HMMptac_2_mix(pf06130) PDU1AB
23335 HMM15953pduput_1_pdu1(pf15953) PDU1AB EP_NTERM
23340 P_babyPink PDU1AB
23345 -pf00005 -
23350 -pf03150pf00034pf14376 -
23355 -pf02613 -
23360 -pf00384pf18364pf01568 -
23365 -pf03264 -
23370 HMMreg00072_2_pdugrm(pf00072pf00486) EUT2E
23375 -pf00532 -
23380 HMMreg00486_1_mix(pf00672pf02518pf00512) CsomeBCy
23385 - -
23390 -pf05689pf05688 -
23395 - -
23400 -pf11924 -
23405 -pf00419 -
BMC-P
BMC-H
BMC-T
Aldehyde dehydrogenase
PTAC
Signature enzyme
Alcohol dehydrogenase
Regulator

Salmonella enterica subsp. enterica serovar Bonn. Bacteria; Proteobacteria; Gammaproteobacteria; Enterobacterales;

EWK23__locus_1

PDU1AB

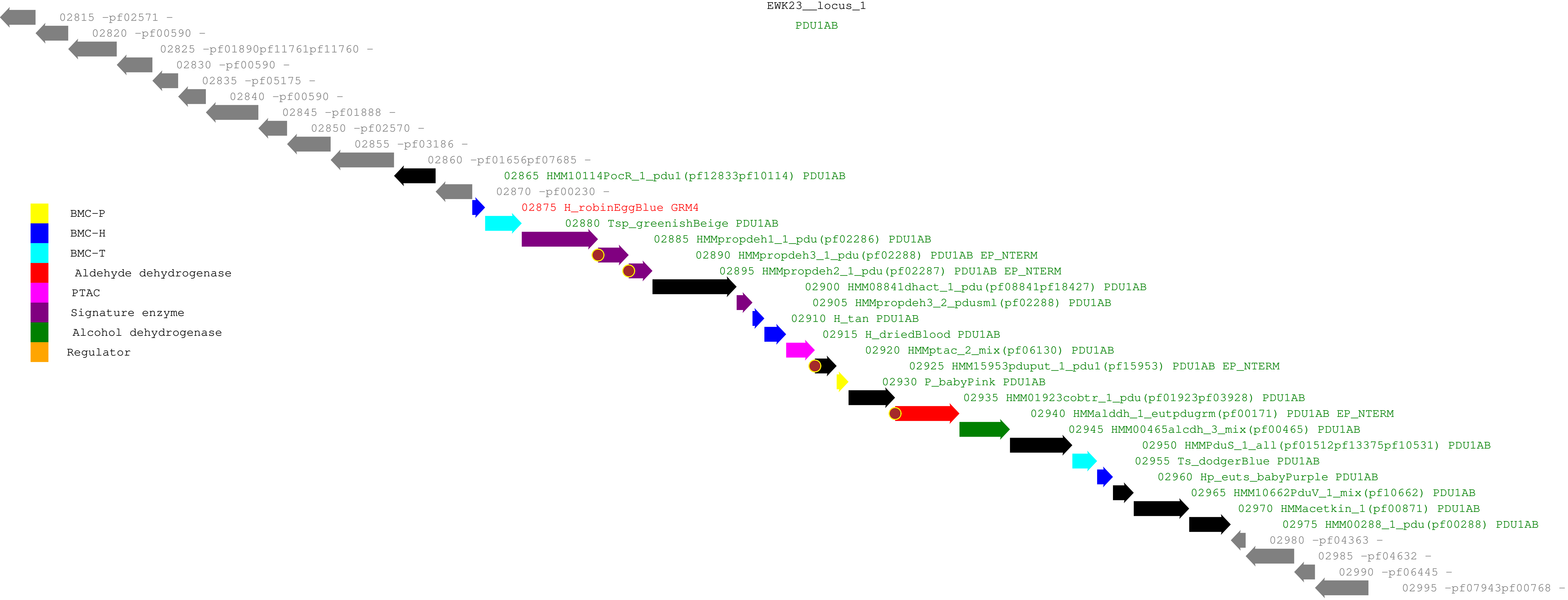

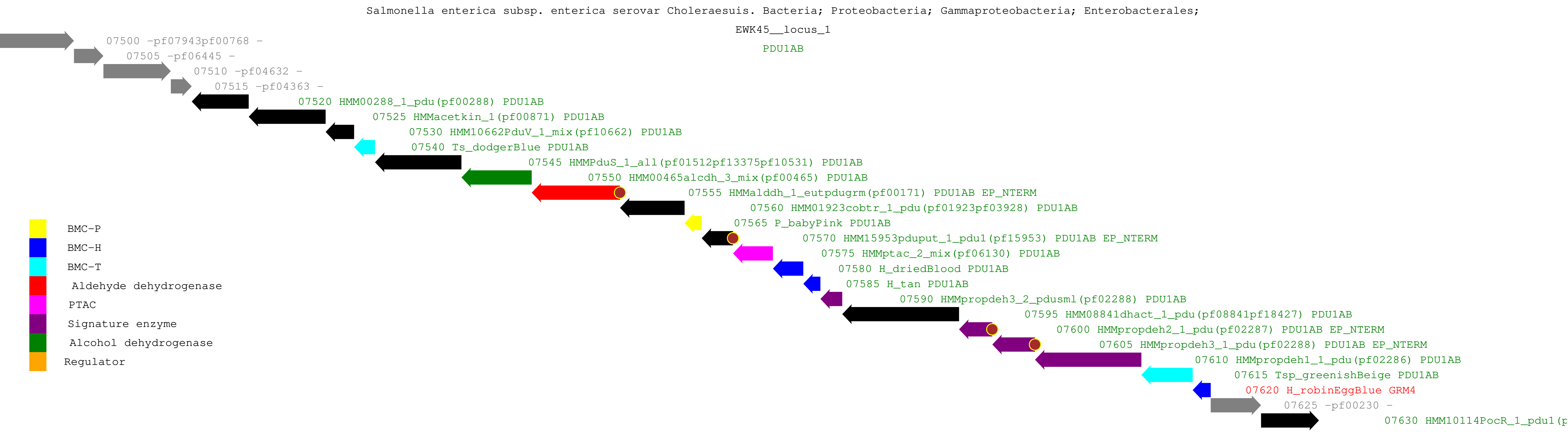

Salmonella enterica subsp. enterica serovar Choleraesuis. Bacteria; Proteobacteria; Gammaproteobacteria; Enterobacterales;
EWK45__locus_1
PDU1AB
07500 -pf07943pf00768 -
07505 -pf06445 -
07510 -pf04632 -
07515 -pf04363 -
07520 HMM00288_1_pdu(pf00288) PDU1AB
07525 HMMacetkin_1(pf00871) PDU1AB
07530 HMM10662PduV_1_mix(pf10662) PDU1AB
07540 Ts_dodgerBlue PDU1AB
07545 HMMPduS_1_all(pf01512pf13375pf10531) PDU1AB
07550 HMM00465alcdh_3_mix(pf00465) PDU1AB
07555 HMMalddh_1_eutpdugrm(pf00171) PDU1AB EP_NTERM
07560 HMM01923cobtr_1_pdu(pf01923pf03928) PDU1AB
07565 P_babyPink PDU1AB
07570 HMM15953pduput_1_pdu1(pf15953) PDU1AB EP_NTERM
07575 HMMptac_2_mix(pf06130) PDU1AB
07580 H_driedBlood PDU1AB
07585 H_tan PDU1AB
07590 HMMpropdeh3_2_pdusml(pf02288) PDU1AB
07595 HMM08841dhact_1_pdu(pf08841pf18427) PDU1AB
07600 HMMpropdeh2_1_pdu(pf02287) PDU1AB EP_NTERM
07605 HMMpropdeh3_1_pdu(pf02288) PDU1AB EP_NTERM
07610 HMMpropdeh1_1_pdu(pf02286) PDU1AB
07615 Tsp_greenishBeige PDU1AB
07620 H_robinEggBlue GRM4
07625 -pf00230 -
07630 HMM10114PocR_1_pdu1(p
BMC-P
BMC-H
BMC-T
Aldehyde dehydrogenase
PTAC
Signature enzyme
Alcohol dehydrogenase
Regulator

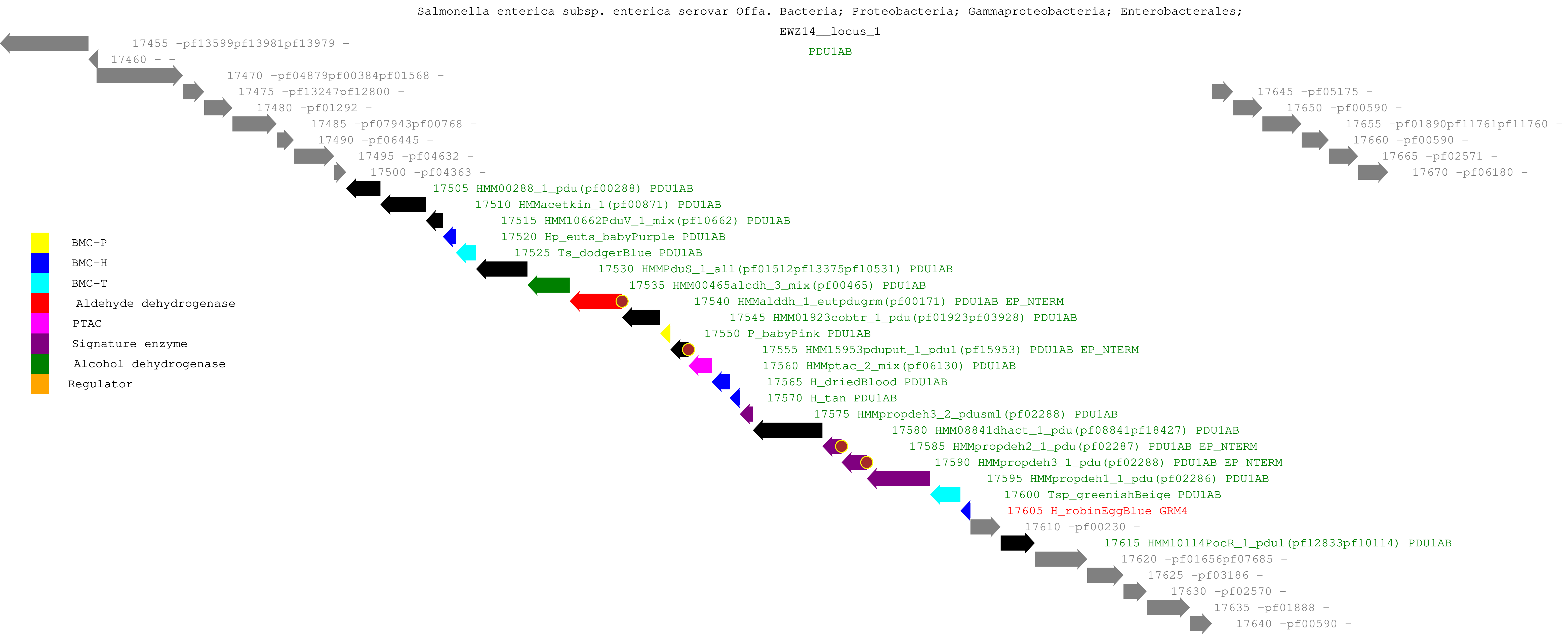
Salmonella enterica subsp. enterica serovar Offa. Bacteria; Proteobacteria; Gammaproteobacteria; Enterobacterales;
EWZ14__locus_1
PDU1AB
17455 -pf13599pf13981pf13979 -
17460 - -
17470 -pf04879pf00384pf01568 -
17475 -pf13247pf12800 -
17480 -pf01292 -
17485 -pf07943pf00768 -
17490 -pf06445 -
17495 -pf04632 -
17500 -pf04363 -
17505 HMM00288_1_pdu(pf00288) PDU1AB
17510 HMMacetkin_1(pf00871) PDU1AB
17515 HMM10662PduV_1_mix(pf10662) PDU1AB
17520 Hp_euts_babyPurple PDU1AB
17525 Ts_dodgerBlue PDU1AB
17530 HMMPduS_1_all(pf01512pf13375pf10531) PDU1AB
17535 HMM00465alcdh_3_mix(pf00465) PDU1AB
17540 HMMalddh_1_eutpdugrm(pf00171) PDU1AB EP_NTERM
17545 HMM01923cobtr_1_pdu(pf01923pf03928) PDU1AB
17550 P_babyPink PDU1AB
17555 HMM15953pduput_1_pdu1(pf15953) PDU1AB EP_NTERM
17560 HMMptac_2_mix(pf06130) PDU1AB
17565 H_driedBlood PDU1AB
17570 H_tan PDU1AB
17575 HMMpropdeh3_2_pdusml(pf02288) PDU1AB
17580 HMM08841dhact_1_pdu(pf08841pf18427) PDU1AB
17585 HMMpropdeh2_1_pdu(pf02287) PDU1AB EP_NTERM
17590 HMMpropdeh3_1_pdu(pf02288) PDU1AB EP_NTERM
17595 HMMpropdeh1_1_pdu(pf02286) PDU1AB
17600 Tsp_greenishBeige PDU1AB
17605 H_robinEggBlue GRM4
17610 -pf00230 -
17615 HMM10114PocR_1_pdu1(pf12833pf10114) PDU1AB
17620 -pf01656pf07685 -
17625 -pf03186 -
17630 -pf02570 -
17635 -pf01888 -
17640 -pf00590 -
17645 -pf05175 -
17650 -pf00590 -
17655 -pf01890pf11761pf11760 -
17660 -pf00590 -
17665 -pf02571 -
17670 -pf06180 -
BMC-P
BMC-H
BMC-T
Aldehyde dehydrogenase
PTAC
Signature enzyme
Alcohol dehydrogenase
Regulator

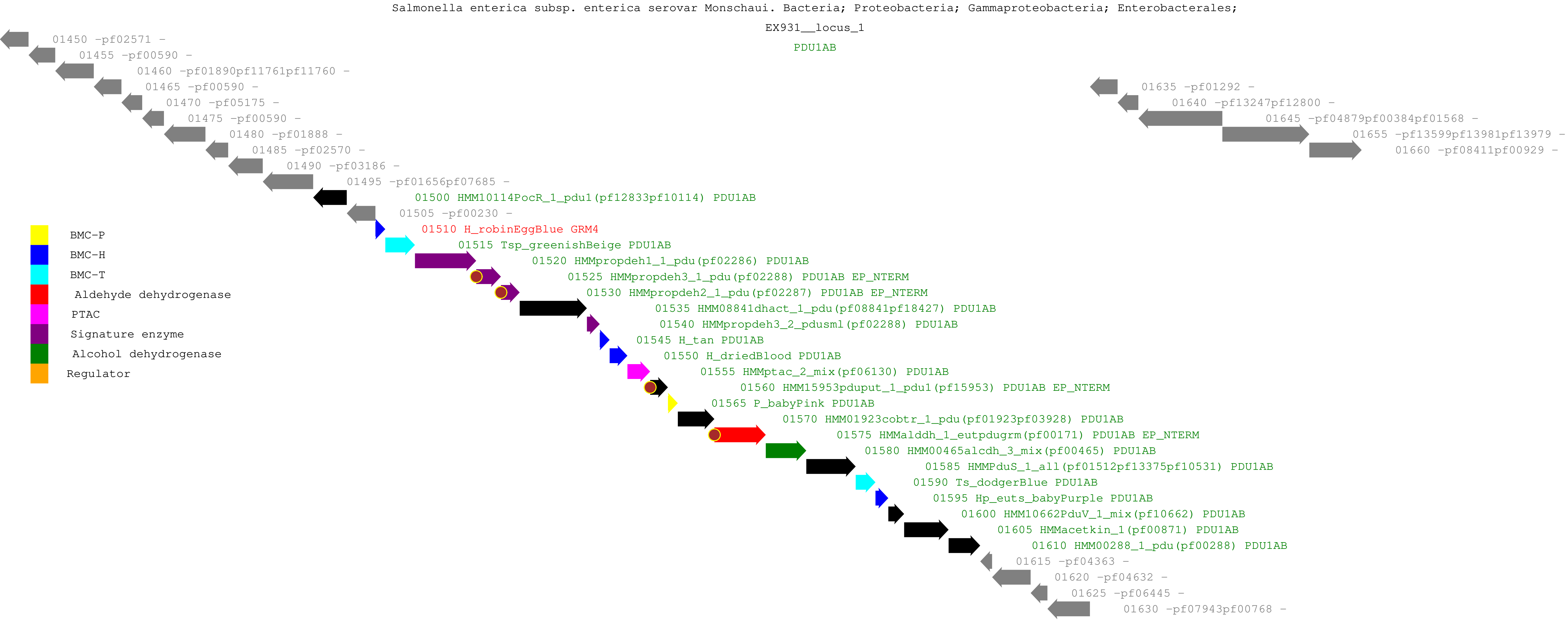
Salmonella enterica subsp. enterica serovar Monschaui. Bacteria; Proteobacteria; Gammaproteobacteria; Enterobacterales;
EX931__locus_1
PDU1AB
01450 -pf02571 -
01455 -pf00590 -
01460 -pf01890pf11761pf11760 -
01465 -pf00590 -
01470 -pf05175 -
01475 -pf00590 -
01480 -pf01888 -
01485 -pf02570 -
01490 -pf03186 -
01495 -pf01656pf07685 -
01500 HMM10114PocR_1_pdu1(pf12833pf10114) PDU1AB
01505 -pf00230 -
01510 H_robinEggBlue GRM4
01515 Tsp_greenishBeige PDU1AB
01520 HMMpropdeh1_1_pdu(pf02286) PDU1AB
01525 HMMpropdeh3_1_pdu(pf02288) PDU1AB EP_NTERM
01530 HMMpropdeh2_1_pdu(pf02287) PDU1AB EP_NTERM
01535 HMM08841dhact_1_pdu(pf08841pf18427) PDU1AB
01540 HMMpropdeh3_2_pdusml(pf02288) PDU1AB
01545 H_tan PDU1AB
01550 H_driedBlood PDU1AB
01555 HMMptac_2_mix(pf06130) PDU1AB
01560 HMM15953pduput_1_pdu1(pf15953) PDU1AB EP_NTERM
01565 P_babyPink PDU1AB
01570 HMM01923cobtr_1_pdu(pf01923pf03928) PDU1AB
01575 HMMalddh_1_eutpdugrm(pf00171) PDU1AB EP_NTERM
01580 HMM00465alcdh_3_mix(pf00465) PDU1AB
01585 HMMPduS_1_all(pf01512pf13375pf10531) PDU1AB
01590 Ts_dodgerBlue PDU1AB
01595 Hp_euts_babyPurple PDU1AB
01600 HMM10662PduV_1_mix(pf10662) PDU1AB
01605 HMMacetkin_1(pf00871) PDU1AB
01610 HMM00288_1_pdu(pf00288) PDU1AB
01615 -pf04363 -
01620 -pf04632 -
01625 -pf06445 -
01630 -pf07943pf00768 -
01635 -pf01292 -
01640 -pf13247pf12800 -
01645 -pf04879pf00384pf01568 -
01655 -pf13599pf13981pf13979 -
01660 -pf08411pf00929 -
BMC-P
BMC-H
BMC-T
Aldehyde dehydrogenase
PTAC
Signature enzyme
Alcohol dehydrogenase
Regulator

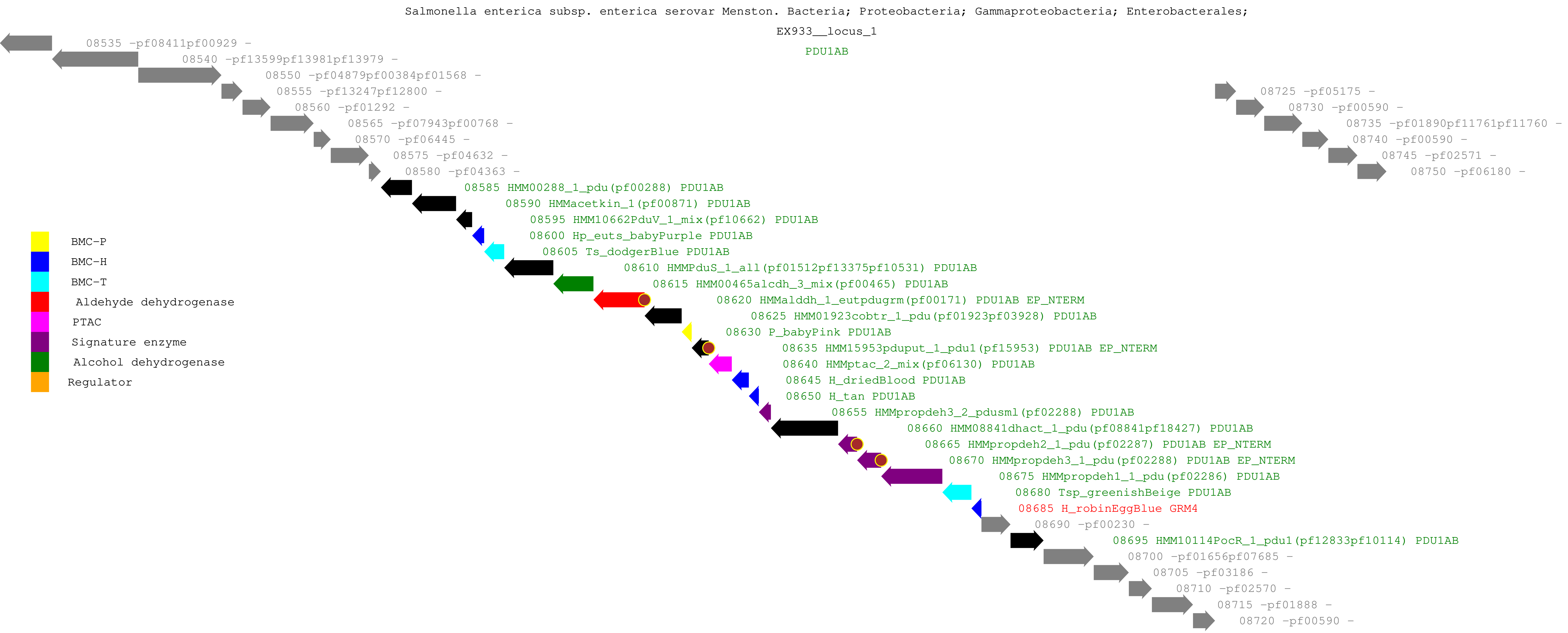
Salmonella enterica subsp. enterica serovar Menston. Bacteria; Proteobacteria; Gammaproteobacteria; Enterobacterales;
EX933__locus_1
PDU1AB
08535 -pf08411pf00929 -
08540 -pf13599pf13981pf13979 -
08550 -pf04879pf00384pf01568 -
08555 -pf13247pf12800 -
08560 -pf01292 -
08565 -pf07943pf00768 -
08570 -pf06445 -
08575 -pf04632 -
08580 -pf04363 -
08585 HMM00288_1_pdu(pf00288) PDU1AB
08590 HMMacetkin_1(pf00871) PDU1AB
08595 HMM10662PduV_1_mix(pf10662) PDU1AB
08600 Hp_euts_babyPurple PDU1AB
08605 Ts_dodgerBlue PDU1AB
08610 HMMPduS_1_all(pf01512pf13375pf10531) PDU1AB
08615 HMM00465alcdh_3_mix(pf00465) PDU1AB
08620 HMMalddh_1_eutpdugrm(pf00171) PDU1AB EP_NTERM
08625 HMM01923cobtr_1_pdu(pf01923pf03928) PDU1AB
08630 P_babyPink PDU1AB
08635 HMM15953pduput_1_pdu1(pf15953) PDU1AB EP_NTERM
08640 HMMptac_2_mix(pf06130) PDU1AB
08645 H_driedBlood PDU1AB
08650 H_tan PDU1AB
08655 HMMpropdeh3_2_pdusml(pf02288) PDU1AB
08660 HMM08841dhact_1_pdu(pf08841pf18427) PDU1AB
08665 HMMpropdeh2_1_pdu(pf02287) PDU1AB EP_NTERM
08670 HMMpropdeh3_1_pdu(pf02288) PDU1AB EP_NTERM
08675 HMMpropdeh1_1_pdu(pf02286) PDU1AB
08680 Tsp_greenishBeige PDU1AB
08685 H_robinEggBlue GRM4
08690 -pf00230 -
08695 HMM10114PocR_1_pdu1(pf12833pf10114) PDU1AB
08700 -pf01656pf07685 -
08705 -pf03186 -
08710 -pf02570 -
08715 -pf01888 -
08720 -pf00590 -
08725 -pf05175 -
08730 -pf00590 -
08735 -pf01890pf11761pf11760 -
08740 -pf00590 -
08745 -pf02571 -
08750 -pf06180 -
BMC-P
BMC-H
BMC-T
Aldehyde dehydrogenase
PTAC
Signature enzyme
Alcohol dehydrogenase
Regulator

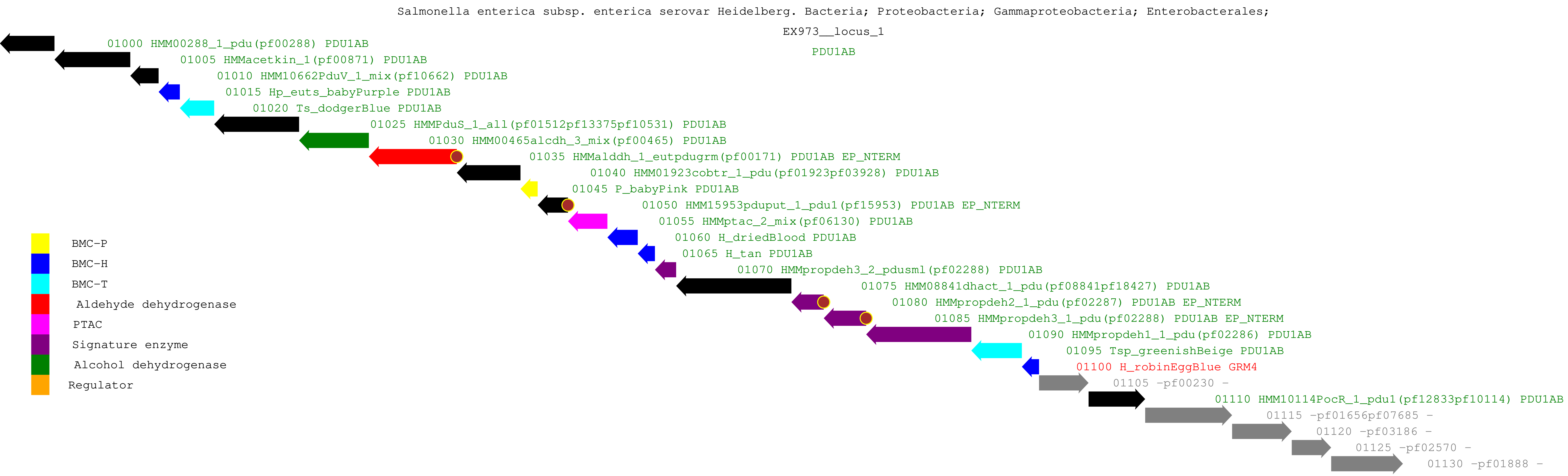
Salmonella enterica subsp. enterica serovar Heidelberg. Bacteria; Proteobacteria; Gammaproteobacteria; Enterobacterales;
EX973__locus_1
PDU1AB
01000 HMM00288_1_pdu(pf00288) PDU1AB
01005 HMMacetkin_1(pf00871) PDU1AB
01010 HMM10662PduV_1_mix(pf10662) PDU1AB
01015 Hp_euts_babyPurple PDU1AB
01020 Ts_dodgerBlue PDU1AB
01025 HMMPduS_1_all(pf01512pf13375pf10531) PDU1AB
01030 HMM00465alcdh_3_mix(pf00465) PDU1AB
01035 HMMalddh_1_eutpdugrm(pf00171) PDU1AB EP_NTERM
01040 HMM01923cobtr_1_pdu(pf01923pf03928) PDU1AB
01045 P_babyPink PDU1AB
01050 HMM15953pduput_1_pdu1(pf15953) PDU1AB EP_NTERM
01055 HMMptac_2_mix(pf06130) PDU1AB
01060 H_driedBlood PDU1AB
01065 H_tan PDU1AB
01070 HMMpropdeh3_2_pdusml(pf02288) PDU1AB
01075 HMM08841dhact_1_pdu(pf08841pf18427) PDU1AB
01080 HMMpropdeh2_1_pdu(pf02287) PDU1AB EP_NTERM
01085 HMMpropdeh3_1_pdu(pf02288) PDU1AB EP_NTERM
01090 HMMpropdeh1_1_pdu(pf02286) PDU1AB
01095 Tsp_greenishBeige PDU1AB
01100 H_robinEggBlue GRM4
01105 -pf00230 -
01110 HMM10114PocR_1_pdu1(pf12833pf10114) PDU1AB
01115 -pf01656pf07685 -
01120 -pf03186 -
01125 -pf02570 -
01130 -pf01888 -
BMC-P
BMC-H
BMC-T
Aldehyde dehydrogenase
PTAC
Signature enzyme
Alcohol dehydrogenase
Regulator

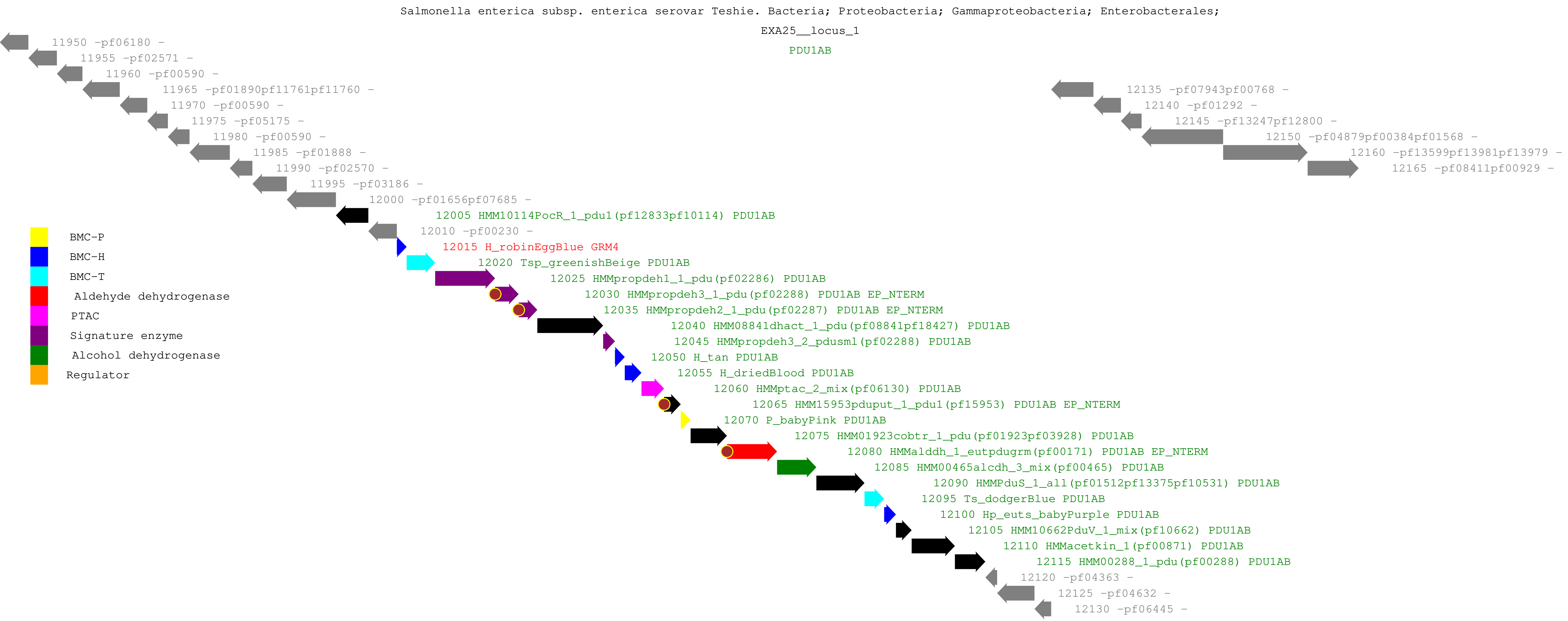
Salmonella enterica subsp. enterica serovar Teshie. Bacteria; Proteobacteria; Gammaproteobacteria; Enterobacterales;
EXA25__locus_1
PDU1AB
11950 -pf06180 -
11955 -pf02571 -
11960 -pf00590 -
11965 -pf01890pf11761pf11760 -
11970 -pf00590 -
11975 -pf05175 -
11980 -pf00590 -
11985 -pf01888 -
11990 -pf02570 -
11995 -pf03186 -
12000 -pf01656pf07685 -
12005 HMM10114PocR_1_pdu1(pf12833pf10114) PDU1AB
12010 -pf00230 -
12015 H_robinEggBlue GRM4
12020 Tsp_greenishBeige PDU1AB
12025 HMMpropdeh1_1_pdu(pf02286) PDU1AB
12030 HMMpropdeh3_1_pdu(pf02288) PDU1AB EP_NTERM
12035 HMMpropdeh2_1_pdu(pf02287) PDU1AB EP_NTERM
12040 HMM08841dhact_1_pdu(pf08841pf18427) PDU1AB
12045 HMMpropdeh3_2_pdusml(pf02288) PDU1AB
12050 H_tan PDU1AB
12055 H_driedBlood PDU1AB
12060 HMMptac_2_mix(pf06130) PDU1AB
12065 HMM15953pduput_1_pdu1(pf15953) PDU1AB EP_NTERM
12070 P_babyPink PDU1AB
12075 HMM01923cobtr_1_pdu(pf01923pf03928) PDU1AB
12080 HMMalddh_1_eutpdugrm(pf00171) PDU1AB EP_NTERM
12085 HMM00465alcdh_3_mix(pf00465) PDU1AB
12090 HMMPduS_1_all(pf01512pf13375pf10531) PDU1AB
12095 Ts_dodgerBlue PDU1AB
12100 Hp_euts_babyPurple PDU1AB
12105 HMM10662PduV_1_mix(pf10662) PDU1AB
12110 HMMacetkin_1(pf00871) PDU1AB
12115 HMM00288_1_pdu(pf00288) PDU1AB
12120 -pf04363 -
12125 -pf04632 -
12130 -pf06445 -
12135 -pf07943pf00768 -
12140 -pf01292 -
12145 -pf13247pf12800 -
12150 -pf04879pf00384pf01568 -
12160 -pf13599pf13981pf13979 -
12165 -pf08411pf00929 -
BMC-P
BMC-H
BMC-T
Aldehyde dehydrogenase
PTAC
Signature enzyme
Alcohol dehydrogenase
Regulator

Salmonella enterica subsp. enterica serovar Korlebu. Bacteria; Proteobacteria; Gammaproteobacteria; Enterobacterales;

EXA36__locus_1

PDU1AB

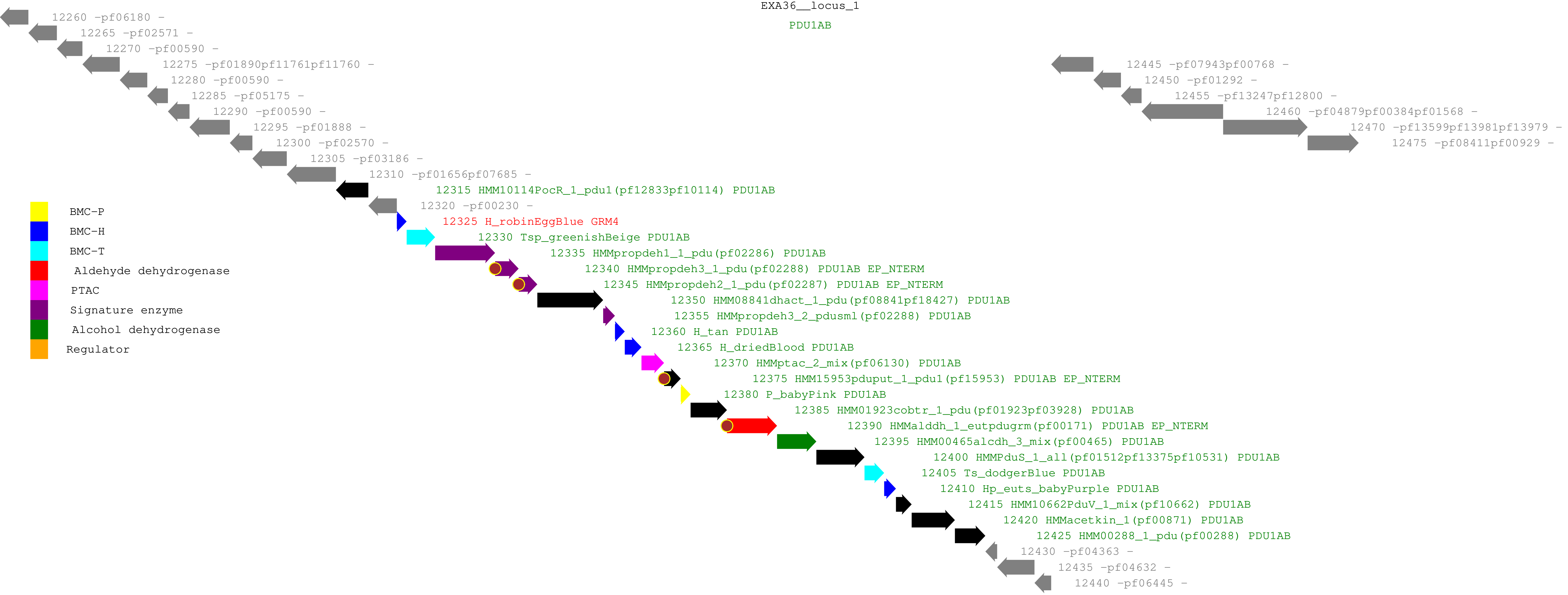

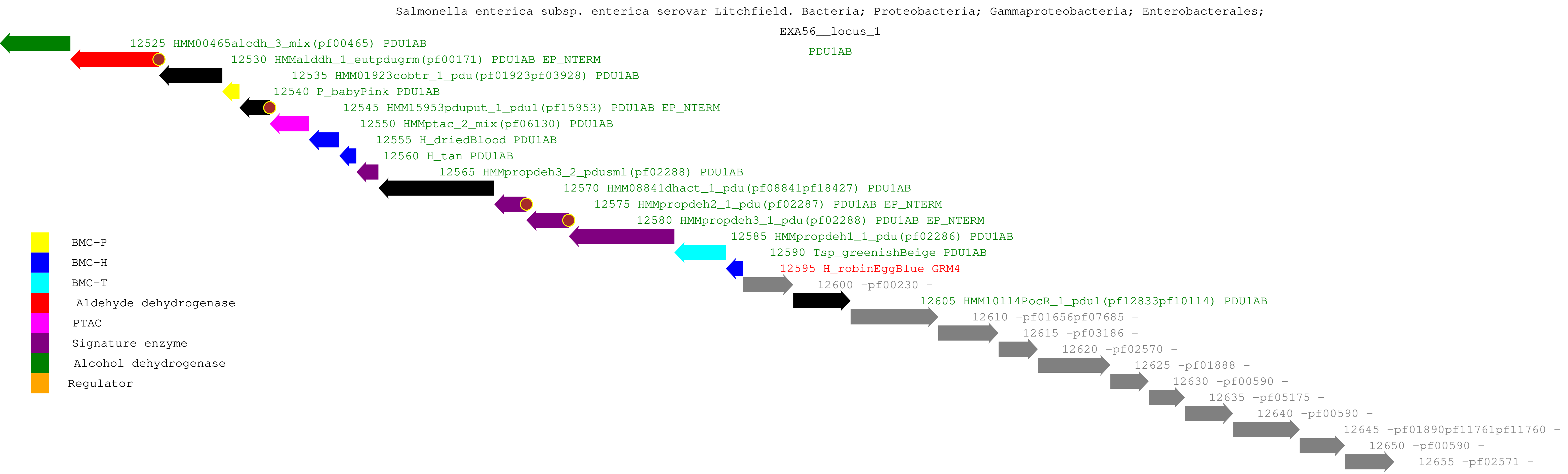

Salmonella enterica subsp. enterica serovar Litchfield. Bacteria; Proteobacteria; Gammaproteobacteria; Enterobacterales;
EXA56__locus_1
PDU1AB
12525 HMM00465alcdh_3_mix(pf00465) PDU1AB
12530 HMMalddh_1_eutpdugrm(pf00171) PDU1AB EP_NTERM
12535 HMM01923cobtr_1_pdu(pf01923pf03928) PDU1AB
12540 P_babyPink PDU1AB
12545 HMM15953pduput_1_pdu1(pf15953) PDU1AB EP_NTERM
12550 HMMptac_2_mix(pf06130) PDU1AB
12555 H_driedBlood PDU1AB
12560 H_tan PDU1AB
12565 HMMpropdeh3_2_pdusml(pf02288) PDU1AB
12570 HMM08841dhact_1_pdu(pf08841pf18427) PDU1AB
12575 HMMpropdeh2_1_pdu(pf02287) PDU1AB EP_NTERM
12580 HMMpropdeh3_1_pdu(pf02288) PDU1AB EP_NTERM
12585 HMMpropdeh1_1_pdu(pf02286) PDU1AB
12590 Tsp_greenishBeige PDU1AB
12595 H_robinEggBlue GRM4
12600 -pf00230 -
12605 HMM10114PocR_1_pdu1(pf12833pf10114) PDU1AB
12610 -pf01656pf07685 -
12615 -pf03186 -
12620 -pf02570 -
12625 -pf01888 -
12630 -pf00590 -
12635 -pf05175 -
12640 -pf00590 -
12645 -pf01890pf11761pf11760 -
12650 -pf00590 -
12655 -pf02571 -
BMC-P
BMC-H
BMC-T
Aldehyde dehydrogenase
PTAC
Signature enzyme
Alcohol dehydrogenase
Regulator

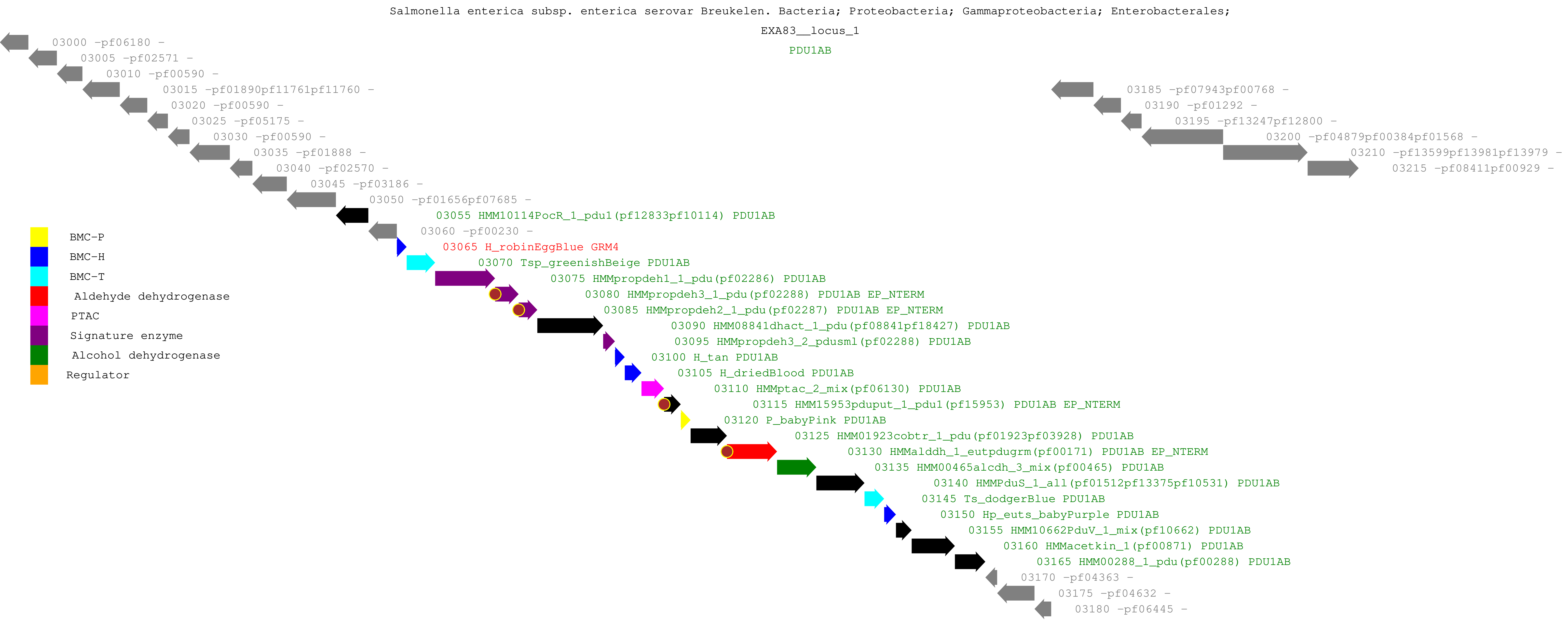
Salmonella enterica subsp. enterica serovar Breukelen. Bacteria; Proteobacteria; Gammaproteobacteria; Enterobacterales;
EXA83__locus_1
PDU1AB
03000 -pf06180 -
03005 -pf02571 -
03010 -pf00590 -
03015 -pf01890pf11761pf11760 -
03020 -pf00590 -
03025 -pf05175 -
03030 -pf00590 -
03035 -pf01888 -
03040 -pf02570 -
03045 -pf03186 -
03050 -pf01656pf07685 -
03055 HMM10114PocR_1_pdu1(pf12833pf10114) PDU1AB
03060 -pf00230 -
03065 H_robinEggBlue GRM4
03070 Tsp_greenishBeige PDU1AB
03075 HMMpropdeh1_1_pdu(pf02286) PDU1AB
03080 HMMpropdeh3_1_pdu(pf02288) PDU1AB EP_NTERM
03085 HMMpropdeh2_1_pdu(pf02287) PDU1AB EP_NTERM
03090 HMM08841dhact_1_pdu(pf08841pf18427) PDU1AB
03095 HMMpropdeh3_2_pdusml(pf02288) PDU1AB
03100 H_tan PDU1AB
03105 H_driedBlood PDU1AB
03110 HMMptac_2_mix(pf06130) PDU1AB
03115 HMM15953pduput_1_pdu1(pf15953) PDU1AB EP_NTERM
03120 P_babyPink PDU1AB
03125 HMM01923cobtr_1_pdu(pf01923pf03928) PDU1AB
03130 HMMaldh_1_eutpdugrm(pf00171) PDU1AB EP_NTERM
03135 HMM00465alcdh_3_mix(pf00465) PDU1AB
03140 HMMPduS_1_all(pf01512pf13375pf10531) PDU1AB
03145 Ts_dodgerBlue PDU1AB
03150 Hp_euts_babyPurple PDU1AB
03155 HMM10662PduV_1_mix(pf10662) PDU1AB
03160 HMMacetkin_1(pf00871) PDU1AB
03165 HMM00288_1_pdu(pf00288) PDU1AB
03170 -pf04363 -
03175 -pf04632 -
03180 -pf06445 -
03185 -pf07943pf00768 -
03190 -pf01292 -
03195 -pf13247pf12800 -
03200 -pf04879pf00384pf01568 -
03210 -pf13599pf13981pf13979 -
03215 -pf08411pf00929 -
BMC-P
BMC-H
BMC-T
Aldehyde dehydrogenase
PTAC
Signature enzyme
Alcohol dehydrogenase
Regulator

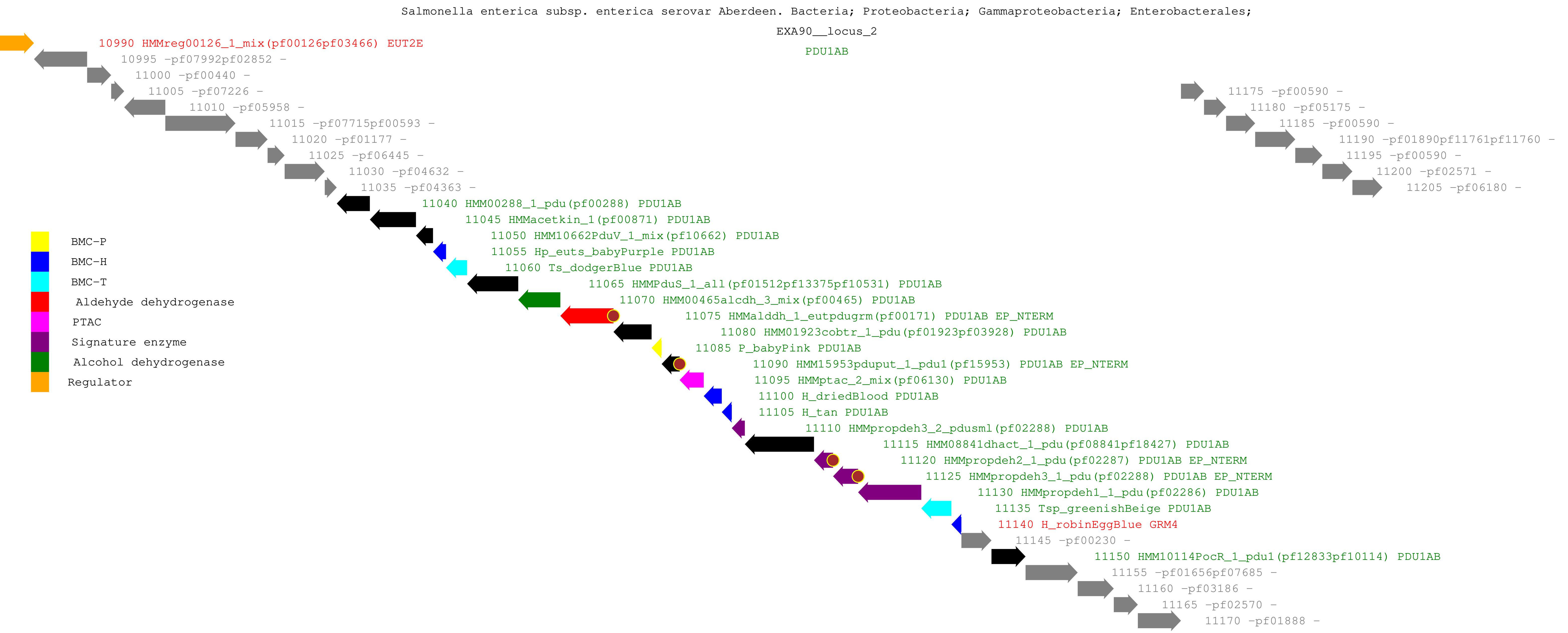

Salmonella enterica subsp. enterica serovar Aberdeen. Bacteria; Proteobacteria; Gammaproteobacteria; Enterobacterales;
EXA90__locus_2
PDU1AB
10990 HMMreg00126_1_mix(pf00126pf03466) EUT2E
10995 -pf07992pf02852 -
11000 -pf00440 -
11005 -pf07226 -
11010 -pf05958 -
11015 -pf07715pf00593 -
11020 -pf01177 -
11025 -pf06445 -
11030 -pf04632 -
11035 -pf04363 -
11040 HMM00288_1_pdu(pf00288) PDU1AB
11045 HMMacetkin_1(pf00871) PDU1AB
11050 HMM10662PduV_1_mix(pf10662) PDU1AB
11055 Hp_euts_babyPurple PDU1AB
11060 Ts_dodgerBlue PDU1AB
11065 HMMPduS_1_all(pf01512pf13375pf10531) PDU1AB
11070 HMM00465alcdh_3_mix(pf00465) PDU1AB
11075 HMMaldh_1_eutpdugrm(pf00171) PDU1AB EP_NTERM
11080 HMM01923cobtr_1_pdu(pf01923pf03928) PDU1AB
11085 P_babyPink PDU1AB
11090 HMM15953pduput_1_pdu1(pf15953) PDU1AB EP_NTERM
11095 HMMptac_2_mix(pf06130) PDU1AB
11100 H_driedBlood PDU1AB
11105 H_tan PDU1AB
11110 HMMpropdeh3_2_pdusml(pf02288) PDU1AB
11115 HMM08841dhact_1_pdu(pf08841pf18427) PDU1AB
11120 HMMpropdeh2_1_pdu(pf02287) PDU1AB EP_NTERM
11125 HMMpropdeh3_1_pdu(pf02288) PDU1AB EP_NTERM
11130 HMMpropdeh1_1_pdu(pf02286) PDU1AB
11135 Tsp_greenishBeige PDU1AB
11140 H_robinEggBlue GRM4
11145 -pf00230 -
11150 HMM10114PocR_1_pdu1(pf12833pf10114) PDU1AB
11155 -pf01656pf07685 -
11160 -pf03186 -
11165 -pf02570 -
11170 -pf01888 -
11175 -pf00590 -
11180 -pf05175 -
11185 -pf00590 -
11190 -pf01890pf11761pf11760 -
11195 -pf00590 -
11200 -pf02571 -
11205 -pf06180 -
BMC-P
BMC-H
BMC-T
Aldehyde dehydrogenase
PTAC
Signature enzyme
Alcohol dehydrogenase
Regulator

Salmonella enterica subsp. enterica serovar Fresno. Bacteria; Proteobacteria; Gammaproteobacteria; Enterobacterales;

EXB02__locus_1

PDU1AB

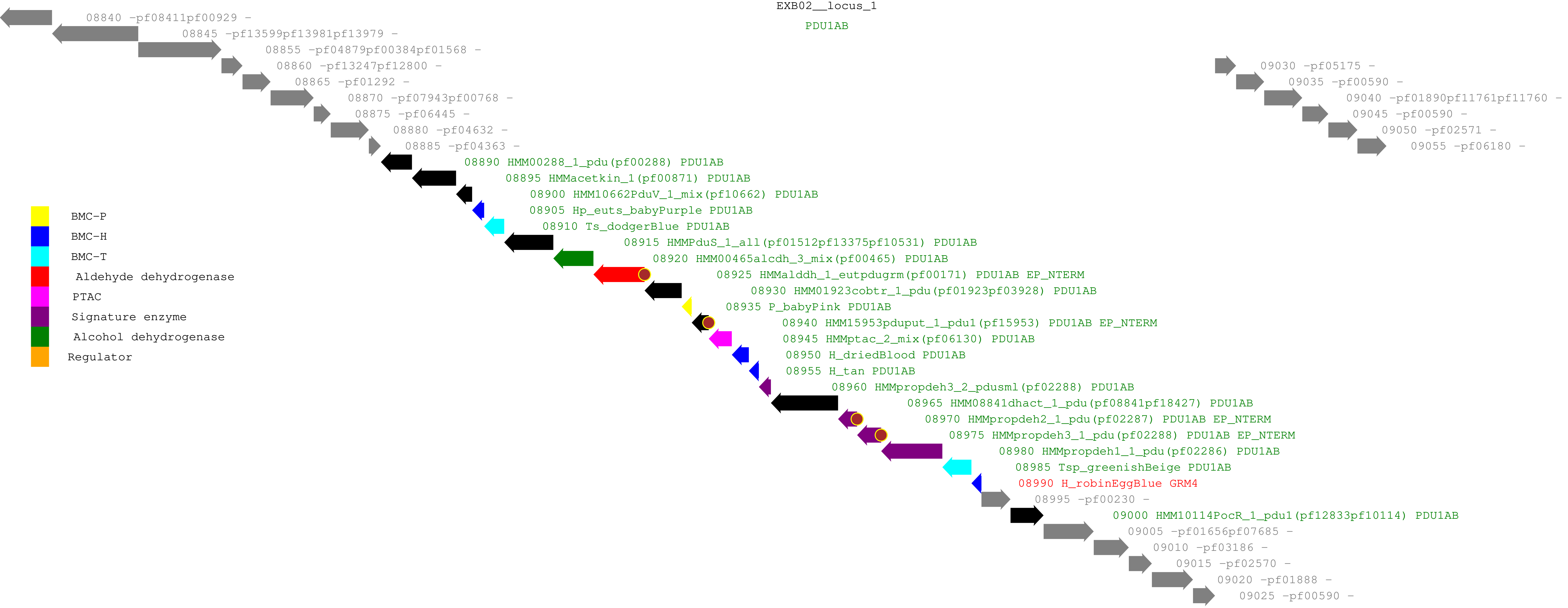

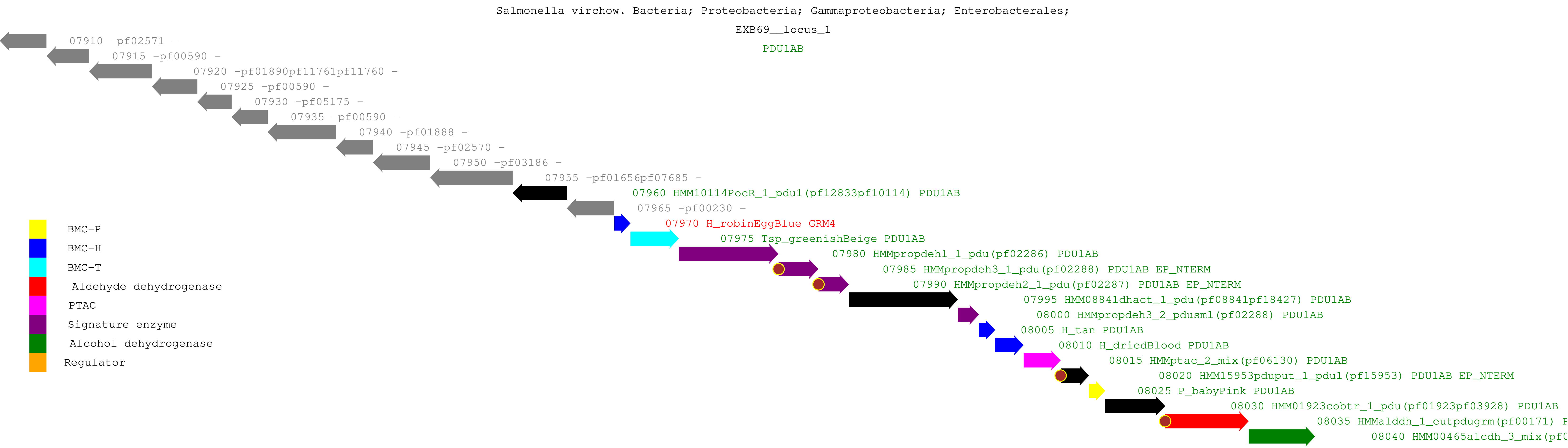
Salmonella virchow. Bacteria; Proteobacteria; Gammaproteobacteria; Enterobacterales;
EXB69__locus_1
PDU1AB
07910 -pf02571 -
07915 -pf00590 -
07920 -pf01890pf11761pf11760 -
07925 -pf00590 -
07930 -pf05175 -
07935 -pf00590 -
07940 -pf01888 -
07945 -pf02570 -
07950 -pf03186 -
07955 -pf01656pf07685 -
07960 HMM10114PocR_1_pdu1(pf12833pf10114) PDU1AB
07965 -pf00230 -
07970 H_robinEggBlue GRM4
07975 Tsp_greenishBeige PDU1AB
07980 HMMpropdeh1_1_pdu(pf02286) PDU1AB
07985 HMMpropdeh3_1_pdu(pf02288) PDU1AB EP_NTERM
07990 HMMpropdeh2_1_pdu(pf02287) PDU1AB EP_NTERM
07995 HMM08841dhact_1_pdu(pf08841pf18427) PDU1AB
08000 HMMpropdeh3_2_pdusml(pf02288) PDU1AB
08005 H_tan PDU1AB
08010 H_driedBlood PDU1AB
08015 HMMptac_2_mix(pf06130) PDU1AB
08020 HMM15953pduput_1_pdu1(pf15953) PDU1AB EP_NTERM
08025 P_babyPink PDU1AB
08030 HMM01923cobtr_1_pdu(pf01923pf03928) PDU1AB
08035 HMMalddh_1_eutpdugrm(pf00171)
08040 HMM00465alcdh_3_mix
BMC-P
BMC-H
BMC-T
Aldehyde dehydrogenase
PTAC
Signature enzyme
Alcohol dehydrogenase
Regulator

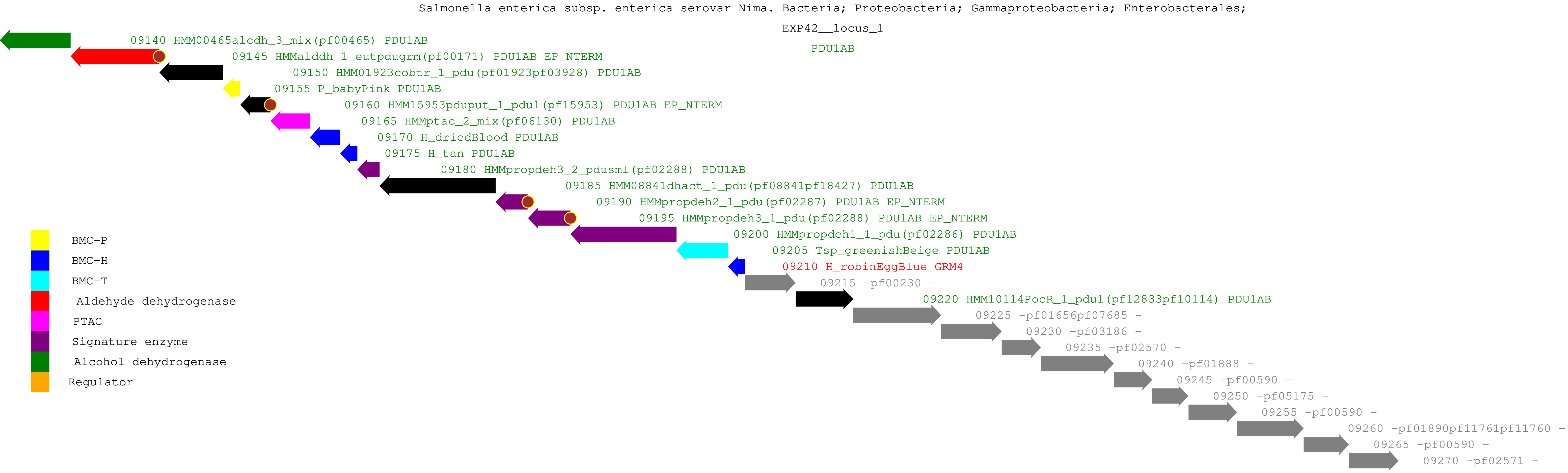

Salmonella enterica subsp. enterica serovar Nima. Bacteria; Proteobacteria; Gammaproteobacteria; Enterobacterales;
EXP42__locus_1
PDU1AB
09140 HMM00465alcdh_3_mix(pf00465) PDU1AB
09145 HMMalddh_1_eutpdugrm(pf00171) PDU1AB EP_NTERM
09150 HMM01923cobtr_1_pdu(pf01923pf03928) PDU1AB
09155 P_babyPink PDU1AB
09160 HMM15953pduput_1_pdu1(pf15953) PDU1AB EP_NTERM
09165 HMMptac_2_mix(pf06130) PDU1AB
09170 H_driedBlood PDU1AB
09175 H_tan PDU1AB
09180 HMMpropdeh3_2_pdusml(pf02288) PDU1AB
09185 HMM08841dhact_1_pdu(pf08841pf18427) PDU1AB
09190 HMMpropdeh2_1_pdu(pf02287) PDU1AB EP_NTERM
09195 HMMpropdeh3_1_pdu(pf02288) PDU1AB EP_NTERM
09200 HMMpropdeh1_1_pdu(pf02286) PDU1AB
09205 Tsp_greenishBeige PDU1AB
09210 H_robinEggBlue GRM4
09215 -pf00230 -
09220 HMM10114PocR_1_pdu1(pf12833pf10114) PDU1AB
09225 -pf01656pf07685 -
09230 -pf03186 -
09235 -pf02570 -
09240 -pf01888 -
09245 -pf00590 -
09250 -pf05175 -
09255 -pf00590 -
09260 -pf01890pf11761pf11760 -
09265 -pf00590 -
09270 -pf02571 -
BMC-P
BMC-H
BMC-T
Aldehyde dehydrogenase
PTAC
Signature enzyme
Alcohol dehydrogenase
Regulator

Salmonella enterica subsp. enterica serovar Anecho. Bacteria; Proteobacteria; Gammaproteobacteria; Enterobacterales;

EXP46__locus_2

PDU1AB

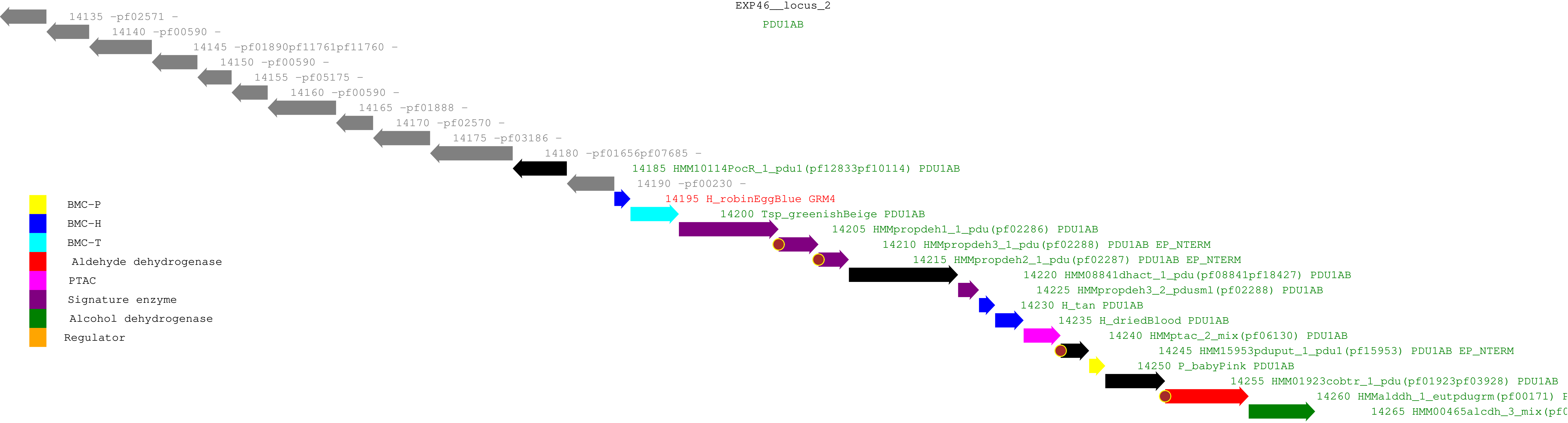

Salmonella enterica subsp. enterica serovar Havana. Bacteria; Proteobacteria; Gammaproteobacteria; Enterobacterales;

EXQ04__locus_1

PDU1AB

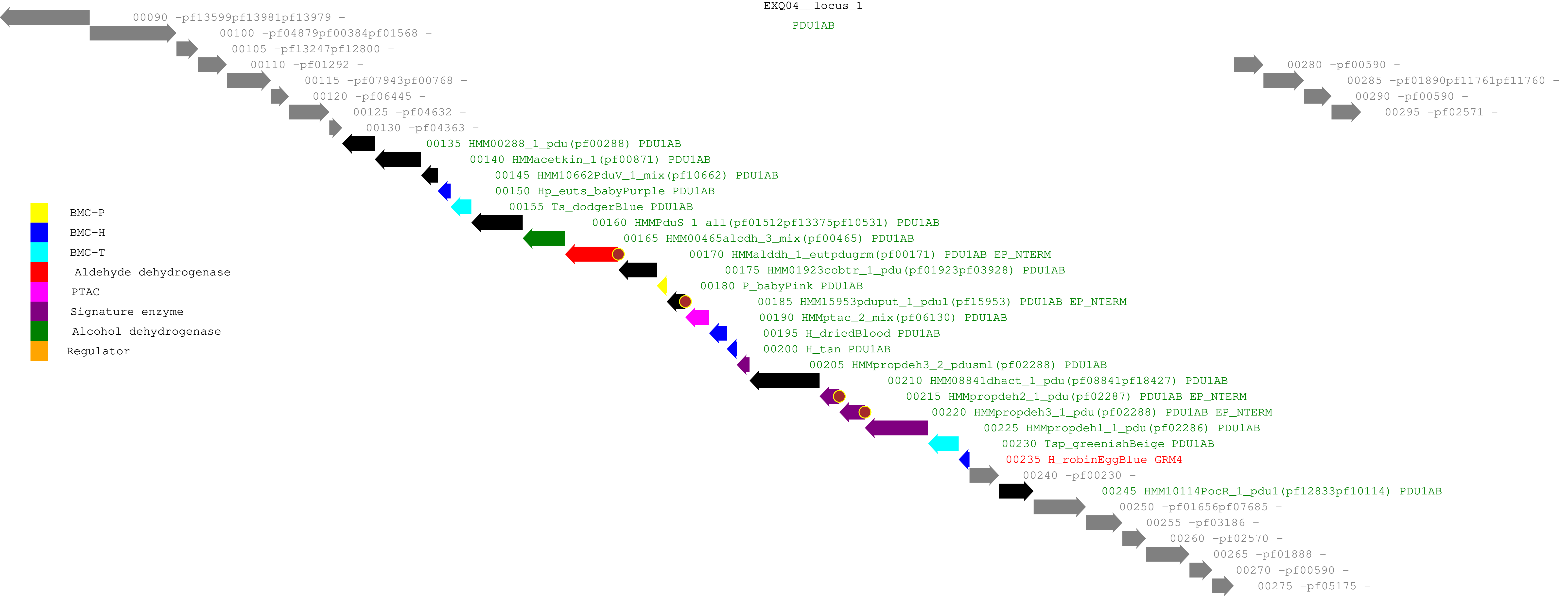

Klebsiella pneumoniae. Bacteria; Proteobacteria; Gammaproteobacteria; Enterobacterales;

EXT45__locus_1

PDU1AB

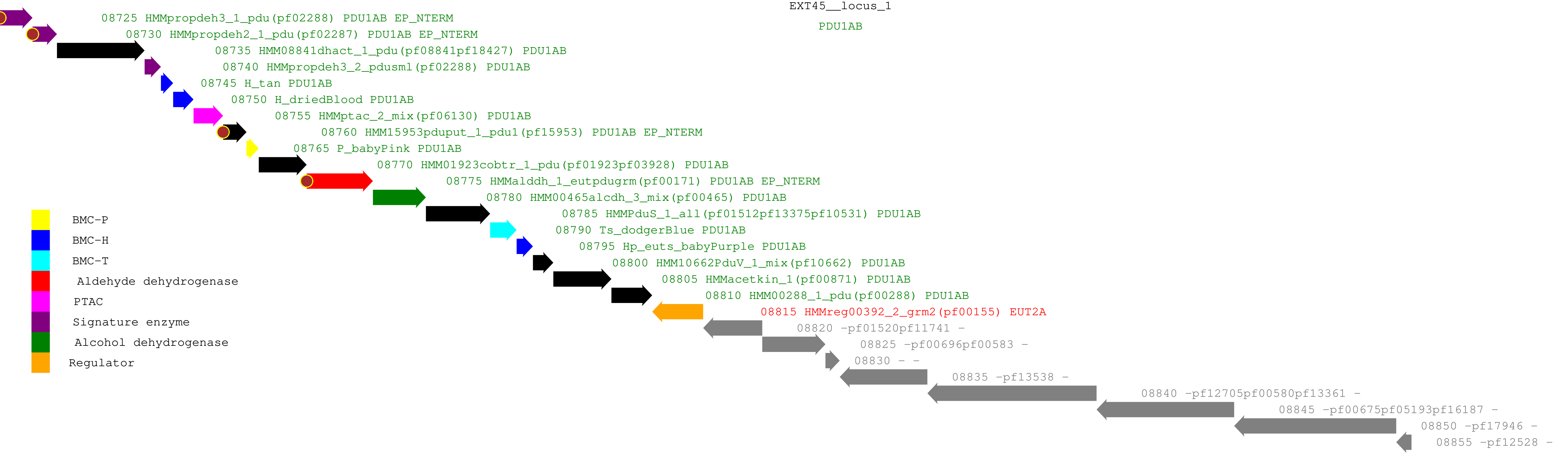

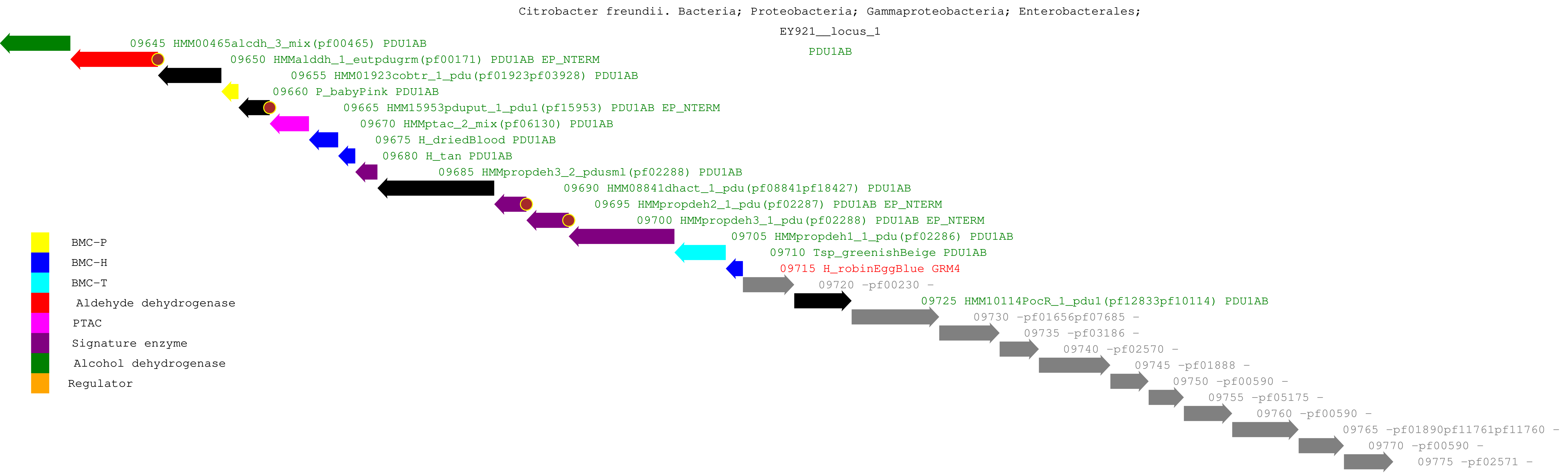
Citrobacter freundii. Bacteria; Proteobacteria; Gammaproteobacteria; Enterobacterales;
EY921__locus_1
PDU1AB
09645 HMM00465alcdh_3_mix(pf00465) PDU1AB
09650 HMMalddh_1_eutpdugrm(pf00171) PDU1AB EP_NTERM
09655 HMM01923cobtr_1_pdu(pf01923pf03928) PDU1AB
09660 P_babyPink PDU1AB
09665 HMM15953pduput_1_pdu1(pf15953) PDU1AB EP_NTERM
09670 HMMptac_2_mix(pf06130) PDU1AB
09675 H_driedBlood PDU1AB
09680 H_tan PDU1AB
09685 HMMpropdeh3_2_pdusml(pf02288) PDU1AB
09690 HMM08841dhact_1_pdu(pf08841pf18427) PDU1AB
09695 HMMpropdeh2_1_pdu(pf02287) PDU1AB EP_NTERM
09700 HMMpropdeh3_1_pdu(pf02288) PDU1AB EP_NTERM
09705 HMMpropdeh1_1_pdu(pf02286) PDU1AB
09710 Tsp_greenishBeige PDU1AB
09715 H_robinEggBlue GRM4
09720 -pf00230 -
09725 HMM10114PocR_1_pdu1(pf12833pf10114) PDU1AB
09730 -pf01656pf07685 -
09735 -pf03186 -
09740 -pf02570 -
09745 -pf01888 -
09750 -pf00590 -
09755 -pf05175 -
09760 -pf00590 -
09765 -pf01890pf11761pf11760 -
09770 -pf00590 -
09775 -pf02571 -
BMC-P
BMC-H
BMC-T
Aldehyde dehydrogenase
PTAC
Signature enzyme
Alcohol dehydrogenase
Regulator

Salmonella enterica subsp. enterica serovar Adjame. Bacteria; Proteobacteria; Gammaproteobacteria; Enterobacterales;

EYA23__locus_2

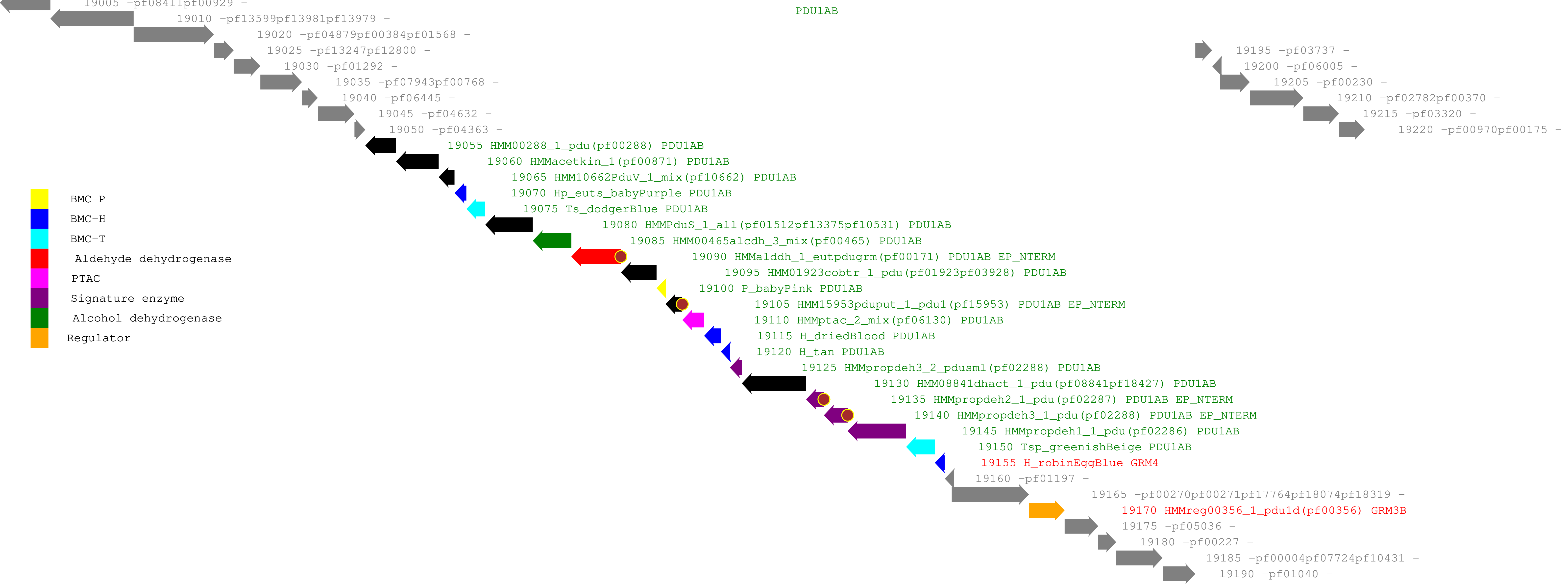

Escherichia albertii. Bacteria; Proteobacteria; Gammaproteobacteria; Enterobacterales;

EYS06__locus_2

PDU1AB

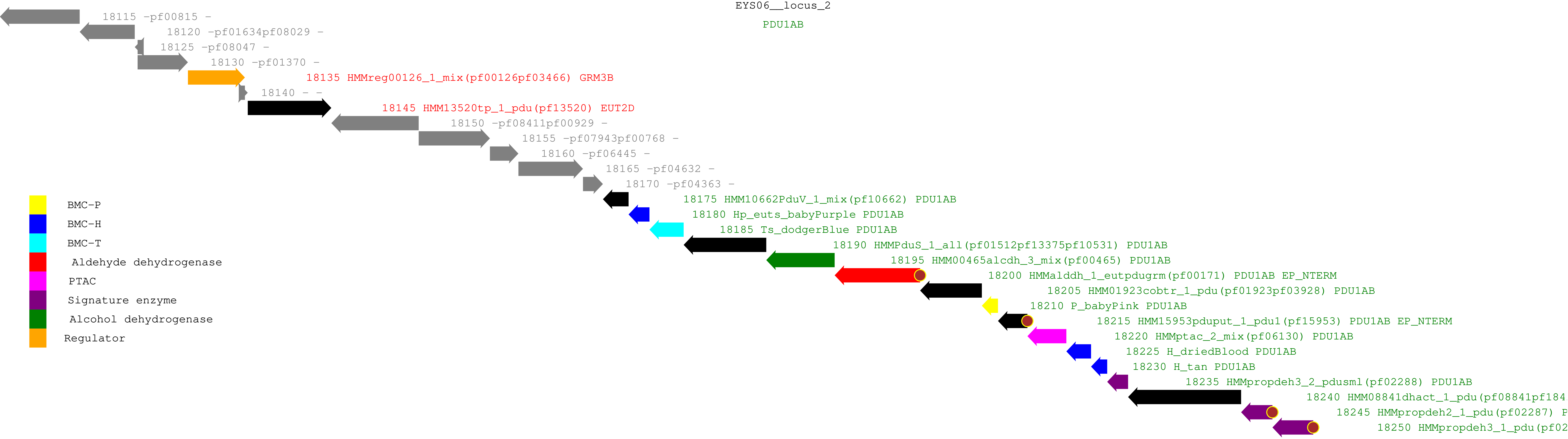

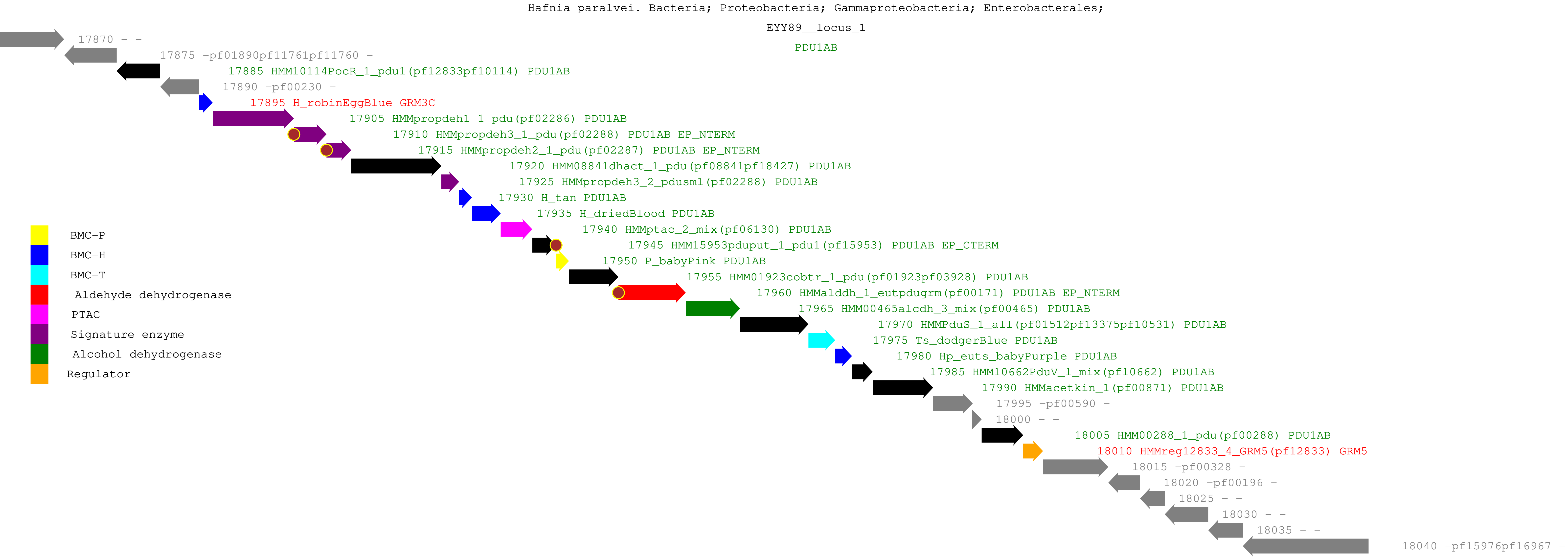

Hafnia paralvei. Bacteria; Proteobacteria; Gammaproteobacteria; Enterobacterales;
EYY89__locus_1
PDU1AB
17870 - -
17875 -pf01890pf11761pf11760 -
17885 HMM10114PocR_1_pdu1(pf12833pf10114) PDU1AB
17890 -pf00230 -
17895 H_robinEggBlue GRM3C
17905 HMMpropdeh1_1_pdu(pf02286) PDU1AB
17910 HMMpropdeh3_1_pdu(pf02288) PDU1AB EP_NTERM
17915 HMMpropdeh2_1_pdu(pf02287) PDU1AB EP_NTERM
17920 HMM08841dhact_1_pdu(pf08841pf18427) PDU1AB
17925 HMMpropdeh3_2_pdusml(pf02288) PDU1AB
17930 H_tan PDU1AB
17935 H_driedBlood PDU1AB
17940 HMMptac_2_mix(pf06130) PDU1AB
17945 HMM15953pduput_1_pdu1(pf15953) PDU1AB EP_CTERM
17950 P_babyPink PDU1AB
17955 HMM01923cobtr_1_pdu(pf01923pf03928) PDU1AB
17960 HMMalddh_1_eutpdugrm(pf00171) PDU1AB EP_NTERM
17965 HMM00465alcdh_3_mix(pf00465) PDU1AB
17970 HMMPduS_1_all(pf01512pf13375pf10531) PDU1AB
17975 Ts_dodgerBlue PDU1AB
17980 Hp_euts_babyPurple PDU1AB
17985 HMM10662PduV_1_mix(pf10662) PDU1AB
17990 HMMacetkin_1(pf00871) PDU1AB
17995 -pf00590 -
18000 - -
18005 HMM00288_1_pdu(pf00288) PDU1AB
18010 HMMreg12833_4_GRM5(pf12833) GRM5
18015 -pf00328 -
18020 -pf00196 -
18025 - -
18030 - -
18035 - -
18040 -pf15976pf16967 -
BMC-P
BMC-H
BMC-T
Aldehyde dehydrogenase
PTAC
Signature enzyme
Alcohol dehydrogenase
Regulator

Salmonella enterica subsp. enterica serovar Idikan. Bacteria; Proteobacteria; Gammaproteobacteria; Enterobacterales;

EZ668__locus_2

PDU1AB

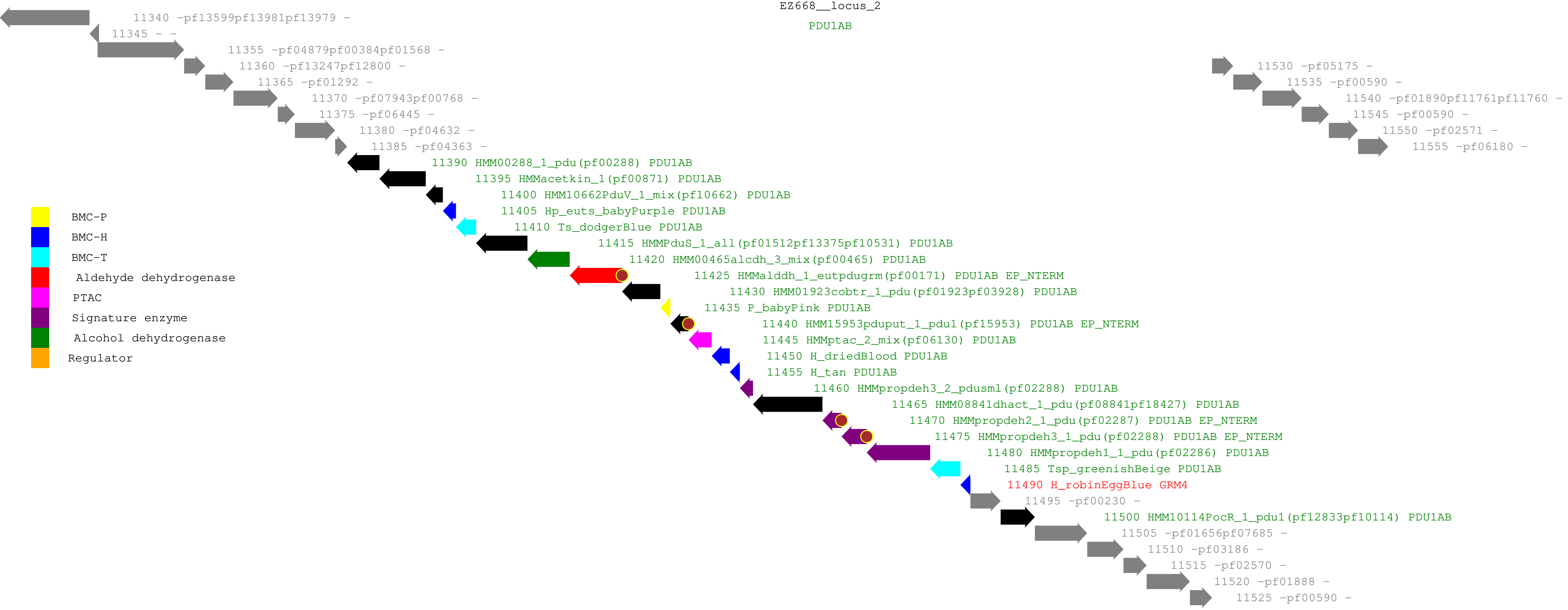

Salmonella enterica subsp. enterica serovar Wien. Bacteria; Proteobacteria; Gammaproteobacteria; Enterobacterales;

EZ751__locus_1

PDU1AB

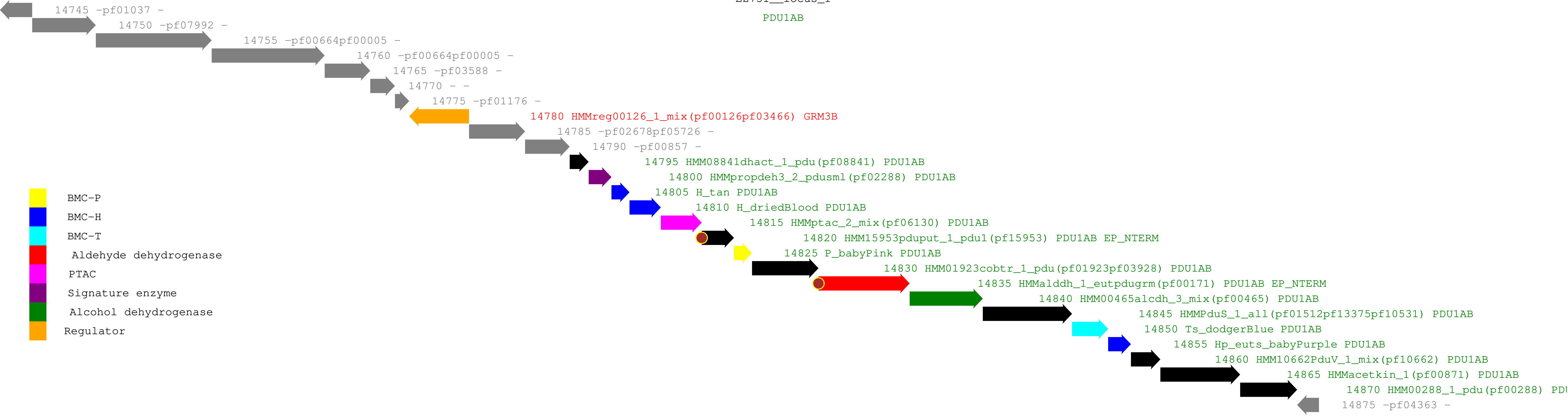

Salmonella enterica subsp. enterica serovar Fulica. Bacteria; Proteobacteria; Gammaproteobacteria; Enterobacterales;

EZ771__locus_2

PDU1AB

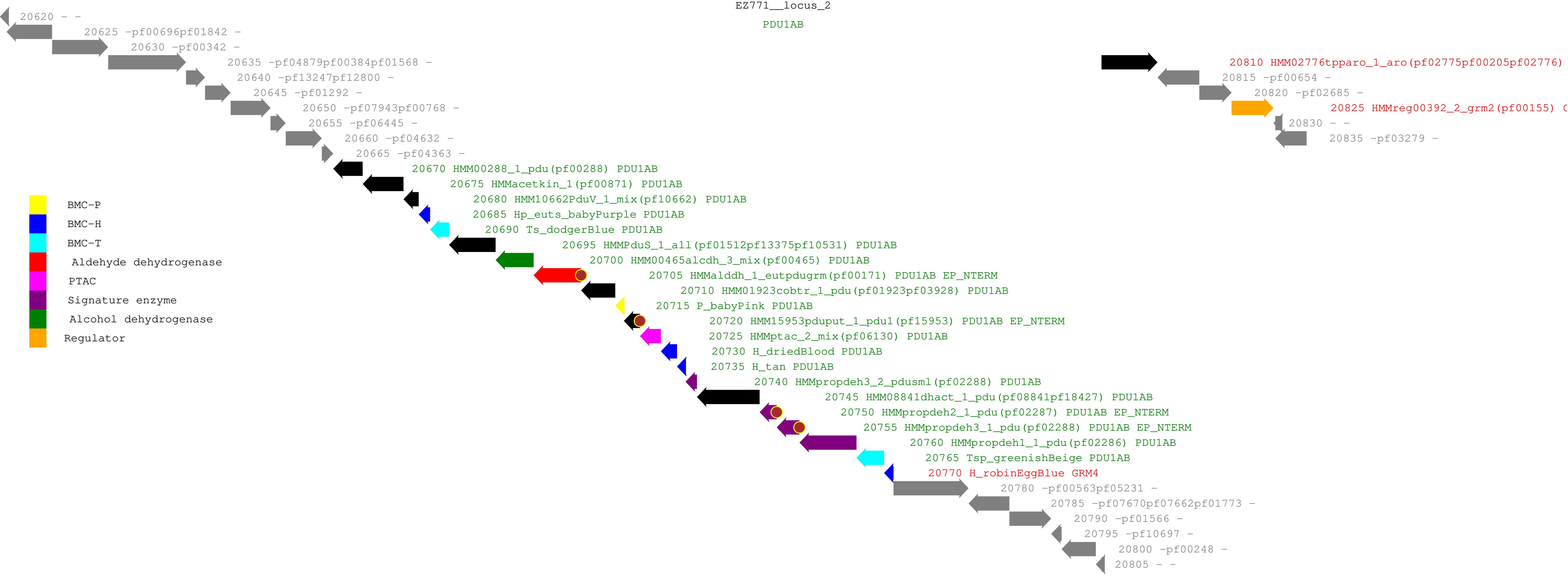

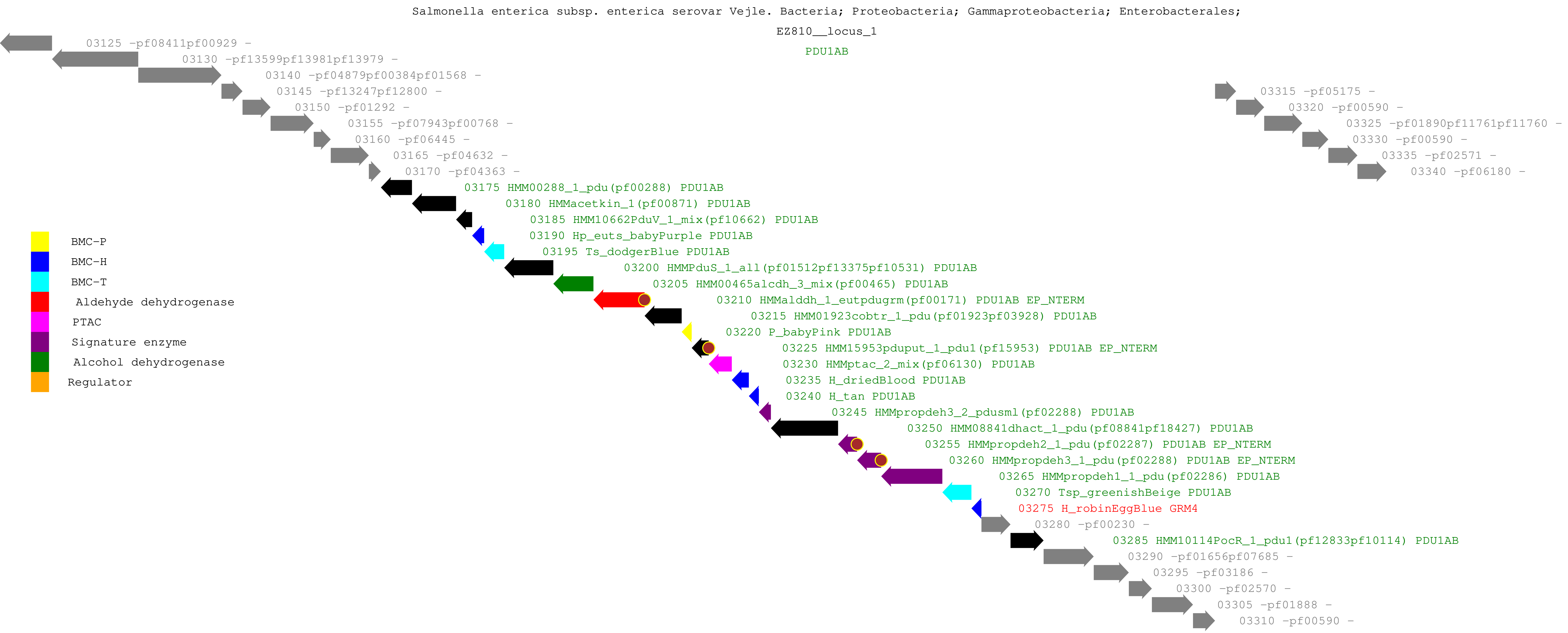

Salmonella enterica subsp. enterica serovar Vejle. Bacteria; Proteobacteria; Gammaproteobacteria; Enterobacterales;
EZ810__locus_1
PDU1AB
03125 -pf08411pf00929 -
03130 -pf13599pf13981pf13979 -
03140 -pf04879pf00384pf01568 -
03145 -pf13247pf12800 -
03150 -pf01292 -
03155 -pf07943pf00768 -
03160 -pf06445 -
03165 -pf04632 -
03170 -pf04363 -
03175 HMM00288_1_pdu(pf00288) PDU1AB
03180 HMMacetkin_1(pf00871) PDU1AB
03185 HMM10662PduV_1_mix(pf10662) PDU1AB
03190 Hp_euts_babyPurple PDU1AB
03195 Ts_dodgerBlue PDU1AB
03200 HMMPduS_1_all(pf01512pf13375pf10531) PDU1AB
03205 HMM00465alcdh_3_mix(pf00465) PDU1AB
03210 HMMalddh_1_eutpdugrm(pf00171) PDU1AB EP_NTERM
03215 HMM01923cobtr_1_pdu(pf01923pf03928) PDU1AB
03220 P_babyPink PDU1AB
03225 HMM15953pduput_1_pdu1(pf15953) PDU1AB EP_NTERM
03230 HMMptac_2_mix(pf06130) PDU1AB
03235 H_driedBlood PDU1AB
03240 H_tan PDU1AB
03245 HMMpropdeh3_2_pdusml(pf02288) PDU1AB
03250 HMM08841dhact_1_pdu(pf08841pf18427) PDU1AB
03255 HMMpropdeh2_1_pdu(pf02287) PDU1AB EP_NTERM
03260 HMMpropdeh3_1_pdu(pf02288) PDU1AB EP_NTERM
03265 HMMpropdeh1_1_pdu(pf02286) PDU1AB
03270 Tsp_greenishBeige PDU1AB
03275 H_robinEggBlue GRM4
03280 -pf00230 -
03285 HMM10114PocR_1_pdu1(pf12833pf10114) PDU1AB
03290 -pf01656pf07685 -
03295 -pf03186 -
03300 -pf02570 -
03305 -pf01888 -
03310 -pf00590 -
03315 -pf05175 -
03320 -pf00590 -
03325 -pf01890pf11761pf11760 -
03330 -pf00590 -
03335 -pf02571 -
03340 -pf06180 -
BMC-P
BMC-H
BMC-T
Aldehyde dehydrogenase
PTAC
Signature enzyme
Alcohol dehydrogenase
Regulator

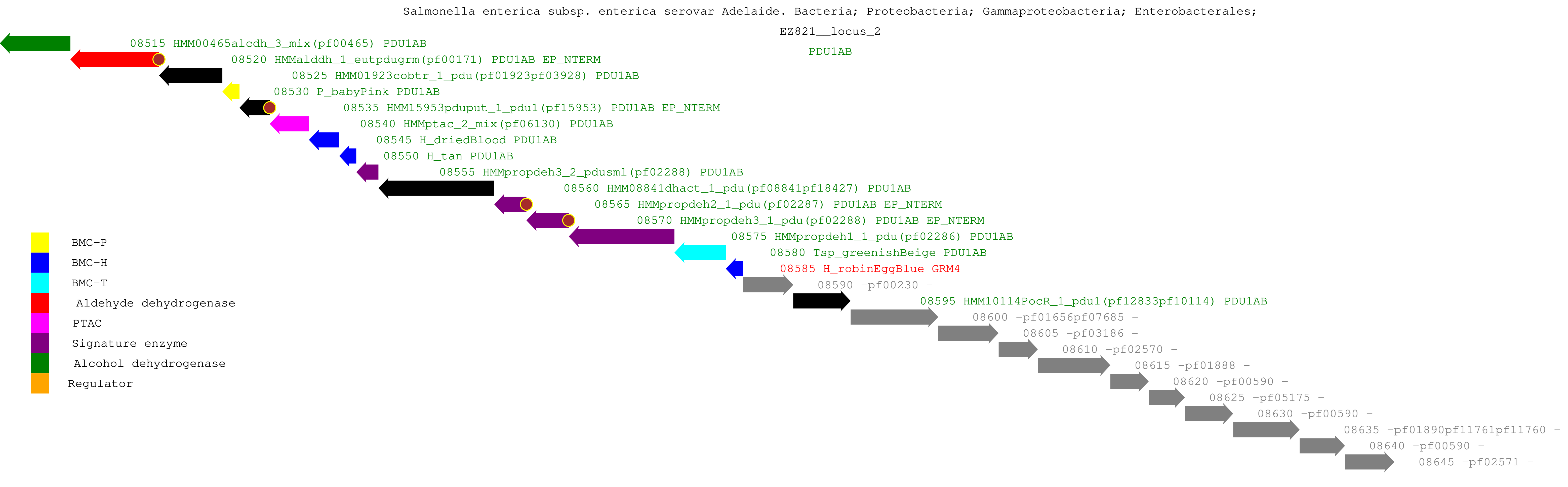
Salmonella enterica subsp. enterica serovar Adelaide. Bacteria; Proteobacteria; Gammaproteobacteria; Enterobacterales;
EZ821__locus_2
PDU1AB
08515 HMM00465alcdh_3_mix(pf00465) PDU1AB
08520 HMMalddh_1_eutpdugrm(pf00171) PDU1AB EP_NTERM
08525 HMM01923cobtr_1_pdu(pf01923pf03928) PDU1AB
08530 P_babyPink PDU1AB
08535 HMM15953pduput_1_pdu1(pf15953) PDU1AB EP_NTERM
08540 HMMptac_2_mix(pf06130) PDU1AB
08545 H_driedBlood PDU1AB
08550 H_tan PDU1AB
08555 HMMpropdeh3_2_pdusml(pf02288) PDU1AB
08560 HMM08841dhact_1_pdu(pf08841pf18427) PDU1AB
08565 HMMpropdeh2_1_pdu(pf02287) PDU1AB EP_NTERM
08570 HMMpropdeh3_1_pdu(pf02288) PDU1AB EP_NTERM
08575 HMMpropdeh1_1_pdu(pf02286) PDU1AB
08580 Tsp_greenishBeige PDU1AB
08585 H_robinEggBlue GRM4
08590 -pf00230 -
08595 HMM10114PocR_1_pdu1(pf12833pf10114) PDU1AB
08600 -pf01656pf07685 -
08605 -pf03186 -
08610 -pf02570 -
08615 -pf01888 -
08620 -pf00590 -
08625 -pf05175 -
08630 -pf00590 -
08635 -pf01890pf11761pf11760 -
08640 -pf00590 -
08645 -pf02571 -
BMC-P
BMC-H
BMC-T
Aldehyde dehydrogenase
PTAC
Signature enzyme
Alcohol dehydrogenase
Regulator

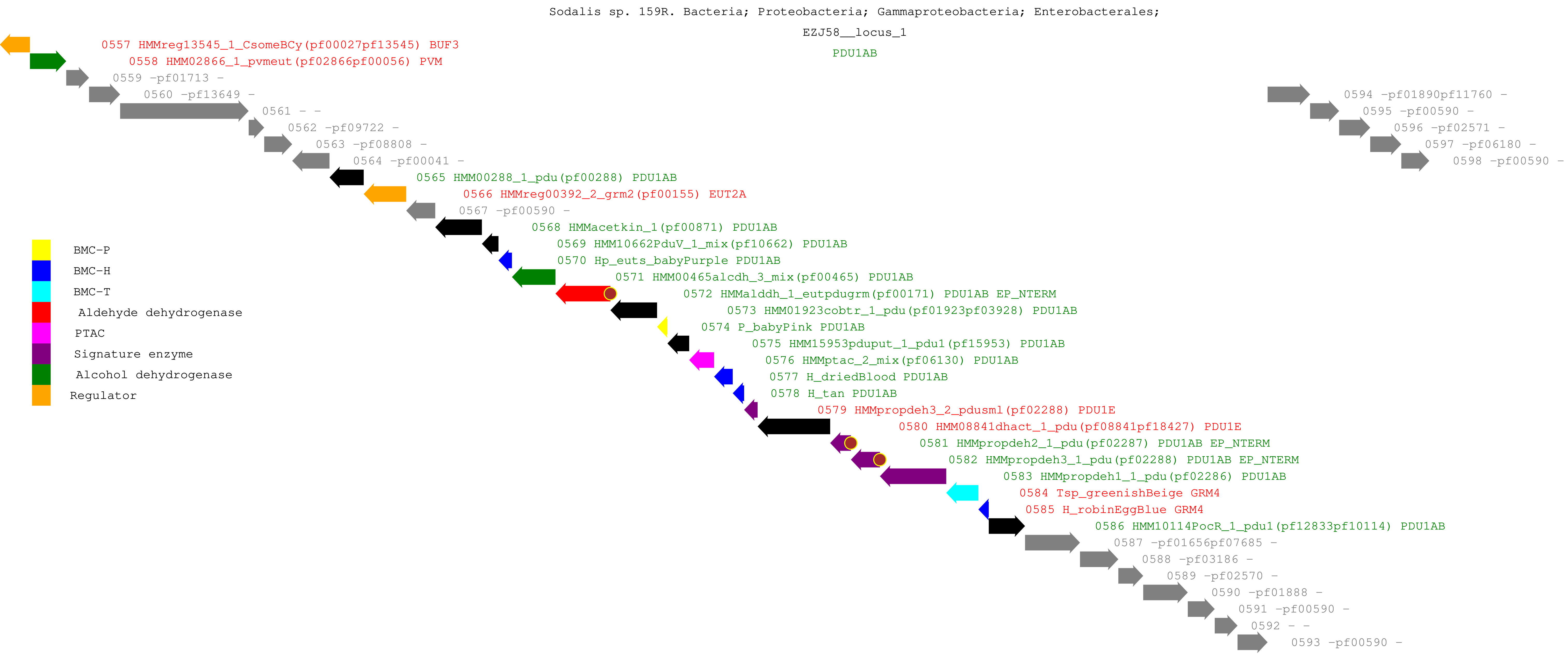
Sodalis sp. 159R. Bacteria; Proteobacteria; Gammaproteobacteria; Enterobacterales;
EZJ58__locus_1
PDU1AB
0557 HMMreg13545_1_CsomeBCy(pf00027pf13545) BUF3
0558 HMM02866_1_pvmeut(pf02866pf00056) PVM
0559 -pf01713 -
0560 -pf13649 -
0561 - -
0562 -pf09722 -
0563 -pf08808 -
0564 -pf00041 -
0565 HMM00288_1_pdu(pf00288) PDU1AB
0566 HMMreg00392_2_grm2(pf00155) EUT2A
0567 -pf00590 -
0568 HMMacetkin_1(pf00871) PDU1AB
0569 HMM10662PduV_1_mix(pf10662) PDU1AB
0570 Hp_euts_babyPurple PDU1AB
0571 HMM00465alcdh_3_mix(pf00465) PDU1AB
0572 HMMalddh_1_eutpdugrm(pf00171) PDU1AB EP_NTERM
0573 HMM01923cobtr_1_pdu(pf01923pf03928) PDU1AB
0574 P_babyPink PDU1AB
0575 HMM15953pduput_1_pdu1(pf15953) PDU1AB
0576 HMMptac_2_mix(pf06130) PDU1AB
0577 H_driedBlood PDU1AB
0578 H_tan PDU1AB
0579 HMMpropdeh3_2_pdusml(pf02288) PDU1E
0580 HMM08841dhact_1_pdu(pf08841pf18427) PDU1E
0581 HMMpropdeh2_1_pdu(pf02287) PDU1AB EP_NTERM
0582 HMMpropdeh3_1_pdu(pf02288) PDU1AB EP_NTERM
0583 HMMpropdeh1_1_pdu(pf02286) PDU1AB
0584 Tsp_greenishBeige GRM4
0585 H_robinEggBlue GRM4
0586 HMM10114PocR_1_pdu1(pf12833pf10114) PDU1AB
0587 -pf01656pf07685 -
0588 -pf03186 -
0589 -pf02570 -
0590 -pf01888 -
0591 -pf00590 -
0592 - -
0593 -pf00590 -
0594 -pf01890pf11760 -
0595 -pf00590 -
0596 -pf02571 -
0597 -pf06180 -
0598 -pf00590 -
BMC-P
BMC-H
BMC-T
Aldehyde dehydrogenase
PTAC
Signature enzyme
Alcohol dehydrogenase
Regulator

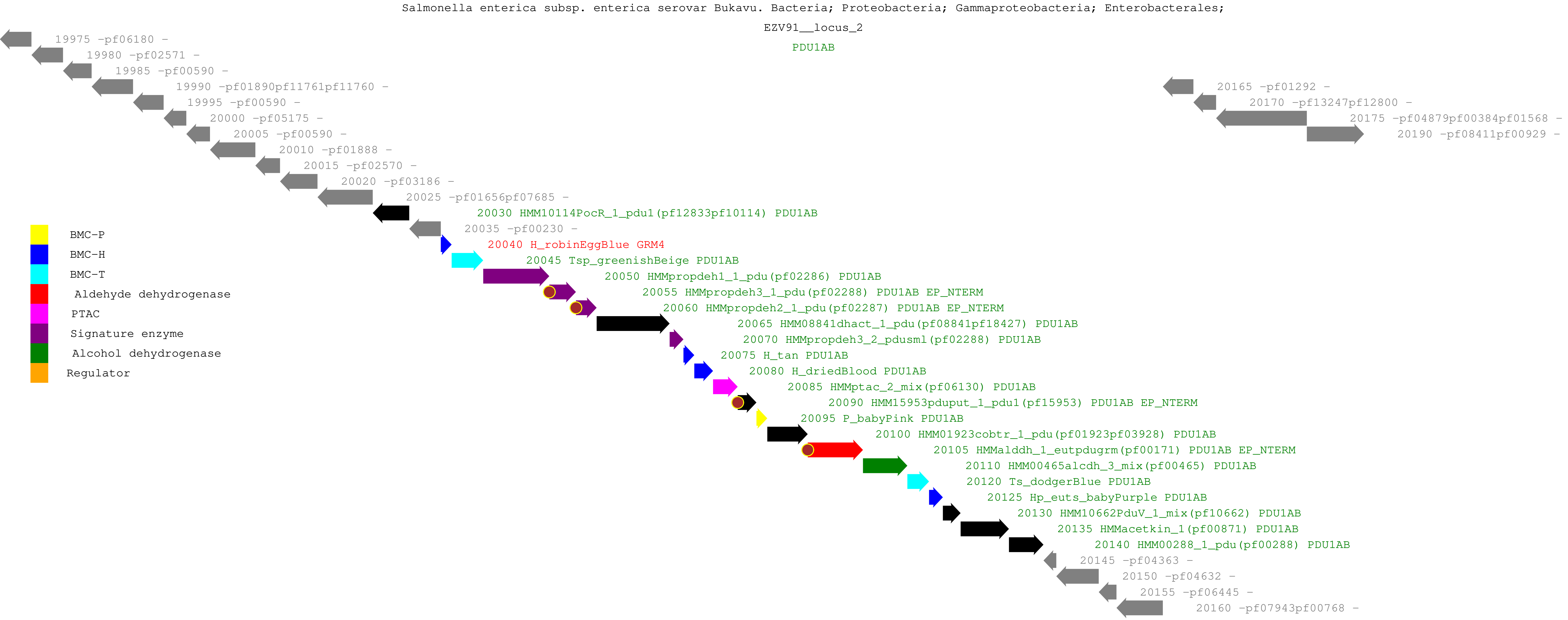

Salmonella enterica subsp. enterica serovar Bukavu. Bacteria; Proteobacteria; Gammaproteobacteria; Enterobacterales;
EZV91__locus_2
PDU1AB
19975 -pf06180 -
19980 -pf02571 -
19985 -pf00590 -
19990 -pf01890pf11761pf11760 -
19995 -pf00590 -
20000 -pf05175 -
20005 -pf00590 -
20010 -pf01888 -
20015 -pf02570 -
20020 -pf03186 -
20025 -pf01656pf07685 -
20030 HMM10114PocR_1_pdu1(pf12833pf10114) PDU1AB
20035 -pf00230 -
20040 H_robinEggBlue GRM4
20045 Tsp_greenishBeige PDU1AB
20050 HMMpropdeh1_1_pdu(pf02286) PDU1AB
20055 HMMpropdeh3_1_pdu(pf02288) PDU1AB EP_NTERM
20060 HMMpropdeh2_1_pdu(pf02287) PDU1AB EP_NTERM
20065 HMM08841dhact_1_pdu(pf08841pf18427) PDU1AB
20070 HMMpropdeh3_2_pdusml(pf02288) PDU1AB
20075 H_tan PDU1AB
20080 H_driedBlood PDU1AB
20085 HMMptac_2_mix(pf06130) PDU1AB
20090 HMM15953pduput_1_pdu1(pf15953) PDU1AB EP_NTERM
20095 P_babyPink PDU1AB
20100 HMM01923cobtr_1_pdu(pf01923pf03928) PDU1AB
20105 HMMalddh_1_eutpdugrm(pf00171) PDU1AB EP_NTERM
20110 HMM00465alcdh_3_mix(pf00465) PDU1AB
20120 Ts_dodgerBlue PDU1AB
20125 Hp_euts_babyPurple PDU1AB
20130 HMM10662PduV_1_mix(pf10662) PDU1AB
20135 HMMacetkin_1(pf00871) PDU1AB
20140 HMM00288_1_pdu(pf00288) PDU1AB
20145 -pf04363 -
20150 -pf04632 -
20155 -pf06445 -
20160 -pf07943pf00768 -
20165 -pf01292 -
20170 -pf13247pf12800 -
20175 -pf04879pf00384pf01568 -
20190 -pf08411pf00929 -
BMC-P
BMC-H
BMC-T
Aldehyde dehydrogenase
PTAC
Signature enzyme
Alcohol dehydrogenase
Regulator

Salmonella oranienberg. Bacteria; Proteobacteria; Gammaproteobacteria; Enterobacterales;

EZX63__locus_2

PDU1AB

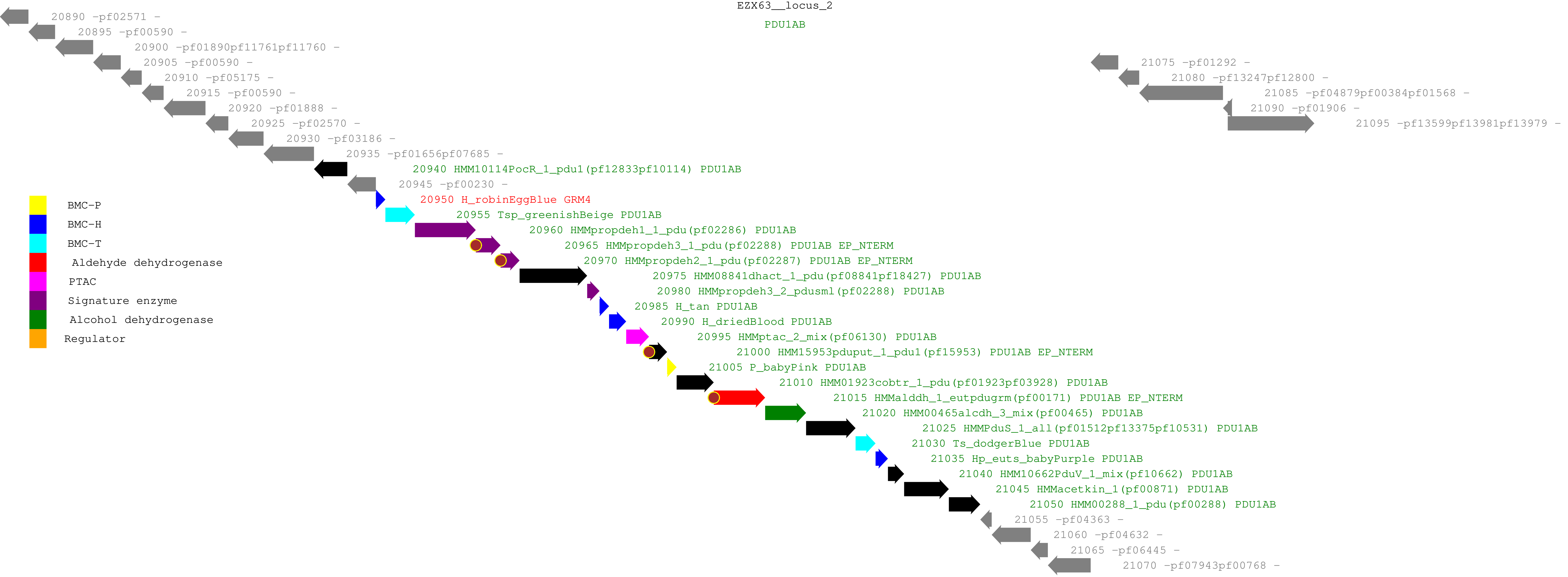

Citrobacter braakii. Bacteria; Proteobacteria; Gammaproteobacteria; Enterobacterales;

F0327__locus_1

PDU1AB

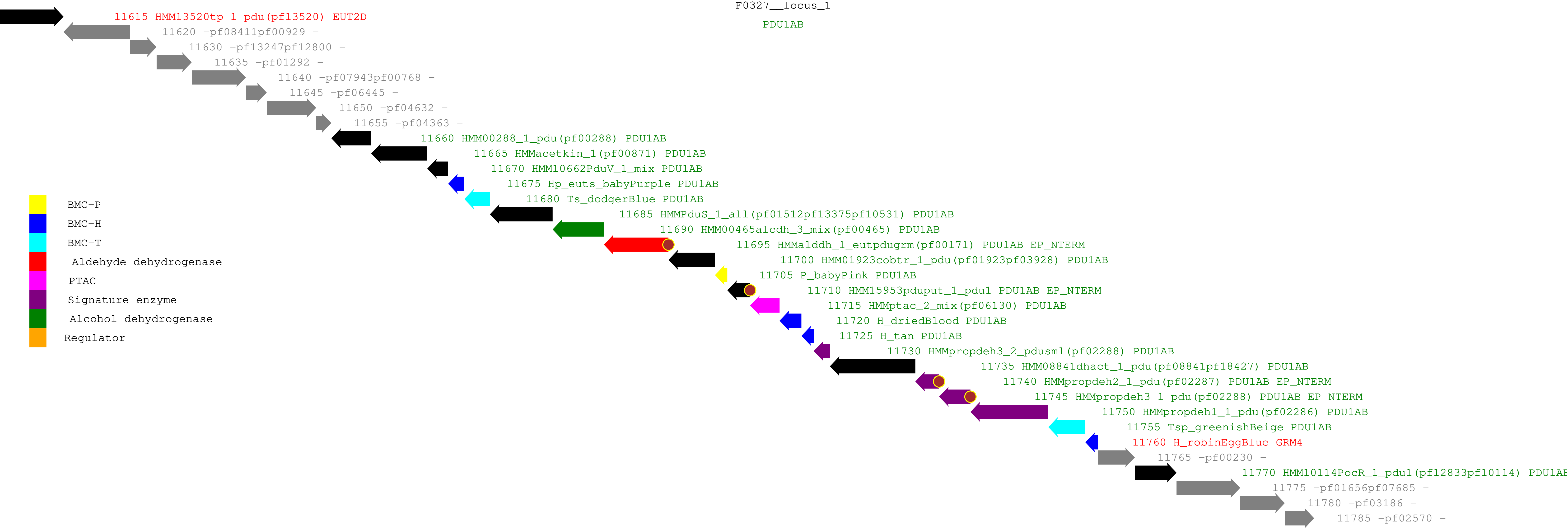

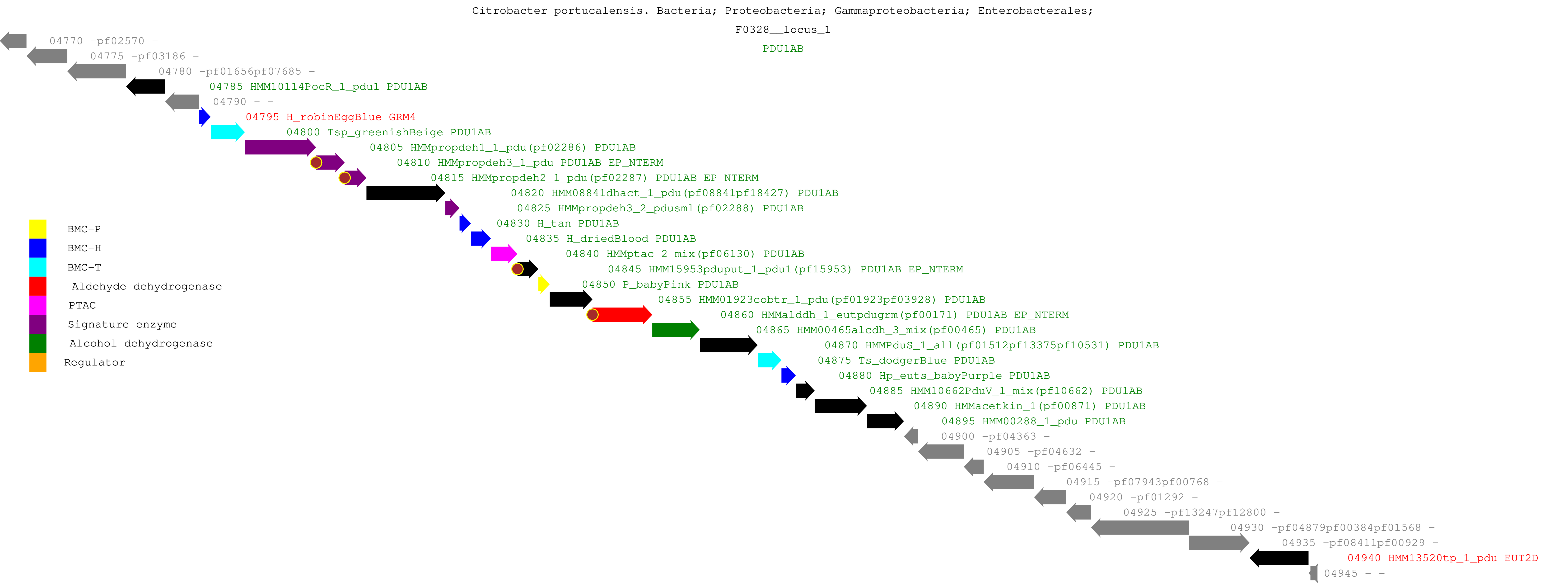

Citrobacter portucalensis. Bacteria; Proteobacteria; Gammaproteobacteria; Enterobacterales;
F0328__locus_1
PDU1AB
04770 -pf02570 -
04775 -pf03186 -
04780 -pf01656pf07685 -
04785 HMM10114PocR_1_pdu1 PDU1AB
04790 - -
04795 H_robinEggBlue GRM4
04800 Tsp_greenishBeige PDU1AB
04805 HMMpropdeh1_1_pdu(pf02286) PDU1AB
04810 HMMpropdeh3_1_pdu PDU1AB EP_NTERM
04815 HMMpropdeh2_1_pdu(pf02287) PDU1AB EP_NTERM
04820 HMM08841dhact_1_pdu(pf08841pf18427) PDU1AB
04825 HMMpropdeh3_2_pdusml(pf02288) PDU1AB
04830 H_tan PDU1AB
04835 H_driedBlood PDU1AB
04840 HMMptac_2_mix(pf06130) PDU1AB
04845 HMM15953pduput_1_pdu1(pf15953) PDU1AB EP_NTERM
04850 P_babyPink PDU1AB
04855 HMM01923cobtr_1_pdu(pf01923pf03928) PDU1AB
04860 HMMalddh_1_eutpdugrm(pf00171) PDU1AB EP_NTERM
04865 HMM00465alcdh_3_mix(pf00465) PDU1AB
04870 HMMPduS_1_all(pf01512pf13375pf10531) PDU1AB
04875 Ts_dodgerBlue PDU1AB
04880 Hp_euts_babyPurple PDU1AB
04885 HMM10662PduV_1_mix(pf10662) PDU1AB
04890 HMMacetkin_1(pf00871) PDU1AB
04895 HMM00288_1_pdu PDU1AB
04900 -pf04363 -
04905 -pf04632 -
04910 -pf06445 -
04915 -pf07943pf00768 -
04920 -pf01292 -
04925 -pf13247pf12800 -
04930 -pf04879pf00384pf01568 -
04935 -pf08411pf00929 -
04940 HMM13520tp_1_pdu EUT2D
04945 - -
BMC-P
BMC-H
BMC-T
Aldehyde dehydrogenase
PTAC
Signature enzyme
Alcohol dehydrogenase
Regulator

Citrobacter werkmanii. Bacteria; Proteobacteria; Gammaproteobacteria; Enterobacterales;

F0329__locus_1

PDU1AB

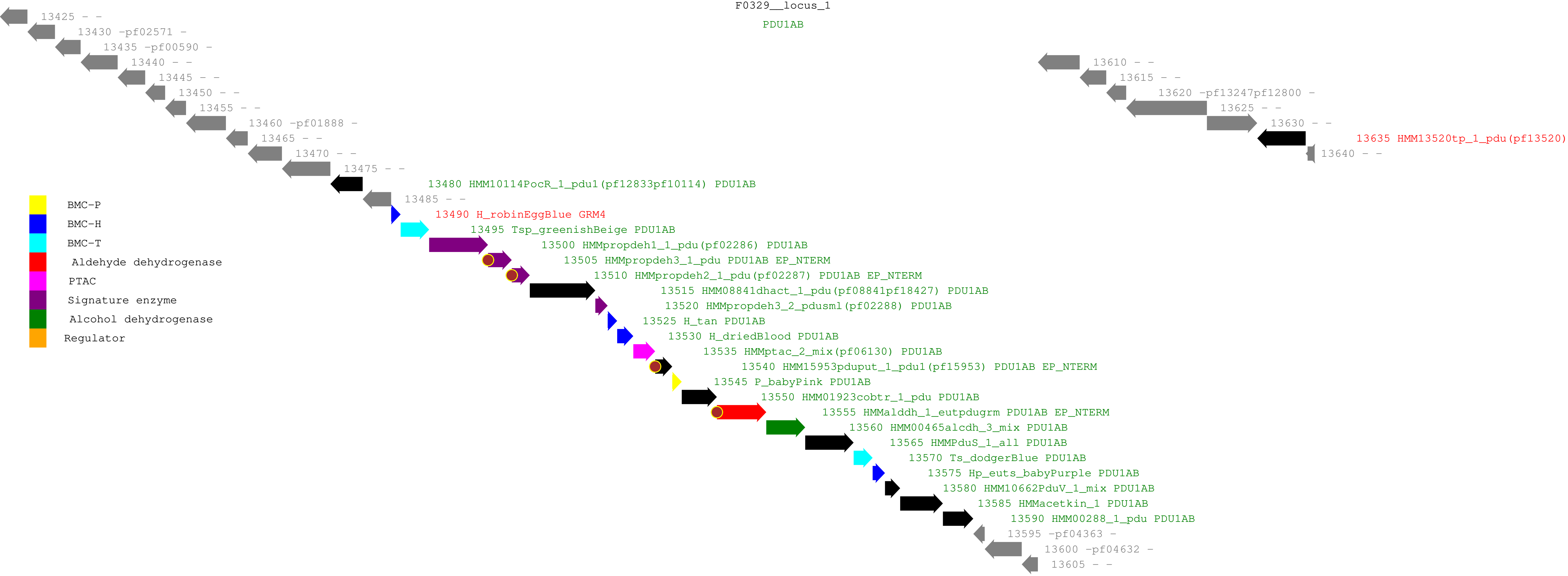

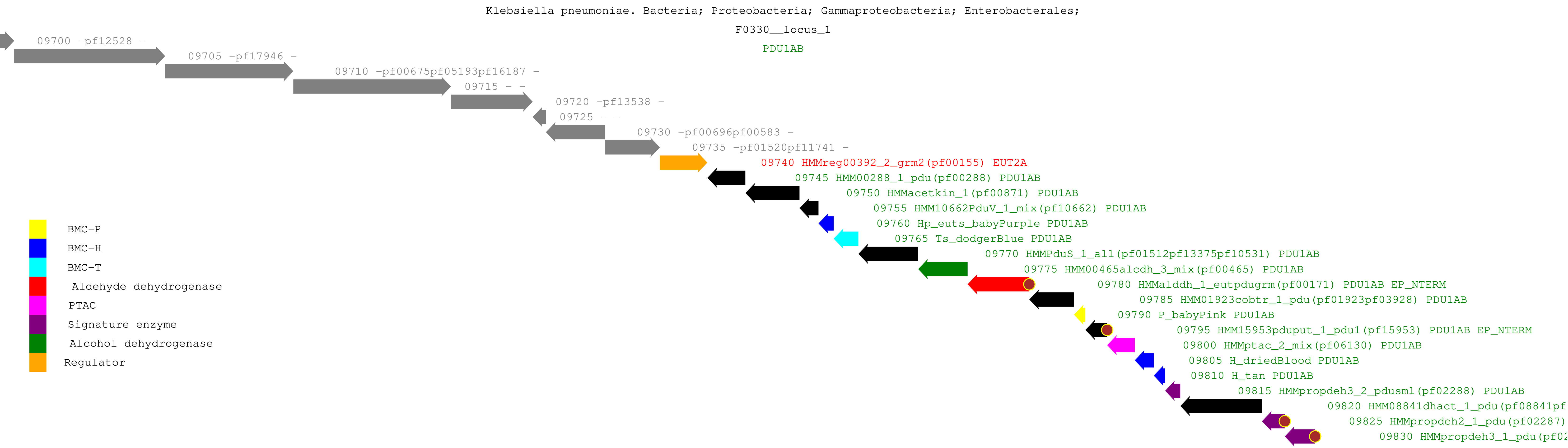
Klebsiella pneumoniae. Bacteria; Proteobacteria; Gammaproteobacteria; Enterobacterales;
F0330__locus_1
PDU1AB
09700 -pf12528 -
09705 -pf17946 -
09710 -pf00675pf05193pf16187 -
09715 - -
09720 -pf13538 -
09725 - -
09730 -pf00696pf00583 -
09735 -pf01520pf11741 -
09740 HMMreg00392_2_grm2(pf00155) EUT2A
09745 HMM00288_1_pdu(pf00288) PDU1AB
09750 HMMacetkin_1(pf00871) PDU1AB
09755 HMM10662PduV_1_mix(pf10662) PDU1AB
09760 Hp_euts_babyPurple PDU1AB
09765 Ts_dodgerBlue PDU1AB
09770 HMMPduS_1_all(pf01512pf13375pf10531) PDU1AB
09775 HMM00465alcdh_3_mix(pf00465) PDU1AB
09780 HMMalddh_1_eutpdugrm(pf00171) PDU1AB EP_NTERM
09785 HMM01923cobtr_1_pdu(pf01923pf03928) PDU1AB
09790 P_babyPink PDU1AB
09795 HMM15953pduput_1_pdu1(pf15953) PDU1AB EP_NTERM
09800 HMMptac_2_mix(pf06130) PDU1AB
09805 H_driedBlood PDU1AB
09810 H_tan PDU1AB
09815 HMMpropdeh3_2_pdusml(pf02288) PDU1AB
09820 HMM08841dhact_1_pdu(pf08841pf
09825 HMMpropdeh2_1_pdu(pf02287)
09830 HMMpropdeh3_1_pdu(pf0
BMC-P
BMC-H
BMC-T
Aldehyde dehydrogenase
PTAC
Signature enzyme
Alcohol dehydrogenase
Regulator

Pseudocitrobacter sp. 73. Bacteria; Proteobacteria; Gammaproteobacteria; Enterobacterales;

F0Q32__locus_2

PDU1AB

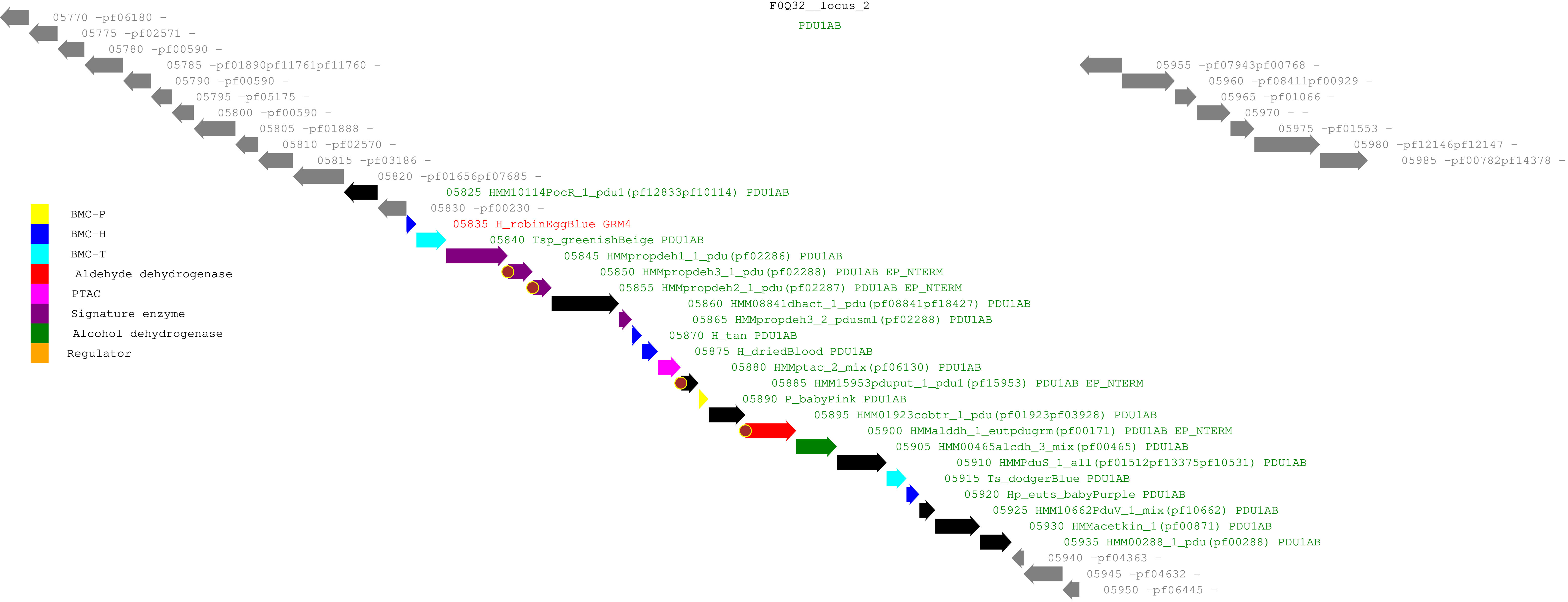

Salmonella enterica (Salmonella choleraesuis). Bacteria; Proteobacteria; Gammaproteobacteria; Enterobacterales;

F1527__locus_1

PDU1AB

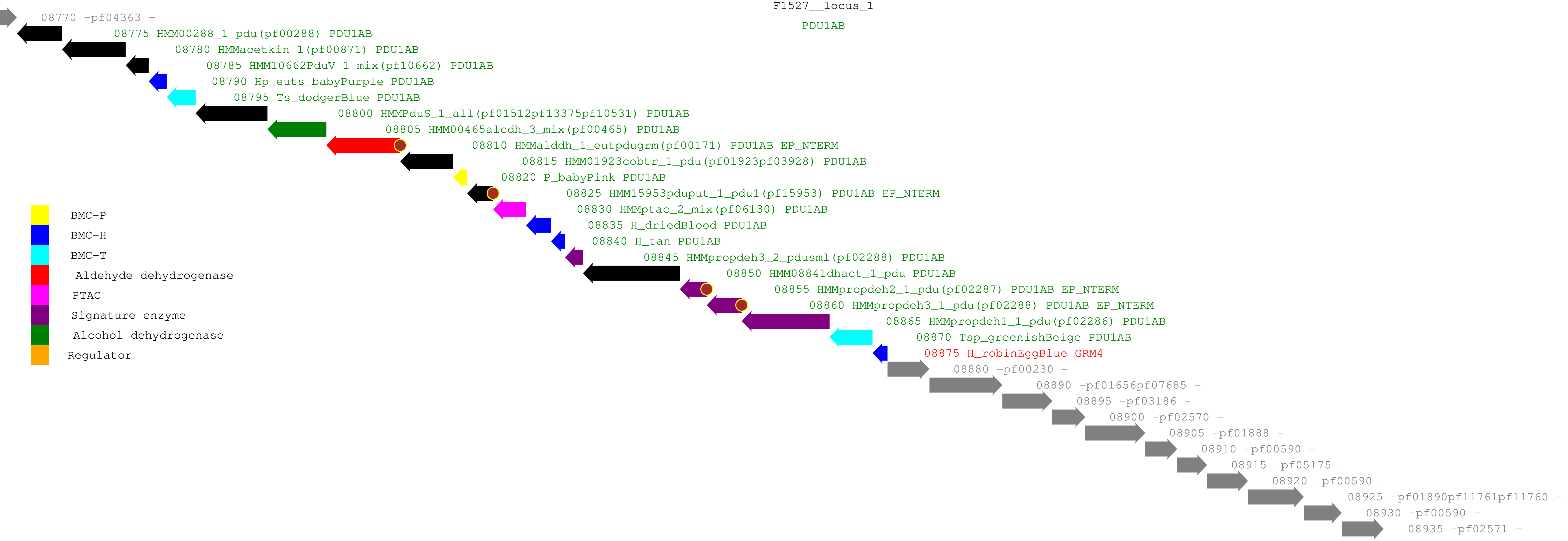

Klebsiella pneumoniae. Bacteria; Proteobacteria; Gammaproteobacteria; Enterobacterales;

F1D54__locus_1

PDU1AB

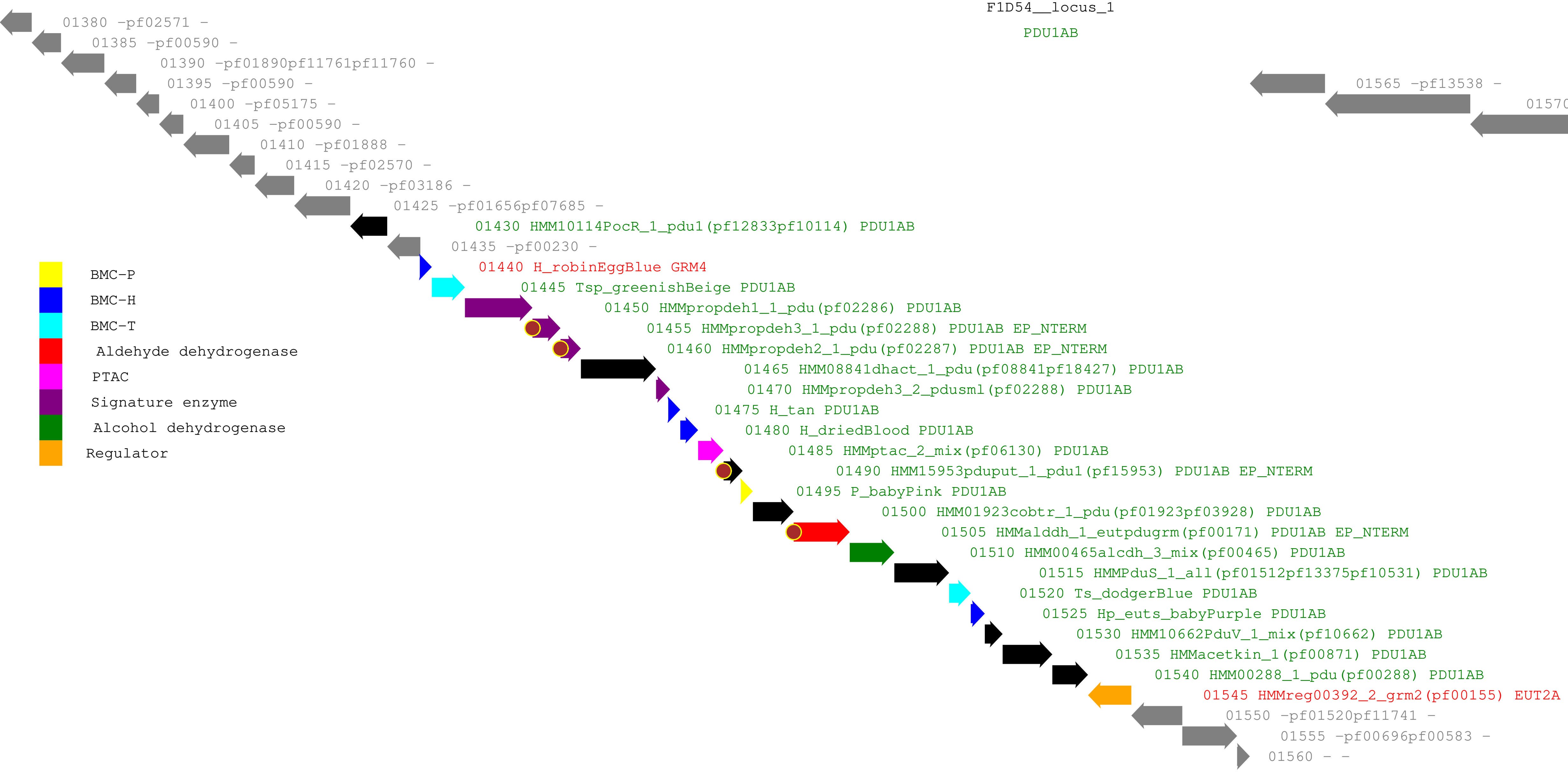

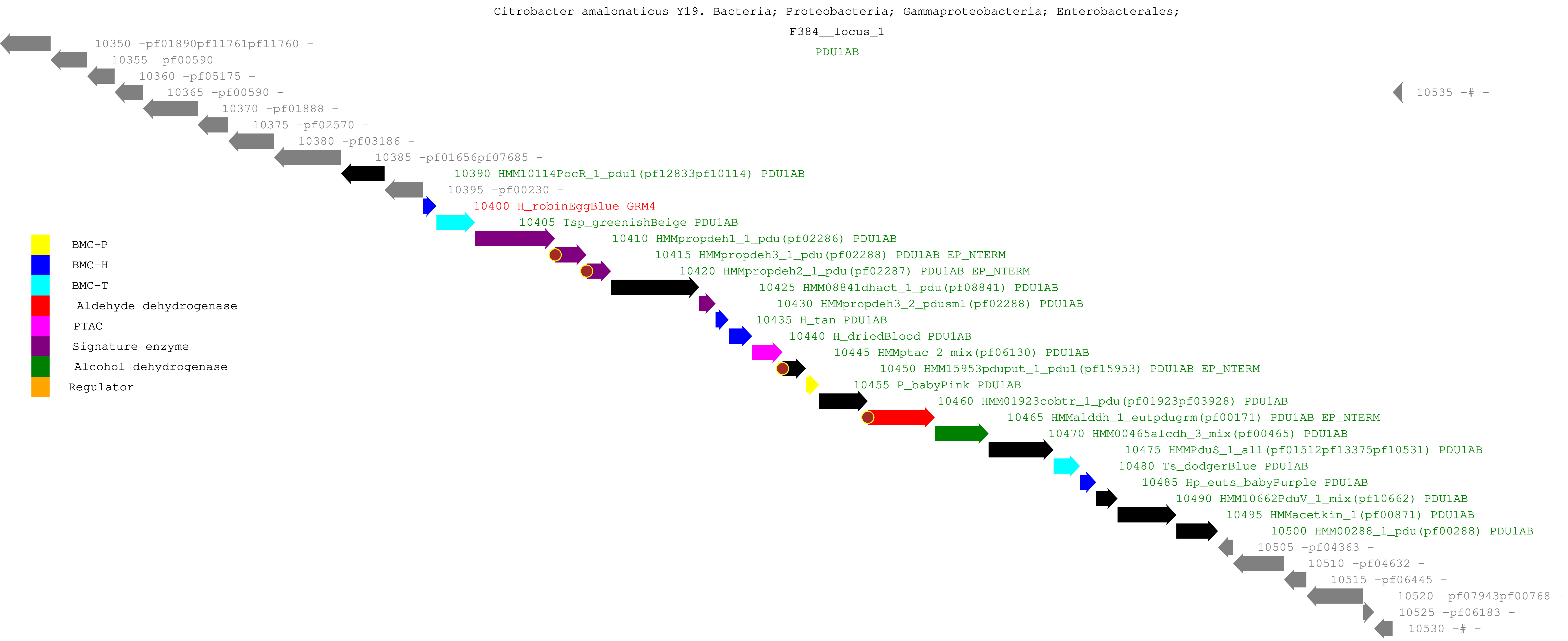
Citrobacter amalonaticus Y19. Bacteria; Proteobacteria; Gammaproteobacteria; Enterobacterales;
F384__locus_1
PDU1AB
10350 -pf01890pf11761pf11760 -
10355 -pf00590 -
10360 -pf05175 -
10365 -pf00590 -
10370 -pf01888 -
10375 -pf02570 -
10380 -pf03186 -
10385 -pf01656pf07685 -
10390 HMM10114PocR_1_pdu1(pf12833pf10114) PDU1AB
10395 -pf00230 -
10400 H_robinEggBlue GRM4
10405 Tsp_greenishBeige PDU1AB
10410 HMMpropdeh1_1_pdu(pf02286) PDU1AB
10415 HMMpropdeh3_1_pdu(pf02288) PDU1AB EP_NTERM
10420 HMMpropdeh2_1_pdu(pf02287) PDU1AB EP_NTERM
10425 HMM08841dhact_1_pdu(pf08841) PDU1AB
10430 HMMpropdeh3_2_pdusml(pf02288) PDU1AB
10435 H_tan PDU1AB
10440 H_driedBlood PDU1AB
10445 HMMptac_2_mix(pf06130) PDU1AB
10450 HMM15953pduput_1_pdu1(pf15953) PDU1AB EP_NTERM
10455 P_babyPink PDU1AB
10460 HMM01923cobtr_1_pdu(pf01923pf03928) PDU1AB
10465 HMMalddh_1_eutpdugrm(pf00171) PDU1AB EP_NTERM
10470 HMM00465alcdh_3_mix(pf00465) PDU1AB
10475 HMMPduS_1_all(pf01512pf13375pf10531) PDU1AB
10480 Ts_dodgerBlue PDU1AB
10485 Hp_euts_babyPurple PDU1AB
10490 HMM10662PduV_1_mix(pf10662) PDU1AB
10495 HMMacetkin_1(pf00871) PDU1AB
10500 HMM00288_1_pdu(pf00288) PDU1AB
10505 -pf04363 -
10510 -pf04632 -
10515 -pf06445 -
10520 -pf07943pf00768 -
10525 -pf06183 -
10530 -# -
10535 -# -
BMC-P
BMC-H
BMC-T
Aldehyde dehydrogenase
PTAC
Signature enzyme
Alcohol dehydrogenase
Regulator

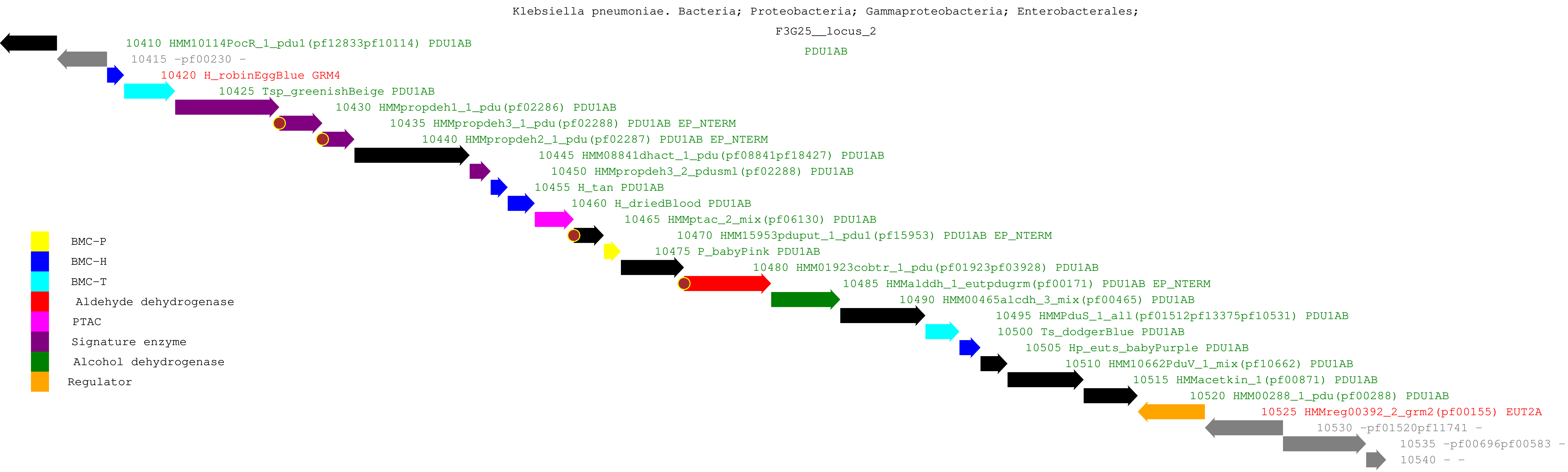
Klebsiella pneumoniae. Bacteria; Proteobacteria; Gammaproteobacteria; Enterobacterales;
F3G25__locus_2
PDU1AB
10410 HMM10114PocR_1_pdu1(pf12833pf10114) PDU1AB
10415 -pf00230 -
10420 H_robinEggBlue GRM4
10425 Tsp_greenishBeige PDU1AB
10430 HMMpropdeh1_1_pdu(pf02286) PDU1AB
10435 HMMpropdeh3_1_pdu(pf02288) PDU1AB EP_NTERM
10440 HMMpropdeh2_1_pdu(pf02287) PDU1AB EP_NTERM
10445 HMM08841dhact_1_pdu(pf08841pf18427) PDU1AB
10450 HMMpropdeh3_2_pdusml(pf02288) PDU1AB
10455 H_tan PDU1AB
10460 H_driedBlood PDU1AB
10465 HMMptac_2_mix(pf06130) PDU1AB
10470 HMM15953pduput_1_pdu1(pf15953) PDU1AB EP_NTERM
10475 P_babyPink PDU1AB
10480 HMM01923cobtr_1_pdu(pf01923pf03928) PDU1AB
10485 HMMalddh_1_eutpdugrm(pf00171) PDU1AB EP_NTERM
10490 HMM00465alcdh_3_mix(pf00465) PDU1AB
10495 HMMPduS_1_all(pf01512pf13375pf10531) PDU1AB
10500 Ts_dodgerBlue PDU1AB
10505 Hp_euts_babyPurple PDU1AB
10510 HMM10662PduV_1_mix(pf10662) PDU1AB
10515 HMMacetkin_1(pf00871) PDU1AB
10520 HMM00288_1_pdu(pf00288) PDU1AB
10525 HMMreg00392_2_grm2(pf00155) EUT2A
10530 -pf01520pf11741 -
10535 -pf00696pf00583 -
10540 - -
BMC-P
BMC-H
BMC-T
Aldehyde dehydrogenase
PTAC
Signature enzyme
Alcohol dehydrogenase
Regulator

Enterobacteriaceae bacterium bta3-1. Bacteria; Proteobacteria; Gammaproteobacteria; Enterobacterales;

F652__locus_1

PDU1AB

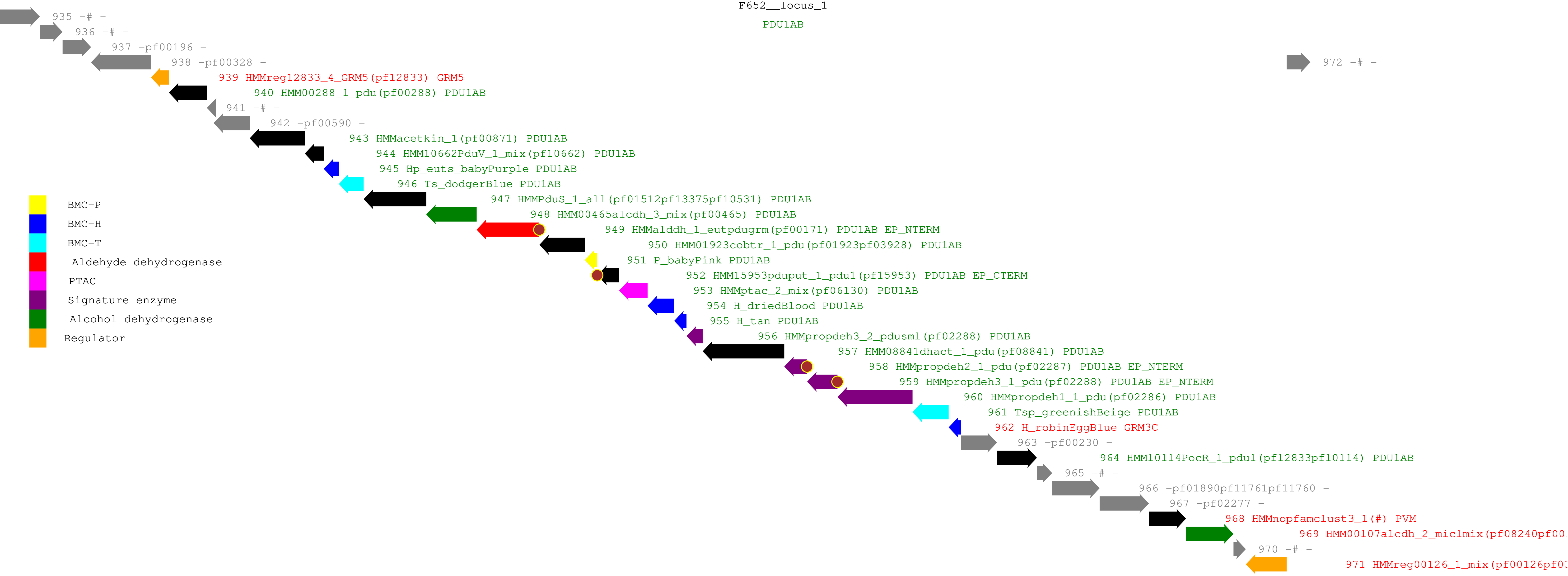

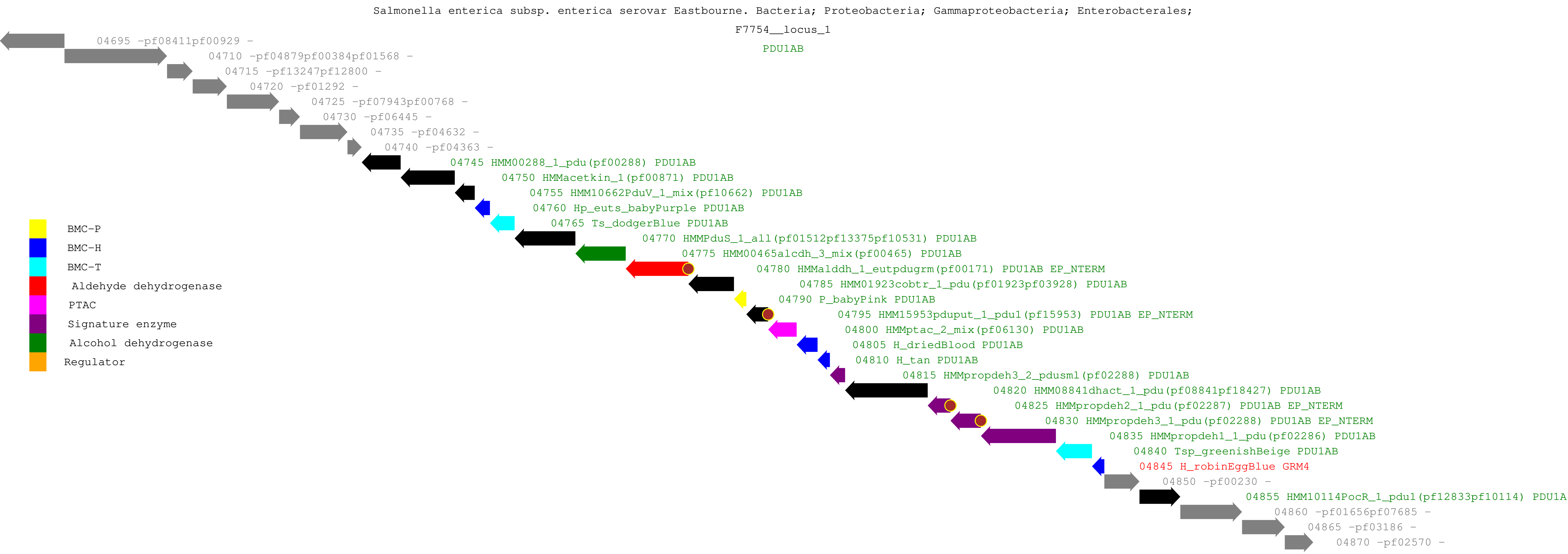
Salmonella enterica subsp. enterica serovar Eastbourne. Bacteria; Proteobacteria; Gammaproteobacteria; Enterobacterales;
F7754__locus_1
PDU1AB
04695 -pf08411pf00929 -
04710 -pf04879pf00384pf01568 -
04715 -pf13247pf12800 -
04720 -pf01292 -
04725 -pf07943pf00768 -
04730 -pf06445 -
04735 -pf04632 -
04740 -pf04363 -
04745 HMM00288_1_pdu(pf00288) PDU1AB
04750 HMMacetkin_1(pf00871) PDU1AB
04755 HMM10662PduV_1_mix(pf10662) PDU1AB
04760 Hp_euts_babyPurple PDU1AB
04765 Ts_dodgerBlue PDU1AB
04770 HMMPduS_1_all(pf01512pf13375pf10531) PDU1AB
04775 HMM00465alcdh_3_mix(pf00465) PDU1AB
04780 HMMalddh_1_eutpdugrm(pf00171) PDU1AB EP_NTERM
04785 HMM01923cobtr_1_pdu(pf01923pf03928) PDU1AB
04790 P_babyPink PDU1AB
04795 HMM15953pduput_1_pdu1(pf15953) PDU1AB EP_NTERM
04800 HMMptac_2_mix(pf06130) PDU1AB
04805 H_driedBlood PDU1AB
04810 H_tan PDU1AB
04815 HMMpropdeh3_2_pdusml(pf02288) PDU1AB
04820 HMM08841dhact_1_pdu(pf08841pf18427) PDU1AB
04825 HMMpropdeh2_1_pdu(pf02287) PDU1AB EP_NTERM
04830 HMMpropdeh3_1_pdu(pf02288) PDU1AB EP_NTERM
04835 HMMpropdeh1_1_pdu(pf02286) PDU1AB
04840 Tsp_greenishBeige PDU1AB
04845 H_robinEggBlue GRM4
04850 -pf00230 -
04855 HMM10114PocR_1_pdu1(pf12833pf10114) PDU1A
04860 -pf01656pf07685 -
04865 -pf03186 -
04870 -pf02570 -
BMC-P
BMC-H
BMC-T
Aldehyde dehydrogenase
PTAC
Signature enzyme
Alcohol dehydrogenase
Regulator

Salmonella enterica subsp. enterica serovar Nchanga. Bacteria; Proteobacteria; Gammaproteobacteria; Enterobacterales;

FA839__locus_1

PDU1AB

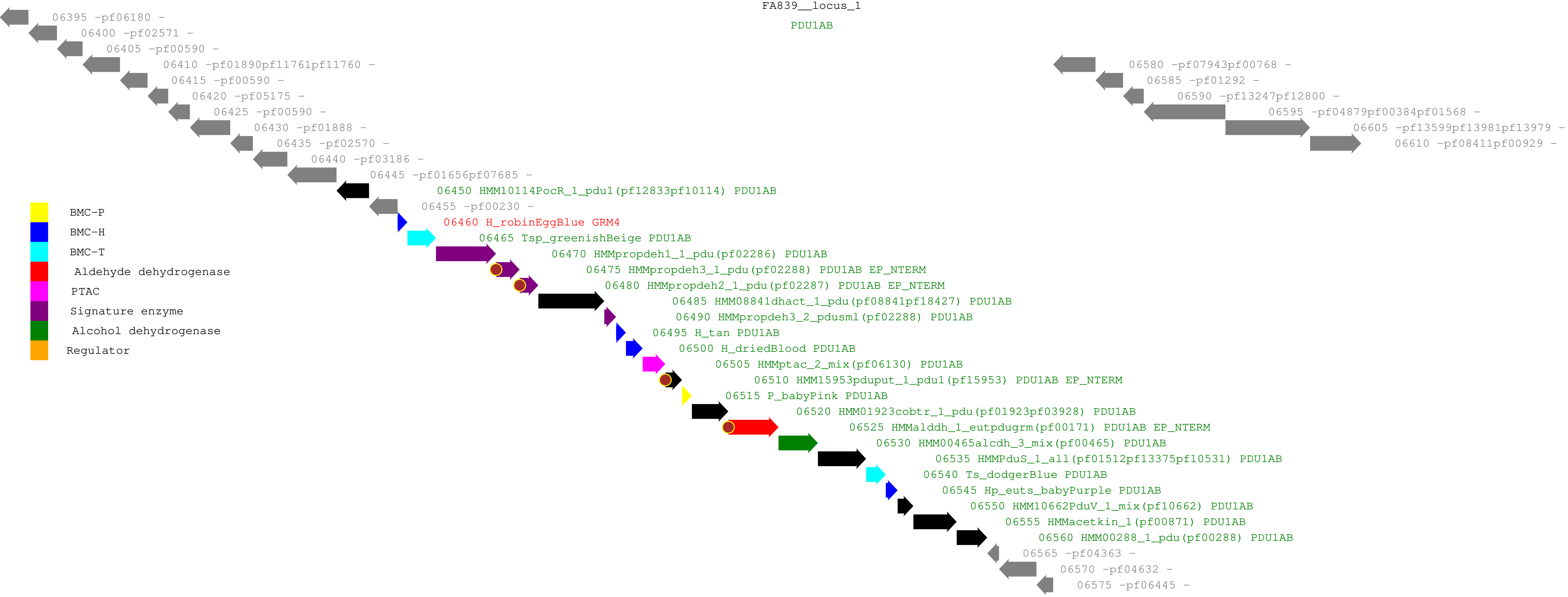

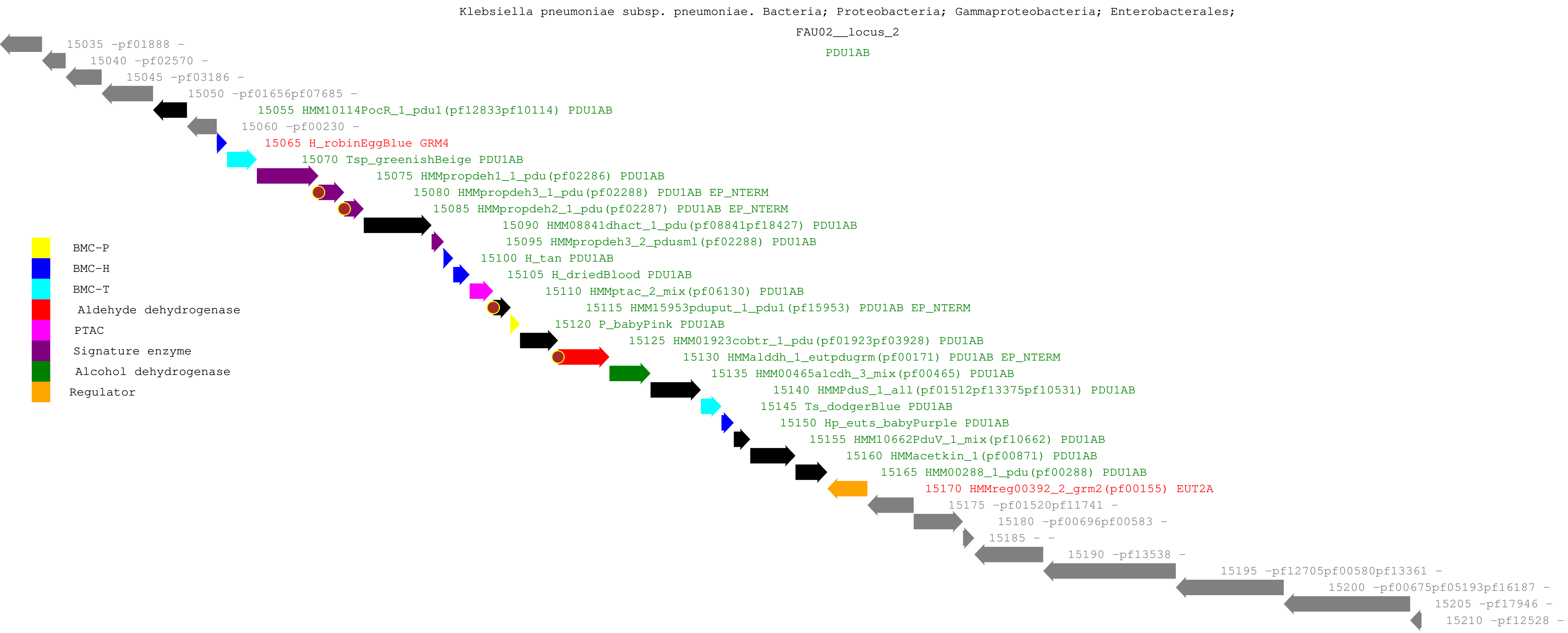
Klebsiella pneumoniae subsp. pneumoniae. Bacteria; Proteobacteria; Gammaproteobacteria; Enterobacterales;
FAU02__locus_2
PDU1AB
15035 -pf01888 -
15040 -pf02570 -
15045 -pf03186 -
15050 -pf01656pf07685 -
15055 HMM10114PocR_1_pdu1(pf12833pf10114) PDU1AB
15060 -pf00230 -
15065 H_robinEggBlue GRM4
15070 Tsp_greenishBeige PDU1AB
15075 HMMpropdeh1_1_pdu(pf02286) PDU1AB
15080 HMMpropdeh3_1_pdu(pf02288) PDU1AB EP_NTERM
15085 HMMpropdeh2_1_pdu(pf02287) PDU1AB EP_NTERM
15090 HMM08841dhact_1_pdu(pf08841pf18427) PDU1AB
15095 HMMpropdeh3_2_pdusml(pf02288) PDU1AB
15100 H_tan PDU1AB
15105 H_driedBlood PDU1AB
15110 HMMptac_2_mix(pf06130) PDU1AB
15115 HMM15953pduput_1_pdu1(pf15953) PDU1AB EP_NTERM
15120 P_babyPink PDU1AB
15125 HMM01923cobtr_1_pdu(pf01923pf03928) PDU1AB
15130 HMMaldh_1_eutpdugrm(pf00171) PDU1AB EP_NTERM
15135 HMM00465alcdh_3_mix(pf00465) PDU1AB
15140 HMMPduS_1_all(pf01512pf13375pf10531) PDU1AB
15145 Ts_dodgerBlue PDU1AB
15150 Hp_euts_babyPurple PDU1AB
15155 HMM10662PduV_1_mix(pf10662) PDU1AB
15160 HMMacetkin_1(pf00871) PDU1AB
15165 HMM00288_1_pdu(pf00288) PDU1AB
15170 HMMreg00392_2_grm2(pf00155) EUT2A
15175 -pf01520pf11741 -
15180 -pf00696pf00583 -
15185 - -
15190 -pf13538 -
15195 -pf12705pf00580pf13361 -
15200 -pf00675pf05193pf16187 -
15205 -pf17946 -
15210 -pf12528 -
BMC-P
BMC-H
BMC-T
Aldehyde dehydrogenase
PTAC
Signature enzyme
Alcohol dehydrogenase
Regulator

Biostraticola sp. BGMRC 2031. Bacteria; Proteobacteria; Gammaproteobacteria; Enterobacterales;

FCN80__locus_1

PDU1AB

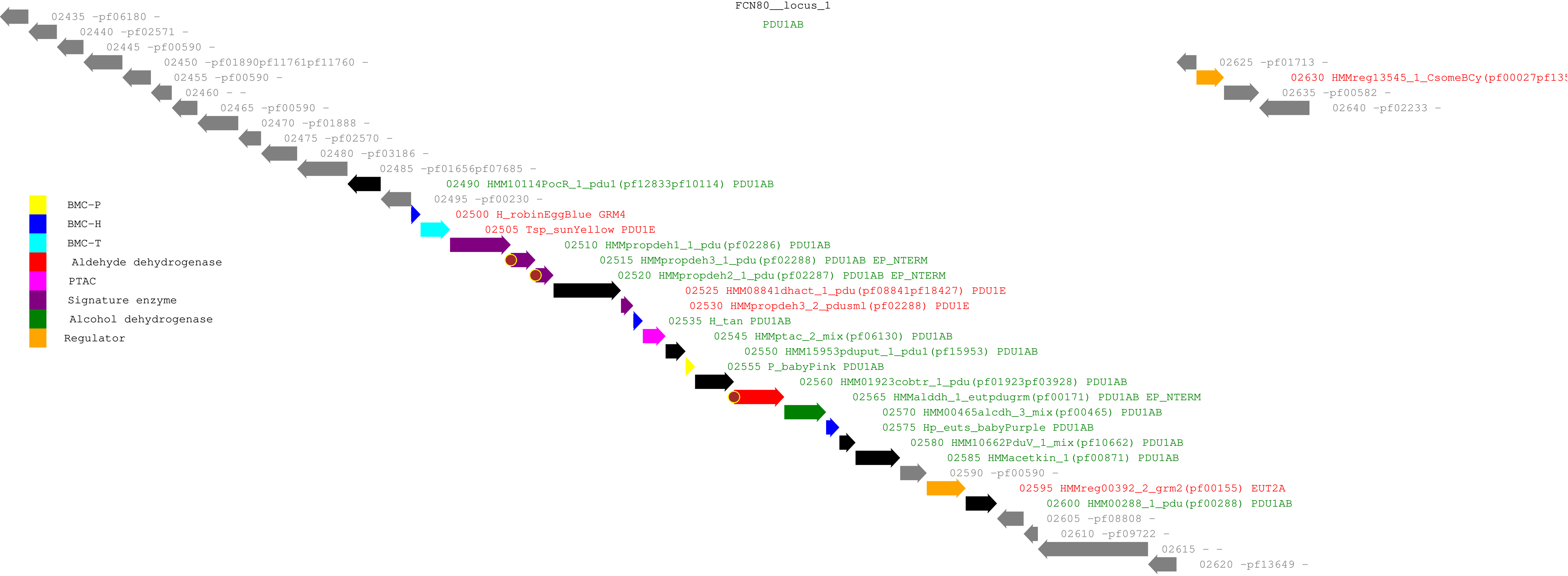

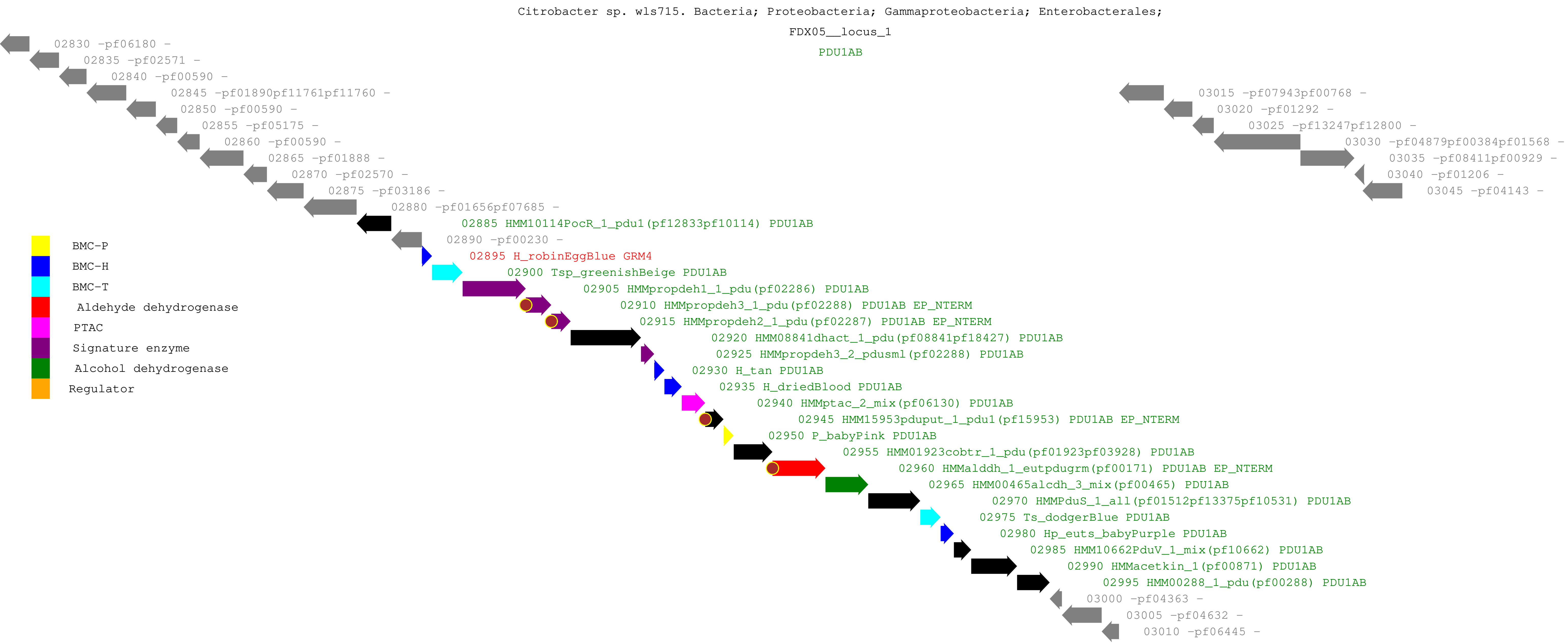
Citrobacter sp. wls715. Bacteria; Proteobacteria; Gammaproteobacteria; Enterobacterales;
FDX05__locus_1
PDU1AB
02830 -pf06180 -
02835 -pf02571 -
02840 -pf00590 -
02845 -pf01890pf11761pf11760 -
02850 -pf00590 -
02855 -pf05175 -
02860 -pf00590 -
02865 -pf01888 -
02870 -pf02570 -
02875 -pf03186 -
02880 -pf01656pf07685 -
02885 HMM10114PocR_1_pdu1(pf12833pf10114) PDU1AB
02890 -pf00230 -
02895 H_robinEggBlue GRM4
02900 Tsp_greenishBeige PDU1AB
02905 HMMpropdeh1_1_pdu(pf02286) PDU1AB
02910 HMMpropdeh3_1_pdu(pf02288) PDU1AB EP_NTERM
02915 HMMpropdeh2_1_pdu(pf02287) PDU1AB EP_NTERM
02920 HMM08841dhact_1_pdu(pf08841pf18427) PDU1AB
02925 HMMpropdeh3_2_pdusml(pf02288) PDU1AB
02930 H_tan PDU1AB
02935 H_driedBlood PDU1AB
02940 HMMptac_2_mix(pf06130) PDU1AB
02945 HMM15953pduput_1_pdu1(pf15953) PDU1AB EP_NTERM
02950 P_babyPink PDU1AB
02955 HMM01923cobtr_1_pdu(pf01923pf03928) PDU1AB
02960 HMMalddh_1_eutpdugrm(pf00171) PDU1AB EP_NTERM
02965 HMM00465alcdh_3_mix(pf00465) PDU1AB
02970 HMMPduS_1_all(pf01512pf13375pf10531) PDU1AB
02975 Ts_dodgerBlue PDU1AB
02980 Hp_euts_babyPurple PDU1AB
02985 HMM10662PduV_1_mix(pf10662) PDU1AB
02990 HMMacetkin_1(pf00871) PDU1AB
02995 HMM00288_1_pdu(pf00288) PDU1AB
03000 -pf04363 -
03005 -pf04632 -
03010 -pf06445 -
03015 -pf07943pf00768 -
03020 -pf01292 -
03025 -pf13247pf12800 -
03030 -pf04879pf00384pf01568 -
03035 -pf08411pf00929 -
03040 -pf01206 -
03045 -pf04143 -
BMC-P
BMC-H
BMC-T
Aldehyde dehydrogenase
PTAC
Signature enzyme
Alcohol dehydrogenase
Regulator

Citrobacter sp. wls714. Bacteria; Proteobacteria; Gammaproteobacteria; Enterobacterales;

FDX11__locus_1

PDU1AB

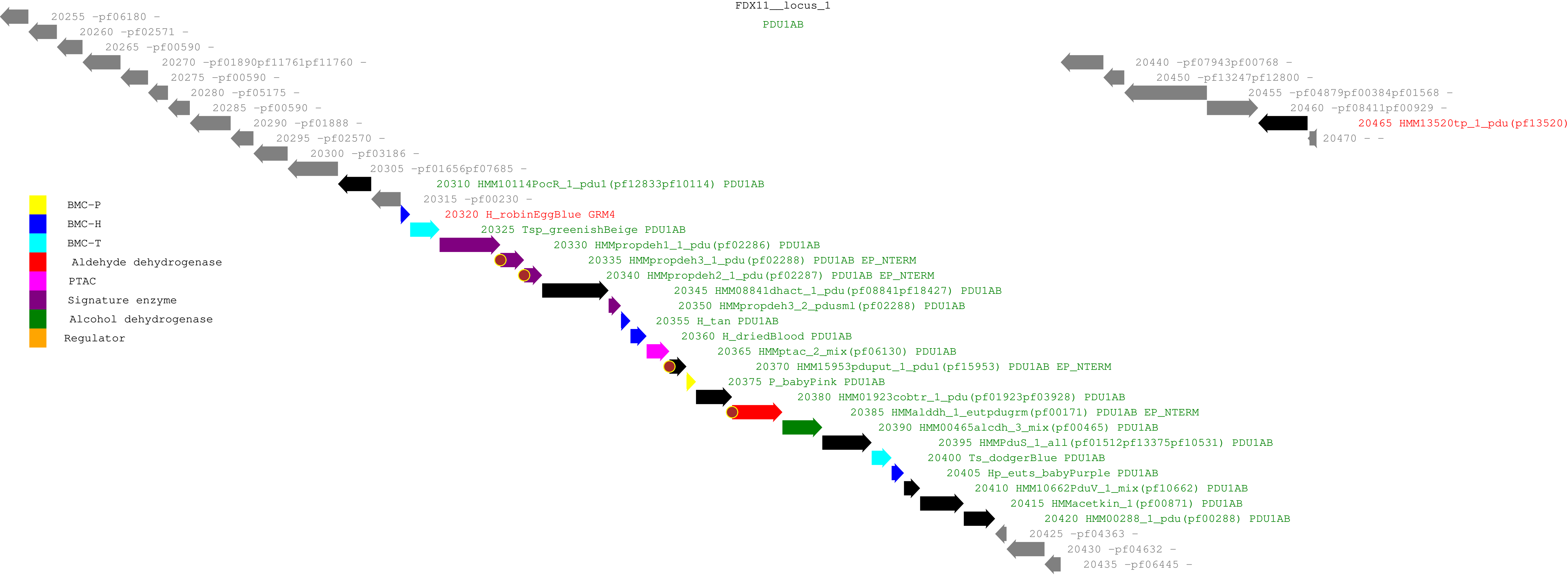

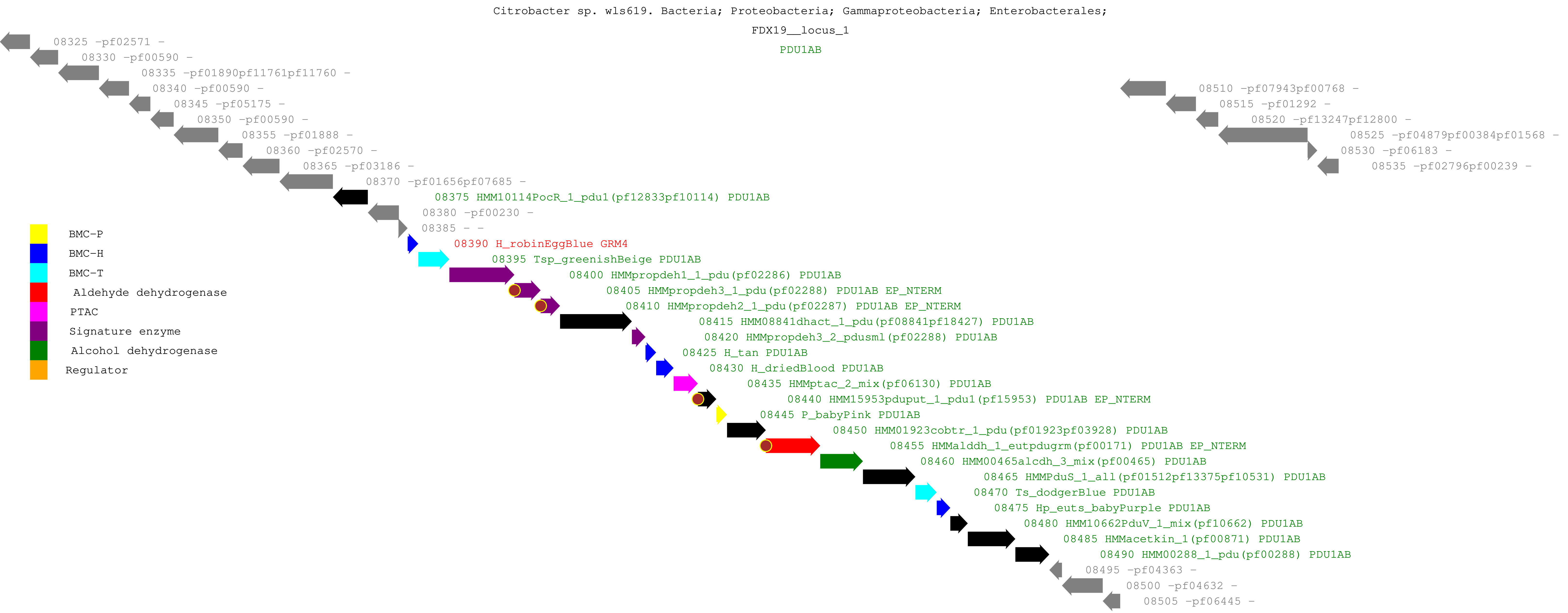

Citrobacter sp. wls619. Bacteria; Proteobacteria; Gammaproteobacteria; Enterobacterales;
FDX19__locus_1
PDU1AB
08325 -pf02571 -
08330 -pf00590 -
08335 -pf01890pf11761pf11760 -
08340 -pf00590 -
08345 -pf05175 -
08350 -pf00590 -
08355 -pf01888 -
08360 -pf02570 -
08365 -pf03186 -
08370 -pf01656pf07685 -
08375 HMM10114PocR_1_pdu1(pf12833pf10114) PDU1AB
08380 -pf00230 -
08385 - -
08390 H_robinEggBlue GRM4
08395 Tsp_greenishBeige PDU1AB
08400 HMMpropdeh1_1_pdu(pf02286) PDU1AB
08405 HMMpropdeh3_1_pdu(pf02288) PDU1AB EP_NTERM
08410 HMMpropdeh2_1_pdu(pf02287) PDU1AB EP_NTERM
08415 HMM08841dhact_1_pdu(pf08841pf18427) PDU1AB
08420 HMMpropdeh3_2_pdusml(pf02288) PDU1AB
08425 H_tan PDU1AB
08430 H_driedBlood PDU1AB
08435 HMMptac_2_mix(pf06130) PDU1AB
08440 HMM15953pduput_1_pdu1(pf15953) PDU1AB EP_NTERM
08445 P_babyPink PDU1AB
08450 HMM01923cobtr_1_pdu(pf01923pf03928) PDU1AB
08455 HMMalddh_1_eutpdugrm(pf00171) PDU1AB EP_NTERM
08460 HMM00465alcdh_3_mix(pf00465) PDU1AB
08465 HMMPduS_1_all(pf01512pf13375pf10531) PDU1AB
08470 Ts_dodgerBlue PDU1AB
08475 Hp_euts_babyPurple PDU1AB
08480 HMM10662PduV_1_mix(pf10662) PDU1AB
08485 HMMacetkin_1(pf00871) PDU1AB
08490 HMM00288_1_pdu(pf00288) PDU1AB
08495 -pf04363 -
08500 -pf04632 -
08505 -pf06445 -
08510 -pf07943pf00768 -
08515 -pf01292 -
08520 -pf13247pf12800 -
08525 -pf04879pf00384pf01568 -
08530 -pf06183 -
08535 -pf02796pf00239 -
BMC-P
BMC-H
BMC-T
Aldehyde dehydrogenase
PTAC
Signature enzyme
Alcohol dehydrogenase
Regulator

Citrobacter sp. TBCS-14. Bacteria; Proteobacteria; Gammaproteobacteria; Enterobacterales;

FDX22__locus_2

PDU1AB

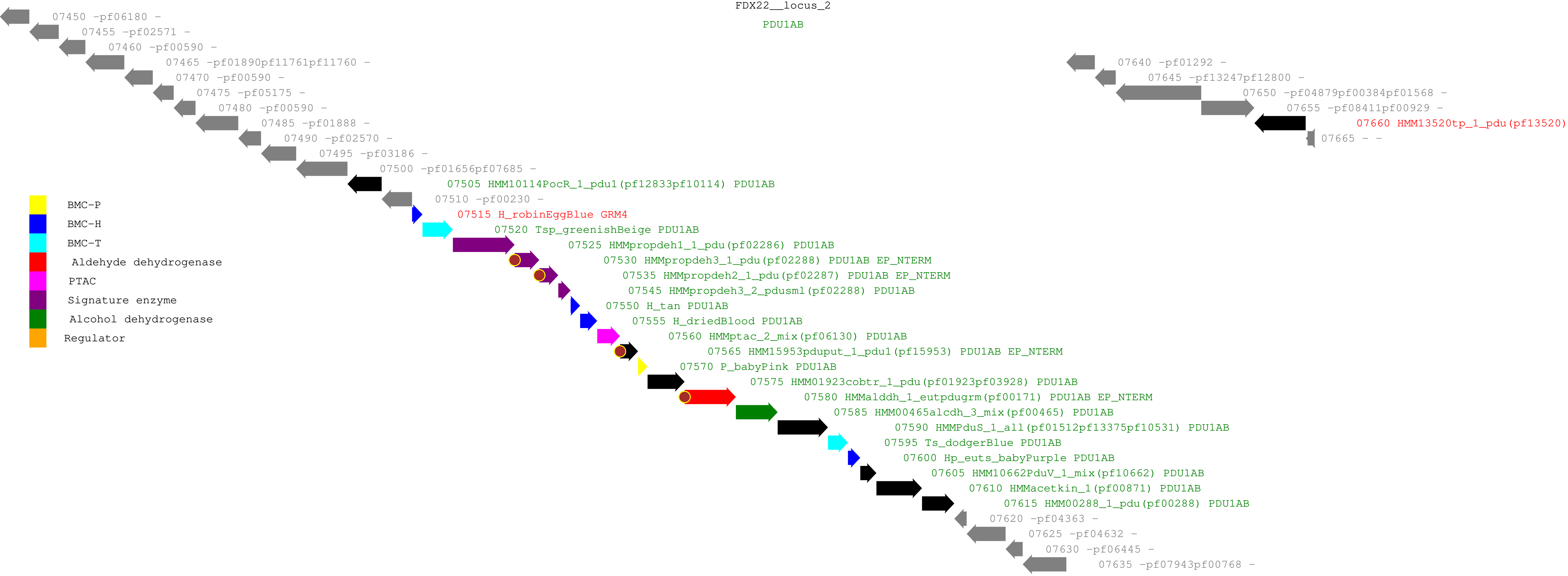

Salmonella enterica subsp. enterica serovar Ibadan. Bacteria; Proteobacteria; Gammaproteobacteria; Enterobacterales;

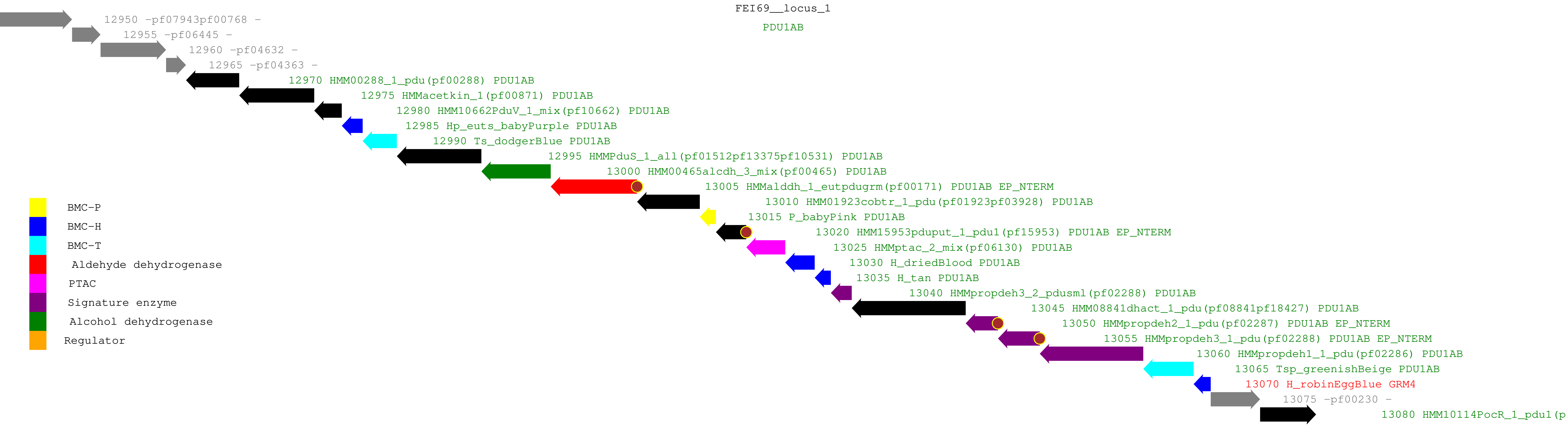

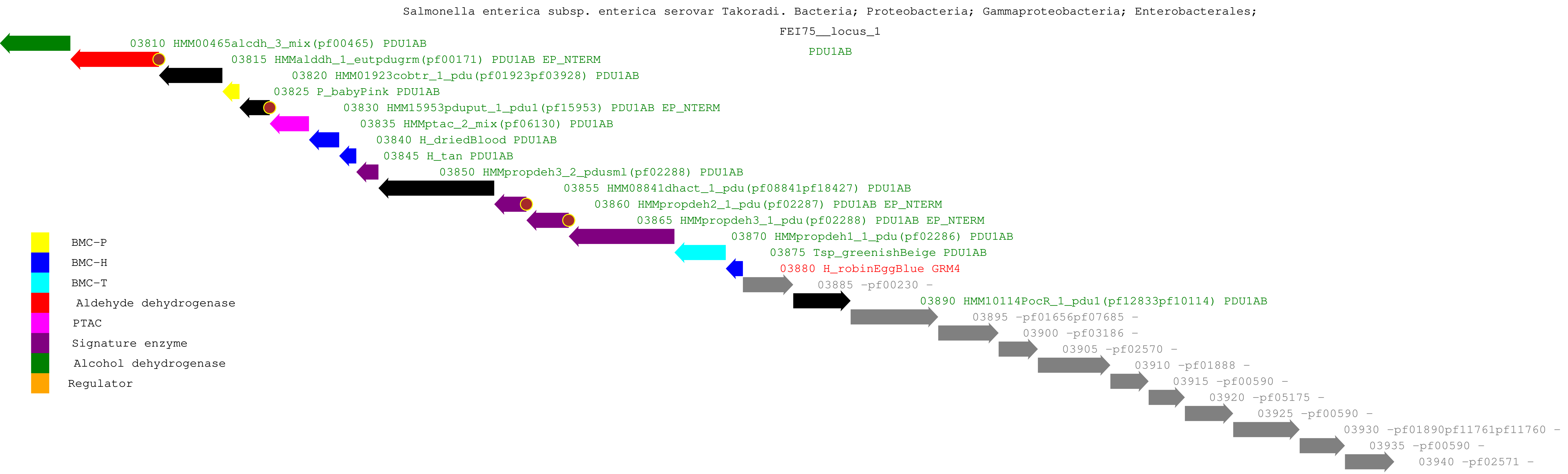
Salmonella enterica subsp. enterica serovar Takoradi. Bacteria; Proteobacteria; Gammaproteobacteria; Enterobacterales;
FEI75__locus_1
PDU1AB
03810 HMM00465alcdh_3_mix(pf00465) PDU1AB
03815 HMMalddh_1_eutpdugrm(pf00171) PDU1AB EP_NTERM
03820 HMM01923cobtr_1_pdu(pf01923pf03928) PDU1AB
03825 P_babyPink PDU1AB
03830 HMM15953pduput_1_pdu1(pf15953) PDU1AB EP_NTERM
03835 HMMptac_2_mix(pf06130) PDU1AB
03840 H_driedBlood PDU1AB
03845 H_tan PDU1AB
03850 HMMpropdeh3_2_pdusml(pf02288) PDU1AB
03855 HMM08841dhact_1_pdu(pf08841pf18427) PDU1AB
03860 HMMpropdeh2_1_pdu(pf02287) PDU1AB EP_NTERM
03865 HMMpropdeh3_1_pdu(pf02288) PDU1AB EP_NTERM
03870 HMMpropdeh1_1_pdu(pf02286) PDU1AB
03875 Tsp_greenishBeige PDU1AB
03880 H_robinEggBlue GRM4
03885 -pf00230 -
03890 HMM10114PocR_1_pdu1(pf12833pf10114) PDU1AB
03895 -pf01656pf07685 -
03900 -pf03186 -
03905 -pf02570 -
03910 -pf01888 -
03915 -pf00590 -
03920 -pf05175 -
03925 -pf00590 -
03930 -pf01890pf11761pf11760 -
03935 -pf00590 -
03940 -pf02571 -
BMC-P
BMC-H
BMC-T
Aldehyde dehydrogenase
PTAC
Signature enzyme
Alcohol dehydrogenase
Regulator

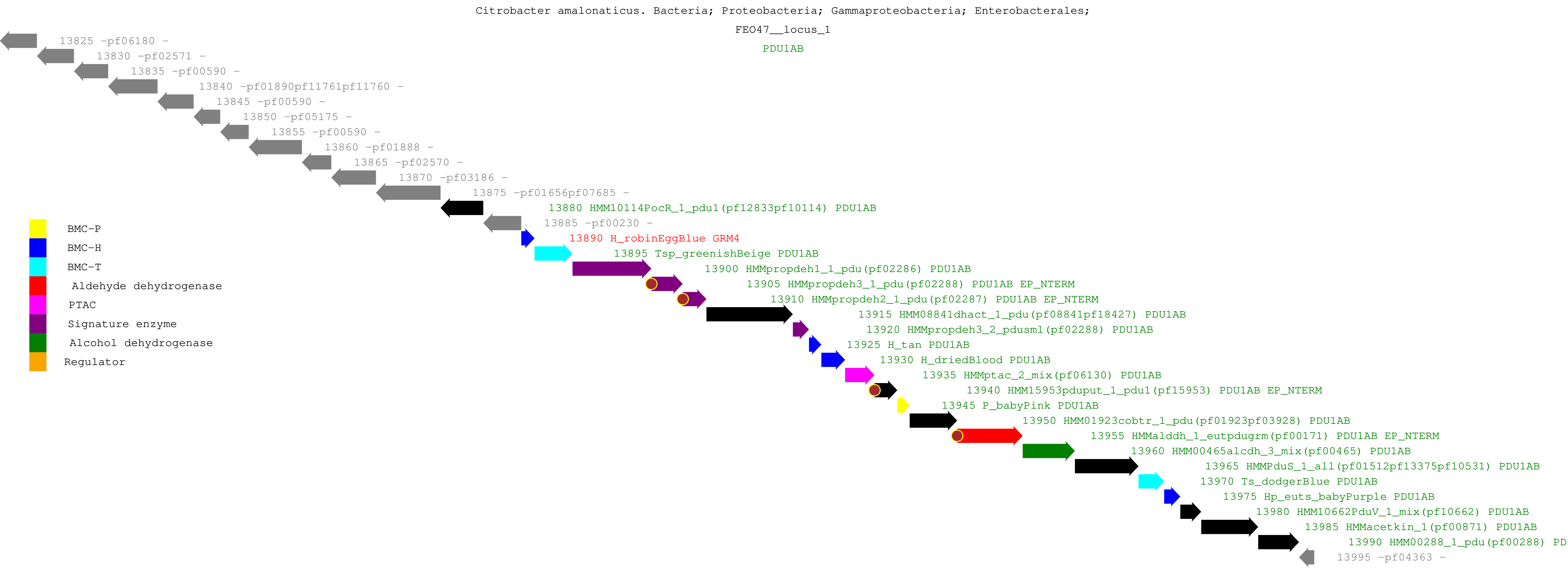

Citrobacter amalonaticus. Bacteria; Proteobacteria; Gammaproteobacteria; Enterobacterales;
FEO47__locus_1
PDU1AB
13825 -pf06180 -
13830 -pf02571 -
13835 -pf00590 -
13840 -pf01890pf11761pf11760 -
13845 -pf00590 -
13850 -pf05175 -
13855 -pf00590 -
13860 -pf01888 -
13865 -pf02570 -
13870 -pf03186 -
13875 -pf01656pf07685 -
13880 HMM10114PocR_1_pdu1(pf12833pf10114) PDU1AB
13885 -pf00230 -
13890 H_robinEggBlue GRM4
13895 Tsp_greenishBeige PDU1AB
13900 HMMpropdeh1_1_pdu(pf02286) PDU1AB
13905 HMMpropdeh3_1_pdu(pf02288) PDU1AB EP_NTERM
13910 HMMpropdeh2_1_pdu(pf02287) PDU1AB EP_NTERM
13915 HMM08841dhact_1_pdu(pf08841pf18427) PDU1AB
13920 HMMpropdeh3_2_pdusml(pf02288) PDU1AB
13925 H_tan PDU1AB
13930 H_driedBlood PDU1AB
13935 HMMptac_2_mix(pf06130) PDU1AB
13940 HMM15953pduput_1_pdu1(pf15953) PDU1AB EP_NTERM
13945 P_babyPink PDU1AB
13950 HMM01923cobtr_1_pdu(pf01923pf03928) PDU1AB
13955 HMMalddh_1_eutpdugrm(pf00171) PDU1AB EP_NTERM
13960 HMM00465alcdh_3_mix(pf00465) PDU1AB
13965 HMMPduS_1_all(pf01512pf13375pf10531) PDU1AB
13970 Ts_dodgerBlue PDU1AB
13975 Hp_euts_babyPurple PDU1AB
13980 HMM10662PduV_1_mix(pf10662) PDU1AB
13985 HMMacetkin_1(pf00871) PDU1AB
13990 HMM00288_1_pdu(pf00288) PD
13995 -pf04363 -
BMC-P
BMC-H
BMC-T
Aldehyde dehydrogenase
PTAC
Signature enzyme
Alcohol dehydrogenase
Regulator

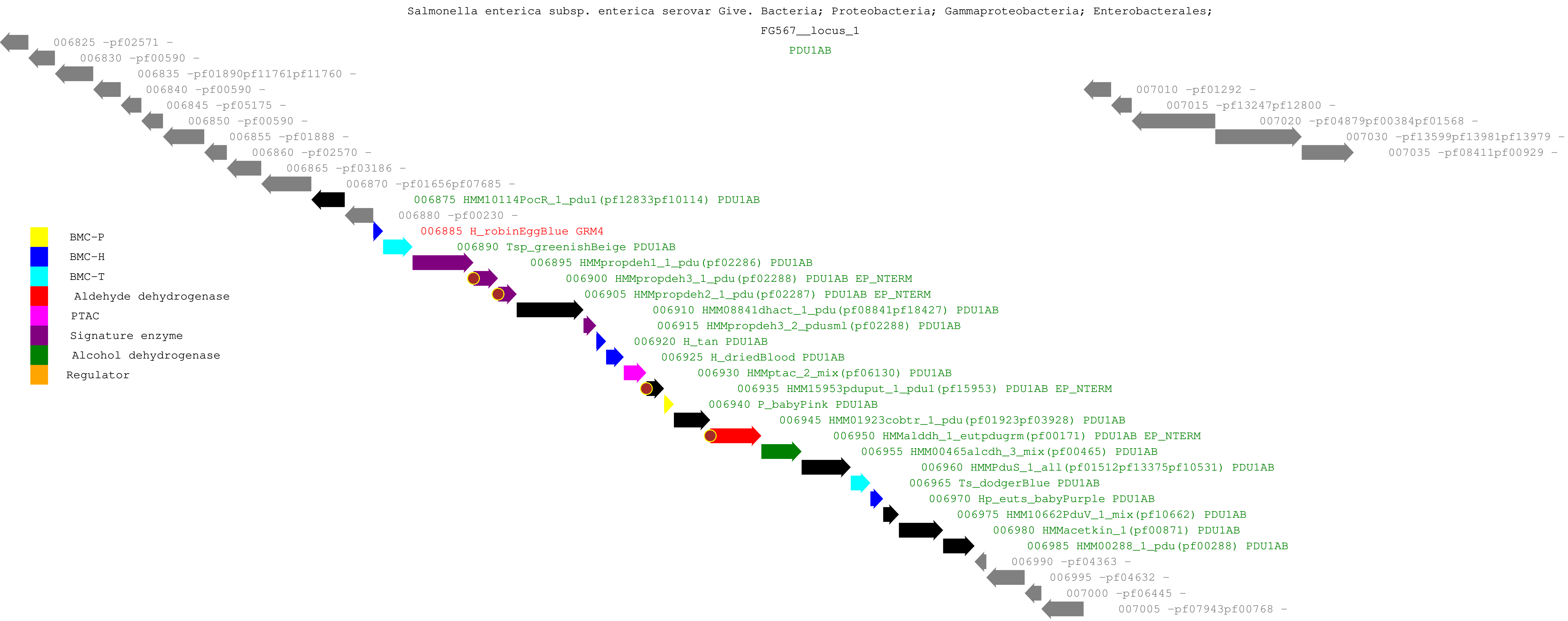
Salmonella enterica subsp. enterica serovar Give. Bacteria; Proteobacteria; Gammaproteobacteria; Enterobacterales;
FG567__locus_1
PDU1AB
006825 -pf02571 -
006830 -pf00590 -
006835 -pf01890pf11761pf11760 -
006840 -pf00590 -
006845 -pf05175 -
006850 -pf00590 -
006855 -pf01888 -
006860 -pf02570 -
006865 -pf03186 -
006870 -pf01656pf07685 -
006875 HMM10114PocR_1_pdu1(pf12833pf10114) PDU1AB
006880 -pf00230 -
006885 H_robinEggBlue GRM4
006890 Tsp_greenishBeige PDU1AB
006895 HMMpropdeh1_1_pdu(pf02286) PDU1AB
006900 HMMpropdeh3_1_pdu(pf02288) PDU1AB EP_NTERM
006905 HMMpropdeh2_1_pdu(pf02287) PDU1AB EP_NTERM
006910 HMM08841dhact_1_pdu(pf08841pf18427) PDU1AB
006915 HMMpropdeh3_2_pdusml(pf02288) PDU1AB
006920 H_tan PDU1AB
006925 H_driedBlood PDU1AB
006930 HMMptac_2_mix(pf06130) PDU1AB
006935 HMM15953pduput_1_pdu1(pf15953) PDU1AB EP_NTERM
006940 P_babyPink PDU1AB
006945 HMM01923cobtr_1_pdu(pf01923pf03928) PDU1AB
006950 HMMalddh_1_eutpdugrm(pf00171) PDU1AB EP_NTERM
006955 HMM00465alcdh_3_mix(pf00465) PDU1AB
006960 HMMPduS_1_all(pf01512pf13375pf10531) PDU1AB
006965 Ts_dodgerBlue PDU1AB
006970 Hp_euts_babyPurple PDU1AB
006975 HMM10662PduV_1_mix(pf10662) PDU1AB
006980 HMMacetkin_1(pf00871) PDU1AB
006985 HMM00288_1_pdu(pf00288) PDU1AB
006990 -pf04363 -
006995 -pf04632 -
007000 -pf06445 -
007005 -pf07943pf00768 -
007010 -pf01292 -
007015 -pf13247pf12800 -
007020 -pf04879pf00384pf01568 -
007030 -pf13599pf13981pf13979 -
007035 -pf08411pf00929 -
BMC-P
BMC-H
BMC-T
Aldehyde dehydrogenase
PTAC
Signature enzyme
Alcohol dehydrogenase
Regulator

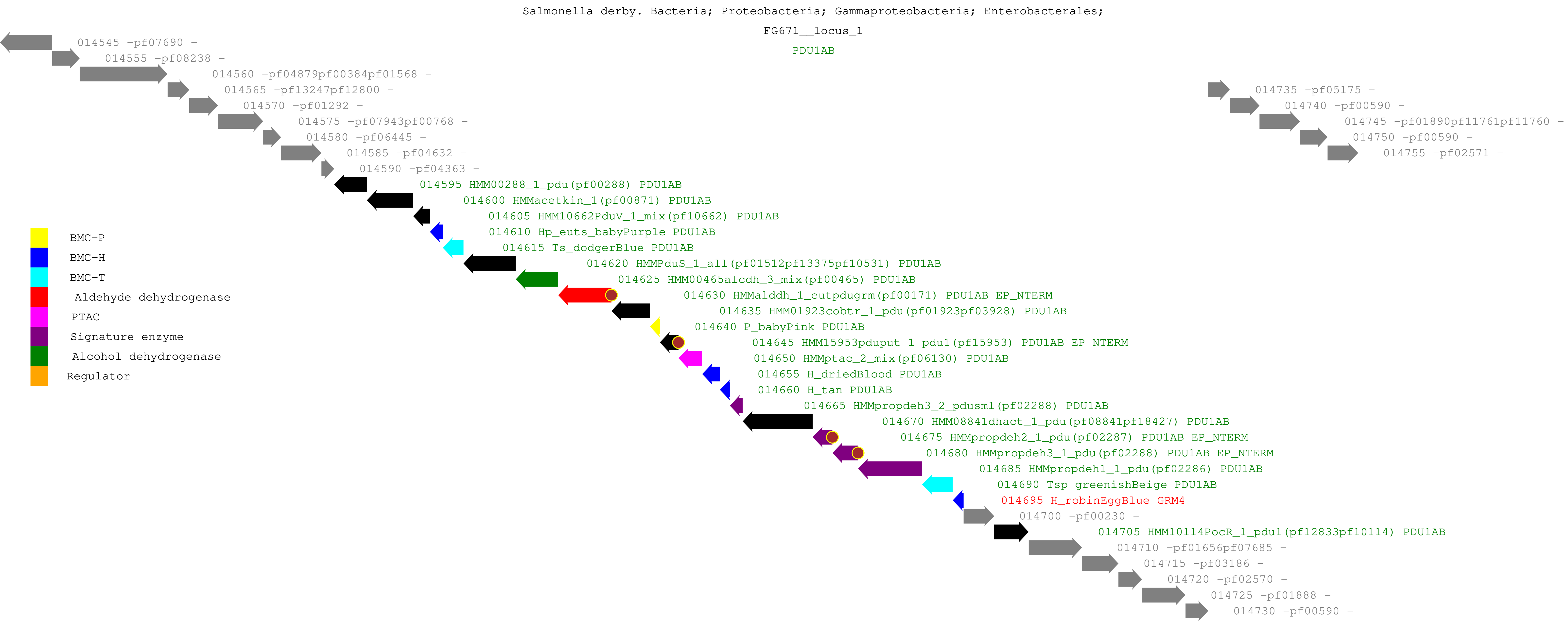
Salmonella derby. Bacteria; Proteobacteria; Gammaproteobacteria; Enterobacterales;
FG671__locus_1
PDU1AB
014545 -pf07690 -
014555 -pf08238 -
014560 -pf04879pf00384pf01568 -
014565 -pf13247pf12800 -
014570 -pf01292 -
014575 -pf07943pf00768 -
014580 -pf06445 -
014585 -pf04632 -
014590 -pf04363 -
014595 HMM00288_1_pdu(pf00288) PDU1AB
014600 HMMacetkin_1(pf00871) PDU1AB
014605 HMM10662PduV_1_mix(pf10662) PDU1AB
014610 Hp_euts_babyPurple PDU1AB
014615 Ts_dodgerBlue PDU1AB
014620 HMMPduS_1_all(pf01512pf13375pf10531) PDU1AB
014625 HMM00465alcdh_3_mix(pf00465) PDU1AB
014630 HMMaldh_1_eutpdugrm(pf00171) PDU1AB EP_NTERM
014635 HMM01923cobtr_1_pdu(pf01923pf03928) PDU1AB
014640 P_babyPink PDU1AB
014645 HMM15953pduput_1_pdu1(pf15953) PDU1AB EP_NTERM
014650 HMMptac_2_mix(pf06130) PDU1AB
014655 H_driedBlood PDU1AB
014660 H_tan PDU1AB
014665 HMMpropdeh3_2_pdusml(pf02288) PDU1AB
014670 HMM08841dhact_1_pdu(pf08841pf18427) PDU1AB
014675 HMMpropdeh2_1_pdu(pf02287) PDU1AB EP_NTERM
014680 HMMpropdeh3_1_pdu(pf02288) PDU1AB EP_NTERM
014685 HMMpropdeh1_1_pdu(pf02286) PDU1AB
014690 Tsp_greenishBeige PDU1AB
014695 H_robinEggBlue GRM4
014700 -pf00230 -
014705 HMM10114PocR_1_pdu1(pf12833pf10114) PDU1AB
014710 -pf01656pf07685 -
014715 -pf03186 -
014720 -pf02570 -
014725 -pf01888 -
014730 -pf00590 -
014735 -pf05175 -
014740 -pf00590 -
014745 -pf01890pf11761pf11760 -
014750 -pf00590 -
014755 -pf02571 -
BMC-P
BMC-H
BMC-T
Aldehyde dehydrogenase
PTAC
Signature enzyme
Alcohol dehydrogenase
Regulator

Salmonella enterica subsp. enterica serovar 1,4,5,12:i:-. Bacteria; Proteobacteria; Gammaproteobacteria; Enterobacterales;

FGF59__locus_1

PDU1AB

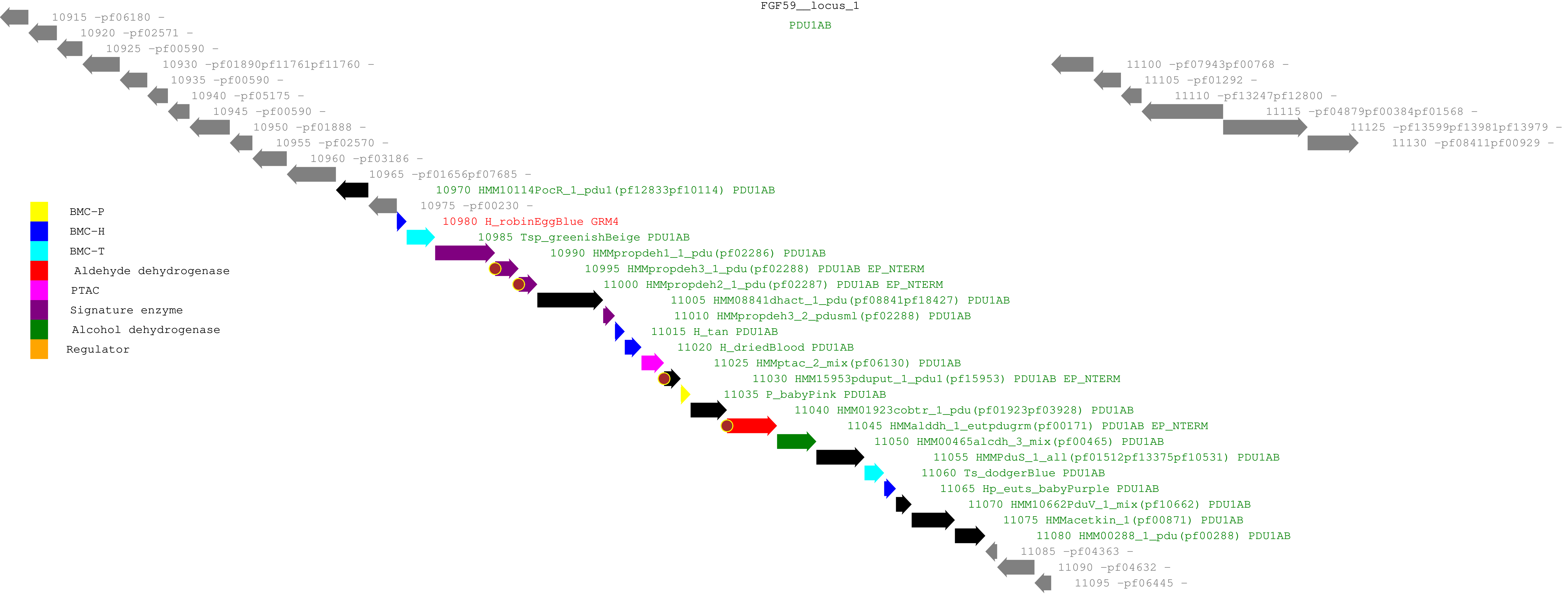

Salmonella enterica subsp. enterica serovar Eko. Bacteria; Proteobacteria; Gammaproteobacteria; Enterobacterales;

FGJ70__locus_2

PDU1AB

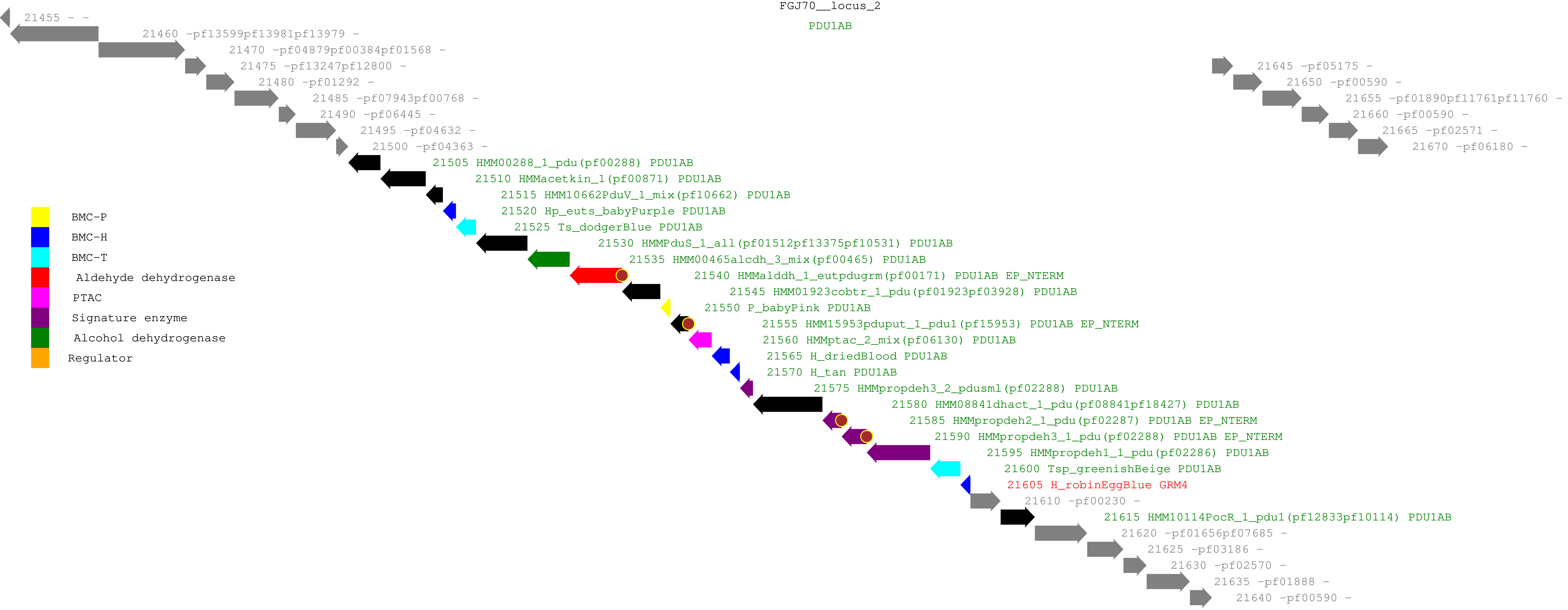

Serratia fonticola. Bacteria; Proteobacteria; Gammaproteobacteria; Enterobacterales;

FHU10__locus_3

PDU1AB

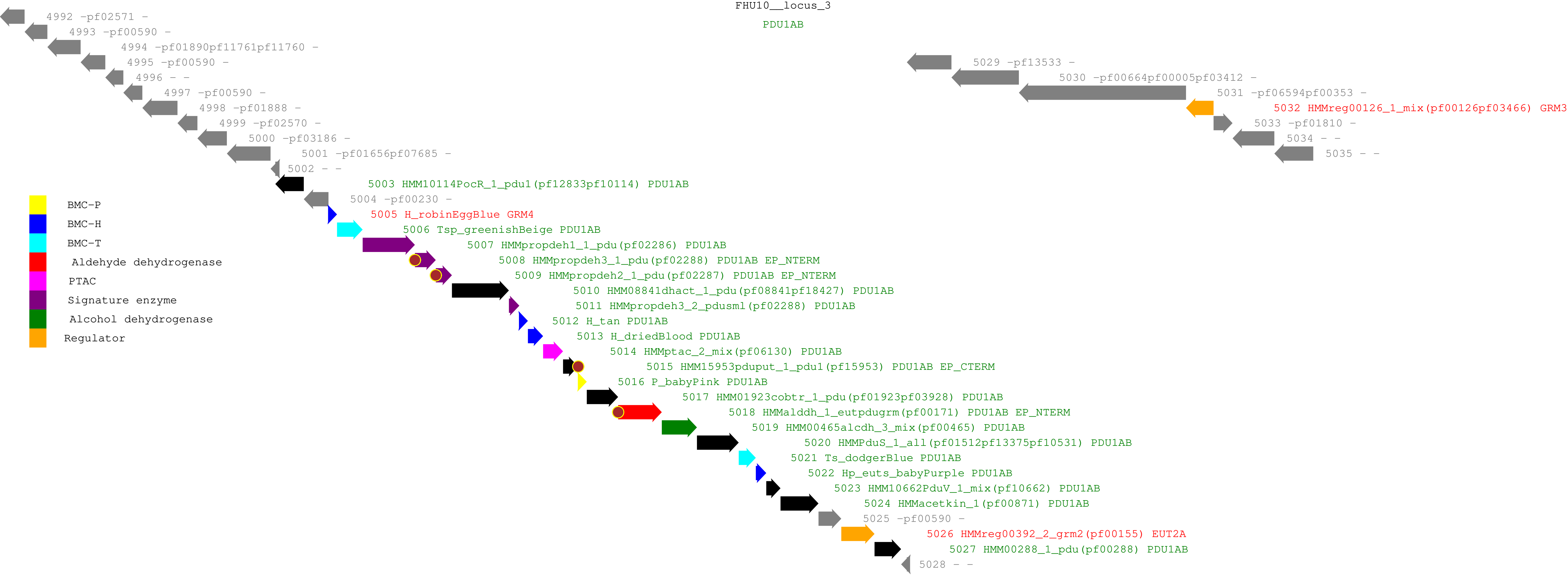

Salmonella enterica (Salmonella choleraesuis). Bacteria; Proteobacteria; Gammaproteobacteria; Enterobacterales;

FJR63__locus_1

PDU1AB

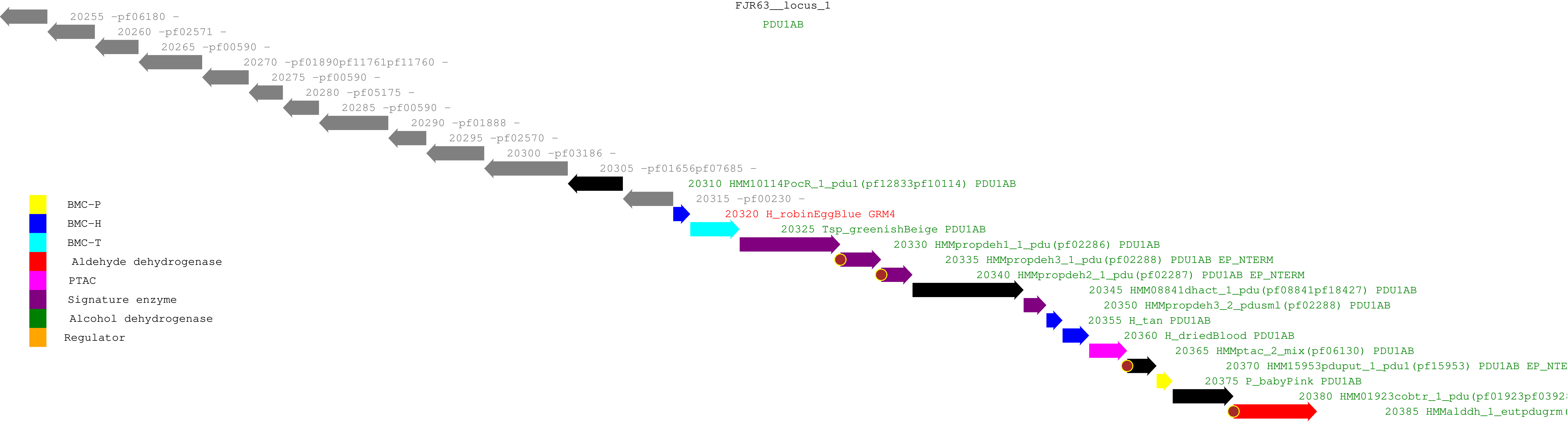

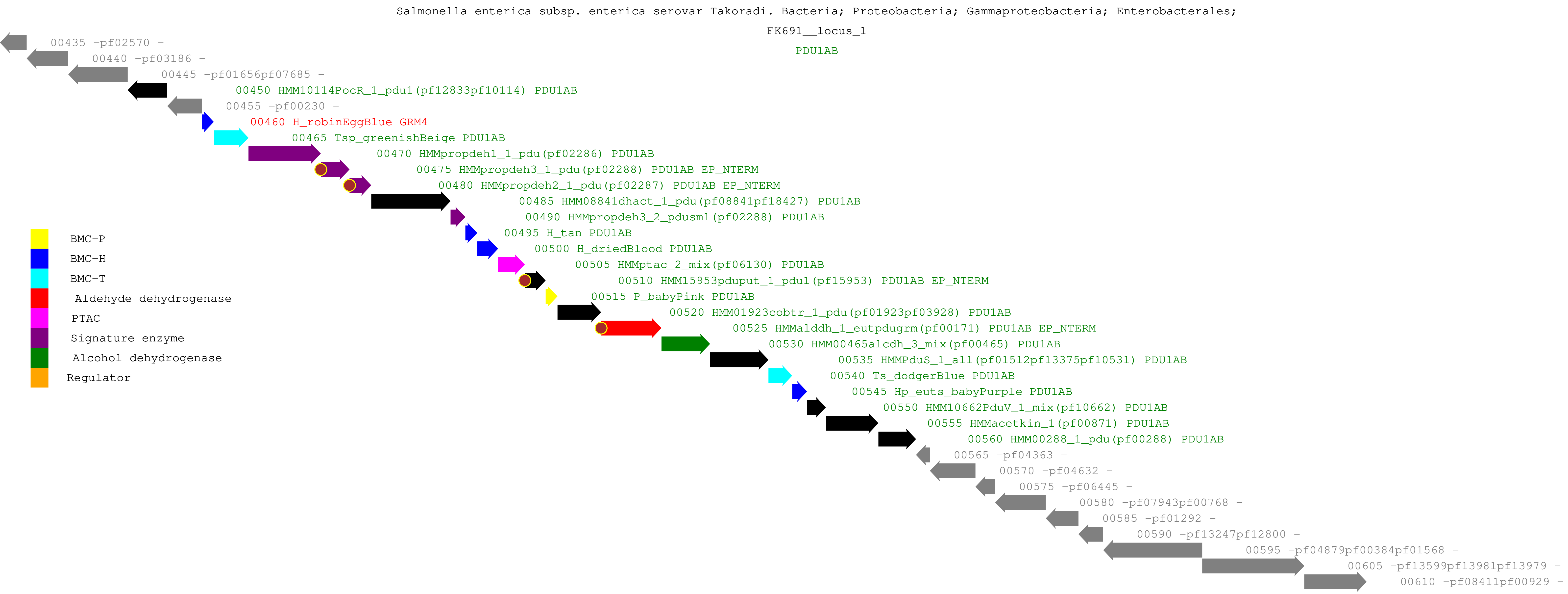
Salmonella enterica subsp. enterica serovar Takoradi. Bacteria; Proteobacteria; Gammaproteobacteria; Enterobacterales;
FK691__locus_1
PDU1AB
00435 -pf02570 -
00440 -pf03186 -
00445 -pf01656pf07685 -
00450 HMM10114PocR_1_pdu1(pf12833pf10114) PDU1AB
00455 -pf00230 -
00460 H_robinEggBlue GRM4
00465 Tsp_greenishBeige PDU1AB
00470 HMMpropdeh1_1_pdu(pf02286) PDU1AB
00475 HMMpropdeh3_1_pdu(pf02288) PDU1AB EP_NTERM
00480 HMMpropdeh2_1_pdu(pf02287) PDU1AB EP_NTERM
00485 HMM08841dhact_1_pdu(pf08841pf18427) PDU1AB
00490 HMMpropdeh3_2_pdusml(pf02288) PDU1AB
00495 H_tan PDU1AB
00500 H_driedBlood PDU1AB
00505 HMMptac_2_mix(pf06130) PDU1AB
00510 HMM15953pduput_1_pdu1(pf15953) PDU1AB EP_NTERM
00515 P_babyPink PDU1AB
00520 HMM01923cobtr_1_pdu(pf01923pf03928) PDU1AB
00525 HMMalddh_1_eutpdugrm(pf00171) PDU1AB EP_NTERM
00530 HMM00465alcdh_3_mix(pf00465) PDU1AB
00535 HMMPduS_1_all(pf01512pf13375pf10531) PDU1AB
00540 Ts_dodgerBlue PDU1AB
00545 Hp_euts_babyPurple PDU1AB
00550 HMM10662PduV_1_mix(pf10662) PDU1AB
00555 HMMacetkin_1(pf00871) PDU1AB
00560 HMM00288_1_pdu(pf00288) PDU1AB
00565 -pf04363 -
00570 -pf04632 -
00575 -pf06445 -
00580 -pf07943pf00768 -
00585 -pf01292 -
00590 -pf13247pf12800 -
00595 -pf04879pf00384pf01568 -
00605 -pf13599pf13981pf13979 -
00610 -pf08411pf00929 -
BMC-P
BMC-H
BMC-T
Aldehyde dehydrogenase
PTAC
Signature enzyme
Alcohol dehydrogenase
Regulator

Salmonella enterica subsp. enterica serovar Bahati. Bacteria; Proteobacteria; Gammaproteobacteria; Enterobacterales;

FK693__locus_2

PDU1AB

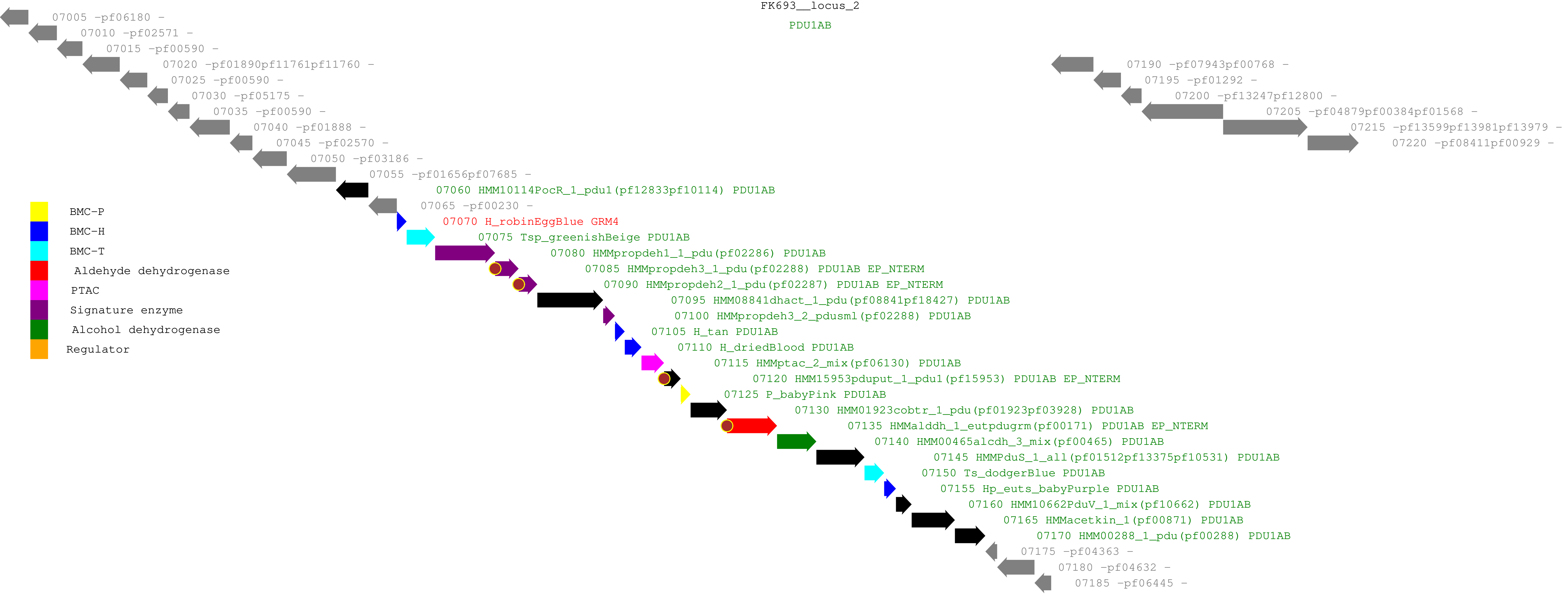

Salmonella enterica subsp. enterica serovar Mikawasima. Bacteria; Proteobacteria; Gammaproteobacteria; Enterobacterales;

FKX61__locus_1

PDU1AB

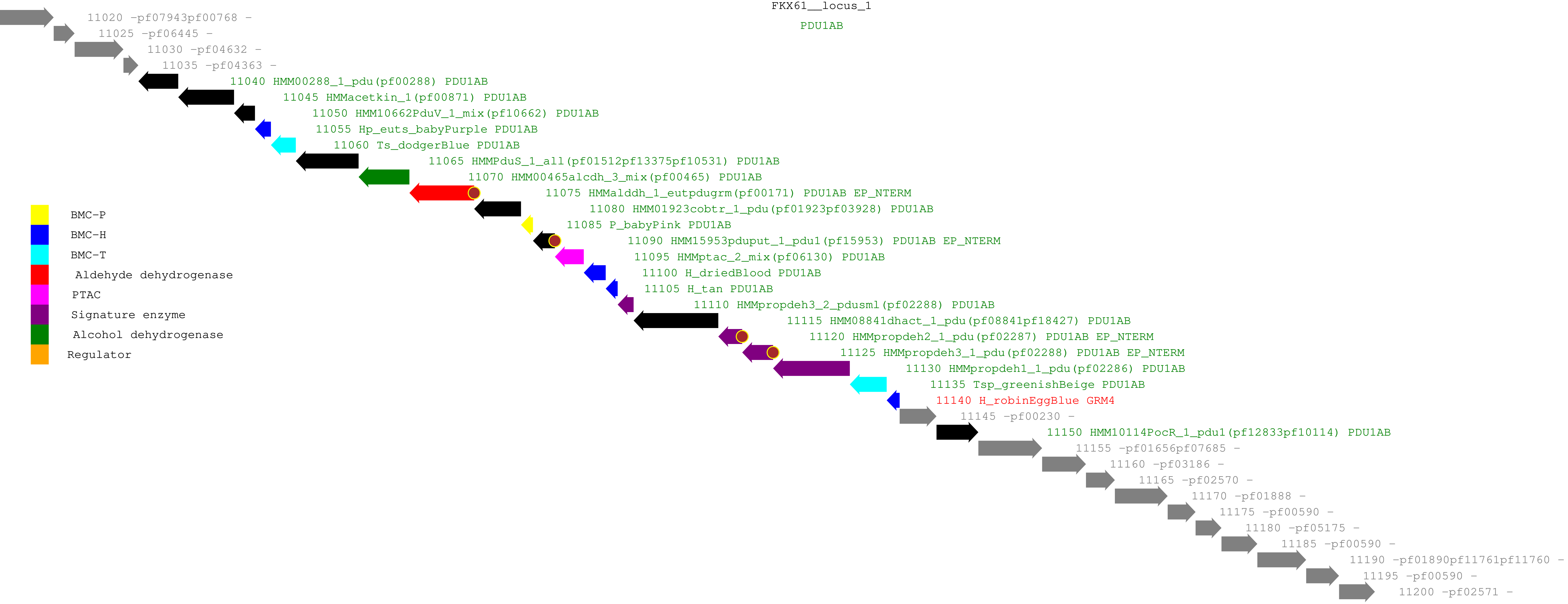

Salmonella enterica subsp. enterica serovar Bahati. Bacteria; Proteobacteria; Gammaproteobacteria; Enterobacterales;

FKX64__locus_1

PDU1AB

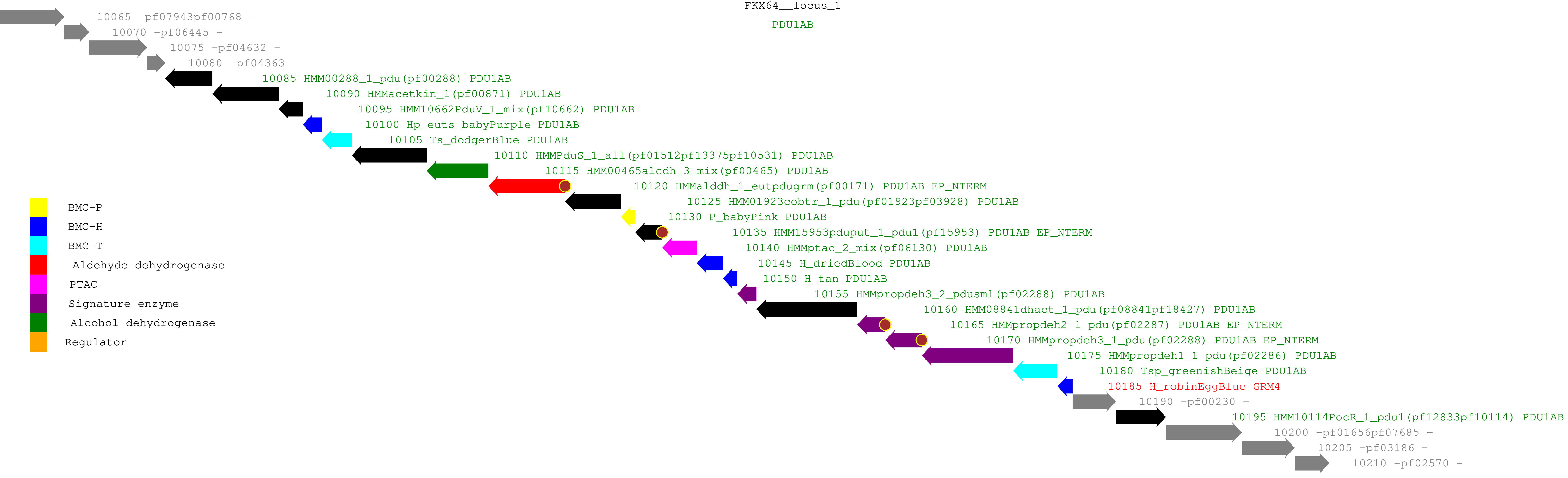

Salmonella enterica subsp. enterica serovar Singapore. Bacteria; Proteobacteria; Gammaproteobacteria; Enterobacterales;

FLK40__locus_1

PDU1AB

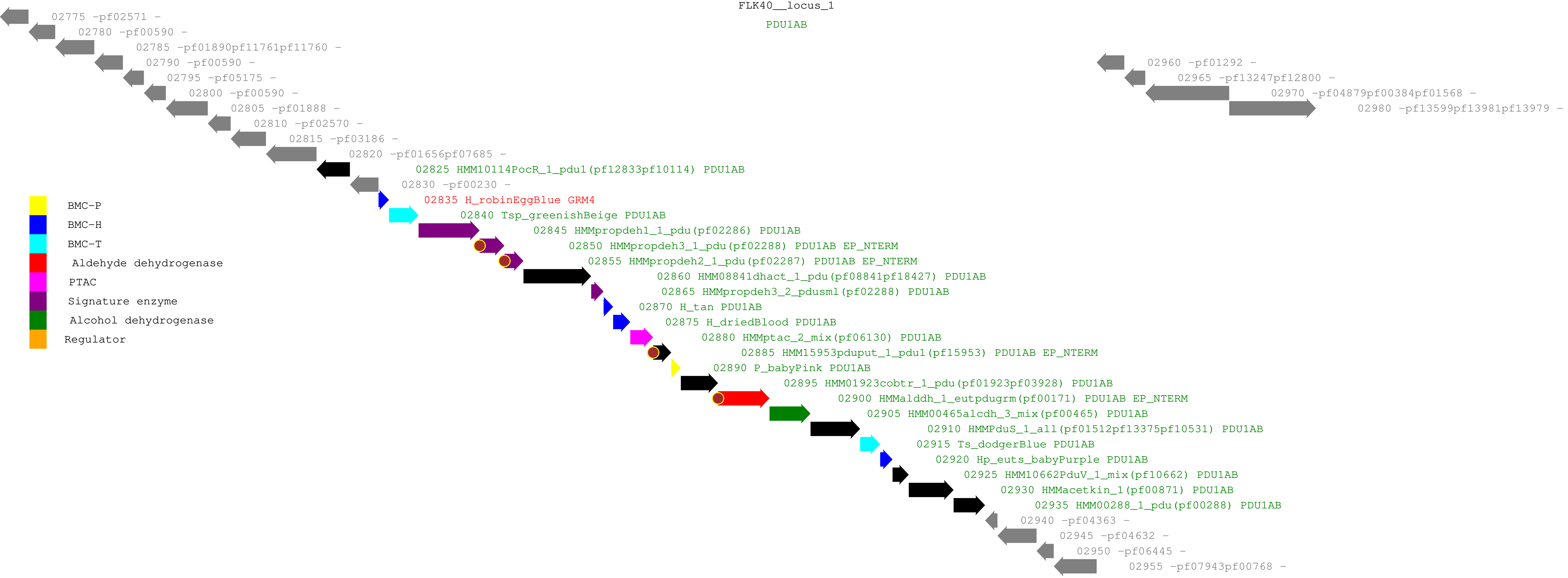

Citrobacter youngae. Bacteria; Proteobacteria; Gammaproteobacteria; Enterobacterales;

FMM65__locus_2

PDU1AB

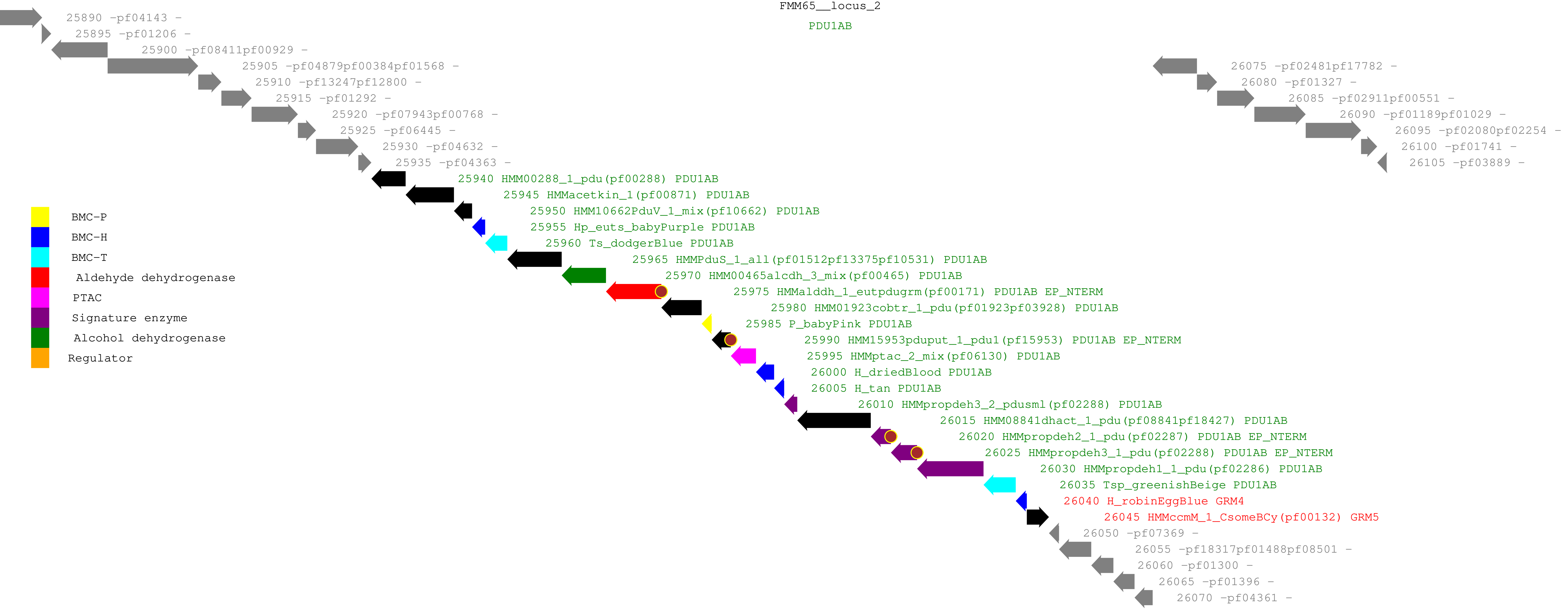

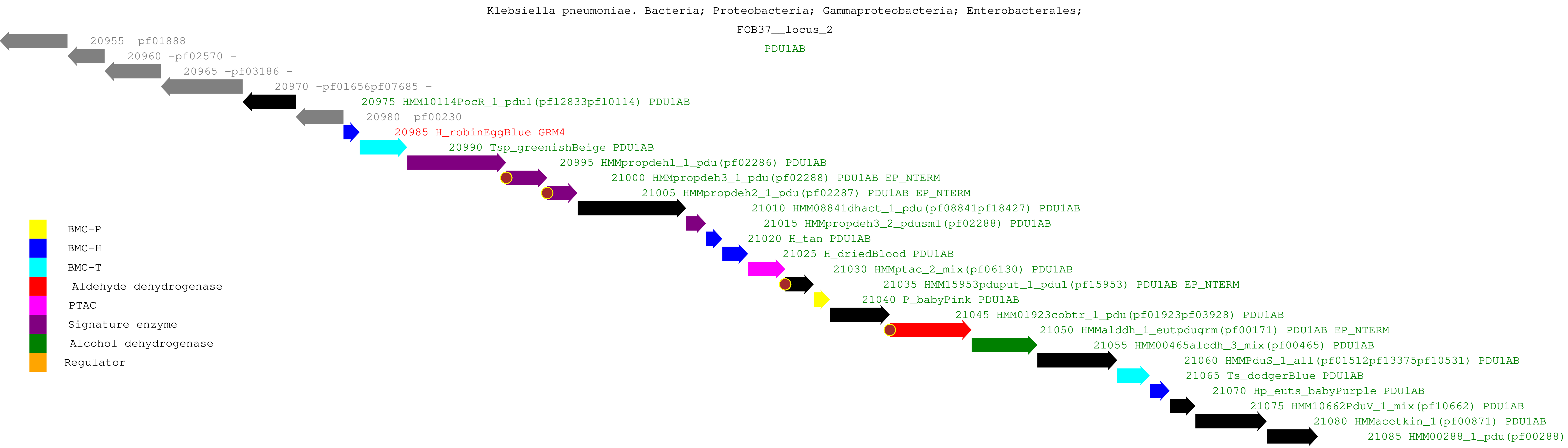
Klebsiella pneumoniae. Bacteria; Proteobacteria; Gammaproteobacteria; Enterobacterales;
FOB37__locus_2
PDU1AB
20955 -pf01888 -
20960 -pf02570 -
20965 -pf03186 -
20970 -pf01656pf07685 -
20975 HMM10114PocR_1_pdu1(pf12833pf10114) PDU1AB
20980 -pf00230 -
20985 H_robinEggBlue GRM4
20990 Tsp_greenishBeige PDU1AB
20995 HMMpropdeh1_1_pdu(pf02286) PDU1AB
21000 HMMpropdeh3_1_pdu(pf02288) PDU1AB EP_NTERM
21005 HMMpropdeh2_1_pdu(pf02287) PDU1AB EP_NTERM
21010 HMM08841dhact_1_pdu(pf08841pf18427) PDU1AB
21015 HMMpropdeh3_2_pdusml(pf02288) PDU1AB
21020 H_tan PDU1AB
21025 H_driedBlood PDU1AB
21030 HMMptac_2_mix(pf06130) PDU1AB
21035 HMM15953pduput_1_pdu1(pf15953) PDU1AB EP_NTERM
21040 P_babyPink PDU1AB
21045 HMM01923cobtr_1_pdu(pf01923pf03928) PDU1AB
21050 HMMalddh_1_eutpdugrm(pf00171) PDU1AB EP_NTERM
21055 HMM00465alcdh_3_mix(pf00465) PDU1AB
21060 HMMPduS_1_all(pf01512pf13375pf10531) PDU1AB
21065 Ts_dodgerBlue PDU1AB
21070 Hp_euts_babyPurple PDU1AB
21075 HMM10662PduV_1_mix(pf10662) PDU1AB
21080 HMMacetkin_1(pf00871) PDU1AB
21085 HMM00288_1_pdu(pf00288)
BMC-P
BMC-H
BMC-T
Aldehyde dehydrogenase
PTAC
Signature enzyme
Alcohol dehydrogenase
Regulator

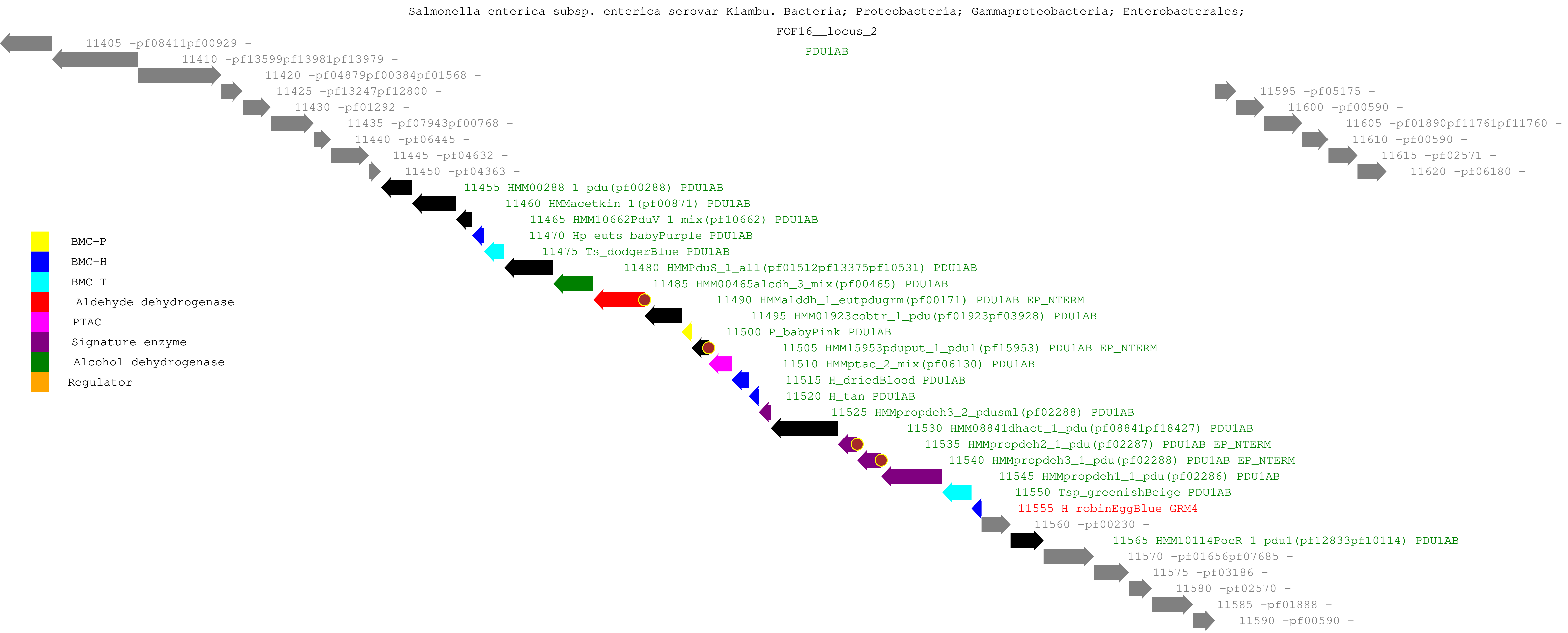
Salmonella enterica subsp. enterica serovar Kiambu. Bacteria; Proteobacteria; Gammaproteobacteria; Enterobacterales;
FOF16__locus_2
PDU1AB
11405 -pf08411pf00929 -
11410 -pf13599pf13981pf13979 -
11420 -pf04879pf00384pf01568 -
11425 -pf13247pf12800 -
11430 -pf01292 -
11435 -pf07943pf00768 -
11440 -pf06445 -
11445 -pf04632 -
11450 -pf04363 -
11455 HMM00288_1_pdu(pf00288) PDU1AB
11460 HMMacetkin_1(pf00871) PDU1AB
11465 HMM10662PduV_1_mix(pf10662) PDU1AB
11470 Hp_euts_babyPurple PDU1AB
11475 Ts_dodgerBlue PDU1AB
11480 HMMPduS_1_all(pf01512pf13375pf10531) PDU1AB
11485 HMM00465alcdh_3_mix(pf00465) PDU1AB
11490 HMMaldh_1_eutpdugrm(pf00171) PDU1AB EP_NTERM
11495 HMM01923cobtr_1_pdu(pf01923pf03928) PDU1AB
11500 P_babyPink PDU1AB
11505 HMM15953pduput_1_pdu1(pf15953) PDU1AB EP_NTERM
11510 HMMptac_2_mix(pf06130) PDU1AB
11515 H_driedBlood PDU1AB
11520 H_tan PDU1AB
11525 HMMpropdeh3_2_pdusml(pf02288) PDU1AB
11530 HMM08841dhact_1_pdu(pf08841pf18427) PDU1AB
11535 HMMpropdeh2_1_pdu(pf02287) PDU1AB EP_NTERM
11540 HMMpropdeh3_1_pdu(pf02288) PDU1AB EP_NTERM
11545 HMMpropdeh1_1_pdu(pf02286) PDU1AB
11550 Tsp_greenishBeige PDU1AB
11555 H_robinEggBlue GRM4
11560 -pf00230 -
11565 HMM10114PocR_1_pdu1(pf12833pf10114) PDU1AB
11570 -pf01656pf07685 -
11575 -pf03186 -
11580 -pf02570 -
11585 -pf01888 -
11590 -pf00590 -
11595 -pf05175 -
11600 -pf00590 -
11605 -pf01890pf11761pf11760 -
11610 -pf00590 -
11615 -pf02571 -
11620 -pf06180 -
BMC-P
BMC-H
BMC-T
Aldehyde dehydrogenase
PTAC
Signature enzyme
Alcohol dehydrogenase
Regulator

Salmonella enteritidis. Bacteria; Proteobacteria; Gammaproteobacteria; Enterobacterales;

FORC52__locus_1

PDU1AB

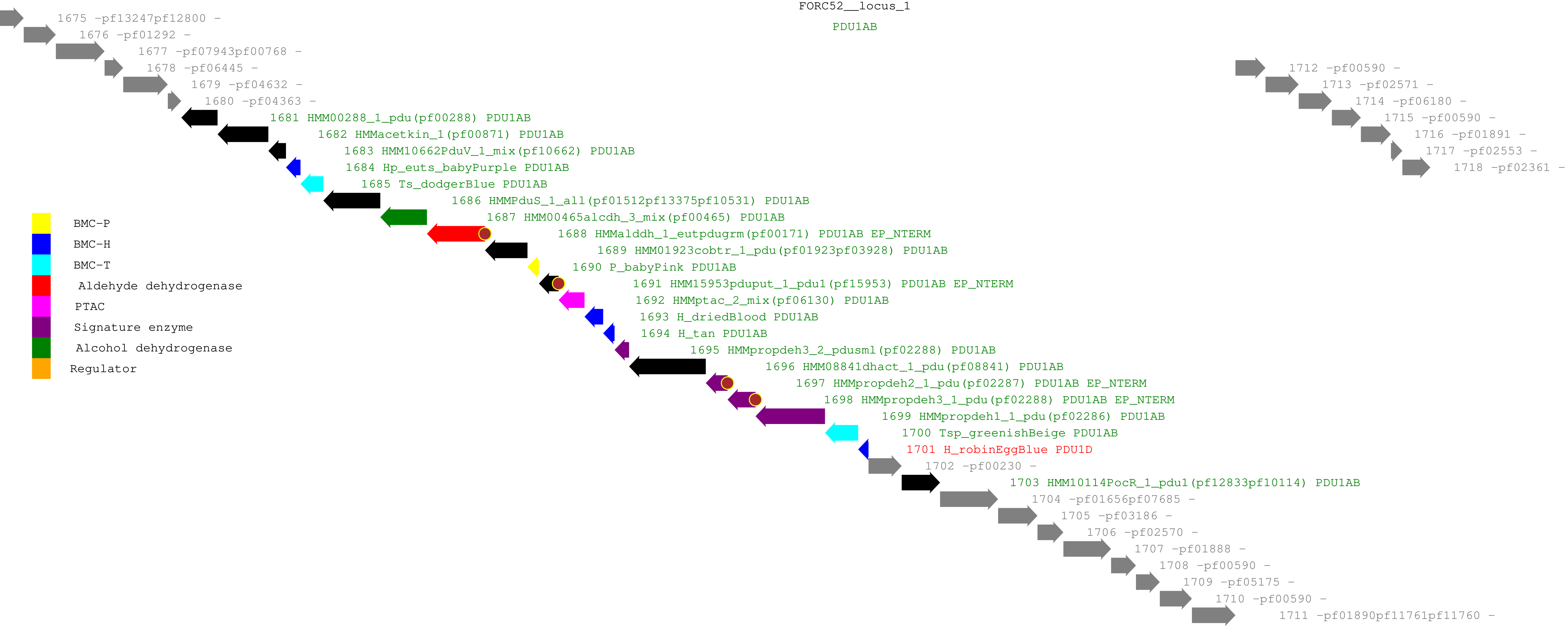

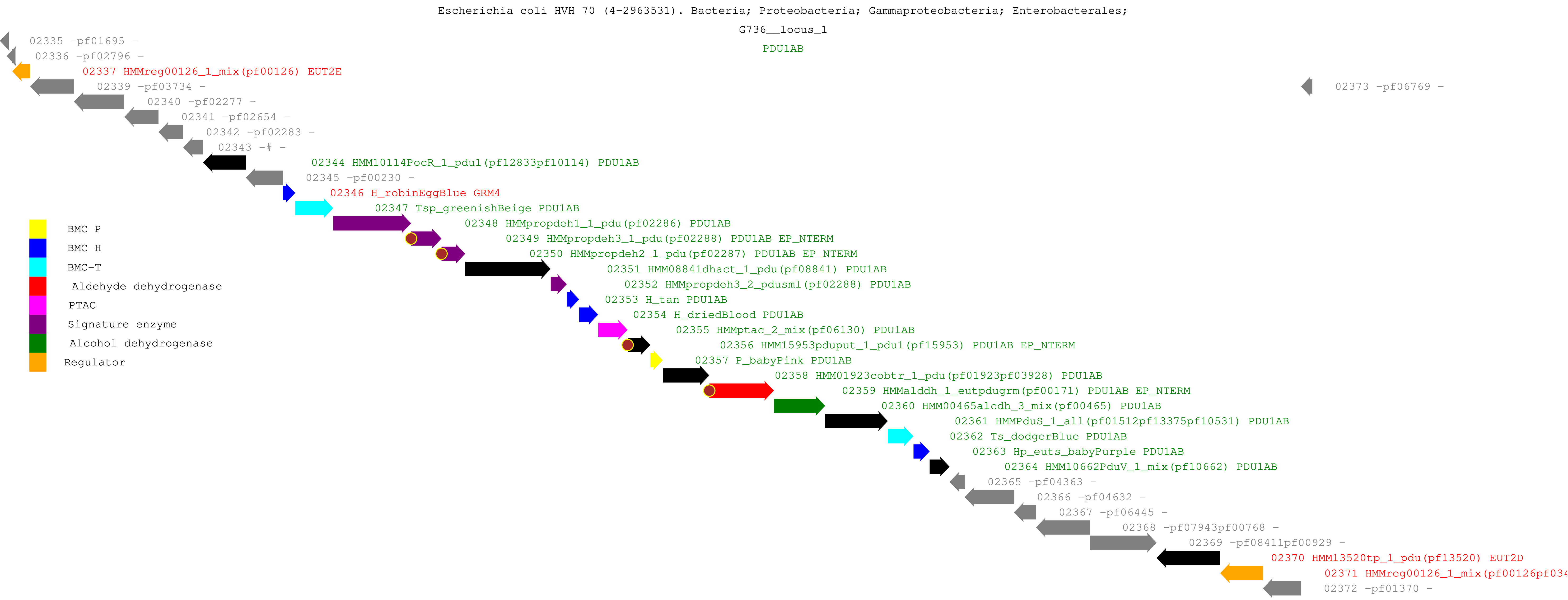

Escherichia coli HVH 70 (4-2963531). Bacteria; Proteobacteria; Gammaproteobacteria; Enterobacterales;
G736__locus_1
PDU1AB
02335 -pf01695 -
02336 -pf02796 -
02337 HMMreg00126_1_mix(pf00126) EUT2E
02339 -pf03734 -
02340 -pf02277 -
02341 -pf02654 -
02342 -pf02283 -
02343 -# -
02344 HMM10114PocR_1_pdu1(pf12833pf10114) PDU1AB
02345 -pf00230 -
02346 H_robinEggBlue GRM4
02347 Tsp_greenishBeige PDU1AB
02348 HMMpropdeh1_1_pdu(pf02286) PDU1AB
02349 HMMpropdeh3_1_pdu(pf02288) PDU1AB EP_NTERM
02350 HMMpropdeh2_1_pdu(pf02287) PDU1AB EP_NTERM
02351 HMM08841dhact_1_pdu(pf08841) PDU1AB
02352 HMMpropdeh3_2_pdusml(pf02288) PDU1AB
02353 H_tan PDU1AB
02354 H_driedBlood PDU1AB
02355 HMMptac_2_mix(pf06130) PDU1AB
02356 HMM15953pduput_1_pdu1(pf15953) PDU1AB EP_NTERM
02357 P_babyPink PDU1AB
02358 HMM01923cobtr_1_pdu(pf01923pf03928) PDU1AB
02359 HMMalddh_1_eutpdugrm(pf00171) PDU1AB EP_NTERM
02360 HMM00465alcdh_3_mix(pf00465) PDU1AB
02361 HMMPduS_1_all(pf01512pf13375pf10531) PDU1AB
02362 Ts_dodgerBlue PDU1AB
02363 Hp_euts_babyPurple PDU1AB
02364 HMM10662PduV_1_mix(pf10662) PDU1AB
02365 -pf04363 -
02366 -pf04632 -
02367 -pf06445 -
02368 -pf07943pf00768 -
02369 -pf08411pf00929 -
02370 HMM13520tp_1_pdu(pf13520) EUT2D
02371 HMMreg00126_1_mix(pf00126pf034
02372 -pf01370 -
02373 -pf06769 -
BMC-P
BMC-H
BMC-T
Aldehyde dehydrogenase
PTAC
Signature enzyme
Alcohol dehydrogenase
Regulator

Escherichia coli KOEGE 71 (186a). Bacteria; Proteobacteria; Gammaproteobacteria; Enterobacterales;

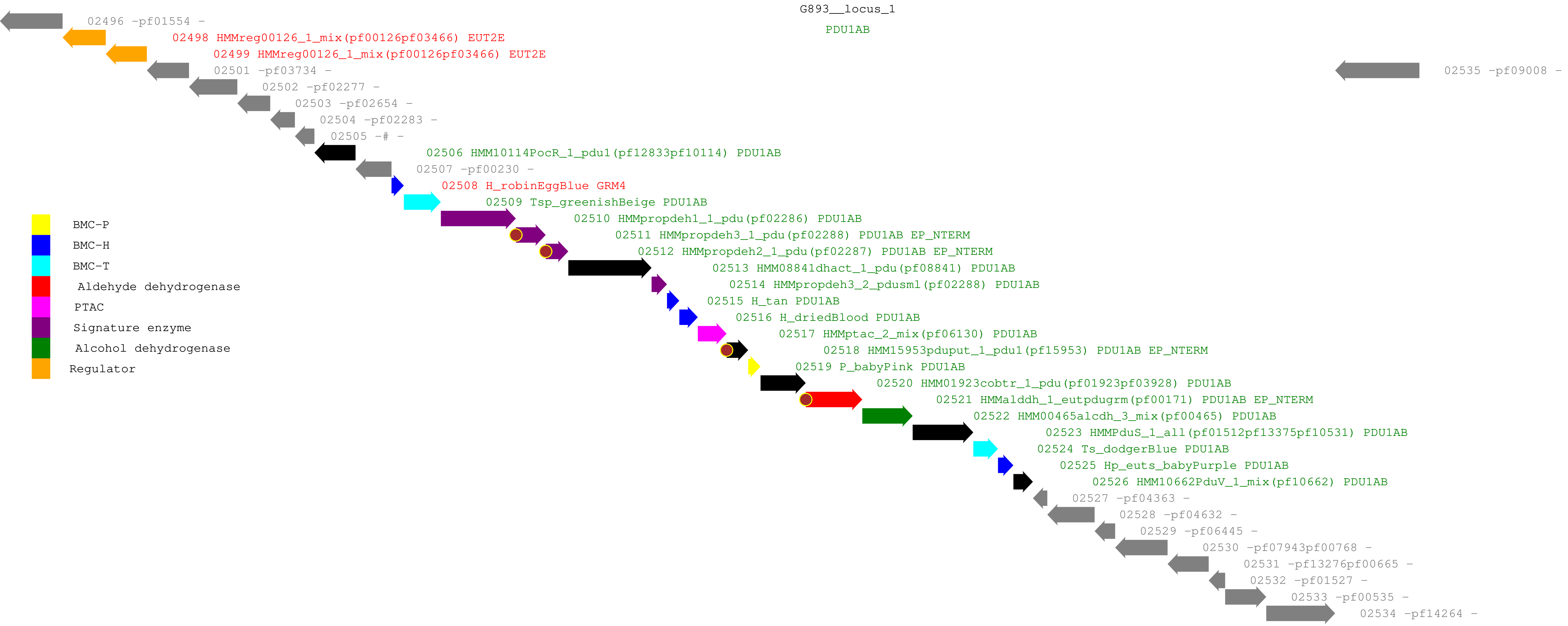

Escherichia coli UMEA 3200-1. Bacteria; Proteobacteria; Gammaproteobacteria; Enterobacterales;

G938__locus_1

PDU1AB

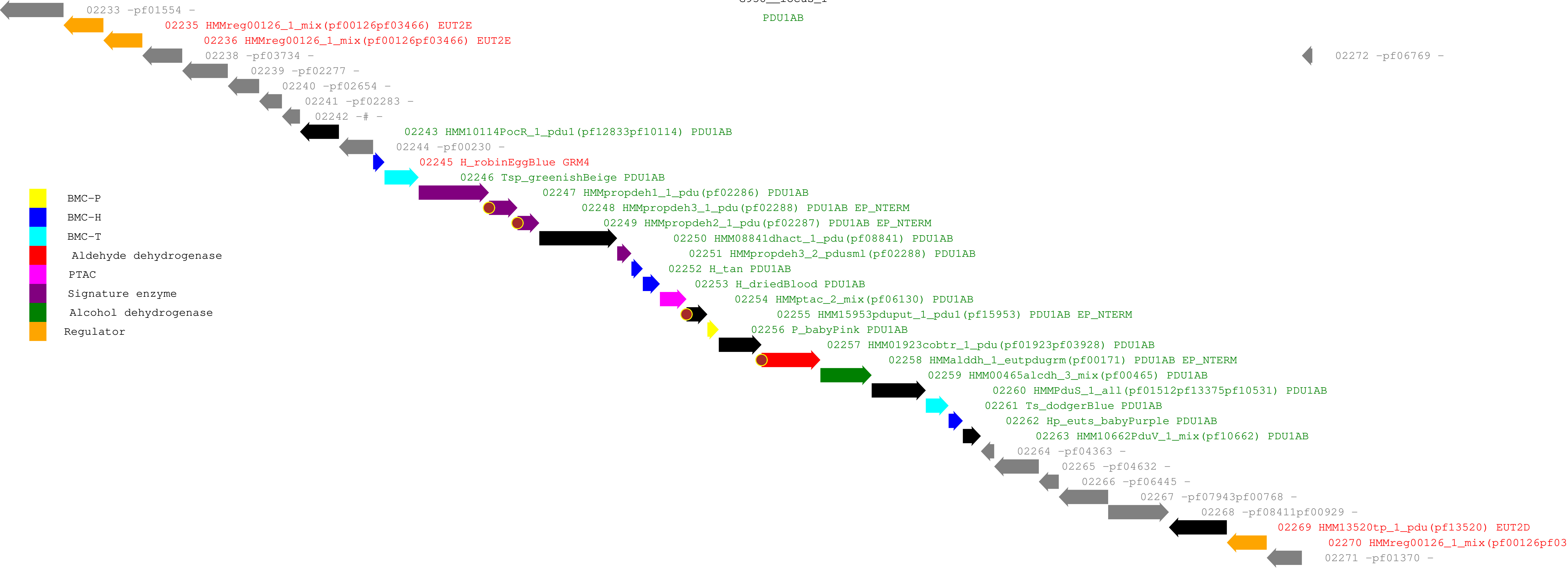

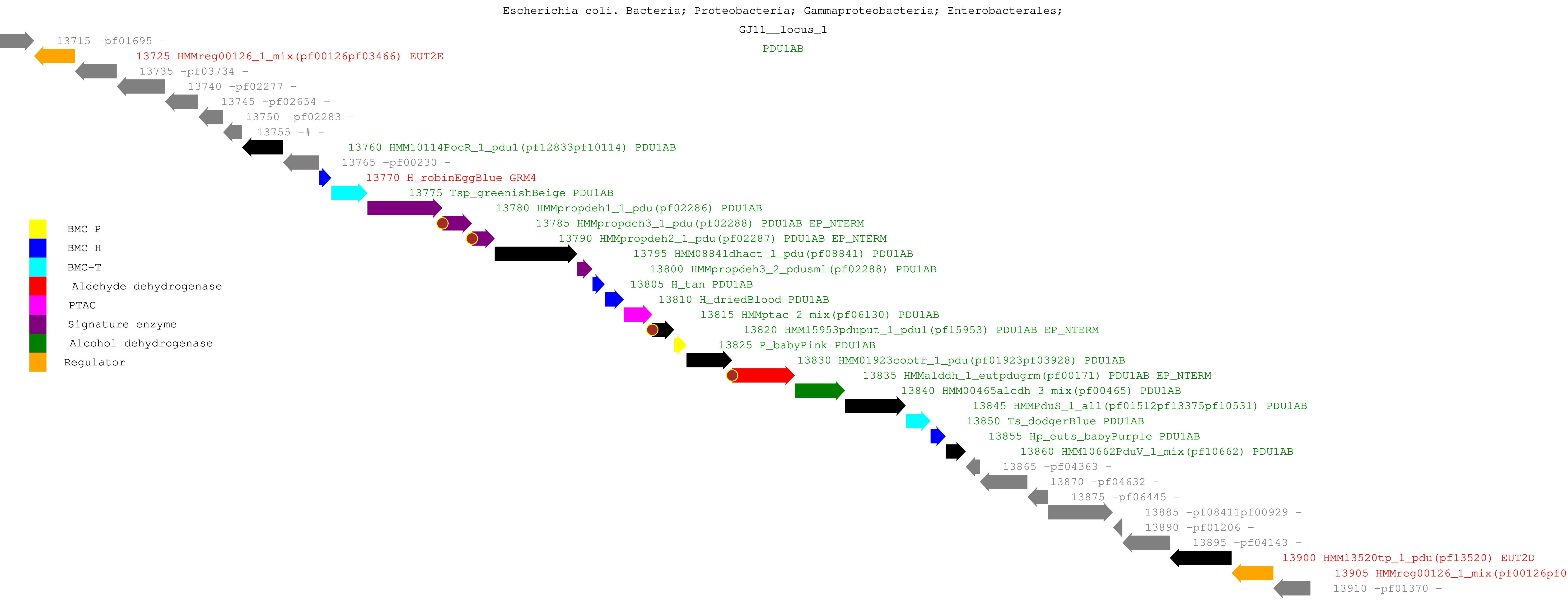
Escherichia coli. Bacteria; Proteobacteria; Gammaproteobacteria; Enterobacterales;
GJ11__locus_1
PDU1AB
13715 -pf01695 -
13725 HMMreg00126_1_mix(pf00126pf03466) EUT2E
13735 -pf03734 -
13740 -pf02277 -
13745 -pf02654 -
13750 -pf02283 -
13755 -# -
13760 HMM10114PocR_1_pdu1(pf12833pf10114) PDU1AB
13765 -pf00230 -
13770 H_robinEggBlue GRM4
13775 Tsp_greenishBeige PDU1AB
13780 HMMpropdeh1_1_pdu(pf02286) PDU1AB
13785 HMMpropdeh3_1_pdu(pf02288) PDU1AB EP_NTERM
13790 HMMpropdeh2_1_pdu(pf02287) PDU1AB EP_NTERM
13795 HMM08841dhact_1_pdu(pf08841) PDU1AB
13800 HMMpropdeh3_2_pdusml(pf02288) PDU1AB
13805 H_tan PDU1AB
13810 H_driedBlood PDU1AB
13815 HMMptac_2_mix(pf06130) PDU1AB
13820 HMM15953pduput_1_pdu1(pf15953) PDU1AB EP_NTERM
13825 P_babyPink PDU1AB
13830 HMM01923cobtr_1_pdu(pf01923pf03928) PDU1AB
13835 HMMalddh_1_eutpdugrm(pf00171) PDU1AB EP_NTERM
13840 HMM00465alcdh_3_mix(pf00465) PDU1AB
13845 HMMPduS_1_all(pf01512pf13375pf10531) PDU1AB
13850 Ts_dodgerBlue PDU1AB
13855 Hp_euts_babyPurple PDU1AB
13860 HMM10662PduV_1_mix(pf10662) PDU1AB
13865 -pf04363 -
13870 -pf04632 -
13875 -pf06445 -
13885 -pf08411pf00929 -
13890 -pf01206 -
13895 -pf04143 -
13900 HMM13520tp_1_pdu(pf13520) EUT2D
13905 HMMreg00126_1_mix(pf00126pf0
13910 -pf01370 -
BMC-P
BMC-H
BMC-T
Aldehyde dehydrogenase
PTAC
Signature enzyme
Alcohol dehydrogenase
Regulator

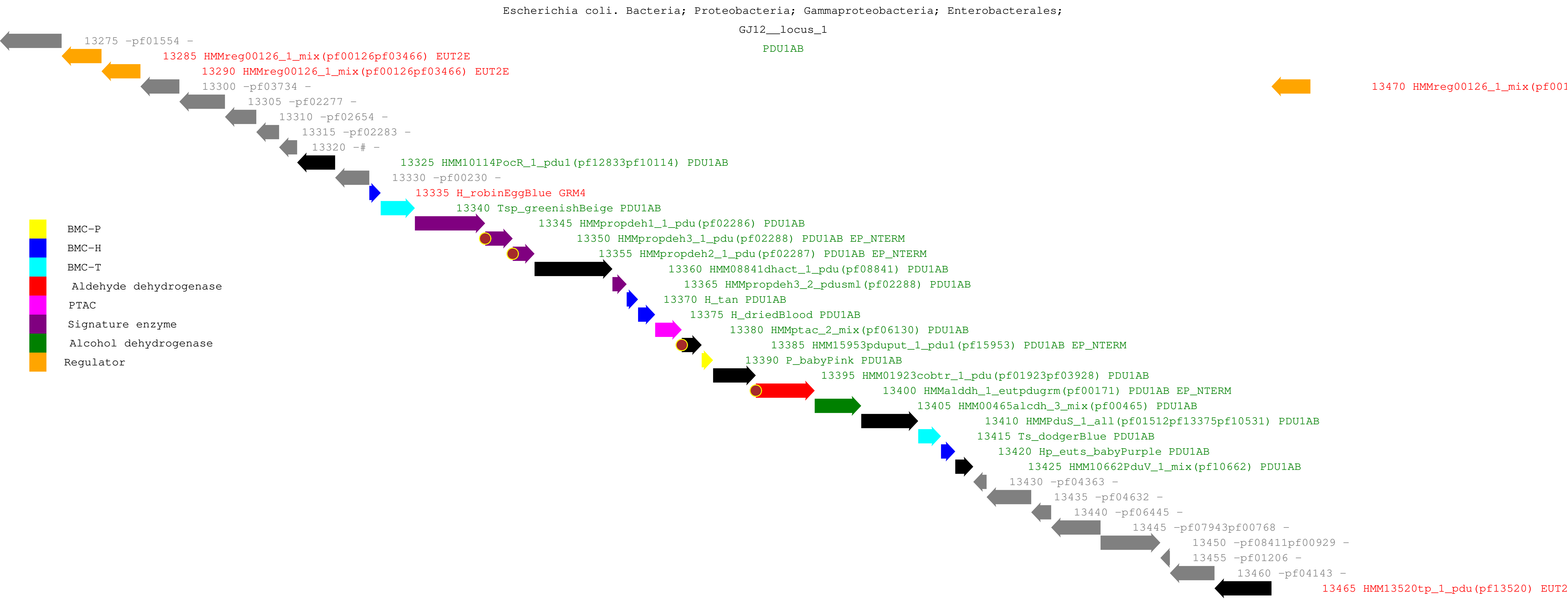
Escherichia coli. Bacteria; Proteobacteria; Gammaproteobacteria; Enterobacterales;
GJ12__locus_1
PDU1AB
13275 -pf01554 -
13285 HMMreg00126_1_mix(pf00126pf03466) EUT2E
13290 HMMreg00126_1_mix(pf00126pf03466) EUT2E
13300 -pf03734 -
13305 -pf02277 -
13310 -pf02654 -
13315 -pf02283 -
13320 -# -
13325 HMM10114PocR_1_pdu1(pf12833pf10114) PDU1AB
13330 -pf00230 -
13335 H_robinEggBlue GRM4
13340 Tsp_greenishBeige PDU1AB
13345 HMMpropdeh1_1_pdu(pf02286) PDU1AB
13350 HMMpropdeh3_1_pdu(pf02288) PDU1AB EP_NTERM
13355 HMMpropdeh2_1_pdu(pf02287) PDU1AB EP_NTERM
13360 HMM08841dhact_1_pdu(pf08841) PDU1AB
13365 HMMpropdeh3_2_pdusml(pf02288) PDU1AB
13370 H_tan PDU1AB
13375 H_driedBlood PDU1AB
13380 HMMptac_2_mix(pf06130) PDU1AB
13385 HMM15953pduput_1_pdu1(pf15953) PDU1AB EP_NTERM
13390 P_babyPink PDU1AB
13395 HMM01923cobtr_1_pdu(pf01923pf03928) PDU1AB
13400 HMMaldh_1_eutpdugrm(pf00171) PDU1AB EP_NTERM
13405 HMM00465alcdh_3_mix(pf00465) PDU1AB
13410 HMMPduS_1_all(pf01512pf13375pf10531) PDU1AB
13415 Ts_dodgerBlue PDU1AB
13420 Hp_euts_babyPurple PDU1AB
13425 HMM10662PduV_1_mix(pf10662) PDU1AB
13430 -pf04363 -
13435 -pf04632 -
13440 -pf06445 -
13445 -pf07943pf00768 -
13450 -pf08411pf00929 -
13455 -pf01206 -
13460 -pf04143 -
13465 HMM13520tp_1_pdu(pf13520) EUT2
13470 HMMreg00126_1_mix(pf001
BMC-P
BMC-H
BMC-T
Aldehyde dehydrogenase
PTAC
Signature enzyme
Alcohol dehydrogenase
Regulator

Hafnia alvei ATCC 51873. Bacteria; Proteobacteria; Gammaproteobacteria; Enterobacterales;

HMPREF0454__locus_1

PDU1AB

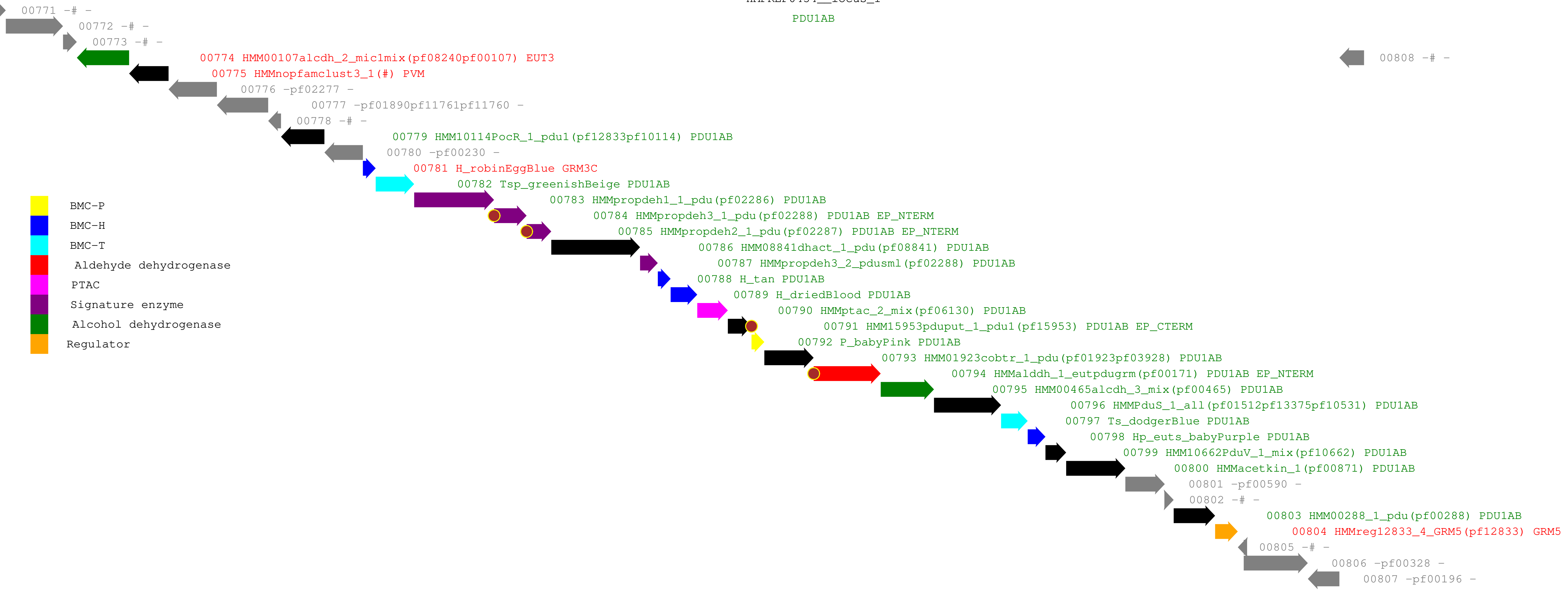

Klebsiella pneumoniae subsp. rhinoscleromatis ATCC 13884. Bacteria; Proteobacteria; Gammaproteobacteria; Enterobacterales;

HMPREF0484__locus_1

PDU1AB

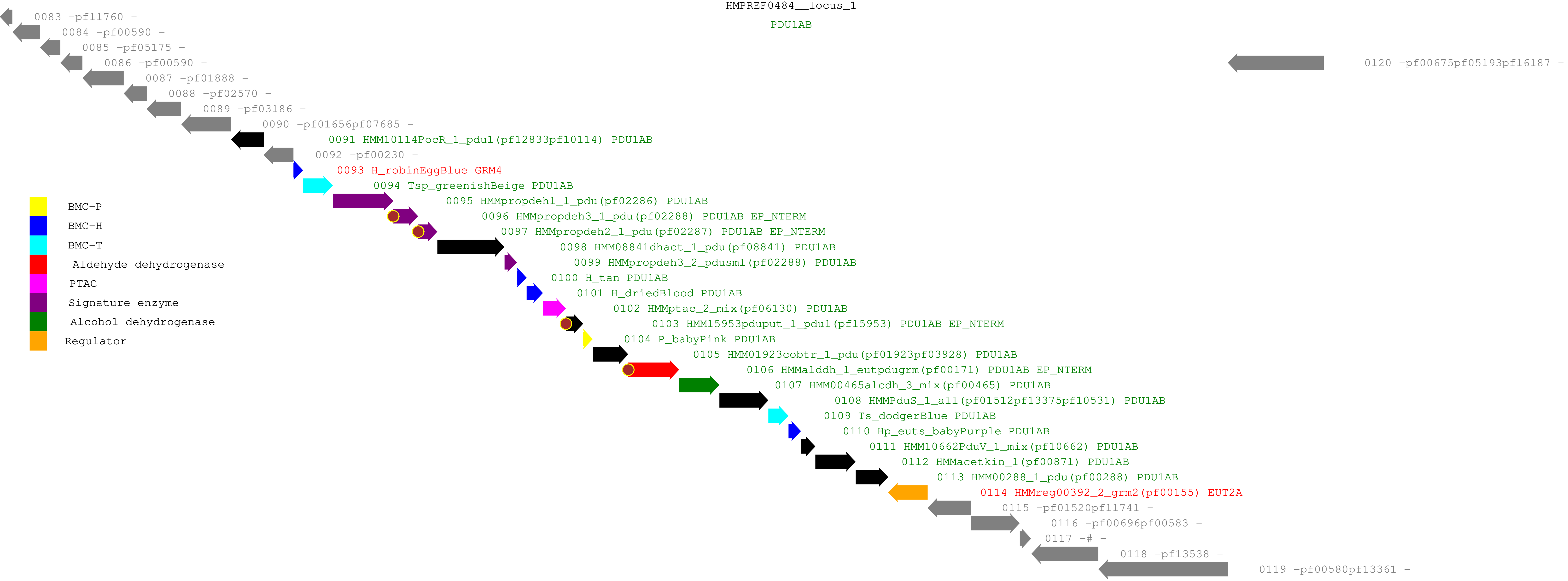

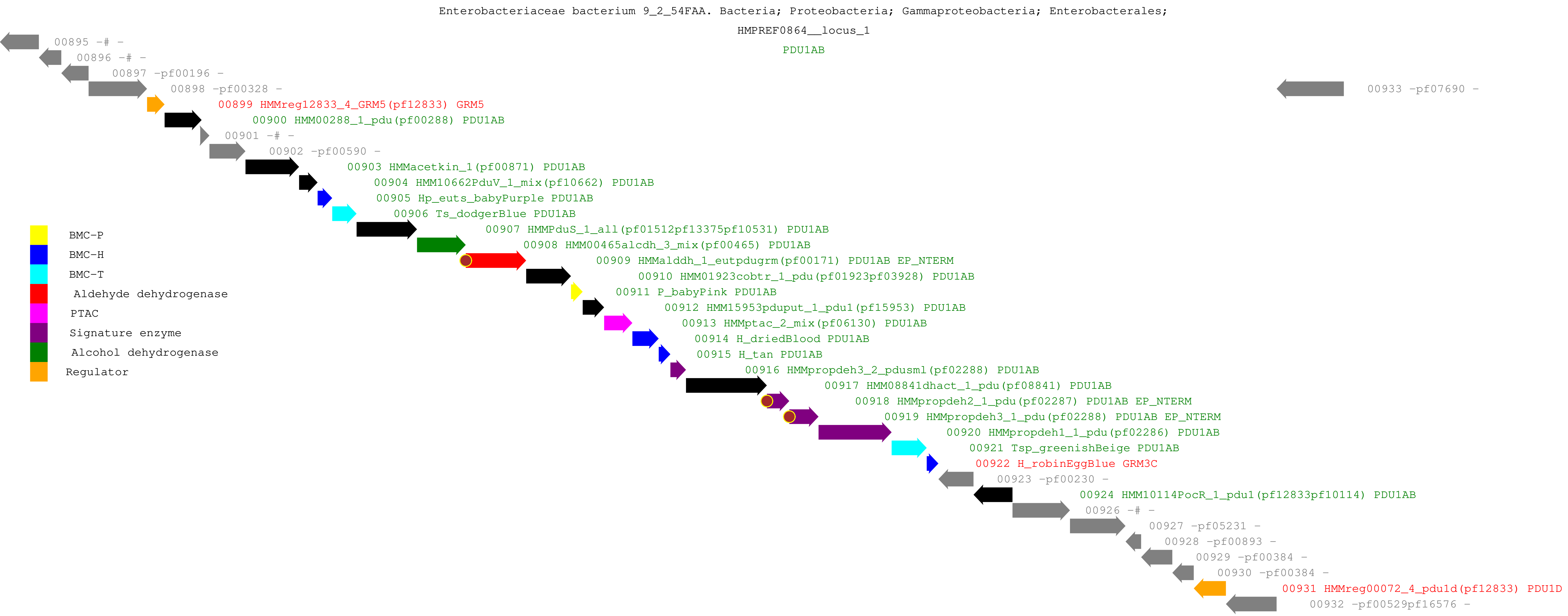

Enterobacteriaceae bacterium 9_2_54FAA. Bacteria; Proteobacteria; Gammaproteobacteria; Enterobacterales;
HMPREF0864__locus_1
PDU1AB
00895 -# -
00896 -# -
00897 -pf00196 -
00898 -pf00328 -
00899 HMMreg12833_4_GRM5(pf12833) GRM5
00900 HMM00288_1_pdu(pf00288) PDU1AB
00901 -# -
00902 -pf00590 -
00903 HMMacetkin_1(pf00871) PDU1AB
00904 HMM10662PduV_1_mix(pf10662) PDU1AB
00905 Hp_euts_babyPurple PDU1AB
00906 Ts_dodgerBlue PDU1AB
00907 HMMPduS_1_all(pf01512pf13375pf10531) PDU1AB
00908 HMM00465alcdh_3_mix(pf00465) PDU1AB
00909 HMMalddh_1_eutpdugrm(pf00171) PDU1AB EP_NTERM
00910 HMM01923cobtr_1_pdu(pf01923pf03928) PDU1AB
00911 P_babyPink PDU1AB
00912 HMM15953pduput_1_pdu1(pf15953) PDU1AB
00913 HMMptac_2_mix(pf06130) PDU1AB
00914 H_driedBlood PDU1AB
00915 H_tan PDU1AB
00916 HMMpropdeh3_2_pdusml(pf02288) PDU1AB
00917 HMM08841dhact_1_pdu(pf08841) PDU1AB
00918 HMMpropdeh2_1_pdu(pf02287) PDU1AB EP_NTERM
00919 HMMpropdeh3_1_pdu(pf02288) PDU1AB EP_NTERM
00920 HMMpropdeh1_1_pdu(pf02286) PDU1AB
00921 Tsp_greenishBeige PDU1AB
00922 H_robinEggBlue GRM3C
00923 -pf00230 -
00924 HMM10114PocR_1_pdu1(pf12833pf10114) PDU1AB
00926 -# -
00927 -pf05231 -
00928 -pf00893 -
00929 -pf00384 -
00930 -pf00384 -
00931 HMMreg00072_4_pdu1d(pf12833) PDU1D
00932 -pf00529pf16576 -
00933 -pf07690 -
BMC-P
BMC-H
BMC-T
Aldehyde dehydrogenase
PTAC
Signature enzyme
Alcohol dehydrogenase
Regulator

Escherichia coli 4_1_47FAA. Bacteria; Proteobacteria; Gammaproteobacteria; Enterobacterales;

HMPREF0986__locus_2

PDU1AB

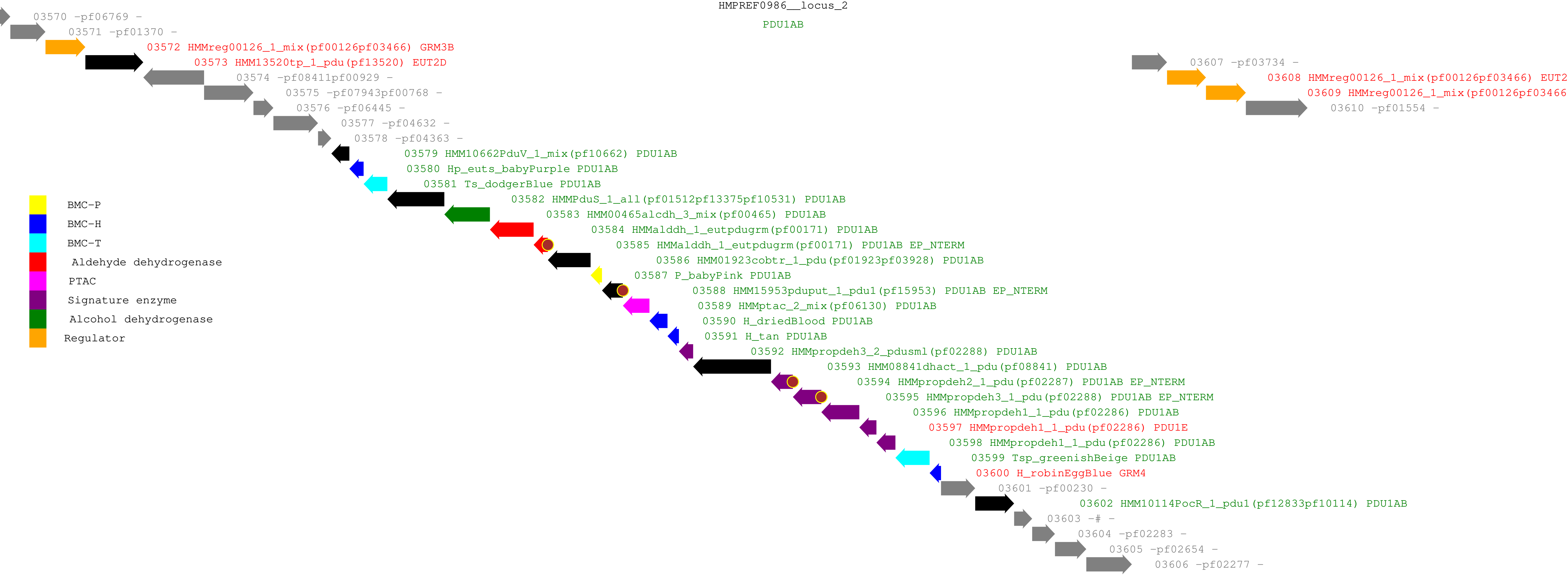

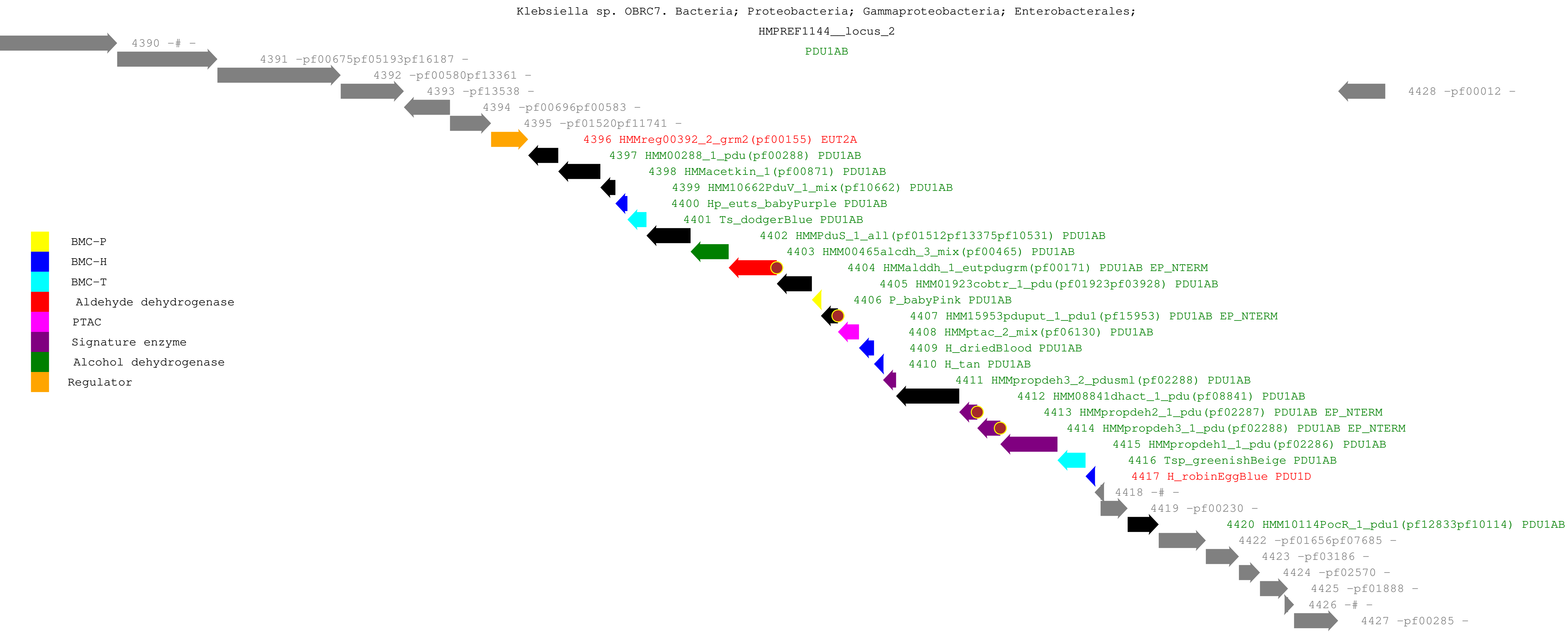
Klebsiella sp. OBRC7. Bacteria; Proteobacteria; Gammaproteobacteria; Enterobacterales;
HMPREF1144__locus_2
PDU1AB
4390 -# -
4391 -pf00675pf05193pf16187 -
4392 -pf00580pf13361 -
4393 -pf13538 -
4394 -pf00696pf00583 -
4395 -pf01520pf11741 -
4396 HMMreg00392_2_grm2(pf00155) EUT2A
4397 HMM00288_1_pdu(pf00288) PDU1AB
4398 HMMacetkin_1(pf00871) PDU1AB
4399 HMM10662PduV_1_mix(pf10662) PDU1AB
4400 Hp_euts_babyPurple PDU1AB
4401 Ts_dodgerBlue PDU1AB
4402 HMMPduS_1_all(pf01512pf13375pf10531) PDU1AB
4403 HMM00465alcdh_3_mix(pf00465) PDU1AB
4404 HMMalddh_1_eutpdugrm(pf00171) PDU1AB EP_NTERM
4405 HMM01923cobtr_1_pdu(pf01923pf03928) PDU1AB
4406 P_babyPink PDU1AB
4407 HMM15953pduput_1_pdu1(pf15953) PDU1AB EP_NTERM
4408 HMMptac_2_mix(pf06130) PDU1AB
4409 H_driedBlood PDU1AB
4410 H_tan PDU1AB
4411 HMMpropdeh3_2_pdusml(pf02288) PDU1AB
4412 HMM08841dhact_1_pdu(pf08841) PDU1AB
4413 HMMpropdeh2_1_pdu(pf02287) PDU1AB EP_NTERM
4414 HMMpropdeh3_1_pdu(pf02288) PDU1AB EP_NTERM
4415 HMMpropdeh1_1_pdu(pf02286) PDU1AB
4416 Tsp_greenishBeige PDU1AB
4417 H_robinEggBlue PDU1D
4418 -# -
4419 -pf00230 -
4420 HMM10114PocR_1_pdu1(pf12833pf10114) PDU1AB
4422 -pf01656pf07685 -
4423 -pf03186 -
4424 -pf02570 -
4425 -pf01888 -
4426 -# -
4427 -pf00285 -
4428 -pf00012 -
BMC-P
BMC-H
BMC-T
Aldehyde dehydrogenase
PTAC
Signature enzyme
Alcohol dehydrogenase
Regulator

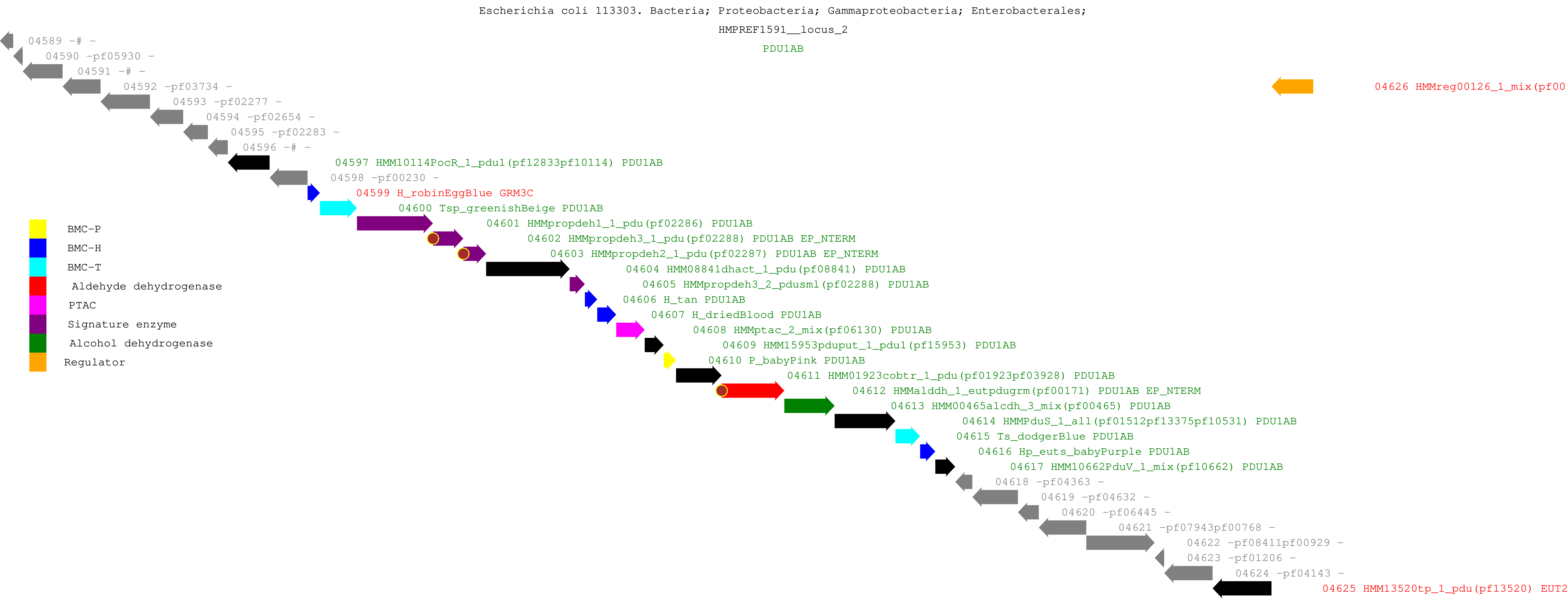

Escherichia coli 113303. Bacteria; Proteobacteria; Gammaproteobacteria; Enterobacterales;
HMPREF1591__locus_2
PDU1AB
04589 -# -
04590 -pf05930 -
04591 -# -
04592 -pf03734 -
04593 -pf02277 -
04594 -pf02654 -
04595 -pf02283 -
04596 -# -
04597 HMM10114PocR_1_pdu1(pf12833pf10114) PDU1AB
04598 -pf00230 -
04599 H_robinEggBlue GRM3C
04600 Tsp_greenishBeige PDU1AB
04601 HMMpropdeh1_1_pdu(pf02286) PDU1AB
04602 HMMpropdeh3_1_pdu(pf02288) PDU1AB EP_NTERM
04603 HMMpropdeh2_1_pdu(pf02287) PDU1AB EP_NTERM
04604 HMM08841dhact_1_pdu(pf08841) PDU1AB
04605 HMMpropdeh3_2_pdusml(pf02288) PDU1AB
04606 H_tan PDU1AB
04607 H_driedBlood PDU1AB
04608 HMMptac_2_mix(pf06130) PDU1AB
04609 HMM15953pduput_1_pdu1(pf15953) PDU1AB
04610 P_babyPink PDU1AB
04611 HMM01923cobtr_1_pdu(pf01923pf03928) PDU1AB
04612 HMMalddh_1_eutpdugrm(pf00171) PDU1AB EP_NTERM
04613 HMM00465alcdh_3_mix(pf00465) PDU1AB
04614 HMMPduS_1_all(pf01512pf13375pf10531) PDU1AB
04615 Ts_dodgerBlue PDU1AB
04616 Hp_euts_babyPurple PDU1AB
04617 HMM10662PduV_1_mix(pf10662) PDU1AB
04618 -pf04363 -
04619 -pf04632 -
04620 -pf06445 -
04621 -pf07943pf00768 -
04622 -pf08411pf00929 -
04623 -pf01206 -
04624 -pf04143 -
04625 HMM13520tp_1_pdu(pf13520) EUT2
04626 HMMreg00126_1_mix(pf00
BMC-P
BMC-H
BMC-T
Aldehyde dehydrogenase
PTAC
Signature enzyme
Alcohol dehydrogenase
Regulator

Escherichia coli 907357. Bacteria; Proteobacteria; Gammaproteobacteria; Enterobacterales;

HMPREF1592__locus_1

PDU1AB

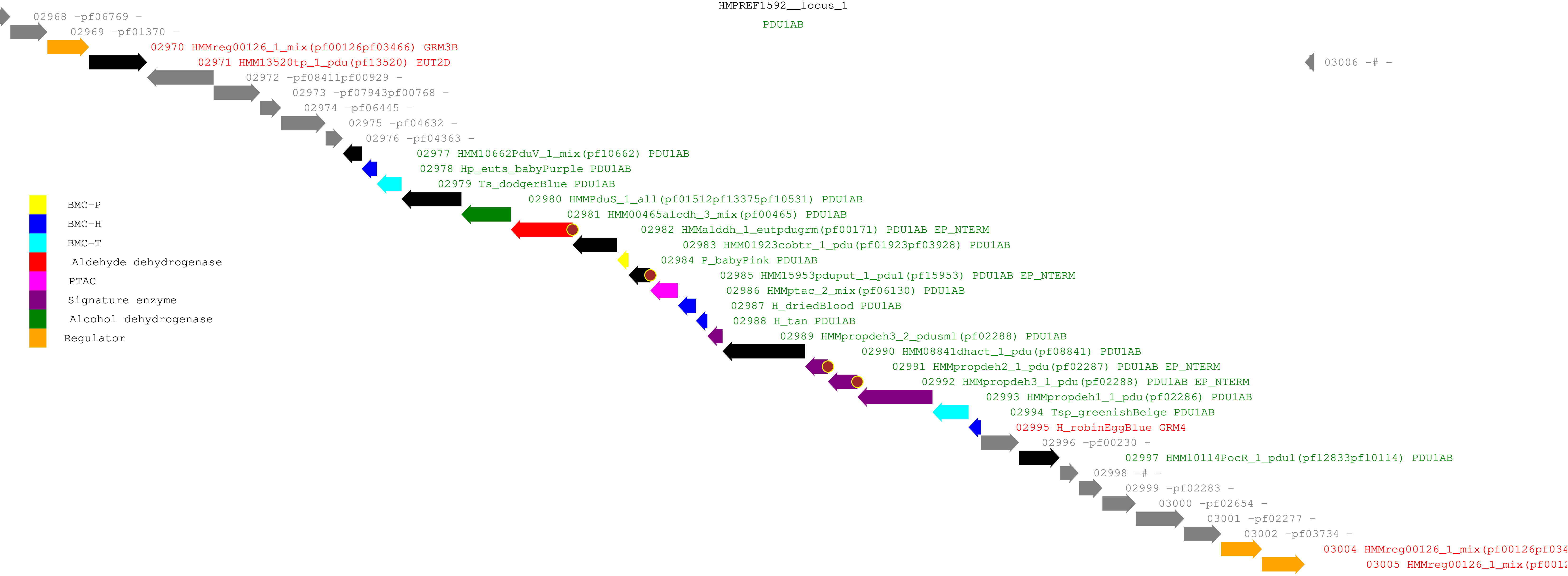

Escherichia coli 907713. Bacteria; Proteobacteria; Gammaproteobacteria; Enterobacterales;

HMPREF1599__locus_2

PDU1AB

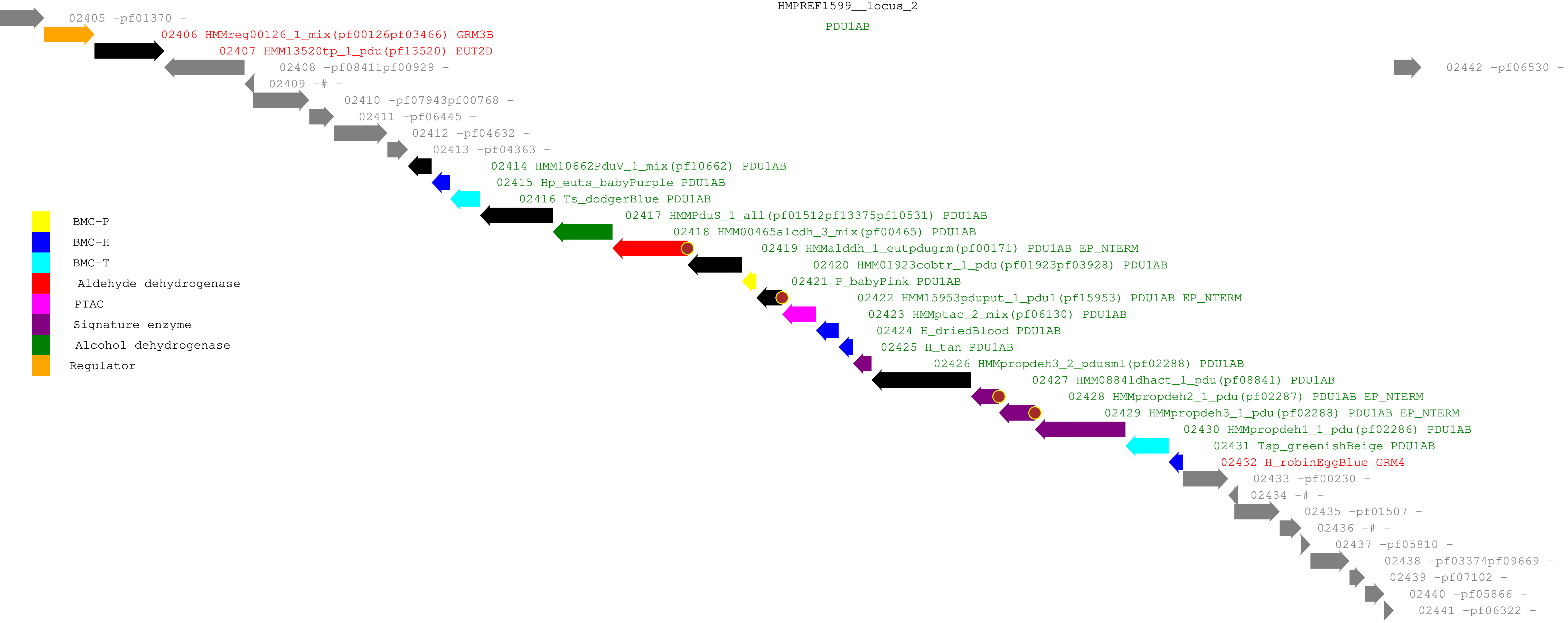

Escherichia coli 908519. Bacteria; Proteobacteria; Gammaproteobacteria; Enterobacterales;

HMPREF1604__locus_2

PDU1AB

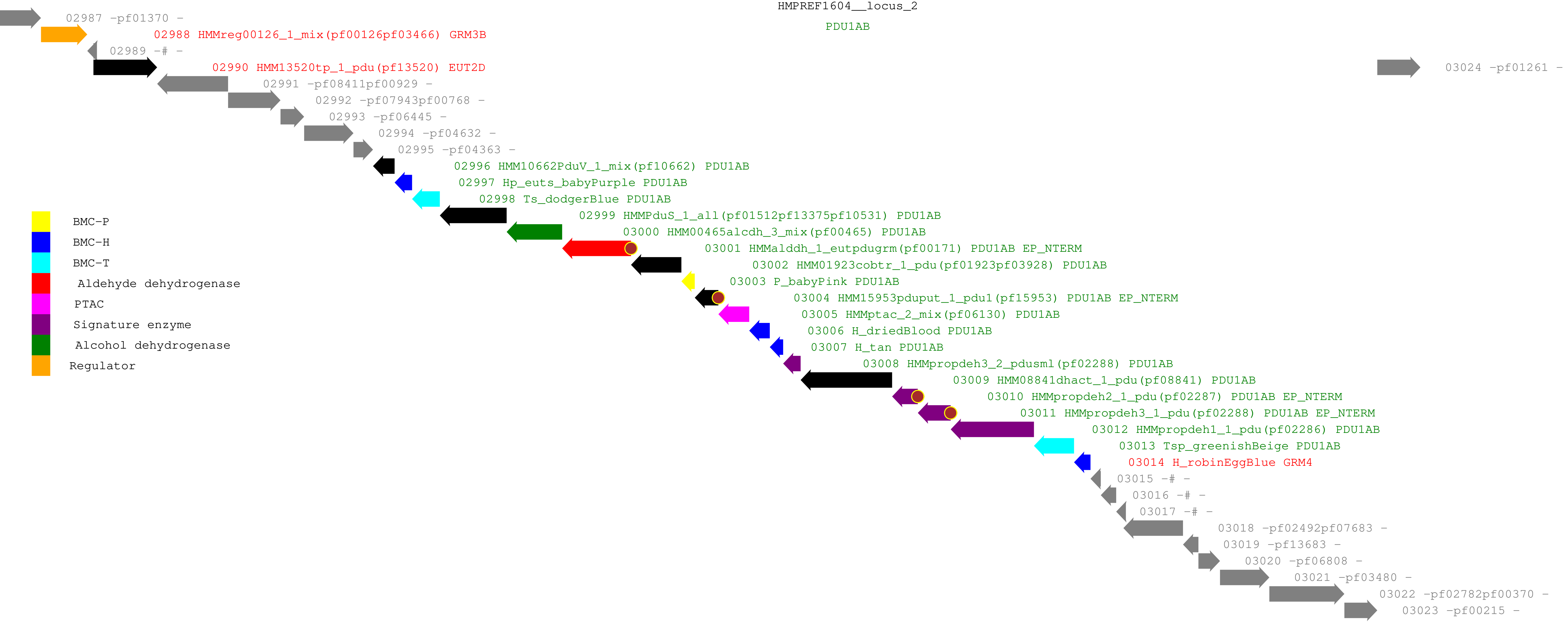

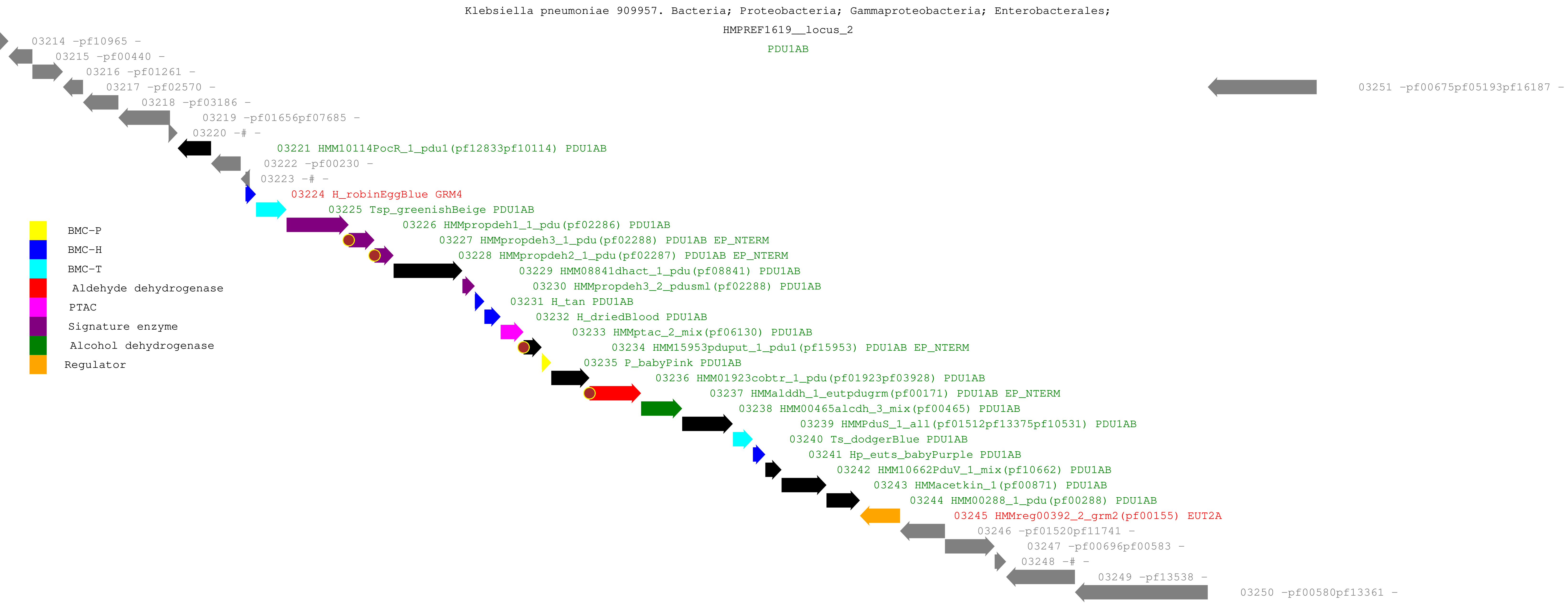
Klebsiella pneumoniae 909957. Bacteria; Proteobacteria; Gammaproteobacteria; Enterobacterales;
HMPREF1619__locus_2
PDU1AB
03214 -pf10965 -
03215 -pf00440 -
03216 -pf01261 -
03217 -pf02570 -
03218 -pf03186 -
03219 -pf01656pf07685 -
03220 -# -
03221 HMM10114PocR_1_pdu1(pf12833pf10114) PDU1AB
03222 -pf00230 -
03223 -# -
03224 H_robinEggBlue GRM4
03225 Tsp_greenishBeige PDU1AB
03226 HMMpropdeh1_1_pdu(pf02286) PDU1AB
03227 HMMpropdeh3_1_pdu(pf02288) PDU1AB EP_NTERM
03228 HMMpropdeh2_1_pdu(pf02287) PDU1AB EP_NTERM
03229 HMM08841dhact_1_pdu(pf08841) PDU1AB
03230 HMMpropdeh3_2_pdusml(pf02288) PDU1AB
03231 H_tan PDU1AB
03232 H_driedBlood PDU1AB
03233 HMMptac_2_mix(pf06130) PDU1AB
03234 HMM15953pduput_1_pdu1(pf15953) PDU1AB EP_NTERM
03235 P_babyPink PDU1AB
03236 HMM01923cobtr_1_pdu(pf01923pf03928) PDU1AB
03237 HMMalddh_1_eutpdugrm(pf00171) PDU1AB EP_NTERM
03238 HMM00465alcdh_3_mix(pf00465) PDU1AB
03239 HMMPduS_1_all(pf01512pf13375pf10531) PDU1AB
03240 Ts_dodgerBlue PDU1AB
03241 Hp_euts_babyPurple PDU1AB
03242 HMM10662PduV_1_mix(pf10662) PDU1AB
03243 HMMacetkin_1(pf00871) PDU1AB
03244 HMM00288_1_pdu(pf00288) PDU1AB
03245 HMMreg00392_2_grm2(pf00155) EUT2A
03246 -pf01520pf11741 -
03247 -pf00696pf00583 -
03248 -# -
03249 -pf13538 -
03250 -pf00580pf13361 -
03251 -pf00675pf05193pf16187 -
BMC-P
BMC-H
BMC-T
Aldehyde dehydrogenase
PTAC
Signature enzyme
Alcohol dehydrogenase
Regulator

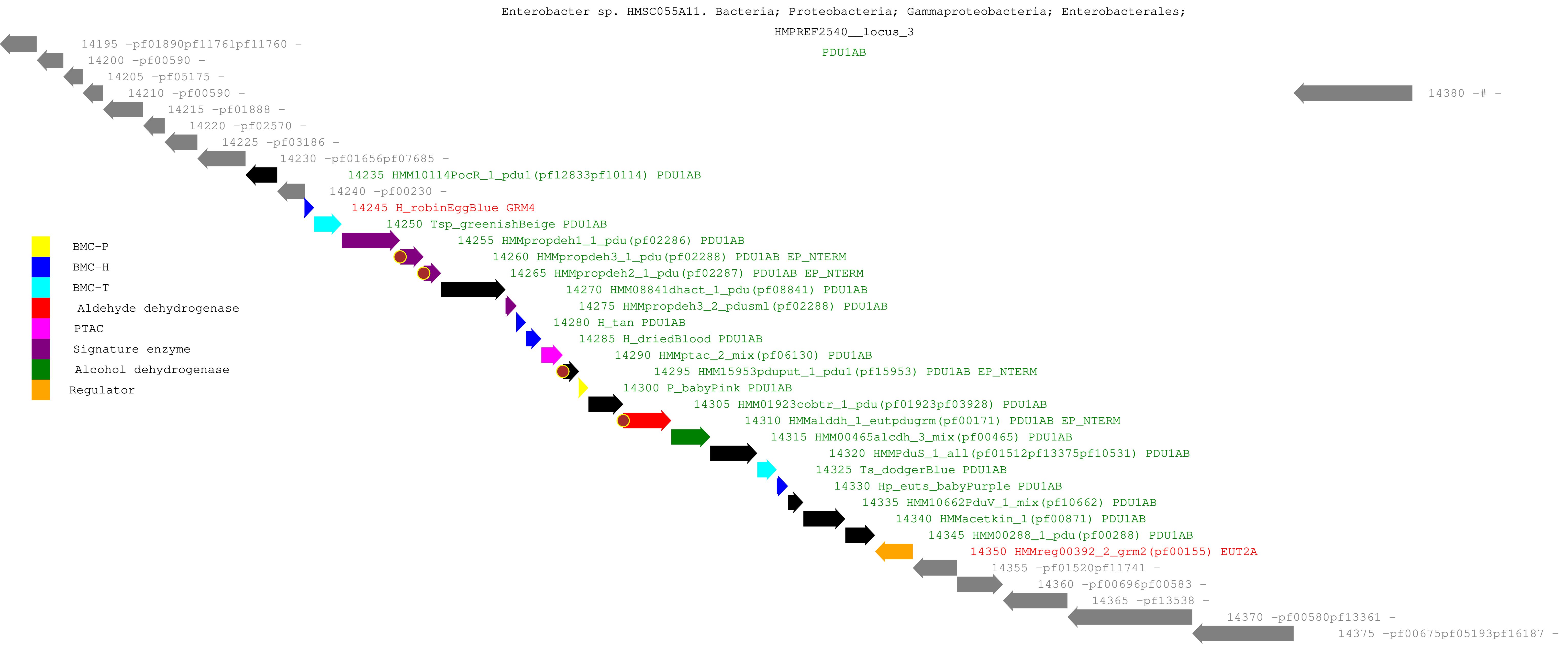
Enterobacter sp. HMSC055A11. Bacteria; Proteobacteria; Gammaproteobacteria; Enterobacterales;
HMPREF2540__locus_3
PDU1AB
14195 -pf01890pf11761pf11760 -
14200 -pf00590 -
14205 -pf05175 -
14210 -pf00590 -
14215 -pf01888 -
14220 -pf02570 -
14225 -pf03186 -
14230 -pf01656pf07685 -
14235 HMM10114PocR_1_pdu1(pf12833pf10114) PDU1AB
14240 -pf00230 -
14245 H_robinEggBlue GRM4
14250 Tsp_greenishBeige PDU1AB
14255 HMMpropdeh1_1_pdu(pf02286) PDU1AB
14260 HMMpropdeh3_1_pdu(pf02288) PDU1AB EP_NTERM
14265 HMMpropdeh2_1_pdu(pf02287) PDU1AB EP_NTERM
14270 HMM08841dhact_1_pdu(pf08841) PDU1AB
14275 HMMpropdeh3_2_pdusml(pf02288) PDU1AB
14280 H_tan PDU1AB
14285 H_driedBlood PDU1AB
14290 HMMptac_2_mix(pf06130) PDU1AB
14295 HMM15953pduput_1_pdu1(pf15953) PDU1AB EP_NTERM
14300 P_babyPink PDU1AB
14305 HMM01923cobtr_1_pdu(pf01923pf03928) PDU1AB
14310 HMMalddh_1_eutpdugrm(pf00171) PDU1AB EP_NTERM
14315 HMM00465alcdh_3_mix(pf00465) PDU1AB
14320 HMMPduS_1_all(pf01512pf13375pf10531) PDU1AB
14325 Ts_dodgerBlue PDU1AB
14330 Hp_euts_babyPurple PDU1AB
14335 HMM10662PduV_1_mix(pf10662) PDU1AB
14340 HMMacetkin_1(pf00871) PDU1AB
14345 HMM00288_1_pdu(pf00288) PDU1AB
14350 HMMreg00392_2_grm2(pf00155) EUT2A
14355 -pf01520pf11741 -
14360 -pf00696pf00583 -
14365 -pf13538 -
14370 -pf00580pf13361 -
14375 -pf00675pf05193pf16187 -
14380 -# -
BMC-P
BMC-H
BMC-T
Aldehyde dehydrogenase
PTAC
Signature enzyme
Alcohol dehydrogenase
Regulator

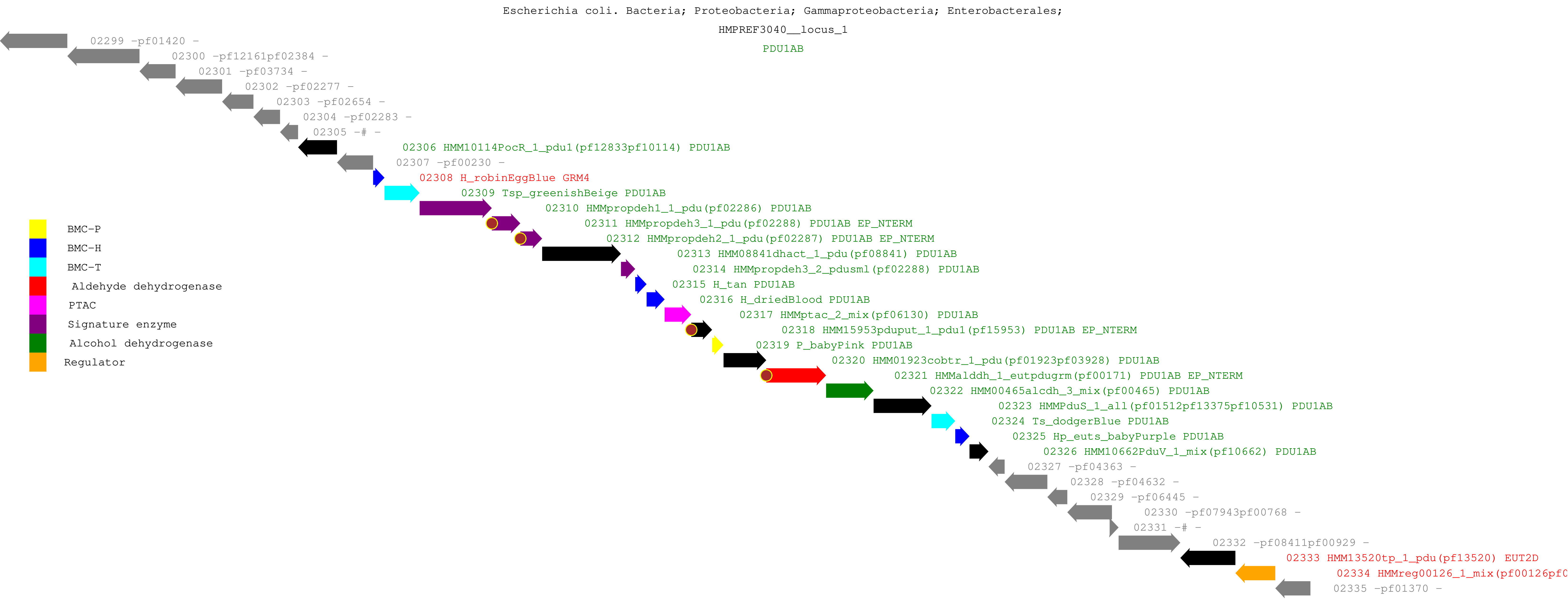

Escherichia coli. Bacteria; Proteobacteria; Gammaproteobacteria; Enterobacterales;
HMPREF3040__locus_1
PDU1AB
02299 -pf01420 -
02300 -pf12161pf02384 -
02301 -pf03734 -
02302 -pf02277 -
02303 -pf02654 -
02304 -pf02283 -
02305 -# -
02306 HMM10114PocR_1_pdu1(pf12833pf10114) PDU1AB
02307 -pf00230 -
02308 H_robinEggBlue GRM4
02309 Tsp_greenishBeige PDU1AB
02310 HMMpropdeh1_1_pdu(pf02286) PDU1AB
02311 HMMpropdeh3_1_pdu(pf02288) PDU1AB EP_NTERM
02312 HMMpropdeh2_1_pdu(pf02287) PDU1AB EP_NTERM
02313 HMM08841dhact_1_pdu(pf08841) PDU1AB
02314 HMMpropdeh3_2_pdusml(pf02288) PDU1AB
02315 H_tan PDU1AB
02316 H_driedBlood PDU1AB
02317 HMMptac_2_mix(pf06130) PDU1AB
02318 HMM15953pduput_1_pdu1(pf15953) PDU1AB EP_NTERM
02319 P_babyPink PDU1AB
02320 HMM01923cobtr_1_pdu(pf01923pf03928) PDU1AB
02321 HMMalddh_1_eutpdugrm(pf00171) PDU1AB EP_NTERM
02322 HMM00465alcdh_3_mix(pf00465) PDU1AB
02323 HMMPduS_1_all(pf01512pf13375pf10531) PDU1AB
02324 Ts_dodgerBlue PDU1AB
02325 Hp_euts_babyPurple PDU1AB
02326 HMM10662PduV_1_mix(pf10662) PDU1AB
02327 -pf04363 -
02328 -pf04632 -
02329 -pf06445 -
02330 -pf07943pf00768 -
02331 -# -
02332 -pf08411pf00929 -
02333 HMM13520tp_1_pdu(pf13520) EUT2D
02334 HMMreg00126_1_mix(pf00126pf0
02335 -pf01370 -
BMC-P
BMC-H
BMC-T
Aldehyde dehydrogenase
PTAC
Signature enzyme
Alcohol dehydrogenase
Regulator

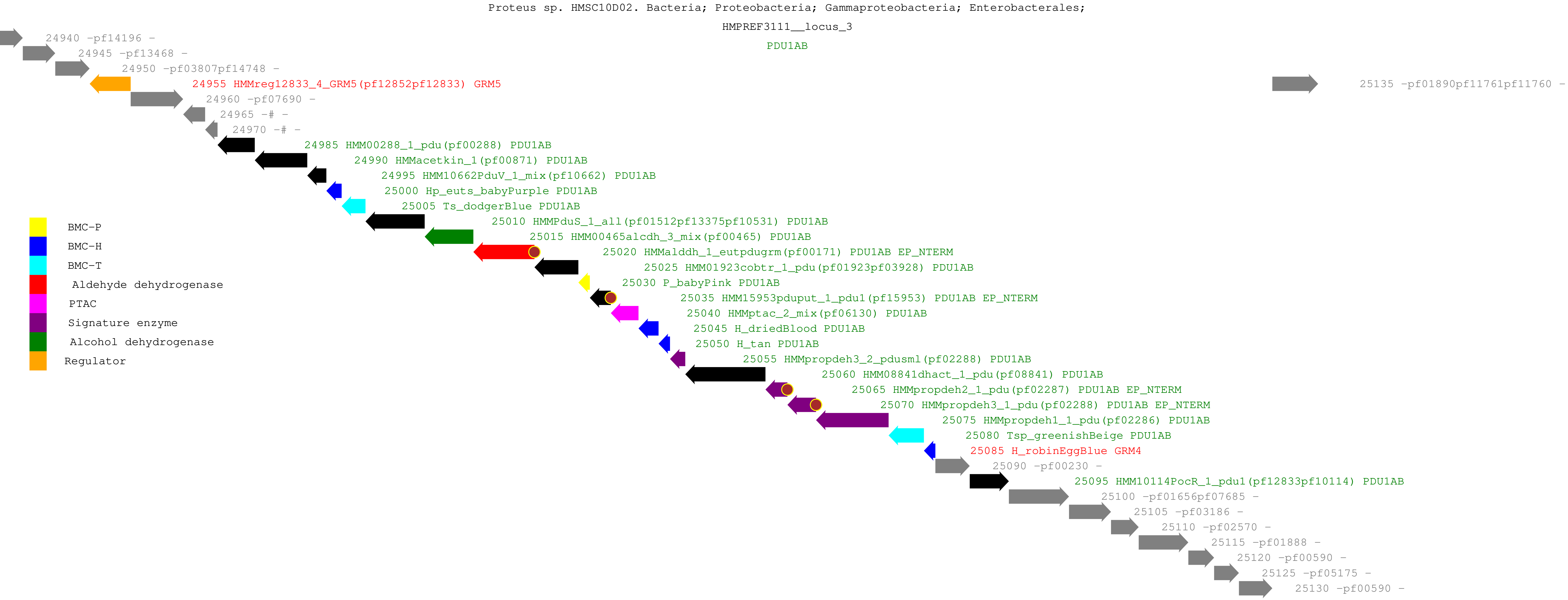

Proteus sp. HMSC10D02. Bacteria; Proteobacteria; Gammaproteobacteria; Enterobacterales;
HMPREF3111__locus_3
PDU1AB
24940 -pf14196 -
24945 -pf13468 -
24950 -pf03807pf14748 -
24955 HMMreg12833_4_GRM5(pf12852pf12833) GRM5
25135 -pf01890pf11761pf11760 -
24960 -pf07690 -
24965 -# -
24970 -# -
24985 HMM00288_1_pdu(pf00288) PDU1AB
24990 HMMacetkin_1(pf00871) PDU1AB
24995 HMM10662PduV_1_mix(pf10662) PDU1AB
25000 Hp_euts_babyPurple PDU1AB
25005 Ts_dodgerBlue PDU1AB
25010 HMMPduS_1_all(pf01512pf13375pf10531) PDU1AB
25015 HMM00465alcdh_3_mix(pf00465) PDU1AB
25020 HMMalddh_1_eutpdugrm(pf00171) PDU1AB EP_NTERM
25025 HMM01923cobtr_1_pdu(pf01923pf03928) PDU1AB
25030 P_babyPink PDU1AB
25035 HMM15953pduput_1_pdu1(pf15953) PDU1AB EP_NTERM
25040 HMMptac_2_mix(pf06130) PDU1AB
25045 H_driedBlood PDU1AB
25050 H_tan PDU1AB
25055 HMMpropdeh3_2_pdusml(pf02288) PDU1AB
25060 HMM08841dhact_1_pdu(pf08841) PDU1AB
25065 HMMpropdeh2_1_pdu(pf02287) PDU1AB EP_NTERM
25070 HMMpropdeh3_1_pdu(pf02288) PDU1AB EP_NTERM
25075 HMMpropdeh1_1_pdu(pf02286) PDU1AB
25080 Tsp_greenishBeige PDU1AB
25085 H_robinEggBlue GRM4
25090 -pf00230 -
25095 HMM10114PocR_1_pdu1(pf12833pf10114) PDU1AB
25100 -pf01656pf07685 -
25105 -pf03186 -
25110 -pf02570 -
25115 -pf01888 -
25120 -pf00590 -
25125 -pf05175 -
25130 -pf00590 -
BMC-P
BMC-H
BMC-T
Aldehyde dehydrogenase
PTAC
Signature enzyme
Alcohol dehydrogenase
Regulator

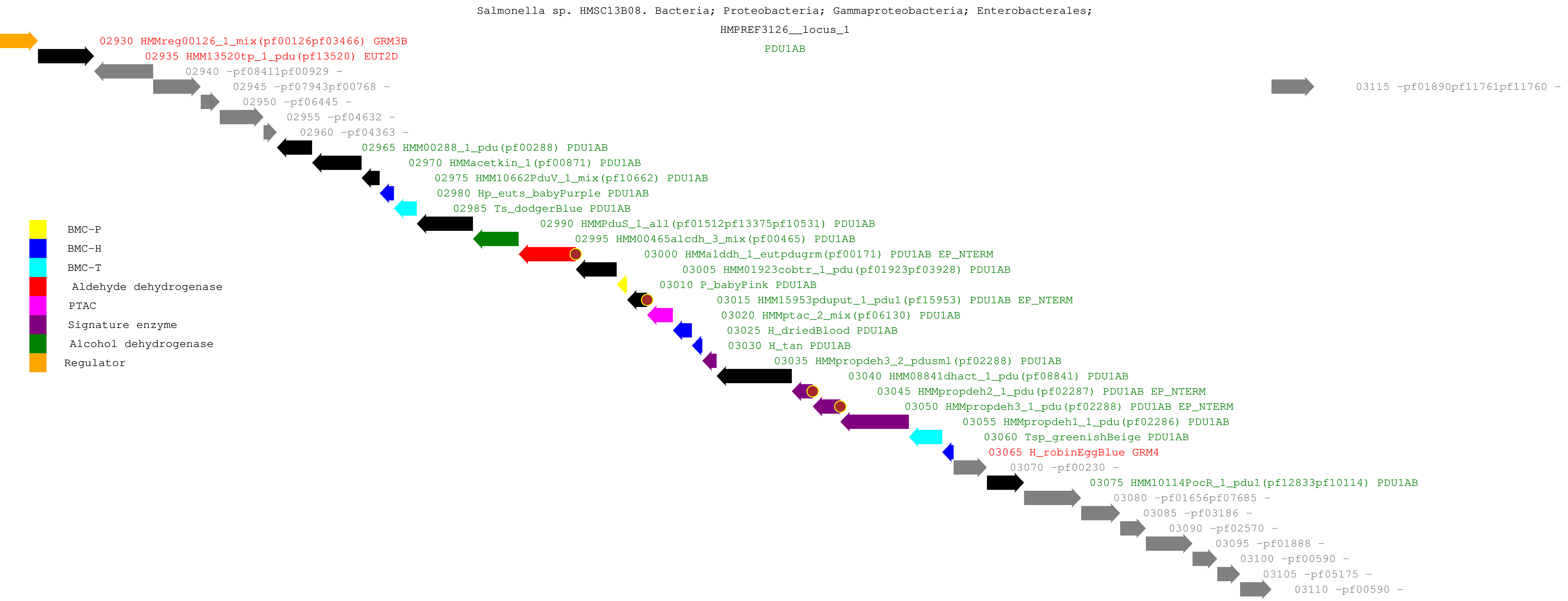

Salmonella sp. HMSC13B08. Bacteria; Proteobacteria; Gammaproteobacteria; Enterobacterales;
HMPREF3126__locus_1
PDU1AB
02930 HMMreg00126_1_mix(pf00126pf03466) GRM3B
02935 HMM13520tp_1_pdu(pf13520) EUT2D
02940 -pf08411pf00929 -
02945 -pf07943pf00768 -
02950 -pf06445 -
02955 -pf04632 -
02960 -pf04363 -
02965 HMM00288_1_pdu(pf00288) PDU1AB
02970 HMMacetkin_1(pf00871) PDU1AB
02975 HMM10662PduV_1_mix(pf10662) PDU1AB
02980 Hp_euts_babyPurple PDU1AB
02985 Ts_dodgerBlue PDU1AB
02990 HMMPduS_1_all(pf01512pf13375pf10531) PDU1AB
02995 HMM00465alcdh_3_mix(pf00465) PDU1AB
03000 HMMalddh_1_eutpdugrm(pf00171) PDU1AB EP_NTERM
03005 HMM01923cobtr_1_pdu(pf01923pf03928) PDU1AB
03010 P_babyPink PDU1AB
03015 HMM15953pduput_1_pdu1(pf15953) PDU1AB EP_NTERM
03020 HMMptac_2_mix(pf06130) PDU1AB
03025 H_driedBlood PDU1AB
03030 H_tan PDU1AB
03035 HMMpropdeh3_2_pdusml(pf02288) PDU1AB
03040 HMM08841dhact_1_pdu(pf08841) PDU1AB
03045 HMMpropdeh2_1_pdu(pf02287) PDU1AB EP_NTERM
03050 HMMpropdeh3_1_pdu(pf02288) PDU1AB EP_NTERM
03055 HMMpropdeh1_1_pdu(pf02286) PDU1AB
03060 Tsp_greenishBeige PDU1AB
03065 H_robinEggBlue GRM4
03070 -pf00230 -
03075 HMM10114PocR_1_pdu1(pf12833pf10114) PDU1AB
03080 -pf01656pf07685 -
03085 -pf03186 -
03090 -pf02570 -
03095 -pf01888 -
03100 -pf00590 -
03105 -pf05175 -
03110 -pf00590 -
03115 -pf01890pf11761pf11760 -
BMC-P
BMC-H
BMC-T
Aldehyde dehydrogenase
PTAC
Signature enzyme
Alcohol dehydrogenase
Regulator

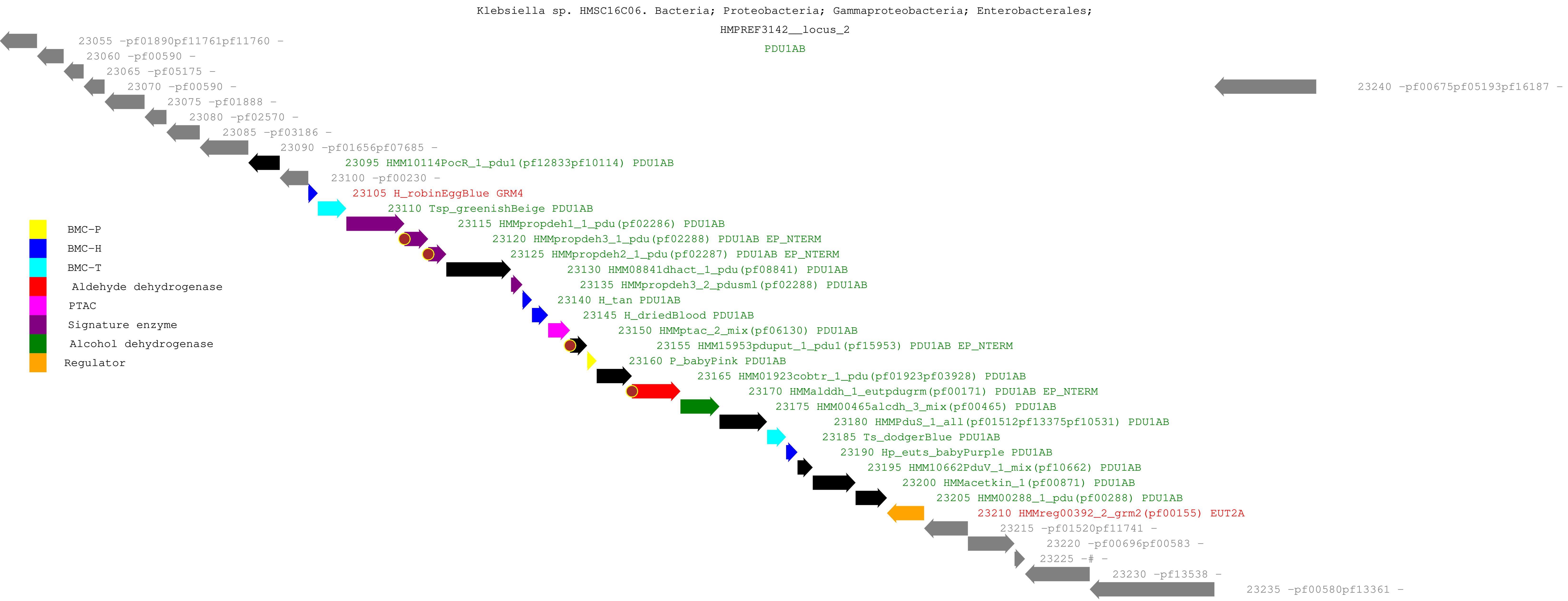

Klebsiella sp. HMSC16C06. Bacteria; Proteobacteria; Gammaproteobacteria; Enterobacterales;
HMPREF3142__locus_2
PDU1AB
23055 -pf01890pf11761pf11760 -
23060 -pf00590 -
23065 -pf05175 -
23070 -pf00590 -
23075 -pf01888 -
23080 -pf02570 -
23085 -pf03186 -
23090 -pf01656pf07685 -
23095 HMM10114PocR_1_pdu1(pf12833pf10114) PDU1AB
23100 -pf00230 -
23105 H_robinEggBlue GRM4
23110 Tsp_greenishBeige PDU1AB
23115 HMMpropdeh1_1_pdu(pf02286) PDU1AB
23120 HMMpropdeh3_1_pdu(pf02288) PDU1AB EP_NTERM
23125 HMMpropdeh2_1_pdu(pf02287) PDU1AB EP_NTERM
23130 HMM08841dhact_1_pdu(pf08841) PDU1AB
23135 HMMpropdeh3_2_pdusml(pf02288) PDU1AB
23140 H_tan PDU1AB
23145 H_driedBlood PDU1AB
23150 HMMptac_2_mix(pf06130) PDU1AB
23155 HMM15953pduput_1_pdu1(pf15953) PDU1AB EP_NTERM
23160 P_babyPink PDU1AB
23165 HMM01923cobtr_1_pdu(pf01923pf03928) PDU1AB
23170 HMMalddh_1_eutpdugrm(pf00171) PDU1AB EP_NTERM
23175 HMM00465alcdh_3_mix(pf00465) PDU1AB
23180 HMMPduS_1_all(pf01512pf13375pf10531) PDU1AB
23185 Ts_dodgerBlue PDU1AB
23190 Hp_euts_babyPurple PDU1AB
23195 HMM10662PduV_1_mix(pf10662) PDU1AB
23200 HMMacetkin_1(pf00871) PDU1AB
23205 HMM00288_1_pdu(pf00288) PDU1AB
23210 HMMreg00392_2_grm2(pf00155) EUT2A
23215 -pf01520pf11741 -
23220 -pf00696pf00583 -
23225 -# -
23230 -pf13538 -
23235 -pf00580pf13361 -
23240 -pf00675pf05193pf16187 -
BMC-P
BMC-H
BMC-T
Aldehyde dehydrogenase
PTAC
Signature enzyme
Alcohol dehydrogenase
Regulator

Citrobacter freundii. Bacteria; Proteobacteria; Gammaproteobacteria; Enterobacterales;

HMPREF3212__locus_1

PDU1AB

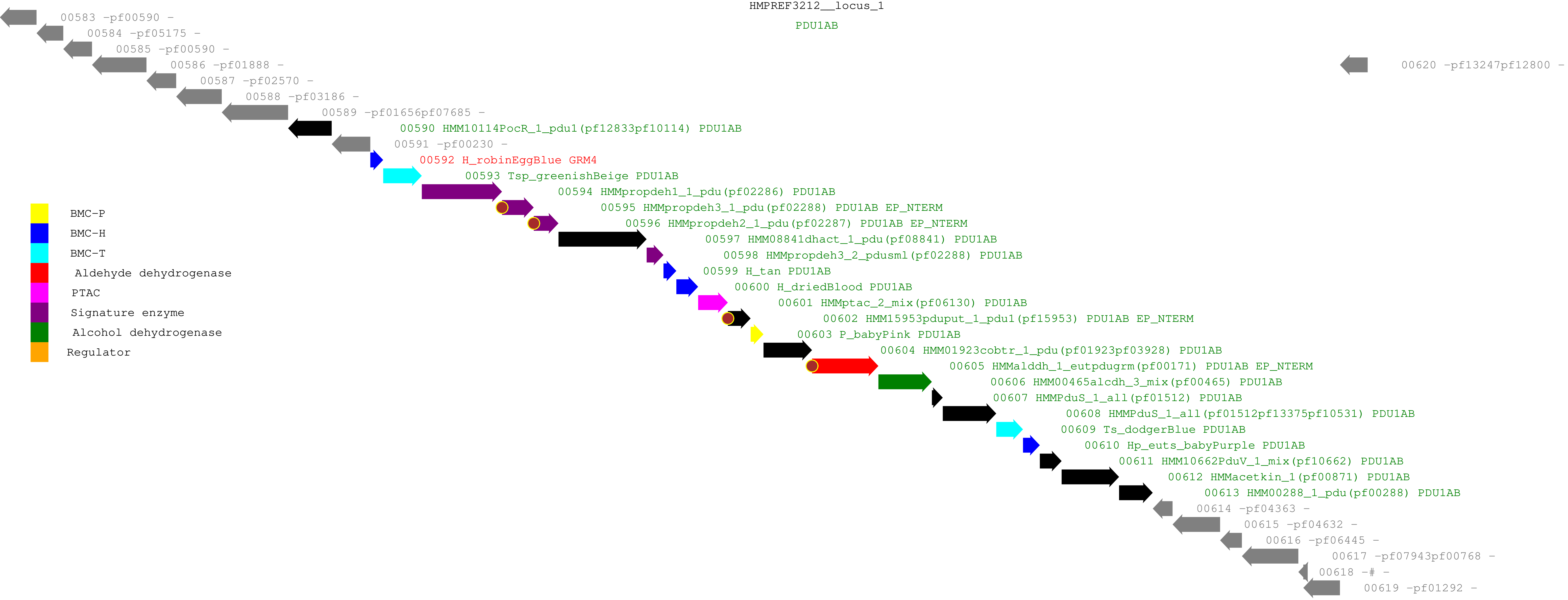

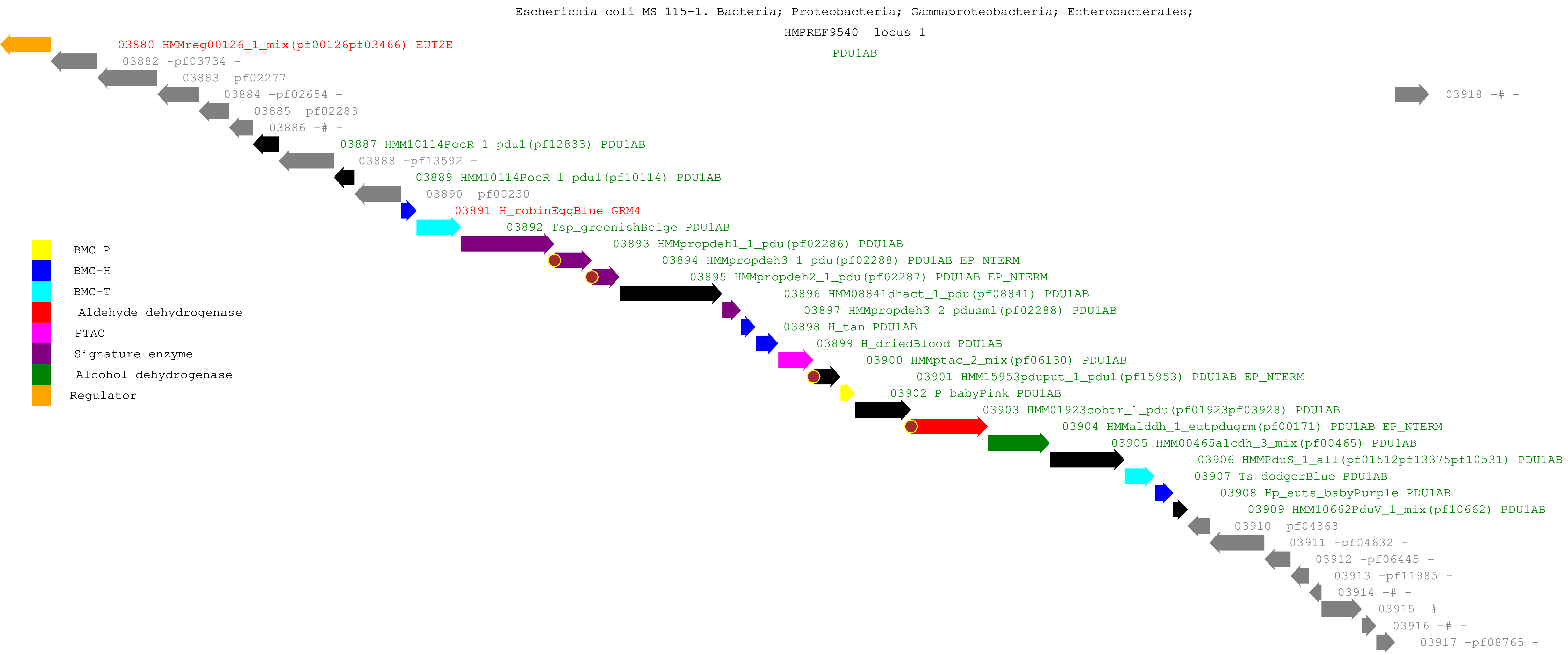

Escherichia coli MS 115-1. Bacteria; Proteobacteria; Gammaproteobacteria; Enterobacterales;
HMPREF9540__locus_1
PDU1AB
03880 HMMreg00126_1_mix(pf00126pf03466) EUT2E
03882 -pf03734 -
03883 -pf02277 -
03884 -pf02654 -
03885 -pf02283 -
03886 -# -
03887 HMM10114PocR_1_pdu1(pf12833) PDU1AB
03888 -pf13592 -
03889 HMM10114PocR_1_pdu1(pf10114) PDU1AB
03890 -pf00230 -
03891 H_robinEggBlue GRM4
03892 Tsp_greenishBeige PDU1AB
03893 HMMpropdeh1_1_pdu(pf02286) PDU1AB
03894 HMMpropdeh3_1_pdu(pf02288) PDU1AB EP_NTERM
03895 HMMpropdeh2_1_pdu(pf02287) PDU1AB EP_NTERM
03896 HMM08841dhact_1_pdu(pf08841) PDU1AB
03897 HMMpropdeh3_2_pdusml(pf02288) PDU1AB
03898 H_tan PDU1AB
03899 H_driedBlood PDU1AB
03900 HMMptac_2_mix(pf06130) PDU1AB
03901 HMM15953pduput_1_pdu1(pf15953) PDU1AB EP_NTERM
03902 P_babyPink PDU1AB
03903 HMM01923cobtr_1_pdu(pf01923pf03928) PDU1AB
03904 HMMalddh_1_eutpdugrm(pf00171) PDU1AB EP_NTERM
03905 HMM00465alcdh_3_mix(pf00465) PDU1AB
03906 HMMPduS_1_all(pf01512pf13375pf10531) PDU1AB
03907 Ts_dodgerBlue PDU1AB
03908 Hp_euts_babyPurple PDU1AB
03909 HMM10662PduV_1_mix(pf10662) PDU1AB
03910 -pf04363 -
03911 -pf04632 -
03912 -pf06445 -
03913 -pf11985 -
03914 -# -
03915 -# -
03916 -# -
03917 -pf08765 -
03918 -# -
BMC-P
BMC-H
BMC-T
Aldehyde dehydrogenase
PTAC
Signature enzyme
Alcohol dehydrogenase
Regulator

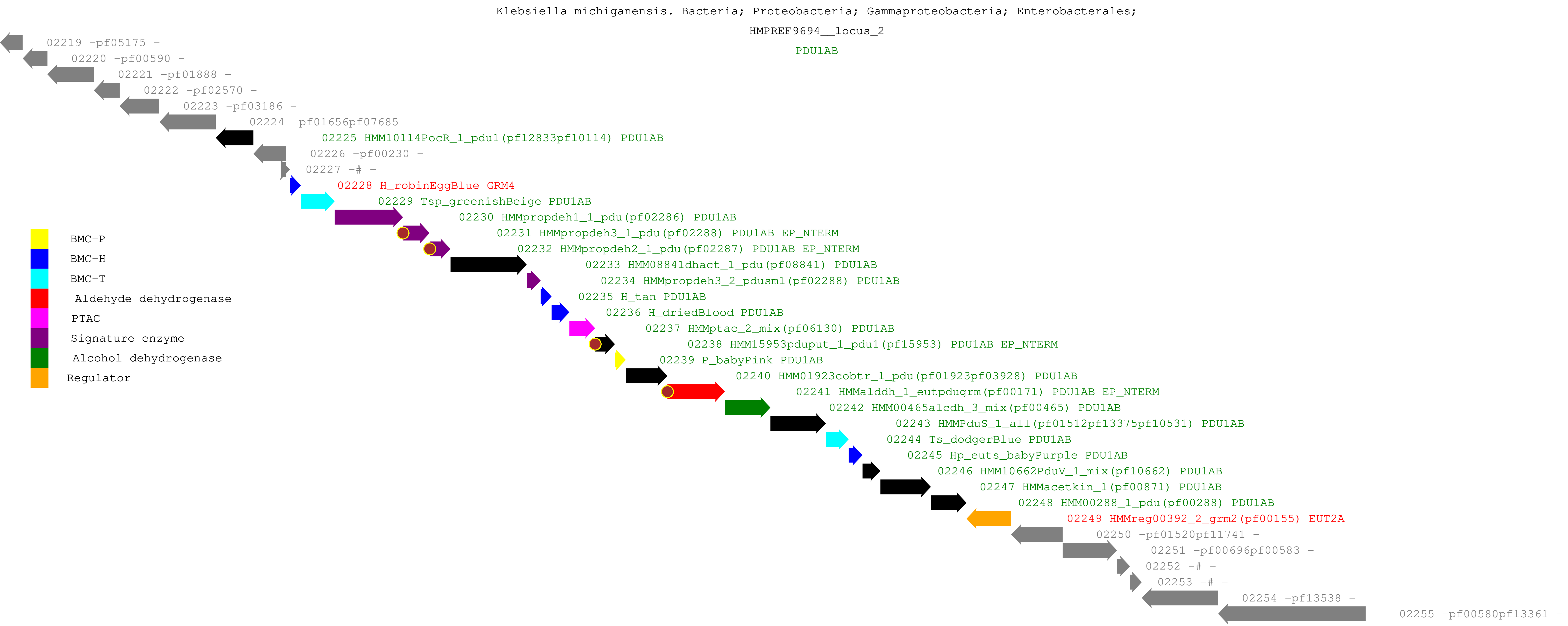
Klebsiella michiganensis. Bacteria; Proteobacteria; Gammaproteobacteria; Enterobacterales;
HMPREF9694__locus_2
PDU1AB
02219 -pf05175 -
02220 -pf00590 -
02221 -pf01888 -
02222 -pf02570 -
02223 -pf03186 -
02224 -pf01656pf07685 -
02225 HMM10114PocR_1_pdu1(pf12833pf10114) PDU1AB
02226 -pf00230 -
02227 -# -
02228 H_robinEggBlue GRM4
02229 Tsp_greenishBeige PDU1AB
02230 HMMpropdeh1_1_pdu(pf02286) PDU1AB
02231 HMMpropdeh3_1_pdu(pf02288) PDU1AB EP_NTERM
02232 HMMpropdeh2_1_pdu(pf02287) PDU1AB EP_NTERM
02233 HMM08841dhact_1_pdu(pf08841) PDU1AB
02234 HMMpropdeh3_2_pdusml(pf02288) PDU1AB
02235 H_tan PDU1AB
02236 H_driedBlood PDU1AB
02237 HMMptac_2_mix(pf06130) PDU1AB
02238 HMM15953pduput_1_pdu1(pf15953) PDU1AB EP_NTERM
02239 P_babyPink PDU1AB
02240 HMM01923cobtr_1_pdu(pf01923pf03928) PDU1AB
02241 HMMalddh_1_eutpdugrm(pf00171) PDU1AB EP_NTERM
02242 HMM00465alcdh_3_mix(pf00465) PDU1AB
02243 HMMPduS_1_all(pf01512pf13375pf10531) PDU1AB
02244 Ts_dodgerBlue PDU1AB
02245 Hp_euts_babyPurple PDU1AB
02246 HMM10662PduV_1_mix(pf10662) PDU1AB
02247 HMMacetkin_1(pf00871) PDU1AB
02248 HMM00288_1_pdu(pf00288) PDU1AB
02249 HMMreg00392_2_grm2(pf00155) EUT2A
02250 -pf01520pf11741 -
02251 -pf00696pf00583 -
02252 -# -
02253 -# -
02254 -pf13538 -
02255 -pf00580pf13361 -
BMC-P
BMC-H
BMC-T
Aldehyde dehydrogenase
PTAC
Signature enzyme
Alcohol dehydrogenase
Regulator

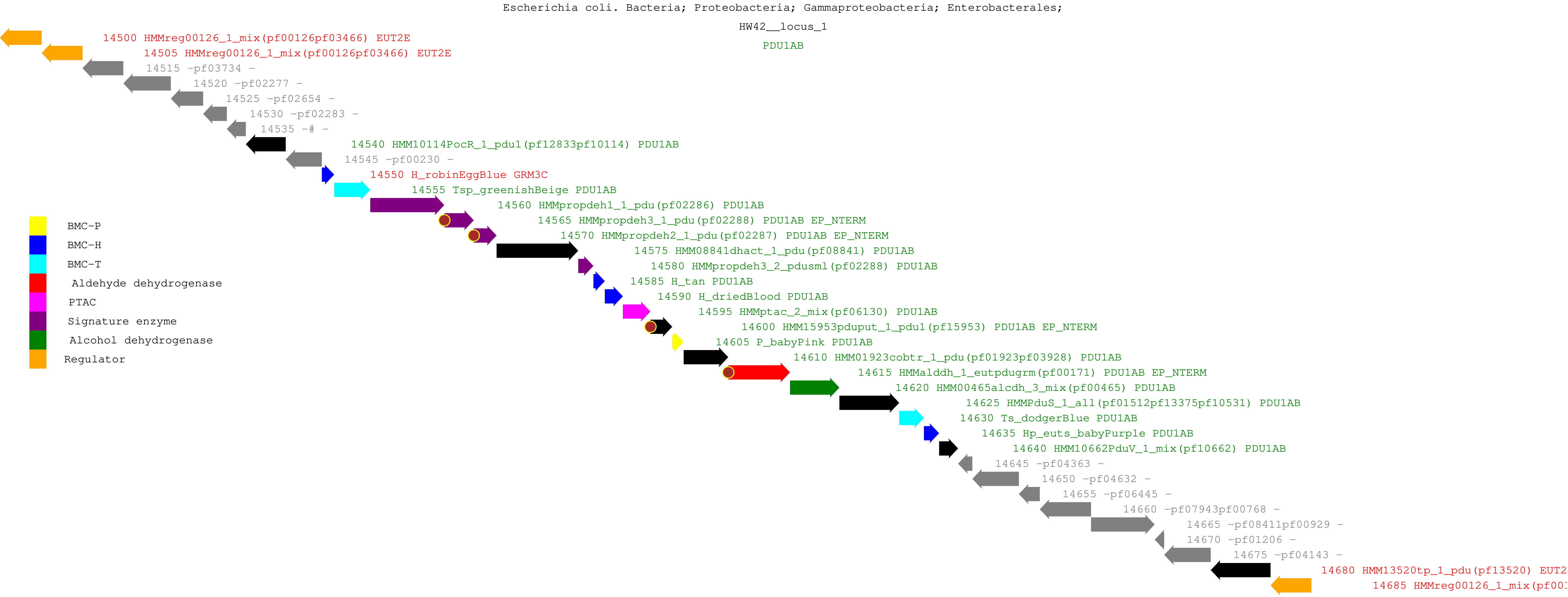

Escherichia coli. Bacteria; Proteobacteria; Gammaproteobacteria; Enterobacterales;
HW42__locus_1
PDU1AB
14500 HMMreg00126_1_mix(pf00126pf03466) EUT2E
14505 HMMreg00126_1_mix(pf00126pf03466) EUT2E
14515 -pf03734 -
14520 -pf02277 -
14525 -pf02654 -
14530 -pf02283 -
14535 -# -
14540 HMM10114PocR_1_pdu1(pf12833pf10114) PDU1AB
14545 -pf00230 -
14550 H_robinEggBlue GRM3C
14555 Tsp_greenishBeige PDU1AB
14560 HMMpropdeh1_1_pdu(pf02286) PDU1AB
14565 HMMpropdeh3_1_pdu(pf02288) PDU1AB EP_NTERM
14570 HMMpropdeh2_1_pdu(pf02287) PDU1AB EP_NTERM
14575 HMM08841dhact_1_pdu(pf08841) PDU1AB
14580 HMMpropdeh3_2_pdusml(pf02288) PDU1AB
14585 H_tan PDU1AB
14590 H_driedBlood PDU1AB
14595 HMMptac_2_mix(pf06130) PDU1AB
14600 HMM15953pduput_1_pdu1(pf15953) PDU1AB EP_NTERM
14605 P_babyPink PDU1AB
14610 HMM01923cobtr_1_pdu(pf01923pf03928) PDU1AB
14615 HMMalddh_1_eutpdugrm(pf00171) PDU1AB EP_NTERM
14620 HMM00465alcdh_3_mix(pf00465) PDU1AB
14625 HMMPduS_1_all(pf01512pf13375pf10531) PDU1AB
14630 Ts_dodgerBlue PDU1AB
14635 Hp_euts_babyPurple PDU1AB
14640 HMM10662PduV_1_mix(pf10662) PDU1AB
14645 -pf04363 -
14650 -pf04632 -
14655 -pf06445 -
14660 -pf07943pf00768 -
14665 -pf08411pf00929 -
14670 -pf01206 -
14675 -pf04143 -
14680 HMM13520tp_1_pdu(pf13520) EUT2
14685 HMMreg00126_1_mix(pf001
BMC-P
BMC-H
BMC-T
Aldehyde dehydrogenase
PTAC
Signature enzyme
Alcohol dehydrogenase
Regulator

Escherichia coli. Bacteria; Proteobacteria; Gammaproteobacteria; Enterobacterales;

HW43__locus_1

PDU1AB

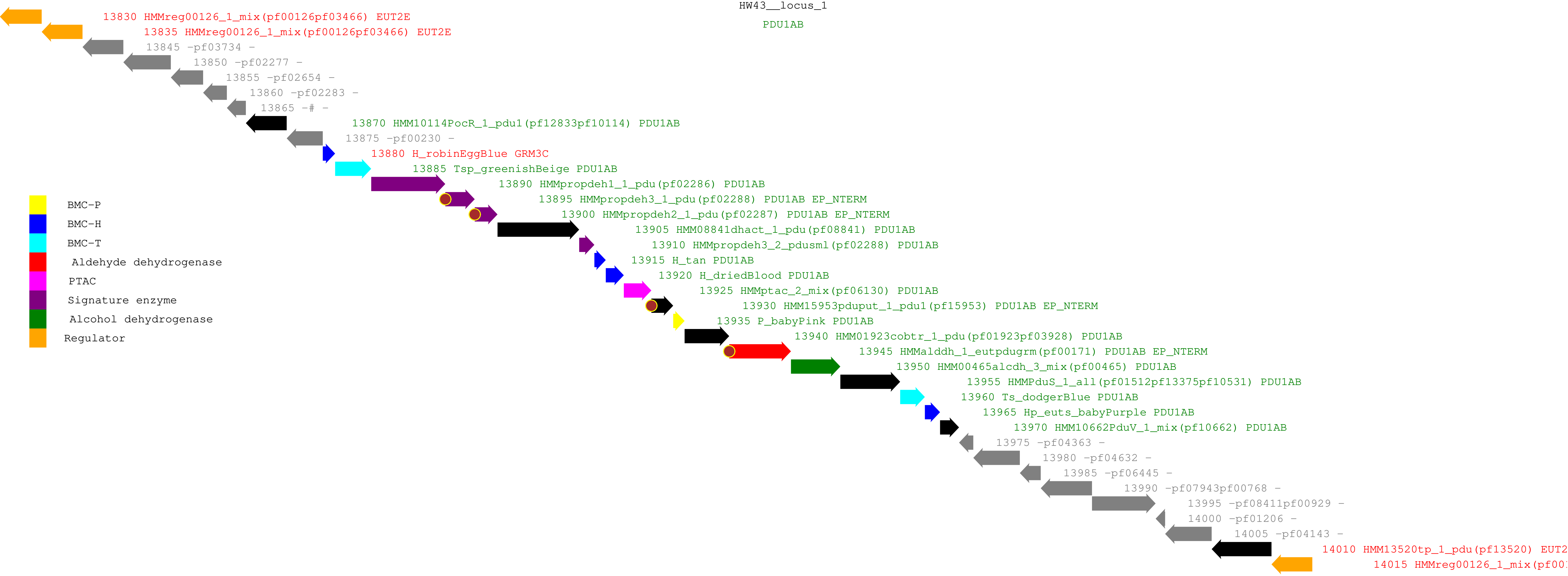

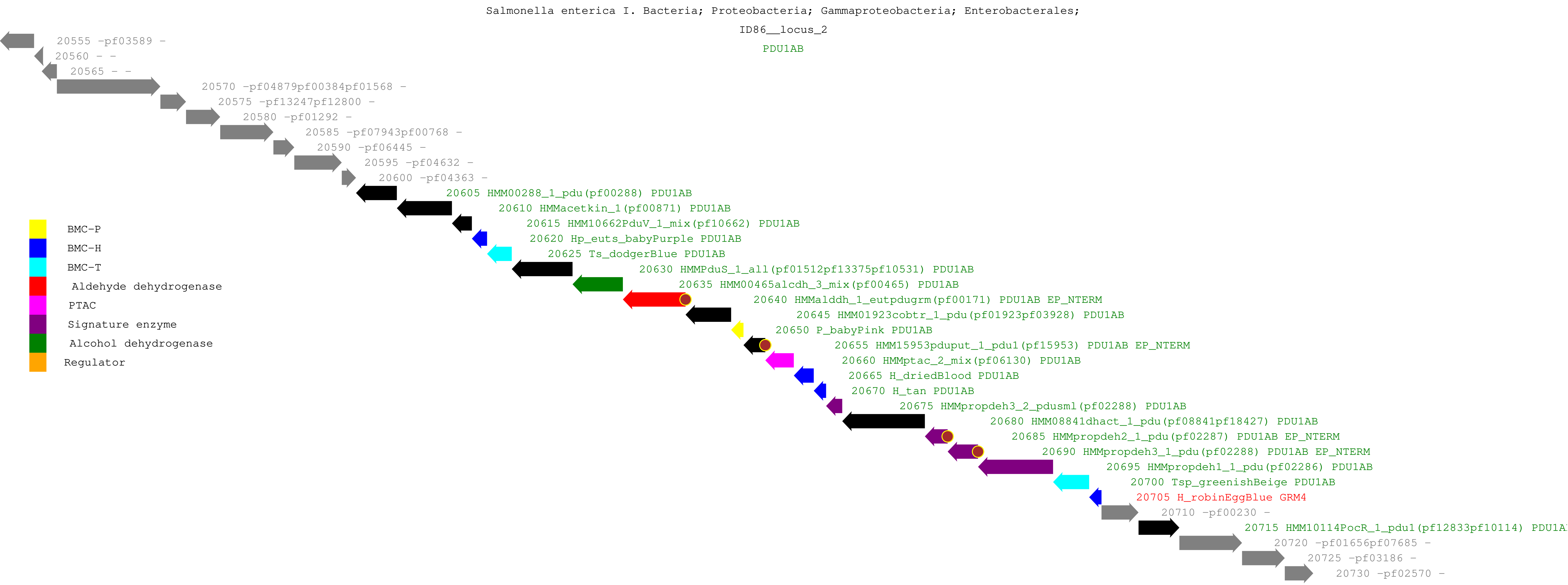
Salmonella enterica I. Bacteria; Proteobacteria; Gammaproteobacteria; Enterobacterales;
ID86__locus_2
PDU1AB
20555 -pf03589 -
20560 - -
20565 - -
20570 -pf04879pf00384pf01568 -
20575 -pf13247pf12800 -
20580 -pf01292 -
20585 -pf07943pf00768 -
20590 -pf06445 -
20595 -pf04632 -
20600 -pf04363 -
20605 HMM00288_1_pdu(pf00288) PDU1AB
20610 HMMacetkin_1(pf00871) PDU1AB
20615 HMM10662PduV_1_mix(pf10662) PDU1AB
20620 Hp_euts_babyPurple PDU1AB
20625 Ts_dodgerBlue PDU1AB
20630 HMMPduS_1_all(pf01512pf13375pf10531) PDU1AB
20635 HMM00465alcdh_3_mix(pf00465) PDU1AB
20640 HMMalddh_1_eutpdugrm(pf00171) PDU1AB EP_NTERM
20645 HMM01923cobtr_1_pdu(pf01923pf03928) PDU1AB
20650 P_babyPink PDU1AB
20655 HMM15953pduput_1_pdu1(pf15953) PDU1AB EP_NTERM
20660 HMMptac_2_mix(pf06130) PDU1AB
20665 H_driedBlood PDU1AB
20670 H_tan PDU1AB
20675 HMMpropdeh3_2_pdusml(pf02288) PDU1AB
20680 HMM08841dhact_1_pdu(pf08841pf18427) PDU1AB
20685 HMMpropdeh2_1_pdu(pf02287) PDU1AB EP_NTERM
20690 HMMpropdeh3_1_pdu(pf02288) PDU1AB EP_NTERM
20695 HMMpropdeh1_1_pdu(pf02286) PDU1AB
20700 Tsp_greenishBeige PDU1AB
20705 H_robinEggBlue GRM4
20710 -pf00230 -
20715 HMM10114PocR_1_pdu1(pf12833pf10114) PDU1A
20720 -pf01656pf07685 -
20725 -pf03186 -
20730 -pf02570 -
BMC-P
BMC-H
BMC-T
Aldehyde dehydrogenase
PTAC
Signature enzyme
Alcohol dehydrogenase
Regulator

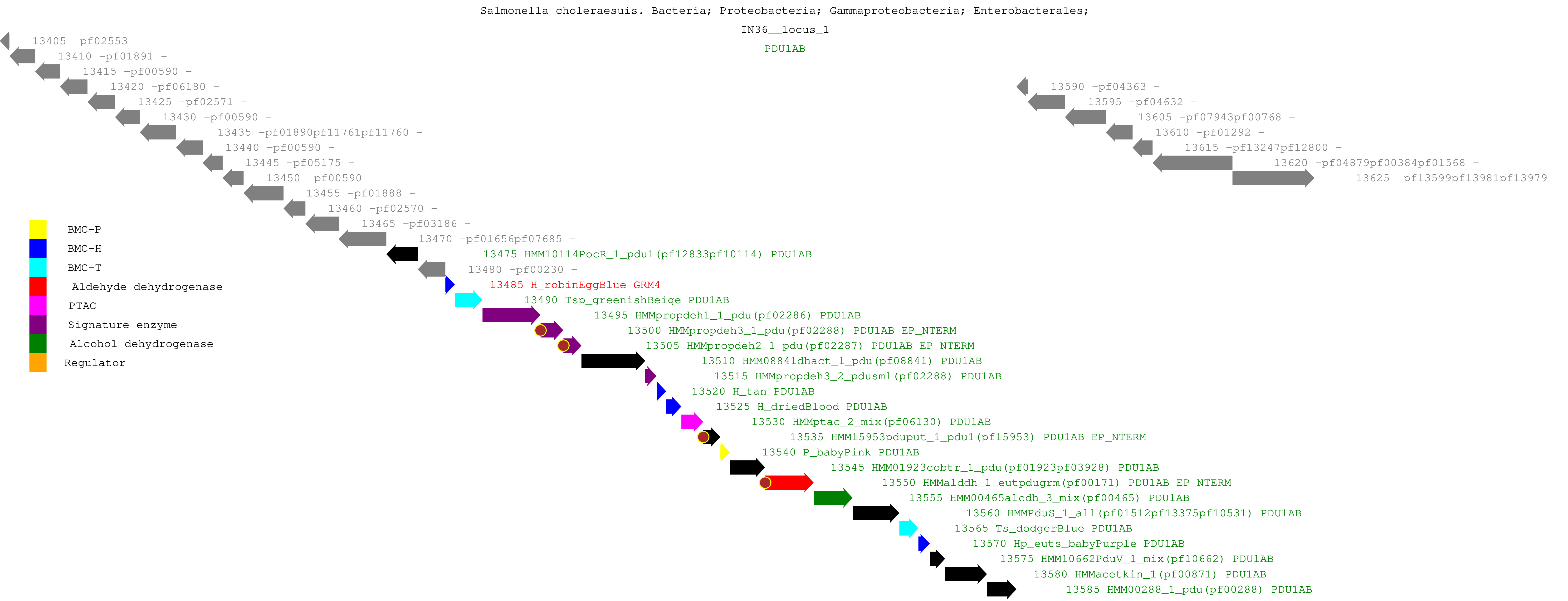
Salmonella choleraesuis. Bacteria; Proteobacteria; Gammaproteobacteria; Enterobacterales;
IN36__locus_1
PDU1AB
13405 -pf02553 -
13410 -pf01891 -
13415 -pf00590 -
13420 -pf06180 -
13425 -pf02571 -
13430 -pf00590 -
13435 -pf01890pf11761pf11760 -
13440 -pf00590 -
13445 -pf05175 -
13450 -pf00590 -
13455 -pf01888 -
13460 -pf02570 -
13465 -pf03186 -
13470 -pf01656pf07685 -
13475 HMM10114PocR_1_pdu1(pf12833pf10114) PDU1AB
13480 -pf00230 -
13485 H_robinEggBlue GRM4
13490 Tsp_greenishBeige PDU1AB
13495 HMMpropdeh1_1_pdu(pf02286) PDU1AB
13500 HMMpropdeh3_1_pdu(pf02288) PDU1AB EP_NTERM
13505 HMMpropdeh2_1_pdu(pf02287) PDU1AB EP_NTERM
13510 HMM08841dhact_1_pdu(pf08841) PDU1AB
13515 HMMpropdeh3_2_pdusml(pf02288) PDU1AB
13520 H_tan PDU1AB
13525 H_driedBlood PDU1AB
13530 HMMptac_2_mix(pf06130) PDU1AB
13535 HMM15953pduput_1_pdu1(pf15953) PDU1AB EP_NTERM
13540 P_babyPink PDU1AB
13545 HMM01923cobtr_1_pdu(pf01923pf03928) PDU1AB
13550 HMMalddh_1_eutpdugrm(pf00171) PDU1AB EP_NTERM
13555 HMM00465alcdh_3_mix(pf00465) PDU1AB
13560 HMMPduS_1_all(pf01512pf13375pf10531) PDU1AB
13565 Ts_dodgerBlue PDU1AB
13570 Hp_euts_babyPurple PDU1AB
13575 HMM10662PduV_1_mix(pf10662) PDU1AB
13580 HMMacetkin_1(pf00871) PDU1AB
13585 HMM00288_1_pdu(pf00288) PDU1AB
13590 -pf04363 -
13595 -pf04632 -
13605 -pf07943pf00768 -
13610 -pf01292 -
13615 -pf13247pf12800 -
13620 -pf04879pf00384pf01568 -
13625 -pf13599pf13981pf13979 -
BMC-P
BMC-H
BMC-T
Aldehyde dehydrogenase
PTAC
Signature enzyme
Alcohol dehydrogenase
Regulator

Salmonella choleraesuis. Bacteria; Proteobacteria; Gammaproteobacteria; Enterobacterales;

IN69__locus_1

PDU1AB

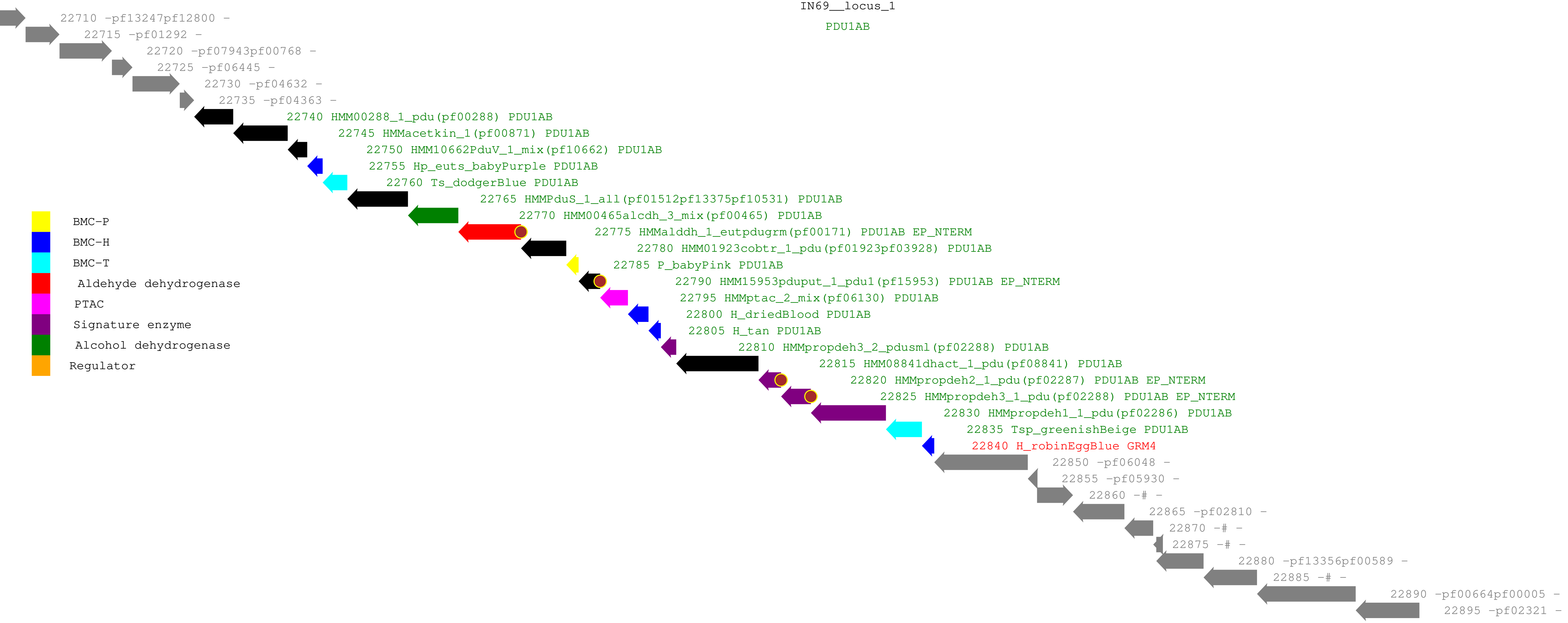

Escherichia coli. Bacteria; Proteobacteria; Gammaproteobacteria; Enterobacterales;

JD73__locus_1

PDU1AB

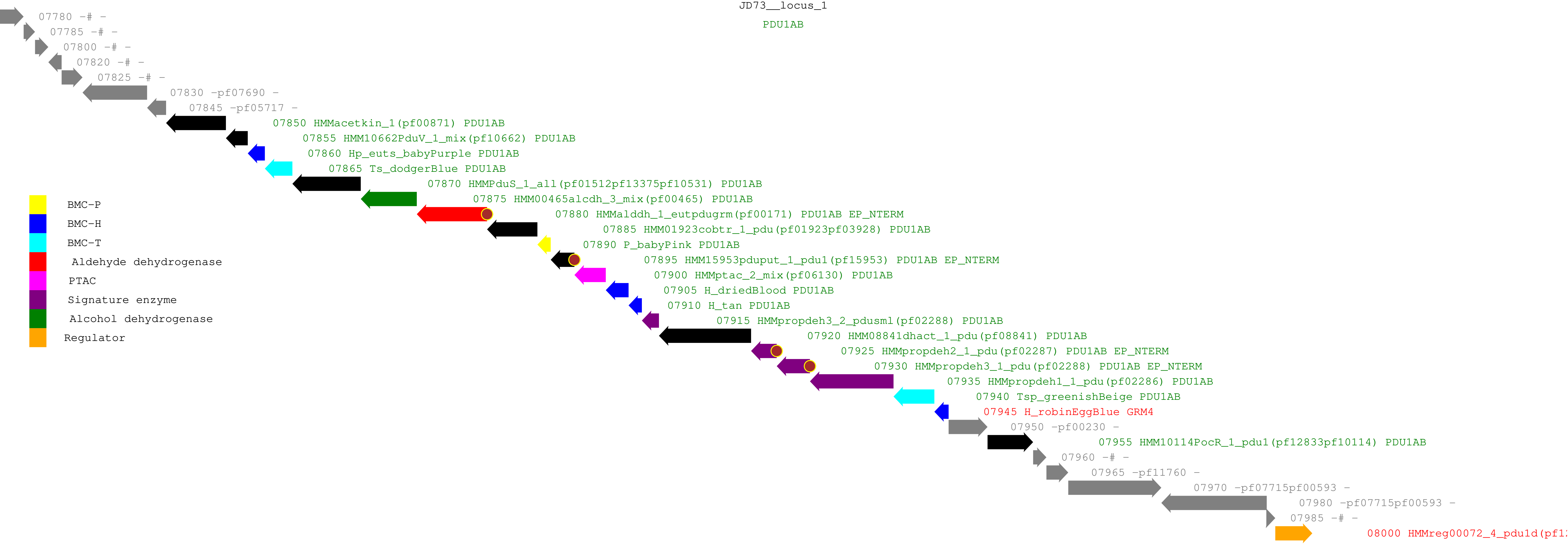

uncultured Citrobacter sp. Bacteria; Proteobacteria; Gammaproteobacteria; Enterobacterales;

KL86CIT2__locus_1

PDU1AB

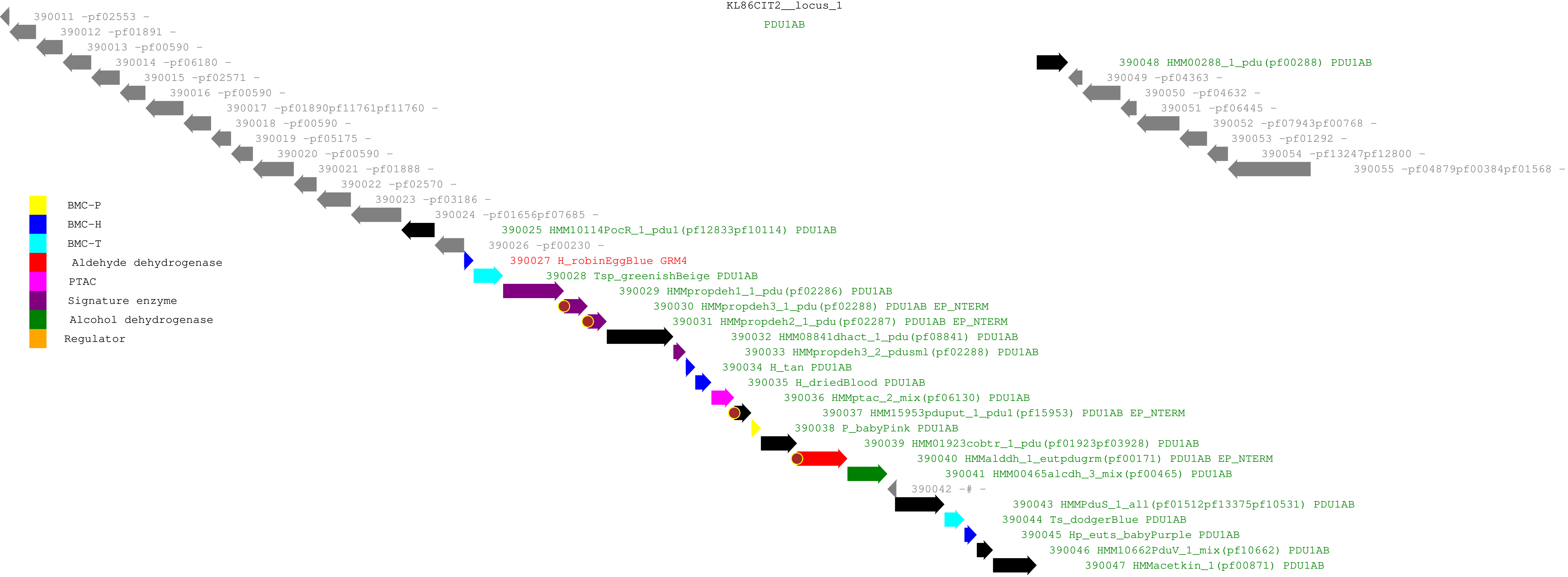

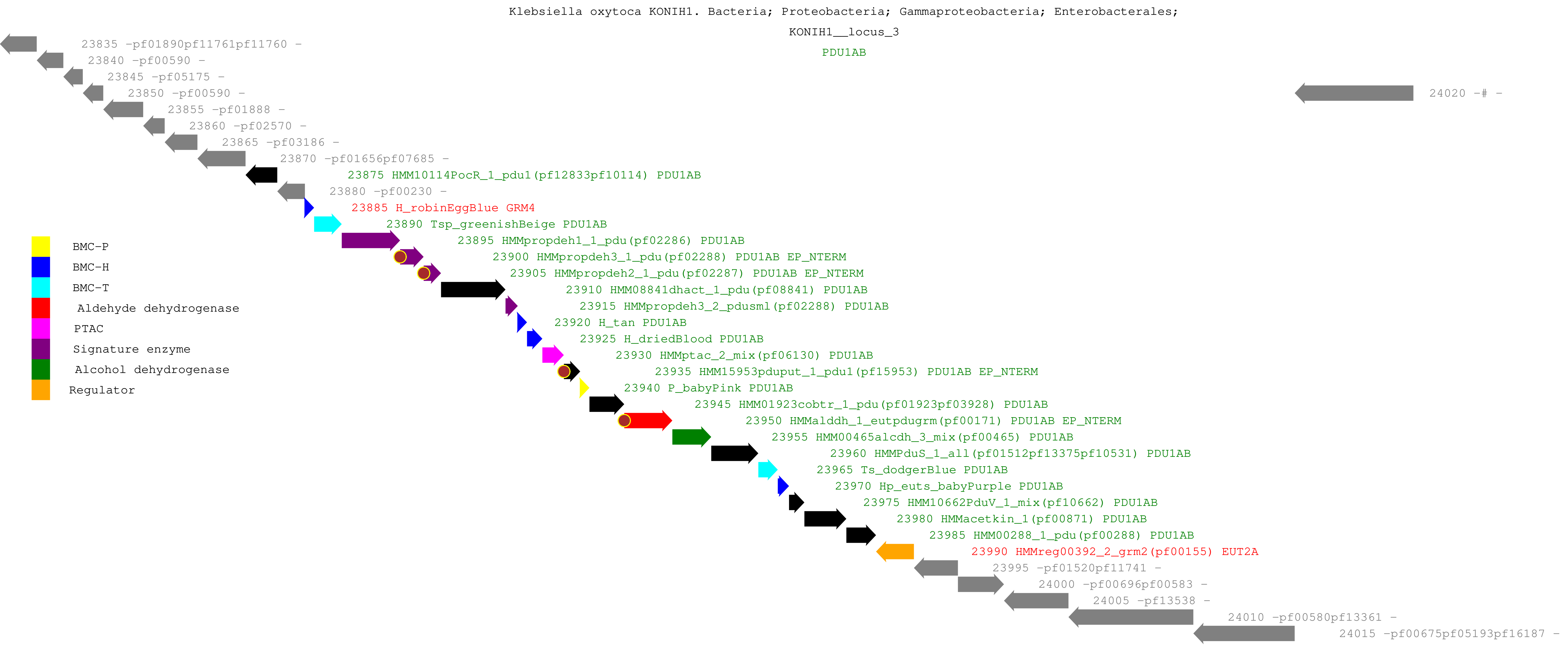

Klebsiella oxytoca KONIH1. Bacteria; Proteobacteria; Gammaproteobacteria; Enterobacterales;
KONIH1__locus_3
PDU1AB
23835 -pf01890pf11761pf11760 -
23840 -pf00590 -
23845 -pf05175 -
23850 -pf00590 -
23855 -pf01888 -
23860 -pf02570 -
23865 -pf03186 -
23870 -pf01656pf07685 -
23875 HMM10114PocR_1_pdu1(pf12833pf10114) PDU1AB
23880 -pf00230 -
23885 H_robinEggBlue GRM4
23890 Tsp_greenishBeige PDU1AB
23895 HMMpropdeh1_1_pdu(pf02286) PDU1AB
23900 HMMpropdeh3_1_pdu(pf02288) PDU1AB EP_NTERM
23905 HMMpropdeh2_1_pdu(pf02287) PDU1AB EP_NTERM
23910 HMM08841dhact_1_pdu(pf08841) PDU1AB
23915 HMMpropdeh3_2_pdusml(pf02288) PDU1AB
23920 H_tan PDU1AB
23925 H_driedBlood PDU1AB
23930 HMMptac_2_mix(pf06130) PDU1AB
23935 HMM15953pduput_1_pdu1(pf15953) PDU1AB EP_NTERM
23940 P_babyPink PDU1AB
23945 HMM01923cobtr_1_pdu(pf01923pf03928) PDU1AB
23950 HMMalddh_1_eutpdugrm(pf00171) PDU1AB EP_NTERM
23955 HMM00465alcdh_3_mix(pf00465) PDU1AB
23960 HMMPduS_1_all(pf01512pf13375pf10531) PDU1AB
23965 Ts_dodgerBlue PDU1AB
23970 Hp_euts_babyPurple PDU1AB
23975 HMM10662PduV_1_mix(pf10662) PDU1AB
23980 HMMacetkin_1(pf00871) PDU1AB
23985 HMM00288_1_pdu(pf00288) PDU1AB
23990 HMMreg00392_2_grm2(pf00155) EUT2A
23995 -pf01520pf11741 -
24000 -pf00696pf00583 -
24005 -pf13538 -
24010 -pf00580pf13361 -
24015 -pf00675pf05193pf16187 -
24020 -# -
BMC-P
BMC-H
BMC-T
Aldehyde dehydrogenase
PTAC
Signature enzyme
Alcohol dehydrogenase
Regulator

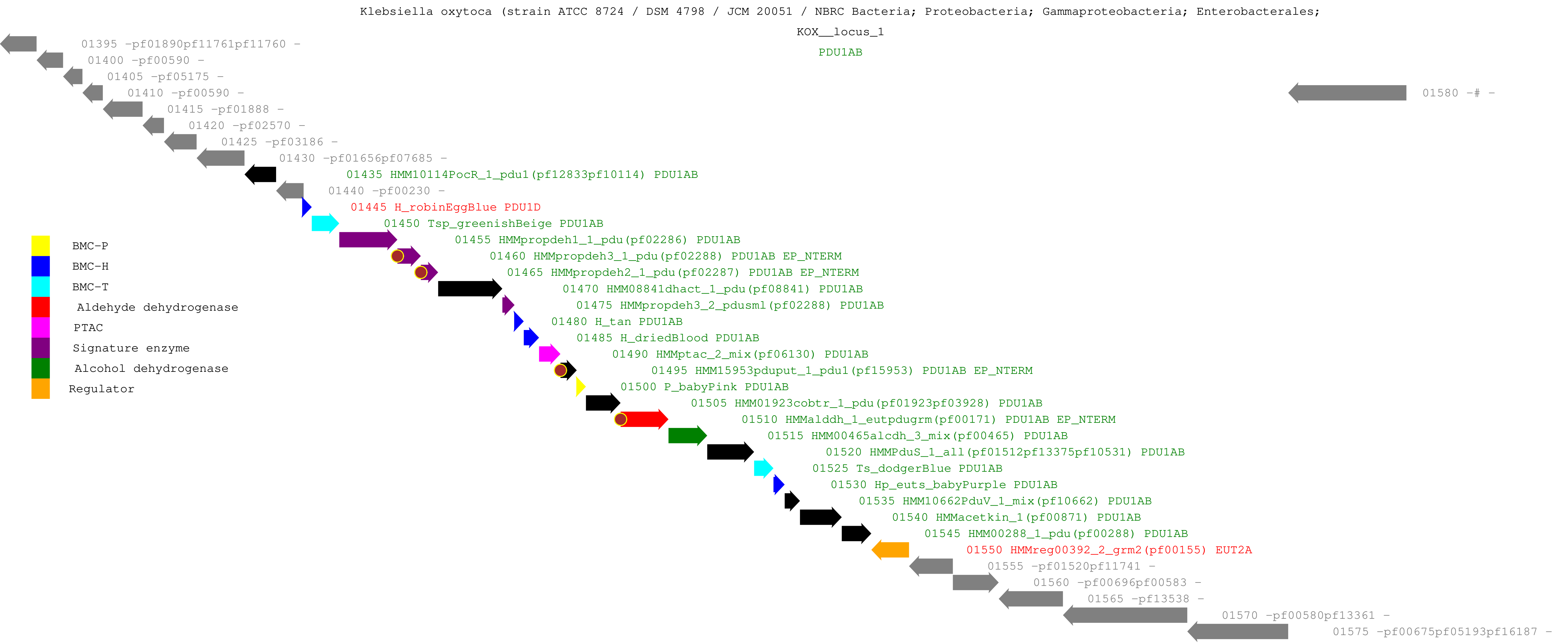

Klebsiella oxytoca (strain ATCC 8724 / DSM 4798 / JCM 20051 / NBRC Bacteria; Proteobacteria; Gammaproteobacteria; Enterobacterales;
KOX__locus_1
PDU1AB
01395 -pf01890pf11761pf11760 -
01400 -pf00590 -
01405 -pf05175 -
01410 -pf00590 -
01415 -pf01888 -
01420 -pf02570 -
01425 -pf03186 -
01430 -pf01656pf07685 -
01435 HMM10114PocR_1_pdu1(pf12833pf10114) PDU1AB
01440 -pf00230 -
01445 H_robinEggBlue PDU1D
01450 Tsp_greenishBeige PDU1AB
01455 HMMpropdeh1_1_pdu(pf02286) PDU1AB
01460 HMMpropdeh3_1_pdu(pf02288) PDU1AB EP_NTERM
01465 HMMpropdeh2_1_pdu(pf02287) PDU1AB EP_NTERM
01470 HMM08841dhact_1_pdu(pf08841) PDU1AB
01475 HMMpropdeh3_2_pdusml(pf02288) PDU1AB
01480 H_tan PDU1AB
01485 H_driedBlood PDU1AB
01490 HMMptac_2_mix(pf06130) PDU1AB
01495 HMM15953pduput_1_pdu1(pf15953) PDU1AB EP_NTERM
01500 P_babyPink PDU1AB
01505 HMM01923cobtr_1_pdu(pf01923pf03928) PDU1AB
01510 HMMalddh_1_eutpdugrm(pf00171) PDU1AB EP_NTERM
01515 HMM00465alcdh_3_mix(pf00465) PDU1AB
01520 HMMPduS_1_all(pf01512pf13375pf10531) PDU1AB
01525 Ts_dodgerBlue PDU1AB
01530 Hp_euts_babyPurple PDU1AB
01535 HMM10662PduV_1_mix(pf10662) PDU1AB
01540 HMMacetkin_1(pf00871) PDU1AB
01545 HMM00288_1_pdu(pf00288) PDU1AB
01550 HMMreg00392_2_grm2(pf00155) EUT2A
01555 -pf01520pf11741 -
01560 -pf00696pf00583 -
01565 -pf13538 -
01570 -pf00580pf13361 -
01575 -pf00675pf05193pf16187 -
01580 -# -
BMC-P
BMC-H
BMC-T
Aldehyde dehydrogenase
PTAC
Signature enzyme
Alcohol dehydrogenase
Regulator

Klebsiella pneumoniae subsp. pneumoniae (strain HS11286). Bacteria; Proteobacteria; Gammaproteobacteria; Enterobacterales;

KPHS__locus_2

PDU1AB

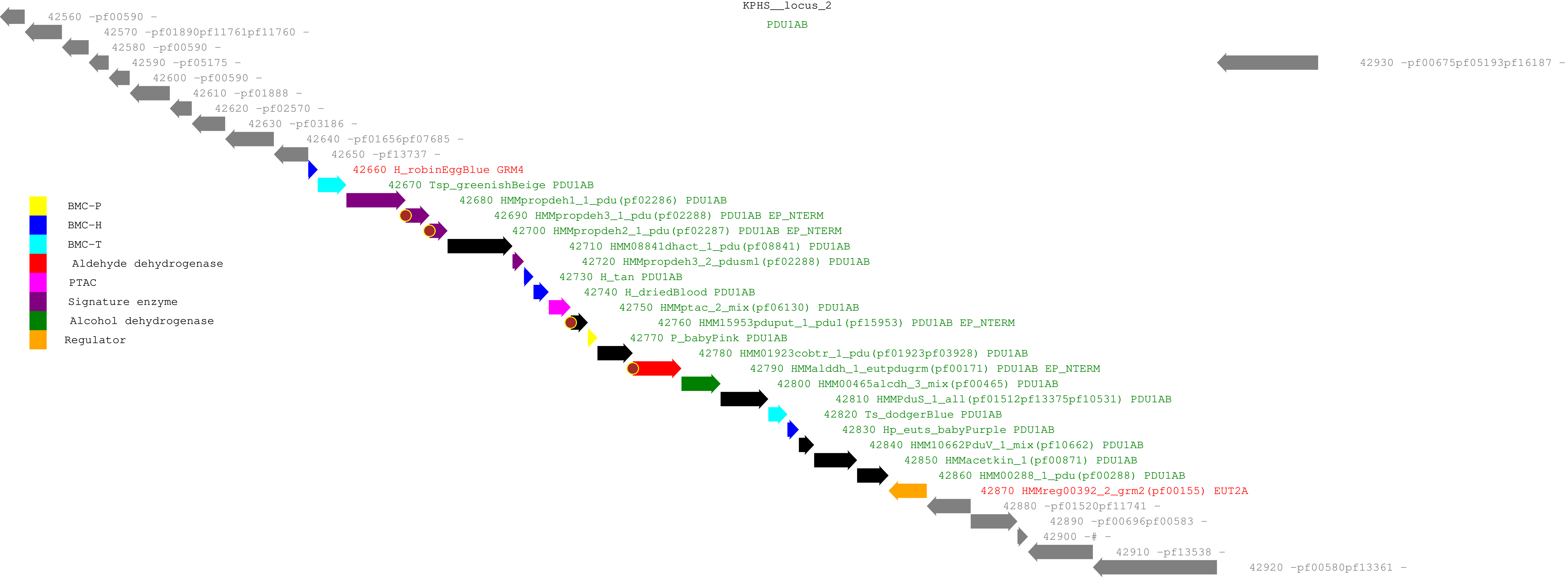

Klebsiella pneumoniae (strain 342). Bacteria; Proteobacteria; Gammaproteobacteria; Enterobacterales;

KPK__locus_1

PDU1AB

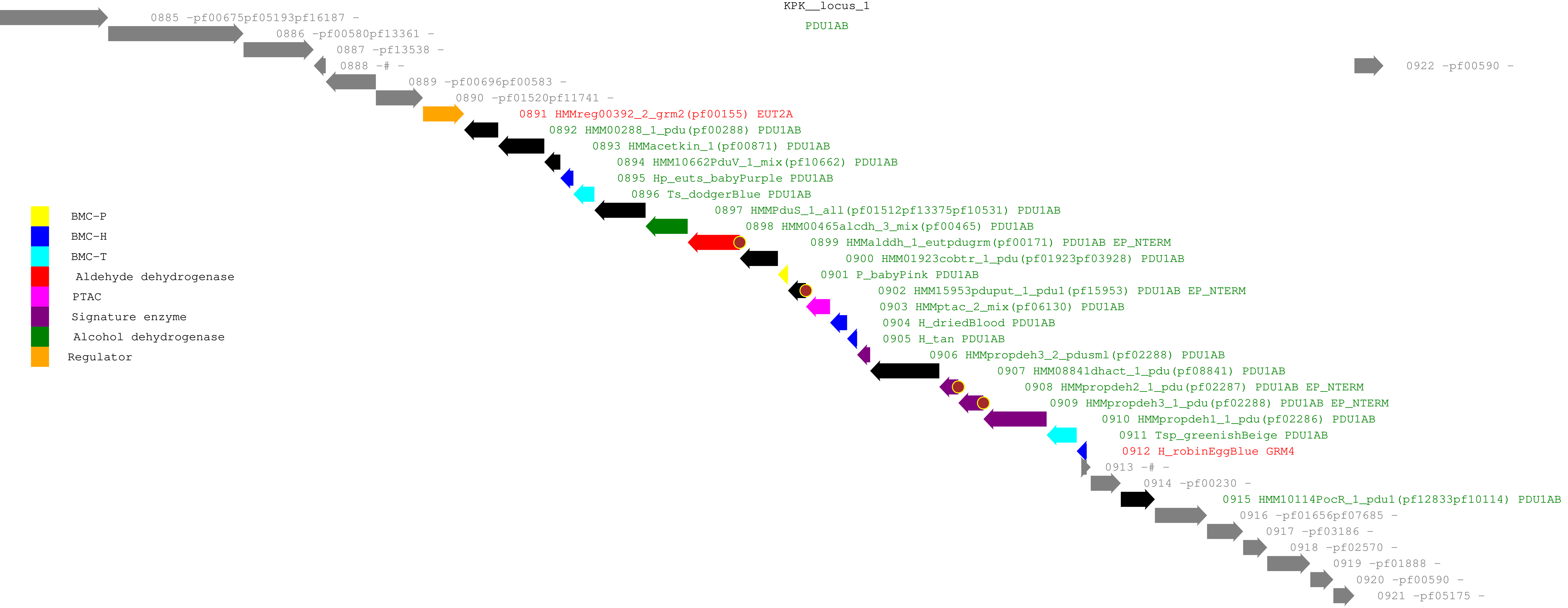

Klebsiella pneumoniae subsp. pneumoniae. Bacteria; Proteobacteria; Gammaproteobacteria; Enterobacterales;

KPNIH32__locus_2

PDU1AB

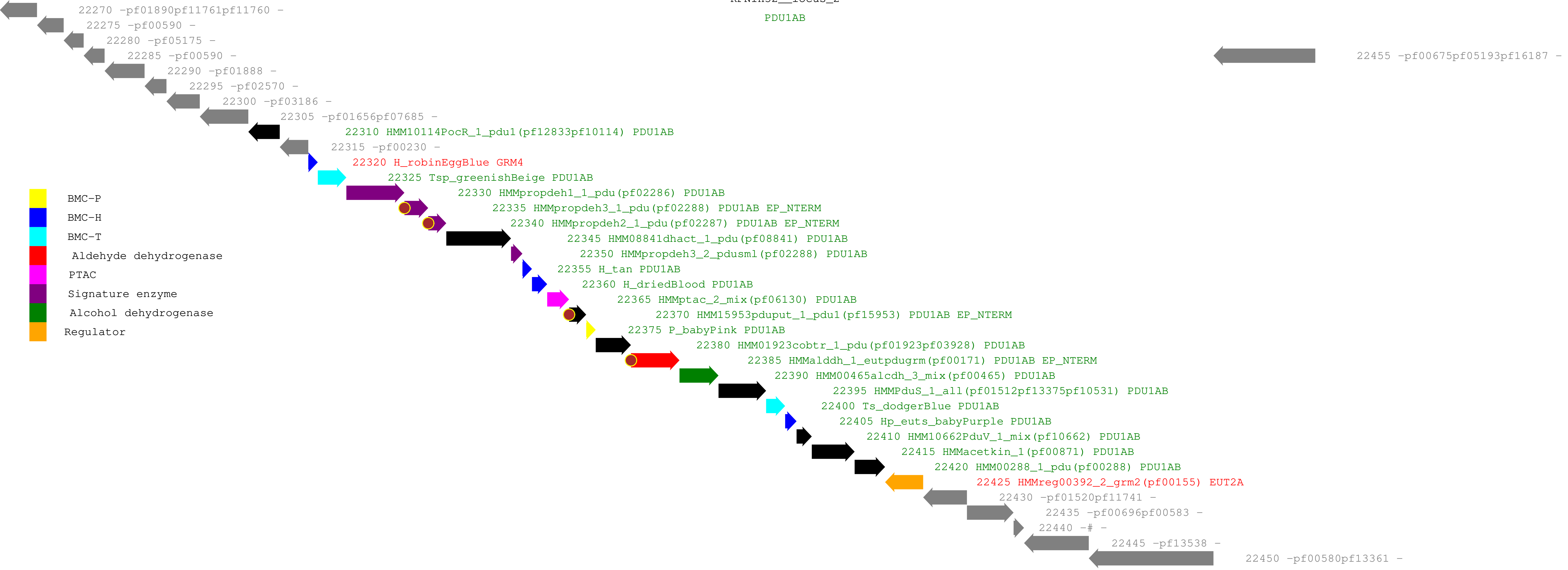

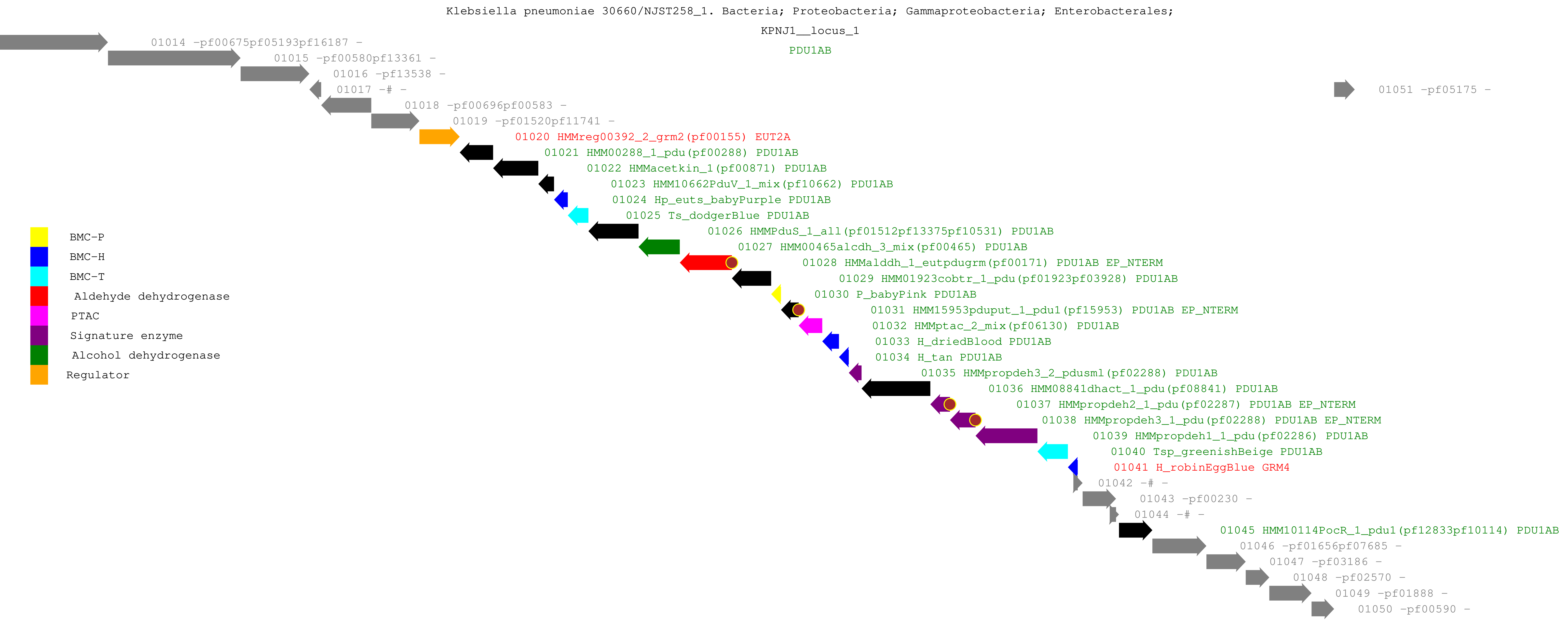
Klebsiella pneumoniae 30660/NJST258_1. Bacteria; Proteobacteria; Gammaproteobacteria; Enterobacterales;
KPNJ1__locus_1
PDU1AB
01014 -pf00675pf05193pf16187 -
01015 -pf00580pf13361 -
01016 -pf13538 -
01017 -# -
01018 -pf00696pf00583 -
01019 -pf01520pf11741 -
01020 HMMreg00392_2_grm2(pf00155) EUT2A
01021 HMM00288_1_pdu(pf00288) PDU1AB
01022 HMMacetkin_1(pf00871) PDU1AB
01023 HMM10662PduV_1_mix(pf10662) PDU1AB
01024 Hp_euts_babyPurple PDU1AB
01025 Ts_dodgerBlue PDU1AB
01026 HMMPduS_1_all(pf01512pf13375pf10531) PDU1AB
01027 HMM00465alcdh_3_mix(pf00465) PDU1AB
01028 HMMalddh_1_eutpdugrm(pf00171) PDU1AB EP_NTERM
01029 HMM01923cobtr_1_pdu(pf01923pf03928) PDU1AB
01030 P_babyPink PDU1AB
01031 HMM15953pduput_1_pdu1(pf15953) PDU1AB EP_NTERM
01032 HMMptac_2_mix(pf06130) PDU1AB
01033 H_driedBlood PDU1AB
01034 H_tan PDU1AB
01035 HMMpropdeh3_2_pdusml(pf02288) PDU1AB
01036 HMM08841dhact_1_pdu(pf08841) PDU1AB
01037 HMMpropdeh2_1_pdu(pf02287) PDU1AB EP_NTERM
01038 HMMpropdeh3_1_pdu(pf02288) PDU1AB EP_NTERM
01039 HMMpropdeh1_1_pdu(pf02286) PDU1AB
01040 Tsp_greenishBeige PDU1AB
01041 H_robinEggBlue GRM4
01042 -# -
01043 -pf00230 -
01044 -# -
01045 HMM10114PocR_1_pdu1(pf12833pf10114) PDU1AB
01046 -pf01656pf07685 -
01047 -pf03186 -
01048 -pf02570 -
01049 -pf01888 -
01050 -pf00590 -
01051 -pf05175 -
BMC-P
BMC-H
BMC-T
Aldehyde dehydrogenase
PTAC
Signature enzyme
Alcohol dehydrogenase
Regulator

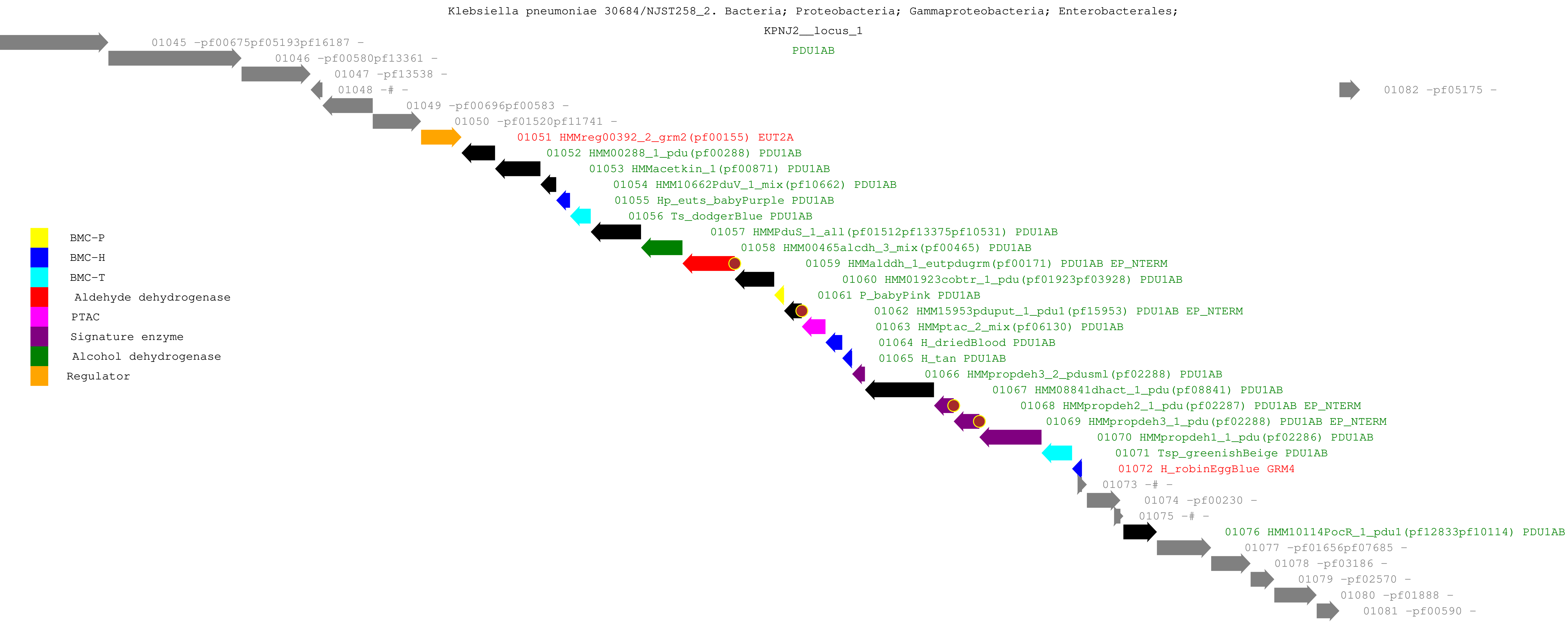
Klebsiella pneumoniae 30684/NJST258_2. Bacteria; Proteobacteria; Gammaproteobacteria; Enterobacterales;
KPNJ2__locus_1
PDU1AB
01045 -pf00675pf05193pf16187 -
01046 -pf00580pf13361 -
01047 -pf13538 -
01048 -# -
01049 -pf00696pf00583 -
01050 -pf01520pf11741 -
01051 HMMreg00392_2_grm2(pf00155) EUT2A
01052 HMM00288_1_pdu(pf00288) PDU1AB
01053 HMMacetkin_1(pf00871) PDU1AB
01054 HMM10662PduV_1_mix(pf10662) PDU1AB
01055 Hp_euts_babyPurple PDU1AB
01056 Ts_dodgerBlue PDU1AB
01057 HMMPduS_1_all(pf01512pf13375pf10531) PDU1AB
01058 HMM00465alcdh_3_mix(pf00465) PDU1AB
01059 HMMalddh_1_eutpdugrm(pf00171) PDU1AB EP_NTERM
01060 HMM01923cobtr_1_pdu(pf01923pf03928) PDU1AB
01061 P_babyPink PDU1AB
01062 HMM15953pduput_1_pdu1(pf15953) PDU1AB EP_NTERM
01063 HMMptac_2_mix(pf06130) PDU1AB
01064 H_driedBlood PDU1AB
01065 H_tan PDU1AB
01066 HMMpropdeh3_2_pdusml(pf02288) PDU1AB
01067 HMM08841dhact_1_pdu(pf08841) PDU1AB
01068 HMMpropdeh2_1_pdu(pf02287) PDU1AB EP_NTERM
01069 HMMpropdeh3_1_pdu(pf02288) PDU1AB EP_NTERM
01070 HMMpropdeh1_1_pdu(pf02286) PDU1AB
01071 Tsp_greenishBeige PDU1AB
01072 H_robinEggBlue GRM4
01073 -# -
01074 -pf00230 -
01075 -# -
01076 HMM10114PocR_1_pdu1(pf12833pf10114) PDU1AB
01077 -pf01656pf07685 -
01078 -pf03186 -
01079 -pf02570 -
01080 -pf01888 -
01081 -pf00590 -
01082 -pf05175 -
BMC-P
BMC-H
BMC-T
Aldehyde dehydrogenase
PTAC
Signature enzyme
Alcohol dehydrogenase
Regulator

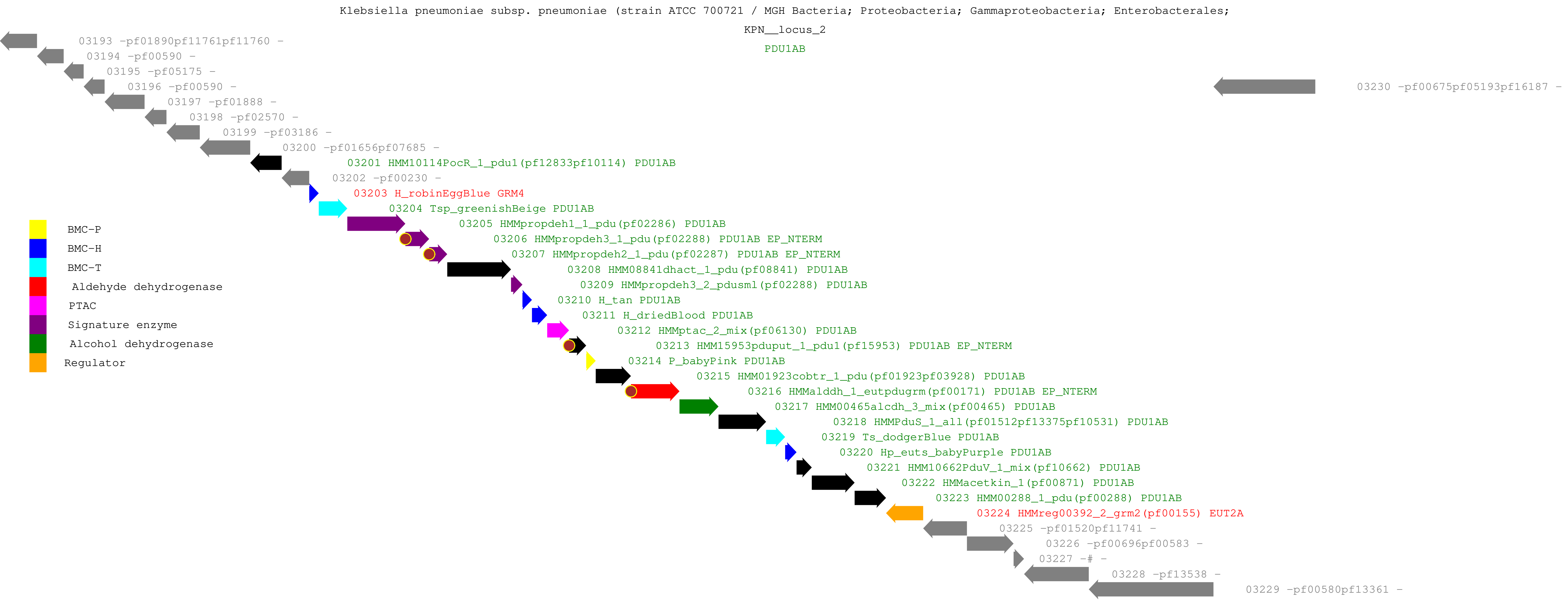
Klebsiella pneumoniae subsp. pneumoniae (strain ATCC 700721 / MGH Bacteria; Proteobacteria; Gammaproteobacteria; Enterobacterales;
KPN__locus_2
PDU1AB
03193 -pf01890pf11761pf11760 -
03194 -pf00590 -
03195 -pf05175 -
03196 -pf00590 -
03197 -pf01888 -
03198 -pf02570 -
03199 -pf03186 -
03200 -pf01656pf07685 -
03201 HMM10114PocR_1_pdu1(pf12833pf10114) PDU1AB
03202 -pf00230 -
03203 H_robinEggBlue GRM4
03204 Tsp_greenishBeige PDU1AB
03205 HMMpropdeh1_1_pdu(pf02286) PDU1AB
03206 HMMpropdeh3_1_pdu(pf02288) PDU1AB EP_NTERM
03207 HMMpropdeh2_1_pdu(pf02287) PDU1AB EP_NTERM
03208 HMM08841dhact_1_pdu(pf08841) PDU1AB
03209 HMMpropdeh3_2_pdusml(pf02288) PDU1AB
03210 H_tan PDU1AB
03211 H_driedBlood PDU1AB
03212 HMMptac_2_mix(pf06130) PDU1AB
03213 HMM15953pduput_1_pdu1(pf15953) PDU1AB EP_NTERM
03214 P_babyPink PDU1AB
03215 HMM01923cobtr_1_pdu(pf01923pf03928) PDU1AB
03216 HMMalddh_1_eutpdugrm(pf00171) PDU1AB EP_NTERM
03217 HMM00465alcdh_3_mix(pf00465) PDU1AB
03218 HMMPduS_1_all(pf01512pf13375pf10531) PDU1AB
03219 Ts_dodgerBlue PDU1AB
03220 Hp_euts_babyPurple PDU1AB
03221 HMM10662PduV_1_mix(pf10662) PDU1AB
03222 HMMacetkin_1(pf00871) PDU1AB
03223 HMM00288_1_pdu(pf00288) PDU1AB
03224 HMMreg00392_2_grm2(pf00155) EUT2A
03225 -pf01520pf11741 -
03226 -pf00696pf00583 -
03227 -# -
03228 -pf13538 -
03229 -pf00580pf13361 -
03230 -pf00675pf05193pf16187 -
BMC-P
BMC-H
BMC-T
Aldehyde dehydrogenase
PTAC
Signature enzyme
Alcohol dehydrogenase
Regulator

Klebsiella pneumoniae subsp. rhinoscleromatis SB3432. Bacteria; Proteobacteria; Gammaproteobacteria; Enterobacterales;

KPR__locus_2

PDU1AB

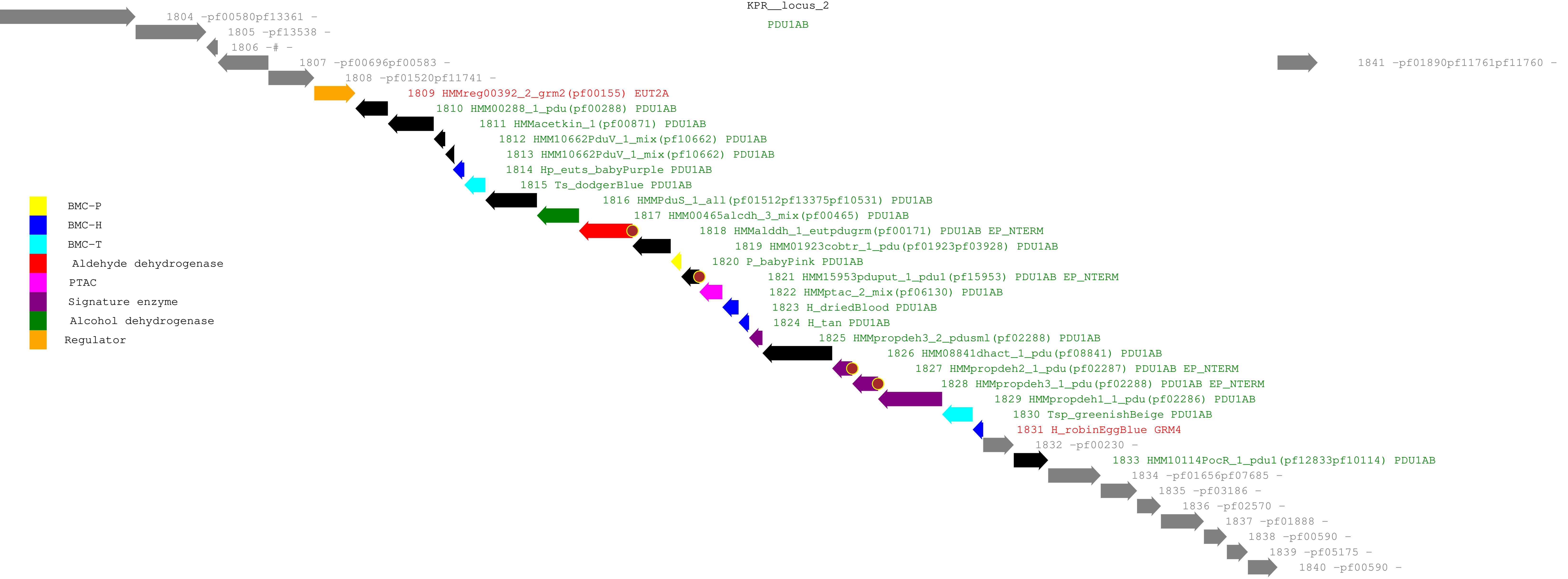

Klebsiella quasipneumoniae subsp. quasipneumoniae. Bacteria; Proteobacteria; Gammaproteobacteria; Enterobacterales;

KQQSB11__locus_1

PDU1AB

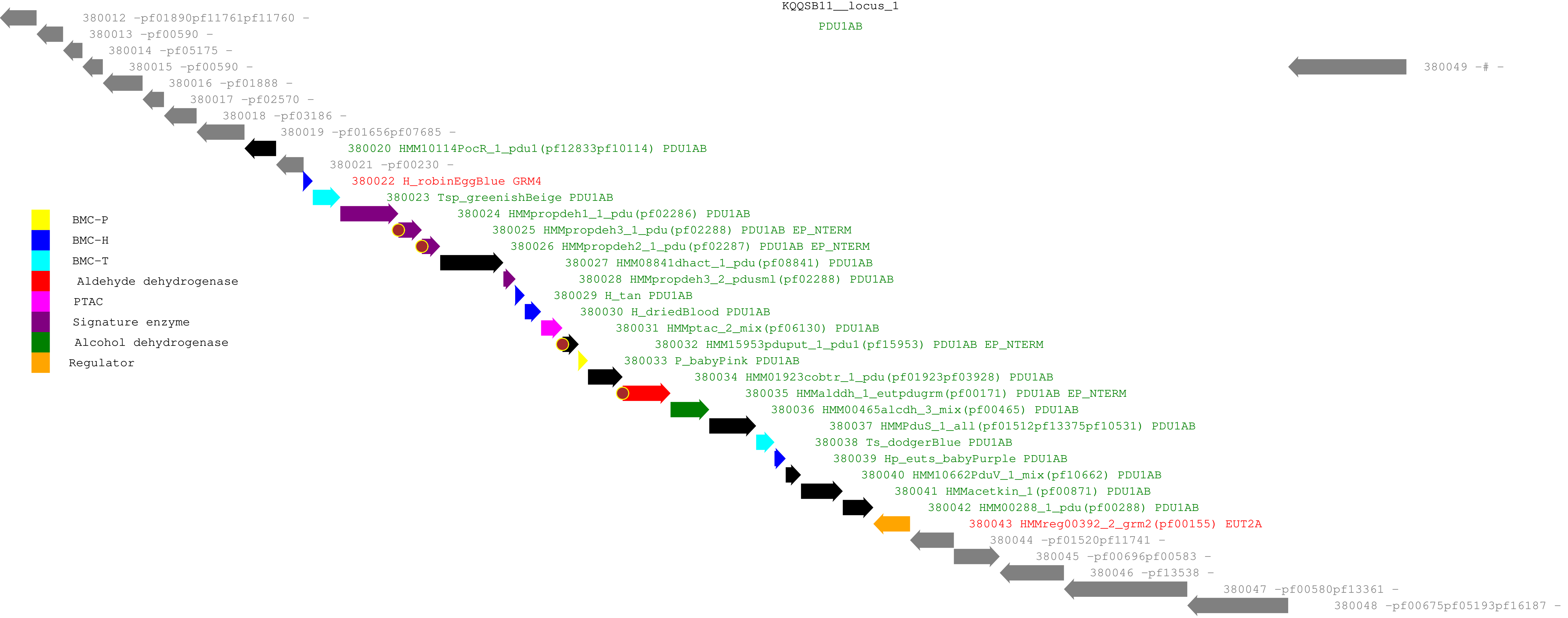

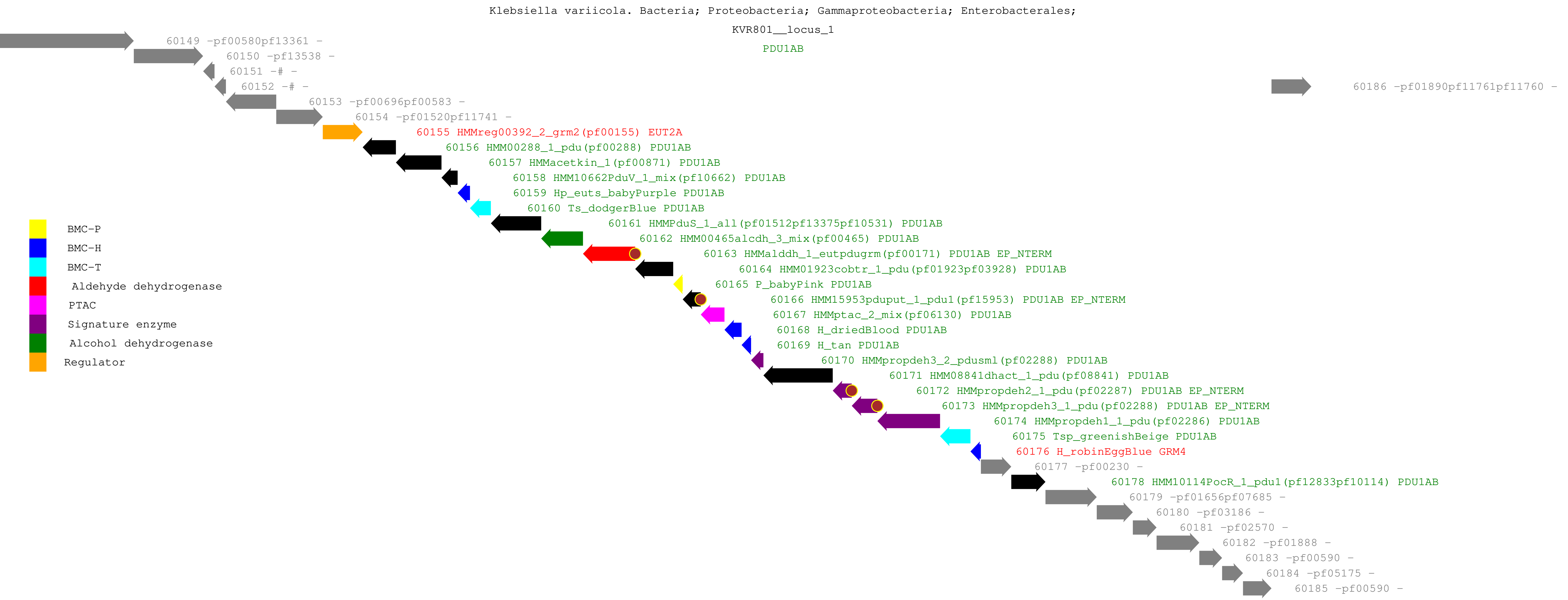
Klebsiella variicola. Bacteria; Proteobacteria; Gammaproteobacteria; Enterobacterales;
KVR801__locus_1
PDU1AB
60149 -pf00580pf13361 -
60150 -pf13538 -
60151 -# -
60152 -# -
60153 -pf00696pf00583 -
60154 -pf01520pf11741 -
60155 HMMreg00392_2_grm2(pf00155) EUT2A
60156 HMM00288_1_pdu(pf00288) PDU1AB
60157 HMMacetkin_1(pf00871) PDU1AB
60158 HMM10662PduV_1_mix(pf10662) PDU1AB
60159 Hp_euts_babyPurple PDU1AB
60160 Ts_dodgerBlue PDU1AB
60161 HMMPduS_1_all(pf01512pf13375pf10531) PDU1AB
60162 HMM00465alcdh_3_mix(pf00465) PDU1AB
60163 HMMalddh_1_eutpdugrm(pf00171) PDU1AB EP_NTERM
60164 HMM01923cobtr_1_pdu(pf01923pf03928) PDU1AB
60165 P_babyPink PDU1AB
60166 HMM15953pduput_1_pdu1(pf15953) PDU1AB EP_NTERM
60167 HMMptac_2_mix(pf06130) PDU1AB
60168 H_driedBlood PDU1AB
60169 H_tan PDU1AB
60170 HMMpropdeh3_2_pdusml(pf02288) PDU1AB
60171 HMM08841dhact_1_pdu(pf08841) PDU1AB
60172 HMMpropdeh2_1_pdu(pf02287) PDU1AB EP_NTERM
60173 HMMpropdeh3_1_pdu(pf02288) PDU1AB EP_NTERM
60174 HMMpropdeh1_1_pdu(pf02286) PDU1AB
60175 Tsp_greenishBeige PDU1AB
60176 H_robinEggBlue GRM4
60177 -pf00230 -
60178 HMM10114PocR_1_pdu1(pf12833pf10114) PDU1AB
60179 -pf01656pf07685 -
60180 -pf03186 -
60181 -pf02570 -
60182 -pf01888 -
60183 -pf00590 -
60184 -pf05175 -
60185 -pf00590 -
60186 -pf01890pf11761pf11760 -
BMC-P
BMC-H
BMC-T
Aldehyde dehydrogenase
PTAC
Signature enzyme
Alcohol dehydrogenase
Regulator

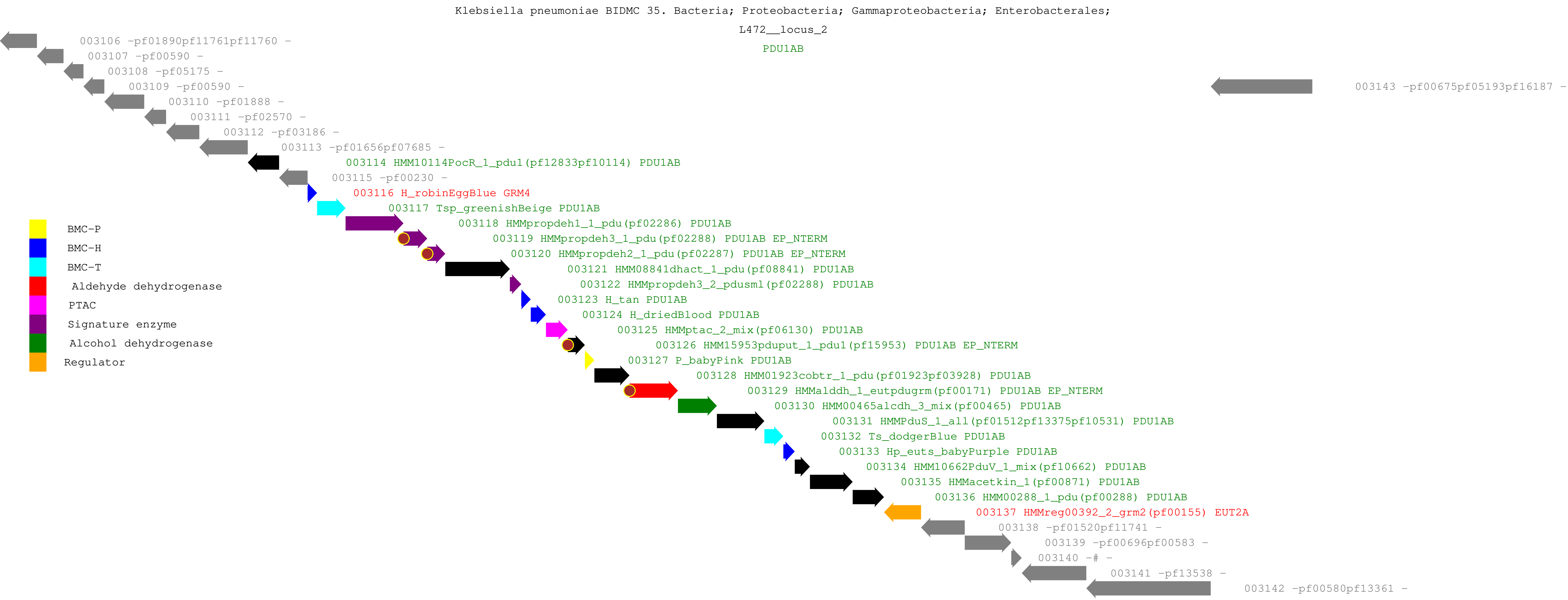
Klebsiella pneumoniae BIDMC 35. Bacteria; Proteobacteria; Gammaproteobacteria; Enterobacterales;
L472__locus_2
PDU1AB
003106 -pf01890pf11761pf11760 -
003107 -pf00590 -
003108 -pf05175 -
003109 -pf00590 -
003110 -pf01888 -
003111 -pf02570 -
003112 -pf03186 -
003113 -pf01656pf07685 -
003114 HMM10114PocR_1_pdu1(pf12833pf10114) PDU1AB
003115 -pf00230 -
003116 H_robinEggBlue GRM4
003117 Tsp_greenishBeige PDU1AB
003118 HMMpropdeh1_1_pdu(pf02286) PDU1AB
003119 HMMpropdeh3_1_pdu(pf02288) PDU1AB EP_NTERM
003120 HMMpropdeh2_1_pdu(pf02287) PDU1AB EP_NTERM
003121 HMM08841dhact_1_pdu(pf08841) PDU1AB
003122 HMMpropdeh3_2_pdusml(pf02288) PDU1AB
003123 H_tan PDU1AB
003124 H_driedBlood PDU1AB
003125 HMMptac_2_mix(pf06130) PDU1AB
003126 HMM15953pduput_1_pdu1(pf15953) PDU1AB EP_NTERM
003127 P_babyPink PDU1AB
003128 HMM01923cobtr_1_pdu(pf01923pf03928) PDU1AB
003129 HMMalddh_1_eutpdugrm(pf00171) PDU1AB EP_NTERM
003130 HMM00465alcdh_3_mix(pf00465) PDU1AB
003131 HMMPduS_1_all(pf01512pf13375pf10531) PDU1AB
003132 Ts_dodgerBlue PDU1AB
003133 Hp_euts_babyPurple PDU1AB
003134 HMM10662PduV_1_mix(pf10662) PDU1AB
003135 HMMacetkin_1(pf00871) PDU1AB
003136 HMM00288_1_pdu(pf00288) PDU1AB
003137 HMMreg00392_2_grm2(pf00155) EUT2A
003138 -pf01520pf11741 -
003139 -pf00696pf00583 -
003140 -# -
003141 -pf13538 -
003142 -pf00580pf13361 -
003143 -pf00675pf05193pf16187 -
BMC-P
BMC-H
BMC-T
Aldehyde dehydrogenase
PTAC
Signature enzyme
Alcohol dehydrogenase
Regulator

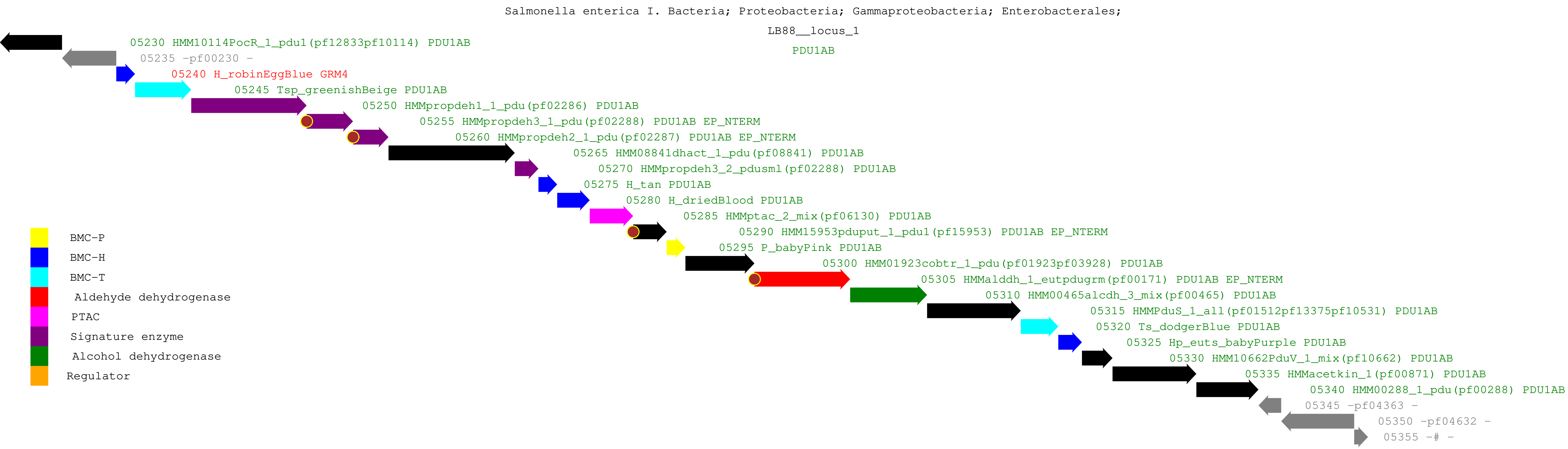
Salmonella enterica I. Bacteria; Proteobacteria; Gammaproteobacteria; Enterobacterales;
LB88__locus_1
PDU1AB
05230 HMM10114PocR_1_pdu1(pf12833pf10114) PDU1AB
05235 -pf00230 -
05240 H_robinEggBlue GRM4
05245 Tsp_greenishBeige PDU1AB
05250 HMMpropdeh1_1_pdu(pf02286) PDU1AB
05255 HMMpropdeh3_1_pdu(pf02288) PDU1AB EP_NTERM
05260 HMMpropdeh2_1_pdu(pf02287) PDU1AB EP_NTERM
05265 HMM08841dhact_1_pdu(pf08841) PDU1AB
05270 HMMpropdeh3_2_pdusml(pf02288) PDU1AB
05275 H_tan PDU1AB
05280 H_driedBlood PDU1AB
05285 HMMptac_2_mix(pf06130) PDU1AB
05290 HMM15953pduput_1_pdu1(pf15953) PDU1AB EP_NTERM
05295 P_babyPink PDU1AB
05300 HMM01923cobtr_1_pdu(pf01923pf03928) PDU1AB
05305 HMMaldh_1_eutpdugrm(pf00171) PDU1AB EP_NTERM
05310 HMM00465alcdh_3_mix(pf00465) PDU1AB
05315 HMMPduS_1_all(pf01512pf13375pf10531) PDU1AB
05320 Ts_dodgerBlue PDU1AB
05325 Hp_euts_babyPurple PDU1AB
05330 HMM10662PduV_1_mix(pf10662) PDU1AB
05335 HMMacetkin_1(pf00871) PDU1AB
05340 HMM00288_1_pdu(pf00288) PDU1AB
05345 -pf04363 -
05350 -pf04632 -
05355 -# -
BMC-P
BMC-H
BMC-T
Aldehyde dehydrogenase
PTAC
Signature enzyme
Alcohol dehydrogenase
Regulator

Salmonella enterica subsp. enterica serovar Gaminara str. SA20063285. Bacteria; Proteobacteria; Gammaproteobacteria; Enterobacterales;

LFZ12__locus_2

PDU1AB

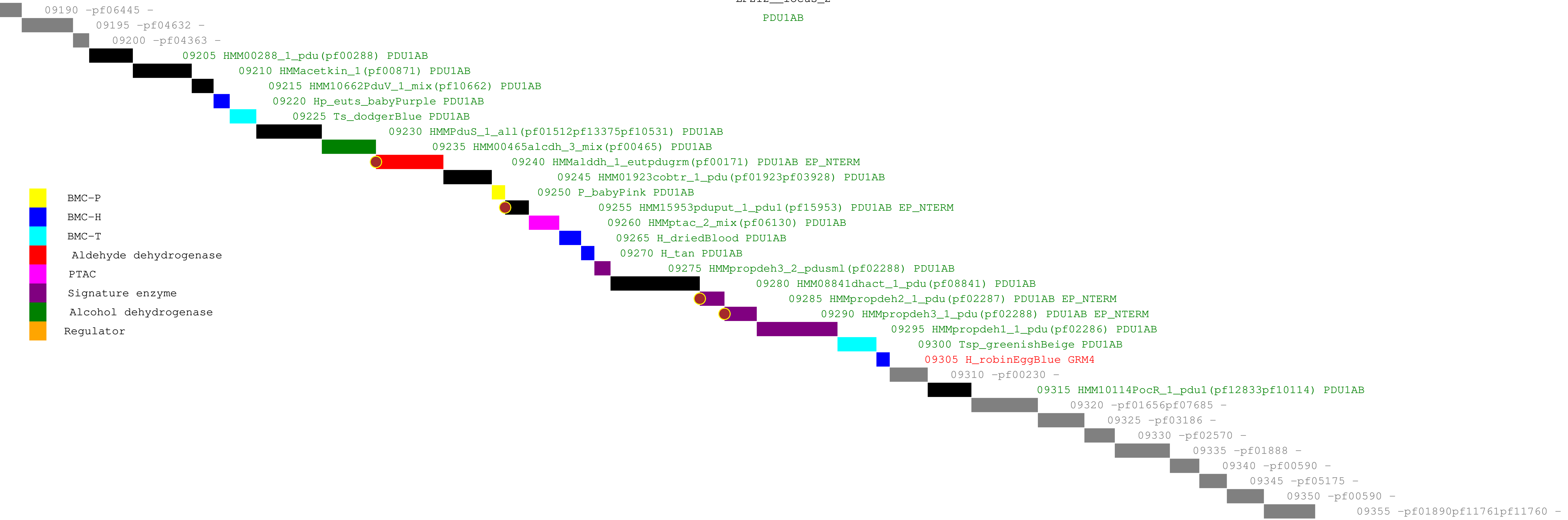

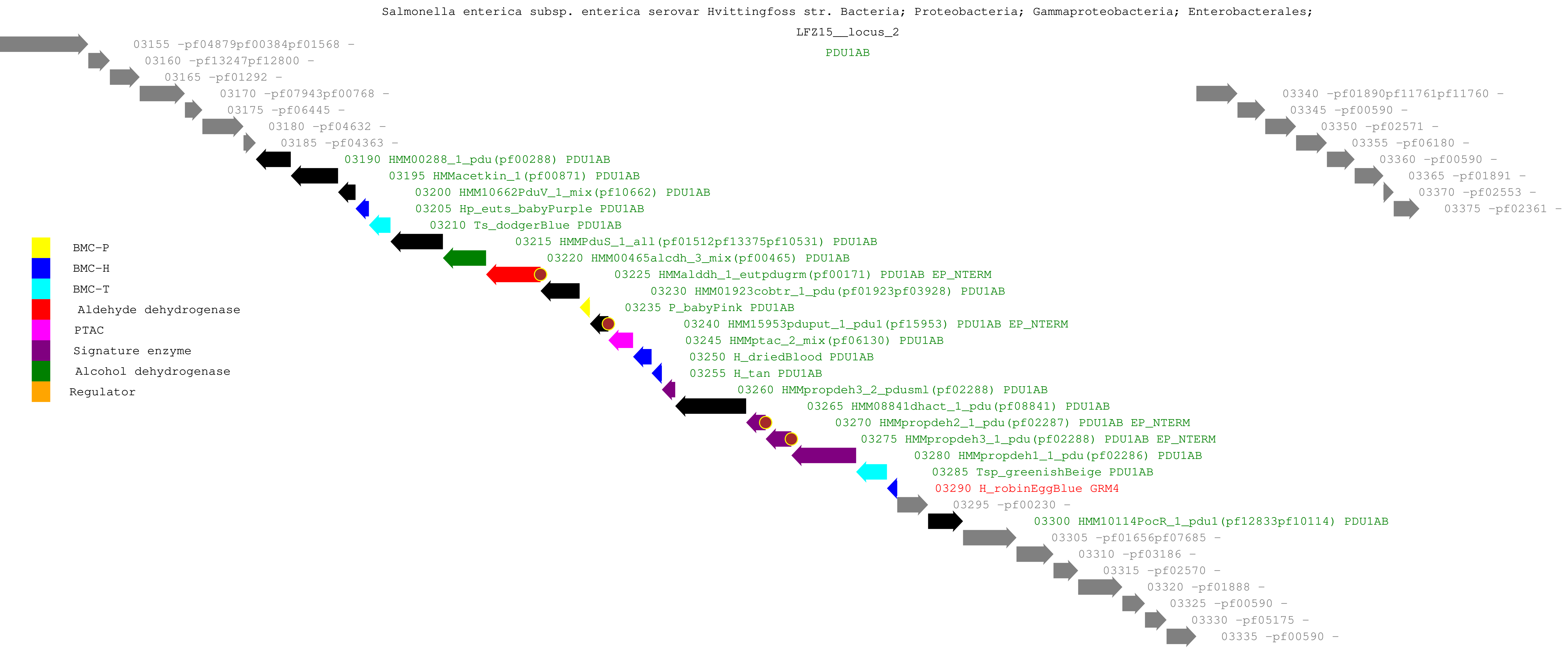
Salmonella enterica subsp. enterica serovar Hvittingfoss str. Bacteria; Proteobacteria; Gammaproteobacteria; Enterobacterales;
LFZ15__locus_2
PDU1AB
03155 -pf04879pf00384pf01568 -
03160 -pf13247pf12800 -
03165 -pf01292 -
03170 -pf07943pf00768 -
03175 -pf06445 -
03180 -pf04632 -
03185 -pf04363 -
03190 HMM00288_1_pdu(pf00288) PDU1AB
03195 HMMacetkin_1(pf00871) PDU1AB
03200 HMM10662PduV_1_mix(pf10662) PDU1AB
03205 Hp_euts_babyPurple PDU1AB
03210 Ts_dodgerBlue PDU1AB
03215 HMMPduS_1_all(pf01512pf13375pf10531) PDU1AB
03220 HMM00465alcdh_3_mix(pf00465) PDU1AB
03225 HMMaldh_1_eutpdugrm(pf00171) PDU1AB EP_NTERM
03230 HMM01923cobtr_1_pdu(pf01923pf03928) PDU1AB
03235 P_babyPink PDU1AB
03240 HMM15953pduput_1_pdu1(pf15953) PDU1AB EP_NTERM
03245 HMMptac_2_mix(pf06130) PDU1AB
03250 H_driedBlood PDU1AB
03255 H_tan PDU1AB
03260 HMMpropdeh3_2_pdusml(pf02288) PDU1AB
03265 HMM08841dhact_1_pdu(pf08841) PDU1AB
03270 HMMpropdeh2_1_pdu(pf02287) PDU1AB EP_NTERM
03275 HMMpropdeh3_1_pdu(pf02288) PDU1AB EP_NTERM
03280 HMMpropdeh1_1_pdu(pf02286) PDU1AB
03285 Tsp_greenishBeige PDU1AB
03290 H_robinEggBlue GRM4
03295 -pf00230 -
03300 HMM10114PocR_1_pdu1(pf12833pf10114) PDU1AB
03305 -pf01656pf07685 -
03310 -pf03186 -
03315 -pf02570 -
03320 -pf01888 -
03325 -pf00590 -
03330 -pf05175 -
03335 -pf00590 -
03340 -pf01890pf11761pf11760 -
03345 -pf00590 -
03350 -pf02571 -
03355 -pf06180 -
03360 -pf00590 -
03365 -pf01891 -
03370 -pf02553 -
03375 -pf02361 -
BMC-P
BMC-H
BMC-T
Aldehyde dehydrogenase
PTAC
Signature enzyme
Alcohol dehydrogenase
Regulator

Salmonella enterica subsp. enterica serovar India str. SA20085604. Bacteria; Proteobacteria; Gammaproteobacteria; Enterobacterales;

LFZ16__locus_2

PDU1AB

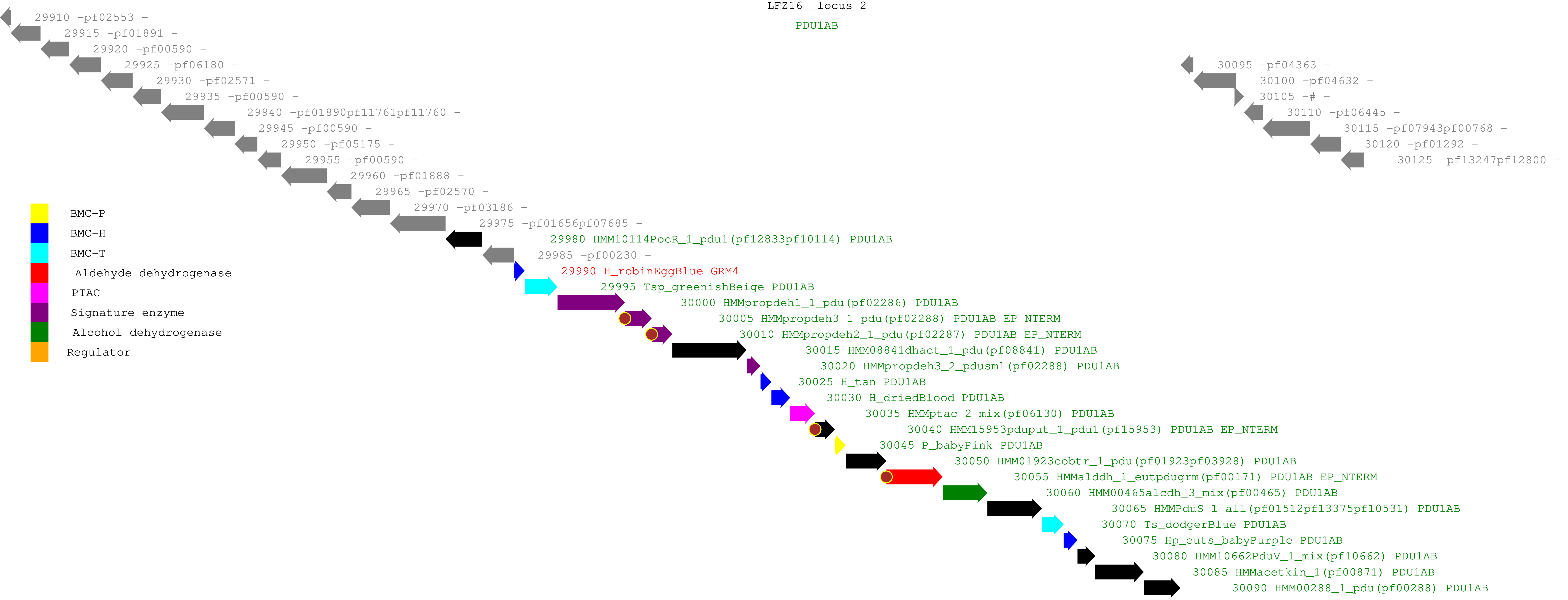

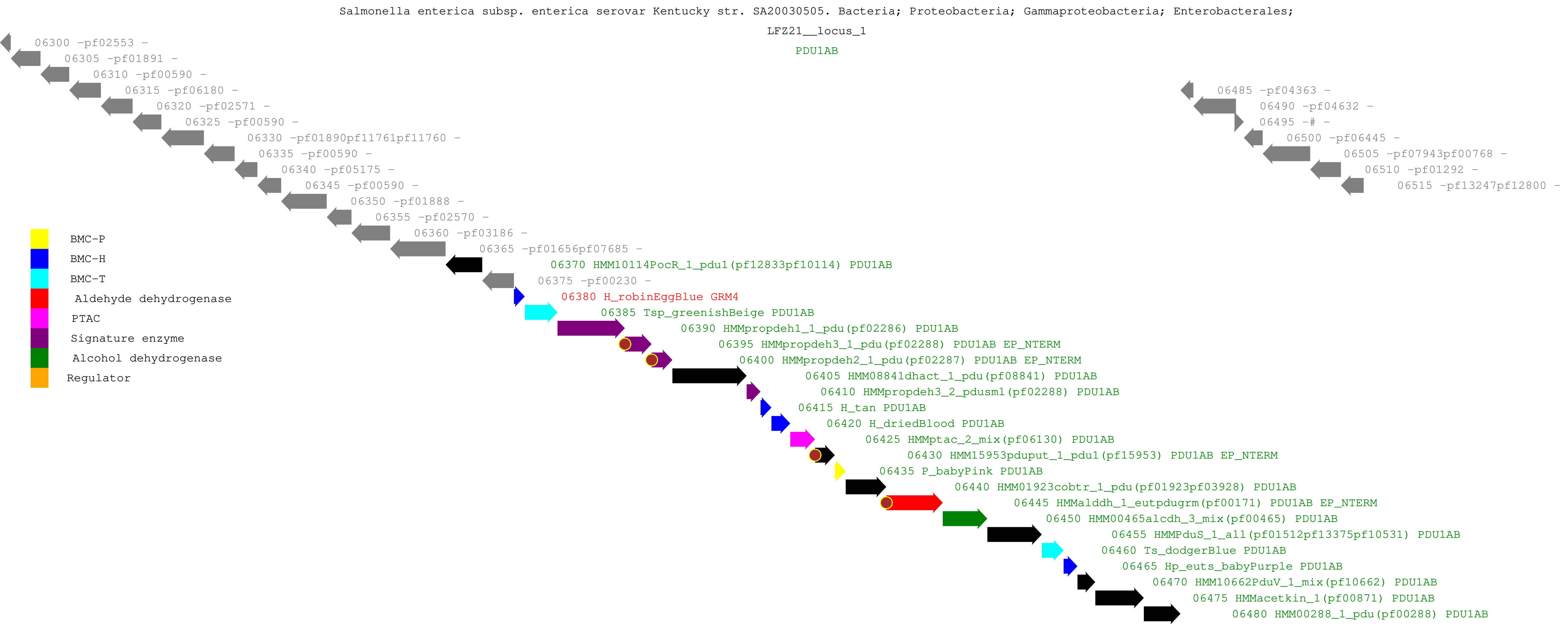
Salmonella enterica subsp. enterica serovar Kentucky str. SA20030505. Bacteria; Proteobacteria; Gammaproteobacteria; Enterobacterales;
LFZ21__locus_1
PDU1AB
06300 -pf02553 -
06305 -pf01891 -
06310 -pf00590 -
06315 -pf06180 -
06320 -pf02571 -
06325 -pf00590 -
06330 -pf01890pf11761pf11760 -
06335 -pf00590 -
06340 -pf05175 -
06345 -pf00590 -
06350 -pf01888 -
06355 -pf02570 -
06360 -pf03186 -
06365 -pf01656pf07685 -
06370 HMM10114PocR_1_pdu1(pf12833pf10114) PDU1AB
06375 -pf00230 -
06380 H_robinEggBlue GRM4
06385 Tsp_greenishBeige PDU1AB
06390 HMMpropdeh1_1_pdu(pf02286) PDU1AB
06395 HMMpropdeh3_1_pdu(pf02288) PDU1AB EP_NTERM
06400 HMMpropdeh2_1_pdu(pf02287) PDU1AB EP_NTERM
06405 HMM08841dhact_1_pdu(pf08841) PDU1AB
06410 HMMpropdeh3_2_pdusml(pf02288) PDU1AB
06415 H_tan PDU1AB
06420 H_driedBlood PDU1AB
06425 HMMptac_2_mix(pf06130) PDU1AB
06430 HMM15953pduput_1_pdu1(pf15953) PDU1AB EP_NTERM
06435 P_babyPink PDU1AB
06440 HMM01923cobtr_1_pdu(pf01923pf03928) PDU1AB
06445 HMMalddh_1_eutpdugrm(pf00171) PDU1AB EP_NTERM
06450 HMM00465alcdh_3_mix(pf00465) PDU1AB
06455 HMMPduS_1_all(pf01512pf13375pf10531) PDU1AB
06460 Ts_dodgerBlue PDU1AB
06465 Hp_euts_babyPurple PDU1AB
06470 HMM10662PduV_1_mix(pf10662) PDU1AB
06475 HMMacetkin_1(pf00871) PDU1AB
06480 HMM00288_1_pdu(pf00288) PDU1AB
06485 -pf04363 -
06490 -pf04632 -
06495 -# -
06500 -pf06445 -
06505 -pf07943pf00768 -
06510 -pf01292 -
06515 -pf13247pf12800 -
BMC-P
BMC-H
BMC-T
Aldehyde dehydrogenase
PTAC
Signature enzyme
Alcohol dehydrogenase
Regulator

Salmonella enterica subsp. enterica serovar Krefeld str. SA20030536. Bacteria; Proteobacteria; Gammaproteobacteria; Enterobacterales;

LFZ24__locus_1

PDU1AB

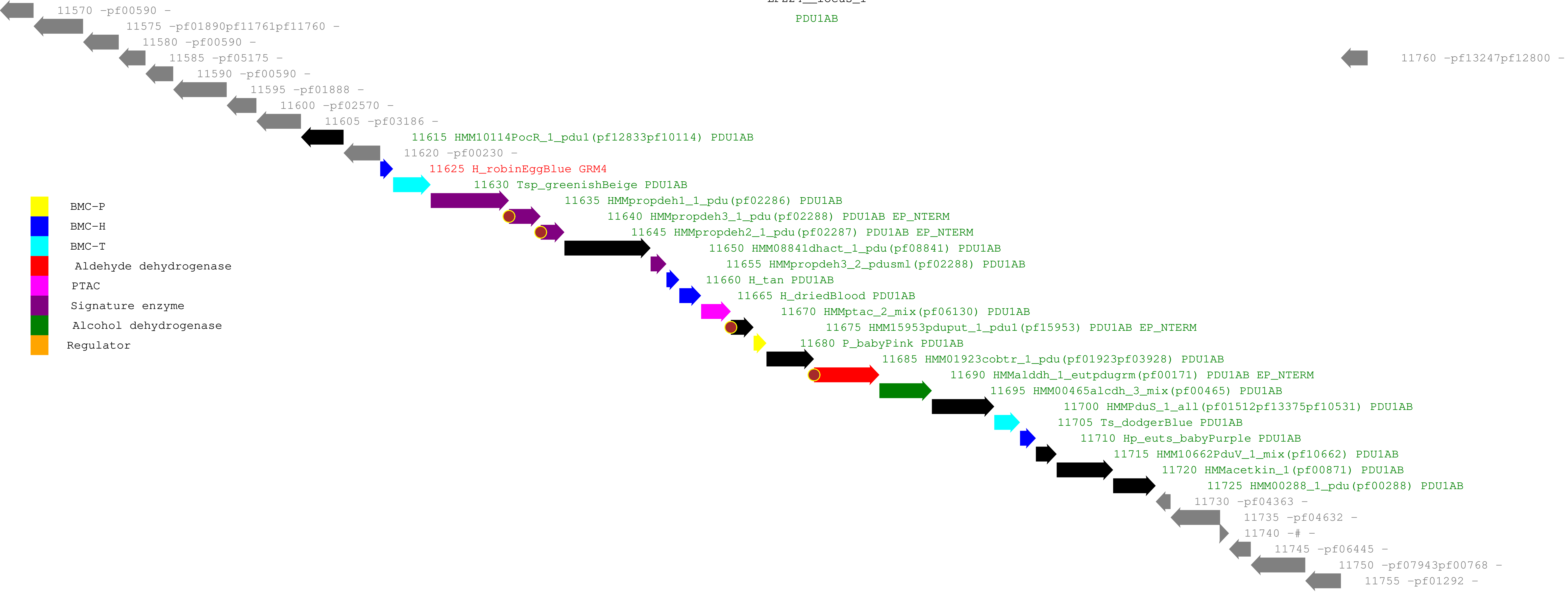

Salmonella enterica subsp. enterica serovar Macclesfield str. S-1643. Bacteria; Proteobacteria; Gammaproteobacteria; Enterobacterales;

LFZ25__locus_2

PDU1AB

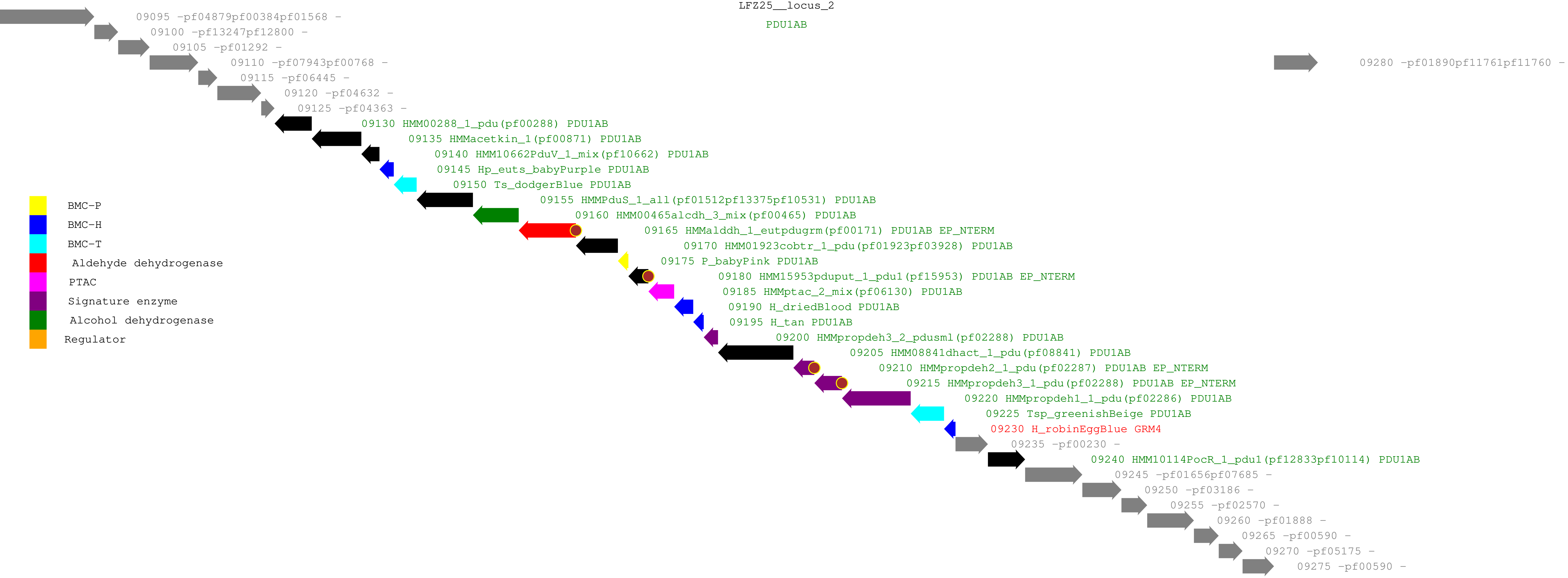

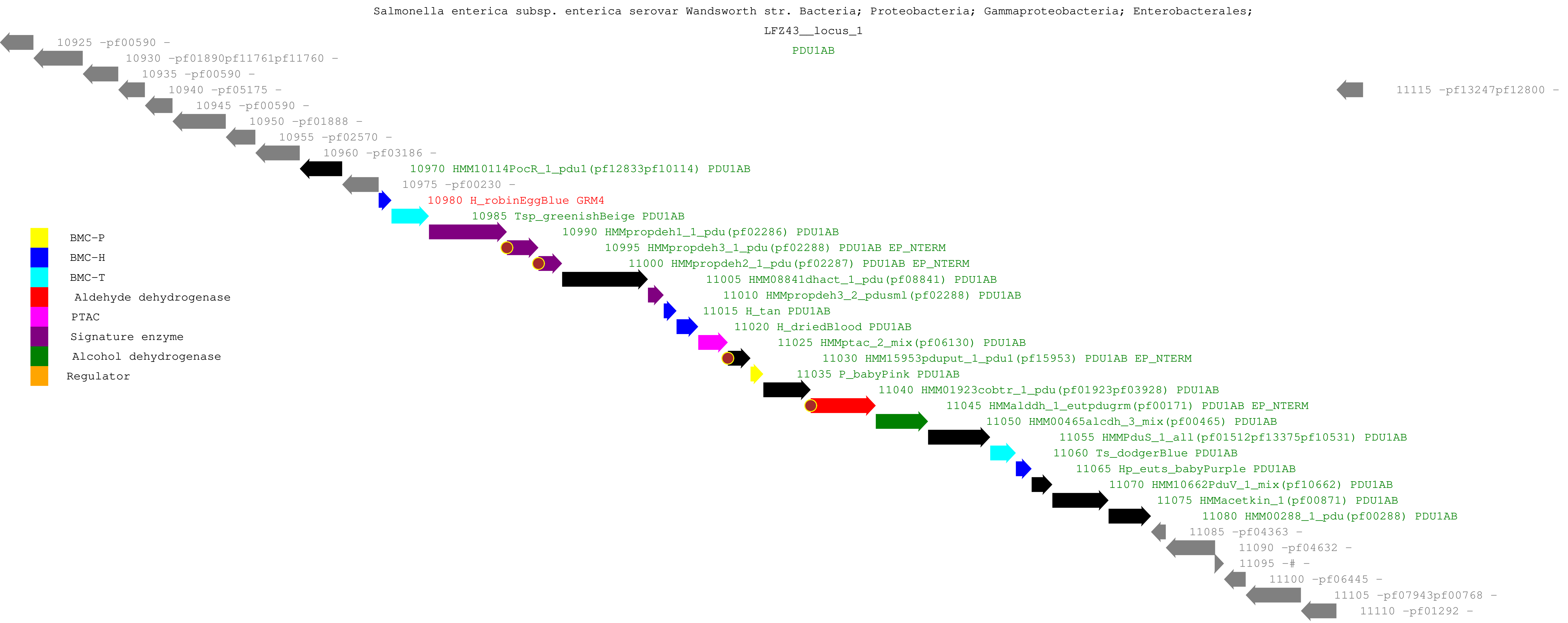

Salmonella enterica subsp. enterica serovar Wandsworth str. Bacteria; Proteobacteria; Gammaproteobacteria; Enterobacterales;
LFZ43__locus_1
PDU1AB
10925 -pf00590 -
10930 -pf01890pf11761pf11760 -
10935 -pf00590 -
10940 -pf05175 -
10945 -pf00590 -
10950 -pf01888 -
10955 -pf02570 -
10960 -pf03186 -
10970 HMM10114PocR_1_pdu1(pf12833pf10114) PDU1AB
10975 -pf00230 -
10980 H_robinEggBlue GRM4
10985 Tsp_greenishBeige PDU1AB
10990 HMMpropdeh1_1_pdu(pf02286) PDU1AB
10995 HMMpropdeh3_1_pdu(pf02288) PDU1AB EP_NTERM
11000 HMMpropdeh2_1_pdu(pf02287) PDU1AB EP_NTERM
11005 HMM08841dhact_1_pdu(pf08841) PDU1AB
11010 HMMpropdeh3_2_pdusml(pf02288) PDU1AB
11015 H_tan PDU1AB
11020 H_driedBlood PDU1AB
11025 HMMptac_2_mix(pf06130) PDU1AB
11030 HMM15953pduput_1_pdu1(pf15953) PDU1AB EP_NTERM
11035 P_babyPink PDU1AB
11040 HMM01923cobtr_1_pdu(pf01923pf03928) PDU1AB
11045 HMMalddh_1_eutpdugrm(pf00171) PDU1AB EP_NTERM
11050 HMM00465alcdh_3_mix(pf00465) PDU1AB
11055 HMMPduS_1_all(pf01512pf13375pf10531) PDU1AB
11060 Ts_dodgerBlue PDU1AB
11065 Hp_euts_babyPurple PDU1AB
11070 HMM10662PduV_1_mix(pf10662) PDU1AB
11075 HMMacetkin_1(pf00871) PDU1AB
11080 HMM00288_1_pdu(pf00288) PDU1AB
11085 -pf04363 -
11090 -pf04632 -
11095 -# -
11100 -pf06445 -
11105 -pf07943pf00768 -
11110 -pf01292 -
11115 -pf13247pf12800 -
BMC-P
BMC-H
BMC-T
Aldehyde dehydrogenase
PTAC
Signature enzyme
Alcohol dehydrogenase
Regulator

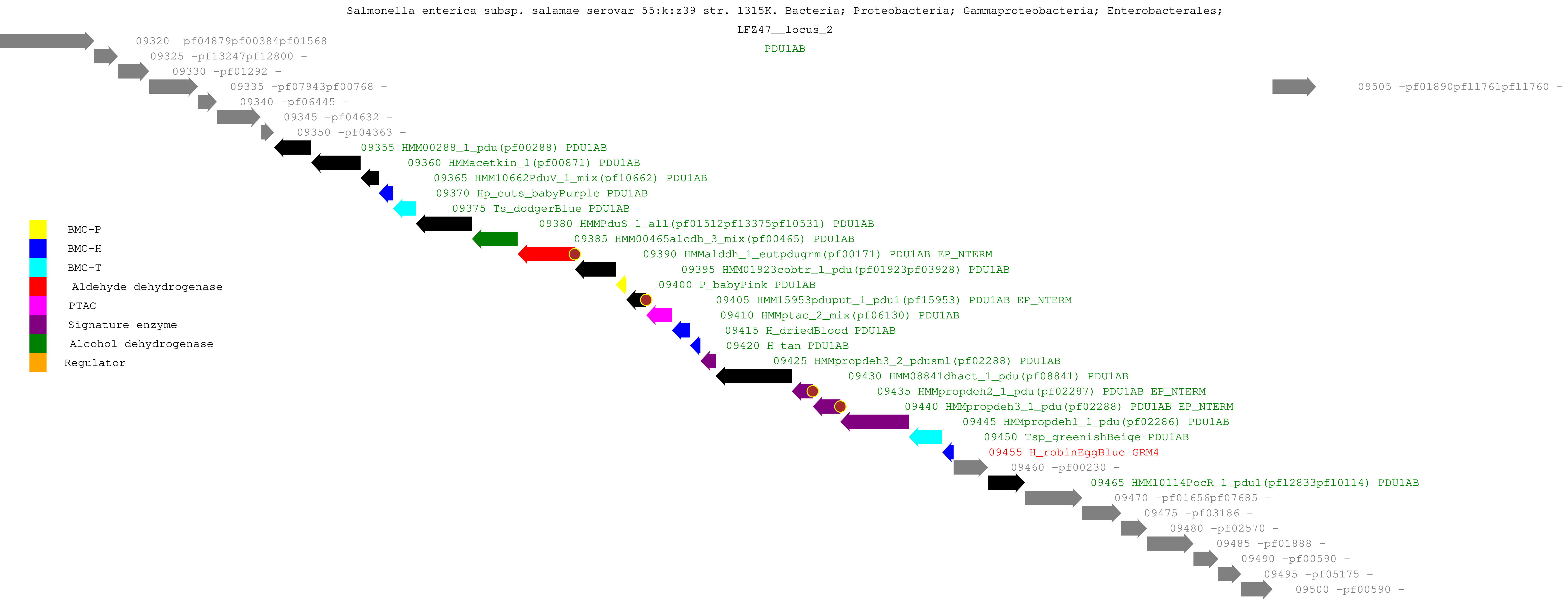
Salmonella enterica subsp. salamae serovar 55:k:z39 str. 1315K. Bacteria; Proteobacteria; Gammaproteobacteria; Enterobacterales;
LFZ47__locus_2
PDU1AB
09320 -pf04879pf00384pf01568 -
09325 -pf13247pf12800 -
09330 -pf01292 -
09335 -pf07943pf00768 -
09340 -pf06445 -
09345 -pf04632 -
09350 -pf04363 -
09355 HMM00288_1_pdu(pf00288) PDU1AB
09360 HMMacetkin_1(pf00871) PDU1AB
09365 HMM10662PduV_1_mix(pf10662) PDU1AB
09370 Hp_euts_babyPurple PDU1AB
09375 Ts_dodgerBlue PDU1AB
09380 HMMPduS_1_all(pf01512pf13375pf10531) PDU1AB
09385 HMM00465alcdh_3_mix(pf00465) PDU1AB
09390 HMMalddh_1_eutpdugrm(pf00171) PDU1AB EP_NTERM
09395 HMM01923cobtr_1_pdu(pf01923pf03928) PDU1AB
09400 P_babyPink PDU1AB
09405 HMM15953pduput_1_pdu1(pf15953) PDU1AB EP_NTERM
09410 HMMptac_2_mix(pf06130) PDU1AB
09415 H_driedBlood PDU1AB
09420 H_tan PDU1AB
09425 HMMpropdeh3_2_pdusml(pf02288) PDU1AB
09430 HMM08841dhact_1_pdu(pf08841) PDU1AB
09435 HMMpropdeh2_1_pdu(pf02287) PDU1AB EP_NTERM
09440 HMMpropdeh3_1_pdu(pf02288) PDU1AB EP_NTERM
09445 HMMpropdeh1_1_pdu(pf02286) PDU1AB
09450 Tsp_greenishBeige PDU1AB
09455 H_robinEggBlue GRM4
09460 -pf00230 -
09465 HMM10114PocR_1_pdu1(pf12833pf10114) PDU1AB
09470 -pf01656pf07685 -
09475 -pf03186 -
09480 -pf02570 -
09485 -pf01888 -
09490 -pf00590 -
09495 -pf05175 -
09500 -pf00590 -
09505 -pf01890pf11761pf11760 -
BMC-P
BMC-H
BMC-T
Aldehyde dehydrogenase
PTAC
Signature enzyme
Alcohol dehydrogenase
Regulator

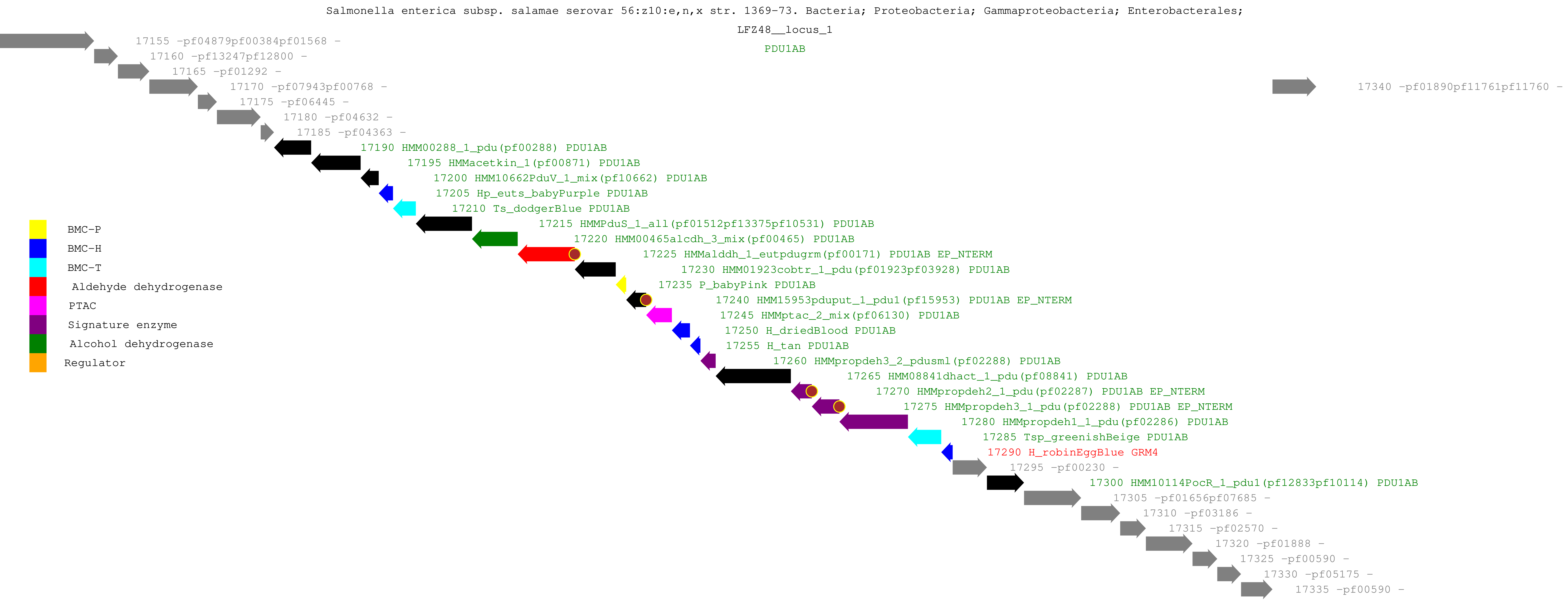
Salmonella enterica subsp. salamae serovar 56:z10:e,n,x str. 1369-73. Bacteria; Proteobacteria; Gammaproteobacteria; Enterobacterales;
LFZ48__locus_1
PDU1AB
17155 -pf04879pf00384pf01568 -
17160 -pf13247pf12800 -
17165 -pf01292 -
17170 -pf07943pf00768 -
17175 -pf06445 -
17180 -pf04632 -
17185 -pf04363 -
17190 HMM00288_1_pdu(pf00288) PDU1AB
17195 HMMacetkin_1(pf00871) PDU1AB
17200 HMM10662PduV_1_mix(pf10662) PDU1AB
17205 Hp_euts_babyPurple PDU1AB
17210 Ts_dodgerBlue PDU1AB
17215 HMMPduS_1_all(pf01512pf13375pf10531) PDU1AB
17220 HMM00465alcdh_3_mix(pf00465) PDU1AB
17225 HMMalddh_1_eutpdugrm(pf00171) PDU1AB EP_NTERM
17230 HMM01923cobtr_1_pdu(pf01923pf03928) PDU1AB
17235 P_babyPink PDU1AB
17240 HMM15953pduput_1_pdu1(pf15953) PDU1AB EP_NTERM
17245 HMMptac_2_mix(pf06130) PDU1AB
17250 H_driedBlood PDU1AB
17255 H_tan PDU1AB
17260 HMMpropdeh3_2_pdusml(pf02288) PDU1AB
17265 HMM08841dhact_1_pdu(pf08841) PDU1AB
17270 HMMpropdeh2_1_pdu(pf02287) PDU1AB EP_NTERM
17275 HMMpropdeh3_1_pdu(pf02288) PDU1AB EP_NTERM
17280 HMMpropdeh1_1_pdu(pf02286) PDU1AB
17285 Tsp_greenishBeige PDU1AB
17290 H_robinEggBlue GRM4
17295 -pf00230 -
17300 HMM10114PocR_1_pdu1(pf12833pf10114) PDU1AB
17305 -pf01656pf07685 -
17310 -pf03186 -
17315 -pf02570 -
17320 -pf01888 -
17325 -pf00590 -
17330 -pf05175 -
17335 -pf00590 -
17340 -pf01890pf11761pf11760 -
BMC-P
BMC-H
BMC-T
Aldehyde dehydrogenase
PTAC
Signature enzyme
Alcohol dehydrogenase
Regulator

Salmonella enterica I. Bacteria; Proteobacteria; Gammaproteobacteria; Enterobacterales;

LFZ50__locus_2

PDU1AB

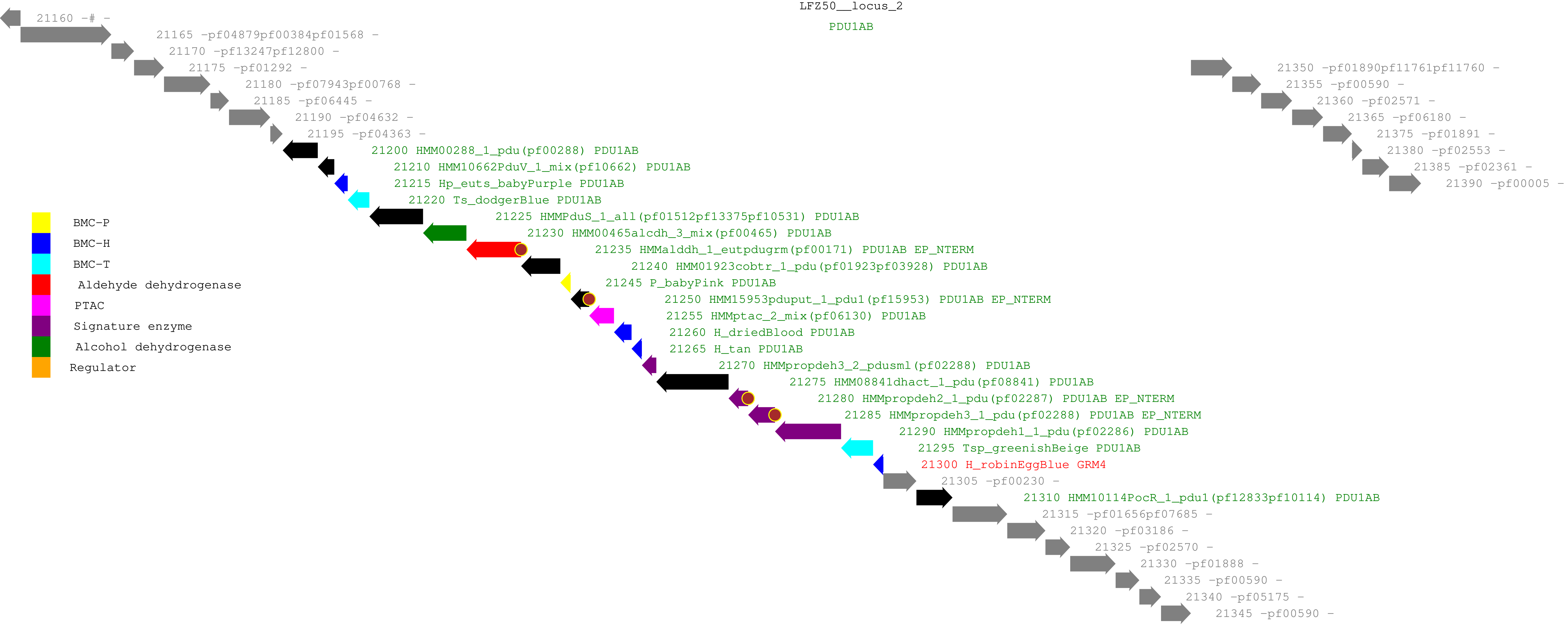

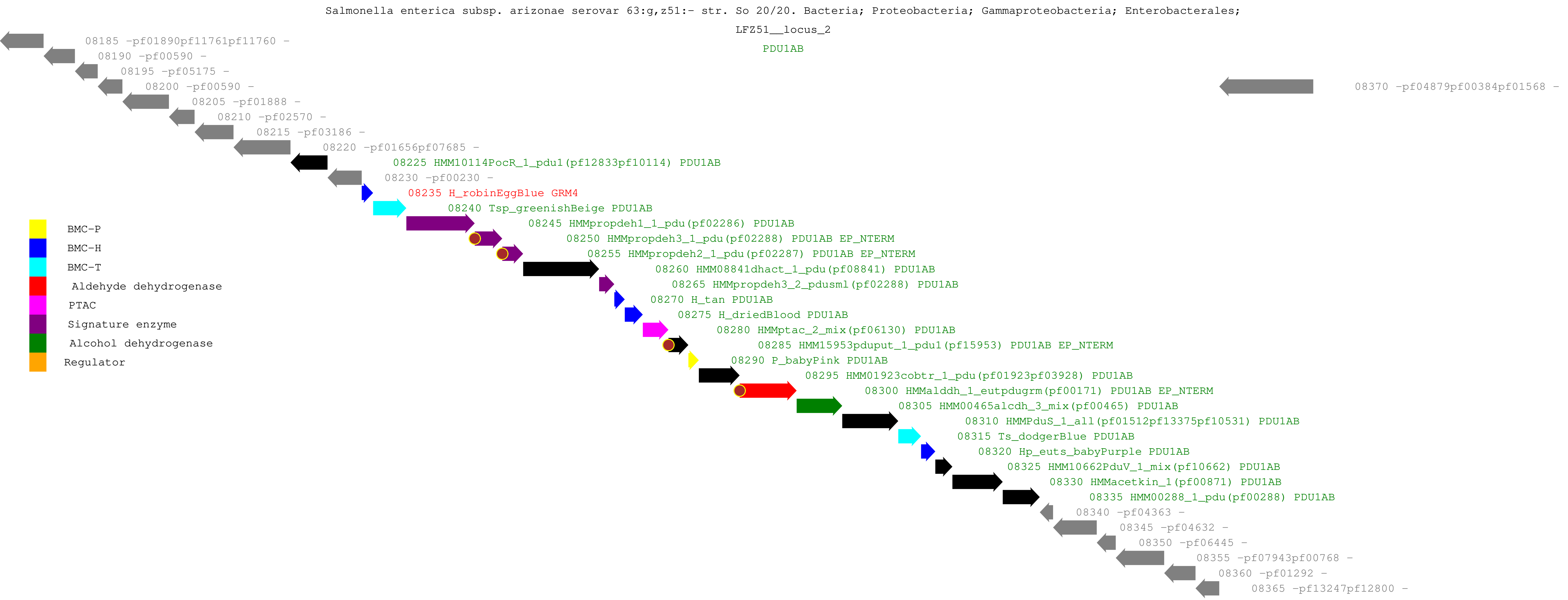

Salmonella enterica subsp. arizonae serovar 63:g,z51:- str. So 20/20. Bacteria; Proteobacteria; Gammaproteobacteria; Enterobacterales;
LFZ51__locus_2
PDU1AB
08185 -pf01890pf11761pf11760 -
08190 -pf00590 -
08195 -pf05175 -
08200 -pf00590 -
08205 -pf01888 -
08210 -pf02570 -
08215 -pf03186 -
08220 -pf01656pf07685 -
08225 HMM10114PocR_1_pdu1(pf12833pf10114) PDU1AB
08230 -pf00230 -
08235 H_robinEggBlue GRM4
08240 Tsp_greenishBeige PDU1AB
08245 HMMpropdeh1_1_pdu(pf02286) PDU1AB
08250 HMMpropdeh3_1_pdu(pf02288) PDU1AB EP_NTERM
08255 HMMpropdeh2_1_pdu(pf02287) PDU1AB EP_NTERM
08260 HMM08841dhact_1_pdu(pf08841) PDU1AB
08265 HMMpropdeh3_2_pdusml(pf02288) PDU1AB
08270 H_tan PDU1AB
08275 H_driedBlood PDU1AB
08280 HMMptac_2_mix(pf06130) PDU1AB
08285 HMM15953pduput_1_pdu1(pf15953) PDU1AB EP_NTERM
08290 P_babyPink PDU1AB
08295 HMM01923cobtr_1_pdu(pf01923pf03928) PDU1AB
08300 HMMalddh_1_eutpdugrm(pf00171) PDU1AB EP_NTERM
08305 HMM00465alcdh_3_mix(pf00465) PDU1AB
08310 HMMPduS_1_all(pf01512pf13375pf10531) PDU1AB
08315 Ts_dodgerBlue PDU1AB
08320 Hp_euts_babyPurple PDU1AB
08325 HMM10662PduV_1_mix(pf10662) PDU1AB
08330 HMMacetkin_1(pf00871) PDU1AB
08335 HMM00288_1_pdu(pf00288) PDU1AB
08340 -pf04363 -
08345 -pf04632 -
08350 -pf06445 -
08355 -pf07943pf00768 -
08360 -pf01292 -
08365 -pf13247pf12800 -
08370 -pf04879pf00384pf01568 -
BMC-P
BMC-H
BMC-T
Aldehyde dehydrogenase
PTAC
Signature enzyme
Alcohol dehydrogenase
Regulator

Salmonella enterica subsp. diarizonae serovar 50:k:z str. MZ0080. Bacteria; Proteobacteria; Gammaproteobacteria; Enterobacterales;

LFZ53__locus_2

PDU1AB

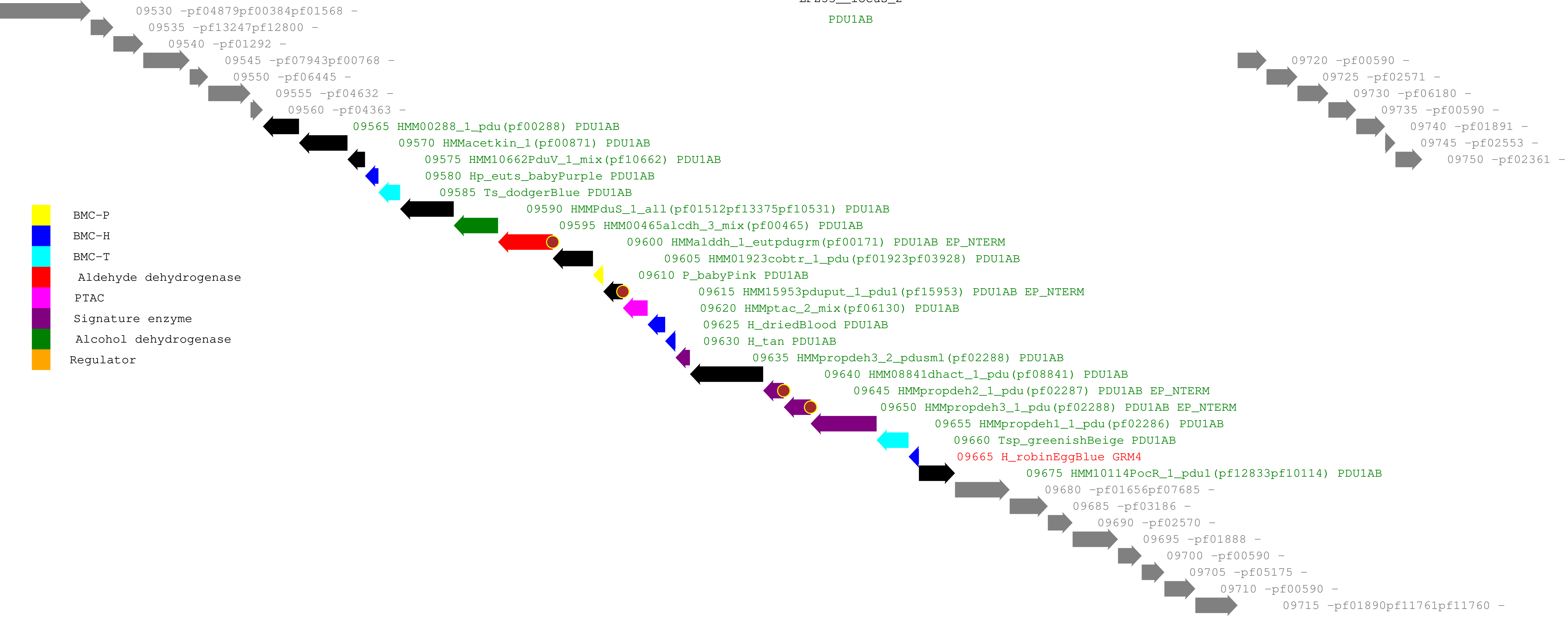

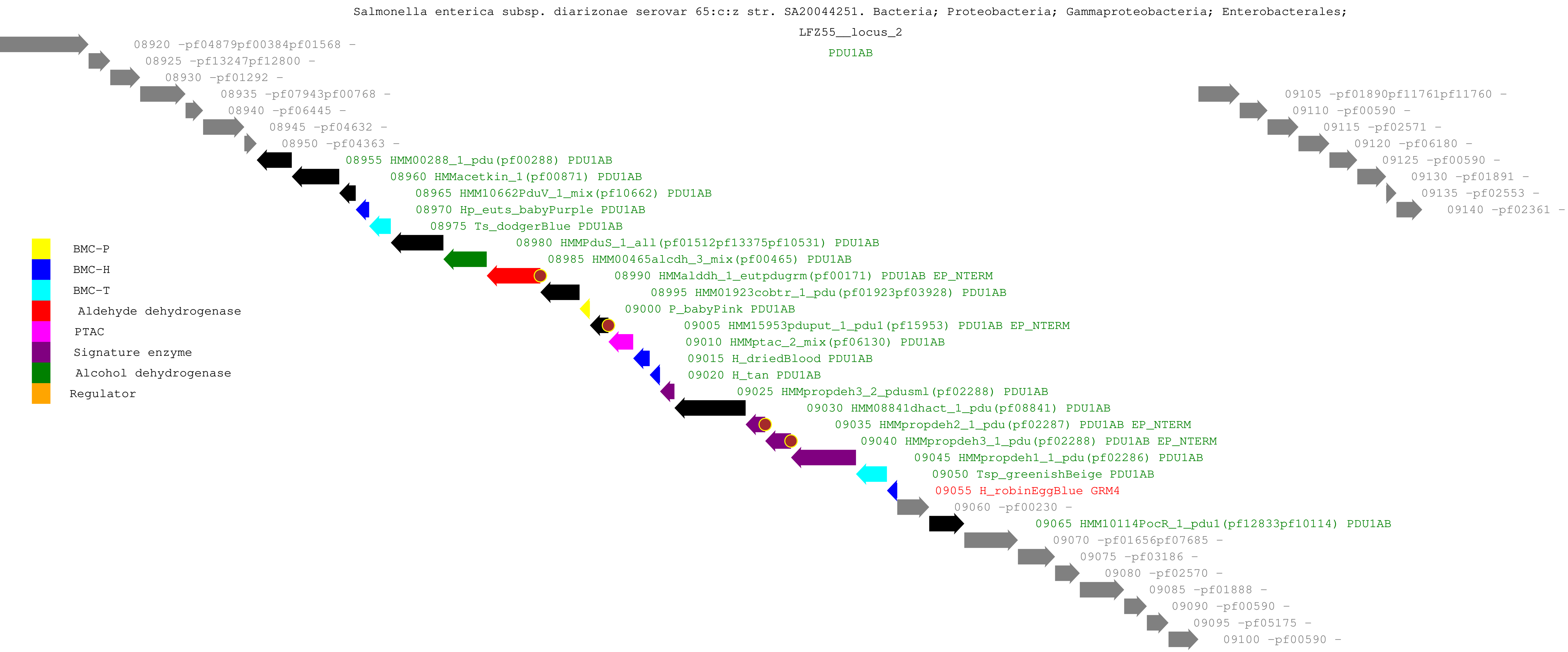

Salmonella enterica subsp. diarizonae serovar 65:c:z str. SA20044251. Bacteria; Proteobacteria; Gammaproteobacteria; Enterobacterales;
LFZ55__locus_2
PDU1AB
08920 -pf04879pf00384pf01568 -
08925 -pf13247pf12800 -
08930 -pf01292 -
08935 -pf07943pf00768 -
08940 -pf06445 -
08945 -pf04632 -
08950 -pf04363 -
08955 HMM00288_1_pdu(pf00288) PDU1AB
08960 HMMacetkin_1(pf00871) PDU1AB
08965 HMM10662PduV_1_mix(pf10662) PDU1AB
08970 Hp_euts_babyPurple PDU1AB
08975 Ts_dodgerBlue PDU1AB
08980 HMMPduS_1_all(pf01512pf13375pf10531) PDU1AB
08985 HMM00465alcdh_3_mix(pf00465) PDU1AB
08990 HMMaldh_1_eutpdugrm(pf00171) PDU1AB EP_NTERM
08995 HMM01923cobtr_1_pdu(pf01923pf03928) PDU1AB
09000 P_babyPink PDU1AB
09005 HMM15953pduput_1_pdu1(pf15953) PDU1AB EP_NTERM
09010 HMMptac_2_mix(pf06130) PDU1AB
09015 H_driedBlood PDU1AB
09020 H_tan PDU1AB
09025 HMMpropdeh3_2_pdusml(pf02288) PDU1AB
09030 HMM08841dhact_1_pdu(pf08841) PDU1AB
09035 HMMpropdeh2_1_pdu(pf02287) PDU1AB EP_NTERM
09040 HMMpropdeh3_1_pdu(pf02288) PDU1AB EP_NTERM
09045 HMMpropdeh1_1_pdu(pf02286) PDU1AB
09050 Tsp_greenishBeige PDU1AB
09055 H_robinEggBlue GRM4
09060 -pf00230 -
09065 HMM10114PocR_1_pdu1(pf12833pf10114) PDU1AB
09070 -pf01656pf07685 -
09075 -pf03186 -
09080 -pf02570 -
09085 -pf01888 -
09090 -pf00590 -
09095 -pf05175 -
09100 -pf00590 -
09105 -pf01890pf11761pf11760 -
09110 -pf00590 -
09115 -pf02571 -
09120 -pf06180 -
09125 -pf00590 -
09130 -pf01891 -
09135 -pf02553 -
09140 -pf02361 -
BMC-P
BMC-H
BMC-T
Aldehyde dehydrogenase
PTAC
Signature enzyme
Alcohol dehydrogenase
Regulator

Salmonella enterica subsp. salamae serovar 57:z29:z42. Bacteria; Proteobacteria; Gammaproteobacteria; Enterobacterales;

LFZ92__locus_2

PDU1AB

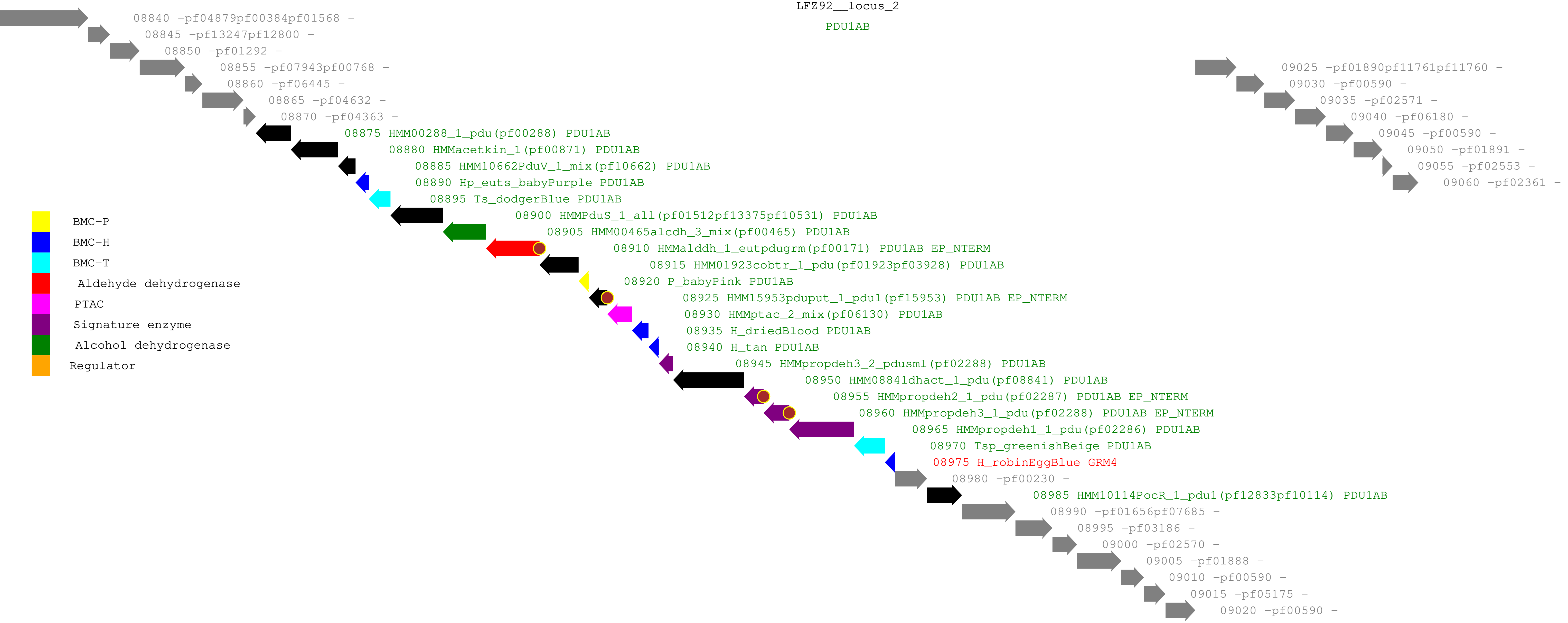

Salmonella enterica subsp. enterica serovar Adelaide str. A4-669. Bacteria; Proteobacteria; Gammaproteobacteria; Enterobacterales;

LTSEADE__locus_1

PDU1AB

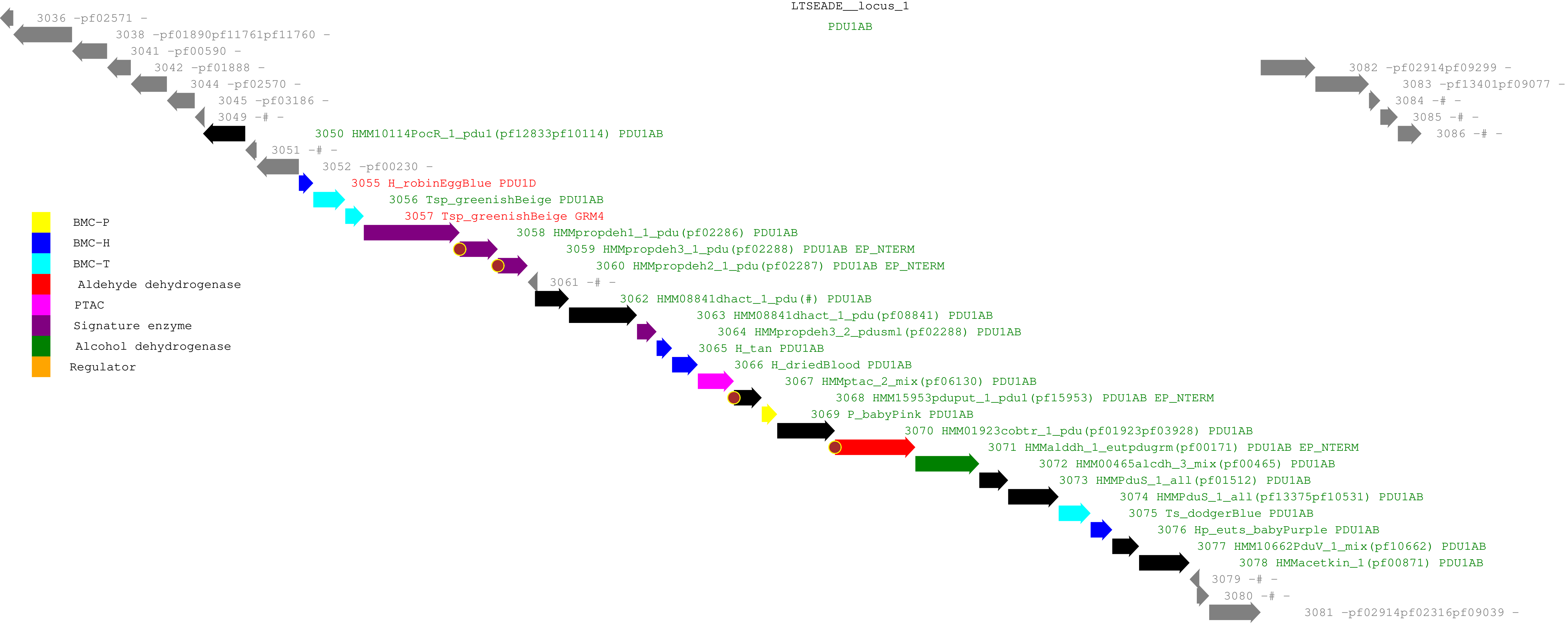

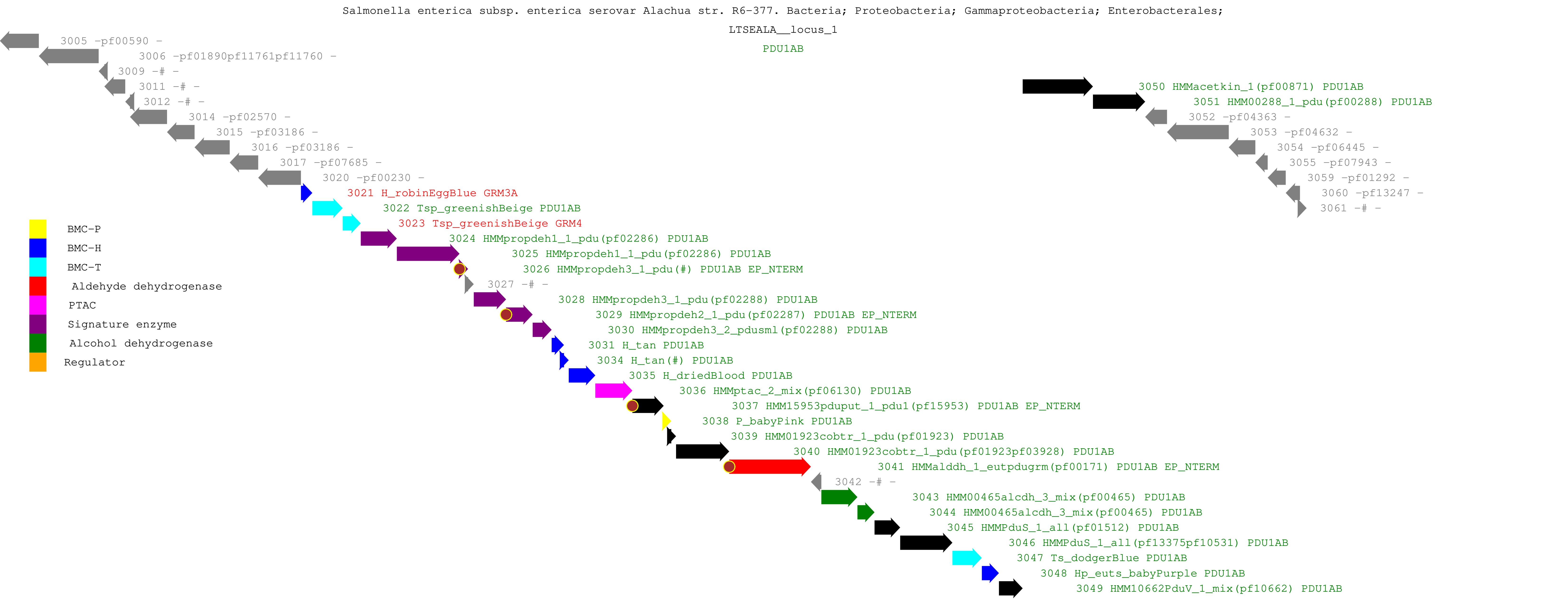

Salmonella enterica subsp. enterica serovar Alachua str. R6-377. Bacteria; Proteobacteria; Gammaproteobacteria; Enterobacterales;
LTSEALA__locus_1
PDU1AB
3005 -pf00590 -
3006 -pf01890pf11761pf11760 -
3009 -# -
3011 -# -
3012 -# -
3014 -pf02570 -
3015 -pf03186 -
3016 -pf03186 -
3017 -pf07685 -
3020 -pf00230 -
3021 H_robinEggBlue GRM3A
3022 Tsp_greenishBeige PDU1AB
3023 Tsp_greenishBeige GRM4
3024 HMMpropdeh1_1_pdu(pf02286) PDU1AB
3025 HMMpropdeh1_1_pdu(pf02286) PDU1AB
3026 HMMpropdeh3_1_pdu(#) PDU1AB EP_NTERM
3027 -# -
3028 HMMpropdeh3_1_pdu(pf02288) PDU1AB
3029 HMMpropdeh2_1_pdu(pf02287) PDU1AB EP_NTERM
3030 HMMpropdeh3_2_pdusml(pf02288) PDU1AB
3031 H_tan PDU1AB
3034 H_tan(#) PDU1AB
3035 H_driedBlood PDU1AB
3036 HMMptac_2_mix(pf06130) PDU1AB
3037 HMM15953pduput_1_pdu1(pf15953) PDU1AB EP_NTERM
3038 P_babyPink PDU1AB
3039 HMM01923cobtr_1_pdu(pf01923) PDU1AB
3040 HMM01923cobtr_1_pdu(pf01923pf03928) PDU1AB
3041 HMMaldh_1_eutpdugrm(pf00171) PDU1AB EP_NTERM
3042 -# -
3043 HMM00465alcdh_3_mix(pf00465) PDU1AB
3044 HMM00465alcdh_3_mix(pf00465) PDU1AB
3045 HMMPduS_1_all(pf01512) PDU1AB
3046 HMMPduS_1_all(pf13375pf10531) PDU1AB
3047 Ts_dodgerBlue PDU1AB
3048 Hp_euts_babyPurple PDU1AB
3049 HMM10662PduV_1_mix(pf10662) PDU1AB
3050 HMMacetkin_1(pf00871) PDU1AB
3051 HMM00288_1_pdu(pf00288) PDU1AB
3052 -pf04363 -
3053 -pf04632 -
3054 -pf06445 -
3055 -pf07943 -
3059 -pf01292 -
3060 -pf13247 -
3061 -# -
BMC-P
BMC-H
BMC-T
Aldehyde dehydrogenase
PTAC
Signature enzyme
Alcohol dehydrogenase
Regulator

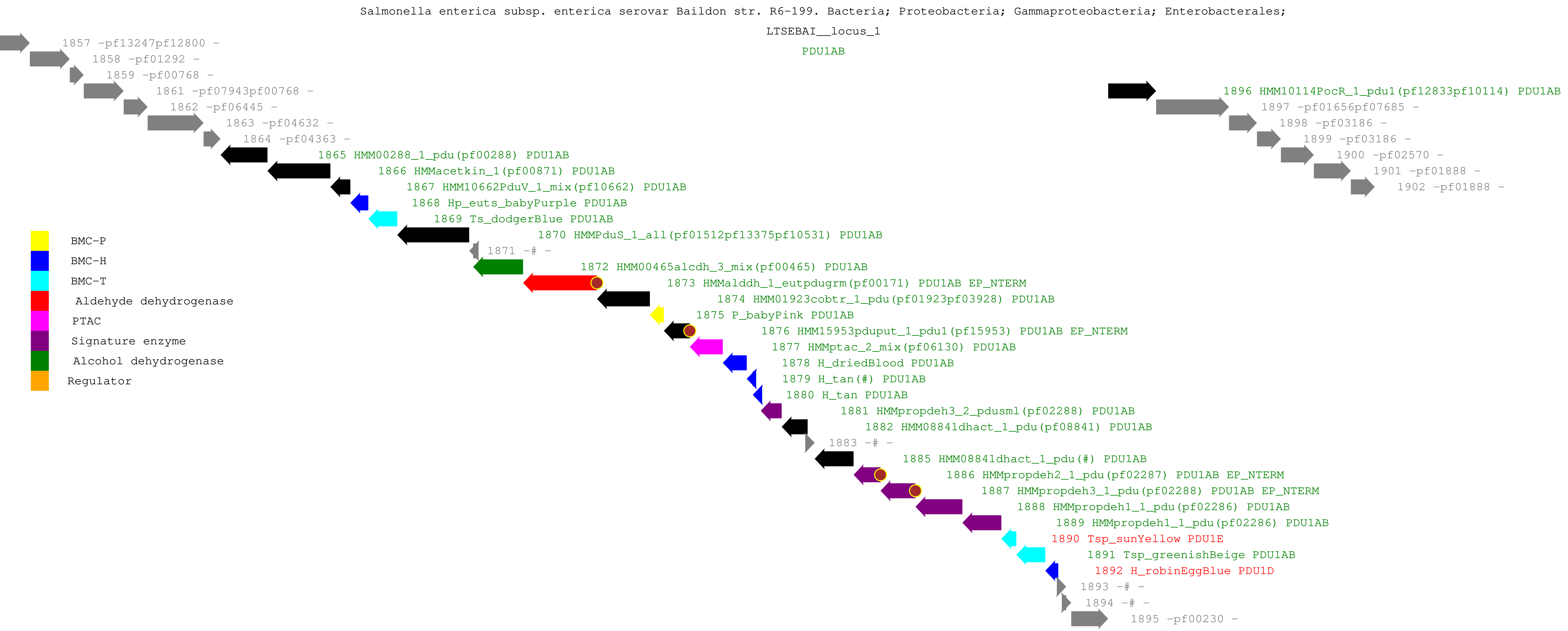
Salmonella enterica subsp. enterica serovar Baildon str. R6-199. Bacteria; Proteobacteria; Gammaproteobacteria; Enterobacterales;
LTSEBAI__locus_1
PDU1AB
1857 -pf13247pf12800 -
1858 -pf01292 -
1859 -pf00768 -
1861 -pf07943pf00768 -
1862 -pf06445 -
1863 -pf04632 -
1864 -pf04363 -
1865 HMM00288_1_pdu(pf00288) PDU1AB
1866 HMMacetkin_1(pf00871) PDU1AB
1867 HMM10662PduV_1_mix(pf10662) PDU1AB
1868 Hp_euts_babyPurple PDU1AB
1869 Ts_dodgerBlue PDU1AB
1870 HMMPduS_1_all(pf01512pf13375pf10531) PDU1AB
1871 -# -
1872 HMM00465alcdh_3_mix(pf00465) PDU1AB
1873 HMMalddh_1_eutpdugrm(pf00171) PDU1AB EP_NTERM
1874 HMM01923cobtr_1_pdu(pf01923pf03928) PDU1AB
1875 P_babyPink PDU1AB
1876 HMM15953pduput_1_pdu1(pf15953) PDU1AB EP_NTERM
1877 HMMptac_2_mix(pf06130) PDU1AB
1878 H_driedBlood PDU1AB
1879 H_tan(#) PDU1AB
1880 H_tan PDU1AB
1881 HMMpropdeh3_2_pdusml(pf02288) PDU1AB
1882 HMM08841dhact_1_pdu(pf08841) PDU1AB
1883 -# -
1885 HMM08841dhact_1_pdu(#) PDU1AB
1886 HMMpropdeh2_1_pdu(pf02287) PDU1AB EP_NTERM
1887 HMMpropdeh3_1_pdu(pf02288) PDU1AB EP_NTERM
1888 HMMpropdeh1_1_pdu(pf02286) PDU1AB
1889 HMMpropdeh1_1_pdu(pf02286) PDU1AB
1890 Tsp_sunYellow PDU1E
1891 Tsp_greenishBeige PDU1AB
1892 H_robinEggBlue PDU1D
1893 -# -
1894 -# -
1895 -pf00230 -
1896 HMM10114PocR_1_pdu1(pf12833pf10114) PDU1AB
1897 -pf01656pf07685 -
1898 -pf03186 -
1899 -pf03186 -
1900 -pf02570 -
1901 -pf01888 -
1902 -pf01888 -
BMC-P
BMC-H
BMC-T
Aldehyde dehydrogenase
PTAC
Signature enzyme
Alcohol dehydrogenase
Regulator

Salmonella enterica subsp. enterica serovar Give str. S5-487. Bacteria; Proteobacteria; Gammaproteobacteria; Enterobacterales;

LTSEGIV__locus_2

PDU1AB

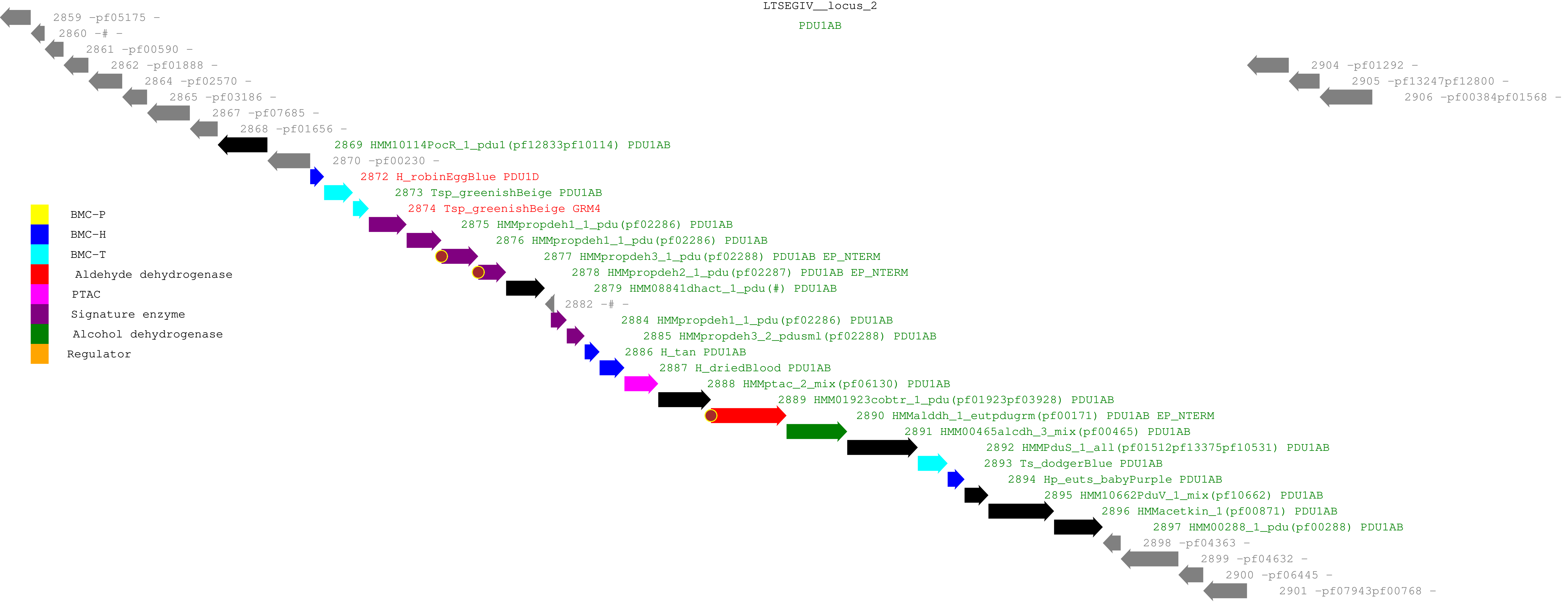

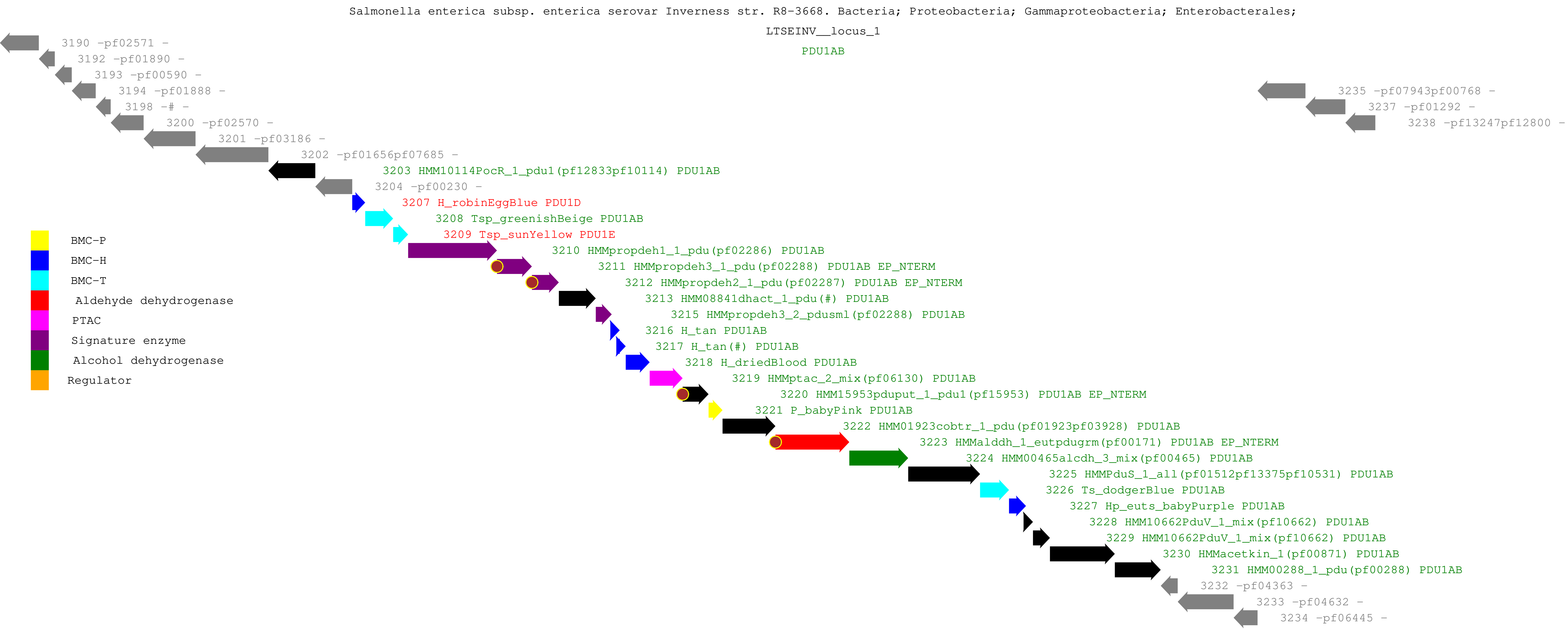
Salmonella enterica subsp. enterica serovar Inverness str. R8-3668. Bacteria; Proteobacteria; Gammaproteobacteria; Enterobacterales;
LTSEINV__locus_1
PDU1AB
3190 -pf02571 -
3192 -pf01890 -
3193 -pf00590 -
3194 -pf01888 -
3198 -# -
3200 -pf02570 -
3201 -pf03186 -
3202 -pf01656pf07685 -
3203 HMM10114PocR_1_pdu1(pf12833pf10114) PDU1AB
3204 -pf00230 -
3207 H_robinEggBlue PDU1D
3208 Tsp_greenishBeige PDU1AB
3209 Tsp_sunYellow PDU1E
3210 HMMpropdeh1_1_pdu(pf02286) PDU1AB
3211 HMMpropdeh3_1_pdu(pf02288) PDU1AB EP_NTERM
3212 HMMpropdeh2_1_pdu(pf02287) PDU1AB EP_NTERM
3213 HMM08841dhact_1_pdu(#) PDU1AB
3215 HMMpropdeh3_2_pdusml(pf02288) PDU1AB
3216 H_tan PDU1AB
3217 H_tan(#) PDU1AB
3218 H_driedBlood PDU1AB
3219 HMMptac_2_mix(pf06130) PDU1AB
3220 HMM15953pduput_1_pdu1(pf15953) PDU1AB EP_NTERM
3221 P_babyPink PDU1AB
3222 HMM01923cobtr_1_pdu(pf01923pf03928) PDU1AB
3223 HMMalddh_1_eutpdugrm(pf00171) PDU1AB EP_NTERM
3224 HMM00465alcdh_3_mix(pf00465) PDU1AB
3225 HMMPduS_1_all(pf01512pf13375pf10531) PDU1AB
3226 Ts_dodgerBlue PDU1AB
3227 Hp_euts_babyPurple PDU1AB
3228 HMM10662PduV_1_mix(pf10662) PDU1AB
3229 HMM10662PduV_1_mix(pf10662) PDU1AB
3230 HMMacetkin_1(pf00871) PDU1AB
3231 HMM00288_1_pdu(pf00288) PDU1AB
3232 -pf04363 -
3233 -pf04632 -
3234 -pf06445 -
3235 -pf07943pf00768 -
3237 -pf01292 -
3238 -pf13247pf12800 -
BMC-P
BMC-H
BMC-T
Aldehyde dehydrogenase
PTAC
Signature enzyme
Alcohol dehydrogenase
Regulator

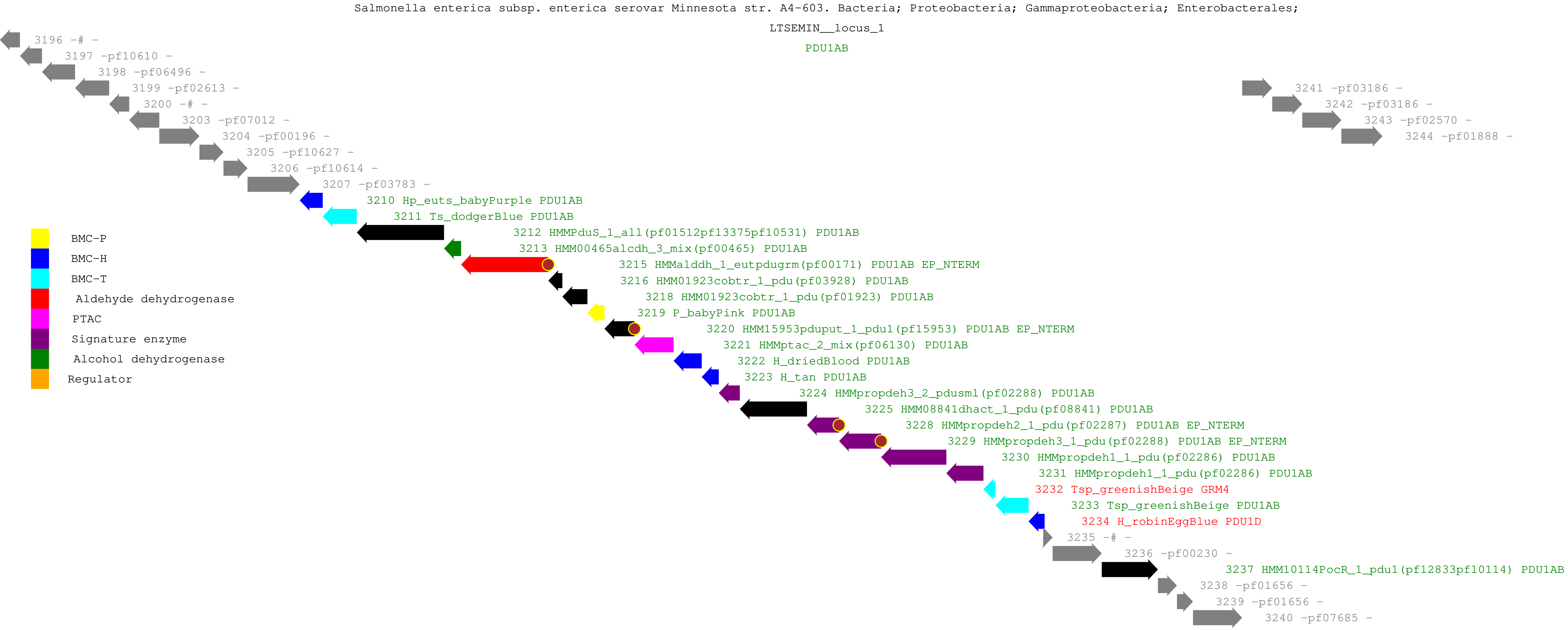

Salmonella enterica subsp. enterica serovar Minnesota str. A4-603. Bacteria; Proteobacteria; Gammaproteobacteria; Enterobacterales;
LTSEMIN__locus_1
PDU1AB
3196 -# -
3197 -pf10610 -
3198 -pf06496 -
3199 -pf02613 -
3200 -# -
3203 -pf07012 -
3204 -pf00196 -
3205 -pf10627 -
3206 -pf10614 -
3207 -pf03783 -
3210 Hp_euts_babyPurple PDU1AB
3211 Ts_dodgerBlue PDU1AB
3212 HMMPduS_1_all(pf01512pf13375pf10531) PDU1AB
3213 HMM00465alcdh_3_mix(pf00465) PDU1AB
3215 HMMalddh_1_eutpdugrm(pf00171) PDU1AB EP_NTERM
3216 HMM01923cobtr_1_pdu(pf03928) PDU1AB
3218 HMM01923cobtr_1_pdu(pf01923) PDU1AB
3219 P_babyPink PDU1AB
3220 HMM15953pduput_1_pdu1(pf15953) PDU1AB EP_NTERM
3221 HMMptac_2_mix(pf06130) PDU1AB
3222 H_driedBlood PDU1AB
3223 H_tan PDU1AB
3224 HMMpropdeh3_2_pdusml(pf02288) PDU1AB
3225 HMM08841dhact_1_pdu(pf08841) PDU1AB
3228 HMMpropdeh2_1_pdu(pf02287) PDU1AB EP_NTERM
3229 HMMpropdeh3_1_pdu(pf02288) PDU1AB EP_NTERM
3230 HMMpropdeh1_1_pdu(pf02286) PDU1AB
3231 HMMpropdeh1_1_pdu(pf02286) PDU1AB
3232 Tsp_greenishBeige GRM4
3233 Tsp_greenishBeige PDU1AB
3234 H_robinEggBlue PDU1D
3235 -# -
3236 -pf00230 -
3237 HMM10114PocR_1_pdu1(pf12833pf10114) PDU1AB
3238 -pf01656 -
3239 -pf01656 -
3240 -pf07685 -
3241 -pf03186 -
3242 -pf03186 -
3243 -pf02570 -
3244 -pf01888 -
BMC-P
BMC-H
BMC-T
Aldehyde dehydrogenase
PTAC
Signature enzyme
Alcohol dehydrogenase
Regulator

Salmonella enterica subsp. enterica serovar Montevideo str. S5-403. Bacteria; Proteobacteria; Gammaproteobacteria; Enterobacterales;

LTSEMON__locus_1

PDU1AB

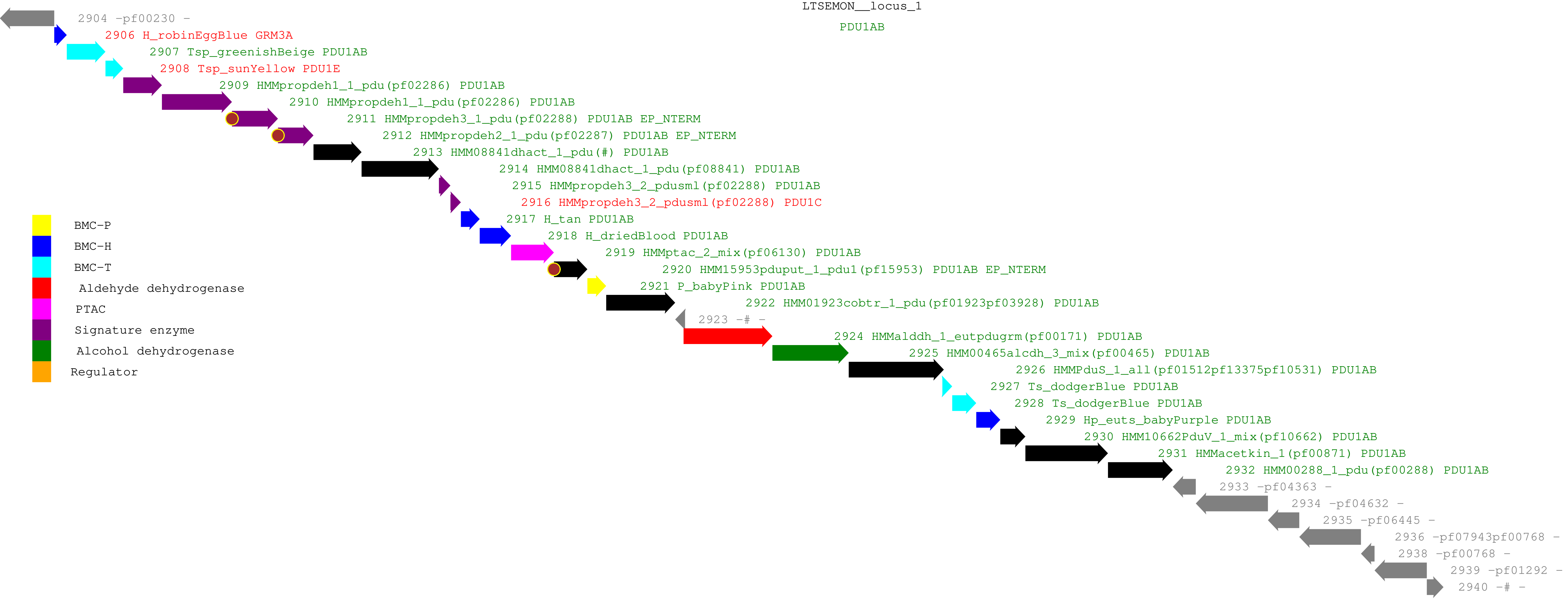

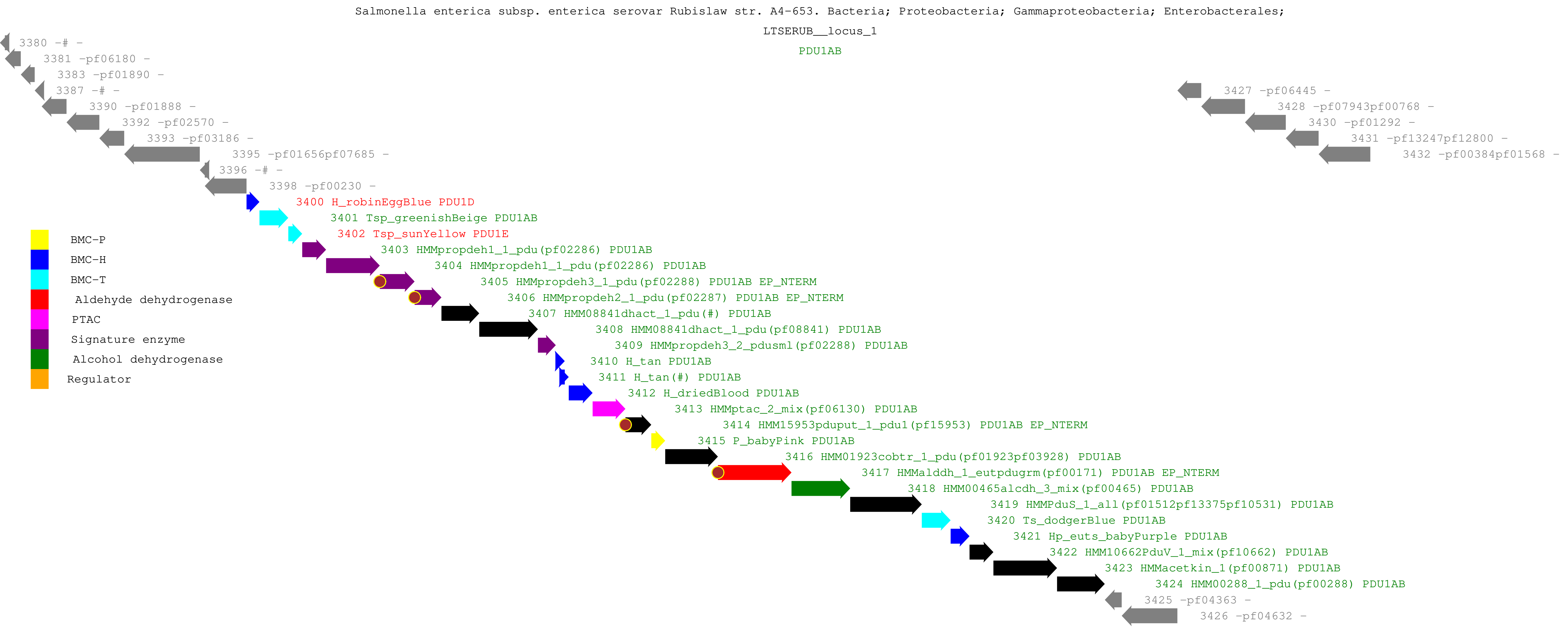
Salmonella enterica subsp. enterica serovar Rubislaw str. A4-653. Bacteria; Proteobacteria; Gammaproteobacteria; Enterobacterales;
LTSERUB__locus_1
PDU1AB
3380 -# -
3381 -pf06180 -
3383 -pf01890 -
3387 -# -
3390 -pf01888 -
3392 -pf02570 -
3393 -pf03186 -
3395 -pf01656pf07685 -
3396 -# -
3398 -pf00230 -
3400 H_robinEggBlue PDU1D
3401 Tsp_greenishBeige PDU1AB
3402 Tsp_sunYellow PDU1E
3403 HMMpropdeh1_1_pdu(pf02286) PDU1AB
3404 HMMpropdeh1_1_pdu(pf02286) PDU1AB
3405 HMMpropdeh3_1_pdu(pf02288) PDU1AB EP_NTERM
3406 HMMpropdeh2_1_pdu(pf02287) PDU1AB EP_NTERM
3407 HMM08841dhact_1_pdu(#) PDU1AB
3408 HMM08841dhact_1_pdu(pf08841) PDU1AB
3409 HMMpropdeh3_2_pdusml(pf02288) PDU1AB
3410 H_tan PDU1AB
3411 H_tan(#) PDU1AB
3412 H_driedBlood PDU1AB
3413 HMMptac_2_mix(pf06130) PDU1AB
3414 HMM15953pduput_1_pdu1(pf15953) PDU1AB EP_NTERM
3415 P_babyPink PDU1AB
3416 HMM01923cobtr_1_pdu(pf01923pf03928) PDU1AB
3417 HMMalddh_1_eutpdugrm(pf00171) PDU1AB EP_NTERM
3418 HMM00465alcdh_3_mix(pf00465) PDU1AB
3419 HMMPduS_1_all(pf01512pf13375pf10531) PDU1AB
3420 Ts_dodgerBlue PDU1AB
3421 Hp_euts_babyPurple PDU1AB
3422 HMM10662PduV_1_mix(pf10662) PDU1AB
3423 HMMacetkin_1(pf00871) PDU1AB
3424 HMM00288_1_pdu(pf00288) PDU1AB
3425 -pf04363 -
3426 -pf04632 -
3427 -pf06445 -
3428 -pf07943pf00768 -
3430 -pf01292 -
3431 -pf13247pf12800 -
3432 -pf00384pf01568 -
BMC-P
BMC-H
BMC-T
Aldehyde dehydrogenase
PTAC
Signature enzyme
Alcohol dehydrogenase
Regulator

Salmonella enterica subsp. enterica serovar Senftenberg str. A4-543. Bacteria; Proteobacteria; Gammaproteobacteria; Enterobacterales;

LTSESEN__locus_1

PDU1AB

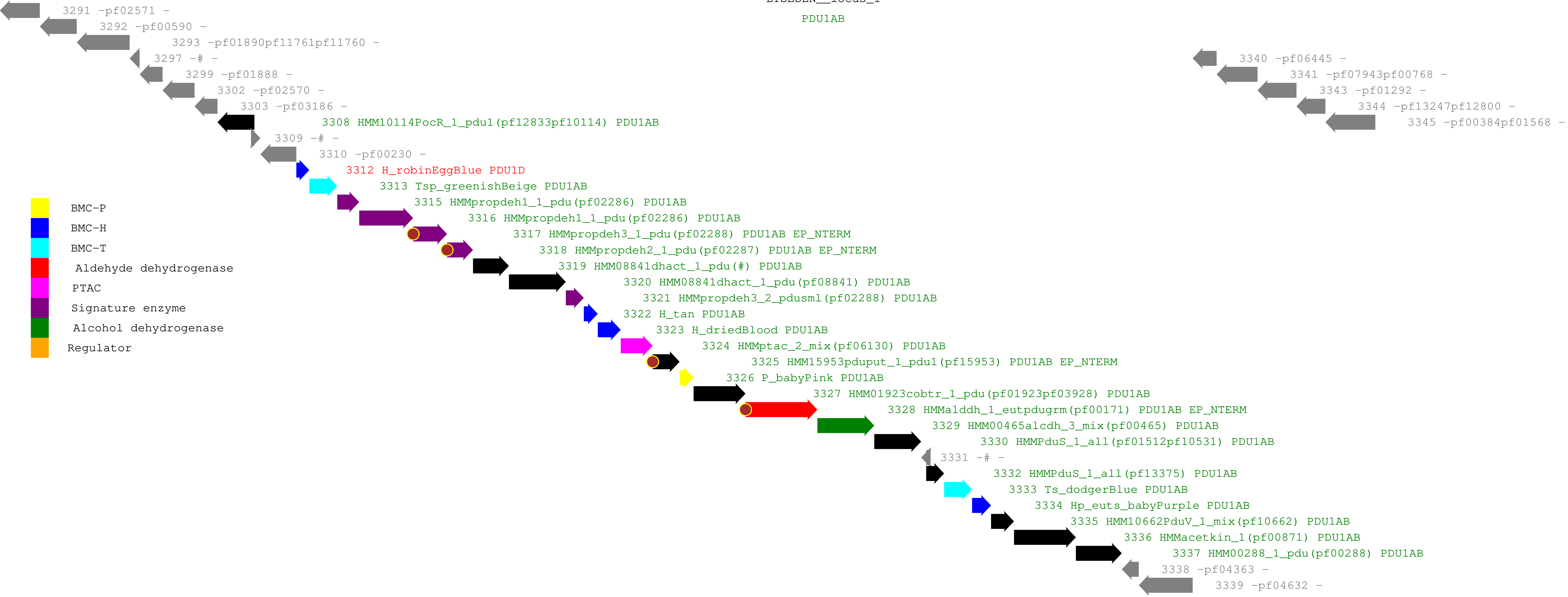

Salmonella enterica subsp. enterica serovar Uganda str. R8-3404. Bacteria; Proteobacteria; Gammaproteobacteria; Enterobacterales;

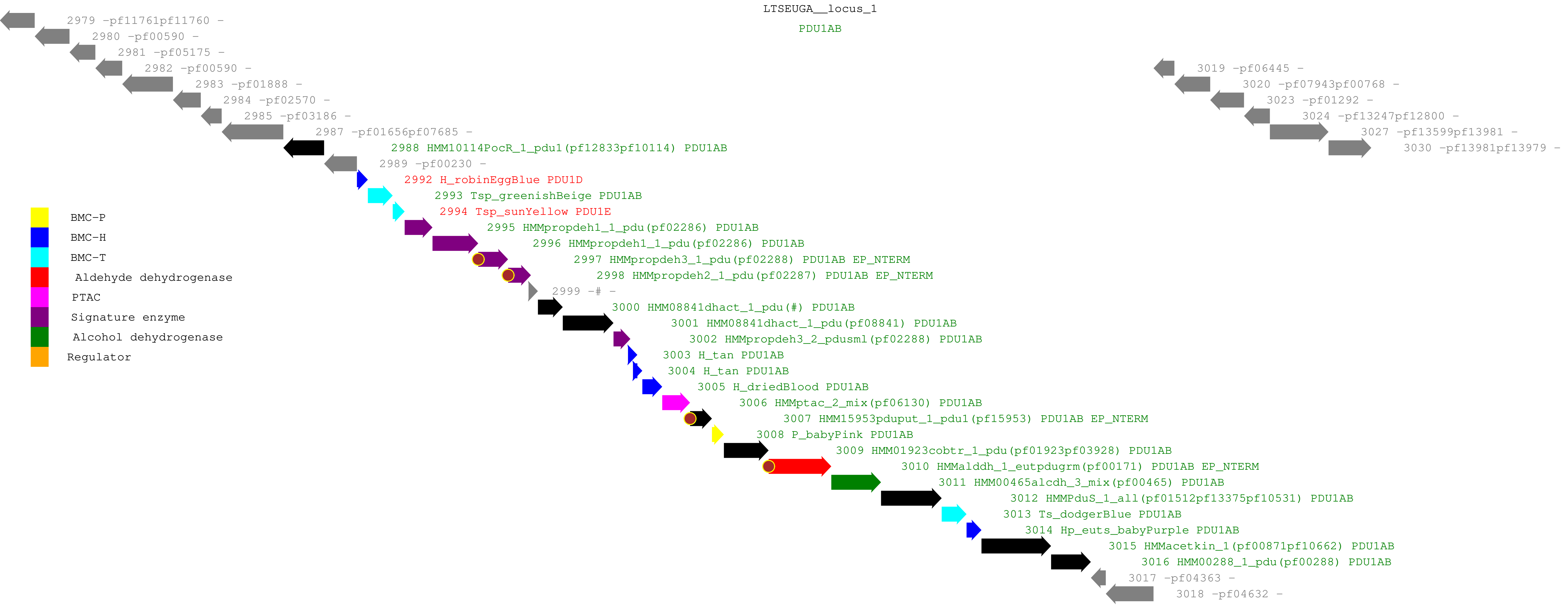

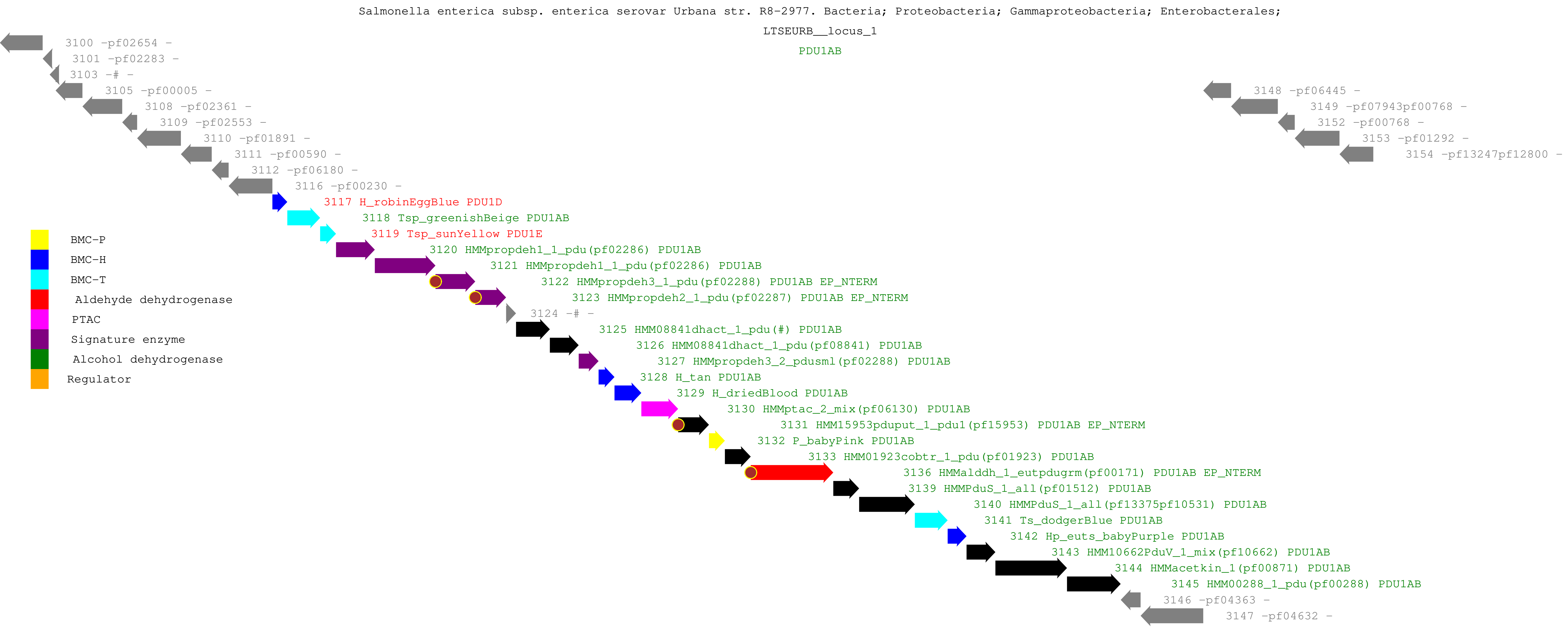

Salmonella enterica subsp. enterica serovar Urbana str. R8-2977. Bacteria; Proteobacteria; Gammaproteobacteria; Enterobacterales;
LTSEURB__locus_1
PDU1AB
3100 -pf02654 -
3101 -pf02283 -
3103 -# -
3105 -pf00005 -
3108 -pf02361 -
3109 -pf02553 -
3110 -pf01891 -
3111 -pf00590 -
3112 -pf06180 -
3116 -pf00230 -
3117 H_robinEggBlue PDU1D
3118 Tsp_greenishBeige PDU1AB
3119 Tsp_sunYellow PDU1E
3120 HMMpropdeh1_1_pdu(pf02286) PDU1AB
3121 HMMpropdeh1_1_pdu(pf02286) PDU1AB
3122 HMMpropdeh3_1_pdu(pf02288) PDU1AB EP_NTERM
3123 HMMpropdeh2_1_pdu(pf02287) PDU1AB EP_NTERM
3124 -# -
3125 HMM08841dhact_1_pdu(#) PDU1AB
3126 HMM08841dhact_1_pdu(pf08841) PDU1AB
3127 HMMpropdeh3_2_pdusml(pf02288) PDU1AB
3128 H_tan PDU1AB
3129 H_driedBlood PDU1AB
3130 HMMptac_2_mix(pf06130) PDU1AB
3131 HMM15953pduput_1_pdu1(pf15953) PDU1AB EP_NTERM
3132 P_babyPink PDU1AB
3133 HMM01923cobtr_1_pdu(pf01923) PDU1AB
3136 HMMalddh_1_eutpdugrm(pf00171) PDU1AB EP_NTERM
3139 HMMPduS_1_all(pf01512) PDU1AB
3140 HMMPduS_1_all(pf13375pf10531) PDU1AB
3141 Ts_dodgerBlue PDU1AB
3142 Hp_euts_babyPurple PDU1AB
3143 HMM10662PduV_1_mix(pf10662) PDU1AB
3144 HMMacetkin_1(pf00871) PDU1AB
3145 HMM00288_1_pdu(pf00288) PDU1AB
3146 -pf04363 -
3147 -pf04632 -
3148 -pf06445 -
3149 -pf07943pf00768 -
3152 -pf00768 -
3153 -pf01292 -
3154 -pf13247pf12800 -
BMC-P
BMC-H
BMC-T
Aldehyde dehydrogenase
PTAC
Signature enzyme
Alcohol dehydrogenase
Regulator

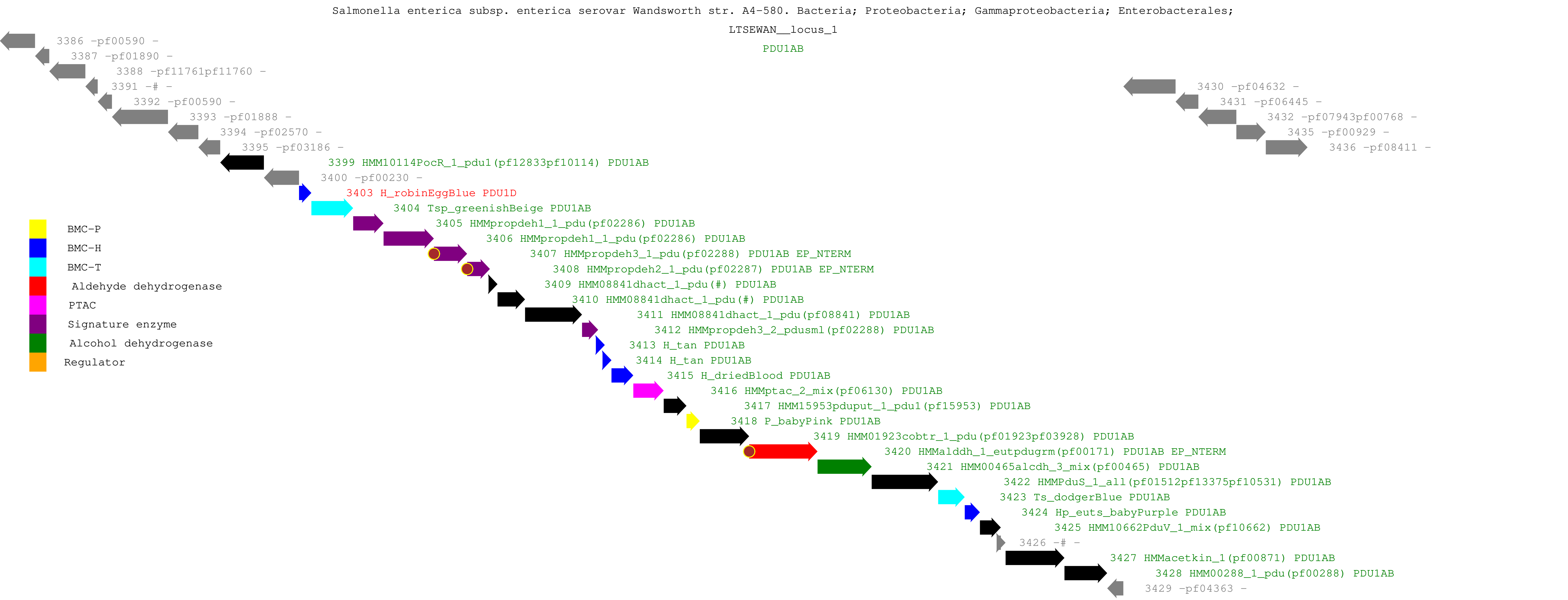
Salmonella enterica subsp. enterica serovar Wandsworth str. A4-580. Bacteria; Proteobacteria; Gammaproteobacteria; Enterobacterales;
LTSEWAN__locus_1
PDU1AB
3386 -pf00590 -
3387 -pf01890 -
3388 -pf11761pf11760 -
3391 -# -
3392 -pf00590 -
3393 -pf01888 -
3394 -pf02570 -
3395 -pf03186 -
3399 HMM10114PocR_1_pdu1(pf12833pf10114) PDU1AB
3400 -pf00230 -
3403 H_robinEggBlue PDU1D
3404 Tsp_greenishBeige PDU1AB
3405 HMMpropdeh1_1_pdu(pf02286) PDU1AB
3406 HMMpropdeh1_1_pdu(pf02286) PDU1AB
3407 HMMpropdeh3_1_pdu(pf02288) PDU1AB EP_NTERM
3408 HMMpropdeh2_1_pdu(pf02287) PDU1AB EP_NTERM
3409 HMM08841dhact_1_pdu(#) PDU1AB
3410 HMM08841dhact_1_pdu(#) PDU1AB
3411 HMM08841dhact_1_pdu(pf08841) PDU1AB
3412 HMMpropdeh3_2_pdusml(pf02288) PDU1AB
3413 H_tan PDU1AB
3414 H_tan PDU1AB
3415 H_driedBlood PDU1AB
3416 HMMptac_2_mix(pf06130) PDU1AB
3417 HMM15953pduput_1_pdu1(pf15953) PDU1AB
3418 P_babyPink PDU1AB
3419 HMM01923cobtr_1_pdu(pf01923pf03928) PDU1AB
3420 HMMalddh_1_eutpdugrm(pf00171) PDU1AB EP_NTERM
3421 HMM00465alcdh_3_mix(pf00465) PDU1AB
3422 HMMPduS_1_all(pf01512pf13375pf10531) PDU1AB
3423 Ts_dodgerBlue PDU1AB
3424 Hp_euts_babyPurple PDU1AB
3425 HMM10662PduV_1_mix(pf10662) PDU1AB
3426 -# -
3427 HMMacetkin_1(pf00871) PDU1AB
3428 HMM00288_1_pdu(pf00288) PDU1AB
3429 -pf04363 -
3430 -pf04632 -
3431 -pf06445 -
3432 -pf07943pf00768 -
3435 -pf00929 -
3436 -pf08411 -
BMC-P
BMC-H
BMC-T
Aldehyde dehydrogenase
PTAC
Signature enzyme
Alcohol dehydrogenase
Regulator

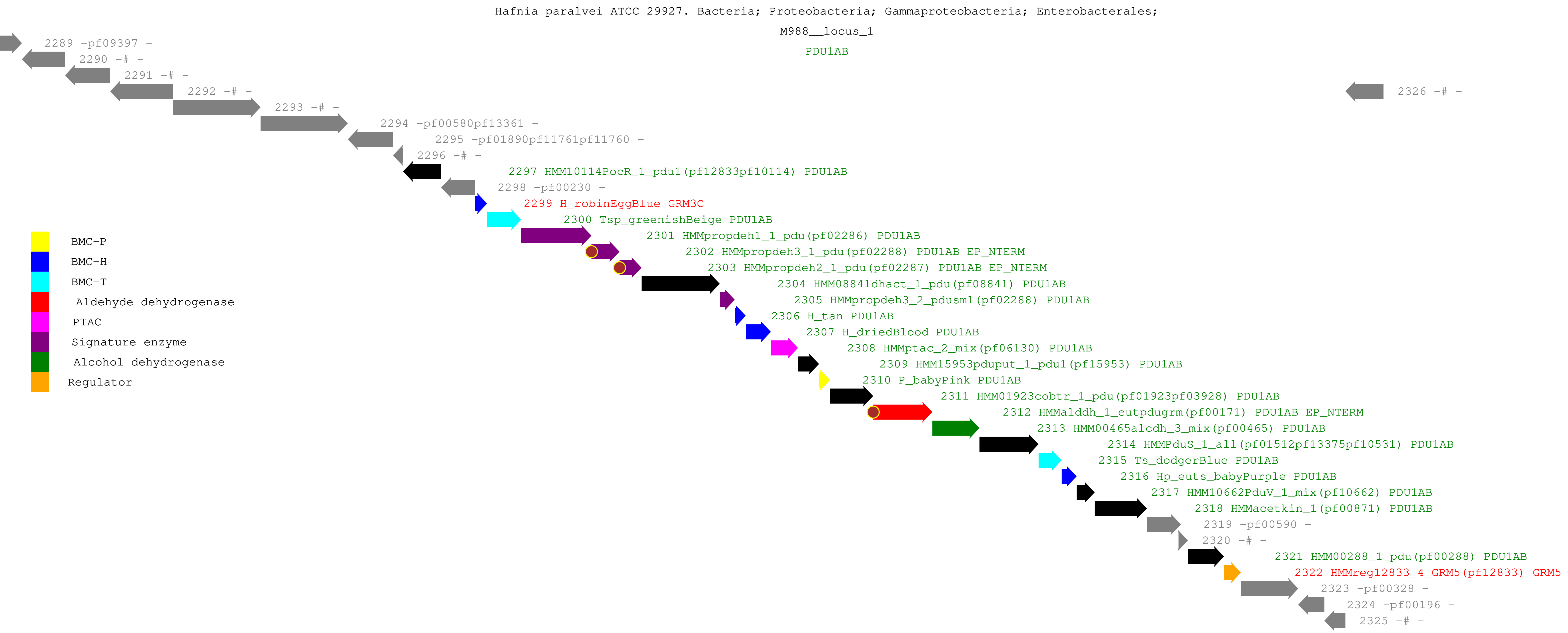
Hafnia paralvei ATCC 29927. Bacteria; Proteobacteria; Gammaproteobacteria; Enterobacterales;
M988__locus_1
PDU1AB
2289 -pf09397 -
2290 -# -
2291 -# -
2292 -# -
2293 -# -
2294 -pf00580pf13361 -
2295 -pf01890pf11761pf11760 -
2296 -# -
2297 HMM10114PocR_1_pdu1(pf12833pf10114) PDU1AB
2298 -pf00230 -
2299 H_robinEggBlue GRM3C
2300 Tsp_greenishBeige PDU1AB
2301 HMMpropdeh1_1_pdu(pf02286) PDU1AB
2302 HMMpropdeh3_1_pdu(pf02288) PDU1AB EP_NTERM
2303 HMMpropdeh2_1_pdu(pf02287) PDU1AB EP_NTERM
2304 HMM08841dhact_1_pdu(pf08841) PDU1AB
2305 HMMpropdeh3_2_pdusml(pf02288) PDU1AB
2306 H_tan PDU1AB
2307 H_driedBlood PDU1AB
2308 HMMptac_2_mix(pf06130) PDU1AB
2309 HMM15953pduput_1_pdu1(pf15953) PDU1AB
2310 P_babyPink PDU1AB
2311 HMM01923cobtr_1_pdu(pf01923pf03928) PDU1AB
2312 HMMaldh_1_eutpdugrm(pf00171) PDU1AB EP_NTERM
2313 HMM00465alcdh_3_mix(pf00465) PDU1AB
2314 HMMPduS_1_all(pf01512pf13375pf10531) PDU1AB
2315 Ts_dodgerBlue PDU1AB
2316 Hp_euts_babyPurple PDU1AB
2317 HMM10662PduV_1_mix(pf10662) PDU1AB
2318 HMMacetkin_1(pf00871) PDU1AB
2319 -pf00590 -
2320 -# -
2321 HMM00288_1_pdu(pf00288) PDU1AB
2322 HMMreg12833_4_GRM5(pf12833) GRM5
2323 -pf00328 -
2324 -pf00196 -
2325 -# -
2326 -# -
BMC-P
BMC-H
BMC-T
Aldehyde dehydrogenase
PTAC
Signature enzyme
Alcohol dehydrogenase
Regulator

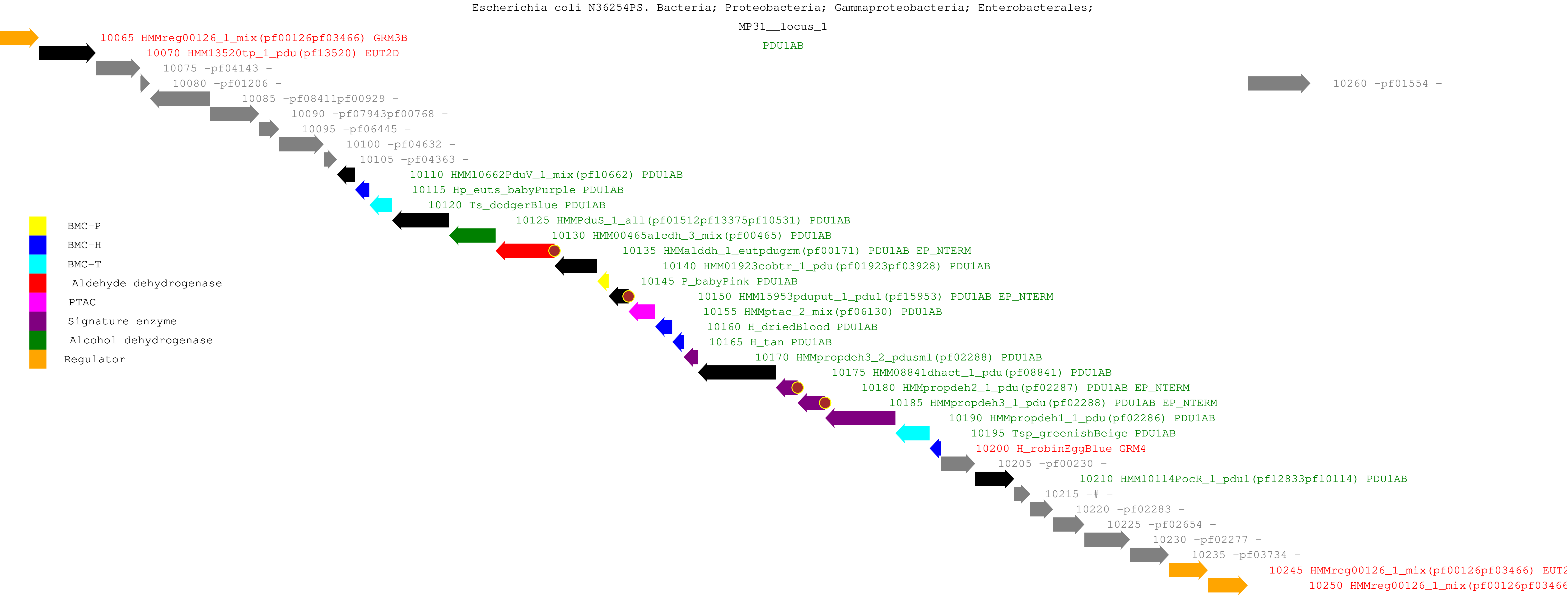
Escherichia coli N36254PS. Bacteria; Proteobacteria; Gammaproteobacteria; Enterobacterales;
MP31__locus_1
PDU1AB
10065 HMMreg00126_1_mix(pf00126pf03466) GRM3B
10070 HMM13520tp_1_pdu(pf13520) EUT2D
10075 -pf04143 -
10080 -pf01206 -
10085 -pf08411pf00929 -
10090 -pf07943pf00768 -
10095 -pf06445 -
10100 -pf04632 -
10105 -pf04363 -
10110 HMM10662PduV_1_mix(pf10662) PDU1AB
10115 Hp_euts_babyPurple PDU1AB
10120 Ts_dodgerBlue PDU1AB
10125 HMMPduS_1_all(pf01512pf13375pf10531) PDU1AB
10130 HMM00465alcdh_3_mix(pf00465) PDU1AB
10135 HMMalddh_1_eutpdugrm(pf00171) PDU1AB EP_NTERM
10140 HMM01923cobtr_1_pdu(pf01923pf03928) PDU1AB
10145 P_babyPink PDU1AB
10150 HMM15953pduput_1_pdu1(pf15953) PDU1AB EP_NTERM
10155 HMMptac_2_mix(pf06130) PDU1AB
10160 H_driedBlood PDU1AB
10165 H_tan PDU1AB
10170 HMMpropdeh3_2_pdusml(pf02288) PDU1AB
10175 HMM08841dhact_1_pdu(pf08841) PDU1AB
10180 HMMpropdeh2_1_pdu(pf02287) PDU1AB EP_NTERM
10185 HMMpropdeh3_1_pdu(pf02288) PDU1AB EP_NTERM
10190 HMMpropdeh1_1_pdu(pf02286) PDU1AB
10195 Tsp_greenishBeige PDU1AB
10200 H_robinEggBlue GRM4
10205 -pf00230 -
10210 HMM10114PocR_1_pdu1(pf12833pf10114) PDU1AB
10215 -# -
10220 -pf02283 -
10225 -pf02654 -
10230 -pf02277 -
10235 -pf03734 -
10245 HMMreg00126_1_mix(pf00126pf03466) EUT2
10250 HMMreg00126_1_mix(pf00126pf03466
10260 -pf01554 -
BMC-P
BMC-H
BMC-T
Aldehyde dehydrogenase
PTAC
Signature enzyme
Alcohol dehydrogenase
Regulator

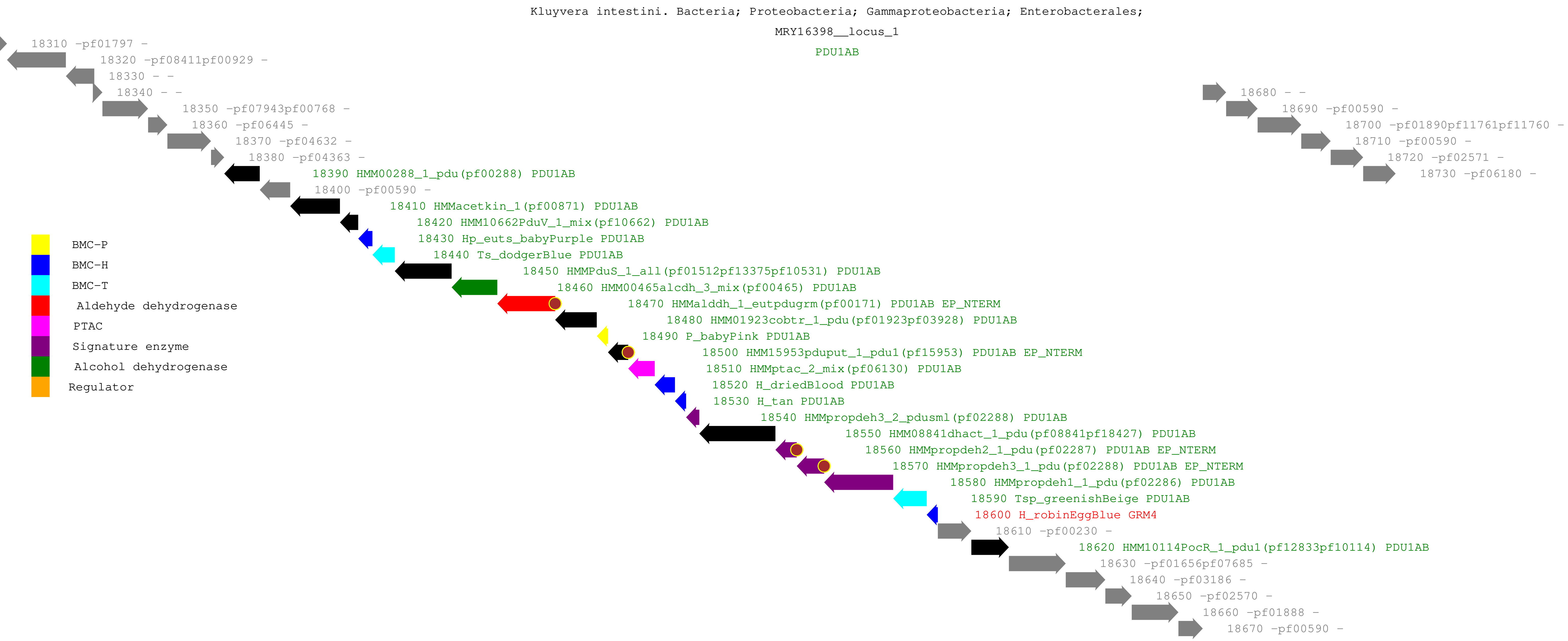
Kluyvera intestini. Bacteria; Proteobacteria; Gammaproteobacteria; Enterobacterales;
MRY16398__locus_1
PDU1AB
18310 -pf01797 -
18320 -pf08411pf00929 -
18330 - -
18340 - -
18350 -pf07943pf00768 -
18360 -pf06445 -
18370 -pf04632 -
18380 -pf04363 -
18390 HMM00288_1_pdu(pf00288) PDU1AB
18400 -pf00590 -
18410 HMMacetkin_1(pf00871) PDU1AB
18420 HMM10662PduV_1_mix(pf10662) PDU1AB
18430 Hp_euts_babyPurple PDU1AB
18440 Ts_dodgerBlue PDU1AB
18450 HMMPduS_1_all(pf01512pf13375pf10531) PDU1AB
18460 HMM00465alcdh_3_mix(pf00465) PDU1AB
18470 HMMalddh_1_eutpdugrm(pf00171) PDU1AB EP_NTERM
18480 HMM01923cobtr_1_pdu(pf01923pf03928) PDU1AB
18490 P_babyPink PDU1AB
18500 HMM15953pduput_1_pdu1(pf15953) PDU1AB EP_NTERM
18510 HMMptac_2_mix(pf06130) PDU1AB
18520 H_driedBlood PDU1AB
18530 H_tan PDU1AB
18540 HMMpropdeh3_2_pdusml(pf02288) PDU1AB
18550 HMM08841dhact_1_pdu(pf08841pf18427) PDU1AB
18560 HMMpropdeh2_1_pdu(pf02287) PDU1AB EP_NTERM
18570 HMMpropdeh3_1_pdu(pf02288) PDU1AB EP_NTERM
18580 HMMpropdeh1_1_pdu(pf02286) PDU1AB
18590 Tsp_greenishBeige PDU1AB
18600 H_robinEggBlue GRM4
18610 -pf00230 -
18620 HMM10114PocR_1_pdu1(pf12833pf10114) PDU1AB
18630 -pf01656pf07685 -
18640 -pf03186 -
18650 -pf02570 -
18660 -pf01888 -
18670 -pf00590 -
18680 - -
18690 -pf00590 -
18700 -pf01890pf11761pf11760 -
18710 -pf00590 -
18720 -pf02571 -
18730 -pf06180 -
BMC-P
BMC-H
BMC-T
Aldehyde dehydrogenase
PTAC
Signature enzyme
Alcohol dehydrogenase
Regulator

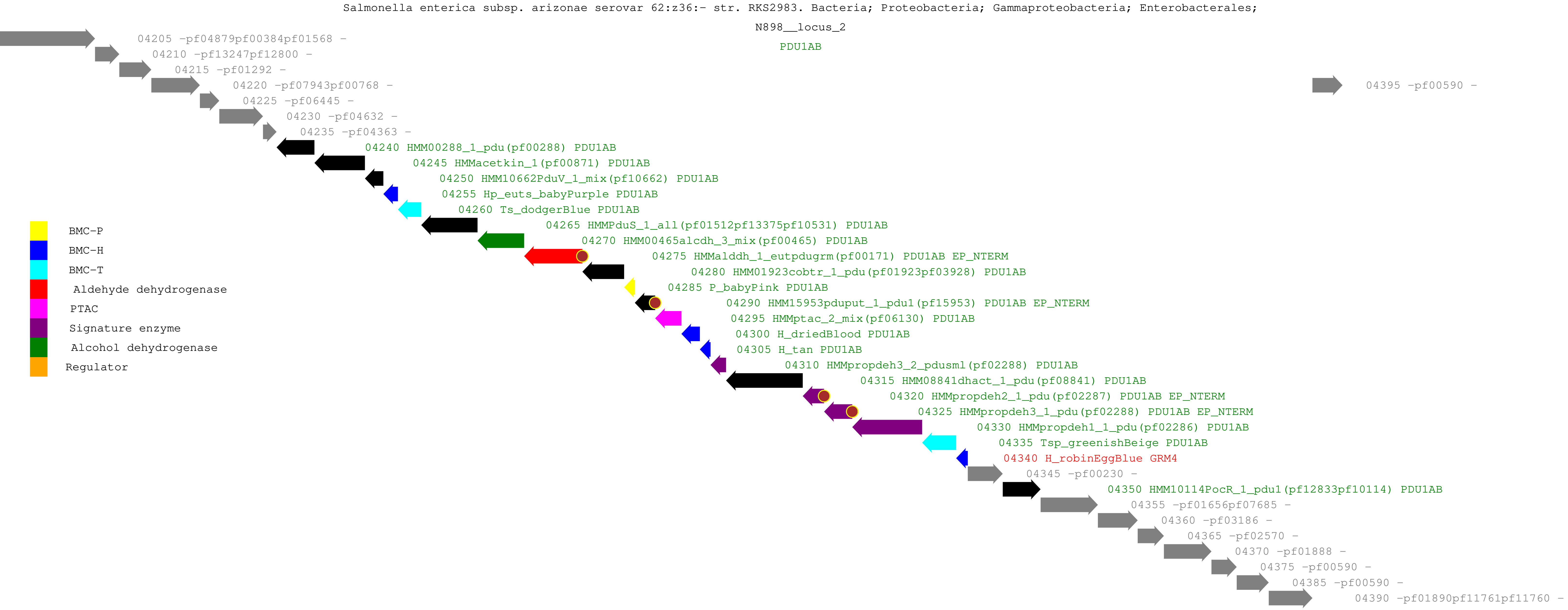

Salmonella enterica subsp. arizonae serovar 62:z36:- str. RKS2983. Bacteria; Proteobacteria; Gammaproteobacteria; Enterobacterales;
N898__locus_2
PDU1AB
04205 -pf04879pf00384pf01568 -
04210 -pf13247pf12800 -
04215 -pf01292 -
04220 -pf07943pf00768 -
04225 -pf06445 -
04230 -pf04632 -
04235 -pf04363 -
04240 HMM00288_1_pdu(pf00288) PDU1AB
04245 HMMacetkin_1(pf00871) PDU1AB
04250 HMM10662PduV_1_mix(pf10662) PDU1AB
04255 Hp_euts_babyPurple PDU1AB
04260 Ts_dodgerBlue PDU1AB
04265 HMMPduS_1_all(pf01512pf13375pf10531) PDU1AB
04270 HMM00465alcdh_3_mix(pf00465) PDU1AB
04275 HMMalddh_1_eutpdugrm(pf00171) PDU1AB EP_NTERM
04280 HMM01923cobtr_1_pdu(pf01923pf03928) PDU1AB
04285 P_babyPink PDU1AB
04290 HMM15953pduput_1_pdu1(pf15953) PDU1AB EP_NTERM
04295 HMMptac_2_mix(pf06130) PDU1AB
04300 H_driedBlood PDU1AB
04305 H_tan PDU1AB
04310 HMMpropdeh3_2_pdusml(pf02288) PDU1AB
04315 HMM08841dhact_1_pdu(pf08841) PDU1AB
04320 HMMpropdeh2_1_pdu(pf02287) PDU1AB EP_NTERM
04325 HMMpropdeh3_1_pdu(pf02288) PDU1AB EP_NTERM
04330 HMMpropdeh1_1_pdu(pf02286) PDU1AB
04335 Tsp_greenishBeige PDU1AB
04340 H_robinEggBlue GRM4
04345 -pf00230 -
04350 HMM10114PocR_1_pdu1(pf12833pf10114) PDU1AB
04355 -pf01656pf07685 -
04360 -pf03186 -
04365 -pf02570 -
04370 -pf01888 -
04375 -pf00590 -
04385 -pf00590 -
04390 -pf01890pf11761pf11760 -
04395 -pf00590 -
BMC-P
BMC-H
BMC-T
Aldehyde dehydrogenase
PTAC
Signature enzyme
Alcohol dehydrogenase
Regulator

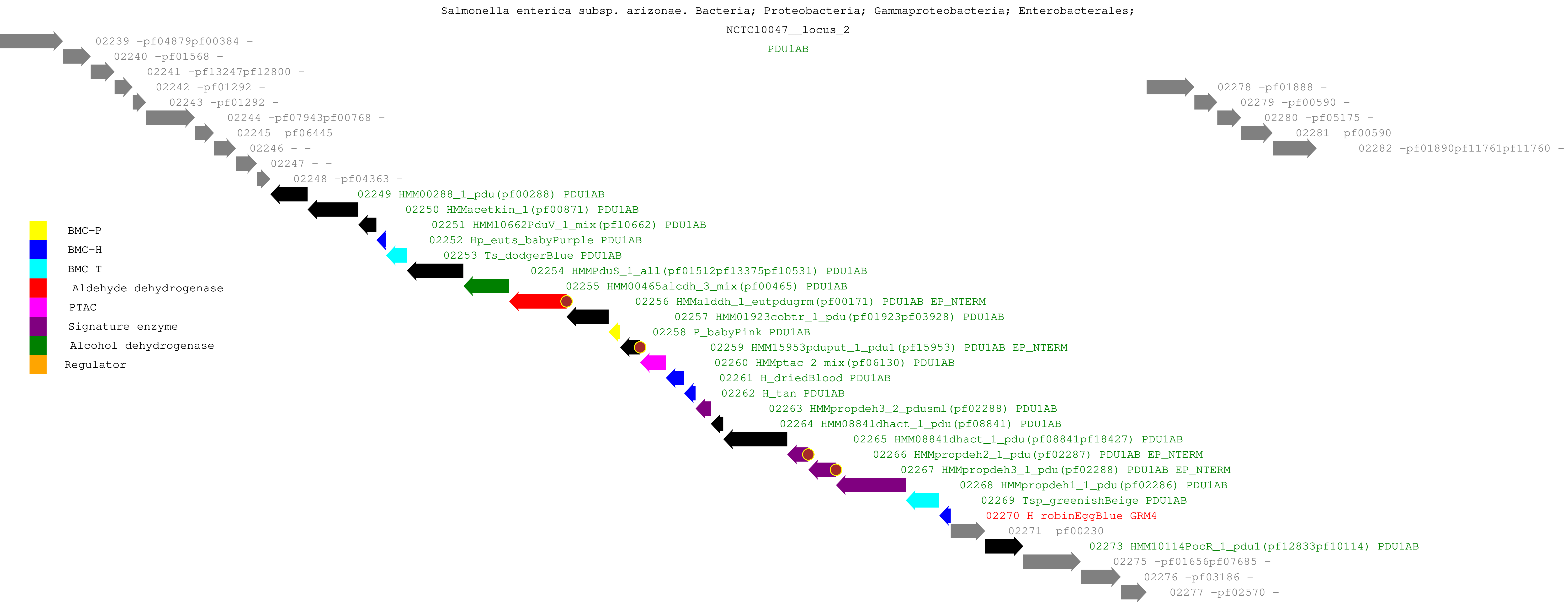
Salmonella enterica subsp. arizonae. Bacteria; Proteobacteria; Gammaproteobacteria; Enterobacterales;
NCTC10047__locus_2
PDU1AB
02239 -pf04879pf00384 -
02240 -pf01568 -
02241 -pf13247pf12800 -
02242 -pf01292 -
02243 -pf01292 -
02244 -pf07943pf00768 -
02245 -pf06445 -
02246 - -
02247 - -
02248 -pf04363 -
02249 HMM00288_1_pdu(pf00288) PDU1AB
02250 HMMacetkin_1(pf00871) PDU1AB
02251 HMM10662PduV_1_mix(pf10662) PDU1AB
02252 Hp_euts_babyPurple PDU1AB
02253 Ts_dodgerBlue PDU1AB
02254 HMMPduS_1_all(pf01512pf13375pf10531) PDU1AB
02255 HMM00465alcdh_3_mix(pf00465) PDU1AB
02256 HMMalddh_1_eutpdugrm(pf00171) PDU1AB EP_NTERM
02257 HMM01923cobtr_1_pdu(pf01923pf03928) PDU1AB
02258 P_babyPink PDU1AB
02259 HMM15953pduput_1_pdu1(pf15953) PDU1AB EP_NTERM
02260 HMMptac_2_mix(pf06130) PDU1AB
02261 H_driedBlood PDU1AB
02262 H_tan PDU1AB
02263 HMMpropdeh3_2_pdusml(pf02288) PDU1AB
02264 HMM08841dhact_1_pdu(pf08841) PDU1AB
02265 HMM08841dhact_1_pdu(pf08841pf18427) PDU1AB
02266 HMMpropdeh2_1_pdu(pf02287) PDU1AB EP_NTERM
02267 HMMpropdeh3_1_pdu(pf02288) PDU1AB EP_NTERM
02268 HMMpropdeh1_1_pdu(pf02286) PDU1AB
02269 Tsp_greenishBeige PDU1AB
02270 H_robinEggBlue GRM4
02271 -pf00230 -
02273 HMM10114PocR_1_pdu1(pf12833pf10114) PDU1AB
02275 -pf01656pf07685 -
02276 -pf03186 -
02277 -pf02570 -
02278 -pf01888 -
02279 -pf00590 -
02280 -pf05175 -
02281 -pf00590 -
02282 -pf01890pf11761pf11760 -
BMC-P
BMC-H
BMC-T
Aldehyde dehydrogenase
PTAC
Signature enzyme
Alcohol dehydrogenase
Regulator

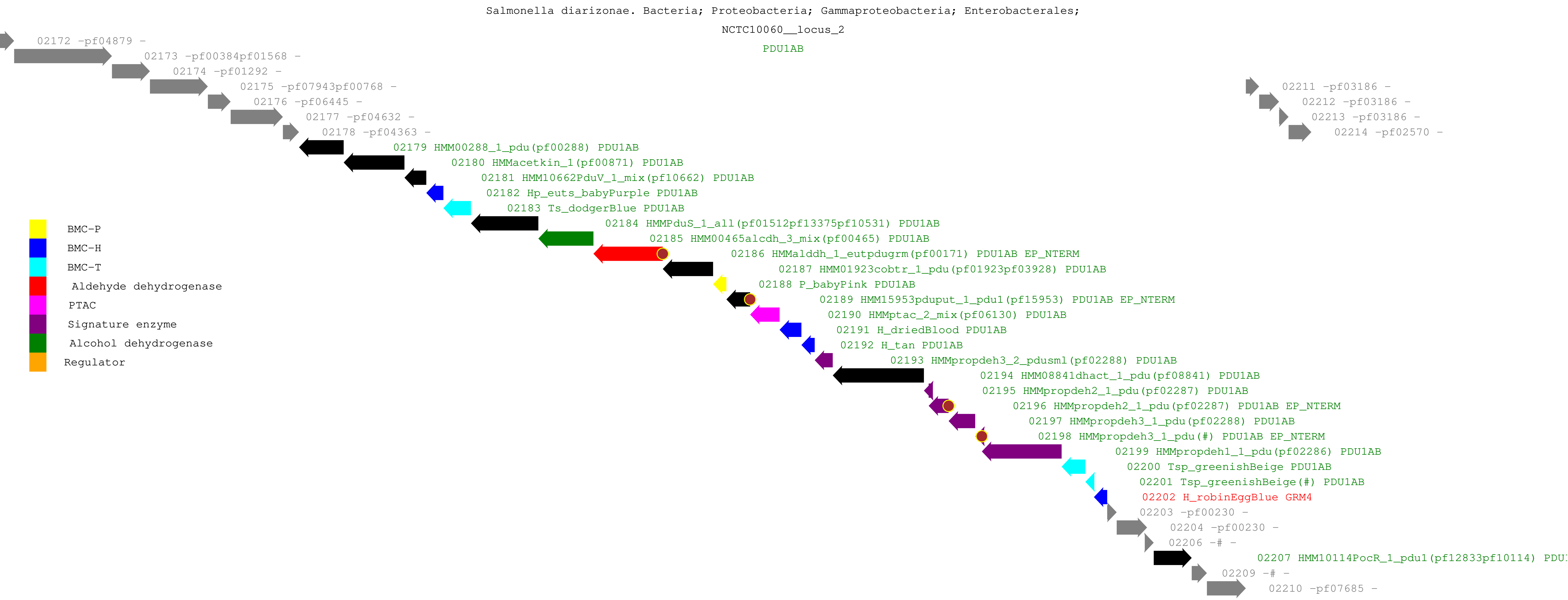

Salmonella diarizonae. Bacteria; Proteobacteria; Gammaproteobacteria; Enterobacterales;
NCTC10060__locus_2
PDU1AB
02172 -pf04879 -
02173 -pf00384pf01568 -
02174 -pf01292 -
02175 -pf07943pf00768 -
02176 -pf06445 -
02177 -pf04632 -
02178 -pf04363 -
02179 HMM00288_1_pdu(pf00288) PDU1AB
02180 HMMacetkin_1(pf00871) PDU1AB
02181 HMM10662PduV_1_mix(pf10662) PDU1AB
02182 Hp_euts_babyPurple PDU1AB
02183 Ts_dodgerBlue PDU1AB
02184 HMMPduS_1_all(pf01512pf13375pf10531) PDU1AB
02185 HMM00465alcdh_3_mix(pf00465) PDU1AB
02186 HMMalddh_1_eutpdugrm(pf00171) PDU1AB EP_NTERM
02187 HMM01923cobtr_1_pdu(pf01923pf03928) PDU1AB
02188 P_babyPink PDU1AB
02189 HMM15953pduput_1_pdu1(pf15953) PDU1AB EP_NTERM
02190 HMMptac_2_mix(pf06130) PDU1AB
02191 H_driedBlood PDU1AB
02192 H_tan PDU1AB
02193 HMMpropdeh3_2_pdusml(pf02288) PDU1AB
02194 HMM08841dhact_1_pdu(pf08841) PDU1AB
02195 HMMpropdeh2_1_pdu(pf02287) PDU1AB
02196 HMMpropdeh2_1_pdu(pf02287) PDU1AB EP_NTERM
02197 HMMpropdeh3_1_pdu(pf02288) PDU1AB
02198 HMMpropdeh3_1_pdu(#) PDU1AB EP_NTERM
02199 HMMpropdeh1_1_pdu(pf02286) PDU1AB
02200 Tsp_greenishBeige PDU1AB
02201 Tsp_greenishBeige(#) PDU1AB
02202 H_robinEggBlue GRM4
02203 -pf00230 -
02204 -pf00230 -
02206 -# -
02207 HMM10114PocR_1_pdu1(pf12833pf10114) PDU
02209 -# -
02210 -pf07685 -
02211 -pf03186 -
02212 -pf03186 -
02213 -pf03186 -
02214 -pf02570 -
BMC-P
BMC-H
BMC-T
Aldehyde dehydrogenase
PTAC
Signature enzyme
Alcohol dehydrogenase
Regulator

Salmonella choleraesuis. Bacteria; Proteobacteria; Gammaproteobacteria; Enterobacterales;

NCTC10252__locus_2

PDU1AB

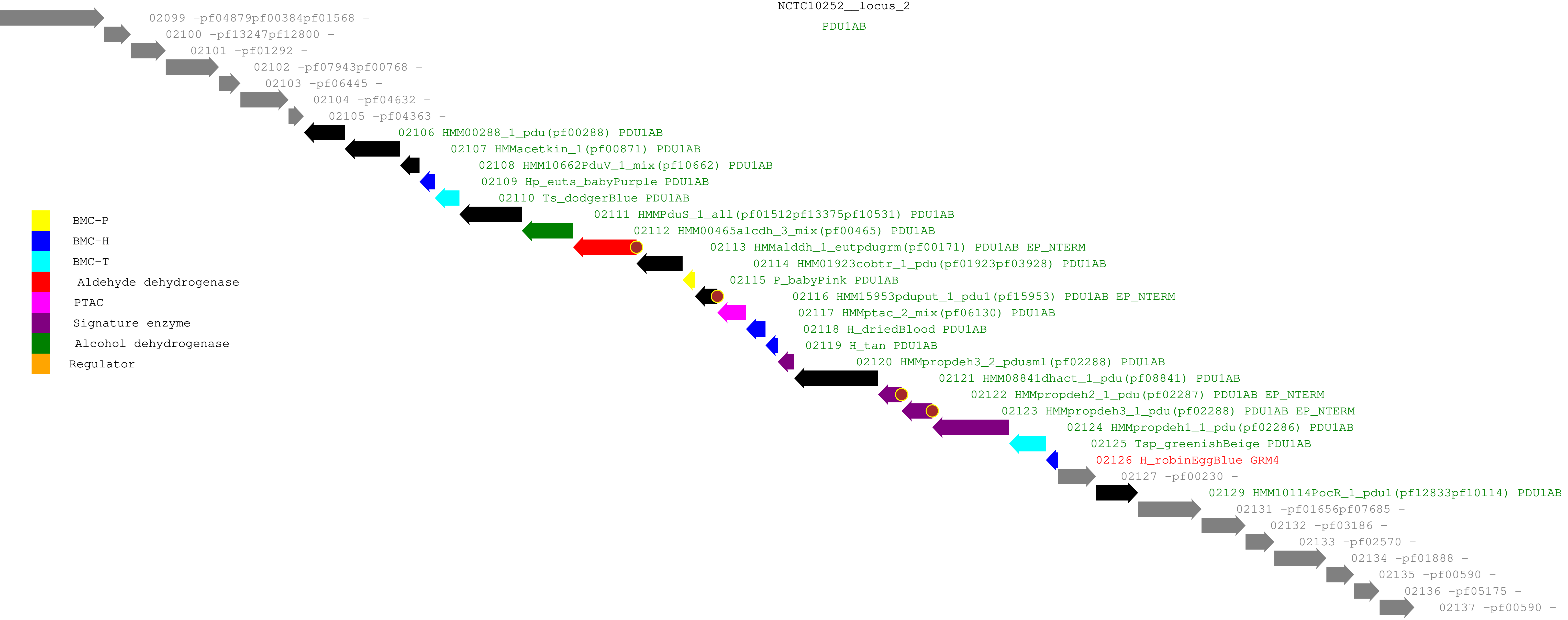

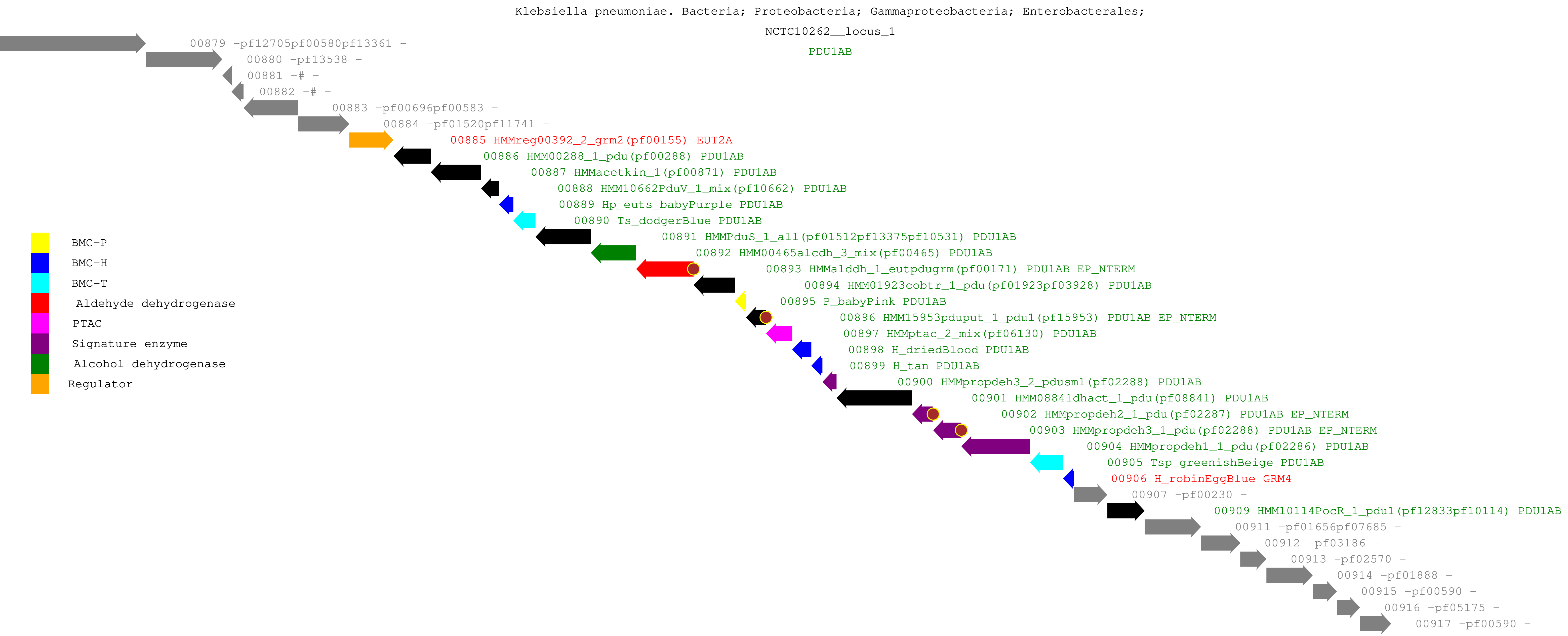
Klebsiella pneumoniae. Bacteria; Proteobacteria; Gammaproteobacteria; Enterobacterales;
NCTC10262__locus_1
PDU1AB
00879 -pf12705pf00580pf13361 -
00880 -pf13538 -
00881 -# -
00882 -# -
00883 -pf00696pf00583 -
00884 -pf01520pf11741 -
00885 HMMreg00392_2_grm2(pf00155) EUT2A
00886 HMM00288_1_pdu(pf00288) PDU1AB
00887 HMMacetkin_1(pf00871) PDU1AB
00888 HMM10662PduV_1_mix(pf10662) PDU1AB
00889 Hp_euts_babyPurple PDU1AB
00890 Ts_dodgerBlue PDU1AB
00891 HMMPduS_1_all(pf01512pf13375pf10531) PDU1AB
00892 HMM00465alcdh_3_mix(pf00465) PDU1AB
00893 HMMalddh_1_eutpdugrm(pf00171) PDU1AB EP_NTERM
00894 HMM01923cobtr_1_pdu(pf01923pf03928) PDU1AB
00895 P_babyPink PDU1AB
00896 HMM15953pduput_1_pdu1(pf15953) PDU1AB EP_NTERM
00897 HMMptac_2_mix(pf06130) PDU1AB
00898 H_driedBlood PDU1AB
00899 H_tan PDU1AB
00900 HMMpropdeh3_2_pdusml(pf02288) PDU1AB
00901 HMM08841dhact_1_pdu(pf08841) PDU1AB
00902 HMMpropdeh2_1_pdu(pf02287) PDU1AB EP_NTERM
00903 HMMpropdeh3_1_pdu(pf02288) PDU1AB EP_NTERM
00904 HMMpropdeh1_1_pdu(pf02286) PDU1AB
00905 Tsp_greenishBeige PDU1AB
00906 H_robinEggBlue GRM4
00907 -pf00230 -
00909 HMM10114PocR_1_pdu1(pf12833pf10114) PDU1AB
00911 -pf01656pf07685 -
00912 -pf03186 -
00913 -pf02570 -
00914 -pf01888 -
00915 -pf00590 -
00916 -pf05175 -
00917 -pf00590 -
BMC-P
BMC-H
BMC-T
Aldehyde dehydrogenase
PTAC
Signature enzyme
Alcohol dehydrogenase
Regulator

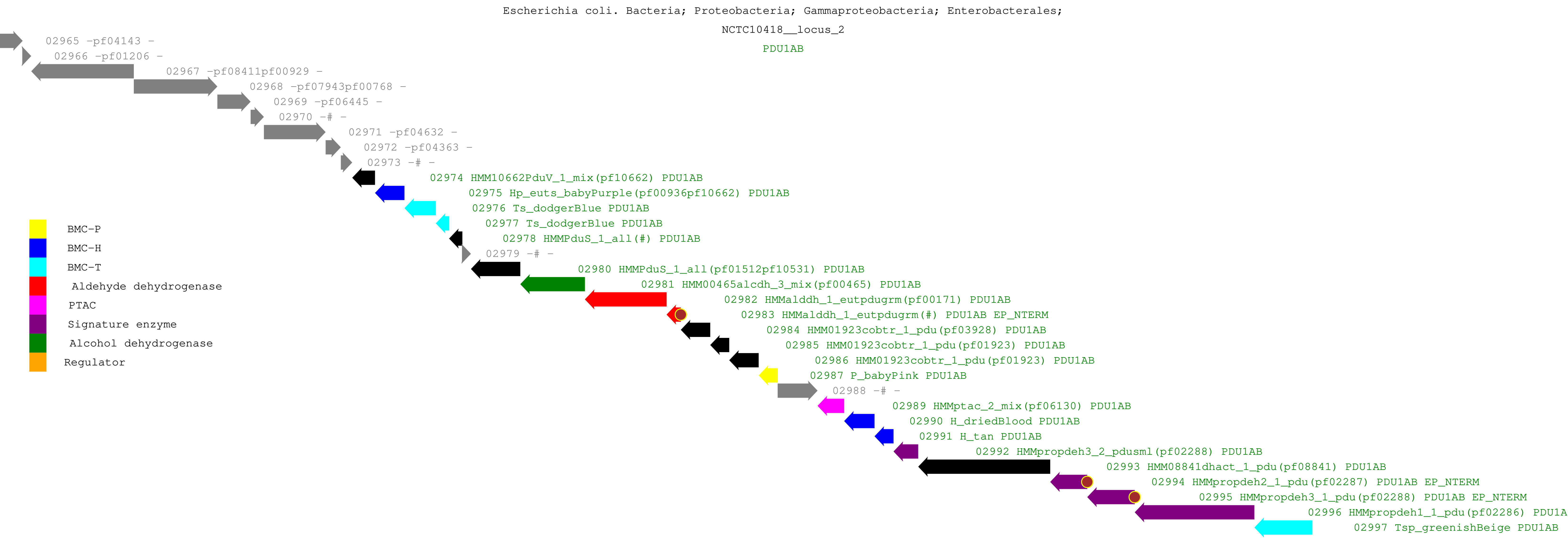
Escherichia coli. Bacteria; Proteobacteria; Gammaproteobacteria; Enterobacterales;
NCTC10418__locus_2
PDU1AB
02965 -pf04143 -
02966 -pf01206 -
02967 -pf08411pf00929 -
02968 -pf07943pf00768 -
02969 -pf06445 -
02970 -# -
02971 -pf04632 -
02972 -pf04363 -
02973 -# -
02974 HMM10662PduV_1_mix(pf10662) PDU1AB
02975 Hp_euts_babyPurple(pf00936pf10662) PDU1AB
02976 Ts_dodgerBlue PDU1AB
02977 Ts_dodgerBlue PDU1AB
02978 HMMPduS_1_all(#) PDU1AB
02979 -# -
02980 HMMPduS_1_all(pf01512pf10531) PDU1AB
02981 HMM00465alcdh_3_mix(pf00465) PDU1AB
02982 HMMalddh_1_eutpdugrm(pf00171) PDU1AB
02983 HMMalddh_1_eutpdugrm(#) PDU1AB EP_NTERM
02984 HMM01923cobtr_1_pdu(pf03928) PDU1AB
02985 HMM01923cobtr_1_pdu(pf01923) PDU1AB
02986 HMM01923cobtr_1_pdu(pf01923) PDU1AB
02987 P_babyPink PDU1AB
02988 -# -
02989 HMMptac_2_mix(pf06130) PDU1AB
02990 H_driedBlood PDU1AB
02991 H_tan PDU1AB
02992 HMMpropdeh3_2_pdusml(pf02288) PDU1AB
02993 HMM08841dhact_1_pdu(pf08841) PDU1AB
02994 HMMpropdeh2_1_pdu(pf02287) PDU1AB EP_NTERM
02995 HMMpropdeh3_1_pdu(pf02288) PDU1AB EP_NTERM
02996 HMMpropdeh1_1_pdu(pf02286) PDU1A
02997 Tsp_greenishBeige PDU1AB
BMC-P
BMC-H
BMC-T
Aldehyde dehydrogenase
PTAC
Signature enzyme
Alcohol dehydrogenase
Regulator

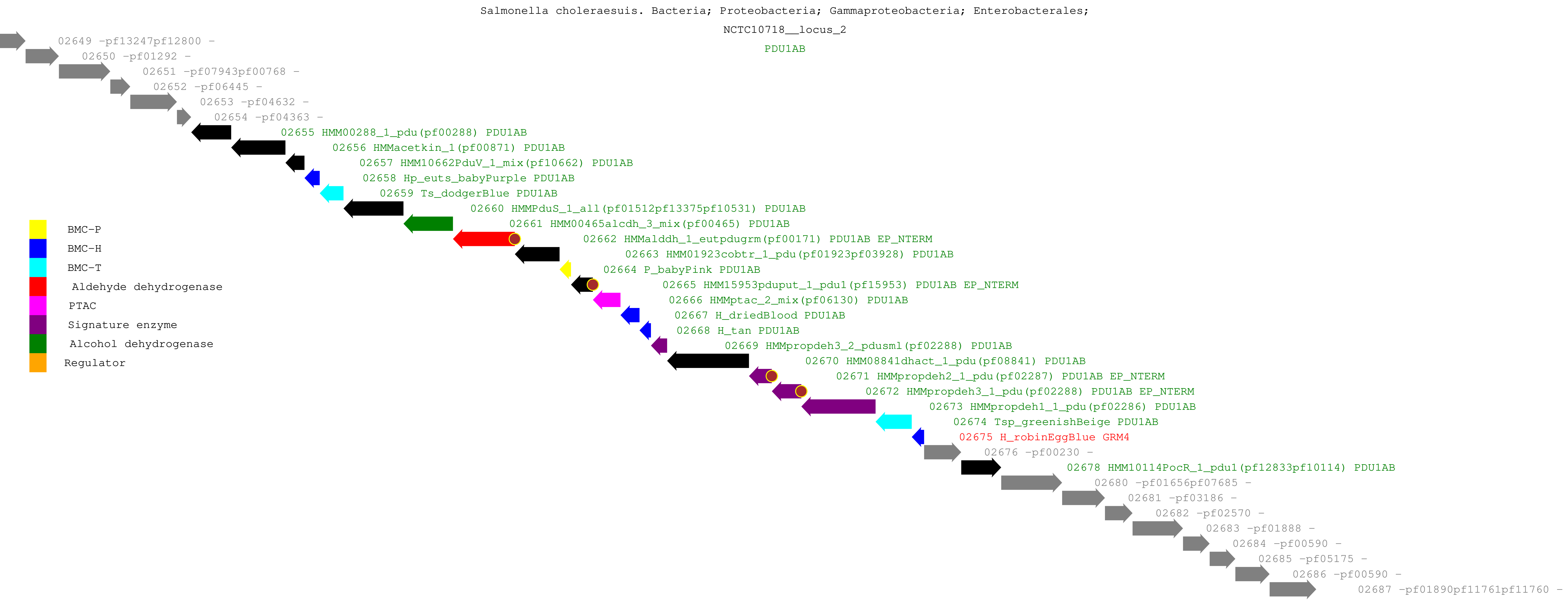
Salmonella choleraesuis. Bacteria; Proteobacteria; Gammaproteobacteria; Enterobacterales;
NCTC10718__locus_2
PDU1AB
02649 -pf13247pf12800 -
02650 -pf01292 -
02651 -pf07943pf00768 -
02652 -pf06445 -
02653 -pf04632 -
02654 -pf04363 -
02655 HMM00288_1_pdu(pf00288) PDU1AB
02656 HMMacetkin_1(pf00871) PDU1AB
02657 HMM10662PduV_1_mix(pf10662) PDU1AB
02658 Hp_euts_babyPurple PDU1AB
02659 Ts_dodgerBlue PDU1AB
02660 HMMPduS_1_all(pf01512pf13375pf10531) PDU1AB
02661 HMM00465alcdh_3_mix(pf00465) PDU1AB
02662 HMMaldh_1_eutpdugrm(pf00171) PDU1AB EP_NTERM
02663 HMM01923cobtr_1_pdu(pf01923pf03928) PDU1AB
02664 P_babyPink PDU1AB
02665 HMM15953pduput_1_pdu1(pf15953) PDU1AB EP_NTERM
02666 HMMptac_2_mix(pf06130) PDU1AB
02667 H_driedBlood PDU1AB
02668 H_tan PDU1AB
02669 HMMpropdeh3_2_pdusml(pf02288) PDU1AB
02670 HMM08841dhact_1_pdu(pf08841) PDU1AB
02671 HMMpropdeh2_1_pdu(pf02287) PDU1AB EP_NTERM
02672 HMMpropdeh3_1_pdu(pf02288) PDU1AB EP_NTERM
02673 HMMpropdeh1_1_pdu(pf02286) PDU1AB
02674 Tsp_greenishBeige PDU1AB
02675 H_robinEggBlue GRM4
02676 -pf00230 -
02678 HMM10114PocR_1_pdu1(pf12833pf10114) PDU1AB
02680 -pf01656pf07685 -
02681 -pf03186 -
02682 -pf02570 -
02683 -pf01888 -
02684 -pf00590 -
02685 -pf05175 -
02686 -pf00590 -
02687 -pf01890pf11761pf11760 -
BMC-P
BMC-H
BMC-T
Aldehyde dehydrogenase
PTAC
Signature enzyme
Alcohol dehydrogenase
Regulator

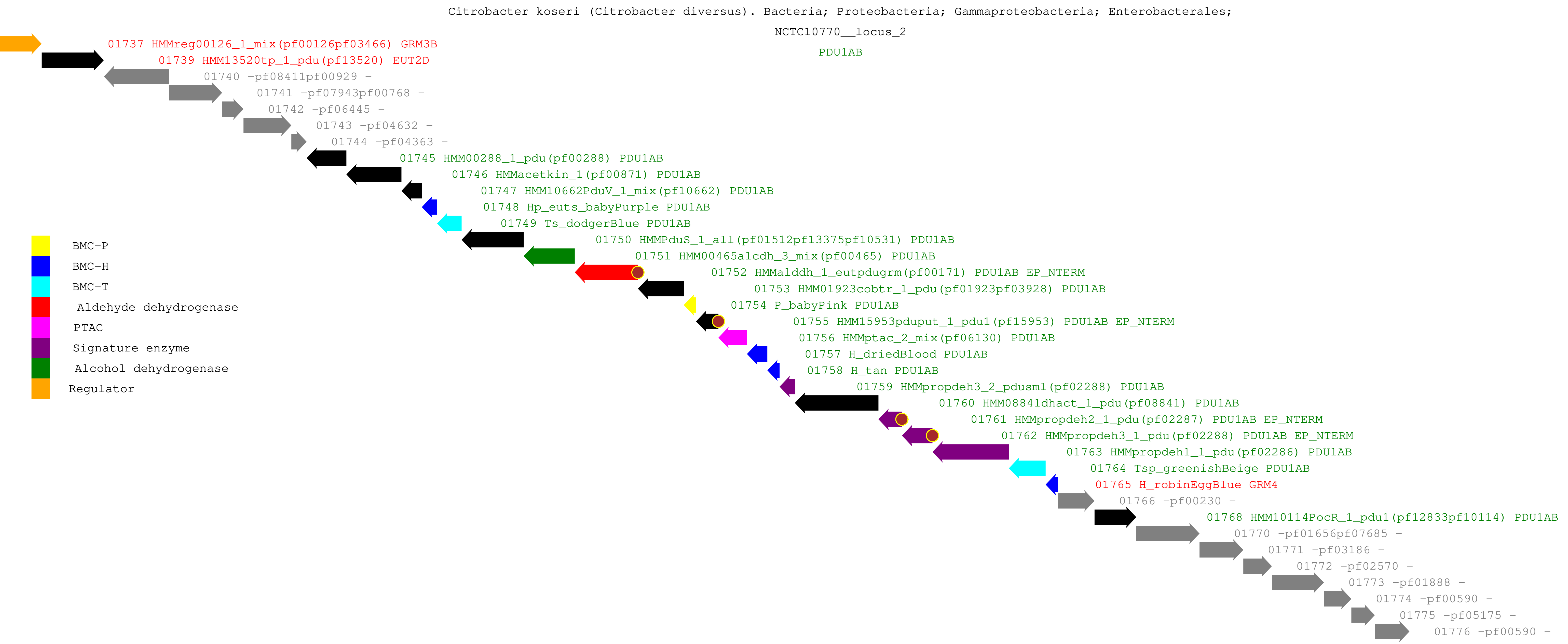

Citrobacter koseri (Citrobacter diversus). Bacteria; Proteobacteria; Gammaproteobacteria; Enterobacterales;
NCTC10770__locus_2
PDU1AB
01737 HMMreg00126_1_mix(pf00126pf03466) GRM3B
01739 HMM13520tp_1_pdu(pf13520) EUT2D
01740 -pf08411pf00929 -
01741 -pf07943pf00768 -
01742 -pf06445 -
01743 -pf04632 -
01744 -pf04363 -
01745 HMM00288_1_pdu(pf00288) PDU1AB
01746 HMMacetkin_1(pf00871) PDU1AB
01747 HMM10662PduV_1_mix(pf10662) PDU1AB
01748 Hp_euts_babyPurple PDU1AB
01749 Ts_dodgerBlue PDU1AB
01750 HMMPduS_1_all(pf01512pf13375pf10531) PDU1AB
01751 HMM00465alcdh_3_mix(pf00465) PDU1AB
01752 HMMalddh_1_eutpdugrm(pf00171) PDU1AB EP_NTERM
01753 HMM01923cobtr_1_pdu(pf01923pf03928) PDU1AB
01754 P_babyPink PDU1AB
01755 HMM15953pduput_1_pdu1(pf15953) PDU1AB EP_NTERM
01756 HMMptac_2_mix(pf06130) PDU1AB
01757 H_driedBlood PDU1AB
01758 H_tan PDU1AB
01759 HMMpropdeh3_2_pdusml(pf02288) PDU1AB
01760 HMM08841dhact_1_pdu(pf08841) PDU1AB
01761 HMMpropdeh2_1_pdu(pf02287) PDU1AB EP_NTERM
01762 HMMpropdeh3_1_pdu(pf02288) PDU1AB EP_NTERM
01763 HMMpropdeh1_1_pdu(pf02286) PDU1AB
01764 Tsp_greenishBeige PDU1AB
01765 H_robinEggBlue GRM4
01766 -pf00230 -
01768 HMM10114PocR_1_pdu1(pf12833pf10114) PDU1AB
01770 -pf01656pf07685 -
01771 -pf03186 -
01772 -pf02570 -
01773 -pf01888 -
01774 -pf00590 -
01775 -pf05175 -
01776 -pf00590 -
BMC-P
BMC-H
BMC-T
Aldehyde dehydrogenase
PTAC
Signature enzyme
Alcohol dehydrogenase
Regulator

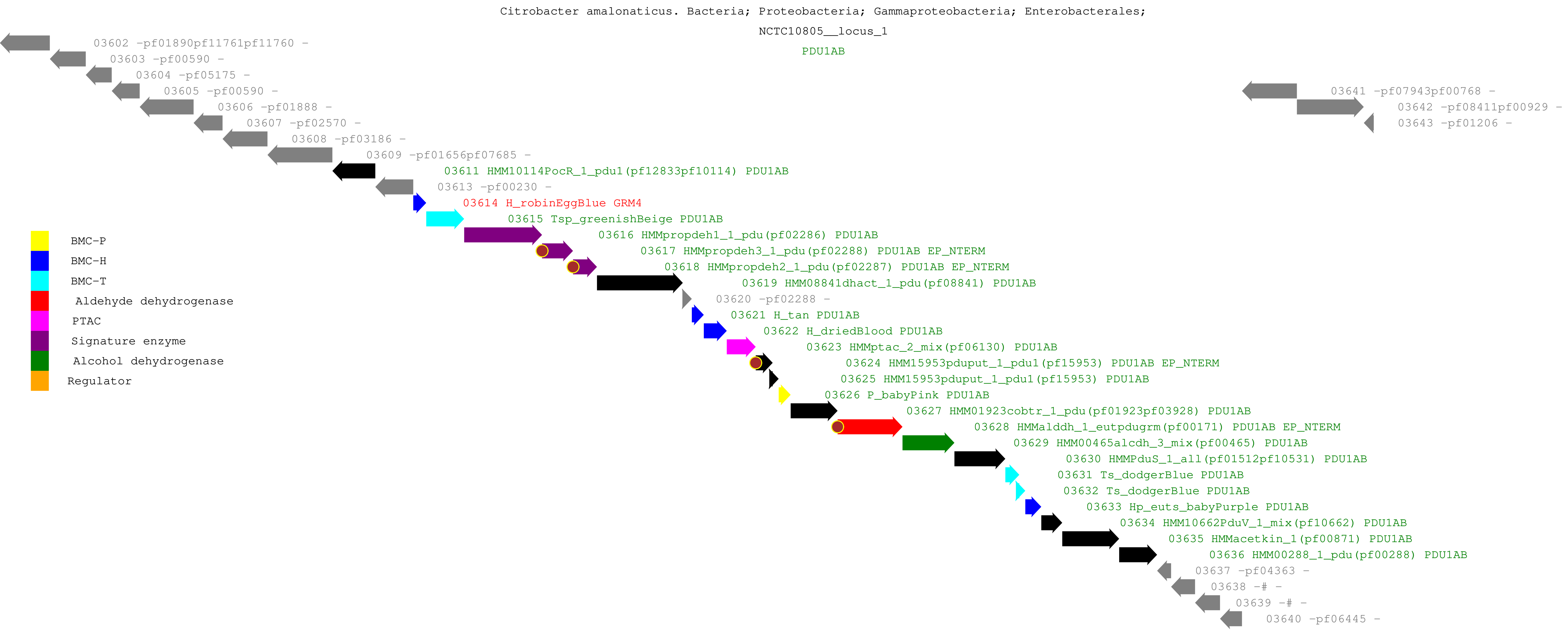

Citrobacter amalonaticus. Bacteria; Proteobacteria; Gammaproteobacteria; Enterobacterales;
NCTC10805__locus_1
PDU1AB
03602 -pf01890pf11761pf11760 -
03603 -pf00590 -
03604 -pf05175 -
03605 -pf00590 -
03606 -pf01888 -
03607 -pf02570 -
03608 -pf03186 -
03609 -pf01656pf07685 -
03611 HMM10114PocR_1_pdu1(pf12833pf10114) PDU1AB
03613 -pf00230 -
03614 H_robinEggBlue GRM4
03615 Tsp_greenishBeige PDU1AB
03616 HMMpropdeh1_1_pdu(pf02286) PDU1AB
03617 HMMpropdeh3_1_pdu(pf02288) PDU1AB EP_NTERM
03618 HMMpropdeh2_1_pdu(pf02287) PDU1AB EP_NTERM
03619 HMM08841dhact_1_pdu(pf08841) PDU1AB
03620 -pf02288 -
03621 H_tan PDU1AB
03622 H_driedBlood PDU1AB
03623 HMMptac_2_mix(pf06130) PDU1AB
03624 HMM15953pduput_1_pdu1(pf15953) PDU1AB EP_NTERM
03625 HMM15953pduput_1_pdu1(pf15953) PDU1AB
03626 P_babyPink PDU1AB
03627 HMM01923cobtr_1_pdu(pf01923pf03928) PDU1AB
03628 HMMalddh_1_eutpdugrm(pf00171) PDU1AB EP_NTERM
03629 HMM00465alcdh_3_mix(pf00465) PDU1AB
03630 HMMPduS_1_all(pf01512pf10531) PDU1AB
03631 Ts_dodgerBlue PDU1AB
03632 Ts_dodgerBlue PDU1AB
03633 Hp_euts_babyPurple PDU1AB
03634 HMM10662PduV_1_mix(pf10662) PDU1AB
03635 HMMacetkin_1(pf00871) PDU1AB
03636 HMM00288_1_pdu(pf00288) PDU1AB
03637 -pf04363 -
03638 -# -
03639 -# -
03640 -pf06445 -
03641 -pf07943pf00768 -
03642 -pf08411pf00929 -
03643 -pf01206 -
BMC-P
BMC-H
BMC-T
Aldehyde dehydrogenase
PTAC
Signature enzyme
Alcohol dehydrogenase
Regulator

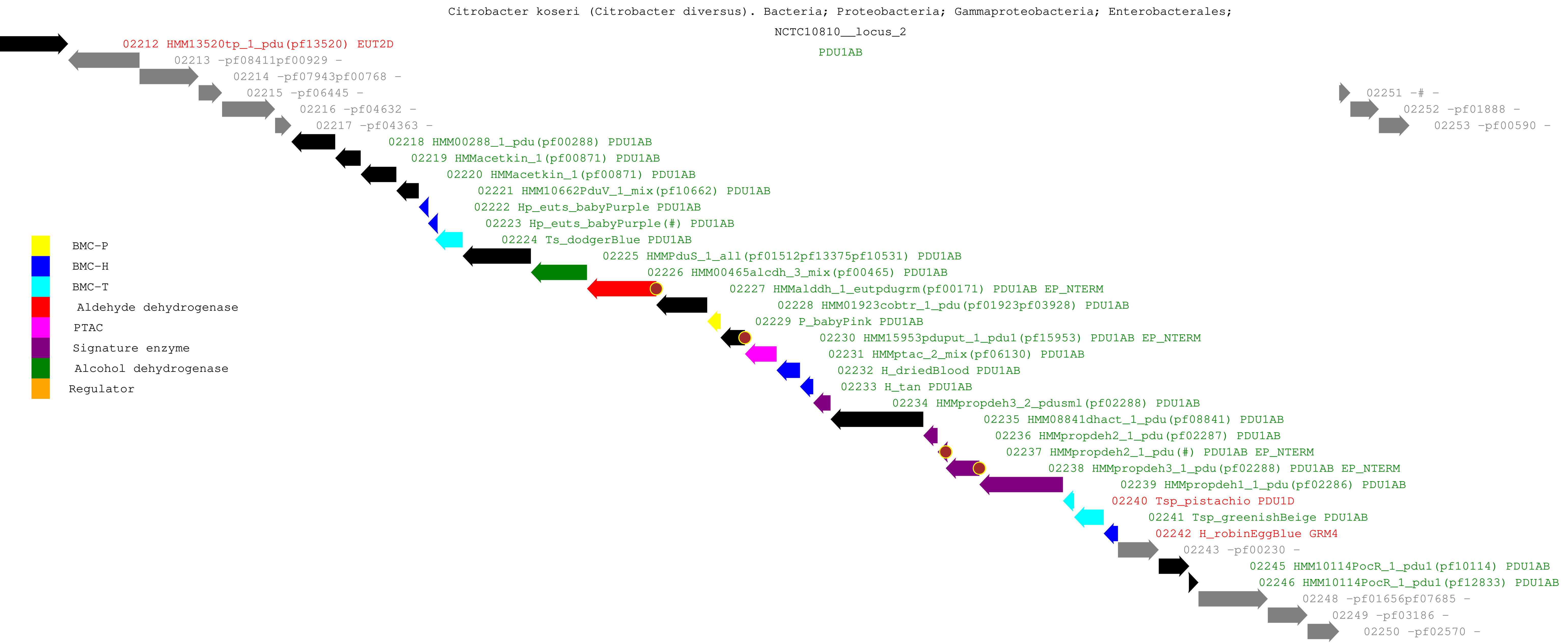

Citrobacter koseri (Citrobacter diversus). Bacteria; Proteobacteria; Gammaproteobacteria; Enterobacterales;
NCTC10810__locus_2
PDU1AB
02212 HMM13520tp_1_pdu(pf13520) EUT2D
02213 -pf08411pf00929 -
02214 -pf07943pf00768 -
02215 -pf06445 -
02216 -pf04632 -
02217 -pf04363 -
02218 HMM00288_1_pdu(pf00288) PDU1AB
02219 HMMacetkin_1(pf00871) PDU1AB
02220 HMMacetkin_1(pf00871) PDU1AB
02221 HMM10662PduV_1_mix(pf10662) PDU1AB
02222 Hp_euts_babyPurple PDU1AB
02223 Hp_euts_babyPurple(#) PDU1AB
02224 Ts_dodgerBlue PDU1AB
02225 HMMPduS_1_all(pf01512pf13375pf10531) PDU1AB
02226 HMM00465alcdh_3_mix(pf00465) PDU1AB
02227 HMMalddh_1_eutpdugrm(pf00171) PDU1AB EP_NTERM
02228 HMM01923cobtr_1_pdu(pf01923pf03928) PDU1AB
02229 P_babyPink PDU1AB
02230 HMM15953pduput_1_pdu1(pf15953) PDU1AB EP_NTERM
02231 HMMptac_2_mix(pf06130) PDU1AB
02232 H_driedBlood PDU1AB
02233 H_tan PDU1AB
02234 HMMpropdeh3_2_pdusml(pf02288) PDU1AB
02235 HMM08841dhact_1_pdu(pf08841) PDU1AB
02236 HMMpropdeh2_1_pdu(pf02287) PDU1AB
02237 HMMpropdeh2_1_pdu(#) PDU1AB EP_NTERM
02238 HMMpropdeh3_1_pdu(pf02288) PDU1AB EP_NTERM
02239 HMMpropdeh1_1_pdu(pf02286) PDU1AB
02240 Tsp_pistachio PDU1D
02241 Tsp_greenishBeige PDU1AB
02242 H_robinEggBlue GRM4
02243 -pf00230 -
02245 HMM10114PocR_1_pdu1(pf10114) PDU1AB
02246 HMM10114PocR_1_pdu1(pf12833) PDU1AB
02248 -pf01656pf07685 -
02249 -pf03186 -
02250 -pf02570 -
02251 -# -
02252 -pf01888 -
02253 -pf00590 -
BMC-P
BMC-H
BMC-T
Aldehyde dehydrogenase
PTAC
Signature enzyme
Alcohol dehydrogenase
Regulator

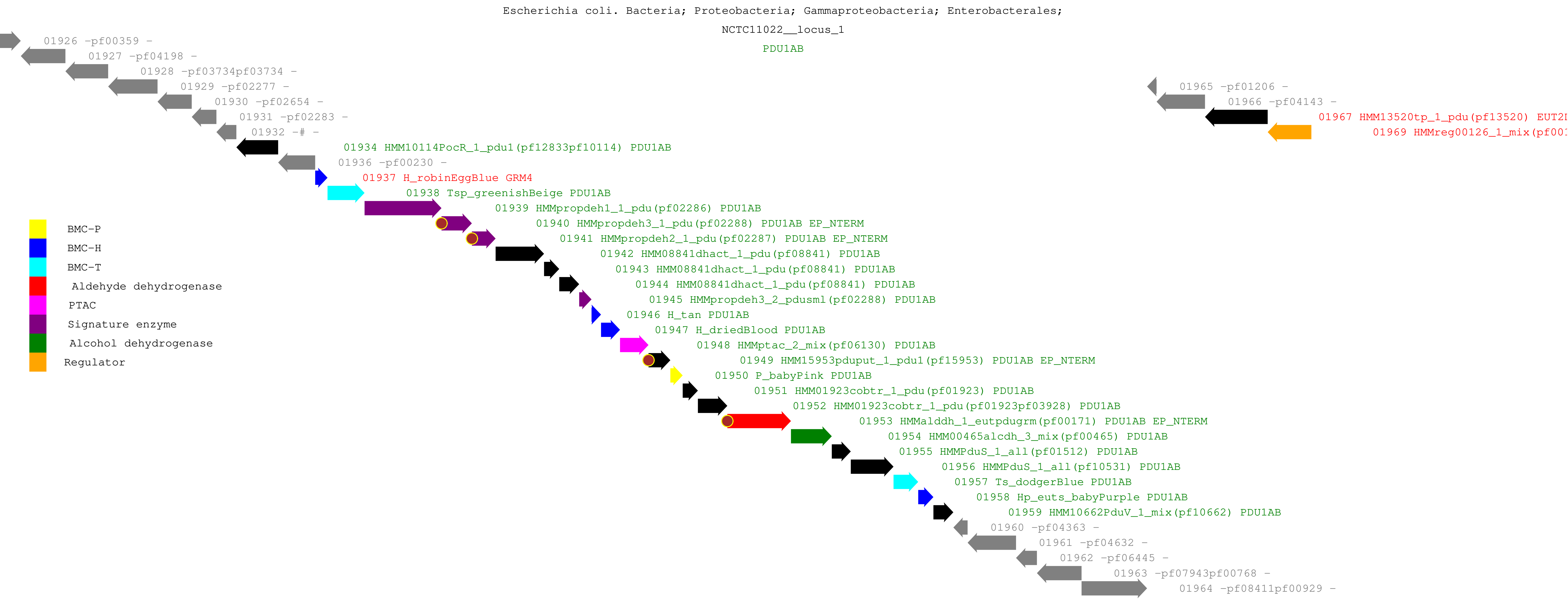
Escherichia coli. Bacteria; Proteobacteria; Gammaproteobacteria; Enterobacterales;
NCTC11022__locus_1
PDU1AB
01926 -pf00359 -
01927 -pf04198 -
01928 -pf03734pf03734 -
01929 -pf02277 -
01930 -pf02654 -
01931 -pf02283 -
01932 -# -
01934 HMM10114PocR_1_pdu1(pf12833pf10114) PDU1AB
01936 -pf00230 -
01937 H_robinEggBlue GRM4
01938 Tsp_greenishBeige PDU1AB
01939 HMMpropdeh1_1_pdu(pf02286) PDU1AB
01940 HMMpropdeh3_1_pdu(pf02288) PDU1AB EP_NTERM
01941 HMMpropdeh2_1_pdu(pf02287) PDU1AB EP_NTERM
01942 HMM08841dhact_1_pdu(pf08841) PDU1AB
01943 HMM08841dhact_1_pdu(pf08841) PDU1AB
01944 HMM08841dhact_1_pdu(pf08841) PDU1AB
01945 HMMpropdeh3_2_pdusml(pf02288) PDU1AB
01946 H_tan PDU1AB
01947 H_driedBlood PDU1AB
01948 HMMptac_2_mix(pf06130) PDU1AB
01949 HMM15953pduput_1_pdu1(pf15953) PDU1AB EP_NTERM
01950 P_babyPink PDU1AB
01951 HMM01923cobtr_1_pdu(pf01923) PDU1AB
01952 HMM01923cobtr_1_pdu(pf01923pf03928) PDU1AB
01953 HMMalddh_1_eutpdugrm(pf00171) PDU1AB EP_NTERM
01954 HMM00465alcdh_3_mix(pf00465) PDU1AB
01955 HMMPduS_1_all(pf01512) PDU1AB
01956 HMMPduS_1_all(pf10531) PDU1AB
01957 Ts_dodgerBlue PDU1AB
01958 Hp_euts_babyPurple PDU1AB
01959 HMM10662PduV_1_mix(pf10662) PDU1AB
01960 -pf04363 -
01961 -pf04632 -
01962 -pf06445 -
01963 -pf07943pf00768 -
01964 -pf08411pf00929 -
01965 -pf01206 -
01966 -pf04143 -
01967 HMM13520tp_1_pdu(pf13520) EUT2I
01969 HMMreg00126_1_mix(pf001
BMC-P
BMC-H
BMC-T
Aldehyde dehydrogenase
PTAC
Signature enzyme
Alcohol dehydrogenase
Regulator

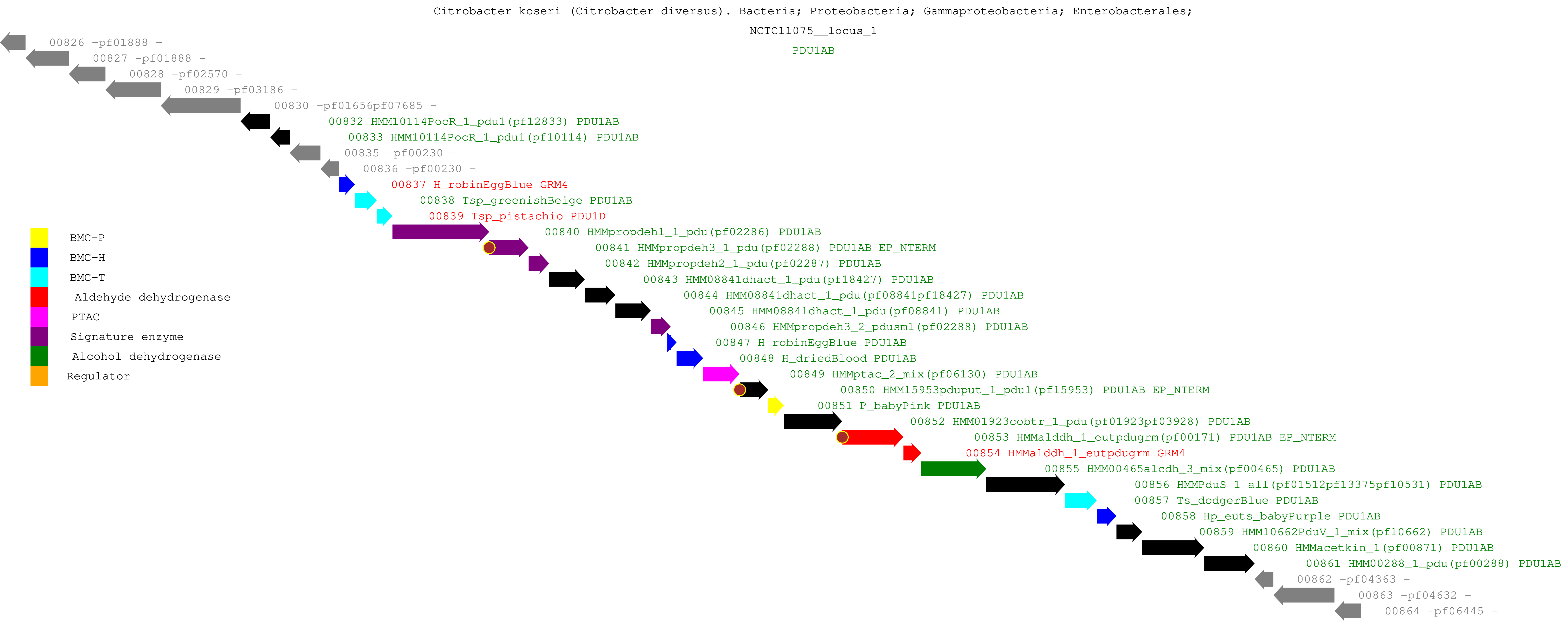
Citrobacter koseri (Citrobacter diversus). Bacteria; Proteobacteria; Gammaproteobacteria; Enterobacterales;
NCTC11075__locus_1
PDU1AB
00826 -pf01888 -
00827 -pf01888 -
00828 -pf02570 -
00829 -pf03186 -
00830 -pf01656pf07685 -
00832 HMM10114PocR_1_pdu1(pf12833) PDU1AB
00833 HMM10114PocR_1_pdu1(pf10114) PDU1AB
00835 -pf00230 -
00836 -pf00230 -
00837 H_robinEggBlue GRM4
00838 Tsp_greenishBeige PDU1AB
00839 Tsp_pistachio PDU1D
00840 HMMpropdeh1_1_pdu(pf02286) PDU1AB
00841 HMMpropdeh3_1_pdu(pf02288) PDU1AB EP_NTERM
00842 HMMpropdeh2_1_pdu(pf02287) PDU1AB
00843 HMM08841dhact_1_pdu(pf18427) PDU1AB
00844 HMM08841dhact_1_pdu(pf08841pf18427) PDU1AB
00845 HMM08841dhact_1_pdu(pf08841) PDU1AB
00846 HMMpropdeh3_2_pdusml(pf02288) PDU1AB
00847 H_robinEggBlue PDU1AB
00848 H_driedBlood PDU1AB
00849 HMMptac_2_mix(pf06130) PDU1AB
00850 HMM15953pduput_1_pdu1(pf15953) PDU1AB EP_NTERM
00851 P_babyPink PDU1AB
00852 HMM01923cobtr_1_pdu(pf01923pf03928) PDU1AB
00853 HMMalddh_1_eutpdugrm(pf00171) PDU1AB EP_NTERM
00854 HMMalddh_1_eutpdugrm GRM4
00855 HMM00465alcdh_3_mix(pf00465) PDU1AB
00856 HMMPduS_1_all(pf01512pf13375pf10531) PDU1AB
00857 Ts_dodgerBlue PDU1AB
00858 Hp_euts_babyPurple PDU1AB
00859 HMM10662PduV_1_mix(pf10662) PDU1AB
00860 HMMacetkin_1(pf00871) PDU1AB
00861 HMM00288_1_pdu(pf00288) PDU1AB
00862 -pf04363 -
00863 -pf04632 -
00864 -pf06445 -
BMC-P
BMC-H
BMC-T
Aldehyde dehydrogenase
PTAC
Signature enzyme
Alcohol dehydrogenase
Regulator

Klebsiella oxytoca. Bacteria; Proteobacteria; Gammaproteobacteria; Enterobacterales;

NCTC11355__locus_2

PDU1AB

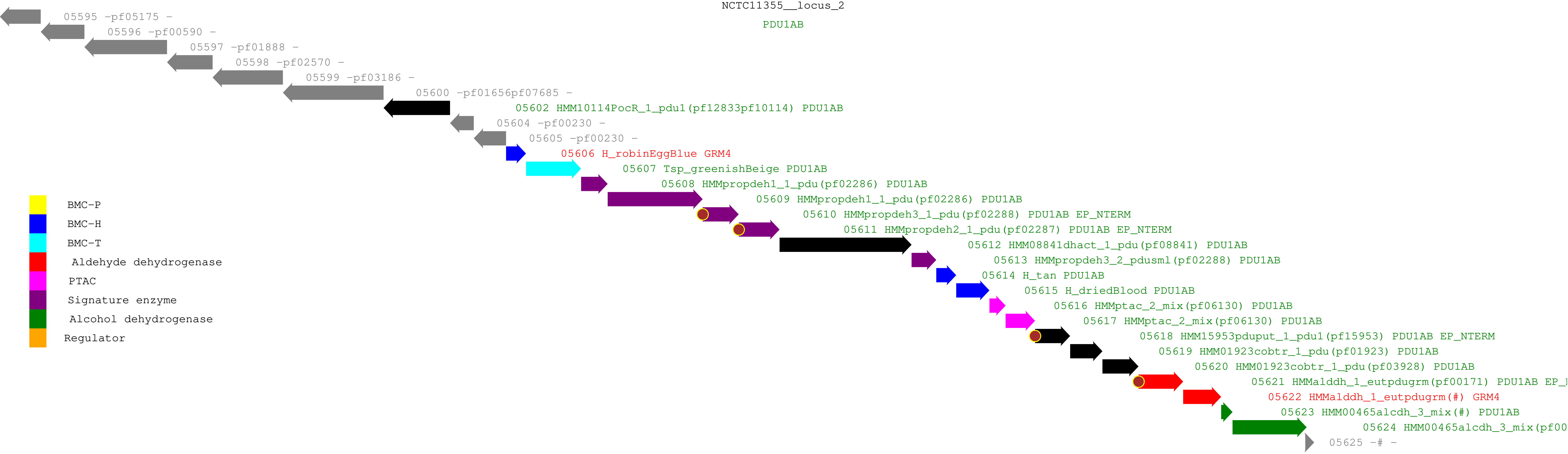

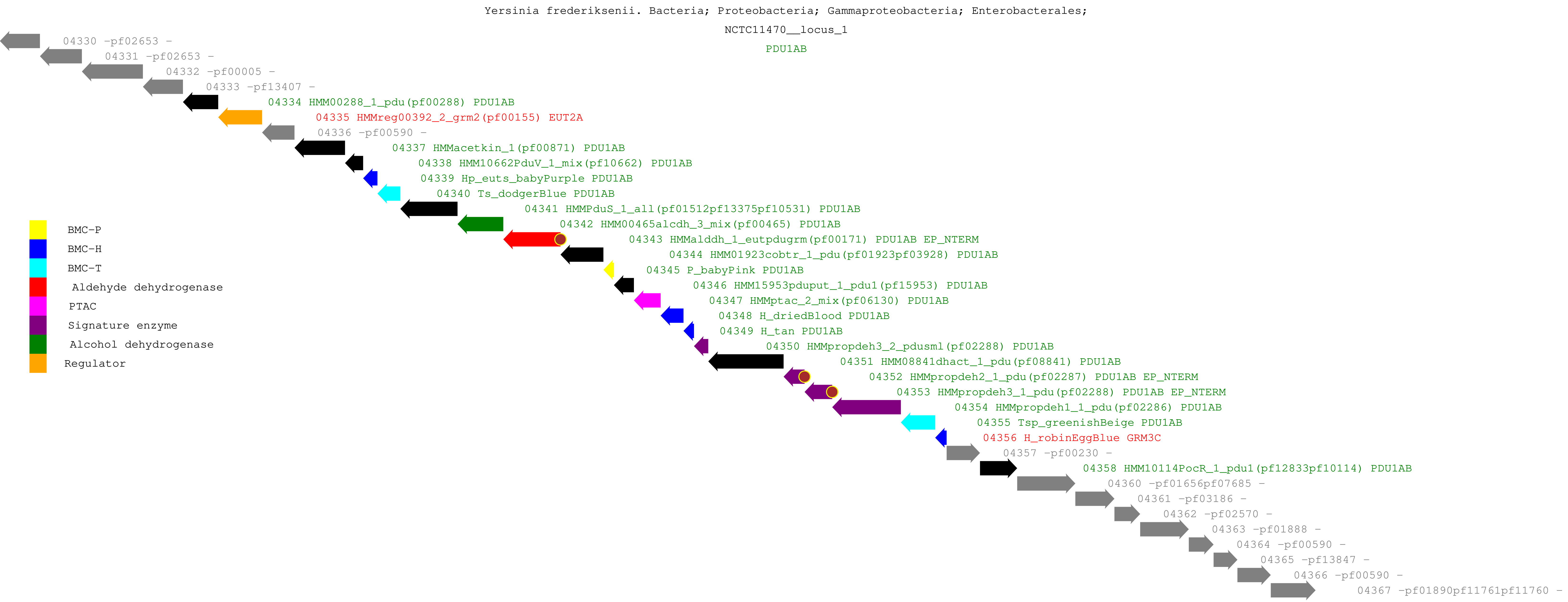
Yersinia frederiksenii. Bacteria; Proteobacteria; Gammaproteobacteria; Enterobacterales;
NCTC11470__locus_1
PDU1AB
04330 -pf02653 -
04331 -pf02653 -
04332 -pf00005 -
04333 -pf13407 -
04334 HMM00288_1_pdu(pf00288) PDU1AB
04335 HMMreg00392_2_grm2(pf00155) EUT2A
04336 -pf00590 -
04337 HMMacetkin_1(pf00871) PDU1AB
04338 HMM10662PduV_1_mix(pf10662) PDU1AB
04339 Hp_euts_babyPurple PDU1AB
04340 Ts_dodgerBlue PDU1AB
04341 HMMPduS_1_all(pf01512pf13375pf10531) PDU1AB
04342 HMM00465alcdh_3_mix(pf00465) PDU1AB
04343 HMMalddh_1_eutpdugrm(pf00171) PDU1AB EP_NTERM
04344 HMM01923cobtr_1_pdu(pf01923pf03928) PDU1AB
04345 P_babyPink PDU1AB
04346 HMM15953pduput_1_pdu1(pf15953) PDU1AB
04347 HMMptac_2_mix(pf06130) PDU1AB
04348 H_driedBlood PDU1AB
04349 H_tan PDU1AB
04350 HMMpropdeh3_2_pdusml(pf02288) PDU1AB
04351 HMM08841dhact_1_pdu(pf08841) PDU1AB
04352 HMMpropdeh2_1_pdu(pf02287) PDU1AB EP_NTERM
04353 HMMpropdeh3_1_pdu(pf02288) PDU1AB EP_NTERM
04354 HMMpropdeh1_1_pdu(pf02286) PDU1AB
04355 Tsp_greenishBeige PDU1AB
04356 H_robinEggBlue GRM3C
04357 -pf00230 -
04358 HMM10114PocR_1_pdu1(pf12833pf10114) PDU1AB
04360 -pf01656pf07685 -
04361 -pf03186 -
04362 -pf02570 -
04363 -pf01888 -
04364 -pf00590 -
04365 -pf13847 -
04366 -pf00590 -
04367 -pf01890pf11761pf11760 -
BMC-P
BMC-H
BMC-T
Aldehyde dehydrogenase
PTAC
Signature enzyme
Alcohol dehydrogenase
Regulator

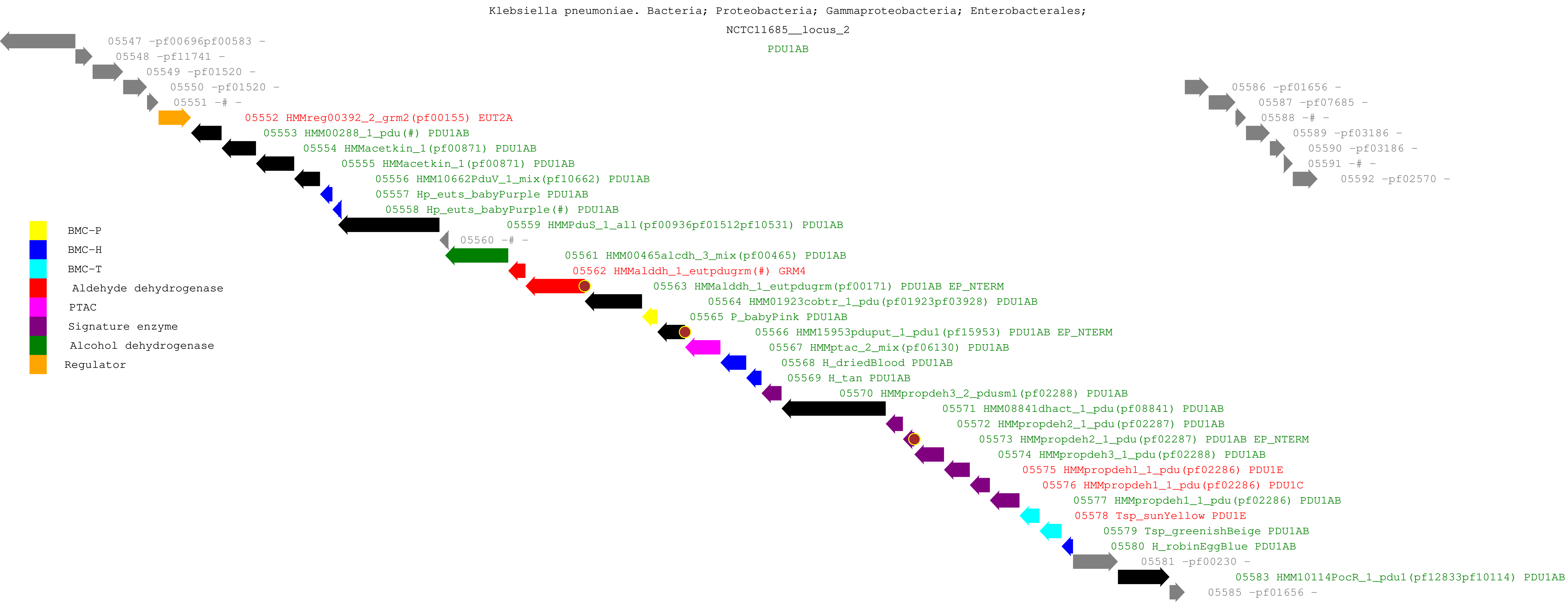
Klebsiella pneumoniae. Bacteria; Proteobacteria; Gammaproteobacteria; Enterobacterales;
NCTC11685__locus_2
PDU1AB
05547 -pf00696pf00583 -
05548 -pf11741 -
05549 -pf01520 -
05550 -pf01520 -
05551 -# -
05552 HMMreg00392_2_grm2(pf00155) EUT2A
05553 HMM00288_1_pdu(#) PDU1AB
05554 HMMacetkin_1(pf00871) PDU1AB
05555 HMMacetkin_1(pf00871) PDU1AB
05556 HMM10662PduV_1_mix(pf10662) PDU1AB
05557 Hp_euts_babyPurple PDU1AB
05558 Hp_euts_babyPurple(#) PDU1AB
05559 HMMPduS_1_all(pf00936pf01512pf10531) PDU1AB
05560 -# -
05561 HMM00465alcdh_3_mix(pf00465) PDU1AB
05562 HMMalddh_1_eutpdugrm(#) GRM4
05563 HMMalddh_1_eutpdugrm(pf00171) PDU1AB EP_NTERM
05564 HMM01923cobtr_1_pdu(pf01923pf03928) PDU1AB
05565 P_babyPink PDU1AB
05566 HMM15953pduput_1_pdu1(pf15953) PDU1AB EP_NTERM
05567 HMMptac_2_mix(pf06130) PDU1AB
05568 H_driedBlood PDU1AB
05569 H_tan PDU1AB
05570 HMMpropdeh3_2_pdusml(pf02288) PDU1AB
05571 HMM08841dhact_1_pdu(pf08841) PDU1AB
05572 HMMpropdeh2_1_pdu(pf02287) PDU1AB
05573 HMMpropdeh2_1_pdu(pf02287) PDU1AB EP_NTERM
05574 HMMpropdeh3_1_pdu(pf02288) PDU1AB
05575 HMMpropdeh1_1_pdu(pf02286) PDU1E
05576 HMMpropdeh1_1_pdu(pf02286) PDU1C
05577 HMMpropdeh1_1_pdu(pf02286) PDU1AB
05578 Tsp_sunYellow PDU1E
05579 Tsp_greenishBeige PDU1AB
05580 H_robinEggBlue PDU1AB
05581 -pf00230 -
05583 HMM10114PocR_1_pdu1(pf12833pf10114) PDU1AB
05585 -pf01656 -
05586 -pf01656 -
05587 -pf07685 -
05588 -# -
05589 -pf03186 -
05590 -pf03186 -
05591 -# -
05592 -pf02570 -
BMC-P
BMC-H
BMC-T
Aldehyde dehydrogenase
PTAC
Signature enzyme
Alcohol dehydrogenase
Regulator

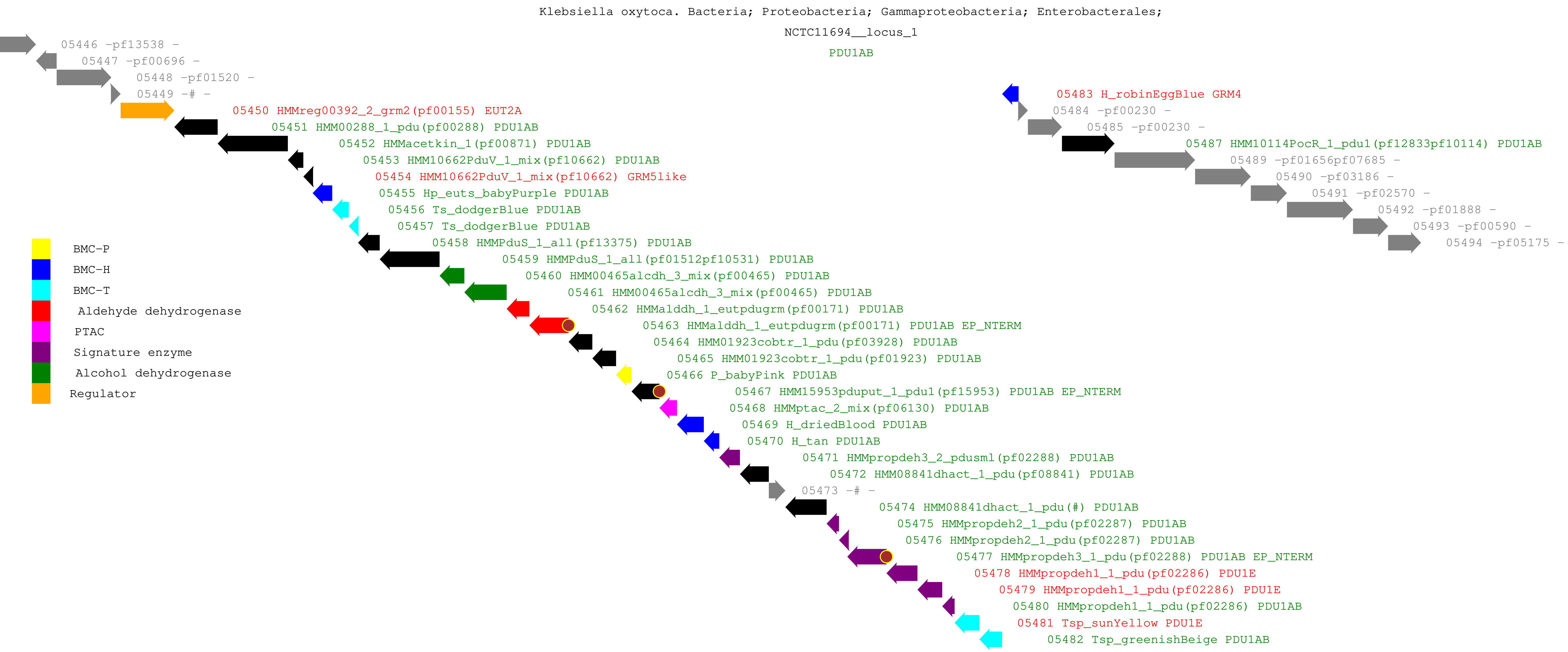

Klebsiella oxytoca. Bacteria; Proteobacteria; Gammaproteobacteria; Enterobacterales;
NCTC11694__locus_1
PDU1AB
05446 -pf13538 -
05447 -pf00696 -
05448 -pf01520 -
05449 -# -
05450 HMMreg00392_2_grm2(pf00155) EUT2A
05451 HMM00288_1_pdu(pf00288) PDU1AB
05452 HMMacetkin_1(pf00871) PDU1AB
05453 HMM10662PduV_1_mix(pf10662) PDU1AB
05454 HMM10662PduV_1_mix(pf10662) GRM5like
05455 Hp_euts_babyPurple PDU1AB
05456 Ts_dodgerBlue PDU1AB
05457 Ts_dodgerBlue PDU1AB
05458 HMMPduS_1_all(pf13375) PDU1AB
05459 HMMPduS_1_all(pf01512pf10531) PDU1AB
05460 HMM00465alcdh_3_mix(pf00465) PDU1AB
05461 HMM00465alcdh_3_mix(pf00465) PDU1AB
05462 HMMalddh_1_eutpdugrm(pf00171) PDU1AB
05463 HMMalddh_1_eutpdugrm(pf00171) PDU1AB EP_NTERM
05464 HMM01923cobtr_1_pdu(pf03928) PDU1AB
05465 HMM01923cobtr_1_pdu(pf01923) PDU1AB
05466 P_babyPink PDU1AB
05467 HMM15953pduput_1_pdu1(pf15953) PDU1AB EP_NTERM
05468 HMMptac_2_mix(pf06130) PDU1AB
05469 H_driedBlood PDU1AB
05470 H_tan PDU1AB
05471 HMMpropdeh3_2_pdusml(pf02288) PDU1AB
05472 HMM08841dhact_1_pdu(pf08841) PDU1AB
05473 -# -
05474 HMM08841dhact_1_pdu(#) PDU1AB
05475 HMMpropdeh2_1_pdu(pf02287) PDU1AB
05476 HMMpropdeh2_1_pdu(pf02287) PDU1AB
05477 HMMpropdeh3_1_pdu(pf02288) PDU1AB EP_NTERM
05478 HMMpropdeh1_1_pdu(pf02286) PDU1E
05479 HMMpropdeh1_1_pdu(pf02286) PDU1E
05480 HMMpropdeh1_1_pdu(pf02286) PDU1AB
05481 Tsp_sunYellow PDU1E
05482 Tsp_greenishBeige PDU1AB
05483 H_robinEggBlue GRM4
05484 -pf00230 -
05485 -pf00230 -
05487 HMM10114PocR_1_pdu1(pf12833pf10114) PDU1AB
05489 -pf01656pf07685 -
05490 -pf03186 -
05491 -pf02570 -
05492 -pf01888 -
05493 -pf00590 -
05494 -pf05175 -
BMC-P
BMC-H
BMC-T
Aldehyde dehydrogenase
PTAC
Signature enzyme
Alcohol dehydrogenase
Regulator

Salmonella sp. NCTC 11881. Bacteria; Proteobacteria; Gammaproteobacteria; Enterobacterales;

NCTC11881__locus_2

PDU1AB

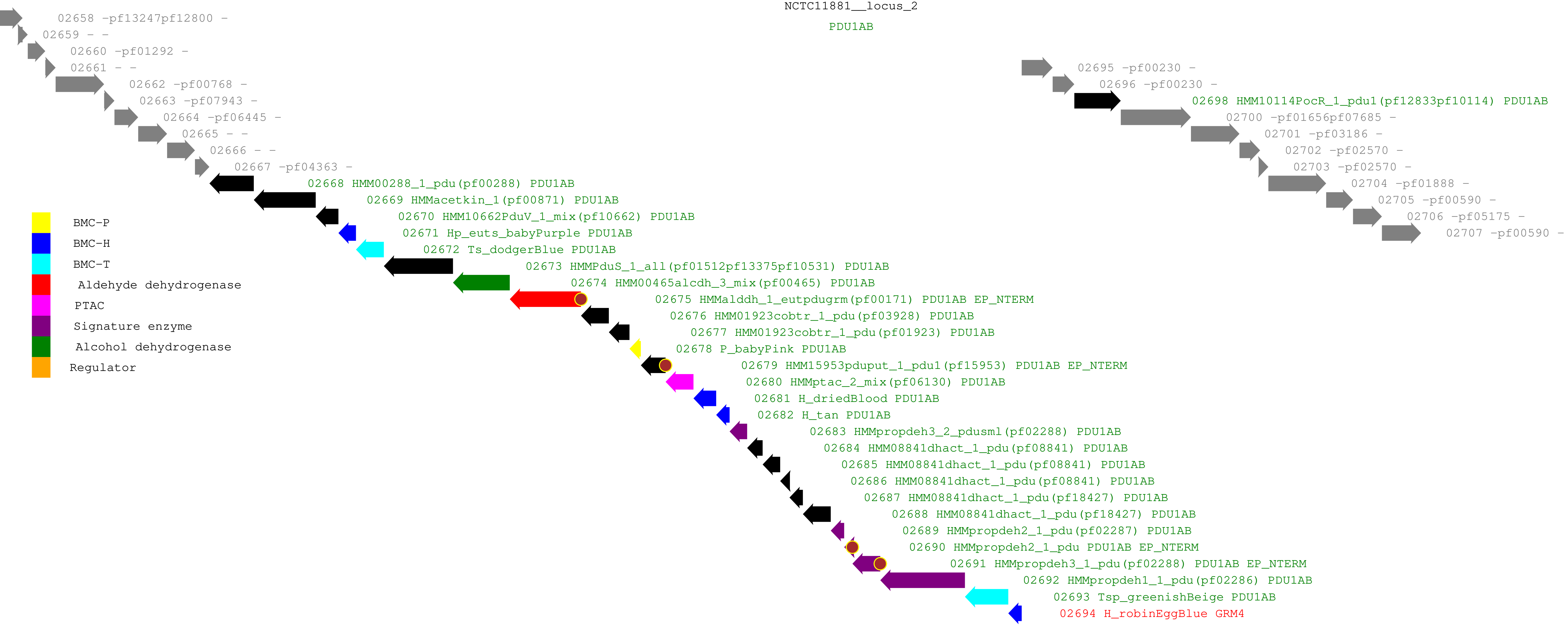

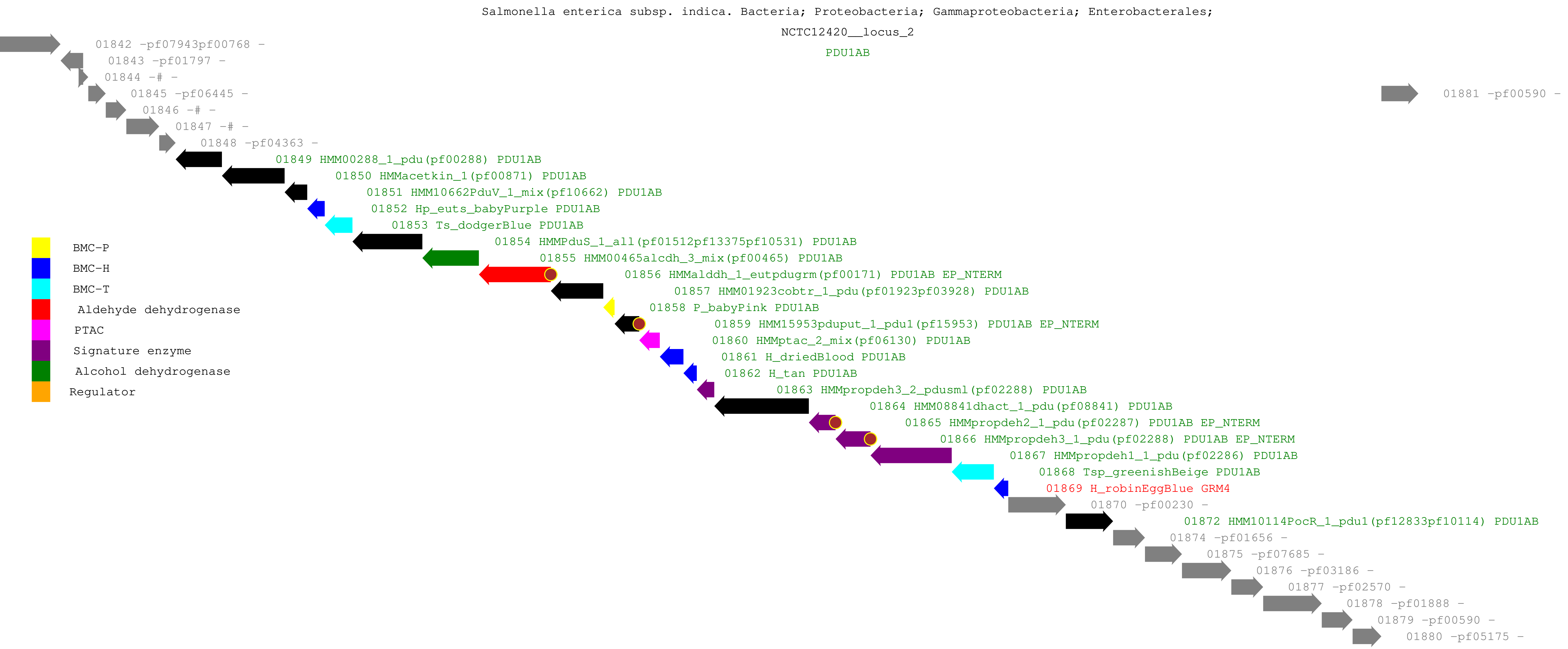
Salmonella enterica subsp. indica. Bacteria; Proteobacteria; Gammaproteobacteria; Enterobacterales;
NCTC12420__locus_2
PDU1AB
01842 -pf07943pf00768 -
01843 -pf01797 -
01844 -# -
01845 -pf06445 -
01846 -# -
01847 -# -
01848 -pf04363 -
01849 HMM00288_1_pdu(pf00288) PDU1AB
01850 HMMacetkin_1(pf00871) PDU1AB
01851 HMM10662PduV_1_mix(pf10662) PDU1AB
01852 Hp_euts_babyPurple PDU1AB
01853 Ts_dodgerBlue PDU1AB
01854 HMMPduS_1_all(pf01512pf13375pf10531) PDU1AB
01855 HMM00465alcdh_3_mix(pf00465) PDU1AB
01856 HMMalddh_1_eutpdugrm(pf00171) PDU1AB EP_NTERM
01857 HMM01923cobtr_1_pdu(pf01923pf03928) PDU1AB
01858 P_babyPink PDU1AB
01859 HMM15953pduput_1_pdu1(pf15953) PDU1AB EP_NTERM
01860 HMMptac_2_mix(pf06130) PDU1AB
01861 H_driedBlood PDU1AB
01862 H_tan PDU1AB
01863 HMMpropdeh3_2_pdusml(pf02288) PDU1AB
01864 HMM08841dhact_1_pdu(pf08841) PDU1AB
01865 HMMpropdeh2_1_pdu(pf02287) PDU1AB EP_NTERM
01866 HMMpropdeh3_1_pdu(pf02288) PDU1AB EP_NTERM
01867 HMMpropdeh1_1_pdu(pf02286) PDU1AB
01868 Tsp_greenishBeige PDU1AB
01869 H_robinEggBlue GRM4
01870 -pf00230 -
01872 HMM10114PocR_1_pdu1(pf12833pf10114) PDU1AB
01874 -pf01656 -
01875 -pf07685 -
01876 -pf03186 -
01877 -pf02570 -
01878 -pf01888 -
01879 -pf00590 -
01880 -pf05175 -
01881 -pf00590 -
BMC-P
BMC-H
BMC-T
Aldehyde dehydrogenase
PTAC
Signature enzyme
Alcohol dehydrogenase
Regulator

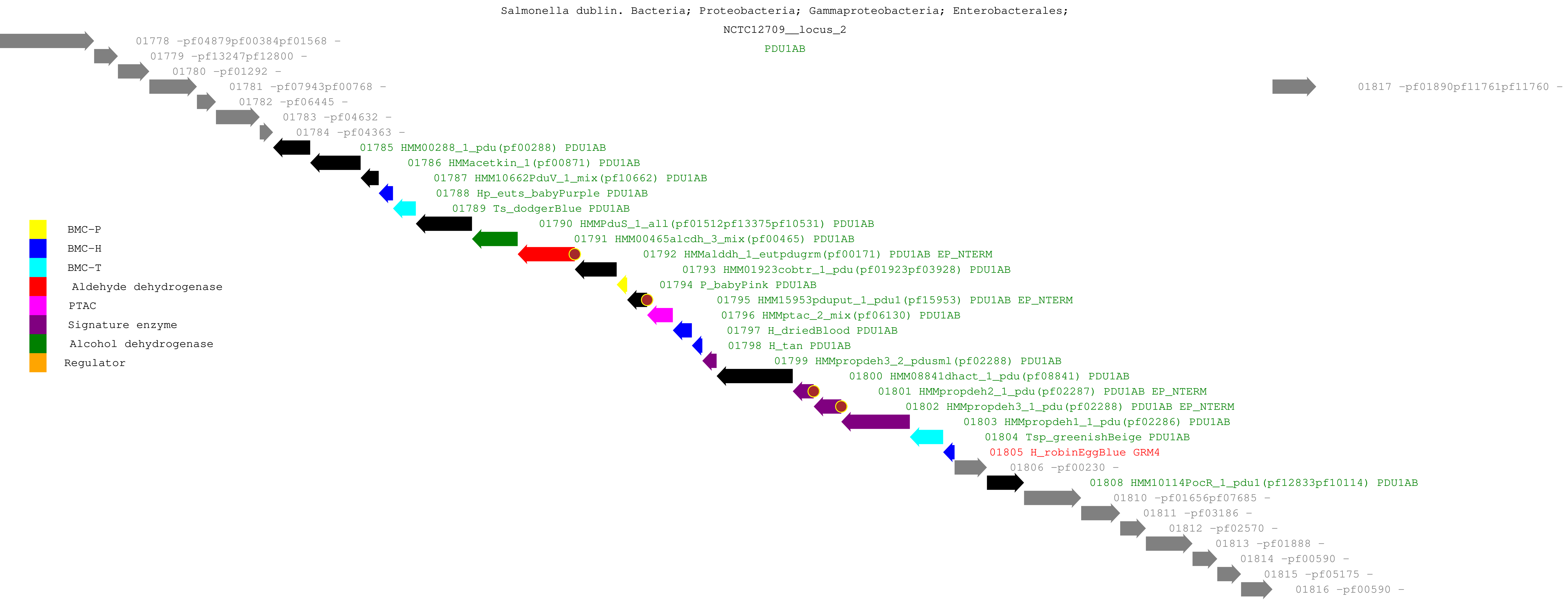
Salmonella dublin. Bacteria; Proteobacteria; Gammaproteobacteria; Enterobacterales;
NCTC12709__locus_2
PDU1AB
01778 -pf04879pf00384pf01568 -
01779 -pf13247pf12800 -
01780 -pf01292 -
01781 -pf07943pf00768 -
01782 -pf06445 -
01783 -pf04632 -
01784 -pf04363 -
01785 HMM00288_1_pdu(pf00288) PDU1AB
01786 HMMacetkin_1(pf00871) PDU1AB
01787 HMM10662PduV_1_mix(pf10662) PDU1AB
01788 Hp_euts_babyPurple PDU1AB
01789 Ts_dodgerBlue PDU1AB
01790 HMMPduS_1_all(pf01512pf13375pf10531) PDU1AB
01791 HMM00465alcdh_3_mix(pf00465) PDU1AB
01792 HMMalddh_1_eutpdugrm(pf00171) PDU1AB EP_NTERM
01793 HMM01923cobtr_1_pdu(pf01923pf03928) PDU1AB
01794 P_babyPink PDU1AB
01795 HMM15953pduput_1_pdu1(pf15953) PDU1AB EP_NTERM
01796 HMMptac_2_mix(pf06130) PDU1AB
01797 H_driedBlood PDU1AB
01798 H_tan PDU1AB
01799 HMMpropdeh3_2_pdusml(pf02288) PDU1AB
01800 HMM08841dhact_1_pdu(pf08841) PDU1AB
01801 HMMpropdeh2_1_pdu(pf02287) PDU1AB EP_NTERM
01802 HMMpropdeh3_1_pdu(pf02288) PDU1AB EP_NTERM
01803 HMMpropdeh1_1_pdu(pf02286) PDU1AB
01804 Tsp_greenishBeige PDU1AB
01805 H_robinEggBlue GRM4
01806 -pf00230 -
01808 HMM10114PocR_1_pdu1(pf12833pf10114) PDU1AB
01810 -pf01656pf07685 -
01811 -pf03186 -
01812 -pf02570 -
01813 -pf01888 -
01814 -pf00590 -
01815 -pf05175 -
01816 -pf00590 -
01817 -pf01890pf11761pf11760 -
BMC-P
BMC-H
BMC-T
Aldehyde dehydrogenase
PTAC
Signature enzyme
Alcohol dehydrogenase
Regulator

Serratia liquefaciens. Bacteria; Proteobacteria; Gammaproteobacteria; Enterobacterales;

NCTC12962__locus_1

PDU1AB

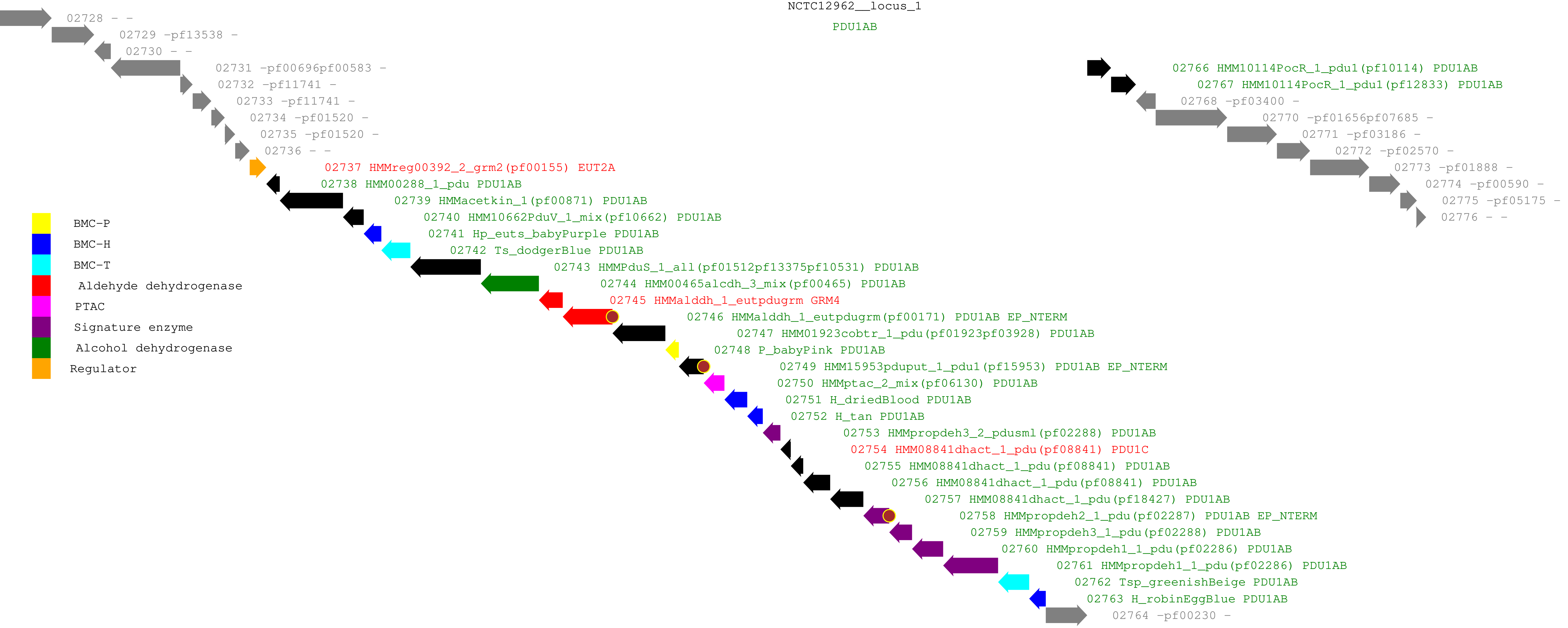

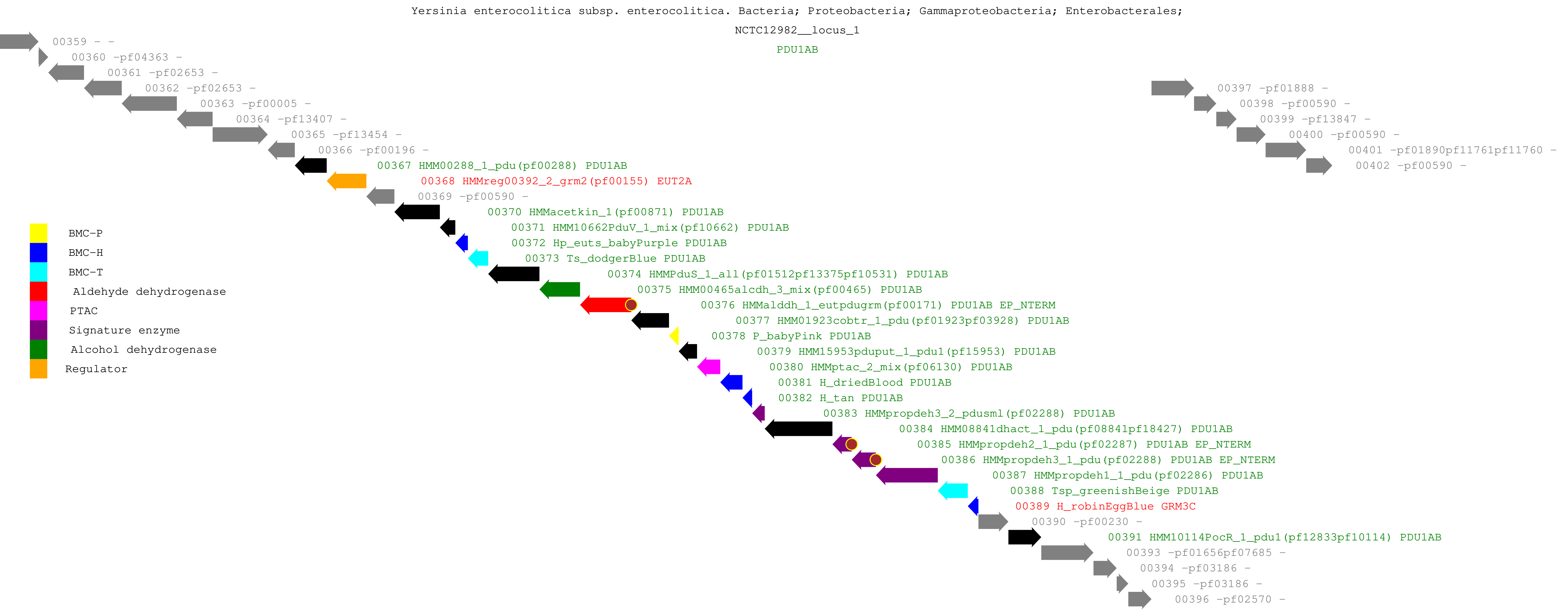
Yersinia enterocolitica subsp. enterocolitica. Bacteria; Proteobacteria; Gammaproteobacteria; Enterobacterales;
NCTC12982__locus_1
PDU1AB
00359 - -
00360 -pf04363 -
00361 -pf02653 -
00362 -pf02653 -
00363 -pf00005 -
00364 -pf13407 -
00365 -pf13454 -
00366 -pf00196 -
00367 HMM00288_1_pdu(pf00288) PDU1AB
00368 HMMreg00392_2_grm2(pf00155) EUT2A
00369 -pf00590 -
00370 HMMacetkin_1(pf00871) PDU1AB
00371 HMM10662PduV_1_mix(pf10662) PDU1AB
00372 Hp_euts_babyPurple PDU1AB
00373 Ts_dodgerBlue PDU1AB
00374 HMMPduS_1_all(pf01512pf13375pf10531) PDU1AB
00375 HMM00465alcdh_3_mix(pf00465) PDU1AB
00376 HMMalddh_1_eutpdugrm(pf00171) PDU1AB EP_NTERM
00377 HMM01923cobtr_1_pdu(pf01923pf03928) PDU1AB
00378 P_babyPink PDU1AB
00379 HMM15953pduput_1_pdu1(pf15953) PDU1AB
00380 HMMptac_2_mix(pf06130) PDU1AB
00381 H_driedBlood PDU1AB
00382 H_tan PDU1AB
00383 HMMpropdeh3_2_pdusml(pf02288) PDU1AB
00384 HMM08841dhact_1_pdu(pf08841pf18427) PDU1AB
00385 HMMpropdeh2_1_pdu(pf02287) PDU1AB EP_NTERM
00386 HMMpropdeh3_1_pdu(pf02288) PDU1AB EP_NTERM
00387 HMMpropdeh1_1_pdu(pf02286) PDU1AB
00388 Tsp_greenishBeige PDU1AB
00389 H_robinEggBlue GRM3C
00390 -pf00230 -
00391 HMM10114PocR_1_pdu1(pf12833pf10114) PDU1AB
00393 -pf01656pf07685 -
00394 -pf03186 -
00395 -pf03186 -
00396 -pf02570 -
00397 -pf01888 -
00398 -pf00590 -
00399 -pf13847 -
00400 -pf00590 -
00401 -pf01890pf11761pf11760 -
00402 -pf00590 -
BMC-P
BMC-H
BMC-T
Aldehyde dehydrogenase
PTAC
Signature enzyme
Alcohol dehydrogenase
Regulator

Salmonella enterica I. Bacteria; Proteobacteria; Gammaproteobacteria; Enterobacterales;

NCTC129__locus_1

PDU1AB

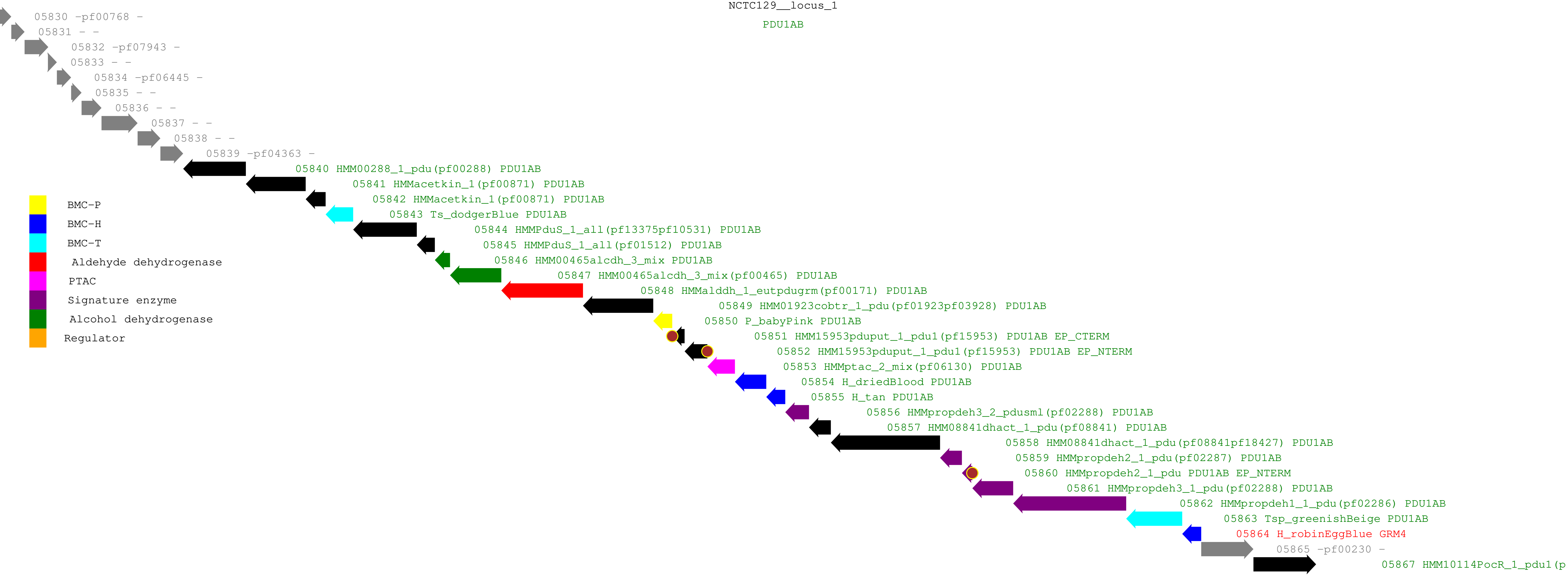

Salmonella typhimurium. Bacteria; Proteobacteria; Gammaproteobacteria; Enterobacterales;

NCTC13348__locus_1

PDU1AB

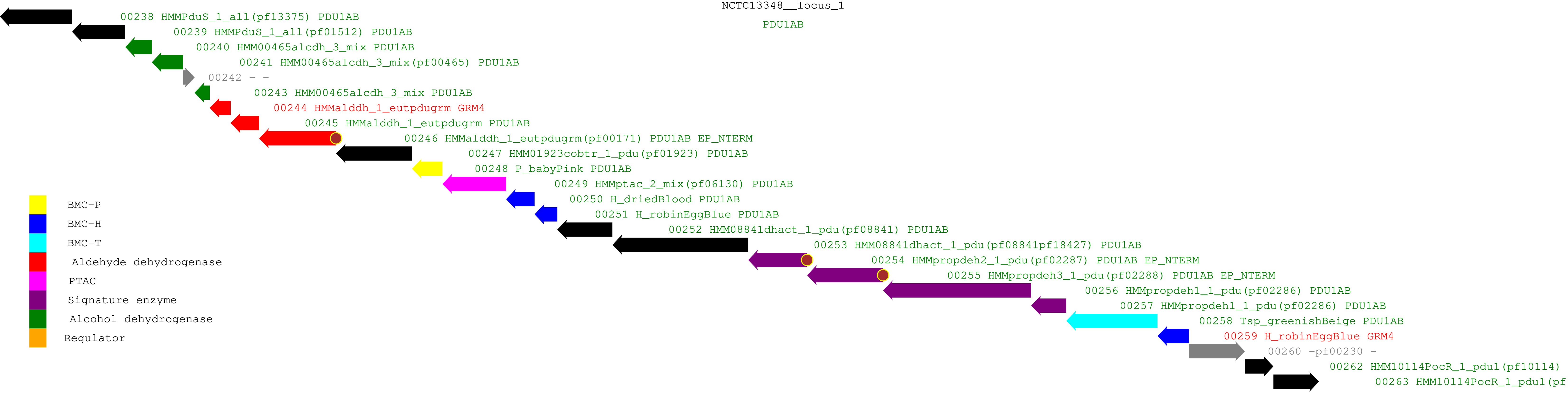

Klebsiella pneumoniae. Bacteria; Proteobacteria; Gammaproteobacteria; Enterobacterales;

NCTC13443__locus_2

PDU1AB

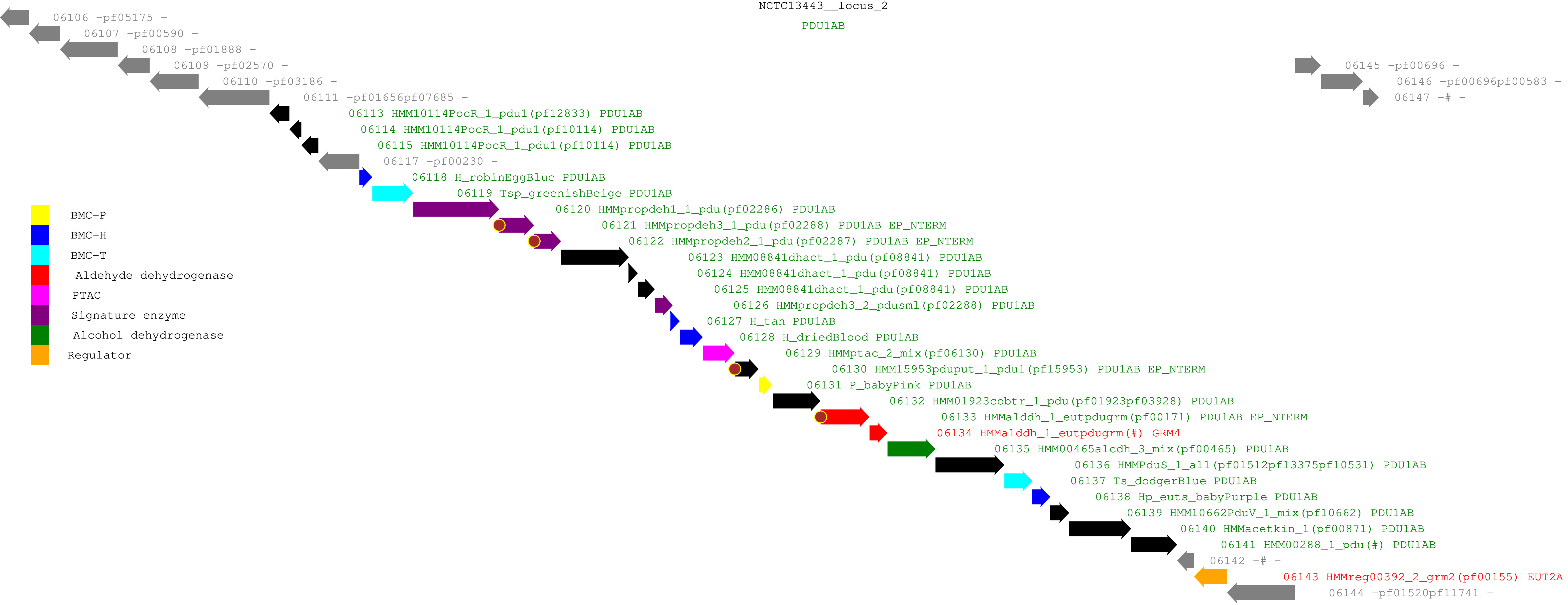

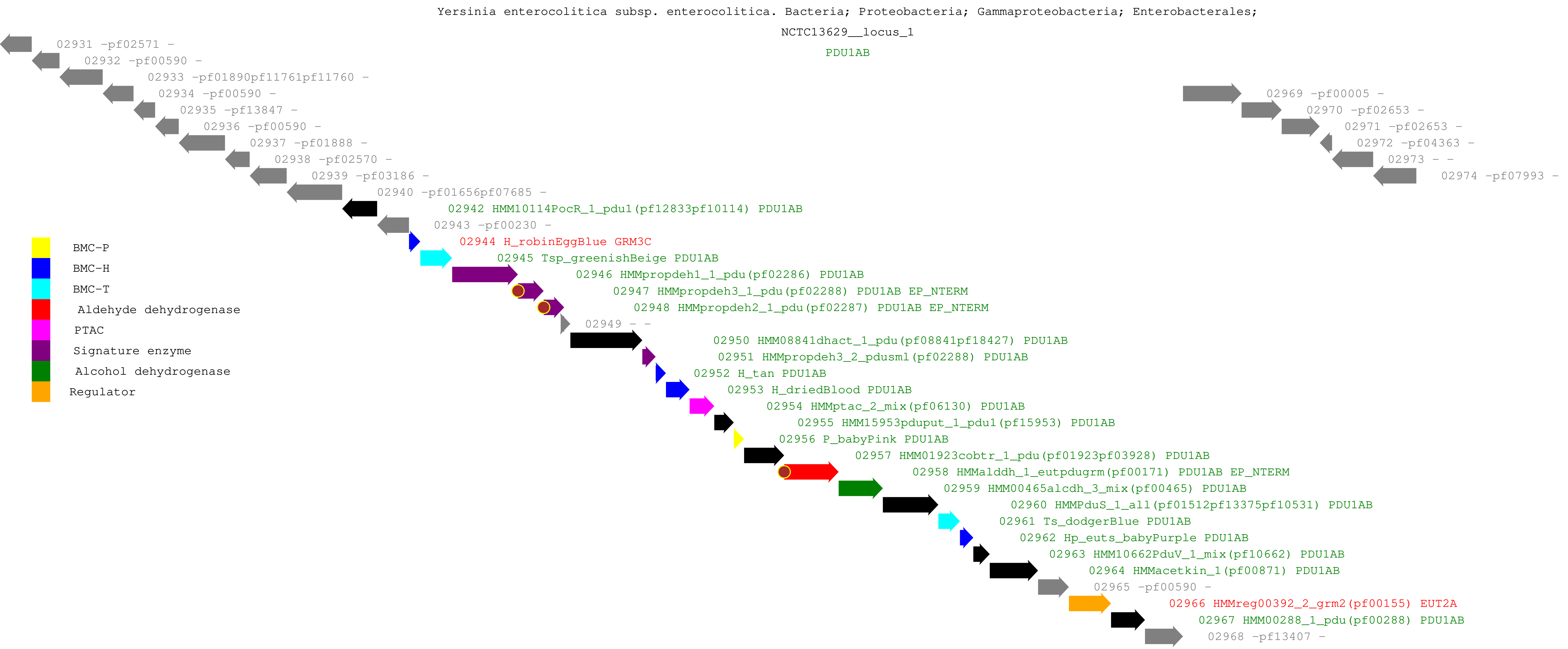

Yersinia enterocolitica subsp. enterocolitica. Bacteria; Proteobacteria; Gammaproteobacteria; Enterobacterales;
NCTC13629__locus_1
PDU1AB
02931 -pf02571 -
02932 -pf00590 -
02933 -pf01890pf11761pf11760 -
02934 -pf00590 -
02935 -pf13847 -
02936 -pf00590 -
02937 -pf01888 -
02938 -pf02570 -
02939 -pf03186 -
02940 -pf01656pf07685 -
02942 HMM10114PocR_1_pdu1(pf12833pf10114) PDU1AB
02943 -pf00230 -
02944 H_robinEggBlue GRM3C
02945 Tsp_greenishBeige PDU1AB
02946 HMMpropdeh1_1_pdu(pf02286) PDU1AB
02947 HMMpropdeh3_1_pdu(pf02288) PDU1AB EP_NTERM
02948 HMMpropdeh2_1_pdu(pf02287) PDU1AB EP_NTERM
02949 - -
02950 HMM08841dhact_1_pdu(pf08841pf18427) PDU1AB
02951 HMMpropdeh3_2_pdusml(pf02288) PDU1AB
02952 H_tan PDU1AB
02953 H_driedBlood PDU1AB
02954 HMMptac_2_mix(pf06130) PDU1AB
02955 HMM15953pduput_1_pdu1(pf15953) PDU1AB
02956 P_babyPink PDU1AB
02957 HMM01923cobtr_1_pdu(pf01923pf03928) PDU1AB
02958 HMMalddh_1_eutpdugrm(pf00171) PDU1AB EP_NTERM
02959 HMM00465alcdh_3_mix(pf00465) PDU1AB
02960 HMMPduS_1_all(pf01512pf13375pf10531) PDU1AB
02961 Ts_dodgerBlue PDU1AB
02962 Hp_euts_babyPurple PDU1AB
02963 HMM10662PduV_1_mix(pf10662) PDU1AB
02964 HMMacetkin_1(pf00871) PDU1AB
02965 -pf00590 -
02966 HMMreg00392_2_grm2(pf00155) EUT2A
02967 HMM00288_1_pdu(pf00288) PDU1AB
02968 -pf13407 -
02969 -pf00005 -
02970 -pf02653 -
02971 -pf02653 -
02972 -pf04363 -
02973 - -
02974 -pf07993 -
BMC-P
BMC-H
BMC-T
Aldehyde dehydrogenase
PTAC
Signature enzyme
Alcohol dehydrogenase
Regulator

Klebsiella pneumoniae. Bacteria; Proteobacteria; Gammaproteobacteria; Enterobacterales;

NCTC13635__locus_2

PDU1AB

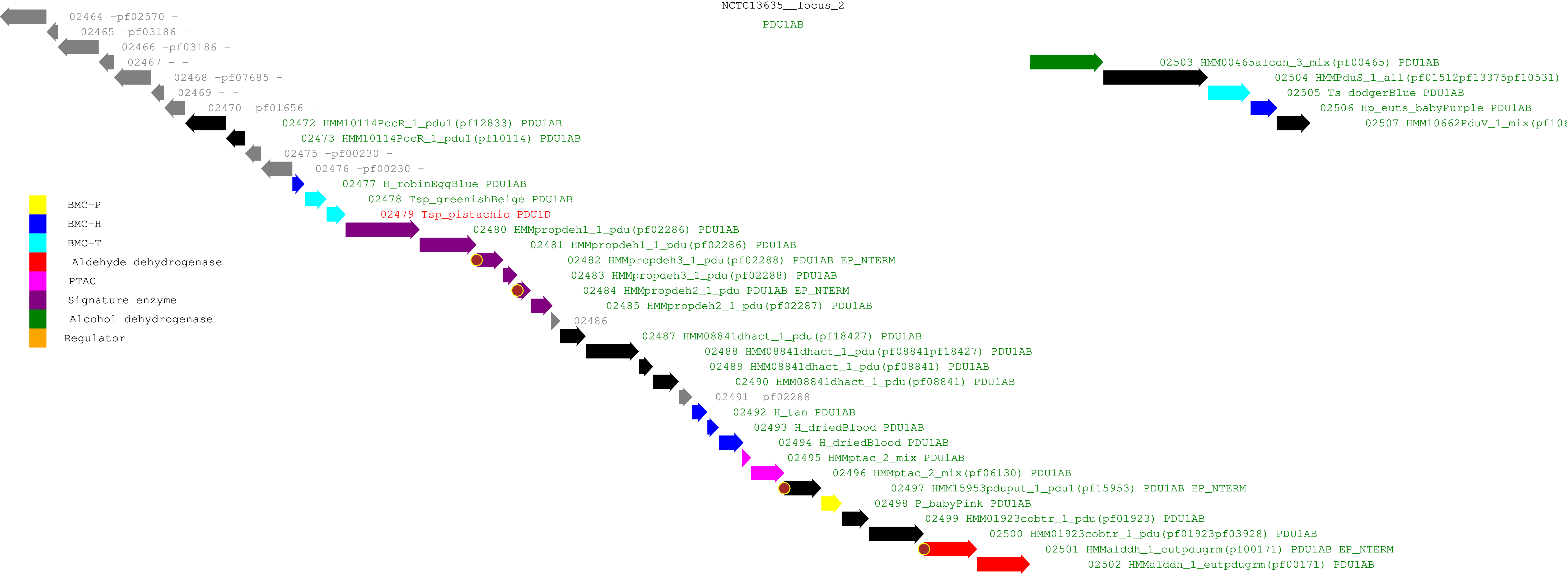

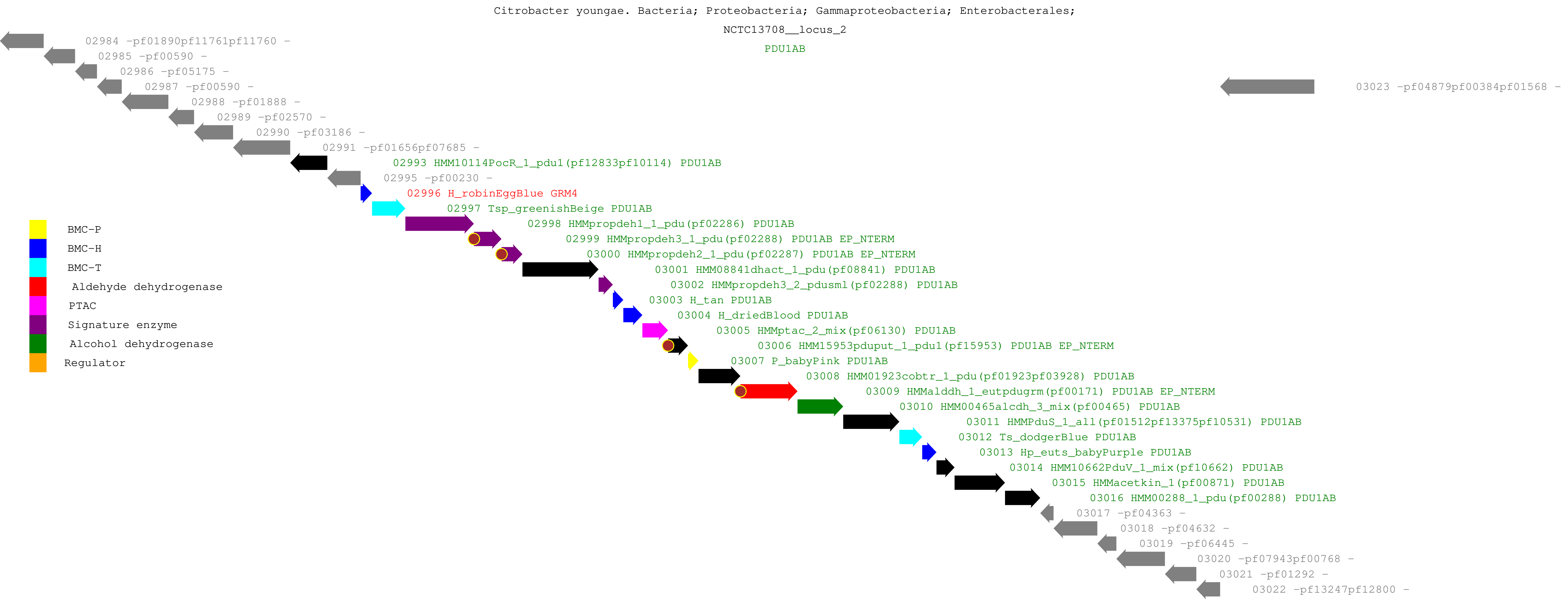
Citrobacter youngae. Bacteria; Proteobacteria; Gammaproteobacteria; Enterobacterales;
NCTC13708__locus_2
PDU1AB
02984 -pf01890pf11761pf11760 -
02985 -pf00590 -
02986 -pf05175 -
02987 -pf00590 -
02988 -pf01888 -
02989 -pf02570 -
02990 -pf03186 -
02991 -pf01656pf07685 -
02993 HMM10114PocR_1_pdu1(pf12833pf10114) PDU1AB
02995 -pf00230 -
02996 H_robinEggBlue GRM4
02997 Tsp_greenishBeige PDU1AB
02998 HMMpropdeh1_1_pdu(pf02286) PDU1AB
02999 HMMpropdeh3_1_pdu(pf02288) PDU1AB EP_NTERM
03000 HMMpropdeh2_1_pdu(pf02287) PDU1AB EP_NTERM
03001 HMM08841dhact_1_pdu(pf08841) PDU1AB
03002 HMMpropdeh3_2_pdusml(pf02288) PDU1AB
03003 H_tan PDU1AB
03004 H_driedBlood PDU1AB
03005 HMMptac_2_mix(pf06130) PDU1AB
03006 HMM15953pduput_1_pdu1(pf15953) PDU1AB EP_NTERM
03007 P_babyPink PDU1AB
03008 HMM01923cobtr_1_pdu(pf01923pf03928) PDU1AB
03009 HMMalddh_1_eutpdugrm(pf00171) PDU1AB EP_NTERM
03010 HMM00465alcdh_3_mix(pf00465) PDU1AB
03011 HMMPduS_1_all(pf01512pf13375pf10531) PDU1AB
03012 Ts_dodgerBlue PDU1AB
03013 Hp_euts_babyPurple PDU1AB
03014 HMM10662PduV_1_mix(pf10662) PDU1AB
03015 HMMacetkin_1(pf00871) PDU1AB
03016 HMM00288_1_pdu(pf00288) PDU1AB
03017 -pf04363 -
03018 -pf04632 -
03019 -pf06445 -
03020 -pf07943pf00768 -
03021 -pf01292 -
03022 -pf13247pf12800 -
03023 -pf04879pf00384pf01568 -
BMC-P
BMC-H
BMC-T
Aldehyde dehydrogenase
PTAC
Signature enzyme
Alcohol dehydrogenase
Regulator

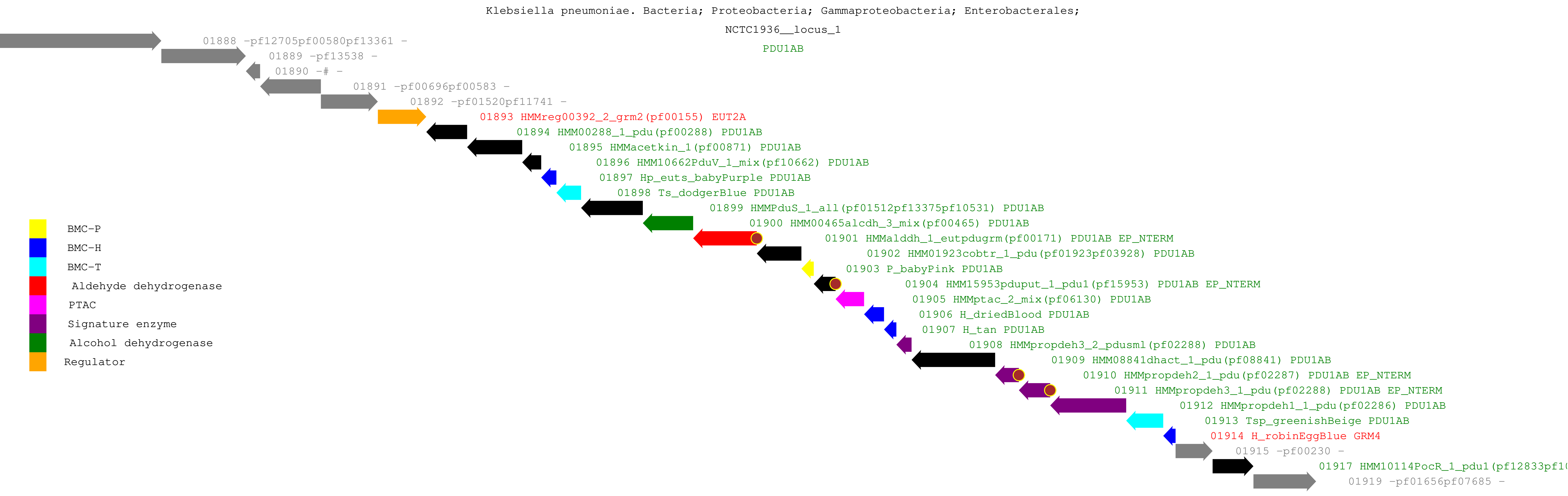
Klebsiella pneumoniae. Bacteria; Proteobacteria; Gammaproteobacteria; Enterobacterales;
NCTC1936__locus_1
PDU1AB
01888 -pf12705pf00580pf13361 -
01889 -pf13538 -
01890 -# -
01891 -pf00696pf00583 -
01892 -pf01520pf11741 -
01893 HMMreg00392_2_grm2(pf00155) EUT2A
01894 HMM00288_1_pdu(pf00288) PDU1AB
01895 HMMacetkin_1(pf00871) PDU1AB
01896 HMM10662PduV_1_mix(pf10662) PDU1AB
01897 Hp_euts_babyPurple PDU1AB
01898 Ts_dodgerBlue PDU1AB
01899 HMMPduS_1_all(pf01512pf13375pf10531) PDU1AB
01900 HMM00465alcdh_3_mix(pf00465) PDU1AB
01901 HMMalddh_1_eutpdugrm(pf00171) PDU1AB EP_NTERM
01902 HMM01923cobtr_1_pdu(pf01923pf03928) PDU1AB
01903 P_babyPink PDU1AB
01904 HMM15953pduput_1_pdu1(pf15953) PDU1AB EP_NTERM
01905 HMMptac_2_mix(pf06130) PDU1AB
01906 H_driedBlood PDU1AB
01907 H_tan PDU1AB
01908 HMMpropdeh3_2_pdusml(pf02288) PDU1AB
01909 HMM08841dhact_1_pdu(pf08841) PDU1AB
01910 HMMpropdeh2_1_pdu(pf02287) PDU1AB EP_NTERM
01911 HMMpropdeh3_1_pdu(pf02288) PDU1AB EP_NTERM
01912 HMMpropdeh1_1_pdu(pf02286) PDU1AB
01913 Tsp_greenishBeige PDU1AB
01914 H_robinEggBlue GRM4
01915 -pf00230 -
01917 HMM10114PocR_1_pdu1(pf12833pf1
01919 -pf01656pf07685 -
BMC-P
BMC-H
BMC-T
Aldehyde dehydrogenase
PTAC
Signature enzyme
Alcohol dehydrogenase
Regulator

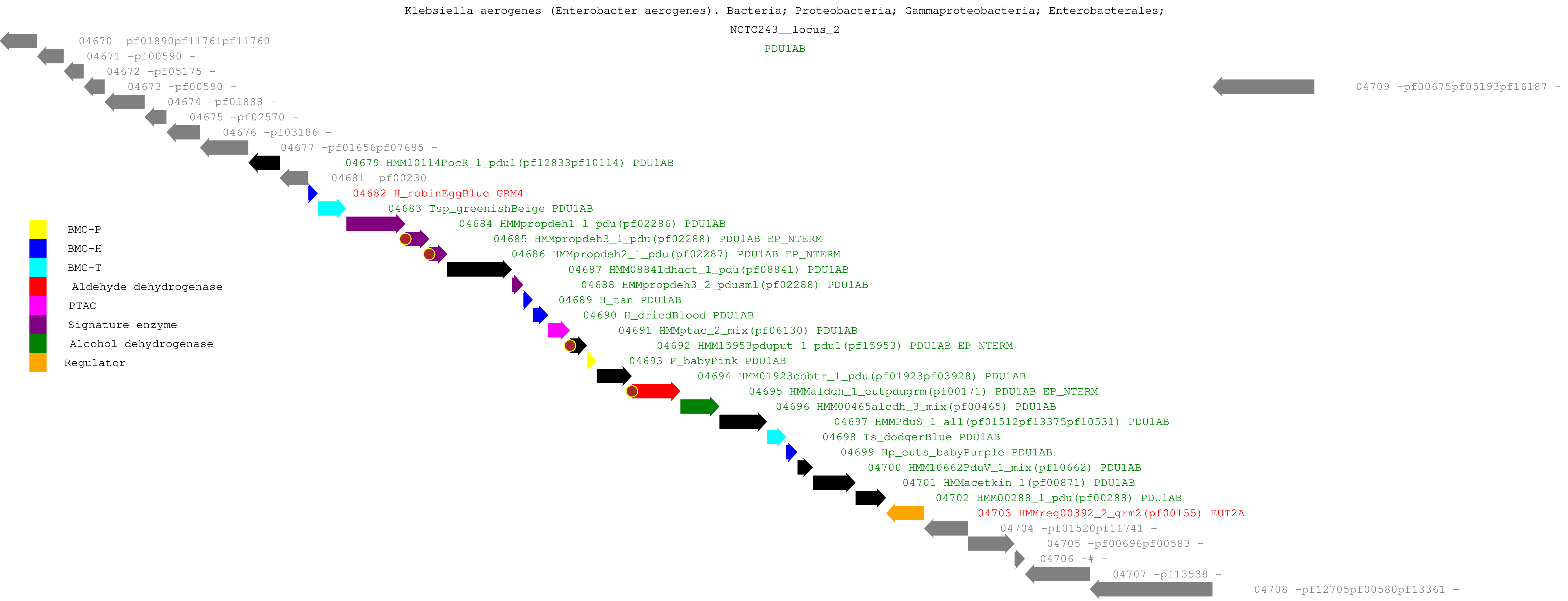
Klebsiella aerogenes (Enterobacter aerogenes). Bacteria; Proteobacteria; Gammaproteobacteria; Enterobacterales;
NCTC243__locus_2
PDU1AB
04670 -pf01890pf11761pf11760 -
04671 -pf00590 -
04672 -pf05175 -
04673 -pf00590 -
04674 -pf01888 -
04675 -pf02570 -
04676 -pf03186 -
04677 -pf01656pf07685 -
04679 HMM10114PocR_1_pdu1(pf12833pf10114) PDU1AB
04681 -pf00230 -
04682 H_robinEggBlue GRM4
04683 Tsp_greenishBeige PDU1AB
04684 HMMpropdeh1_1_pdu(pf02286) PDU1AB
04685 HMMpropdeh3_1_pdu(pf02288) PDU1AB EP_NTERM
04686 HMMpropdeh2_1_pdu(pf02287) PDU1AB EP_NTERM
04687 HMM08841dhact_1_pdu(pf08841) PDU1AB
04688 HMMpropdeh3_2_pdusml(pf02288) PDU1AB
04689 H_tan PDU1AB
04690 H_driedBlood PDU1AB
04691 HMMptac_2_mix(pf06130) PDU1AB
04692 HMM15953pduput_1_pdu1(pf15953) PDU1AB EP_NTERM
04693 P_babyPink PDU1AB
04694 HMM01923cobtr_1_pdu(pf01923pf03928) PDU1AB
04695 HMMalddh_1_eutpdugrm(pf00171) PDU1AB EP_NTERM
04696 HMM00465alcdh_3_mix(pf00465) PDU1AB
04697 HMMPduS_1_all(pf01512pf13375pf10531) PDU1AB
04698 Ts_dodgerBlue PDU1AB
04699 Hp_euts_babyPurple PDU1AB
04700 HMM10662PduV_1_mix(pf10662) PDU1AB
04701 HMMacetkin_1(pf00871) PDU1AB
04702 HMM00288_1_pdu(pf00288) PDU1AB
04703 HMMreg00392_2_grm2(pf00155) EUT2A
04704 -pf01520pf11741 -
04705 -pf00696pf00583 -
04706 -# -
04707 -pf13538 -
04708 -pf12705pf00580pf13361 -
04709 -pf00675pf05193pf16187 -
BMC-P
BMC-H
BMC-T
Aldehyde dehydrogenase
PTAC
Signature enzyme
Alcohol dehydrogenase
Regulator

Salmonella sp. NCTC 3046. Bacteria; Proteobacteria; Gammaproteobacteria; Enterobacterales;

NCTC3046__locus_2

PDU1AB

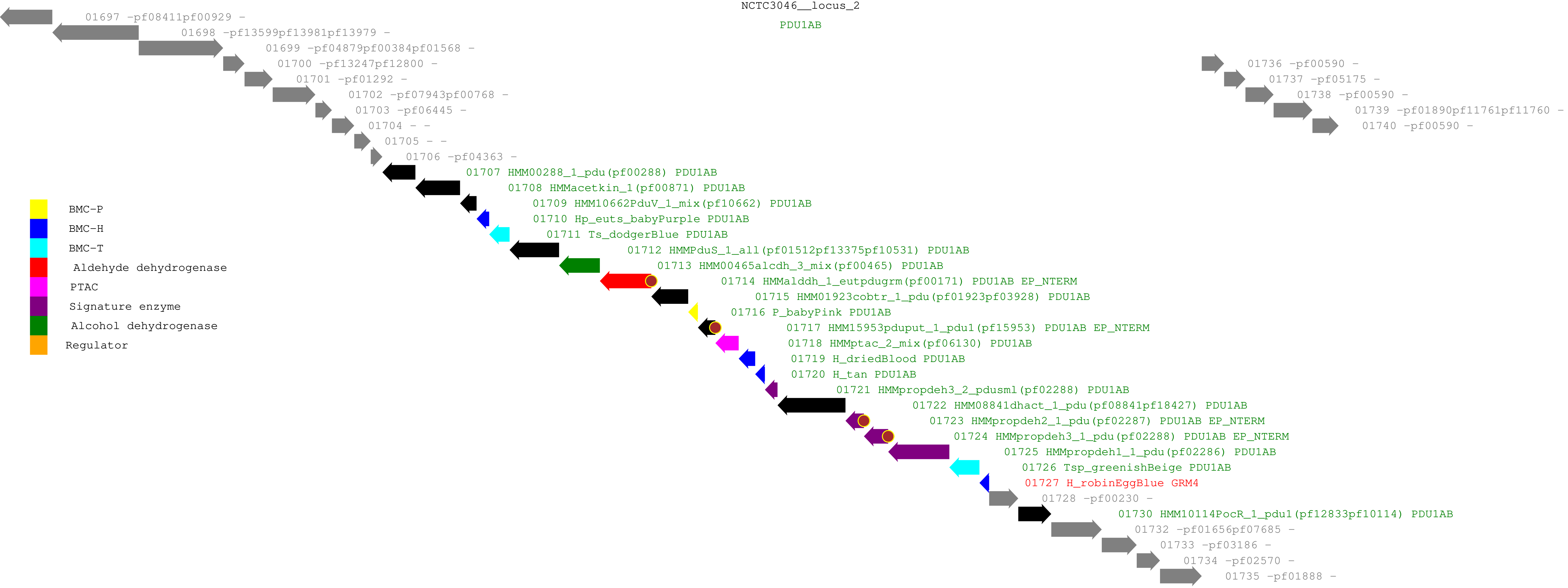

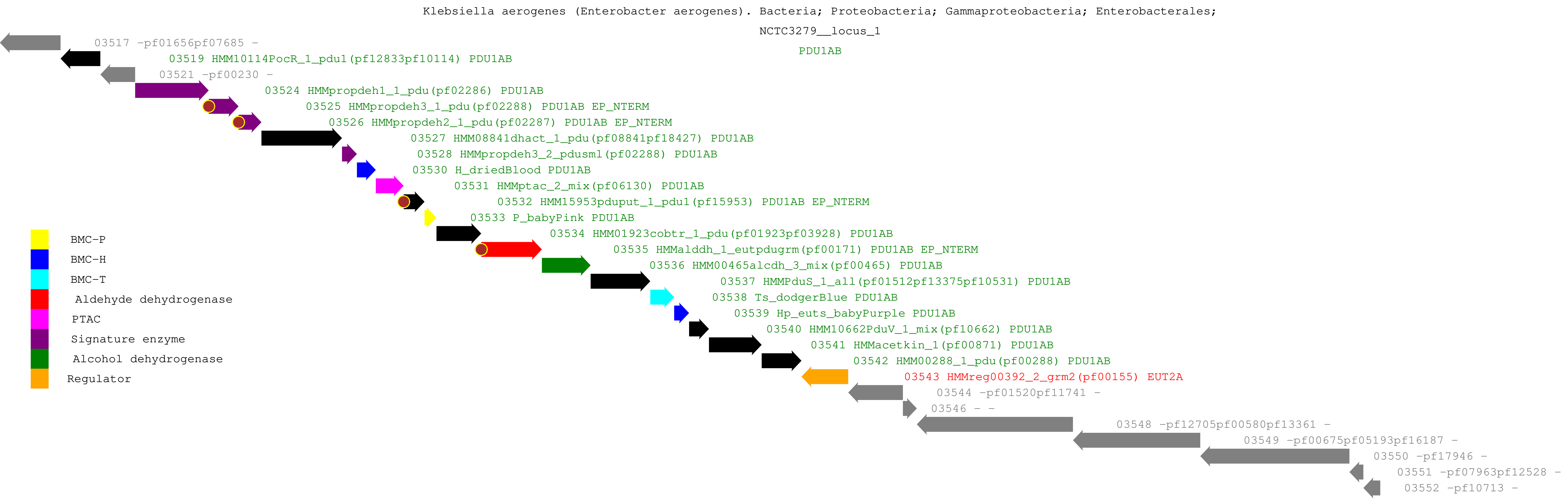
Klebsiella aerogenes (Enterobacter aerogenes). Bacteria; Proteobacteria; Gammaproteobacteria; Enterobacterales;
NCTC3279__locus_1
PDU1AB
03517 -pf01656pf07685 -
03519 HMM10114PocR_1_pdu1(pf12833pf10114) PDU1AB
03521 -pf00230 -
03524 HMMpropdeh1_1_pdu(pf02286) PDU1AB
03525 HMMpropdeh3_1_pdu(pf02288) PDU1AB EP_NTERM
03526 HMMpropdeh2_1_pdu(pf02287) PDU1AB EP_NTERM
03527 HMM08841dhact_1_pdu(pf08841pf18427) PDU1AB
03528 HMMpropdeh3_2_pdusml(pf02288) PDU1AB
03530 H_driedBlood PDU1AB
03531 HMMptac_2_mix(pf06130) PDU1AB
03532 HMM15953pduput_1_pdu1(pf15953) PDU1AB EP_NTERM
03533 P_babyPink PDU1AB
03534 HMM01923cobtr_1_pdu(pf01923pf03928) PDU1AB
03535 HMMalddh_1_eutpdugrm(pf00171) PDU1AB EP_NTERM
03536 HMM00465alcdh_3_mix(pf00465) PDU1AB
03537 HMMPduS_1_all(pf01512pf13375pf10531) PDU1AB
03538 Ts_dodgerBlue PDU1AB
03539 Hp_euts_babyPurple PDU1AB
03540 HMM10662PduV_1_mix(pf10662) PDU1AB
03541 HMMacetkin_1(pf00871) PDU1AB
03542 HMM00288_1_pdu(pf00288) PDU1AB
03543 HMMreg00392_2_grm2(pf00155) EUT2A
03544 -pf01520pf11741 -
03546 - -
03548 -pf12705pf00580pf13361 -
03549 -pf00675pf05193pf16187 -
03550 -pf17946 -
03551 -pf07963pf12528 -
03552 -pf10713 -
BMC-P
BMC-H
BMC-T
Aldehyde dehydrogenase
PTAC
Signature enzyme
Alcohol dehydrogenase
Regulator

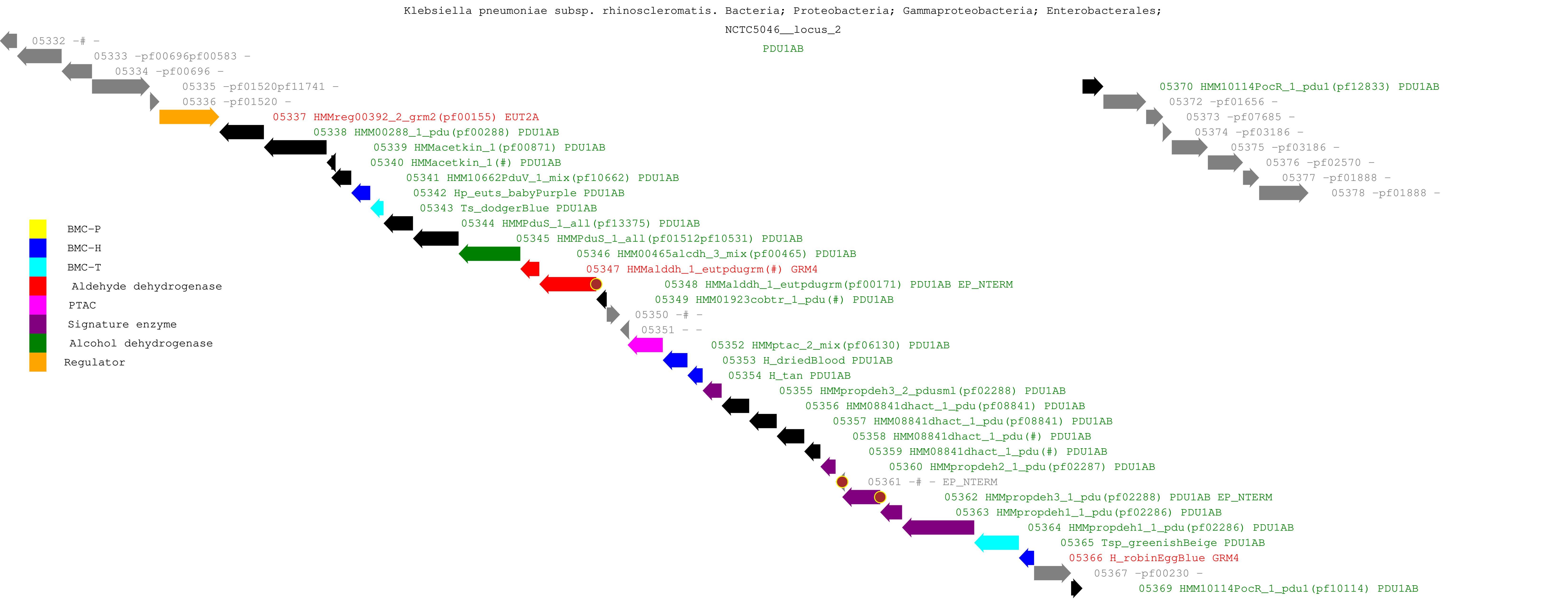
Klebsiella pneumoniae subsp. rhinoscleromatis. Bacteria; Proteobacteria; Gammaproteobacteria; Enterobacterales;
NCTC5046__locus_2
PDU1AB
05332 -# -
05333 -pf00696pf00583 -
05334 -pf00696 -
05335 -pf01520pf11741 -
05336 -pf01520 -
05337 HMMreg00392_2_grm2(pf00155) EUT2A
05338 HMM00288_1_pdu(pf00288) PDU1AB
05339 HMMacetkin_1(pf00871) PDU1AB
05340 HMMacetkin_1(#) PDU1AB
05341 HMM10662PduV_1_mix(pf10662) PDU1AB
05342 Hp_euts_babyPurple PDU1AB
05343 Ts_dodgerBlue PDU1AB
05344 HMMPduS_1_all(pf13375) PDU1AB
05345 HMMPduS_1_all(pf01512pf10531) PDU1AB
05346 HMM00465alcdh_3_mix(pf00465) PDU1AB
05347 HMMaldh_1_eutpdugrm(#) GRM4
05348 HMMaldh_1_eutpdugrm(pf00171) PDU1AB EP_NTERM
05349 HMM01923cobtr_1_pdu(#) PDU1AB
05350 -# -
05351 - -
05352 HMMptac_2_mix(pf06130) PDU1AB
05353 H_driedBlood PDU1AB
05354 H_tan PDU1AB
05355 HMMpropdeh3_2_pdusml(pf02288) PDU1AB
05356 HMM08841dhact_1_pdu(pf08841) PDU1AB
05357 HMM08841dhact_1_pdu(pf08841) PDU1AB
05358 HMM08841dhact_1_pdu(#) PDU1AB
05359 HMM08841dhact_1_pdu(#) PDU1AB
05360 HMMpropdeh2_1_pdu(pf02287) PDU1AB
05361 -# - EP_NTERM
05362 HMMpropdeh3_1_pdu(pf02288) PDU1AB EP_NTERM
05363 HMMpropdeh1_1_pdu(pf02286) PDU1AB
05364 HMMpropdeh1_1_pdu(pf02286) PDU1AB
05365 Tsp_greenishBeige PDU1AB
05366 H_robinEggBlue GRM4
05367 -pf00230 -
05369 HMM10114PocR_1_pdu1(pf10114) PDU1AB
05370 HMM10114PocR_1_pdu1(pf12833) PDU1AB
05372 -pf01656 -
05373 -pf07685 -
05374 -pf03186 -
05375 -pf03186 -
05376 -pf02570 -
05377 -pf01888 -
05378 -pf01888 -
BMC-P
BMC-H
BMC-T
Aldehyde dehydrogenase
PTAC
Signature enzyme
Alcohol dehydrogenase
Regulator

Klebsiella pneumoniae. Bacteria; Proteobacteria; Gammaproteobacteria; Enterobacterales;

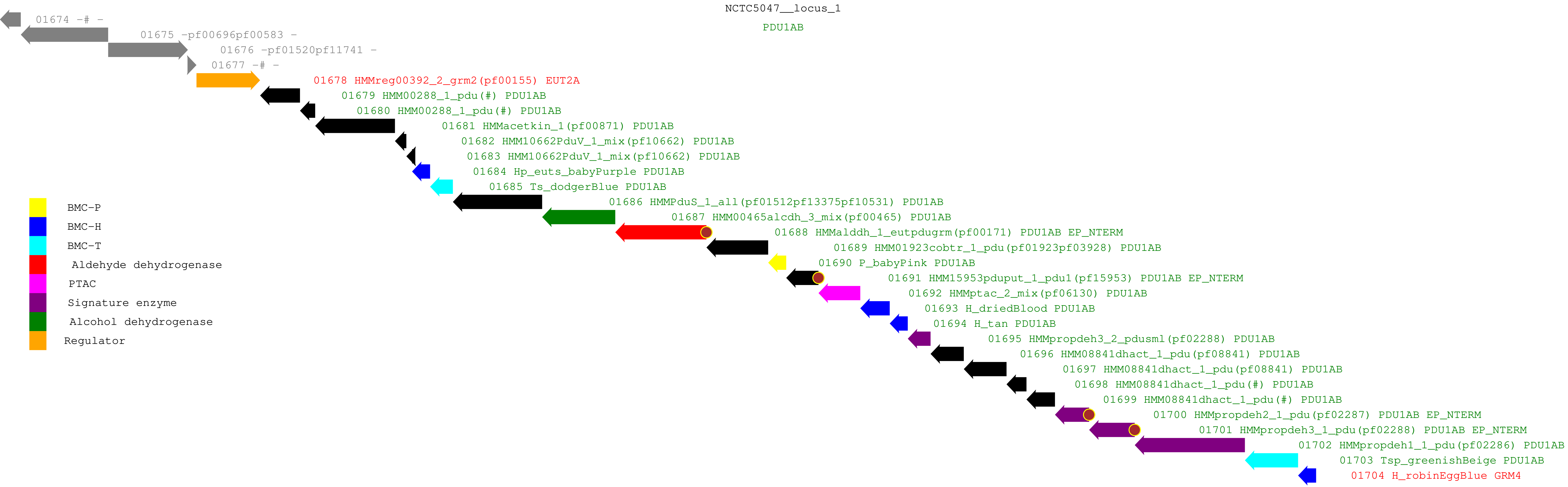

Klebsiella pneumoniae subsp. ozaenae. Bacteria; Proteobacteria; Gammaproteobacteria; Enterobacterales;

NCTC5050__locus_1

PDU1AB

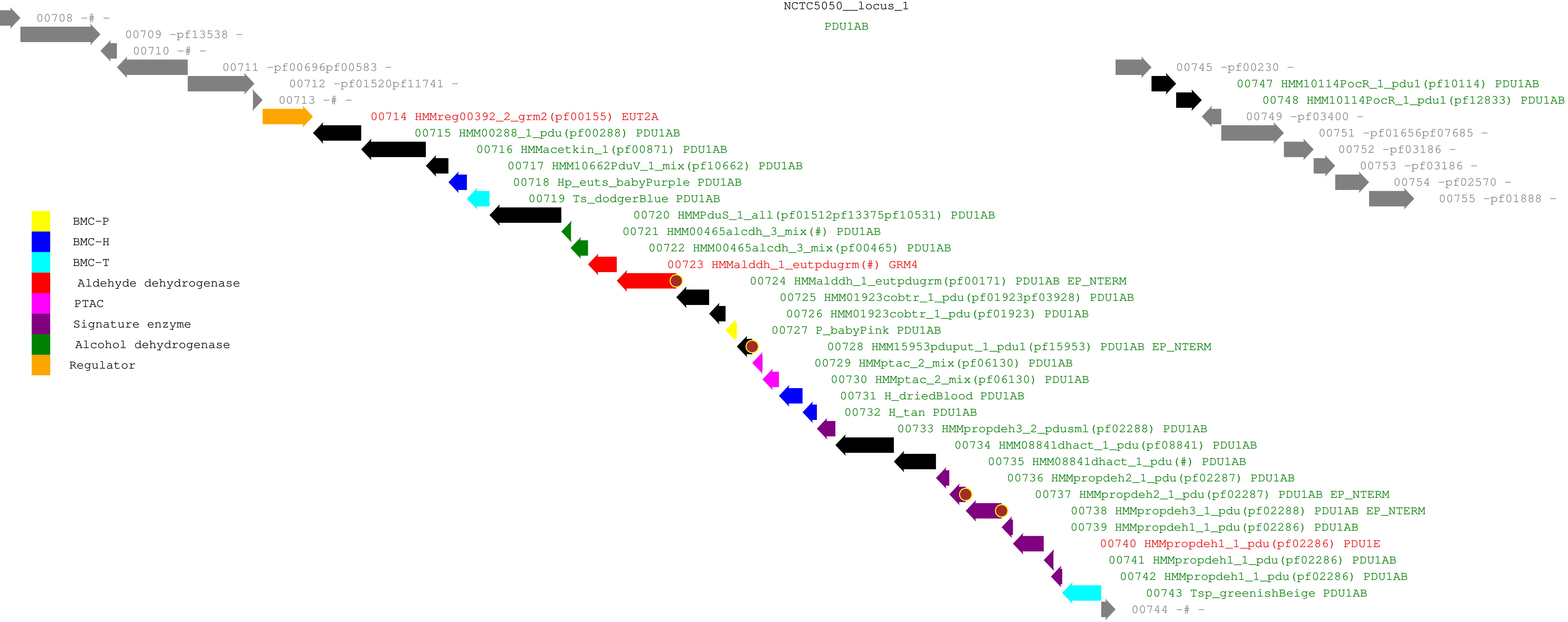

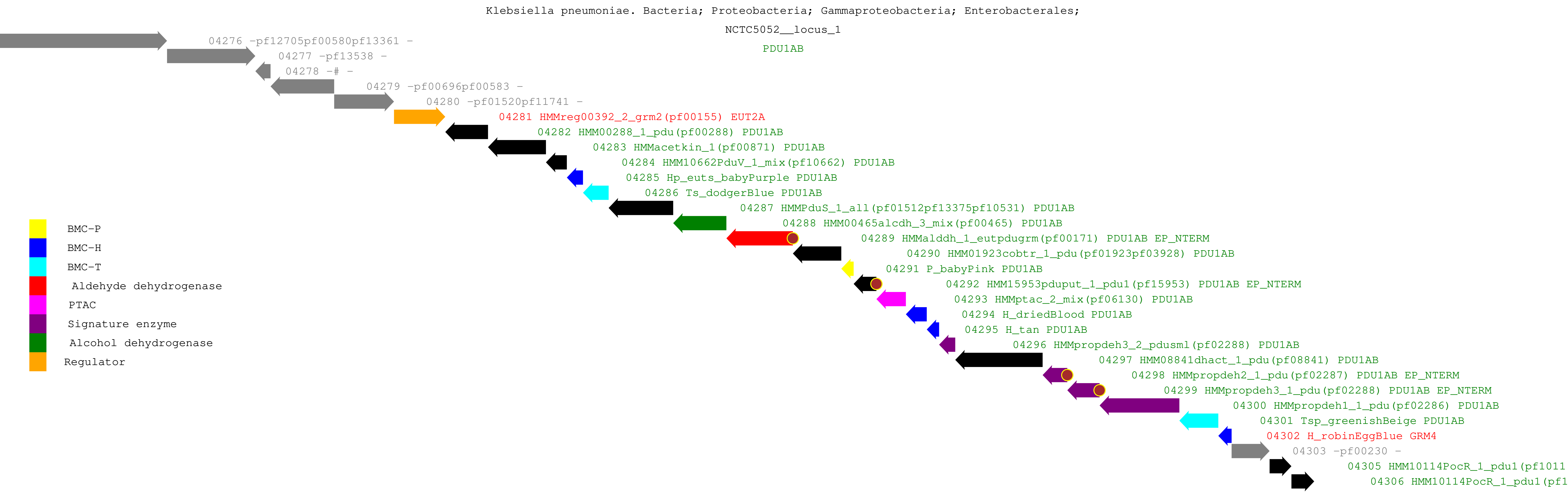
Klebsiella pneumoniae. Bacteria; Proteobacteria; Gammaproteobacteria; Enterobacterales;
NCTC5052__locus_1
PDU1AB
04276 -pf12705pf00580pf13361 -
04277 -pf13538 -
04278 -# -
04279 -pf00696pf00583 -
04280 -pf01520pf11741 -
04281 HMMreg00392_2_grm2(pf00155) EUT2A
04282 HMM00288_1_pdu(pf00288) PDU1AB
04283 HMMacetkin_1(pf00871) PDU1AB
04284 HMM10662PduV_1_mix(pf10662) PDU1AB
04285 Hp_euts_babyPurple PDU1AB
04286 Ts_dodgerBlue PDU1AB
04287 HMMPduS_1_all(pf01512pf13375pf10531) PDU1AB
04288 HMM00465alcdh_3_mix(pf00465) PDU1AB
04289 HMMalddh_1_eutpdugrm(pf00171) PDU1AB EP_NTERM
04290 HMM01923cobtr_1_pdu(pf01923pf03928) PDU1AB
04291 P_babyPink PDU1AB
04292 HMM15953pduput_1_pdu1(pf15953) PDU1AB EP_NTERM
04293 HMMptac_2_mix(pf06130) PDU1AB
04294 H_driedBlood PDU1AB
04295 H_tan PDU1AB
04296 HMMpropdeh3_2_pdusml(pf02288) PDU1AB
04297 HMM08841dhact_1_pdu(pf08841) PDU1AB
04298 HMMpropdeh2_1_pdu(pf02287) PDU1AB EP_NTERM
04299 HMMpropdeh3_1_pdu(pf02288) PDU1AB EP_NTERM
04300 HMMpropdeh1_1_pdu(pf02286) PDU1AB
04301 Tsp_greenishBeige PDU1AB
04302 H_robinEggBlue GRM4
04303 -pf00230 -
04305 HMM10114PocR_1_pdu1(pf1011
04306 HMM10114PocR_1_pdu1(pf1
BMC-P
BMC-H
BMC-T
Aldehyde dehydrogenase
PTAC
Signature enzyme
Alcohol dehydrogenase
Regulator

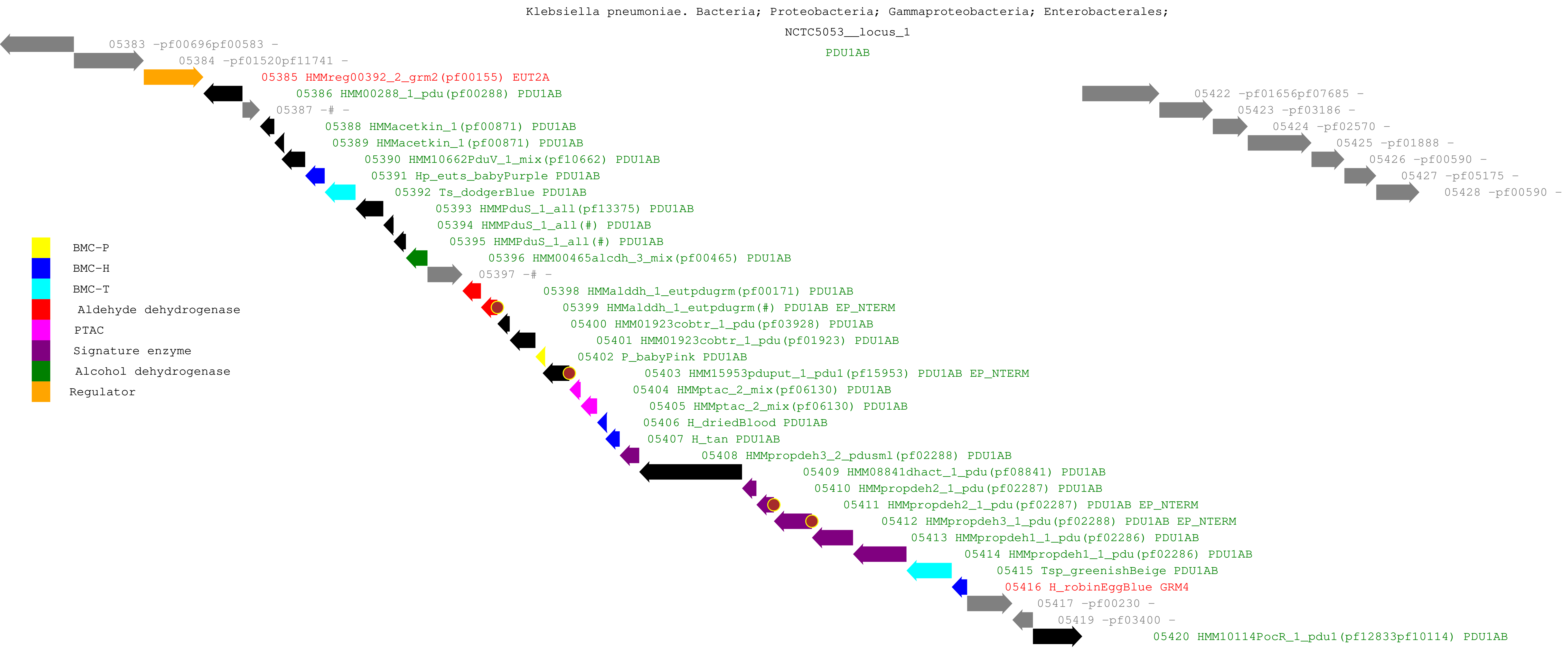
Klebsiella pneumoniae. Bacteria; Proteobacteria; Gammaproteobacteria; Enterobacterales;
NCTC5053__locus_1
PDU1AB
05383 -pf00696pf00583 -
05384 -pf01520pf11741 -
05385 HMMreg00392_2_grm2(pf00155) EUT2A
05386 HMM00288_1_pdu(pf00288) PDU1AB
05387 -# -
05388 HMMacetkin_1(pf00871) PDU1AB
05389 HMMacetkin_1(pf00871) PDU1AB
05390 HMM10662PduV_1_mix(pf10662) PDU1AB
05391 Hp_euts_babyPurple PDU1AB
05392 Ts_dodgerBlue PDU1AB
05393 HMMPduS_1_all(pf13375) PDU1AB
05394 HMMPduS_1_all(#) PDU1AB
05395 HMMPduS_1_all(#) PDU1AB
05396 HMM00465alcdh_3_mix(pf00465) PDU1AB
05397 -# -
05398 HMMaldddh_1_eutpdugrm(pf00171) PDU1AB
05399 HMMaldddh_1_eutpdugrm(#) PDU1AB EP_NTERM
05400 HMM01923cobtr_1_pdu(pf03928) PDU1AB
05401 HMM01923cobtr_1_pdu(pf01923) PDU1AB
05402 P_babyPink PDU1AB
05403 HMM15953pduput_1_pdu1(pf15953) PDU1AB EP_NTERM
05404 HMMptac_2_mix(pf06130) PDU1AB
05405 HMMptac_2_mix(pf06130) PDU1AB
05406 H_driedBlood PDU1AB
05407 H_tan PDU1AB
05408 HMMpropdeh3_2_pdusml(pf02288) PDU1AB
05409 HMM08841dhact_1_pdu(pf08841) PDU1AB
05410 HMMpropdeh2_1_pdu(pf02287) PDU1AB
05411 HMMpropdeh2_1_pdu(pf02287) PDU1AB EP_NTERM
05412 HMMpropdeh3_1_pdu(pf02288) PDU1AB EP_NTERM
05413 HMMpropdeh1_1_pdu(pf02286) PDU1AB
05414 HMMpropdeh1_1_pdu(pf02286) PDU1AB
05415 Tsp_greenishBeige PDU1AB
05416 H_robinEggBlue GRM4
05417 -pf00230 -
05419 -pf03400 -
05420 HMM10114PocR_1_pdu1(pf12833pf10114) PDU1AB
05422 -pf01656pf07685 -
05423 -pf03186 -
05424 -pf02570 -
05425 -pf01888 -
05426 -pf00590 -
05427 -pf05175 -
05428 -pf00590 -
BMC-P
BMC-H
BMC-T
Aldehyde dehydrogenase
PTAC
Signature enzyme
Alcohol dehydrogenase
Regulator

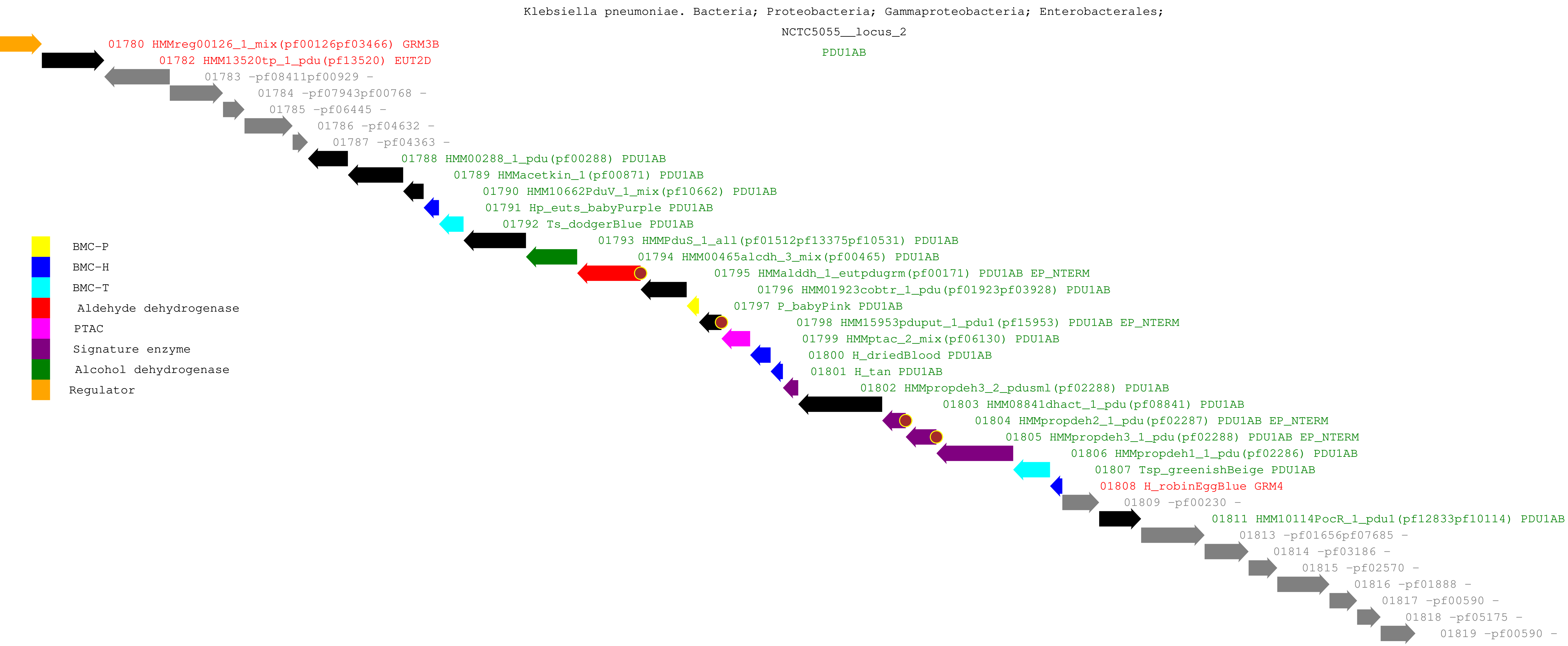

Klebsiella pneumoniae. Bacteria; Proteobacteria; Gammaproteobacteria; Enterobacterales;
NCTC5055__locus_2
PDU1AB
01780 HMMreg00126_1_mix(pf00126pf03466) GRM3B
01782 HMM13520tp_1_pdu(pf13520) EUT2D
01783 -pf08411pf00929 -
01784 -pf07943pf00768 -
01785 -pf06445 -
01786 -pf04632 -
01787 -pf04363 -
01788 HMM00288_1_pdu(pf00288) PDU1AB
01789 HMMacetkin_1(pf00871) PDU1AB
01790 HMM10662PduV_1_mix(pf10662) PDU1AB
01791 Hp_euts_babyPurple PDU1AB
01792 Ts_dodgerBlue PDU1AB
01793 HMMPduS_1_all(pf01512pf13375pf10531) PDU1AB
01794 HMM00465alcdh_3_mix(pf00465) PDU1AB
01795 HMMalddh_1_eutpdugrm(pf00171) PDU1AB EP_NTERM
01796 HMM01923cobtr_1_pdu(pf01923pf03928) PDU1AB
01797 P_babyPink PDU1AB
01798 HMM15953pduput_1_pdu1(pf15953) PDU1AB EP_NTERM
01799 HMMptac_2_mix(pf06130) PDU1AB
01800 H_driedBlood PDU1AB
01801 H_tan PDU1AB
01802 HMMpropdeh3_2_pdusml(pf02288) PDU1AB
01803 HMM08841dhact_1_pdu(pf08841) PDU1AB
01804 HMMpropdeh2_1_pdu(pf02287) PDU1AB EP_NTERM
01805 HMMpropdeh3_1_pdu(pf02288) PDU1AB EP_NTERM
01806 HMMpropdeh1_1_pdu(pf02286) PDU1AB
01807 Tsp_greenishBeige PDU1AB
01808 H_robinEggBlue GRM4
01809 -pf00230 -
01811 HMM10114PocR_1_pdu1(pf12833pf10114) PDU1AB
01813 -pf01656pf07685 -
01814 -pf03186 -
01815 -pf02570 -
01816 -pf01888 -
01817 -pf00590 -
01818 -pf05175 -
01819 -pf00590 -
BMC-P
BMC-H
BMC-T
Aldehyde dehydrogenase
PTAC
Signature enzyme
Alcohol dehydrogenase
Regulator

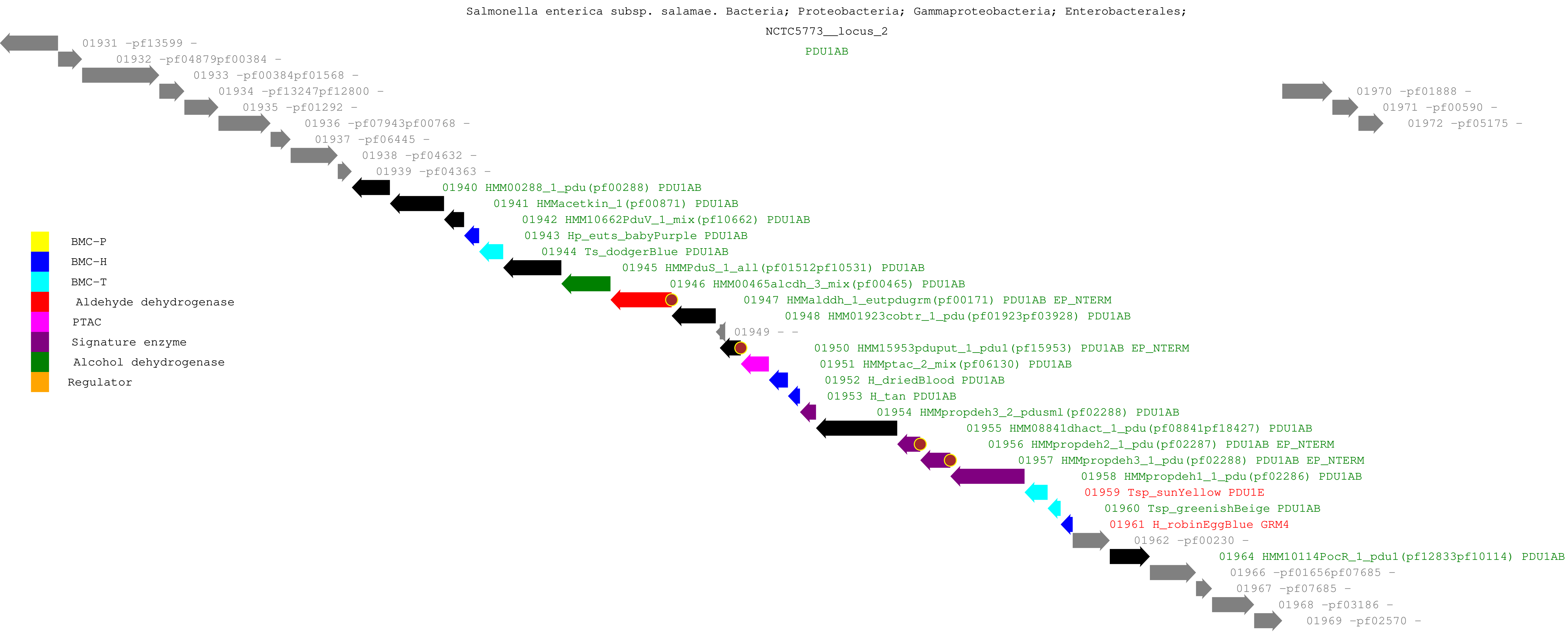
Salmonella enterica subsp. salamae. Bacteria; Proteobacteria; Gammaproteobacteria; Enterobacterales;
NCTC5773__locus_2
PDU1AB
01931 -pf13599 -
01932 -pf04879pf00384 -
01933 -pf00384pf01568 -
01934 -pf13247pf12800 -
01935 -pf01292 -
01936 -pf07943pf00768 -
01937 -pf06445 -
01938 -pf04632 -
01939 -pf04363 -
01940 HMM00288_1_pdu(pf00288) PDU1AB
01941 HMMacetkin_1(pf00871) PDU1AB
01942 HMM10662PduV_1_mix(pf10662) PDU1AB
01943 Hp_euts_babyPurple PDU1AB
01944 Ts_dodgerBlue PDU1AB
01945 HMMPduS_1_all(pf01512pf10531) PDU1AB
01946 HMM00465alcdh_3_mix(pf00465) PDU1AB
01947 HMMalddh_1_eutpdugrm(pf00171) PDU1AB EP_NTERM
01948 HMM01923cobtr_1_pdu(pf01923pf03928) PDU1AB
01949 - -
01950 HMM15953pduput_1_pdu1(pf15953) PDU1AB EP_NTERM
01951 HMMptac_2_mix(pf06130) PDU1AB
01952 H_driedBlood PDU1AB
01953 H_tan PDU1AB
01954 HMMpropdeh3_2_pdusml(pf02288) PDU1AB
01955 HMM08841dhact_1_pdu(pf08841pf18427) PDU1AB
01956 HMMpropdeh2_1_pdu(pf02287) PDU1AB EP_NTERM
01957 HMMpropdeh3_1_pdu(pf02288) PDU1AB EP_NTERM
01958 HMMpropdeh1_1_pdu(pf02286) PDU1AB
01959 Tsp_sunYellow PDU1E
01960 Tsp_greenishBeige PDU1AB
01961 H_robinEggBlue GRM4
01962 -pf00230 -
01964 HMM10114PocR_1_pdu1(pf12833pf10114) PDU1AB
01966 -pf01656pf07685 -
01967 -pf07685 -
01968 -pf03186 -
01969 -pf02570 -
01970 -pf01888 -
01971 -pf00590 -
01972 -pf05175 -
BMC-P
BMC-H
BMC-T
Aldehyde dehydrogenase
PTAC
Signature enzyme
Alcohol dehydrogenase
Regulator

Salmonella enterica I. Bacteria; Proteobacteria; Gammaproteobacteria; Enterobacterales;

NCTC5798__locus_1

PDU1AB

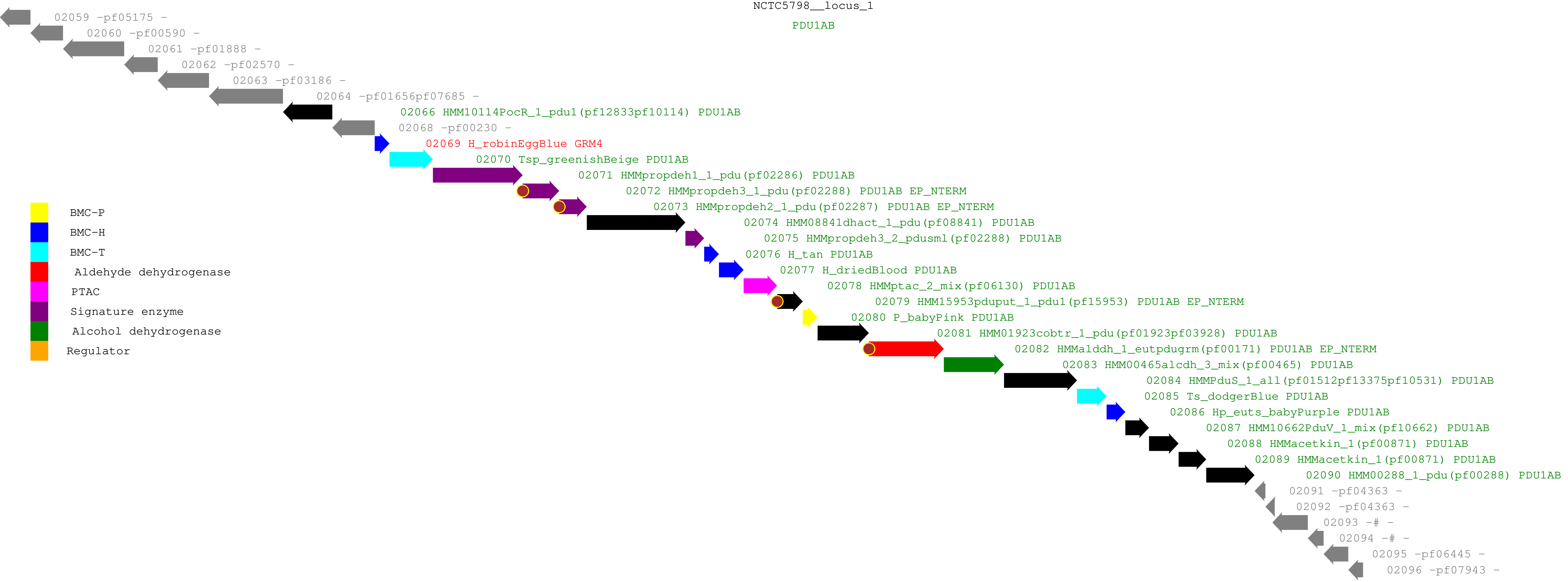

Citrobacter freundii. Bacteria; Proteobacteria; Gammaproteobacteria; Enterobacterales;

NCTC6266__locus_1

PDU1AB

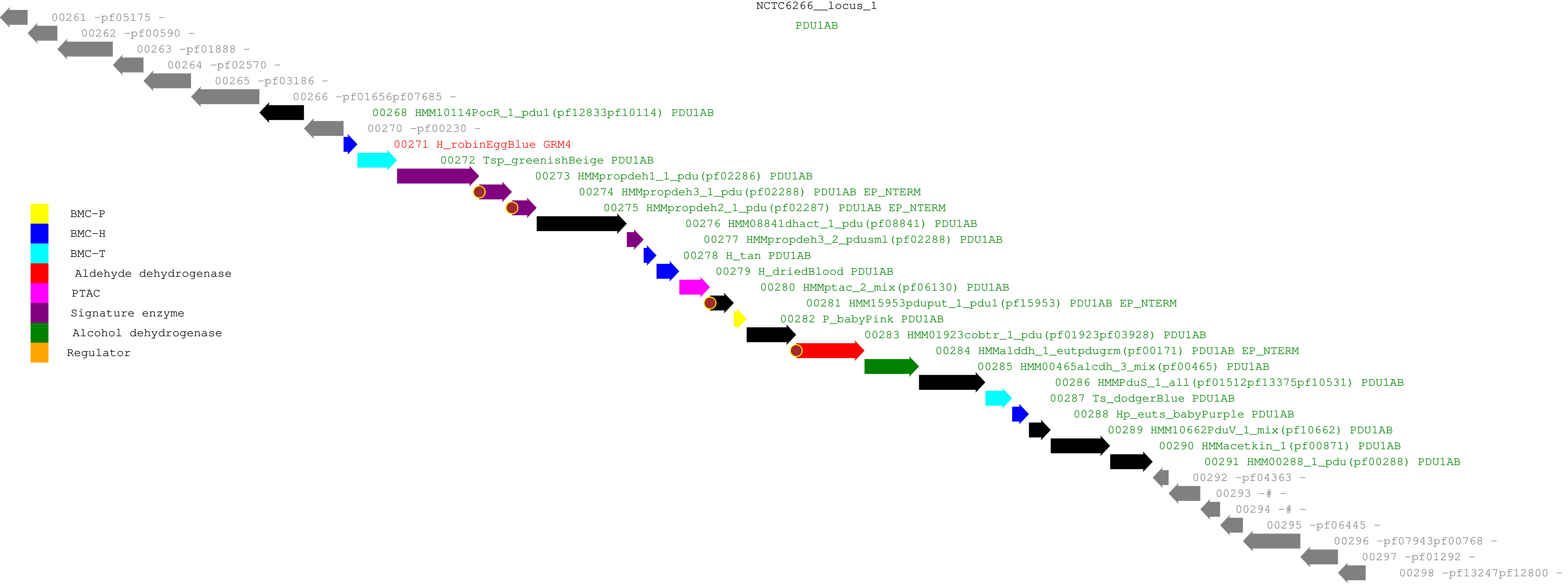

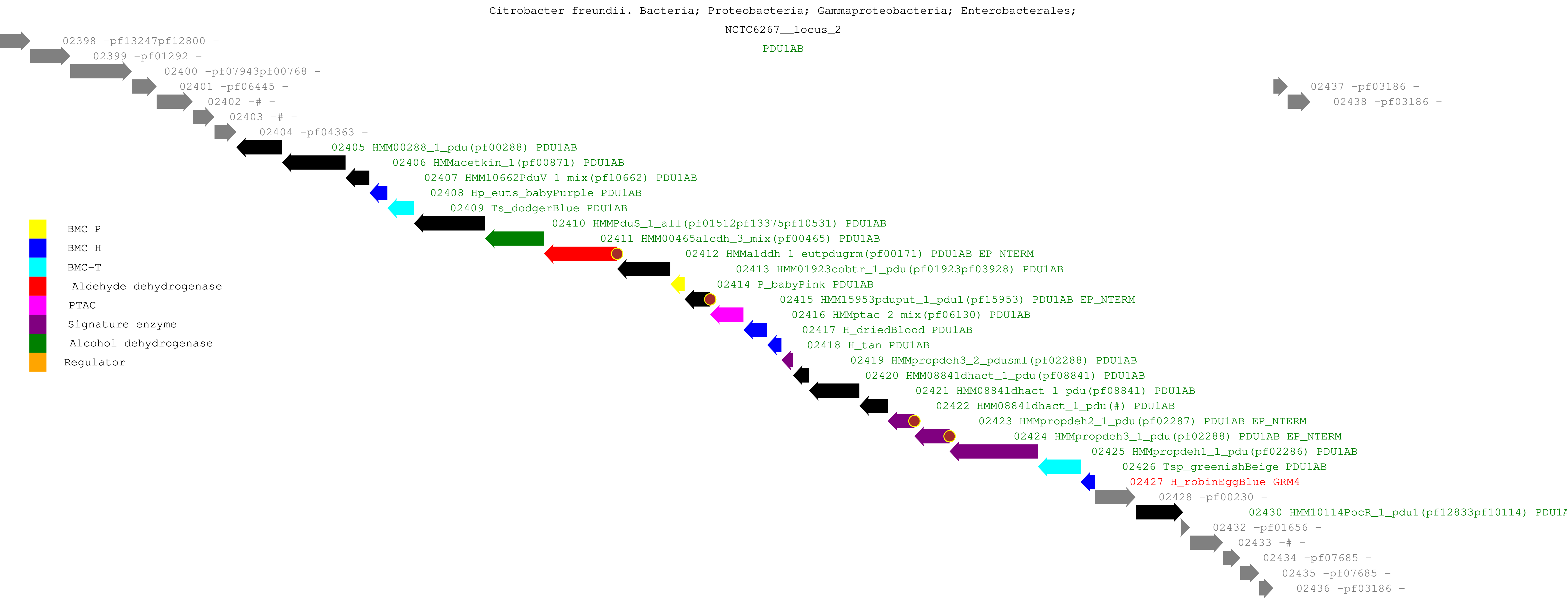
Citrobacter freundii. Bacteria; Proteobacteria; Gammaproteobacteria; Enterobacterales;
NCTC6267__locus_2
PDU1AB
02398 -pf13247pf12800 -
02399 -pf01292 -
02400 -pf07943pf00768 -
02401 -pf06445 -
02402 -# -
02403 -# -
02404 -pf04363 -
02405 HMM00288_1_pdu(pf00288) PDU1AB
02406 HMMacetkin_1(pf00871) PDU1AB
02407 HMM10662PduV_1_mix(pf10662) PDU1AB
02408 Hp_euts_babyPurple PDU1AB
02409 Ts_dodgerBlue PDU1AB
02410 HMMPduS_1_all(pf01512pf13375pf10531) PDU1AB
02411 HMM00465alcdh_3_mix(pf00465) PDU1AB
02412 HMMalddh_1_eutpdugrm(pf00171) PDU1AB EP_NTERM
02413 HMM01923cobtr_1_pdu(pf01923pf03928) PDU1AB
02414 P_babyPink PDU1AB
02415 HMM15953pduput_1_pdu1(pf15953) PDU1AB EP_NTERM
02416 HMMptac_2_mix(pf06130) PDU1AB
02417 H_driedBlood PDU1AB
02418 H_tan PDU1AB
02419 HMMpropdeh3_2_pdusml(pf02288) PDU1AB
02420 HMM08841dhact_1_pdu(pf08841) PDU1AB
02421 HMM08841dhact_1_pdu(pf08841) PDU1AB
02422 HMM08841dhact_1_pdu(#) PDU1AB
02423 HMMpropdeh2_1_pdu(pf02287) PDU1AB EP_NTERM
02424 HMMpropdeh3_1_pdu(pf02288) PDU1AB EP_NTERM
02425 HMMpropdeh1_1_pdu(pf02286) PDU1AB
02426 Tsp_greenishBeige PDU1AB
02427 H_robinEggBlue GRM4
02428 -pf00230 -
02430 HMM10114PocR_1_pdu1(pf12833pf10114) PDU1A
02432 -pf01656 -
02433 -# -
02434 -pf07685 -
02435 -pf07685 -
02436 -pf03186 -
02437 -pf03186 -
02438 -pf03186 -
BMC-P
BMC-H
BMC-T
Aldehyde dehydrogenase
PTAC
Signature enzyme
Alcohol dehydrogenase
Regulator

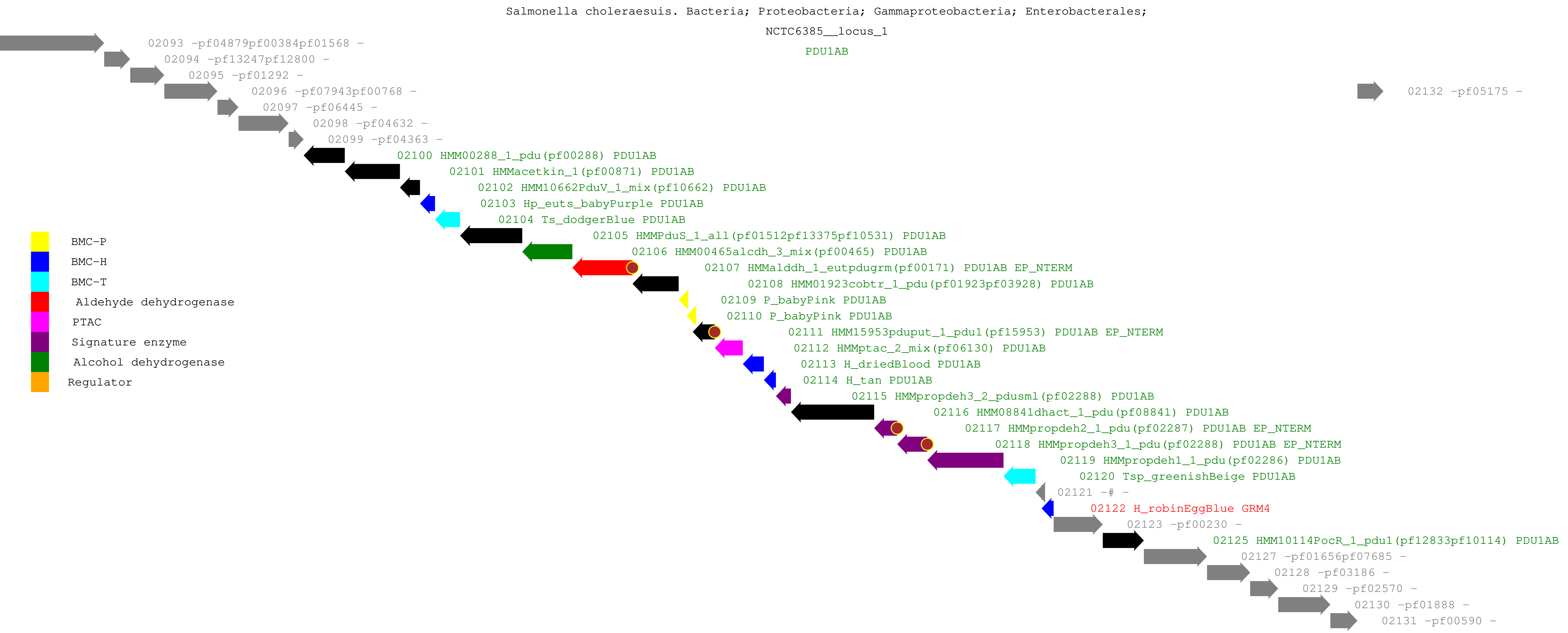
Salmonella choleraesuis. Bacteria; Proteobacteria; Gammaproteobacteria; Enterobacterales;
NCTC6385__locus_1
PDU1AB
02093 -pf04879pf00384pf01568 -
02094 -pf13247pf12800 -
02095 -pf01292 -
02096 -pf07943pf00768 -
02097 -pf06445 -
02098 -pf04632 -
02099 -pf04363 -
02100 HMM00288_1_pdu(pf00288) PDU1AB
02101 HMMacetkin_1(pf00871) PDU1AB
02102 HMM10662PduV_1_mix(pf10662) PDU1AB
02103 Hp_euts_babyPurple PDU1AB
02104 Ts_dodgerBlue PDU1AB
02105 HMMPduS_1_all(pf01512pf13375pf10531) PDU1AB
02106 HMM00465alcdh_3_mix(pf00465) PDU1AB
02107 HMMalddh_1_eutpdugrm(pf00171) PDU1AB EP_NTERM
02108 HMM01923cobtr_1_pdu(pf01923pf03928) PDU1AB
02109 P_babyPink PDU1AB
02110 P_babyPink PDU1AB
02111 HMM15953pduput_1_pdu1(pf15953) PDU1AB EP_NTERM
02112 HMMptac_2_mix(pf06130) PDU1AB
02113 H_driedBlood PDU1AB
02114 H_tan PDU1AB
02115 HMMpropdeh3_2_pdusml(pf02288) PDU1AB
02116 HMM08841dhact_1_pdu(pf08841) PDU1AB
02117 HMMpropdeh2_1_pdu(pf02287) PDU1AB EP_NTERM
02118 HMMpropdeh3_1_pdu(pf02288) PDU1AB EP_NTERM
02119 HMMpropdeh1_1_pdu(pf02286) PDU1AB
02120 Tsp_greenishBeige PDU1AB
02121 -# -
02122 H_robinEggBlue GRM4
02123 -pf00230 -
02125 HMM10114PocR_1_pdu1(pf12833pf10114) PDU1AB
02127 -pf01656pf07685 -
02128 -pf03186 -
02129 -pf02570 -
02130 -pf01888 -
02131 -pf00590 -
02132 -pf05175 -
BMC-P
BMC-H
BMC-T
Aldehyde dehydrogenase
PTAC
Signature enzyme
Alcohol dehydrogenase
Regulator

Salmonella enterica I. Bacteria; Proteobacteria; Gammaproteobacteria; Enterobacterales;

NCTC6754__locus_2

PDU1AB

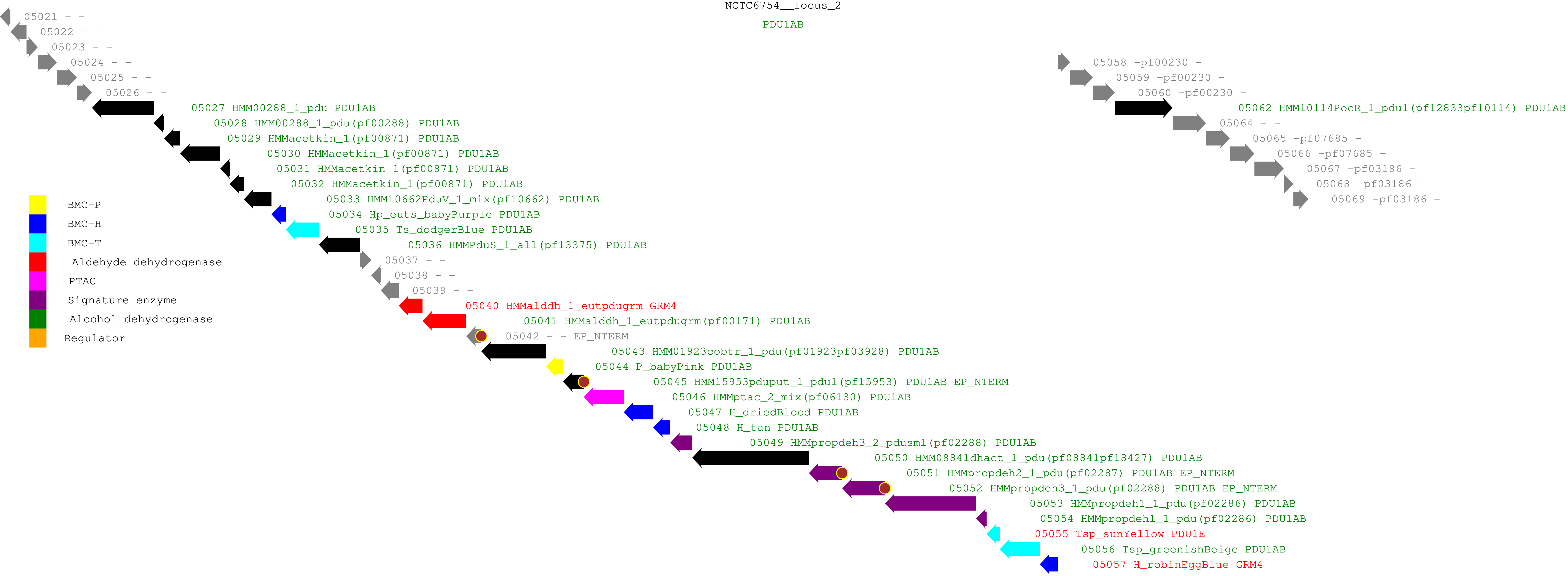

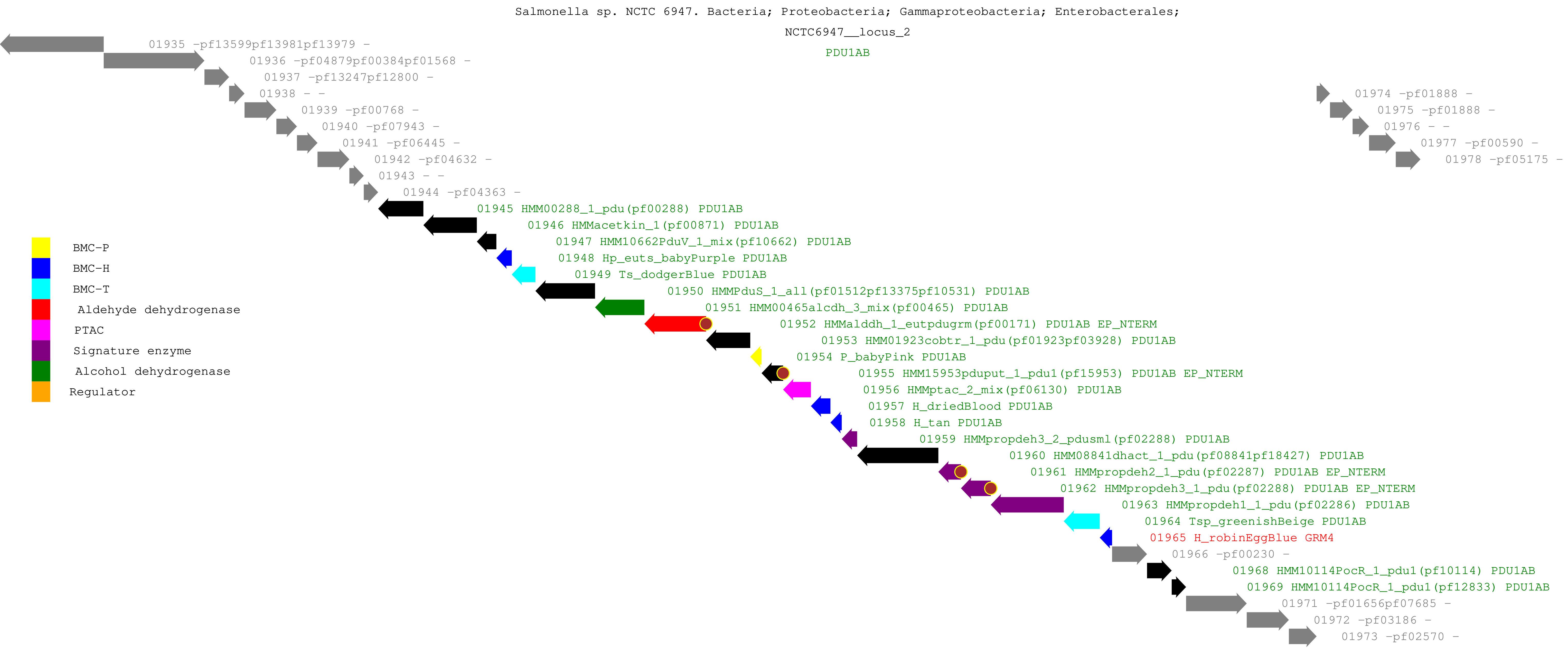
Salmonella sp. NCTC 6947. Bacteria; Proteobacteria; Gammaproteobacteria; Enterobacterales;
NCTC6947__locus_2
PDU1AB
01935 -pf13599pf13981pf13979 -
01936 -pf04879pf00384pf01568 -
01937 -pf13247pf12800 -
01938 - -
01939 -pf00768 -
01940 -pf07943 -
01941 -pf06445 -
01942 -pf04632 -
01943 - -
01944 -pf04363 -
01945 HMM00288_1_pdu(pf00288) PDU1AB
01946 HMMacetkin_1(pf00871) PDU1AB
01947 HMM10662PduV_1_mix(pf10662) PDU1AB
01948 Hp_euts_babyPurple PDU1AB
01949 Ts_dodgerBlue PDU1AB
01950 HMMPduS_1_all(pf01512pf13375pf10531) PDU1AB
01951 HMM00465alcdh_3_mix(pf00465) PDU1AB
01952 HMMalddh_1_eutpdugrm(pf00171) PDU1AB EP_NTERM
01953 HMM01923cobtr_1_pdu(pf01923pf03928) PDU1AB
01954 P_babyPink PDU1AB
01955 HMM15953pduput_1_pdu1(pf15953) PDU1AB EP_NTERM
01956 HMMptac_2_mix(pf06130) PDU1AB
01957 H_driedBlood PDU1AB
01958 H_tan PDU1AB
01959 HMMpropdeh3_2_pdusml(pf02288) PDU1AB
01960 HMM08841dhact_1_pdu(pf08841pf18427) PDU1AB
01961 HMMpropdeh2_1_pdu(pf02287) PDU1AB EP_NTERM
01962 HMMpropdeh3_1_pdu(pf02288) PDU1AB EP_NTERM
01963 HMMpropdeh1_1_pdu(pf02286) PDU1AB
01964 Tsp_greenishBeige PDU1AB
01965 H_robinEggBlue GRM4
01966 -pf00230 -
01968 HMM10114PocR_1_pdu1(pf10114) PDU1AB
01969 HMM10114PocR_1_pdu1(pf12833) PDU1AB
01971 -pf01656pf07685 -
01972 -pf03186 -
01973 -pf02570 -
01974 -pf01888 -
01975 -pf01888 -
01976 - -
01977 -pf00590 -
01978 -pf05175 -
BMC-P
BMC-H
BMC-T
Aldehyde dehydrogenase
PTAC
Signature enzyme
Alcohol dehydrogenase
Regulator

Salmonella enterica subsp. enterica serovar Daytona. Bacteria; Proteobacteria; Gammaproteobacteria; Enterobacterales;

NCTC7102__locus_2

PDU1AB

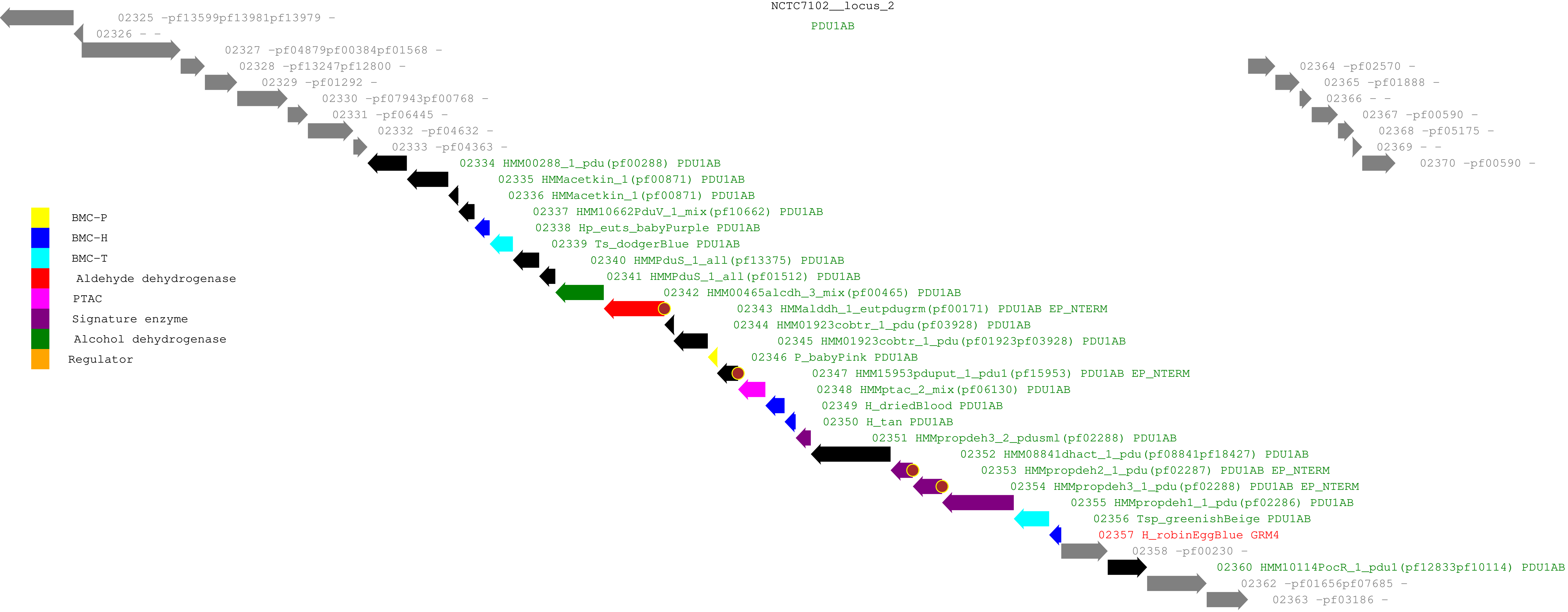

Salmonella enterica subsp. arizonae. Bacteria; Proteobacteria; Gammaproteobacteria; Enterobacterales;

NCTC7295__locus_2

PDU1AB

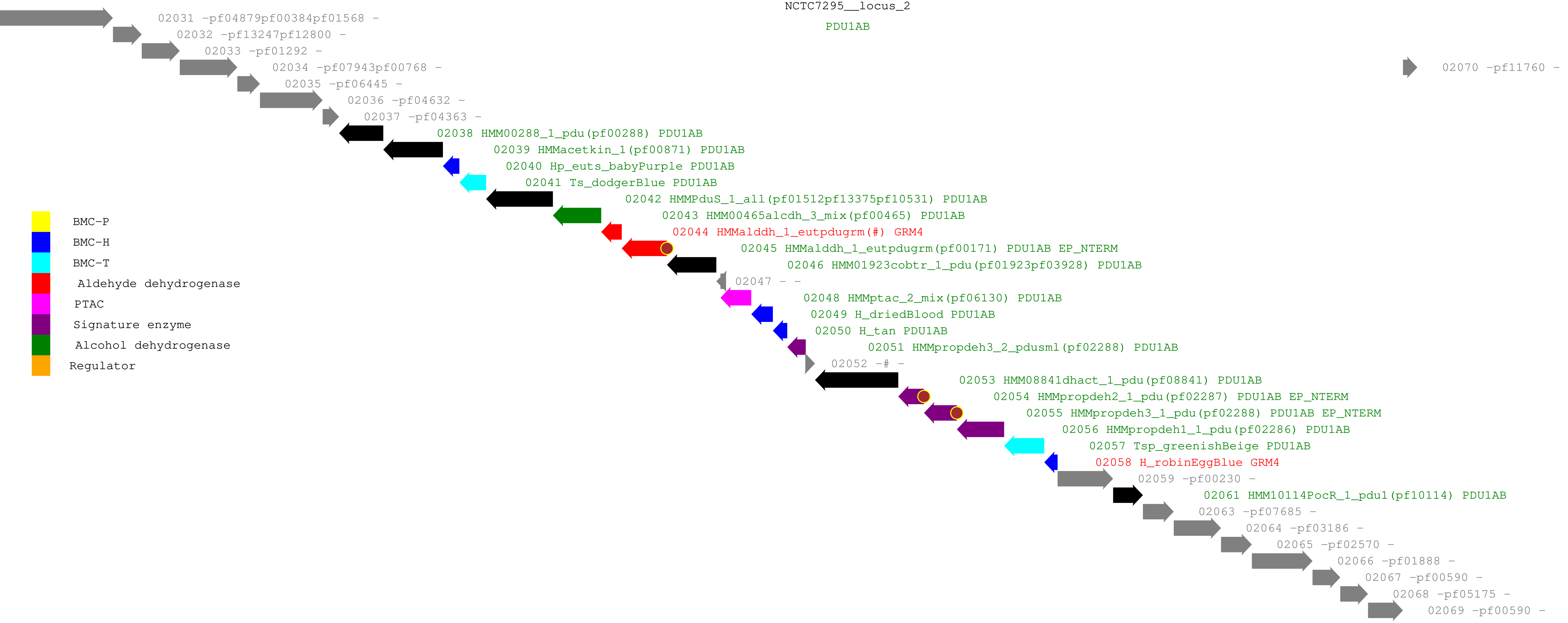

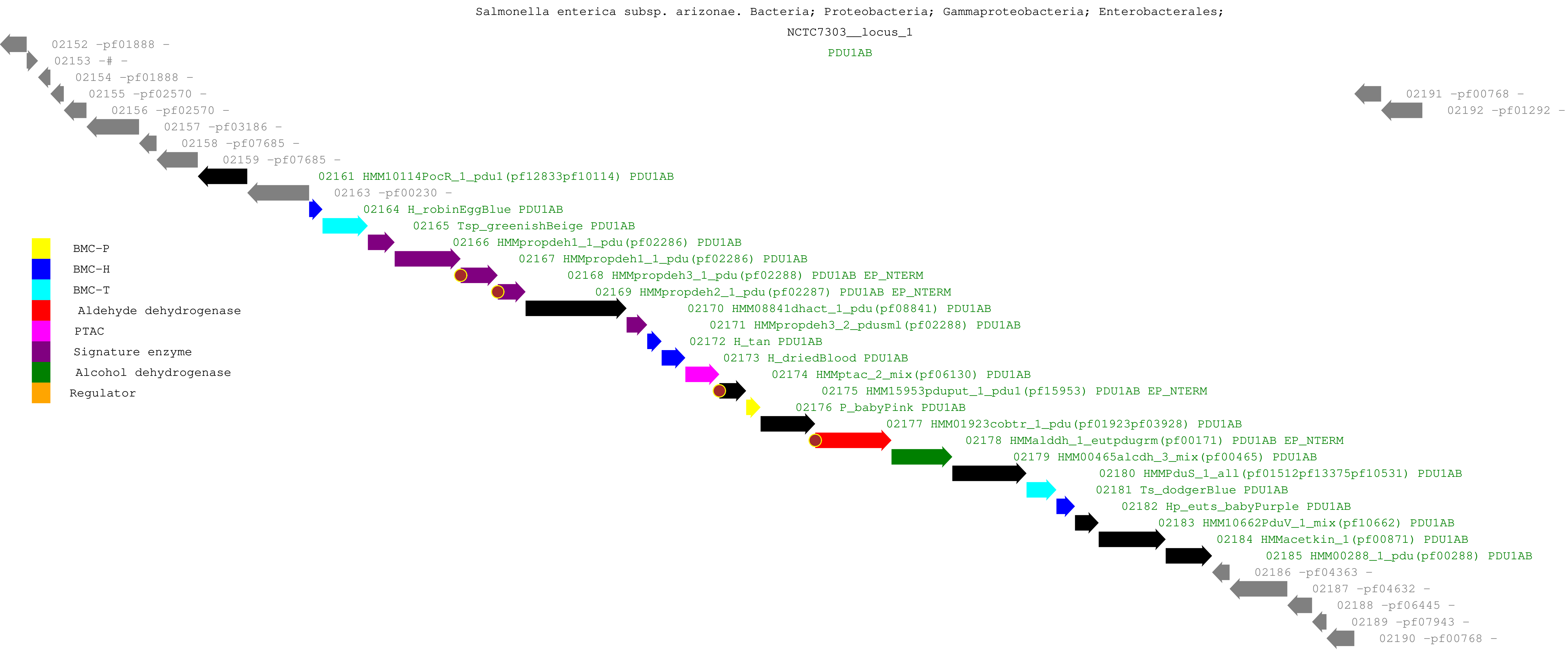
Salmonella enterica subsp. arizonae. Bacteria; Proteobacteria; Gammaproteobacteria; Enterobacterales;
NCTC7303__locus_1
PDU1AB
02152 -pf01888 -
02153 -# -
02154 -pf01888 -
02155 -pf02570 -
02156 -pf02570 -
02157 -pf03186 -
02158 -pf07685 -
02159 -pf07685 -
02161 HMM10114PocR_1_pdu1(pf12833pf10114) PDU1AB
02163 -pf00230 -
02164 H_robinEggBlue PDU1AB
02165 Tsp_greenishBeige PDU1AB
02166 HMMpropdeh1_1_pdu(pf02286) PDU1AB
02167 HMMpropdeh1_1_pdu(pf02286) PDU1AB
02168 HMMpropdeh3_1_pdu(pf02288) PDU1AB EP_NTERM
02169 HMMpropdeh2_1_pdu(pf02287) PDU1AB EP_NTERM
02170 HMM08841dhact_1_pdu(pf08841) PDU1AB
02171 HMMpropdeh3_2_pdusml(pf02288) PDU1AB
02172 H_tan PDU1AB
02173 H_driedBlood PDU1AB
02174 HMMptac_2_mix(pf06130) PDU1AB
02175 HMM15953pduput_1_pdu1(pf15953) PDU1AB EP_NTERM
02176 P_babyPink PDU1AB
02177 HMM01923cobtr_1_pdu(pf01923pf03928) PDU1AB
02178 HMMaldh_1_eutpdugrm(pf00171) PDU1AB EP_NTERM
02179 HMM00465alcdh_3_mix(pf00465) PDU1AB
02180 HMMPduS_1_all(pf01512pf13375pf10531) PDU1AB
02181 Ts_dodgerBlue PDU1AB
02182 Hp_euts_babyPurple PDU1AB
02183 HMM10662PduV_1_mix(pf10662) PDU1AB
02184 HMMacetkin_1(pf00871) PDU1AB
02185 HMM00288_1_pdu(pf00288) PDU1AB
02186 -pf04363 -
02187 -pf04632 -
02188 -pf06445 -
02189 -pf07943 -
02190 -pf00768 -
02191 -pf00768 -
02192 -pf01292 -
BMC-P
BMC-H
BMC-T
Aldehyde dehydrogenase
PTAC
Signature enzyme
Alcohol dehydrogenase
Regulator

Salmonella enterica subsp. arizonae. Bacteria; Proteobacteria; Gammaproteobacteria; Enterobacterales;

NCTC7304__locus_2

PDU1AB

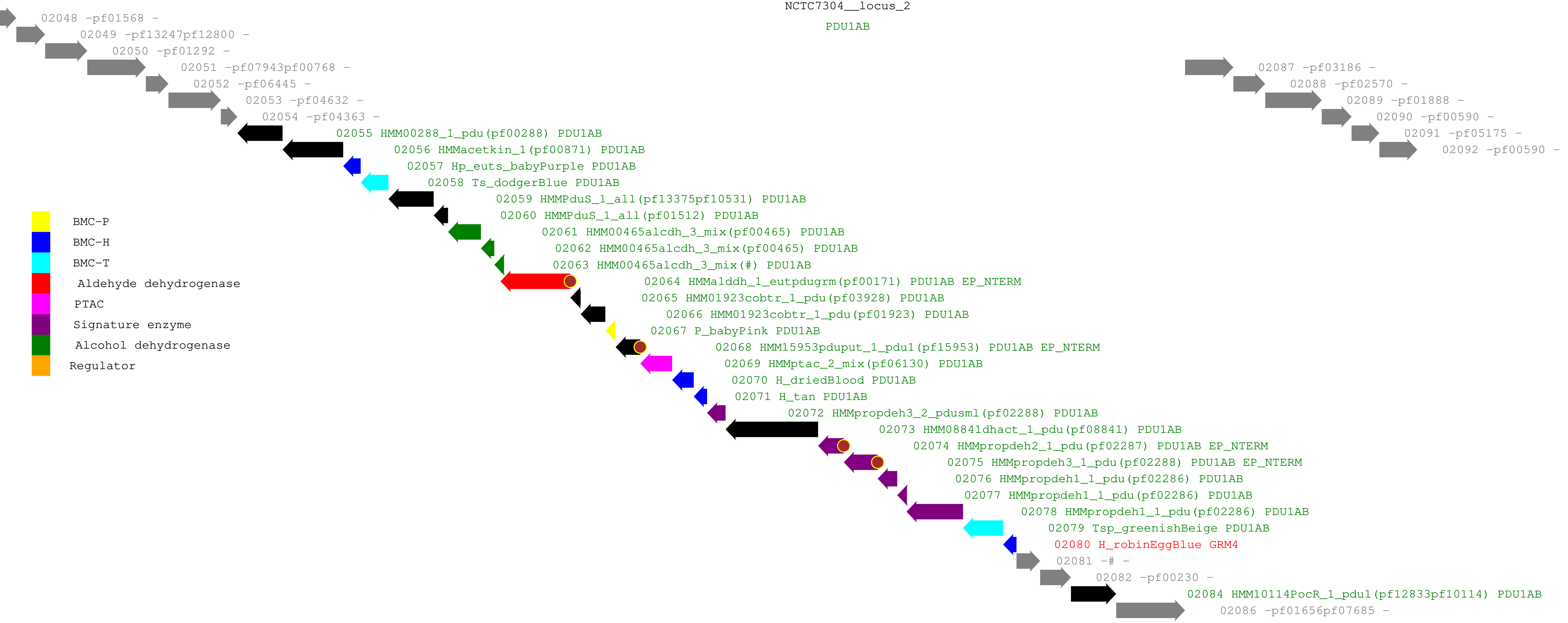

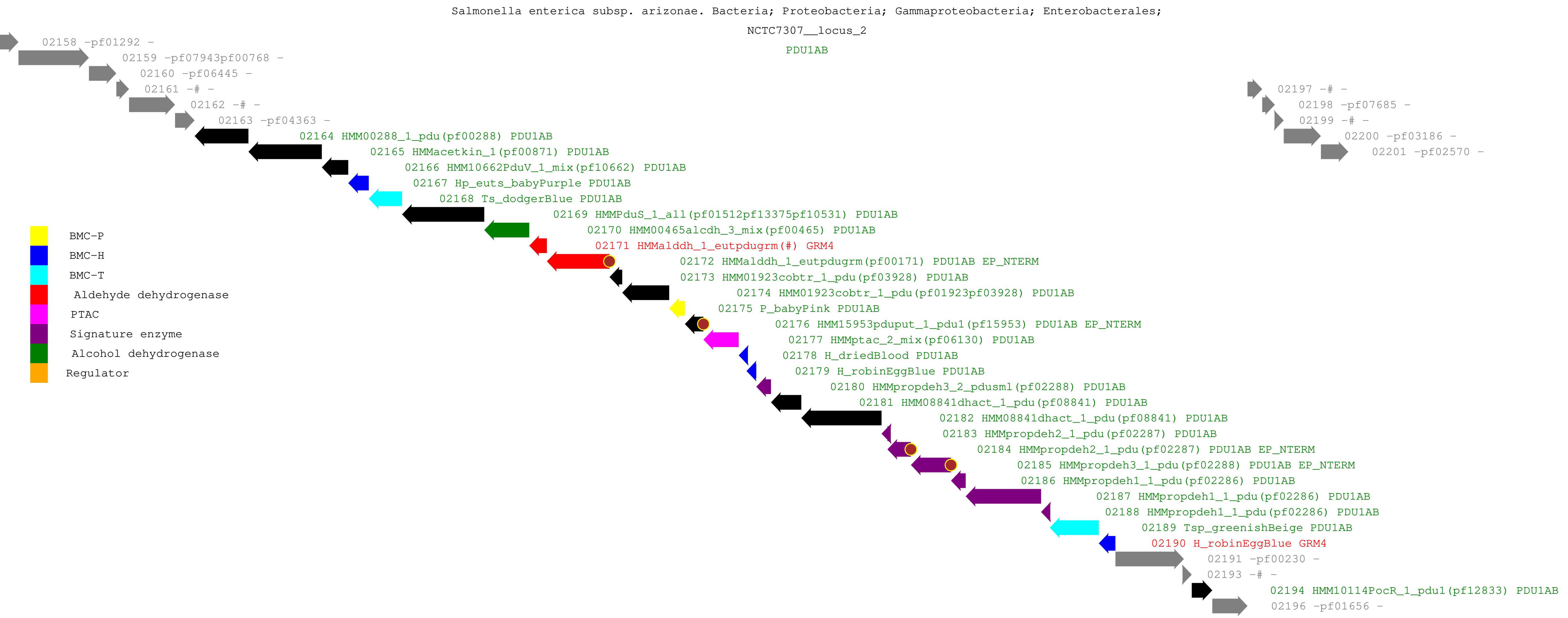
Salmonella enterica subsp. arizonae. Bacteria; Proteobacteria; Gammaproteobacteria; Enterobacterales;
NCTC7307__locus_2
PDU1AB
02158 -pf01292 -
02159 -pf07943pf00768 -
02160 -pf06445 -
02161 -# -
02162 -# -
02163 -pf04363 -
02164 HMM00288_1_pdu(pf00288) PDU1AB
02165 HMMacetkin_1(pf00871) PDU1AB
02166 HMM10662PduV_1_mix(pf10662) PDU1AB
02167 Hp_euts_babyPurple PDU1AB
02168 Ts_dodgerBlue PDU1AB
02169 HMMPduS_1_all(pf01512pf13375pf10531) PDU1AB
02170 HMM00465alcdh_3_mix(pf00465) PDU1AB
02171 HMMalddh_1_eutpdugrm(#) GRM4
02172 HMMalddh_1_eutpdugrm(pf00171) PDU1AB EP_NTERM
02173 HMM01923cobtr_1_pdu(pf03928) PDU1AB
02174 HMM01923cobtr_1_pdu(pf01923pf03928) PDU1AB
02175 P_babyPink PDU1AB
02176 HMM15953pduput_1_pdu1(pf15953) PDU1AB EP_NTERM
02177 HMMptac_2_mix(pf06130) PDU1AB
02178 H_driedBlood PDU1AB
02179 H_robinEggBlue PDU1AB
02180 HMMpropdeh3_2_pdusml(pf02288) PDU1AB
02181 HMM08841dhact_1_pdu(pf08841) PDU1AB
02182 HMM08841dhact_1_pdu(pf08841) PDU1AB
02183 HMMpropdeh2_1_pdu(pf02287) PDU1AB
02184 HMMpropdeh2_1_pdu(pf02287) PDU1AB EP_NTERM
02185 HMMpropdeh3_1_pdu(pf02288) PDU1AB EP_NTERM
02186 HMMpropdeh1_1_pdu(pf02286) PDU1AB
02187 HMMpropdeh1_1_pdu(pf02286) PDU1AB
02188 HMMpropdeh1_1_pdu(pf02286) PDU1AB
02189 Tsp_greenishBeige PDU1AB
02190 H_robinEggBlue GRM4
02191 -pf00230 -
02193 -# -
02194 HMM10114PocR_1_pdu1(pf12833) PDU1AB
02196 -pf01656 -
02197 -# -
02198 -pf07685 -
02199 -# -
02200 -pf03186 -
02201 -pf02570 -
BMC-P
BMC-H
BMC-T
Aldehyde dehydrogenase
PTAC
Signature enzyme
Alcohol dehydrogenase
Regulator

Salmonella choleraesuis. Bacteria; Proteobacteria; Gammaproteobacteria; Enterobacterales;

NCTC7346__locus_2

PDU1AB

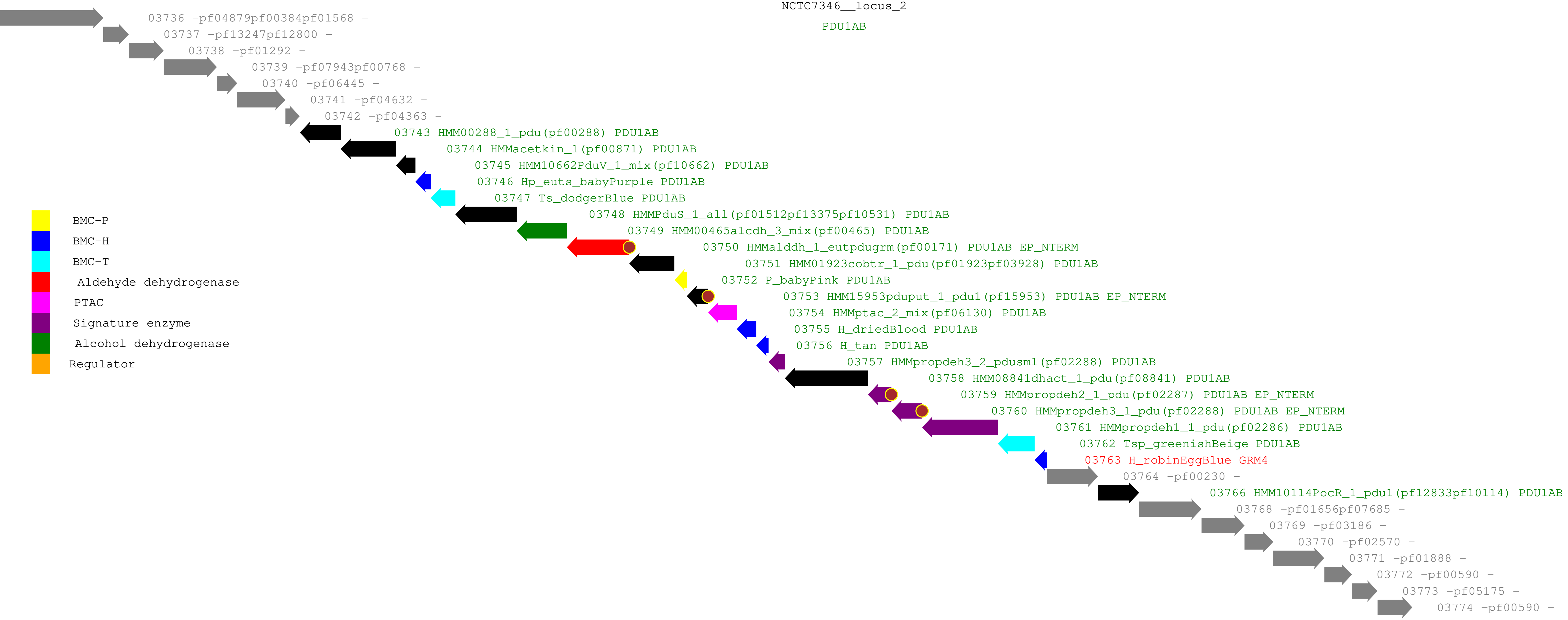

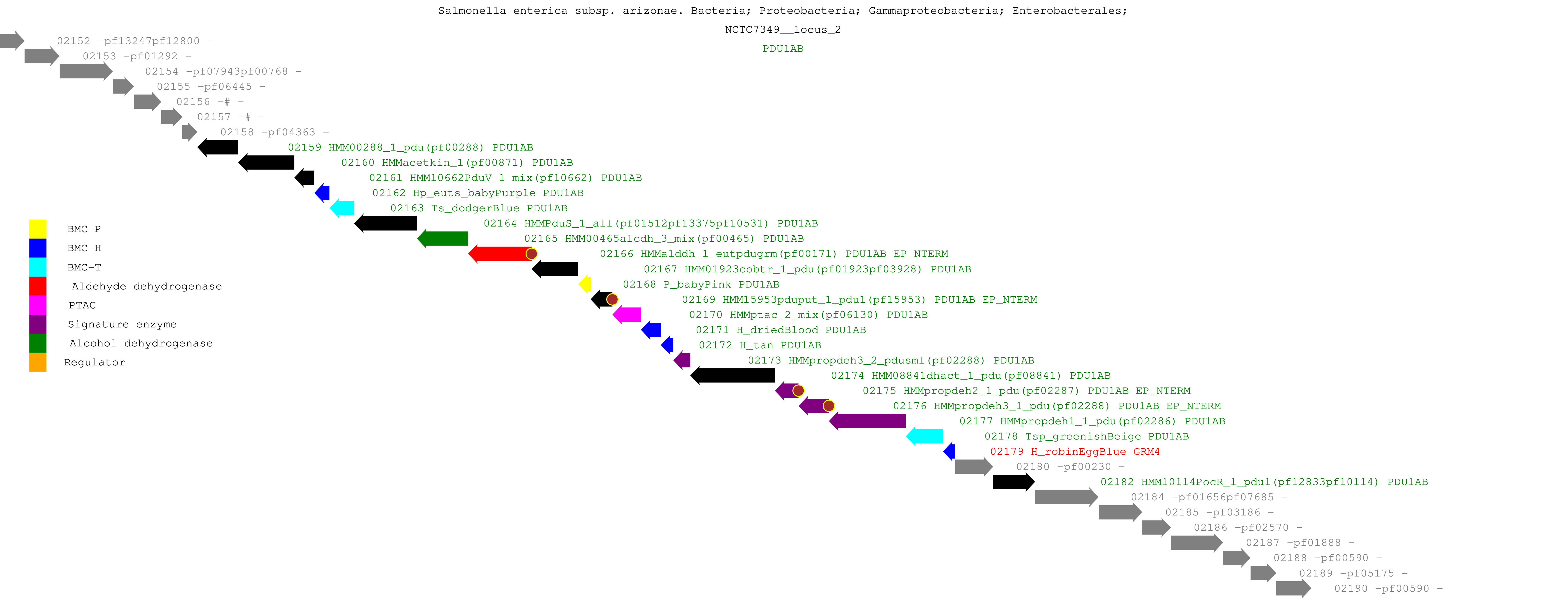
Salmonella enterica subsp. arizonae. Bacteria; Proteobacteria; Gammaproteobacteria; Enterobacterales;
NCTC7349__locus_2
PDU1AB
02152 -pf13247pf12800 -
02153 -pf01292 -
02154 -pf07943pf00768 -
02155 -pf06445 -
02156 -# -
02157 -# -
02158 -pf04363 -
02159 HMM00288_1_pdu(pf00288) PDU1AB
02160 HMMacetkin_1(pf00871) PDU1AB
02161 HMM10662PduV_1_mix(pf10662) PDU1AB
02162 Hp_euts_babyPurple PDU1AB
02163 Ts_dodgerBlue PDU1AB
02164 HMMPduS_1_all(pf01512pf13375pf10531) PDU1AB
02165 HMM00465alcdh_3_mix(pf00465) PDU1AB
02166 HMMalddh_1_eutpdugrm(pf00171) PDU1AB EP_NTERM
02167 HMM01923cobtr_1_pdu(pf01923pf03928) PDU1AB
02168 P_babyPink PDU1AB
02169 HMM15953pduput_1_pdu1(pf15953) PDU1AB EP_NTERM
02170 HMMptac_2_mix(pf06130) PDU1AB
02171 H_driedBlood PDU1AB
02172 H_tan PDU1AB
02173 HMMpropdeh3_2_pdusml(pf02288) PDU1AB
02174 HMM08841dhact_1_pdu(pf08841) PDU1AB
02175 HMMpropdeh2_1_pdu(pf02287) PDU1AB EP_NTERM
02176 HMMpropdeh3_1_pdu(pf02288) PDU1AB EP_NTERM
02177 HMMpropdeh1_1_pdu(pf02286) PDU1AB
02178 Tsp_greenishBeige PDU1AB
02179 H_robinEggBlue GRM4
02180 -pf00230 -
02182 HMM10114PocR_1_pdu1(pf12833pf10114) PDU1AB
02184 -pf01656pf07685 -
02185 -pf03186 -
02186 -pf02570 -
02187 -pf01888 -
02188 -pf00590 -
02189 -pf05175 -
02190 -pf00590 -
BMC-P
BMC-H
BMC-T
Aldehyde dehydrogenase
PTAC
Signature enzyme
Alcohol dehydrogenase
Regulator

Salmonella enterica subsp. enterica serovar Sanjuan. Bacteria; Proteobacteria; Gammaproteobacteria; Enterobacterales;

NCTC7406__locus_2

PDU1AB

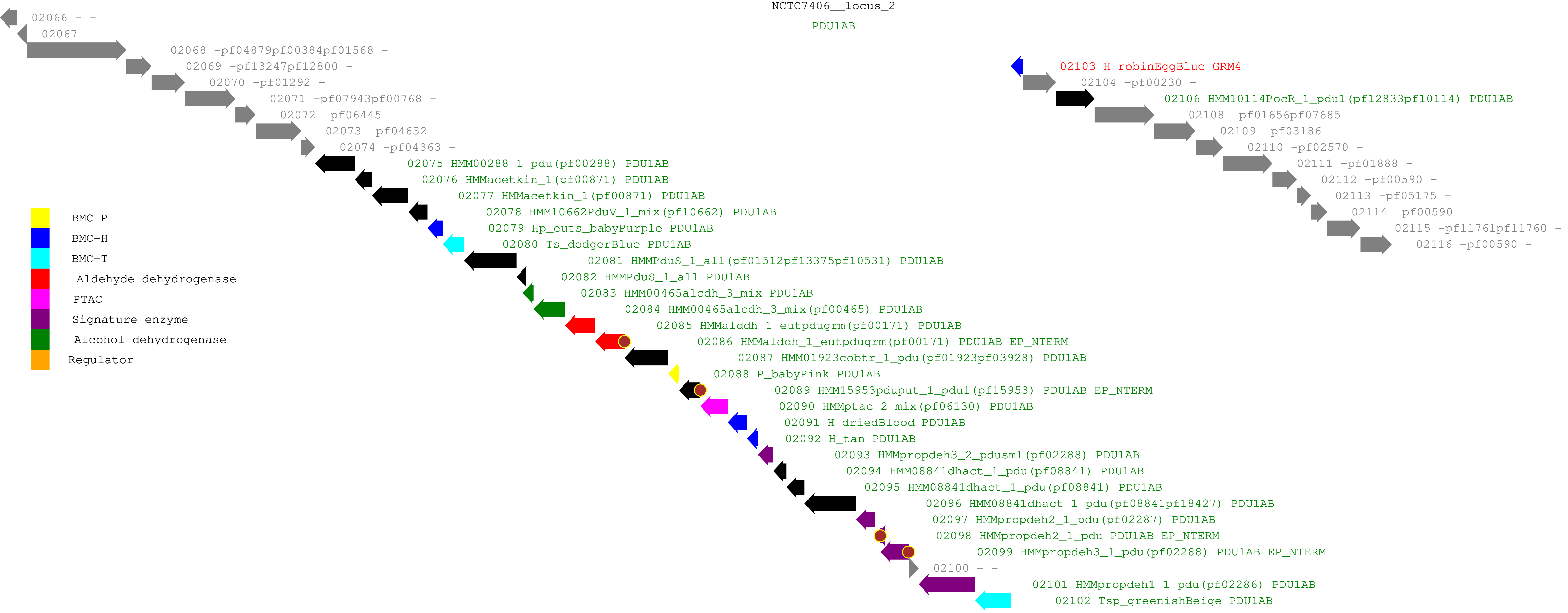

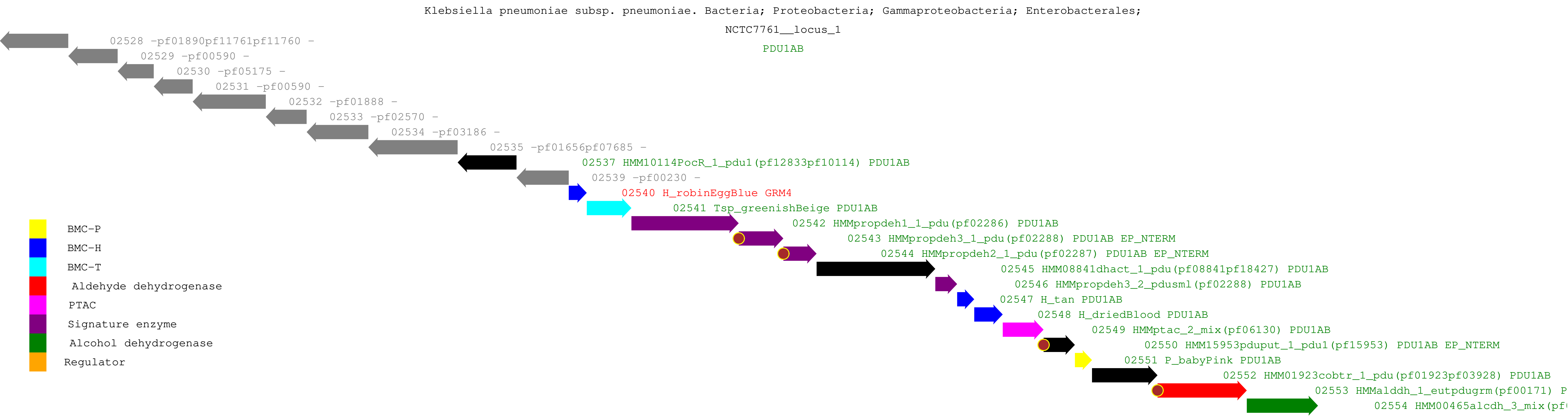
Klebsiella pneumoniae subsp. pneumoniae. Bacteria; Proteobacteria; Gammaproteobacteria; Enterobacterales;
NCTC7761__locus_1
PDU1AB
02528 -pf01890pf11761pf11760 -
02529 -pf00590 -
02530 -pf05175 -
02531 -pf00590 -
02532 -pf01888 -
02533 -pf02570 -
02534 -pf03186 -
02535 -pf01656pf07685 -
02537 HMM10114PocR_1_pdu1(pf12833pf10114) PDU1AB
02539 -pf00230 -
02540 H_robinEggBlue GRM4
02541 Tsp_greenishBeige PDU1AB
02542 HMMpropdeh1_1_pdu(pf02286) PDU1AB
02543 HMMpropdeh3_1_pdu(pf02288) PDU1AB EP_NTERM
02544 HMMpropdeh2_1_pdu(pf02287) PDU1AB EP_NTERM
02545 HMM08841dhact_1_pdu(pf08841pf18427) PDU1AB
02546 HMMpropdeh3_2_pdusml(pf02288) PDU1AB
02547 H_tan PDU1AB
02548 H_driedBlood PDU1AB
02549 HMMptac_2_mix(pf06130) PDU1AB
02550 HMM15953pduput_1_pdu1(pf15953) PDU1AB EP_NTERM
02551 P_babyPink PDU1AB
02552 HMM01923cobtr_1_pdu(pf01923pf03928) PDU1AB
02553 HMMalddh_1_eutpdugrm(pf00171) P
02554 HMM00465alcdh_3_mix(pf
BMC-P
BMC-H
BMC-T
Aldehyde dehydrogenase
PTAC
Signature enzyme
Alcohol dehydrogenase
Regulator

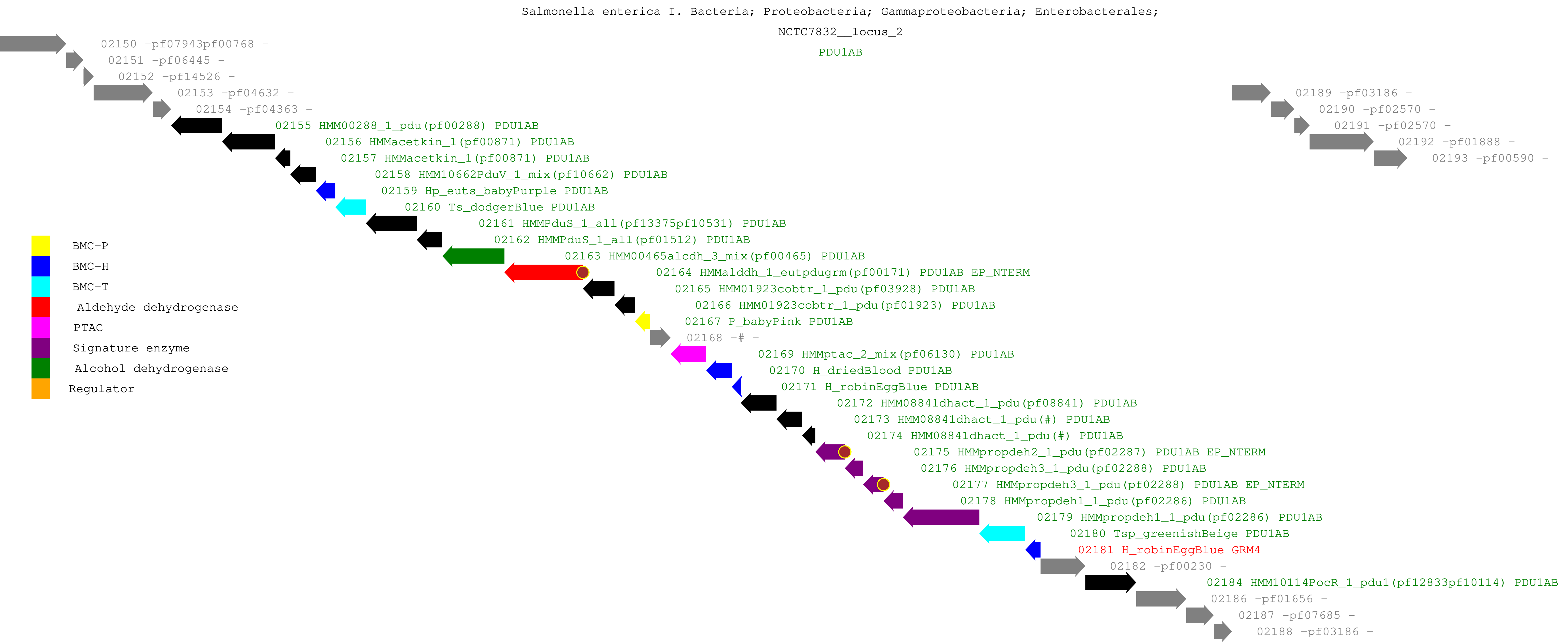
Salmonella enterica I. Bacteria; Proteobacteria; Gammaproteobacteria; Enterobacterales;
NCTC7832__locus_2
PDU1AB
02150 -pf07943pf00768 -
02151 -pf06445 -
02152 -pf14526 -
02153 -pf04632 -
02154 -pf04363 -
02155 HMM00288_1_pdu(pf00288) PDU1AB
02156 HMMacetkin_1(pf00871) PDU1AB
02157 HMMacetkin_1(pf00871) PDU1AB
02158 HMM10662PduV_1_mix(pf10662) PDU1AB
02159 Hp_euts_babyPurple PDU1AB
02160 Ts_dodgerBlue PDU1AB
02161 HMMPduS_1_all(pf13375pf10531) PDU1AB
02162 HMMPduS_1_all(pf01512) PDU1AB
02163 HMM00465alcdh_3_mix(pf00465) PDU1AB
02164 HMMalddh_1_eutpdugrm(pf00171) PDU1AB EP_NTERM
02165 HMM01923cobtr_1_pdu(pf03928) PDU1AB
02166 HMM01923cobtr_1_pdu(pf01923) PDU1AB
02167 P_babyPink PDU1AB
02168 -# -
02169 HMMptac_2_mix(pf06130) PDU1AB
02170 H_driedBlood PDU1AB
02171 H_robinEggBlue PDU1AB
02172 HMM08841dhact_1_pdu(pf08841) PDU1AB
02173 HMM08841dhact_1_pdu(#) PDU1AB
02174 HMM08841dhact_1_pdu(#) PDU1AB
02175 HMMpropdeh2_1_pdu(pf02287) PDU1AB EP_NTERM
02176 HMMpropdeh3_1_pdu(pf02288) PDU1AB
02177 HMMpropdeh3_1_pdu(pf02288) PDU1AB EP_NTERM
02178 HMMpropdeh1_1_pdu(pf02286) PDU1AB
02179 HMMpropdeh1_1_pdu(pf02286) PDU1AB
02180 Tsp_greenishBeige PDU1AB
02181 H_robinEggBlue GRM4
02182 -pf00230 -
02184 HMM10114PocR_1_pdu1(pf12833pf10114) PDU1AB
02186 -pf01656 -
02187 -pf07685 -
02188 -pf03186 -
02189 -pf03186 -
02190 -pf02570 -
02191 -pf02570 -
02192 -pf01888 -
02193 -pf00590 -
BMC-P
BMC-H
BMC-T
Aldehyde dehydrogenase
PTAC
Signature enzyme
Alcohol dehydrogenase
Regulator

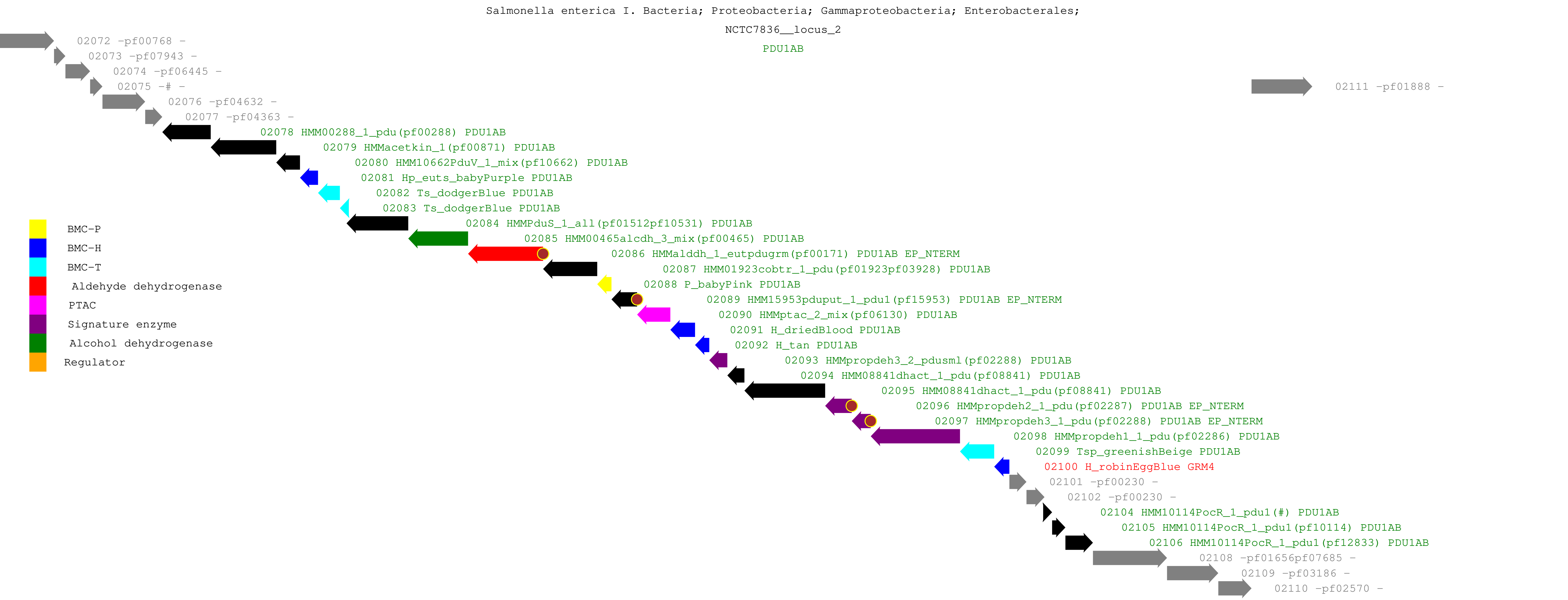

Salmonella enterica I. Bacteria; Proteobacteria; Gammaproteobacteria; Enterobacterales;
NCTC7836__locus_2
PDU1AB
02072 -pf00768 -
02073 -pf07943 -
02074 -pf06445 -
02075 -# -
02076 -pf04632 -
02077 -pf04363 -
02078 HMM00288_1_pdu(pf00288) PDU1AB
02079 HMMacetkin_1(pf00871) PDU1AB
02080 HMM10662PduV_1_mix(pf10662) PDU1AB
02081 Hp_euts_babyPurple PDU1AB
02082 Ts_dodgerBlue PDU1AB
02083 Ts_dodgerBlue PDU1AB
02084 HMMPduS_1_all(pf01512pf10531) PDU1AB
02085 HMM00465alcdh_3_mix(pf00465) PDU1AB
02086 HMMalddh_1_eutpdugrm(pf00171) PDU1AB EP_NTERM
02087 HMM01923cobtr_1_pdu(pf01923pf03928) PDU1AB
02088 P_babyPink PDU1AB
02089 HMM15953pduput_1_pdu1(pf15953) PDU1AB EP_NTERM
02090 HMMptac_2_mix(pf06130) PDU1AB
02091 H_driedBlood PDU1AB
02092 H_tan PDU1AB
02093 HMMpropdeh3_2_pdusml(pf02288) PDU1AB
02094 HMM08841dhact_1_pdu(pf08841) PDU1AB
02095 HMM08841dhact_1_pdu(pf08841) PDU1AB
02096 HMMpropdeh2_1_pdu(pf02287) PDU1AB EP_NTERM
02097 HMMpropdeh3_1_pdu(pf02288) PDU1AB EP_NTERM
02098 HMMpropdeh1_1_pdu(pf02286) PDU1AB
02099 Tsp_greenishBeige PDU1AB
02100 H_robinEggBlue GRM4
02101 -pf00230 -
02102 -pf00230 -
02104 HMM10114PocR_1_pdu1(#) PDU1AB
02105 HMM10114PocR_1_pdu1(pf10114) PDU1AB
02106 HMM10114PocR_1_pdu1(pf12833) PDU1AB
02108 -pf01656pf07685 -
02109 -pf03186 -
02110 -pf02570 -
02111 -pf01888 -
BMC-P
BMC-H
BMC-T
Aldehyde dehydrogenase
PTAC
Signature enzyme
Alcohol dehydrogenase
Regulator

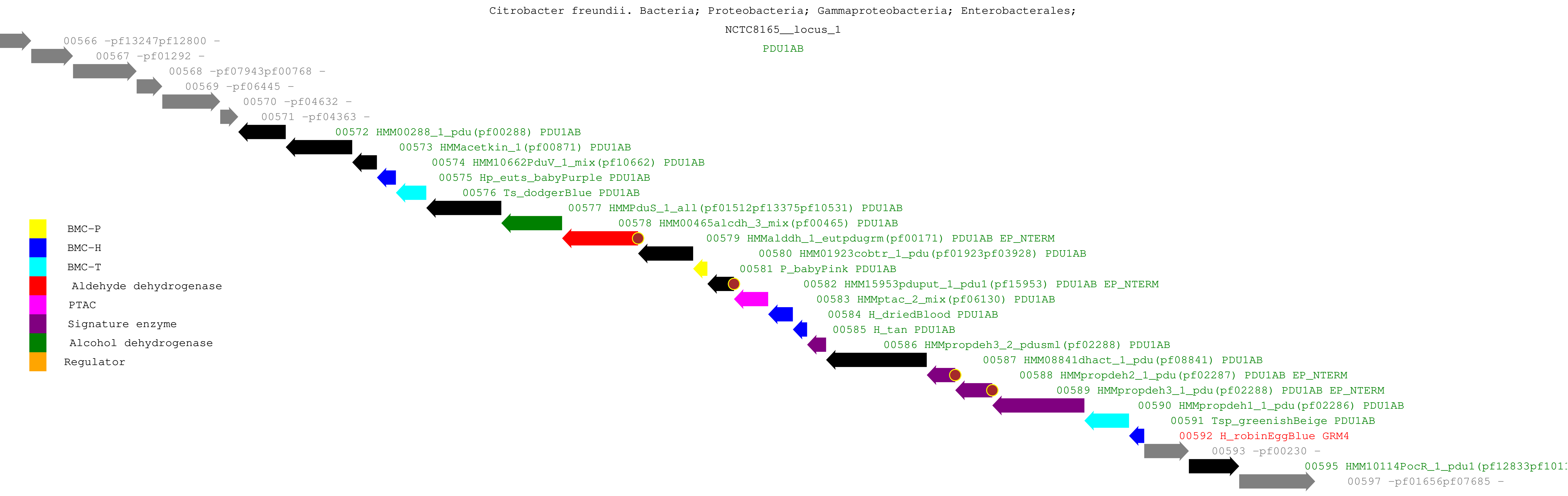
Citrobacter freundii. Bacteria; Proteobacteria; Gammaproteobacteria; Enterobacterales;
NCTC8165__locus_1
PDU1AB
00566 -pf13247pf12800 -
00567 -pf01292 -
00568 -pf07943pf00768 -
00569 -pf06445 -
00570 -pf04632 -
00571 -pf04363 -
00572 HMM00288_1_pdu(pf00288) PDU1AB
00573 HMMacetkin_1(pf00871) PDU1AB
00574 HMM10662PduV_1_mix(pf10662) PDU1AB
00575 Hp_euts_babyPurple PDU1AB
00576 Ts_dodgerBlue PDU1AB
00577 HMMPduS_1_all(pf01512pf13375pf10531) PDU1AB
00578 HMM00465alcdh_3_mix(pf00465) PDU1AB
00579 HMMalddh_1_eutpdugrm(pf00171) PDU1AB EP_NTERM
00580 HMM01923cobtr_1_pdu(pf01923pf03928) PDU1AB
00581 P_babyPink PDU1AB
00582 HMM15953pduput_1_pdu1(pf15953) PDU1AB EP_NTERM
00583 HMMptac_2_mix(pf06130) PDU1AB
00584 H_driedBlood PDU1AB
00585 H_tan PDU1AB
00586 HMMpropdeh3_2_pdusml(pf02288) PDU1AB
00587 HMM08841dhact_1_pdu(pf08841) PDU1AB
00588 HMMpropdeh2_1_pdu(pf02287) PDU1AB EP_NTERM
00589 HMMpropdeh3_1_pdu(pf02288) PDU1AB EP_NTERM
00590 HMMpropdeh1_1_pdu(pf02286) PDU1AB
00591 Tsp_greenishBeige PDU1AB
00592 H_robinEggBlue GRM4
00593 -pf00230 -
00595 HMM10114PocR_1_pdu1(pf12833pf101
00597 -pf01656pf07685 -
BMC-P
BMC-H
BMC-T
Aldehyde dehydrogenase
PTAC
Signature enzyme
Alcohol dehydrogenase
Regulator

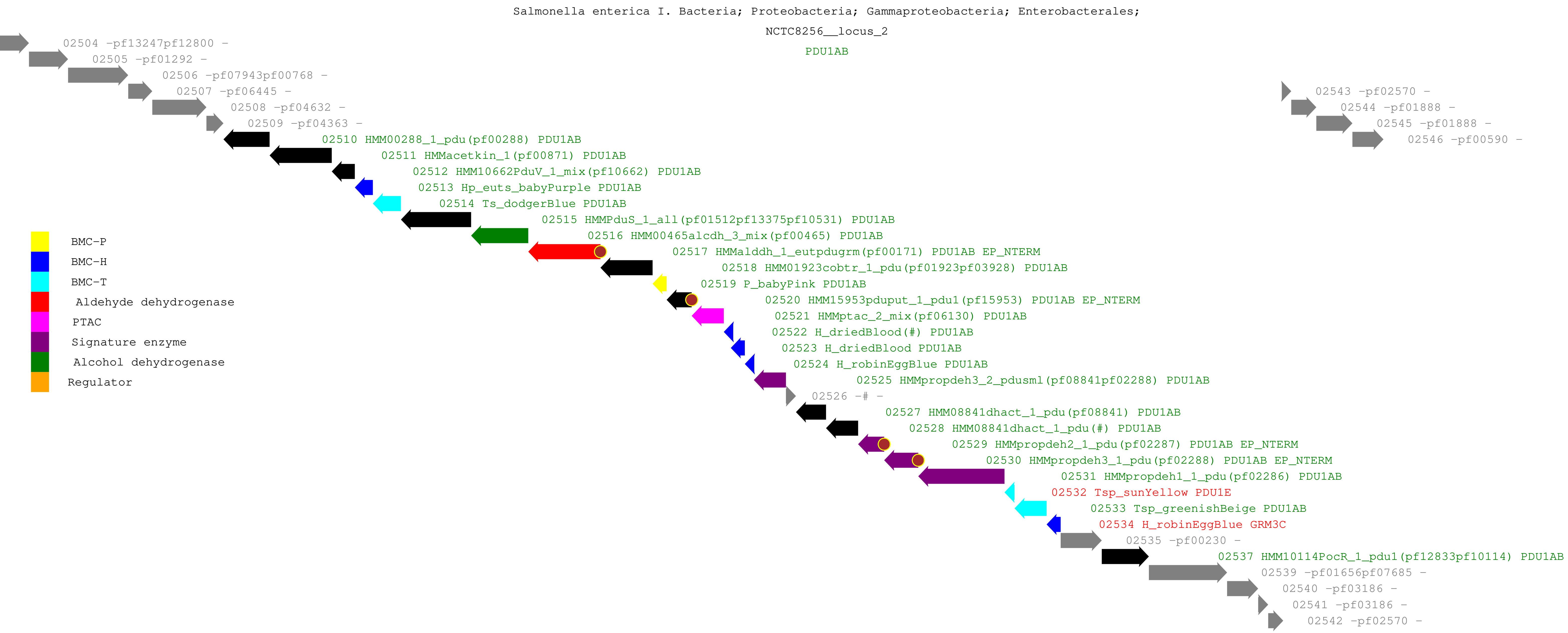

Salmonella enterica I. Bacteria; Proteobacteria; Gammaproteobacteria; Enterobacterales;
NCTC8256__locus_2
PDU1AB
02504 -pf13247pf12800 -
02505 -pf01292 -
02506 -pf07943pf00768 -
02507 -pf06445 -
02508 -pf04632 -
02509 -pf04363 -
02510 HMM00288_1_pdu(pf00288) PDU1AB
02511 HMMacetkin_1(pf00871) PDU1AB
02512 HMM10662PduV_1_mix(pf10662) PDU1AB
02513 Hp_euts_babyPurple PDU1AB
02514 Ts_dodgerBlue PDU1AB
02515 HMMPduS_1_all(pf01512pf13375pf10531) PDU1AB
02516 HMM00465alcdh_3_mix(pf00465) PDU1AB
02517 HMMaldh_1_eutpdugrm(pf00171) PDU1AB EP_NTERM
02518 HMM01923cobtr_1_pdu(pf01923pf03928) PDU1AB
02519 P_babyPink PDU1AB
02520 HMM15953pduput_1_pdu1(pf15953) PDU1AB EP_NTERM
02521 HMMptac_2_mix(pf06130) PDU1AB
02522 H_driedBlood(#) PDU1AB
02523 H_driedBlood PDU1AB
02524 H_robinEggBlue PDU1AB
02525 HMMpropdeh3_2_pdusml(pf08841pf02288) PDU1AB
02526 -# -
02527 HMM08841dhact_1_pdu(pf08841) PDU1AB
02528 HMM08841dhact_1_pdu(#) PDU1AB
02529 HMMpropdeh2_1_pdu(pf02287) PDU1AB EP_NTERM
02530 HMMpropdeh3_1_pdu(pf02288) PDU1AB EP_NTERM
02531 HMMpropdeh1_1_pdu(pf02286) PDU1AB
02532 Tsp_sunYellow PDU1E
02533 Tsp_greenishBeige PDU1AB
02534 H_robinEggBlue GRM3C
02535 -pf00230 -
02537 HMM10114PocR_1_pdu1(pf12833pf10114) PDU1AB
02539 -pf01656pf07685 -
02540 -pf03186 -
02541 -pf03186 -
02542 -pf02570 -
02543 -pf02570 -
02544 -pf01888 -
02545 -pf01888 -
02546 -pf00590 -
BMC-P
BMC-H
BMC-T
Aldehyde dehydrogenase
PTAC
Signature enzyme
Alcohol dehydrogenase
Regulator

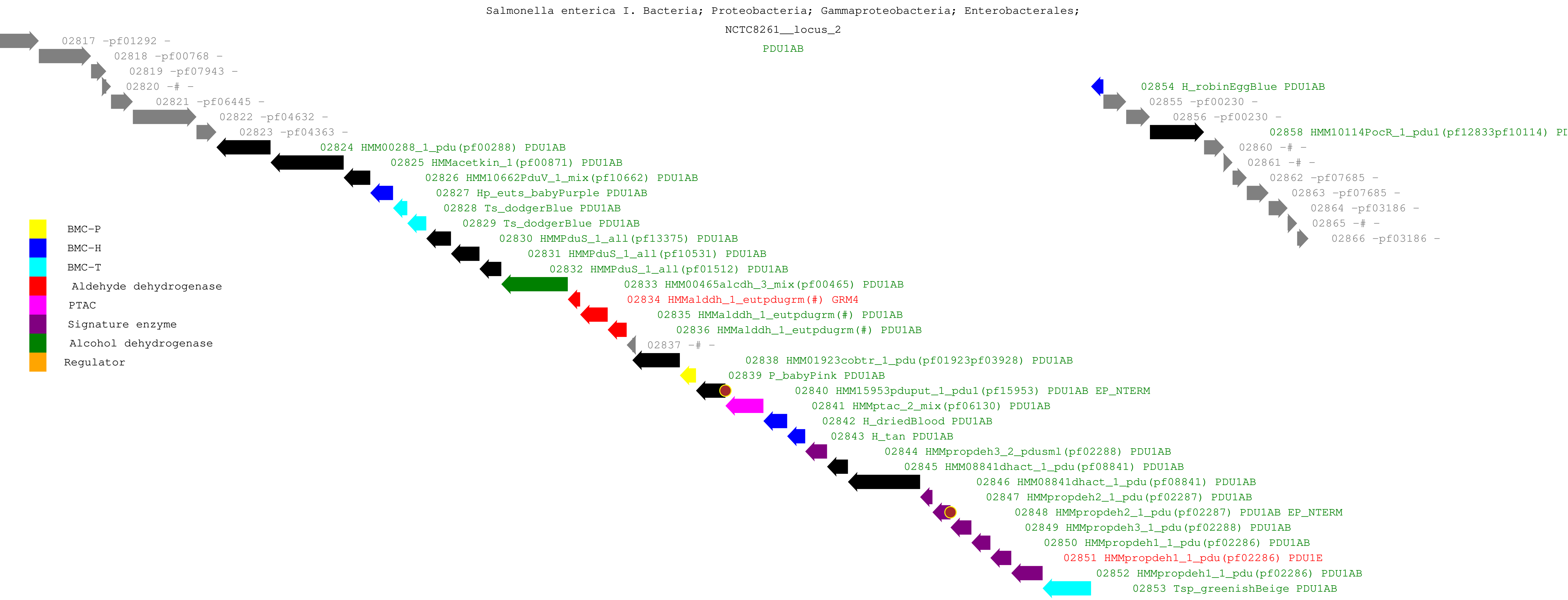

Salmonella enterica I. Bacteria; Proteobacteria; Gammaproteobacteria; Enterobacterales;
NCTC8261__locus_2
PDU1AB
02817 -pf01292 -
02818 -pf00768 -
02819 -pf07943 -
02820 -# -
02821 -pf06445 -
02822 -pf04632 -
02823 -pf04363 -
02824 HMM00288_1_pdu(pf00288) PDU1AB
02825 HMMacetkin_1(pf00871) PDU1AB
02826 HMM10662PduV_1_mix(pf10662) PDU1AB
02827 Hp_euts_babyPurple PDU1AB
02828 Ts_dodgerBlue PDU1AB
02829 Ts_dodgerBlue PDU1AB
02830 HMMPduS_1_all(pf13375) PDU1AB
02831 HMMPduS_1_all(pf10531) PDU1AB
02832 HMMPduS_1_all(pf01512) PDU1AB
02833 HMM00465alcdh_3_mix(pf00465) PDU1AB
02834 HMMalddh_1_eutpdugrm(#) GRM4
02835 HMMalddh_1_eutpdugrm(#) PDU1AB
02836 HMMalddh_1_eutpdugrm(#) PDU1AB
02837 -# -
02838 HMM01923cobtr_1_pdu(pf01923pf03928) PDU1AB
02839 P_babyPink PDU1AB
02840 HMM15953pduput_1_pdu1(pf15953) PDU1AB EP_NTERM
02841 HMMptac_2_mix(pf06130) PDU1AB
02842 H_driedBlood PDU1AB
02843 H_tan PDU1AB
02844 HMMpropdeh3_2_pdusml(pf02288) PDU1AB
02845 HMM08841dhact_1_pdu(pf08841) PDU1AB
02846 HMM08841dhact_1_pdu(pf08841) PDU1AB
02847 HMMpropdeh2_1_pdu(pf02287) PDU1AB
02848 HMMpropdeh2_1_pdu(pf02287) PDU1AB EP_NTERM
02849 HMMpropdeh3_1_pdu(pf02288) PDU1AB
02850 HMMpropdeh1_1_pdu(pf02286) PDU1AB
02851 HMMpropdeh1_1_pdu(pf02286) PDU1E
02852 HMMpropdeh1_1_pdu(pf02286) PDU1AB
02853 Tsp_greenishBeige PDU1AB
02854 H_robinEggBlue PDU1AB
02855 -pf00230 -
02856 -pf00230 -
02858 HMM10114PocR_1_pdu1(pf12833pf10114) PD
02860 -# -
02861 -# -
02862 -pf07685 -
02863 -pf07685 -
02864 -pf03186 -
02865 -# -
02866 -pf03186 -
BMC-P
BMC-H
BMC-T
Aldehyde dehydrogenase
PTAC
Signature enzyme
Alcohol dehydrogenase
Regulator

Salmonella enterica I. Bacteria; Proteobacteria; Gammaproteobacteria; Enterobacterales;

NCTC8271__locus_2

PDU1AB

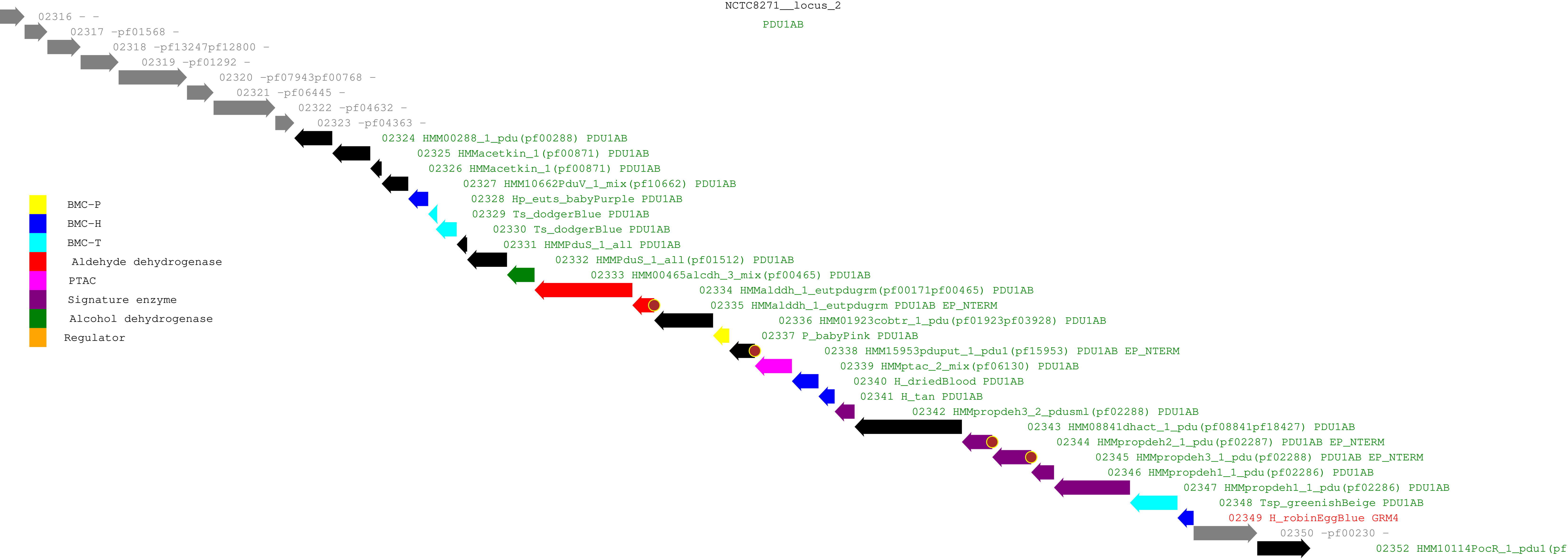

Salmonella enterica I. Bacteria; Proteobacteria; Gammaproteobacteria; Enterobacterales;

NCTC8272__locus_2

PDU1AB

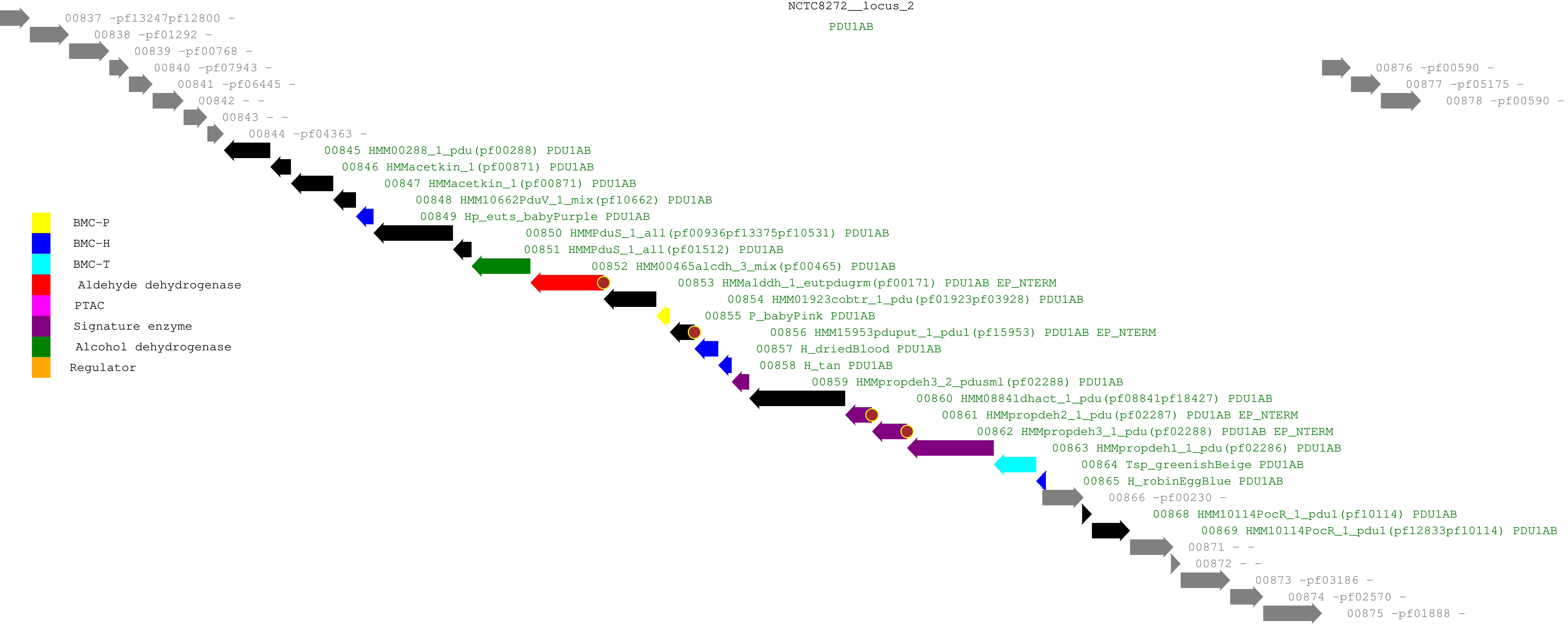

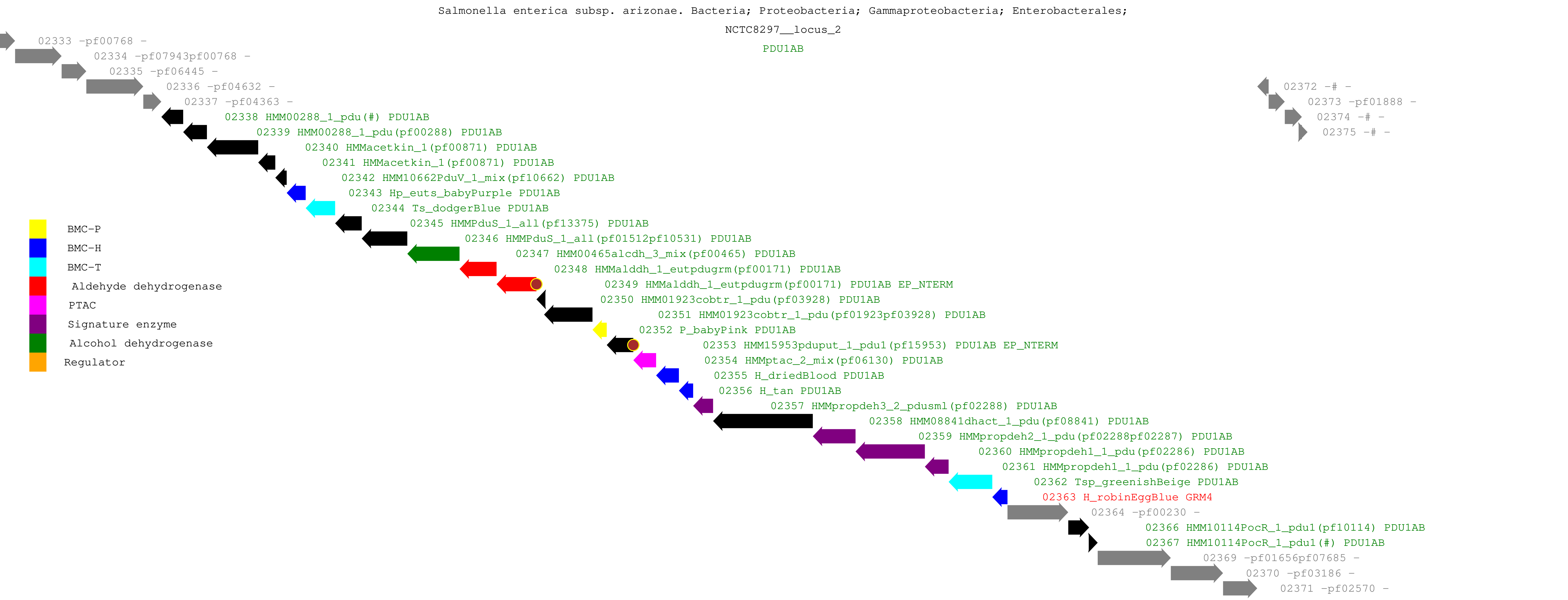
Salmonella enterica subsp. arizonae. Bacteria; Proteobacteria; Gammaproteobacteria; Enterobacterales;
NCTC8297__locus_2
PDU1AB
02333 -pf00768 -
02334 -pf07943pf00768 -
02335 -pf06445 -
02336 -pf04632 -
02337 -pf04363 -
02338 HMM00288_1_pdu(#) PDU1AB
02339 HMM00288_1_pdu(pf00288) PDU1AB
02340 HMMacetkin_1(pf00871) PDU1AB
02341 HMMacetkin_1(pf00871) PDU1AB
02342 HMM10662PduV_1_mix(pf10662) PDU1AB
02343 Hp_euts_babyPurple PDU1AB
02344 Ts_dodgerBlue PDU1AB
02345 HMMPduS_1_all(pf13375) PDU1AB
02346 HMMPduS_1_all(pf01512pf10531) PDU1AB
02347 HMM00465alcdh_3_mix(pf00465) PDU1AB
02348 HMMalddh_1_eutpdugrm(pf00171) PDU1AB
02349 HMMalddh_1_eutpdugrm(pf00171) PDU1AB EP_NTERM
02350 HMM01923cobtr_1_pdu(pf03928) PDU1AB
02351 HMM01923cobtr_1_pdu(pf01923pf03928) PDU1AB
02352 P_babyPink PDU1AB
02353 HMM15953pduput_1_pdu1(pf15953) PDU1AB EP_NTERM
02354 HMMptac_2_mix(pf06130) PDU1AB
02355 H_driedBlood PDU1AB
02356 H_tan PDU1AB
02357 HMMpropdeh3_2_pdusml(pf02288) PDU1AB
02358 HMM08841dhact_1_pdu(pf08841) PDU1AB
02359 HMMpropdeh2_1_pdu(pf02288pf02287) PDU1AB
02360 HMMpropdeh1_1_pdu(pf02286) PDU1AB
02361 HMMpropdeh1_1_pdu(pf02286) PDU1AB
02362 Tsp_greenishBeige PDU1AB
02363 H_robinEggBlue GRM4
02364 -pf00230 -
02366 HMM10114PocR_1_pdu1(pf10114) PDU1AB
02367 HMM10114PocR_1_pdu1(#) PDU1AB
02369 -pf01656pf07685 -
02370 -pf03186 -
02371 -pf02570 -
02372 -# -
02373 -pf01888 -
02374 -# -
02375 -# -
BMC-P
BMC-H
BMC-T
Aldehyde dehydrogenase
PTAC
Signature enzyme
Alcohol dehydrogenase
Regulator

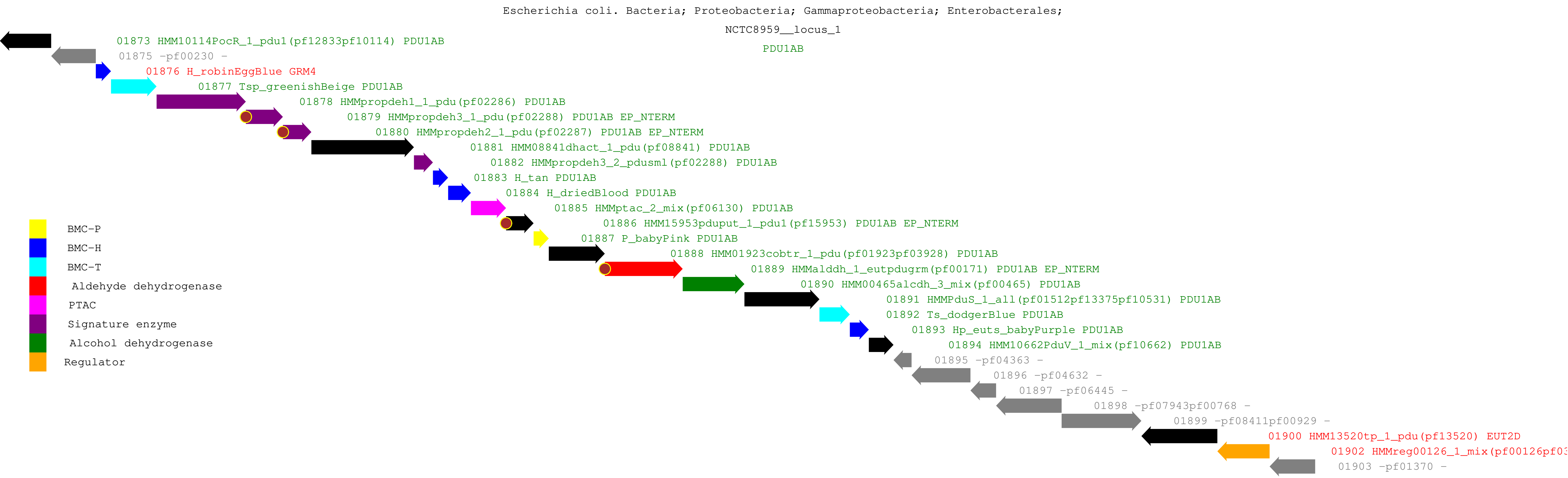
Escherichia coli. Bacteria; Proteobacteria; Gammaproteobacteria; Enterobacterales;
NCTC8959__locus_1
PDU1AB
01873 HMM10114PocR_1_pdu1(pf12833pf10114) PDU1AB
01875 -pf00230 -
01876 H_robinEggBlue GRM4
01877 Tsp_greenishBeige PDU1AB
01878 HMMpropdeh1_1_pdu(pf02286) PDU1AB
01879 HMMpropdeh3_1_pdu(pf02288) PDU1AB EP_NTERM
01880 HMMpropdeh2_1_pdu(pf02287) PDU1AB EP_NTERM
01881 HMM08841dhact_1_pdu(pf08841) PDU1AB
01882 HMMpropdeh3_2_pdusml(pf02288) PDU1AB
01883 H_tan PDU1AB
01884 H_driedBlood PDU1AB
01885 HMMptac_2_mix(pf06130) PDU1AB
01886 HMM15953pduput_1_pdu1(pf15953) PDU1AB EP_NTERM
01887 P_babyPink PDU1AB
01888 HMM01923cobtr_1_pdu(pf01923pf03928) PDU1AB
01889 HMMalddh_1_eutpdugrm(pf00171) PDU1AB EP_NTERM
01890 HMM00465alcdh_3_mix(pf00465) PDU1AB
01891 HMMPduS_1_all(pf01512pf13375pf10531) PDU1AB
01892 Ts_dodgerBlue PDU1AB
01893 Hp_euts_babyPurple PDU1AB
01894 HMM10662PduV_1_mix(pf10662) PDU1AB
01895 -pf04363 -
01896 -pf04632 -
01897 -pf06445 -
01898 -pf07943pf00768 -
01899 -pf08411pf00929 -
01900 HMM13520tp_1_pdu(pf13520) EUT2D
01902 HMMreg00126_1_mix(pf00126pf03
01903 -pf01370 -
BMC-P
BMC-H
BMC-T
Aldehyde dehydrogenase
PTAC
Signature enzyme
Alcohol dehydrogenase
Regulator

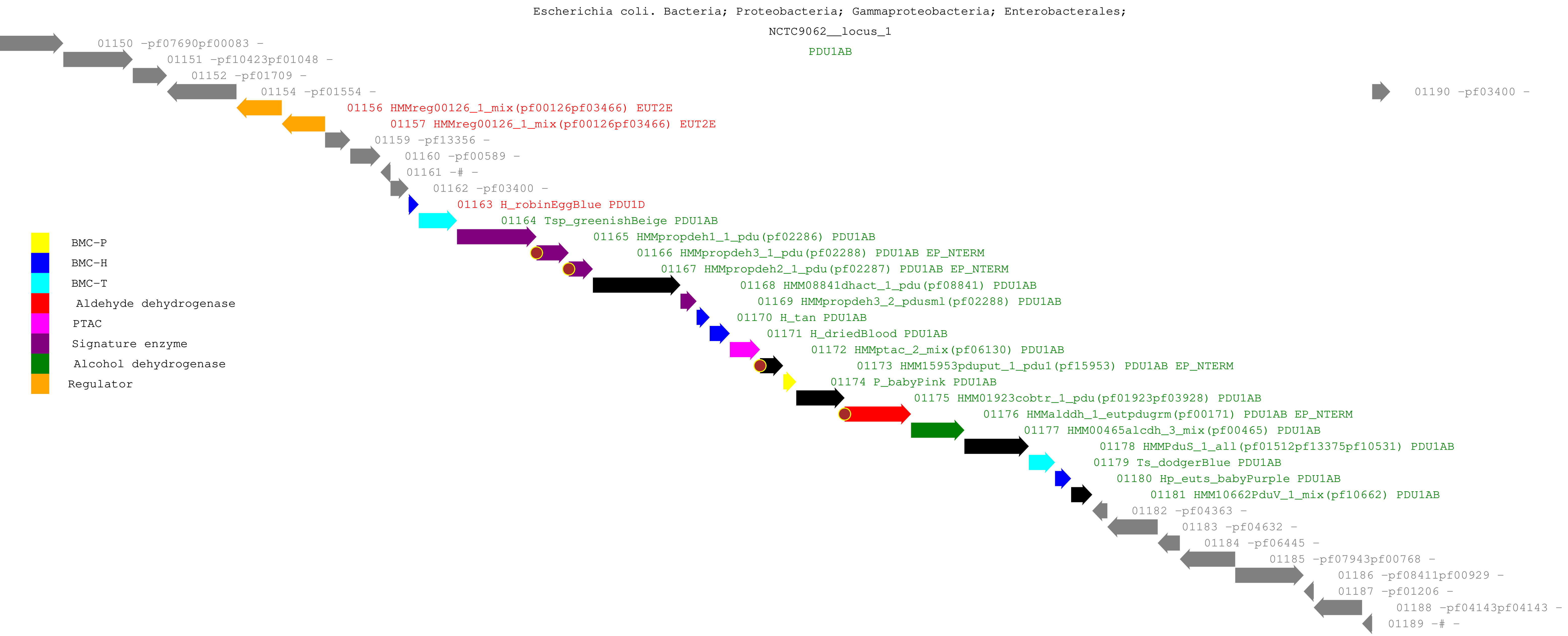
Escherichia coli. Bacteria; Proteobacteria; Gammaproteobacteria; Enterobacterales;
NCTC9062__locus_1
PDU1AB
01150 -pf07690pf00083 -
01151 -pf10423pf01048 -
01152 -pf01709 -
01154 -pf01554 -
01156 HMMreg00126_1_mix(pf00126pf03466) EUT2E
01157 HMMreg00126_1_mix(pf00126pf03466) EUT2E
01159 -pf13356 -
01160 -pf00589 -
01161 -# -
01162 -pf03400 -
01163 H_robinEggBlue PDU1D
01164 Tsp_greenishBeige PDU1AB
01165 HMMpropdeh1_1_pdu(pf02286) PDU1AB
01166 HMMpropdeh3_1_pdu(pf02288) PDU1AB EP_NTERM
01167 HMMpropdeh2_1_pdu(pf02287) PDU1AB EP_NTERM
01168 HMM08841dhact_1_pdu(pf08841) PDU1AB
01169 HMMpropdeh3_2_pdusml(pf02288) PDU1AB
01170 H_tan PDU1AB
01171 H_driedBlood PDU1AB
01172 HMMptac_2_mix(pf06130) PDU1AB
01173 HMM15953pduput_1_pdu1(pf15953) PDU1AB EP_NTERM
01174 P_babyPink PDU1AB
01175 HMM01923cobtr_1_pdu(pf01923pf03928) PDU1AB
01176 HMMalddh_1_eutpdugrm(pf00171) PDU1AB EP_NTERM
01177 HMM00465alcdh_3_mix(pf00465) PDU1AB
01178 HMMPduS_1_all(pf01512pf13375pf10531) PDU1AB
01179 Ts_dodgerBlue PDU1AB
01180 Hp_euts_babyPurple PDU1AB
01181 HMM10662PduV_1_mix(pf10662) PDU1AB
01182 -pf04363 -
01183 -pf04632 -
01184 -pf06445 -
01185 -pf07943pf00768 -
01186 -pf08411pf00929 -
01187 -pf01206 -
01188 -pf04143pf04143 -
01189 -# -
01190 -pf03400 -
BMC-P
BMC-H
BMC-T
Aldehyde dehydrogenase
PTAC
Signature enzyme
Alcohol dehydrogenase
Regulator

Escherichia coli. Bacteria; Proteobacteria; Gammaproteobacteria; Enterobacterales;

NCTC9067__locus_2

PDU1AB

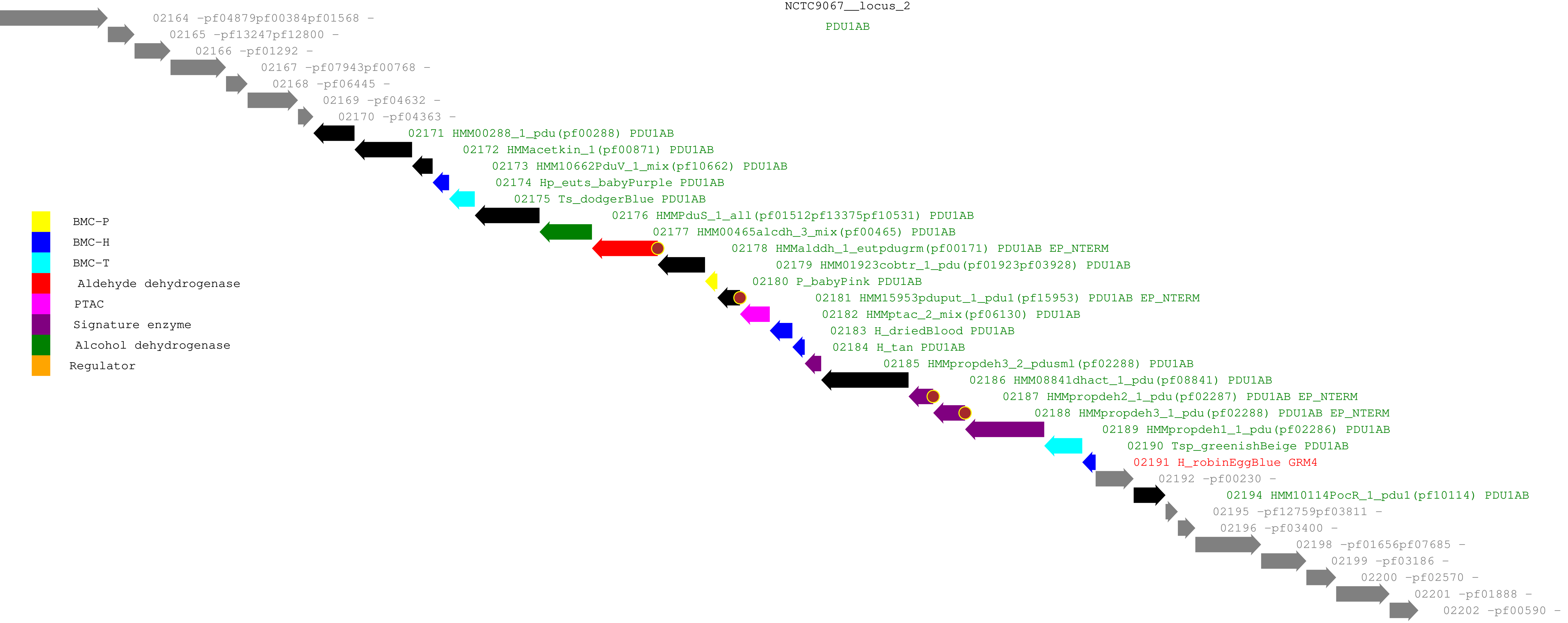

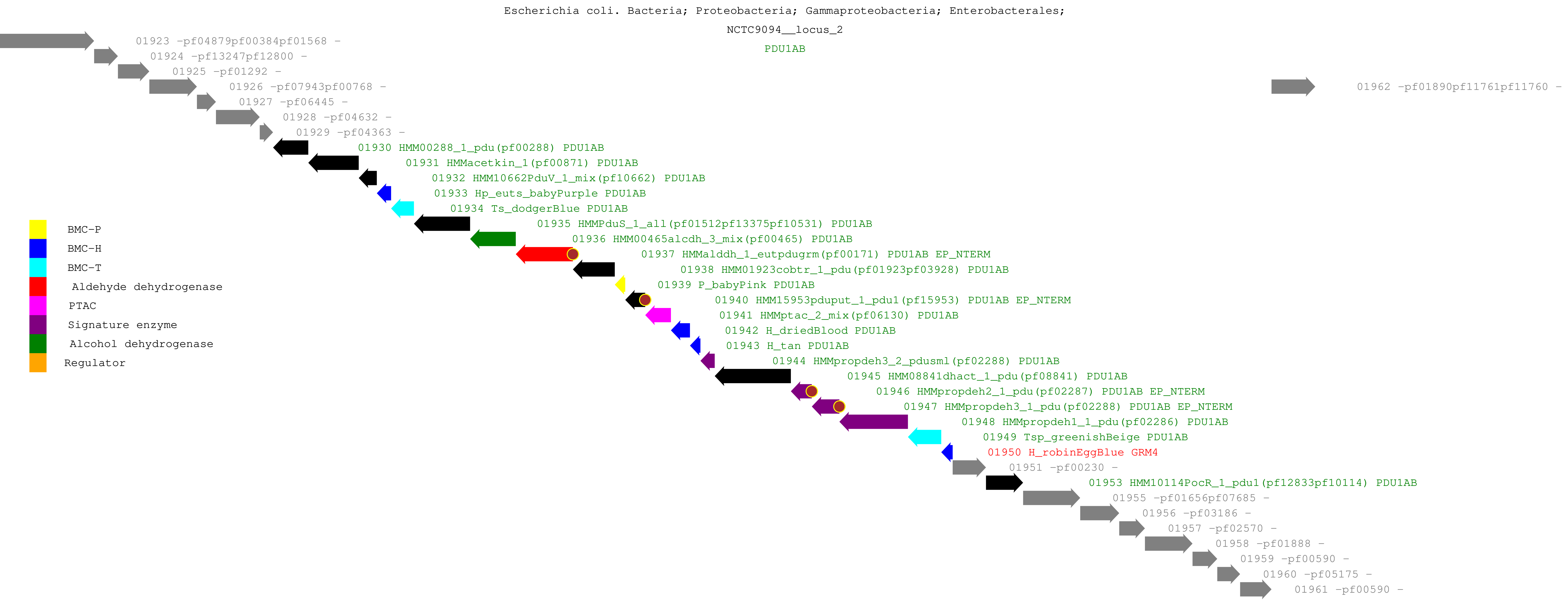

Escherichia coli. Bacteria; Proteobacteria; Gammaproteobacteria; Enterobacterales;
NCTC9094__locus_2
PDU1AB
01923 -pf04879pf00384pf01568 -
01924 -pf13247pf12800 -
01925 -pf01292 -
01926 -pf07943pf00768 -
01927 -pf06445 -
01928 -pf04632 -
01929 -pf04363 -
01930 HMM00288_1_pdu(pf00288) PDU1AB
01931 HMMacetkin_1(pf00871) PDU1AB
01932 HMM10662PduV_1_mix(pf10662) PDU1AB
01933 Hp_euts_babyPurple PDU1AB
01934 Ts_dodgerBlue PDU1AB
01935 HMMPduS_1_all(pf01512pf13375pf10531) PDU1AB
01936 HMM00465alcdh_3_mix(pf00465) PDU1AB
01937 HMMalddh_1_eutpdugrm(pf00171) PDU1AB EP_NTERM
01938 HMM01923cobtr_1_pdu(pf01923pf03928) PDU1AB
01939 P_babyPink PDU1AB
01940 HMM15953pduput_1_pdu1(pf15953) PDU1AB EP_NTERM
01941 HMMptac_2_mix(pf06130) PDU1AB
01942 H_driedBlood PDU1AB
01943 H_tan PDU1AB
01944 HMMpropdeh3_2_pdusml(pf02288) PDU1AB
01945 HMM08841dhact_1_pdu(pf08841) PDU1AB
01946 HMMpropdeh2_1_pdu(pf02287) PDU1AB EP_NTERM
01947 HMMpropdeh3_1_pdu(pf02288) PDU1AB EP_NTERM
01948 HMMpropdeh1_1_pdu(pf02286) PDU1AB
01949 Tsp_greenishBeige PDU1AB
01950 H_robinEggBlue GRM4
01951 -pf00230 -
01953 HMM10114PocR_1_pdu1(pf12833pf10114) PDU1AB
01955 -pf01656pf07685 -
01956 -pf03186 -
01957 -pf02570 -
01958 -pf01888 -
01959 -pf00590 -
01960 -pf05175 -
01961 -pf00590 -
01962 -pf01890pf11761pf11760 -
BMC-P
BMC-H
BMC-T
Aldehyde dehydrogenase
PTAC
Signature enzyme
Alcohol dehydrogenase
Regulator

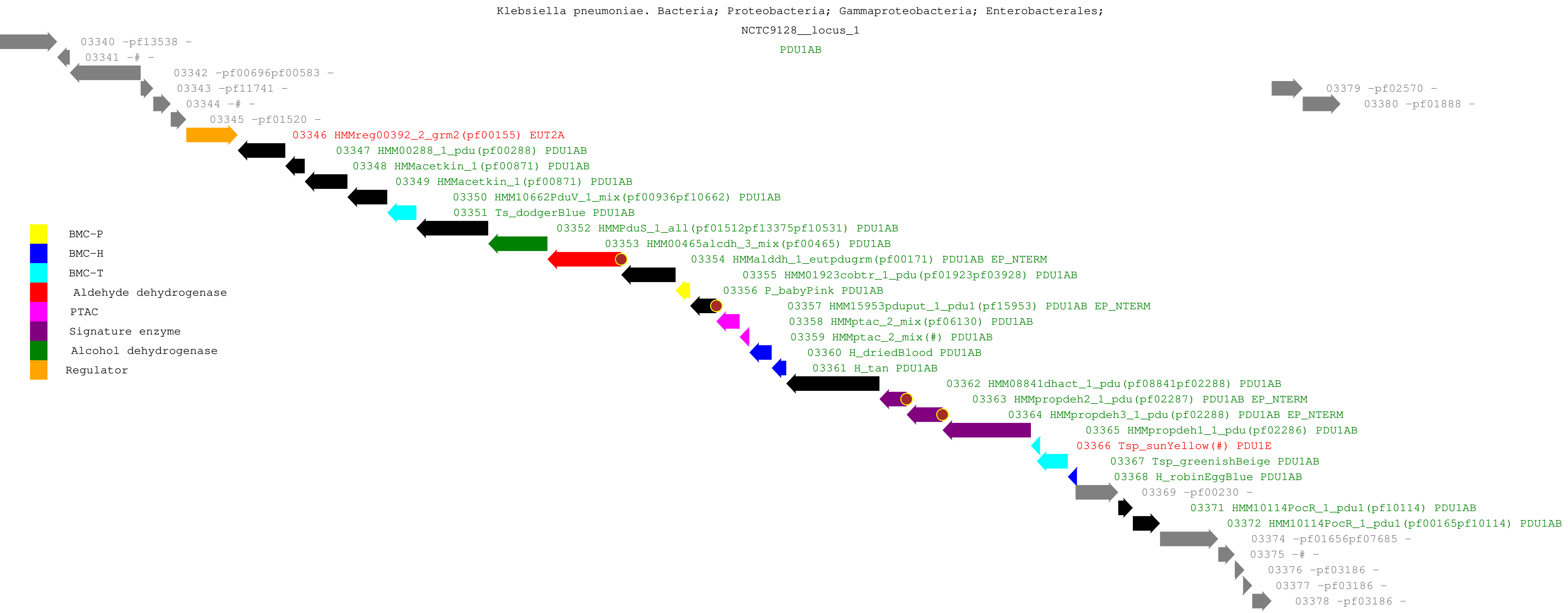
Klebsiella pneumoniae. Bacteria; Proteobacteria; Gammaproteobacteria; Enterobacterales;
NCTC9128__locus_1
PDU1AB
03340 -pf13538 -
03341 -# -
03342 -pf00696pf00583 -
03343 -pf11741 -
03344 -# -
03345 -pf01520 -
03346 HMMreg00392_2_grm2(pf00155) EUT2A
03347 HMM00288_1_pdu(pf00288) PDU1AB
03348 HMMacetkin_1(pf00871) PDU1AB
03349 HMMacetkin_1(pf00871) PDU1AB
03350 HMM10662PduV_1_mix(pf00936pf10662) PDU1AB
03351 Ts_dodgerBlue PDU1AB
03352 HMMPduS_1_all(pf01512pf13375pf10531) PDU1AB
03353 HMM00465alcdh_3_mix(pf00465) PDU1AB
03354 HMMaldh_1_eutpdugrm(pf00171) PDU1AB EP_NTERM
03355 HMM01923cobtr_1_pdu(pf01923pf03928) PDU1AB
03356 P_babyPink PDU1AB
03357 HMM15953pduput_1_pdu1(pf15953) PDU1AB EP_NTERM
03358 HMMptac_2_mix(pf06130) PDU1AB
03359 HMMptac_2_mix(#) PDU1AB
03360 H_driedBlood PDU1AB
03361 H_tan PDU1AB
03362 HMM08841dhact_1_pdu(pf08841pf02288) PDU1AB
03363 HMMpropdeh2_1_pdu(pf02287) PDU1AB EP_NTERM
03364 HMMpropdeh3_1_pdu(pf02288) PDU1AB EP_NTERM
03365 HMMpropdeh1_1_pdu(pf02286) PDU1AB
03366 Tsp_sunYellow(#) PDU1E
03367 Tsp_greenishBeige PDU1AB
03368 H_robinEggBlue PDU1AB
03369 -pf00230 -
03371 HMM10114PocR_1_pdu1(pf10114) PDU1AB
03372 HMM10114PocR_1_pdu1(pf00165pf10114) PDU1AB
03374 -pf01656pf07685 -
03375 -# -
03376 -pf03186 -
03377 -pf03186 -
03378 -pf03186 -
03379 -pf02570 -
03380 -pf01888 -
BMC-P
BMC-H
BMC-T
Aldehyde dehydrogenase
PTAC
Signature enzyme
Alcohol dehydrogenase
Regulator

Klebsiella oxytoca. Bacteria; Proteobacteria; Gammaproteobacteria; Enterobacterales;

NCTC9149__locus_1

PDU1AB

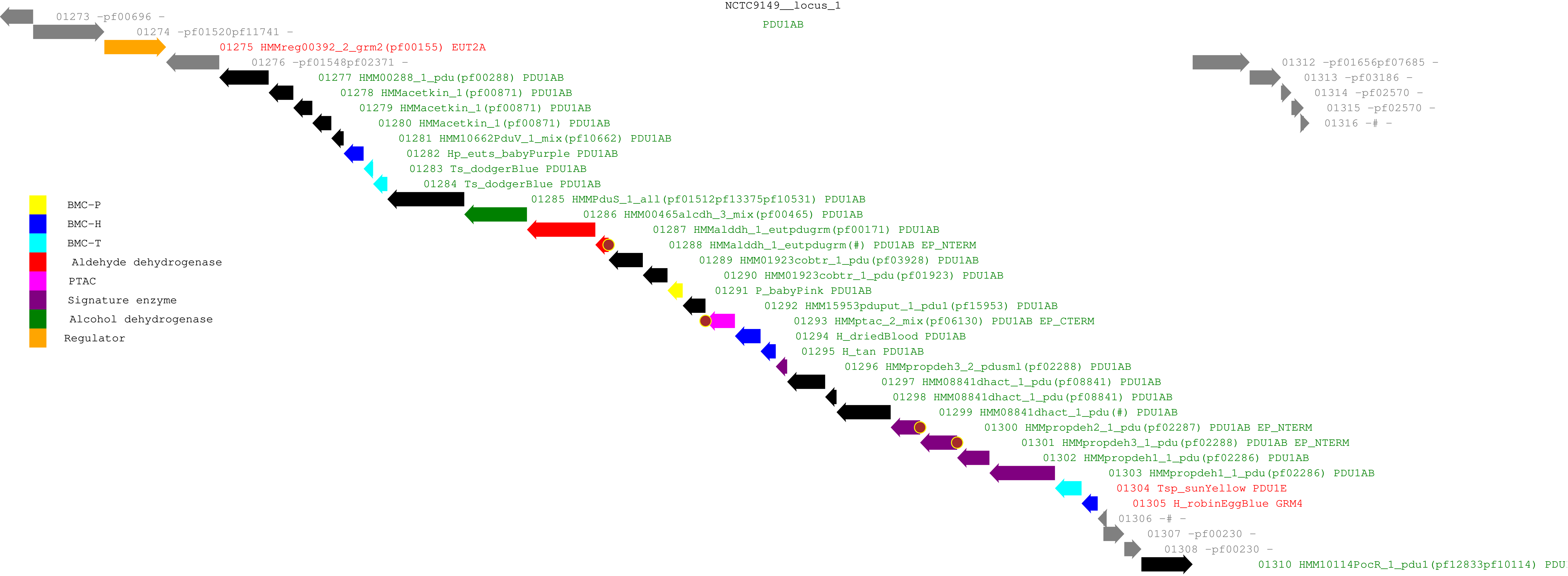

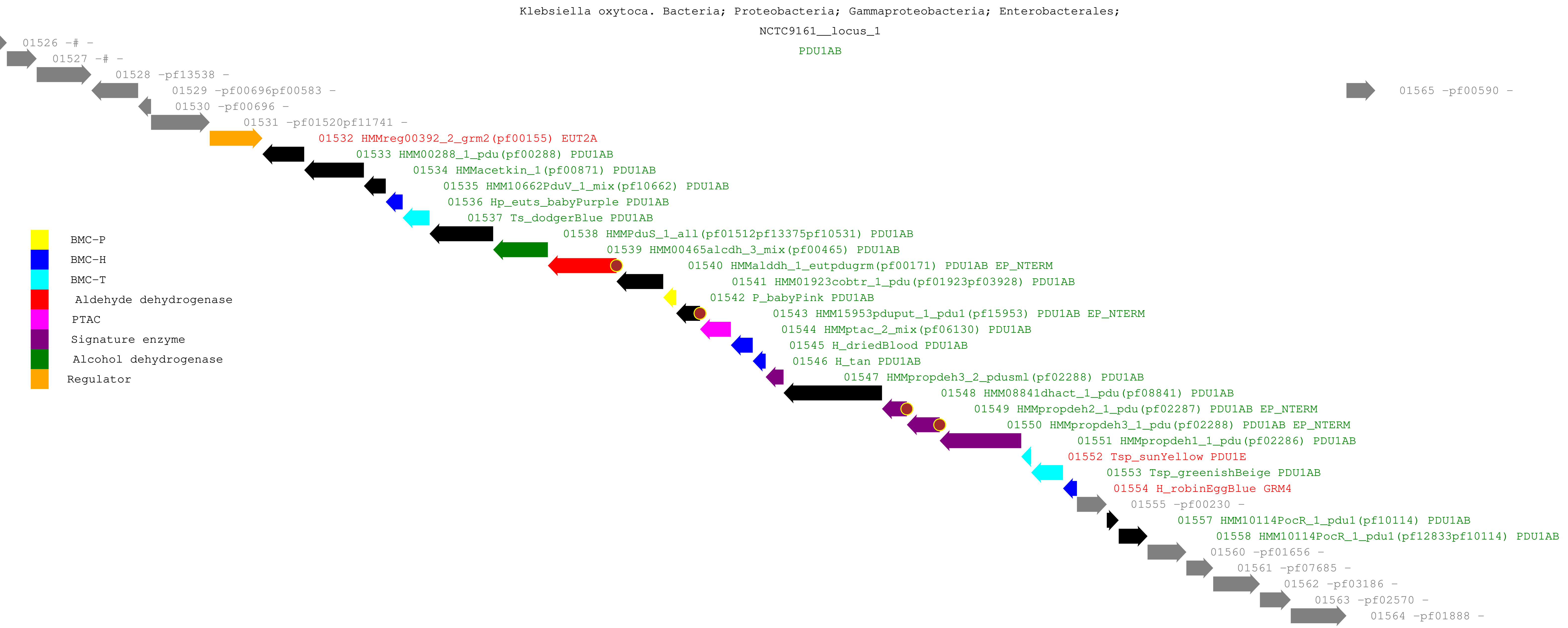
Klebsiella oxytoca. Bacteria; Proteobacteria; Gammaproteobacteria; Enterobacterales;
NCTC9161__locus_1
PDU1AB
01526 -# -
01527 -# -
01528 -pf13538 -
01529 -pf00696pf00583 -
01565 -pf00590 -
01530 -pf00696 -
01531 -pf01520pf11741 -
01532 HMMreg00392_2_grm2(pf00155) EUT2A
01533 HMM00288_1_pdu(pf00288) PDU1AB
01534 HMMacetkin_1(pf00871) PDU1AB
01535 HMM10662PduV_1_mix(pf10662) PDU1AB
01536 Hp_euts_babyPurple PDU1AB
01537 Ts_dodgerBlue PDU1AB
01538 HMMPduS_1_all(pf01512pf13375pf10531) PDU1AB
01539 HMM00465alcdh_3_mix(pf00465) PDU1AB
01540 HMMalddh_1_eutpdugrm(pf00171) PDU1AB EP_NTERM
01541 HMM01923cobtr_1_pdu(pf01923pf03928) PDU1AB
01542 P_babyPink PDU1AB
01543 HMM15953pduput_1_pdu1(pf15953) PDU1AB EP_NTERM
01544 HMMptac_2_mix(pf06130) PDU1AB
01545 H_driedBlood PDU1AB
01546 H_tan PDU1AB
01547 HMMpropdeh3_2_pdusml(pf02288) PDU1AB
01548 HMM08841dhact_1_pdu(pf08841) PDU1AB
01549 HMMpropdeh2_1_pdu(pf02287) PDU1AB EP_NTERM
01550 HMMpropdeh3_1_pdu(pf02288) PDU1AB EP_NTERM
01551 HMMpropdeh1_1_pdu(pf02286) PDU1AB
01552 Tsp_sunYellow PDU1E
01553 Tsp_greenishBeige PDU1AB
01554 H_robinEggBlue GRM4
01555 -pf00230 -
01557 HMM10114PocR_1_pdu1(pf10114) PDU1AB
01558 HMM10114PocR_1_pdu1(pf12833pf10114) PDU1AB
01560 -pf01656 -
01561 -pf07685 -
01562 -pf03186 -
01563 -pf02570 -
01564 -pf01888 -
BMC-P
BMC-H
BMC-T
Aldehyde dehydrogenase
PTAC
Signature enzyme
Alcohol dehydrogenase
Regulator

Klebsiella pneumoniae. Bacteria; Proteobacteria; Gammaproteobacteria; Enterobacterales;

NCTC9177__locus_3

PDU1AB

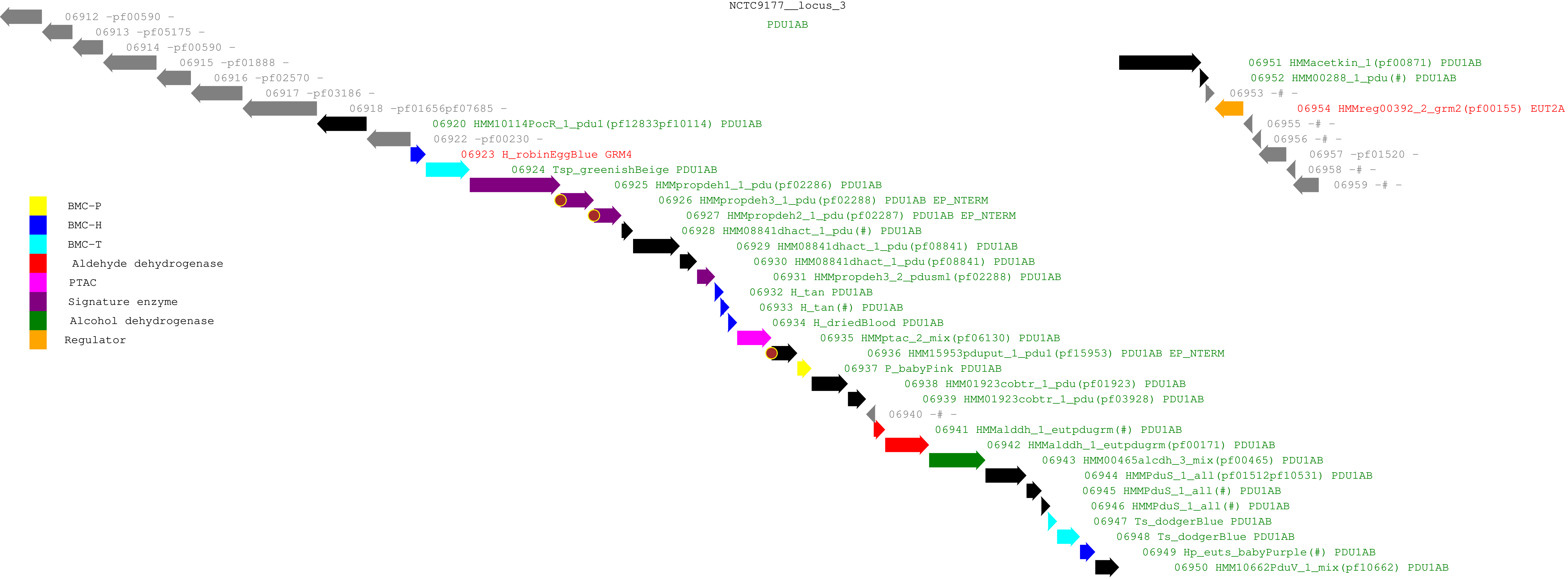

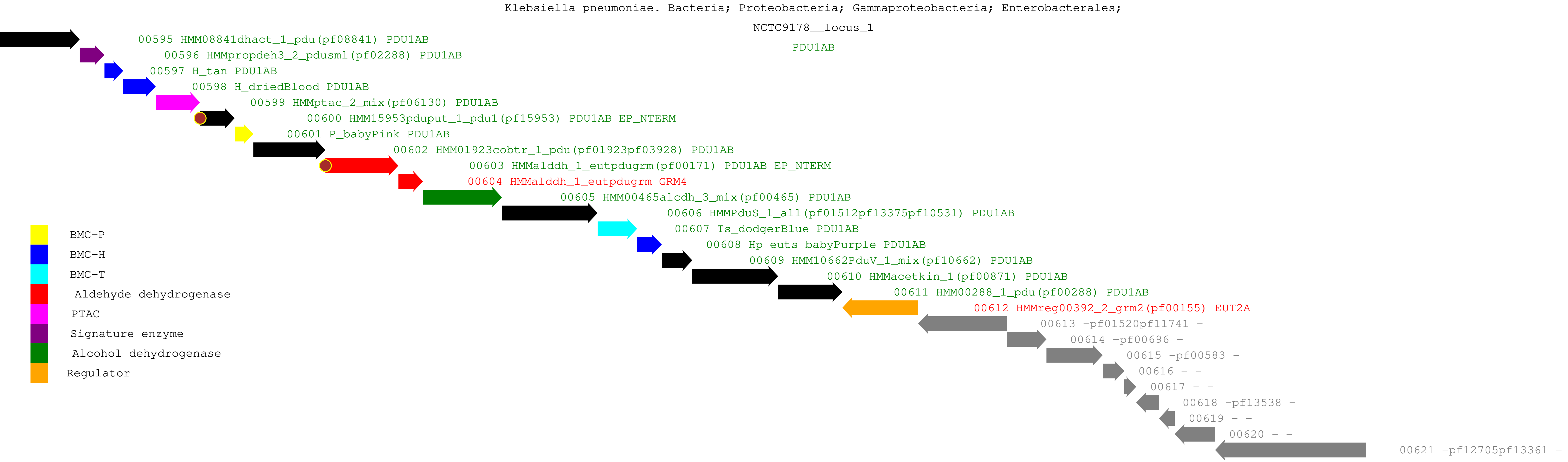
Klebsiella pneumoniae. Bacteria; Proteobacteria; Gammaproteobacteria; Enterobacterales;
NCTC9178__locus_1
PDU1AB
00595 HMM08841dhact_1_pdu(pf08841) PDU1AB
00596 HMMpropdeh3_2_pdusml(pf02288) PDU1AB
00597 H_tan PDU1AB
00598 H_driedBlood PDU1AB
00599 HMMptac_2_mix(pf06130) PDU1AB
00600 HMM15953pduput_1_pdu1(pf15953) PDU1AB EP_NTERM
00601 P_babyPink PDU1AB
00602 HMM01923cobtr_1_pdu(pf01923pf03928) PDU1AB
00603 HMMalddh_1_eutpdugrm(pf00171) PDU1AB EP_NTERM
00604 HMMalddh_1_eutpdugrm GRM4
00605 HMM00465alcdh_3_mix(pf00465) PDU1AB
00606 HMMPduS_1_all(pf01512pf13375pf10531) PDU1AB
00607 Ts_dodgerBlue PDU1AB
00608 Hp_euts_babyPurple PDU1AB
00609 HMM10662PduV_1_mix(pf10662) PDU1AB
00610 HMMacetkin_1(pf00871) PDU1AB
00611 HMM00288_1_pdu(pf00288) PDU1AB
00612 HMMreg00392_2_grm2(pf00155) EUT2A
00613 -pf01520pf11741 -
00614 -pf00696 -
00615 -pf00583 -
00616 - -
00617 - -
00618 -pf13538 -
00619 - -
00620 - -
00621 -pf12705pf13361 -
BMC-P
BMC-H
BMC-T
Aldehyde dehydrogenase
PTAC
Signature enzyme
Alcohol dehydrogenase
Regulator

Klebsiella oxytoca. Bacteria; Proteobacteria; Gammaproteobacteria; Enterobacterales;

NCTC9186__locus_1

PDU1AB

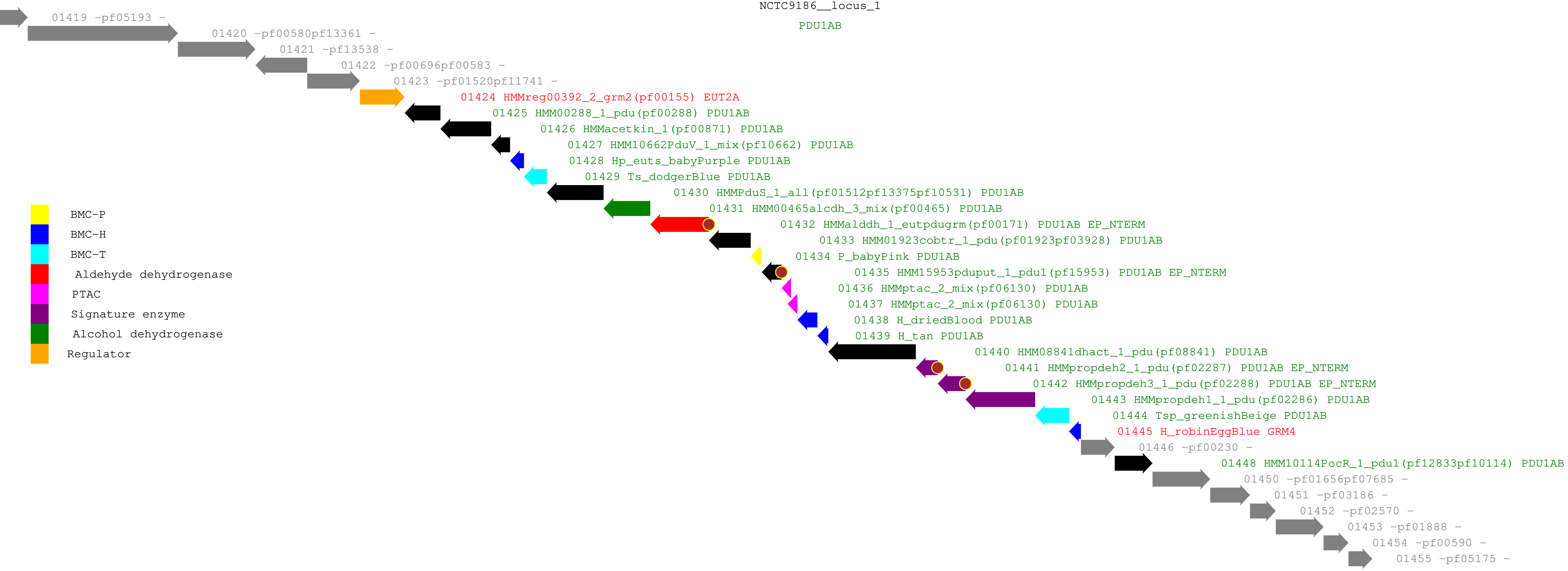

Klebsiella pneumoniae subsp. pneumoniae. Bacteria; Proteobacteria; Gammaproteobacteria; Enterobacterales;

NCTC9494__locus_1

PDU1AB

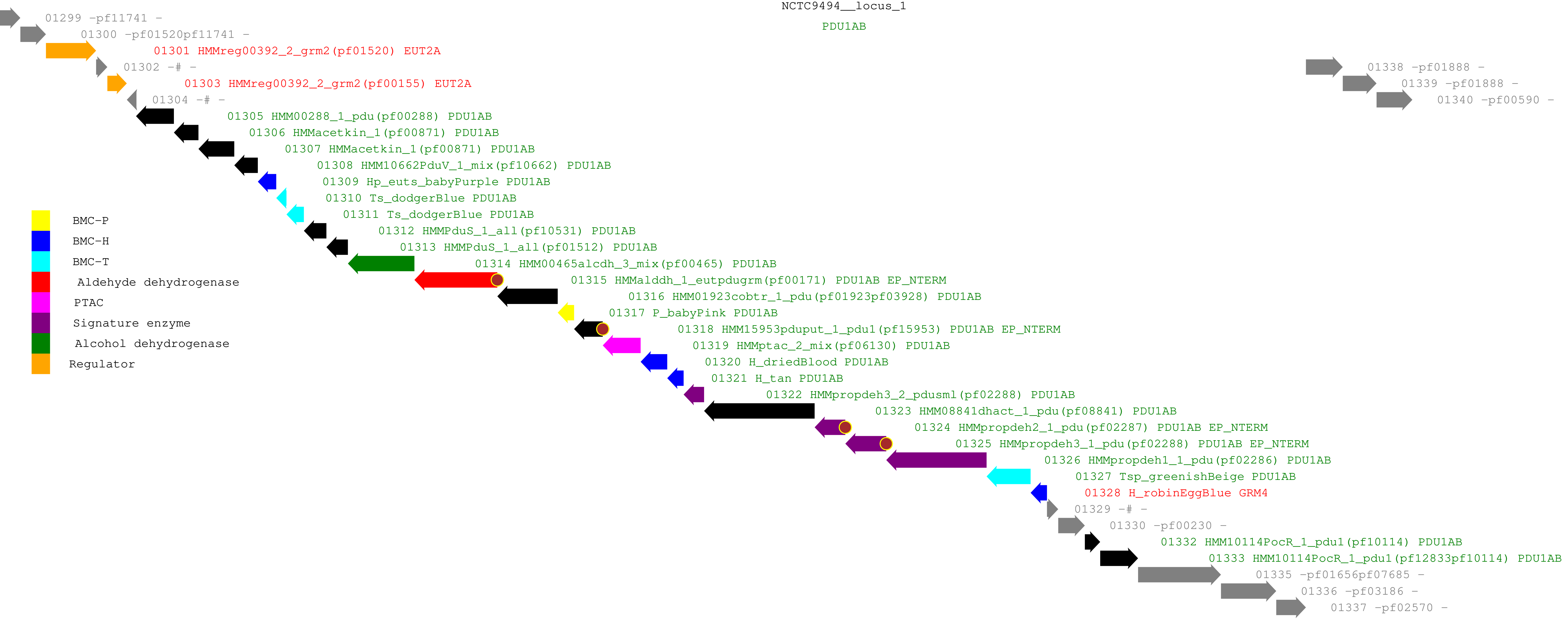

Klebsiella aerogenes (Enterobacter aerogenes). Bacteria; Proteobacteria; Gammaproteobacteria; Enterobacterales;

NCTC9499__locus_2

PDU1AB

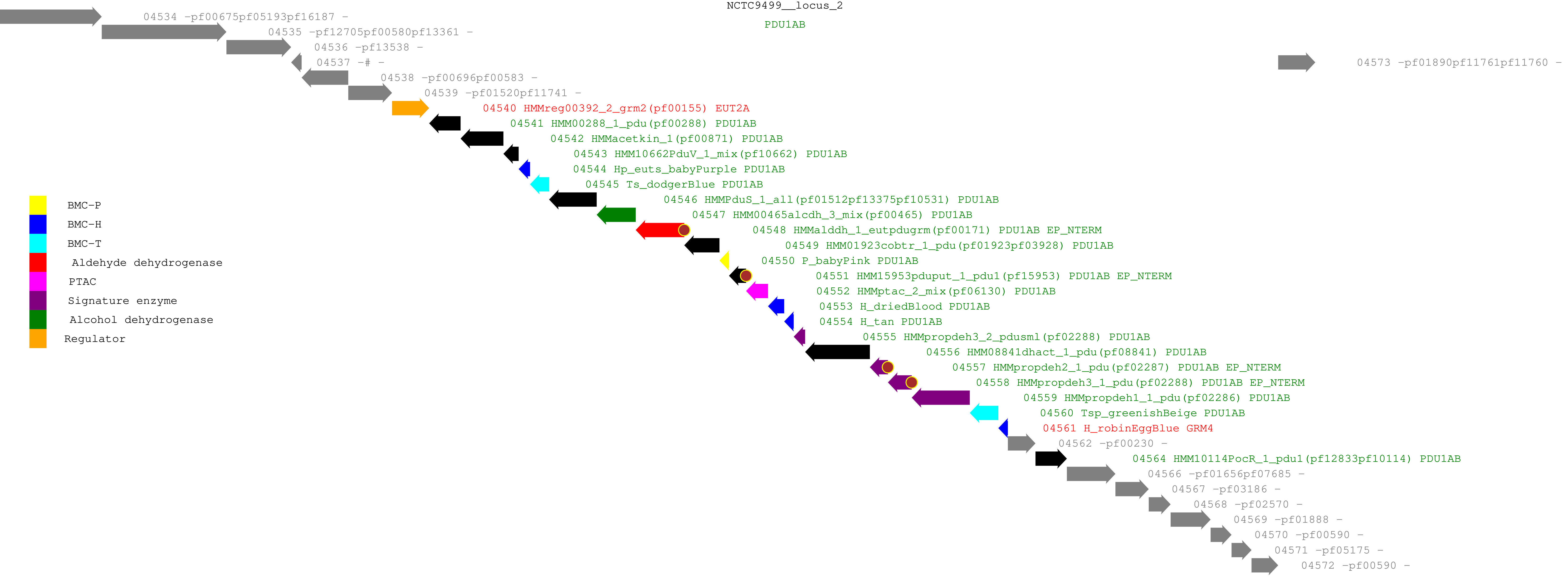

Klebsiella pneumoniae subsp. pneumoniae. Bacteria; Proteobacteria; Gammaproteobacteria; Enterobacterales;

NCTC9504__locus_1

PDU1AB

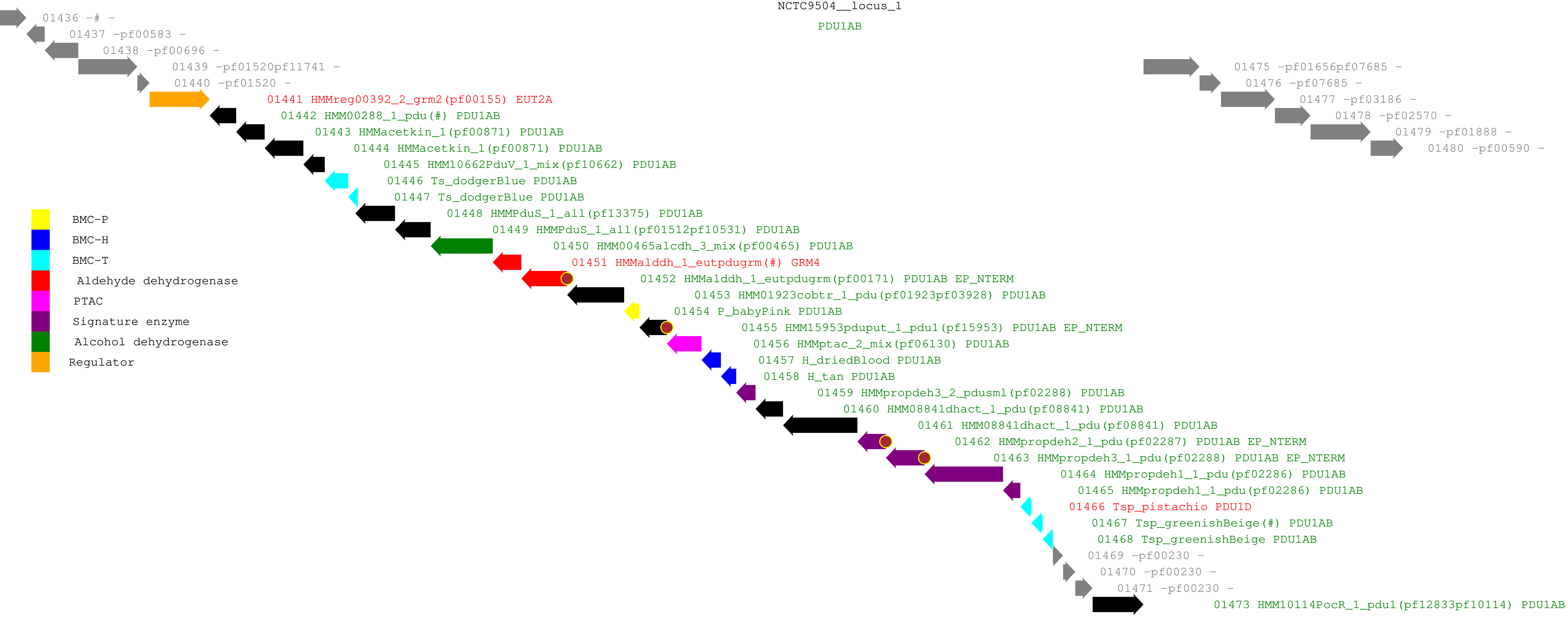

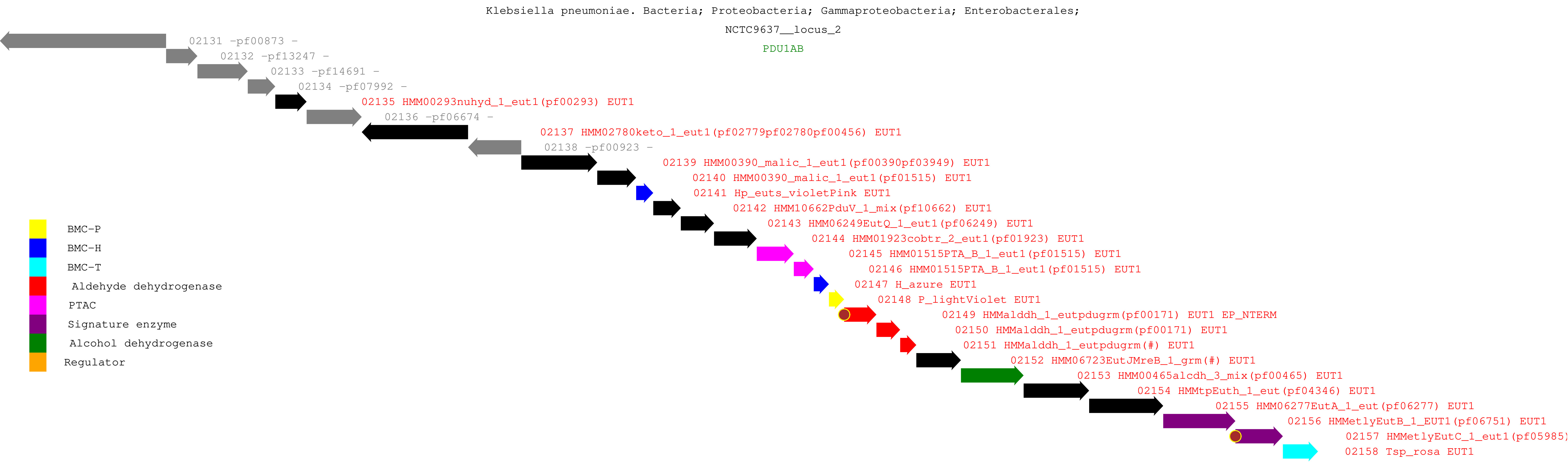

Klebsiella pneumoniae. Bacteria; Proteobacteria; Gammaproteobacteria; Enterobacterales;
NCTC9637__locus_2
PDU1AB
02131 -pf00873 -
02132 -pf13247 -
02133 -pf14691 -
02134 -pf07992 -
02135 HMM00293nuhyd_1_eut1(pf00293) EUT1
02136 -pf06674 -
02137 HMM02780keto_1_eut1(pf02779pf02780pf00456) EUT1
02138 -pf00923 -
02139 HMM00390_malic_1_eut1(pf00390pf03949) EUT1
02140 HMM00390_malic_1_eut1(pf01515) EUT1
02141 Hp_euts_violetPink EUT1
02142 HMM10662PduV_1_mix(pf10662) EUT1
02143 HMM06249EutQ_1_eut1(pf06249) EUT1
02144 HMM01923cobtr_2_eut1(pf01923) EUT1
02145 HMM01515PTA_B_1_eut1(pf01515) EUT1
02146 HMM01515PTA_B_1_eut1(pf01515) EUT1
02147 H_azure EUT1
02148 P_lightViolet EUT1
02149 HMMalddh_1_eutpdugrm(pf00171) EUT1 EP_NTERM
02150 HMMalddh_1_eutpdugrm(pf00171) EUT1
02151 HMMalddh_1_eutpdugrm(#) EUT1
02152 HMM06723EutJMreB_1_grm(#) EUT1
02153 HMM00465alcdh_3_mix(pf00465) EUT1
02154 HMMtpEuth_1_eut(pf04346) EUT1
02155 HMM06277EutA_1_eut(pf06277) EUT1
02156 HMMetlyEutB_1_EUT1(pf06751) EUT1
02157 HMMetlyEutC_1_eut1(pf05985)
02158 Tsp_rosa EUT1
BMC-P
BMC-H
BMC-T
Aldehyde dehydrogenase
PTAC
Signature enzyme
Alcohol dehydrogenase
Regulator

Klebsiella pneumoniae. Bacteria; Proteobacteria; Gammaproteobacteria; Enterobacterales;

NCTC9645__locus_2

PDU1AB

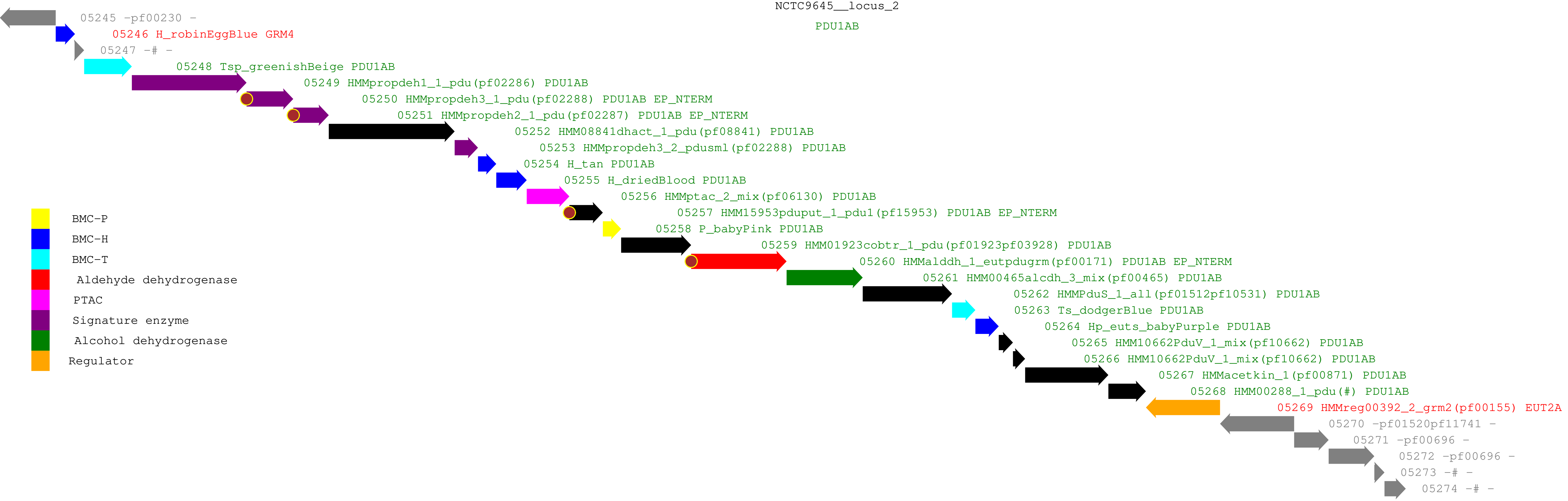

Klebsiella aerogenes (Enterobacter aerogenes). Bacteria; Proteobacteria; Gammaproteobacteria; Enterobacterales;

NCTC9652__locus_1

PDU1AB

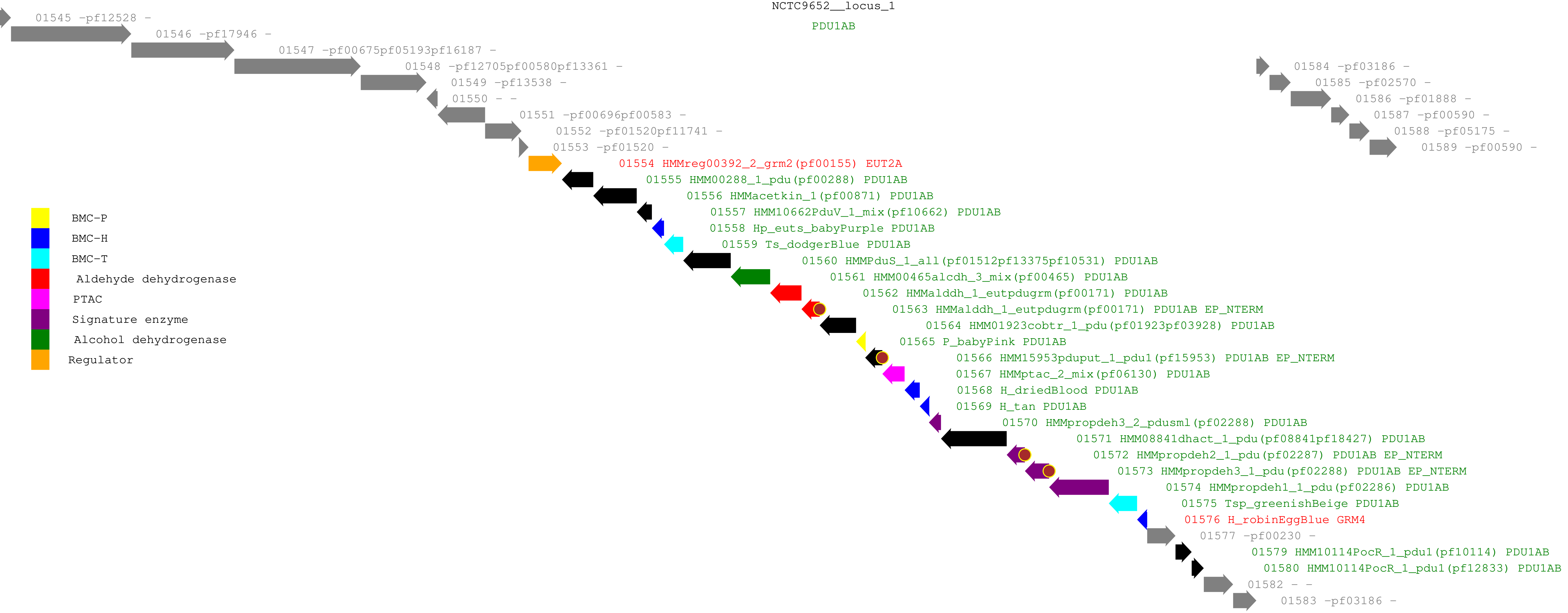

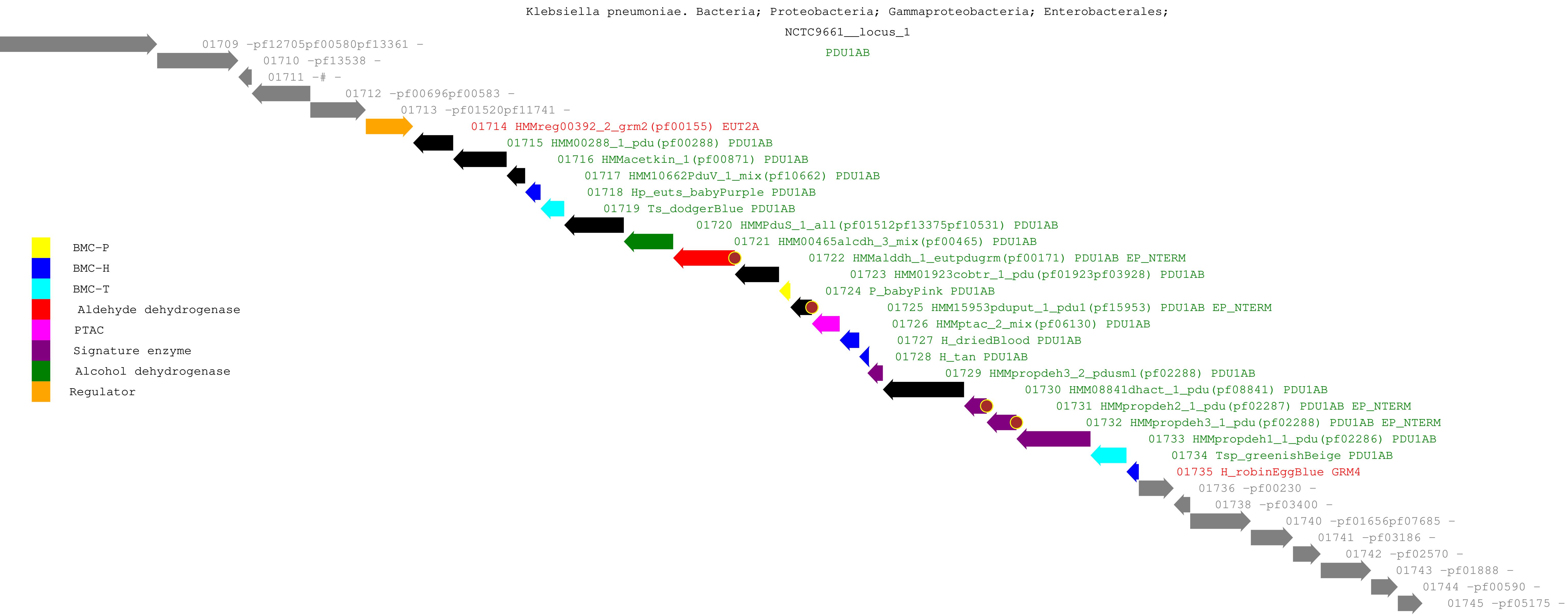

Klebsiella pneumoniae. Bacteria; Proteobacteria; Gammaproteobacteria; Enterobacterales;
NCTC9661__locus_1
PDU1AB
01709 -pf12705pf00580pf13361 -
01710 -pf13538 -
01711 -# -
01712 -pf00696pf00583 -
01713 -pf01520pf11741 -
01714 HMMreg00392_2_grm2(pf00155) EUT2A
01715 HMM00288_1_pdu(pf00288) PDU1AB
01716 HMMacetkin_1(pf00871) PDU1AB
01717 HMM10662PduV_1_mix(pf10662) PDU1AB
01718 Hp_euts_babyPurple PDU1AB
01719 Ts_dodgerBlue PDU1AB
01720 HMMPduS_1_all(pf01512pf13375pf10531) PDU1AB
01721 HMM00465alcdh_3_mix(pf00465) PDU1AB
01722 HMMaldh_1_eutpdugrm(pf00171) PDU1AB EP_NTERM
01723 HMM01923cobtr_1_pdu(pf01923pf03928) PDU1AB
01724 P_babyPink PDU1AB
01725 HMM15953pduput_1_pdu1(pf15953) PDU1AB EP_NTERM
01726 HMMptac_2_mix(pf06130) PDU1AB
01727 H_driedBlood PDU1AB
01728 H_tan PDU1AB
01729 HMMpropdeh3_2_pdusml(pf02288) PDU1AB
01730 HMM08841dhact_1_pdu(pf08841) PDU1AB
01731 HMMpropdeh2_1_pdu(pf02287) PDU1AB EP_NTERM
01732 HMMpropdeh3_1_pdu(pf02288) PDU1AB EP_NTERM
01733 HMMpropdeh1_1_pdu(pf02286) PDU1AB
01734 Tsp_greenishBeige PDU1AB
01735 H_robinEggBlue GRM4
01736 -pf00230 -
01738 -pf03400 -
01740 -pf01656pf07685 -
01741 -pf03186 -
01742 -pf02570 -
01743 -pf01888 -
01744 -pf00590 -
01745 -pf05175 -
BMC-P
BMC-H
BMC-T
Aldehyde dehydrogenase
PTAC
Signature enzyme
Alcohol dehydrogenase
Regulator

Salmonella choleraesuis. Bacteria; Proteobacteria; Gammaproteobacteria; Enterobacterales;

NCTC9854__locus_2

PDU1AB

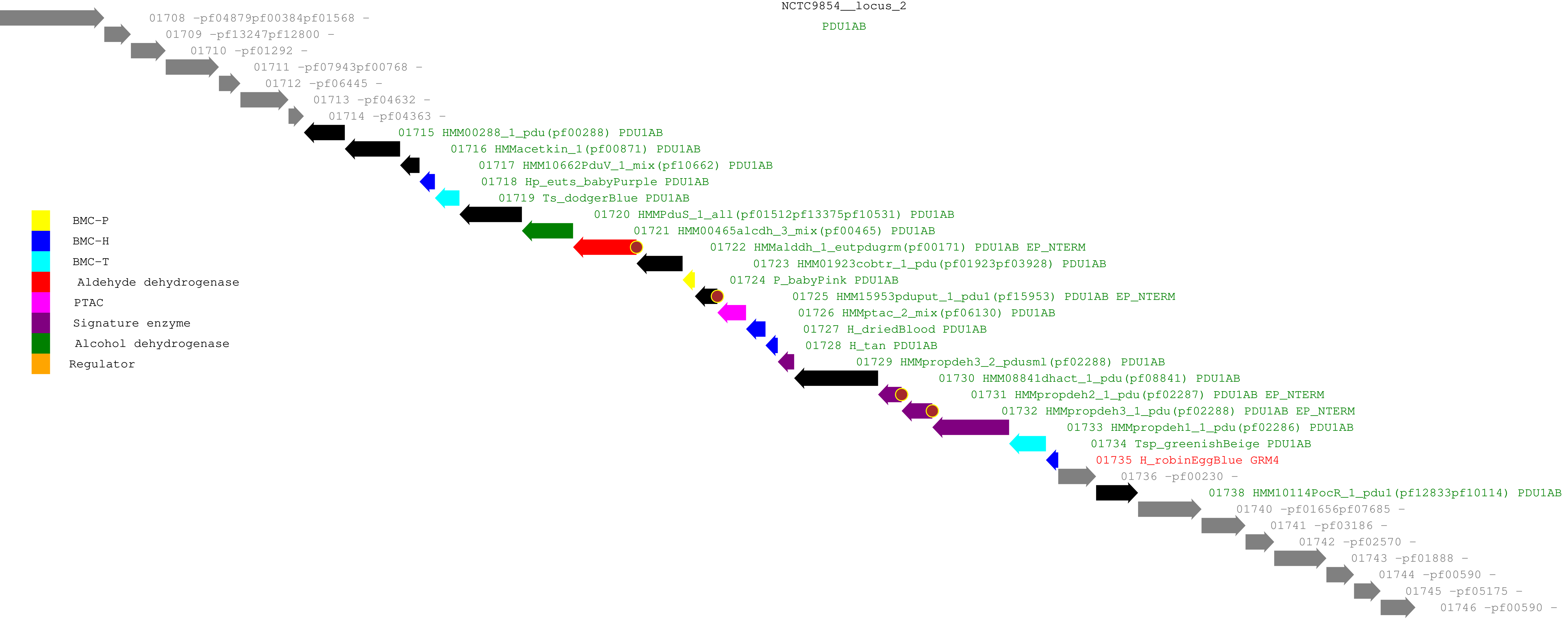

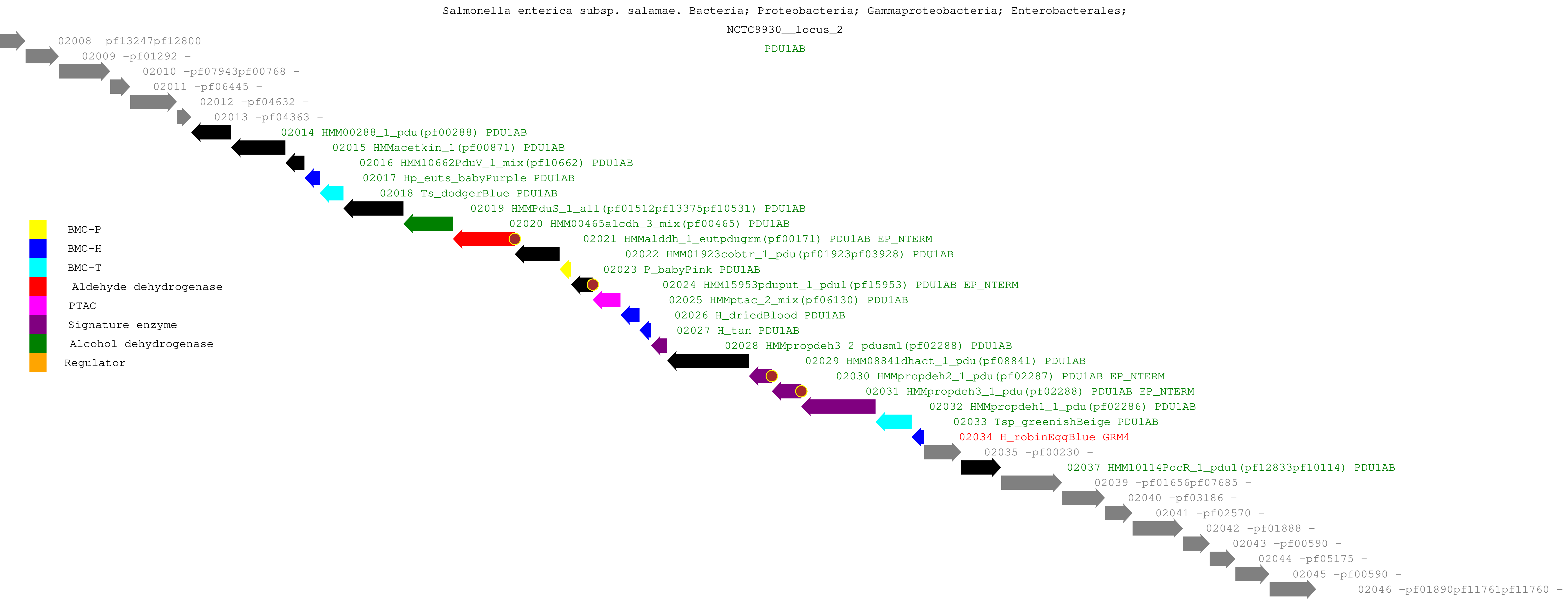
Salmonella enterica subsp. salamae. Bacteria; Proteobacteria; Gammaproteobacteria; Enterobacterales;
NCTC9930__locus_2
PDU1AB
02008 -pf13247pf12800 -
02009 -pf01292 -
02010 -pf07943pf00768 -
02011 -pf06445 -
02012 -pf04632 -
02013 -pf04363 -
02014 HMM00288_1_pdu(pf00288) PDU1AB
02015 HMMacetkin_1(pf00871) PDU1AB
02016 HMM10662PduV_1_mix(pf10662) PDU1AB
02017 Hp_euts_babyPurple PDU1AB
02018 Ts_dodgerBlue PDU1AB
02019 HMMPduS_1_all(pf01512pf13375pf10531) PDU1AB
02020 HMM00465alcdh_3_mix(pf00465) PDU1AB
02021 HMMalddh_1_eutpdugrm(pf00171) PDU1AB EP_NTERM
02022 HMM01923cobtr_1_pdu(pf01923pf03928) PDU1AB
02023 P_babyPink PDU1AB
02024 HMM15953pduput_1_pdu1(pf15953) PDU1AB EP_NTERM
02025 HMMptac_2_mix(pf06130) PDU1AB
02026 H_driedBlood PDU1AB
02027 H_tan PDU1AB
02028 HMMpropdeh3_2_pdusml(pf02288) PDU1AB
02029 HMM08841dhact_1_pdu(pf08841) PDU1AB
02030 HMMpropdeh2_1_pdu(pf02287) PDU1AB EP_NTERM
02031 HMMpropdeh3_1_pdu(pf02288) PDU1AB EP_NTERM
02032 HMMpropdeh1_1_pdu(pf02286) PDU1AB
02033 Tsp_greenishBeige PDU1AB
02034 H_robinEggBlue GRM4
02035 -pf00230 -
02037 HMM10114PocR_1_pdu1(pf12833pf10114) PDU1AB
02039 -pf01656pf07685 -
02040 -pf03186 -
02041 -pf02570 -
02042 -pf01888 -
02043 -pf00590 -
02044 -pf05175 -
02045 -pf00590 -
02046 -pf01890pf11761pf11760 -
BMC-P
BMC-H
BMC-T
Aldehyde dehydrogenase
PTAC
Signature enzyme
Alcohol dehydrogenase
Regulator

Salmonella choleraesuis. Bacteria; Proteobacteria; Gammaproteobacteria; Enterobacterales;

NGUA18__locus_2

PDU1AB

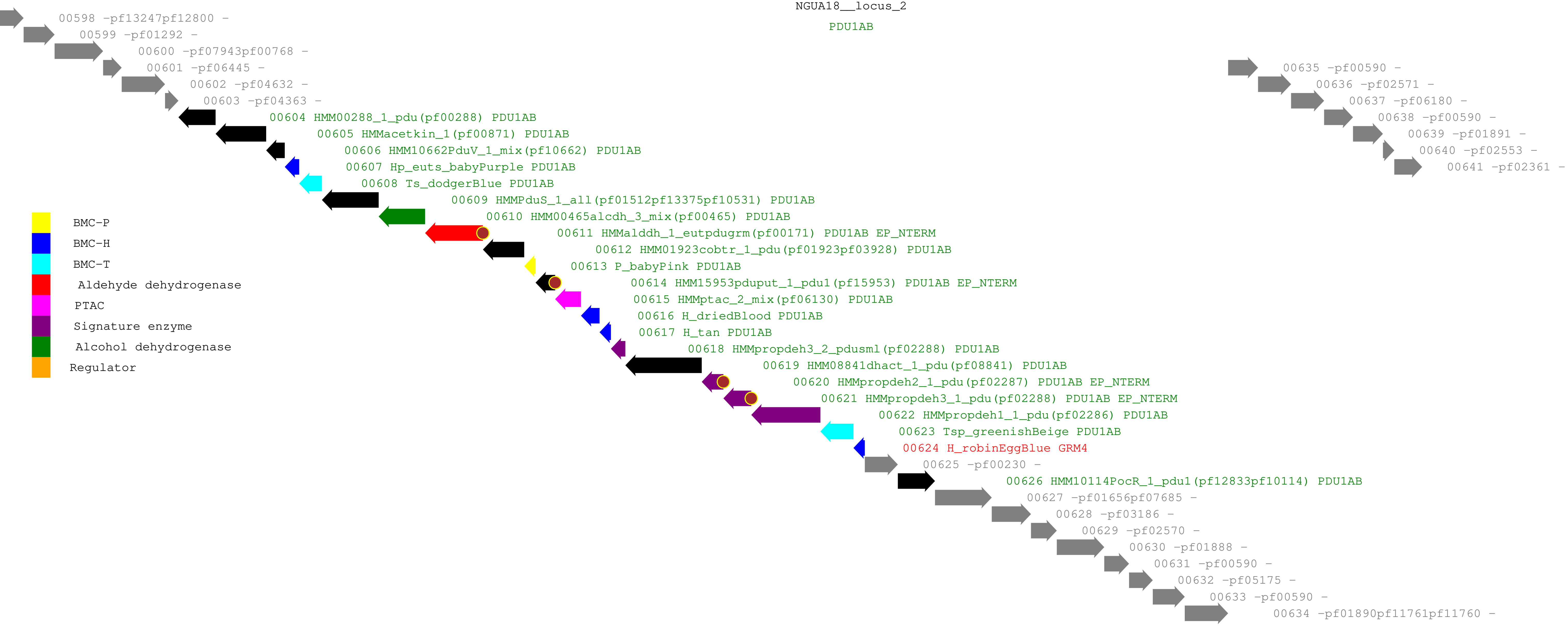

Salmonella choleraesuis. Bacteria; Proteobacteria; Gammaproteobacteria; Enterobacterales;

NGUA41__locus_1

PDU1AB

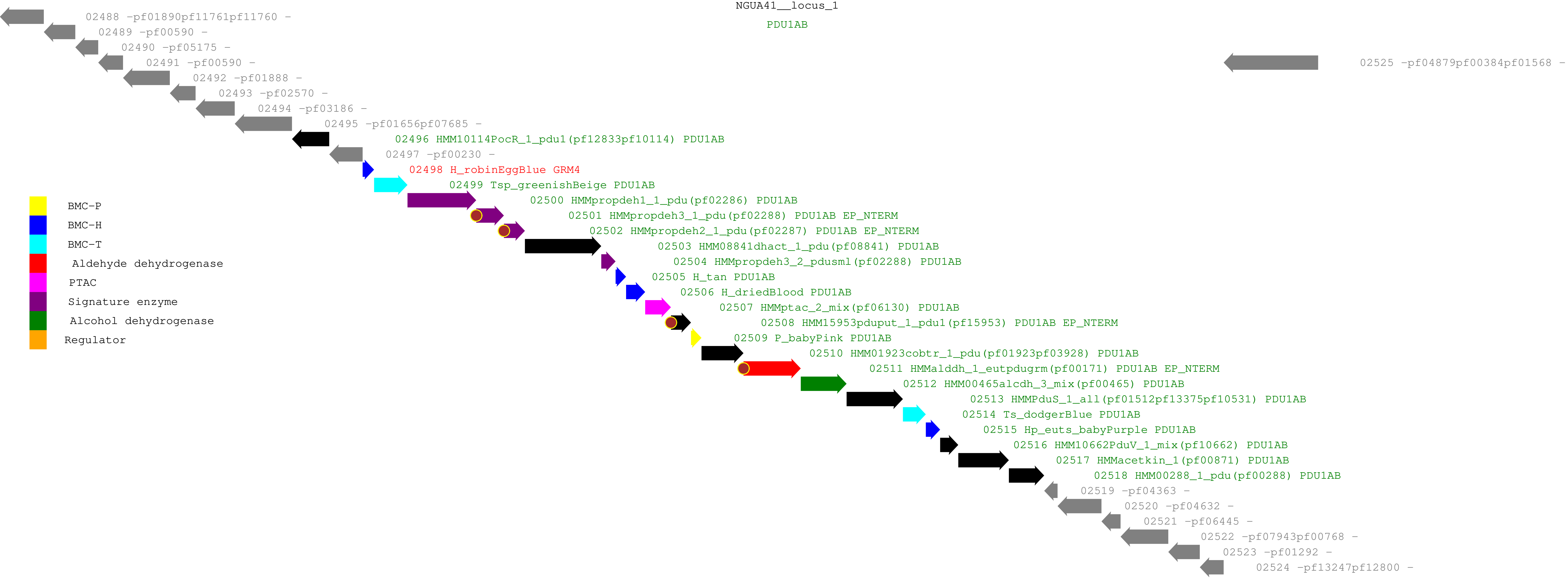

Salmonella enterica subsp. enterica serovar Heidelberg. Bacteria; Proteobacteria; Gammaproteobacteria; Enterobacterales;

NJ03__locus_1

PDU1AB

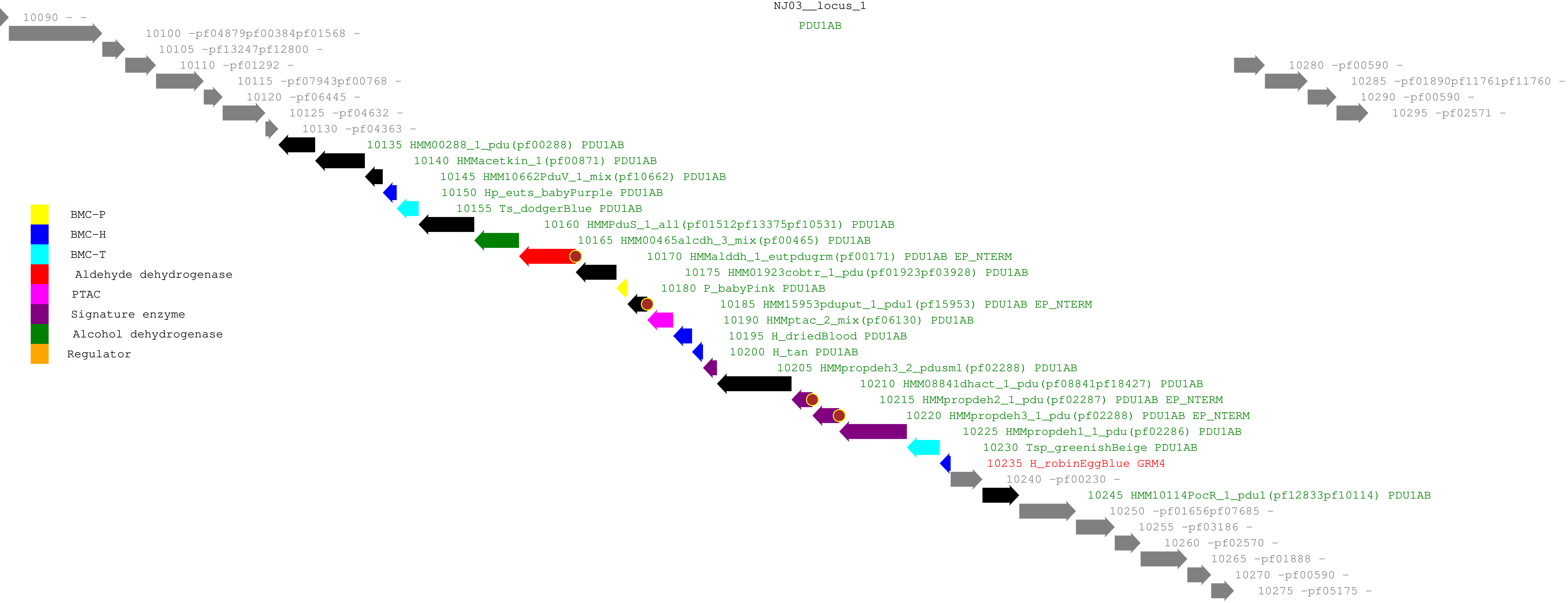

Escherichia coli O83:H1 (strain NRG 857C / AIEC). Bacteria; Proteobacteria; Gammaproteobacteria; Enterobacterales;

NRG857__locus_1

PDU1AB

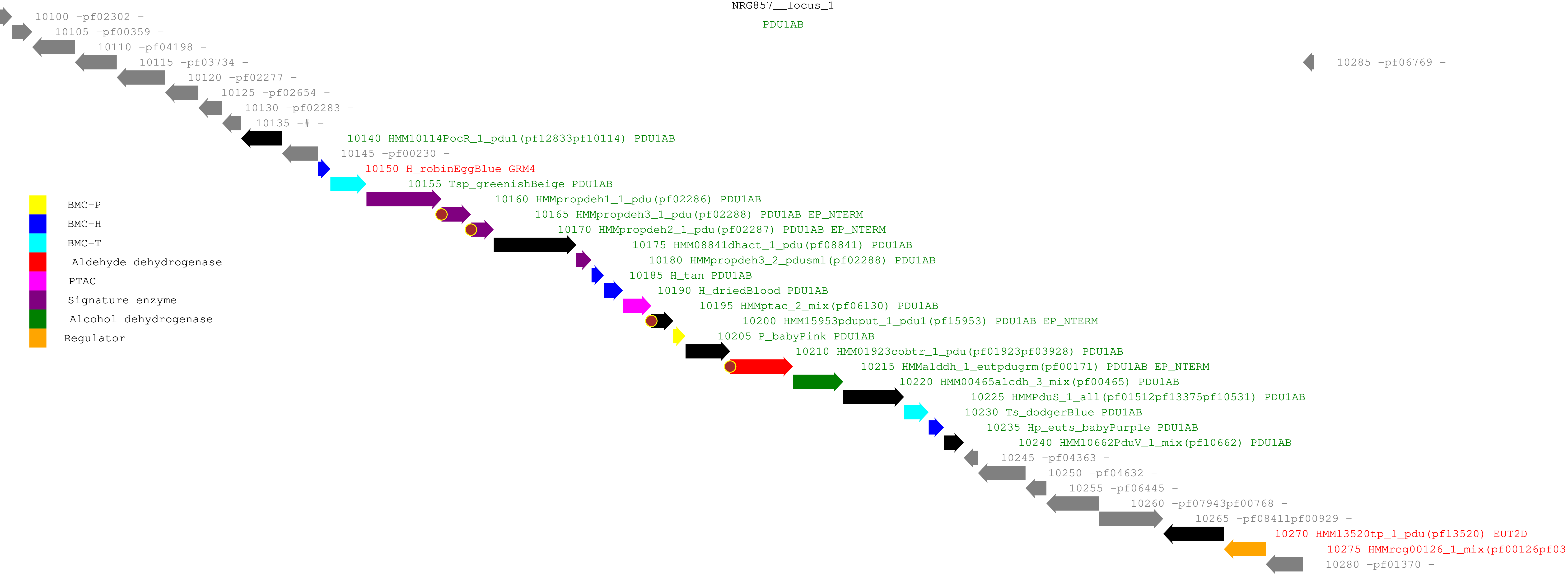

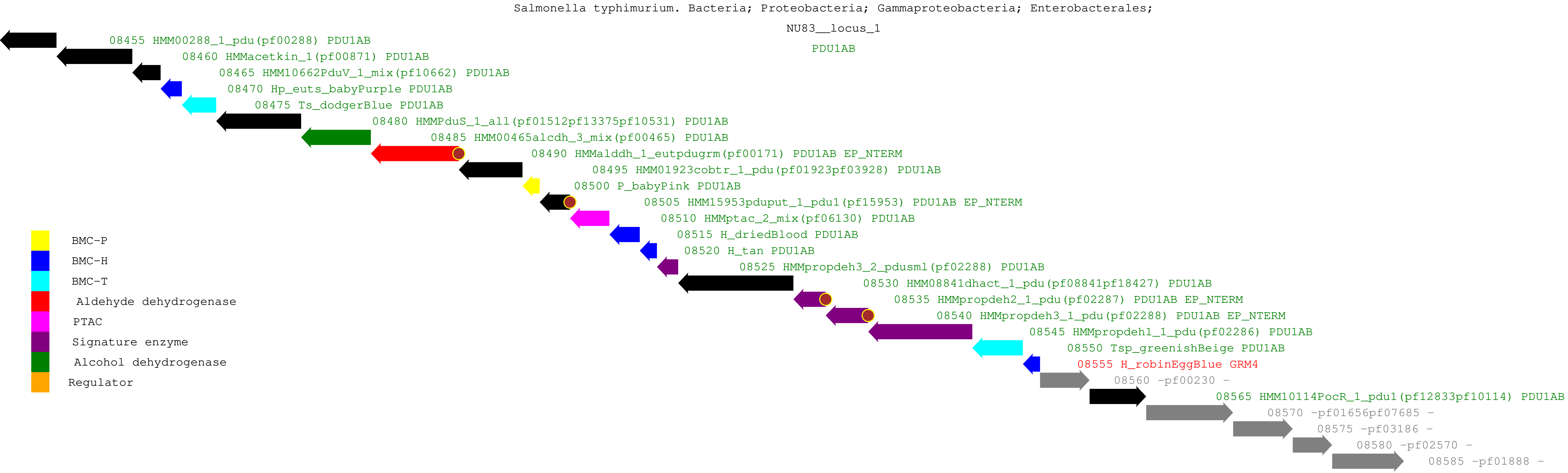
Salmonella typhimurium. Bacteria; Proteobacteria; Gammaproteobacteria; Enterobacterales;
NU83__locus_1
PDU1AB
08455 HMM00288_1_pdu(pf00288) PDU1AB
08460 HMMacetkin_1(pf00871) PDU1AB
08465 HMM10662PduV_1_mix(pf10662) PDU1AB
08470 Hp_euts_babyPurple PDU1AB
08475 Ts_dodgerBlue PDU1AB
08480 HMMPduS_1_all(pf01512pf13375pf10531) PDU1AB
08485 HMM00465alcdh_3_mix(pf00465) PDU1AB
08490 HMMalddh_1_eutpdugrm(pf00171) PDU1AB EP_NTERM
08495 HMM01923cobtr_1_pdu(pf01923pf03928) PDU1AB
08500 P_babyPink PDU1AB
08505 HMM15953pduput_1_pdu1(pf15953) PDU1AB EP_NTERM
08510 HMMptac_2_mix(pf06130) PDU1AB
08515 H_driedBlood PDU1AB
08520 H_tan PDU1AB
08525 HMMpropdeh3_2_pdusml(pf02288) PDU1AB
08530 HMM08841dhact_1_pdu(pf08841pf18427) PDU1AB
08535 HMMpropdeh2_1_pdu(pf02287) PDU1AB EP_NTERM
08540 HMMpropdeh3_1_pdu(pf02288) PDU1AB EP_NTERM
08545 HMMpropdeh1_1_pdu(pf02286) PDU1AB
08550 Tsp_greenishBeige PDU1AB
08555 H_robinEggBlue GRM4
08560 -pf00230 -
08565 HMM10114PocR_1_pdu1(pf12833pf10114) PDU1AB
08570 -pf01656pf07685 -
08575 -pf03186 -
08580 -pf02570 -
08585 -pf01888 -
BMC-P
BMC-H
BMC-T
Aldehyde dehydrogenase
PTAC
Signature enzyme
Alcohol dehydrogenase
Regulator

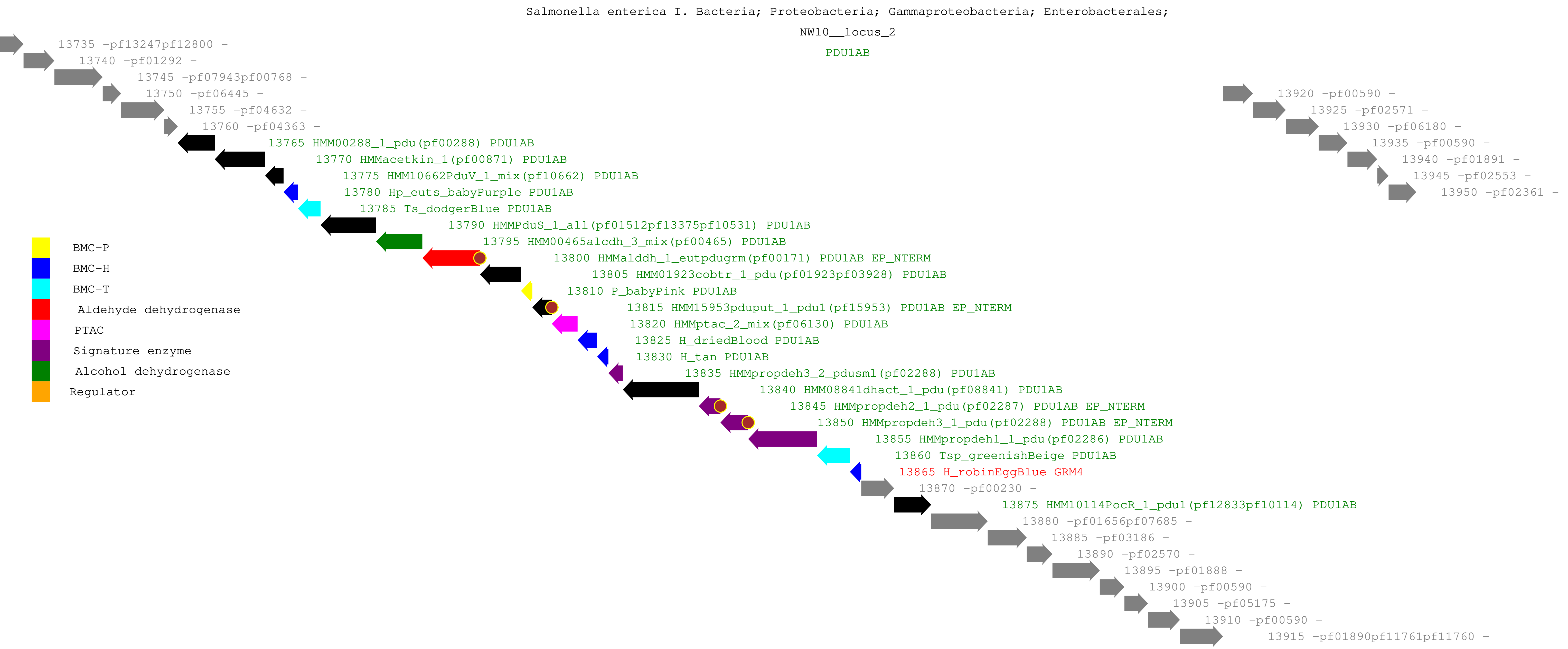
Salmonella enterica I. Bacteria; Proteobacteria; Gammaproteobacteria; Enterobacterales;
NW10__locus_2
PDU1AB
13735 -pf13247pf12800 -
13740 -pf01292 -
13745 -pf07943pf00768 -
13750 -pf06445 -
13755 -pf04632 -
13760 -pf04363 -
13765 HMM00288_1_pdu(pf00288) PDU1AB
13770 HMMacetkin_1(pf00871) PDU1AB
13775 HMM10662PduV_1_mix(pf10662) PDU1AB
13780 Hp_euts_babyPurple PDU1AB
13785 Ts_dodgerBlue PDU1AB
13790 HMMPduS_1_all(pf01512pf13375pf10531) PDU1AB
13795 HMM00465alcdh_3_mix(pf00465) PDU1AB
13800 HMMalddh_1_eutpdugrm(pf00171) PDU1AB EP_NTERM
13805 HMM01923cobtr_1_pdu(pf01923pf03928) PDU1AB
13810 P_babyPink PDU1AB
13815 HMM15953pduput_1_pdu1(pf15953) PDU1AB EP_NTERM
13820 HMMptac_2_mix(pf06130) PDU1AB
13825 H_driedBlood PDU1AB
13830 H_tan PDU1AB
13835 HMMpropdeh3_2_pdusml(pf02288) PDU1AB
13840 HMM08841dhact_1_pdu(pf08841) PDU1AB
13845 HMMpropdeh2_1_pdu(pf02287) PDU1AB EP_NTERM
13850 HMMpropdeh3_1_pdu(pf02288) PDU1AB EP_NTERM
13855 HMMpropdeh1_1_pdu(pf02286) PDU1AB
13860 Tsp_greenishBeige PDU1AB
13865 H_robinEggBlue GRM4
13870 -pf00230 -
13875 HMM10114PocR_1_pdu1(pf12833pf10114) PDU1AB
13880 -pf01656pf07685 -
13885 -pf03186 -
13890 -pf02570 -
13895 -pf01888 -
13900 -pf00590 -
13905 -pf05175 -
13910 -pf00590 -
13915 -pf01890pf11761pf11760 -
13920 -pf00590 -
13925 -pf02571 -
13930 -pf06180 -
13935 -pf00590 -
13940 -pf01891 -
13945 -pf02553 -
13950 -pf02361 -
BMC-P
BMC-H
BMC-T
Aldehyde dehydrogenase
PTAC
Signature enzyme
Alcohol dehydrogenase
Regulator

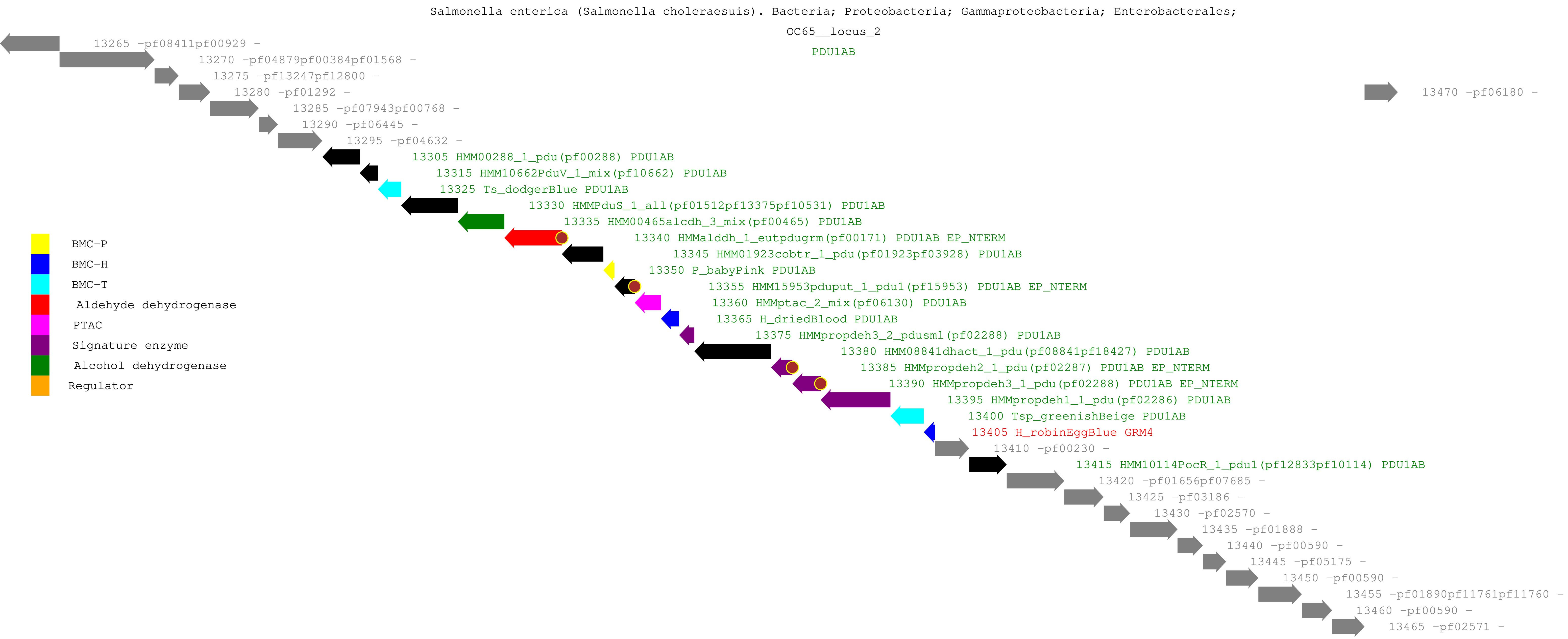
Salmonella enterica (Salmonella choleraesuis). Bacteria; Proteobacteria; Gammaproteobacteria; Enterobacterales;
OC65__locus_2
PDU1AB
13265 -pf08411pf00929 -
13270 -pf04879pf00384pf01568 -
13275 -pf13247pf12800 -
13280 -pf01292 -
13470 -pf06180 -
13285 -pf07943pf00768 -
13290 -pf06445 -
13295 -pf04632 -
13305 HMM00288_1_pdu(pf00288) PDU1AB
13315 HMM10662PduV_1_mix(pf10662) PDU1AB
13325 Ts_dodgerBlue PDU1AB
13330 HMMPduS_1_all(pf01512pf13375pf10531) PDU1AB
13335 HMM00465alcdh_3_mix(pf00465) PDU1AB
BMC-P
BMC-H
BMC-T
Aldehyde dehydrogenase
PTAC
Signature enzyme
Alcohol dehydrogenase
Regulator
13340 HMMalddh_1_eutpdugrm(pf00171) PDU1AB EP_NTERM
13345 HMM01923cobtr_1_pdu(pf01923pf03928) PDU1AB
13350 P_babyPink PDU1AB
13355 HMM15953pduput_1_pdu1(pf15953) PDU1AB EP_NTERM
13360 HMMptac_2_mix(pf06130) PDU1AB
13365 H_driedBlood PDU1AB
13375 HMMpropdeh3_2_pdusml(pf02288) PDU1AB
13380 HMM08841dhact_1_pdu(pf08841pf18427) PDU1AB
13385 HMMpropdeh2_1_pdu(pf02287) PDU1AB EP_NTERM
13390 HMMpropdeh3_1_pdu(pf02288) PDU1AB EP_NTERM
13395 HMMpropdeh1_1_pdu(pf02286) PDU1AB
13400 Tsp_greenishBeige PDU1AB
13405 H_robinEggBlue GRM4
13410 -pf00230 -
13415 HMM10114PocR_1_pdu1(pf12833pf10114) PDU1AB
13420 -pf01656pf07685 -
13425 -pf03186 -
13430 -pf02570 -
13435 -pf01888 -
13440 -pf00590 -
13445 -pf05175 -
13450 -pf00590 -
13455 -pf01890pf11761pf11760 -
13460 -pf00590 -
13465 -pf02571 -

Salmonella enterica subsp. arizonae serovar 18:z4,z23:- str. CVM Bacteria; Proteobacteria; Gammaproteobacteria; Enterobacterales;

P287__locus_1

PDU1AB

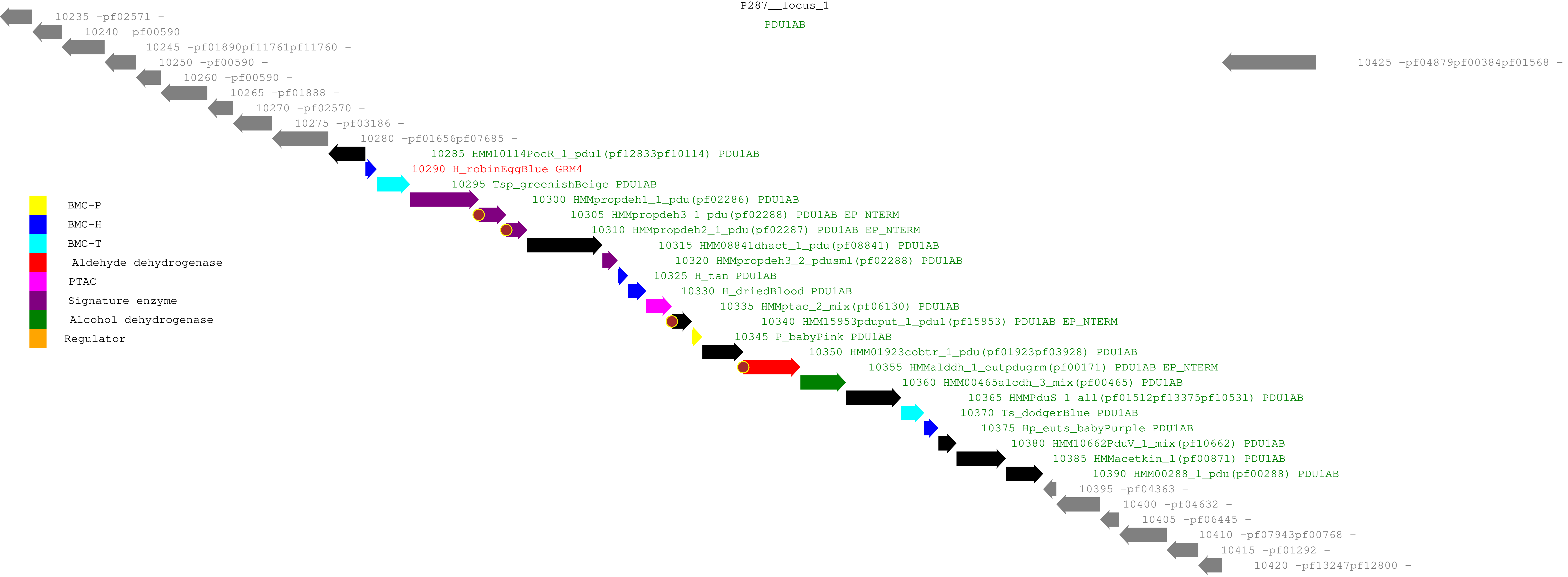

Salmonella enterica subsp. arizonae serovar 18:z4,z23:- str. CVM N26626. Bacteria; Proteobacteria; Gammaproteobacteria; Enterobacterales;

P298__locus_2

PDU1AB

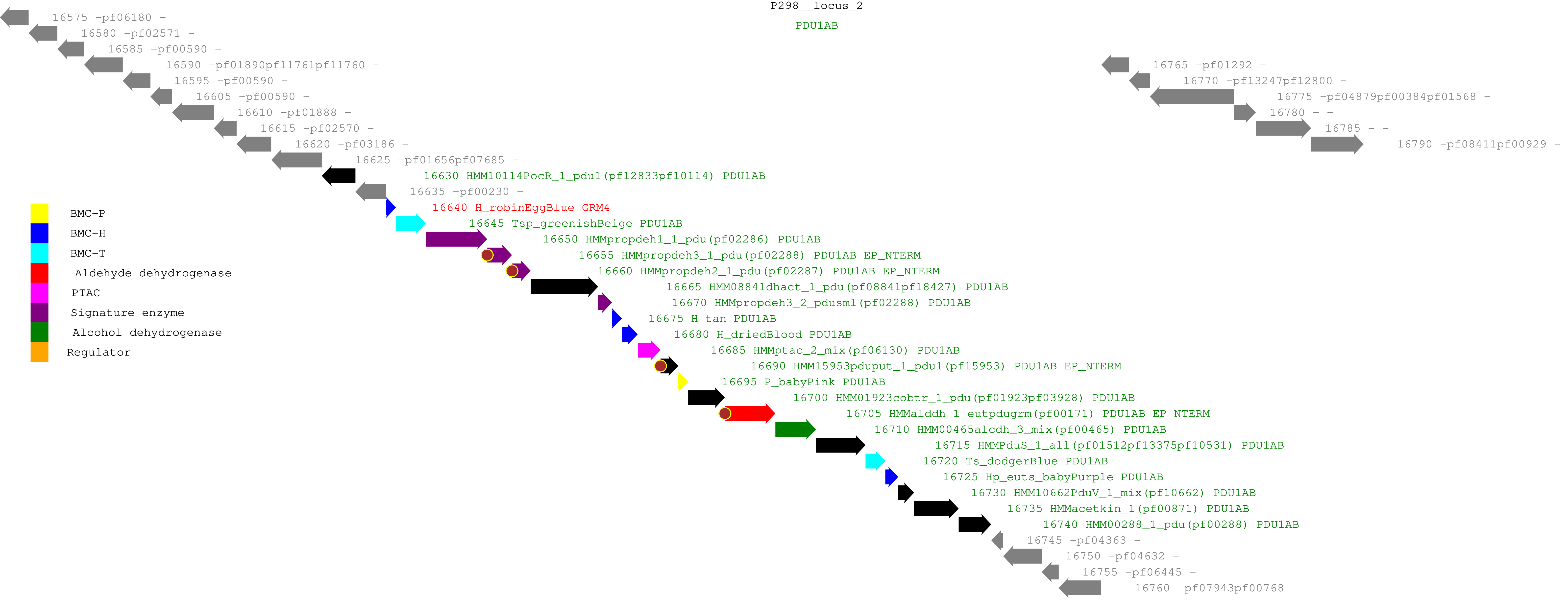

Salmonella enterica subsp. enterica serovar Newport str. CVM80_2288. Bacteria; Proteobacteria; Gammaproteobacteria; Enterobacterales;

P701__locus_1

PDU1AB

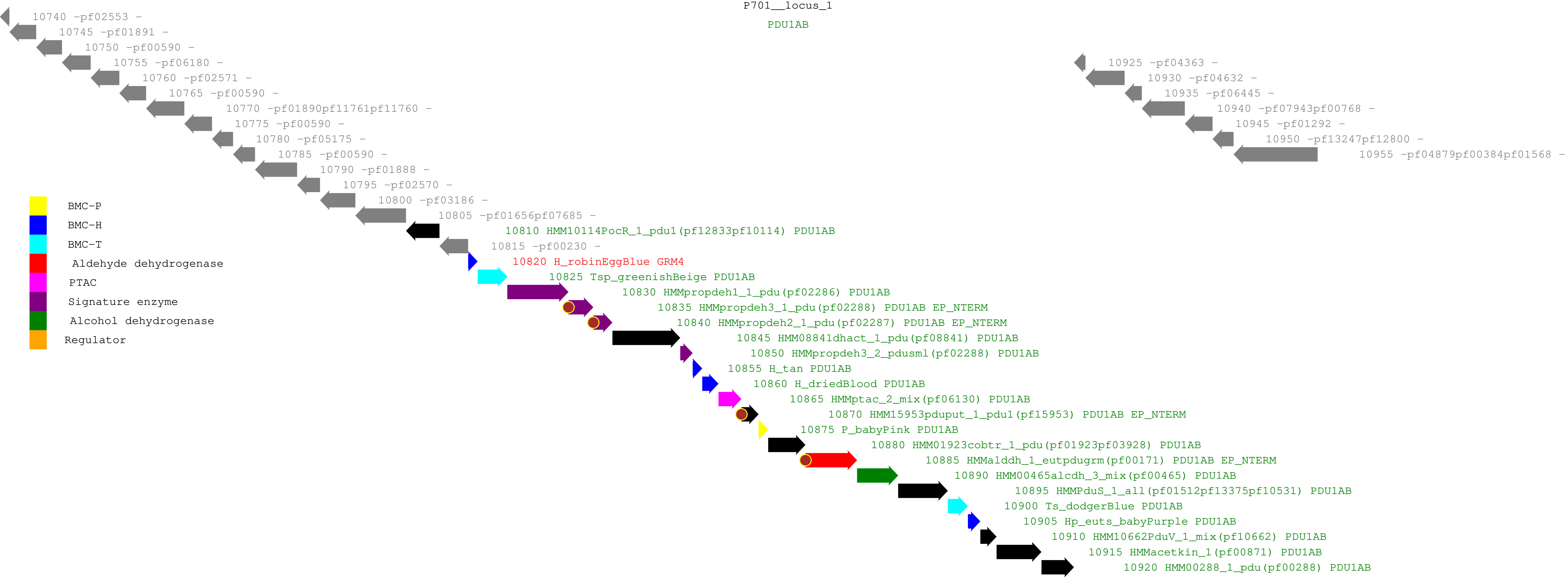

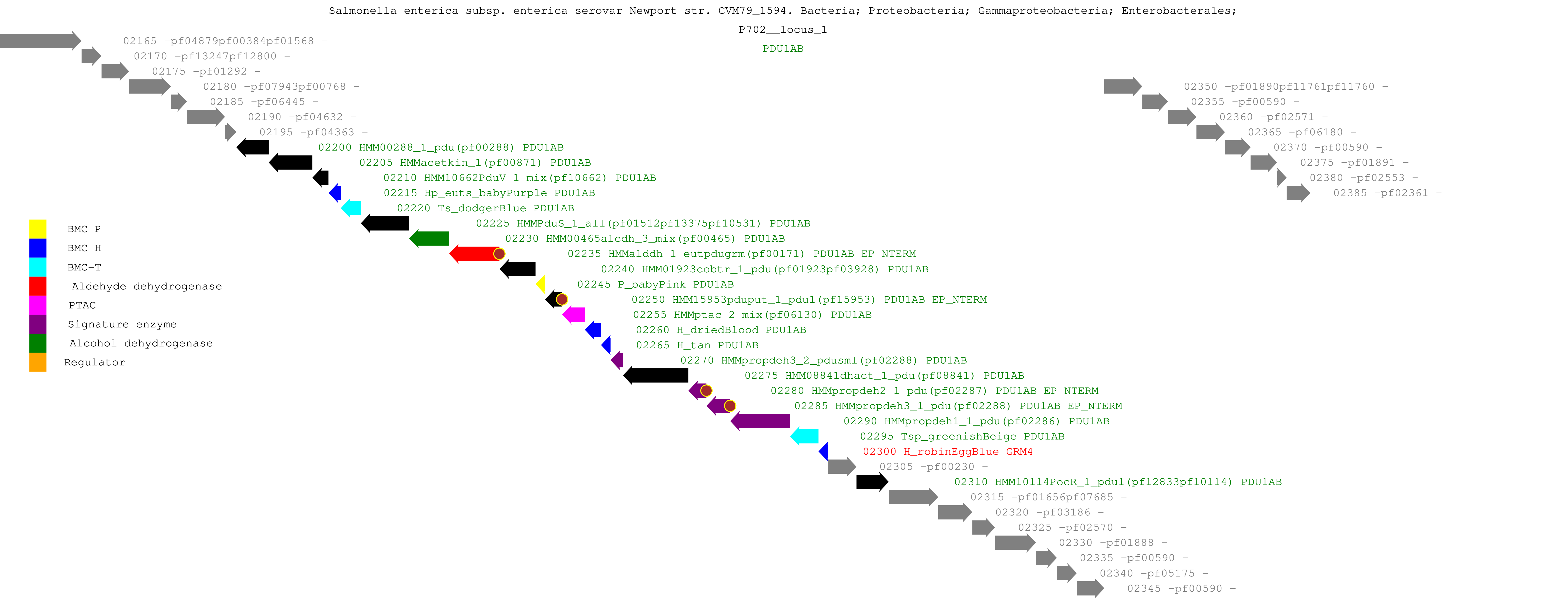

Salmonella enterica subsp. enterica serovar Newport str. CVM79_1594. Bacteria; Proteobacteria; Gammaproteobacteria; Enterobacterales;
P702__locus_1
PDU1AB
02165 -pf04879pf00384pf01568 -
02170 -pf13247pf12800 -
02175 -pf01292 -
02180 -pf07943pf00768 -
02185 -pf06445 -
02190 -pf04632 -
02195 -pf04363 -
02200 HMM00288_1_pdu(pf00288) PDU1AB
02205 HMMacetkin_1(pf00871) PDU1AB
02210 HMM10662PduV_1_mix(pf10662) PDU1AB
02215 Hp_euts_babyPurple PDU1AB
02220 Ts_dodgerBlue PDU1AB
02225 HMMPduS_1_all(pf01512pf13375pf10531) PDU1AB
02230 HMM00465alcdh_3_mix(pf00465) PDU1AB
02235 HMMalddh_1_eutpdugrm(pf00171) PDU1AB EP_NTERM
02240 HMM01923cobtr_1_pdu(pf01923pf03928) PDU1AB
02245 P_babyPink PDU1AB
02250 HMM15953pduput_1_pdu1(pf15953) PDU1AB EP_NTERM
02255 HMMptac_2_mix(pf06130) PDU1AB
02260 H_driedBlood PDU1AB
02265 H_tan PDU1AB
02270 HMMpropdeh3_2_pdusml(pf02288) PDU1AB
02275 HMM08841dhact_1_pdu(pf08841) PDU1AB
02280 HMMpropdeh2_1_pdu(pf02287) PDU1AB EP_NTERM
02285 HMMpropdeh3_1_pdu(pf02288) PDU1AB EP_NTERM
02290 HMMpropdeh1_1_pdu(pf02286) PDU1AB
02295 Tsp_greenishBeige PDU1AB
02300 H_robinEggBlue GRM4
02305 -pf00230 -
02310 HMM10114PocR_1_pdu1(pf12833pf10114) PDU1AB
02315 -pf01656pf07685 -
02320 -pf03186 -
02325 -pf02570 -
02330 -pf01888 -
02335 -pf00590 -
02340 -pf05175 -
02345 -pf00590 -
02350 -pf01890pf11761pf11760 -
02355 -pf00590 -
02360 -pf02571 -
02365 -pf06180 -
02370 -pf00590 -
02375 -pf01891 -
02380 -pf02553 -
02385 -pf02361 -
BMC-P
BMC-H
BMC-T
Aldehyde dehydrogenase
PTAC
Signature enzyme
Alcohol dehydrogenase
Regulator

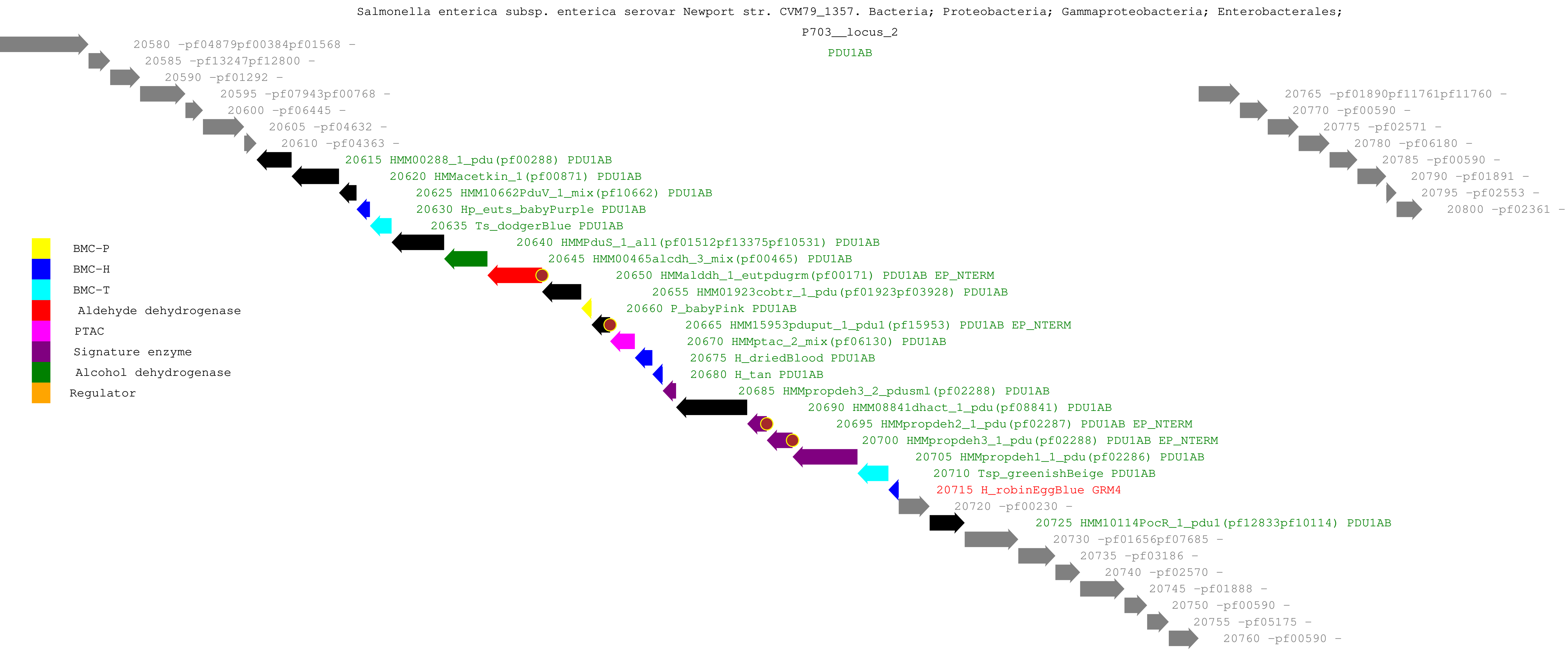
Salmonella enterica subsp. enterica serovar Newport str. CVM79_1357. Bacteria; Proteobacteria; Gammaproteobacteria; Enterobacterales;
P703__locus_2
PDU1AB
20580 -pf04879pf00384pf01568 -
20585 -pf13247pf12800 -
20590 -pf01292 -
20595 -pf07943pf00768 -
20600 -pf06445 -
20605 -pf04632 -
20610 -pf04363 -
20615 HMM00288_1_pdu(pf00288) PDU1AB
20620 HMMacetkin_1(pf00871) PDU1AB
20625 HMM10662PduV_1_mix(pf10662) PDU1AB
20630 Hp_euts_babyPurple PDU1AB
20635 Ts_dodgerBlue PDU1AB
20640 HMMPduS_1_all(pf01512pf13375pf10531) PDU1AB
20645 HMM00465alcdh_3_mix(pf00465) PDU1AB
20650 HMMalddh_1_eutpdugrm(pf00171) PDU1AB EP_NTERM
20655 HMM01923cobtr_1_pdu(pf01923pf03928) PDU1AB
20660 P_babyPink PDU1AB
20665 HMM15953pduput_1_pdu1(pf15953) PDU1AB EP_NTERM
20670 HMMptac_2_mix(pf06130) PDU1AB
20675 H_driedBlood PDU1AB
20680 H_tan PDU1AB
20685 HMMpropdeh3_2_pdusml(pf02288) PDU1AB
20690 HMM08841dhact_1_pdu(pf08841) PDU1AB
20695 HMMpropdeh2_1_pdu(pf02287) PDU1AB EP_NTERM
20700 HMMpropdeh3_1_pdu(pf02288) PDU1AB EP_NTERM
20705 HMMpropdeh1_1_pdu(pf02286) PDU1AB
20710 Tsp_greenishBeige PDU1AB
20715 H_robinEggBlue GRM4
20720 -pf00230 -
20725 HMM10114PocR_1_pdu1(pf12833pf10114) PDU1AB
20730 -pf01656pf07685 -
20735 -pf03186 -
20740 -pf02570 -
20745 -pf01888 -
20750 -pf00590 -
20755 -pf05175 -
20760 -pf00590 -
20765 -pf01890pf11761pf11760 -
20770 -pf00590 -
20775 -pf02571 -
20780 -pf06180 -
20785 -pf00590 -
20790 -pf01891 -
20795 -pf02553 -
20800 -pf02361 -
BMC-P
BMC-H
BMC-T
Aldehyde dehydrogenase
PTAC
Signature enzyme
Alcohol dehydrogenase
Regulator

Salmonella enterica subsp. enterica serovar Newport str. CVM21537. Bacteria; Proteobacteria; Gammaproteobacteria; Enterobacterales;

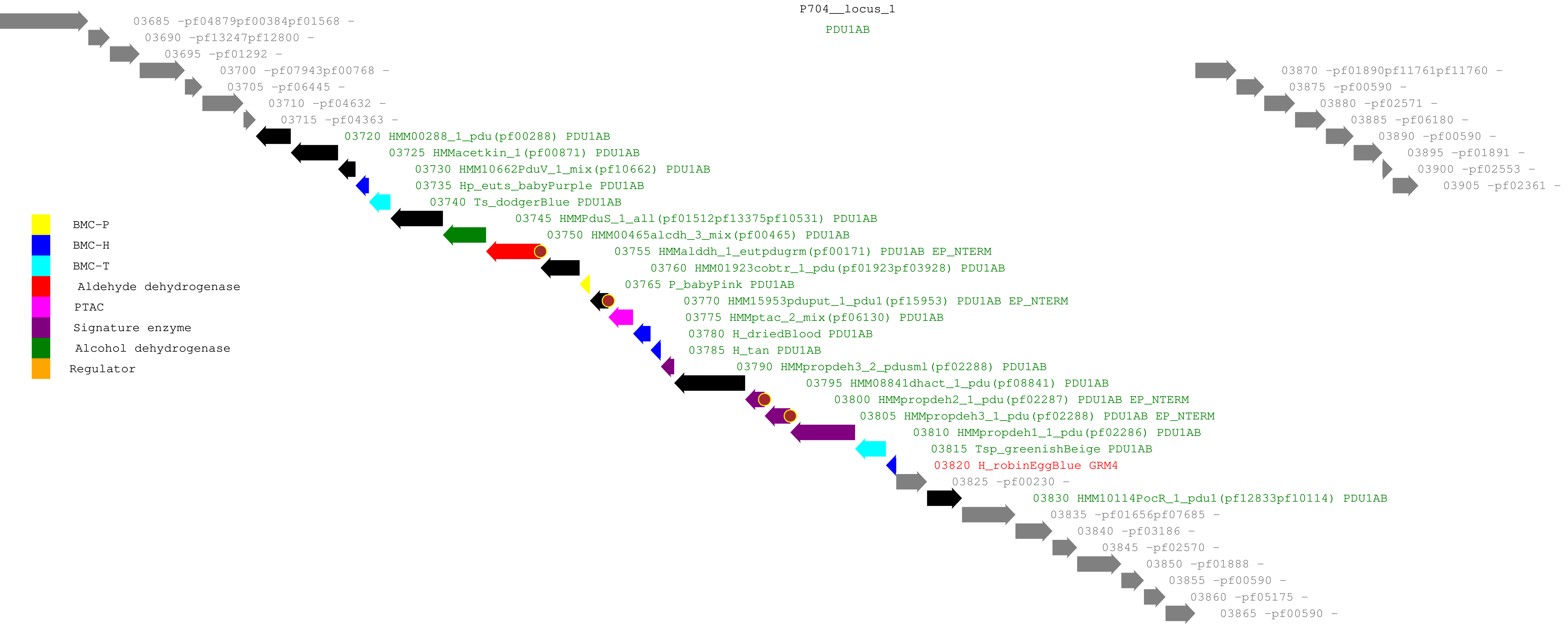

Salmonella enterica subsp. enterica serovar Newport str. CVM18000. Bacteria; Proteobacteria; Gammaproteobacteria; Enterobacterales;

P705__locus_2

PDU1AB

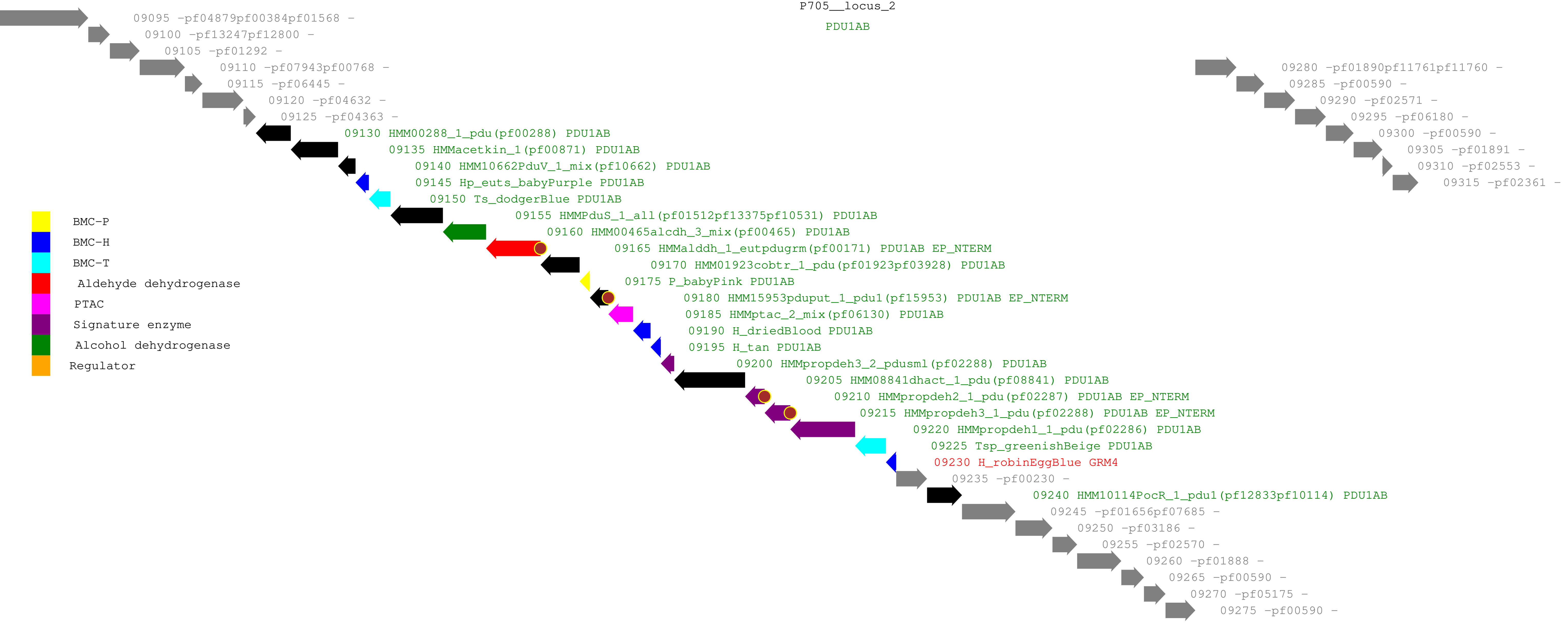

Salmonella enterica subsp. enterica serovar Newport str. CVM20776. Bacteria; Proteobacteria; Gammaproteobacteria; Enterobacterales;

P706__locus_2

PDU1AB

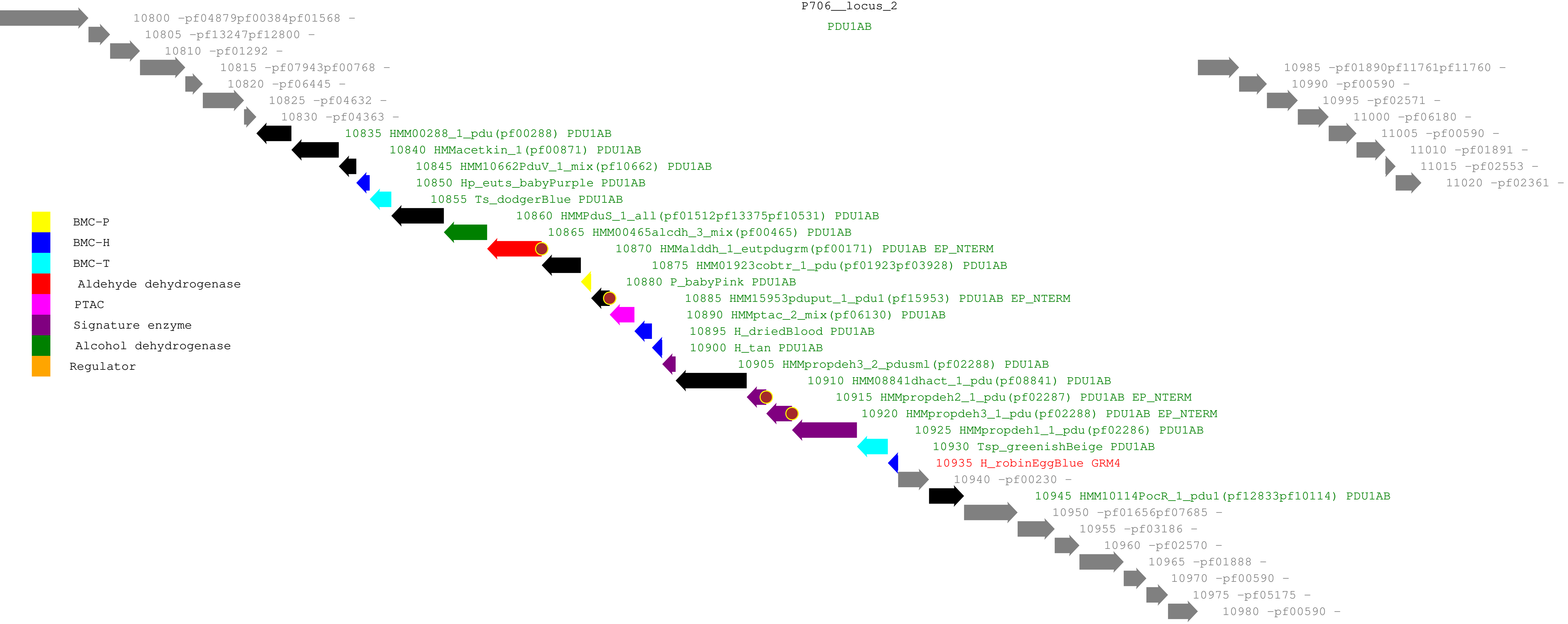

Salmonella enterica subsp. enterica serovar Newport str. CVM22707. Bacteria; Proteobacteria; Gammaproteobacteria; Enterobacterales;

P707__locus_2

PDU1AB

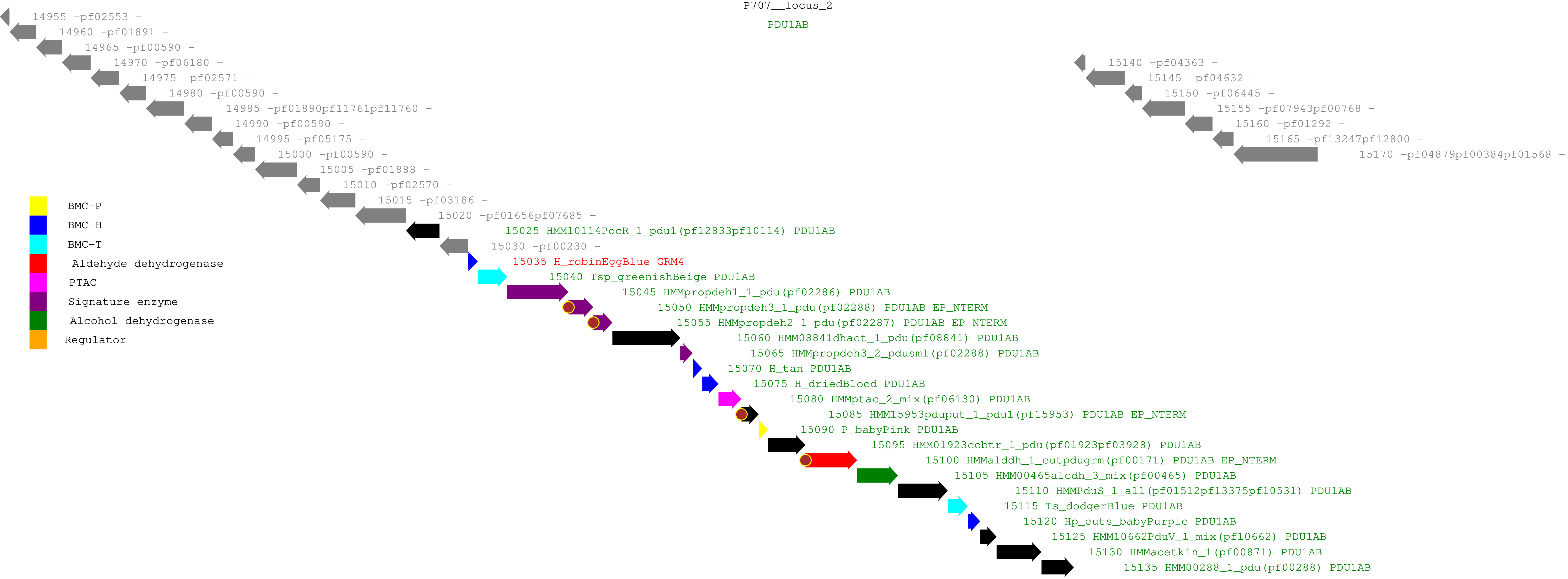

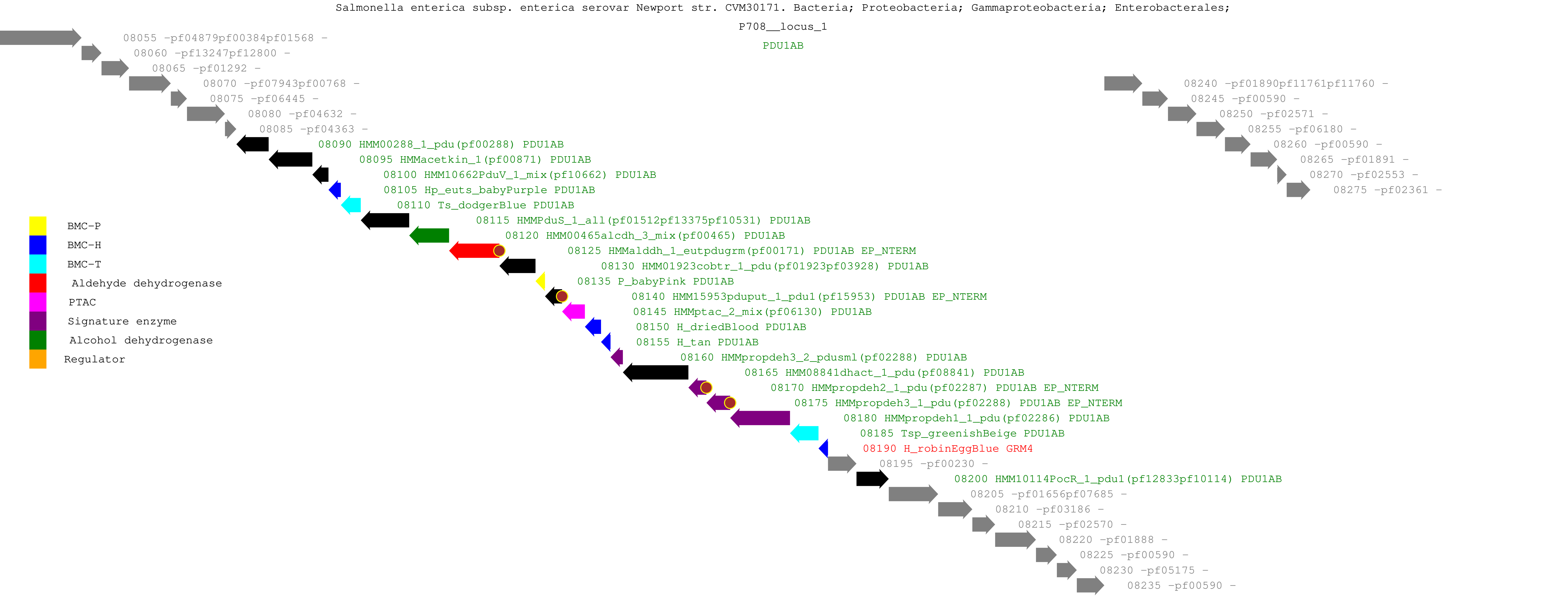
Salmonella enterica subsp. enterica serovar Newport str. CVM30171. Bacteria; Proteobacteria; Gammaproteobacteria; Enterobacterales;
P708__locus_1
PDU1AB
08055 -pf04879pf00384pf01568 -
08060 -pf13247pf12800 -
08065 -pf01292 -
08070 -pf07943pf00768 -
08075 -pf06445 -
08080 -pf04632 -
08085 -pf04363 -
08090 HMM00288_1_pdu(pf00288) PDU1AB
08095 HMMacetkin_1(pf00871) PDU1AB
08100 HMM10662PduV_1_mix(pf10662) PDU1AB
08105 Hp_euts_babyPurple PDU1AB
08110 Ts_dodgerBlue PDU1AB
08115 HMMPduS_1_all(pf01512pf13375pf10531) PDU1AB
08120 HMM00465alcdh_3_mix(pf00465) PDU1AB
08125 HMMaldh_1_eutpdugrm(pf00171) PDU1AB EP_NTERM
08130 HMM01923cobtr_1_pdu(pf01923pf03928) PDU1AB
08135 P_babyPink PDU1AB
08140 HMM15953pduput_1_pdu1(pf15953) PDU1AB EP_NTERM
08145 HMMptac_2_mix(pf06130) PDU1AB
08150 H_driedBlood PDU1AB
08155 H_tan PDU1AB
08160 HMMpropdeh3_2_pdusml(pf02288) PDU1AB
08165 HMM08841dhact_1_pdu(pf08841) PDU1AB
08170 HMMpropdeh2_1_pdu(pf02287) PDU1AB EP_NTERM
08175 HMMpropdeh3_1_pdu(pf02288) PDU1AB EP_NTERM
08180 HMMpropdeh1_1_pdu(pf02286) PDU1AB
08185 Tsp_greenishBeige PDU1AB
08190 H_robinEggBlue GRM4
08195 -pf00230 -
08200 HMM10114PocR_1_pdu1(pf12833pf10114) PDU1AB
08205 -pf01656pf07685 -
08210 -pf03186 -
08215 -pf02570 -
08220 -pf01888 -
08225 -pf00590 -
08230 -pf05175 -
08235 -pf00590 -
08240 -pf01890pf11761pf11760 -
08245 -pf00590 -
08250 -pf02571 -
08255 -pf06180 -
08260 -pf00590 -
08265 -pf01891 -
08270 -pf02553 -
08275 -pf02361 -
BMC-P
BMC-H
BMC-T
Aldehyde dehydrogenase
PTAC
Signature enzyme
Alcohol dehydrogenase
Regulator

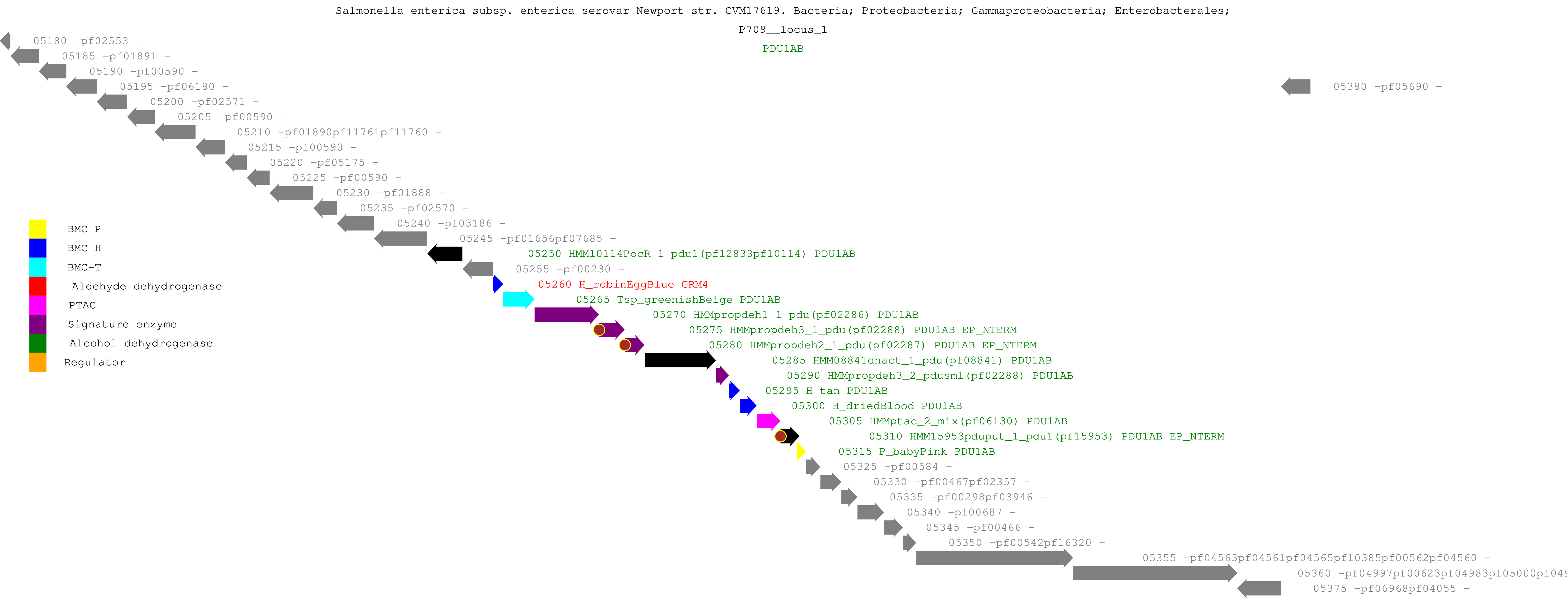
Salmonella enterica subsp. enterica serovar Newport str. CVM17619. Bacteria; Proteobacteria; Gammaproteobacteria; Enterobacterales;
P709__locus_1
PDU1AB
05180 -pf02553 -
05185 -pf01891 -
05190 -pf00590 -
05195 -pf06180 -
05200 -pf02571 -
05205 -pf00590 -
05210 -pf01890pf11761pf11760 -
05215 -pf00590 -
05220 -pf05175 -
05225 -pf00590 -
05230 -pf01888 -
05235 -pf02570 -
05240 -pf03186 -
05245 -pf01656pf07685 -
05250 HMM10114PocR_1_pdu1(pf12833pf10114) PDU1AB
05255 -pf00230 -
05260 H_robinEggBlue GRM4
05265 Tsp_greenishBeige PDU1AB
05270 HMMpropdeh1_1_pdu(pf02286) PDU1AB
05275 HMMpropdeh3_1_pdu(pf02288) PDU1AB EP_NTERM
05280 HMMpropdeh2_1_pdu(pf02287) PDU1AB EP_NTERM
05285 HMM08841dhact_1_pdu(pf08841) PDU1AB
05290 HMMpropdeh3_2_pdusml(pf02288) PDU1AB
05295 H_tan PDU1AB
05300 H_driedBlood PDU1AB
05305 HMMptac_2_mix(pf06130) PDU1AB
05310 HMM15953pduput_1_pdu1(pf15953) PDU1AB EP_NTERM
05315 P_babyPink PDU1AB
05325 -pf00584 -
05330 -pf00467pf02357 -
05335 -pf00298pf03946 -
05340 -pf00687 -
05345 -pf00466 -
05350 -pf00542pf16320 -
05355 -pf04563pf04561pf04565pf10385pf00562pf04560 -
05360 -pf04997pf00623pf04983pf05000pf04
05375 -pf06968pf04055 -
05380 -pf05690 -
BMC-P
BMC-H
BMC-T
Aldehyde dehydrogenase
PTAC
Signature enzyme
Alcohol dehydrogenase
Regulator

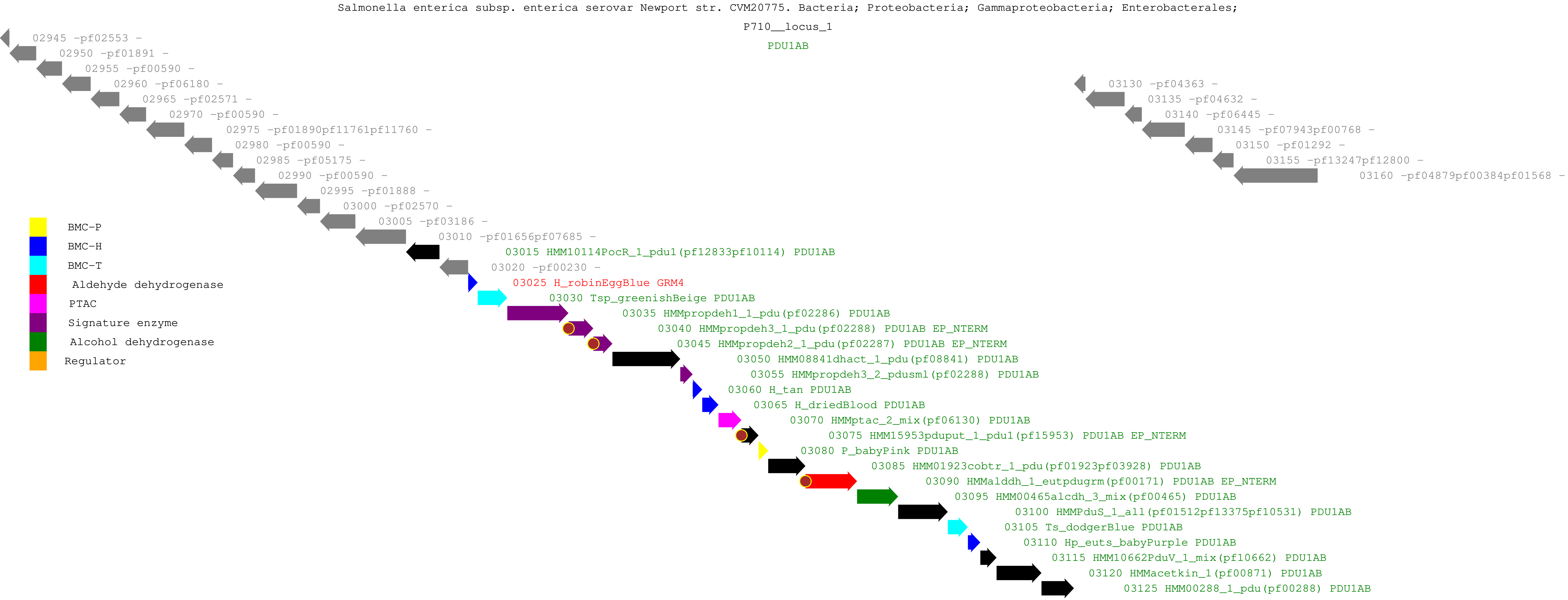
Salmonella enterica subsp. enterica serovar Newport str. CVM20775. Bacteria; Proteobacteria; Gammaproteobacteria; Enterobacterales;
P710__locus_1
PDU1AB
02945 -pf02553 -
02950 -pf01891 -
02955 -pf00590 -
02960 -pf06180 -
02965 -pf02571 -
02970 -pf00590 -
02975 -pf01890pf11761pf11760 -
02980 -pf00590 -
02985 -pf05175 -
02990 -pf00590 -
02995 -pf01888 -
03000 -pf02570 -
03005 -pf03186 -
03010 -pf01656pf07685 -
03015 HMM10114PocR_1_pdu1(pf12833pf10114) PDU1AB
03020 -pf00230 -
03025 H_robinEggBlue GRM4
03030 Tsp_greenishBeige PDU1AB
03035 HMMpropdeh1_1_pdu(pf02286) PDU1AB
03040 HMMpropdeh3_1_pdu(pf02288) PDU1AB EP_NTERM
03045 HMMpropdeh2_1_pdu(pf02287) PDU1AB EP_NTERM
03050 HMM08841dhact_1_pdu(pf08841) PDU1AB
03055 HMMpropdeh3_2_pdusml(pf02288) PDU1AB
03060 H_tan PDU1AB
03065 H_driedBlood PDU1AB
03070 HMMptac_2_mix(pf06130) PDU1AB
03075 HMM15953pduput_1_pdu1(pf15953) PDU1AB EP_NTERM
03080 P_babyPink PDU1AB
03085 HMM01923cobtr_1_pdu(pf01923pf03928) PDU1AB
03090 HMMalddh_1_eutpdugrm(pf00171) PDU1AB EP_NTERM
03095 HMM00465alcdh_3_mix(pf00465) PDU1AB
03100 HMMPduS_1_all(pf01512pf13375pf10531) PDU1AB
03105 Ts_dodgerBlue PDU1AB
03110 Hp_euts_babyPurple PDU1AB
03115 HMM10662PduV_1_mix(pf10662) PDU1AB
03120 HMMacetkin_1(pf00871) PDU1AB
03125 HMM00288_1_pdu(pf00288) PDU1AB
03130 -pf04363 -
03135 -pf04632 -
03140 -pf06445 -
03145 -pf07943pf00768 -
03150 -pf01292 -
03155 -pf13247pf12800 -
03160 -pf04879pf00384pf01568 -
BMC-P
BMC-H
BMC-T
Aldehyde dehydrogenase
PTAC
Signature enzyme
Alcohol dehydrogenase
Regulator

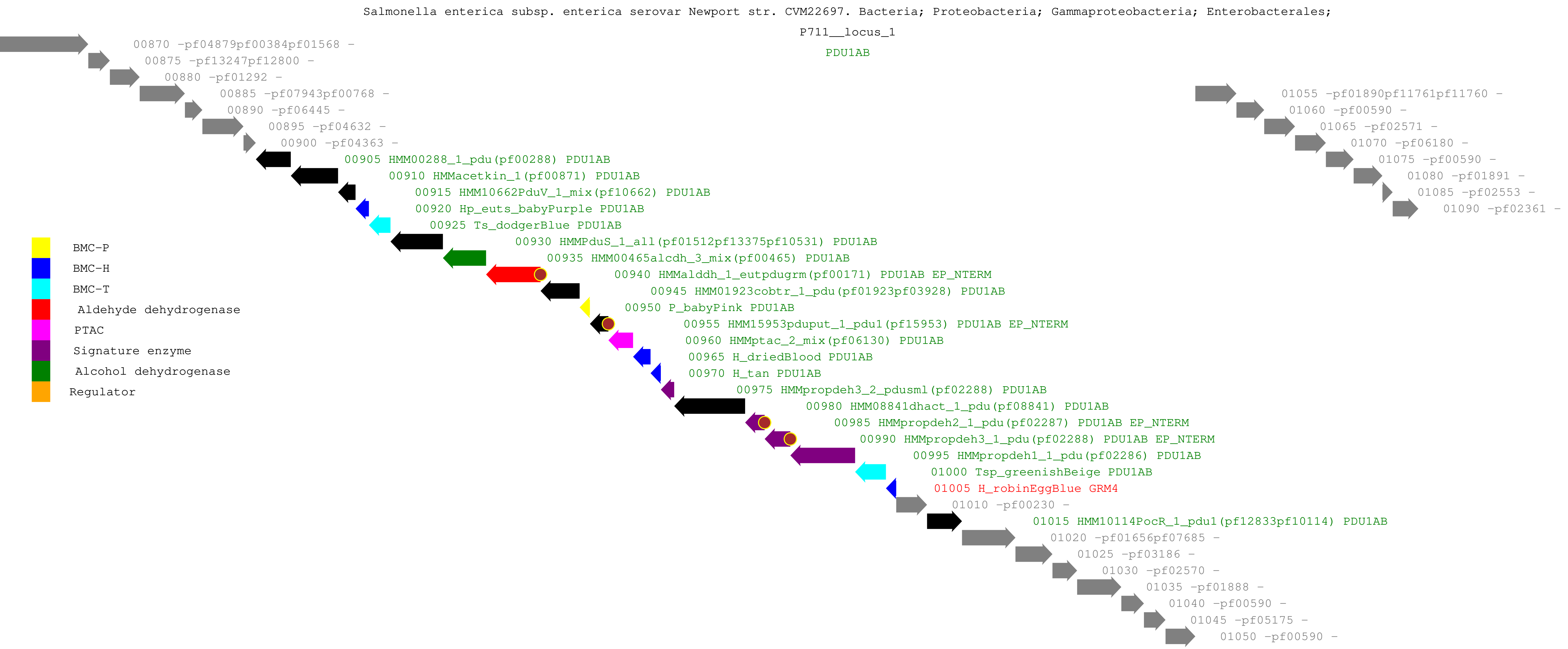

Salmonella enterica subsp. enterica serovar Newport str. CVM22697. Bacteria; Proteobacteria; Gammaproteobacteria; Enterobacterales;
P711__locus_1
PDU1AB
00870 -pf04879pf00384pf01568 -
00875 -pf13247pf12800 -
00880 -pf01292 -
00885 -pf07943pf00768 -
00890 -pf06445 -
00895 -pf04632 -
00900 -pf04363 -
00905 HMM00288_1_pdu(pf00288) PDU1AB
00910 HMMacetkin_1(pf00871) PDU1AB
00915 HMM10662PduV_1_mix(pf10662) PDU1AB
00920 Hp_euts_babyPurple PDU1AB
00925 Ts_dodgerBlue PDU1AB
00930 HMMPduS_1_all(pf01512pf13375pf10531) PDU1AB
00935 HMM00465alcdh_3_mix(pf00465) PDU1AB
00940 HMMalddh_1_eutpdugrm(pf00171) PDU1AB EP_NTERM
00945 HMM01923cobtr_1_pdu(pf01923pf03928) PDU1AB
00950 P_babyPink PDU1AB
00955 HMM15953pduput_1_pdu1(pf15953) PDU1AB EP_NTERM
00960 HMMptac_2_mix(pf06130) PDU1AB
00965 H_driedBlood PDU1AB
00970 H_tan PDU1AB
00975 HMMpropdeh3_2_pdusml(pf02288) PDU1AB
00980 HMM08841dhact_1_pdu(pf08841) PDU1AB
00985 HMMpropdeh2_1_pdu(pf02287) PDU1AB EP_NTERM
00990 HMMpropdeh3_1_pdu(pf02288) PDU1AB EP_NTERM
00995 HMMpropdeh1_1_pdu(pf02286) PDU1AB
01000 Tsp_greenishBeige PDU1AB
01005 H_robinEggBlue GRM4
01010 -pf00230 -
01015 HMM10114PocR_1_pdu1(pf12833pf10114) PDU1AB
01020 -pf01656pf07685 -
01025 -pf03186 -
01030 -pf02570 -
01035 -pf01888 -
01040 -pf00590 -
01045 -pf05175 -
01050 -pf00590 -
01055 -pf01890pf11761pf11760 -
01060 -pf00590 -
01065 -pf02571 -
01070 -pf06180 -
01075 -pf00590 -
01080 -pf01891 -
01085 -pf02553 -
01090 -pf02361 -
BMC-P
BMC-H
BMC-T
Aldehyde dehydrogenase
PTAC
Signature enzyme
Alcohol dehydrogenase
Regulator

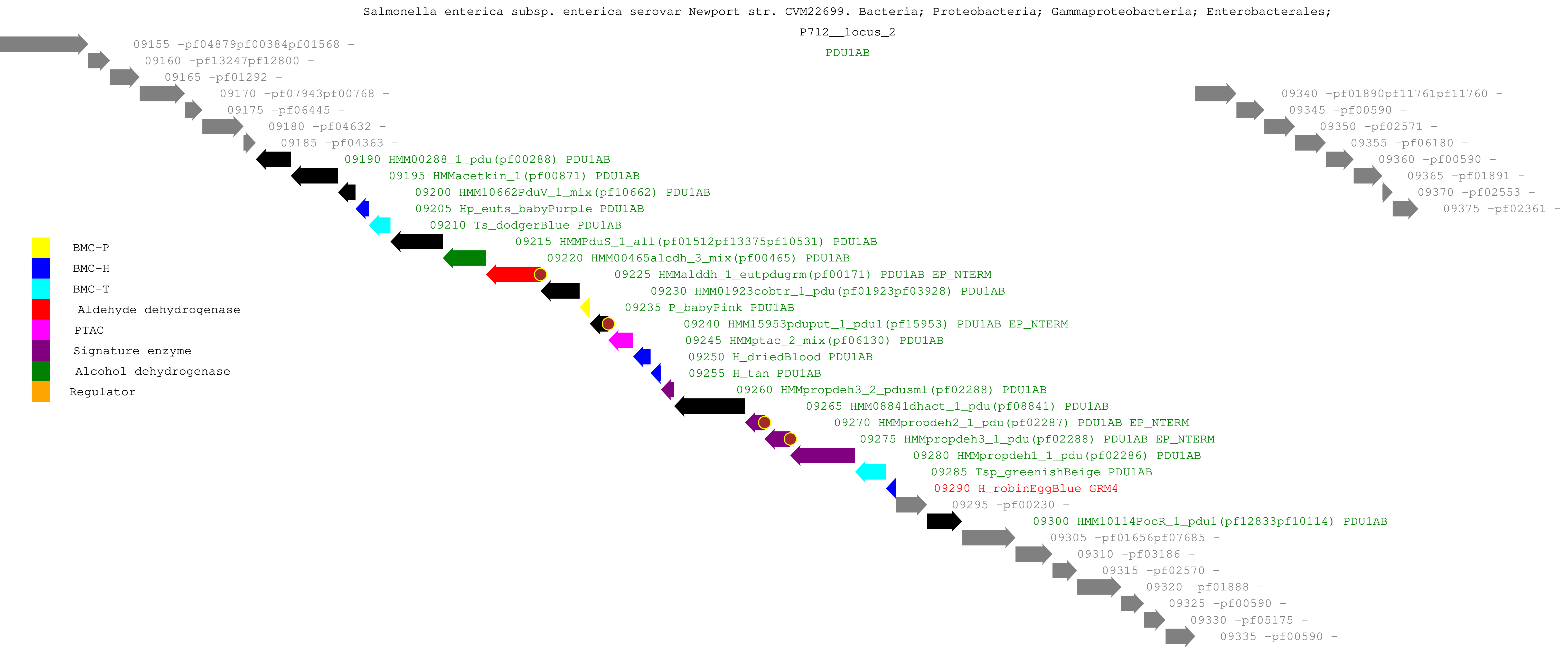
Salmonella enterica subsp. enterica serovar Newport str. CVM22699. Bacteria; Proteobacteria; Gammaproteobacteria; Enterobacterales;
P712__locus_2
PDU1AB
09155 -pf04879pf00384pf01568 -
09160 -pf13247pf12800 -
09165 -pf01292 -
09170 -pf07943pf00768 -
09175 -pf06445 -
09180 -pf04632 -
09185 -pf04363 -
09190 HMM00288_1_pdu(pf00288) PDU1AB
09195 HMMacetkin_1(pf00871) PDU1AB
09200 HMM10662PduV_1_mix(pf10662) PDU1AB
09205 Hp_euts_babyPurple PDU1AB
09210 Ts_dodgerBlue PDU1AB
09215 HMMPduS_1_all(pf01512pf13375pf10531) PDU1AB
09220 HMM00465alcdh_3_mix(pf00465) PDU1AB
09225 HMMalddh_1_eutpdugrm(pf00171) PDU1AB EP_NTERM
09230 HMM01923cobtr_1_pdu(pf01923pf03928) PDU1AB
09235 P_babyPink PDU1AB
09240 HMM15953pduput_1_pdu1(pf15953) PDU1AB EP_NTERM
09245 HMMptac_2_mix(pf06130) PDU1AB
09250 H_driedBlood PDU1AB
09255 H_tan PDU1AB
09260 HMMpropdeh3_2_pdusml(pf02288) PDU1AB
09265 HMM08841dhact_1_pdu(pf08841) PDU1AB
09270 HMMpropdeh2_1_pdu(pf02287) PDU1AB EP_NTERM
09275 HMMpropdeh3_1_pdu(pf02288) PDU1AB EP_NTERM
09280 HMMpropdeh1_1_pdu(pf02286) PDU1AB
09285 Tsp_greenishBeige PDU1AB
09290 H_robinEggBlue GRM4
09295 -pf00230 -
09300 HMM10114PocR_1_pdu1(pf12833pf10114) PDU1AB
09305 -pf01656pf07685 -
09310 -pf03186 -
09315 -pf02570 -
09320 -pf01888 -
09325 -pf00590 -
09330 -pf05175 -
09335 -pf00590 -
09340 -pf01890pf11761pf11760 -
09345 -pf00590 -
09350 -pf02571 -
09355 -pf06180 -
09360 -pf00590 -
09365 -pf01891 -
09370 -pf02553 -
09375 -pf02361 -
BMC-P
BMC-H
BMC-T
Aldehyde dehydrogenase
PTAC
Signature enzyme
Alcohol dehydrogenase
Regulator

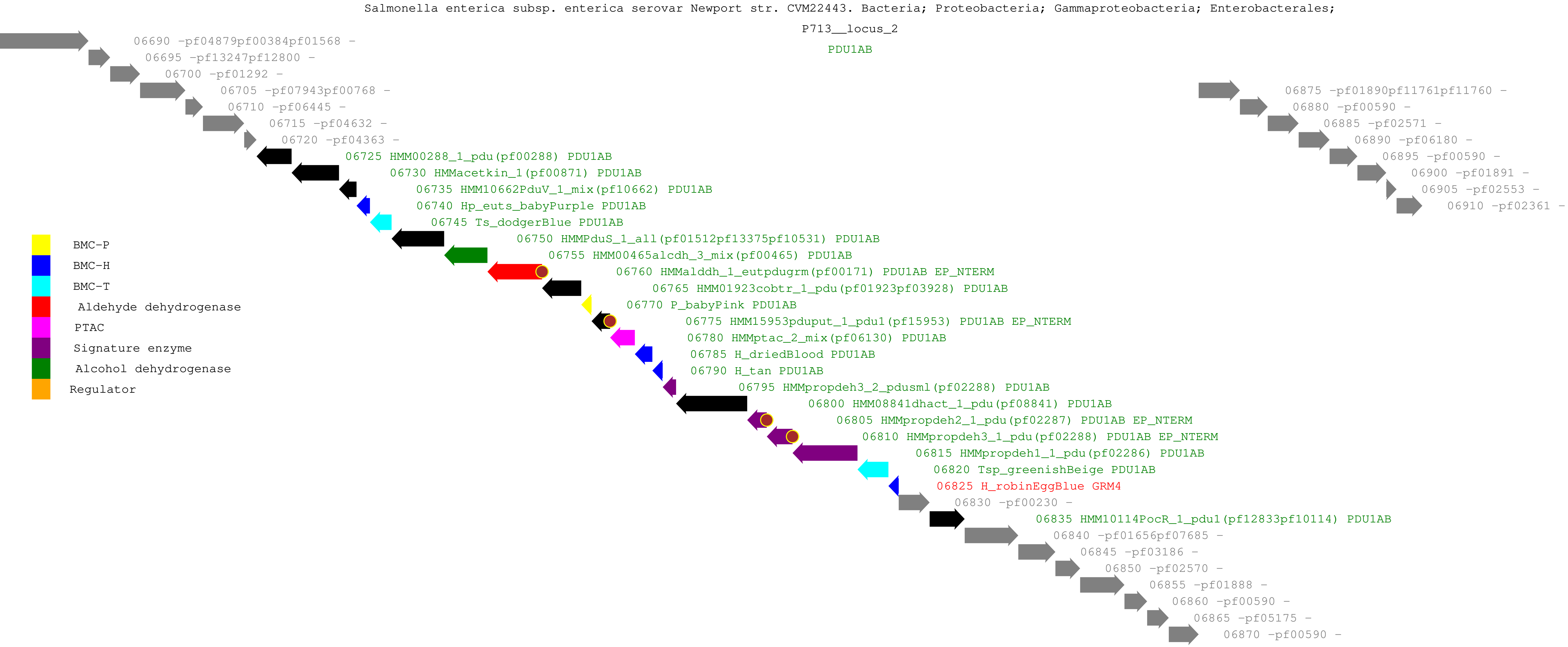

Salmonella enterica subsp. enterica serovar Newport str. CVM22443. Bacteria; Proteobacteria; Gammaproteobacteria; Enterobacterales;
P713__locus_2
PDU1AB
06690 -pf04879pf00384pf01568 -
06695 -pf13247pf12800 -
06700 -pf01292 -
06705 -pf07943pf00768 -
06710 -pf06445 -
06715 -pf04632 -
06720 -pf04363 -
06725 HMM00288_1_pdu(pf00288) PDU1AB
06730 HMMacetkin_1(pf00871) PDU1AB
06735 HMM10662PduV_1_mix(pf10662) PDU1AB
06740 Hp_euts_babyPurple PDU1AB
06745 Ts_dodgerBlue PDU1AB
06750 HMMPduS_1_all(pf01512pf13375pf10531) PDU1AB
06755 HMM00465alcdh_3_mix(pf00465) PDU1AB
06760 HMMalddh_1_eutpdugrm(pf00171) PDU1AB EP_NTERM
06765 HMM01923cobtr_1_pdu(pf01923pf03928) PDU1AB
06770 P_babyPink PDU1AB
06775 HMM15953pduput_1_pdu1(pf15953) PDU1AB EP_NTERM
06780 HMMptac_2_mix(pf06130) PDU1AB
06785 H_driedBlood PDU1AB
06790 H_tan PDU1AB
06795 HMMpropdeh3_2_pdusml(pf02288) PDU1AB
06800 HMM08841dhact_1_pdu(pf08841) PDU1AB
06805 HMMpropdeh2_1_pdu(pf02287) PDU1AB EP_NTERM
06810 HMMpropdeh3_1_pdu(pf02288) PDU1AB EP_NTERM
06815 HMMpropdeh1_1_pdu(pf02286) PDU1AB
06820 Tsp_greenishBeige PDU1AB
06825 H_robinEggBlue GRM4
06830 -pf00230 -
06835 HMM10114PocR_1_pdu1(pf12833pf10114) PDU1AB
06840 -pf01656pf07685 -
06845 -pf03186 -
06850 -pf02570 -
06855 -pf01888 -
06860 -pf00590 -
06865 -pf05175 -
06870 -pf00590 -
06875 -pf01890pf11761pf11760 -
06880 -pf00590 -
06885 -pf02571 -
06890 -pf06180 -
06895 -pf00590 -
06900 -pf01891 -
06905 -pf02553 -
06910 -pf02361 -
BMC-P
BMC-H
BMC-T
Aldehyde dehydrogenase
PTAC
Signature enzyme
Alcohol dehydrogenase
Regulator

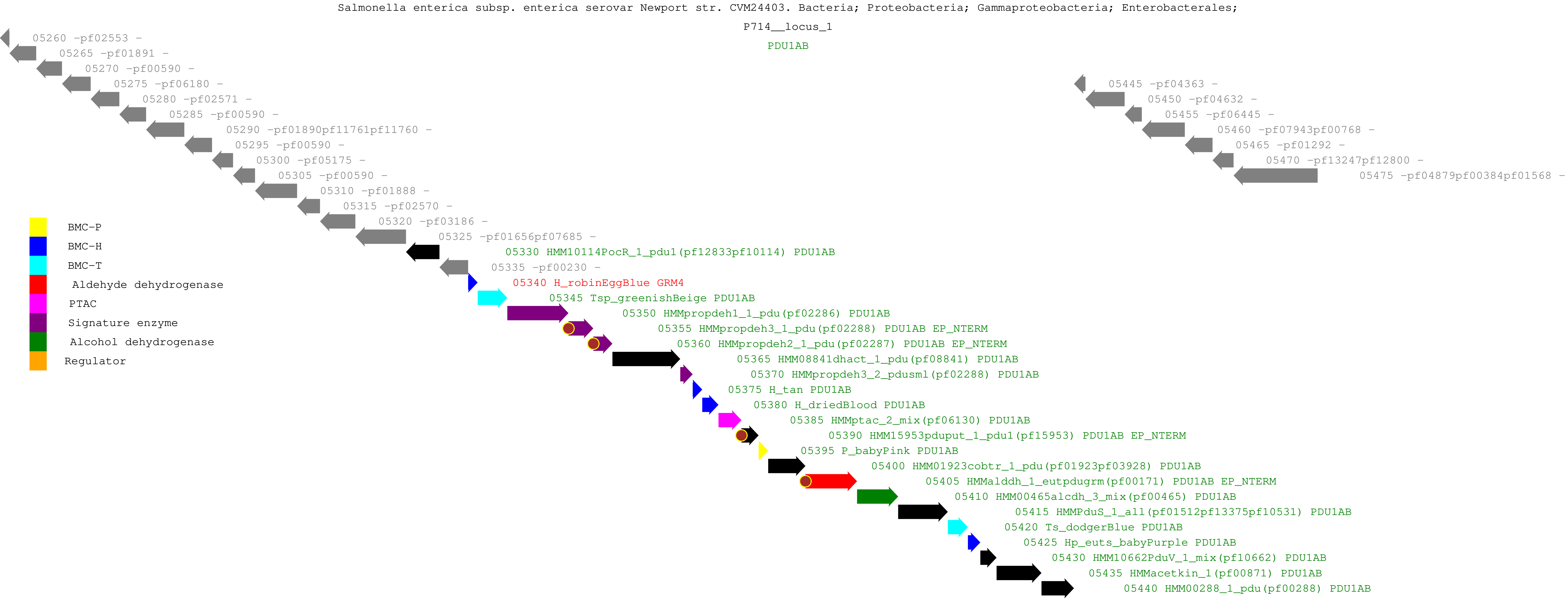

Salmonella enterica subsp. enterica serovar Newport str. CVM24403. Bacteria; Proteobacteria; Gammaproteobacteria; Enterobacterales;
P714__locus_1
PDU1AB
05260 -pf02553 -
05265 -pf01891 -
05270 -pf00590 -
05275 -pf06180 -
05280 -pf02571 -
05285 -pf00590 -
05290 -pf01890pf11761pf11760 -
05295 -pf00590 -
05300 -pf05175 -
05305 -pf00590 -
05310 -pf01888 -
05315 -pf02570 -
05320 -pf03186 -
05325 -pf01656pf07685 -
05330 HMM10114PocR_1_pdu1(pf12833pf10114) PDU1AB
05335 -pf00230 -
05340 H_robinEggBlue GRM4
05345 Tsp_greenishBeige PDU1AB
05350 HMMpropdeh1_1_pdu(pf02286) PDU1AB
05355 HMMpropdeh3_1_pdu(pf02288) PDU1AB EP_NTERM
05360 HMMpropdeh2_1_pdu(pf02287) PDU1AB EP_NTERM
05365 HMM08841dhact_1_pdu(pf08841) PDU1AB
05370 HMMpropdeh3_2_pdusml(pf02288) PDU1AB
05375 H_tan PDU1AB
05380 H_driedBlood PDU1AB
05385 HMMptac_2_mix(pf06130) PDU1AB
05390 HMM15953pduput_1_pdu1(pf15953) PDU1AB EP_NTERM
05395 P_babyPink PDU1AB
05400 HMM01923cobtr_1_pdu(pf01923pf03928) PDU1AB
05405 HMMaldh_1_eutpdugrm(pf00171) PDU1AB EP_NTERM
05410 HMM00465alcdh_3_mix(pf00465) PDU1AB
05415 HMMPduS_1_all(pf01512pf13375pf10531) PDU1AB
05420 Ts_dodgerBlue PDU1AB
05425 Hp_euts_babyPurple PDU1AB
05430 HMM10662PduV_1_mix(pf10662) PDU1AB
05435 HMMacetkin_1(pf00871) PDU1AB
05440 HMM00288_1_pdu(pf00288) PDU1AB
05445 -pf04363 -
05450 -pf04632 -
05455 -pf06445 -
05460 -pf07943pf00768 -
05465 -pf01292 -
05470 -pf13247pf12800 -
05475 -pf04879pf00384pf01568 -
BMC-P
BMC-H
BMC-T
Aldehyde dehydrogenase
PTAC
Signature enzyme
Alcohol dehydrogenase
Regulator

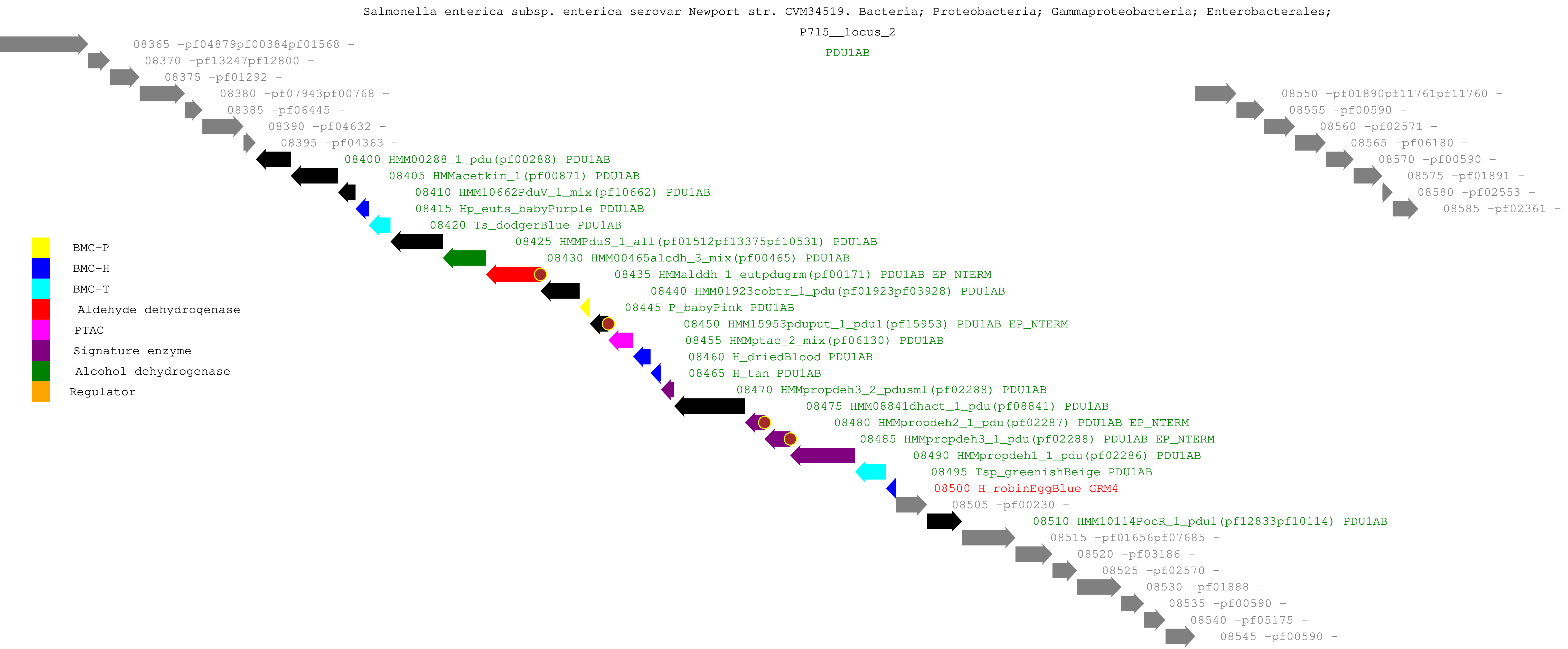
Salmonella enterica subsp. enterica serovar Newport str. CVM34519. Bacteria; Proteobacteria; Gammaproteobacteria; Enterobacterales;
P715__locus_2
PDU1AB
08365 -pf04879pf00384pf01568 -
08370 -pf13247pf12800 -
08375 -pf01292 -
08380 -pf07943pf00768 -
08385 -pf06445 -
08390 -pf04632 -
08395 -pf04363 -
08400 HMM00288_1_pdu(pf00288) PDU1AB
08405 HMMacetkin_1(pf00871) PDU1AB
08410 HMM10662PduV_1_mix(pf10662) PDU1AB
08415 Hp_euts_babyPurple PDU1AB
08420 Ts_dodgerBlue PDU1AB
08425 HMMPduS_1_all(pf01512pf13375pf10531) PDU1AB
08430 HMM00465alcdh_3_mix(pf00465) PDU1AB
08435 HMMalddh_1_eutpdugrm(pf00171) PDU1AB EP_NTERM
08440 HMM01923cobtr_1_pdu(pf01923pf03928) PDU1AB
08445 P_babyPink PDU1AB
08450 HMM15953pduput_1_pdu1(pf15953) PDU1AB EP_NTERM
08455 HMMptac_2_mix(pf06130) PDU1AB
08460 H_driedBlood PDU1AB
08465 H_tan PDU1AB
08470 HMMpropdeh3_2_pdusml(pf02288) PDU1AB
08475 HMM08841dhact_1_pdu(pf08841) PDU1AB
08480 HMMpropdeh2_1_pdu(pf02287) PDU1AB EP_NTERM
08485 HMMpropdeh3_1_pdu(pf02288) PDU1AB EP_NTERM
08490 HMMpropdeh1_1_pdu(pf02286) PDU1AB
08495 Tsp_greenishBeige PDU1AB
08500 H_robinEggBlue GRM4
08505 -pf00230 -
08510 HMM10114PocR_1_pdu1(pf12833pf10114) PDU1AB
08515 -pf01656pf07685 -
08520 -pf03186 -
08525 -pf02570 -
08530 -pf01888 -
08535 -pf00590 -
08540 -pf05175 -
08545 -pf00590 -
08550 -pf01890pf11761pf11760 -
08555 -pf00590 -
08560 -pf02571 -
08565 -pf06180 -
08570 -pf00590 -
08575 -pf01891 -
08580 -pf02553 -
08585 -pf02361 -
BMC-P
BMC-H
BMC-T
Aldehyde dehydrogenase
PTAC
Signature enzyme
Alcohol dehydrogenase
Regulator

Salmonella enterica subsp. enterica serovar Newport str. CVM18005. Bacteria; Proteobacteria; Gammaproteobacteria; Enterobacterales;

P716__locus_2

PDU1AB

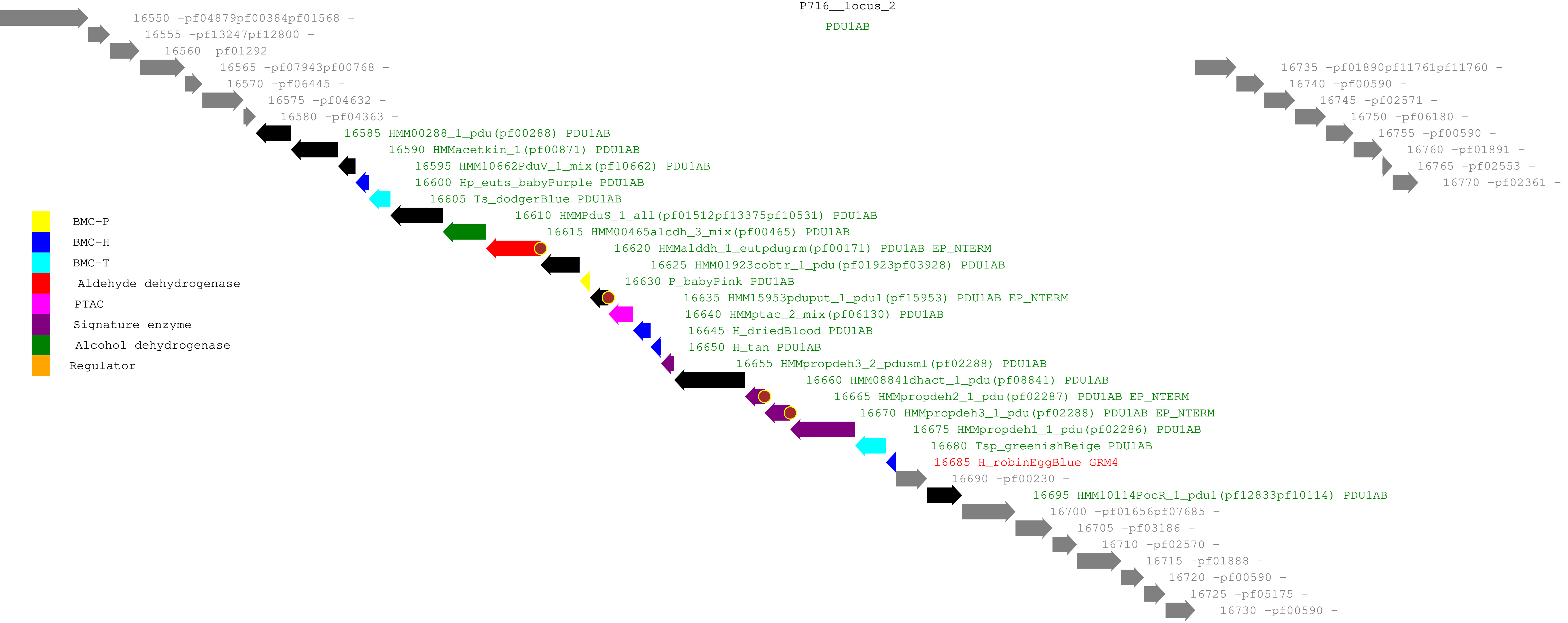

Salmonella enterica subsp. enterica serovar Enteritidis str. SHSE001. Bacteria; Proteobacteria; Gammaproteobacteria; Enterobacterales;

P727__locus_2

PDU1AB

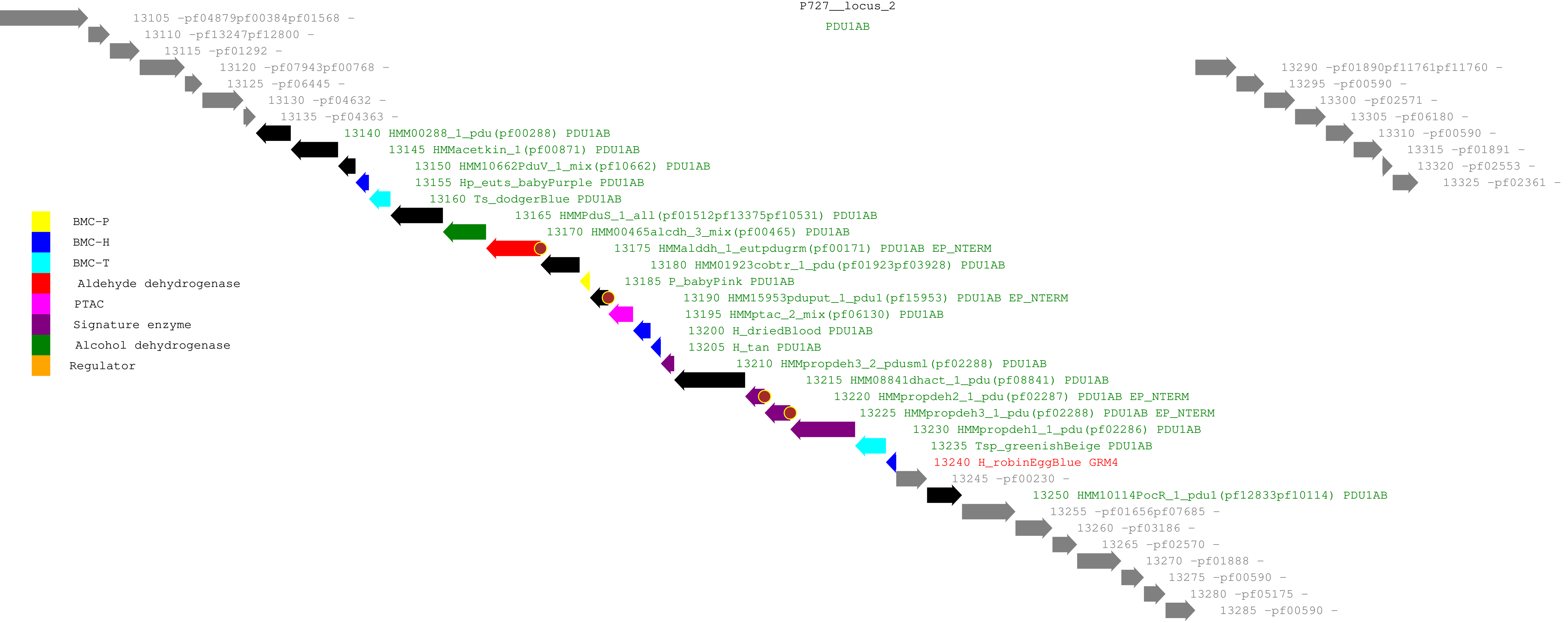

Salmonella enterica subsp. enterica serovar Enteritidis str. SHSE002. Bacteria; Proteobacteria; Gammaproteobacteria; Enterobacterales;

P728__locus_1

PDU1AB

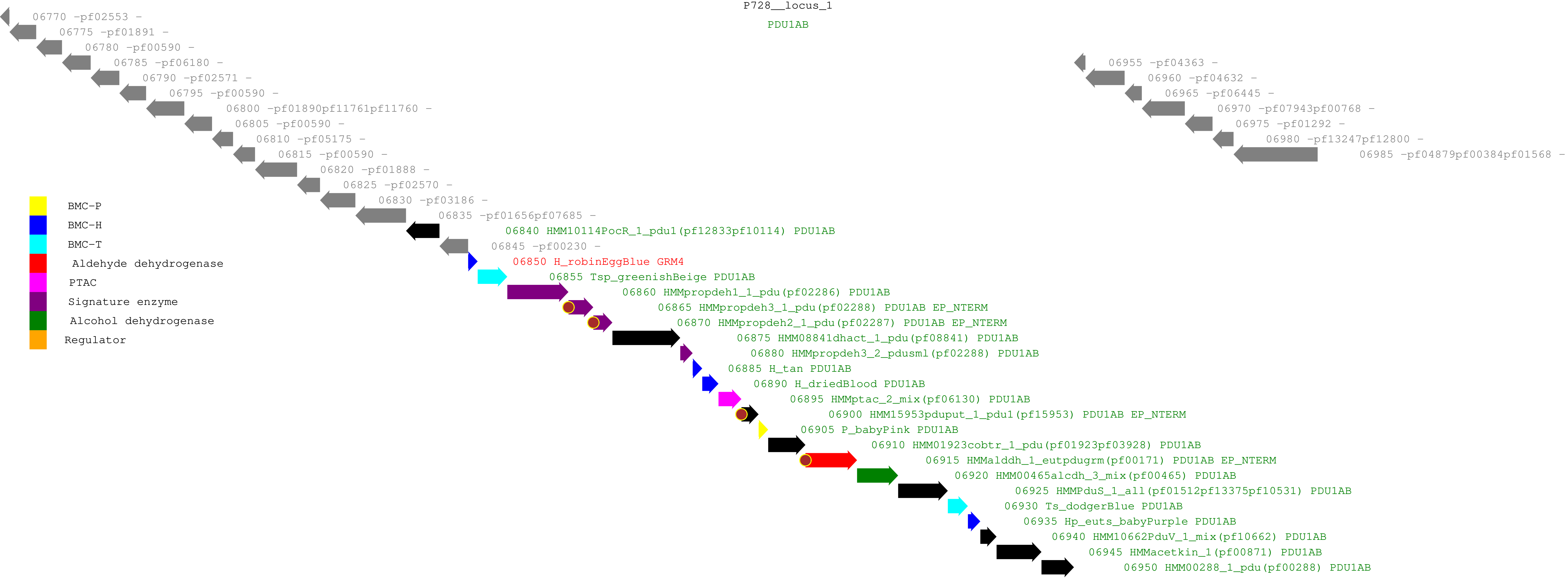

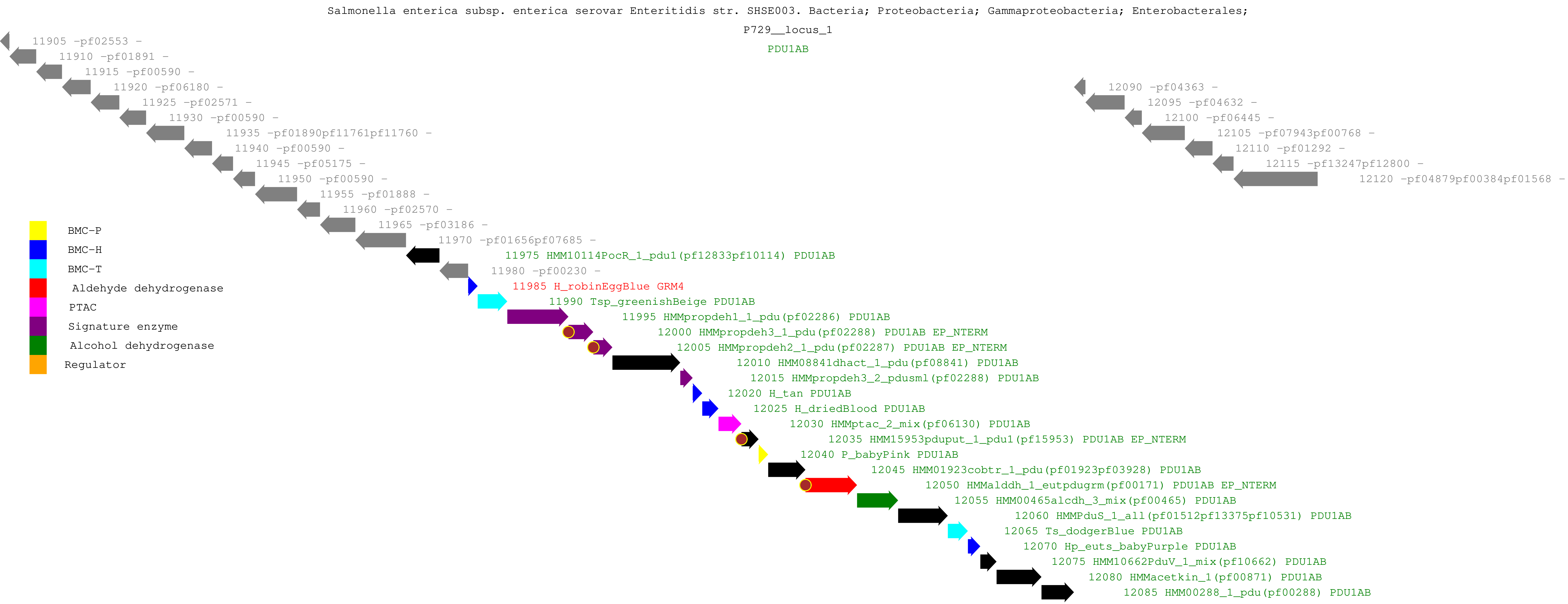
Salmonella enterica subsp. enterica serovar Enteritidis str. SHSE003. Bacteria; Proteobacteria; Gammaproteobacteria; Enterobacterales;
P729__locus_1
PDU1AB
11905 -pf02553 -
11910 -pf01891 -
11915 -pf00590 -
11920 -pf06180 -
11925 -pf02571 -
11930 -pf00590 -
11935 -pf01890pf11761pf11760 -
11940 -pf00590 -
11945 -pf05175 -
11950 -pf00590 -
11955 -pf01888 -
11960 -pf02570 -
11965 -pf03186 -
11970 -pf01656pf07685 -
11975 HMM10114PocR_1_pdu1(pf12833pf10114) PDU1AB
11980 -pf00230 -
11985 H_robinEggBlue GRM4
11990 Tsp_greenishBeige PDU1AB
11995 HMMpropdeh1_1_pdu(pf02286) PDU1AB
12000 HMMpropdeh3_1_pdu(pf02288) PDU1AB EP_NTERM
12005 HMMpropdeh2_1_pdu(pf02287) PDU1AB EP_NTERM
12010 HMM08841dhact_1_pdu(pf08841) PDU1AB
12015 HMMpropdeh3_2_pdusml(pf02288) PDU1AB
12020 H_tan PDU1AB
12025 H_driedBlood PDU1AB
12030 HMMptac_2_mix(pf06130) PDU1AB
12035 HMM15953pduput_1_pdu1(pf15953) PDU1AB EP_NTERM
12040 P_babyPink PDU1AB
12045 HMM01923cobtr_1_pdu(pf01923pf03928) PDU1AB
12050 HMMaldh_1_eutpdugrm(pf00171) PDU1AB EP_NTERM
12055 HMM00465alcdh_3_mix(pf00465) PDU1AB
12060 HMMPduS_1_all(pf01512pf13375pf10531) PDU1AB
12065 Ts_dodgerBlue PDU1AB
12070 Hp_euts_babyPurple PDU1AB
12075 HMM10662PduV_1_mix(pf10662) PDU1AB
12080 HMMacetkin_1(pf00871) PDU1AB
12085 HMM00288_1_pdu(pf00288) PDU1AB
12090 -pf04363 -
12095 -pf04632 -
12100 -pf06445 -
12105 -pf07943pf00768 -
12110 -pf01292 -
12115 -pf13247pf12800 -
12120 -pf04879pf00384pf01568 -
BMC-P
BMC-H
BMC-T
Aldehyde dehydrogenase
PTAC
Signature enzyme
Alcohol dehydrogenase
Regulator

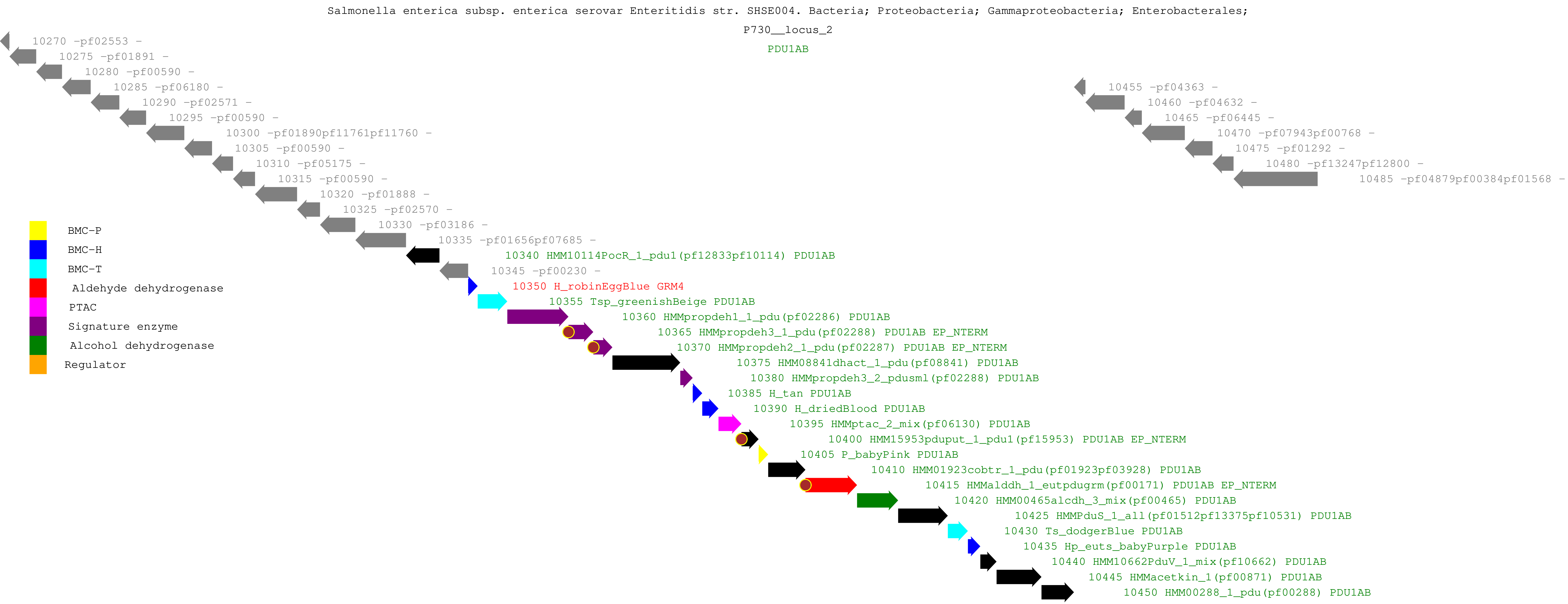
Salmonella enterica subsp. enterica serovar Enteritidis str. SHSE004. Bacteria; Proteobacteria; Gammaproteobacteria; Enterobacterales;
P730__locus_2
PDU1AB
10270 -pf02553 -
10275 -pf01891 -
10280 -pf00590 -
10285 -pf06180 -
10290 -pf02571 -
10295 -pf00590 -
10300 -pf01890pf11761pf11760 -
10305 -pf00590 -
10310 -pf05175 -
10315 -pf00590 -
10320 -pf01888 -
10325 -pf02570 -
10330 -pf03186 -
10335 -pf01656pf07685 -
10340 HMM10114PocR_1_pdu1(pf12833pf10114) PDU1AB
10345 -pf00230 -
10350 H_robinEggBlue GRM4
10355 Tsp_greenishBeige PDU1AB
10360 HMMpropdeh1_1_pdu(pf02286) PDU1AB
10365 HMMpropdeh3_1_pdu(pf02288) PDU1AB EP_NTERM
10370 HMMpropdeh2_1_pdu(pf02287) PDU1AB EP_NTERM
10375 HMM08841dhact_1_pdu(pf08841) PDU1AB
10380 HMMpropdeh3_2_pdusml(pf02288) PDU1AB
10385 H_tan PDU1AB
10390 H_driedBlood PDU1AB
10395 HMMptac_2_mix(pf06130) PDU1AB
10400 HMM15953pduput_1_pdu1(pf15953) PDU1AB EP_NTERM
10405 P_babyPink PDU1AB
10410 HMM01923cobtr_1_pdu(pf01923pf03928) PDU1AB
10415 HMMalddh_1_eutpdugrm(pf00171) PDU1AB EP_NTERM
10420 HMM00465alcdh_3_mix(pf00465) PDU1AB
10425 HMMPduS_1_all(pf01512pf13375pf10531) PDU1AB
10430 Ts_dodgerBlue PDU1AB
10435 Hp_euts_babyPurple PDU1AB
10440 HMM10662PduV_1_mix(pf10662) PDU1AB
10445 HMMacetkin_1(pf00871) PDU1AB
10450 HMM00288_1_pdu(pf00288) PDU1AB
10455 -pf04363 -
10460 -pf04632 -
10465 -pf06445 -
10470 -pf07943pf00768 -
10475 -pf01292 -
10480 -pf13247pf12800 -
10485 -pf04879pf00384pf01568 -
BMC-P
BMC-H
BMC-T
Aldehyde dehydrogenase
PTAC
Signature enzyme
Alcohol dehydrogenase
Regulator

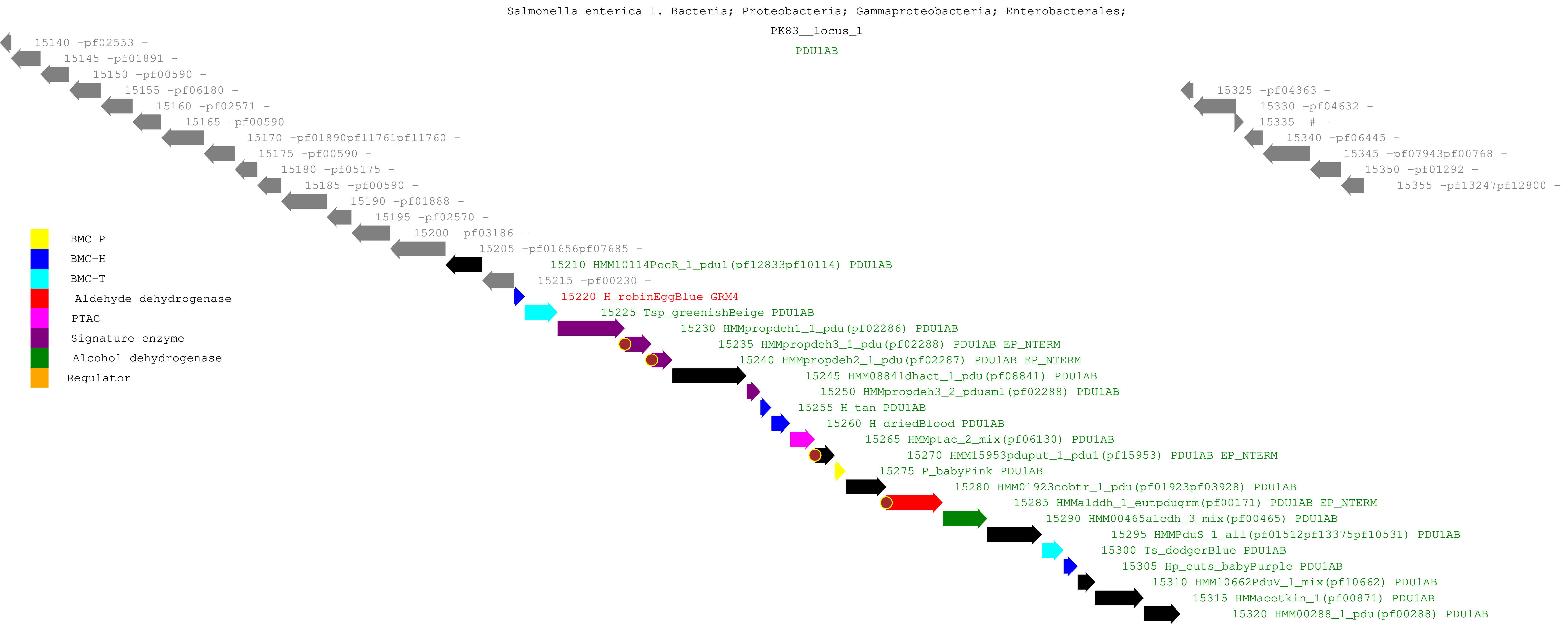

Salmonella enterica I. Bacteria; Proteobacteria; Gammaproteobacteria; Enterobacterales;
PK83__locus_1
PDU1AB
15140 -pf02553 -
15145 -pf01891 -
15150 -pf00590 -
15155 -pf06180 -
15160 -pf02571 -
15165 -pf00590 -
15170 -pf01890pf11761pf11760 -
15175 -pf00590 -
15180 -pf05175 -
15185 -pf00590 -
15190 -pf01888 -
15195 -pf02570 -
15200 -pf03186 -
15205 -pf01656pf07685 -
15210 HMM10114PocR_1_pdu1(pf12833pf10114) PDU1AB
15215 -pf00230 -
15220 H_robinEggBlue GRM4
15225 Tsp_greenishBeige PDU1AB
15230 HMMpropdeh1_1_pdu(pf02286) PDU1AB
15235 HMMpropdeh3_1_pdu(pf02288) PDU1AB EP_NTERM
15240 HMMpropdeh2_1_pdu(pf02287) PDU1AB EP_NTERM
15245 HMM08841dhact_1_pdu(pf08841) PDU1AB
15250 HMMpropdeh3_2_pdusml(pf02288) PDU1AB
15255 H_tan PDU1AB
15260 H_driedBlood PDU1AB
15265 HMMptac_2_mix(pf06130) PDU1AB
15270 HMM15953pduput_1_pdu1(pf15953) PDU1AB EP_NTERM
15275 P_babyPink PDU1AB
15280 HMM01923cobtr_1_pdu(pf01923pf03928) PDU1AB
15285 HMMalddh_1_eutpdugrm(pf00171) PDU1AB EP_NTERM
15290 HMM00465alcdh_3_mix(pf00465) PDU1AB
15295 HMMPduS_1_all(pf01512pf13375pf10531) PDU1AB
15300 Ts_dodgerBlue PDU1AB
15305 Hp_euts_babyPurple PDU1AB
15310 HMM10662PduV_1_mix(pf10662) PDU1AB
15315 HMMacetkin_1(pf00871) PDU1AB
15320 HMM00288_1_pdu(pf00288) PDU1AB
15325 -pf04363 -
15330 -pf04632 -
15335 -# -
15340 -pf06445 -
15345 -pf07943pf00768 -
15350 -pf01292 -
15355 -pf13247pf12800 -
BMC-P
BMC-H
BMC-T
Aldehyde dehydrogenase
PTAC
Signature enzyme
Alcohol dehydrogenase
Regulator

Salmonella enterica I. Bacteria; Proteobacteria; Gammaproteobacteria; Enterobacterales;

PK95__locus_1

PDU1AB

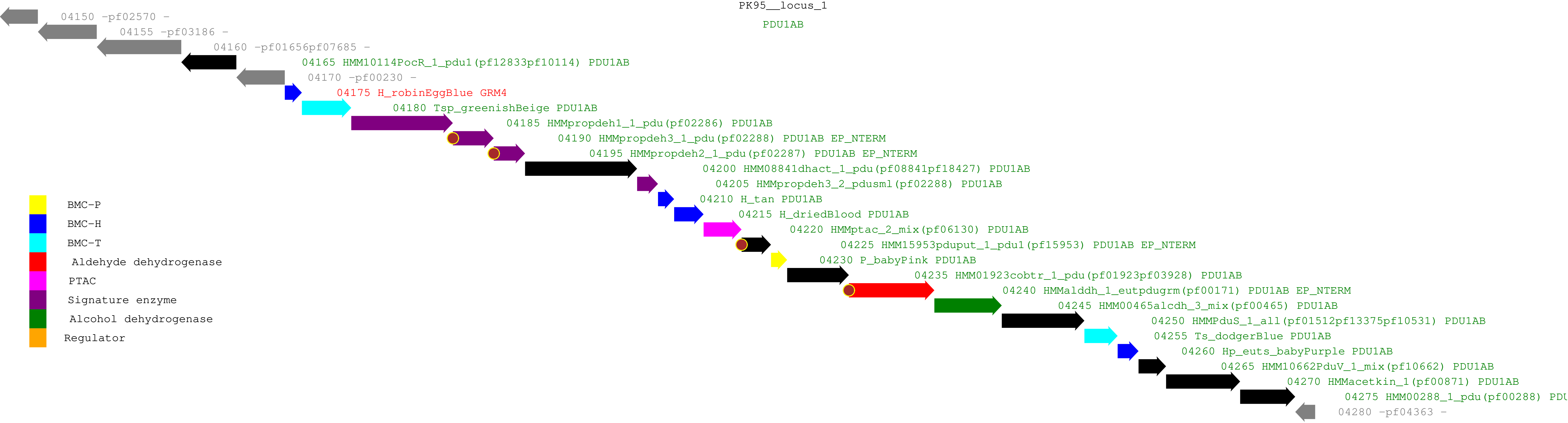

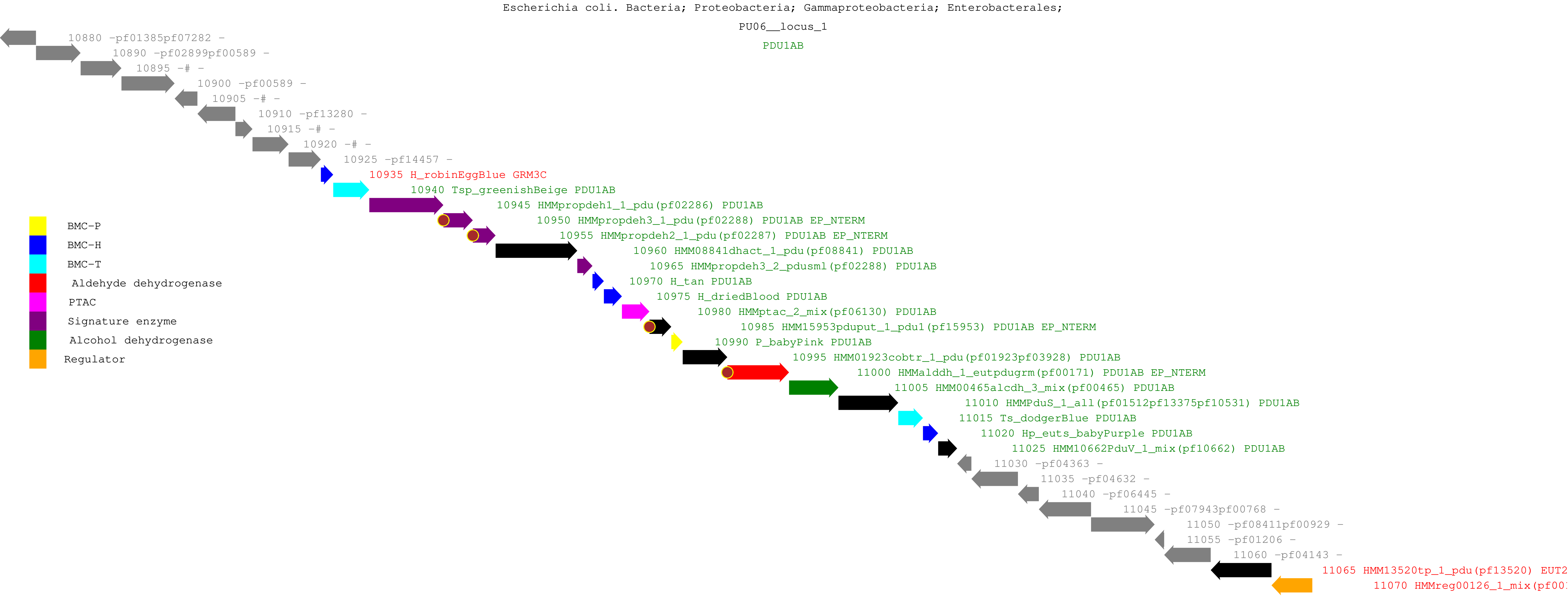

Escherichia coli. Bacteria; Proteobacteria; Gammaproteobacteria; Enterobacterales;
PU06__locus_1
PDU1AB
10880 -pf01385pf07282 -
10890 -pf02899pf00589 -
10895 -# -
10900 -pf00589 -
10905 -# -
10910 -pf13280 -
10915 -# -
10920 -# -
10925 -pf14457 -
10935 H_robinEggBlue GRM3C
10940 Tsp_greenishBeige PDU1AB
10945 HMMpropdeh1_1_pdu(pf02286) PDU1AB
10950 HMMpropdeh3_1_pdu(pf02288) PDU1AB EP_NTERM
10955 HMMpropdeh2_1_pdu(pf02287) PDU1AB EP_NTERM
10960 HMM08841dhact_1_pdu(pf08841) PDU1AB
10965 HMMpropdeh3_2_pdusml(pf02288) PDU1AB
10970 H_tan PDU1AB
10975 H_driedBlood PDU1AB
10980 HMMptac_2_mix(pf06130) PDU1AB
10985 HMM15953pduput_1_pdu1(pf15953) PDU1AB EP_NTERM
10990 P_babyPink PDU1AB
10995 HMM01923cobtr_1_pdu(pf01923pf03928) PDU1AB
11000 HMMalddh_1_eutpdugrm(pf00171) PDU1AB EP_NTERM
11005 HMM00465alcdh_3_mix(pf00465) PDU1AB
11010 HMMPduS_1_all(pf01512pf13375pf10531) PDU1AB
11015 Ts_dodgerBlue PDU1AB
11020 Hp_euts_babyPurple PDU1AB
11025 HMM10662PduV_1_mix(pf10662) PDU1AB
11030 -pf04363 -
11035 -pf04632 -
11040 -pf06445 -
11045 -pf07943pf00768 -
11050 -pf08411pf00929 -
11055 -pf01206 -
11060 -pf04143 -
11065 HMM13520tp_1_pdu(pf13520) EUT2
11070 HMMreg00126_1_mix(pf00
BMC-P
BMC-H
BMC-T
Aldehyde dehydrogenase
PTAC
Signature enzyme
Alcohol dehydrogenase
Regulator

Phytobacter ursingii. Bacteria; Proteobacteria; Gammaproteobacteria; Enterobacterales;

PUATCC27989T__locus_1

PDU1AB

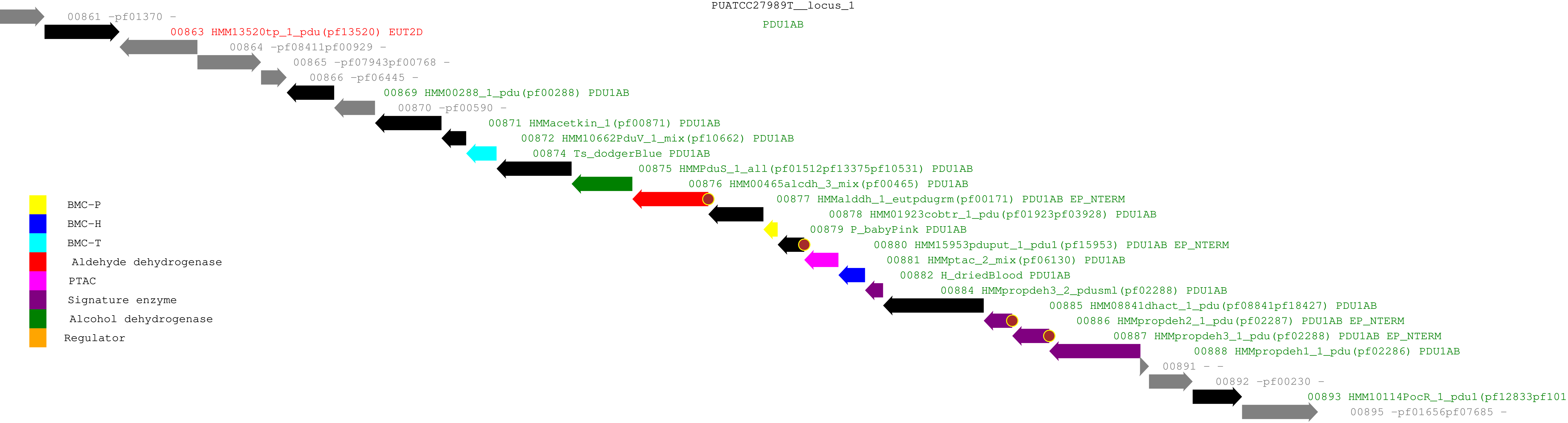

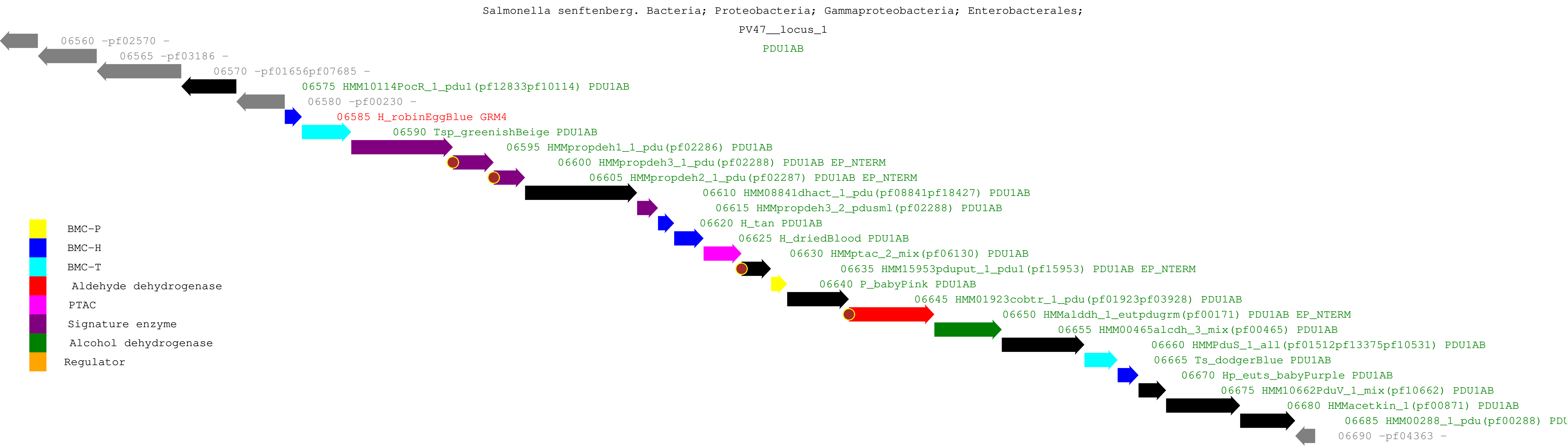

Salmonella senftenberg. Bacteria; Proteobacteria; Gammaproteobacteria; Enterobacterales;
PV47__locus_1
PDU1AB
06560 -pf02570 -
06565 -pf03186 -
06570 -pf01656pf07685 -
06575 HMM10114PocR_1_pdu1(pf12833pf10114) PDU1AB
06580 -pf00230 -
06585 H_robinEggBlue GRM4
06590 Tsp_greenishBeige PDU1AB
06595 HMMpropdeh1_1_pdu(pf02286) PDU1AB
06600 HMMpropdeh3_1_pdu(pf02288) PDU1AB EP_NTERM
06605 HMMpropdeh2_1_pdu(pf02287) PDU1AB EP_NTERM
06610 HMM08841dhact_1_pdu(pf08841pf18427) PDU1AB
06615 HMMpropdeh3_2_pdusml(pf02288) PDU1AB
06620 H_tan PDU1AB
06625 H_driedBlood PDU1AB
06630 HMMptac_2_mix(pf06130) PDU1AB
06635 HMM15953pduput_1_pdu1(pf15953) PDU1AB EP_NTERM
06640 P_babyPink PDU1AB
06645 HMM01923cobtr_1_pdu(pf01923pf03928) PDU1AB
06650 HMMalddh_1_eutpdugrm(pf00171) PDU1AB EP_NTERM
06655 HMM00465alcdh_3_mix(pf00465) PDU1AB
06660 HMMPduS_1_all(pf01512pf13375pf10531) PDU1AB
06665 Ts_dodgerBlue PDU1AB
06670 Hp_euts_babyPurple PDU1AB
06675 HMM10662PduV_1_mix(pf10662) PDU1AB
06680 HMMacetkin_1(pf00871) PDU1AB
06685 HMM00288_1_pdu(pf00288) PDU
06690 -pf04363 -
BMC-P
BMC-H
BMC-T
Aldehyde dehydrogenase
PTAC
Signature enzyme
Alcohol dehydrogenase
Regulator

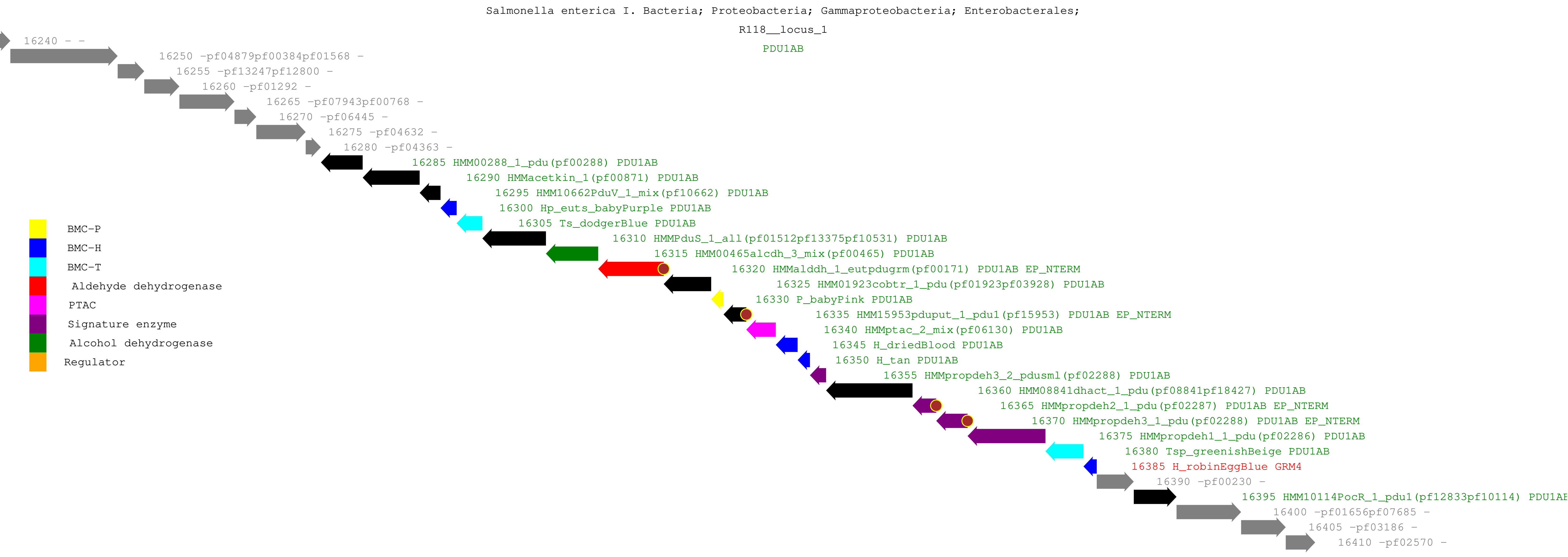
Salmonella enterica I. Bacteria; Proteobacteria; Gammaproteobacteria; Enterobacterales;
R118__locus_1
PDU1AB
16240 - -
16250 -pf04879pf00384pf01568 -
16255 -pf13247pf12800 -
16260 -pf01292 -
16265 -pf07943pf00768 -
16270 -pf06445 -
16275 -pf04632 -
16280 -pf04363 -
16285 HMM00288_1_pdu(pf00288) PDU1AB
16290 HMMacetkin_1(pf00871) PDU1AB
16295 HMM10662PduV_1_mix(pf10662) PDU1AB
16300 Hp_euts_babyPurple PDU1AB
16305 Ts_dodgerBlue PDU1AB
16310 HMMPduS_1_all(pf01512pf13375pf10531) PDU1AB
16315 HMM00465alcdh_3_mix(pf00465) PDU1AB
16320 HMMalddh_1_eutpdugrm(pf00171) PDU1AB EP_NTERM
16325 HMM01923cobtr_1_pdu(pf01923pf03928) PDU1AB
16330 P_babyPink PDU1AB
16335 HMM15953pduput_1_pdu1(pf15953) PDU1AB EP_NTERM
16340 HMMptac_2_mix(pf06130) PDU1AB
16345 H_driedBlood PDU1AB
16350 H_tan PDU1AB
16355 HMMpropdeh3_2_pdusml(pf02288) PDU1AB
16360 HMM08841dhact_1_pdu(pf08841pf18427) PDU1AB
16365 HMMpropdeh2_1_pdu(pf02287) PDU1AB EP_NTERM
16370 HMMpropdeh3_1_pdu(pf02288) PDU1AB EP_NTERM
16375 HMMpropdeh1_1_pdu(pf02286) PDU1AB
16380 Tsp_greenishBeige PDU1AB
16385 H_robinEggBlue GRM4
16390 -pf00230 -
16395 HMM10114PocR_1_pdu1(pf12833pf10114) PDU1AB
16400 -pf01656pf07685 -
16405 -pf03186 -
16410 -pf02570 -
BMC-P
BMC-H
BMC-T
Aldehyde dehydrogenase
PTAC
Signature enzyme
Alcohol dehydrogenase
Regulator

Salmonella enterica subsp. arizonae serovar 41:z4,z23:-. Bacteria; Proteobacteria; Gammaproteobacteria; Enterobacterales;

R515__locus_2

PDU1AB

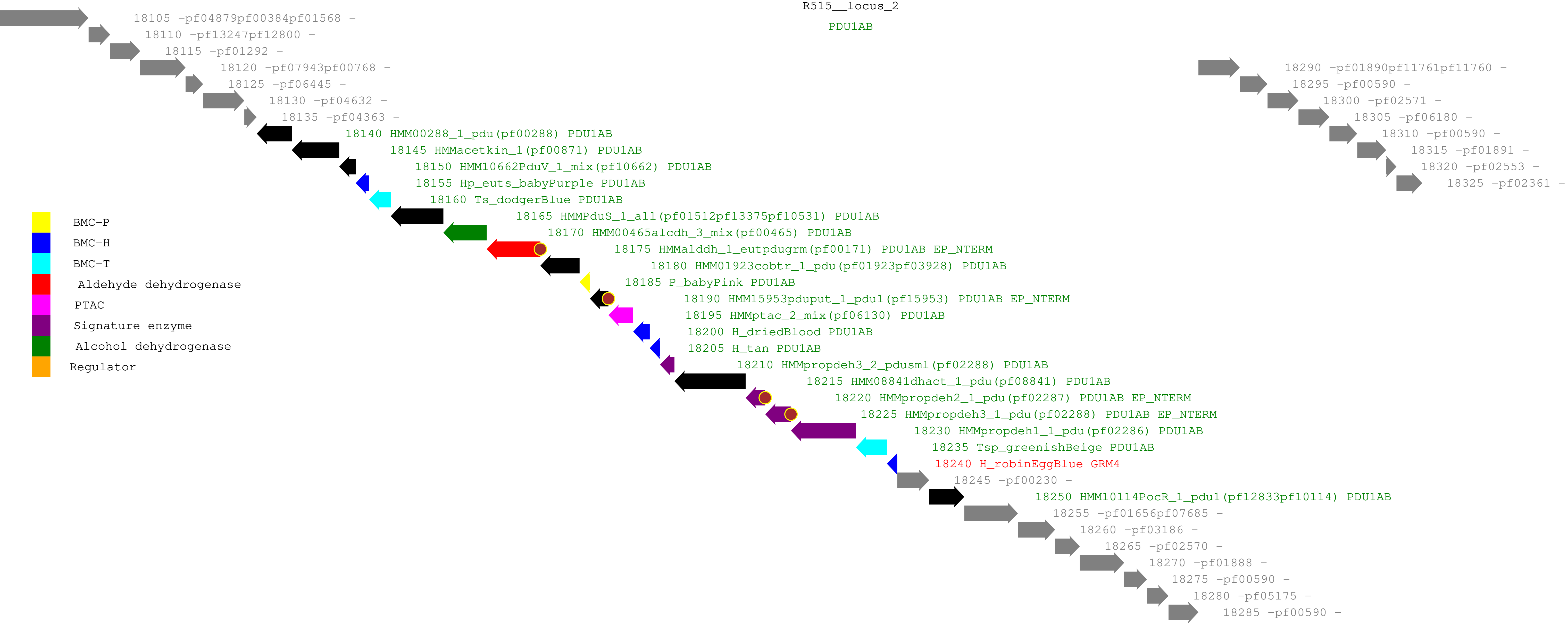

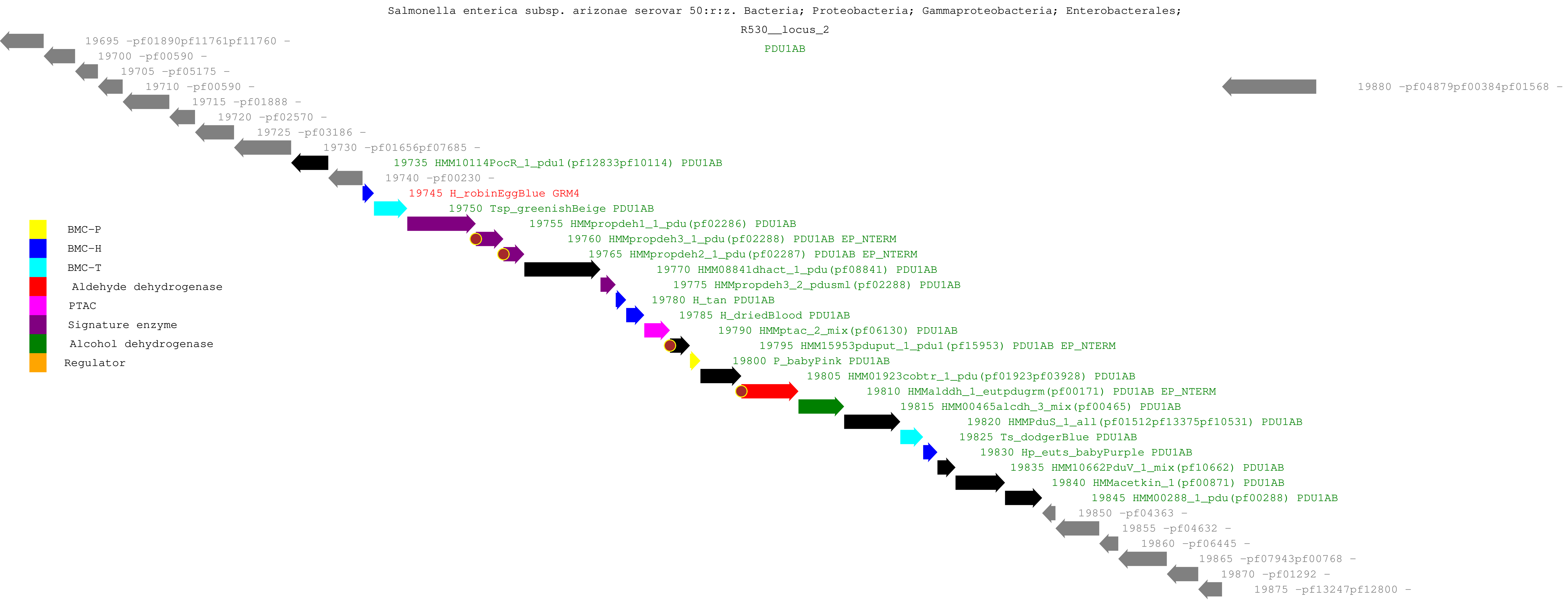
Salmonella enterica subsp. arizonae serovar 50:r:z. Bacteria; Proteobacteria; Gammaproteobacteria; Enterobacterales;
R530__locus_2
PDU1AB
19695 -pf01890pf11761pf11760 -
19700 -pf00590 -
19705 -pf05175 -
19710 -pf00590 -
19715 -pf01888 -
19720 -pf02570 -
19725 -pf03186 -
19730 -pf01656pf07685 -
19735 HMM10114PocR_1_pdu1(pf12833pf10114) PDU1AB
19740 -pf00230 -
19745 H_robinEggBlue GRM4
19750 Tsp_greenishBeige PDU1AB
19755 HMMpropdeh1_1_pdu(pf02286) PDU1AB
19760 HMMpropdeh3_1_pdu(pf02288) PDU1AB EP_NTERM
19765 HMMpropdeh2_1_pdu(pf02287) PDU1AB EP_NTERM
19770 HMM08841dhact_1_pdu(pf08841) PDU1AB
19775 HMMpropdeh3_2_pdusml(pf02288) PDU1AB
19780 H_tan PDU1AB
19785 H_driedBlood PDU1AB
19790 HMMptac_2_mix(pf06130) PDU1AB
19795 HMM15953pduput_1_pdu1(pf15953) PDU1AB EP_NTERM
19800 P_babyPink PDU1AB
19805 HMM01923cobtr_1_pdu(pf01923pf03928) PDU1AB
19810 HMMalddh_1_eutpdugrm(pf00171) PDU1AB EP_NTERM
19815 HMM00465alcdh_3_mix(pf00465) PDU1AB
19820 HMMPduS_1_all(pf01512pf13375pf10531) PDU1AB
19825 Ts_dodgerBlue PDU1AB
19830 Hp_euts_babyPurple PDU1AB
19835 HMM10662PduV_1_mix(pf10662) PDU1AB
19840 HMMacetkin_1(pf00871) PDU1AB
19845 HMM00288_1_pdu(pf00288) PDU1AB
19850 -pf04363 -
19855 -pf04632 -
19860 -pf06445 -
19865 -pf07943pf00768 -
19870 -pf01292 -
19875 -pf13247pf12800 -
19880 -pf04879pf00384pf01568 -
BMC-P
BMC-H
BMC-T
Aldehyde dehydrogenase
PTAC
Signature enzyme
Alcohol dehydrogenase
Regulator

Salmonella enterica subsp. enterica serovar Rough O:d:1,7. Bacteria; Proteobacteria; Gammaproteobacteria; Enterobacterales;

R537__locus_1

PDU1AB

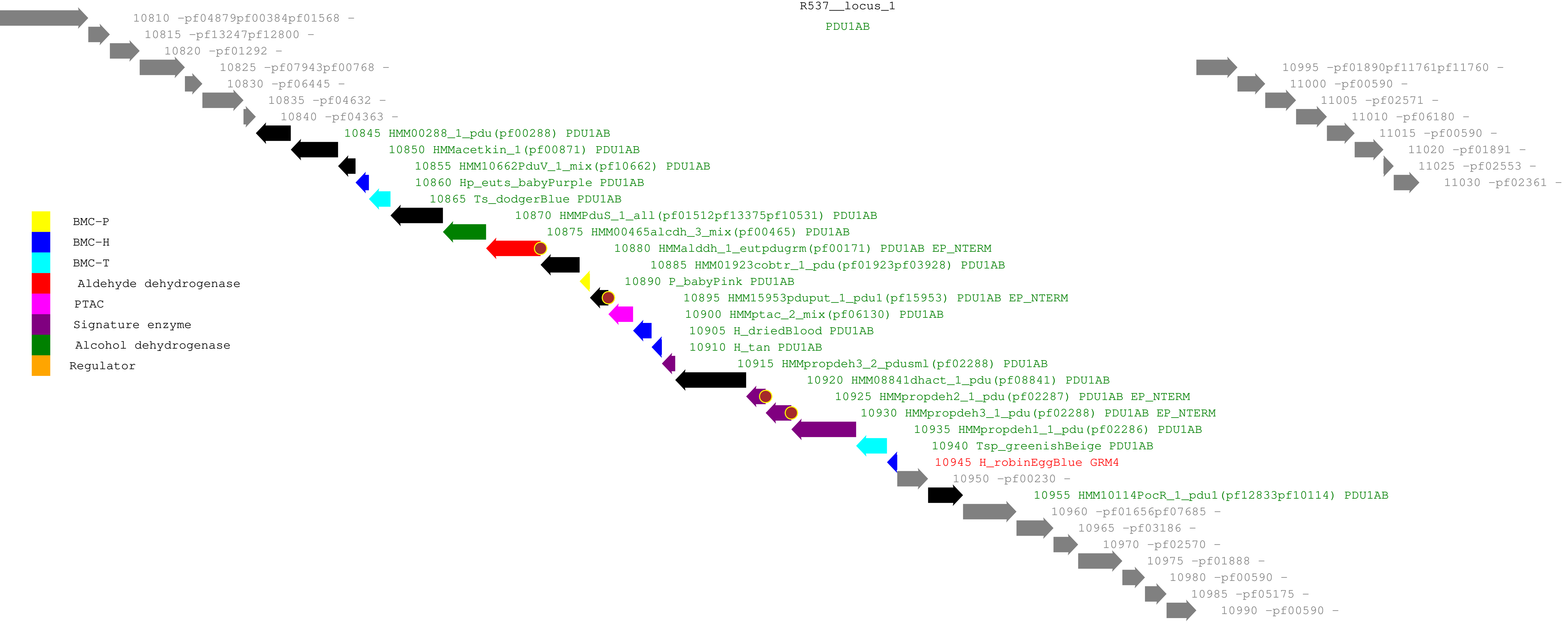

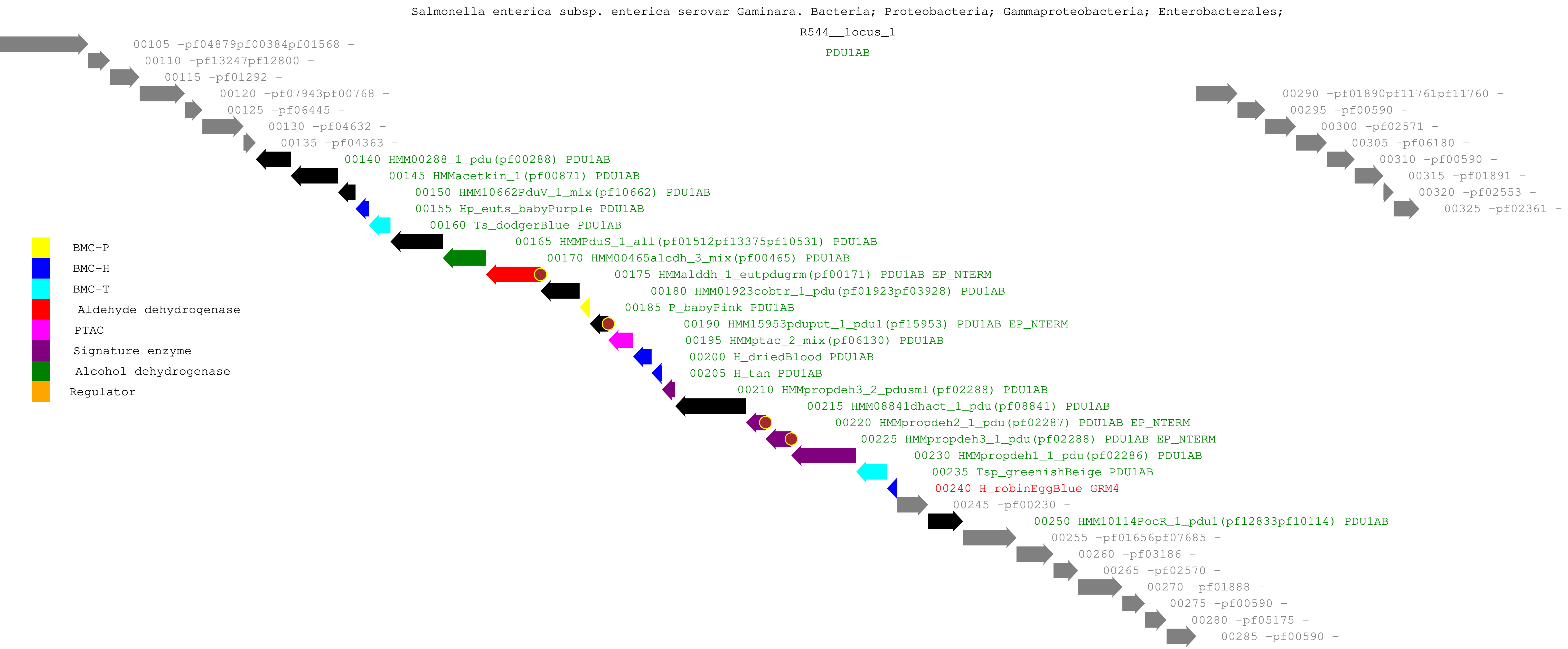

Salmonella enterica subsp. enterica serovar Gaminara. Bacteria; Proteobacteria; Gammaproteobacteria; Enterobacterales;
R544__locus_1
PDU1AB
00105 -pf04879pf00384pf01568 -
00110 -pf13247pf12800 -
00115 -pf01292 -
00120 -pf07943pf00768 -
00125 -pf06445 -
00130 -pf04632 -
00135 -pf04363 -
00140 HMM00288_1_pdu(pf00288) PDU1AB
00145 HMMacetkin_1(pf00871) PDU1AB
00150 HMM10662PduV_1_mix(pf10662) PDU1AB
00155 Hp_euts_babyPurple PDU1AB
00160 Ts_dodgerBlue PDU1AB
00165 HMMPduS_1_all(pf01512pf13375pf10531) PDU1AB
00170 HMM00465alcdh_3_mix(pf00465) PDU1AB
00175 HMMalddh_1_eutpdugrm(pf00171) PDU1AB EP_NTERM
00180 HMM01923cobtr_1_pdu(pf01923pf03928) PDU1AB
00185 P_babyPink PDU1AB
00190 HMM15953pduput_1_pdu1(pf15953) PDU1AB EP_NTERM
00195 HMMptac_2_mix(pf06130) PDU1AB
00200 H_driedBlood PDU1AB
00205 H_tan PDU1AB
00210 HMMpropdeh3_2_pdusml(pf02288) PDU1AB
00215 HMM08841dhact_1_pdu(pf08841) PDU1AB
00220 HMMpropdeh2_1_pdu(pf02287) PDU1AB EP_NTERM
00225 HMMpropdeh3_1_pdu(pf02288) PDU1AB EP_NTERM
00230 HMMpropdeh1_1_pdu(pf02286) PDU1AB
00235 Tsp_greenishBeige PDU1AB
00240 H_robinEggBlue GRM4
00245 -pf00230 -
00250 HMM10114PocR_1_pdu1(pf12833pf10114) PDU1AB
00255 -pf01656pf07685 -
00260 -pf03186 -
00265 -pf02570 -
00270 -pf01888 -
00275 -pf00590 -
00280 -pf05175 -
00285 -pf00590 -
00290 -pf01890pf11761pf11760 -
00295 -pf00590 -
00300 -pf02571 -
00305 -pf06180 -
00310 -pf00590 -
00315 -pf01891 -
00320 -pf02553 -
00325 -pf02361 -
BMC-P
BMC-H
BMC-T
Aldehyde dehydrogenase
PTAC
Signature enzyme
Alcohol dehydrogenase
Regulator

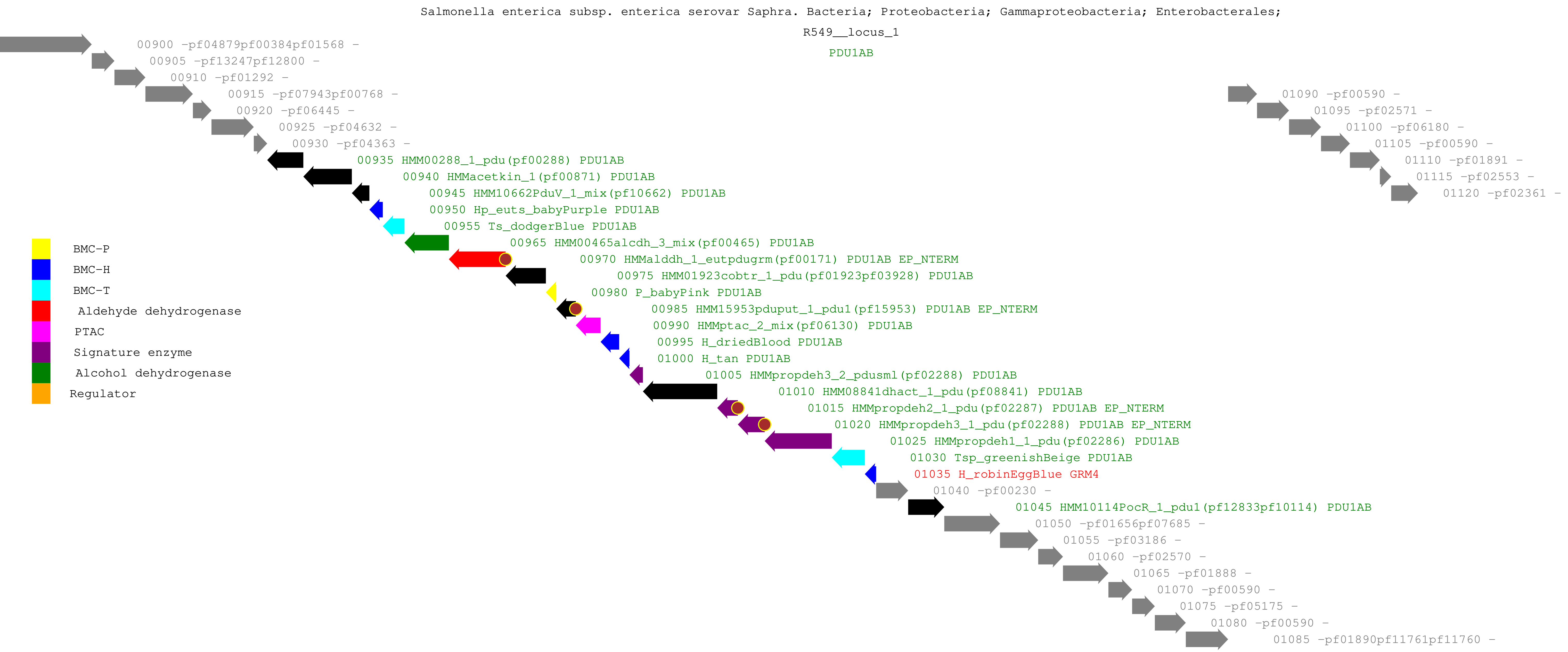
Salmonella enterica subsp. enterica serovar Saphra. Bacteria; Proteobacteria; Gammaproteobacteria; Enterobacterales;
R549__locus_1
PDU1AB
00900 -pf04879pf00384pf01568 -
00905 -pf13247pf12800 -
00910 -pf01292 -
00915 -pf07943pf00768 -
00920 -pf06445 -
00925 -pf04632 -
00930 -pf04363 -
00935 HMM00288_1_pdu(pf00288) PDU1AB
00940 HMMacetkin_1(pf00871) PDU1AB
00945 HMM10662PduV_1_mix(pf10662) PDU1AB
00950 Hp_euts_babyPurple PDU1AB
00955 Ts_dodgerBlue PDU1AB
00965 HMM00465alcdh_3_mix(pf00465) PDU1AB
00970 HMMaldh_1_eutpdugrm(pf00171) PDU1AB EP_NTERM
00975 HMM01923cobtr_1_pdu(pf01923pf03928) PDU1AB
00980 P_babyPink PDU1AB
00985 HMM15953pduput_1_pdu1(pf15953) PDU1AB EP_NTERM
00990 HMMptac_2_mix(pf06130) PDU1AB
00995 H_driedBlood PDU1AB
01000 H_tan PDU1AB
01005 HMMpropdeh3_2_pdusml(pf02288) PDU1AB
01010 HMM08841dhact_1_pdu(pf08841) PDU1AB
01015 HMMpropdeh2_1_pdu(pf02287) PDU1AB EP_NTERM
01020 HMMpropdeh3_1_pdu(pf02288) PDU1AB EP_NTERM
01025 HMMpropdeh1_1_pdu(pf02286) PDU1AB
01030 Tsp_greenishBeige PDU1AB
01035 H_robinEggBlue GRM4
01040 -pf00230 -
01045 HMM10114PocR_1_pdu1(pf12833pf10114) PDU1AB
01050 -pf01656pf07685 -
01055 -pf03186 -
01060 -pf02570 -
01065 -pf01888 -
01070 -pf00590 -
01075 -pf05175 -
01080 -pf00590 -
01085 -pf01890pf11761pf11760 -
01090 -pf00590 -
01095 -pf02571 -
01100 -pf06180 -
01105 -pf00590 -
01110 -pf01891 -
01115 -pf02553 -
01120 -pf02361 -
BMC-P
BMC-H
BMC-T
Aldehyde dehydrogenase
PTAC
Signature enzyme
Alcohol dehydrogenase
Regulator

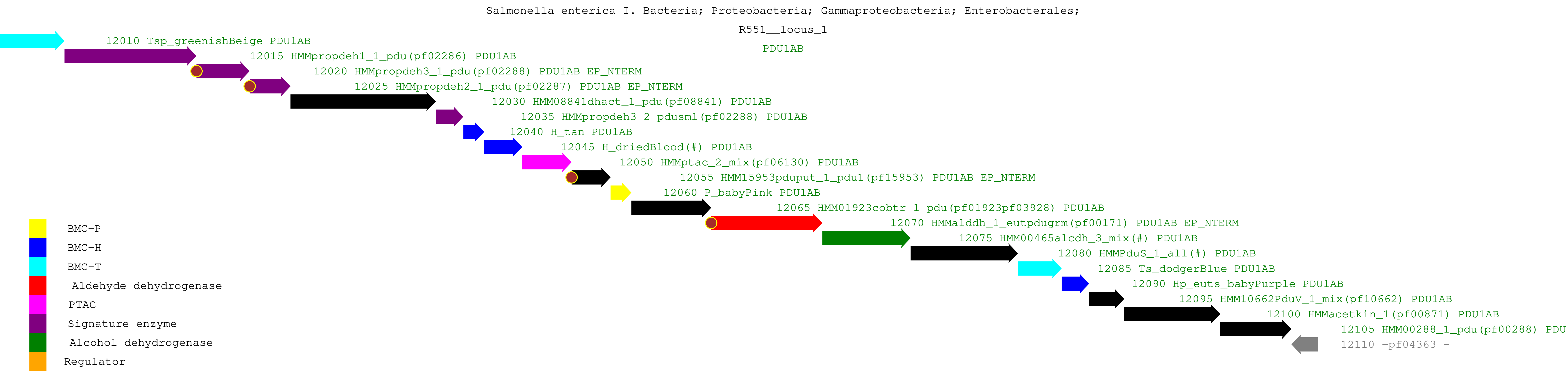
Salmonella enterica I. Bacteria; Proteobacteria; Gammaproteobacteria; Enterobacterales;
R551__locus_1
PDU1AB
12010 Tsp_greenishBeige PDU1AB
12015 HMMpropdeh1_1_pdu(pf02286) PDU1AB
12020 HMMpropdeh3_1_pdu(pf02288) PDU1AB EP_NTERM
12025 HMMpropdeh2_1_pdu(pf02287) PDU1AB EP_NTERM
12030 HMM08841dhact_1_pdu(pf08841) PDU1AB
12035 HMMpropdeh3_2_pdusml(pf02288) PDU1AB
12040 H_tan PDU1AB
12045 H_driedBlood(#) PDU1AB
12050 HMMptac_2_mix(pf06130) PDU1AB
12055 HMM15953pduput_1_pdu1(pf15953) PDU1AB EP_NTERM
12060 P_babyPink PDU1AB
12065 HMM01923cobtr_1_pdu(pf01923pf03928) PDU1AB
12070 HMMalddh_1_eutpdugrm(pf00171) PDU1AB EP_NTERM
12075 HMM00465alcdh_3_mix(#) PDU1AB
12080 HMMPduS_1_all(#) PDU1AB
12085 Ts_dodgerBlue PDU1AB
12090 Hp_euts_babyPurple PDU1AB
12095 HMM10662PduV_1_mix(pf10662) PDU1AB
12100 HMMacetkin_1(pf00871) PDU1AB
12105 HMM00288_1_pdu(pf00288) PDU
12110 -pf04363 -
BMC-P
BMC-H
BMC-T
Aldehyde dehydrogenase
PTAC
Signature enzyme
Alcohol dehydrogenase
Regulator

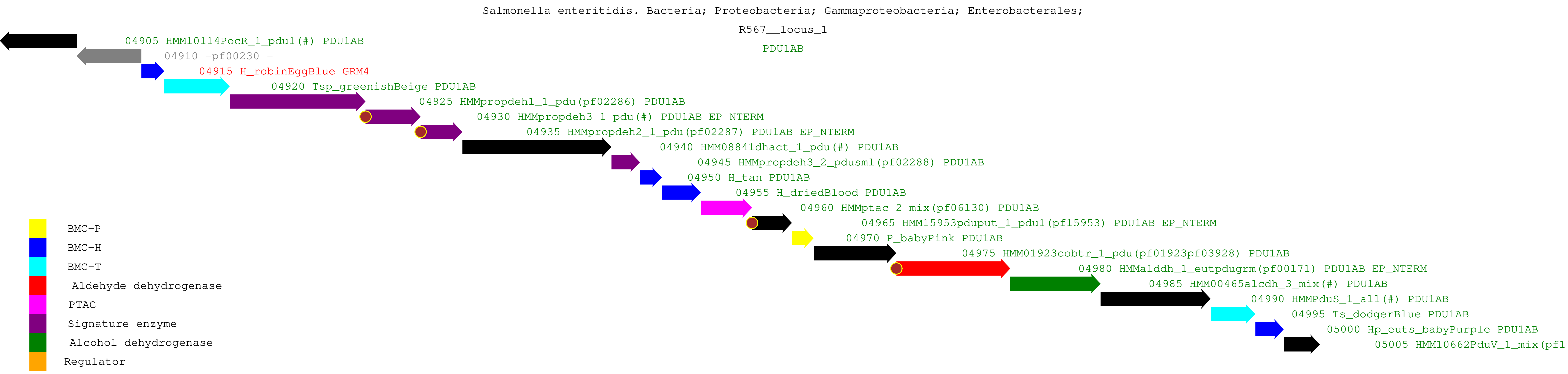
Salmonella enteritidis. Bacteria; Proteobacteria; Gammaproteobacteria; Enterobacterales;
R567__locus_1
PDU1AB
04905 HMM10114PocR_1_pdu1(#) PDU1AB
04910 -pf00230 -
04915 H_robinEggBlue GRM4
04920 Tsp_greenishBeige PDU1AB
04925 HMMpropdeh1_1_pdu(pf02286) PDU1AB
04930 HMMpropdeh3_1_pdu(#) PDU1AB EP_NTERM
04935 HMMpropdeh2_1_pdu(pf02287) PDU1AB EP_NTERM
04940 HMM08841dhact_1_pdu(#) PDU1AB
04945 HMMpropdeh3_2_pdusml(pf02288) PDU1AB
04950 H_tan PDU1AB
04955 H_driedBlood PDU1AB
04960 HMMptac_2_mix(pf06130) PDU1AB
04965 HMM15953pduput_1_pdu1(pf15953) PDU1AB EP_NTERM
04970 P_babyPink PDU1AB
04975 HMM01923cobtr_1_pdu(pf01923pf03928) PDU1AB
04980 HMMalddh_1_eutpdugrm(pf00171) PDU1AB EP_NTERM
04985 HMM00465alcdh_3_mix(#) PDU1AB
04990 HMMPduS_1_all(#) PDU1AB
04995 Ts_dodgerBlue PDU1AB
05000 Hp_euts_babyPurple PDU1AB
05005 HMM10662PduV_1_mix(pf1
BMC-P
BMC-H
BMC-T
Aldehyde dehydrogenase
PTAC
Signature enzyme
Alcohol dehydrogenase
Regulator

Salmonella enterica subsp. enterica serovar Seftenburg. Bacteria; Proteobacteria; Gammaproteobacteria; Enterobacterales;

R583__locus_1

PDU1AB

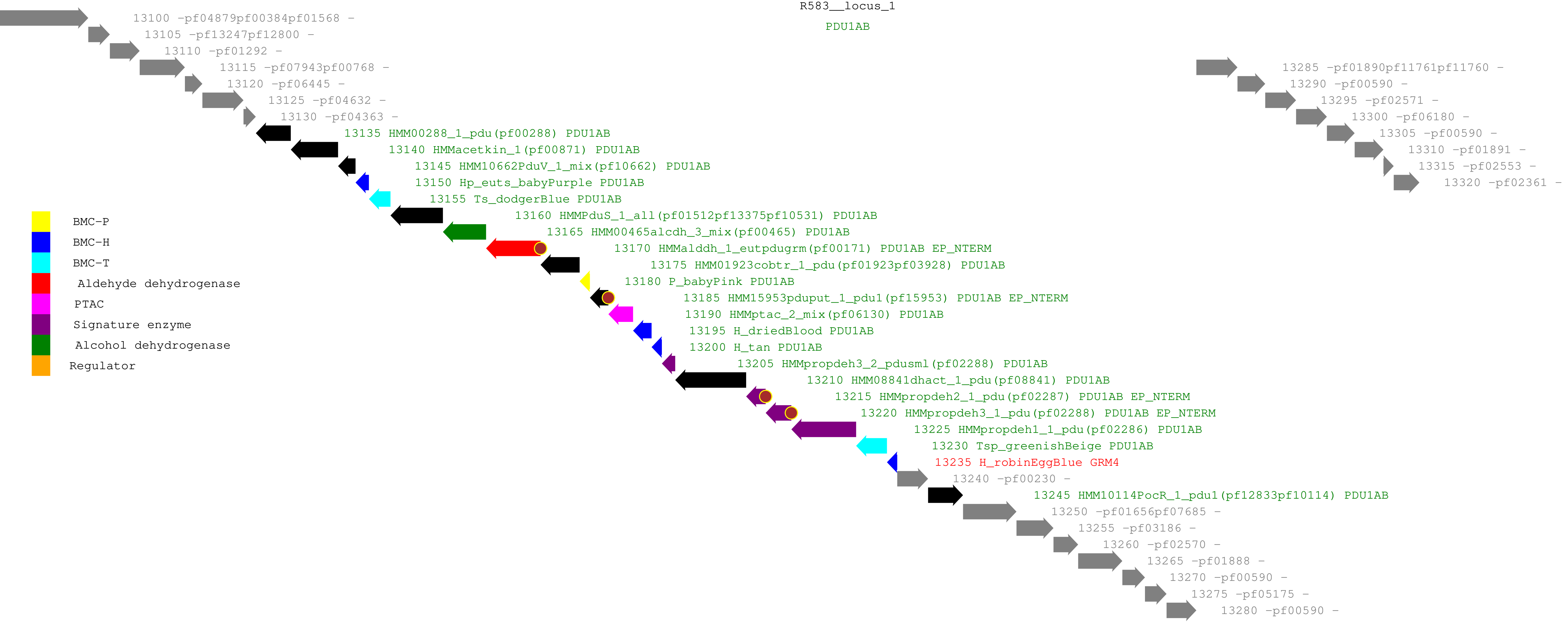

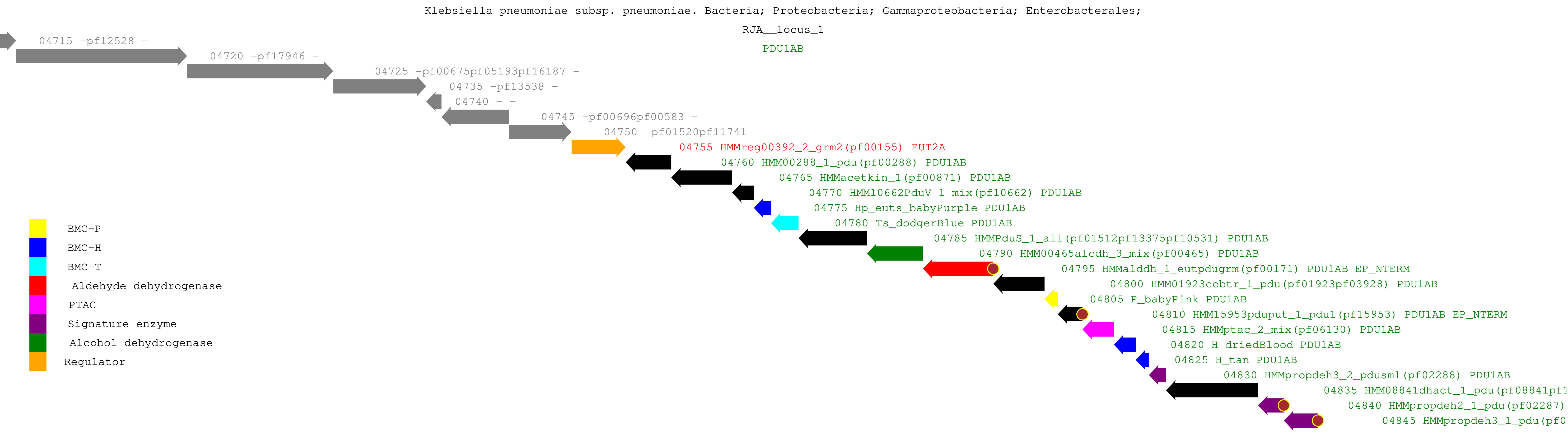

Klebsiella pneumoniae subsp. pneumoniae. Bacteria; Proteobacteria; Gammaproteobacteria; Enterobacterales;
RJA__locus_1
PDU1AB
04715 -pf12528 -
04720 -pf17946 -
04725 -pf00675pf05193pf16187 -
04735 -pf13538 -
04740 - -
04745 -pf00696pf00583 -
04750 -pf01520pf11741 -
04755 HMMreg00392_2_grm2(pf00155) EUT2A
04760 HMM00288_1_pdu(pf00288) PDU1AB
04765 HMMacetkin_1(pf00871) PDU1AB
04770 HMM10662PduV_1_mix(pf10662) PDU1AB
04775 Hp_euts_babyPurple PDU1AB
04780 Ts_dodgerBlue PDU1AB
04785 HMMPduS_1_all(pf01512pf13375pf10531) PDU1AB
04790 HMM00465alcdh_3_mix(pf00465) PDU1AB
04795 HMMalddh_1_eutpdugrm(pf00171) PDU1AB EP_NTERM
04800 HMM01923cobtr_1_pdu(pf01923pf03928) PDU1AB
04805 P_babyPink PDU1AB
04810 HMM15953pduput_1_pdu1(pf15953) PDU1AB EP_NTERM
04815 HMMptac_2_mix(pf06130) PDU1AB
04820 H_driedBlood PDU1AB
04825 H_tan PDU1AB
04830 HMMpropdeh3_2_pdusml(pf02288) PDU1AB
04835 HMM08841dhact_1_pdu(pf08841pf1
04840 HMMpropdeh2_1_pdu(pf02287)
04845 HMMpropdeh3_1_pdu(pf0
BMC-P
BMC-H
BMC-T
Aldehyde dehydrogenase
PTAC
Signature enzyme
Alcohol dehydrogenase
Regulator

Salmonella enterica subsp. enterica serovar Paratyphi B. Bacteria; Proteobacteria; Gammaproteobacteria; Enterobacterales;

RL96__locus_2

PDU1AB

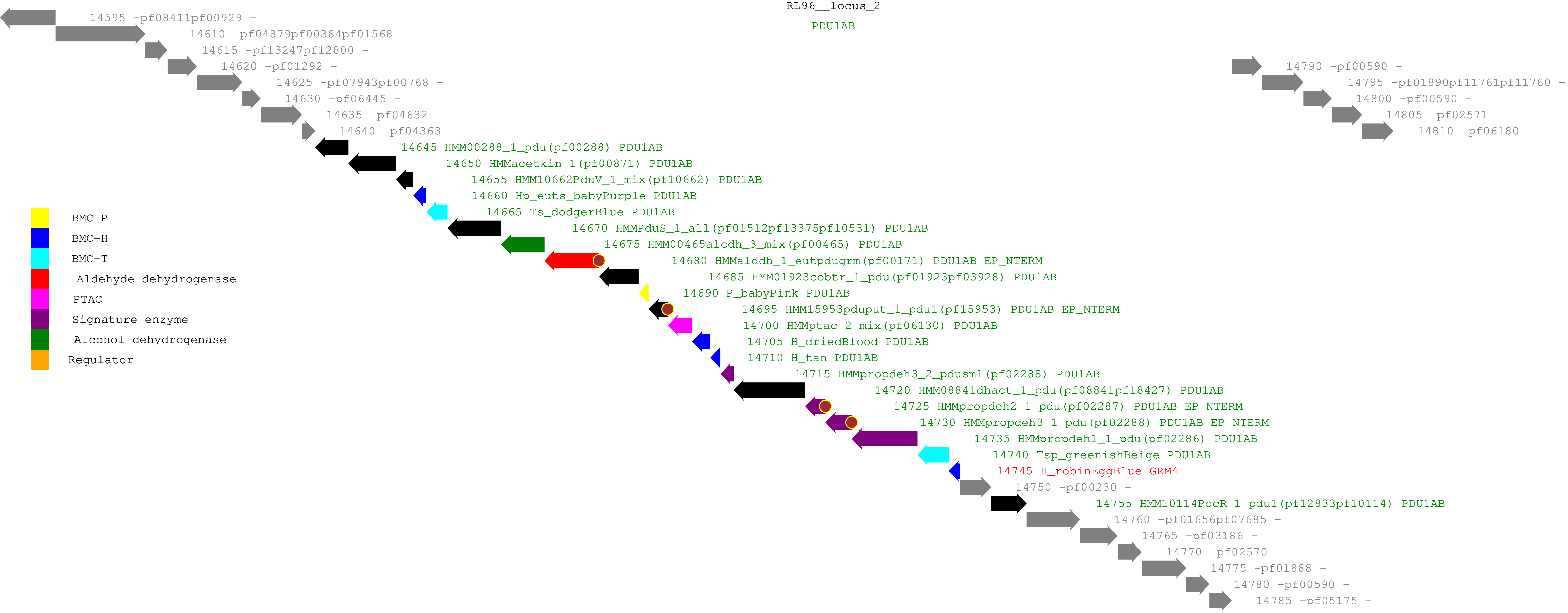

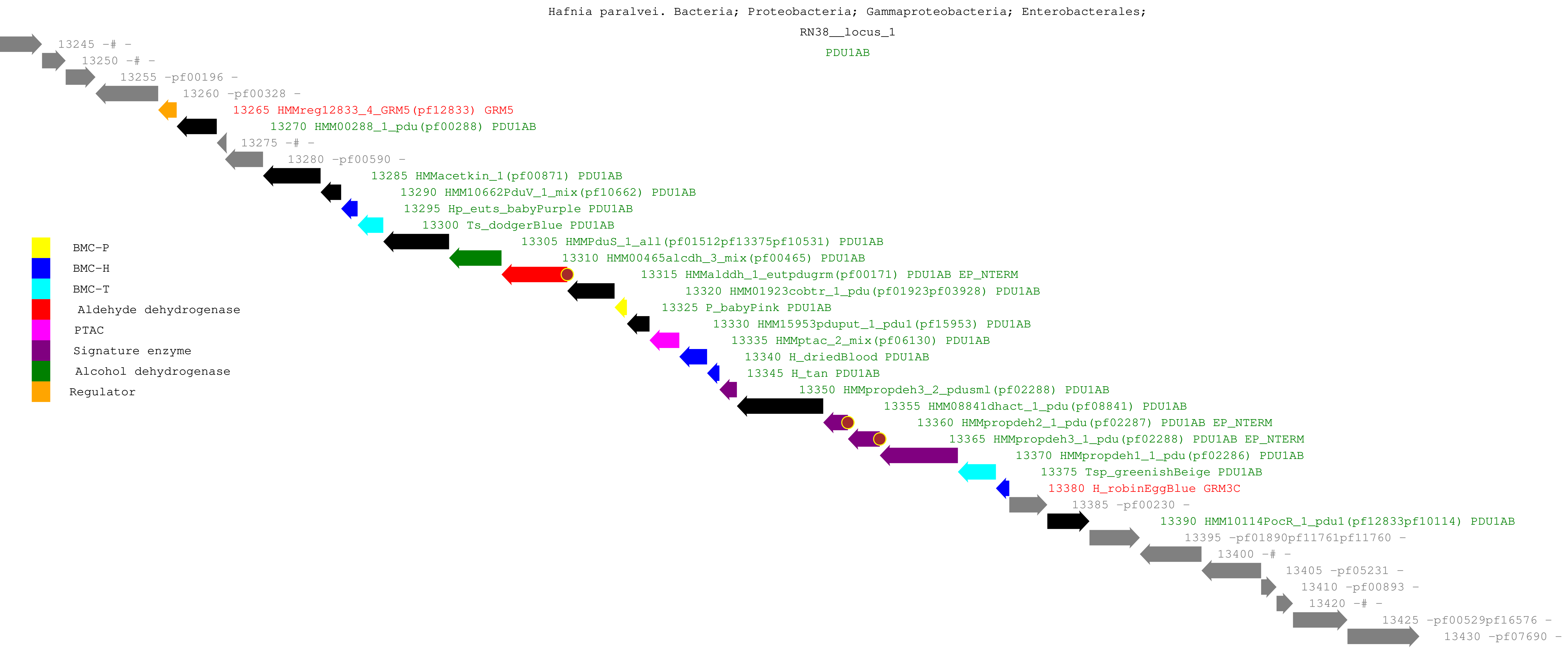
Hafnia paralvei. Bacteria; Proteobacteria; Gammaproteobacteria; Enterobacterales;
RN38__locus_1
PDU1AB
13245 -# -
13250 -# -
13255 -pf00196 -
13260 -pf00328 -
13265 HMMreg12833_4_GRM5(pf12833) GRM5
13270 HMM00288_1_pdu(pf00288) PDU1AB
13275 -# -
13280 -pf00590 -
13285 HMMacetkin_1(pf00871) PDU1AB
13290 HMM10662PduV_1_mix(pf10662) PDU1AB
13295 Hp_euts_babyPurple PDU1AB
13300 Ts_dodgerBlue PDU1AB
13305 HMMPduS_1_all(pf01512pf13375pf10531) PDU1AB
13310 HMM00465alcdh_3_mix(pf00465) PDU1AB
13315 HMMaldDh_1_eutpdugrm(pf00171) PDU1AB EP_NTERM
13320 HMM01923cobtr_1_pdu(pf01923pf03928) PDU1AB
13325 P_babyPink PDU1AB
13330 HMM15953pduput_1_pdu1(pf15953) PDU1AB
13335 HMMptac_2_mix(pf06130) PDU1AB
13340 H_driedBlood PDU1AB
13345 H_tan PDU1AB
13350 HMMpropdeh3_2_pdusml(pf02288) PDU1AB
13355 HMM08841dhact_1_pdu(pf08841) PDU1AB
13360 HMMpropdeh2_1_pdu(pf02287) PDU1AB EP_NTERM
13365 HMMpropdeh3_1_pdu(pf02288) PDU1AB EP_NTERM
13370 HMMpropdeh1_1_pdu(pf02286) PDU1AB
13375 Tsp_greenishBeige PDU1AB
13380 H_robinEggBlue GRM3C
13385 -pf00230 -
13390 HMM10114PocR_1_pdu1(pf12833pf10114) PDU1AB
13395 -pf01890pf11761pf11760 -
13400 -# -
13405 -pf05231 -
13410 -pf00893 -
13420 -# -
13425 -pf00529pf16576 -
13430 -pf07690 -
BMC-P
BMC-H
BMC-T
Aldehyde dehydrogenase
PTAC
Signature enzyme
Alcohol dehydrogenase
Regulator

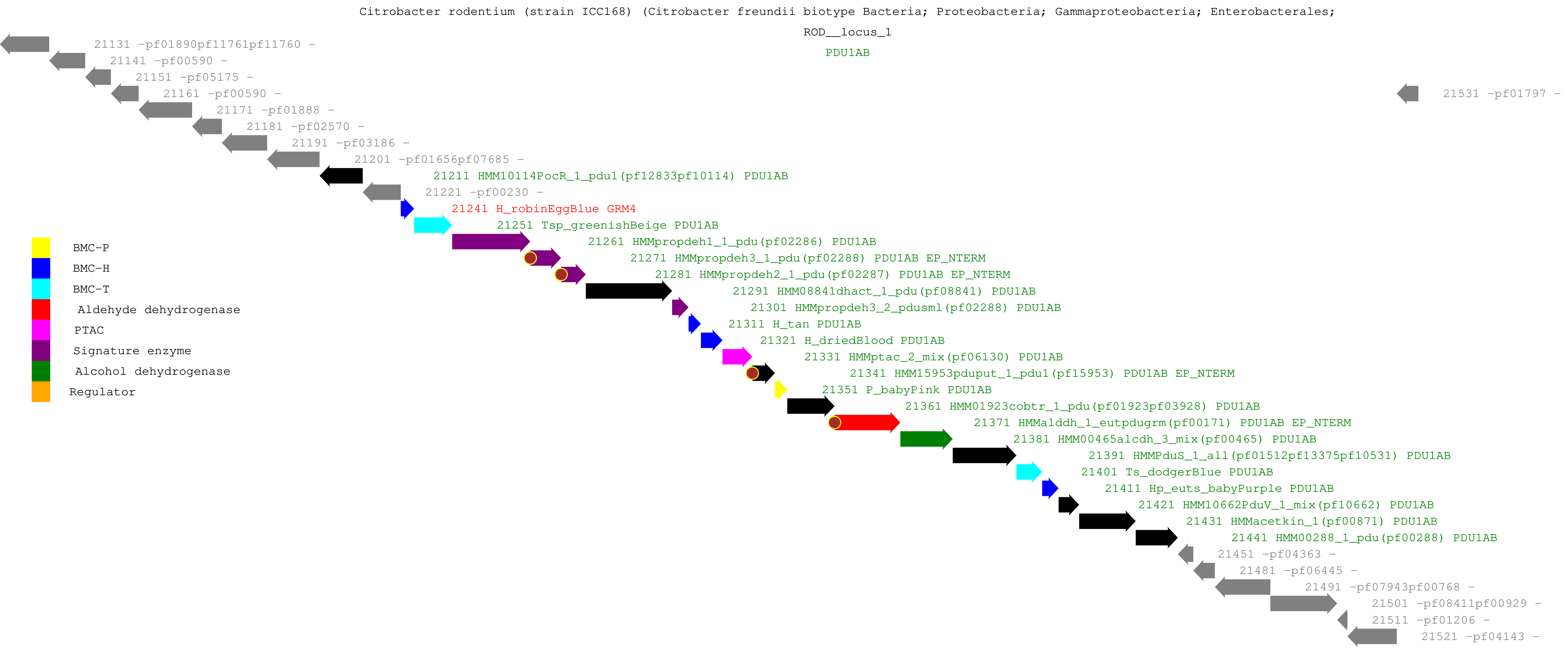
Citrobacter rodentium (strain ICC168) (Citrobacter freundii biotype Bacteria; Proteobacteria; Gammaproteobacteria; Enterobacterales;
ROD__locus_1
PDU1AB
21131 -pf01890pf11761pf11760 -
21141 -pf00590 -
21151 -pf05175 -
21161 -pf00590 -
21171 -pf01888 -
21181 -pf02570 -
21191 -pf03186 -
21201 -pf01656pf07685 -
21211 HMM10114PocR_1_pdu1(pf12833pf10114) PDU1AB
21221 -pf00230 -
21241 H_robinEggBlue GRM4
21251 Tsp_greenishBeige PDU1AB
21261 HMMpropdeh1_1_pdu(pf02286) PDU1AB
21271 HMMpropdeh3_1_pdu(pf02288) PDU1AB EP_NTERM
21281 HMMpropdeh2_1_pdu(pf02287) PDU1AB EP_NTERM
21291 HMM08841dhact_1_pdu(pf08841) PDU1AB
21301 HMMpropdeh3_2_pdusml(pf02288) PDU1AB
21311 H_tan PDU1AB
21321 H_driedBlood PDU1AB
21331 HMMptac_2_mix(pf06130) PDU1AB
21341 HMM15953pduput_1_pdu1(pf15953) PDU1AB EP_NTERM
21351 P_babyPink PDU1AB
21361 HMM01923cobtr_1_pdu(pf01923pf03928) PDU1AB
21371 HMMalddh_1_eutpdugrm(pf00171) PDU1AB EP_NTERM
21381 HMM00465alcdh_3_mix(pf00465) PDU1AB
21391 HMMPduS_1_all(pf01512pf13375pf10531) PDU1AB
21401 Ts_dodgerBlue PDU1AB
21411 Hp_euts_babyPurple PDU1AB
21421 HMM10662PduV_1_mix(pf10662) PDU1AB
21431 HMMacetkin_1(pf00871) PDU1AB
21441 HMM00288_1_pdu(pf00288) PDU1AB
21451 -pf04363 -
21481 -pf06445 -
21491 -pf07943pf00768 -
21501 -pf08411pf00929 -
21511 -pf01206 -
21521 -pf04143 -
21531 -pf01797 -
BMC-P
BMC-H
BMC-T
Aldehyde dehydrogenase
PTAC
Signature enzyme
Alcohol dehydrogenase
Regulator

Salmonella enterica I. Bacteria; Proteobacteria; Gammaproteobacteria; Enterobacterales;

RU17__locus_1

PDU1AB

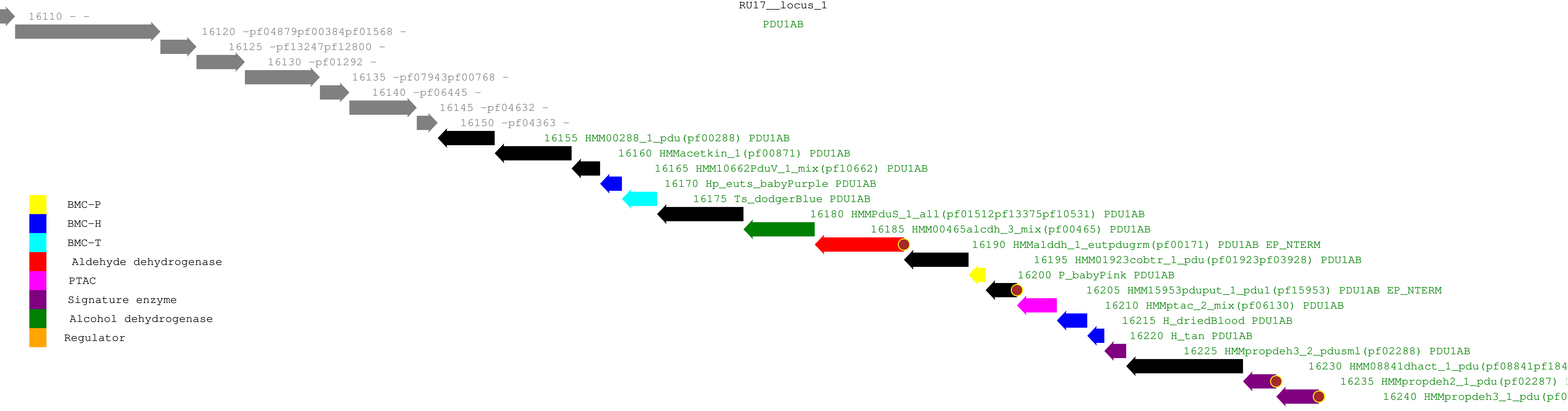

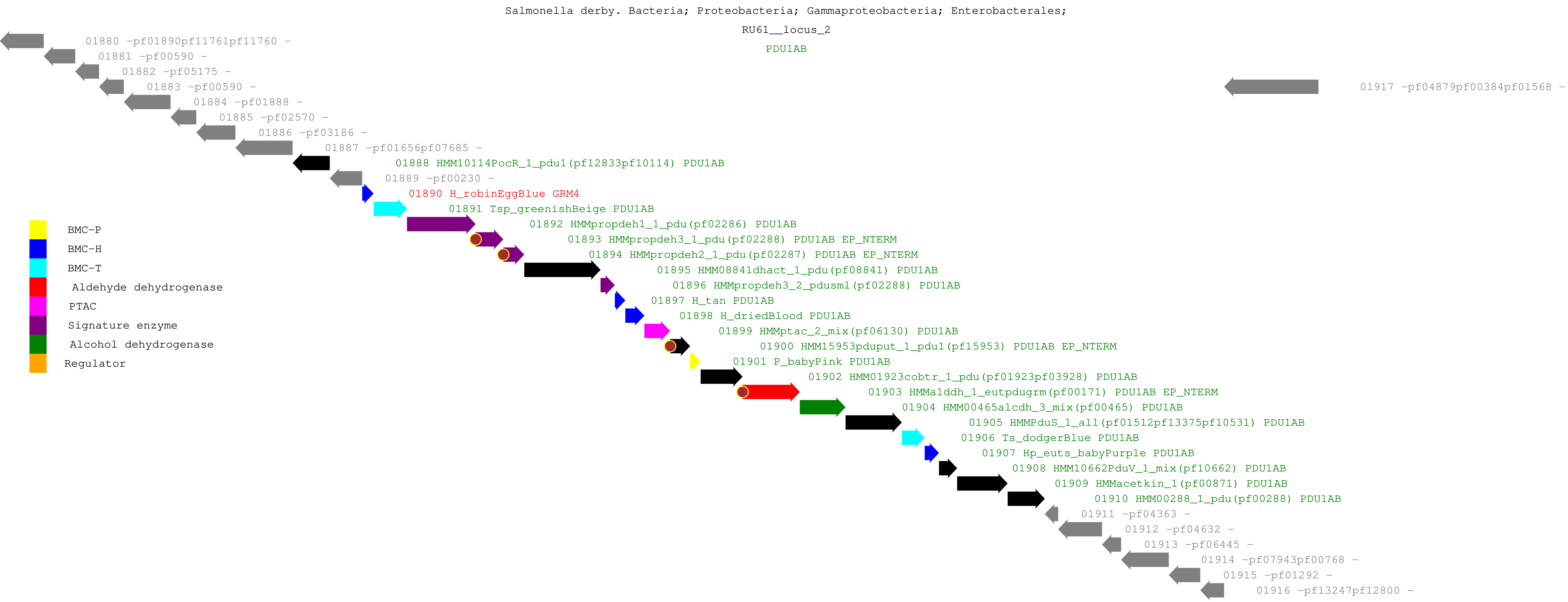
Salmonella derby. Bacteria; Proteobacteria; Gammaproteobacteria; Enterobacterales;
RU61__locus_2
PDU1AB
01880 -pf01890pf11761pf11760 -
01881 -pf00590 -
01882 -pf05175 -
01883 -pf00590 -
01884 -pf01888 -
01885 -pf02570 -
01886 -pf03186 -
01887 -pf01656pf07685 -
01888 HMM10114PocR_1_pdu1(pf12833pf10114) PDU1AB
01889 -pf00230 -
01890 H_robinEggBlue GRM4
01891 Tsp_greenishBeige PDU1AB
01892 HMMpropdeh1_1_pdu(pf02286) PDU1AB
01893 HMMpropdeh3_1_pdu(pf02288) PDU1AB EP_NTERM
01894 HMMpropdeh2_1_pdu(pf02287) PDU1AB EP_NTERM
01895 HMM08841dhact_1_pdu(pf08841) PDU1AB
01896 HMMpropdeh3_2_pdusml(pf02288) PDU1AB
01897 H_tan PDU1AB
01898 H_driedBlood PDU1AB
01899 HMMptac_2_mix(pf06130) PDU1AB
01900 HMM15953pduput_1_pdu1(pf15953) PDU1AB EP_NTERM
01901 P_babyPink PDU1AB
01902 HMM01923cobtr_1_pdu(pf01923pf03928) PDU1AB
01903 HMMalddh_1_eutpdugrm(pf00171) PDU1AB EP_NTERM
01904 HMM00465alcdh_3_mix(pf00465) PDU1AB
01905 HMMPduS_1_all(pf01512pf13375pf10531) PDU1AB
01906 Ts_dodgerBlue PDU1AB
01907 Hp_euts_babyPurple PDU1AB
01908 HMM10662PduV_1_mix(pf10662) PDU1AB
01909 HMMacetkin_1(pf00871) PDU1AB
01910 HMM00288_1_pdu(pf00288) PDU1AB
01911 -pf04363 -
01912 -pf04632 -
01913 -pf06445 -
01914 -pf07943pf00768 -
01915 -pf01292 -
01916 -pf13247pf12800 -
01917 -pf04879pf00384pf01568 -
BMC-P
BMC-H
BMC-T
Aldehyde dehydrogenase
PTAC
Signature enzyme
Alcohol dehydrogenase
Regulator

Salmonella enterica I. Bacteria; Proteobacteria; Gammaproteobacteria; Enterobacterales;

S021__locus_1

PDU1AB

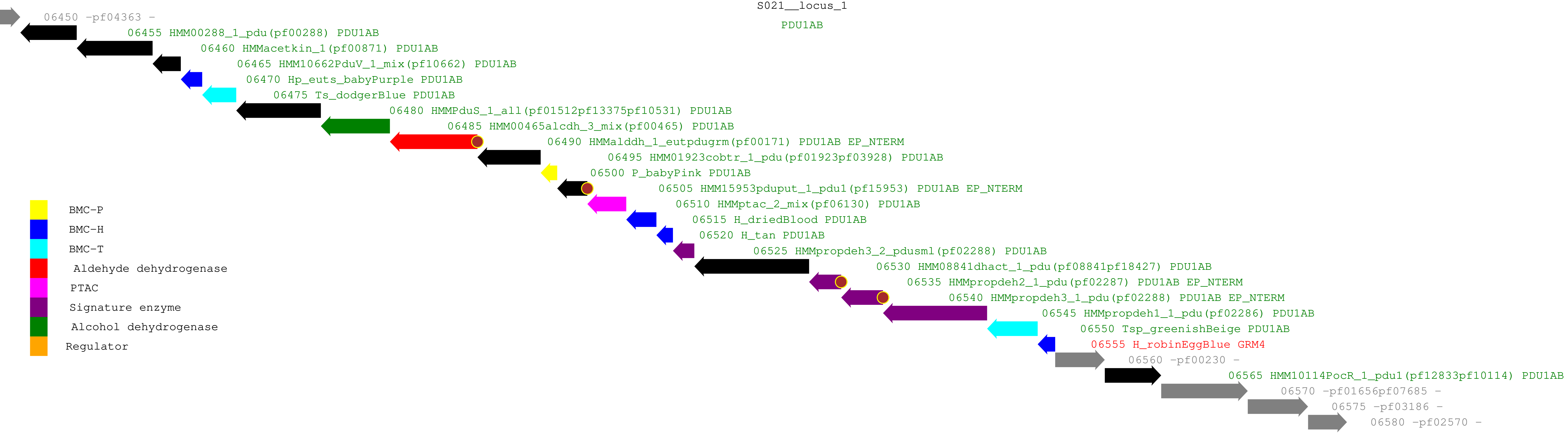

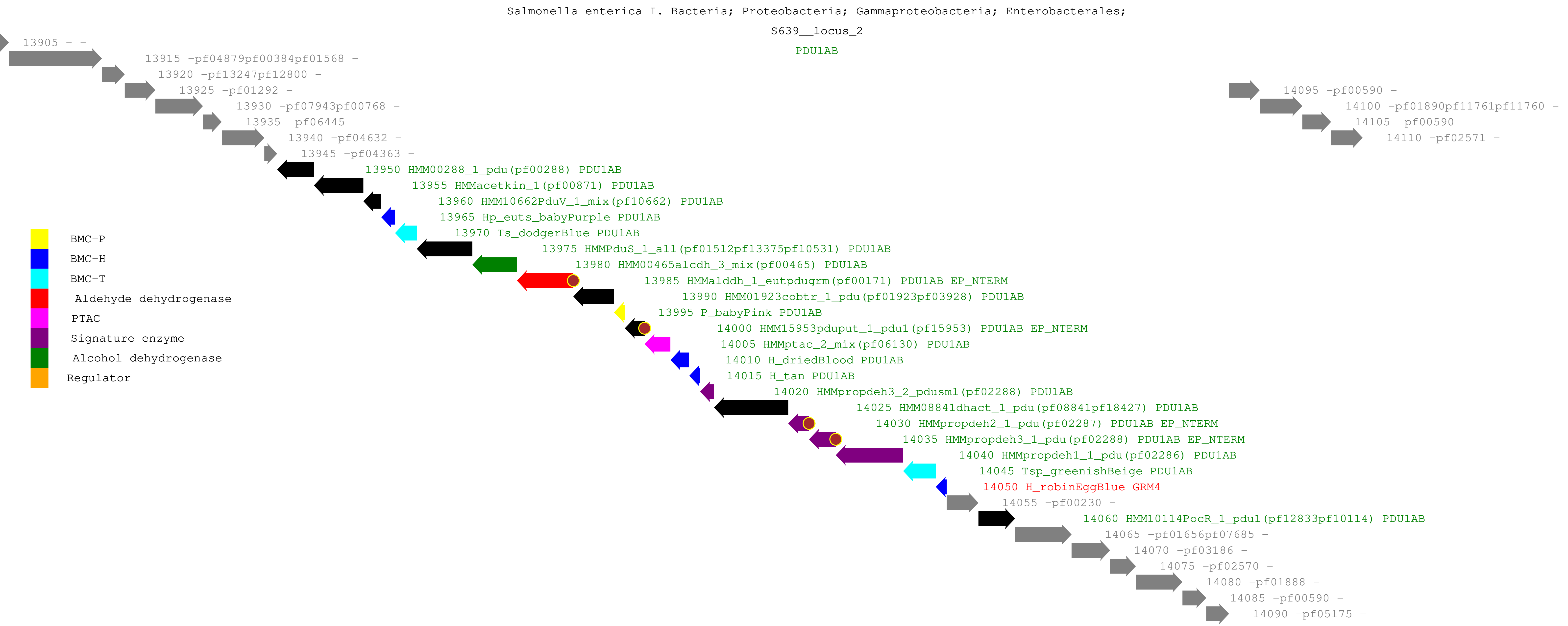
Salmonella enterica I. Bacteria; Proteobacteria; Gammaproteobacteria; Enterobacterales;
S639__locus_2
PDU1AB
13905 - -
13915 -pf04879pf00384pf01568 -
13920 -pf13247pf12800 -
13925 -pf01292 -
13930 -pf07943pf00768 -
13935 -pf06445 -
13940 -pf04632 -
13945 -pf04363 -
13950 HMM00288_1_pdu(pf00288) PDU1AB
13955 HMMacetkin_1(pf00871) PDU1AB
13960 HMM10662PduV_1_mix(pf10662) PDU1AB
13965 Hp_euts_babyPurple PDU1AB
13970 Ts_dodgerBlue PDU1AB
13975 HMMPduS_1_all(pf01512pf13375pf10531) PDU1AB
13980 HMM00465alcdh_3_mix(pf00465) PDU1AB
13985 HMMalddh_1_eutpdugrm(pf00171) PDU1AB EP_NTERM
13990 HMM01923cobtr_1_pdu(pf01923pf03928) PDU1AB
13995 P_babyPink PDU1AB
14000 HMM15953pduput_1_pdu1(pf15953) PDU1AB EP_NTERM
14005 HMMptac_2_mix(pf06130) PDU1AB
14010 H_driedBlood PDU1AB
14015 H_tan PDU1AB
14020 HMMpropdeh3_2_pdusml(pf02288) PDU1AB
14025 HMM08841dhact_1_pdu(pf08841pf18427) PDU1AB
14030 HMMpropdeh2_1_pdu(pf02287) PDU1AB EP_NTERM
14035 HMMpropdeh3_1_pdu(pf02288) PDU1AB EP_NTERM
14040 HMMpropdeh1_1_pdu(pf02286) PDU1AB
14045 Tsp_greenishBeige PDU1AB
14050 H_robinEggBlue GRM4
14055 -pf00230 -
14060 HMM10114PocR_1_pdu1(pf12833pf10114) PDU1AB
14065 -pf01656pf07685 -
14070 -pf03186 -
14075 -pf02570 -
14080 -pf01888 -
14085 -pf00590 -
14090 -pf05175 -
14095 -pf00590 -
14100 -pf01890pf11761pf11760 -
14105 -pf00590 -
14110 -pf02571 -
BMC-P
BMC-H
BMC-T
Aldehyde dehydrogenase
PTAC
Signature enzyme
Alcohol dehydrogenase
Regulator

Klebsiella pneumoniae. Bacteria; Proteobacteria; Gammaproteobacteria; Enterobacterales;

SAMEA104567797__locus_1

PDU1AB

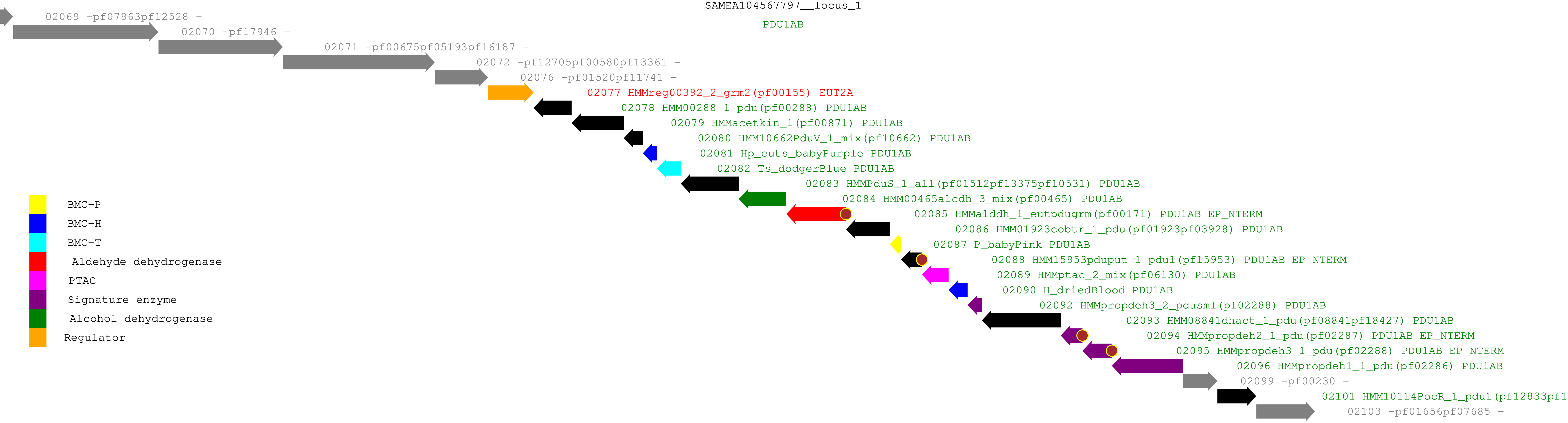

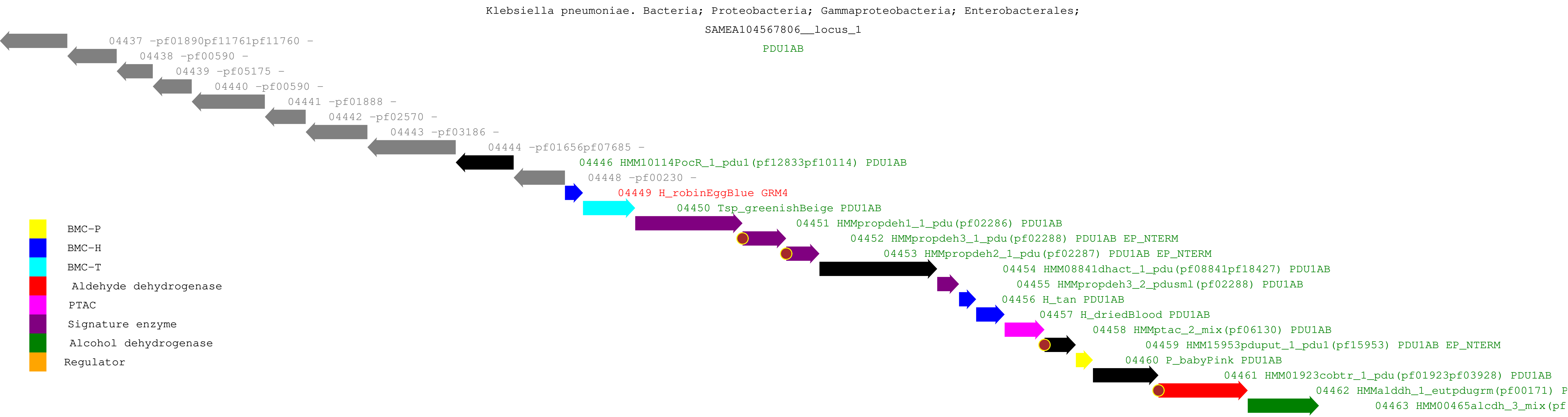

Klebsiella pneumoniae. Bacteria; Proteobacteria; Gammaproteobacteria; Enterobacterales;
SAMEA104567806__locus_1
PDU1AB
04437 -pf01890pf11761pf11760 -
04438 -pf00590 -
04439 -pf05175 -
04440 -pf00590 -
04441 -pf01888 -
04442 -pf02570 -
04443 -pf03186 -
04444 -pf01656pf07685 -
04446 HMM10114PocR_1_pdu1(pf12833pf10114) PDU1AB
04448 -pf00230 -
04449 H_robinEggBlue GRM4
04450 Tsp_greenishBeige PDU1AB
04451 HMMpropdeh1_1_pdu(pf02286) PDU1AB
04452 HMMpropdeh3_1_pdu(pf02288) PDU1AB EP_NTERM
04453 HMMpropdeh2_1_pdu(pf02287) PDU1AB EP_NTERM
04454 HMM08841dhact_1_pdu(pf08841pf18427) PDU1AB
04455 HMMpropdeh3_2_pdusml(pf02288) PDU1AB
04456 H_tan PDU1AB
04457 H_driedBlood PDU1AB
04458 HMMptac_2_mix(pf06130) PDU1AB
04459 HMM15953pduput_1_pdu1(pf15953) PDU1AB EP_NTERM
04460 P_babyPink PDU1AB
04461 HMM01923cobtr_1_pdu(pf01923pf03928) PDU1AB
04462 HMMalddh_1_eutpdugrm(pf00171) P
04463 HMM00465alcdh_3_mix(pf
BMC-P
BMC-H
BMC-T
Aldehyde dehydrogenase
PTAC
Signature enzyme
Alcohol dehydrogenase
Regulator

Klebsiella pneumoniae. Bacteria; Proteobacteria; Gammaproteobacteria; Enterobacterales;

SAMEA104567857__locus_1

PDU1AB

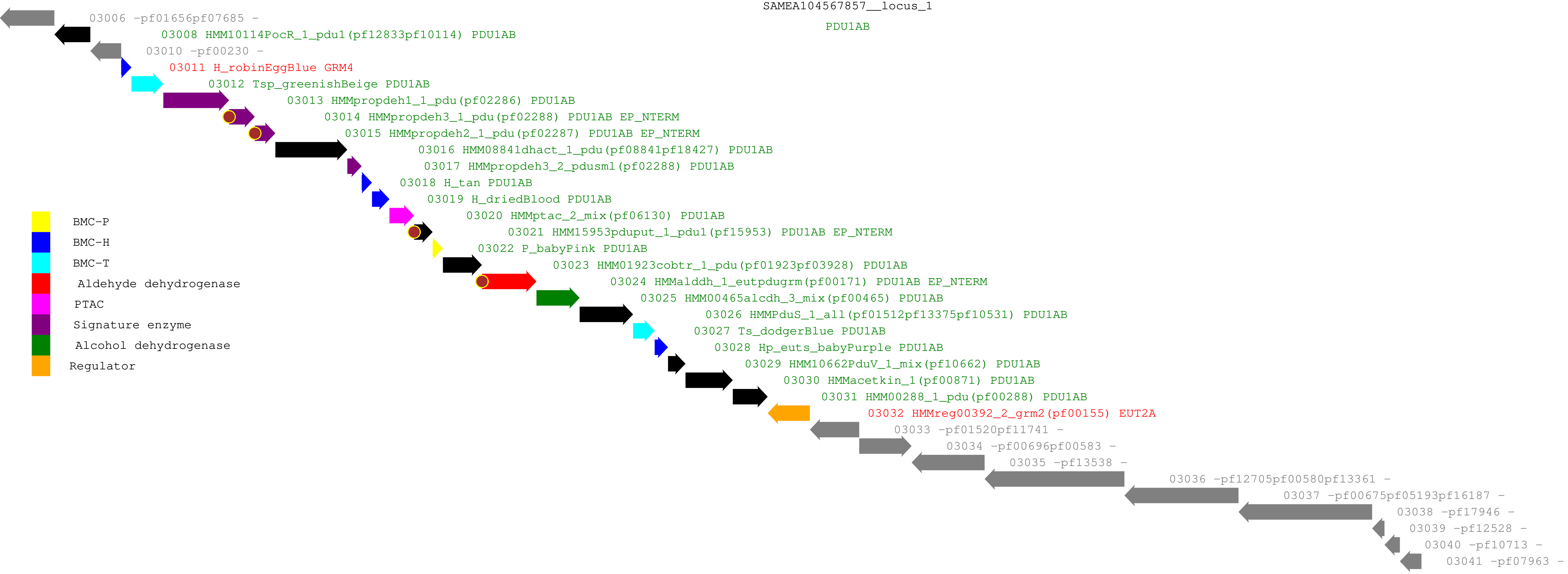

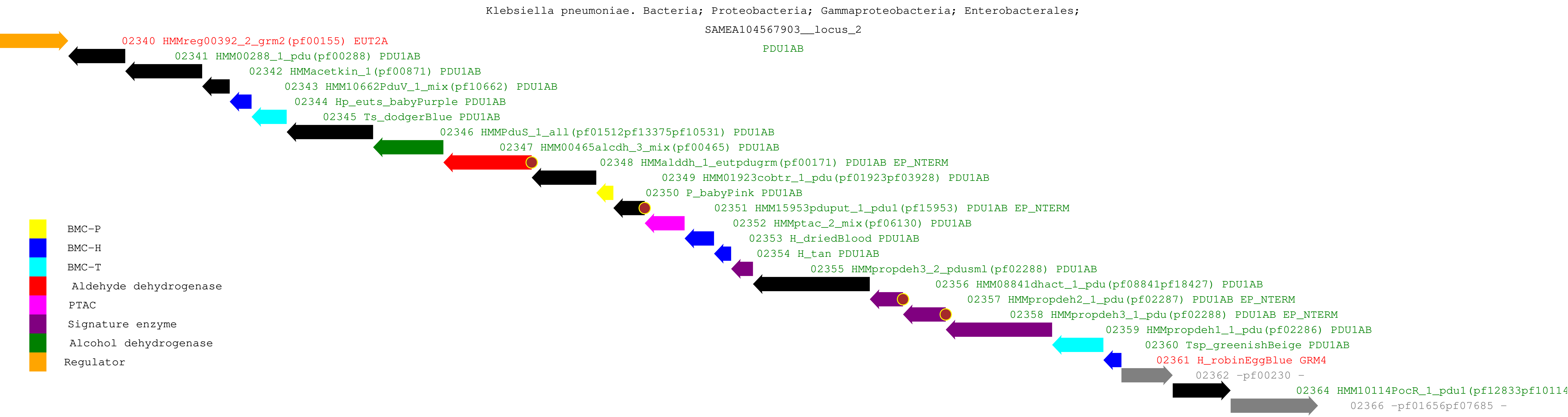
Klebsiella pneumoniae. Bacteria; Proteobacteria; Gammaproteobacteria; Enterobacterales;
SAMEA104567903__locus_2
PDU1AB
02340 HMMreg00392_2_grm2(pf00155) EUT2A
02341 HMM00288_1_pdu(pf00288) PDU1AB
02342 HMMacetkin_1(pf00871) PDU1AB
02343 HMM10662PduV_1_mix(pf10662) PDU1AB
02344 Hp_euts_babyPurple PDU1AB
02345 Ts_dodgerBlue PDU1AB
02346 HMMPduS_1_all(pf01512pf13375pf10531) PDU1AB
02347 HMM00465alcdh_3_mix(pf00465) PDU1AB
02348 HMMaldh_1_eutpdugrm(pf00171) PDU1AB EP_NTERM
02349 HMM01923cobtr_1_pdu(pf01923pf03928) PDU1AB
02350 P_babyPink PDU1AB
02351 HMM15953pduput_1_pdu1(pf15953) PDU1AB EP_NTERM
02352 HMMptac_2_mix(pf06130) PDU1AB
02353 H_driedBlood PDU1AB
02354 H_tan PDU1AB
02355 HMMpropdeh3_2_pdusml(pf02288) PDU1AB
02356 HMM08841dhact_1_pdu(pf08841pf18427) PDU1AB
02357 HMMpropdeh2_1_pdu(pf02287) PDU1AB EP_NTERM
02358 HMMpropdeh3_1_pdu(pf02288) PDU1AB EP_NTERM
02359 HMMpropdeh1_1_pdu(pf02286) PDU1AB
02360 Tsp_greenishBeige PDU1AB
02361 H_robinEggBlue GRM4
02362 -pf00230 -
02364 HMM10114PocR_1_pdu1(pf12833pf10114
02366 -pf01656pf07685 -
BMC-P
BMC-H
BMC-T
Aldehyde dehydrogenase
PTAC
Signature enzyme
Alcohol dehydrogenase
Regulator

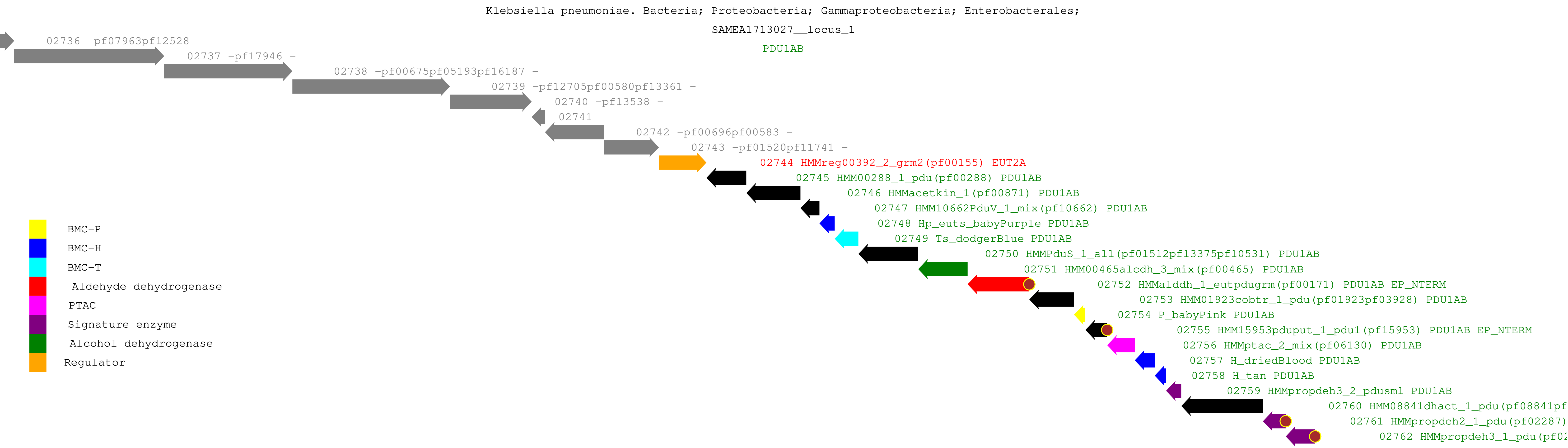

Klebsiella pneumoniae. Bacteria; Proteobacteria; Gammaproteobacteria; Enterobacterales;
SAMEA1713027__locus_1
PDU1AB
02736 -pf07963pf12528 -
02737 -pf17946 -
02738 -pf00675pf05193pf16187 -
02739 -pf12705pf00580pf13361 -
02740 -pf13538 -
02741 - -
02742 -pf00696pf00583 -
02743 -pf01520pf11741 -
02744 HMMreg00392_2_grm2(pf00155) EUT2A
02745 HMM00288_1_pdu(pf00288) PDU1AB
02746 HMMacetkin_1(pf00871) PDU1AB
02747 HMM10662PduV_1_mix(pf10662) PDU1AB
02748 Hp_euts_babyPurple PDU1AB
02749 Ts_dodgerBlue PDU1AB
02750 HMMPduS_1_all(pf01512pf13375pf10531) PDU1AB
02751 HMM00465alcdh_3_mix(pf00465) PDU1AB
02752 HMMalddh_1_eutpdugrm(pf00171) PDU1AB EP_NTERM
02753 HMM01923cobtr_1_pdu(pf01923pf03928) PDU1AB
02754 P_babyPink PDU1AB
02755 HMM15953pduput_1_pdu1(pf15953) PDU1AB EP_NTERM
02756 HMMptac_2_mix(pf06130) PDU1AB
02757 H_driedBlood PDU1AB
02758 H_tan PDU1AB
02759 HMMpropdeh3_2_pdusml PDU1AB
02760 HMM08841dhact_1_pdu(pf08841pf
02761 HMMpropdeh2_1_pdu(pf02287)
02762 HMMpropdeh3_1_pdu(pf0
BMC-P
BMC-H
BMC-T
Aldehyde dehydrogenase
PTAC
Signature enzyme
Alcohol dehydrogenase
Regulator

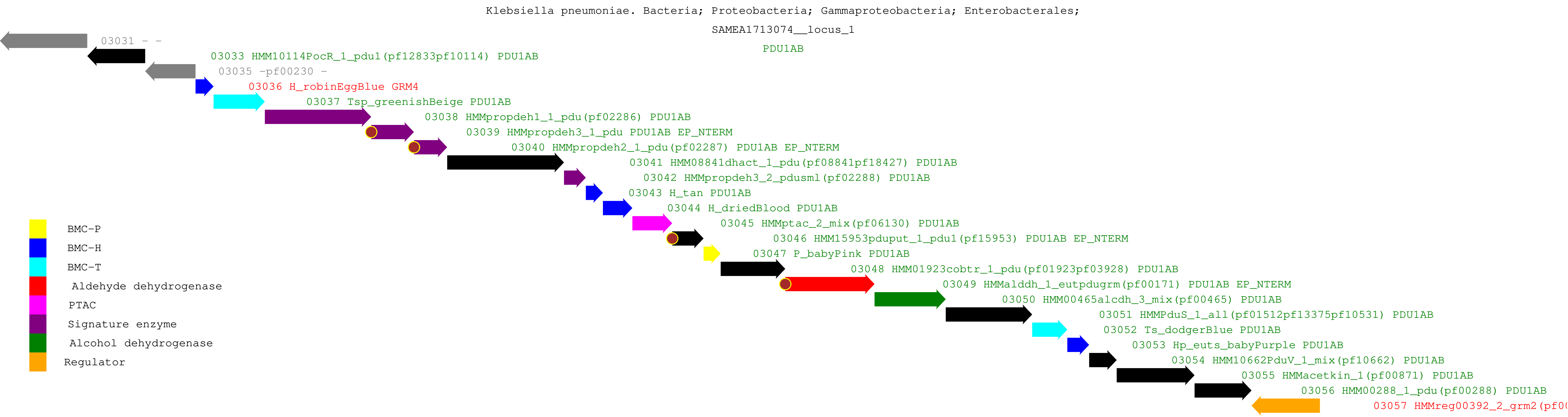

Klebsiella pneumoniae. Bacteria; Proteobacteria; Gammaproteobacteria; Enterobacterales;
SAMEA1713074__locus_1
PDU1AB
03031 - -
03033 HMM10114PocR_1_pdu1(pf12833pf10114) PDU1AB
03035 -pf00230 -
03036 H_robinEggBlue GRM4
03037 Tsp_greenishBeige PDU1AB
03038 HMMpropdeh1_1_pdu(pf02286) PDU1AB
03039 HMMpropdeh3_1_pdu PDU1AB EP_NTERM
03040 HMMpropdeh2_1_pdu(pf02287) PDU1AB EP_NTERM
03041 HMM08841dhact_1_pdu(pf08841pf18427) PDU1AB
03042 HMMpropdeh3_2_pdusml(pf02288) PDU1AB
03043 H_tan PDU1AB
03044 H_driedBlood PDU1AB
03045 HMMptac_2_mix(pf06130) PDU1AB
03046 HMM15953pduput_1_pdu1(pf15953) PDU1AB EP_NTERM
03047 P_babyPink PDU1AB
03048 HMM01923cobtr_1_pdu(pf01923pf03928) PDU1AB
03049 HMMalddh_1_eutpdugrm(pf00171) PDU1AB EP_NTERM
03050 HMM00465alcdh_3_mix(pf00465) PDU1AB
03051 HMMPduS_1_all(pf01512pf13375pf10531) PDU1AB
03052 Ts_dodgerBlue PDU1AB
03053 Hp_euts_babyPurple PDU1AB
03054 HMM10662PduV_1_mix(pf10662) PDU1AB
03055 HMMacetkin_1(pf00871) PDU1AB
03056 HMM00288_1_pdu(pf00288) PDU1AB
03057 HMMreg00392_2_grm2(pf0
BMC-P
BMC-H
BMC-T
Aldehyde dehydrogenase
PTAC
Signature enzyme
Alcohol dehydrogenase
Regulator

Klebsiella pneumoniae. Bacteria; Proteobacteria; Gammaproteobacteria; Enterobacterales;

SAMEA1713115__locus_2

PDU1AB

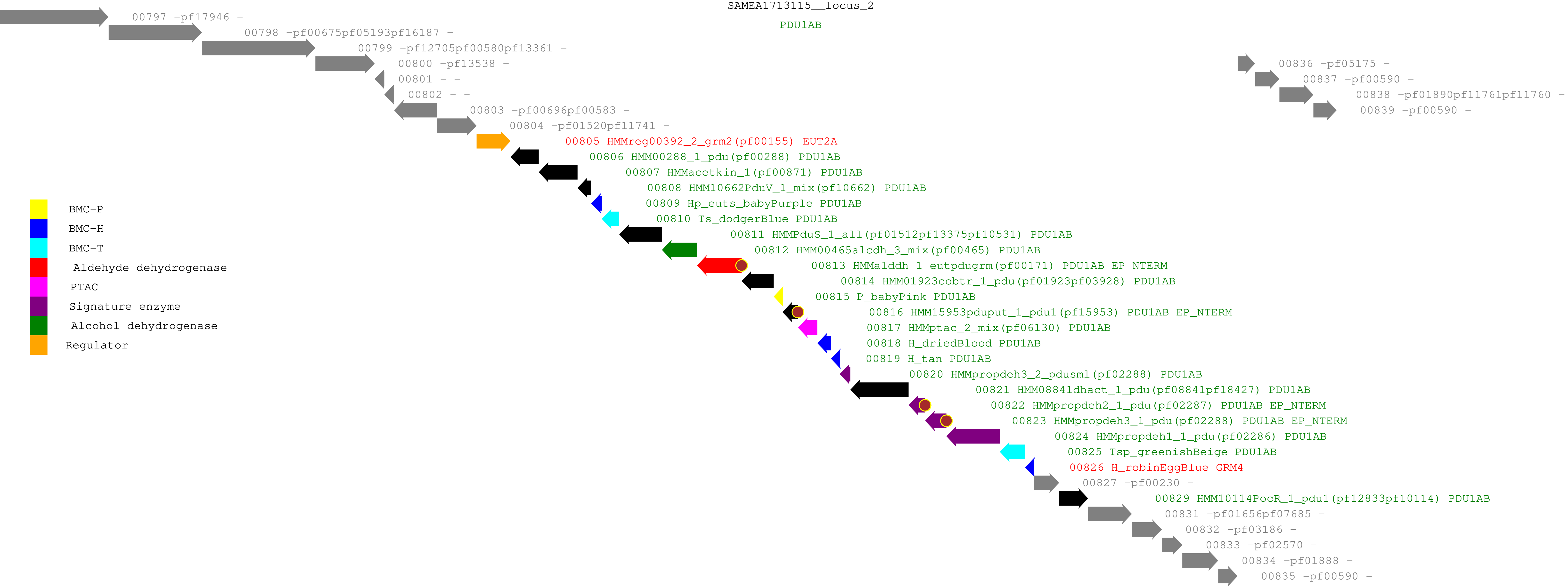

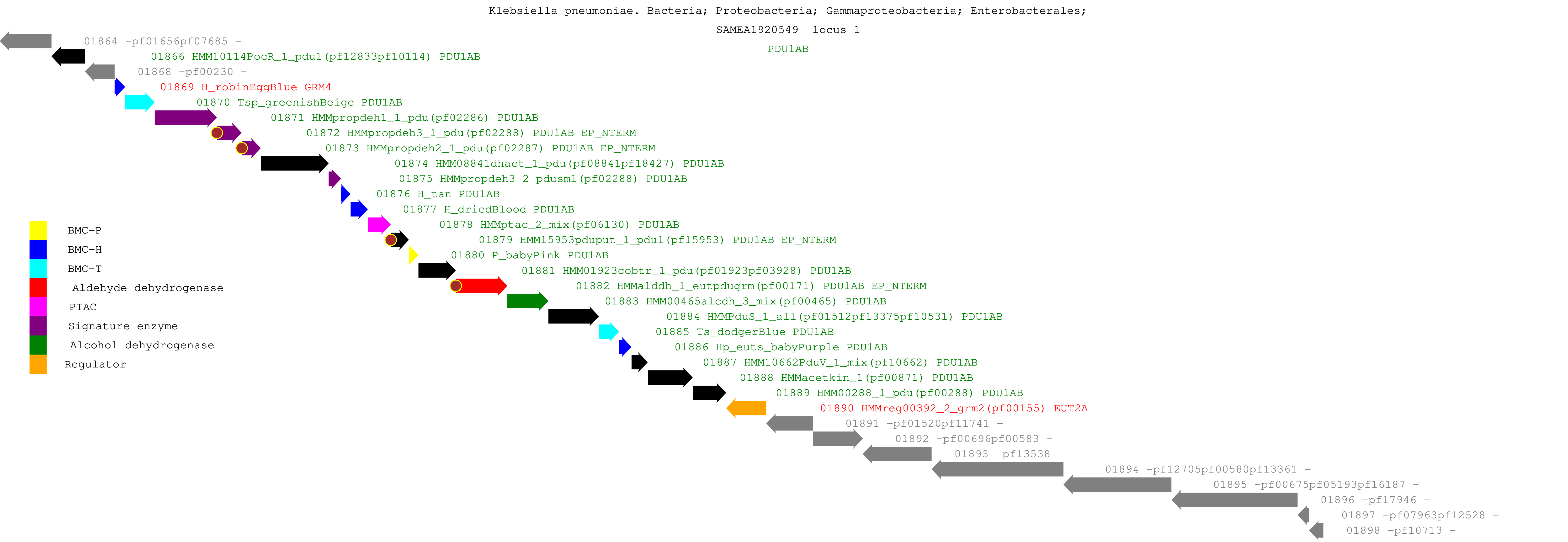

Klebsiella pneumoniae. Bacteria; Proteobacteria; Gammaproteobacteria; Enterobacterales;
SAMEA1920549__locus_1
PDU1AB
01864 -pf01656pf07685 -
01866 HMM10114PocR_1_pdu1(pf12833pf10114) PDU1AB
01868 -pf00230 -
01869 H_robinEggBlue GRM4
01870 Tsp_greenishBeige PDU1AB
01871 HMMpropdeh1_1_pdu(pf02286) PDU1AB
01872 HMMpropdeh3_1_pdu(pf02288) PDU1AB EP_NTERM
01873 HMMpropdeh2_1_pdu(pf02287) PDU1AB EP_NTERM
01874 HMM08841dhact_1_pdu(pf08841pf18427) PDU1AB
01875 HMMpropdeh3_2_pdusml(pf02288) PDU1AB
01876 H_tan PDU1AB
01877 H_driedBlood PDU1AB
01878 HMMptac_2_mix(pf06130) PDU1AB
01879 HMM15953pduput_1_pdu1(pf15953) PDU1AB EP_NTERM
01880 P_babyPink PDU1AB
01881 HMM01923cobtr_1_pdu(pf01923pf03928) PDU1AB
01882 HMMalddh_1_eutpdugrm(pf00171) PDU1AB EP_NTERM
01883 HMM00465alcdh_3_mix(pf00465) PDU1AB
01884 HMMPduS_1_all(pf01512pf13375pf10531) PDU1AB
01885 Ts_dodgerBlue PDU1AB
01886 Hp_euts_babyPurple PDU1AB
01887 HMM10662PduV_1_mix(pf10662) PDU1AB
01888 HMMacetkin_1(pf00871) PDU1AB
01889 HMM00288_1_pdu(pf00288) PDU1AB
01890 HMMreg00392_2_grm2(pf00155) EUT2A
01891 -pf01520pf11741 -
01892 -pf00696pf00583 -
01893 -pf13538 -
01894 -pf12705pf00580pf13361 -
01895 -pf00675pf05193pf16187 -
01896 -pf17946 -
01897 -pf07963pf12528 -
01898 -pf10713 -
BMC-P
BMC-H
BMC-T
Aldehyde dehydrogenase
PTAC
Signature enzyme
Alcohol dehydrogenase
Regulator

Klebsiella pneumoniae. Bacteria; Proteobacteria; Gammaproteobacteria; Enterobacterales;

SAMEA2053874__locus_1

PDU1AB

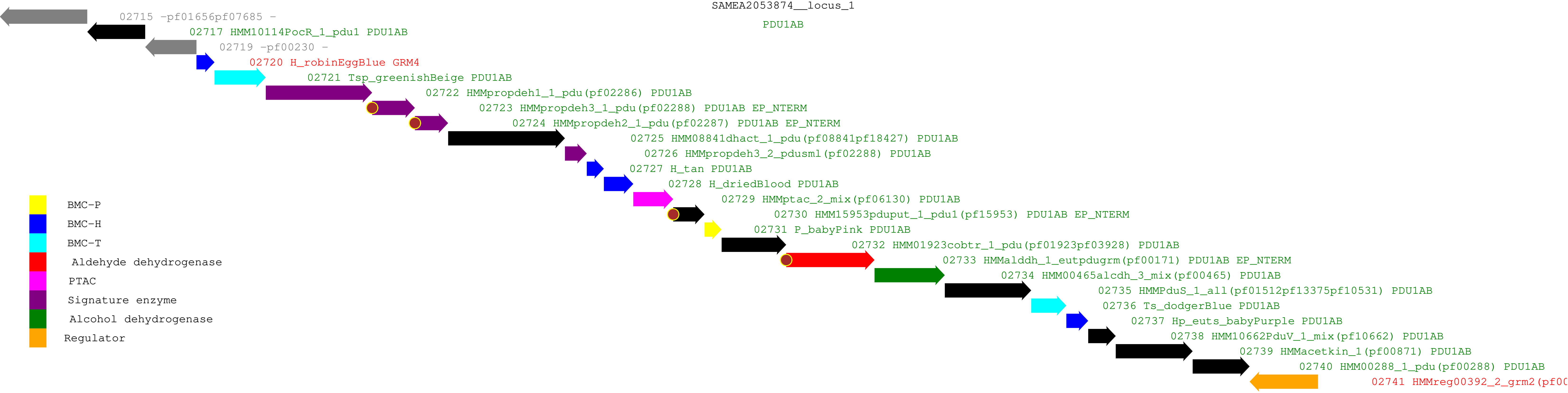

Klebsiella pneumoniae. Bacteria; Proteobacteria; Gammaproteobacteria; Enterobacterales;

SAMEA2053876__locus_1

PDU1AB

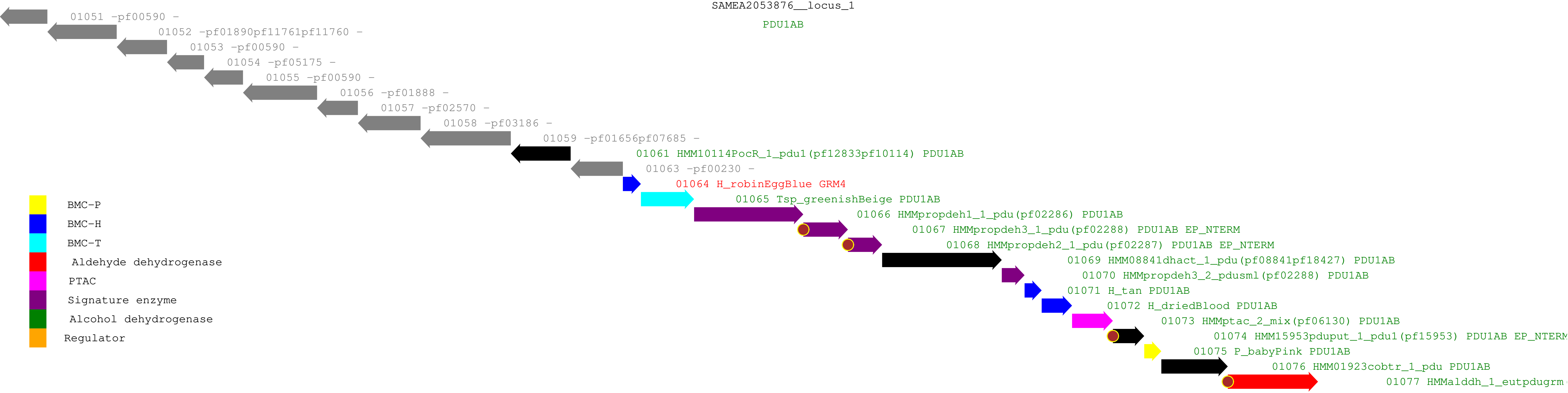

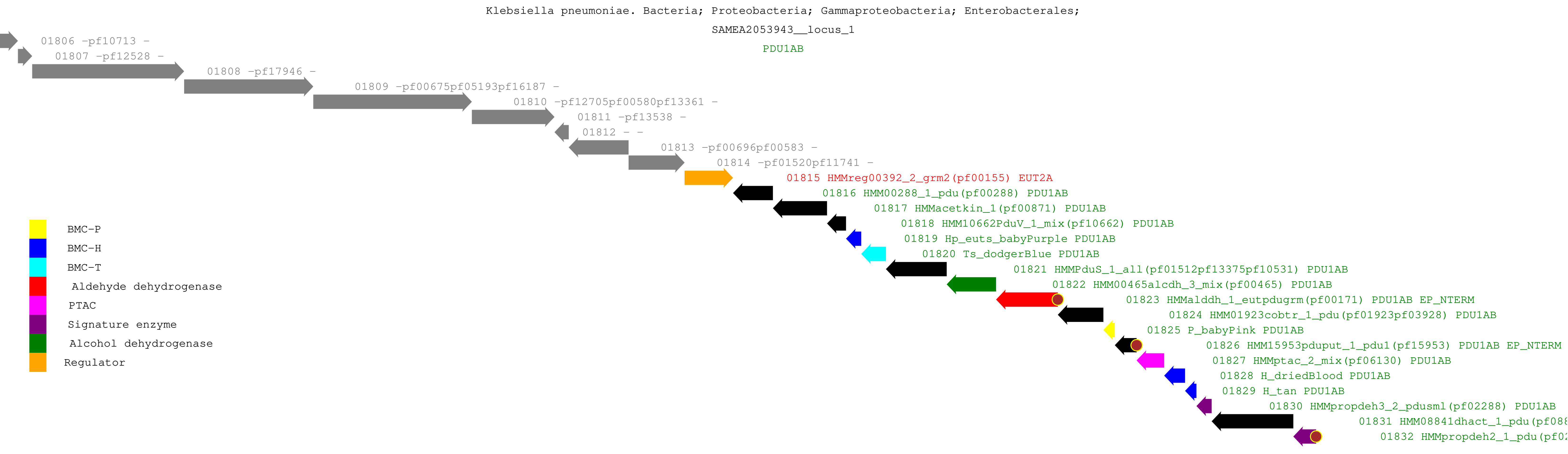

Klebsiella pneumoniae. Bacteria; Proteobacteria; Gammaproteobacteria; Enterobacterales;
SAMEA2053943__locus_1
PDU1AB
01806 -pf10713 -
01807 -pf12528 -
01808 -pf17946 -
01809 -pf00675pf05193pf16187 -
01810 -pf12705pf00580pf13361 -
01811 -pf13538 -
01812 - -
01813 -pf00696pf00583 -
01814 -pf01520pf11741 -
01815 HMMreg00392_2_grm2(pf00155) EUT2A
01816 HMM00288_1_pdu(pf00288) PDU1AB
01817 HMMacetkin_1(pf00871) PDU1AB
01818 HMM10662PduV_1_mix(pf10662) PDU1AB
01819 Hp_euts_babyPurple PDU1AB
01820 Ts_dodgerBlue PDU1AB
01821 HMMPduS_1_all(pf01512pf13375pf10531) PDU1AB
01822 HMM00465alcdh_3_mix(pf00465) PDU1AB
01823 HMMaldh_1_eutpdugrm(pf00171) PDU1AB EP_NTERM
01824 HMM01923cobtr_1_pdu(pf01923pf03928) PDU1AB
01825 P_babyPink PDU1AB
01826 HMM15953pduput_1_pdu1(pf15953) PDU1AB EP_NTERM
01827 HMMptac_2_mix(pf06130) PDU1AB
01828 H_driedBlood PDU1AB
01829 H_tan PDU1AB
01830 HMMpropdeh3_2_pdusml(pf02288) PDU1AB
01831 HMM08841dhact_1_pdu(pf08
01832 HMMpropdeh2_1_pdu(pf0
BMC-P
BMC-H
BMC-T
Aldehyde dehydrogenase
PTAC
Signature enzyme
Alcohol dehydrogenase
Regulator

Klebsiella pneumoniae. Bacteria; Proteobacteria; Gammaproteobacteria; Enterobacterales;

SAMEA2054826__locus_1

PDU1AB

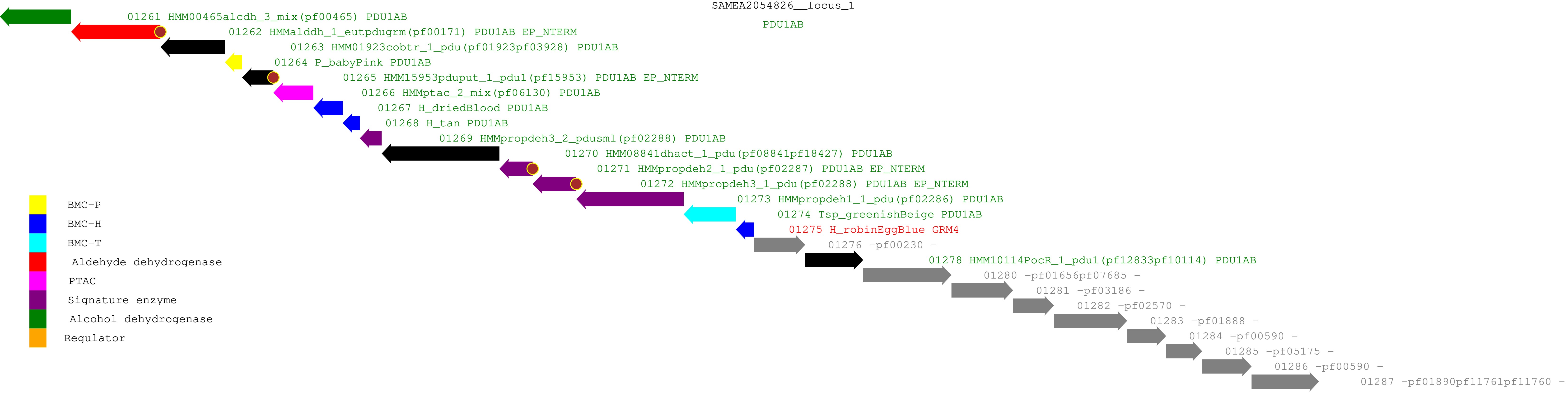

Klebsiella pneumoniae. Bacteria; Proteobacteria; Gammaproteobacteria; Enterobacterales;

SAMEA2055014__locus_1

PDU1AB

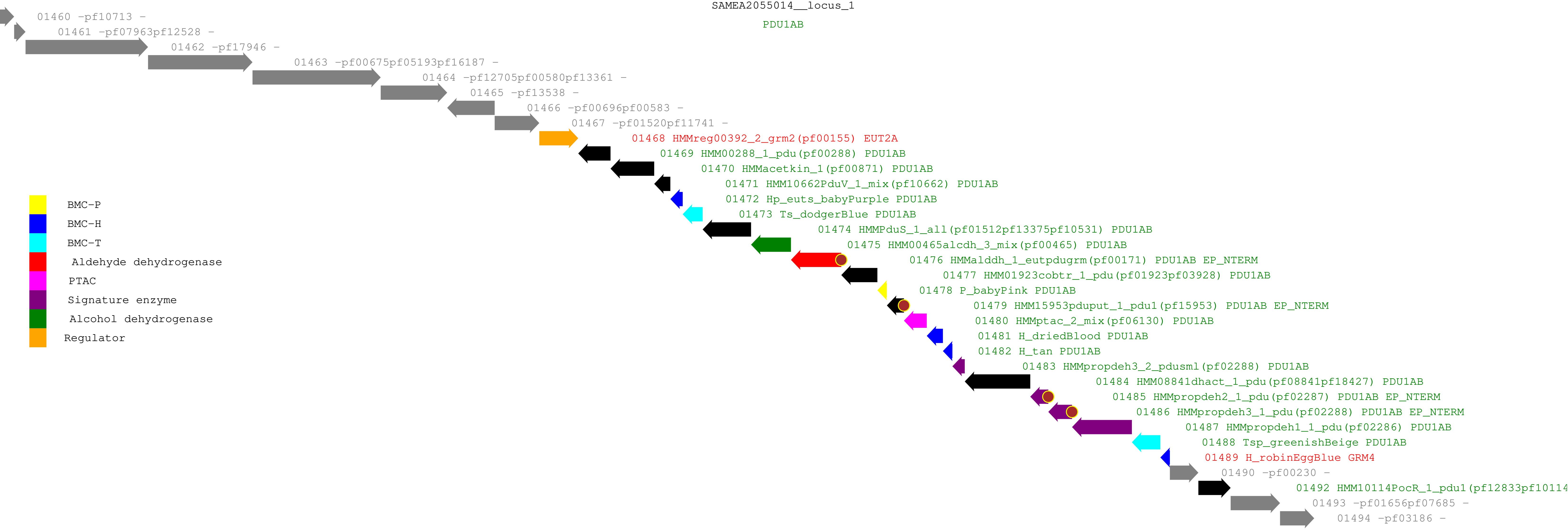

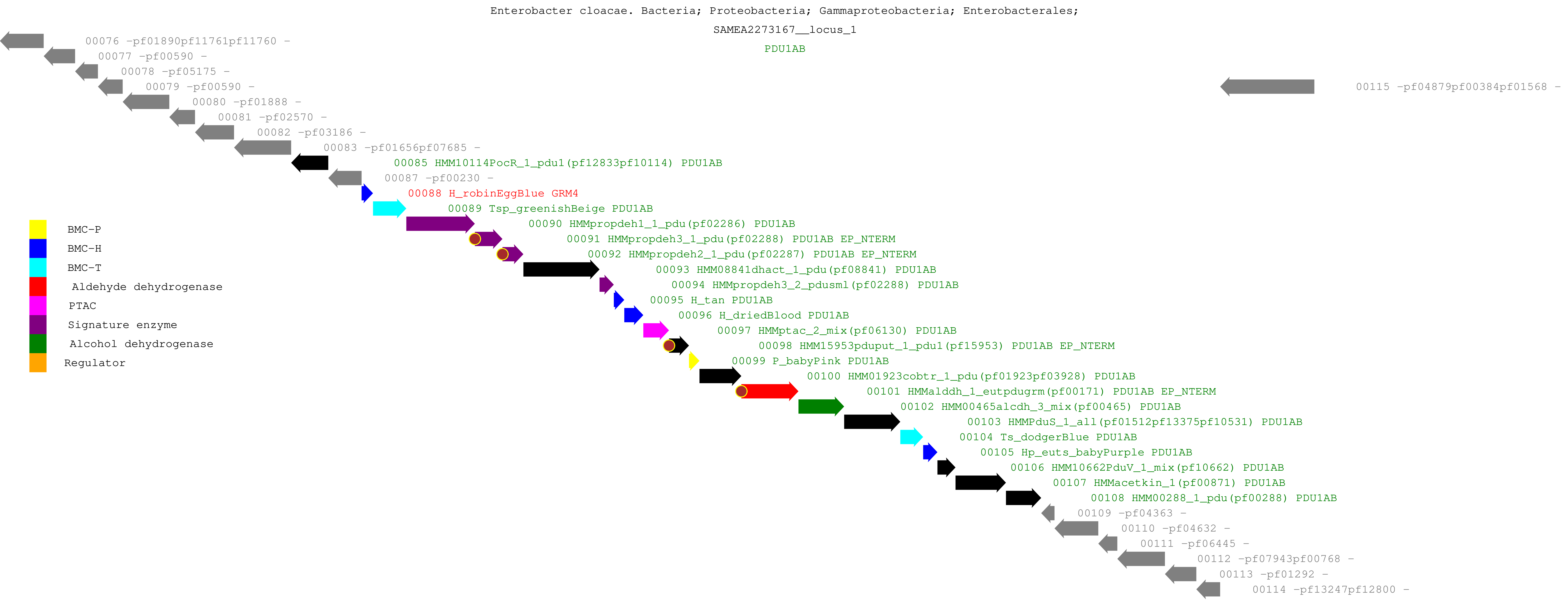
Enterobacter cloacae. Bacteria; Proteobacteria; Gammaproteobacteria; Enterobacterales;
SAMEA2273167__locus_1
PDU1AB
00076 -pf01890pf11761pf11760 -
00077 -pf00590 -
00078 -pf05175 -
00079 -pf00590 -
00080 -pf01888 -
00081 -pf02570 -
00082 -pf03186 -
00083 -pf01656pf07685 -
00085 HMM10114PocR_1_pdu1(pf12833pf10114) PDU1AB
00087 -pf00230 -
00088 H_robinEggBlue GRM4
00089 Tsp_greenishBeige PDU1AB
00090 HMMpropdeh1_1_pdu(pf02286) PDU1AB
00091 HMMpropdeh3_1_pdu(pf02288) PDU1AB EP_NTERM
00092 HMMpropdeh2_1_pdu(pf02287) PDU1AB EP_NTERM
00093 HMM08841dhact_1_pdu(pf08841) PDU1AB
00094 HMMpropdeh3_2_pdusml(pf02288) PDU1AB
00095 H_tan PDU1AB
00096 H_driedBlood PDU1AB
00097 HMMptac_2_mix(pf06130) PDU1AB
00098 HMM15953pduput_1_pdu1(pf15953) PDU1AB EP_NTERM
00099 P_babyPink PDU1AB
00100 HMM01923cobtr_1_pdu(pf01923pf03928) PDU1AB
00101 HMMalddh_1_eutpdugrm(pf00171) PDU1AB EP_NTERM
00102 HMM00465alcdh_3_mix(pf00465) PDU1AB
00103 HMMPduS_1_all(pf01512pf13375pf10531) PDU1AB
00104 Ts_dodgerBlue PDU1AB
00105 Hp_euts_babyPurple PDU1AB
00106 HMM10662PduV_1_mix(pf10662) PDU1AB
00107 HMMacetkin_1(pf00871) PDU1AB
00108 HMM00288_1_pdu(pf00288) PDU1AB
00109 -pf04363 -
00110 -pf04632 -
00111 -pf06445 -
00112 -pf07943pf00768 -
00113 -pf01292 -
00114 -pf13247pf12800 -
00115 -pf04879pf00384pf01568 -
BMC-P
BMC-H
BMC-T
Aldehyde dehydrogenase
PTAC
Signature enzyme
Alcohol dehydrogenase
Regulator

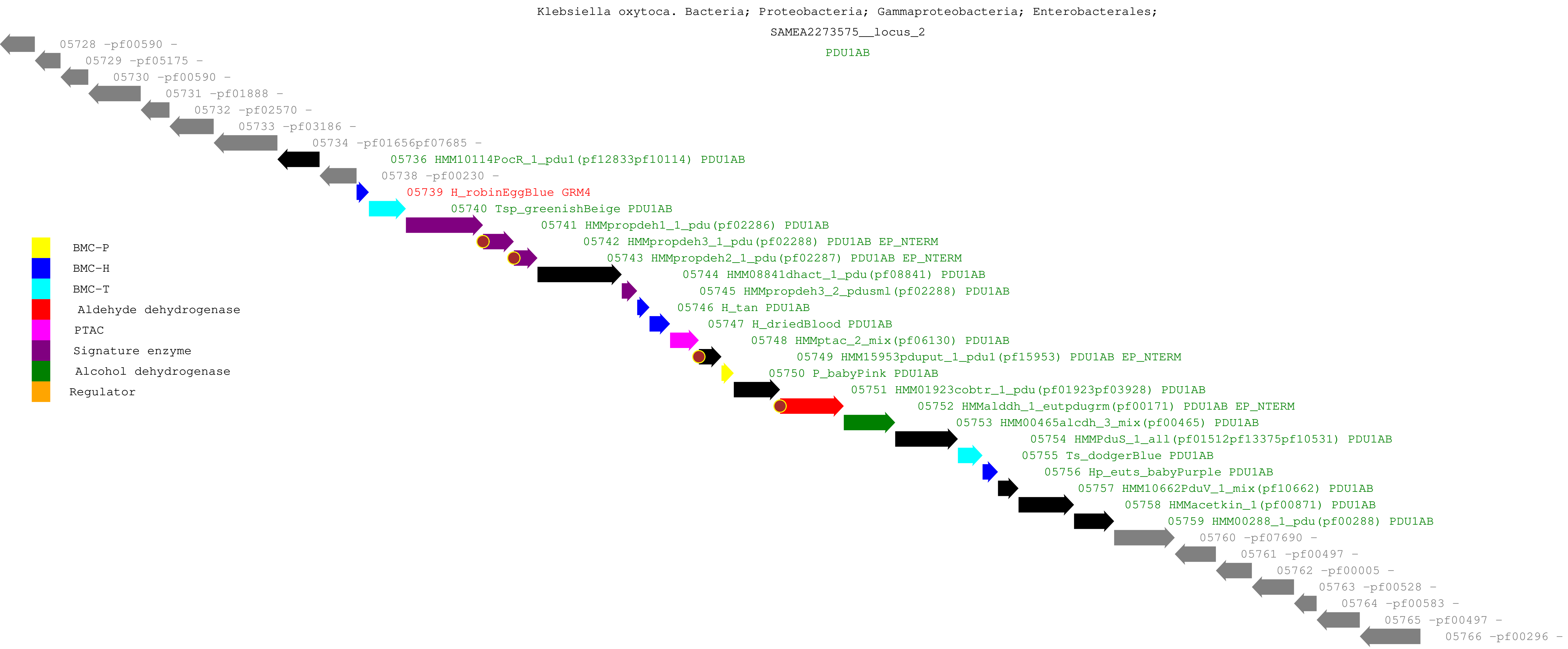
Klebsiella oxytoca. Bacteria; Proteobacteria; Gammaproteobacteria; Enterobacterales;
SAMEA2273575__locus_2
PDU1AB
05728 -pf00590 -
05729 -pf05175 -
05730 -pf00590 -
05731 -pf01888 -
05732 -pf02570 -
05733 -pf03186 -
05734 -pf01656pf07685 -
05736 HMM10114PocR_1_pdu1(pf12833pf10114) PDU1AB
05738 -pf00230 -
05739 H_robinEggBlue GRM4
05740 Tsp_greenishBeige PDU1AB
05741 HMMpropdeh1_1_pdu(pf02286) PDU1AB
05742 HMMpropdeh3_1_pdu(pf02288) PDU1AB EP_NTERM
05743 HMMpropdeh2_1_pdu(pf02287) PDU1AB EP_NTERM
05744 HMM08841dhact_1_pdu(pf08841) PDU1AB
05745 HMMpropdeh3_2_pdusml(pf02288) PDU1AB
05746 H_tan PDU1AB
05747 H_driedBlood PDU1AB
05748 HMMptac_2_mix(pf06130) PDU1AB
05749 HMM15953pduput_1_pdu1(pf15953) PDU1AB EP_NTERM
05750 P_babyPink PDU1AB
05751 HMM01923cobtr_1_pdu(pf01923pf03928) PDU1AB
05752 HMMalddh_1_eutpdugrm(pf00171) PDU1AB EP_NTERM
05753 HMM00465alcdh_3_mix(pf00465) PDU1AB
05754 HMMPduS_1_all(pf01512pf13375pf10531) PDU1AB
05755 Ts_dodgerBlue PDU1AB
05756 Hp_euts_babyPurple PDU1AB
05757 HMM10662PduV_1_mix(pf10662) PDU1AB
05758 HMMacetkin_1(pf00871) PDU1AB
05759 HMM00288_1_pdu(pf00288) PDU1AB
05760 -pf07690 -
05761 -pf00497 -
05762 -pf00005 -
05763 -pf00528 -
05764 -pf00583 -
05765 -pf00497 -
05766 -pf00296 -
BMC-P
BMC-H
BMC-T
Aldehyde dehydrogenase
PTAC
Signature enzyme
Alcohol dehydrogenase
Regulator

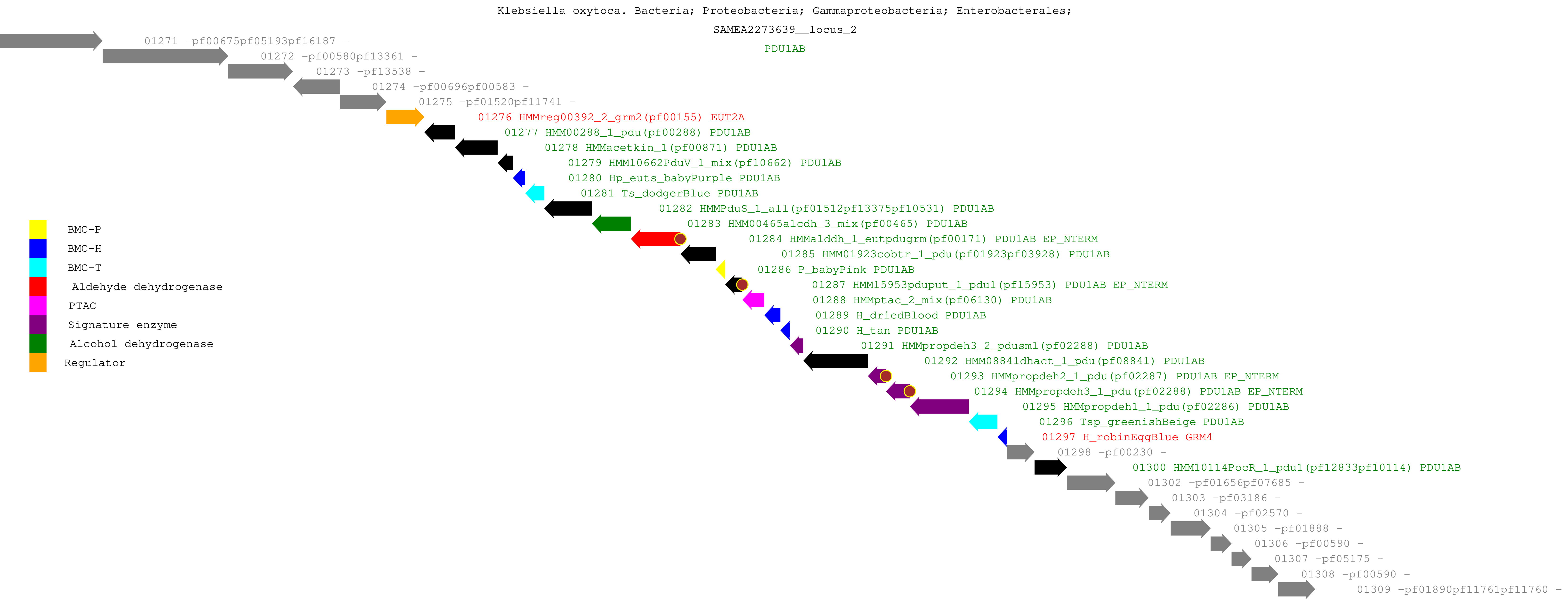
Klebsiella oxytoca. Bacteria; Proteobacteria; Gammaproteobacteria; Enterobacterales;
SAMEA2273639__locus_2
PDU1AB
01271 -pf00675pf05193pf16187 -
01272 -pf00580pf13361 -
01273 -pf13538 -
01274 -pf00696pf00583 -
01275 -pf01520pf11741 -
01276 HMMreg00392_2_grm2(pf00155) EUT2A
01277 HMM00288_1_pdu(pf00288) PDU1AB
01278 HMMacetkin_1(pf00871) PDU1AB
01279 HMM10662PduV_1_mix(pf10662) PDU1AB
01280 Hp_euts_babyPurple PDU1AB
01281 Ts_dodgerBlue PDU1AB
01282 HMMPduS_1_all(pf01512pf13375pf10531) PDU1AB
01283 HMM00465alcdh_3_mix(pf00465) PDU1AB
01284 HMMalddh_1_eutpdugrm(pf00171) PDU1AB EP_NTERM
01285 HMM01923cobtr_1_pdu(pf01923pf03928) PDU1AB
01286 P_babyPink PDU1AB
01287 HMM15953pduput_1_pdu1(pf15953) PDU1AB EP_NTERM
01288 HMMptac_2_mix(pf06130) PDU1AB
01289 H_driedBlood PDU1AB
01290 H_tan PDU1AB
01291 HMMpropdeh3_2_pdusml(pf02288) PDU1AB
01292 HMM08841dhact_1_pdu(pf08841) PDU1AB
01293 HMMpropdeh2_1_pdu(pf02287) PDU1AB EP_NTERM
01294 HMMpropdeh3_1_pdu(pf02288) PDU1AB EP_NTERM
01295 HMMpropdeh1_1_pdu(pf02286) PDU1AB
01296 Tsp_greenishBeige PDU1AB
01297 H_robinEggBlue GRM4
01298 -pf00230 -
01300 HMM10114PocR_1_pdu1(pf12833pf10114) PDU1AB
01302 -pf01656pf07685 -
01303 -pf03186 -
01304 -pf02570 -
01305 -pf01888 -
01306 -pf00590 -
01307 -pf05175 -
01308 -pf00590 -
01309 -pf01890pf11761pf11760 -
BMC-P
BMC-H
BMC-T
Aldehyde dehydrogenase
PTAC
Signature enzyme
Alcohol dehydrogenase
Regulator

Klebsiella oxytoca. Bacteria; Proteobacteria; Gammaproteobacteria; Enterobacterales;

SAMEA2273697__locus_2

PDU1AB

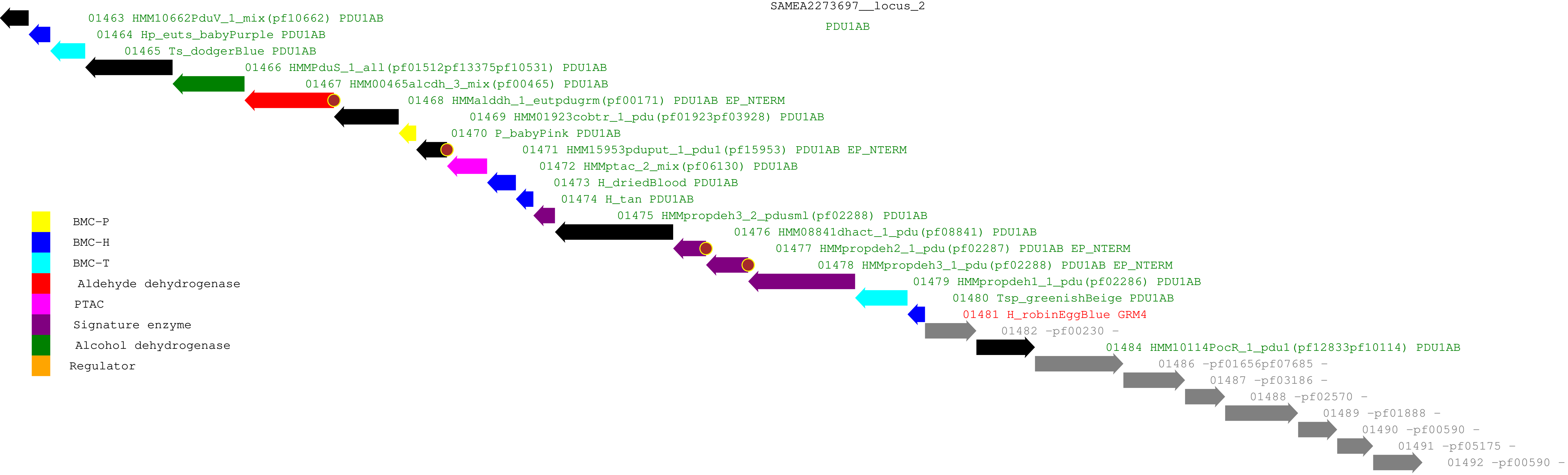

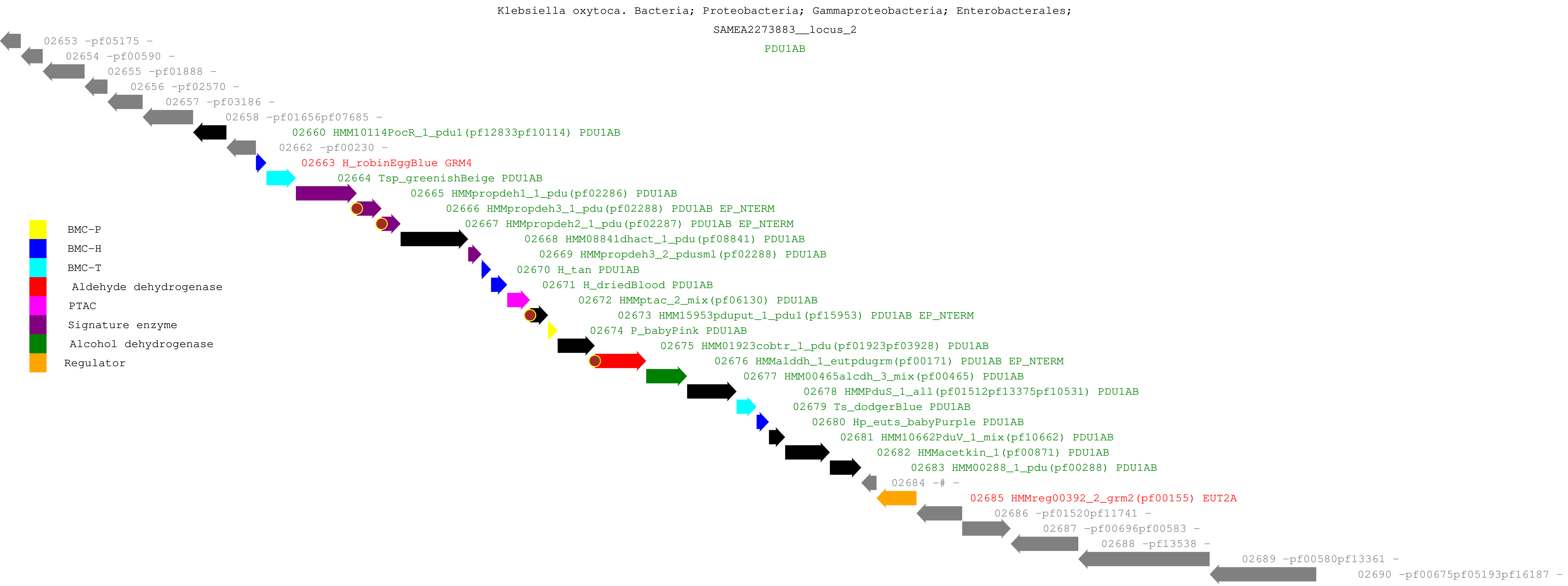
Klebsiella oxytoca. Bacteria; Proteobacteria; Gammaproteobacteria; Enterobacterales;
SAMEA2273883__locus_2
PDU1AB
02653 -pf05175 -
02654 -pf00590 -
02655 -pf01888 -
02656 -pf02570 -
02657 -pf03186 -
02658 -pf01656pf07685 -
02660 HMM10114PocR_1_pdu1(pf12833pf10114) PDU1AB
02662 -pf00230 -
02663 H_robinEggBlue GRM4
02664 Tsp_greenishBeige PDU1AB
02665 HMMpropdeh1_1_pdu(pf02286) PDU1AB
02666 HMMpropdeh3_1_pdu(pf02288) PDU1AB EP_NTERM
02667 HMMpropdeh2_1_pdu(pf02287) PDU1AB EP_NTERM
02668 HMM08841dhact_1_pdu(pf08841) PDU1AB
02669 HMMpropdeh3_2_pdusml(pf02288) PDU1AB
02670 H_tan PDU1AB
02671 H_driedBlood PDU1AB
02672 HMMptac_2_mix(pf06130) PDU1AB
02673 HMM15953pduput_1_pdu1(pf15953) PDU1AB EP_NTERM
02674 P_babyPink PDU1AB
02675 HMM01923cobtr_1_pdu(pf01923pf03928) PDU1AB
02676 HMMalddh_1_eutpdugrm(pf00171) PDU1AB EP_NTERM
02677 HMM00465alcdh_3_mix(pf00465) PDU1AB
02678 HMMPduS_1_all(pf01512pf13375pf10531) PDU1AB
02679 Ts_dodgerBlue PDU1AB
02680 Hp_euts_babyPurple PDU1AB
02681 HMM10662PduV_1_mix(pf10662) PDU1AB
02682 HMMacetkin_1(pf00871) PDU1AB
02683 HMM00288_1_pdu(pf00288) PDU1AB
02684 -# -
02685 HMMreg00392_2_grm2(pf00155) EUT2A
02686 -pf01520pf11741 -
02687 -pf00696pf00583 -
02688 -pf13538 -
02689 -pf00580pf13361 -
02690 -pf00675pf05193pf16187 -
BMC-P
BMC-H
BMC-T
Aldehyde dehydrogenase
PTAC
Signature enzyme
Alcohol dehydrogenase
Regulator

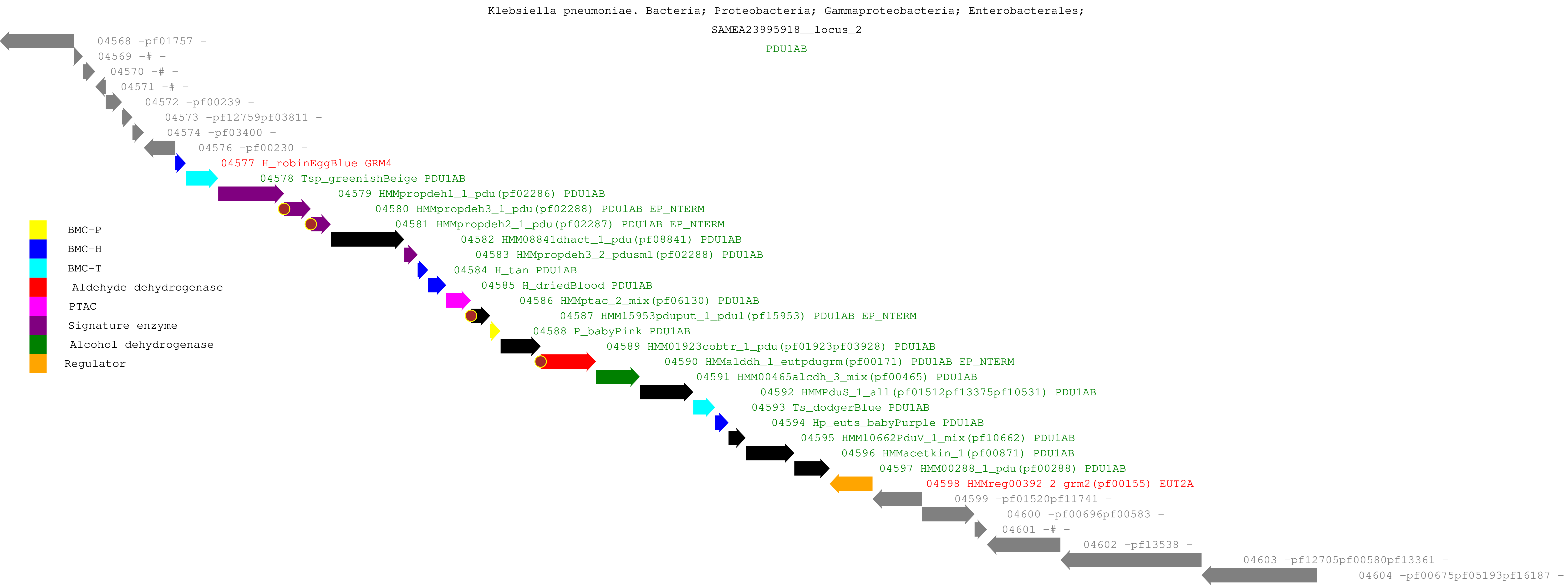

Klebsiella pneumoniae. Bacteria; Proteobacteria; Gammaproteobacteria; Enterobacterales;
SAMEA23995918__locus_2
PDU1AB
04568 -pf01757 -
04569 -# -
04570 -# -
04571 -# -
04572 -pf00239 -
04573 -pf12759pf03811 -
04574 -pf03400 -
04576 -pf00230 -
04577 H_robinEggBlue GRM4
04578 Tsp_greenishBeige PDU1AB
04579 HMMpropdeh1_1_pdu(pf02286) PDU1AB
04580 HMMpropdeh3_1_pdu(pf02288) PDU1AB EP_NTERM
04581 HMMpropdeh2_1_pdu(pf02287) PDU1AB EP_NTERM
04582 HMM08841dhact_1_pdu(pf08841) PDU1AB
04583 HMMpropdeh3_2_pdusml(pf02288) PDU1AB
04584 H_tan PDU1AB
04585 H_driedBlood PDU1AB
04586 HMMptac_2_mix(pf06130) PDU1AB
04587 HMM15953pduput_1_pdu1(pf15953) PDU1AB EP_NTERM
04588 P_babyPink PDU1AB
04589 HMM01923cobtr_1_pdu(pf01923pf03928) PDU1AB
04590 HMMalddh_1_eutpdugrm(pf00171) PDU1AB EP_NTERM
04591 HMM00465alcdh_3_mix(pf00465) PDU1AB
04592 HMMPduS_1_all(pf01512pf13375pf10531) PDU1AB
04593 Ts_dodgerBlue PDU1AB
04594 Hp_euts_babyPurple PDU1AB
04595 HMM10662PduV_1_mix(pf10662) PDU1AB
04596 HMMacetkin_1(pf00871) PDU1AB
04597 HMM00288_1_pdu(pf00288) PDU1AB
04598 HMMreg00392_2_grm2(pf00155) EUT2A
04599 -pf01520pf11741 -
04600 -pf00696pf00583 -
04601 -# -
04602 -pf13538 -
04603 -pf12705pf00580pf13361 -
04604 -pf00675pf05193pf16187 -
BMC-P
BMC-H
BMC-T
Aldehyde dehydrogenase
PTAC
Signature enzyme
Alcohol dehydrogenase
Regulator

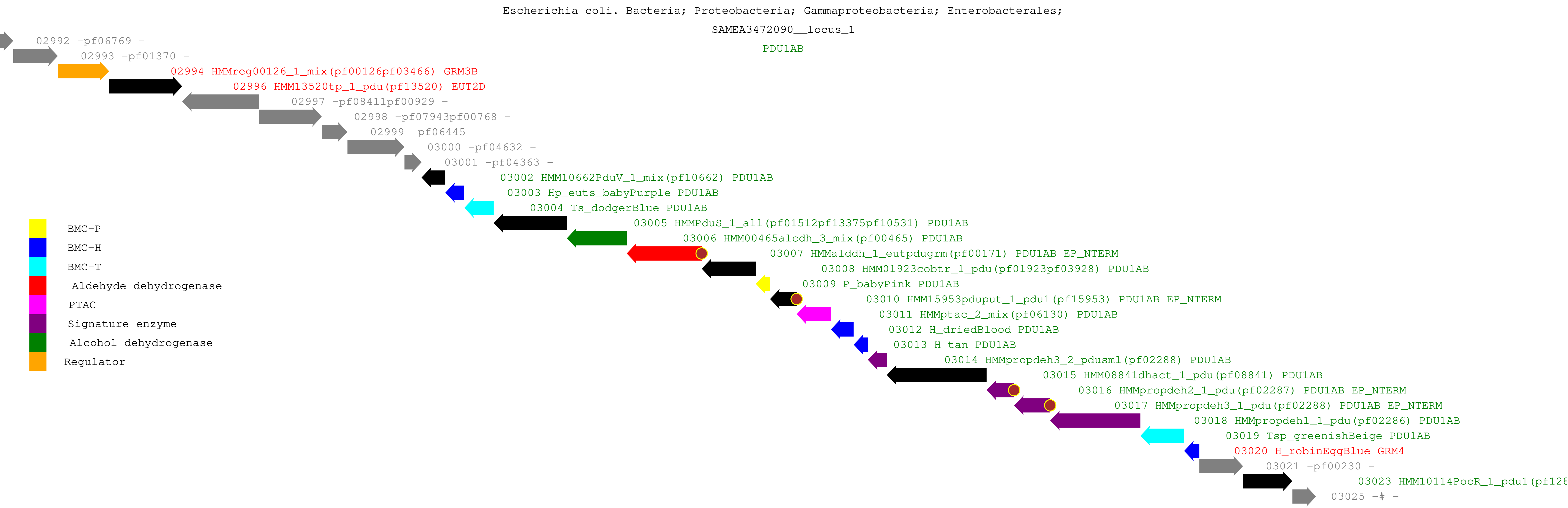
Escherichia coli. Bacteria; Proteobacteria; Gammaproteobacteria; Enterobacterales;
SAMEA3472090__locus_1
PDU1AB
02992 -pf06769 -
02993 -pf01370 -
02994 HMMreg00126_1_mix(pf00126pf03466) GRM3B
02996 HMM13520tp_1_pdu(pf13520) EUT2D
02997 -pf08411pf00929 -
02998 -pf07943pf00768 -
02999 -pf06445 -
03000 -pf04632 -
03001 -pf04363 -
03002 HMM10662PduV_1_mix(pf10662) PDU1AB
03003 Hp_euts_babyPurple PDU1AB
03004 Ts_dodgerBlue PDU1AB
03005 HMMPduS_1_all(pf01512pf13375pf10531) PDU1AB
03006 HMM00465alcdh_3_mix(pf00465) PDU1AB
03007 HMMalddh_1_eutpdugrm(pf00171) PDU1AB EP_NTERM
03008 HMM01923cobtr_1_pdu(pf01923pf03928) PDU1AB
03009 P_babyPink PDU1AB
03010 HMM15953pduput_1_pdu1(pf15953) PDU1AB EP_NTERM
03011 HMMptac_2_mix(pf06130) PDU1AB
03012 H_driedBlood PDU1AB
03013 H_tan PDU1AB
03014 HMMpropdeh3_2_pdusml(pf02288) PDU1AB
03015 HMM08841dhact_1_pdu(pf08841) PDU1AB
03016 HMMpropdeh2_1_pdu(pf02287) PDU1AB EP_NTERM
03017 HMMpropdeh3_1_pdu(pf02288) PDU1AB EP_NTERM
03018 HMMpropdeh1_1_pdu(pf02286) PDU1AB
03019 Tsp_greenishBeige PDU1AB
03020 H_robinEggBlue GRM4
03021 -pf00230 -
03023 HMM10114PocR_1_pdu1(pf128
03025 -# -
BMC-P
BMC-H
BMC-T
Aldehyde dehydrogenase
PTAC
Signature enzyme
Alcohol dehydrogenase
Regulator

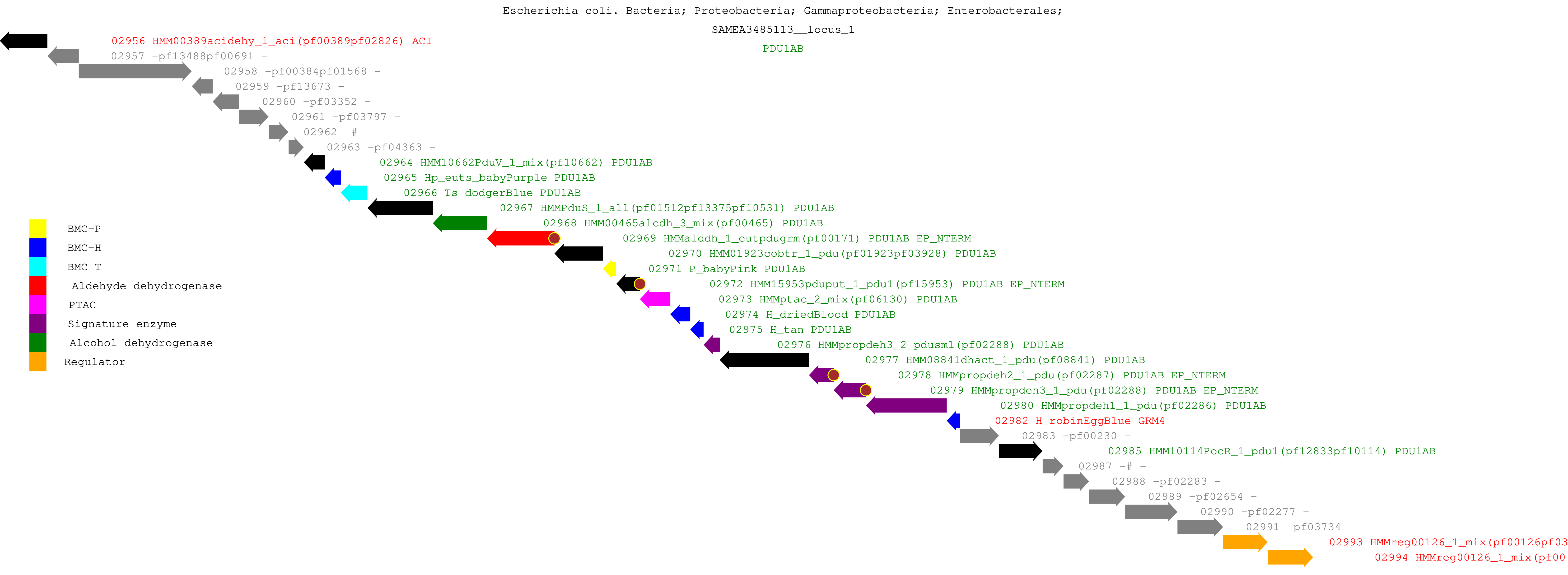
Escherichia coli. Bacteria; Proteobacteria; Gammaproteobacteria; Enterobacterales;
SAMEA3485113__locus_1
PDU1AB
02956 HMM00389acidehy_1_aci(pf00389pf02826) ACI
02957 -pf13488pf00691 -
02958 -pf00384pf01568 -
02959 -pf13673 -
02960 -pf03352 -
02961 -pf03797 -
02962 -# -
02963 -pf04363 -
02964 HMM10662PduV_1_mix(pf10662) PDU1AB
02965 Hp_euts_babyPurple PDU1AB
02966 Ts_dodgerBlue PDU1AB
02967 HMMPduS_1_all(pf01512pf13375pf10531) PDU1AB
02968 HMM00465alcdh_3_mix(pf00465) PDU1AB
02969 HMMalddh_1_eutpdugrm(pf00171) PDU1AB EP_NTERM
02970 HMM01923cobtr_1_pdu(pf01923pf03928) PDU1AB
02971 P_babyPink PDU1AB
02972 HMM15953pduput_1_pdu1(pf15953) PDU1AB EP_NTERM
02973 HMMptac_2_mix(pf06130) PDU1AB
02974 H_driedBlood PDU1AB
02975 H_tan PDU1AB
02976 HMMpropdeh3_2_pdusml(pf02288) PDU1AB
02977 HMM08841dhact_1_pdu(pf08841) PDU1AB
02978 HMMpropdeh2_1_pdu(pf02287) PDU1AB EP_NTERM
02979 HMMpropdeh3_1_pdu(pf02288) PDU1AB EP_NTERM
02980 HMMpropdeh1_1_pdu(pf02286) PDU1AB
02982 H_robinEggBlue GRM4
02983 -pf00230 -
02985 HMM10114PocR_1_pdu1(pf12833pf10114) PDU1AB
02987 -# -
02988 -pf02283 -
02989 -pf02654 -
02990 -pf02277 -
02991 -pf03734 -
02993 HMMreg00126_1_mix(pf00126pf03
02994 HMMreg00126_1_mix(pf00
BMC-P
BMC-H
BMC-T
Aldehyde dehydrogenase
PTAC
Signature enzyme
Alcohol dehydrogenase
Regulator

Klebsiella variicola. Bacteria; Proteobacteria; Gammaproteobacteria; Enterobacterales;

SAMEA3515120__locus_1

PDU1AB

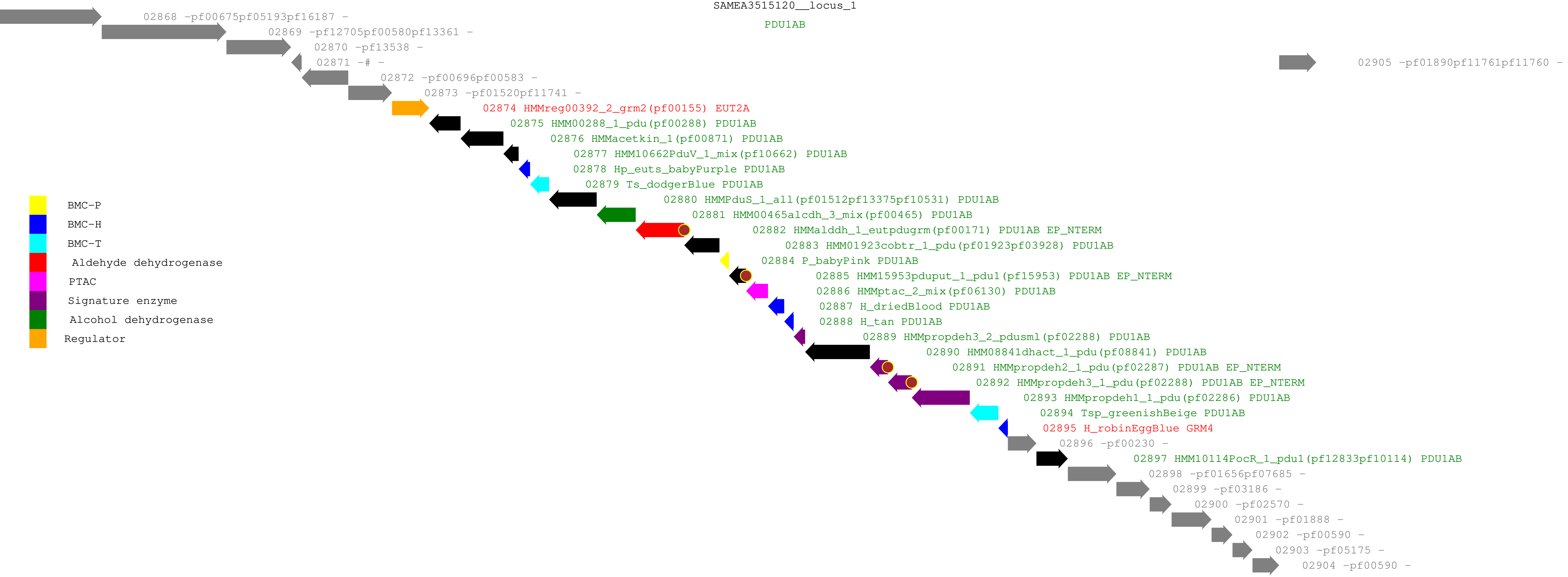

Klebsiella pneumoniae. Bacteria; Proteobacteria; Gammaproteobacteria; Enterobacterales;

SAMEA3531778__locus_1

PDU1AB

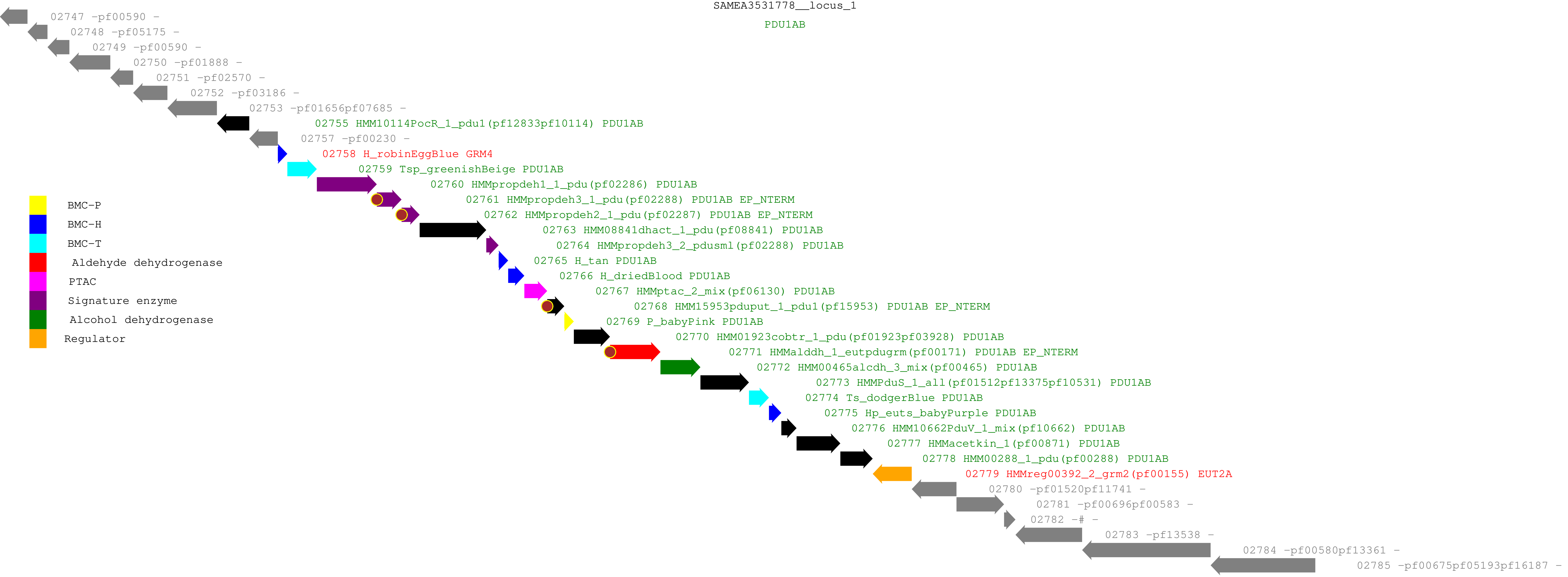

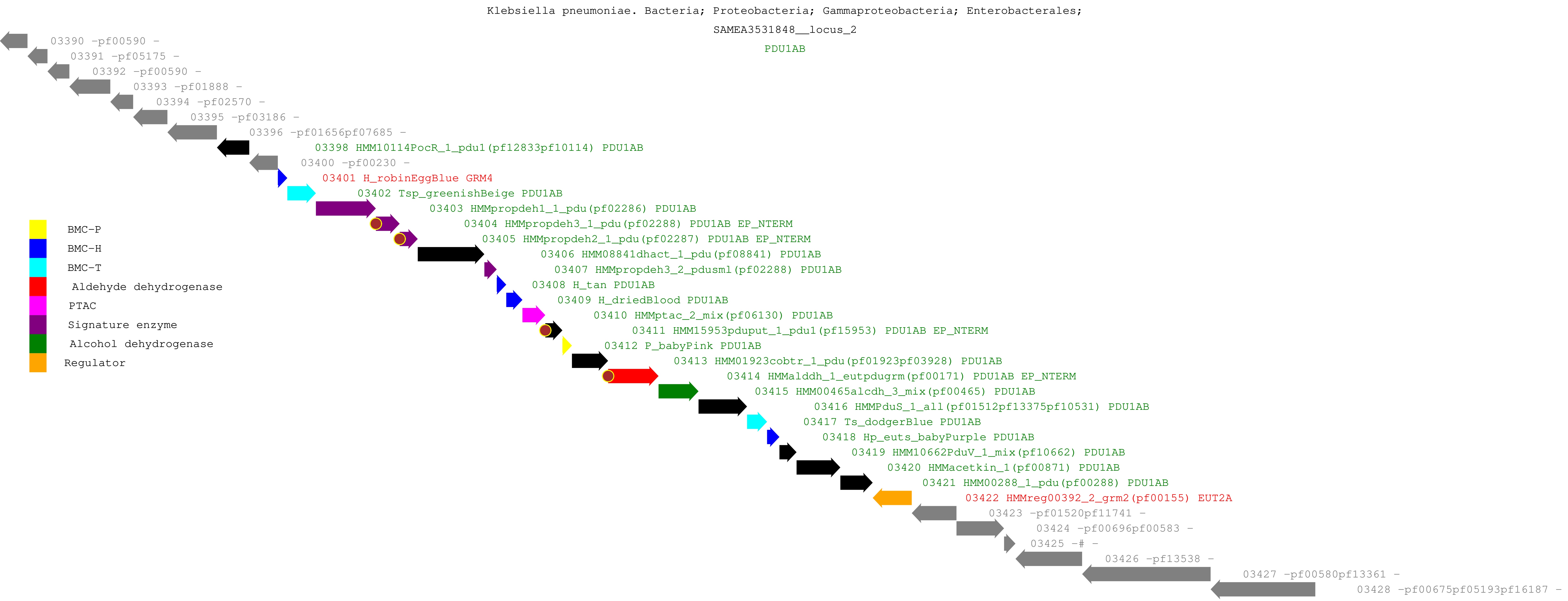
Klebsiella pneumoniae. Bacteria; Proteobacteria; Gammaproteobacteria; Enterobacterales;
SAMEA3531848__locus_2
PDU1AB
03390 -pf00590 -
03391 -pf05175 -
03392 -pf00590 -
03393 -pf01888 -
03394 -pf02570 -
03395 -pf03186 -
03396 -pf01656pf07685 -
03398 HMM10114PocR_1_pdu1(pf12833pf10114) PDU1AB
03400 -pf00230 -
03401 H_robinEggBlue GRM4
03402 Tsp_greenishBeige PDU1AB
03403 HMMpropdeh1_1_pdu(pf02286) PDU1AB
03404 HMMpropdeh3_1_pdu(pf02288) PDU1AB EP_NTERM
03405 HMMpropdeh2_1_pdu(pf02287) PDU1AB EP_NTERM
03406 HMM08841dhact_1_pdu(pf08841) PDU1AB
03407 HMMpropdeh3_2_pdusml(pf02288) PDU1AB
03408 H_tan PDU1AB
03409 H_driedBlood PDU1AB
03410 HMMptac_2_mix(pf06130) PDU1AB
03411 HMM15953pduput_1_pdu1(pf15953) PDU1AB EP_NTERM
03412 P_babyPink PDU1AB
03413 HMM01923cobtr_1_pdu(pf01923pf03928) PDU1AB
03414 HMMalddh_1_eutpdugrm(pf00171) PDU1AB EP_NTERM
03415 HMM00465alcdh_3_mix(pf00465) PDU1AB
03416 HMMPduS_1_all(pf01512pf13375pf10531) PDU1AB
03417 Ts_dodgerBlue PDU1AB
03418 Hp_euts_babyPurple PDU1AB
03419 HMM10662PduV_1_mix(pf10662) PDU1AB
03420 HMMacetkin_1(pf00871) PDU1AB
03421 HMM00288_1_pdu(pf00288) PDU1AB
03422 HMMreg00392_2_grm2(pf00155) EUT2A
03423 -pf01520pf11741 -
03424 -pf00696pf00583 -
03425 -# -
03426 -pf13538 -
03427 -pf00580pf13361 -
03428 -pf00675pf05193pf16187 -
BMC-P
BMC-H
BMC-T
Aldehyde dehydrogenase
PTAC
Signature enzyme
Alcohol dehydrogenase
Regulator

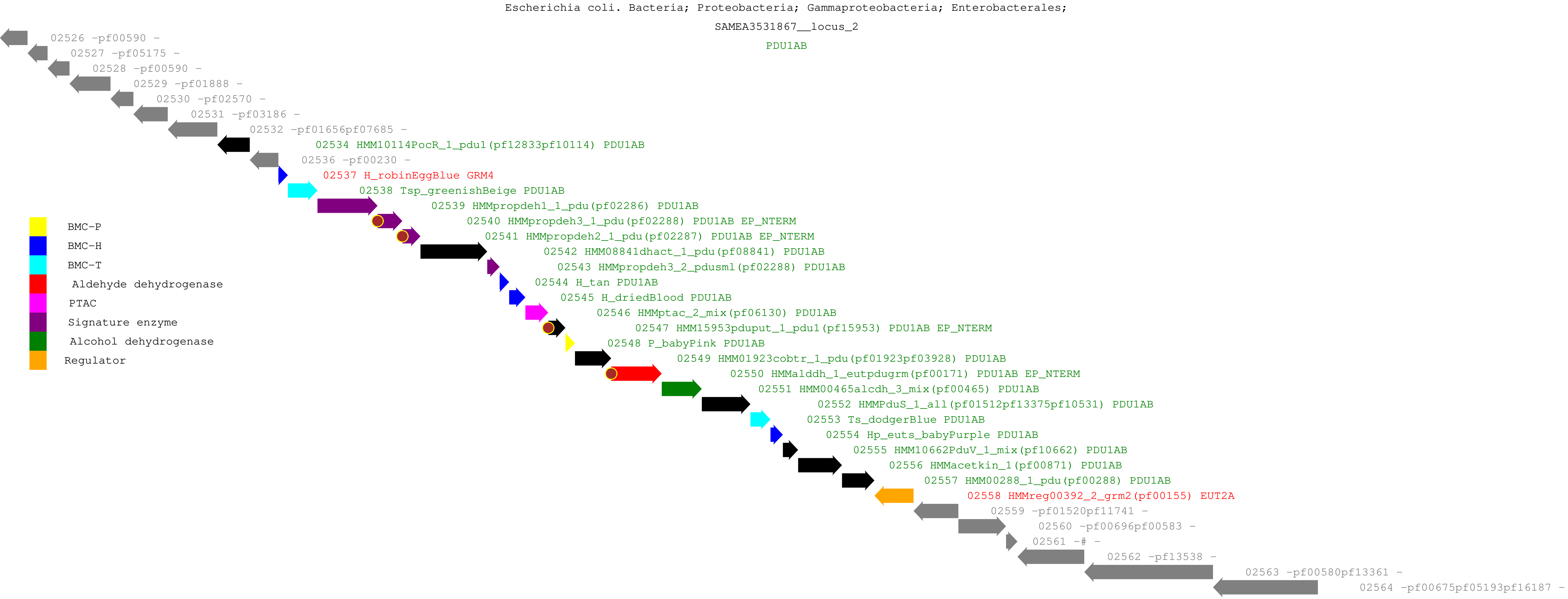

Escherichia coli. Bacteria; Proteobacteria; Gammaproteobacteria; Enterobacterales;
SAMEA3531867__locus_2
PDU1AB
02526 -pf00590 -
02527 -pf05175 -
02528 -pf00590 -
02529 -pf01888 -
02530 -pf02570 -
02531 -pf03186 -
02532 -pf01656pf07685 -
02534 HMM10114PocR_1_pdu1(pf12833pf10114) PDU1AB
02536 -pf00230 -
02537 H_robinEggBlue GRM4
02538 Tsp_greenishBeige PDU1AB
02539 HMMpropdeh1_1_pdu(pf02286) PDU1AB
02540 HMMpropdeh3_1_pdu(pf02288) PDU1AB EP_NTERM
02541 HMMpropdeh2_1_pdu(pf02287) PDU1AB EP_NTERM
02542 HMM08841dhact_1_pdu(pf08841) PDU1AB
02543 HMMpropdeh3_2_pdusml(pf02288) PDU1AB
02544 H_tan PDU1AB
02545 H_driedBlood PDU1AB
02546 HMMptac_2_mix(pf06130) PDU1AB
02547 HMM15953pduput_1_pdu1(pf15953) PDU1AB EP_NTERM
02548 P_babyPink PDU1AB
02549 HMM01923cobtr_1_pdu(pf01923pf03928) PDU1AB
02550 HMMaldh_1_eutpdugrm(pf00171) PDU1AB EP_NTERM
02551 HMM00465alcdh_3_mix(pf00465) PDU1AB
02552 HMMPduS_1_all(pf01512pf13375pf10531) PDU1AB
02553 Ts_dodgerBlue PDU1AB
02554 Hp_euts_babyPurple PDU1AB
02555 HMM10662PduV_1_mix(pf10662) PDU1AB
02556 HMMacetkin_1(pf00871) PDU1AB
02557 HMM00288_1_pdu(pf00288) PDU1AB
02558 HMMreg00392_2_grm2(pf00155) EUT2A
02559 -pf01520pf11741 -
02560 -pf00696pf00583 -
02561 -# -
02562 -pf13538 -
02563 -pf00580pf13361 -
02564 -pf00675pf05193pf16187 -
BMC-P
BMC-H
BMC-T
Aldehyde dehydrogenase
PTAC
Signature enzyme
Alcohol dehydrogenase
Regulator

Klebsiella quasivariicola. Bacteria; Proteobacteria; Gammaproteobacteria; Enterobacterales;

SAMEA3538468__locus_2

PDU1AB

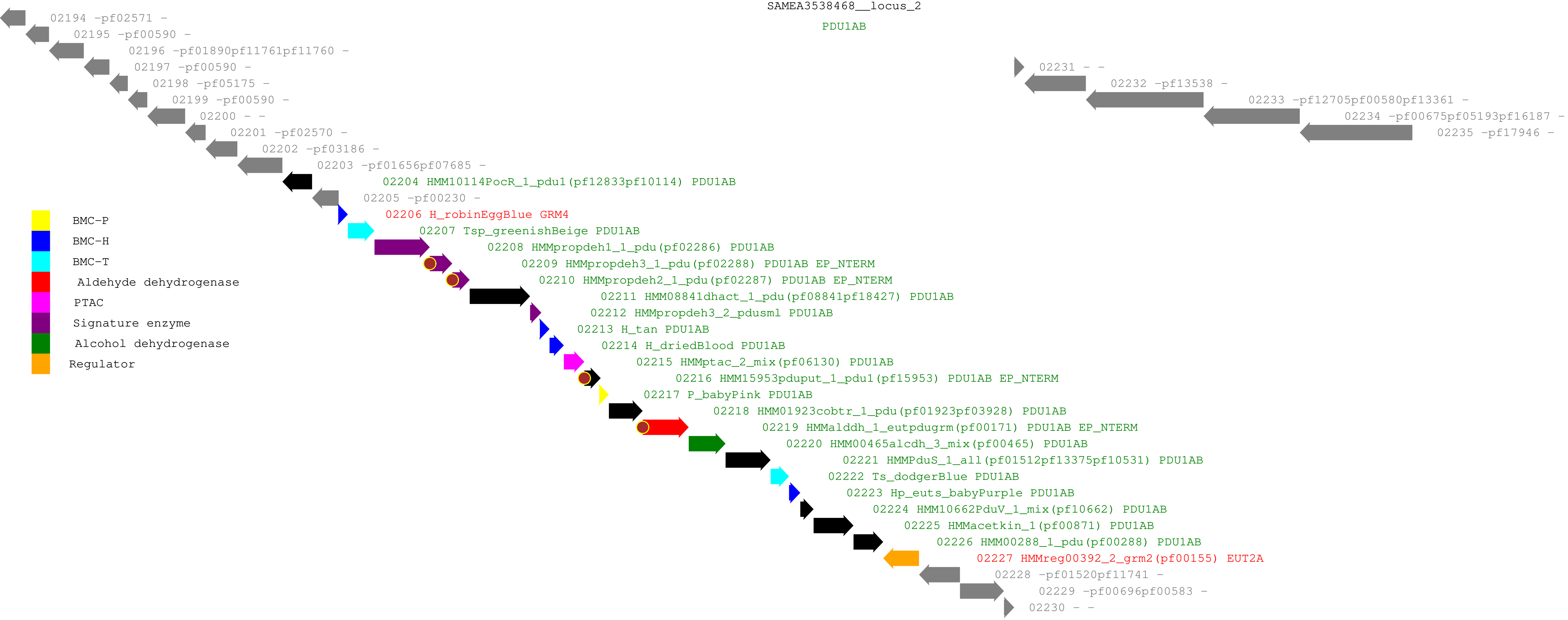

Klebsiella quasipneumoniae. Bacteria; Proteobacteria; Gammaproteobacteria; Enterobacterales;

SAMEA3649790__locus_1

PDU1AB

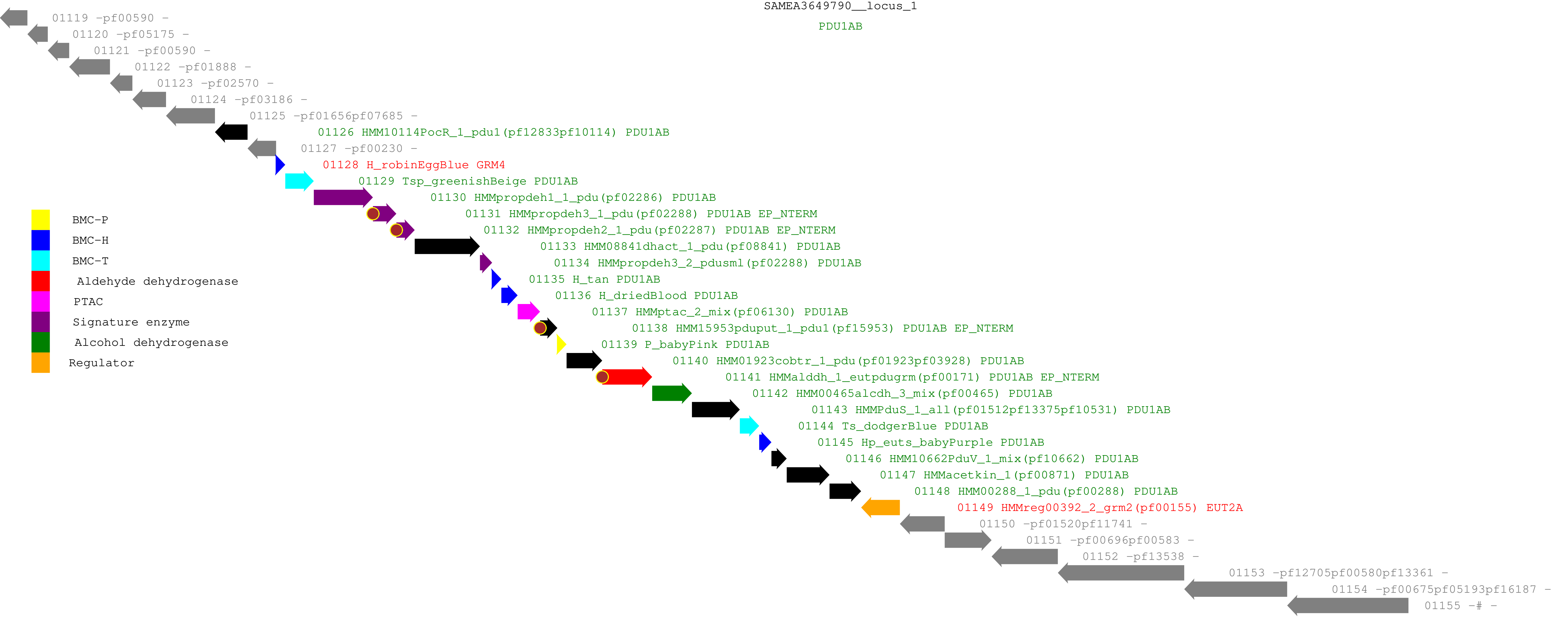

Klebsiella pneumoniae. Bacteria; Proteobacteria; Gammaproteobacteria; Enterobacterales;

SAMEA3729690__locus_2

PDU1AB

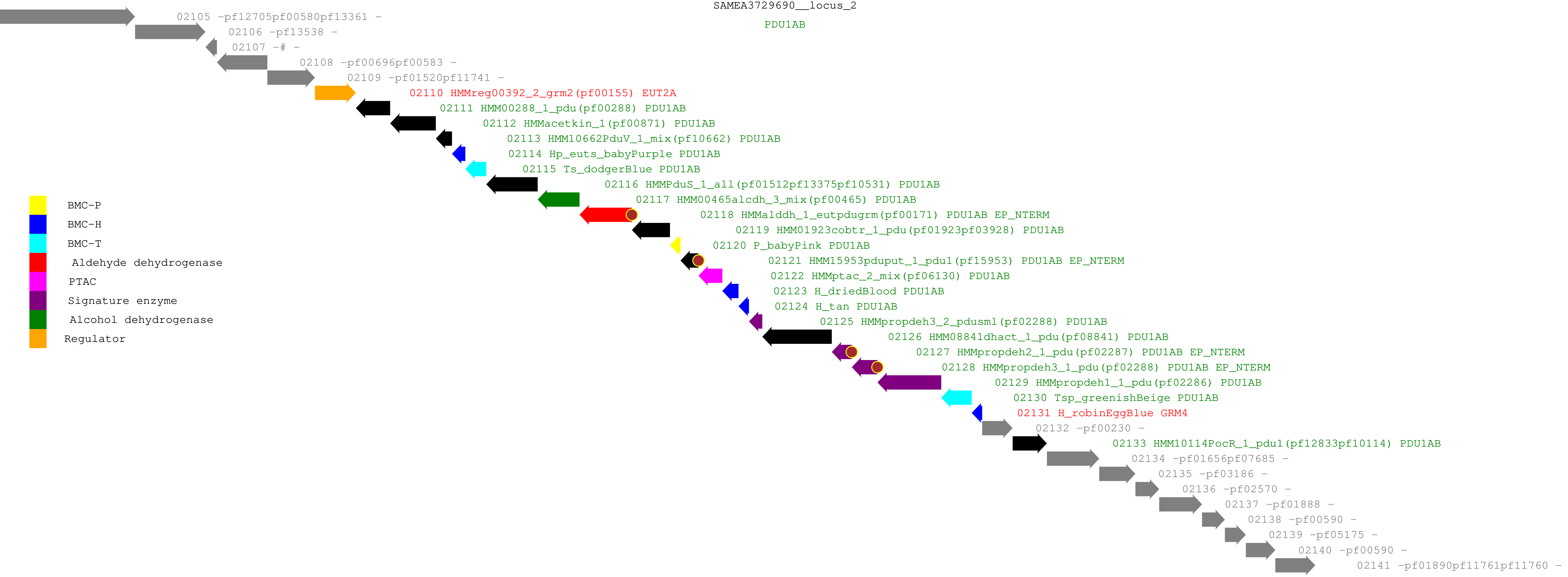

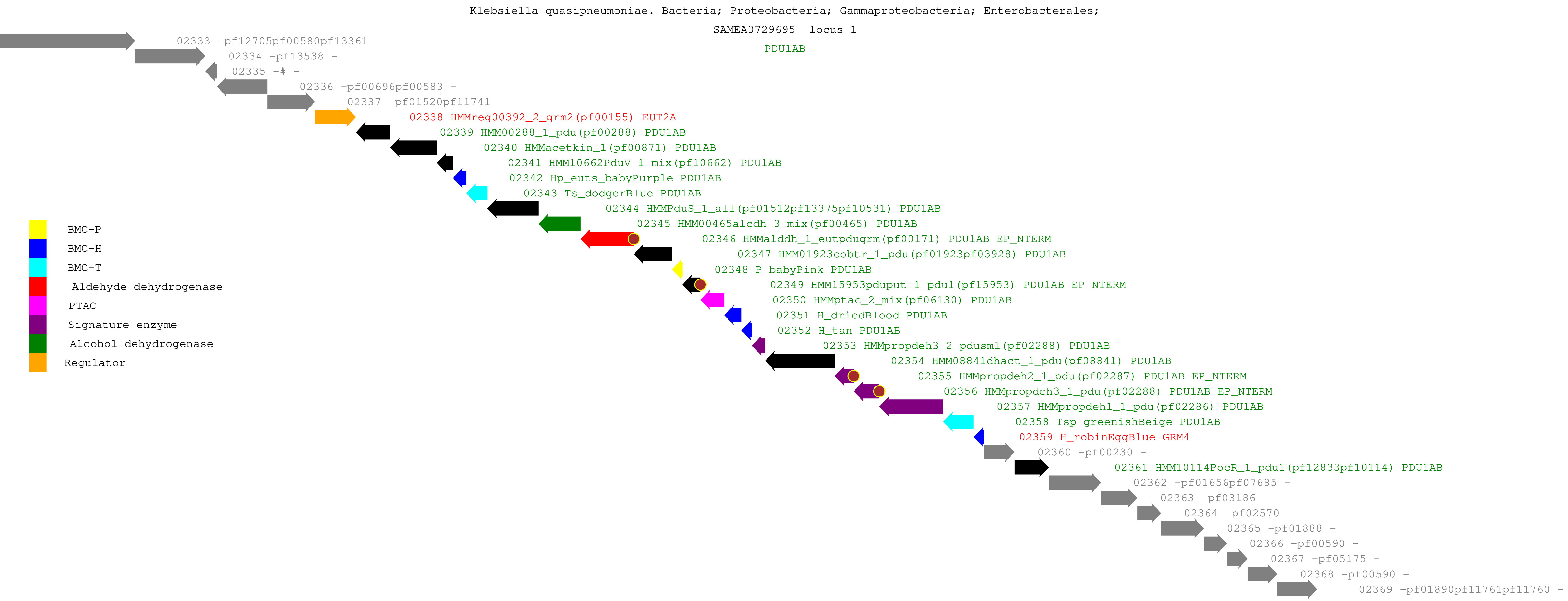
Klebsiella quasipneumoniae. Bacteria; Proteobacteria; Gammaproteobacteria; Enterobacterales;
SAMEA3729695__locus_1
PDU1AB
02333 -pf12705pf00580pf13361 -
02334 -pf13538 -
02335 -# -
02336 -pf00696pf00583 -
02337 -pf01520pf11741 -
02338 HMMreg00392_2_grm2(pf00155) EUT2A
02339 HMM00288_1_pdu(pf00288) PDU1AB
02340 HMMacetkin_1(pf00871) PDU1AB
02341 HMM10662PduV_1_mix(pf10662) PDU1AB
02342 Hp_euts_babyPurple PDU1AB
02343 Ts_dodgerBlue PDU1AB
02344 HMMPduS_1_all(pf01512pf13375pf10531) PDU1AB
02345 HMM00465alcdh_3_mix(pf00465) PDU1AB
02346 HMMalddh_1_eutpdugrm(pf00171) PDU1AB EP_NTERM
02347 HMM01923cobtr_1_pdu(pf01923pf03928) PDU1AB
02348 P_babyPink PDU1AB
02349 HMM15953pduput_1_pdu1(pf15953) PDU1AB EP_NTERM
02350 HMMptac_2_mix(pf06130) PDU1AB
02351 H_driedBlood PDU1AB
02352 H_tan PDU1AB
02353 HMMpropdeh3_2_pdusml(pf02288) PDU1AB
02354 HMM08841dhact_1_pdu(pf08841) PDU1AB
02355 HMMpropdeh2_1_pdu(pf02287) PDU1AB EP_NTERM
02356 HMMpropdeh3_1_pdu(pf02288) PDU1AB EP_NTERM
02357 HMMpropdeh1_1_pdu(pf02286) PDU1AB
02358 Tsp_greenishBeige PDU1AB
02359 H_robinEggBlue GRM4
02360 -pf00230 -
02361 HMM10114PocR_1_pdu1(pf12833pf10114) PDU1AB
02362 -pf01656pf07685 -
02363 -pf03186 -
02364 -pf02570 -
02365 -pf01888 -
02366 -pf00590 -
02367 -pf05175 -
02368 -pf00590 -
02369 -pf01890pf11761pf11760 -
BMC-P
BMC-H
BMC-T
Aldehyde dehydrogenase
PTAC
Signature enzyme
Alcohol dehydrogenase
Regulator

Escherichia coli. Bacteria; Proteobacteria; Gammaproteobacteria; Enterobacterales;

SAMEA3753137__locus_1

PDU1AB

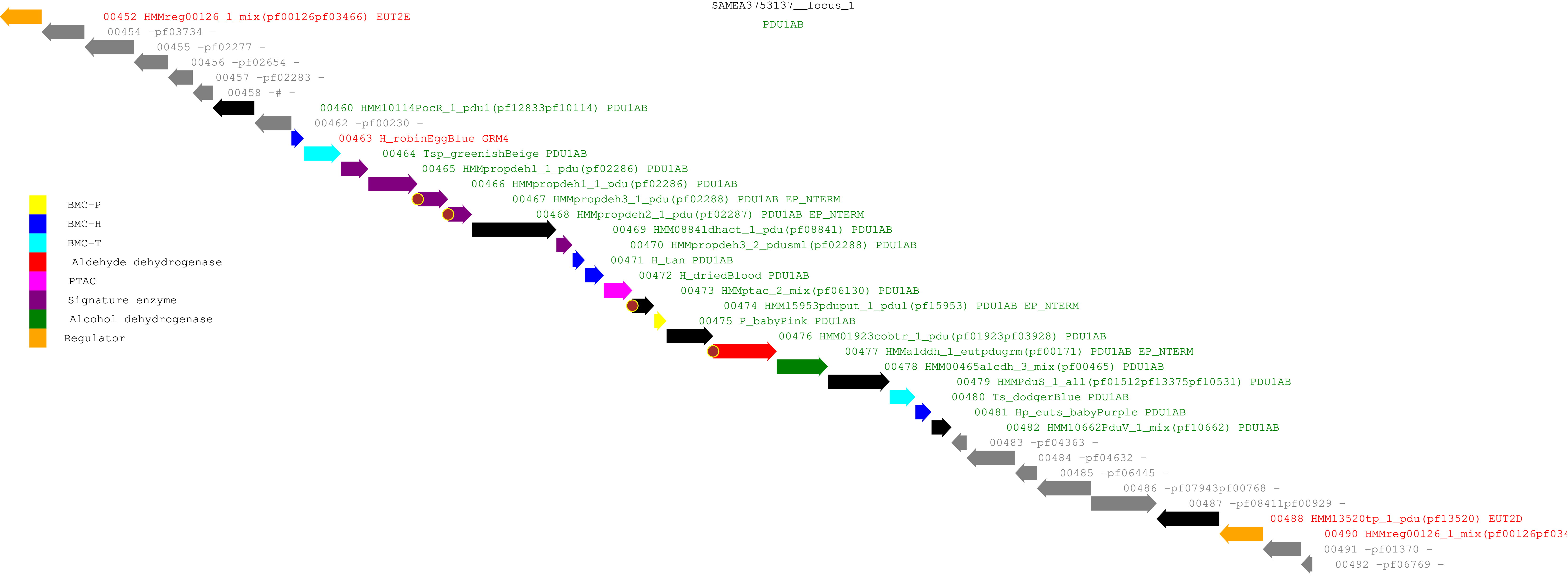

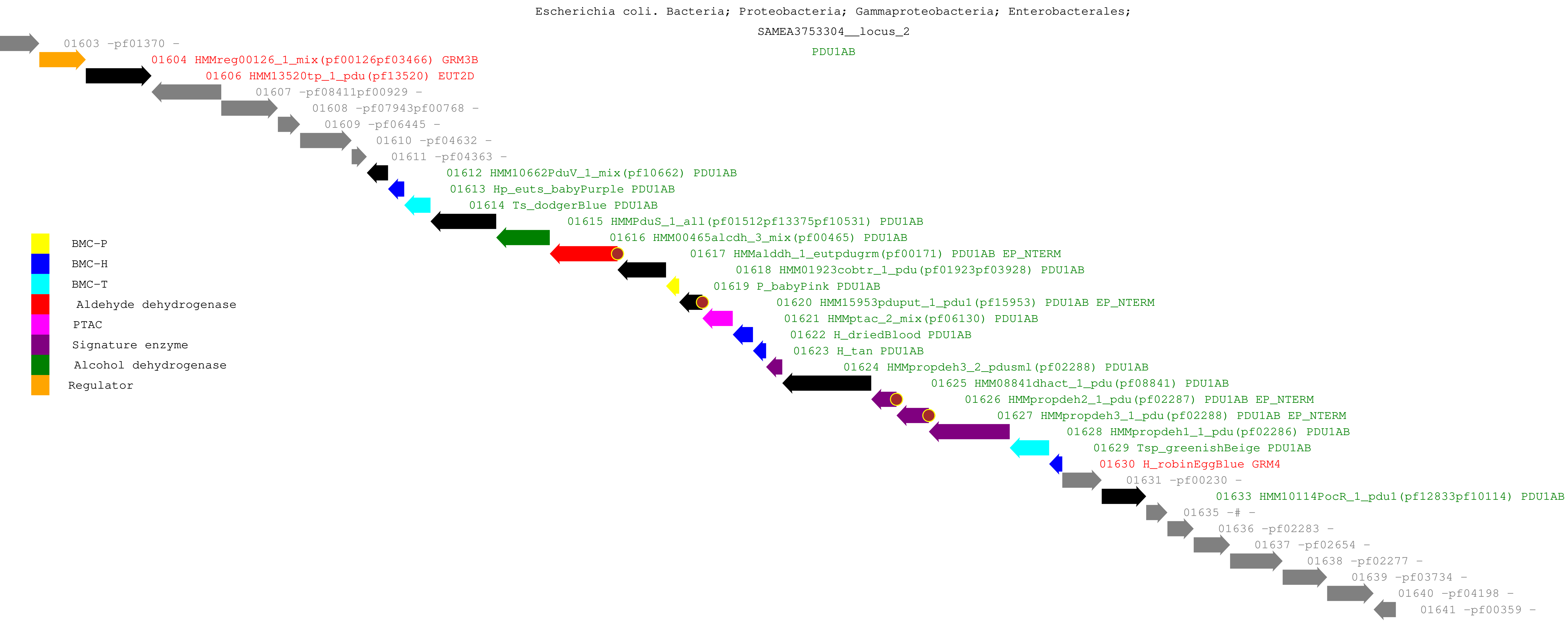

Escherichia coli. Bacteria; Proteobacteria; Gammaproteobacteria; Enterobacterales;
SAMEA3753304__locus_2
PDU1AB
01603 -pf01370 -
01604 HMMreg00126_1_mix(pf00126pf03466) GRM3B
01606 HMM13520tp_1_pdu(pf13520) EUT2D
01607 -pf08411pf00929 -
01608 -pf07943pf00768 -
01609 -pf06445 -
01610 -pf04632 -
01611 -pf04363 -
01612 HMM10662PduV_1_mix(pf10662) PDU1AB
01613 Hp_euts_babyPurple PDU1AB
01614 Ts_dodgerBlue PDU1AB
01615 HMMPduS_1_all(pf01512pf13375pf10531) PDU1AB
01616 HMM00465alcdh_3_mix(pf00465) PDU1AB
01617 HMMalddh_1_eutpdugrm(pf00171) PDU1AB EP_NTERM
01618 HMM01923cobtr_1_pdu(pf01923pf03928) PDU1AB
01619 P_babyPink PDU1AB
01620 HMM15953pduput_1_pdu1(pf15953) PDU1AB EP_NTERM
01621 HMMptac_2_mix(pf06130) PDU1AB
01622 H_driedBlood PDU1AB
01623 H_tan PDU1AB
01624 HMMpropdeh3_2_pdusml(pf02288) PDU1AB
01625 HMM08841dhact_1_pdu(pf08841) PDU1AB
01626 HMMpropdeh2_1_pdu(pf02287) PDU1AB EP_NTERM
01627 HMMpropdeh3_1_pdu(pf02288) PDU1AB EP_NTERM
01628 HMMpropdeh1_1_pdu(pf02286) PDU1AB
01629 Tsp_greenishBeige PDU1AB
01630 H_robinEggBlue GRM4
01631 -pf00230 -
01633 HMM10114PocR_1_pdu1(pf12833pf10114) PDU1AB
01635 -# -
01636 -pf02283 -
01637 -pf02654 -
01638 -pf02277 -
01639 -pf03734 -
01640 -pf04198 -
01641 -pf00359 -
BMC-P
BMC-H
BMC-T
Aldehyde dehydrogenase
PTAC
Signature enzyme
Alcohol dehydrogenase
Regulator

Klebsiella pneumoniae. Bacteria; Proteobacteria; Gammaproteobacteria; Enterobacterales;

SAMEA4362597__locus_1

PDU1AB

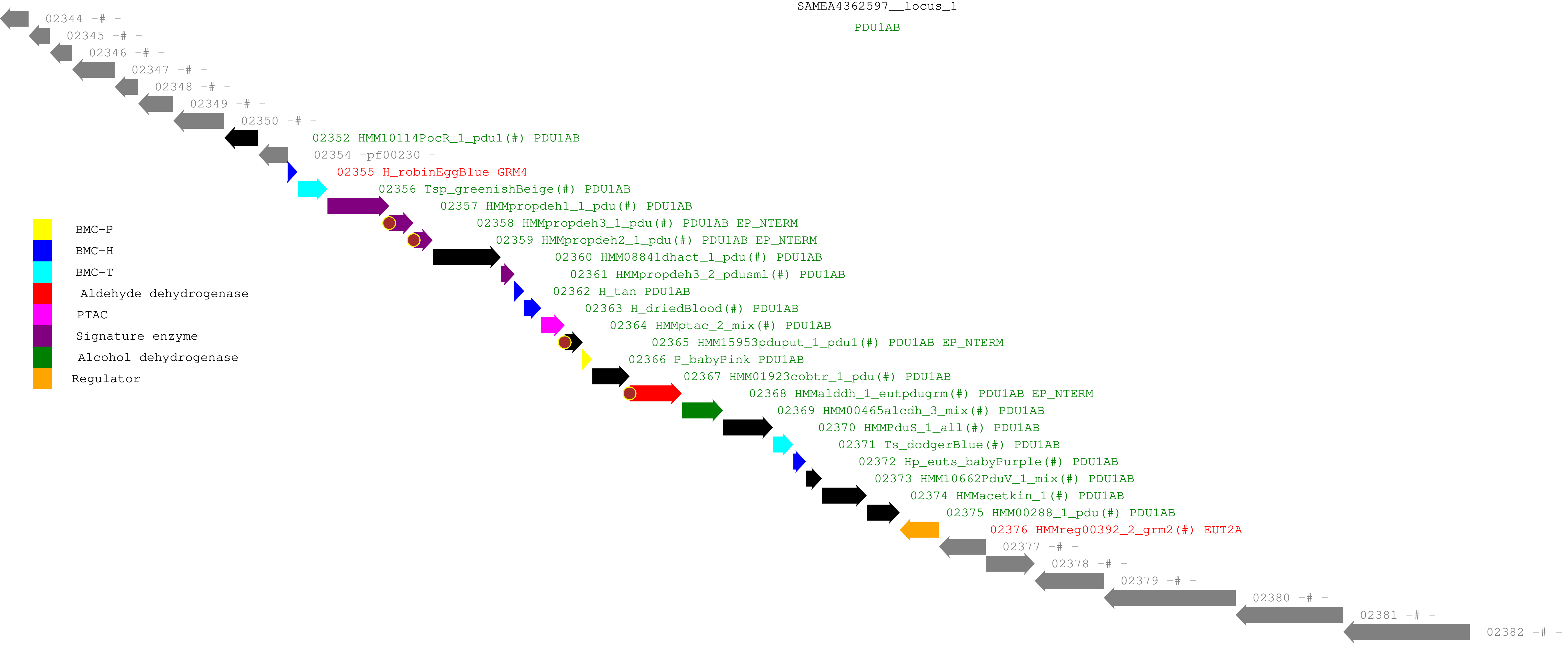

Klebsiella pneumoniae. Bacteria; Proteobacteria; Gammaproteobacteria; Enterobacterales;

SAMEA4362680__locus_2

PDU1AB

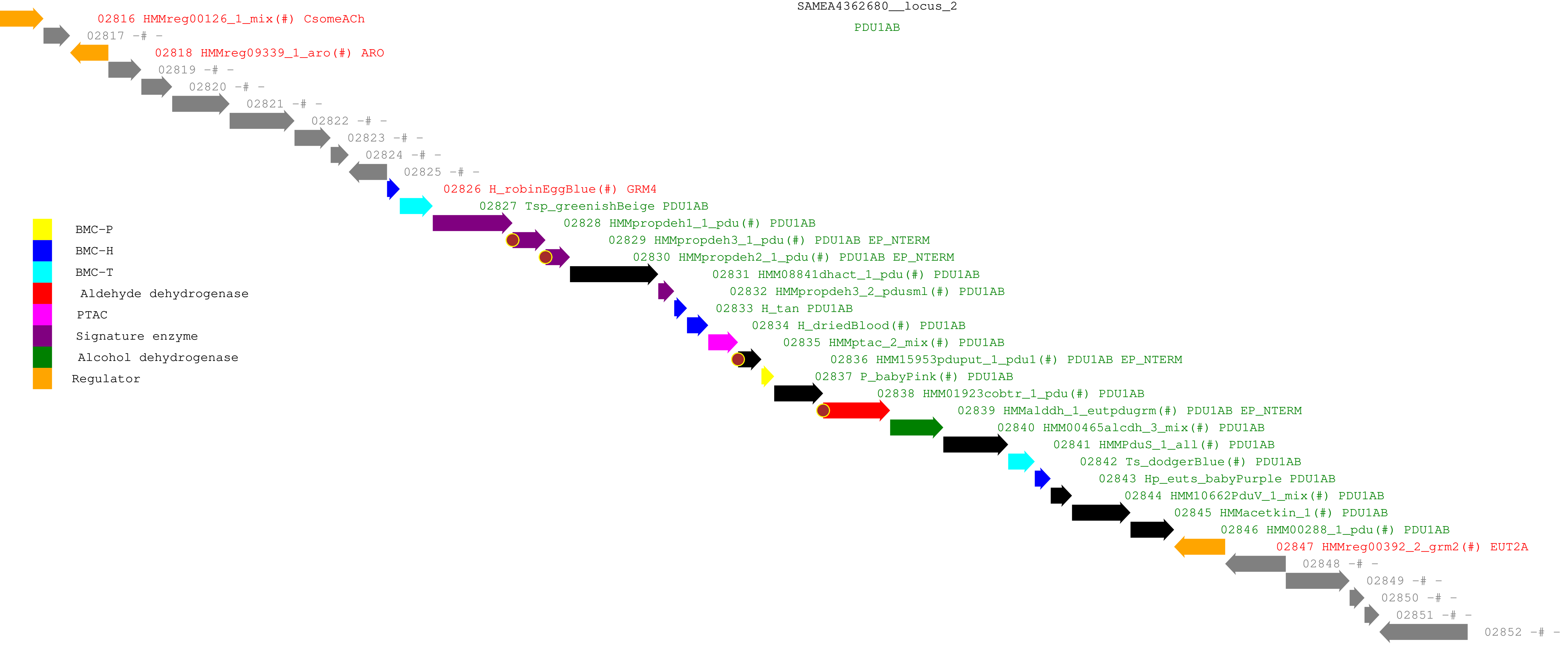

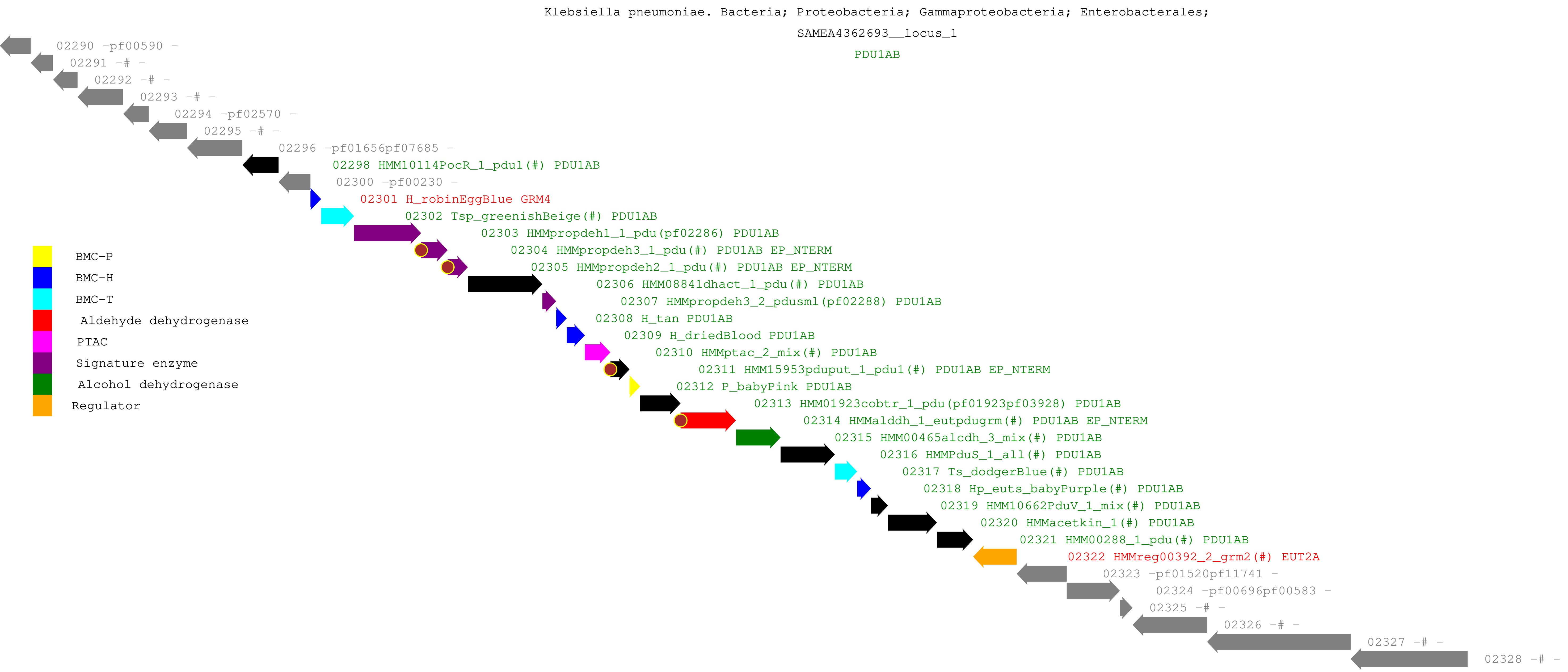

Klebsiella pneumoniae. Bacteria; Proteobacteria; Gammaproteobacteria; Enterobacterales;
SAMEA4362693__locus_1
PDU1AB
02290 -pf00590 -
02291 -# -
02292 -# -
02293 -# -
02294 -pf02570 -
02295 -# -
02296 -pf01656pf07685 -
02298 HMM10114PocR_1_pdu1(#) PDU1AB
02300 -pf00230 -
02301 H_robinEggBlue GRM4
02302 Tsp_greenishBeige(#) PDU1AB
02303 HMMpropdeh1_1_pdu(pf02286) PDU1AB
02304 HMMpropdeh3_1_pdu(#) PDU1AB EP_NTERM
02305 HMMpropdeh2_1_pdu(#) PDU1AB EP_NTERM
02306 HMM08841dhact_1_pdu(#) PDU1AB
02307 HMMpropdeh3_2_pdusml(pf02288) PDU1AB
02308 H_tan PDU1AB
02309 H_driedBlood PDU1AB
02310 HMMptac_2_mix(#) PDU1AB
02311 HMM15953pduput_1_pdu1(#) PDU1AB EP_NTERM
02312 P_babyPink PDU1AB
02313 HMM01923cobtr_1_pdu(pf01923pf03928) PDU1AB
02314 HMMalddh_1_eutpdugrm(#) PDU1AB EP_NTERM
02315 HMM00465alcdh_3_mix(#) PDU1AB
02316 HMMPduS_1_all(#) PDU1AB
02317 Ts_dodgerBlue(#) PDU1AB
02318 Hp_euts_babyPurple(#) PDU1AB
02319 HMM10662PduV_1_mix(#) PDU1AB
02320 HMMacetkin_1(#) PDU1AB
02321 HMM00288_1_pdu(#) PDU1AB
02322 HMMreg00392_2_grm2(#) EUT2A
02323 -pf01520pf11741 -
02324 -pf00696pf00583 -
02325 -# -
02326 -# -
02327 -# -
02328 -# -
BMC-P
BMC-H
BMC-T
Aldehyde dehydrogenase
PTAC
Signature enzyme
Alcohol dehydrogenase
Regulator

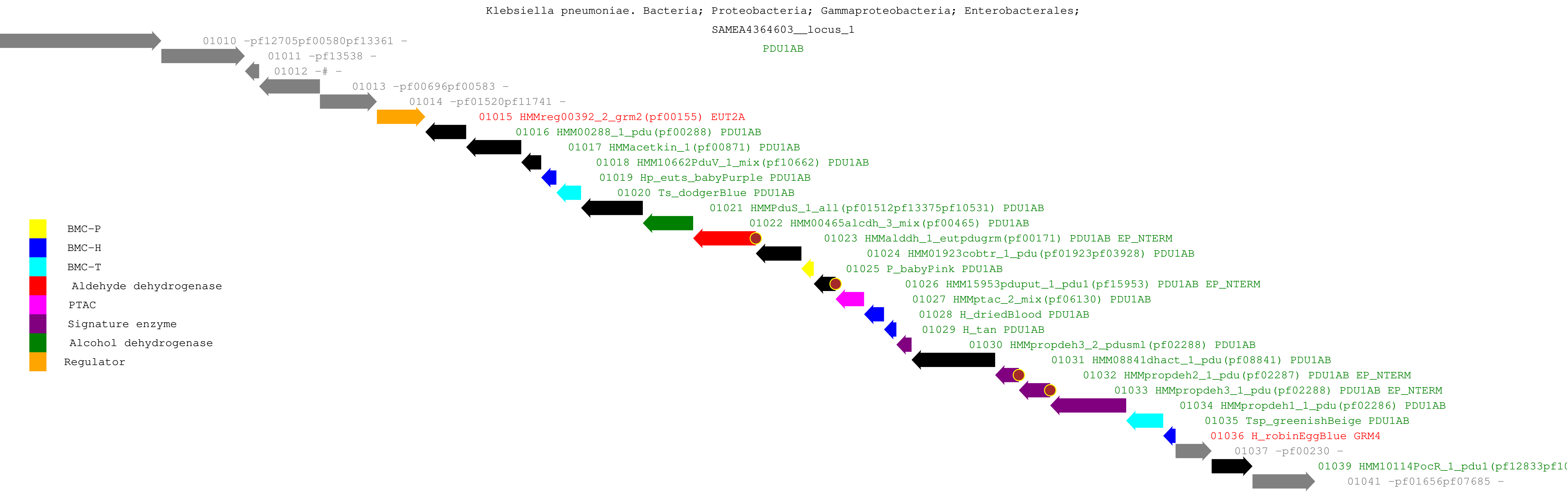
Klebsiella pneumoniae. Bacteria; Proteobacteria; Gammaproteobacteria; Enterobacterales;
SAMEA4364603__locus_1
PDU1AB
01010 -pf12705pf00580pf13361 -
01011 -pf13538 -
01012 -# -
01013 -pf00696pf00583 -
01014 -pf01520pf11741 -
01015 HMMreg00392_2_grm2(pf00155) EUT2A
01016 HMM00288_1_pdu(pf00288) PDU1AB
01017 HMMacetkin_1(pf00871) PDU1AB
01018 HMM10662PduV_1_mix(pf10662) PDU1AB
01019 Hp_euts_babyPurple PDU1AB
01020 Ts_dodgerBlue PDU1AB
01021 HMMPduS_1_all(pf01512pf13375pf10531) PDU1AB
01022 HMM00465alcdh_3_mix(pf00465) PDU1AB
01023 HMMalddh_1_eutpdugrm(pf00171) PDU1AB EP_NTERM
01024 HMM01923cobtr_1_pdu(pf01923pf03928) PDU1AB
01025 P_babyPink PDU1AB
01026 HMM15953pduput_1_pdu1(pf15953) PDU1AB EP_NTERM
01027 HMMptac_2_mix(pf06130) PDU1AB
01028 H_driedBlood PDU1AB
01029 H_tan PDU1AB
01030 HMMpropdeh3_2_pdusml(pf02288) PDU1AB
01031 HMM08841dhact_1_pdu(pf08841) PDU1AB
01032 HMMpropdeh2_1_pdu(pf02287) PDU1AB EP_NTERM
01033 HMMpropdeh3_1_pdu(pf02288) PDU1AB EP_NTERM
01034 HMMpropdeh1_1_pdu(pf02286) PDU1AB
01035 Tsp_greenishBeige PDU1AB
01036 H_robinEggBlue GRM4
01037 -pf00230 -
01039 HMM10114PocR_1_pdu1(pf12833pf1
01041 -pf01656pf07685 -
BMC-P
BMC-H
BMC-T
Aldehyde dehydrogenase
PTAC
Signature enzyme
Alcohol dehydrogenase
Regulator

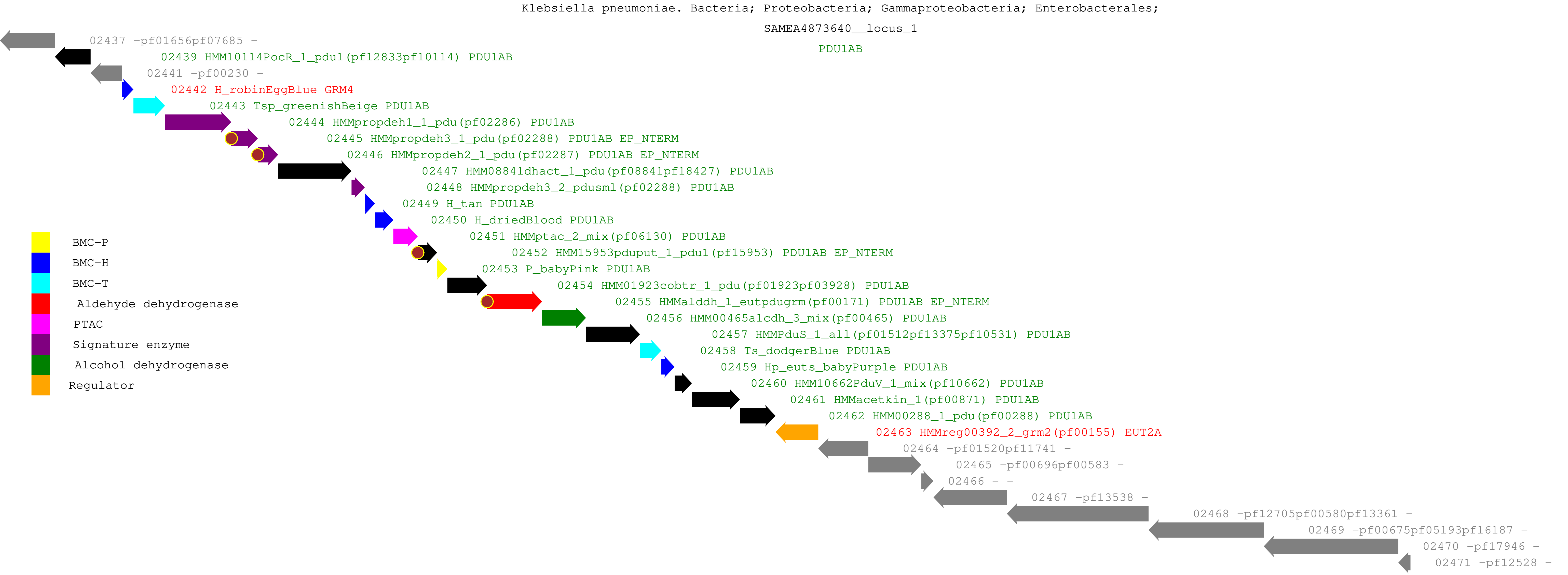

Klebsiella pneumoniae. Bacteria; Proteobacteria; Gammaproteobacteria; Enterobacterales;
SAMEA4873640__locus_1
PDU1AB
02437 -pf01656pf07685 -
02439 HMM10114PocR_1_pdu1(pf12833pf10114) PDU1AB
02441 -pf00230 -
02442 H_robinEggBlue GRM4
02443 Tsp_greenishBeige PDU1AB
02444 HMMpropdeh1_1_pdu(pf02286) PDU1AB
02445 HMMpropdeh3_1_pdu(pf02288) PDU1AB EP_NTERM
02446 HMMpropdeh2_1_pdu(pf02287) PDU1AB EP_NTERM
02447 HMM08841dhact_1_pdu(pf08841pf18427) PDU1AB
02448 HMMpropdeh3_2_pdusml(pf02288) PDU1AB
02449 H_tan PDU1AB
02450 H_driedBlood PDU1AB
02451 HMMptac_2_mix(pf06130) PDU1AB
02452 HMM15953pduput_1_pdu1(pf15953) PDU1AB EP_NTERM
02453 P_babyPink PDU1AB
02454 HMM01923cobtr_1_pdu(pf01923pf03928) PDU1AB
02455 HMMaldh_1_eutpdugrm(pf00171) PDU1AB EP_NTERM
02456 HMM00465alcdh_3_mix(pf00465) PDU1AB
02457 HMMPduS_1_all(pf01512pf13375pf10531) PDU1AB
02458 Ts_dodgerBlue PDU1AB
02459 Hp_euts_babyPurple PDU1AB
02460 HMM10662PduV_1_mix(pf10662) PDU1AB
02461 HMMacetkin_1(pf00871) PDU1AB
02462 HMM00288_1_pdu(pf00288) PDU1AB
02463 HMMreg00392_2_grm2(pf00155) EUT2A
02464 -pf01520pf11741 -
02465 -pf00696pf00583 -
02466 - -
02467 -pf13538 -
02468 -pf12705pf00580pf13361 -
02469 -pf00675pf05193pf16187 -
02470 -pf17946 -
02471 -pf12528 -
BMC-P
BMC-H
BMC-T
Aldehyde dehydrogenase
PTAC
Signature enzyme
Alcohol dehydrogenase
Regulator

Klebsiella sp. NFIX38. Bacteria; Proteobacteria; Gammaproteobacteria; Enterobacterales;

SAMN03159330__locus_1

PDU1AB

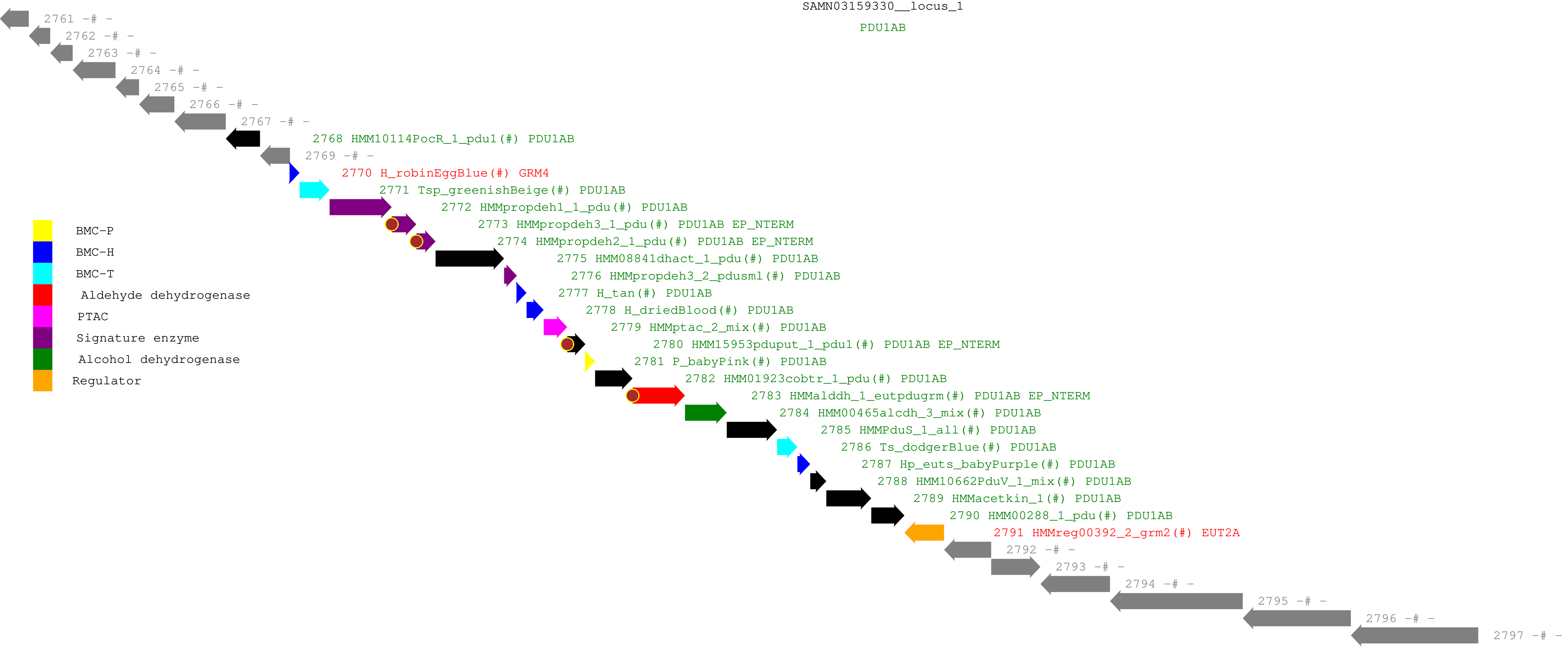

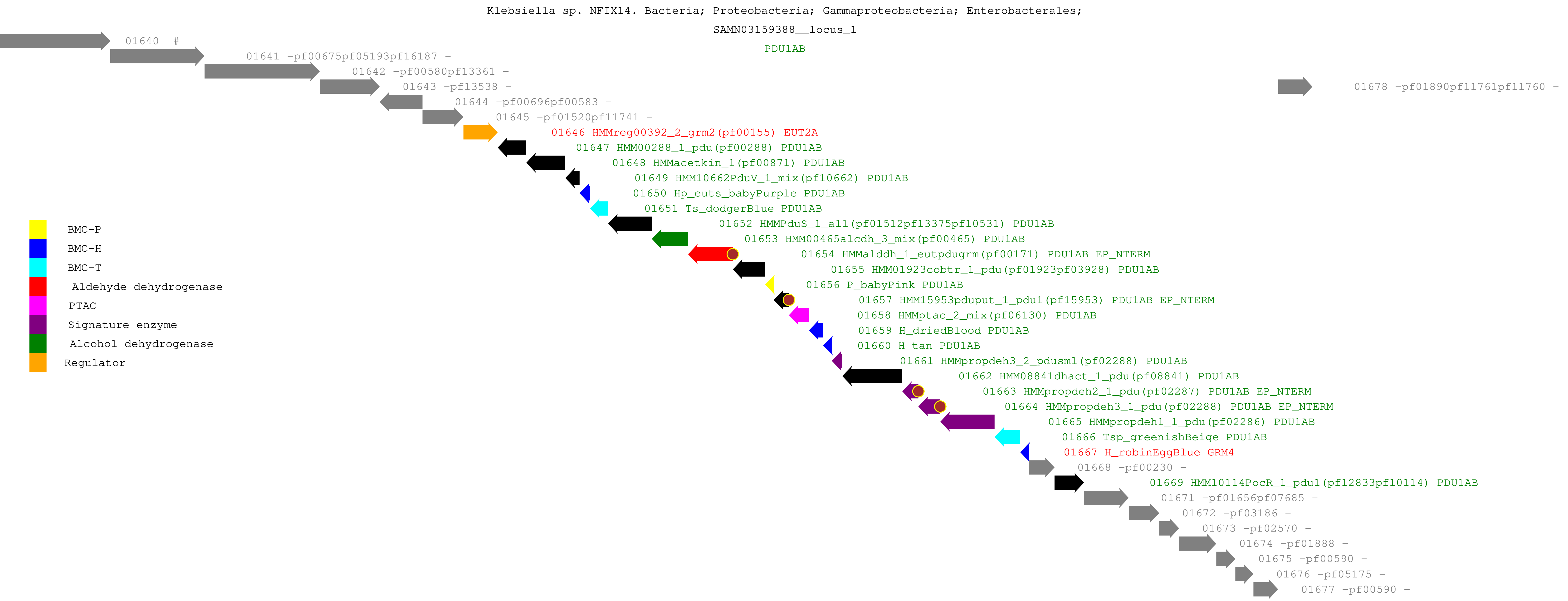
Klebsiella sp. NFIX14. Bacteria; Proteobacteria; Gammaproteobacteria; Enterobacterales;
SAMN03159388__locus_1
PDU1AB
01640 -# -
01641 -pf00675pf05193pf16187 -
01642 -pf00580pf13361 -
01643 -pf13538 -
01644 -pf00696pf00583 -
01645 -pf01520pf11741 -
01646 HMMreg00392_2_grm2(pf00155) EUT2A
01647 HMM00288_1_pdu(pf00288) PDU1AB
01648 HMMacetkin_1(pf00871) PDU1AB
01649 HMM10662PduV_1_mix(pf10662) PDU1AB
01650 Hp_euts_babyPurple PDU1AB
01651 Ts_dodgerBlue PDU1AB
01652 HMMPduS_1_all(pf01512pf13375pf10531) PDU1AB
01653 HMM00465alcdh_3_mix(pf00465) PDU1AB
01654 HMMalddh_1_eutpdugrm(pf00171) PDU1AB EP_NTERM
01655 HMM01923cobtr_1_pdu(pf01923pf03928) PDU1AB
01656 P_babyPink PDU1AB
01657 HMM15953pduput_1_pdu1(pf15953) PDU1AB EP_NTERM
01658 HMMptac_2_mix(pf06130) PDU1AB
01659 H_driedBlood PDU1AB
01660 H_tan PDU1AB
01661 HMMpropdeh3_2_pdusml(pf02288) PDU1AB
01662 HMM08841dhact_1_pdu(pf08841) PDU1AB
01663 HMMpropdeh2_1_pdu(pf02287) PDU1AB EP_NTERM
01664 HMMpropdeh3_1_pdu(pf02288) PDU1AB EP_NTERM
01665 HMMpropdeh1_1_pdu(pf02286) PDU1AB
01666 Tsp_greenishBeige PDU1AB
01667 H_robinEggBlue GRM4
01668 -pf00230 -
01669 HMM10114PocR_1_pdu1(pf12833pf10114) PDU1AB
01671 -pf01656pf07685 -
01672 -pf03186 -
01673 -pf02570 -
01674 -pf01888 -
01675 -pf00590 -
01676 -pf05175 -
01677 -pf00590 -
01678 -pf01890pf11761pf11760 -
BMC-P
BMC-H
BMC-T
Aldehyde dehydrogenase
PTAC
Signature enzyme
Alcohol dehydrogenase
Regulator

Enterobacter sp. NFR05. Bacteria; Proteobacteria; Gammaproteobacteria; Enterobacterales;

SAMN03159434__locus_1

PDU1AB

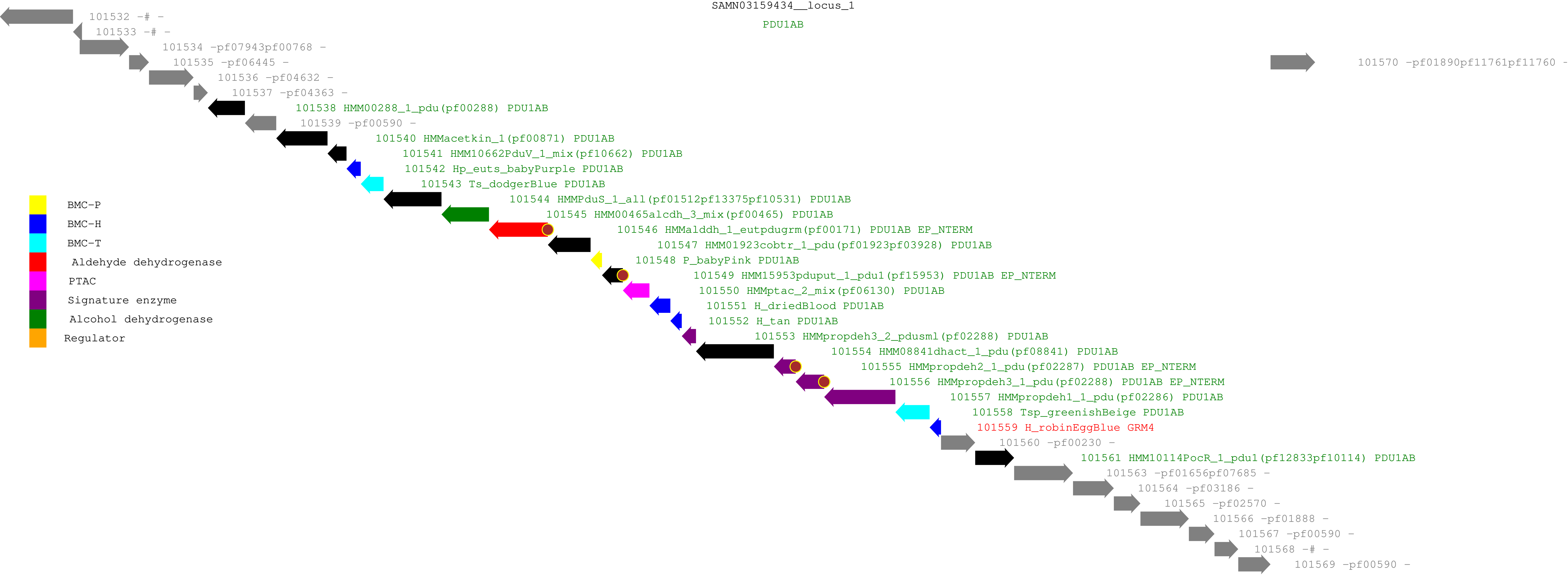

Vibrio xiamenensis. Bacteria; Proteobacteria; Gammaproteobacteria; Vibrionales;

SAMN04488136__locus_1_main_sat

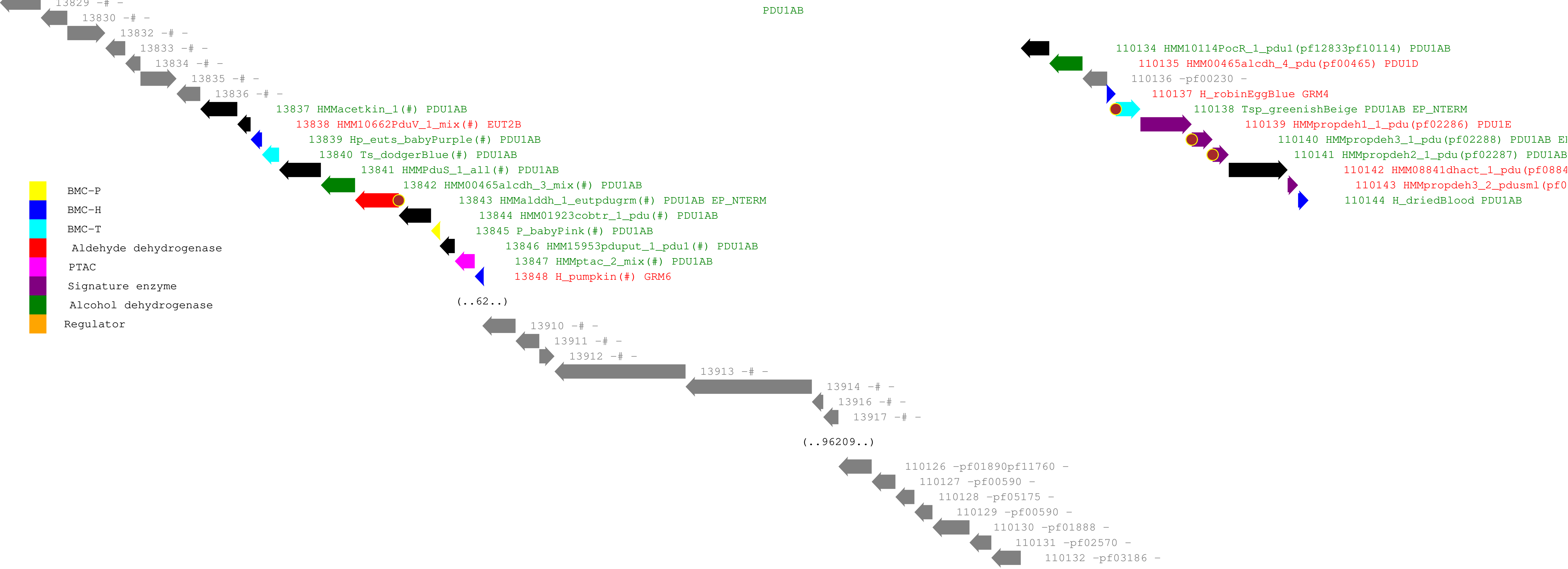

Citrobacter sp. NLAE-zl-C269. Bacteria; Proteobacteria; Gammaproteobacteria; Enterobacterales;

SAMN05216502__locus_1

PDU1AB

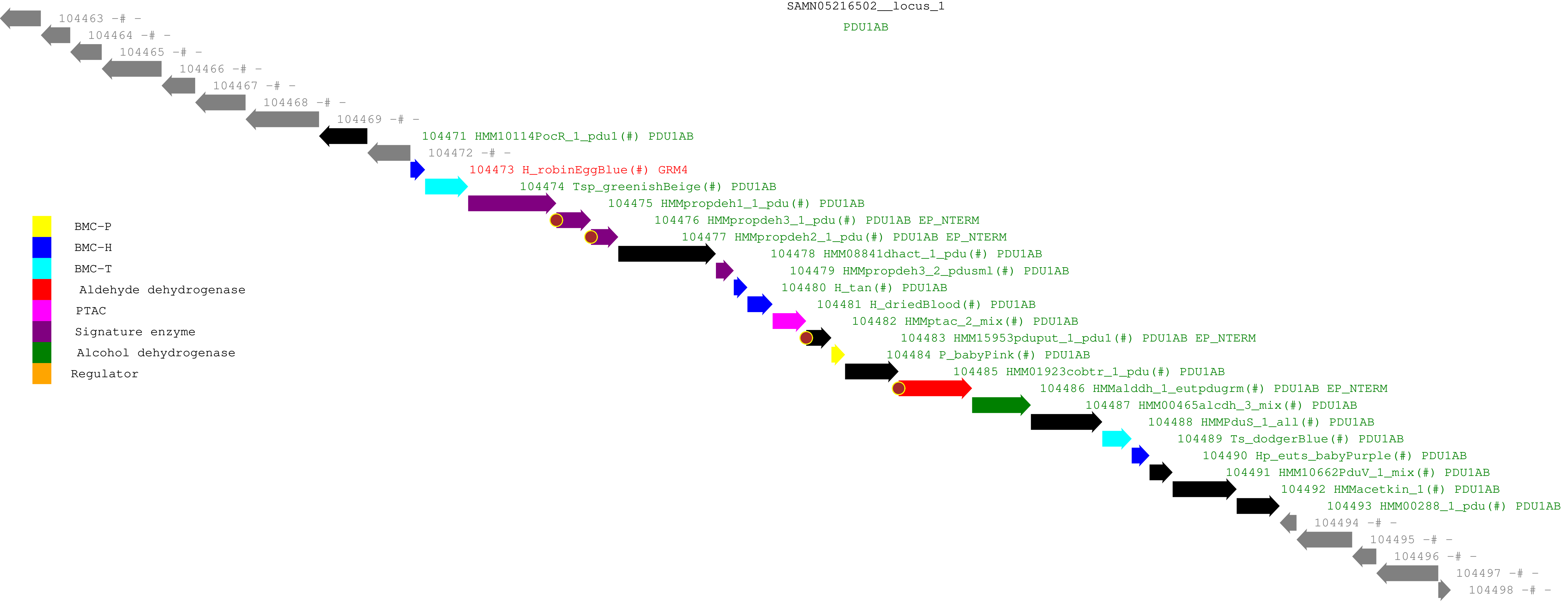

Kosakonia sp. S29. Bacteria; Proteobacteria; Gammaproteobacteria; Enterobacterales;

SAMN05216563__locus_1

PDU1AB

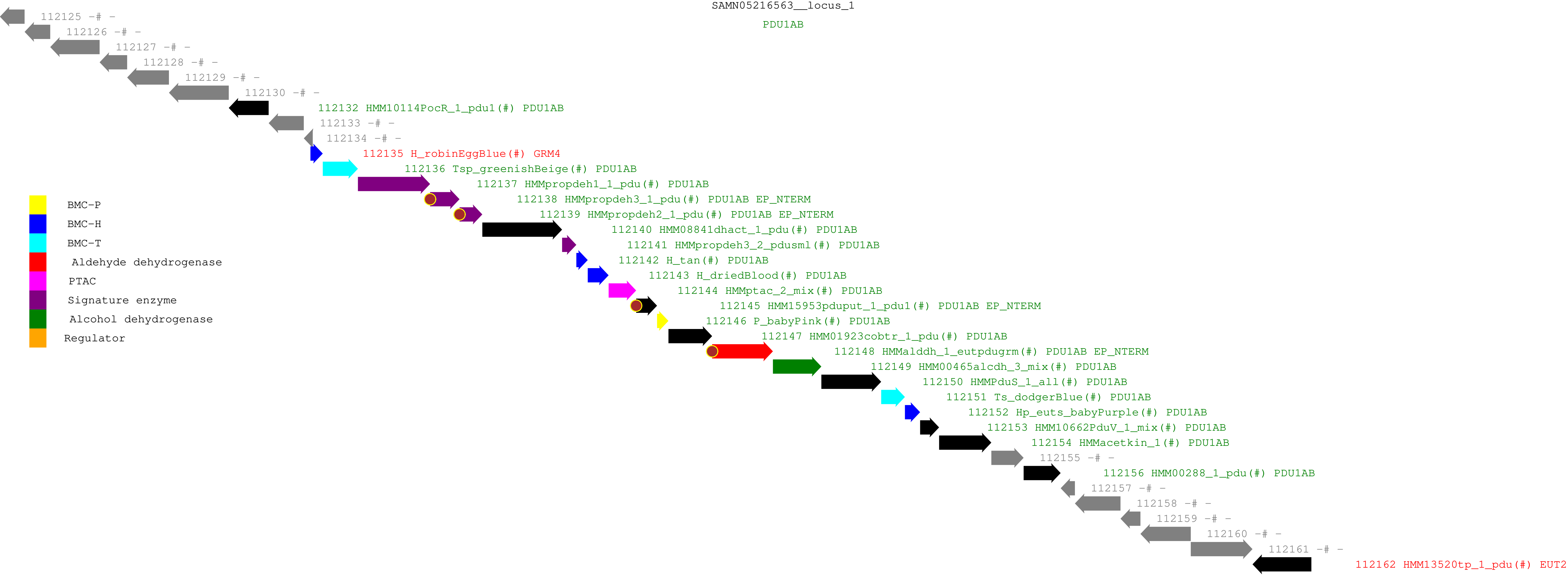

Aeromonas sp. RU39B. Bacteria; Proteobacteria; Gammaproteobacteria; Aeromonadales;

SAMN05880558__locus_1

PDU1AB

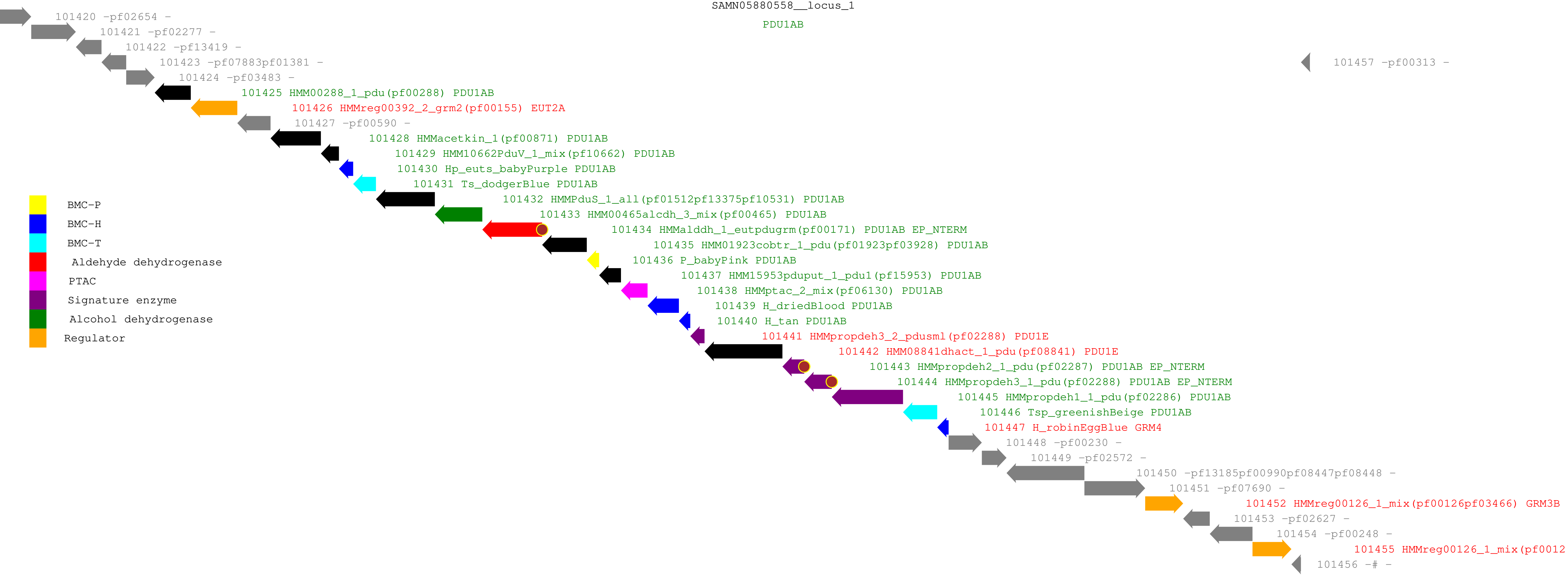

Salmonella arizonae (strain ATCC BAA-731 / CDC346-86 / RSK2980). Bacteria; Proteobacteria; Gammaproteobacteria; Enterobacterales;

SARI__locus_2

PDU1AB

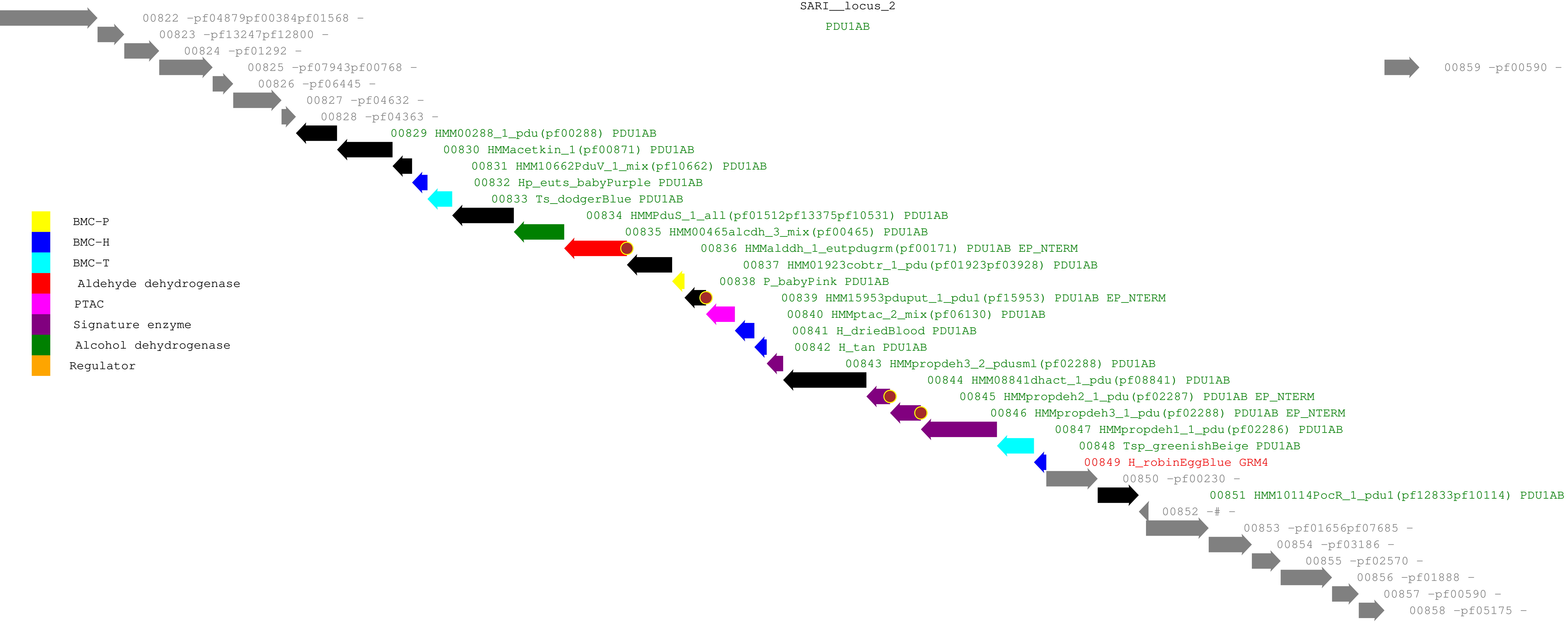

Klebsiella michiganensis. Bacteria; Proteobacteria; Gammaproteobacteria; Enterobacterales;

SB6408__locus_1

PDU1AB

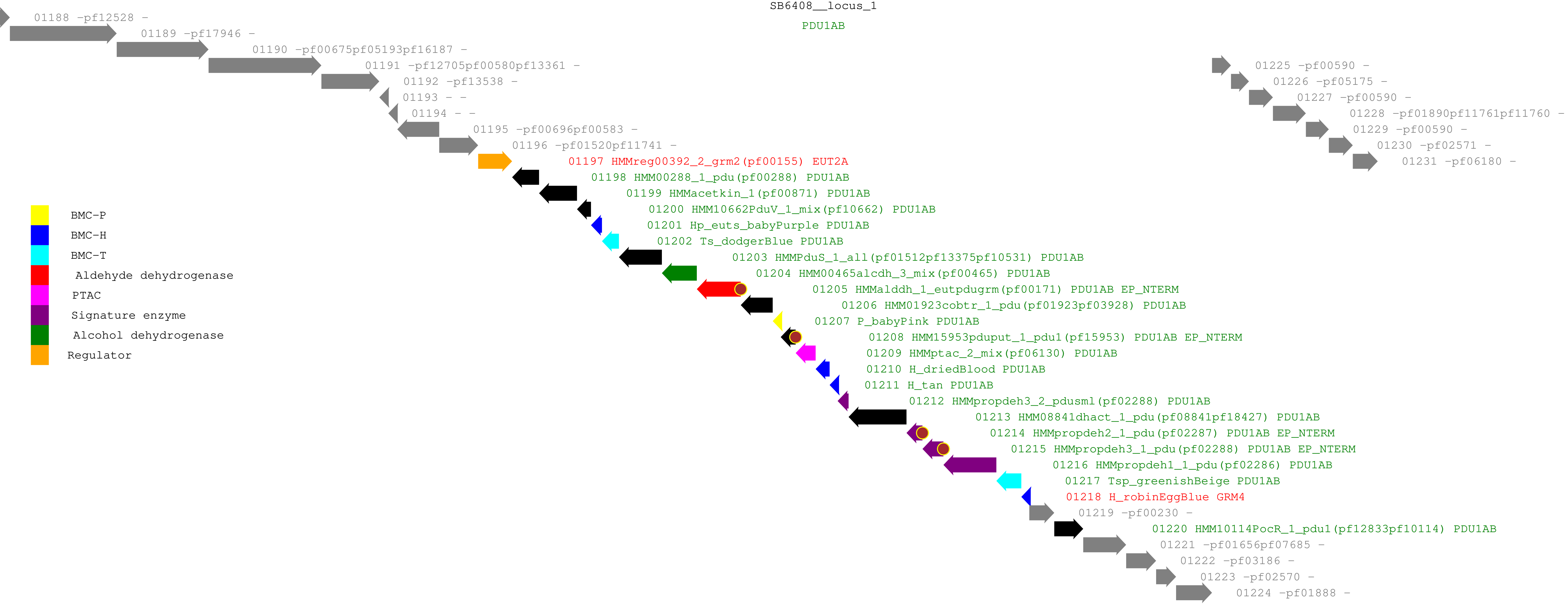

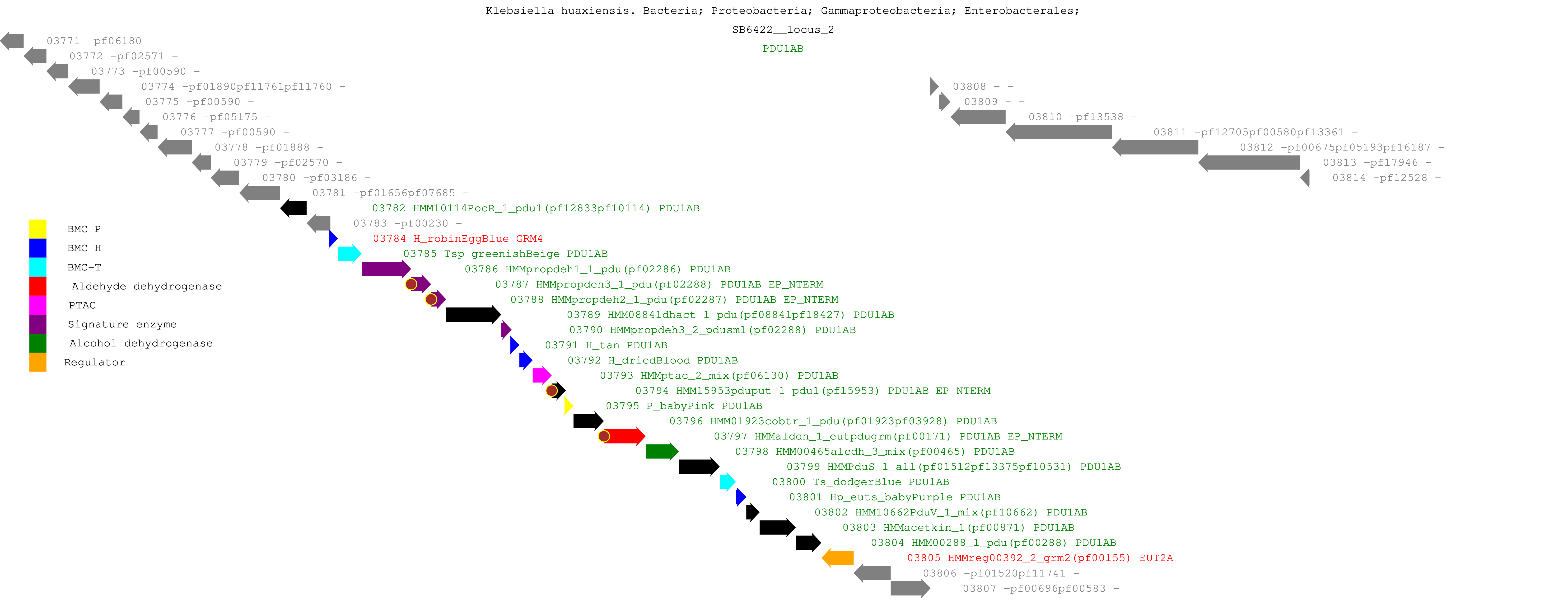
Klebsiella huaxiensis. Bacteria; Proteobacteria; Gammaproteobacteria; Enterobacterales;
SB6422__locus_2
PDU1AB
03771 -pf06180 -
03772 -pf02571 -
03773 -pf00590 -
03774 -pf01890pf11761pf11760 -
03775 -pf00590 -
03776 -pf05175 -
03777 -pf00590 -
03778 -pf01888 -
03779 -pf02570 -
03780 -pf03186 -
03781 -pf01656pf07685 -
03782 HMM10114PocR_1_pdu1(pf12833pf10114) PDU1AB
03783 -pf00230 -
03784 H_robinEggBlue GRM4
03785 Tsp_greenishBeige PDU1AB
03786 HMMpropdeh1_1_pdu(pf02286) PDU1AB
03787 HMMpropdeh3_1_pdu(pf02288) PDU1AB EP_NTERM
03788 HMMpropdeh2_1_pdu(pf02287) PDU1AB EP_NTERM
03789 HMM08841dhact_1_pdu(pf08841pf18427) PDU1AB
03790 HMMpropdeh3_2_pdusml(pf02288) PDU1AB
03791 H_tan PDU1AB
03792 H_driedBlood PDU1AB
03793 HMMptac_2_mix(pf06130) PDU1AB
03794 HMM15953pduput_1_pdu1(pf15953) PDU1AB EP_NTERM
03795 P_babyPink PDU1AB
03796 HMM01923cobtr_1_pdu(pf01923pf03928) PDU1AB
03797 HMMalddh_1_eutpdugrm(pf00171) PDU1AB EP_NTERM
03798 HMM00465alcdh_3_mix(pf00465) PDU1AB
03799 HMMPduS_1_all(pf01512pf13375pf10531) PDU1AB
03800 Ts_dodgerBlue PDU1AB
03801 Hp_euts_babyPurple PDU1AB
03802 HMM10662PduV_1_mix(pf10662) PDU1AB
03803 HMMacetkin_1(pf00871) PDU1AB
03804 HMM00288_1_pdu(pf00288) PDU1AB
03805 HMMreg00392_2_grm2(pf00155) EUT2A
03806 -pf01520pf11741 -
03807 -pf00696pf00583 -
03808 - -
03809 - -
03810 -pf13538 -
03811 -pf12705pf00580pf13361 -
03812 -pf00675pf05193pf16187 -
03813 -pf17946 -
03814 -pf12528 -
BMC-P
BMC-H
BMC-T
Aldehyde dehydrogenase
PTAC
Signature enzyme
Alcohol dehydrogenase
Regulator

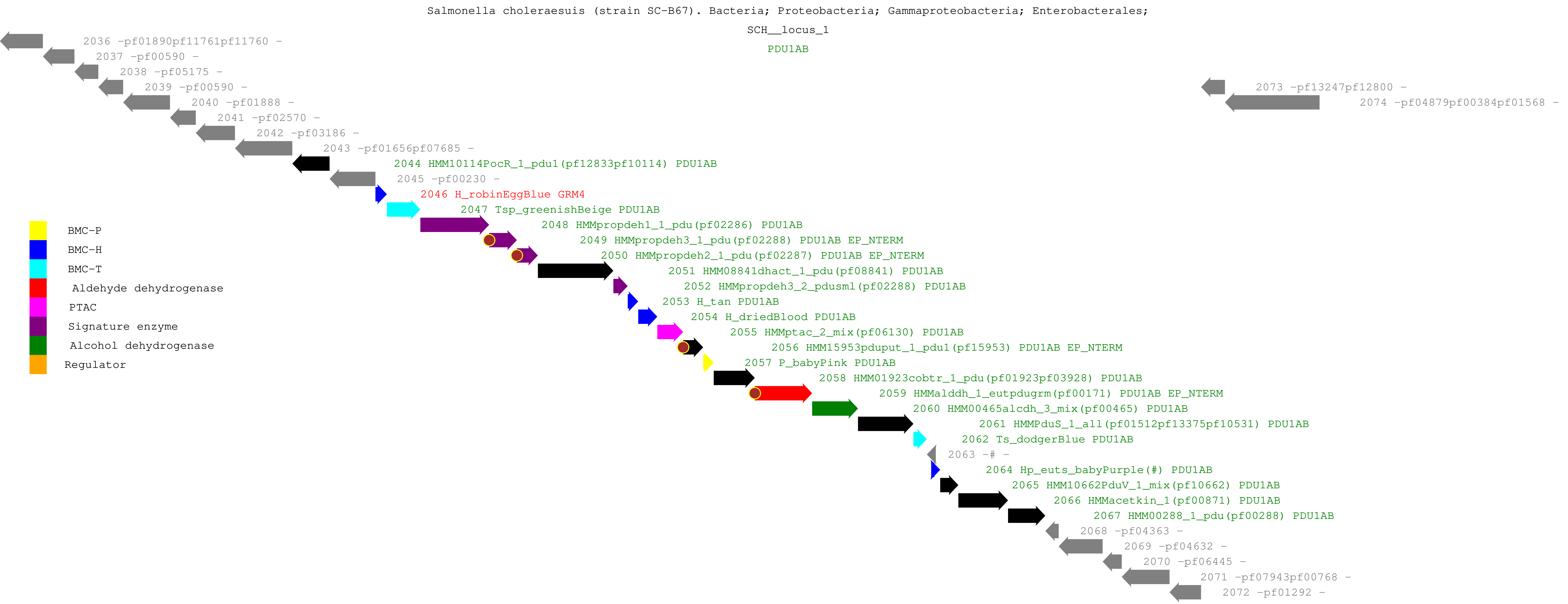
Salmonella choleraesuis (strain SC-B67). Bacteria; Proteobacteria; Gammaproteobacteria; Enterobacterales;
SCH__locus_1
PDU1AB
2036 -pf01890pf11761pf11760 -
2037 -pf00590 -
2038 -pf05175 -
2039 -pf00590 -
2040 -pf01888 -
2041 -pf02570 -
2042 -pf03186 -
2043 -pf01656pf07685 -
2044 HMM10114PocR_1_pdu1(pf12833pf10114) PDU1AB
2045 -pf00230 -
2046 H_robinEggBlue GRM4
2047 Tsp_greenishBeige PDU1AB
2048 HMMpropdeh1_1_pdu(pf02286) PDU1AB
2049 HMMpropdeh3_1_pdu(pf02288) PDU1AB EP_NTERM
2050 HMMpropdeh2_1_pdu(pf02287) PDU1AB EP_NTERM
2051 HMM08841dhact_1_pdu(pf08841) PDU1AB
2052 HMMpropdeh3_2_pdusml(pf02288) PDU1AB
2053 H_tan PDU1AB
2054 H_driedBlood PDU1AB
2055 HMMptac_2_mix(pf06130) PDU1AB
2056 HMM15953pduput_1_pdu1(pf15953) PDU1AB EP_NTERM
2057 P_babyPink PDU1AB
2058 HMM01923cobtr_1_pdu(pf01923pf03928) PDU1AB
2059 HMMalddh_1_eutpdugrm(pf00171) PDU1AB EP_NTERM
2060 HMM00465alcdh_3_mix(pf00465) PDU1AB
2061 HMMPduS_1_all(pf01512pf13375pf10531) PDU1AB
2062 Ts_dodgerBlue PDU1AB
2063 -# -
2064 Hp_euts_babyPurple(#) PDU1AB
2065 HMM10662PduV_1_mix(pf10662) PDU1AB
2066 HMMacetkin_1(pf00871) PDU1AB
2067 HMM00288_1_pdu(pf00288) PDU1AB
2068 -pf04363 -
2069 -pf04632 -
2070 -pf06445 -
2071 -pf07943pf00768 -
2072 -pf01292 -
2073 -pf13247pf12800 -
2074 -pf04879pf00384pf01568 -
BMC-P
BMC-H
BMC-T
Aldehyde dehydrogenase
PTAC
Signature enzyme
Alcohol dehydrogenase
Regulator

Salmonella enterica I. Bacteria; Proteobacteria; Gammaproteobacteria; Enterobacterales;

SE14__locus_1

PDU1AB

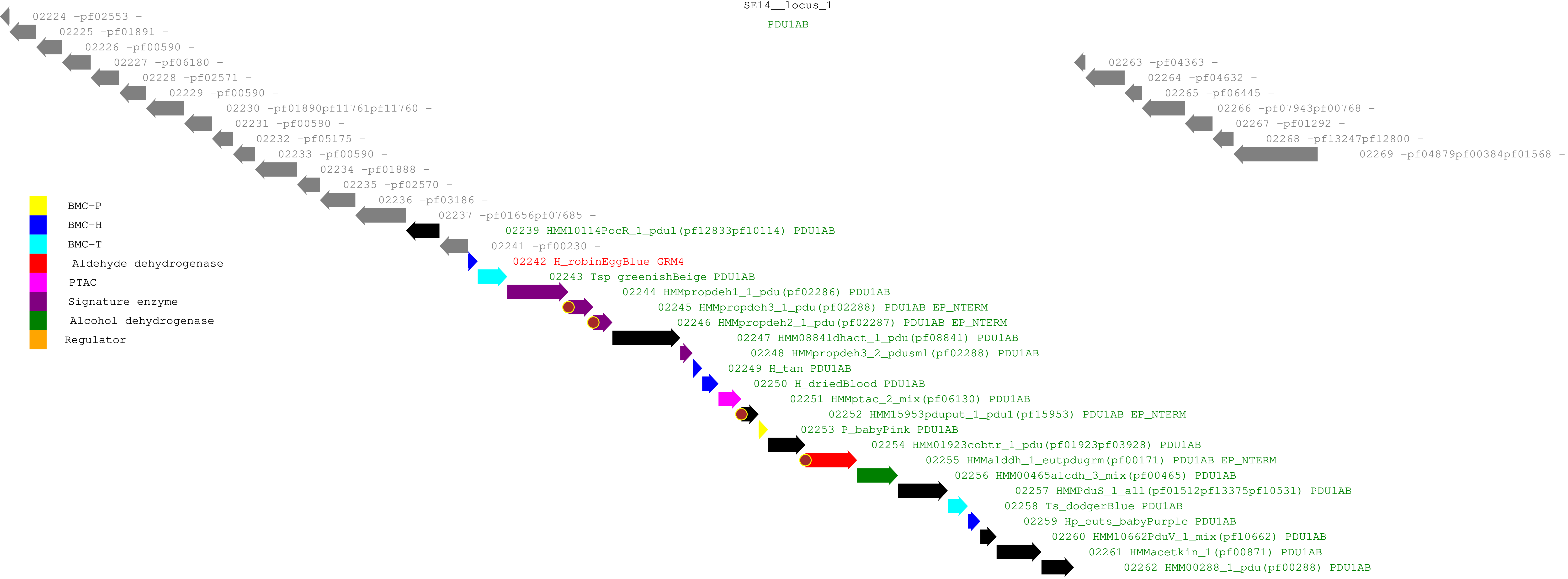

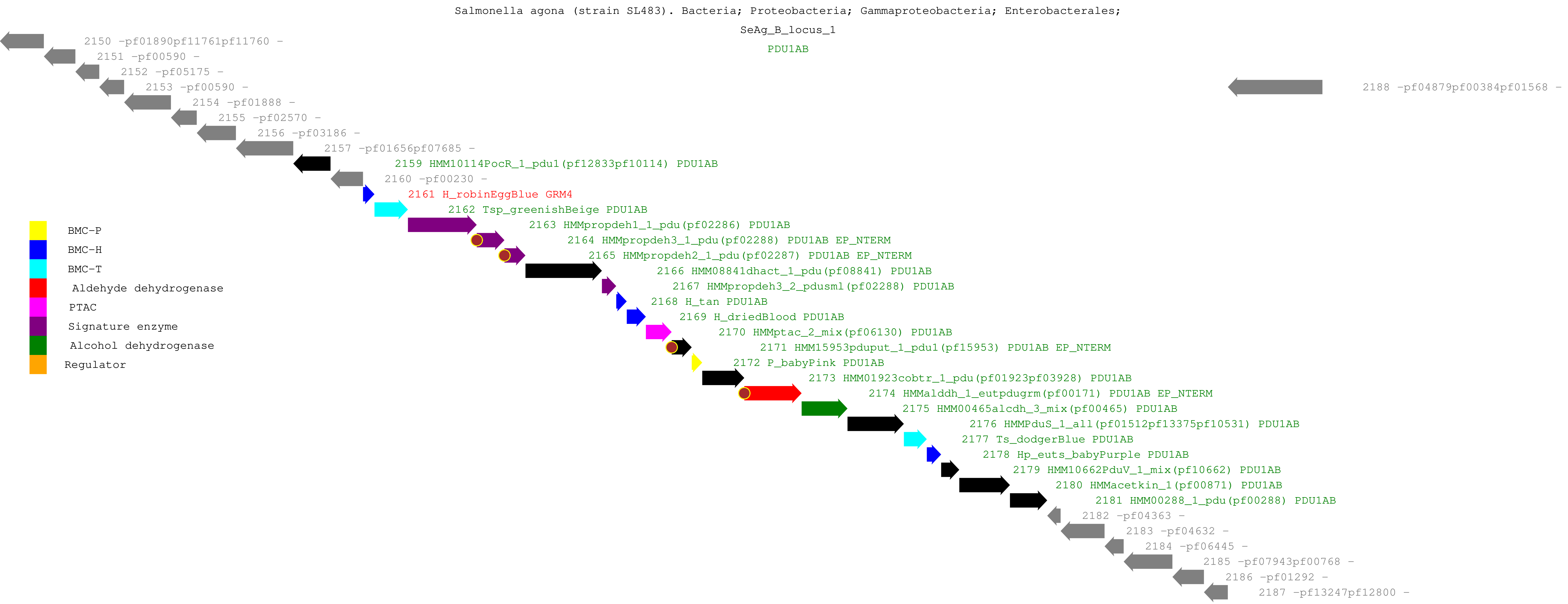
Salmonella agona (strain SL483). Bacteria; Proteobacteria; Gammaproteobacteria; Enterobacterales;
SeAg_B_locus_1
PDU1AB
2150 -pf01890pf11761pf11760 -
2151 -pf00590 -
2152 -pf05175 -
2153 -pf00590 -
2154 -pf01888 -
2155 -pf02570 -
2156 -pf03186 -
2157 -pf01656pf07685 -
2159 HMM10114PocR_1_pdu1(pf12833pf10114) PDU1AB
2160 -pf00230 -
2161 H_robinEggBlue GRM4
2162 Tsp_greenishBeige PDU1AB
2163 HMMpropdeh1_1_pdu(pf02286) PDU1AB
2164 HMMpropdeh3_1_pdu(pf02288) PDU1AB EP_NTERM
2165 HMMpropdeh2_1_pdu(pf02287) PDU1AB EP_NTERM
2166 HMM08841dhact_1_pdu(pf08841) PDU1AB
2167 HMMpropdeh3_2_pdusml(pf02288) PDU1AB
2168 H_tan PDU1AB
2169 H_driedBlood PDU1AB
2170 HMMptac_2_mix(pf06130) PDU1AB
2171 HMM15953pduput_1_pdu1(pf15953) PDU1AB EP_NTERM
2172 P_babyPink PDU1AB
2173 HMM01923cobtr_1_pdu(pf01923pf03928) PDU1AB
2174 HMMalddh_1_eutpdugrm(pf00171) PDU1AB EP_NTERM
2175 HMM00465alcdh_3_mix(pf00465) PDU1AB
2176 HMMPduS_1_all(pf01512pf13375pf10531) PDU1AB
2177 Ts_dodgerBlue PDU1AB
2178 Hp_euts_babyPurple PDU1AB
2179 HMM10662PduV_1_mix(pf10662) PDU1AB
2180 HMMacetkin_1(pf00871) PDU1AB
2181 HMM00288_1_pdu(pf00288) PDU1AB
2182 -pf04363 -
2183 -pf04632 -
2184 -pf06445 -
2185 -pf07943pf00768 -
2186 -pf01292 -
2187 -pf13247pf12800 -
2188 -pf04879pf00384pf01568 -
BMC-P
BMC-H
BMC-T
Aldehyde dehydrogenase
PTAC
Signature enzyme
Alcohol dehydrogenase
Regulator

Salmonella enterica subsp. enterica serovar Chester str. ATCC 11997. Bacteria; Proteobacteria; Gammaproteobacteria; Enterobacterales;

SEECH997__locus_2

PDU1AB

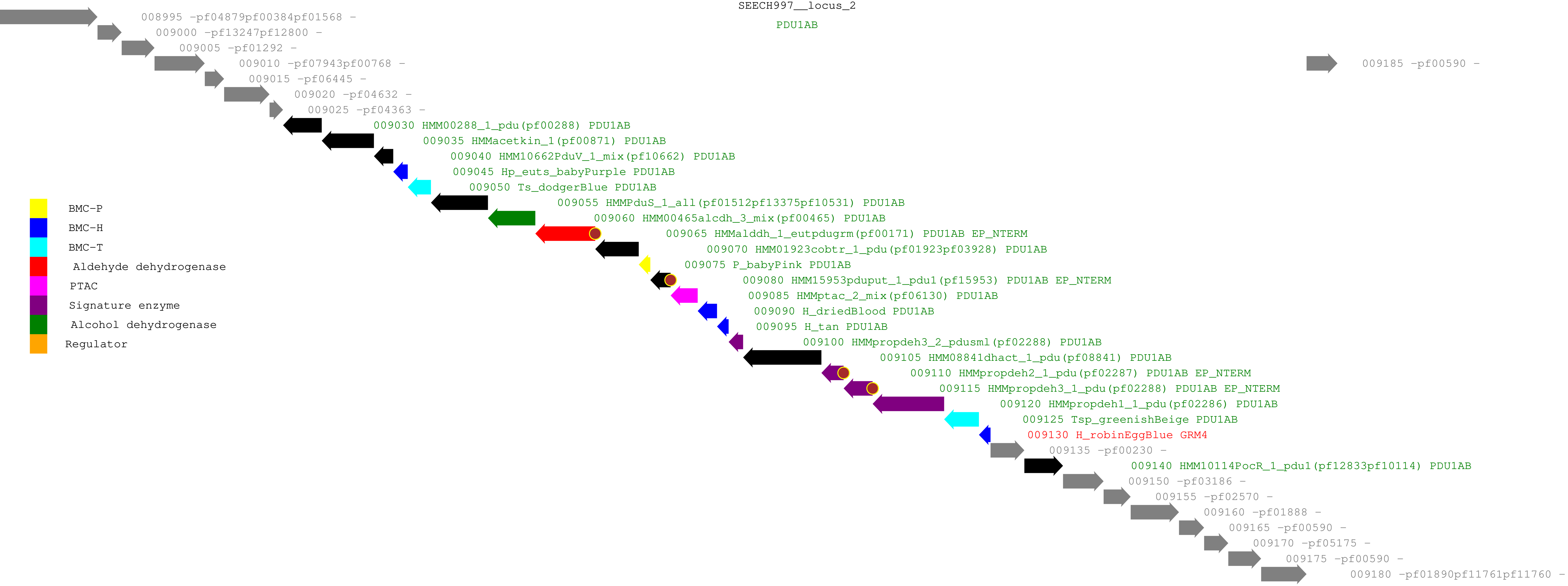

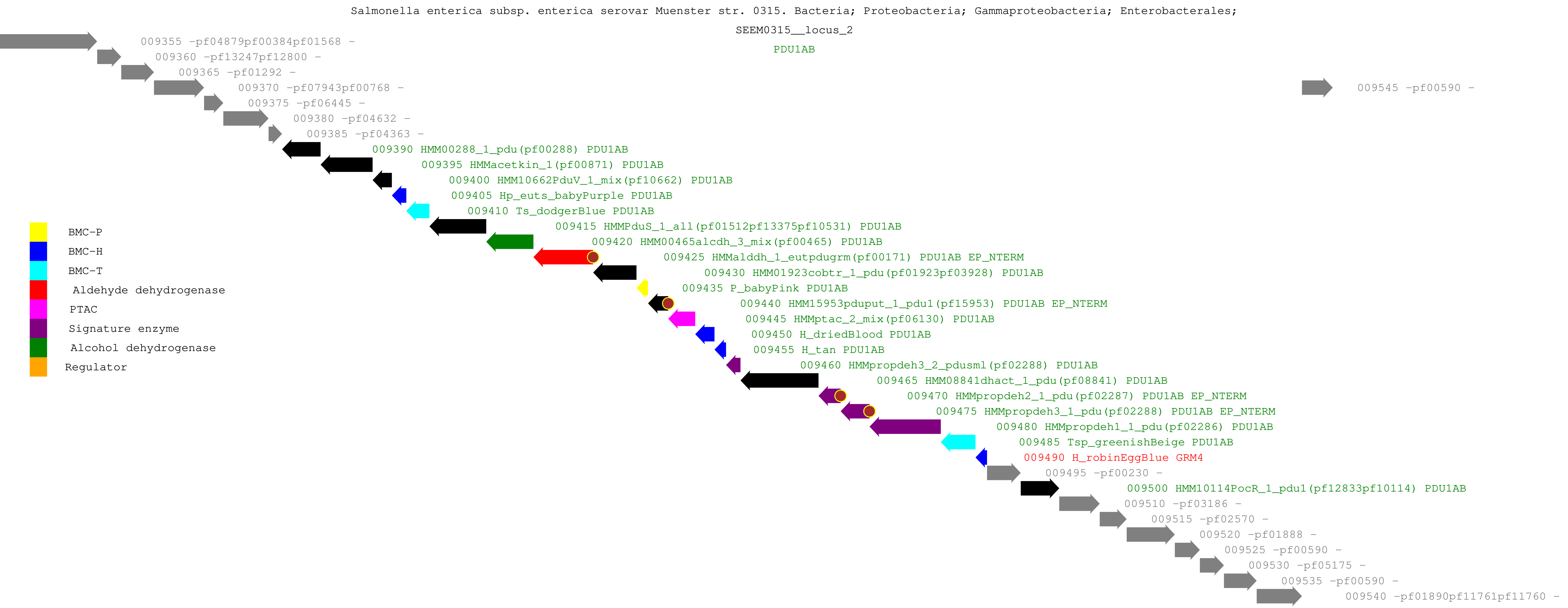
Salmonella enterica subsp. enterica serovar Muenster str. 0315. Bacteria; Proteobacteria; Gammaproteobacteria; Enterobacterales;
SEEM0315__locus_2
PDU1AB
009355 -pf04879pf00384pf01568 -
009360 -pf13247pf12800 -
009365 -pf01292 -
009370 -pf07943pf00768 -
009375 -pf06445 -
009380 -pf04632 -
009385 -pf04363 -
009390 HMM00288_1_pdu(pf00288) PDU1AB
009395 HMMacetkin_1(pf00871) PDU1AB
009400 HMM10662PduV_1_mix(pf10662) PDU1AB
009405 Hp_euts_babyPurple PDU1AB
009410 Ts_dodgerBlue PDU1AB
009415 HMMPduS_1_all(pf01512pf13375pf10531) PDU1AB
009420 HMM00465alcdh_3_mix(pf00465) PDU1AB
009425 HMMalddh_1_eutpdugrm(pf00171) PDU1AB EP_NTERM
009430 HMM01923cobtr_1_pdu(pf01923pf03928) PDU1AB
009435 P_babyPink PDU1AB
009440 HMM15953pduput_1_pdu1(pf15953) PDU1AB EP_NTERM
009445 HMMptac_2_mix(pf06130) PDU1AB
009450 H_driedBlood PDU1AB
009455 H_tan PDU1AB
009460 HMMpropdeh3_2_pdusml(pf02288) PDU1AB
009465 HMM08841dhact_1_pdu(pf08841) PDU1AB
009470 HMMpropdeh2_1_pdu(pf02287) PDU1AB EP_NTERM
009475 HMMpropdeh3_1_pdu(pf02288) PDU1AB EP_NTERM
009480 HMMpropdeh1_1_pdu(pf02286) PDU1AB
009485 Tsp_greenishBeige PDU1AB
009490 H_robinEggBlue GRM4
009495 -pf00230 -
009500 HMM10114PocR_1_pdu1(pf12833pf10114) PDU1AB
009510 -pf03186 -
009515 -pf02570 -
009520 -pf01888 -
009525 -pf00590 -
009530 -pf05175 -
009535 -pf00590 -
009540 -pf01890pf11761pf11760 -
009545 -pf00590 -
BMC-P
BMC-H
BMC-T
Aldehyde dehydrogenase
PTAC
Signature enzyme
Alcohol dehydrogenase
Regulator

Salmonella enterica subsp. enterica serovar Muenchen str. baa1594. Bacteria; Proteobacteria; Gammaproteobacteria; Enterobacterales;

SEEM1594__locus_1

PDU1AB

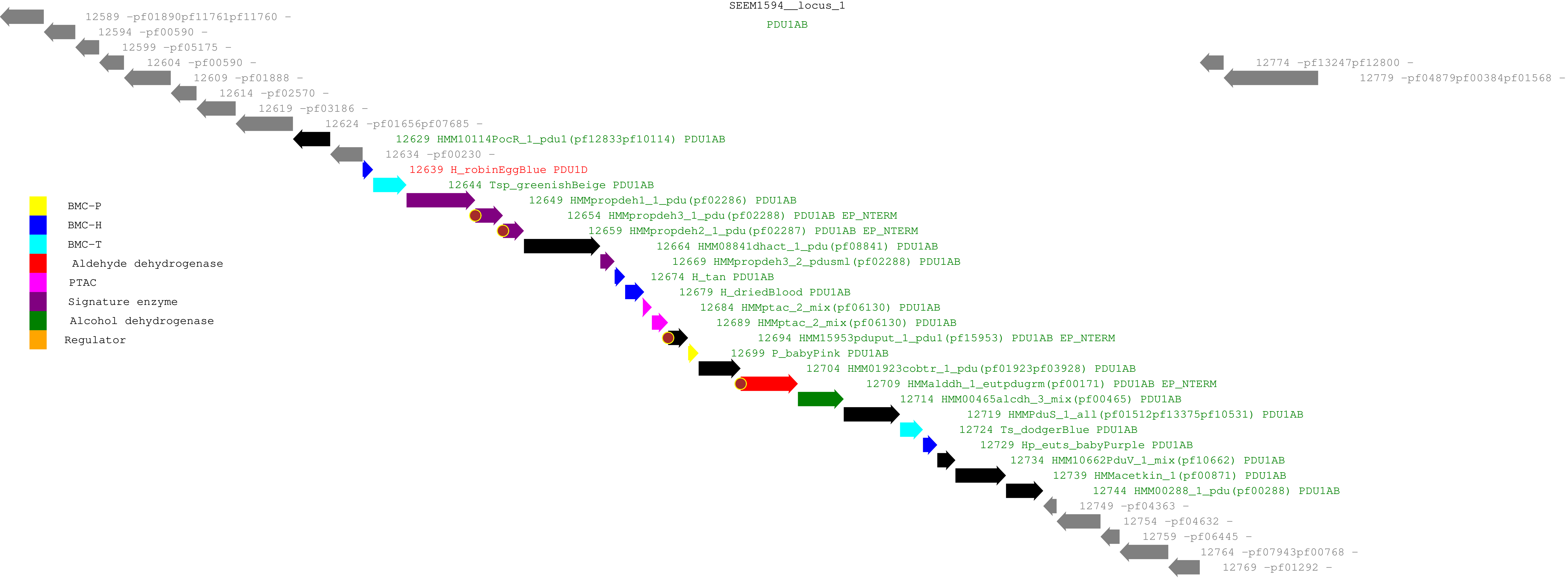

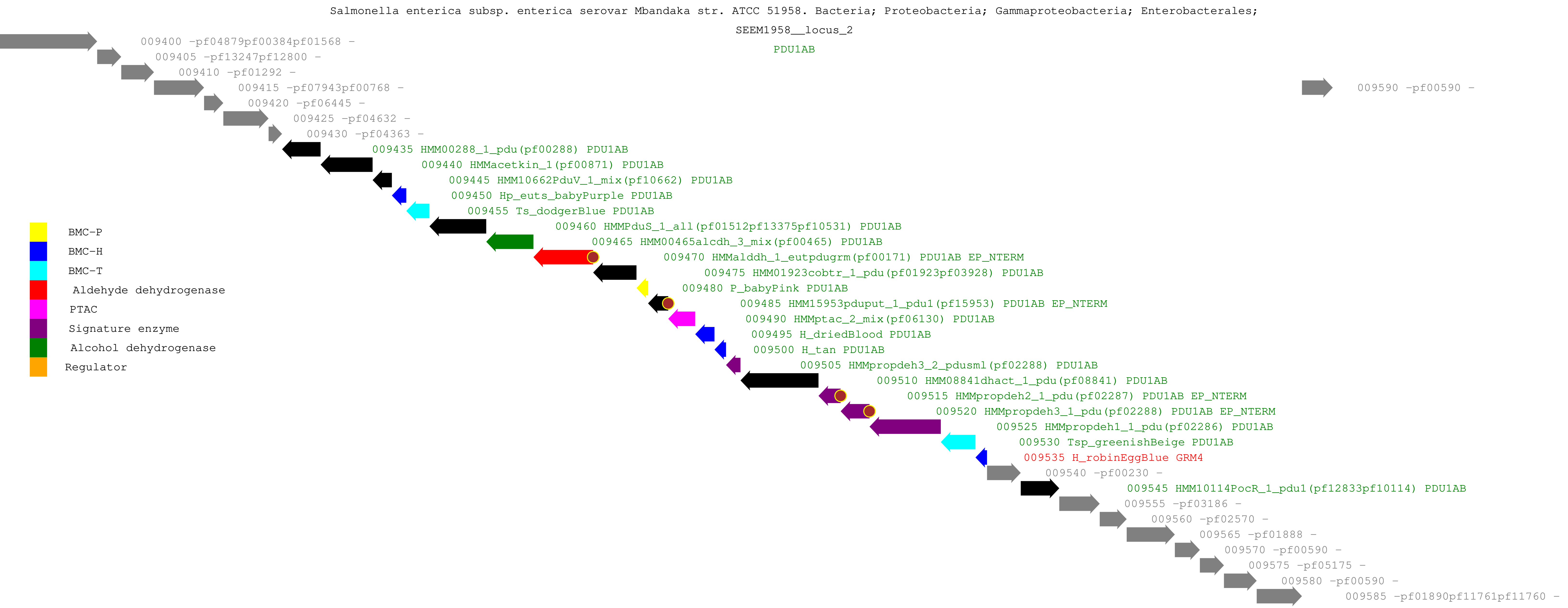
Salmonella enterica subsp. enterica serovar Mbandaka str. ATCC 51958. Bacteria; Proteobacteria; Gammaproteobacteria; Enterobacterales;
SEEM1958__locus_2
PDU1AB
009400 -pf04879pf00384pf01568 -
009405 -pf13247pf12800 -
009410 -pf01292 -
009415 -pf07943pf00768 -
009420 -pf06445 -
009425 -pf04632 -
009430 -pf04363 -
009435 HMM00288_1_pdu(pf00288) PDU1AB
009440 HMMacetkin_1(pf00871) PDU1AB
009445 HMM10662PduV_1_mix(pf10662) PDU1AB
009450 Hp_euts_babyPurple PDU1AB
009455 Ts_dodgerBlue PDU1AB
009460 HMMPduS_1_all(pf01512pf13375pf10531) PDU1AB
009465 HMM00465alcdh_3_mix(pf00465) PDU1AB
009470 HMMalddh_1_eutpdugrm(pf00171) PDU1AB EP_NTERM
009475 HMM01923cobtr_1_pdu(pf01923pf03928) PDU1AB
009480 P_babyPink PDU1AB
009485 HMM15953pduput_1_pdu1(pf15953) PDU1AB EP_NTERM
009490 HMMptac_2_mix(pf06130) PDU1AB
009495 H_driedBlood PDU1AB
009500 H_tan PDU1AB
009505 HMMpropdeh3_2_pdusml(pf02288) PDU1AB
009510 HMM08841dhact_1_pdu(pf08841) PDU1AB
009515 HMMpropdeh2_1_pdu(pf02287) PDU1AB EP_NTERM
009520 HMMpropdeh3_1_pdu(pf02288) PDU1AB EP_NTERM
009525 HMMpropdeh1_1_pdu(pf02286) PDU1AB
009530 Tsp_greenishBeige PDU1AB
009535 H_robinEggBlue GRM4
009540 -pf00230 -
009545 HMM10114PocR_1_pdu1(pf12833pf10114) PDU1AB
009555 -pf03186 -
009560 -pf02570 -
009565 -pf01888 -
009570 -pf00590 -
009575 -pf05175 -
009580 -pf00590 -
009585 -pf01890pf11761pf11760 -
009590 -pf00590 -
BMC-P
BMC-H
BMC-T
Aldehyde dehydrogenase
PTAC
Signature enzyme
Alcohol dehydrogenase
Regulator

Salmonella enterica subsp. enterica serovar Montevideo str. 42N. Bacteria; Proteobacteria; Gammaproteobacteria; Enterobacterales;

SEEM42N__locus_2

PDU1AB

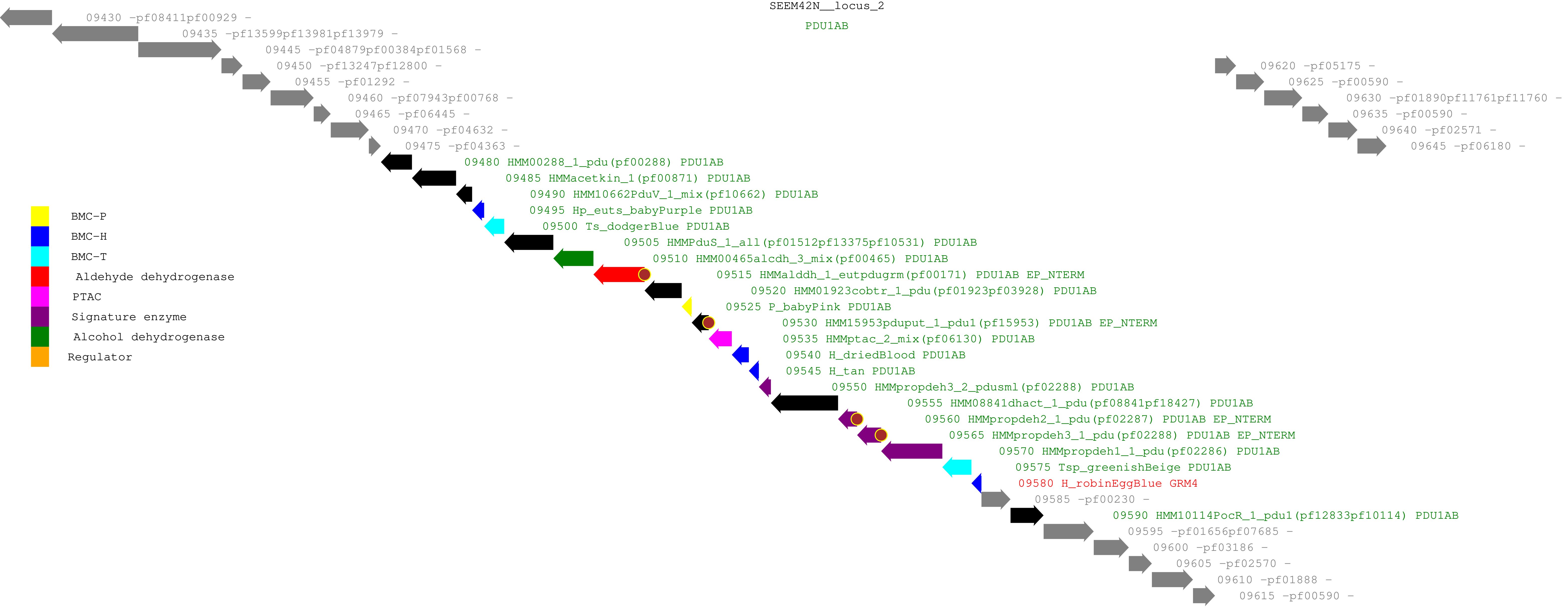

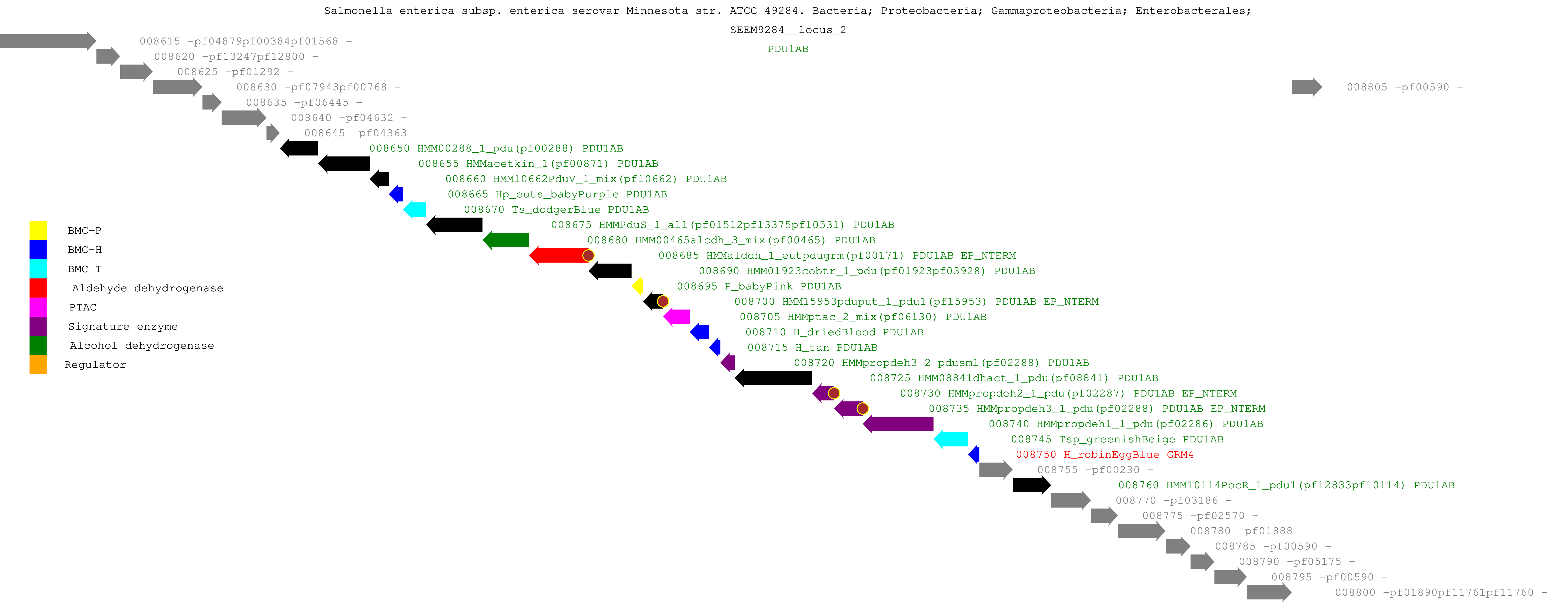
Salmonella enterica subsp. enterica serovar Minnesota str. ATCC 49284. Bacteria; Proteobacteria; Gammaproteobacteria; Enterobacterales;
SEEM9284__locus_2
PDU1AB
008615 -pf04879pf00384pf01568 -
008620 -pf13247pf12800 -
008625 -pf01292 -
008630 -pf07943pf00768 -
008635 -pf06445 -
008640 -pf04632 -
008645 -pf04363 -
008650 HMM00288_1_pdu(pf00288) PDU1AB
008655 HMMacetkin_1(pf00871) PDU1AB
008660 HMM10662PduV_1_mix(pf10662) PDU1AB
008665 Hp_euts_babyPurple PDU1AB
008670 Ts_dodgerBlue PDU1AB
008675 HMMPduS_1_all(pf01512pf13375pf10531) PDU1AB
008680 HMM00465alcdh_3_mix(pf00465) PDU1AB
008685 HMMalddh_1_eutpdugrm(pf00171) PDU1AB EP_NTERM
008690 HMM01923cobtr_1_pdu(pf01923pf03928) PDU1AB
008695 P_babyPink PDU1AB
008700 HMM15953pduput_1_pdu1(pf15953) PDU1AB EP_NTERM
008705 HMMptac_2_mix(pf06130) PDU1AB
008710 H_driedBlood PDU1AB
008715 H_tan PDU1AB
008720 HMMpropdeh3_2_pdusml(pf02288) PDU1AB
008725 HMM08841dhact_1_pdu(pf08841) PDU1AB
008730 HMMpropdeh2_1_pdu(pf02287) PDU1AB EP_NTERM
008735 HMMpropdeh3_1_pdu(pf02288) PDU1AB EP_NTERM
008740 HMMpropdeh1_1_pdu(pf02286) PDU1AB
008745 Tsp_greenishBeige PDU1AB
008750 H_robinEggBlue GRM4
008755 -pf00230 -
008760 HMM10114PocR_1_pdu1(pf12833pf10114) PDU1AB
008770 -pf03186 -
008775 -pf02570 -
008780 -pf01888 -
008785 -pf00590 -
008790 -pf05175 -
008795 -pf00590 -
008800 -pf01890pf11761pf11760 -
008805 -pf00590 -
BMC-P
BMC-H
BMC-T
Aldehyde dehydrogenase
PTAC
Signature enzyme
Alcohol dehydrogenase
Regulator

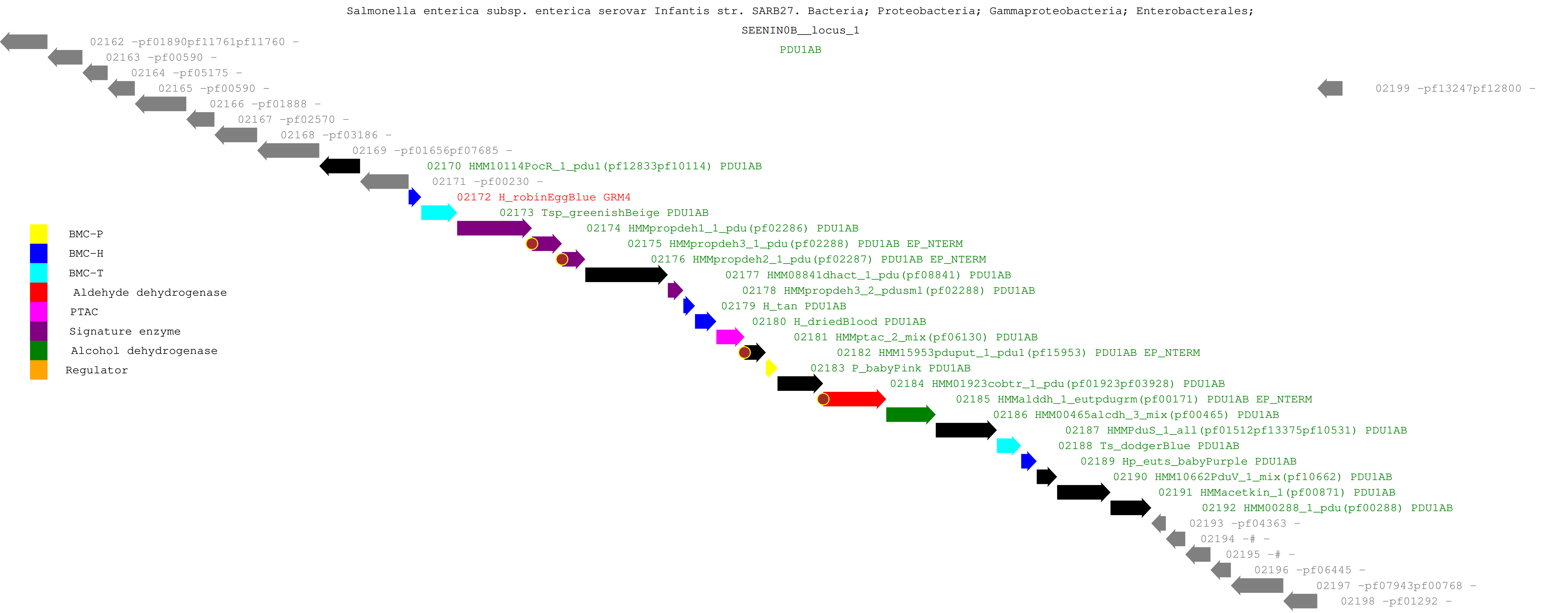
Salmonella enterica subsp. enterica serovar Infantis str. SARB27. Bacteria; Proteobacteria; Gammaproteobacteria; Enterobacterales;
SEENIN0B__locus_1
PDU1AB
02162 -pf01890pf11761pf11760 -
02163 -pf00590 -
02164 -pf05175 -
02165 -pf00590 -
02166 -pf01888 -
02167 -pf02570 -
02168 -pf03186 -
02169 -pf01656pf07685 -
02170 HMM10114PocR_1_pdu1(pf12833pf10114) PDU1AB
02171 -pf00230 -
02172 H_robinEggBlue GRM4
02173 Tsp_greenishBeige PDU1AB
02174 HMMpropdeh1_1_pdu(pf02286) PDU1AB
02175 HMMpropdeh3_1_pdu(pf02288) PDU1AB EP_NTERM
02176 HMMpropdeh2_1_pdu(pf02287) PDU1AB EP_NTERM
02177 HMM08841dhact_1_pdu(pf08841) PDU1AB
02178 HMMpropdeh3_2_pdusml(pf02288) PDU1AB
02179 H_tan PDU1AB
02180 H_driedBlood PDU1AB
02181 HMMptac_2_mix(pf06130) PDU1AB
02182 HMM15953pduput_1_pdu1(pf15953) PDU1AB EP_NTERM
02183 P_babyPink PDU1AB
02184 HMM01923cobtr_1_pdu(pf01923pf03928) PDU1AB
02185 HMMalddh_1_eutpdugrm(pf00171) PDU1AB EP_NTERM
02186 HMM00465alcdh_3_mix(pf00465) PDU1AB
02187 HMMPduS_1_all(pf01512pf13375pf10531) PDU1AB
02188 Ts_dodgerBlue PDU1AB
02189 Hp_euts_babyPurple PDU1AB
02190 HMM10662PduV_1_mix(pf10662) PDU1AB
02191 HMMacetkin_1(pf00871) PDU1AB
02192 HMM00288_1_pdu(pf00288) PDU1AB
02193 -pf04363 -
02194 -# -
02195 -# -
02196 -pf06445 -
02197 -pf07943pf00768 -
02198 -pf01292 -
02199 -pf13247pf12800 -
BMC-P
BMC-H
BMC-T
Aldehyde dehydrogenase
PTAC
Signature enzyme
Alcohol dehydrogenase
Regulator

Salmonella enterica subsp. enterica serovar Rubislaw str. ATCC 10717. Bacteria; Proteobacteria; Gammaproteobacteria; Enterobacterales;

SEERU717__locus_2

PDU1AB

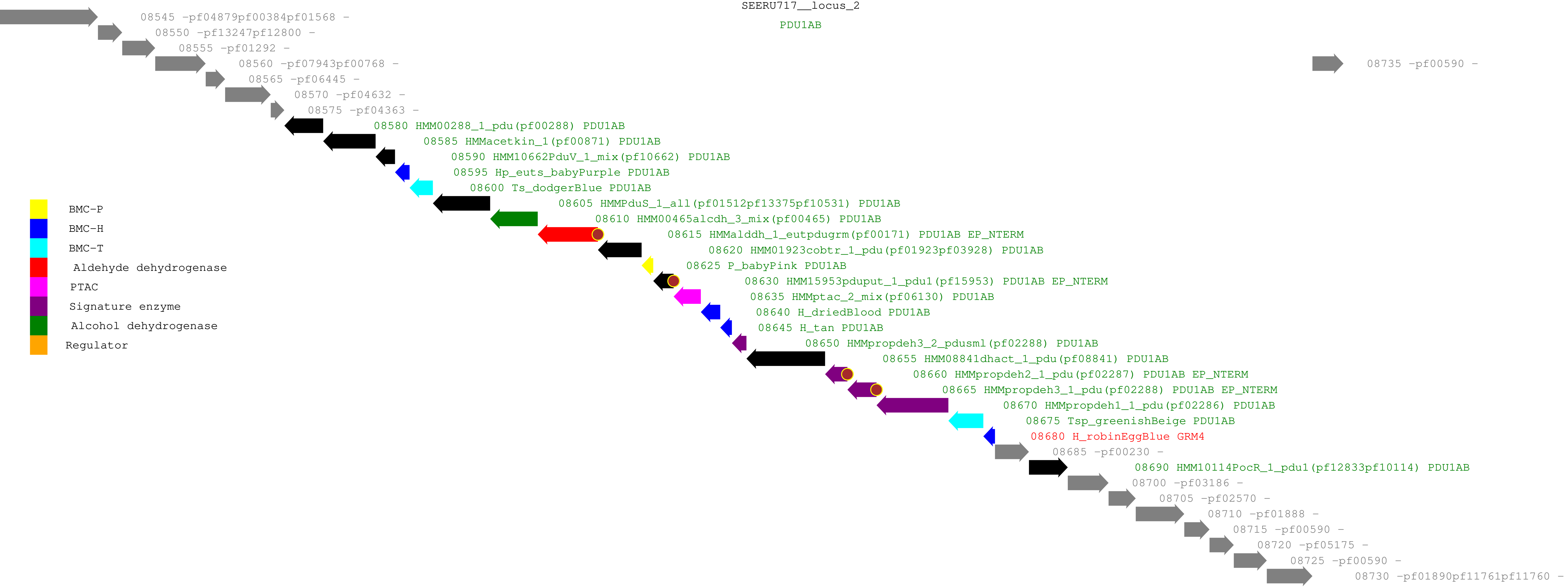

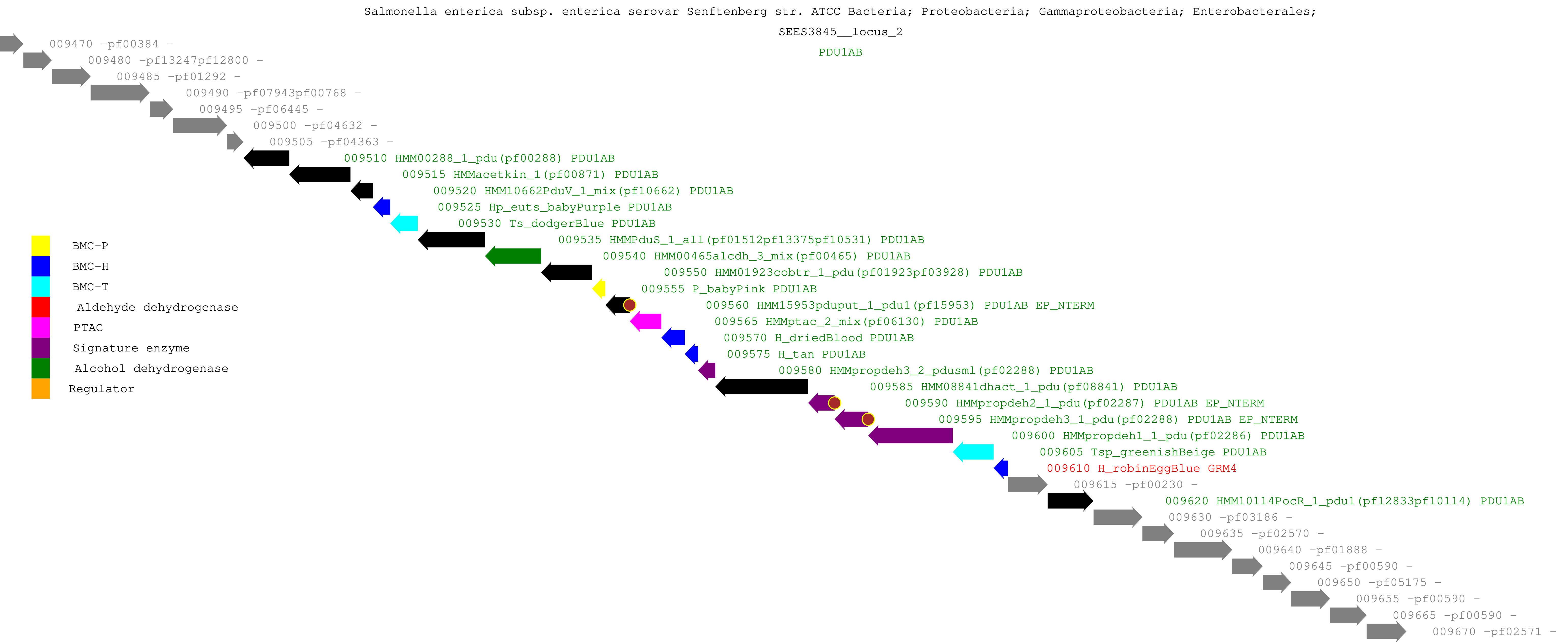
Salmonella enterica subsp. enterica serovar Senftenberg str. ATCC Bacteria; Proteobacteria; Gammaproteobacteria; Enterobacterales;
SEES3845__locus_2
PDU1AB
009470 -pf00384 -
009480 -pf13247pf12800 -
009485 -pf01292 -
009490 -pf07943pf00768 -
009495 -pf06445 -
009500 -pf04632 -
009505 -pf04363 -
009510 HMM00288_1_pdu(pf00288) PDU1AB
009515 HMMacetkin_1(pf00871) PDU1AB
009520 HMM10662PduV_1_mix(pf10662) PDU1AB
009525 Hp_euts_babyPurple PDU1AB
009530 Ts_dodgerBlue PDU1AB
009535 HMMPduS_1_all(pf01512pf13375pf10531) PDU1AB
009540 HMM00465alcdh_3_mix(pf00465) PDU1AB
009550 HMM01923cobtr_1_pdu(pf01923pf03928) PDU1AB
009555 P_babyPink PDU1AB
009560 HMM15953pduput_1_pdu1(pf15953) PDU1AB EP_NTERM
009565 HMMptac_2_mix(pf06130) PDU1AB
009570 H_driedBlood PDU1AB
009575 H_tan PDU1AB
009580 HMMpropdeh3_2_pdusml(pf02288) PDU1AB
009585 HMM08841dhact_1_pdu(pf08841) PDU1AB
009590 HMMpropdeh2_1_pdu(pf02287) PDU1AB EP_NTERM
009595 HMMpropdeh3_1_pdu(pf02288) PDU1AB EP_NTERM
009600 HMMpropdeh1_1_pdu(pf02286) PDU1AB
009605 Tsp_greenishBeige PDU1AB
009610 H_robinEggBlue GRM4
009615 -pf00230 -
009620 HMM10114PocR_1_pdu1(pf12833pf10114) PDU1AB
009630 -pf03186 -
009635 -pf02570 -
009640 -pf01888 -
009645 -pf00590 -
009650 -pf05175 -
009655 -pf00590 -
009665 -pf00590 -
009670 -pf02571 -
BMC-P
BMC-H
BMC-T
Aldehyde dehydrogenase
PTAC
Signature enzyme
Alcohol dehydrogenase
Regulator

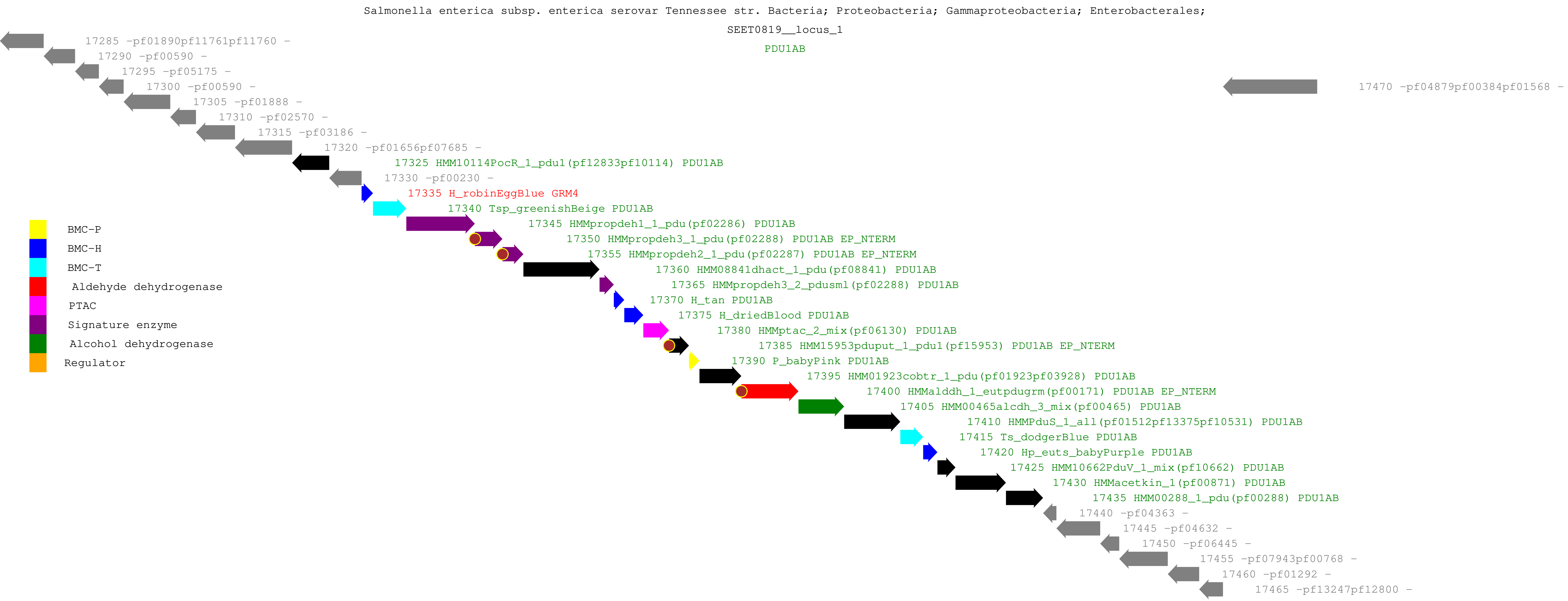
Salmonella enterica subsp. enterica serovar Tennessee str. Bacteria; Proteobacteria; Gammaproteobacteria; Enterobacterales;
SEET0819__locus_1
PDU1AB
17285 -pf01890pf11761pf11760 -
17290 -pf00590 -
17295 -pf05175 -
17300 -pf00590 -
17305 -pf01888 -
17310 -pf02570 -
17315 -pf03186 -
17320 -pf01656pf07685 -
17325 HMM10114PocR_1_pdu1(pf12833pf10114) PDU1AB
17330 -pf00230 -
17335 H_robinEggBlue GRM4
17340 Tsp_greenishBeige PDU1AB
17345 HMMpropdeh1_1_pdu(pf02286) PDU1AB
17350 HMMpropdeh3_1_pdu(pf02288) PDU1AB EP_NTERM
17355 HMMpropdeh2_1_pdu(pf02287) PDU1AB EP_NTERM
17360 HMM08841dhact_1_pdu(pf08841) PDU1AB
17365 HMMpropdeh3_2_pdusml(pf02288) PDU1AB
17370 H_tan PDU1AB
17375 H_driedBlood PDU1AB
17380 HMMptac_2_mix(pf06130) PDU1AB
17385 HMM15953pduput_1_pdu1(pf15953) PDU1AB EP_NTERM
17390 P_babyPink PDU1AB
17395 HMM01923cobtr_1_pdu(pf01923pf03928) PDU1AB
17400 HMMaldh_1_eutpdugrm(pf00171) PDU1AB EP_NTERM
17405 HMM00465alcdh_3_mix(pf00465) PDU1AB
17410 HMMPduS_1_all(pf01512pf13375pf10531) PDU1AB
17415 Ts_dodgerBlue PDU1AB
17420 Hp_euts_babyPurple PDU1AB
17425 HMM10662PduV_1_mix(pf10662) PDU1AB
17430 HMMacetkin_1(pf00871) PDU1AB
17435 HMM00288_1_pdu(pf00288) PDU1AB
17440 -pf04363 -
17445 -pf04632 -
17450 -pf06445 -
17455 -pf07943pf00768 -
17460 -pf01292 -
17465 -pf13247pf12800 -
17470 -pf04879pf00384pf01568 -
BMC-P
BMC-H
BMC-T
Aldehyde dehydrogenase
PTAC
Signature enzyme
Alcohol dehydrogenase
Regulator

Salmonella enterica subsp. indica serovar 6,14,25:z10:1,(2),7 str. Bacteria; Proteobacteria; Gammaproteobacteria; Enterobacterales;

SEI61121__locus_1

PDU1AB

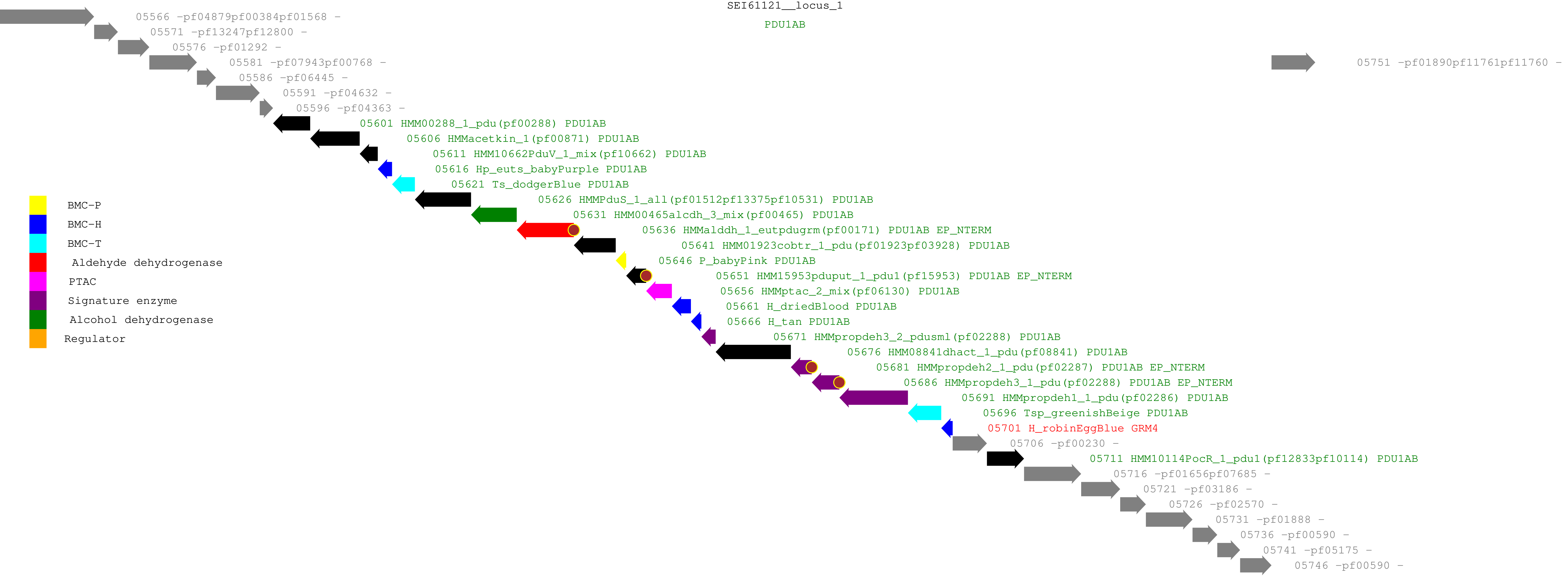

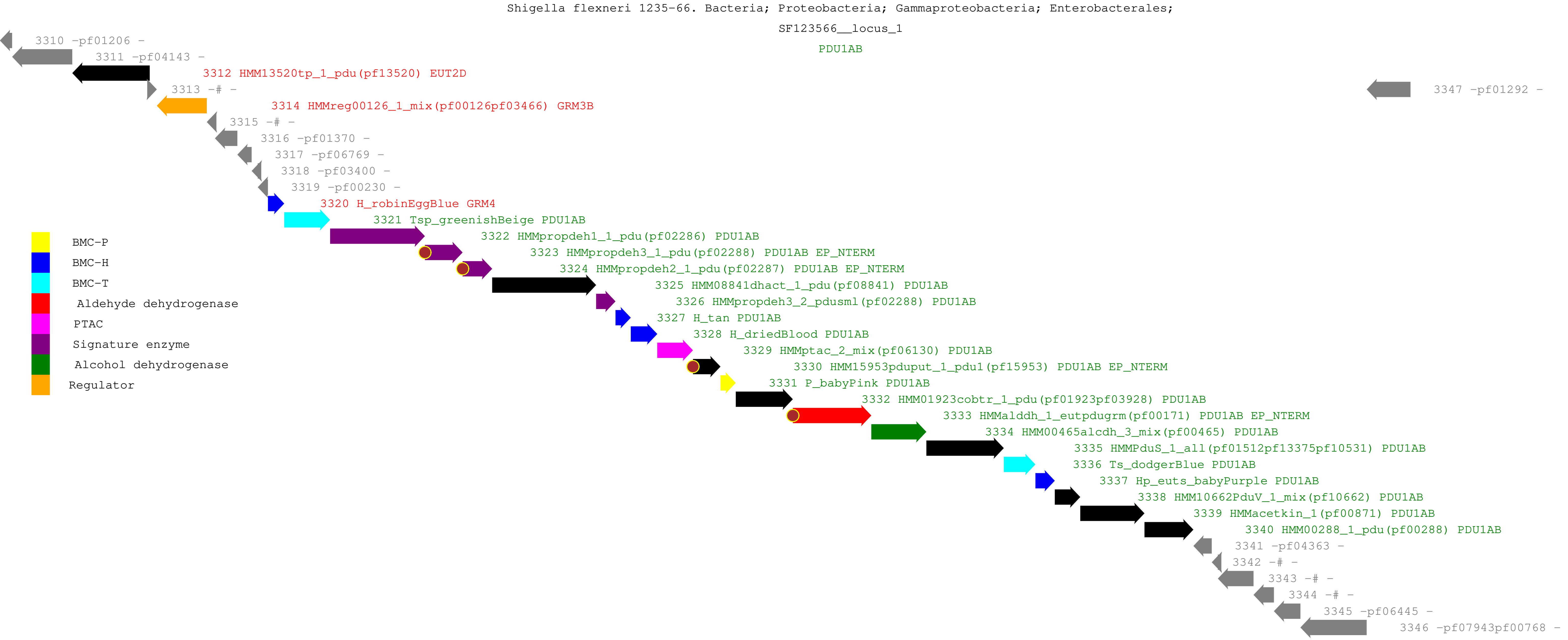
Shigella flexneri 1235-66. Bacteria; Proteobacteria; Gammaproteobacteria; Enterobacterales;
SF123566__locus_1
PDU1AB
3310 -pf01206 -
3311 -pf04143 -
3312 HMM13520tp_1_pdu(pf13520) EUT2D
3313 -# -
3314 HMMreg00126_1_mix(pf00126pf03466) GRM3B
3315 -# -
3316 -pf01370 -
3317 -pf06769 -
3318 -pf03400 -
3319 -pf00230 -
3320 H_robinEggBlue GRM4
3321 Tsp_greenishBeige PDU1AB
3322 HMMpropdeh1_1_pdu(pf02286) PDU1AB
3323 HMMpropdeh3_1_pdu(pf02288) PDU1AB EP_NTERM
3324 HMMpropdeh2_1_pdu(pf02287) PDU1AB EP_NTERM
3325 HMM08841dhact_1_pdu(pf08841) PDU1AB
3326 HMMpropdeh3_2_pdusml(pf02288) PDU1AB
3327 H_tan PDU1AB
3328 H_driedBlood PDU1AB
3329 HMMptac_2_mix(pf06130) PDU1AB
3330 HMM15953pduput_1_pdu1(pf15953) PDU1AB EP_NTERM
3331 P_babyPink PDU1AB
3332 HMM01923cobtr_1_pdu(pf01923pf03928) PDU1AB
3333 HMMalddh_1_eutpdugrm(pf00171) PDU1AB EP_NTERM
3334 HMM00465alcdh_3_mix(pf00465) PDU1AB
3335 HMMPduS_1_all(pf01512pf13375pf10531) PDU1AB
3336 Ts_dodgerBlue PDU1AB
3337 Hp_euts_babyPurple PDU1AB
3338 HMM10662PduV_1_mix(pf10662) PDU1AB
3339 HMMacetkin_1(pf00871) PDU1AB
3340 HMM00288_1_pdu(pf00288) PDU1AB
3341 -pf04363 -
3342 -# -
3343 -# -
3344 -# -
3345 -pf06445 -
3346 -pf07943pf00768 -
3347 -pf01292 -
BMC-P
BMC-H
BMC-T
Aldehyde dehydrogenase
PTAC
Signature enzyme
Alcohol dehydrogenase
Regulator

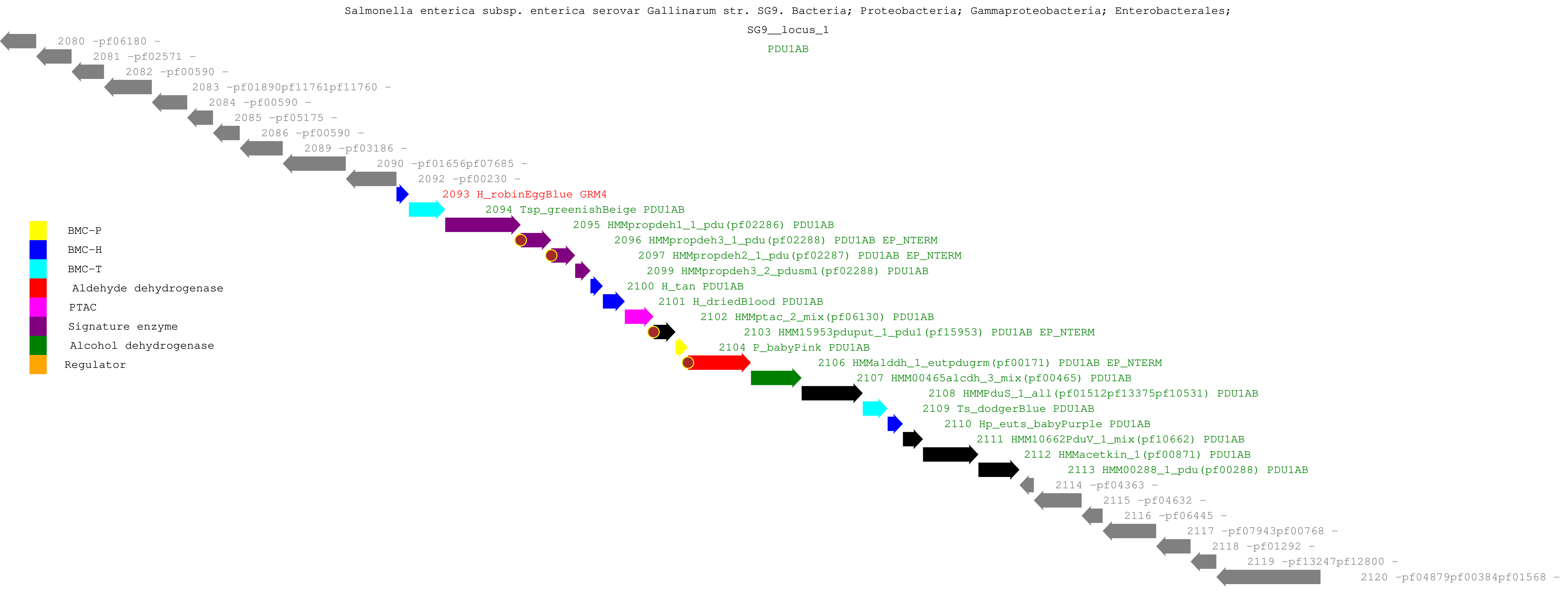

Salmonella enterica subsp. enterica serovar Gallinarum str. SG9. Bacteria; Proteobacteria; Gammaproteobacteria; Enterobacterales;
SG9__locus_1
PDU1AB
2080 -pf06180 -
2081 -pf02571 -
2082 -pf00590 -
2083 -pf01890pf11761pf11760 -
2084 -pf00590 -
2085 -pf05175 -
2086 -pf00590 -
2089 -pf03186 -
2090 -pf01656pf07685 -
2092 -pf00230 -
2093 H_robinEggBlue GRM4
2094 Tsp_greenishBeige PDU1AB
2095 HMMpropdeh1_1_pdu(pf02286) PDU1AB
2096 HMMpropdeh3_1_pdu(pf02288) PDU1AB EP_NTERM
2097 HMMpropdeh2_1_pdu(pf02287) PDU1AB EP_NTERM
2099 HMMpropdeh3_2_pdusml(pf02288) PDU1AB
2100 H_tan PDU1AB
2101 H_driedBlood PDU1AB
2102 HMMptac_2_mix(pf06130) PDU1AB
2103 HMM15953pduput_1_pdu1(pf15953) PDU1AB EP_NTERM
2104 P_babyPink PDU1AB
2106 HMMaldd h_1_eutpdugrm(pf00171) PDU1AB EP_NTERM
2107 HMM00465alcdh_3_mix(pf00465) PDU1AB
2108 HMMPduS_1_all(pf01512pf13375pf10531) PDU1AB
2109 Ts_dodgerBlue PDU1AB
2110 Hp_euts_babyPurple PDU1AB
2111 HMM10662PduV_1_mix(pf10662) PDU1AB
2112 HMMacetkin_1(pf00871) PDU1AB
2113 HMM00288_1_pdu(pf00288) PDU1AB
2114 -pf04363 -
2115 -pf04632 -
2116 -pf06445 -
2117 -pf07943pf00768 -
2118 -pf01292 -
2119 -pf13247pf12800 -
2120 -pf04879pf00384pf01568 -
BMC-P
BMC-H
BMC-T
Aldehyde dehydrogenase
PTAC
Signature enzyme
Alcohol dehydrogenase
Regulator

Citrobacter sp. MGH100. Bacteria; Proteobacteria; Gammaproteobacteria; Enterobacterales;

SK32__locus_2

PDU1AB

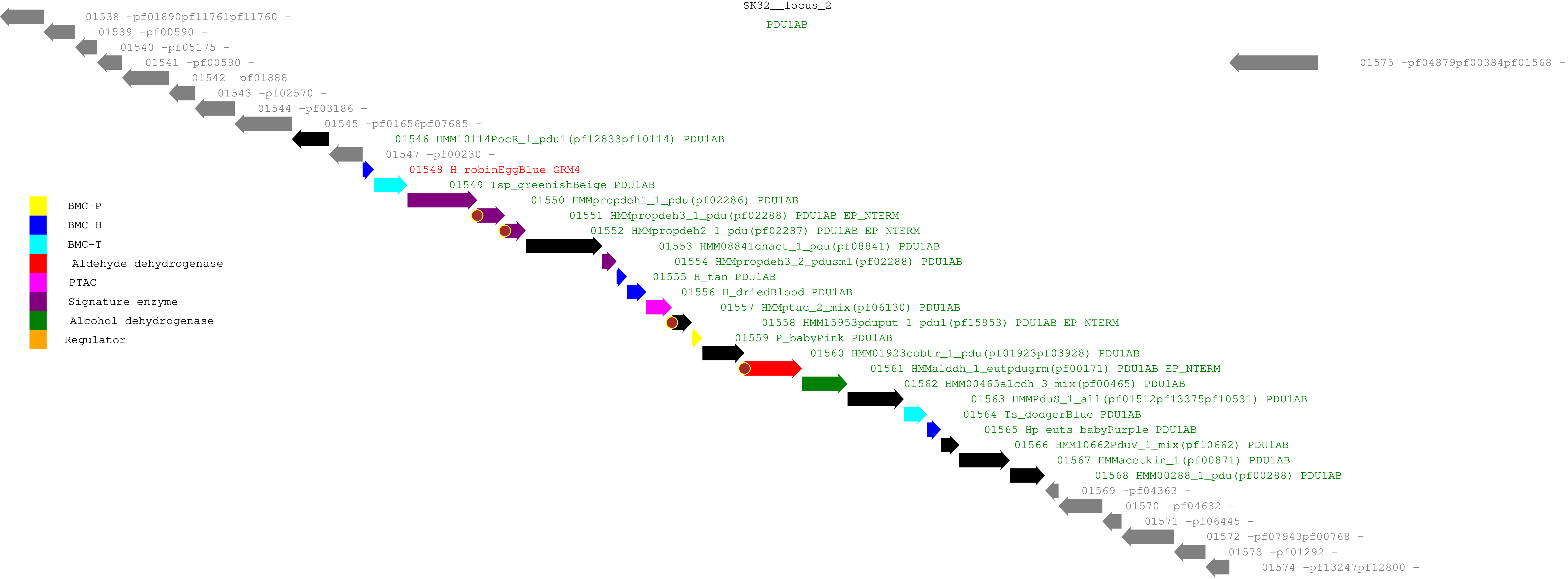

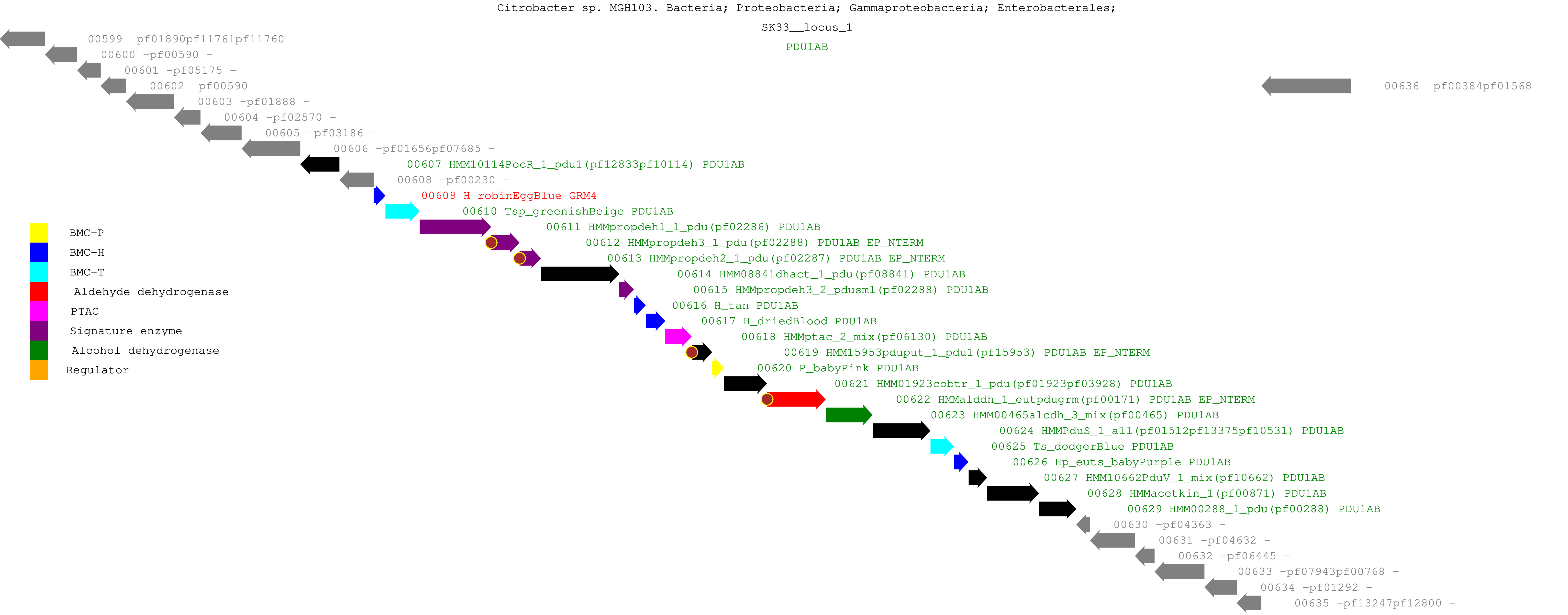
Citrobacter sp. MGH103. Bacteria; Proteobacteria; Gammaproteobacteria; Enterobacterales;
SK33__locus_1
PDU1AB
00599 -pf01890pf11761pf11760 -
00600 -pf00590 -
00601 -pf05175 -
00602 -pf00590 -
00603 -pf01888 -
00604 -pf02570 -
00605 -pf03186 -
00606 -pf01656pf07685 -
00607 HMM10114PocR_1_pdu1(pf12833pf10114) PDU1AB
00608 -pf00230 -
00609 H_robinEggBlue GRM4
00610 Tsp_greenishBeige PDU1AB
00611 HMMpropdeh1_1_pdu(pf02286) PDU1AB
00612 HMMpropdeh3_1_pdu(pf02288) PDU1AB EP_NTERM
00613 HMMpropdeh2_1_pdu(pf02287) PDU1AB EP_NTERM
00614 HMM08841dhact_1_pdu(pf08841) PDU1AB
00615 HMMpropdeh3_2_pdusml(pf02288) PDU1AB
00616 H_tan PDU1AB
00617 H_driedBlood PDU1AB
00618 HMMptac_2_mix(pf06130) PDU1AB
00619 HMM15953pduput_1_pdu1(pf15953) PDU1AB EP_NTERM
00620 P_babyPink PDU1AB
00621 HMM01923cobtr_1_pdu(pf01923pf03928) PDU1AB
00622 HMMalddh_1_eutpdugrm(pf00171) PDU1AB EP_NTERM
00623 HMM00465alcdh_3_mix(pf00465) PDU1AB
00624 HMMPduS_1_all(pf01512pf13375pf10531) PDU1AB
00625 Ts_dodgerBlue PDU1AB
00626 Hp_euts_babyPurple PDU1AB
00627 HMM10662PduV_1_mix(pf10662) PDU1AB
00628 HMMacetkin_1(pf00871) PDU1AB
00629 HMM00288_1_pdu(pf00288) PDU1AB
00630 -pf04363 -
00631 -pf04632 -
00632 -pf06445 -
00633 -pf07943pf00768 -
00634 -pf01292 -
00635 -pf13247pf12800 -
00636 -pf00384pf01568 -
BMC-P
BMC-H
BMC-T
Aldehyde dehydrogenase
PTAC
Signature enzyme
Alcohol dehydrogenase
Regulator

Citrobacter sp. MGH105. Bacteria; Proteobacteria; Gammaproteobacteria; Enterobacterales;

SK35__locus_3

PDU1AB

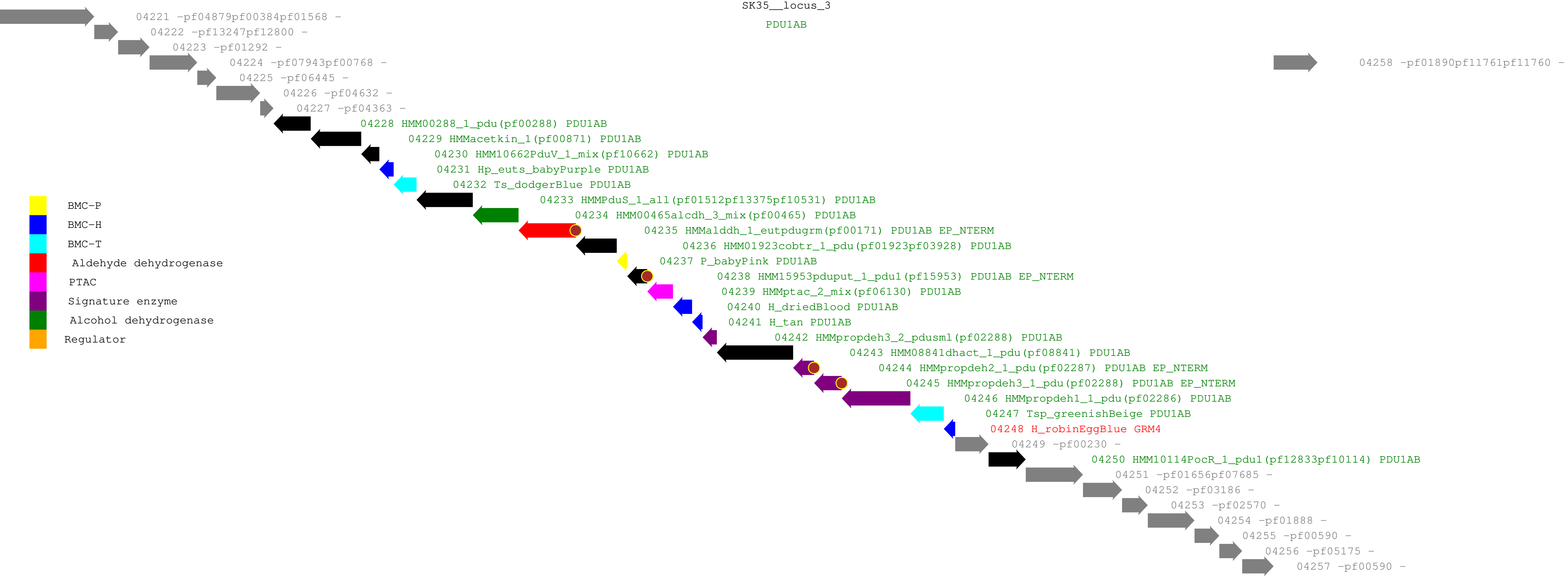

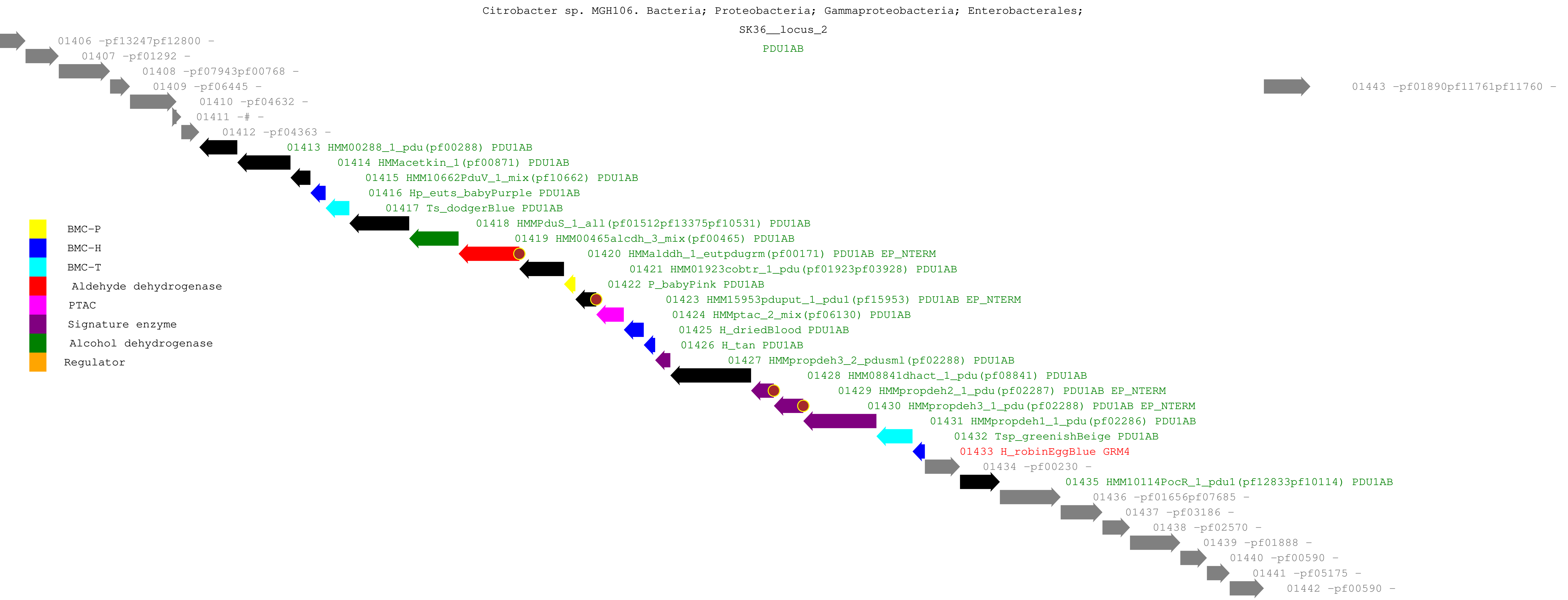

Citrobacter sp. MGH106. Bacteria; Proteobacteria; Gammaproteobacteria; Enterobacterales;
SK36__locus_2
PDU1AB
01406 -pf13247pf12800 -
01407 -pf01292 -
01408 -pf07943pf00768 -
01409 -pf06445 -
01410 -pf04632 -
01411 -# -
01412 -pf04363 -
01413 HMM00288_1_pdu(pf00288) PDU1AB
01414 HMMacetkin_1(pf00871) PDU1AB
01415 HMM10662PduV_1_mix(pf10662) PDU1AB
01416 Hp_euts_babyPurple PDU1AB
01417 Ts_dodgerBlue PDU1AB
01418 HMMPduS_1_all(pf01512pf13375pf10531) PDU1AB
01419 HMM00465alcdh_3_mix(pf00465) PDU1AB
01420 HMMalddh_1_eutpdugrm(pf00171) PDU1AB EP_NTERM
01421 HMM01923cobtr_1_pdu(pf01923pf03928) PDU1AB
01422 P_babyPink PDU1AB
01423 HMM15953pduput_1_pdu1(pf15953) PDU1AB EP_NTERM
01424 HMMptac_2_mix(pf06130) PDU1AB
01425 H_driedBlood PDU1AB
01426 H_tan PDU1AB
01427 HMMpropdeh3_2_pdusml(pf02288) PDU1AB
01428 HMM08841dhact_1_pdu(pf08841) PDU1AB
01429 HMMpropdeh2_1_pdu(pf02287) PDU1AB EP_NTERM
01430 HMMpropdeh3_1_pdu(pf02288) PDU1AB EP_NTERM
01431 HMMpropdeh1_1_pdu(pf02286) PDU1AB
01432 Tsp_greenishBeige PDU1AB
01433 H_robinEggBlue GRM4
01434 -pf00230 -
01435 HMM10114PocR_1_pdu1(pf12833pf10114) PDU1AB
01436 -pf01656pf07685 -
01437 -pf03186 -
01438 -pf02570 -
01439 -pf01888 -
01440 -pf00590 -
01441 -pf05175 -
01442 -pf00590 -
01443 -pf01890pf11761pf11760 -
BMC-P
BMC-H
BMC-T
Aldehyde dehydrogenase
PTAC
Signature enzyme
Alcohol dehydrogenase
Regulator

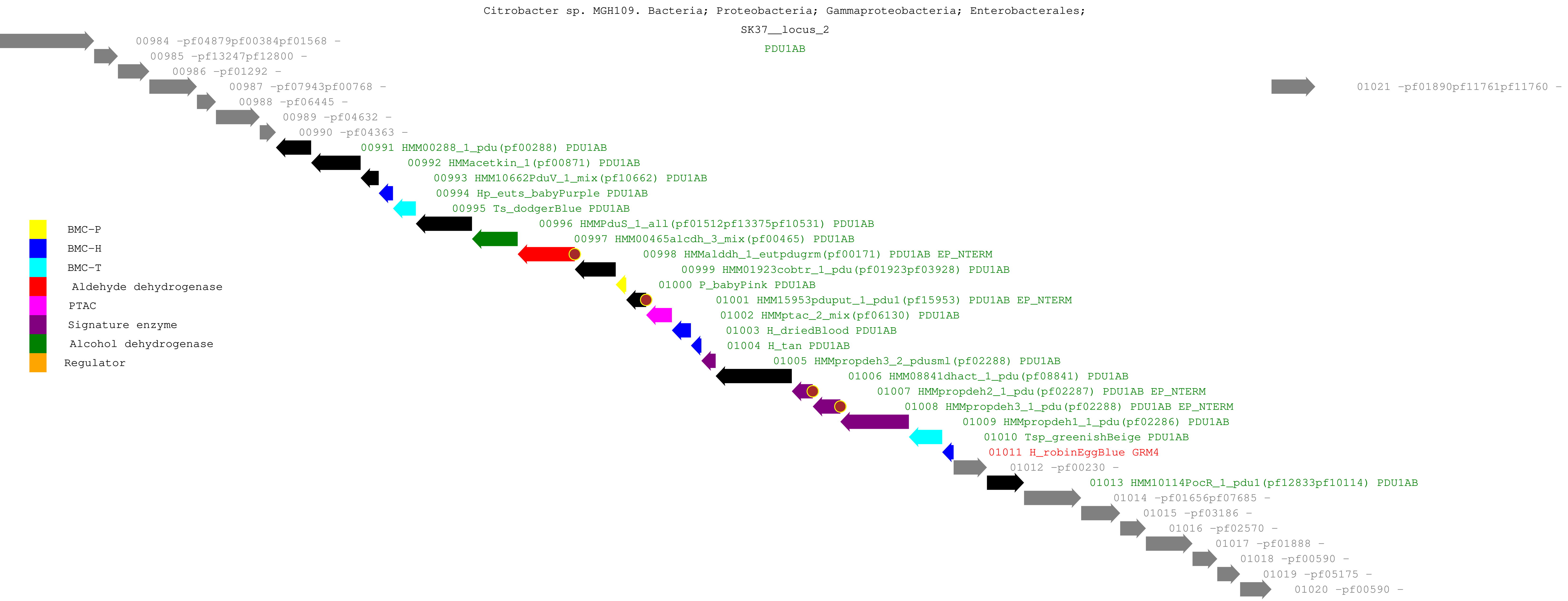
Citrobacter sp. MGH109. Bacteria; Proteobacteria; Gammaproteobacteria; Enterobacterales;
SK37__locus_2
PDU1AB
00984 -pf04879pf00384pf01568 -
00985 -pf13247pf12800 -
00986 -pf01292 -
00987 -pf07943pf00768 -
00988 -pf06445 -
00989 -pf04632 -
00990 -pf04363 -
00991 HMM00288_1_pdu(pf00288) PDU1AB
00992 HMMacetkin_1(pf00871) PDU1AB
00993 HMM10662PduV_1_mix(pf10662) PDU1AB
00994 Hp_euts_babyPurple PDU1AB
00995 Ts_dodgerBlue PDU1AB
00996 HMMPduS_1_all(pf01512pf13375pf10531) PDU1AB
00997 HMM00465alcdh_3_mix(pf00465) PDU1AB
00998 HMMalddh_1_eutpdugrm(pf00171) PDU1AB EP_NTERM
00999 HMM01923cobtr_1_pdu(pf01923pf03928) PDU1AB
01000 P_babyPink PDU1AB
01001 HMM15953pduput_1_pdu1(pf15953) PDU1AB EP_NTERM
01002 HMMptac_2_mix(pf06130) PDU1AB
01003 H_driedBlood PDU1AB
01004 H_tan PDU1AB
01005 HMMpropdeh3_2_pdusml(pf02288) PDU1AB
01006 HMM08841dhact_1_pdu(pf08841) PDU1AB
01007 HMMpropdeh2_1_pdu(pf02287) PDU1AB EP_NTERM
01008 HMMpropdeh3_1_pdu(pf02288) PDU1AB EP_NTERM
01009 HMMpropdeh1_1_pdu(pf02286) PDU1AB
01010 Tsp_greenishBeige PDU1AB
01011 H_robinEggBlue GRM4
01012 -pf00230 -
01013 HMM10114PocR_1_pdu1(pf12833pf10114) PDU1AB
01014 -pf01656pf07685 -
01015 -pf03186 -
01016 -pf02570 -
01017 -pf01888 -
01018 -pf00590 -
01019 -pf05175 -
01020 -pf00590 -
01021 -pf01890pf11761pf11760 -
BMC-P
BMC-H
BMC-T
Aldehyde dehydrogenase
PTAC
Signature enzyme
Alcohol dehydrogenase
Regulator

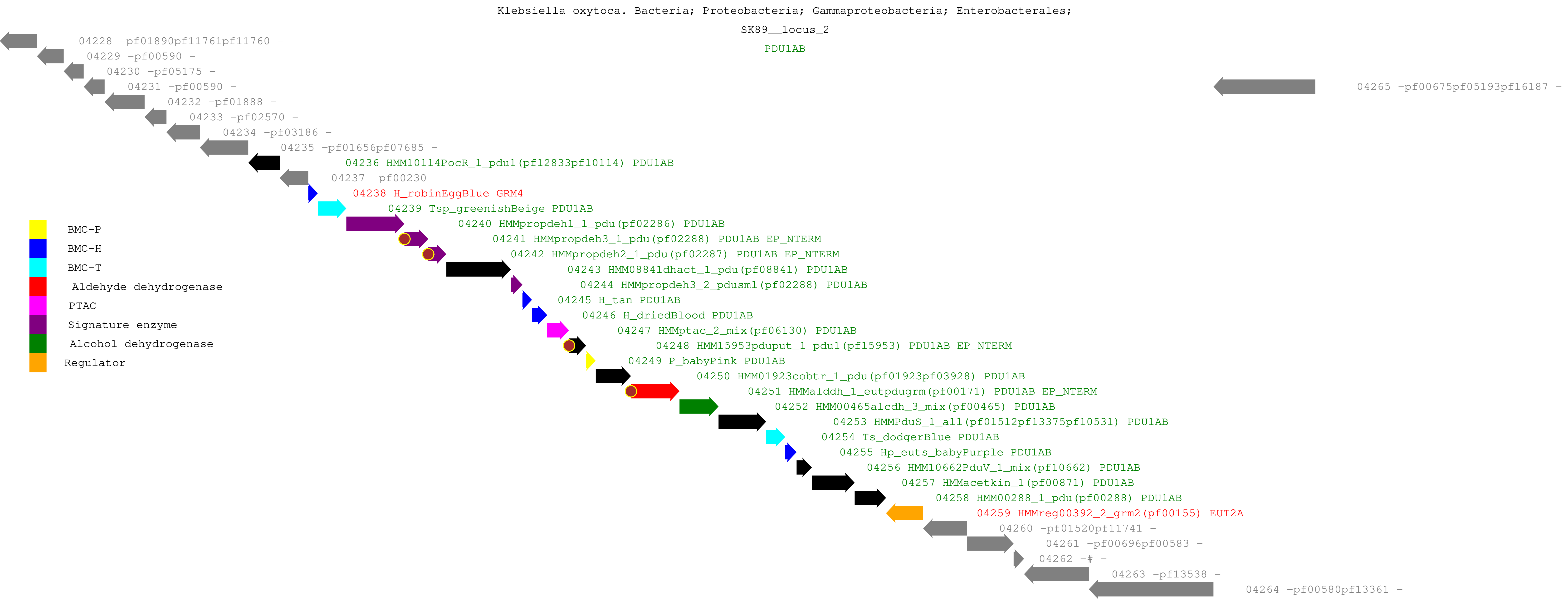
Klebsiella oxytoca. Bacteria; Proteobacteria; Gammaproteobacteria; Enterobacterales;
SK89__locus_2
PDU1AB
04228 -pf01890pf11761pf11760 -
04229 -pf00590 -
04230 -pf05175 -
04231 -pf00590 -
04232 -pf01888 -
04233 -pf02570 -
04234 -pf03186 -
04235 -pf01656pf07685 -
04236 HMM10114PocR_1_pdu1(pf12833pf10114) PDU1AB
04237 -pf00230 -
04238 H_robinEggBlue GRM4
04239 Tsp_greenishBeige PDU1AB
04240 HMMpropdeh1_1_pdu(pf02286) PDU1AB
04241 HMMpropdeh3_1_pdu(pf02288) PDU1AB EP_NTERM
04242 HMMpropdeh2_1_pdu(pf02287) PDU1AB EP_NTERM
04243 HMM08841dhact_1_pdu(pf08841) PDU1AB
04244 HMMpropdeh3_2_pdusml(pf02288) PDU1AB
04245 H_tan PDU1AB
04246 H_driedBlood PDU1AB
04247 HMMptac_2_mix(pf06130) PDU1AB
04248 HMM15953pduput_1_pdu1(pf15953) PDU1AB EP_NTERM
04249 P_babyPink PDU1AB
04250 HMM01923cobtr_1_pdu(pf01923pf03928) PDU1AB
04251 HMMalddh_1_eutpdugrm(pf00171) PDU1AB EP_NTERM
04252 HMM00465alcdh_3_mix(pf00465) PDU1AB
04253 HMMPduS_1_all(pf01512pf13375pf10531) PDU1AB
04254 Ts_dodgerBlue PDU1AB
04255 Hp_euts_babyPurple PDU1AB
04256 HMM10662PduV_1_mix(pf10662) PDU1AB
04257 HMMacetkin_1(pf00871) PDU1AB
04258 HMM00288_1_pdu(pf00288) PDU1AB
04259 HMMreg00392_2_grm2(pf00155) EUT2A
04260 -pf01520pf11741 -
04261 -pf00696pf00583 -
04262 -# -
04263 -pf13538 -
04264 -pf00580pf13361 -
04265 -pf00675pf05193pf16187 -
BMC-P
BMC-H
BMC-T
Aldehyde dehydrogenase
PTAC
Signature enzyme
Alcohol dehydrogenase
Regulator

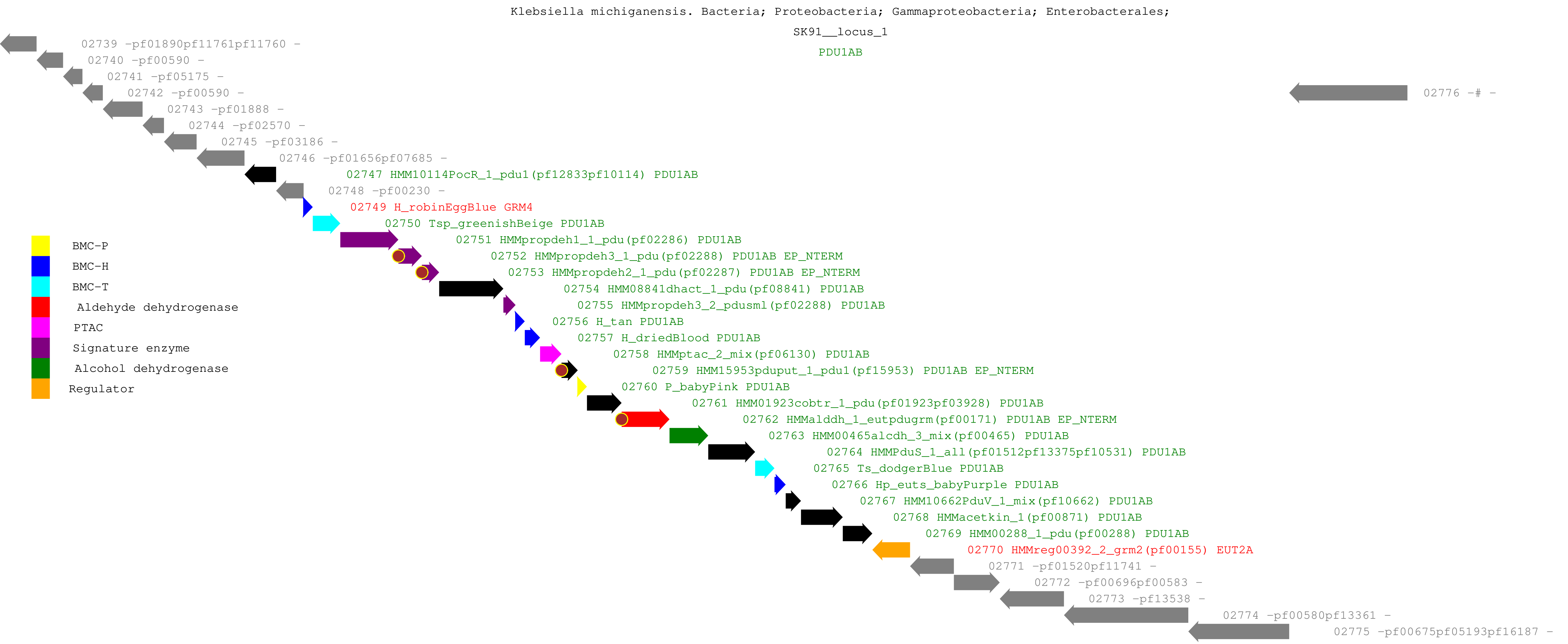

Klebsiella michiganensis. Bacteria; Proteobacteria; Gammaproteobacteria; Enterobacterales;
SK91__locus_1
PDU1AB
02739 -pf01890pf11761pf11760 -
02740 -pf00590 -
02741 -pf05175 -
02742 -pf00590 -
02743 -pf01888 -
02744 -pf02570 -
02745 -pf03186 -
02746 -pf01656pf07685 -
02747 HMM10114PocR_1_pdu1(pf12833pf10114) PDU1AB
02748 -pf00230 -
02749 H_robinEggBlue GRM4
02750 Tsp_greenishBeige PDU1AB
02751 HMMpropdeh1_1_pdu(pf02286) PDU1AB
02752 HMMpropdeh3_1_pdu(pf02288) PDU1AB EP_NTERM
02753 HMMpropdeh2_1_pdu(pf02287) PDU1AB EP_NTERM
02754 HMM08841dhact_1_pdu(pf08841) PDU1AB
02755 HMMpropdeh3_2_pdusml(pf02288) PDU1AB
02756 H_tan PDU1AB
02757 H_driedBlood PDU1AB
02758 HMMptac_2_mix(pf06130) PDU1AB
02759 HMM15953pduput_1_pdu1(pf15953) PDU1AB EP_NTERM
02760 P_babyPink PDU1AB
02761 HMM01923cobtr_1_pdu(pf01923pf03928) PDU1AB
02762 HMMalddh_1_eutpdugrm(pf00171) PDU1AB EP_NTERM
02763 HMM00465alcdh_3_mix(pf00465) PDU1AB
02764 HMMPduS_1_all(pf01512pf13375pf10531) PDU1AB
02765 Ts_dodgerBlue PDU1AB
02766 Hp_euts_babyPurple PDU1AB
02767 HMM10662PduV_1_mix(pf10662) PDU1AB
02768 HMMacetkin_1(pf00871) PDU1AB
02769 HMM00288_1_pdu(pf00288) PDU1AB
02770 HMMreg00392_2_grm2(pf00155) EUT2A
02771 -pf01520pf11741 -
02772 -pf00696pf00583 -
02773 -pf13538 -
02774 -pf00580pf13361 -
02775 -pf00675pf05193pf16187 -
02776 -# -
BMC-P
BMC-H
BMC-T
Aldehyde dehydrogenase
PTAC
Signature enzyme
Alcohol dehydrogenase
Regulator

Salmonella typhimurium (strain SL1344). Bacteria; Proteobacteria; Gammaproteobacteria; Enterobacterales;

SL1344__locus_1

PDU1AB

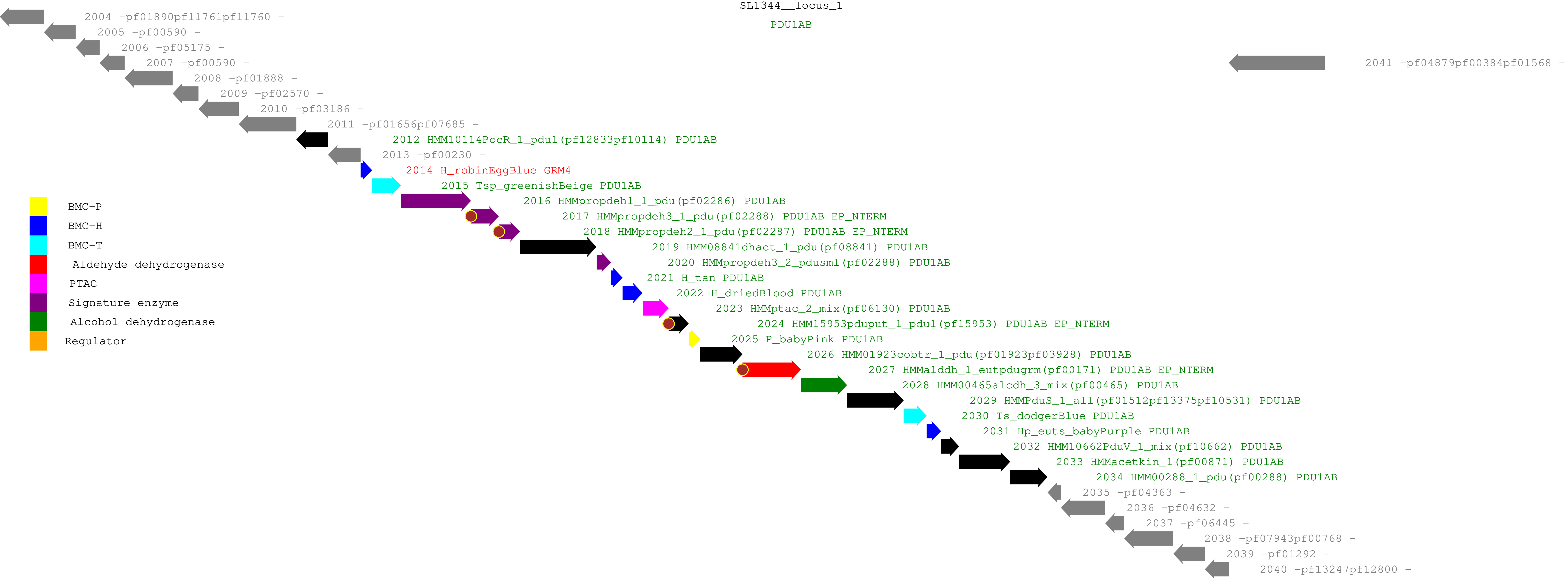

Klebsiella pneumoniae. Bacteria; Proteobacteria; Gammaproteobacteria; Enterobacterales;

SM87__locus_3

PDU1AB

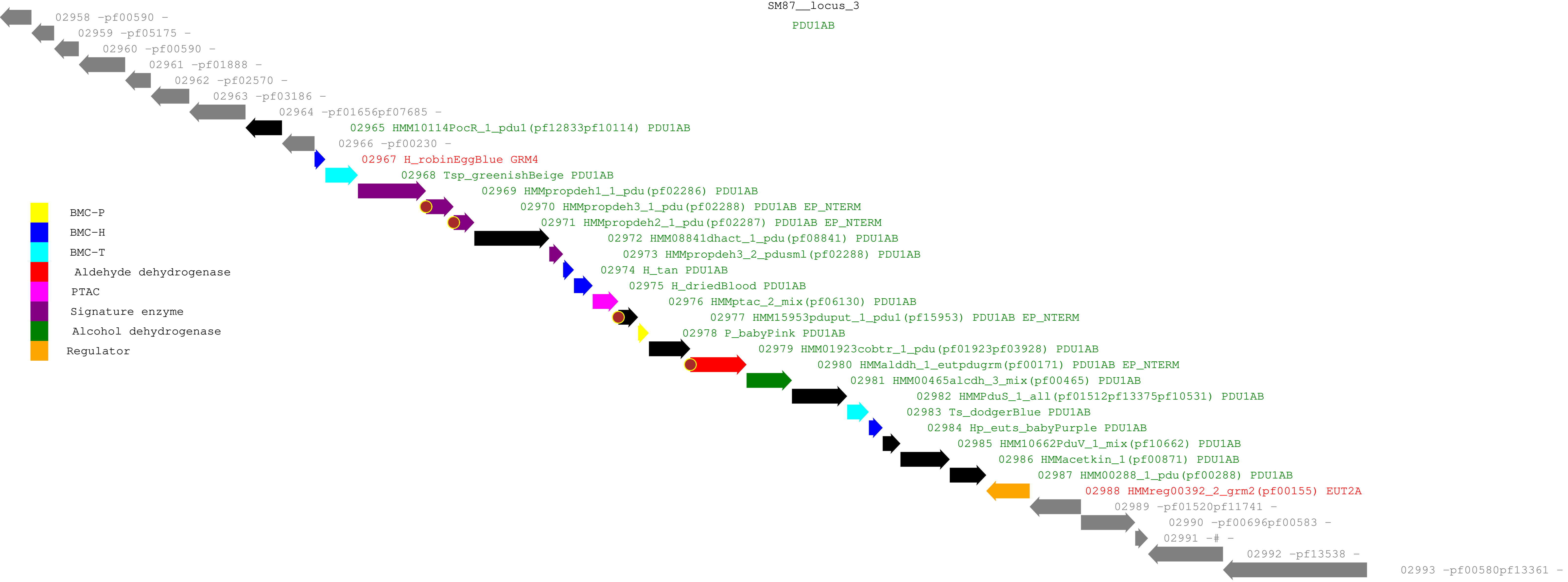

Salmonella newport (strain SL254). Bacteria; Proteobacteria; Gammaproteobacteria; Enterobacterales;

SNSL254_A_locus_1

PDU1AB

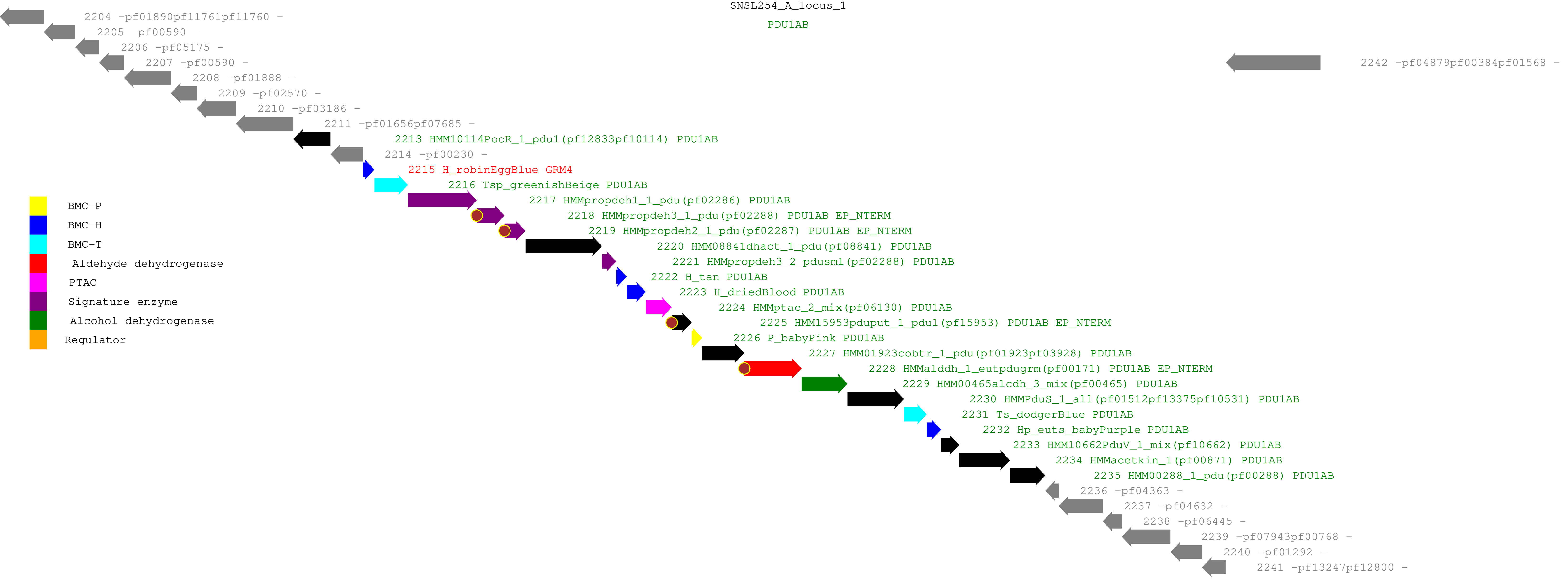

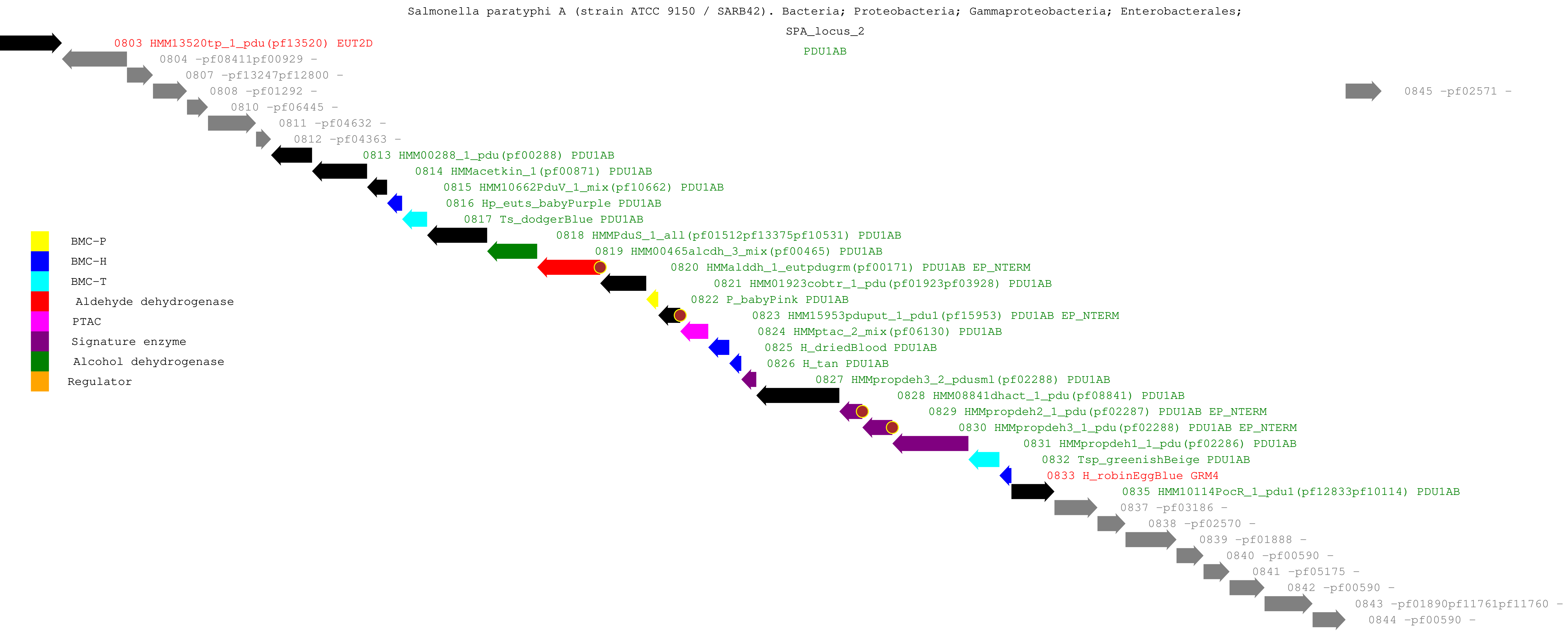
Salmonella paratyphi A (strain ATCC 9150 / SARB42). Bacteria; Proteobacteria; Gammaproteobacteria; Enterobacterales;
SPA_locus_2
PDU1AB
0803 HMM13520tp_1_pdu(pf13520) EUT2D
0804 -pf08411pf00929 -
0807 -pf13247pf12800 -
0808 -pf01292 -
0810 -pf06445 -
0811 -pf04632 -
0812 -pf04363 -
0813 HMM00288_1_pdu(pf00288) PDU1AB
0814 HMMacetkin_1(pf00871) PDU1AB
0815 HMM10662PduV_1_mix(pf10662) PDU1AB
0816 Hp_euts_babyPurple PDU1AB
0817 Ts_dodgerBlue PDU1AB
0818 HMMPduS_1_all(pf01512pf13375pf10531) PDU1AB
0819 HMM00465alcdh_3_mix(pf00465) PDU1AB
0820 HMMalddh_1_eutpdugrm(pf00171) PDU1AB EP_NTERM
0821 HMM01923cobtr_1_pdu(pf01923pf03928) PDU1AB
0822 P_babyPink PDU1AB
0823 HMM15953pduput_1_pdu1(pf15953) PDU1AB EP_NTERM
0824 HMMptac_2_mix(pf06130) PDU1AB
0825 H_driedBlood PDU1AB
0826 H_tan PDU1AB
0827 HMMpropdeh3_2_pdusml(pf02288) PDU1AB
0828 HMM08841dhact_1_pdu(pf08841) PDU1AB
0829 HMMpropdeh2_1_pdu(pf02287) PDU1AB EP_NTERM
0830 HMMpropdeh3_1_pdu(pf02288) PDU1AB EP_NTERM
0831 HMMpropdeh1_1_pdu(pf02286) PDU1AB
0832 Tsp_greenishBeige PDU1AB
0833 H_robinEggBlue GRM4
0835 HMM10114PocR_1_pdu1(pf12833pf10114) PDU1AB
0837 -pf03186 -
0838 -pf02570 -
0839 -pf01888 -
0840 -pf00590 -
0841 -pf05175 -
0842 -pf00590 -
0843 -pf01890pf11761pf11760 -
0844 -pf00590 -
0845 -pf02571 -
BMC-P
BMC-H
BMC-T
Aldehyde dehydrogenase
PTAC
Signature enzyme
Alcohol dehydrogenase
Regulator

Shigella sonnei str. Moseley. Bacteria; Proteobacteria; Gammaproteobacteria; Enterobacterales;

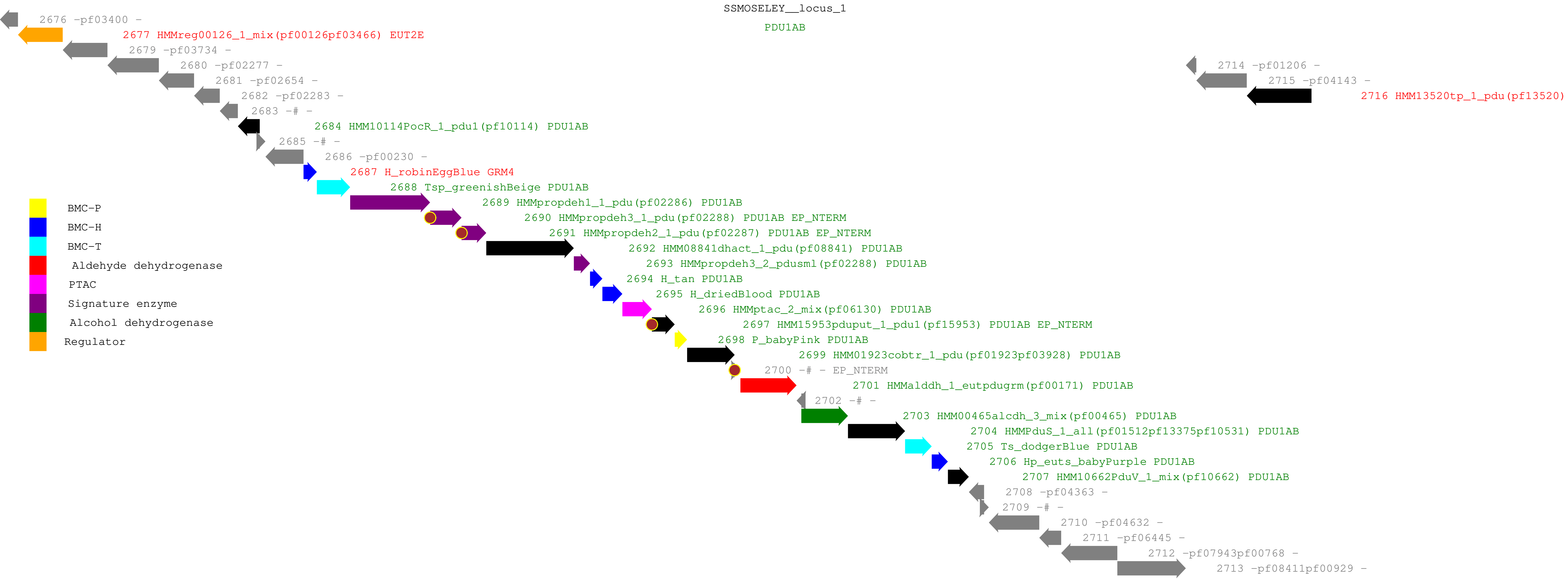

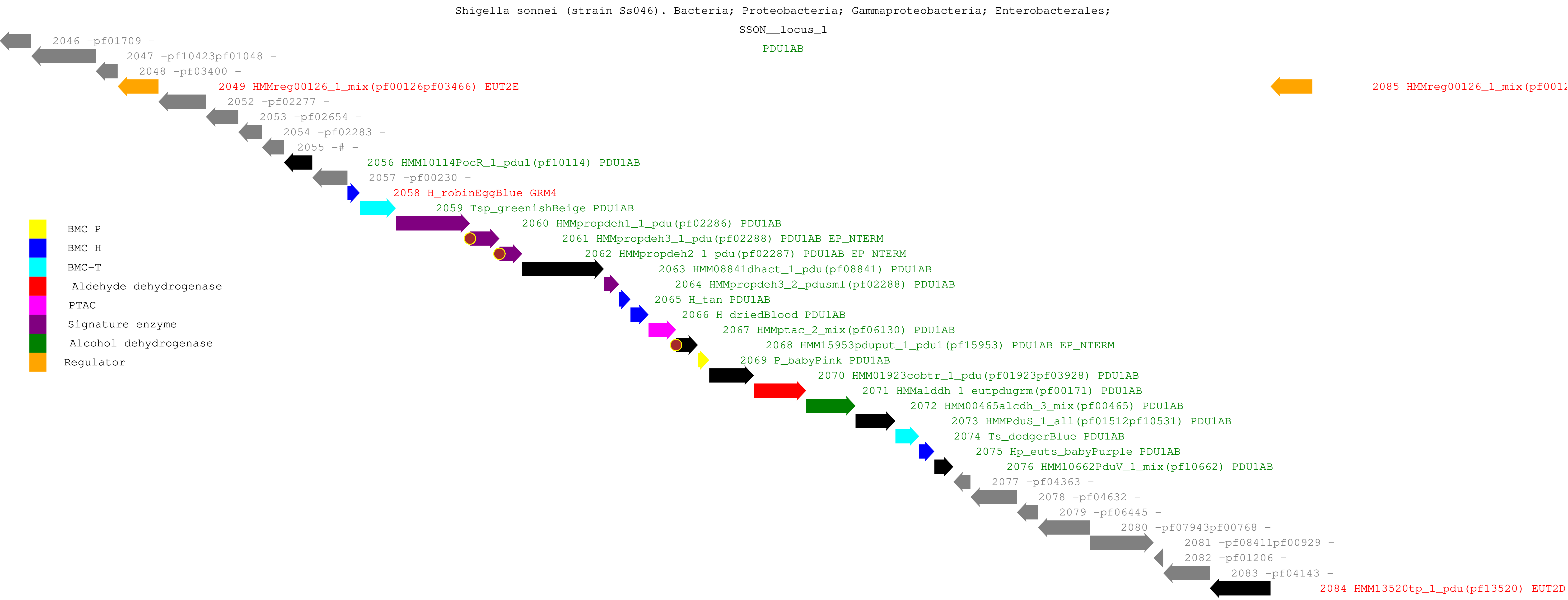
Shigella sonnei (strain Ss046). Bacteria; Proteobacteria; Gammaproteobacteria; Enterobacterales;
SSON__locus_1
PDU1AB
2046 -pf01709 -
2047 -pf10423pf01048 -
2048 -pf03400 -
2049 HMMreg00126_1_mix(pf00126pf03466) EUT2E
2085 HMMreg00126_1_mix(pf0012
2052 -pf02277 -
2053 -pf02654 -
2054 -pf02283 -
2055 -# -
2056 HMM10114PocR_1_pdu1(pf10114) PDU1AB
2057 -pf00230 -
2058 H_robinEggBlue GRM4
2059 Tsp_greenishBeige PDU1AB
2060 HMMpropdeh1_1_pdu(pf02286) PDU1AB
2061 HMMpropdeh3_1_pdu(pf02288) PDU1AB EP_NTERM
2062 HMMpropdeh2_1_pdu(pf02287) PDU1AB EP_NTERM
2063 HMM08841dhact_1_pdu(pf08841) PDU1AB
2064 HMMpropdeh3_2_pdusml(pf02288) PDU1AB
2065 H_tan PDU1AB
2066 H_driedBlood PDU1AB
2067 HMMptac_2_mix(pf06130) PDU1AB
2068 HMM15953pduput_1_pdu1(pf15953) PDU1AB EP_NTERM
2069 P_babyPink PDU1AB
2070 HMM01923cobtr_1_pdu(pf01923pf03928) PDU1AB
2071 HMMalddh_1_eutpdugrm(pf00171) PDU1AB
2072 HMM00465alcdh_3_mix(pf00465) PDU1AB
2073 HMMPduS_1_all(pf01512pf10531) PDU1AB
2074 Ts_dodgerBlue PDU1AB
2075 Hp_euts_babyPurple PDU1AB
2076 HMM10662PduV_1_mix(pf10662) PDU1AB
2077 -pf04363 -
2078 -pf04632 -
2079 -pf06445 -
2080 -pf07943pf00768 -
2081 -pf08411pf00929 -
2082 -pf01206 -
2083 -pf04143 -
2084 HMM13520tp_1_pdu(pf13520) EUT2D
BMC-P
BMC-H
BMC-T
Aldehyde dehydrogenase
PTAC
Signature enzyme
Alcohol dehydrogenase
Regulator

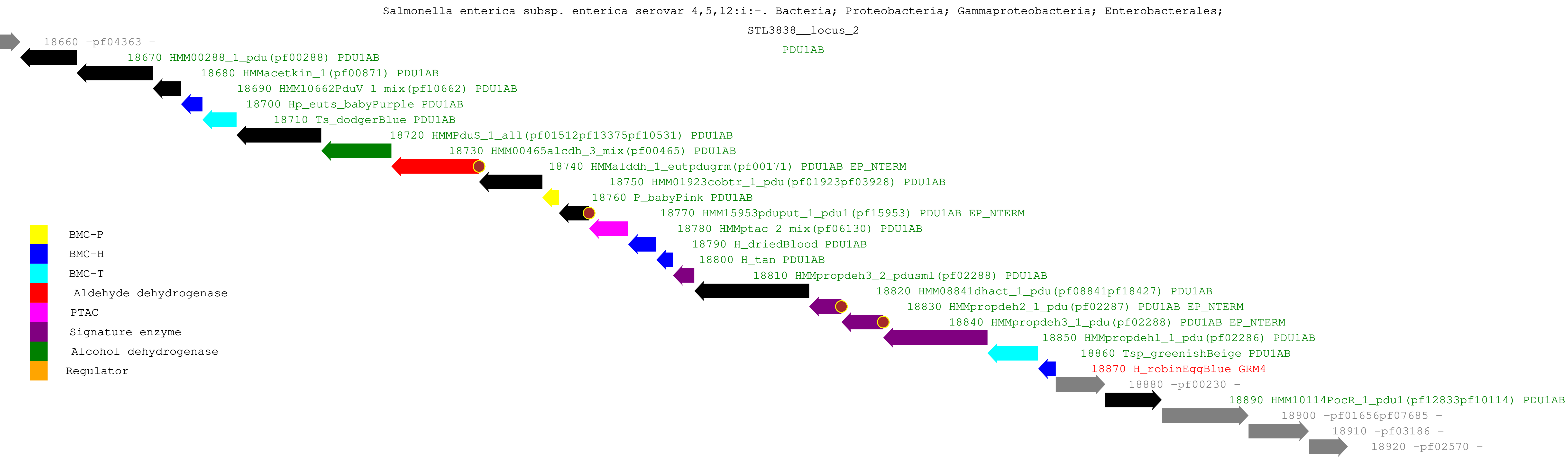
Salmonella enterica subsp. enterica serovar 4,5,12:i:-. Bacteria; Proteobacteria; Gammaproteobacteria; Enterobacterales;
STL3838__locus_2
PDU1AB
18660 -pf04363 -
18670 HMM00288_1_pdu(pf00288) PDU1AB
18680 HMMacetkin_1(pf00871) PDU1AB
18690 HMM10662PduV_1_mix(pf10662) PDU1AB
18700 Hp_euts_babyPurple PDU1AB
18710 Ts_dodgerBlue PDU1AB
18720 HMMPduS_1_all(pf01512pf13375pf10531) PDU1AB
18730 HMM00465alcdh_3_mix(pf00465) PDU1AB
18740 HMMalddh_1_eutpdugrm(pf00171) PDU1AB EP_NTERM
18750 HMM01923cobtr_1_pdu(pf01923pf03928) PDU1AB
18760 P_babyPink PDU1AB
18770 HMM15953pduput_1_pdu1(pf15953) PDU1AB EP_NTERM
18780 HMMptac_2_mix(pf06130) PDU1AB
18790 H_driedBlood PDU1AB
18800 H_tan PDU1AB
18810 HMMpropdeh3_2_pdusml(pf02288) PDU1AB
18820 HMM08841dhact_1_pdu(pf08841pf18427) PDU1AB
18830 HMMpropdeh2_1_pdu(pf02287) PDU1AB EP_NTERM
18840 HMMpropdeh3_1_pdu(pf02288) PDU1AB EP_NTERM
18850 HMMpropdeh1_1_pdu(pf02286) PDU1AB
18860 Tsp_greenishBeige PDU1AB
18870 H_robinEggBlue GRM4
18880 -pf00230 -
18890 HMM10114PocR_1_pdu1(pf12833pf10114) PDU1AB
18900 -pf01656pf07685 -
18910 -pf03186 -
18920 -pf02570 -
BMC-P
BMC-H
BMC-T
Aldehyde dehydrogenase
PTAC
Signature enzyme
Alcohol dehydrogenase
Regulator

Salmonella typhimurium (strain 14028s / SGSC 2262). Bacteria; Proteobacteria; Gammaproteobacteria; Enterobacterales;

STM14__locus_1

PDU1AB

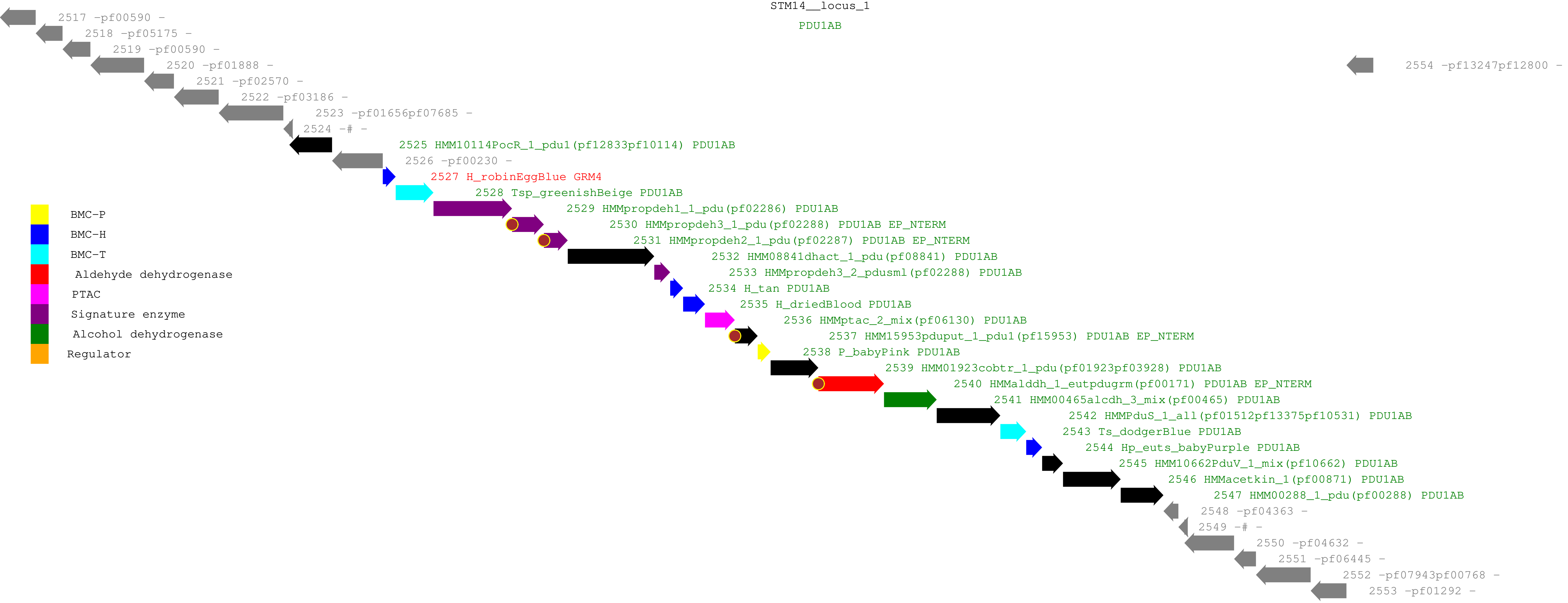

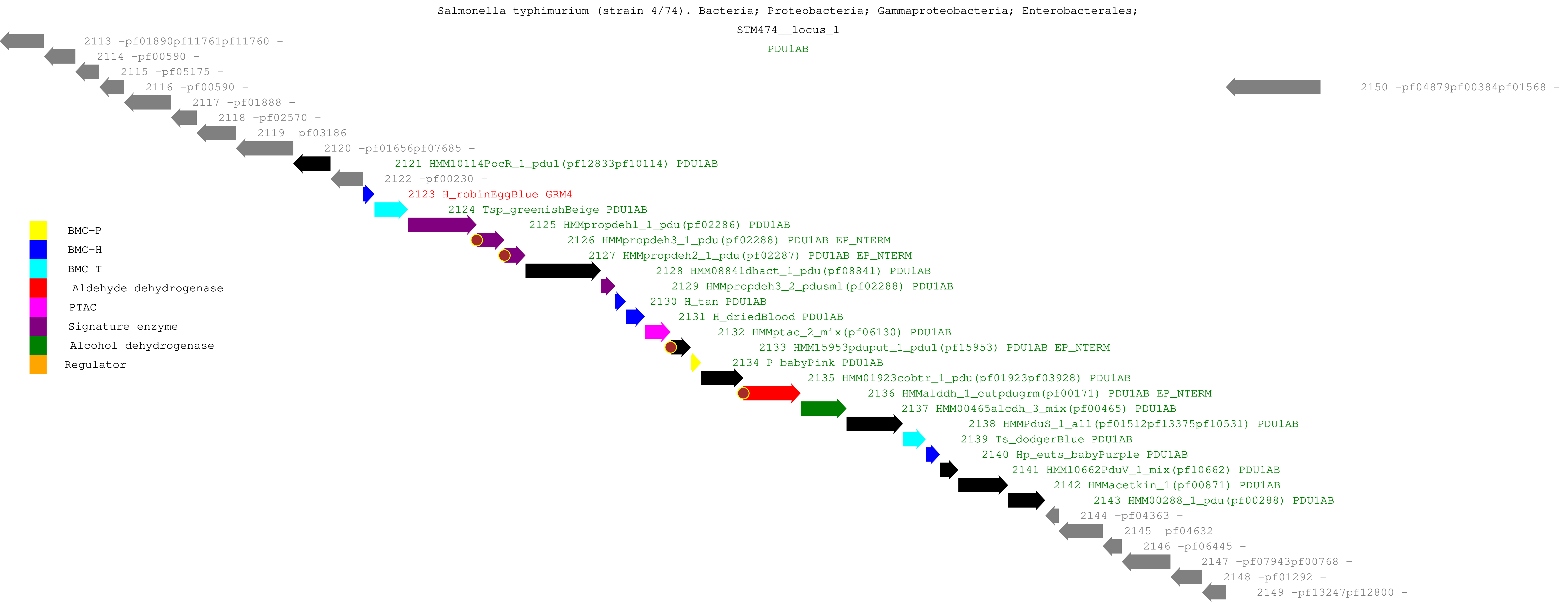

Salmonella typhimurium (strain 4/74). Bacteria; Proteobacteria; Gammaproteobacteria; Enterobacterales;
STM474__locus_1
PDU1AB
2113 -pf01890pf11761pf11760 -
2114 -pf00590 -
2115 -pf05175 -
2116 -pf00590 -
2117 -pf01888 -
2118 -pf02570 -
2119 -pf03186 -
2120 -pf01656pf07685 -
2121 HMM10114PocR_1_pdu1(pf12833pf10114) PDU1AB
2122 -pf00230 -
2123 H_robinEggBlue GRM4
2124 Tsp_greenishBeige PDU1AB
2125 HMMpropdeh1_1_pdu(pf02286) PDU1AB
2126 HMMpropdeh3_1_pdu(pf02288) PDU1AB EP_NTERM
2127 HMMpropdeh2_1_pdu(pf02287) PDU1AB EP_NTERM
2128 HMM08841dhact_1_pdu(pf08841) PDU1AB
2129 HMMpropdeh3_2_pdusml(pf02288) PDU1AB
2130 H_tan PDU1AB
2131 H_driedBlood PDU1AB
2132 HMMptac_2_mix(pf06130) PDU1AB
2133 HMM15953pduput_1_pdu1(pf15953) PDU1AB EP_NTERM
2134 P_babyPink PDU1AB
2135 HMM01923cobtr_1_pdu(pf01923pf03928) PDU1AB
2136 HMMalddh_1_eutpdugrm(pf00171) PDU1AB EP_NTERM
2137 HMM00465alcdh_3_mix(pf00465) PDU1AB
2138 HMMPduS_1_all(pf01512pf13375pf10531) PDU1AB
2139 Ts_dodgerBlue PDU1AB
2140 Hp_euts_babyPurple PDU1AB
2141 HMM10662PduV_1_mix(pf10662) PDU1AB
2142 HMMacetkin_1(pf00871) PDU1AB
2143 HMM00288_1_pdu(pf00288) PDU1AB
2144 -pf04363 -
2145 -pf04632 -
2146 -pf06445 -
2147 -pf07943pf00768 -
2148 -pf01292 -
2149 -pf13247pf12800 -
2150 -pf04879pf00384pf01568 -
BMC-P
BMC-H
BMC-T
Aldehyde dehydrogenase
PTAC
Signature enzyme
Alcohol dehydrogenase
Regulator

Salmonella typhimurium (strain LT2 / SGSC1412 / ATCC 700720). Bacteria; Proteobacteria; Gammaproteobacteria; Enterobacterales;

STM_locus_1

PDU1AB

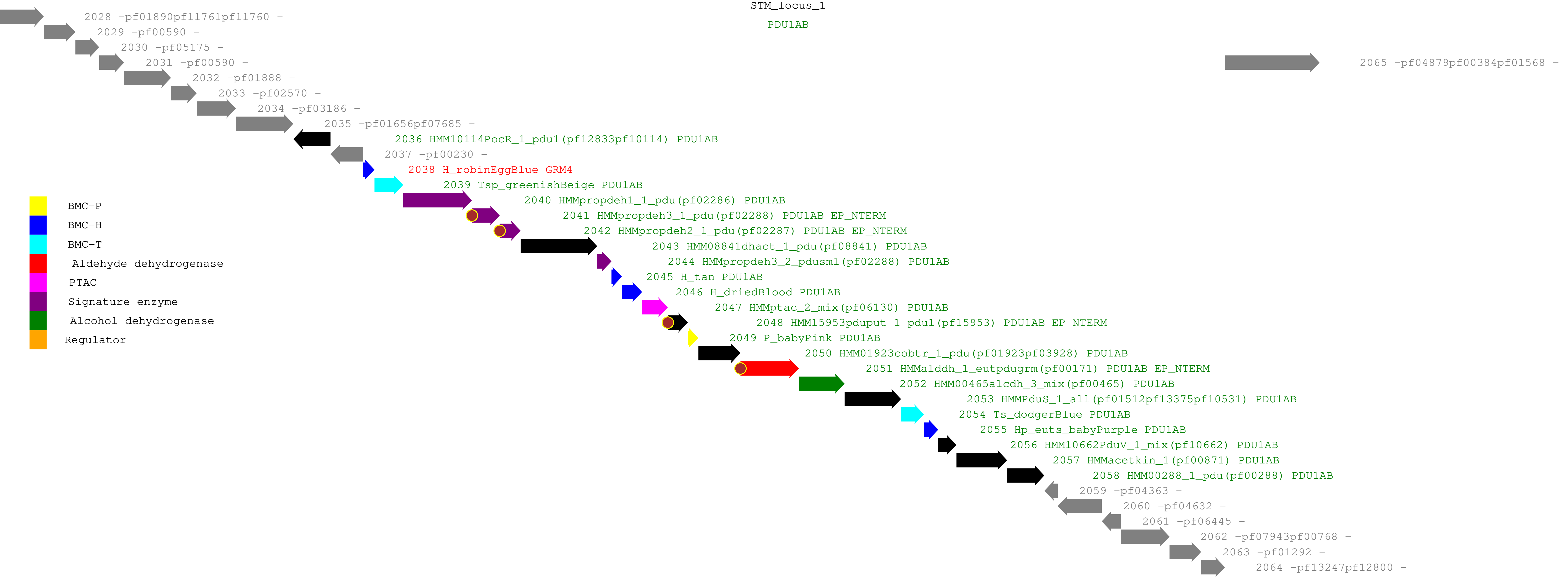

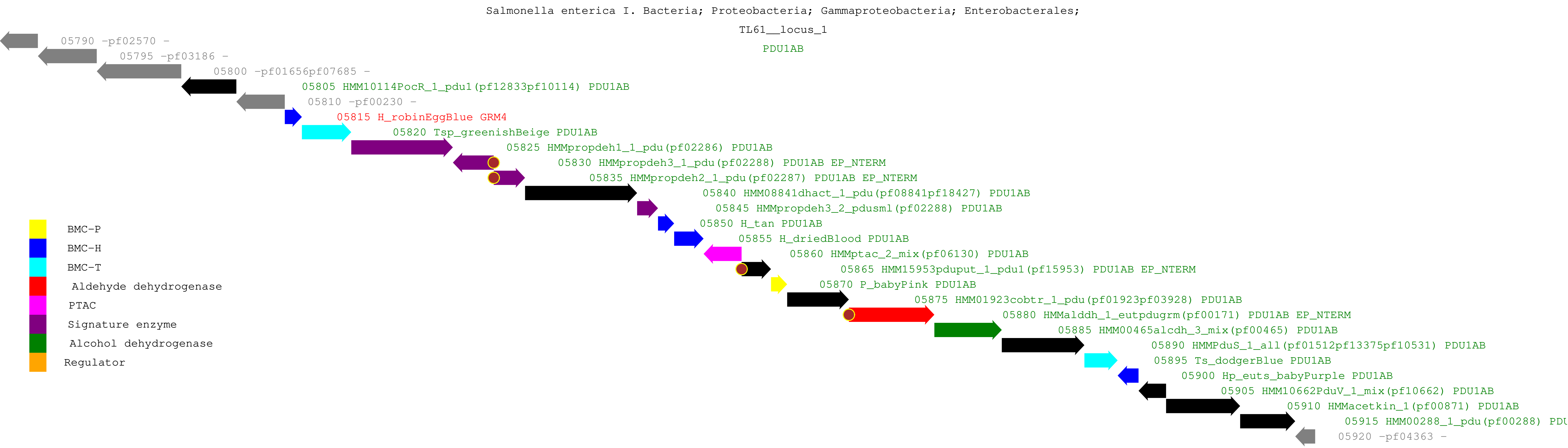
Salmonella enterica I. Bacteria; Proteobacteria; Gammaproteobacteria; Enterobacterales;
TL61__locus_1
PDU1AB
05790 -pf02570 -
05795 -pf03186 -
05800 -pf01656pf07685 -
05805 HMM10114PocR_1_pdu1(pf12833pf10114) PDU1AB
05810 -pf00230 -
05815 H_robinEggBlue GRM4
05820 Tsp_greenishBeige PDU1AB
05825 HMMpropdeh1_1_pdu(pf02286) PDU1AB
05830 HMMpropdeh3_1_pdu(pf02288) PDU1AB EP_NTERM
05835 HMMpropdeh2_1_pdu(pf02287) PDU1AB EP_NTERM
05840 HMM08841dhact_1_pdu(pf08841pf18427) PDU1AB
05845 HMMpropdeh3_2_pdusml(pf02288) PDU1AB
05850 H_tan PDU1AB
05855 H_driedBlood PDU1AB
05860 HMMptac_2_mix(pf06130) PDU1AB
05865 HMM15953pduput_1_pdu1(pf15953) PDU1AB EP_NTERM
05870 P_babyPink PDU1AB
05875 HMM01923cobtr_1_pdu(pf01923pf03928) PDU1AB
05880 HMMalddh_1_eutpdugrm(pf00171) PDU1AB EP_NTERM
05885 HMM00465alcdh_3_mix(pf00465) PDU1AB
05890 HMMPduS_1_all(pf01512pf13375pf10531) PDU1AB
05895 Ts_dodgerBlue PDU1AB
05900 Hp_euts_babyPurple PDU1AB
05905 HMM10662PduV_1_mix(pf10662) PDU1AB
05910 HMMacetkin_1(pf00871) PDU1AB
05915 HMM00288_1_pdu(pf00288) PDU
05920 -pf04363 -
BMC-P
BMC-H
BMC-T
Aldehyde dehydrogenase
PTAC
Signature enzyme
Alcohol dehydrogenase
Regulator

Salmonella enterica subsp. salamae. Bacteria; Proteobacteria; Gammaproteobacteria; Enterobacterales;

TM63__locus_2

PDU1AB

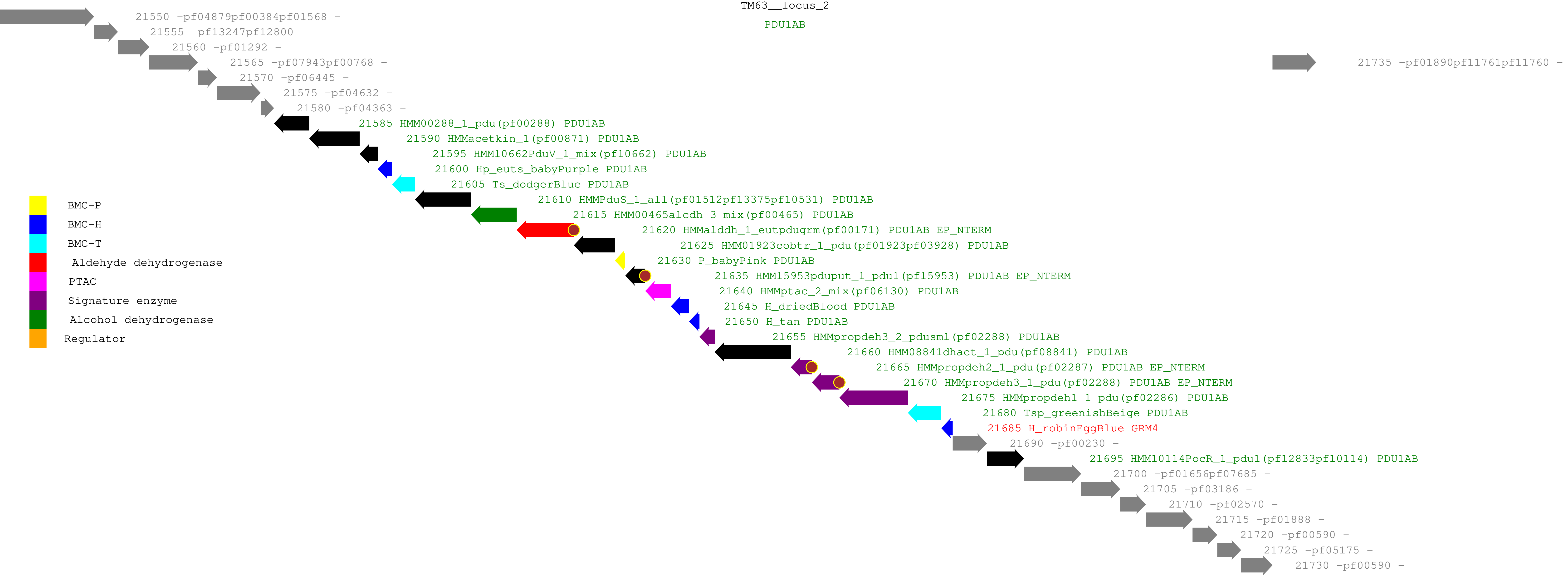

Citrobacter freundii. Bacteria; Proteobacteria; Gammaproteobacteria; Enterobacterales;

TO64__locus_1

PDU1AB

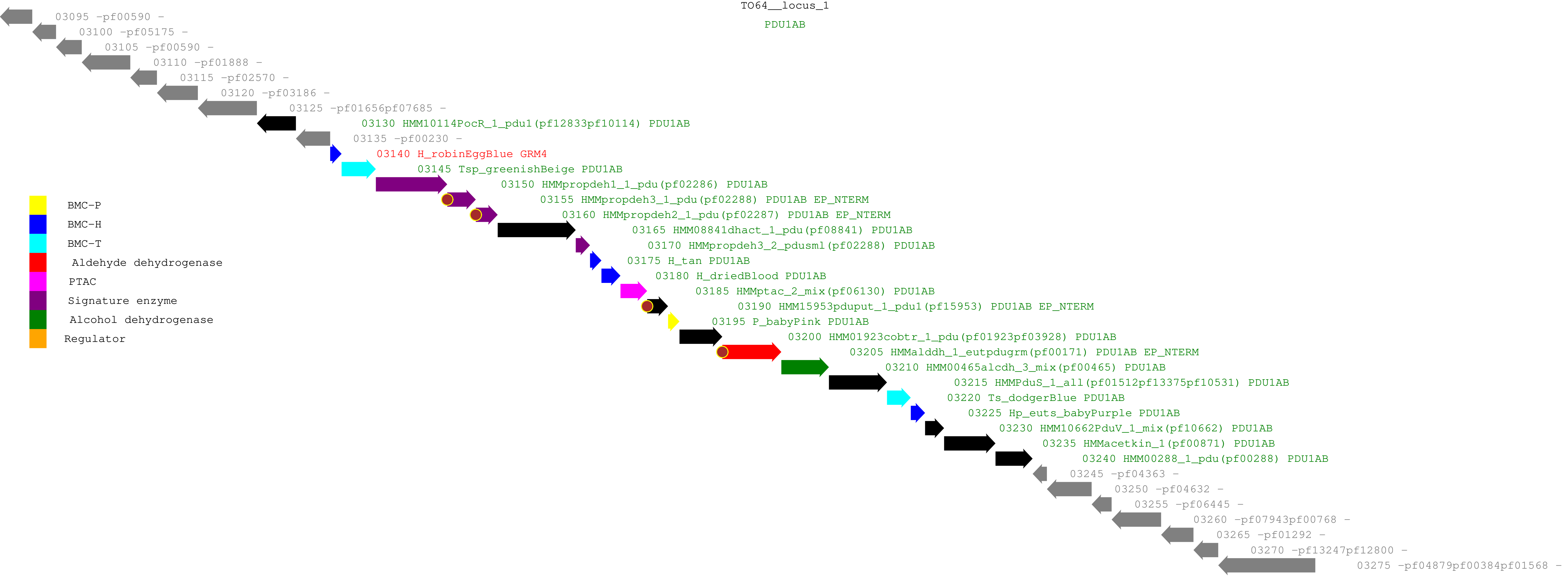

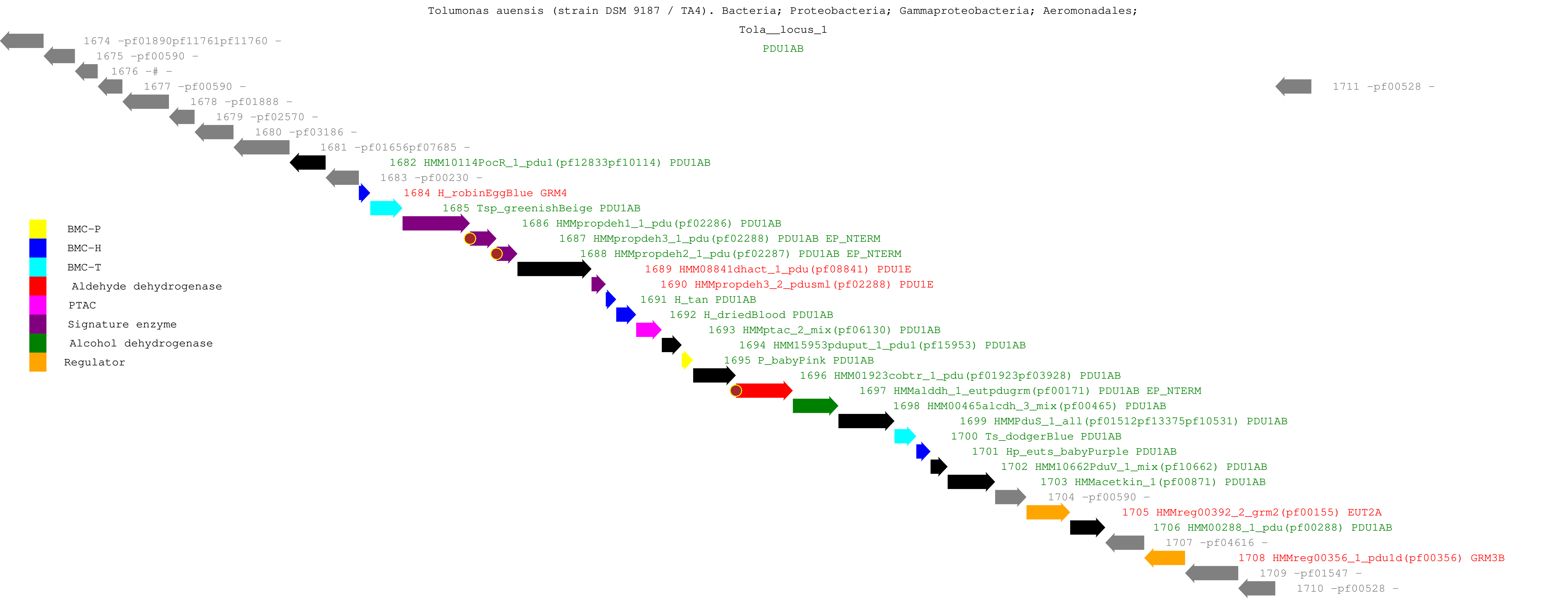
Tolumonas auensis (strain DSM 9187 / TA4). Bacteria; Proteobacteria; Gammaproteobacteria; Aeromonadales;
Tola__locus_1
PDU1AB
1674 -pf01890pf11761pf11760 -
1675 -pf00590 -
1676 -# -
1677 -pf00590 -
1678 -pf01888 -
1679 -pf02570 -
1680 -pf03186 -
1681 -pf01656pf07685 -
1682 HMM10114PocR_1_pdu1(pf12833pf10114) PDU1AB
1683 -pf00230 -
1684 H_robinEggBlue GRM4
1685 Tsp_greenishBeige PDU1AB
1686 HMMpropdeh1_1_pdu(pf02286) PDU1AB
1687 HMMpropdeh3_1_pdu(pf02288) PDU1AB EP_NTERM
1688 HMMpropdeh2_1_pdu(pf02287) PDU1AB EP_NTERM
1689 HMM08841dhact_1_pdu(pf08841) PDU1E
1690 HMMpropdeh3_2_pdusml(pf02288) PDU1E
1691 H_tan PDU1AB
1692 H_driedBlood PDU1AB
1693 HMMptac_2_mix(pf06130) PDU1AB
1694 HMM15953pduput_1_pdu1(pf15953) PDU1AB
1695 P_babyPink PDU1AB
1696 HMM01923cobtr_1_pdu(pf01923pf03928) PDU1AB
1697 HMMalddh_1_eutpdugrm(pf00171) PDU1AB EP_NTERM
1698 HMM00465alcdh_3_mix(pf00465) PDU1AB
1699 HMMPduS_1_all(pf01512pf13375pf10531) PDU1AB
1700 Ts_dodgerBlue PDU1AB
1701 Hp_euts_babyPurple PDU1AB
1702 HMM10662PduV_1_mix(pf10662) PDU1AB
1703 HMMacetkin_1(pf00871) PDU1AB
1704 -pf00590 -
1705 HMMreg00392_2_grm2(pf00155) EUT2A
1706 HMM00288_1_pdu(pf00288) PDU1AB
1707 -pf04616 -
1708 HMMreg00356_1_pdu1d(pf00356) GRM3B
1709 -pf01547 -
1710 -pf00528 -
1711 -pf00528 -
BMC-P
BMC-H
BMC-T
Aldehyde dehydrogenase
PTAC
Signature enzyme
Alcohol dehydrogenase
Regulator

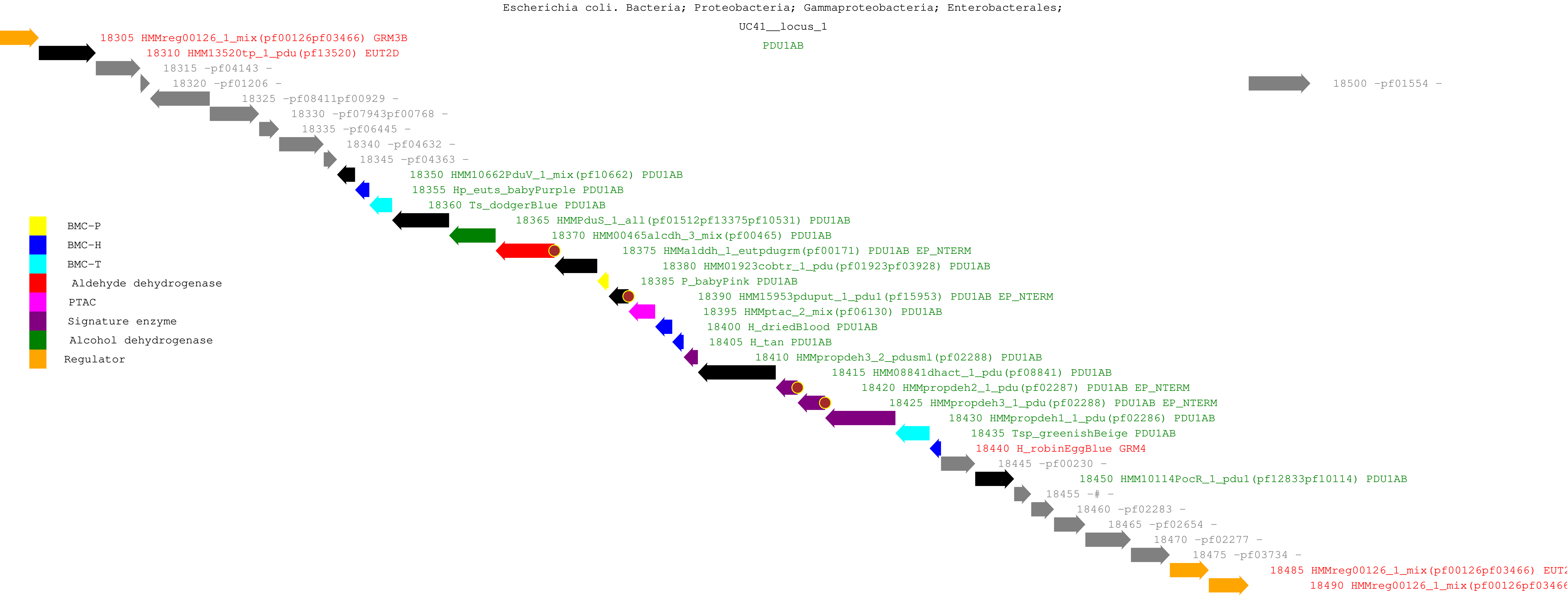
Escherichia coli. Bacteria; Proteobacteria; Gammaproteobacteria; Enterobacterales;
UC41__locus_1
PDU1AB
18305 HMMreg00126_1_mix(pf00126pf03466) GRM3B
18310 HMM13520tp_1_pdu(pf13520) EUT2D
18315 -pf04143 -
18320 -pf01206 -
18325 -pf08411pf00929 -
18330 -pf07943pf00768 -
18335 -pf06445 -
18340 -pf04632 -
18345 -pf04363 -
18350 HMM10662PduV_1_mix(pf10662) PDU1AB
18355 Hp_euts_babyPurple PDU1AB
18360 Ts_dodgerBlue PDU1AB
18365 HMMPduS_1_all(pf01512pf13375pf10531) PDU1AB
18370 HMM00465alcdh_3_mix(pf00465) PDU1AB
18375 HMMalddh_1_eutpdugrm(pf00171) PDU1AB EP_NTERM
18380 HMM01923cobtr_1_pdu(pf01923pf03928) PDU1AB
18385 P_babyPink PDU1AB
18390 HMM15953pduput_1_pdu1(pf15953) PDU1AB EP_NTERM
18395 HMMptac_2_mix(pf06130) PDU1AB
18400 H_driedBlood PDU1AB
18405 H_tan PDU1AB
18410 HMMpropdeh3_2_pdusml(pf02288) PDU1AB
18415 HMM08841dhact_1_pdu(pf08841) PDU1AB
18420 HMMpropdeh2_1_pdu(pf02287) PDU1AB EP_NTERM
18425 HMMpropdeh3_1_pdu(pf02288) PDU1AB EP_NTERM
18430 HMMpropdeh1_1_pdu(pf02286) PDU1AB
18435 Tsp_greenishBeige PDU1AB
18440 H_robinEggBlue GRM4
18445 -pf00230 -
18450 HMM10114PocR_1_pdu1(pf12833pf10114) PDU1AB
18455 -# -
18460 -pf02283 -
18465 -pf02654 -
18470 -pf02277 -
18475 -pf03734 -
18485 HMMreg00126_1_mix(pf00126pf03466) EUT2
18490 HMMreg00126_1_mix(pf00126pf03466
18500 -pf01554 -
BMC-P
BMC-H
BMC-T
Aldehyde dehydrogenase
PTAC
Signature enzyme
Alcohol dehydrogenase
Regulator

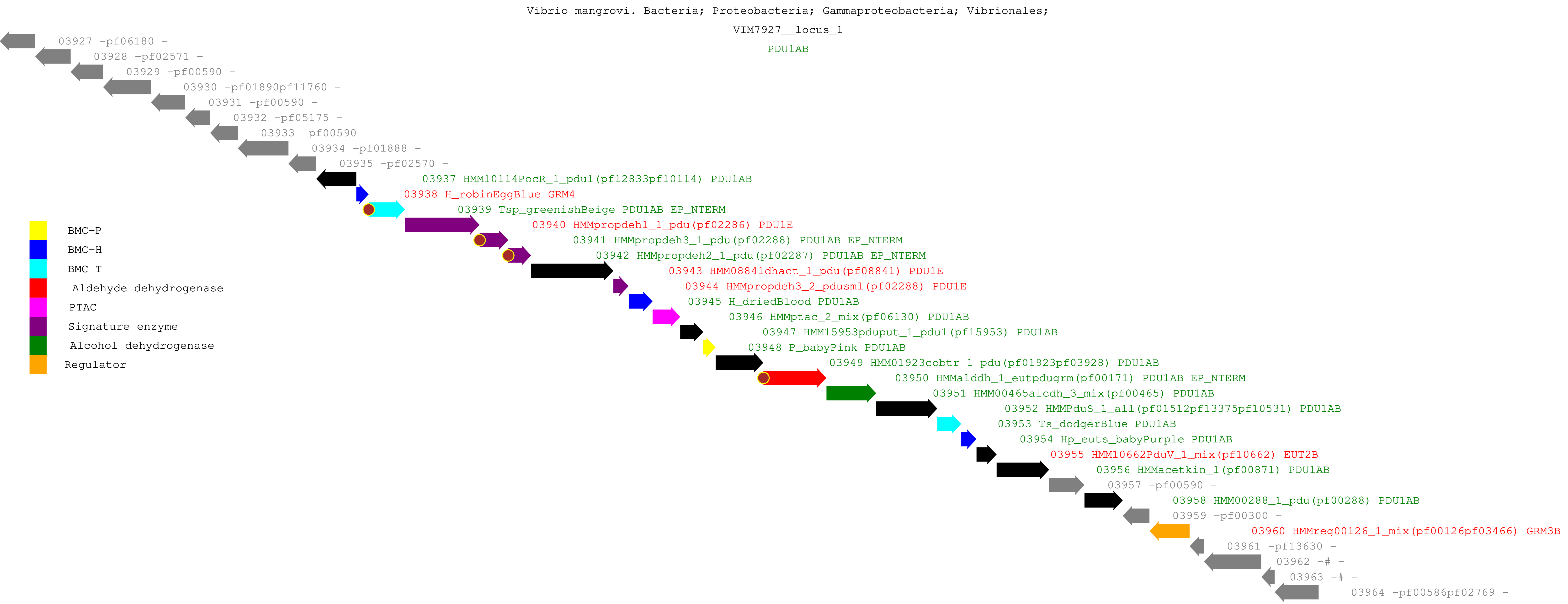
Vibrio mangrovi. Bacteria; Proteobacteria; Gammaproteobacteria; Vibrionales;
VIM7927__locus_1
PDU1AB
03927 -pf06180 -
03928 -pf02571 -
03929 -pf00590 -
03930 -pf01890pf11760 -
03931 -pf00590 -
03932 -pf05175 -
03933 -pf00590 -
03934 -pf01888 -
03935 -pf02570 -
03937 HMM10114PocR_1_pdu1(pf12833pf10114) PDU1AB
03938 H_robinEggBlue GRM4
03939 Tsp_greenishBeige PDU1AB EP_NTERM
03940 HMMpropdeh1_1_pdu(pf02286) PDU1E
03941 HMMpropdeh3_1_pdu(pf02288) PDU1AB EP_NTERM
03942 HMMpropdeh2_1_pdu(pf02287) PDU1AB EP_NTERM
03943 HMM08841dhact_1_pdu(pf08841) PDU1E
03944 HMMpropdeh3_2_pdusml(pf02288) PDU1E
03945 H_driedBlood PDU1AB
03946 HMMptac_2_mix(pf06130) PDU1AB
03947 HMM15953pduput_1_pdu1(pf15953) PDU1AB
03948 P_babyPink PDU1AB
03949 HMM01923cobtr_1_pdu(pf01923pf03928) PDU1AB
03950 HMMalddh_1_eutpdugrm(pf00171) PDU1AB EP_NTERM
03951 HMM00465alcdh_3_mix(pf00465) PDU1AB
03952 HMMPduS_1_all(pf01512pf13375pf10531) PDU1AB
03953 Ts_dodgerBlue PDU1AB
03954 Hp_euts_babyPurple PDU1AB
03955 HMM10662PduV_1_mix(pf10662) EUT2B
03956 HMMacetkin_1(pf00871) PDU1AB
03957 -pf00590 -
03958 HMM00288_1_pdu(pf00288) PDU1AB
03959 -pf00300 -
03960 HMMreg00126_1_mix(pf00126pf03466) GRM3B
03961 -pf13630 -
03962 -# -
03963 -# -
03964 -pf00586pf02769 -
BMC-P
BMC-H
BMC-T
Aldehyde dehydrogenase
PTAC
Signature enzyme
Alcohol dehydrogenase
Regulator

Escherichia sp. KTE114. Bacteria; Proteobacteria; Gammaproteobacteria; Enterobacterales;

WC5__locus_1

PDU1AB

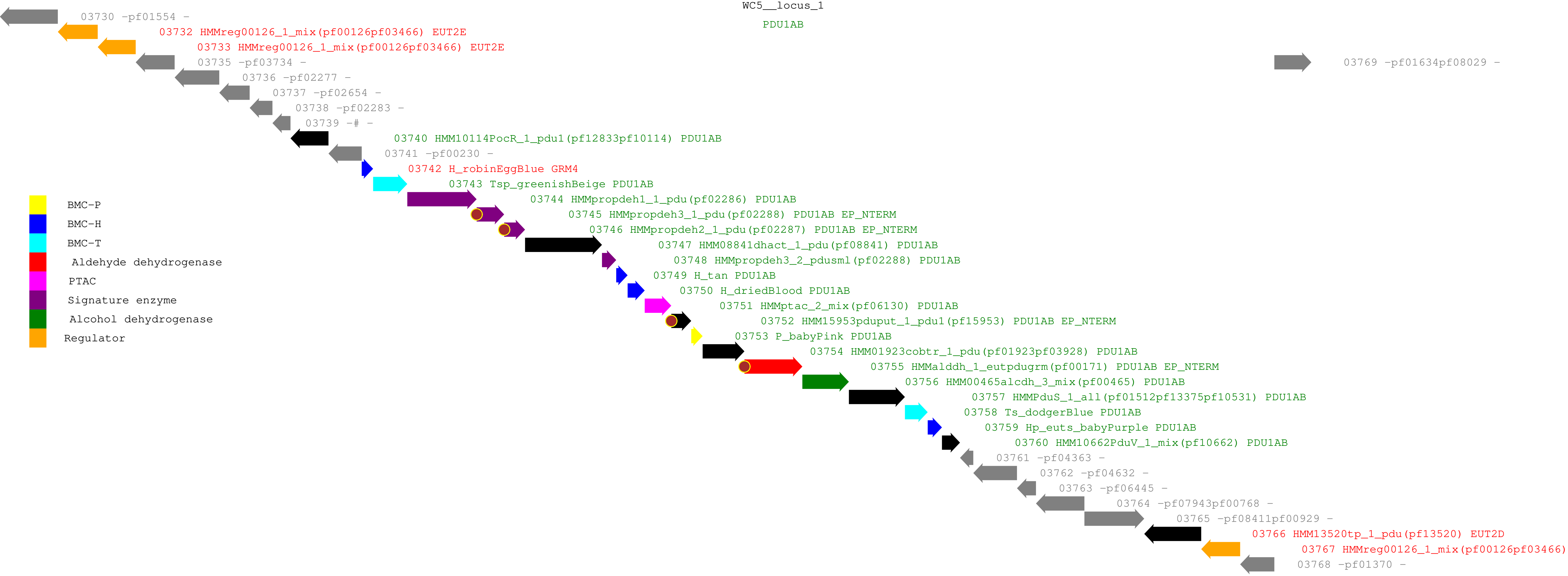

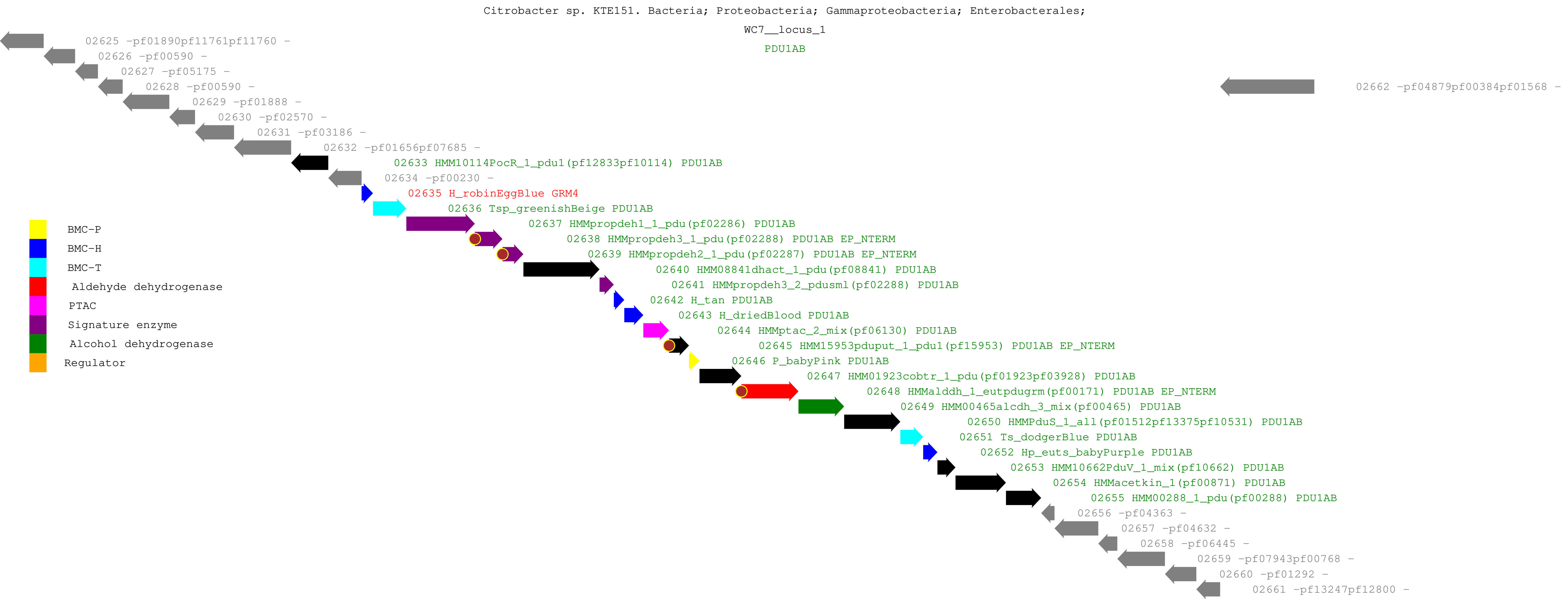
Citrobacter sp. KTE151. Bacteria; Proteobacteria; Gammaproteobacteria; Enterobacterales;
WC7__locus_1
PDU1AB
02625 -pf01890pf11761pf11760 -
02626 -pf00590 -
02627 -pf05175 -
02628 -pf00590 -
02629 -pf01888 -
02630 -pf02570 -
02631 -pf03186 -
02632 -pf01656pf07685 -
02633 HMM10114PocR_1_pdu1(pf12833pf10114) PDU1AB
02634 -pf00230 -
02635 H_robinEggBlue GRM4
02636 Tsp_greenishBeige PDU1AB
02637 HMMpropdeh1_1_pdu(pf02286) PDU1AB
02638 HMMpropdeh3_1_pdu(pf02288) PDU1AB EP_NTERM
02639 HMMpropdeh2_1_pdu(pf02287) PDU1AB EP_NTERM
02640 HMM08841dhact_1_pdu(pf08841) PDU1AB
02641 HMMpropdeh3_2_pdusml(pf02288) PDU1AB
02642 H_tan PDU1AB
02643 H_driedBlood PDU1AB
02644 HMMptac_2_mix(pf06130) PDU1AB
02645 HMM15953pduput_1_pdu1(pf15953) PDU1AB EP_NTERM
02646 P_babyPink PDU1AB
02647 HMM01923cobtr_1_pdu(pf01923pf03928) PDU1AB
02648 HMMalddh_1_eutpdugrm(pf00171) PDU1AB EP_NTERM
02649 HMM00465alcdh_3_mix(pf00465) PDU1AB
02650 HMMPduS_1_all(pf01512pf13375pf10531) PDU1AB
02651 Ts_dodgerBlue PDU1AB
02652 Hp_euts_babyPurple PDU1AB
02653 HMM10662PduV_1_mix(pf10662) PDU1AB
02654 HMMacetkin_1(pf00871) PDU1AB
02655 HMM00288_1_pdu(pf00288) PDU1AB
02656 -pf04363 -
02657 -pf04632 -
02658 -pf06445 -
02659 -pf07943pf00768 -
02660 -pf01292 -
02661 -pf13247pf12800 -
02662 -pf04879pf00384pf01568 -
BMC-P
BMC-H
BMC-T
Aldehyde dehydrogenase
PTAC
Signature enzyme
Alcohol dehydrogenase
Regulator

Escherichia sp. KTE11. Bacteria; Proteobacteria; Gammaproteobacteria; Enterobacterales;

WCO__locus_1

PDU1AB

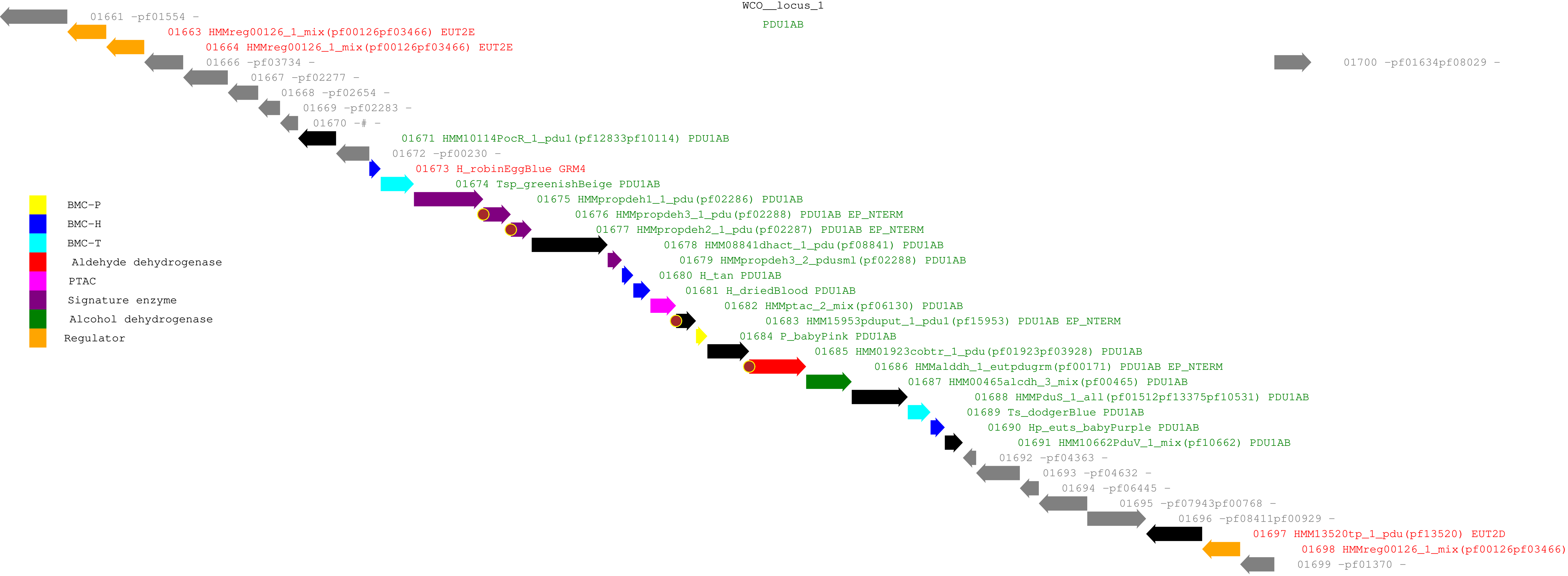

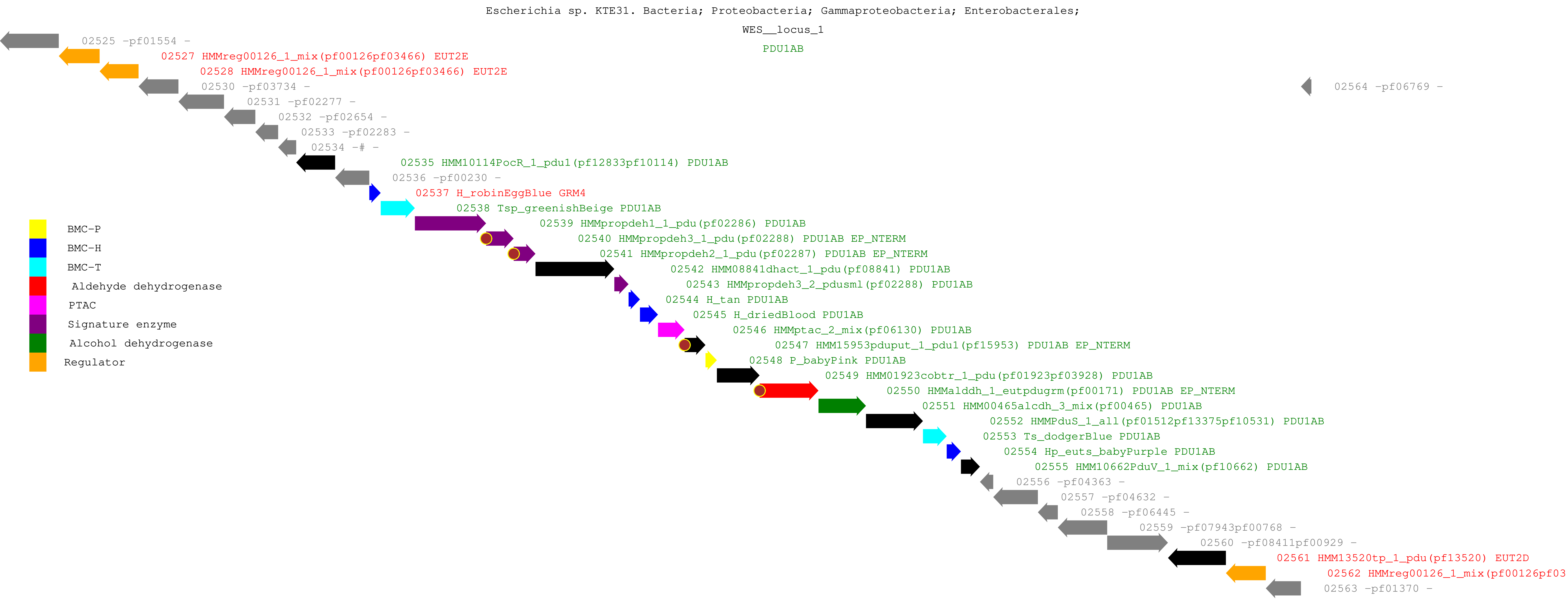

Escherichia sp. KTE31. Bacteria; Proteobacteria; Gammaproteobacteria; Enterobacterales;
WES__locus_1
PDU1AB
02525 -pf01554 -
02527 HMMreg00126_1_mix(pf00126pf03466) EUT2E
02528 HMMreg00126_1_mix(pf00126pf03466) EUT2E
02530 -pf03734 -
02531 -pf02277 -
02532 -pf02654 -
02533 -pf02283 -
02534 -# -
02535 HMM10114PocR_1_pdu1(pf12833pf10114) PDU1AB
02536 -pf00230 -
02537 H_robinEggBlue GRM4
02538 Tsp_greenishBeige PDU1AB
02539 HMMpropdeh1_1_pdu(pf02286) PDU1AB
02540 HMMpropdeh3_1_pdu(pf02288) PDU1AB EP_NTERM
02541 HMMpropdeh2_1_pdu(pf02287) PDU1AB EP_NTERM
02542 HMM08841dhact_1_pdu(pf08841) PDU1AB
02543 HMMpropdeh3_2_pdusml(pf02288) PDU1AB
02544 H_tan PDU1AB
02545 H_driedBlood PDU1AB
02546 HMMptac_2_mix(pf06130) PDU1AB
02547 HMM15953pduput_1_pdu1(pf15953) PDU1AB EP_NTERM
02548 P_babyPink PDU1AB
02549 HMM01923cobtr_1_pdu(pf01923pf03928) PDU1AB
02550 HMMalddh_1_eutpdugrm(pf00171) PDU1AB EP_NTERM
02551 HMM00465alcdh_3_mix(pf00465) PDU1AB
02552 HMMPduS_1_all(pf01512pf13375pf10531) PDU1AB
02553 Ts_dodgerBlue PDU1AB
02554 Hp_euts_babyPurple PDU1AB
02555 HMM10662PduV_1_mix(pf10662) PDU1AB
02556 -pf04363 -
02557 -pf04632 -
02558 -pf06445 -
02559 -pf07943pf00768 -
02560 -pf08411pf00929 -
02561 HMM13520tp_1_pdu(pf13520) EUT2D
02562 HMMreg00126_1_mix(pf00126pf03
02563 -pf01370 -
02564 -pf06769 -
BMC-P
BMC-H
BMC-T
Aldehyde dehydrogenase
PTAC
Signature enzyme
Alcohol dehydrogenase
Regulator

Escherichia coli KTE33. Bacteria; Proteobacteria; Gammaproteobacteria; Enterobacterales;

WEW__locus_1

PDU1AB

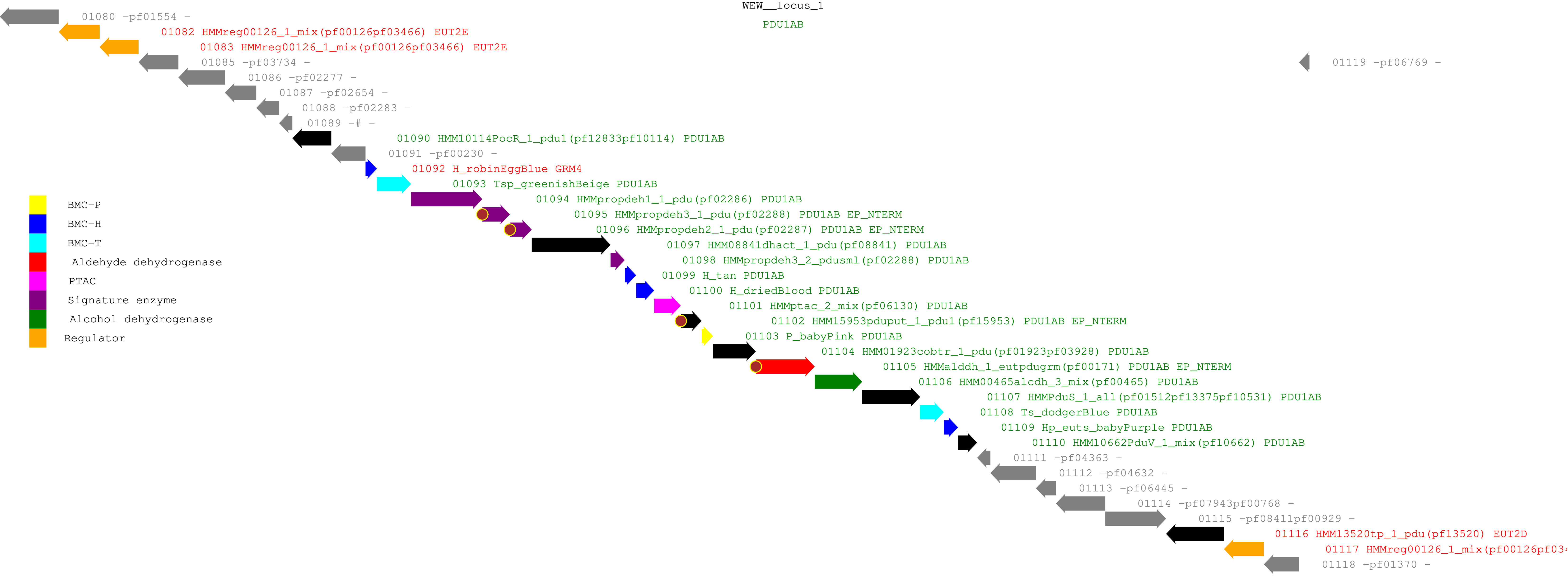

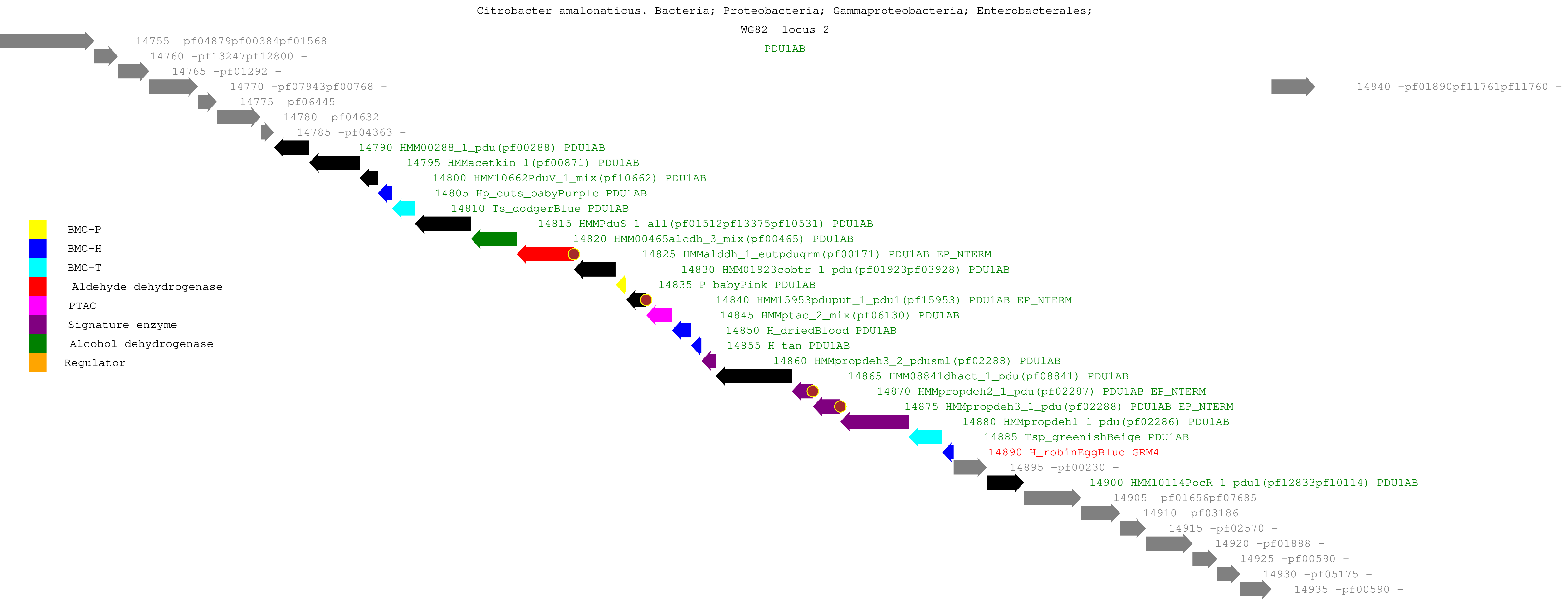
Citrobacter amalonaticus. Bacteria; Proteobacteria; Gammaproteobacteria; Enterobacterales;
WG82__locus_2
PDU1AB
14755 -pf04879pf00384pf01568 -
14760 -pf13247pf12800 -
14765 -pf01292 -
14770 -pf07943pf00768 -
14775 -pf06445 -
14780 -pf04632 -
14785 -pf04363 -
14790 HMM00288_1_pdu(pf00288) PDU1AB
14795 HMMacetkin_1(pf00871) PDU1AB
14800 HMM10662PduV_1_mix(pf10662) PDU1AB
14805 Hp_euts_babyPurple PDU1AB
14810 Ts_dodgerBlue PDU1AB
14815 HMMPduS_1_all(pf01512pf13375pf10531) PDU1AB
14820 HMM00465alcdh_3_mix(pf00465) PDU1AB
14825 HMMalddh_1_eutpdugrm(pf00171) PDU1AB EP_NTERM
14830 HMM01923cobtr_1_pdu(pf01923pf03928) PDU1AB
14835 P_babyPink PDU1AB
14840 HMM15953pduput_1_pdu1(pf15953) PDU1AB EP_NTERM
14845 HMMptac_2_mix(pf06130) PDU1AB
14850 H_driedBlood PDU1AB
14855 H_tan PDU1AB
14860 HMMpropdeh3_2_pdusml(pf02288) PDU1AB
14865 HMM08841dhact_1_pdu(pf08841) PDU1AB
14870 HMMpropdeh2_1_pdu(pf02287) PDU1AB EP_NTERM
14875 HMMpropdeh3_1_pdu(pf02288) PDU1AB EP_NTERM
14880 HMMpropdeh1_1_pdu(pf02286) PDU1AB
14885 Tsp_greenishBeige PDU1AB
14890 H_robinEggBlue GRM4
14895 -pf00230 -
14900 HMM10114PocR_1_pdu1(pf12833pf10114) PDU1AB
14905 -pf01656pf07685 -
14910 -pf03186 -
14915 -pf02570 -
14920 -pf01888 -
14925 -pf00590 -
14930 -pf05175 -
14935 -pf00590 -
14940 -pf01890pf11761pf11760 -
BMC-P
BMC-H
BMC-T
Aldehyde dehydrogenase
PTAC
Signature enzyme
Alcohol dehydrogenase
Regulator

Citrobacter freundii complex sp. CFNIH2. Bacteria; Proteobacteria; Gammaproteobacteria; Enterobacterales;

WM46__locus_2

PDU1AB

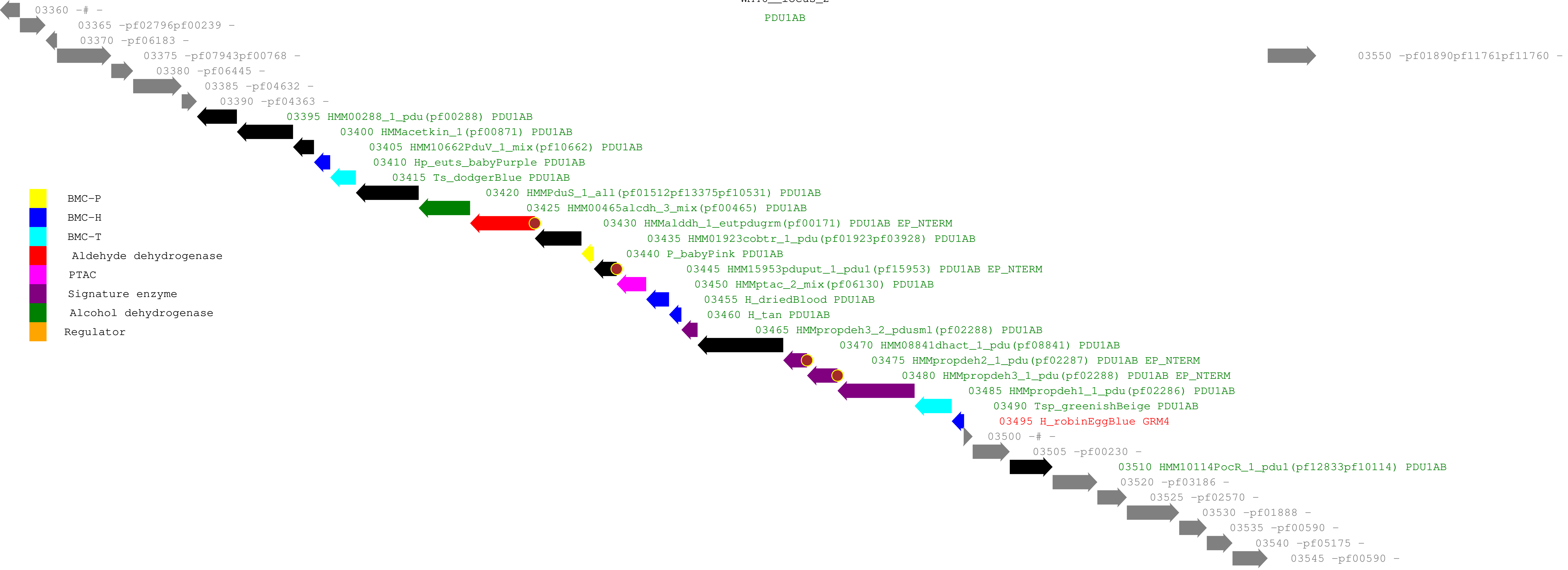

Salmonella enterica I. Bacteria; Proteobacteria; Gammaproteobacteria; Enterobacterales;

XJ41__locus_1_main_sat

PDU1AB

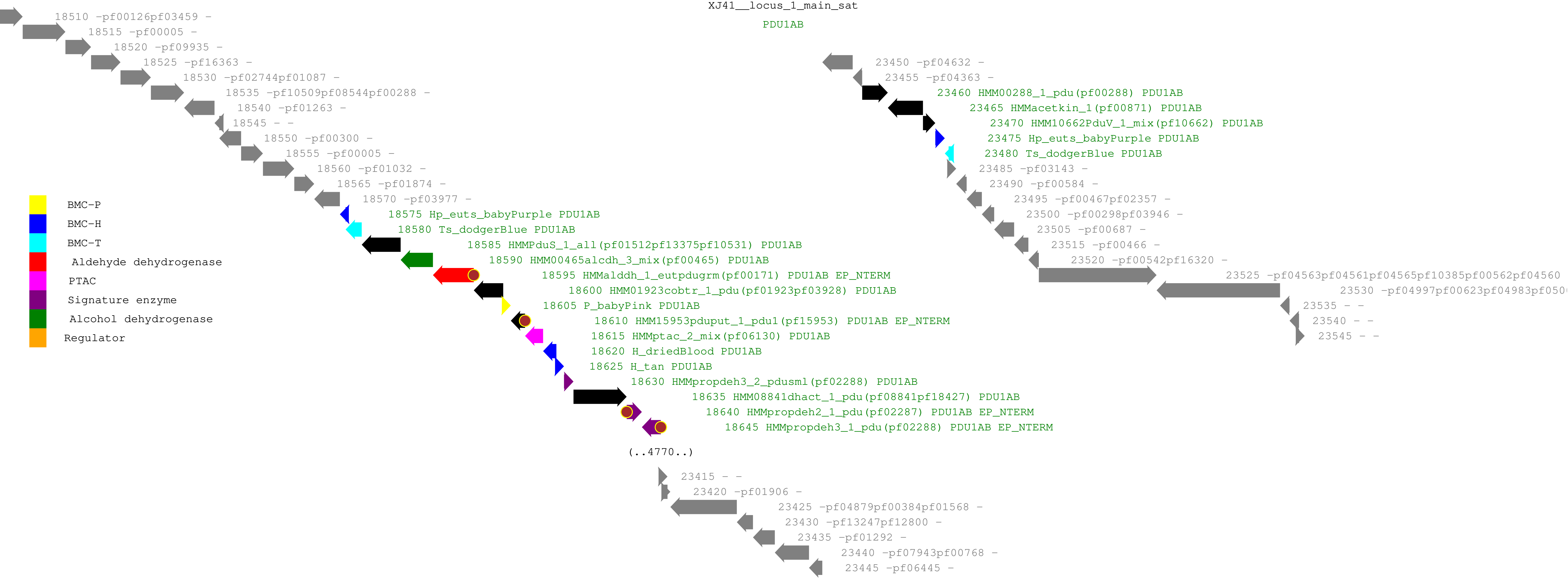

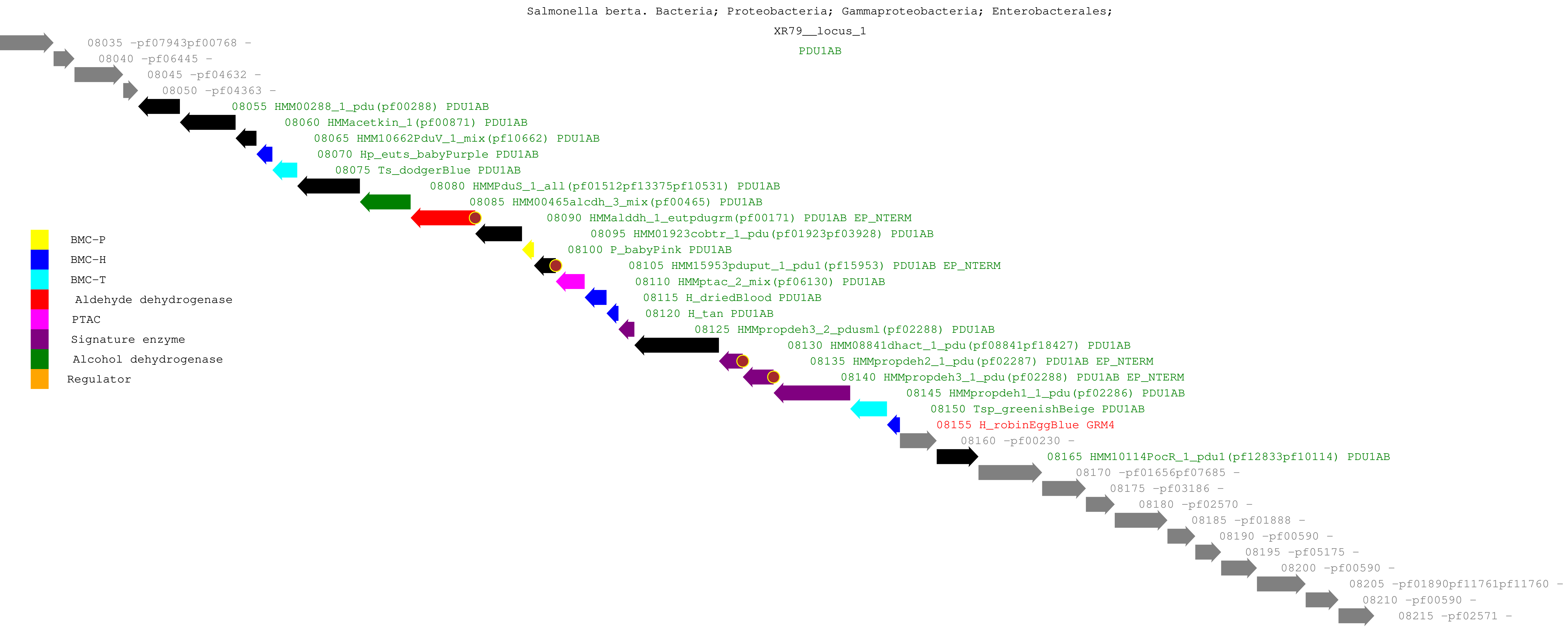
Salmonella berta. Bacteria; Proteobacteria; Gammaproteobacteria; Enterobacterales;
XR79__locus_1
PDU1AB
08035 -pf07943pf00768 -
08040 -pf06445 -
08045 -pf04632 -
08050 -pf04363 -
08055 HMM00288_1_pdu(pf00288) PDU1AB
08060 HMMacetkin_1(pf00871) PDU1AB
08065 HMM10662PduV_1_mix(pf10662) PDU1AB
08070 Hp_euts_babyPurple PDU1AB
08075 Ts_dodgerBlue PDU1AB
08080 HMMPduS_1_all(pf01512pf13375pf10531) PDU1AB
08085 HMM00465alcdh_3_mix(pf00465) PDU1AB
08090 HMMalddh_1_eutpdugrm(pf00171) PDU1AB EP_NTERM
08095 HMM01923cobtr_1_pdu(pf01923pf03928) PDU1AB
08100 P_babyPink PDU1AB
08105 HMM15953pduput_1_pdu1(pf15953) PDU1AB EP_NTERM
08110 HMMptac_2_mix(pf06130) PDU1AB
08115 H_driedBlood PDU1AB
08120 H_tan PDU1AB
08125 HMMpropdeh3_2_pdusml(pf02288) PDU1AB
08130 HMM08841dhact_1_pdu(pf08841pf18427) PDU1AB
08135 HMMpropdeh2_1_pdu(pf02287) PDU1AB EP_NTERM
08140 HMMpropdeh3_1_pdu(pf02288) PDU1AB EP_NTERM
08145 HMMpropdeh1_1_pdu(pf02286) PDU1AB
08150 Tsp_greenishBeige PDU1AB
08155 H_robinEggBlue GRM4
08160 -pf00230 -
08165 HMM10114PocR_1_pdu1(pf12833pf10114) PDU1AB
08170 -pf01656pf07685 -
08175 -pf03186 -
08180 -pf02570 -
08185 -pf01888 -
08190 -pf00590 -
08195 -pf05175 -
08200 -pf00590 -
08205 -pf01890pf11761pf11760 -
08210 -pf00590 -
08215 -pf02571 -
BMC-P
BMC-H
BMC-T
Aldehyde dehydrogenase
PTAC
Signature enzyme
Alcohol dehydrogenase
Regulator

Salmonella enterica subsp. salamae. Bacteria; Proteobacteria; Gammaproteobacteria; Enterobacterales;

XT66__locus_1

PDU1AB

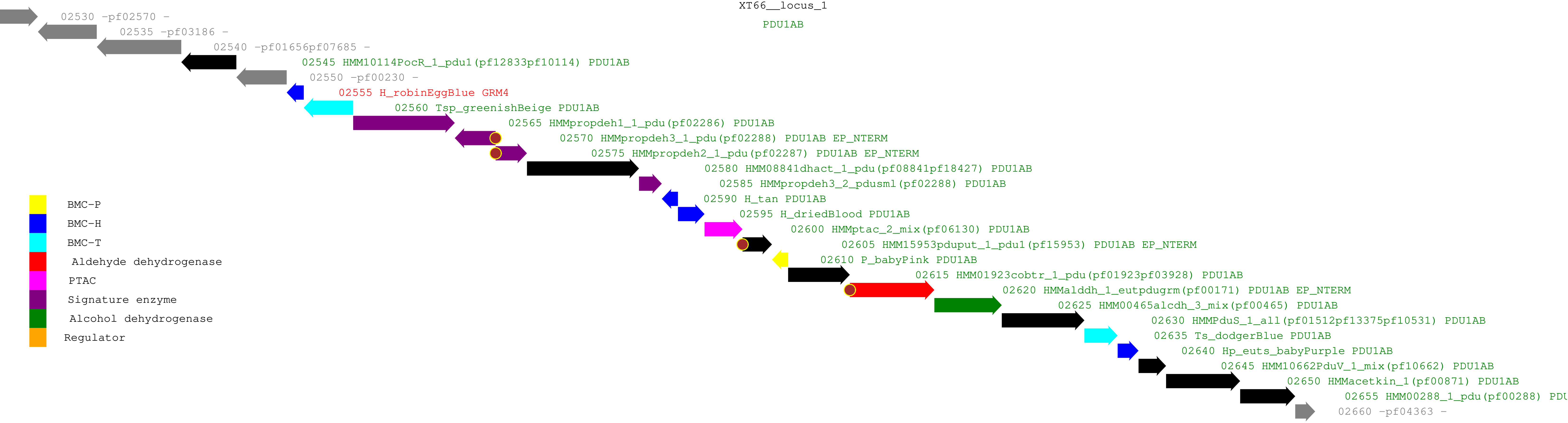

Salmonella enterica subsp. enterica serovar Infantis str. CVM Bacteria; Proteobacteria; Gammaproteobacteria; Enterobacterales;

Y082__locus_2

PDU1AB

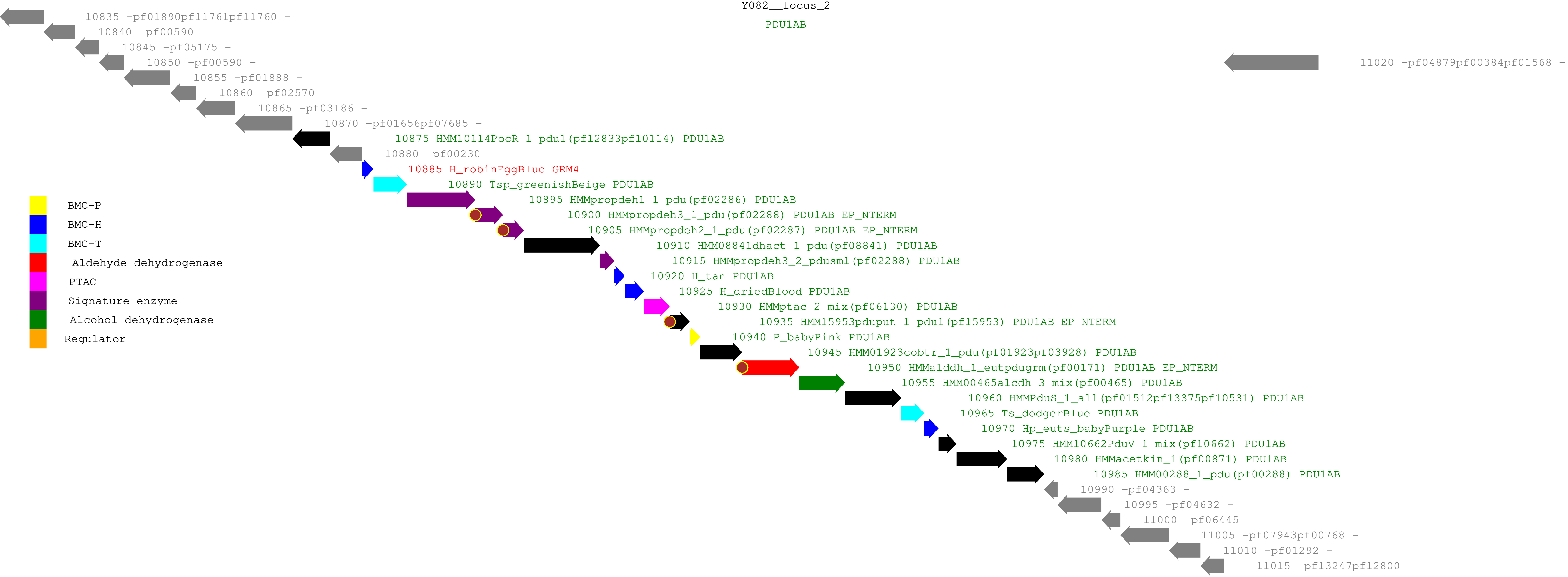

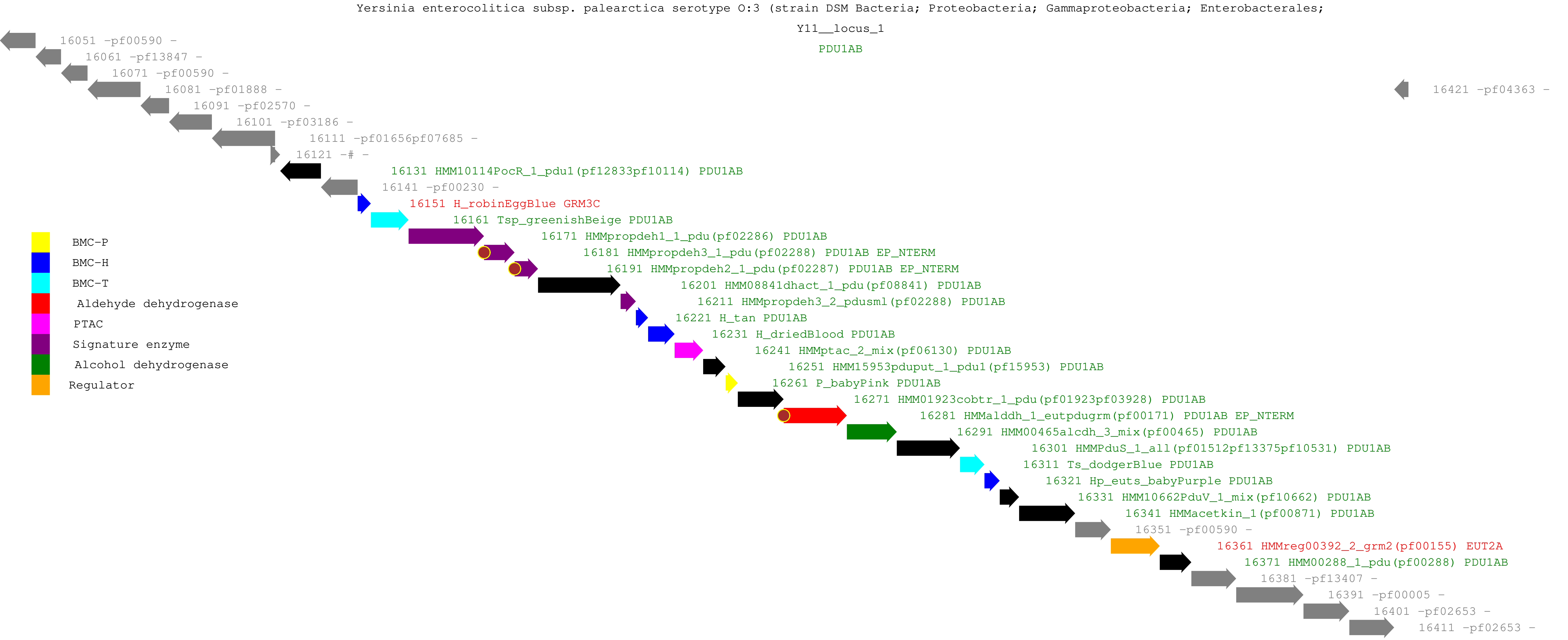
Yersinia enterocolitica subsp. palearctica serotype O:3 (strain DSM Bacteria; Proteobacteria; Gammaproteobacteria; Enterobacterales;
Y11__locus_1
PDU1AB
16051 -pf00590 -
16061 -pf13847 -
16071 -pf00590 -
16081 -pf01888 -
16091 -pf02570 -
16101 -pf03186 -
16111 -pf01656pf07685 -
16121 -# -
16131 HMM10114PocR_1_pdu1(pf12833pf10114) PDU1AB
16141 -pf00230 -
16151 H_robinEggBlue GRM3C
16161 Tsp_greenishBeige PDU1AB
16171 HMMpropdeh1_1_pdu(pf02286) PDU1AB
16181 HMMpropdeh3_1_pdu(pf02288) PDU1AB EP_NTERM
16191 HMMpropdeh2_1_pdu(pf02287) PDU1AB EP_NTERM
16201 HMM08841dhact_1_pdu(pf08841) PDU1AB
16211 HMMpropdeh3_2_pdusml(pf02288) PDU1AB
16221 H_tan PDU1AB
16231 H_driedBlood PDU1AB
16241 HMMptac_2_mix(pf06130) PDU1AB
16251 HMM15953pduput_1_pdu1(pf15953) PDU1AB
16261 P_babyPink PDU1AB
16271 HMM01923cobtr_1_pdu(pf01923pf03928) PDU1AB
16281 HMMalddh_1_eutpdugrm(pf00171) PDU1AB EP_NTERM
16291 HMM00465alcdh_3_mix(pf00465) PDU1AB
16301 HMMPduS_1_all(pf01512pf13375pf10531) PDU1AB
16311 Ts_dodgerBlue PDU1AB
16321 Hp_euts_babyPurple PDU1AB
16331 HMM10662PduV_1_mix(pf10662) PDU1AB
16341 HMMacetkin_1(pf00871) PDU1AB
16351 -pf00590 -
16361 HMMreg00392_2_grm2(pf00155) EUT2A
16371 HMM00288_1_pdu(pf00288) PDU1AB
16381 -pf13407 -
16391 -pf00005 -
16401 -pf02653 -
16411 -pf02653 -
16421 -pf04363 -
BMC-P
BMC-H
BMC-T
Aldehyde dehydrogenase
PTAC
Signature enzyme
Alcohol dehydrogenase
Regulator

Yersinia enterocolitica serotype O:8 / biotype 1B (strain NCTC 13174 / Bacteria; Proteobacteria; Gammaproteobacteria; Enterobacterales;

YE_locus_1

PDU1AB

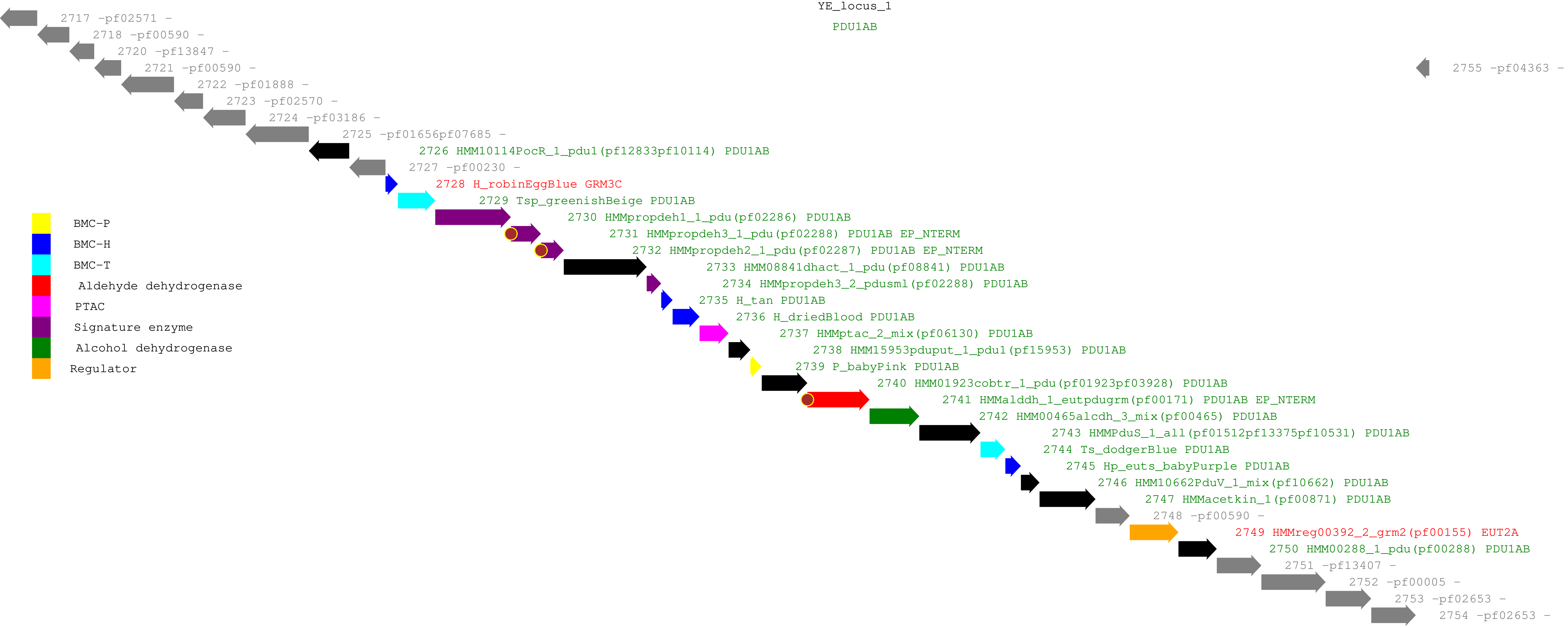

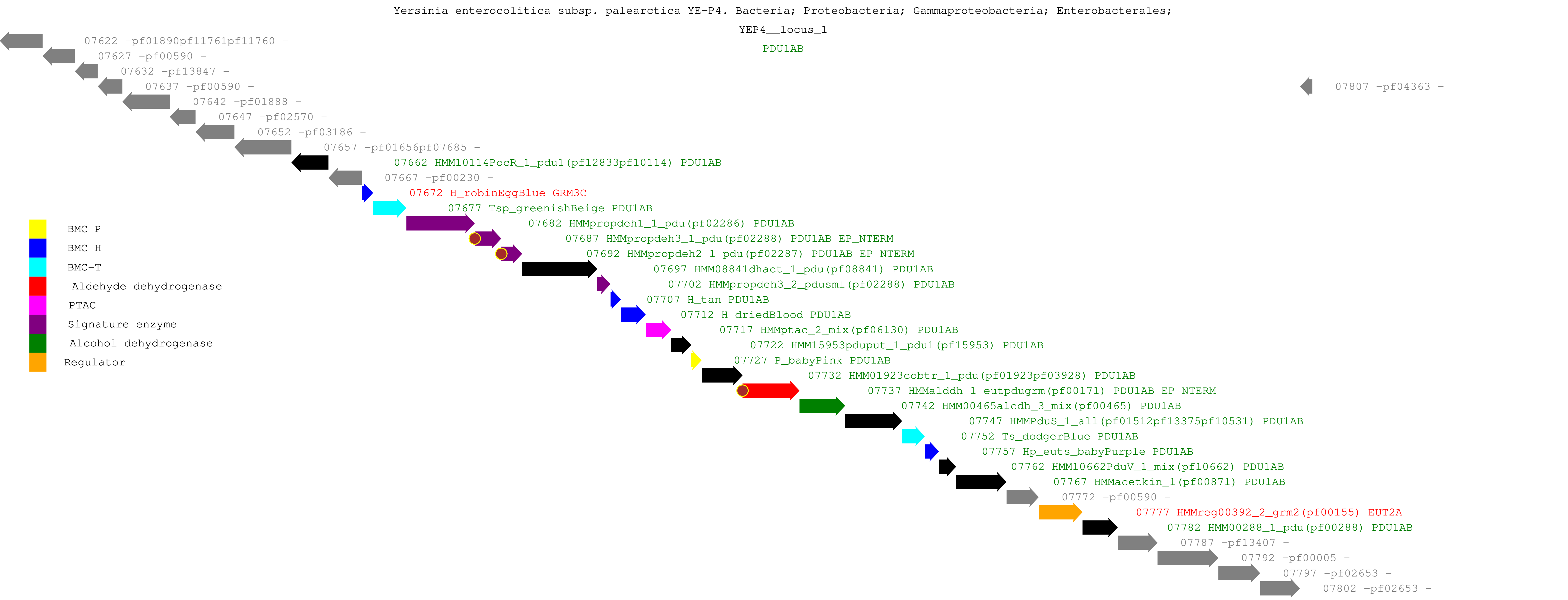
Yersinia enterocolitica subsp. palearctica YE-P4. Bacteria; Proteobacteria; Gammaproteobacteria; Enterobacterales;
YEP4__locus_1
PDU1AB
07622 -pf01890pf11761pf11760 -
07627 -pf00590 -
07632 -pf13847 -
07637 -pf00590 -
07642 -pf01888 -
07647 -pf02570 -
07652 -pf03186 -
07657 -pf01656pf07685 -
07662 HMM10114PocR_1_pdu1(pf12833pf10114) PDU1AB
07667 -pf00230 -
07672 H_robinEggBlue GRM3C
07677 Tsp_greenishBeige PDU1AB
07682 HMMpropdeh1_1_pdu(pf02286) PDU1AB
07687 HMMpropdeh3_1_pdu(pf02288) PDU1AB EP_NTERM
07692 HMMpropdeh2_1_pdu(pf02287) PDU1AB EP_NTERM
07697 HMM08841dhact_1_pdu(pf08841) PDU1AB
07702 HMMpropdeh3_2_pdusml(pf02288) PDU1AB
07707 H_tan PDU1AB
07712 H_driedBlood PDU1AB
07717 HMMptac_2_mix(pf06130) PDU1AB
07722 HMM15953pduput_1_pdu1(pf15953) PDU1AB
07727 P_babyPink PDU1AB
07732 HMM01923cobtr_1_pdu(pf01923pf03928) PDU1AB
07737 HMMalddh_1_eutpdugrm(pf00171) PDU1AB EP_NTERM
07742 HMM00465alcdh_3_mix(pf00465) PDU1AB
07747 HMMPduS_1_all(pf01512pf13375pf10531) PDU1AB
07752 Ts_dodgerBlue PDU1AB
07757 Hp_euts_babyPurple PDU1AB
07762 HMM10662PduV_1_mix(pf10662) PDU1AB
07767 HMMacetkin_1(pf00871) PDU1AB
07772 -pf00590 -
07777 HMMreg00392_2_grm2(pf00155) EUT2A
07782 HMM00288_1_pdu(pf00288) PDU1AB
07787 -pf13407 -
07792 -pf00005 -
07797 -pf02653 -
07802 -pf02653 -
07807 -pf04363 -
BMC-P
BMC-H
BMC-T
Aldehyde dehydrogenase
PTAC
Signature enzyme
Alcohol dehydrogenase
Regulator

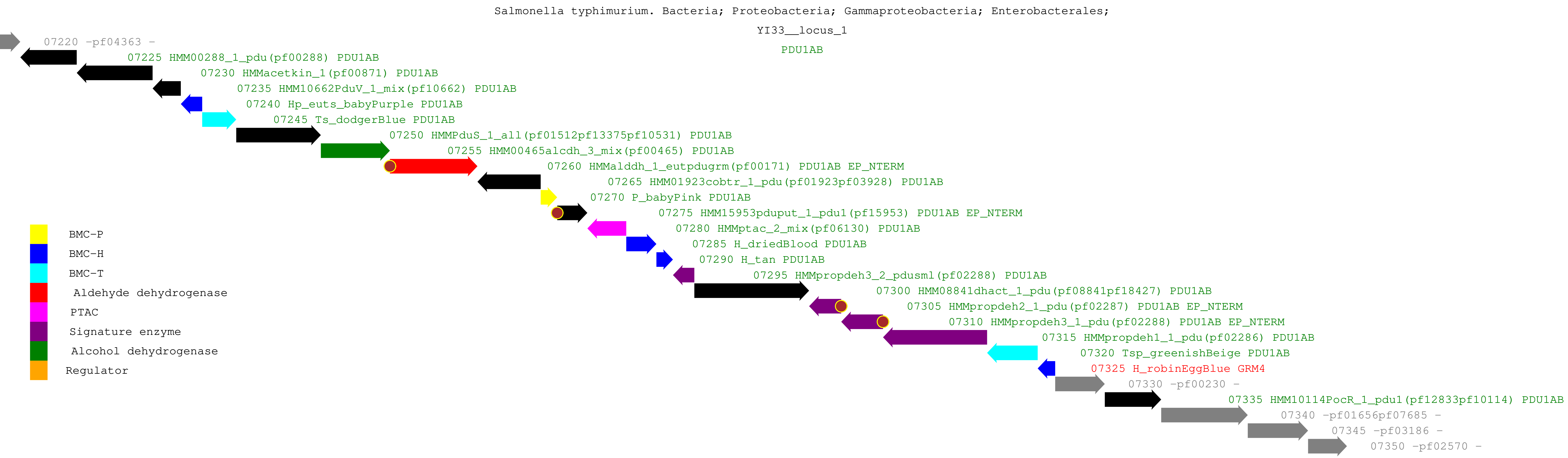
Salmonella typhimurium. Bacteria; Proteobacteria; Gammaproteobacteria; Enterobacterales;
YI33__locus_1
PDU1AB
07220 -pf04363 -
07225 HMM00288_1_pdu(pf00288) PDU1AB
07230 HMMacetkin_1(pf00871) PDU1AB
07235 HMM10662PduV_1_mix(pf10662) PDU1AB
07240 Hp_euts_babyPurple PDU1AB
07245 Ts_dodgerBlue PDU1AB
07250 HMMPduS_1_all(pf01512pf13375pf10531) PDU1AB
07255 HMM00465alcdh_3_mix(pf00465) PDU1AB
07260 HMMalddh_1_eutpdugrm(pf00171) PDU1AB EP_NTERM
07265 HMM01923cobtr_1_pdu(pf01923pf03928) PDU1AB
07270 P_babyPink PDU1AB
07275 HMM15953pduput_1_pdu1(pf15953) PDU1AB EP_NTERM
07280 HMMptac_2_mix(pf06130) PDU1AB
07285 H_driedBlood PDU1AB
07290 H_tan PDU1AB
07295 HMMpropdeh3_2_pdusml(pf02288) PDU1AB
07300 HMM08841dhact_1_pdu(pf08841pf18427) PDU1AB
07305 HMMpropdeh2_1_pdu(pf02287) PDU1AB EP_NTERM
07310 HMMpropdeh3_1_pdu(pf02288) PDU1AB EP_NTERM
07315 HMMpropdeh1_1_pdu(pf02286) PDU1AB
07320 Tsp_greenishBeige PDU1AB
07325 H_robinEggBlue GRM4
07330 -pf00230 -
07335 HMM10114PocR_1_pdu1(pf12833pf10114) PDU1AB
07340 -pf01656pf07685 -
07345 -pf03186 -
07350 -pf02570 -
BMC-P
BMC-H
BMC-T
Aldehyde dehydrogenase
PTAC
Signature enzyme
Alcohol dehydrogenase
Regulator

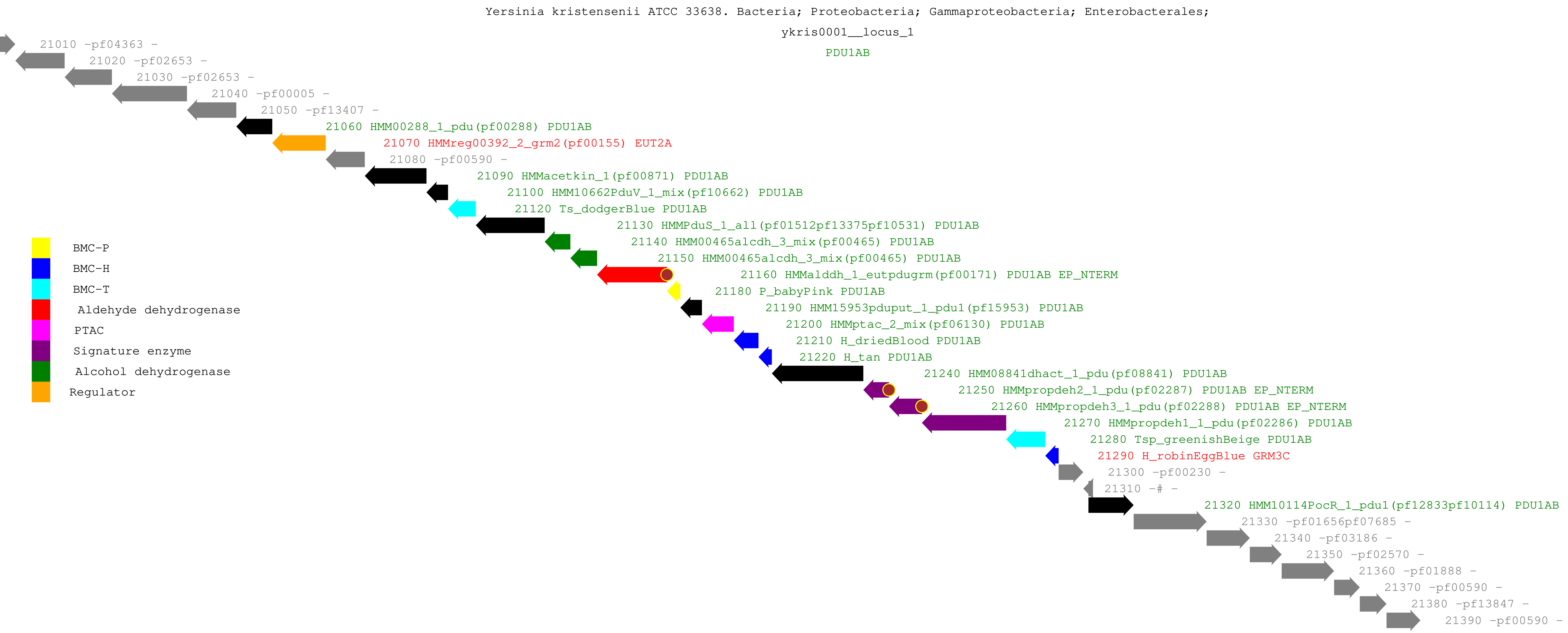

Yersinia kristensenii ATCC 33638. Bacteria; Proteobacteria; Gammaproteobacteria; Enterobacterales;
ykris0001__locus_1
PDU1AB
21010 -pf04363 -
21020 -pf02653 -
21030 -pf02653 -
21040 -pf00005 -
21050 -pf13407 -
21060 HMM00288_1_pdu(pf00288) PDU1AB
21070 HMMreg00392_2_grm2(pf00155) EUT2A
21080 -pf00590 -
21090 HMMacetkin_1(pf00871) PDU1AB
21100 HMM10662PduV_1_mix(pf10662) PDU1AB
21120 Ts_dodgerBlue PDU1AB
21130 HMMPduS_1_all(pf01512pf13375pf10531) PDU1AB
21140 HMM00465alcdh_3_mix(pf00465) PDU1AB
21150 HMM00465alcdh_3_mix(pf00465) PDU1AB
21160 HMMalddh_1_eutpdugrm(pf00171) PDU1AB EP_NTERM
21180 P_babyPink PDU1AB
21190 HMM15953pduput_1_pdu1(pf15953) PDU1AB
21200 HMMptac_2_mix(pf06130) PDU1AB
21210 H_driedBlood PDU1AB
21220 H_tan PDU1AB
21240 HMM08841dhact_1_pdu(pf08841) PDU1AB
21250 HMMpropdeh2_1_pdu(pf02287) PDU1AB EP_NTERM
21260 HMMpropdeh3_1_pdu(pf02288) PDU1AB EP_NTERM
21270 HMMpropdeh1_1_pdu(pf02286) PDU1AB
21280 Tsp_greenishBeige PDU1AB
21290 H_robinEggBlue GRM3C
21300 -pf00230 -
21310 -# -
21320 HMM10114PocR_1_pdu1(pf12833pf10114) PDU1AB
21330 -pf01656pf07685 -
21340 -pf03186 -
21350 -pf02570 -
21360 -pf01888 -
21370 -pf00590 -
21380 -pf13847 -
21390 -pf00590 -
BMC-P
BMC-H
BMC-T
Aldehyde dehydrogenase
PTAC
Signature enzyme
Alcohol dehydrogenase
Regulator

Yersinia mollaretii ATCC 43969. Bacteria; Proteobacteria; Gammaproteobacteria; Enterobacterales;

ymoll0001__locus_1

PDU1AB

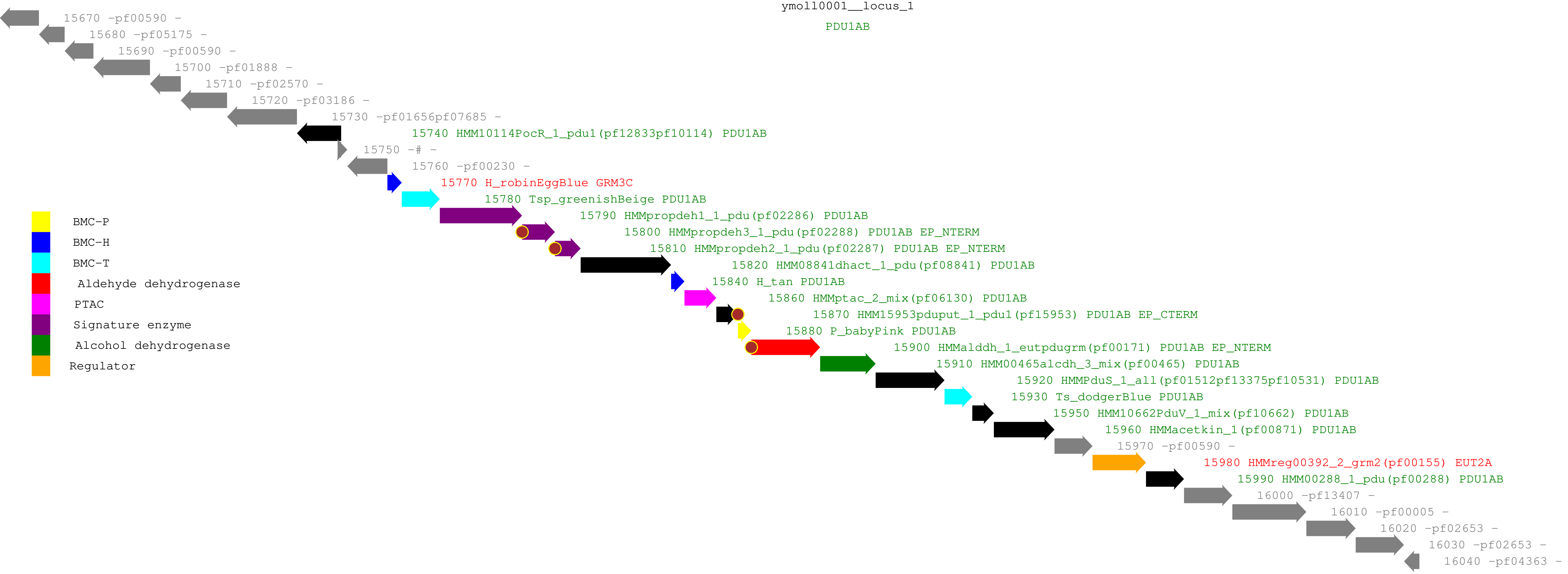

Salmonella enteritidis. Bacteria; Proteobacteria; Gammaproteobacteria; Enterobacterales;

YQ47__locus_1

PDU1AB

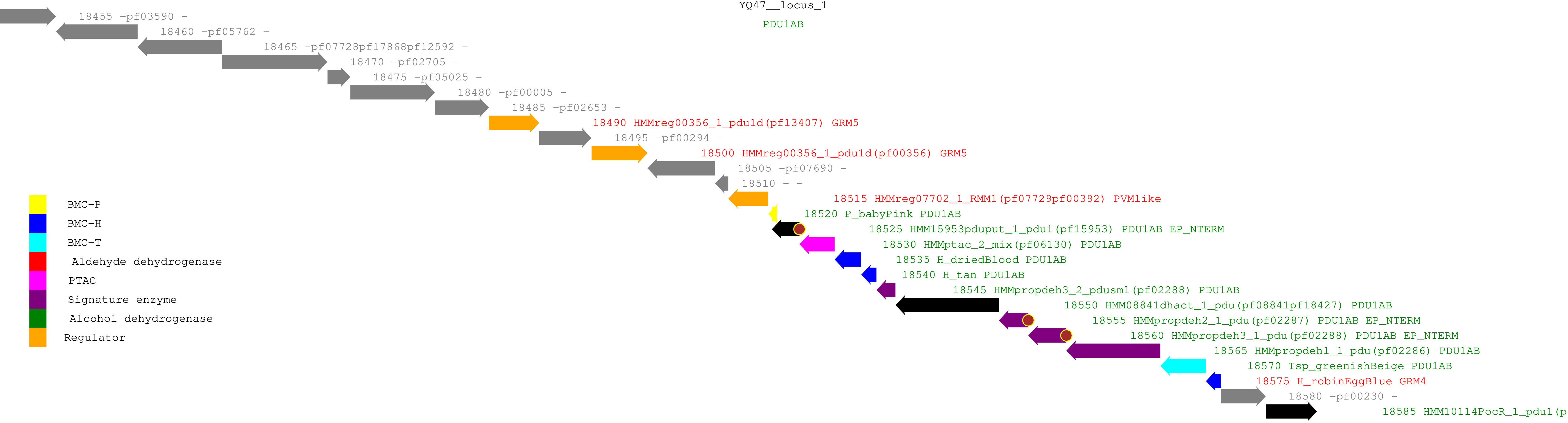

Salmonella enteritidis. Bacteria; Proteobacteria; Gammaproteobacteria; Enterobacterales;

YQ47__locus_2

PDU1AB

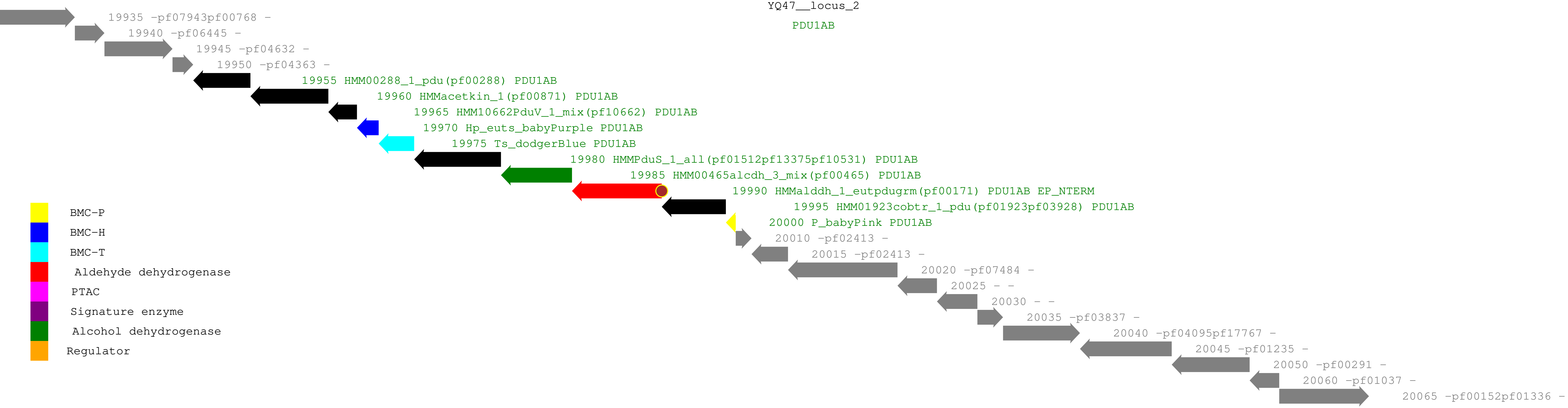

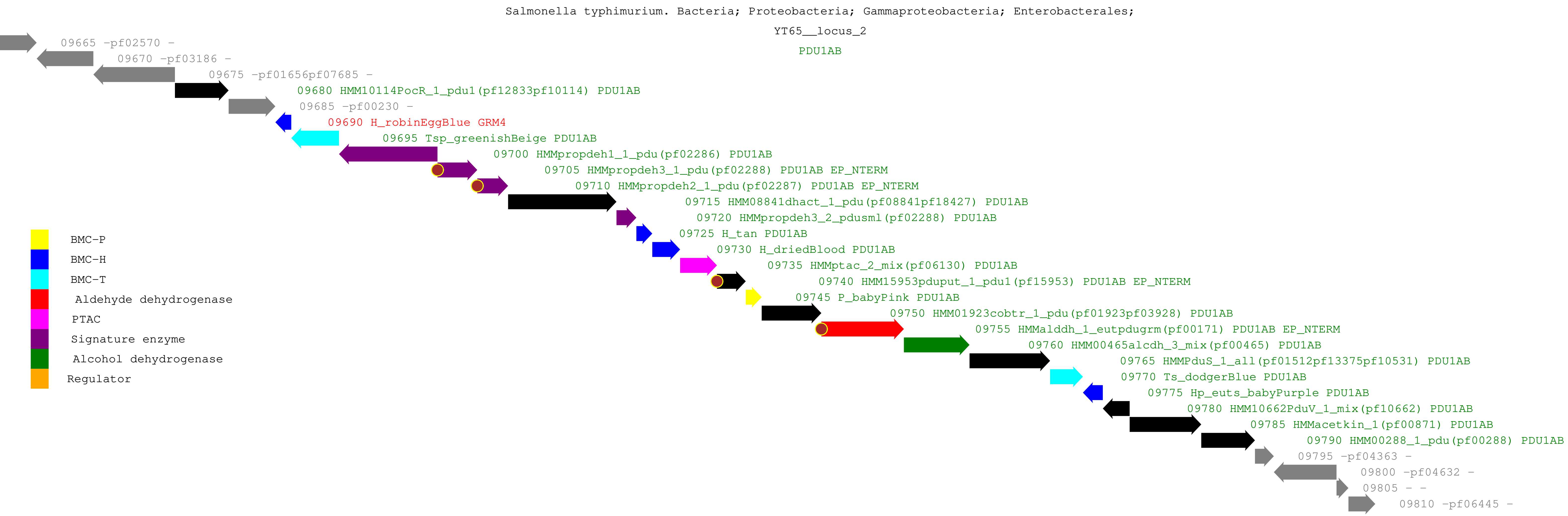
Salmonella typhimurium. Bacteria; Proteobacteria; Gammaproteobacteria; Enterobacterales;
YT65__locus_2
PDU1AB
09665 -pf02570 -
09670 -pf03186 -
09675 -pf01656pf07685 -
09680 HMM10114PocR_1_pdu1(pf12833pf10114) PDU1AB
09685 -pf00230 -
09690 H_robinEggBlue GRM4
09695 Tsp_greenishBeige PDU1AB
09700 HMMpropdeh1_1_pdu(pf02286) PDU1AB
09705 HMMpropdeh3_1_pdu(pf02288) PDU1AB EP_NTERM
09710 HMMpropdeh2_1_pdu(pf02287) PDU1AB EP_NTERM
09715 HMM08841dhact_1_pdu(pf08841pf18427) PDU1AB
09720 HMMpropdeh3_2_pdusml(pf02288) PDU1AB
09725 H_tan PDU1AB
09730 H_driedBlood PDU1AB
09735 HMMptac_2_mix(pf06130) PDU1AB
09740 HMM15953pduput_1_pdu1(pf15953) PDU1AB EP_NTERM
09745 P_babyPink PDU1AB
09750 HMM01923cobtr_1_pdu(pf01923pf03928) PDU1AB
09755 HMMalddh_1_eutpdugrm(pf00171) PDU1AB EP_NTERM
09760 HMM00465alcdh_3_mix(pf00465) PDU1AB
09765 HMMPduS_1_all(pf01512pf13375pf10531) PDU1AB
09770 Ts_dodgerBlue PDU1AB
09775 Hp_euts_babyPurple PDU1AB
09780 HMM10662PduV_1_mix(pf10662) PDU1AB
09785 HMMacetkin_1(pf00871) PDU1AB
09790 HMM00288_1_pdu(pf00288) PDU1AB
09795 -pf04363 -
09800 -pf04632 -
09805 - -
09810 -pf06445 -
BMC-P
BMC-H
BMC-T
Aldehyde dehydrogenase
PTAC
Signature enzyme
Alcohol dehydrogenase
Regulator

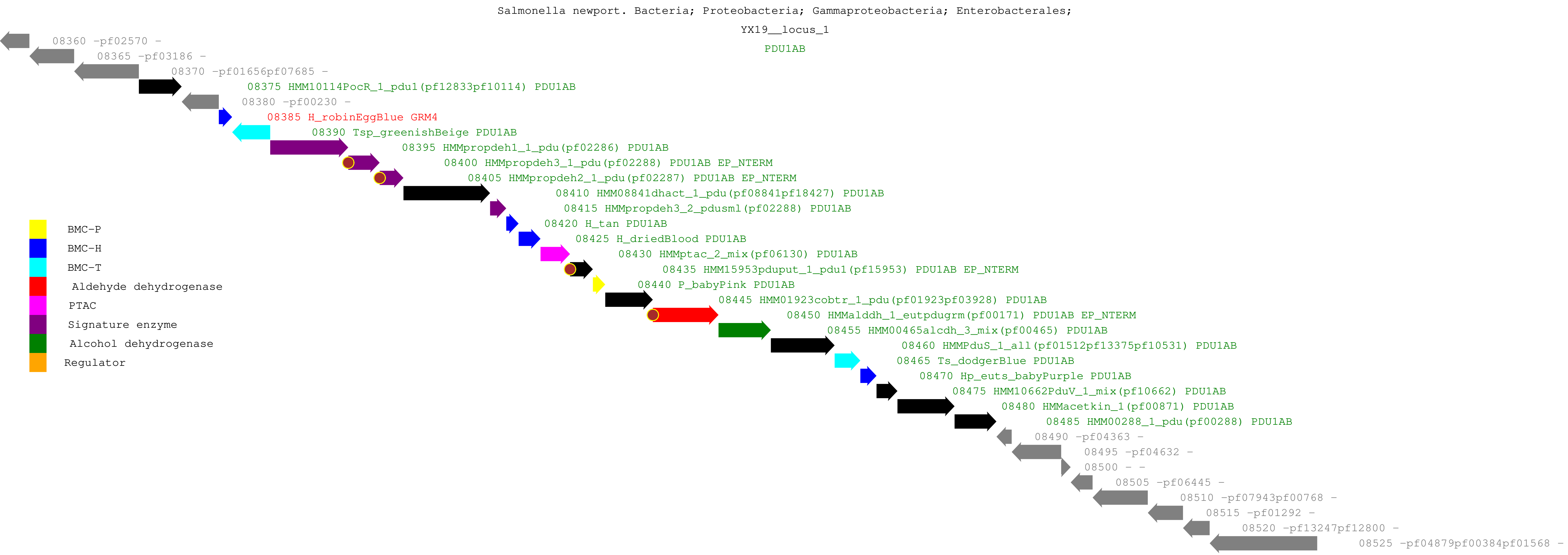
Salmonella newport. Bacteria; Proteobacteria; Gammaproteobacteria; Enterobacterales;
YX19__locus_1
PDU1AB
08360 -pf02570 -
08365 -pf03186 -
08370 -pf01656pf07685 -
08375 HMM10114PocR_1_pdu1(pf12833pf10114) PDU1AB
08380 -pf00230 -
08385 H_robinEggBlue GRM4
08390 Tsp_greenishBeige PDU1AB
08395 HMMpropdeh1_1_pdu(pf02286) PDU1AB
08400 HMMpropdeh3_1_pdu(pf02288) PDU1AB EP_NTERM
08405 HMMpropdeh2_1_pdu(pf02287) PDU1AB EP_NTERM
08410 HMM08841dhact_1_pdu(pf08841pf18427) PDU1AB
08415 HMMpropdeh3_2_pdusml(pf02288) PDU1AB
08420 H_tan PDU1AB
08425 H_driedBlood PDU1AB
08430 HMMptac_2_mix(pf06130) PDU1AB
08435 HMM15953pduput_1_pdu1(pf15953) PDU1AB EP_NTERM
08440 P_babyPink PDU1AB
08445 HMM01923cobtr_1_pdu(pf01923pf03928) PDU1AB
08450 HMMalddh_1_eutpdugrm(pf00171) PDU1AB EP_NTERM
08455 HMM00465alcdh_3_mix(pf00465) PDU1AB
08460 HMMPduS_1_all(pf01512pf13375pf10531) PDU1AB
08465 Ts_dodgerBlue PDU1AB
08470 Hp_euts_babyPurple PDU1AB
08475 HMM10662PduV_1_mix(pf10662) PDU1AB
08480 HMMacetkin_1(pf00871) PDU1AB
08485 HMM00288_1_pdu(pf00288) PDU1AB
08490 -pf04363 -
08495 -pf04632 -
08500 - -
08505 -pf06445 -
08510 -pf07943pf00768 -
08515 -pf01292 -
08520 -pf13247pf12800 -
08525 -pf04879pf00384pf01568 -
BMC-P
BMC-H
BMC-T
Aldehyde dehydrogenase
PTAC
Signature enzyme
Alcohol dehydrogenase
Regulator

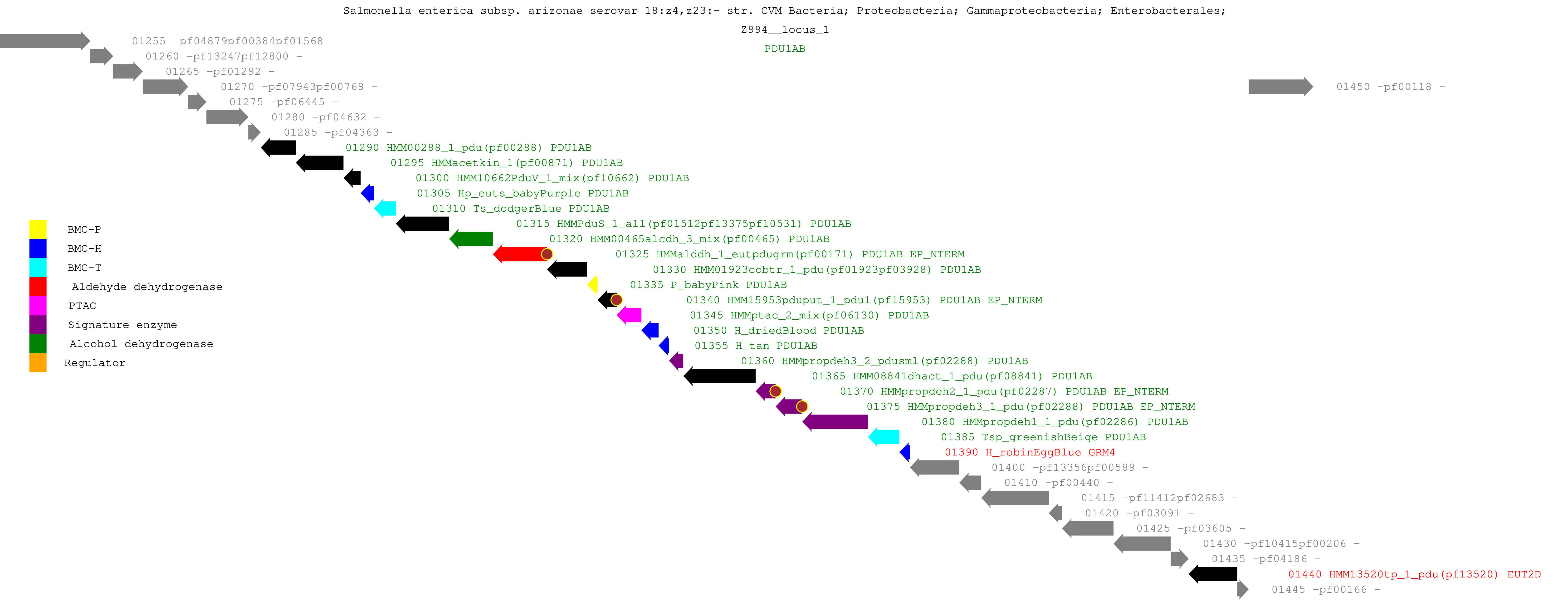

Salmonella enterica subsp. arizonae serovar 18:z4,z23:- str. CVM Bacteria; Proteobacteria; Gammaproteobacteria; Enterobacterales;
Z994__locus_1
PDU1AB
01255 -pf04879pf00384pf01568 -
01260 -pf13247pf12800 -
01265 -pf01292 -
01270 -pf07943pf00768 -
01275 -pf06445 -
01280 -pf04632 -
01285 -pf04363 -
01290 HMM00288_1_pdu(pf00288) PDU1AB
01295 HMMacetkin_1(pf00871) PDU1AB
01300 HMM10662PduV_1_mix(pf10662) PDU1AB
01305 Hp_euts_babyPurple PDU1AB
01310 Ts_dodgerBlue PDU1AB
01315 HMMPduS_1_all(pf01512pf13375pf10531) PDU1AB
01320 HMM00465alcdh_3_mix(pf00465) PDU1AB
01325 HMMalddh_1_eutpdugrm(pf00171) PDU1AB EP_NTERM
01330 HMM01923cobtr_1_pdu(pf01923pf03928) PDU1AB
01335 P_babyPink PDU1AB
01340 HMM15953pduput_1_pdu1(pf15953) PDU1AB EP_NTERM
01345 HMMptac_2_mix(pf06130) PDU1AB
01350 H_driedBlood PDU1AB
01355 H_tan PDU1AB
01360 HMMpropdeh3_2_pdusml(pf02288) PDU1AB
01365 HMM08841dhact_1_pdu(pf08841) PDU1AB
01370 HMMpropdeh2_1_pdu(pf02287) PDU1AB EP_NTERM
01375 HMMpropdeh3_1_pdu(pf02288) PDU1AB EP_NTERM
01380 HMMpropdeh1_1_pdu(pf02286) PDU1AB
01385 Tsp_greenishBeige PDU1AB
01390 H_robinEggBlue GRM4
01400 -pf13356pf00589 -
01410 -pf00440 -
01415 -pf11412pf02683 -
01420 -pf03091 -
01425 -pf03605 -
01430 -pf10415pf00206 -
01435 -pf04186 -
01440 HMM13520tp_1_pdu(pf13520) EUT2D
01445 -pf00166 -
01450 -pf00118 -
BMC-P
BMC-H
BMC-T
Aldehyde dehydrogenase
PTAC
Signature enzyme
Alcohol dehydrogenase
Regulator

Salmonella enterica I. Bacteria; Proteobacteria; Gammaproteobacteria; Enterobacterales;

ZA28__locus_1

PDU1AB

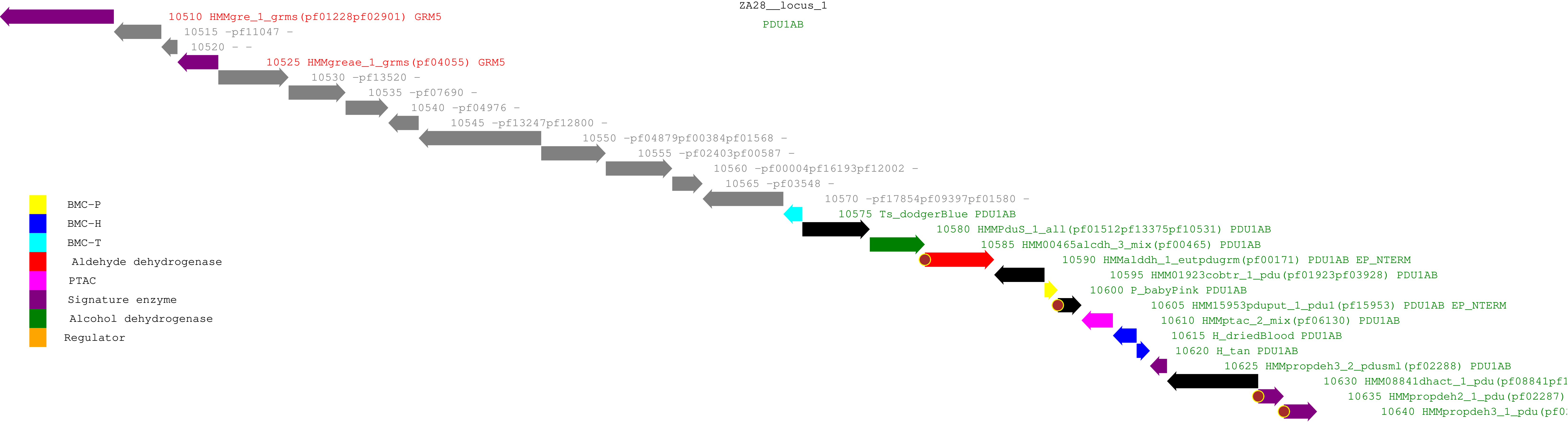

Salmonella typhimurium. Bacteria; Proteobacteria; Gammaproteobacteria; Enterobacterales;

ZA53__locus_1

PDU1AB

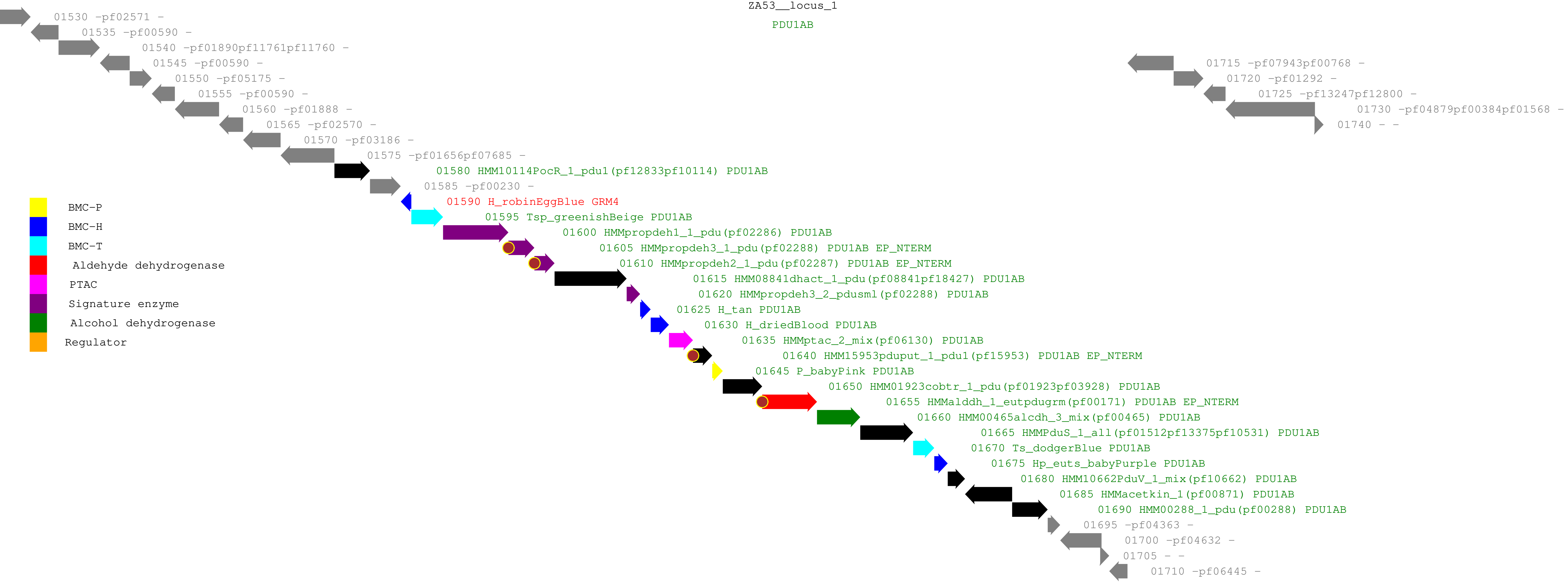

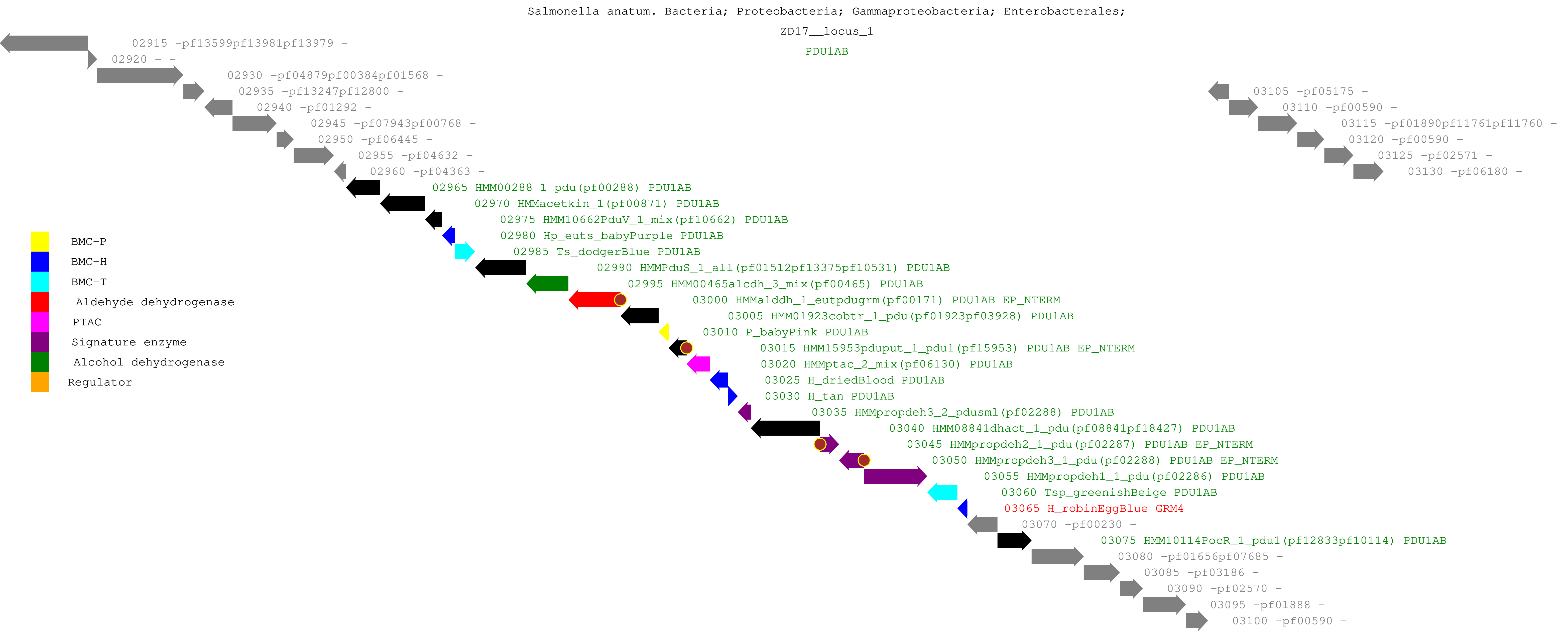
Salmonella anatum. Bacteria; Proteobacteria; Gammaproteobacteria; Enterobacterales;
ZD17__locus_1
PDU1AB
02915 -pf13599pf13981pf13979 -
02920 - -
02930 -pf04879pf00384pf01568 -
02935 -pf13247pf12800 -
02940 -pf01292 -
02945 -pf07943pf00768 -
02950 -pf06445 -
02955 -pf04632 -
02960 -pf04363 -
02965 HMM00288_1_pdu(pf00288) PDU1AB
02970 HMMacetkin_1(pf00871) PDU1AB
02975 HMM10662PduV_1_mix(pf10662) PDU1AB
02980 Hp_euts_babyPurple PDU1AB
02985 Ts_dodgerBlue PDU1AB
02990 HMMPduS_1_all(pf01512pf13375pf10531) PDU1AB
02995 HMM00465alcdh_3_mix(pf00465) PDU1AB
03000 HMMalddh_1_eutpdugrm(pf00171) PDU1AB EP_NTERM
03005 HMM01923cobtr_1_pdu(pf01923pf03928) PDU1AB
03010 P_babyPink PDU1AB
03015 HMM15953pduput_1_pdu1(pf15953) PDU1AB EP_NTERM
03020 HMMptac_2_mix(pf06130) PDU1AB
03025 H_driedBlood PDU1AB
03030 H_tan PDU1AB
03035 HMMpropdeh3_2_pdusml(pf02288) PDU1AB
03040 HMM08841dhact_1_pdu(pf08841pf18427) PDU1AB
03045 HMMpropdeh2_1_pdu(pf02287) PDU1AB EP_NTERM
03050 HMMpropdeh3_1_pdu(pf02288) PDU1AB EP_NTERM
03055 HMMpropdeh1_1_pdu(pf02286) PDU1AB
03060 Tsp_greenishBeige PDU1AB
03065 H_robinEggBlue GRM4
03070 -pf00230 -
03075 HMM10114PocR_1_pdu1(pf12833pf10114) PDU1AB
03080 -pf01656pf07685 -
03085 -pf03186 -
03090 -pf02570 -
03095 -pf01888 -
03100 -pf00590 -
03105 -pf05175 -
03110 -pf00590 -
03115 -pf01890pf11761pf11760 -
03120 -pf00590 -
03125 -pf02571 -
03130 -pf06180 -
BMC-P
BMC-H
BMC-T
Aldehyde dehydrogenase
PTAC
Signature enzyme
Alcohol dehydrogenase
Regulator

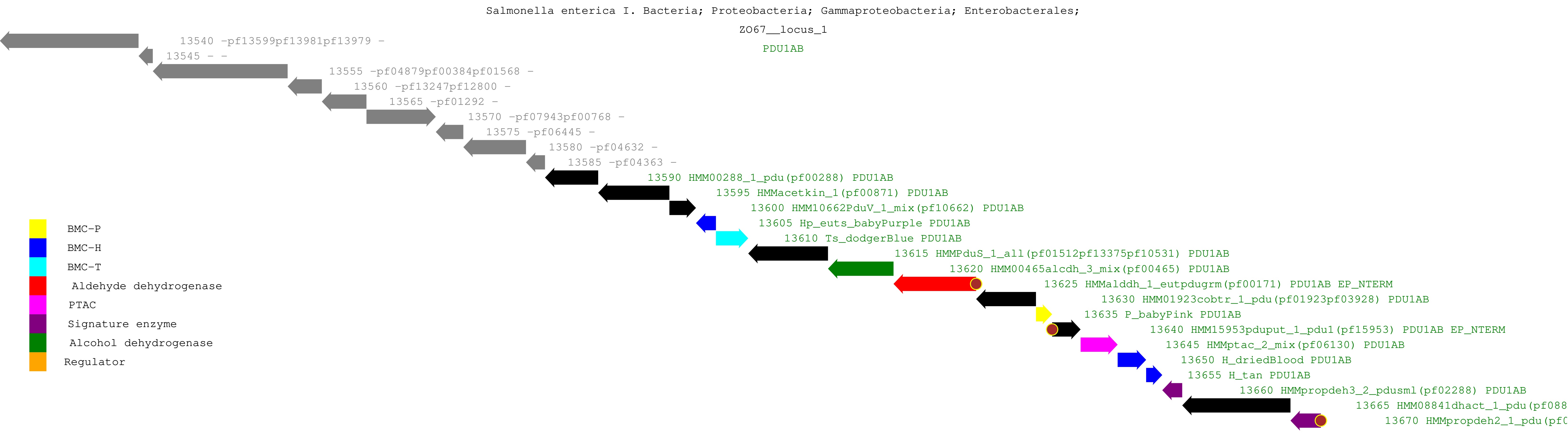
Salmonella enterica I. Bacteria; Proteobacteria; Gammaproteobacteria; Enterobacterales;
Z067__locus_1
PDU1AB
13540 -pf13599pf13981pf13979 -
13545 - -
13555 -pf04879pf00384pf01568 -
13560 -pf13247pf12800 -
13565 -pf01292 -
13570 -pf07943pf00768 -
13575 -pf06445 -
13580 -pf04632 -
13585 -pf04363 -
13590 HMM00288_1_pdu(pf00288) PDU1AB
13595 HMMacetkin_1(pf00871) PDU1AB
13600 HMM10662PduV_1_mix(pf10662) PDU1AB
13605 Hp_euts_babyPurple PDU1AB
13610 Ts_dodgerBlue PDU1AB
13615 HMMPduS_1_all(pf01512pf13375pf10531) PDU1AB
13620 HMM00465alcdh_3_mix(pf00465) PDU1AB
13625 HMMalddh_1_eutpdugrm(pf00171) PDU1AB EP_NTERM
13630 HMM01923cobtr_1_pdu(pf01923pf03928) PDU1AB
13635 P_babyPink PDU1AB
13640 HMM15953pduput_1_pdu1(pf15953) PDU1AB EP_NTERM
13645 HMMptac_2_mix(pf06130) PDU1AB
13650 H_driedBlood PDU1AB
13655 H_tan PDU1AB
13660 HMMpropdeh3_2_pdusml(pf02288) PDU1AB
13665 HMM08841dhact_1_pdu(pf088
13670 HMMpropdeh2_1_pdu(pf0
BMC-P
BMC-H
BMC-T
Aldehyde dehydrogenase
PTAC
Signature enzyme
Alcohol dehydrogenase
Regulator

Salmonella oranienberg. Bacteria; Proteobacteria; Gammaproteobacteria; Enterobacterales;

ZU40__locus_1

PDU1AB

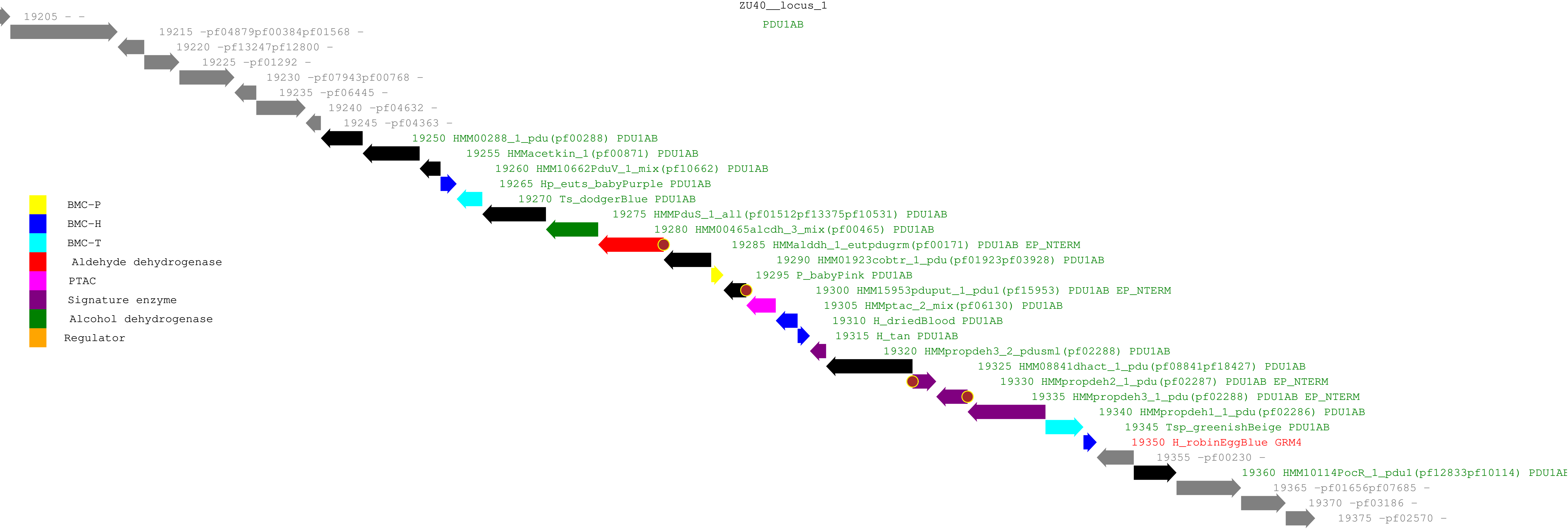

Salmonella enterica subsp. enterica serovar Corvallis. Bacteria; Proteobacteria; Gammaproteobacteria; Enterobacterales;

ZU68__locus_1

PDU1AB

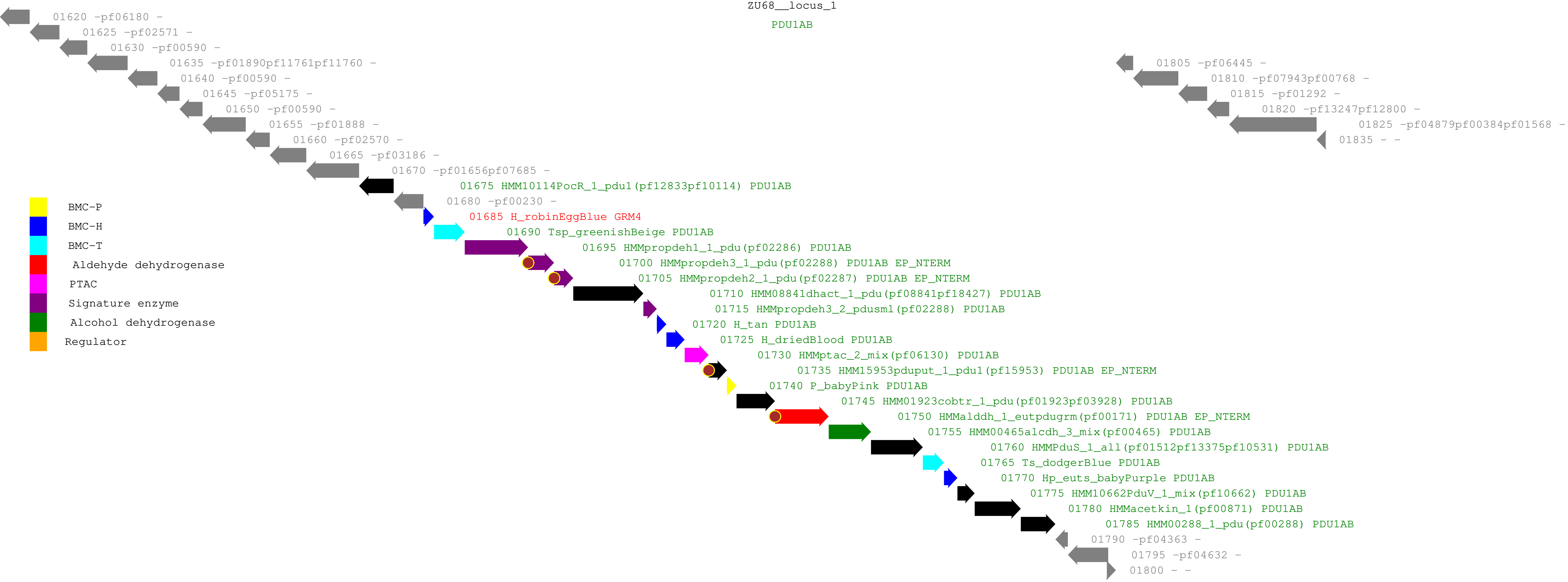

Lactobacillus reuteri. Bacteria; Firmicutes; Bacilli; Lactobacillales; Lactobacillaceae;

A4V07__locus_1

PDU1C

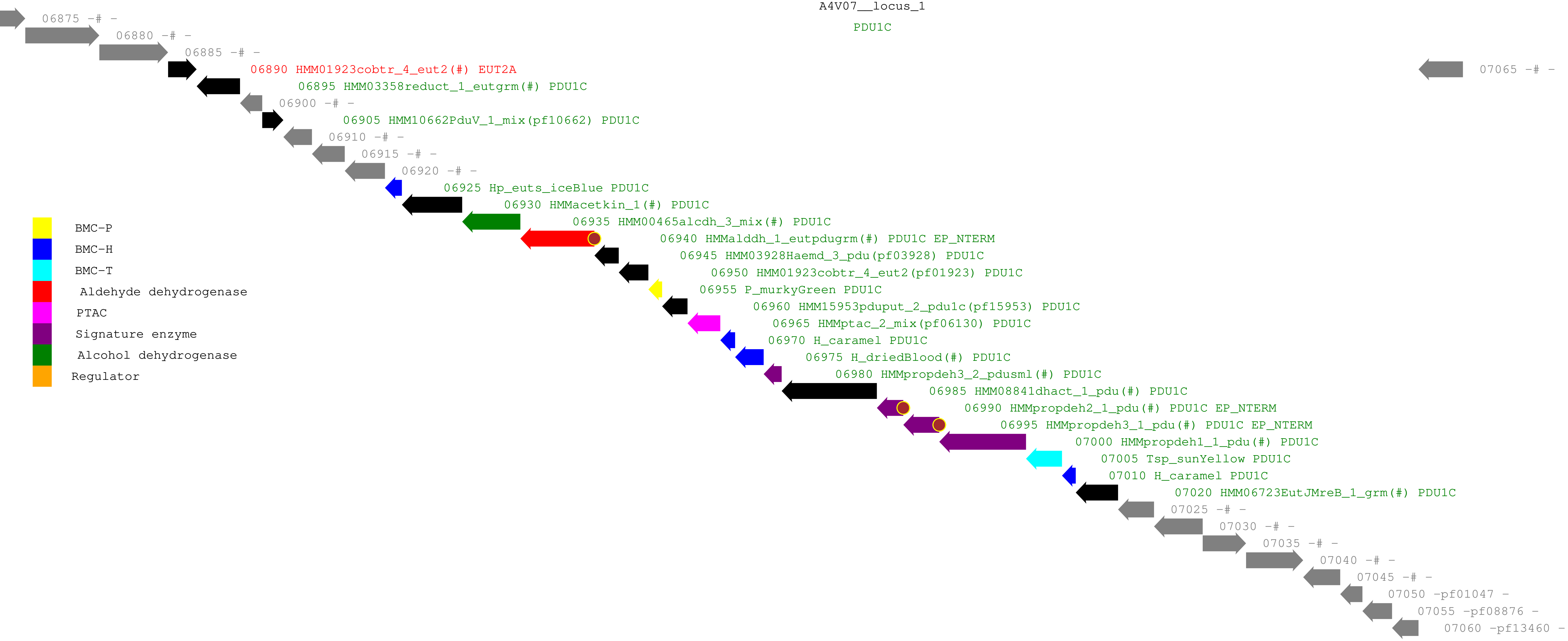

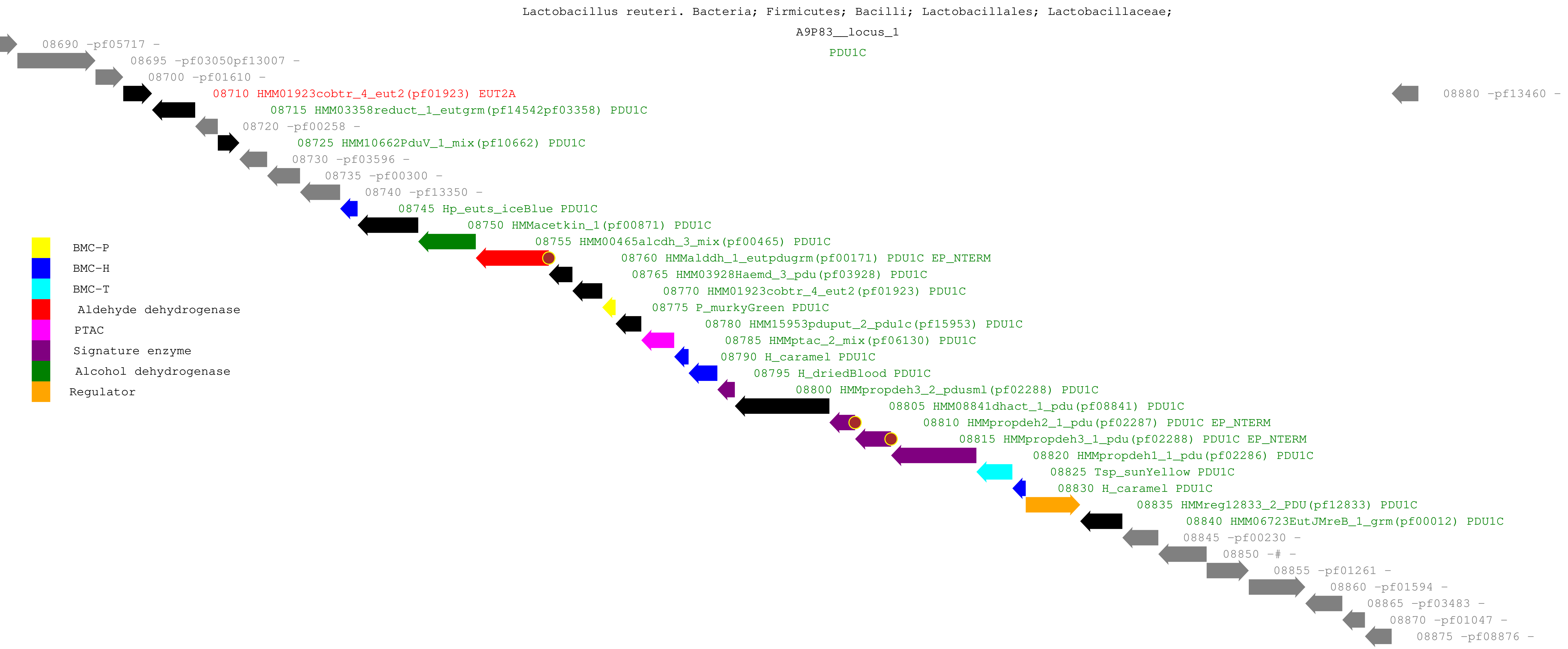
Lactobacillus reuteri. Bacteria; Firmicutes; Bacilli; Lactobacillales; Lactobacillaceae;
A9P83__locus_1
PDU1C
08690 -pf05717 -
08695 -pf03050pf13007 -
08700 -pf01610 -
08710 HMM01923cobtr_4_eut2(pf01923) EUT2A
08715 HMM03358reduct_1_eutgrm(pf14542pf03358) PDU1C
08720 -pf00258 -
08725 HMM10662PduV_1_mix(pf10662) PDU1C
08730 -pf03596 -
08735 -pf00300 -
08740 -pf13350 -
08745 Hp_euts_iceBlue PDU1C
08750 HMMacetkin_1(pf00871) PDU1C
08755 HMM00465alcdh_3_mix(pf00465) PDU1C
08760 HMMalddh_1_eutpdugrm(pf00171) PDU1C EP_NTERM
08765 HMM03928Haemd_3_pdu(pf03928) PDU1C
08770 HMM01923cobtr_4_eut2(pf01923) PDU1C
08775 P_murkyGreen PDU1C
08780 HMM15953pduput_2_pdu1c(pf15953) PDU1C
08785 HMMptac_2_mix(pf06130) PDU1C
08790 H_caramel PDU1C
08795 H_driedBlood PDU1C
08800 HMMpropdeh3_2_pdusml(pf02288) PDU1C
08805 HMM08841dhact_1_pdu(pf08841) PDU1C
08810 HMMpropdeh2_1_pdu(pf02287) PDU1C EP_NTERM
08815 HMMpropdeh3_1_pdu(pf02288) PDU1C EP_NTERM
08820 HMMpropdeh1_1_pdu(pf02286) PDU1C
08825 Tsp_sunYellow PDU1C
08830 H_caramel PDU1C
08835 HMMreg12833_2_PDU(pf12833) PDU1C
08840 HMM06723EutJMreB_1_grm(pf00012) PDU1C
08845 -pf00230 -
08850 -# -
08855 -pf01261 -
08860 -pf01594 -
08865 -pf03483 -
08870 -pf01047 -
08875 -pf08876 -
08880 -pf13460 -
BMC-P
BMC-H
BMC-T
Aldehyde dehydrogenase
PTAC
Signature enzyme
Alcohol dehydrogenase
Regulator

Lactobacillus ginsenosidimutans. Bacteria; Firmicutes; Bacilli; Lactobacillales; Lactobacillaceae;

ABM34__locus_1

PDU1C

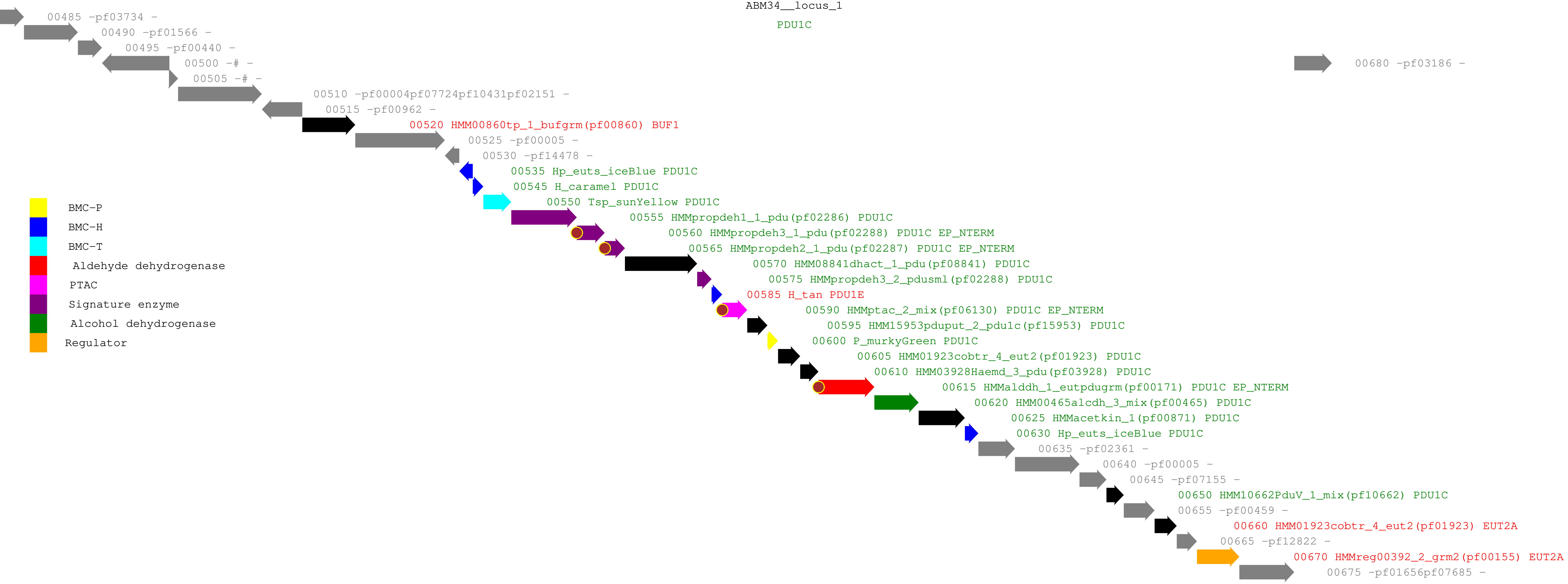

Lactobacillus reuteri. Bacteria; Firmicutes; Bacilli; Lactobacillales; Lactobacillaceae;

ADV92__locus_1

PDU1C

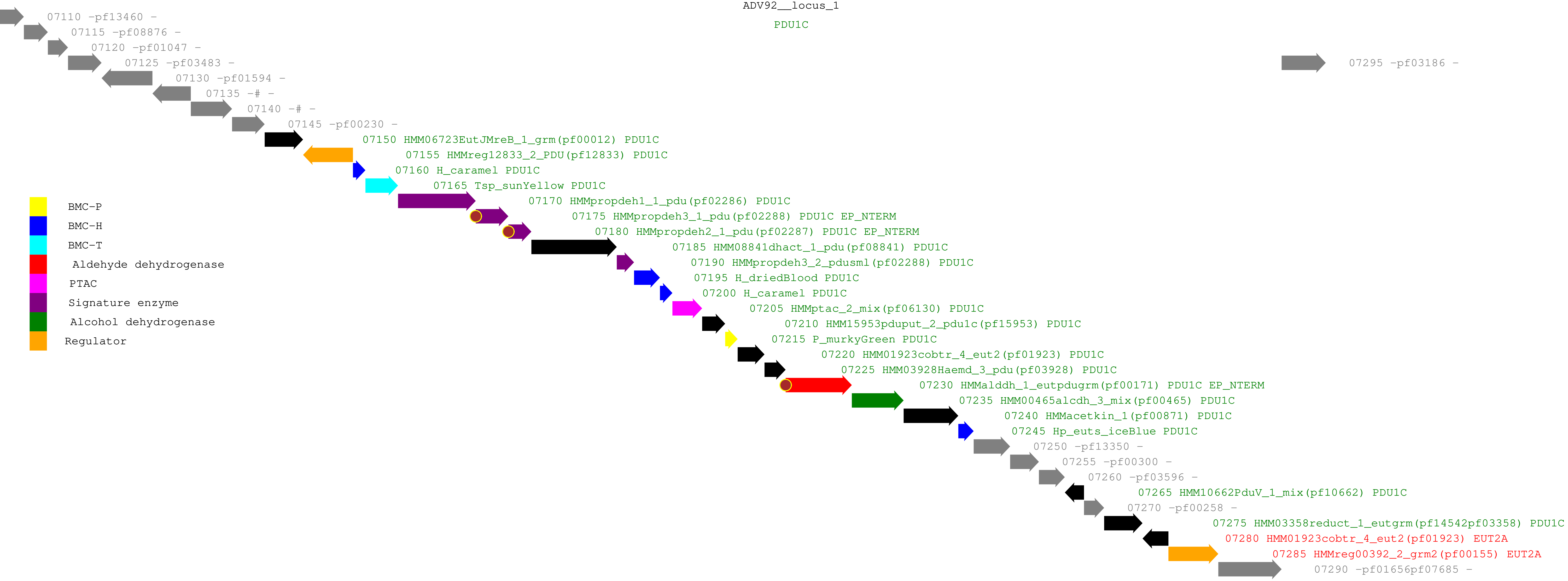

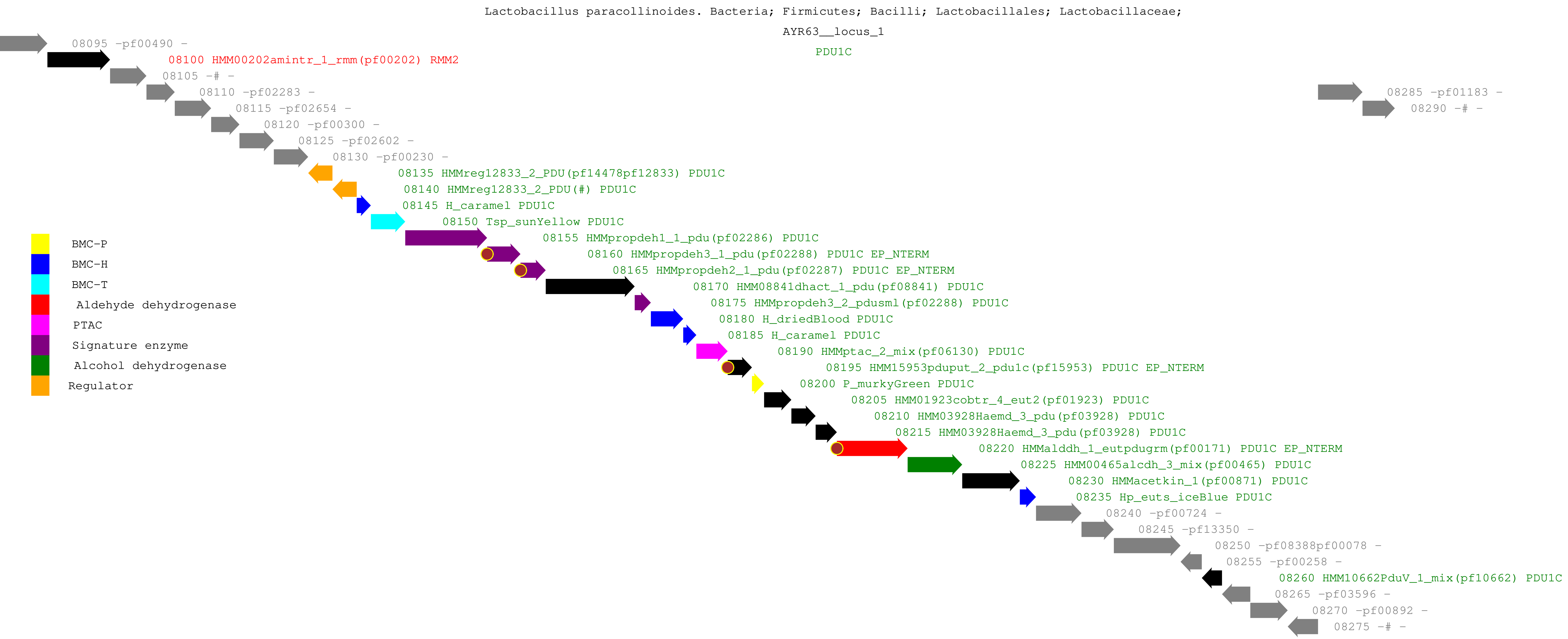
Lactobacillus paracollinoides. Bacteria; Firmicutes; Bacilli; Lactobacillales; Lactobacillaceae;
AYR63__locus_1
PDU1C
08095 -pf00490 -
08100 HMM00202amintr_1_rmm(pf00202) RMM2
08105 -# -
08110 -pf02283 -
08115 -pf02654 -
08120 -pf00300 -
08125 -pf02602 -
08130 -pf00230 -
08135 HMMreg12833_2_PDU(pf14478pf12833) PDU1C
08140 HMMreg12833_2_PDU(#) PDU1C
08145 H_caramel PDU1C
08150 Tsp_sunYellow PDU1C
08155 HMMpropdeh1_1_pdu(pf02286) PDU1C
08160 HMMpropdeh3_1_pdu(pf02288) PDU1C EP_NTERM
08165 HMMpropdeh2_1_pdu(pf02287) PDU1C EP_NTERM
08170 HMM08841dhact_1_pdu(pf08841) PDU1C
08175 HMMpropdeh3_2_pdusml(pf02288) PDU1C
08180 H_driedBlood PDU1C
08185 H_caramel PDU1C
08190 HMMptac_2_mix(pf06130) PDU1C
08195 HMM15953pduput_2_pdu1c(pf15953) PDU1C EP_NTERM
08200 P_murkyGreen PDU1C
08205 HMM01923cobtr_4_eut2(pf01923) PDU1C
08210 HMM03928Haemd_3_pdu(pf03928) PDU1C
08215 HMM03928Haemd_3_pdu(pf03928) PDU1C
08220 HMMalddh_1_eutpdugrm(pf00171) PDU1C EP_NTERM
08225 HMM00465alcdh_3_mix(pf00465) PDU1C
08230 HMMacetkin_1(pf00871) PDU1C
08235 Hp_euts_iceBlue PDU1C
08240 -pf00724 -
08245 -pf13350 -
08250 -pf08388pf00078 -
08255 -pf00258 -
08260 HMM10662PduV_1_mix(pf10662) PDU1C
08265 -pf03596 -
08270 -pf00892 -
08275 -# -
08285 -pf01183 -
08290 -# -
BMC-P
BMC-H
BMC-T
Aldehyde dehydrogenase
PTAC
Signature enzyme
Alcohol dehydrogenase
Regulator

Lactobacillus brevis. Bacteria; Firmicutes; Bacilli; Lactobacillales; Lactobacillaceae;

AZI09__locus_1_main_sat

PDU1C

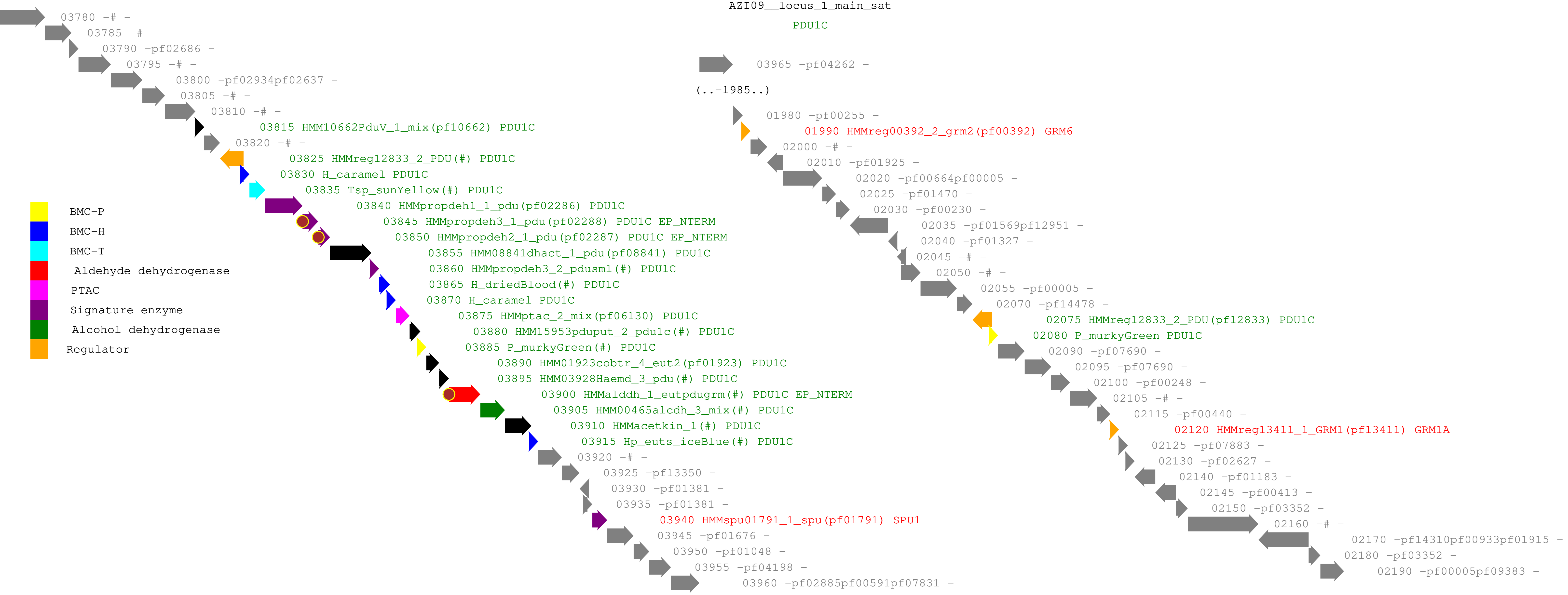

Lactobacillus brevis. Bacteria; Firmicutes; Bacilli; Lactobacillales; Lactobacillaceae;

AZI11__locus_1_main_sat

PDU1C

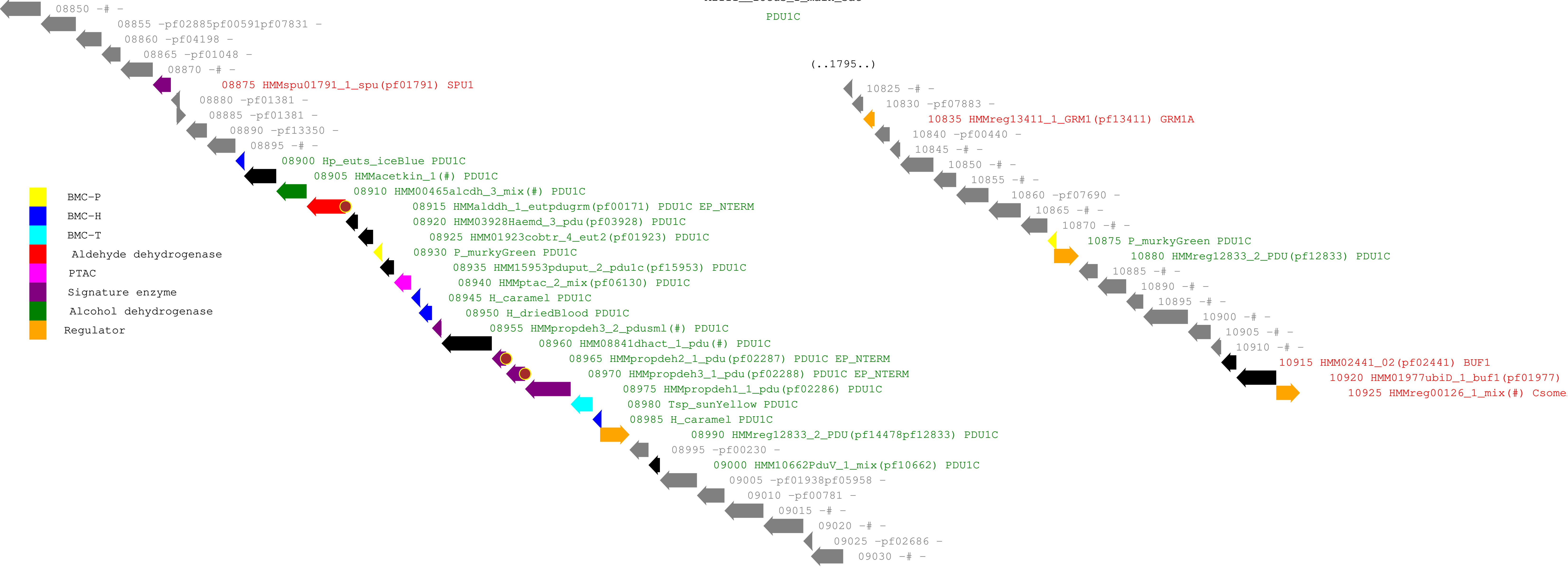

Lactobacillus reuteri. Bacteria; Firmicutes; Bacilli; Lactobacillales; Lactobacillaceae;

B5F04__locus_1

PDU1C

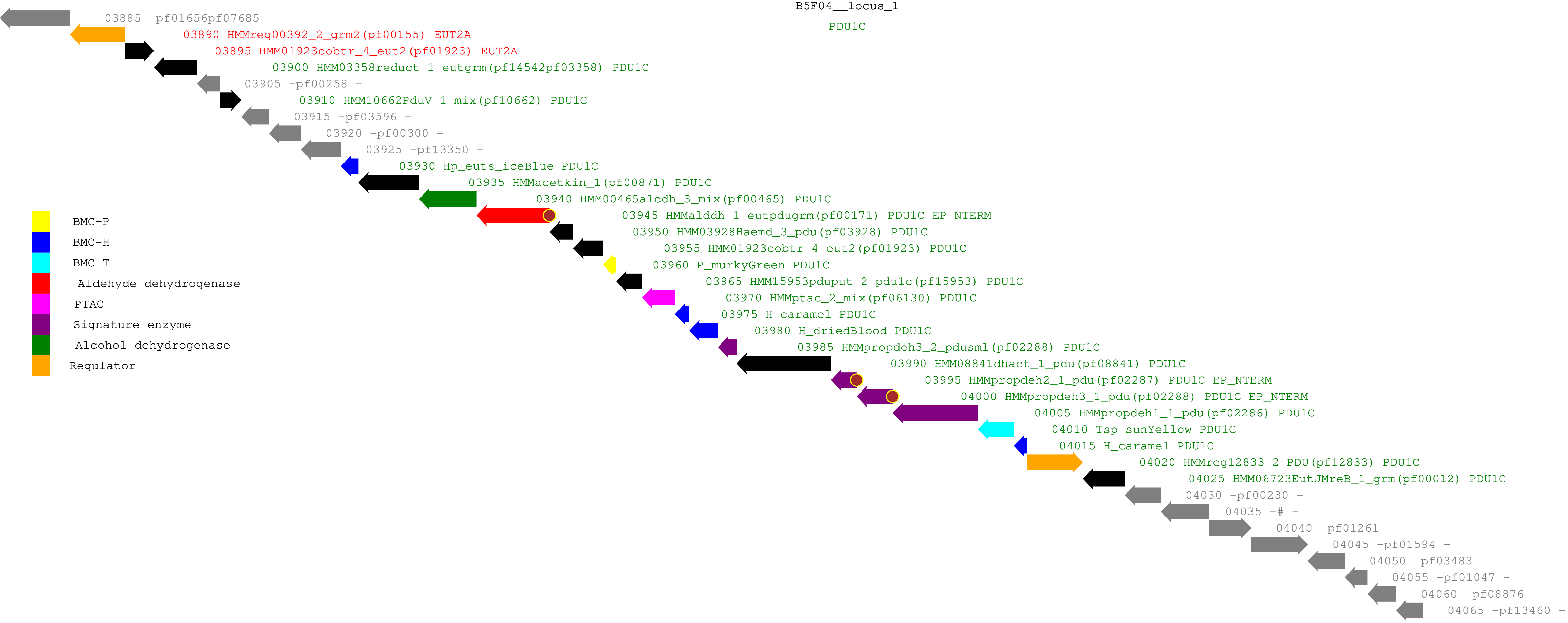

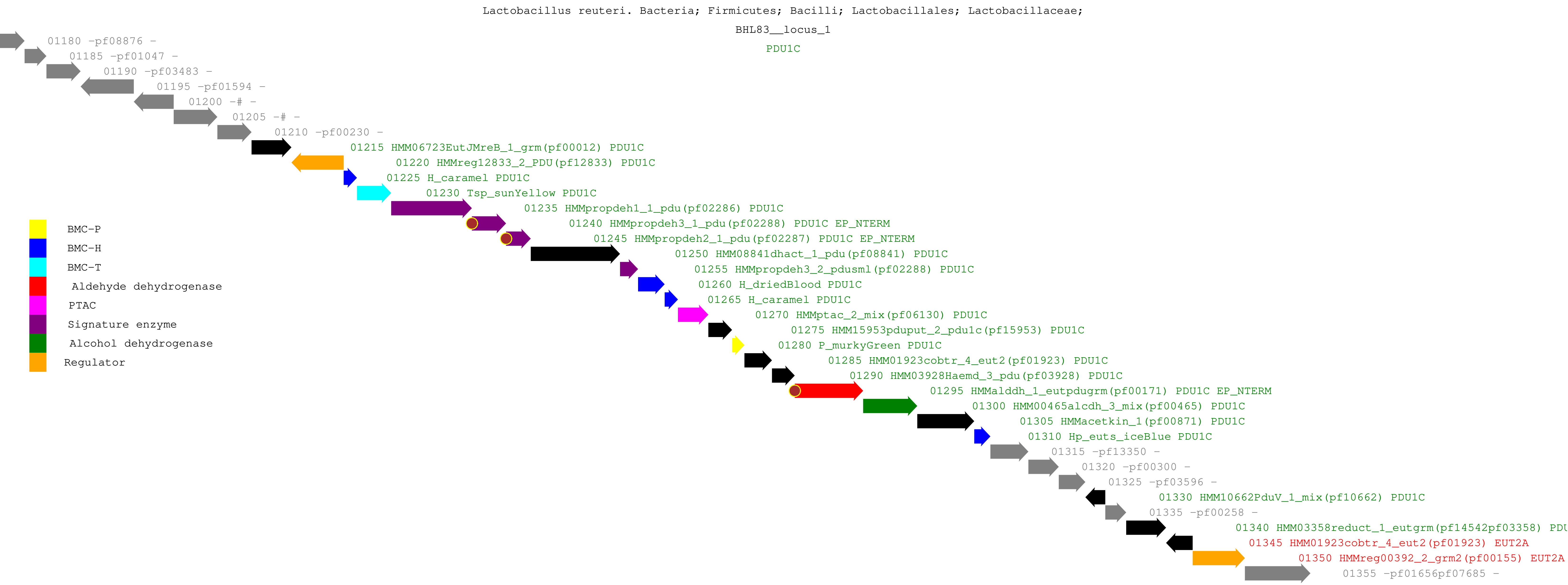
Lactobacillus reuteri. Bacteria; Firmicutes; Bacilli; Lactobacillales; Lactobacillaceae;
BHL83__locus_1
PDU1C
01180 -pf08876 -
01185 -pf01047 -
01190 -pf03483 -
01195 -pf01594 -
01200 -# -
01205 -# -
01210 -pf00230 -
01215 HMM06723EutJMreB_1_grm(pf00012) PDU1C
01220 HMMreg12833_2_PDU(pf12833) PDU1C
01225 H_caramel PDU1C
01230 Tsp_sunYellow PDU1C
01235 HMMpropdeh1_1_pdu(pf02286) PDU1C
01240 HMMpropdeh3_1_pdu(pf02288) PDU1C EP_NTERM
01245 HMMpropdeh2_1_pdu(pf02287) PDU1C EP_NTERM
01250 HMM08841dhact_1_pdu(pf08841) PDU1C
01255 HMMpropdeh3_2_pdusml(pf02288) PDU1C
01260 H_driedBlood PDU1C
01265 H_caramel PDU1C
01270 HMMptac_2_mix(pf06130) PDU1C
01275 HMM15953pduput_2_pdu1c(pf15953) PDU1C
01280 P_murkyGreen PDU1C
01285 HMM01923cobtr_4_eut2(pf01923) PDU1C
01290 HMM03928Haemd_3_pdu(pf03928) PDU1C
01295 HMMalddh_1_eutpdugrm(pf00171) PDU1C EP_NTERM
01300 HMM00465alcdh_3_mix(pf00465) PDU1C
01305 HMMacetkin_1(pf00871) PDU1C
01310 Hp_euts_iceBlue PDU1C
01315 -pf13350 -
01320 -pf00300 -
01325 -pf03596 -
01330 HMM10662PduV_1_mix(pf10662) PDU1C
01335 -pf00258 -
01340 HMM03358reduct_1_eutgrm(pf14542pf03358) PDU
01345 HMM01923cobtr_4_eut2(pf01923) EUT2A
01350 HMMreg00392_2_grm2(pf00155) EUT2A
01355 -pf01656pf07685 -
BMC-P
BMC-H
BMC-T
Aldehyde dehydrogenase
PTAC
Signature enzyme
Alcohol dehydrogenase
Regulator

Pediococcus acidilactici. Bacteria; Firmicutes; Bacilli; Lactobacillales; Lactobacillaceae;

BTW26__locus_1

PDU1C

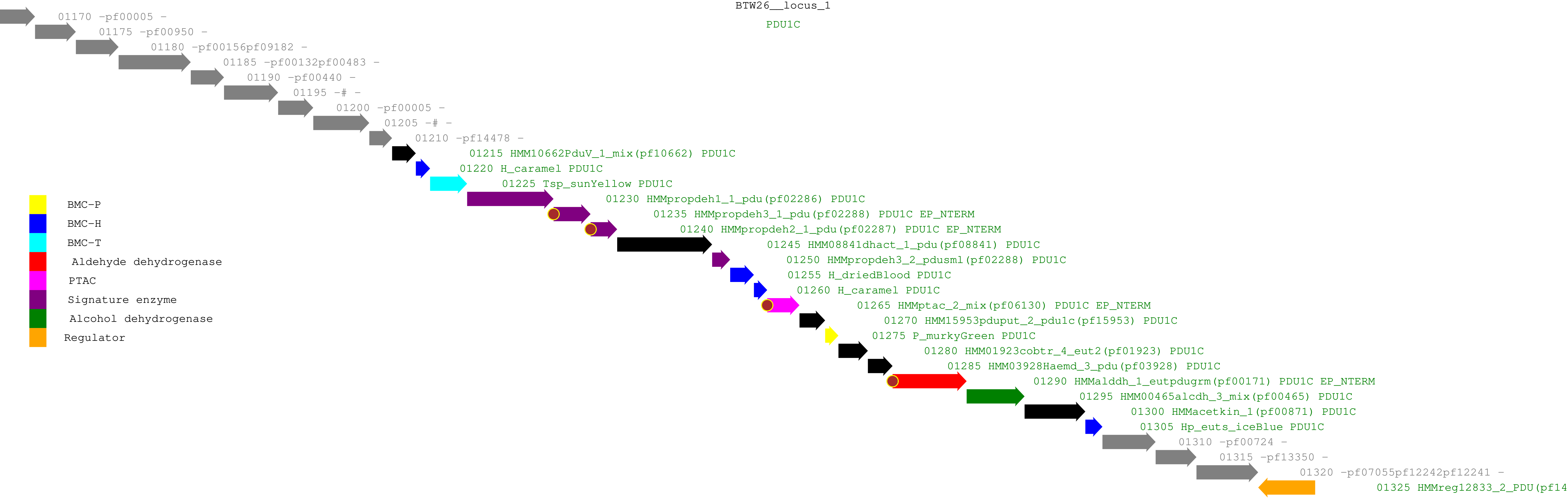

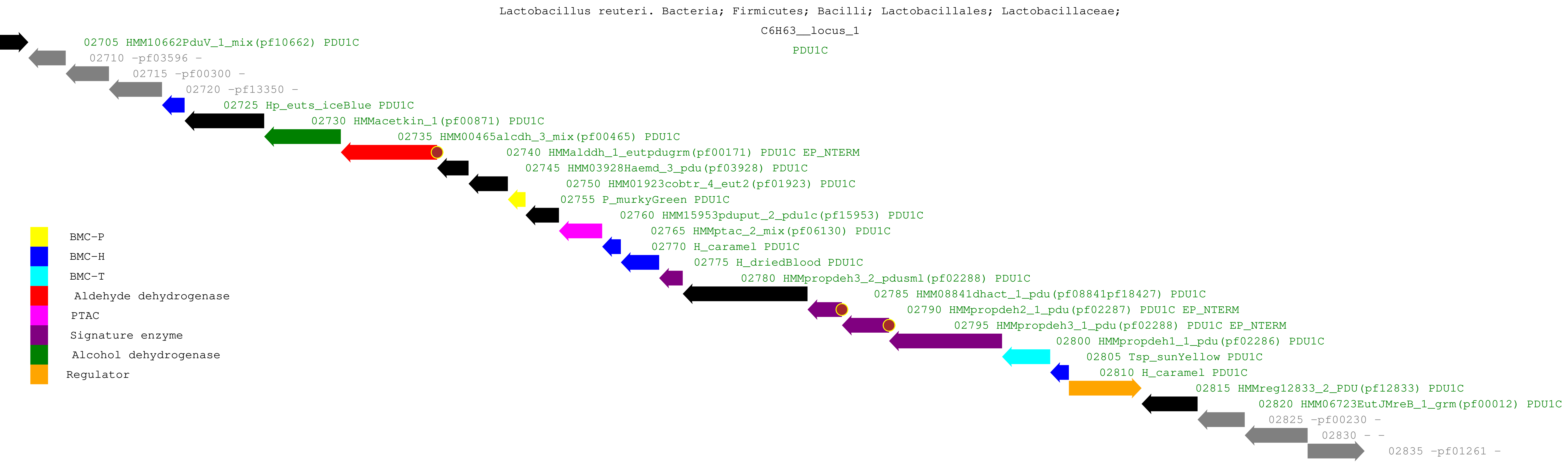
Lactobacillus reuteri. Bacteria; Firmicutes; Bacilli; Lactobacillales; Lactobacillaceae;
C6H63__locus_1
PDU1C
02705 HMM10662PduV_1_mix(pf10662) PDU1C
02710 -pf03596 -
02715 -pf00300 -
02720 -pf13350 -
02725 Hp_euts_iceBlue PDU1C
02730 HMMacetkin_1(pf00871) PDU1C
02735 HMM00465alcdh_3_mix(pf00465) PDU1C
02740 HMMalddh_1_eutpdugrm(pf00171) PDU1C EP_NTERM
02745 HMM03928Haemd_3_pdu(pf03928) PDU1C
02750 HMM01923cobtr_4_eut2(pf01923) PDU1C
02755 P_murkyGreen PDU1C
02760 HMM15953pduput_2_pdu1c(pf15953) PDU1C
02765 HMMptac_2_mix(pf06130) PDU1C
02770 H_caramel PDU1C
02775 H_driedBlood PDU1C
02780 HMMpropdeh3_2_pdusml(pf02288) PDU1C
02785 HMM08841dhact_1_pdu(pf08841pf18427) PDU1C
02790 HMMpropdeh2_1_pdu(pf02287) PDU1C EP_NTERM
02795 HMMpropdeh3_1_pdu(pf02288) PDU1C EP_NTERM
02800 HMMpropdeh1_1_pdu(pf02286) PDU1C
02805 Tsp_sunYellow PDU1C
02810 H_caramel PDU1C
02815 HMMreg12833_2_PDU(pf12833) PDU1C
02820 HMM06723EutJMreB_1_grm(pf00012) PDU1C
02825 -pf00230 -
02830 - -
02835 -pf01261 -
BMC-P
BMC-H
BMC-T
Aldehyde dehydrogenase
PTAC
Signature enzyme
Alcohol dehydrogenase
Regulator

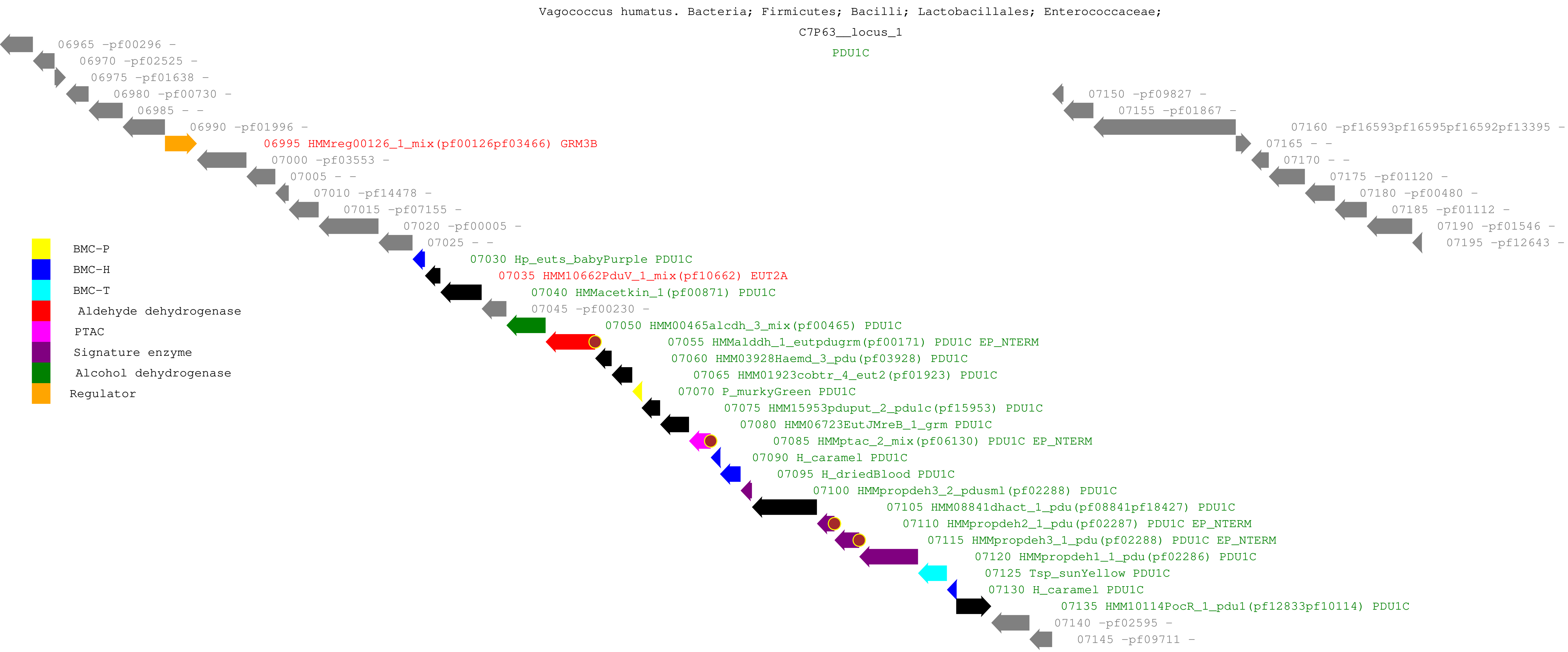
Vagococcus humatus. Bacteria; Firmicutes; Bacilli; Lactobacillales; Enterococcaceae;
C7P63__locus_1
PDU1C
06965 -pf00296 -
06970 -pf02525 -
06975 -pf01638 -
06980 -pf00730 -
06985 - -
06990 -pf01996 -
06995 HMMreg00126_1_mix(pf00126pf03466) GRM3B
07000 -pf03553 -
07005 - -
07010 -pf14478 -
07015 -pf07155 -
07020 -pf00005 -
07025 - -
07030 Hp_euts_babyPurple PDU1C
07035 HMM10662PduV_1_mix(pf10662) EUT2A
07040 HMMacetkin_1(pf00871) PDU1C
07045 -pf00230 -
07050 HMM00465alcdh_3_mix(pf00465) PDU1C
07055 HMMalddh_1_eutpdugrm(pf00171) PDU1C EP_NTERM
07060 HMM03928Haemd_3_pdu(pf03928) PDU1C
07065 HMM01923cobtr_4_eut2(pf01923) PDU1C
07070 P_murkyGreen PDU1C
07075 HMM15953pduput_2_pdu1c(pf15953) PDU1C
07080 HMM06723EutJMreB_1_grm PDU1C
07085 HMMptac_2_mix(pf06130) PDU1C EP_NTERM
07090 H_caramel PDU1C
07095 H_driedBlood PDU1C
07100 HMMpropdeh3_2_pdusml(pf02288) PDU1C
07105 HMM08841dhact_1_pdu(pf08841pf18427) PDU1C
07110 HMMpropdeh2_1_pdu(pf02287) PDU1C EP_NTERM
07115 HMMpropdeh3_1_pdu(pf02288) PDU1C EP_NTERM
07120 HMMpropdeh1_1_pdu(pf02286) PDU1C
07125 Tsp_sunYellow PDU1C
07130 H_caramel PDU1C
07135 HMM10114PocR_1_pdu1(pf12833pf10114) PDU1C
07140 -pf02595 -
07145 -pf09711 -
07150 -pf09827 -
07155 -pf01867 -
07160 -pf16593pf16595pf16592pf13395 -
07165 - -
07170 - -
07175 -pf01120 -
07180 -pf00480 -
07185 -pf01112 -
07190 -pf01546 -
07195 -pf12643 -
BMC-P
BMC-H
BMC-T
Aldehyde dehydrogenase
PTAC
Signature enzyme
Alcohol dehydrogenase
Regulator

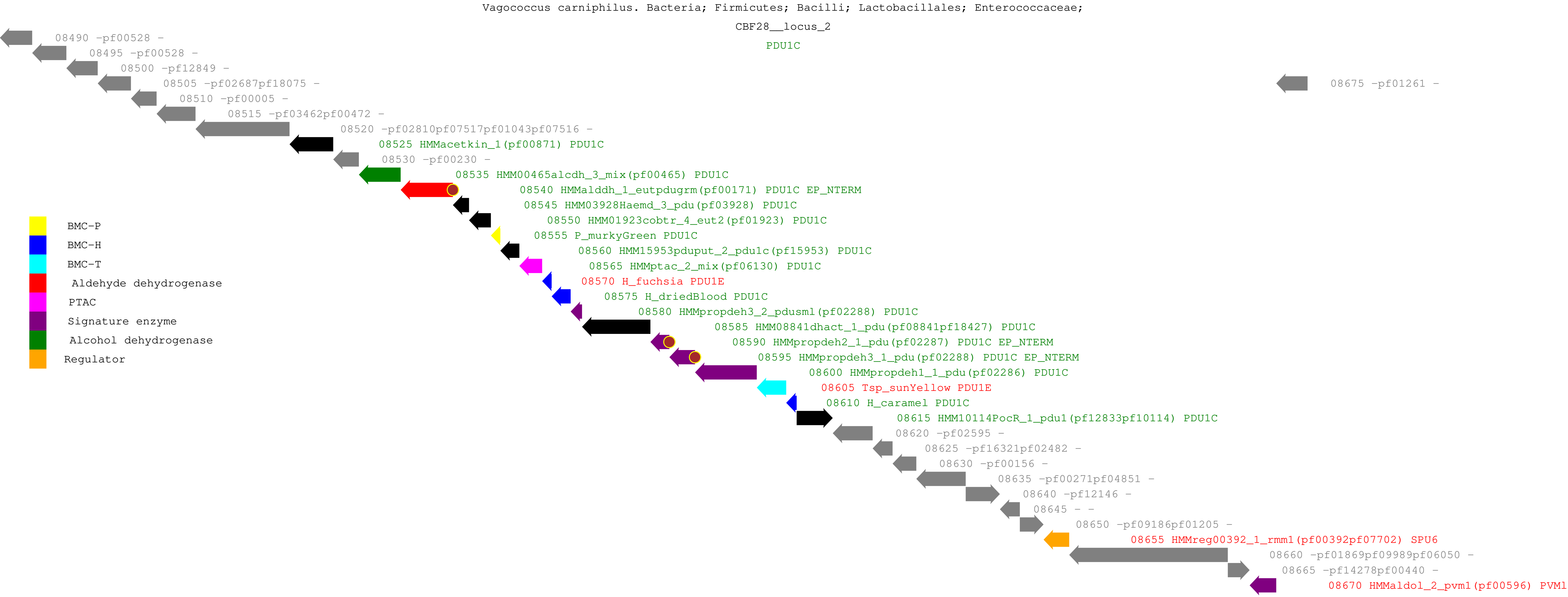
Vagococcus carniphilus. Bacteria; Firmicutes; Bacilli; Lactobacillales; Enterococcaceae;
CBF28__locus_2
PDU1C
08490 -pf00528 -
08495 -pf00528 -
08500 -pf12849 -
08505 -pf02687pf18075 -
08510 -pf00005 -
08515 -pf03462pf00472 -
08520 -pf02810pf07517pf01043pf07516 -
08525 HMMacetkin_1(pf00871) PDU1C
08530 -pf00230 -
08535 HMM00465alcdh_3_mix(pf00465) PDU1C
08540 HMMalddh_1_eutpdugrm(pf00171) PDU1C EP_NTERM
08545 HMM03928Haemd_3_pdu(pf03928) PDU1C
08550 HMM01923cobtr_4_eut2(pf01923) PDU1C
08555 P_murkyGreen PDU1C
08560 HMM15953pduput_2_pdu1c(pf15953) PDU1C
08565 HMMptac_2_mix(pf06130) PDU1C
08570 H_fuchsia PDU1E
08575 H_driedBlood PDU1C
08580 HMMpropdeh3_2_pdusml(pf02288) PDU1C
08585 HMM08841dhact_1_pdu(pf08841pf18427) PDU1C
08590 HMMpropdeh2_1_pdu(pf02287) PDU1C EP_NTERM
08595 HMMpropdeh3_1_pdu(pf02288) PDU1C EP_NTERM
08600 HMMpropdeh1_1_pdu(pf02286) PDU1C
08605 Tsp_sunYellow PDU1E
08610 H_caramel PDU1C
08615 HMM10114PocR_1_pdu1(pf12833pf10114) PDU1C
08620 -pf02595 -
08625 -pf16321pf02482 -
08630 -pf00156 -
08635 -pf00271pf04851 -
08640 -pf12146 -
08645 - -
08650 -pf09186pf01205 -
08655 HMMreg00392_1_rmm1(pf00392pf07702) SPU6
08660 -pf01869pf09989pf06050 -
08665 -pf14278pf00440 -
08670 HMMaldol_2_pvm1(pf00596) PVM1
08675 -pf01261 -
BMC-P
BMC-H
BMC-T
Aldehyde dehydrogenase
PTAC
Signature enzyme
Alcohol dehydrogenase
Regulator

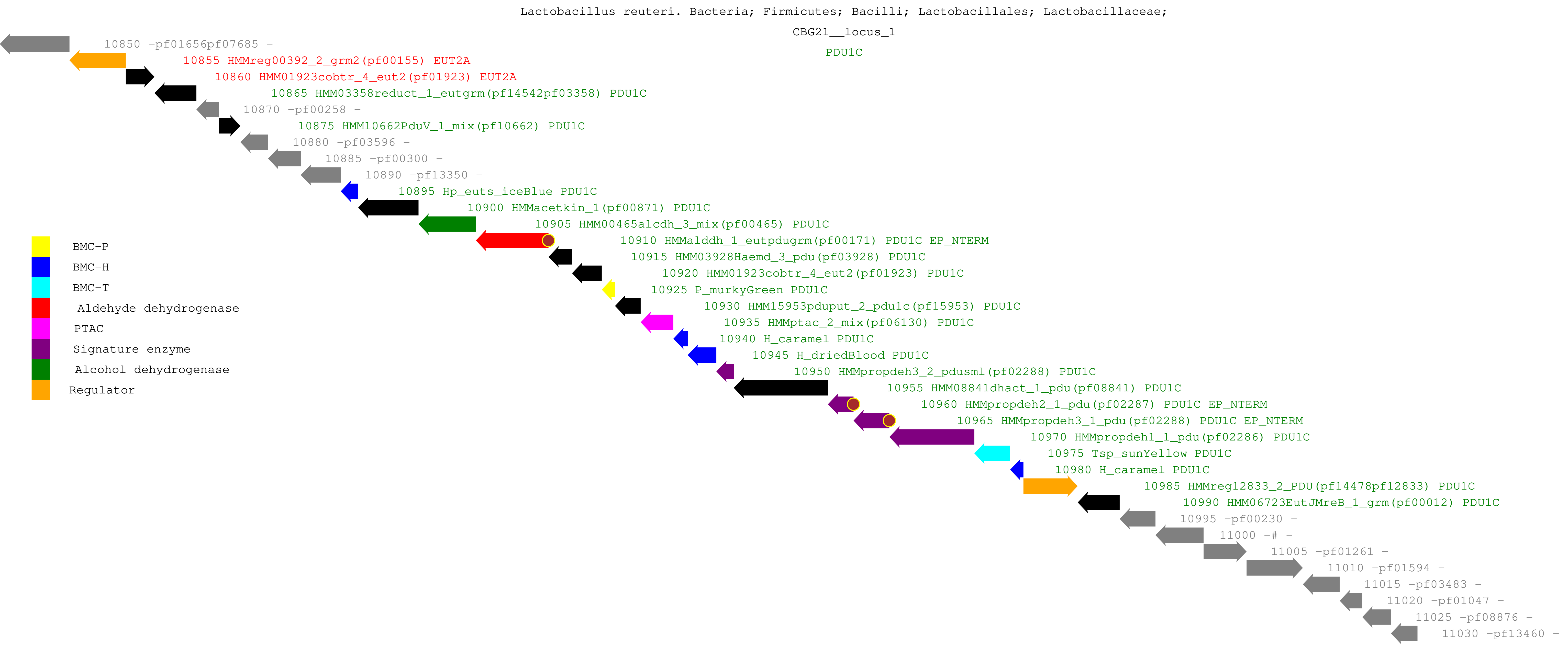
Lactobacillus reuteri. Bacteria; Firmicutes; Bacilli; Lactobacillales; Lactobacillaceae;
CBG21__locus_1
PDU1C
10850 -pf01656pf07685 -
10855 HMMreg00392_2_grm2(pf00155) EUT2A
10860 HMM01923cobtr_4_eut2(pf01923) EUT2A
10865 HMM03358reduct_1_eutgrm(pf14542pf03358) PDU1C
10870 -pf00258 -
10875 HMM10662PduV_1_mix(pf10662) PDU1C
10880 -pf03596 -
10885 -pf00300 -
10890 -pf13350 -
10895 Hp_euts_iceBlue PDU1C
10900 HMMacetkin_1(pf00871) PDU1C
10905 HMM00465alcdh_3_mix(pf00465) PDU1C
10910 HMMalddh_1_eutpdugrm(pf00171) PDU1C EP_NTERM
10915 HMM03928Haemd_3_pdu(pf03928) PDU1C
10920 HMM01923cobtr_4_eut2(pf01923) PDU1C
10925 P_murkyGreen PDU1C
10930 HMM15953pduput_2_pdu1c(pf15953) PDU1C
10935 HMMptac_2_mix(pf06130) PDU1C
10940 H_caramel PDU1C
10945 H_driedBlood PDU1C
10950 HMMpropdeh3_2_pdusml(pf02288) PDU1C
10955 HMM08841dhact_1_pdu(pf08841) PDU1C
10960 HMMpropdeh2_1_pdu(pf02287) PDU1C EP_NTERM
10965 HMMpropdeh3_1_pdu(pf02288) PDU1C EP_NTERM
10970 HMMpropdeh1_1_pdu(pf02286) PDU1C
10975 Tsp_sunYellow PDU1C
10980 H_caramel PDU1C
10985 HMMreg12833_2_PDU(pf14478pf12833) PDU1C
10990 HMM06723EutJMreB_1_grm(pf00012) PDU1C
10995 -pf00230 -
11000 -# -
11005 -pf01261 -
11010 -pf01594 -
11015 -pf03483 -
11020 -pf01047 -
11025 -pf08876 -
11030 -pf13460 -
BMC-P
BMC-H
BMC-T
Aldehyde dehydrogenase
PTAC
Signature enzyme
Alcohol dehydrogenase
Regulator

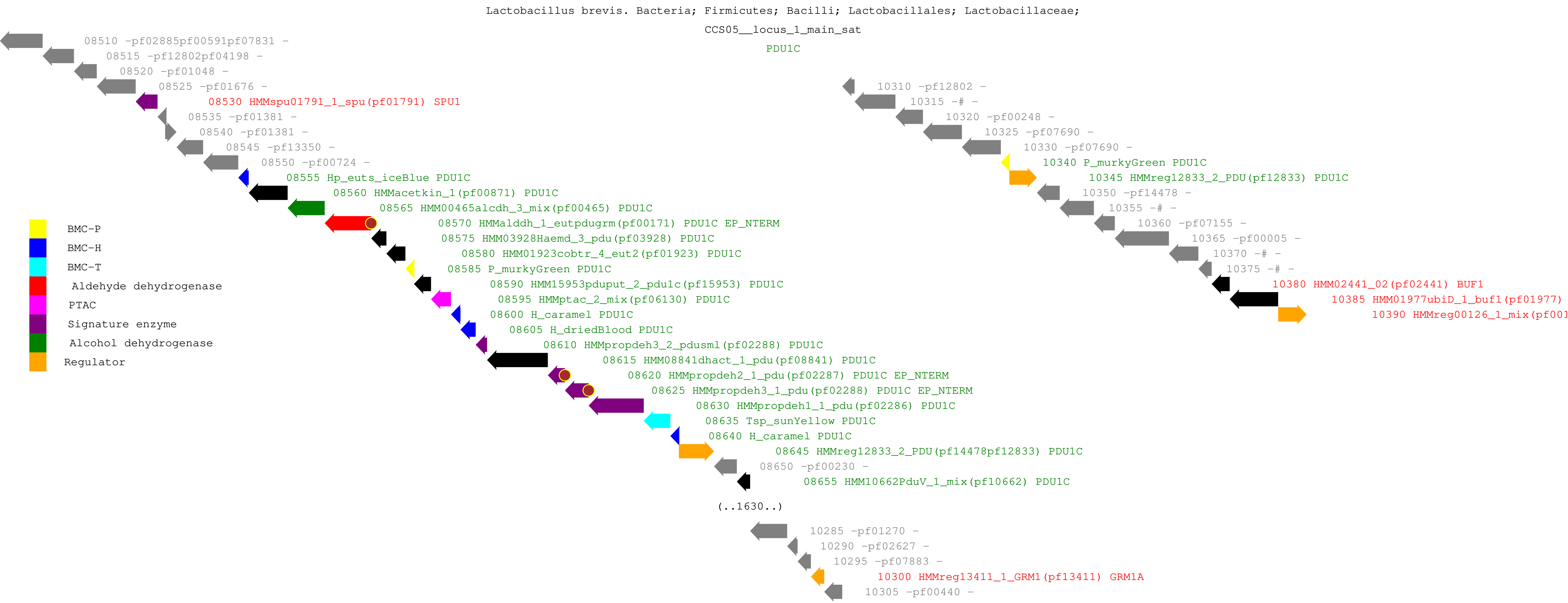
Lactobacillus brevis. Bacteria; Firmicutes; Bacilli; Lactobacillales; Lactobacillaceae;
CCS05__locus_1_main_sat
PDU1C
08510 -pf02885pf00591pf07831 -
08515 -pf12802pf04198 -
08520 -pf01048 -
08525 -pf01676 -
08530 HMMspu01791_1_spu(pf01791) SPU1
08535 -pf01381 -
08540 -pf01381 -
08545 -pf13350 -
08550 -pf00724 -
08555 Hp_euts_iceBlue PDU1C
08560 HMMacetkin_1(pf00871) PDU1C
08565 HMM00465alcdh_3_mix(pf00465) PDU1C
08570 HMMalddh_1_eutpdugrm(pf00171) PDU1C EP_NTERM
08575 HMM03928Haemd_3_pdu(pf03928) PDU1C
08580 HMM01923cobtr_4_eut2(pf01923) PDU1C
08585 P_murkyGreen PDU1C
08590 HMM15953pduput_2_pdu1c(pf15953) PDU1C
08595 HMMptac_2_mix(pf06130) PDU1C
08600 H_caramel PDU1C
08605 H_driedBlood PDU1C
08610 HMMpropdeh3_2_pdusml(pf02288) PDU1C
08615 HMM08841dhact_1_pdu(pf08841) PDU1C
08620 HMMpropdeh2_1_pdu(pf02287) PDU1C EP_NTERM
08625 HMMpropdeh3_1_pdu(pf02288) PDU1C EP_NTERM
08630 HMMpropdeh1_1_pdu(pf02286) PDU1C
08635 Tsp_sunYellow PDU1C
08640 H_caramel PDU1C
08645 HMMreg12833_2_PDU(pf14478pf12833) PDU1C
08650 -pf00230 -
08655 HMM10662PduV_1_mix(pf10662) PDU1C
(..1630..)
10285 -pf01270 -
10290 -pf02627 -
10295 -pf07883 -
10300 HMMreg13411_1_GRM1(pf13411) GRM1A
10305 -pf00440 -
10310 -pf12802 -
10315 -# -
10320 -pf00248 -
10325 -pf07690 -
10330 -pf07690 -
10340 P_murkyGreen PDU1C
10345 HMMreg12833_2_PDU(pf12833) PDU1C
10350 -pf14478 -
10355 -# -
10360 -pf07155 -
10365 -pf00005 -
10370 -# -
10375 -# -
10380 HMM02441_02(pf02441) BUF1
10385 HMM01977ubiD_1_buf1(pf01977)
10390 HMMreg00126_1_mix(pf001
BMC-P
BMC-H
BMC-T
Aldehyde dehydrogenase
PTAC
Signature enzyme
Alcohol dehydrogenase
Regulator

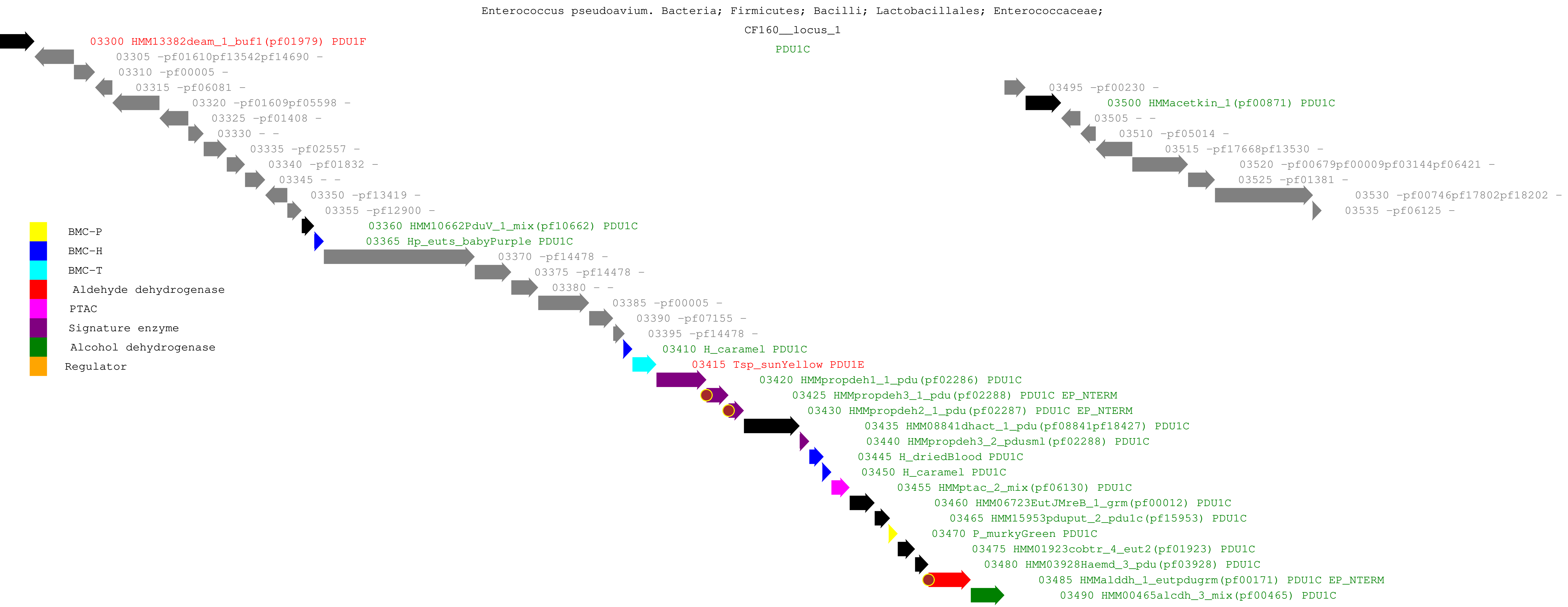
Enterococcus pseudoavium. Bacteria; Firmicutes; Bacilli; Lactobacillales; Enterococcaceae;
CF160__locus_1
PDU1C
03300 HMM13382deam_1_buf1(pf01979) PDU1F
03305 -pf01610pf13542pf14690 -
03310 -pf00005 -
03315 -pf06081 -
03320 -pf01609pf05598 -
03325 -pf01408 -
03330 - -
03335 -pf02557 -
03340 -pf01832 -
03345 - -
03350 -pf13419 -
03355 -pf12900 -
03360 HMM10662PduV_1_mix(pf10662) PDU1C
03365 Hp_euts_babyPurple PDU1C
03370 -pf14478 -
03375 -pf14478 -
03380 - -
03385 -pf00005 -
03390 -pf07155 -
03395 -pf14478 -
03410 H_caramel PDU1C
03415 Tsp_sunYellow PDU1E
03420 HMMpropdeh1_1_pdu(pf02286) PDU1C
03425 HMMpropdeh3_1_pdu(pf02288) PDU1C EP_NTERM
03430 HMMpropdeh2_1_pdu(pf02287) PDU1C EP_NTERM
03435 HMM08841dhact_1_pdu(pf08841pf18427) PDU1C
03440 HMMpropdeh3_2_pdusml(pf02288) PDU1C
03445 H_driedBlood PDU1C
03450 H_caramel PDU1C
03455 HMMptac_2_mix(pf06130) PDU1C
03460 HMM06723EutJMreB_1_grm(pf00012) PDU1C
03465 HMM15953pduput_2_pdu1c(pf15953) PDU1C
03470 P_murkyGreen PDU1C
03475 HMM01923cobtr_4_eut2(pf01923) PDU1C
03480 HMM03928Haemd_3_pdu(pf03928) PDU1C
03485 HMMalddh_1_eutpdugrm(pf00171) PDU1C EP_NTERM
03490 HMM00465alcdh_3_mix(pf00465) PDU1C
03495 -pf00230 -
03500 HMMacetkin_1(pf00871) PDU1C
03505 - -
03510 -pf05014 -
03515 -pf17668pf13530 -
03520 -pf00679pf00009pf03144pf06421 -
03525 -pf01381 -
03530 -pf00746pf17802pf18202 -
03535 -pf06125 -
BMC-P
BMC-H
BMC-T
Aldehyde dehydrogenase
PTAC
Signature enzyme
Alcohol dehydrogenase
Regulator

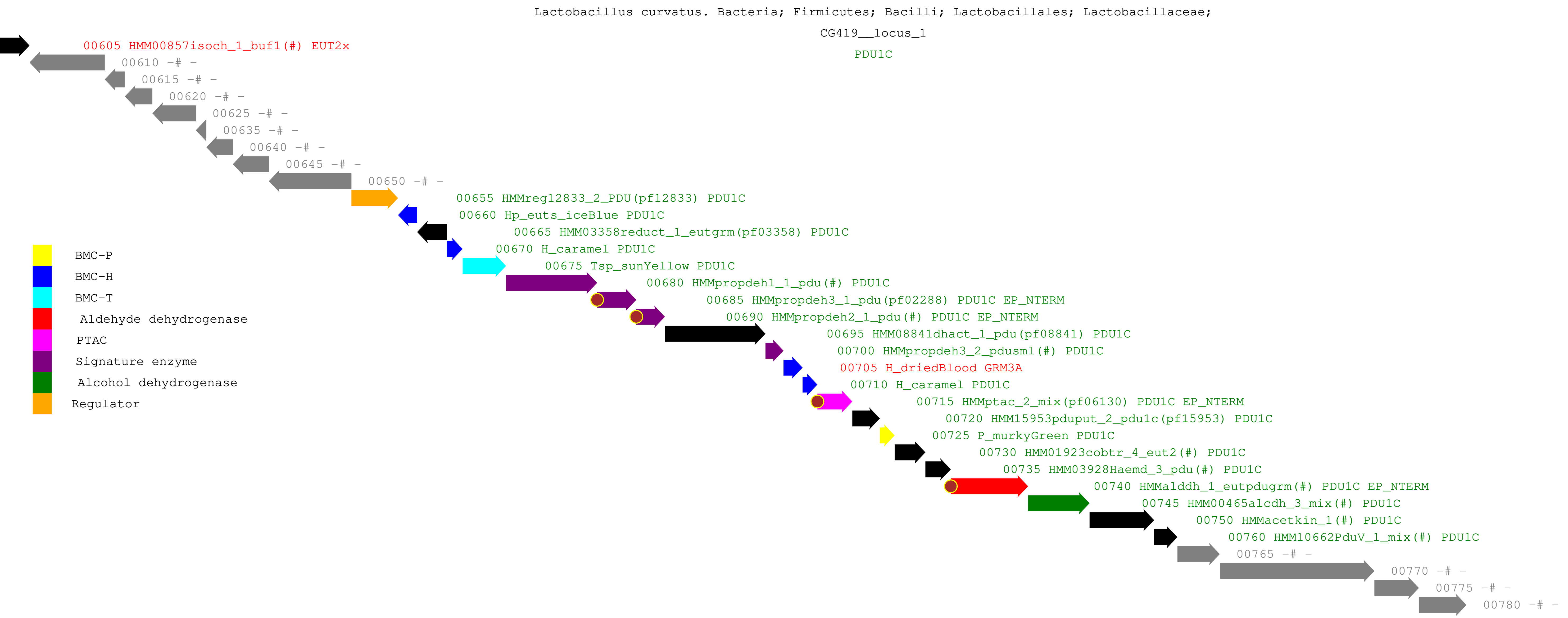
Lactobacillus curvatus. Bacteria; Firmicutes; Bacilli; Lactobacillales; Lactobacillaceae;
CG419__locus_1
PDU1C
00605 HMM00857isoch_1_buf1(#) EUT2x
00610 -# -
00615 -# -
00620 -# -
00625 -# -
00635 -# -
00640 -# -
00645 -# -
00650 -# -
00655 HMMreg12833_2_PDU(pf12833) PDU1C
00660 Hp_euts_iceBlue PDU1C
00665 HMM03358reduct_1_eutgrm(pf03358) PDU1C
00670 H_caramel PDU1C
00675 Tsp_sunYellow PDU1C
00680 HMMpropdeh1_1_pdu(#) PDU1C
00685 HMMpropdeh3_1_pdu(pf02288) PDU1C EP_NTERM
00690 HMMpropdeh2_1_pdu(#) PDU1C EP_NTERM
00695 HMM08841dhact_1_pdu(pf08841) PDU1C
00700 HMMpropdeh3_2_pdusml(#) PDU1C
00705 H_driedBlood GRM3A
00710 H_caramel PDU1C
00715 HMMptac_2_mix(pf06130) PDU1C EP_NTERM
00720 HMM15953pduput_2_pdu1c(pf15953) PDU1C
00725 P_murkyGreen PDU1C
00730 HMM01923cobtr_4_eut2(#) PDU1C
00735 HMM03928Haemd_3_pdu(#) PDU1C
00740 HMMalddh_1_eutpdugrm(#) PDU1C EP_NTERM
00745 HMM00465alcdh_3_mix(#) PDU1C
00750 HMMacetkin_1(#) PDU1C
00760 HMM10662PduV_1_mix(#) PDU1C
00765 -# -
00770 -# -
00775 -# -
00780 -# -
BMC-P
BMC-H
BMC-T
Aldehyde dehydrogenase
PTAC
Signature enzyme
Alcohol dehydrogenase
Regulator

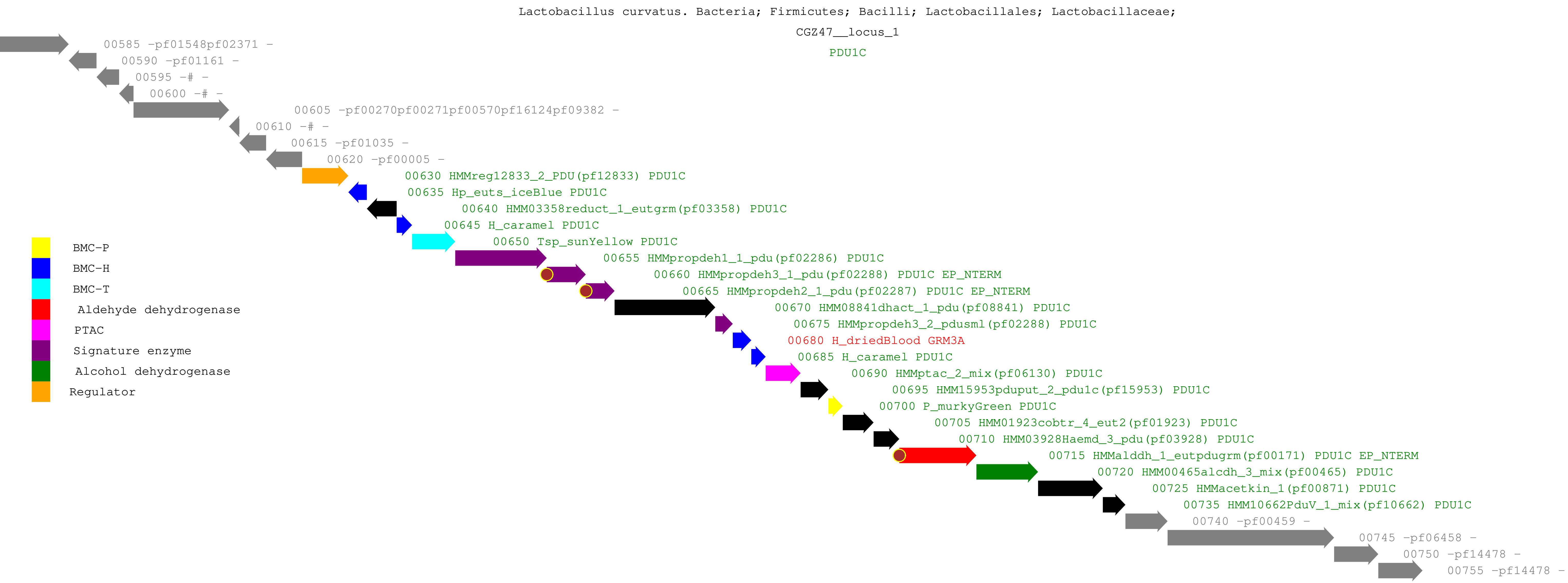
Lactobacillus curvatus. Bacteria; Firmicutes; Bacilli; Lactobacillales; Lactobacillaceae;
CGZ47__locus_1
PDU1C
00585 -pf01548pf02371 -
00590 -pf01161 -
00595 -# -
00600 -# -
00605 -pf00270pf00271pf00570pf16124pf09382 -
00610 -# -
00615 -pf01035 -
00620 -pf00005 -
00630 HMMreg12833_2_PDU(pf12833) PDU1C
00635 Hp_euts_iceBlue PDU1C
00640 HMM03358reduct_1_eutgrm(pf03358) PDU1C
00645 H_caramel PDU1C
00650 Tsp_sunYellow PDU1C
00655 HMMpropdeh1_1_pdu(pf02286) PDU1C
00660 HMMpropdeh3_1_pdu(pf02288) PDU1C EP_NTERM
00665 HMMpropdeh2_1_pdu(pf02287) PDU1C EP_NTERM
00670 HMM08841dhact_1_pdu(pf08841) PDU1C
00675 HMMpropdeh3_2_pdusml(pf02288) PDU1C
00680 H_driedBlood GRM3A
00685 H_caramel PDU1C
00690 HMMptac_2_mix(pf06130) PDU1C
00695 HMM15953pduput_2_pdu1c(pf15953) PDU1C
00700 P_murkyGreen PDU1C
00705 HMM01923cobtr_4_eut2(pf01923) PDU1C
00710 HMM03928Haemd_3_pdu(pf03928) PDU1C
00715 HMMalddh_1_eutpdugrm(pf00171) PDU1C EP_NTERM
00720 HMM00465alcdh_3_mix(pf00465) PDU1C
00725 HMMacetkin_1(pf00871) PDU1C
00735 HMM10662PduV_1_mix(pf10662) PDU1C
00740 -pf00459 -
00745 -pf06458 -
00750 -pf14478 -
00755 -pf14478 -
BMC-P
BMC-H
BMC-T
Aldehyde dehydrogenase
PTAC
Signature enzyme
Alcohol dehydrogenase
Regulator

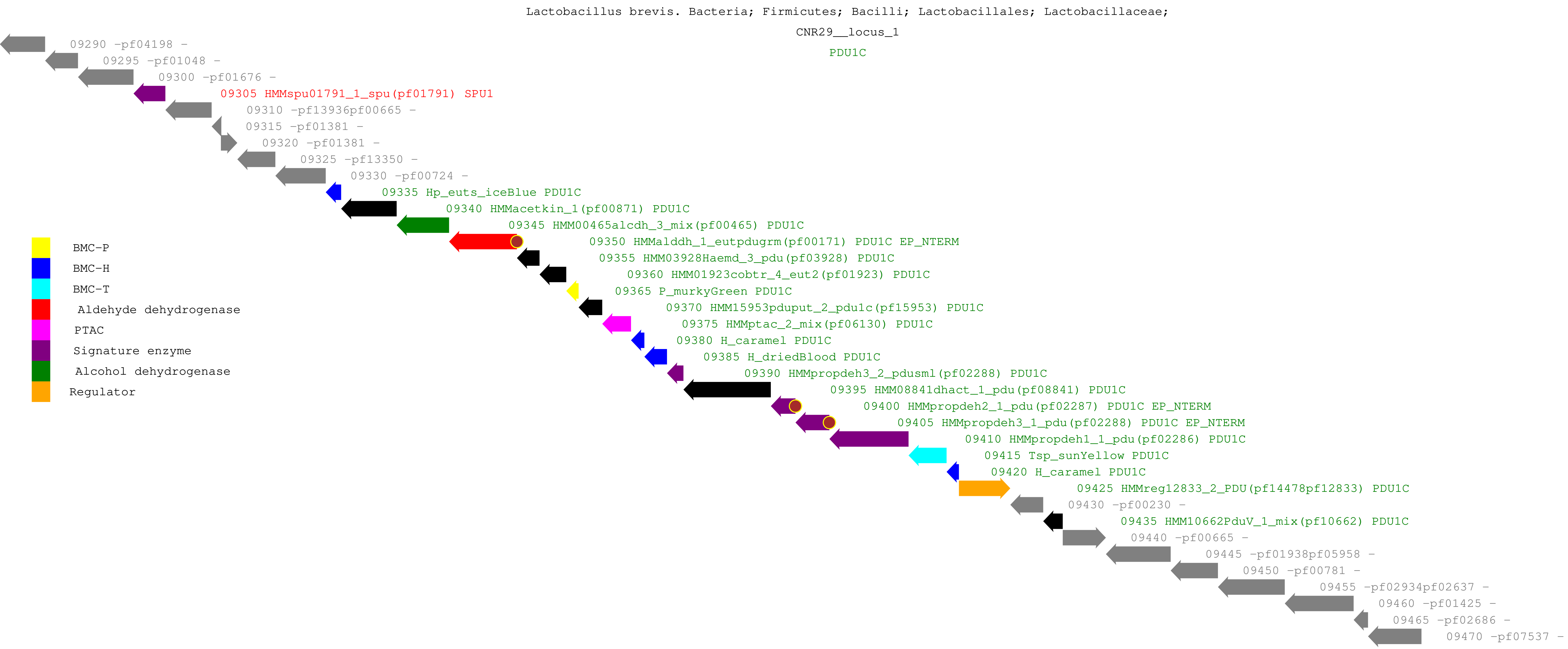

Lactobacillus brevis. Bacteria; Firmicutes; Bacilli; Lactobacillales; Lactobacillaceae;
CNR29__locus_1
PDU1C
09290 -pf04198 -
09295 -pf01048 -
09300 -pf01676 -
09305 HMMspu01791_1_spu(pf01791) SPU1
09310 -pf13936pf00665 -
09315 -pf01381 -
09320 -pf01381 -
09325 -pf13350 -
09330 -pf00724 -
09335 Hp_euts_iceBlue PDU1C
09340 HMMacetkin_1(pf00871) PDU1C
09345 HMM00465alcdh_3_mix(pf00465) PDU1C
09350 HMMalddh_1_eutpdugrm(pf00171) PDU1C EP_NTERM
09355 HMM03928Haemd_3_pdu(pf03928) PDU1C
09360 HMM01923cobtr_4_eut2(pf01923) PDU1C
09365 P_murkyGreen PDU1C
09370 HMM15953pduput_2_pdu1c(pf15953) PDU1C
09375 HMMptac_2_mix(pf06130) PDU1C
09380 H_caramel PDU1C
09385 H_driedBlood PDU1C
09390 HMMpropdeh3_2_pdusml(pf02288) PDU1C
09395 HMM08841dhact_1_pdu(pf08841) PDU1C
09400 HMMpropdeh2_1_pdu(pf02287) PDU1C EP_NTERM
09405 HMMpropdeh3_1_pdu(pf02288) PDU1C EP_NTERM
09410 HMMpropdeh1_1_pdu(pf02286) PDU1C
09415 Tsp_sunYellow PDU1C
09420 H_caramel PDU1C
09425 HMMreg12833_2_PDU(pf14478pf12833) PDU1C
09430 -pf00230 -
09435 HMM10662PduV_1_mix(pf10662) PDU1C
09440 -pf00665 -
09445 -pf01938pf05958 -
09450 -pf00781 -
09455 -pf02934pf02637 -
09460 -pf01425 -
09465 -pf02686 -
09470 -pf07537 -
BMC-P
BMC-H
BMC-T
Aldehyde dehydrogenase
PTAC
Signature enzyme
Alcohol dehydrogenase
Regulator

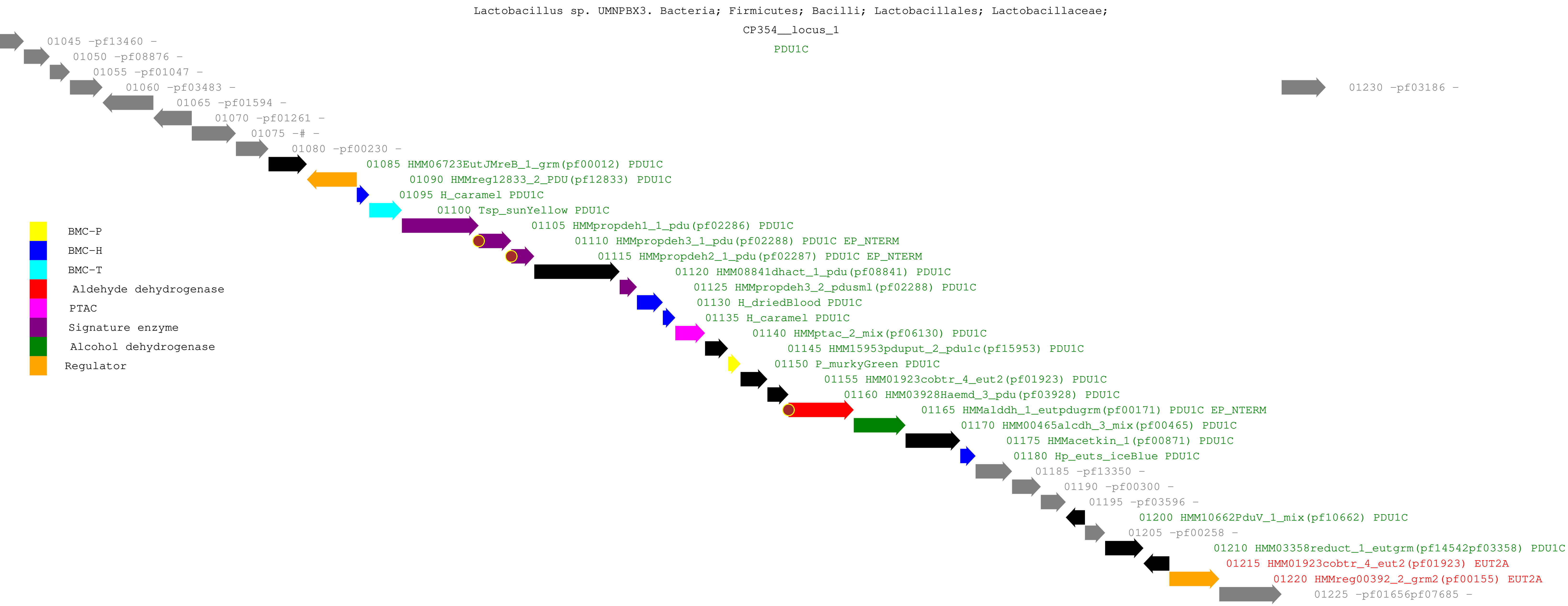
Lactobacillus sp. UMNPBX3. Bacteria; Firmicutes; Bacilli; Lactobacillales; Lactobacillaceae;
CP354__locus_1
PDU1C
01045 -pf13460 -
01050 -pf08876 -
01055 -pf01047 -
01060 -pf03483 -
01065 -pf01594 -
01070 -pf01261 -
01075 -# -
01080 -pf00230 -
01085 HMM06723EutJMreB_1_grm(pf00012) PDU1C
01090 HMMreg12833_2_PDU(pf12833) PDU1C
01095 H_caramel PDU1C
01100 Tsp_sunYellow PDU1C
01105 HMMpropdeh1_1_pdu(pf02286) PDU1C
01110 HMMpropdeh3_1_pdu(pf02288) PDU1C EP_NTERM
01115 HMMpropdeh2_1_pdu(pf02287) PDU1C EP_NTERM
01120 HMM08841dhact_1_pdu(pf08841) PDU1C
01125 HMMpropdeh3_2_pdusml(pf02288) PDU1C
01130 H_driedBlood PDU1C
01135 H_caramel PDU1C
01140 HMMptac_2_mix(pf06130) PDU1C
01145 HMM15953pduput_2_pdu1c(pf15953) PDU1C
01150 P_murkyGreen PDU1C
01155 HMM01923cobtr_4_eut2(pf01923) PDU1C
01160 HMM03928Haemd_3_pdu(pf03928) PDU1C
01165 HMMaldh_1_eutpdugrm(pf00171) PDU1C EP_NTERM
01170 HMM00465alcdh_3_mix(pf00465) PDU1C
01175 HMMacetkin_1(pf00871) PDU1C
01180 Hp_euts_iceBlue PDU1C
01185 -pf13350 -
01190 -pf00300 -
01195 -pf03596 -
01200 HMM10662PduV_1_mix(pf10662) PDU1C
01205 -pf00258 -
01210 HMM03358reduct_1_eutgrm(pf14542pf03358) PDU1C
01215 HMM01923cobtr_4_eut2(pf01923) EUT2A
01220 HMMreg00392_2_grm2(pf00155) EUT2A
01225 -pf01656pf07685 -
01230 -pf03186 -
BMC-P
BMC-H
BMC-T
Aldehyde dehydrogenase
PTAC
Signature enzyme
Alcohol dehydrogenase
Regulator

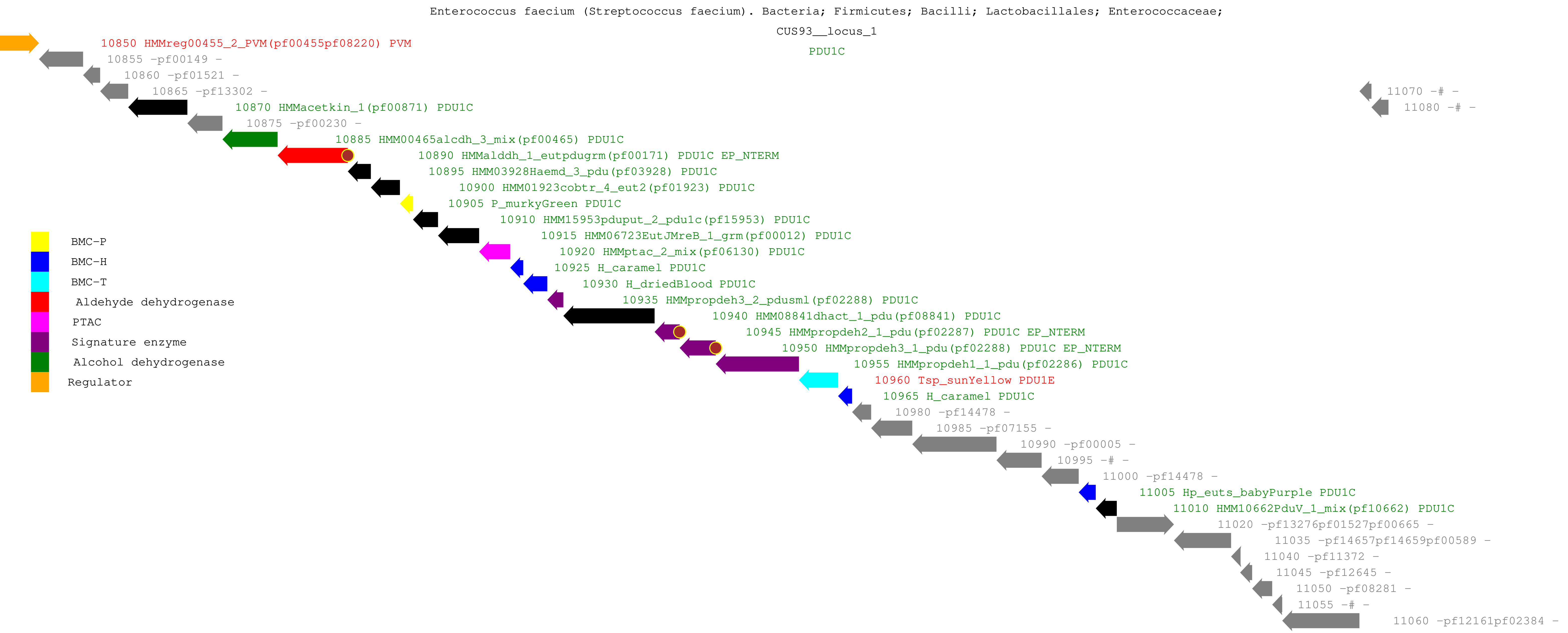
Enterococcus faecium (Streptococcus faecium). Bacteria; Firmicutes; Bacilli; Lactobacillales; Enterococcaceae;
CUS93__locus_1
PDU1C
10850 HMMreg00455_2_PVM(pf00455pf08220) PVM
10855 -pf00149 -
10860 -pf01521 -
10865 -pf13302 -
10870 HMMacetkin_1(pf00871) PDU1C
10875 -pf00230 -
10885 HMM00465alcdh_3_mix(pf00465) PDU1C
10890 HMMalddh_1_eutpdugrm(pf00171) PDU1C EP_NTERM
10895 HMM03928Haemd_3_pdu(pf03928) PDU1C
10900 HMM01923cobtr_4_eut2(pf01923) PDU1C
10905 P_murkyGreen PDU1C
10910 HMM15953pduput_2_pdu1c(pf15953) PDU1C
10915 HMM06723EutJMreB_1_grm(pf00012) PDU1C
10920 HMMptac_2_mix(pf06130) PDU1C
10925 H_caramel PDU1C
10930 H_driedBlood PDU1C
10935 HMMpropdeh3_2_pdusml(pf02288) PDU1C
10940 HMM08841dhact_1_pdu(pf08841) PDU1C
10945 HMMpropdeh2_1_pdu(pf02287) PDU1C EP_NTERM
10950 HMMpropdeh3_1_pdu(pf02288) PDU1C EP_NTERM
10955 HMMpropdeh1_1_pdu(pf02286) PDU1C
10960 Tsp_sunYellow PDU1E
10965 H_caramel PDU1C
10980 -pf14478 -
10985 -pf07155 -
10990 -pf00005 -
10995 -# -
11000 -pf14478 -
11005 Hp_euts_babyPurple PDU1C
11010 HMM10662PduV_1_mix(pf10662) PDU1C
11020 -pf13276pf01527pf00665 -
11035 -pf14657pf14659pf00589 -
11040 -pf11372 -
11045 -pf12645 -
11050 -pf08281 -
11055 -# -
11060 -pf12161pf02384 -
11070 -# -
11080 -# -
BMC-P
BMC-H
BMC-T
Aldehyde dehydrogenase
PTAC
Signature enzyme
Alcohol dehydrogenase
Regulator

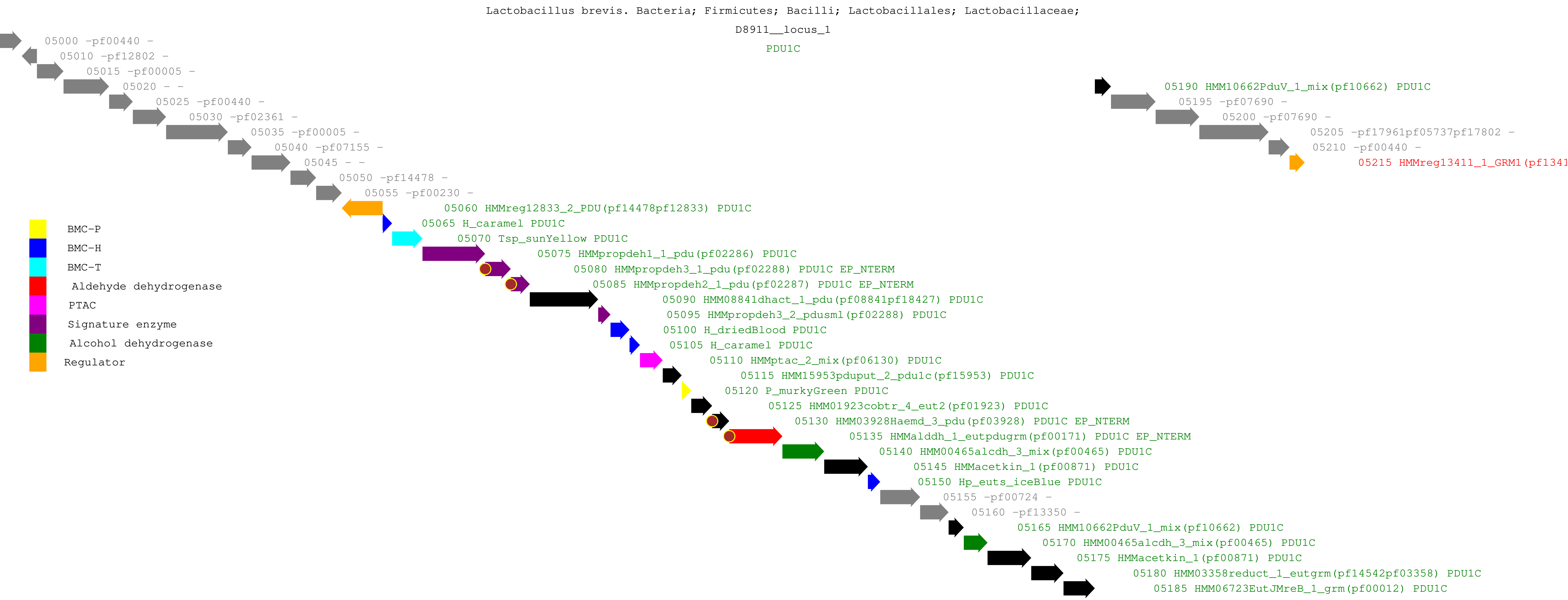

Lactobacillus brevis. Bacteria; Firmicutes; Bacilli; Lactobacillales; Lactobacillaceae;
D8911__locus_1
PDU1C
05000 -pf00440 -
05010 -pf12802 -
05015 -pf00005 -
05020 - -
05025 -pf00440 -
05030 -pf02361 -
05035 -pf00005 -
05040 -pf07155 -
05045 - -
05050 -pf14478 -
05055 -pf00230 -
05060 HMMreg12833_2_PDU(pf14478pf12833) PDU1C
05065 H_caramel PDU1C
05070 Tsp_sunYellow PDU1C
05075 HMMpropdeh1_1_pdu(pf02286) PDU1C
05080 HMMpropdeh3_1_pdu(pf02288) PDU1C EP_NTERM
05085 HMMpropdeh2_1_pdu(pf02287) PDU1C EP_NTERM
05090 HMM08841dhact_1_pdu(pf08841pf18427) PDU1C
05095 HMMpropdeh3_2_pdusml(pf02288) PDU1C
05100 H_driedBlood PDU1C
05105 H_caramel PDU1C
05110 HMMptac_2_mix(pf06130) PDU1C
05115 HMM15953pduput_2_pdu1c(pf15953) PDU1C
05120 P_murkyGreen PDU1C
05125 HMM01923cobtr_4_eut2(pf01923) PDU1C
05130 HMM03928Haemd_3_pdu(pf03928) PDU1C EP_NTERM
05135 HMMalddh_1_eutpdugrm(pf00171) PDU1C EP_NTERM
05140 HMM00465alcdh_3_mix(pf00465) PDU1C
05145 HMMacetkin_1(pf00871) PDU1C
05150 Hp_euts_iceBlue PDU1C
05155 -pf00724 -
05160 -pf13350 -
05165 HMM10662PduV_1_mix(pf10662) PDU1C
05170 HMM00465alcdh_3_mix(pf00465) PDU1C
05175 HMMacetkin_1(pf00871) PDU1C
05180 HMM03358reduct_1_eutgrm(pf14542pf03358) PDU1C
05185 HMM06723EutJMreB_1_grm(pf00012) PDU1C
05190 HMM10662PduV_1_mix(pf10662) PDU1C
05195 -pf07690 -
05200 -pf07690 -
05205 -pf17961pf05737pf17802 -
05210 -pf00440 -
05215 HMMreg13411_1_GRM1(pf1341
BMC-P
BMC-H
BMC-T
Aldehyde dehydrogenase
PTAC
Signature enzyme
Alcohol dehydrogenase
Regulator

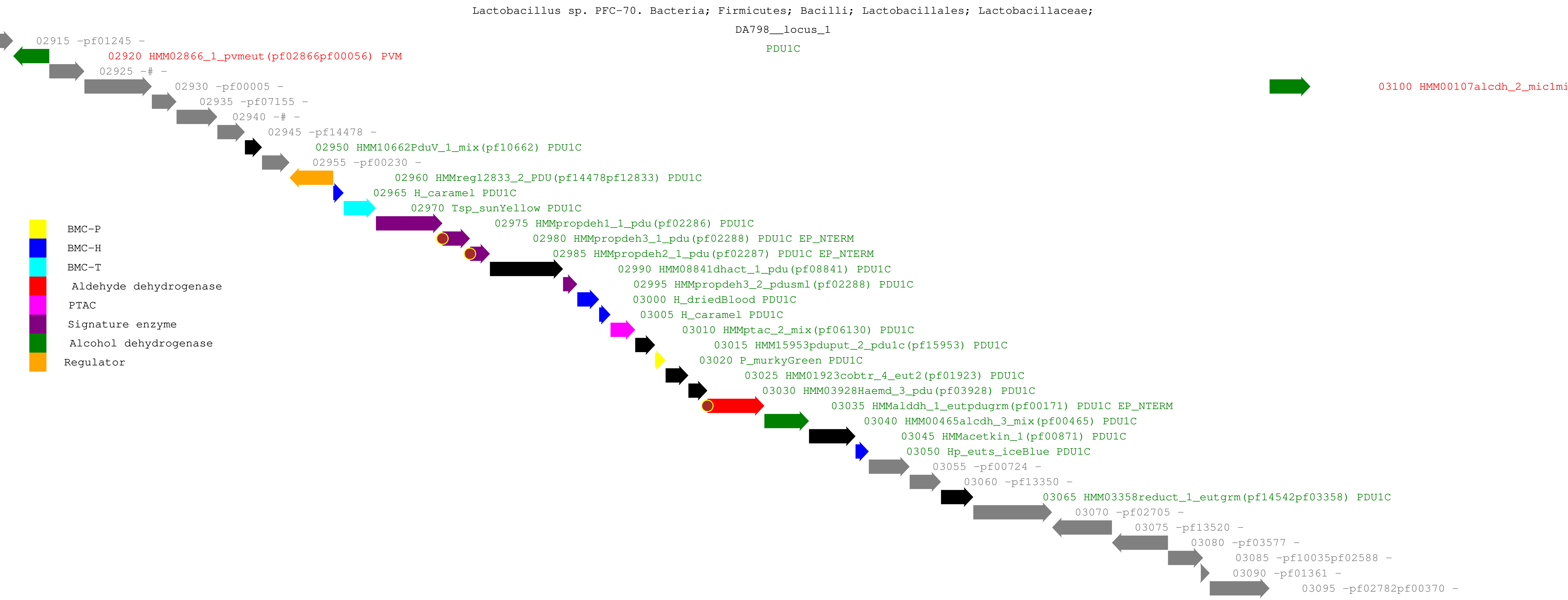
Lactobacillus sp. PFC-70. Bacteria; Firmicutes; Bacilli; Lactobacillales; Lactobacillaceae;
DA798__locus_1
PDU1C
02915 -pf01245 -
02920 HMM02866_1_pvmeut(pf02866pf00056) PVM
02925 -# -
02930 -pf00005 -
02935 -pf07155 -
02940 -# -
02945 -pf14478 -
02950 HMM10662PduV_1_mix(pf10662) PDU1C
02955 -pf00230 -
02960 HMMreg12833_2_PDU(pf14478pf12833) PDU1C
02965 H_caramel PDU1C
02970 Tsp_sunYellow PDU1C
02975 HMMpropdeh1_1_pdu(pf02286) PDU1C
02980 HMMpropdeh3_1_pdu(pf02288) PDU1C EP_NTERM
02985 HMMpropdeh2_1_pdu(pf02287) PDU1C EP_NTERM
02990 HMM08841dhact_1_pdu(pf08841) PDU1C
02995 HMMpropdeh3_2_pdusml(pf02288) PDU1C
03000 H_driedBlood PDU1C
03005 H_caramel PDU1C
03010 HMMptac_2_mix(pf06130) PDU1C
03015 HMM15953pduput_2_pdu1c(pf15953) PDU1C
03020 P_murkyGreen PDU1C
03025 HMM01923cobtr_4_eut2(pf01923) PDU1C
03030 HMM03928Haemd_3_pdu(pf03928) PDU1C
03035 HMMalddh_1_eutpdugrm(pf00171) PDU1C EP_NTERM
03040 HMM00465alcdh_3_mix(pf00465) PDU1C
03045 HMMacetkin_1(pf00871) PDU1C
03050 Hp_euts_iceBlue PDU1C
03055 -pf00724 -
03060 -pf13350 -
03065 HMM03358reduct_1_eutgrm(pf14542pf03358) PDU1C
03070 -pf02705 -
03075 -pf13520 -
03080 -pf03577 -
03085 -pf10035pf02588 -
03090 -pf01361 -
03095 -pf02782pf00370 -
03100 HMM00107alcdh_2_mic1mi
BMC-P
BMC-H
BMC-T
Aldehyde dehydrogenase
PTAC
Signature enzyme
Alcohol dehydrogenase
Regulator

Lactobacillus reuteri. Bacteria; Firmicutes; Bacilli; Lactobacillales; Lactobacillaceae;

DBQ34__locus_1

PDU1C

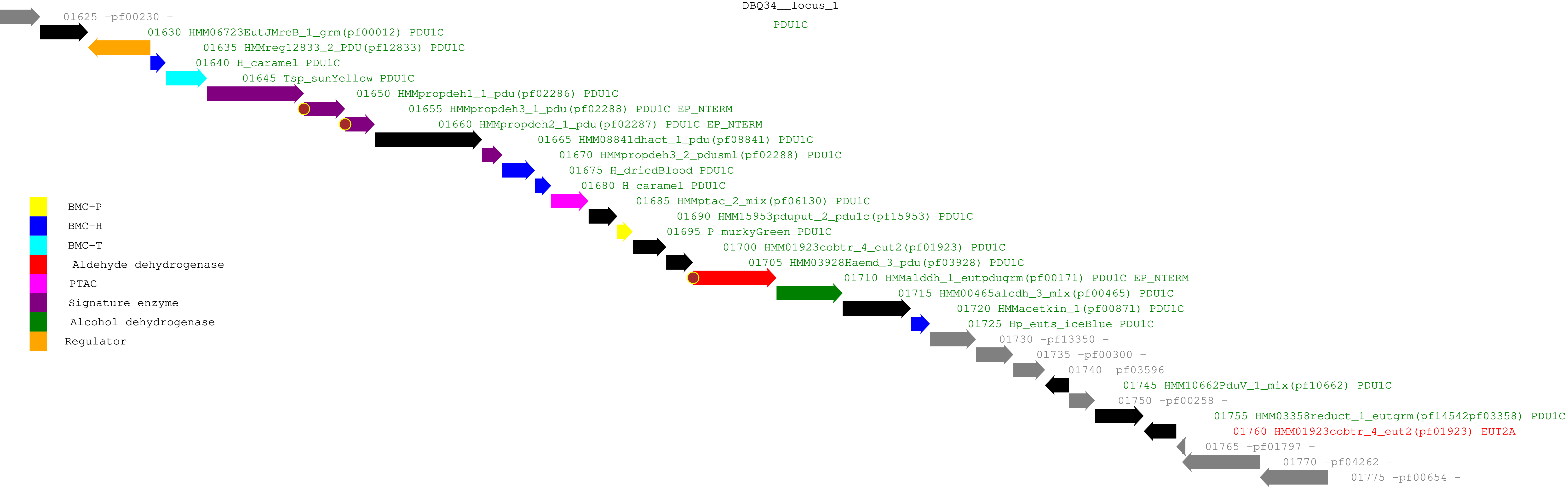

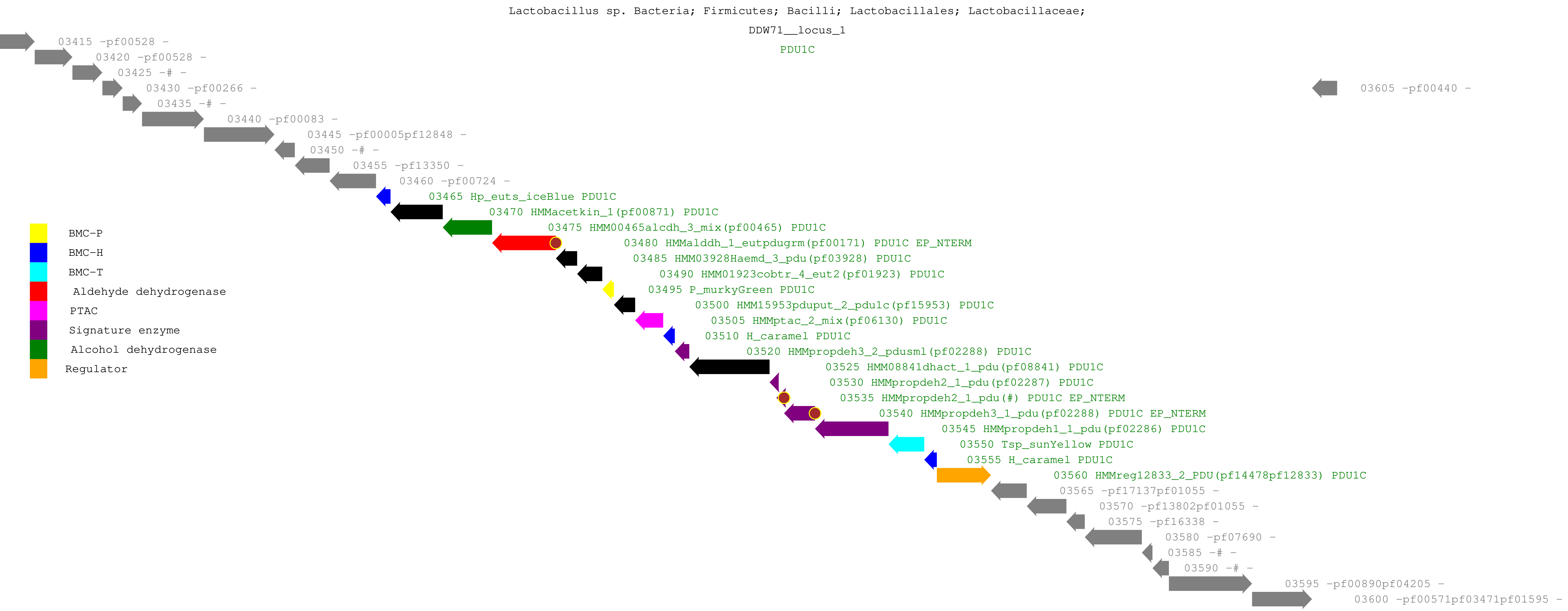
Lactobacillus sp. Bacteria; Firmicutes; Bacilli; Lactobacillales; Lactobacillaceae;
DDW71__locus_1
PDU1C
03415 -pf00528 -
03420 -pf00528 -
03425 -# -
03430 -pf00266 -
03435 -# -
03440 -pf00083 -
03445 -pf00005pf12848 -
03450 -# -
03455 -pf13350 -
03460 -pf00724 -
03465 Hp_euts_iceBlue PDU1C
03470 HMMacetkin_1(pf00871) PDU1C
03475 HMM00465alcdh_3_mix(pf00465) PDU1C
03480 HMMalddh_1_eutpdugrm(pf00171) PDU1C EP_NTERM
03485 HMM03928Haemd_3_pdu(pf03928) PDU1C
03490 HMM01923cobtr_4_eut2(pf01923) PDU1C
03495 P_murkyGreen PDU1C
03500 HMM15953pduput_2_pdu1c(pf15953) PDU1C
03505 HMMptac_2_mix(pf06130) PDU1C
03510 H_caramel PDU1C
03520 HMMpropdeh3_2_pdusml(pf02288) PDU1C
03525 HMM08841dhact_1_pdu(pf08841) PDU1C
03530 HMMpropdeh2_1_pdu(pf02287) PDU1C
03535 HMMpropdeh2_1_pdu(#) PDU1C EP_NTERM
03540 HMMpropdeh3_1_pdu(pf02288) PDU1C EP_NTERM
03545 HMMpropdeh1_1_pdu(pf02286) PDU1C
03550 Tsp_sunYellow PDU1C
03555 H_caramel PDU1C
03560 HMMreg12833_2_PDU(pf14478pf12833) PDU1C
03565 -pf17137pf01055 -
03570 -pf13802pf01055 -
03575 -pf16338 -
03580 -pf07690 -
03585 -# -
03590 -# -
03595 -pf00890pf04205 -
03600 -pf00571pf03471pf01595 -
03605 -pf00440 -
BMC-P
BMC-H
BMC-T
Aldehyde dehydrogenase
PTAC
Signature enzyme
Alcohol dehydrogenase
Regulator

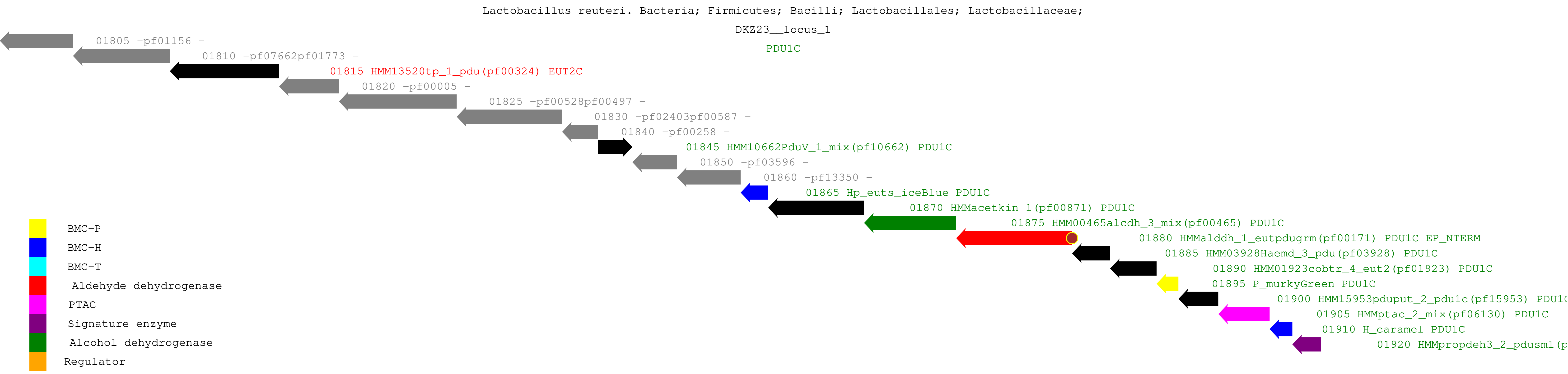
Lactobacillus reuteri. Bacteria; Firmicutes; Bacilli; Lactobacillales; Lactobacillaceae;
DKZ23__locus_1
PDU1C
01805 -pf01156 -
01810 -pf07662pf01773 -
01815 HMM13520tp_1_pdu(pf00324) EUT2C
01820 -pf00005 -
01825 -pf00528pf00497 -
01830 -pf02403pf00587 -
01840 -pf00258 -
01845 HMM10662PduV_1_mix(pf10662) PDU1C
01850 -pf03596 -
01860 -pf13350 -
01865 Hp_euts_iceBlue PDU1C
01870 HMMacetkin_1(pf00871) PDU1C
01875 HMM00465alcdh_3_mix(pf00465) PDU1C
01880 HMMaldh_1_eutpdugrm(pf00171) PDU1C EP_NTERM
01885 HMM03928Haemd_3_pdu(pf03928) PDU1C
01890 HMM01923cobtr_4_eut2(pf01923) PDU1C
01895 P_murkyGreen PDU1C
01900 HMM15953pduput_2_pdu1c(pf15953) PDU1
01905 HMMptac_2_mix(pf06130) PDU1C
01910 H_caramel PDU1C
01920 HMMpropdeh3_2_pdusml(p
BMC-P
BMC-H
BMC-T
Aldehyde dehydrogenase
PTAC
Signature enzyme
Alcohol dehydrogenase
Regulator

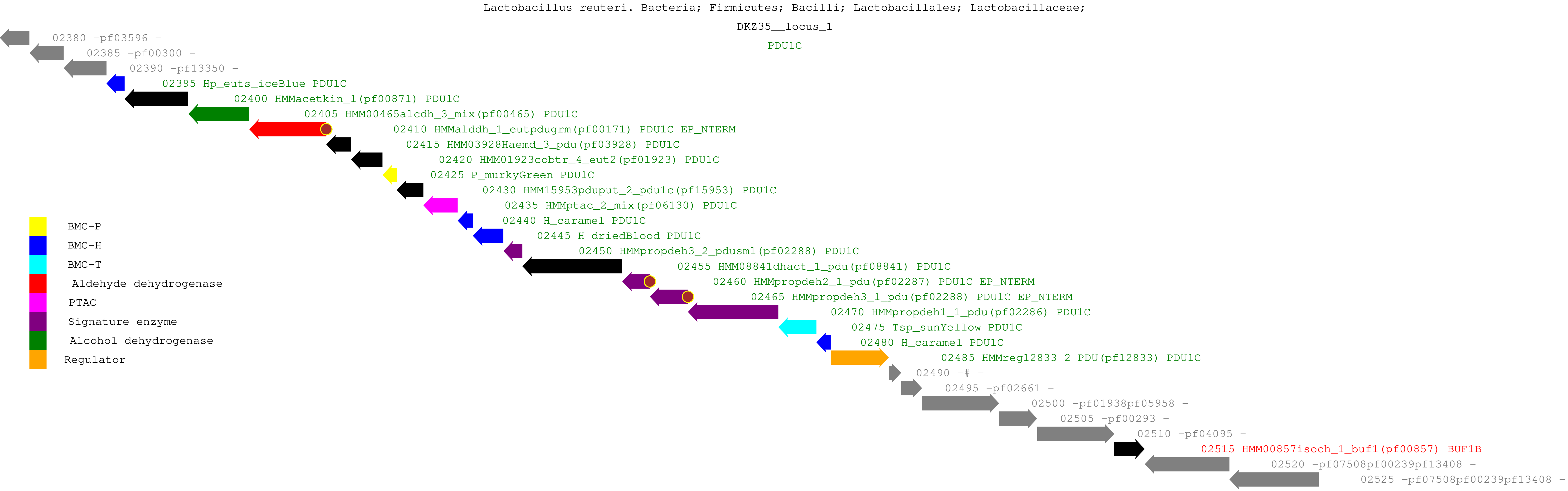
Lactobacillus reuteri. Bacteria; Firmicutes; Bacilli; Lactobacillales; Lactobacillaceae;
DKZ35__locus_1
PDU1C
02380 -pf03596 -
02385 -pf00300 -
02390 -pf13350 -
02395 Hp_euts_iceBlue PDU1C
02400 HMMacetkin_1(pf00871) PDU1C
02405 HMM00465alcdh_3_mix(pf00465) PDU1C
02410 HMMalddh_1_eutpdugrm(pf00171) PDU1C EP_NTERM
02415 HMM03928Haemd_3_pdu(pf03928) PDU1C
02420 HMM01923cobtr_4_eut2(pf01923) PDU1C
02425 P_murkyGreen PDU1C
02430 HMM15953pduput_2_pdu1c(pf15953) PDU1C
02435 HMMptac_2_mix(pf06130) PDU1C
02440 H_caramel PDU1C
02445 H_driedBlood PDU1C
02450 HMMpropdeh3_2_pdusml(pf02288) PDU1C
02455 HMM08841dhact_1_pdu(pf08841) PDU1C
02460 HMMpropdeh2_1_pdu(pf02287) PDU1C EP_NTERM
02465 HMMpropdeh3_1_pdu(pf02288) PDU1C EP_NTERM
02470 HMMpropdeh1_1_pdu(pf02286) PDU1C
02475 Tsp_sunYellow PDU1C
02480 H_caramel PDU1C
02485 HMMreg12833_2_PDU(pf12833) PDU1C
02490 -# -
02495 -pf02661 -
02500 -pf01938pf05958 -
02505 -pf00293 -
02510 -pf04095 -
02515 HMM00857isoch_1_buf1(pf00857) BUF1B
02520 -pf07508pf00239pf13408 -
02525 -pf07508pf00239pf13408 -
BMC-P
BMC-H
BMC-T
Aldehyde dehydrogenase
PTAC
Signature enzyme
Alcohol dehydrogenase
Regulator

Lactobacillus sp. Bacteria; Firmicutes; Bacilli; Lactobacillales; Lactobacillaceae;

DUD28__locus_1_main_sat

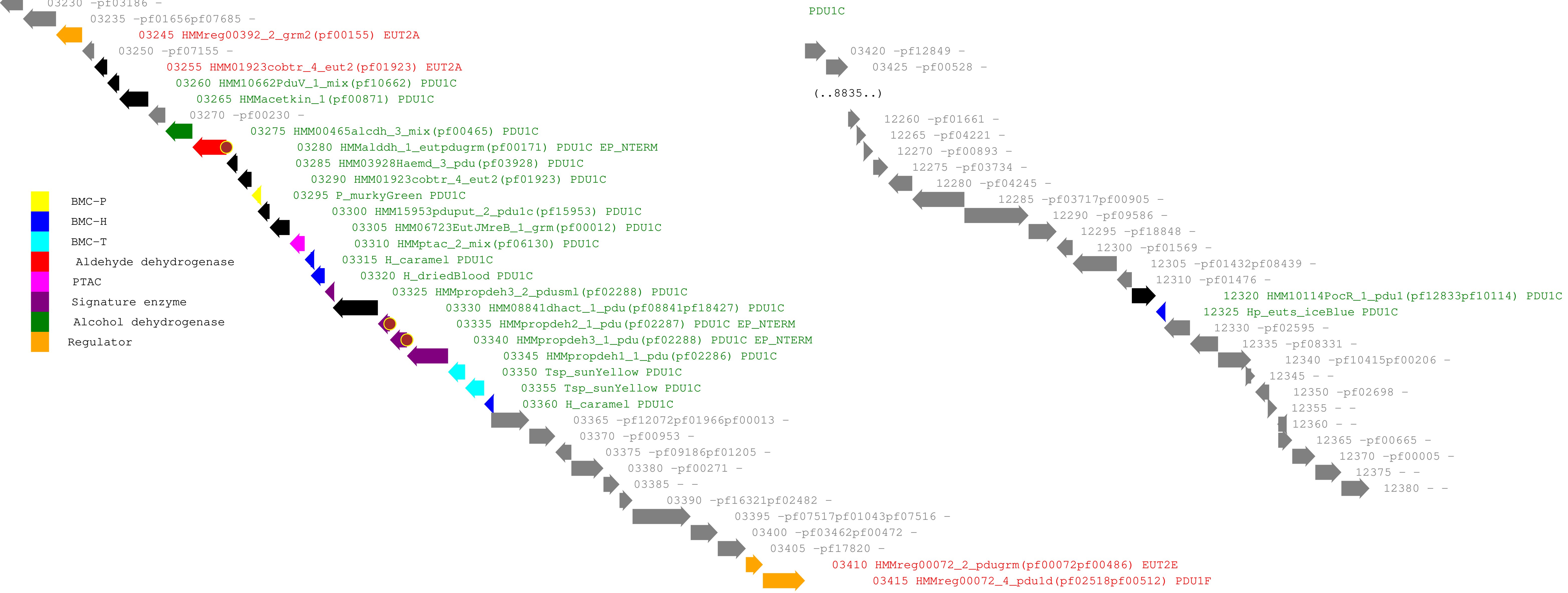

Lactobacillus sp. Bacteria; Firmicutes; Bacilli; Lactobacillales; Lactobacillaceae;

DUD34__locus_2_main_sat

PDU1C

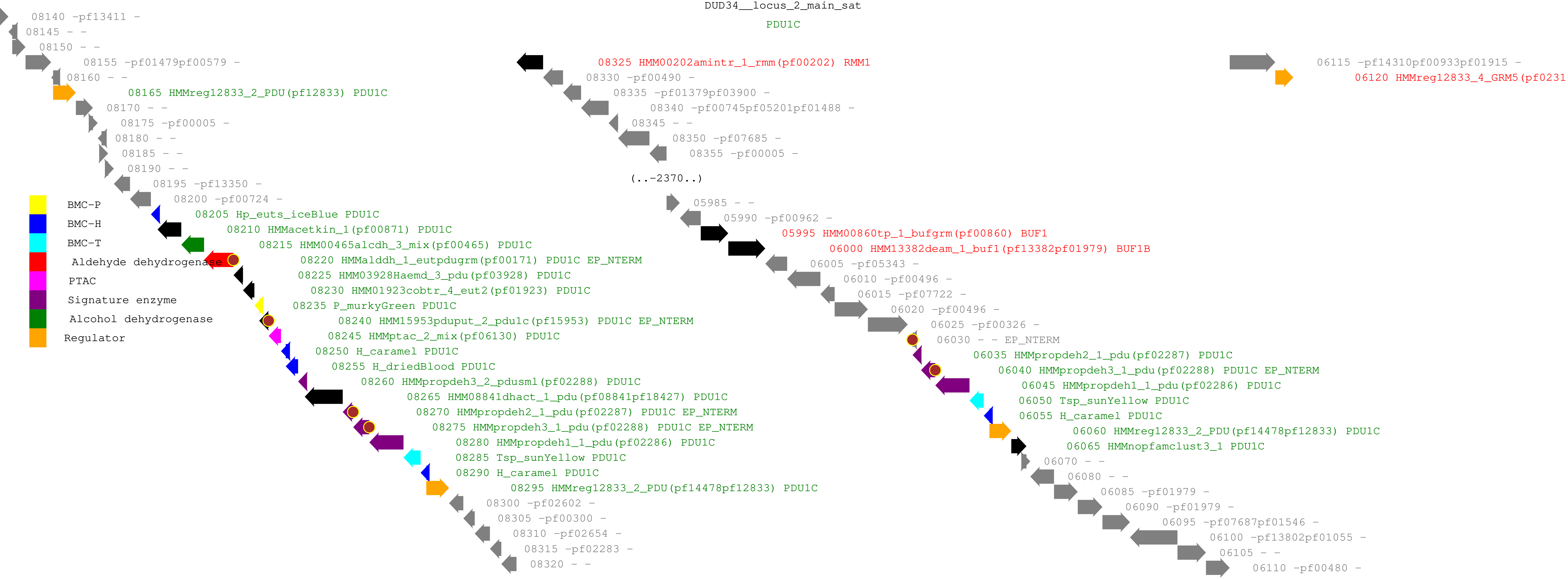

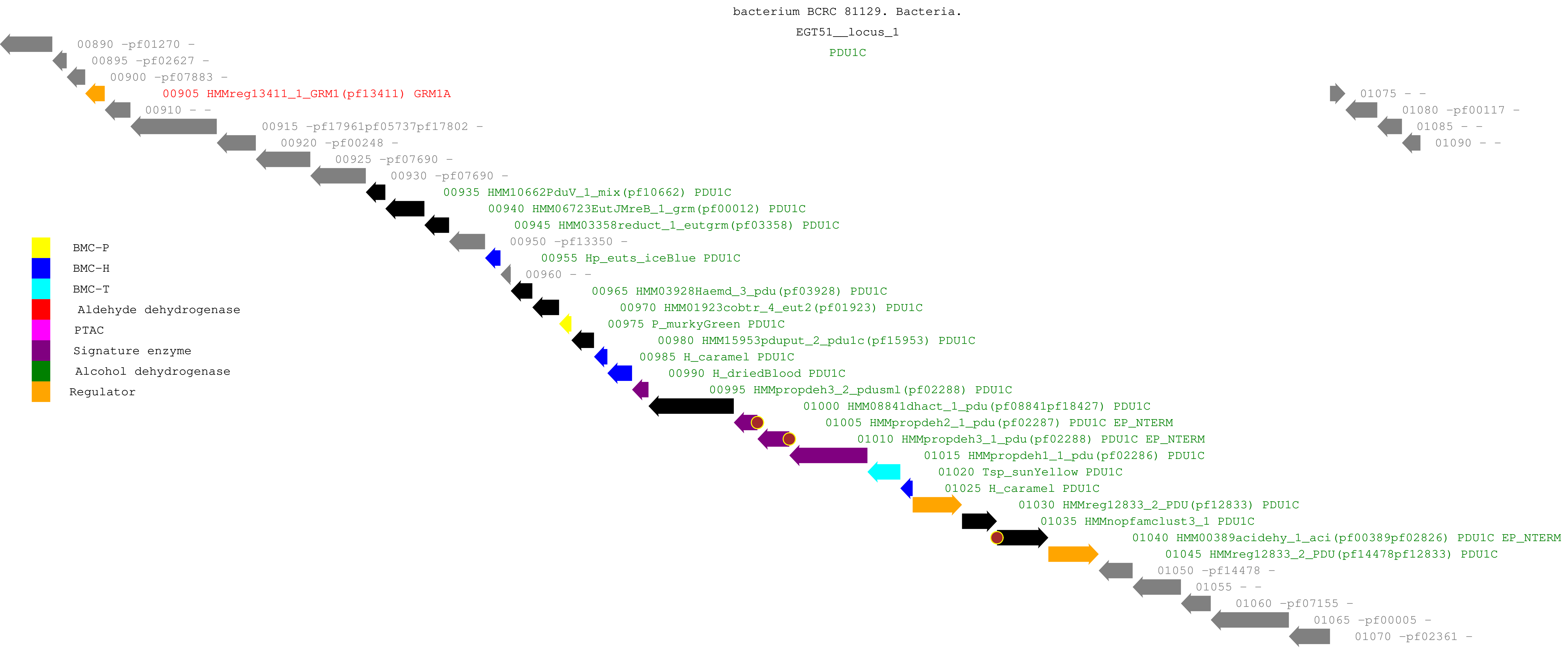

bacterium BCRC 81129. Bacteria.
EGT51__locus_1
PDU1C
00890 -pf01270 -
00895 -pf02627 -
00900 -pf07883 -
00905 HMMreg13411_1_GRM1(pf13411) GRM1A
00910 - -
00915 -pf17961pf05737pf17802 -
00920 -pf00248 -
00925 -pf07690 -
00930 -pf07690 -
00935 HMM10662PduV_1_mix(pf10662) PDU1C
00940 HMM06723EutJMreB_1_grm(pf00012) PDU1C
00945 HMM03358reduct_1_eutgrm(pf03358) PDU1C
00950 -pf13350 -
00955 Hp_euts_iceBlue PDU1C
00960 - -
00965 HMM03928Haemd_3_pdu(pf03928) PDU1C
00970 HMM01923cobtr_4_eut2(pf01923) PDU1C
00975 P_murkyGreen PDU1C
00980 HMM15953pduput_2_pdu1c(pf15953) PDU1C
00985 H_caramel PDU1C
00990 H_driedBlood PDU1C
00995 HMMpropdeh3_2_pdusml(pf02288) PDU1C
01000 HMM08841dhact_1_pdu(pf08841pf18427) PDU1C
01005 HMMpropdeh2_1_pdu(pf02287) PDU1C EP_NTERM
01010 HMMpropdeh3_1_pdu(pf02288) PDU1C EP_NTERM
01015 HMMpropdeh1_1_pdu(pf02286) PDU1C
01020 Tsp_sunYellow PDU1C
01025 H_caramel PDU1C
01030 HMMreg12833_2_PDU(pf12833) PDU1C
01035 HMMnopfamclust3_1 PDU1C
01040 HMM00389acidehy_1_aci(pf00389pf02826) PDU1C EP_NTERM
01045 HMMreg12833_2_PDU(pf14478pf12833) PDU1C
01050 -pf14478 -
01055 - -
01060 -pf07155 -
01065 -pf00005 -
01070 -pf02361 -
01075 - -
01080 -pf00117 -
01085 - -
01090 - -
BMC-P
BMC-H
BMC-T
Aldehyde dehydrogenase
PTAC
Signature enzyme
Alcohol dehydrogenase
Regulator

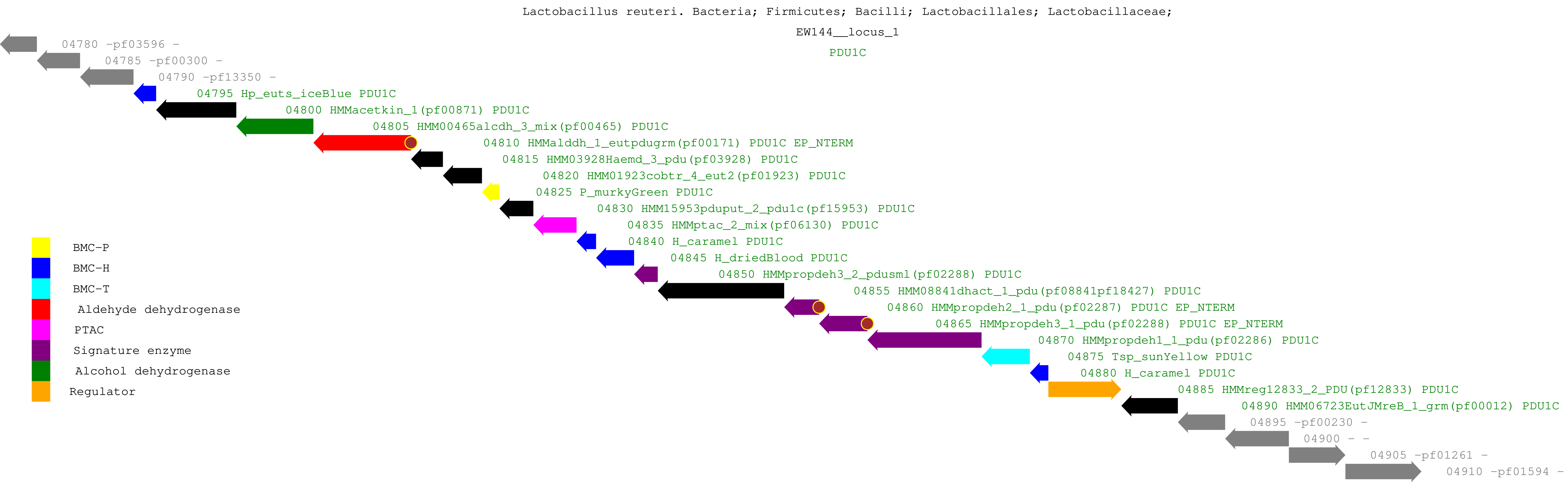

Lactobacillus reuteri. Bacteria; Firmicutes; Bacilli; Lactobacillales; Lactobacillaceae;
EW144__locus_1
PDU1C
04780 -pf03596 -
04785 -pf00300 -
04790 -pf13350 -
04795 Hp_euts_iceBlue PDU1C
04800 HMMacetkin_1(pf00871) PDU1C
04805 HMM00465alcdh_3_mix(pf00465) PDU1C
04810 HMMalddh_1_eutpdugrm(pf00171) PDU1C EP_NTERM
04815 HMM03928Haemd_3_pdu(pf03928) PDU1C
04820 HMM01923cobtr_4_eut2(pf01923) PDU1C
04825 P_murkyGreen PDU1C
04830 HMM15953pduput_2_pdu1c(pf15953) PDU1C
04835 HMMptac_2_mix(pf06130) PDU1C
04840 H_caramel PDU1C
04845 H_driedBlood PDU1C
04850 HMMpropdeh3_2_pdusml(pf02288) PDU1C
04855 HMM08841dhact_1_pdu(pf08841pf18427) PDU1C
04860 HMMpropdeh2_1_pdu(pf02287) PDU1C EP_NTERM
04865 HMMpropdeh3_1_pdu(pf02288) PDU1C EP_NTERM
04870 HMMpropdeh1_1_pdu(pf02286) PDU1C
04875 Tsp_sunYellow PDU1C
04880 H_caramel PDU1C
04885 HMMreg12833_2_PDU(pf12833) PDU1C
04890 HMM06723EutJMreB_1_grm(pf00012) PDU1C
04895 -pf00230 -
04900 - -
04905 -pf01261 -
04910 -pf01594 -
BMC-P
BMC-H
BMC-T
Aldehyde dehydrogenase
PTAC
Signature enzyme
Alcohol dehydrogenase
Regulator

Lactobacillus coryniformis subsp. torquens DSM 20004 = KCTC 3535. Bacteria; Firmicutes; Bacilli; Lactobacillales; Lactobacillaceae;

FC16_GL_locus_1

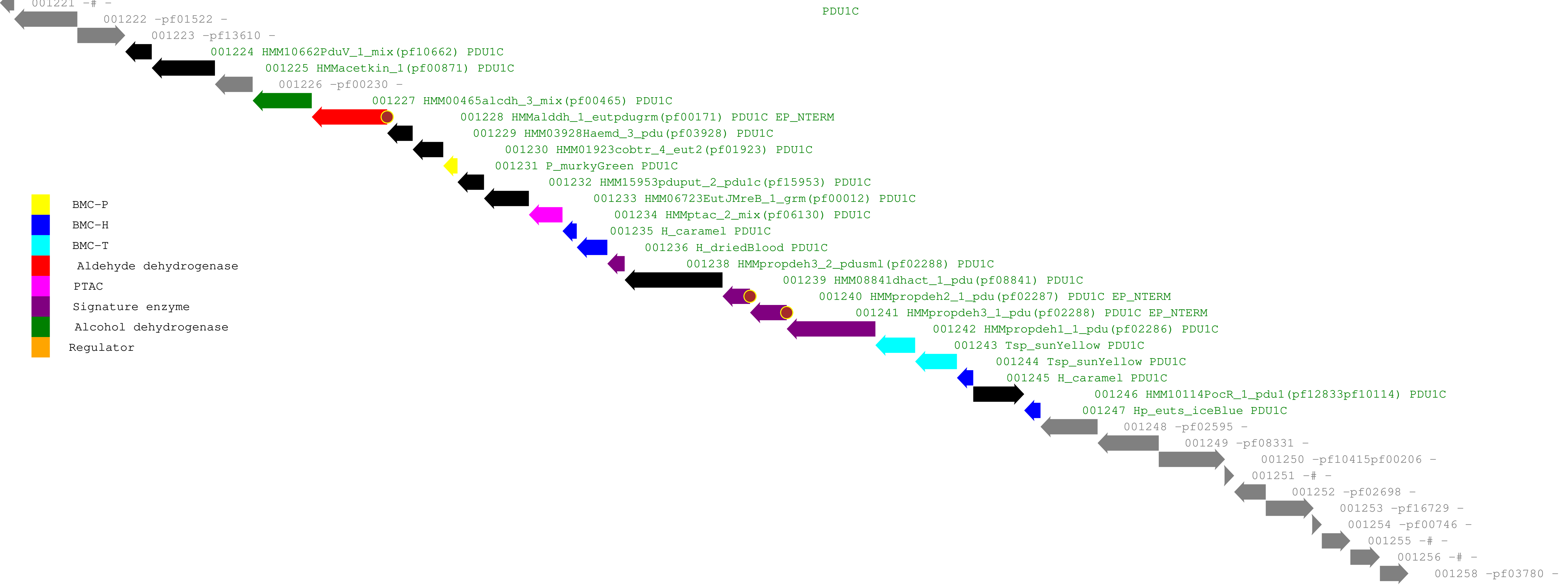

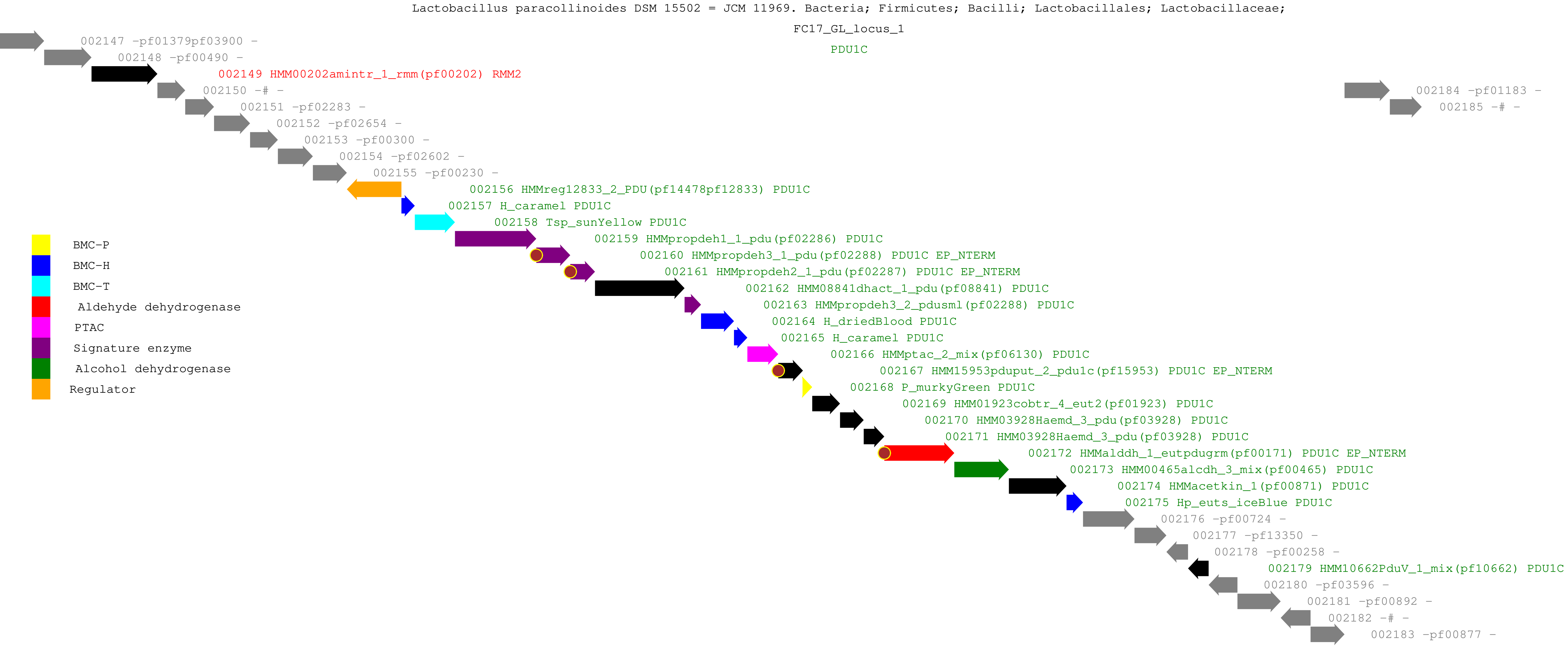

Lactobacillus paracollinoides DSM 15502 = JCM 11969. Bacteria; Firmicutes; Bacilli; Lactobacillales; Lactobacillaceae;
FC17_GL_locus_1
PDU1C
002147 -pf01379pf03900 -
002148 -pf00490 -
002149 HMM00202amintr_1_rmm(pf00202) RMM2
002150 -# -
002151 -pf02283 -
002152 -pf02654 -
002153 -pf00300 -
002154 -pf02602 -
002155 -pf00230 -
002156 HMMreg12833_2_PDU(pf14478pf12833) PDU1C
002157 H_caramel PDU1C
002158 Tsp_sunYellow PDU1C
002159 HMMpropdeh1_1_pdu(pf02286) PDU1C
002160 HMMpropdeh3_1_pdu(pf02288) PDU1C EP_NTERM
002161 HMMpropdeh2_1_pdu(pf02287) PDU1C EP_NTERM
002162 HMM08841dhact_1_pdu(pf08841) PDU1C
002163 HMMpropdeh3_2_pdusml(pf02288) PDU1C
002164 H_driedBlood PDU1C
002165 H_caramel PDU1C
002166 HMMptac_2_mix(pf06130) PDU1C
002167 HMM15953pduput_2_pdu1c(pf15953) PDU1C EP_NTERM
002168 P_murkyGreen PDU1C
002169 HMM01923cobtr_4_eut2(pf01923) PDU1C
002170 HMM03928Haemd_3_pdu(pf03928) PDU1C
002171 HMM03928Haemd_3_pdu(pf03928) PDU1C
002172 HMMalddh_1_eutpdugrm(pf00171) PDU1C EP_NTERM
002173 HMM00465alcdh_3_mix(pf00465) PDU1C
002174 HMMacetkin_1(pf00871) PDU1C
002175 Hp_euts_iceBlue PDU1C
002176 -pf00724 -
002177 -pf13350 -
002178 -pf00258 -
002179 HMM10662PduV_1_mix(pf10662) PDU1C
002180 -pf03596 -
002181 -pf00892 -
002182 -# -
002183 -pf00877 -
002184 -pf01183 -
002185 -# -
BMC-P
BMC-H
BMC-T
Aldehyde dehydrogenase
PTAC
Signature enzyme
Alcohol dehydrogenase
Regulator

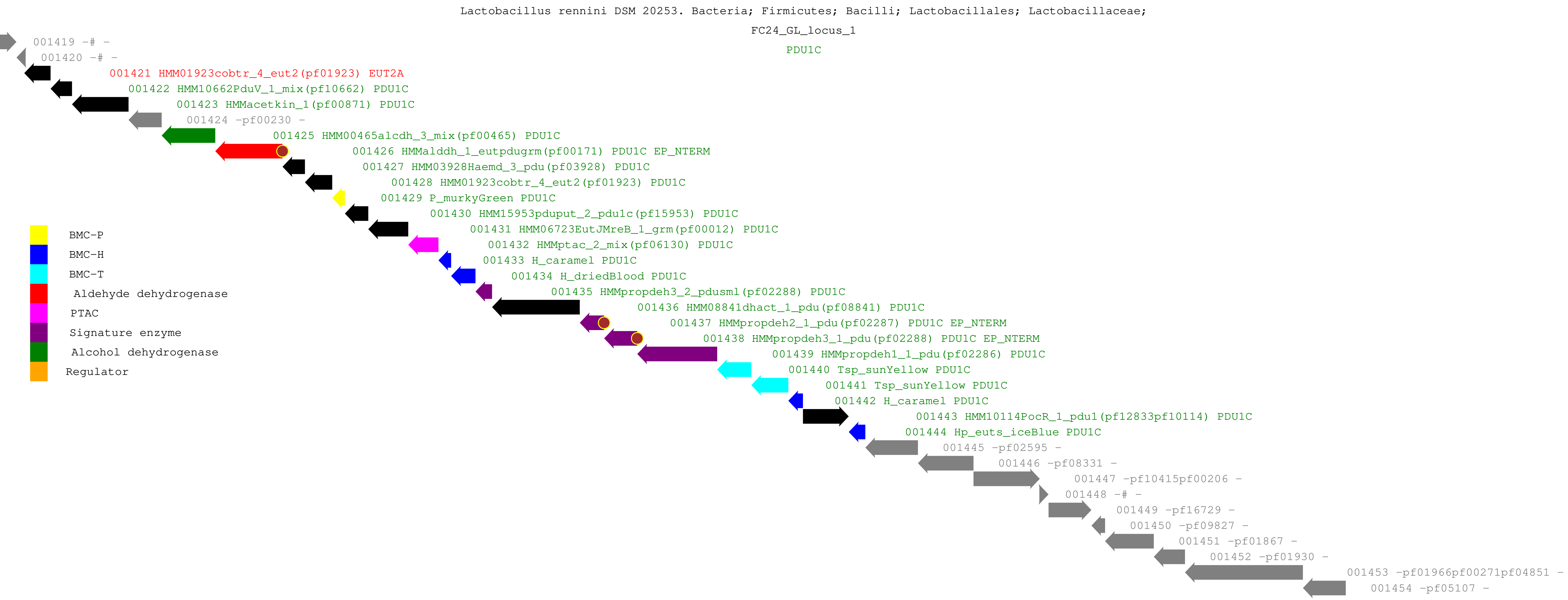
Lactobacillus rennini DSM 20253. Bacteria; Firmicutes; Bacilli; Lactobacillales; Lactobacillaceae;
FC24_GL_locus_1
PDU1C
001419 -# -
001420 -# -
001421 HMM01923cobtr_4_eut2(pf01923) EUT2A
001422 HMM10662PduV_1_mix(pf10662) PDU1C
001423 HMMacetkin_1(pf00871) PDU1C
001424 -pf00230 -
001425 HMM00465alcdh_3_mix(pf00465) PDU1C
001426 HMMalddh_1_eutpdugrm(pf00171) PDU1C EP_NTERM
001427 HMM03928Haemd_3_pdu(pf03928) PDU1C
001428 HMM01923cobtr_4_eut2(pf01923) PDU1C
001429 P_murkyGreen PDU1C
001430 HMM15953pduput_2_pdu1c(pf15953) PDU1C
001431 HMM06723EutJMreB_1_grm(pf00012) PDU1C
001432 HMMptac_2_mix(pf06130) PDU1C
001433 H_caramel PDU1C
001434 H_driedBlood PDU1C
001435 HMMpropdeh3_2_pdusml(pf02288) PDU1C
001436 HMM08841dhact_1_pdu(pf08841) PDU1C
001437 HMMpropdeh2_1_pdu(pf02287) PDU1C EP_NTERM
001438 HMMpropdeh3_1_pdu(pf02288) PDU1C EP_NTERM
001439 HMMpropdeh1_1_pdu(pf02286) PDU1C
001440 Tsp_sunYellow PDU1C
001441 Tsp_sunYellow PDU1C
001442 H_caramel PDU1C
001443 HMM10114PocR_1_pdu1(pf12833pf10114) PDU1C
001444 Hp_euts_iceBlue PDU1C
001445 -pf02595 -
001446 -pf08331 -
001447 -pf10415pf00206 -
001448 -# -
001449 -pf16729 -
001450 -pf09827 -
001451 -pf01867 -
001452 -pf01930 -
001453 -pf01966pf00271pf04851 -
001454 -pf05107 -
BMC-P
BMC-H
BMC-T
Aldehyde dehydrogenase
PTAC
Signature enzyme
Alcohol dehydrogenase
Regulator

Lactobacillus versmoldensis DSM 14857 = KCTC 3814. Bacteria; Firmicutes; Bacilli; Lactobacillales; Lactobacillaceae;

FC27_GL_locus_1

PDU1C

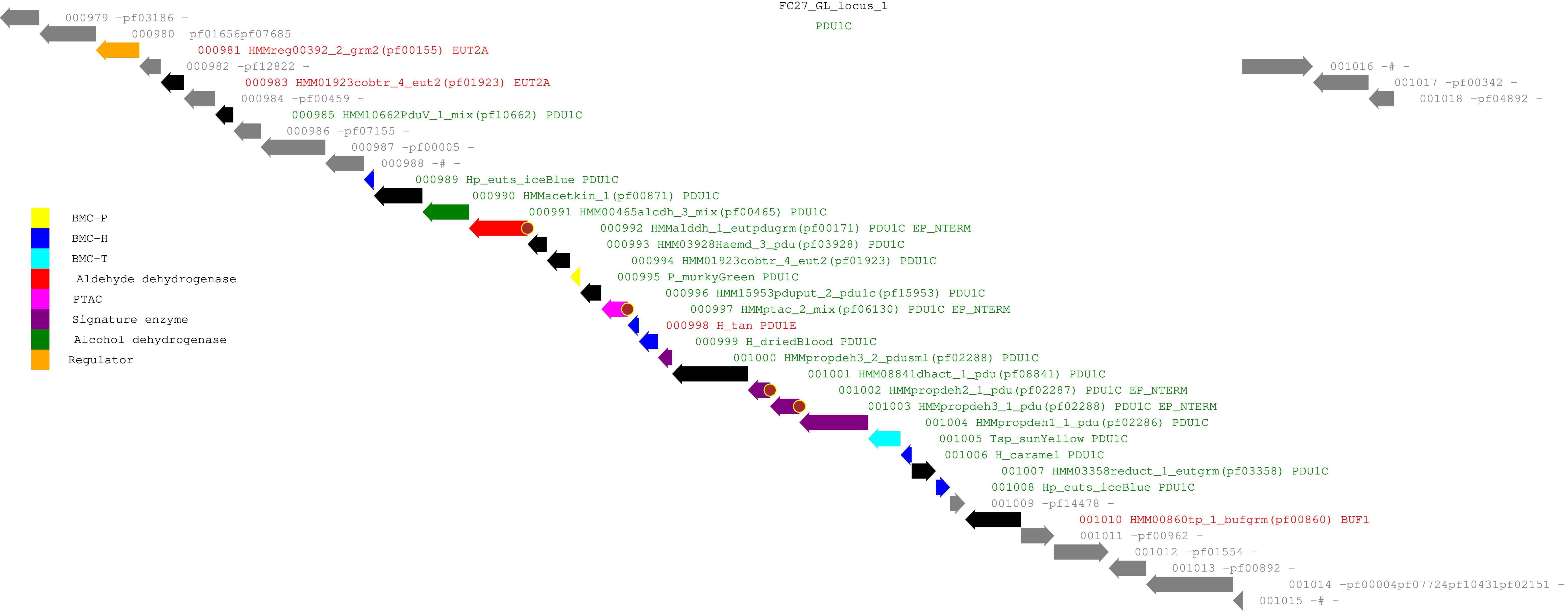

Lactobacillus fuchuensis DSM 14340 = JCM 11249. Bacteria; Firmicutes; Bacilli; Lactobacillales; Lactobacillaceae;

FC69_GL_locus_1

PDU1C

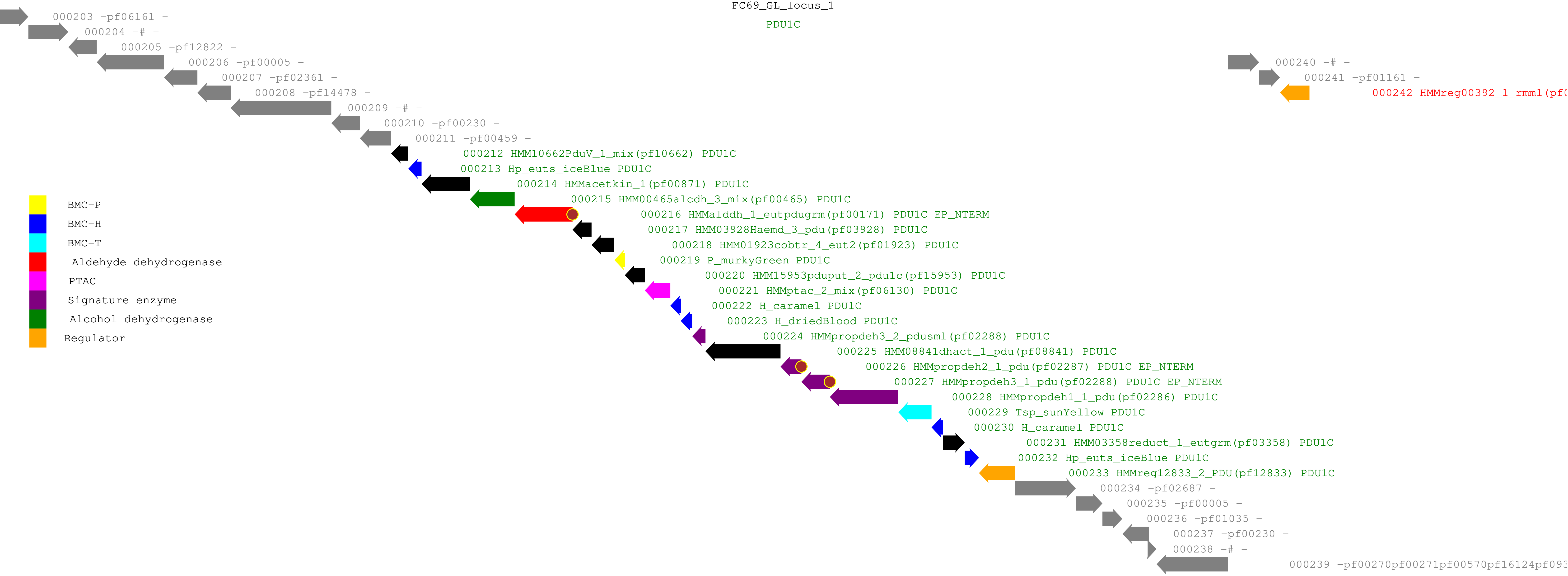

Lactobacillus collinoides DSM 20515 = JCM 1123. Bacteria; Firmicutes; Bacilli; Lactobacillales; Lactobacillaceae;

FC82_GL_locus_1

PDU1C

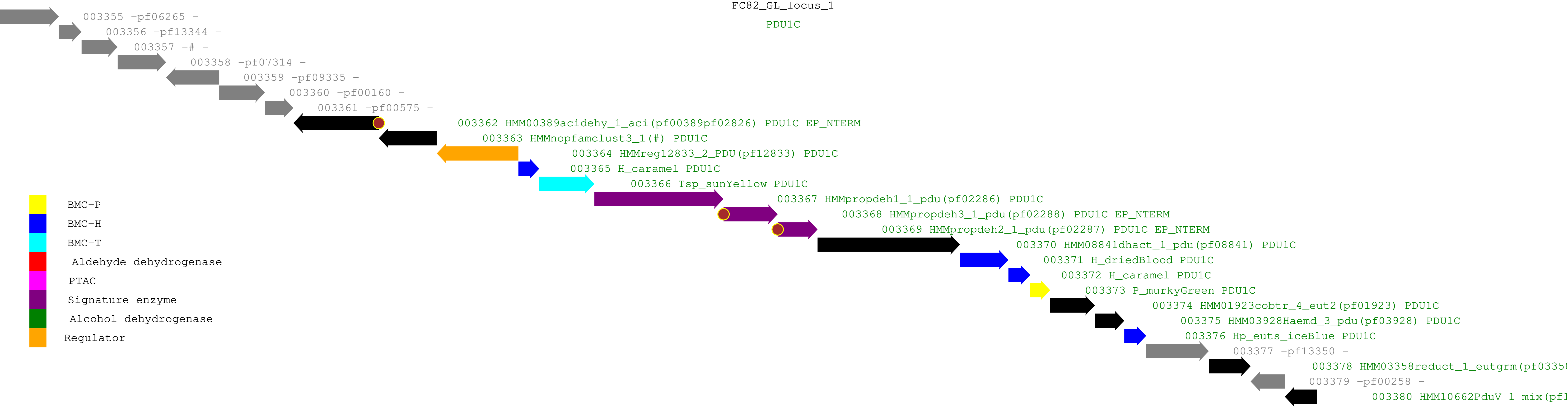

Lactobacillus diolivorans DSM 14421. Bacteria; Firmicutes; Bacilli; Lactobacillales; Lactobacillaceae;

FC85_GL_locus_1

PDU1C

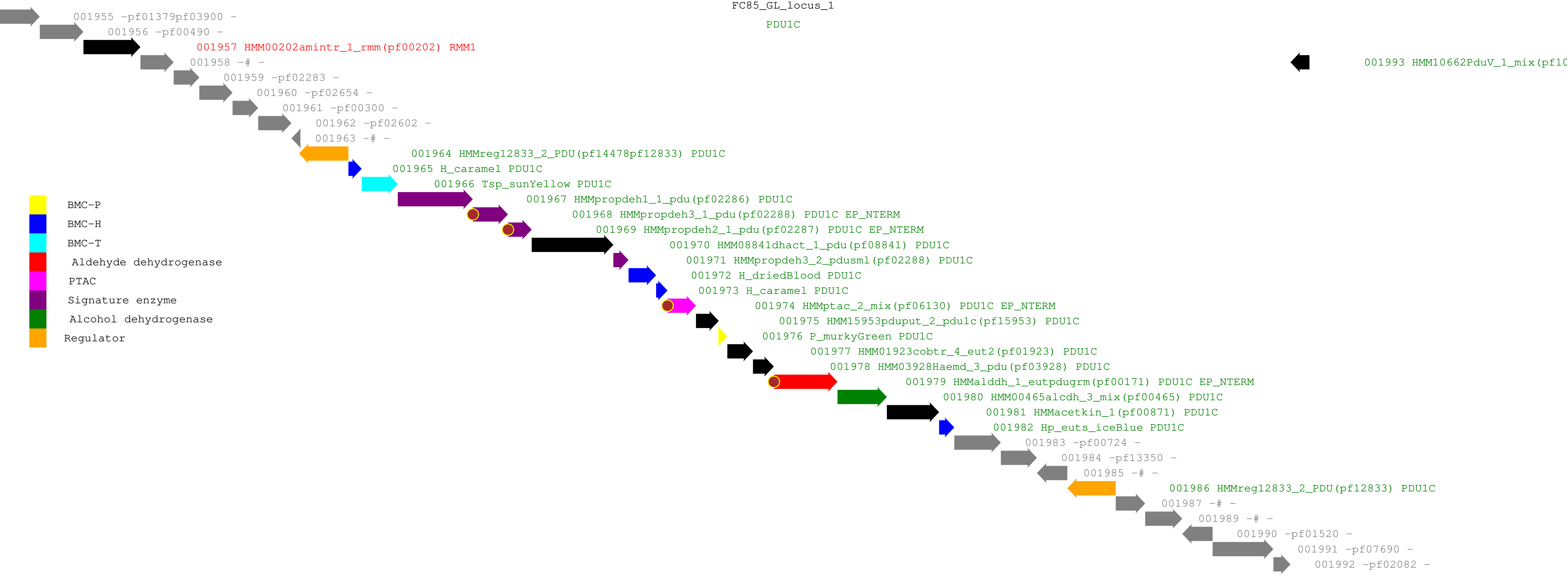

Lactobacillus graminis DSM 20719. Bacteria; Firmicutes; Bacilli; Lactobacillales; Lactobacillaceae;

FC90_GL_locus_1

PDU1C

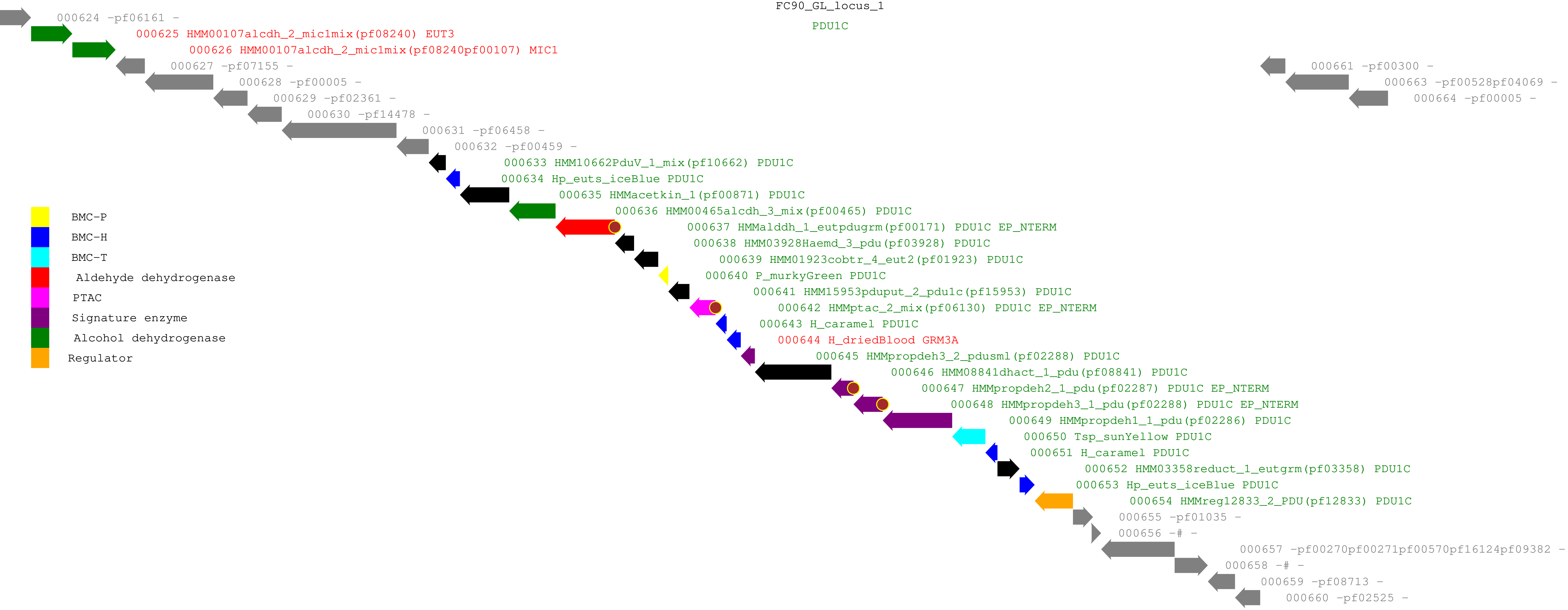

Lactobacillus kimchicus JCM 15530. Bacteria; Firmicutes; Bacilli; Lactobacillales; Lactobacillaceae;

FC96_GL_locus_1

PDU1C

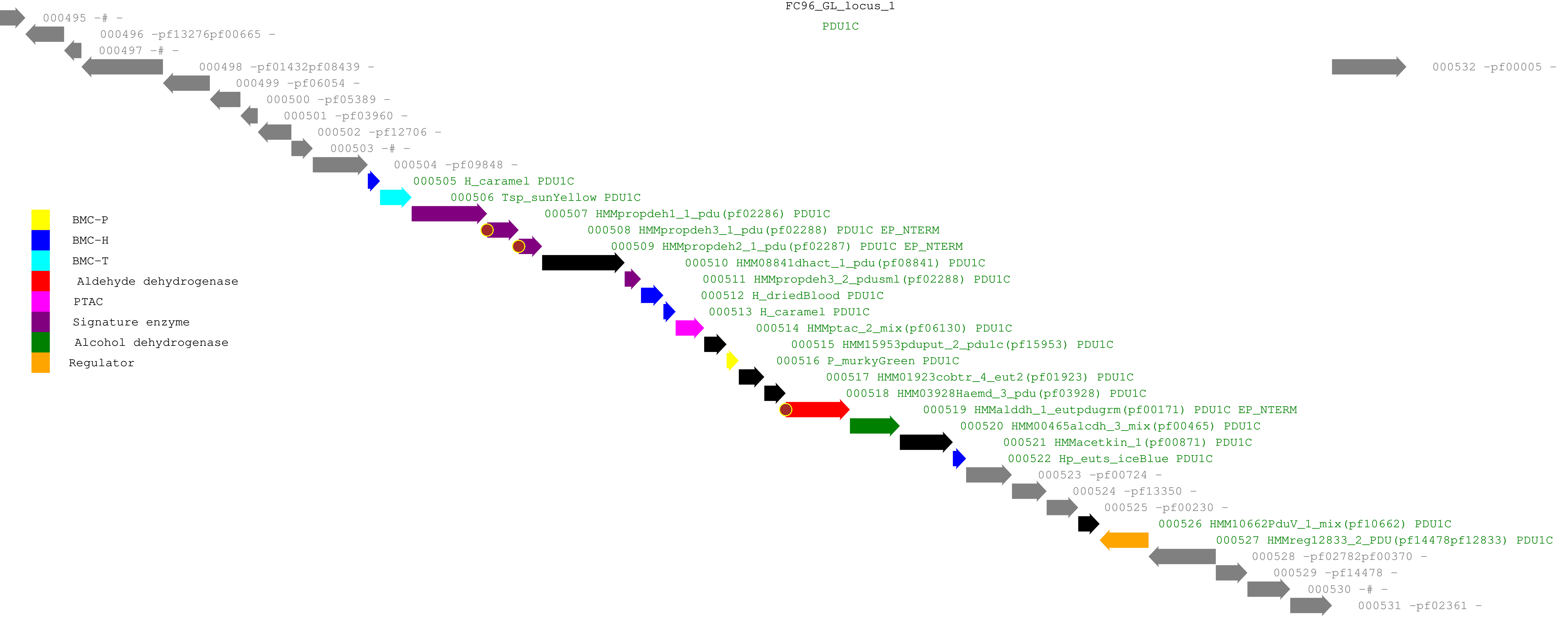

Lactobacillus kisonensis DSM 19906 = JCM 15041. Bacteria; Firmicutes; Bacilli; Lactobacillales; Lactobacillaceae;

FC98_GL_locus_1

PDU1C

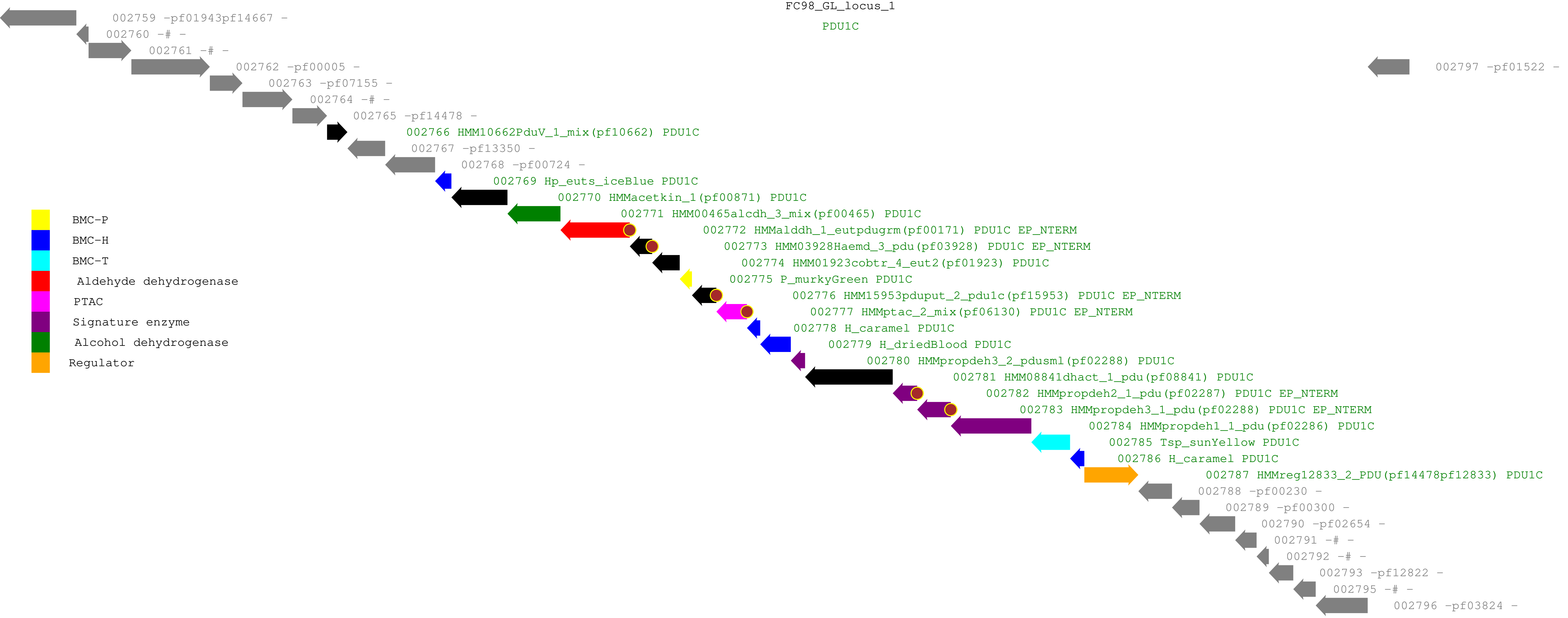

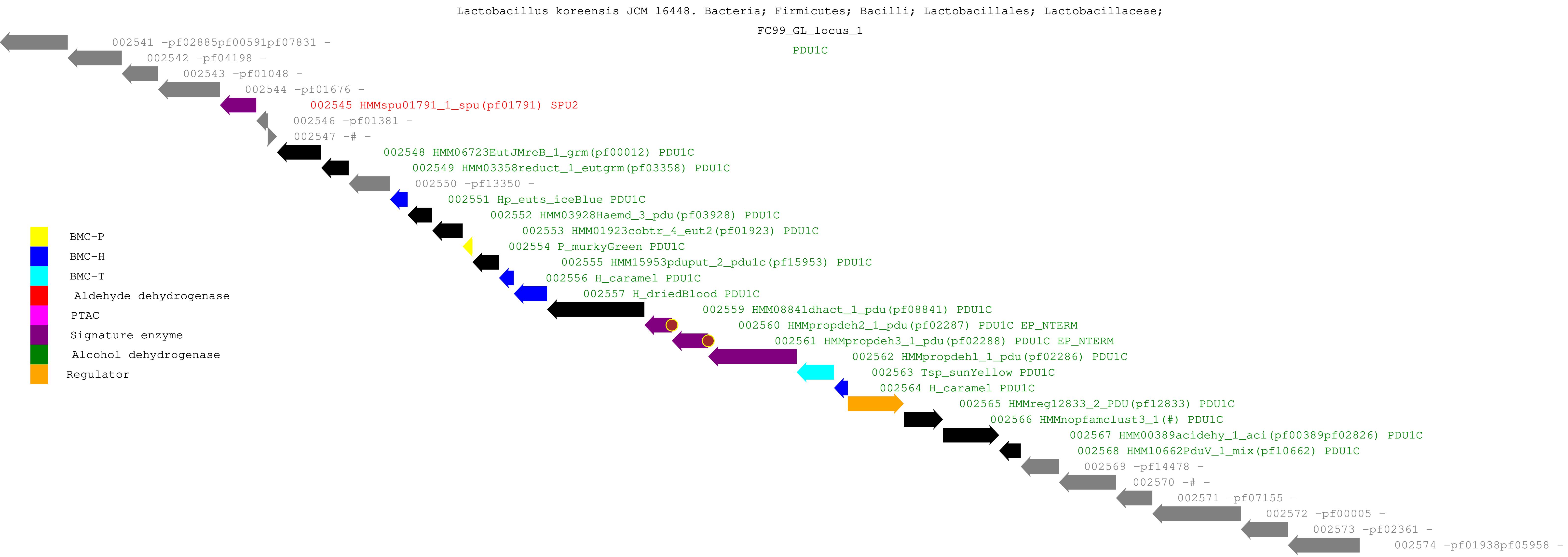
Lactobacillus koreensis JCM 16448. Bacteria; Firmicutes; Bacilli; Lactobacillales; Lactobacillaceae;
FC99_GL_locus_1
PDU1C
002541 -pf02885pf00591pf07831 -
002542 -pf04198 -
002543 -pf01048 -
002544 -pf01676 -
002545 HMMspu01791_1_spu(pf01791) SPU2
002546 -pf01381 -
002547 -# -
002548 HMM06723EutJMreB_1_grm(pf00012) PDU1C
002549 HMM03358reduct_1_eutgrm(pf03358) PDU1C
002550 -pf13350 -
002551 Hp_euts_iceBlue PDU1C
002552 HMM03928Haemd_3_pdu(pf03928) PDU1C
002553 HMM01923cobtr_4_eut2(pf01923) PDU1C
002554 P_murkyGreen PDU1C
002555 HMM15953pduput_2_pdu1c(pf15953) PDU1C
002556 H_caramel PDU1C
002557 H_driedBlood PDU1C
002559 HMM08841dhact_1_pdu(pf08841) PDU1C
002560 HMMpropdeh2_1_pdu(pf02287) PDU1C EP_NTERM
002561 HMMpropdeh3_1_pdu(pf02288) PDU1C EP_NTERM
002562 HMMpropdeh1_1_pdu(pf02286) PDU1C
002563 Tsp_sunYellow PDU1C
002564 H_caramel PDU1C
002565 HMMreg12833_2_PDU(pf12833) PDU1C
002566 HMMnopfamclust3_1(#) PDU1C
002567 HMM00389acidehy_1_aci(pf00389pf02826) PDU1C
002568 HMM10662PduV_1_mix(pf10662) PDU1C
002569 -pf14478 -
002570 -# -
002571 -pf07155 -
002572 -pf00005 -
002573 -pf02361 -
002574 -pf01938pf05958 -
BMC-P
BMC-H
BMC-T
Aldehyde dehydrogenase
PTAC
Signature enzyme
Alcohol dehydrogenase
Regulator

Lactobacillus rapi DSM 19907 = JCM 15042. Bacteria; Firmicutes; Bacilli; Lactobacillales; Lactobacillaceae;

FD12_GL_locus_1

PDU1C

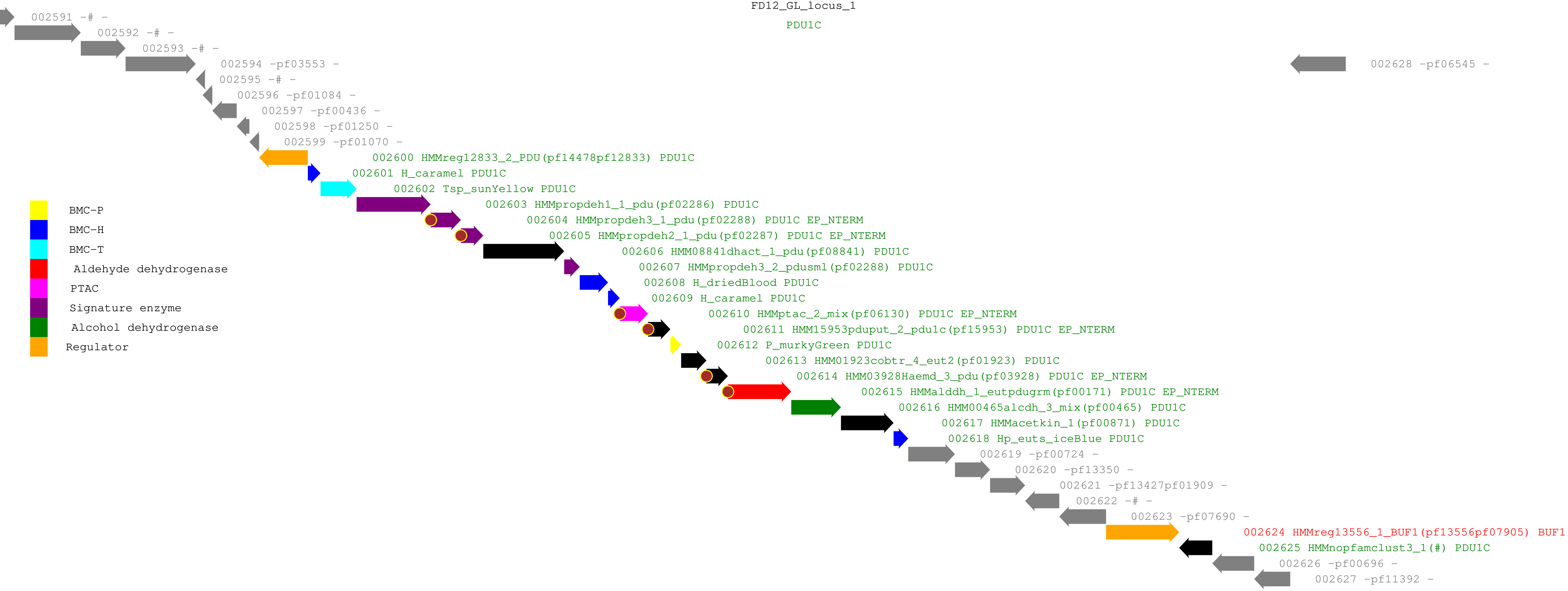

Lactobacillus similis DSM 23365 = JCM 2765. Bacteria; Firmicutes; Bacilli; Lactobacillales; Lactobacillaceae;

FD14_GL_locus_2

PDU1C

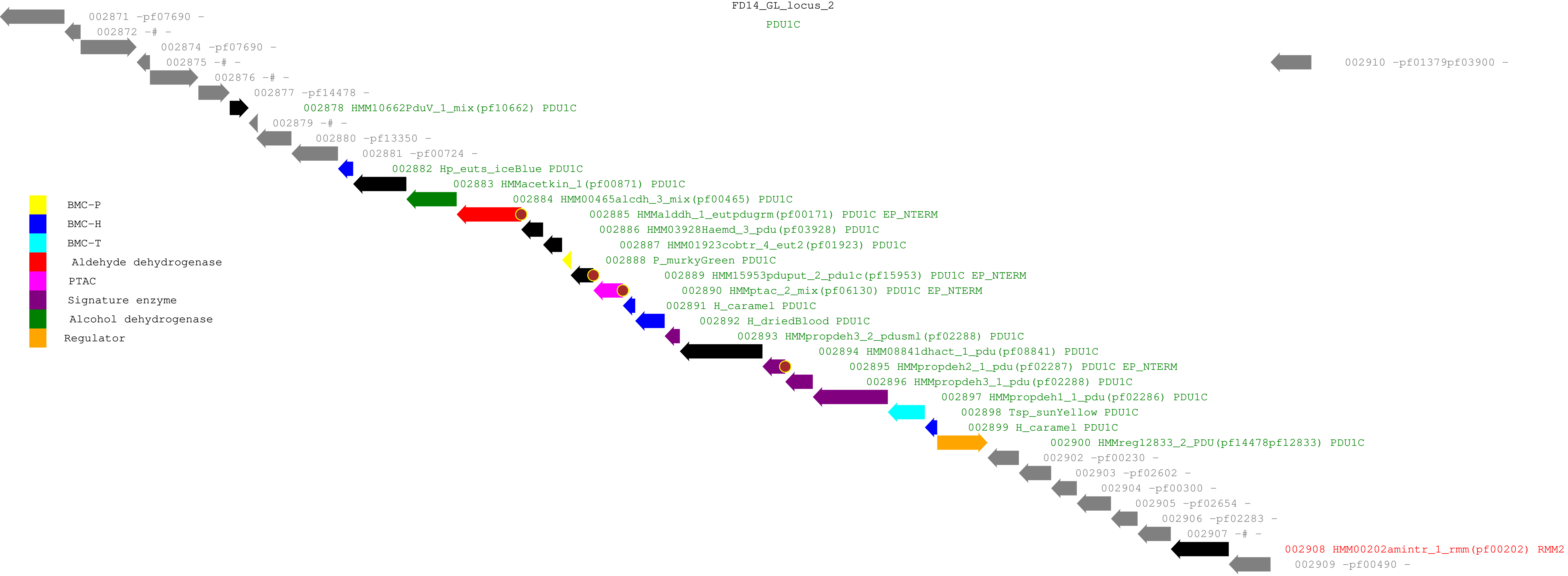

Lactobacillus coryniformis subsp. coryniformis KCTC 3167 = DSM 20001. Bacteria; Firmicutes; Bacilli; Lactobacillales; Lactobacillaceae;

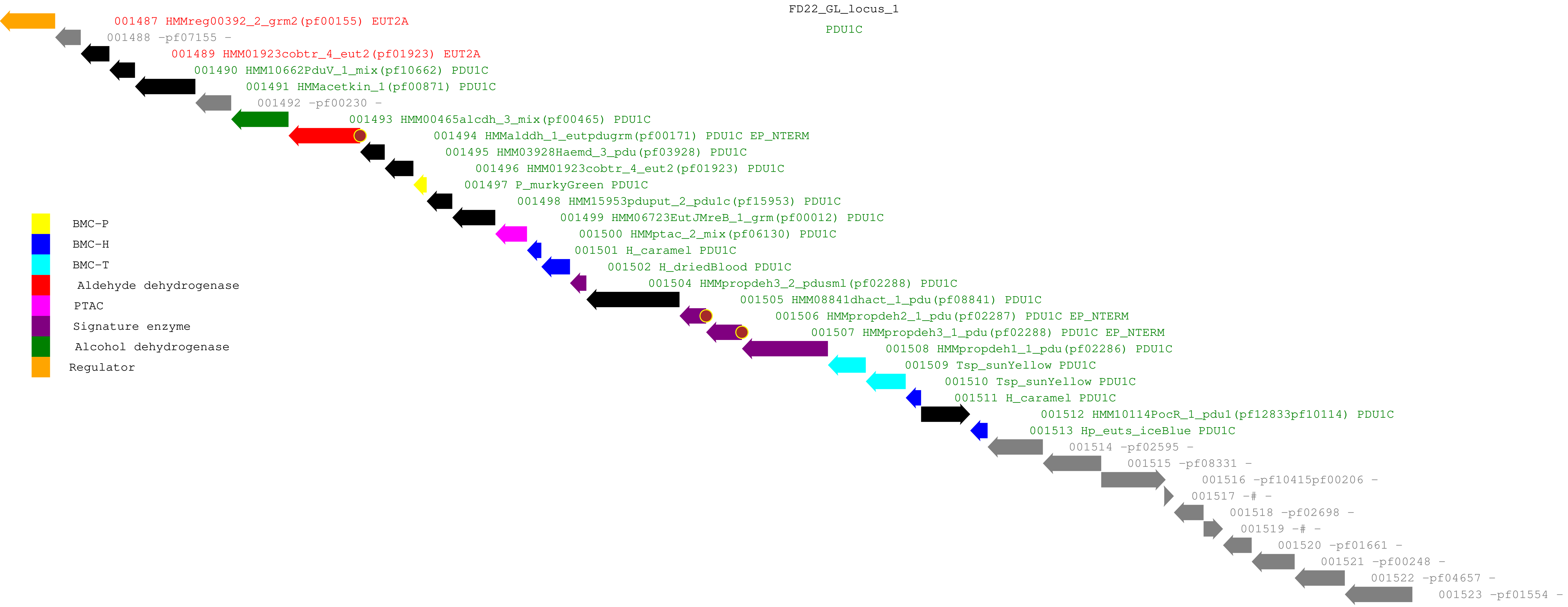

Lactobacillus namurensis DSM 19117. Bacteria; Firmicutes; Bacilli; Lactobacillales; Lactobacillaceae;

FD30_GL_locus_1

PDU1C

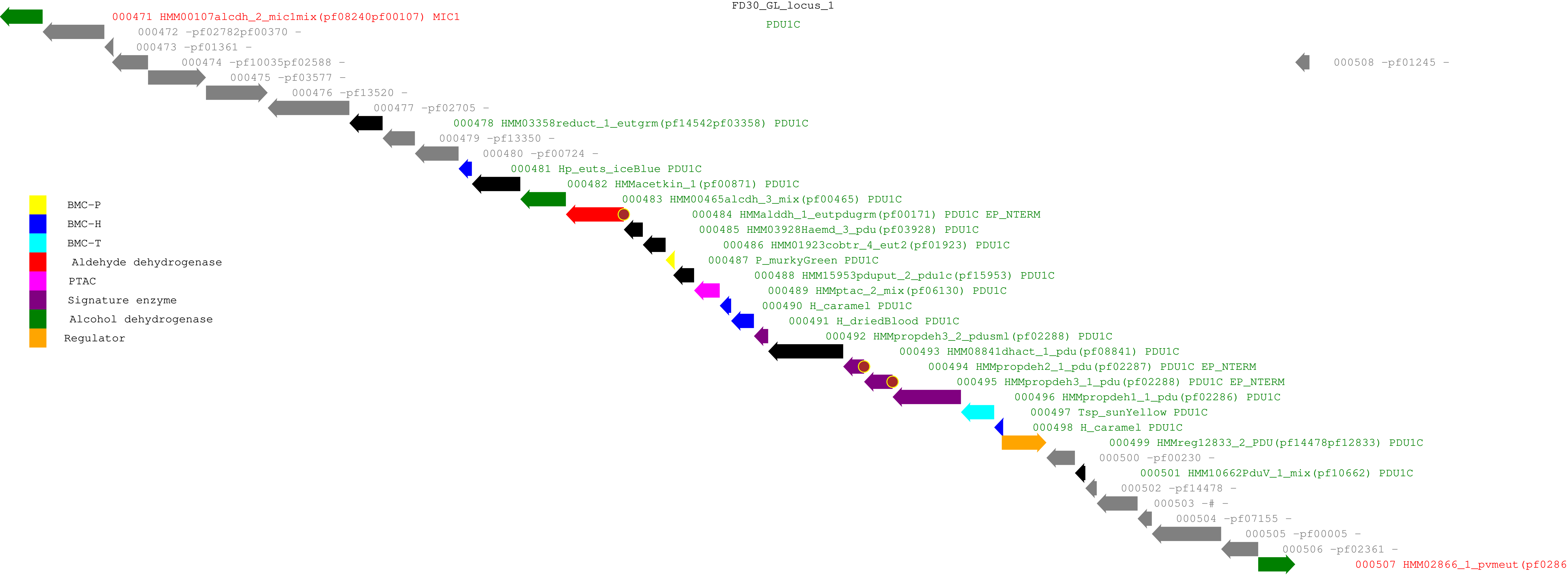

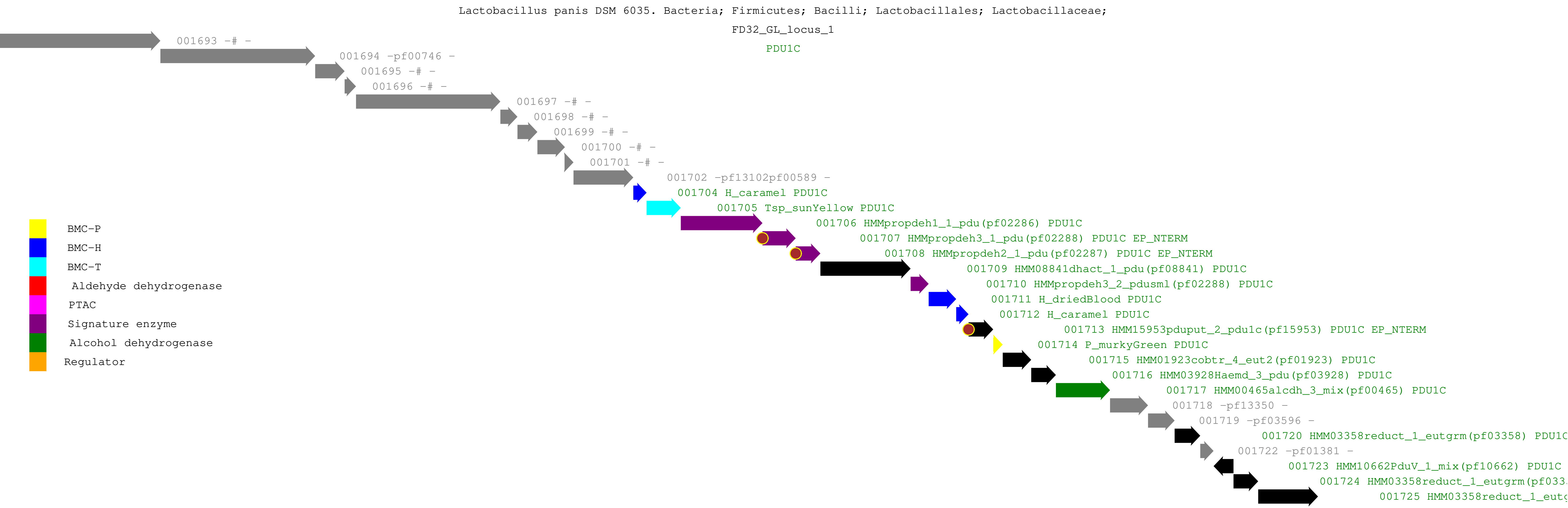

Lactobacillus panis DSM 6035. Bacteria; Firmicutes; Bacilli; Lactobacillales; Lactobacillaceae;
FD32_GL_locus_1
PDU1C
001693 -# -
001694 -pf00746 -
001695 -# -
001696 -# -
001697 -# -
001698 -# -
001699 -# -
001700 -# -
001701 -# -
001702 -pf13102pf00589 -
001704 H_caramel PDU1C
001705 Tsp_sunYellow PDU1C
001706 HMMpropdeh1_1_pdu(pf02286) PDU1C
001707 HMMpropdeh3_1_pdu(pf02288) PDU1C EP_NTERM
001708 HMMpropdeh2_1_pdu(pf02287) PDU1C EP_NTERM
001709 HMM08841dhact_1_pdu(pf08841) PDU1C
001710 HMMpropdeh3_2_pdusml(pf02288) PDU1C
001711 H_driedBlood PDU1C
001712 H_caramel PDU1C
001713 HMM15953pduput_2_pdu1c(pf15953) PDU1C EP_NTERM
001714 P_murkyGreen PDU1C
001715 HMM01923cobtr_4_eut2(pf01923) PDU1C
001716 HMM03928Haemd_3_pdu(pf03928) PDU1C
001717 HMM00465alcdh_3_mix(pf00465) PDU1C
001718 -pf13350 -
001719 -pf03596 -
001720 HMM03358reduct_1_eutgrm(pf03358) PDU1C
001722 -pf01381 -
001723 HMM10662PduV_1_mix(pf10662) PDU1C
001724 HMM03358reduct_1_eutgrm(pf033
001725 HMM03358reduct_1_eutg
BMC-P
BMC-H
BMC-T
Aldehyde dehydrogenase
PTAC
Signature enzyme
Alcohol dehydrogenase
Regulator

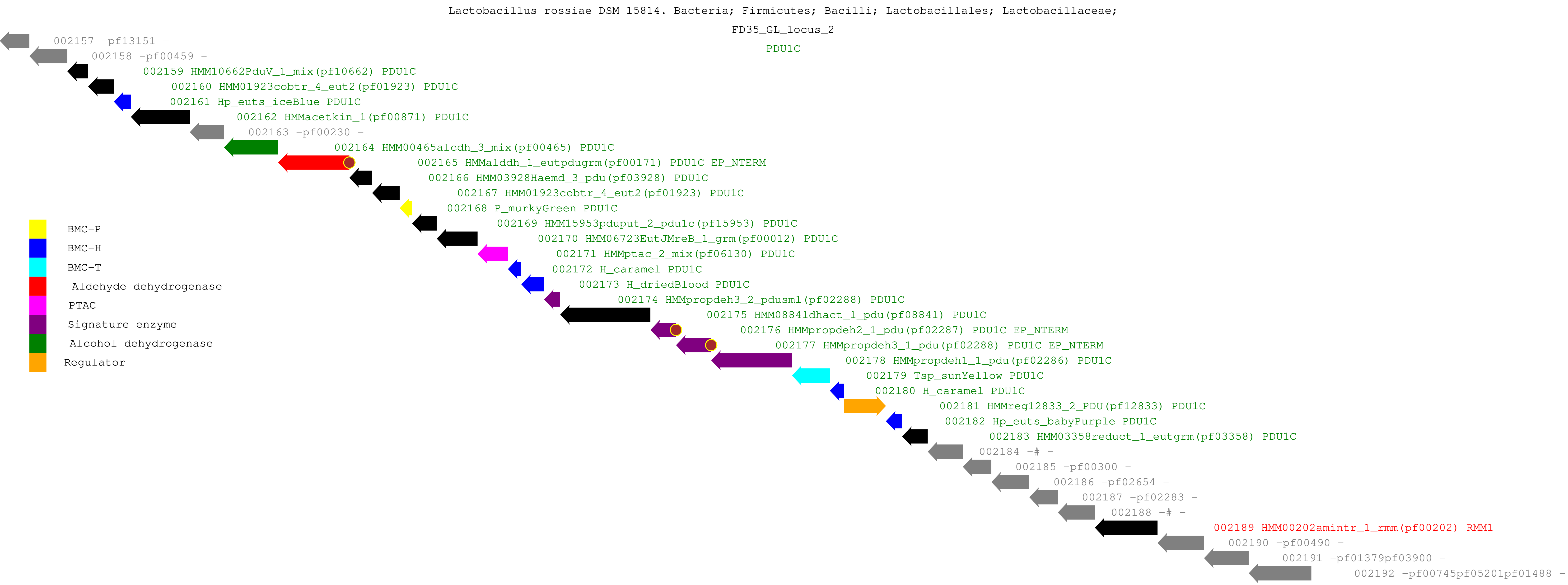

Lactobacillus rossiae DSM 15814. Bacteria; Firmicutes; Bacilli; Lactobacillales; Lactobacillaceae;
FD35_GL_locus_2
PDU1C
002157 -pf13151 -
002158 -pf00459 -
002159 HMM10662PduV_1_mix(pf10662) PDU1C
002160 HMM01923cobtr_4_eut2(pf01923) PDU1C
002161 Hp_euts_iceBlue PDU1C
002162 HMMacetkin_1(pf00871) PDU1C
002163 -pf00230 -
002164 HMM00465alcdh_3_mix(pf00465) PDU1C
002165 HMMalddh_1_eutpdugrm(pf00171) PDU1C EP_NTERM
002166 HMM03928Haemd_3_pdu(pf03928) PDU1C
002167 HMM01923cobtr_4_eut2(pf01923) PDU1C
002168 P_murkyGreen PDU1C
002169 HMM15953pduput_2_pdu1c(pf15953) PDU1C
002170 HMM06723EutJMreB_1_grm(pf00012) PDU1C
002171 HMMptac_2_mix(pf06130) PDU1C
002172 H_caramel PDU1C
002173 H_driedBlood PDU1C
002174 HMMpropdeh3_2_pdusml(pf02288) PDU1C
002175 HMM08841dhact_1_pdu(pf08841) PDU1C
002176 HMMpropdeh2_1_pdu(pf02287) PDU1C EP_NTERM
002177 HMMpropdeh3_1_pdu(pf02288) PDU1C EP_NTERM
002178 HMMpropdeh1_1_pdu(pf02286) PDU1C
002179 Tsp_sunYellow PDU1C
002180 H_caramel PDU1C
002181 HMMreg12833_2_PDU(pf12833) PDU1C
002182 Hp_euts_babyPurple PDU1C
002183 HMM03358reduct_1_eutgrm(pf03358) PDU1C
002184 -# -
002185 -pf00300 -
002186 -pf02654 -
002187 -pf02283 -
002188 -# -
002189 HMM00202amintr_1_rmm(pf00202) RMM1
002190 -pf00490 -
002191 -pf01379pf03900 -
002192 -pf00745pf05201pf01488 -
BMC-P
BMC-H
BMC-T
Aldehyde dehydrogenase
PTAC
Signature enzyme
Alcohol dehydrogenase
Regulator

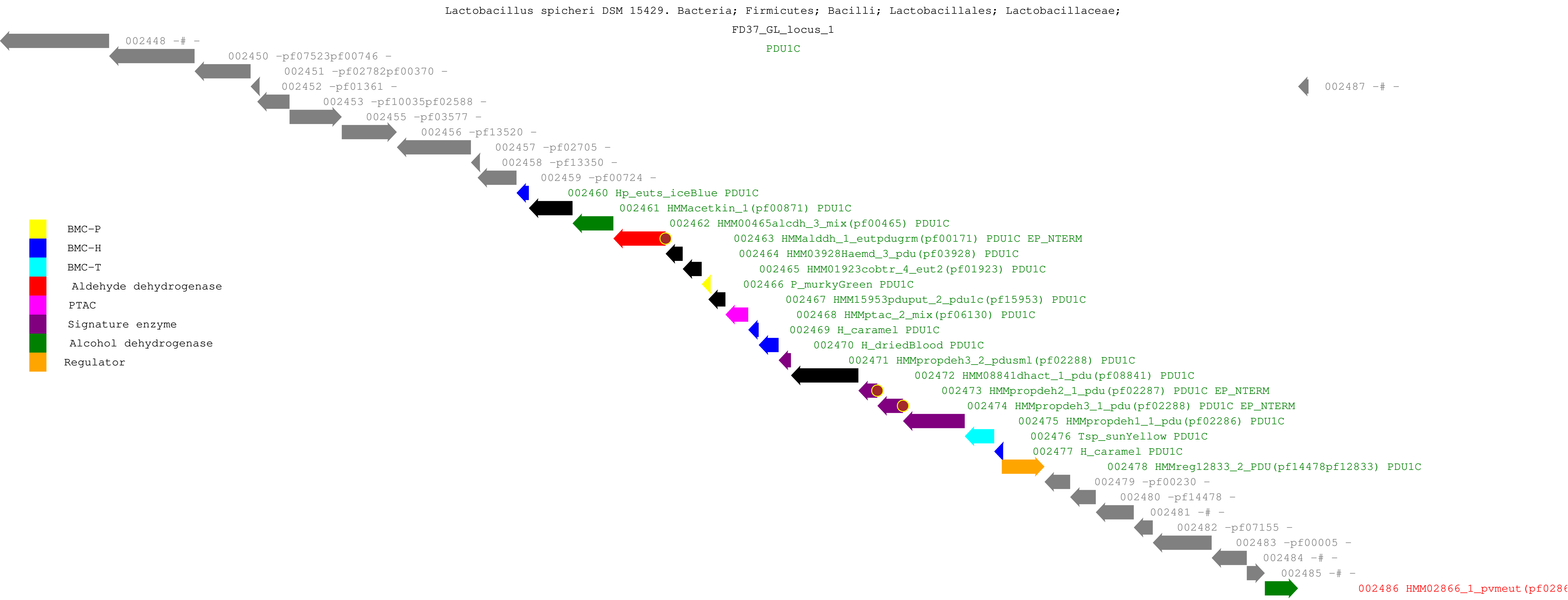
Lactobacillus spicheri DSM 15429. Bacteria; Firmicutes; Bacilli; Lactobacillales; Lactobacillaceae;
FD37_GL_locus_1
PDU1C
002448 -# -
002450 -pf07523pf00746 -
002451 -pf02782pf00370 -
002452 -pf01361 -
002453 -pf10035pf02588 -
002455 -pf03577 -
002456 -pf13520 -
002457 -pf02705 -
002458 -pf13350 -
002459 -pf00724 -
002460 Hp_euts_iceBlue PDU1C
002461 HMMacetkin_1(pf00871) PDU1C
002462 HMM00465alcdh_3_mix(pf00465) PDU1C
002463 HMMalddh_1_eutpdugrm(pf00171) PDU1C EP_NTERM
002464 HMM03928Haemd_3_pdu(pf03928) PDU1C
002465 HMM01923cobtr_4_eut2(pf01923) PDU1C
002466 P_murkyGreen PDU1C
002467 HMM15953pduput_2_pdu1c(pf15953) PDU1C
002468 HMMptac_2_mix(pf06130) PDU1C
002469 H_caramel PDU1C
002470 H_driedBlood PDU1C
002471 HMMpropdeh3_2_pdusml(pf02288) PDU1C
002472 HMM08841dhact_1_pdu(pf08841) PDU1C
002473 HMMpropdeh2_1_pdu(pf02287) PDU1C EP_NTERM
002474 HMMpropdeh3_1_pdu(pf02288) PDU1C EP_NTERM
002475 HMMpropdeh1_1_pdu(pf02286) PDU1C
002476 Tsp_sunYellow PDU1C
002477 H_caramel PDU1C
002478 HMMreg12833_2_PDU(pf14478pf12833) PDU1C
002479 -pf00230 -
002480 -pf14478 -
002481 -# -
002482 -pf07155 -
002483 -pf00005 -
002484 -# -
002485 -# -
002486 HMM02866_1_pvmeut(pf0286
002487 -# -
BMC-P
BMC-H
BMC-T
Aldehyde dehydrogenase
PTAC
Signature enzyme
Alcohol dehydrogenase
Regulator

Lactobacillus zymae DSM 19395. Bacteria; Firmicutes; Bacilli; Lactobacillales; Lactobacillaceae;

FD38_GL_locus_1

PDU1C

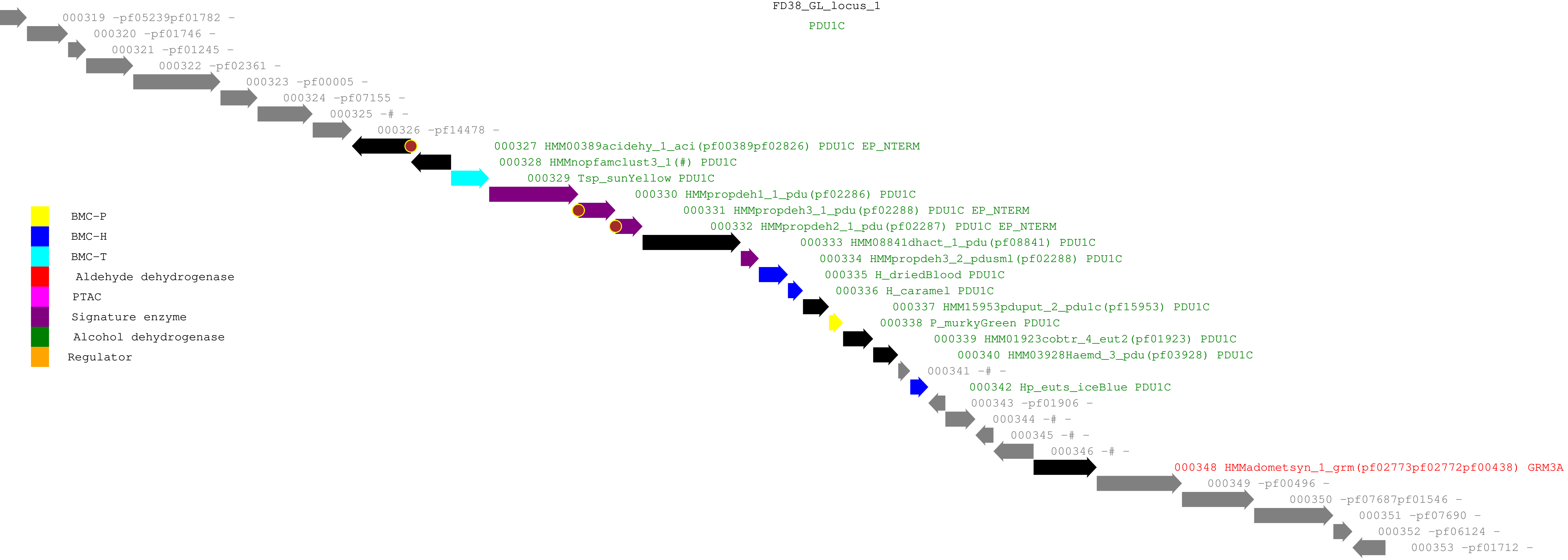

Lactobacillus parafarraginis DSM 18390 = JCM 14109. Bacteria; Firmicutes; Bacilli; Lactobacillales; Lactobacillaceae;

FD47_GL_locus_1

PDU1C

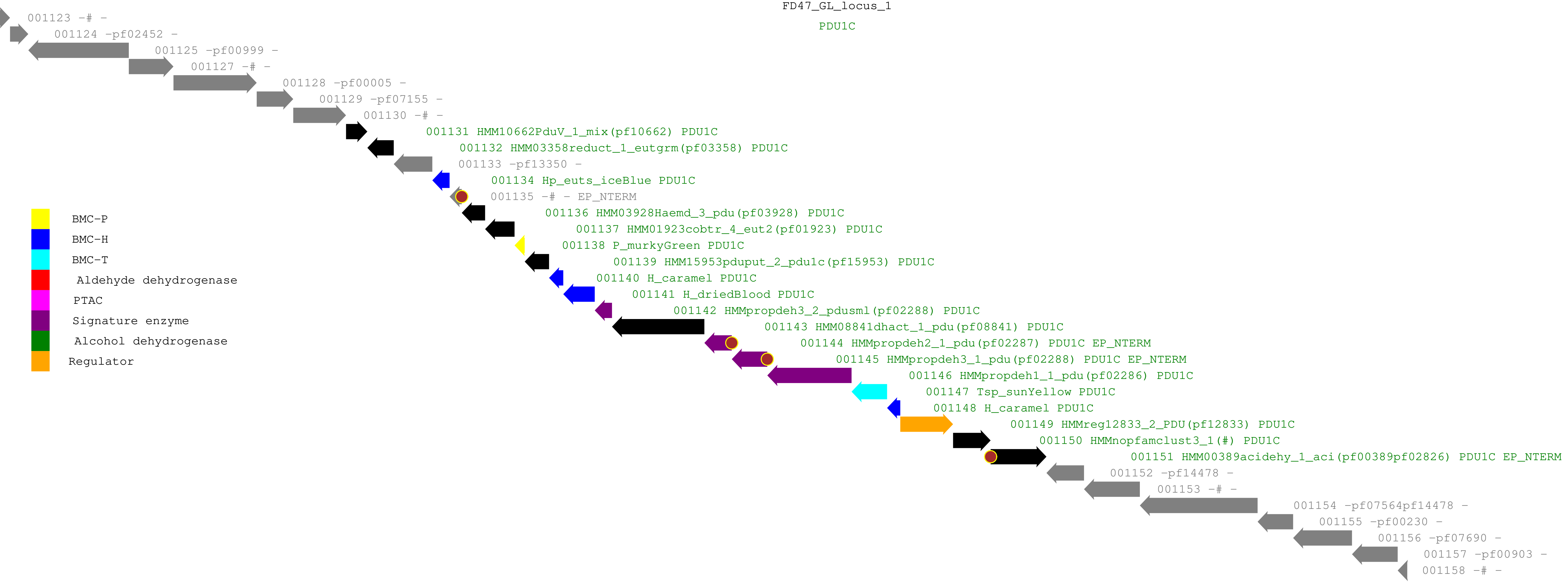

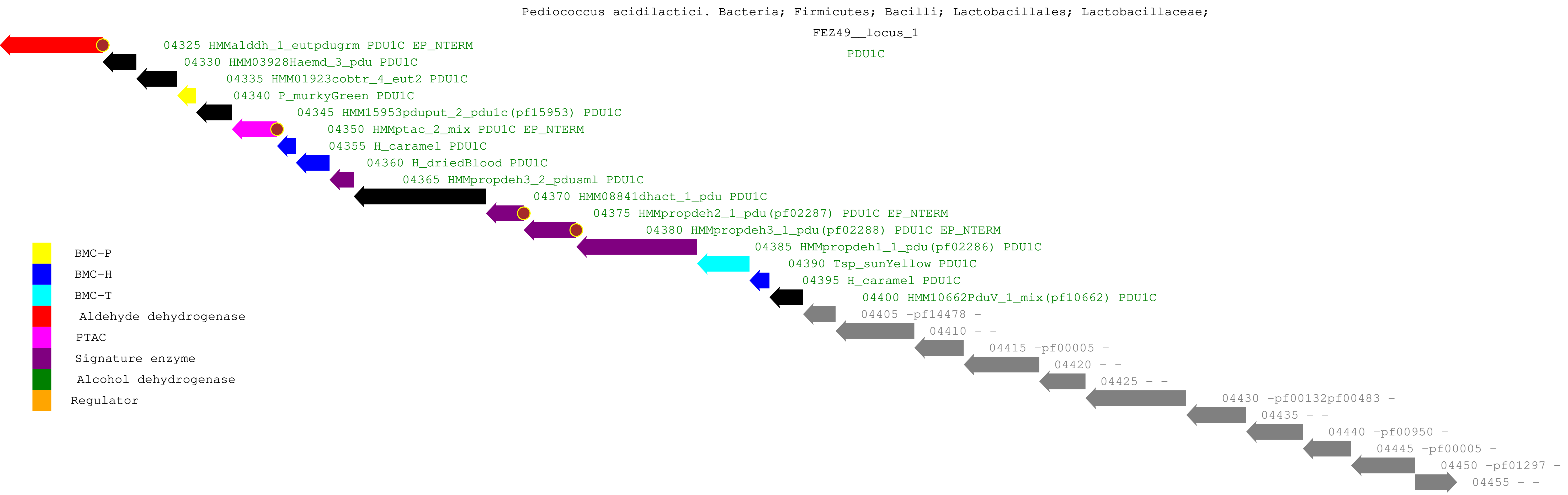
Pediococcus acidilactici. Bacteria; Firmicutes; Bacilli; Lactobacillales; Lactobacillaceae;
FEZ49__locus_1
PDU1C
04325 HMMalddh_1_eutpdugrm PDU1C EP_NTERM
04330 HMM03928Haemd_3_pdu PDU1C
04335 HMM01923cobtr_4_eut2 PDU1C
04340 P_murkyGreen PDU1C
04345 HMM15953pduput_2_pdu1c(pf15953) PDU1C
04350 HMMptac_2_mix PDU1C EP_NTERM
04355 H_caramel PDU1C
04360 H_driedBlood PDU1C
04365 HMMpropdeh3_2_pdusml PDU1C
04370 HMM08841dhact_1_pdu PDU1C
04375 HMMpropdeh2_1_pdu(pf02287) PDU1C EP_NTERM
04380 HMMpropdeh3_1_pdu(pf02288) PDU1C EP_NTERM
04385 HMMpropdeh1_1_pdu(pf02286) PDU1C
04390 Tsp_sunYellow PDU1C
04395 H_caramel PDU1C
04400 HMM10662PduV_1_mix(pf10662) PDU1C
04405 -pf14478 -
04410 - -
04415 -pf00005 -
04420 - -
04425 - -
04430 -pf00132pf00483 -
04435 - -
04440 -pf00950 -
04445 -pf00005 -
04450 -pf01297 -
04455 - -
BMC-P
BMC-H
BMC-T
Aldehyde dehydrogenase
PTAC
Signature enzyme
Alcohol dehydrogenase
Regulator

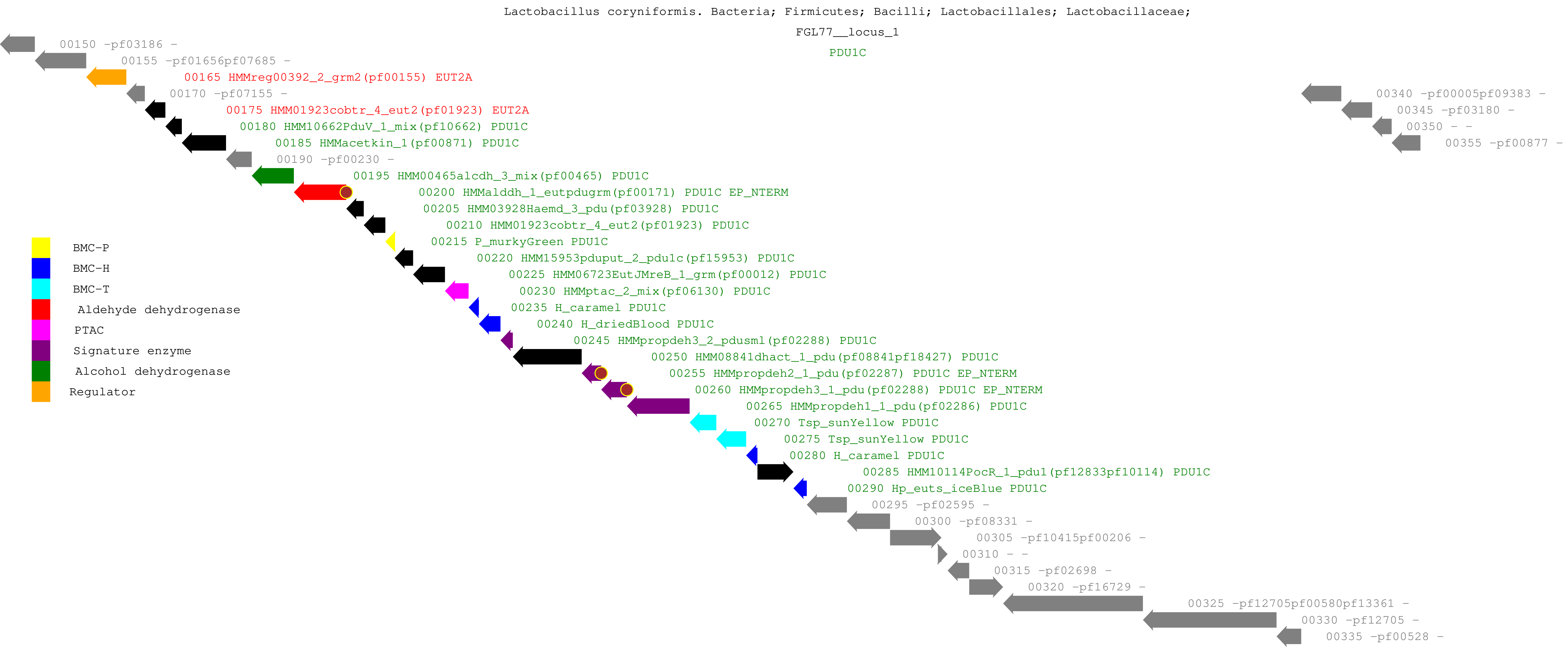

Lactobacillus coryniformis. Bacteria; Firmicutes; Bacilli; Lactobacillales; Lactobacillaceae;
FGL77__locus_1
PDU1C
00150 -pf03186 -
00155 -pf01656pf07685 -
00165 HMMreg00392_2_grm2(pf00155) EUT2A
00170 -pf07155 -
00175 HMM01923cobtr_4_eut2(pf01923) EUT2A
00180 HMM10662PduV_1_mix(pf10662) PDU1C
00185 HMMacetkin_1(pf00871) PDU1C
00190 -pf00230 -
00195 HMM00465alcdh_3_mix(pf00465) PDU1C
00200 HMMaldh_1_eutpdugrm(pf00171) PDU1C EP_NTERM
00205 HMM03928Haemd_3_pdu(pf03928) PDU1C
00210 HMM01923cobtr_4_eut2(pf01923) PDU1C
00215 P_murkyGreen PDU1C
00220 HMM15953pduput_2_pdu1c(pf15953) PDU1C
00225 HMM06723EutJMreB_1_grm(pf00012) PDU1C
00230 HMMptac_2_mix(pf06130) PDU1C
00235 H_caramel PDU1C
00240 H_driedBlood PDU1C
00245 HMMpropdeh3_2_pdusml(pf02288) PDU1C
00250 HMM08841dhact_1_pdu(pf08841pf18427) PDU1C
00255 HMMpropdeh2_1_pdu(pf02287) PDU1C EP_NTERM
00260 HMMpropdeh3_1_pdu(pf02288) PDU1C EP_NTERM
00265 HMMpropdeh1_1_pdu(pf02286) PDU1C
00270 Tsp_sunYellow PDU1C
00275 Tsp_sunYellow PDU1C
00280 H_caramel PDU1C
00285 HMM10114PocR_1_pdu1(pf12833pf10114) PDU1C
00290 Hp_euts_iceBlue PDU1C
00295 -pf02595 -
00300 -pf08331 -
00305 -pf10415pf00206 -
00310 - -
00315 -pf02698 -
00320 -pf16729 -
00325 -pf12705pf00580pf13361 -
00330 -pf12705 -
00335 -pf00528 -
00340 -pf00005pf09383 -
00345 -pf03180 -
00350 - -
00355 -pf00877 -
BMC-P
BMC-H
BMC-T
Aldehyde dehydrogenase
PTAC
Signature enzyme
Alcohol dehydrogenase
Regulator

Vagococcus fluvialis bH819. Bacteria; Firmicutes; Bacilli; Lactobacillales; Enterococcaceae;

FM121__locus_2

PDU1C

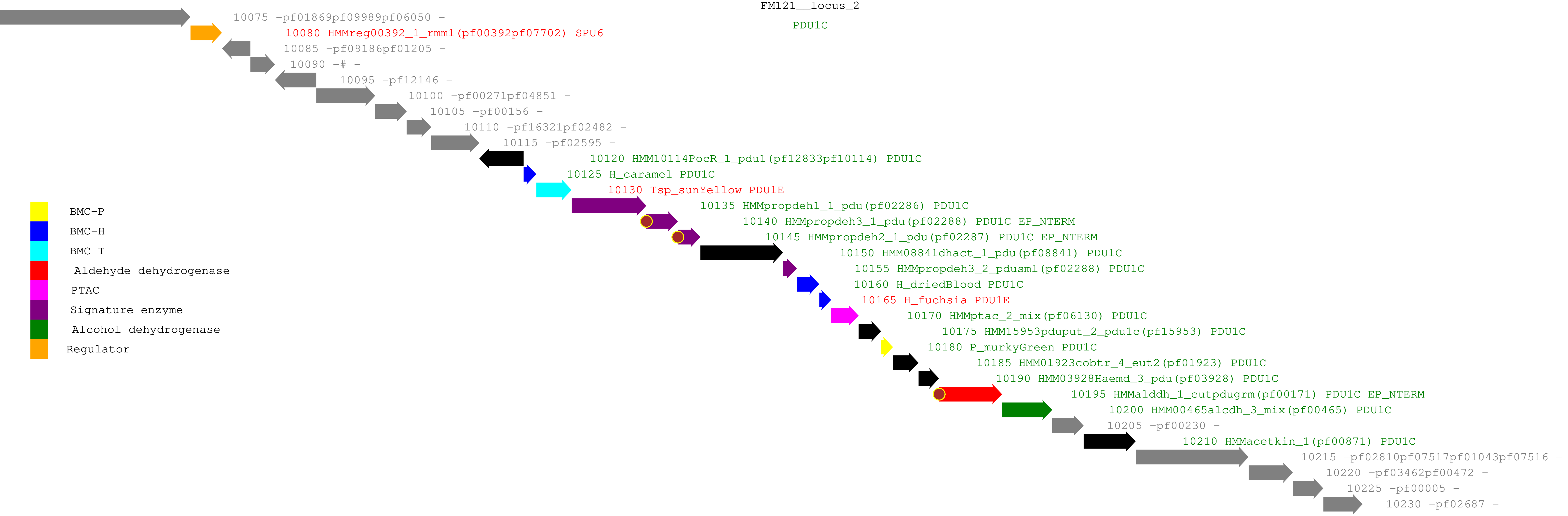

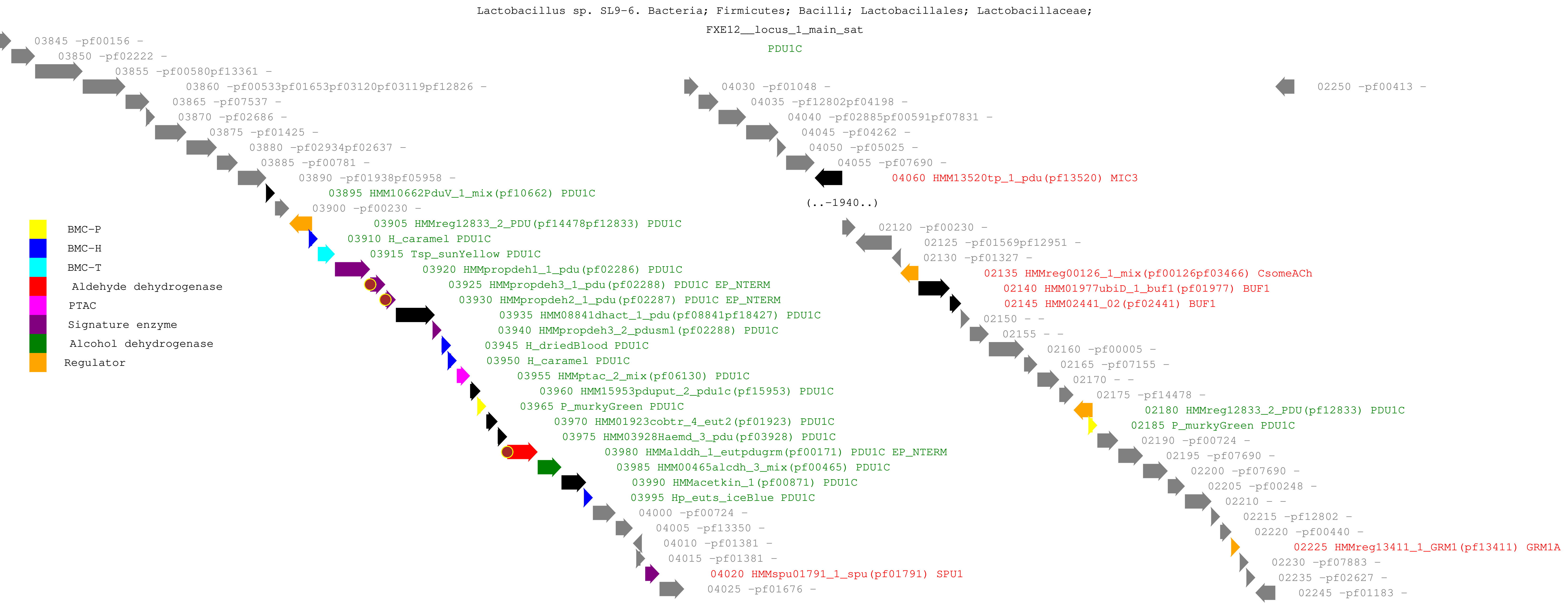
Lactobacillus sp. SL9-6. Bacteria; Firmicutes; Bacilli; Lactobacillales; Lactobacillaceae;
FXE12__locus_1_main_sat
PDU1C
03845 -pf00156 -
03850 -pf02222 -
03855 -pf00580pf13361 -
03860 -pf00533pf01653pf03120pf03119pf12826 -
03865 -pf07537 -
03870 -pf02686 -
03875 -pf01425 -
03880 -pf02934pf02637 -
03885 -pf00781 -
03890 -pf01938pf05958 -
03895 HMM10662PduV_1_mix(pf10662) PDU1C
03900 -pf00230 -
03905 HMMreg12833_2_PDU(pf14478pf12833) PDU1C
03910 H_caramel PDU1C
03915 Tsp_sunYellow PDU1C
03920 HMMpropdeh1_1_pdu(pf02286) PDU1C
03925 HMMpropdeh3_1_pdu(pf02288) PDU1C EP_NTERM
03930 HMMpropdeh2_1_pdu(pf02287) PDU1C EP_NTERM
03935 HMM08841dhact_1_pdu(pf08841pf18427) PDU1C
03940 HMMpropdeh3_2_pdusml(pf02288) PDU1C
03945 H_driedBlood PDU1C
03950 H_caramel PDU1C
03955 HMMptac_2_mix(pf06130) PDU1C
03960 HMM15953pduput_2_pdu1c(pf15953) PDU1C
03965 P_murkyGreen PDU1C
03970 HMM01923cobtr_4_eut2(pf01923) PDU1C
03975 HMM03928Haemd_3_pdu(pf03928) PDU1C
03980 HMMaldh_1_eutpdugrm(pf00171) PDU1C EP_NTERM
03985 HMM00465alcdh_3_mix(pf00465) PDU1C
03990 HMMacetkin_1(pf00871) PDU1C
03995 Hp_euts_iceBlue PDU1C
04000 -pf00724 -
04005 -pf13350 -
04010 -pf01381 -
04015 -pf01381 -
04020 HMMspu01791_1_spu(pf01791) SPU1
04025 -pf01676 -
04030 -pf01048 -
04035 -pf12802pf04198 -
04040 -pf02885pf00591pf07831 -
04045 -pf04262 -
04050 -pf05025 -
04055 -pf07690 -
04060 HMM13520tp_1_pdu(pf13520) MIC3
(..-1940..)
02120 -pf00230 -
02125 -pf01569pf12951 -
02130 -pf01327 -
02135 HMMreg00126_1_mix(pf00126pf03466) CsomeACh
02140 HMM01977ubiD_1_buf1(pf01977) BUF1
02145 HMM02441_02(pf02441) BUF1
02150 - -
02155 - -
02160 -pf00005 -
02165 -pf07155 -
02170 - -
02175 -pf14478 -
02180 HMMreg12833_2_PDU(pf12833) PDU1C
02185 P_murkyGreen PDU1C
02190 -pf00724 -
02195 -pf07690 -
02200 -pf07690 -
02205 -pf00248 -
02210 - -
02215 -pf12802 -
02220 -pf00440 -
02225 HMMreg13411_1_GRM1(pf13411) GRM1A
02230 -pf07883 -
02235 -pf02627 -
02245 -pf01183 -
02250 -pf00413 -
BMC-P
BMC-H
BMC-T
Aldehyde dehydrogenase
PTAC
Signature enzyme
Alcohol dehydrogenase
Regulator

Lactobacillus brevis ATCC 14869 = DSM 20054. Bacteria; Firmicutes; Bacilli; Lactobacillales; Lactobacillaceae;

HMPREF0495__locus_2

PDU1C

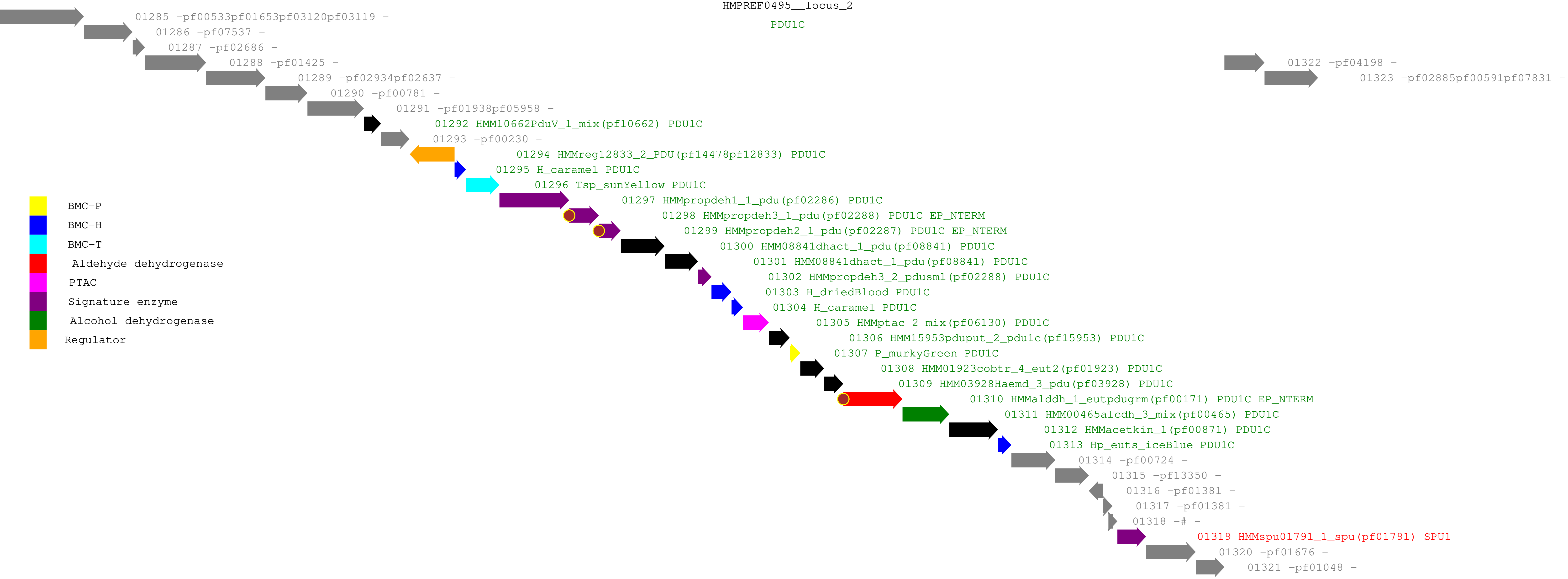

Lactobacillus reuteri MM2-3. Bacteria; Firmicutes; Bacilli; Lactobacillales; Lactobacillaceae;

HMPREF0535__locus_1

PDU1C

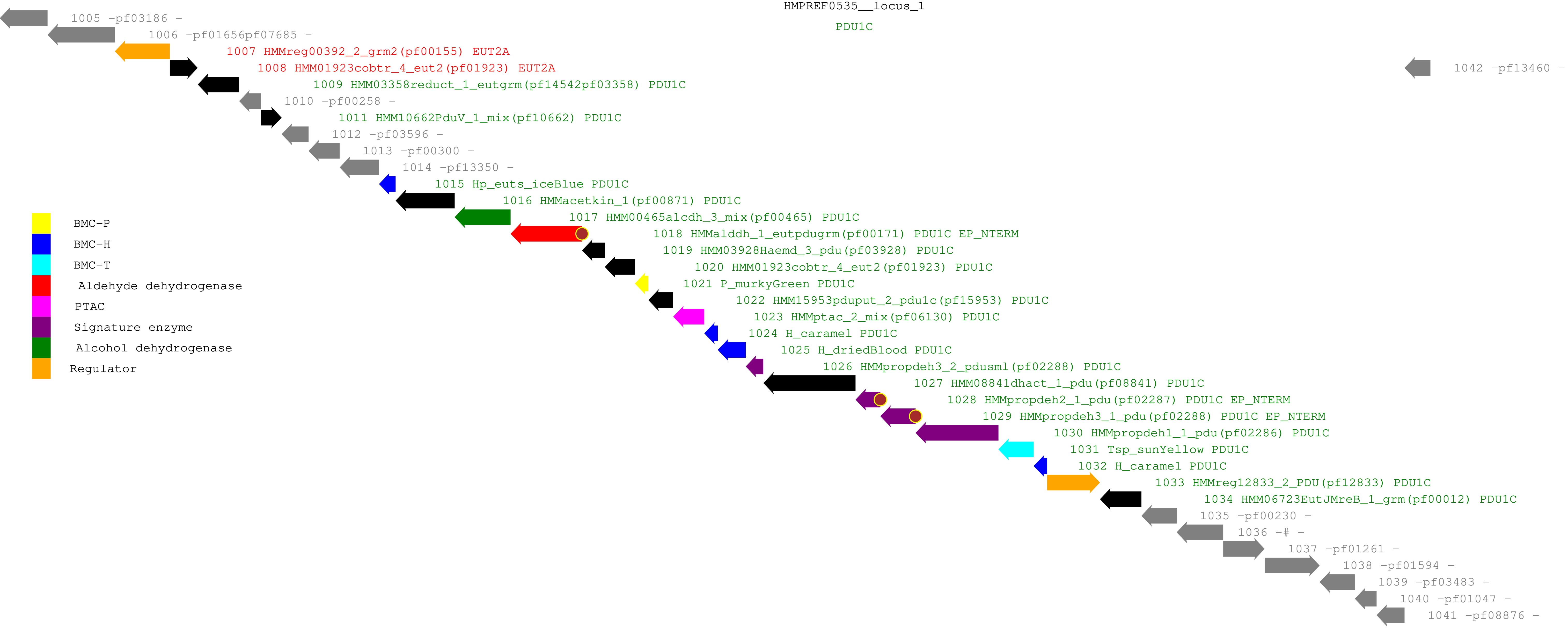

Lactobacillus reuteri (strain ATCC 55730 / SD2112). Bacteria; Firmicutes; Bacilli; Lactobacillales; Lactobacillaceae;

HMPREF0538__locus_1

PDU1C

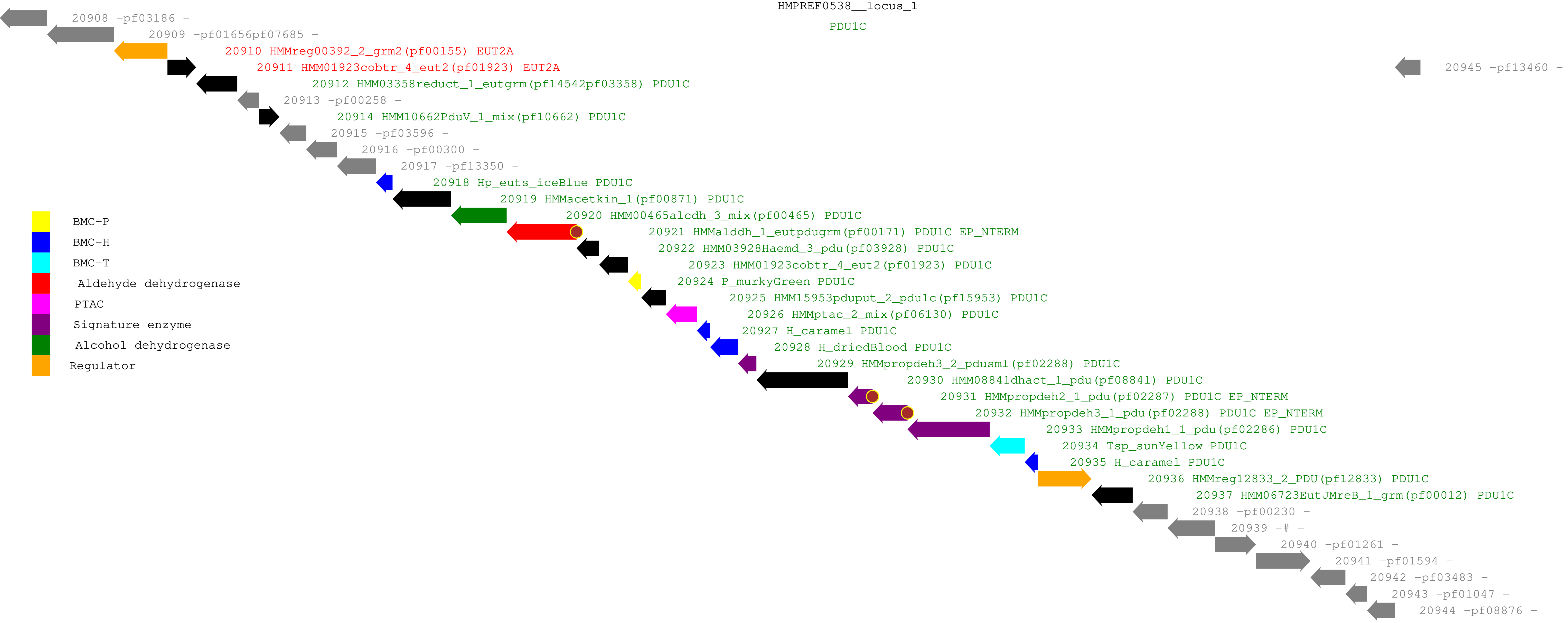

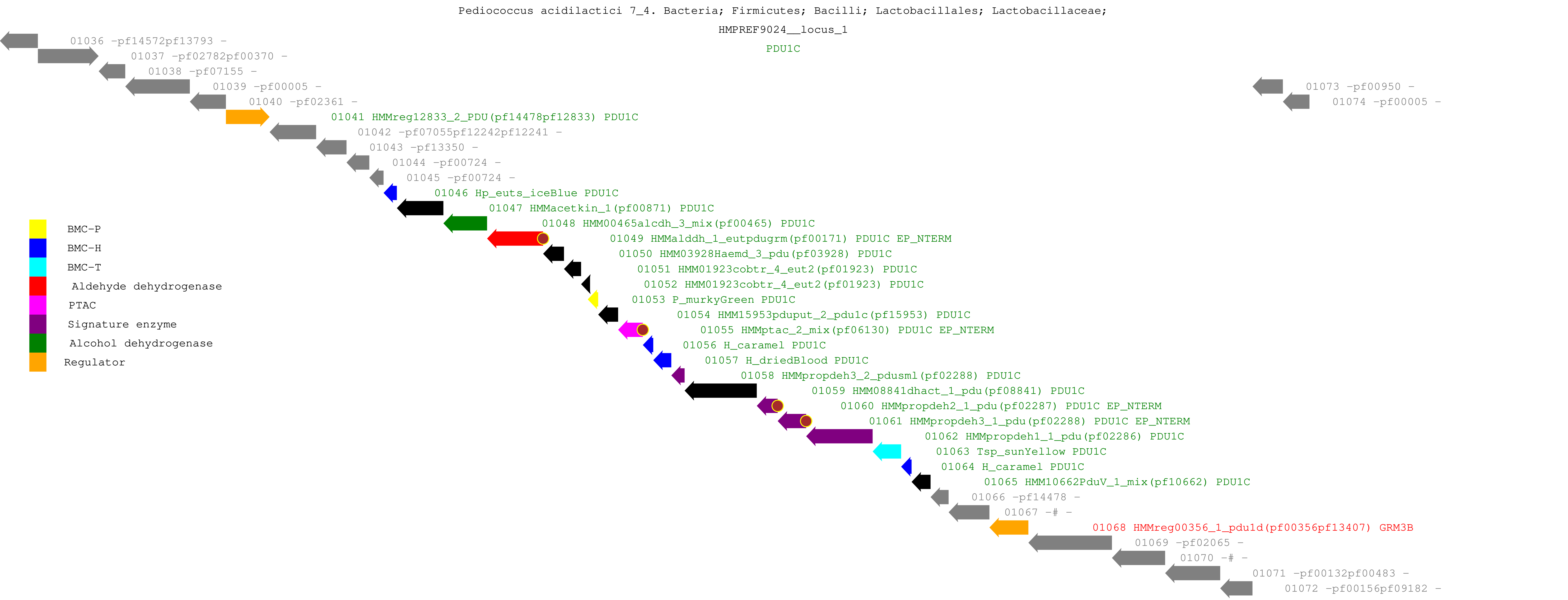
Pediococcus acidilactici 7_4. Bacteria; Firmicutes; Bacilli; Lactobacillales; Lactobacillaceae;
HMPREF9024__locus_1
PDU1C
01036 -pf14572pf13793 -
01037 -pf02782pf00370 -
01038 -pf07155 -
01039 -pf00005 -
01040 -pf02361 -
01041 HMMreg12833_2_PDU(pf14478pf12833) PDU1C
01042 -pf07055pf12242pf12241 -
01043 -pf13350 -
01044 -pf00724 -
01045 -pf00724 -
01046 Hp_euts_iceBlue PDU1C
01047 HMMacetkin_1(pf00871) PDU1C
01048 HMM00465alcdh_3_mix(pf00465) PDU1C
01049 HMMaldd h_1_eutpdugrm(pf00171) PDU1C EP_NTERM
01050 HMM03928Haemd_3_pdu(pf03928) PDU1C
01051 HMM01923cobtr_4_eut2(pf01923) PDU1C
01052 HMM01923cobtr_4_eut2(pf01923) PDU1C
01053 P_murkyGreen PDU1C
01054 HMM15953pduput_2_pdu1c(pf15953) PDU1C
01055 HMMptac_2_mix(pf06130) PDU1C EP_NTERM
01056 H_caramel PDU1C
01057 H_driedBlood PDU1C
01058 HMMpropdeh3_2_pdusml(pf02288) PDU1C
01059 HMM08841dhact_1_pdu(pf08841) PDU1C
01060 HMMpropdeh2_1_pdu(pf02287) PDU1C EP_NTERM
01061 HMMpropdeh3_1_pdu(pf02288) PDU1C EP_NTERM
01062 HMMpropdeh1_1_pdu(pf02286) PDU1C
01063 Tsp_sunYellow PDU1C
01064 H_caramel PDU1C
01065 HMM10662PduV_1_mix(pf10662) PDU1C
01066 -pf14478 -
01067 -# -
01068 HMMreg00356_1_pdu1d(pf00356pf13407) GRM3B
01069 -pf02065 -
01070 -# -
01071 -pf00132pf00483 -
01072 -pf00156pf09182 -
01073 -pf00950 -
01074 -pf00005 -
BMC-P
BMC-H
BMC-T
Aldehyde dehydrogenase
PTAC
Signature enzyme
Alcohol dehydrogenase
Regulator

Lactobacillus kisonensis F0435. Bacteria; Firmicutes; Bacilli; Lactobacillales; Lactobacillaceae;

HMPREF9104__locus_1

PDU1C

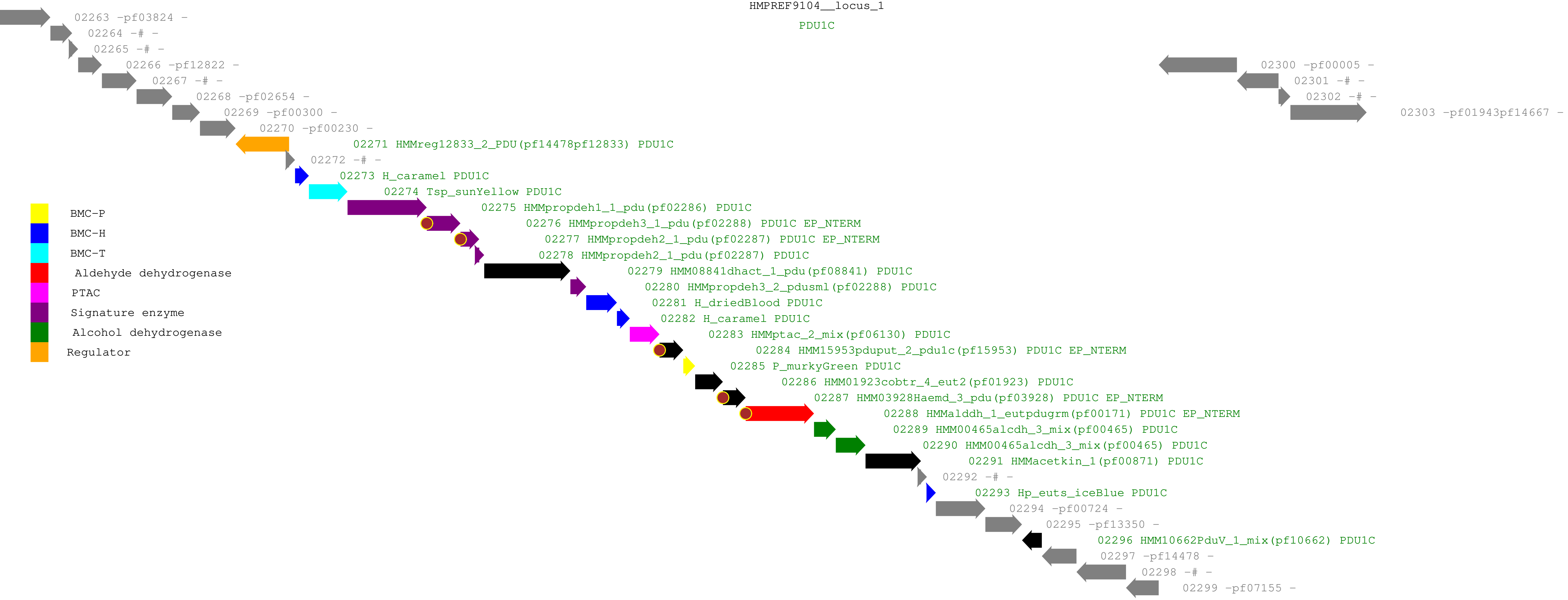

Lactobacillus siliginis. Bacteria; Firmicutes; Bacilli; Lactobacillales; Lactobacillaceae;

IV55_GL_locus_1

PDU1C

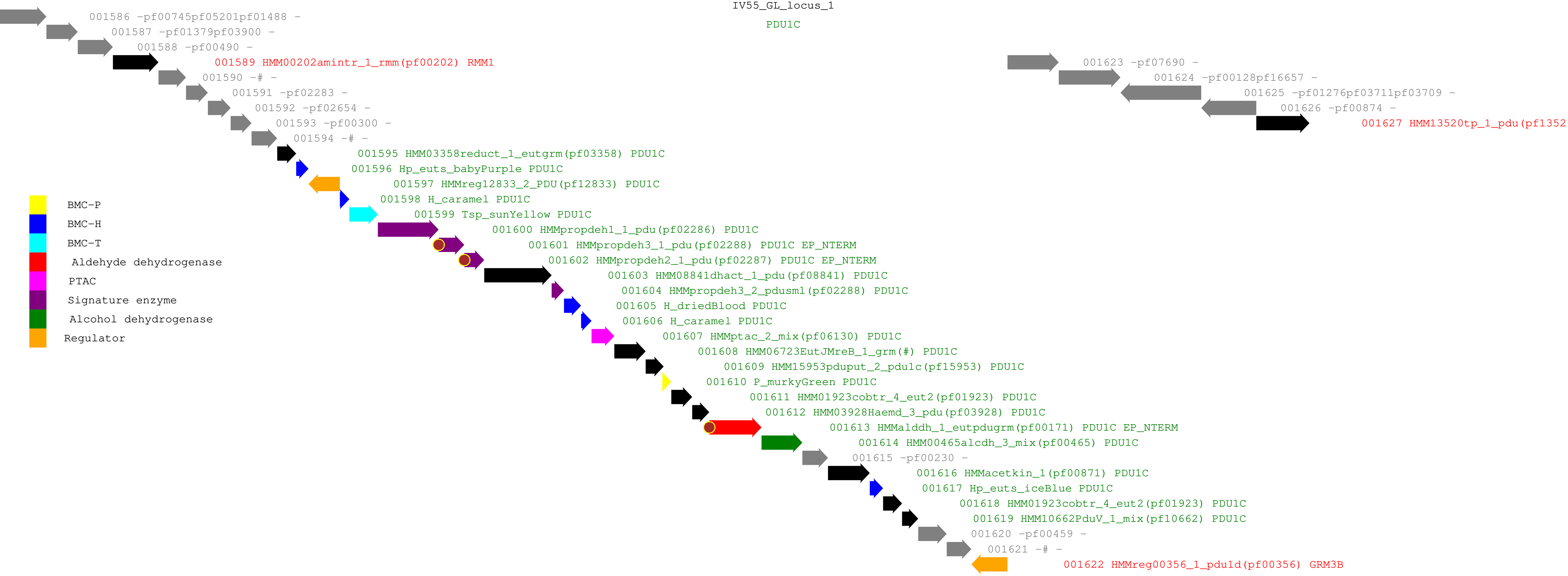

Lactobacillus silagei JCM 19001. Bacteria; Firmicutes; Bacilli; Lactobacillales; Lactobacillaceae;

IWT126__locus_1

PDU1C

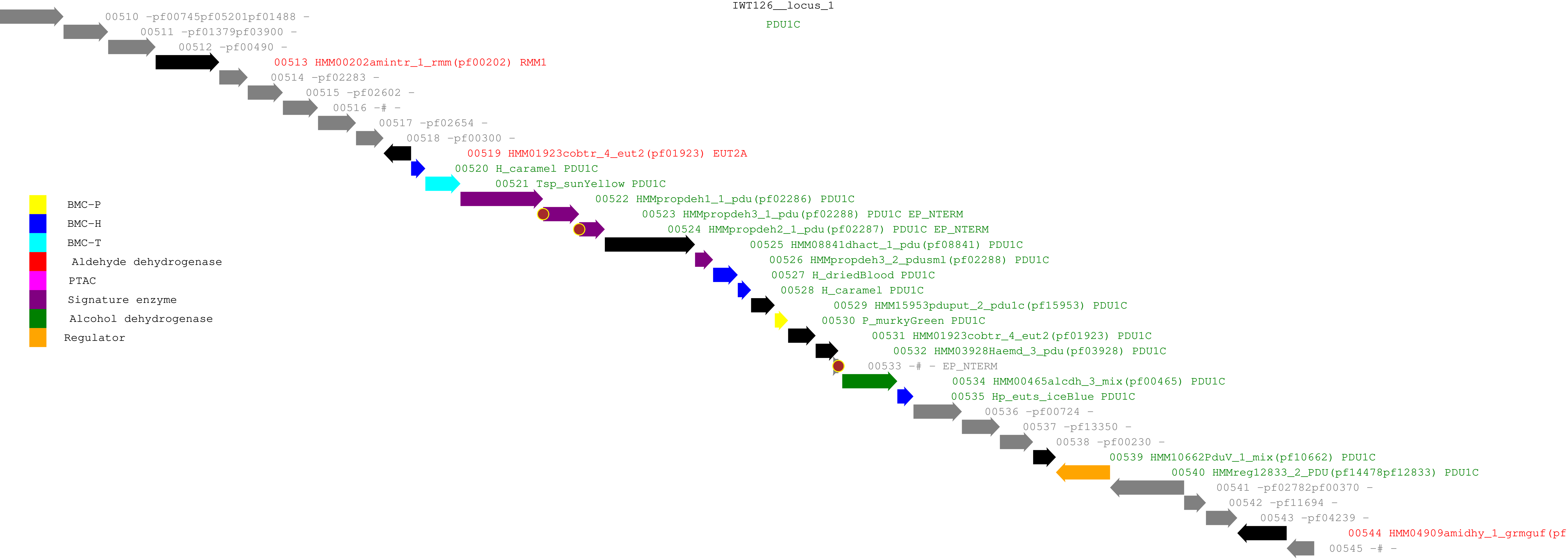

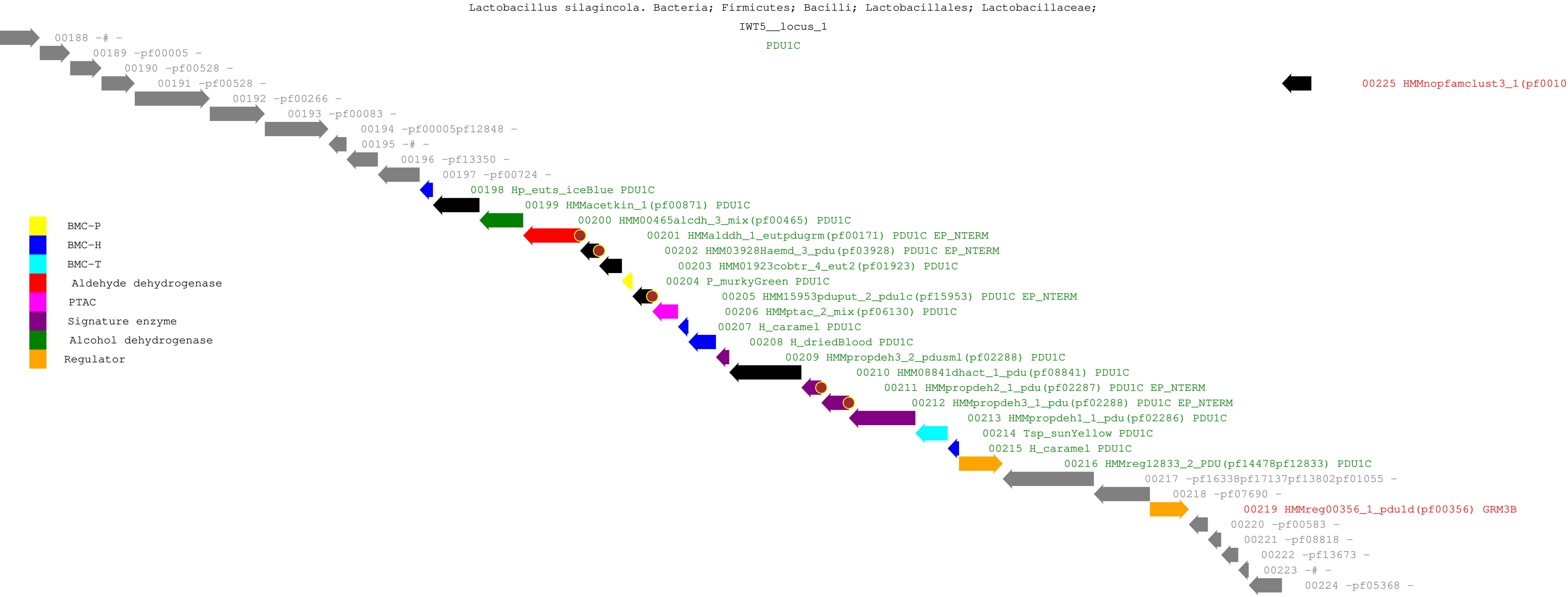
Lactobacillus silagincola. Bacteria; Firmicutes; Bacilli; Lactobacillales; Lactobacillaceae;
IWT5__locus_1
PDU1C
00188 -# -
00189 -pf00005 -
00190 -pf00528 -
00191 -pf00528 -
00192 -pf00266 -
00193 -pf00083 -
00194 -pf00005pf12848 -
00195 -# -
00196 -pf13350 -
00197 -pf00724 -
00198 Hp_euts_iceBlue PDU1C
00199 HMMacetkin_1(pf00871) PDU1C
00200 HMM00465alcdh_3_mix(pf00465) PDU1C
00201 HMMaldh_1_eutpdugrm(pf00171) PDU1C EP_NTERM
00202 HMM03928Haemd_3_pdu(pf03928) PDU1C EP_NTERM
00203 HMM01923cobtr_4_eut2(pf01923) PDU1C
00204 P_murkyGreen PDU1C
00205 HMM15953pduput_2_pdu1c(pf15953) PDU1C EP_NTERM
00206 HMMptac_2_mix(pf06130) PDU1C
00207 H_caramel PDU1C
00208 H_driedBlood PDU1C
00209 HMMpropdeh3_2_pdusml(pf02288) PDU1C
00210 HMM08841dhact_1_pdu(pf08841) PDU1C
00211 HMMpropdeh2_1_pdu(pf02287) PDU1C EP_NTERM
00212 HMMpropdeh3_1_pdu(pf02288) PDU1C EP_NTERM
00213 HMMpropdeh1_1_pdu(pf02286) PDU1C
00214 Tsp_sunYellow PDU1C
00215 H_caramel PDU1C
00216 HMMreg12833_2_PDU(pf14478pf12833) PDU1C
00217 -pf16338pf17137pf13802pf01055 -
00218 -pf07690 -
00219 HMMreg00356_1_pdu1d(pf00356) GRM3B
00220 -pf00583 -
00221 -pf08818 -
00222 -pf13673 -
00223 -# -
00224 -pf05368 -
00225 HMMnopfamclust3_1(pf0010
BMC-P
BMC-H
BMC-T
Aldehyde dehydrogenase
PTAC
Signature enzyme
Alcohol dehydrogenase
Regulator

Lactobacillus mellifer. Bacteria; Firmicutes; Bacilli; Lactobacillales; Lactobacillaceae;

JG30__locus_1

PDU1C

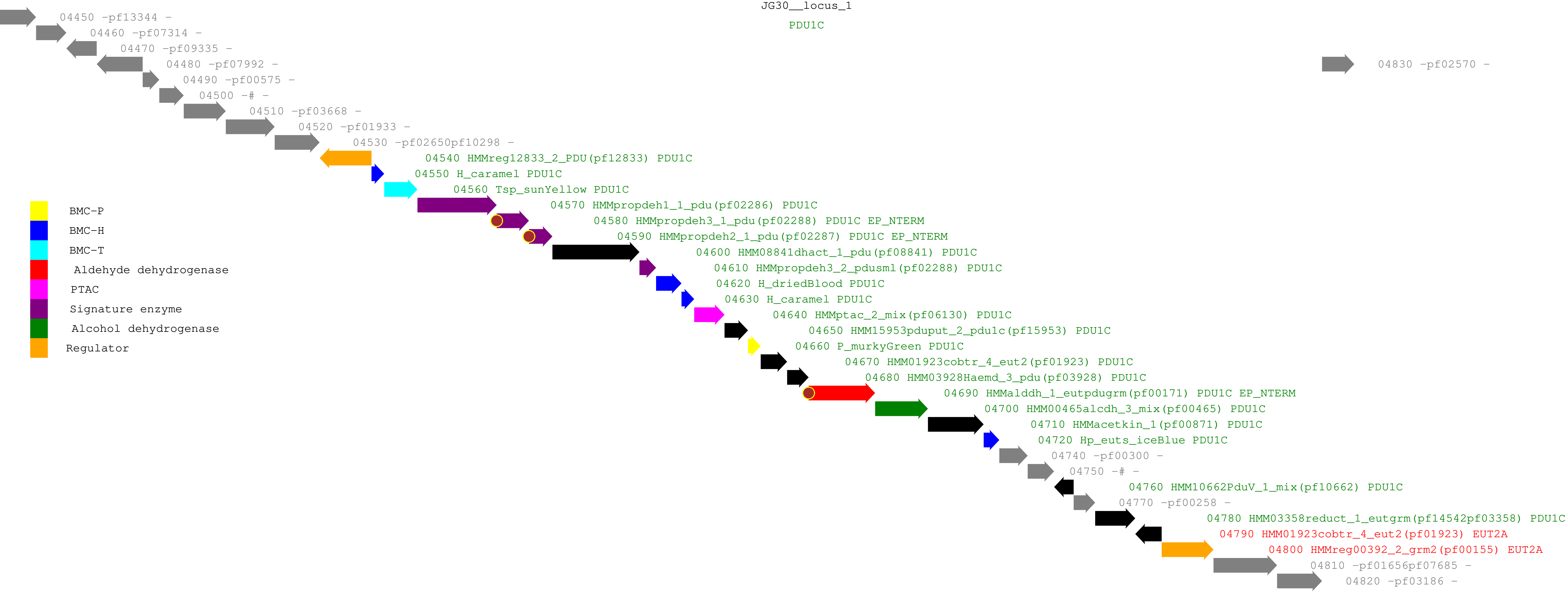

Lactobacillus brevis BSO 464. Bacteria; Firmicutes; Bacilli; Lactobacillales; Lactobacillaceae;

L747__locus_1_main_sat

PDU1C

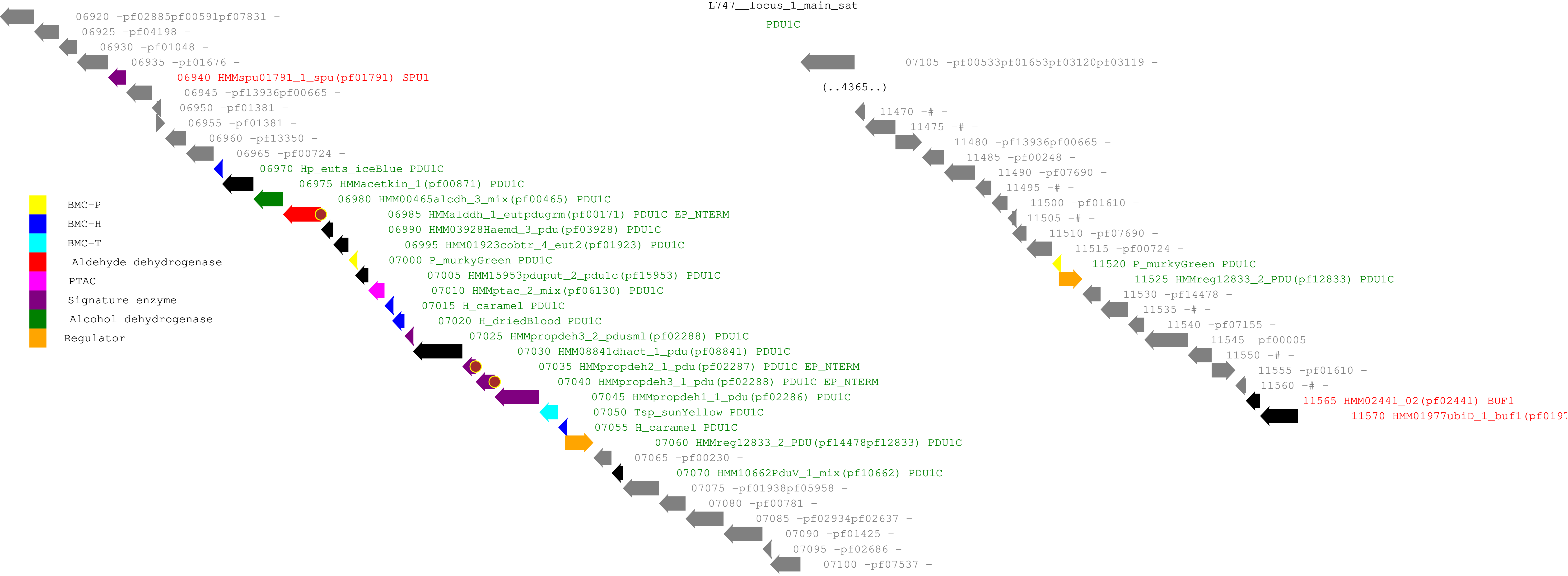

Lactobacillus brevis. Bacteria; Firmicutes; Bacilli; Lactobacillales; Lactobacillaceae;

LbDm2__locus_1_main_sat

PDU1C

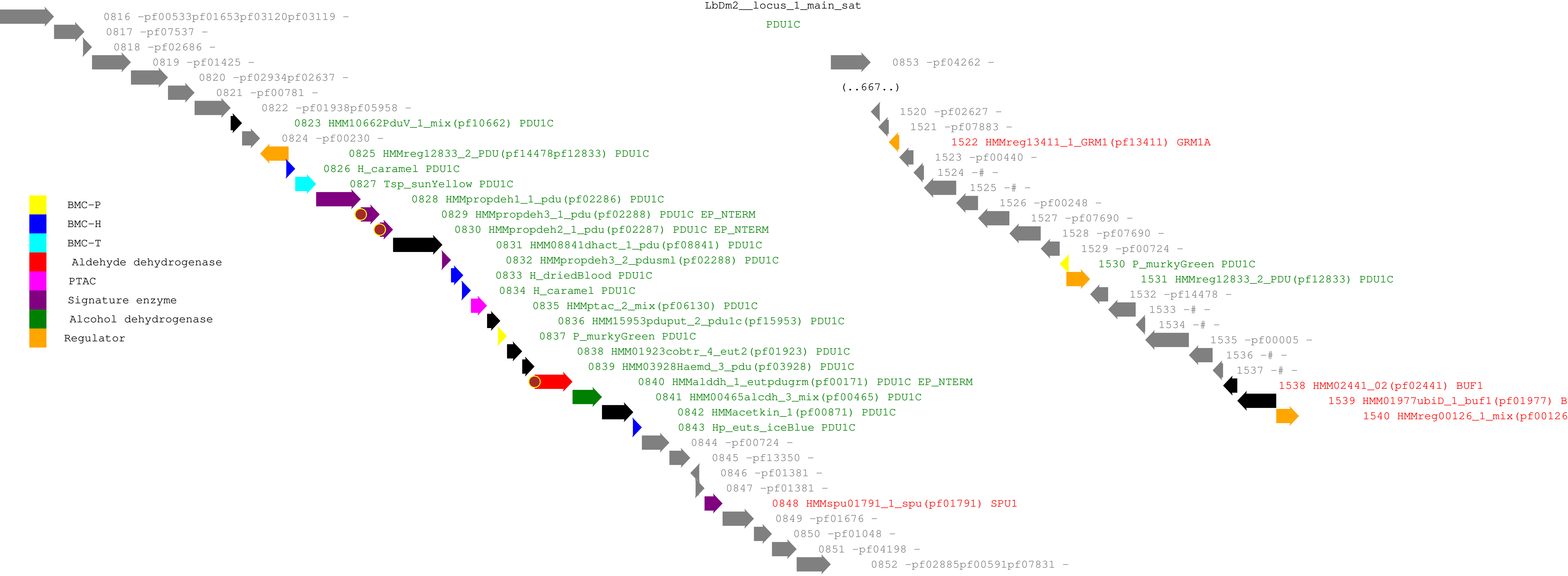

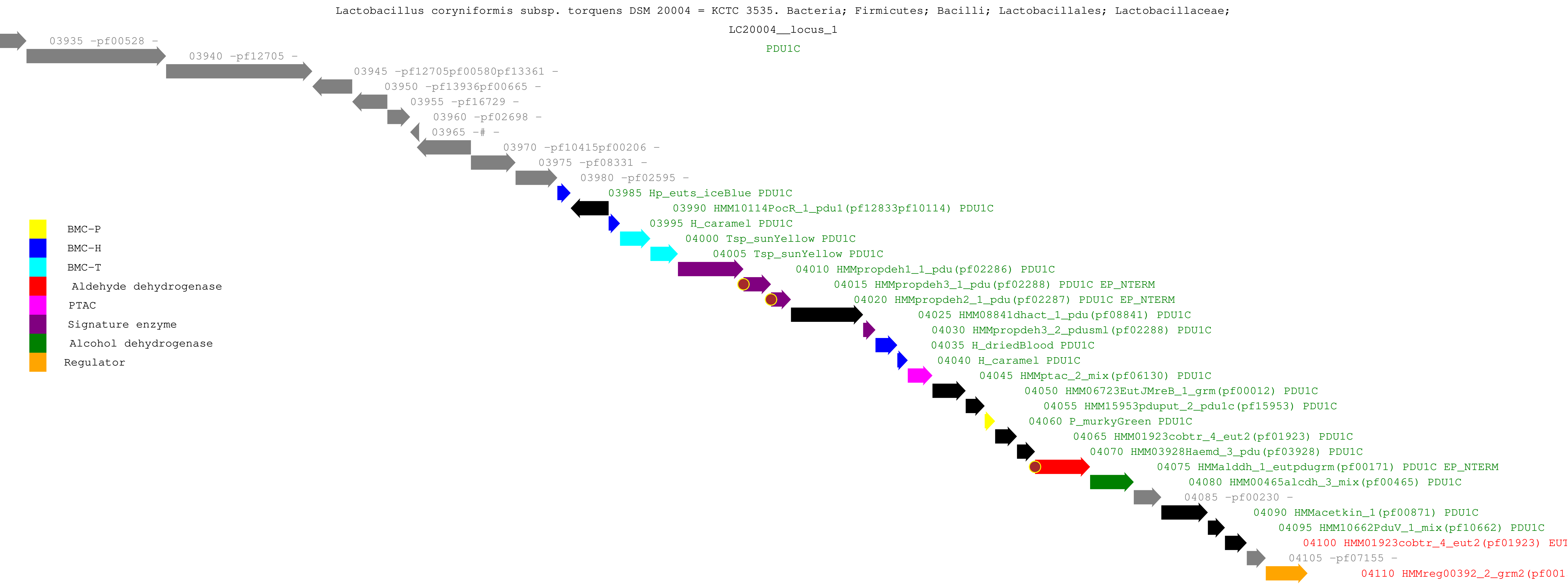
Lactobacillus coryniformis subsp. torquens DSM 20004 = KCTC 3535. Bacteria; Firmicutes; Bacilli; Lactobacillales; Lactobacillaceae;
LC20004__locus_1
PDU1C
03935 -pf00528 -
03940 -pf12705 -
03945 -pf12705pf00580pf13361 -
03950 -pf13936pf00665 -
03955 -pf16729 -
03960 -pf02698 -
03965 -# -
03970 -pf10415pf00206 -
03975 -pf08331 -
03980 -pf02595 -
03985 Hp_euts_iceBlue PDU1C
03990 HMM10114PocR_1_pdu1(pf12833pf10114) PDU1C
03995 H_caramel PDU1C
04000 Tsp_sunYellow PDU1C
04005 Tsp_sunYellow PDU1C
04010 HMMpropdeh1_1_pdu(pf02286) PDU1C
04015 HMMpropdeh3_1_pdu(pf02288) PDU1C EP_NTERM
04020 HMMpropdeh2_1_pdu(pf02287) PDU1C EP_NTERM
04025 HMM08841dhact_1_pdu(pf08841) PDU1C
04030 HMMpropdeh3_2_pdusml(pf02288) PDU1C
04035 H_driedBlood PDU1C
04040 H_caramel PDU1C
04045 HMMptac_2_mix(pf06130) PDU1C
04050 HMM06723EutJMreB_1_grm(pf00012) PDU1C
04055 HMM15953pduput_2_pdu1c(pf15953) PDU1C
04060 P_murkyGreen PDU1C
04065 HMM01923cobtr_4_eut2(pf01923) PDU1C
04070 HMM03928Haemd_3_pdu(pf03928) PDU1C
04075 HMMalddh_1_eutpdugrm(pf00171) PDU1C EP_NTERM
04080 HMM00465alcdh_3_mix(pf00465) PDU1C
04085 -pf00230 -
04090 HMMacetkin_1(pf00871) PDU1C
04095 HMM10662PduV_1_mix(pf10662) PDU1C
04100 HMM01923cobtr_4_eut2(pf01923) EU1
04105 -pf07155 -
04110 HMMreg00392_2_grm2(pf001
BMC-P
BMC-H
BMC-T
Aldehyde dehydrogenase
PTAC
Signature enzyme
Alcohol dehydrogenase
Regulator

Lactobacillus curvatus. Bacteria; Firmicutes; Bacilli; Lactobacillales; Lactobacillaceae;

LCUFL03__locus_1

PDU1C

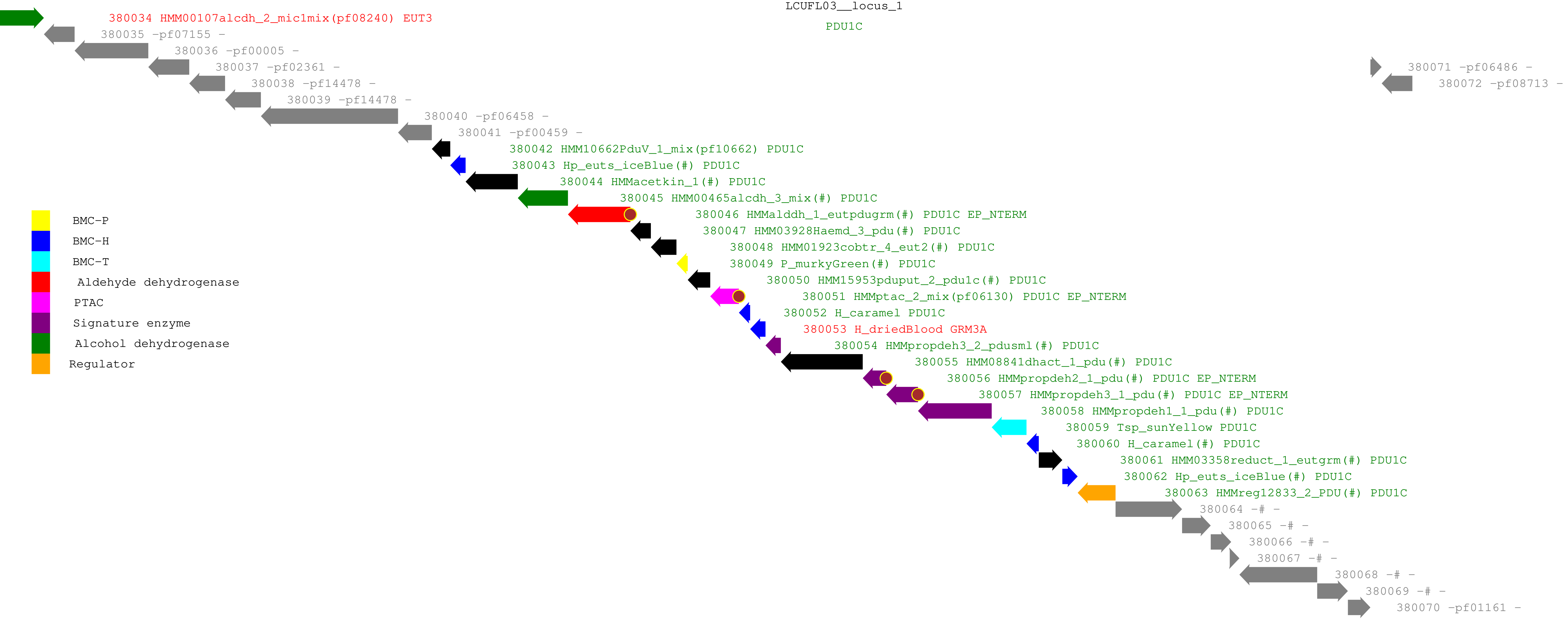

Lactobacillus fuchuensis. Bacteria; Firmicutes; Bacilli; Lactobacillales; Lactobacillaceae;

LFUMFP__locus_1

PDU1C

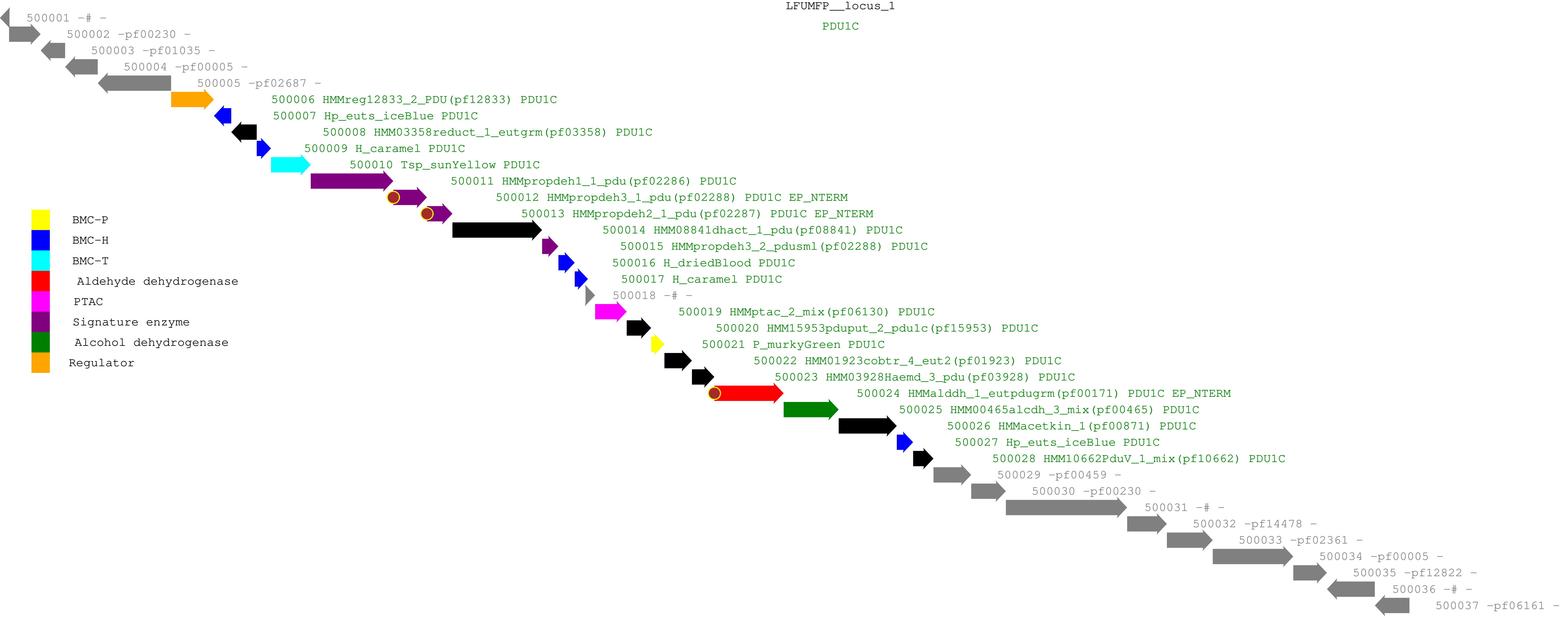

Lactobacillus graminis. Bacteria; Firmicutes; Bacilli; Lactobacillales; Lactobacillaceae;

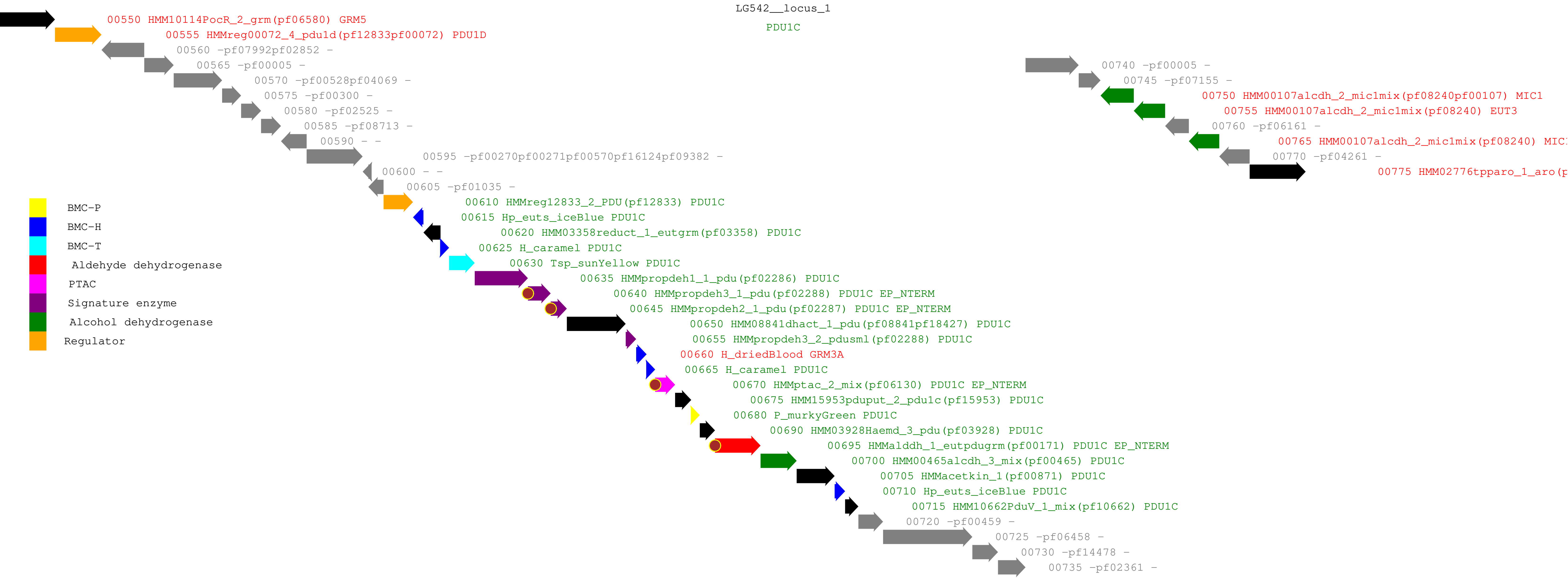

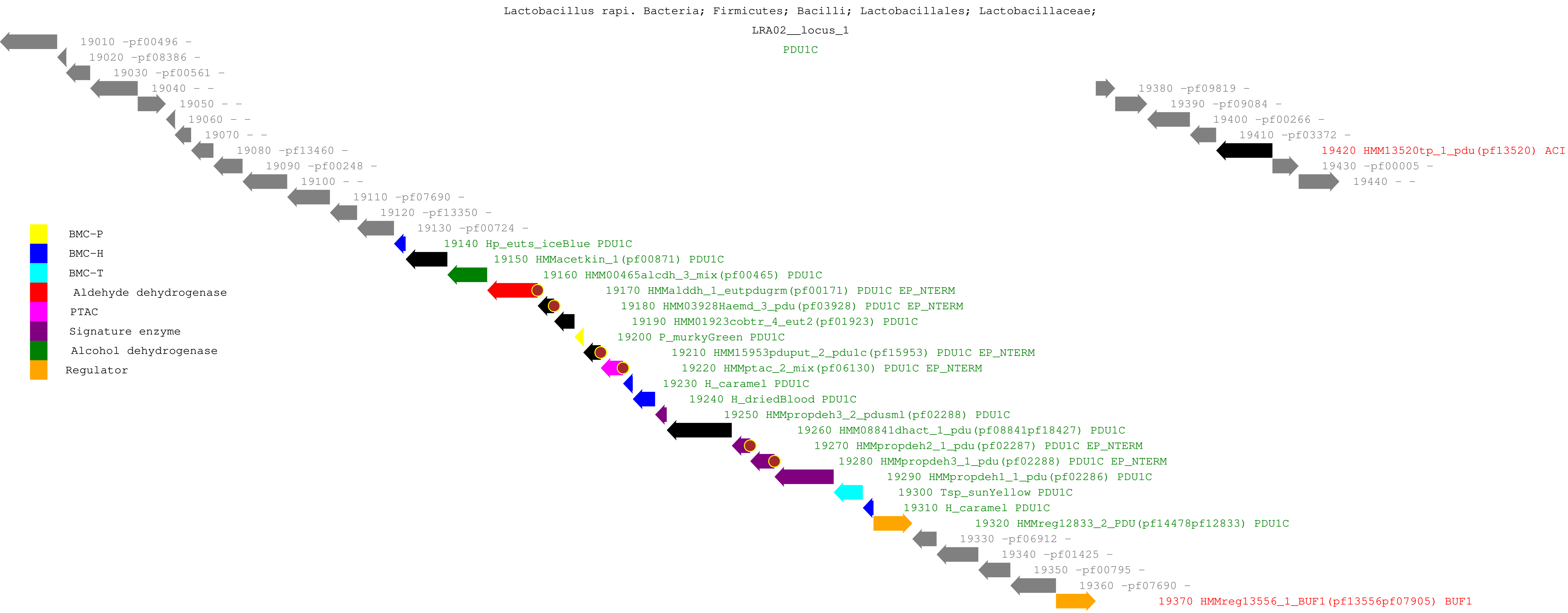
Lactobacillus rapi. Bacteria; Firmicutes; Bacilli; Lactobacillales; Lactobacillaceae;
LRA02__locus_1
PDU1C
19010 -pf00496 -
19020 -pf08386 -
19030 -pf00561 -
19040 - -
19050 - -
19060 - -
19070 - -
19080 -pf13460 -
19090 -pf00248 -
19100 - -
19110 -pf07690 -
19120 -pf13350 -
19130 -pf00724 -
19140 Hp_euts_iceBlue PDU1C
19150 HMMacetkin_1(pf00871) PDU1C
19160 HMM00465alcdh_3_mix(pf00465) PDU1C
19170 HMMalddh_1_eutpdugrm(pf00171) PDU1C EP_NTERM
19180 HMM03928Haemd_3_pdu(pf03928) PDU1C EP_NTERM
19190 HMM01923cobtr_4_eut2(pf01923) PDU1C
19200 P_murkyGreen PDU1C
19210 HMM15953pduput_2_pdu1c(pf15953) PDU1C EP_NTERM
19220 HMMptac_2_mix(pf06130) PDU1C EP_NTERM
19230 H_caramel PDU1C
19240 H_driedBlood PDU1C
19250 HMMpropdeh3_2_pdusml(pf02288) PDU1C
19260 HMM08841dhact_1_pdu(pf08841pf18427) PDU1C
19270 HMMpropdeh2_1_pdu(pf02287) PDU1C EP_NTERM
19280 HMMpropdeh3_1_pdu(pf02288) PDU1C EP_NTERM
19290 HMMpropdeh1_1_pdu(pf02286) PDU1C
19300 Tsp_sunYellow PDU1C
19310 H_caramel PDU1C
19320 HMMreg12833_2_PDU(pf14478pf12833) PDU1C
19330 -pf06912 -
19340 -pf01425 -
19350 -pf00795 -
19360 -pf07690 -
19370 HMMreg13556_1_BUF1(pf13556pf07905) BUF1
19380 -pf09819 -
19390 -pf09084 -
19400 -pf00266 -
19410 -pf03372 -
19420 HMM13520tp_1_pdu(pf13520) ACI
19430 -pf00005 -
19440 - -
BMC-P
BMC-H
BMC-T
Aldehyde dehydrogenase
PTAC
Signature enzyme
Alcohol dehydrogenase
Regulator

Lactobacillus reuteri ATCC 53608. Bacteria; Firmicutes; Bacilli; Lactobacillales; Lactobacillaceae;

LRATCC53608__locus_1_main_sat

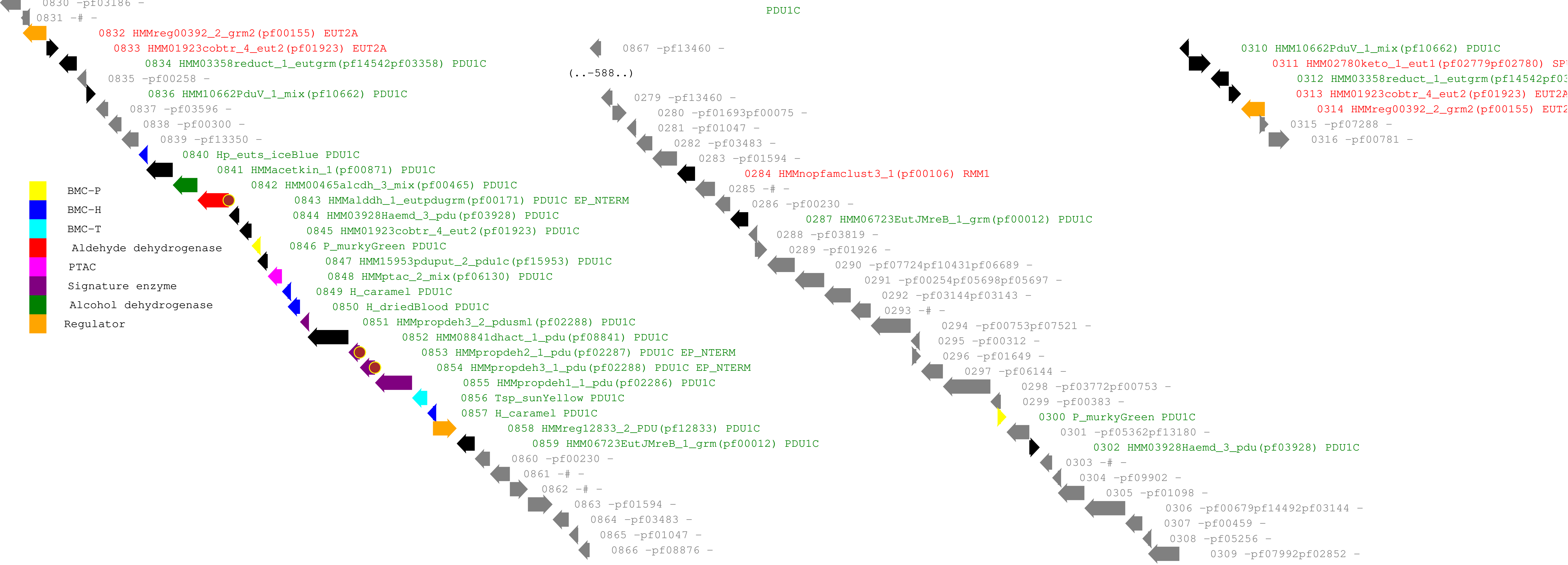

Lactobacillus rennini. Bacteria; Firmicutes; Bacilli; Lactobacillales; Lactobacillaceae;

LREN565__locus_1

PDU1C

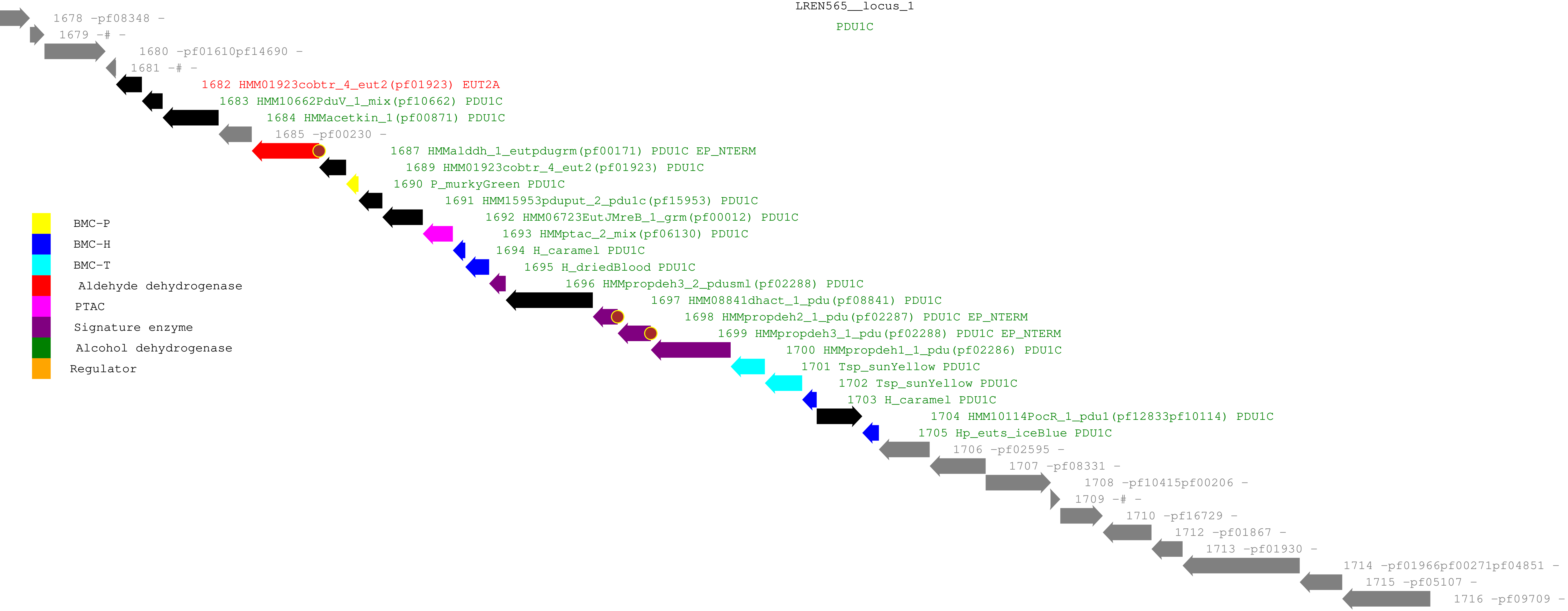

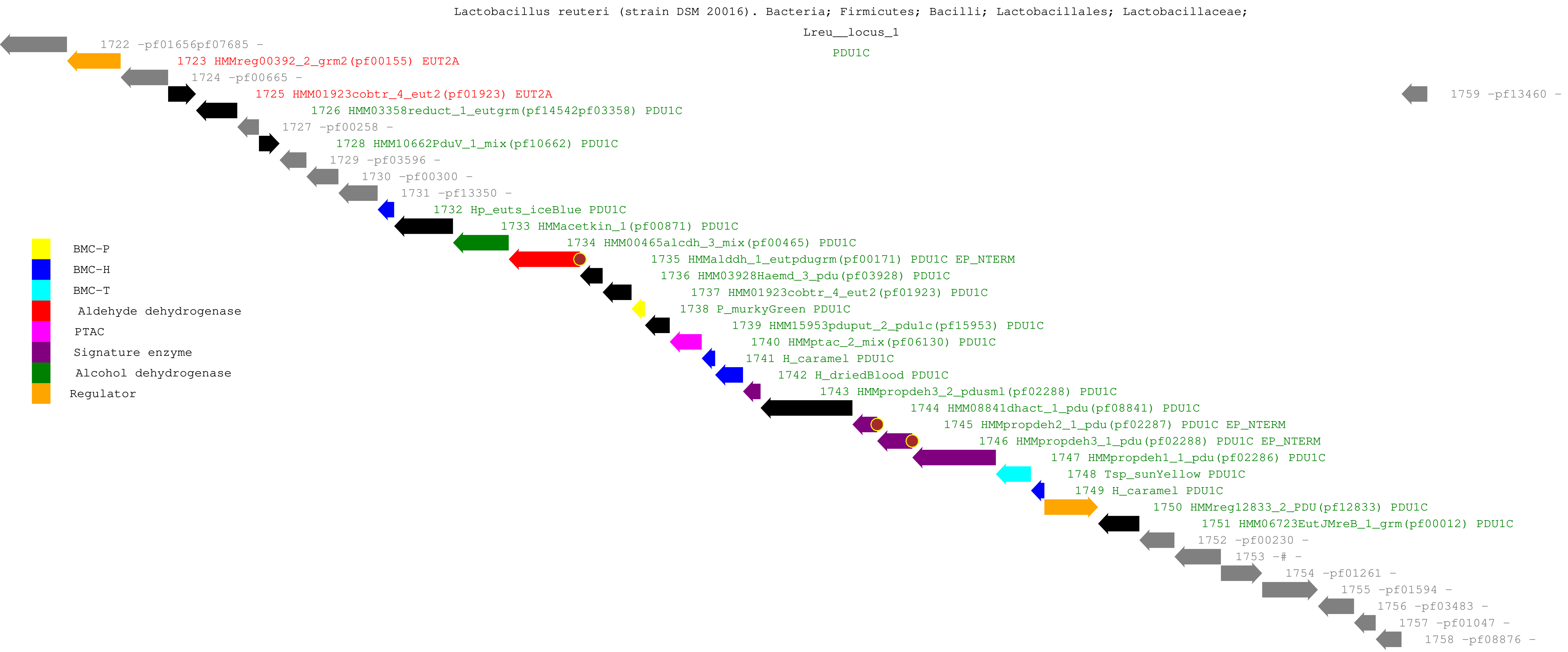
Lactobacillus reuteri (strain DSM 20016). Bacteria; Firmicutes; Bacilli; Lactobacillales; Lactobacillaceae;
Lreu__locus_1
PDU1C
1722 -pf01656pf07685 -
1723 HMMreg00392_2_grm2(pf00155) EUT2A
1724 -pf00665 -
1725 HMM01923cobtr_4_eut2(pf01923) EUT2A
1726 HMM03358reduct_1_eutgrm(pf14542pf03358) PDU1C
1727 -pf00258 -
1728 HMM10662PduV_1_mix(pf10662) PDU1C
1729 -pf03596 -
1730 -pf00300 -
1731 -pf13350 -
1732 Hp_euts_iceBlue PDU1C
1733 HMMacetkin_1(pf00871) PDU1C
1734 HMM00465alcdh_3_mix(pf00465) PDU1C
1735 HMMaldddh_1_eutpdugrm(pf00171) PDU1C EP_NTERM
1736 HMM03928Haemd_3_pdu(pf03928) PDU1C
1737 HMM01923cobtr_4_eut2(pf01923) PDU1C
1738 P_murkyGreen PDU1C
1739 HMM15953pduput_2_pdu1c(pf15953) PDU1C
1740 HMMptac_2_mix(pf06130) PDU1C
1741 H_caramel PDU1C
1742 H_driedBlood PDU1C
1743 HMMpropdeh3_2_pdusml(pf02288) PDU1C
1744 HMM08841dhact_1_pdu(pf08841) PDU1C
1745 HMMpropdeh2_1_pdu(pf02287) PDU1C EP_NTERM
1746 HMMpropdeh3_1_pdu(pf02288) PDU1C EP_NTERM
1747 HMMpropdeh1_1_pdu(pf02286) PDU1C
1748 Tsp_sunYellow PDU1C
1749 H_caramel PDU1C
1750 HMMreg12833_2_PDU(pf12833) PDU1C
1751 HMM06723EutJMreB_1_grm(pf00012) PDU1C
1752 -pf00230 -
1753 -# -
1754 -pf01261 -
1755 -pf01594 -
1756 -pf03483 -
1757 -pf01047 -
1758 -pf08876 -
1759 -pf13460 -
BMC-P
BMC-H
BMC-T
Aldehyde dehydrogenase
PTAC
Signature enzyme
Alcohol dehydrogenase
Regulator

Lactobacillus reuteri. Bacteria; Firmicutes; Bacilli; Lactobacillales; Lactobacillaceae;

lr_locus_1

PDU1C

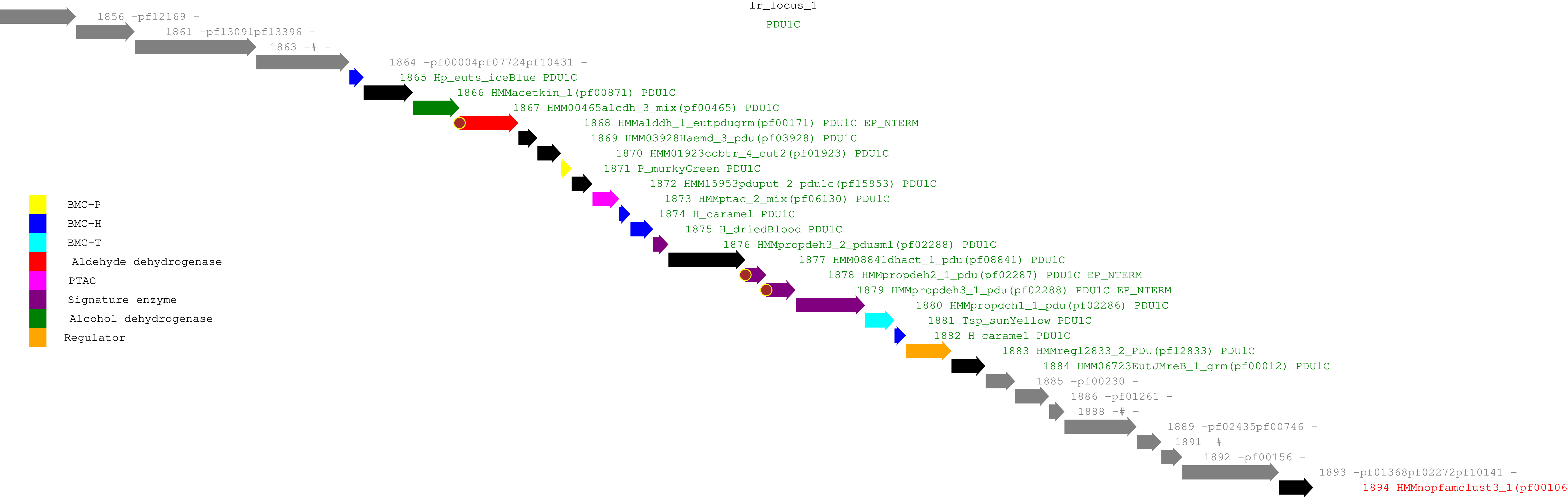

Lactobacillus siliginis. Bacteria; Firmicutes; Bacilli; Lactobacillales; Lactobacillaceae;

LSI01__locus_1

PDU1C

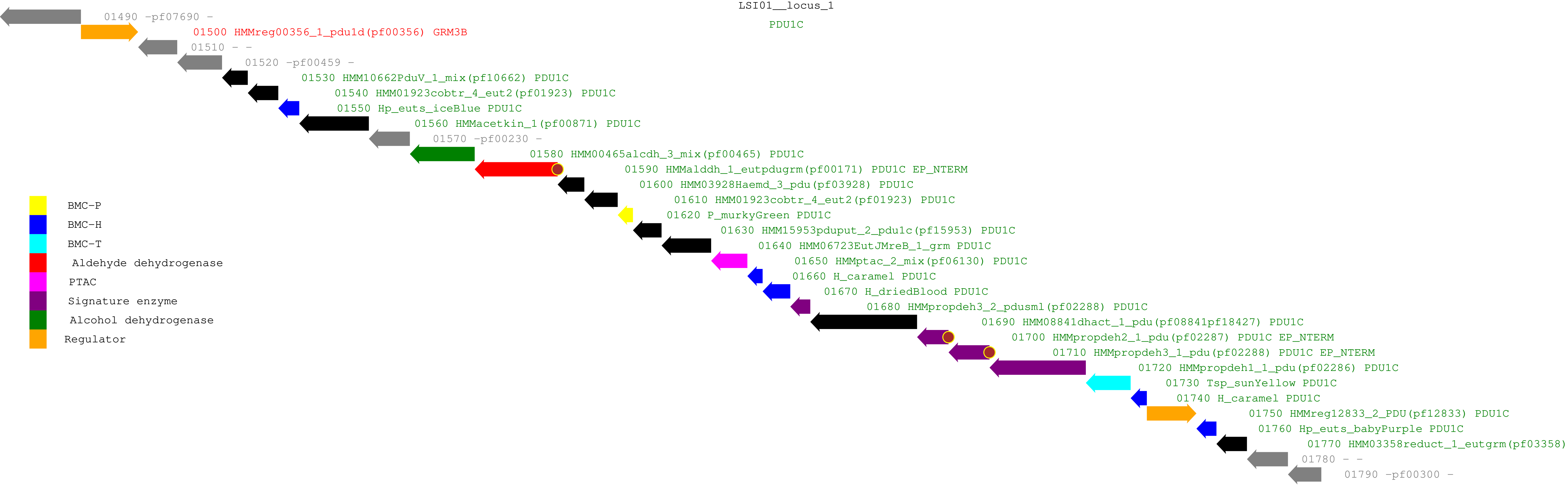

Lactobacillus brevis KB290. Bacteria; Firmicutes; Bacilli; Lactobacillales; Lactobacillaceae;

LVISKB__locus_1_main_sat

PDU1C

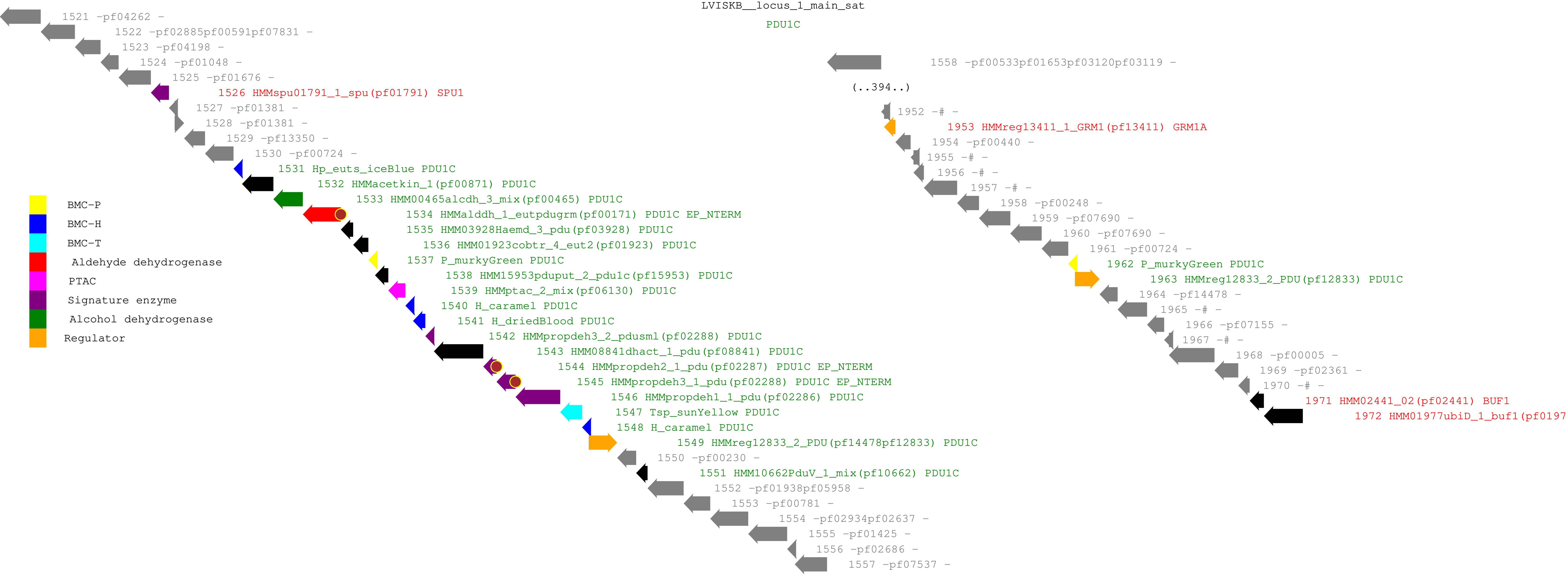

Lactobacillus brevis (strain ATCC 367 / JCM 1170). Bacteria; Firmicutes; Bacilli; Lactobacillales; Lactobacillaceae;

LVIS__locus_1_main_sat

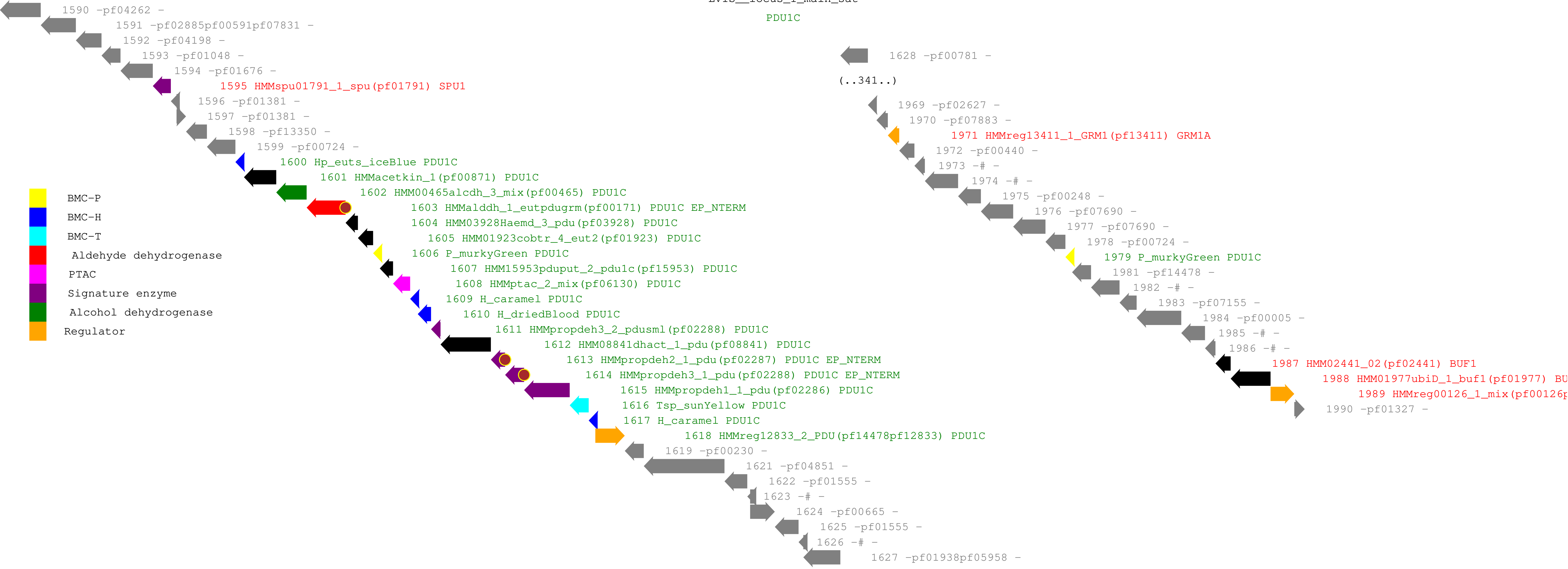

Lactobacillus brevis. Bacteria; Firmicutes; Bacilli; Lactobacillales; Lactobacillaceae;

N624__locus_1

PDU1C

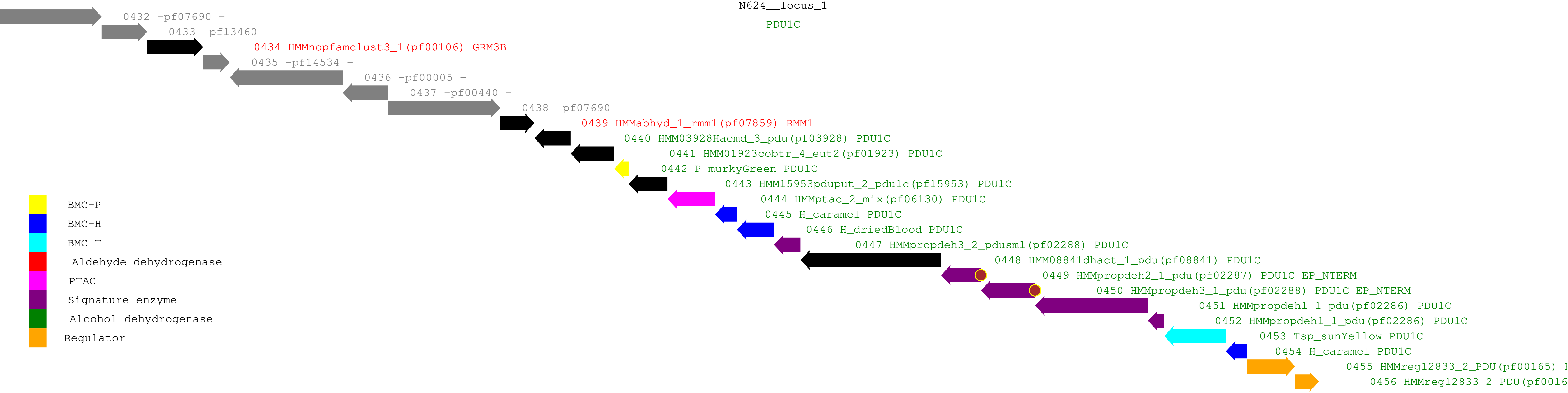

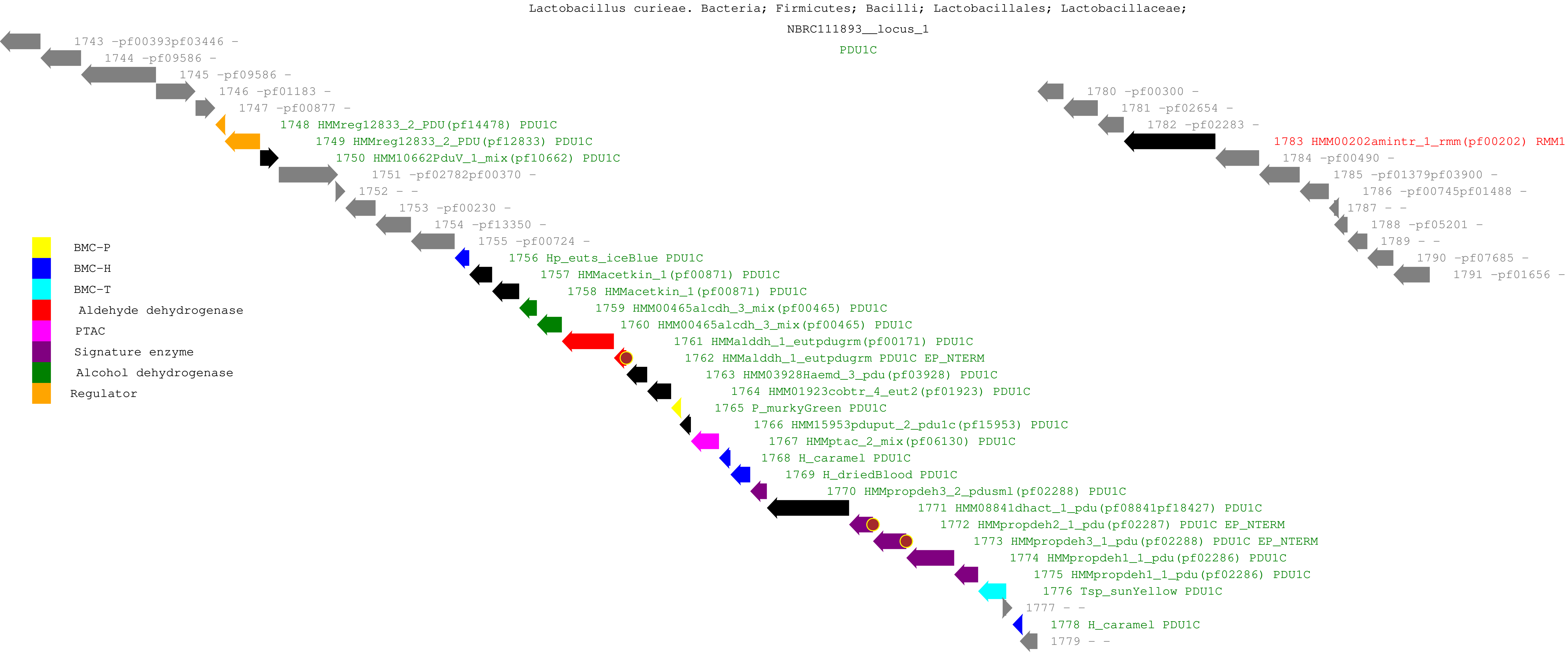
Lactobacillus curieae. Bacteria; Firmicutes; Bacilli; Lactobacillales; Lactobacillaceae;
NBRC111893__locus_1
PDU1C
1743 -pf00393pf03446 -
1744 -pf09586 -
1745 -pf09586 -
1746 -pf01183 -
1747 -pf00877 -
1748 HMMreg12833_2_PDU(pf14478) PDU1C
1749 HMMreg12833_2_PDU(pf12833) PDU1C
1750 HMM10662PduV_1_mix(pf10662) PDU1C
1751 -pf02782pf00370 -
1752 - -
1753 -pf00230 -
1754 -pf13350 -
1755 -pf00724 -
1756 Hp_euts_iceBlue PDU1C
1757 HMMacetkin_1(pf00871) PDU1C
1758 HMMacetkin_1(pf00871) PDU1C
1759 HMM00465alcdh_3_mix(pf00465) PDU1C
1760 HMM00465alcdh_3_mix(pf00465) PDU1C
1761 HMMalddh_1_eutpdugrm(pf00171) PDU1C
1762 HMMalddh_1_eutpdugrm PDU1C EP_NTERM
1763 HMM03928Haemd_3_pdu(pf03928) PDU1C
1764 HMM01923cobtr_4_eut2(pf01923) PDU1C
1765 P_murkyGreen PDU1C
1766 HMM15953pduput_2_pdu1c(pf15953) PDU1C
1767 HMMptac_2_mix(pf06130) PDU1C
1768 H_caramel PDU1C
1769 H_driedBlood PDU1C
1770 HMMpropdeh3_2_pdusml(pf02288) PDU1C
1771 HMM08841dhact_1_pdu(pf08841pf18427) PDU1C
1772 HMMpropdeh2_1_pdu(pf02287) PDU1C EP_NTERM
1773 HMMpropdeh3_1_pdu(pf02288) PDU1C EP_NTERM
1774 HMMpropdeh1_1_pdu(pf02286) PDU1C
1775 HMMpropdeh1_1_pdu(pf02286) PDU1C
1776 Tsp_sunYellow PDU1C
1777 - -
1778 H_caramel PDU1C
1779 - -
1780 -pf00300 -
1781 -pf02654 -
1782 -pf02283 -
1783 HMM00202amintr_1_rmm(pf00202) RMM1
1784 -pf00490 -
1785 -pf01379pf03900 -
1786 -pf00745pf01488 -
1787 - -
1788 -pf05201 -
1789 - -
1790 -pf07685 -
1791 -pf01656 -
BMC-P
BMC-H
BMC-T
Aldehyde dehydrogenase
PTAC
Signature enzyme
Alcohol dehydrogenase
Regulator

Lactobacillus brevis. Bacteria; Firmicutes; Bacilli; Lactobacillales; Lactobacillaceae;

NCTC13386__locus_1

PDU1C

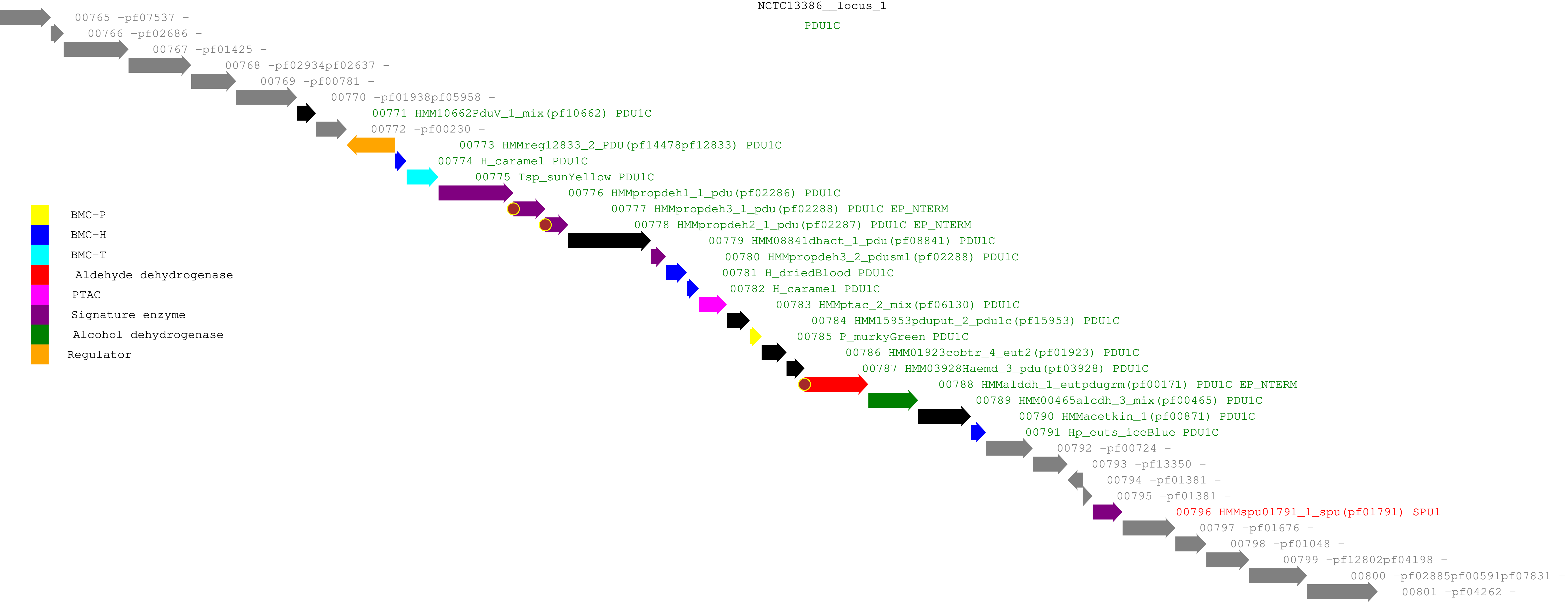

Pediococcus pentosaceus. Bacteria; Firmicutes; Bacilli; Lactobacillales; Lactobacillaceae;

NCTC8066__locus_1

PDU1C

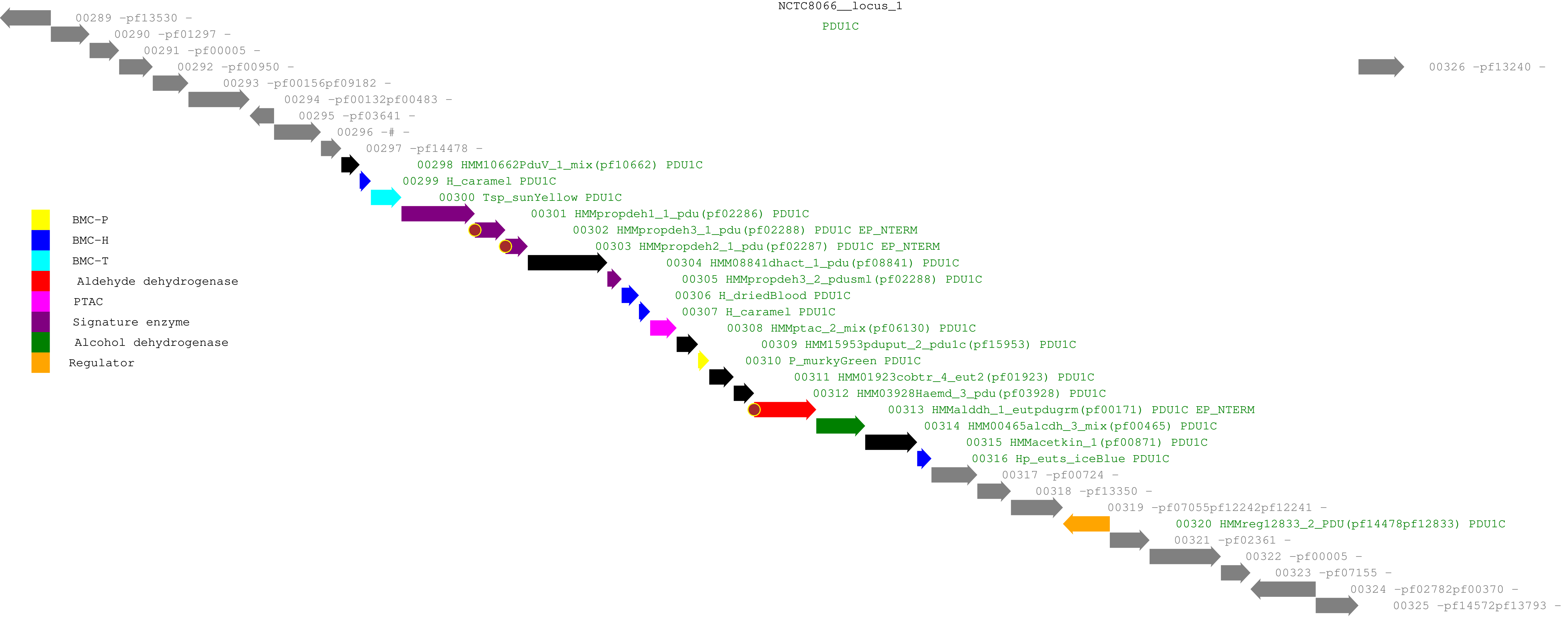

Lactobacillus curvatus. Bacteria; Firmicutes; Bacilli; Lactobacillales; Lactobacillaceae;

OA78__locus_1

PDU1C

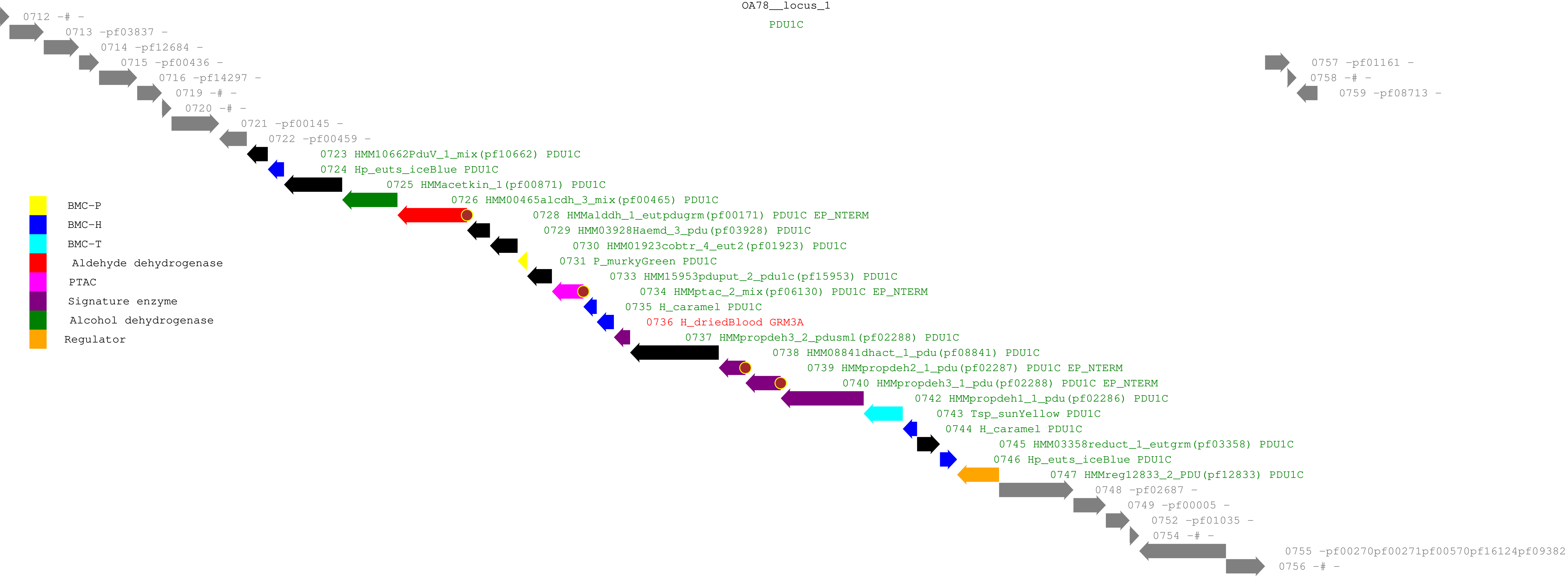

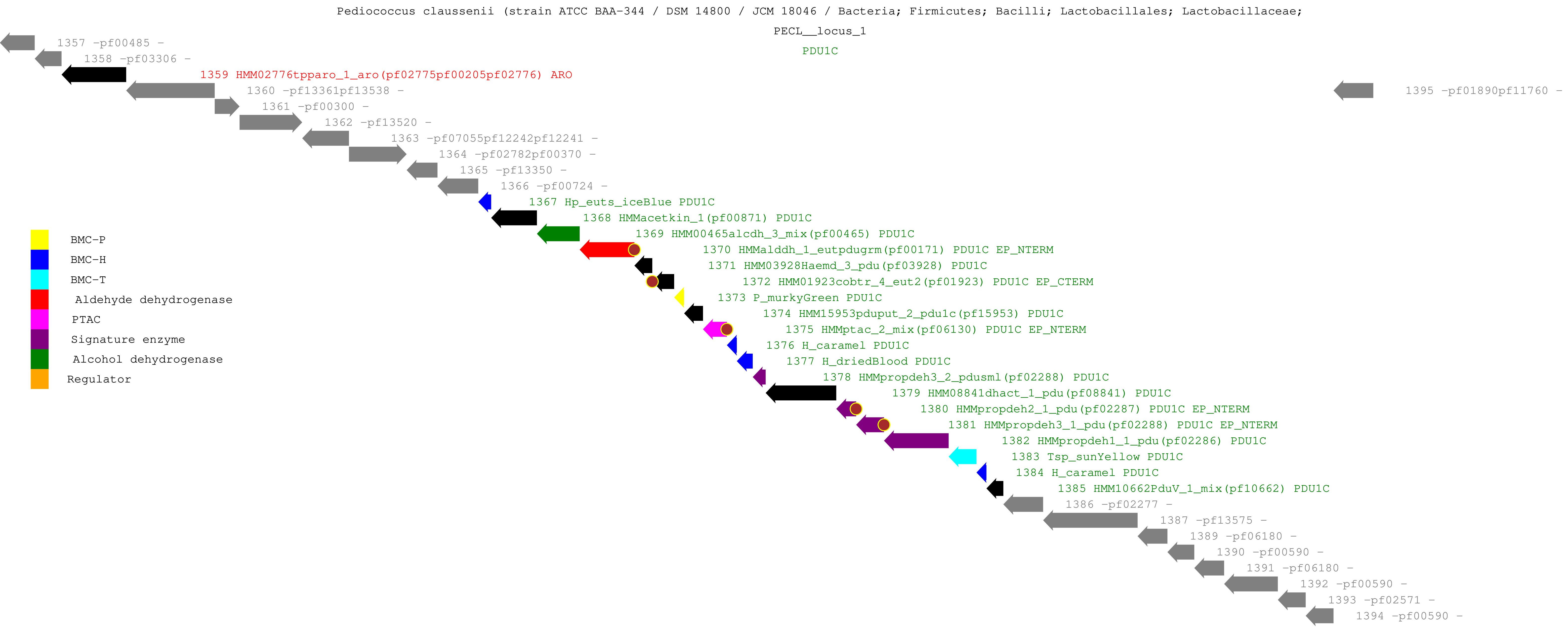
Pediococcus claussenii (strain ATCC BAA-344 / DSM 14800 / JCM 18046 / Bacteria; Firmicutes; Bacilli; Lactobacillales; Lactobacillaceae;
PECL__locus_1
PDU1C
1357 -pf00485 -
1358 -pf03306 -
1359 HMM02776tpparo_1_aro(pf02775pf00205pf02776) ARO
1360 -pf13361pf13538 -
1361 -pf00300 -
1362 -pf13520 -
1363 -pf07055pf12242pf12241 -
1364 -pf02782pf00370 -
1365 -pf13350 -
1366 -pf00724 -
1367 Hp_euts_iceBlue PDU1C
1368 HMMacetkin_1(pf00871) PDU1C
1369 HMM00465alcdh_3_mix(pf00465) PDU1C
1370 HMMalddh_1_eutpdugrm(pf00171) PDU1C EP_NTERM
1371 HMM03928Haemd_3_pdu(pf03928) PDU1C
1372 HMM01923cobtr_4_eut2(pf01923) PDU1C EP_CTERM
1373 P_murkyGreen PDU1C
1374 HMM15953pduput_2_pdu1c(pf15953) PDU1C
1375 HMMptac_2_mix(pf06130) PDU1C EP_NTERM
1376 H_caramel PDU1C
1377 H_driedBlood PDU1C
1378 HMMpropdeh3_2_pdusml(pf02288) PDU1C
1379 HMM08841dhact_1_pdu(pf08841) PDU1C
1380 HMMpropdeh2_1_pdu(pf02287) PDU1C EP_NTERM
1381 HMMpropdeh3_1_pdu(pf02288) PDU1C EP_NTERM
1382 HMMpropdeh1_1_pdu(pf02286) PDU1C
1383 Tsp_sunYellow PDU1C
1384 H_caramel PDU1C
1385 HMM10662PduV_1_mix(pf10662) PDU1C
1386 -pf02277 -
1387 -pf13575 -
1389 -pf06180 -
1390 -pf00590 -
1391 -pf06180 -
1392 -pf00590 -
1393 -pf02571 -
1394 -pf00590 -
1395 -pf01890pf11760 -
BMC-P
BMC-H
BMC-T
Aldehyde dehydrogenase
PTAC
Signature enzyme
Alcohol dehydrogenase
Regulator

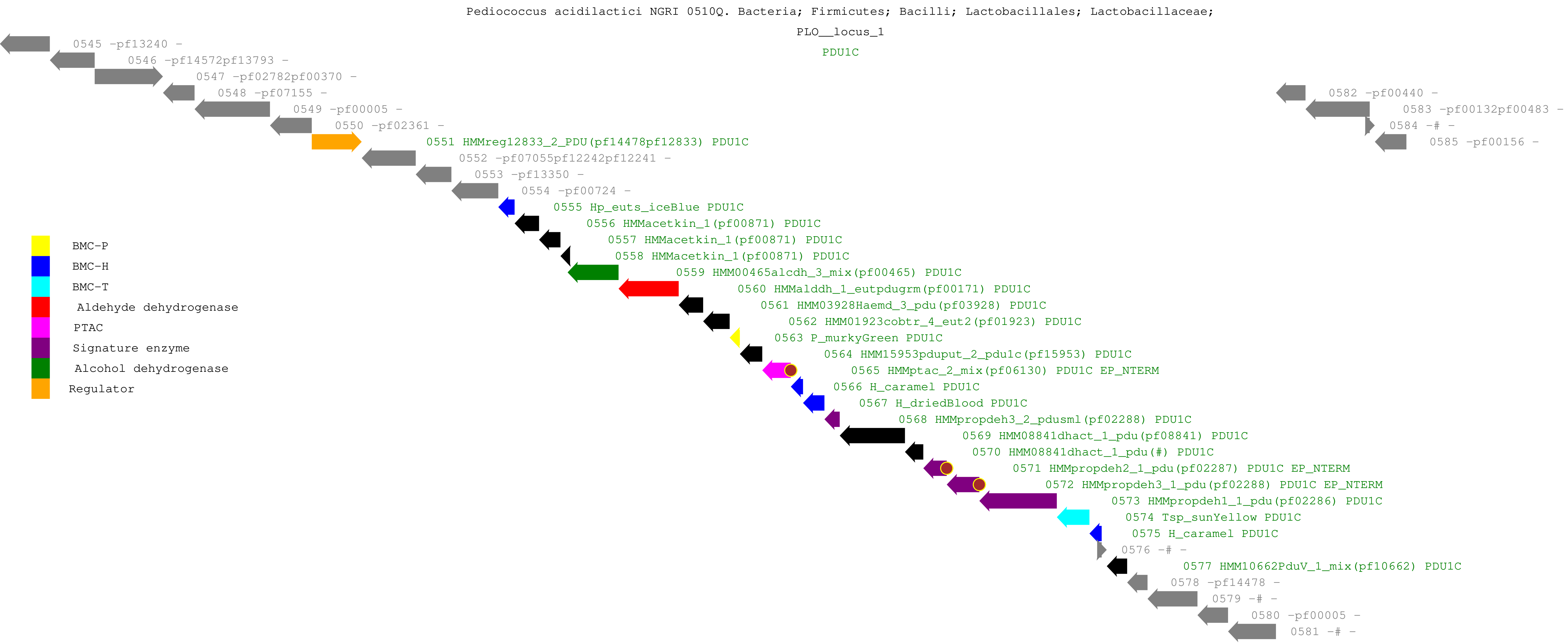
Pediococcus acidilactici NGRI 0510Q. Bacteria; Firmicutes; Bacilli; Lactobacillales; Lactobacillaceae;
PLO__locus_1
PDU1C
0545 -pf13240 -
0546 -pf14572pf13793 -
0547 -pf02782pf00370 -
0548 -pf07155 -
0549 -pf00005 -
0550 -pf02361 -
0551 HMMreg12833_2_PDU(pf14478pf12833) PDU1C
0552 -pf07055pf12242pf12241 -
0553 -pf13350 -
0554 -pf00724 -
0555 Hp_euts_iceBlue PDU1C
0556 HMMacetkin_1(pf00871) PDU1C
0557 HMMacetkin_1(pf00871) PDU1C
0558 HMMacetkin_1(pf00871) PDU1C
0559 HMM00465alcdh_3_mix(pf00465) PDU1C
0560 HMMalddh_1_eutpdugrm(pf00171) PDU1C
0561 HMM03928Haemd_3_pdu(pf03928) PDU1C
0562 HMM01923cobtr_4_eut2(pf01923) PDU1C
0563 P_murkyGreen PDU1C
0564 HMM15953pduput_2_pdu1c(pf15953) PDU1C
0565 HMMptac_2_mix(pf06130) PDU1C EP_NTERM
0566 H_caramel PDU1C
0567 H_driedBlood PDU1C
0568 HMMpropdeh3_2_pdusml(pf02288) PDU1C
0569 HMM08841dhact_1_pdu(pf08841) PDU1C
0570 HMM08841dhact_1_pdu(#) PDU1C
0571 HMMpropdeh2_1_pdu(pf02287) PDU1C EP_NTERM
0572 HMMpropdeh3_1_pdu(pf02288) PDU1C EP_NTERM
0573 HMMpropdeh1_1_pdu(pf02286) PDU1C
0574 Tsp_sunYellow PDU1C
0575 H_caramel PDU1C
0576 -# -
0577 HMM10662PduV_1_mix(pf10662) PDU1C
0578 -pf14478 -
0579 -# -
0580 -pf00005 -
0581 -# -
0582 -pf00440 -
0583 -pf00132pf00483 -
0584 -# -
0585 -pf00156 -
BMC-P
BMC-H
BMC-T
Aldehyde dehydrogenase
PTAC
Signature enzyme
Alcohol dehydrogenase
Regulator

Pediococcus acidilactici. Bacteria; Firmicutes; Bacilli; Lactobacillales; Lactobacillaceae;

SRCM100320__locus_1

PDU1C

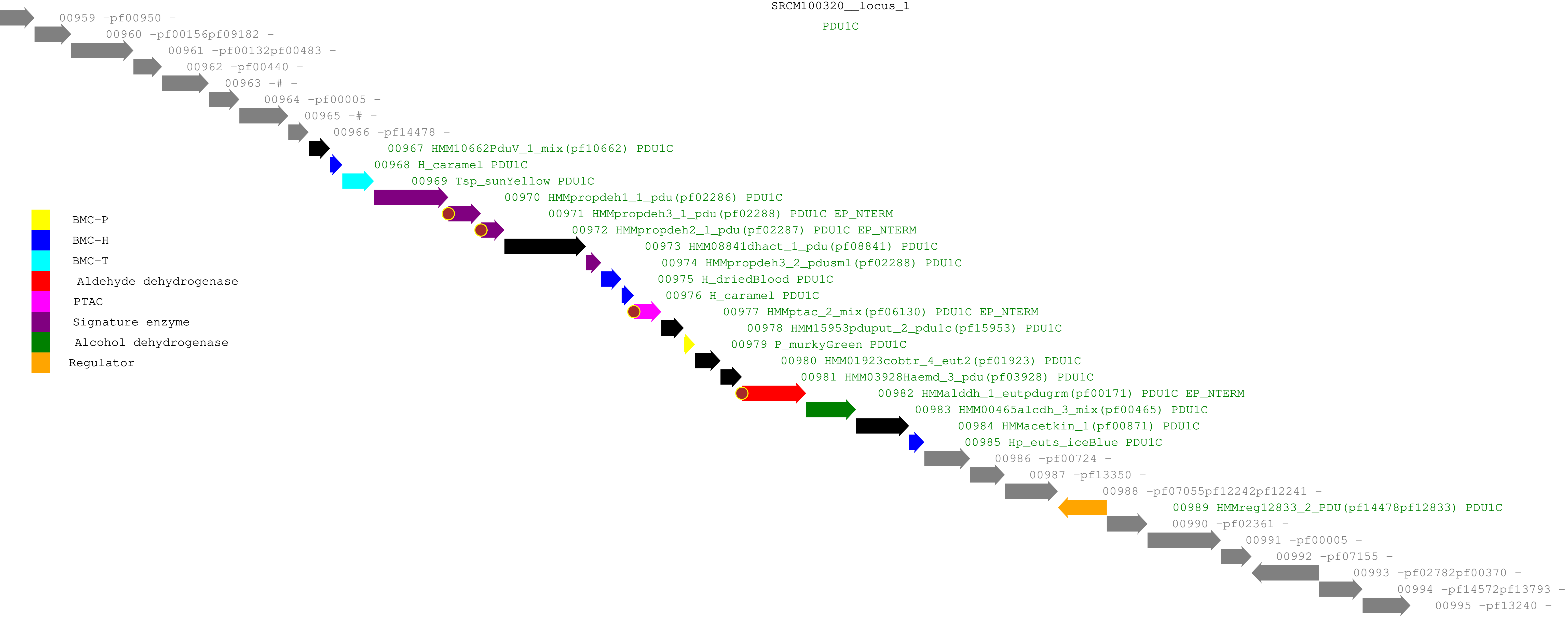

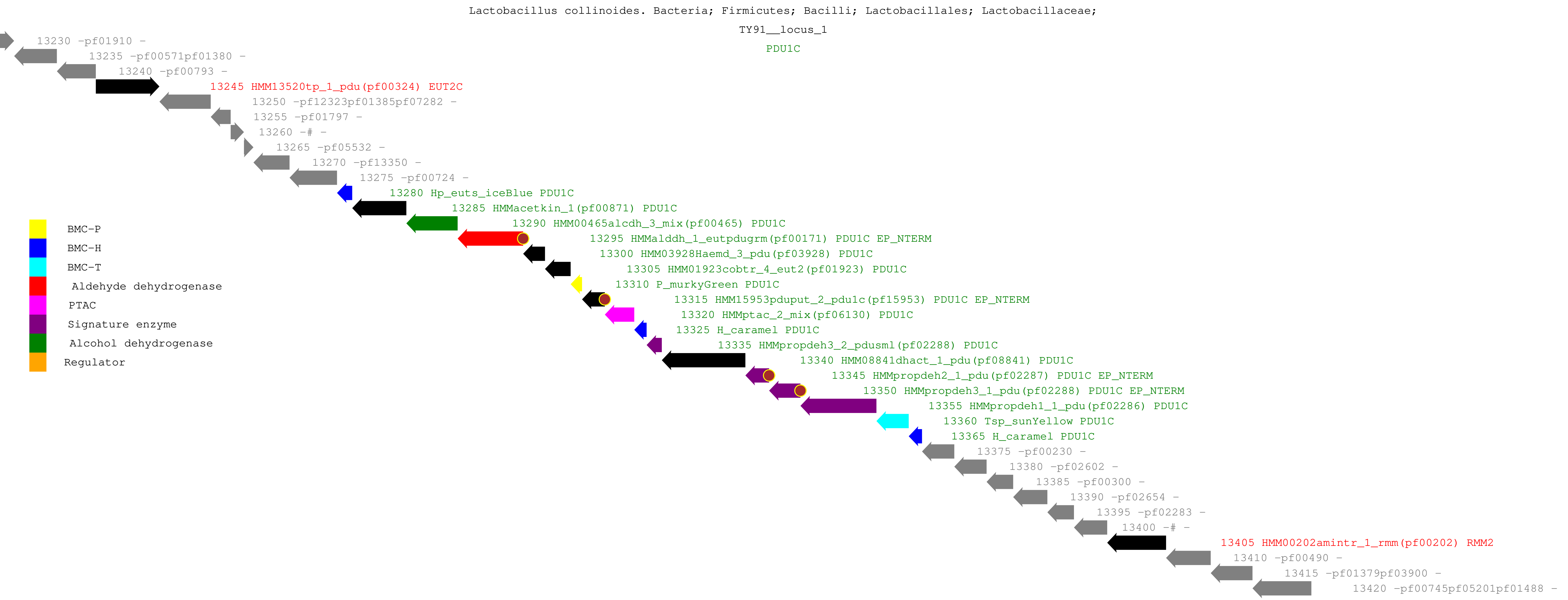
Lactobacillus collinoides. Bacteria; Firmicutes; Bacilli; Lactobacillales; Lactobacillaceae;
TY91__locus_1
PDU1C
13230 -pf01910 -
13235 -pf00571pf01380 -
13240 -pf00793 -
13245 HMM13520tp_1_pdu(pf00324) EUT2C
13250 -pf12323pf01385pf07282 -
13255 -pf01797 -
13260 -# -
13265 -pf05532 -
13270 -pf13350 -
13275 -pf00724 -
13280 Hp_euts_iceBlue PDU1C
13285 HMMacetkin_1(pf00871) PDU1C
13290 HMM00465alcdh_3_mix(pf00465) PDU1C
13295 HMMalddh_1_eutpdugrm(pf00171) PDU1C EP_NTERM
13300 HMM03928Haemd_3_pdu(pf03928) PDU1C
13305 HMM01923cobtr_4_eut2(pf01923) PDU1C
13310 P_murkyGreen PDU1C
13315 HMM15953pduput_2_pdu1c(pf15953) PDU1C EP_NTERM
13320 HMMptac_2_mix(pf06130) PDU1C
13325 H_caramel PDU1C
13335 HMMpropdeh3_2_pdusml(pf02288) PDU1C
13340 HMM08841dhact_1_pdu(pf08841) PDU1C
13345 HMMpropdeh2_1_pdu(pf02287) PDU1C EP_NTERM
13350 HMMpropdeh3_1_pdu(pf02288) PDU1C EP_NTERM
13355 HMMpropdeh1_1_pdu(pf02286) PDU1C
13360 Tsp_sunYellow PDU1C
13365 H_caramel PDU1C
13375 -pf00230 -
13380 -pf02602 -
13385 -pf00300 -
13390 -pf02654 -
13395 -pf02283 -
13400 -# -
13405 HMM00202amintr_1_rmm(pf00202) RMM2
13410 -pf00490 -
13415 -pf01379pf03900 -
13420 -pf00745pf05201pf01488 -
BMC-P
BMC-H
BMC-T
Aldehyde dehydrogenase
PTAC
Signature enzyme
Alcohol dehydrogenase
Regulator

Lactobacillus brevis. Bacteria; Firmicutes; Bacilli; Lactobacillales; Lactobacillaceae;

UCCLBBS449__locus_1

PDU1C

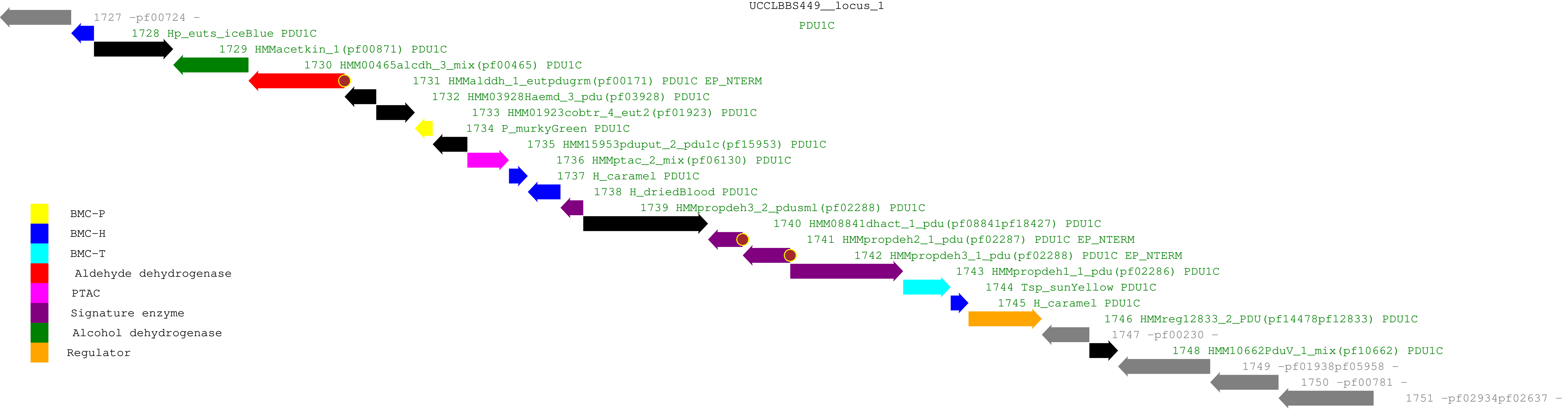

Lactobacillus spicheri. Bacteria; Firmicutes; Bacilli; Lactobacillales; Lactobacillaceae;

VC81__locus_1

PDU1C

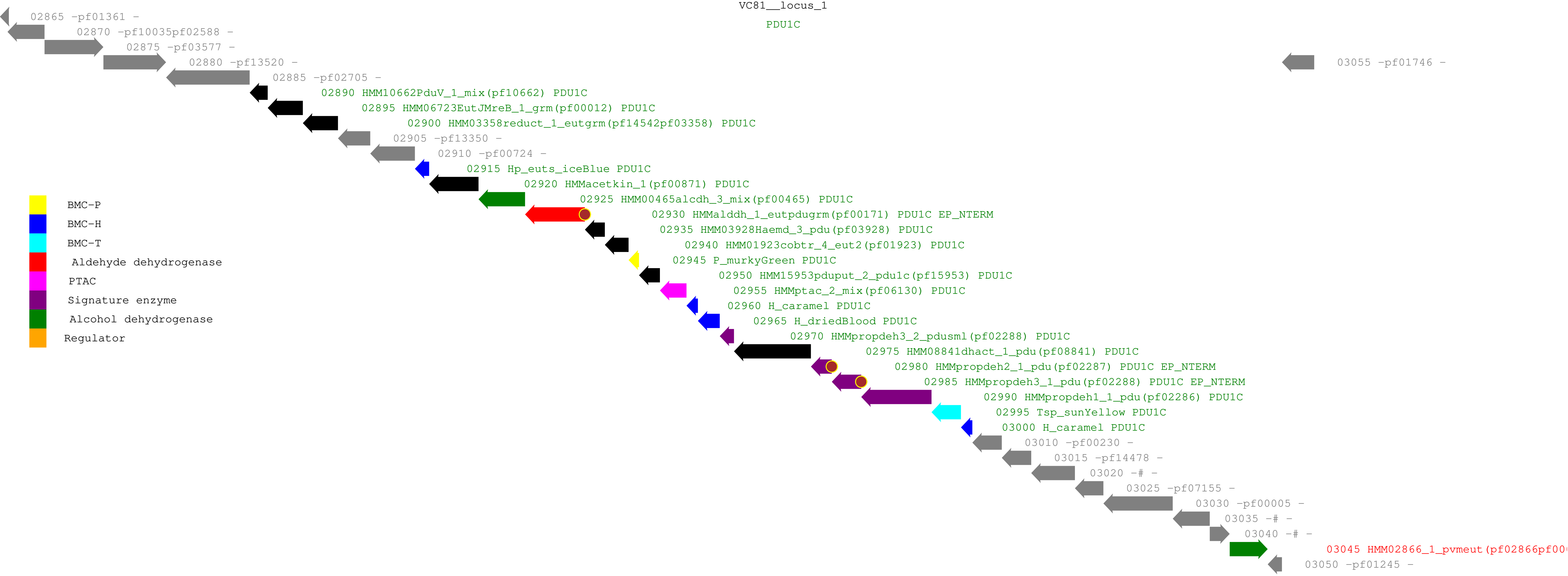

Syntrophobacterales bacterium GWC2_56_13. Bacteria; Proteobacteria; Deltaproteobacteria; Syntrophobacterales.

A2X96__locus_1

PDU1D

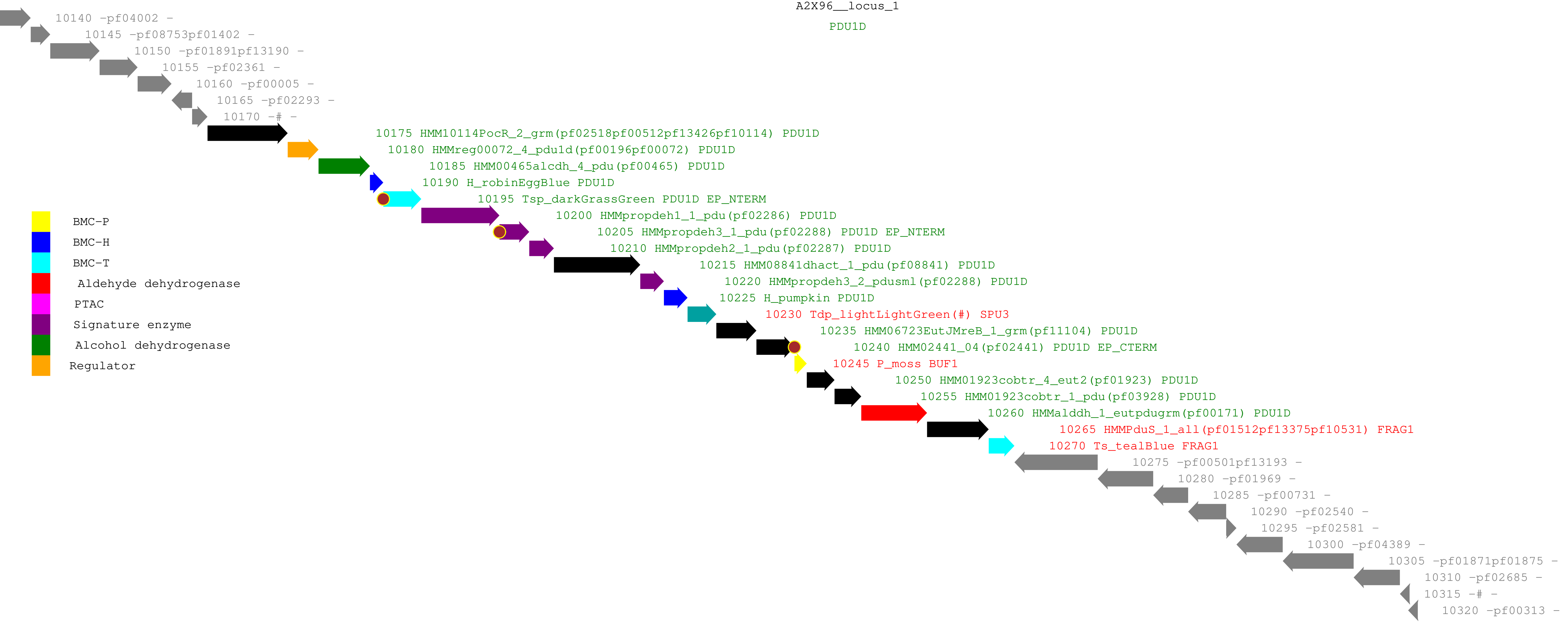

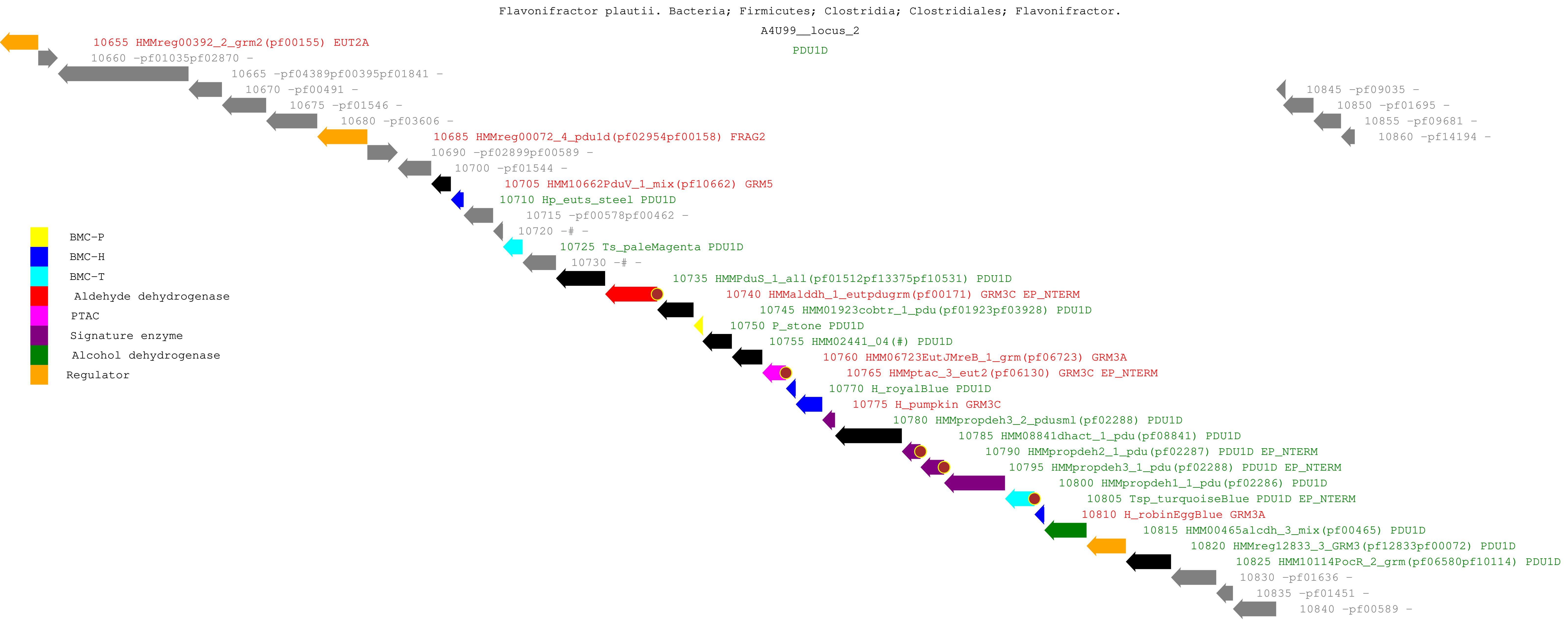
Flavonifractor plautii. Bacteria; Firmicutes; Clostridia; Clostridiales; Flavonifractor.
A4U99__locus_2
PDU1D
10655 HMMreg00392_2_grm2(pf00155) EUT2A
10660 -pf01035pf02870 -
10665 -pf04389pf00395pf01841 -
10670 -pf00491 -
10675 -pf01546 -
10680 -pf03606 -
10685 HMMreg00072_4_pdu1d(pf02954pf00158) FRAG2
10690 -pf02899pf00589 -
10700 -pf01544 -
10705 HMM10662PduV_1_mix(pf10662) GRM5
10710 Hp_euts_steel PDU1D
10715 -pf00578pf00462 -
10720 -# -
10725 Ts_paleMagenta PDU1D
10730 -# -
10735 HMMPduS_1_all(pf01512pf13375pf10531) PDU1D
10740 HMMaldh_1_eutpdugrm(pf00171) GRM3C EP_NTERM
10745 HMM01923cobtr_1_pdu(pf01923pf03928) PDU1D
10750 P_stone PDU1D
10755 HMM02441_04(#) PDU1D
10760 HMM06723EutJMreB_1_grm(pf06723) GRM3A
10765 HMMptac_3_eut2(pf06130) GRM3C EP_NTERM
10770 H_royalBlue PDU1D
10775 H_pumpkin GRM3C
10780 HMMpropdeh3_2_pdusml(pf02288) PDU1D
10785 HMM08841dhact_1_pdu(pf08841) PDU1D
10790 HMMpropdeh2_1_pdu(pf02287) PDU1D EP_NTERM
10795 HMMpropdeh3_1_pdu(pf02288) PDU1D EP_NTERM
10800 HMMpropdeh1_1_pdu(pf02286) PDU1D
10805 Tsp_turquoiseBlue PDU1D EP_NTERM
10810 H_robinEggBlue GRM3A
10815 HMM00465alcdh_3_mix(pf00465) PDU1D
10820 HMMreg12833_3_GRM3(pf12833pf00072) PDU1D
10825 HMM10114PocR_2_grm(pf06580pf10114) PDU1D
10830 -pf01636 -
10835 -pf01451 -
10840 -pf00589 -
10845 -pf09035 -
10850 -pf01695 -
10855 -pf09681 -
10860 -pf14194 -
BMC-P
BMC-H
BMC-T
Aldehyde dehydrogenase
PTAC
Signature enzyme
Alcohol dehydrogenase
Regulator

Clostridium estertheticum subsp. estertheticum. Bacteria; Firmicutes; Clostridia; Clostridiales; Clostridiaceae;

A7L45__locus_1

PDU1D

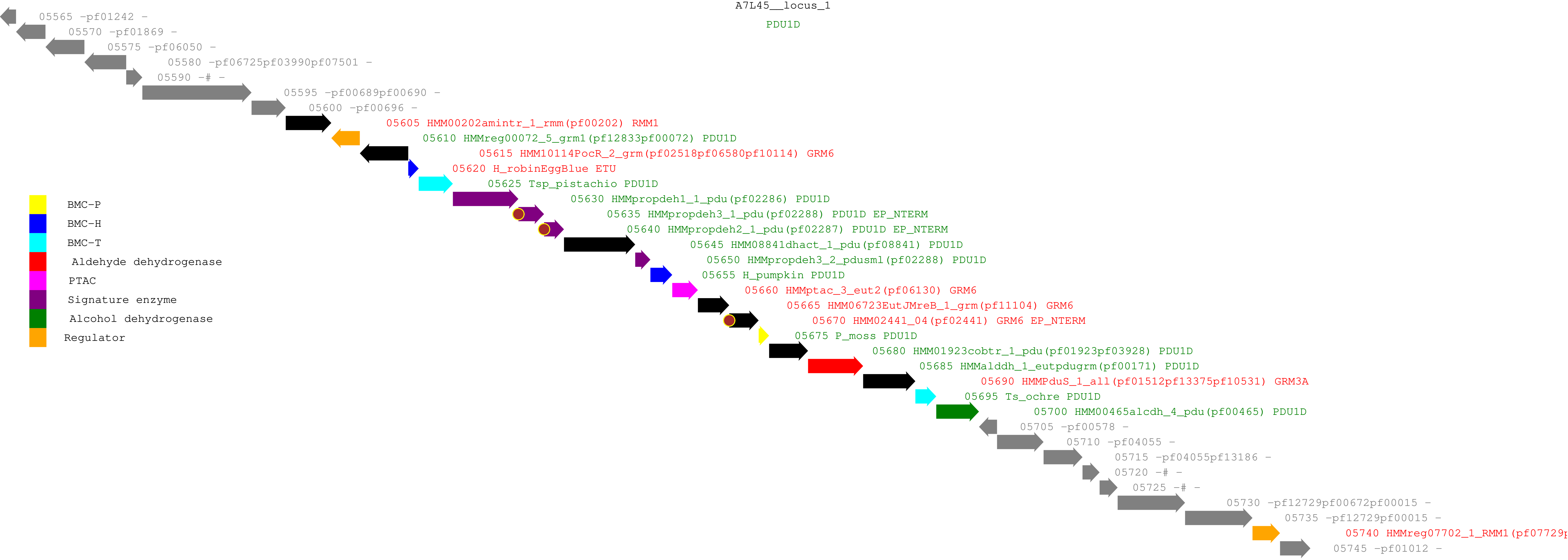

Brachyspira hampsonii. Bacteria; Spirochaetes; Brachyspirales; Brachyspiraceae; Brachyspira.

A9495__locus_1

PDU1D

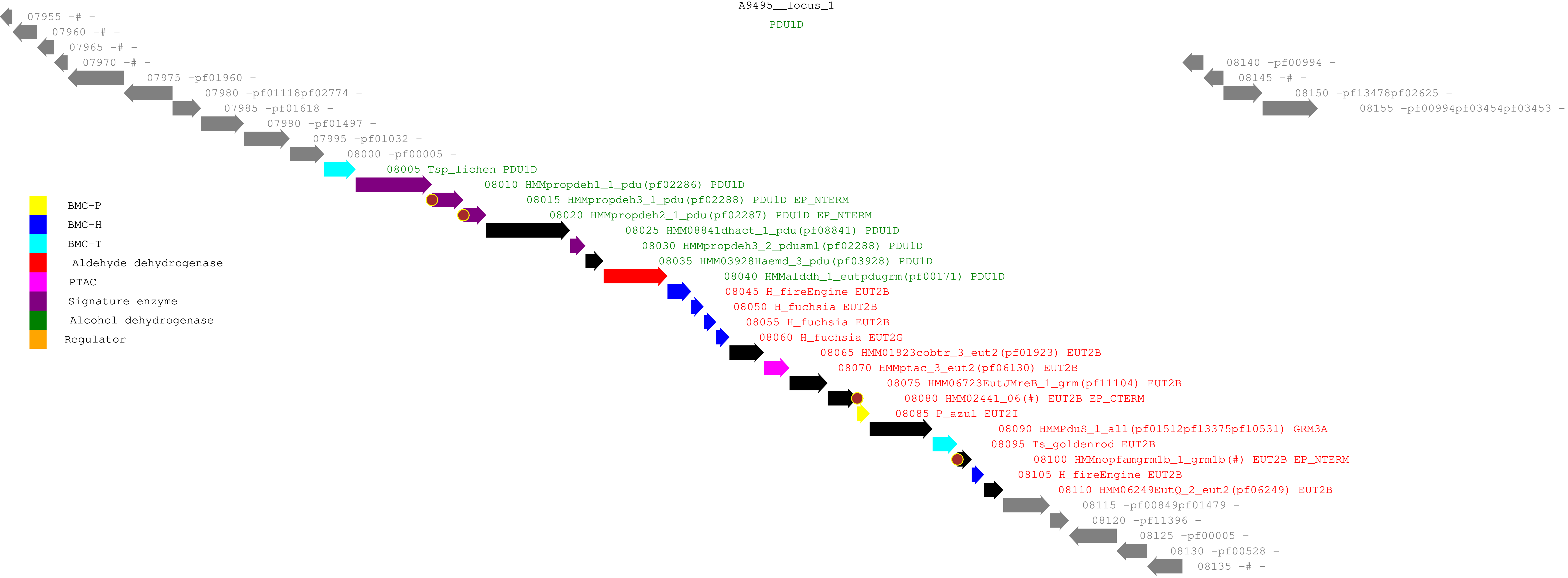

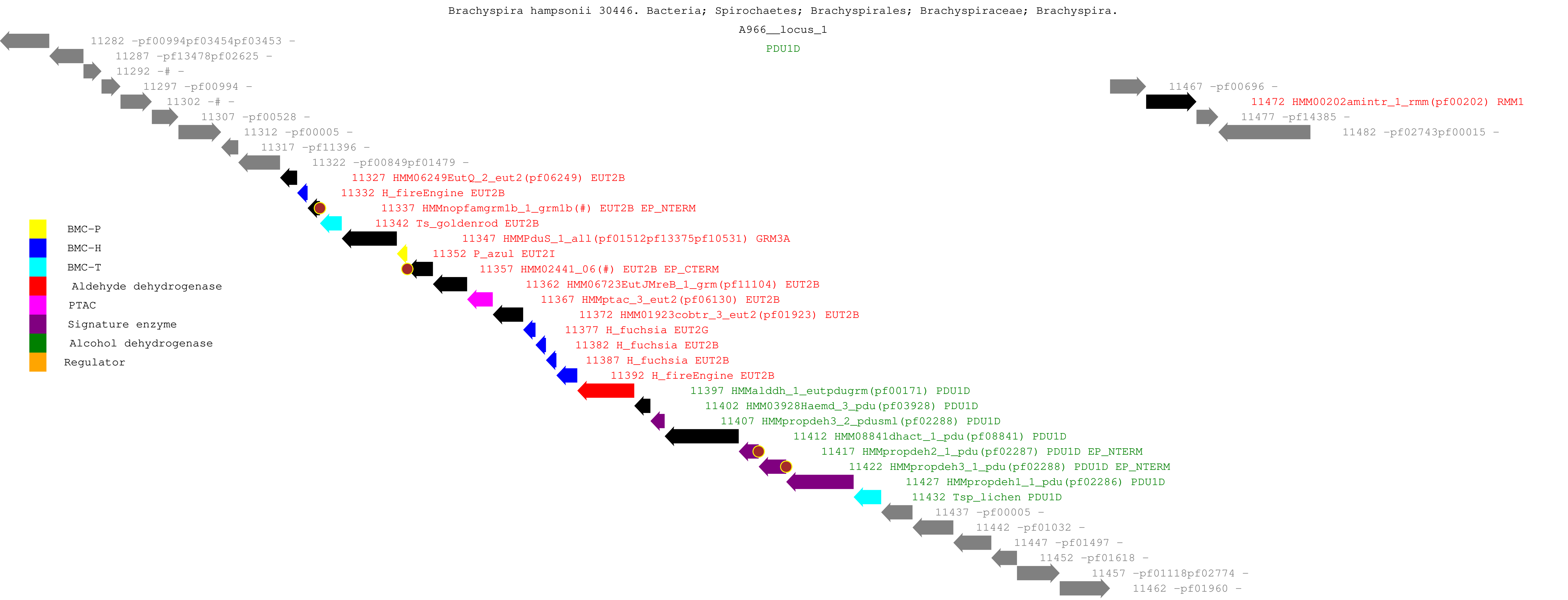

Brachyspira hampsonii 30446. Bacteria; Spirochaetes; Brachyspirales; Brachyspiraceae; Brachyspira.
A966__locus_1
PDU1D
11282 -pf00994pf03454pf03453 -
11287 -pf13478pf02625 -
11292 -# -
11297 -pf00994 -
11302 -# -
11307 -pf00528 -
11312 -pf00005 -
11317 -pf11396 -
11322 -pf00849pf01479 -
11327 HMM06249EutQ_2_eut2(pf06249) EUT2B
11332 H_fireEngine EUT2B
11337 HMMnopfamgrm1b_1_grm1b(#) EUT2B EP_NTERM
11342 Ts_goldenrod EUT2B
11347 HMMPduS_1_all(pf01512pf13375pf10531) GRM3A
11352 P_azul EUT2I
11357 HMM02441_06(#) EUT2B EP_CTERM
11362 HMM06723EutJMreB_1_grm(pf11104) EUT2B
11367 HMMptac_3_eut2(pf06130) EUT2B
11372 HMM01923cobtr_3_eut2(pf01923) EUT2B
11377 H_fuchsia EUT2G
11382 H_fuchsia EUT2B
11387 H_fuchsia EUT2B
11392 H_fireEngine EUT2B
11397 HMMalddh_1_eutpdugrm(pf00171) PDU1D
11402 HMM03928Haemd_3_pdu(pf03928) PDU1D
11407 HMMpropdeh3_2_pdusml(pf02288) PDU1D
11412 HMM08841dhact_1_pdu(pf08841) PDU1D
11417 HMMpropdeh2_1_pdu(pf02287) PDU1D EP_NTERM
11422 HMMpropdeh3_1_pdu(pf02288) PDU1D EP_NTERM
11427 HMMpropdeh1_1_pdu(pf02286) PDU1D
11432 Tsp_lichen PDU1D
11437 -pf00005 -
11442 -pf01032 -
11447 -pf01497 -
11452 -pf01618 -
11457 -pf01118pf02774 -
11462 -pf01960 -
11467 -pf00696 -
11472 HMM00202amintr_1_rmm(pf00202) RMM1
11477 -pf14385 -
11482 -pf02743pf00015 -
BMC-P
BMC-H
BMC-T
Aldehyde dehydrogenase
PTAC
Signature enzyme
Alcohol dehydrogenase
Regulator

Eubacterium limosum. Bacteria; Firmicutes; Clostridia; Clostridiales; Eubacteriaceae;

ACH52__locus_1_main_sat

PDU1D

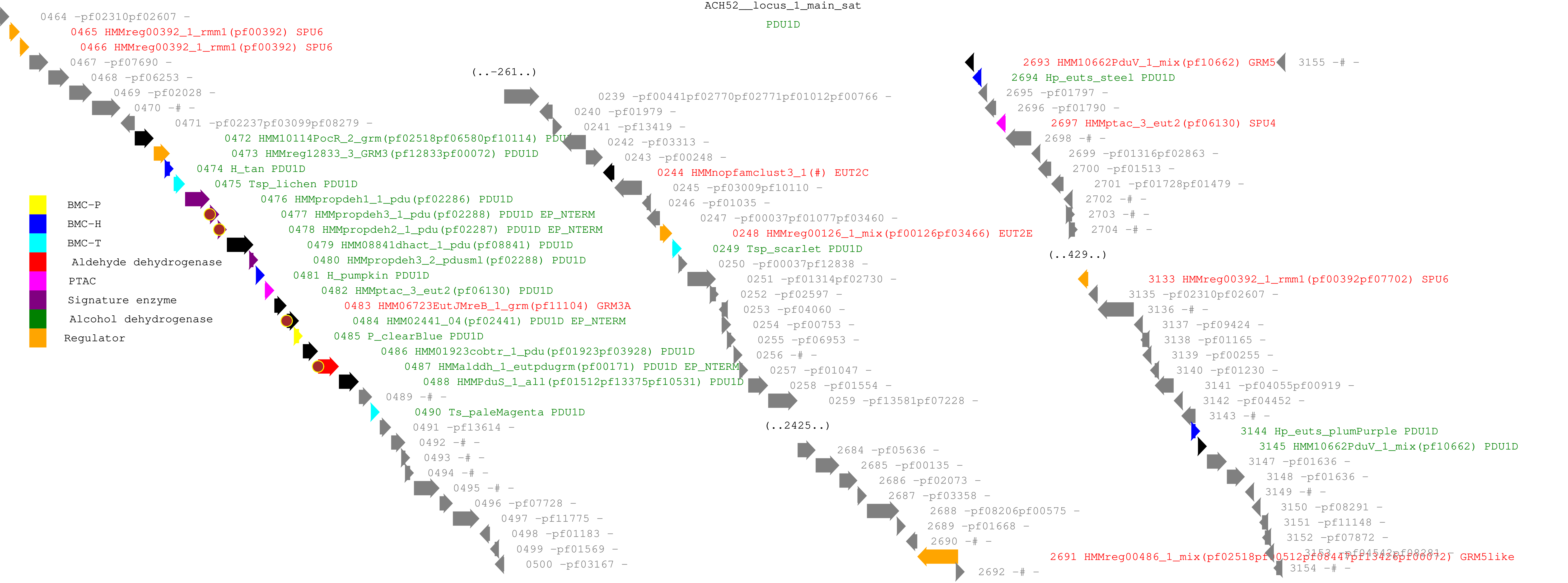

Acetobacterium wieringae. Bacteria; Firmicutes; Clostridia; Clostridiales; Eubacteriaceae;

ACWI__locus_1_main_sat

PDU1D

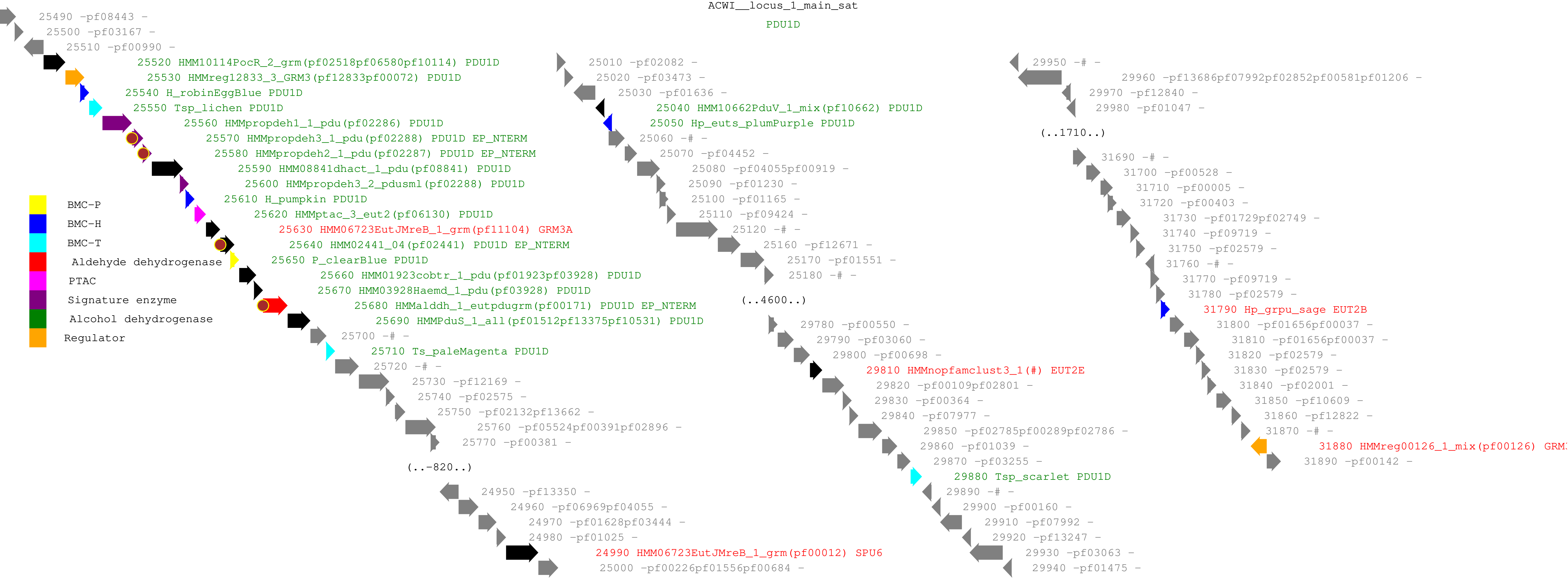

Acetobacterium bakii. Bacteria; Firmicutes; Clostridia; Clostridiales; Eubacteriaceae;

AKG39__locus_2

PDU1D

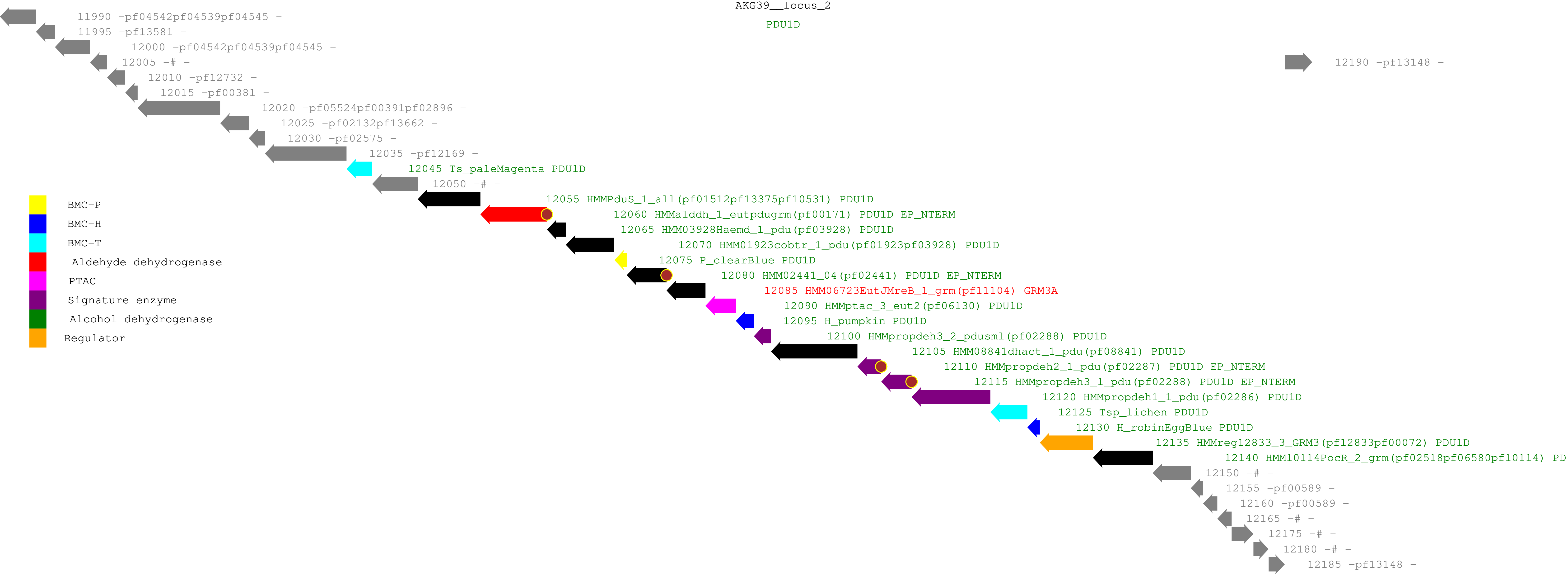

Acetonema longum DSM 6540. Bacteria; Firmicutes; Negativicutes; Selenomonadales; Sporomusaceae;

ALO__locus_1

PDU1D

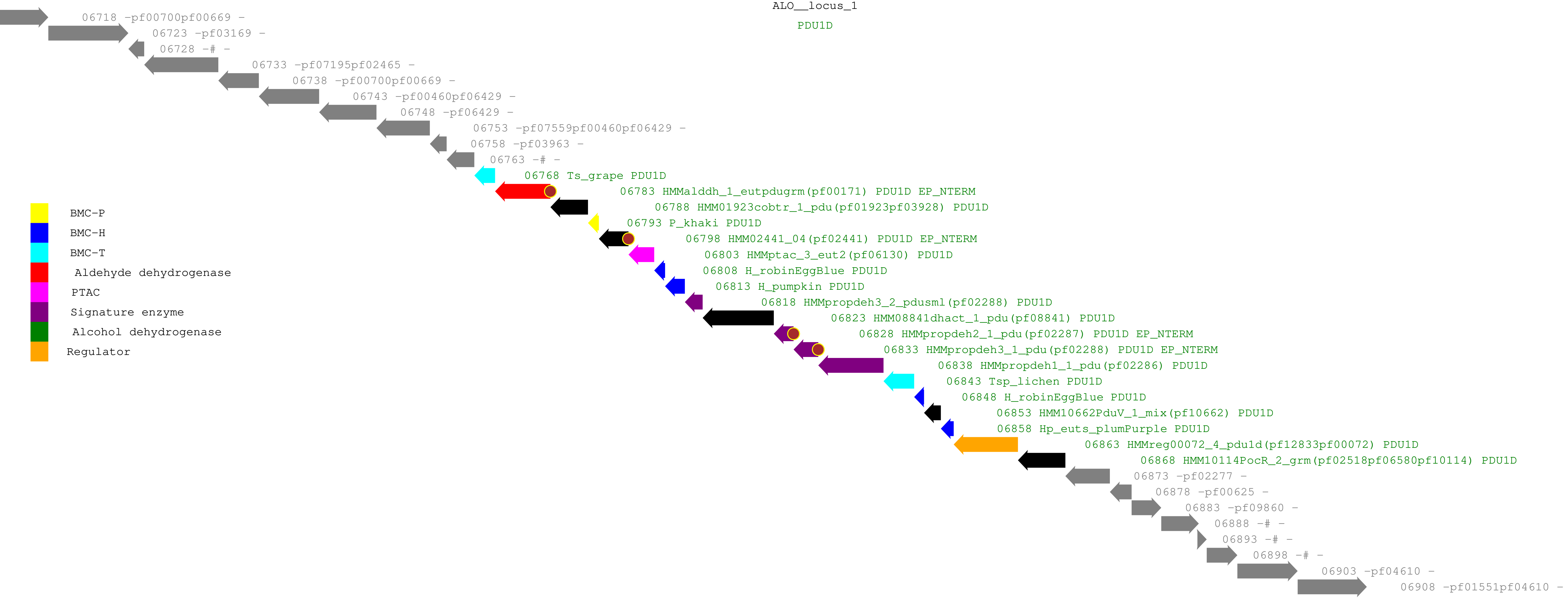

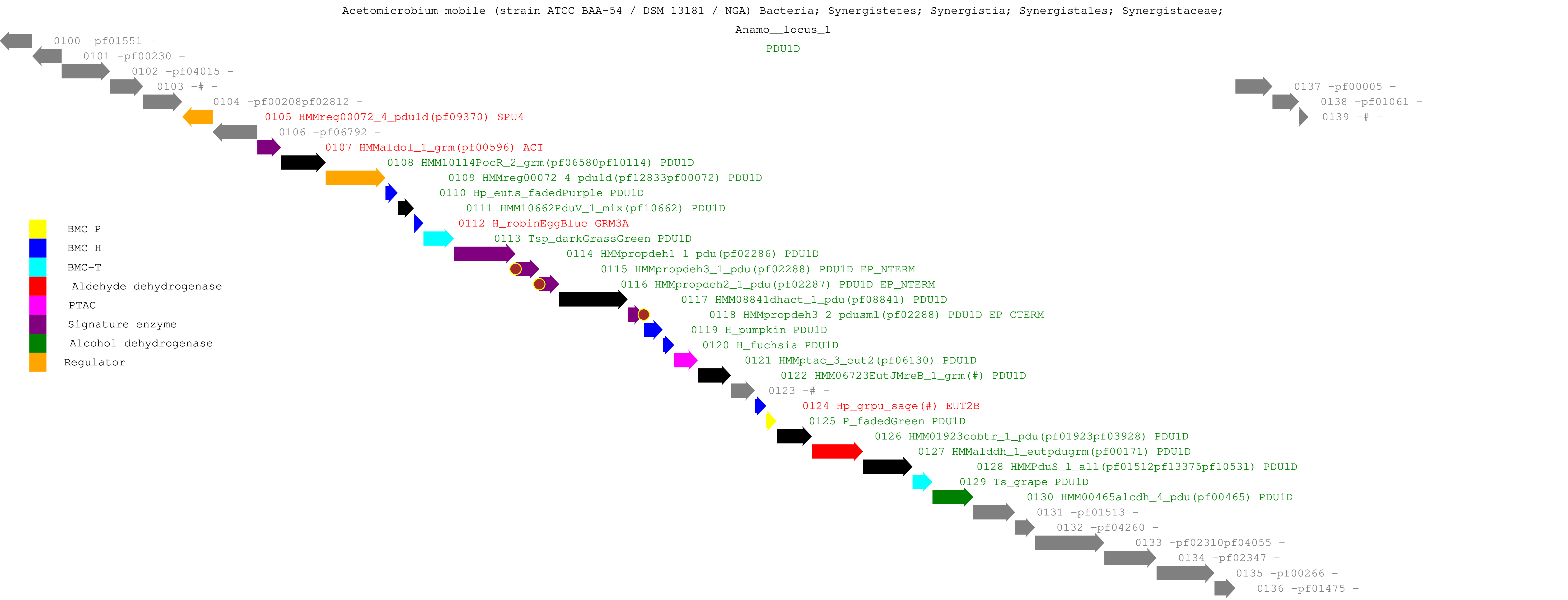
Acetomicrobium mobile (strain ATCC BAA-54 / DSM 13181 / NGA) Bacteria; Synergistetes; Synergistia; Synergistales; Synergistaceae;
Anamo__locus_1
PDU1D
0100 -pf01551 -
0101 -pf00230 -
0102 -pf04015 -
0103 -# -
0104 -pf00208pf02812 -
0105 HMMreg00072_4_pdu1d(pf09370) SPU4
0106 -pf06792 -
0107 HMMaldol_1_grm(pf00596) ACI
0108 HMM10114PocR_2_grm(pf06580pf10114) PDU1D
0109 HMMreg00072_4_pdu1d(pf12833pf00072) PDU1D
0110 Hp_euts_fadedPurple PDU1D
0111 HMM10662PduV_1_mix(pf10662) PDU1D
0112 H_robinEggBlue GRM3A
0113 Tsp_darkGrassGreen PDU1D
0114 HMMpropdeh1_1_pdu(pf02286) PDU1D
0115 HMMpropdeh3_1_pdu(pf02288) PDU1D EP_NTERM
0116 HMMpropdeh2_1_pdu(pf02287) PDU1D EP_NTERM
0117 HMM08841dhact_1_pdu(pf08841) PDU1D
0118 HMMpropdeh3_2_pdusml(pf02288) PDU1D EP_CTERM
0119 H_pumpkin PDU1D
0120 H_fuchsia PDU1D
0121 HMMptac_3_eut2(pf06130) PDU1D
0122 HMM06723EutJMreB_1_grm(#) PDU1D
0123 -# -
0124 Hp_grpu_sage(#) EUT2B
0125 P_fadedGreen PDU1D
0126 HMM01923cobtr_1_pdu(pf01923pf03928) PDU1D
0127 HMMalddh_1_eutpdugrm(pf00171) PDU1D
0128 HMMPduS_1_all(pf01512pf13375pf10531) PDU1D
0129 Ts_grape PDU1D
0130 HMM00465alcdh_4_pdu(pf00465) PDU1D
0131 -pf01513 -
0132 -pf04260 -
0133 -pf02310pf04055 -
0134 -pf02347 -
0135 -pf00266 -
0136 -pf01475 -
0137 -pf00005 -
0138 -pf01061 -
0139 -# -
BMC-P
BMC-H
BMC-T
Aldehyde dehydrogenase
PTAC
Signature enzyme
Alcohol dehydrogenase
Regulator

Aminomonas paucivorans DSM 12260. Bacteria; Synergistetes; Synergistia; Synergistales; Synergistaceae;

Apau__locus_1

PDU1D

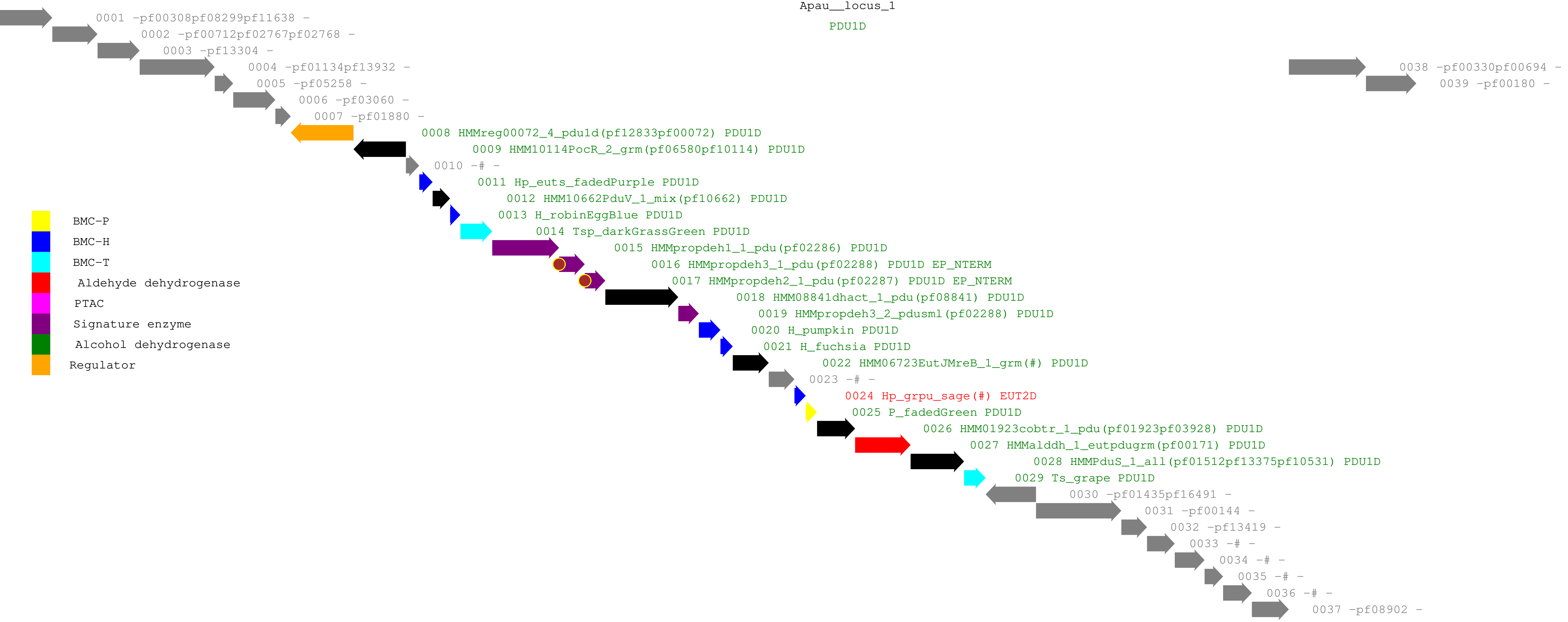

Halanaerobium sp. T82-1. Bacteria; Firmicutes; Clostridia; Halanaerobiales; Halanaerobiaceae;

AWL62__locus_2

PDU1D

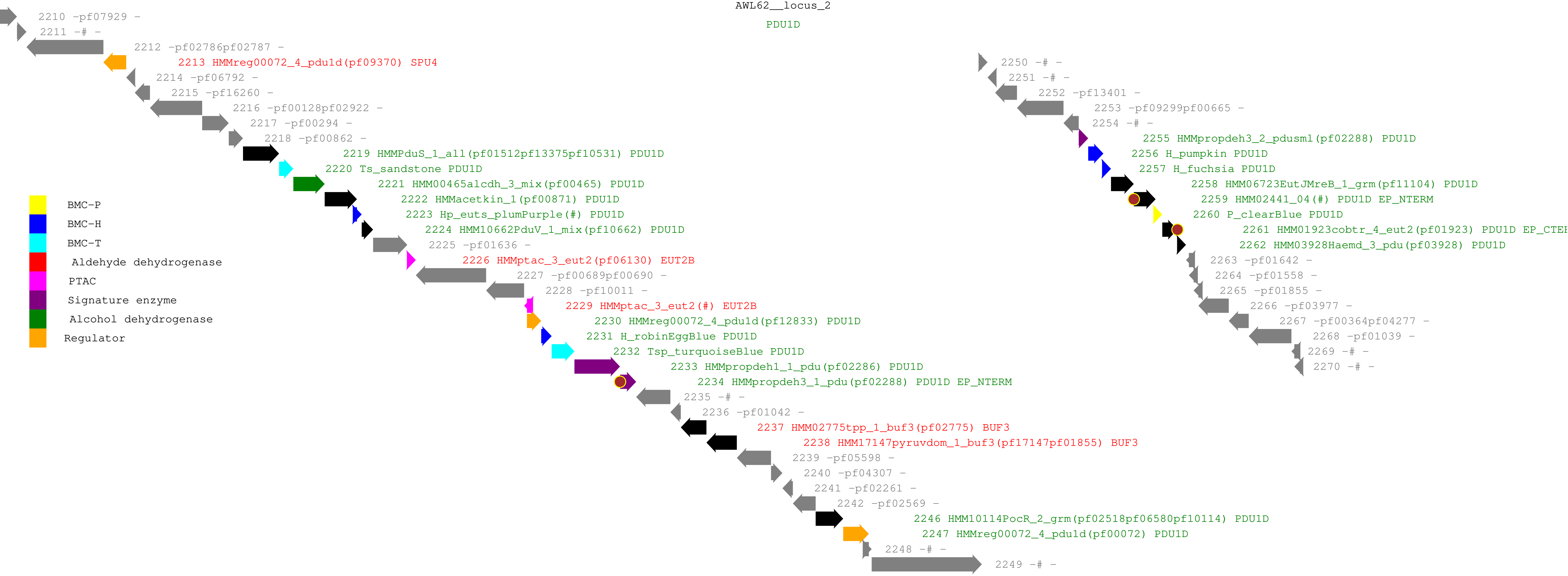

Acetobacterium woodii (strain ATCC 29683 / DSM 1030 / JCM 2381 / KCTC Bacteria; Firmicutes; Clostridia; Clostridiales; Eubacteriaceae;

Awo_c_locus_1_main_sat

PDU1D

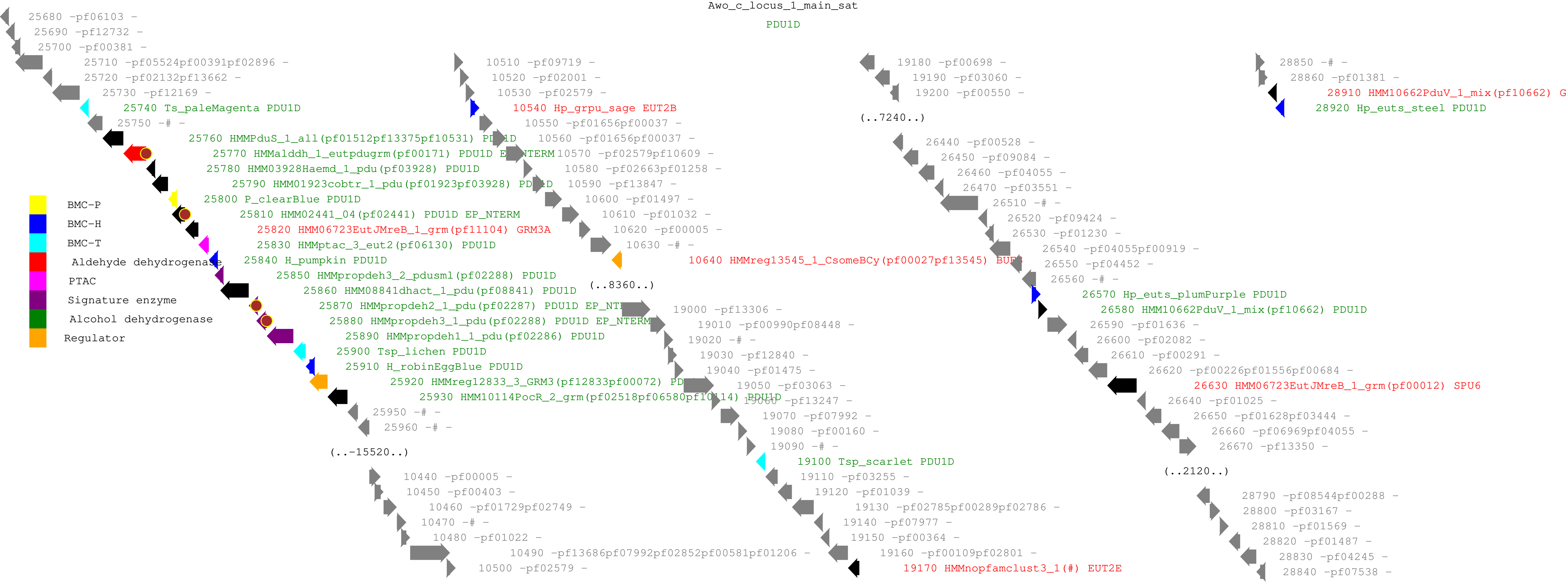

Eubacterium limosum. Bacteria; Firmicutes; Clostridia; Clostridiales; Eubacteriaceae;

B2M23__locus_1_main_sat

PDU1D

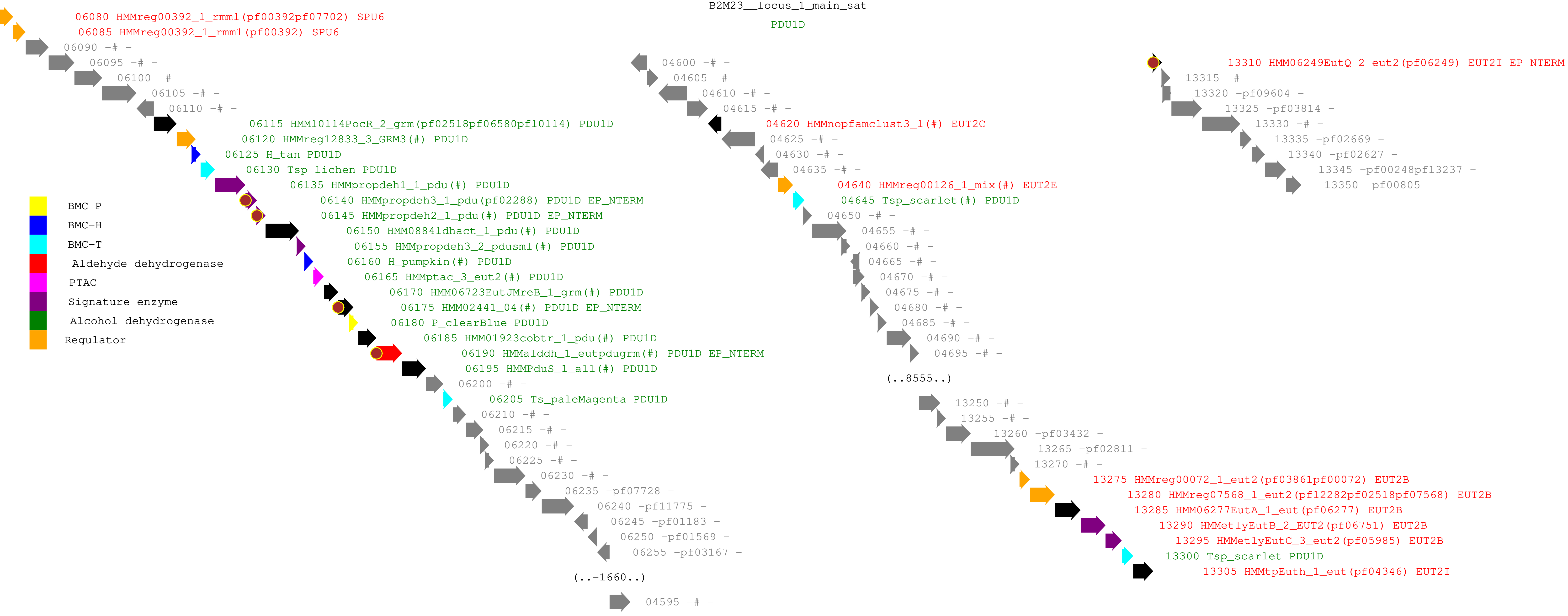

Anoxybacillus flavithermus. Bacteria; Firmicutes; Bacilli; Bacillales; Bacillaceae; Anoxybacillus.

B4168__locus_1

PDU1D

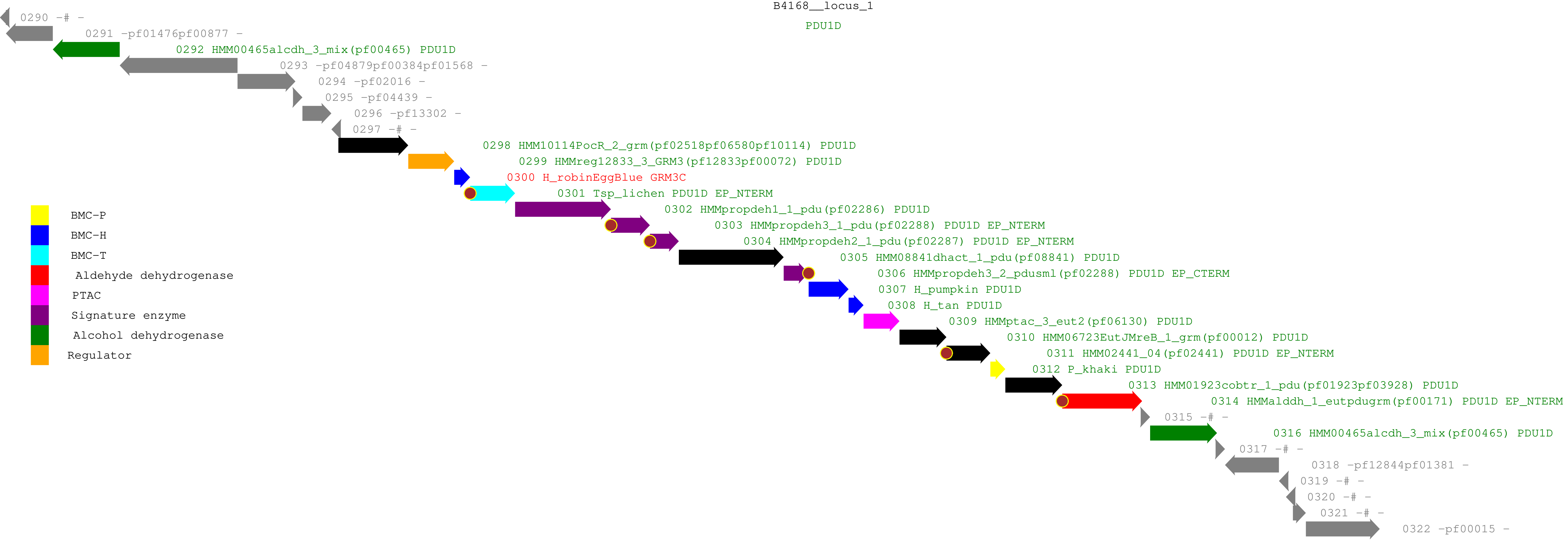

Eubacterium sp. An11. Bacteria; Firmicutes; Clostridia; Clostridiales; Eubacteriaceae;

B5E53__locus_1_main_sat

PDU1D

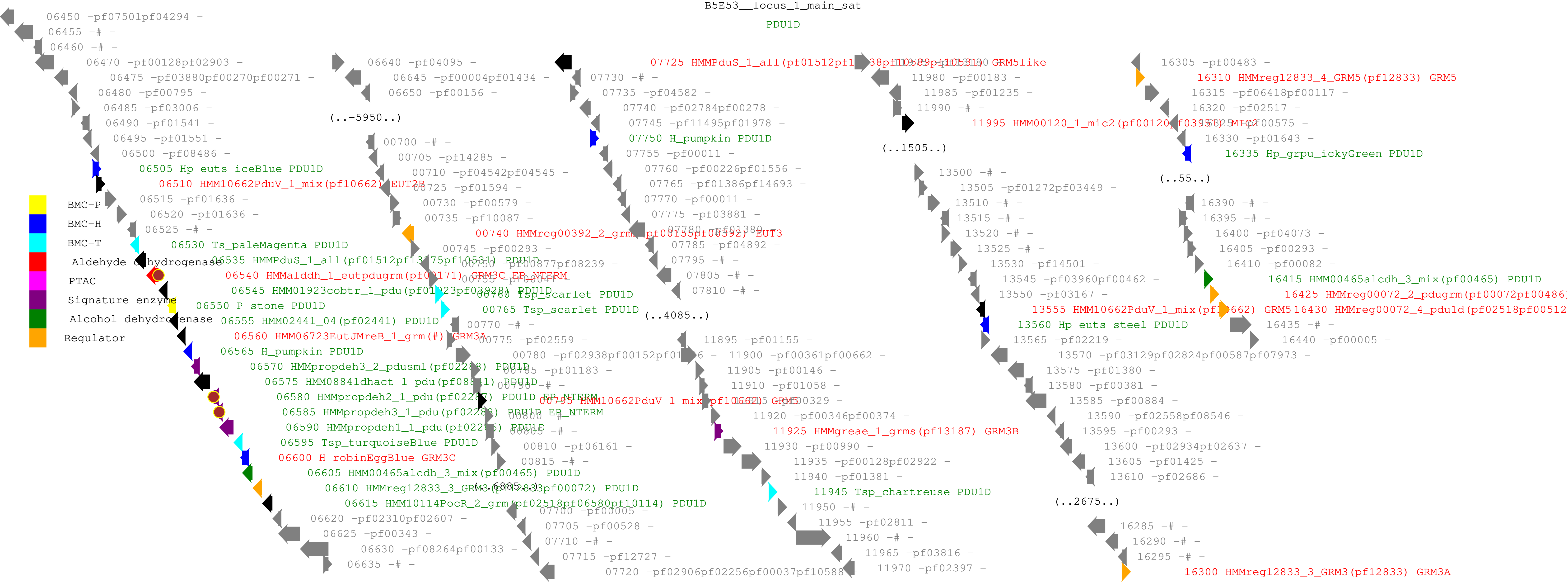

Flavonifractor plautii. Bacteria; Firmicutes; Clostridia; Clostridiales; Flavonifractor.

B5F52__locus_2

PDU1D

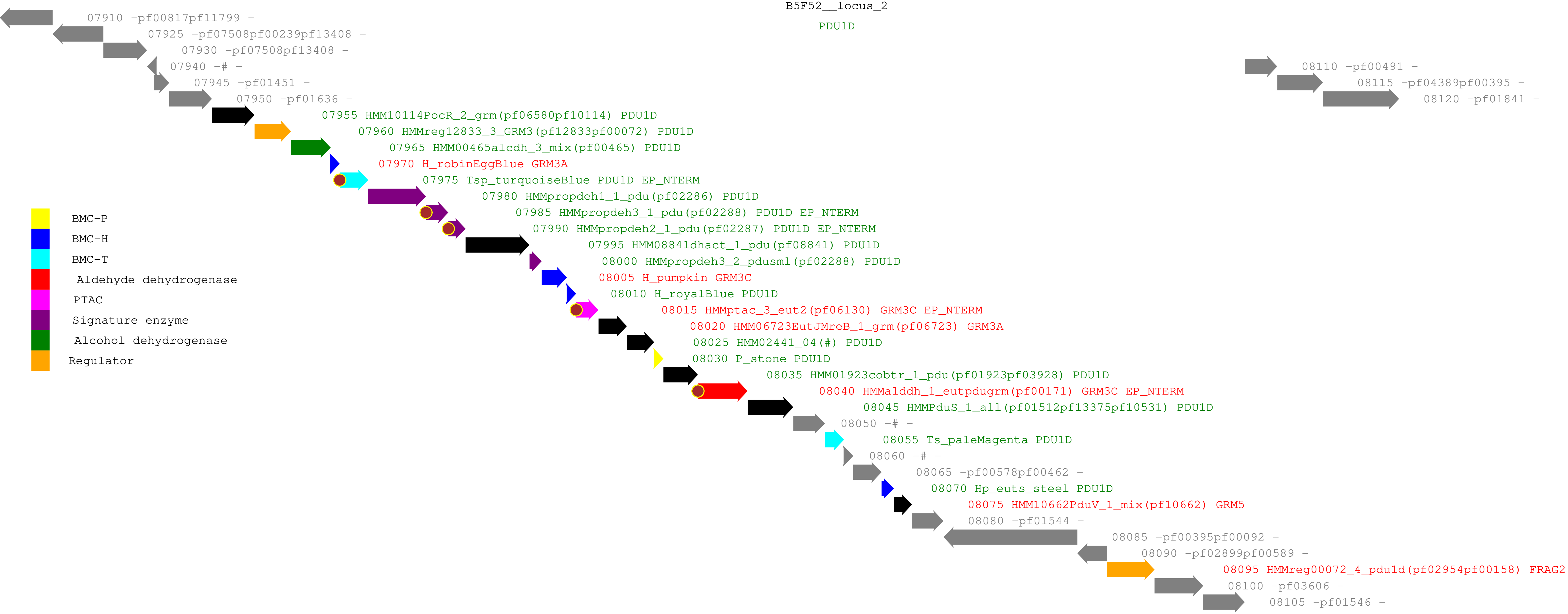

Eubacterium sp. An3. Bacteria; Firmicutes; Clostridia; Clostridiales; Eubacteriaceae;

B5F87__locus_1_main_sat

PDU1D

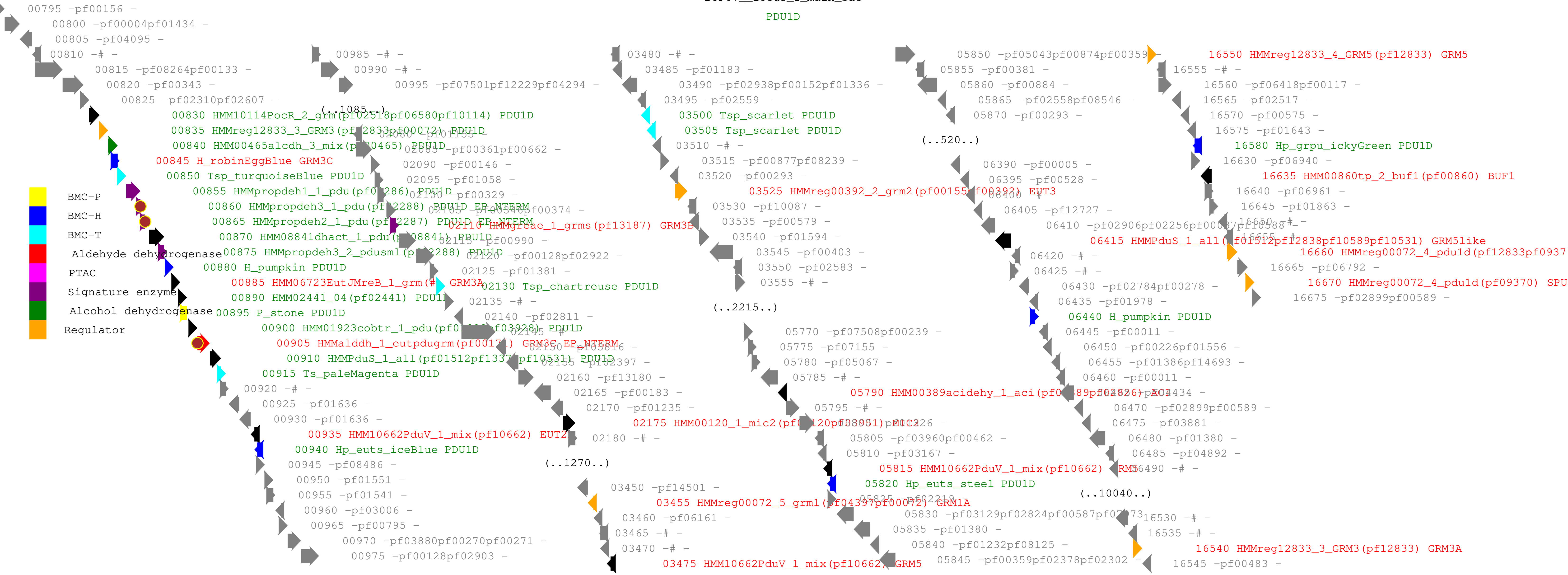

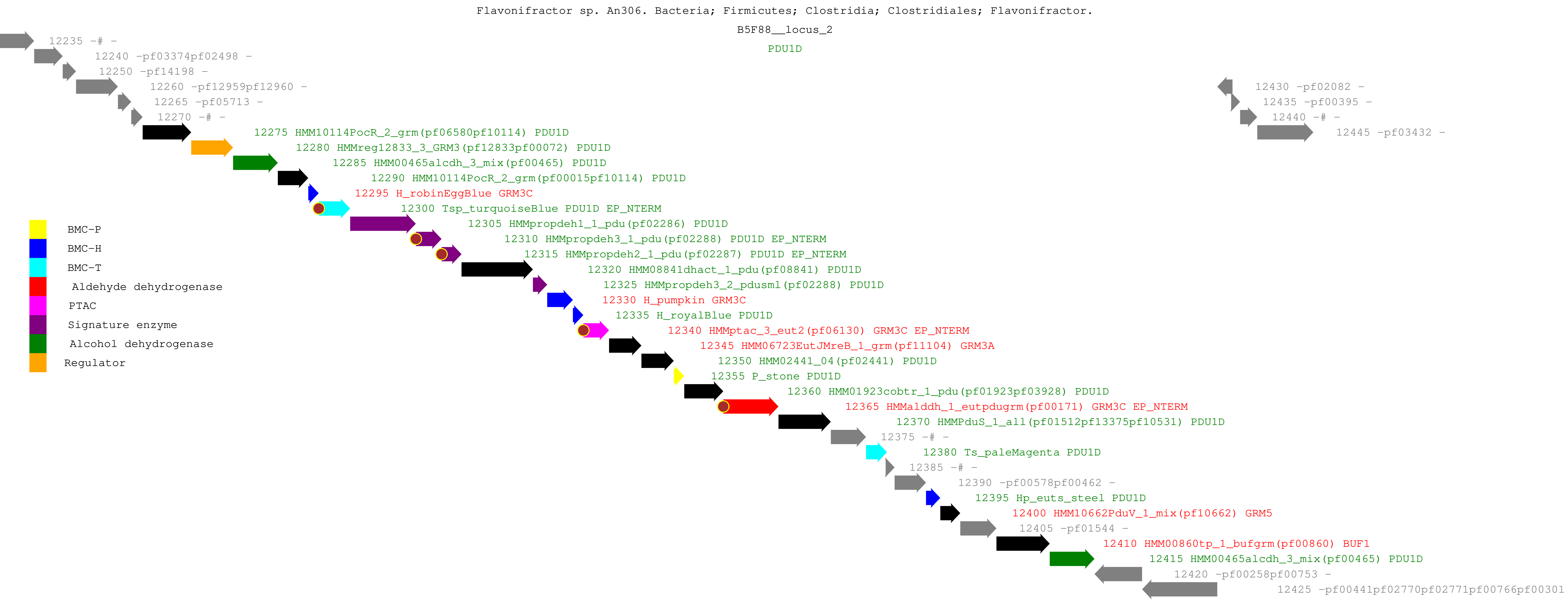

Flavonifractor sp. An306. Bacteria; Firmicutes; Clostridia; Clostridiales; Flavonifractor.
B5F88__locus_2
PDU1D
12235 -# -
12240 -pf03374pf02498 -
12250 -pf14198 -
12260 -pf12959pf12960 -
12265 -pf05713 -
12270 -# -
12275 HMM10114PocR_2_grm(pf06580pf10114) PDU1D
12280 HMMreg12833_3_GRM3(pf12833pf00072) PDU1D
12285 HMM00465alcdh_3_mix(pf00465) PDU1D
12290 HMM10114PocR_2_grm(pf00015pf10114) PDU1D
12295 H_robinEggBlue GRM3C
12300 Tsp_turquoiseBlue PDU1D EP_NTERM
12305 HMMpropdeh1_1_pdu(pf02286) PDU1D
12310 HMMpropdeh3_1_pdu(pf02288) PDU1D EP_NTERM
12315 HMMpropdeh2_1_pdu(pf02287) PDU1D EP_NTERM
12320 HMM08841dhact_1_pdu(pf08841) PDU1D
12325 HMMpropdeh3_2_pdusml(pf02288) PDU1D
12330 H_pumpkin GRM3C
12335 H_royalBlue PDU1D
12340 HMMptac_3_eut2(pf06130) GRM3C EP_NTERM
12345 HMM06723EutJMreB_1_grm(pf11104) GRM3A
12350 HMM02441_04(pf02441) PDU1D
12355 P_stone PDU1D
12360 HMM01923cobtr_1_pdu(pf01923pf03928) PDU1D
12365 HMMalddh_1_eutpdugrm(pf00171) GRM3C EP_NTERM
12370 HMMPduS_1_all(pf01512pf13375pf10531) PDU1D
12375 -# -
12380 Ts_paleMagenta PDU1D
12385 -# -
12390 -pf00578pf00462 -
12395 Hp_euts_steel PDU1D
12400 HMM10662PduV_1_mix(pf10662) GRM5
12405 -pf01544 -
12410 HMM00860tp_1_bufgrm(pf00860) BUF1
12415 HMM00465alcdh_3_mix(pf00465) PDU1D
12420 -pf00258pf00753 -
12425 -pf00441pf02770pf02771pf00766pf00301
12430 -pf02082 -
12435 -pf00395 -
12440 -# -
12445 -pf03432 -
BMC-P
BMC-H
BMC-T
Aldehyde dehydrogenase
PTAC
Signature enzyme
Alcohol dehydrogenase
Regulator

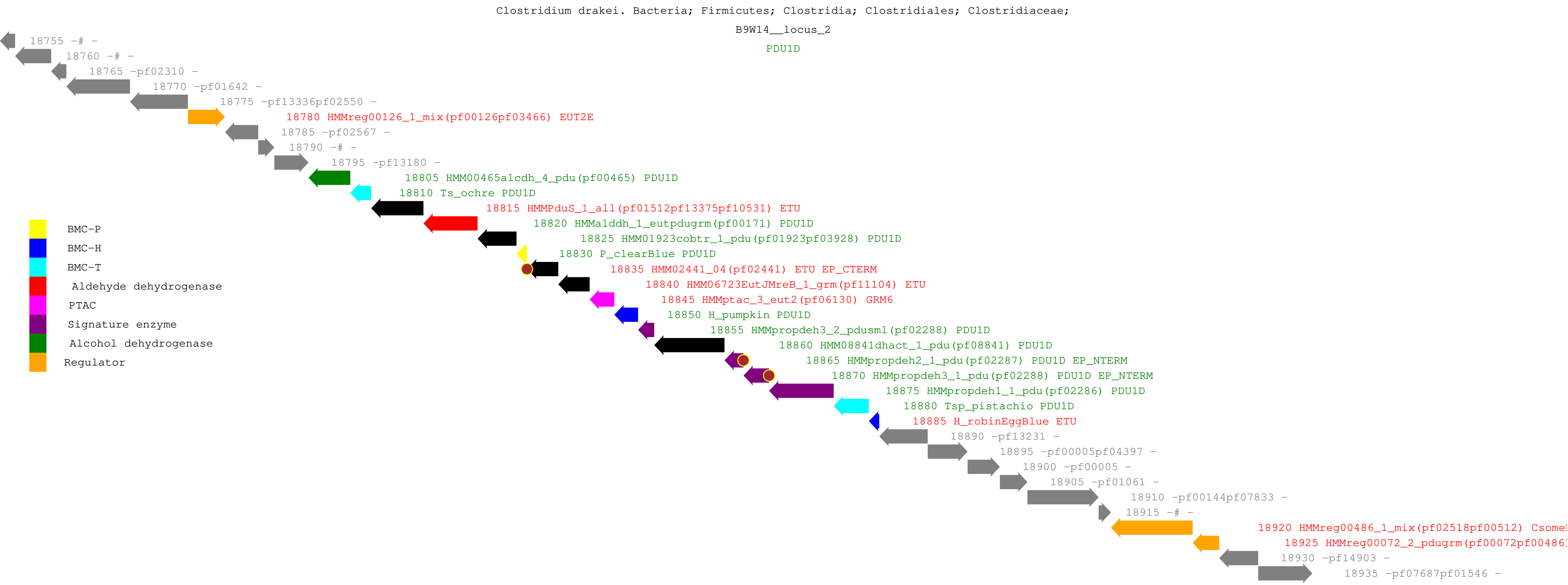
Clostridium drakei. Bacteria; Firmicutes; Clostridia; Clostridiales; Clostridiaceae;
B9W14__locus_2
PDU1D
18755 -# -
18760 -# -
18765 -pf02310 -
18770 -pf01642 -
18775 -pf13336pf02550 -
18780 HMMreg00126_1_mix(pf00126pf03466) EUT2E
18785 -pf02567 -
18790 -# -
18795 -pf13180 -
18805 HMM00465alcdh_4_pdu(pf00465) PDU1D
18810 Ts_ochre PDU1D
18815 HMMPduS_1_all(pf01512pf13375pf10531) ETU
18820 HMMaldh_1_eutpdugrm(pf00171) PDU1D
18825 HMM01923cobtr_1_pdu(pf01923pf03928) PDU1D
18830 P_clearBlue PDU1D
18835 HMM02441_04(pf02441) ETU EP_CTERM
18840 HMM06723EutJMreB_1_grm(pf11104) ETU
18845 HMMptac_3_eut2(pf06130) GRM6
18850 H_pumpkin PDU1D
18855 HMMpropdeh3_2_pdusml(pf02288) PDU1D
18860 HMM08841dhact_1_pdu(pf08841) PDU1D
18865 HMMpropdeh2_1_pdu(pf02287) PDU1D EP_NTERM
18870 HMMpropdeh3_1_pdu(pf02288) PDU1D EP_NTERM
18875 HMMpropdeh1_1_pdu(pf02286) PDU1D
18880 Tsp_pistachio PDU1D
18885 H_robinEggBlue ETU
18890 -pf13231 -
18895 -pf00005pf04397 -
18900 -pf00005 -
18905 -pf01061 -
18910 -pf00144pf07833 -
18915 -# -
18920 HMMreg00486_1_mix(pf02518pf00512) Csome
18925 HMMreg00072_2_pdugrm(pf00072pf00486)
18930 -pf14903 -
18935 -pf07687pf01546 -
BMC-P
BMC-H
BMC-T
Aldehyde dehydrogenase
PTAC
Signature enzyme
Alcohol dehydrogenase
Regulator

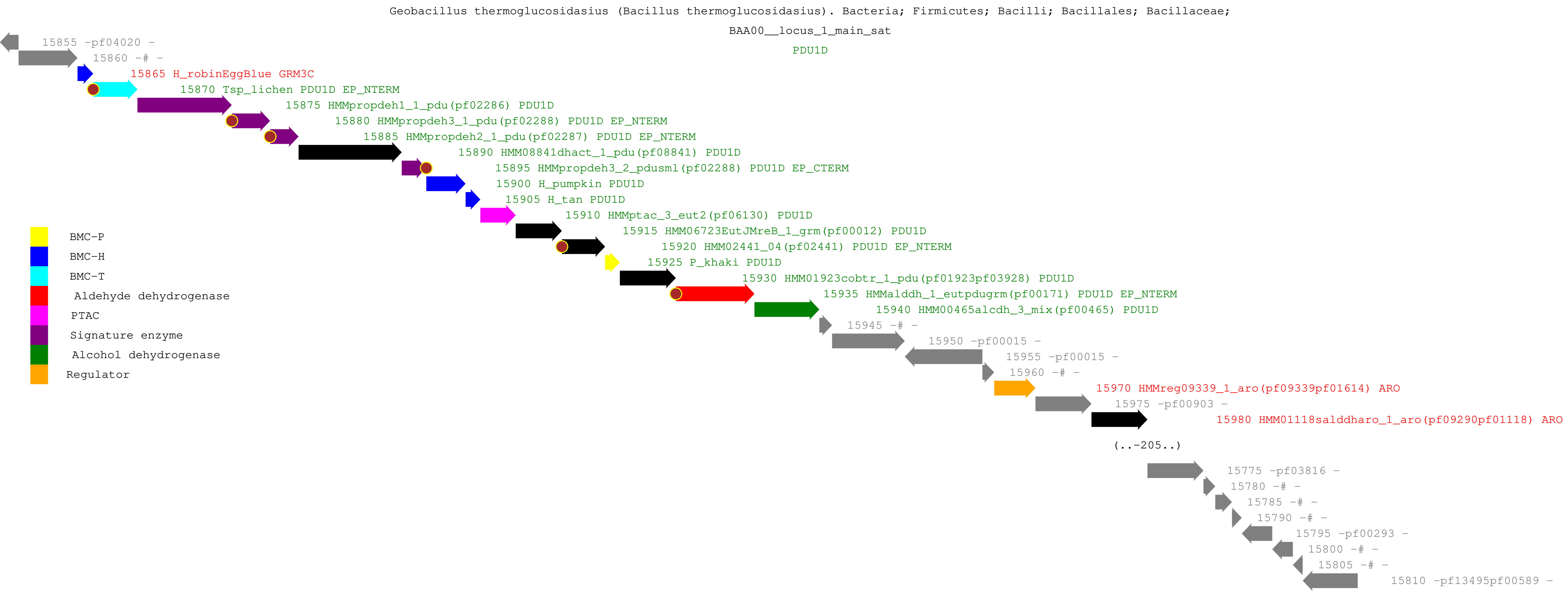

Geobacillus thermoglucosidasius (Bacillus thermoglucosidasius). Bacteria; Firmicutes; Bacilli; Bacillales; Bacillaceae;
BAA00__locus_1_main_sat
PDU1D
15855 -pf04020 -
15860 -# -
15865 H_robinEggBlue GRM3C
15870 Tsp_lichen PDU1D EP_NTERM
15875 HMMpropdeh1_1_pdu(pf02286) PDU1D
15880 HMMpropdeh3_1_pdu(pf02288) PDU1D EP_NTERM
15885 HMMpropdeh2_1_pdu(pf02287) PDU1D EP_NTERM
15890 HMM08841dhact_1_pdu(pf08841) PDU1D
15895 HMMpropdeh3_2_pdusml(pf02288) PDU1D EP_CTERM
15900 H_pumpkin PDU1D
15905 H_tan PDU1D
15910 HMMptac_3_eut2(pf06130) PDU1D
15915 HMM06723EutJMreB_1_grm(pf00012) PDU1D
15920 HMM02441_04(pf02441) PDU1D EP_NTERM
15925 P_khaki PDU1D
15930 HMM01923cobtr_1_pdu(pf01923pf03928) PDU1D
15935 HMMalddh_1_eutpdugrm(pf00171) PDU1D EP_NTERM
15940 HMM00465alcdh_3_mix(pf00465) PDU1D
15945 -# -
15950 -pf00015 -
15955 -pf00015 -
15960 -# -
15970 HMMreg09339_1_aro(pf09339pf01614) ARO
15975 -pf00903 -
15980 HMM01118salddharo_1_aro(pf09290pf01118) ARO
(..-205..)
15775 -pf03816 -
15780 -# -
15785 -# -
15790 -# -
15795 -pf00293 -
15800 -# -
15805 -# -
15810 -pf13495pf00589 -
BMC-P
BMC-H
BMC-T
Aldehyde dehydrogenase
PTAC
Signature enzyme
Alcohol dehydrogenase
Regulator

Bacillus azotoformans LMG 9581. Bacteria; Firmicutes; Bacilli; Bacillales; Bacillaceae; Bacillus.

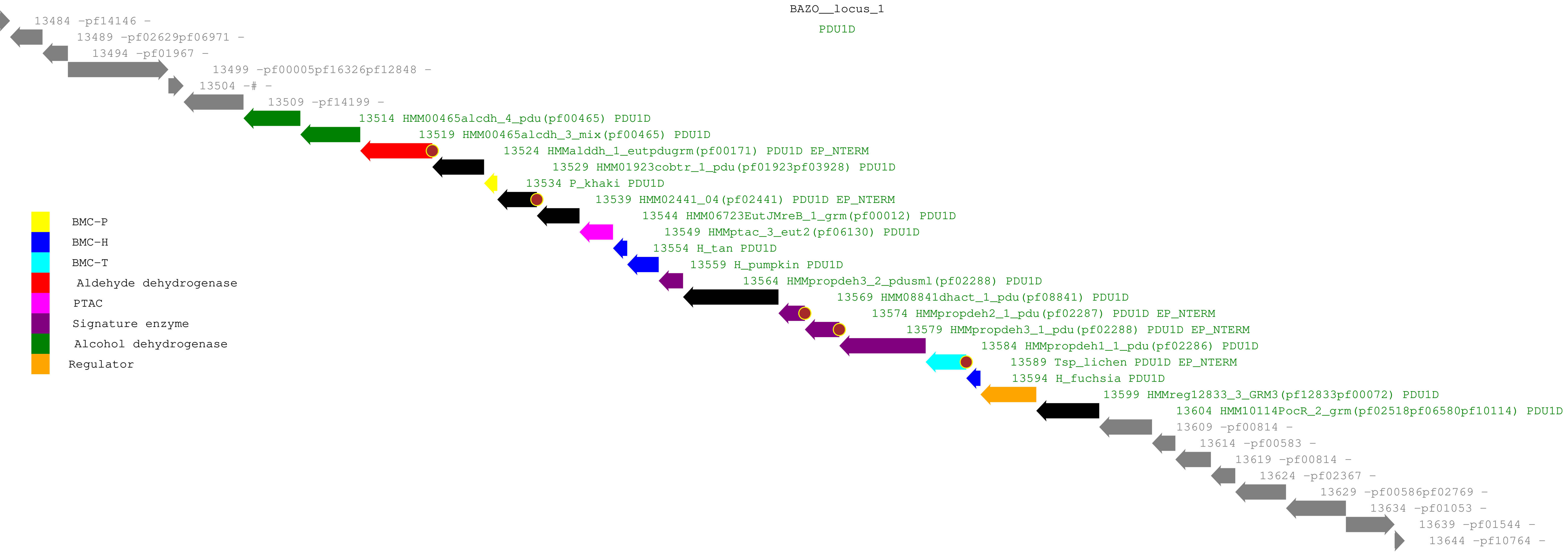

Cloacibacillus porcorum. Bacteria; Synergistetes; Synergistia; Synergistales; Synergistaceae;

BED41__locus_2

PDU1D

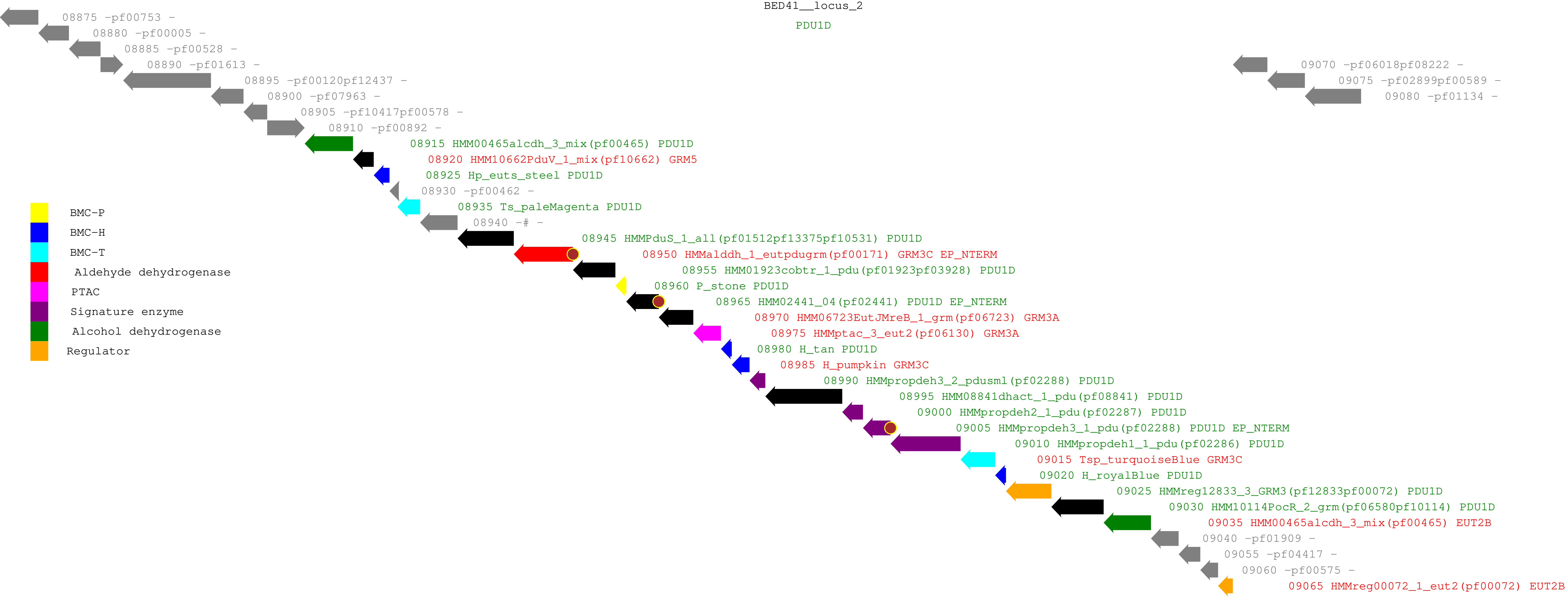

Thermohalobacter berrensis. Bacteria; Firmicutes; Clostridia; Clostridiales; Clostridiaceae;

BET03__locus_1

PDU1D

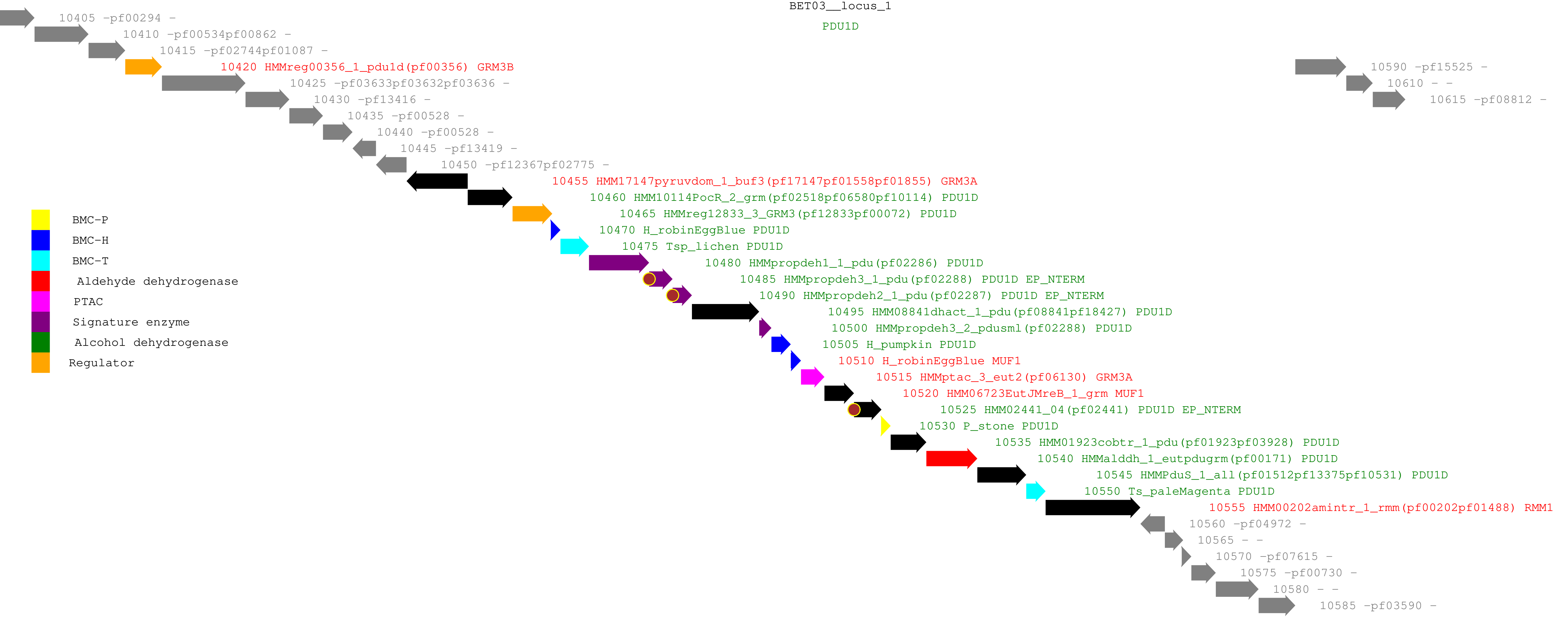

Eubacterium sp. 38_16. Bacteria; Firmicutes; Clostridia; Clostridiales; Eubacteriaceae;

BHW19__locus_1_main_sat

PDU1D

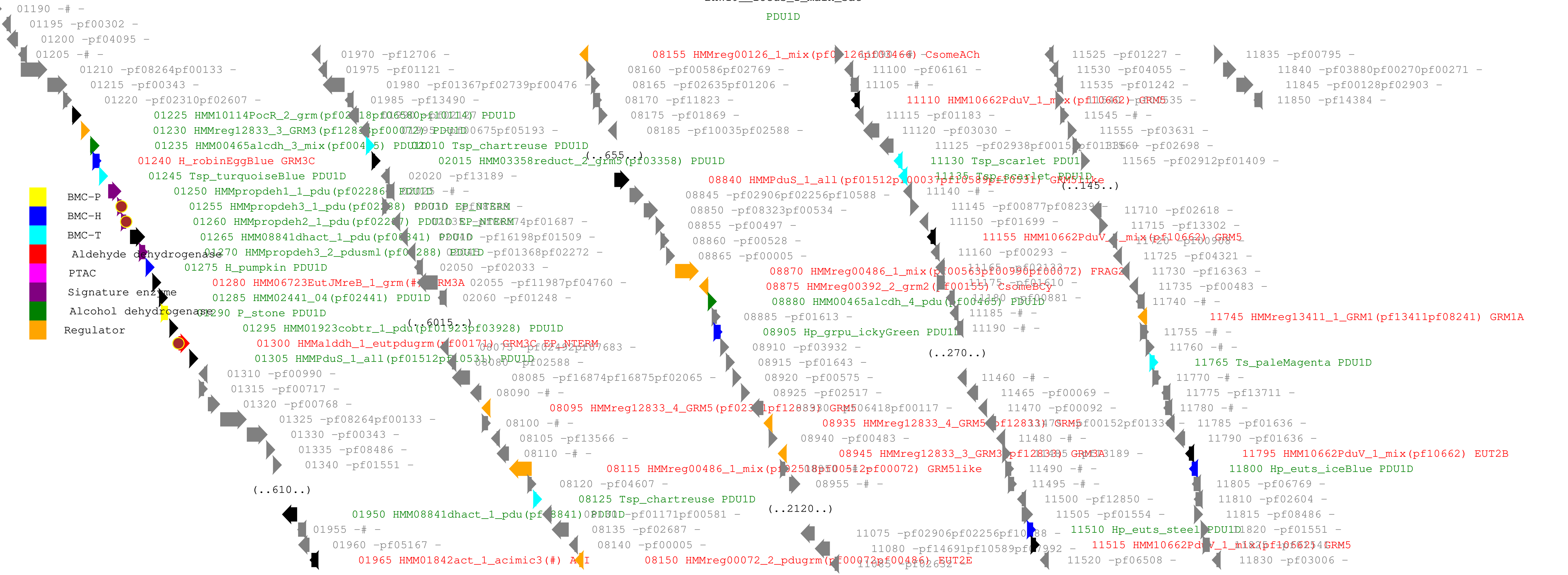

Acetobacterium sp. MES1. Bacteria; Firmicutes; Clostridia; Clostridiales; Eubacteriaceae;

BI182__locus_1_main_sat

PDU1D

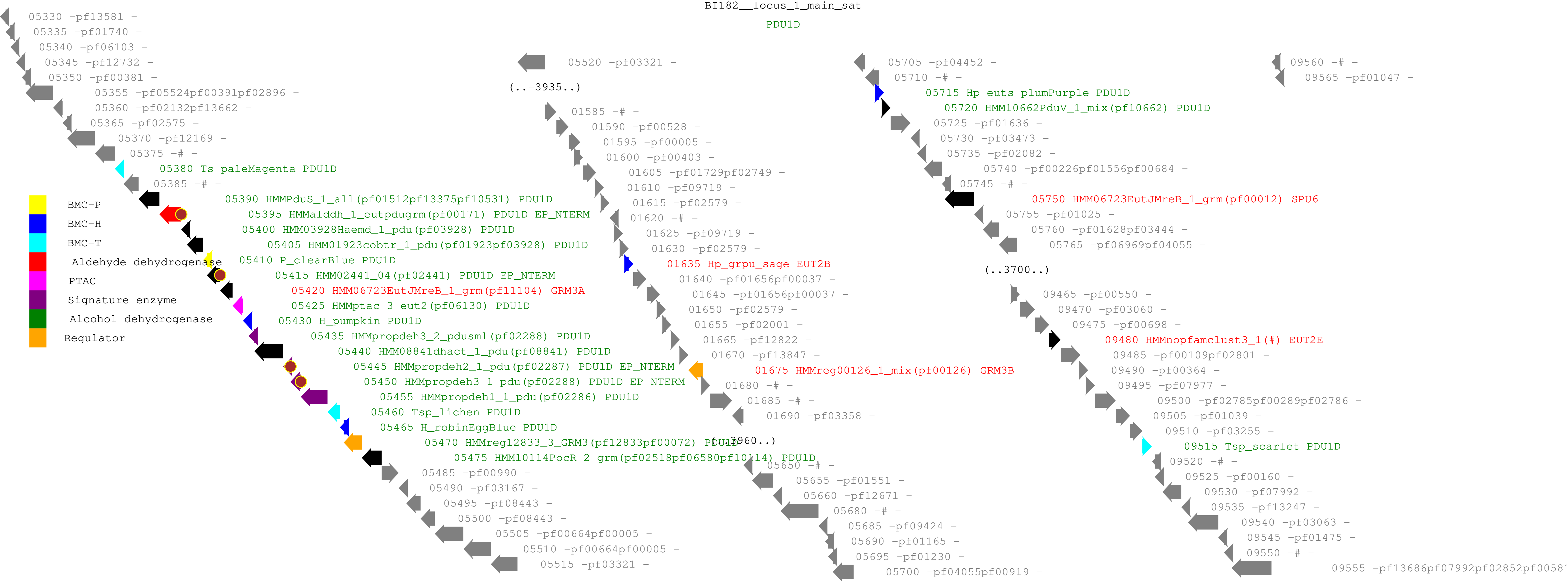

Eubacteriaceae bacterium CHKCI004. Bacteria; Firmicutes; Clostridia; Clostridiales; Eubacteriaceae.

BN3660__locus_1_main_sat

PDU1D

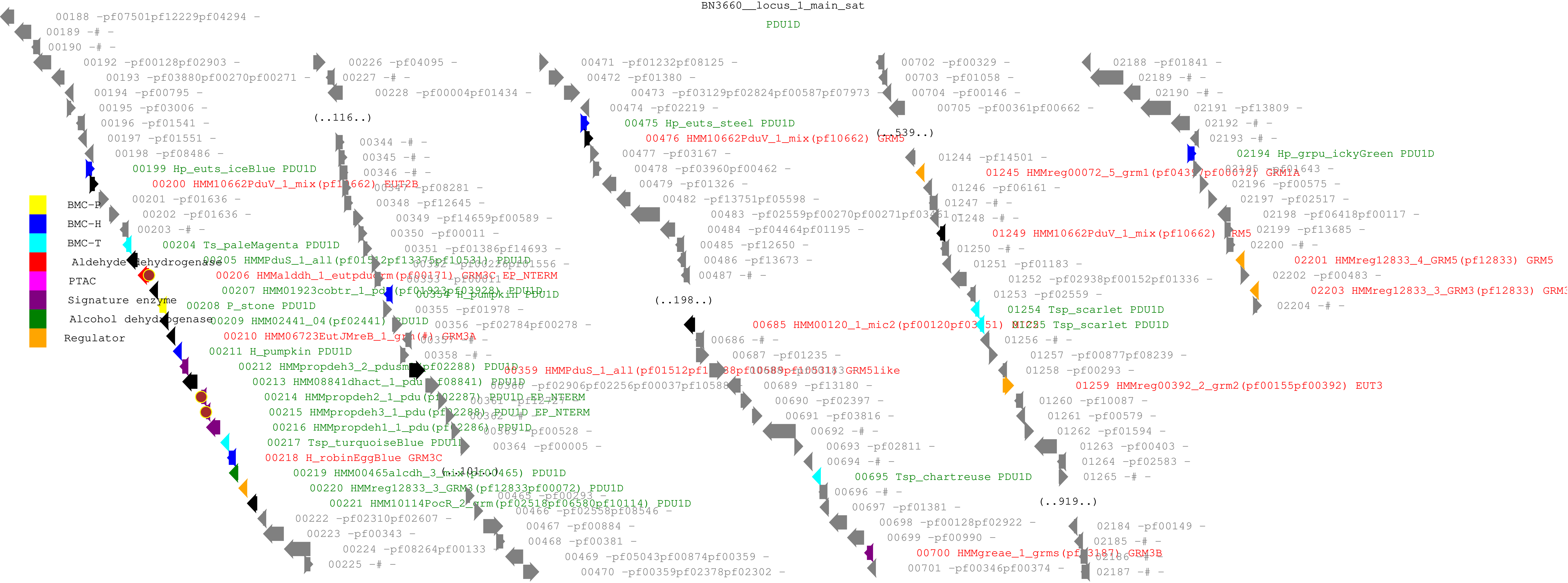

Eubacterium hallii CAG:12. Bacteria; Firmicutes; Clostridia; Clostridiales; Eubacteriaceae;

BN476__locus_1_main_sat

PDU1D

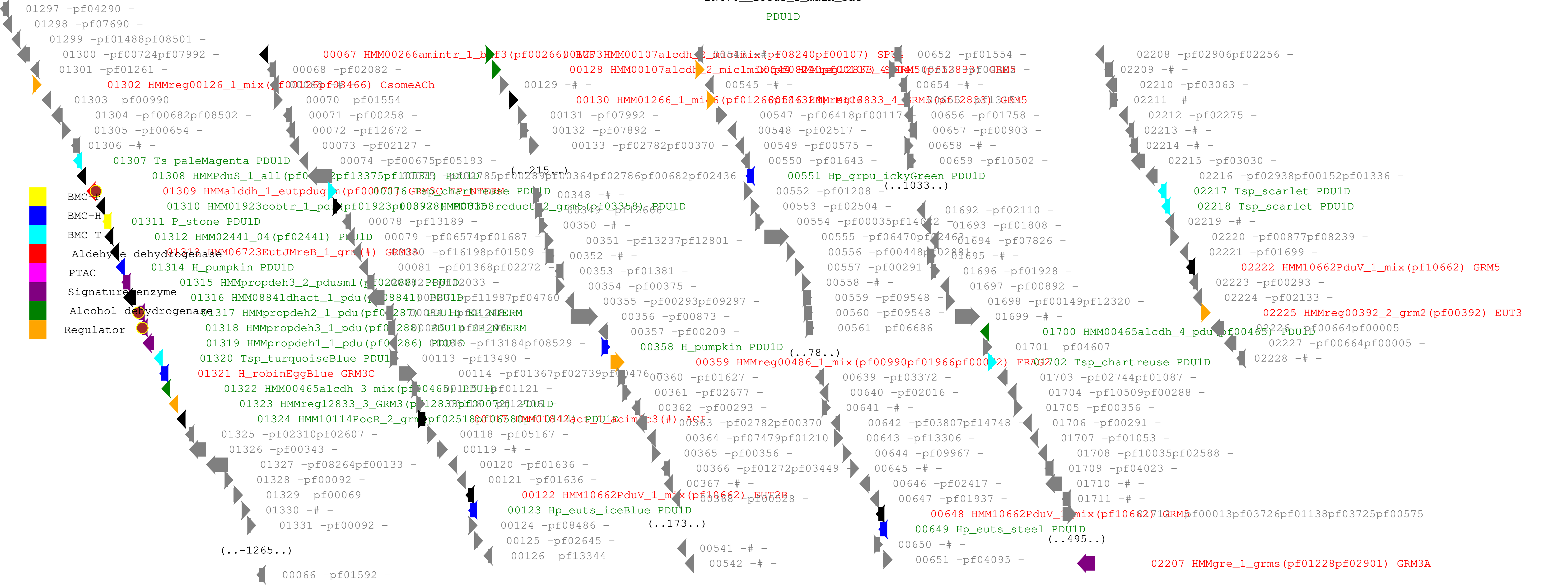

Eubacterium sp. CAG:146. Bacteria; Firmicutes; Clostridia; Clostridiales; Eubacteriaceae;

BN498__locus_1_main_sat

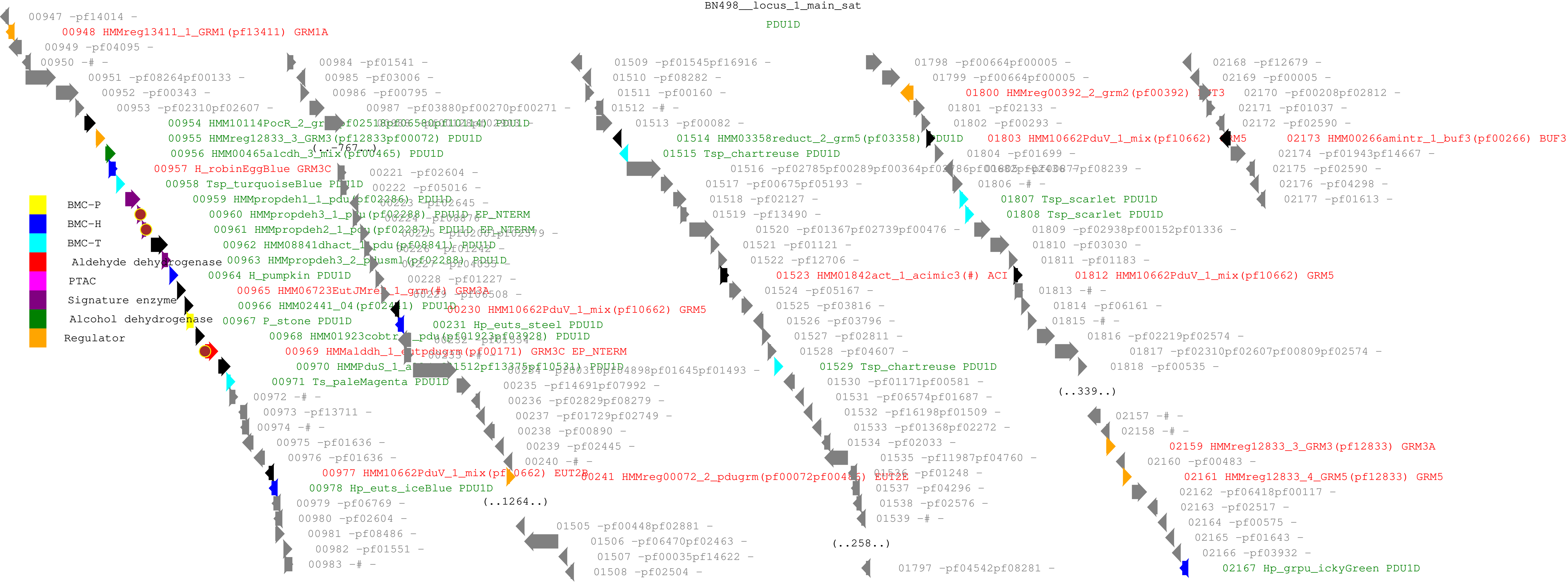

Butyribacterium methylotrophicum. Bacteria; Firmicutes; Clostridia; Clostridiales; Clostridiaceae;

BUME__locus_1_main_sat

PDU1D

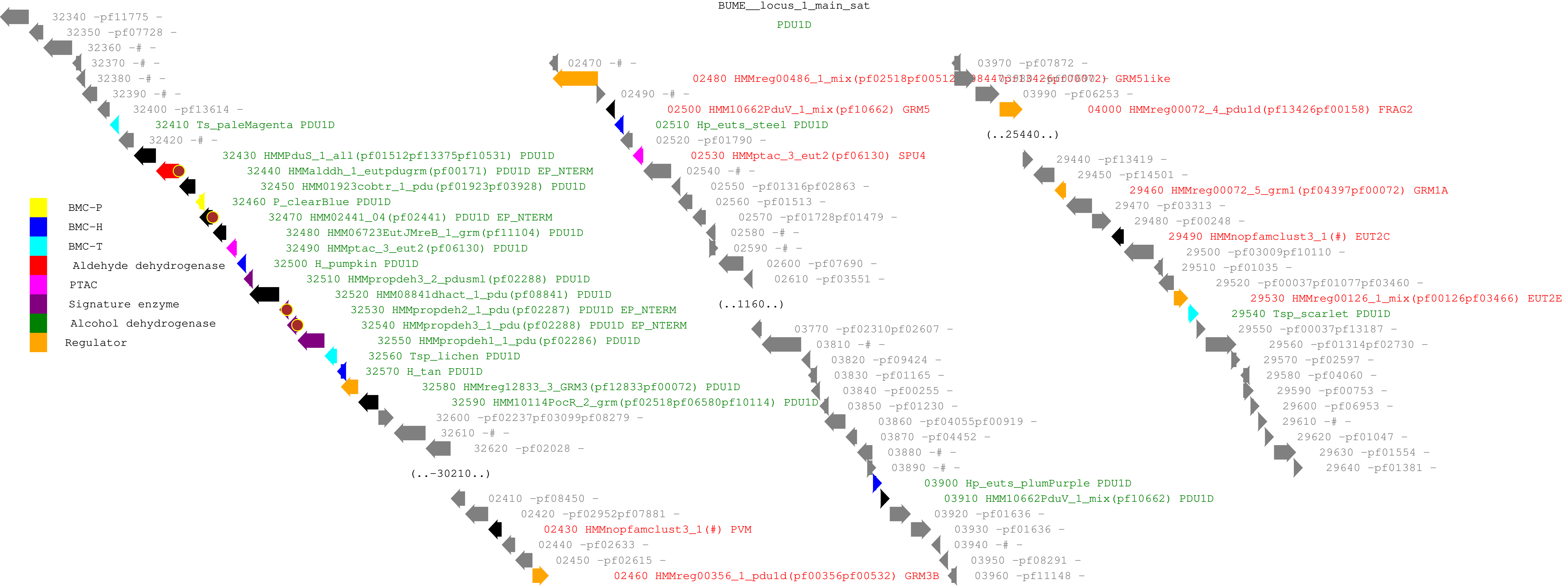

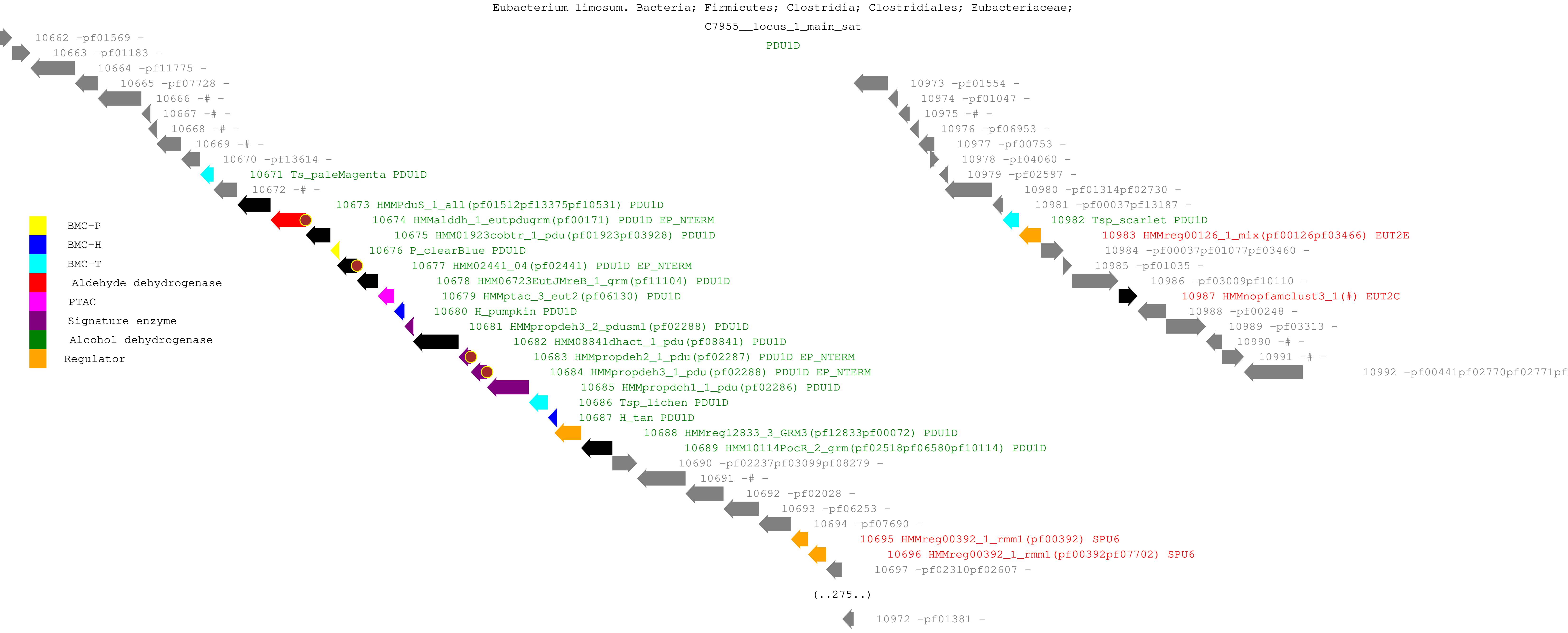
Eubacterium limosum. Bacteria; Firmicutes; Clostridia; Clostridiales; Eubacteriaceae;
C7955__locus_1_main_sat
PDU1D
10662 -pf01569 -
10663 -pf01183 -
10664 -pf11775 -
10665 -pf07728 -
10666 -# -
10667 -# -
10668 -# -
10669 -# -
10670 -pf13614 -
10671 Ts_paleMagenta PDU1D
10672 -# -
10673 HMMPduS_1_all(pf01512pf13375pf10531) PDU1D
10674 HMMalddh_1_eutpdugrm(pf00171) PDU1D EP_NTERM
10675 HMM01923cobtr_1_pdu(pf01923pf03928) PDU1D
10676 P_clearBlue PDU1D
10677 HMM02441_04(pf02441) PDU1D EP_NTERM
10678 HMM06723EutJMreB_1_grm(pf11104) PDU1D
10679 HMMptac_3_eut2(pf06130) PDU1D
10680 H_pumpkin PDU1D
10681 HMMpropdeh3_2_pdusml(pf02288) PDU1D
10682 HMM08841dhact_1_pdu(pf08841) PDU1D
10683 HMMpropdeh2_1_pdu(pf02287) PDU1D EP_NTERM
10684 HMMpropdeh3_1_pdu(pf02288) PDU1D EP_NTERM
10685 HMMpropdeh1_1_pdu(pf02286) PDU1D
10686 Tsp_lichen PDU1D
10687 H_tan PDU1D
10688 HMMreg12833_3_GRM3(pf12833pf00072) PDU1D
10689 HMM10114PocR_2_grm(pf02518pf06580pf10114) PDU1D
10690 -pf02237pf03099pf08279 -
10691 -# -
10692 -pf02028 -
10693 -pf06253 -
10694 -pf07690 -
10695 HMMreg00392_1_rmm1(pf00392) SPU6
10696 HMMreg00392_1_rmm1(pf00392pf07702) SPU6
10697 -pf02310pf02607 -
(..275..)
10972 -pf01381 -
10973 -pf01554 -
10974 -pf01047 -
10975 -# -
10976 -pf06953 -
10977 -pf00753 -
10978 -pf04060 -
10979 -pf02597 -
10980 -pf01314pf02730 -
10981 -pf00037pf13187 -
10982 Tsp_scarlet PDU1D
10983 HMMreg00126_1_mix(pf00126pf03466) EUT2E
10984 -pf00037pf01077pf03460 -
10985 -pf01035 -
10986 -pf03009pf10110 -
10987 HMMnopfamclust3_1(#) EUT2C
10988 -pf00248 -
10989 -pf03313 -
10990 -# -
10991 -# -
10992 -pf00441pf02770pf02771pf
BMC-P
BMC-H
BMC-T
Aldehyde dehydrogenase
PTAC
Signature enzyme
Alcohol dehydrogenase
Regulator

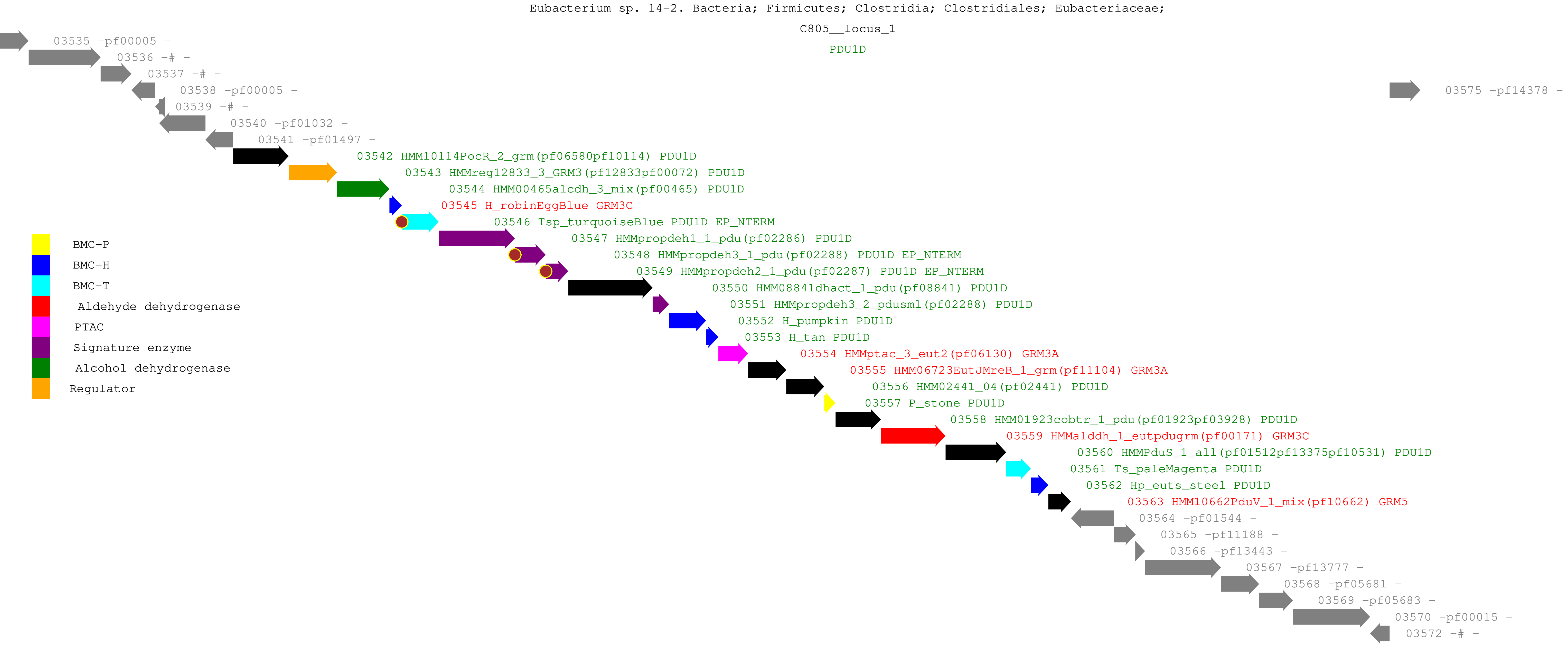
Eubacterium sp. 14-2. Bacteria; Firmicutes; Clostridia; Clostridiales; Eubacteriaceae;
C805__locus_1
PDU1D
03535 -pf00005 -
03536 -# -
03537 -# -
03538 -pf00005 -
03539 -# -
03540 -pf01032 -
03541 -pf01497 -
03542 HMM10114PocR_2_grm(pf06580pf10114) PDU1D
03543 HMMreg12833_3_GRM3(pf12833pf00072) PDU1D
03544 HMM00465alcdh_3_mix(pf00465) PDU1D
03545 H_robinEggBlue GRM3C
03546 Tsp_turquoiseBlue PDU1D EP_NTERM
03547 HMMpropdeh1_1_pdu(pf02286) PDU1D
03548 HMMpropdeh3_1_pdu(pf02288) PDU1D EP_NTERM
03549 HMMpropdeh2_1_pdu(pf02287) PDU1D EP_NTERM
03550 HMM08841dhact_1_pdu(pf08841) PDU1D
03551 HMMpropdeh3_2_pdusml(pf02288) PDU1D
03552 H_pumpkin PDU1D
03553 H_tan PDU1D
03554 HMMptac_3_eut2(pf06130) GRM3A
03555 HMM06723EutJMreB_1_grm(pf11104) GRM3A
03556 HMM02441_04(pf02441) PDU1D
03557 P_stone PDU1D
03558 HMM01923cobtr_1_pdu(pf01923pf03928) PDU1D
03559 HMMalddh_1_eutpdugrm(pf00171) GRM3C
03560 HMMPduS_1_all(pf01512pf13375pf10531) PDU1D
03561 Ts_paleMagenta PDU1D
03562 Hp_euts_steel PDU1D
03563 HMM10662PduV_1_mix(pf10662) GRM5
03564 -pf01544 -
03565 -pf11188 -
03566 -pf13443 -
03567 -pf13777 -
03568 -pf05681 -
03569 -pf05683 -
03570 -pf00015 -
03572 -# -
03575 -pf14378 -
BMC-P
BMC-H
BMC-T
Aldehyde dehydrogenase
PTAC
Signature enzyme
Alcohol dehydrogenase
Regulator

Lachnospiraceae bacterium 3-1. Bacteria; Firmicutes; Clostridia; Clostridiales; Lachnospiraceae.

C806__locus_1

PDU1D

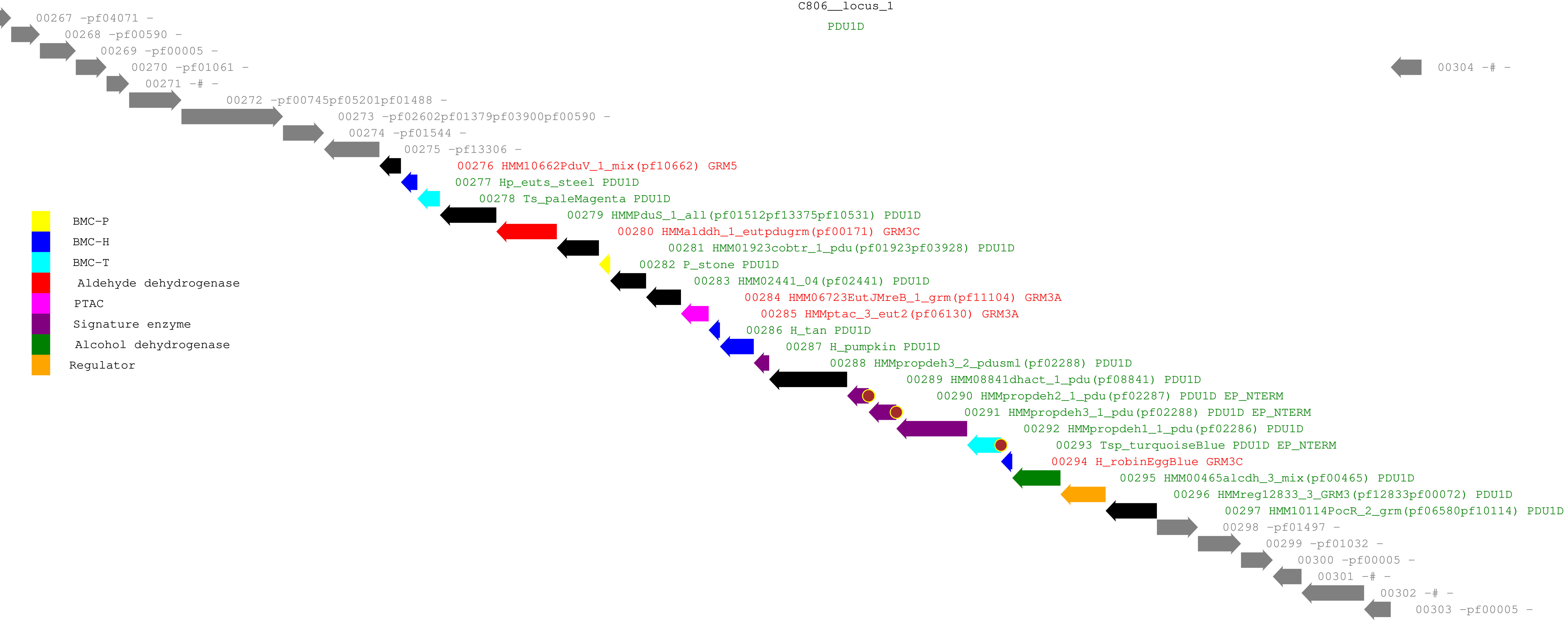

Lachnospiraceae bacterium A2. Bacteria; Firmicutes; Clostridia; Clostridiales; Lachnospiraceae.

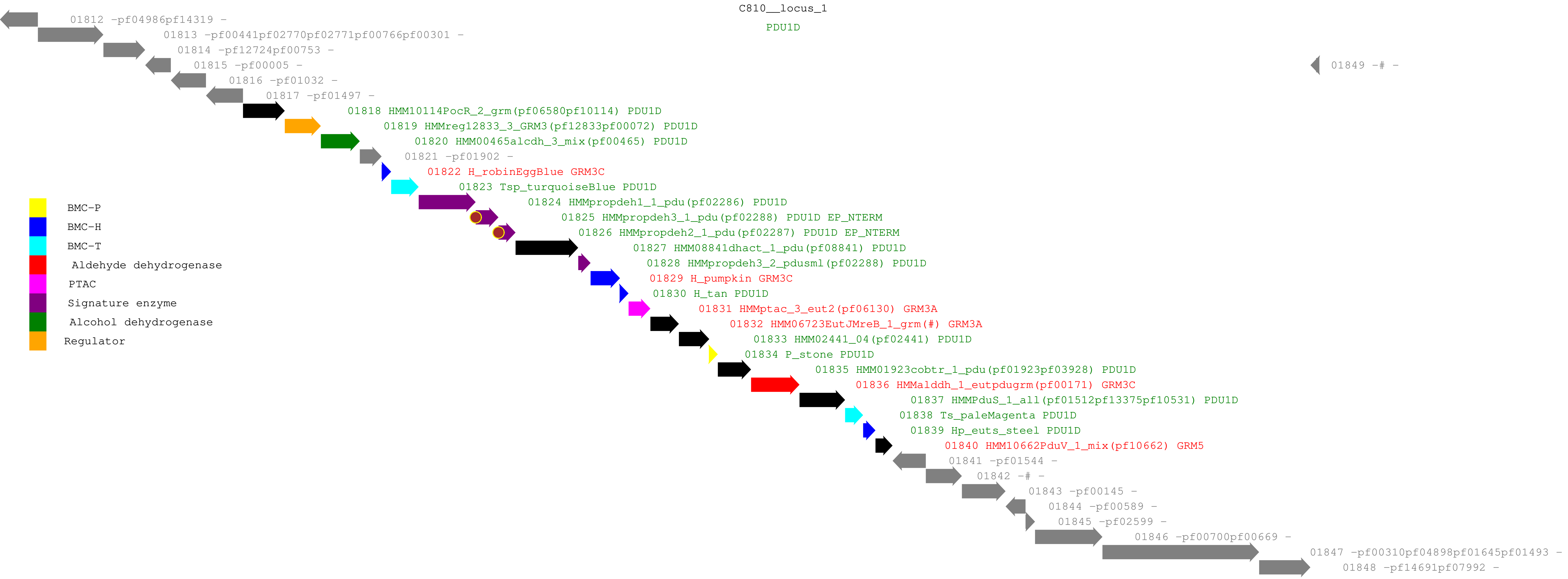

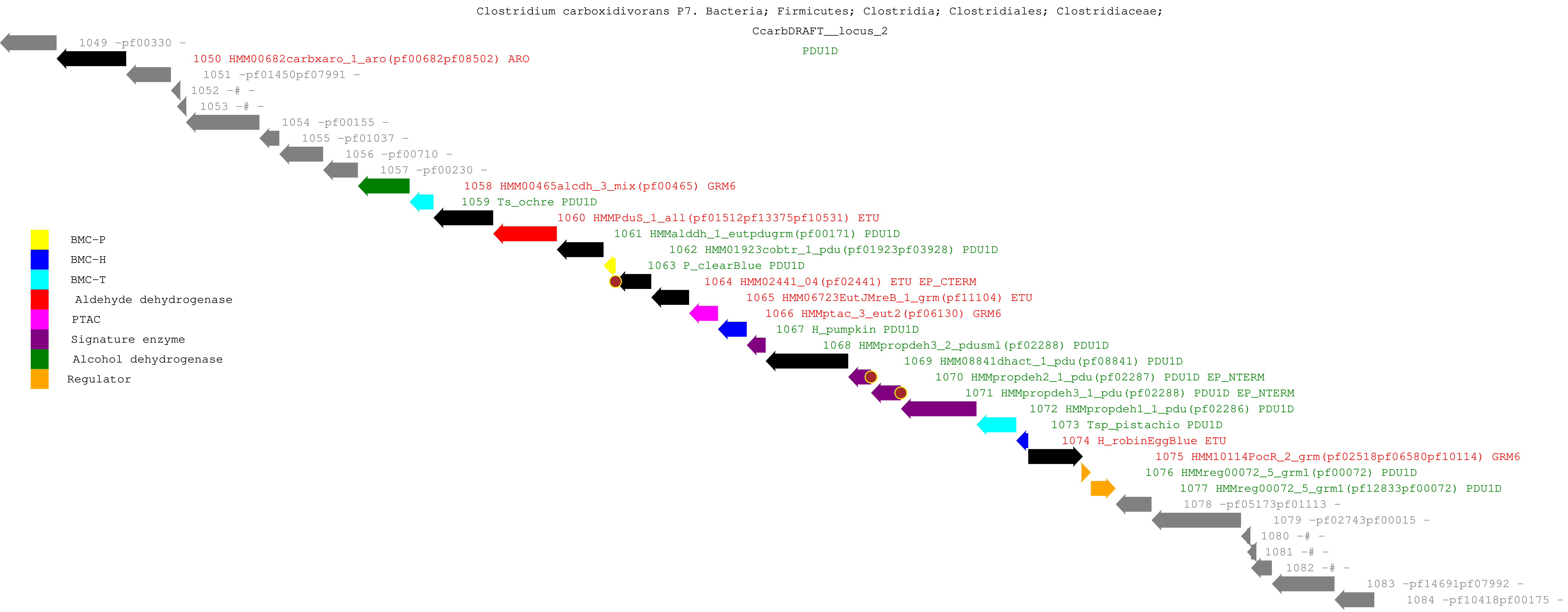
Clostridium carboxidivorans P7. Bacteria; Firmicutes; Clostridia; Clostridiales; Clostridiaceae;
CcarbDRAFT__locus_2
PDU1D
1049 -pf00330 -
1050 HMM00682carbxaro_1_aro(pf00682pf08502) ARO
1051 -pf01450pf07991 -
1052 -# -
1053 -# -
1054 -pf00155 -
1055 -pf01037 -
1056 -pf00710 -
1057 -pf00230 -
1058 HMM00465alcdh_3_mix(pf00465) GRM6
1059 Ts_ochre PDU1D
1060 HMMPduS_1_all(pf01512pf13375pf10531) ETU
1061 HMMalddh_1_eutpdugrm(pf00171) PDU1D
1062 HMM01923cobtr_1_pdu(pf01923pf03928) PDU1D
1063 P_clearBlue PDU1D
1064 HMM02441_04(pf02441) ETU EP_CTERM
1065 HMM06723EutJMreB_1_grm(pf11104) ETU
1066 HMMptac_3_eut2(pf06130) GRM6
1067 H_pumpkin PDU1D
1068 HMMpropdeh3_2_pdusml(pf02288) PDU1D
1069 HMM08841dhact_1_pdu(pf08841) PDU1D
1070 HMMpropdeh2_1_pdu(pf02287) PDU1D EP_NTERM
1071 HMMpropdeh3_1_pdu(pf02288) PDU1D EP_NTERM
1072 HMMpropdeh1_1_pdu(pf02286) PDU1D
1073 Tsp_pistachio PDU1D
1074 H_robinEggBlue ETU
1075 HMM10114PocR_2_grm(pf02518pf06580pf10114) GRM6
1076 HMMreg00072_5_grm1(pf00072) PDU1D
1077 HMMreg00072_5_grm1(pf12833pf00072) PDU1D
1078 -pf05173pf01113 -
1079 -pf02743pf00015 -
1080 -# -
1081 -# -
1082 -# -
1083 -pf14691pf07992 -
1084 -pf10418pf00175 -
BMC-P
BMC-H
BMC-T
Aldehyde dehydrogenase
PTAC
Signature enzyme
Alcohol dehydrogenase
Regulator

Clostridium carboxidivorans P7. Bacteria; Firmicutes; Clostridia; Clostridiales; Clostridiaceae;

CcarbDRAFT__locus_3

PDU1D

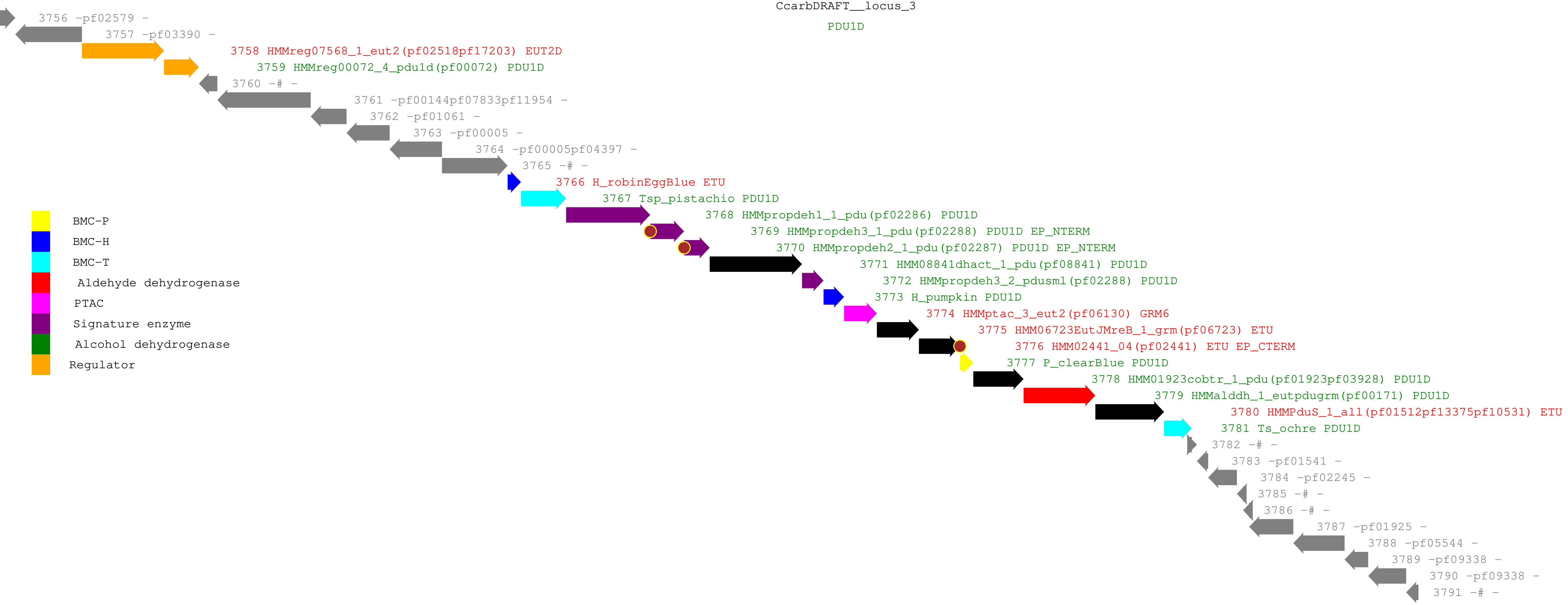

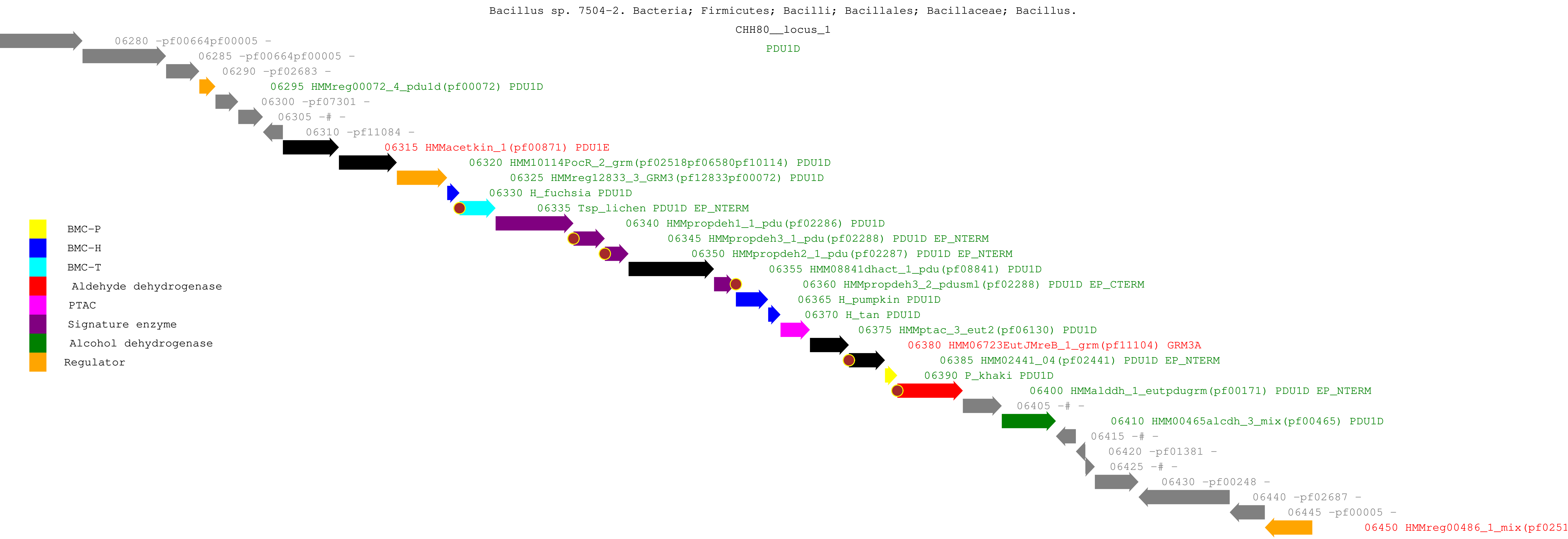
Bacillus sp. 7504-2. Bacteria; Firmicutes; Bacilli; Bacillales; Bacillaceae; Bacillus.
CHH80__locus_1
PDU1D
06280 -pf00664pf00005 -
06285 -pf00664pf00005 -
06290 -pf02683 -
06295 HMMreg00072_4_pdu1d(pf00072) PDU1D
06300 -pf07301 -
06305 -# -
06310 -pf11084 -
06315 HMMacetkin_1(pf00871) PDU1E
06320 HMM10114PocR_2_grm(pf02518pf06580pf10114) PDU1D
06325 HMMreg12833_3_GRM3(pf12833pf00072) PDU1D
06330 H_fuchsia PDU1D
06335 Tsp_lichen PDU1D EP_NTERM
06340 HMMpropdeh1_1_pdu(pf02286) PDU1D
06345 HMMpropdeh3_1_pdu(pf02288) PDU1D EP_NTERM
06350 HMMpropdeh2_1_pdu(pf02287) PDU1D EP_NTERM
06355 HMM08841dhact_1_pdu(pf08841) PDU1D
06360 HMMpropdeh3_2_pdusml(pf02288) PDU1D EP_CTERM
06365 H_pumpkin PDU1D
06370 H_tan PDU1D
06375 HMMptac_3_eut2(pf06130) PDU1D
06380 HMM06723EutJMreB_1_grm(pf11104) GRM3A
06385 HMM02441_04(pf02441) PDU1D EP_NTERM
06390 P_khaki PDU1D
06400 HMMalddh_1_eutpdugrm(pf00171) PDU1D EP_NTERM
06405 -# -
06410 HMM00465alcdh_3_mix(pf00465) PDU1D
06415 -# -
06420 -pf01381 -
06425 -# -
06430 -pf00248 -
06440 -pf02687 -
06445 -pf00005 -
06450 HMMreg00486_1_mix(pf0251
BMC-P
BMC-H
BMC-T
Aldehyde dehydrogenase
PTAC
Signature enzyme
Alcohol dehydrogenase
Regulator

Bacillus sp. PK3_68. Bacteria; Firmicutes; Bacilli; Bacillales; Bacillaceae; Bacillus.

CJ483__locus_2

PDU1D

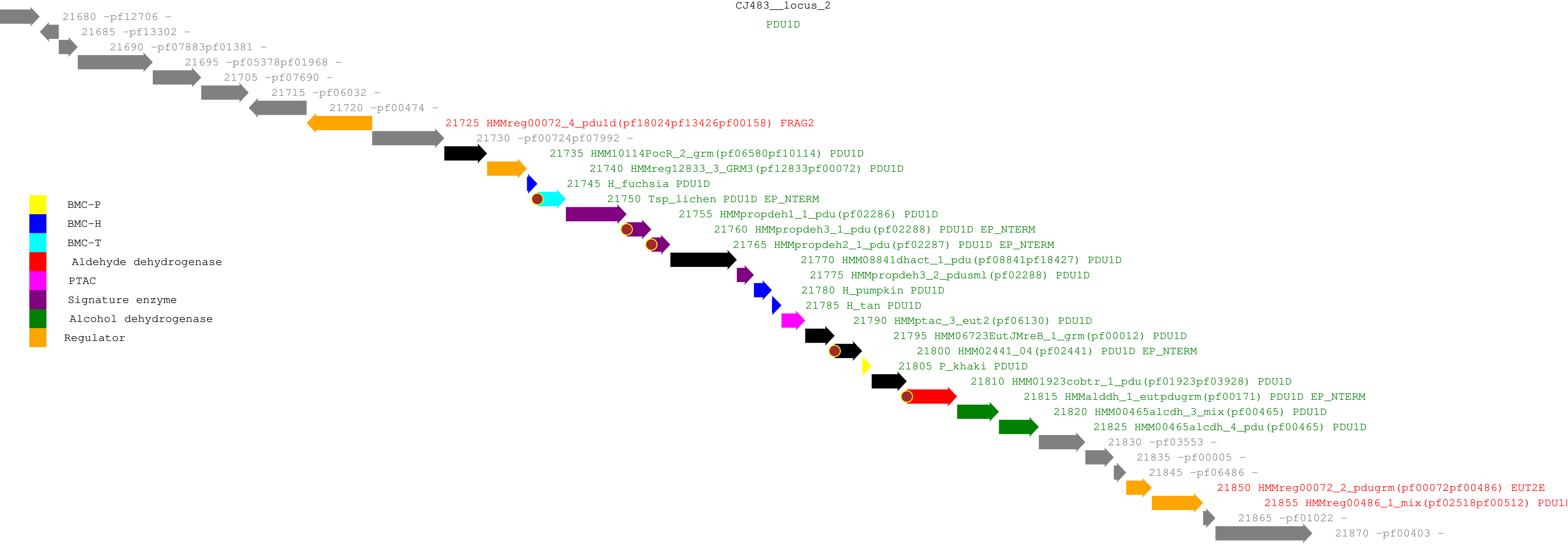

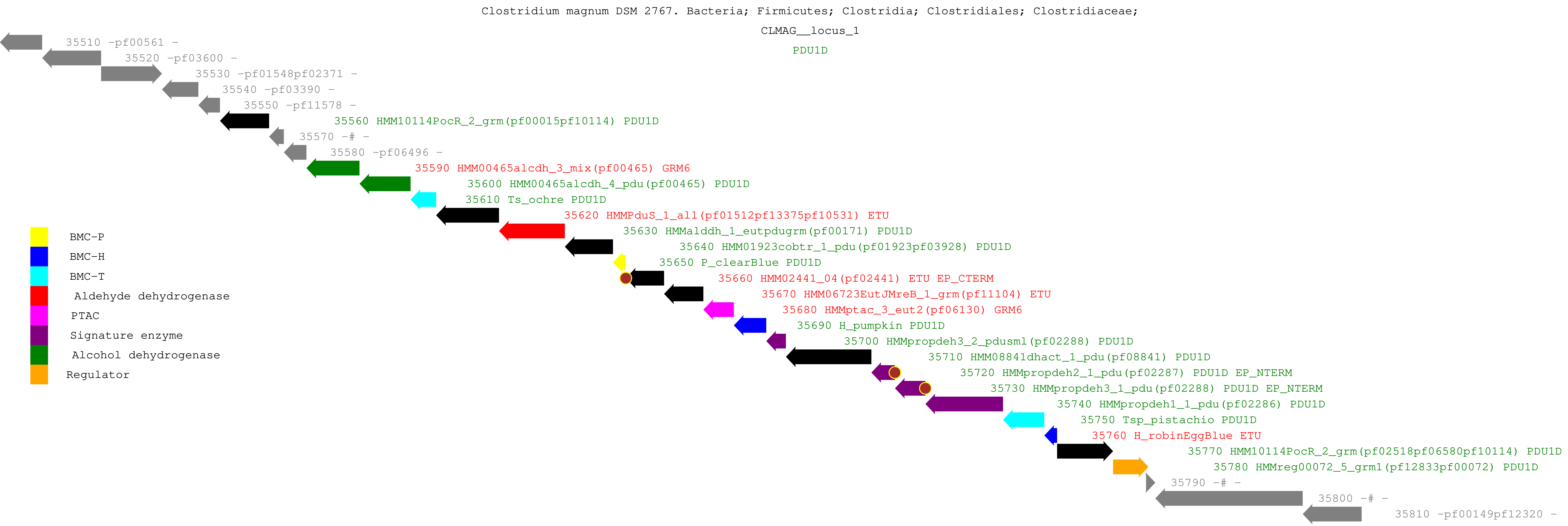

Clostridium magnum DSM 2767. Bacteria; Firmicutes; Clostridia; Clostridiales; Clostridiaceae;
CLMAG__locus_1
PDU1D
35510 -pf00561 -
35520 -pf03600 -
35530 -pf01548pf02371 -
35540 -pf03390 -
35550 -pf11578 -
35560 HMM10114PocR_2_grm(pf00015pf10114) PDU1D
35570 -# -
35580 -pf06496 -
35590 HMM00465alcdh_3_mix(pf00465) GRM6
35600 HMM00465alcdh_4_pdu(pf00465) PDU1D
35610 Ts_ochre PDU1D
35620 HMMPduS_1_all(pf01512pf13375pf10531) ETU
35630 HMMalddh_1_eutpdugrm(pf00171) PDU1D
35640 HMM01923cobtr_1_pdu(pf01923pf03928) PDU1D
35650 P_clearBlue PDU1D
35660 HMM02441_04(pf02441) ETU EP_CTERM
35670 HMM06723EutJMreB_1_grm(pf11104) ETU
35680 HMMptac_3_eut2(pf06130) GRM6
35690 H_pumpkin PDU1D
35700 HMMpropdeh3_2_pdusml(pf02288) PDU1D
35710 HMM08841dhact_1_pdu(pf08841) PDU1D
35720 HMMpropdeh2_1_pdu(pf02287) PDU1D EP_NTERM
35730 HMMpropdeh3_1_pdu(pf02288) PDU1D EP_NTERM
35740 HMMpropdeh1_1_pdu(pf02286) PDU1D
35750 Tsp_pistachio PDU1D
35760 H_robinEggBlue ETU
35770 HMM10114PocR_2_grm(pf02518pf06580pf10114) PDU1D
35780 HMMreg00072_5_grm1(pf12833pf00072) PDU1D
35790 -# -
35800 -# -
35810 -pf00149pf12320 -
BMC-P
BMC-H
BMC-T
Aldehyde dehydrogenase
PTAC
Signature enzyme
Alcohol dehydrogenase
Regulator

Eubacterium sp. YI. Bacteria; Firmicutes; Clostridia; Clostridiales; Eubacteriaceae;

CPZ25__locus_1_main_sat

PDU1D

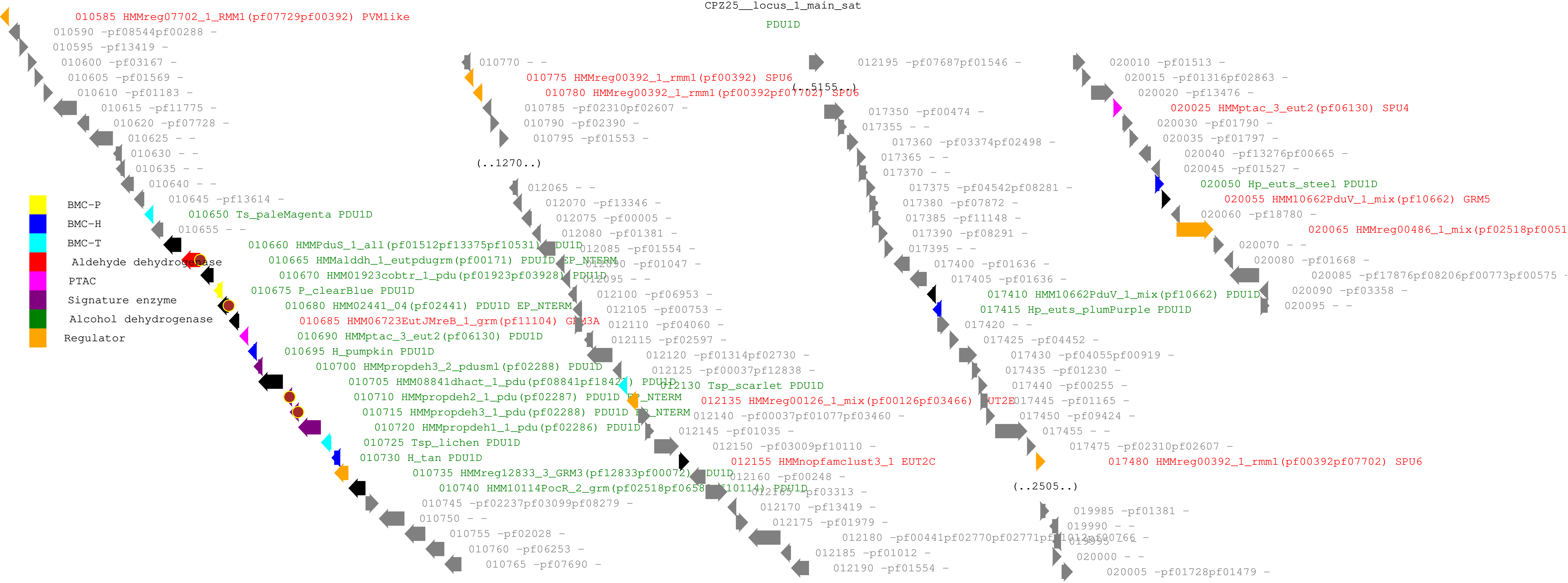

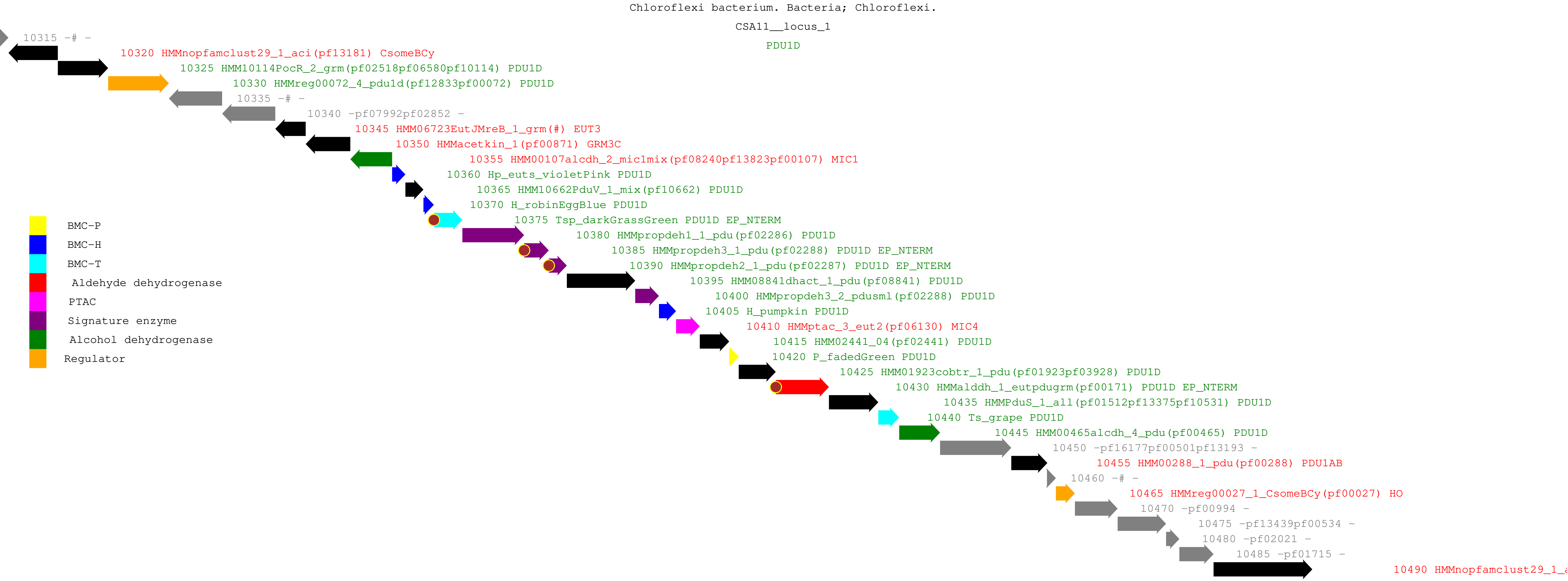

Chloroflexi bacterium. Bacteria; Chloroflexi.
CSA11__locus_1
PDU1D
10315 -# -
10320 HMMnopfamclust29_1_aci(pf13181) CsomeBCy
10325 HMM10114PocR_2_grm(pf02518pf06580pf10114) PDU1D
10330 HMMreg00072_4_pdu1d(pf12833pf00072) PDU1D
10335 -# -
10340 -pf07992pf02852 -
10345 HMM06723EutJMreB_1_grm(#) EUT3
10350 HMMacetkin_1(pf00871) GRM3C
10355 HMM00107alcdh_2_mic1mix(pf08240pf13823pf00107) MIC1
10360 Hp_euts_violetPink PDU1D
10365 HMM10662PduV_1_mix(pf10662) PDU1D
10370 H_robinEggBlue PDU1D
10375 Tsp_darkGrassGreen PDU1D EP_NTERM
10380 HMMpropdeh1_1_pdu(pf02286) PDU1D
10385 HMMpropdeh3_1_pdu(pf02288) PDU1D EP_NTERM
10390 HMMpropdeh2_1_pdu(pf02287) PDU1D EP_NTERM
10395 HMM08841dhact_1_pdu(pf08841) PDU1D
10400 HMMpropdeh3_2_pdusml(pf02288) PDU1D
10405 H_pumpkin PDU1D
10410 HMMptac_3_eut2(pf06130) MIC4
10415 HMM02441_04(pf02441) PDU1D
10420 P_fadedGreen PDU1D
10425 HMM01923cobtr_1_pdu(pf01923pf03928) PDU1D
10430 HMMalddh_1_eutpdugrm(pf00171) PDU1D EP_NTERM
10435 HMMPduS_1_all(pf01512pf13375pf10531) PDU1D
10440 Ts_grape PDU1D
10445 HMM00465alcdh_4_pdu(pf00465) PDU1D
10450 -pf16177pf00501pf13193 -
10455 HMM00288_1_pdu(pf00288) PDU1AB
10460 -# -
10465 HMMreg00027_1_CsomeBCy(pf00027) HO
10470 -pf00994 -
10475 -pf13439pf00534 -
10480 -pf02021 -
10485 -pf01715 -
10490 HMMnopfamclust29_1_a
BMC-P
BMC-H
BMC-T
Aldehyde dehydrogenase
PTAC
Signature enzyme
Alcohol dehydrogenase
Regulator

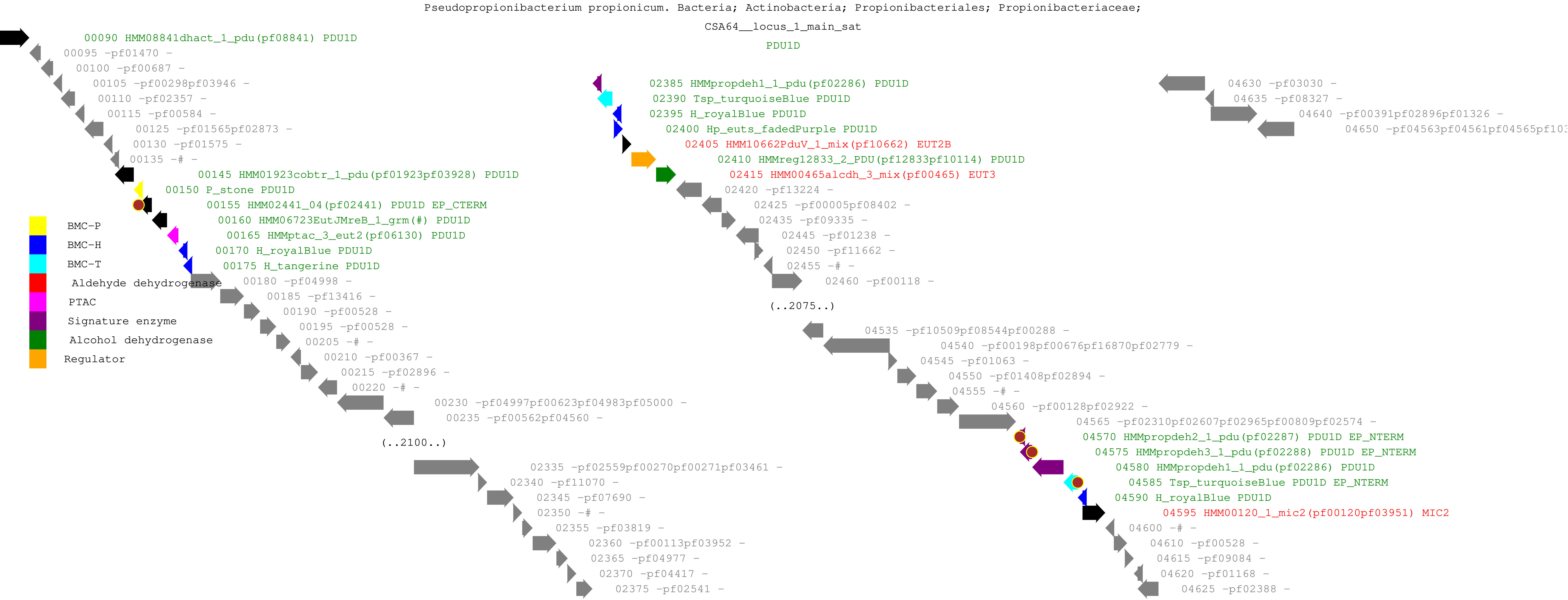
Pseudopropionibacterium propionicum. Bacteria; Actinobacteria; Propionibacteriales; Propionibacteriaceae;
CSA64__locus_1_main_sat
PDU1D
00090 HMM08841dhact_1_pdu(pf08841) PDU1D
00095 -pf01470 -
00100 -pf00687 -
00105 -pf00298pf03946 -
00110 -pf02357 -
00115 -pf00584 -
00125 -pf01565pf02873 -
00130 -pf01575 -
00135 -# -
00145 HMM01923cobtr_1_pdu(pf01923pf03928) PDU1D
00150 P_stone PDU1D
00155 HMM02441_04(pf02441) PDU1D EP_CTERM
00160 HMM06723EutJMreB_1_grm(#) PDU1D
00165 HMMptac_3_eut2(pf06130) PDU1D
00170 H_royalBlue PDU1D
00175 H_tangerine PDU1D
00180 -pf04998 -
00185 -pf13416 -
00190 -pf00528 -
00195 -pf00528 -
00205 -# -
00210 -pf00367 -
00215 -pf02896 -
00220 -# -
00230 -pf04997pf00623pf04983pf05000 -
00235 -pf00562pf04560 -
(..2100..)
02335 -pf02559pf00270pf00271pf03461 -
02340 -pf11070 -
02345 -pf07690 -
02350 -# -
02355 -pf03819 -
02360 -pf00113pf03952 -
02365 -pf04977 -
02370 -pf04417 -
02375 -pf02541 -
BMC-P
BMC-H
BMC-T
Aldehyde dehydrogenase
PTAC
Signature enzyme
Alcohol dehydrogenase
Regulator
02385 HMMpropdeh1_1_pdu(pf02286) PDU1D
02390 Tsp_turquoiseBlue PDU1D
02395 H_royalBlue PDU1D
02400 Hp_euts_fadedPurple PDU1D
02405 HMM10662PduV_1_mix(pf10662) EUT2B
02410 HMMreg12833_2_PDU(pf12833pf10114) PDU1D
02415 HMM00465alcdh_3_mix(pf00465) EUT3
02420 -pf13224 -
02425 -pf00005pf08402 -
02435 -pf09335 -
02445 -pf01238 -
02450 -pf11662 -
02455 -# -
02460 -pf00118 -
(..2075..)
04535 -pf10509pf08544pf00288 -
04540 -pf00198pf00676pf16870pf02779 -
04545 -pf01063 -
04550 -pf01408pf02894 -
04555 -# -
04560 -pf00128pf02922 -
04565 -pf02310pf02607pf02965pf00809pf02574 -
04570 HMMpropdeh2_1_pdu(pf02287) PDU1D EP_NTERM
04575 HMMpropdeh3_1_pdu(pf02288) PDU1D EP_NTERM
04580 HMMpropdeh1_1_pdu(pf02286) PDU1D
04585 Tsp_turquoiseBlue PDU1D EP_NTERM
04590 H_royalBlue PDU1D
04595 HMM00120_1_mic2(pf00120pf03951) MIC2
04600 -# -
04610 -pf00528 -
04615 -pf09084 -
04620 -pf01168 -
04625 -pf02388 -
04630 -pf03030 -
04635 -pf08327 -
04640 -pf00391pf02896pf01326 -
04650 -pf04563pf04561pf04565pf103

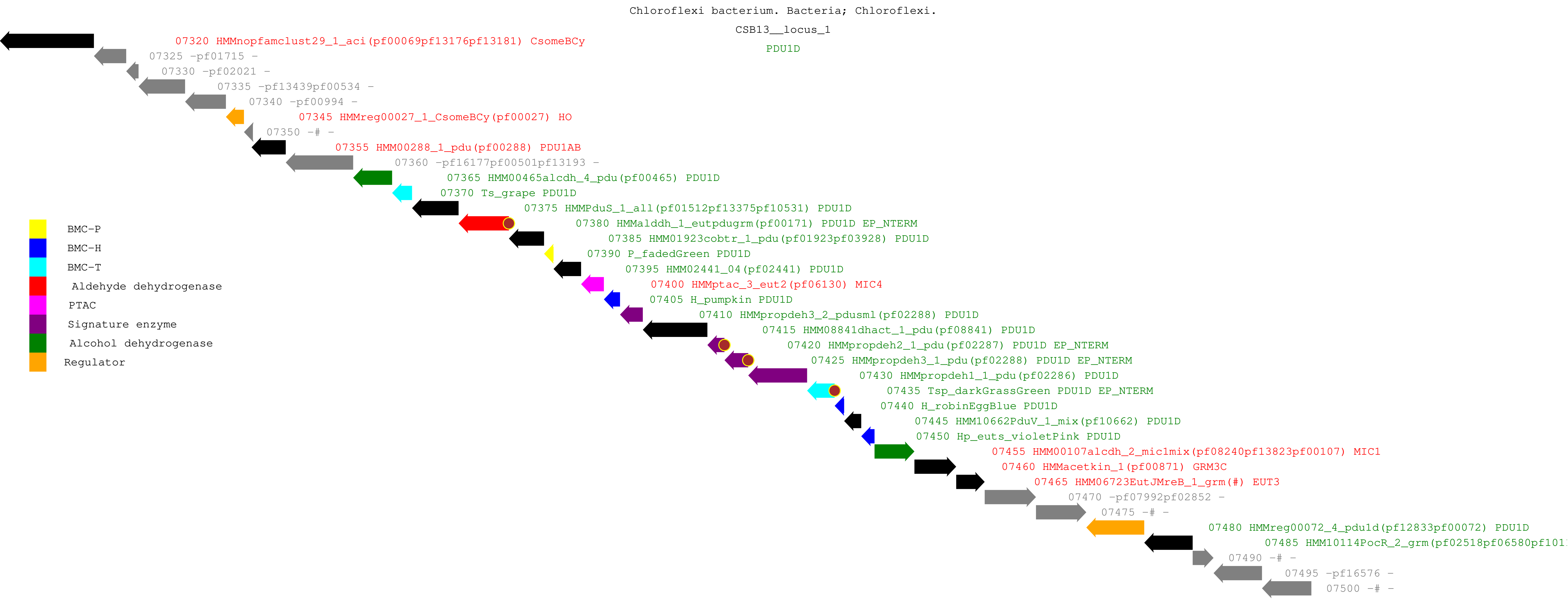
Chloroflexi bacterium. Bacteria; Chloroflexi.
CSB13__locus_1
PDU1D
07320 HMMnopfamclust29_1_aci(pf00069pf13176pf13181) CsomeBCy
07325 -pf01715 -
07330 -pf02021 -
07335 -pf13439pf00534 -
07340 -pf00994 -
07345 HMMreg00027_1_CsomeBCy(pf00027) HO
07350 -# -
07355 HMM00288_1_pdu(pf00288) PDU1AB
07360 -pf16177pf00501pf13193 -
07365 HMM00465alcdh_4_pdu(pf00465) PDU1D
07370 Ts_grape PDU1D
07375 HMMPduS_1_all(pf01512pf13375pf10531) PDU1D
07380 HMMalddh_1_eutpdugrm(pf00171) PDU1D EP_NTERM
07385 HMM01923cobtr_1_pdu(pf01923pf03928) PDU1D
07390 P_fadedGreen PDU1D
07395 HMM02441_04(pf02441) PDU1D
07400 HMMptac_3_eut2(pf06130) MIC4
07405 H_pumpkin PDU1D
07410 HMMpropdeh3_2_pdusml(pf02288) PDU1D
07415 HMM08841dhact_1_pdu(pf08841) PDU1D
07420 HMMpropdeh2_1_pdu(pf02287) PDU1D EP_NTERM
07425 HMMpropdeh3_1_pdu(pf02288) PDU1D EP_NTERM
07430 HMMpropdeh1_1_pdu(pf02286) PDU1D
07435 Tsp_darkGrassGreen PDU1D EP_NTERM
07440 H_robinEggBlue PDU1D
07445 HMM10662PduV_1_mix(pf10662) PDU1D
07450 Hp_euts_violetPink PDU1D
07455 HMM00107alcdh_2_mic1mix(pf08240pf13823pf00107) MIC1
07460 HMMacetkin_1(pf00871) GRM3C
07465 HMM06723EutJMreB_1_grm(#) EUT3
07470 -pf07992pf02852 -
07475 -# -
07480 HMMreg00072_4_pdu1d(pf12833pf00072) PDU1D
07485 HMM10114PocR_2_grm(pf02518pf06580pf101
07490 -# -
07495 -pf16576 -
07500 -# -
BMC-P
BMC-H
BMC-T
Aldehyde dehydrogenase
PTAC
Signature enzyme
Alcohol dehydrogenase
Regulator

Clostridium scatologenes. Bacteria; Firmicutes; Clostridia; Clostridiales; Clostridiaceae;

CSCA__locus_3

PDU1D

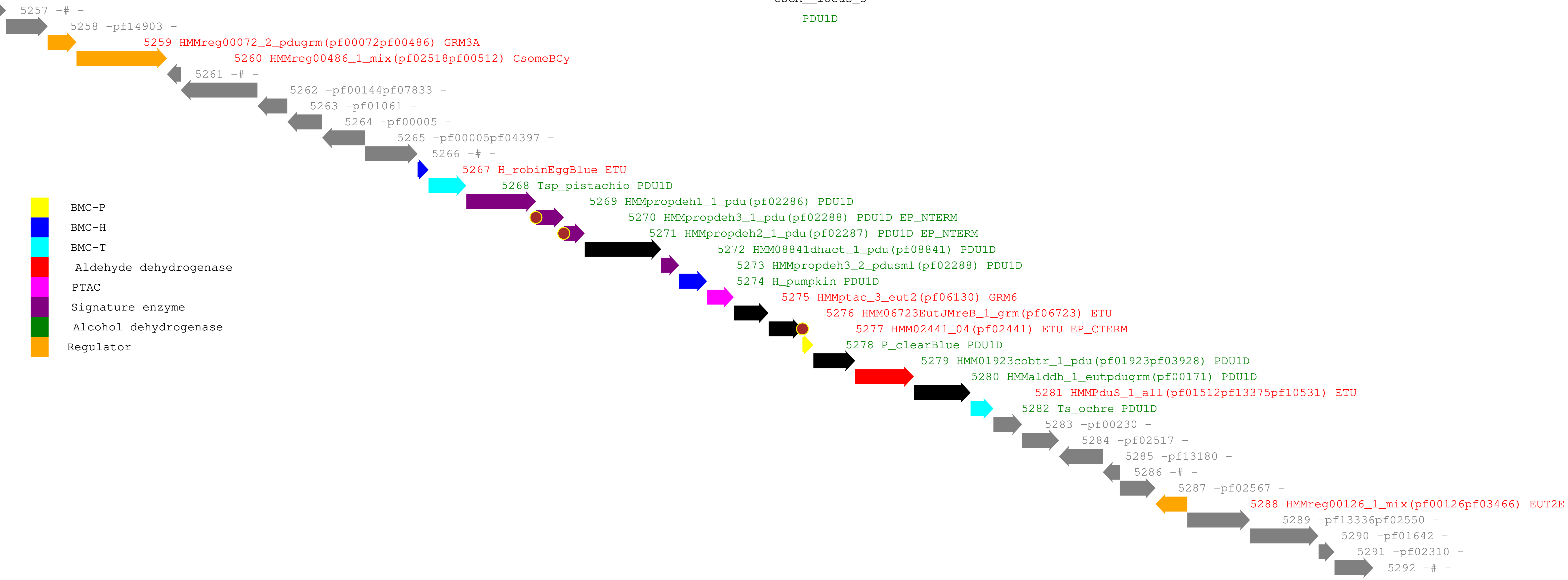

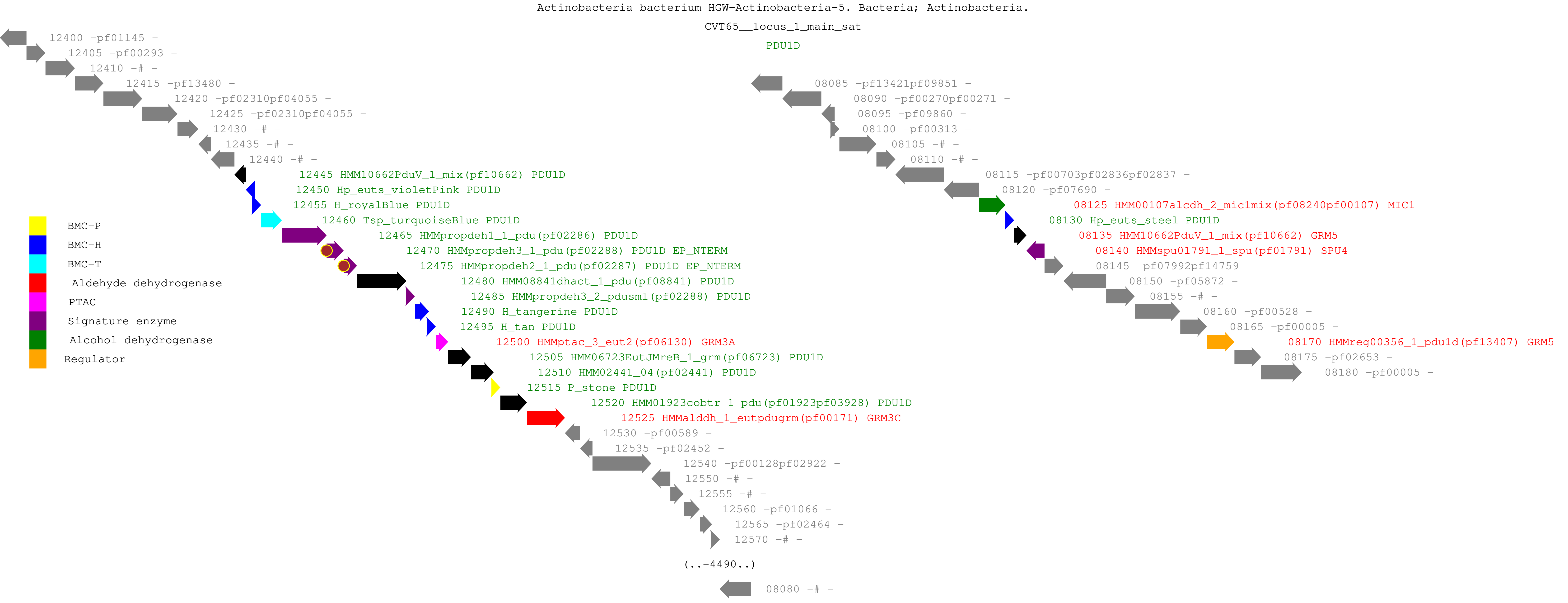

Actinobacteria bacterium HGW-Actinobacteria-5. Bacteria; Actinobacteria.
CVT65__locus_1_main_sat
PDU1D
12400 -pf01145 -
12405 -pf00293 -
12410 -# -
12415 -pf13480 -
12420 -pf02310pf04055 -
12425 -pf02310pf04055 -
12430 -# -
12435 -# -
12440 -# -
12445 HMM10662PduV_1_mix(pf10662) PDU1D
12450 Hp_euts_violetPink PDU1D
12455 H_royalBlue PDU1D
12460 Tsp_turquoiseBlue PDU1D
12465 HMMpropdeh1_1_pdu(pf02286) PDU1D
12470 HMMpropdeh3_1_pdu(pf02288) PDU1D EP_NTERM
12475 HMMpropdeh2_1_pdu(pf02287) PDU1D EP_NTERM
12480 HMM08841dhact_1_pdu(pf08841) PDU1D
12485 HMMpropdeh3_2_pdusml(pf02288) PDU1D
12490 H_tangerine PDU1D
12495 H_tan PDU1D
12500 HMMptac_3_eut2(pf06130) GRM3A
12505 HMM06723EutJMreB_1_grm(pf06723) PDU1D
12510 HMM02441_04(pf02441) PDU1D
12515 P_stone PDU1D
12520 HMM01923cobtr_1_pdu(pf01923pf03928) PDU1D
12525 HMMalddh_1_eutpdugrm(pf00171) GRM3C
12530 -pf00589 -
12535 -pf02452 -
12540 -pf00128pf02922 -
12550 -# -
12555 -# -
12560 -pf01066 -
12565 -pf02464 -
12570 -# -
(..-4490..)
08080 -# -
08085 -pf13421pf09851 -
08090 -pf00270pf00271 -
08095 -pf09860 -
08100 -pf00313 -
08105 -# -
08110 -# -
08115 -pf00703pf02836pf02837 -
08120 -pf07690 -
08125 HMM00107alcdh_2_mic1mix(pf08240pf00107) MIC1
08130 Hp_euts_steel PDU1D
08135 HMM10662PduV_1_mix(pf10662) GRM5
08140 HMMspu01791_1_spu(pf01791) SPU4
08145 -pf07992pf14759 -
08150 -pf05872 -
08155 -# -
08160 -pf00528 -
08165 -pf00005 -
08170 HMMreg00356_1_pdu1d(pf13407) GRM5
08175 -pf02653 -
08180 -pf00005 -
BMC-P
BMC-H
BMC-T
Aldehyde dehydrogenase
PTAC
Signature enzyme
Alcohol dehydrogenase
Regulator

Firmicutes bacterium HGW-Firmicutes-1. Bacteria; Firmicutes.

CVU84__locus_1_main_sat

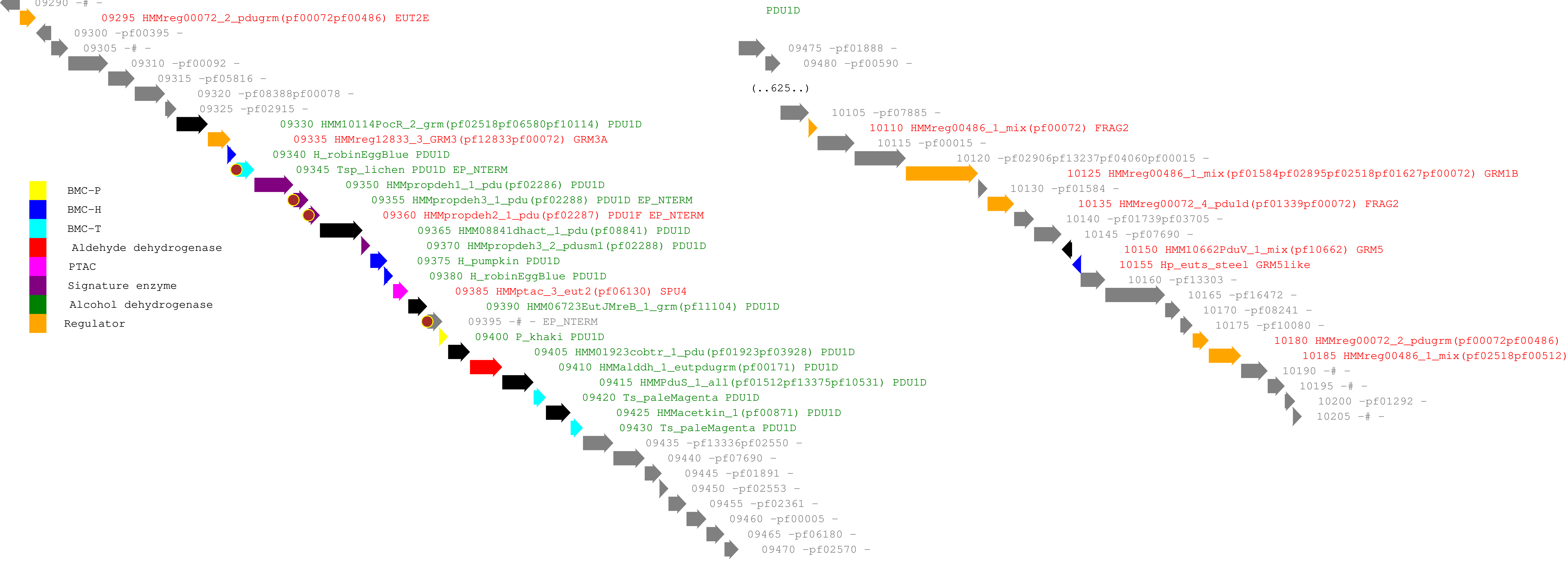

Firmicutes bacterium HGW-Firmicutes-4. Bacteria; Firmicutes.

CVU99__locus_2_main_sat

PDU1D

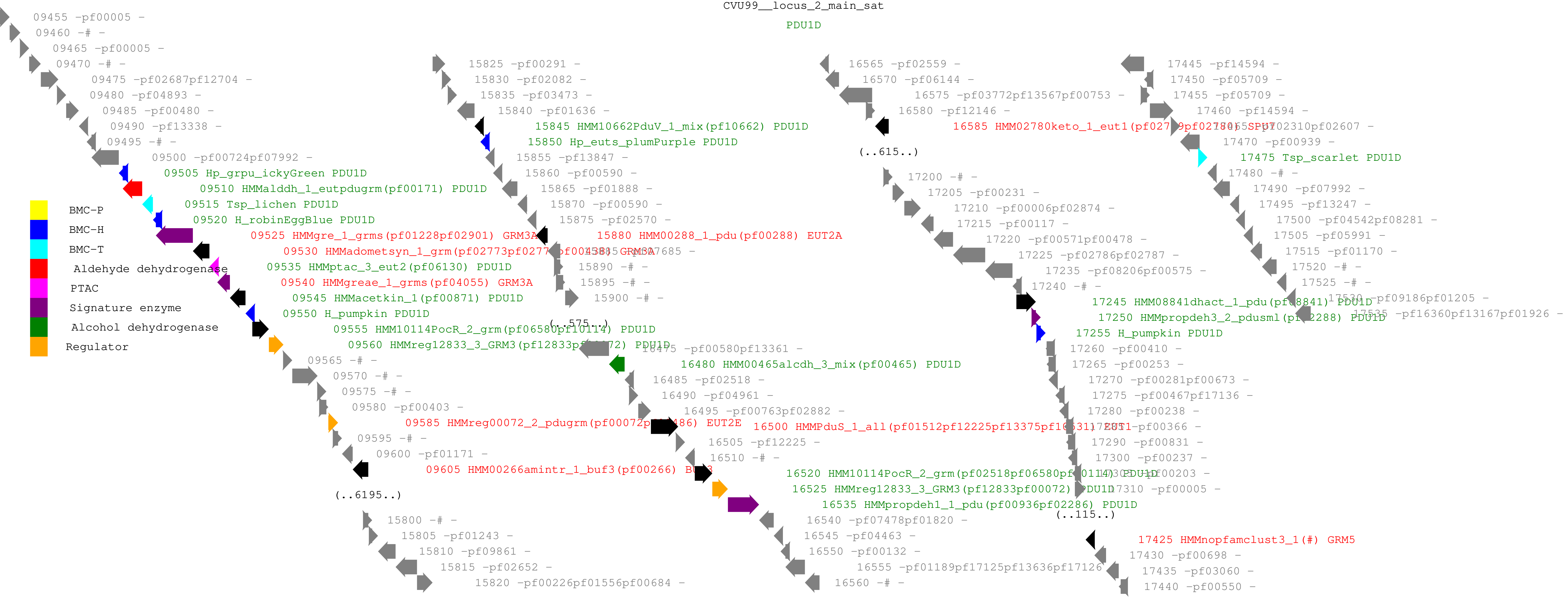

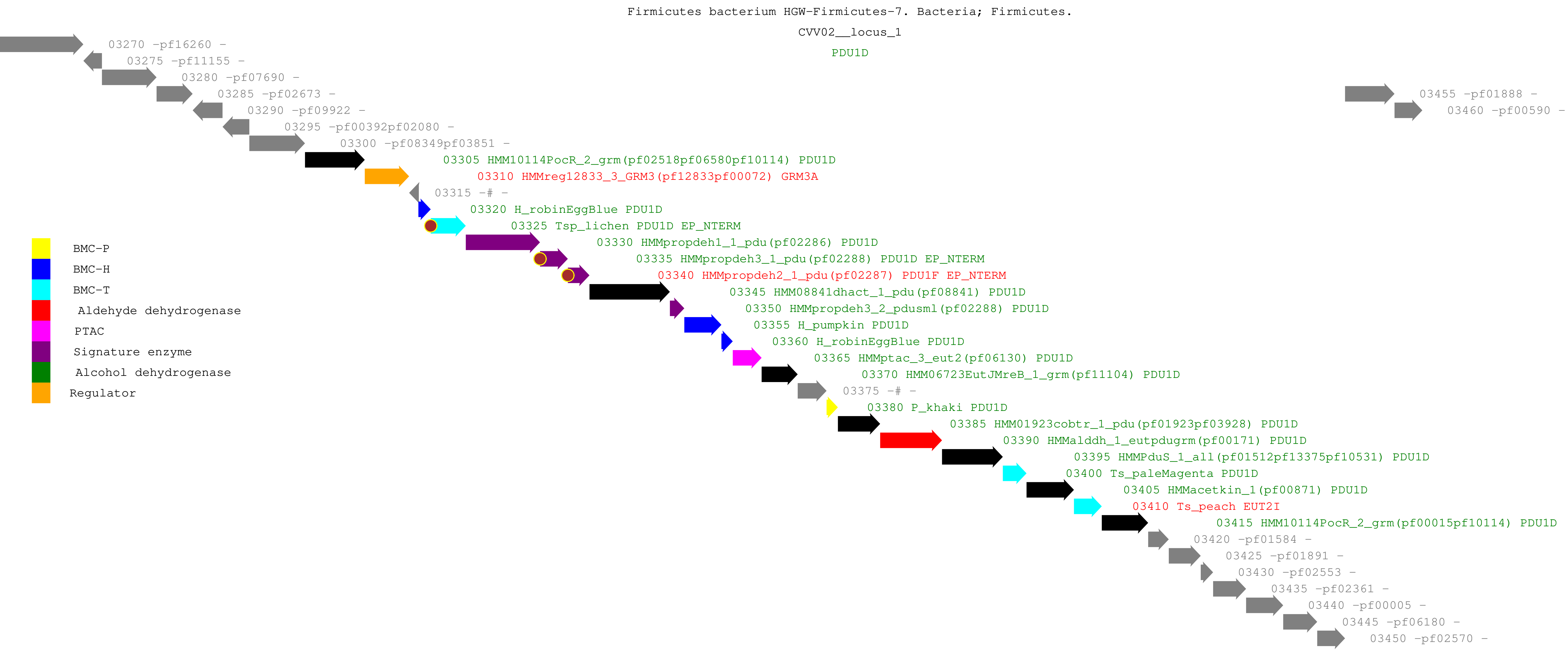

Firmicutes bacterium HGW-Firmicutes-7. Bacteria; Firmicutes.
CVV02__locus_1
PDU1D
03270 -pf16260 -
03275 -pf11155 -
03280 -pf07690 -
03285 -pf02673 -
03290 -pf09922 -
03295 -pf00392pf02080 -
03300 -pf08349pf03851 -
03305 HMM10114PocR_2_grm(pf02518pf06580pf10114) PDU1D
03310 HMMreg12833_3_GRM3(pf12833pf00072) GRM3A
03315 -# -
03320 H_robinEggBlue PDU1D
03325 Tsp_lichen PDU1D EP_NTERM
03330 HMMpropdeh1_1_pdu(pf02286) PDU1D
03335 HMMpropdeh3_1_pdu(pf02288) PDU1D EP_NTERM
03340 HMMpropdeh2_1_pdu(pf02287) PDU1F EP_NTERM
03345 HMM08841dhact_1_pdu(pf08841) PDU1D
03350 HMMpropdeh3_2_pdusml(pf02288) PDU1D
03355 H_pumpkin PDU1D
03360 H_robinEggBlue PDU1D
03365 HMMptac_3_eut2(pf06130) PDU1D
03370 HMM06723EutJMreB_1_grm(pf11104) PDU1D
03375 -# -
03380 P_khaki PDU1D
03385 HMM01923cobtr_1_pdu(pf01923pf03928) PDU1D
03390 HMMalddh_1_eutpdugrm(pf00171) PDU1D
03395 HMMPduS_1_all(pf01512pf13375pf10531) PDU1D
03400 Ts_paleMagenta PDU1D
03405 HMMacetkin_1(pf00871) PDU1D
03410 Ts_peach EUT2I
03415 HMM10114PocR_2_grm(pf00015pf10114) PDU1D
03420 -pf01584 -
03425 -pf01891 -
03430 -pf02553 -
03435 -pf02361 -
03440 -pf00005 -
03445 -pf06180 -
03450 -pf02570 -
03455 -pf01888 -
03460 -pf00590 -
BMC-P
BMC-H
BMC-T
Aldehyde dehydrogenase
PTAC
Signature enzyme
Alcohol dehydrogenase
Regulator

Telmatospirillum siberiense. Bacteria; Proteobacteria; Alphaproteobacteria; Rhodospirillales;

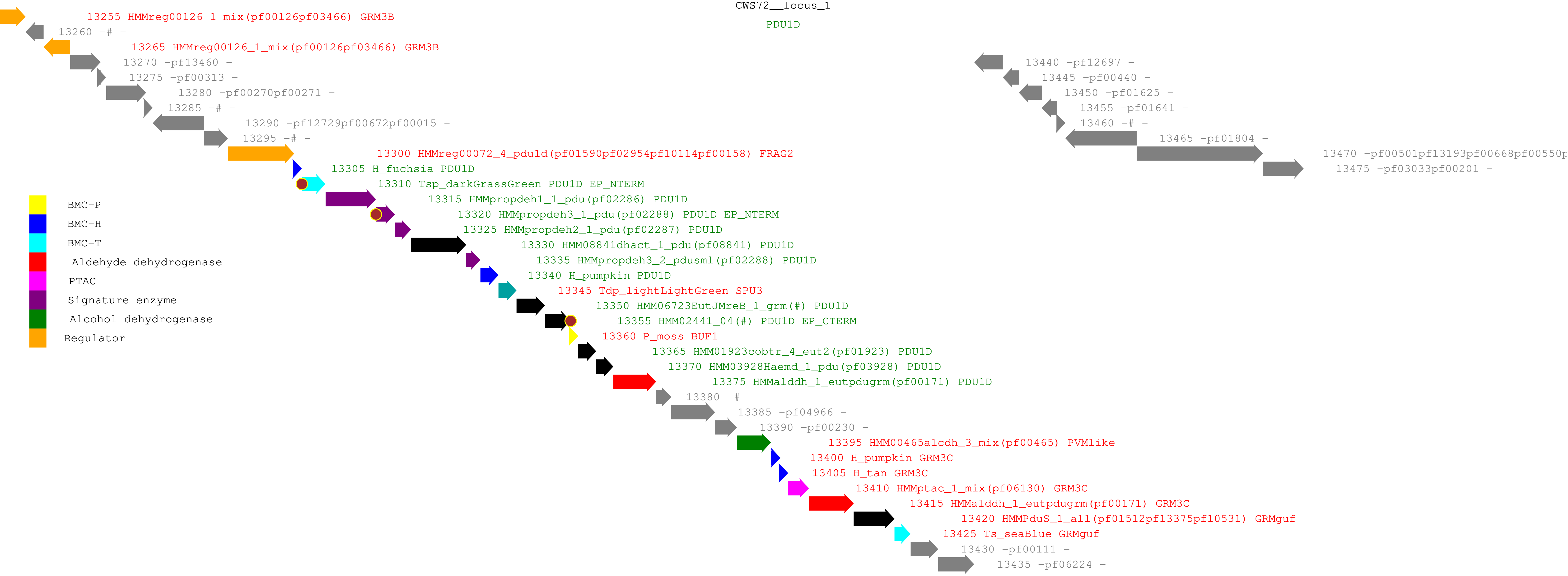

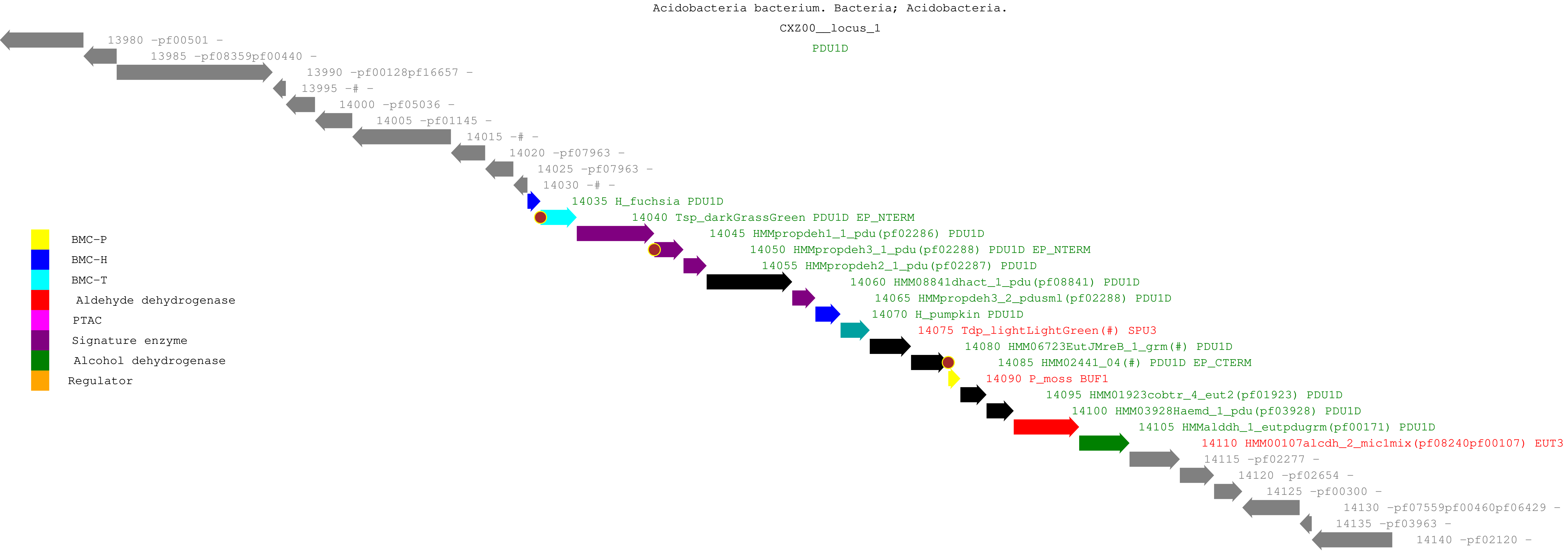

Acidobacteria bacterium. Bacteria; Acidobacteria.
CXZ00__locus_1
PDU1D
13980 -pf00501 -
13985 -pf08359pf00440 -
13990 -pf00128pf16657 -
13995 -# -
14000 -pf05036 -
14005 -pf01145 -
14015 -# -
14020 -pf07963 -
14025 -pf07963 -
14030 -# -
14035 H_fuchsia PDU1D
14040 Tsp_darkGrassGreen PDU1D EP_NTERM
14045 HMMpropdeh1_1_pdu(pf02286) PDU1D
14050 HMMpropdeh3_1_pdu(pf02288) PDU1D EP_NTERM
14055 HMMpropdeh2_1_pdu(pf02287) PDU1D
14060 HMM08841dhact_1_pdu(pf08841) PDU1D
14065 HMMpropdeh3_2_pdusml(pf02288) PDU1D
14070 H_pumpkin PDU1D
14075 Tdp_lightLightGreen(#) SPU3
14080 HMM06723EutJMreB_1_grm(#) PDU1D
14085 HMM02441_04(#) PDU1D EP_CTERM
14090 P_moss BUF1
14095 HMM01923cobtr_4_eut2(pf01923) PDU1D
14100 HMM03928Haemd_1_pdu(pf03928) PDU1D
14105 HMMalddh_1_eutpdugrm(pf00171) PDU1D
14110 HMM00107alcdh_2_mic1mix(pf08240pf00107) EUT3
14115 -pf02277 -
14120 -pf02654 -
14125 -pf00300 -
14130 -pf07559pf00460pf06429 -
14135 -pf03963 -
14140 -pf02120 -
BMC-P
BMC-H
BMC-T
Aldehyde dehydrogenase
PTAC
Signature enzyme
Alcohol dehydrogenase
Regulator

Halanaerobium sp. MSAO_Bac5. Bacteria; Firmicutes; Clostridia; Halanaerobiales; Halanaerobiaceae;

D5S01__locus_1_main_sat

PDU1D

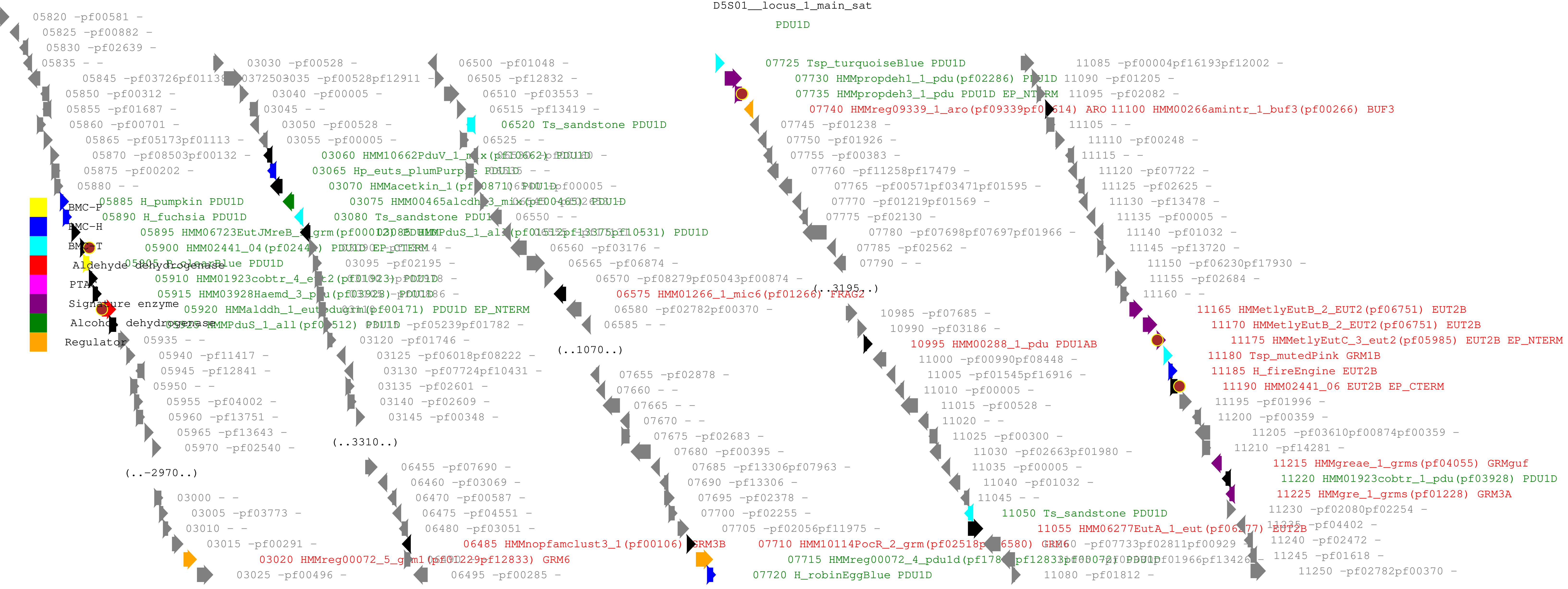

Desulfofundulus salinum. Bacteria; Firmicutes; Clostridia; Clostridiales; Peptococcaceae;

D7024__locus_1

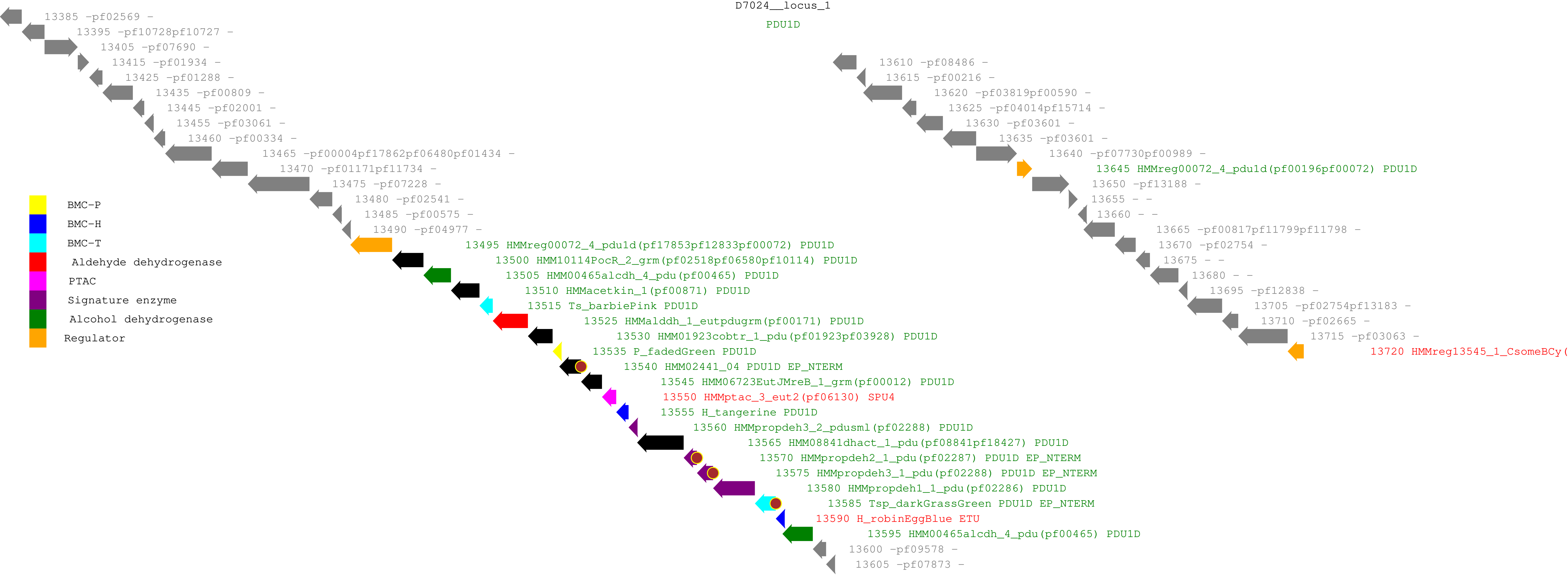

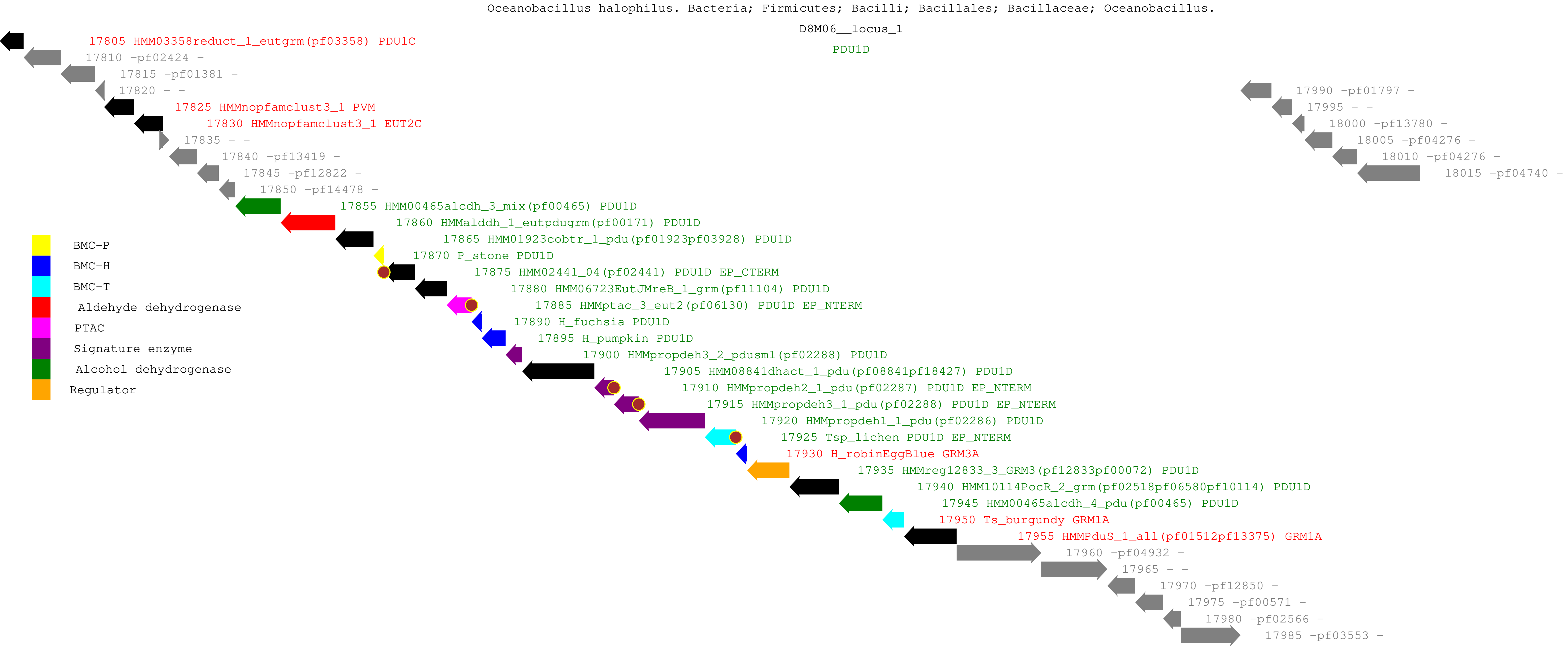
Oceanobacillus halophilus. Bacteria; Firmicutes; Bacilli; Bacillales; Bacillaceae; Oceanobacillus.
D8M06__locus_1
PDU1D
17805 HMM03358reduct_1_eutgrm(pf03358) PDU1C
17810 -pf02424 -
17815 -pf01381 -
17820 - -
17825 HMMnopfamclust3_1 PVM
17830 HMMnopfamclust3_1 EUT2C
17835 - -
17840 -pf13419 -
17845 -pf12822 -
17850 -pf14478 -
17855 HMM00465alcdh_3_mix(pf00465) PDU1D
17860 HMMalddh_1_eutpdugrm(pf00171) PDU1D
17865 HMM01923cobtr_1_pdu(pf01923pf03928) PDU1D
17870 P_stone PDU1D
17875 HMM02441_04(pf02441) PDU1D EP_CTERM
17880 HMM06723EutJMreB_1_grm(pf11104) PDU1D
17885 HMMptac_3_eut2(pf06130) PDU1D EP_NTERM
17890 H_fuchsia PDU1D
17895 H_pumpkin PDU1D
17900 HMMpropdeh3_2_pdusml(pf02288) PDU1D
17905 HMM08841dhact_1_pdu(pf08841pf18427) PDU1D
17910 HMMpropdeh2_1_pdu(pf02287) PDU1D EP_NTERM
17915 HMMpropdeh3_1_pdu(pf02288) PDU1D EP_NTERM
17920 HMMpropdeh1_1_pdu(pf02286) PDU1D
17925 Tsp_lichen PDU1D EP_NTERM
17930 H_robinEggBlue GRM3A
17935 HMMreg12833_3_GRM3(pf12833pf00072) PDU1D
17940 HMM10114PocR_2_grm(pf02518pf06580pf10114) PDU1D
17945 HMM00465alcdh_4_pdu(pf00465) PDU1D
17950 Ts_burgundy GRM1A
17955 HMMPduS_1_all(pf01512pf13375) GRM1A
17960 -pf04932 -
17965 - -
17970 -pf12850 -
17975 -pf00571 -
17980 -pf02566 -
17985 -pf03553 -
17990 -pf01797 -
17995 - -
18000 -pf13780 -
18005 -pf04276 -
18010 -pf04276 -
18015 -pf04740 -
BMC-P
BMC-H
BMC-T
Aldehyde dehydrogenase
PTAC
Signature enzyme
Alcohol dehydrogenase
Regulator

Desulfatibacillum alkenivorans (strain AK-01). Bacteria; Proteobacteria; Deltaproteobacteria; Desulfobacterales;

Dalk__locus_1

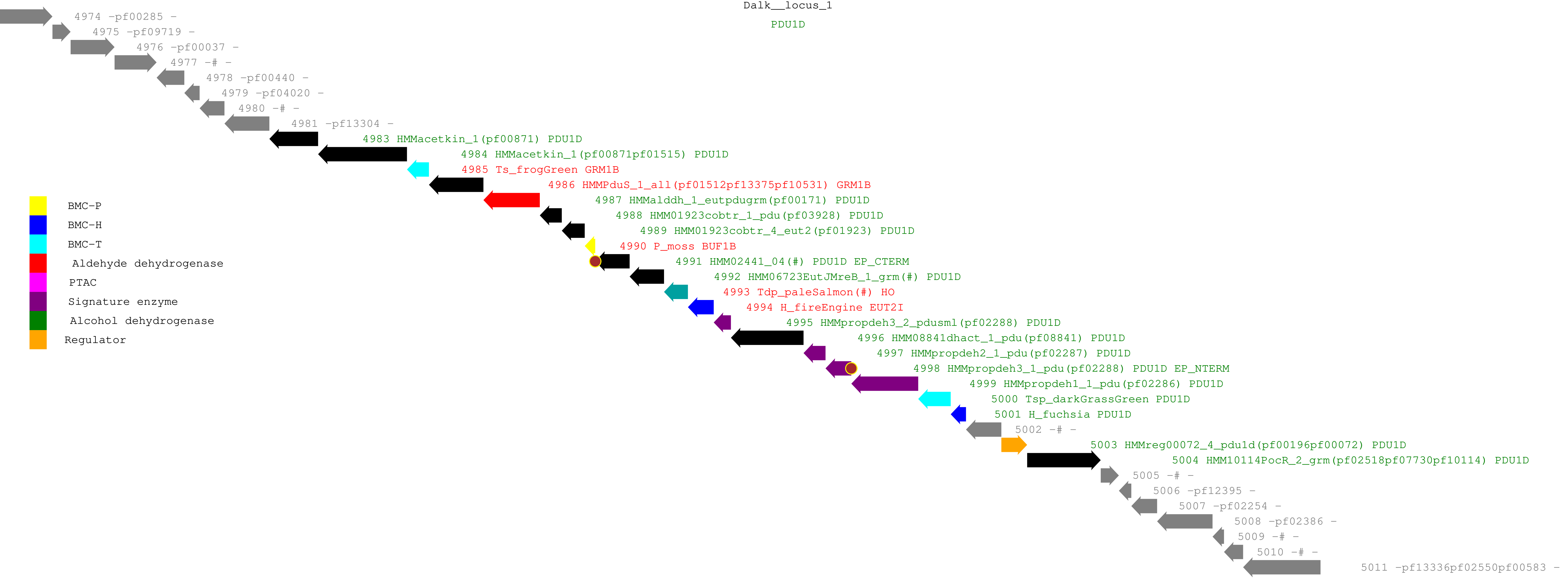

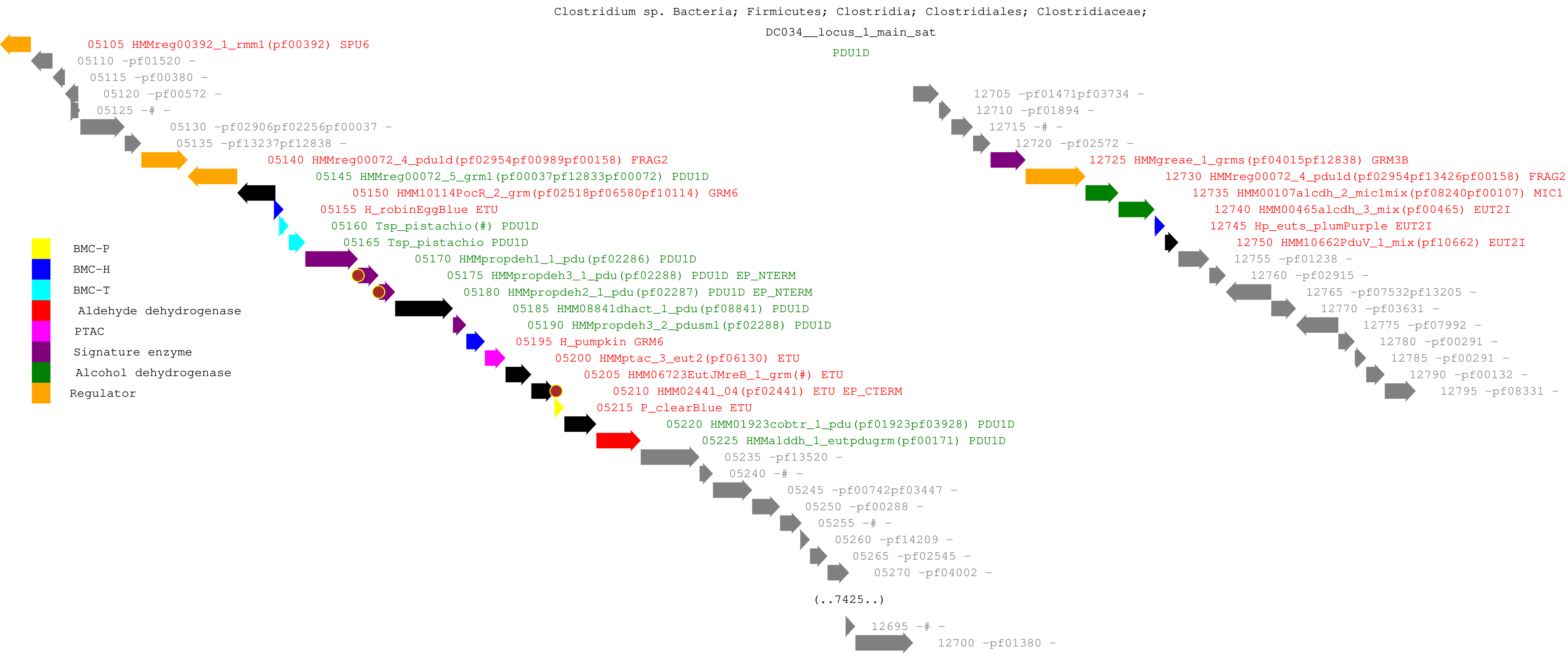
Clostridium sp. Bacteria; Firmicutes; Clostridia; Clostridiales; Clostridiaceae;
DC034__locus_1_main_sat
PDU1D
05105 HMMreg00392_1_rmm1(pf00392) SPU6
05110 -pf01520 -
05115 -pf00380 -
05120 -pf00572 -
05125 -# -
05130 -pf02906pf02256pf00037 -
05135 -pf13237pf12838 -
05140 HMMreg00072_4_pdu1d(pf02954pf00989pf00158) FRAG2
05145 HMMreg00072_5_grm1(pf00037pf12833pf00072) PDU1D
05150 HMM10114PocR_2_grm(pf02518pf06580pf10114) GRM6
05155 H_robinEggBlue ETU
05160 Tsp_pistachio(#) PDU1D
05165 Tsp_pistachio PDU1D
05170 HMMpropdeh1_1_pdu(pf02286) PDU1D
05175 HMMpropdeh3_1_pdu(pf02288) PDU1D EP_NTERM
05180 HMMpropdeh2_1_pdu(pf02287) PDU1D EP_NTERM
05185 HMM08841dhact_1_pdu(pf08841) PDU1D
05190 HMMpropdeh3_2_pdusml(pf02288) PDU1D
05195 H_pumpkin GRM6
05200 HMMptac_3_eut2(pf06130) ETU
05205 HMM06723EutJMreB_1_grm(#) ETU
05210 HMM02441_04(pf02441) ETU EP_CTERM
05215 P_clearBlue ETU
05220 HMM01923cobtr_1_pdu(pf01923pf03928) PDU1D
05225 HMMalddh_1_eutpdugrm(pf00171) PDU1D
05235 -pf13520 -
05240 -# -
05245 -pf00742pf03447 -
05250 -pf00288 -
05255 -# -
05260 -pf14209 -
05265 -pf02545 -
05270 -pf04002 -
(..7425..)
12695 -# -
12700 -pf01380 -
12705 -pf01471pf03734 -
12710 -pf01894 -
12715 -# -
12720 -pf02572 -
12725 HMMgreae_1_grms(pf04015pf12838) GRM3B
12730 HMMreg00072_4_pdu1d(pf02954pf13426pf00158) FRAG2
12735 HMM00107alcdh_2_mic1mix(pf08240pf00107) MIC1
12740 HMM00465alcdh_3_mix(pf00465) EUT2I
12745 Hp_euts_plumPurple EUT2I
12750 HMM10662PduV_1_mix(pf10662) EUT2I
12755 -pf01238 -
12760 -pf02915 -
12765 -pf07532pf13205 -
12770 -pf03631 -
12775 -pf07992 -
12780 -pf00291 -
12785 -pf00291 -
12790 -pf00132 -
12795 -pf08331 -
BMC-P
BMC-H
BMC-T
Aldehyde dehydrogenase
PTAC
Signature enzyme
Alcohol dehydrogenase
Regulator

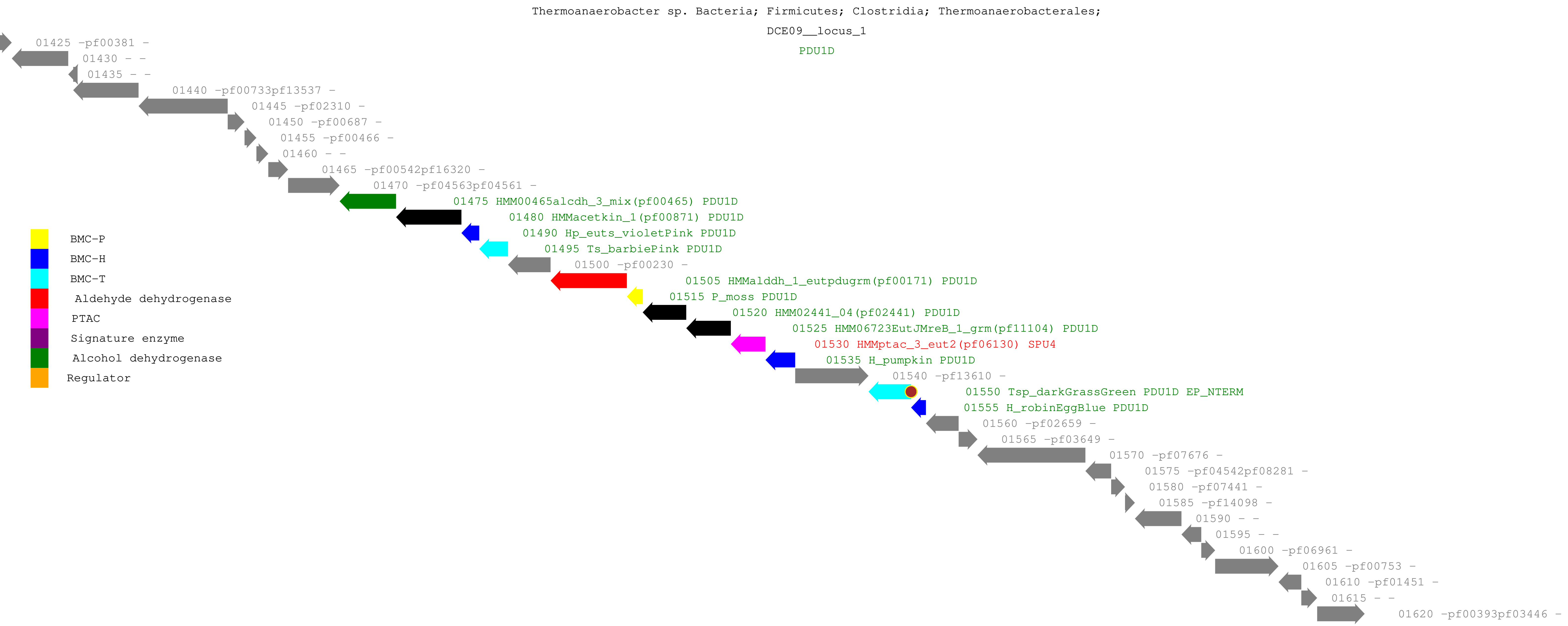
Thermoanaerobacter sp. Bacteria; Firmicutes; Clostridia; Thermoanaerobacterales;
DCE09__locus_1
PDU1D
01425 -pf00381 -
01430 - -
01435 - -
01440 -pf00733pf13537 -
01445 -pf02310 -
01450 -pf00687 -
01455 -pf00466 -
01460 - -
01465 -pf00542pf16320 -
01470 -pf04563pf04561 -
01475 HMM00465alcdh_3_mix(pf00465) PDU1D
01480 HMMacetkin_1(pf00871) PDU1D
01490 Hp_euts_violetPink PDU1D
01495 Ts_barbiePink PDU1D
01500 -pf00230 -
01505 HMMaldh_1_eutpdugrm(pf00171) PDU1D
01515 P_moss PDU1D
01520 HMM02441_04(pf02441) PDU1D
01525 HMM06723EutJMreB_1_grm(pf11104) PDU1D
01530 HMMptac_3_eut2(pf06130) SPU4
01535 H_pumpkin PDU1D
01540 -pf13610 -
01550 Tsp_darkGrassGreen PDU1D EP_NTERM
01555 H_robinEggBlue PDU1D
01560 -pf02659 -
01565 -pf03649 -
01570 -pf07676 -
01575 -pf04542pf08281 -
01580 -pf07441 -
01585 -pf14098 -
01590 - -
01595 - -
01600 -pf06961 -
01605 -pf00753 -
01610 -pf01451 -
01615 - -
01620 -pf00393pf03446 -
BMC-P
BMC-H
BMC-T
Aldehyde dehydrogenase
PTAC
Signature enzyme
Alcohol dehydrogenase
Regulator

Peptococcaceae bacterium DCMF. Bacteria; Firmicutes; Clostridia; Clostridiales; Peptococcaceae.

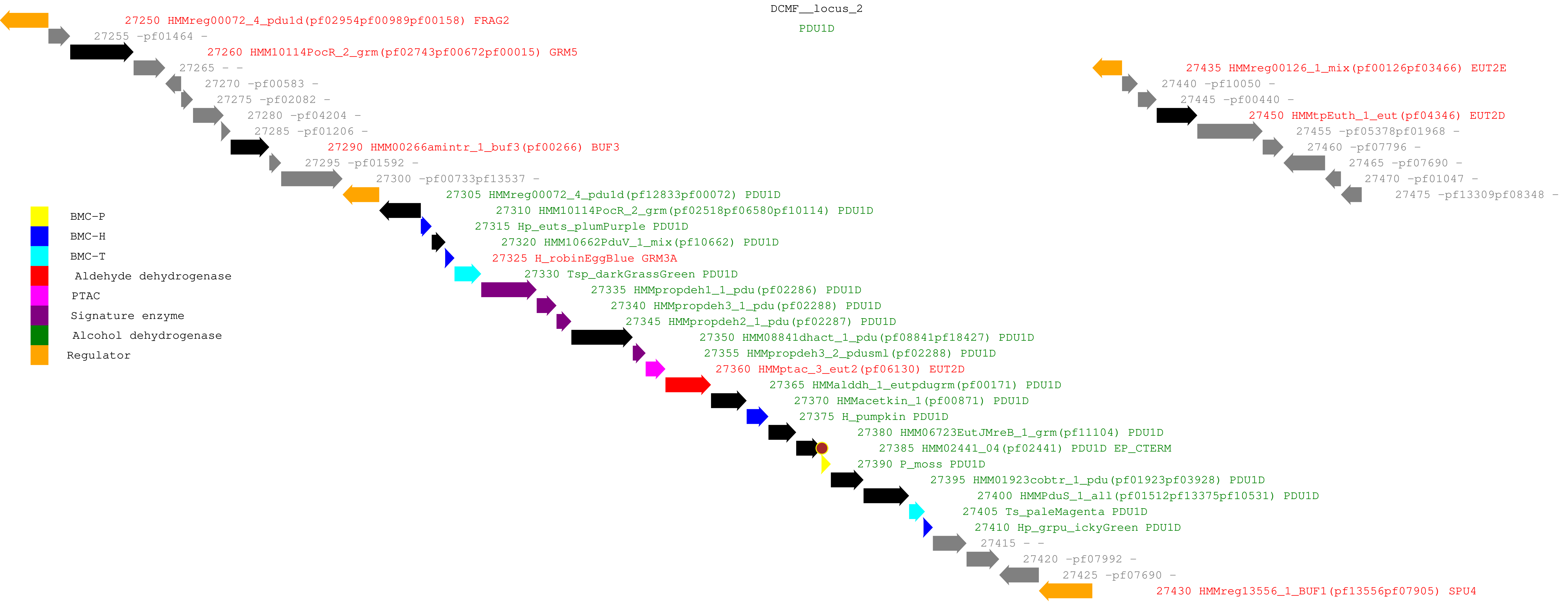

Sporomusaceae bacterium. Bacteria; Firmicutes; Negativicutes; Selenomonadales; Sporomusaceae;

DCP36__locus_1

PDU1D

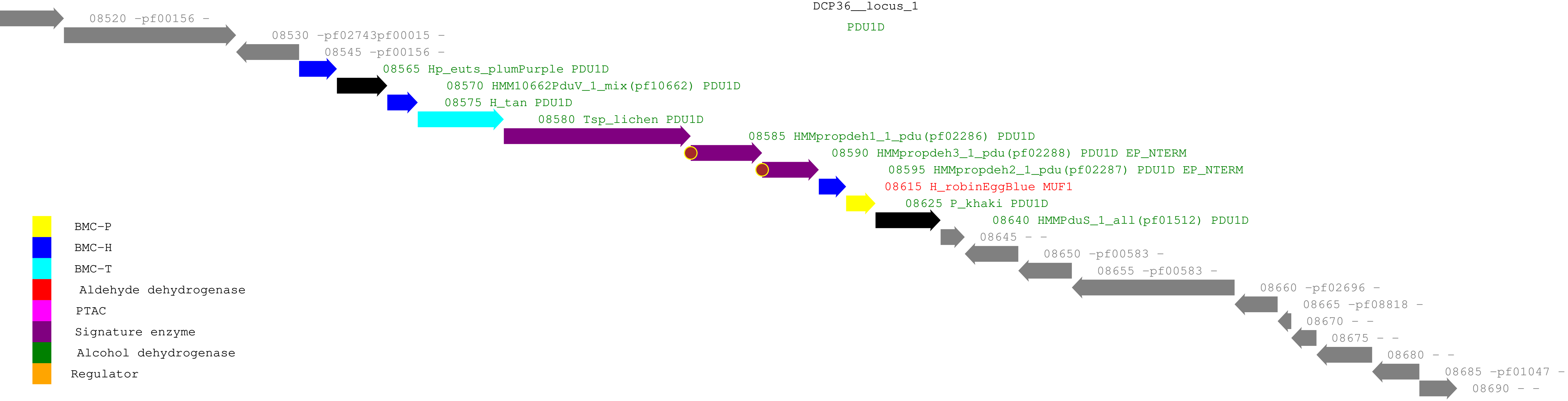

Acetobacterium sp. Bacteria; Firmicutes; Clostridia; Clostridiales; Eubacteriaceae;

DCY58__locus_1_main_sat

PDU1D

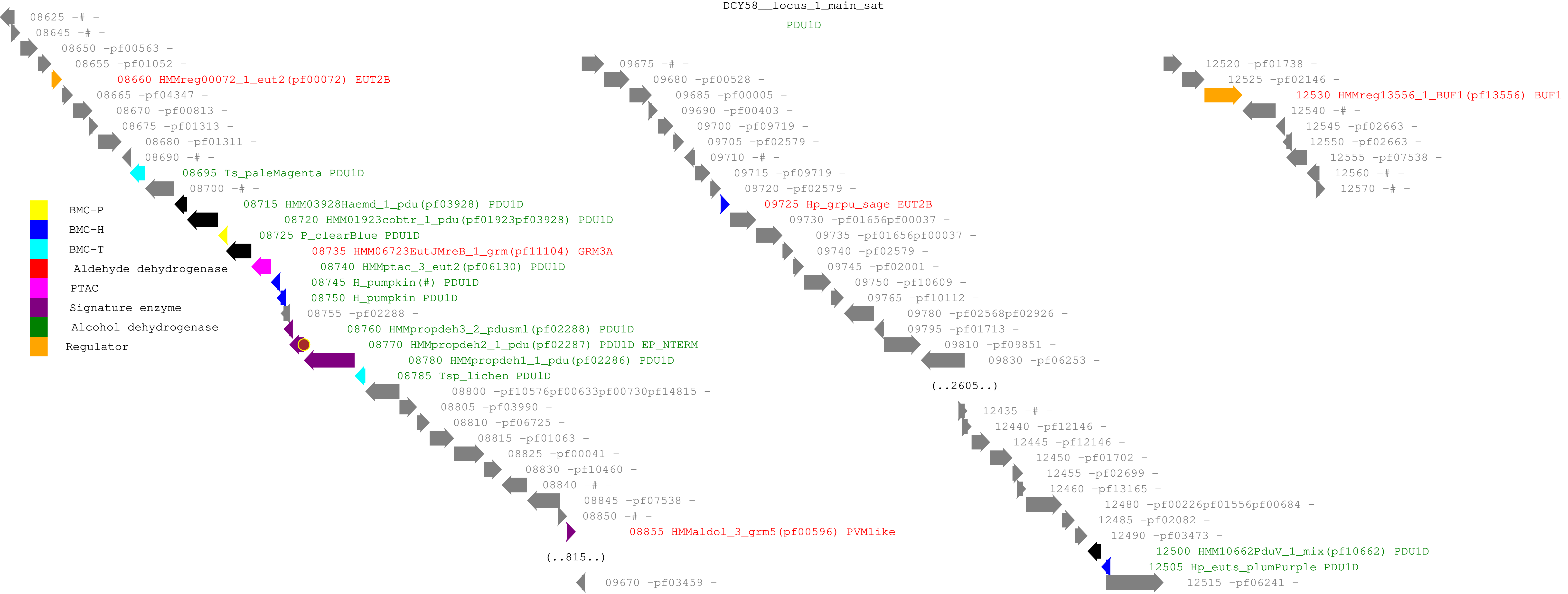

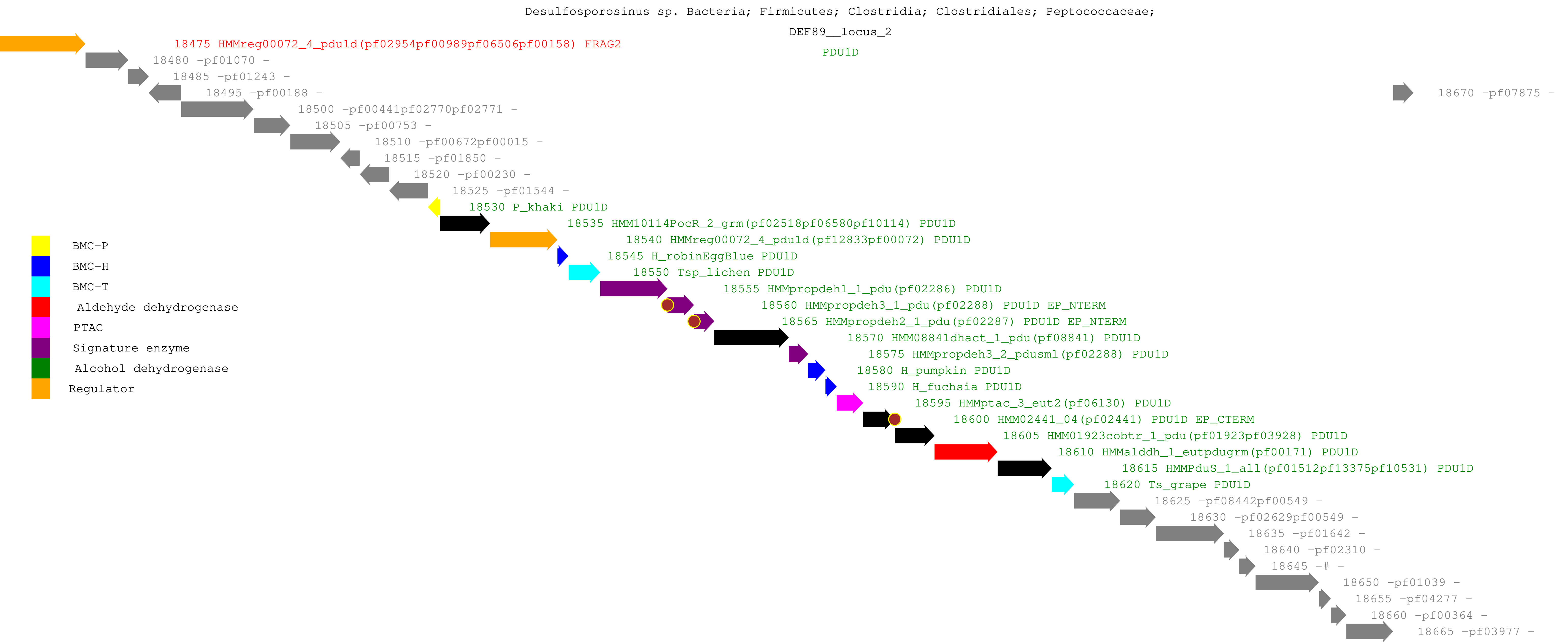

Desulfosporosinus sp. Bacteria; Firmicutes; Clostridia; Clostridiales; Peptococcaceae;
DEF89__locus_2
PDU1D
18475 HMMreg00072_4_pdu1d(pf02954pf00989pf06506pf00158) FRAG2
18480 -pf01070 -
18485 -pf01243 -
18495 -pf00188 -
18500 -pf00441pf02770pf02771 -
18505 -pf00753 -
18510 -pf00672pf00015 -
18515 -pf01850 -
18520 -pf00230 -
18525 -pf01544 -
18530 P_khaki PDU1D
18535 HMM10114PocR_2_grm(pf02518pf06580pf10114) PDU1D
18540 HMMreg00072_4_pdu1d(pf12833pf00072) PDU1D
18545 H_robinEggBlue PDU1D
18550 Tsp_lichen PDU1D
18555 HMMpropdeh1_1_pdu(pf02286) PDU1D
18560 HMMpropdeh3_1_pdu(pf02288) PDU1D EP_NTERM
18565 HMMpropdeh2_1_pdu(pf02287) PDU1D EP_NTERM
18570 HMM08841dhact_1_pdu(pf08841) PDU1D
18575 HMMpropdeh3_2_pdusml(pf02288) PDU1D
18580 H_pumpkin PDU1D
18590 H_fuchsia PDU1D
18595 HMMptac_3_eut2(pf06130) PDU1D
18600 HMM02441_04(pf02441) PDU1D EP_CTERM
18605 HMM01923cobtr_1_pdu(pf01923pf03928) PDU1D
18610 HMMalddh_1_eutpdugrm(pf00171) PDU1D
18615 HMMPduS_1_all(pf01512pf13375pf10531) PDU1D
18620 Ts_grape PDU1D
18625 -pf08442pf00549 -
18630 -pf02629pf00549 -
18635 -pf01642 -
18640 -pf02310 -
18645 -# -
18650 -pf01039 -
18655 -pf04277 -
18660 -pf00364 -
18665 -pf03977 -
18670 -pf07875 -
BMC-P
BMC-H
BMC-T
Aldehyde dehydrogenase
PTAC
Signature enzyme
Alcohol dehydrogenase
Regulator

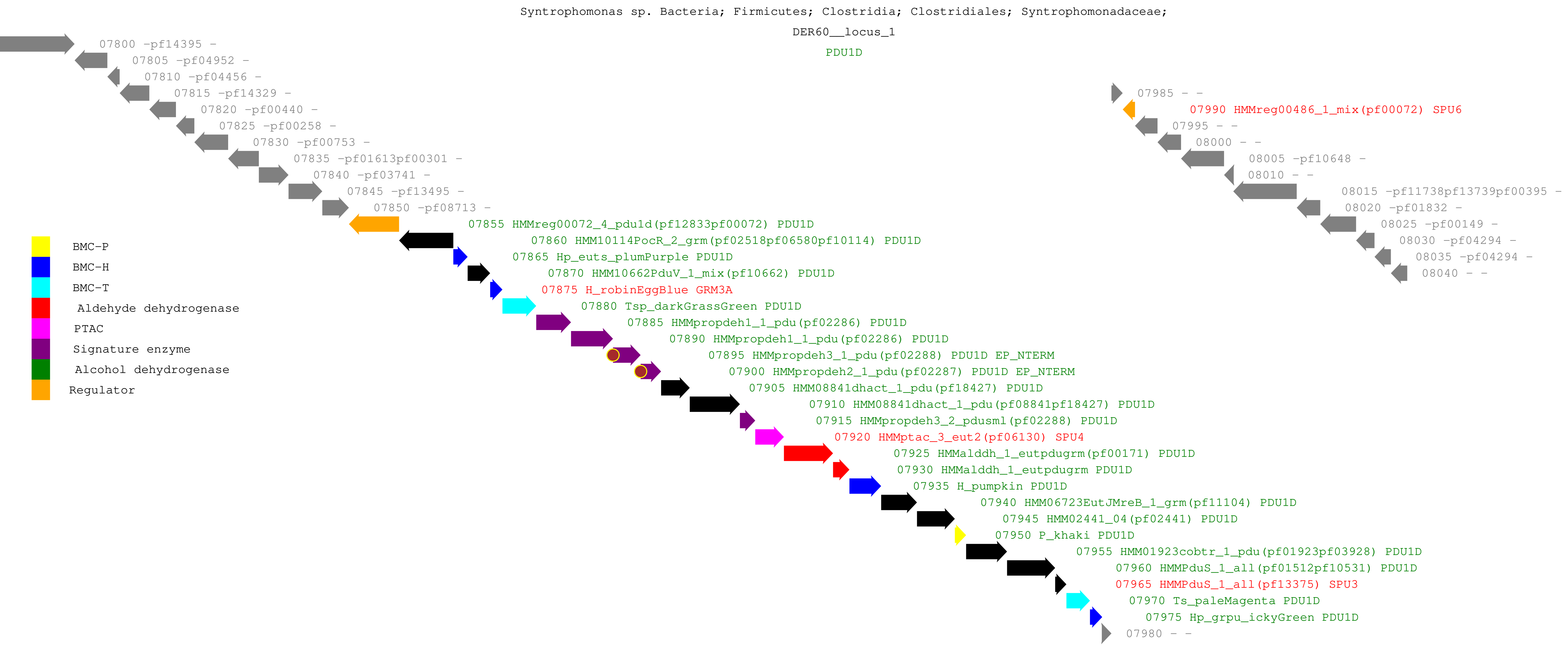

Syntrophomonas sp. Bacteria; Firmicutes; Clostridia; Clostridiales; Syntrophomonadaceae;
DER60__locus_1
PDU1D
07800 -pf14395 -
07805 -pf04952 -
07810 -pf04456 -
07815 -pf14329 -
07820 -pf00440 -
07825 -pf00258 -
07830 -pf00753 -
07835 -pf01613pf00301 -
07840 -pf03741 -
07845 -pf13495 -
07850 -pf08713 -
07855 HMMreg00072_4_pdu1d(pf12833pf00072) PDU1D
07860 HMM10114PocR_2_grm(pf02518pf06580pf10114) PDU1D
07865 Hp_euts_plumPurple PDU1D
07870 HMM10662PduV_1_mix(pf10662) PDU1D
07875 H_robinEggBlue GRM3A
07880 Tsp_darkGrassGreen PDU1D
07885 HMMpropdeh1_1_pdu(pf02286) PDU1D
07890 HMMpropdeh1_1_pdu(pf02286) PDU1D
07895 HMMpropdeh3_1_pdu(pf02288) PDU1D EP_NTERM
07900 HMMpropdeh2_1_pdu(pf02287) PDU1D EP_NTERM
07905 HMM08841dhact_1_pdu(pf18427) PDU1D
07910 HMM08841dhact_1_pdu(pf08841pf18427) PDU1D
07915 HMMpropdeh3_2_pdusml(pf02288) PDU1D
07920 HMMptac_3_eut2(pf06130) SPU4
07925 HMMaldd h_1_eutpdugrm(pf00171) PDU1D
07930 HMMaldd h_1_eutpdugrm PDU1D
07935 H_pumpkin PDU1D
07940 HMM06723EutJMreB_1_grm(pf11104) PDU1D
07945 HMM02441_04(pf02441) PDU1D
07950 P_khaki PDU1D
07955 HMM01923cobtr_1_pdu(pf01923pf03928) PDU1D
07960 HMMPduS_1_all(pf01512pf10531) PDU1D
07965 HMMPduS_1_all(pf13375) SPU3
07970 Ts_paleMagenta PDU1D
07975 Hp_grpu_ickyGreen PDU1D
07980 - -
07985 - -
07990 HMMreg00486_1_mix(pf00072) SPU6
07995 - -
08000 - -
08005 -pf10648 -
08010 - -
08015 -pf11738pf13739pf00395 -
08020 -pf01832 -
08025 -pf00149 -
08030 -pf04294 -
08035 -pf04294 -
08040 - -
BMC-P
BMC-H
BMC-T
Aldehyde dehydrogenase
PTAC
Signature enzyme
Alcohol dehydrogenase
Regulator

Halanaerobium sp. DL-01. Bacteria; Firmicutes; Clostridia; Halanaerobiales; Halanaerobiaceae;

DER71__locus_1_main_sat

PDU1D

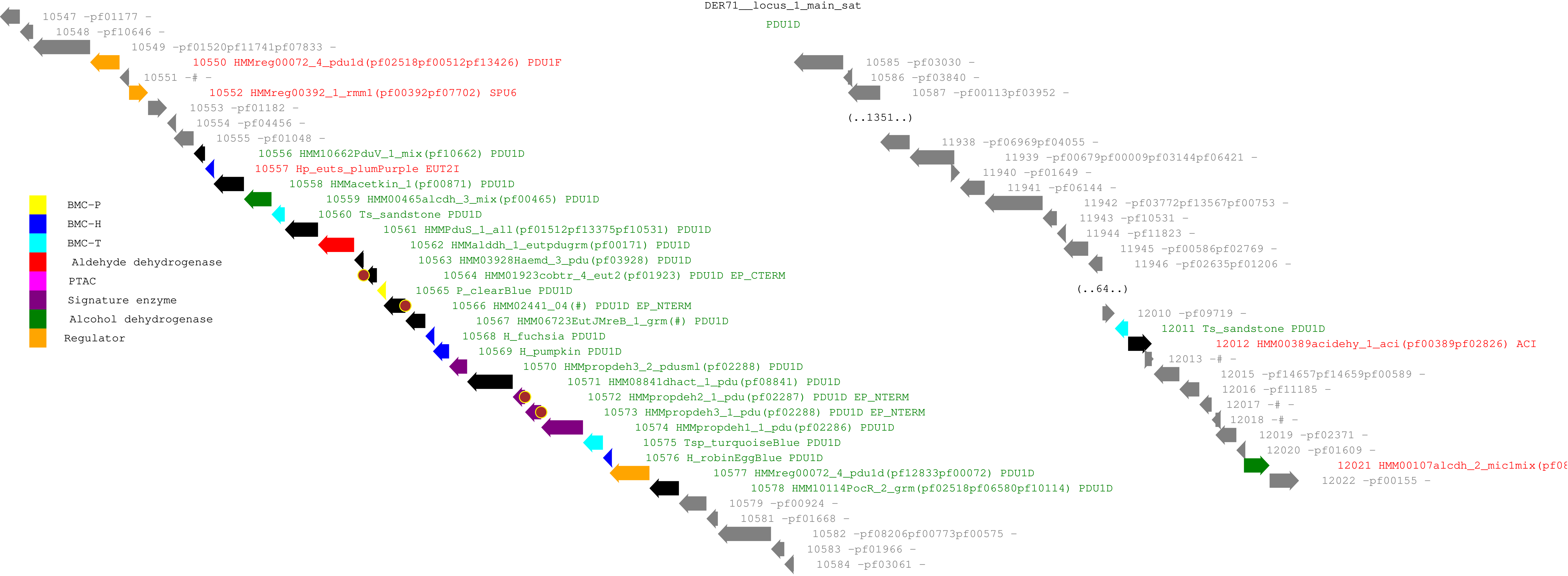

Clostridiales bacterium. Bacteria; Firmicutes; Clostridia; Clostridiales.

DHM42__locus_fragmented

PDU1D

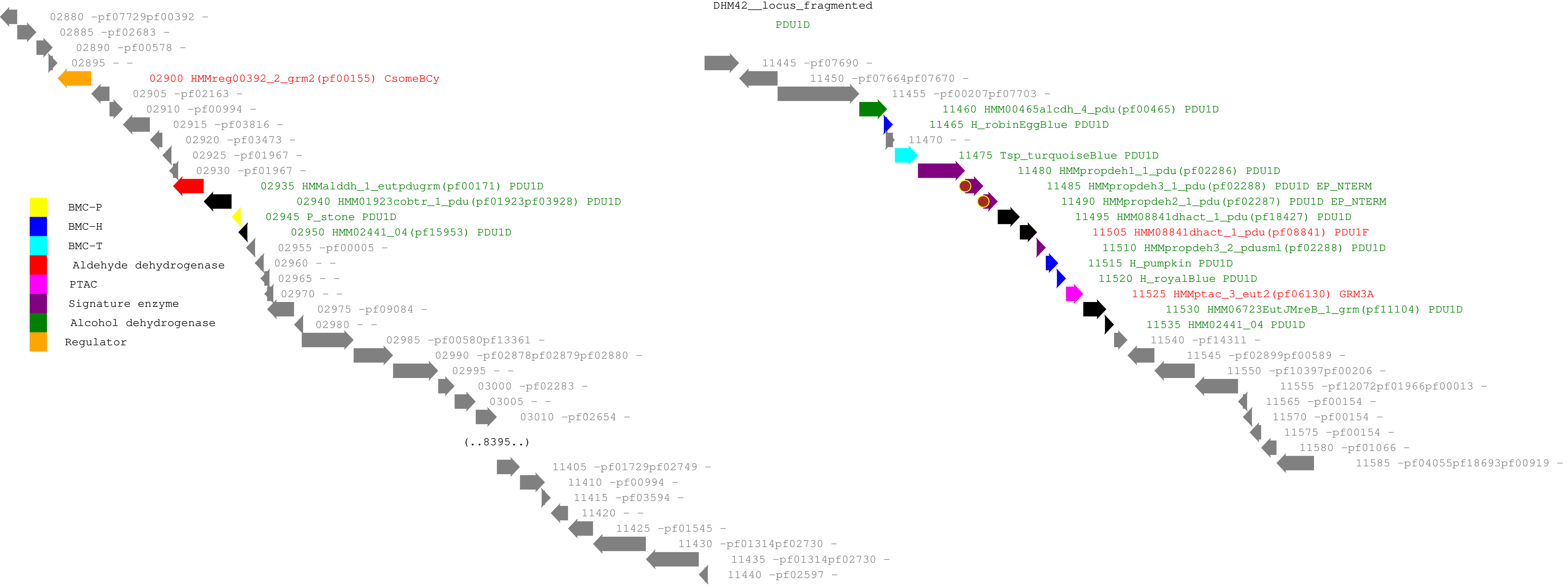

Firmicutes bacterium. Bacteria; Firmicutes.

DIU66__locus_fragmented

PDU1D

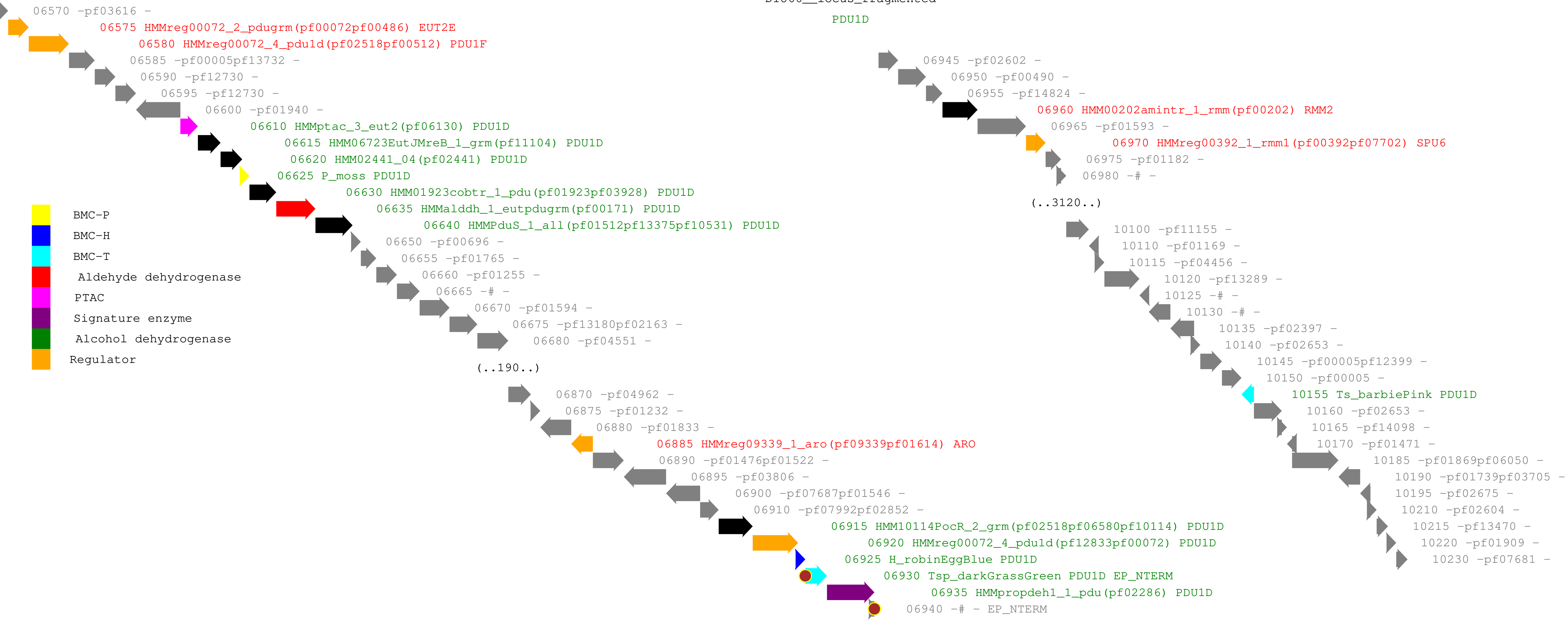

Desulfosporosinus sp. OT. Bacteria; Firmicutes; Clostridia; Clostridiales; Peptococcaceae;

DOT__locus_3

PDU1D

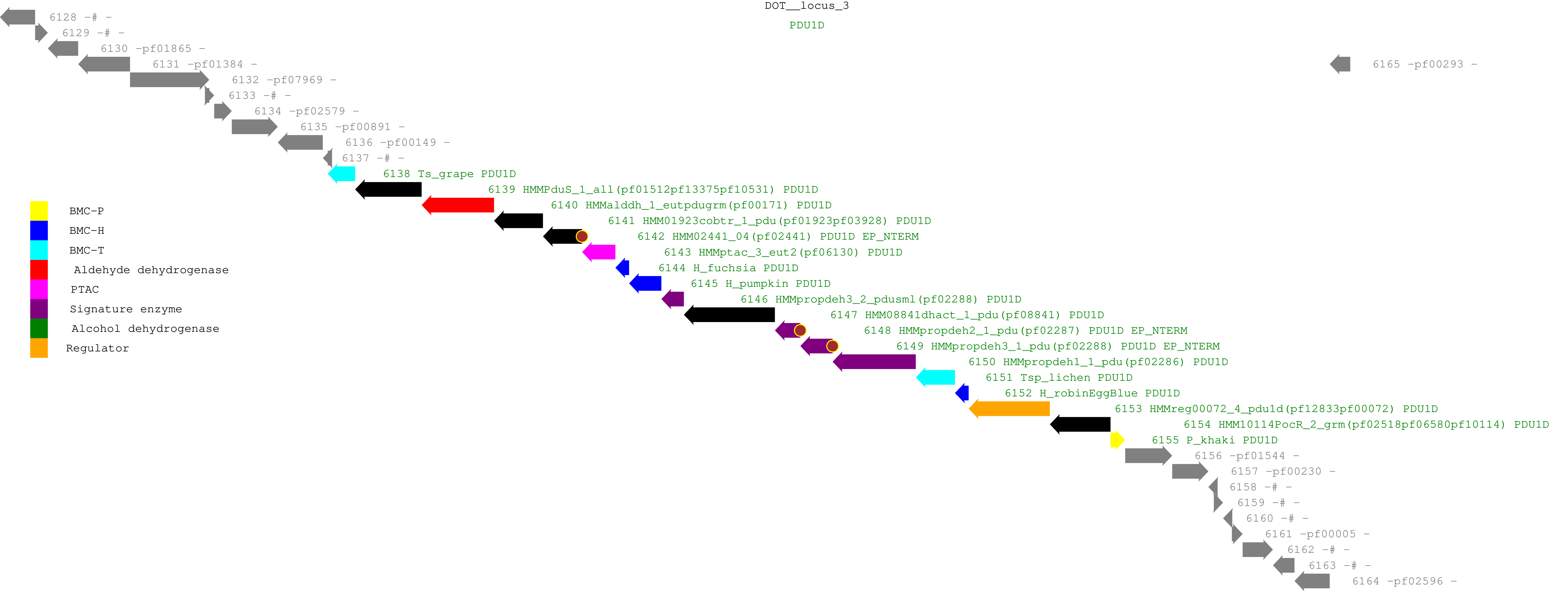

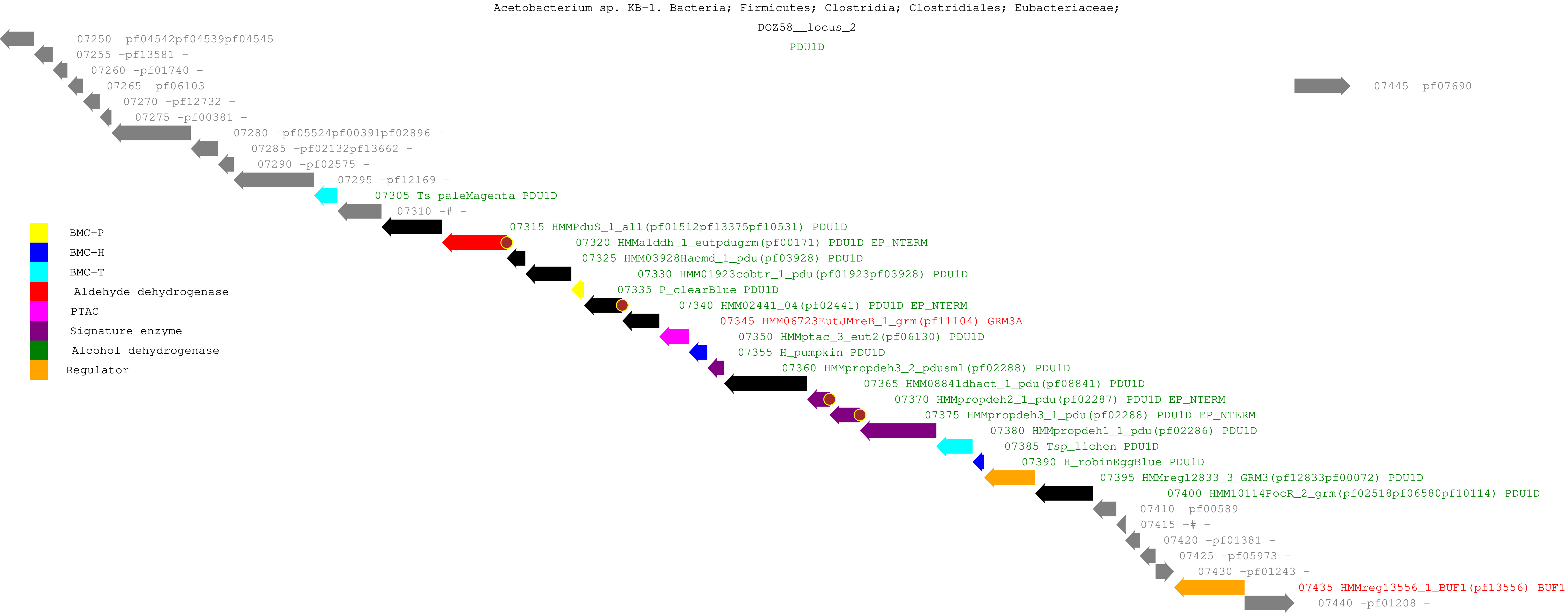

Acetobacterium sp. KB-1. Bacteria; Firmicutes; Clostridia; Clostridiales; Eubacteriaceae;
DOZ58__locus_2
PDU1D
07250 -pf04542pf04539pf04545 -
07255 -pf13581 -
07260 -pf01740 -
07265 -pf06103 -
07270 -pf12732 -
07275 -pf00381 -
07280 -pf05524pf00391pf02896 -
07285 -pf02132pf13662 -
07290 -pf02575 -
07295 -pf12169 -
07305 Ts_paleMagenta PDU1D
07310 -# -
07315 HMMPduS_1_all(pf01512pf13375pf10531) PDU1D
07320 HMMaldh_1_eutpdugrm(pf00171) PDU1D EP_NTERM
07325 HMM03928Haemd_1_pdu(pf03928) PDU1D
07330 HMM01923cobtr_1_pdu(pf01923pf03928) PDU1D
07335 P_clearBlue PDU1D
07340 HMM02441_04(pf02441) PDU1D EP_NTERM
07345 HMM06723EutJMreB_1_grm(pf11104) GRM3A
07350 HMMptac_3_eut2(pf06130) PDU1D
07355 H_pumpkin PDU1D
07360 HMMpropdeh3_2_pdusml(pf02288) PDU1D
07365 HMM08841dhact_1_pdu(pf08841) PDU1D
07370 HMMpropdeh2_1_pdu(pf02287) PDU1D EP_NTERM
07375 HMMpropdeh3_1_pdu(pf02288) PDU1D EP_NTERM
07380 HMMpropdeh1_1_pdu(pf02286) PDU1D
07385 Tsp_lichen PDU1D
07390 H_robinEggBlue PDU1D
07395 HMMreg12833_3_GRM3(pf12833pf00072) PDU1D
07400 HMM10114PocR_2_grm(pf02518pf06580pf10114) PDU1D
07410 -pf00589 -
07415 -# -
07420 -pf01381 -
07425 -pf05973 -
07430 -pf01243 -
07435 HMMreg13556_1_BUF1(pf13556) BUF1
07440 -pf01208 -
07445 -pf07690 -
BMC-P
BMC-H
BMC-T
Aldehyde dehydrogenase
PTAC
Signature enzyme
Alcohol dehydrogenase
Regulator

Clostridium sp. HMP27. Bacteria; Firmicutes; Clostridia; Clostridiales; Clostridiaceae;

DP68__locus_1

PDU1D

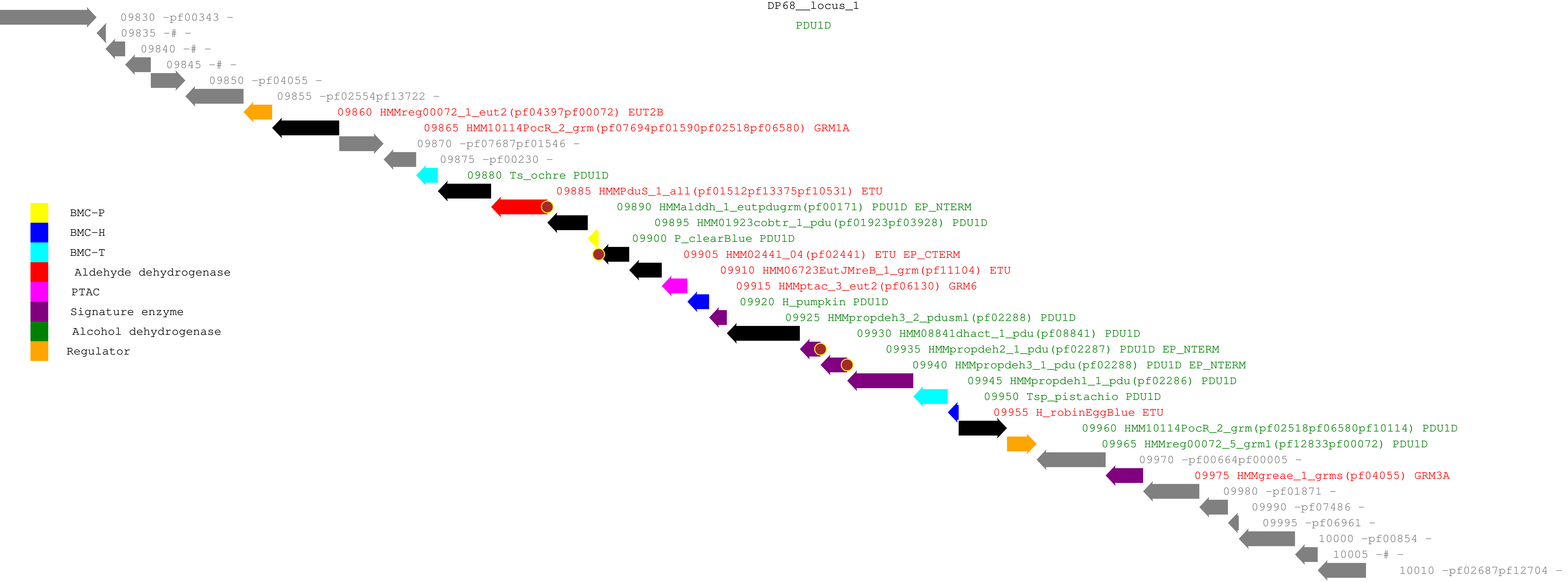

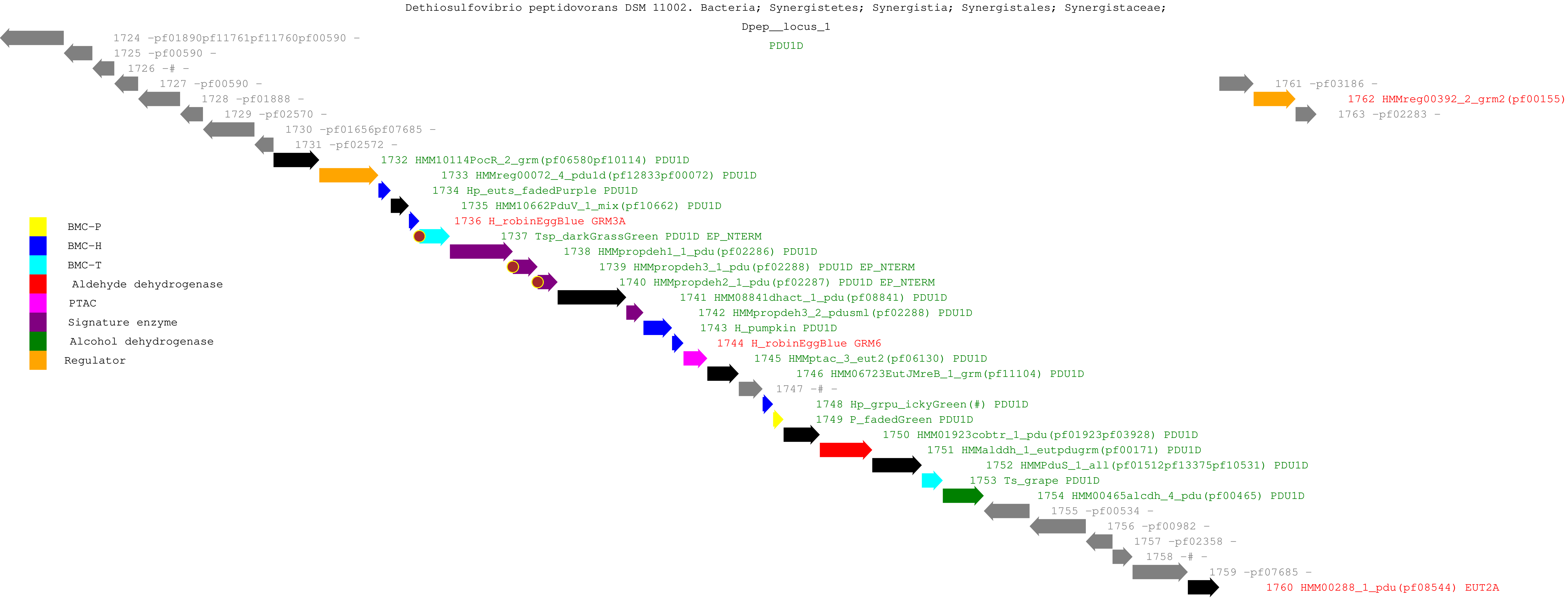
Dethiosulfovibrio peptidovorans DSM 11002. Bacteria; Synergistetes; Synergistia; Synergistales; Synergistaceae;
Dpep__locus_1
PDU1D
1724 -pf01890pf11761pf11760pf00590 -
1725 -pf00590 -
1726 -# -
1727 -pf00590 -
1728 -pf01888 -
1729 -pf02570 -
1730 -pf01656pf07685 -
1731 -pf02572 -
1732 HMM10114PocR_2_grm(pf06580pf10114) PDU1D
1733 HMMreg00072_4_pdu1d(pf12833pf00072) PDU1D
1734 Hp_euts_fadedPurple PDU1D
1735 HMM10662PduV_1_mix(pf10662) PDU1D
1736 H_robinEggBlue GRM3A
1737 Tsp_darkGrassGreen PDU1D EP_NTERM
1738 HMMpropdeh1_1_pdu(pf02286) PDU1D
1739 HMMpropdeh3_1_pdu(pf02288) PDU1D EP_NTERM
1740 HMMpropdeh2_1_pdu(pf02287) PDU1D EP_NTERM
1741 HMM08841dhact_1_pdu(pf08841) PDU1D
1742 HMMpropdeh3_2_pdusml(pf02288) PDU1D
1743 H_pumpkin PDU1D
1744 H_robinEggBlue GRM6
1745 HMMptac_3_eut2(pf06130) PDU1D
1746 HMM06723EutJMreB_1_grm(pf11104) PDU1D
1747 -# -
1748 Hp_grpu_ickyGreen(#) PDU1D
1749 P_fadedGreen PDU1D
1750 HMM01923cobtr_1_pdu(pf01923pf03928) PDU1D
1751 HMMalddh_1_eutpdugrm(pf00171) PDU1D
1752 HMMPduS_1_all(pf01512pf13375pf10531) PDU1D
1753 Ts_grape PDU1D
1754 HMM00465alcdh_4_pdu(pf00465) PDU1D
1755 -pf00534 -
1756 -pf00982 -
1757 -pf02358 -
1758 -# -
1759 -pf07685 -
1760 HMM00288_1_pdu(pf08544) EUT2A
1761 -pf03186 -
1762 HMMreg00392_2_grm2(pf00155)
1763 -pf02283 -
BMC-P
BMC-H
BMC-T
Aldehyde dehydrogenase
PTAC
Signature enzyme
Alcohol dehydrogenase
Regulator

Eubacteriaceae bacterium. Bacteria; Firmicutes; Clostridia; Clostridiales; Eubacteriaceae.

DUD26__locus_1_main_sat

PDU1D

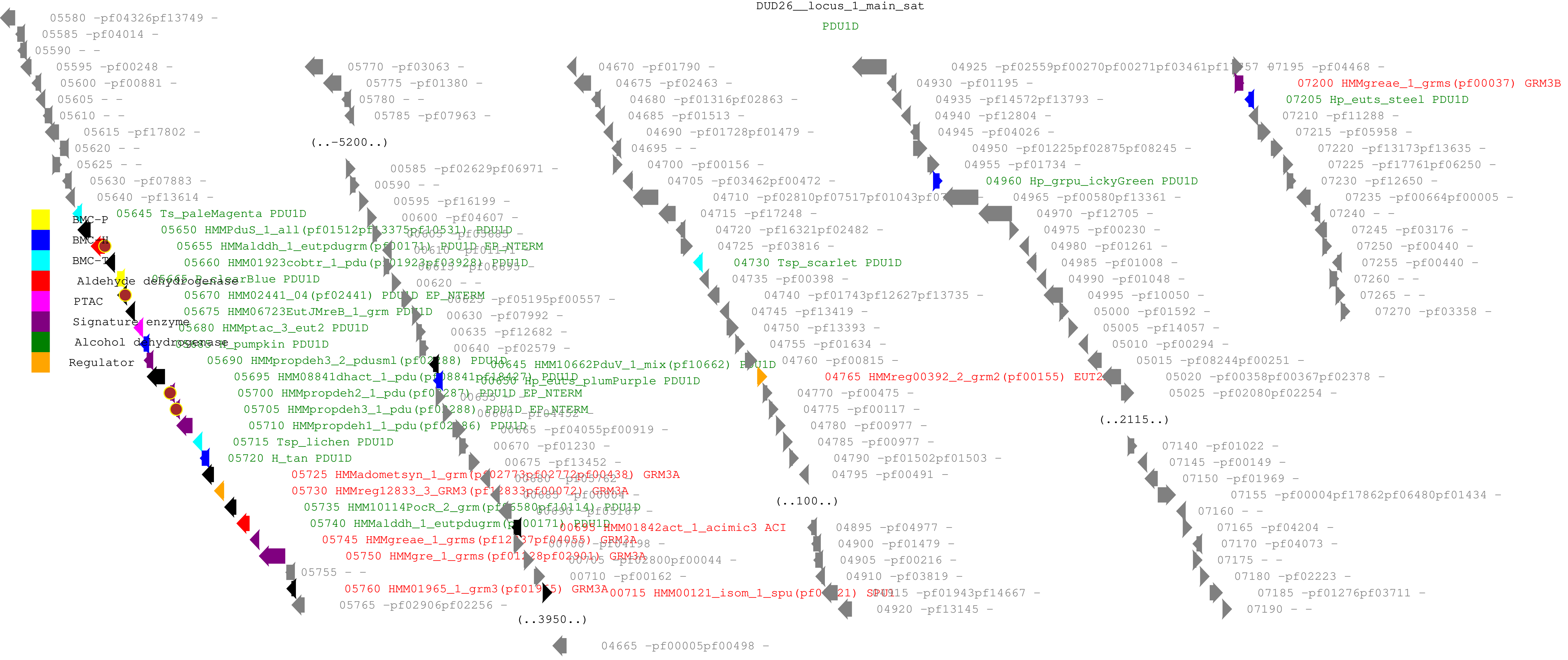

Anaerobutyricum hallii. Bacteria; Firmicutes; Clostridia; Clostridiales; Lachnospiraceae;

DW068__locus_1_main_sat

PDU1D

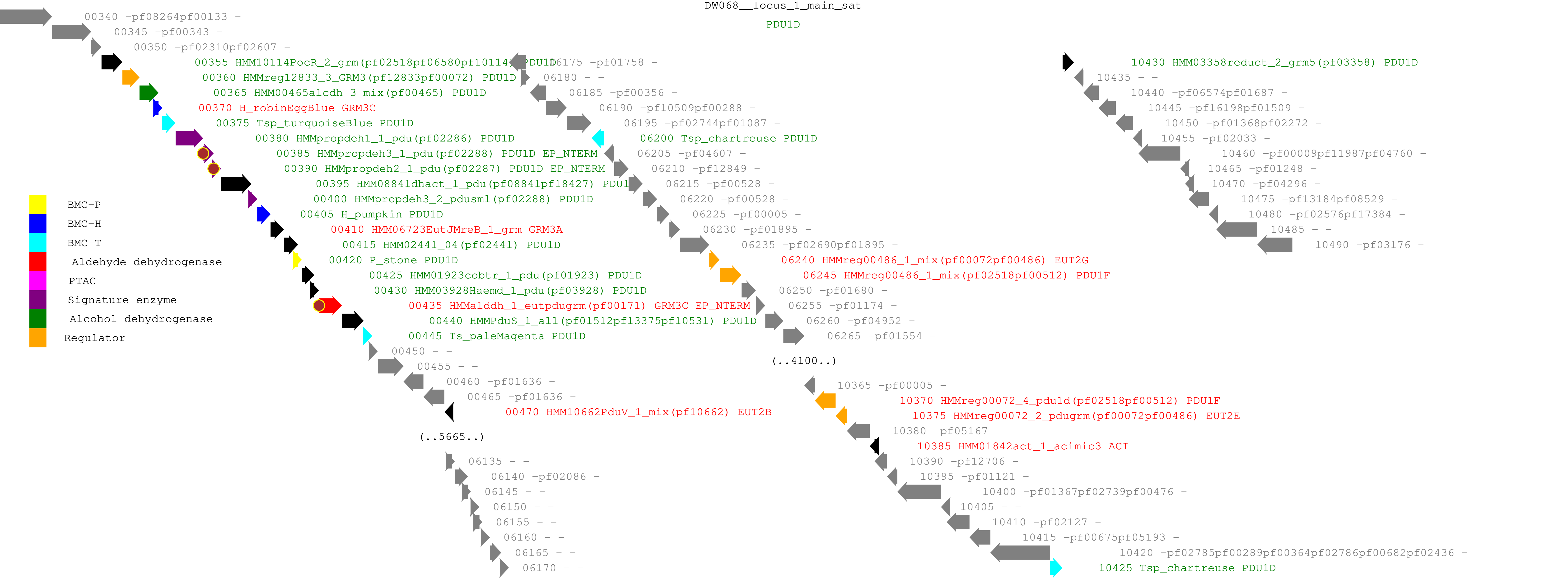

Eubacterium sp. AM05-23. Bacteria; Firmicutes; Clostridia; Clostridiales; Eubacteriaceae;

DW091__locus_1_main_sat

PDU1D

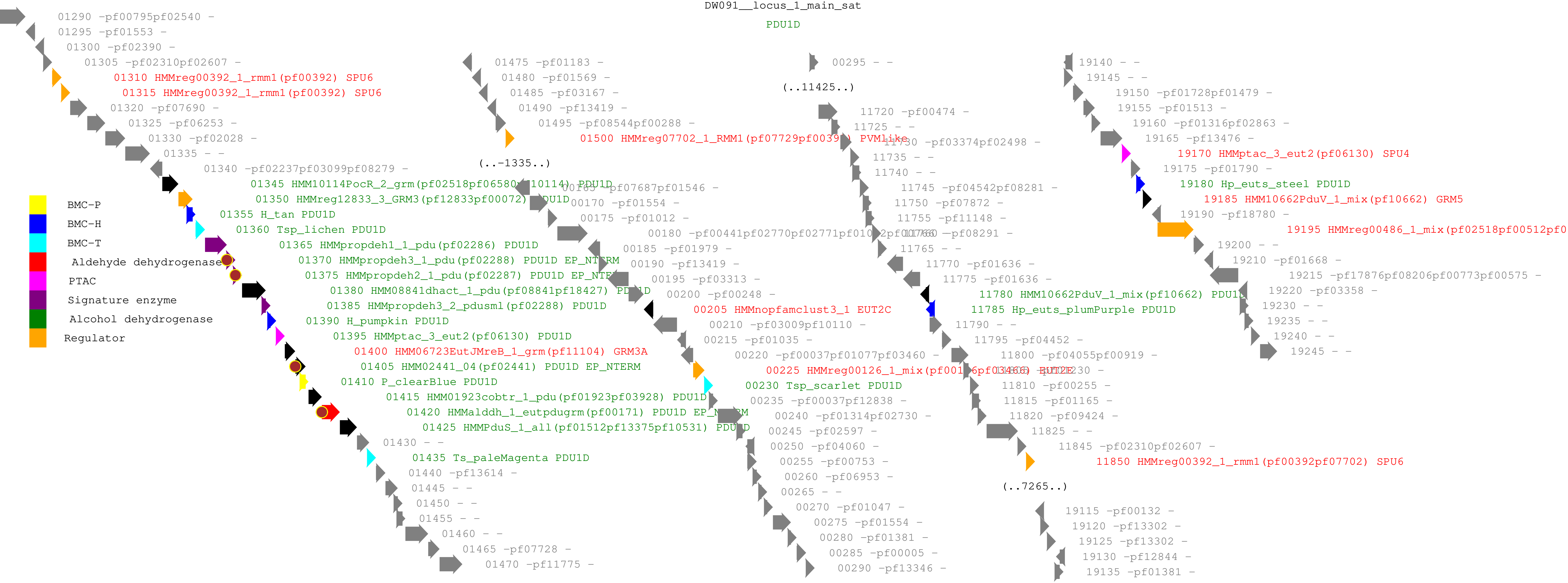

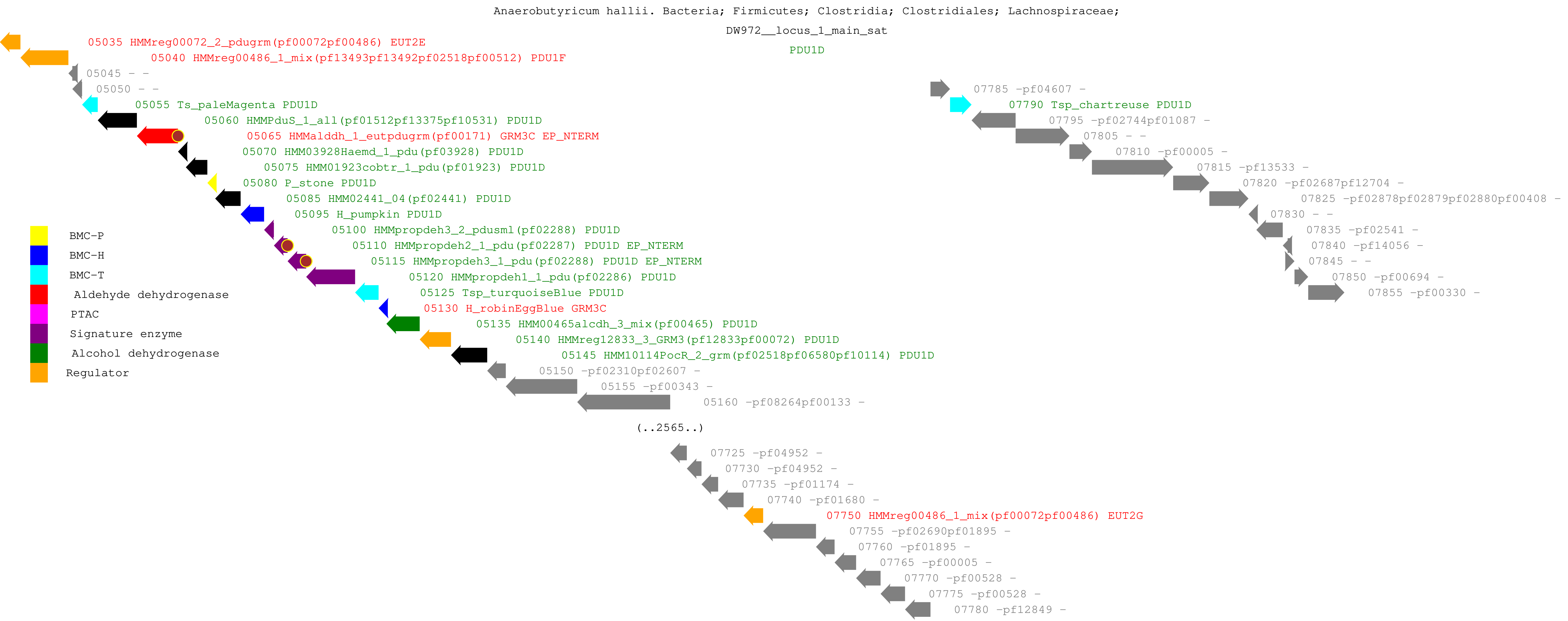
Anaerobutyricum hallii. Bacteria; Firmicutes; Clostridia; Clostridiales; Lachnospiraceae;
DW972__locus_1_main_sat
PDU1D
05035 HMMreg00072_2_pdugrm(pf00072pf00486) EUT2E
05040 HMMreg00486_1_mix(pf13493pf13492pf02518pf00512) PDU1F
05045 - -
05050 - -
05055 Ts_paleMagenta PDU1D
05060 HMMPduS_1_all(pf01512pf13375pf10531) PDU1D
05065 HMMalddh_1_eutpdugrm(pf00171) GRM3C EP_NTERM
05070 HMM03928Haemd_1_pdu(pf03928) PDU1D
05075 HMM01923cobtr_1_pdu(pf01923) PDU1D
05080 P_stone PDU1D
05085 HMM02441_04(pf02441) PDU1D
05095 H_pumpkin PDU1D
05100 HMMpropdeh3_2_pdusml(pf02288) PDU1D
05110 HMMpropdeh2_1_pdu(pf02287) PDU1D EP_NTERM
05115 HMMpropdeh3_1_pdu(pf02288) PDU1D EP_NTERM
05120 HMMpropdeh1_1_pdu(pf02286) PDU1D
05125 Tsp_turquoiseBlue PDU1D
05130 H_robinEggBlue GRM3C
05135 HMM00465alcdh_3_mix(pf00465) PDU1D
05140 HMMreg12833_3_GRM3(pf12833pf00072) PDU1D
05145 HMM10114PocR_2_grm(pf02518pf06580pf10114) PDU1D
05150 -pf02310pf02607 -
05155 -pf00343 -
05160 -pf08264pf00133 -
(..2565..)
07725 -pf04952 -
07730 -pf04952 -
07735 -pf01174 -
07740 -pf01680 -
07750 HMMreg00486_1_mix(pf00072pf00486) EUT2G
07755 -pf02690pf01895 -
07760 -pf01895 -
07765 -pf00005 -
07770 -pf00528 -
07775 -pf00528 -
07780 -pf12849 -
07785 -pf04607 -
07790 Tsp_chartreuse PDU1D
07795 -pf02744pf01087 -
07805 - -
07810 -pf00005 -
07815 -pf13533 -
07820 -pf02687pf12704 -
07825 -pf02878pf02879pf02880pf00408 -
07830 - -
07835 -pf02541 -
07840 -pf14056 -
07845 - -
07850 -pf00694 -
07855 -pf00330 -
BMC-P
BMC-H
BMC-T
Aldehyde dehydrogenase
PTAC
Signature enzyme
Alcohol dehydrogenase
Regulator

Anaerobutyricum hallii. Bacteria; Firmicutes; Clostridia; Clostridiales; Lachnospiraceae;

DWZ29__locus_1_main_sat

PDU1D

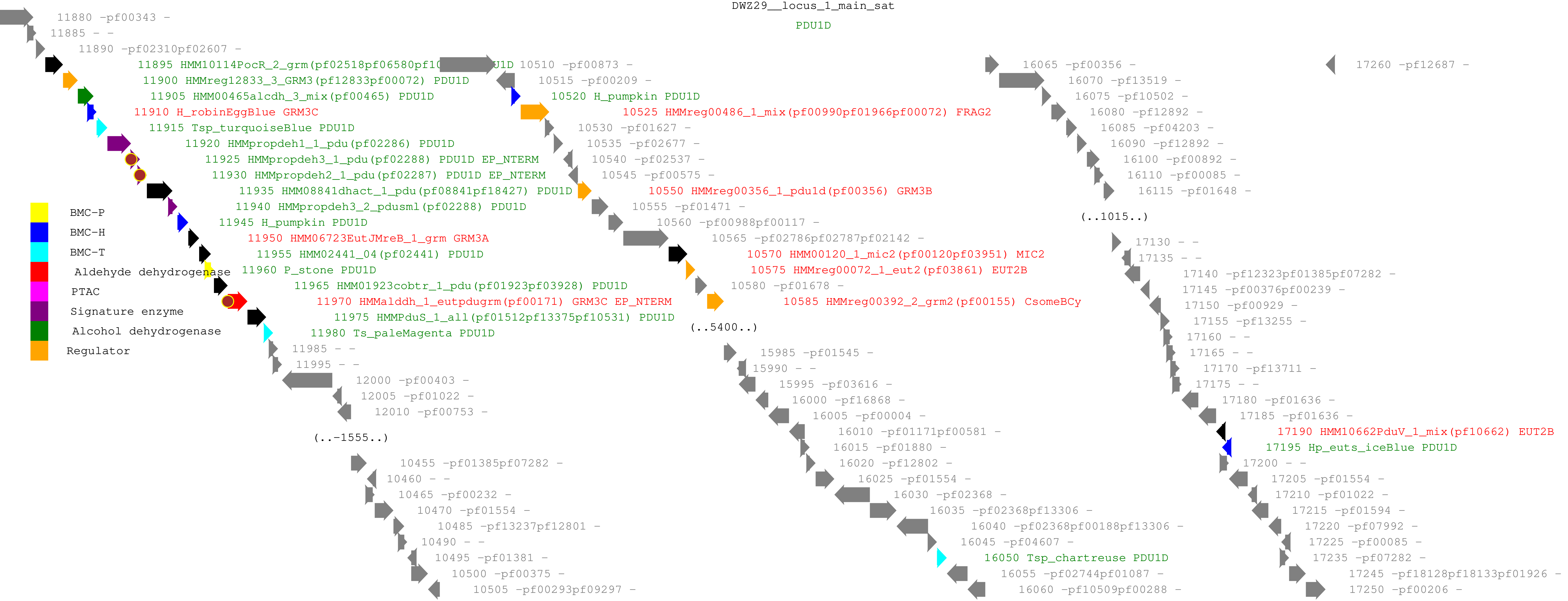

Desulfosporosinus fructosivorans. Bacteria; Firmicutes; Clostridia; Clostridiales; Peptococcaceae;

E4K67__locus_2

PDU1D

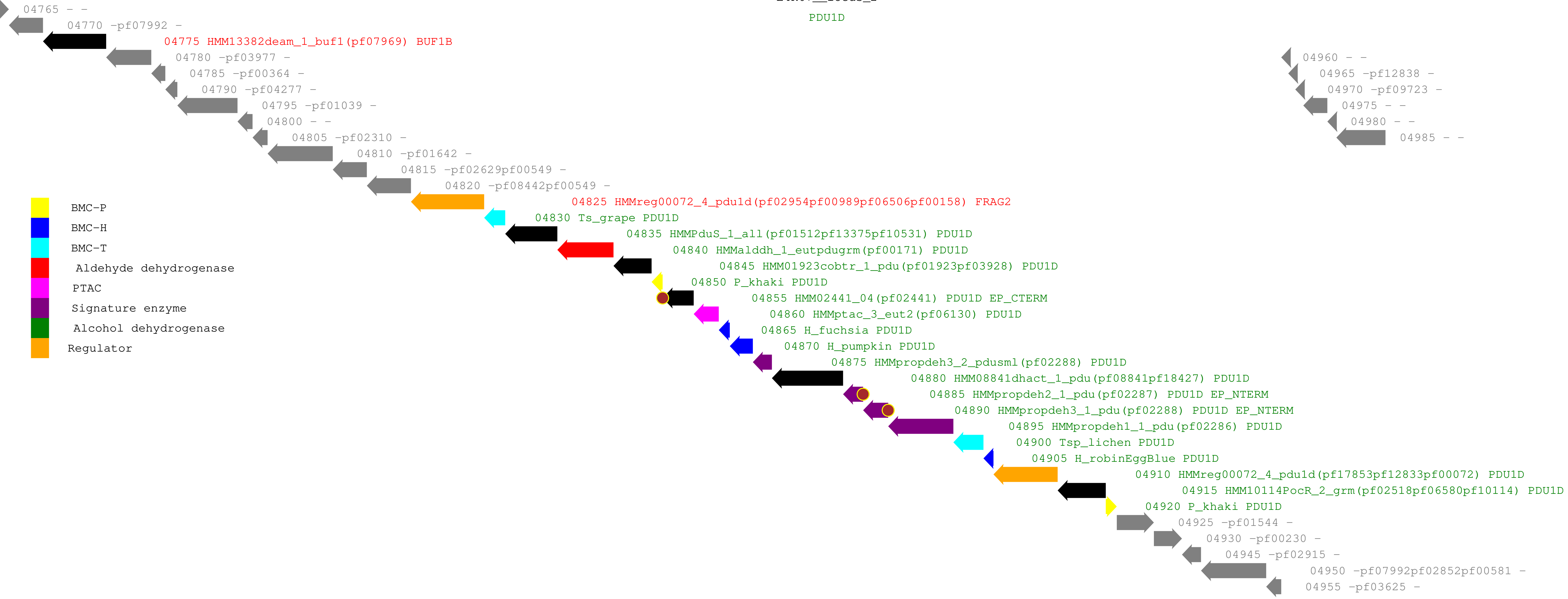

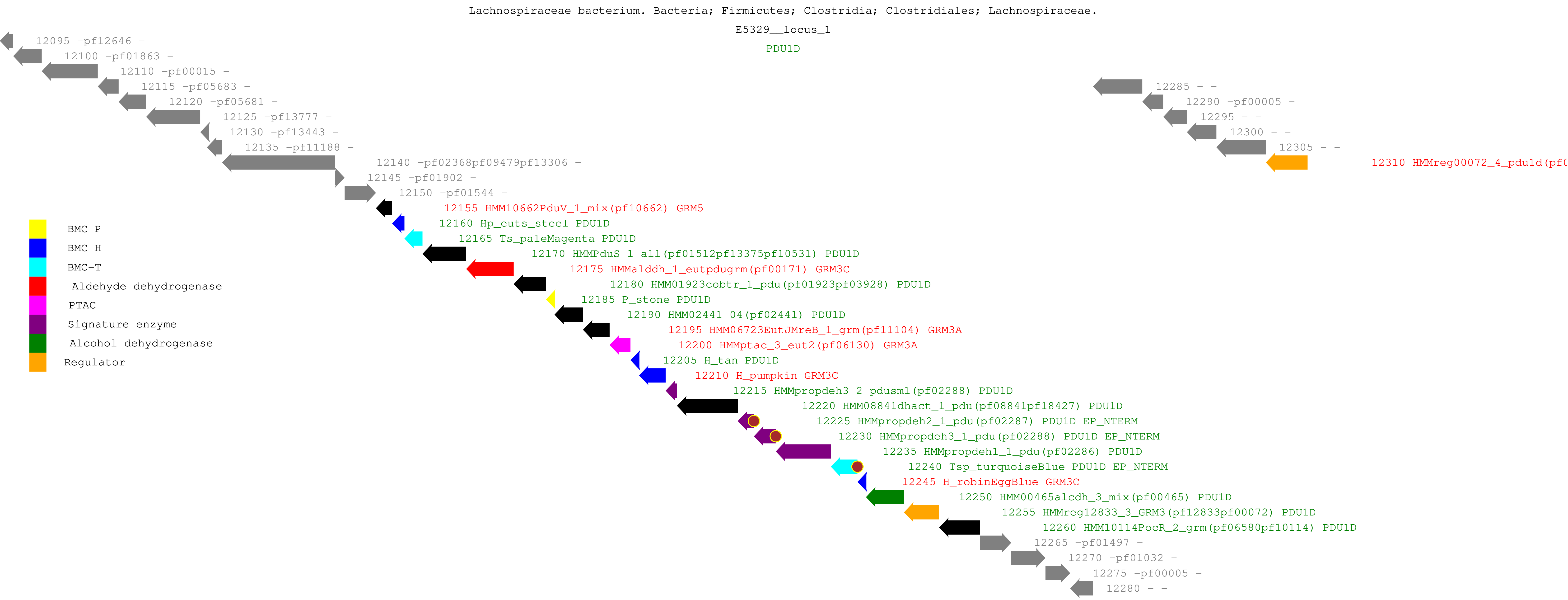
Lachnospiraceae bacterium. Bacteria; Firmicutes; Clostridia; Clostridiales; Lachnospiraceae.
E5329__locus_1
PDU1D
12095 -pf12646 -
12100 -pf01863 -
12110 -pf00015 -
12115 -pf05683 -
12120 -pf05681 -
12125 -pf13777 -
12130 -pf13443 -
12135 -pf11188 -
12140 -pf02368pf09479pf13306 -
12145 -pf01902 -
12150 -pf01544 -
12155 HMM10662PduV_1_mix(pf10662) GRM5
12160 Hp_euts_steel PDU1D
12165 Ts_paleMagenta PDU1D
12170 HMMPduS_1_all(pf01512pf13375pf10531) PDU1D
12175 HMMaldh_1_eutpdugrm(pf00171) GRM3C
12180 HMM01923cobtr_1_pdu(pf01923pf03928) PDU1D
12185 P_stone PDU1D
12190 HMM02441_04(pf02441) PDU1D
12195 HMM06723EutJMreB_1_grm(pf11104) GRM3A
12200 HMMptac_3_eut2(pf06130) GRM3A
12205 H_tan PDU1D
12210 H_pumpkin GRM3C
12215 HMMpropdeh3_2_pdusml(pf02288) PDU1D
12220 HMM08841dhact_1_pdu(pf08841pf18427) PDU1D
12225 HMMpropdeh2_1_pdu(pf02287) PDU1D EP_NTERM
12230 HMMpropdeh3_1_pdu(pf02288) PDU1D EP_NTERM
12235 HMMpropdeh1_1_pdu(pf02286) PDU1D
12240 Tsp_turquoiseBlue PDU1D EP_NTERM
12245 H_robinEggBlue GRM3C
12250 HMM00465alcdh_3_mix(pf00465) PDU1D
12255 HMMreg12833_3_GRM3(pf12833pf00072) PDU1D
12260 HMM10114PocR_2_grm(pf06580pf10114) PDU1D
12265 -pf01497 -
12270 -pf01032 -
12275 -pf00005 -
12280 - -
12285 - -
12290 -pf00005 -
12295 - -
12300 - -
12305 - -
12310 HMMreg00072_4_pdu1d(pf0
BMC-P
BMC-H
BMC-T
Aldehyde dehydrogenase
PTAC
Signature enzyme
Alcohol dehydrogenase
Regulator

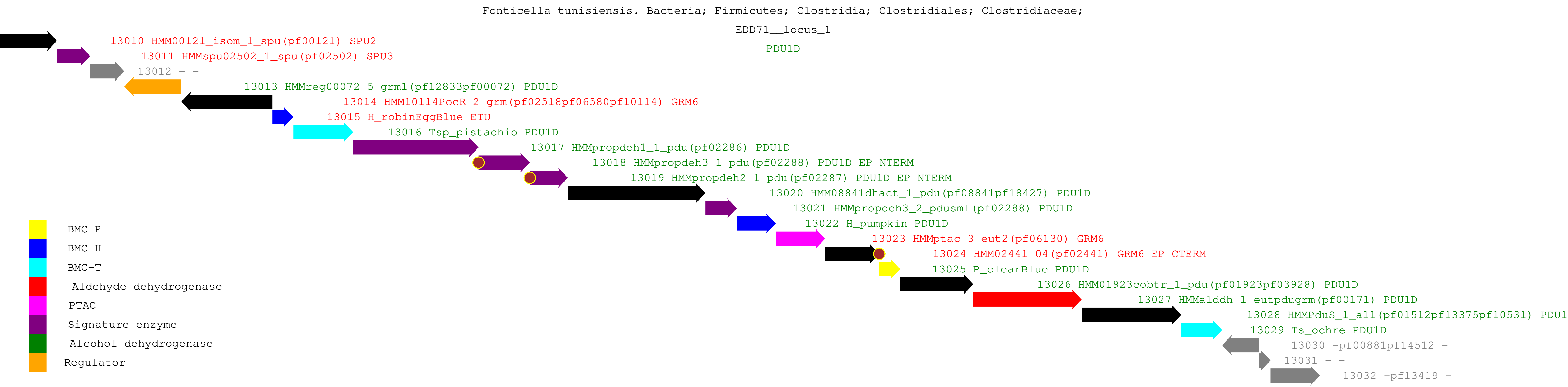
Fonticella tunisiensis. Bacteria; Firmicutes; Clostridia; Clostridiales; Clostridiaceae;
EDD71__locus_1
PDU1D
13010 HMM00121_isom_1_spu(pf00121) SPU2
13011 HMMspu02502_1_spu(pf02502) SPU3
13012 - -
13013 HMMreg00072_5_grm1(pf12833pf00072) PDU1D
13014 HMM10114PocR_2_grm(pf02518pf06580pf10114) GRM6
13015 H_robinEggBlue ETU
13016 Tsp_pistachio PDU1D
13017 HMMpropdeh1_1_pdu(pf02286) PDU1D
13018 HMMpropdeh3_1_pdu(pf02288) PDU1D EP_NTERM
13019 HMMpropdeh2_1_pdu(pf02287) PDU1D EP_NTERM
13020 HMM08841dhact_1_pdu(pf08841pf18427) PDU1D
13021 HMMpropdeh3_2_pdusml(pf02288) PDU1D
13022 H_pumpkin PDU1D
13023 HMMptac_3_eut2(pf06130) GRM6
13024 HMM02441_04(pf02441) GRM6 EP_CTERM
13025 P_clearBlue PDU1D
13026 HMM01923cobtr_1_pdu(pf01923pf03928) PDU1D
13027 HMMalddh_1_eutpdugrm(pf00171) PDU1D
13028 HMMPduS_1_all(pf01512pf13375pf10531) PDU1
13029 Ts_ochre PDU1D
13030 -pf00881pf14512 -
13031 - -
13032 -pf13419 -
BMC-P
BMC-H
BMC-T
Aldehyde dehydrogenase
PTAC
Signature enzyme
Alcohol dehydrogenase
Regulator

Propionibacterium acidifaciens. Bacteria; Actinobacteria; Propionibacteriales; Propionibacteriaceae;

EGX94__locus_1

PDU1D

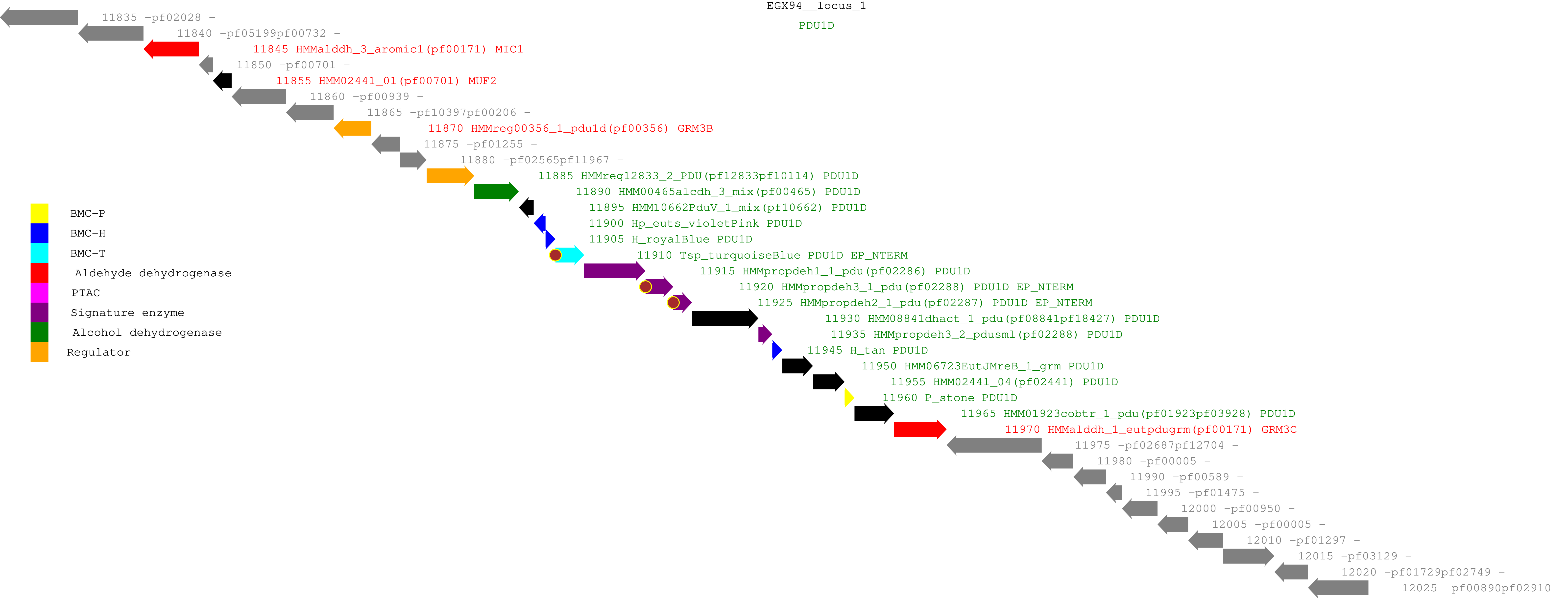

Eubacterium hallii. Bacteria; Firmicutes; Clostridia; Clostridiales; Eubacteriaceae;

EHLA__locus_1_main_sat

PDU1D

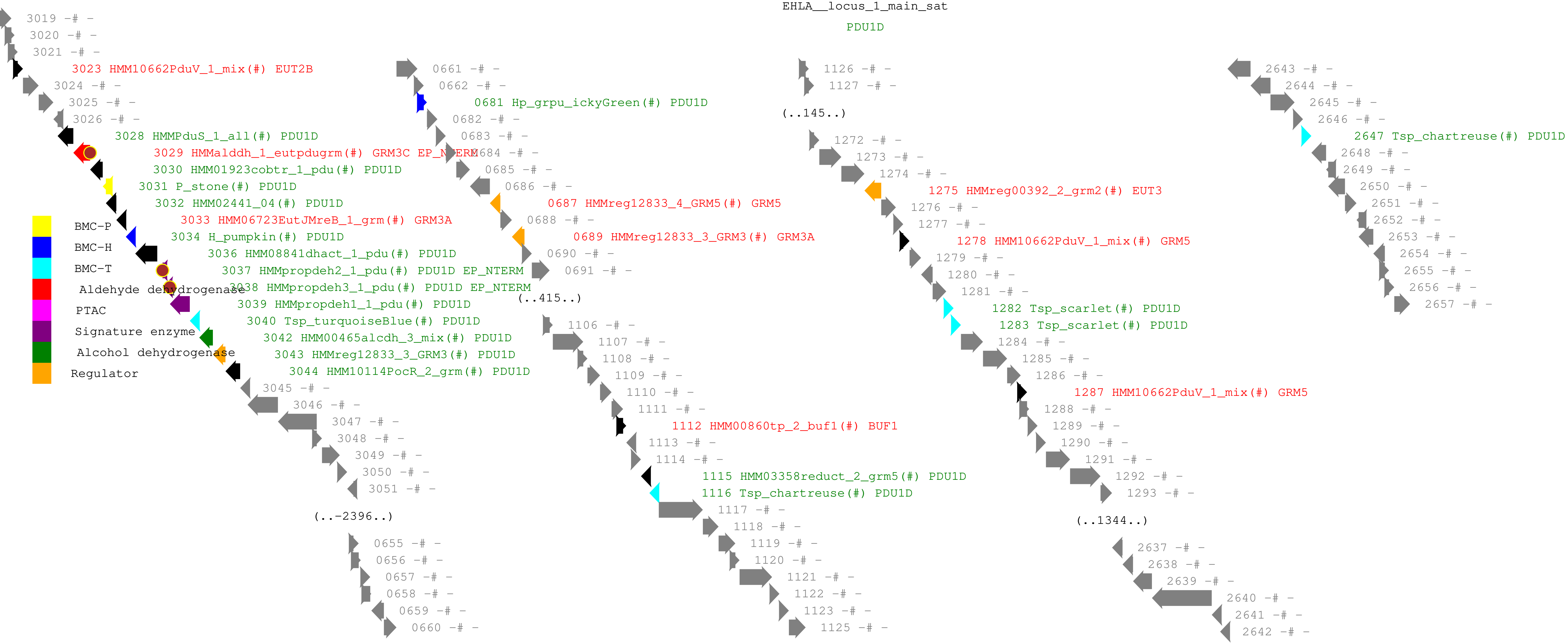

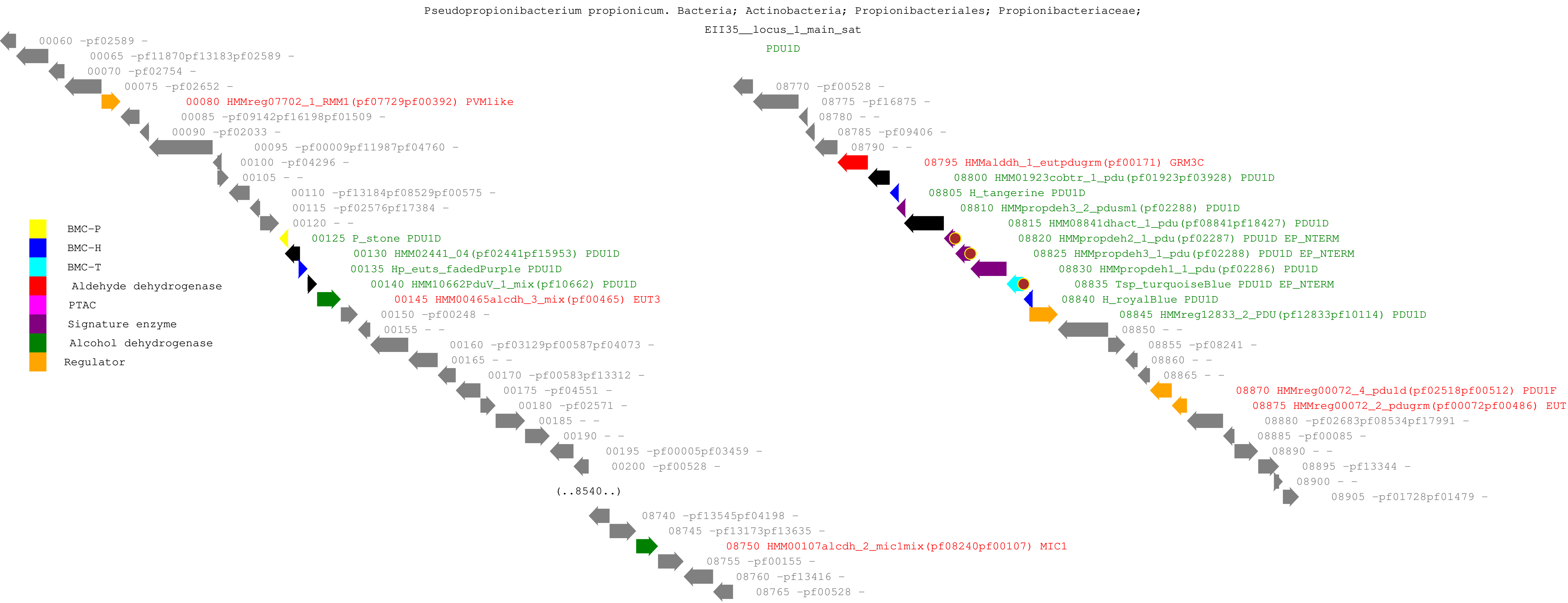
Pseudopropionibacterium propionicum. Bacteria; Actinobacteria; Propionibacteriales; Propionibacteriaceae;
EII35__locus_1_main_sat
PDU1D
00060 -pf02589 -
00065 -pf11870pf13183pf02589 -
00070 -pf02754 -
00075 -pf02652 -
00080 HMMreg07702_1_RMM1(pf07729pf00392) PVMlike
00085 -pf09142pf16198pf01509 -
00090 -pf02033 -
00095 -pf00009pf11987pf04760 -
00100 -pf04296 -
00105 - -
00110 -pf13184pf08529pf00575 -
00115 -pf02576pf17384 -
00120 - -
00125 P_stone PDU1D
00130 HMM02441_04(pf02441pf15953) PDU1D
00135 Hp_euts_fadedPurple PDU1D
00140 HMM10662PduV_1_mix(pf10662) PDU1D
00145 HMM00465alcdh_3_mix(pf00465) EUT3
00150 -pf00248 -
00155 - -
00160 -pf03129pf00587pf04073 -
00165 - -
00170 -pf00583pf13312 -
00175 -pf04551 -
00180 -pf02571 -
00185 - -
00190 - -
00195 -pf00005pf03459 -
00200 -pf00528 -
(..8540..)
08740 -pf13545pf04198 -
08745 -pf13173pf13635 -
08750 HMM00107alcdh_2_mic1mix(pf08240pf00107) MIC1
08755 -pf00155 -
08760 -pf13416 -
08765 -pf00528 -
08770 -pf00528 -
08775 -pf16875 -
08780 - -
08785 -pf09406 -
08790 - -
08795 HMMaldh_1_eutpdugrm(pf00171) GRM3C
08800 HMM01923cobtr_1_pdu(pf01923pf03928) PDU1D
08805 H_tangerine PDU1D
08810 HMMpropdeh3_2_pdusml(pf02288) PDU1D
08815 HMM08841dhact_1_pdu(pf08841pf18427) PDU1D
08820 HMMpropdeh2_1_pdu(pf02287) PDU1D EP_NTERM
08825 HMMpropdeh3_1_pdu(pf02288) PDU1D EP_NTERM
08830 HMMpropdeh1_1_pdu(pf02286) PDU1D
08835 Tsp_turquoiseBlue PDU1D EP_NTERM
08840 H_royalBlue PDU1D
08845 HMMreg12833_2_PDU(pf12833pf10114) PDU1D
08850 - -
08855 -pf08241 -
08860 - -
08865 - -
08870 HMMreg00072_4_pdu1d(pf02518pf00512) PDU1F
08875 HMMreg00072_2_pdugrm(pf00072pf00486) EUT
08880 -pf02683pf08534pf17991 -
08885 -pf00085 -
08890 - -
08895 -pf13344 -
08900 - -
08905 -pf01728pf01479 -
BMC-P
BMC-H
BMC-T
Aldehyde dehydrogenase
PTAC
Signature enzyme
Alcohol dehydrogenase
Regulator

Tessaracoccus sp. OH4464_COT-324. Bacteria; Actinobacteria; Propionibacteriales; Propionibacteriaceae;

EII42__locus_1

PDU1D

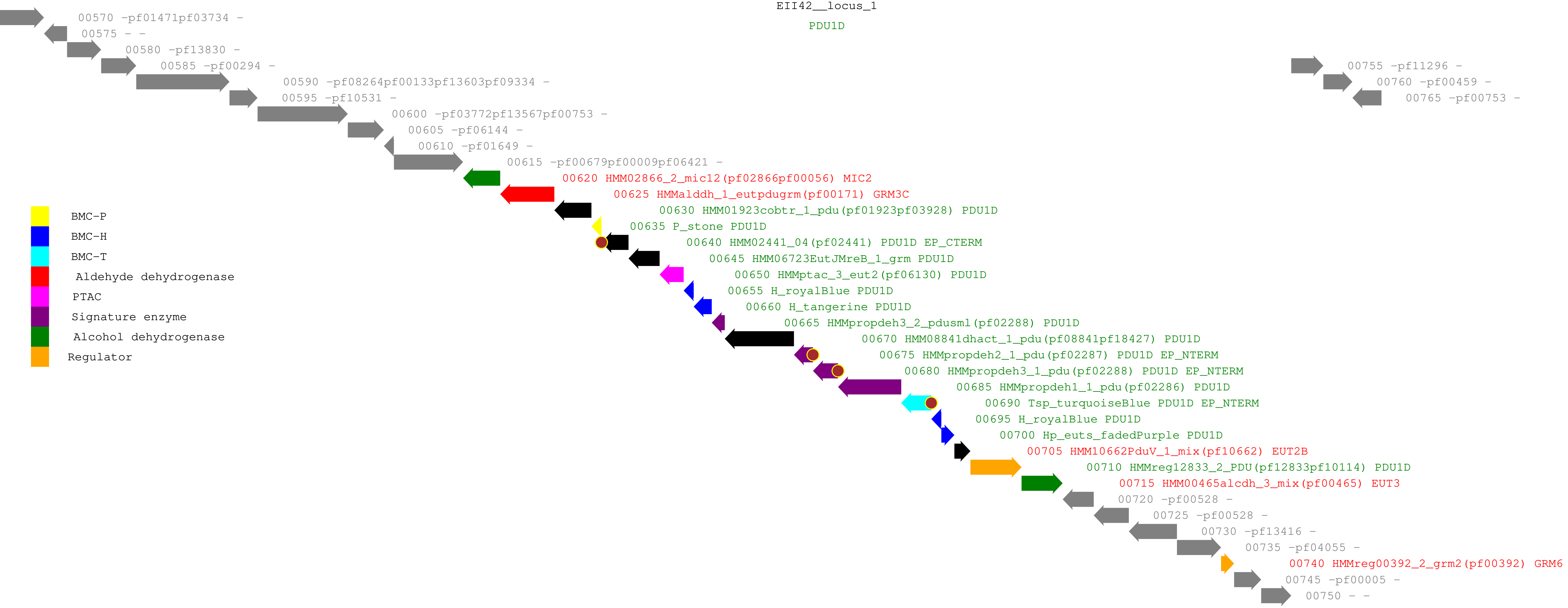

Eubacterium limosum (strain KIST612). Bacteria; Firmicutes; Clostridia; Clostridiales; Eubacteriaceae;

ELI__locus_1_main_sat

PDU1D

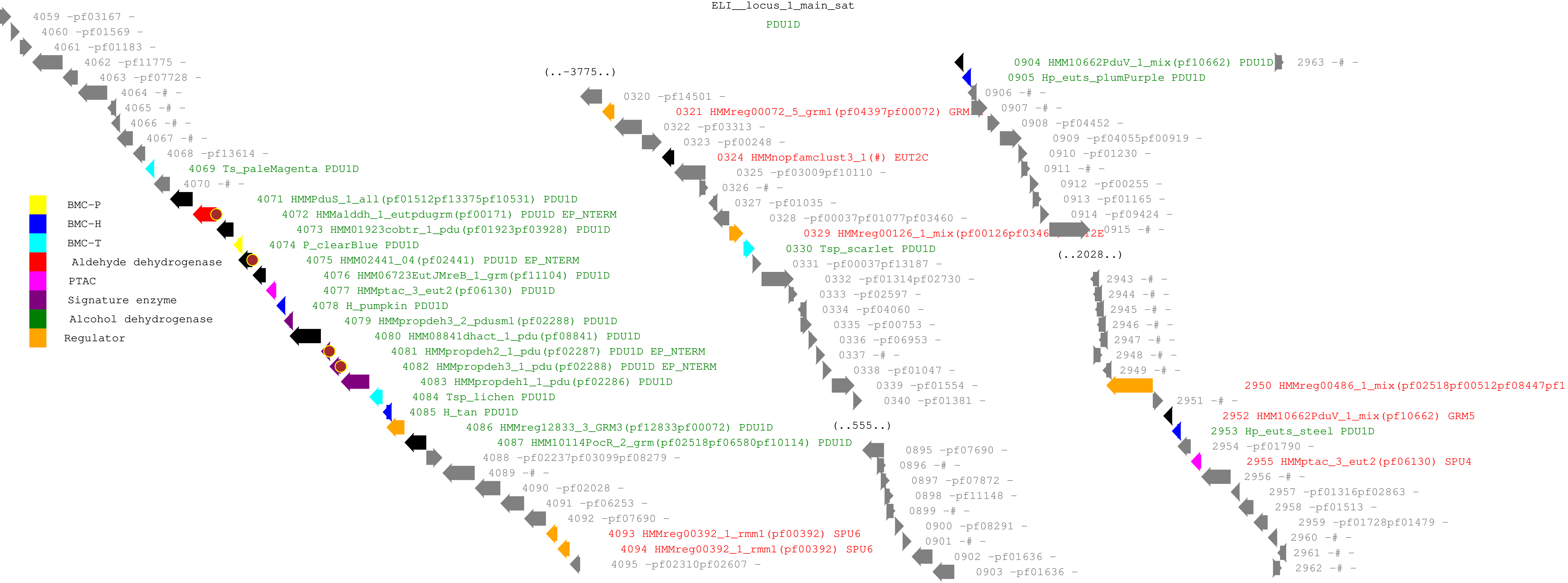

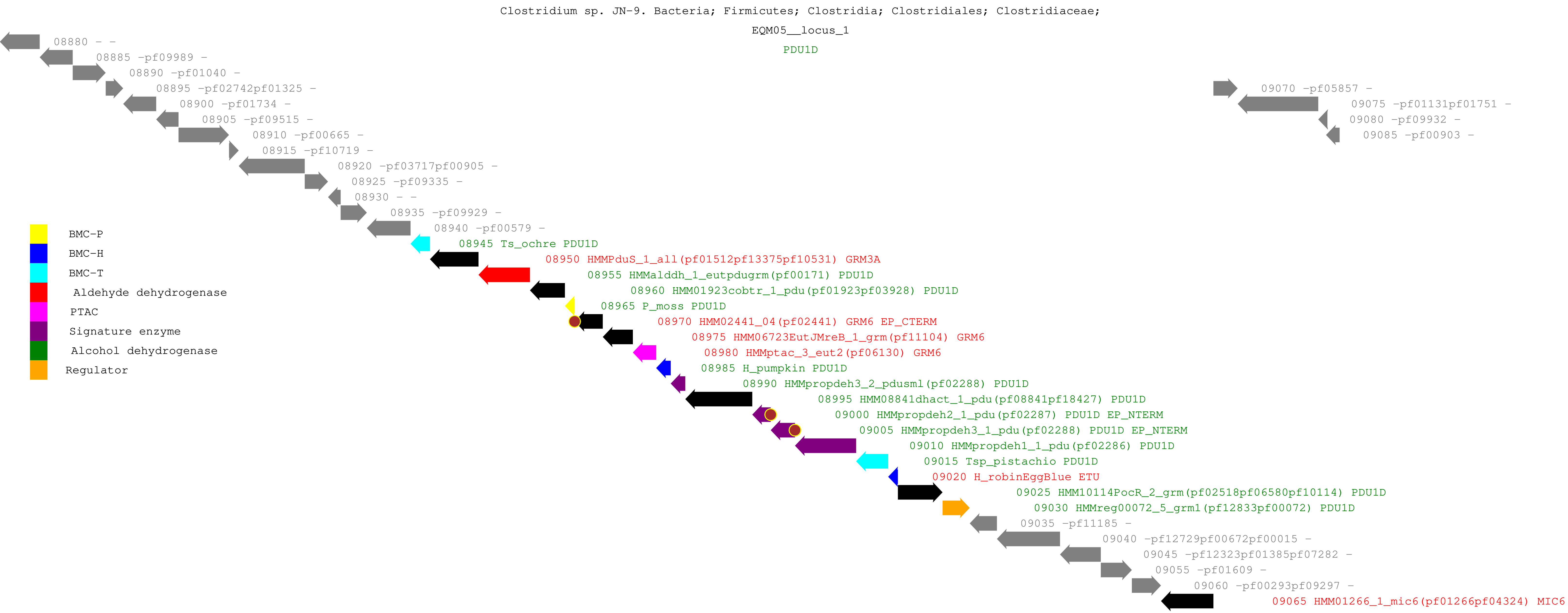
Clostridium sp. JN-9. Bacteria; Firmicutes; Clostridia; Clostridiales; Clostridiaceae;
EQM05__locus_1
PDU1D
08880 - -
08885 -pf09989 -
08890 -pf01040 -
08895 -pf02742pf01325 -
08900 -pf01734 -
08905 -pf09515 -
08910 -pf00665 -
08915 -pf10719 -
08920 -pf03717pf00905 -
08925 -pf09335 -
08930 - -
08935 -pf09929 -
08940 -pf00579 -
08945 Ts_ochre PDU1D
08950 HMMPduS_1_all(pf01512pf13375pf10531) GRM3A
08955 HMMalddh_1_eutpdugrm(pf00171) PDU1D
08960 HMM01923cobtr_1_pdu(pf01923pf03928) PDU1D
08965 P_moss PDU1D
08970 HMM02441_04(pf02441) GRM6 EP_CTERM
08975 HMM06723EutJMreB_1_grm(pf11104) GRM6
08980 HMMptac_3_eut2(pf06130) GRM6
08985 H_pumpkin PDU1D
08990 HMMpropdeh3_2_pdusml(pf02288) PDU1D
08995 HMM08841dhact_1_pdu(pf08841pf18427) PDU1D
09000 HMMpropdeh2_1_pdu(pf02287) PDU1D EP_NTERM
09005 HMMpropdeh3_1_pdu(pf02288) PDU1D EP_NTERM
09010 HMMpropdeh1_1_pdu(pf02286) PDU1D
09015 Tsp_pistachio PDU1D
09020 H_robinEggBlue ETU
09025 HMM10114PocR_2_grm(pf02518pf06580pf10114) PDU1D
09030 HMMreg00072_5_grm1(pf12833pf00072) PDU1D
09035 -pf11185 -
09040 -pf12729pf00672pf00015 -
09045 -pf12323pf01385pf07282 -
09055 -pf01609 -
09060 -pf00293pf09297 -
09065 HMM01266_1_mic6(pf01266pf04324) MIC6
09070 -pf05857 -
09075 -pf01131pf01751 -
09080 -pf09932 -
09085 -pf00903 -
BMC-P
BMC-H
BMC-T
Aldehyde dehydrogenase
PTAC
Signature enzyme
Alcohol dehydrogenase
Regulator

Eubacterium hallii. Bacteria; Firmicutes; Clostridia; Clostridiales; Eubacteriaceae;

ERS852450__locus_1_main_sat

PDU1D

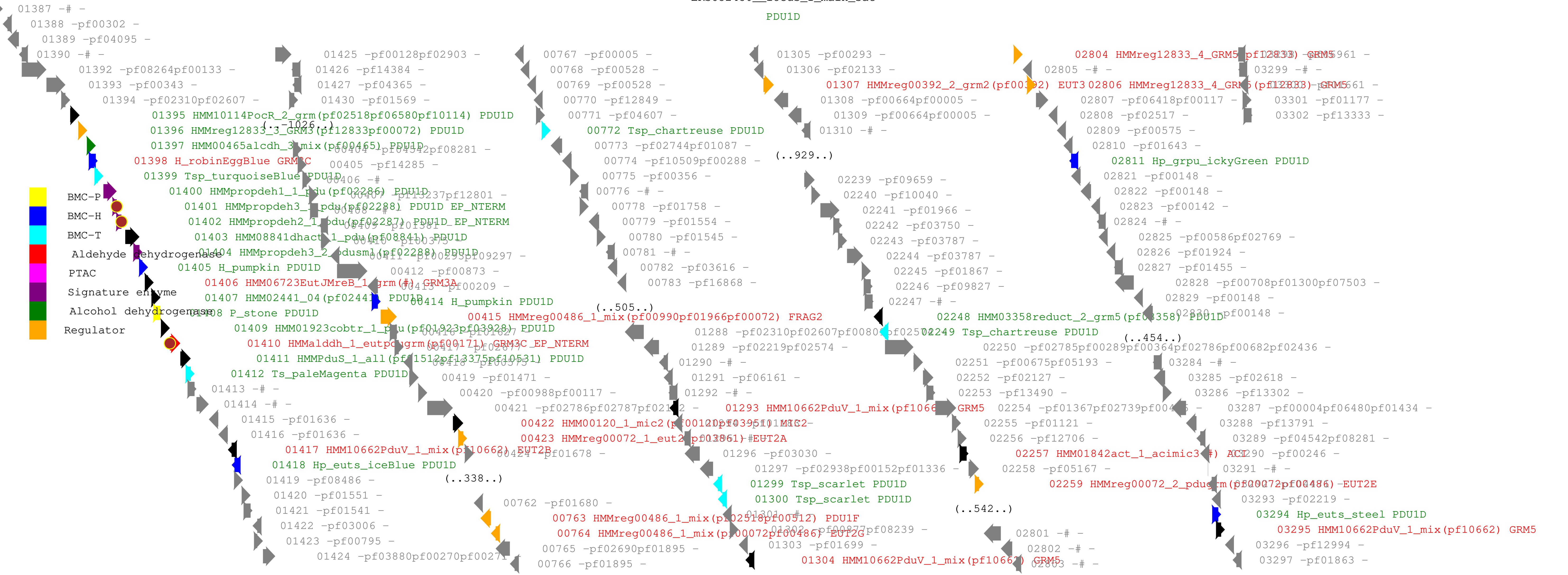

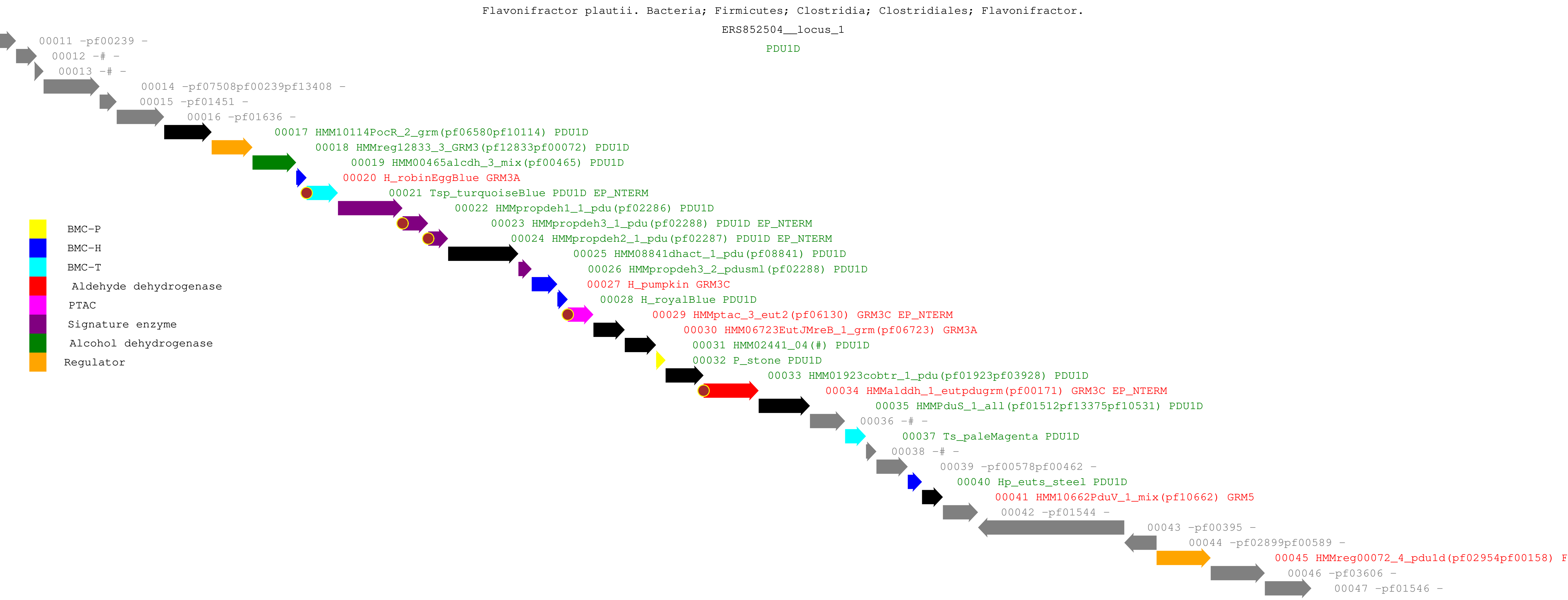
Flavonifractor plautii. Bacteria; Firmicutes; Clostridia; Clostridiales; Flavonifractor.
ERS852504__locus_1
PDU1D
00011 -pf00239 -
00012 -# -
00013 -# -
00014 -pf07508pf00239pf13408 -
00015 -pf01451 -
00016 -pf01636 -
00017 HMM10114PocR_2_grm(pf06580pf10114) PDU1D
00018 HMMreg12833_3_GRM3(pf12833pf00072) PDU1D
00019 HMM00465alcdh_3_mix(pf00465) PDU1D
00020 H_robinEggBlue GRM3A
00021 Tsp_turquoiseBlue PDU1D EP_NTERM
00022 HMMpropdeh1_1_pdu(pf02286) PDU1D
00023 HMMpropdeh3_1_pdu(pf02288) PDU1D EP_NTERM
00024 HMMpropdeh2_1_pdu(pf02287) PDU1D EP_NTERM
00025 HMM08841dhact_1_pdu(pf08841) PDU1D
00026 HMMpropdeh3_2_pdusml(pf02288) PDU1D
00027 H_pumpkin GRM3C
00028 H_royalBlue PDU1D
00029 HMMptac_3_eut2(pf06130) GRM3C EP_NTERM
00030 HMM06723EutJMreB_1_grm(pf06723) GRM3A
00031 HMM02441_04(#) PDU1D
00032 P_stone PDU1D
00033 HMM01923cobtr_1_pdu(pf01923pf03928) PDU1D
00034 HMMalddh_1_eutpdugrm(pf00171) GRM3C EP_NTERM
00035 HMMPduS_1_all(pf01512pf13375pf10531) PDU1D
00036 -# -
00037 Ts_paleMagenta PDU1D
00038 -# -
00039 -pf00578pf00462 -
00040 Hp_euts_steel PDU1D
00041 HMM10662PduV_1_mix(pf10662) GRM5
00042 -pf01544 -
00043 -pf00395 -
00044 -pf02899pf00589 -
00045 HMMreg00072_4_pdu1d(pf02954pf00158) F
00046 -pf03606 -
00047 -pf01546 -
BMC-P
BMC-H
BMC-T
Aldehyde dehydrogenase
PTAC
Signature enzyme
Alcohol dehydrogenase
Regulator

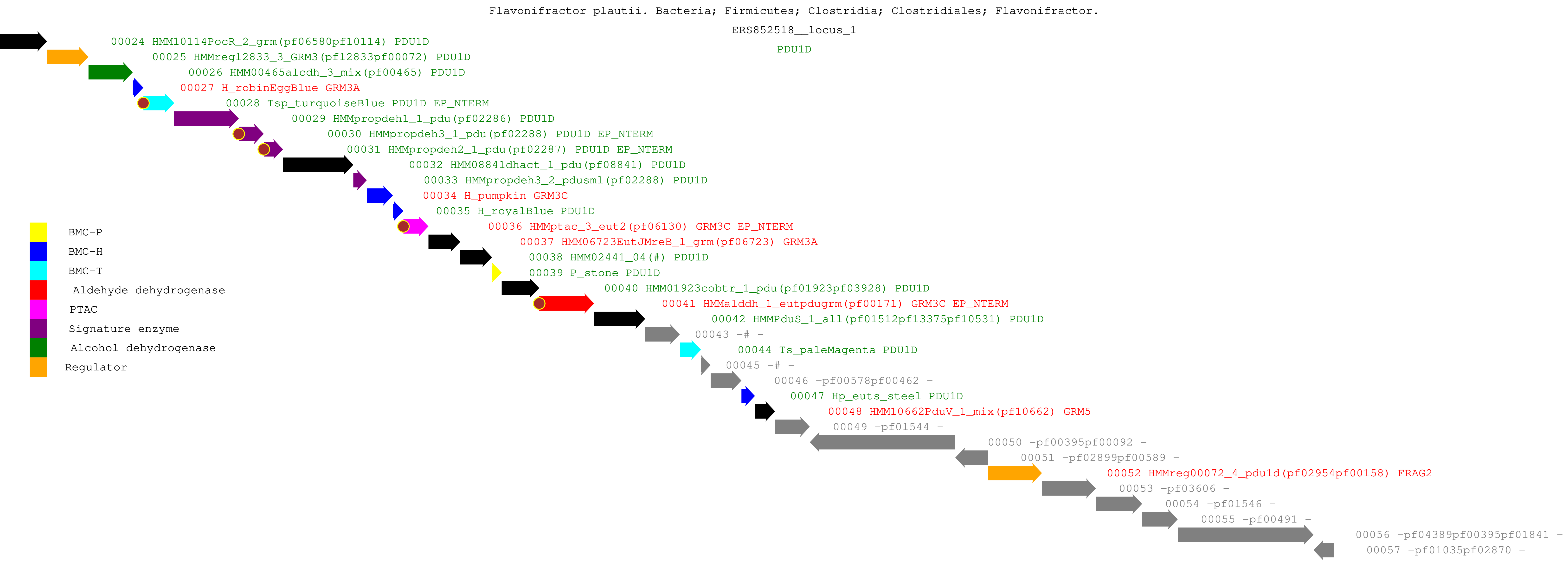
Flavonifractor plautii. Bacteria; Firmicutes; Clostridia; Clostridiales; Flavonifractor.
ERS852518__locus_1
PDU1D
00024 HMM10114PocR_2_grm(pf06580pf10114) PDU1D
00025 HMMreg12833_3_GRM3(pf12833pf00072) PDU1D
00026 HMM00465alcdh_3_mix(pf00465) PDU1D
00027 H_robinEggBlue GRM3A
00028 Tsp_turquoiseBlue PDU1D EP_NTERM
00029 HMMpropdeh1_1_pdu(pf02286) PDU1D
00030 HMMpropdeh3_1_pdu(pf02288) PDU1D EP_NTERM
00031 HMMpropdeh2_1_pdu(pf02287) PDU1D EP_NTERM
00032 HMM08841dhact_1_pdu(pf08841) PDU1D
00033 HMMpropdeh3_2_pdusml(pf02288) PDU1D
00034 H_pumpkin GRM3C
00035 H_royalBlue PDU1D
00036 HMMptac_3_eut2(pf06130) GRM3C EP_NTERM
00037 HMM06723EutJMreB_1_grm(pf06723) GRM3A
00038 HMM02441_04(#) PDU1D
00039 P_stone PDU1D
00040 HMM01923cobtr_1_pdu(pf01923pf03928) PDU1D
00041 HMMalddh_1_eutpdugrm(pf00171) GRM3C EP_NTERM
00042 HMMPduS_1_all(pf01512pf13375pf10531) PDU1D
00043 -# -
00044 Ts_paleMagenta PDU1D
00045 -# -
00046 -pf00578pf00462 -
00047 Hp_euts_steel PDU1D
00048 HMM10662PduV_1_mix(pf10662) GRM5
00049 -pf01544 -
00050 -pf00395pf00092 -
00051 -pf02899pf00589 -
00052 HMMreg00072_4_pdu1d(pf02954pf00158) FRAG2
00053 -pf03606 -
00054 -pf01546 -
00055 -pf00491 -
00056 -pf04389pf00395pf01841 -
00057 -pf01035pf02870 -
BMC-P
BMC-H
BMC-T
Aldehyde dehydrogenase
PTAC
Signature enzyme
Alcohol dehydrogenase
Regulator

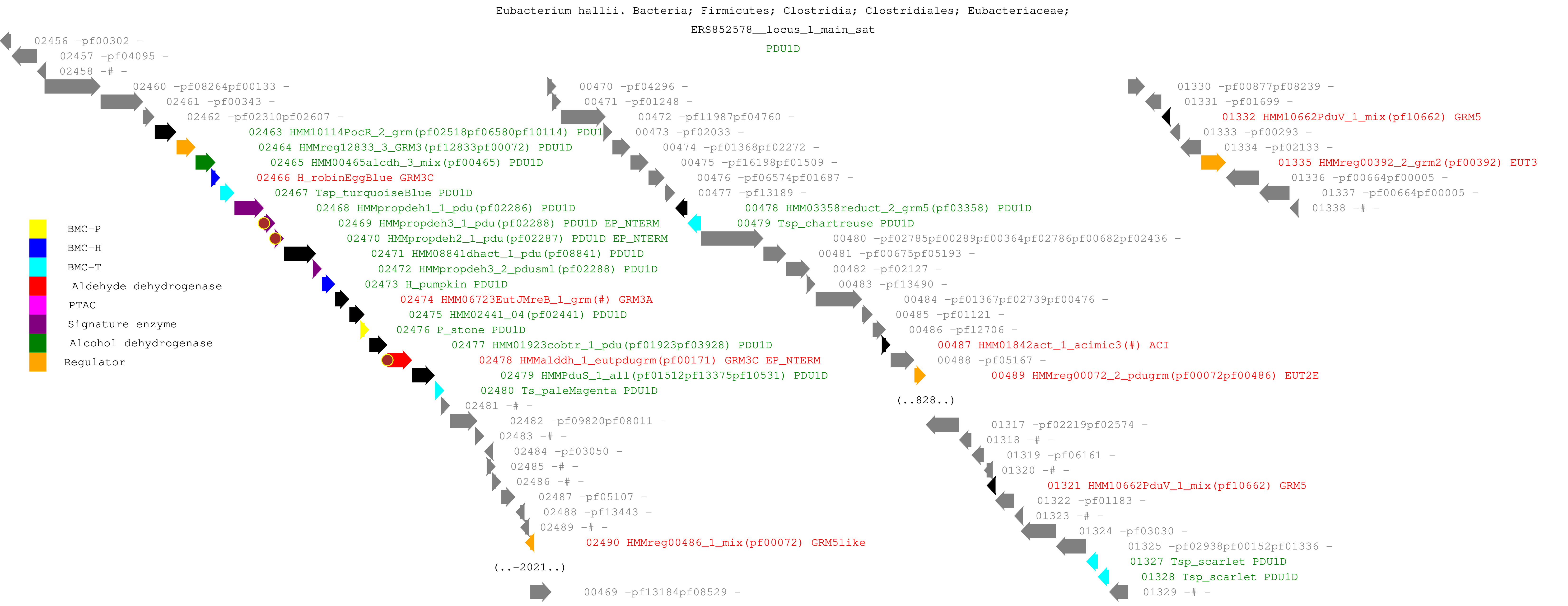
Eubacterium hallii. Bacteria; Firmicutes; Clostridia; Clostridiales; Eubacteriaceae;
ERS852578__locus_1_main_sat
PDU1D
02456 -pf00302 -
02457 -pf04095 -
02458 -# -
02460 -pf08264pf00133 -
02461 -pf00343 -
02462 -pf02310pf02607 -
02463 HMM10114PocR_2_grm(pf02518pf06580pf10114) PDU1
02464 HMMreg12833_3_GRM3(pf12833pf00072) PDU1D
02465 HMM00465alcdh_3_mix(pf00465) PDU1D
02466 H_robinEggBlue GRM3C
02467 Tsp_turquoiseBlue PDU1D
02468 HMMpropdeh1_1_pdu(pf02286) PDU1D
02469 HMMpropdeh3_1_pdu(pf02288) PDU1D EP_NTERM
02470 HMMpropdeh2_1_pdu(pf02287) PDU1D EP_NTERM
02471 HMM08841dhact_1_pdu(pf08841) PDU1D
02472 HMMpropdeh3_2_pdusml(pf02288) PDU1D
02473 H_pumpkin PDU1D
02474 HMM06723EutJMreB_1_grm(#) GRM3A
02475 HMM02441_04(pf02441) PDU1D
02476 P_stone PDU1D
02477 HMM01923cobtr_1_pdu(pf01923pf03928) PDU1D
02478 HMMalddh_1_eutpdugrm(pf00171) GRM3C EP_NTERM
02479 HMMPduS_1_all(pf01512pf13375pf10531) PDU1D
02480 Ts_paleMagenta PDU1D
02481 -# -
02482 -pf09820pf08011 -
02483 -# -
02484 -pf03050 -
02485 -# -
02486 -# -
02487 -pf05107 -
02488 -pf13443 -
02489 -# -
02490 HMMreg00486_1_mix(pf00072) GRM5like
(..-2021..)
00469 -pf13184pf08529 -
00470 -pf04296 -
00471 -pf01248 -
00472 -pf11987pf04760 -
00473 -pf02033 -
00474 -pf01368pf02272 -
00475 -pf16198pf01509 -
00476 -pf06574pf01687 -
00477 -pf13189 -
00478 HMM03358reduct_2_grm5(pf03358) PDU1D
00479 Tsp_chartreuse PDU1D
00480 -pf02785pf00289pf00364pf02786pf00682pf02436 -
00481 -pf00675pf05193 -
00482 -pf02127 -
00483 -pf13490 -
00484 -pf01367pf02739pf00476 -
00485 -pf01121 -
00486 -pf12706 -
00487 HMM01842act_1_acimic3(#) ACI
00488 -pf05167 -
00489 HMMreg00072_2_pdugrm(pf00072pf00486) EUT2E
(..828..)
01317 -pf02219pf02574 -
01318 -# -
01319 -pf06161 -
01320 -# -
01321 HMM10662PduV_1_mix(pf10662) GRM5
01322 -pf01183 -
01323 -# -
01324 -pf03030 -
01325 -pf02938pf00152pf01336 -
01327 Tsp_scarlet PDU1D
01328 Tsp_scarlet PDU1D
01329 -# -
01330 -pf00877pf08239 -
01331 -pf01699 -
01332 HMM10662PduV_1_mix(pf10662) GRM5
01333 -pf00293 -
01334 -pf02133 -
01335 HMMreg00392_2_grm2(pf00392) EUT3
01336 -pf00664pf00005 -
01337 -pf00664pf00005 -
01338 -# -
BMC-P
BMC-H
BMC-T
Aldehyde dehydrogenase
PTAC
Signature enzyme
Alcohol dehydrogenase
Regulator

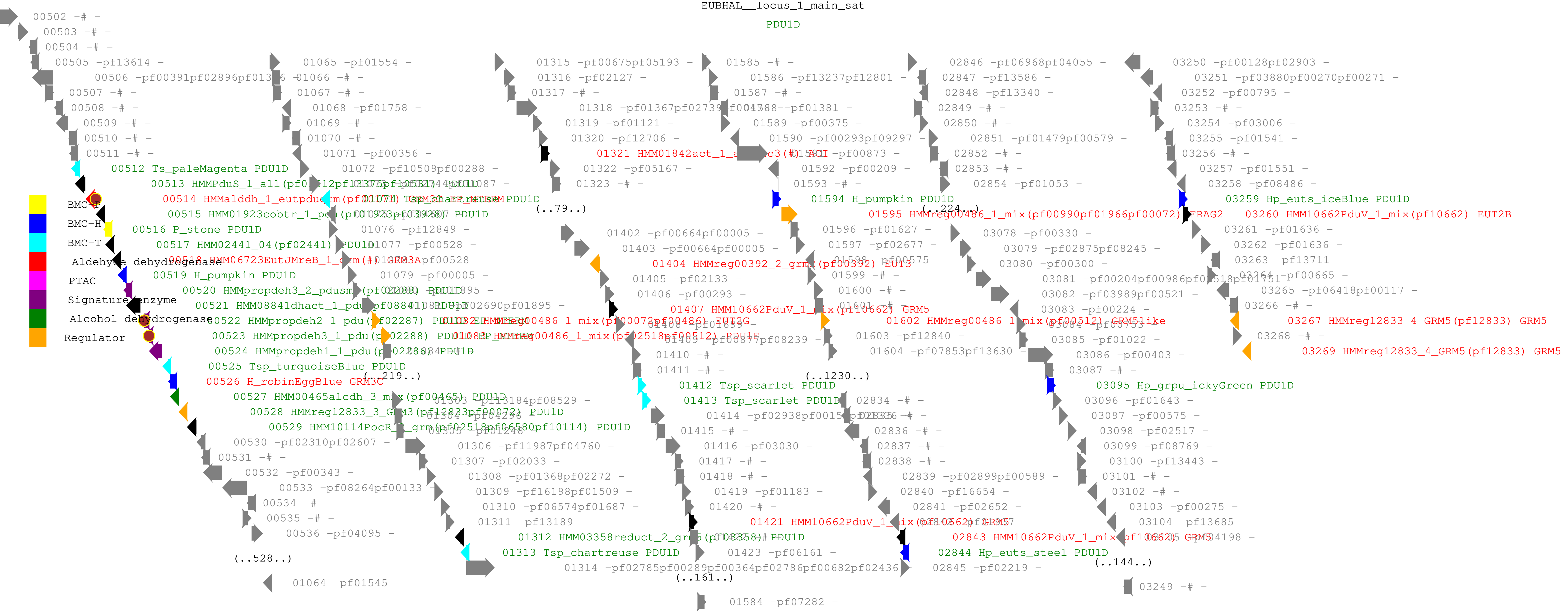

Eubacterium hallii DSM 3353. Bacteria; Firmicutes; Clostridia; Clostridiales; Eubacteriaceae;
EUBHAL__locus_1_main_sat
PDU1D
BMC-P
BMC-H
BMC-T
Aldehyde dehydrogenase
PTAC
Signature enzyme
Alcohol dehydrogenase
Regulator
00512 Ts_paleMagenta PDU1D
00513 HMMPduS_1_all PDU1D
00514 HMMaldh_1_eutpduccm
00515 HMM01923cobtr_1_pdu PDU1D
00516 P_stone PDU1D
00517 HMM02441_04(pf02441) PDU1D
00519 H_pumpkin PDU1D
00520 HMMpropdeh3_2_pdusm PDU1D
00521 HMM08841dhact_1_pdu PDU1D
00522 HMMpropdeh2_1_pdu(pf02287) PDU1D
00523 HMMpropdeh3_1_pdu(pf02288) PDU1D
00524 HMMpropdeh1_1_pdu PDU1D
00525 Tsp_turquoiseBlue PDU1D
00526 H_robinEggBlue GRM3C
00527 HMM00465alcdh_3_mix(pf00465) PDU1D
00528 HMMreg12833_3_GRM3(pf12833pf00072) PDU1D
00529 HMM10114PocR_1_grm(pf02518pf06580pf10114) PDU1D
01312 HMM03358reduct_2_grm5 PDU1D
01313 Tsp_chartreuse PDU1D
01321 HMM01842act_1_a
01404 HMMreg00392_2_grm(pf00392) EUT3
01407 HMM10662PduV_1_mix(pf10662) GRM5
01412 Tsp_scarlet PDU1D
01413 Tsp_scarlet PDU1D
01421 HMM10662PduV_1_mix(pf10662) GRM5
01594 H_pumpkin PDU1D
01595 HMMreg00486_1_mix(pf00990pf01966pf00072) FRAG2
01602 HMMreg00486_1_mix(pf00512) GRM5like
02843 HMM10662PduV_1_mix(pf10662) GRM5
02844 Hp_euts_steel PDU1D
03095 Hp_grpu_ickyGreen PDU1D
03259 Hp_euts_iceBlue PDU1D
03260 HMM10662PduV_1_mix(pf10662) EUT2B
03267 HMMreg12833_4_GRM5(pf12833) GRM5
03269 HMMreg12833_4_GRM5(pf12833) GRM5
(..528..)
(..219..)
(..79..)
(..161..)
(..1230..)
(..224..)
(..144..)

Flavonifractor plautii DSM 6740. Bacteria; Firmicutes; Clostridia; Clostridiales; Ruminococcaceae;

EV206__locus_3

PDU1D

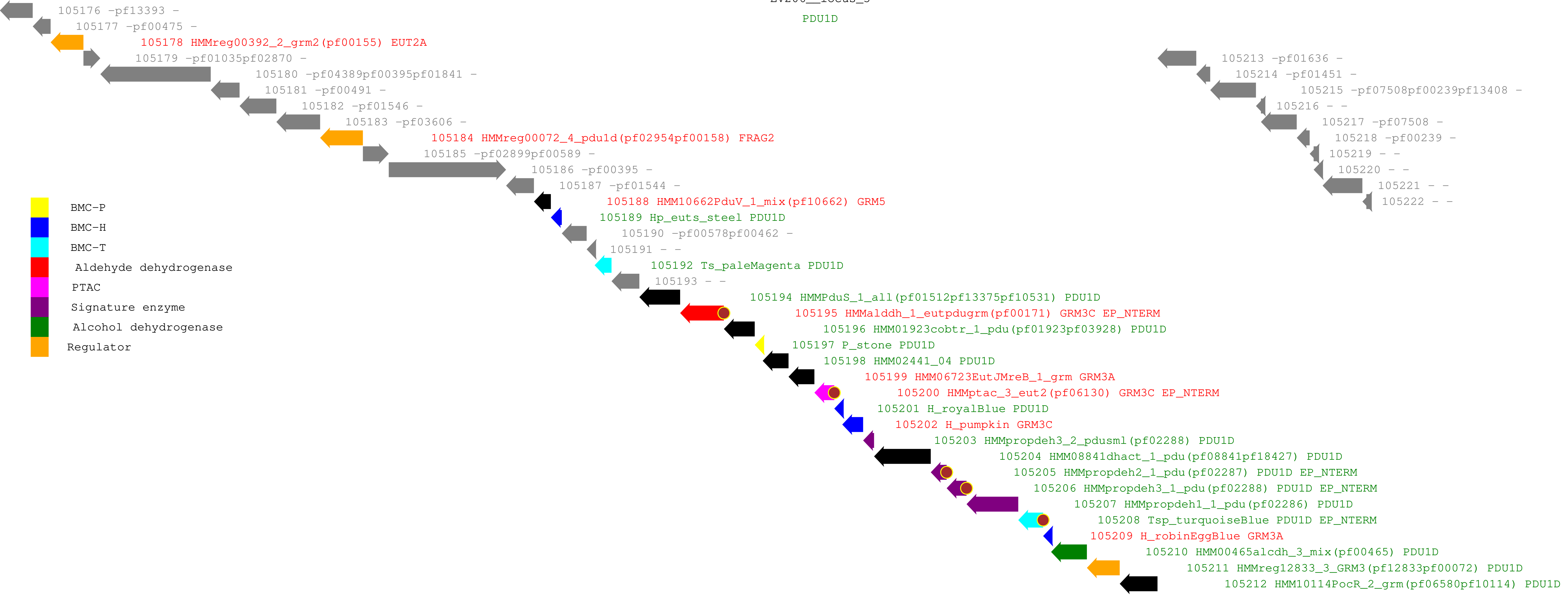

Anaerospora hongkongensis. Bacteria; Firmicutes; Negativicutes; Selenomonadales; Sporomusaceae;

EV210__locus_1

PDU1D

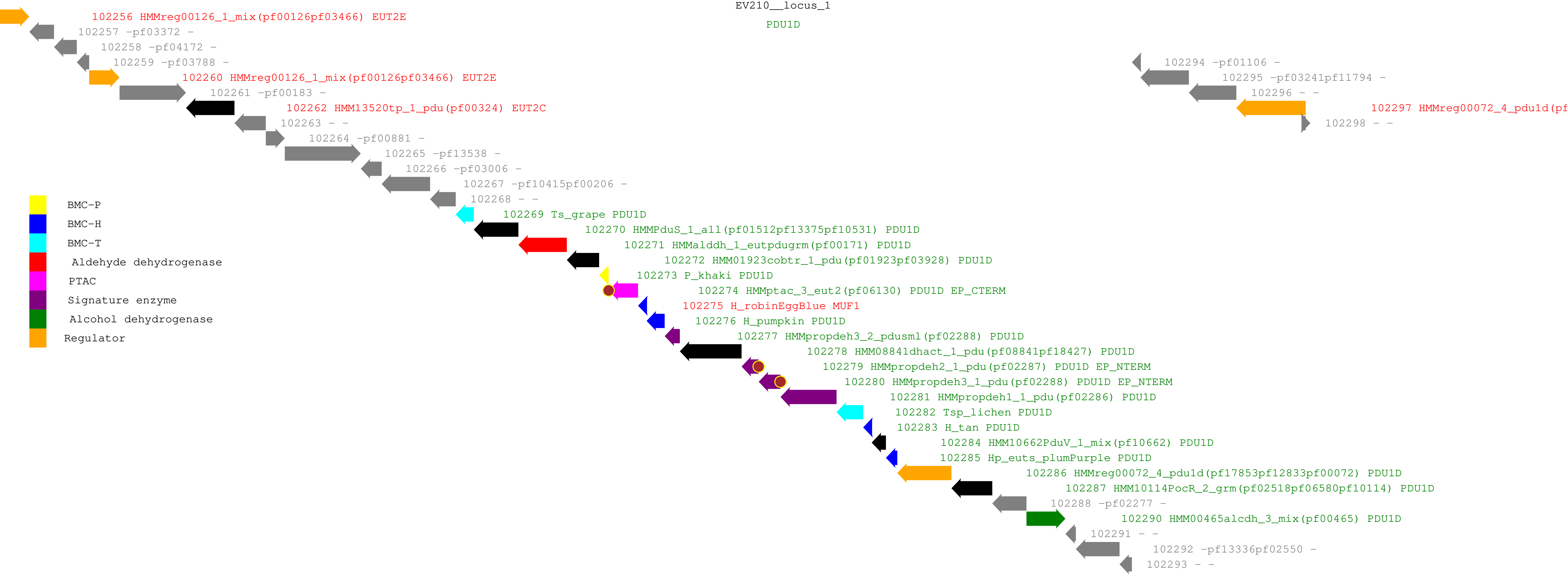

Marinisporobacter balticus. Bacteria; Firmicutes; Clostridia; Clostridiales; Clostridiaceae;

EV214__locus_1

PDU1D

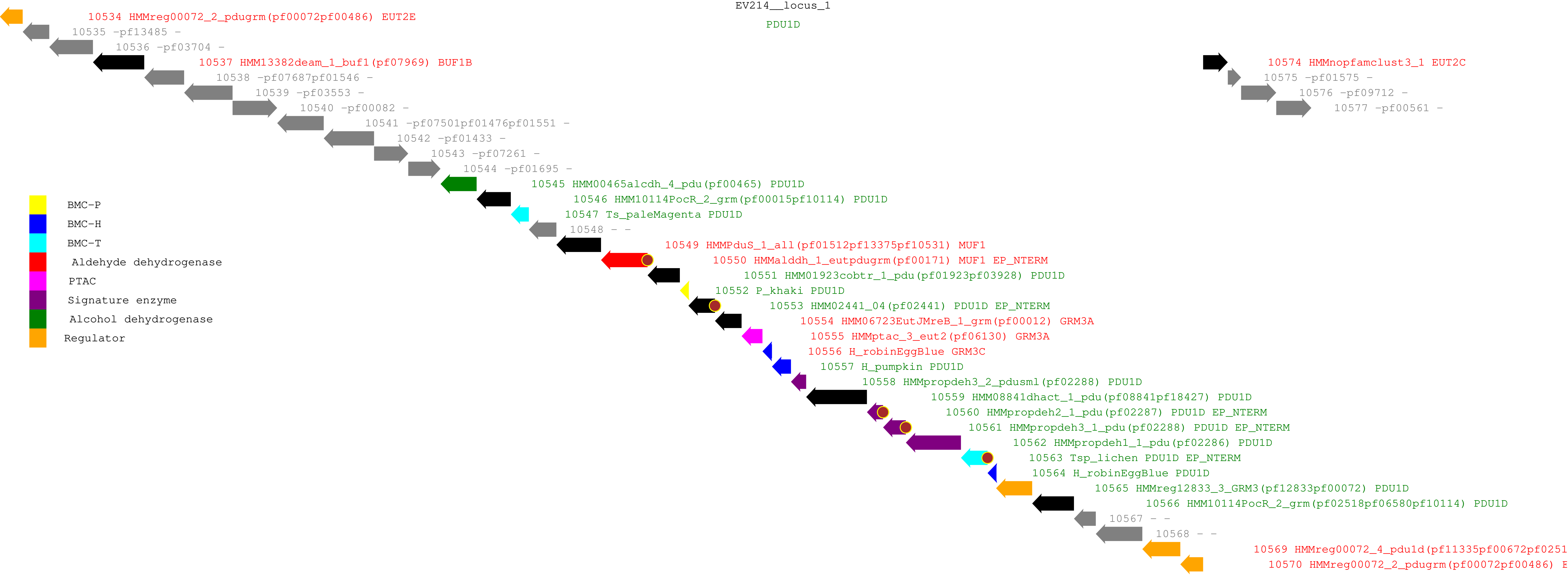

Siculibacillus lacustris. Bacteria; Proteobacteria; Alphaproteobacteria; Rhizobiales;

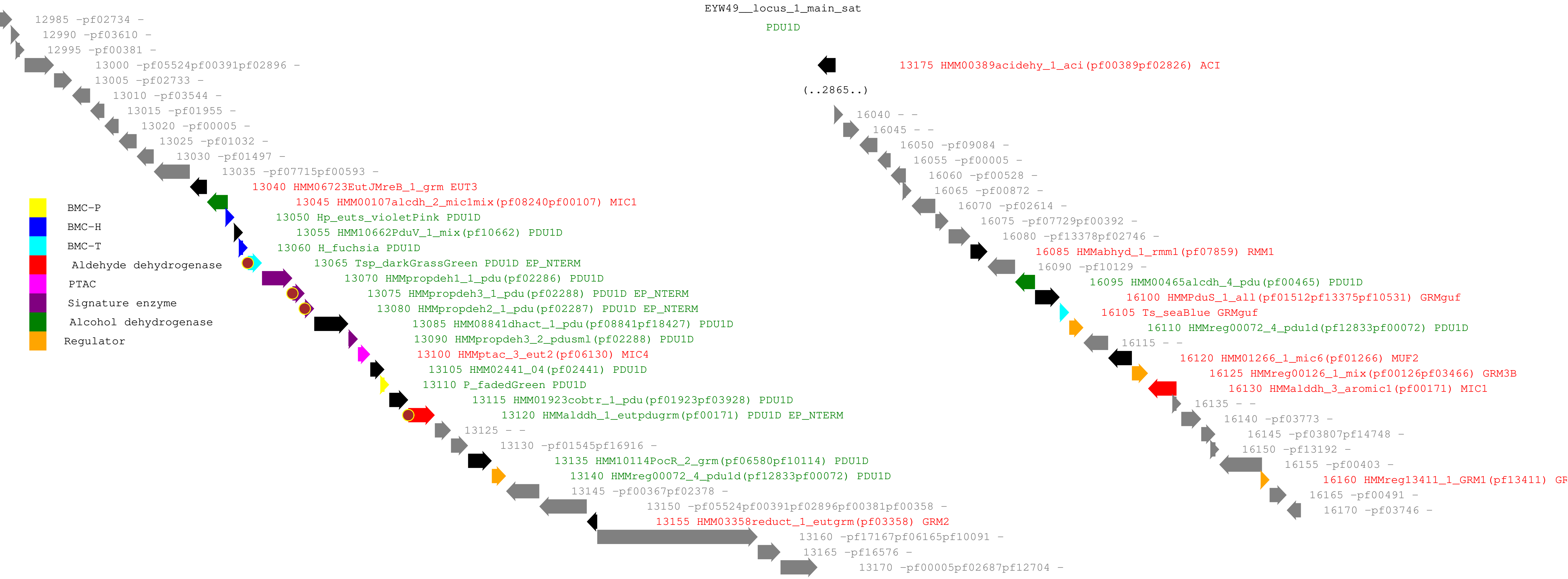

Pelosinus fermentans B4. Bacteria; Firmicutes; Negativicutes; Selenomonadales; Sporomusaceae;

FB4__locus_1

PDU1D

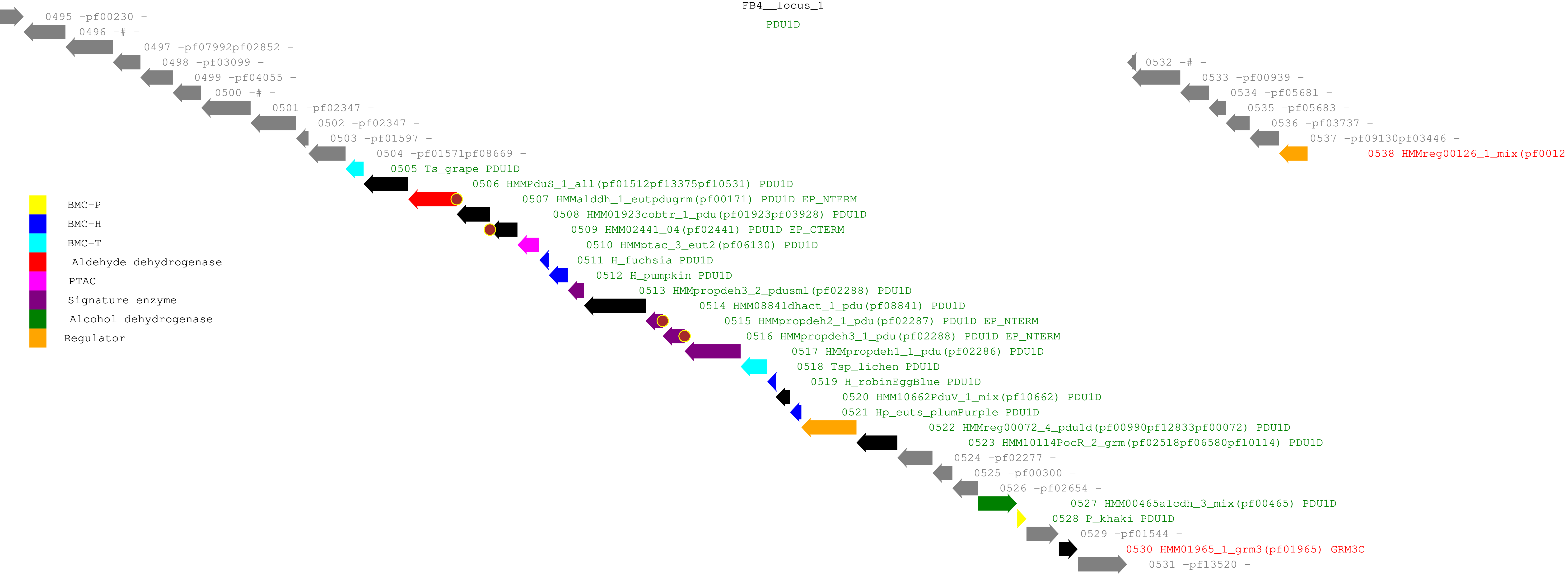

Pseudopropionibacterium propionicum. Bacteria; Actinobacteria; Propionibacteriales; Propionibacteriaceae;

FBF34__locus_1

PDU1D

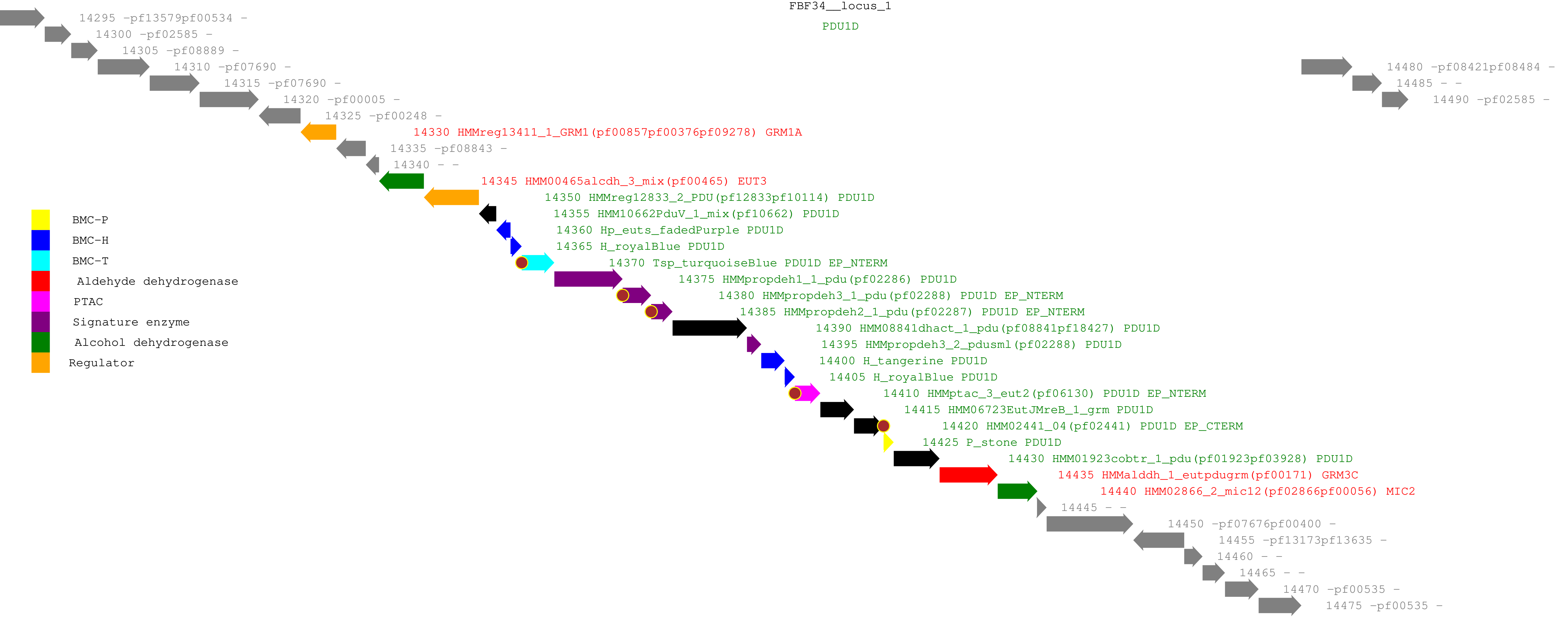

Caloramator sp. E03. Bacteria; Firmicutes; Clostridia; Clostridiales; Clostridiaceae;

FDN13__locus_2

PDU1D

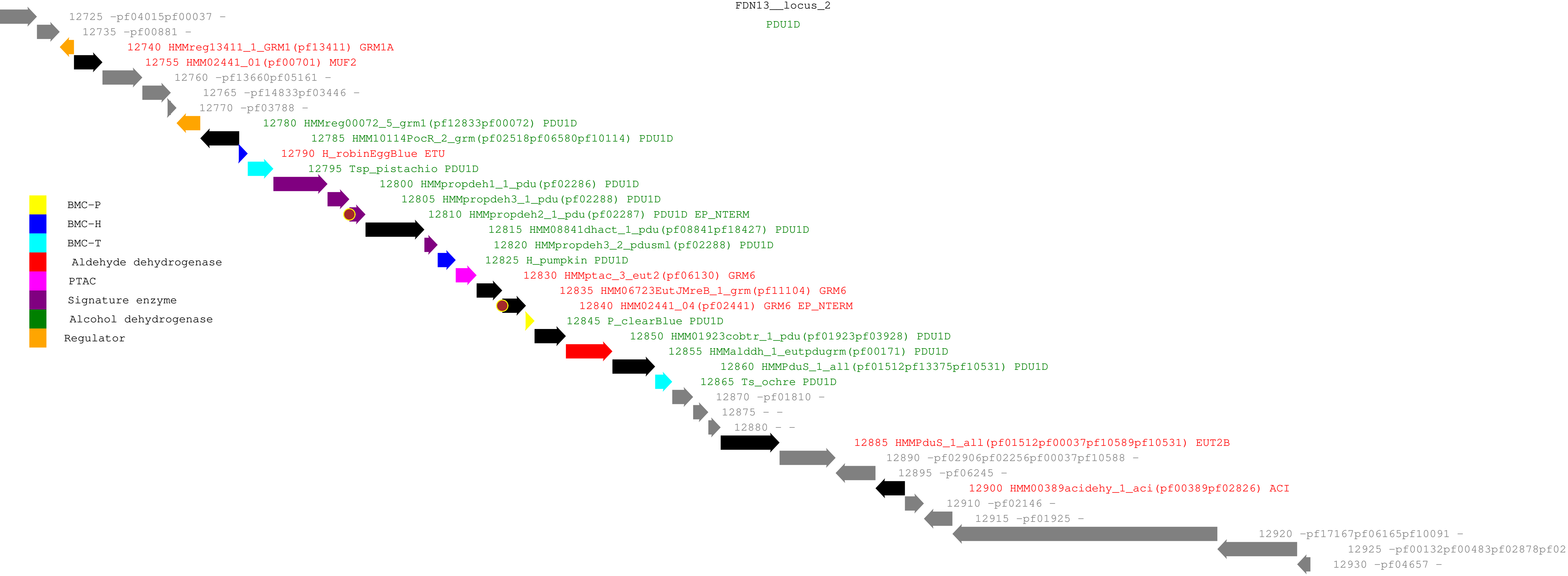

Romboutsia sp. Frifi. Bacteria; Firmicutes; Clostridia; Clostridiales;

FRIFI__locus_1

PDU1D

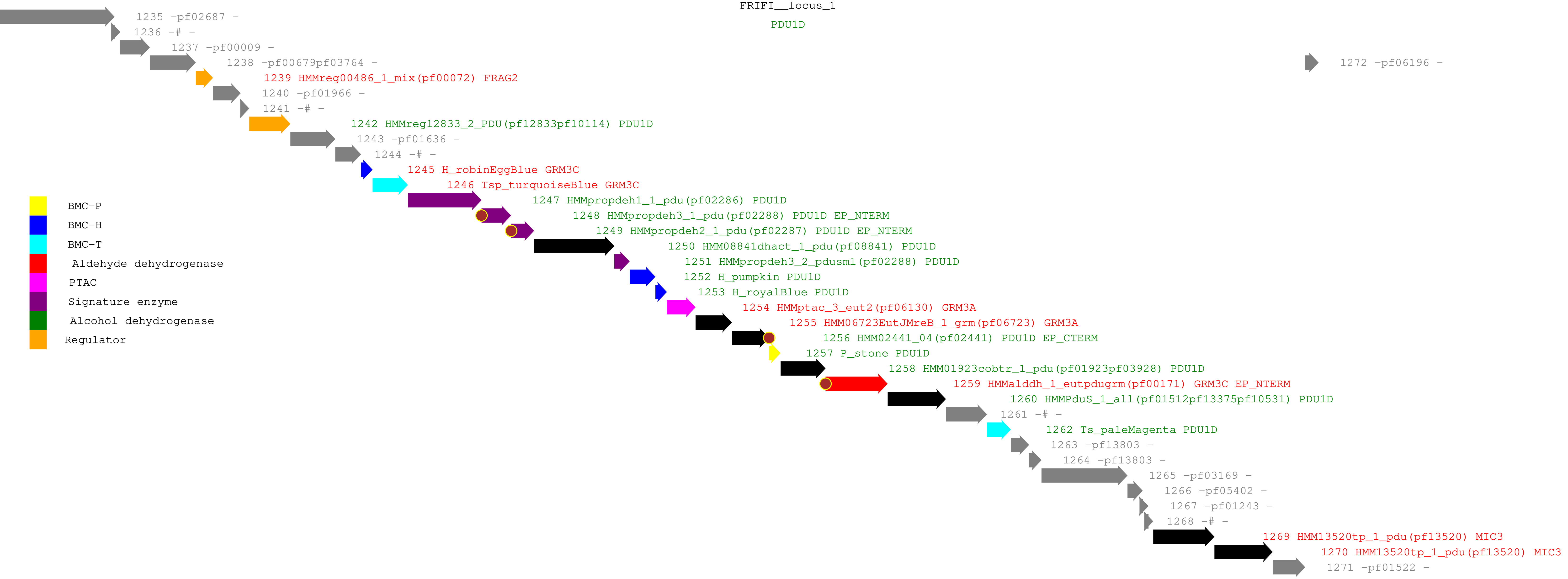

Thermoanaerobacteraceae bacterium A05MB. Bacteria; Firmicutes; Clostridia; Thermoanaerobacterales;

FWJ32__locus_1

PDU1D

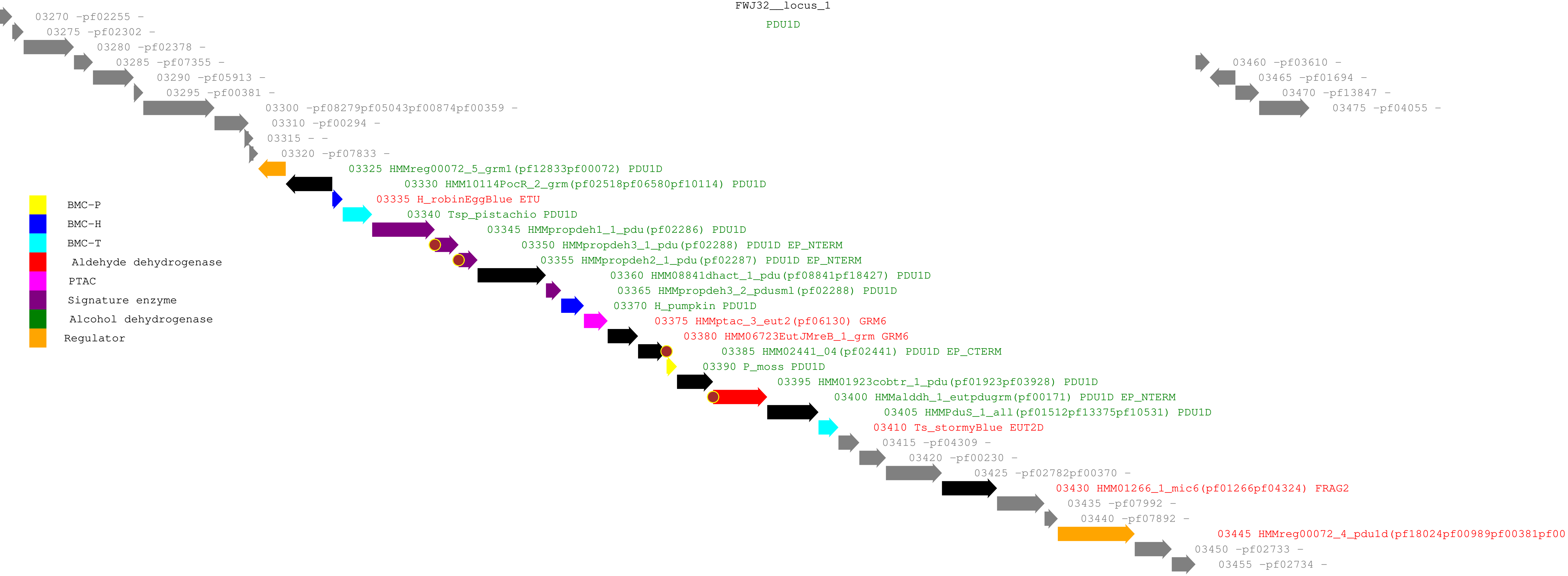

Acetobacterium wieringae. Bacteria; Firmicutes; Clostridia; Clostridiales; Eubacteriaceae;

FXB42__locus_fragmented

PDU1D

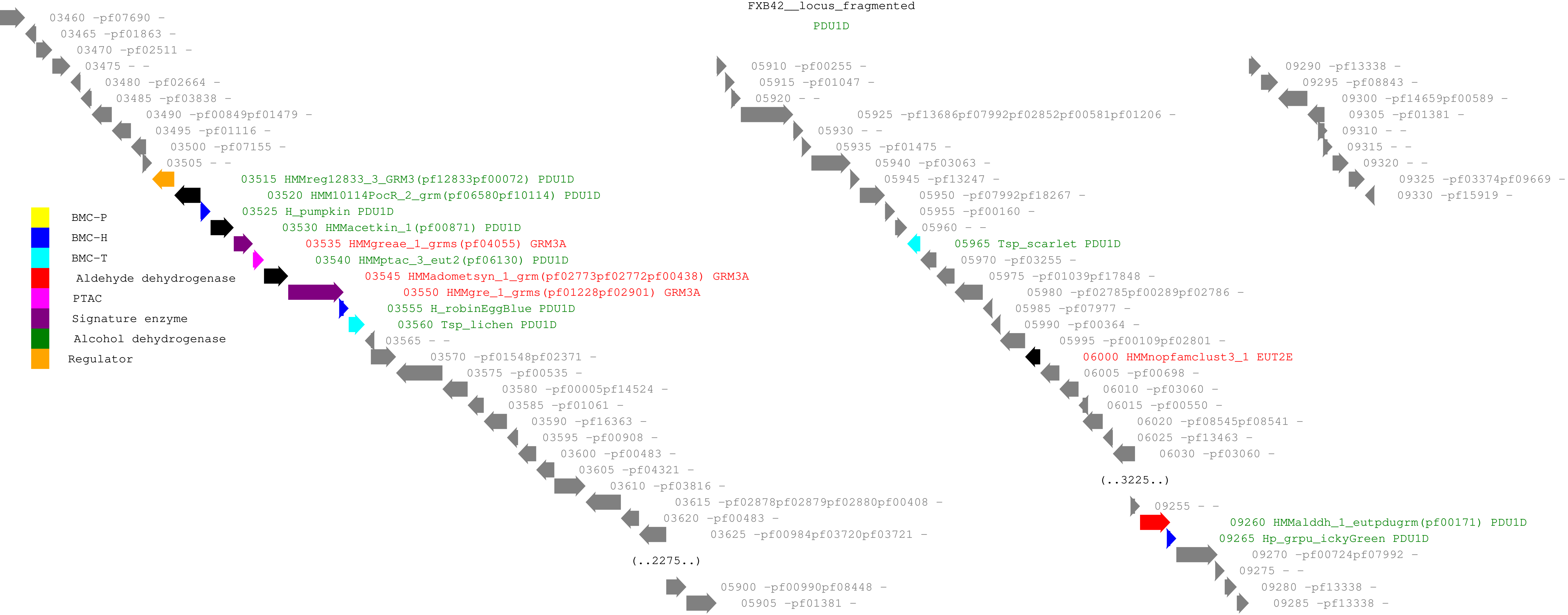

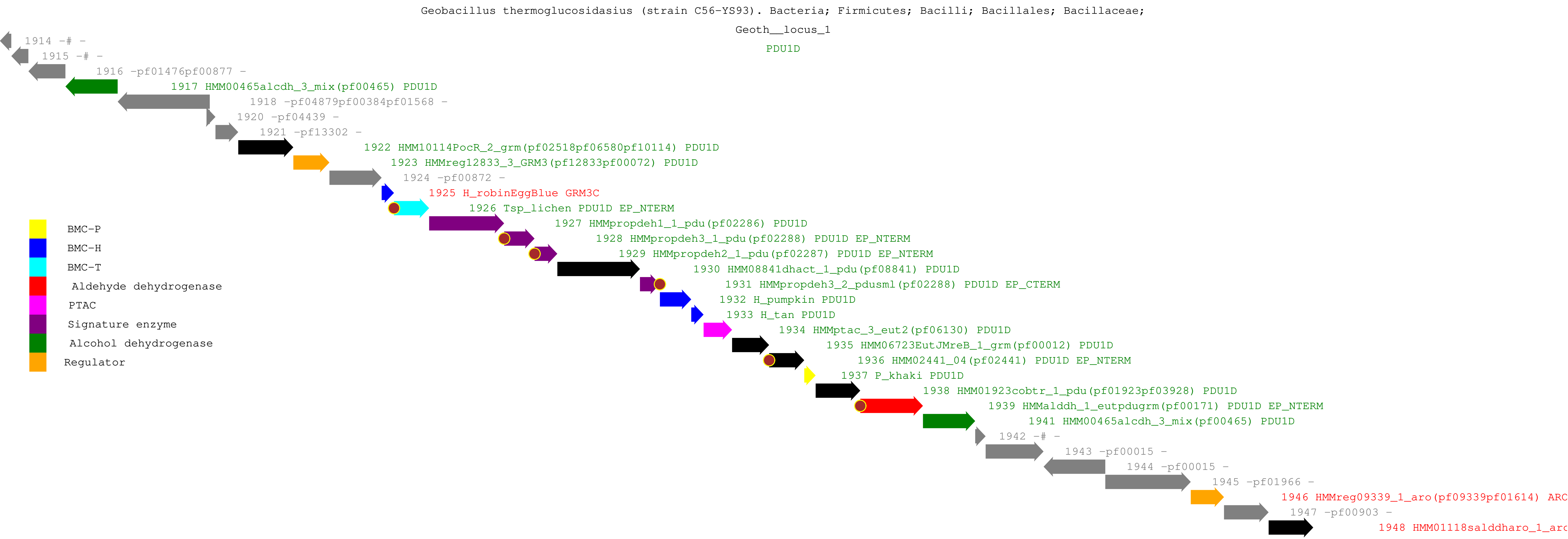
Geobacillus thermoglucosidasius (strain C56-YS93). Bacteria; Firmicutes; Bacilli; Bacillales; Bacillaceae;
Geoth__locus_1
PDU1D
1914 -# -
1915 -# -
1916 -pf01476pf00877 -
1917 HMM00465alcdh_3_mix(pf00465) PDU1D
1918 -pf04879pf00384pf01568 -
1920 -pf04439 -
1921 -pf13302 -
1922 HMM10114PocR_2_grm(pf02518pf06580pf10114) PDU1D
1923 HMMreg12833_3_GRM3(pf12833pf00072) PDU1D
1924 -pf00872 -
1925 H_robinEggBlue GRM3C
1926 Tsp_lichen PDU1D EP_NTERM
1927 HMMpropdeh1_1_pdu(pf02286) PDU1D
1928 HMMpropdeh3_1_pdu(pf02288) PDU1D EP_NTERM
1929 HMMpropdeh2_1_pdu(pf02287) PDU1D EP_NTERM
1930 HMM08841dhact_1_pdu(pf08841) PDU1D
1931 HMMpropdeh3_2_pdusml(pf02288) PDU1D EP_CTERM
1932 H_pumpkin PDU1D
1933 H_tan PDU1D
1934 HMMptac_3_eut2(pf06130) PDU1D
1935 HMM06723EutJMreB_1_grm(pf00012) PDU1D
1936 HMM02441_04(pf02441) PDU1D EP_NTERM
1937 P_khaki PDU1D
1938 HMM01923cobtr_1_pdu(pf01923pf03928) PDU1D
1939 HMMaldh_1_eutpdugrm(pf00171) PDU1D EP_NTERM
1941 HMM00465alcdh_3_mix(pf00465) PDU1D
1942 -# -
1943 -pf00015 -
1944 -pf00015 -
1945 -pf01966 -
1946 HMMreg09339_1_aro(pf09339pf01614) ARC
1947 -pf00903 -
1948 HMM01118salddharo_1_arc
BMC-P
BMC-H
BMC-T
Aldehyde dehydrogenase
PTAC
Signature enzyme
Alcohol dehydrogenase
Regulator

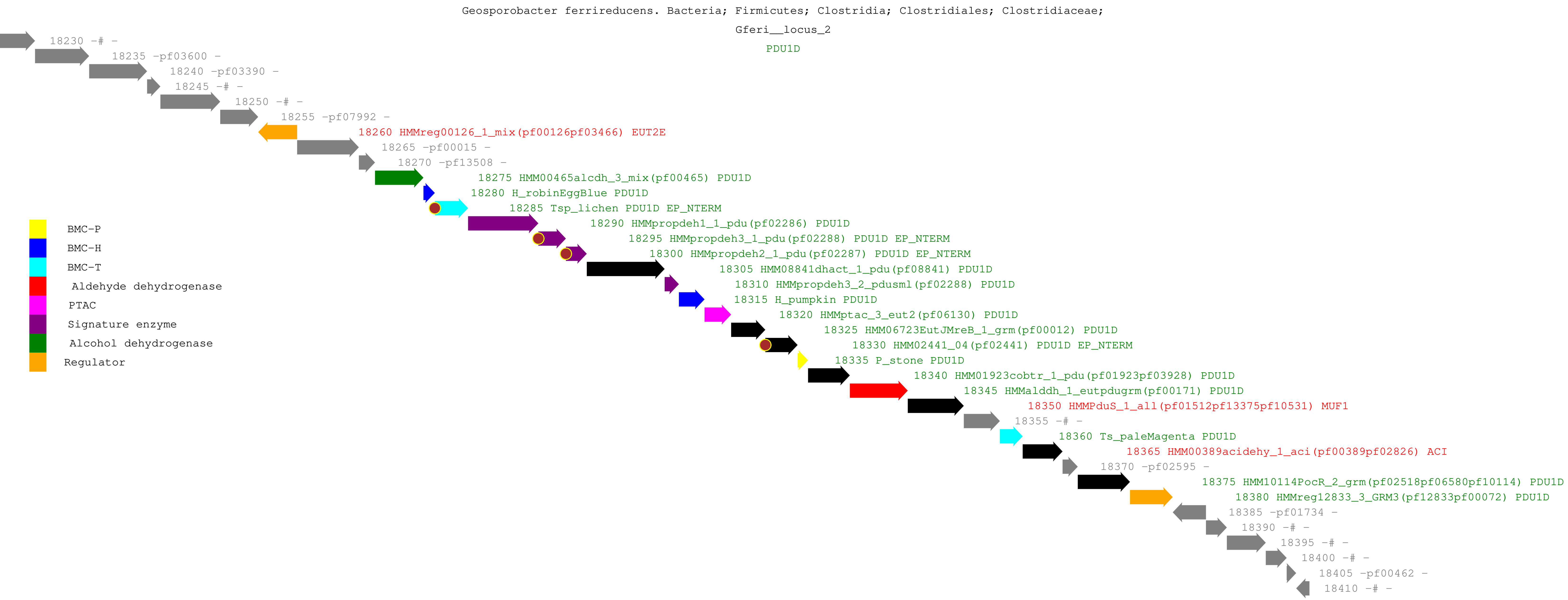
Geosporobacter ferrireducens. Bacteria; Firmicutes; Clostridia; Clostridiales; Clostridiaceae;
Gferi__locus_2
PDU1D
18230 -# -
18235 -pf03600 -
18240 -pf03390 -
18245 -# -
18250 -# -
18255 -pf07992 -
18260 HMMreg00126_1_mix(pf00126pf03466) EUT2E
18265 -pf00015 -
18270 -pf13508 -
18275 HMM00465alcdh_3_mix(pf00465) PDU1D
18280 H_robinEggBlue PDU1D
18285 Tsp_lichen PDU1D EP_NTERM
18290 HMMpropdeh1_1_pdu(pf02286) PDU1D
18295 HMMpropdeh3_1_pdu(pf02288) PDU1D EP_NTERM
18300 HMMpropdeh2_1_pdu(pf02287) PDU1D EP_NTERM
18305 HMM08841dhact_1_pdu(pf08841) PDU1D
18310 HMMpropdeh3_2_pdusml(pf02288) PDU1D
18315 H_pumpkin PDU1D
18320 HMMptac_3_eut2(pf06130) PDU1D
18325 HMM06723EutJMreB_1_grm(pf00012) PDU1D
18330 HMM02441_04(pf02441) PDU1D EP_NTERM
18335 P_stone PDU1D
18340 HMM01923cobtr_1_pdu(pf01923pf03928) PDU1D
18345 HMMaldдh_1_eutpdugrm(pf00171) PDU1D
18350 HMMPduS_1_all(pf01512pf13375pf10531) MUF1
18355 -# -
18360 Ts_paleMagenta PDU1D
18365 HMM00389acidehy_1_aci(pf00389pf02826) ACI
18370 -pf02595 -
18375 HMM10114PocR_2_grm(pf02518pf06580pf10114) PDU1D
18380 HMMreg12833_3_GRM3(pf12833pf00072) PDU1D
18385 -pf01734 -
18390 -# -
18395 -# -
18400 -# -
18405 -pf00462 -
18410 -# -
BMC-P
BMC-H
BMC-T
Aldehyde dehydrogenase
PTAC
Signature enzyme
Alcohol dehydrogenase
Regulator

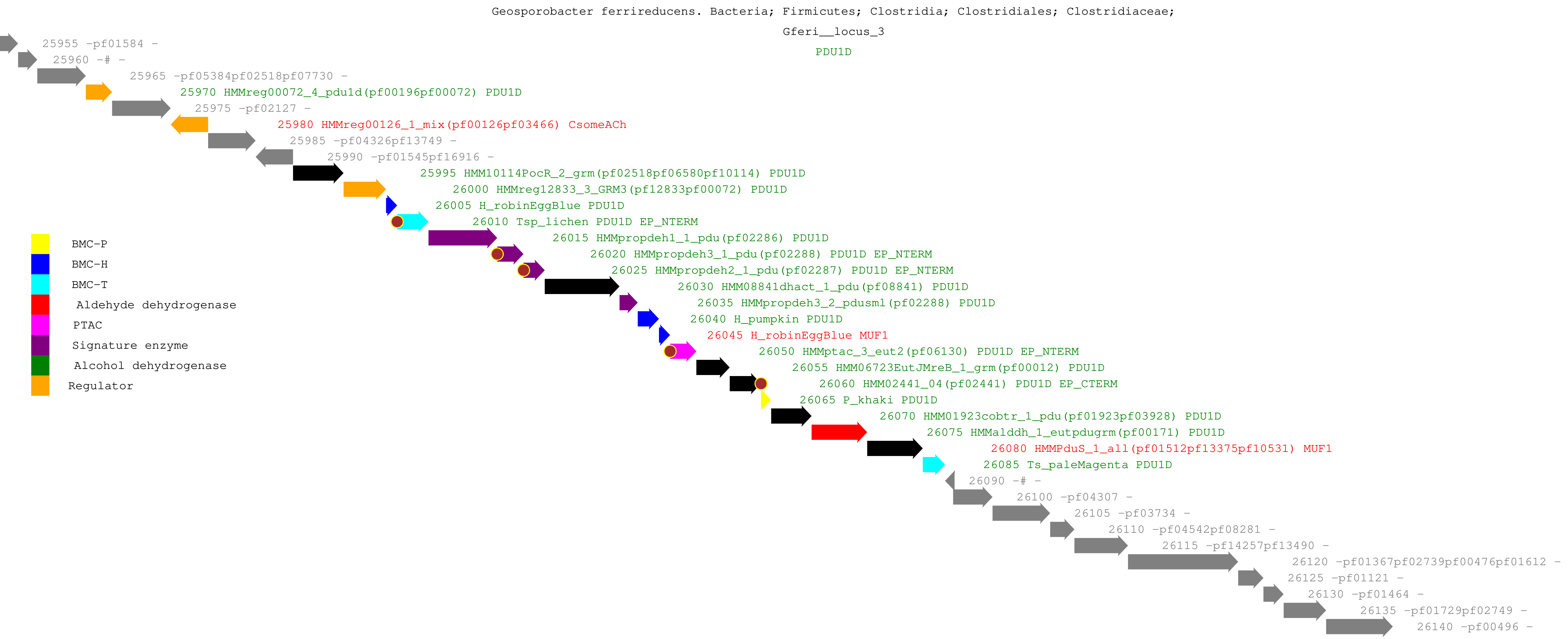

Geosporobacter ferrireducens. Bacteria; Firmicutes; Clostridia; Clostridiales; Clostridiaceae;
Gferi__locus_3
PDU1D
25955 -pf01584 -
25960 -# -
25965 -pf05384pf02518pf07730 -
25970 HMMreg00072_4_pdu1d(pf00196pf00072) PDU1D
25975 -pf02127 -
25980 HMMreg00126_1_mix(pf00126pf03466) CsomeACh
25985 -pf04326pf13749 -
25990 -pf01545pf16916 -
25995 HMM10114PocR_2_grm(pf02518pf06580pf10114) PDU1D
26000 HMMreg12833_3_GRM3(pf12833pf00072) PDU1D
26005 H_robinEggBlue PDU1D
26010 Tsp_lichen PDU1D EP_NTERM
26015 HMMpropdeh1_1_pdu(pf02286) PDU1D
26020 HMMpropdeh3_1_pdu(pf02288) PDU1D EP_NTERM
26025 HMMpropdeh2_1_pdu(pf02287) PDU1D EP_NTERM
26030 HMM08841dhact_1_pdu(pf08841) PDU1D
26035 HMMpropdeh3_2_pdusml(pf02288) PDU1D
26040 H_pumpkin PDU1D
26045 H_robinEggBlue MUF1
26050 HMMptac_3_eut2(pf06130) PDU1D EP_NTERM
26055 HMM06723EutJMreB_1_grm(pf00012) PDU1D
26060 HMM02441_04(pf02441) PDU1D EP_CTERM
26065 P_khaki PDU1D
26070 HMM01923cobtr_1_pdu(pf01923pf03928) PDU1D
26075 HMMalddh_1_eutpdugrm(pf00171) PDU1D
26080 HMMPduS_1_all(pf01512pf13375pf10531) MUF1
26085 Ts_paleMagenta PDU1D
26090 -# -
26100 -pf04307 -
26105 -pf03734 -
26110 -pf04542pf08281 -
26115 -pf14257pf13490 -
26120 -pf01367pf02739pf00476pf01612 -
26125 -pf01121 -
26130 -pf01464 -
26135 -pf01729pf02749 -
26140 -pf00496 -
BMC-P
BMC-H
BMC-T
Aldehyde dehydrogenase
PTAC
Signature enzyme
Alcohol dehydrogenase
Regulator

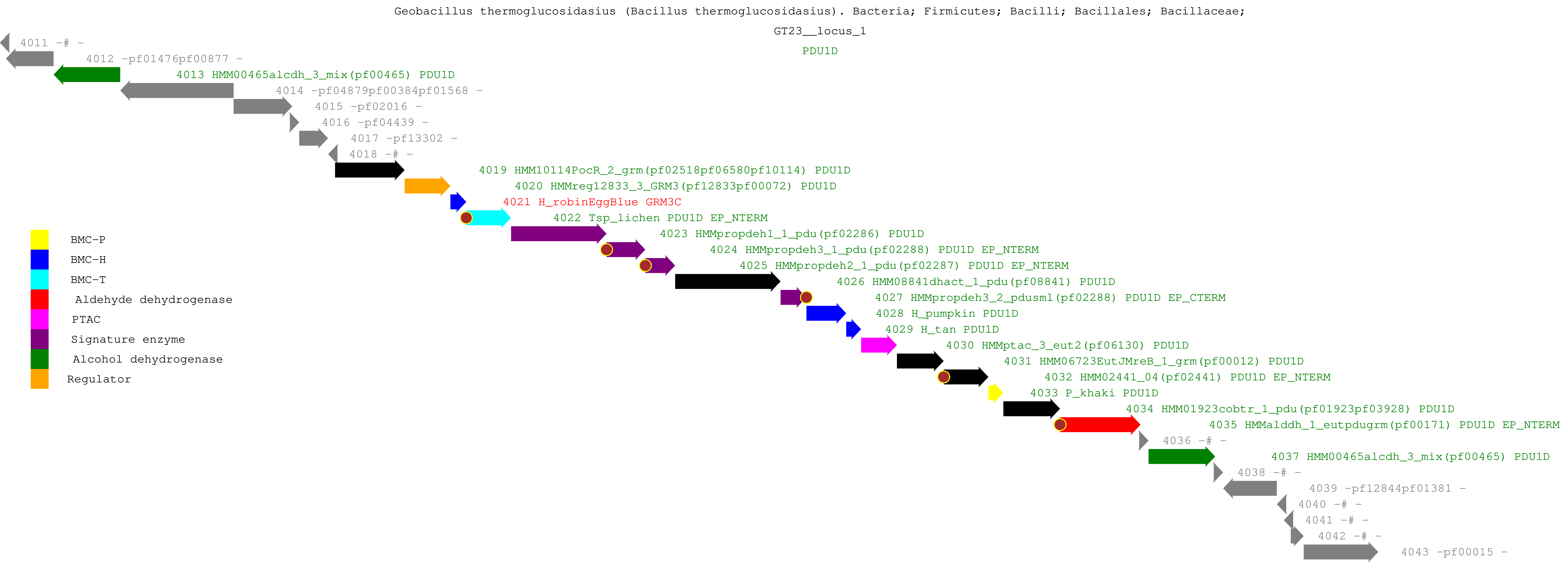
Geobacillus thermoglucosidasius (Bacillus thermoglucosidasius). Bacteria; Firmicutes; Bacilli; Bacillales; Bacillaceae;
GT23__locus_1
PDU1D
4011 -# -
4012 -pf01476pf00877 -
4013 HMM00465alcdh_3_mix(pf00465) PDU1D
4014 -pf04879pf00384pf01568 -
4015 -pf02016 -
4016 -pf04439 -
4017 -pf13302 -
4018 -# -
4019 HMM10114PocR_2_grm(pf02518pf06580pf10114) PDU1D
4020 HMMreg12833_3_GRM3(pf12833pf00072) PDU1D
4021 H_robinEggBlue GRM3C
4022 Tsp_lichen PDU1D EP_NTERM
4023 HMMpropdeh1_1_pdu(pf02286) PDU1D
4024 HMMpropdeh3_1_pdu(pf02288) PDU1D EP_NTERM
4025 HMMpropdeh2_1_pdu(pf02287) PDU1D EP_NTERM
4026 HMM08841dhact_1_pdu(pf08841) PDU1D
4027 HMMpropdeh3_2_pdusml(pf02288) PDU1D EP_CTERM
4028 H_pumpkin PDU1D
4029 H_tan PDU1D
4030 HMMptac_3_eut2(pf06130) PDU1D
4031 HMM06723EutJMreB_1_grm(pf00012) PDU1D
4032 HMM02441_04(pf02441) PDU1D EP_NTERM
4033 P_khaki PDU1D
4034 HMM01923cobtr_1_pdu(pf01923pf03928) PDU1D
4035 HMMalddh_1_eutpdugrm(pf00171) PDU1D EP_NTERM
4036 -# -
4037 HMM00465alcdh_3_mix(pf00465) PDU1D
4038 -# -
4039 -pf12844pf01381 -
4040 -# -
4041 -# -
4042 -# -
4043 -pf00015 -
BMC-P
BMC-H
BMC-T
Aldehyde dehydrogenase
PTAC
Signature enzyme
Alcohol dehydrogenase
Regulator

Geobacillus sp. (strain Y4.1MC1). Bacteria; Firmicutes; Bacilli; Bacillales; Bacillaceae; Geobacillus.

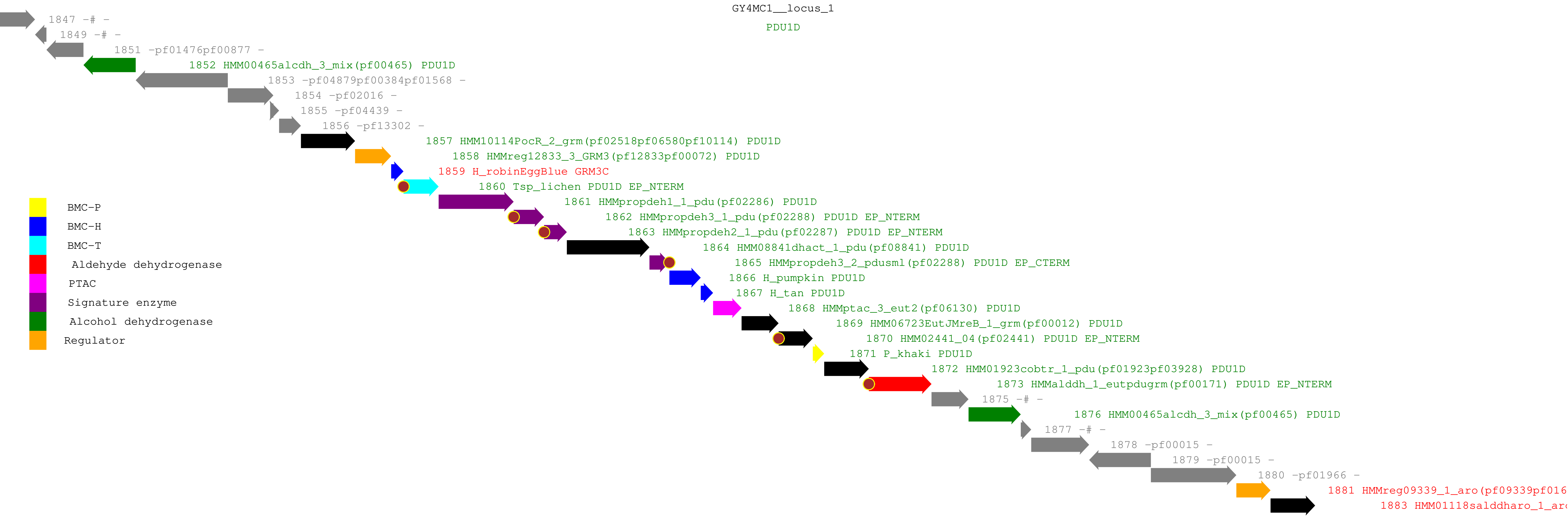

Halanaerobium hydrogeniformans (Halanaerobium sp. (strain Bacteria; Firmicutes; Clostridia; Halanaerobiales; Halanaerobiaceae;

Halsa__locus_1

PDU1D

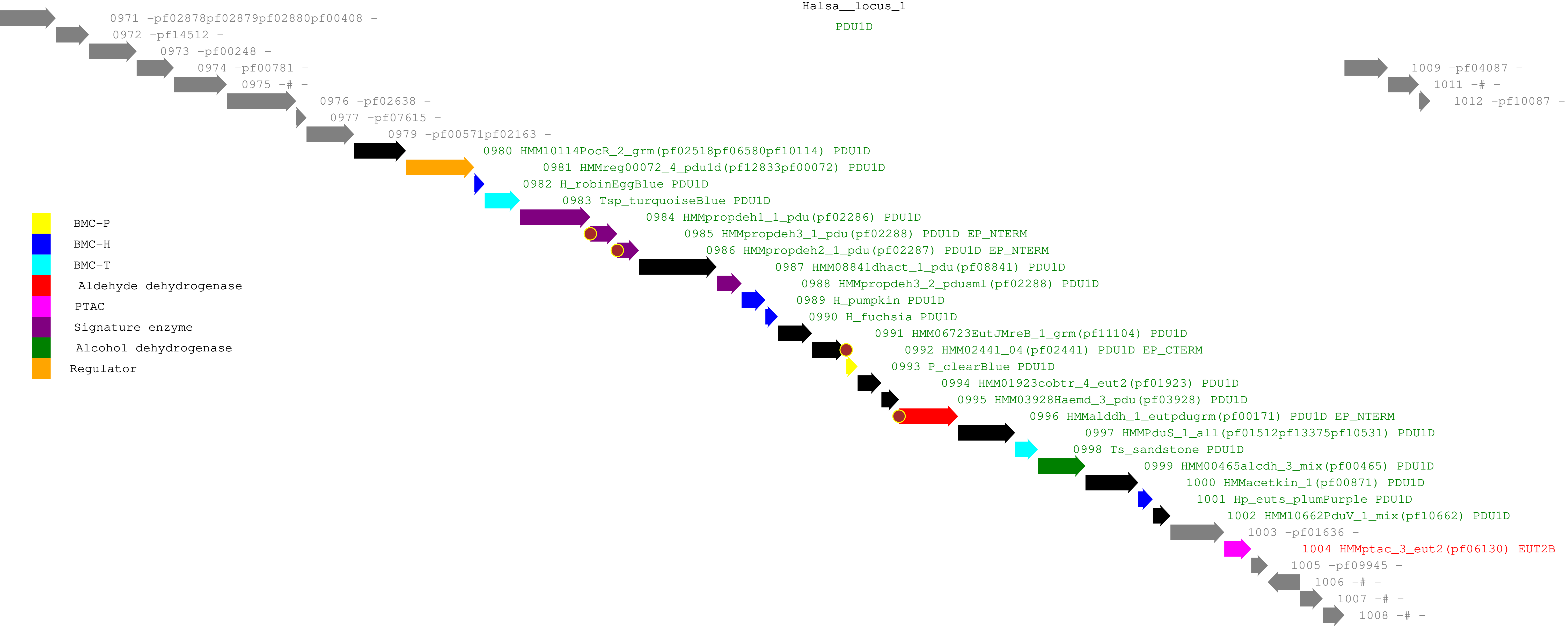

Pseudoramibacter alactolyticus ATCC 23263. Bacteria; Firmicutes; Clostridia; Clostridiales; Eubacteriaceae;

HMP0721__locus_2

PDU1D

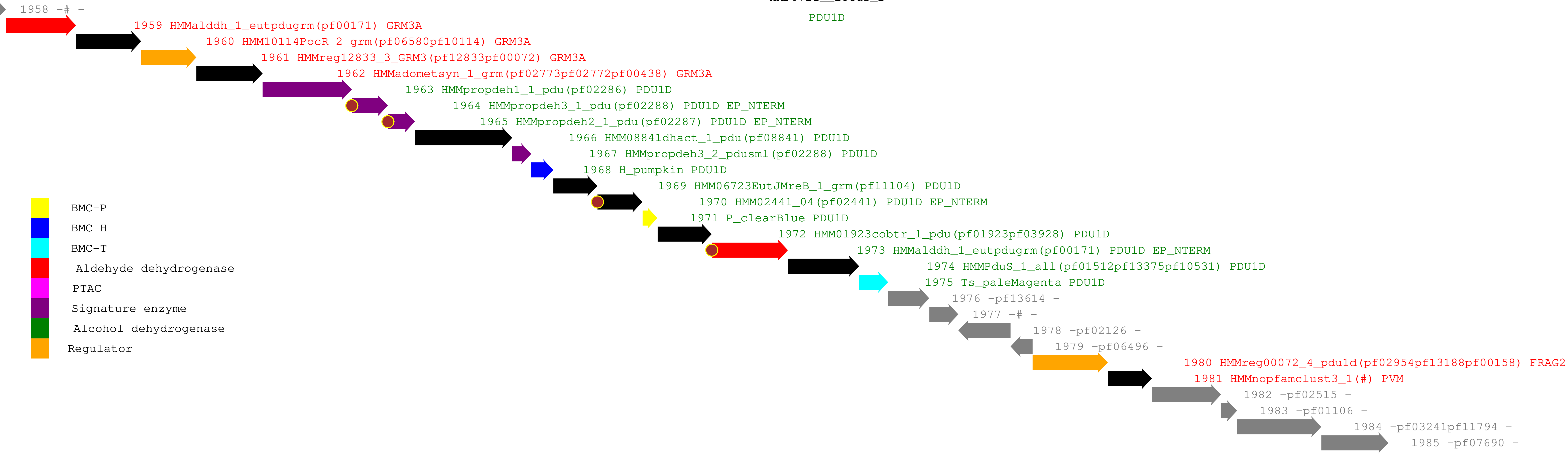

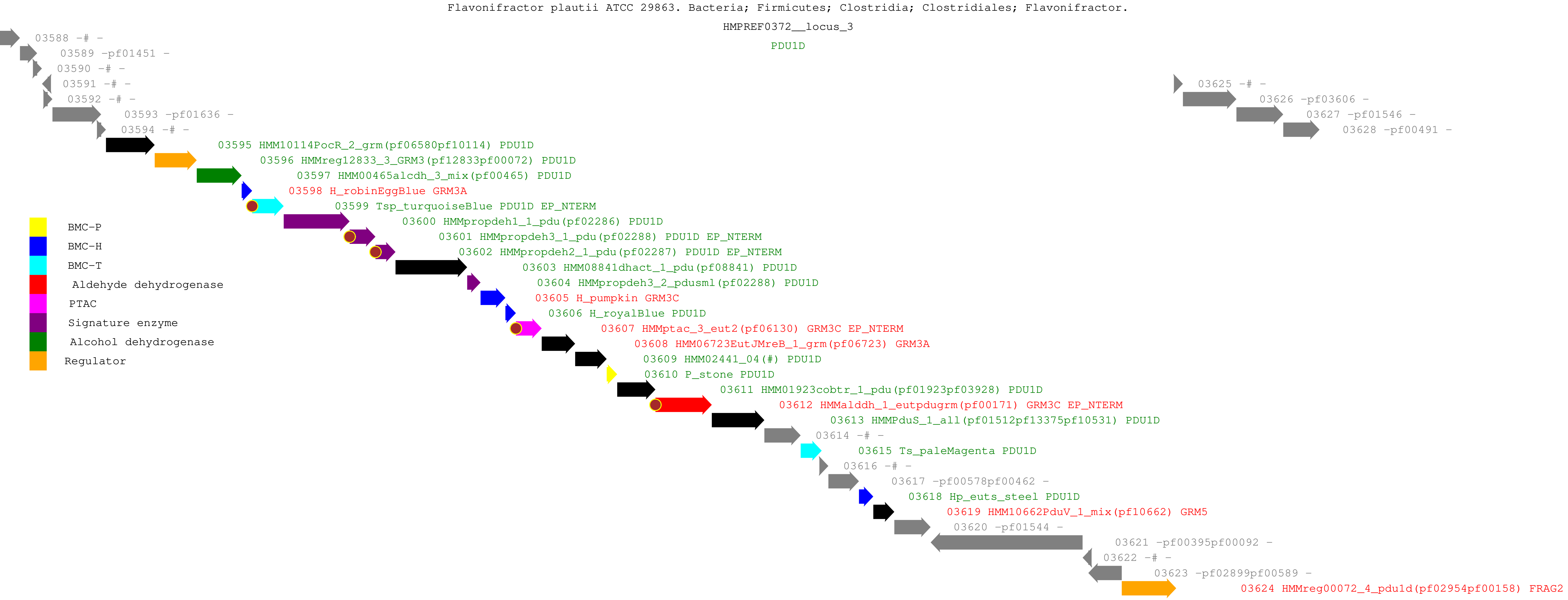
Flavonifractor plautii ATCC 29863. Bacteria; Firmicutes; Clostridia; Clostridiales; Flavonifractor.
HMPREF0372__locus_3
PDU1D
03588 -# -
03589 -pf01451 -
03590 -# -
03591 -# -
03592 -# -
03593 -pf01636 -
03594 -# -
03595 HMM10114PocR_2_grm(pf06580pf10114) PDU1D
03596 HMMreg12833_3_GRM3(pf12833pf00072) PDU1D
03597 HMM00465alcdh_3_mix(pf00465) PDU1D
03598 H_robinEggBlue GRM3A
03599 Tsp_turquoiseBlue PDU1D EP_NTERM
03600 HMMpropdeh1_1_pdu(pf02286) PDU1D
03601 HMMpropdeh3_1_pdu(pf02288) PDU1D EP_NTERM
03602 HMMpropdeh2_1_pdu(pf02287) PDU1D EP_NTERM
03603 HMM08841dhact_1_pdu(pf08841) PDU1D
03604 HMMpropdeh3_2_pdusml(pf02288) PDU1D
03605 H_pumpkin GRM3C
03606 H_royalBlue PDU1D
03607 HMMptac_3_eut2(pf06130) GRM3C EP_NTERM
03608 HMM06723EutJMreB_1_grm(pf06723) GRM3A
03609 HMM02441_04(#) PDU1D
03610 P_stone PDU1D
03611 HMM01923cobtr_1_pdu(pf01923pf03928) PDU1D
03612 HMMalddh_1_eutpdugrm(pf00171) GRM3C EP_NTERM
03613 HMMPduS_1_all(pf01512pf13375pf10531) PDU1D
03614 -# -
03615 Ts_paleMagenta PDU1D
03616 -# -
03617 -pf00578pf00462 -
03618 Hp_euts_steel PDU1D
03619 HMM10662PduV_1_mix(pf10662) GRM5
03620 -pf01544 -
03621 -pf00395pf00092 -
03622 -# -
03623 -pf02899pf00589 -
03624 HMMreg00072_4_pdu1d(pf02954pf00158) FRAG2
03625 -# -
03626 -pf03606 -
03627 -pf01546 -
03628 -pf00491 -
BMC-P
BMC-H
BMC-T
Aldehyde dehydrogenase
PTAC
Signature enzyme
Alcohol dehydrogenase
Regulator

Propionibacterium acidifaciens F0233. Bacteria; Actinobacteria; Propionibacteriales; Propionibacteriaceae;

HMPREF0682__locus_1_main_sat

PDU1D

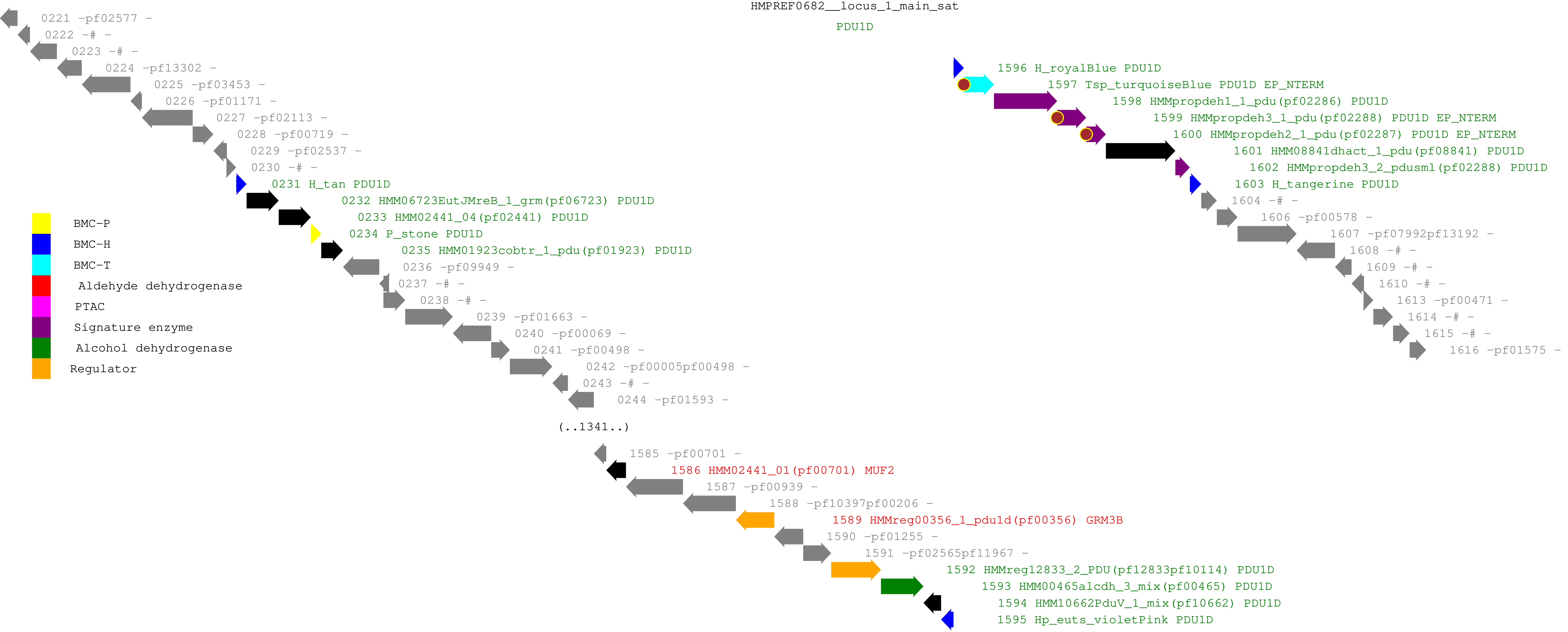

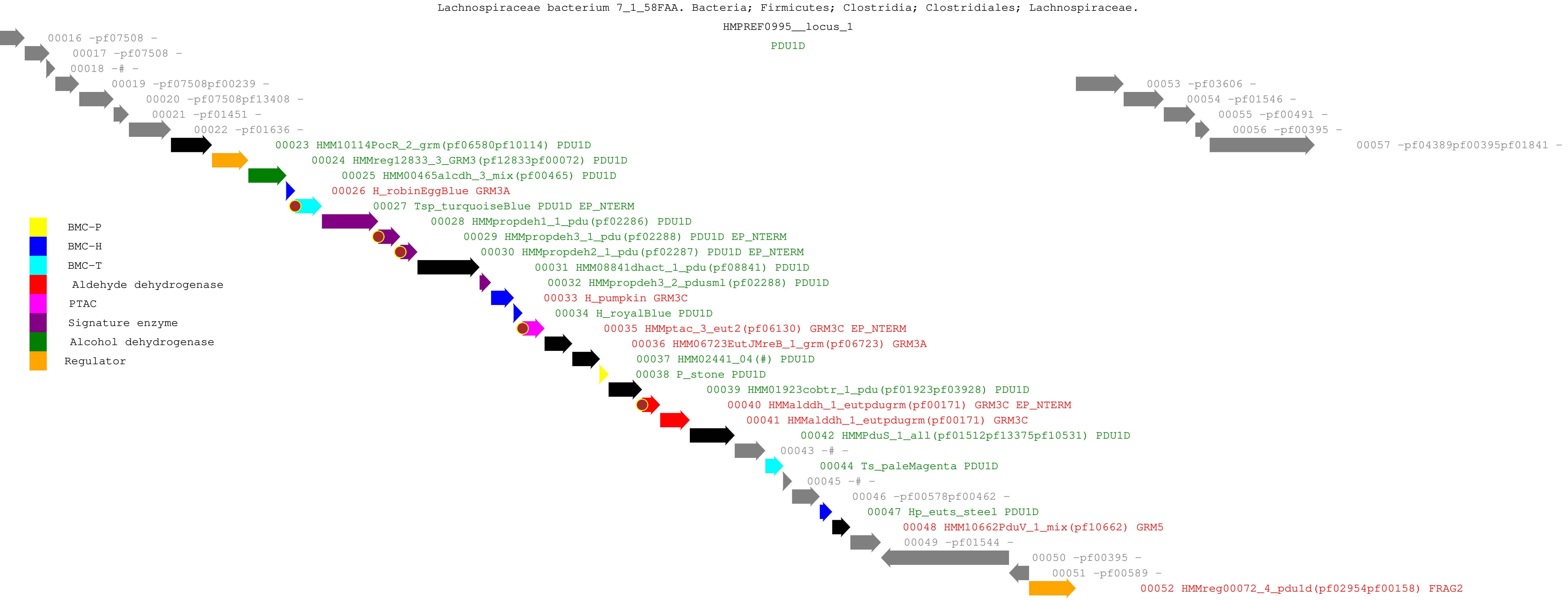

Lachnospiraceae bacterium 7_1_58FAA. Bacteria; Firmicutes; Clostridia; Clostridiales; Lachnospiraceae.
HMPREF0995__locus_1
PDU1D
00016 -pf07508 -
00017 -pf07508 -
00018 -# -
00019 -pf07508pf00239 -
00020 -pf07508pf13408 -
00021 -pf01451 -
00022 -pf01636 -
00023 HMM10114PocR_2_grm(pf06580pf10114) PDU1D
00024 HMMreg12833_3_GRM3(pf12833pf00072) PDU1D
00025 HMM00465alcdh_3_mix(pf00465) PDU1D
00026 H_robinEggBlue GRM3A
00027 Tsp_turquoiseBlue PDU1D EP_NTERM
00028 HMMpropdeh1_1_pdu(pf02286) PDU1D
00029 HMMpropdeh3_1_pdu(pf02288) PDU1D EP_NTERM
00030 HMMpropdeh2_1_pdu(pf02287) PDU1D EP_NTERM
00031 HMM08841dhact_1_pdu(pf08841) PDU1D
00032 HMMpropdeh3_2_pdusml(pf02288) PDU1D
00033 H_pumpkin GRM3C
00034 H_royalBlue PDU1D
00035 HMMptac_3_eut2(pf06130) GRM3C EP_NTERM
00036 HMM06723EutJMreB_1_grm(pf06723) GRM3A
00037 HMM02441_04(#) PDU1D
00038 P_stone PDU1D
00039 HMM01923cobtr_1_pdu(pf01923pf03928) PDU1D
00040 HMMalddh_1_eutpdugrm(pf00171) GRM3C EP_NTERM
00041 HMMalddh_1_eutpdugrm(pf00171) GRM3C
00042 HMMPduS_1_all(pf01512pf13375pf10531) PDU1D
00043 -# -
00044 Ts_paleMagenta PDU1D
00045 -# -
00046 -pf00578pf00462 -
00047 Hp_euts_steel PDU1D
00048 HMM10662PduV_1_mix(pf10662) GRM5
00049 -pf01544 -
00050 -pf00395 -
00051 -pf00589 -
00052 HMMreg00072_4_pdu1d(pf02954pf00158) FRAG2
00053 -pf03606 -
00054 -pf01546 -
00055 -pf00491 -
00056 -pf00395 -
00057 -pf04389pf00395pf01841 -
BMC-P
BMC-H
BMC-T
Aldehyde dehydrogenase
PTAC
Signature enzyme
Alcohol dehydrogenase
Regulator

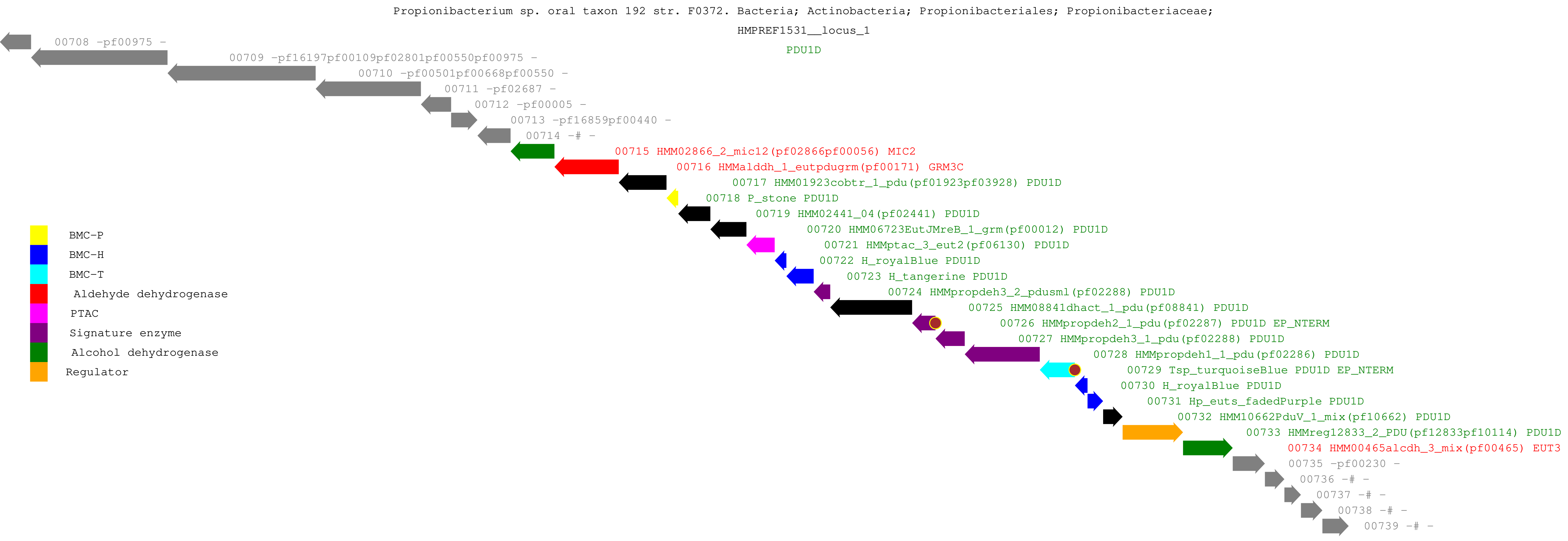
Propionibacterium sp. oral taxon 192 str. F0372. Bacteria; Actinobacteria; Propionibacteriales; Propionibacteriaceae;
HMPREF1531__locus_1
PDU1D
00708 -pf00975 -
00709 -pf16197pf00109pf02801pf00550pf00975 -
00710 -pf00501pf00668pf00550 -
00711 -pf02687 -
00712 -pf00005 -
00713 -pf16859pf00440 -
00714 -# -
00715 HMM02866_2_mic12(pf02866pf00056) MIC2
00716 HMMalddh_1_eutpdugrm(pf00171) GRM3C
00717 HMM01923cobtr_1_pdu(pf01923pf03928) PDU1D
00718 P_stone PDU1D
00719 HMM02441_04(pf02441) PDU1D
00720 HMM06723EutJMreB_1_grm(pf00012) PDU1D
00721 HMMptac_3_eut2(pf06130) PDU1D
00722 H_royalBlue PDU1D
00723 H_tangerine PDU1D
00724 HMMpropdeh3_2_pdusml(pf02288) PDU1D
00725 HMM08841dhact_1_pdu(pf08841) PDU1D
00726 HMMpropdeh2_1_pdu(pf02287) PDU1D EP_NTERM
00727 HMMpropdeh3_1_pdu(pf02288) PDU1D
00728 HMMpropdeh1_1_pdu(pf02286) PDU1D
00729 Tsp_turquoiseBlue PDU1D EP_NTERM
00730 H_royalBlue PDU1D
00731 Hp_euts_fadedPurple PDU1D
00732 HMM10662PduV_1_mix(pf10662) PDU1D
00733 HMMreg12833_2_PDU(pf12833pf10114) PDU1D
00734 HMM00465alcdh_3_mix(pf00465) EUT3
00735 -pf00230 -
00736 -# -
00737 -# -
00738 -# -
00739 -# -
BMC-P
BMC-H
BMC-T
Aldehyde dehydrogenase
PTAC
Signature enzyme
Alcohol dehydrogenase
Regulator

Tissierellia bacterium S5-A11. Bacteria; Firmicutes; Tissierellia.

HMPREF1633__locus_2

PDU1D

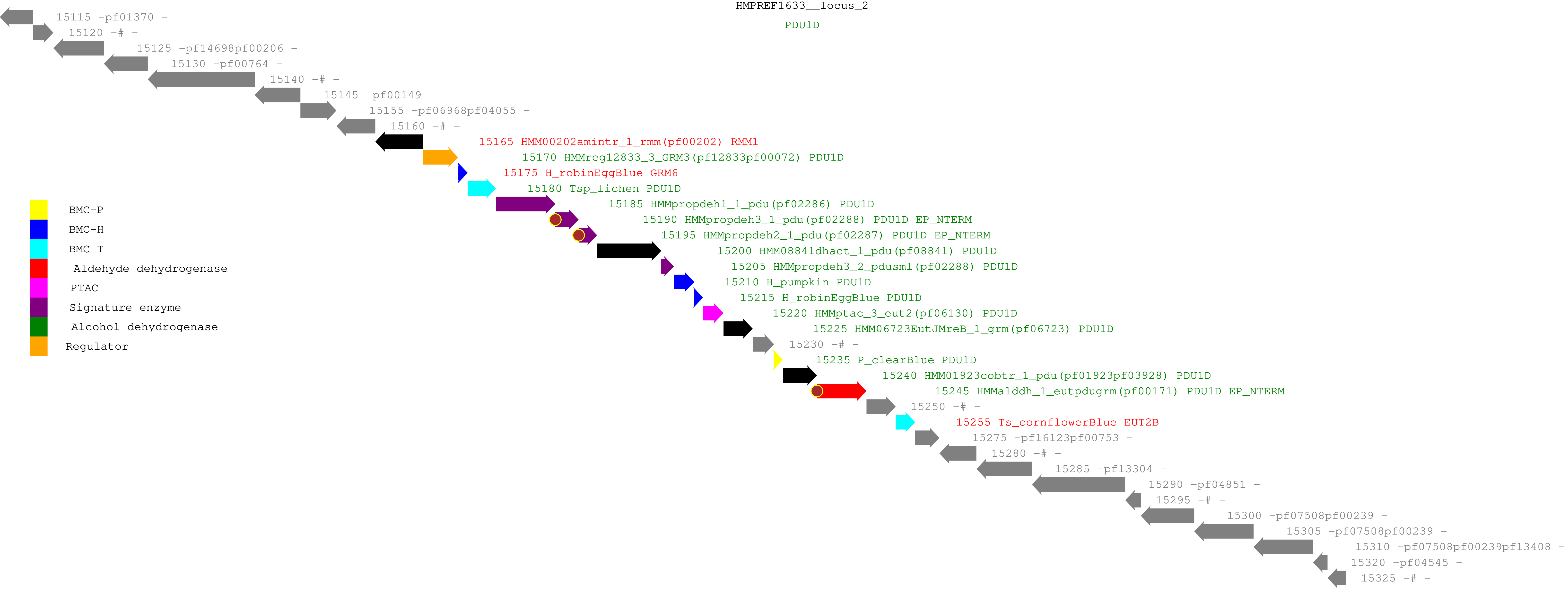

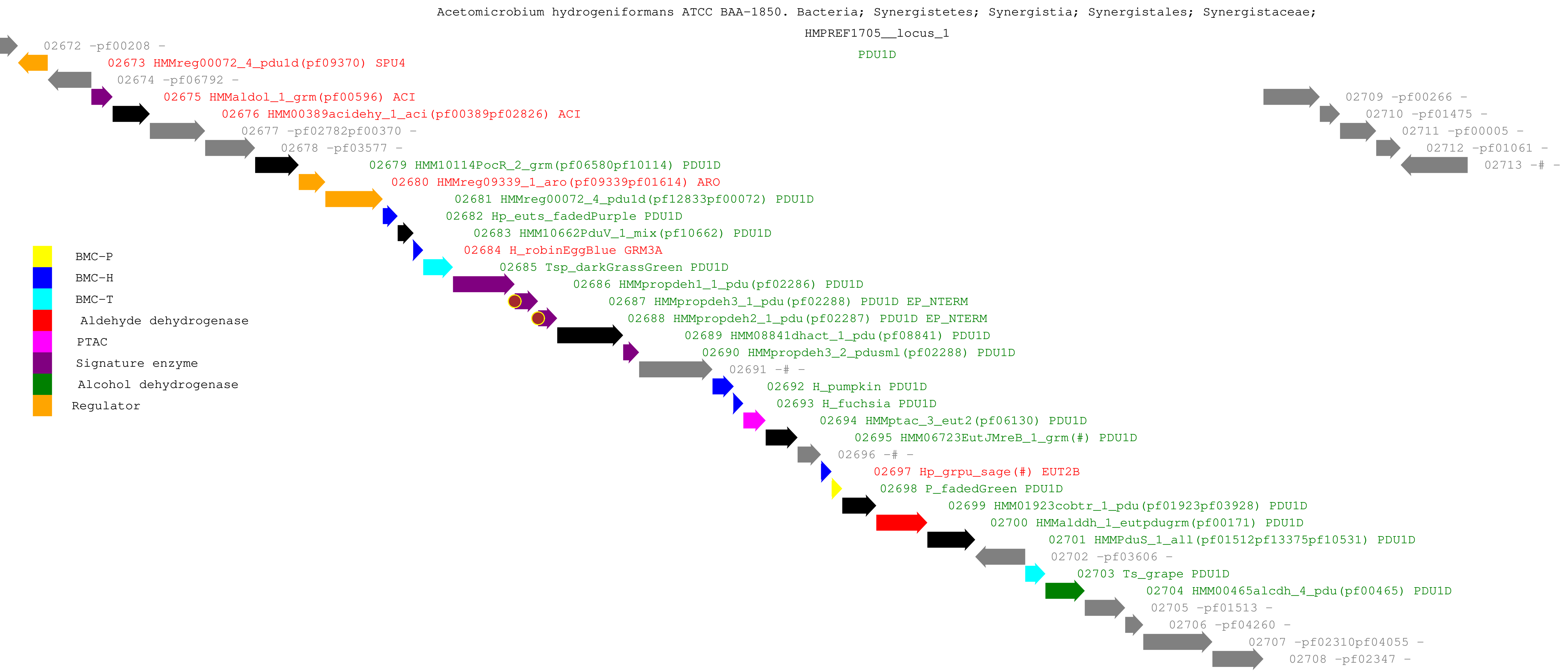
Acetomicrobium hydrogeniformans ATCC BAA-1850. Bacteria; Synergistetes; Synergistia; Synergistales; Synergistaceae;
HMPREF1705__locus_1
PDU1D
02672 -pf00208 -
02673 HMMreg00072_4_pdu1d(pf09370) SPU4
02674 -pf06792 -
02675 HMMaldol_1_grm(pf00596) ACI
02676 HMM00389acidehy_1_aci(pf00389pf02826) ACI
02677 -pf02782pf00370 -
02678 -pf03577 -
02679 HMM10114PocR_2_grm(pf06580pf10114) PDU1D
02680 HMMreg09339_1_aro(pf09339pf01614) ARO
02681 HMMreg00072_4_pdu1d(pf12833pf00072) PDU1D
02682 Hp_euts_fadedPurple PDU1D
02683 HMM10662PduV_1_mix(pf10662) PDU1D
02684 H_robinEggBlue GRM3A
02685 Tsp_darkGrassGreen PDU1D
02686 HMMpropdeh1_1_pdu(pf02286) PDU1D
02687 HMMpropdeh3_1_pdu(pf02288) PDU1D EP_NTERM
02688 HMMpropdeh2_1_pdu(pf02287) PDU1D EP_NTERM
02689 HMM08841dhact_1_pdu(pf08841) PDU1D
02690 HMMpropdeh3_2_pdusml(pf02288) PDU1D
02691 -# -
02692 H_pumpkin PDU1D
02693 H_fuchsia PDU1D
02694 HMMptac_3_eut2(pf06130) PDU1D
02695 HMM06723EutJMreB_1_grm(#) PDU1D
02696 -# -
02697 Hp_grpu_sage(#) EUT2B
02698 P_fadedGreen PDU1D
02699 HMM01923cobtr_1_pdu(pf01923pf03928) PDU1D
02700 HMMalddh_1_eutpdugrm(pf00171) PDU1D
02701 HMMPduS_1_all(pf01512pf13375pf10531) PDU1D
02702 -pf03606 -
02703 Ts_grape PDU1D
02704 HMM00465alcdh_4_pdu(pf00465) PDU1D
02705 -pf01513 -
02706 -pf04260 -
02707 -pf02310pf04055 -
02708 -pf02347 -
02709 -pf00266 -
02710 -pf01475 -
02711 -pf00005 -
02712 -pf01061 -
02713 -# -
BMC-P
BMC-H
BMC-T
Aldehyde dehydrogenase
PTAC
Signature enzyme
Alcohol dehydrogenase
Regulator

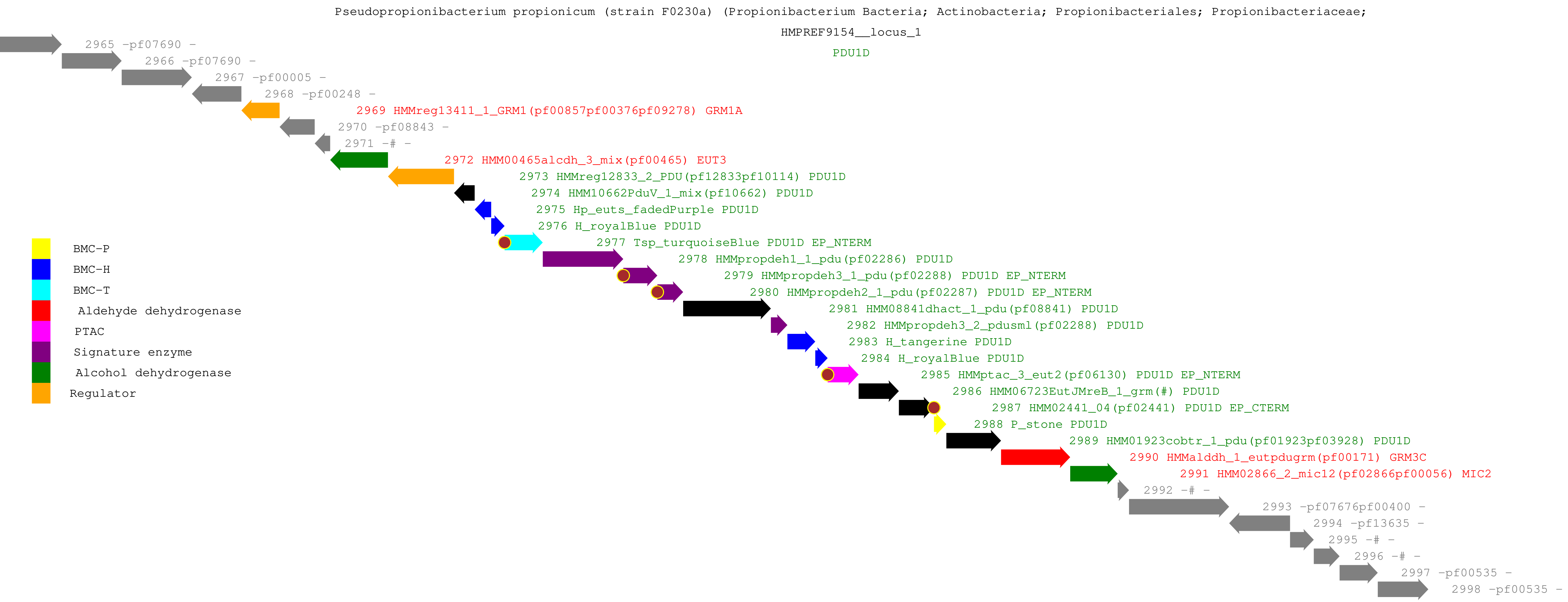
Pseudopropionibacterium propionicum (strain F0230a) (Propionibacterium Bacteria; Actinobacteria; Propionibacteriales; Propionibacteriaceae;
HMPREF9154__locus_1
PDU1D
2965 -pf07690 -
2966 -pf07690 -
2967 -pf00005 -
2968 -pf00248 -
2969 HMMreg13411_1_GRM1(pf00857pf00376pf09278) GRM1A
2970 -pf08843 -
2971 -# -
2972 HMM00465alcdh_3_mix(pf00465) EUT3
2973 HMMreg12833_2_PDU(pf12833pf10114) PDU1D
2974 HMM10662PduV_1_mix(pf10662) PDU1D
2975 Hp_euts_fadedPurple PDU1D
2976 H_royalBlue PDU1D
2977 Tsp_turquoiseBlue PDU1D EP_NTERM
2978 HMMpropdeh1_1_pdu(pf02286) PDU1D
2979 HMMpropdeh3_1_pdu(pf02288) PDU1D EP_NTERM
2980 HMMpropdeh2_1_pdu(pf02287) PDU1D EP_NTERM
2981 HMM08841dhact_1_pdu(pf08841) PDU1D
2982 HMMpropdeh3_2_pdusml(pf02288) PDU1D
2983 H_tangerine PDU1D
2984 H_royalBlue PDU1D
2985 HMMptac_3_eut2(pf06130) PDU1D EP_NTERM
2986 HMM06723EutJMreB_1_grm(#) PDU1D
2987 HMM02441_04(pf02441) PDU1D EP_CTERM
2988 P_stone PDU1D
2989 HMM01923cobtr_1_pdu(pf01923pf03928) PDU1D
2990 HMMaldh_1_eutpdugrm(pf00171) GRM3C
2991 HMM02866_2_mic12(pf02866pf00056) MIC2
2992 -# -
2993 -pf07676pf00400 -
2994 -pf13635 -
2995 -# -
2996 -# -
2997 -pf00535 -
2998 -pf00535 -
BMC-P
BMC-H
BMC-T
Aldehyde dehydrogenase
PTAC
Signature enzyme
Alcohol dehydrogenase
Regulator

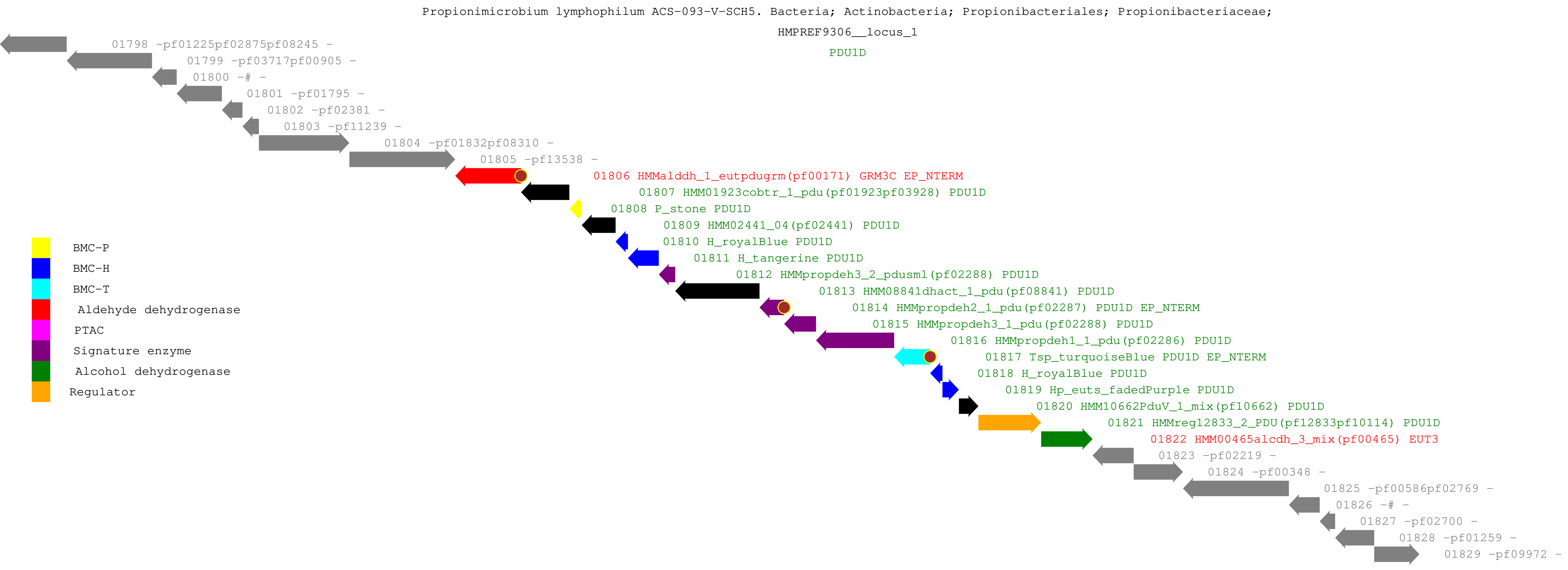
Propionimicrobium lymphophilum ACS-093-V-SCH5. Bacteria; Actinobacteria; Propionibacteriales; Propionibacteriaceae;
HMPREF9306__locus_1
PDU1D
01798 -pf01225pf02875pf08245 -
01799 -pf03717pf00905 -
01800 -# -
01801 -pf01795 -
01802 -pf02381 -
01803 -pf11239 -
01804 -pf01832pf08310 -
01805 -pf13538 -
01806 HMMalddh_1_eutpdugrm(pf00171) GRM3C EP_NTERM
01807 HMM01923cobtr_1_pdu(pf01923pf03928) PDU1D
01808 P_stone PDU1D
01809 HMM02441_04(pf02441) PDU1D
01810 H_royalBlue PDU1D
01811 H_tangerine PDU1D
01812 HMMpropdeh3_2_pdusml(pf02288) PDU1D
01813 HMM08841dhact_1_pdu(pf08841) PDU1D
01814 HMMpropdeh2_1_pdu(pf02287) PDU1D EP_NTERM
01815 HMMpropdeh3_1_pdu(pf02288) PDU1D
01816 HMMpropdeh1_1_pdu(pf02286) PDU1D
01817 Tsp_turquoiseBlue PDU1D EP_NTERM
01818 H_royalBlue PDU1D
01819 Hp_euts_fadedPurple PDU1D
01820 HMM10662PduV_1_mix(pf10662) PDU1D
01821 HMMreg12833_2_PDU(pf12833pf10114) PDU1D
01822 HMM00465alcdh_3_mix(pf00465) EUT3
01823 -pf02219 -
01824 -pf00348 -
01825 -pf00586pf02769 -
01826 -# -
01827 -pf02700 -
01828 -pf01259 -
01829 -pf09972 -
BMC-P
BMC-H
BMC-T
Aldehyde dehydrogenase
PTAC
Signature enzyme
Alcohol dehydrogenase
Regulator

Clostridium orbiscindens 1_3_50AFAA. Bacteria; Firmicutes; Clostridia; Clostridiales; Flavonifractor.

HMPREF9460__locus_1

PDU1D

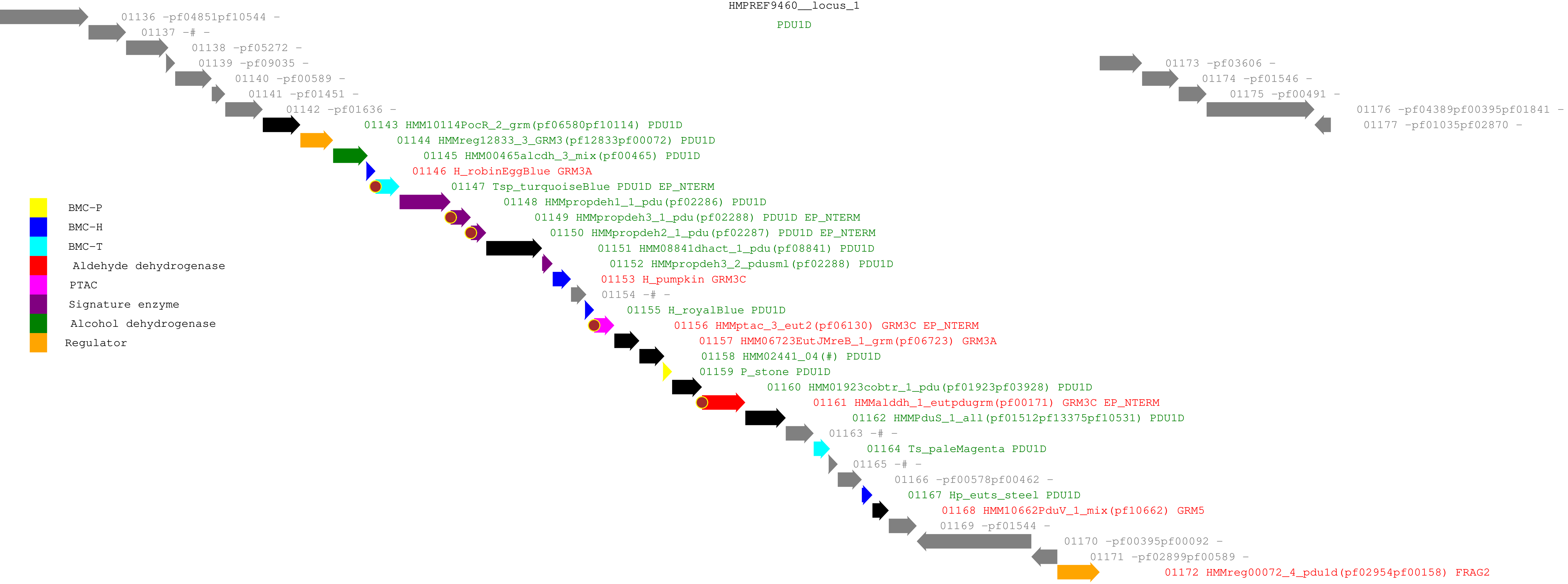

Halanaerobium saccharolyticum subsp. saccharolyticum DSM 6643. Bacteria; Firmicutes; Clostridia; Halanaerobiales; Halanaerobiaceae;

HSACCH__locus_1_main_sat

PDU1D

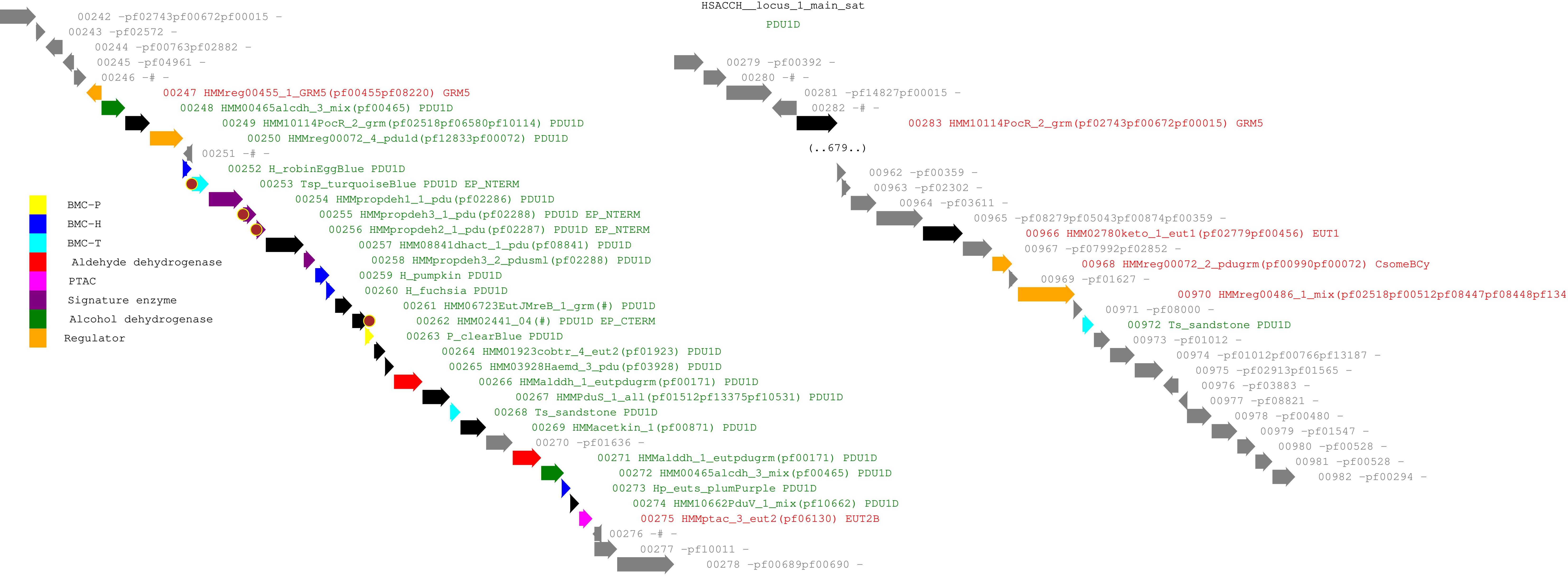

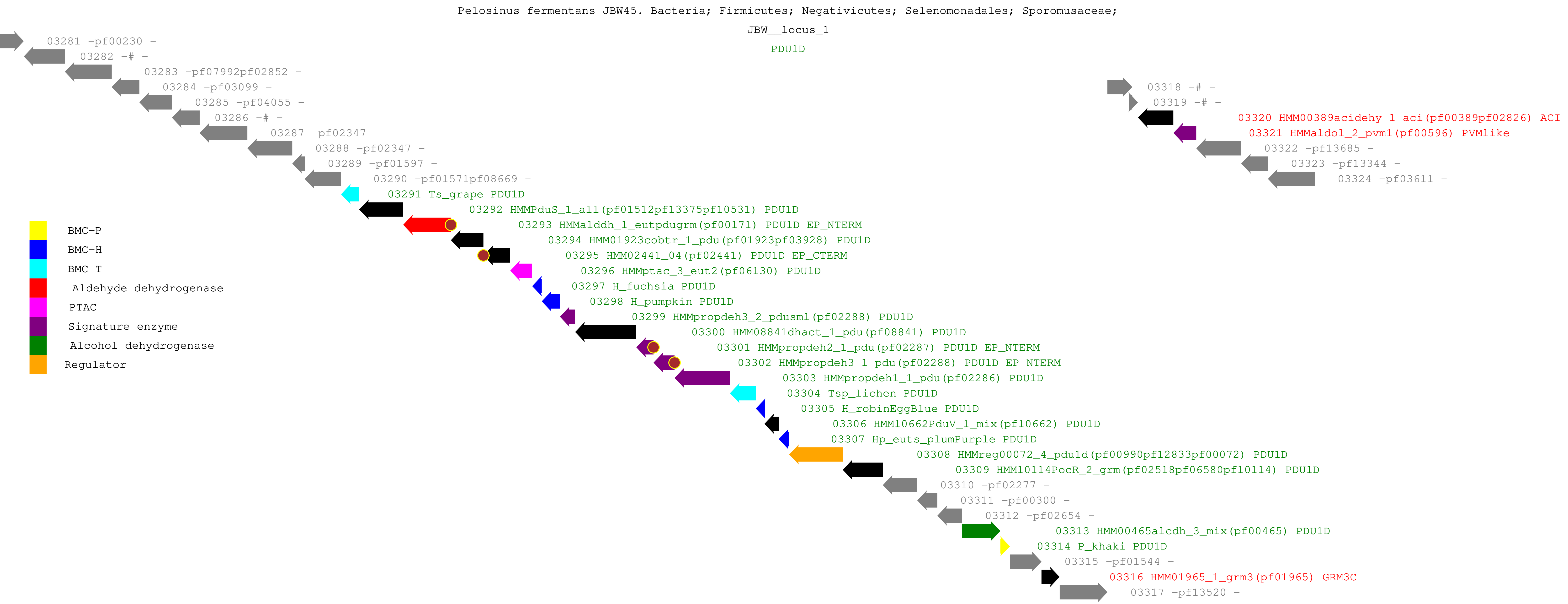
Pelosinus fermentans JBW45. Bacteria; Firmicutes; Negativicutes; Selenomonadales; Sporomusaceae;
JBW__locus_1
PDU1D
03281 -pf00230 -
03282 -# -
03283 -pf07992pf02852 -
03284 -pf03099 -
03285 -pf04055 -
03286 -# -
03287 -pf02347 -
03288 -pf02347 -
03289 -pf01597 -
03290 -pf01571pf08669 -
03291 Ts_grape PDU1D
03292 HMMPduS_1_all(pf01512pf13375pf10531) PDU1D
03293 HMMalddh_1_eutpdugrm(pf00171) PDU1D EP_NTERM
03294 HMM01923cobtr_1_pdu(pf01923pf03928) PDU1D
03295 HMM02441_04(pf02441) PDU1D EP_CTERM
03296 HMMptac_3_eut2(pf06130) PDU1D
03297 H_fuchsia PDU1D
03298 H_pumpkin PDU1D
03299 HMMpropdeh3_2_pdusml(pf02288) PDU1D
03300 HMM08841dhact_1_pdu(pf08841) PDU1D
03301 HMMpropdeh2_1_pdu(pf02287) PDU1D EP_NTERM
03302 HMMpropdeh3_1_pdu(pf02288) PDU1D EP_NTERM
03303 HMMpropdeh1_1_pdu(pf02286) PDU1D
03304 Tsp_lichen PDU1D
03305 H_robinEggBlue PDU1D
03306 HMM10662PduV_1_mix(pf10662) PDU1D
03307 Hp_euts_plumPurple PDU1D
03308 HMMreg00072_4_pdu1d(pf00990pf12833pf00072) PDU1D
03309 HMM10114PocR_2_grm(pf02518pf06580pf10114) PDU1D
03310 -pf02277 -
03311 -pf00300 -
03312 -pf02654 -
03313 HMM00465alcdh_3_mix(pf00465) PDU1D
03314 P_khaki PDU1D
03315 -pf01544 -
03316 HMM01965_1_grm3(pf01965) GRM3C
03317 -pf13520 -
03318 -# -
03319 -# -
03320 HMM00389acidehy_1_aci(pf00389pf02826) ACI
03321 HMMaldol_2_pvm1(pf00596) PVMlike
03322 -pf13685 -
03323 -pf13344 -
03324 -pf03611 -
BMC-P
BMC-H
BMC-T
Aldehyde dehydrogenase
PTAC
Signature enzyme
Alcohol dehydrogenase
Regulator

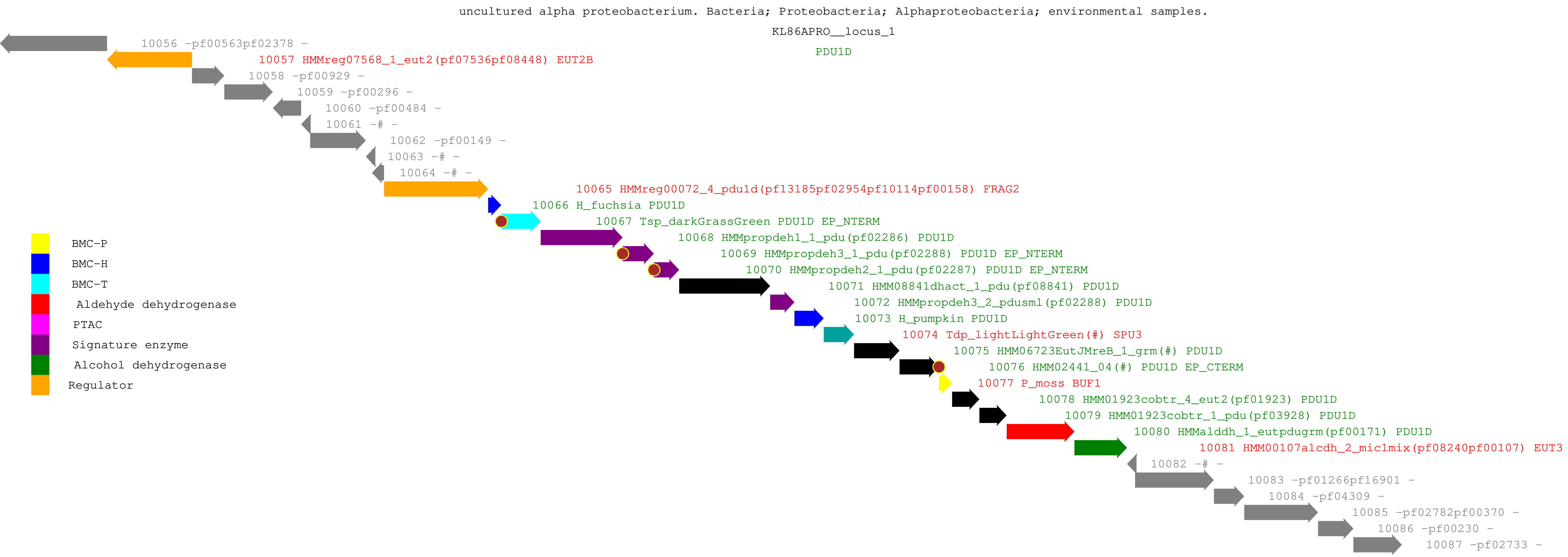
uncultured alpha proteobacterium. Bacteria; Proteobacteria; Alphaproteobacteria; environmental samples.
KL86APRO__locus_1
PDU1D
10056 -pf00563pf02378 -
10057 HMMreg07568_1_eut2(pf07536pf08448) EUT2B
10058 -pf00929 -
10059 -pf00296 -
10060 -pf00484 -
10061 -# -
10062 -pf00149 -
10063 -# -
10064 -# -
10065 HMMreg00072_4_pdu1d(pf13185pf02954pf10114pf00158) FRAG2
10066 H_fuchsia PDU1D
10067 Tsp_darkGrassGreen PDU1D EP_NTERM
10068 HMMpropdeh1_1_pdu(pf02286) PDU1D
10069 HMMpropdeh3_1_pdu(pf02288) PDU1D EP_NTERM
10070 HMMpropdeh2_1_pdu(pf02287) PDU1D EP_NTERM
10071 HMM08841dhact_1_pdu(pf08841) PDU1D
10072 HMMpropdeh3_2_pdusml(pf02288) PDU1D
10073 H_pumpkin PDU1D
10074 Tdp_lightLightGreen(#) SPU3
10075 HMM06723EutJMreB_1_grm(#) PDU1D
10076 HMM02441_04(#) PDU1D EP_CTERM
10077 P_moss BUF1
10078 HMM01923cobtr_4_eut2(pf01923) PDU1D
10079 HMM01923cobtr_1_pdu(pf03928) PDU1D
10080 HMMaldddh_1_eutpdugrm(pf00171) PDU1D
10081 HMM00107alcdh_2_mic1mix(pf08240pf00107) EUT3
10082 -# -
10083 -pf01266pf16901 -
10084 -pf04309 -
10085 -pf02782pf00370 -
10086 -pf00230 -
10087 -pf02733 -
BMC-P
BMC-H
BMC-T
Aldehyde dehydrogenase
PTAC
Signature enzyme
Alcohol dehydrogenase
Regulator

uncultured Sporomusa sp. Bacteria; Firmicutes; Negativicutes; Selenomonadales; Sporomusaceae;

KL86SPO__locus_1

PDU1D

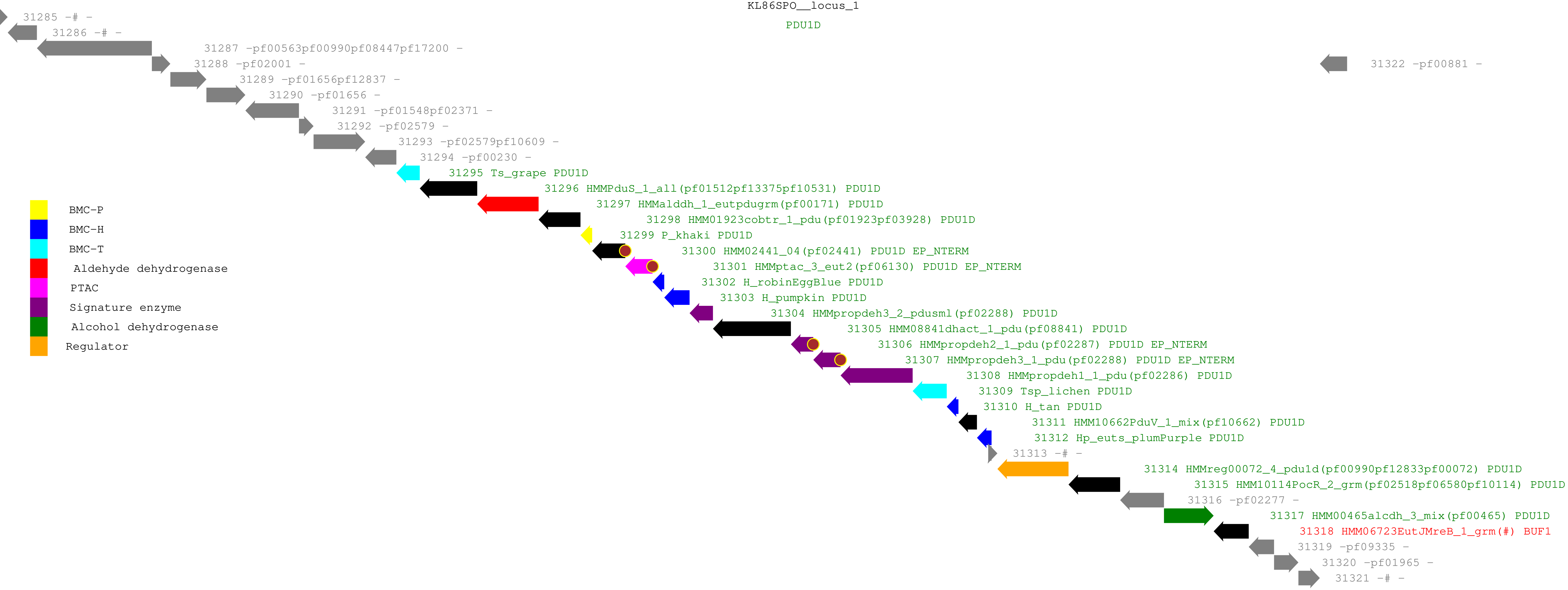

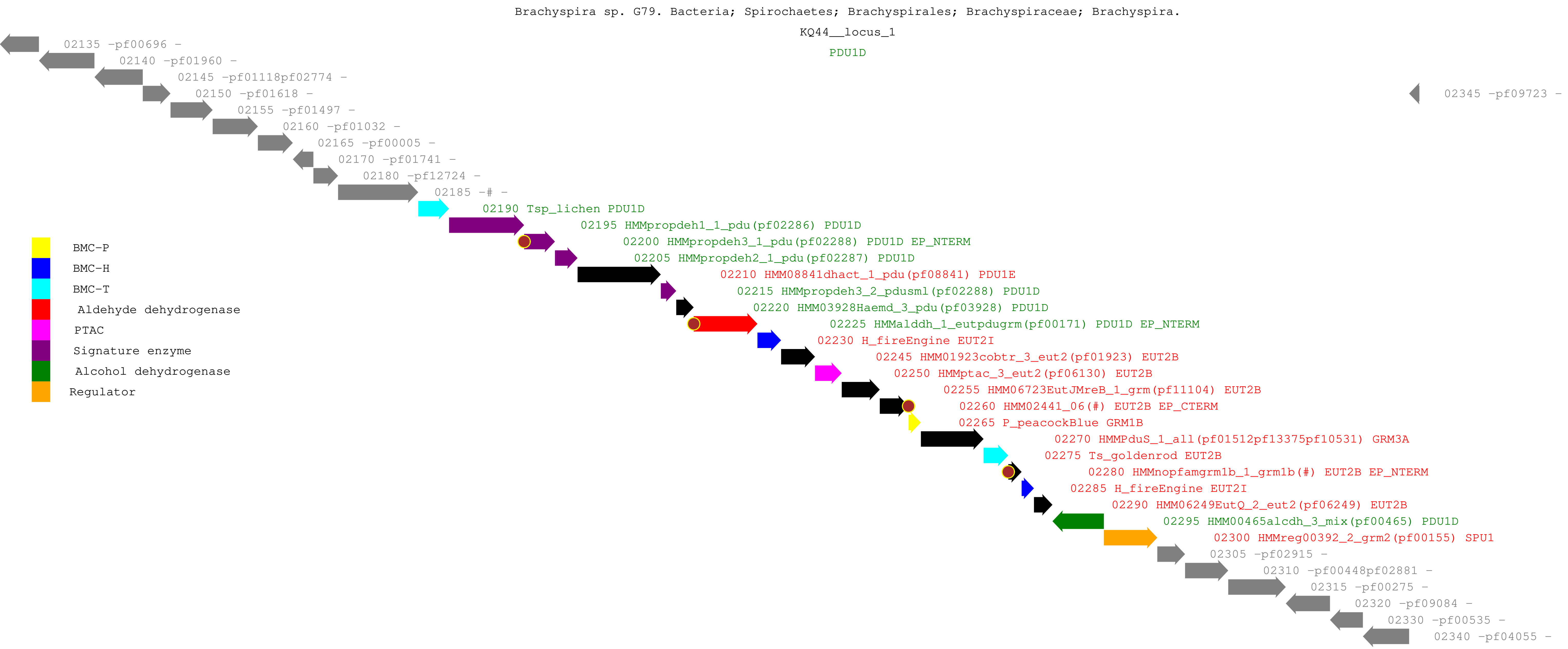

Brachyspira sp. G79. Bacteria; Spirochaetes; Brachyspirales; Brachyspiraceae; Brachyspira.
KQ44__locus_1
PDU1D
02135 -pf00696 -
02140 -pf01960 -
02145 -pf01118pf02774 -
02150 -pf01618 -
02155 -pf01497 -
02160 -pf01032 -
02165 -pf00005 -
02170 -pf01741 -
02180 -pf12724 -
02185 -# -
02190 Tsp_lichen PDU1D
02195 HMMpropdeh1_1_pdu(pf02286) PDU1D
02200 HMMpropdeh3_1_pdu(pf02288) PDU1D EP_NTERM
02205 HMMpropdeh2_1_pdu(pf02287) PDU1D
02210 HMM08841dhact_1_pdu(pf08841) PDU1E
02215 HMMpropdeh3_2_pdusml(pf02288) PDU1D
02220 HMM03928Haemd_3_pdu(pf03928) PDU1D
02225 HMMalddh_1_eutpdugrm(pf00171) PDU1D EP_NTERM
02230 H_fireEngine EUT2I
02245 HMM01923cobtr_3_eut2(pf01923) EUT2B
02250 HMMptac_3_eut2(pf06130) EUT2B
02255 HMM06723EutJMreB_1_grm(pf11104) EUT2B
02260 HMM02441_06(#) EUT2B EP_CTERM
02265 P_peacockBlue GRM1B
02270 HMMPduS_1_all(pf01512pf13375pf10531) GRM3A
02275 Ts_goldenrod EUT2B
02280 HMMnopfamgrm1b_1_grm1b(#) EUT2B EP_NTERM
02285 H_fireEngine EUT2I
02290 HMM06249EutQ_2_eut2(pf06249) EUT2B
02295 HMM00465alcdh_3_mix(pf00465) PDU1D
02300 HMMreg00392_2_grm2(pf00155) SPU1
02305 -pf02915 -
02310 -pf00448pf02881 -
02315 -pf00275 -
02320 -pf09084 -
02330 -pf00535 -
02340 -pf04055 -
02345 -pf09723 -
BMC-P
BMC-H
BMC-T
Aldehyde dehydrogenase
PTAC
Signature enzyme
Alcohol dehydrogenase
Regulator

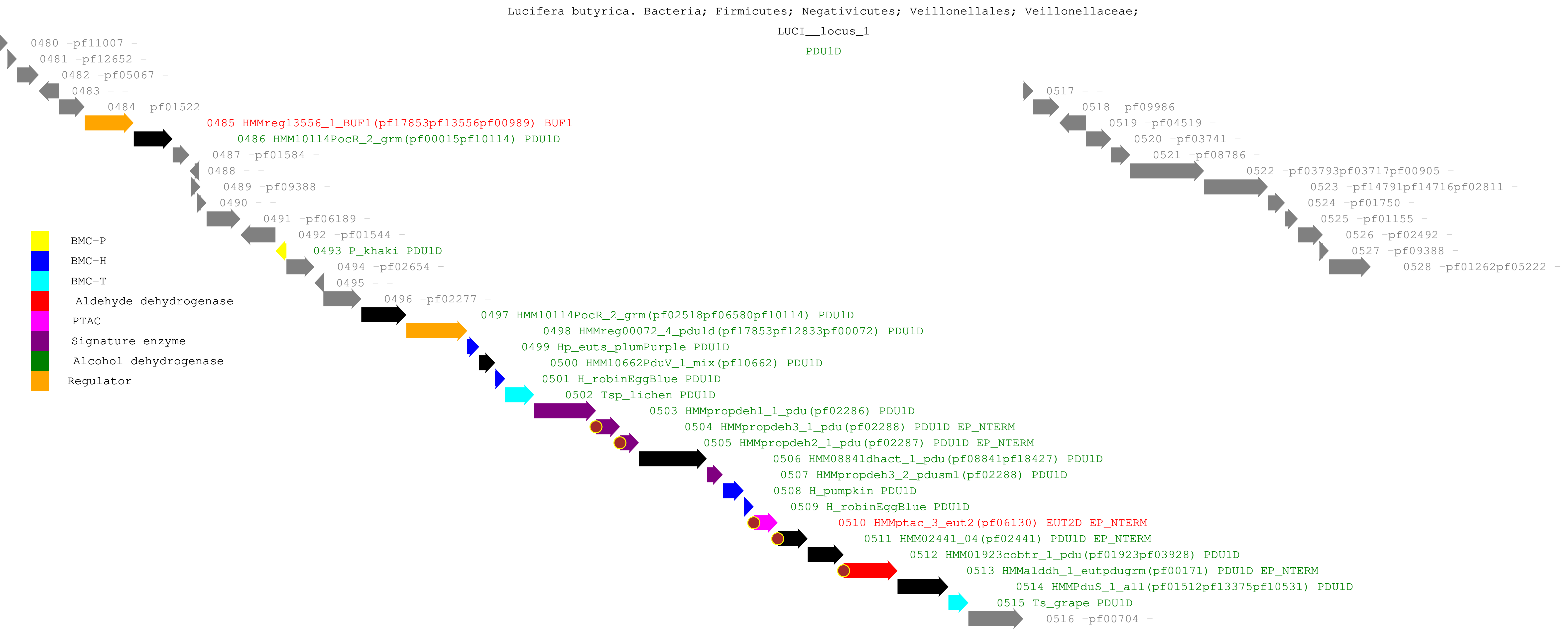
Lucifera butyrica. Bacteria; Firmicutes; Negativicutes; Veillonellales; Veillonellaceae;
LUCI__locus_1
PDU1D
0480 -pf11007 -
0481 -pf12652 -
0482 -pf05067 -
0483 - -
0484 -pf01522 -
0485 HMMreg13556_1_BUF1(pf17853pf13556pf00989) BUF1
0486 HMM10114PocR_2_grm(pf00015pf10114) PDU1D
0487 -pf01584 -
0488 - -
0489 -pf09388 -
0490 - -
0491 -pf06189 -
0492 -pf01544 -
0493 P_khaki PDU1D
0494 -pf02654 -
0495 - -
0496 -pf02277 -
0497 HMM10114PocR_2_grm(pf02518pf06580pf10114) PDU1D
0498 HMMreg00072_4_pdu1d(pf17853pf12833pf00072) PDU1D
0499 Hp_euts_plumPurple PDU1D
0500 HMM10662PduV_1_mix(pf10662) PDU1D
0501 H_robinEggBlue PDU1D
0502 Tsp_lichen PDU1D
0503 HMMpropdeh1_1_pdu(pf02286) PDU1D
0504 HMMpropdeh3_1_pdu(pf02288) PDU1D EP_NTERM
0505 HMMpropdeh2_1_pdu(pf02287) PDU1D EP_NTERM
0506 HMM08841dhact_1_pdu(pf08841pf18427) PDU1D
0507 HMMpropdeh3_2_pdusml(pf02288) PDU1D
0508 H_pumpkin PDU1D
0509 H_robinEggBlue PDU1D
0510 HMMptac_3_eut2(pf06130) EUT2D EP_NTERM
0511 HMM02441_04(pf02441) PDU1D EP_NTERM
0512 HMM01923cobtr_1_pdu(pf01923pf03928) PDU1D
0513 HMMalddh_1_eutpdugrm(pf00171) PDU1D EP_NTERM
0514 HMMPduS_1_all(pf01512pf13375pf10531) PDU1D
0515 Ts_grape PDU1D
0516 -pf00704 -
0517 - -
0518 -pf09986 -
0519 -pf04519 -
0520 -pf03741 -
0521 -pf08786 -
0522 -pf03793pf03717pf00905 -
0523 -pf14791pf14716pf02811 -
0524 -pf01750 -
0525 -pf01155 -
0526 -pf02492 -
0527 -pf09388 -
0528 -pf01262pf05222 -
BMC-P
BMC-H
BMC-T
Aldehyde dehydrogenase
PTAC
Signature enzyme
Alcohol dehydrogenase
Regulator

Bacillus azotoformans MEV2011. Bacteria; Firmicutes; Bacilli; Bacillales; Bacillaceae; Bacillus.

M670__locus_2

PDU1D

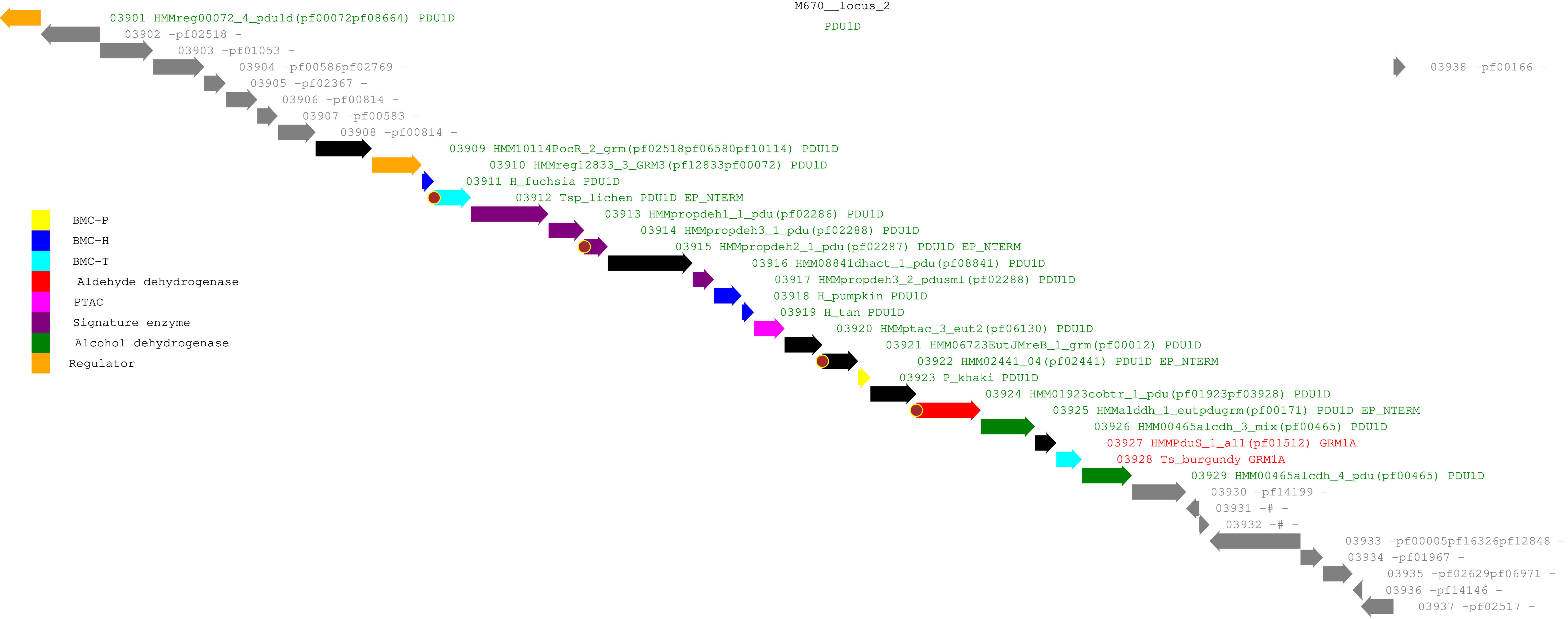

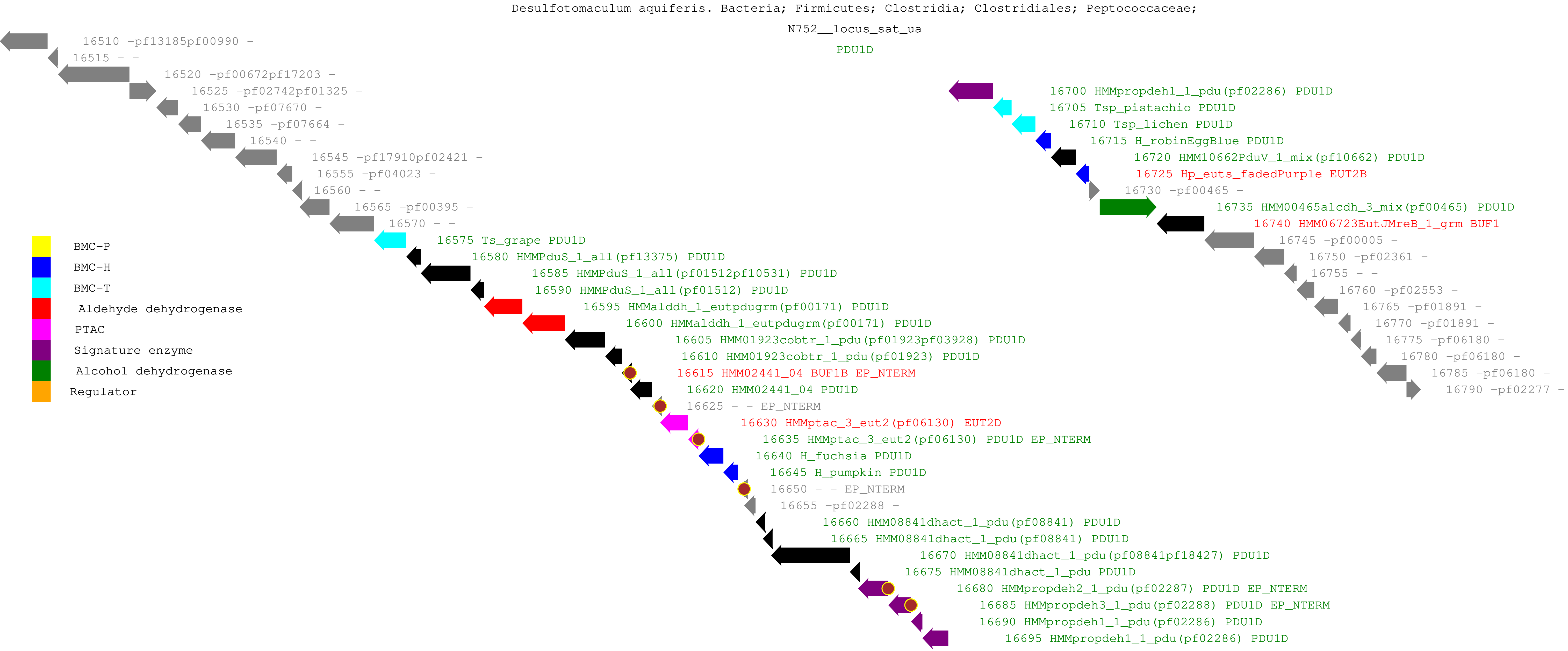
Desulfotomaculum aquiferis. Bacteria; Firmicutes; Clostridia; Clostridiales; Peptococcaceae;
N752__locus_sat_ua
PDU1D
16510 -pf13185pf00990 -
16515 - -
16520 -pf00672pf17203 -
16525 -pf02742pf01325 -
16530 -pf07670 -
16535 -pf07664 -
16540 - -
16545 -pf17910pf02421 -
16555 -pf04023 -
16560 - -
16565 -pf00395 -
16570 - -
16575 Ts_grape PDU1D
16580 HMMPduS_1_all(pf13375) PDU1D
16585 HMMPduS_1_all(pf01512pf10531) PDU1D
16590 HMMPduS_1_all(pf01512) PDU1D
16595 HMMaldd h_1_eutpdugrm(pf00171) PDU1D
16600 HMMaldd h_1_eutpdugrm(pf00171) PDU1D
16605 HMM01923cobtr_1_pdu(pf01923pf03928) PDU1D
16610 HMM01923cobtr_1_pdu(pf01923) PDU1D
16615 HMM02441_04 BUF1B EP_NTERM
16620 HMM02441_04 PDU1D
16625 - - EP_NTERM
16630 HMMptac_3_eut2(pf06130) EUT2D
16635 HMMptac_3_eut2(pf06130) PDU1D EP_NTERM
16640 H_fuchsia PDU1D
16645 H_pumpkin PDU1D
16650 - - EP_NTERM
16655 -pf02288 -
16660 HMM08841dhact_1_pdu(pf08841) PDU1D
16665 HMM08841dhact_1_pdu(pf08841) PDU1D
16670 HMM08841dhact_1_pdu(pf08841pf18427) PDU1D
16675 HMM08841dhact_1_pdu PDU1D
16680 HMMpropdeh2_1_pdu(pf02287) PDU1D EP_NTERM
16685 HMMpropdeh3_1_pdu(pf02288) PDU1D EP_NTERM
16690 HMMpropdeh1_1_pdu(pf02286) PDU1D
16695 HMMpropdeh1_1_pdu(pf02286) PDU1D
16700 HMMpropdeh1_1_pdu(pf02286) PDU1D
16705 Tsp_pistachio PDU1D
16710 Tsp_lichen PDU1D
16715 H_robinEggBlue PDU1D
16720 HMM10662PduV_1_mix(pf10662) PDU1D
16725 Hp_euts_fadedPurple EUT2B
16730 -pf00465 -
16735 HMM00465alcdh_3_mix(pf00465) PDU1D
16740 HMM06723EutJMreB_1_grm BUF1
16745 -pf00005 -
16750 -pf02361 -
16755 - -
16760 -pf02553 -
16765 -pf01891 -
16770 -pf01891 -
16775 -pf06180 -
16780 -pf06180 -
16785 -pf06180 -
16790 -pf02277 -
BMC-P
BMC-H
BMC-T
Aldehyde dehydrogenase
PTAC
Signature enzyme
Alcohol dehydrogenase
Regulator

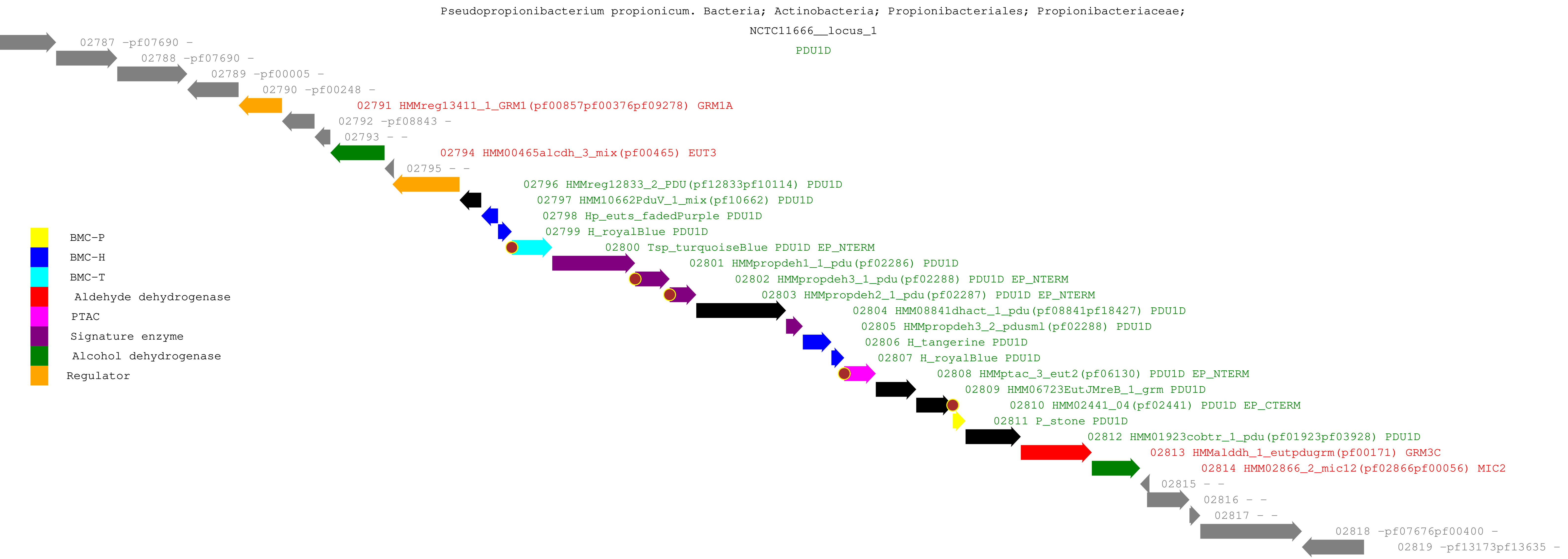

Pseudopropionibacterium propionicum. Bacteria; Actinobacteria; Propionibacteriales; Propionibacteriaceae;
NCTC11666__locus_1
PDU1D
02787 -pf07690 -
02788 -pf07690 -
02789 -pf00005 -
02790 -pf00248 -
02791 HMMreg13411_1_GRM1(pf00857pf00376pf09278) GRM1A
02792 -pf08843 -
02793 - -
02794 HMM00465alcdh_3_mix(pf00465) EUT3
02795 - -
02796 HMMreg12833_2_PDU(pf12833pf10114) PDU1D
02797 HMM10662PduV_1_mix(pf10662) PDU1D
02798 Hp_euts_fadedPurple PDU1D
02799 H_royalBlue PDU1D
02800 Tsp_turquoiseBlue PDU1D EP_NTERM
02801 HMMpropdeh1_1_pdu(pf02286) PDU1D
02802 HMMpropdeh3_1_pdu(pf02288) PDU1D EP_NTERM
02803 HMMpropdeh2_1_pdu(pf02287) PDU1D EP_NTERM
02804 HMM08841dhact_1_pdu(pf08841pf18427) PDU1D
02805 HMMpropdeh3_2_pdusml(pf02288) PDU1D
02806 H_tangerine PDU1D
02807 H_royalBlue PDU1D
02808 HMMptac_3_eut2(pf06130) PDU1D EP_NTERM
02809 HMM06723EutJMreB_1_grm PDU1D
02810 HMM02441_04(pf02441) PDU1D EP_CTERM
02811 P_stone PDU1D
02812 HMM01923cobtr_1_pdu(pf01923pf03928) PDU1D
02813 HMMalddh_1_eutpdugrm(pf00171) GRM3C
02814 HMM02866_2_mic12(pf02866pf00056) MIC2
02815 - -
02816 - -
02817 - -
02818 -pf07676pf00400 -
02819 -pf13173pf13635 -
BMC-P
BMC-H
BMC-T
Aldehyde dehydrogenase
PTAC
Signature enzyme
Alcohol dehydrogenase
Regulator

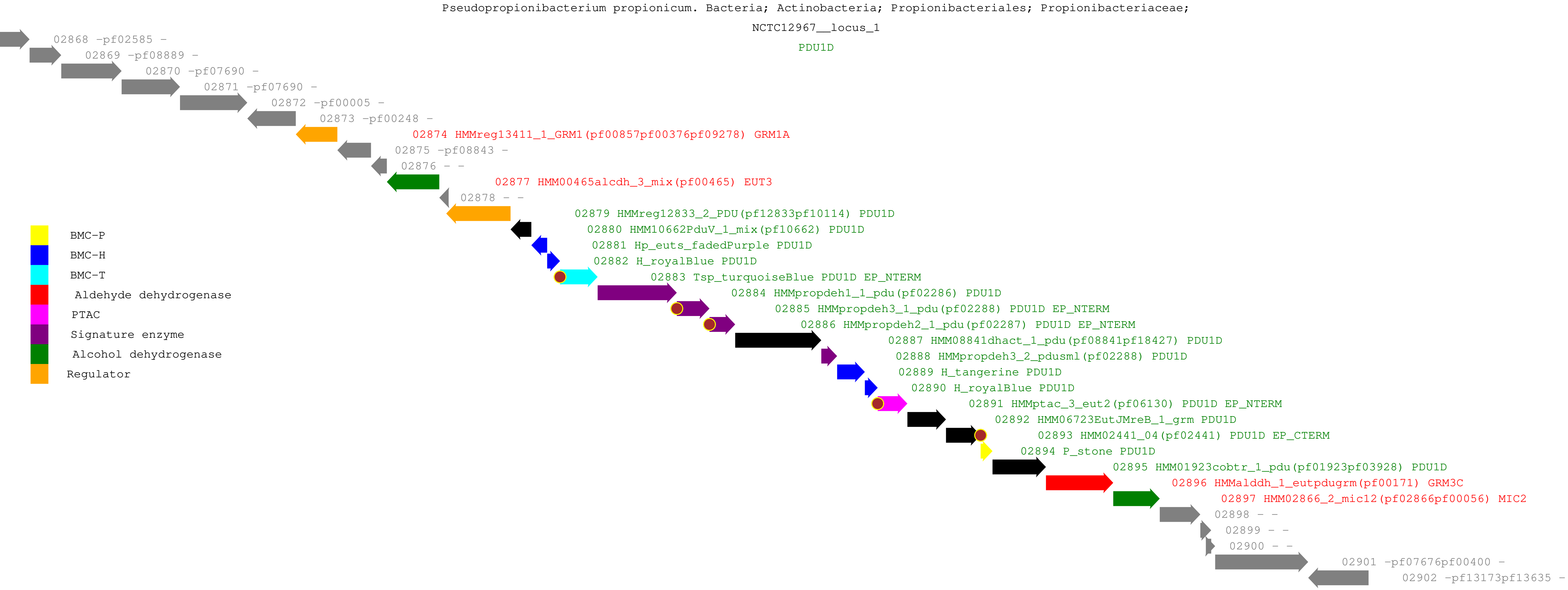
Pseudopropionibacterium propionicum. Bacteria; Actinobacteria; Propionibacteriales; Propionibacteriaceae;
NCTC12967__locus_1
PDU1D
02868 -pf02585 -
02869 -pf08889 -
02870 -pf07690 -
02871 -pf07690 -
02872 -pf00005 -
02873 -pf00248 -
02874 HMMreg13411_1_GRM1(pf00857pf00376pf09278) GRM1A
02875 -pf08843 -
02876 - -
02877 HMM00465alcdh_3_mix(pf00465) EUT3
02878 - -
02879 HMMreg12833_2_PDU(pf12833pf10114) PDU1D
02880 HMM10662PduV_1_mix(pf10662) PDU1D
02881 Hp_euts_fadedPurple PDU1D
02882 H_royalBlue PDU1D
02883 Tsp_turquoiseBlue PDU1D EP_NTERM
02884 HMMpropdeh1_1_pdu(pf02286) PDU1D
02885 HMMpropdeh3_1_pdu(pf02288) PDU1D EP_NTERM
02886 HMMpropdeh2_1_pdu(pf02287) PDU1D EP_NTERM
02887 HMM08841dhact_1_pdu(pf08841pf18427) PDU1D
02888 HMMpropdeh3_2_pdusml(pf02288) PDU1D
02889 H_tangerine PDU1D
02890 H_royalBlue PDU1D
02891 HMMptac_3_eut2(pf06130) PDU1D EP_NTERM
02892 HMM06723EutJMreB_1_grm PDU1D
02893 HMM02441_04(pf02441) PDU1D EP_CTERM
02894 P_stone PDU1D
02895 HMM01923cobtr_1_pdu(pf01923pf03928) PDU1D
02896 HMMalddh_1_eutpdugrm(pf00171) GRM3C
02897 HMM02866_2_mic12(pf02866pf00056) MIC2
02898 - -
02899 - -
02900 - -
02901 -pf07676pf00400 -
02902 -pf13173pf13635 -
BMC-P
BMC-H
BMC-T
Aldehyde dehydrogenase
PTAC
Signature enzyme
Alcohol dehydrogenase
Regulator

Anaerococcus prevotii. Bacteria; Firmicutes; Tissierellia; Tissierellales; Peptoniphilaceae;

NCTC13150__locus_1

PDU1D

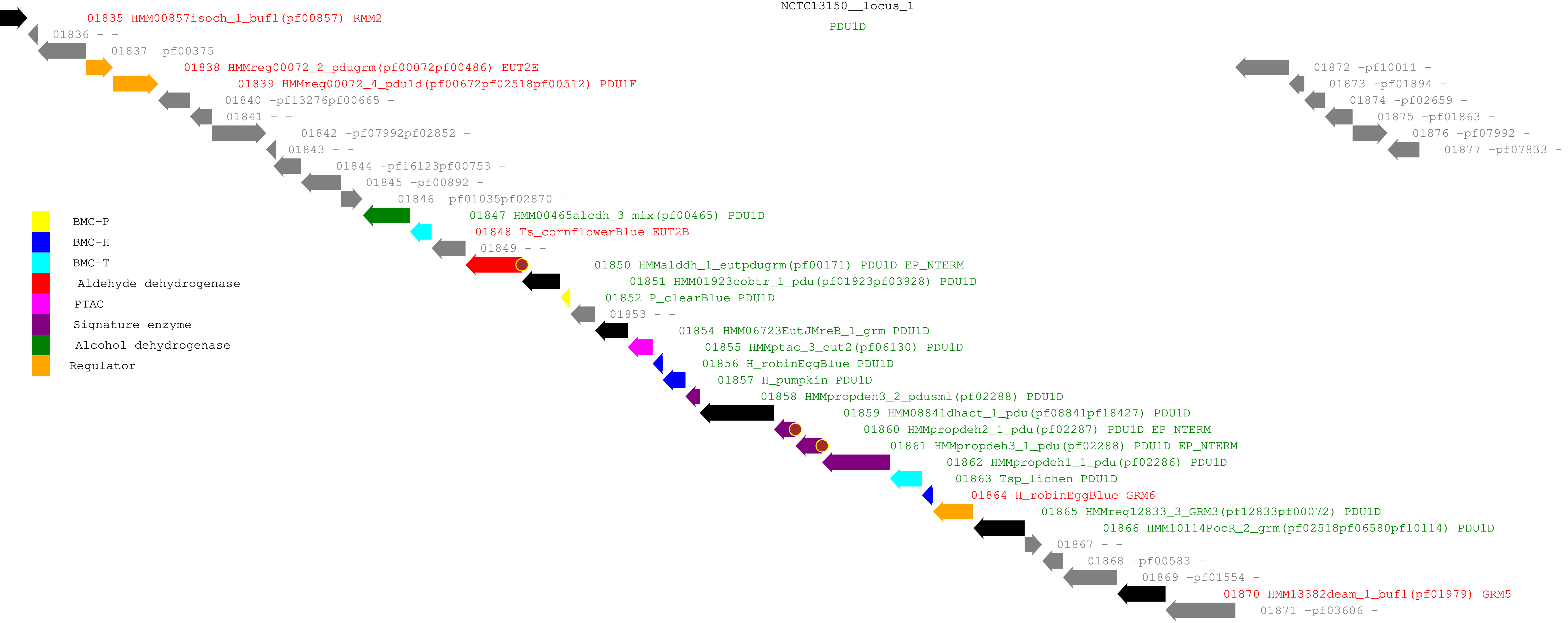

Propionispora sp. 2/2-37. Bacteria; Firmicutes; Negativicutes; Selenomonadales; Sporomusaceae;

P22__locus_1

PDU1D

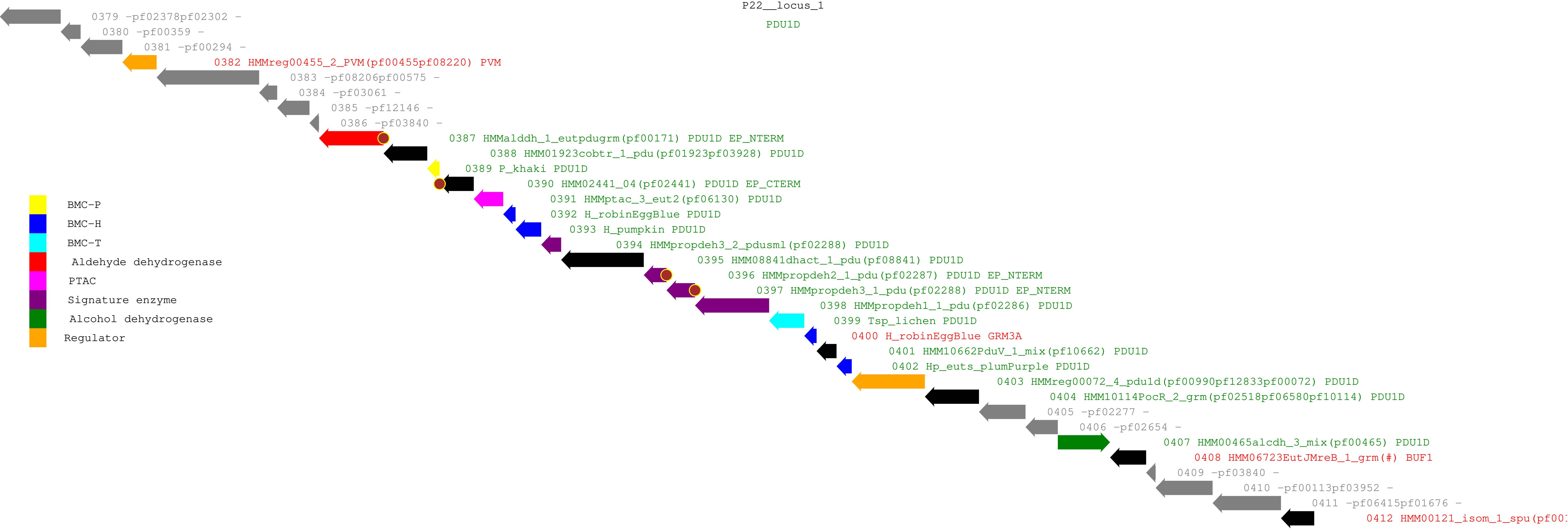

Propionibacterium freudenreichii. Bacteria; Actinobacteria; Propionibacteriales; Propionibacteriaceae;

PFCIRM512__locus_1

PDU1D

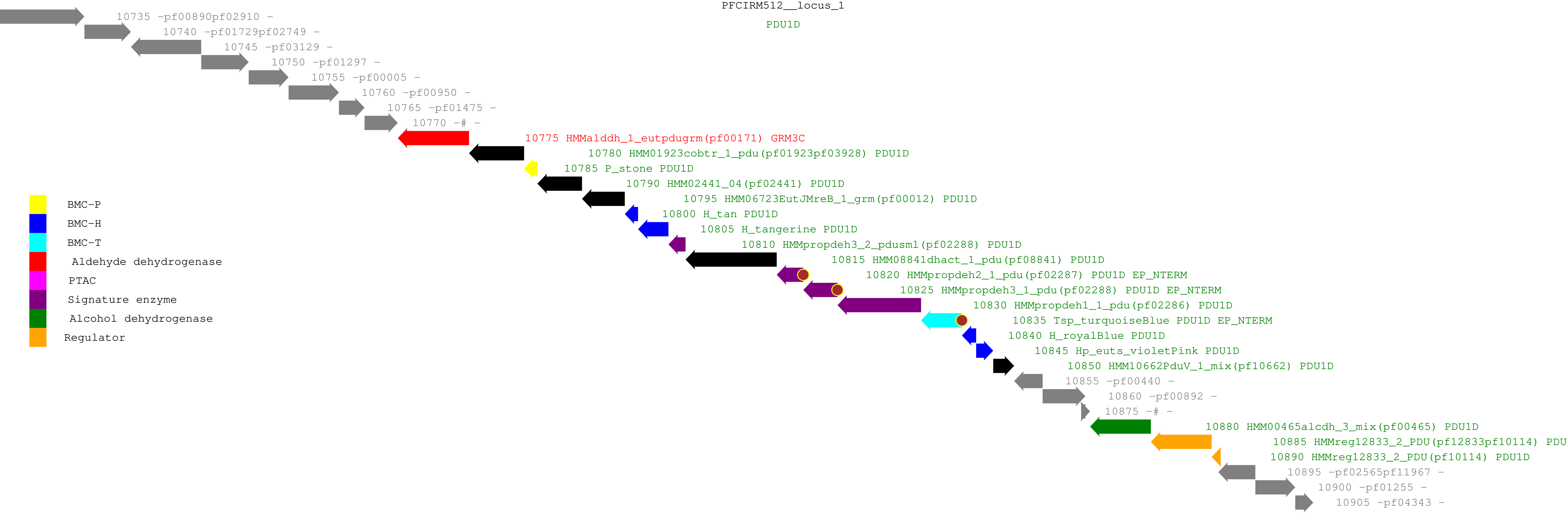

Propionibacterium freudenreichii subsp. shermanii. Bacteria; Actinobacteria; Propionibacteriales; Propionibacteriaceae;

PFREUDJS001__locus_1

PDU1D

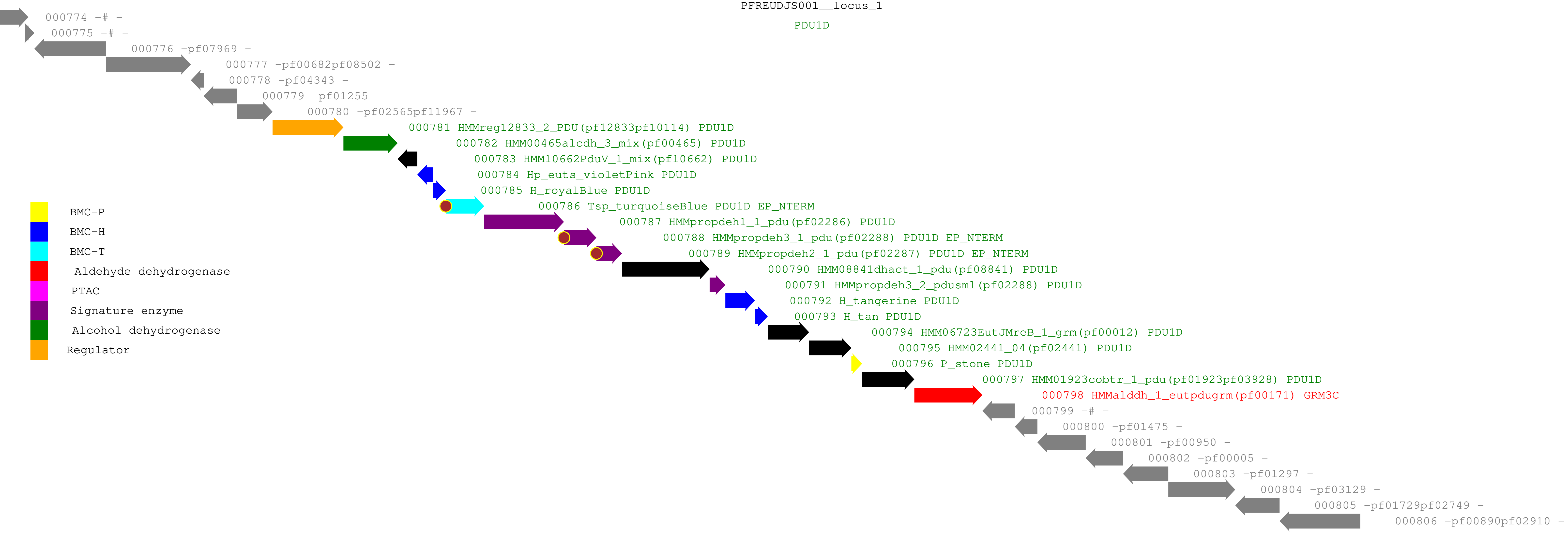

Propionibacterium freudenreichii subsp. shermanii (strain ATCC 9614 / Bacteria; Actinobacteria; Propionibacteriales; Propionibacteriaceae;

PFREUD__locus_1

PDU1D

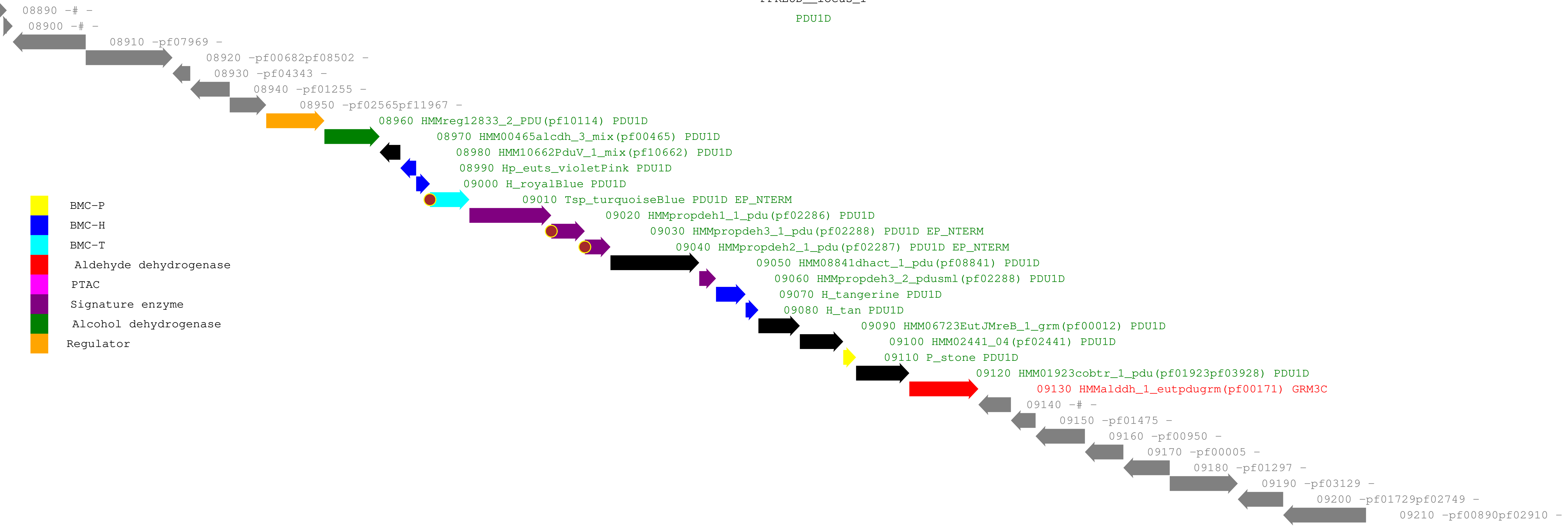

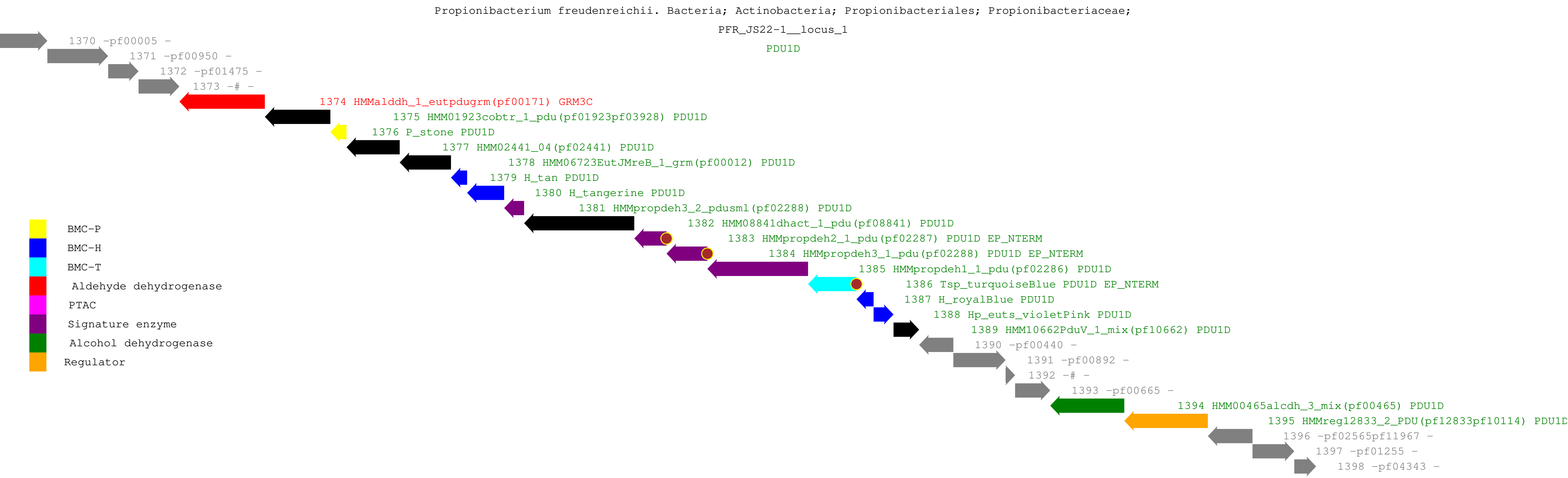
Propionibacterium freudenreichii. Bacteria; Actinobacteria; Propionibacteriales; Propionibacteriaceae;
PFR_JS22-1__locus_1
PDU1D
1370 -pf00005 -
1371 -pf00950 -
1372 -pf01475 -
1373 -# -
1374 HMMalddh_1_eutpdugrm(pf00171) GRM3C
1375 HMM01923cobtr_1_pdu(pf01923pf03928) PDU1D
1376 P_stone PDU1D
1377 HMM02441_04(pf02441) PDU1D
1378 HMM06723EutJMreB_1_grm(pf00012) PDU1D
1379 H_tan PDU1D
1380 H_tangerine PDU1D
1381 HMMpropdeh3_2_pdusml(pf02288) PDU1D
1382 HMM08841dhact_1_pdu(pf08841) PDU1D
1383 HMMpropdeh2_1_pdu(pf02287) PDU1D EP_NTERM
1384 HMMpropdeh3_1_pdu(pf02288) PDU1D EP_NTERM
1385 HMMpropdeh1_1_pdu(pf02286) PDU1D
1386 Tsp_turquoiseBlue PDU1D EP_NTERM
1387 H_royalBlue PDU1D
1388 Hp_euts_violetPink PDU1D
1389 HMM10662PduV_1_mix(pf10662) PDU1D
1390 -pf00440 -
1391 -pf00892 -
1392 -# -
1393 -pf00665 -
1394 HMM00465alcdh_3_mix(pf00465) PDU1D
1395 HMMreg12833_2_PDU(pf12833pf10114) PDU1D
1396 -pf02565pf11967 -
1397 -pf01255 -
1398 -pf04343 -
BMC-P
BMC-H
BMC-T
Aldehyde dehydrogenase
PTAC
Signature enzyme
Alcohol dehydrogenase
Regulator

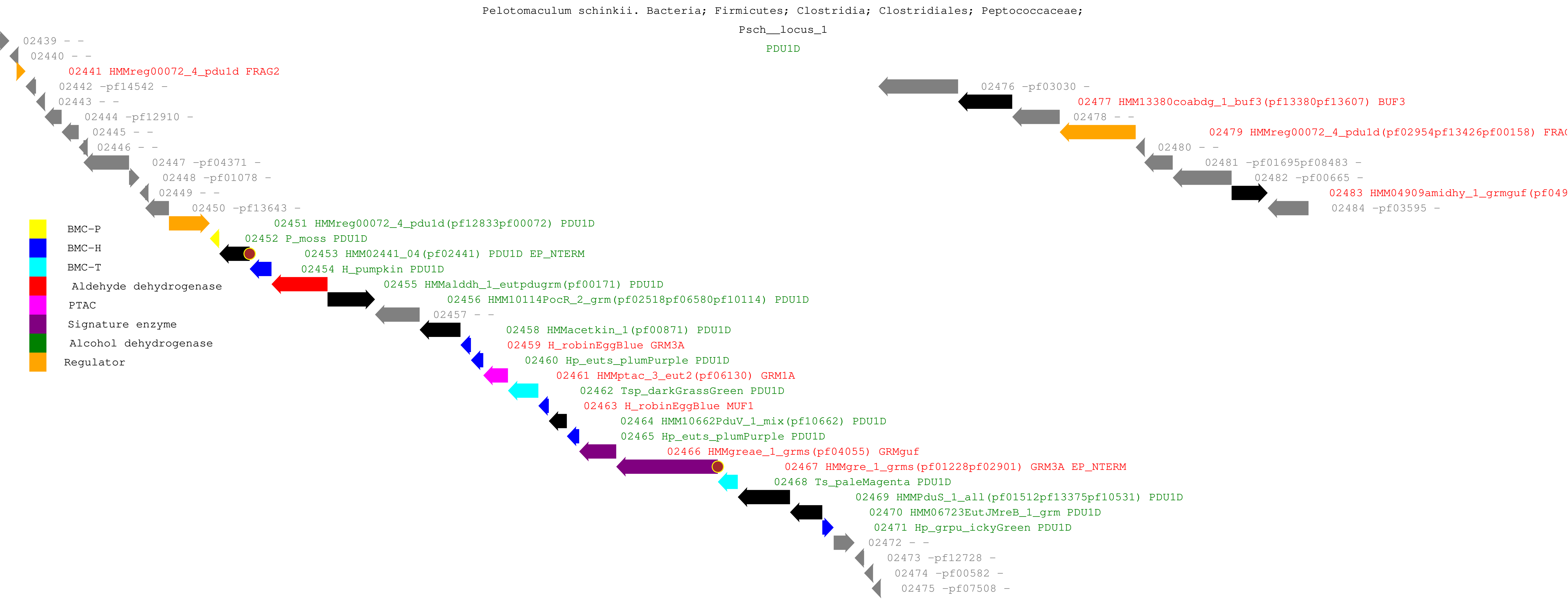

Pelotomaculum schinkii. Bacteria; Firmicutes; Clostridia; Clostridiales; Peptococcaceae;
Psch__locus_1
PDU1D
02439 - -
02440 - -
02441 HMMreg00072_4_pdu1d FRAG2
02442 -pf14542 -
02443 - -
02444 -pf12910 -
02445 - -
02446 - -
02447 -pf04371 -
02448 -pf01078 -
02449 - -
02450 -pf13643 -
02451 HMMreg00072_4_pdu1d(pf12833pf00072) PDU1D
02452 P_moss PDU1D
02453 HMM02441_04(pf02441) PDU1D EP_NTERM
02454 H_pumpkin PDU1D
02455 HMMalddh_1_eutpdugrm(pf00171) PDU1D
02456 HMM10114PocR_2_grm(pf02518pf06580pf10114) PDU1D
02457 - -
02458 HMMacetkin_1(pf00871) PDU1D
02459 H_robinEggBlue GRM3A
02460 Hp_euts_plumPurple PDU1D
02461 HMMptac_3_eut2(pf06130) GRM1A
02462 Tsp_darkGrassGreen PDU1D
02463 H_robinEggBlue MUF1
02464 HMM10662PduV_1_mix(pf10662) PDU1D
02465 Hp_euts_plumPurple PDU1D
02466 HMMgreae_1_grms(pf04055) GRMguf
02467 HMMgre_1_grms(pf01228pf02901) GRM3A EP_NTERM
02468 Ts_paleMagenta PDU1D
02469 HMMPduS_1_all(pf01512pf13375pf10531) PDU1D
02470 HMM06723EutJMreB_1_grm PDU1D
02471 Hp_grpu_ickyGreen PDU1D
02472 - -
02473 -pf12728 -
02474 -pf00582 -
02475 -pf07508 -
02476 -pf03030 -
02477 HMM13380coabdg_1_buf3(pf13380pf13607) BUF3
02478 - -
02479 HMMreg00072_4_pdu1d(pf02954pf13426pf00158) FRA
02480 - -
02481 -pf01695pf08483 -
02482 -pf00665 -
02483 HMM04909amidhy_1_grmguf(pf049
02484 -pf03595 -
BMC-P
BMC-H
BMC-T
Aldehyde dehydrogenase
PTAC
Signature enzyme
Alcohol dehydrogenase
Regulator

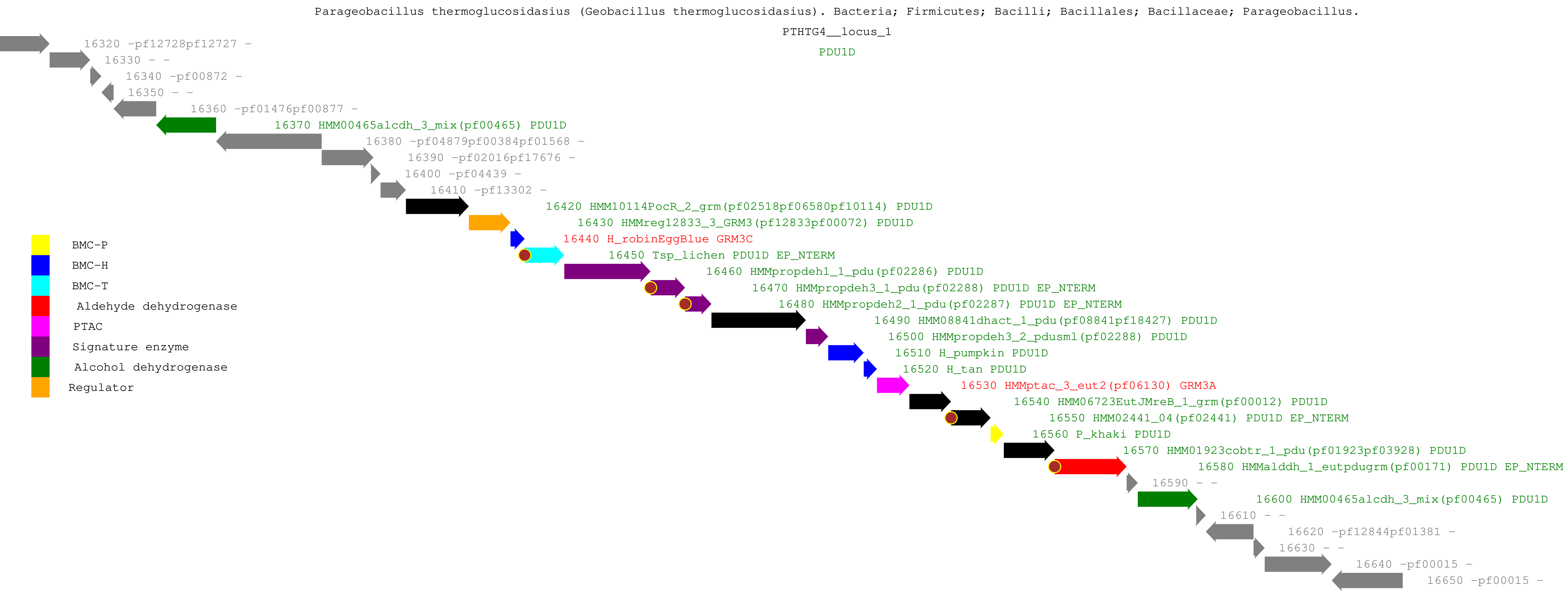
Parageobacillus thermoglucosidasius (Geobacillus thermoglucosidasius). Bacteria; Firmicutes; Bacilli; Bacillales; Bacillaceae; Parageobacillus.
PTHTG4__locus_1
PDU1D
16320 -pf12728pf12727 -
16330 - -
16340 -pf00872 -
16350 - -
16360 -pf01476pf00877 -
16370 HMM00465alcdh_3_mix(pf00465) PDU1D
16380 -pf04879pf00384pf01568 -
16390 -pf02016pf17676 -
16400 -pf04439 -
16410 -pf13302 -
16420 HMM10114PocR_2_grm(pf02518pf06580pf10114) PDU1D
16430 HMMreg12833_3_GRM3(pf12833pf00072) PDU1D
16440 H_robinEggBlue GRM3C
16450 Tsp_lichen PDU1D EP_NTERM
16460 HMMpropdeh1_1_pdu(pf02286) PDU1D
16470 HMMpropdeh3_1_pdu(pf02288) PDU1D EP_NTERM
16480 HMMpropdeh2_1_pdu(pf02287) PDU1D EP_NTERM
16490 HMM08841dhact_1_pdu(pf08841pf18427) PDU1D
16500 HMMpropdeh3_2_pdusml(pf02288) PDU1D
16510 H_pumpkin PDU1D
16520 H_tan PDU1D
16530 HMMptac_3_eut2(pf06130) GRM3A
16540 HMM06723EutJMreB_1_grm(pf00012) PDU1D
16550 HMM02441_04(pf02441) PDU1D EP_NTERM
16560 P_khaki PDU1D
16570 HMM01923cobtr_1_pdu(pf01923pf03928) PDU1D
16580 HMMaldh_1_eutpdugrm(pf00171) PDU1D EP_NTERM
16590 - -
16600 HMM00465alcdh_3_mix(pf00465) PDU1D
16610 - -
16620 -pf12844pf01381 -
16630 - -
16640 -pf00015 -
16650 -pf00015 -
BMC-P
BMC-H
BMC-T
Aldehyde dehydrogenase
PTAC
Signature enzyme
Alcohol dehydrogenase
Regulator

Bacillaceae bacterium MTCC 8252. Bacteria; Firmicutes; Bacilli; Bacillales; Bacillaceae;

QY95__locus_1

PDU1D

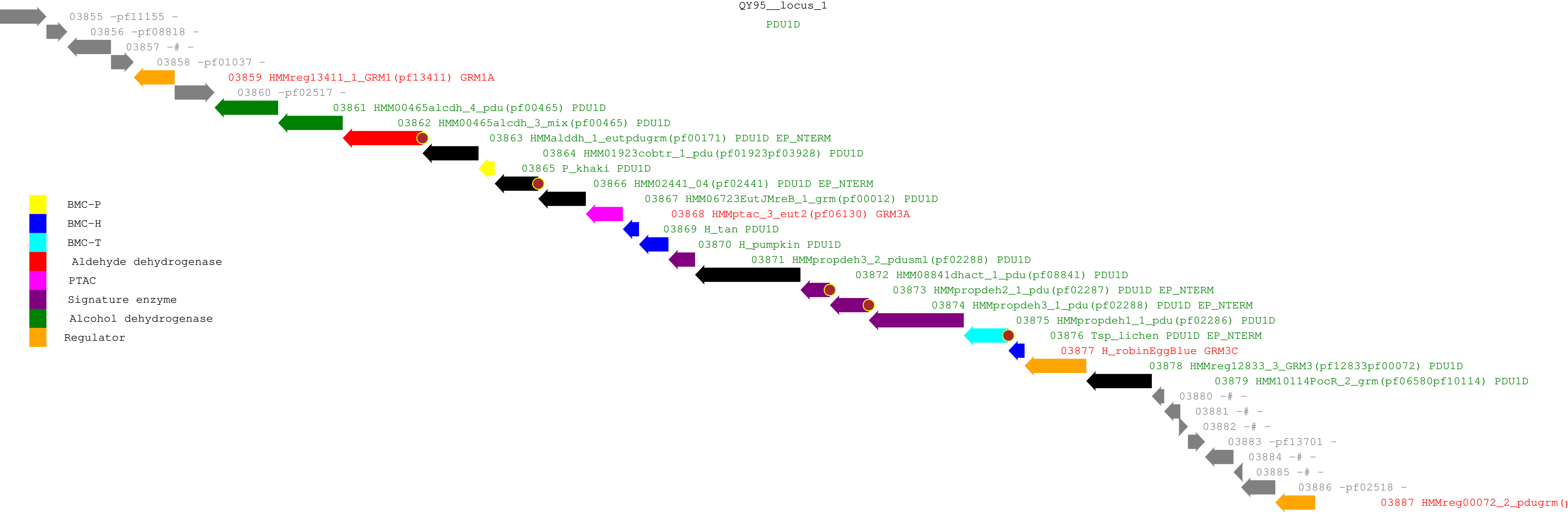

Bacillus thermotolerans. Bacteria; Firmicutes; Bacilli; Bacillales; Bacillaceae; Bacillus.

QY97__locus_1

PDU1D

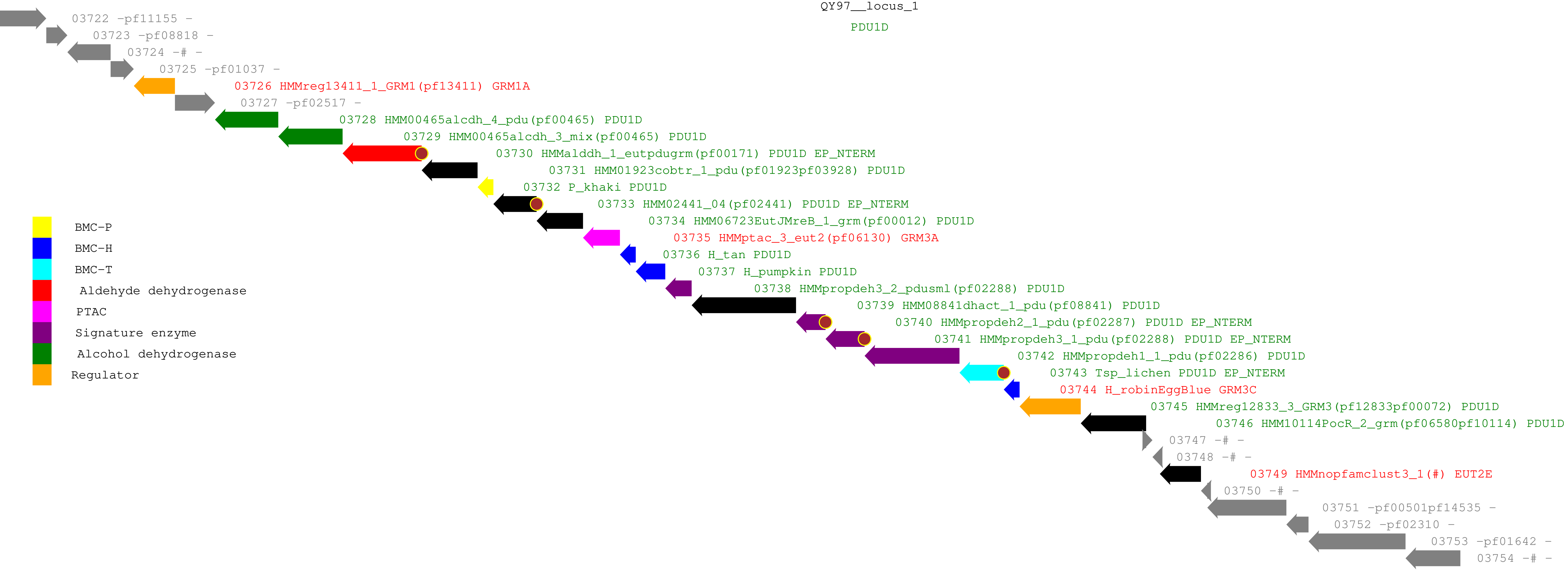

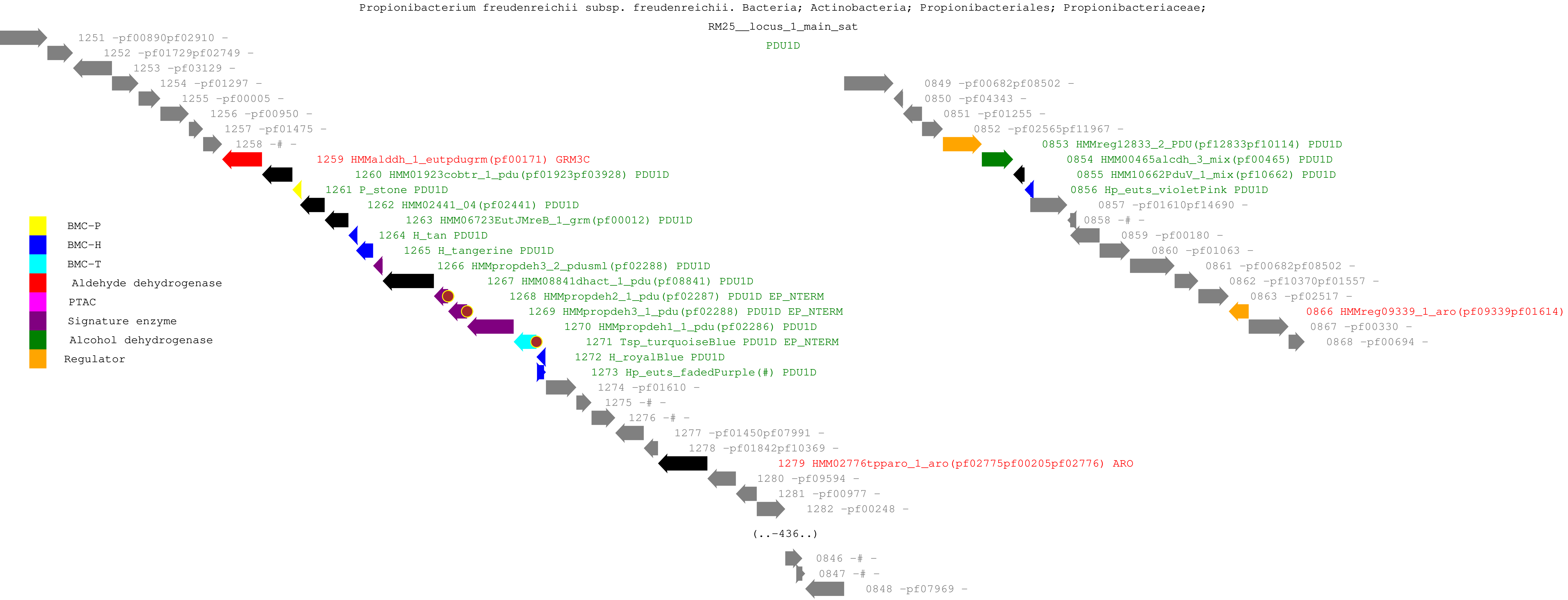
Propionibacterium freudenreichii subsp. freudenreichii. Bacteria; Actinobacteria; Propionibacteriales; Propionibacteriaceae;
RM25__locus_1_main_sat
PDU1D
1251 -pf00890pf02910 -
1252 -pf01729pf02749 -
1253 -pf03129 -
1254 -pf01297 -
1255 -pf00005 -
1256 -pf00950 -
1257 -pf01475 -
1258 -# -
1259 HMMalddh_1_eutpdugrm(pf00171) GRM3C
1260 HMM01923cobtr_1_pdu(pf01923pf03928) PDU1D
1261 P_stone PDU1D
1262 HMM02441_04(pf02441) PDU1D
1263 HMM06723EutJMreB_1_grm(pf00012) PDU1D
1264 H_tan PDU1D
1265 H_tangerine PDU1D
1266 HMMpropdeh3_2_pdusml(pf02288) PDU1D
1267 HMM08841dhact_1_pdu(pf08841) PDU1D
1268 HMMpropdeh2_1_pdu(pf02287) PDU1D EP_NTERM
1269 HMMpropdeh3_1_pdu(pf02288) PDU1D EP_NTERM
1270 HMMpropdeh1_1_pdu(pf02286) PDU1D
1271 Tsp_turquoiseBlue PDU1D EP_NTERM
1272 H_royalBlue PDU1D
1273 Hp_euts_fadedPurple(#) PDU1D
1274 -pf01610 -
1275 -# -
1276 -# -
1277 -pf01450pf07991 -
1278 -pf01842pf10369 -
1279 HMM02776tpparo_1_aro(pf02775pf00205pf02776) ARO
1280 -pf09594 -
1281 -pf00977 -
1282 -pf00248 -
(..-436..)
0846 -# -
0847 -# -
0848 -pf07969 -
0849 -pf00682pf08502 -
0850 -pf04343 -
0851 -pf01255 -
0852 -pf02565pf11967 -
0853 HMMreg12833_2_PDU(pf12833pf10114) PDU1D
0854 HMM00465alcdh_3_mix(pf00465) PDU1D
0855 HMM10662PduV_1_mix(pf10662) PDU1D
0856 Hp_euts_violetPink PDU1D
0857 -pf01610pf14690 -
0858 -# -
0859 -pf00180 -
0860 -pf01063 -
0861 -pf00682pf08502 -
0862 -pf10370pf01557 -
0863 -pf02517 -
0866 HMMreg09339_1_aro(pf09339pf01614)
0867 -pf00330 -
0868 -pf00694 -
BMC-P
BMC-H
BMC-T
Aldehyde dehydrogenase
PTAC
Signature enzyme
Alcohol dehydrogenase
Regulator

uncultured Eubacterium sp. Bacteria; Firmicutes; Clostridia; Clostridiales; Eubacteriaceae;

SAMEA3545253__locus_1_main_sat

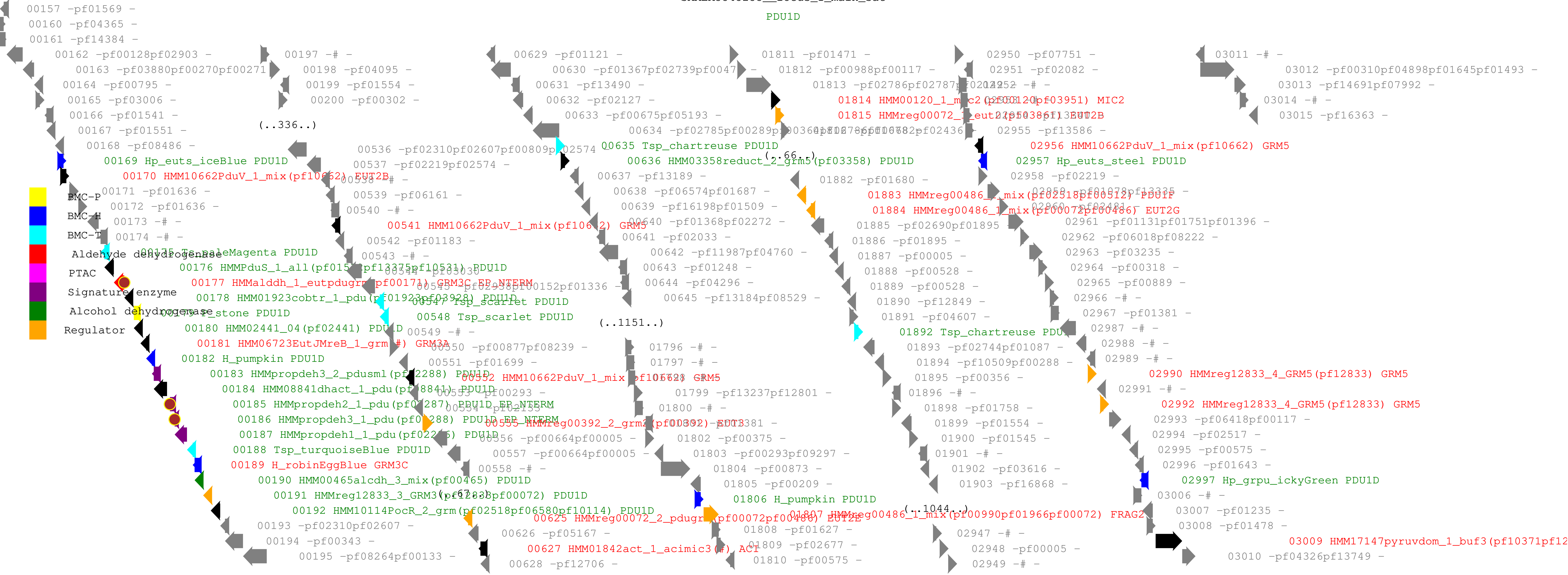

uncultured Clostridium sp. Bacteria; Firmicutes; Clostridia; Clostridiales; Clostridiaceae;

SAMEA3545325__locus_1_main_sat

PDU1D

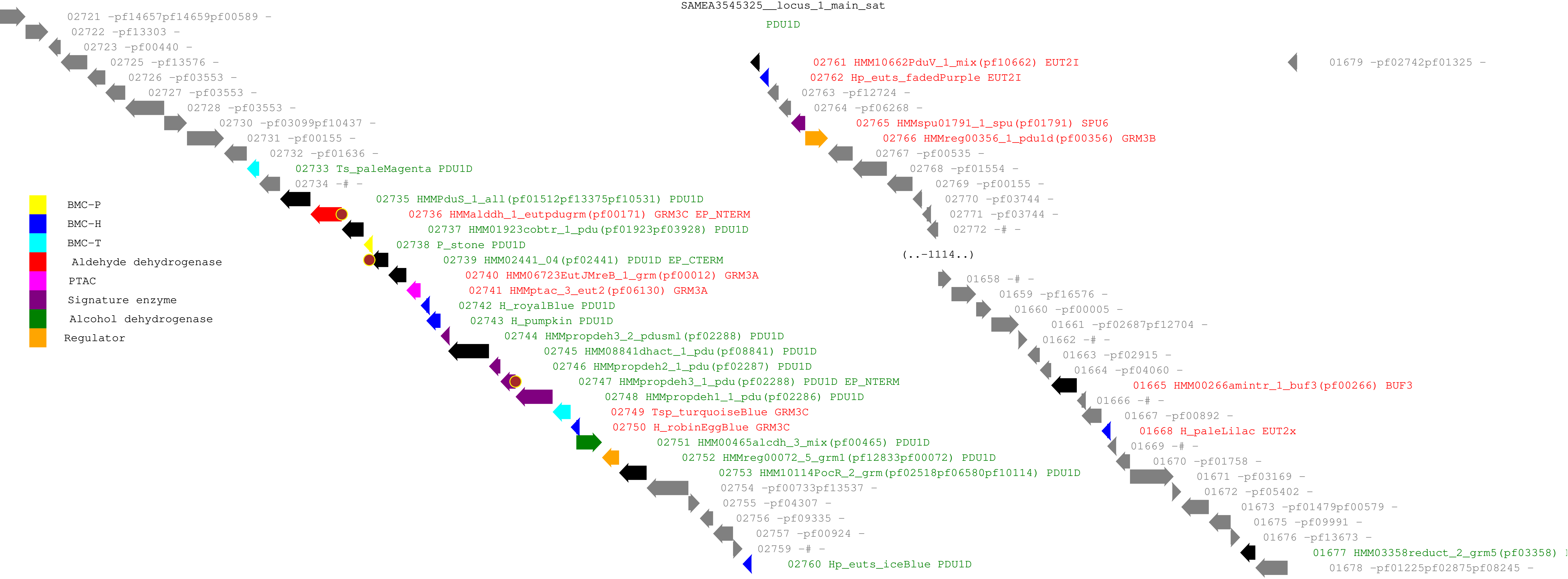

uncultured Eubacterium sp. Bacteria; Firmicutes; Clostridia; Clostridiales; Eubacteriaceae;

SAMEA3545427__locus_1_main_sat

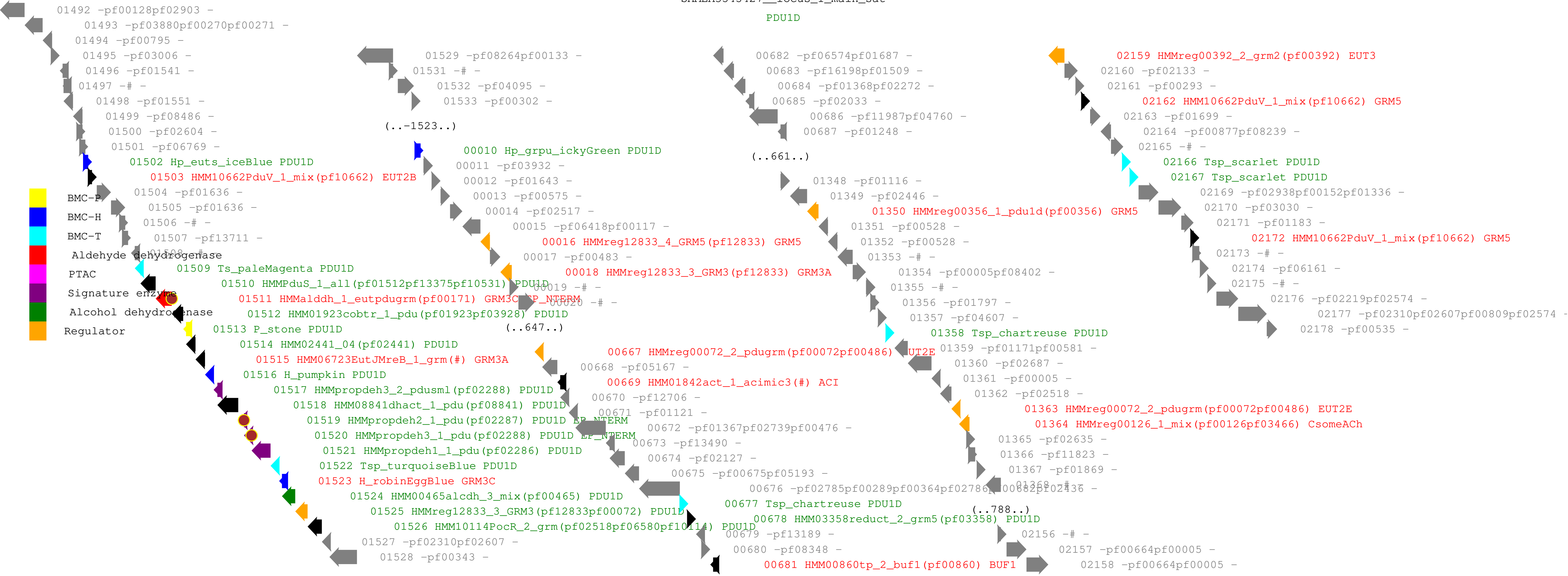

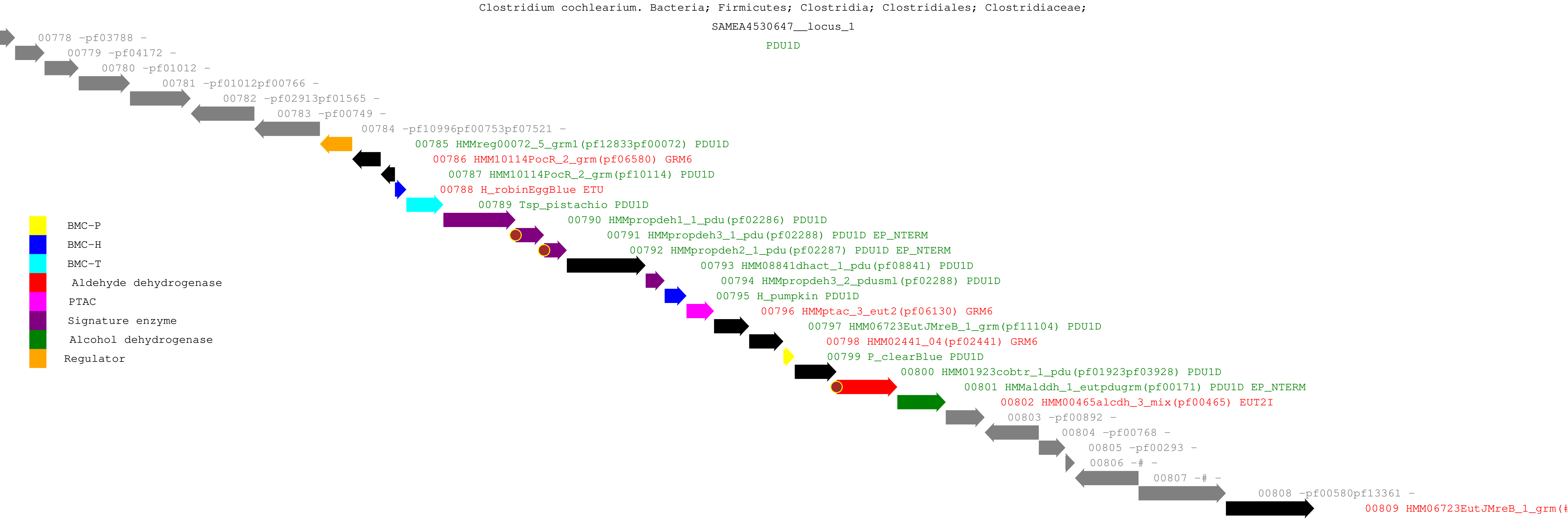
Clostridium cochlearium. Bacteria; Firmicutes; Clostridia; Clostridiales; Clostridiaceae;
SAMEA4530647__locus_1
PDU1D
00778 -pf03788 -
00779 -pf04172 -
00780 -pf01012 -
00781 -pf01012pf00766 -
00782 -pf02913pf01565 -
00783 -pf00749 -
00784 -pf10996pf00753pf07521 -
00785 HMMreg00072_5_grm1(pf12833pf00072) PDU1D
00786 HMM10114PocR_2_grm(pf06580) GRM6
00787 HMM10114PocR_2_grm(pf10114) PDU1D
00788 H_robinEggBlue ETU
00789 Tsp_pistachio PDU1D
00790 HMMpropdeh1_1_pdu(pf02286) PDU1D
00791 HMMpropdeh3_1_pdu(pf02288) PDU1D EP_NTERM
00792 HMMpropdeh2_1_pdu(pf02287) PDU1D EP_NTERM
00793 HMM08841dhact_1_pdu(pf08841) PDU1D
00794 HMMpropdeh3_2_pdusml(pf02288) PDU1D
00795 H_pumpkin PDU1D
00796 HMMptac_3_eut2(pf06130) GRM6
00797 HMM06723EutJMreB_1_grm(pf11104) PDU1D
00798 HMM02441_04(pf02441) GRM6
00799 P_clearBlue PDU1D
00800 HMM01923cobtr_1_pdu(pf01923pf03928) PDU1D
00801 HMMalddh_1_eutpdugrm(pf00171) PDU1D EP_NTERM
00802 HMM00465alcdh_3_mix(pf00465) EUT2I
00803 -pf00892 -
00804 -pf00768 -
00805 -pf00293 -
00806 -# -
00807 -# -
00808 -pf00580pf13361 -
00809 HMM06723EutJMreB_1_grm(#
BMC-P
BMC-H
BMC-T
Aldehyde dehydrogenase
PTAC
Signature enzyme
Alcohol dehydrogenase
Regulator

Maledivibacter halophilus. Bacteria; Firmicutes; Clostridia; Clostridiales; Clostridiaceae;

SAMN02194393__locus_6

PDU1D

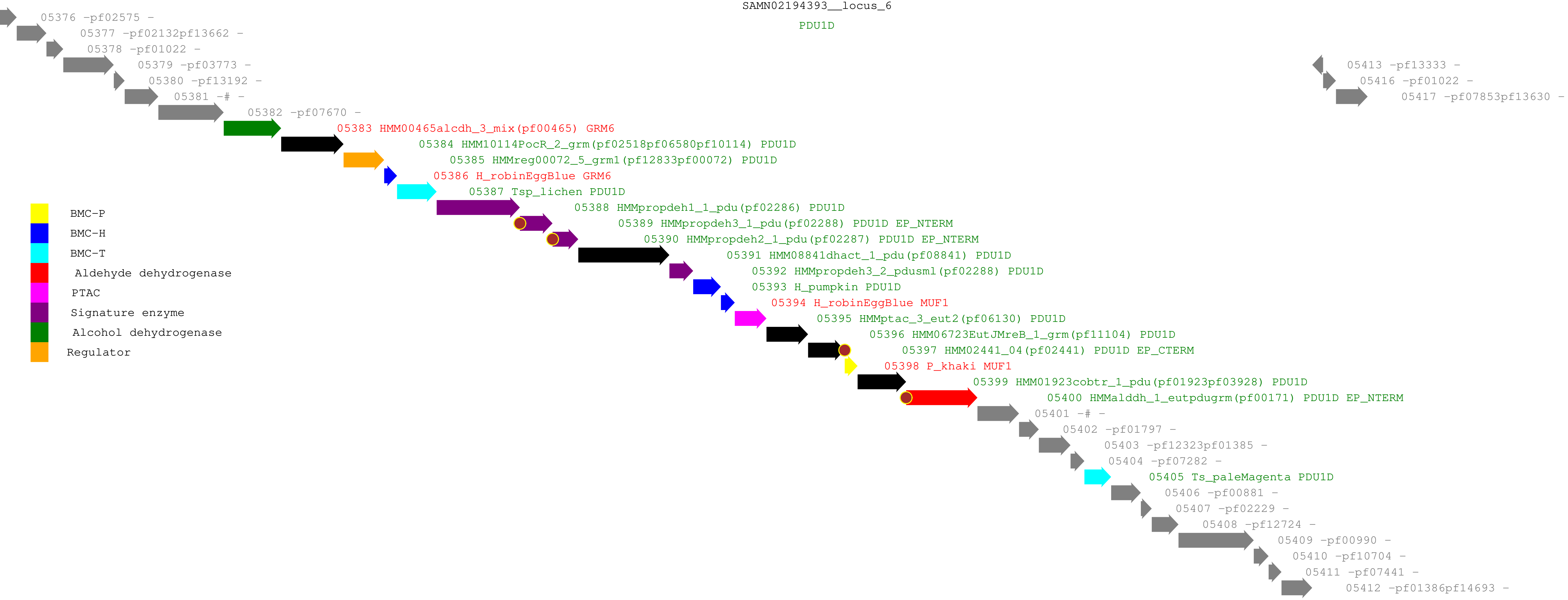

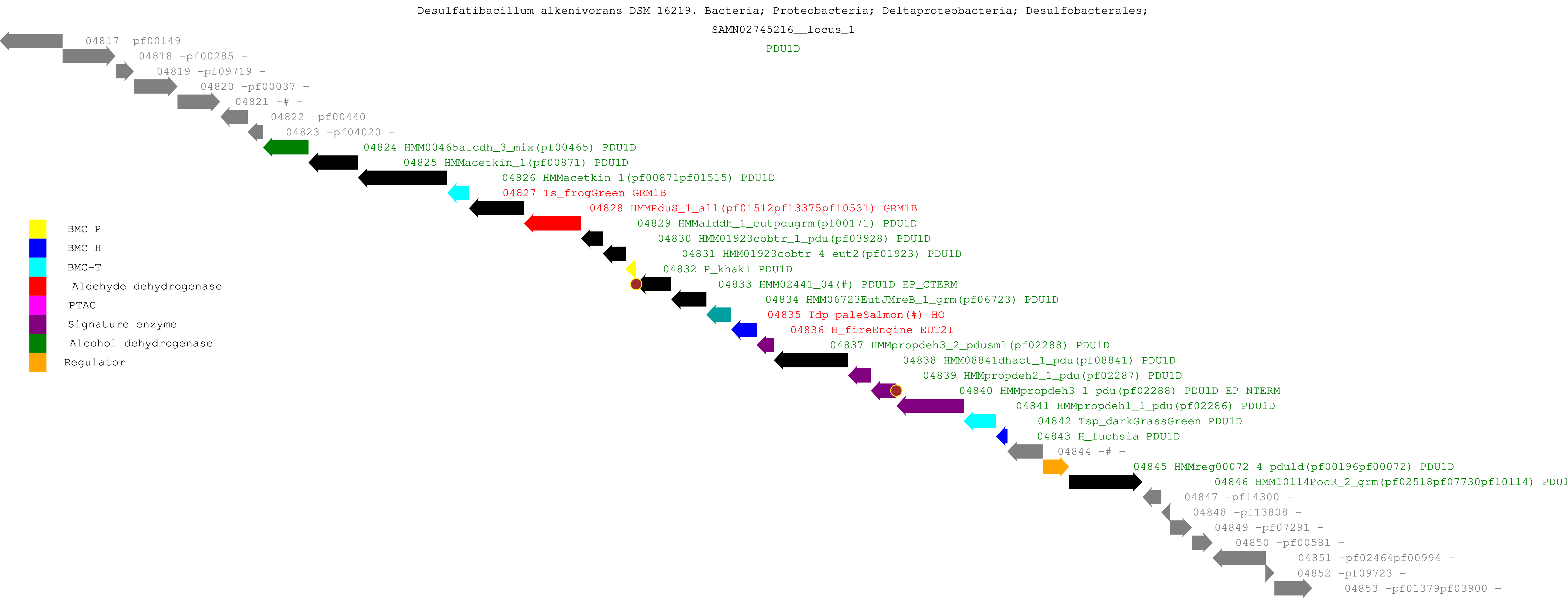
Desulfatibacillum alkenivorans DSM 16219. Bacteria; Proteobacteria; Deltaproteobacteria; Desulfobacterales;
SAMN02745216__locus_1
PDU1D
04817 -pf00149 -
04818 -pf00285 -
04819 -pf09719 -
04820 -pf00037 -
04821 -# -
04822 -pf00440 -
04823 -pf04020 -
04824 HMM00465alcdh_3_mix(pf00465) PDU1D
04825 HMMacetkin_1(pf00871) PDU1D
04826 HMMacetkin_1(pf00871pf01515) PDU1D
04827 Ts_frogGreen GRM1B
04828 HMMPduS_1_all(pf01512pf13375pf10531) GRM1B
04829 HMMalddh_1_eutpdugrm(pf00171) PDU1D
04830 HMM01923cobtr_1_pdu(pf03928) PDU1D
04831 HMM01923cobtr_4_eut2(pf01923) PDU1D
04832 P_khaki PDU1D
04833 HMM02441_04(#) PDU1D EP_CTERM
04834 HMM06723EutJMreB_1_grm(pf06723) PDU1D
04835 Tdp_paleSalmon(#) HO
04836 H_fireEngine EUT2I
04837 HMMpropdeh3_2_pdusml(pf02288) PDU1D
04838 HMM08841dhact_1_pdu(pf08841) PDU1D
04839 HMMpropdeh2_1_pdu(pf02287) PDU1D
04840 HMMpropdeh3_1_pdu(pf02288) PDU1D EP_NTERM
04841 HMMpropdeh1_1_pdu(pf02286) PDU1D
04842 Tsp_darkGrassGreen PDU1D
04843 H_fuchsia PDU1D
04844 -# -
04845 HMMreg00072_4_pdu1d(pf00196pf00072) PDU1D
04846 HMM10114PocR_2_grm(pf02518pf07730pf10114) PDU1
04847 -pf14300 -
04848 -pf13808 -
04849 -pf07291 -
04850 -pf00581 -
04851 -pf02464pf00994 -
04852 -pf09723 -
04853 -pf01379pf03900 -
BMC-P
BMC-H
BMC-T
Aldehyde dehydrogenase
PTAC
Signature enzyme
Alcohol dehydrogenase
Regulator

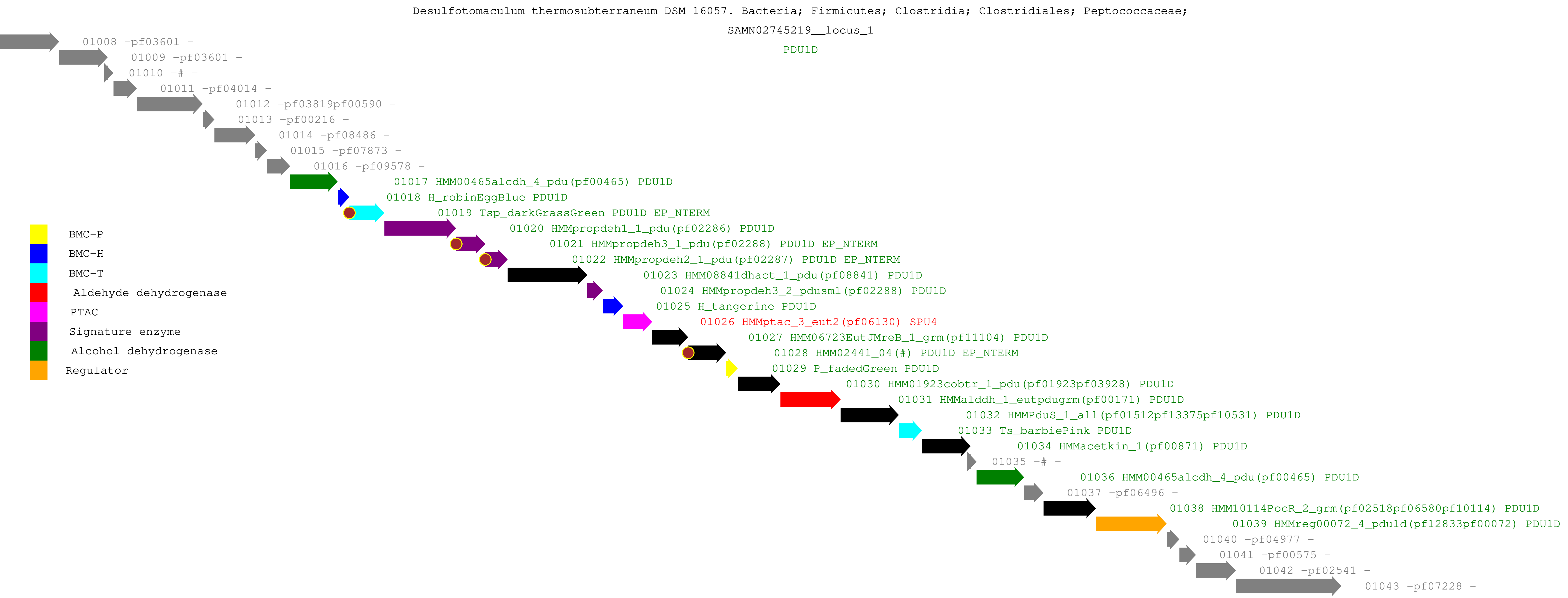

Desulfotomaculum thermosubterraneum DSM 16057. Bacteria; Firmicutes; Clostridia; Clostridiales; Peptococcaceae;
SAMN02745219__locus_1
PDU1D
01008 -pf03601 -
01009 -pf03601 -
01010 -# -
01011 -pf04014 -
01012 -pf03819pf00590 -
01013 -pf00216 -
01014 -pf08486 -
01015 -pf07873 -
01016 -pf09578 -
01017 HMM00465alcdh_4_pdu(pf00465) PDU1D
01018 H_robinEggBlue PDU1D
01019 Tsp_darkGrassGreen PDU1D EP_NTERM
01020 HMMpropdeh1_1_pdu(pf02286) PDU1D
01021 HMMpropdeh3_1_pdu(pf02288) PDU1D EP_NTERM
01022 HMMpropdeh2_1_pdu(pf02287) PDU1D EP_NTERM
01023 HMM08841dhact_1_pdu(pf08841) PDU1D
01024 HMMpropdeh3_2_pdusml(pf02288) PDU1D
01025 H_tangerine PDU1D
01026 HMMptac_3_eut2(pf06130) SPU4
01027 HMM06723EutJMreB_1_grm(pf11104) PDU1D
01028 HMM02441_04(#) PDU1D EP_NTERM
01029 P_fadedGreen PDU1D
01030 HMM01923cobtr_1_pdu(pf01923pf03928) PDU1D
01031 HMMalddh_1_eutpdugrm(pf00171) PDU1D
01032 HMMPduS_1_all(pf01512pf13375pf10531) PDU1D
01033 Ts_barbiePink PDU1D
01034 HMMacetkin_1(pf00871) PDU1D
01035 -# -
01036 HMM00465alcdh_4_pdu(pf00465) PDU1D
01037 -pf06496 -
01038 HMM10114PocR_2_grm(pf02518pf06580pf10114) PDU1D
01039 HMMreg00072_4_pdu1d(pf12833pf00072) PDU1D
01040 -pf04977 -
01041 -pf00575 -
01042 -pf02541 -
01043 -pf07228 -
BMC-P
BMC-H
BMC-T
Aldehyde dehydrogenase
PTAC
Signature enzyme
Alcohol dehydrogenase
Regulator

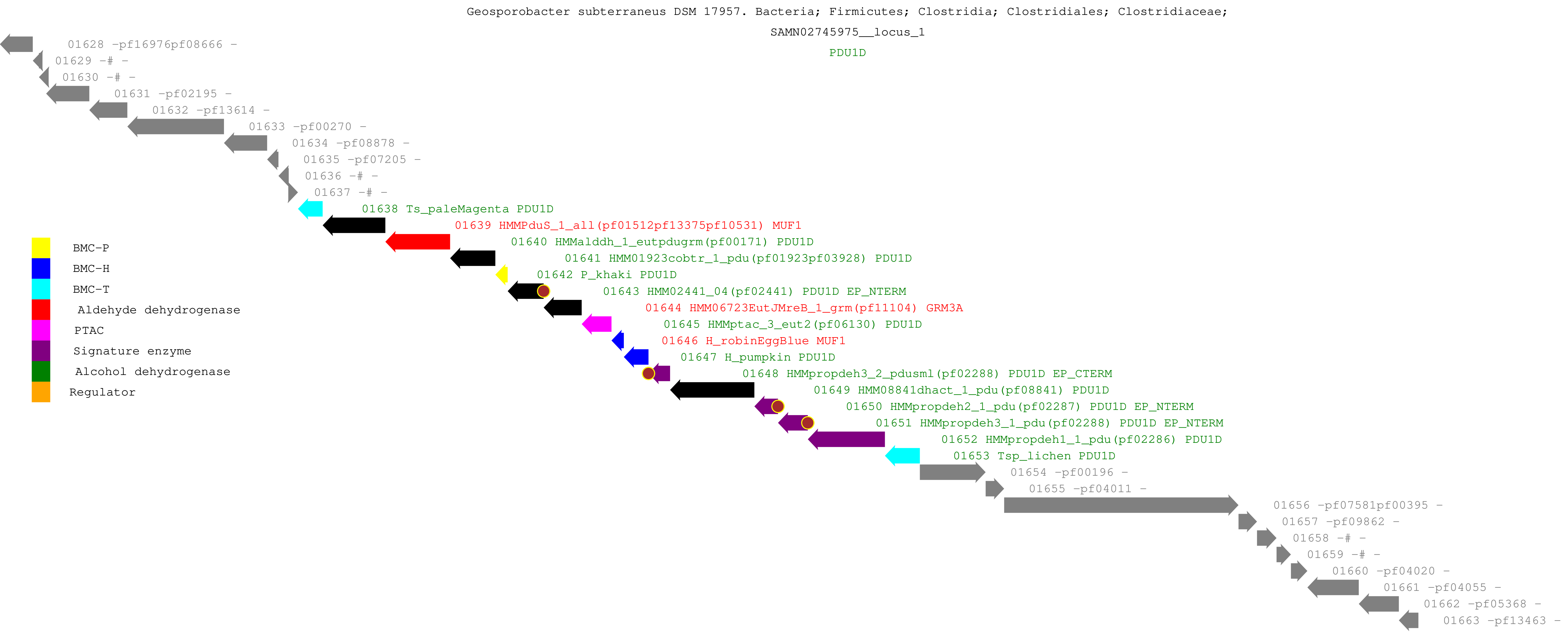
Geosporobacter subterraneus DSM 17957. Bacteria; Firmicutes; Clostridia; Clostridiales; Clostridiaceae;
SAMN02745975__locus_1
PDU1D
01628 -pf16976pf08666 -
01629 -# -
01630 -# -
01631 -pf02195 -
01632 -pf13614 -
01633 -pf00270 -
01634 -pf08878 -
01635 -pf07205 -
01636 -# -
01637 -# -
01638 Ts_paleMagenta PDU1D
01639 HMMPduS_1_all(pf01512pf13375pf10531) MUF1
01640 HMMalddh_1_eutpdugrm(pf00171) PDU1D
01641 HMM01923cobtr_1_pdu(pf01923pf03928) PDU1D
01642 P_khaki PDU1D
01643 HMM02441_04(pf02441) PDU1D EP_NTERM
01644 HMM06723EutJMreB_1_grm(pf11104) GRM3A
01645 HMMptac_3_eut2(pf06130) PDU1D
01646 H_robinEggBlue MUF1
01647 H_pumpkin PDU1D
01648 HMMpropdeh3_2_pdusml(pf02288) PDU1D EP_CTERM
01649 HMM08841dhact_1_pdu(pf08841) PDU1D
01650 HMMpropdeh2_1_pdu(pf02287) PDU1D EP_NTERM
01651 HMMpropdeh3_1_pdu(pf02288) PDU1D EP_NTERM
01652 HMMpropdeh1_1_pdu(pf02286) PDU1D
01653 Tsp_lichen PDU1D
01654 -pf00196 -
01655 -pf04011 -
01656 -pf07581pf00395 -
01657 -pf09862 -
01658 -# -
01659 -# -
01660 -pf04020 -
01661 -pf04055 -
01662 -pf05368 -
01663 -pf13463 -
BMC-P
BMC-H
BMC-T
Aldehyde dehydrogenase
PTAC
Signature enzyme
Alcohol dehydrogenase
Regulator

Geosporobacter subterraneus DSM 17957. Bacteria; Firmicutes; Clostridia; Clostridiales; Clostridiaceae;

SAMN02745975__locus_3

PDU1D

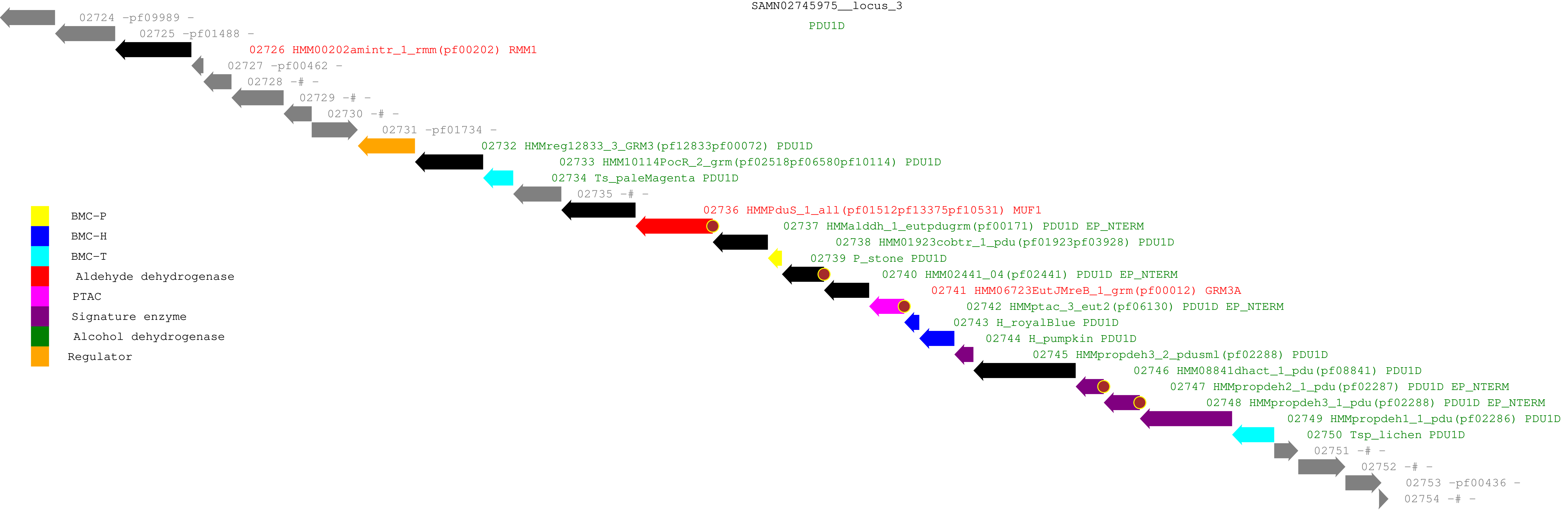

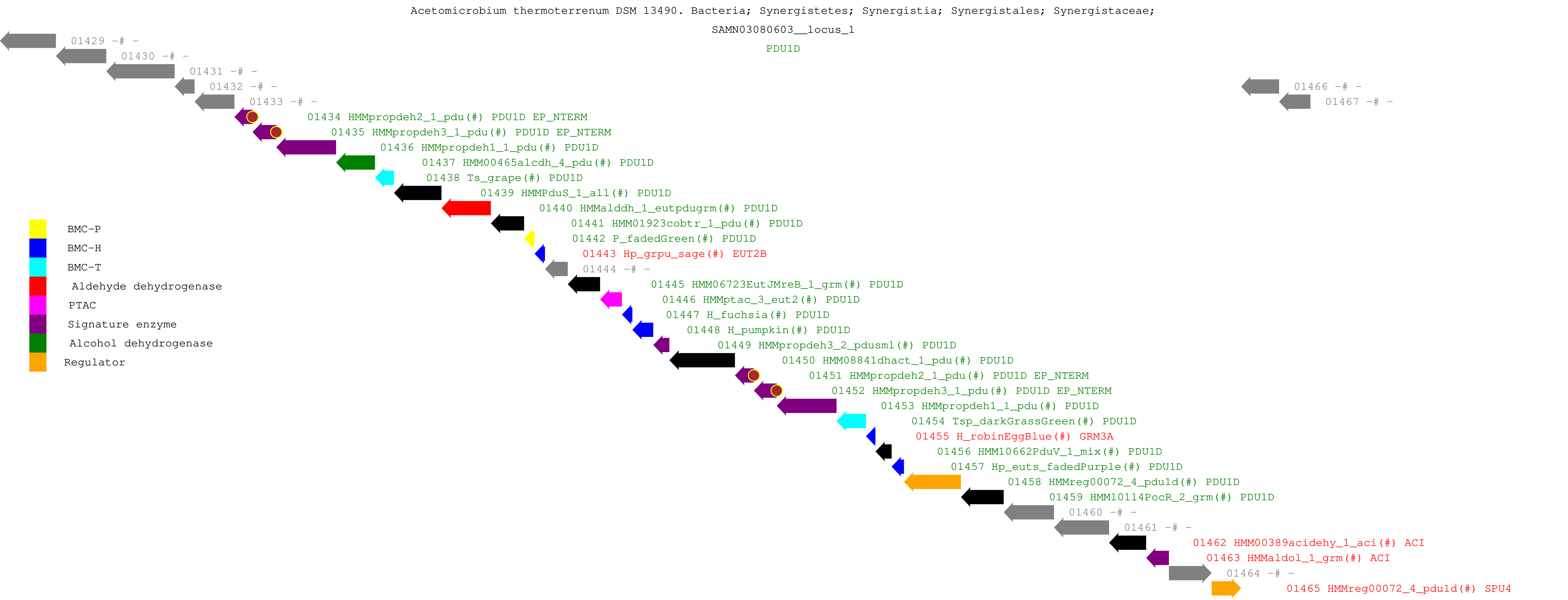
Acetomicrobium thermoterrenum DSM 13490. Bacteria; Synergistetes; Synergistia; Synergistales; Synergistaceae;
SAMN03080603__locus_1
PDU1D
01429 -# -
01430 -# -
01431 -# -
01432 -# -
01433 -# -
01434 HMMpropdeh2_1_pdu(#) PDU1D EP_NTERM
01435 HMMpropdeh3_1_pdu(#) PDU1D EP_NTERM
01436 HMMpropdeh1_1_pdu(#) PDU1D
01437 HMM00465alcdh_4_pdu(#) PDU1D
01438 Ts_grape(#) PDU1D
01439 HMMPduS_1_all(#) PDU1D
01440 HMMalddh_1_eutpdugrm(#) PDU1D
01441 HMM01923cobtr_1_pdu(#) PDU1D
01442 P_fadedGreen(#) PDU1D
01443 Hp_grpu_sage(#) EUT2B
01444 -# -
01445 HMM06723EutJMreB_1_grm(#) PDU1D
01446 HMMptac_3_eut2(#) PDU1D
01447 H_fuchsia(#) PDU1D
01448 H_pumpkin(#) PDU1D
01449 HMMpropdeh3_2_pdusml(#) PDU1D
01450 HMM08841dhact_1_pdu(#) PDU1D
01451 HMMpropdeh2_1_pdu(#) PDU1D EP_NTERM
01452 HMMpropdeh3_1_pdu(#) PDU1D EP_NTERM
01453 HMMpropdeh1_1_pdu(#) PDU1D
01454 Tsp_darkGrassGreen(#) PDU1D
01455 H_robinEggBlue(#) GRM3A
01456 HMM10662PduV_1_mix(#) PDU1D
01457 Hp_euts_fadedPurple(#) PDU1D
01458 HMMreg00072_4_pdu1d(#) PDU1D
01459 HMM10114PocR_2_grm(#) PDU1D
01460 -# -
01461 -# -
01462 HMM00389acidehy_1_aci(#) ACI
01463 HMMaldol_1_grm(#) ACI
01464 -# -
01465 HMMreg00072_4_pdu1d(#) SPU4
01466 -# -
01467 -# -
BMC-P
BMC-H
BMC-T
Aldehyde dehydrogenase
PTAC
Signature enzyme
Alcohol dehydrogenase
Regulator

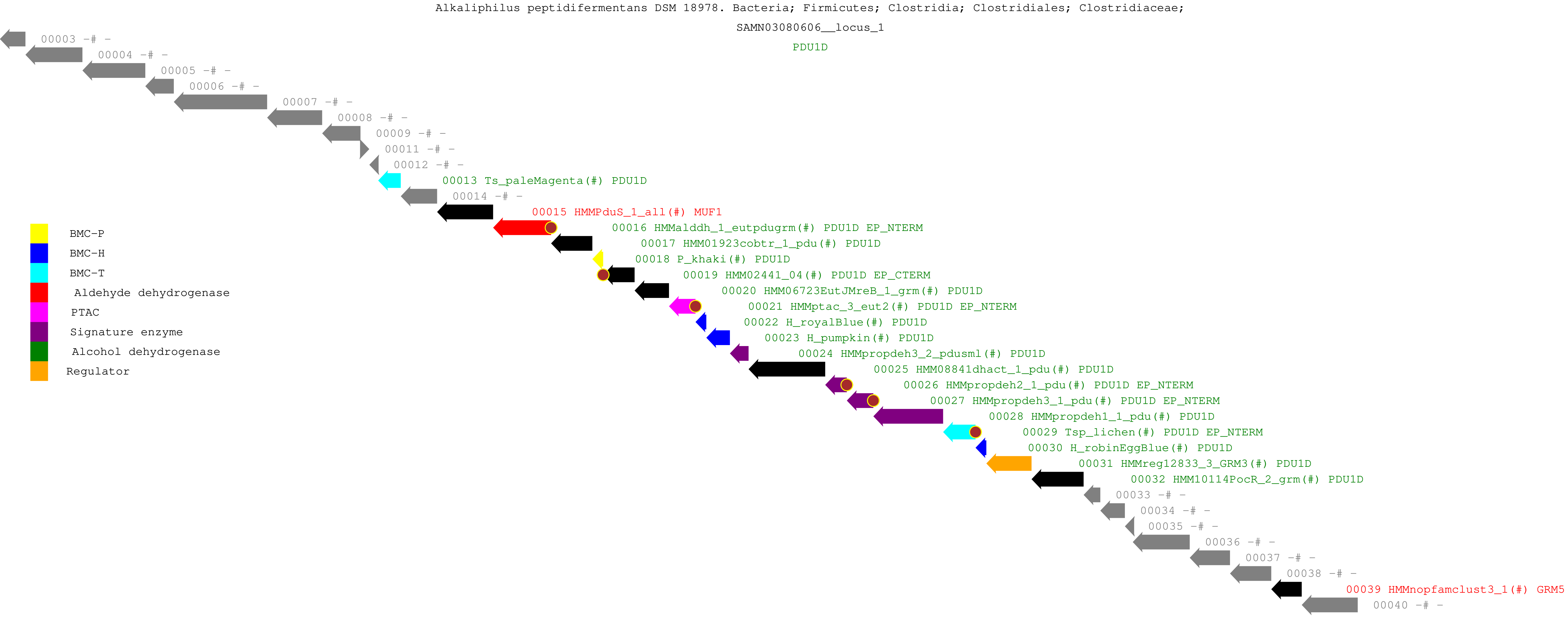

Alkaliphilus peptidifermentans DSM 18978. Bacteria; Firmicutes; Clostridia; Clostridiales; Clostridiaceae;
SAMN03080606__locus_1
PDU1D
00003 -# -
00004 -# -
00005 -# -
00006 -# -
00007 -# -
00008 -# -
00009 -# -
00011 -# -
00012 -# -
00013 Ts_paleMagenta(#) PDU1D
00014 -# -
00015 HMMPduS_1_all(#) MUF1
00016 HMMaldh_1_eutpdugrm(#) PDU1D EP_NTERM
00017 HMM01923cobtr_1_pdu(#) PDU1D
00018 P_khaki(#) PDU1D
00019 HMM02441_04(#) PDU1D EP_CTERM
00020 HMM06723EutJMreB_1_grm(#) PDU1D
00021 HMMptac_3_eut2(#) PDU1D EP_NTERM
00022 H_royalBlue(#) PDU1D
00023 H_pumpkin(#) PDU1D
00024 HMMpropdeh3_2_pdusml(#) PDU1D
00025 HMM08841dhact_1_pdu(#) PDU1D
00026 HMMpropdeh2_1_pdu(#) PDU1D EP_NTERM
00027 HMMpropdeh3_1_pdu(#) PDU1D EP_NTERM
00028 HMMpropdeh1_1_pdu(#) PDU1D
00029 Tsp_lichen(#) PDU1D EP_NTERM
00030 H_robinEggBlue(#) PDU1D
00031 HMMreg12833_3_GRM3(#) PDU1D
00032 HMM10114PocR_2_grm(#) PDU1D
00033 -# -
00034 -# -
00035 -# -
00036 -# -
00037 -# -
00038 -# -
00039 HMMnopfamclust3_1(#) GRM5
00040 -# -
BMC-P
BMC-H
BMC-T
Aldehyde dehydrogenase
PTAC
Signature enzyme
Alcohol dehydrogenase
Regulator

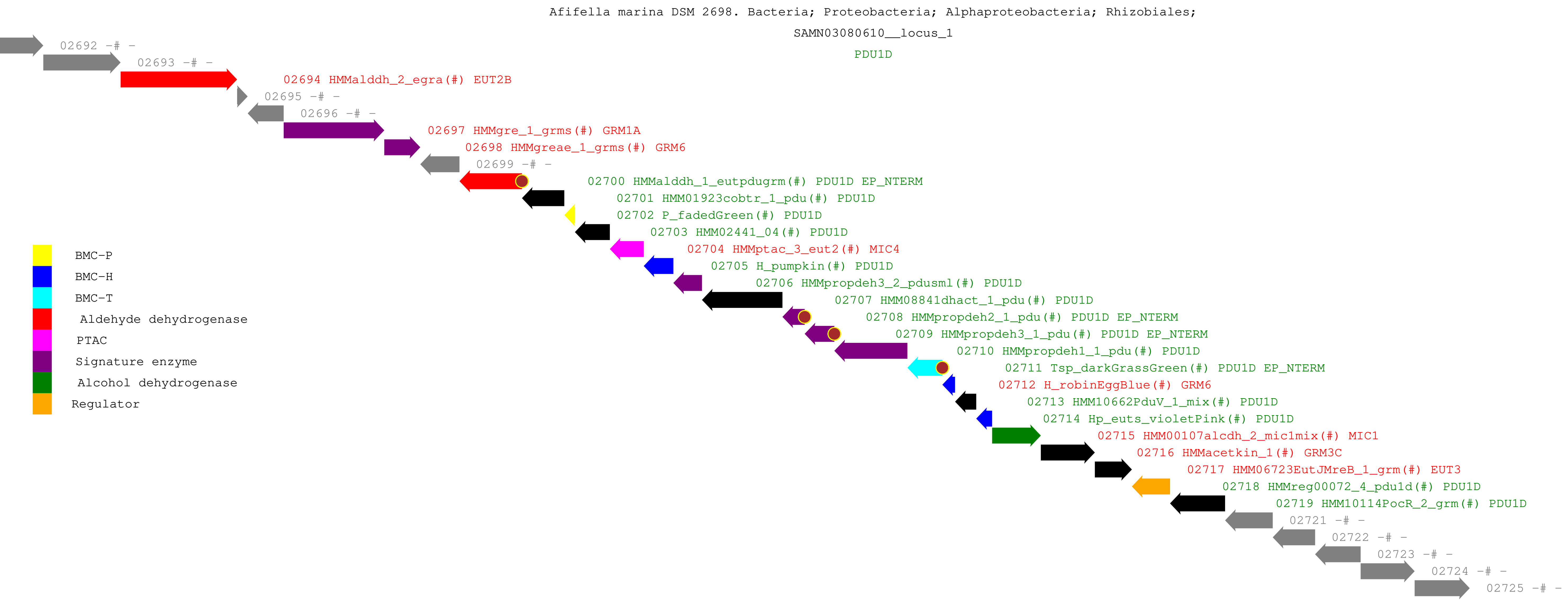
Afifella marina DSM 2698. Bacteria; Proteobacteria; Alphaproteobacteria; Rhizobiales;
SAMN03080610__locus_1
PDU1D
02692 -# -
02693 -# -
02694 HMMalddh_2_egra(#) EUT2B
02695 -# -
02696 -# -
02697 HMMgre_1_grms(#) GRM1A
02698 HMMgreae_1_grms(#) GRM6
02699 -# -
02700 HMMalddh_1_eutpdugrm(#) PDU1D EP_NTERM
02701 HMM01923cobtr_1_pdu(#) PDU1D
02702 P_fadedGreen(#) PDU1D
02703 HMM02441_04(#) PDU1D
02704 HMMptac_3_eut2(#) MIC4
02705 H_pumpkin(#) PDU1D
02706 HMMpropdeh3_2_pdusml(#) PDU1D
02707 HMM08841dhact_1_pdu(#) PDU1D
02708 HMMpropdeh2_1_pdu(#) PDU1D EP_NTERM
02709 HMMpropdeh3_1_pdu(#) PDU1D EP_NTERM
02710 HMMpropdeh1_1_pdu(#) PDU1D
02711 Tsp_darkGrassGreen(#) PDU1D EP_NTERM
02712 H_robinEggBlue(#) GRM6
02713 HMM10662PduV_1_mix(#) PDU1D
02714 Hp_euts_violetPink(#) PDU1D
02715 HMM00107alcdh_2_mic1mix(#) MIC1
02716 HMMacetkin_1(#) GRM3C
02717 HMM06723EutJMreB_1_grm(#) EUT3
02718 HMMreg00072_4_pdu1d(#) PDU1D
02719 HMM10114PocR_2_grm(#) PDU1D
02721 -# -
02722 -# -
02723 -# -
02724 -# -
02725 -# -
BMC-P
BMC-H
BMC-T
Aldehyde dehydrogenase
PTAC
Signature enzyme
Alcohol dehydrogenase
Regulator

Eubacterium callanderi. Bacteria; Firmicutes; Clostridia; Clostridiales; Eubacteriaceae;

SAMN04487888__locus_1_main_sat

PDU1D

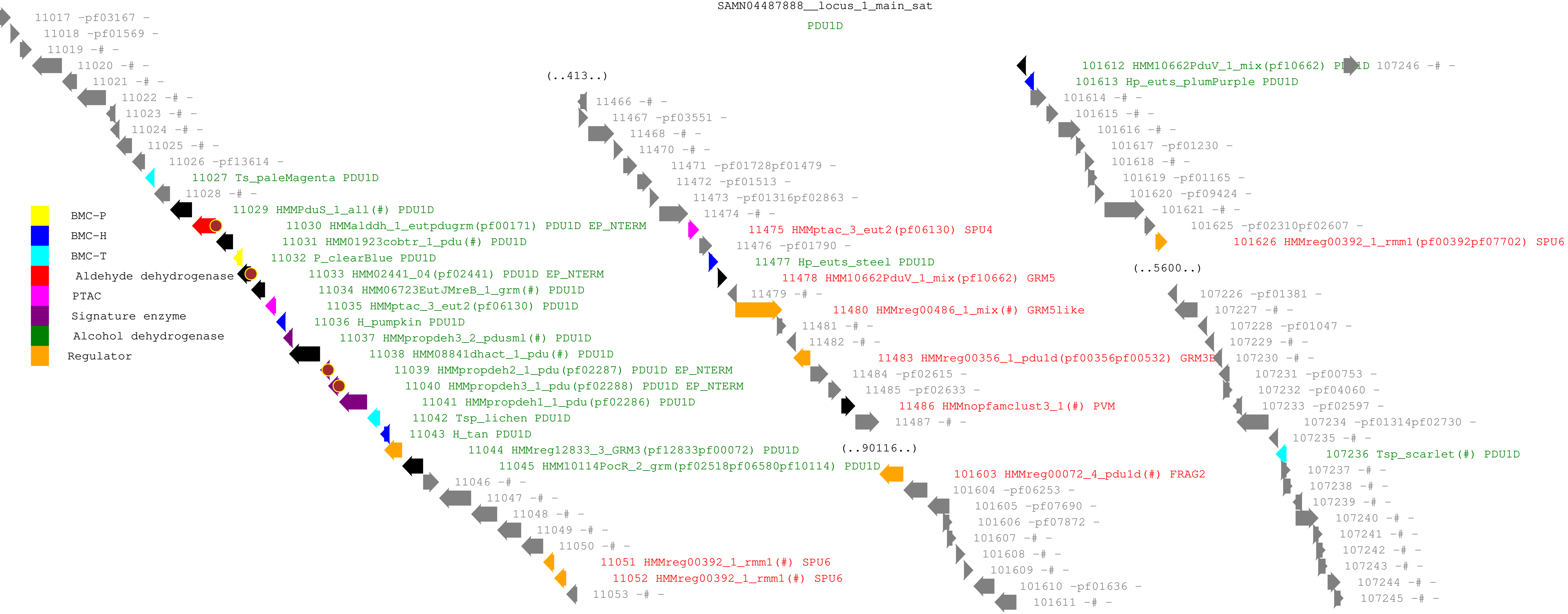

Sporomusa acidovorans. Bacteria; Firmicutes; Negativicutes; Selenomonadales; Sporomusaceae;

SAMN04488499__locus_2

PDU1D

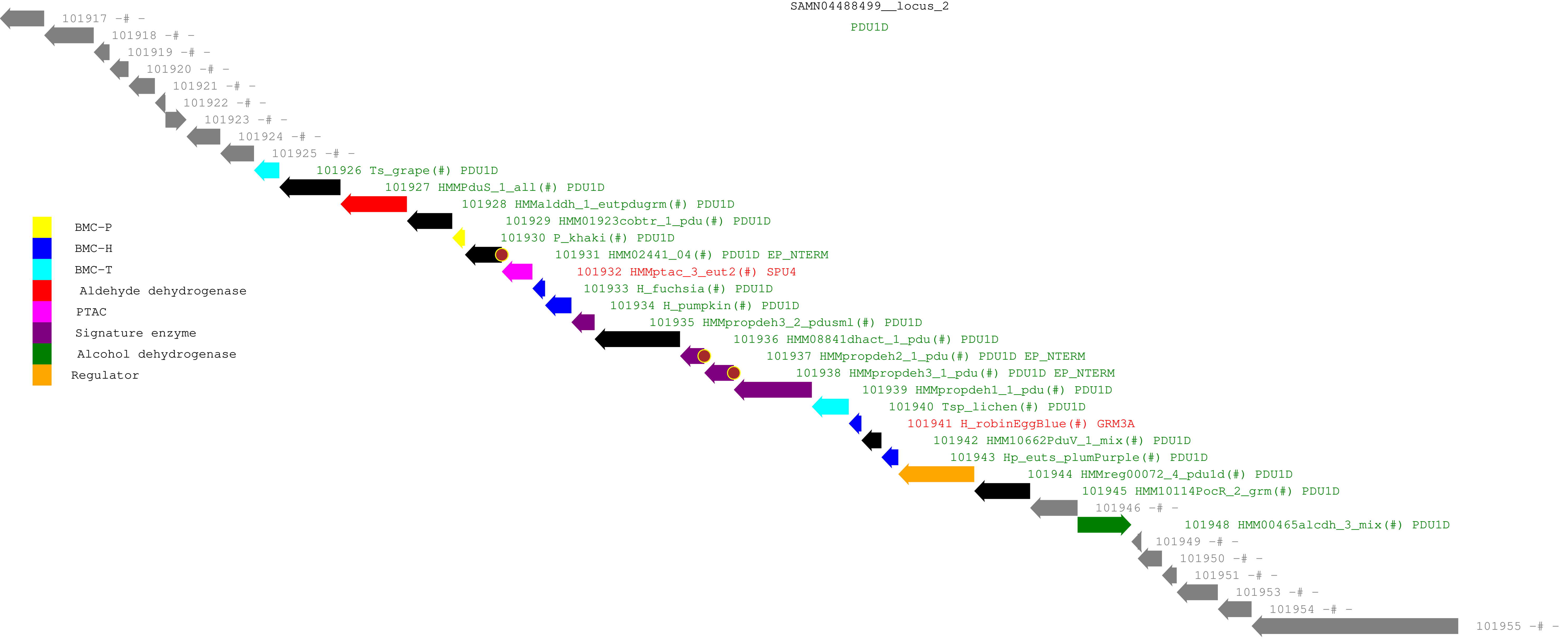

Sporomusa malonica. Bacteria; Firmicutes; Negativicutes; Selenomonadales; Sporomusaceae;

SAMN04488500__locus_3

PDU1D

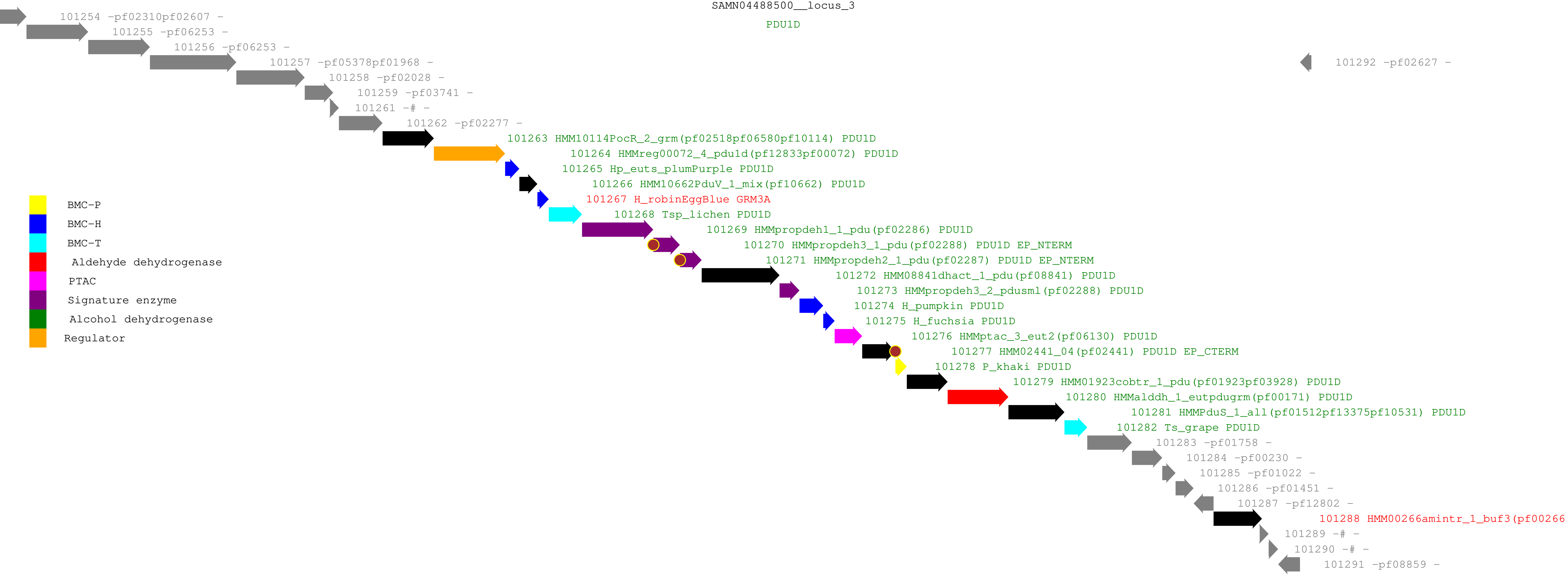

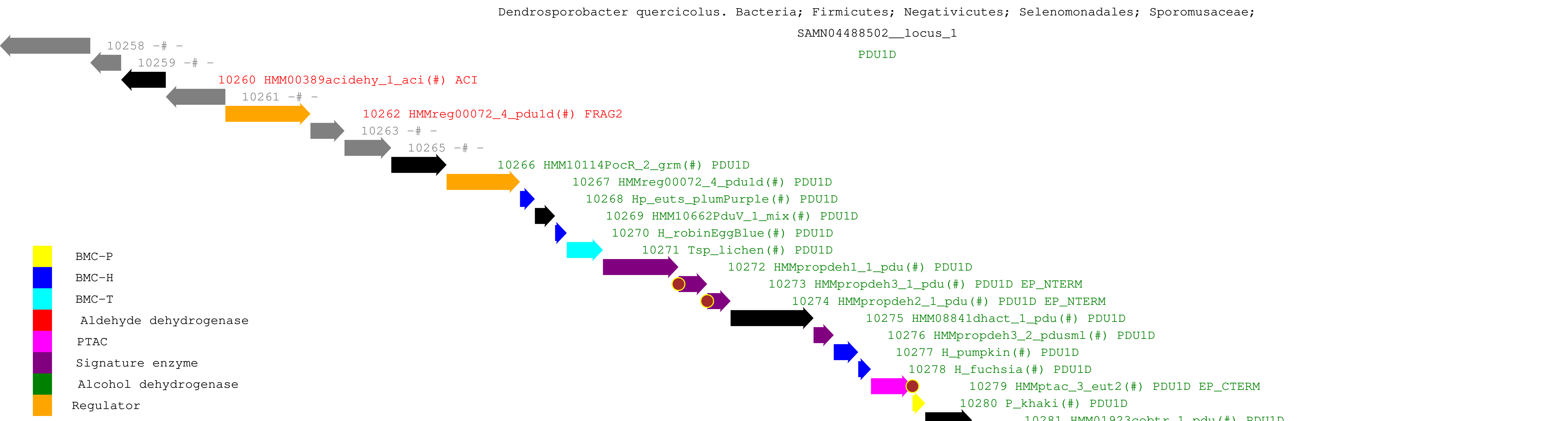
Dendrosporobacter quercicolus. Bacteria; Firmicutes; Negativicutes; Selenomonadales; Sporomusaceae;
SAMN04488502__locus_1
PDU1D
10258 -# -
10259 -# -
10260 HMM00389acidehy_1_aci(#) ACI
10261 -# -
10262 HMMreg00072_4_pdu1d(#) FRAG2
10263 -# -
10265 -# -
10266 HMM10114PocR_2_grm(#) PDU1D
10267 HMMreg00072_4_pdu1d(#) PDU1D
10268 Hp_euts_plumPurple(#) PDU1D
10269 HMM10662PduV_1_mix(#) PDU1D
10270 H_robinEggBlue(#) PDU1D
10271 Tsp_lichen(#) PDU1D
10272 HMMpropdeh1_1_pdu(#) PDU1D
10273 HMMpropdeh3_1_pdu(#) PDU1D EP_NTERM
10274 HMMpropdeh2_1_pdu(#) PDU1D EP_NTERM
10275 HMM08841dhact_1_pdu(#) PDU1D
10276 HMMpropdeh3_2_pdusml(#) PDU1D
10277 H_pumpkin(#) PDU1D
10278 H_fuchsia(#) PDU1D
10279 HMMptac_3_eut2(#) PDU1D EP_CTERM
10280 P_khaki(#) PDU1D
10281 HMM01923cobtr_1_pdu(#) PDU1D
10282 HMMalddh_1_eutpdugrm(#) PDU1D EP_NTERM
10283 -# -
10284 HMM02776tpparo_1_aro(#) ARO
10285 -# -
10286 -# -
10287 -# -
10288 -# -
10289 -# -
10290 -# -
BMC-P
BMC-H
BMC-T
Aldehyde dehydrogenase
PTAC
Signature enzyme
Alcohol dehydrogenase
Regulator

Eubacterium barkeri (Clostridium barkeri). Bacteria; Firmicutes; Clostridia; Clostridiales; Eubacteriaceae;

SAMN04488579__locus_2

PDU1D

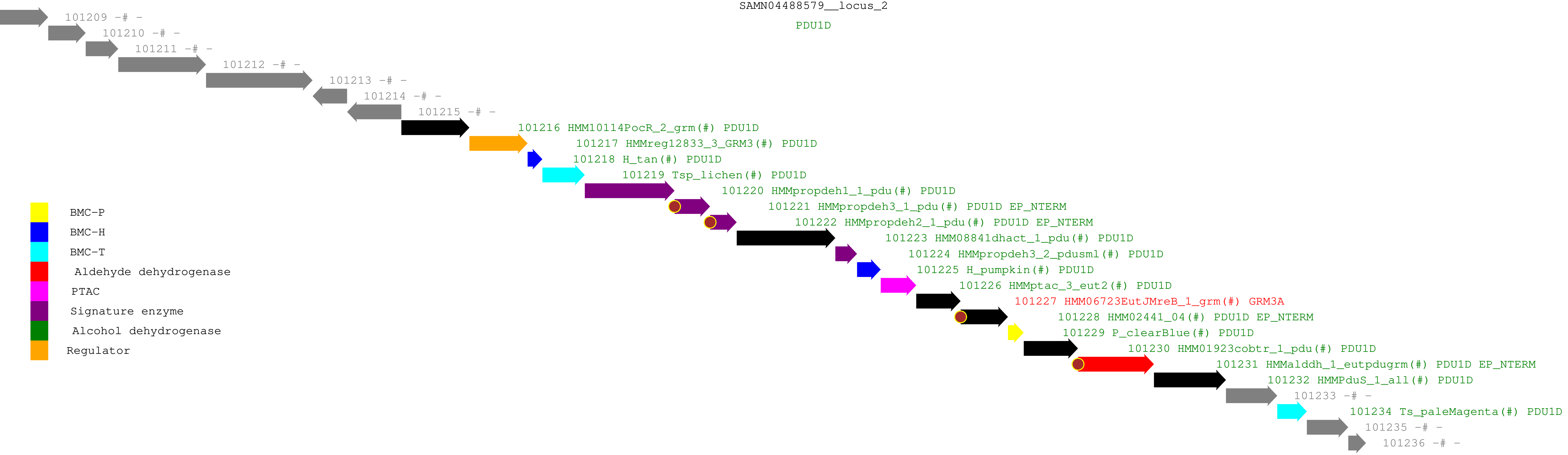

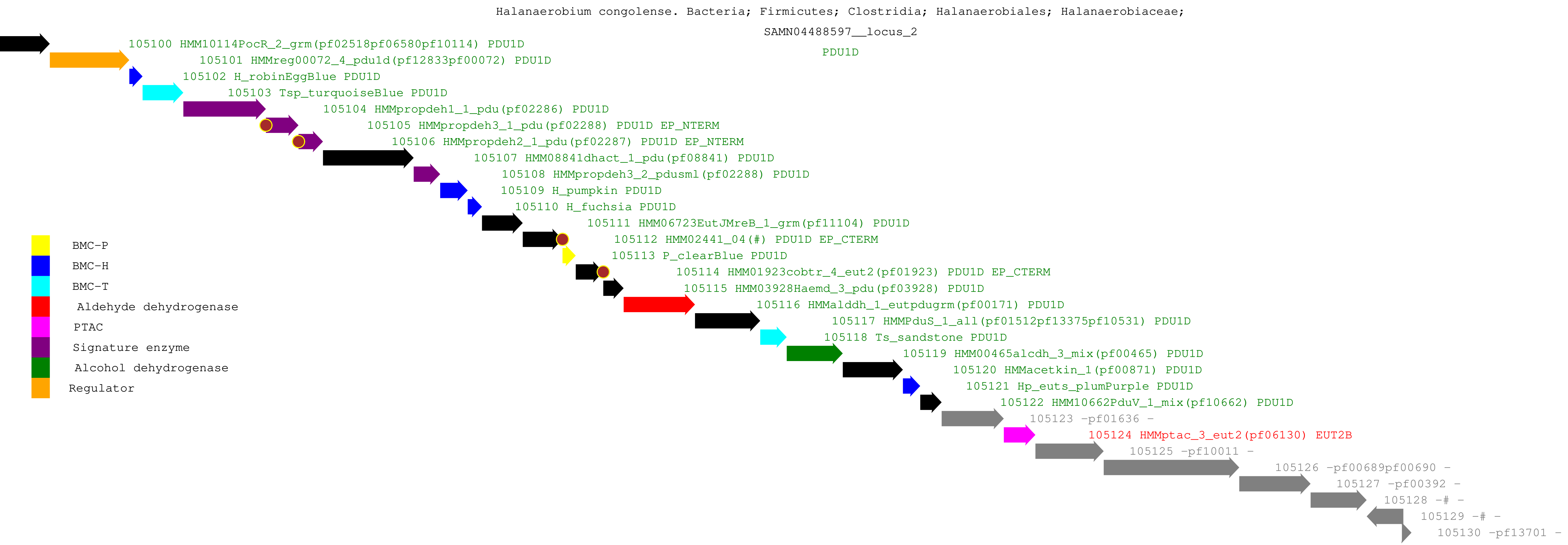
Halanaerobium congolense. Bacteria; Firmicutes; Clostridia; Halanaerobiales; Halanaerobiaceae;
SAMN04488597__locus_2
PDU1D
105100 HMM10114PocR_2_grm(pf02518pf06580pf10114) PDU1D
105101 HMMreg00072_4_pdu1d(pf12833pf00072) PDU1D
105102 H_robinEggBlue PDU1D
105103 Tsp_turquoiseBlue PDU1D
105104 HMMpropdeh1_1_pdu(pf02286) PDU1D
105105 HMMpropdeh3_1_pdu(pf02288) PDU1D EP_NTERM
105106 HMMpropdeh2_1_pdu(pf02287) PDU1D EP_NTERM
105107 HMM08841dhact_1_pdu(pf08841) PDU1D
105108 HMMpropdeh3_2_pdusml(pf02288) PDU1D
105109 H_pumpkin PDU1D
105110 H_fuchsia PDU1D
105111 HMM06723EutJMreB_1_grm(pf11104) PDU1D
105112 HMM02441_04(#) PDU1D EP_CTERM
105113 P_clearBlue PDU1D
105114 HMM01923cobtr_4_eut2(pf01923) PDU1D EP_CTERM
105115 HMM03928Haemd_3_pdu(pf03928) PDU1D
105116 HMMalddh_1_eutpdugrm(pf00171) PDU1D
105117 HMMPduS_1_all(pf01512pf13375pf10531) PDU1D
105118 Ts_sandstone PDU1D
105119 HMM00465alcdh_3_mix(pf00465) PDU1D
105120 HMMacetkin_1(pf00871) PDU1D
105121 Hp_euts_plumPurple PDU1D
105122 HMM10662PduV_1_mix(pf10662) PDU1D
105123 -pf01636 -
105124 HMMptac_3_eut2(pf06130) EUT2B
105125 -pf10011 -
105126 -pf00689pf00690 -
105127 -pf00392 -
105128 -# -
105129 -# -
105130 -pf13701 -
BMC-P
BMC-H
BMC-T
Aldehyde dehydrogenase
PTAC
Signature enzyme
Alcohol dehydrogenase
Regulator

Halanaerobium congolense. Bacteria; Firmicutes; Clostridia; Halanaerobiales; Halanaerobiaceae;

SAMN04488599__locus_1

PDU1D

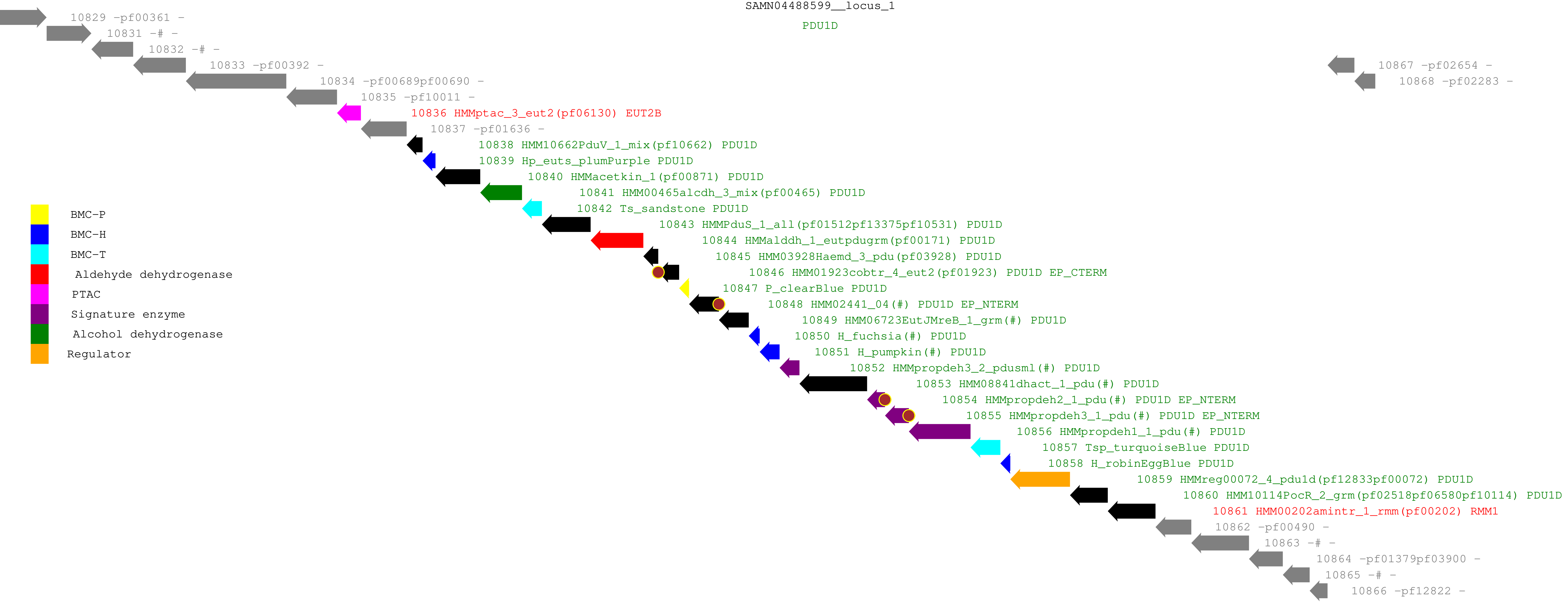

Propionispora vibrioides. Bacteria; Firmicutes; Negativicutes; Selenomonadales; Sporomusaceae;

SAMN04490178__locus_1

PDU1D

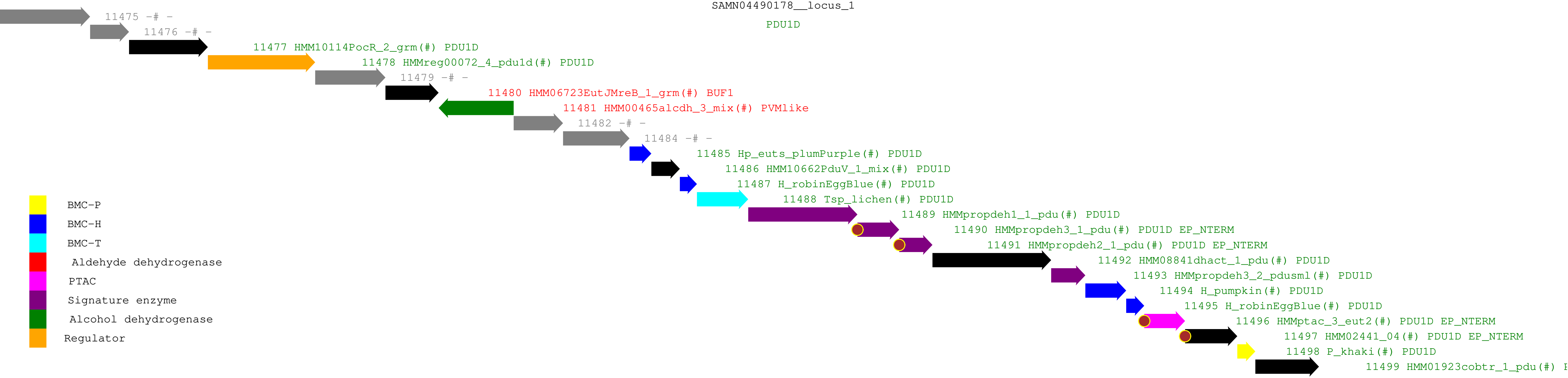

Pelosinus propionicus DSM 13327. Bacteria; Firmicutes; Negativicutes; Selenomonadales; Sporomusaceae;

SAMN04490355__locus_1

PDU1D

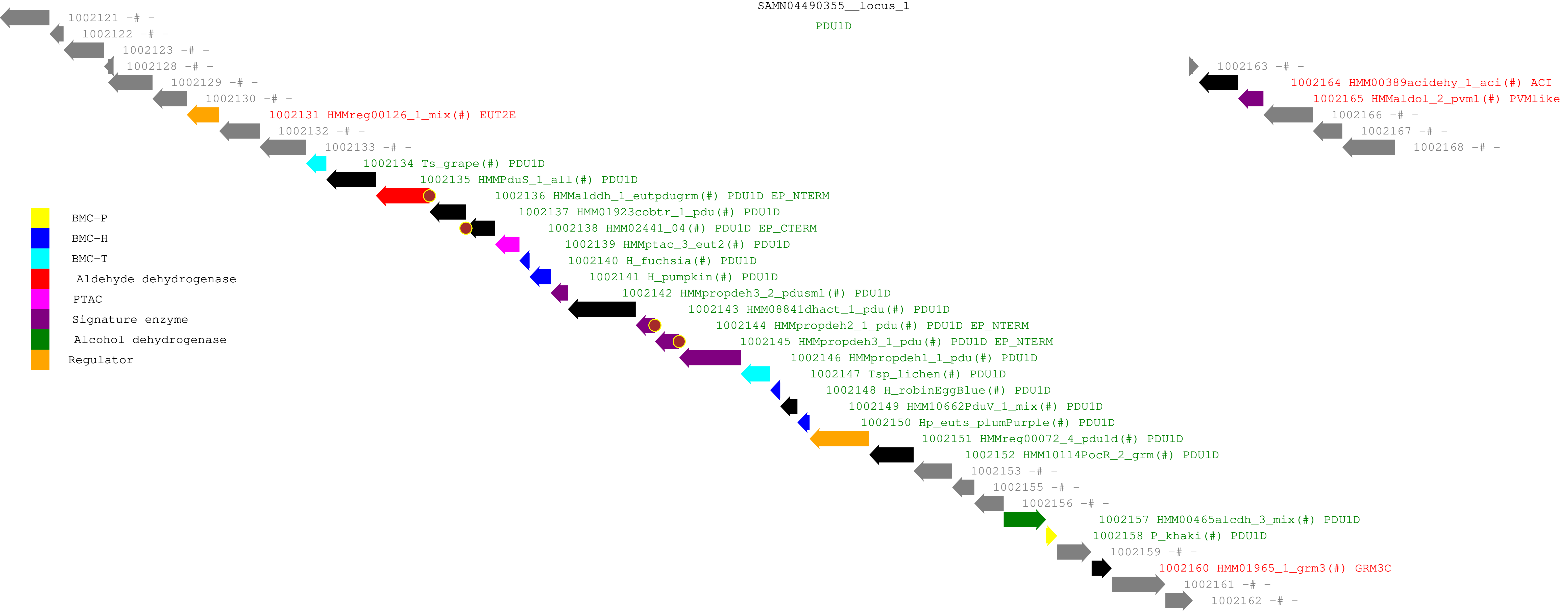

Eubacterium callanderi. Bacteria; Firmicutes; Clostridia; Clostridiales; Eubacteriaceae;

SAMN04515649__locus_1_main_sat

PDU1D

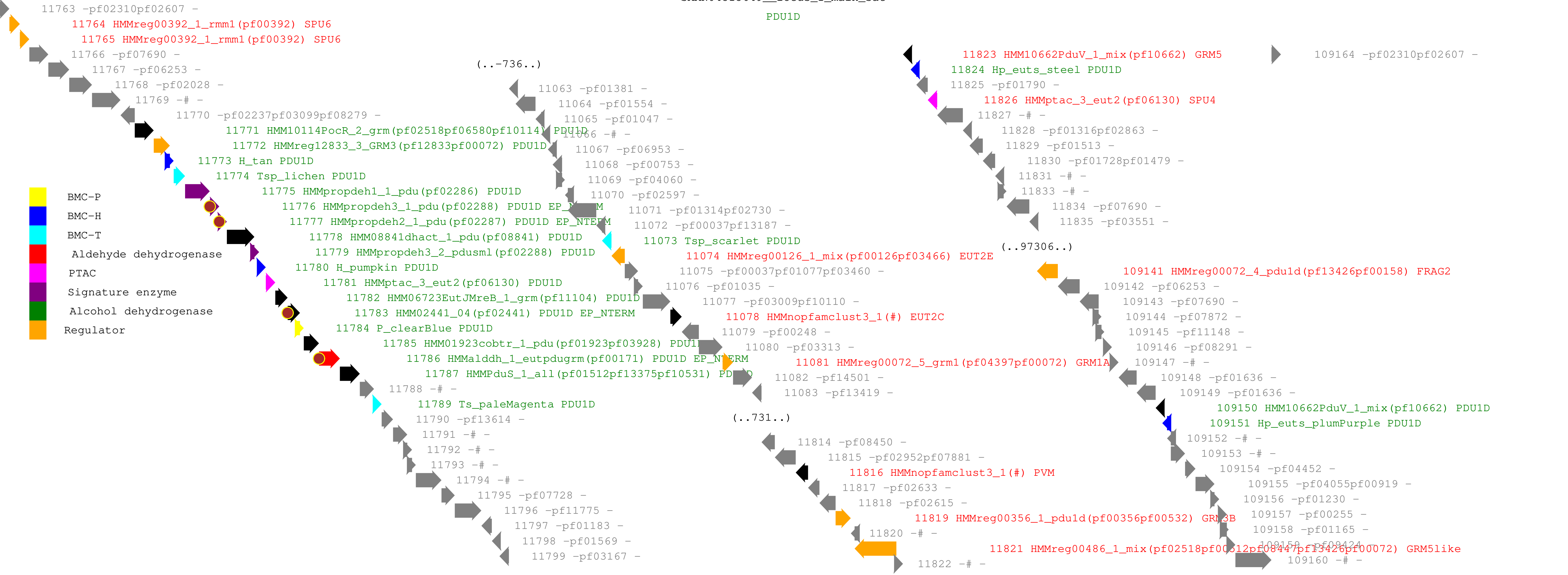

Halanaerobium congolense. Bacteria; Firmicutes; Clostridia; Halanaerobiales; Halanaerobiaceae;

SAMN04515654__locus_1

PDU1D

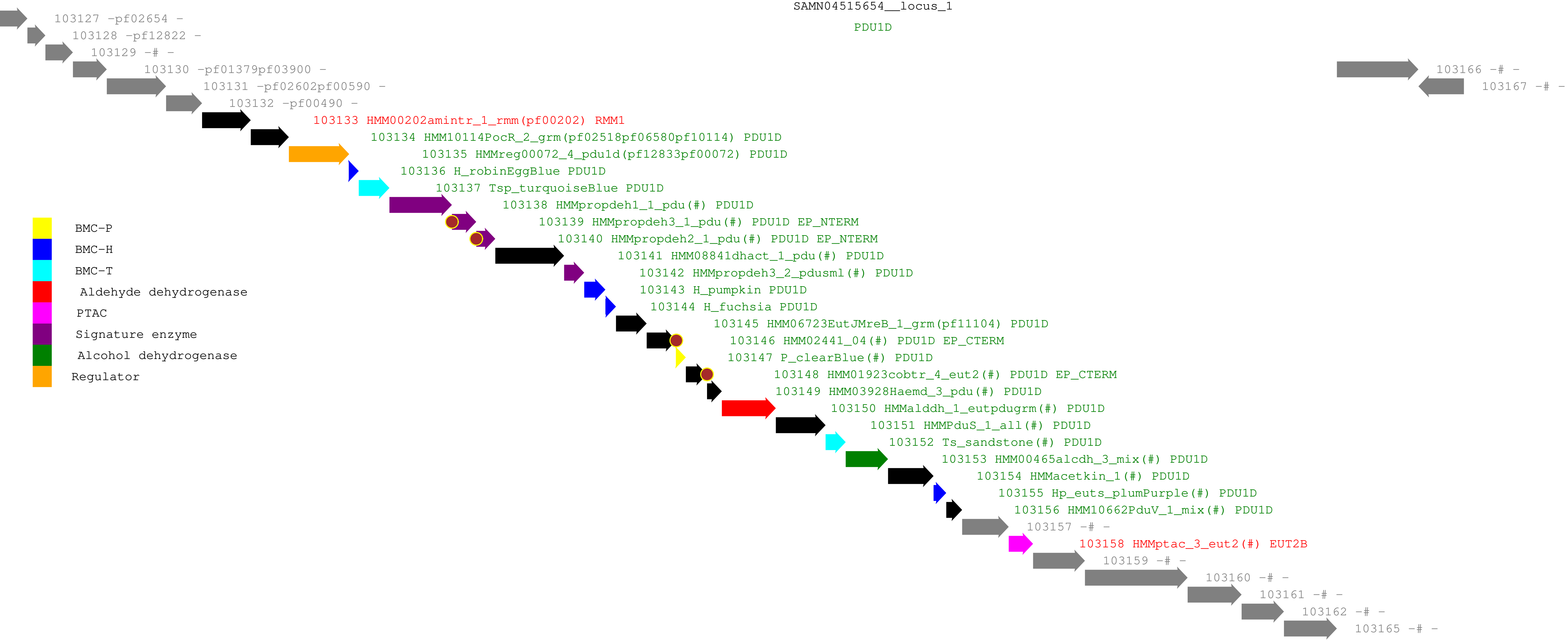

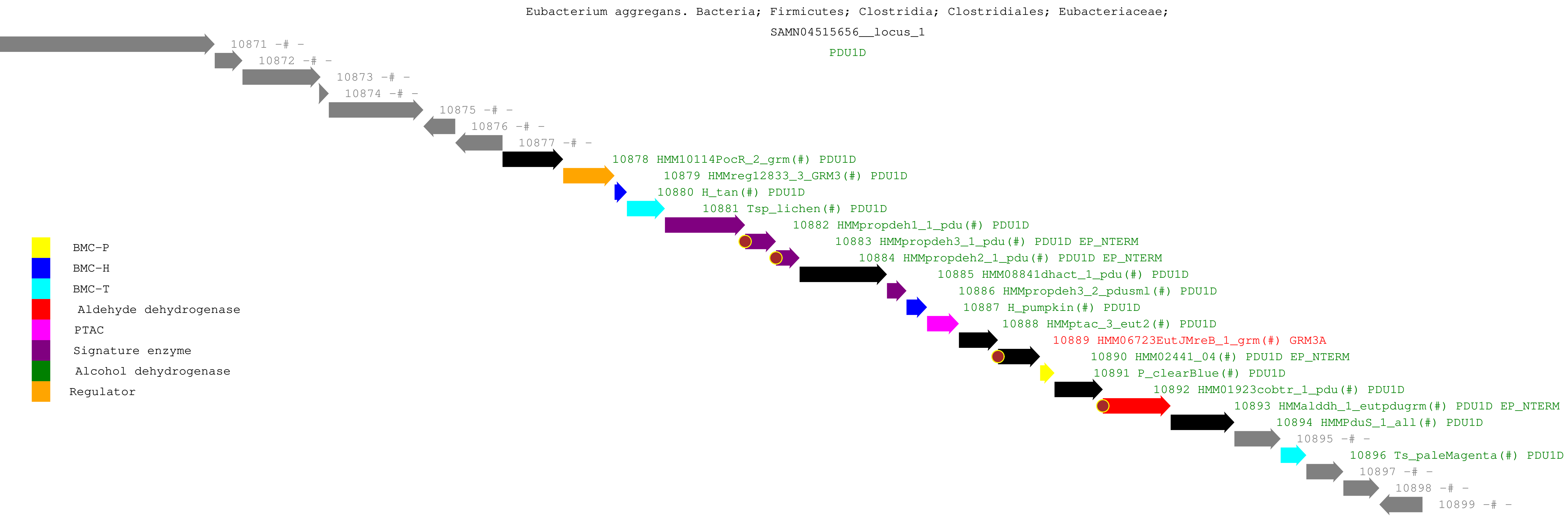
Eubacterium aggregans. Bacteria; Firmicutes; Clostridia; Clostridiales; Eubacteriaceae;
SAMN04515656__locus_1
PDU1D
10871 -# -
10872 -# -
10873 -# -
10874 -# -
10875 -# -
10876 -# -
10877 -# -
10878 HMM10114PocR_2_grm(#) PDU1D
10879 HMMreg12833_3_GRM3(#) PDU1D
10880 H_tan(#) PDU1D
10881 Tsp_lichen(#) PDU1D
10882 HMMpropdeh1_1_pdu(#) PDU1D
10883 HMMpropdeh3_1_pdu(#) PDU1D EP_NTERM
10884 HMMpropdeh2_1_pdu(#) PDU1D EP_NTERM
10885 HMM08841dhact_1_pdu(#) PDU1D
10886 HMMpropdeh3_2_pdusml(#) PDU1D
10887 H_pumpkin(#) PDU1D
10888 HMMptac_3_eut2(#) PDU1D
10889 HMM06723EutJMreB_1_grm(#) GRM3A
10890 HMM02441_04(#) PDU1D EP_NTERM
10891 P_clearBlue(#) PDU1D
10892 HMM01923cobtr_1_pdu(#) PDU1D
10893 HMMalddh_1_eutpdugrm(#) PDU1D EP_NTERM
10894 HMMPduS_1_all(#) PDU1D
10895 -# -
10896 Ts_paleMagenta(#) PDU1D
10897 -# -
10898 -# -
10899 -# -
BMC-P
BMC-H
BMC-T
Aldehyde dehydrogenase
PTAC
Signature enzyme
Alcohol dehydrogenase
Regulator

Sporolituus thermophilus DSM 23256. Bacteria; Firmicutes; Negativicutes; Selenomonadales; Sporomusaceae;

SAMN05660235__locus_1

PDU1D

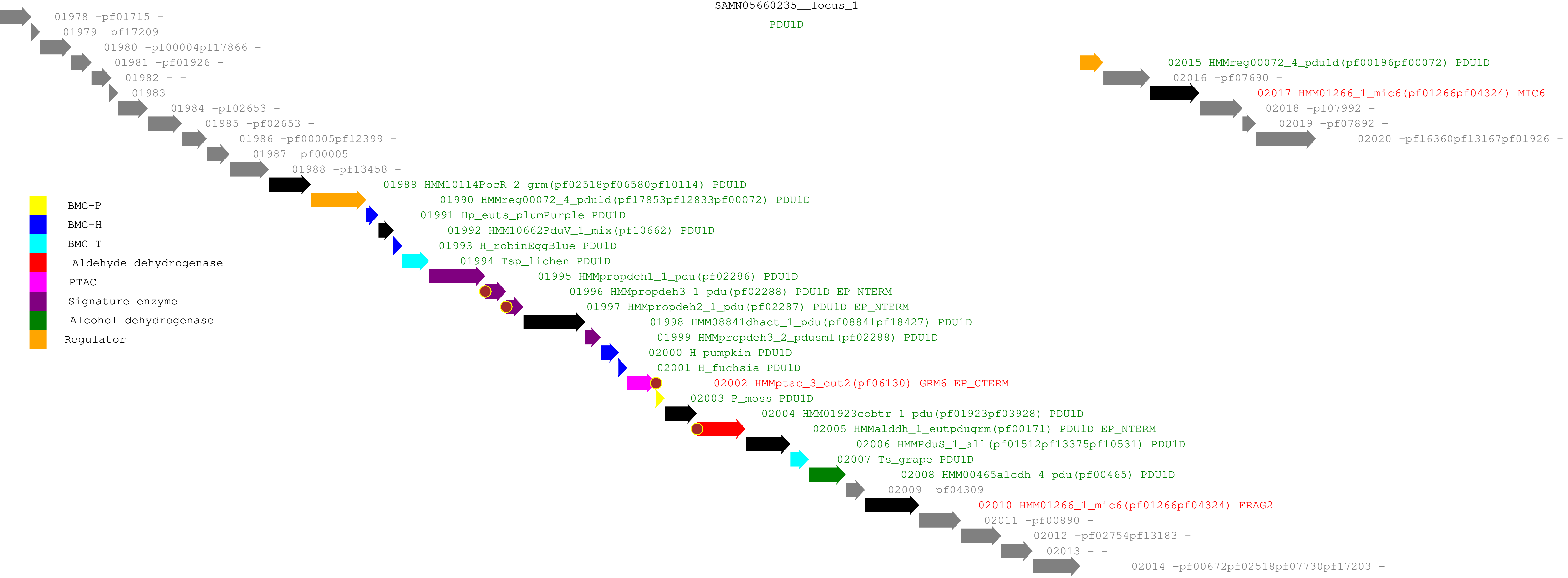

Thermoanaerobacterium sp. RBIITD. Bacteria; Firmicutes; Clostridia; Thermoanaerobacterales;

SAMN05660242__locus_1

PDU1D

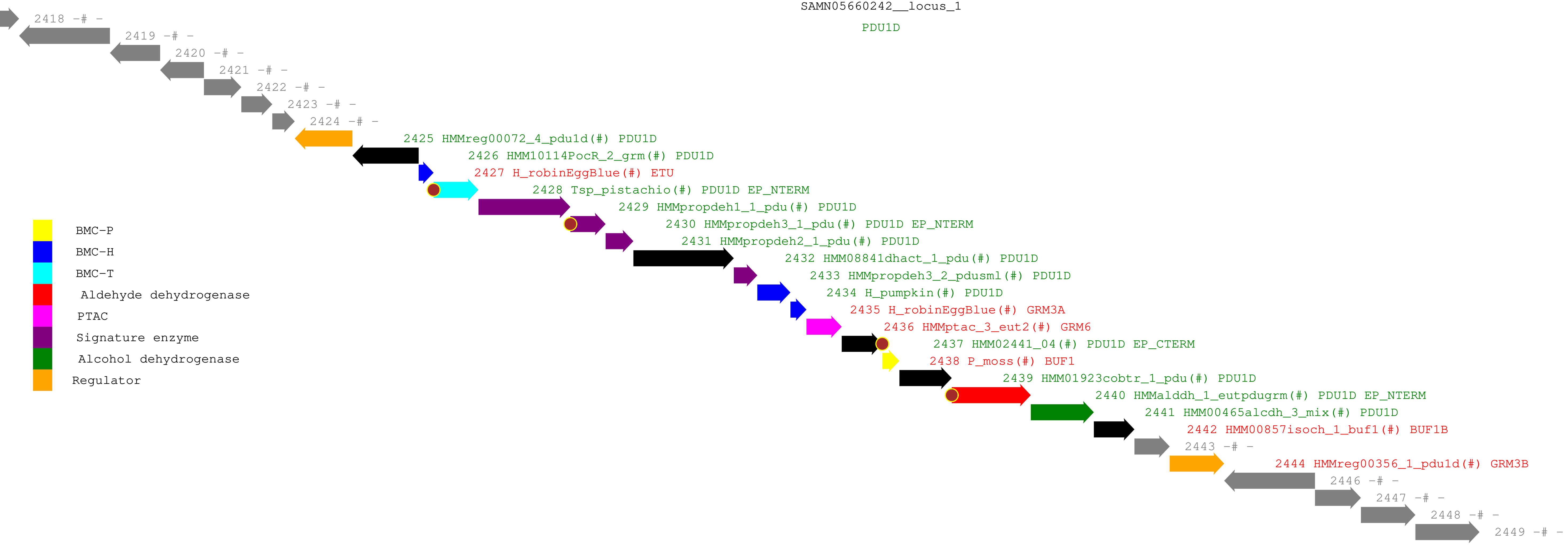

Caldanaerovirga acetigignens. Bacteria; Firmicutes; Clostridia; Thermoanaerobacterales;

SAMN05660826__locus_1

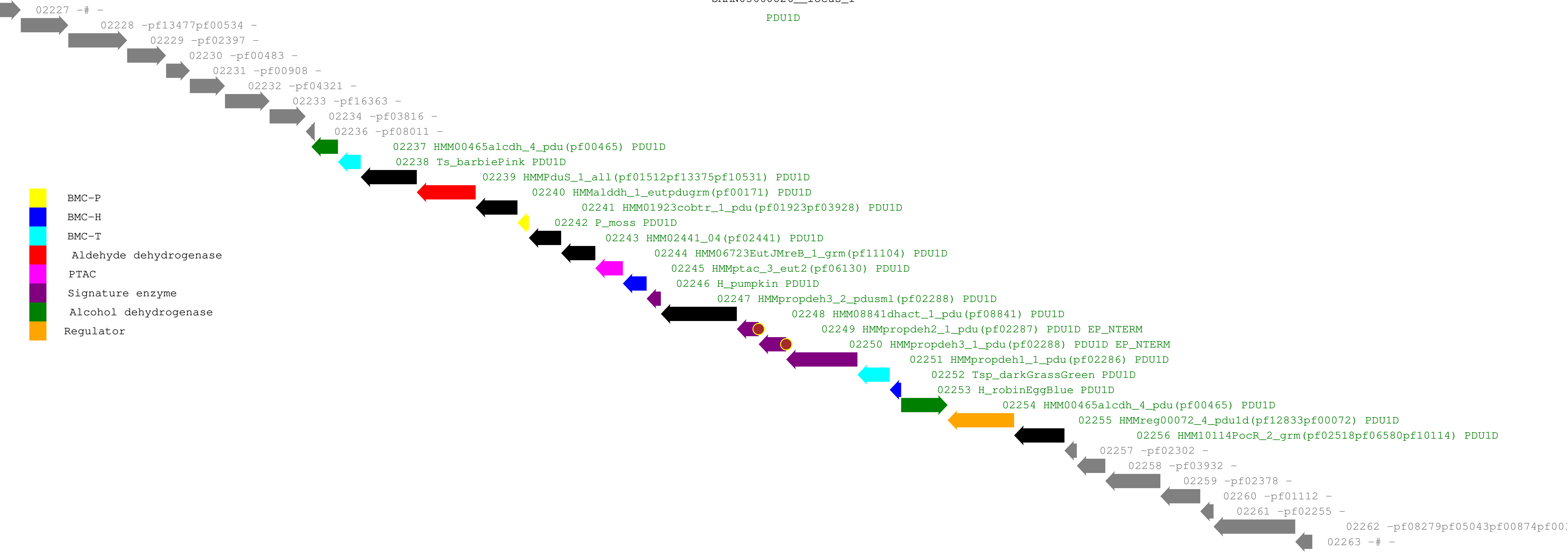

Sporomusa acidovorans DSM 3132. Bacteria; Firmicutes; Negativicutes; Selenomonadales; Sporomusaceae;

SPACI__locus_3

PDU1D

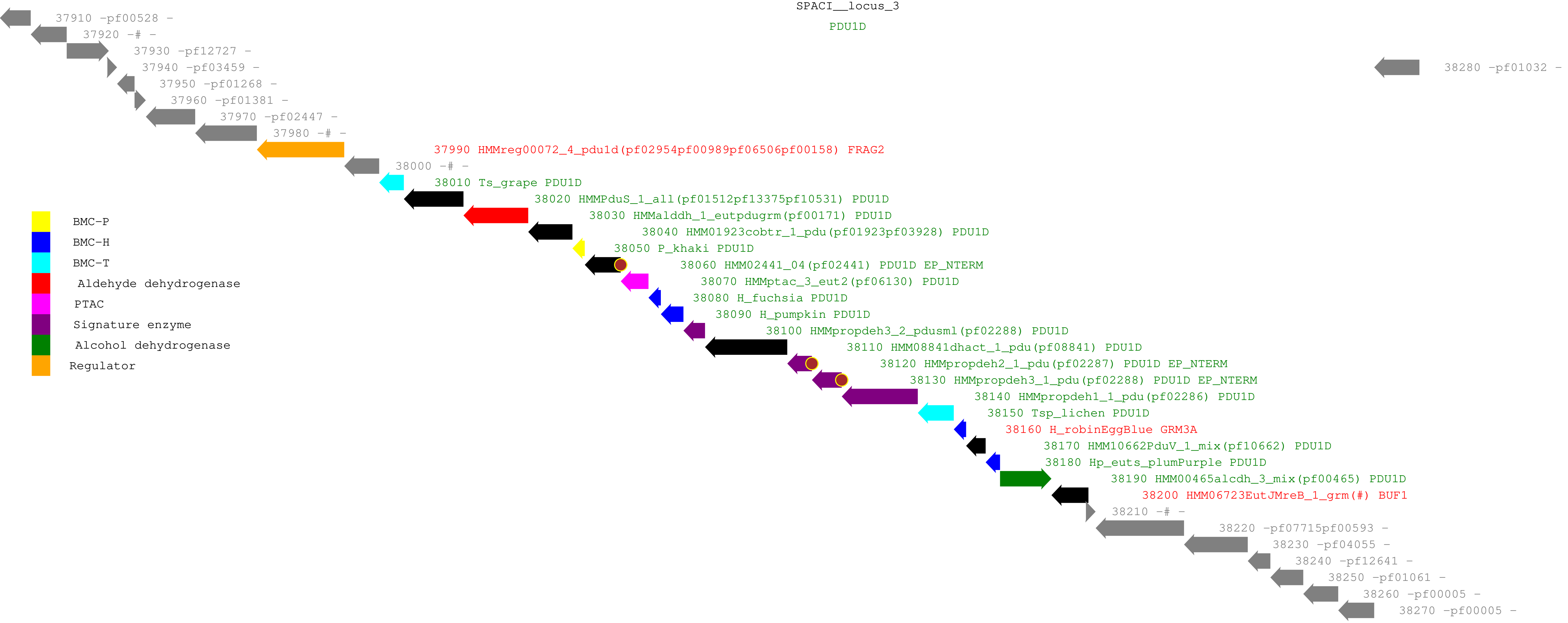

Sporomusa acidovorans DSM 3132. Bacteria; Firmicutes; Negativicutes; Selenomonadales; Sporomusaceae;

SPACI__locus_4

PDU1D

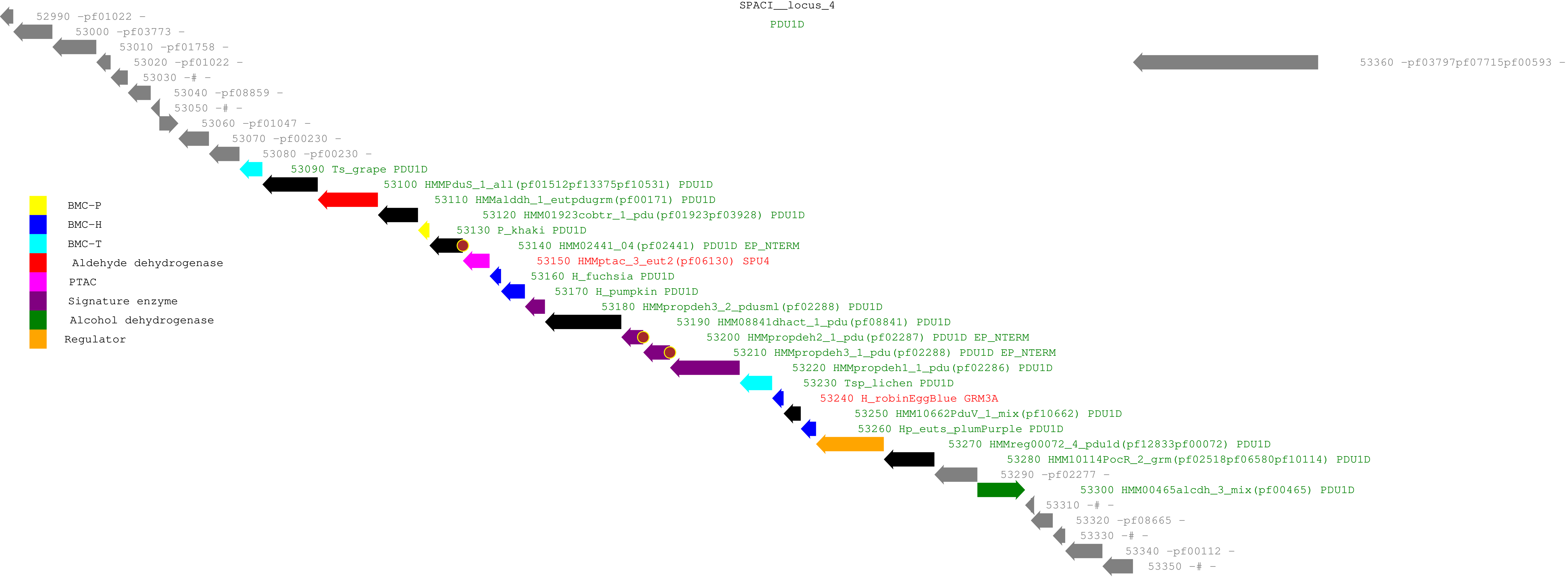

Sporomusaceae bacterium. Bacteria; Firmicutes; Negativicutes; Selenomonadales; Sporomusaceae;

SPFL3102__locus_1

PDU1D

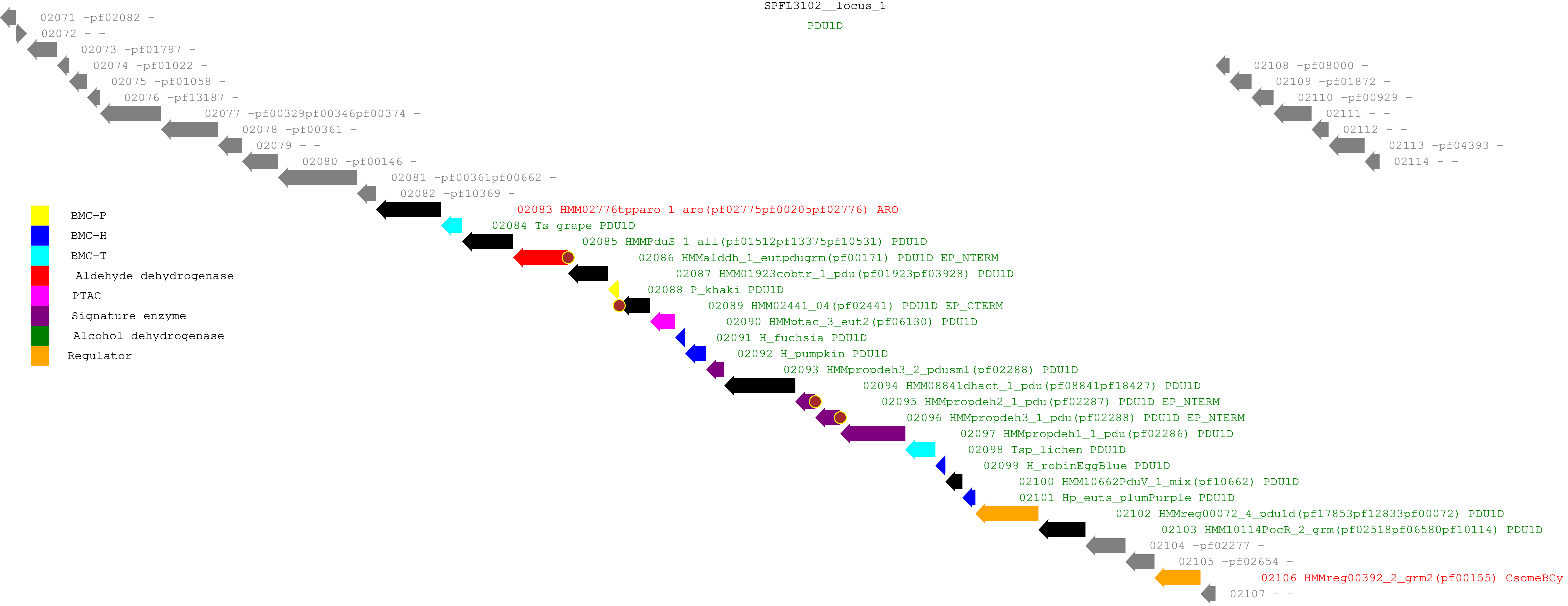

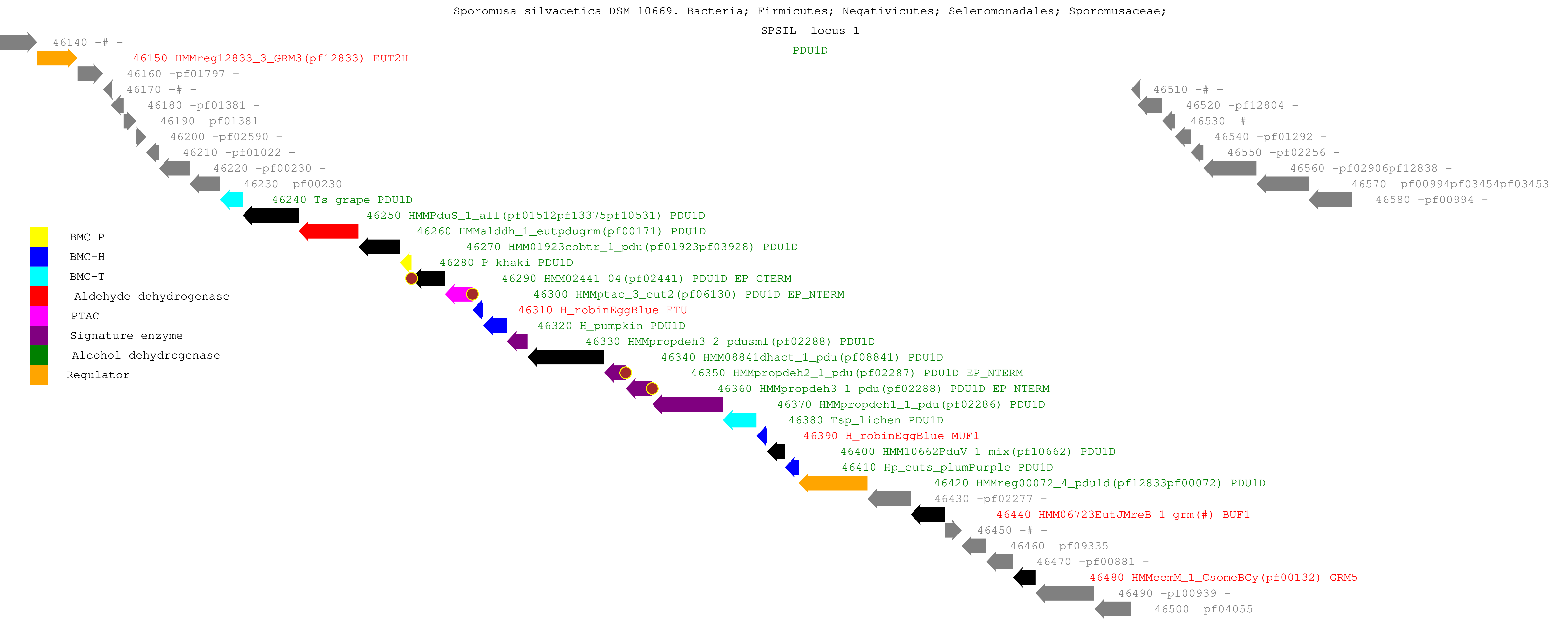
Sporomusa silvacetica DSM 10669. Bacteria; Firmicutes; Negativicutes; Selenomonadales; Sporomusaceae;
SPSIL__locus_1
PDU1D
46140 -# -
46150 HMMreg12833_3_GRM3(pf12833) EUT2H
46160 -pf01797 -
46170 -# -
46180 -pf01381 -
46190 -pf01381 -
46200 -pf02590 -
46210 -pf01022 -
46220 -pf00230 -
46230 -pf00230 -
46240 Ts_grape PDU1D
46250 HMMPduS_1_all(pf01512pf13375pf10531) PDU1D
46260 HMMalddh_1_eutpdugrm(pf00171) PDU1D
46270 HMM01923cobtr_1_pdu(pf01923pf03928) PDU1D
46280 P_khaki PDU1D
46290 HMM02441_04(pf02441) PDU1D EP_CTERM
46300 HMMptac_3_eut2(pf06130) PDU1D EP_NTERM
46310 H_robinEggBlue ETU
46320 H_pumpkin PDU1D
46330 HMMpropdeh3_2_pdusml(pf02288) PDU1D
46340 HMM08841dhact_1_pdu(pf08841) PDU1D
46350 HMMpropdeh2_1_pdu(pf02287) PDU1D EP_NTERM
46360 HMMpropdeh3_1_pdu(pf02288) PDU1D EP_NTERM
46370 HMMpropdeh1_1_pdu(pf02286) PDU1D
46380 Tsp_lichen PDU1D
46390 H_robinEggBlue MUF1
46400 HMM10662PduV_1_mix(pf10662) PDU1D
46410 Hp_euts_plumPurple PDU1D
46420 HMMreg00072_4_pdu1d(pf12833pf00072) PDU1D
46430 -pf02277 -
46440 HMM06723EutJMreB_1_grm(#) BUF1
46450 -# -
46460 -pf09335 -
46470 -pf00881 -
46480 HMMccmM_1_CsomeBCy(pf00132) GRM5
46490 -pf00939 -
46500 -pf04055 -
46510 -# -
46520 -pf12804 -
46530 -# -
46540 -pf01292 -
46550 -pf02256 -
46560 -pf02906pf12838 -
46570 -pf00994pf03454pf03453 -
46580 -pf00994 -
BMC-P
BMC-H
BMC-T
Aldehyde dehydrogenase
PTAC
Signature enzyme
Alcohol dehydrogenase
Regulator

Sporomusa sphaeroides DSM 2875. Bacteria; Firmicutes; Negativicutes; Selenomonadales; Sporomusaceae;

SPSPH__locus_1

PDU1D

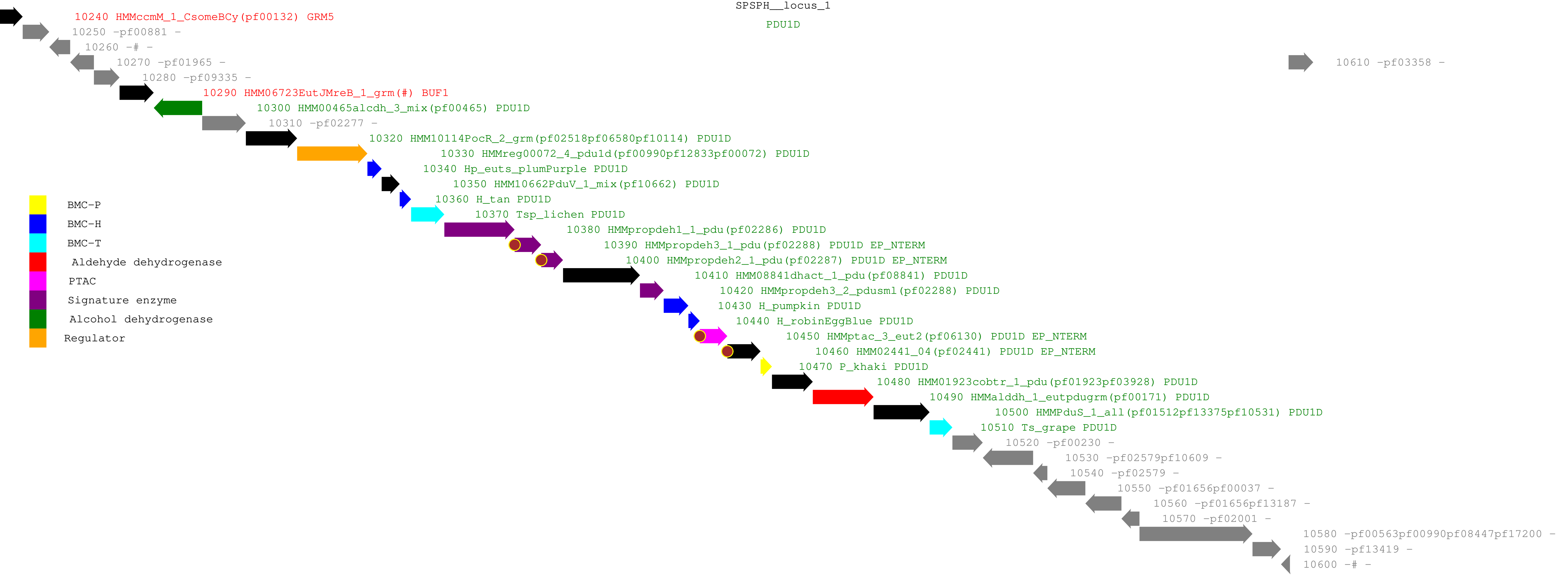

Clostridiales bacterium PH28_bin88. Bacteria; Firmicutes; Clostridia; Clostridiales.

SY88__locus_1

PDU1D

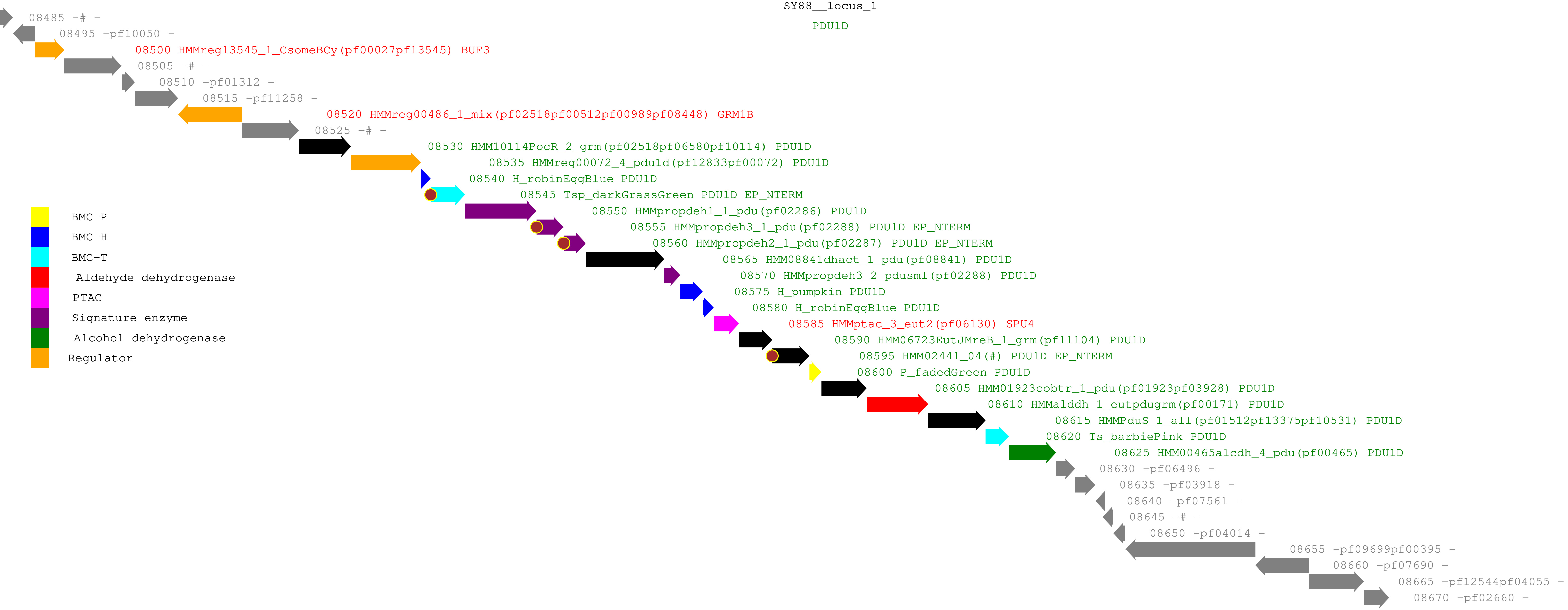

Thermoanaerobacter sp. (strain X514). Bacteria; Firmicutes; Clostridia; Thermoanaerobacterales;

Teth514__locus_1

PDU1D

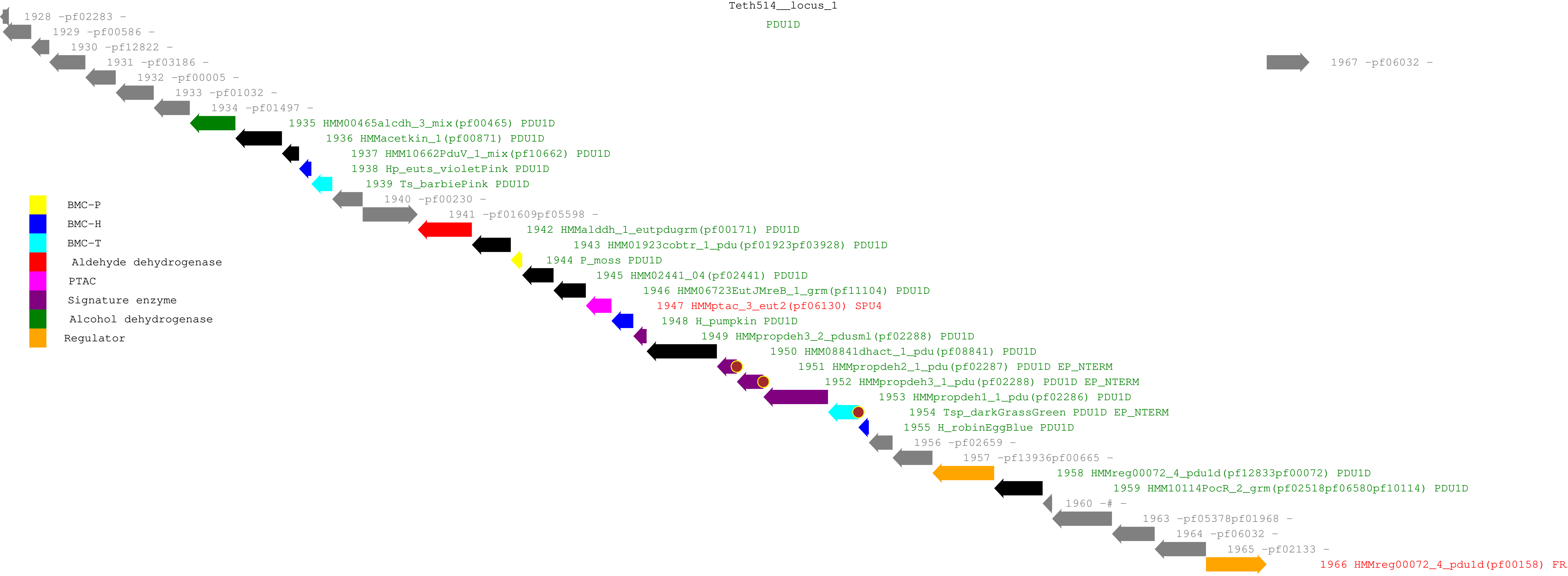

Thermoanaerobacter sp. X561. Bacteria; Firmicutes; Clostridia; Thermoanaerobacterales;

Teth561_PD_locus_1

PDU1D

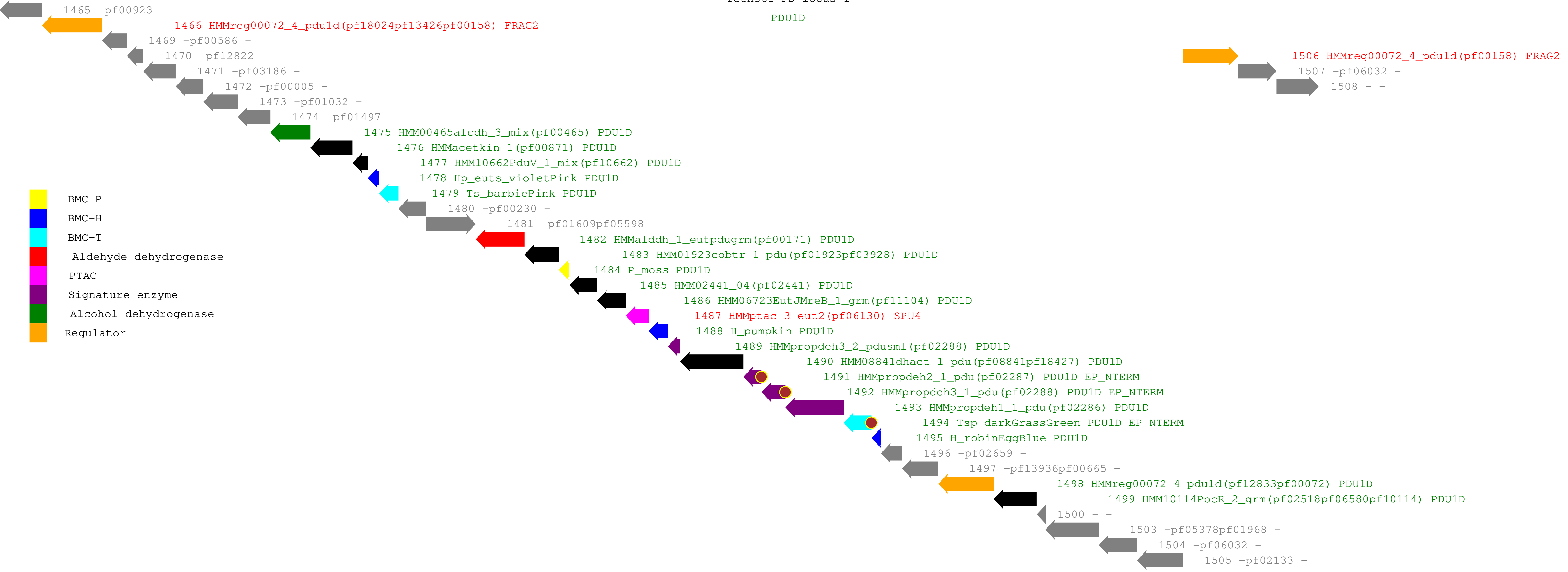

Thermincola ferriacetica. Bacteria; Firmicutes; Clostridia; Clostridiales; Peptococcaceae;

Tfer__locus_1

PDU1D

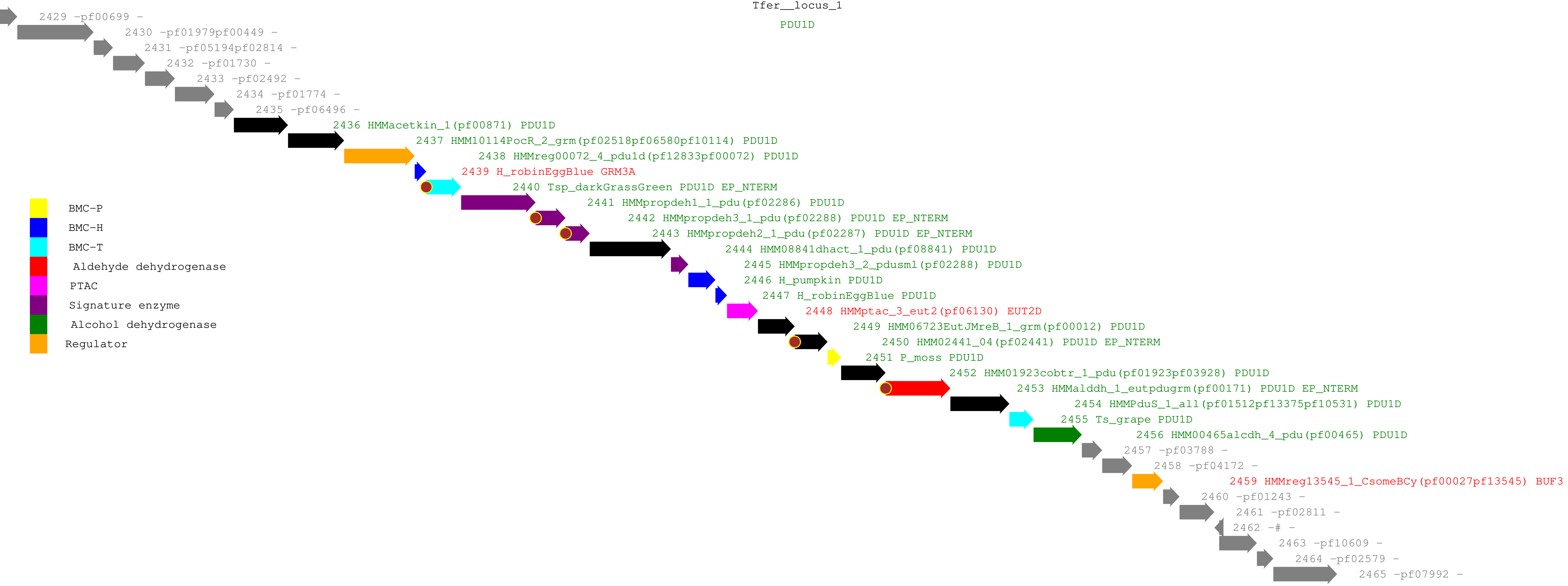

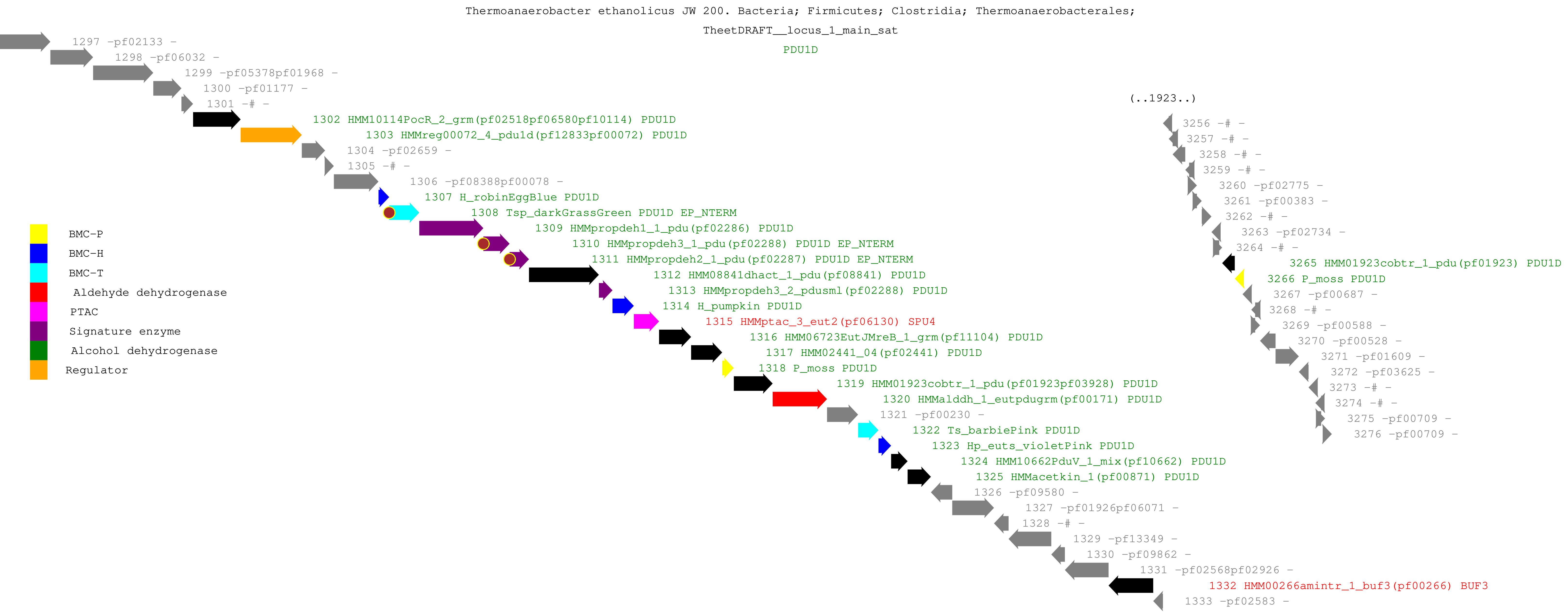
Thermoanaerobacter ethanolicus JW 200. Bacteria; Firmicutes; Clostridia; Thermoanaerobacterales;
TheetDRAFT__locus_1_main_sat
PDU1D
1297 -pf02133 -
1298 -pf06032 -
1299 -pf05378pf01968 -
1300 -pf01177 -
1301 -# -
1302 HMM10114PocR_2_grm(pf02518pf06580pf10114) PDU1D
1303 HMMreg00072_4_pdu1d(pf12833pf00072) PDU1D
1304 -pf02659 -
1305 -# -
1306 -pf08388pf00078 -
1307 H_robinEggBlue PDU1D
1308 Tsp_darkGrassGreen PDU1D EP_NTERM
1309 HMMpropdeh1_1_pdu(pf02286) PDU1D
1310 HMMpropdeh3_1_pdu(pf02288) PDU1D EP_NTERM
1311 HMMpropdeh2_1_pdu(pf02287) PDU1D EP_NTERM
1312 HMM08841dhact_1_pdu(pf08841) PDU1D
1313 HMMpropdeh3_2_pdusml(pf02288) PDU1D
1314 H_pumpkin PDU1D
1315 HMMptac_3_eut2(pf06130) SPU4
1316 HMM06723EutJMreB_1_grm(pf11104) PDU1D
1317 HMM02441_04(pf02441) PDU1D
1318 P_moss PDU1D
1319 HMM01923cobtr_1_pdu(pf01923pf03928) PDU1D
1320 HMMalddh_1_eutpdugrm(pf00171) PDU1D
1321 -pf00230 -
1322 Ts_barbiePink PDU1D
1323 Hp_euts_violetPink PDU1D
1324 HMM10662PduV_1_mix(pf10662) PDU1D
1325 HMMacetkin_1(pf00871) PDU1D
1326 -pf09580 -
1327 -pf01926pf06071 -
1328 -# -
1329 -pf13349 -
1330 -pf09862 -
1331 -pf02568pf02926 -
1332 HMM00266amintr_1_buf3(pf00266) BUF3
1333 -pf02583 -
(..1923..)
3256 -# -
3257 -# -
3258 -# -
3259 -# -
3260 -pf02775 -
3261 -pf00383 -
3262 -# -
3263 -pf02734 -
3264 -# -
3265 HMM01923cobtr_1_pdu(pf01923) PDU1D
3266 P_moss PDU1D
3267 -pf00687 -
3268 -# -
3269 -pf00588 -
3270 -pf00528 -
3271 -pf01609 -
3272 -pf03625 -
3273 -# -
3274 -# -
3275 -pf00709 -
3276 -pf00709 -
BMC-P
BMC-H
BMC-T
Aldehyde dehydrogenase
PTAC
Signature enzyme
Alcohol dehydrogenase
Regulator

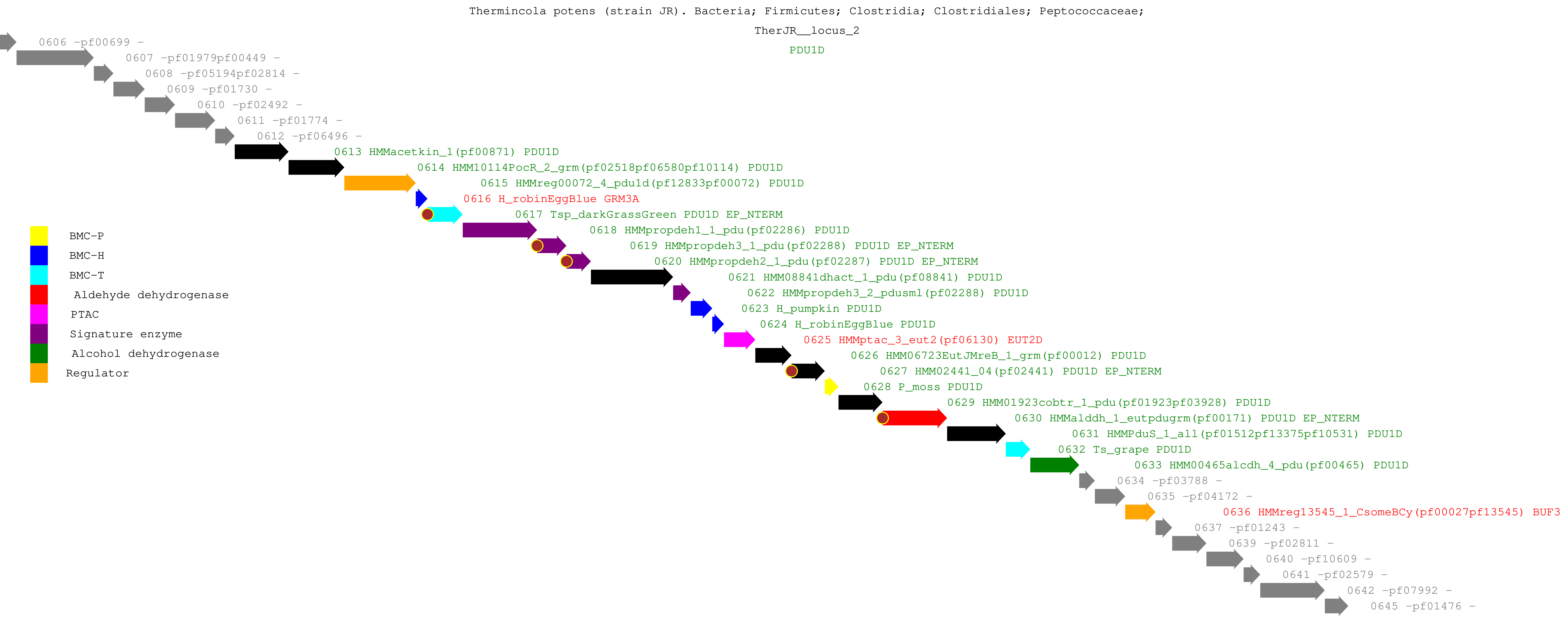
Thermincola potens (strain JR). Bacteria; Firmicutes; Clostridia; Clostridiales; Peptococcaceae;
TherJR__locus_2
PDU1D
0606 -pf00699 -
0607 -pf01979pf00449 -
0608 -pf05194pf02814 -
0609 -pf01730 -
0610 -pf02492 -
0611 -pf01774 -
0612 -pf06496 -
0613 HMMacetkin_1(pf00871) PDU1D
0614 HMM10114PocR_2_grm(pf02518pf06580pf10114) PDU1D
0615 HMMreg00072_4_pdu1d(pf12833pf00072) PDU1D
0616 H_robinEggBlue GRM3A
0617 Tsp_darkGrassGreen PDU1D EP_NTERM
0618 HMMpropdeh1_1_pdu(pf02286) PDU1D
0619 HMMpropdeh3_1_pdu(pf02288) PDU1D EP_NTERM
0620 HMMpropdeh2_1_pdu(pf02287) PDU1D EP_NTERM
0621 HMM08841dhact_1_pdu(pf08841) PDU1D
0622 HMMpropdeh3_2_pdusml(pf02288) PDU1D
0623 H_pumpkin PDU1D
0624 H_robinEggBlue PDU1D
0625 HMMptac_3_eut2(pf06130) EUT2D
0626 HMM06723EutJMreB_1_grm(pf00012) PDU1D
0627 HMM02441_04(pf02441) PDU1D EP_NTERM
0628 P_moss PDU1D
0629 HMM01923cobtr_1_pdu(pf01923pf03928) PDU1D
0630 HMMalddh_1_eutpdugrm(pf00171) PDU1D EP_NTERM
0631 HMMPduS_1_all(pf01512pf13375pf10531) PDU1D
0632 Ts_grape PDU1D
0633 HMM00465alcdh_4_pdu(pf00465) PDU1D
0634 -pf03788 -
0635 -pf04172 -
0636 HMMreg13545_1_CsomeBCy(pf00027pf13545) BUF3
0637 -pf01243 -
0639 -pf02811 -
0640 -pf10609 -
0641 -pf02579 -
0642 -pf07992 -
0645 -pf01476 -
BMC-P
BMC-H
BMC-T
Aldehyde dehydrogenase
PTAC
Signature enzyme
Alcohol dehydrogenase
Regulator

Thermoanaerobacterium thermosaccharolyticum (Clostridium Bacteria; Firmicutes; Clostridia; Thermoanaerobacterales;

Thert__locus_1

PDU1D

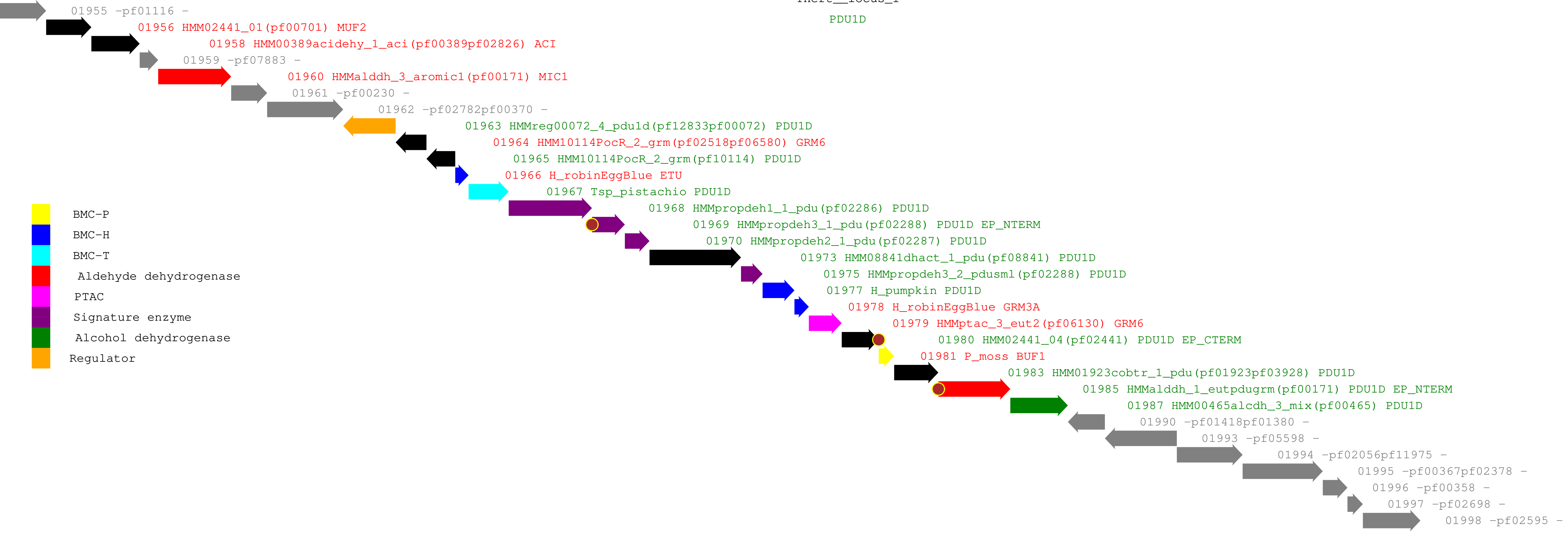

Thermoanaerobacterium thermosaccharolyticum M0795. Bacteria; Firmicutes; Clostridia; Thermoanaerobacterales;

Thethe__locus_1

PDU1D

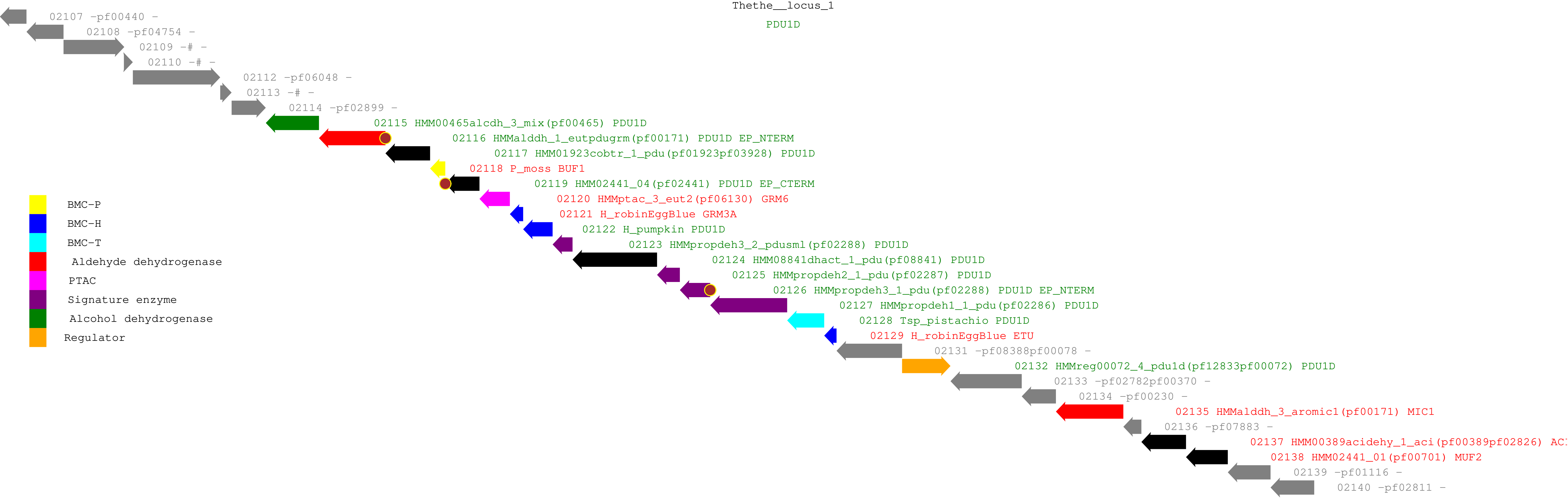

Thermovirga lienii (strain ATCC BAA-1197 / DSM 17291 / Cas60314). Bacteria; Synergistetes; Synergistia; Synergistales; Synergistaceae;

Tlie__locus_1

PDU1D

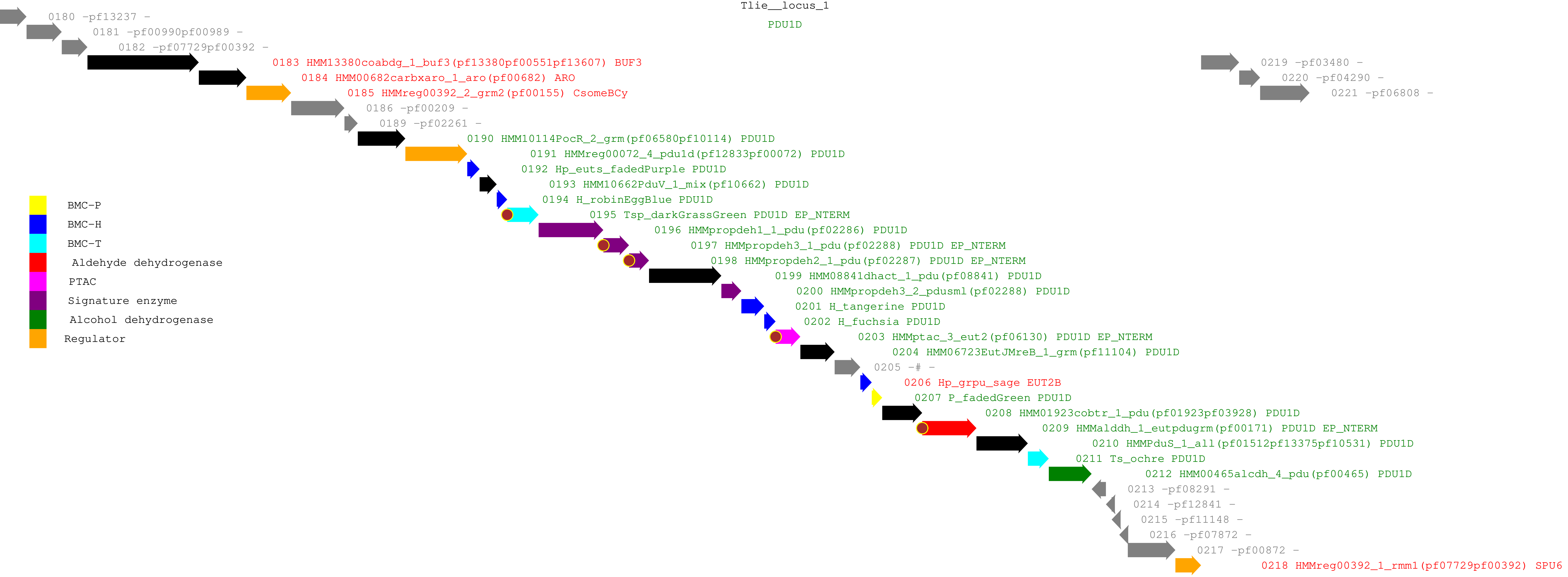

Thermosediminibacter oceani (strain ATCC BAA-1034 / DSM 16646 / Bacteria; Firmicutes; Clostridia; Thermoanaerobacterales;

Toce__locus_1

PDU1D

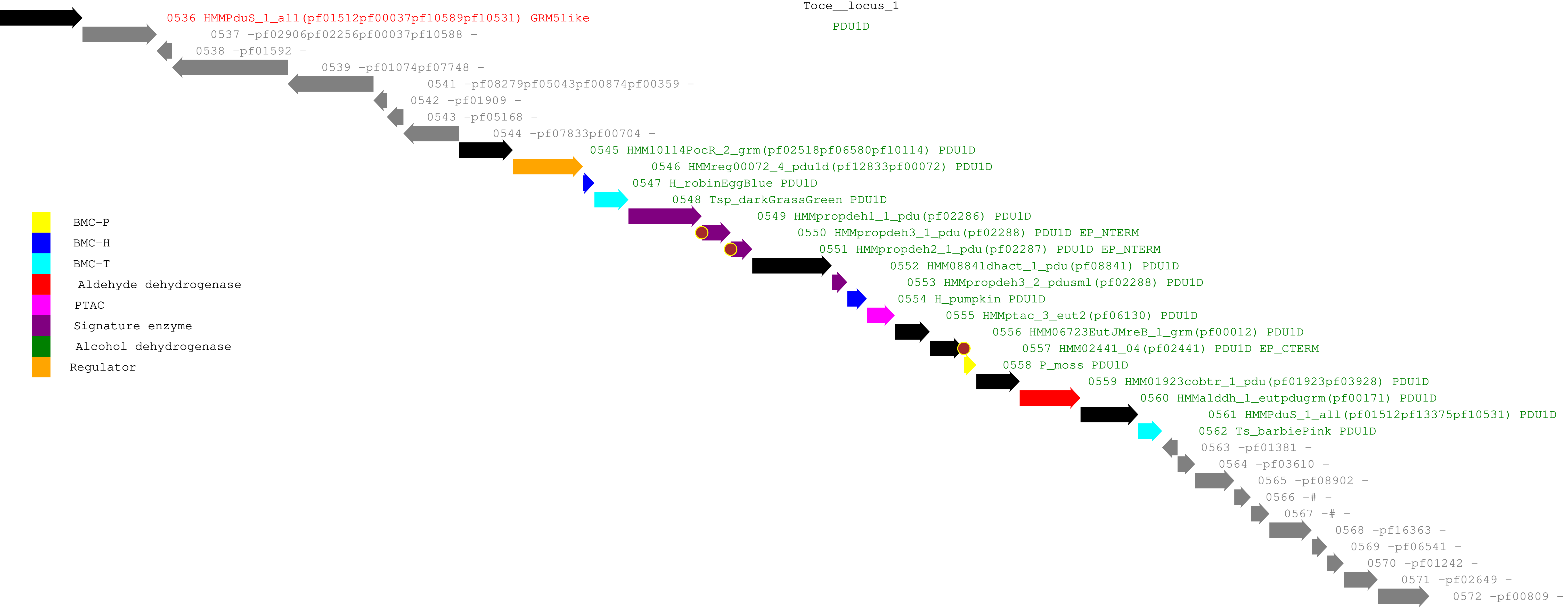

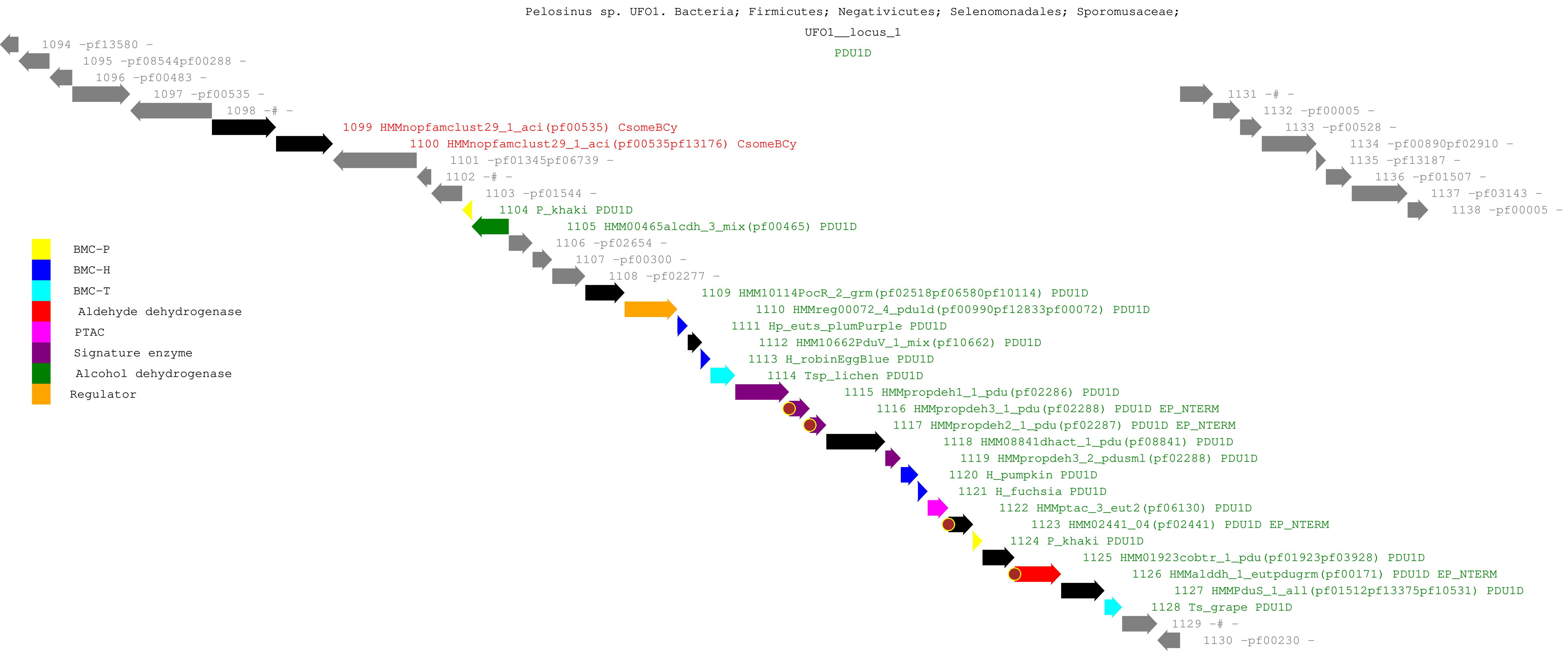
Pelosinus sp. UFO1. Bacteria; Firmicutes; Negativicutes; Selenomonadales; Sporomusaceae;
UFO1__locus_1
PDU1D
1094 -pf13580 -
1095 -pf08544pf00288 -
1096 -pf00483 -
1097 -pf00535 -
1098 -# -
1099 HMMnopfamclust29_1_aci(pf00535) CsomeBCy
1100 HMMnopfamclust29_1_aci(pf00535pf13176) CsomeBCy
1101 -pf01345pf06739 -
1102 -# -
1103 -pf01544 -
1104 P_khaki PDU1D
1105 HMM00465alcdh_3_mix(pf00465) PDU1D
1106 -pf02654 -
1107 -pf00300 -
1108 -pf02277 -
1109 HMM10114PocR_2_grm(pf02518pf06580pf10114) PDU1D
1110 HMMreg00072_4_pdu1d(pf00990pf12833pf00072) PDU1D
1111 Hp_euts_plumPurple PDU1D
1112 HMM10662PduV_1_mix(pf10662) PDU1D
1113 H_robinEggBlue PDU1D
1114 Tsp_lichen PDU1D
1115 HMMpropdeh1_1_pdu(pf02286) PDU1D
1116 HMMpropdeh3_1_pdu(pf02288) PDU1D EP_NTERM
1117 HMMpropdeh2_1_pdu(pf02287) PDU1D EP_NTERM
1118 HMM08841dhact_1_pdu(pf08841) PDU1D
1119 HMMpropdeh3_2_pdusml(pf02288) PDU1D
1120 H_pumpkin PDU1D
1121 H_fuchsia PDU1D
1122 HMMptac_3_eut2(pf06130) PDU1D
1123 HMM02441_04(pf02441) PDU1D EP_NTERM
1124 P_khaki PDU1D
1125 HMM01923cobtr_1_pdu(pf01923pf03928) PDU1D
1126 HMMalddh_1_eutpdugrm(pf00171) PDU1D EP_NTERM
1127 HMMPduS_1_all(pf01512pf13375pf10531) PDU1D
1128 Ts_grape PDU1D
1129 -# -
1130 -pf00230 -
1131 -# -
1132 -pf00005 -
1133 -pf00528 -
1134 -pf00890pf02910 -
1135 -pf13187 -
1136 -pf01507 -
1137 -pf03143 -
1138 -pf00005 -
BMC-P
BMC-H
BMC-T
Aldehyde dehydrogenase
PTAC
Signature enzyme
Alcohol dehydrogenase
Regulator

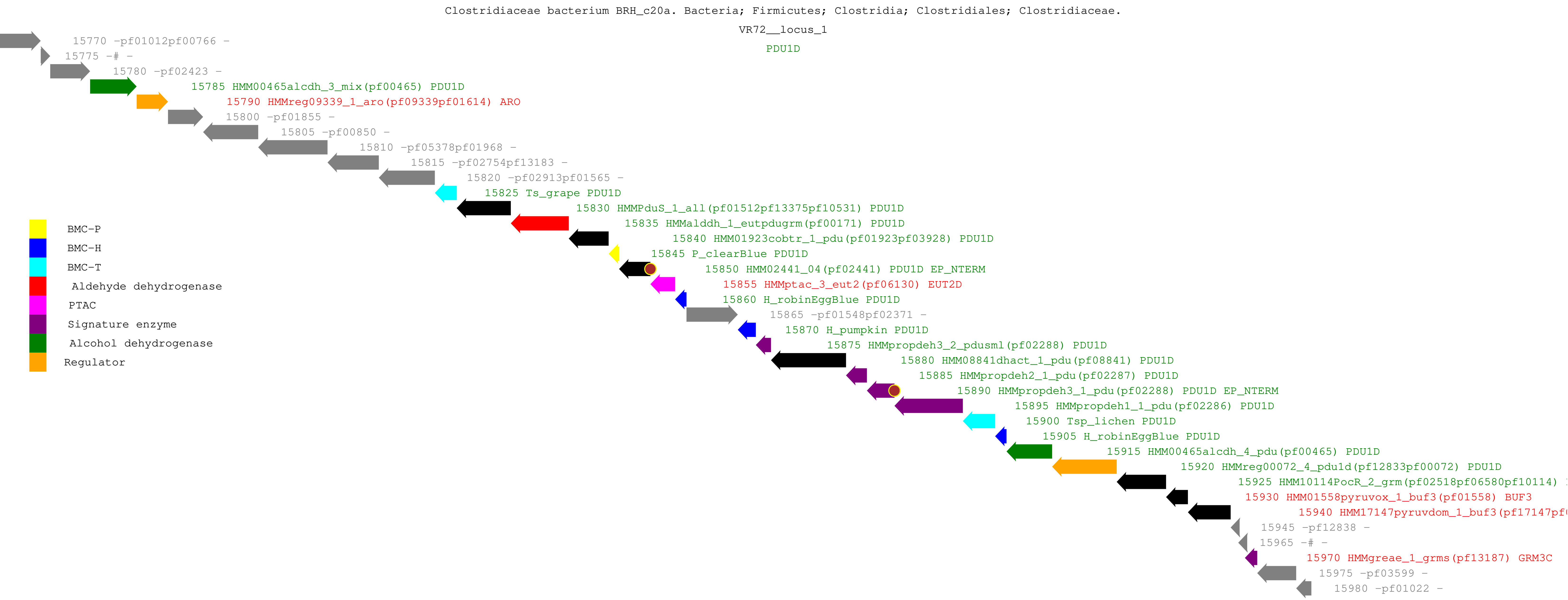
Clostridiaceae bacterium BRH_c20a. Bacteria; Firmicutes; Clostridia; Clostridiales; Clostridiaceae.
VR72__locus_1
PDU1D
15770 -pf01012pf00766 -
15775 -# -
15780 -pf02423 -
15785 HMM00465alcdh_3_mix(pf00465) PDU1D
15790 HMMreg09339_1_aro(pf09339pf01614) ARO
15800 -pf01855 -
15805 -pf00850 -
15810 -pf05378pf01968 -
15815 -pf02754pf13183 -
15820 -pf02913pf01565 -
15825 Ts_grape PDU1D
15830 HMMPduS_1_all(pf01512pf13375pf10531) PDU1D
15835 HMMalddh_1_eutpdugrm(pf00171) PDU1D
15840 HMM01923cobtr_1_pdu(pf01923pf03928) PDU1D
15845 P_clearBlue PDU1D
15850 HMM02441_04(pf02441) PDU1D EP_NTERM
15855 HMMptac_3_eut2(pf06130) EUT2D
15860 H_robinEggBlue PDU1D
15865 -pf01548pf02371 -
15870 H_pumpkin PDU1D
15875 HMMpropdeh3_2_pdusml(pf02288) PDU1D
15880 HMM08841dhact_1_pdu(pf08841) PDU1D
15885 HMMpropdeh2_1_pdu(pf02287) PDU1D
15890 HMMpropdeh3_1_pdu(pf02288) PDU1D EP_NTERM
15895 HMMpropdeh1_1_pdu(pf02286) PDU1D
15900 Tsp_lichen PDU1D
15905 H_robinEggBlue PDU1D
15915 HMM00465alcdh_4_pdu(pf00465) PDU1D
15920 HMMreg00072_4_pdu1d(pf12833pf00072) PDU1D
15925 HMM10114PocR_2_grm(pf02518pf06580pf10114)
15930 HMM01558pyruvox_1_buf3(pf01558) BUF3
15940 HMM17147pyruvdom_1_buf3(pf17147pf
15945 -pf12838 -
15965 -# -
15970 HMMgreae_1_grms(pf13187) GRM3C
15975 -pf03599 -
15980 -pf01022 -
BMC-P
BMC-H
BMC-T
Aldehyde dehydrogenase
PTAC
Signature enzyme
Alcohol dehydrogenase
Regulator

Caldanaerobacter subterraneus. Bacteria; Firmicutes; Clostridia; Thermoanaerobacterales;

XD49__locus_fragmented

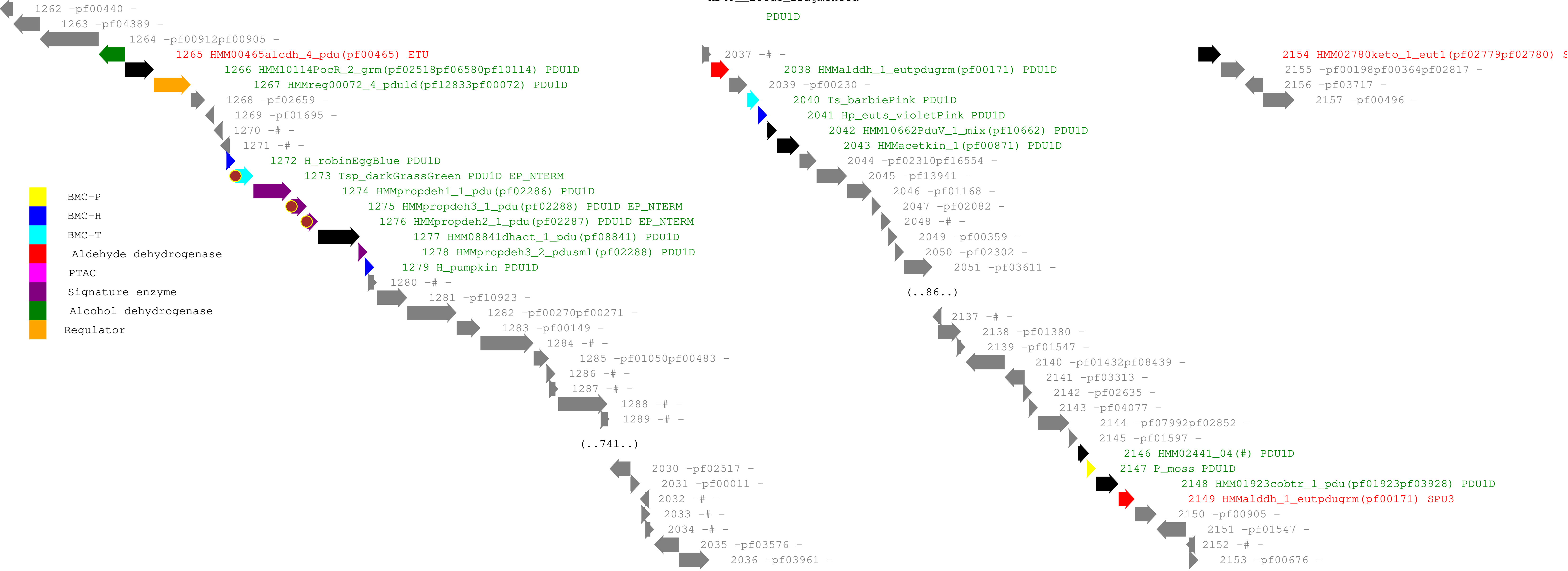

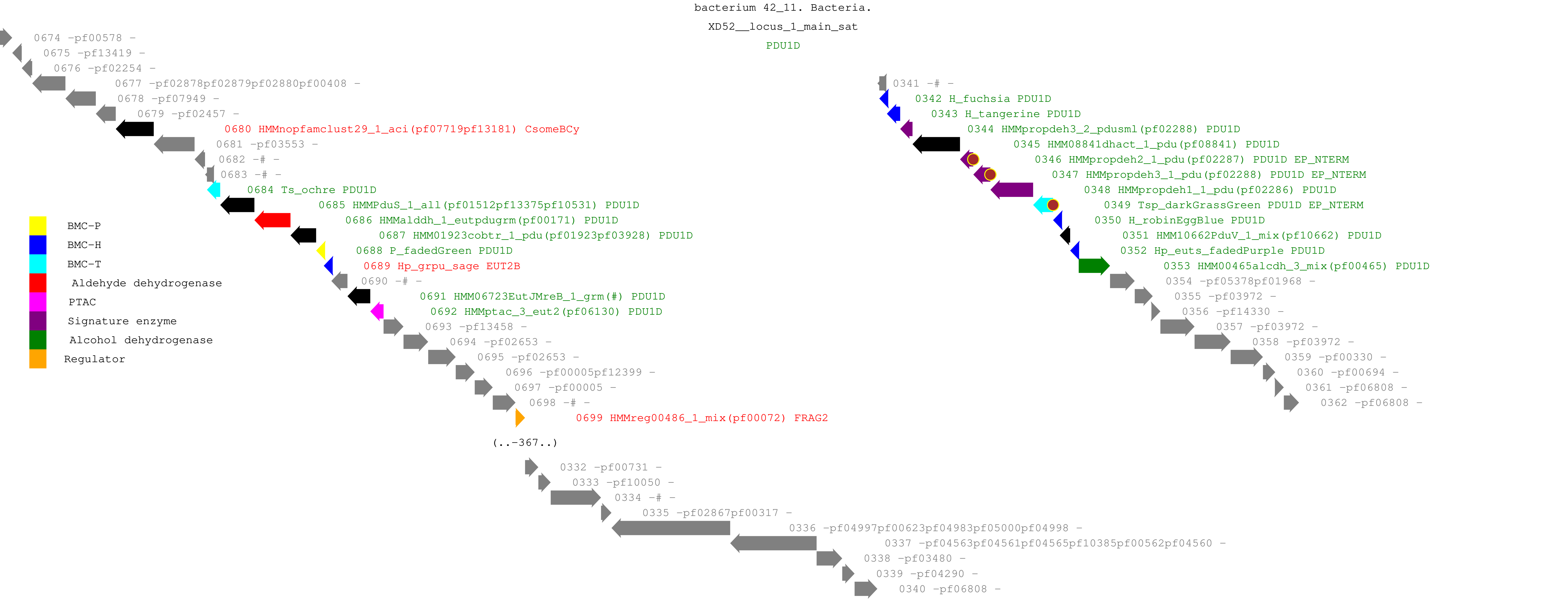

bacterium 42_11. Bacteria.
XD52__locus_1_main_sat
PDU1D
0674 -pf00578 -
0675 -pf13419 -
0676 -pf02254 -
0677 -pf02878pf02879pf02880pf00408 -
0678 -pf07949 -
0679 -pf02457 -
0680 HMMnopfamclust29_1_aci(pf07719pf13181) CsomeBCy
0681 -pf03553 -
0682 -# -
0683 -# -
0684 Ts_ochre PDU1D
0685 HMMPduS_1_all(pf01512pf13375pf10531) PDU1D
0686 HMMalddh_1_eutpdugrm(pf00171) PDU1D
0687 HMM01923cobtr_1_pdu(pf01923pf03928) PDU1D
0688 P_fadedGreen PDU1D
0689 Hp_grpu_sage EUT2B
0690 -# -
0691 HMM06723EutJMreB_1_grm(#) PDU1D
0692 HMMptac_3_eut2(pf06130) PDU1D
0693 -pf13458 -
0694 -pf02653 -
0695 -pf02653 -
0696 -pf00005pf12399 -
0697 -pf00005 -
0698 -# -
0699 HMMreg00486_1_mix(pf00072) FRAG2
(..-367..)
0332 -pf00731 -
0333 -pf10050 -
0334 -# -
0335 -pf02867pf00317 -
0336 -pf04997pf00623pf04983pf05000pf04998 -
0337 -pf04563pf04561pf04565pf10385pf00562pf04560 -
0338 -pf03480 -
0339 -pf04290 -
0340 -pf06808 -
0341 -# -
0342 H_fuchsia PDU1D
0343 H_tangerine PDU1D
0344 HMMpropdeh3_2_pdusml(pf02288) PDU1D
0345 HMM08841dhact_1_pdu(pf08841) PDU1D
0346 HMMpropdeh2_1_pdu(pf02287) PDU1D EP_NTERM
0347 HMMpropdeh3_1_pdu(pf02288) PDU1D EP_NTERM
0348 HMMpropdeh1_1_pdu(pf02286) PDU1D
0349 Tsp_darkGrassGreen PDU1D EP_NTERM
0350 H_robinEggBlue PDU1D
0351 HMM10662PduV_1_mix(pf10662) PDU1D
0352 Hp_euts_fadedPurple PDU1D
0353 HMM00465alcdh_3_mix(pf00465) PDU1D
0354 -pf05378pf01968 -
0355 -pf03972 -
0356 -pf14330 -
0357 -pf03972 -
0358 -pf03972 -
0359 -pf00330 -
0360 -pf00694 -
0361 -pf06808 -
0362 -pf06808 -
BMC-P
BMC-H
BMC-T
Aldehyde dehydrogenase
PTAC
Signature enzyme
Alcohol dehydrogenase
Regulator

Thermoanaerobacterales bacterium 50_218. Bacteria; Firmicutes; Clostridia; Thermoanaerobacterales.

XD63__locus_fragmented

PDU1D

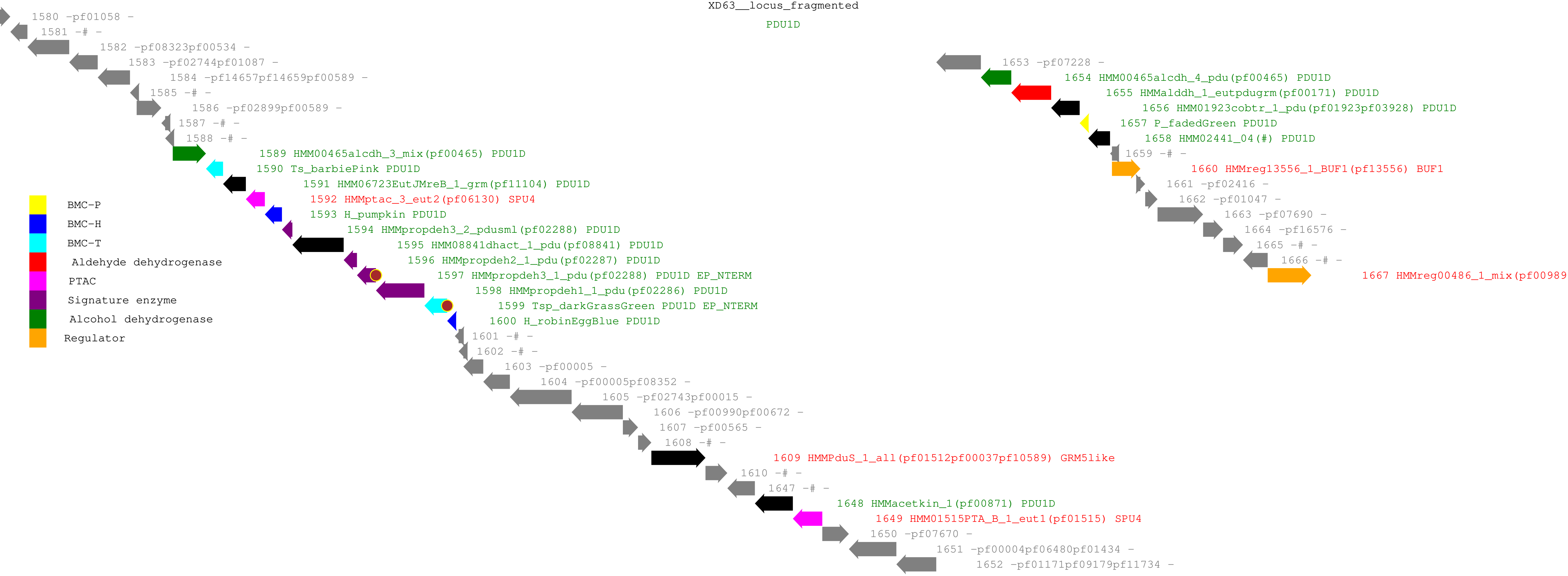

Enterococcus mundtii. Bacteria; Firmicutes; Bacilli; Lactobacillales; Enterococcaceae;

A5802__locus_1

PDU1E

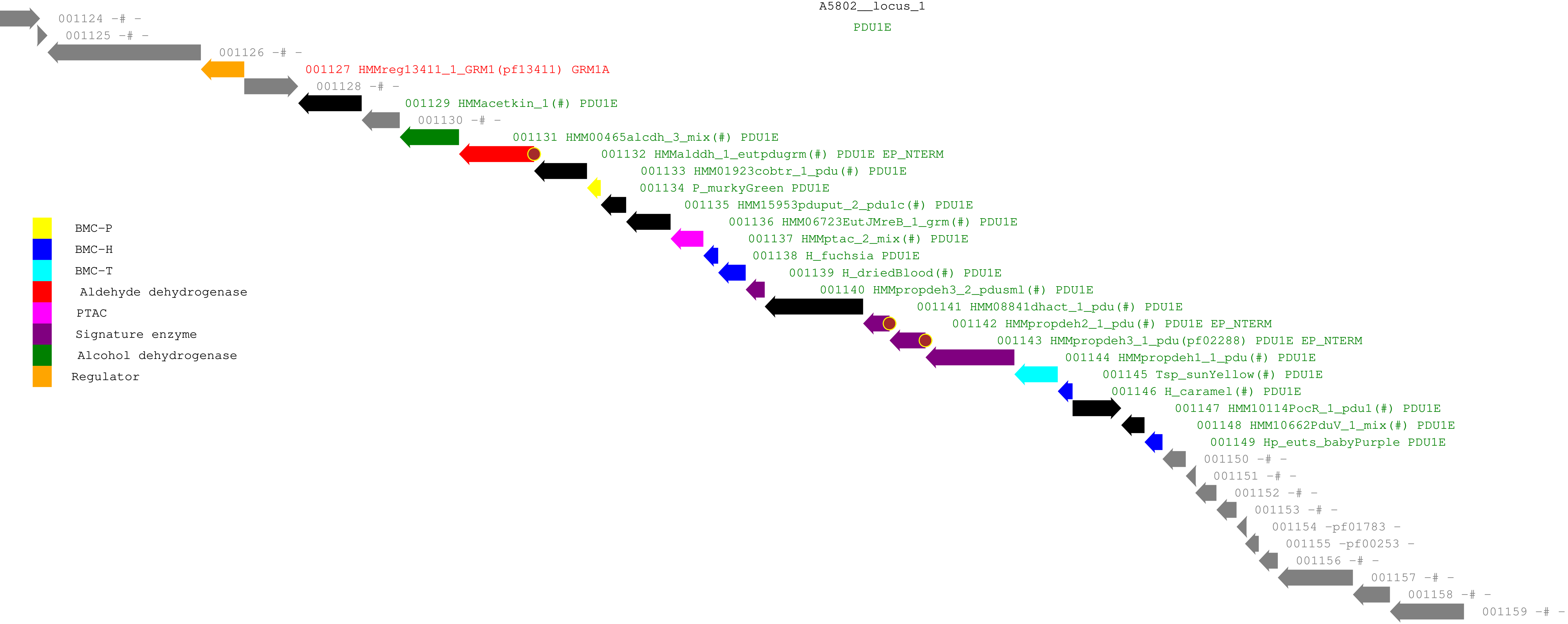

Enterococcus sp. 3H8_DIV0648. Bacteria; Firmicutes; Bacilli; Lactobacillales; Enterococcaceae;

A5875__locus_1

PDU1E

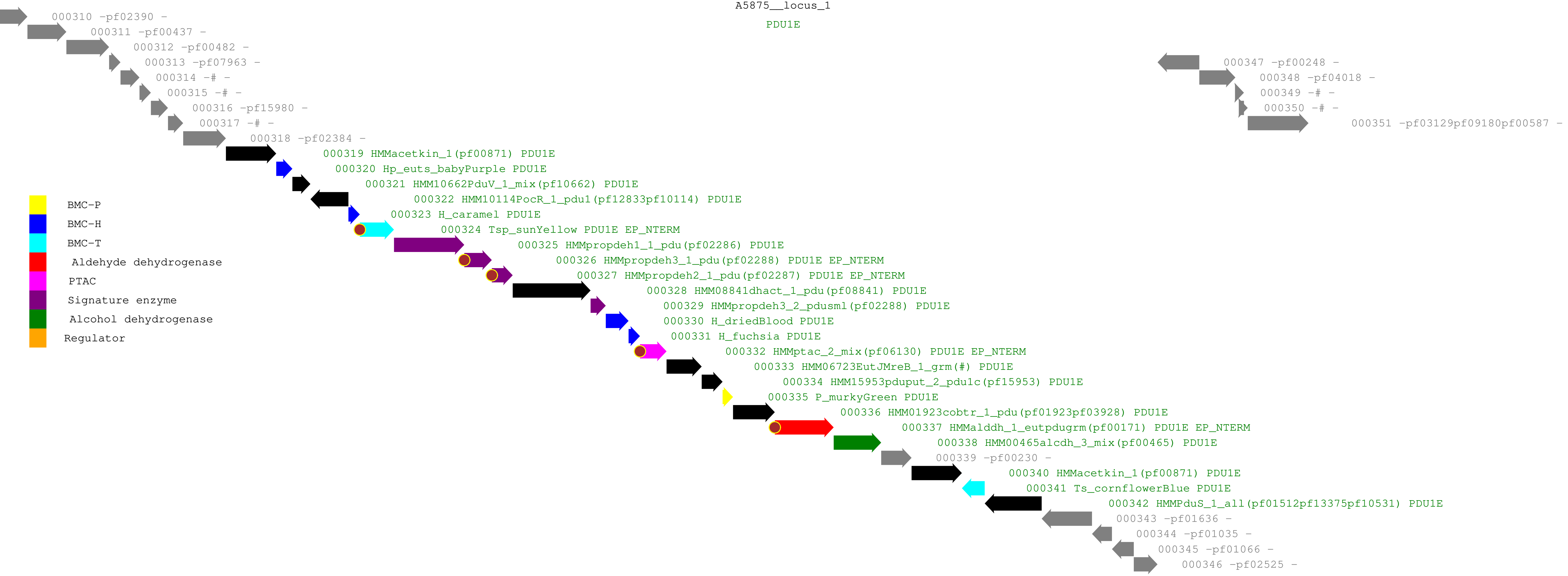

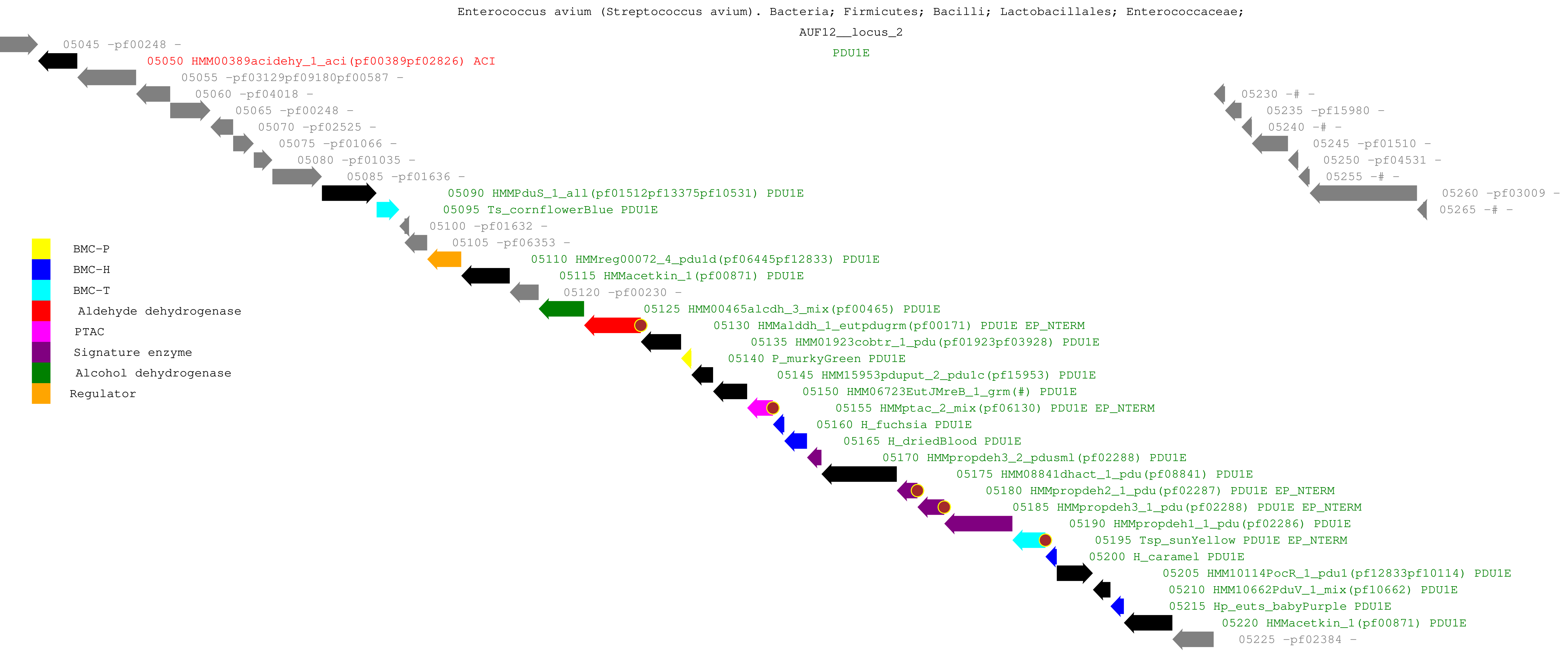
Enterococcus avium (Streptococcus avium). Bacteria; Firmicutes; Bacilli; Lactobacillales; Enterococcaceae;
AUF12__locus_2
PDU1E
05045 -pf00248 -
05050 HMM00389acidehy_1_aci(pf00389pf02826) ACI
05055 -pf03129pf09180pf00587 -
05060 -pf04018 -
05065 -pf00248 -
05070 -pf02525 -
05075 -pf01066 -
05080 -pf01035 -
05085 -pf01636 -
05090 HMMPduS_1_all(pf01512pf13375pf10531) PDU1E
05095 Ts_cornflowerBlue PDU1E
05100 -pf01632 -
05105 -pf06353 -
05110 HMMreg00072_4_pdu1d(pf06445pf12833) PDU1E
05115 HMMacetkin_1(pf00871) PDU1E
05120 -pf00230 -
05125 HMM00465alcdh_3_mix(pf00465) PDU1E
05130 HMMalddh_1_eutpdugrm(pf00171) PDU1E EP_NTERM
05135 HMM01923cobtr_1_pdu(pf01923pf03928) PDU1E
05140 P_murkyGreen PDU1E
05145 HMM15953pduput_2_pdu1c(pf15953) PDU1E
05150 HMM06723EutJMreB_1_grm(#) PDU1E
05155 HMMptac_2_mix(pf06130) PDU1E EP_NTERM
05160 H_fuchsia PDU1E
05165 H_driedBlood PDU1E
05170 HMMpropdeh3_2_pdusml(pf02288) PDU1E
05175 HMM08841dhact_1_pdu(pf08841) PDU1E
05180 HMMpropdeh2_1_pdu(pf02287) PDU1E EP_NTERM
05185 HMMpropdeh3_1_pdu(pf02288) PDU1E EP_NTERM
05190 HMMpropdeh1_1_pdu(pf02286) PDU1E
05195 Tsp_sunYellow PDU1E EP_NTERM
05200 H_caramel PDU1E
05205 HMM10114PocR_1_pdu1(pf12833pf10114) PDU1E
05210 HMM10662PduV_1_mix(pf10662) PDU1E
05215 Hp_euts_babyPurple PDU1E
05220 HMMacetkin_1(pf00871) PDU1E
05225 -pf02384 -
05230 -# -
05235 -pf15980 -
05240 -# -
05245 -pf01510 -
05250 -pf04531 -
05255 -# -
05260 -pf03009 -
05265 -# -
BMC-P
BMC-H
BMC-T
Aldehyde dehydrogenase
PTAC
Signature enzyme
Alcohol dehydrogenase
Regulator

Enterococcus avium (Streptococcus avium). Bacteria; Firmicutes; Bacilli; Lactobacillales; Enterococcaceae;

AUF16__locus_2

PDU1E

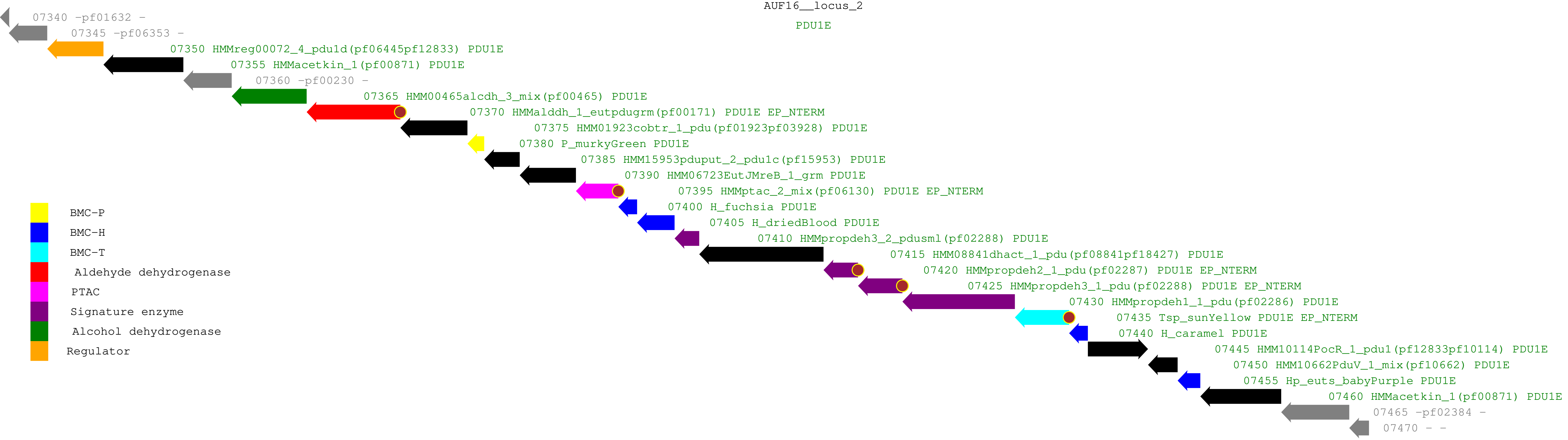

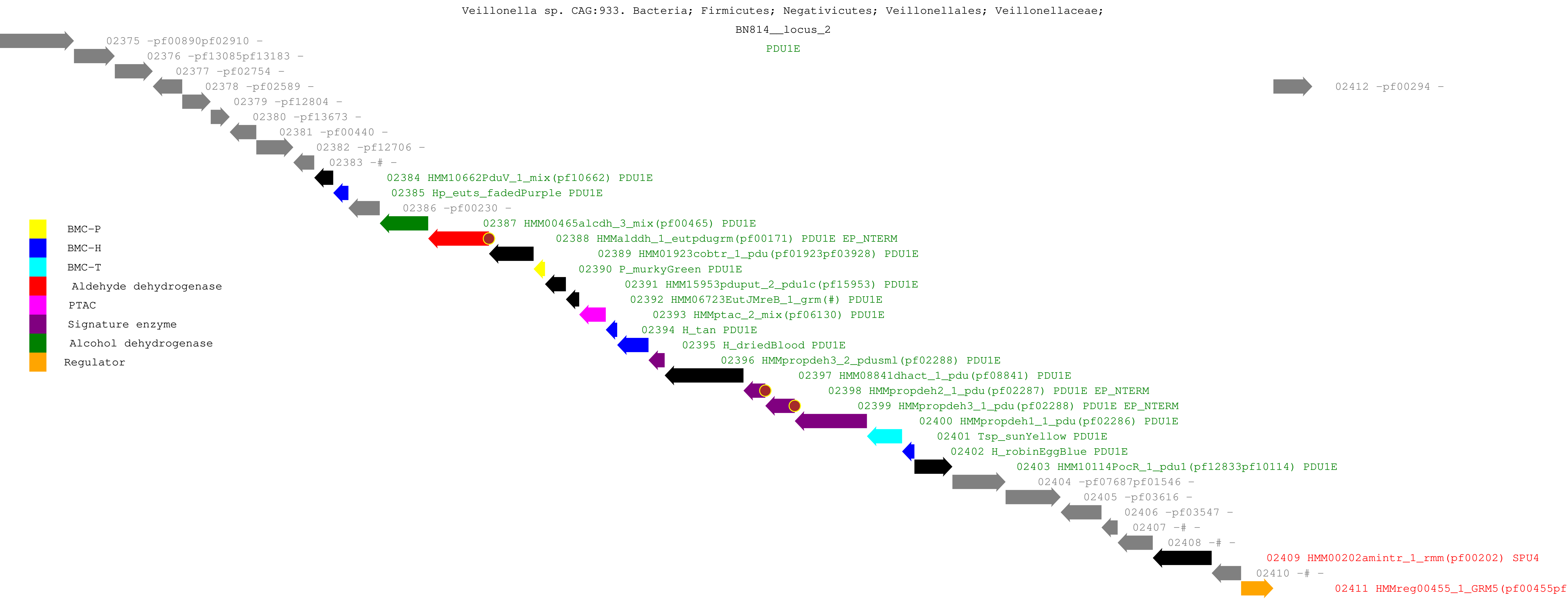
Veillonella sp. CAG:933. Bacteria; Firmicutes; Negativicutes; Veillonellales; Veillonellaceae;
BN814__locus_2
PDU1E
02375 -pf00890pf02910 -
02376 -pf13085pf13183 -
02377 -pf02754 -
02378 -pf02589 -
02379 -pf12804 -
02380 -pf13673 -
02381 -pf00440 -
02382 -pf12706 -
02383 -# -
02384 HMM10662PduV_1_mix(pf10662) PDU1E
02385 Hp_euts_fadedPurple PDU1E
02386 -pf00230 -
02387 HMM00465alcdh_3_mix(pf00465) PDU1E
02388 HMMalddh_1_eutpdugrm(pf00171) PDU1E EP_NTERM
02389 HMM01923cobtr_1_pdu(pf01923pf03928) PDU1E
02390 P_murkyGreen PDU1E
02391 HMM15953pduput_2_pdu1c(pf15953) PDU1E
02392 HMM06723EutJMreB_1_grm(#) PDU1E
02393 HMMptac_2_mix(pf06130) PDU1E
02394 H_tan PDU1E
02395 H_driedBlood PDU1E
02396 HMMpropdeh3_2_pdusml(pf02288) PDU1E
02397 HMM08841dhact_1_pdu(pf08841) PDU1E
02398 HMMpropdeh2_1_pdu(pf02287) PDU1E EP_NTERM
02399 HMMpropdeh3_1_pdu(pf02288) PDU1E EP_NTERM
02400 HMMpropdeh1_1_pdu(pf02286) PDU1E
02401 Tsp_sunYellow PDU1E
02402 H_robinEggBlue PDU1E
02403 HMM10114PocR_1_pdu1(pf12833pf10114) PDU1E
02404 -pf07687pf01546 -
02405 -pf03616 -
02406 -pf03547 -
02407 -# -
02408 -# -
02409 HMM00202amintr_1_rmm(pf00202) SPU4
02410 -# -
02411 HMMreg00455_1_GRM5(pf00455pf
02412 -pf00294 -
BMC-P
BMC-H
BMC-T
Aldehyde dehydrogenase
PTAC
Signature enzyme
Alcohol dehydrogenase
Regulator

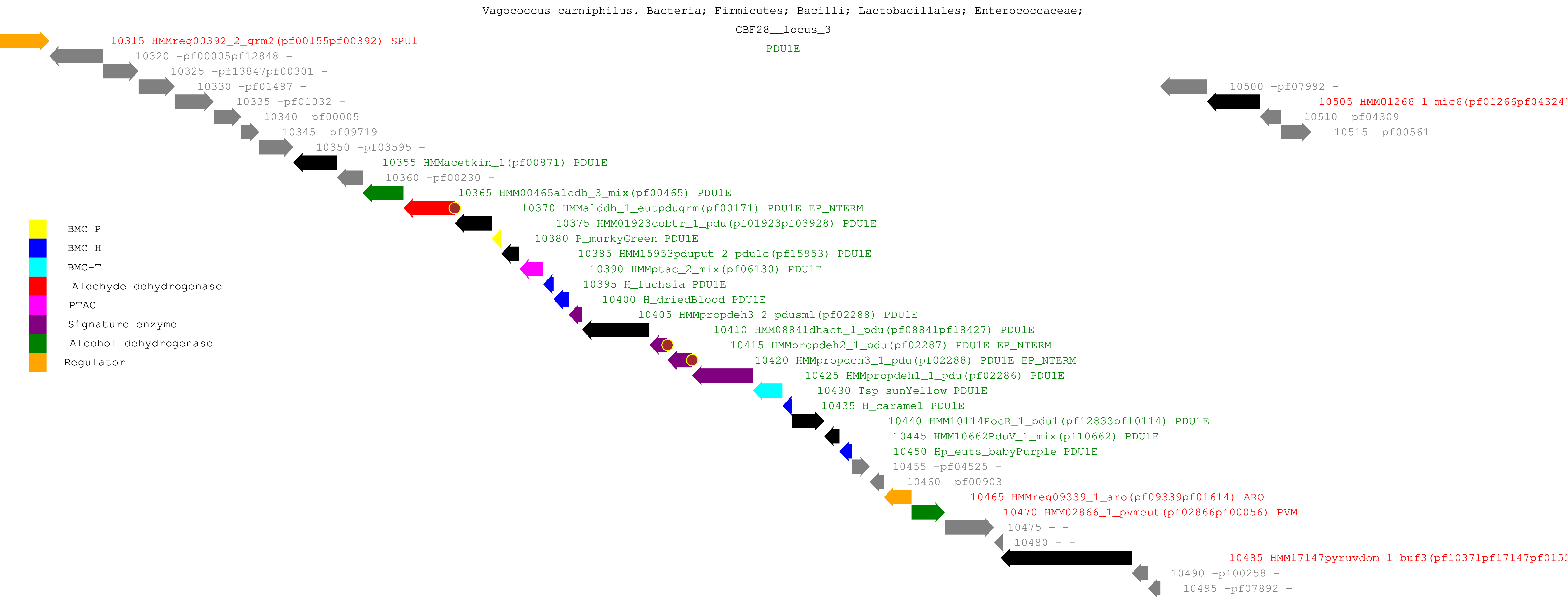
Vagococcus carniphilus. Bacteria; Firmicutes; Bacilli; Lactobacillales; Enterococcaceae;
CBF28__locus_3
PDU1E
10315 HMMreg00392_2_grm2(pf00155pf00392) SPU1
10320 -pf00005pf12848 -
10325 -pf13847pf00301 -
10330 -pf01497 -
10335 -pf01032 -
10340 -pf00005 -
10345 -pf09719 -
10350 -pf03595 -
10355 HMMacetkin_1(pf00871) PDU1E
10360 -pf00230 -
10365 HMM00465alcdh_3_mix(pf00465) PDU1E
10370 HMMaldh_1_eutpdugrm(pf00171) PDU1E EP_NTERM
10375 HMM01923cobtr_1_pdu(pf01923pf03928) PDU1E
10380 P_murkyGreen PDU1E
10385 HMM15953pduput_2_pdu1c(pf15953) PDU1E
10390 HMMptac_2_mix(pf06130) PDU1E
10395 H_fuchsia PDU1E
10400 H_driedBlood PDU1E
10405 HMMpropdeh3_2_pdusml(pf02288) PDU1E
10410 HMM08841dhact_1_pdu(pf08841pf18427) PDU1E
10415 HMMpropdeh2_1_pdu(pf02287) PDU1E EP_NTERM
10420 HMMpropdeh3_1_pdu(pf02288) PDU1E EP_NTERM
10425 HMMpropdeh1_1_pdu(pf02286) PDU1E
10430 Tsp_sunYellow PDU1E
10435 H_caramel PDU1E
10440 HMM10114PocR_1_pdu1(pf12833pf10114) PDU1E
10445 HMM10662PduV_1_mix(pf10662) PDU1E
10450 Hp_euts_babyPurple PDU1E
10455 -pf04525 -
10460 -pf00903 -
10465 HMMreg09339_1_aro(pf09339pf01614) ARO
10470 HMM02866_1_pvmeut(pf02866pf00056) PVM
10475 - -
10480 - -
10485 HMM17147pyruvdom_1_buf3(pf10371pf17147pf015
10490 -pf00258 -
10495 -pf07892 -
10500 -pf07992 -
10505 HMM01266_1_mic6(pf01266pf04324
10510 -pf04309 -
10515 -pf00561 -
BMC-P
BMC-H
BMC-T
Aldehyde dehydrogenase
PTAC
Signature enzyme
Alcohol dehydrogenase
Regulator

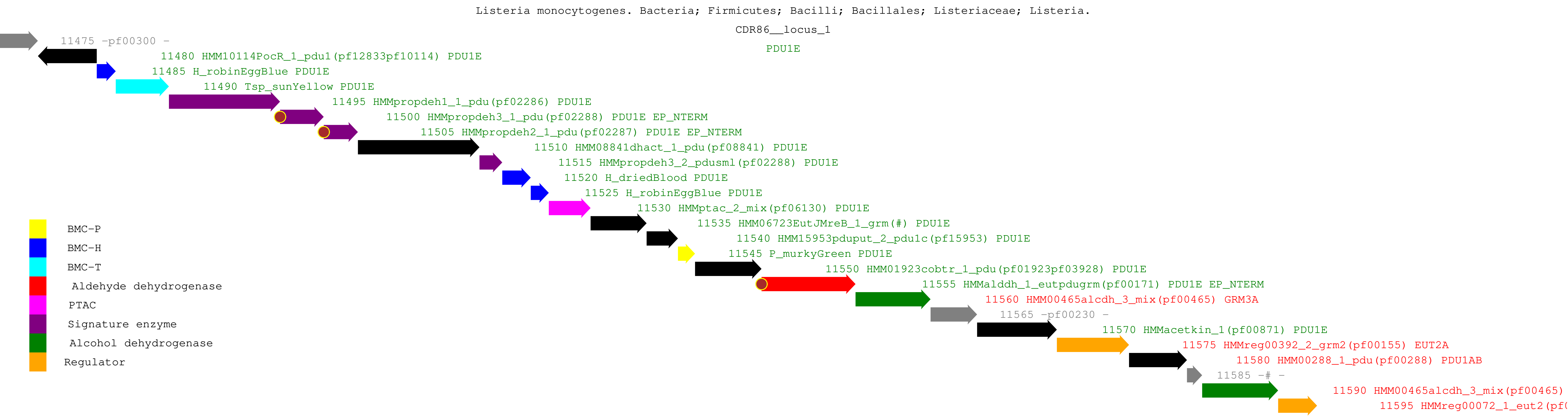
Listeria monocytogenes. Bacteria; Firmicutes; Bacilli; Bacillales; Listeriaceae; Listeria.
CDR86__locus_1
PDU1E
11475 -pf00300 -
11480 HMM10114PocR_1_pdu1(pf12833pf10114) PDU1E
11485 H_robinEggBlue PDU1E
11490 Tsp_sunYellow PDU1E
11495 HMMpropdeh1_1_pdu(pf02286) PDU1E
11500 HMMpropdeh3_1_pdu(pf02288) PDU1E EP_NTERM
11505 HMMpropdeh2_1_pdu(pf02287) PDU1E EP_NTERM
11510 HMM08841dhact_1_pdu(pf08841) PDU1E
11515 HMMpropdeh3_2_pdusml(pf02288) PDU1E
11520 H_driedBlood PDU1E
11525 H_robinEggBlue PDU1E
11530 HMMptac_2_mix(pf06130) PDU1E
11535 HMM06723EutJMreB_1_grm(#) PDU1E
11540 HMM15953pduput_2_pdu1c(pf15953) PDU1E
11545 P_murkyGreen PDU1E
11550 HMM01923cobtr_1_pdu(pf01923pf03928) PDU1E
11555 HMMalddh_1_eutpdugrm(pf00171) PDU1E EP_NTERM
11560 HMM00465alcdh_3_mix(pf00465) GRM3A
11565 -pf00230 -
11570 HMMacetkin_1(pf00871) PDU1E
11575 HMMreg00392_2_grm2(pf00155) EUT2A
11580 HMM00288_1_pdu(pf00288) PDU1AB
11585 -# -
11590 HMM00465alcdh_3_mix(pf00465)
11595 HMMreg00072_1_eut2(pf
BMC-P
BMC-H
BMC-T
Aldehyde dehydrogenase
PTAC
Signature enzyme
Alcohol dehydrogenase
Regulator

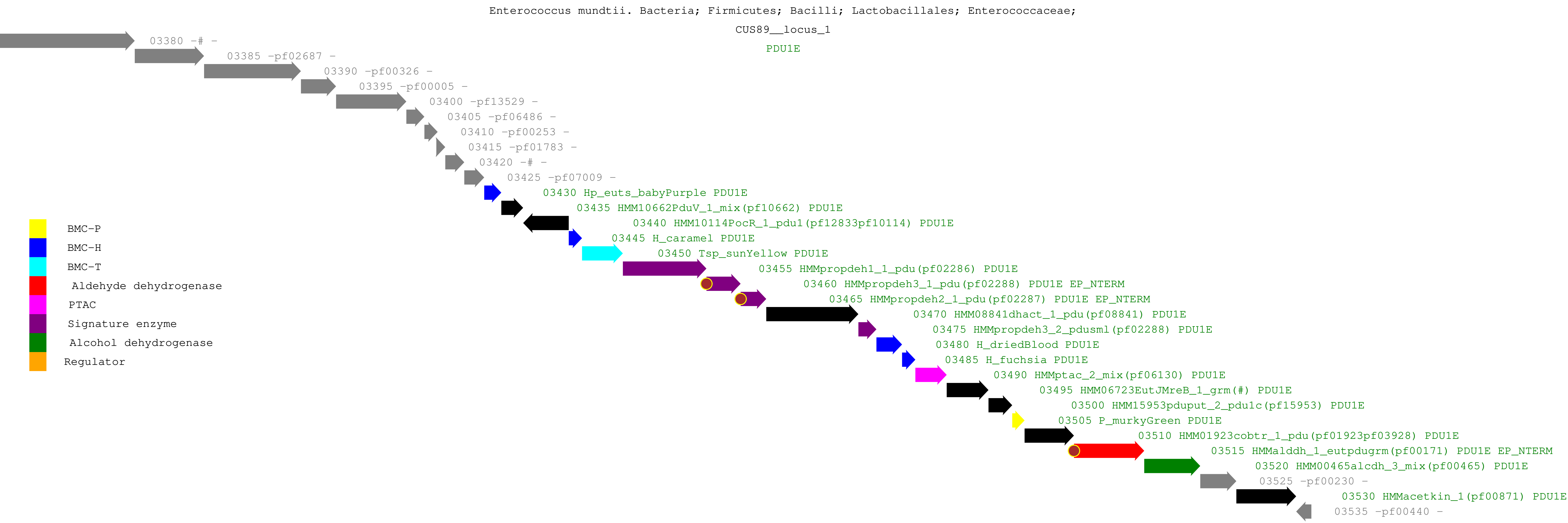
Enterococcus mundtii. Bacteria; Firmicutes; Bacilli; Lactobacillales; Enterococcaceae;
CUS89__locus_1
PDU1E
03380 -# -
03385 -pf02687 -
03390 -pf00326 -
03395 -pf00005 -
03400 -pf13529 -
03405 -pf06486 -
03410 -pf00253 -
03415 -pf01783 -
03420 -# -
03425 -pf07009 -
03430 Hp_euts_babyPurple PDU1E
03435 HMM10662PduV_1_mix(pf10662) PDU1E
03440 HMM10114PocR_1_pdu1(pf12833pf10114) PDU1E
03445 H_caramel PDU1E
03450 Tsp_sunYellow PDU1E
03455 HMMpropdeh1_1_pdu(pf02286) PDU1E
03460 HMMpropdeh3_1_pdu(pf02288) PDU1E EP_NTERM
03465 HMMpropdeh2_1_pdu(pf02287) PDU1E EP_NTERM
03470 HMM08841dhact_1_pdu(pf08841) PDU1E
03475 HMMpropdeh3_2_pdusml(pf02288) PDU1E
03480 H_driedBlood PDU1E
03485 H_fuchsia PDU1E
03490 HMMptac_2_mix(pf06130) PDU1E
03495 HMM06723EutJMreB_1_grm(#) PDU1E
03500 HMM15953pduput_2_pdu1c(pf15953) PDU1E
03505 P_murkyGreen PDU1E
03510 HMM01923cobtr_1_pdu(pf01923pf03928) PDU1E
03515 HMMalddh_1_eutpdugrm(pf00171) PDU1E EP_NTERM
03520 HMM00465alcdh_3_mix(pf00465) PDU1E
03525 -pf00230 -
03530 HMMacetkin_1(pf00871) PDU1E
03535 -pf00440 -
BMC-P
BMC-H
BMC-T
Aldehyde dehydrogenase
PTAC
Signature enzyme
Alcohol dehydrogenase
Regulator

Veillonella parvula. Bacteria; Firmicutes; Negativicutes; Veillonellales; Veillonellaceae;

D3219__locus_1

PDU1E

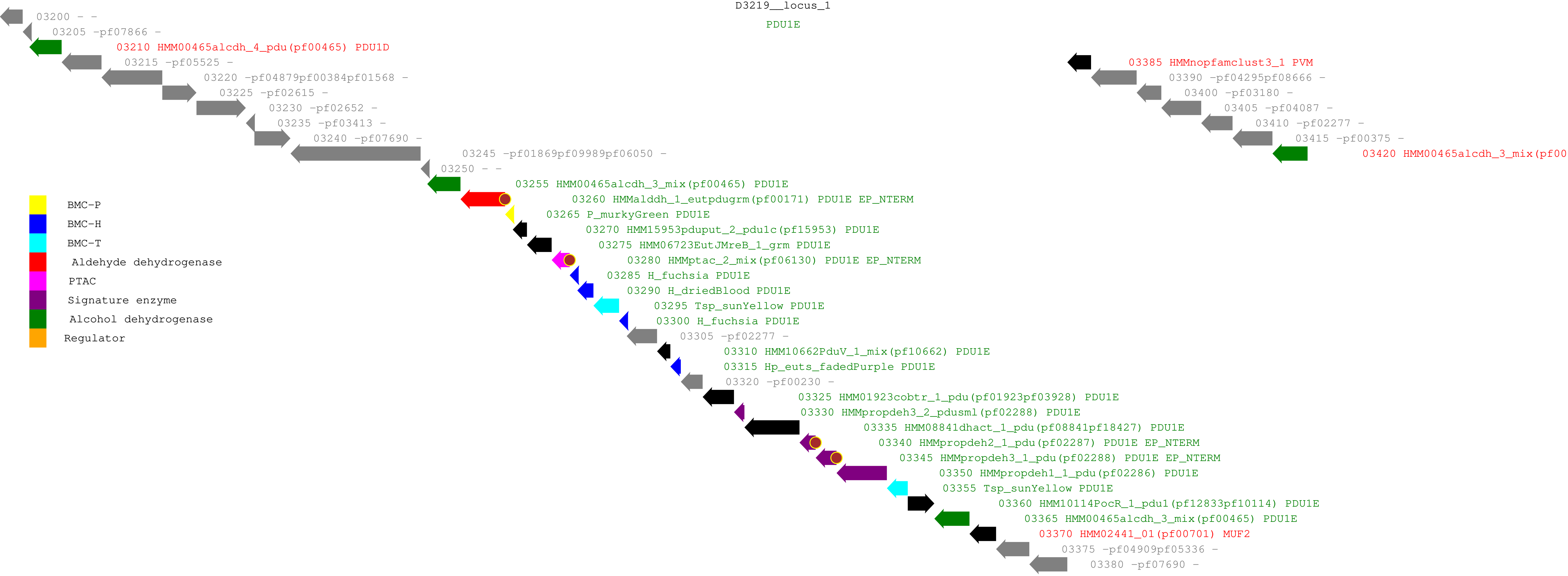

Enterococcus faecium (Streptococcus faecium). Bacteria; Firmicutes; Bacilli; Lactobacillales; Enterococcaceae;

DTPHA__locus_1

PDU1E

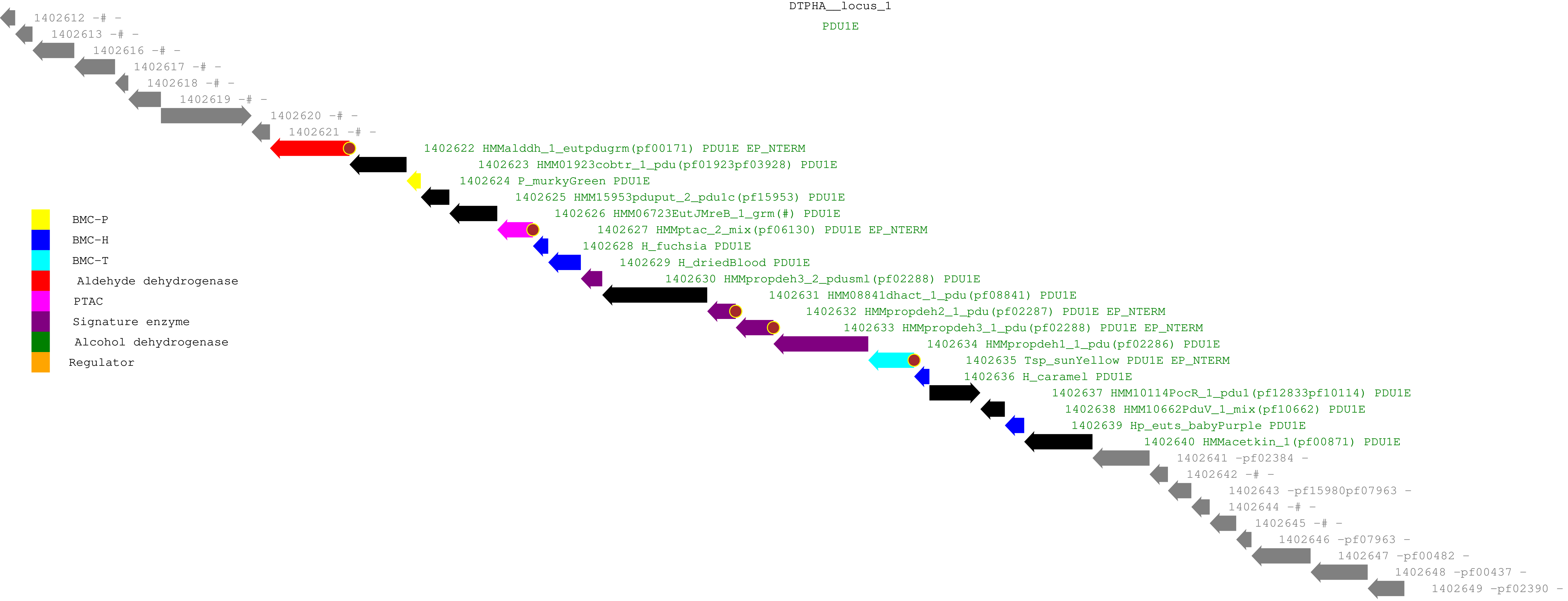

Veillonella sp. AF42-16. Bacteria; Firmicutes; Negativicutes; Veillonellales; Veillonellaceae;

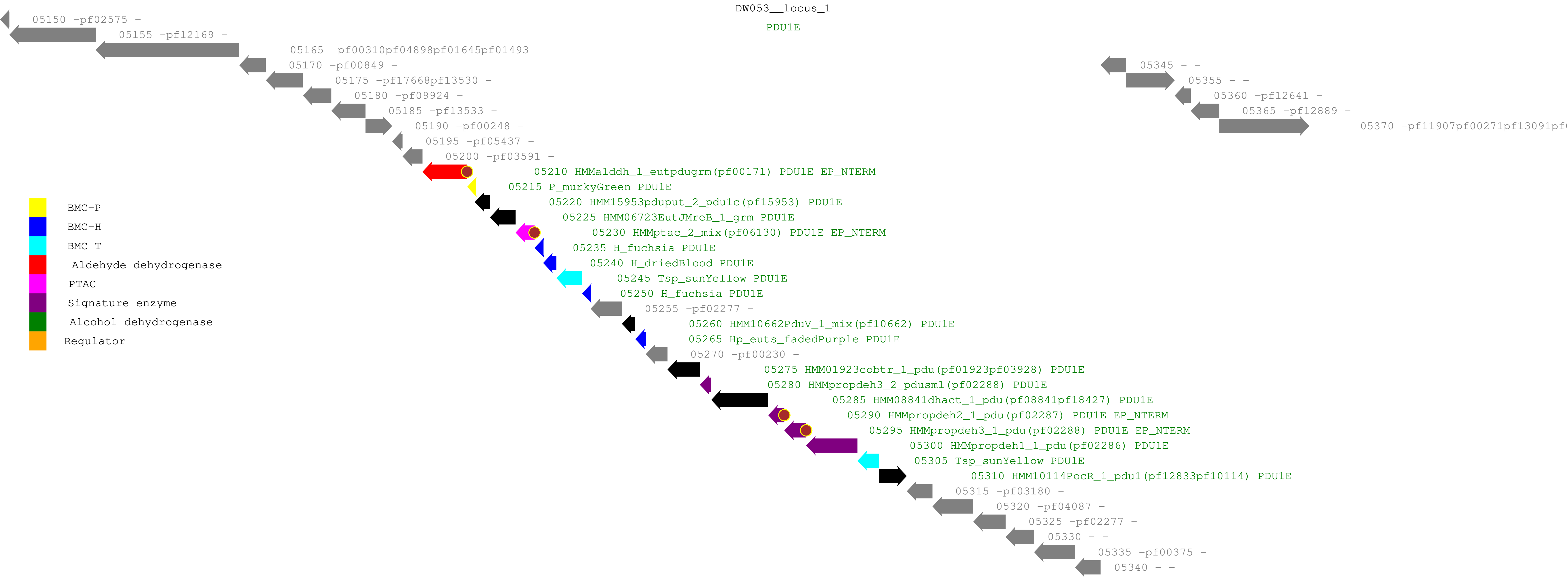

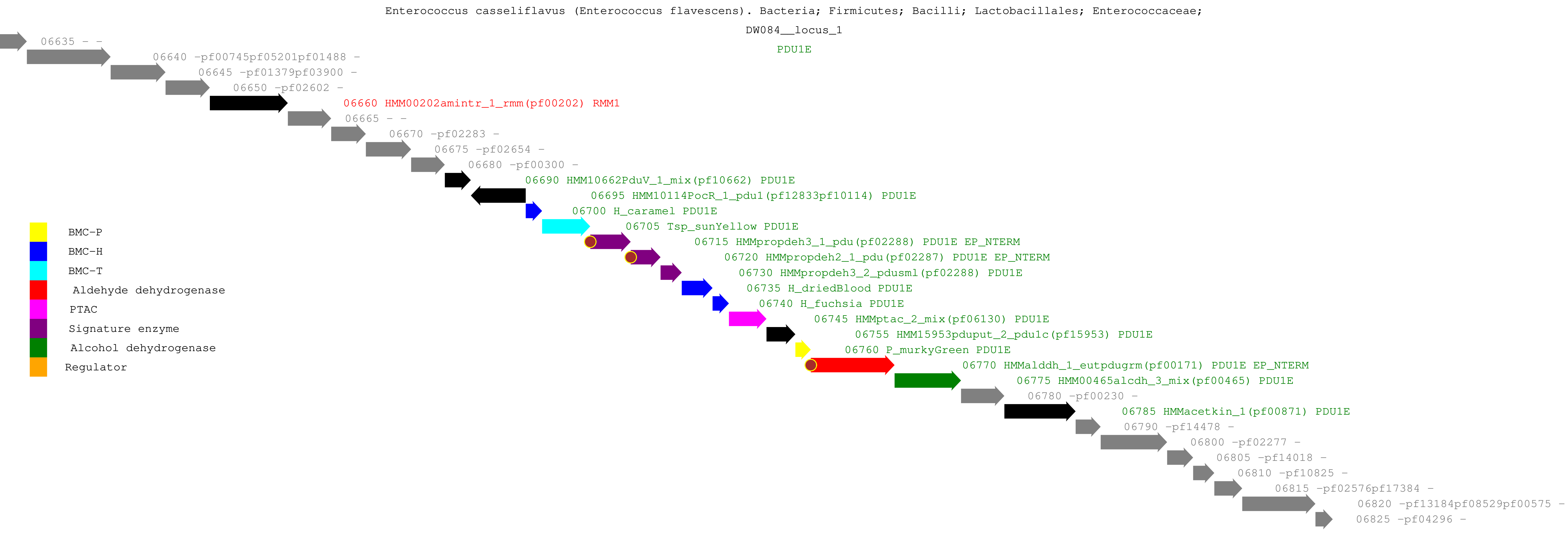
Enterococcus casseliflavus (Enterococcus flavescens). Bacteria; Firmicutes; Bacilli; Lactobacillales; Enterococcaceae;
DW084__locus_1
PDU1E
06635 - -
06640 -pf00745pf05201pf01488 -
06645 -pf01379pf03900 -
06650 -pf02602 -
06660 HMM00202amintr_1_rmm(pf00202) RMM1
06665 - -
06670 -pf02283 -
06675 -pf02654 -
06680 -pf00300 -
06690 HMM10662PduV_1_mix(pf10662) PDU1E
06695 HMM10114PocR_1_pdu1(pf12833pf10114) PDU1E
06700 H_caramel PDU1E
06705 Tsp_sunYellow PDU1E
06715 HMMpropdeh3_1_pdu(pf02288) PDU1E EP_NTERM
06720 HMMpropdeh2_1_pdu(pf02287) PDU1E EP_NTERM
06730 HMMpropdeh3_2_pdusml(pf02288) PDU1E
06735 H_driedBlood PDU1E
06740 H_fuchsia PDU1E
06745 HMMptac_2_mix(pf06130) PDU1E
06755 HMM15953pduput_2_pdu1c(pf15953) PDU1E
06760 P_murkyGreen PDU1E
06770 HMMalddh_1_eutpdugrm(pf00171) PDU1E EP_NTERM
06775 HMM00465alcdh_3_mix(pf00465) PDU1E
06780 -pf00230 -
06785 HMMacetkin_1(pf00871) PDU1E
06790 -pf14478 -
06800 -pf02277 -
06805 -pf14018 -
06810 -pf10825 -
06815 -pf02576pf17384 -
06820 -pf13184pf08529pf00575 -
06825 -pf04296 -
BMC-P
BMC-H
BMC-T
Aldehyde dehydrogenase
PTAC
Signature enzyme
Alcohol dehydrogenase
Regulator

Veillonella dispar. Bacteria; Firmicutes; Negativicutes; Veillonellales; Veillonellaceae;

EAI97__locus_1

PDU1E

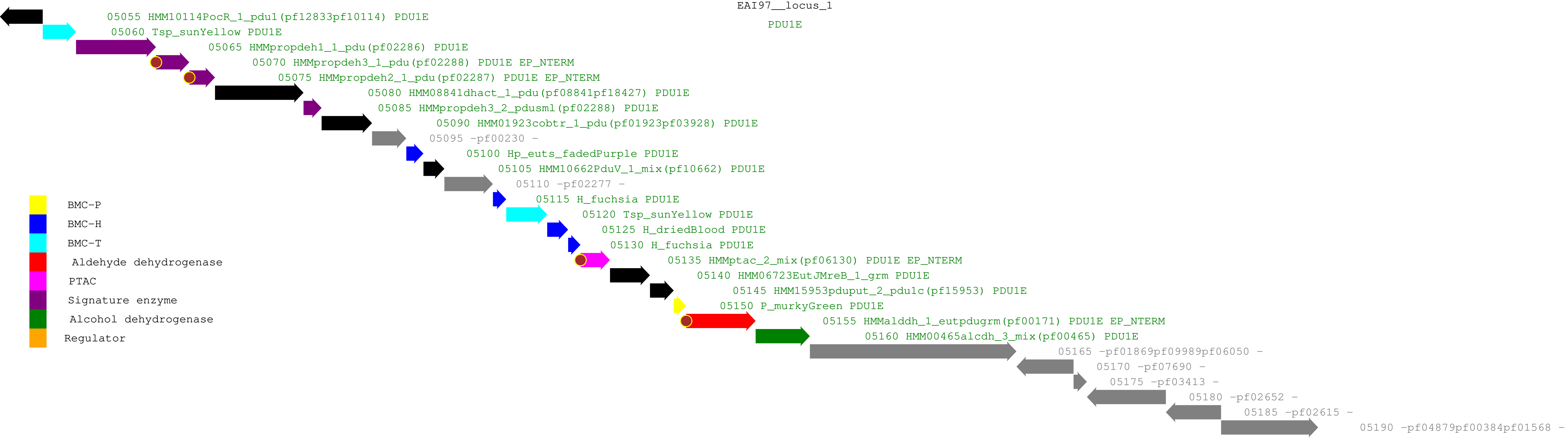

Enterococcus avium (Streptococcus avium). Bacteria; Firmicutes; Bacilli; Lactobacillales; Enterococcaceae;

EH197__locus_2

PDU1E

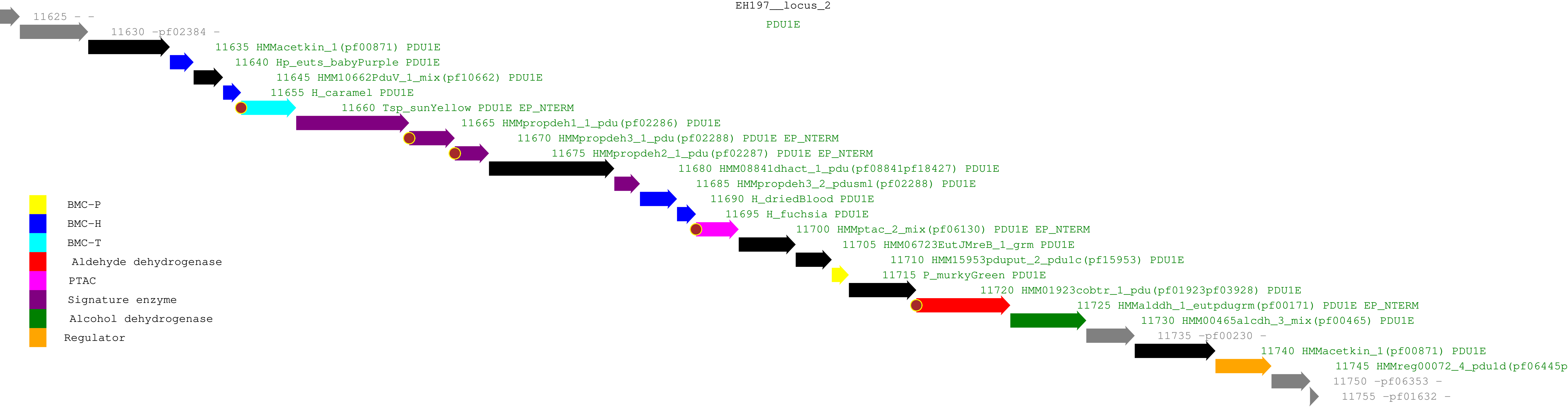

Enterococcus avium (Streptococcus avium). Bacteria; Firmicutes; Bacilli; Lactobacillales; Enterococcaceae;

EK398__locus_2

PDU1E

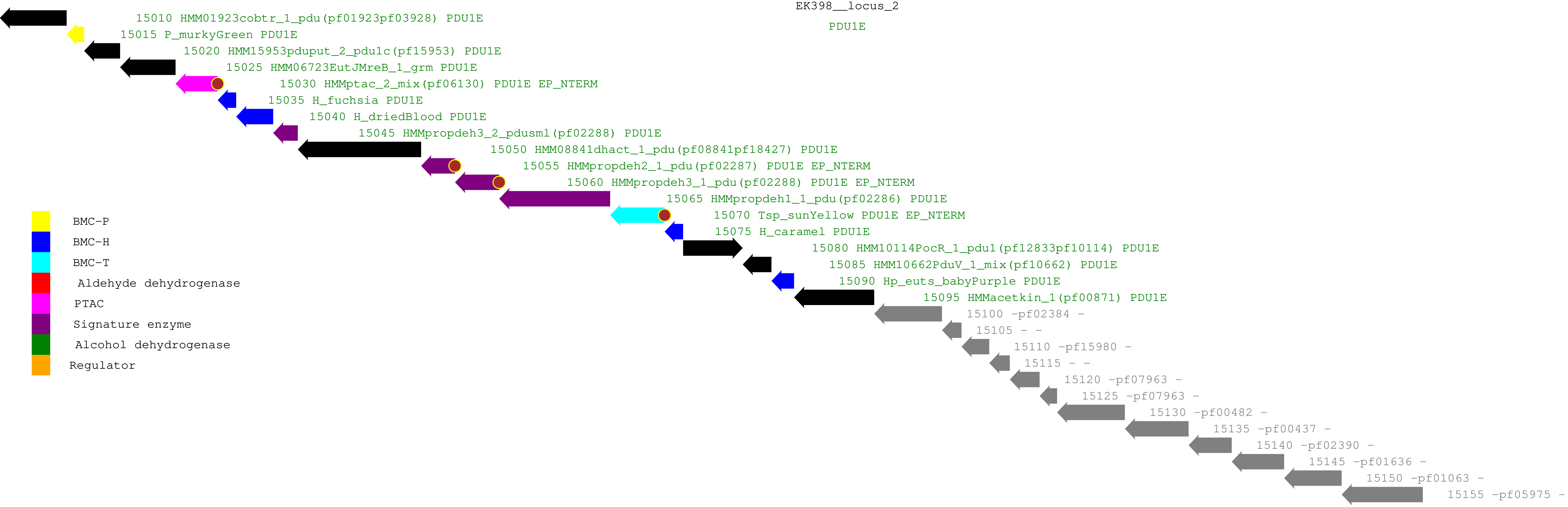

Enterococcus mundtii. Bacteria; Firmicutes; Bacilli; Lactobacillales; Enterococcaceae;

EM4838__locus_1

PDU1E

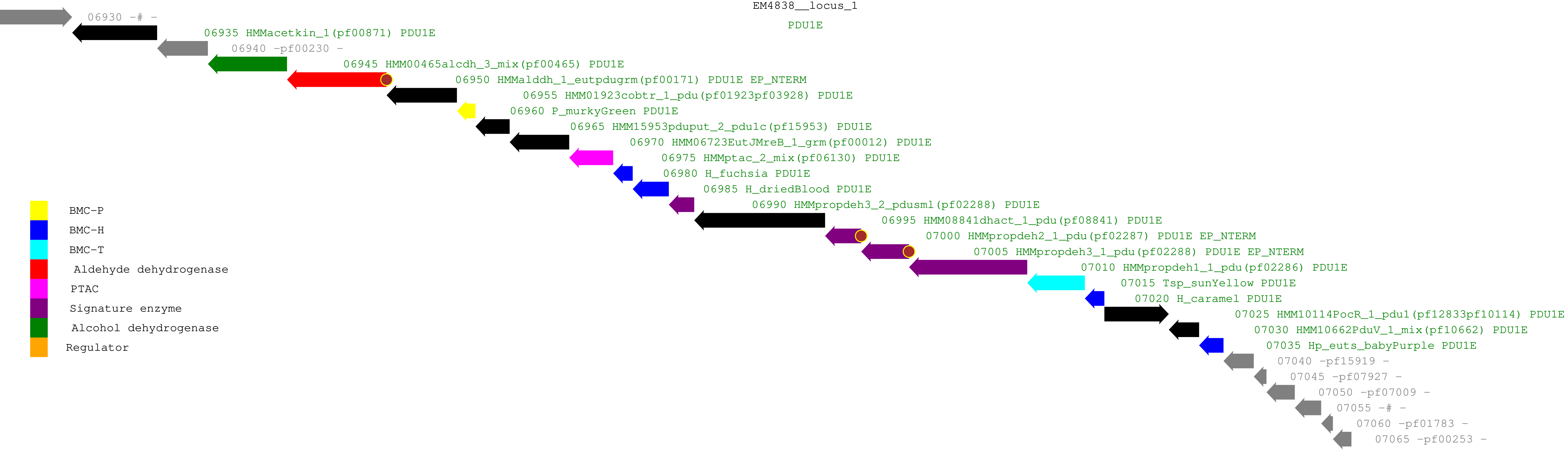

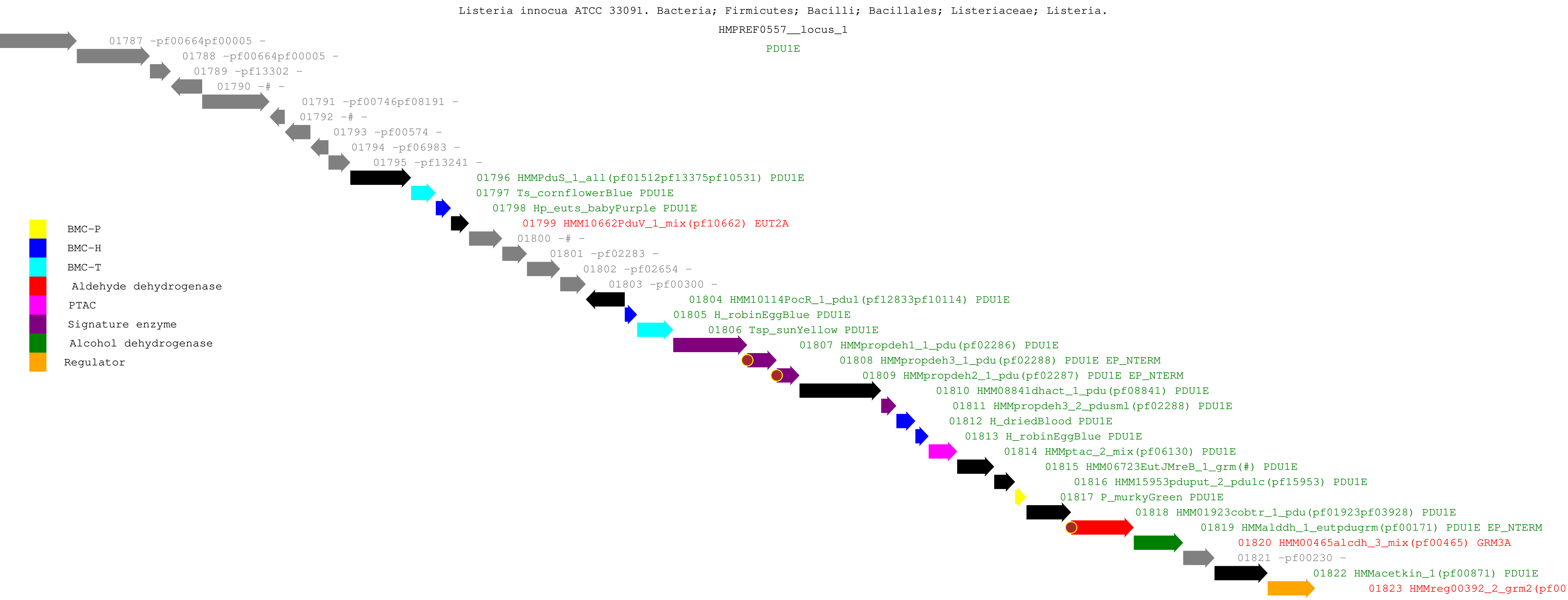

Listeria innocua ATCC 33091. Bacteria; Firmicutes; Bacilli; Bacillales; Listeriaceae; Listeria.
HMPREF0557__locus_1
PDU1E
01787 -pf00664pf00005 -
01788 -pf00664pf00005 -
01789 -pf13302 -
01790 -# -
01791 -pf00746pf08191 -
01792 -# -
01793 -pf00574 -
01794 -pf06983 -
01795 -pf13241 -
01796 HMMPduS_1_all(pf01512pf13375pf10531) PDU1E
01797 Ts_cornflowerBlue PDU1E
01798 Hp_euts_babyPurple PDU1E
01799 HMM10662PduV_1_mix(pf10662) EUT2A
01800 -# -
01801 -pf02283 -
01802 -pf02654 -
01803 -pf00300 -
01804 HMM10114PocR_1_pdu1(pf12833pf10114) PDU1E
01805 H_robinEggBlue PDU1E
01806 Tsp_sunYellow PDU1E
01807 HMMpropdeh1_1_pdu(pf02286) PDU1E
01808 HMMpropdeh3_1_pdu(pf02288) PDU1E EP_NTERM
01809 HMMpropdeh2_1_pdu(pf02287) PDU1E EP_NTERM
01810 HMM08841dhact_1_pdu(pf08841) PDU1E
01811 HMMpropdeh3_2_pdusml(pf02288) PDU1E
01812 H_driedBlood PDU1E
01813 H_robinEggBlue PDU1E
01814 HMMptac_2_mix(pf06130) PDU1E
01815 HMM06723EutJMreB_1_grm(#) PDU1E
01816 HMM15953pduput_2_pdu1c(pf15953) PDU1E
01817 P_murkyGreen PDU1E
01818 HMM01923cobtr_1_pdu(pf01923pf03928) PDU1E
01819 HMMalddh_1_eutpdugrm(pf00171) PDU1E EP_NTERM
01820 HMM00465alcdh_3_mix(pf00465) GRM3A
01821 -pf00230 -
01822 HMMacetkin_1(pf00871) PDU1E
01823 HMMreg00392_2_grm2(pf00
BMC-P
BMC-H
BMC-T
Aldehyde dehydrogenase
PTAC
Signature enzyme
Alcohol dehydrogenase
Regulator

Veillonella sp. 6_1_27. Bacteria; Firmicutes; Negativicutes; Veillonellales; Veillonellaceae;

HMPREF0874__locus_1

PDU1E

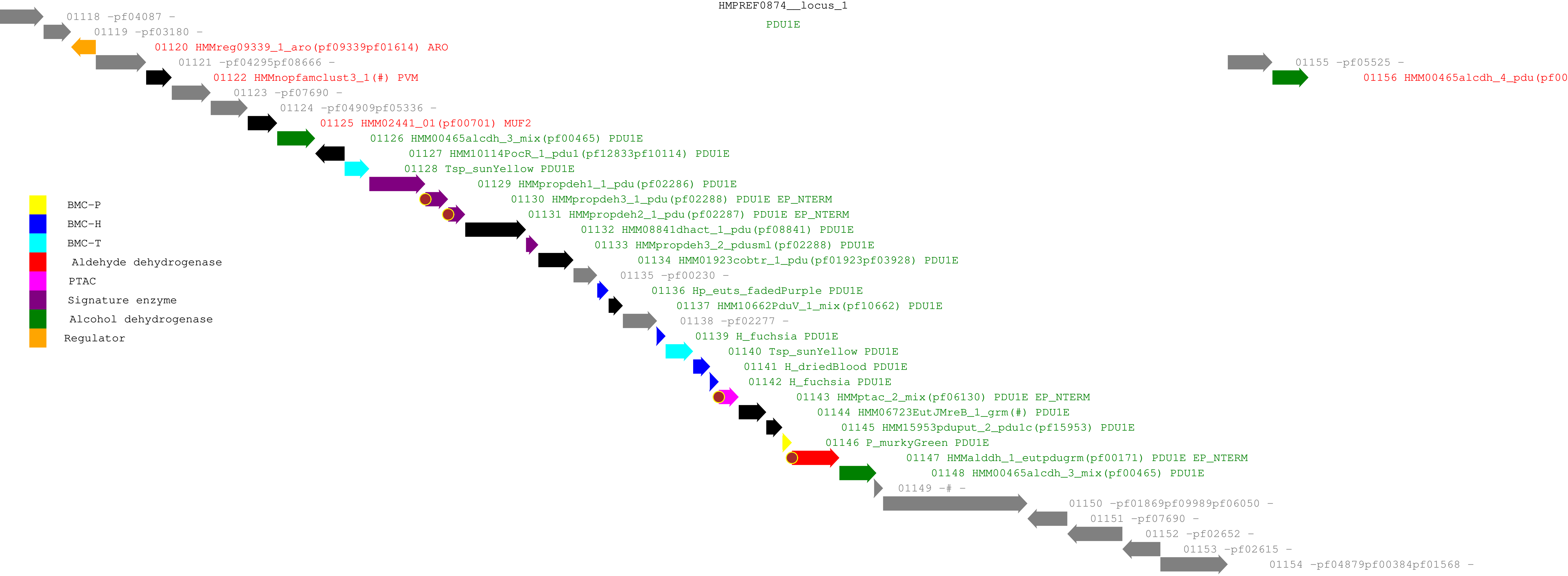

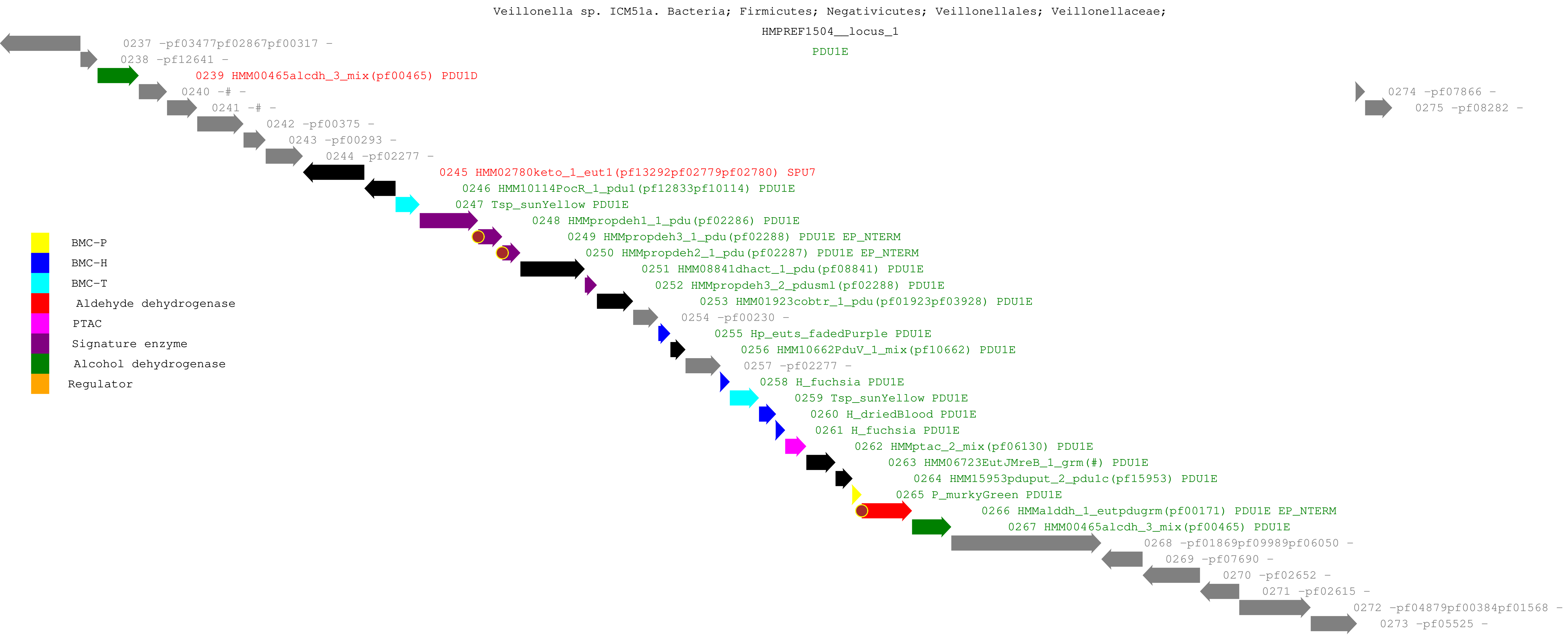

Veillonella sp. ICM51a. Bacteria; Firmicutes; Negativicutes; Veillonellales; Veillonellaceae;
HMPREF1504__locus_1
PDU1E
0237 -pf03477pf02867pf00317 -
0238 -pf12641 -
0239 HMM00465alcdh_3_mix(pf00465) PDU1D
0240 -# -
0241 -# -
0242 -pf00375 -
0243 -pf00293 -
0244 -pf02277 -
0245 HMM02780keto_1_eut1(pf13292pf02779pf02780) SPU7
0246 HMM10114PocR_1_pdu1(pf12833pf10114) PDU1E
0247 Tsp_sunYellow PDU1E
0248 HMMpropdeh1_1_pdu(pf02286) PDU1E
0249 HMMpropdeh3_1_pdu(pf02288) PDU1E EP_NTERM
0250 HMMpropdeh2_1_pdu(pf02287) PDU1E EP_NTERM
0251 HMM08841dhact_1_pdu(pf08841) PDU1E
0252 HMMpropdeh3_2_pdusml(pf02288) PDU1E
0253 HMM01923cobtr_1_pdu(pf01923pf03928) PDU1E
0254 -pf00230 -
0255 Hp_euts_fadedPurple PDU1E
0256 HMM10662PduV_1_mix(pf10662) PDU1E
0257 -pf02277 -
0258 H_fuchsia PDU1E
0259 Tsp_sunYellow PDU1E
0260 H_driedBlood PDU1E
0261 H_fuchsia PDU1E
0262 HMMptac_2_mix(pf06130) PDU1E
0263 HMM06723EutJMreB_1_grm(#) PDU1E
0264 HMM15953pduput_2_pdu1c(pf15953) PDU1E
0265 P_murkyGreen PDU1E
0266 HMMaldDh_1_eutpdugrm(pf00171) PDU1E EP_NTERM
0267 HMM00465alcdh_3_mix(pf00465) PDU1E
0268 -pf01869pf09989pf06050 -
0269 -pf07690 -
0270 -pf02652 -
0271 -pf02615 -
0272 -pf04879pf00384pf01568 -
0273 -pf05525 -
0274 -pf07866 -
0275 -pf08282 -
BMC-P
BMC-H
BMC-T
Aldehyde dehydrogenase
PTAC
Signature enzyme
Alcohol dehydrogenase
Regulator

Veillonella sp. AS16. Bacteria; Firmicutes; Negativicutes; Veillonellales; Veillonellaceae;

HMPREF1521__locus_1

PDU1E

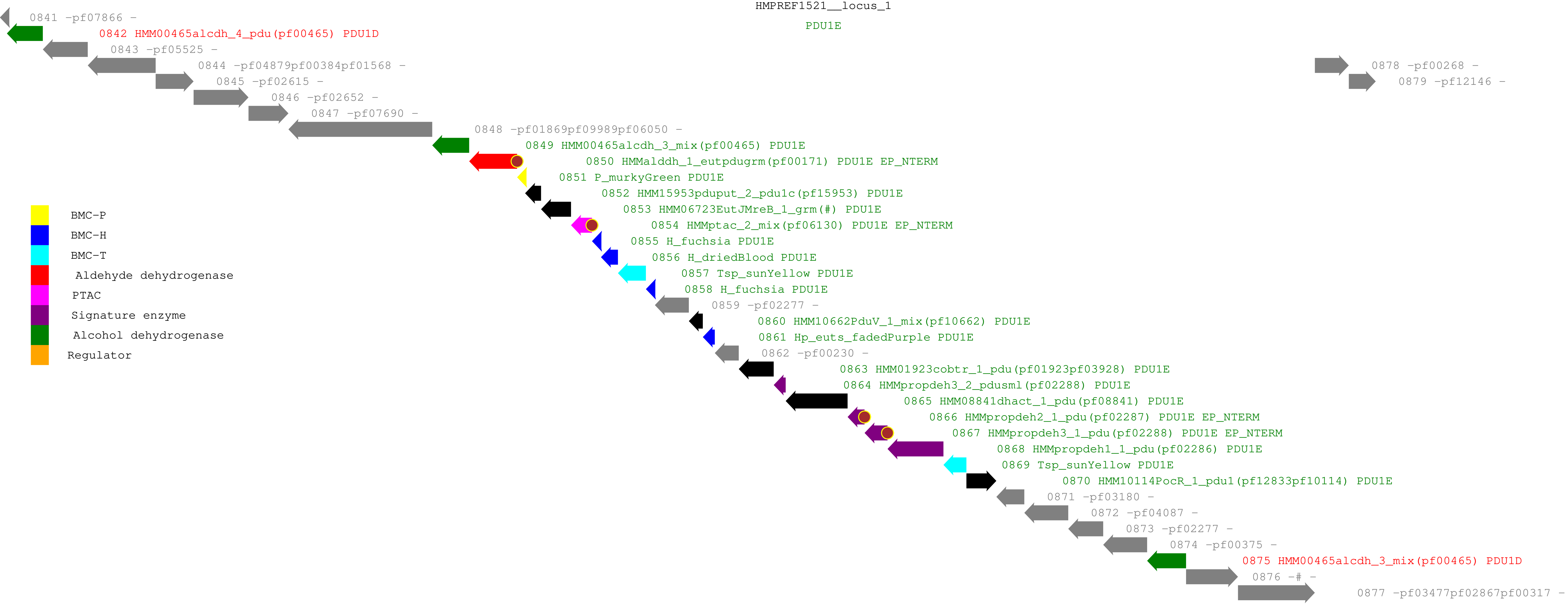

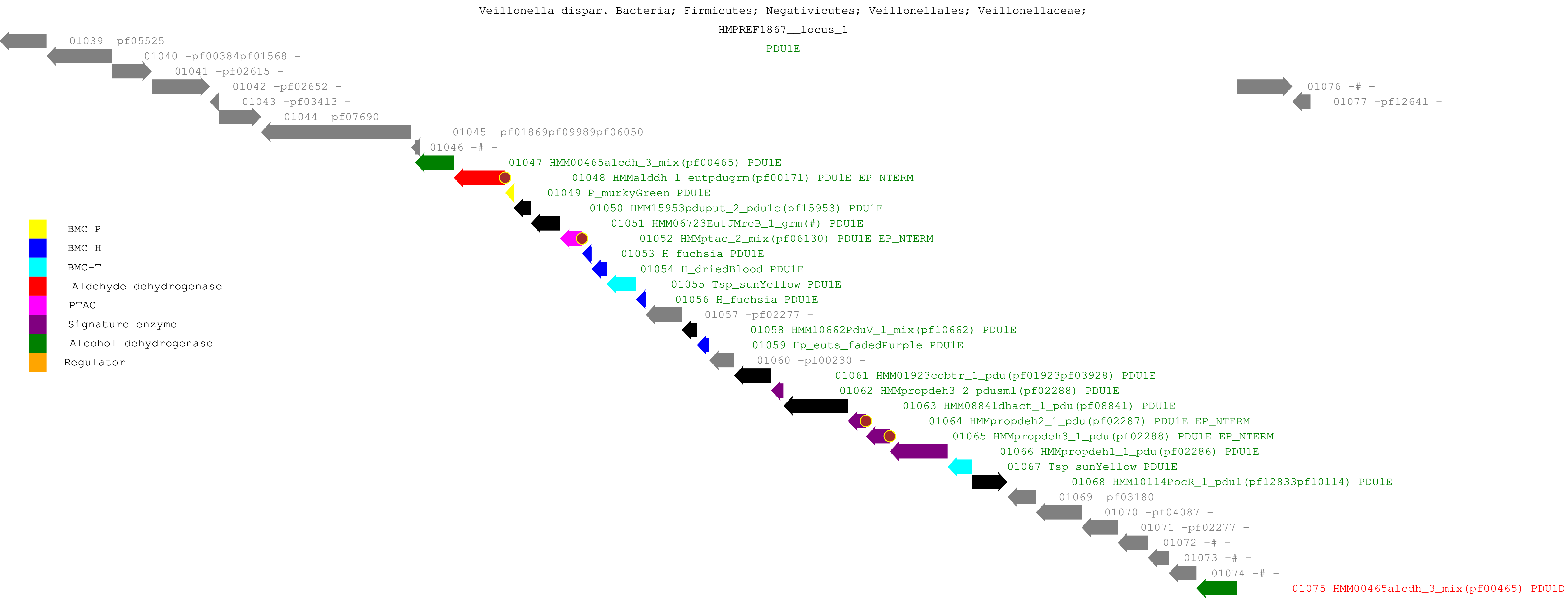
Veillonella dispar. Bacteria; Firmicutes; Negativicutes; Veillonellales; Veillonellaceae;
HMPREF1867__locus_1
PDU1E
01039 -pf05525 -
01040 -pf00384pf01568 -
01041 -pf02615 -
01042 -pf02652 -
01043 -pf03413 -
01044 -pf07690 -
01045 -pf01869pf09989pf06050 -
01046 -# -
01047 HMM00465alcdh_3_mix(pf00465) PDU1E
01048 HMMaldh_1_eutpdugrm(pf00171) PDU1E EP_NTERM
01049 P_murkyGreen PDU1E
01050 HMM15953pduput_2_pdu1c(pf15953) PDU1E
01051 HMM06723EutJMreB_1_grm(#) PDU1E
01052 HMMptac_2_mix(pf06130) PDU1E EP_NTERM
01053 H_fuchsia PDU1E
01054 H_driedBlood PDU1E
01055 Tsp_sunYellow PDU1E
01056 H_fuchsia PDU1E
01057 -pf02277 -
01058 HMM10662PduV_1_mix(pf10662) PDU1E
01059 Hp_euts_fadedPurple PDU1E
01060 -pf00230 -
01061 HMM01923cobtr_1_pdu(pf01923pf03928) PDU1E
01062 HMMpropdeh3_2_pdusml(pf02288) PDU1E
01063 HMM08841dhact_1_pdu(pf08841) PDU1E
01064 HMMpropdeh2_1_pdu(pf02287) PDU1E EP_NTERM
01065 HMMpropdeh3_1_pdu(pf02288) PDU1E EP_NTERM
01066 HMMpropdeh1_1_pdu(pf02286) PDU1E
01067 Tsp_sunYellow PDU1E
01068 HMM10114PocR_1_pdu1(pf12833pf10114) PDU1E
01069 -pf03180 -
01070 -pf04087 -
01071 -pf02277 -
01072 -# -
01073 -# -
01074 -# -
01075 HMM00465alcdh_3_mix(pf00465) PDU1D
01076 -# -
01077 -pf12641 -
BMC-P
BMC-H
BMC-T
Aldehyde dehydrogenase
PTAC
Signature enzyme
Alcohol dehydrogenase
Regulator

Veillonella sp. oral taxon 158 str. F0412. Bacteria; Firmicutes; Negativicutes; Veillonellales; Veillonellaceae;

HMPREF9199__locus_1

PDU1E

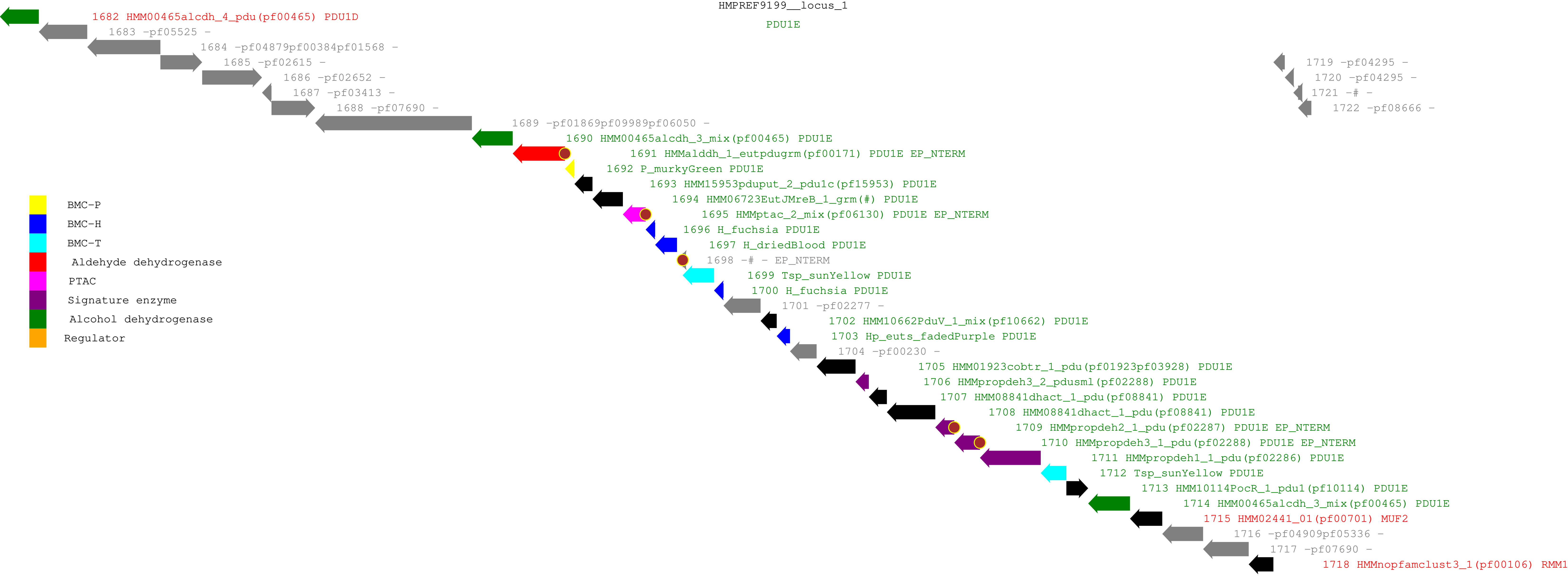

Veillonella seminalis ACS-216-V-Col6b. Bacteria; Firmicutes; Negativicutes; Veillonellales; Veillonellaceae;

HMPREF9282__locus_1

PDU1E

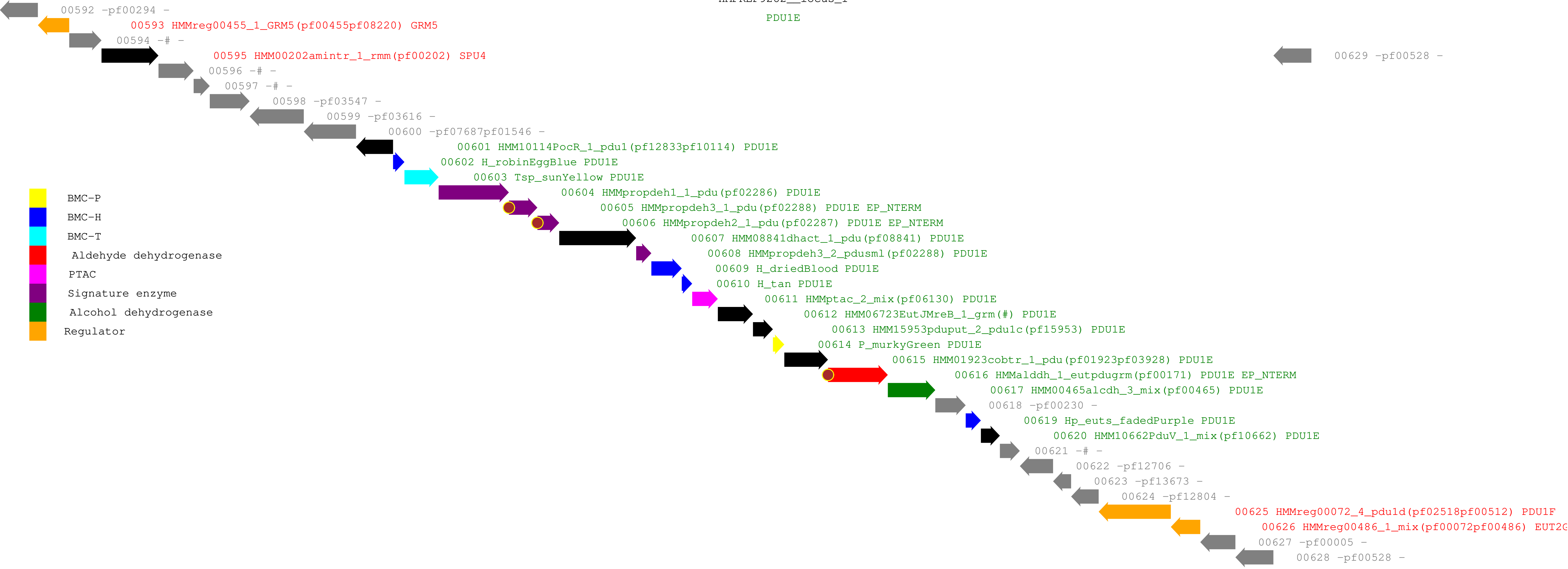

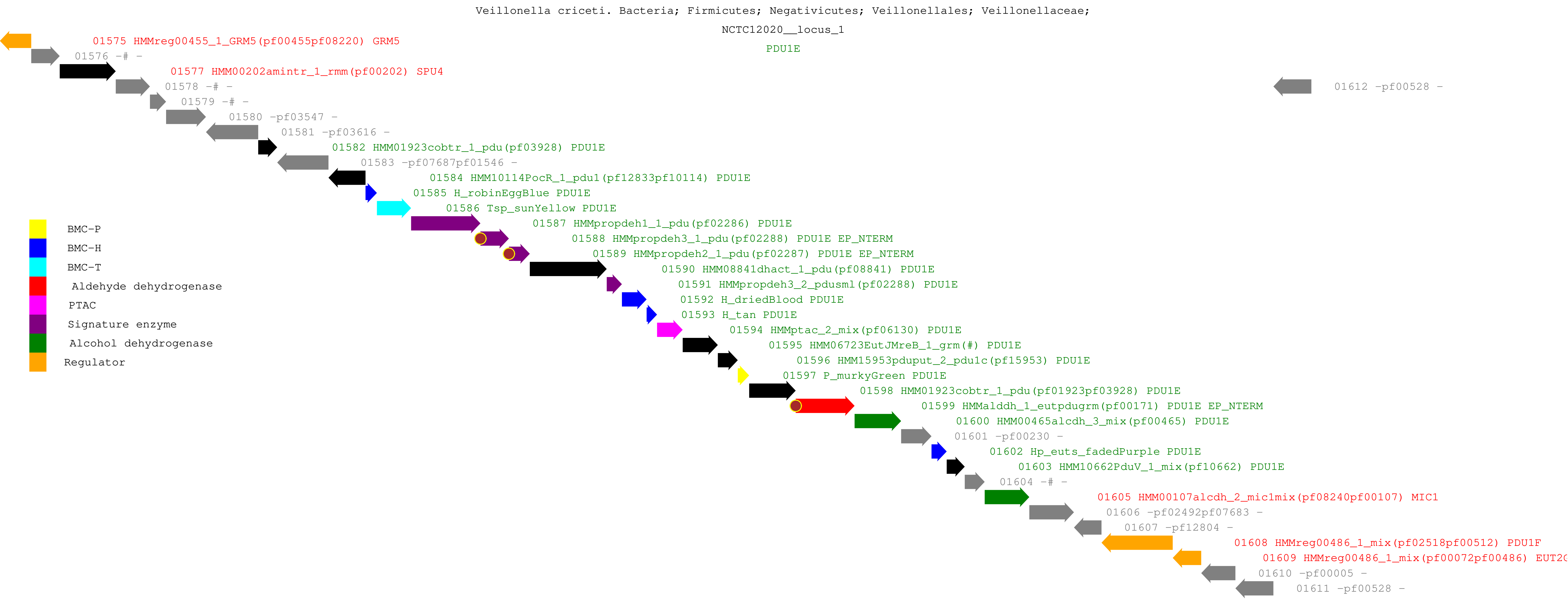
Veillonella criceti. Bacteria; Firmicutes; Negativicutes; Veillonellales; Veillonellaceae;
NCTC12020__locus_1
PDU1E
01575 HMMreg00455_1_GRM5(pf00455pf08220) GRM5
01576 -# -
01577 HMM00202amintr_1_rmm(pf00202) SPU4
01578 -# -
01579 -# -
01580 -pf03547 -
01581 -pf03616 -
01582 HMM01923cobtr_1_pdu(pf03928) PDU1E
01583 -pf07687pf01546 -
01584 HMM10114PocR_1_pdu1(pf12833pf10114) PDU1E
01585 H_robinEggBlue PDU1E
01586 Tsp_sunYellow PDU1E
01587 HMMpropdeh1_1_pdu(pf02286) PDU1E
01588 HMMpropdeh3_1_pdu(pf02288) PDU1E EP_NTERM
01589 HMMpropdeh2_1_pdu(pf02287) PDU1E EP_NTERM
01590 HMM08841dhact_1_pdu(pf08841) PDU1E
01591 HMMpropdeh3_2_pdusml(pf02288) PDU1E
01592 H_driedBlood PDU1E
01593 H_tan PDU1E
01594 HMMptac_2_mix(pf06130) PDU1E
01595 HMM06723EutJMreB_1_grm(#) PDU1E
01596 HMM15953pduput_2_pdu1c(pf15953) PDU1E
01597 P_murkyGreen PDU1E
01598 HMM01923cobtr_1_pdu(pf01923pf03928) PDU1E
01599 HMMalddh_1_eutpdugrm(pf00171) PDU1E EP_NTERM
01600 HMM00465alcdh_3_mix(pf00465) PDU1E
01601 -pf00230 -
01602 Hp_euts_fadedPurple PDU1E
01603 HMM10662PduV_1_mix(pf10662) PDU1E
01604 -# -
01605 HMM00107alcdh_2_mic1mix(pf08240pf00107) MIC1
01606 -pf02492pf07683 -
01607 -pf12804 -
01608 HMMreg00486_1_mix(pf02518pf00512) PDU1F
01609 HMMreg00486_1_mix(pf00072pf00486) EUT2C
01610 -pf00005 -
01611 -pf00528 -
01612 -pf00528 -
BMC-P
BMC-H
BMC-T
Aldehyde dehydrogenase
PTAC
Signature enzyme
Alcohol dehydrogenase
Regulator

Enterococcus florum. Bacteria; Firmicutes; Bacilli; Lactobacillales; Enterococcaceae;

NRIC__locus_2

PDU1E

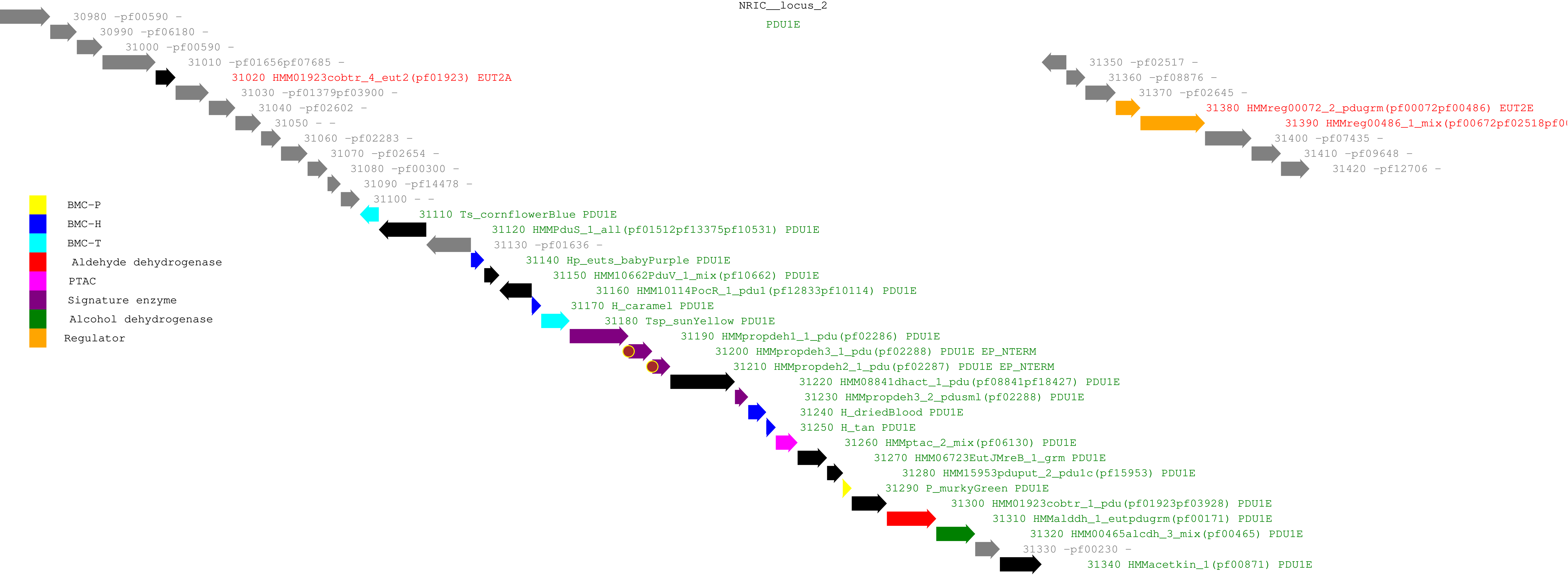

Enterococcus avium ATCC 14025. Bacteria; Firmicutes; Bacilli; Lactobacillales; Enterococcaceae;

OMU__locus_3

PDU1E

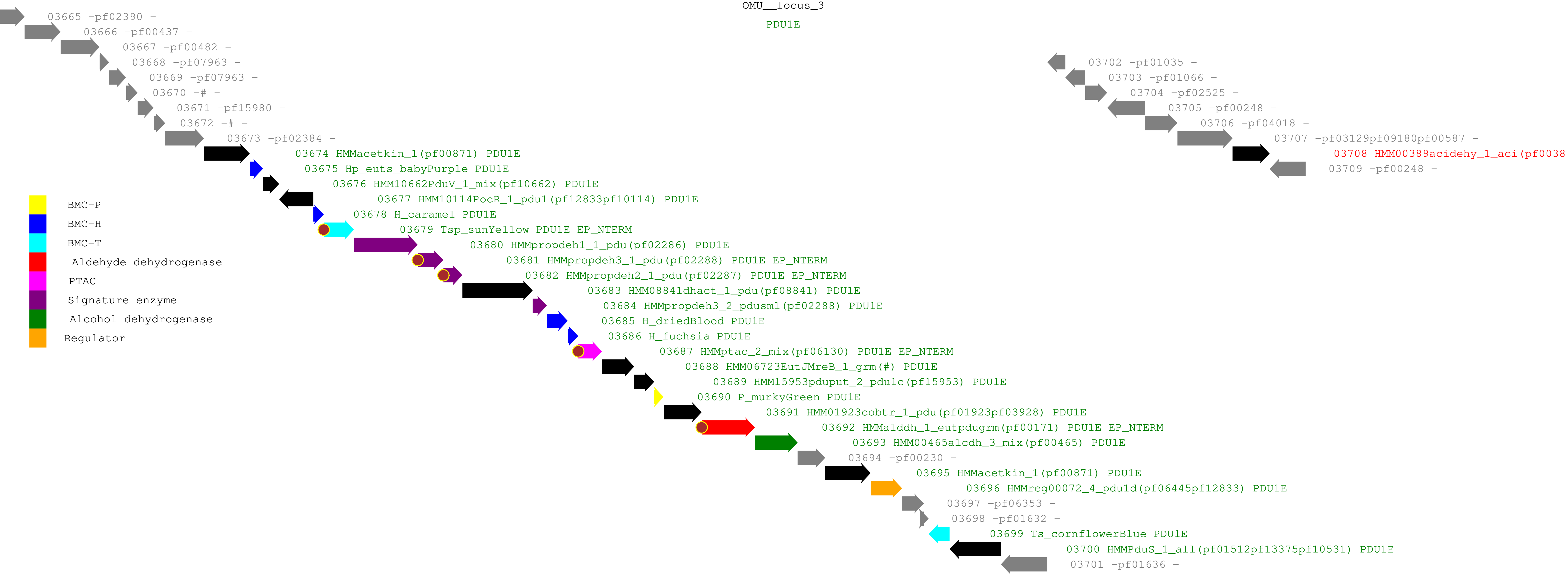

Veillonella tobetsuensis. Bacteria; Firmicutes; Negativicutes; Veillonellales; Veillonellaceae;

PAGU1578__locus_1

PDU1E

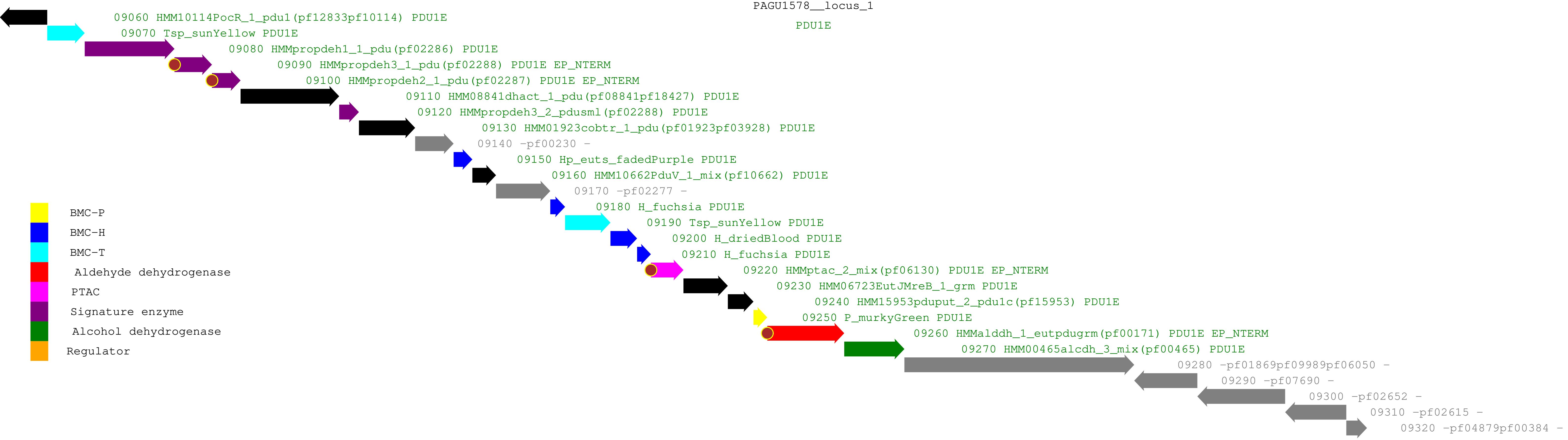

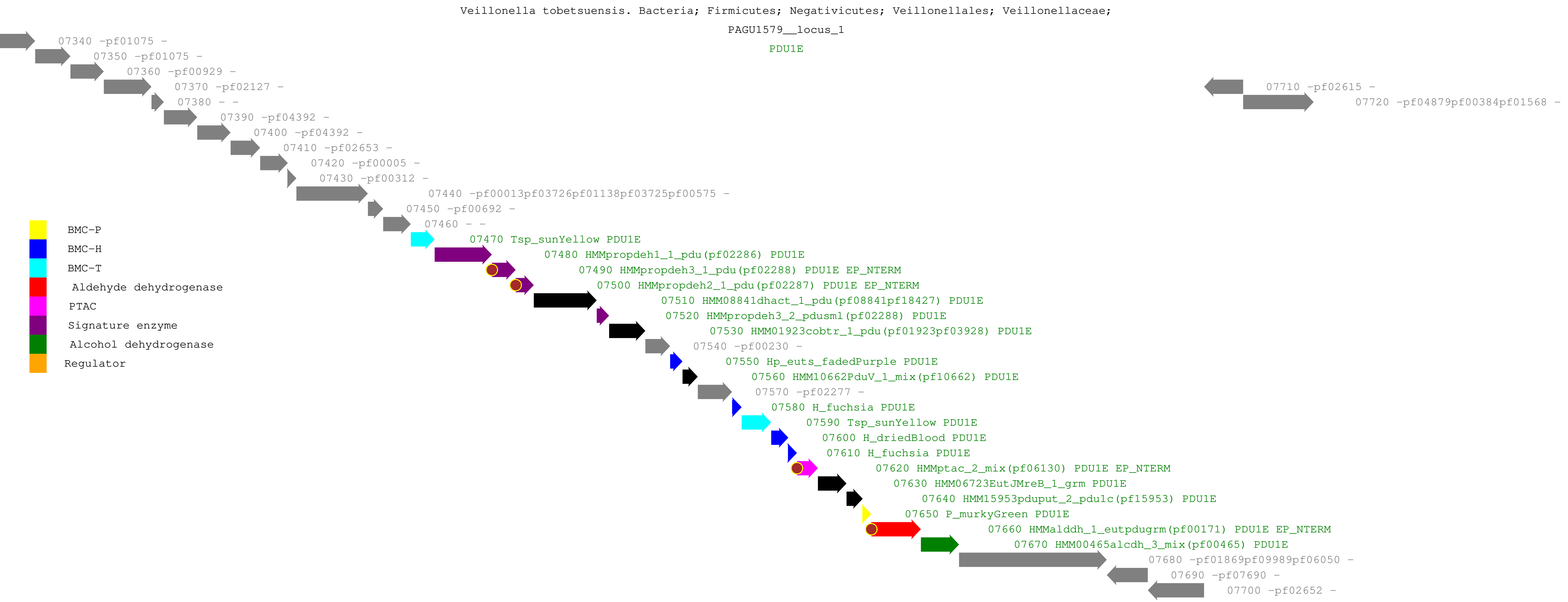
Veillonella tobetsuensis. Bacteria; Firmicutes; Negativicutes; Veillonellales; Veillonellaceae;
PAGU1579__locus_1
PDU1E
07340 -pf01075 -
07350 -pf01075 -
07360 -pf00929 -
07370 -pf02127 -
07380 - -
07390 -pf04392 -
07400 -pf04392 -
07410 -pf02653 -
07420 -pf00005 -
07430 -pf00312 -
07440 -pf00013pf03726pf01138pf03725pf00575 -
07450 -pf00692 -
07460 - -
07470 Tsp_sunYellow PDU1E
07480 HMMpropdeh1_1_pdu(pf02286) PDU1E
07490 HMMpropdeh3_1_pdu(pf02288) PDU1E EP_NTERM
07500 HMMpropdeh2_1_pdu(pf02287) PDU1E EP_NTERM
07510 HMM08841dhact_1_pdu(pf08841pf18427) PDU1E
07520 HMMpropdeh3_2_pdusml(pf02288) PDU1E
07530 HMM01923cobtr_1_pdu(pf01923pf03928) PDU1E
07540 -pf00230 -
07550 Hp_euts_fadedPurple PDU1E
07560 HMM10662PduV_1_mix(pf10662) PDU1E
07570 -pf02277 -
07580 H_fuchsia PDU1E
07590 Tsp_sunYellow PDU1E
07600 H_driedBlood PDU1E
07610 H_fuchsia PDU1E
07620 HMMptac_2_mix(pf06130) PDU1E EP_NTERM
07630 HMM06723EutJMreB_1_grm PDU1E
07640 HMM15953pduput_2_pdu1c(pf15953) PDU1E
07650 P_murkyGreen PDU1E
07660 HMMalddh_1_eutpdugrm(pf00171) PDU1E EP_NTERM
07670 HMM00465alcdh_3_mix(pf00465) PDU1E
07680 -pf01869pf09989pf06050 -
07690 -pf07690 -
07700 -pf02652 -
07710 -pf02615 -
07720 -pf04879pf00384pf01568 -
BMC-P
BMC-H
BMC-T
Aldehyde dehydrogenase
PTAC
Signature enzyme
Alcohol dehydrogenase
Regulator

Veillonella sp. DORA_A_3_16_22. Bacteria; Firmicutes; Negativicutes; Veillonellales; Veillonellaceae;

Q620_VSAC01225G_locus_1

PDU1E

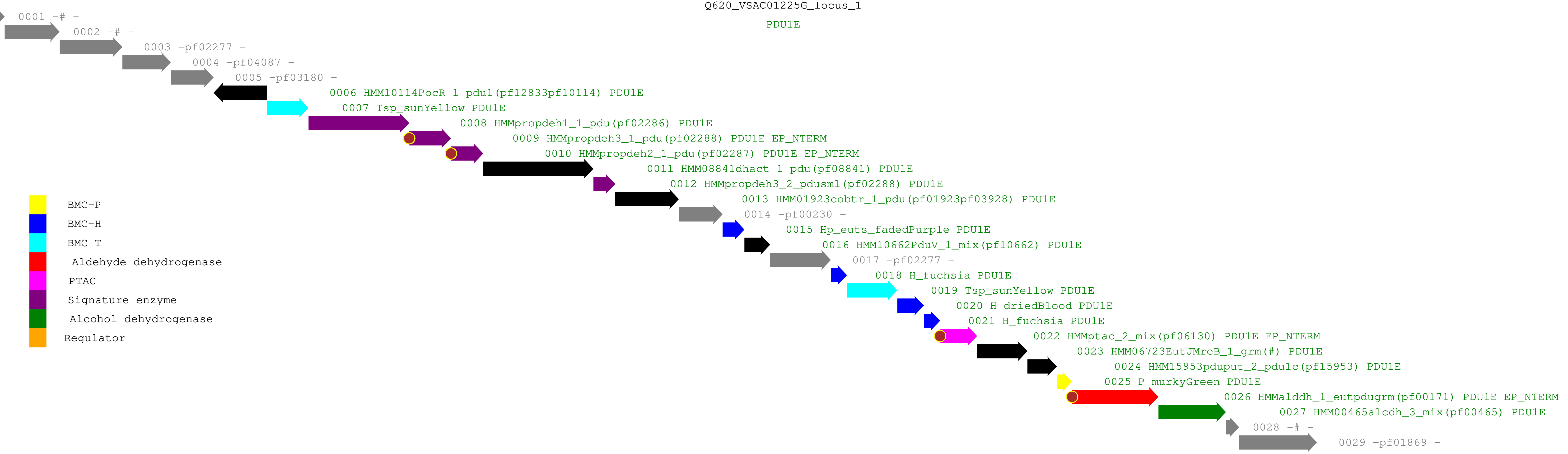

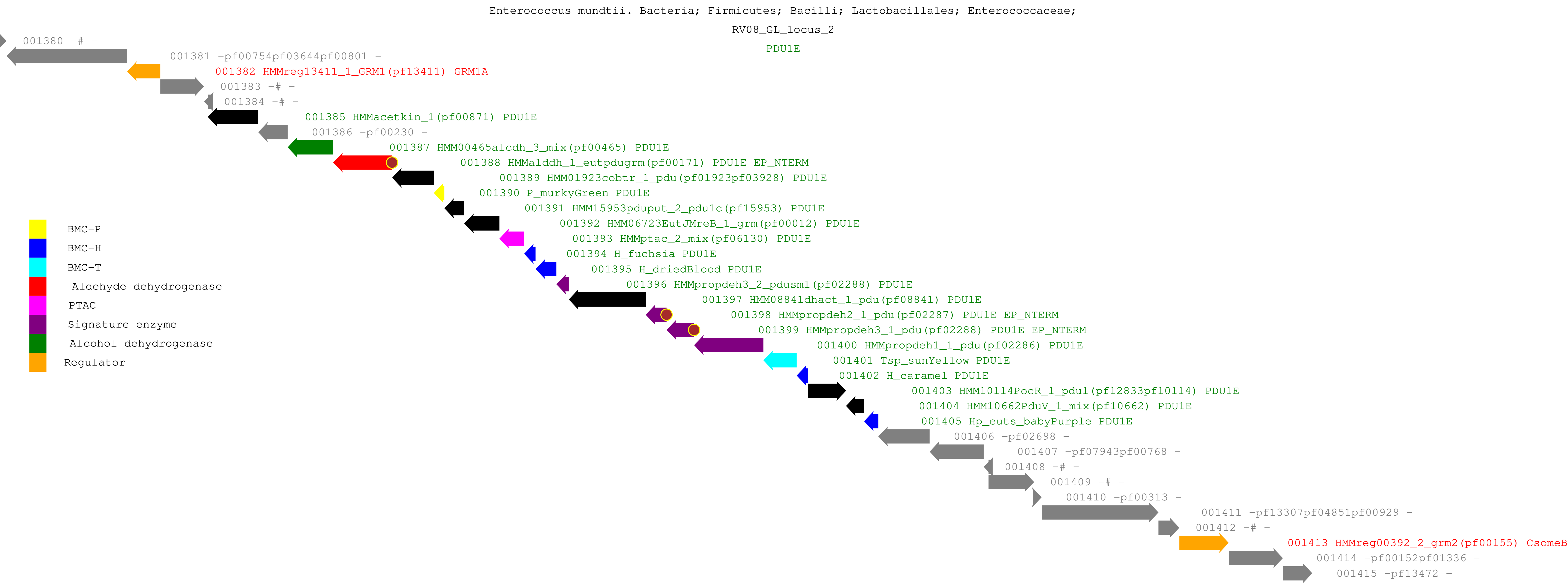
Enterococcus mundtii. Bacteria; Firmicutes; Bacilli; Lactobacillales; Enterococcaceae;
RV08_GL_locus_2
PDU1E
001380 -# -
001381 -pf00754pf03644pf00801 -
001382 HMMreg13411_1_GRM1(pf13411) GRM1A
001383 -# -
001384 -# -
001385 HMMacetkin_1(pf00871) PDU1E
001386 -pf00230 -
001387 HMM00465alcdh_3_mix(pf00465) PDU1E
001388 HMMalddh_1_eutpdugrm(pf00171) PDU1E EP_NTERM
001389 HMM01923cobtr_1_pdu(pf01923pf03928) PDU1E
001390 P_murkyGreen PDU1E
001391 HMM15953pduput_2_pdu1c(pf15953) PDU1E
001392 HMM06723EutJMreB_1_grm(pf00012) PDU1E
001393 HMMptac_2_mix(pf06130) PDU1E
001394 H_fuchsia PDU1E
001395 H_driedBlood PDU1E
001396 HMMpropdeh3_2_pdusml(pf02288) PDU1E
001397 HMM08841dhact_1_pdu(pf08841) PDU1E
001398 HMMpropdeh2_1_pdu(pf02287) PDU1E EP_NTERM
001399 HMMpropdeh3_1_pdu(pf02288) PDU1E EP_NTERM
001400 HMMpropdeh1_1_pdu(pf02286) PDU1E
001401 Tsp_sunYellow PDU1E
001402 H_caramel PDU1E
001403 HMM10114PocR_1_pdu1(pf12833pf10114) PDU1E
001404 HMM10662PduV_1_mix(pf10662) PDU1E
001405 Hp_euts_babyPurple PDU1E
001406 -pf02698 -
001407 -pf07943pf00768 -
001408 -# -
001409 -# -
001410 -pf00313 -
001411 -pf13307pf04851pf00929 -
001412 -# -
001413 HMMreg00392_2_grm2(pf00155) CsomeB
001414 -pf00152pf01336 -
001415 -pf13472 -
BMC-P
BMC-H
BMC-T
Aldehyde dehydrogenase
PTAC
Signature enzyme
Alcohol dehydrogenase
Regulator

Enterococcus raffinosus. Bacteria; Firmicutes; Bacilli; Lactobacillales; Enterococcaceae;

RV13_GL_locus_2

PDU1E

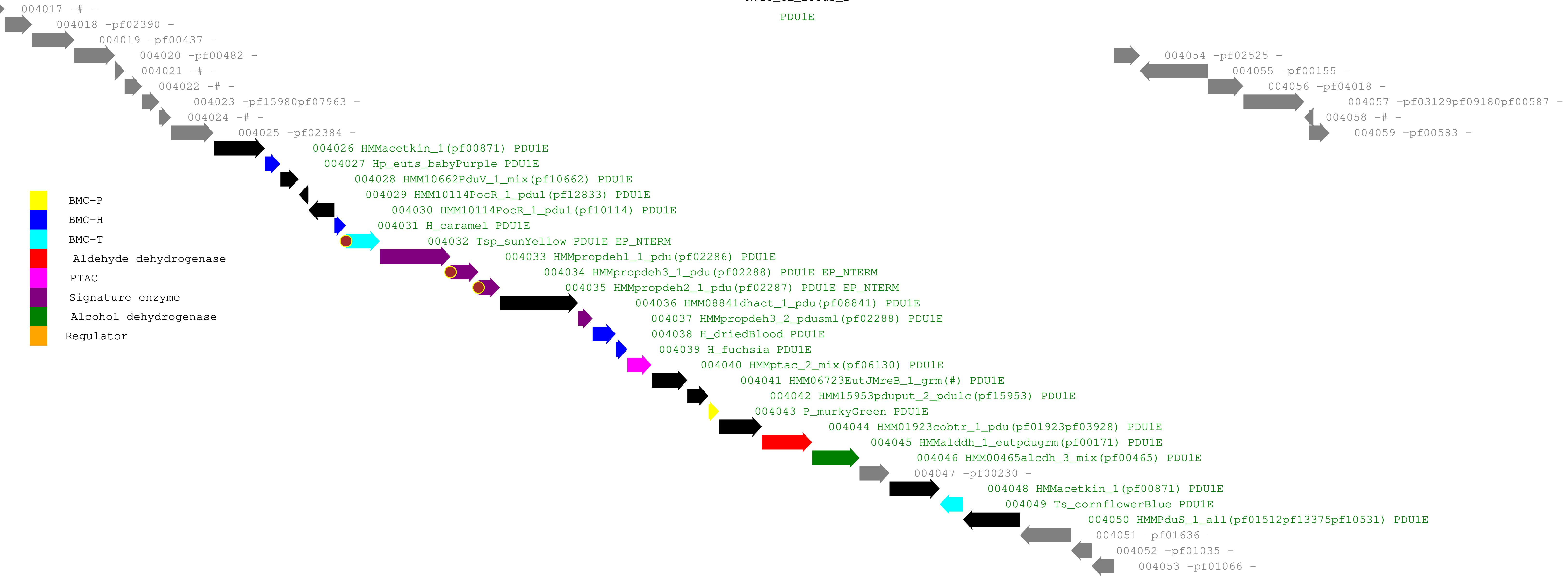

Pilibacter termitis. Bacteria; Firmicutes; Bacilli; Lactobacillales; Enterococcaceae;

SAMN02745116__locus_2_main_sat

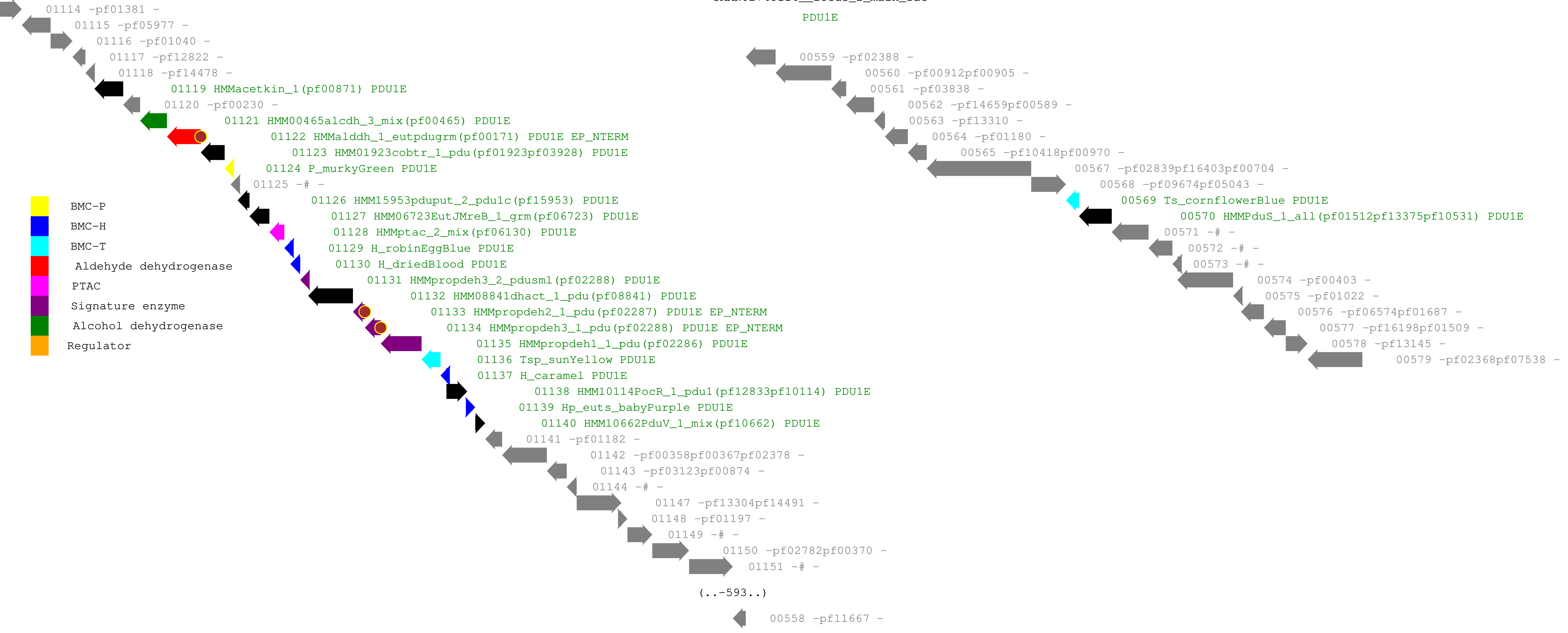

Enterococcus mundtii. Bacteria; Firmicutes; Bacilli; Lactobacillales; Enterococcaceae;

SAMN04487758__locus_2

PDU1E

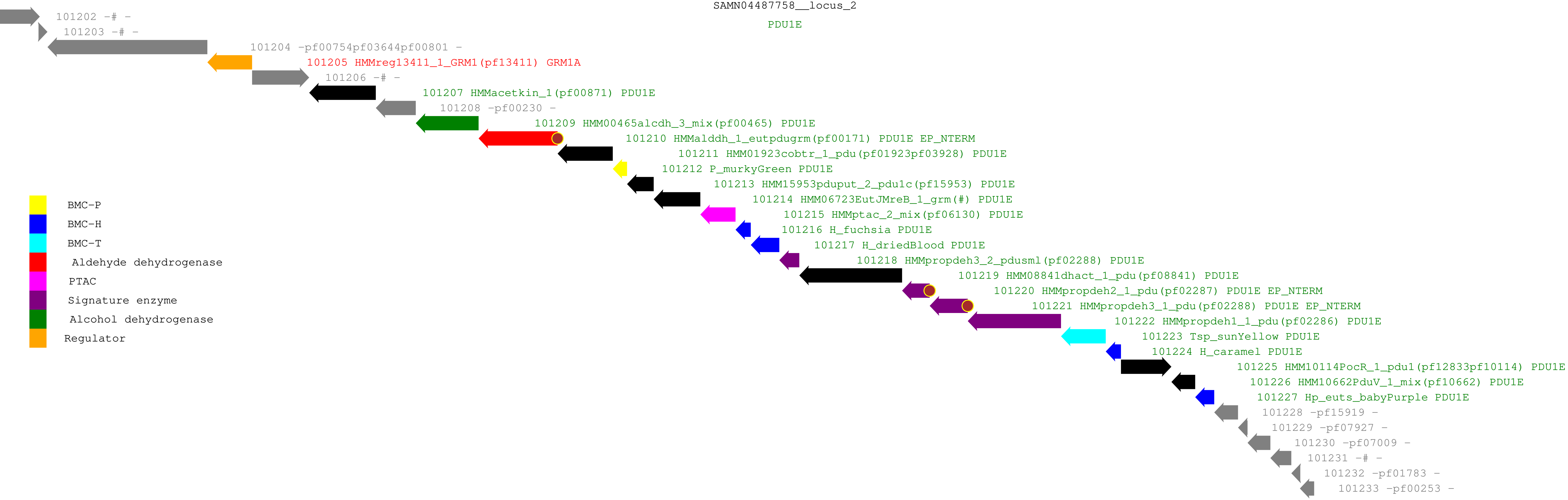

Enterococcus malodoratus ATCC 43197. Bacteria; Firmicutes; Bacilli; Lactobacillales; Enterococcaceae;

UAI__locus_1

PDU1E

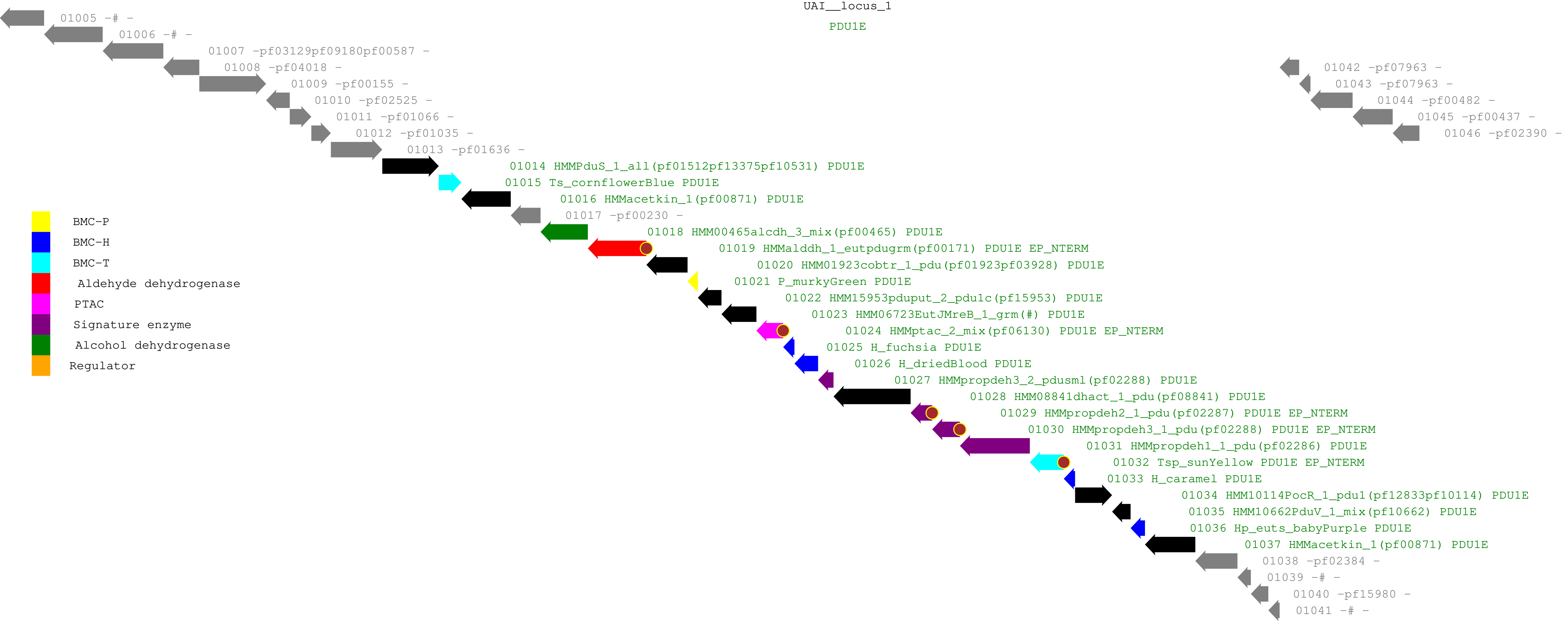

Enterococcus raffinosus ATCC 49464. Bacteria; Firmicutes; Bacilli; Lactobacillales; Enterococcaceae;

UAK__locus_2

PDU1E

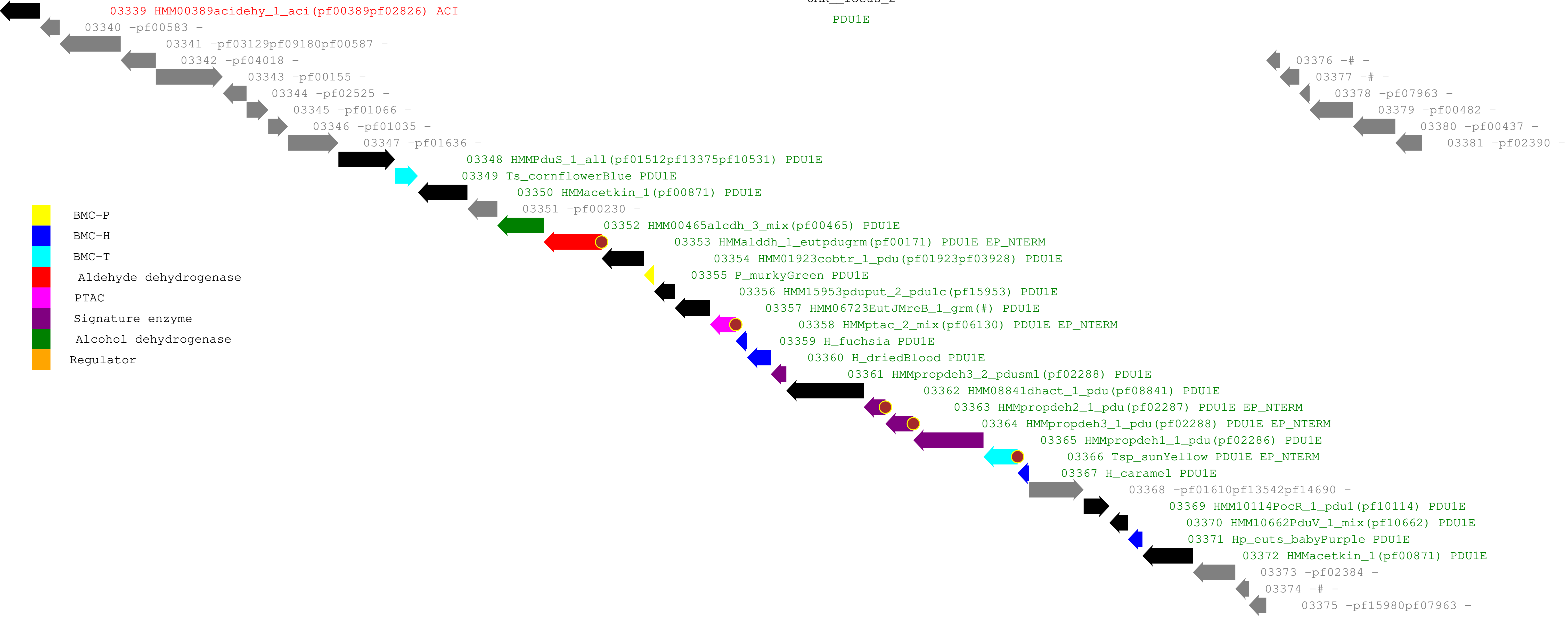

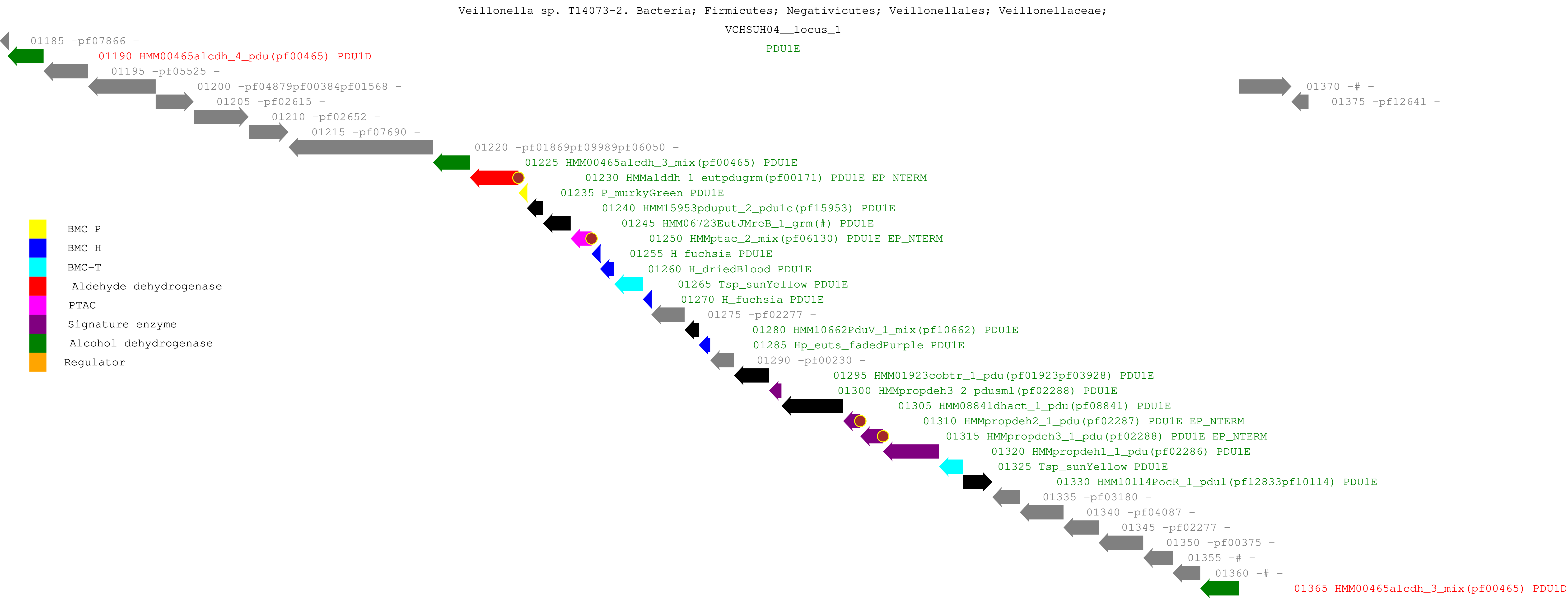

Veillonella sp. T14073-2. Bacteria; Firmicutes; Negativicutes; Veillonellales; Veillonellaceae;
VCHSUH04__locus_1
PDU1E
01185 -pf07866 -
01190 HMM00465alcdh_4_pdu(pf00465) PDU1D
01195 -pf05525 -
01200 -pf04879pf00384pf01568 -
01205 -pf02615 -
01210 -pf02652 -
01215 -pf07690 -
01220 -pf01869pf09989pf06050 -
01225 HMM00465alcdh_3_mix(pf00465) PDU1E
01230 HMMalddh_1_eutpdugrm(pf00171) PDU1E EP_NTERM
01235 P_murkyGreen PDU1E
01240 HMM15953pduput_2_pdu1c(pf15953) PDU1E
01245 HMM06723EutJMreB_1_grm(#) PDU1E
01250 HMMptac_2_mix(pf06130) PDU1E EP_NTERM
01255 H_fuchsia PDU1E
01260 H_driedBlood PDU1E
01265 Tsp_sunYellow PDU1E
01270 H_fuchsia PDU1E
01275 -pf02277 -
01280 HMM10662PduV_1_mix(pf10662) PDU1E
01285 Hp_euts_fadedPurple PDU1E
01290 -pf00230 -
01295 HMM01923cobtr_1_pdu(pf01923pf03928) PDU1E
01300 HMMpropdeh3_2_pdusml(pf02288) PDU1E
01305 HMM08841dhact_1_pdu(pf08841) PDU1E
01310 HMMpropdeh2_1_pdu(pf02287) PDU1E EP_NTERM
01315 HMMpropdeh3_1_pdu(pf02288) PDU1E EP_NTERM
01320 HMMpropdeh1_1_pdu(pf02286) PDU1E
01325 Tsp_sunYellow PDU1E
01330 HMM10114PocR_1_pdu1(pf12833pf10114) PDU1E
01335 -pf03180 -
01340 -pf04087 -
01345 -pf02277 -
01350 -pf00375 -
01355 -# -
01360 -# -
01365 HMM00465alcdh_3_mix(pf00465) PDU1D
01370 -# -
01375 -pf12641 -
BMC-P
BMC-H
BMC-T
Aldehyde dehydrogenase
PTAC
Signature enzyme
Alcohol dehydrogenase
Regulator

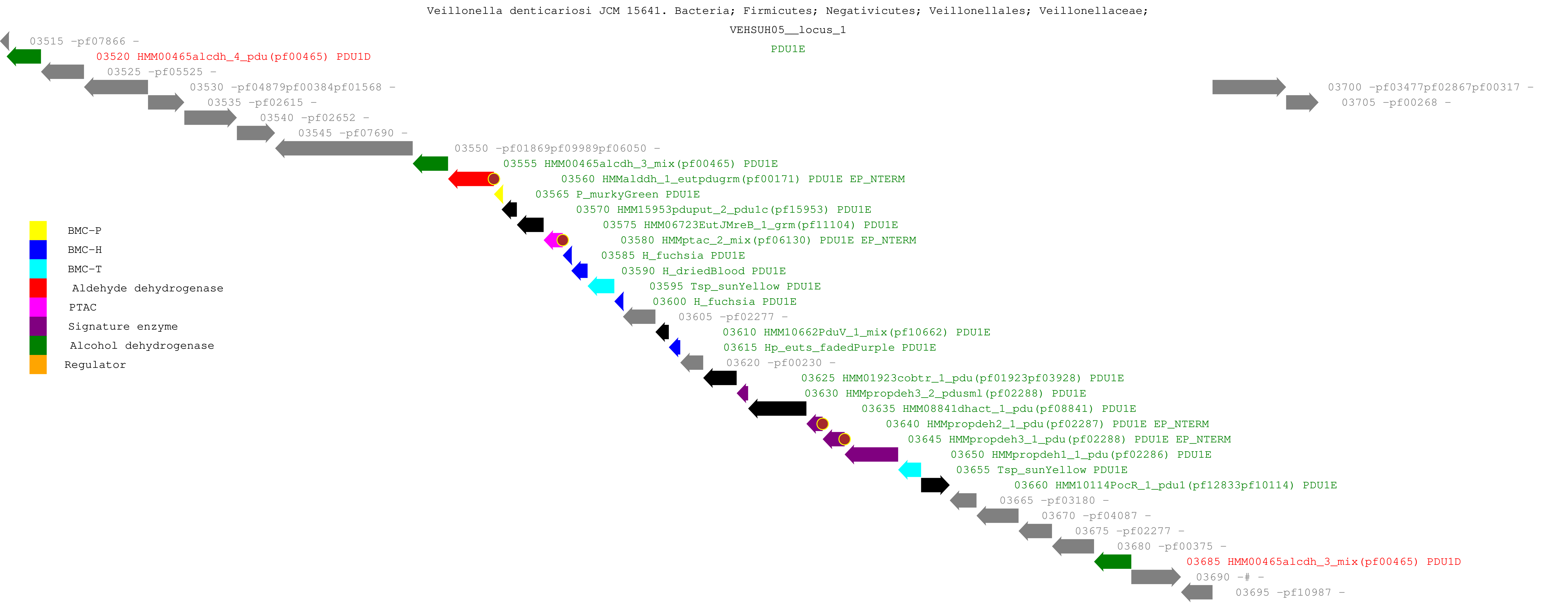
Veillonella denticariosi JCM 15641. Bacteria; Firmicutes; Negativicutes; Veillonellales; Veillonellaceae;
VEHSUH05__locus_1
PDU1E
03515 -pf07866 -
03520 HMM00465alcdh_4_pdu(pf00465) PDU1D
03525 -pf05525 -
03530 -pf04879pf00384pf01568 -
03535 -pf02615 -
03540 -pf02652 -
03545 -pf07690 -
03550 -pf01869pf09989pf06050 -
03555 HMM00465alcdh_3_mix(pf00465) PDU1E
03560 HMMaldh_1_eutpdugrm(pf00171) PDU1E EP_NTERM
03565 P_murkyGreen PDU1E
03570 HMM15953pduput_2_pdu1c(pf15953) PDU1E
03575 HMM06723EutJMreB_1_grm(pf11104) PDU1E
03580 HMMptac_2_mix(pf06130) PDU1E EP_NTERM
03585 H_fuchsia PDU1E
03590 H_driedBlood PDU1E
03595 Tsp_sunYellow PDU1E
03600 H_fuchsia PDU1E
03605 -pf02277 -
03610 HMM10662PduV_1_mix(pf10662) PDU1E
03615 Hp_euts_fadedPurple PDU1E
03620 -pf00230 -
03625 HMM01923cobtr_1_pdu(pf01923pf03928) PDU1E
03630 HMMpropdeh3_2_pdusml(pf02288) PDU1E
03635 HMM08841dhact_1_pdu(pf08841) PDU1E
03640 HMMpropdeh2_1_pdu(pf02287) PDU1E EP_NTERM
03645 HMMpropdeh3_1_pdu(pf02288) PDU1E EP_NTERM
03650 HMMpropdeh1_1_pdu(pf02286) PDU1E
03655 Tsp_sunYellow PDU1E
03660 HMM10114PocR_1_pdu1(pf12833pf10114) PDU1E
03665 -pf03180 -
03670 -pf04087 -
03675 -pf02277 -
03680 -pf00375 -
03685 HMM00465alcdh_3_mix(pf00465) PDU1D
03690 -# -
03695 -pf10987 -
03700 -pf03477pf02867pf00317 -
03705 -pf00268 -
BMC-P
BMC-H
BMC-T
Aldehyde dehydrogenase
PTAC
Signature enzyme
Alcohol dehydrogenase
Regulator

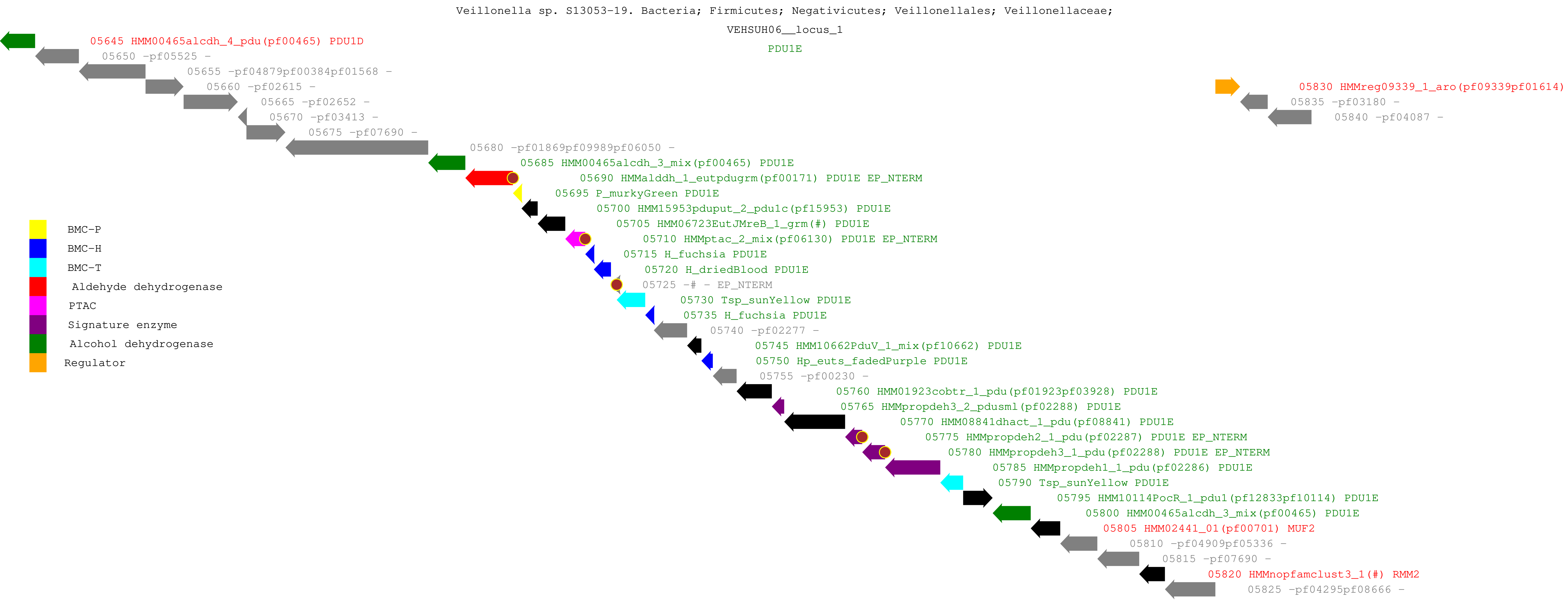
Veillonella sp. S13053-19. Bacteria; Firmicutes; Negativicutes; Veillonellales; Veillonellaceae;
VEHSUH06__locus_1
PDU1E
05645 HMM00465alcdh_4_pdu(pf00465) PDU1D
05650 -pf05525 -
05655 -pf04879pf00384pf01568 -
05660 -pf02615 -
05665 -pf02652 -
05670 -pf03413 -
05675 -pf07690 -
05680 -pf01869pf09989pf06050 -
05685 HMM00465alcdh_3_mix(pf00465) PDU1E
05690 HMMalddh_1_eutpdugrm(pf00171) PDU1E EP_NTERM
05695 P_murkyGreen PDU1E
05700 HMM15953pduput_2_pdu1c(pf15953) PDU1E
05705 HMM06723EutJMreB_1_grm(#) PDU1E
05710 HMMptac_2_mix(pf06130) PDU1E EP_NTERM
05715 H_fuchsia PDU1E
05720 H_driedBlood PDU1E
05725 -# - EP_NTERM
05730 Tsp_sunYellow PDU1E
05735 H_fuchsia PDU1E
05740 -pf02277 -
05745 HMM10662PduV_1_mix(pf10662) PDU1E
05750 Hp_euts_fadedPurple PDU1E
05755 -pf00230 -
05760 HMM01923cobtr_1_pdu(pf01923pf03928) PDU1E
05765 HMMpropdeh3_2_pdusml(pf02288) PDU1E
05770 HMM08841dhact_1_pdu(pf08841) PDU1E
05775 HMMpropdeh2_1_pdu(pf02287) PDU1E EP_NTERM
05780 HMMpropdeh3_1_pdu(pf02288) PDU1E EP_NTERM
05785 HMMpropdeh1_1_pdu(pf02286) PDU1E
05790 Tsp_sunYellow PDU1E
05795 HMM10114PocR_1_pdu1(pf12833pf10114) PDU1E
05800 HMM00465alcdh_3_mix(pf00465) PDU1E
05805 HMM02441_01(pf00701) MUF2
05810 -pf04909pf05336 -
05815 -pf07690 -
05820 HMMnopfamclust3_1(#) RMM2
05825 -pf04295pf08666 -
05830 HMMreg09339_1_aro(pf09339pf01614)
05835 -pf03180 -
05840 -pf04087 -
BMC-P
BMC-H
BMC-T
Aldehyde dehydrogenase
PTAC
Signature enzyme
Alcohol dehydrogenase
Regulator

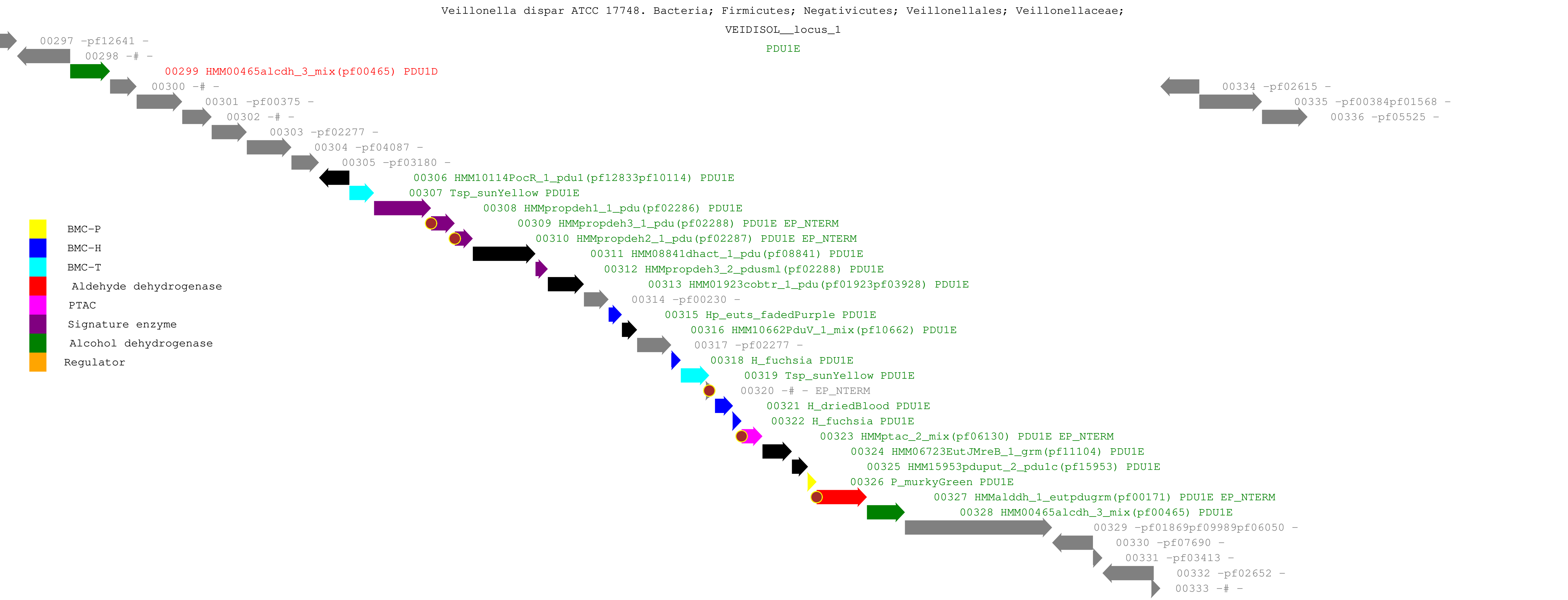
Veillonella dispar ATCC 17748. Bacteria; Firmicutes; Negativicutes; Veillonellales; Veillonellaceae;
VEIDISOL__locus_1
PDU1E
00297 -pf12641 -
00298 -# -
00299 HMM00465alcdh_3_mix(pf00465) PDU1D
00300 -# -
00301 -pf00375 -
00302 -# -
00303 -pf02277 -
00304 -pf04087 -
00305 -pf03180 -
00306 HMM10114PocR_1_pdu1(pf12833pf10114) PDU1E
00307 Tsp_sunYellow PDU1E
00308 HMMpropdeh1_1_pdu(pf02286) PDU1E
00309 HMMpropdeh3_1_pdu(pf02288) PDU1E EP_NTERM
00310 HMMpropdeh2_1_pdu(pf02287) PDU1E EP_NTERM
00311 HMM08841dhact_1_pdu(pf08841) PDU1E
00312 HMMpropdeh3_2_pdusml(pf02288) PDU1E
00313 HMM01923cobtr_1_pdu(pf01923pf03928) PDU1E
00314 -pf00230 -
00315 Hp_euts_fadedPurple PDU1E
00316 HMM10662PduV_1_mix(pf10662) PDU1E
00317 -pf02277 -
00318 H_fuchsia PDU1E
00319 Tsp_sunYellow PDU1E
00320 -# - EP_NTERM
00321 H_driedBlood PDU1E
00322 H_fuchsia PDU1E
00323 HMMptac_2_mix(pf06130) PDU1E EP_NTERM
00324 HMM06723EutJMreB_1_grm(pf11104) PDU1E
00325 HMM15953pduput_2_pdu1c(pf15953) PDU1E
00326 P_murkyGreen PDU1E
00327 HMMalddh_1_eutpdugrm(pf00171) PDU1E EP_NTERM
00328 HMM00465alcdh_3_mix(pf00465) PDU1E
00329 -pf01869pf09989pf06050 -
00330 -pf07690 -
00331 -pf03413 -
00332 -pf02652 -
00333 -# -
00334 -pf02615 -
00335 -pf00384pf01568 -
00336 -pf05525 -
BMC-P
BMC-H
BMC-T
Aldehyde dehydrogenase
PTAC
Signature enzyme
Alcohol dehydrogenase
Regulator

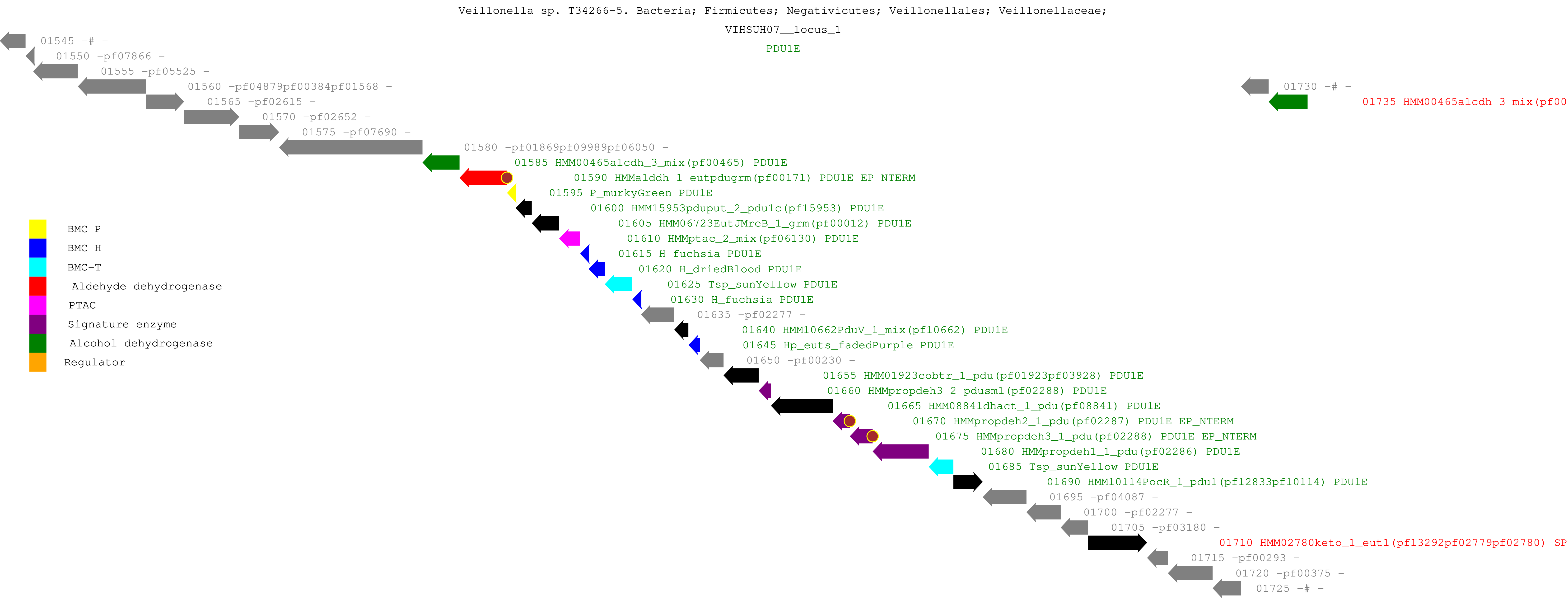

Veillonella sp. T34266-5. Bacteria; Firmicutes; Negativicutes; Veillonellales; Veillonellaceae;
VIHSUH07__locus_1
PDU1E
01545 -# -
01550 -pf07866 -
01555 -pf05525 -
01560 -pf04879pf00384pf01568 -
01565 -pf02615 -
01570 -pf02652 -
01575 -pf07690 -
01580 -pf01869pf09989pf06050 -
01585 HMM00465alcdh_3_mix(pf00465) PDU1E
01590 HMMaldh_1_eutpdugrm(pf00171) PDU1E EP_NTERM
01595 P_murkyGreen PDU1E
01600 HMM15953pduput_2_pdu1c(pf15953) PDU1E
01605 HMM06723EutJMreB_1_grm(pf00012) PDU1E
01610 HMMptac_2_mix(pf06130) PDU1E
01615 H_fuchsia PDU1E
01620 H_driedBlood PDU1E
01625 Tsp_sunYellow PDU1E
01630 H_fuchsia PDU1E
01635 -pf02277 -
01640 HMM10662PduV_1_mix(pf10662) PDU1E
01645 Hp_euts_fadedPurple PDU1E
01650 -pf00230 -
01655 HMM01923cobtr_1_pdu(pf01923pf03928) PDU1E
01660 HMMpropdeh3_2_pdusml(pf02288) PDU1E
01665 HMM08841dhact_1_pdu(pf08841) PDU1E
01670 HMMpropdeh2_1_pdu(pf02287) PDU1E EP_NTERM
01675 HMMpropdeh3_1_pdu(pf02288) PDU1E EP_NTERM
01680 HMMpropdeh1_1_pdu(pf02286) PDU1E
01685 Tsp_sunYellow PDU1E
01690 HMM10114PocR_1_pdu1(pf12833pf10114) PDU1E
01695 -pf04087 -
01700 -pf02277 -
01705 -pf03180 -
01710 HMM02780keto_1_eut1(pf13292pf02779pf02780) SP
01715 -pf00293 -
01720 -pf00375 -
01725 -# -
01730 -# -
01735 HMM00465alcdh_3_mix(pf00
BMC-P
BMC-H
BMC-T
Aldehyde dehydrogenase
PTAC
Signature enzyme
Alcohol dehydrogenase
Regulator

Veillonella rogosae JCM 15642. Bacteria; Firmicutes; Negativicutes; Veillonellales; Veillonellaceae;

VRHSUH09__locus_1

PDU1E

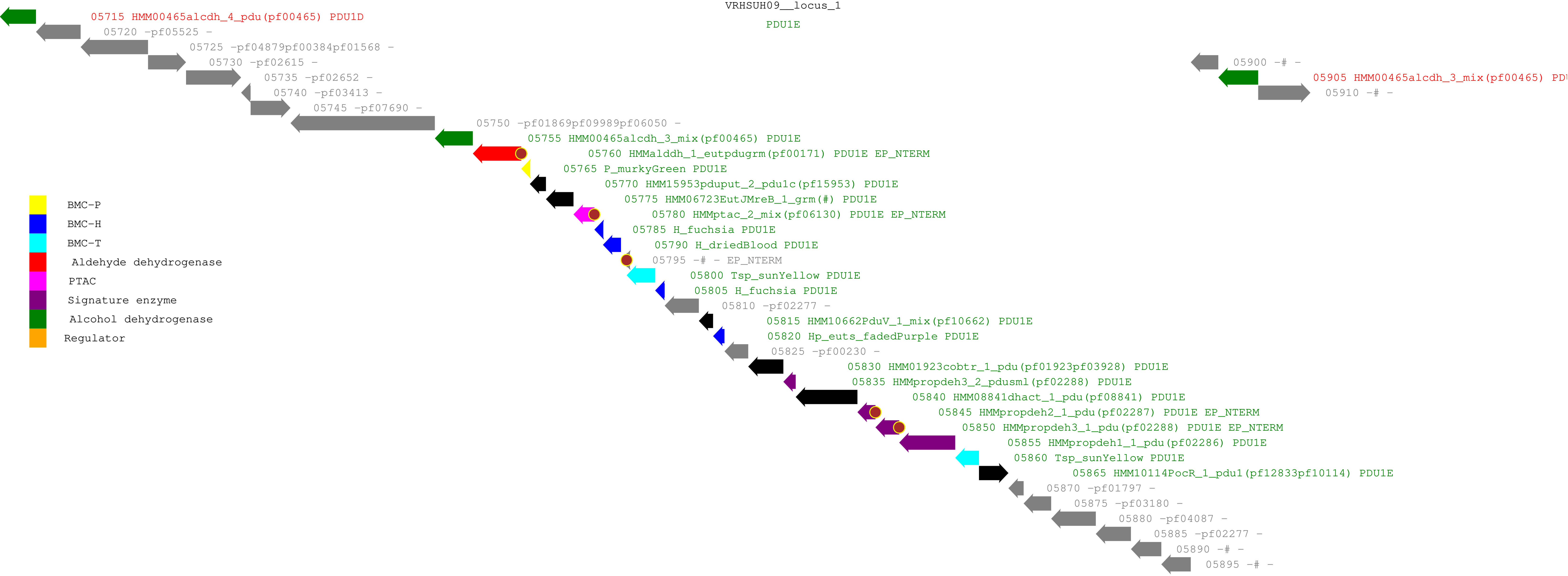

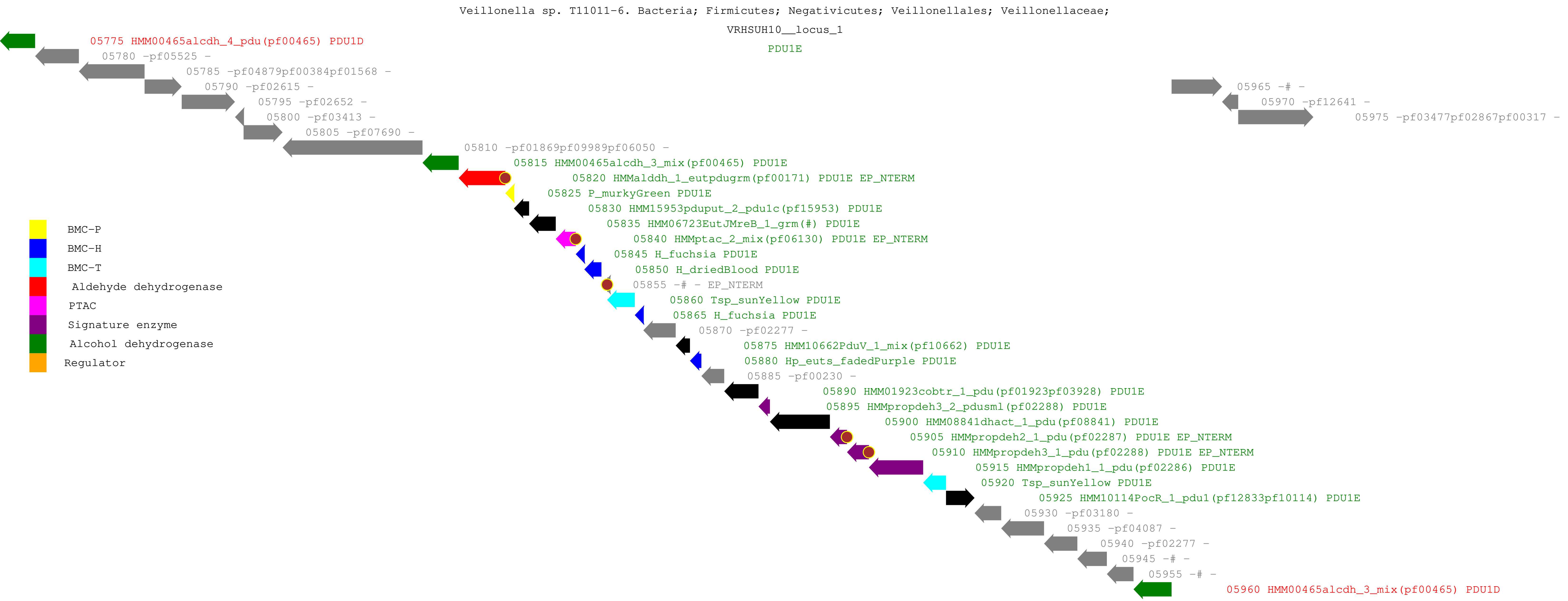

Veillonella sp. T11011-6. Bacteria; Firmicutes; Negativicutes; Veillonellales; Veillonellaceae;
VRHSUH10__locus_1
PDU1E
05775 HMM00465alcdh_4_pdu(pf00465) PDU1D
05780 -pf05525 -
05785 -pf04879pf00384pf01568 -
05790 -pf02615 -
05795 -pf02652 -
05800 -pf03413 -
05805 -pf07690 -
05810 -pf01869pf09989pf06050 -
05815 HMM00465alcdh_3_mix(pf00465) PDU1E
05820 HMMalddh_1_eutpdugrm(pf00171) PDU1E EP_NTERM
05825 P_murkyGreen PDU1E
05830 HMM15953pduput_2_pdu1c(pf15953) PDU1E
05835 HMM06723EutJMreB_1_grm(#) PDU1E
05840 HMMptac_2_mix(pf06130) PDU1E EP_NTERM
05845 H_fuchsia PDU1E
05850 H_driedBlood PDU1E
05855 -# - EP_NTERM
05860 Tsp_sunYellow PDU1E
05865 H_fuchsia PDU1E
05870 -pf02277 -
05875 HMM10662PduV_1_mix(pf10662) PDU1E
05880 Hp_euts_fadedPurple PDU1E
05885 -pf00230 -
05890 HMM01923cobtr_1_pdu(pf01923pf03928) PDU1E
05895 HMMpropdeh3_2_pdusml(pf02288) PDU1E
05900 HMM08841dhact_1_pdu(pf08841) PDU1E
05905 HMMpropdeh2_1_pdu(pf02287) PDU1E EP_NTERM
05910 HMMpropdeh3_1_pdu(pf02288) PDU1E EP_NTERM
05915 HMMpropdeh1_1_pdu(pf02286) PDU1E
05920 Tsp_sunYellow PDU1E
05925 HMM10114PocR_1_pdu1(pf12833pf10114) PDU1E
05930 -pf03180 -
05935 -pf04087 -
05940 -pf02277 -
05945 -# -
05955 -# -
05960 HMM00465alcdh_3_mix(pf00465) PDU1D
05965 -# -
05970 -pf12641 -
05975 -pf03477pf02867pf00317 -
BMC-P
BMC-H
BMC-T
Aldehyde dehydrogenase
PTAC
Signature enzyme
Alcohol dehydrogenase
Regulator

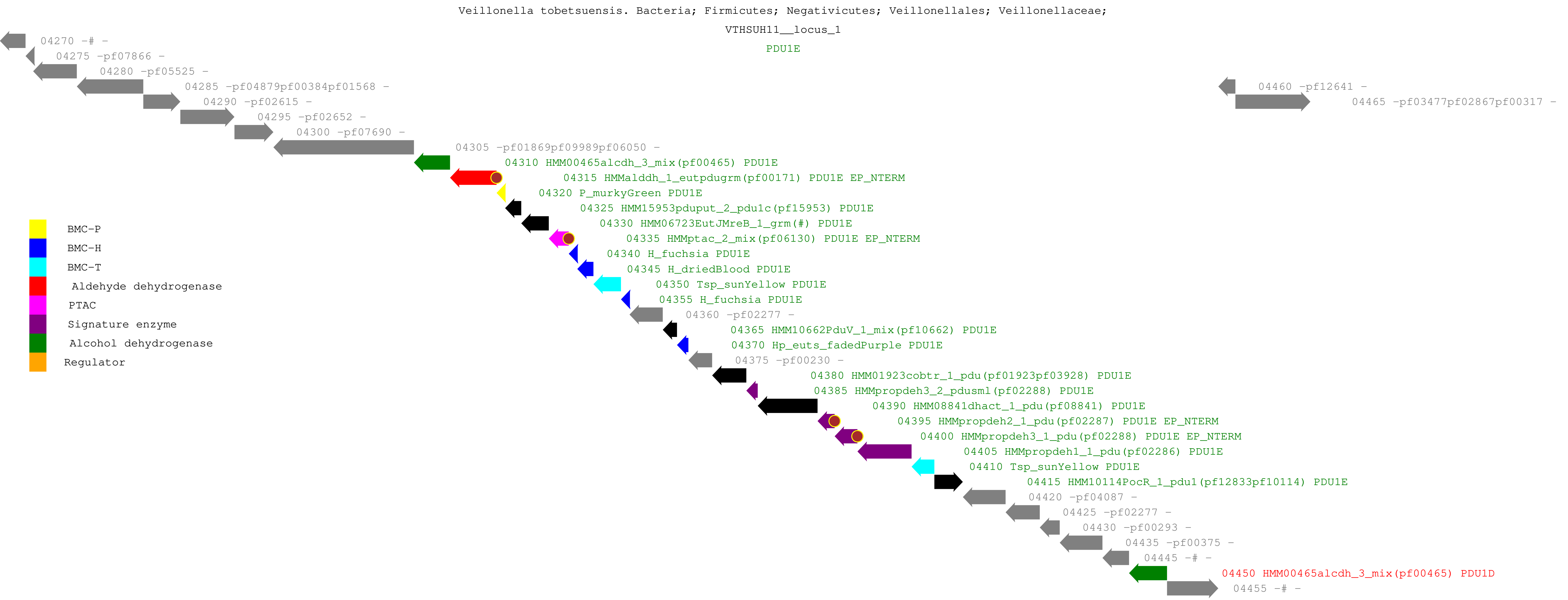

Veillonella tobetsuensis. Bacteria; Firmicutes; Negativicutes; Veillonellales; Veillonellaceae;
VTHSUH11__locus_1
PDU1E
04270 -# -
04275 -pf07866 -
04280 -pf05525 -
04285 -pf04879pf00384pf01568 -
04290 -pf02615 -
04295 -pf02652 -
04300 -pf07690 -
04305 -pf01869pf09989pf06050 -
04310 HMM00465alcdh_3_mix(pf00465) PDU1E
04315 HMMalddh_1_eutpdugrm(pf00171) PDU1E EP_NTERM
04320 P_murkyGreen PDU1E
04325 HMM15953pduput_2_pdu1c(pf15953) PDU1E
04330 HMM06723EutJMreB_1_grm(#) PDU1E
04335 HMMptac_2_mix(pf06130) PDU1E EP_NTERM
04340 H_fuchsia PDU1E
04345 H_driedBlood PDU1E
04350 Tsp_sunYellow PDU1E
04355 H_fuchsia PDU1E
04360 -pf02277 -
04365 HMM10662PduV_1_mix(pf10662) PDU1E
04370 Hp_euts_fadedPurple PDU1E
04375 -pf00230 -
04380 HMM01923cobtr_1_pdu(pf01923pf03928) PDU1E
04385 HMMpropdeh3_2_pdusml(pf02288) PDU1E
04390 HMM08841dhact_1_pdu(pf08841) PDU1E
04395 HMMpropdeh2_1_pdu(pf02287) PDU1E EP_NTERM
04400 HMMpropdeh3_1_pdu(pf02288) PDU1E EP_NTERM
04405 HMMpropdeh1_1_pdu(pf02286) PDU1E
04410 Tsp_sunYellow PDU1E
04415 HMM10114PocR_1_pdu1(pf12833pf10114) PDU1E
04420 -pf04087 -
04425 -pf02277 -
04430 -pf00293 -
04435 -pf00375 -
04445 -# -
04450 HMM00465alcdh_3_mix(pf00465) PDU1D
04455 -# -
04460 -pf12641 -
04465 -pf03477pf02867pf00317 -
BMC-P
BMC-H
BMC-T
Aldehyde dehydrogenase
PTAC
Signature enzyme
Alcohol dehydrogenase
Regulator

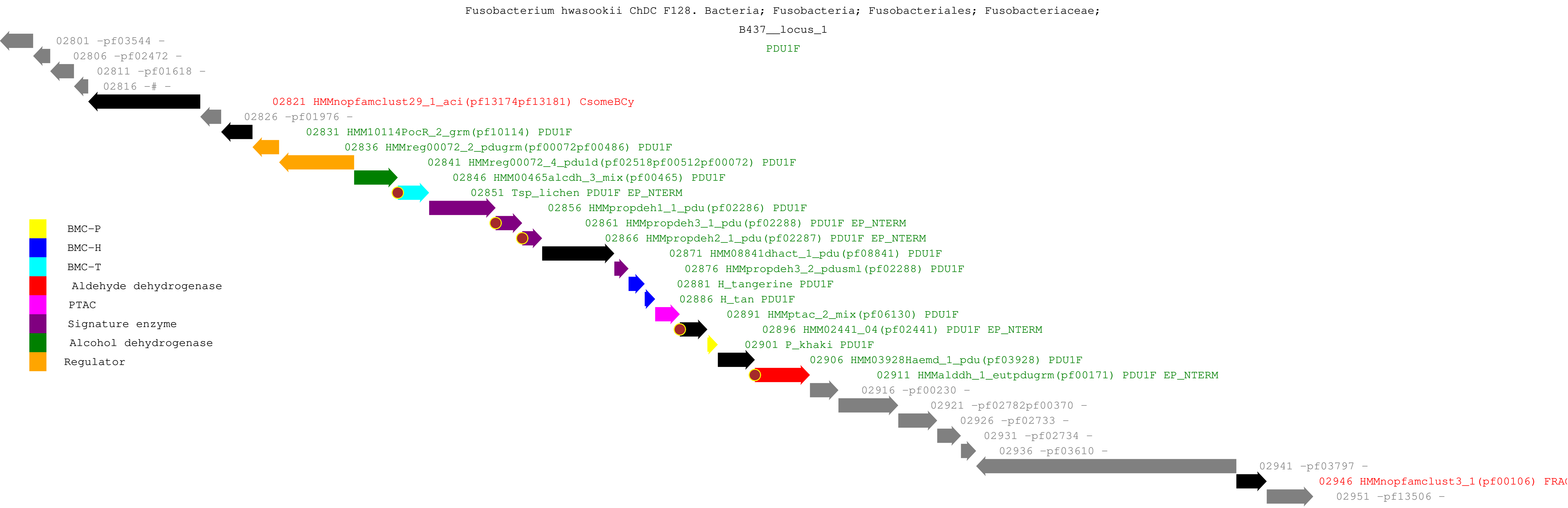
Fusobacterium hwasookii ChDC F128. Bacteria; Fusobacteria; Fusobacteriales; Fusobacteriaceae;
B437__locus_1
PDU1F
02801 -pf03544 -
02806 -pf02472 -
02811 -pf01618 -
02816 -# -
02821 HMMnopfamclust29_1_aci(pf13174pf13181) CsomeBCy
02826 -pf01976 -
02831 HMM10114PocR_2_grm(pf10114) PDU1F
02836 HMMreg00072_2_pdugrm(pf00072pf00486) PDU1F
02841 HMMreg00072_4_pdu1d(pf02518pf00512pf00072) PDU1F
02846 HMM00465alcdh_3_mix(pf00465) PDU1F
02851 Tsp_lichen PDU1F EP_NTERM
02856 HMMpropdeh1_1_pdu(pf02286) PDU1F
02861 HMMpropdeh3_1_pdu(pf02288) PDU1F EP_NTERM
02866 HMMpropdeh2_1_pdu(pf02287) PDU1F EP_NTERM
02871 HMM08841dhact_1_pdu(pf08841) PDU1F
02876 HMMpropdeh3_2_pdusml(pf02288) PDU1F
02881 H_tangerine PDU1F
02886 H_tan PDU1F
02891 HMMptac_2_mix(pf06130) PDU1F
02896 HMM02441_04(pf02441) PDU1F EP_NTERM
02901 P_khaki PDU1F
02906 HMM03928Haemd_1_pdu(pf03928) PDU1F
02911 HMMalddh_1_eutpdugrm(pf00171) PDU1F EP_NTERM
02916 -pf00230 -
02921 -pf02782pf00370 -
02926 -pf02733 -
02931 -pf02734 -
02936 -pf03610 -
02941 -pf03797 -
02946 HMMnopfamclust3_1(pf00106) FRA
02951 -pf13506 -
BMC-P
BMC-H
BMC-T
Aldehyde dehydrogenase
PTAC
Signature enzyme
Alcohol dehydrogenase
Regulator

Fusobacterium ulcerans. Bacteria; Fusobacteria; Fusobacteriales; Fusobacteriaceae;

C4N20__locus_1

PDU1F

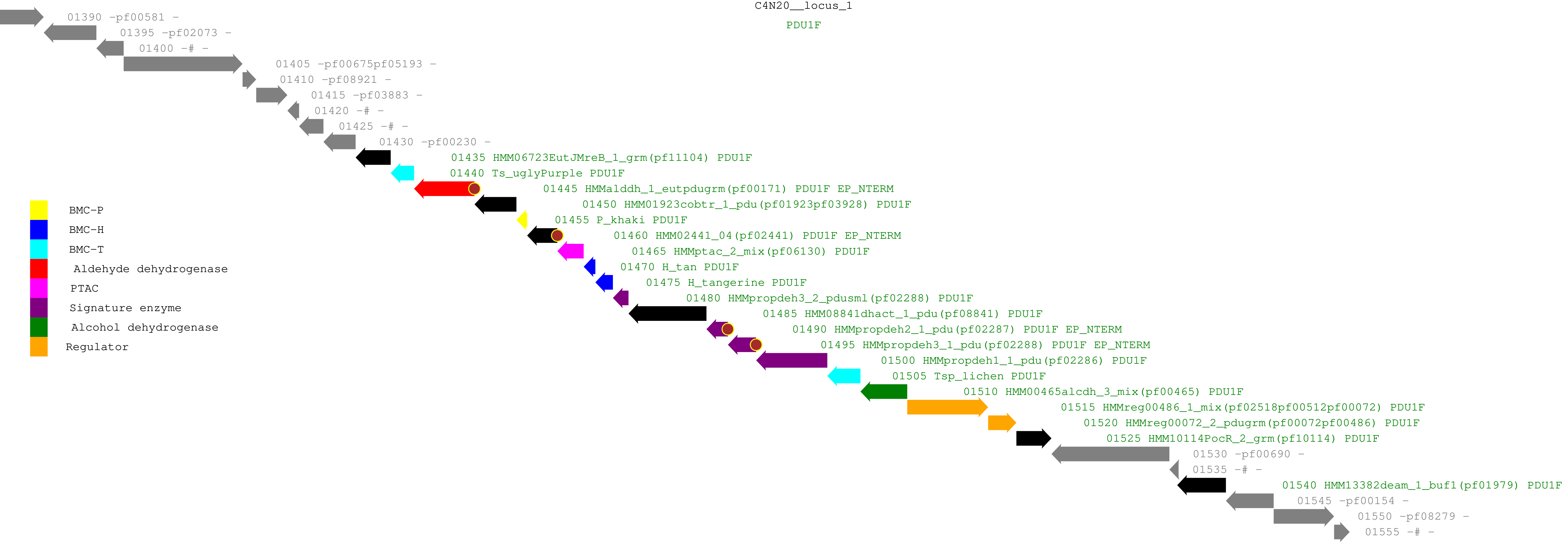

Fusobacterium nucleatum subsp. polymorphum. Bacteria; Fusobacteria; Fusobacteriales; Fusobacteriaceae;

CA839__locus_1

PDU1F

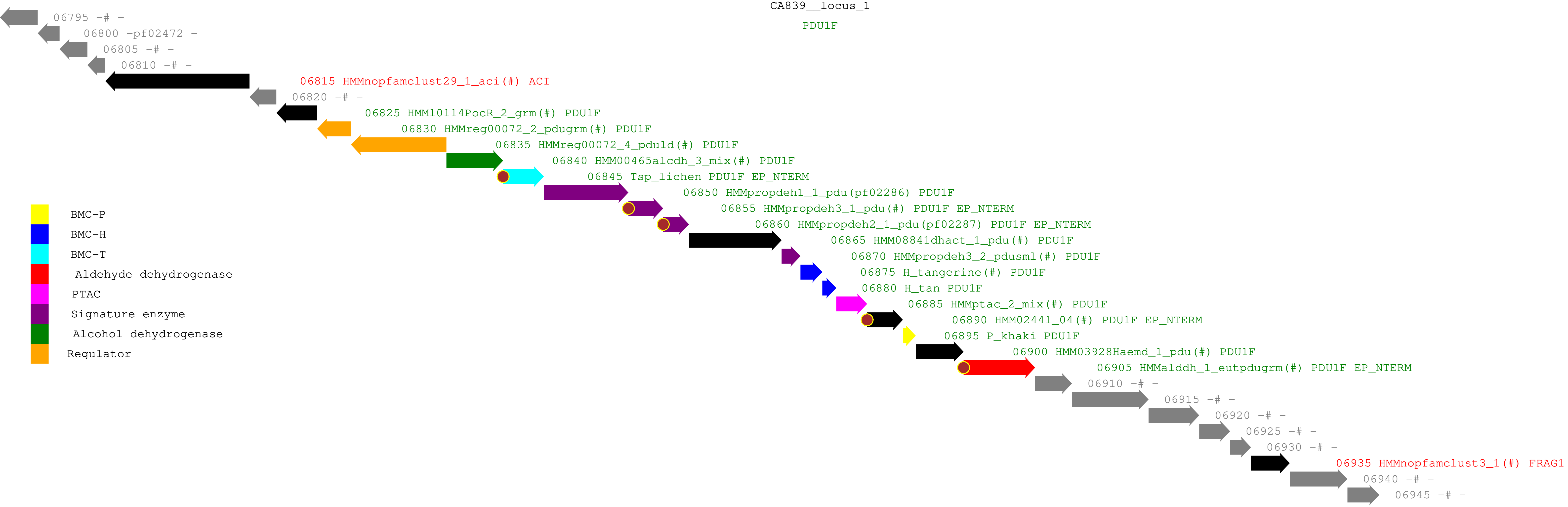

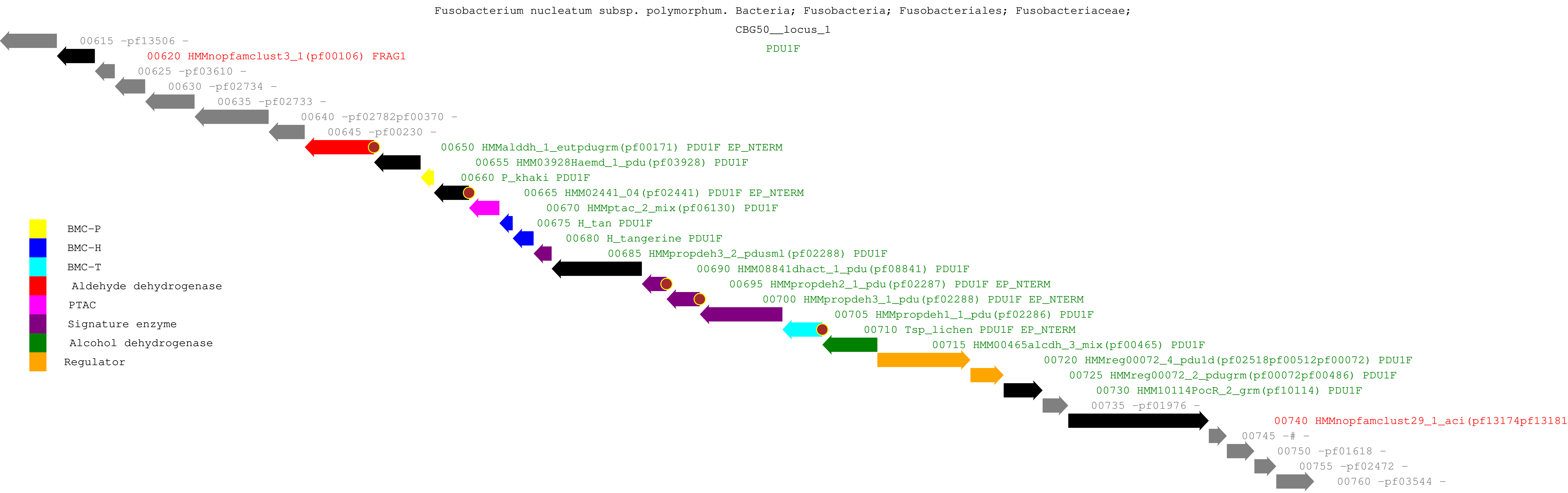

Fusobacterium nucleatum subsp. polymorphum. Bacteria; Fusobacteria; Fusobacteriales; Fusobacteriaceae;
CBG50__locus_1
PDU1F
00615 -pf13506 -
00620 HMMnopfamclust3_1(pf00106) FRAG1
00625 -pf03610 -
00630 -pf02734 -
00635 -pf02733 -
00640 -pf02782pf00370 -
00645 -pf00230 -
00650 HMMalddh_1_eutpdugrm(pf00171) PDU1F EP_NTERM
00655 HMM03928Haemd_1_pdu(pf03928) PDU1F
00660 P_khaki PDU1F
00665 HMM02441_04(pf02441) PDU1F EP_NTERM
00670 HMMptac_2_mix(pf06130) PDU1F
00675 H_tan PDU1F
00680 H_tangerine PDU1F
00685 HMMpropdeh3_2_pdusml(pf02288) PDU1F
00690 HMM08841dhact_1_pdu(pf08841) PDU1F
00695 HMMpropdeh2_1_pdu(pf02287) PDU1F EP_NTERM
00700 HMMpropdeh3_1_pdu(pf02288) PDU1F EP_NTERM
00705 HMMpropdeh1_1_pdu(pf02286) PDU1F
00710 Tsp_lichen PDU1F EP_NTERM
00715 HMM00465alcdh_3_mix(pf00465) PDU1F
00720 HMMreg00072_4_pdu1d(pf02518pf00512pf00072) PDU1F
00725 HMMreg00072_2_pdugrm(pf00072pf00486) PDU1F
00730 HMM10114PocR_2_grm(pf10114) PDU1F
00735 -pf01976 -
00740 HMMnopfamclust29_1_aci(pf13174pf13181
00745 -# -
00750 -pf01618 -
00755 -pf02472 -
00760 -pf03544 -
BMC-P
BMC-H
BMC-T
Aldehyde dehydrogenase
PTAC
Signature enzyme
Alcohol dehydrogenase
Regulator

Fusobacterium nucleatum subsp. polymorphum. Bacteria; Fusobacteria; Fusobacteriales; Fusobacteriaceae;

CBG59__locus_2

PDU1F

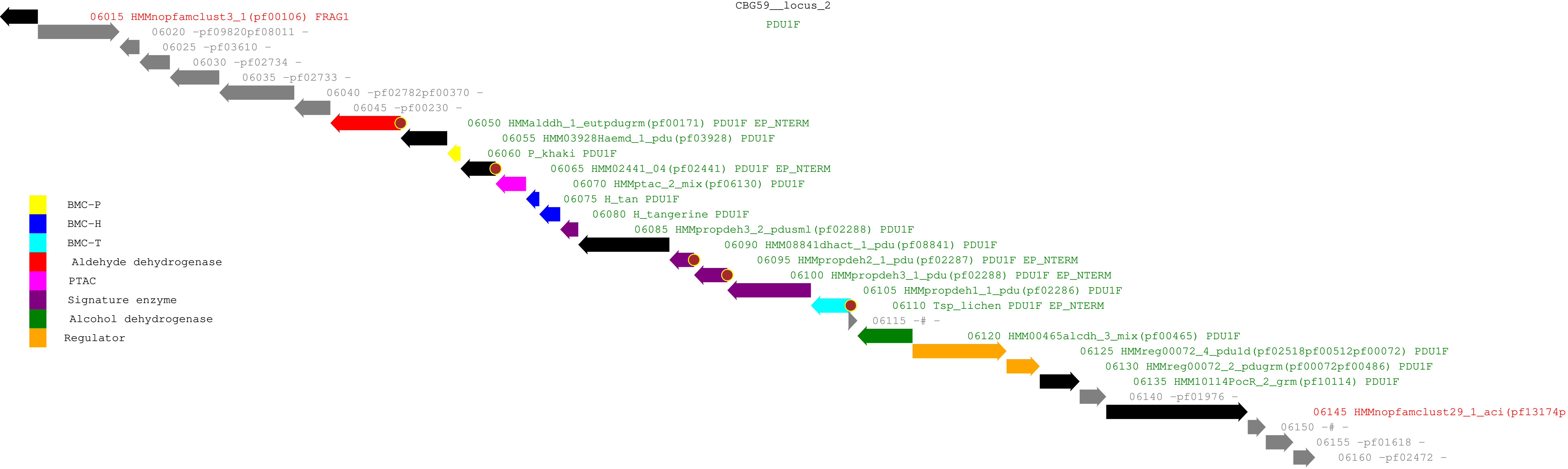

Fusobacterium nucleatum subsp. animalis. Bacteria; Fusobacteria; Fusobacteriales; Fusobacteriaceae;

CI114__locus_1

PDU1F

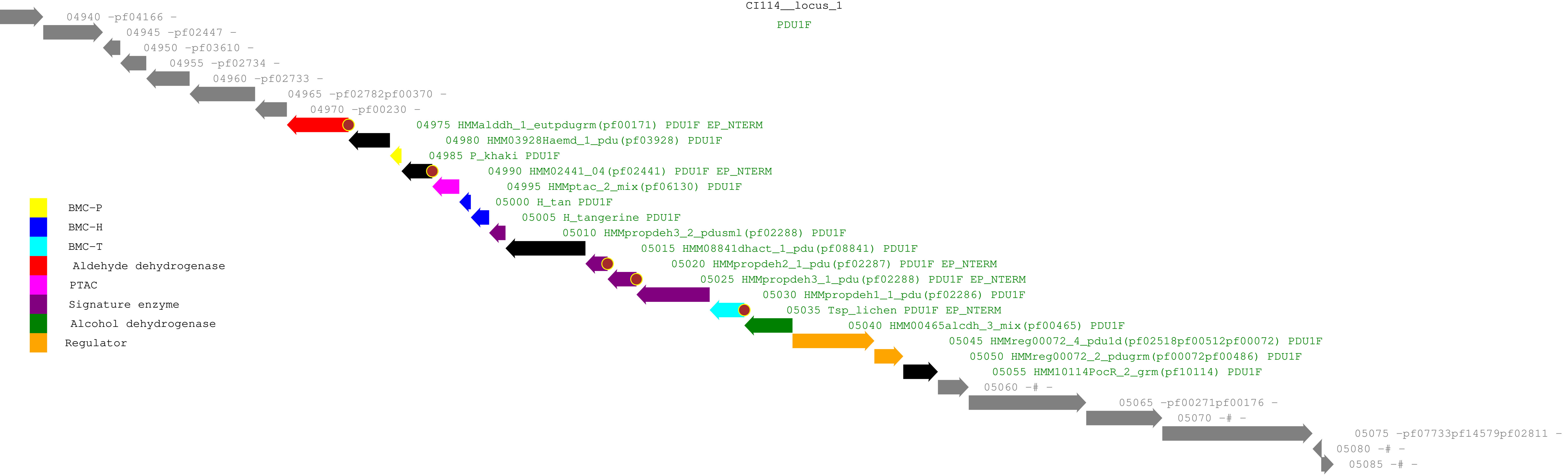

Fusobacterium nucleatum. Bacteria; Fusobacteria; Fusobacteriales; Fusobacteriaceae;

CJ209__locus_1

PDU1F

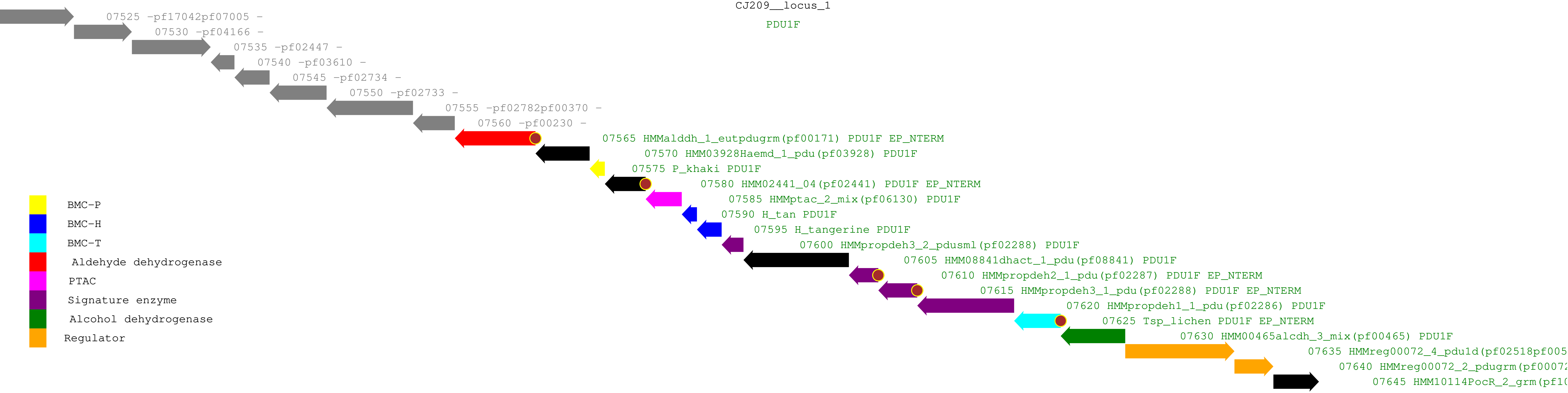

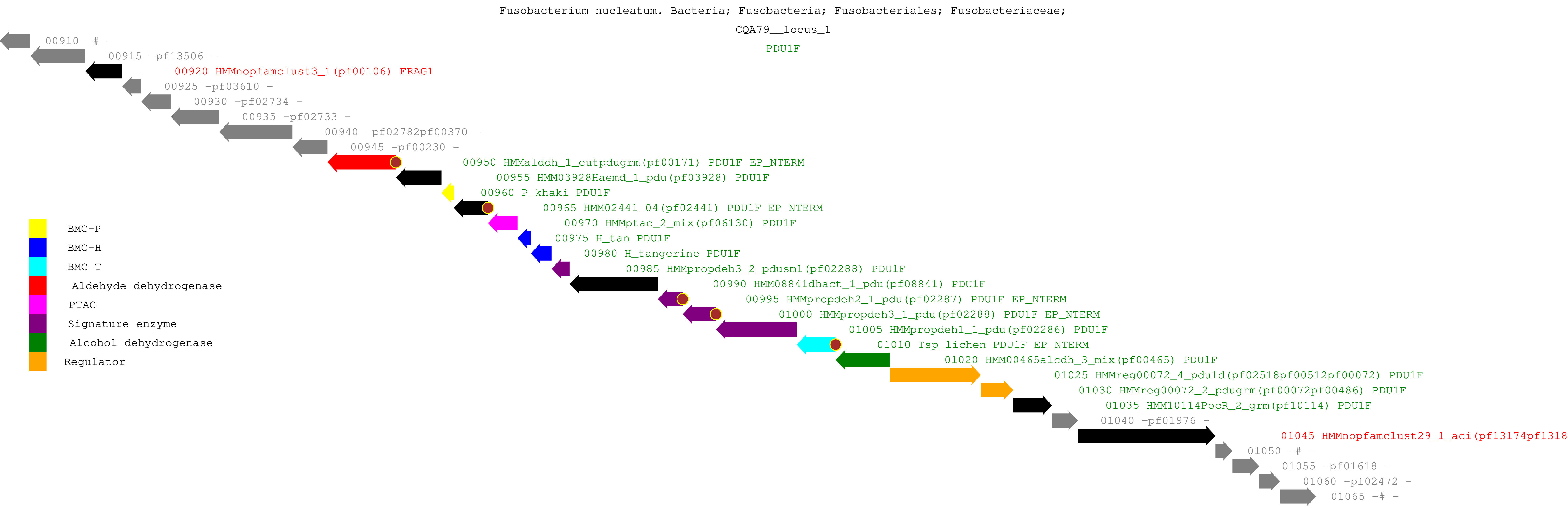

Fusobacterium nucleatum. Bacteria; Fusobacteria; Fusobacteriales; Fusobacteriaceae;
CQA79__locus_1
PDU1F
00910 -# -
00915 -pf13506 -
00920 HMMnopfamclust3_1(pf00106) FRAG1
00925 -pf03610 -
00930 -pf02734 -
00935 -pf02733 -
00940 -pf02782pf00370 -
00945 -pf00230 -
00950 HMMaldh_1_eutpdugrm(pf00171) PDU1F EP_NTERM
00955 HMM03928Haemd_1_pdu(pf03928) PDU1F
00960 P_khaki PDU1F
00965 HMM02441_04(pf02441) PDU1F EP_NTERM
00970 HMMptac_2_mix(pf06130) PDU1F
00975 H_tan PDU1F
00980 H_tangerine PDU1F
00985 HMMpropdeh3_2_pdusml(pf02288) PDU1F
00990 HMM08841dhact_1_pdu(pf08841) PDU1F
00995 HMMpropdeh2_1_pdu(pf02287) PDU1F EP_NTERM
01000 HMMpropdeh3_1_pdu(pf02288) PDU1F EP_NTERM
01005 HMMpropdeh1_1_pdu(pf02286) PDU1F
01010 Tsp_lichen PDU1F EP_NTERM
01020 HMM00465alcdh_3_mix(pf00465) PDU1F
01025 HMMreg00072_4_pdu1d(pf02518pf00512pf00072) PDU1F
01030 HMMreg00072_2_pdugrm(pf00072pf00486) PDU1F
01035 HMM10114PocR_2_grm(pf10114) PDU1F
01040 -pf01976 -
01045 HMMnopfamclust29_1_aci(pf13174pf1318
01050 -# -
01055 -pf01618 -
01060 -pf02472 -
01065 -# -
BMC-P
BMC-H
BMC-T
Aldehyde dehydrogenase
PTAC
Signature enzyme
Alcohol dehydrogenase
Regulator

Fusobacterium periodonticum. Bacteria; Fusobacteria; Fusobacteriales; Fusobacteriaceae;

CTM65__locus_1

PDU1F

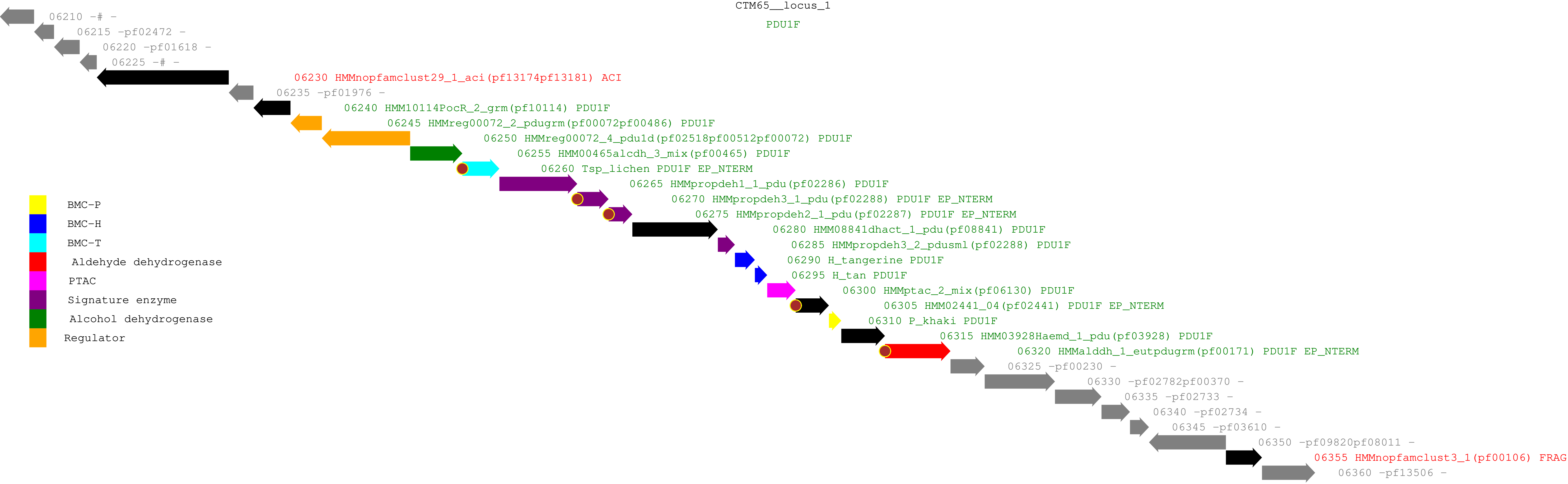

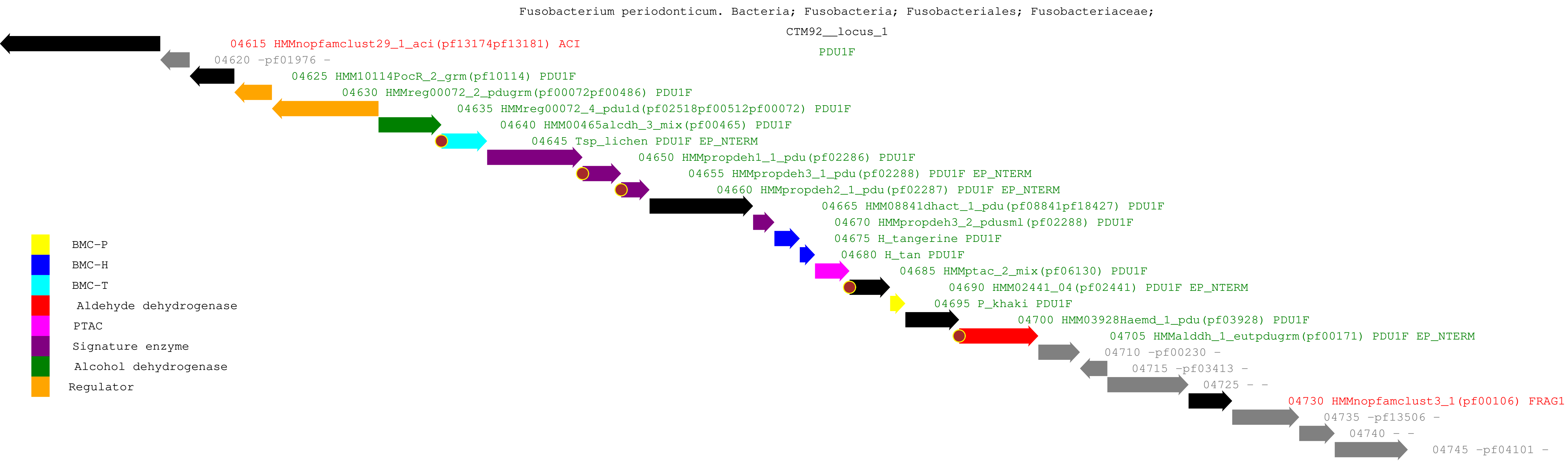
Fusobacterium periodonticum. Bacteria; Fusobacteria; Fusobacteriales; Fusobacteriaceae;
CTM92__locus_1
PDU1F
04615 HMMnopfamclust29_1_aci(pf13174pf13181) ACI
04620 -pf01976 -
04625 HMM10114PocR_2_grm(pf10114) PDU1F
04630 HMMreg00072_2_pdugrm(pf00072pf00486) PDU1F
04635 HMMreg00072_4_pdu1d(pf02518pf00512pf00072) PDU1F
04640 HMM00465alcdh_3_mix(pf00465) PDU1F
04645 Tsp_lichen PDU1F EP_NTERM
04650 HMMpropdeh1_1_pdu(pf02286) PDU1F
04655 HMMpropdeh3_1_pdu(pf02288) PDU1F EP_NTERM
04660 HMMpropdeh2_1_pdu(pf02287) PDU1F EP_NTERM
04665 HMM08841dhact_1_pdu(pf08841pf18427) PDU1F
04670 HMMpropdeh3_2_pdusml(pf02288) PDU1F
04675 H_tangerine PDU1F
04680 H_tan PDU1F
04685 HMMptac_2_mix(pf06130) PDU1F
04690 HMM02441_04(pf02441) PDU1F EP_NTERM
04695 P_khaki PDU1F
04700 HMM03928Haemd_1_pdu(pf03928) PDU1F
04705 HMMalddh_1_eutpdugrm(pf00171) PDU1F EP_NTERM
04710 -pf00230 -
04715 -pf03413 -
04725 - -
04730 HMMnopfamclust3_1(pf00106) FRAG1
04735 -pf13506 -
04740 - -
04745 -pf04101 -
BMC-P
BMC-H
BMC-T
Aldehyde dehydrogenase
PTAC
Signature enzyme
Alcohol dehydrogenase
Regulator

Fusobacterium sp. Bacteria; Fusobacteria; Fusobacteriales; Fusobacteriaceae;

DDY85__locus_1

PDU1F

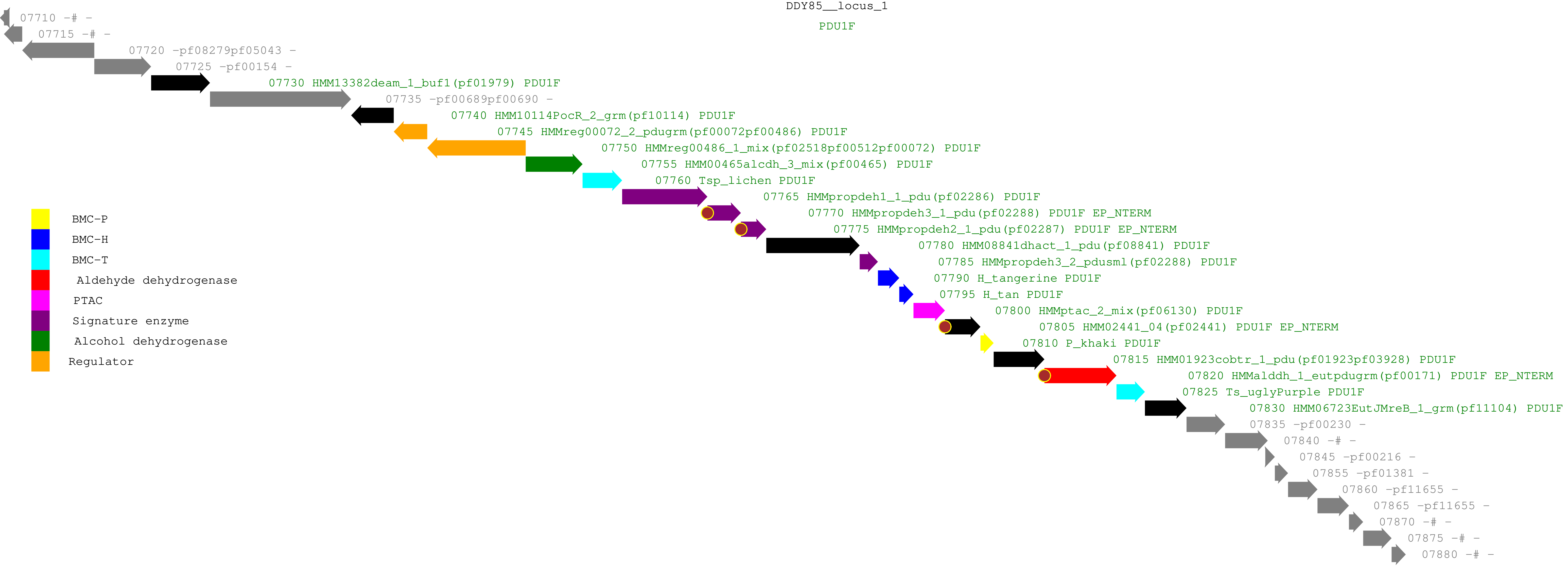

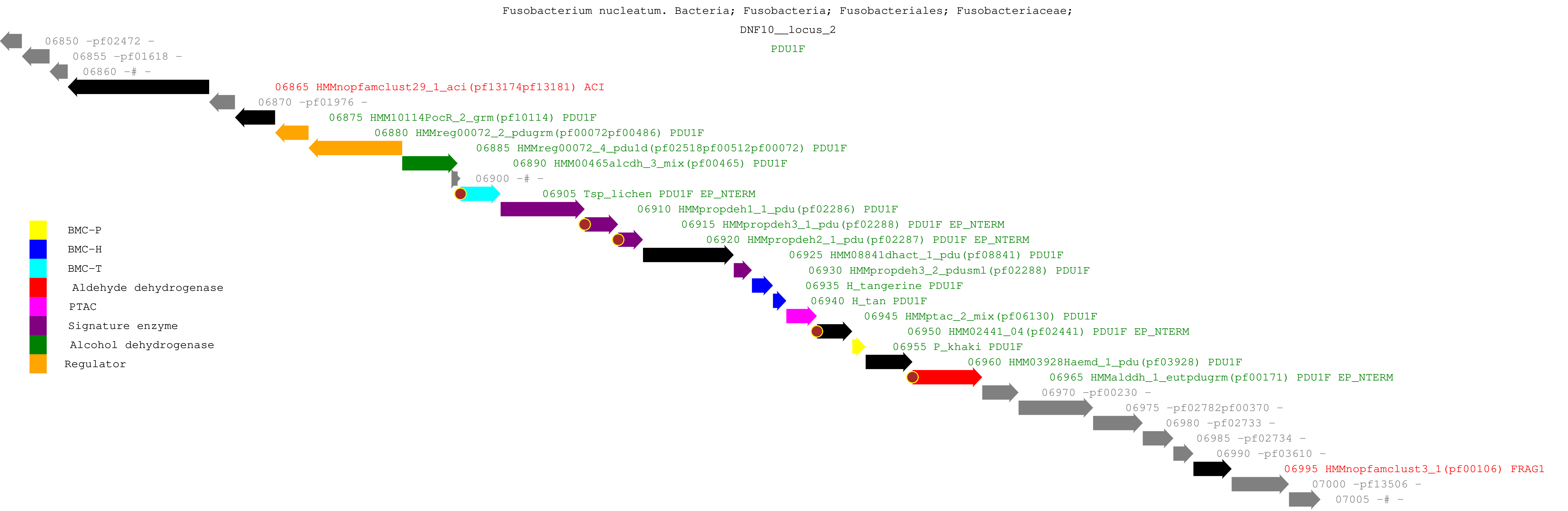
Fusobacterium nucleatum. Bacteria; Fusobacteria; Fusobacteriales; Fusobacteriaceae;
DNF10__locus_2
PDU1F
06850 -pf02472 -
06855 -pf01618 -
06860 -# -
06865 HMMnopfamclust29_1_aci(pf13174pf13181) ACI
06870 -pf01976 -
06875 HMM10114PocR_2_grm(pf10114) PDU1F
06880 HMMreg00072_2_pdugrm(pf00072pf00486) PDU1F
06885 HMMreg00072_4_pdu1d(pf02518pf00512pf00072) PDU1F
06890 HMM00465alcdh_3_mix(pf00465) PDU1F
06900 -# -
06905 Tsp_lichen PDU1F EP_NTERM
06910 HMMpropdeh1_1_pdu(pf02286) PDU1F
06915 HMMpropdeh3_1_pdu(pf02288) PDU1F EP_NTERM
06920 HMMpropdeh2_1_pdu(pf02287) PDU1F EP_NTERM
06925 HMM08841dhact_1_pdu(pf08841) PDU1F
06930 HMMpropdeh3_2_pdusml(pf02288) PDU1F
06935 H_tangerine PDU1F
06940 H_tan PDU1F
06945 HMMptac_2_mix(pf06130) PDU1F
06950 HMM02441_04(pf02441) PDU1F EP_NTERM
06955 P_khaki PDU1F
06960 HMM03928Haemd_1_pdu(pf03928) PDU1F
06965 HMMaldh_1_eutpdugrm(pf00171) PDU1F EP_NTERM
06970 -pf00230 -
06975 -pf02782pf00370 -
06980 -pf02733 -
06985 -pf02734 -
06990 -pf03610 -
06995 HMMnopfamclust3_1(pf00106) FRAG1
07000 -pf13506 -
07005 -# -
BMC-P
BMC-H
BMC-T
Aldehyde dehydrogenase
PTAC
Signature enzyme
Alcohol dehydrogenase
Regulator

Fusobacterium varium. Bacteria; Fusobacteria; Fusobacteriales; Fusobacteriaceae; Fusobacterium.

DW261__locus_1

PDU1F

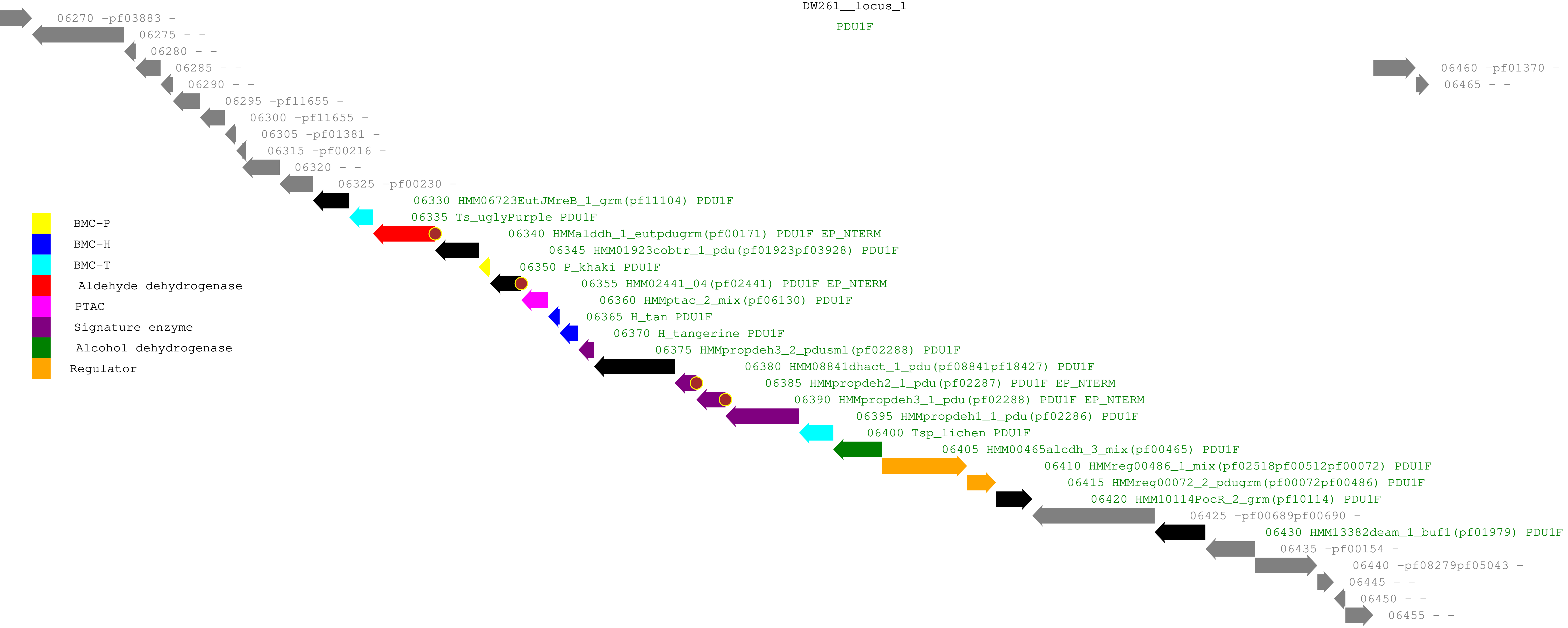

Fusobacterium canifelinum. Bacteria; Fusobacteria; Fusobacteriales; Fusobacteriaceae; Fusobacterium.

EII27__locus_2

PDU1F

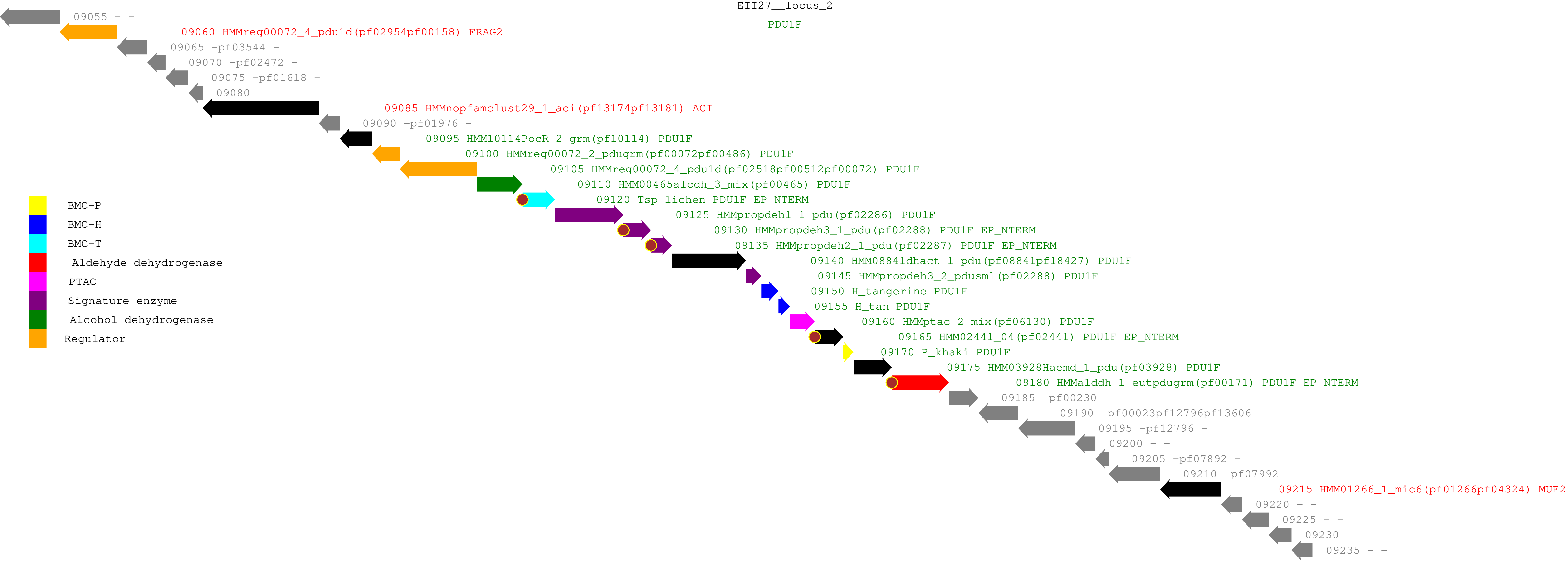

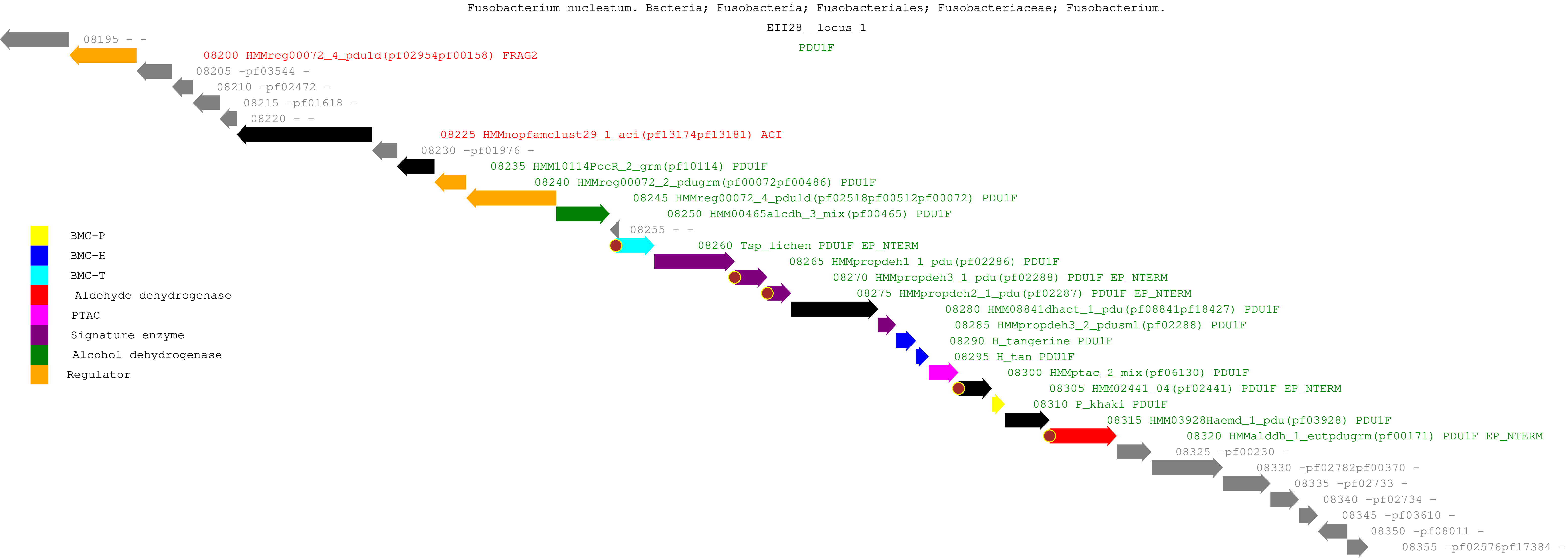
Fusobacterium nucleatum. Bacteria; Fusobacteria; Fusobacteriales; Fusobacteriaceae; Fusobacterium.
EII28__locus_1
PDU1F
08195 - -
08200 HMMreg00072_4_pdu1d(pf02954pf00158) FRAG2
08205 -pf03544 -
08210 -pf02472 -
08215 -pf01618 -
08220 - -
08225 HMMnopfamclust29_1_aci(pf13174pf13181) ACI
08230 -pf01976 -
08235 HMM10114PocR_2_grm(pf10114) PDU1F
08240 HMMreg00072_2_pdugrm(pf00072pf00486) PDU1F
08245 HMMreg00072_4_pdu1d(pf02518pf00512pf00072) PDU1F
08250 HMM00465alcdh_3_mix(pf00465) PDU1F
08255 - -
08260 Tsp_lichen PDU1F EP_NTERM
08265 HMMpropdeh1_1_pdu(pf02286) PDU1F
08270 HMMpropdeh3_1_pdu(pf02288) PDU1F EP_NTERM
08275 HMMpropdeh2_1_pdu(pf02287) PDU1F EP_NTERM
08280 HMM08841dhact_1_pdu(pf08841pf18427) PDU1F
08285 HMMpropdeh3_2_pdusml(pf02288) PDU1F
08290 H_tangerine PDU1F
08295 H_tan PDU1F
08300 HMMptac_2_mix(pf06130) PDU1F
08305 HMM02441_04(pf02441) PDU1F EP_NTERM
08310 P_khaki PDU1F
08315 HMM03928Haemd_1_pdu(pf03928) PDU1F
08320 HMMalddh_1_eutpdugrm(pf00171) PDU1F EP_NTERM
08325 -pf00230 -
08330 -pf02782pf00370 -
08335 -pf02733 -
08340 -pf02734 -
08345 -pf03610 -
08350 -pf08011 -
08355 -pf02576pf17384 -
BMC-P
BMC-H
BMC-T
Aldehyde dehydrogenase
PTAC
Signature enzyme
Alcohol dehydrogenase
Regulator

Fusobacterium nucleatum subsp. polymorphum ATCC 10953. Bacteria; Fusobacteria; Fusobacteriales; Fusobacteriaceae;

FNP__locus_2

PDU1F

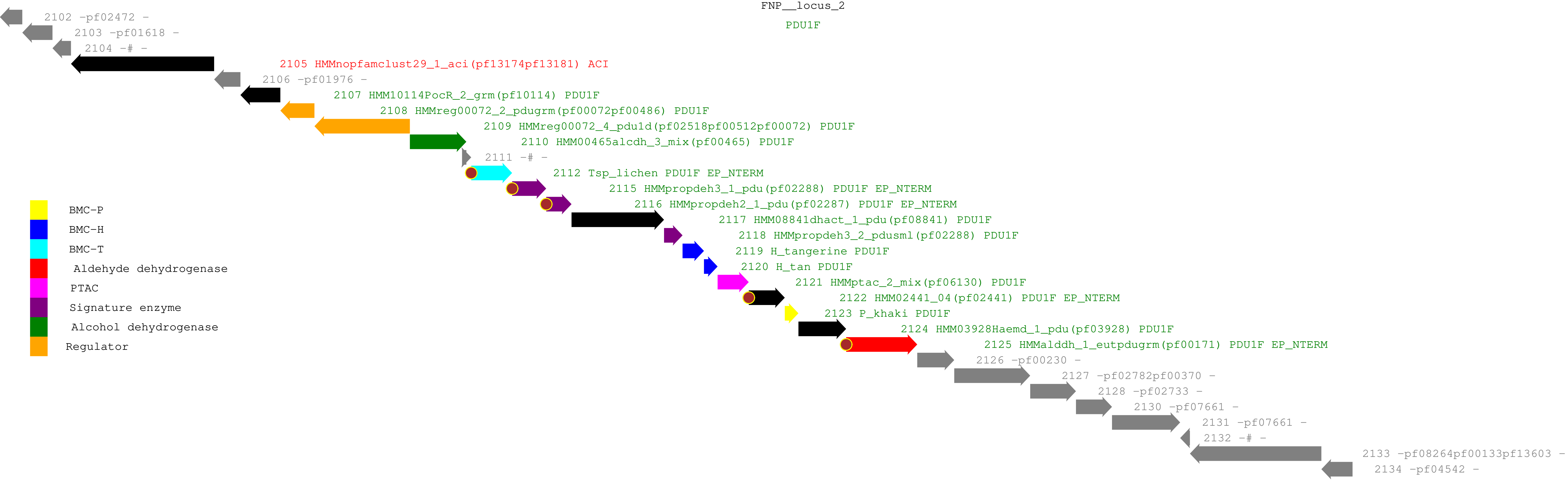

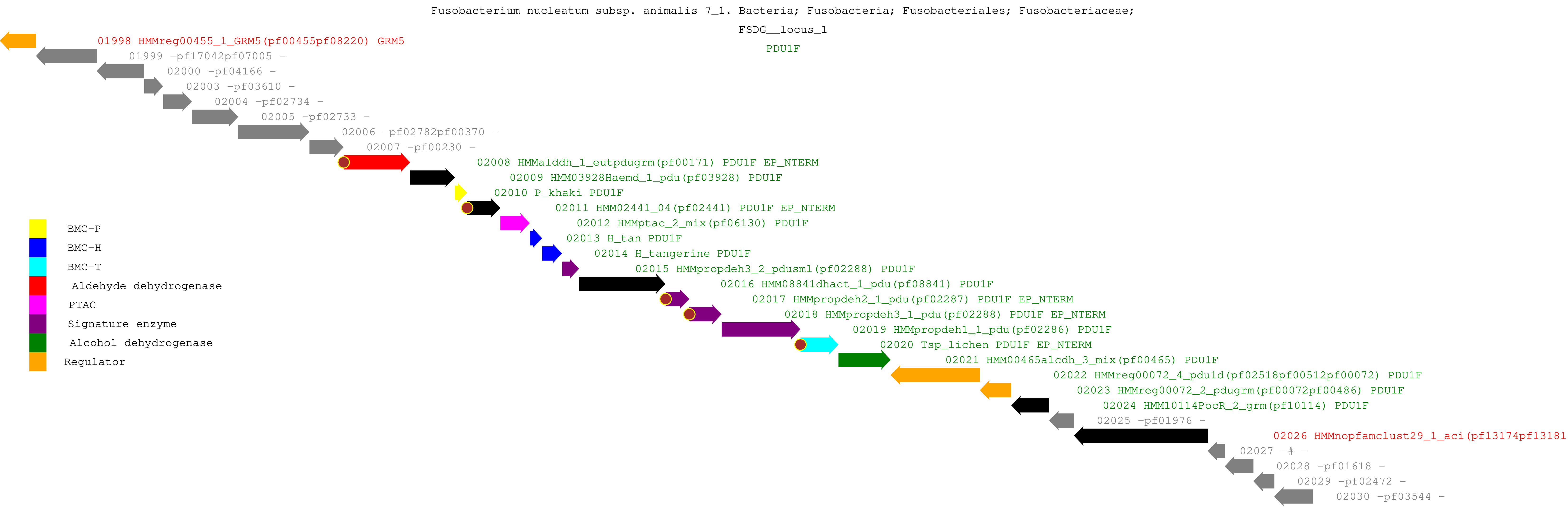

Fusobacterium nucleatum subsp. animalis 7_1. Bacteria; Fusobacteria; Fusobacteriales; Fusobacteriaceae;
FSDG__locus_1
PDU1F
01998 HMMreg00455_1_GRM5(pf00455pf08220) GRM5
01999 -pf17042pf07005 -
02000 -pf04166 -
02003 -pf03610 -
02004 -pf02734 -
02005 -pf02733 -
02006 -pf02782pf00370 -
02007 -pf00230 -
02008 HMMaldh_1_eutpdugrm(pf00171) PDU1F EP_NTERM
02009 HMM03928Haemd_1_pdu(pf03928) PDU1F
02010 P_khaki PDU1F
02011 HMM02441_04(pf02441) PDU1F EP_NTERM
02012 HMMptac_2_mix(pf06130) PDU1F
02013 H_tan PDU1F
02014 H_tangerine PDU1F
02015 HMMpropdeh3_2_pdusml(pf02288) PDU1F
02016 HMM08841dhact_1_pdu(pf08841) PDU1F
02017 HMMpropdeh2_1_pdu(pf02287) PDU1F EP_NTERM
02018 HMMpropdeh3_1_pdu(pf02288) PDU1F EP_NTERM
02019 HMMpropdeh1_1_pdu(pf02286) PDU1F
02020 Tsp_lichen PDU1F EP_NTERM
02021 HMM00465alcdh_3_mix(pf00465) PDU1F
02022 HMMreg00072_4_pdu1d(pf02518pf00512pf00072) PDU1F
02023 HMMreg00072_2_pdugrm(pf00072pf00486) PDU1F
02024 HMM10114PocR_2_grm(pf10114) PDU1F
02025 -pf01976 -
02026 HMMnopfamclust29_1_aci(pf13174pf13181
02027 -# -
02028 -pf01618 -
02029 -pf02472 -
02030 -pf03544 -
BMC-P
BMC-H
BMC-T
Aldehyde dehydrogenase
PTAC
Signature enzyme
Alcohol dehydrogenase
Regulator

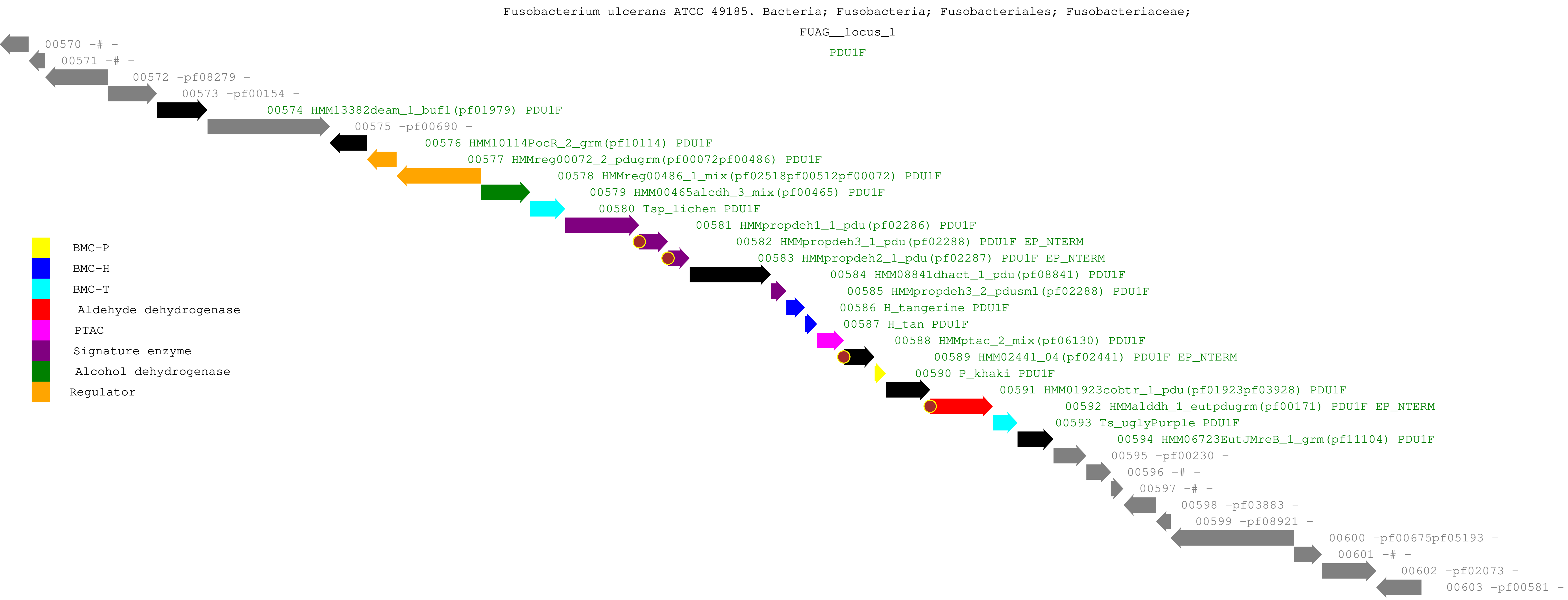

Fusobacterium ulcerans ATCC 49185. Bacteria; Fusobacteria; Fusobacteriales; Fusobacteriaceae;
FUAG__locus_1
PDU1F
00570 -# -
00571 -# -
00572 -pf08279 -
00573 -pf00154 -
00574 HMM13382deam_1_buf1(pf01979) PDU1F
00575 -pf00690 -
00576 HMM10114PocR_2_grm(pf10114) PDU1F
00577 HMMreg00072_2_pdugrm(pf00072pf00486) PDU1F
00578 HMMreg00486_1_mix(pf02518pf00512pf00072) PDU1F
00579 HMM00465alcdh_3_mix(pf00465) PDU1F
00580 Tsp_lichen PDU1F
00581 HMMpropdeh1_1_pdu(pf02286) PDU1F
00582 HMMpropdeh3_1_pdu(pf02288) PDU1F EP_NTERM
00583 HMMpropdeh2_1_pdu(pf02287) PDU1F EP_NTERM
00584 HMM08841dhact_1_pdu(pf08841) PDU1F
00585 HMMpropdeh3_2_pdusml(pf02288) PDU1F
00586 H_tangerine PDU1F
00587 H_tan PDU1F
00588 HMMptac_2_mix(pf06130) PDU1F
00589 HMM02441_04(pf02441) PDU1F EP_NTERM
00590 P_khaki PDU1F
00591 HMM01923cobtr_1_pdu(pf01923pf03928) PDU1F
00592 HMMalddh_1_eutpdugrm(pf00171) PDU1F EP_NTERM
00593 Ts_uglyPurple PDU1F
00594 HMM06723EutJMreB_1_grm(pf11104) PDU1F
00595 -pf00230 -
00596 -# -
00597 -# -
00598 -pf03883 -
00599 -pf08921 -
00600 -pf00675pf05193 -
00601 -# -
00602 -pf02073 -
00603 -pf00581 -
BMC-P
BMC-H
BMC-T
Aldehyde dehydrogenase
PTAC
Signature enzyme
Alcohol dehydrogenase
Regulator

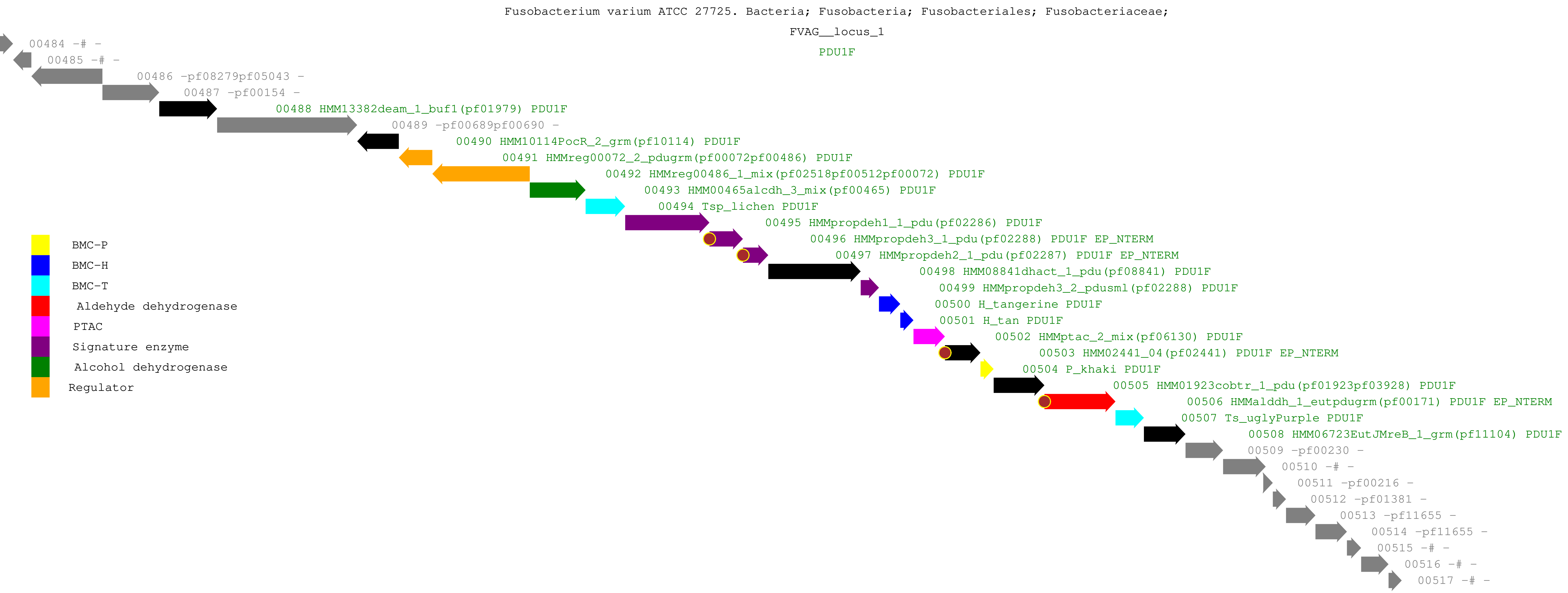
Fusobacterium varium ATCC 27725. Bacteria; Fusobacteria; Fusobacteriales; Fusobacteriaceae;
FVAG__locus_1
PDU1F
00484 -# -
00485 -# -
00486 -pf08279pf05043 -
00487 -pf00154 -
00488 HMM13382deam_1_buf1(pf01979) PDU1F
00489 -pf00689pf00690 -
00490 HMM10114PocR_2_grm(pf10114) PDU1F
00491 HMMreg00072_2_pdugrm(pf00072pf00486) PDU1F
00492 HMMreg00486_1_mix(pf02518pf00512pf00072) PDU1F
00493 HMM00465alcdh_3_mix(pf00465) PDU1F
00494 Tsp_lichen PDU1F
00495 HMMpropdeh1_1_pdu(pf02286) PDU1F
00496 HMMpropdeh3_1_pdu(pf02288) PDU1F EP_NTERM
00497 HMMpropdeh2_1_pdu(pf02287) PDU1F EP_NTERM
00498 HMM08841dhact_1_pdu(pf08841) PDU1F
00499 HMMpropdeh3_2_pdusml(pf02288) PDU1F
00500 H_tangerine PDU1F
00501 H_tan PDU1F
00502 HMMptac_2_mix(pf06130) PDU1F
00503 HMM02441_04(pf02441) PDU1F EP_NTERM
00504 P_khaki PDU1F
00505 HMM01923cobtr_1_pdu(pf01923pf03928) PDU1F
00506 HMMalddh_1_eutpdugrm(pf00171) PDU1F EP_NTERM
00507 Ts_uglyPurple PDU1F
00508 HMM06723EutJMreB_1_grm(pf11104) PDU1F
00509 -pf00230 -
00510 -# -
00511 -pf00216 -
00512 -pf01381 -
00513 -pf11655 -
00514 -pf11655 -
00515 -# -
00516 -# -
00517 -# -
BMC-P
BMC-H
BMC-T
Aldehyde dehydrogenase
PTAC
Signature enzyme
Alcohol dehydrogenase
Regulator

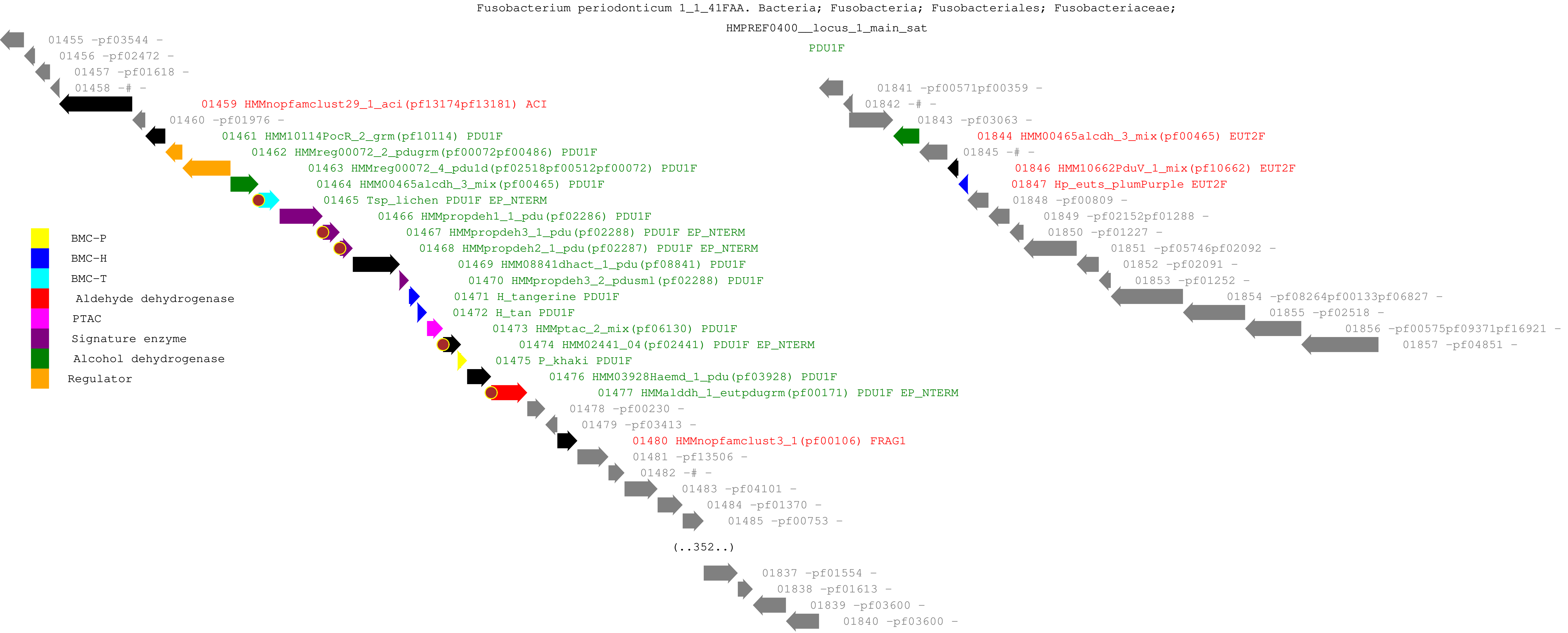
Fusobacterium periodonticum 1_1_41FAA. Bacteria; Fusobacteria; Fusobacteriales; Fusobacteriaceae;
HMPREF0400__locus_1_main_sat
PDU1F
01455 -pf03544 -
01456 -pf02472 -
01457 -pf01618 -
01458 -# -
01459 HMMnopfamclust29_1_aci(pf13174pf13181) ACI
01460 -pf01976 -
01461 HMM10114PocR_2_grm(pf10114) PDU1F
01462 HMMreg00072_2_pdugrm(pf00072pf00486) PDU1F
01463 HMMreg00072_4_pdu1d(pf02518pf00512pf00072) PDU1F
01464 HMM00465alcdh_3_mix(pf00465) PDU1F
01465 Tsp_lichen PDU1F EP_NTERM
01466 HMMpropdeh1_1_pdu(pf02286) PDU1F
01467 HMMpropdeh3_1_pdu(pf02288) PDU1F EP_NTERM
01468 HMMpropdeh2_1_pdu(pf02287) PDU1F EP_NTERM
01469 HMM08841dhact_1_pdu(pf08841) PDU1F
01470 HMMpropdeh3_2_pdusml(pf02288) PDU1F
01471 H_tangerine PDU1F
01472 H_tan PDU1F
01473 HMMptac_2_mix(pf06130) PDU1F
01474 HMM02441_04(pf02441) PDU1F EP_NTERM
01475 P_khaki PDU1F
01476 HMM03928Haemd_1_pdu(pf03928) PDU1F
01477 HMMaldh_1_eutpdugrm(pf00171) PDU1F EP_NTERM
01478 -pf00230 -
01479 -pf03413 -
01480 HMMnopfamclust3_1(pf00106) FRAG1
01481 -pf13506 -
01482 -# -
01483 -pf04101 -
01484 -pf01370 -
01485 -pf00753 -
(..352..)
01837 -pf01554 -
01838 -pf01613 -
01839 -pf03600 -
01840 -pf03600 -
01841 -pf00571pf00359 -
01842 -# -
01843 -pf03063 -
01844 HMM00465alcdh_3_mix(pf00465) EUT2F
01845 -# -
01846 HMM10662PduV_1_mix(pf10662) EUT2F
01847 Hp_euts_plumPurple EUT2F
01848 -pf00809 -
01849 -pf02152pf01288 -
01850 -pf01227 -
01851 -pf05746pf02092 -
01852 -pf02091 -
01853 -pf01252 -
01854 -pf08264pf00133pf06827 -
01855 -pf02518 -
01856 -pf00575pf09371pf16921 -
01857 -pf04851 -
BMC-P
BMC-H
BMC-T
Aldehyde dehydrogenase
PTAC
Signature enzyme
Alcohol dehydrogenase
Regulator

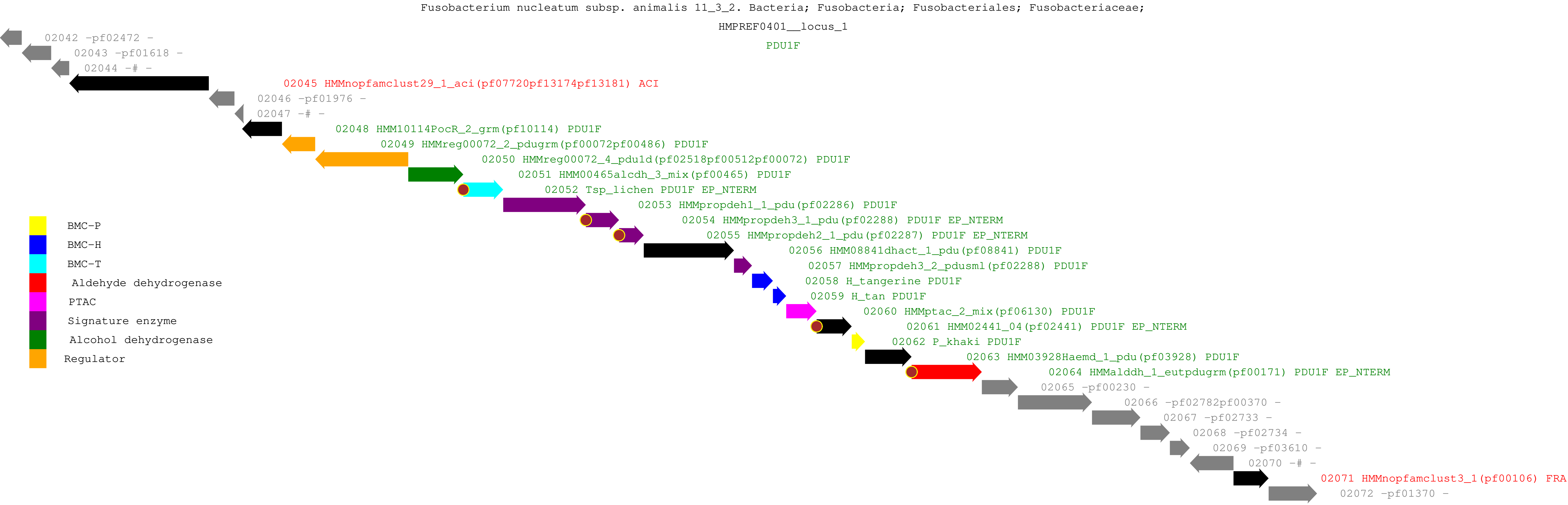

Fusobacterium nucleatum subsp. animalis 11_3_2. Bacteria; Fusobacteria; Fusobacteriales; Fusobacteriaceae;
HMPREF0401__locus_1
PDU1F
02042 -pf02472 -
02043 -pf01618 -
02044 -# -
02045 HMMnopfamclust29_1_aci(pf07720pf13174pf13181) ACI
02046 -pf01976 -
02047 -# -
02048 HMM10114PocR_2_grm(pf10114) PDU1F
02049 HMMreg00072_2_pdugrm(pf00072pf00486) PDU1F
02050 HMMreg00072_4_pdu1d(pf02518pf00512pf00072) PDU1F
02051 HMM00465alcdh_3_mix(pf00465) PDU1F
02052 Tsp_lichen PDU1F EP_NTERM
02053 HMMpropdeh1_1_pdu(pf02286) PDU1F
02054 HMMpropdeh3_1_pdu(pf02288) PDU1F EP_NTERM
02055 HMMpropdeh2_1_pdu(pf02287) PDU1F EP_NTERM
02056 HMM08841dhact_1_pdu(pf08841) PDU1F
02057 HMMpropdeh3_2_pdusml(pf02288) PDU1F
02058 H_tangerine PDU1F
02059 H_tan PDU1F
02060 HMMptac_2_mix(pf06130) PDU1F
02061 HMM02441_04(pf02441) PDU1F EP_NTERM
02062 P_khaki PDU1F
02063 HMM03928Haemd_1_pdu(pf03928) PDU1F
02064 HMMalddh_1_eutpdugrm(pf00171) PDU1F EP_NTERM
02065 -pf00230 -
02066 -pf02782pf00370 -
02067 -pf02733 -
02068 -pf02734 -
02069 -pf03610 -
02070 -# -
02071 HMMnopfamclust3_1(pf00106) FRA
02072 -pf01370 -
BMC-P
BMC-H
BMC-T
Aldehyde dehydrogenase
PTAC
Signature enzyme
Alcohol dehydrogenase
Regulator

Fusobacterium ulcerans 12-1B. Bacteria; Fusobacteria; Fusobacteriales; Fusobacteriaceae;

HMPREF0402__locus_1

PDU1F

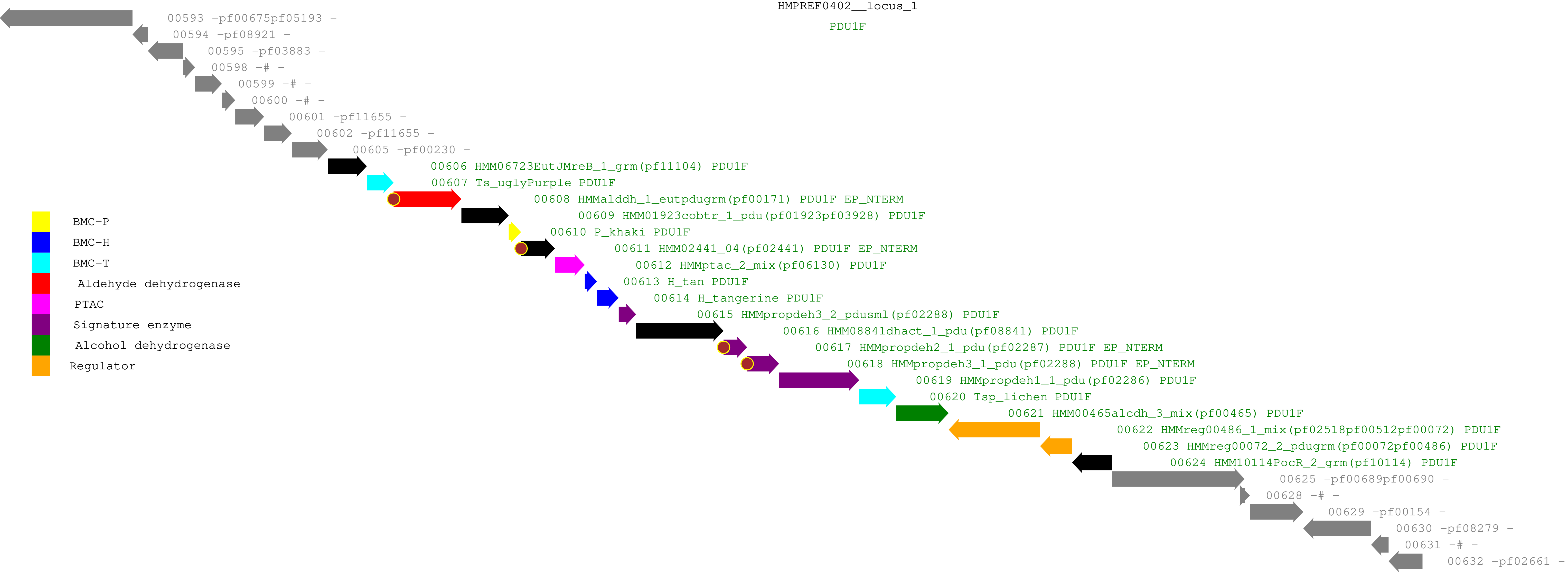

Fusobacterium nucleatum subsp. animalis 4_8. Bacteria; Fusobacteria; Fusobacteriales; Fusobacteriaceae;

HMPREF0409__locus_1

PDU1F

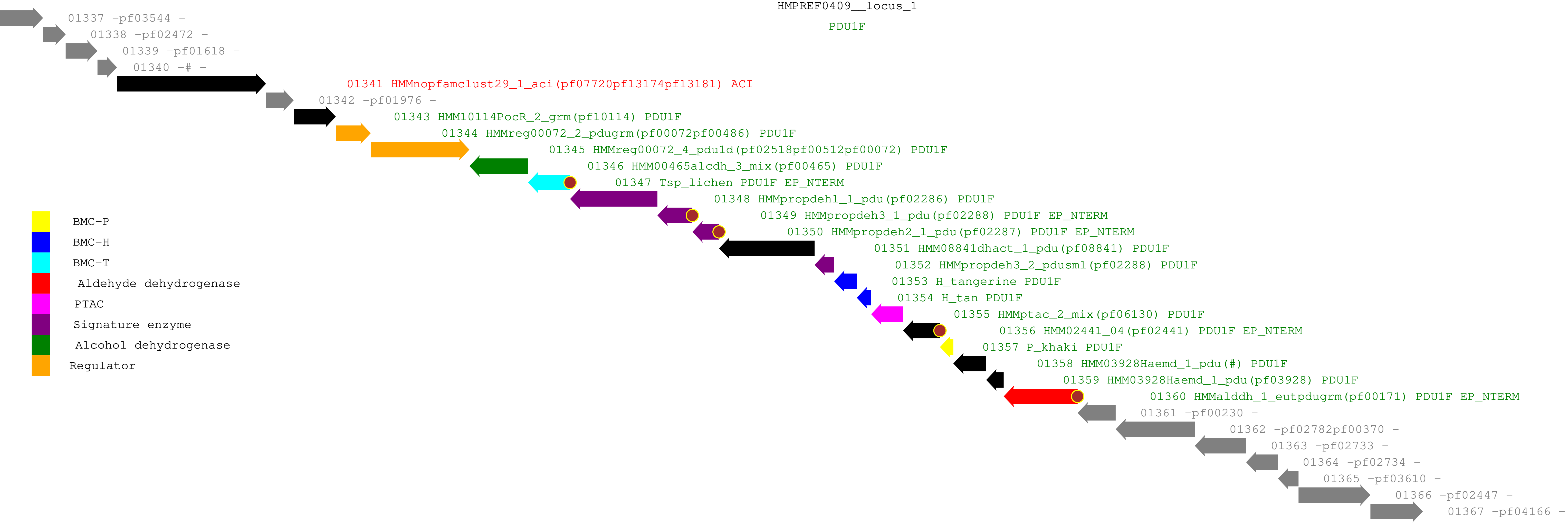

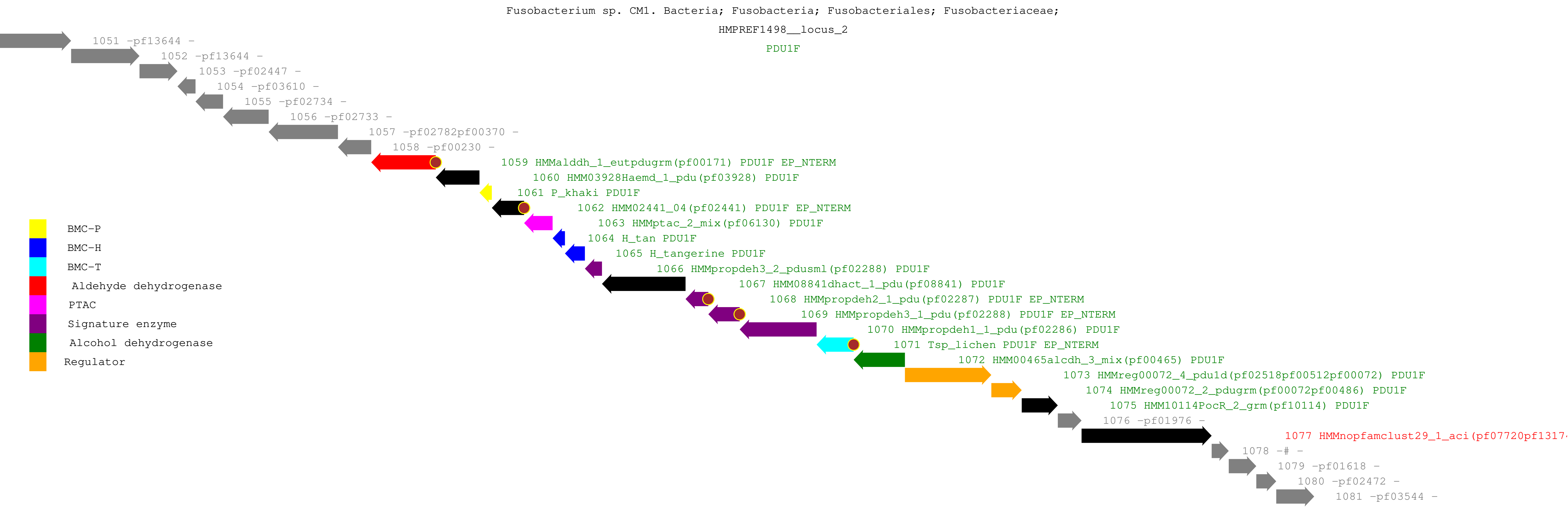
Fusobacterium sp. CM1. Bacteria; Fusobacteria; Fusobacteriales; Fusobacteriaceae;
HMPREF1498__locus_2
PDU1F
1051 -pf13644 -
1052 -pf13644 -
1053 -pf02447 -
1054 -pf03610 -
1055 -pf02734 -
1056 -pf02733 -
1057 -pf02782pf00370 -
1058 -pf00230 -
1059 HMMaldh_1_eutpdugrm(pf00171) PDU1F EP_NTERM
1060 HMM03928Haemd_1_pdu(pf03928) PDU1F
1061 P_khaki PDU1F
1062 HMM02441_04(pf02441) PDU1F EP_NTERM
1063 HMMptac_2_mix(pf06130) PDU1F
1064 H_tan PDU1F
1065 H_tangerine PDU1F
1066 HMMpropdeh3_2_pdusml(pf02288) PDU1F
1067 HMM08841dhact_1_pdu(pf08841) PDU1F
1068 HMMpropdeh2_1_pdu(pf02287) PDU1F EP_NTERM
1069 HMMpropdeh3_1_pdu(pf02288) PDU1F EP_NTERM
1070 HMMpropdeh1_1_pdu(pf02286) PDU1F
1071 Tsp_lichen PDU1F EP_NTERM
1072 HMM00465alcdh_3_mix(pf00465) PDU1F
1073 HMMreg00072_4_pdu1d(pf02518pf00512pf00072) PDU1F
1074 HMMreg00072_2_pdugrm(pf00072pf00486) PDU1F
1075 HMM10114PocR_2_grm(pf10114) PDU1F
1076 -pf01976 -
1077 HMMnopfamclust29_1_aci(pf07720pf1317
1078 -# -
1079 -pf01618 -
1080 -pf02472 -
1081 -pf03544 -
BMC-P
BMC-H
BMC-T
Aldehyde dehydrogenase
PTAC
Signature enzyme
Alcohol dehydrogenase
Regulator

Fusobacterium sp. CM22. Bacteria; Fusobacteria; Fusobacteriales; Fusobacteriaceae;

HMPREF1500__locus_1

PDU1F

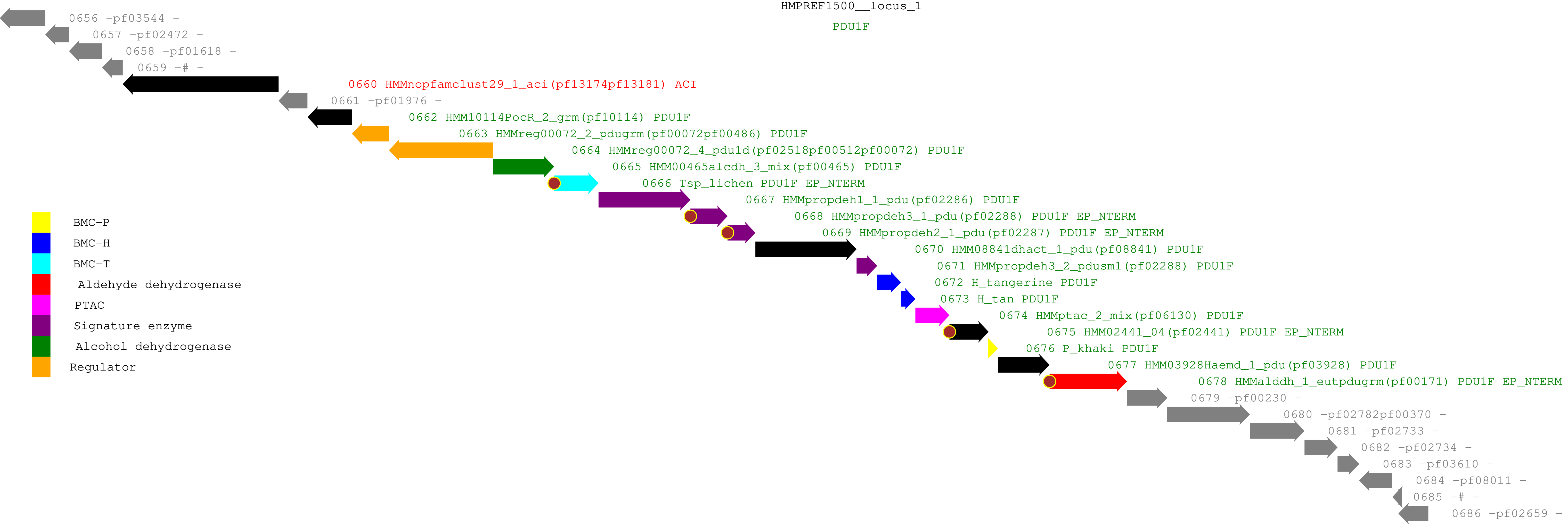

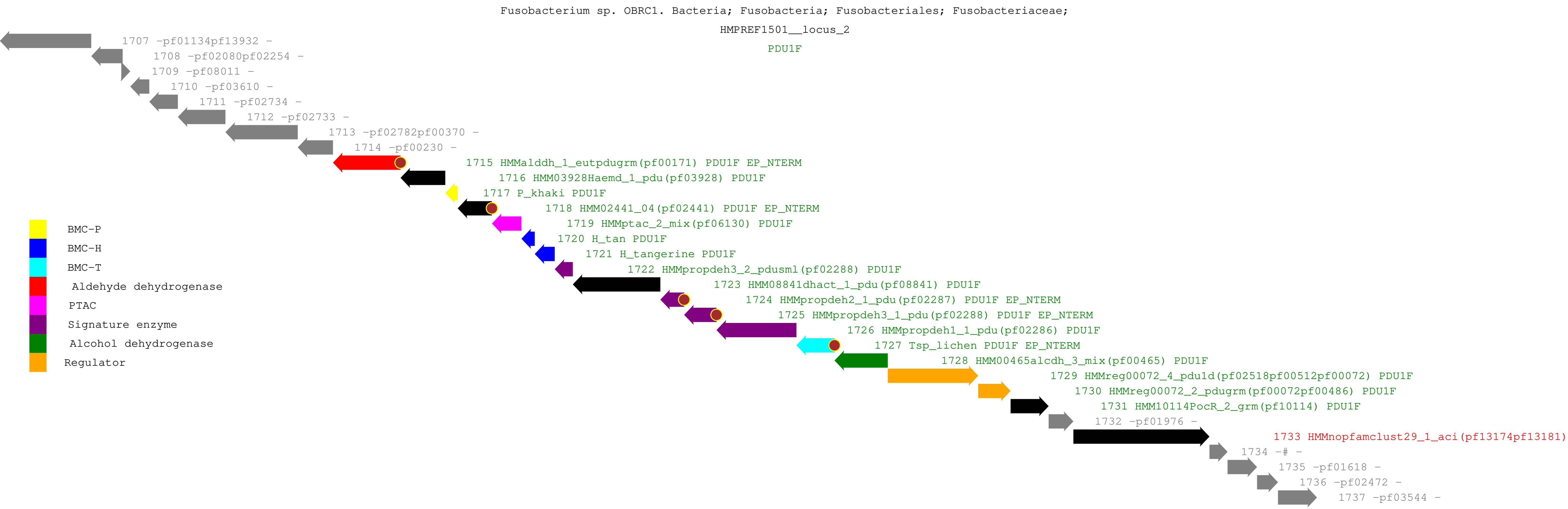
Fusobacterium sp. OBRC1. Bacteria; Fusobacteria; Fusobacteriales; Fusobacteriaceae;
HMPREF1501__locus_2
PDU1F
1707 -pf01134pf13932 -
1708 -pf02080pf02254 -
1709 -pf08011 -
1710 -pf03610 -
1711 -pf02734 -
1712 -pf02733 -
1713 -pf02782pf00370 -
1714 -pf00230 -
1715 HMMaldh_1_eutpdugrm(pf00171) PDU1F EP_NTERM
1716 HMM03928Haemd_1_pdu(pf03928) PDU1F
1717 P_khaki PDU1F
1718 HMM02441_04(pf02441) PDU1F EP_NTERM
1719 HMMptac_2_mix(pf06130) PDU1F
1720 H_tan PDU1F
1721 H_tangerine PDU1F
1722 HMMpropdeh3_2_pdusml(pf02288) PDU1F
1723 HMM08841dhact_1_pdu(pf08841) PDU1F
1724 HMMpropdeh2_1_pdu(pf02287) PDU1F EP_NTERM
1725 HMMpropdeh3_1_pdu(pf02288) PDU1F EP_NTERM
1726 HMMpropdeh1_1_pdu(pf02286) PDU1F
1727 Tsp_lichen PDU1F EP_NTERM
1728 HMM00465alcdh_3_mix(pf00465) PDU1F
1729 HMMreg00072_4_pdu1d(pf02518pf00512pf00072) PDU1F
1730 HMMreg00072_2_pdugrm(pf00072pf00486) PDU1F
1731 HMM10114PocR_2_grm(pf10114) PDU1F
1732 -pf01976 -
1733 HMMnopfamclust29_1_aci(pf13174pf13181)
1734 -# -
1735 -pf01618 -
1736 -pf02472 -
1737 -pf03544 -
BMC-P
BMC-H
BMC-T
Aldehyde dehydrogenase
PTAC
Signature enzyme
Alcohol dehydrogenase
Regulator

Fusobacterium nucleatum CTI-1. Bacteria; Fusobacteria; Fusobacteriales; Fusobacteriaceae;

HMPREF1538__locus_1

PDU1F

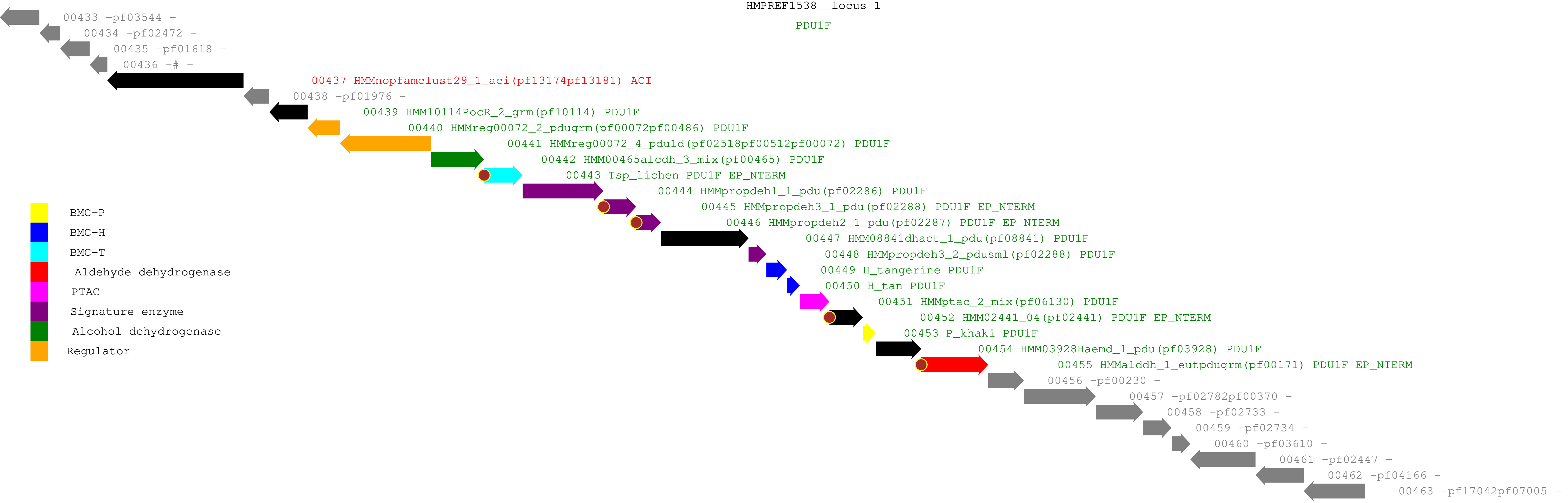

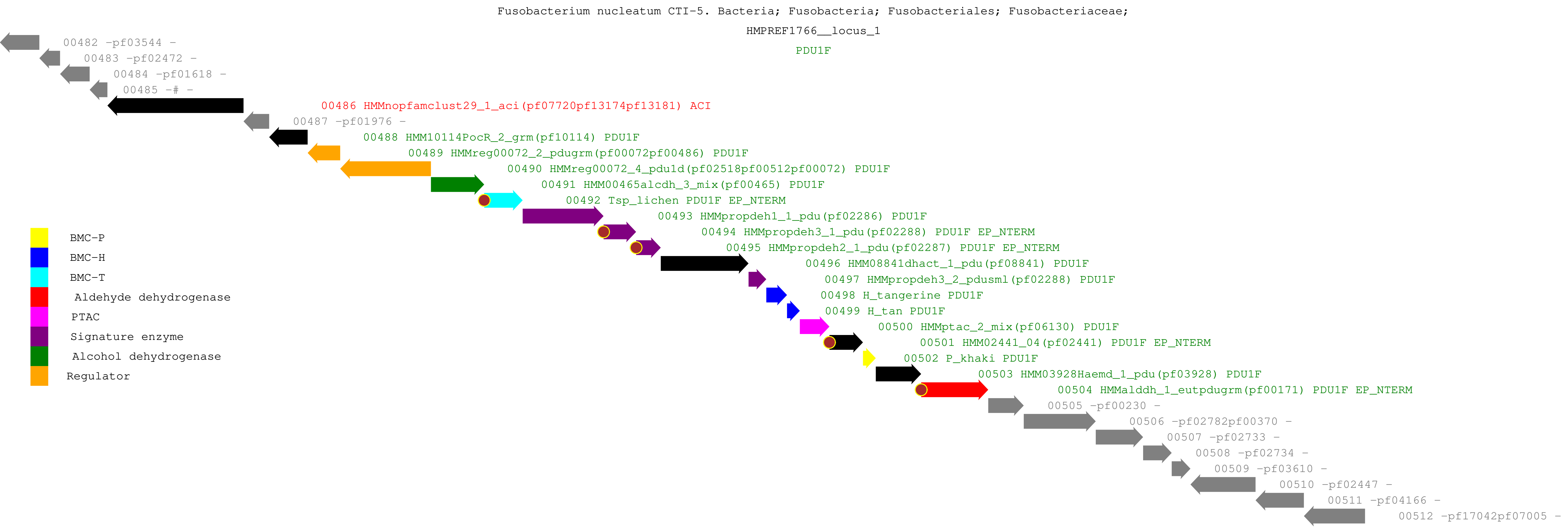
Fusobacterium nucleatum CTI-5. Bacteria; Fusobacteria; Fusobacteriales; Fusobacteriaceae;
HMPREF1766__locus_1
PDU1F
00482 -pf03544 -
00483 -pf02472 -
00484 -pf01618 -
00485 -# -
00486 HMMnopfamclust29_1_aci(pf07720pf13174pf13181) ACI
00487 -pf01976 -
00488 HMM10114PocR_2_grm(pf10114) PDU1F
00489 HMMreg00072_2_pdugrm(pf00072pf00486) PDU1F
00490 HMMreg00072_4_pdu1d(pf02518pf00512pf00072) PDU1F
00491 HMM00465alcdh_3_mix(pf00465) PDU1F
00492 Tsp_lichen PDU1F EP_NTERM
00493 HMMpropdeh1_1_pdu(pf02286) PDU1F
00494 HMMpropdeh3_1_pdu(pf02288) PDU1F EP_NTERM
00495 HMMpropdeh2_1_pdu(pf02287) PDU1F EP_NTERM
00496 HMM08841dhact_1_pdu(pf08841) PDU1F
00497 HMMpropdeh3_2_pdusml(pf02288) PDU1F
00498 H_tangerine PDU1F
00499 H_tan PDU1F
00500 HMMptac_2_mix(pf06130) PDU1F
00501 HMM02441_04(pf02441) PDU1F EP_NTERM
00502 P_khaki PDU1F
00503 HMM03928Haemd_1_pdu(pf03928) PDU1F
00504 HMMalddh_1_eutpdugrm(pf00171) PDU1F EP_NTERM
00505 -pf00230 -
00506 -pf02782pf00370 -
00507 -pf02733 -
00508 -pf02734 -
00509 -pf03610 -
00510 -pf02447 -
00511 -pf04166 -
00512 -pf17042pf07005 -
BMC-P
BMC-H
BMC-T
Aldehyde dehydrogenase
PTAC
Signature enzyme
Alcohol dehydrogenase
Regulator

Fusobacterium nucleatum CTI-6. Bacteria; Fusobacteria; Fusobacteriales; Fusobacteriaceae;

HMPREF1767__locus_1

PDU1F

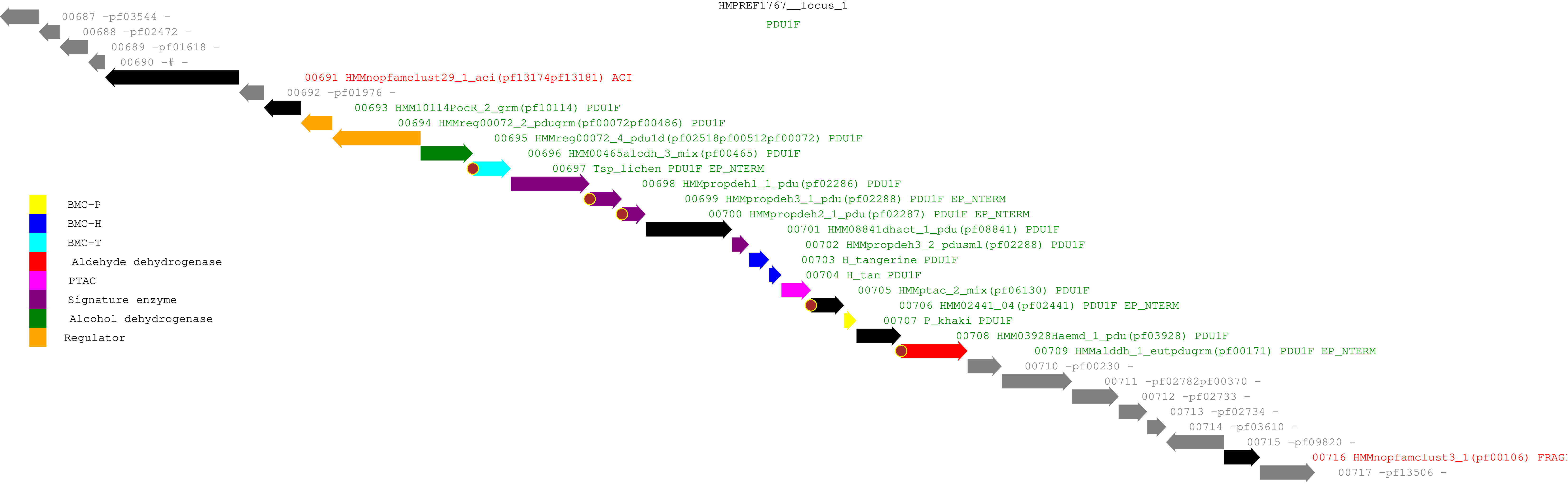

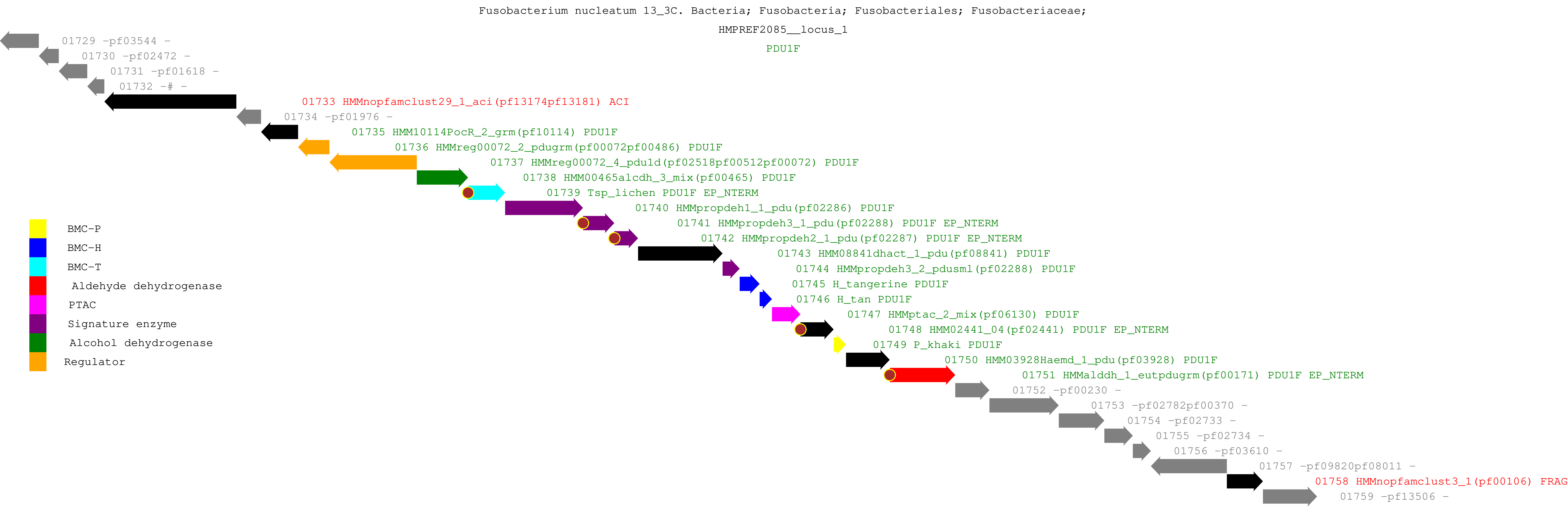
Fusobacterium nucleatum 13_3C. Bacteria; Fusobacteria; Fusobacteriales; Fusobacteriaceae;
HMPREF2085__locus_1
PDU1F
01729 -pf03544 -
01730 -pf02472 -
01731 -pf01618 -
01732 -# -
01733 HMMnopfamclust29_1_aci(pf13174pf13181) ACI
01734 -pf01976 -
01735 HMM10114PocR_2_grm(pf10114) PDU1F
01736 HMMreg00072_2_pdugrm(pf00072pf00486) PDU1F
01737 HMMreg00072_4_pdu1d(pf02518pf00512pf00072) PDU1F
01738 HMM00465alcdh_3_mix(pf00465) PDU1F
01739 Tsp_lichen PDU1F EP_NTERM
01740 HMMpropdeh1_1_pdu(pf02286) PDU1F
01741 HMMpropdeh3_1_pdu(pf02288) PDU1F EP_NTERM
01742 HMMpropdeh2_1_pdu(pf02287) PDU1F EP_NTERM
01743 HMM08841dhact_1_pdu(pf08841) PDU1F
01744 HMMpropdeh3_2_pdusml(pf02288) PDU1F
01745 H_tangerine PDU1F
01746 H_tan PDU1F
01747 HMMptac_2_mix(pf06130) PDU1F
01748 HMM02441_04(pf02441) PDU1F EP_NTERM
01749 P_khaki PDU1F
01750 HMM03928Haemd_1_pdu(pf03928) PDU1F
01751 HMMalddh_1_eutpdugrm(pf00171) PDU1F EP_NTERM
01752 -pf00230 -
01753 -pf02782pf00370 -
01754 -pf02733 -
01755 -pf02734 -
01756 -pf03610 -
01757 -pf09820pf08011 -
01758 HMMnopfamclust3_1(pf00106) FRAG
01759 -pf13506 -
BMC-P
BMC-H
BMC-T
Aldehyde dehydrogenase
PTAC
Signature enzyme
Alcohol dehydrogenase
Regulator

Fusobacterium sp. HMSC073F01. Bacteria; Fusobacteria; Fusobacteriales; Fusobacteriaceae;

HMPREF2747__locus_2

PDU1F

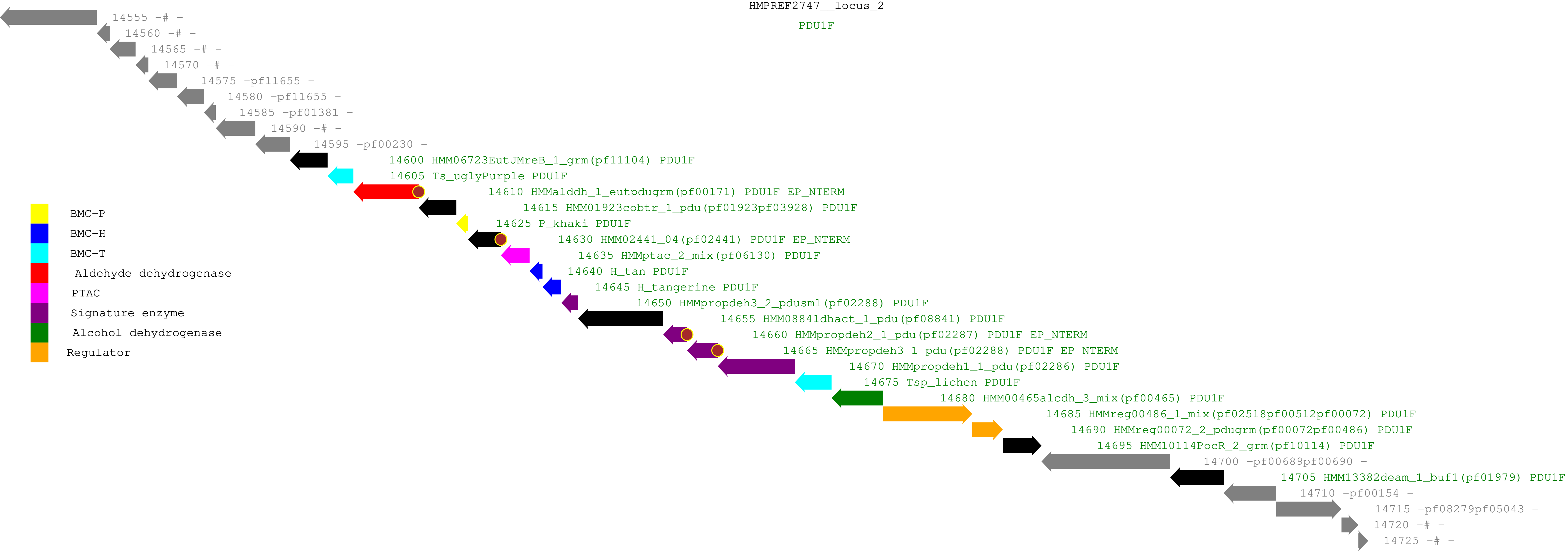

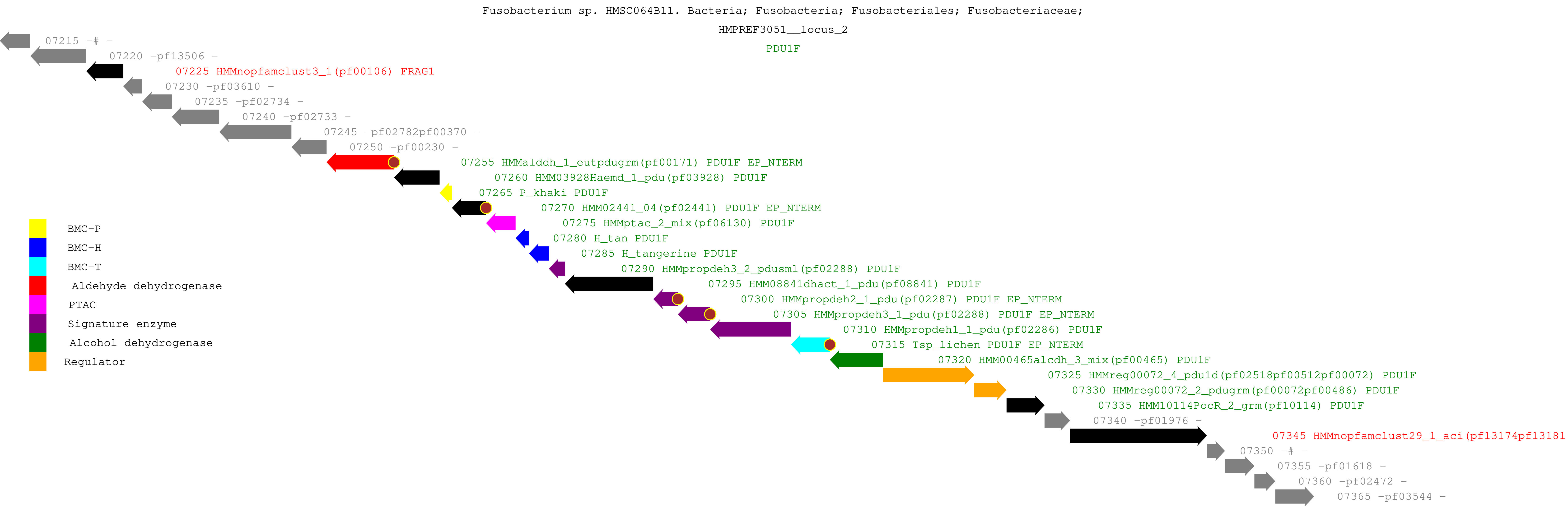
Fusobacterium sp. HMSC064B11. Bacteria; Fusobacteria; Fusobacteriales; Fusobacteriaceae;
HMPREF3051__locus_2
PDU1F
07215 -# -
07220 -pf13506 -
07225 HMMnopfamclust3_1(pf00106) FRAG1
07230 -pf03610 -
07235 -pf02734 -
07240 -pf02733 -
07245 -pf02782pf00370 -
07250 -pf00230 -
07255 HMMalddh_1_eutpdugrm(pf00171) PDU1F EP_NTERM
07260 HMM03928Haemd_1_pdu(pf03928) PDU1F
07265 P_khaki PDU1F
07270 HMM02441_04(pf02441) PDU1F EP_NTERM
07275 HMMptac_2_mix(pf06130) PDU1F
07280 H_tan PDU1F
07285 H_tangerine PDU1F
07290 HMMpropdeh3_2_pdusml(pf02288) PDU1F
07295 HMM08841dhact_1_pdu(pf08841) PDU1F
07300 HMMpropdeh2_1_pdu(pf02287) PDU1F EP_NTERM
07305 HMMpropdeh3_1_pdu(pf02288) PDU1F EP_NTERM
07310 HMMpropdeh1_1_pdu(pf02286) PDU1F
07315 Tsp_lichen PDU1F EP_NTERM
07320 HMM00465alcdh_3_mix(pf00465) PDU1F
07325 HMMreg00072_4_pdu1d(pf02518pf00512pf00072) PDU1F
07330 HMMreg00072_2_pdugrm(pf00072pf00486) PDU1F
07335 HMM10114PocR_2_grm(pf10114) PDU1F
07340 -pf01976 -
07345 HMMnopfamclust29_1_aci(pf13174pf13181
07350 -# -
07355 -pf01618 -
07360 -pf02472 -
07365 -pf03544 -
BMC-P
BMC-H
BMC-T
Aldehyde dehydrogenase
PTAC
Signature enzyme
Alcohol dehydrogenase
Regulator

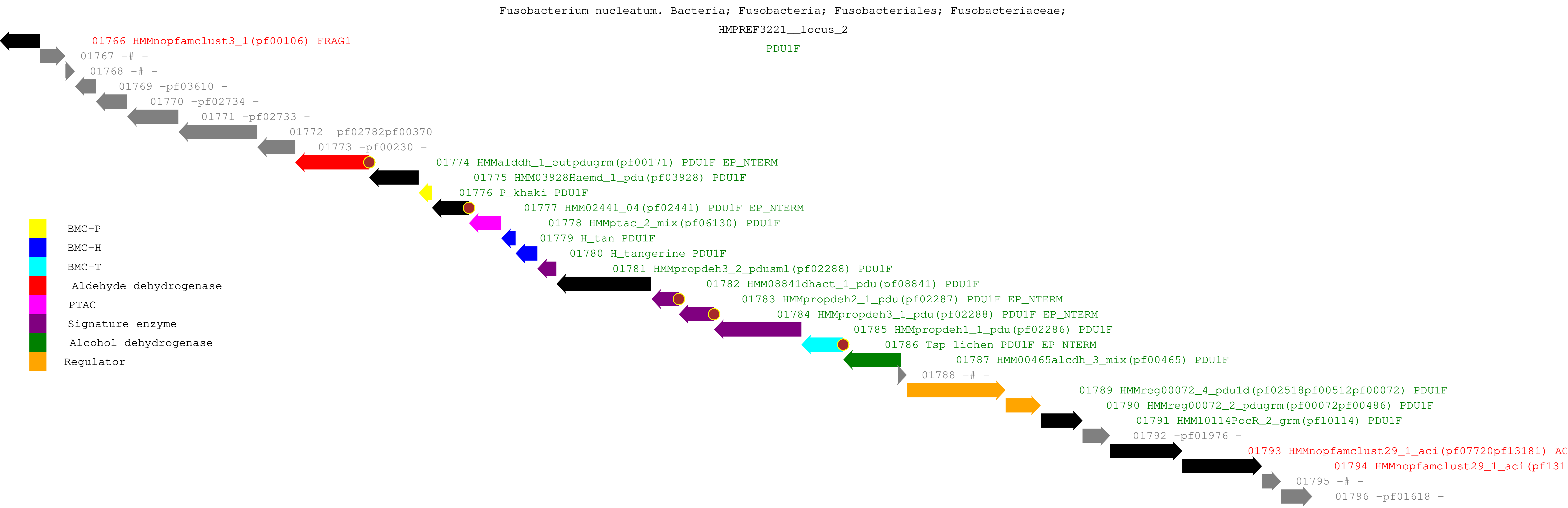
Fusobacterium nucleatum. Bacteria; Fusobacteria; Fusobacteriales; Fusobacteriaceae;
HMPREF3221__locus_2
PDU1F
01766 HMMnopfamclust3_1(pf00106) FRAG1
01767 -# -
01768 -# -
01769 -pf03610 -
01770 -pf02734 -
01771 -pf02733 -
01772 -pf02782pf00370 -
01773 -pf00230 -
01774 HMMaldh_1_eutpdugrm(pf00171) PDU1F EP_NTERM
01775 HMM03928Haemd_1_pdu(pf03928) PDU1F
01776 P_khaki PDU1F
01777 HMM02441_04(pf02441) PDU1F EP_NTERM
01778 HMMptac_2_mix(pf06130) PDU1F
01779 H_tan PDU1F
01780 H_tangerine PDU1F
01781 HMMpropdeh3_2_pdusml(pf02288) PDU1F
01782 HMM08841dhact_1_pdu(pf08841) PDU1F
01783 HMMpropdeh2_1_pdu(pf02287) PDU1F EP_NTERM
01784 HMMpropdeh3_1_pdu(pf02288) PDU1F EP_NTERM
01785 HMMpropdeh1_1_pdu(pf02286) PDU1F
01786 Tsp_lichen PDU1F EP_NTERM
01787 HMM00465alcdh_3_mix(pf00465) PDU1F
01788 -# -
01789 HMMreg00072_4_pdu1d(pf02518pf00512pf00072) PDU1F
01790 HMMreg00072_2_pdugrm(pf00072pf00486) PDU1F
01791 HMM10114PocR_2_grm(pf10114) PDU1F
01792 -pf01976 -
01793 HMMnopfamclust29_1_aci(pf07720pf13181) AC
01794 HMMnopfamclust29_1_aci(pf131
01795 -# -
01796 -pf01618 -
BMC-P
BMC-H
BMC-T
Aldehyde dehydrogenase
PTAC
Signature enzyme
Alcohol dehydrogenase
Regulator

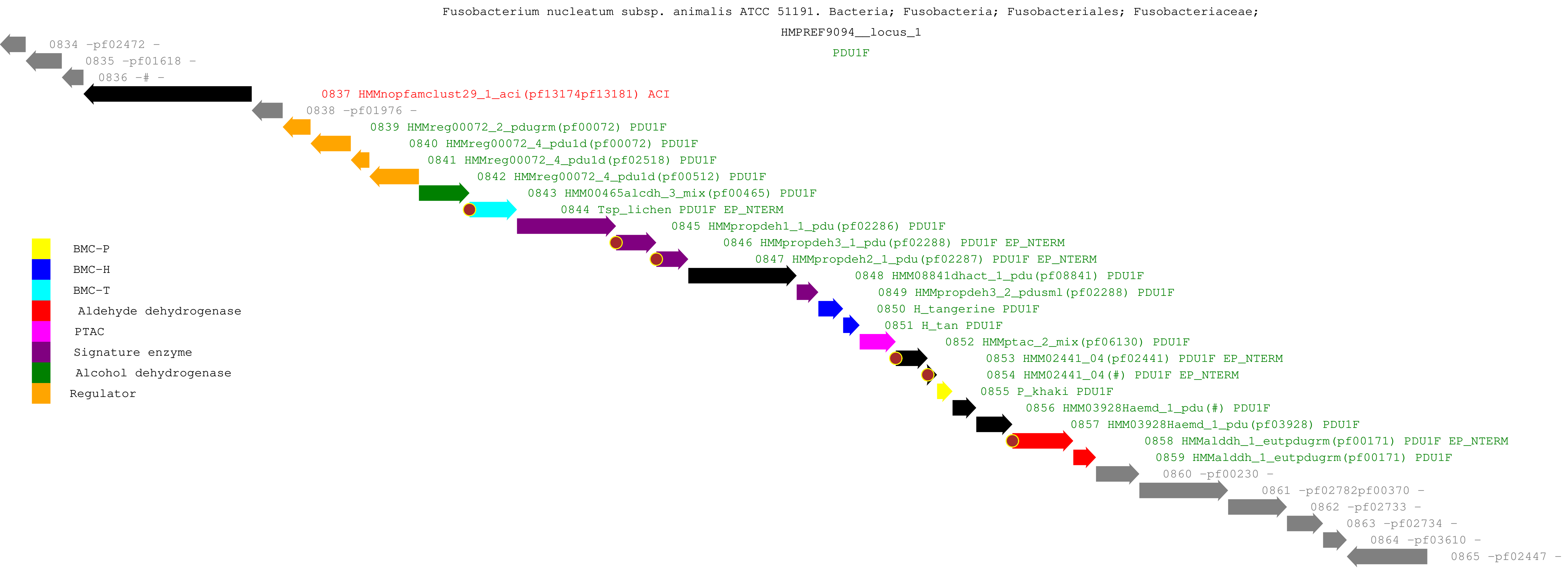
Fusobacterium nucleatum subsp. animalis ATCC 51191. Bacteria; Fusobacteria; Fusobacteriales; Fusobacteriaceae;
HMPREF9094__locus_1
PDU1F
0834 -pf02472 -
0835 -pf01618 -
0836 -# -
0837 HMMnopfamclust29_1_aci(pf13174pf13181) ACI
0838 -pf01976 -
0839 HMMreg00072_2_pdugrm(pf00072) PDU1F
0840 HMMreg00072_4_pdu1d(pf00072) PDU1F
0841 HMMreg00072_4_pdu1d(pf02518) PDU1F
0842 HMMreg00072_4_pdu1d(pf00512) PDU1F
0843 HMM00465alcdh_3_mix(pf00465) PDU1F
0844 Tsp_lichen PDU1F EP_NTERM
0845 HMMpropdeh1_1_pdu(pf02286) PDU1F
0846 HMMpropdeh3_1_pdu(pf02288) PDU1F EP_NTERM
0847 HMMpropdeh2_1_pdu(pf02287) PDU1F EP_NTERM
0848 HMM08841dhact_1_pdu(pf08841) PDU1F
0849 HMMpropdeh3_2_pdusml(pf02288) PDU1F
0850 H_tangerine PDU1F
0851 H_tan PDU1F
0852 HMMptac_2_mix(pf06130) PDU1F
0853 HMM02441_04(pf02441) PDU1F EP_NTERM
0854 HMM02441_04(#) PDU1F EP_NTERM
0855 P_khaki PDU1F
0856 HMM03928Haemd_1_pdu(#) PDU1F
0857 HMM03928Haemd_1_pdu(pf03928) PDU1F
0858 HMMalddh_1_eutpdugrm(pf00171) PDU1F EP_NTERM
0859 HMMalddh_1_eutpdugrm(pf00171) PDU1F
0860 -pf00230 -
0861 -pf02782pf00370 -
0862 -pf02733 -
0863 -pf02734 -
0864 -pf03610 -
0865 -pf02447 -
BMC-P
BMC-H
BMC-T
Aldehyde dehydrogenase
PTAC
Signature enzyme
Alcohol dehydrogenase
Regulator

Fusobacterium nucleatum subsp. animalis F0419. Bacteria; Fusobacteria; Fusobacteriales; Fusobacteriaceae;

HMPREF9942__locus_1

PDU1F

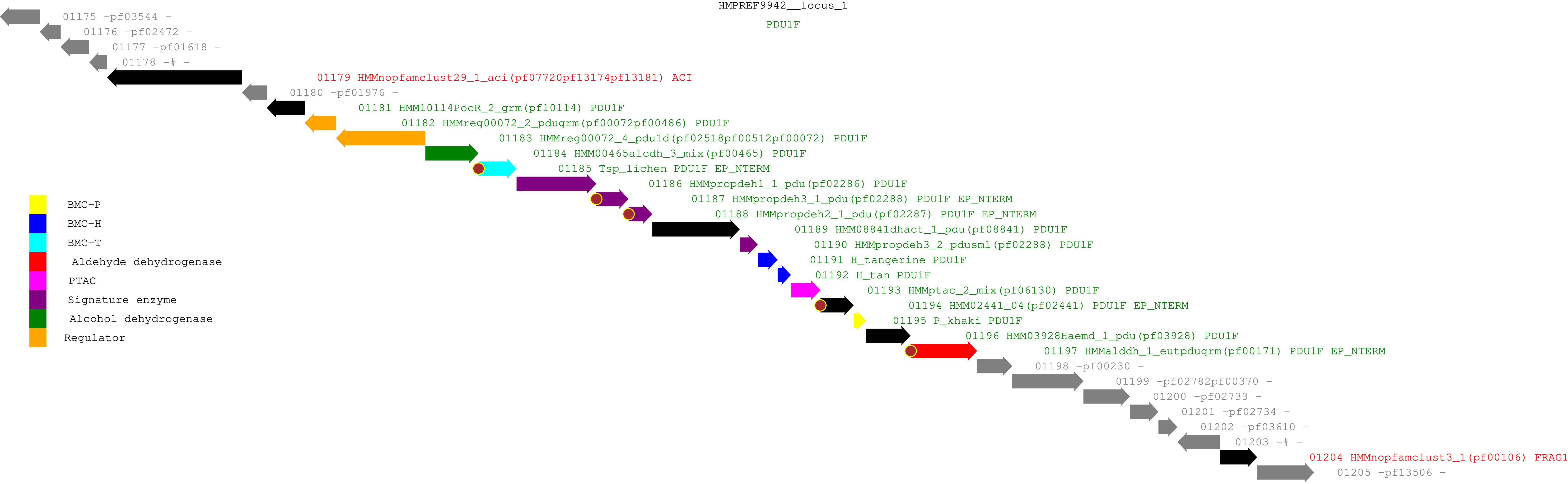

Ilyobacter polytropus (strain DSM 2926 / CuHBu1). Bacteria; Fusobacteria; Fusobacteriales; Fusobacteriaceae; Ilyobacter.

Ilyop__locus_1

PDU1F

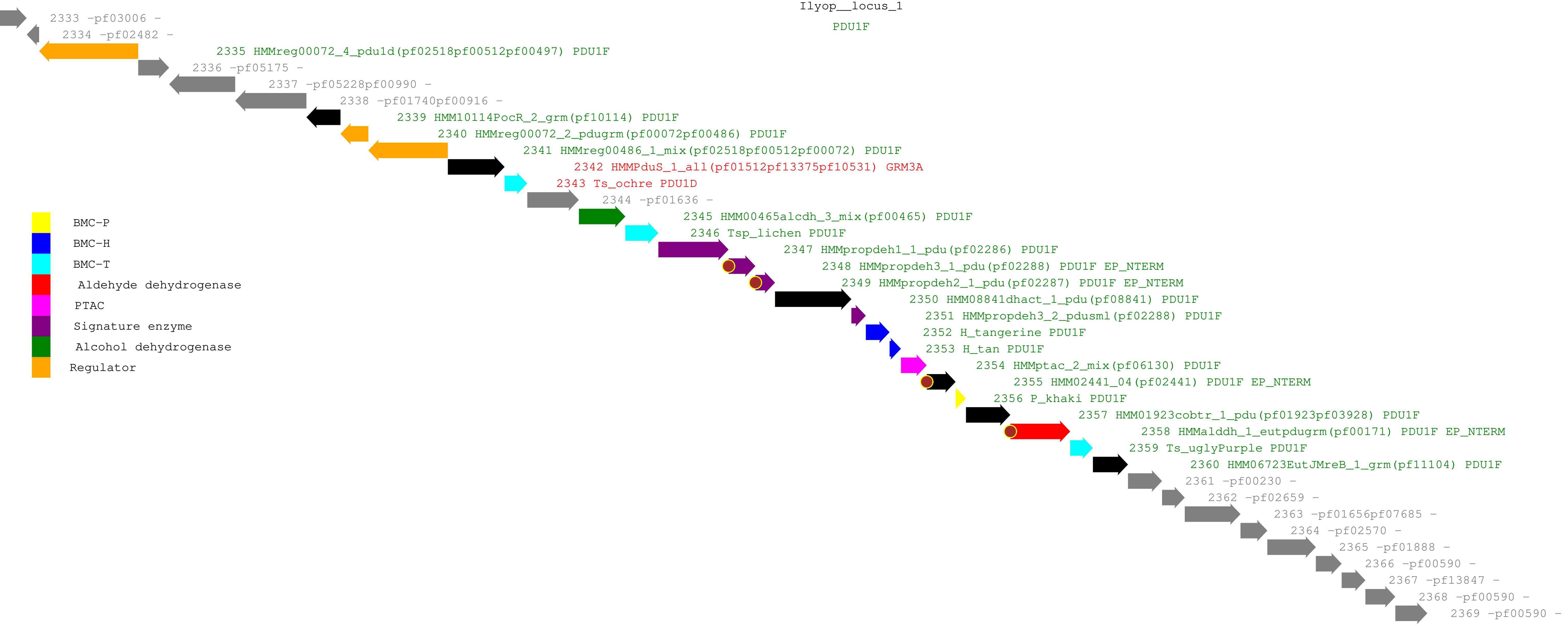

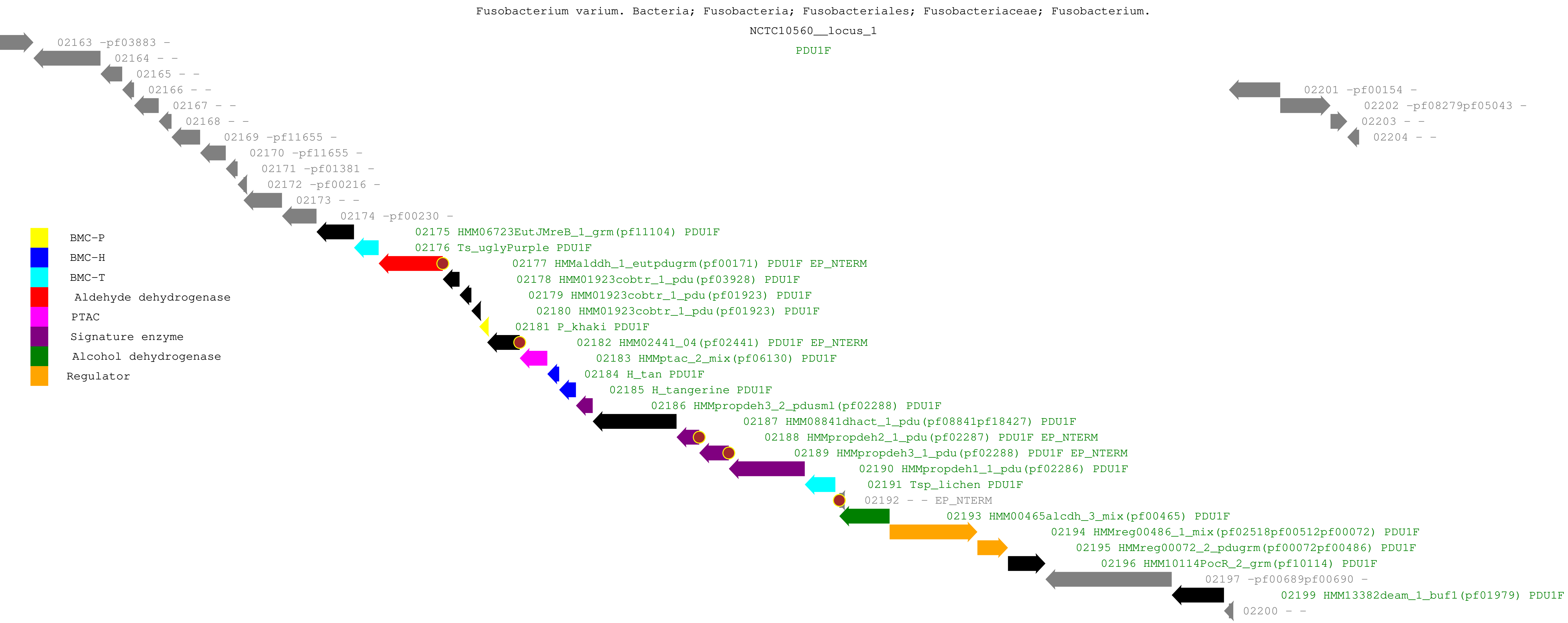
Fusobacterium varium. Bacteria; Fusobacteria; Fusobacteriales; Fusobacteriaceae; Fusobacterium.
NCTC10560__locus_1
PDU1F
02163 -pf03883 -
02164 - -
02165 - -
02166 - -
02167 - -
02168 - -
02169 -pf11655 -
02170 -pf11655 -
02171 -pf01381 -
02172 -pf00216 -
02173 - -
02174 -pf00230 -
02175 HMM06723EutJMreB_1_grm(pf11104) PDU1F
02176 Ts_uglyPurple PDU1F
02177 HMMalddh_1_eutpdugrm(pf00171) PDU1F EP_NTERM
02178 HMM01923cobtr_1_pdu(pf03928) PDU1F
02179 HMM01923cobtr_1_pdu(pf01923) PDU1F
02180 HMM01923cobtr_1_pdu(pf01923) PDU1F
02181 P_khaki PDU1F
02182 HMM02441_04(pf02441) PDU1F EP_NTERM
02183 HMMptac_2_mix(pf06130) PDU1F
02184 H_tan PDU1F
02185 H_tangerine PDU1F
02186 HMMpropdeh3_2_pdusml(pf02288) PDU1F
02187 HMM08841dhact_1_pdu(pf08841pf18427) PDU1F
02188 HMMpropdeh2_1_pdu(pf02287) PDU1F EP_NTERM
02189 HMMpropdeh3_1_pdu(pf02288) PDU1F EP_NTERM
02190 HMMpropdeh1_1_pdu(pf02286) PDU1F
02191 Tsp_lichen PDU1F
02192 - - EP_NTERM
02193 HMM00465alcdh_3_mix(pf00465) PDU1F
02194 HMMreg00486_1_mix(pf02518pf00512pf00072) PDU1F
02195 HMMreg00072_2_pdugrm(pf00072pf00486) PDU1F
02196 HMM10114PocR_2_grm(pf10114) PDU1F
02197 -pf00689pf00690 -
02199 HMM13382deam_1_buf1(pf01979) PDU1F
02200 - -
02201 -pf00154 -
02202 -pf08279pf05043 -
02203 - -
02204 - -
BMC-P
BMC-H
BMC-T
Aldehyde dehydrogenase
PTAC
Signature enzyme
Alcohol dehydrogenase
Regulator

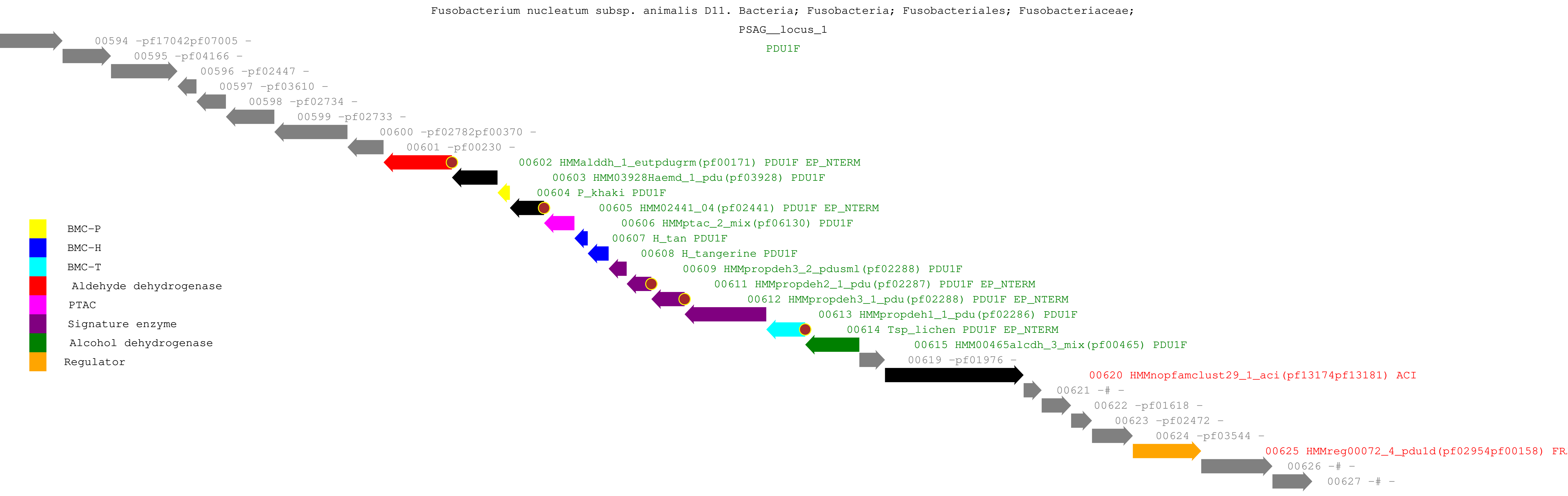
Fusobacterium nucleatum subsp. animalis D11. Bacteria; Fusobacteria; Fusobacteriales; Fusobacteriaceae;
PSAG__locus_1
PDU1F
00594 -pf17042pf07005 -
00595 -pf04166 -
00596 -pf02447 -
00597 -pf03610 -
00598 -pf02734 -
00599 -pf02733 -
00600 -pf02782pf00370 -
00601 -pf00230 -
00602 HMMalddh_1_eutpdugrm(pf00171) PDU1F EP_NTERM
00603 HMM03928Haemd_1_pdu(pf03928) PDU1F
00604 P_khaki PDU1F
00605 HMM02441_04(pf02441) PDU1F EP_NTERM
00606 HMMptac_2_mix(pf06130) PDU1F
00607 H_tan PDU1F
00608 H_tangerine PDU1F
00609 HMMpropdeh3_2_pdusml(pf02288) PDU1F
00611 HMMpropdeh2_1_pdu(pf02287) PDU1F EP_NTERM
00612 HMMpropdeh3_1_pdu(pf02288) PDU1F EP_NTERM
00613 HMMpropdeh1_1_pdu(pf02286) PDU1F
00614 Tsp_lichen PDU1F EP_NTERM
00615 HMM00465alcdh_3_mix(pf00465) PDU1F
00619 -pf01976 -
00620 HMMnopfamclust29_1_aci(pf13174pf13181) ACI
00621 -# -
00622 -pf01618 -
00623 -pf02472 -
00624 -pf03544 -
00625 HMMreg00072_4_pdu1d(pf02954pf00158) FR
00626 -# -
00627 -# -
BMC-P
BMC-H
BMC-T
Aldehyde dehydrogenase
PTAC
Signature enzyme
Alcohol dehydrogenase
Regulator

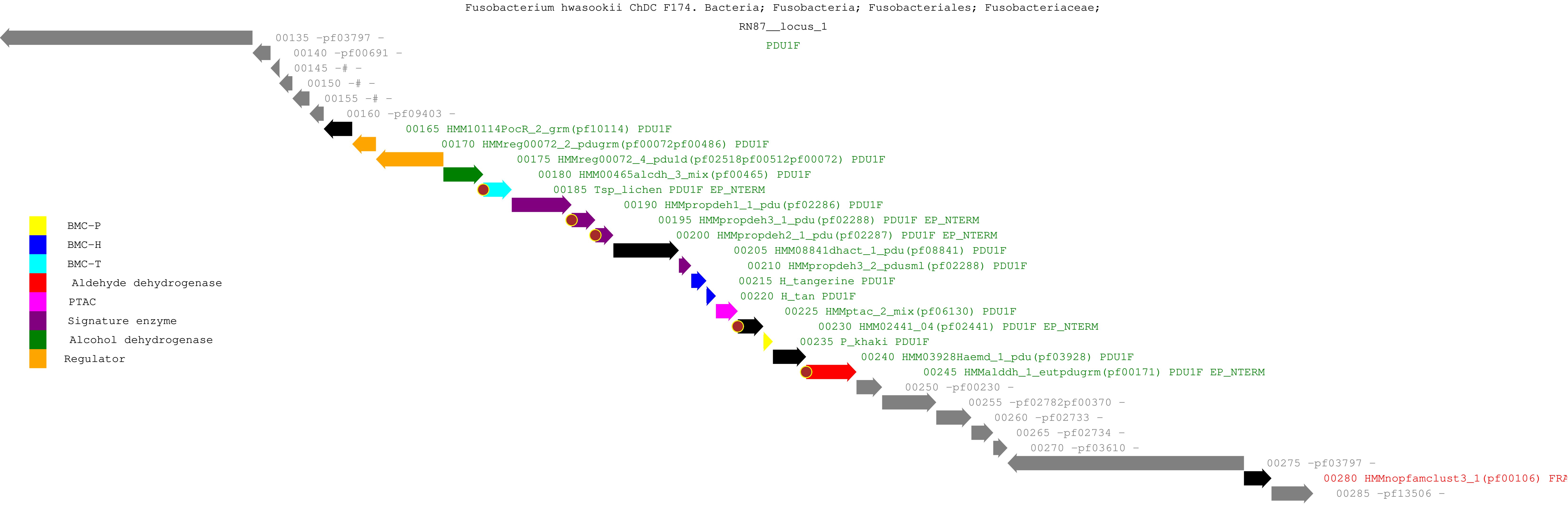
Fusobacterium hwasookii ChDC F174. Bacteria; Fusobacteria; Fusobacteriales; Fusobacteriaceae;
RN87__locus_1
PDU1F
00135 -pf03797 -
00140 -pf00691 -
00145 -# -
00150 -# -
00155 -# -
00160 -pf09403 -
00165 HMM10114PocR_2_grm(pf10114) PDU1F
00170 HMMreg00072_2_pdugrm(pf00072pf00486) PDU1F
00175 HMMreg00072_4_pdu1d(pf02518pf00512pf00072) PDU1F
00180 HMM00465alcdh_3_mix(pf00465) PDU1F
00185 Tsp_lichen PDU1F EP_NTERM
00190 HMMpropdeh1_1_pdu(pf02286) PDU1F
00195 HMMpropdeh3_1_pdu(pf02288) PDU1F EP_NTERM
00200 HMMpropdeh2_1_pdu(pf02287) PDU1F EP_NTERM
00205 HMM08841dhact_1_pdu(pf08841) PDU1F
00210 HMMpropdeh3_2_pdusml(pf02288) PDU1F
00215 H_tangerine PDU1F
00220 H_tan PDU1F
00225 HMMptac_2_mix(pf06130) PDU1F
00230 HMM02441_04(pf02441) PDU1F EP_NTERM
00235 P_khaki PDU1F
00240 HMM03928Haemd_1_pdu(pf03928) PDU1F
00245 HMMalddh_1_eutpdugrm(pf00171) PDU1F EP_NTERM
00250 -pf00230 -
00255 -pf02782pf00370 -
00260 -pf02733 -
00265 -pf02734 -
00270 -pf03610 -
00275 -pf03797 -
00280 HMMnopfamclust3_1(pf00106) FRA
00285 -pf13506 -
BMC-P
BMC-H
BMC-T
Aldehyde dehydrogenase
PTAC
Signature enzyme
Alcohol dehydrogenase
Regulator

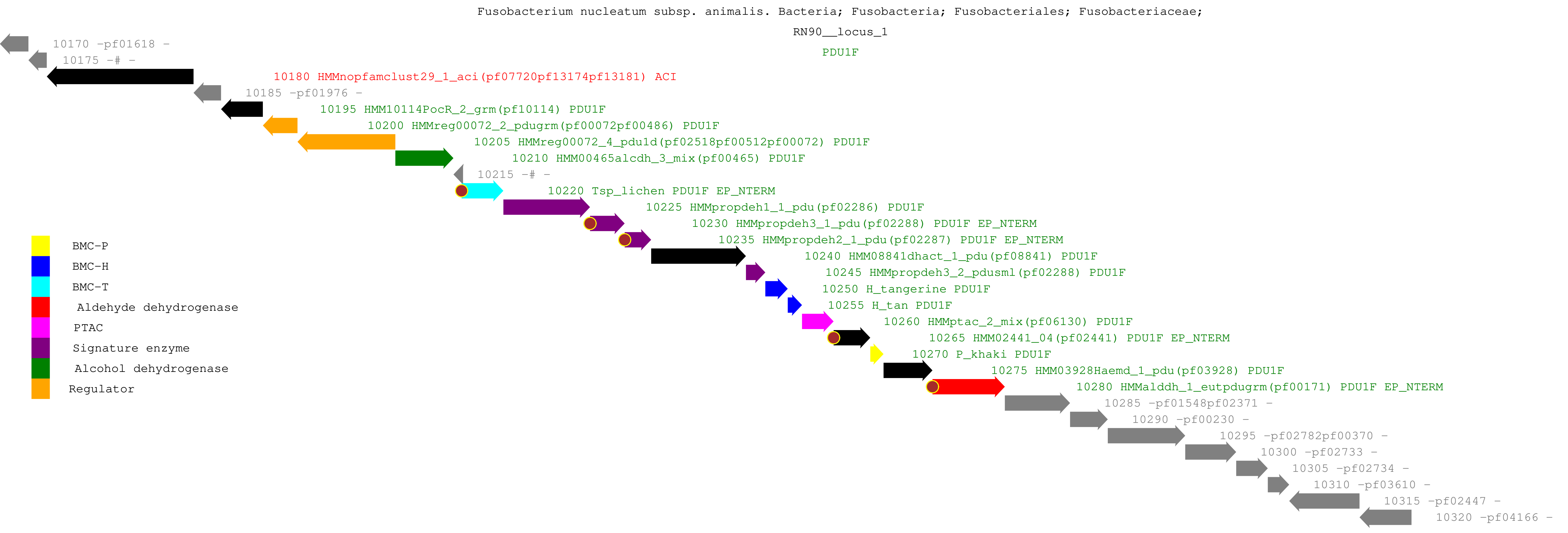
Fusobacterium nucleatum subsp. animalis. Bacteria; Fusobacteria; Fusobacteriales; Fusobacteriaceae;
RN90__locus_1
PDU1F
10170 -pf01618 -
10175 -# -
10180 HMMnopfamclust29_1_aci(pf07720pf13174pf13181) ACI
10185 -pf01976 -
10195 HMM10114PocR_2_grm(pf10114) PDU1F
10200 HMMreg00072_2_pdugrm(pf00072pf00486) PDU1F
10205 HMMreg00072_4_pdu1d(pf02518pf00512pf00072) PDU1F
10210 HMM00465alcdh_3_mix(pf00465) PDU1F
10215 -# -
10220 Tsp_lichen PDU1F EP_NTERM
10225 HMMpropdeh1_1_pdu(pf02286) PDU1F
10230 HMMpropdeh3_1_pdu(pf02288) PDU1F EP_NTERM
10235 HMMpropdeh2_1_pdu(pf02287) PDU1F EP_NTERM
10240 HMM08841dhact_1_pdu(pf08841) PDU1F
10245 HMMpropdeh3_2_pdusml(pf02288) PDU1F
10250 H_tangerine PDU1F
10255 H_tan PDU1F
10260 HMMptac_2_mix(pf06130) PDU1F
10265 HMM02441_04(pf02441) PDU1F EP_NTERM
10270 P_khaki PDU1F
10275 HMM03928Haemd_1_pdu(pf03928) PDU1F
10280 HMMalddh_1_eutpdugrm(pf00171) PDU1F EP_NTERM
10285 -pf01548pf02371 -
10290 -pf00230 -
10295 -pf02782pf00370 -
10300 -pf02733 -
10305 -pf02734 -
10310 -pf03610 -
10315 -pf02447 -
10320 -pf04166 -
BMC-P
BMC-H
BMC-T
Aldehyde dehydrogenase
PTAC
Signature enzyme
Alcohol dehydrogenase
Regulator

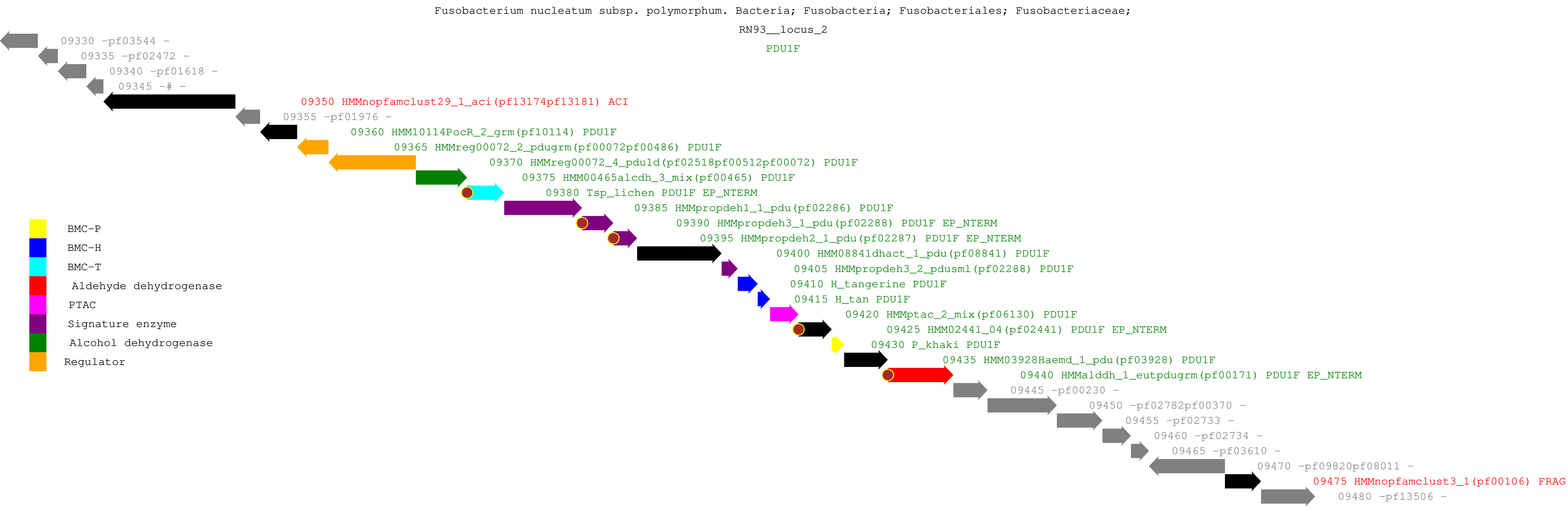
Fusobacterium nucleatum subsp. polymorphum. Bacteria; Fusobacteria; Fusobacteriales; Fusobacteriaceae;
RN93__locus_2
PDU1F
09330 -pf03544 -
09335 -pf02472 -
09340 -pf01618 -
09345 -# -
09350 HMMnopfamclust29_1_aci(pf13174pf13181) ACI
09355 -pf01976 -
09360 HMM10114PocR_2_grm(pf10114) PDU1F
09365 HMMreg00072_2_pdugrm(pf00072pf00486) PDU1F
09370 HMMreg00072_4_pdu1d(pf02518pf00512pf00072) PDU1F
09375 HMM00465alcdh_3_mix(pf00465) PDU1F
09380 Tsp_lichen PDU1F EP_NTERM
09385 HMMpropdeh1_1_pdu(pf02286) PDU1F
09390 HMMpropdeh3_1_pdu(pf02288) PDU1F EP_NTERM
09395 HMMpropdeh2_1_pdu(pf02287) PDU1F EP_NTERM
09400 HMM08841dhact_1_pdu(pf08841) PDU1F
09405 HMMpropdeh3_2_pdusml(pf02288) PDU1F
09410 H_tangerine PDU1F
09415 H_tan PDU1F
09420 HMMptac_2_mix(pf06130) PDU1F
09425 HMM02441_04(pf02441) PDU1F EP_NTERM
09430 P_khaki PDU1F
09435 HMM03928Haemd_1_pdu(pf03928) PDU1F
09440 HMMalddh_1_eutpdugrm(pf00171) PDU1F EP_NTERM
09445 -pf00230 -
09450 -pf02782pf00370 -
09455 -pf02733 -
09460 -pf02734 -
09465 -pf03610 -
09470 -pf09820pf08011 -
09475 HMMnopfamclust3_1(pf00106) FRAG
09480 -pf13506 -
BMC-P
BMC-H
BMC-T
Aldehyde dehydrogenase
PTAC
Signature enzyme
Alcohol dehydrogenase
Regulator

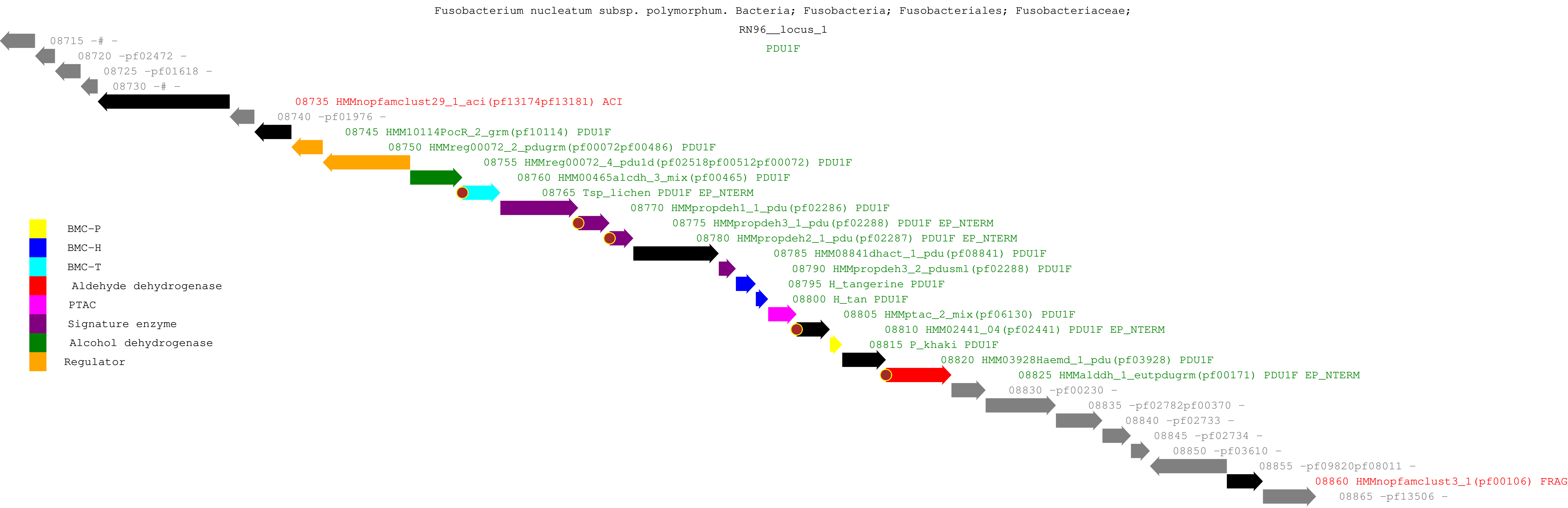
Fusobacterium nucleatum subsp. polymorphum. Bacteria; Fusobacteria; Fusobacteriales; Fusobacteriaceae;
RN96__locus_1
PDU1F
08715 -# -
08720 -pf02472 -
08725 -pf01618 -
08730 -# -
08735 HMMnopfamclust29_1_aci(pf13174pf13181) ACI
08740 -pf01976 -
08745 HMM10114PocR_2_grm(pf10114) PDU1F
08750 HMMreg00072_2_pdugrm(pf00072pf00486) PDU1F
08755 HMMreg00072_4_pdu1d(pf02518pf00512pf00072) PDU1F
08760 HMM00465alcdh_3_mix(pf00465) PDU1F
08765 Tsp_lichen PDU1F EP_NTERM
08770 HMMpropdeh1_1_pdu(pf02286) PDU1F
08775 HMMpropdeh3_1_pdu(pf02288) PDU1F EP_NTERM
08780 HMMpropdeh2_1_pdu(pf02287) PDU1F EP_NTERM
08785 HMM08841dhact_1_pdu(pf08841) PDU1F
08790 HMMpropdeh3_2_pdusml(pf02288) PDU1F
08795 H_tangerine PDU1F
08800 H_tan PDU1F
08805 HMMptac_2_mix(pf06130) PDU1F
08810 HMM02441_04(pf02441) PDU1F EP_NTERM
08815 P_khaki PDU1F
08820 HMM03928Haemd_1_pdu(pf03928) PDU1F
08825 HMMalddh_1_eutpdugrm(pf00171) PDU1F EP_NTERM
08830 -pf00230 -
08835 -pf02782pf00370 -
08840 -pf02733 -
08845 -pf02734 -
08850 -pf03610 -
08855 -pf09820pf08011 -
08860 HMMnopfamclust3_1(pf00106) FRAG
08865 -pf13506 -
BMC-P
BMC-H
BMC-T
Aldehyde dehydrogenase
PTAC
Signature enzyme
Alcohol dehydrogenase
Regulator

Fusobacterium hwasookii ChDC F300. Bacteria; Fusobacteria; Fusobacteriales; Fusobacteriaceae;

RN97__locus_2

PDU1F

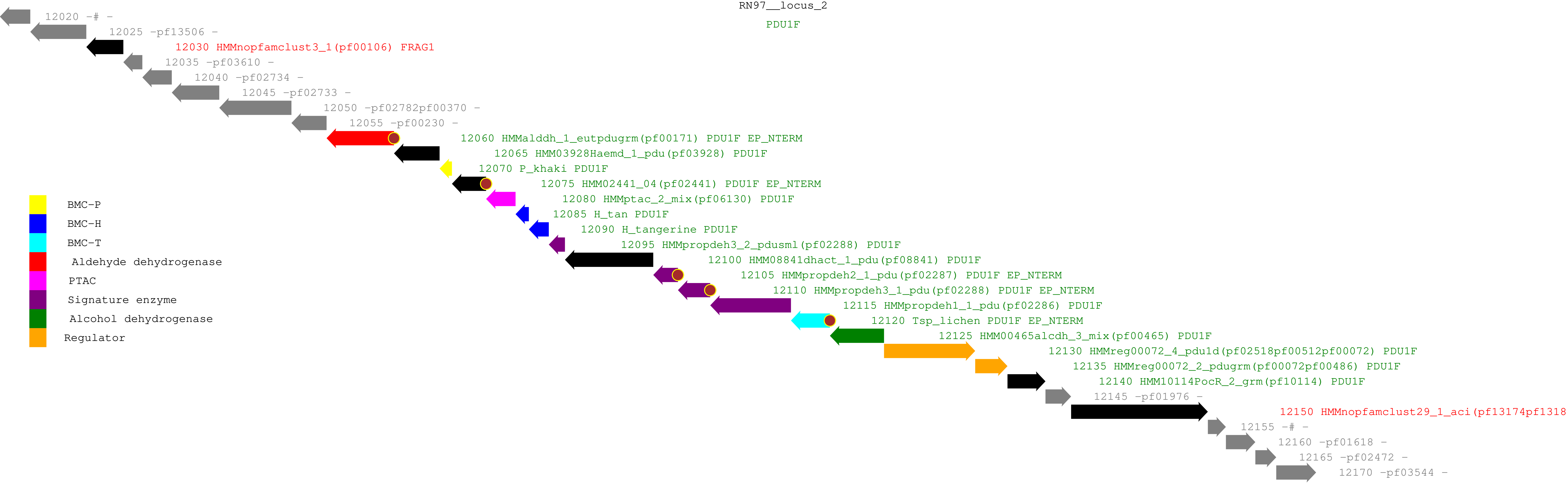

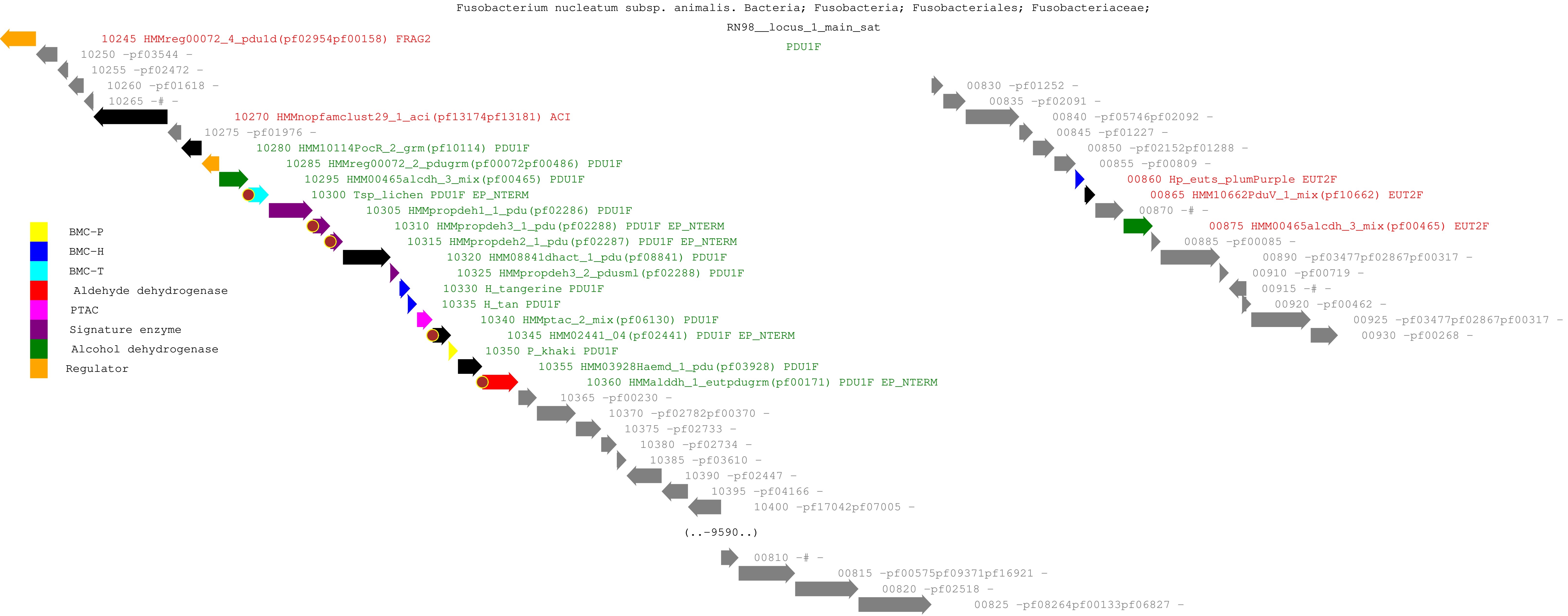

Fusobacterium nucleatum subsp. animalis. Bacteria; Fusobacteria; Fusobacteriales; Fusobacteriaceae;
RN98__locus_1_main_sat
PDU1F
10245 HMMreg00072_4_pdu1d(pf02954pf00158) FRAG2
10250 -pf03544 -
10255 -pf02472 -
10260 -pf01618 -
10265 -# -
10270 HMMnopfamclust29_1_aci(pf13174pf13181) ACI
10275 -pf01976 -
10280 HMM10114PocR_2_grm(pf10114) PDU1F
10285 HMMreg00072_2_pdugrm(pf00072pf00486) PDU1F
10295 HMM00465alcdh_3_mix(pf00465) PDU1F
10300 Tsp_lichen PDU1F EP_NTERM
10305 HMMpropdeh1_1_pdu(pf02286) PDU1F
10310 HMMpropdeh3_1_pdu(pf02288) PDU1F EP_NTERM
10315 HMMpropdeh2_1_pdu(pf02287) PDU1F EP_NTERM
10320 HMM08841dhact_1_pdu(pf08841) PDU1F
10325 HMMpropdeh3_2_pdusml(pf02288) PDU1F
10330 H_tangerine PDU1F
10335 H_tan PDU1F
10340 HMMptac_2_mix(pf06130) PDU1F
10345 HMM02441_04(pf02441) PDU1F EP_NTERM
10350 P_khaki PDU1F
10355 HMM03928Haemd_1_pdu(pf03928) PDU1F
10360 HMMalddh_1_eutpdugrm(pf00171) PDU1F EP_NTERM
10365 -pf00230 -
10370 -pf02782pf00370 -
10375 -pf02733 -
10380 -pf02734 -
10385 -pf03610 -
10390 -pf02447 -
10395 -pf04166 -
10400 -pf17042pf07005 -
(..-9590..)
00810 -# -
00815 -pf00575pf09371pf16921 -
00820 -pf02518 -
00825 -pf08264pf00133pf06827 -
00830 -pf01252 -
00835 -pf02091 -
00840 -pf05746pf02092 -
00845 -pf01227 -
00850 -pf02152pf01288 -
00855 -pf00809 -
00860 Hp_euts_plumPurple EUT2F
00865 HMM10662PduV_1_mix(pf10662) EUT2F
00870 -# -
00875 HMM00465alcdh_3_mix(pf00465) EUT2F
00885 -pf00085 -
00890 -pf03477pf02867pf00317 -
00910 -pf00719 -
00915 -# -
00920 -pf00462 -
00925 -pf03477pf02867pf00317 -
00930 -pf00268 -
BMC-P
BMC-H
BMC-T
Aldehyde dehydrogenase
PTAC
Signature enzyme
Alcohol dehydrogenase
Regulator

Verrucomicrobia bacterium GWF2_62_7. Bacteria; Verrucomicrobia; unclassified Verrucomicrobia.

A2107__locus_1

PVM

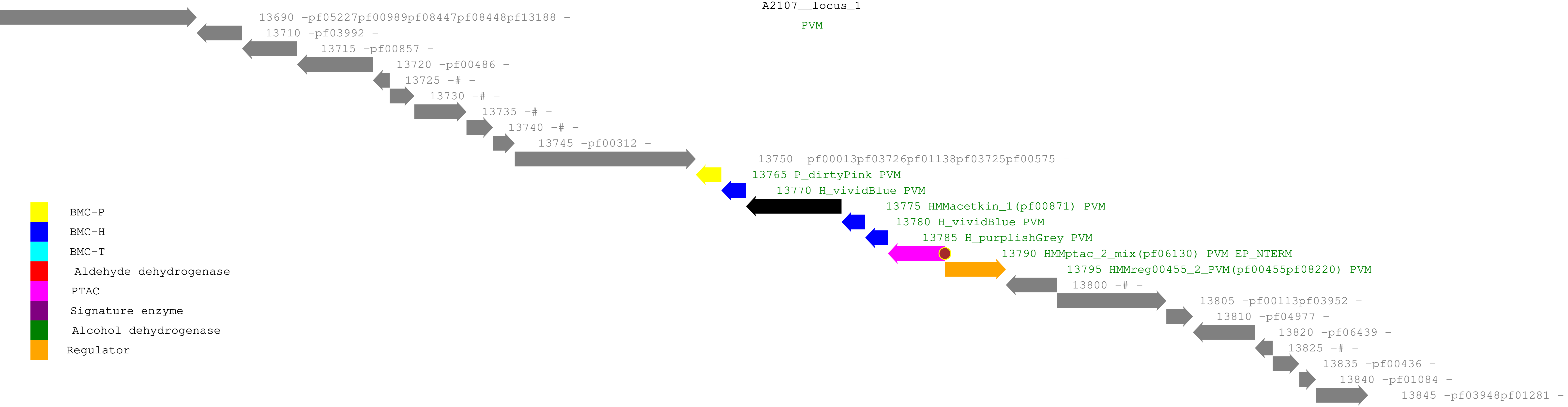

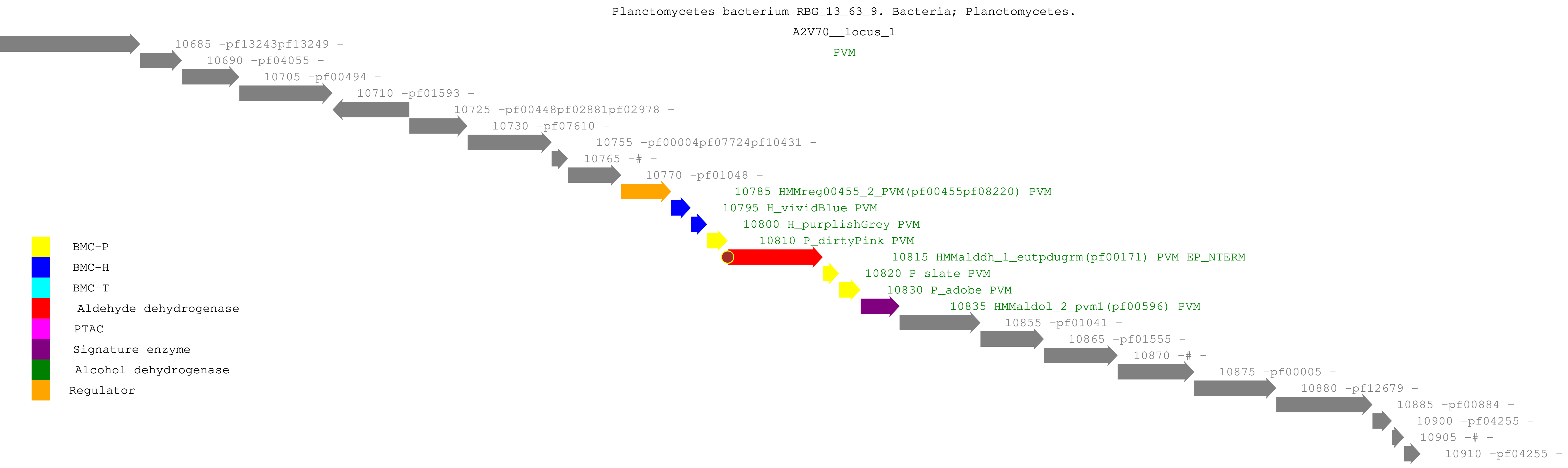
Planctomycetes bacterium RBG_13_63_9. Bacteria; Planctomycetes.
A2V70__locus_1
PVM
10685 -pf13243pf13249 -
10690 -pf04055 -
10705 -pf00494 -
10710 -pf01593 -
10725 -pf00448pf02881pf02978 -
10730 -pf07610 -
10755 -pf00004pf07724pf10431 -
10765 -# -
10770 -pf01048 -
10785 HMMreg00455_2_PVM(pf00455pf08220) PVM
10795 H_vividBlue PVM
10800 H_purplishGrey PVM
10810 P_dirtyPink PVM
10815 HMMalddh_1_eutpdugrm(pf00171) PVM EP_NTERM
10820 P_slate PVM
10830 P_adobe PVM
10835 HMMaldol_2_pvm1(pf00596) PVM
10855 -pf01041 -
10865 -pf01555 -
10870 -# -
10875 -pf00005 -
10880 -pf12679 -
10885 -pf00884 -
10900 -pf04255 -
10905 -# -
10910 -pf04255 -
BMC-P
BMC-H
BMC-T
Aldehyde dehydrogenase
PTAC
Signature enzyme
Alcohol dehydrogenase
Regulator

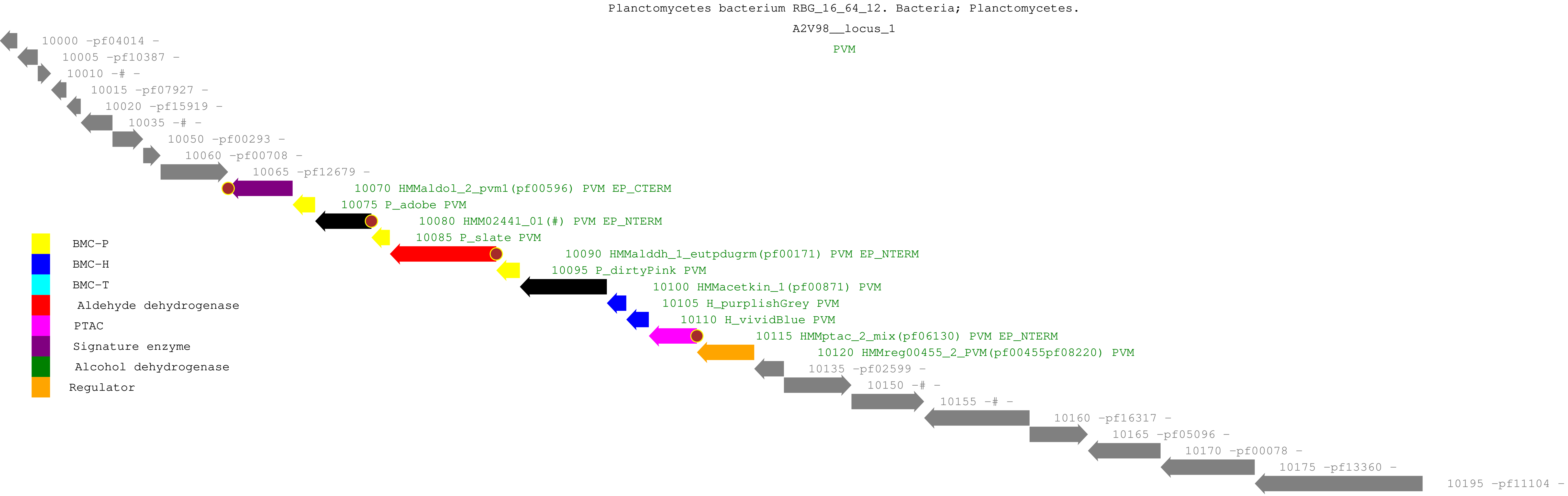

Planctomycetes bacterium RBG_16_64_12. Bacteria; Planctomycetes.
A2V98__locus_1
PVM
10000 -pf04014 -
10005 -pf10387 -
10010 -# -
10015 -pf07927 -
10020 -pf15919 -
10035 -# -
10050 -pf00293 -
10060 -pf00708 -
10065 -pf12679 -
10070 HMMaldol_2_pvm1(pf00596) PVM EP_CTERM
10075 P_adobe PVM
10080 HMM02441_01(#) PVM EP_NTERM
10085 P_slate PVM
10090 HMMalddh_1_eutpdugrm(pf00171) PVM EP_NTERM
10095 P_dirtyPink PVM
10100 HMMacetkin_1(pf00871) PVM
10105 H_purplishGrey PVM
10110 H_vividBlue PVM
10115 HMMptac_2_mix(pf06130) PVM EP_NTERM
10120 HMMreg00455_2_PVM(pf00455pf08220) PVM
10135 -pf02599 -
10150 -# -
10155 -# -
10160 -pf16317 -
10165 -pf05096 -
10170 -pf00078 -
10175 -pf13360 -
10195 -pf11104 -
BMC-P
BMC-H
BMC-T
Aldehyde dehydrogenase
PTAC
Signature enzyme
Alcohol dehydrogenase
Regulator

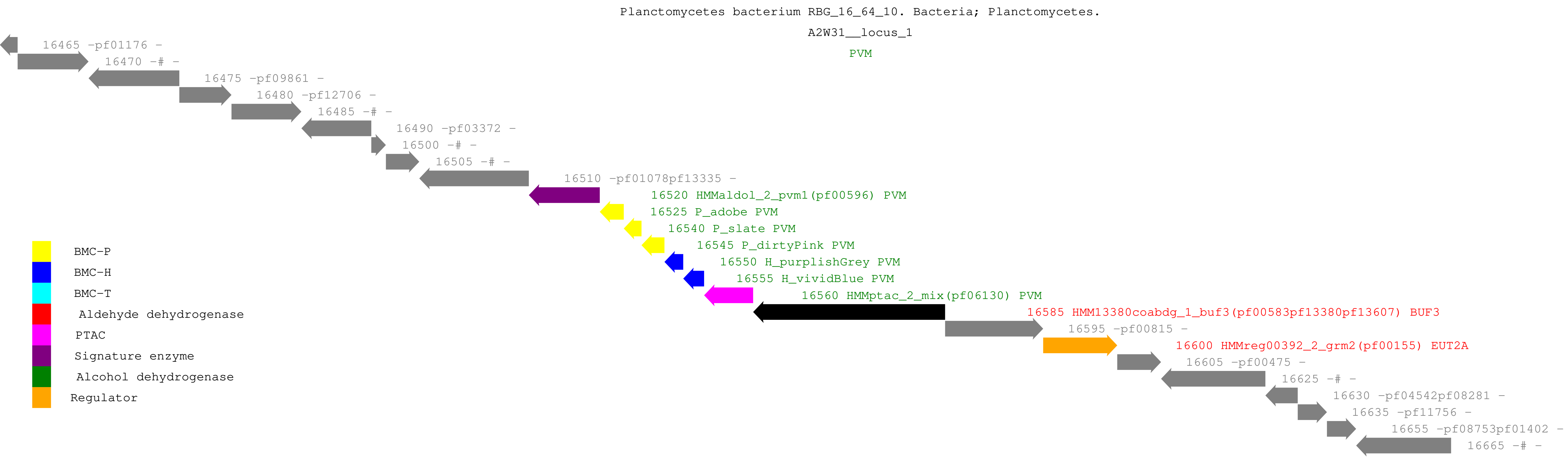
Planctomycetes bacterium RBG_16_64_10. Bacteria; Planctomycetes.
A2W31__locus_1
PVM
16465 -pf01176 -
16470 -# -
16475 -pf09861 -
16480 -pf12706 -
16485 -# -
16490 -pf03372 -
16500 -# -
16505 -# -
16510 -pf01078pf13335 -
16520 HMMaldol_2_pvm1(pf00596) PVM
16525 P_adobe PVM
16540 P_slate PVM
16545 P_dirtyPink PVM
16550 H_purplishGrey PVM
16555 H_vividBlue PVM
16560 HMMptac_2_mix(pf06130) PVM
16585 HMM13380coabdg_1_buf3(pf00583pf13380pf13607) BUF3
16595 -pf00815 -
16600 HMMreg00392_2_grm2(pf00155) EUT2A
16605 -pf00475 -
16625 -# -
16630 -pf04542pf08281 -
16635 -pf11756 -
16655 -pf08753pf01402 -
16665 -# -
BMC-P
BMC-H
BMC-T
Aldehyde dehydrogenase
PTAC
Signature enzyme
Alcohol dehydrogenase
Regulator

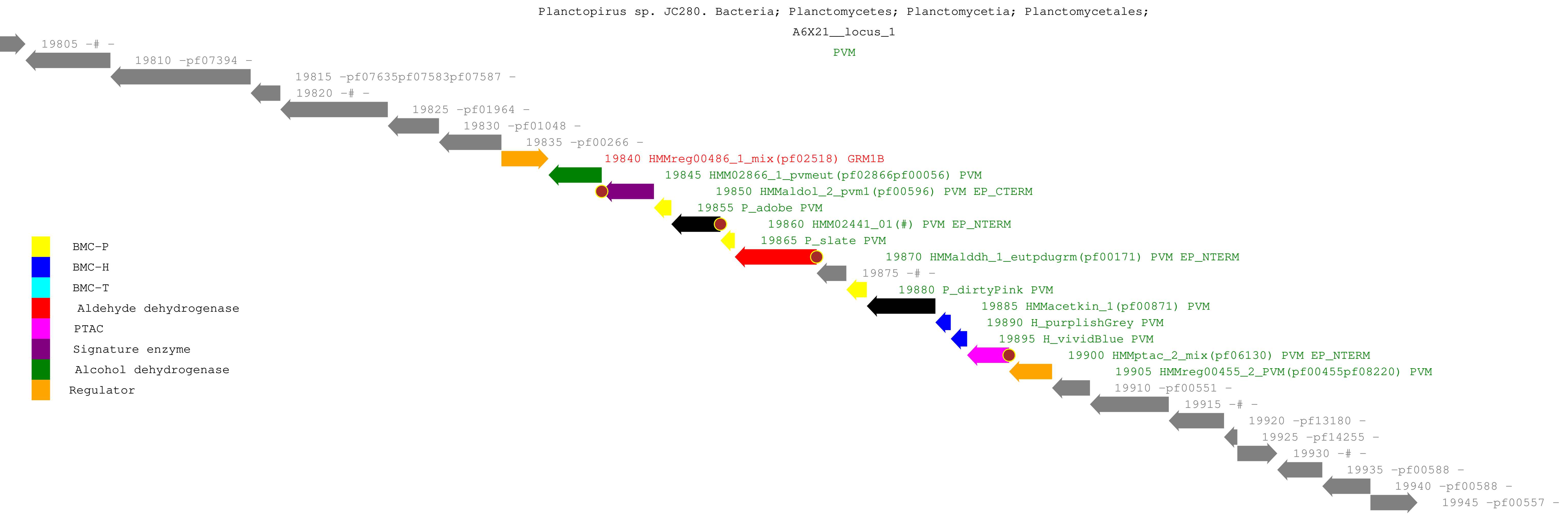
Planctopirus sp. JC280. Bacteria; Planctomycetes; Planctomycetia; Planctomycetales;
A6X21__locus_1
PVM
19805 -# -
19810 -pf07394 -
19815 -pf07635pf07583pf07587 -
19820 -# -
19825 -pf01964 -
19830 -pf01048 -
19835 -pf00266 -
19840 HMMreg00486_1_mix(pf02518) GRM1B
19845 HMM02866_1_pvmeut(pf02866pf00056) PVM
19850 HMMaldol_2_pvm1(pf00596) PVM EP_CTERM
19855 P_adobe PVM
19860 HMM02441_01(#) PVM EP_NTERM
19865 P_slate PVM
19870 HMMalddh_1_eutpdugrm(pf00171) PVM EP_NTERM
19875 -# -
19880 P_dirtyPink PVM
19885 HMMacetkin_1(pf00871) PVM
19890 H_purplishGrey PVM
19895 H_vividBlue PVM
19900 HMMptac_2_mix(pf06130) PVM EP_NTERM
19905 HMMreg00455_2_PVM(pf00455pf08220) PVM
19910 -pf00551 -
19915 -# -
19920 -pf13180 -
19925 -pf14255 -
19930 -# -
19935 -pf00588 -
19940 -pf00588 -
19945 -pf00557 -
BMC-P
BMC-H
BMC-T
Aldehyde dehydrogenase
PTAC
Signature enzyme
Alcohol dehydrogenase
Regulator

Opitutaceae bacterium BACL24 MAG-120322-bin51. Bacteria; Verrucomicrobia; Opitutae; Opitutales; Opitutaceae.

ABS34__locus_1

PVM

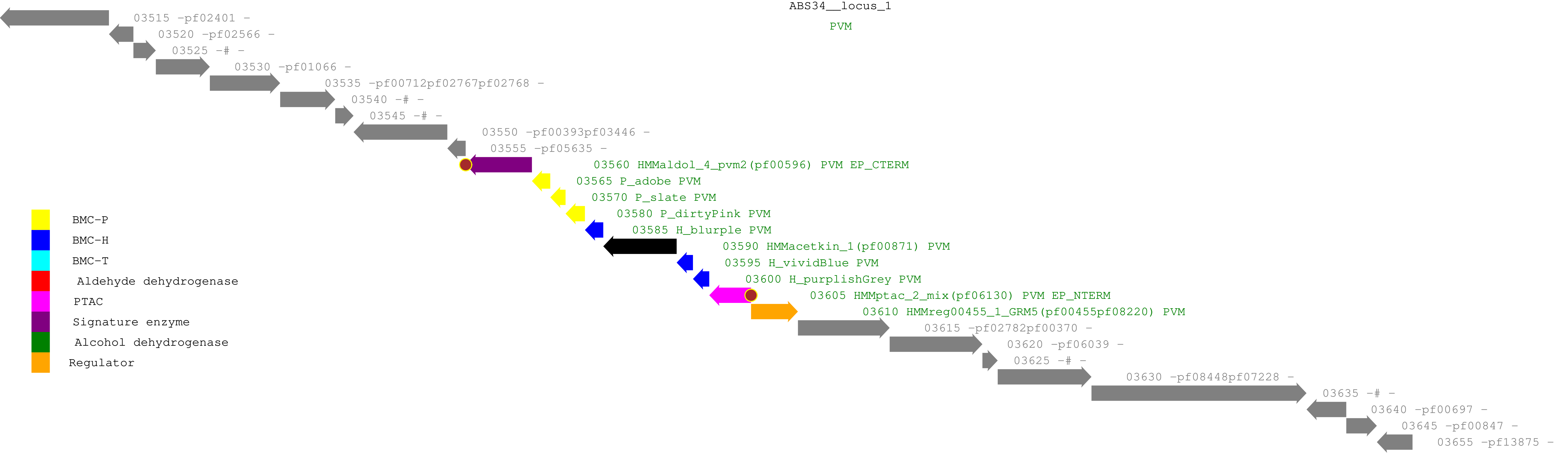

Opitutaceae bacterium TSB47. Bacteria; Verrucomicrobia; Opitutae; Opitutales; Opitutaceae.

AW736__locus_1

PVM

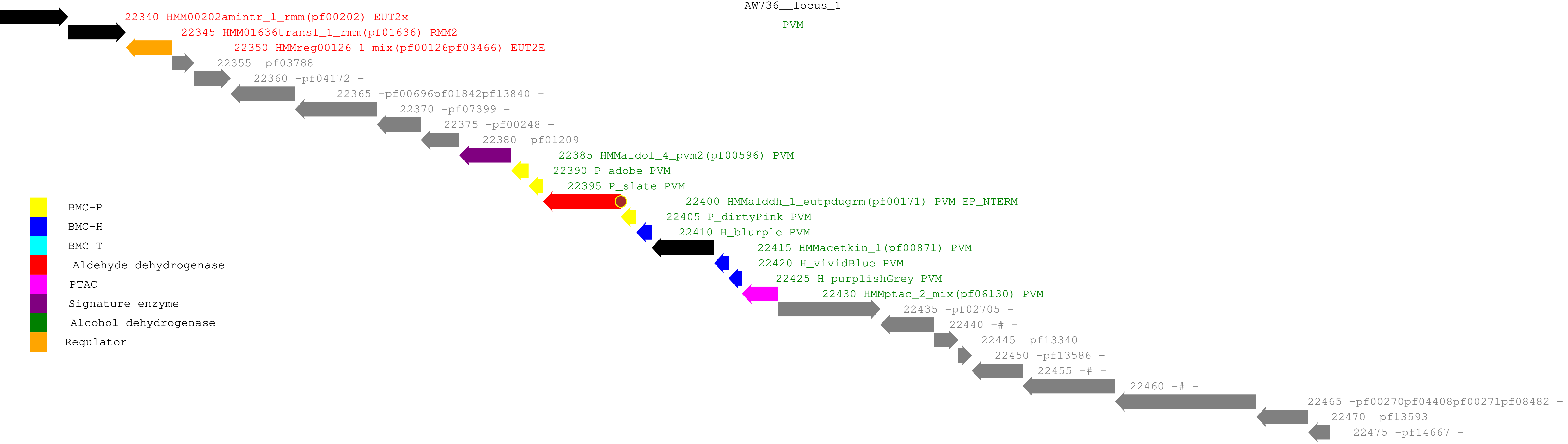

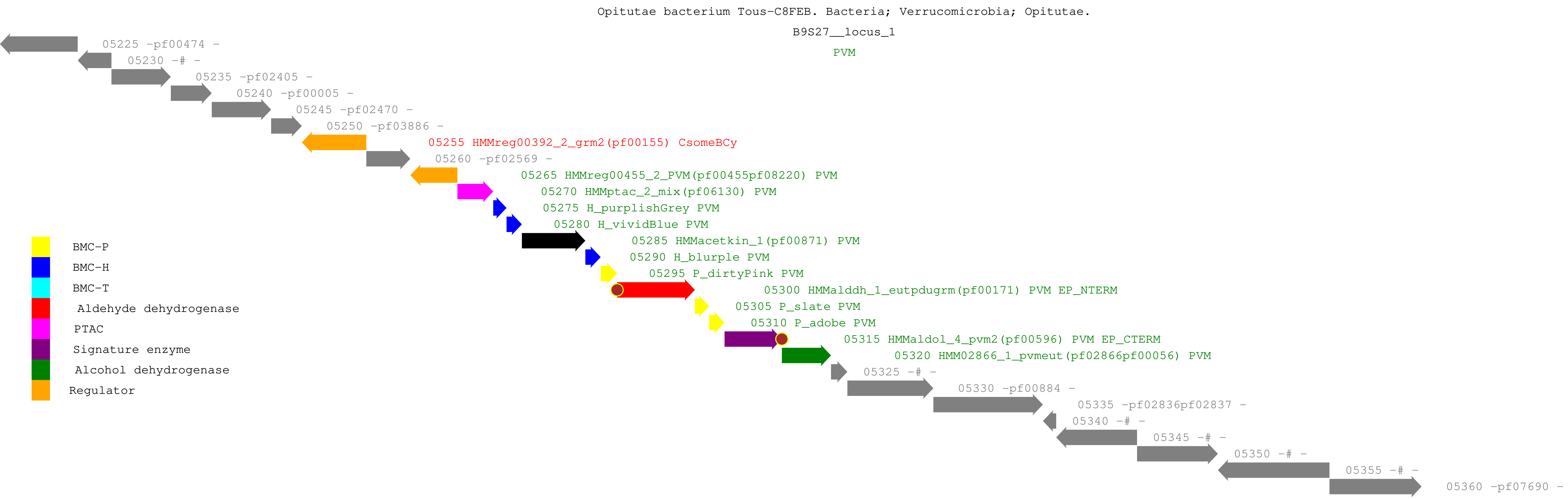
Opitutae bacterium Tous-C8FEB. Bacteria; Verrucomicrobia; Opitutae.
B9S27__locus_1
PVM
05225 -pf00474 -
05230 -# -
05235 -pf02405 -
05240 -pf00005 -
05245 -pf02470 -
05250 -pf03886 -
05255 HMMreg00392_2_grm2(pf00155) CsomeBCy
05260 -pf02569 -
05265 HMMreg00455_2_PVM(pf00455pf08220) PVM
05270 HMMptac_2_mix(pf06130) PVM
05275 H_purplishGrey PVM
05280 H_vividBlue PVM
05285 HMMacetkin_1(pf00871) PVM
05290 H_blurple PVM
05295 P_dirtyPink PVM
05300 HMMaldDH_1_eutpdugrm(pf00171) PVM EP_NTERM
05305 P_slate PVM
05310 P_adobe PVM
05315 HMMaldol_4_pvm2(pf00596) PVM EP_CTERM
05320 HMM02866_1_pvmeut(pf02866pf00056) PVM
05325 -# -
05330 -pf00884 -
05335 -pf02836pf02837 -
05340 -# -
05345 -# -
05350 -# -
05355 -# -
05360 -pf07690 -
BMC-P
BMC-H
BMC-T
Aldehyde dehydrogenase
PTAC
Signature enzyme
Alcohol dehydrogenase
Regulator

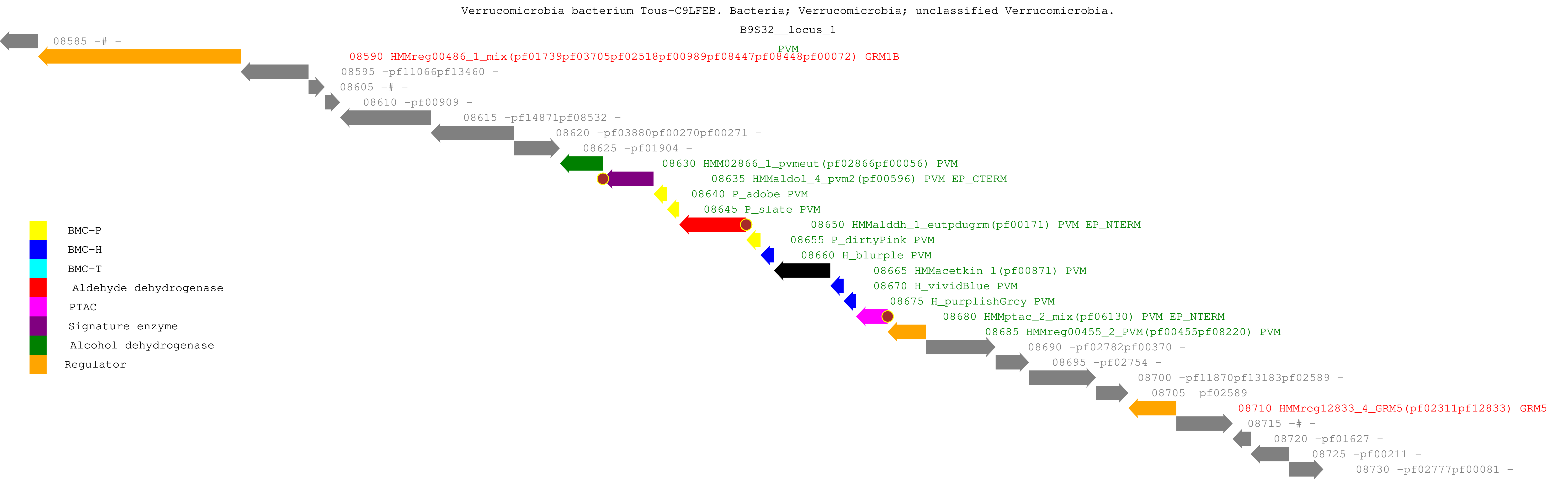
Verrucomicrobia bacterium Tous-C9LFEB. Bacteria; Verrucomicrobia; unclassified Verrucomicrobia.
B9S32__locus_1
PVM
08585 -# -
08590 HMMreg00486_1_mix(pf01739pf03705pf02518pf00989pf08447pf08448pf00072) GRM1B
08595 -pf11066pf13460 -
08605 -# -
08610 -pf00909 -
08615 -pf14871pf08532 -
08620 -pf03880pf00270pf00271 -
08625 -pf01904 -
08630 HMM02866_1_pvmeut(pf02866pf00056) PVM
08635 HMMaldol_4_pvm2(pf00596) PVM EP_CTERM
08640 P_adobe PVM
08645 P_slate PVM
08650 HMMalddh_1_eutpdugrm(pf00171) PVM EP_NTERM
08655 P_dirtyPink PVM
08660 H_blurple PVM
08665 HMMacetkin_1(pf00871) PVM
08670 H_vividBlue PVM
08675 H_purplishGrey PVM
08680 HMMptac_2_mix(pf06130) PVM EP_NTERM
08685 HMMreg00455_2_PVM(pf00455pf08220) PVM
08690 -pf02782pf00370 -
08695 -pf02754 -
08700 -pf11870pf13183pf02589 -
08705 -pf02589 -
08710 HMMreg12833_4_GRM5(pf02311pf12833) GRM5
08715 -# -
08720 -pf01627 -
08725 -pf00211 -
08730 -pf02777pf00081 -
BMC-P
BMC-H
BMC-T
Aldehyde dehydrogenase
PTAC
Signature enzyme
Alcohol dehydrogenase
Regulator

Pedosphaera sp. Tous-C6FEB. Bacteria; Verrucomicrobia; Verrucomicrobiae; Verrucomicrobiales;

B9S33__locus_1

PVM

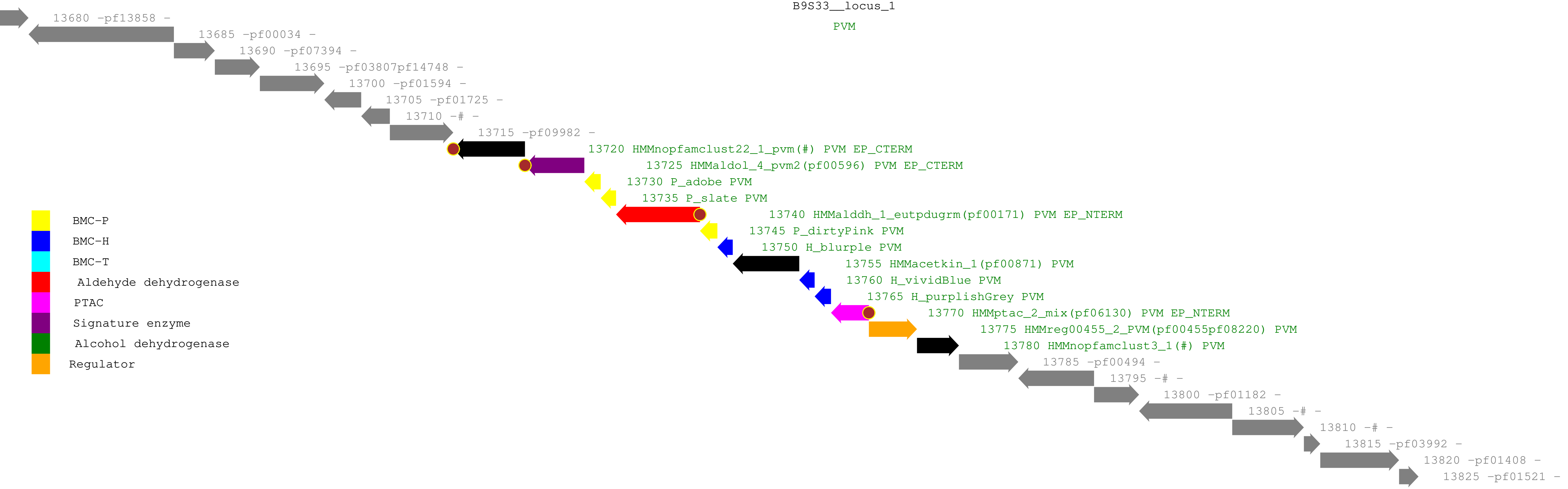

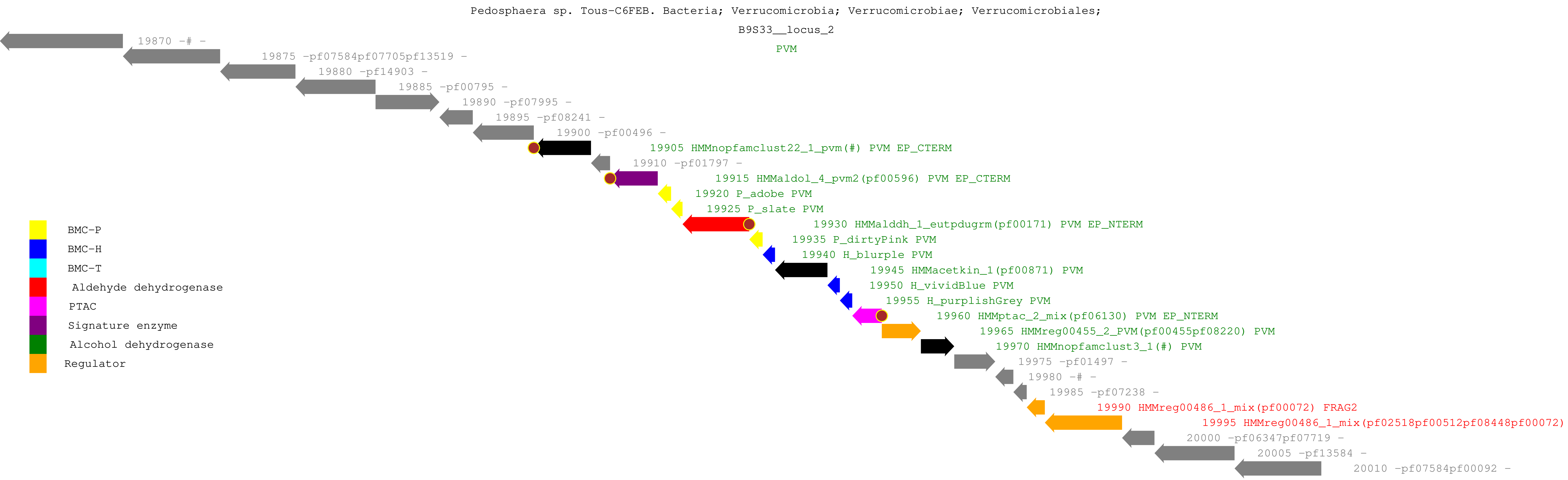
Pedosphaera sp. Tous-C6FEB. Bacteria; Verrucomicrobia; Verrucomicrobiae; Verrucomicrobiales;
B9S33__locus_2
PVM
19870 -# -
19875 -pf07584pf07705pf13519 -
19880 -pf14903 -
19885 -pf00795 -
19890 -pf07995 -
19895 -pf08241 -
19900 -pf00496 -
19905 HMMnopfamclust22_1_pvm(#) PVM EP_CTERM
19910 -pf01797 -
19915 HMMaldol_4_pvm2(pf00596) PVM EP_CTERM
19920 P_adobe PVM
19925 P_slate PVM
19930 HMMaldh_1_eutpdugrm(pf00171) PVM EP_NTERM
19935 P_dirtyPink PVM
19940 H_blurple PVM
19945 HMMacetkin_1(pf00871) PVM
19950 H_vividBlue PVM
19955 H_purplishGrey PVM
19960 HMMptac_2_mix(pf06130) PVM EP_NTERM
19965 HMMreg00455_2_PVM(pf00455pf08220) PVM
19970 HMMnopfamclust3_1(#) PVM
19975 -pf01497 -
19980 -# -
19985 -pf07238 -
19990 HMMreg00486_1_mix(pf00072) FRAG2
19995 HMMreg00486_1_mix(pf02518pf00512pf08448pf00072)
20000 -pf06347pf07719 -
20005 -pf13584 -
20010 -pf07584pf00092 -
BMC-P
BMC-H
BMC-T
Aldehyde dehydrogenase
PTAC
Signature enzyme
Alcohol dehydrogenase
Regulator

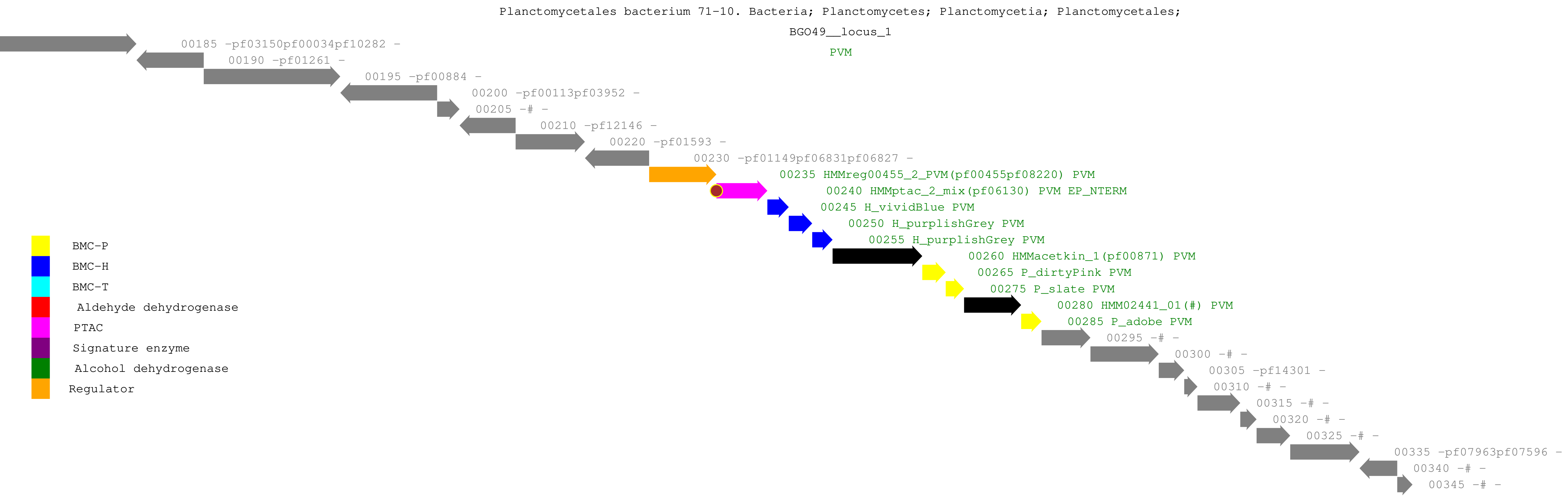
Planctomycetales bacterium 71-10. Bacteria; Planctomycetes; Planctomycetia; Planctomycetales;
BGO49__locus_1
PVM
00185 -pf03150pf00034pf10282 -
00190 -pf01261 -
00195 -pf00884 -
00200 -pf00113pf03952 -
00205 -# -
00210 -pf12146 -
00220 -pf01593 -
00230 -pf01149pf06831pf06827 -
00235 HMMreg00455_2_PVM(pf00455pf08220) PVM
00240 HMMptac_2_mix(pf06130) PVM EP_NTERM
00245 H_vividBlue PVM
00250 H_purplishGrey PVM
00255 H_purplishGrey PVM
00260 HMMacetkin_1(pf00871) PVM
00265 P_dirtyPink PVM
00275 P_slate PVM
00280 HMM02441_01(#) PVM
00285 P_adobe PVM
00295 -# -
00300 -# -
00305 -pf14301 -
00310 -# -
00315 -# -
00320 -# -
00325 -# -
00335 -pf07963pf07596 -
00340 -# -
00345 -# -
BMC-P
BMC-H
BMC-T
Aldehyde dehydrogenase
PTAC
Signature enzyme
Alcohol dehydrogenase
Regulator

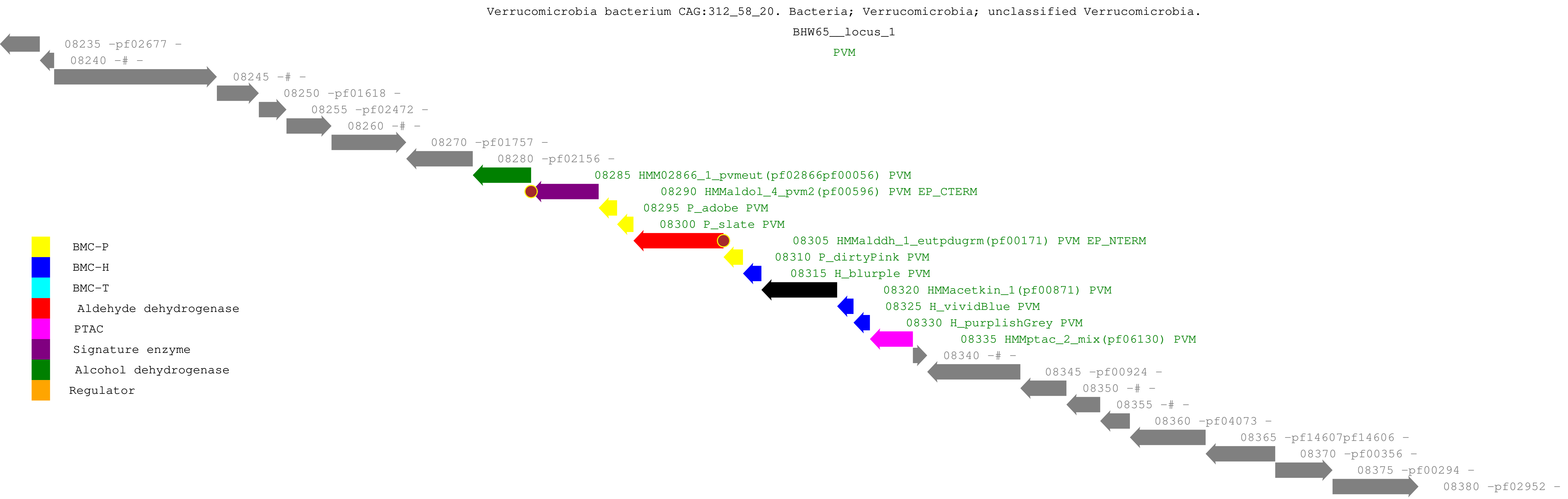
Verrucomicrobia bacterium CAG:312_58_20. Bacteria; Verrucomicrobia; unclassified Verrucomicrobia.
BHW65__locus_1
PVM
08235 -pf02677 -
08240 -# -
08245 -# -
08250 -pf01618 -
08255 -pf02472 -
08260 -# -
08270 -pf01757 -
08280 -pf02156 -
08285 HMM02866_1_pvmeut(pf02866pf00056) PVM
08290 HMMaldol_4_pvm2(pf00596) PVM EP_CTERM
08295 P_adobe PVM
08300 P_slate PVM
08305 HMMalddh_1_eutpdugrm(pf00171) PVM EP_NTERM
08310 P_dirtyPink PVM
08315 H_blurple PVM
08320 HMMacetkin_1(pf00871) PVM
08325 H_vividBlue PVM
08330 H_purplishGrey PVM
08335 HMMptac_2_mix(pf06130) PVM
08340 -# -
08345 -pf00924 -
08350 -# -
08355 -# -
08360 -pf04073 -
08365 -pf14607pf14606 -
08370 -pf00356 -
08375 -pf00294 -
08380 -pf02952 -
BMC-P
BMC-H
BMC-T
Aldehyde dehydrogenase
PTAC
Signature enzyme
Alcohol dehydrogenase
Regulator

Paludisphaera borealis. Bacteria; Planctomycetes; Planctomycetia; Planctomycetales;

BSF38__locus_1

PVM

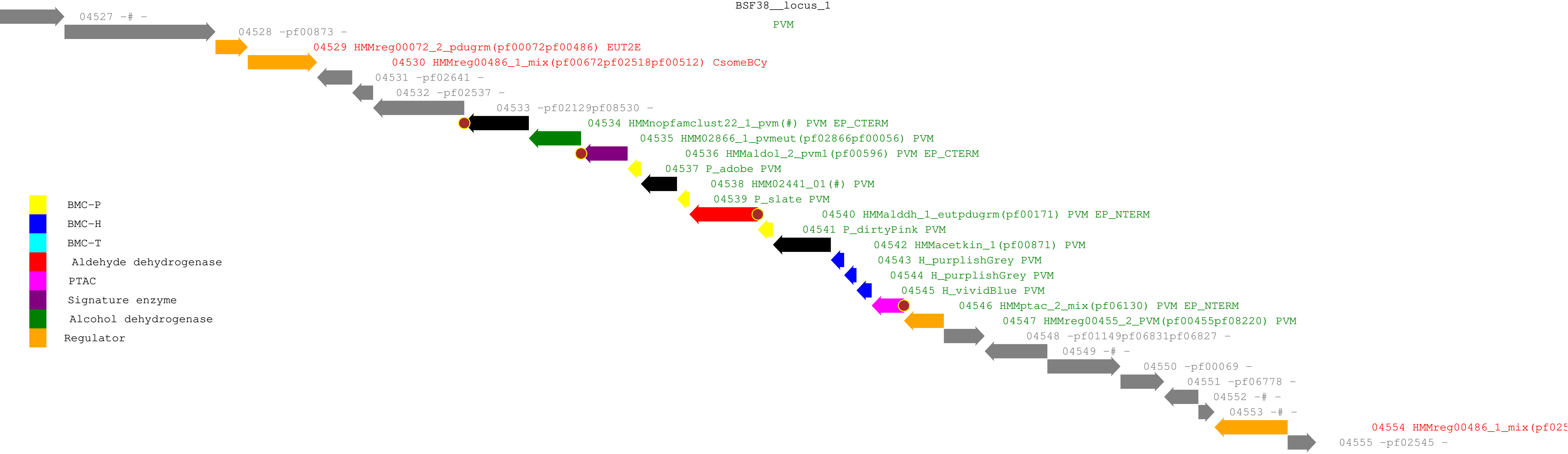

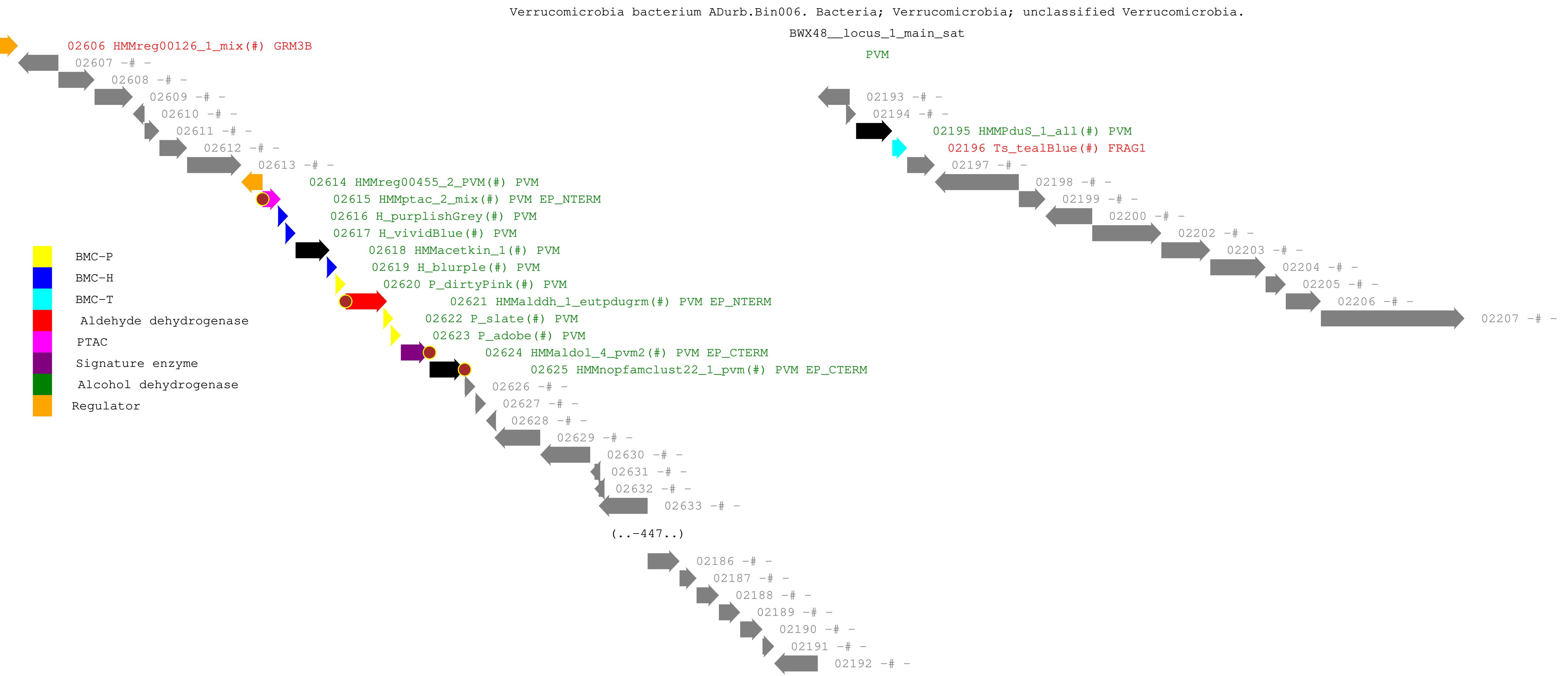

Verrucomicrobia bacterium ADurb.Bin006. Bacteria; Verrucomicrobia; unclassified Verrucomicrobia.
BWX48__locus_1_main_sat
PVM
02606 HMMreg00126_1_mix(#) GRM3B
02607 -# -
02608 -# -
02609 -# -
02610 -# -
02611 -# -
02612 -# -
02613 -# -
02614 HMMreg00455_2_PVM(#) PVM
02615 HMMptac_2_mix(#) PVM EP_NTERM
02616 H_purplishGrey(#) PVM
02617 H_vividBlue(#) PVM
02618 HMMacetkin_1(#) PVM
02619 H_blurple(#) PVM
02620 P_dirtyPink(#) PVM
02621 HMMaldDh_1_eutpdugrm(#) PVM EP_NTERM
02622 P_slate(#) PVM
02623 P_adobe(#) PVM
02624 HMMaldol_4_pvm2(#) PVM EP_CTERM
02625 HMMnopfamclust22_1_pvm(#) PVM EP_CTERM
02626 -# -
02627 -# -
02628 -# -
02629 -# -
02630 -# -
02631 -# -
02632 -# -
02633 -# -
(..-447..)
02186 -# -
02187 -# -
02188 -# -
02189 -# -
02190 -# -
02191 -# -
02192 -# -
02193 -# -
02194 -# -
02195 HMMPduS_1_all(#) PVM
02196 Ts_tealBlue(#) FRAG1
02197 -# -
02198 -# -
02199 -# -
02200 -# -
02202 -# -
02203 -# -
02204 -# -
02205 -# -
02206 -# -
02207 -# -
BMC-P
BMC-H
BMC-T
Aldehyde dehydrogenase
PTAC
Signature enzyme
Alcohol dehydrogenase
Regulator

Verrucomicrobia bacterium ADurb.Bin118. Bacteria; Verrucomicrobia; unclassified Verrucomicrobia.

BWX84__locus_1_main_sat

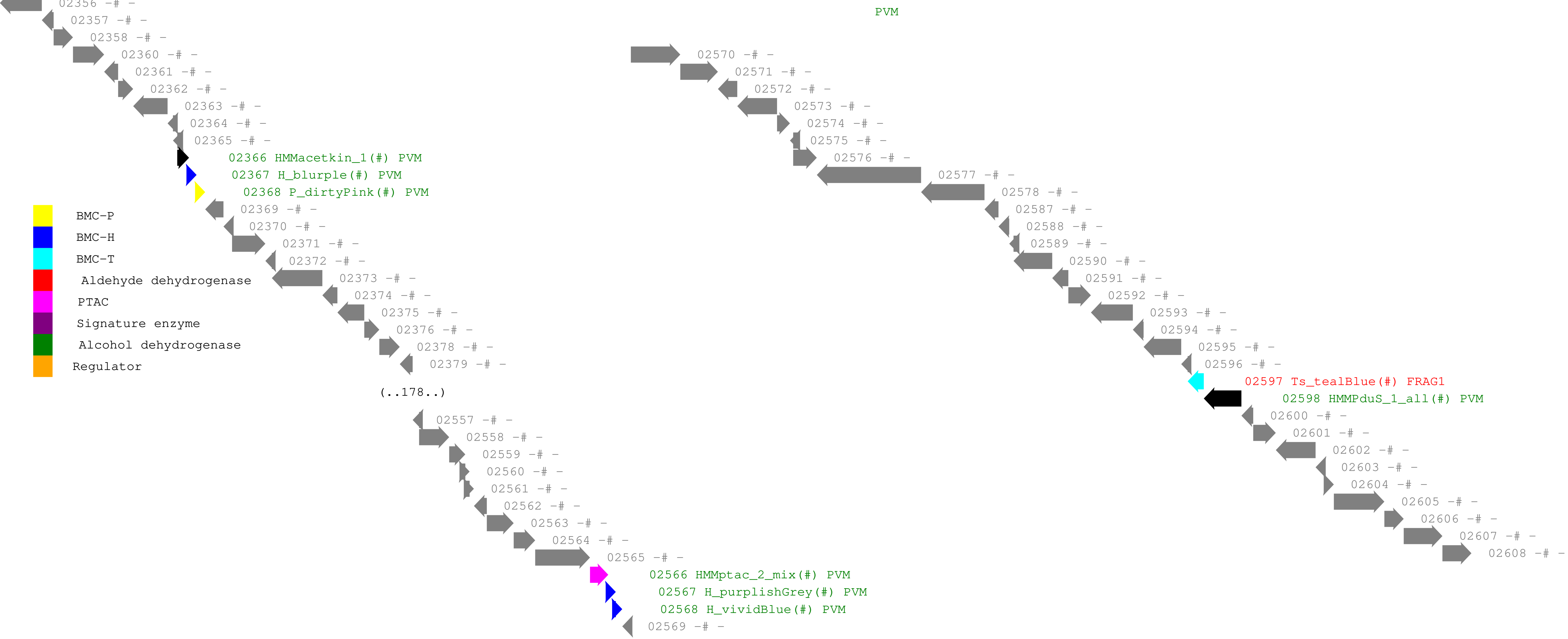

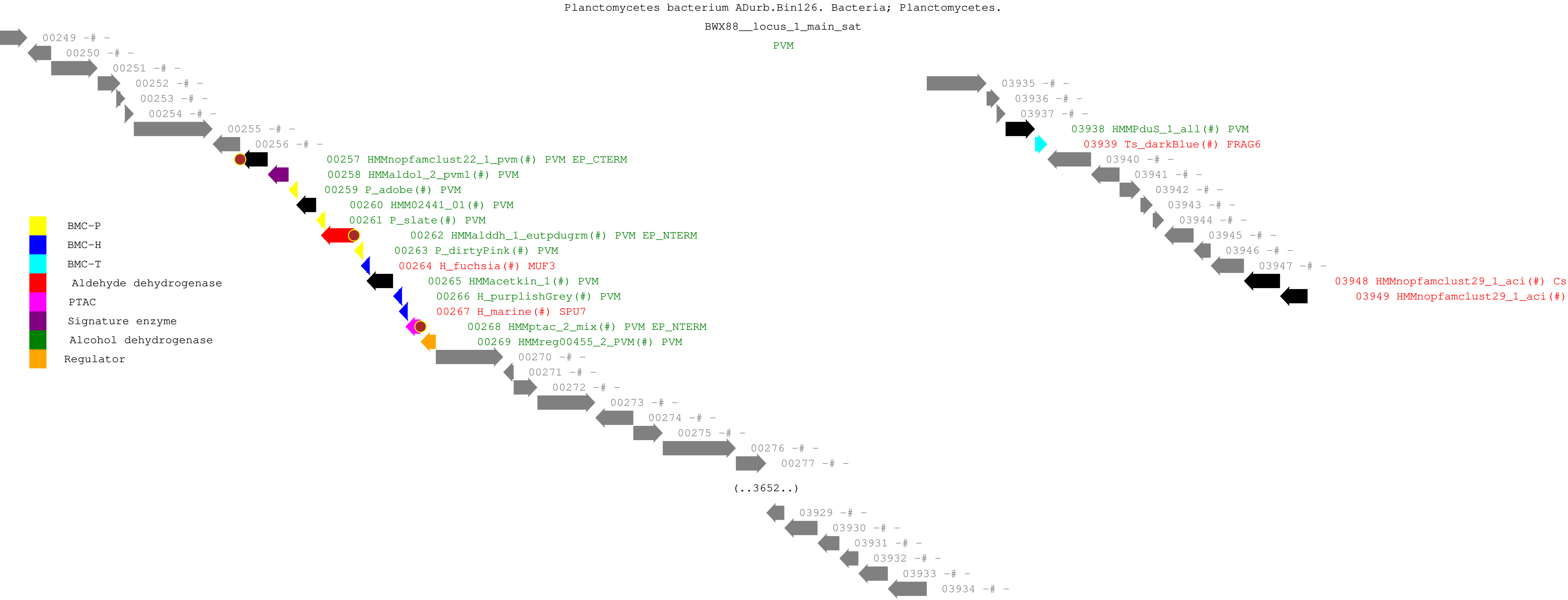

Planctomycetes bacterium ADurb.Bin126. Bacteria; Planctomycetes.
BWX88__locus_1_main_sat
PVM
00249 -# -
00250 -# -
00251 -# -
00252 -# -
00253 -# -
00254 -# -
00255 -# -
00256 -# -
00257 HMMnopfamclust22_1_pvm(#) PVM EP_CTERM
00258 HMMaldol_2_pvm1(#) PVM
00259 P_adobe(#) PVM
00260 HMM02441_01(#) PVM
00261 P_slate(#) PVM
00262 HMMalddh_1_eutpdugrm(#) PVM EP_NTERM
00263 P_dirtyPink(#) PVM
00264 H_fuchsia(#) MUF3
00265 HMMacetkin_1(#) PVM
00266 H_purplishGrey(#) PVM
00267 H_marine(#) SPU7
00268 HMMptac_2_mix(#) PVM EP_NTERM
00269 HMMreg00455_2_PVM(#) PVM
00270 -# -
00271 -# -
00272 -# -
00273 -# -
00274 -# -
00275 -# -
00276 -# -
00277 -# -
(..3652..)
03929 -# -
03930 -# -
03931 -# -
03932 -# -
03933 -# -
03934 -# -
03935 -# -
03936 -# -
03937 -# -
03938 HMMPduS_1_all(#) PVM
03939 Ts_darkBlue(#) FRAG6
03940 -# -
03941 -# -
03942 -# -
03943 -# -
03944 -# -
03945 -# -
03946 -# -
03947 -# -
03948 HMMnopfamclust29_1_aci(#) Cs
03949 HMMnopfamclust29_1_aci(#)
BMC-P
BMC-H
BMC-T
Aldehyde dehydrogenase
PTAC
Signature enzyme
Alcohol dehydrogenase
Regulator

Verrucomicrobia bacterium. Bacteria; Verrucomicrobia; unclassified Verrucomicrobia.

C5B50__locus_1

PVM

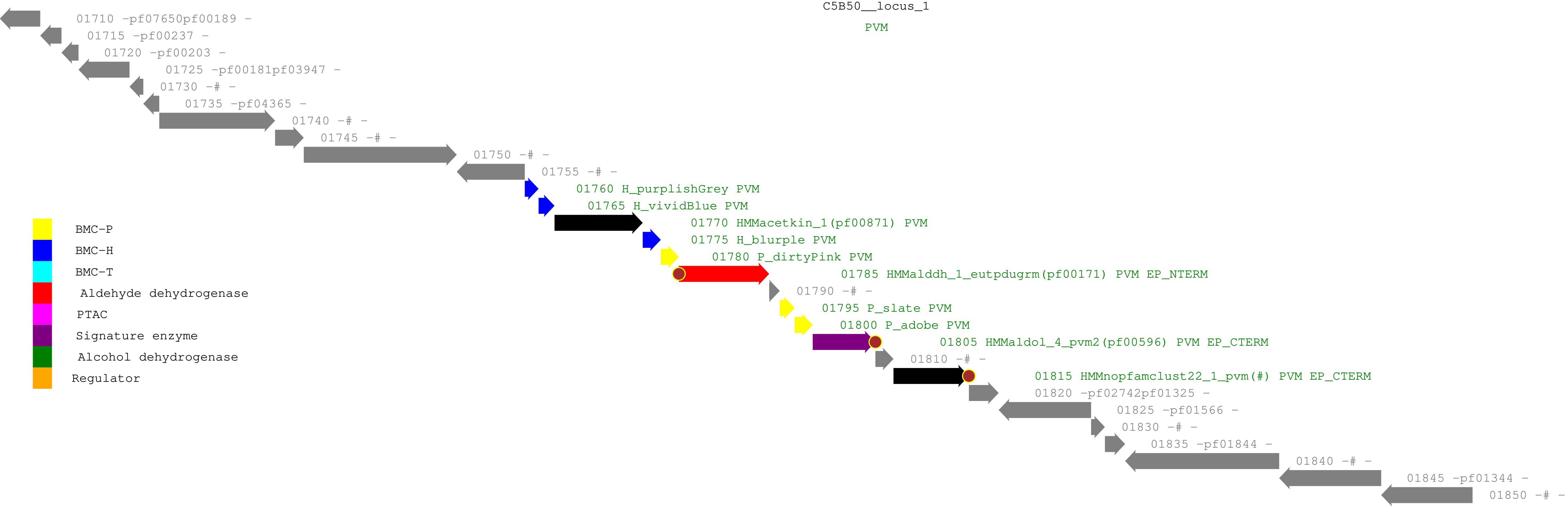

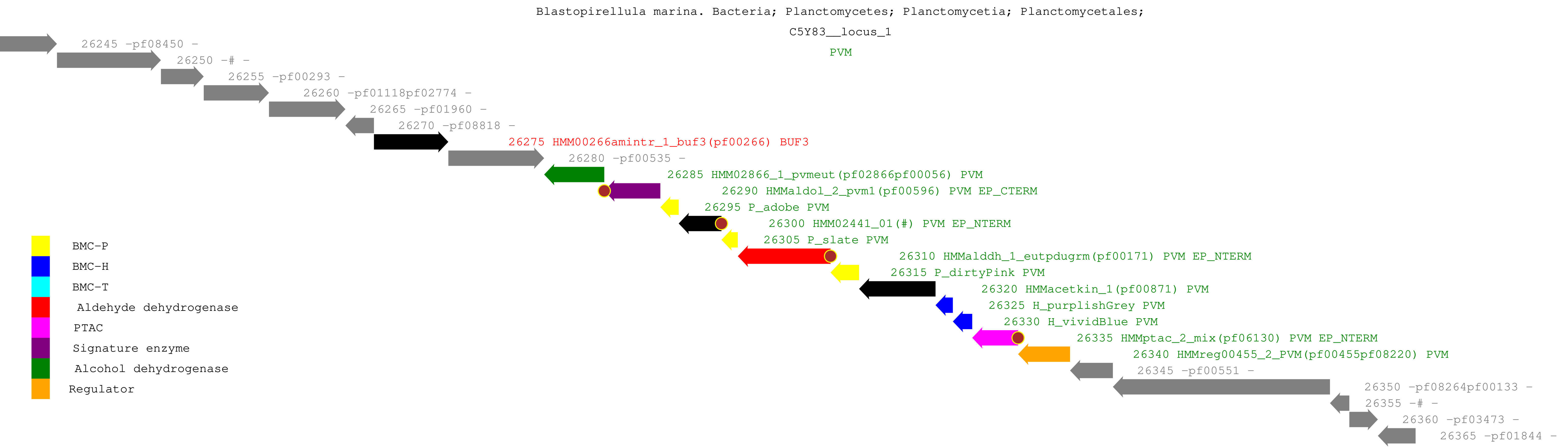
Blastopirellula marina. Bacteria; Planctomycetes; Planctomycetia; Planctomycetales;
C5Y83__locus_1
PVM
26245 -pf08450 -
26250 -# -
26255 -pf00293 -
26260 -pf01118pf02774 -
26265 -pf01960 -
26270 -pf08818 -
26275 HMM00266amintr_1_buf3(pf00266) BUF3
26280 -pf00535 -
26285 HMM02866_1_pvmeut(pf02866pf00056) PVM
26290 HMMaldol_2_pvm1(pf00596) PVM EP_CTERM
26295 P_adobe PVM
26300 HMM02441_01(#) PVM EP_NTERM
26305 P_slate PVM
26310 HMMalddh_1_eutpdugrm(pf00171) PVM EP_NTERM
26315 P_dirtyPink PVM
26320 HMMacetkin_1(pf00871) PVM
26325 H_purplishGrey PVM
26330 H_vividBlue PVM
26335 HMMptac_2_mix(pf06130) PVM EP_NTERM
26340 HMMreg00455_2_PVM(pf00455pf08220) PVM
26345 -pf00551 -
26350 -pf08264pf00133 -
26355 -# -
26360 -pf03473 -
26365 -pf01844 -
BMC-P
BMC-H
BMC-T
Aldehyde dehydrogenase
PTAC
Signature enzyme
Alcohol dehydrogenase
Regulator

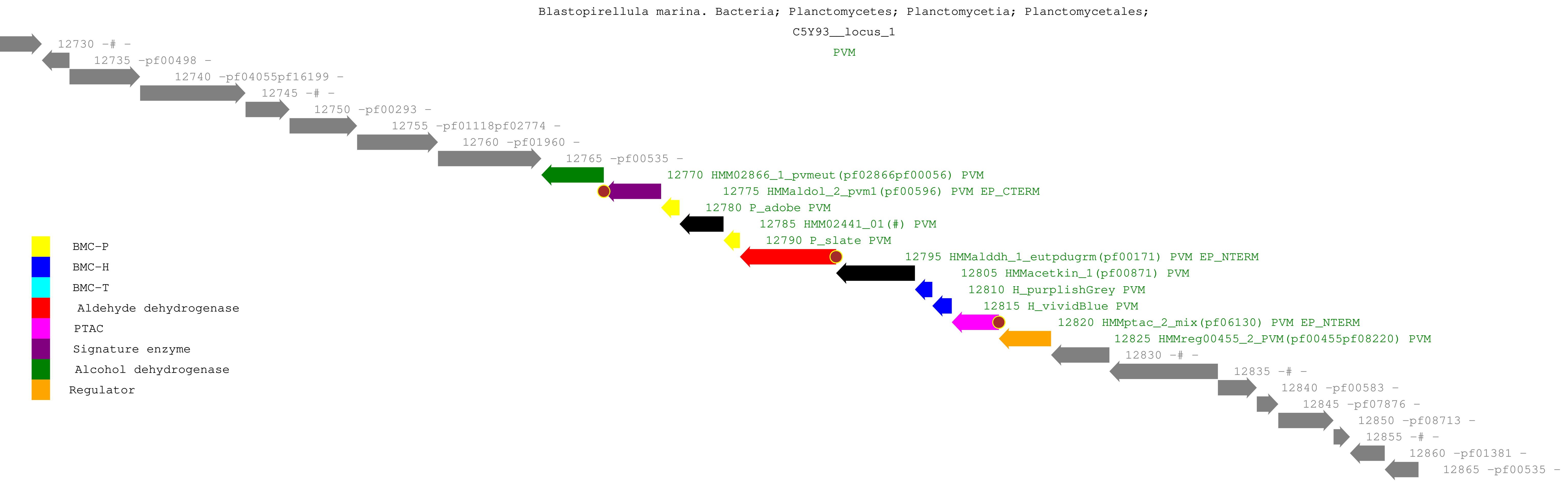
Blastopirellula marina. Bacteria; Planctomycetes; Planctomycetia; Planctomycetales;
C5Y93__locus_1
PVM
12730 -# -
12735 -pf00498 -
12740 -pf04055pf16199 -
12745 -# -
12750 -pf00293 -
12755 -pf01118pf02774 -
12760 -pf01960 -
12765 -pf00535 -
12770 HMM02866_1_pvmeut(pf02866pf00056) PVM
12775 HMMaldol_2_pvm1(pf00596) PVM EP_CTERM
12780 P_adobe PVM
12785 HMM02441_01(#) PVM
12790 P_slate PVM
12795 HMMalddh_1_eutpdugrm(pf00171) PVM EP_NTERM
12805 HMMacetkin_1(pf00871) PVM
12810 H_purplishGrey PVM
12815 H_vividBlue PVM
12820 HMMptac_2_mix(pf06130) PVM EP_NTERM
12825 HMMreg00455_2_PVM(pf00455pf08220) PVM
12830 -# -
12835 -# -
12840 -pf00583 -
12845 -pf07876 -
12850 -pf08713 -
12855 -# -
12860 -pf01381 -
12865 -pf00535 -
BMC-P
BMC-H
BMC-T
Aldehyde dehydrogenase
PTAC
Signature enzyme
Alcohol dehydrogenase
Regulator

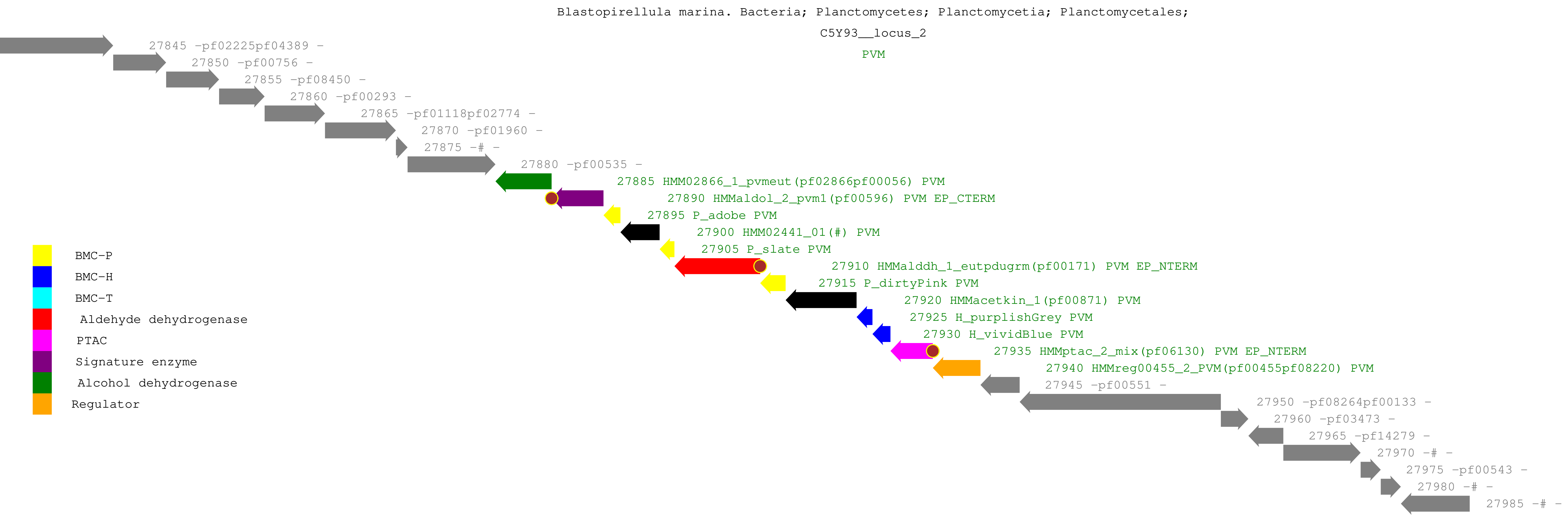
Blastopirellula marina. Bacteria; Planctomycetes; Planctomycetia; Planctomycetales;
C5Y93__locus_2
PVM
27845 -pf02225pf04389 -
27850 -pf00756 -
27855 -pf08450 -
27860 -pf00293 -
27865 -pf01118pf02774 -
27870 -pf01960 -
27875 -# -
27880 -pf00535 -
27885 HMM02866_1_pvmeut(pf02866pf00056) PVM
27890 HMMaldol_2_pvm1(pf00596) PVM EP_CTERM
27895 P_adobe PVM
27900 HMM02441_01(#) PVM
27905 P_slate PVM
27910 HMMalddh_1_eutpdugrm(pf00171) PVM EP_NTERM
27915 P_dirtyPink PVM
27920 HMMacetkin_1(pf00871) PVM
27925 H_purplishGrey PVM
27930 H_vividBlue PVM
27935 HMMptac_2_mix(pf06130) PVM EP_NTERM
27940 HMMreg00455_2_PVM(pf00455pf08220) PVM
27945 -pf00551 -
27950 -pf08264pf00133 -
27960 -pf03473 -
27965 -pf14279 -
27970 -# -
27975 -pf00543 -
27980 -# -
27985 -# -
BMC-P
BMC-H
BMC-T
Aldehyde dehydrogenase
PTAC
Signature enzyme
Alcohol dehydrogenase
Regulator

Blastopirellula marina. Bacteria; Planctomycetes; Planctomycetia; Planctomycetales;

C5Y96__locus_1

PVM

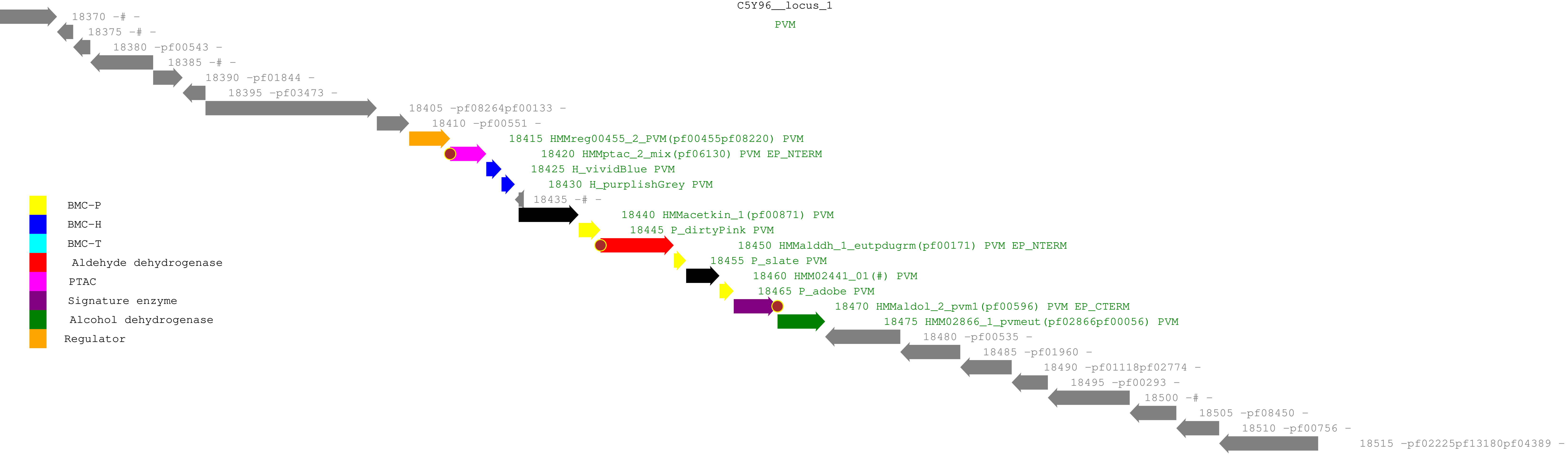

Blastopirellula marina. Bacteria; Planctomycetes; Planctomycetia; Planctomycetales;
C5Y98__locus_1
PVM

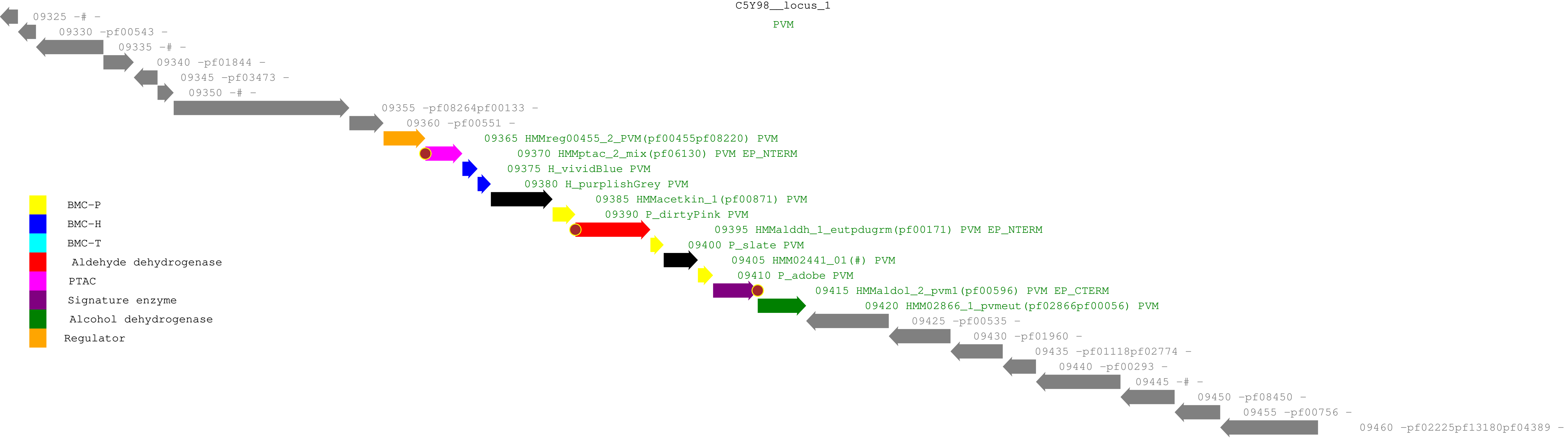

Planctomycetes bacterium CA11. Bacteria; Planctomycetes.

CA11__locus_1

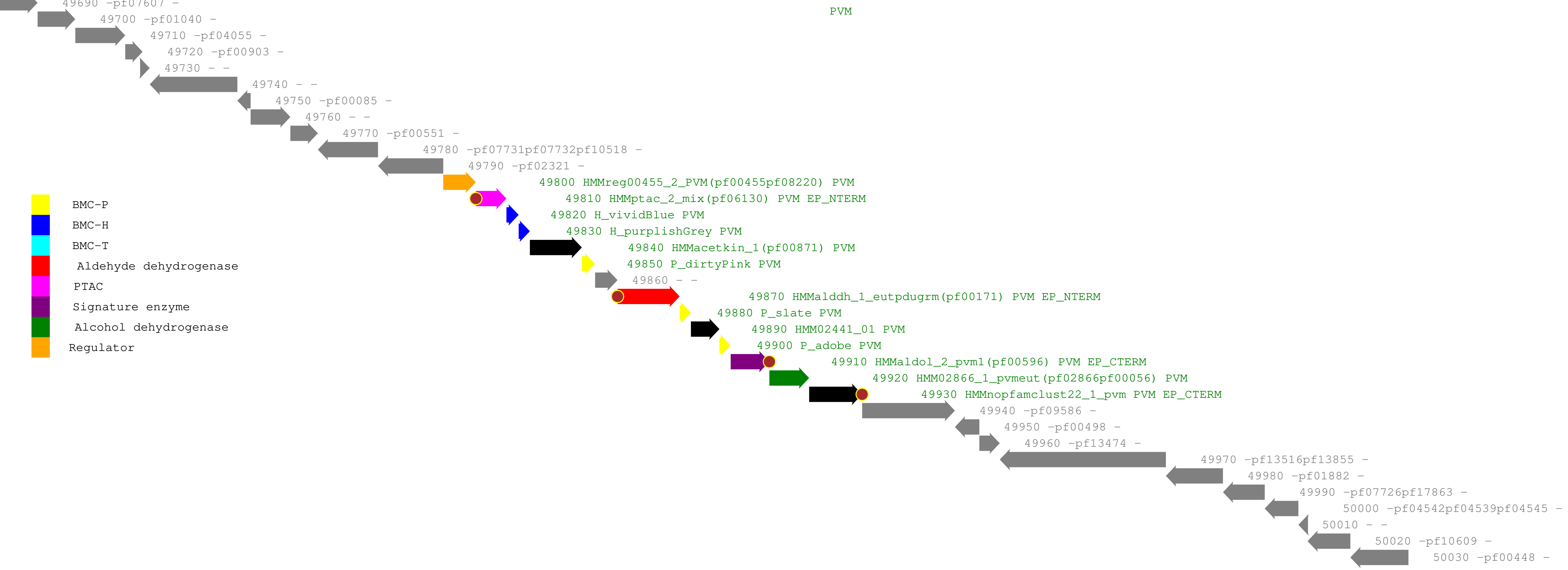

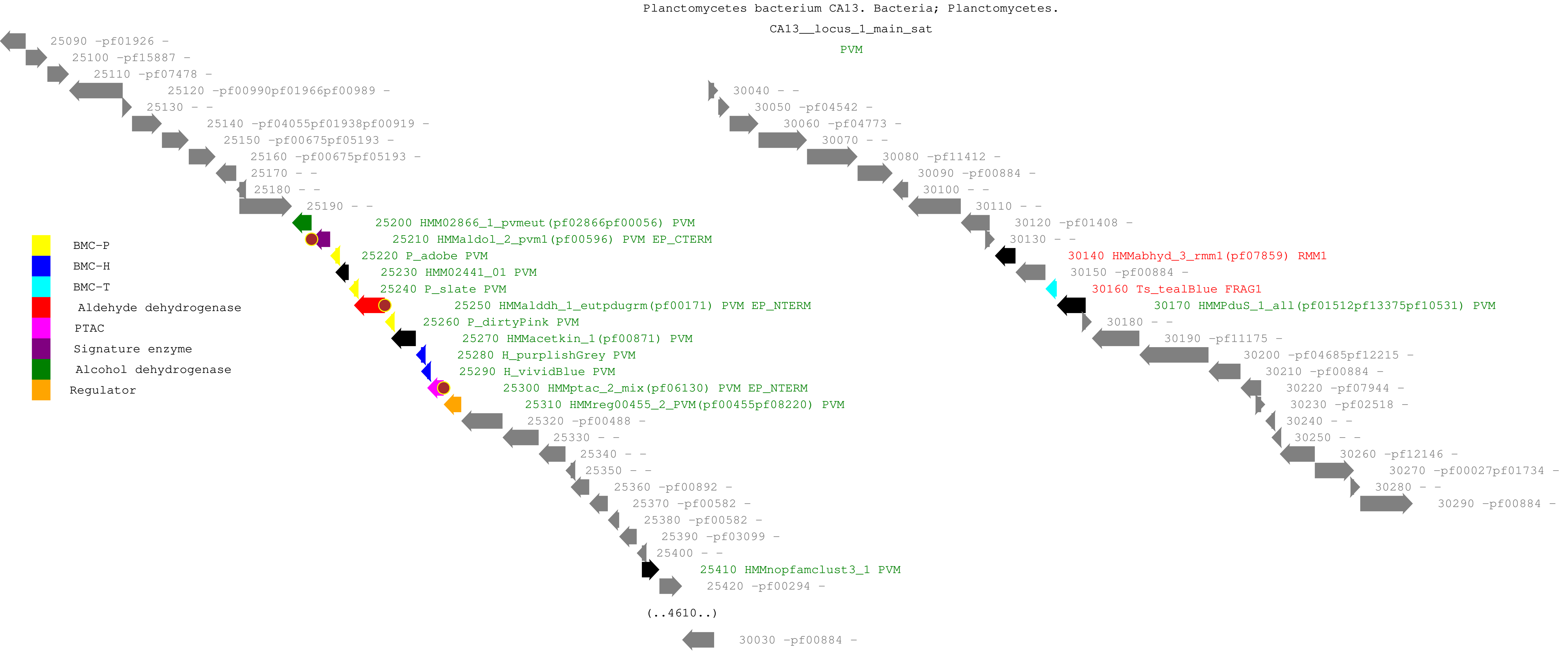

Planctomycetes bacterium CA13. Bacteria; Planctomycetes.
CA13__locus_1_main_sat
PVM
25090 -pf01926 -
25100 -pf15887 -
25110 -pf07478 -
25120 -pf00990pf01966pf00989 -
25130 - -
25140 -pf04055pf01938pf00919 -
25150 -pf00675pf05193 -
25160 -pf00675pf05193 -
25170 - -
25180 - -
25190 - -
25200 HMM02866_1_pvmeut(pf02866pf00056) PVM
25210 HMMaldol_2_pvm1(pf00596) PVM EP_CTERM
25220 P_adobe PVM
25230 HMM02441_01 PVM
25240 P_slate PVM
25250 HMMalddh_1_eutpdugrm(pf00171) PVM EP_NTERM
25260 P_dirtyPink PVM
25270 HMMacetkin_1(pf00871) PVM
25280 H_purplishGrey PVM
25290 H_vividBlue PVM
25300 HMMptac_2_mix(pf06130) PVM EP_NTERM
25310 HMMreg00455_2_PVM(pf00455pf08220) PVM
25320 -pf00488 -
25330 - -
25340 - -
25350 - -
25360 -pf00892 -
25370 -pf00582 -
25380 -pf00582 -
25390 -pf03099 -
25400 - -
25410 HMMnopfamclust3_1 PVM
25420 -pf00294 -
(..4610..)
30030 -pf00884 -
30040 - -
30050 -pf04542 -
30060 -pf04773 -
30070 - -
30080 -pf11412 -
30090 -pf00884 -
30100 - -
30110 - -
30120 -pf01408 -
30130 - -
30140 HMMabhyd_3_rmm1(pf07859) RMM1
30150 -pf00884 -
30160 Ts_tealBlue FRAG1
30170 HMMPduS_1_all(pf01512pf13375pf10531) PVM
30180 - -
30190 -pf11175 -
30200 -pf04685pf12215 -
30210 -pf00884 -
30220 -pf07944 -
30230 -pf02518 -
30240 - -
30250 - -
30260 -pf12146 -
30270 -pf00027pf01734 -
30280 - -
30290 -pf00884 -
BMC-P
BMC-H
BMC-T
Aldehyde dehydrogenase
PTAC
Signature enzyme
Alcohol dehydrogenase
Regulator

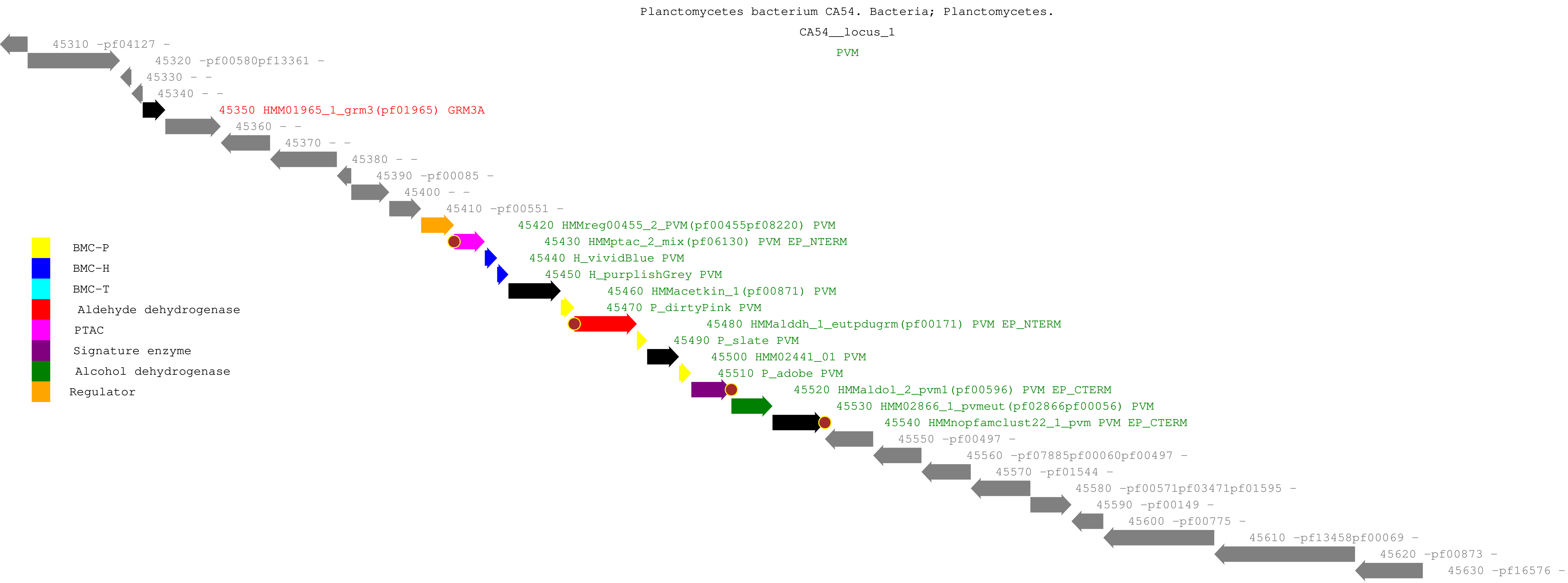

Planctomycetes bacterium CA54. Bacteria; Planctomycetes.
CA54__locus_1
PVM
45310 -pf04127 -
45320 -pf00580pf13361 -
45330 - -
45340 - -
45350 HMM01965_1_grm3(pf01965) GRM3A
45360 - -
45370 - -
45380 - -
45390 -pf00085 -
45400 - -
45410 -pf00551 -
45420 HMMreg00455_2_PVM(pf00455pf08220) PVM
45430 HMMptac_2_mix(pf06130) PVM EP_NTERM
45440 H_vividBlue PVM
45450 H_purplishGrey PVM
45460 HMMacetkin_1(pf00871) PVM
45470 P_dirtyPink PVM
45480 HMMalddh_1_eutpdugrm(pf00171) PVM EP_NTERM
45490 P_slate PVM
45500 HMM02441_01 PVM
45510 P_adobe PVM
45520 HMMaldol_2_pvm1(pf00596) PVM EP_CTERM
45530 HMM02866_1_pvmeut(pf02866pf00056) PVM
45540 HMMnopfamclust22_1_pvm PVM EP_CTERM
45550 -pf00497 -
45560 -pf07885pf00060pf00497 -
45570 -pf01544 -
45580 -pf00571pf03471pf01595 -
45590 -pf00149 -
45600 -pf00775 -
45610 -pf13458pf00069 -
45620 -pf00873 -
45630 -pf16576 -
BMC-P
BMC-H
BMC-T
Aldehyde dehydrogenase
PTAC
Signature enzyme
Alcohol dehydrogenase
Regulator

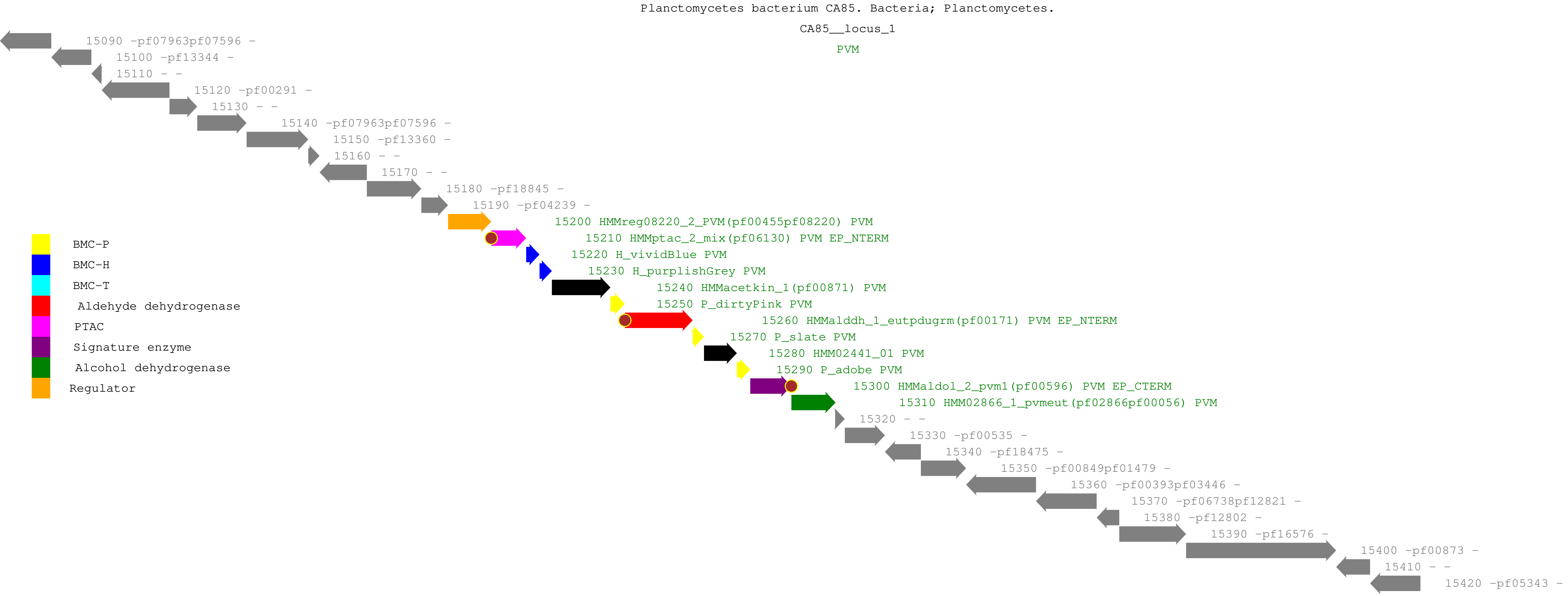

Planctomycetes bacterium CA85. Bacteria; Planctomycetes.
CA85__locus_1
PVM
15090 -pf07963pf07596 -
15100 -pf13344 -
15110 - -
15120 -pf00291 -
15130 - -
15140 -pf07963pf07596 -
15150 -pf13360 -
15160 - -
15170 - -
15180 -pf18845 -
15190 -pf04239 -
15200 HMMreg08220_2_PVM(pf00455pf08220) PVM
15210 HMMptac_2_mix(pf06130) PVM EP_NTERM
15220 H_vividBlue PVM
15230 H_purplishGrey PVM
15240 HMMacetkin_1(pf00871) PVM
15250 P_dirtyPink PVM
15260 HMMalddh_1_eutpdugrm(pf00171) PVM EP_NTERM
15270 P_slate PVM
15280 HMM02441_01 PVM
15290 P_adobe PVM
15300 HMMaldol_2_pvm1(pf00596) PVM EP_CTERM
15310 HMM02866_1_pvmeut(pf02866pf00056) PVM
15320 - -
15330 -pf00535 -
15340 -pf18475 -
15350 -pf00849pf01479 -
15360 -pf00393pf03446 -
15370 -pf06738pf12821 -
15380 -pf12802 -
15390 -pf16576 -
15400 -pf00873 -
15410 - -
15420 -pf05343 -
BMC-P
BMC-H
BMC-T
Aldehyde dehydrogenase
PTAC
Signature enzyme
Alcohol dehydrogenase
Regulator

Spartobacteria bacterium AMD-G5. Bacteria; Verrucomicrobia; Spartobacteria.

CAK85__locus_1

PVM

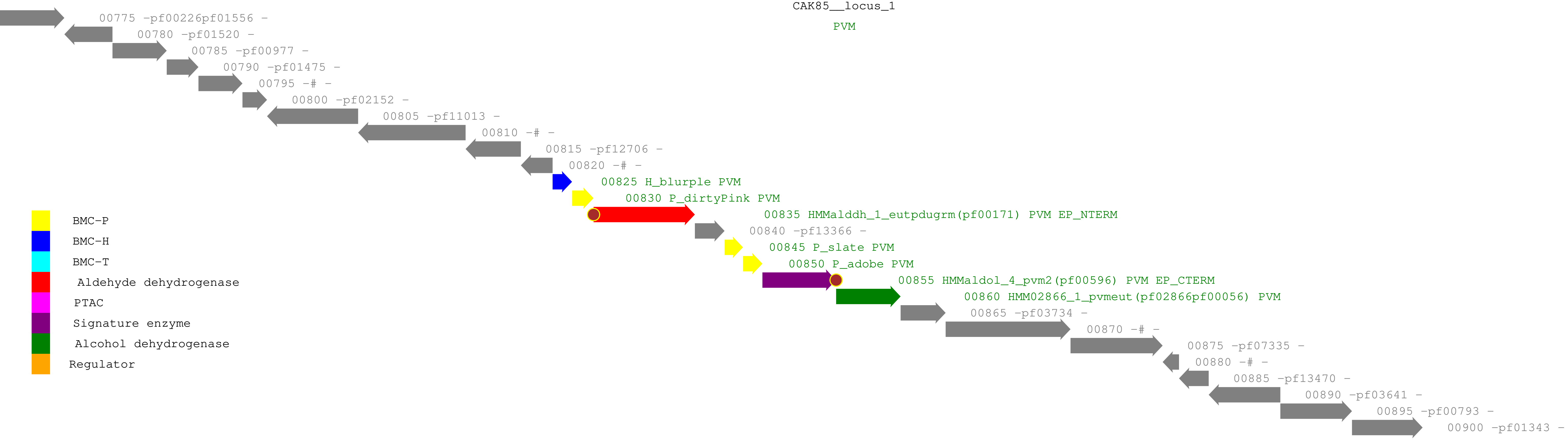

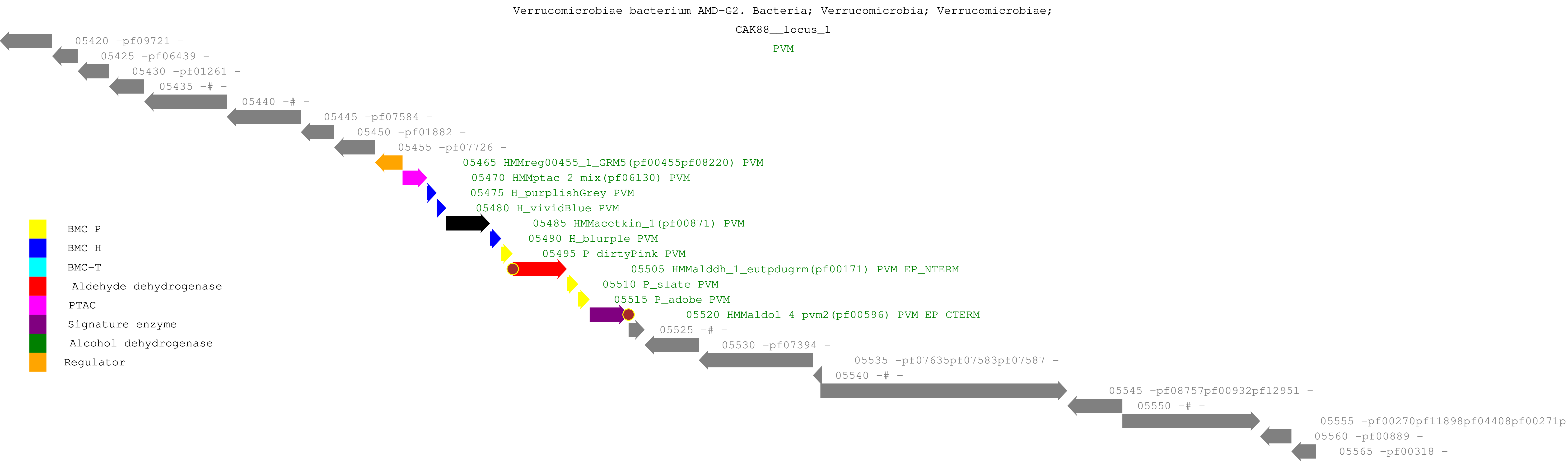
Verrucomicrobiae bacterium AMD-G2. Bacteria; Verrucomicrobia; Verrucomicrobiae;
CAK88__locus_1
PVM
05420 -pf09721 -
05425 -pf06439 -
05430 -pf01261 -
05435 -# -
05440 -# -
05445 -pf07584 -
05450 -pf01882 -
05455 -pf07726 -
05465 HMMreg00455_1_GRM5(pf00455pf08220) PVM
05470 HMMptac_2_mix(pf06130) PVM
05475 H_purplishGrey PVM
05480 H_vividBlue PVM
05485 HMMacetkin_1(pf00871) PVM
05490 H_blurple PVM
05495 P_dirtyPink PVM
05505 HMMalddh_1_eutpdugrm(pf00171) PVM EP_NTERM
05510 P_slate PVM
05515 P_adobe PVM
05520 HMMaldol_4_pvm2(pf00596) PVM EP_CTERM
05525 -# -
05530 -pf07394 -
05535 -pf07635pf07583pf07587 -
05540 -# -
05545 -pf08757pf00932pf12951 -
05550 -# -
05555 -pf00270pf11898pf04408pf00271p
05560 -pf00889 -
05565 -pf00318 -
BMC-P
BMC-H
BMC-T
Aldehyde dehydrogenase
PTAC
Signature enzyme
Alcohol dehydrogenase
Regulator

Planctomycetaceae bacterium TMED10. Bacteria; Planctomycetes; Planctomycetia; Planctomycetales;

CBB70__locus_1

PVM

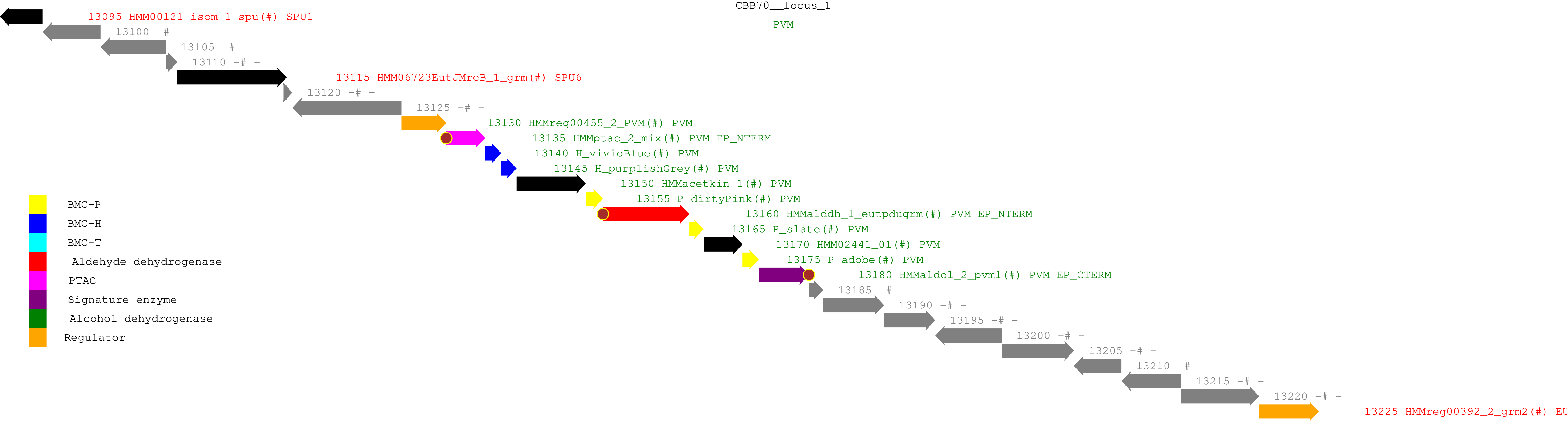

Rhodopirellula sp. TMED11. Bacteria; Planctomycetes; Planctomycetia; Planctomycetales;

CBB71__locus_1

PVM

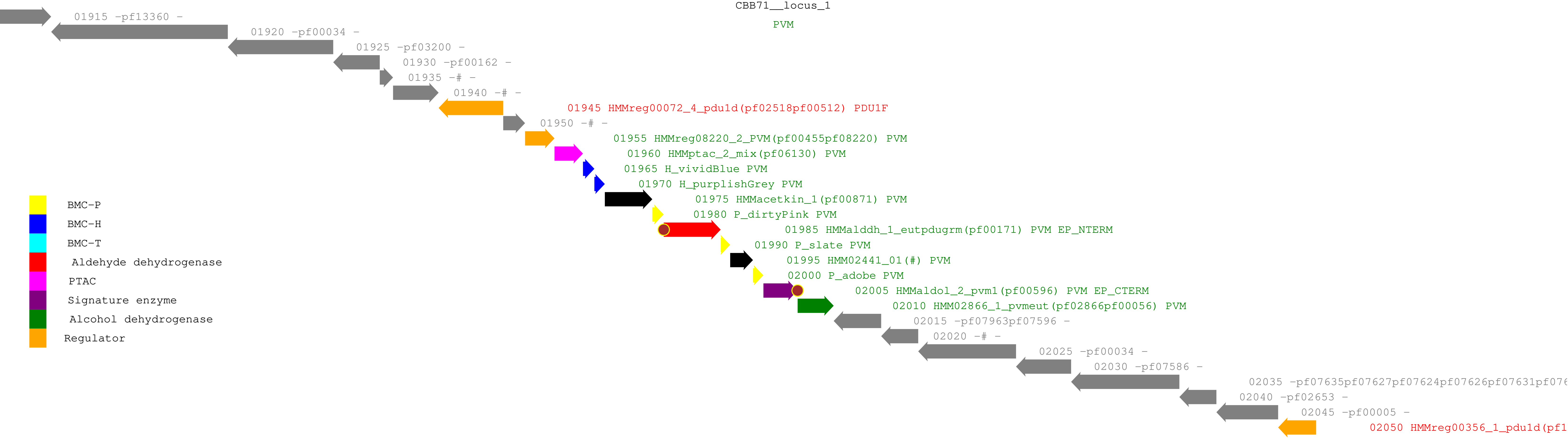

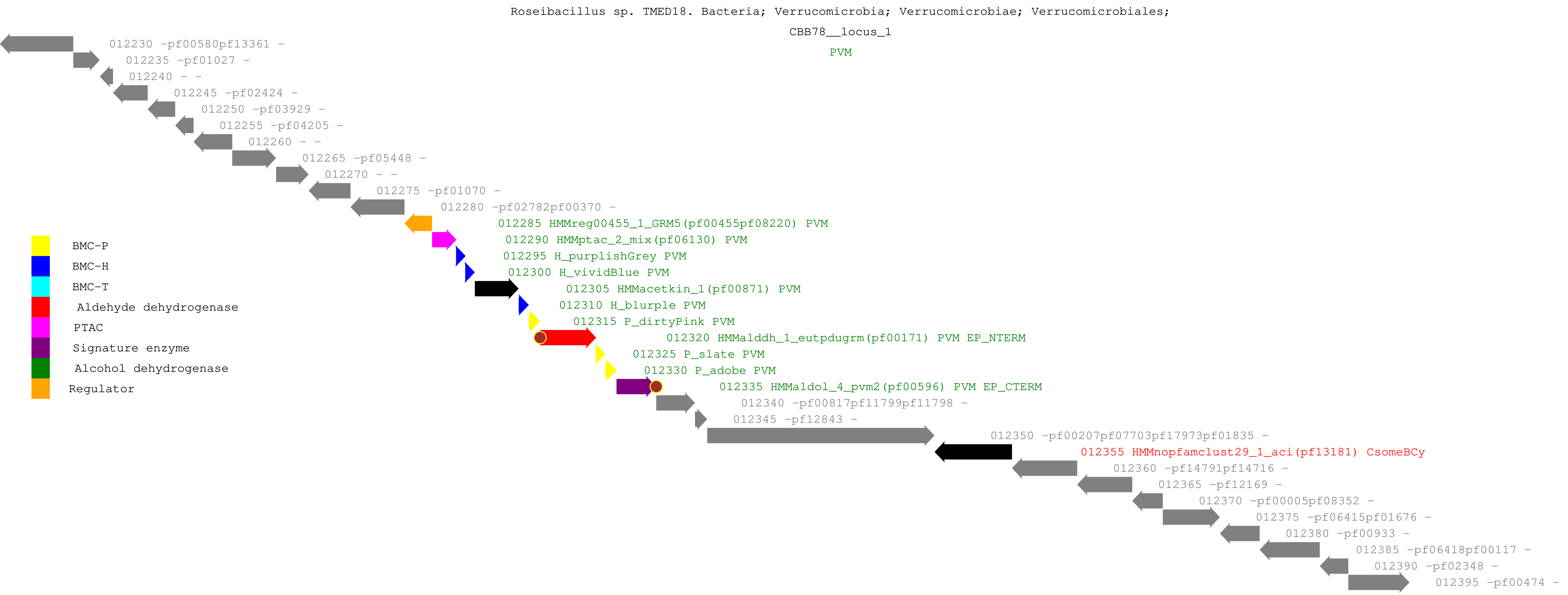
Roseibacillus sp. TMED18. Bacteria; Verrucomicrobia; Verrucomicrobiae; Verrucomicrobiales;
CBB78__locus_1
PVM
012230 -pf00580pf13361 -
012235 -pf01027 -
012240 - -
012245 -pf02424 -
012250 -pf03929 -
012255 -pf04205 -
012260 - -
012265 -pf05448 -
012270 - -
012275 -pf01070 -
012280 -pf02782pf00370 -
012285 HMMreg00455_1_GRM5(pf00455pf08220) PVM
012290 HMMptac_2_mix(pf06130) PVM
012295 H_purplishGrey PVM
012300 H_vividBlue PVM
012305 HMMacetkin_1(pf00871) PVM
012310 H_blurple PVM
012315 P_dirtyPink PVM
012320 HMMalddh_1_eutpdugrm(pf00171) PVM EP_NTERM
012325 P_slate PVM
012330 P_adobe PVM
012335 HMMaldol_4_pvm2(pf00596) PVM EP_CTERM
012340 -pf00817pf11799pf11798 -
012345 -pf12843 -
012350 -pf00207pf07703pf17973pf01835 -
012355 HMMnopfamclust29_1_aci(pf13181) CsomeBCy
012360 -pf14791pf14716 -
012365 -pf12169 -
012370 -pf00005pf08352 -
012375 -pf06415pf01676 -
012380 -pf00933 -
012385 -pf06418pf00117 -
012390 -pf02348 -
012395 -pf00474 -
BMC-P
BMC-H
BMC-T
Aldehyde dehydrogenase
PTAC
Signature enzyme
Alcohol dehydrogenase
Regulator

Opitutae bacterium TMED67. Bacteria; Verrucomicrobia; Opitutae.

CBC27__locus_1

PVM

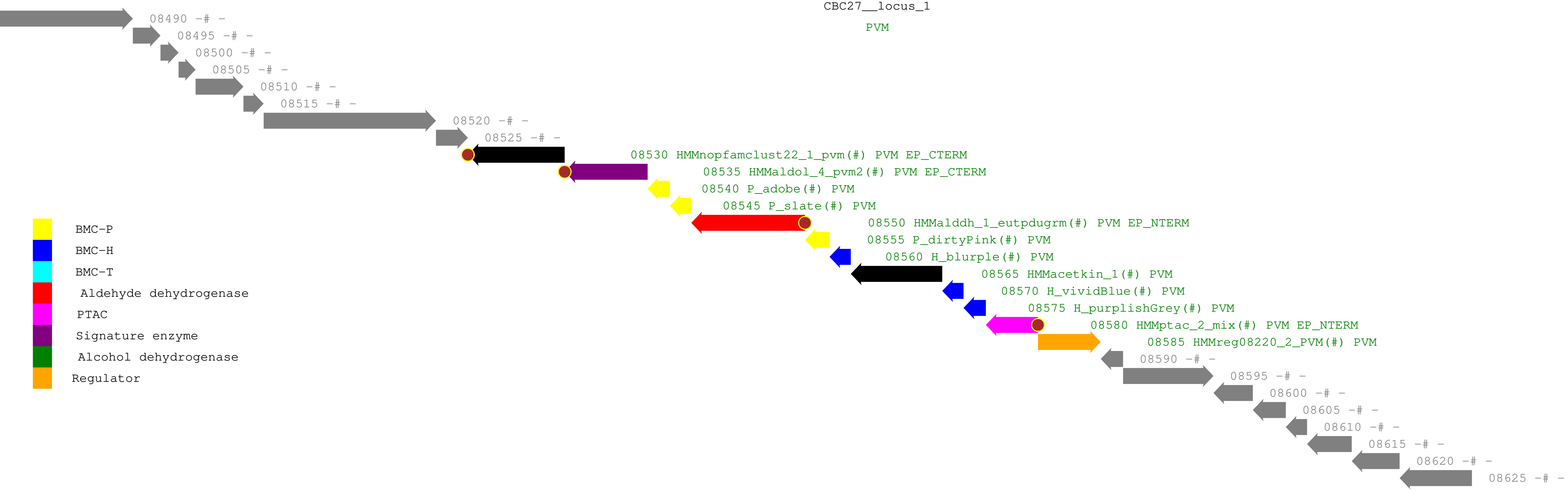

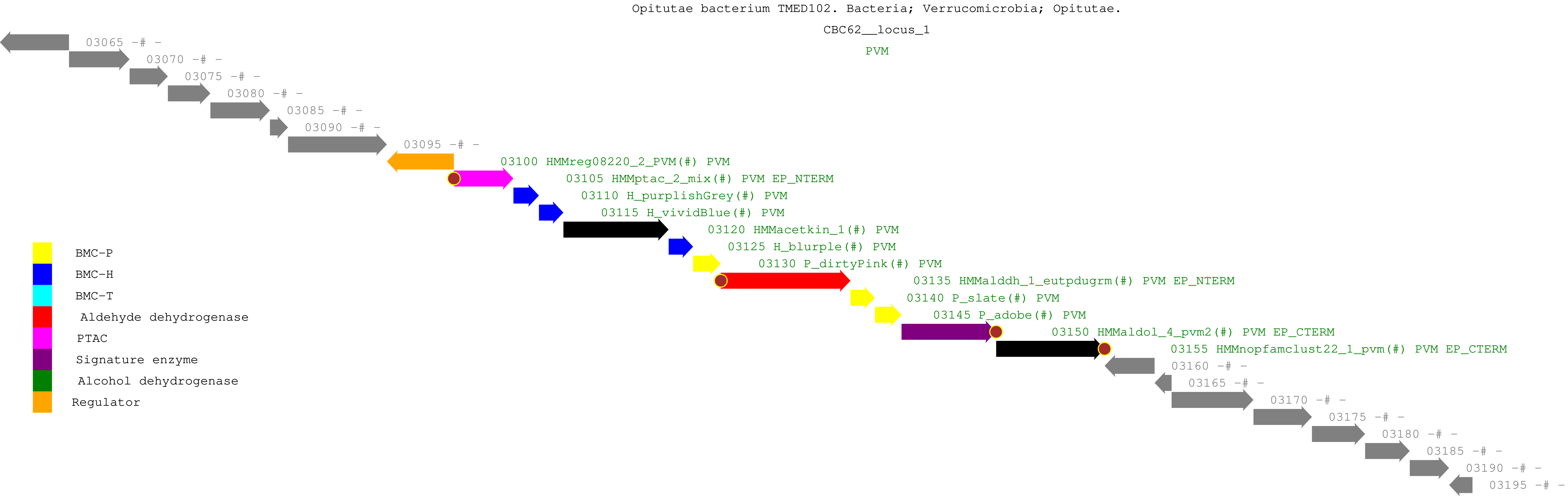
Opitutae bacterium TMED102. Bacteria; Verrucomicrobia; Opitutae.
CBC62__locus_1
PVM
03065 -# -
03070 -# -
03075 -# -
03080 -# -
03085 -# -
03090 -# -
03095 -# -
03100 HMMreg08220_2_PVM(#) PVM
03105 HMMptac_2_mix(#) PVM EP_NTERM
03110 H_purplishGrey(#) PVM
03115 H_vividBlue(#) PVM
03120 HMMacetkin_1(#) PVM
03125 H_blurple(#) PVM
03130 P_dirtyPink(#) PVM
03135 HMMalddh_1_eutpdugrm(#) PVM EP_NTERM
03140 P_slate(#) PVM
03145 P_adobe(#) PVM
03150 HMMaldol_4_pvm2(#) PVM EP_CTERM
03155 HMMnopfamclust22_1_pvm(#) PVM EP_CTERM
03160 -# -
03165 -# -
03170 -# -
03175 -# -
03180 -# -
03185 -# -
03190 -# -
03195 -# -
BMC-P
BMC-H
BMC-T
Aldehyde dehydrogenase
PTAC
Signature enzyme
Alcohol dehydrogenase
Regulator

Saprospirales bacterium TMED214. Bacteria; Bacteroidetes; Saprospiria; Saprospirales.

CBD74__locus_1

PVM

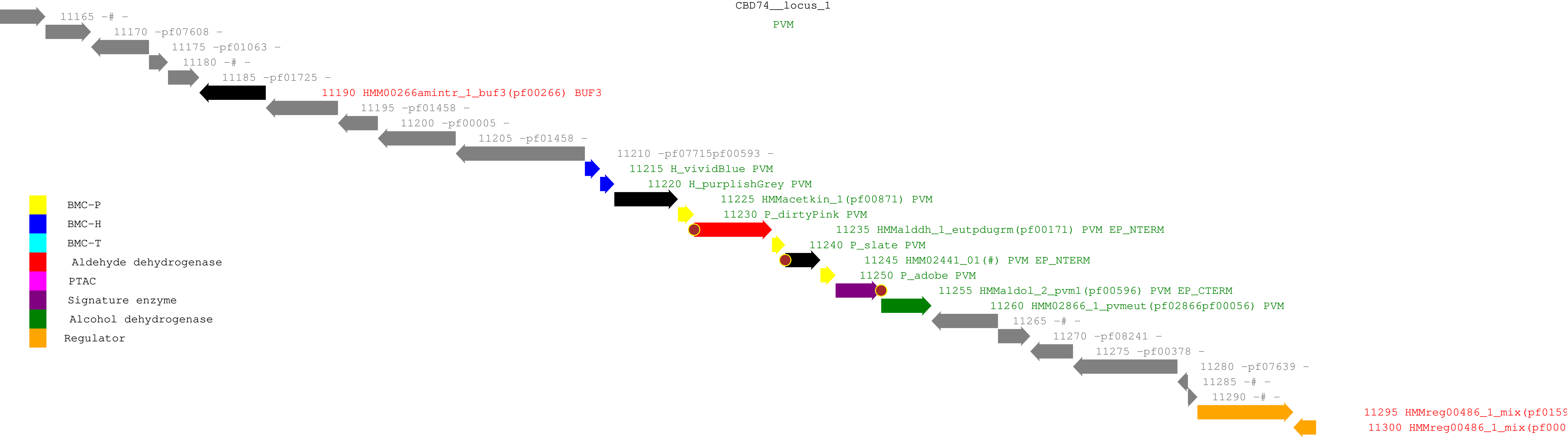

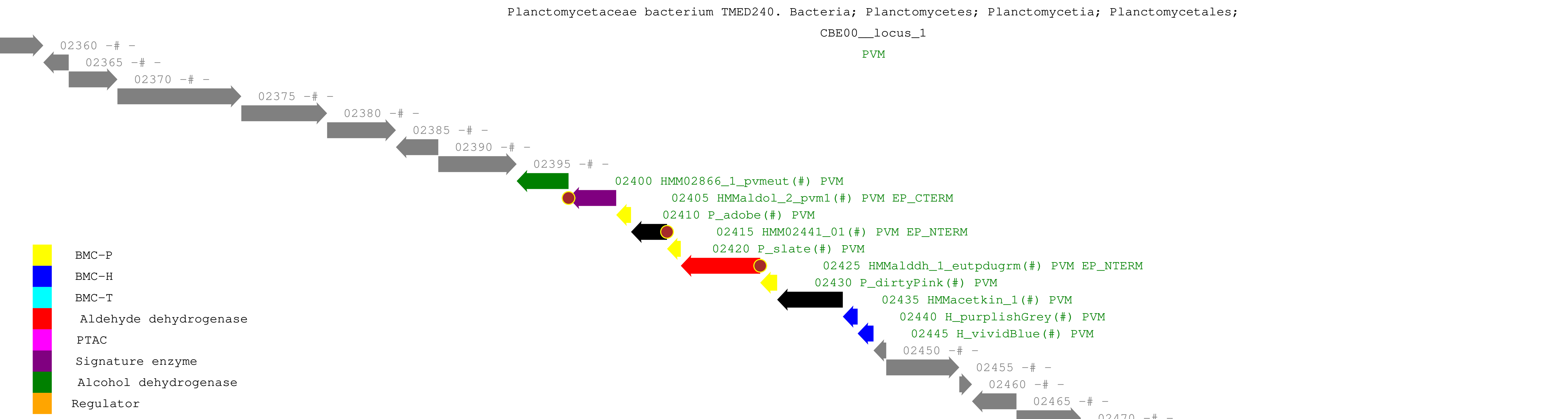
Planctomycetaceae bacterium TMED240. Bacteria; Planctomycetes; Planctomycetia; Planctomycetales;
CBE00__locus_1
PVM
02360 -# -
02365 -# -
02370 -# -
02375 -# -
02380 -# -
02385 -# -
02390 -# -
02395 -# -
02400 HMM02866_1_pvmeut(#) PVM
02405 HMMaldol_2_pvm1(#) PVM EP_CTERM
02410 P_adobe(#) PVM
02415 HMM02441_01(#) PVM EP_NTERM
02420 P_slate(#) PVM
02425 HMMalddh_1_eutpdugrm(#) PVM EP_NTERM
02430 P_dirtyPink(#) PVM
02435 HMMacetkin_1(#) PVM
02440 H_purplishGrey(#) PVM
02445 H_vividBlue(#) PVM
02450 -# -
02455 -# -
02460 -# -
02465 -# -
02470 -# -
02475 -# -
02480 -# -
02485 -# -
02490 -# -
02495 -# -
BMC-P
BMC-H
BMC-T
Aldehyde dehydrogenase
PTAC
Signature enzyme
Alcohol dehydrogenase
Regulator

Rhodopirellula sp. TMED283. Bacteria; Planctomycetes; Planctomycetia; Planctomycetales;

CBE43__locus_1

PVM

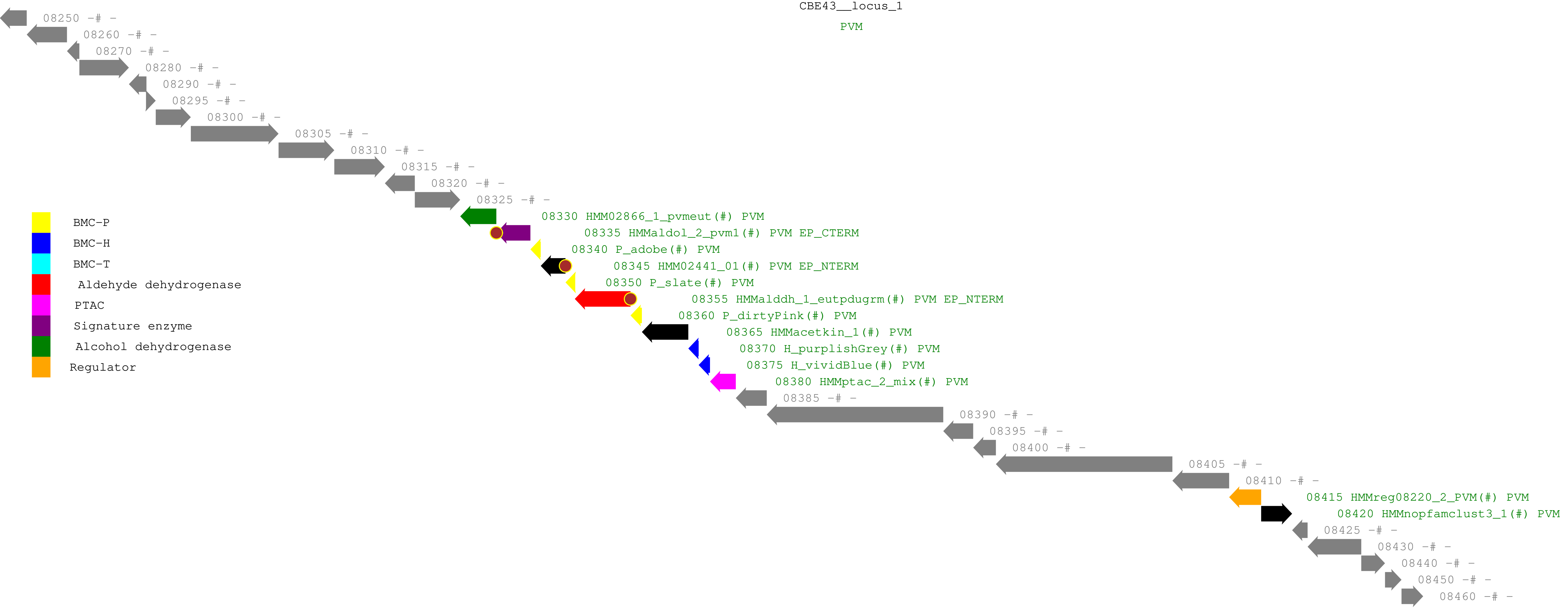

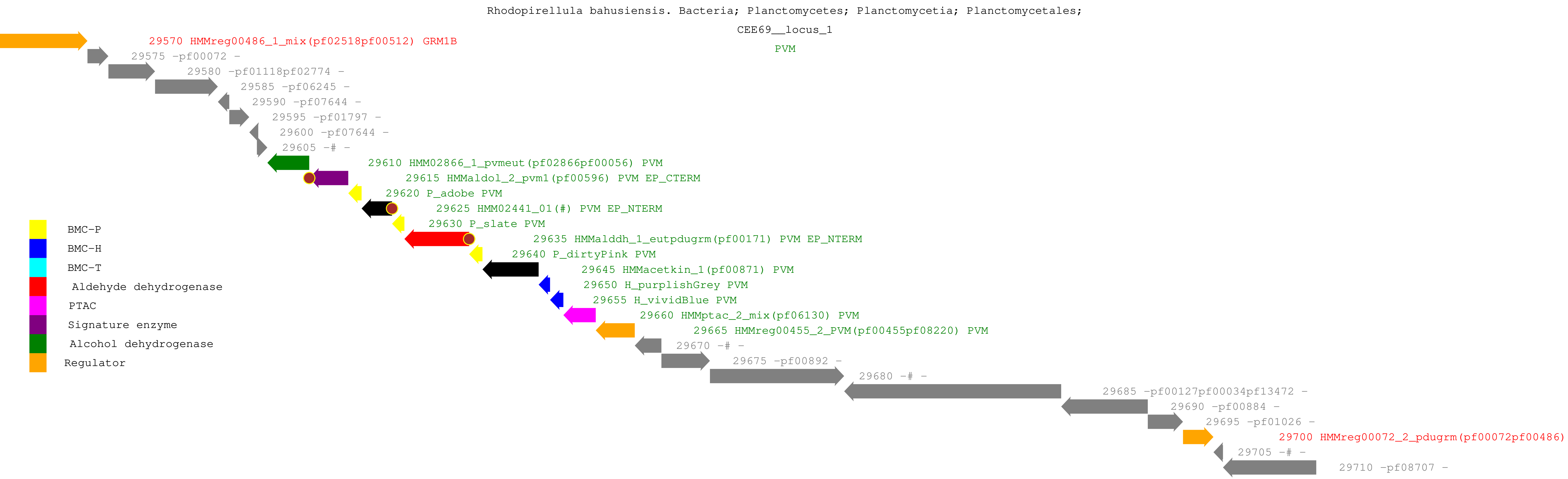

Rhodopirellula bahusiensis. Bacteria; Planctomycetes; Planctomycetia; Planctomycetales;
CEE69__locus_1
PVM
29570 HMMreg00486_1_mix(pf02518pf00512) GRM1B
29575 -pf00072 -
29580 -pf01118pf02774 -
29585 -pf06245 -
29590 -pf07644 -
29595 -pf01797 -
29600 -pf07644 -
29605 -# -
29610 HMM02866_1_pvmeut(pf02866pf00056) PVM
29615 HMMaldol_2_pvm1(pf00596) PVM EP_CTERM
29620 P_adobe PVM
29625 HMM02441_01(#) PVM EP_NTERM
29630 P_slate PVM
29635 HMMalddh_1_eutpdugrm(pf00171) PVM EP_NTERM
29640 P_dirtyPink PVM
29645 HMMacetkin_1(pf00871) PVM
29650 H_purplishGrey PVM
29655 H_vividBlue PVM
29660 HMMptac_2_mix(pf06130) PVM
29665 HMMreg00455_2_PVM(pf00455pf08220) PVM
29670 -# -
29675 -pf00892 -
29680 -# -
29685 -pf00127pf00034pf13472 -
29690 -pf00884 -
29695 -pf01026 -
29700 HMMreg00072_2_pdugrm(pf00072pf00486)
29705 -# -
29710 -pf08707 -
BMC-P
BMC-H
BMC-T
Aldehyde dehydrogenase
PTAC
Signature enzyme
Alcohol dehydrogenase
Regulator

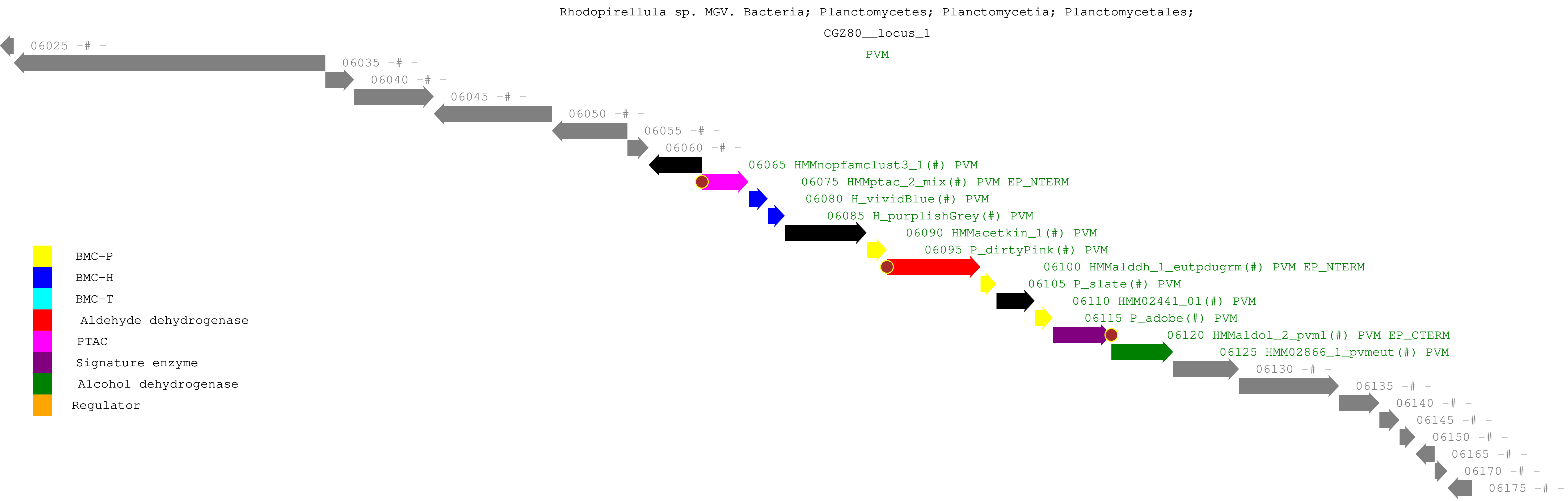
Rhodopirellula sp. MGV. Bacteria; Planctomycetes; Planctomycetia; Planctomycetales;
CGZ80__locus_1
PVM
06025 -# -
06035 -# -
06040 -# -
06045 -# -
06050 -# -
06055 -# -
06060 -# -
06065 HMMnopfamclust3_1(#) PVM
06075 HMMptac_2_mix(#) PVM EP_NTERM
06080 H_vividBlue(#) PVM
06085 H_purplishGrey(#) PVM
06090 HMMacetkin_1(#) PVM
06095 P_dirtyPink(#) PVM
06100 HMMalddh_1_eutpdugrm(#) PVM EP_NTERM
06105 P_slate(#) PVM
06110 HMM02441_01(#) PVM
06115 P_adobe(#) PVM
06120 HMMaldol_2_pvm1(#) PVM EP_CTERM
06125 HMM02866_1_pvmeut(#) PVM
06130 -# -
06135 -# -
06140 -# -
06145 -# -
06150 -# -
06165 -# -
06170 -# -
06175 -# -
BMC-P
BMC-H
BMC-T
Aldehyde dehydrogenase
PTAC
Signature enzyme
Alcohol dehydrogenase
Regulator

Planctomycetaceae bacterium. Bacteria; Planctomycetes; Planctomycetia; Planctomycetales;

CK530__locus_1

PVM

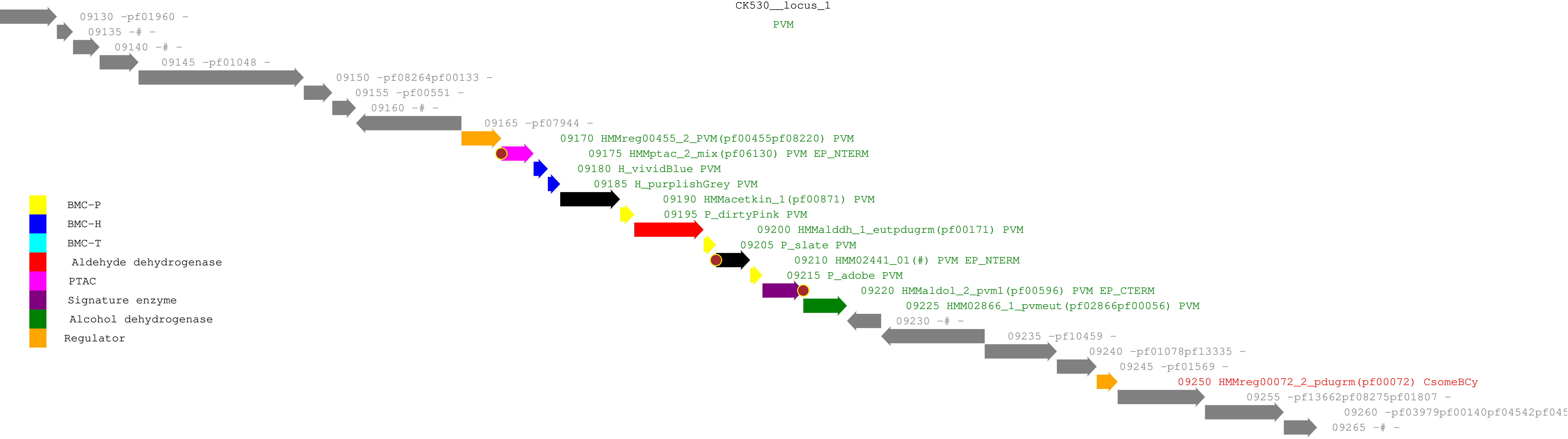

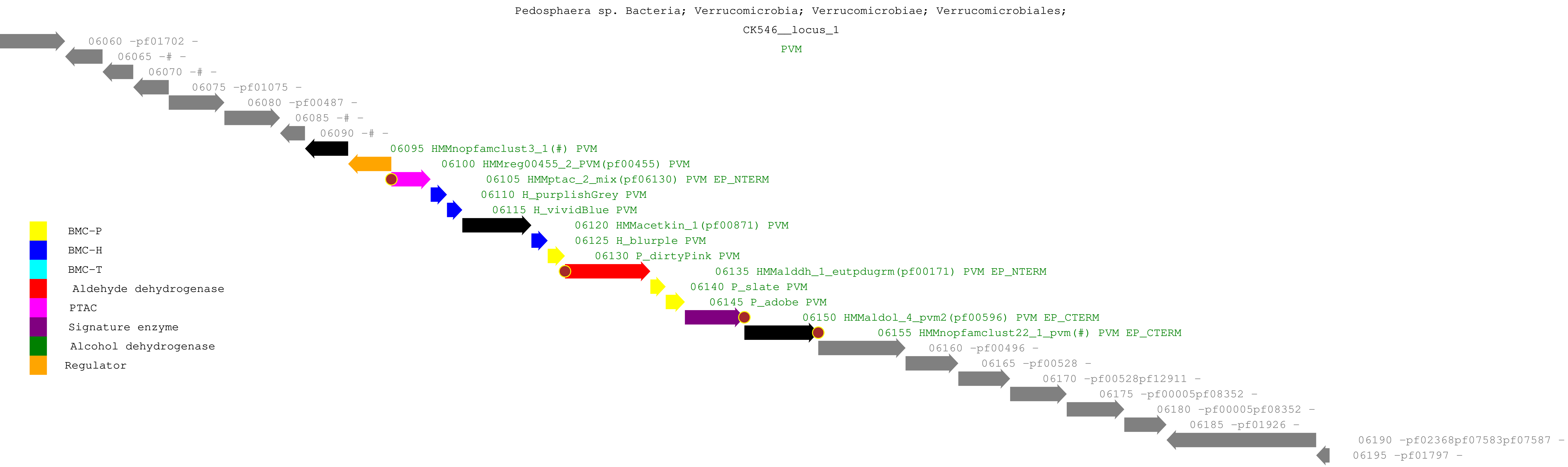

Pedosphaera sp. Bacteria; Verrucomicrobia; Verrucomicrobiae; Verrucomicrobiales;
CK546__locus_1
PVM
06060 -pf01702 -
06065 -# -
06070 -# -
06075 -pf01075 -
06080 -pf00487 -
06085 -# -
06090 -# -
06095 HMMnopfamclust3_1(#) PVM
06100 HMMreg00455_2_PVM(pf00455) PVM
06105 HMMptac_2_mix(pf06130) PVM EP_NTERM
06110 H_purplishGrey PVM
06115 H_vividBlue PVM
06120 HMMacetkin_1(pf00871) PVM
06125 H_blurple PVM
06130 P_dirtyPink PVM
06135 HMMaldh_1_eutpdugrm(pf00171) PVM EP_NTERM
06140 P_slate PVM
06145 P_adobe PVM
06150 HMMaldol_4_pvm2(pf00596) PVM EP_CTERM
06155 HMMnopfamclust22_1_pvm(#) PVM EP_CTERM
06160 -pf00496 -
06165 -pf00528 -
06170 -pf00528pf12911 -
06175 -pf00005pf08352 -
06180 -pf00005pf08352 -
06185 -pf01926 -
06190 -pf02368pf07583pf07587 -
06195 -pf01797 -
BMC-P
BMC-H
BMC-T
Aldehyde dehydrogenase
PTAC
Signature enzyme
Alcohol dehydrogenase
Regulator

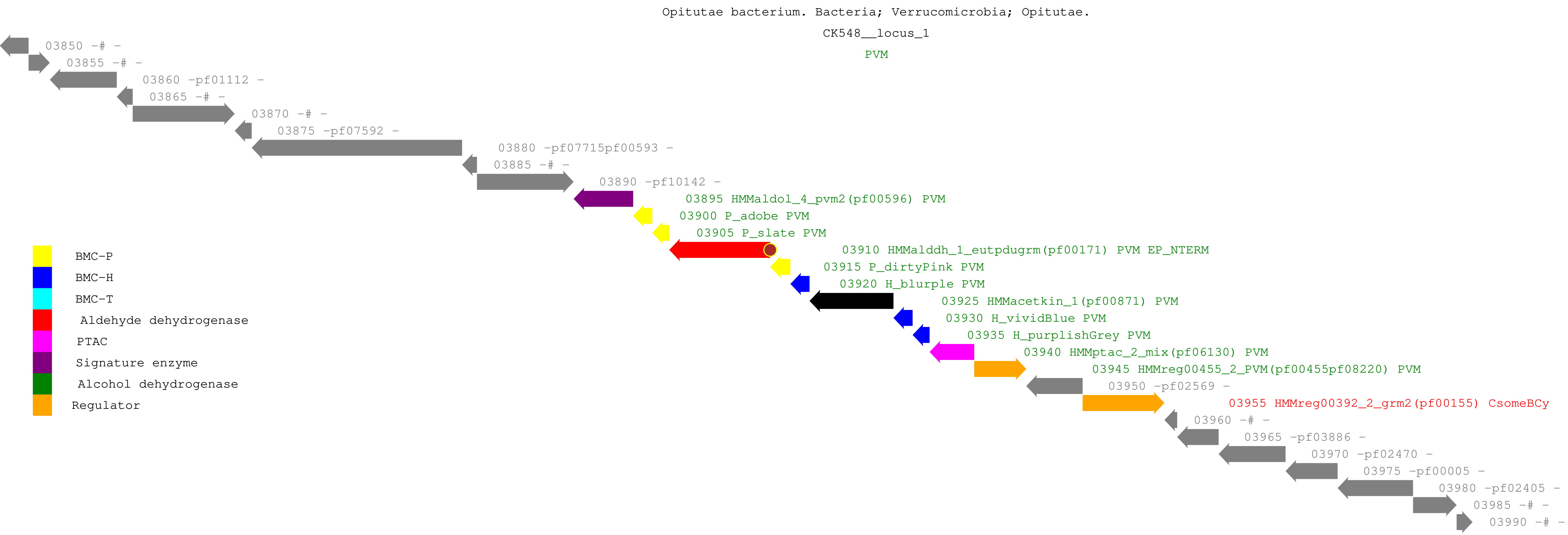
Opitutae bacterium. Bacteria; Verrucomicrobia; Opitutae.
CK548__locus_1
PVM
03850 -# -
03855 -# -
03860 -pf01112 -
03865 -# -
03870 -# -
03875 -pf07592 -
03880 -pf07715pf00593 -
03885 -# -
03890 -pf10142 -
03895 HMMaldol_4_pvm2(pf00596) PVM
03900 P_adobe PVM
03905 P_slate PVM
03910 HMMalddh_1_eutpdugrm(pf00171) PVM EP_NTERM
03915 P_dirtyPink PVM
03920 H_blurple PVM
03925 HMMacetkin_1(pf00871) PVM
03930 H_vividBlue PVM
03935 H_purplishGrey PVM
03940 HMMptac_2_mix(pf06130) PVM
03945 HMMreg00455_2_PVM(pf00455pf08220) PVM
03950 -pf02569 -
03955 HMMreg00392_2_grm2(pf00155) CsomeBCy
03960 -# -
03965 -pf03886 -
03970 -pf02470 -
03975 -pf00005 -
03980 -pf02405 -
03985 -# -
03990 -# -
BMC-P
BMC-H
BMC-T
Aldehyde dehydrogenase
PTAC
Signature enzyme
Alcohol dehydrogenase
Regulator

Planctomycetaceae bacterium. Bacteria; Planctomycetes; Planctomycetia; Planctomycetales;

CK551__locus_1

PVM

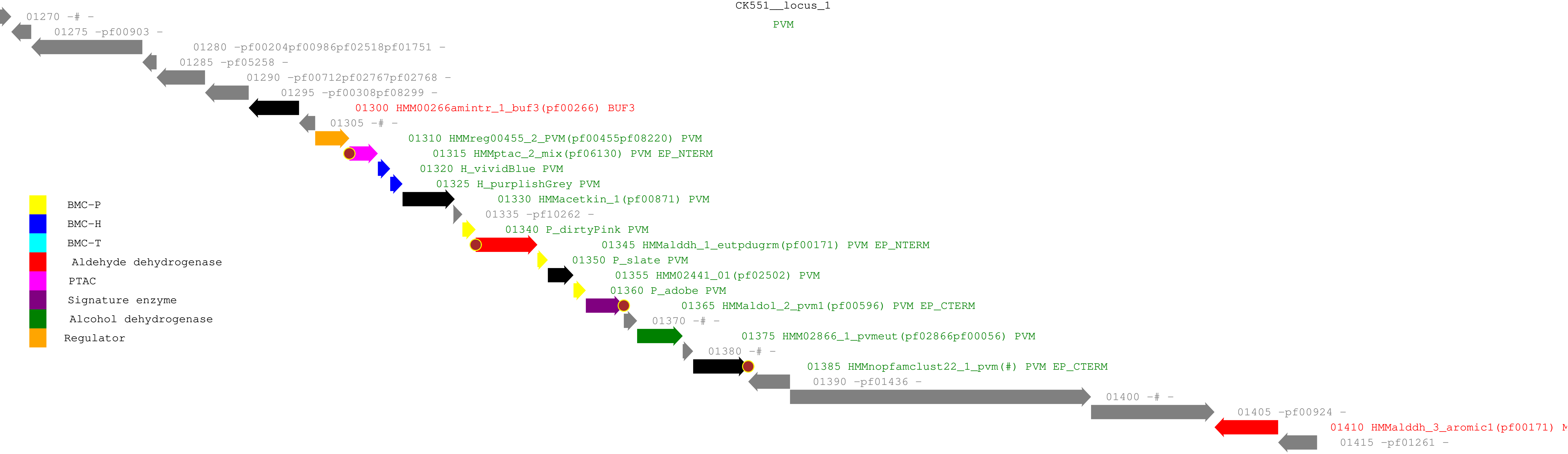

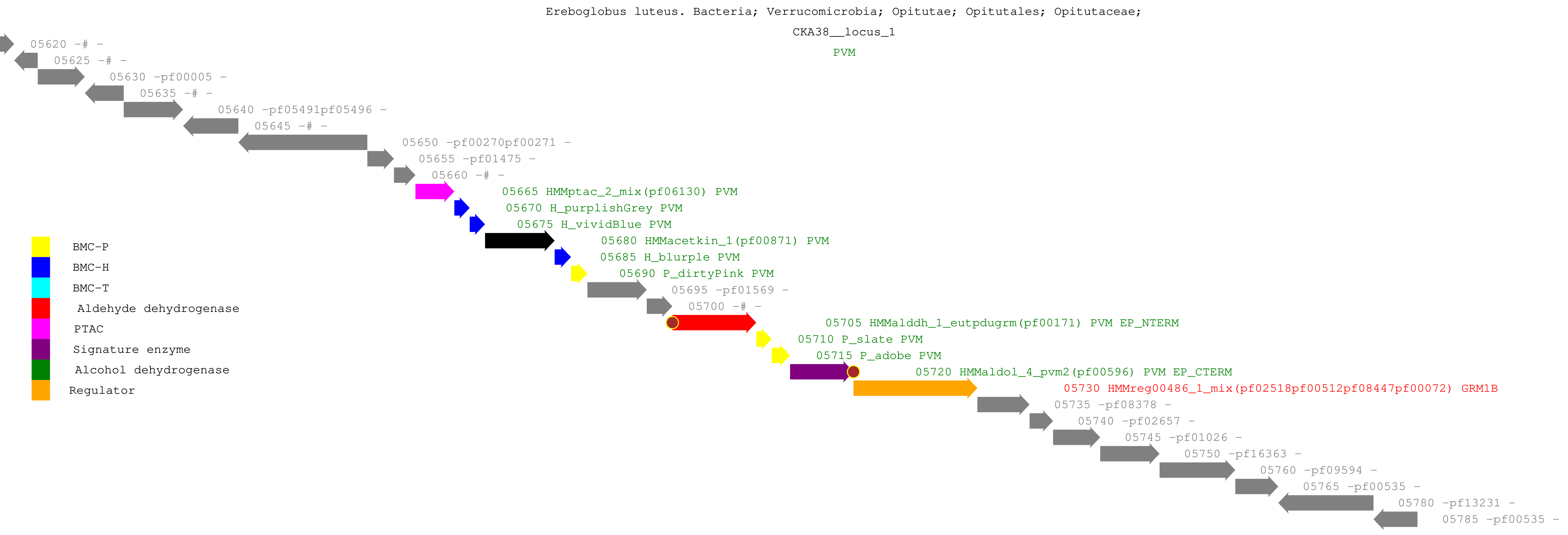
Ereboglobus luteus. Bacteria; Verrucomicrobia; Opitutae; Opitutales; Opitutaceae;
CKA38__locus_1
PVM
05620 -# -
05625 -# -
05630 -pf00005 -
05635 -# -
05640 -pf05491pf05496 -
05645 -# -
05650 -pf00270pf00271 -
05655 -pf01475 -
05660 -# -
05665 HMMptac_2_mix(pf06130) PVM
05670 H_purplishGrey PVM
05675 H_vividBlue PVM
05680 HMMacetkin_1(pf00871) PVM
05685 H_blurple PVM
05690 P_dirtyPink PVM
05695 -pf01569 -
05700 -# -
05705 HMMalddh_1_eutpdugrm(pf00171) PVM EP_NTERM
05710 P_slate PVM
05715 P_adobe PVM
05720 HMMaldol_4_pvm2(pf00596) PVM EP_CTERM
05730 HMMreg00486_1_mix(pf02518pf00512pf08447pf00072) GRM1B
05735 -pf08378 -
05740 -pf02657 -
05745 -pf01026 -
05750 -pf16363 -
05760 -pf09594 -
05765 -pf00535 -
05780 -pf13231 -
05785 -pf00535 -
BMC-P
BMC-H
BMC-T
Aldehyde dehydrogenase
PTAC
Signature enzyme
Alcohol dehydrogenase
Regulator

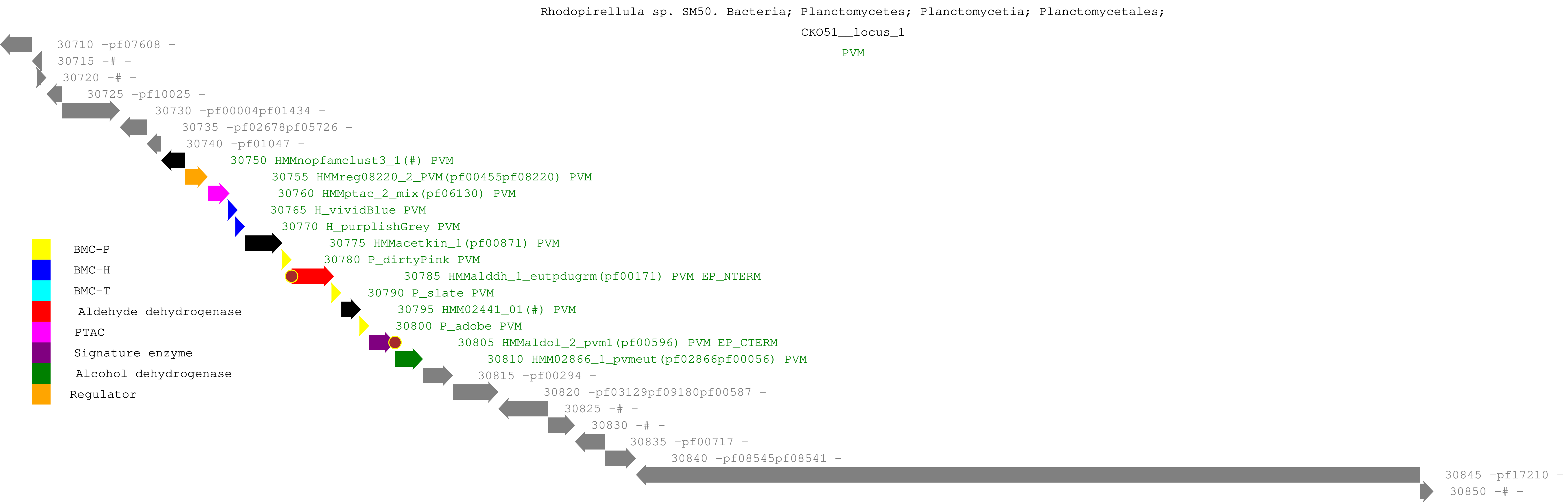
Rhodopirellula sp. SM50. Bacteria; Planctomycetes; Planctomycetia; Planctomycetales;
CKO51__locus_1
PVM
30710 -pf07608 -
30715 -# -
30720 -# -
30725 -pf10025 -
30730 -pf00004pf01434 -
30735 -pf02678pf05726 -
30740 -pf01047 -
30750 HMMnopfamclust3_1(#) PVM
30755 HMMreg08220_2_PVM(pf00455pf08220) PVM
30760 HMMptac_2_mix(pf06130) PVM
30765 H_vividBlue PVM
30770 H_purplishGrey PVM
30775 HMMacetkin_1(pf00871) PVM
30780 P_dirtyPink PVM
30785 HMMalddh_1_eutpdugrm(pf00171) PVM EP_NTERM
30790 P_slate PVM
30795 HMM02441_01(#) PVM
30800 P_adobe PVM
30805 HMMaldol_2_pvm1(pf00596) PVM EP_CTERM
30810 HMM02866_1_pvmeut(pf02866pf00056) PVM
30815 -pf00294 -
30820 -pf03129pf09180pf00587 -
30825 -# -
30830 -# -
30835 -pf00717 -
30840 -pf08545pf08541 -
30845 -pf17210 -
30850 -# -
BMC-P
BMC-H
BMC-T
Aldehyde dehydrogenase
PTAC
Signature enzyme
Alcohol dehydrogenase
Regulator

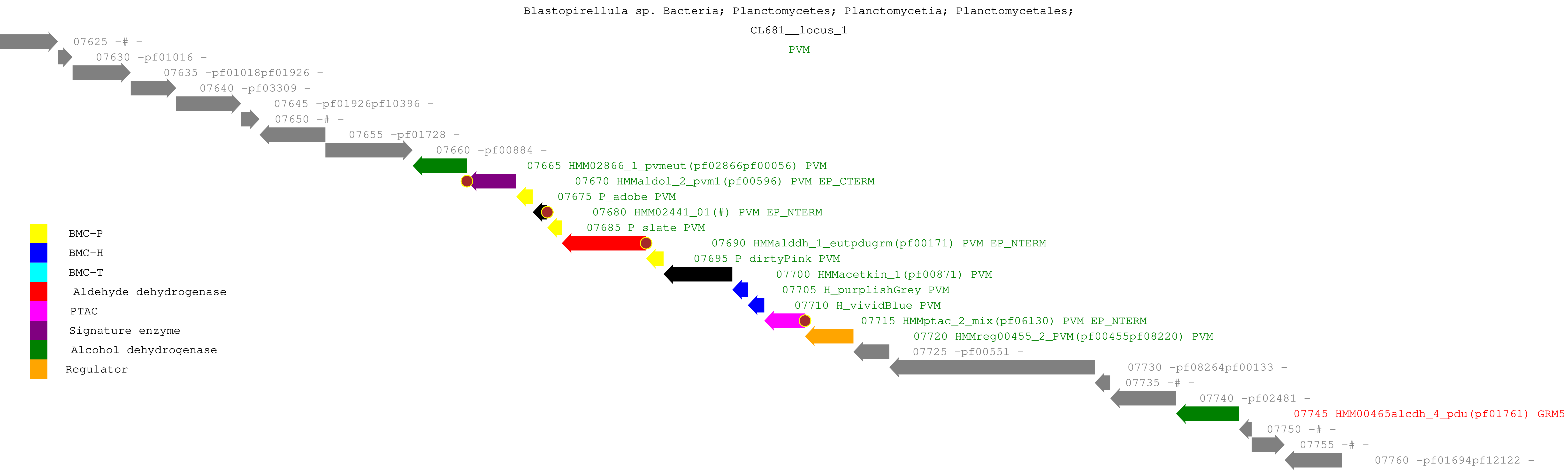

Blastopirellula sp. Bacteria; Planctomycetes; Planctomycetia; Planctomycetales;
CL681__locus_1
PVM
07625 -# -
07630 -pf01016 -
07635 -pf01018pf01926 -
07640 -pf03309 -
07645 -pf01926pf10396 -
07650 -# -
07655 -pf01728 -
07660 -pf00884 -
07665 HMM02866_1_pvmeut(pf02866pf00056) PVM
07670 HMMaldol_2_pvm1(pf00596) PVM EP_CTERM
07675 P_adobe PVM
07680 HMM02441_01(#) PVM EP_NTERM
07685 P_slate PVM
07690 HMMalddh_1_eutpdugrm(pf00171) PVM EP_NTERM
07695 P_dirtyPink PVM
07700 HMMacetkin_1(pf00871) PVM
07705 H_purplishGrey PVM
07710 H_vividBlue PVM
07715 HMMptac_2_mix(pf06130) PVM EP_NTERM
07720 HMMreg00455_2_PVM(pf00455pf08220) PVM
07725 -pf00551 -
07730 -pf08264pf00133 -
07735 -# -
07740 -pf02481 -
07745 HMM00465alcdh_4_pdu(pf01761) GRM5
07750 -# -
07755 -# -
07760 -pf01694pf12122 -
BMC-P
BMC-H
BMC-T
Aldehyde dehydrogenase
PTAC
Signature enzyme
Alcohol dehydrogenase
Regulator

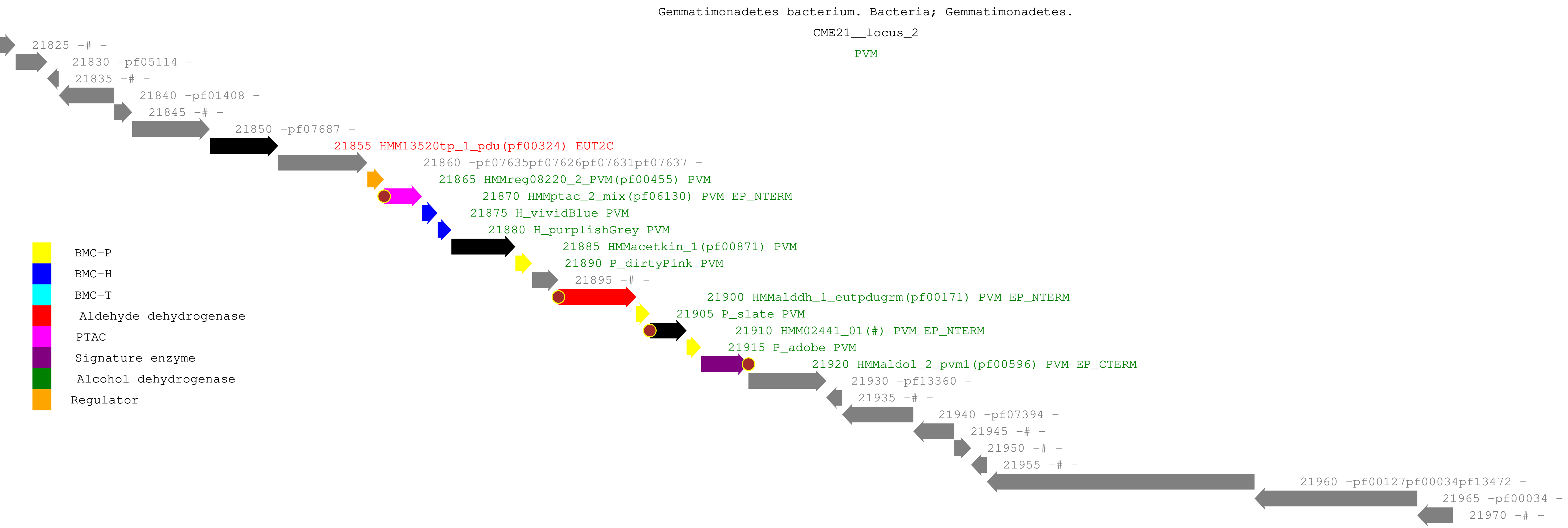
Gemmatimonadetes bacterium. Bacteria; Gemmatimonadetes.
CME21__locus_2
PVM
21825 -# -
21830 -pf05114 -
21835 -# -
21840 -pf01408 -
21845 -# -
21850 -pf07687 -
21855 HMM13520tp_1_pdu(pf00324) EUT2C
21860 -pf07635pf07626pf07631pf07637 -
21865 HMMreg08220_2_PVM(pf00455) PVM
21870 HMMptac_2_mix(pf06130) PVM EP_NTERM
21875 H_vividBlue PVM
21880 H_purplishGrey PVM
21885 HMMacetkin_1(pf00871) PVM
21890 P_dirtyPink PVM
21895 -# -
21900 HMMalddh_1_eutpdugrm(pf00171) PVM EP_NTERM
21905 P_slate PVM
21910 HMM02441_01(#) PVM EP_NTERM
21915 P_adobe PVM
21920 HMMaldol_2_pvm1(pf00596) PVM EP_CTERM
21930 -pf13360 -
21935 -# -
21940 -pf07394 -
21945 -# -
21950 -# -
21955 -# -
21960 -pf00127pf00034pf13472 -
21965 -pf00034 -
21970 -# -
BMC-P
BMC-H
BMC-T
Aldehyde dehydrogenase
PTAC
Signature enzyme
Alcohol dehydrogenase
Regulator

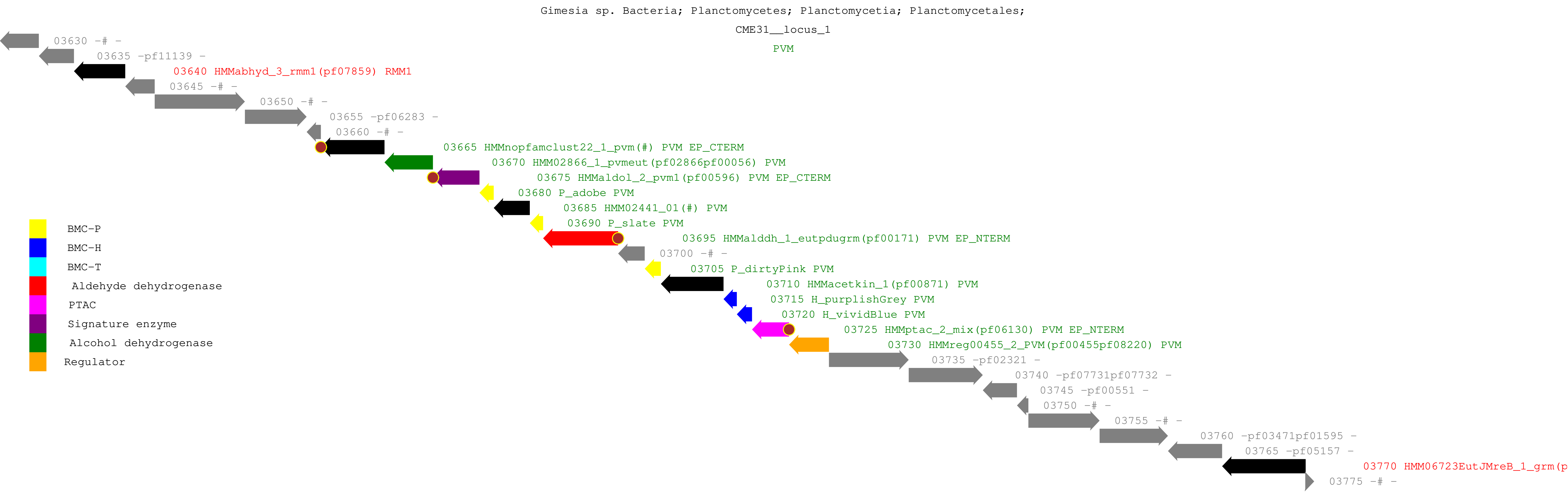
Gimesia sp. Bacteria; Planctomycetes; Planctomycetia; Planctomycetales;
CME31__locus_1
PVM
03630 -# -
03635 -pf11139 -
03640 HMMabhyd_3_rmm1(pf07859) RMM1
03645 -# -
03650 -# -
03655 -pf06283 -
03660 -# -
03665 HMMnopfamclust22_1_pvm(#) PVM EP_CTERM
03670 HMM02866_1_pvmeut(pf02866pf00056) PVM
03675 HMMaldol_2_pvm1(pf00596) PVM EP_CTERM
03680 P_adobe PVM
03685 HMM02441_01(#) PVM
03690 P_slate PVM
03695 HMMalddh_1_eutpdugrm(pf00171) PVM EP_NTERM
03700 -# -
03705 P_dirtyPink PVM
03710 HMMacetkin_1(pf00871) PVM
03715 H_purplishGrey PVM
03720 H_vividBlue PVM
03725 HMMptac_2_mix(pf06130) PVM EP_NTERM
03730 HMMreg00455_2_PVM(pf00455pf08220) PVM
03735 -pf02321 -
03740 -pf07731pf07732 -
03745 -pf00551 -
03750 -# -
03755 -# -
03760 -pf03471pf01595 -
03765 -pf05157 -
03770 HMM06723EutJMreB_1_grm(p
03775 -# -
BMC-P
BMC-H
BMC-T
Aldehyde dehydrogenase
PTAC
Signature enzyme
Alcohol dehydrogenase
Regulator

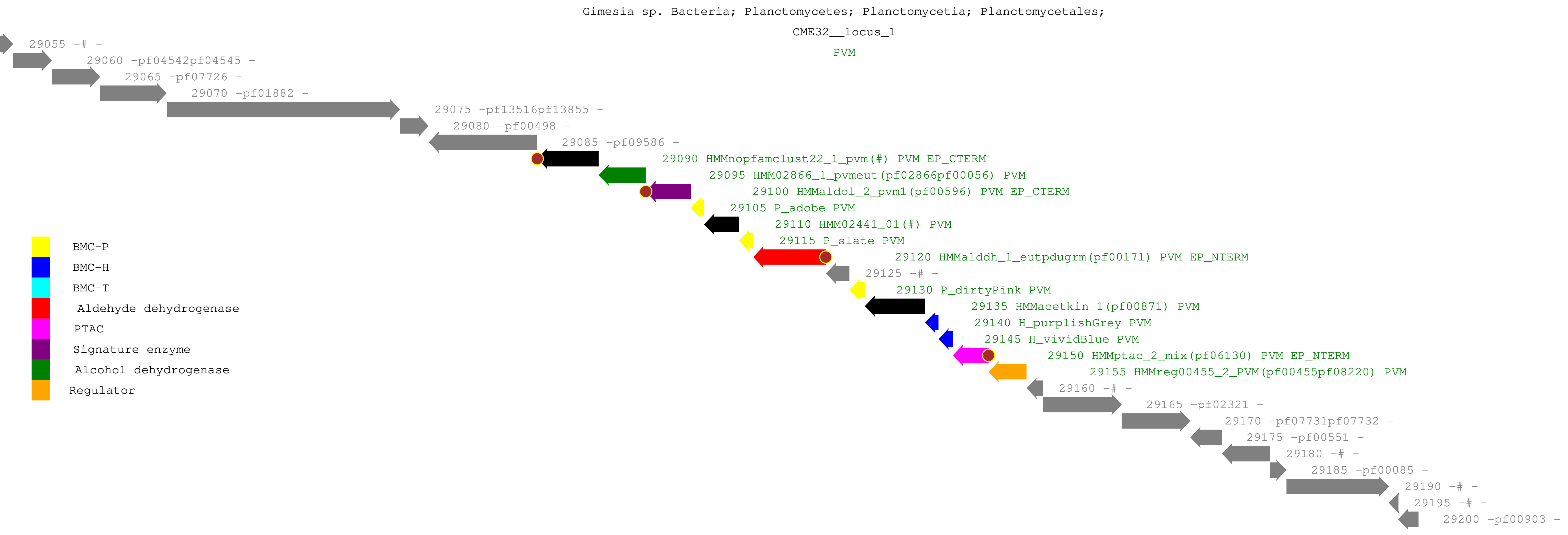
Gimesia sp. Bacteria; Planctomycetes; Planctomycetia; Planctomycetales;
CME32__locus_1
PVM
29055 -# -
29060 -pf04542pf04545 -
29065 -pf07726 -
29070 -pf01882 -
29075 -pf13516pf13855 -
29080 -pf00498 -
29085 -pf09586 -
29090 HMMnopfamclust22_1_pvm(#) PVM EP_CTERM
29095 HMM02866_1_pvmeut(pf02866pf00056) PVM
29100 HMMaldol_2_pvm1(pf00596) PVM EP_CTERM
29105 P_adobe PVM
29110 HMM02441_01(#) PVM
29115 P_slate PVM
29120 HMMalddh_1_eutpdugrm(pf00171) PVM EP_NTERM
29125 -# -
29130 P_dirtyPink PVM
29135 HMMacetkin_1(pf00871) PVM
29140 H_purplishGrey PVM
29145 H_vividBlue PVM
29150 HMMptac_2_mix(pf06130) PVM EP_NTERM
29155 HMMreg00455_2_PVM(pf00455pf08220) PVM
29160 -# -
29165 -pf02321 -
29170 -pf07731pf07732 -
29175 -pf00551 -
29180 -# -
29185 -pf00085 -
29190 -# -
29195 -# -
29200 -pf00903 -
BMC-P
BMC-H
BMC-T
Aldehyde dehydrogenase
PTAC
Signature enzyme
Alcohol dehydrogenase
Regulator

Gimesia sp. Bacteria; Planctomycetes; Planctomycetia; Planctomycetales;

CME33__locus_1

PVM

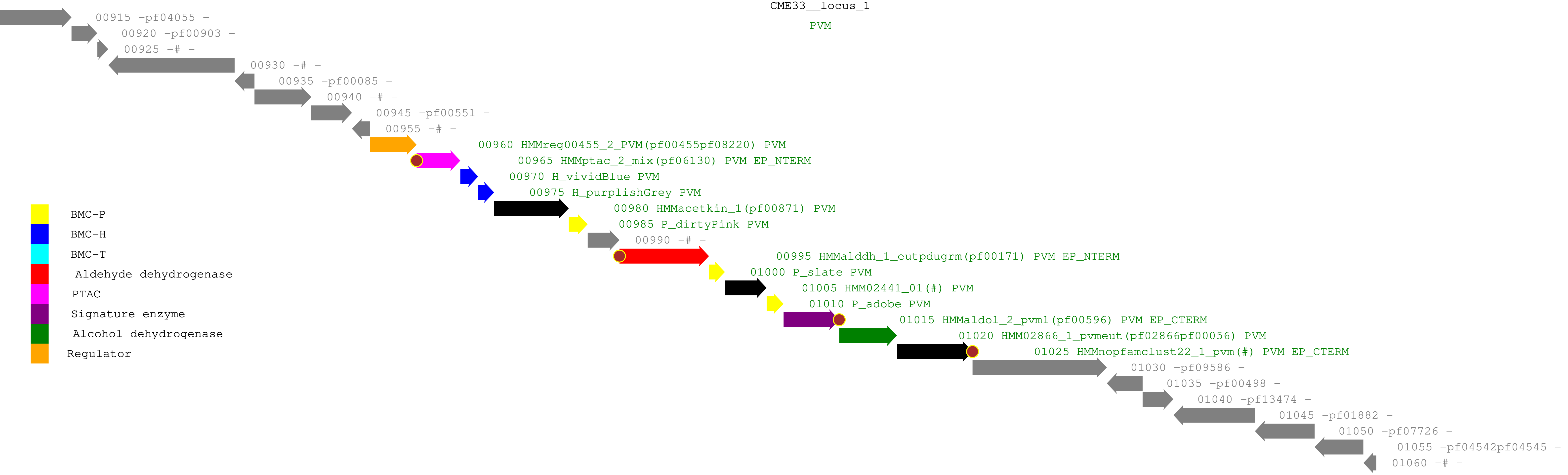

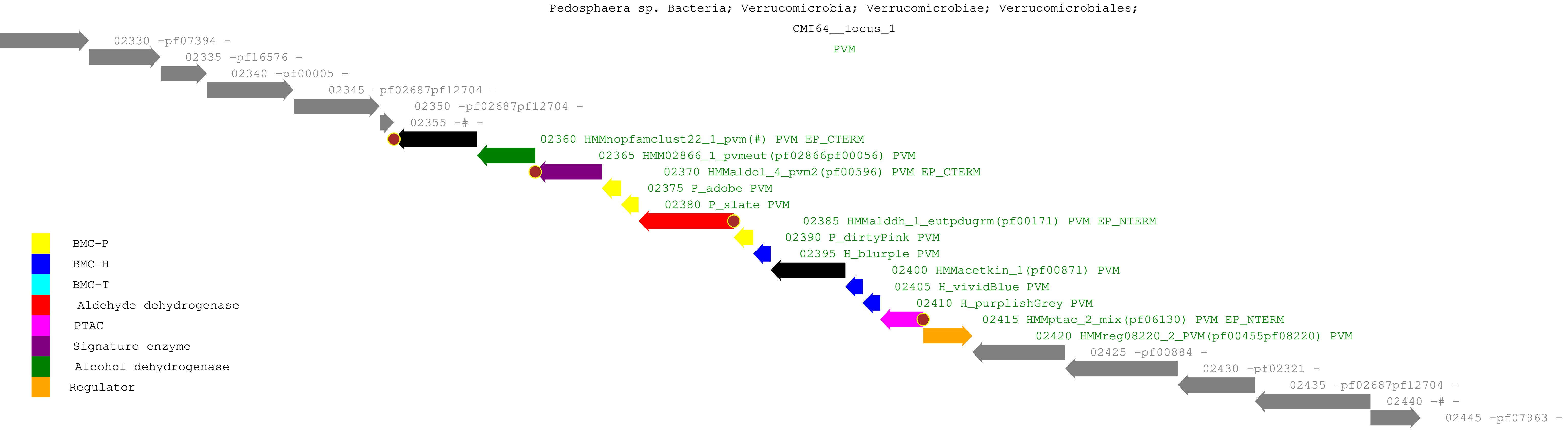
Pedosphaera sp. Bacteria; Verrucomicrobia; Verrucomicrobiae; Verrucomicrobiales;
CMI64__locus_1
PVM
02330 -pf07394 -
02335 -pf16576 -
02340 -pf00005 -
02345 -pf02687pf12704 -
02350 -pf02687pf12704 -
02355 -# -
02360 HMMnopfamclust22_1_pvm(#) PVM EP_CTERM
02365 HMM02866_1_pvmeut(pf02866pf00056) PVM
02370 HMMaldol_4_pvm2(pf00596) PVM EP_CTERM
02375 P_adobe PVM
02380 P_slate PVM
02385 HMMalddh_1_eutpdugrm(pf00171) PVM EP_NTERM
02390 P_dirtyPink PVM
02395 H_blurple PVM
02400 HMMacetkin_1(pf00871) PVM
02405 H_vividBlue PVM
02410 H_purplishGrey PVM
02415 HMMptac_2_mix(pf06130) PVM EP_NTERM
02420 HMMreg08220_2_PVM(pf00455pf08220) PVM
02425 -pf00884 -
02430 -pf02321 -
02435 -pf02687pf12704 -
02440 -# -
02445 -pf07963 -
BMC-P
BMC-H
BMC-T
Aldehyde dehydrogenase
PTAC
Signature enzyme
Alcohol dehydrogenase
Regulator

Pedosphaera sp. Bacteria; Verrucomicrobia; Verrucomicrobiae; Verrucomicrobiales;

CMI65__locus_1

PVM

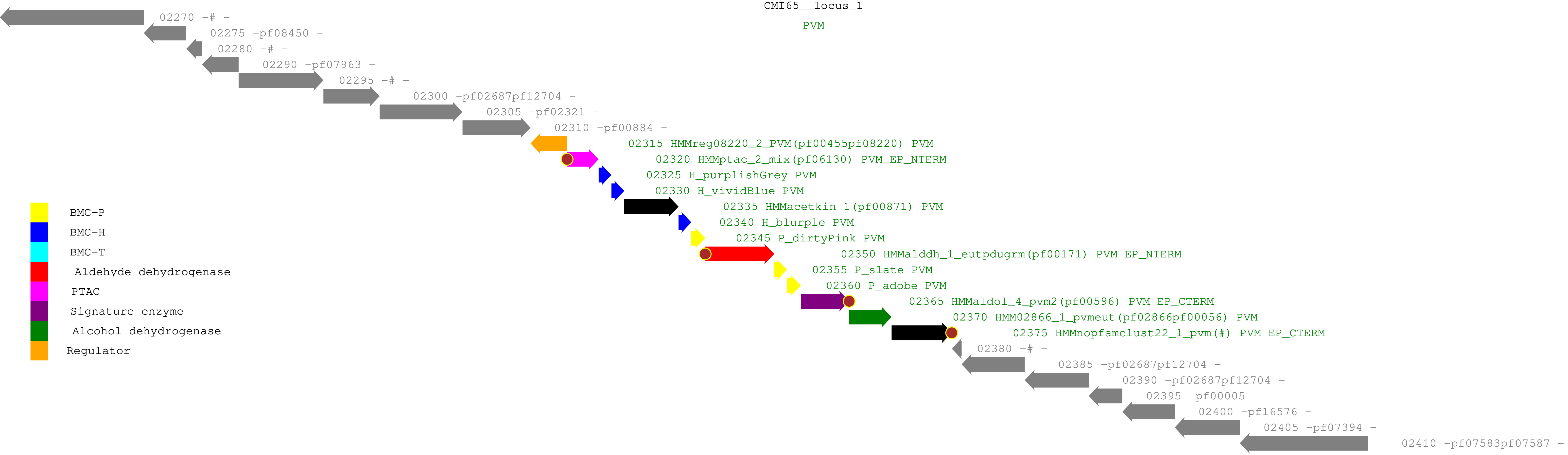

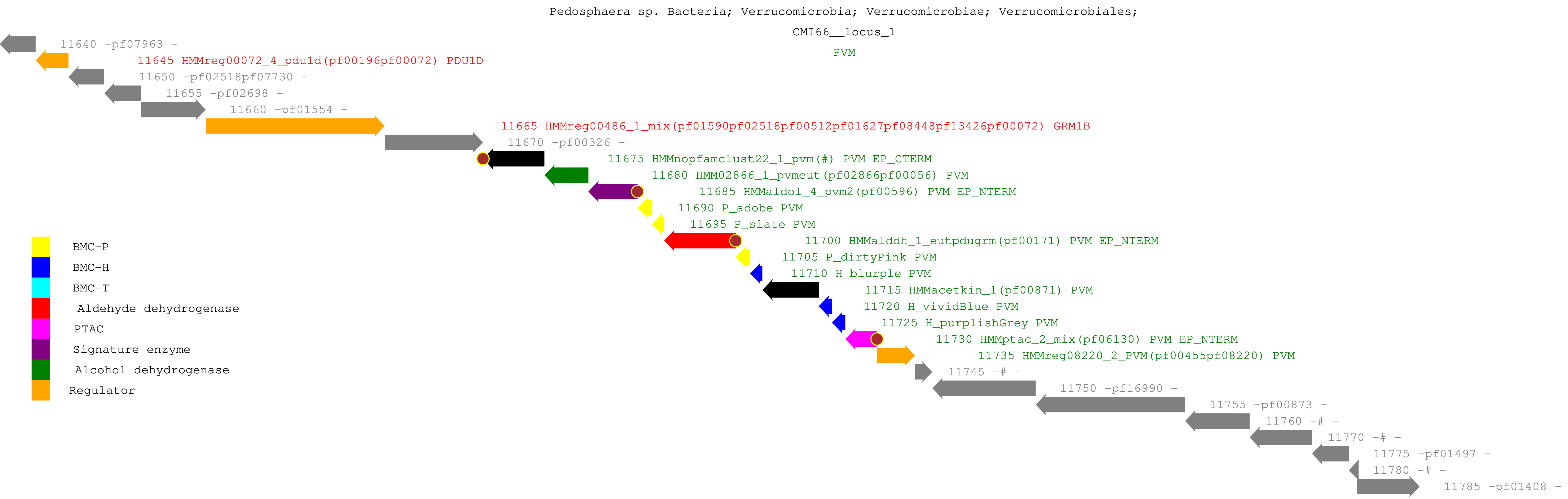
Pedosphaera sp. Bacteria; Verrucomicrobia; Verrucomicrobiae; Verrucomicrobiales;
CMI66__locus_1
PVM
11640 -pf07963 -
11645 HMMreg00072_4_pdu1d(pf00196pf00072) PDU1D
11650 -pf02518pf07730 -
11655 -pf02698 -
11660 -pf01554 -
11665 HMMreg00486_1_mix(pf01590pf02518pf00512pf01627pf08448pf13426pf00072) GRM1B
11670 -pf00326 -
11675 HMMnopfamclust22_1_pvm(#) PVM EP_CTERM
11680 HMM02866_1_pvmeut(pf02866pf00056) PVM
11685 HMMaldol_4_pvm2(pf00596) PVM EP_NTERM
11690 P_adobe PVM
11695 P_slate PVM
11700 HMMalddh_1_eutpdugrm(pf00171) PVM EP_NTERM
11705 P_dirtyPink PVM
11710 H_blurple PVM
11715 HMMacetkin_1(pf00871) PVM
11720 H_vividBlue PVM
11725 H_purplishGrey PVM
11730 HMMptac_2_mix(pf06130) PVM EP_NTERM
11735 HMMreg08220_2_PVM(pf00455pf08220) PVM
11745 -# -
11750 -pf16990 -
11755 -pf00873 -
11760 -# -
11770 -# -
11775 -pf01497 -
11780 -# -
11785 -pf01408 -
BMC-P
BMC-H
BMC-T
Aldehyde dehydrogenase
PTAC
Signature enzyme
Alcohol dehydrogenase
Regulator

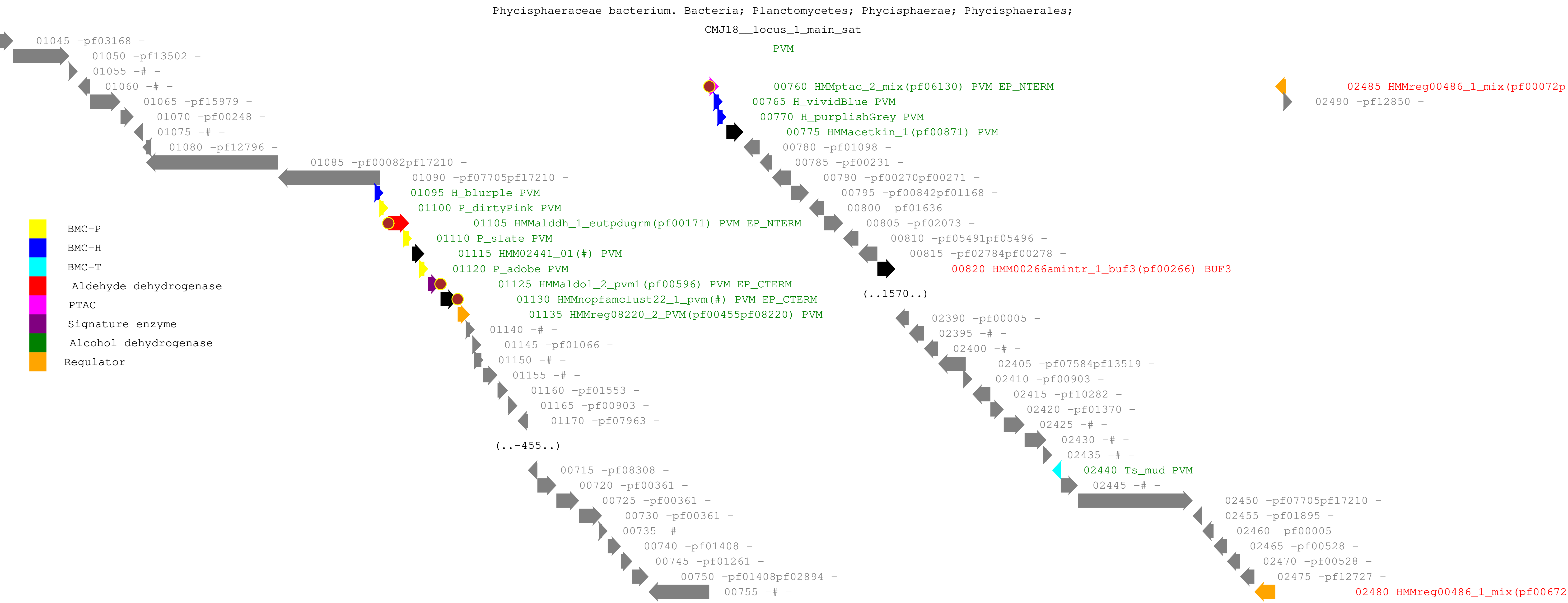
Phycisphaeraceae bacterium. Bacteria; Planctomycetes; Phycisphaerae; Phycisphaerales;
CMJ18__locus_1_main_sat
PVM
01045 -pf03168 -
01050 -pf13502 -
01055 -# -
01060 -# -
01065 -pf15979 -
01070 -pf00248 -
01075 -# -
01080 -pf12796 -
01085 -pf00082pf17210 -
01090 -pf07705pf17210 -
01095 H_blurple PVM
01100 P_dirtyPink PVM
01105 HMMalddh_1_eutpdugrm(pf00171) PVM EP_NTERM
01110 P_slate PVM
01115 HMM02441_01(#) PVM
01120 P_adobe PVM
01125 HMMaldol_2_pvm1(pf00596) PVM EP_CTERM
01130 HMMnopfamclust22_1_pvm(#) PVM EP_CTERM
01135 HMMreg08220_2_PVM(pf00455pf08220) PVM
01140 -# -
01145 -pf01066 -
01150 -# -
01155 -# -
01160 -pf01553 -
01165 -pf00903 -
01170 -pf07963 -
(..-455..)
00715 -pf08308 -
00720 -pf00361 -
00725 -pf00361 -
00730 -pf00361 -
00735 -# -
00740 -pf01408 -
00745 -pf01261 -
00750 -pf01408pf02894 -
00755 -# -
00760 HMMptac_2_mix(pf06130) PVM EP_NTERM
00765 H_vividBlue PVM
00770 H_purplishGrey PVM
00775 HMMacetkin_1(pf00871) PVM
00780 -pf01098 -
00785 -pf00231 -
00790 -pf00270pf00271 -
00795 -pf00842pf01168 -
00800 -pf01636 -
00805 -pf02073 -
00810 -pf05491pf05496 -
00815 -pf02784pf00278 -
00820 HMM00266amintr_1_buf3(pf00266) BUF3
(..1570..)
02390 -pf00005 -
02395 -# -
02400 -# -
02405 -pf07584pf13519 -
02410 -pf00903 -
02415 -pf10282 -
02420 -pf01370 -
02425 -# -
02430 -# -
02435 -# -
02440 Ts_mud PVM
02445 -# -
02450 -pf07705pf17210 -
02455 -pf01895 -
02460 -pf00005 -
02465 -pf00528 -
02470 -pf00528 -
02475 -pf12727 -
02480 HMMreg00486_1_mix(pf00672
02485 HMMreg00486_1_mix(pf00072p
02490 -pf12850 -
BMC-P
BMC-H
BMC-T
Aldehyde dehydrogenase
PTAC
Signature enzyme
Alcohol dehydrogenase
Regulator

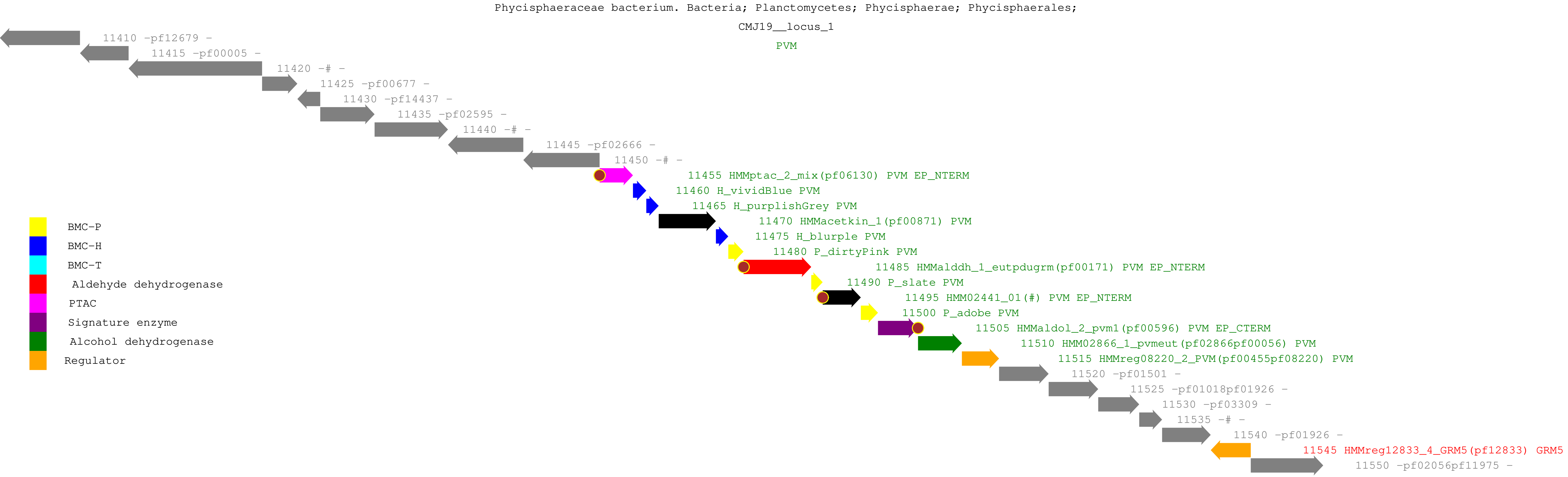

Phycisphaeraceae bacterium. Bacteria; Planctomycetes; Phycisphaerae; Phycisphaerales;
CMJ19__locus_1
PVM
11410 -pf12679 -
11415 -pf00005 -
11420 -# -
11425 -pf00677 -
11430 -pf14437 -
11435 -pf02595 -
11440 -# -
11445 -pf02666 -
11450 -# -
11455 HMMptac_2_mix(pf06130) PVM EP_NTERM
11460 H_vividBlue PVM
11465 H_purplishGrey PVM
11470 HMMacetkin_1(pf00871) PVM
11475 H_blurple PVM
11480 P_dirtyPink PVM
11485 HMMaldDH_1_eutpdugrm(pf00171) PVM EP_NTERM
11490 P_slate PVM
11495 HMM02441_01(#) PVM EP_NTERM
11500 P_adobe PVM
11505 HMMaldol_2_pvm1(pf00596) PVM EP_CTERM
11510 HMM02866_1_pvmeut(pf02866pf00056) PVM
11515 HMMreg08220_2_PVM(pf00455pf08220) PVM
11520 -pf01501 -
11525 -pf01018pf01926 -
11530 -pf03309 -
11535 -# -
11540 -pf01926 -
11545 HMMreg12833_4_GRM5(pf12833) GRM5
11550 -pf02056pf11975 -
BMC-P
BMC-H
BMC-T
Aldehyde dehydrogenase
PTAC
Signature enzyme
Alcohol dehydrogenase
Regulator

Phycisphaeraceae bacterium. Bacteria; Planctomycetes; Phycisphaerae; Phycisphaerales;

CMJ20__locus_1

PVM

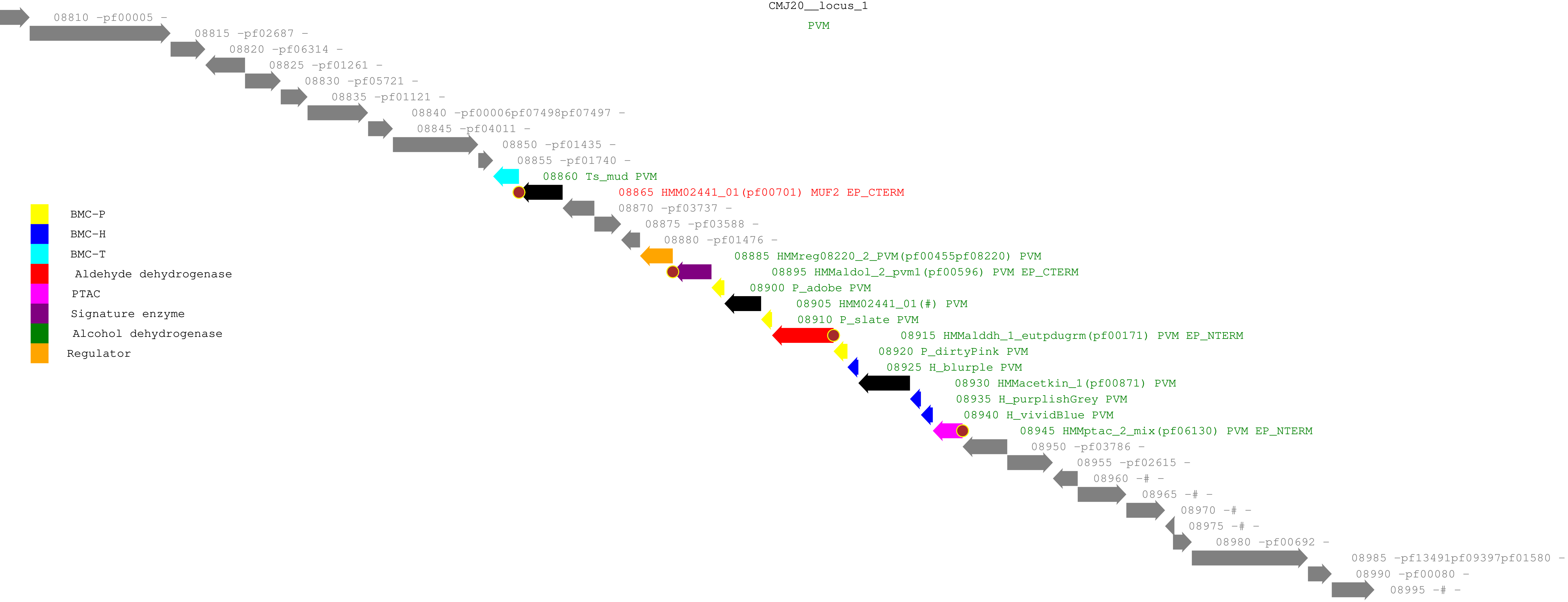

Planctomycetaceae bacterium. Bacteria; Planctomycetes; Planctomycetia; Planctomycetales;

CMJ49__locus_fragmented

PVM

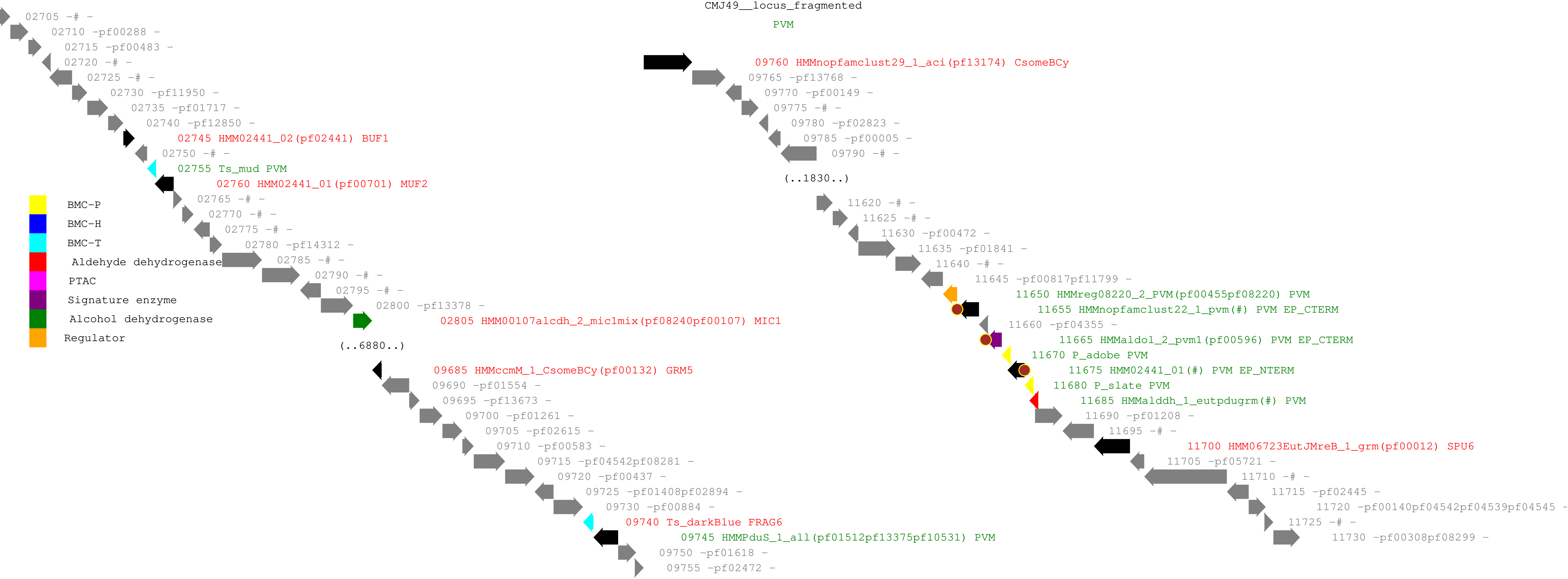

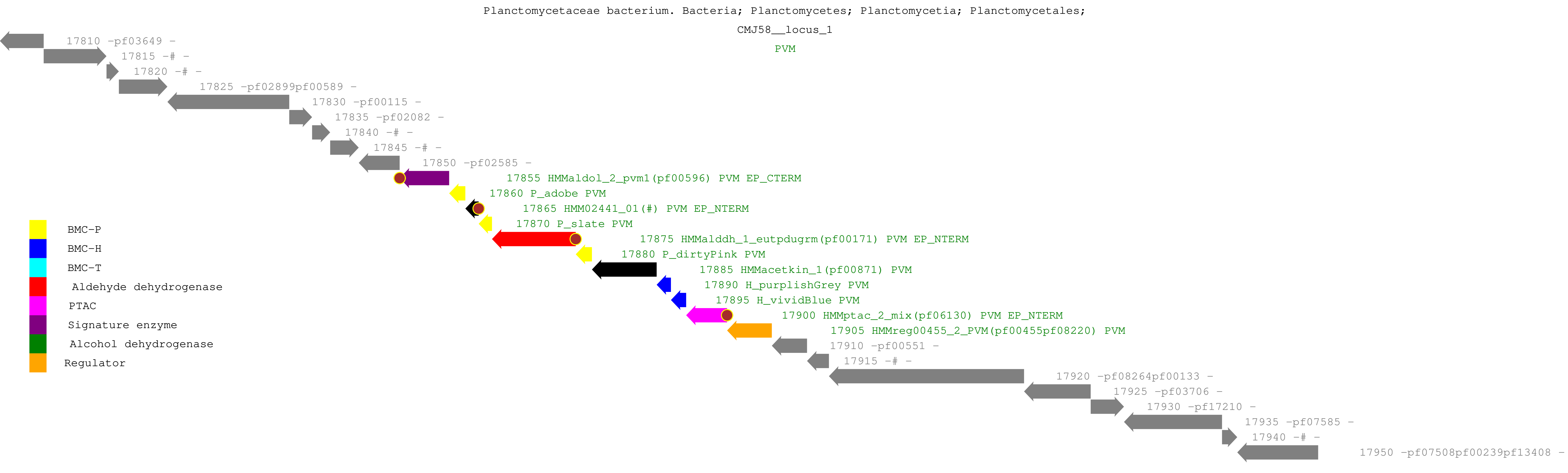
Planctomycetaceae bacterium. Bacteria; Planctomycetes; Planctomycetia; Planctomycetales;
CMJ58__locus_1
PVM
17810 -pf03649 -
17815 -# -
17820 -# -
17825 -pf02899pf00589 -
17830 -pf00115 -
17835 -pf02082 -
17840 -# -
17845 -# -
17850 -pf02585 -
17855 HMMaldol_2_pvm1(pf00596) PVM EP_CTERM
17860 P_adobe PVM
17865 HMM02441_01(#) PVM EP_NTERM
17870 P_slate PVM
17875 HMMalddh_1_eutpdugrm(pf00171) PVM EP_NTERM
17880 P_dirtyPink PVM
17885 HMMacetkin_1(pf00871) PVM
17890 H_purplishGrey PVM
17895 H_vividBlue PVM
17900 HMMptac_2_mix(pf06130) PVM EP_NTERM
17905 HMMreg00455_2_PVM(pf00455pf08220) PVM
17910 -pf00551 -
17915 -# -
17920 -pf08264pf00133 -
17925 -pf03706 -
17930 -pf17210 -
17935 -pf07585 -
17940 -# -
17950 -pf07508pf00239pf13408 -
BMC-P
BMC-H
BMC-T
Aldehyde dehydrogenase
PTAC
Signature enzyme
Alcohol dehydrogenase
Regulator

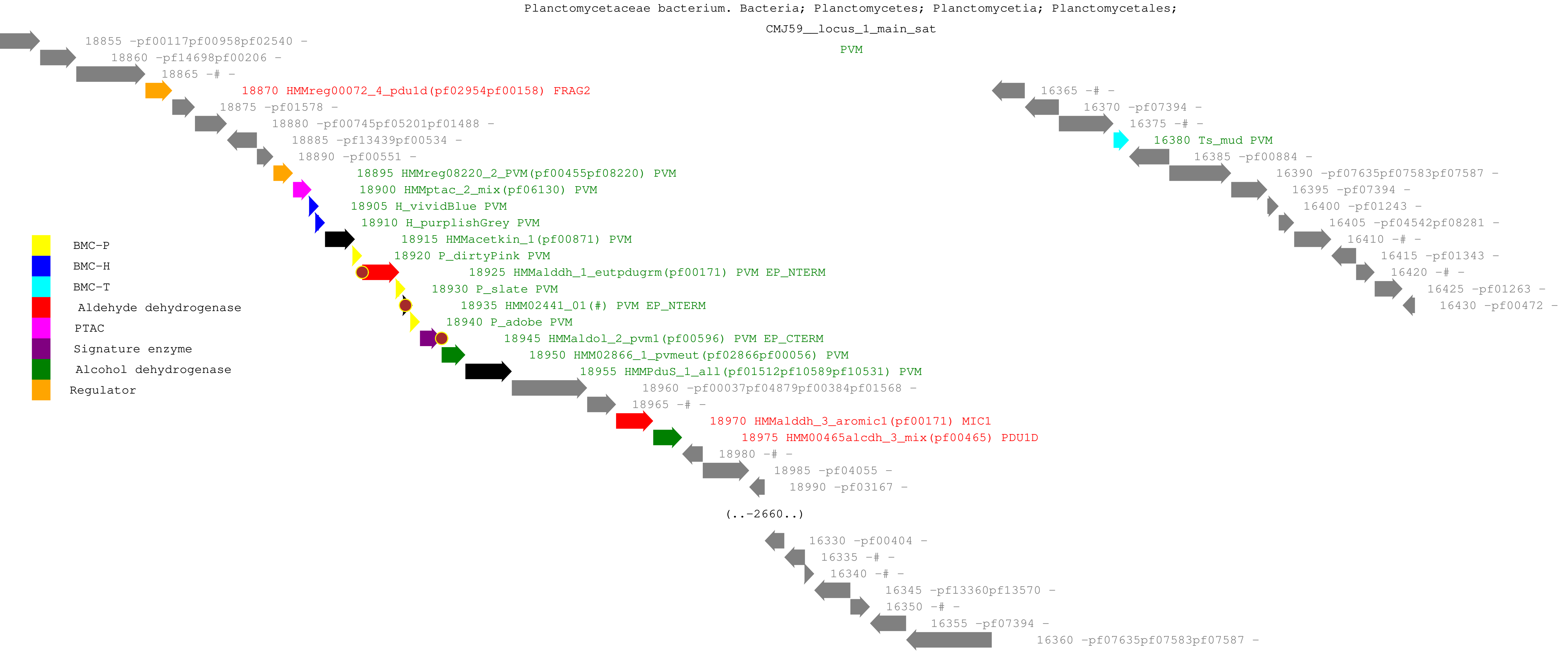
Planctomycetaceae bacterium. Bacteria; Planctomycetes; Planctomycetia; Planctomycetales;
CMJ59__locus_1_main_sat
PVM
18855 -pf00117pf00958pf02540 -
18860 -pf14698pf00206 -
18865 -# -
18870 HMMreg00072_4_pdu1d(pf02954pf00158) FRAG2
18875 -pf01578 -
18880 -pf00745pf05201pf01488 -
18885 -pf13439pf00534 -
18890 -pf00551 -
18895 HMMreg08220_2_PVM(pf00455pf08220) PVM
18900 HMMptac_2_mix(pf06130) PVM
18905 H_vividBlue PVM
18910 H_purplishGrey PVM
18915 HMMacetkin_1(pf00871) PVM
18920 P_dirtyPink PVM
18925 HMMalddh_1_eutpdugrm(pf00171) PVM EP_NTERM
18930 P_slate PVM
18935 HMM02441_01(#) PVM EP_NTERM
18940 P_adobe PVM
18945 HMMaldol_2_pvm1(pf00596) PVM EP_CTERM
18950 HMM02866_1_pvmeut(pf02866pf00056) PVM
18955 HMMPduS_1_all(pf01512pf10589pf10531) PVM
18960 -pf00037pf04879pf00384pf01568 -
18965 -# -
18970 HMMaldDH_3_aromic1(pf00171) MIC1
18975 HMM00465alcdh_3_mix(pf00465) PDU1D
18980 -# -
18985 -pf04055 -
18990 -pf03167 -
(..-2660..)
16330 -pf00404 -
16335 -# -
16340 -# -
16345 -pf13360pf13570 -
16350 -# -
16355 -pf07394 -
16360 -pf07635pf07583pf07587 -
16365 -# -
16370 -pf07394 -
16375 -# -
16380 Ts_mud PVM
16385 -pf00884 -
16390 -pf07635pf07583pf07587 -
16395 -pf07394 -
16400 -pf01243 -
16405 -pf04542pf08281 -
16410 -# -
16415 -pf01343 -
16420 -# -
16425 -pf01263 -
16430 -pf00472 -
BMC-P
BMC-H
BMC-T
Aldehyde dehydrogenase
PTAC
Signature enzyme
Alcohol dehydrogenase
Regulator

Planctomycetaceae bacterium. Bacteria; Planctomycetes; Planctomycetia; Planctomycetales;

CMJ64__locus_1_main_sat

PVM

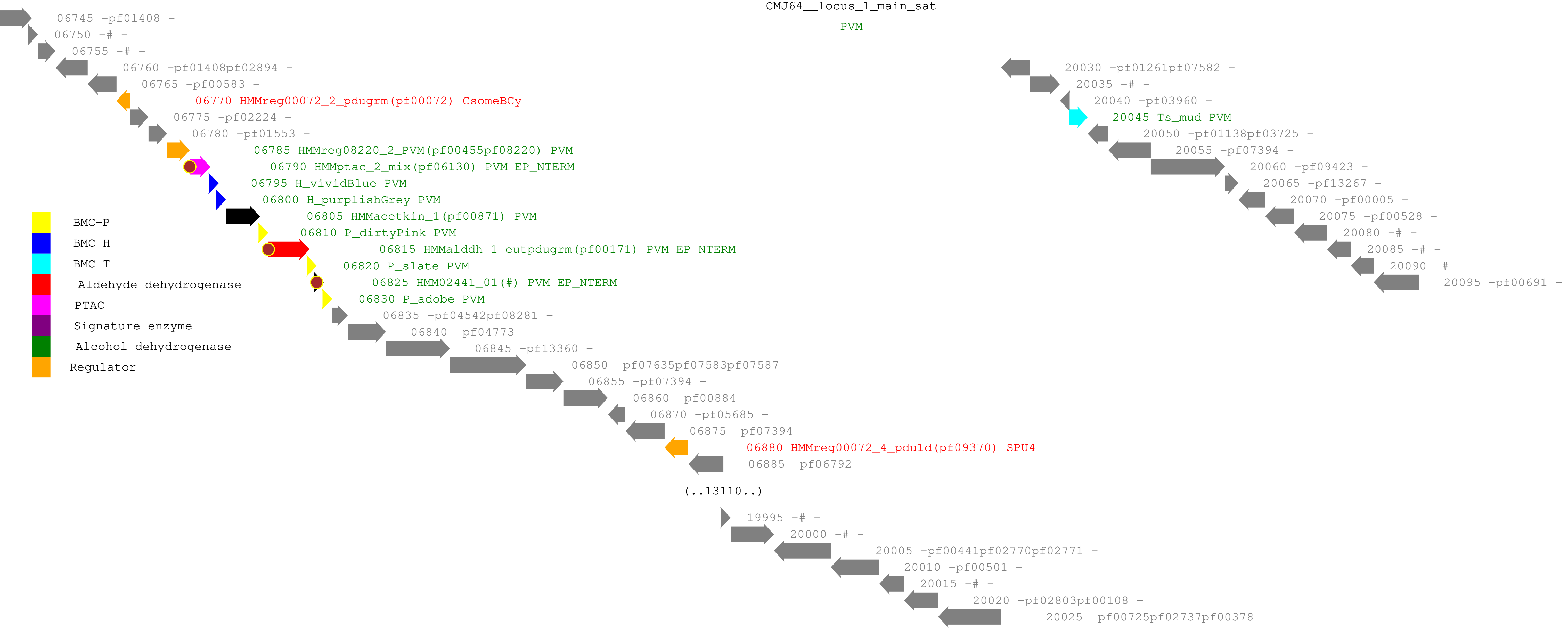

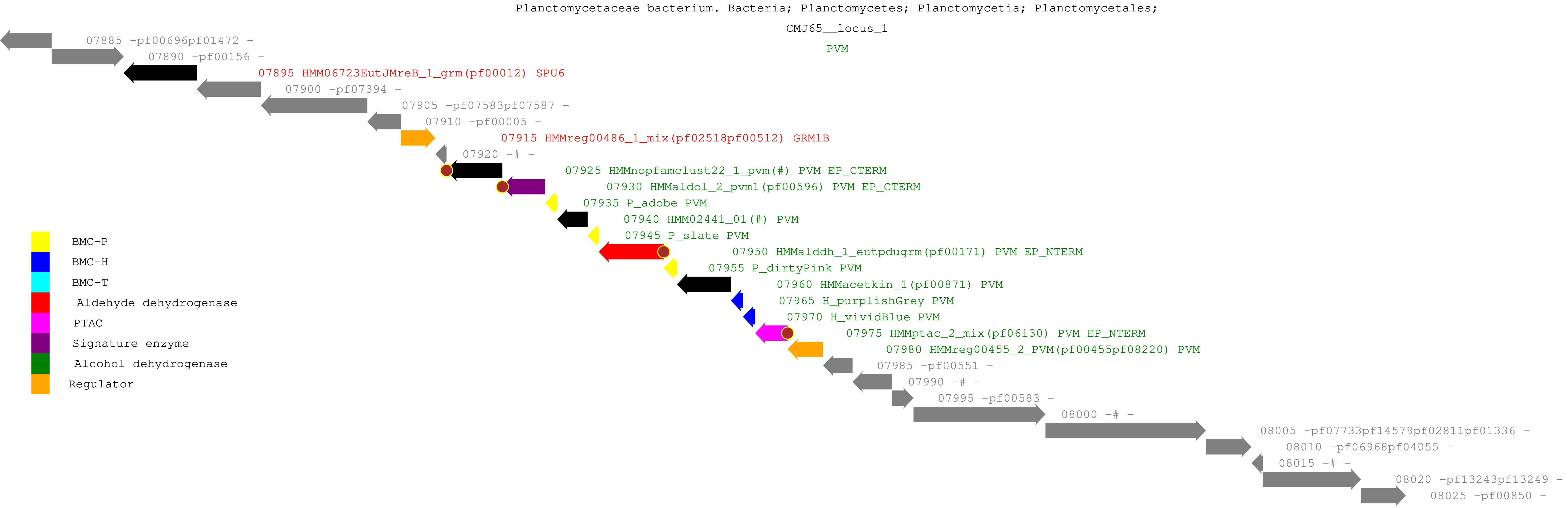

Planctomycetaceae bacterium. Bacteria; Planctomycetes; Planctomycetia; Planctomycetales;
CMJ65__locus_1
PVM
07885 -pf00696pf01472 -
07890 -pf00156 -
07895 HMM06723EutJMreB_1_grm(pf00012) SPU6
07900 -pf07394 -
07905 -pf07583pf07587 -
07910 -pf00005 -
07915 HMMreg00486_1_mix(pf02518pf00512) GRM1B
07920 -# -
07925 HMMnopfamclust22_1_pvm(#) PVM EP_CTERM
07930 HMMaldol_2_pvm1(pf00596) PVM EP_CTERM
07935 P_adobe PVM
07940 HMM02441_01(#) PVM
07945 P_slate PVM
07950 HMMalddh_1_eutpdugrm(pf00171) PVM EP_NTERM
07955 P_dirtyPink PVM
07960 HMMacetkin_1(pf00871) PVM
07965 H_purplishGrey PVM
07970 H_vividBlue PVM
07975 HMMptac_2_mix(pf06130) PVM EP_NTERM
07980 HMMreg00455_2_PVM(pf00455pf08220) PVM
07985 -pf00551 -
07990 -# -
07995 -pf00583 -
08000 -# -
08005 -pf07733pf14579pf02811pf01336 -
08010 -pf06968pf04055 -
08015 -# -
08020 -pf13243pf13249 -
08025 -pf00850 -
BMC-P
BMC-H
BMC-T
Aldehyde dehydrogenase
PTAC
Signature enzyme
Alcohol dehydrogenase
Regulator

Planctomycetaceae bacterium. Bacteria; Planctomycetes; Planctomycetia; Planctomycetales;

CMJ66__locus_1

PVM

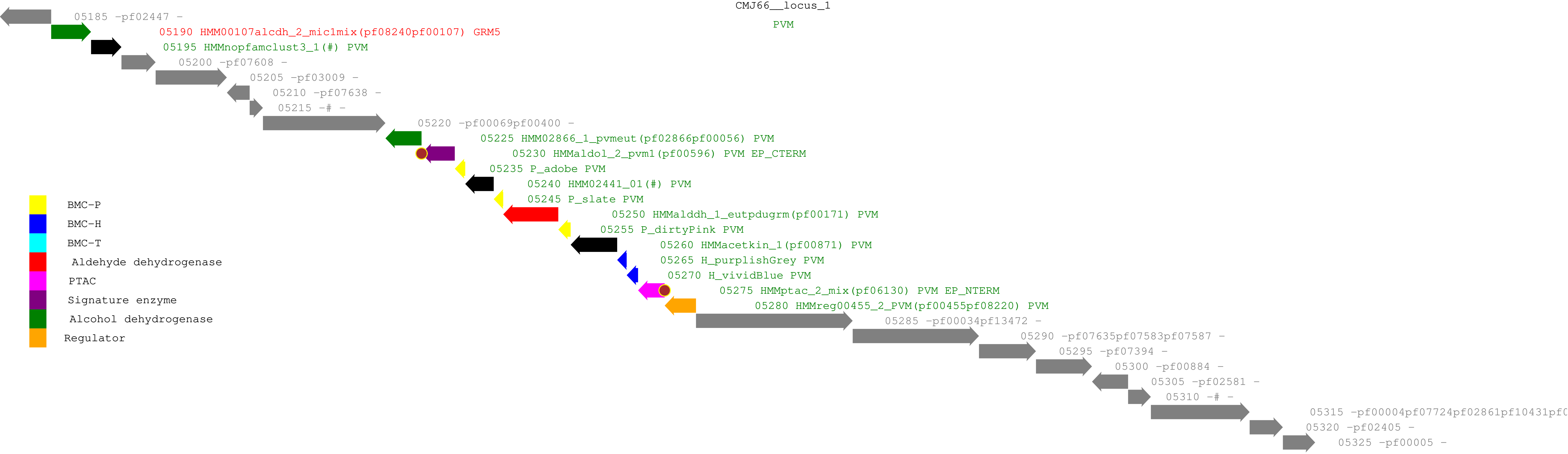

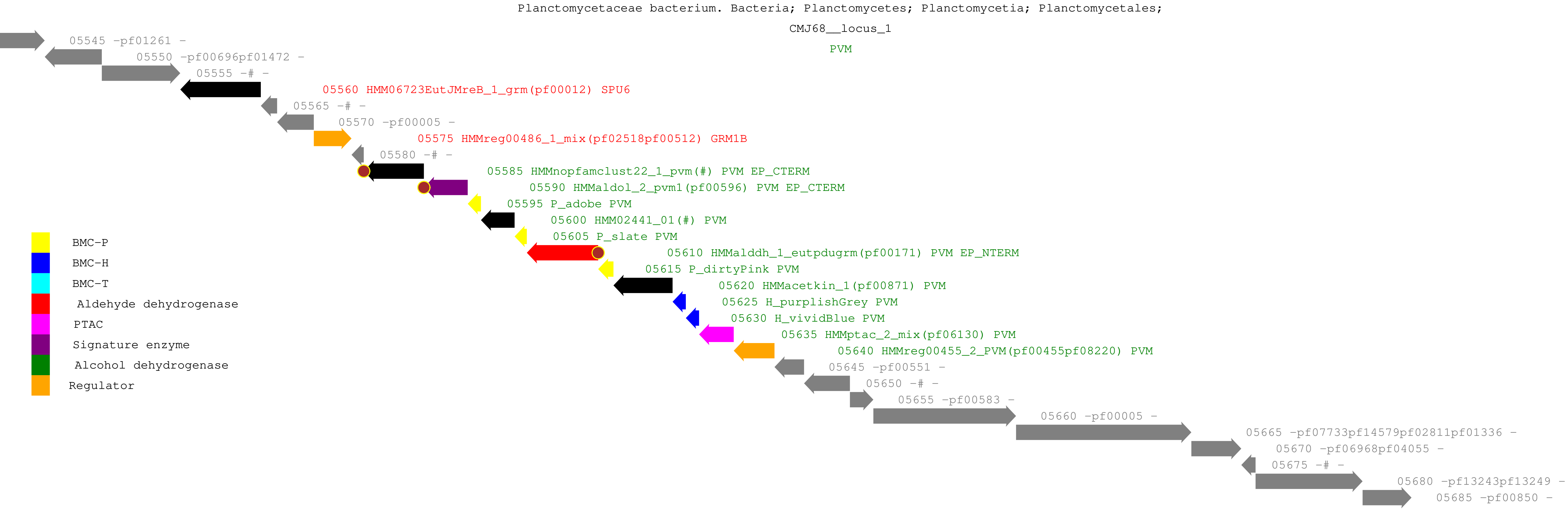
Planctomycetaceae bacterium. Bacteria; Planctomycetes; Planctomycetia; Planctomycetales;
CMJ68__locus_1
PVM
05545 -pf01261 -
05550 -pf00696pf01472 -
05555 -# -
05560 HMM06723EutJMreB_1_grm(pf00012) SPU6
05565 -# -
05570 -pf00005 -
05575 HMMreg00486_1_mix(pf02518pf00512) GRM1B
05580 -# -
05585 HMMnopfamclust22_1_pvm(#) PVM EP_CTERM
05590 HMMaldol_2_pvm1(pf00596) PVM EP_CTERM
05595 P_adobe PVM
05600 HMM02441_01(#) PVM
05605 P_slate PVM
05610 HMMalddh_1_eutpdugrm(pf00171) PVM EP_NTERM
05615 P_dirtyPink PVM
05620 HMMacetkin_1(pf00871) PVM
05625 H_purplishGrey PVM
05630 H_vividBlue PVM
05635 HMMptac_2_mix(pf06130) PVM
05640 HMMreg00455_2_PVM(pf00455pf08220) PVM
05645 -pf00551 -
05650 -# -
05655 -pf00583 -
05660 -pf00005 -
05665 -pf07733pf14579pf02811pf01336 -
05670 -pf06968pf04055 -
05675 -# -
05680 -pf13243pf13249 -
05685 -pf00850 -
BMC-P
BMC-H
BMC-T
Aldehyde dehydrogenase
PTAC
Signature enzyme
Alcohol dehydrogenase
Regulator

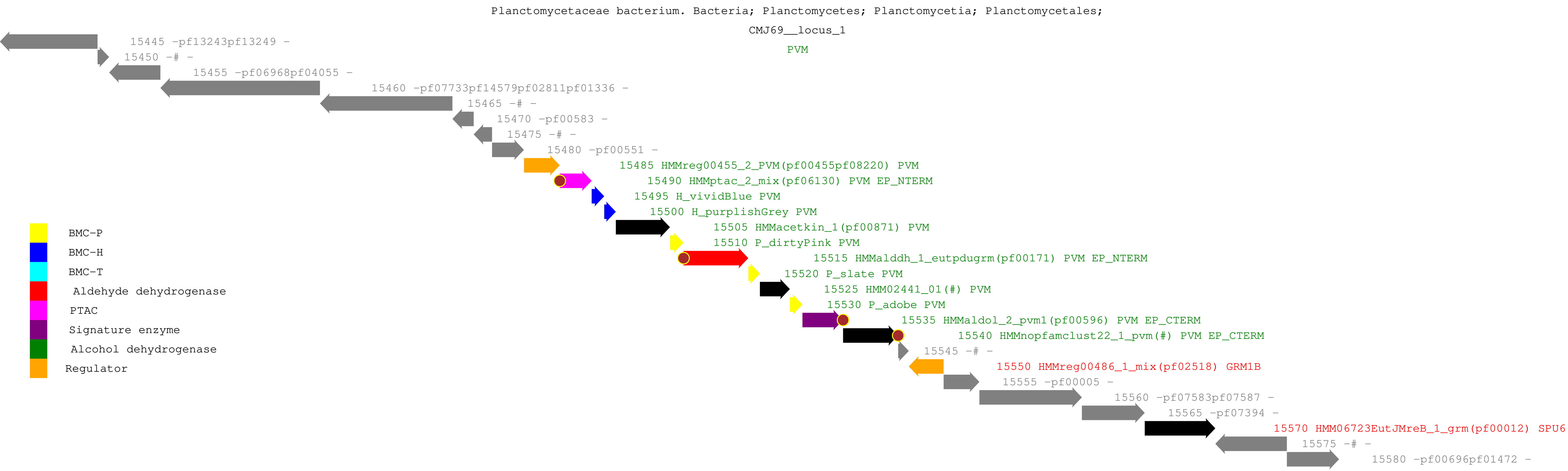

Planctomycetaceae bacterium. Bacteria; Planctomycetes; Planctomycetia; Planctomycetales;
CMJ69__locus_1
PVM
15445 -pf13243pf13249 -
15450 -# -
15455 -pf06968pf04055 -
15460 -pf07733pf14579pf02811pf01336 -
15465 -# -
15470 -pf00583 -
15475 -# -
15480 -pf00551 -
15485 HMMreg00455_2_PVM(pf00455pf08220) PVM
15490 HMMptac_2_mix(pf06130) PVM EP_NTERM
15495 H_vividBlue PVM
15500 H_purplishGrey PVM
15505 HMMacetkin_1(pf00871) PVM
15510 P_dirtyPink PVM
15515 HMMalddh_1_eutpdugrm(pf00171) PVM EP_NTERM
15520 P_slate PVM
15525 HMM02441_01(#) PVM
15530 P_adobe PVM
15535 HMMaldol_2_pvm1(pf00596) PVM EP_CTERM
15540 HMMnopfamclust22_1_pvm(#) PVM EP_CTERM
15545 -# -
15550 HMMreg00486_1_mix(pf02518) GRM1B
15555 -pf00005 -
15560 -pf07583pf07587 -
15565 -pf07394 -
15570 HMM06723EutJMreB_1_grm(pf00012) SPU6
15575 -# -
15580 -pf00696pf01472 -
BMC-P
BMC-H
BMC-T
Aldehyde dehydrogenase
PTAC
Signature enzyme
Alcohol dehydrogenase
Regulator

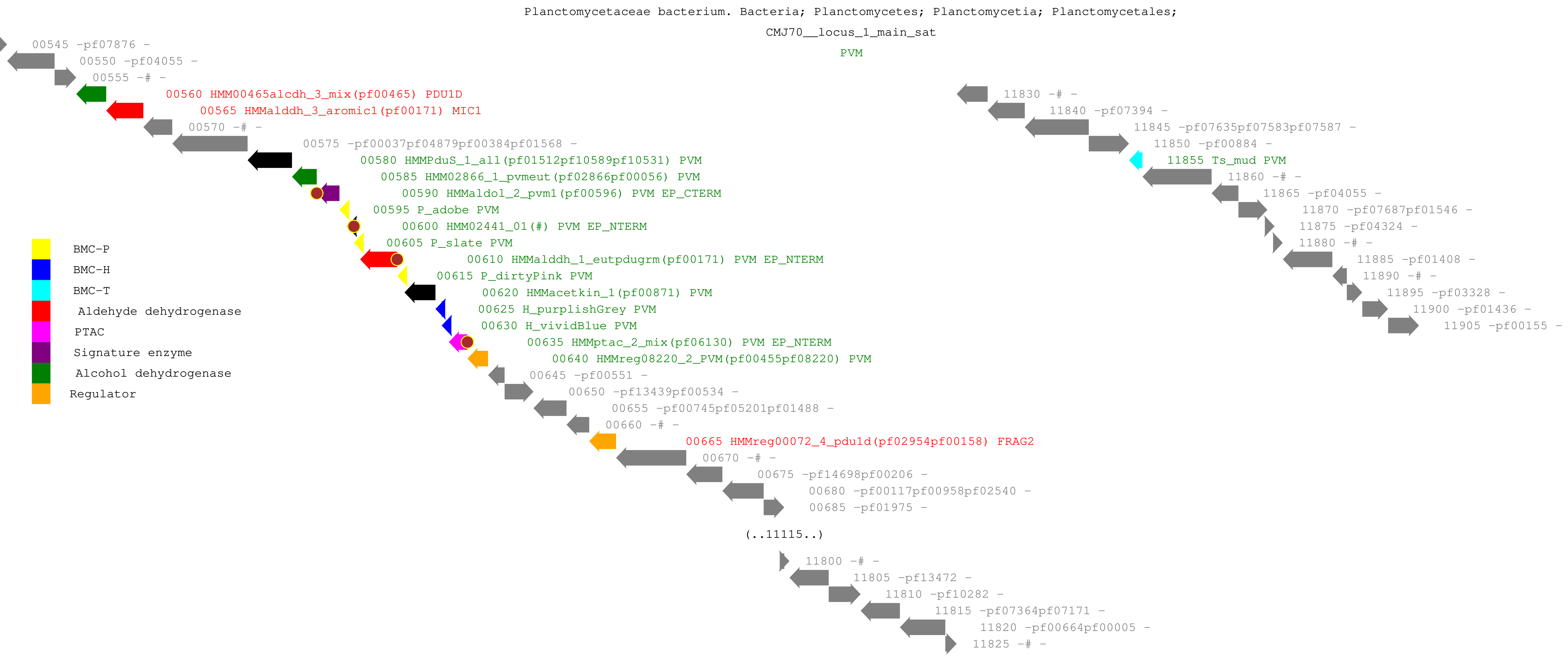
Planctomycetaceae bacterium. Bacteria; Planctomycetes; Planctomycetia; Planctomycetales;
CMJ70__locus_1_main_sat
PVM
00545 -pf07876 -
00550 -pf04055 -
00555 -# -
00560 HMM00465alcdh_3_mix(pf00465) PDU1D
00565 HMMalddh_3_aromic1(pf00171) MIC1
00570 -# -
00575 -pf00037pf04879pf00384pf01568 -
00580 HMMPduS_1_all(pf01512pf10589pf10531) PVM
00585 HMM02866_1_pvmeut(pf02866pf00056) PVM
00590 HMMaldol_2_pvm1(pf00596) PVM EP_CTERM
00595 P_adobe PVM
00600 HMM02441_01(#) PVM EP_NTERM
00605 P_slate PVM
00610 HMMalddh_1_eutpdugrm(pf00171) PVM EP_NTERM
00615 P_dirtyPink PVM
00620 HMMacetkin_1(pf00871) PVM
00625 H_purplishGrey PVM
00630 H_vividBlue PVM
00635 HMMptac_2_mix(pf06130) PVM EP_NTERM
00640 HMMreg08220_2_PVM(pf00455pf08220) PVM
00645 -pf00551 -
00650 -pf13439pf00534 -
00655 -pf00745pf05201pf01488 -
00660 -# -
00665 HMMreg00072_4_pdu1d(pf02954pf00158) FRAG2
00670 -# -
00675 -pf14698pf00206 -
00680 -pf00117pf00958pf02540 -
00685 -pf01975 -
(..11115..)
11800 -# -
11805 -pf13472 -
11810 -pf10282 -
11815 -pf07364pf07171 -
11820 -pf00664pf00005 -
11825 -# -
11830 -# -
11840 -pf07394 -
11845 -pf07635pf07583pf07587 -
11850 -pf00884 -
11855 Ts_mud PVM
11860 -# -
11865 -pf04055 -
11870 -pf07687pf01546 -
11875 -pf04324 -
11880 -# -
11885 -pf01408 -
11890 -# -
11895 -pf03328 -
11900 -pf01436 -
11905 -pf00155 -
BMC-P
BMC-H
BMC-T
Aldehyde dehydrogenase
PTAC
Signature enzyme
Alcohol dehydrogenase
Regulator

Planctomycetaceae bacterium. Bacteria; Planctomycetes; Planctomycetia; Planctomycetales;

CMJ73__locus_1

PVM

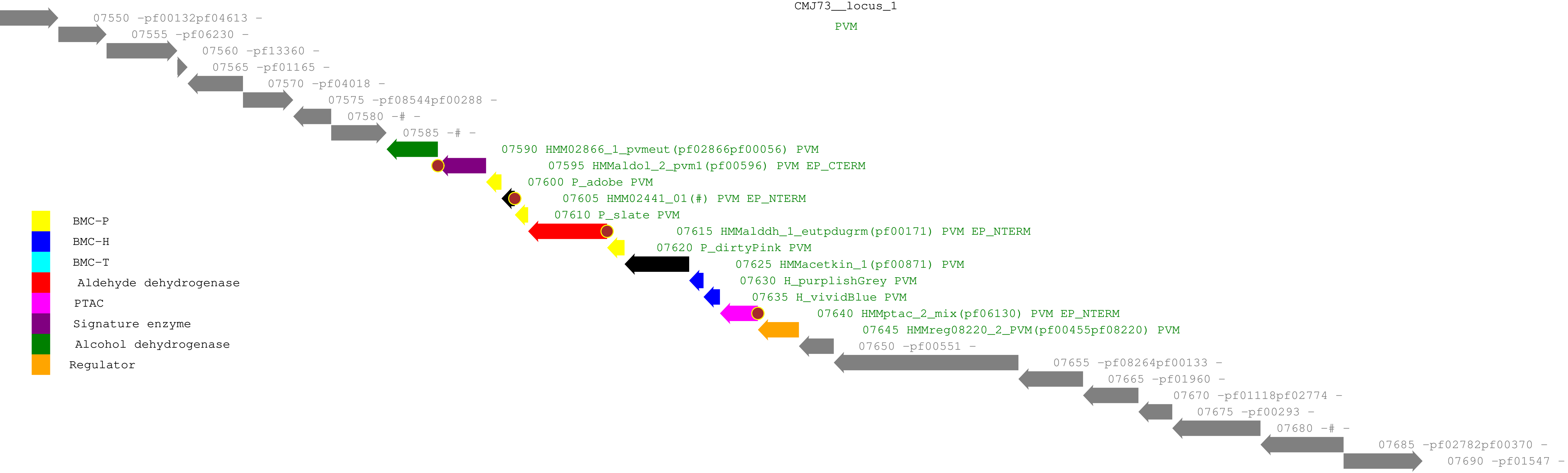

Planctomycetaceae bacterium. Bacteria; Planctomycetes; Planctomycetia; Planctomycetales;

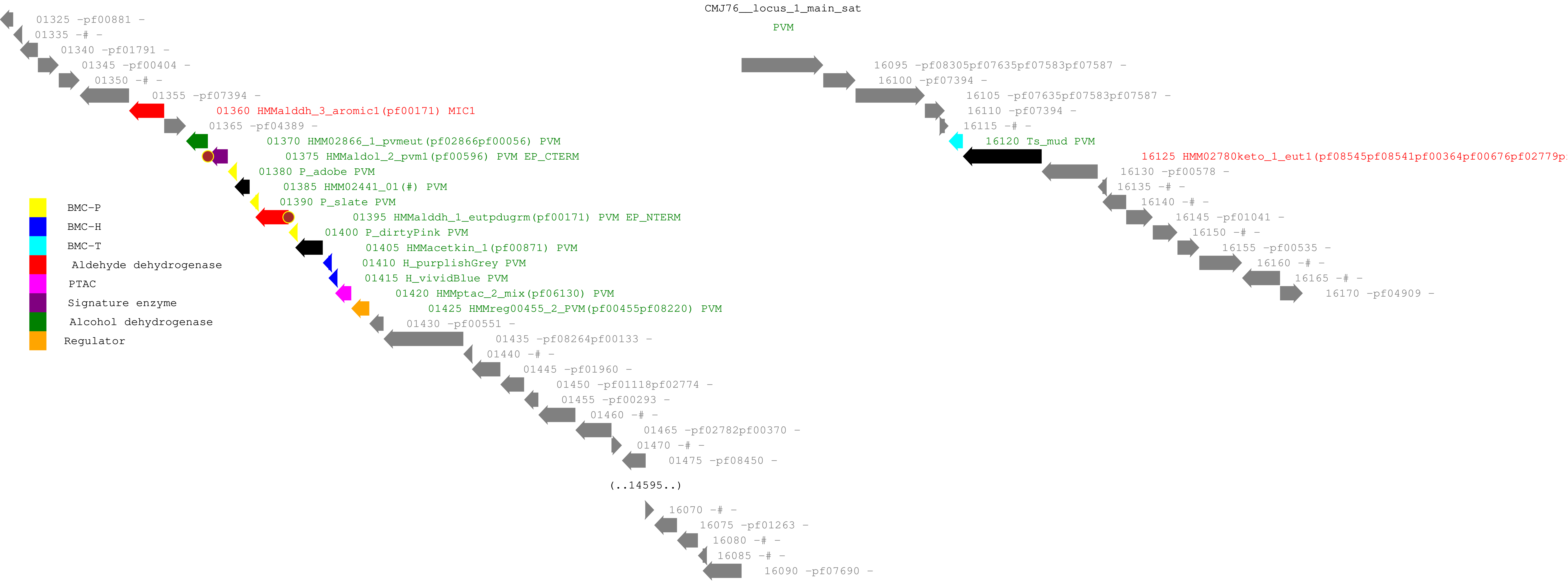

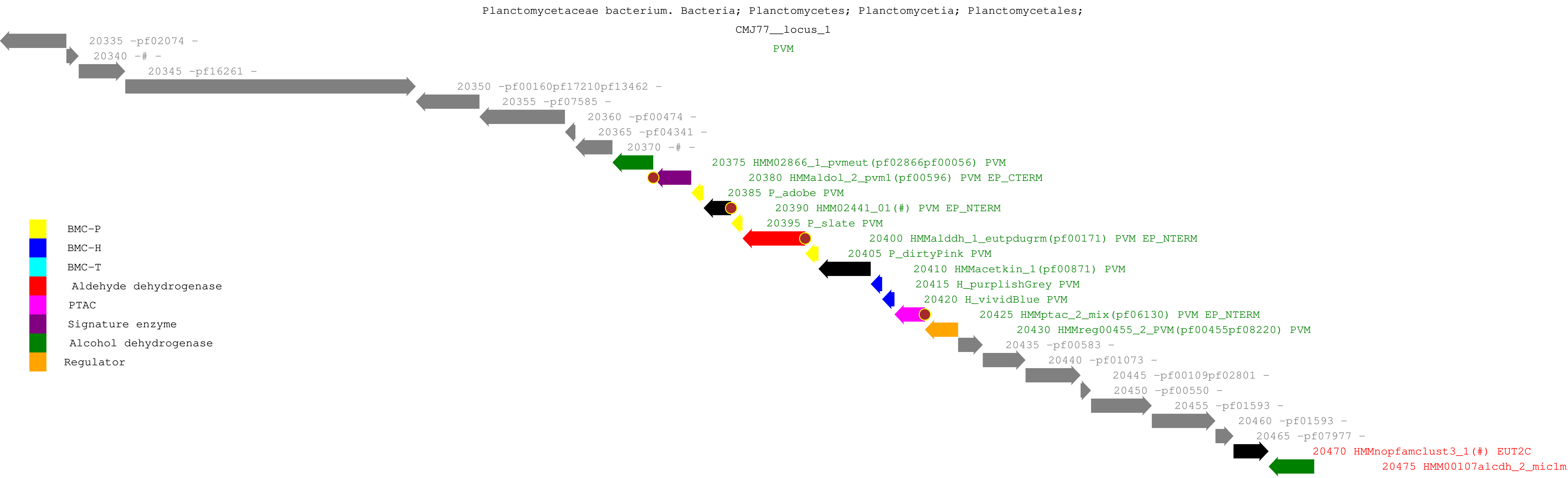
Planctomycetaceae bacterium. Bacteria; Planctomycetes; Planctomycetia; Planctomycetales;
CMJ77__locus_1
PVM
20335 -pf02074 -
20340 -# -
20345 -pf16261 -
20350 -pf00160pf17210pf13462 -
20355 -pf07585 -
20360 -pf00474 -
20365 -pf04341 -
20370 -# -
20375 HMM02866_1_pvmeut(pf02866pf00056) PVM
20380 HMMaldol_2_pvm1(pf00596) PVM EP_CTERM
20385 P_adobe PVM
20390 HMM02441_01(#) PVM EP_NTERM
20395 P_slate PVM
20400 HMMalddh_1_eutpdugrm(pf00171) PVM EP_NTERM
20405 P_dirtyPink PVM
20410 HMMacetkin_1(pf00871) PVM
20415 H_purplishGrey PVM
20420 H_vividBlue PVM
20425 HMMptac_2_mix(pf06130) PVM EP_NTERM
20430 HMMreg00455_2_PVM(pf00455pf08220) PVM
20435 -pf00583 -
20440 -pf01073 -
20445 -pf00109pf02801 -
20450 -pf00550 -
20455 -pf01593 -
20460 -pf01593 -
20465 -pf07977 -
20470 HMMnopfamclust3_1(#) EUT2C
20475 HMM00107alcdh_2_mic1m
BMC-P
BMC-H
BMC-T
Aldehyde dehydrogenase
PTAC
Signature enzyme
Alcohol dehydrogenase
Regulator

Planctomycetaceae bacterium. Bacteria; Planctomycetes; Planctomycetia; Planctomycetales;

CMJ78__locus_1_main_sat

PVM

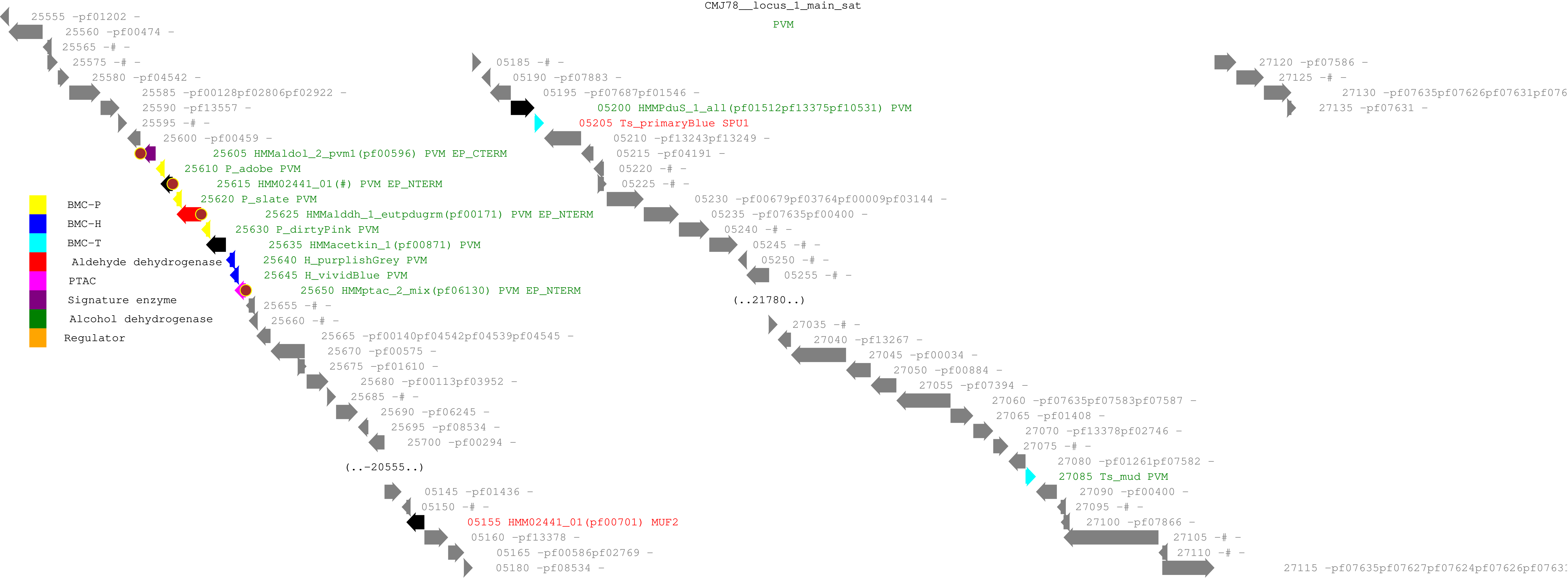

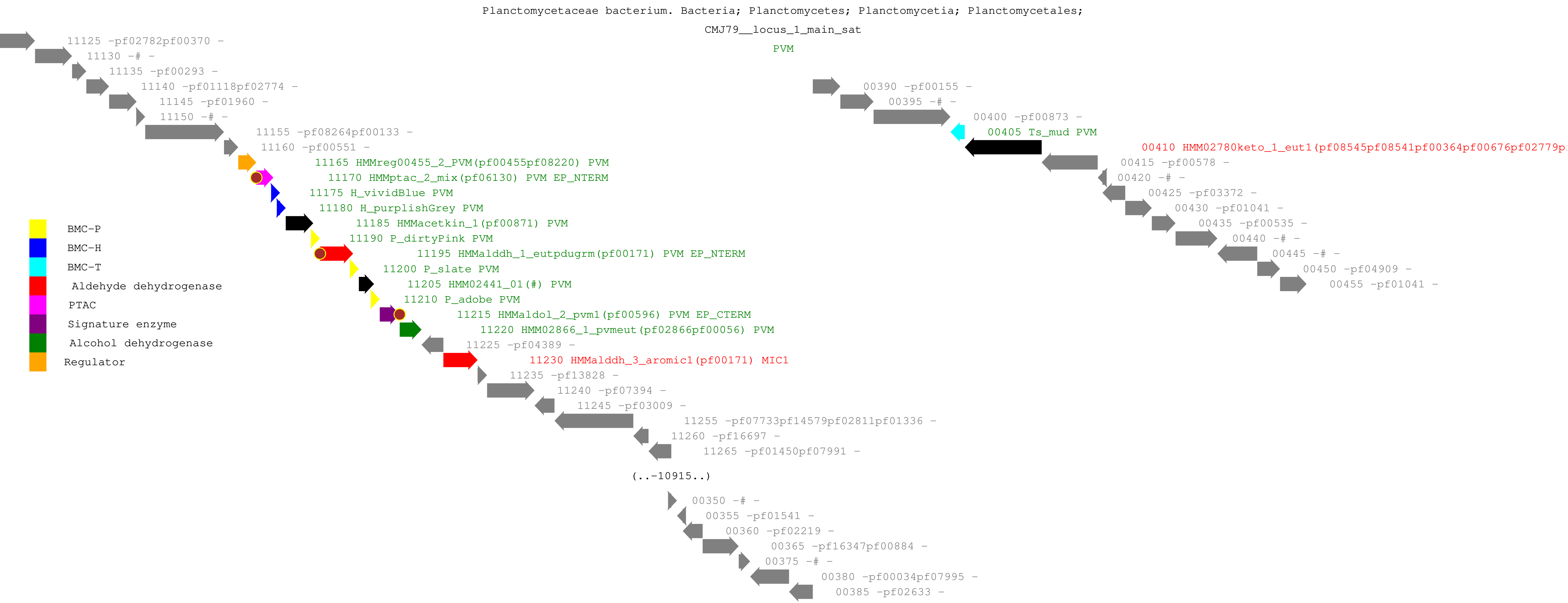

Planctomycetaceae bacterium. Bacteria; Planctomycetes; Planctomycetia; Planctomycetales;
CMJ79__locus_1_main_sat
PVM
11125 -pf02782pf00370 -
11130 -# -
11135 -pf00293 -
11140 -pf01118pf02774 -
11145 -pf01960 -
11150 -# -
11155 -pf08264pf00133 -
11160 -pf00551 -
11165 HMMreg00455_2_PVM(pf00455pf08220) PVM
11170 HMMptac_2_mix(pf06130) PVM EP_NTERM
11175 H_vividBlue PVM
11180 H_purplishGrey PVM
11185 HMMacetkin_1(pf00871) PVM
11190 P_dirtyPink PVM
11195 HMMalddh_1_eutpdugrm(pf00171) PVM EP_NTERM
11200 P_slate PVM
11205 HMM02441_01(#) PVM
11210 P_adobe PVM
11215 HMMaldol_2_pvm1(pf00596) PVM EP_CTERM
11220 HMM02866_1_pvmeut(pf02866pf00056) PVM
11225 -pf04389 -
11230 HMMalddh_3_aromic1(pf00171) MIC1
11235 -pf13828 -
11240 -pf07394 -
11245 -pf03009 -
11255 -pf07733pf14579pf02811pf01336 -
11260 -pf16697 -
11265 -pf01450pf07991 -
(..-10915..)
00350 -# -
00355 -pf01541 -
00360 -pf02219 -
00365 -pf16347pf00884 -
00375 -# -
00380 -pf00034pf07995 -
00385 -pf02633 -
00390 -pf00155 -
00395 -# -
00400 -pf00873 -
00405 Ts_mud PVM
00410 HMM02780keto_1_eut1(pf08545pf08541pf00364pf00676pf02779p
00415 -pf00578 -
00420 -# -
00425 -pf03372 -
00430 -pf01041 -
00435 -pf00535 -
00440 -# -
00445 -# -
00450 -pf04909 -
00455 -pf01041 -
BMC-P
BMC-H
BMC-T
Aldehyde dehydrogenase
PTAC
Signature enzyme
Alcohol dehydrogenase
Regulator

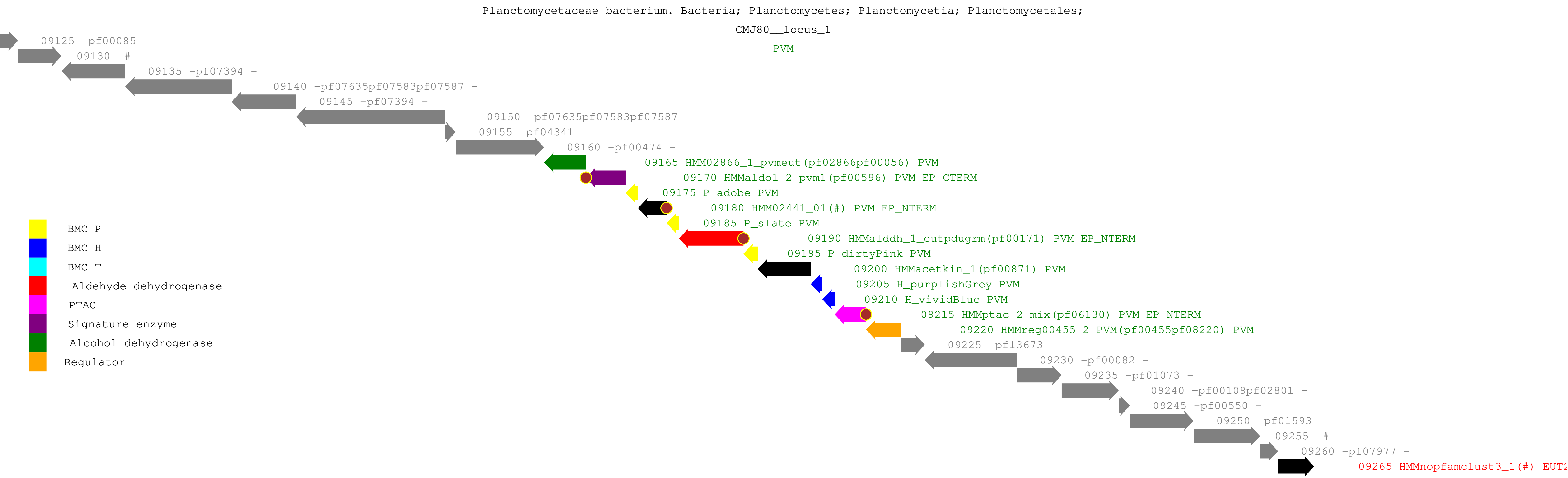

Planctomycetaceae bacterium. Bacteria; Planctomycetes; Planctomycetia; Planctomycetales;
CMJ80__locus_1
PVM
09125 -pf00085 -
09130 -# -
09135 -pf07394 -
09140 -pf07635pf07583pf07587 -
09145 -pf07394 -
09150 -pf07635pf07583pf07587 -
09155 -pf04341 -
09160 -pf00474 -
09165 HMM02866_1_pvmeut(pf02866pf00056) PVM
09170 HMMaldol_2_pvm1(pf00596) PVM EP_CTERM
09175 P_adobe PVM
09180 HMM02441_01(#) PVM EP_NTERM
09185 P_slate PVM
09190 HMMalddh_1_eutpdugrm(pf00171) PVM EP_NTERM
09195 P_dirtyPink PVM
09200 HMMacetkin_1(pf00871) PVM
09205 H_purplishGrey PVM
09210 H_vividBlue PVM
09215 HMMptac_2_mix(pf06130) PVM EP_NTERM
09220 HMMreg00455_2_PVM(pf00455pf08220) PVM
09225 -pf13673 -
09230 -pf00082 -
09235 -pf01073 -
09240 -pf00109pf02801 -
09245 -pf00550 -
09250 -pf01593 -
09255 -# -
09260 -pf07977 -
09265 HMMnopfamclust3_1(#) EUT2
BMC-P
BMC-H
BMC-T
Aldehyde dehydrogenase
PTAC
Signature enzyme
Alcohol dehydrogenase
Regulator

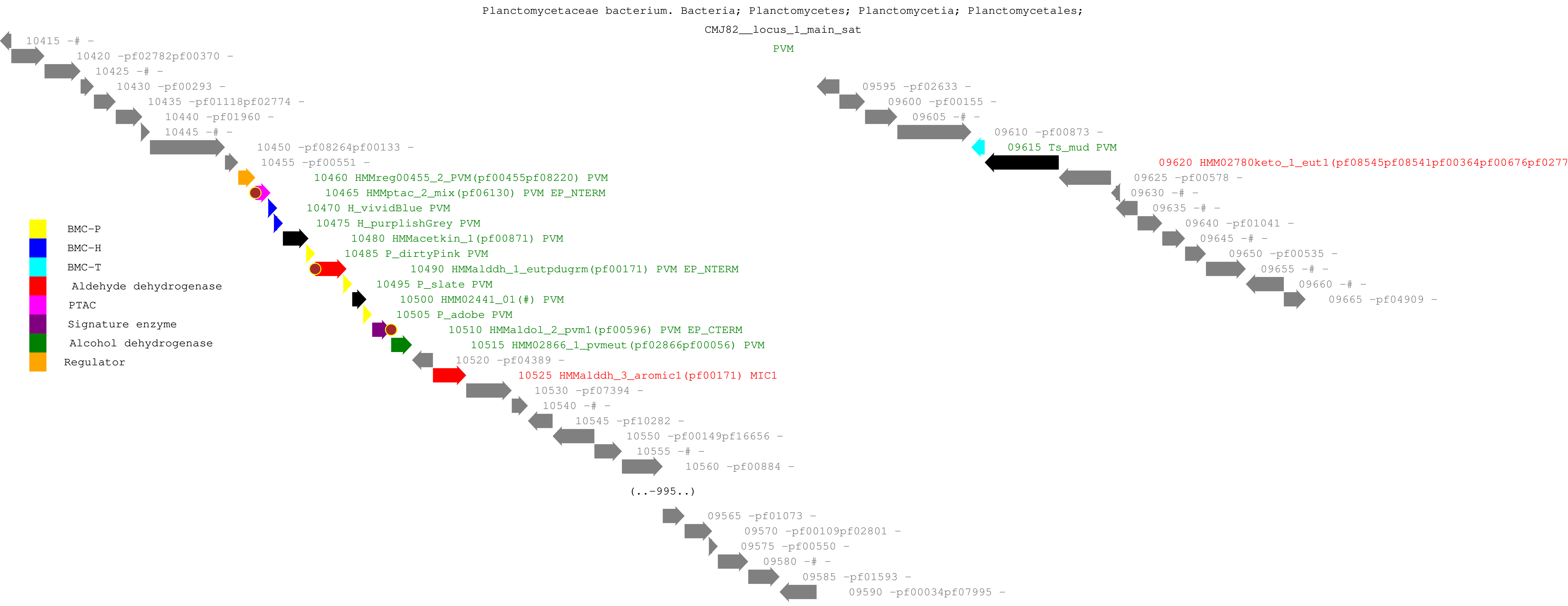
Planctomycetaceae bacterium. Bacteria; Planctomycetes; Planctomycetia; Planctomycetales;
CMJ82__locus_1_main_sat
PVM
10415 -# -
10420 -pf02782pf00370 -
10425 -# -
10430 -pf00293 -
10435 -pf01118pf02774 -
10440 -pf01960 -
10445 -# -
10450 -pf08264pf00133 -
10455 -pf00551 -
10460 HMMreg00455_2_PVM(pf00455pf08220) PVM
10465 HMMptac_2_mix(pf06130) PVM EP_NTERM
10470 H_vividBlue PVM
10475 H_purplishGrey PVM
10480 HMMacetkin_1(pf00871) PVM
10485 P_dirtyPink PVM
10490 HMMalddh_1_eutpdugrm(pf00171) PVM EP_NTERM
10495 P_slate PVM
10500 HMM02441_01(#) PVM
10505 P_adobe PVM
10510 HMMaldol_2_pvm1(pf00596) PVM EP_CTERM
10515 HMM02866_1_pvmeut(pf02866pf00056) PVM
10520 -pf04389 -
10525 HMMalddh_3_aromic1(pf00171) MIC1
10530 -pf07394 -
10540 -# -
10545 -pf10282 -
10550 -pf00149pf16656 -
10555 -# -
10560 -pf00884 -
(..-995..)
09565 -pf01073 -
09570 -pf00109pf02801 -
09575 -pf00550 -
09580 -# -
09585 -pf01593 -
09590 -pf00034pf07995 -
09595 -pf02633 -
09600 -pf00155 -
09605 -# -
09610 -pf00873 -
09615 Ts_mud PVM
09620 HMM02780keto_1_eut1(pf08545pf08541pf00364pf00676pf0277
09625 -pf00578 -
09630 -# -
09635 -# -
09640 -pf01041 -
09645 -# -
09650 -pf00535 -
09655 -# -
09660 -# -
09665 -pf04909 -
BMC-P
BMC-H
BMC-T
Aldehyde dehydrogenase
PTAC
Signature enzyme
Alcohol dehydrogenase
Regulator

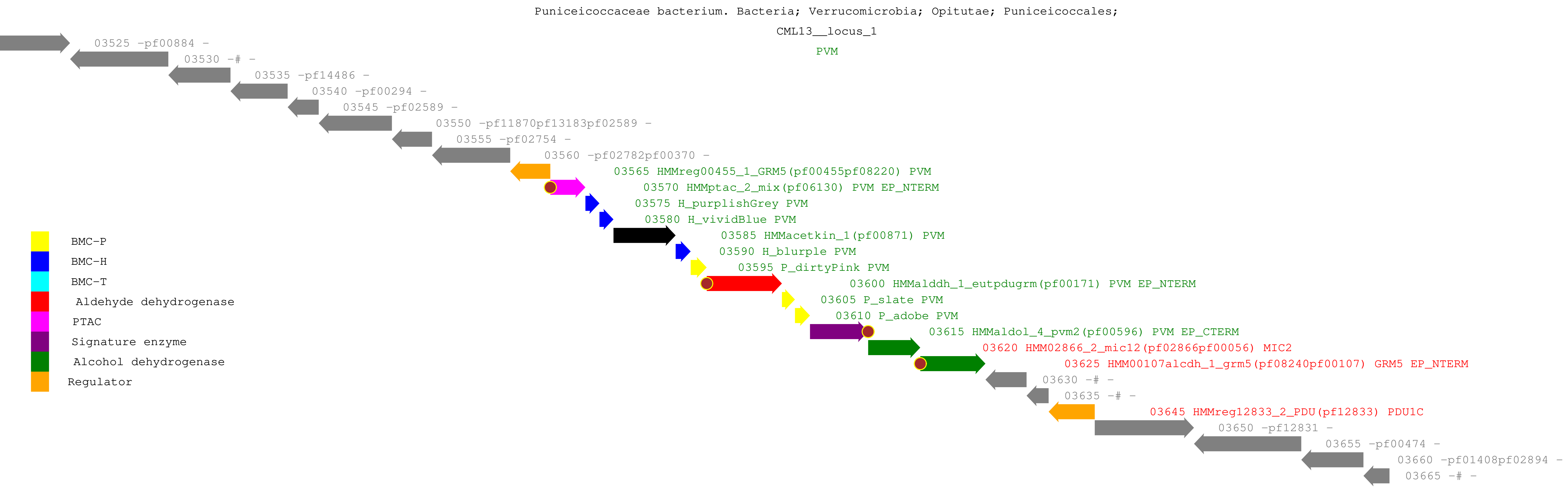
Puniceicoccaceae bacterium. Bacteria; Verrucomicrobia; Opitutae; Puniceicoccales;
CML13__locus_1
PVM
03525 -pf00884 -
03530 -# -
03535 -pf14486 -
03540 -pf00294 -
03545 -pf02589 -
03550 -pf11870pf13183pf02589 -
03555 -pf02754 -
03560 -pf02782pf00370 -
03565 HMMreg00455_1_GRM5(pf00455pf08220) PVM
03570 HMMptac_2_mix(pf06130) PVM EP_NTERM
03575 H_purplishGrey PVM
03580 H_vividBlue PVM
03585 HMMacetkin_1(pf00871) PVM
03590 H_blurple PVM
03595 P_dirtyPink PVM
03600 HMMaldh_1_eutpdugrm(pf00171) PVM EP_NTERM
03605 P_slate PVM
03610 P_adobe PVM
03615 HMMaldol_4_pvm2(pf00596) PVM EP_CTERM
03620 HMM02866_2_mic12(pf02866pf00056) MIC2
03625 HMM00107alcdh_1_grm5(pf08240pf00107) GRM5 EP_NTERM
03630 -# -
03635 -# -
03645 HMMreg12833_2_PDU(pf12833) PDU1C
03650 -pf12831 -
03655 -pf00474 -
03660 -pf01408pf02894 -
03665 -# -
BMC-P
BMC-H
BMC-T
Aldehyde dehydrogenase
PTAC
Signature enzyme
Alcohol dehydrogenase
Regulator

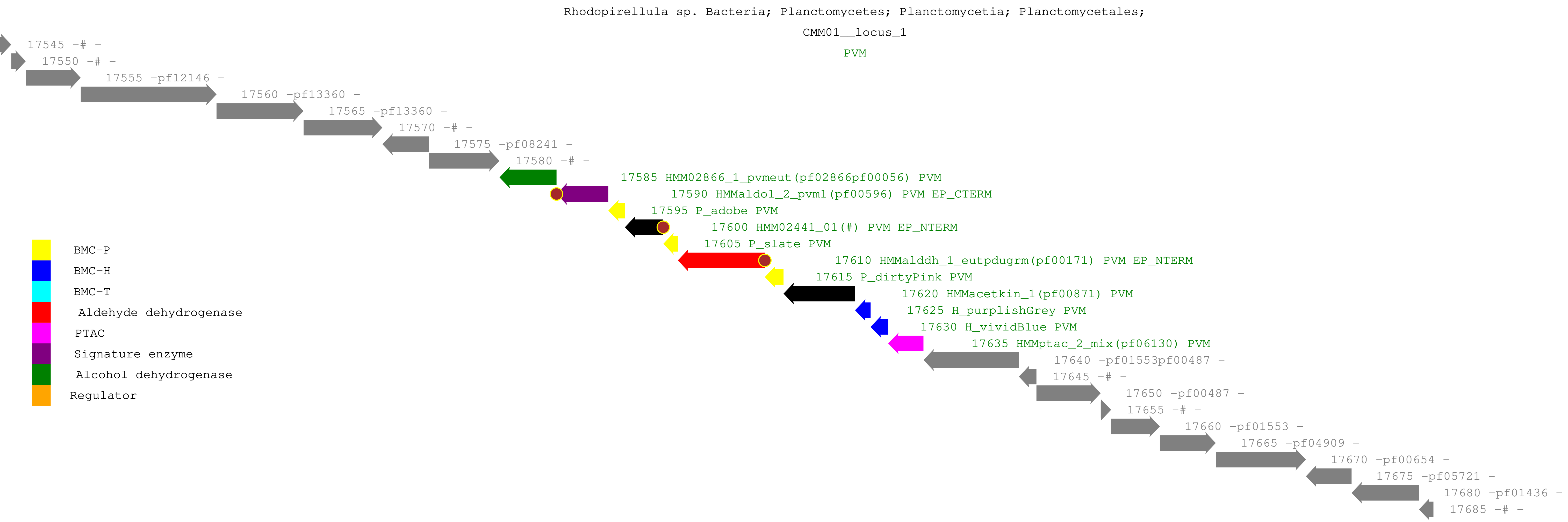
Rhodopirellula sp. Bacteria; Planctomycetes; Planctomycetia; Planctomycetales;
CMM01__locus_1
PVM
17545 -# -
17550 -# -
17555 -pf12146 -
17560 -pf13360 -
17565 -pf13360 -
17570 -# -
17575 -pf08241 -
17580 -# -
17585 HMM02866_1_pvmeut(pf02866pf00056) PVM
17590 HMMaldol_2_pvm1(pf00596) PVM EP_CTERM
17595 P_adobe PVM
17600 HMM02441_01(#) PVM EP_NTERM
17605 P_slate PVM
17610 HMMalddh_1_eutpdugrm(pf00171) PVM EP_NTERM
17615 P_dirtyPink PVM
17620 HMMacetkin_1(pf00871) PVM
17625 H_purplishGrey PVM
17630 H_vividBlue PVM
17635 HMMptac_2_mix(pf06130) PVM
17640 -pf01553pf00487 -
17645 -# -
17650 -pf00487 -
17655 -# -
17660 -pf01553 -
17665 -pf04909 -
17670 -pf00654 -
17675 -pf05721 -
17680 -pf01436 -
17685 -# -
BMC-P
BMC-H
BMC-T
Aldehyde dehydrogenase
PTAC
Signature enzyme
Alcohol dehydrogenase
Regulator

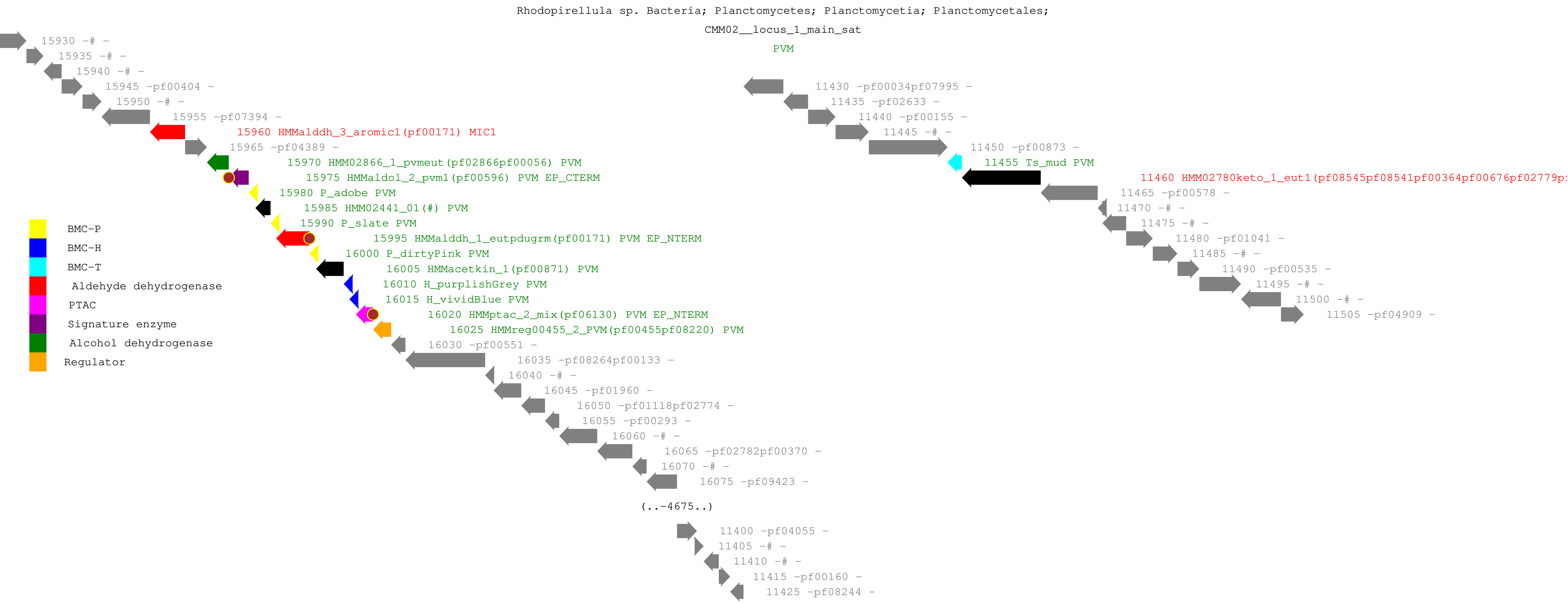
Rhodopirellula sp. Bacteria; Planctomycetes; Planctomycetia; Planctomycetales;
CMM02__locus_1_main_sat
PVM
15930 -# -
15935 -# -
15940 -# -
15945 -pf00404 -
15950 -# -
15955 -pf07394 -
15960 HMMalddh_3_aromic1(pf00171) MIC1
15965 -pf04389 -
15970 HMM02866_1_pvmeut(pf02866pf00056) PVM
15975 HMMaldol_2_pvm1(pf00596) PVM EP_CTERM
15980 P_adobe PVM
15985 HMM02441_01(#) PVM
15990 P_slate PVM
15995 HMMalddh_1_eutpdugrm(pf00171) PVM EP_NTERM
16000 P_dirtyPink PVM
16005 HMMacetkin_1(pf00871) PVM
16010 H_purplishGrey PVM
16015 H_vividBlue PVM
16020 HMMptac_2_mix(pf06130) PVM EP_NTERM
16025 HMMreg00455_2_PVM(pf00455pf08220) PVM
16030 -pf00551 -
16035 -pf08264pf00133 -
16040 -# -
16045 -pf01960 -
16050 -pf01118pf02774 -
16055 -pf00293 -
16060 -# -
16065 -pf02782pf00370 -
16070 -# -
16075 -pf09423 -
(..-4675..)
11400 -pf04055 -
11405 -# -
11410 -# -
11415 -pf00160 -
11425 -pf08244 -
11430 -pf00034pf07995 -
11435 -pf02633 -
11440 -pf00155 -
11445 -# -
11450 -pf00873 -
11455 Ts_mud PVM
11460 HMM02780keto_1_eut1(pf08545pf08541pf00364pf00676pf02779p
11465 -pf00578 -
11470 -# -
11475 -# -
11480 -pf01041 -
11485 -# -
11490 -pf00535 -
11495 -# -
11500 -# -
11505 -pf04909 -
BMC-P
BMC-H
BMC-T
Aldehyde dehydrogenase
PTAC
Signature enzyme
Alcohol dehydrogenase
Regulator

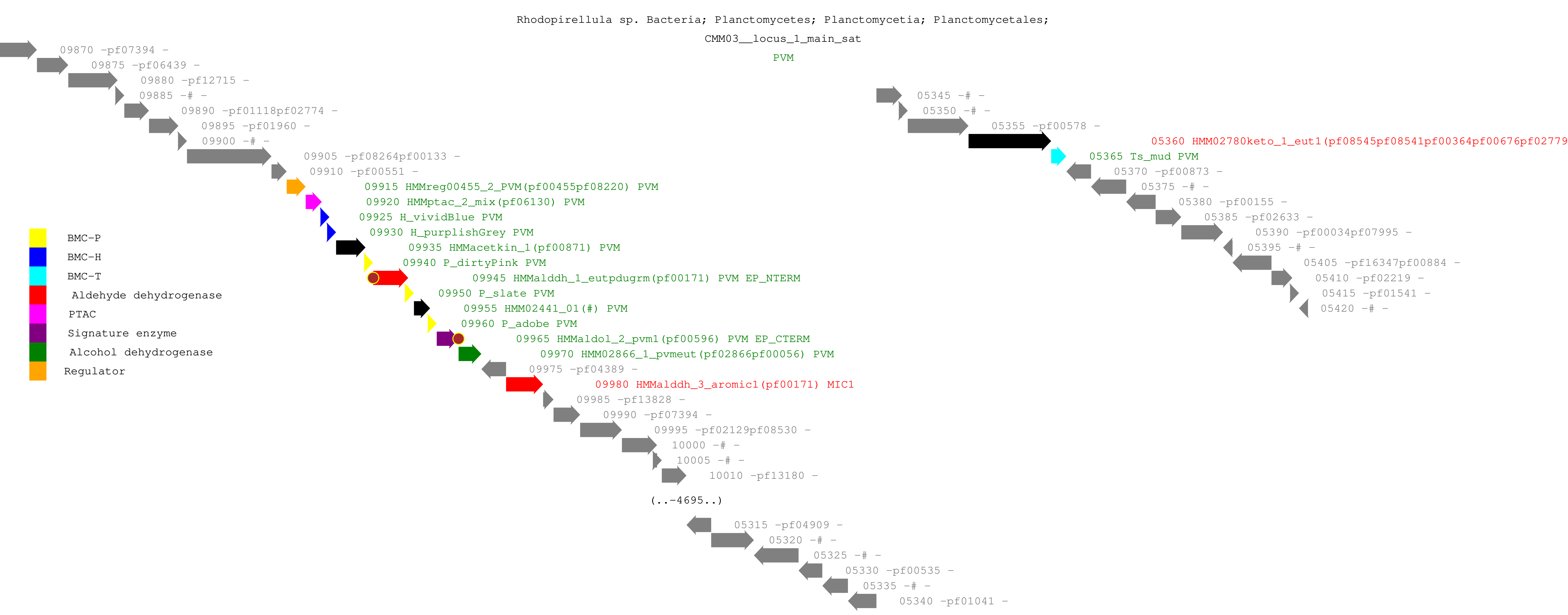
Rhodopirellula sp. Bacteria; Planctomycetes; Planctomycetia; Planctomycetales;
CMM03__locus_1_main_sat
PVM
09870 -pf07394 -
09875 -pf06439 -
09880 -pf12715 -
09885 -# -
09890 -pf01118pf02774 -
09895 -pf01960 -
09900 -# -
09905 -pf08264pf00133 -
09910 -pf00551 -
09915 HMMreg00455_2_PVM(pf00455pf08220) PVM
09920 HMMptac_2_mix(pf06130) PVM
09925 H_vividBlue PVM
09930 H_purplishGrey PVM
09935 HMMacetkin_1(pf00871) PVM
09940 P_dirtyPink PVM
09945 HMMaldh_1_eutpdugrm(pf00171) PVM EP_NTERM
09950 P_slate PVM
09955 HMM02441_01(#) PVM
09960 P_adobe PVM
09965 HMMaldol_2_pvm1(pf00596) PVM EP_CTERM
09970 HMM02866_1_pvmeut(pf02866pf00056) PVM
09975 -pf04389 -
09980 HMMaldh_3_aromic1(pf00171) MIC1
09985 -pf13828 -
09990 -pf07394 -
09995 -pf02129pf08530 -
10000 -# -
10005 -# -
10010 -pf13180 -
(..-4695..)
05315 -pf04909 -
05320 -# -
05325 -# -
05330 -pf00535 -
05335 -# -
05340 -pf01041 -
05345 -# -
05350 -# -
05355 -pf00578 -
05360 HMM02780keto_1_eut1(pf08545pf08541pf00364pf00676pf02779
05365 Ts_mud PVM
05370 -pf00873 -
05375 -# -
05380 -pf00155 -
05385 -pf02633 -
05390 -pf00034pf07995 -
05395 -# -
05405 -pf16347pf00884 -
05410 -pf02219 -
05415 -pf01541 -
05420 -# -
BMC-P
BMC-H
BMC-T
Aldehyde dehydrogenase
PTAC
Signature enzyme
Alcohol dehydrogenase
Regulator

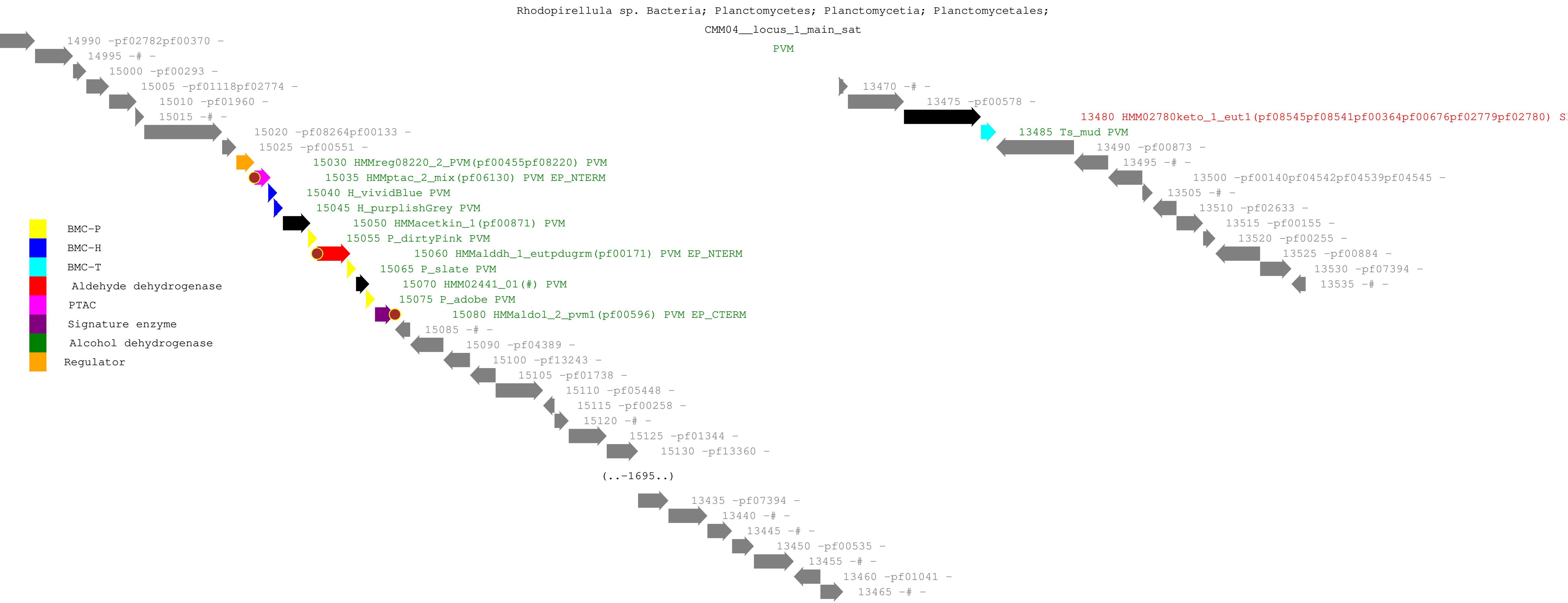
Rhodopirellula sp. Bacteria; Planctomycetes; Planctomycetia; Planctomycetales;
CMM04__locus_1_main_sat
PVM
14990 -pf02782pf00370 -
14995 -# -
15000 -pf00293 -
15005 -pf01118pf02774 -
15010 -pf01960 -
15015 -# -
15020 -pf08264pf00133 -
15025 -pf00551 -
15030 HMMreg08220_2_PVM(pf00455pf08220) PVM
15035 HMMptac_2_mix(pf06130) PVM EP_NTERM
15040 H_vividBlue PVM
15045 H_purplishGrey PVM
15050 HMMacetkin_1(pf00871) PVM
15055 P_dirtyPink PVM
15060 HMMalddh_1_eutpdugrm(pf00171) PVM EP_NTERM
15065 P_slate PVM
15070 HMM02441_01(#) PVM
15075 P_adobe PVM
15080 HMMaldol_2_pvm1(pf00596) PVM EP_CTERM
15085 -# -
15090 -pf04389 -
15100 -pf13243 -
15105 -pf01738 -
15110 -pf05448 -
15115 -pf00258 -
15120 -# -
15125 -pf01344 -
15130 -pf13360 -
(..-1695..)
13435 -pf07394 -
13440 -# -
13445 -# -
13450 -pf00535 -
13455 -# -
13460 -pf01041 -
13465 -# -
13470 -# -
13475 -pf00578 -
13480 HMM02780keto_1_eut1(pf08545pf08541pf00364pf00676pf02779pf02780)
13485 Ts_mud PVM
13490 -pf00873 -
13495 -# -
13500 -pf00140pf04542pf04539pf04545 -
13505 -# -
13510 -pf02633 -
13515 -pf00155 -
13520 -pf00255 -
13525 -pf00884 -
13530 -pf07394 -
13535 -# -
BMC-P
BMC-H
BMC-T
Aldehyde dehydrogenase
PTAC
Signature enzyme
Alcohol dehydrogenase
Regulator

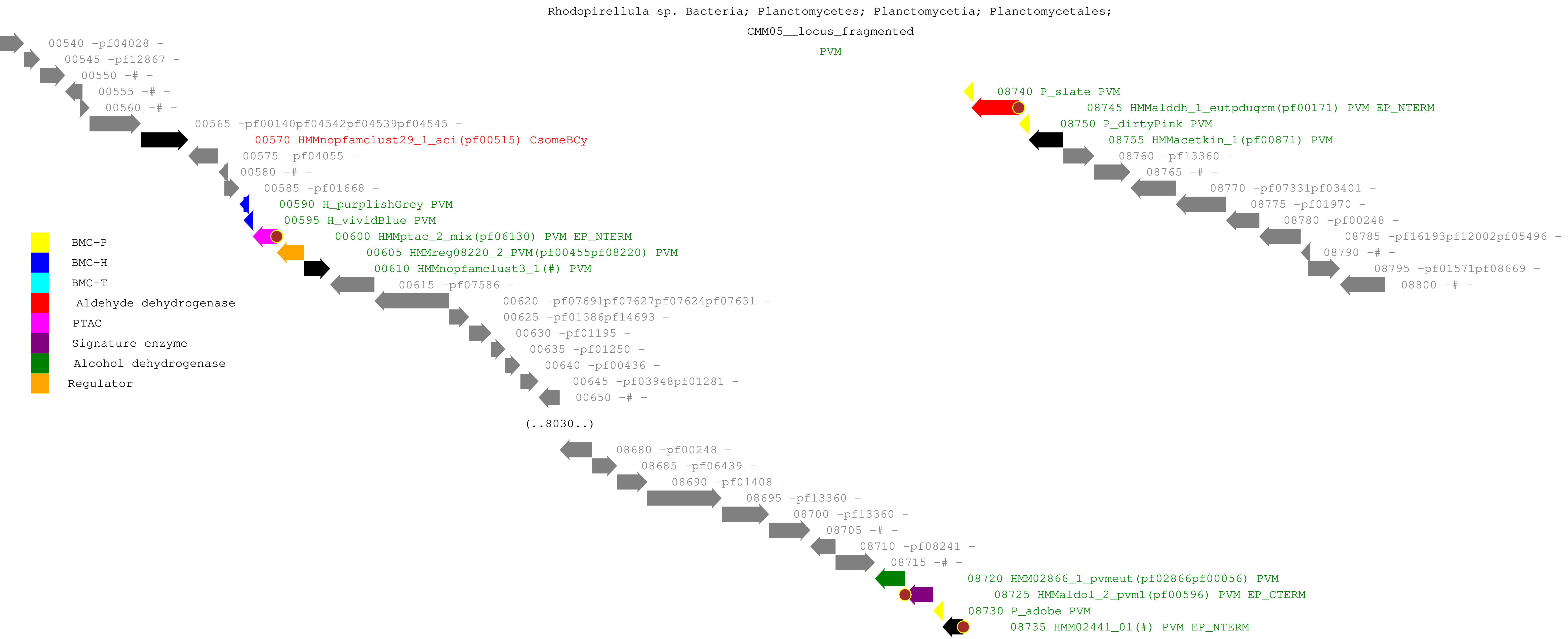
Rhodopirellula sp. Bacteria; Planctomycetes; Planctomycetia; Planctomycetales;
CMM05__locus_fragmented
PVM
00540 -pf04028 -
00545 -pf12867 -
00550 -# -
00555 -# -
00560 -# -
00565 -pf00140pf04542pf04539pf04545 -
00570 HMMnopfamclust29_1_aci(pf00515) CsomeBCy
00575 -pf04055 -
00580 -# -
00585 -pf01668 -
00590 H_purplishGrey PVM
00595 H_vividBlue PVM
00600 HMMptac_2_mix(pf06130) PVM EP_NTERM
00605 HMMreg08220_2_PVM(pf00455pf08220) PVM
00610 HMMnopfamclust3_1(#) PVM
00615 -pf07586 -
00620 -pf07691pf07627pf07624pf07631 -
00625 -pf01386pf14693 -
00630 -pf01195 -
00635 -pf01250 -
00640 -pf00436 -
00645 -pf03948pf01281 -
00650 -# -
(..8030..)
08680 -pf00248 -
08685 -pf06439 -
08690 -pf01408 -
08695 -pf13360 -
08700 -pf13360 -
08705 -# -
08710 -pf08241 -
08715 -# -
08720 HMM02866_1_pvmeut(pf02866pf00056) PVM
08725 HMMaldol_2_pvm1(pf00596) PVM EP_CTERM
08730 P_adobe PVM
08735 HMM02441_01(#) PVM EP_NTERM
08740 P_slate PVM
08745 HMMalddh_1_eutpdugrm(pf00171) PVM EP_NTERM
08750 P_dirtyPink PVM
08755 HMMacetkin_1(pf00871) PVM
08760 -pf13360 -
08765 -# -
08770 -pf07331pf03401 -
08775 -pf01970 -
08780 -pf00248 -
08785 -pf16193pf12002pf05496 -
08790 -# -
08795 -pf01571pf08669 -
08800 -# -
BMC-P
BMC-H
BMC-T
Aldehyde dehydrogenase
PTAC
Signature enzyme
Alcohol dehydrogenase
Regulator

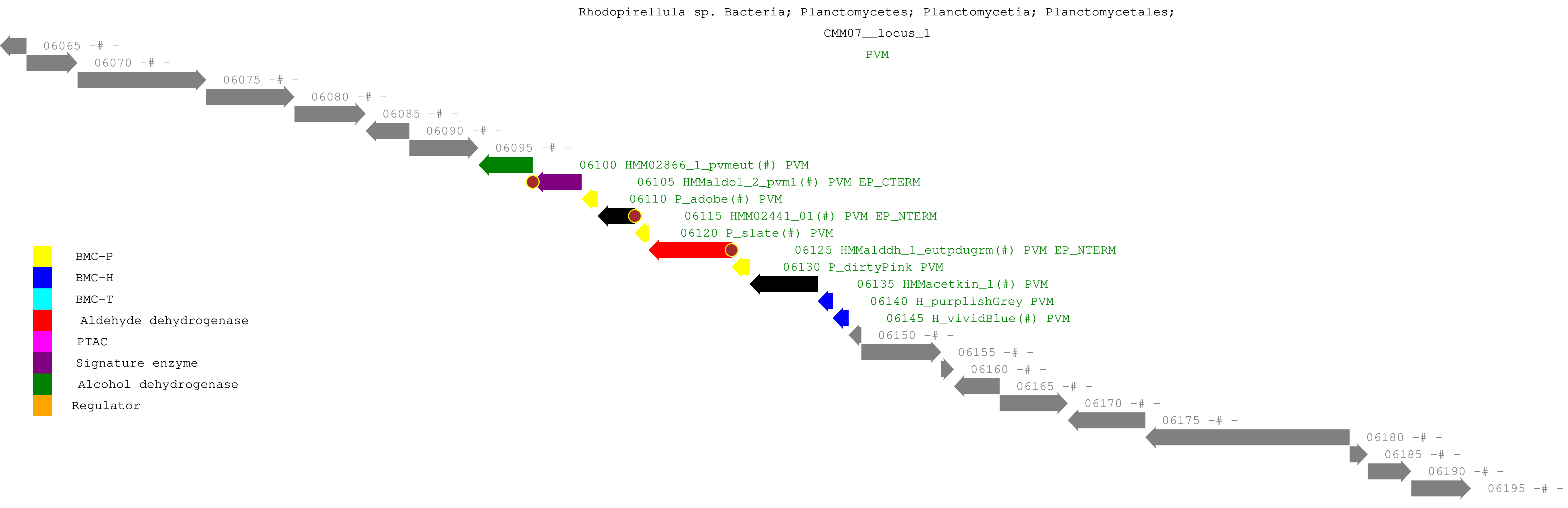

Rhodopirellula sp. Bacteria; Planctomycetes; Planctomycetia; Planctomycetales;
CMM07__locus_1
PVM
06065 -# -
06070 -# -
06075 -# -
06080 -# -
06085 -# -
06090 -# -
06095 -# -
06100 HMM02866_1_pvmeut(#) PVM
06105 HMMaldol_2_pvm1(#) PVM EP_CTERM
06110 P_adobe(#) PVM
06115 HMM02441_01(#) PVM EP_NTERM
06120 P_slate(#) PVM
06125 HMMalddh_1_eutpdugrm(#) PVM EP_NTERM
06130 P_dirtyPink PVM
06135 HMMacetkin_1(#) PVM
06140 H_purplishGrey PVM
06145 H_vividBlue(#) PVM
06150 -# -
06155 -# -
06160 -# -
06165 -# -
06170 -# -
06175 -# -
06180 -# -
06185 -# -
06190 -# -
06195 -# -
BMC-P
BMC-H
BMC-T
Aldehyde dehydrogenase
PTAC
Signature enzyme
Alcohol dehydrogenase
Regulator

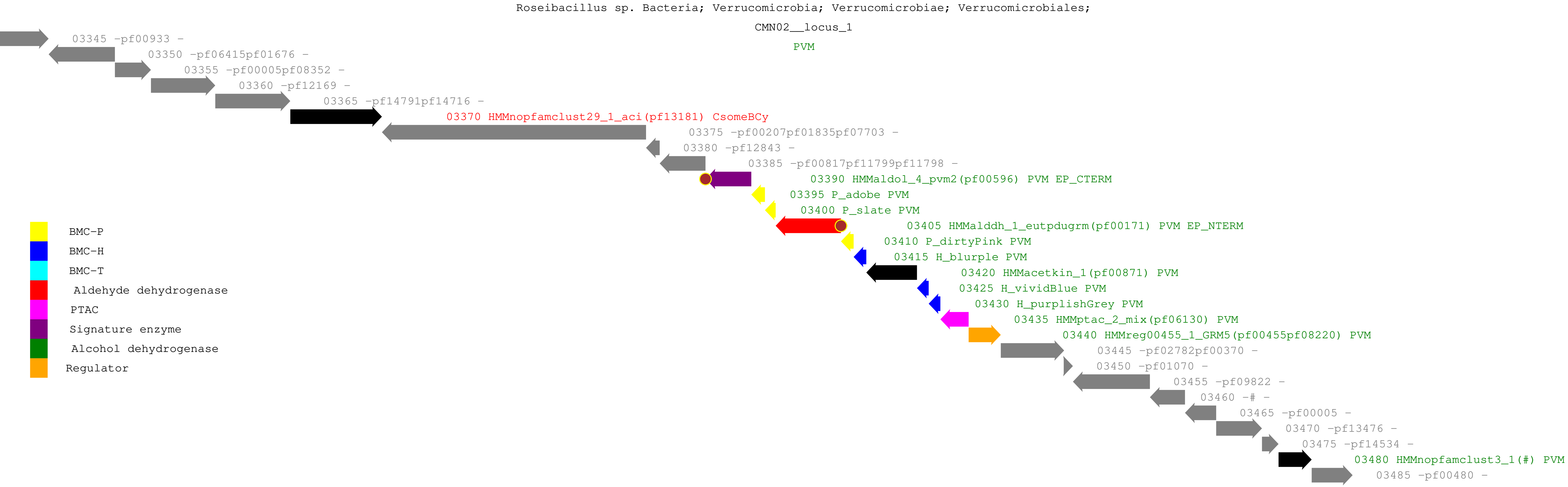
Roseibacillus sp. Bacteria; Verrucomicrobia; Verrucomicrobiae; Verrucomicrobiales;
CMN02__locus_1
PVM
03345 -pf00933 -
03350 -pf06415pf01676 -
03355 -pf00005pf08352 -
03360 -pf12169 -
03365 -pf14791pf14716 -
03370 HMMnopfamclust29_1_aci(pf13181) CsomeBCy
03375 -pf00207pf01835pf07703 -
03380 -pf12843 -
03385 -pf00817pf11799pf11798 -
03390 HMMaldol_4_pvm2(pf00596) PVM EP_CTERM
03395 P_adobe PVM
03400 P_slate PVM
03405 HMMalddh_1_eutpdugrm(pf00171) PVM EP_NTERM
03410 P_dirtyPink PVM
03415 H_blurple PVM
03420 HMMacetkin_1(pf00871) PVM
03425 H_vividBlue PVM
03430 H_purplishGrey PVM
03435 HMMptac_2_mix(pf06130) PVM
03440 HMMreg00455_1_GRM5(pf00455pf08220) PVM
03445 -pf02782pf00370 -
03450 -pf01070 -
03455 -pf09822 -
03460 -# -
03465 -pf00005 -
03470 -pf13476 -
03475 -pf14534 -
03480 HMMnopfamclust3_1(#) PVM
03485 -pf00480 -
BMC-P
BMC-H
BMC-T
Aldehyde dehydrogenase
PTAC
Signature enzyme
Alcohol dehydrogenase
Regulator

Roseibacillus sp. Bacteria; Verrucomicrobia; Verrucomicrobiae; Verrucomicrobiales;

CMN04__locus_1

PVM

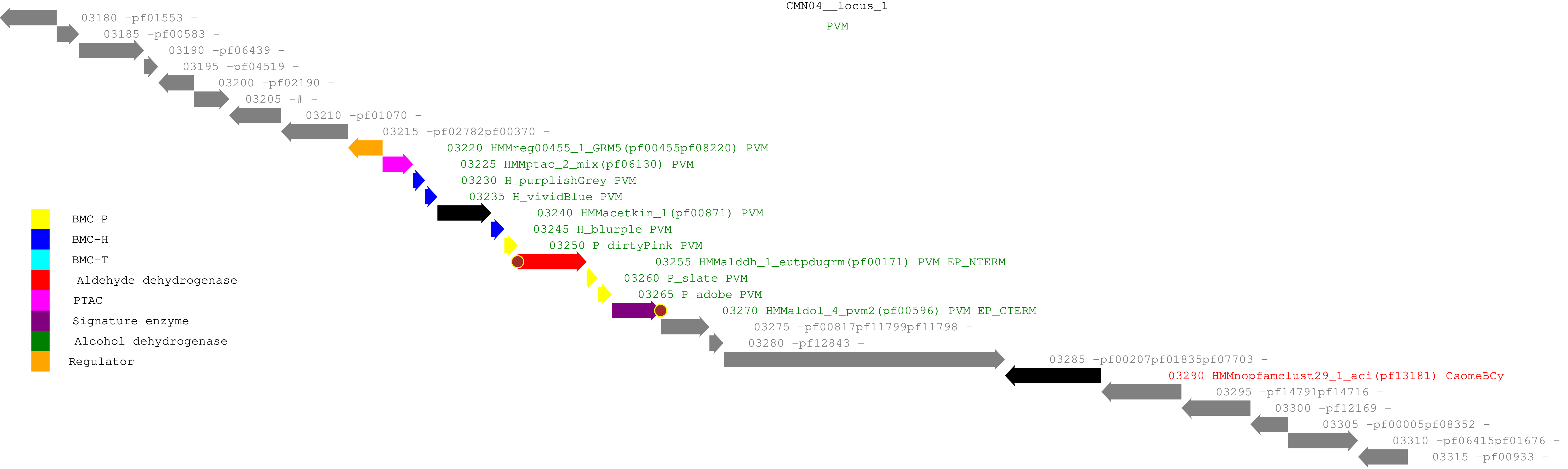

Roseibacillus sp. Bacteria; Verrucomicrobia; Verrucomicrobiae; Verrucomicrobiales;

CMN05__locus_1

PVM

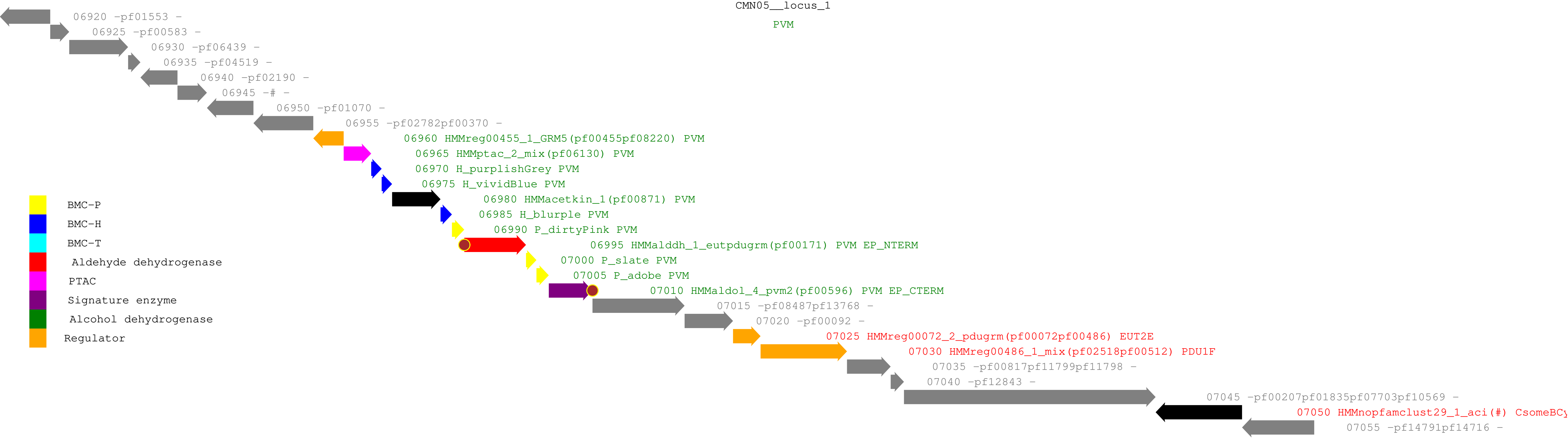

Roseibacillus sp. Bacteria; Verrucomicrobia; Verrucomicrobiae; Verrucomicrobiales;

CMN06__locus_1

PVM

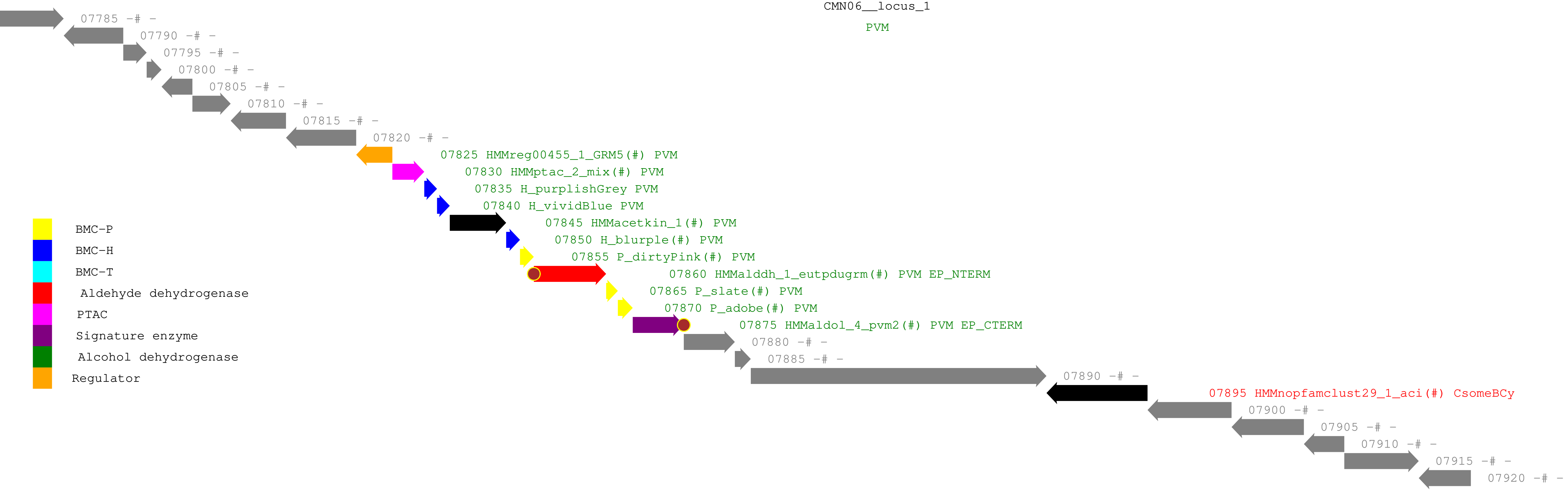

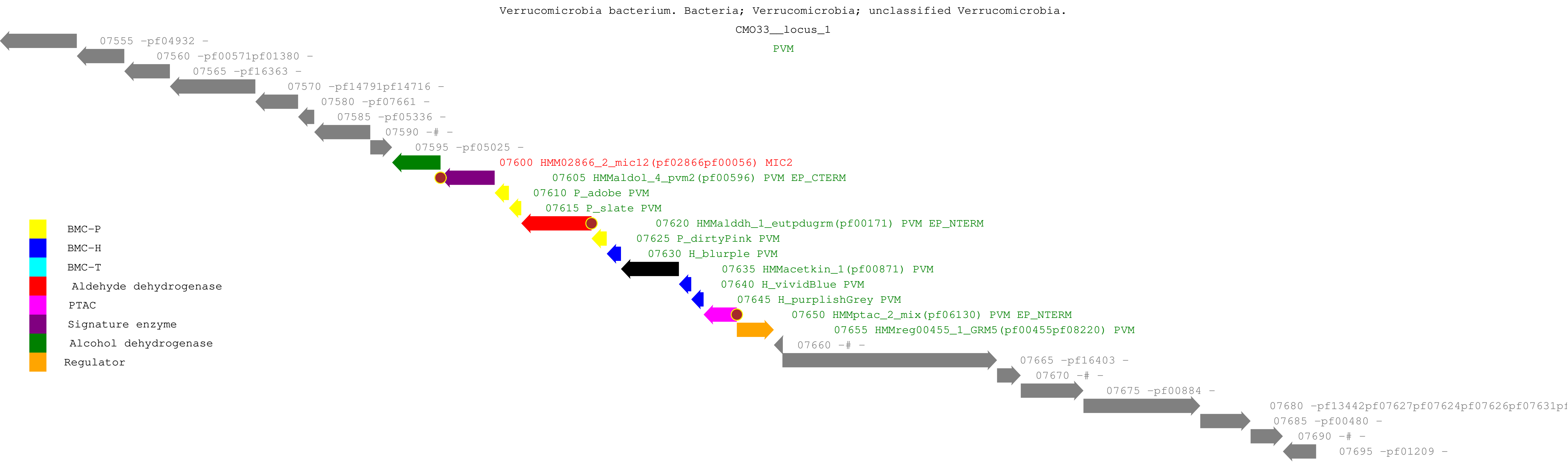
Verrucomicrobia bacterium. Bacteria; Verrucomicrobia; unclassified Verrucomicrobia.
CMO33__locus_1
PVM
07555 -pf04932 -
07560 -pf00571pf01380 -
07565 -pf16363 -
07570 -pf14791pf14716 -
07580 -pf07661 -
07585 -pf05336 -
07590 -# -
07595 -pf05025 -
07600 HMM02866_2_mic12(pf02866pf00056) MIC2
07605 HMMaldol_4_pvm2(pf00596) PVM EP_CTERM
07610 P_adobe PVM
07615 P_slate PVM
07620 HMMalddh_1_eutpdugrm(pf00171) PVM EP_NTERM
07625 P_dirtyPink PVM
07630 H_blurple PVM
07635 HMMacetkin_1(pf00871) PVM
07640 H_vividBlue PVM
07645 H_purplishGrey PVM
07650 HMMptac_2_mix(pf06130) PVM EP_NTERM
07655 HMMreg00455_1_GRM5(pf00455pf08220) PVM
07660 -# -
07665 -pf16403 -
07670 -# -
07675 -pf00884 -
07680 -pf13442pf07627pf07624pf07626pf07631pf
07685 -pf00480 -
07690 -# -
07695 -pf01209 -
BMC-P
BMC-H
BMC-T
Aldehyde dehydrogenase
PTAC
Signature enzyme
Alcohol dehydrogenase
Regulator

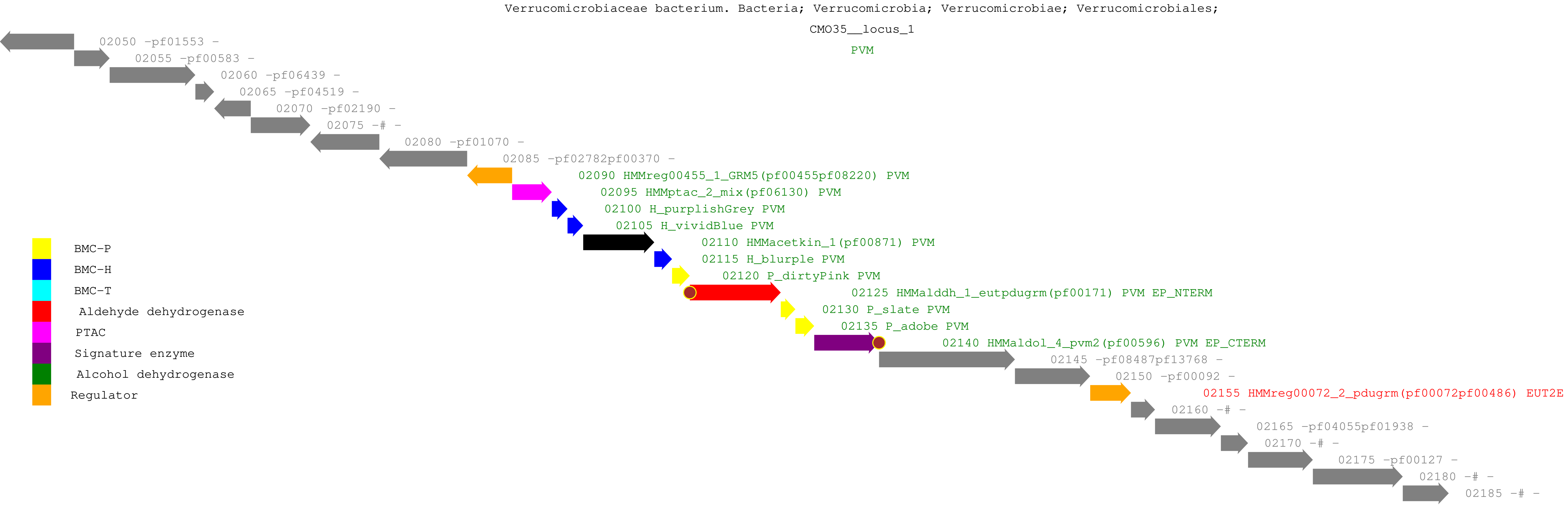

Verrucomicrobiaceae bacterium. Bacteria; Verrucomicrobia; Verrucomicrobiae; Verrucomicrobiales;
CMO35__locus_1
PVM
02050 -pf01553 -
02055 -pf00583 -
02060 -pf06439 -
02065 -pf04519 -
02070 -pf02190 -
02075 -# -
02080 -pf01070 -
02085 -pf02782pf00370 -
02090 HMMreg00455_1_GRM5(pf00455pf08220) PVM
02095 HMMptac_2_mix(pf06130) PVM
02100 H_purplishGrey PVM
02105 H_vividBlue PVM
02110 HMMacetkin_1(pf00871) PVM
02115 H_blurple PVM
02120 P_dirtyPink PVM
02125 HMMalddh_1_eutpdugrm(pf00171) PVM EP_NTERM
02130 P_slate PVM
02135 P_adobe PVM
02140 HMMaldol_4_pvm2(pf00596) PVM EP_CTERM
02145 -pf08487pf13768 -
02150 -pf00092 -
02155 HMMreg00072_2_pdugrm(pf00072pf00486) EUT2E
02160 -# -
02165 -pf04055pf01938 -
02170 -# -
02175 -pf00127 -
02180 -# -
02185 -# -
BMC-P
BMC-H
BMC-T
Aldehyde dehydrogenase
PTAC
Signature enzyme
Alcohol dehydrogenase
Regulator

Verrucomicrobiaceae bacterium. Bacteria; Verrucomicrobia; Verrucomicrobiae; Verrucomicrobiales;

CMO40__locus_1

PVM

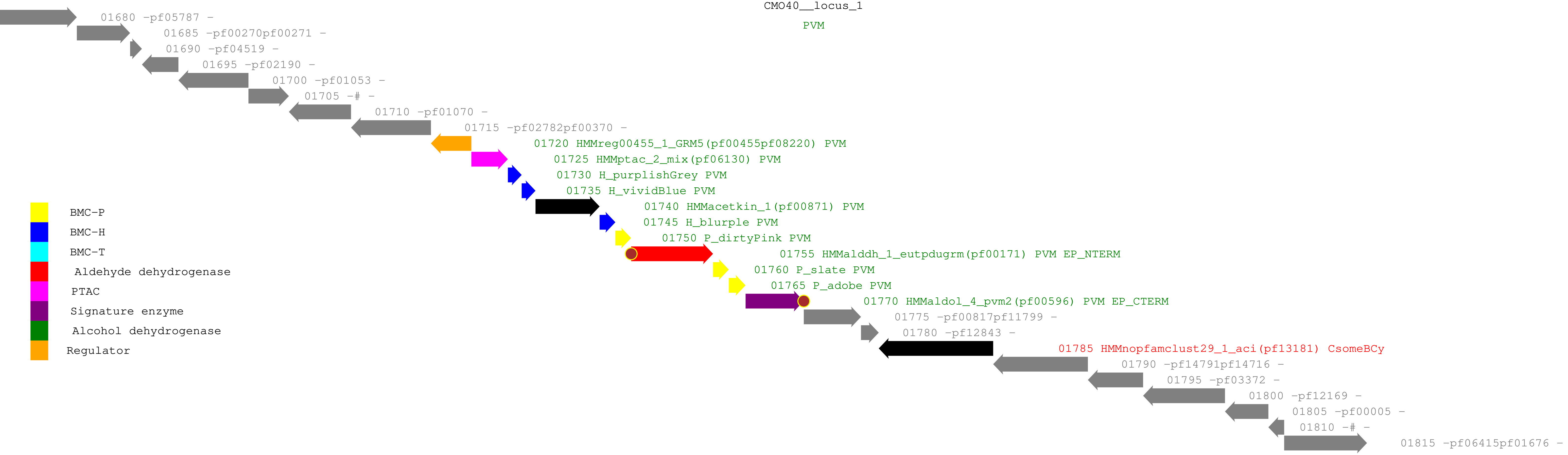

Verrucomicrobiales bacterium. Bacteria; Verrucomicrobia; Verrucomicrobiae; Verrucomicrobiales.

CMO43__locus_1

PVM

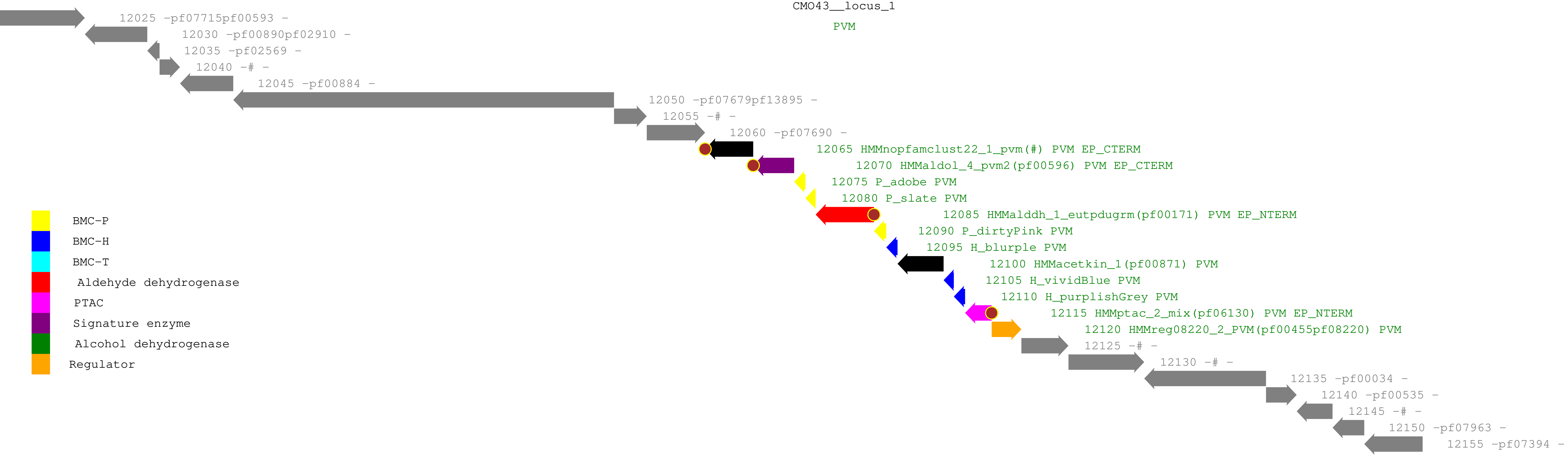

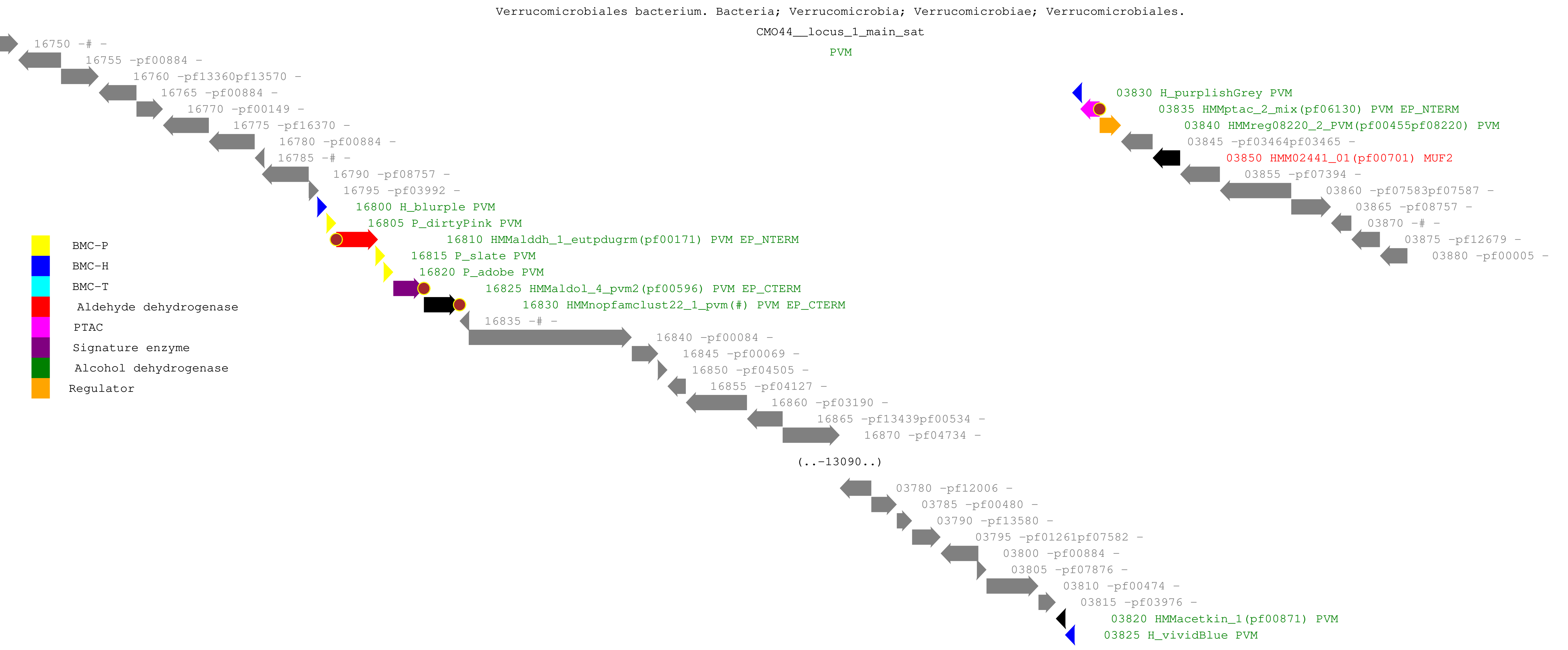
Verrucomicrobiales bacterium. Bacteria; Verrucomicrobia; Verrucomicrobiae; Verrucomicrobiales.
CMO44__locus_1_main_sat
PVM
16750 -# -
16755 -pf00884 -
16760 -pf13360pf13570 -
16765 -pf00884 -
16770 -pf00149 -
16775 -pf16370 -
16780 -pf00884 -
16785 -# -
16790 -pf08757 -
16795 -pf03992 -
16800 H_blurple PVM
16805 P_dirtyPink PVM
16810 HMMalddh_1_eutpdugrm(pf00171) PVM EP_NTERM
16815 P_slate PVM
16820 P_adobe PVM
16825 HMMaldol_4_pvm2(pf00596) PVM EP_CTERM
16830 HMMnopfamclust22_1_pvm(#) PVM EP_CTERM
16835 -# -
16840 -pf00084 -
16845 -pf00069 -
16850 -pf04505 -
16855 -pf04127 -
16860 -pf03190 -
16865 -pf13439pf00534 -
16870 -pf04734 -
(..-13090..)
03780 -pf12006 -
03785 -pf00480 -
03790 -pf13580 -
03795 -pf01261pf07582 -
03800 -pf00884 -
03805 -pf07876 -
03810 -pf00474 -
03815 -pf03976 -
03820 HMMacetkin_1(pf00871) PVM
03825 H_vividBlue PVM
03830 H_purplishGrey PVM
03835 HMMptac_2_mix(pf06130) PVM EP_NTERM
03840 HMMreg08220_2_PVM(pf00455pf08220) PVM
03845 -pf03464pf03465 -
03850 HMM02441_01(pf00701) MUF2
03855 -pf07394 -
03860 -pf07583pf07587 -
03865 -pf08757 -
03870 -# -
03875 -pf12679 -
03880 -pf00005 -
BMC-P
BMC-H
BMC-T
Aldehyde dehydrogenase
PTAC
Signature enzyme
Alcohol dehydrogenase
Regulator

Verrucomicrobiales bacterium. Bacteria; Verrucomicrobia; Verrucomicrobiae; Verrucomicrobiales.

CMO51__locus_1

PVM

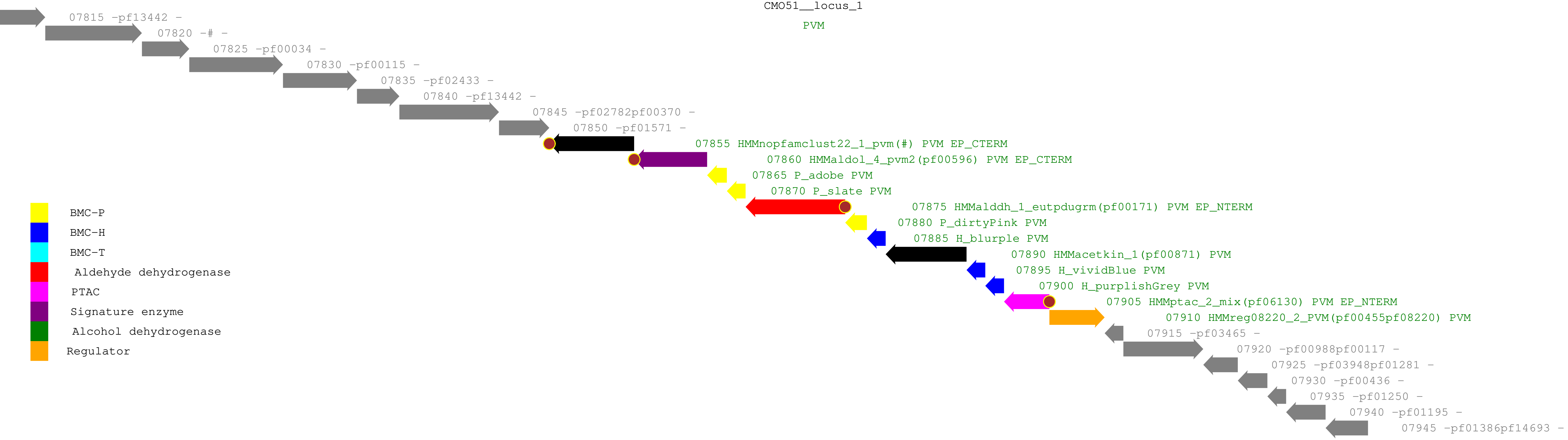

Verrucomicrobiales bacterium. Bacteria; Verrucomicrobia; Verrucomicrobiae; Verrucomicrobiales.

CMO52__locus_1

PVM

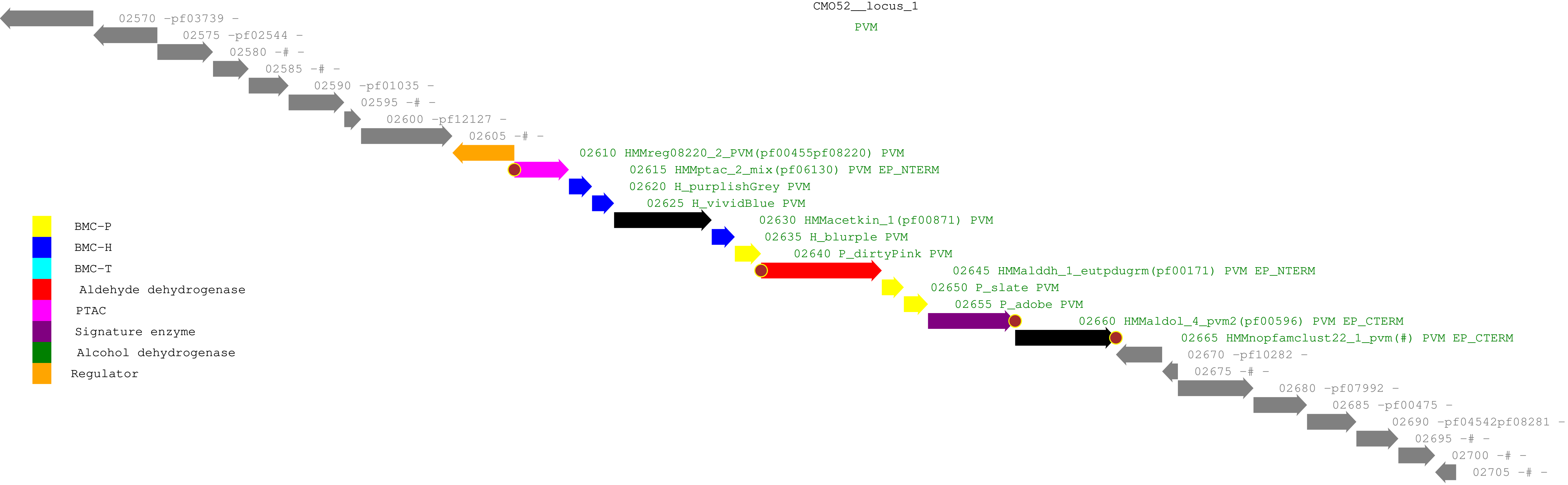

Verrucomicrobiales bacterium. Bacteria; Verrucomicrobia; Verrucomicrobiae; Verrucomicrobiales.

CMO57__locus_1

PVM

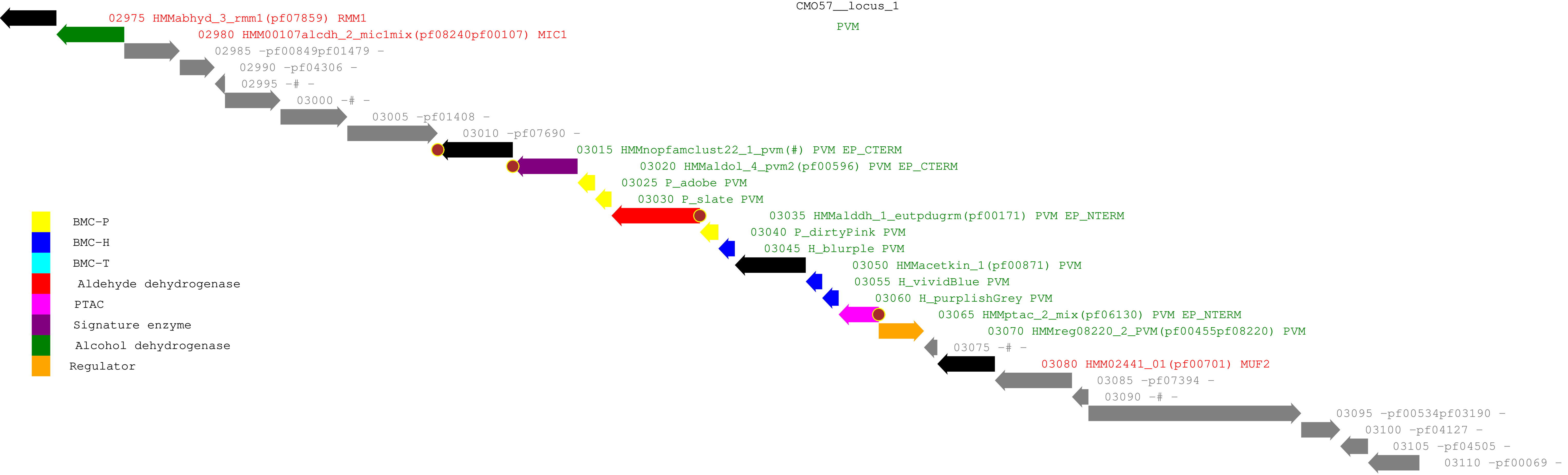

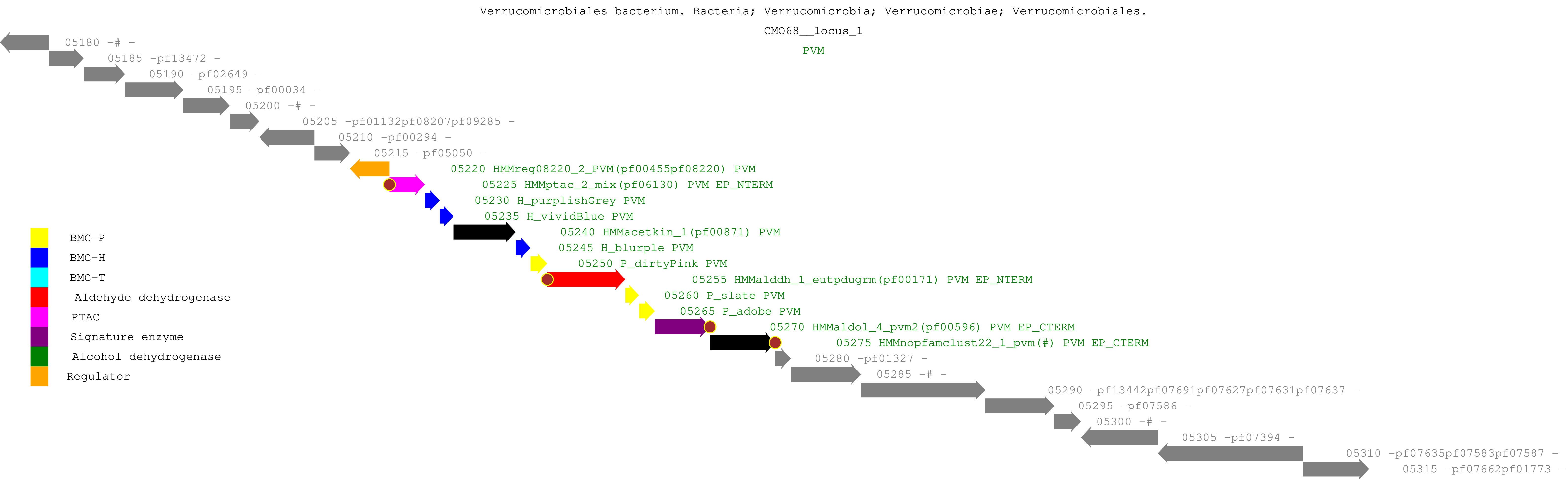

Verrucomicrobiales bacterium. Bacteria; Verrucomicrobia; Verrucomicrobiae; Verrucomicrobiales.
CMO68__locus_1
PVM
05180 -# -
05185 -pf13472 -
05190 -pf02649 -
05195 -pf00034 -
05200 -# -
05205 -pf01132pf08207pf09285 -
05210 -pf00294 -
05215 -pf05050 -
05220 HMMreg08220_2_PVM(pf00455pf08220) PVM
05225 HMMptac_2_mix(pf06130) PVM EP_NTERM
05230 H_purplishGrey PVM
05235 H_vividBlue PVM
05240 HMMacetkin_1(pf00871) PVM
05245 H_blurple PVM
05250 P_dirtyPink PVM
05255 HMMaldDh_1_eutpdugrm(pf00171) PVM EP_NTERM
05260 P_slate PVM
05265 P_adobe PVM
05270 HMMaldol_4_pvm2(pf00596) PVM EP_CTERM
05275 HMMnopfamclust22_1_pvm(#) PVM EP_CTERM
05280 -pf01327 -
05285 -# -
05290 -pf13442pf07691pf07627pf07631pf07637 -
05295 -pf07586 -
05300 -# -
05305 -pf07394 -
05310 -pf07635pf07583pf07587 -
05315 -pf07662pf01773 -
BMC-P
BMC-H
BMC-T
Aldehyde dehydrogenase
PTAC
Signature enzyme
Alcohol dehydrogenase
Regulator

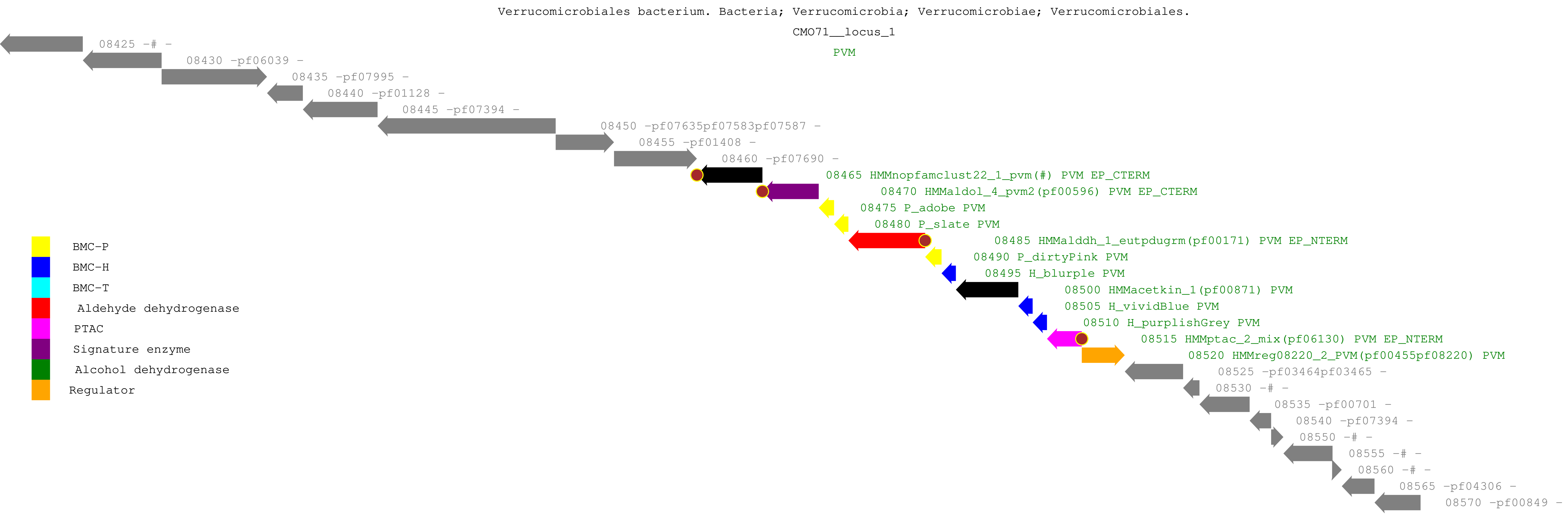

Verrucomicrobiales bacterium. Bacteria; Verrucomicrobia; Verrucomicrobiae; Verrucomicrobiales.
CMO71__locus_1
PVM
08425 -# -
08430 -pf06039 -
08435 -pf07995 -
08440 -pf01128 -
08445 -pf07394 -
08450 -pf07635pf07583pf07587 -
08455 -pf01408 -
08460 -pf07690 -
08465 HMMnopfamclust22_1_pvm(#) PVM EP_CTERM
08470 HMMaldol_4_pvm2(pf00596) PVM EP_CTERM
08475 P_adobe PVM
08480 P_slate PVM
08485 HMMaldh_1_eutpdugrm(pf00171) PVM EP_NTERM
08490 P_dirtyPink PVM
08495 H_blurple PVM
08500 HMMacetkin_1(pf00871) PVM
08505 H_vividBlue PVM
08510 H_purplishGrey PVM
08515 HMMptac_2_mix(pf06130) PVM EP_NTERM
08520 HMMreg08220_2_PVM(pf00455pf08220) PVM
08525 -pf03464pf03465 -
08530 -# -
08535 -pf00701 -
08540 -pf07394 -
08550 -# -
08555 -# -
08560 -# -
08565 -pf04306 -
08570 -pf00849 -
BMC-P
BMC-H
BMC-T
Aldehyde dehydrogenase
PTAC
Signature enzyme
Alcohol dehydrogenase
Regulator

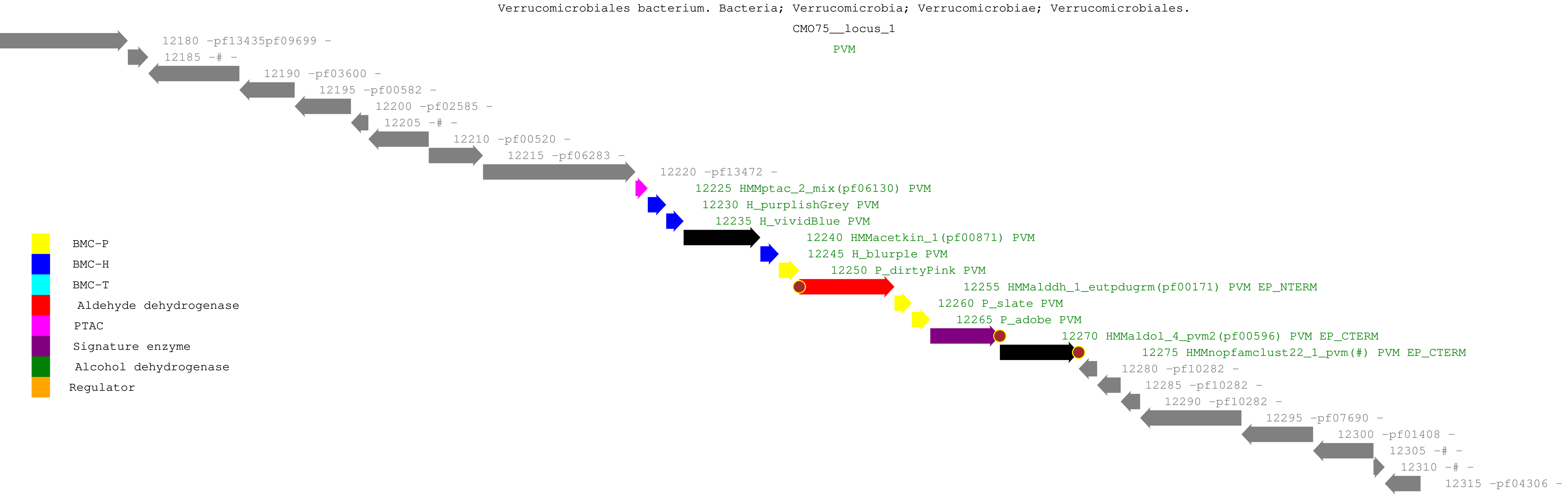
Verrucomicrobiales bacterium. Bacteria; Verrucomicrobia; Verrucomicrobiae; Verrucomicrobiales.
CMO75__locus_1
PVM
12180 -pf13435pf09699 -
12185 -# -
12190 -pf03600 -
12195 -pf00582 -
12200 -pf02585 -
12205 -# -
12210 -pf00520 -
12215 -pf06283 -
12220 -pf13472 -
12225 HMMptac_2_mix(pf06130) PVM
12230 H_purplishGrey PVM
12235 H_vividBlue PVM
12240 HMMacetkin_1(pf00871) PVM
12245 H_blurple PVM
12250 P_dirtyPink PVM
12255 HMMalddh_1_eutpdugrm(pf00171) PVM EP_NTERM
12260 P_slate PVM
12265 P_adobe PVM
12270 HMMaldol_4_pvm2(pf00596) PVM EP_CTERM
12275 HMMnopfamclust22_1_pvm(#) PVM EP_CTERM
12280 -pf10282 -
12285 -pf10282 -
12290 -pf10282 -
12295 -pf07690 -
12300 -pf01408 -
12305 -# -
12310 -# -
12315 -pf04306 -
BMC-P
BMC-H
BMC-T
Aldehyde dehydrogenase
PTAC
Signature enzyme
Alcohol dehydrogenase
Regulator

Verrucomicrobiales bacterium. Bacteria; Verrucomicrobia; Verrucomicrobiae; Verrucomicrobiales.

CMO80__locus_1

PVM

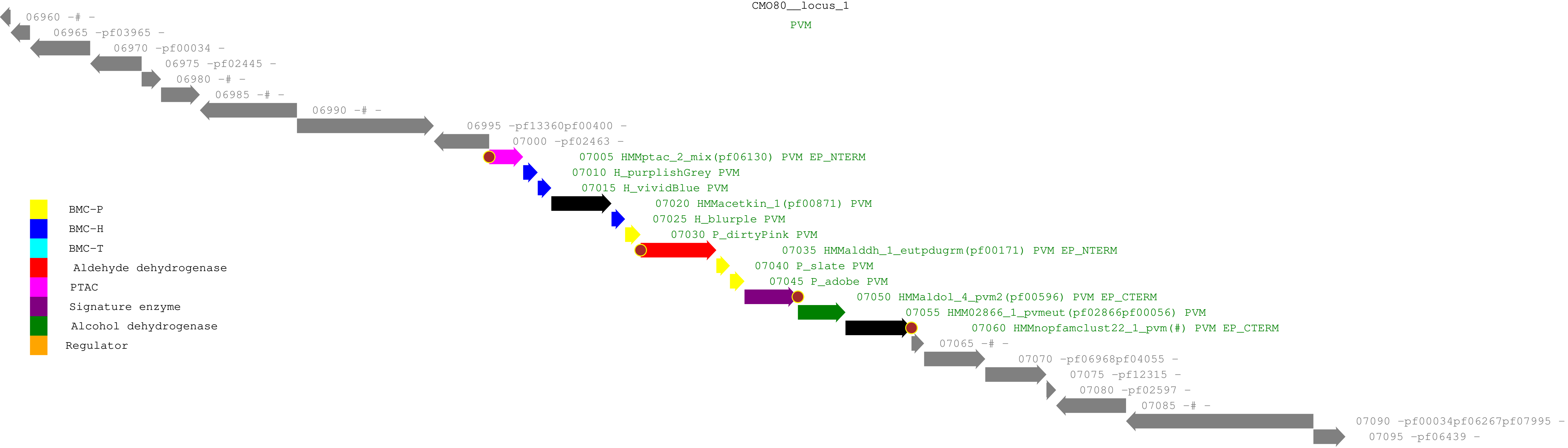

Roseibacillus sp. Bacteria; Verrucomicrobia; Verrucomicrobiae; Verrucomicrobiales;

CMP26__locus_1

PVM

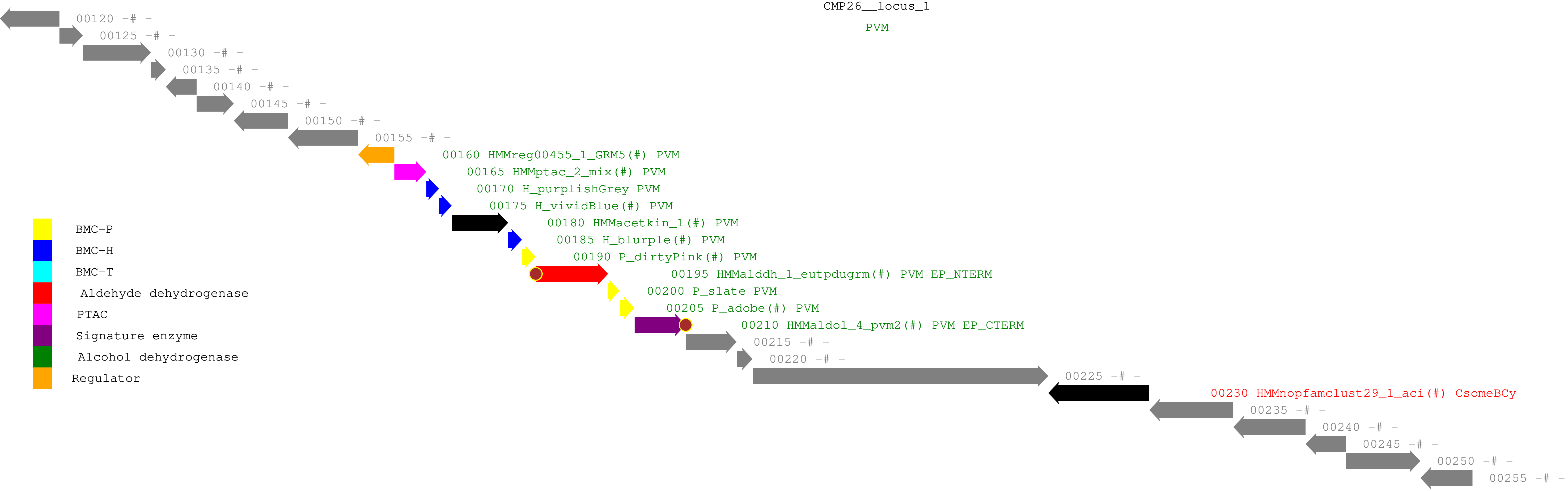

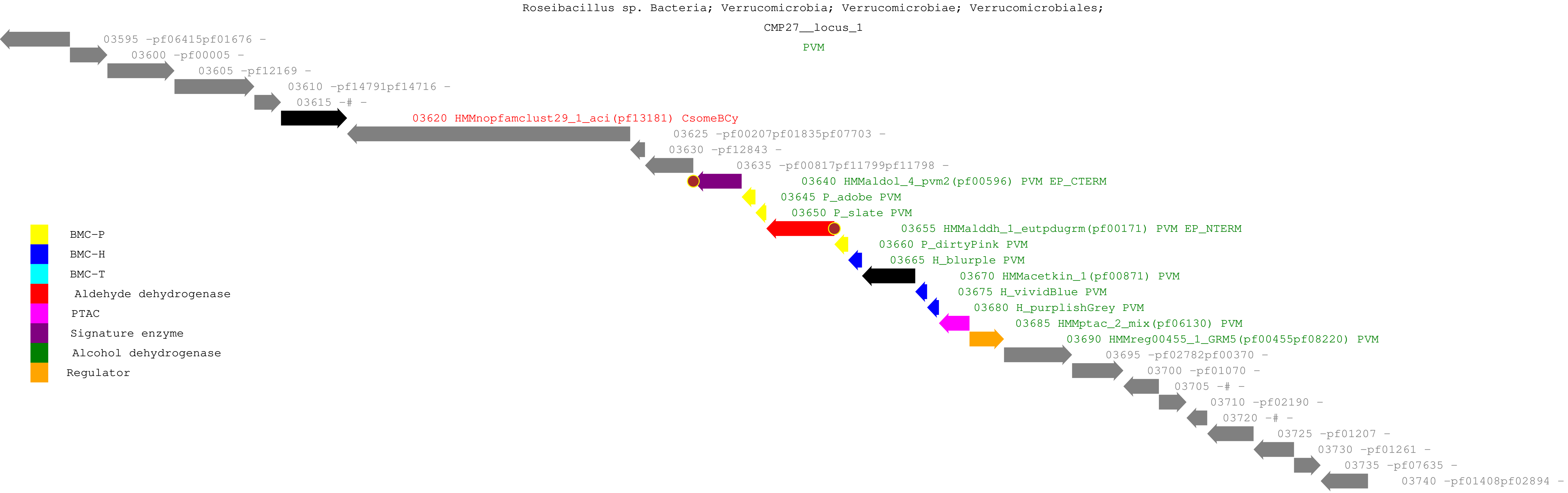
Roseibacillus sp. Bacteria; Verrucomicrobia; Verrucomicrobiae; Verrucomicrobiales;
CMP27__locus_1
PVM
03595 -pf06415pf01676 -
03600 -pf00005 -
03605 -pf12169 -
03610 -pf14791pf14716 -
03615 -# -
03620 HMMnopfamclust29_1_aci(pf13181) CsomeBCy
03625 -pf00207pf01835pf07703 -
03630 -pf12843 -
03635 -pf00817pf11799pf11798 -
03640 HMMaldol_4_pvm2(pf00596) PVM EP_CTERM
03645 P_adobe PVM
03650 P_slate PVM
03655 HMMalddh_1_eutpdugrm(pf00171) PVM EP_NTERM
03660 P_dirtyPink PVM
03665 H_blurple PVM
03670 HMMacetkin_1(pf00871) PVM
03675 H_vividBlue PVM
03680 H_purplishGrey PVM
03685 HMMptac_2_mix(pf06130) PVM
03690 HMMreg00455_1_GRM5(pf00455pf08220) PVM
03695 -pf02782pf00370 -
03700 -pf01070 -
03705 -# -
03710 -pf02190 -
03720 -# -
03725 -pf01207 -
03730 -pf01261 -
03735 -pf07635 -
03740 -pf01408pf02894 -
BMC-P
BMC-H
BMC-T
Aldehyde dehydrogenase
PTAC
Signature enzyme
Alcohol dehydrogenase
Regulator

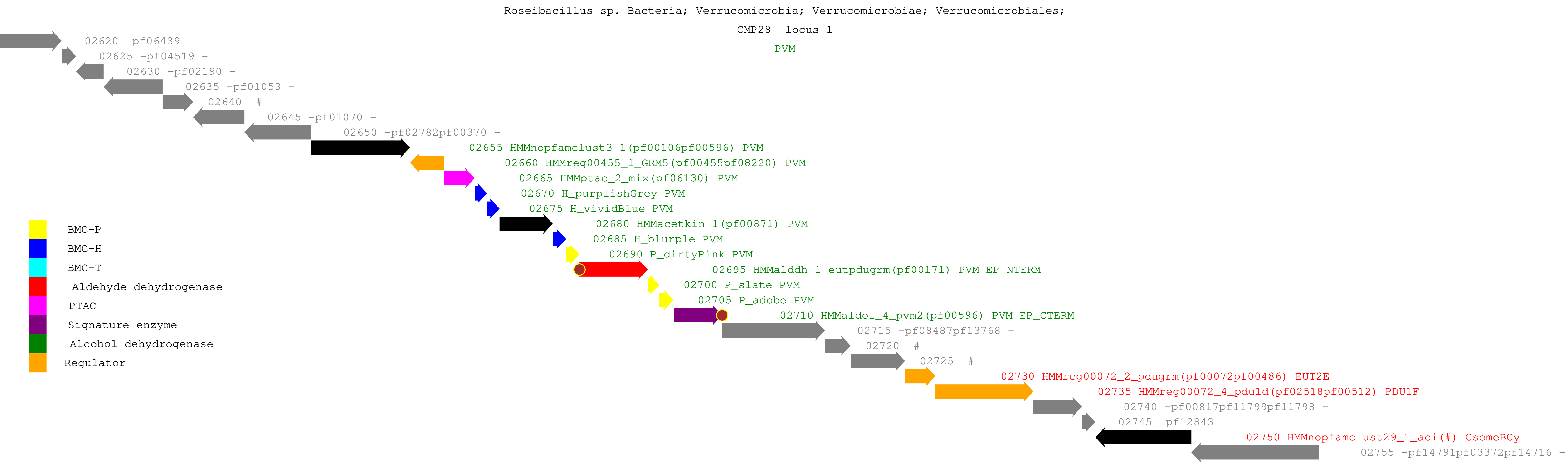
Roseibacillus sp. Bacteria; Verrucomicrobia; Verrucomicrobiae; Verrucomicrobiales;
CMP28__locus_1
PVM
02620 -pf06439 -
02625 -pf04519 -
02630 -pf02190 -
02635 -pf01053 -
02640 -# -
02645 -pf01070 -
02650 -pf02782pf00370 -
02655 HMMnopfamclust3_1(pf00106pf00596) PVM
02660 HMMreg00455_1_GRM5(pf00455pf08220) PVM
02665 HMMptac_2_mix(pf06130) PVM
02670 H_purplishGrey PVM
02675 H_vividBlue PVM
02680 HMMacetkin_1(pf00871) PVM
02685 H_blurple PVM
02690 P_dirtyPink PVM
02695 HMMalddh_1_eutpdugrm(pf00171) PVM EP_NTERM
02700 P_slate PVM
02705 P_adobe PVM
02710 HMMaldol_4_pvm2(pf00596) PVM EP_CTERM
02715 -pf08487pf13768 -
02720 -# -
02725 -# -
02730 HMMreg00072_2_pdugrm(pf00072pf00486) EUT2E
02735 HMMreg00072_4_pdu1d(pf02518pf00512) PDU1F
02740 -pf00817pf11799pf11798 -
02745 -pf12843 -
02750 HMMnopfamclust29_1_aci(#) CsomeBCy
02755 -pf14791pf03372pf14716 -
BMC-P
BMC-H
BMC-T
Aldehyde dehydrogenase
PTAC
Signature enzyme
Alcohol dehydrogenase
Regulator

Roseibacillus sp. Bacteria; Verrucomicrobia; Verrucomicrobiae; Verrucomicrobiales;

CMP30__locus_1

PVM

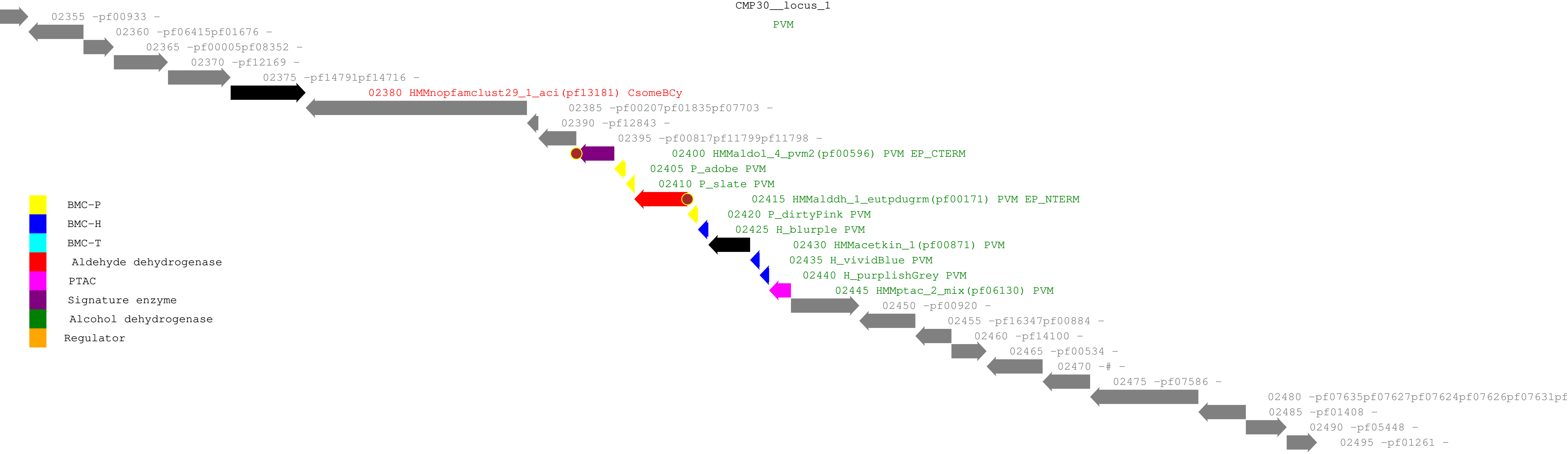

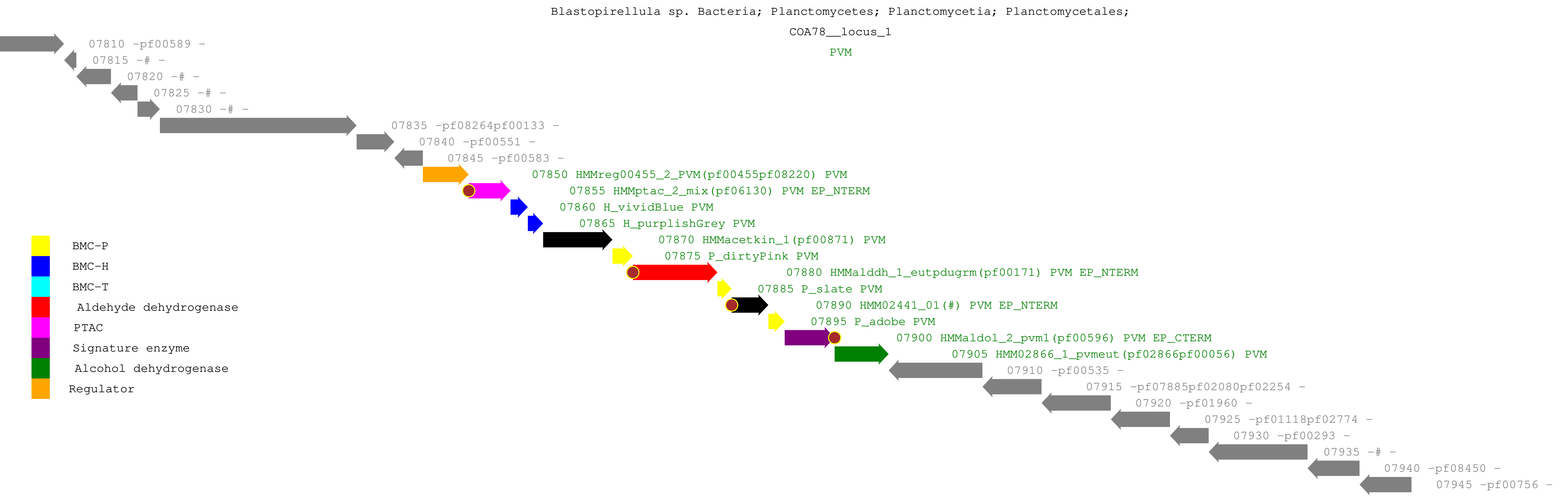
Blastopirellula sp. Bacteria; Planctomycetes; Planctomycetia; Planctomycetales;
COA78__locus_1
PVM
07810 -pf00589 -
07815 -# -
07820 -# -
07825 -# -
07830 -# -
07835 -pf08264pf00133 -
07840 -pf00551 -
07845 -pf00583 -
07850 HMMreg00455_2_PVM(pf00455pf08220) PVM
07855 HMMptac_2_mix(pf06130) PVM EP_NTERM
07860 H_vividBlue PVM
07865 H_purplishGrey PVM
07870 HMMacetkin_1(pf00871) PVM
07875 P_dirtyPink PVM
07880 HMMalddh_1_eutpdugrm(pf00171) PVM EP_NTERM
07885 P_slate PVM
07890 HMM02441_01(#) PVM EP_NTERM
07895 P_adobe PVM
07900 HMMaldol_2_pvm1(pf00596) PVM EP_CTERM
07905 HMM02866_1_pvmeut(pf02866pf00056) PVM
07910 -pf00535 -
07915 -pf07885pf02080pf02254 -
07920 -pf01960 -
07925 -pf01118pf02774 -
07930 -pf00293 -
07935 -# -
07940 -pf08450 -
07945 -pf00756 -
BMC-P
BMC-H
BMC-T
Aldehyde dehydrogenase
PTAC
Signature enzyme
Alcohol dehydrogenase
Regulator

Planctomycetes bacterium. Bacteria; Planctomycetes.

D6725__locus_1

PVM

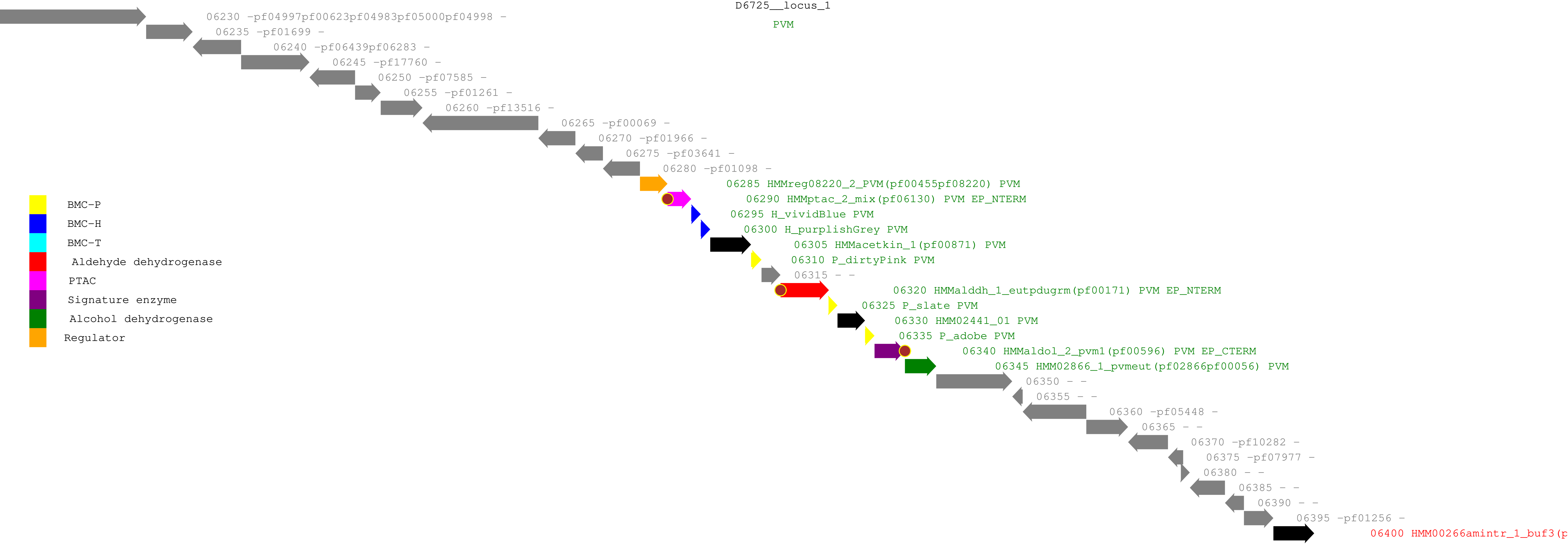

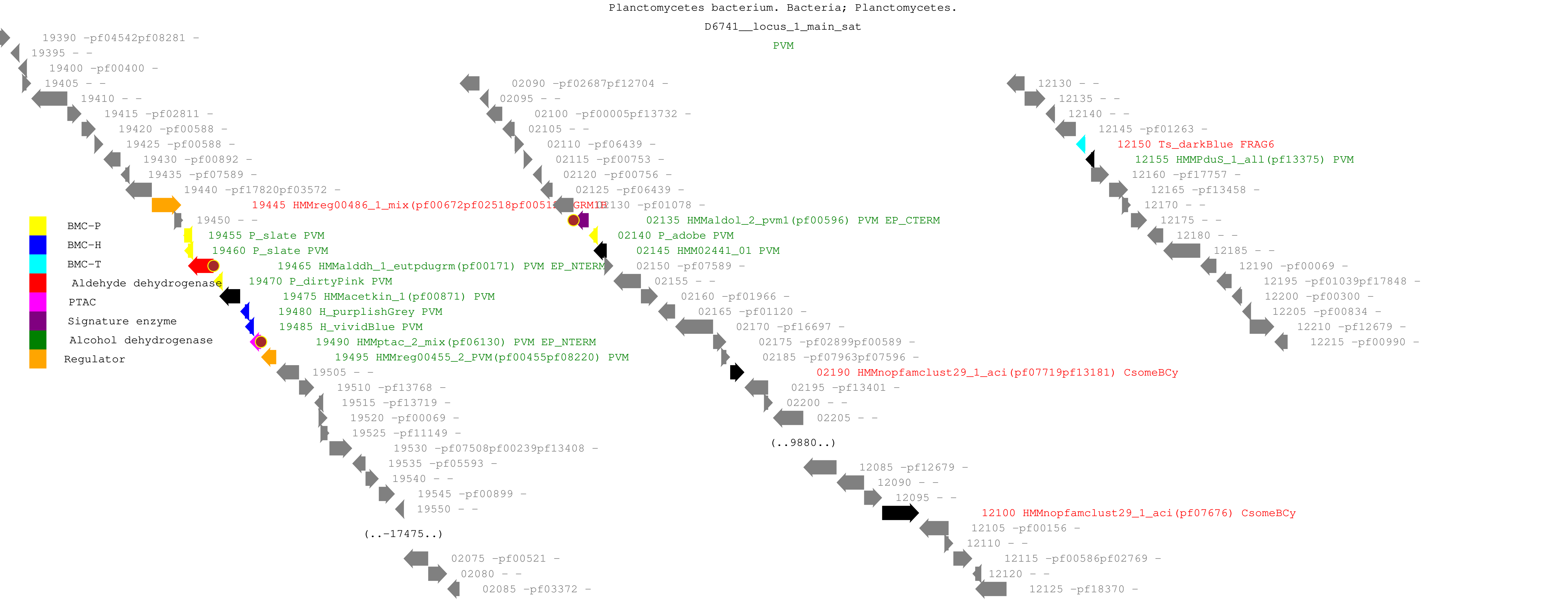

Planctomycetes bacterium. Bacteria; Planctomycetes.
D6741__locus_1_main_sat
PVM
BMC-P
BMC-H
BMC-T
Aldehyde dehydrogenase
PTAC
Signature enzyme
Alcohol dehydrogenase
Regulator
19390 -pf04542pf08281 -
19395 - -
19400 -pf00400 -
19405 - -
19410 - -
19415 -pf02811 -
19420 -pf00588 -
19425 -pf00588 -
19430 -pf00892 -
19435 -pf07589 -
19440 -pf17820pf03572 -
19445 HMMreg00486_1_mix(pf00672pf02518pf00512) GRM1B
19450 - -
19455 P_slate PVM
19460 P_slate PVM
19465 HMMaldDh_1_eutpdugrm(pf00171) PVM EP_NTERM
19470 P_dirtyPink PVM
19475 HMMacetkin_1(pf00871) PVM
19480 H_purplishGrey PVM
19485 H_vividBlue PVM
19490 HMMptac_2_mix(pf06130) PVM EP_NTERM
19495 HMMreg00455_2_PVM(pf00455pf08220) PVM
19505 - -
19510 -pf13768 -
19515 -pf13719 -
19520 -pf00069 -
19525 -pf11149 -
19530 -pf07508pf00239pf13408 -
19535 -pf05593 -
19540 - -
19545 -pf00899 -
19550 - -
(..-17475..)
02075 -pf00521 -
02080 - -
02085 -pf03372 -
02090 -pf02687pf12704 -
02095 - -
02100 -pf00005pf13732 -
02105 - -
02110 -pf06439 -
02115 -pf00753 -
02120 -pf00756 -
02125 -pf06439 -
02130 -pf01078 -
02135 HMMaldol_2_pvm1(pf00596) PVM EP_CTERM
02140 P_adobe PVM
02145 HMM02441_01 PVM
02150 -pf07589 -
02155 - -
02160 -pf01966 -
02165 -pf01120 -
02170 -pf16697 -
02175 -pf02899pf00589 -
02185 -pf07963pf07596 -
02190 HMMnopfamclust29_1_aci(pf07719pf13181) CsomeBCy
02195 -pf13401 -
02200 - -
02205 - -
(..9880..)
12085 -pf12679 -
12090 - -
12095 - -
12100 HMMnopfamclust29_1_aci(pf07676) CsomeBCy
12105 -pf00156 -
12110 - -
12115 -pf00586pf02769 -
12120 - -
12125 -pf18370 -
12130 - -
12135 - -
12140 - -
12145 -pf01263 -
12150 Ts_darkBlue FRAG6
12155 HMMPduS_1_all(pf13375) PVM
12160 -pf17757 -
12165 -pf13458 -
12170 - -
12175 - -
12180 - -
12185 - -
12190 -pf00069 -
12195 -pf01039pf17848 -
12200 -pf00300 -
12205 -pf00834 -
12210 -pf12679 -
12215 -pf00990 -

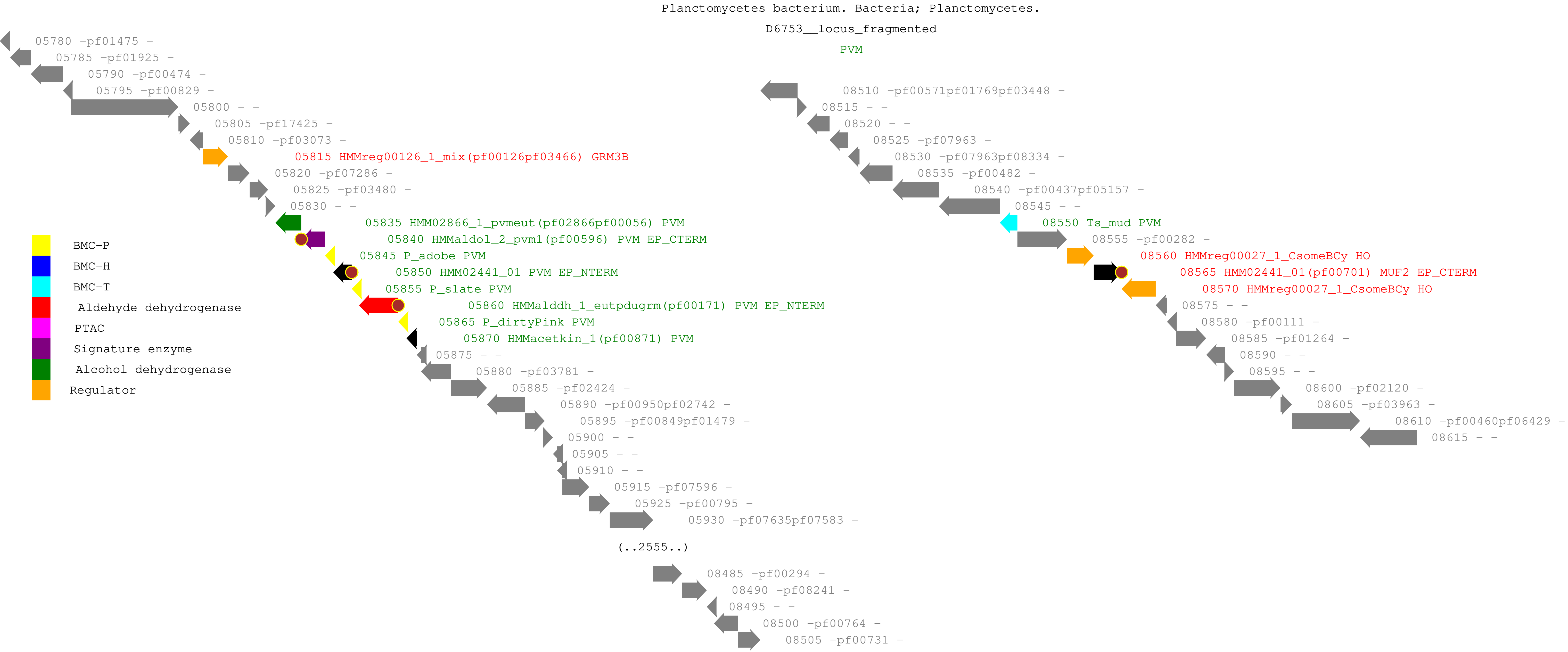
Planctomycetes bacterium. Bacteria; Planctomycetes.
D6753__locus_fragmented
PVM
05780 -pf01475 -
05785 -pf01925 -
05790 -pf00474 -
05795 -pf00829 -
05800 - -
05805 -pf17425 -
05810 -pf03073 -
05815 HMMreg00126_1_mix(pf00126pf03466) GRM3B
05820 -pf07286 -
05825 -pf03480 -
05830 - -
05835 HMM02866_1_pvmeut(pf02866pf00056) PVM
05840 HMMaldol_2_pvm1(pf00596) PVM EP_CTERM
05845 P_adobe PVM
05850 HMM02441_01 PVM EP_NTERM
05855 P_slate PVM
05860 HMMalddh_1_eutpdugrm(pf00171) PVM EP_NTERM
05865 P_dirtyPink PVM
05870 HMMacetkin_1(pf00871) PVM
05875 - -
05880 -pf03781 -
05885 -pf02424 -
05890 -pf00950pf02742 -
05895 -pf00849pf01479 -
05900 - -
05905 - -
05910 - -
05915 -pf07596 -
05925 -pf00795 -
05930 -pf07635pf07583 -
(..2555..)
08485 -pf00294 -
08490 -pf08241 -
08495 - -
08500 -pf00764 -
08505 -pf00731 -
08510 -pf00571pf01769pf03448 -
08515 - -
08520 - -
08525 -pf07963 -
08530 -pf07963pf08334 -
08535 -pf00482 -
08540 -pf00437pf05157 -
08545 - -
08550 Ts_mud PVM
08555 -pf00282 -
08560 HMMreg00027_1_CsomeBCy HO
08565 HMM02441_01(pf00701) MUF2 EP_CTERM
08570 HMMreg00027_1_CsomeBCy HO
08575 - -
08580 -pf00111 -
08585 -pf01264 -
08590 - -
08595 - -
08600 -pf02120 -
08605 -pf03963 -
08610 -pf00460pf06429 -
08615 - -
BMC-P
BMC-H
BMC-T
Aldehyde dehydrogenase
PTAC
Signature enzyme
Alcohol dehydrogenase
Regulator

Verrucomicrobia bacterium. Bacteria; Verrucomicrobia; unclassified Verrucomicrobia.

D6766__locus_1_main_sat

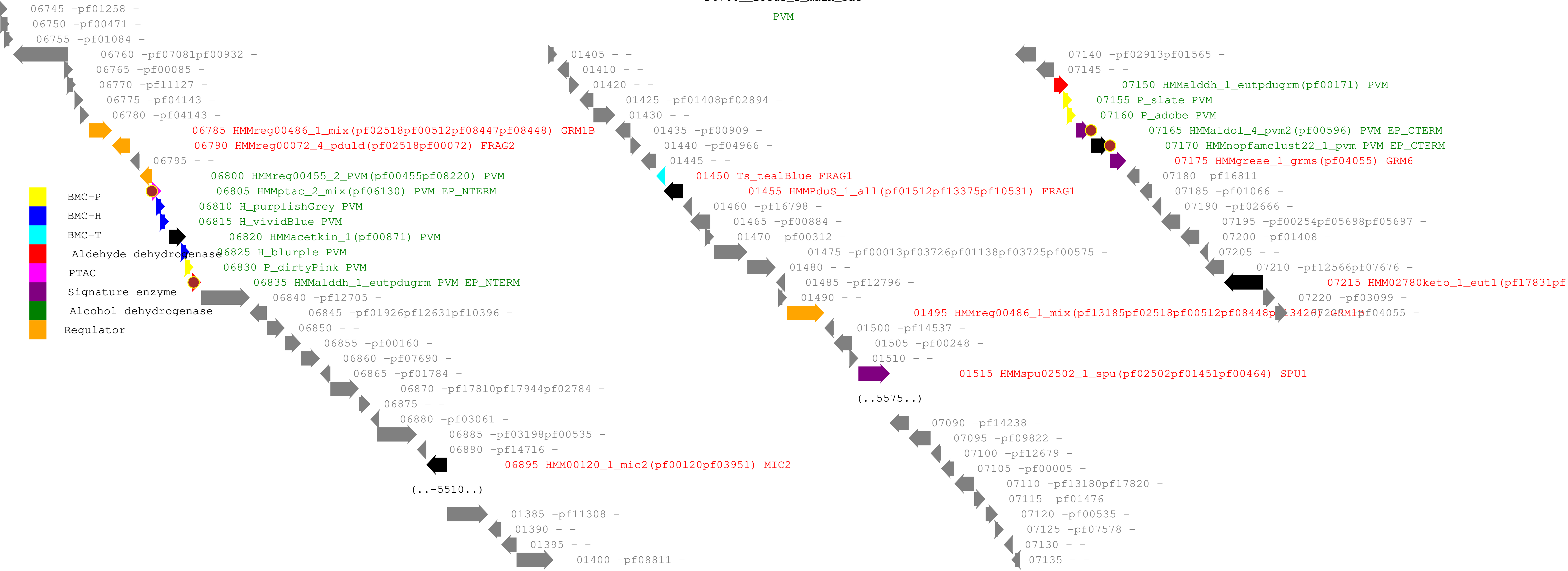

Verrucomicrobia bacterium. Bacteria; Verrucomicrobia; unclassified Verrucomicrobia.

D6781__locus_1

PVM

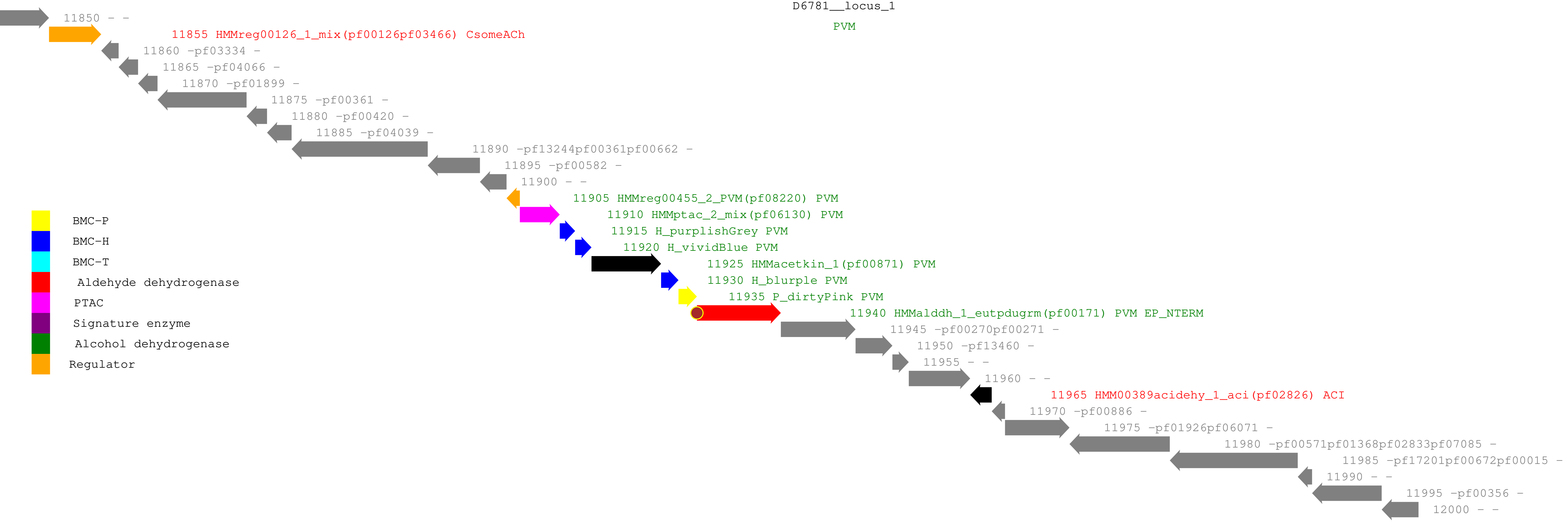

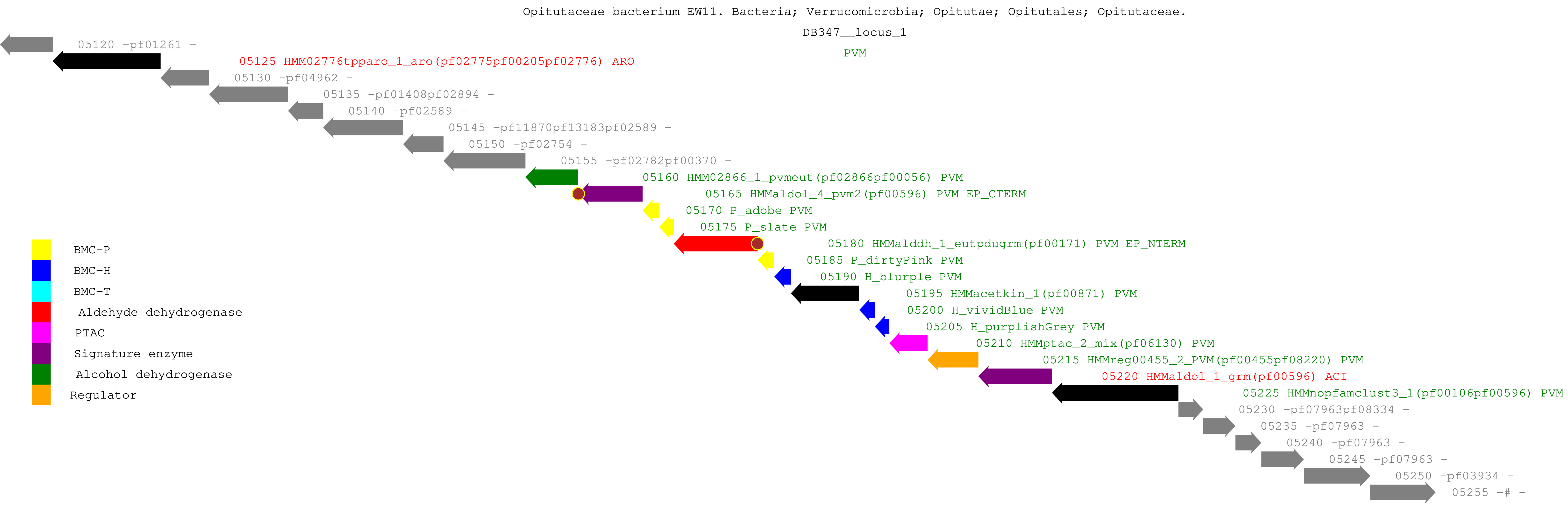
Opitutaceae bacterium EW11. Bacteria; Verrucomicrobia; Opitutae; Opitutales; Opitutaceae.
DB347__locus_1
PVM
05120 -pf01261 -
05125 HMM02776tpparo_1_aro(pf02775pf00205pf02776) ARO
05130 -pf04962 -
05135 -pf01408pf02894 -
05140 -pf02589 -
05145 -pf11870pf13183pf02589 -
05150 -pf02754 -
05155 -pf02782pf00370 -
05160 HMM02866_1_pvmeut(pf02866pf00056) PVM
05165 HMMaldol_4_pvm2(pf00596) PVM EP_CTERM
05170 P_adobe PVM
05175 P_slate PVM
05180 HMMalddh_1_eutpdugrm(pf00171) PVM EP_NTERM
05185 P_dirtyPink PVM
05190 H_blurple PVM
05195 HMMacetkin_1(pf00871) PVM
05200 H_vividBlue PVM
05205 H_purplishGrey PVM
05210 HMMptac_2_mix(pf06130) PVM
05215 HMMreg00455_2_PVM(pf00455pf08220) PVM
05220 HMMaldol_1_grm(pf00596) ACI
05225 HMMnopfamclust3_1(pf00106pf00596) PVM
05230 -pf07963pf08334 -
05235 -pf07963 -
05240 -pf07963 -
05245 -pf07963 -
05250 -pf03934 -
05255 -# -
BMC-P
BMC-H
BMC-T
Aldehyde dehydrogenase
PTAC
Signature enzyme
Alcohol dehydrogenase
Regulator

Opitutus sp. ER46. Bacteria; Verrucomicrobia; Opitutae; Opitutales; Opitutaceae;

DB354__locus_1_main_sat

PVM

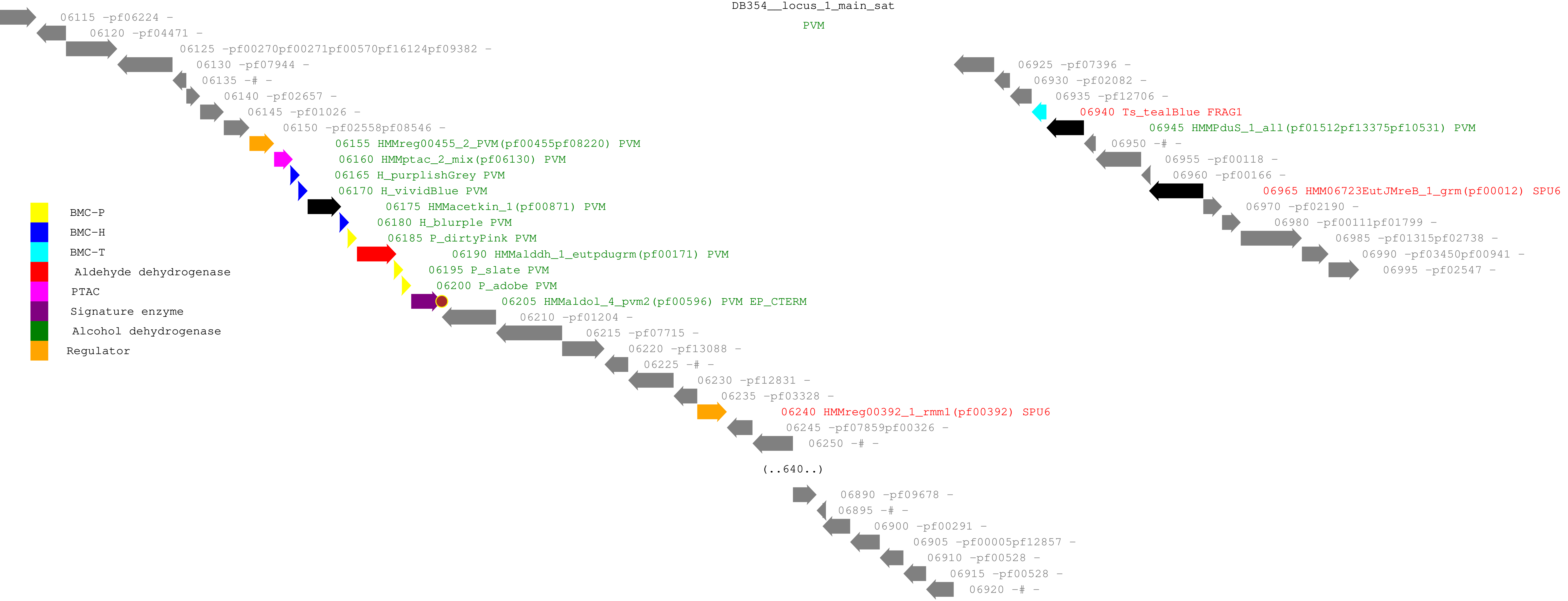

Verrucomicrobia bacterium. Bacteria; Verrucomicrobia; unclassified Verrucomicrobia.

DBX00__locus_1_main_sat

PVM

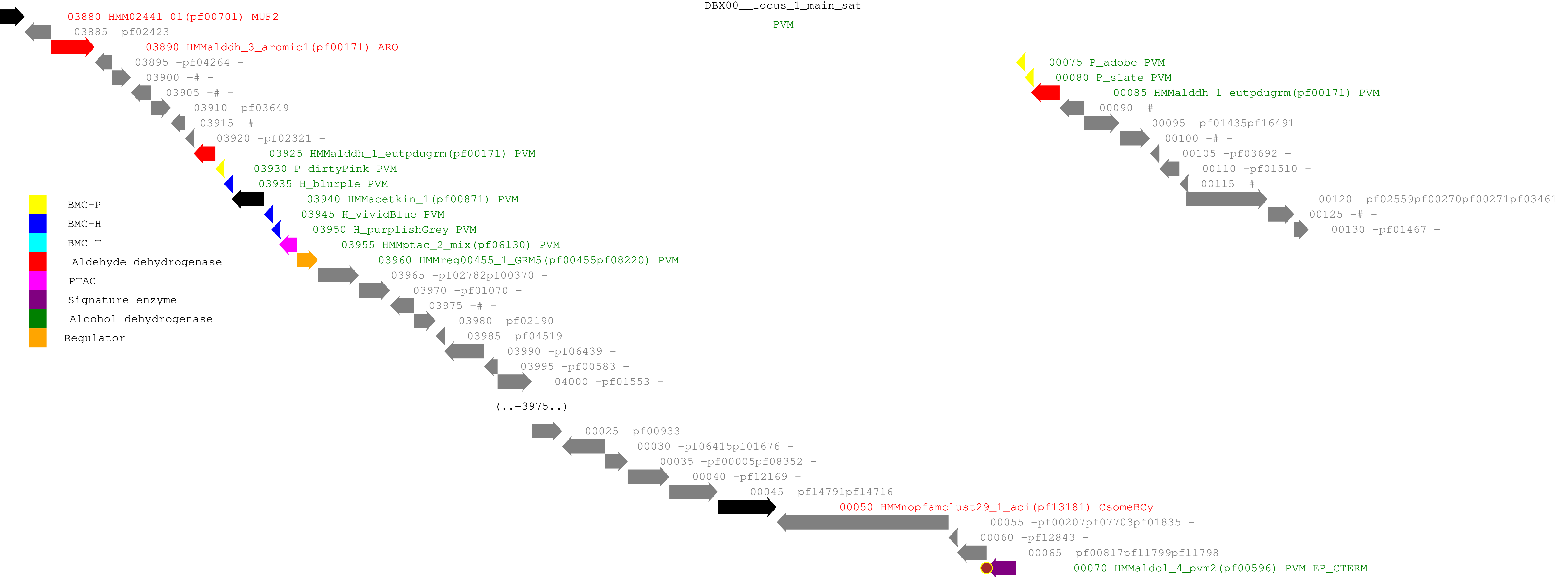

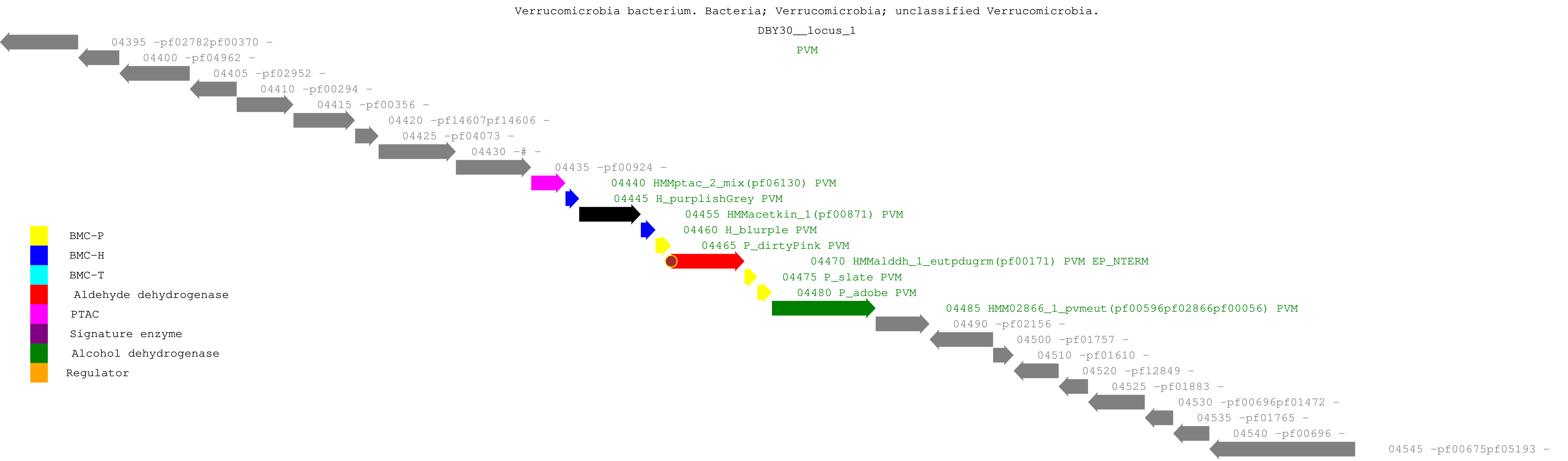

Verrucomicrobia bacterium. Bacteria; Verrucomicrobia; unclassified Verrucomicrobia.
DBY30__locus_1
PVM
04395 -pf02782pf00370 -
04400 -pf04962 -
04405 -pf02952 -
04410 -pf00294 -
04415 -pf00356 -
04420 -pf14607pf14606 -
04425 -pf04073 -
04430 -# -
04435 -pf00924 -
04440 HMMptac_2_mix(pf06130) PVM
04445 H_purplishGrey PVM
04455 HMMacetkin_1(pf00871) PVM
04460 H_blurple PVM
04465 P_dirtyPink PVM
04470 HMMaldd h_1_eutpdugrm(pf00171) PVM EP_NTERM
04475 P_slate PVM
04480 P_adobe PVM
04485 HMM02866_1_pvmeut(pf00596pf02866pf00056) PVM
04490 -pf02156 -
04500 -pf01757 -
04510 -pf01610 -
04520 -pf12849 -
04525 -pf01883 -
04530 -pf00696pf01472 -
04535 -pf01765 -
04540 -pf00696 -
04545 -pf00675pf05193 -
BMC-P
BMC-H
BMC-T
Aldehyde dehydrogenase
PTAC
Signature enzyme
Alcohol dehydrogenase
Regulator

Planctomycetaceae bacterium. Bacteria; Planctomycetes; Planctomycetia; Planctomycetales;

DC048__locus_1_main_sat

PVM

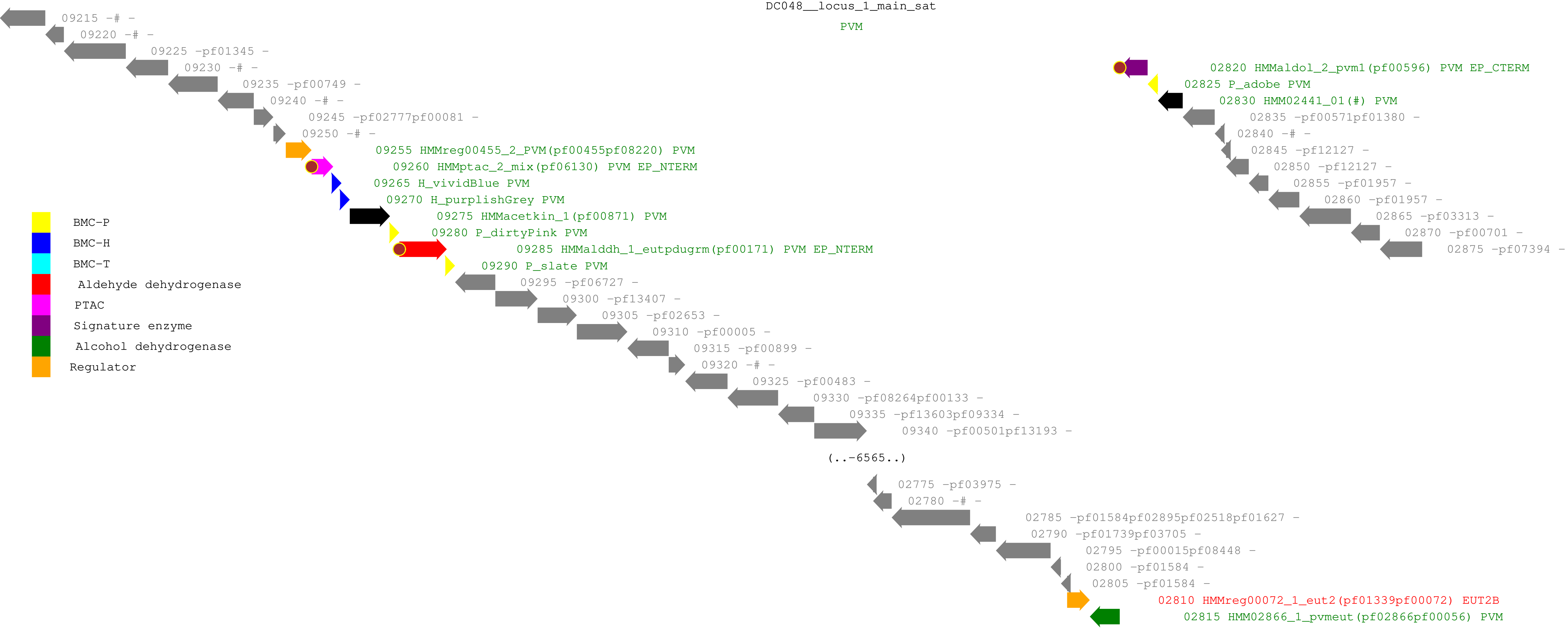

Planctomycetes bacterium. Bacteria; Planctomycetes.

DCC68__locus_fragmented

PVM

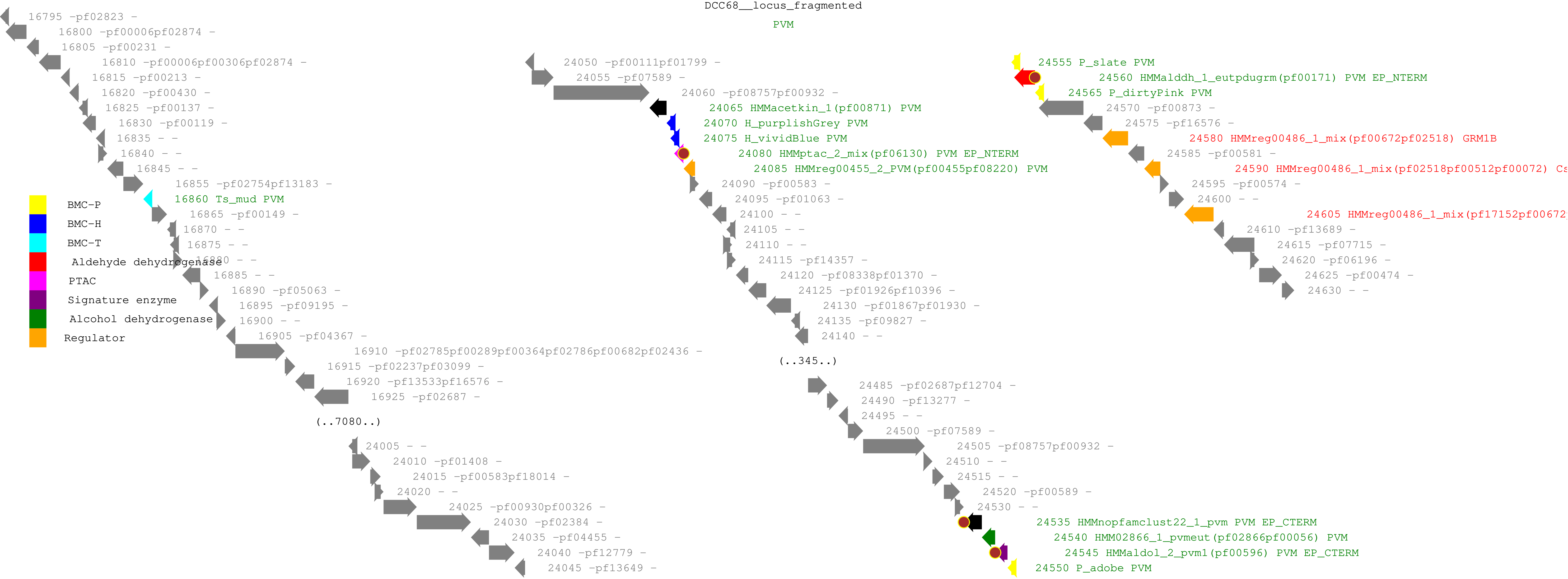

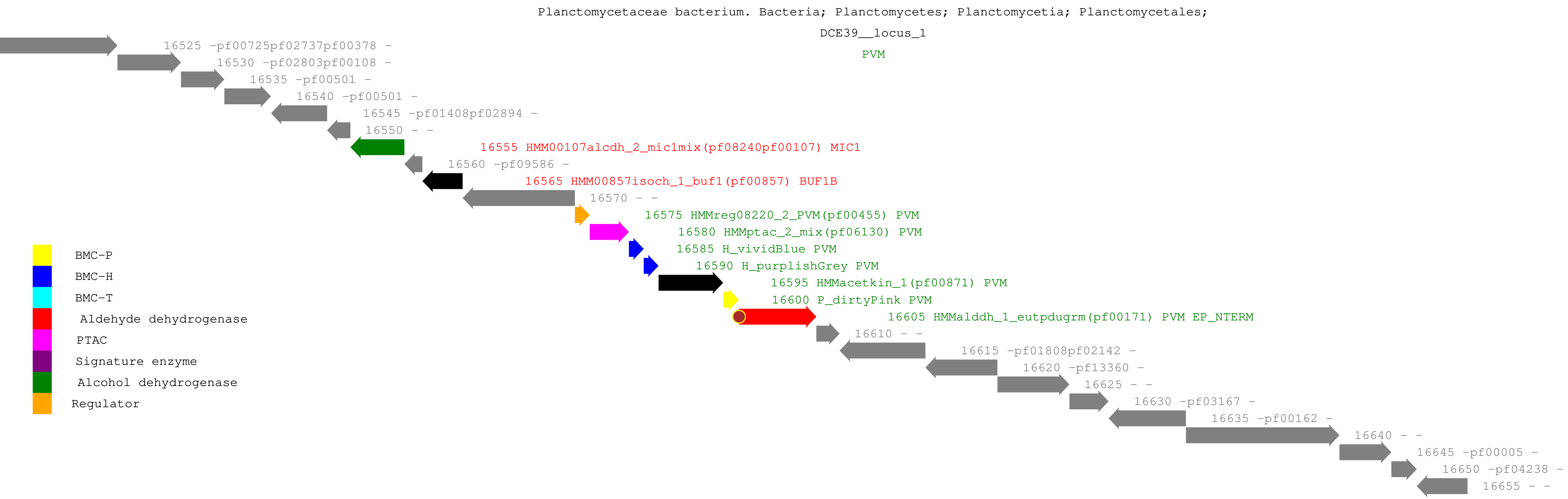
Planctomycetaceae bacterium. Bacteria; Planctomycetes; Planctomycetia; Planctomycetales;
DCE39__locus_1
PVM
16525 -pf00725pf02737pf00378 -
16530 -pf02803pf00108 -
16535 -pf00501 -
16540 -pf00501 -
16545 -pf01408pf02894 -
16550 - -
16555 HMM00107alcdh_2_mic1mix(pf08240pf00107) MIC1
16560 -pf09586 -
16565 HMM00857isoch_1_buf1(pf00857) BUF1B
16570 - -
16575 HMMreg08220_2_PVM(pf00455) PVM
16580 HMMptac_2_mix(pf06130) PVM
16585 H_vividBlue PVM
16590 H_purplishGrey PVM
16595 HMMacetkin_1(pf00871) PVM
16600 P_dirtyPink PVM
16605 HMMaldh_1_eutpdugrm(pf00171) PVM EP_NTERM
16610 - -
16615 -pf01808pf02142 -
16620 -pf13360 -
16625 - -
16630 -pf03167 -
16635 -pf00162 -
16640 - -
16645 -pf00005 -
16650 -pf04238 -
16655 - -
BMC-P
BMC-H
BMC-T
Aldehyde dehydrogenase
PTAC
Signature enzyme
Alcohol dehydrogenase
Regulator

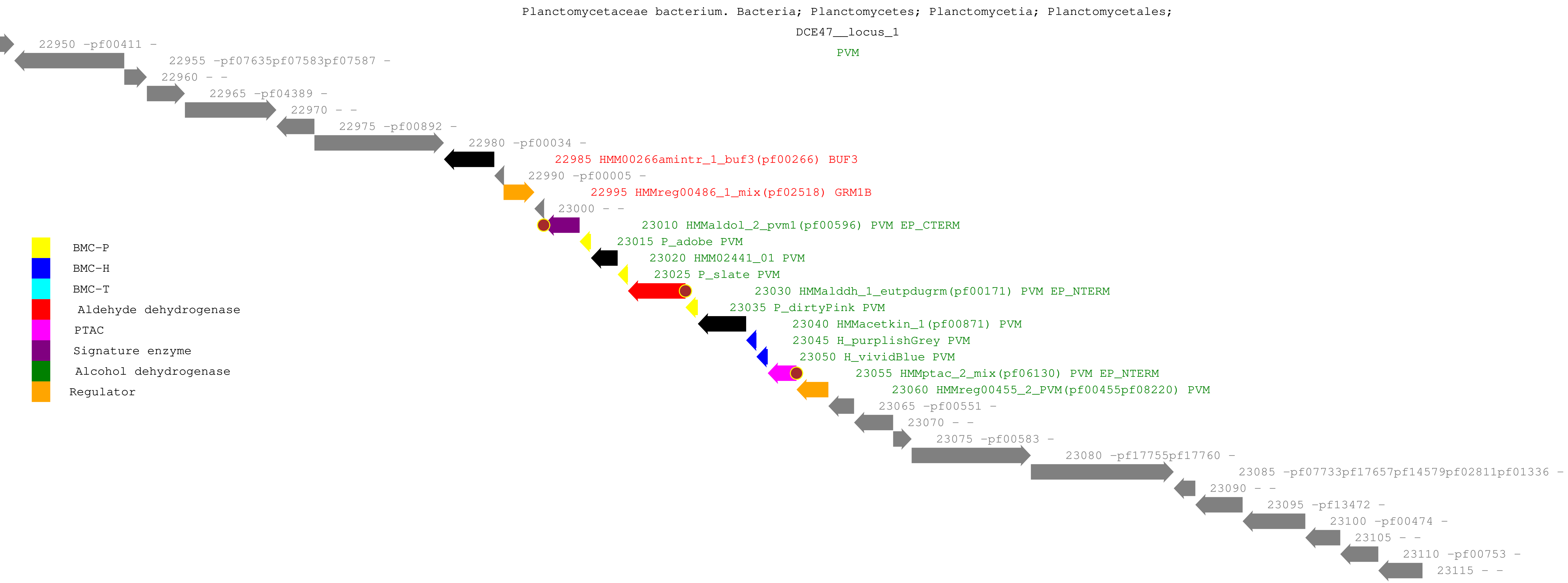

Planctomycetaceae bacterium. Bacteria; Planctomycetes; Planctomycetia; Planctomycetales;
DCE47__locus_1
PVM
22950 -pf00411 -
22955 -pf07635pf07583pf07587 -
22960 - -
22965 -pf04389 -
22970 - -
22975 -pf00892 -
22980 -pf00034 -
22985 HMM00266amintr_1_buf3(pf00266) BUF3
22990 -pf00005 -
22995 HMMreg00486_1_mix(pf02518) GRM1B
23000 - -
23010 HMMaldol_2_pvm1(pf00596) PVM EP_CTERM
23015 P_adobe PVM
23020 HMM02441_01 PVM
23025 P_slate PVM
23030 HMMaldh_1_eutpdugrm(pf00171) PVM EP_NTERM
23035 P_dirtyPink PVM
23040 HMMacetkin_1(pf00871) PVM
23045 H_purplishGrey PVM
23050 H_vividBlue PVM
23055 HMMptac_2_mix(pf06130) PVM EP_NTERM
23060 HMMreg00455_2_PVM(pf00455pf08220) PVM
23065 -pf00551 -
23070 - -
23075 -pf00583 -
23080 -pf17755pf17760 -
23085 -pf07733pf17657pf14579pf02811pf01336 -
23090 - -
23095 -pf13472 -
23100 -pf00474 -
23105 - -
23110 -pf00753 -
23115 - -
BMC-P
BMC-H
BMC-T
Aldehyde dehydrogenase
PTAC
Signature enzyme
Alcohol dehydrogenase
Regulator

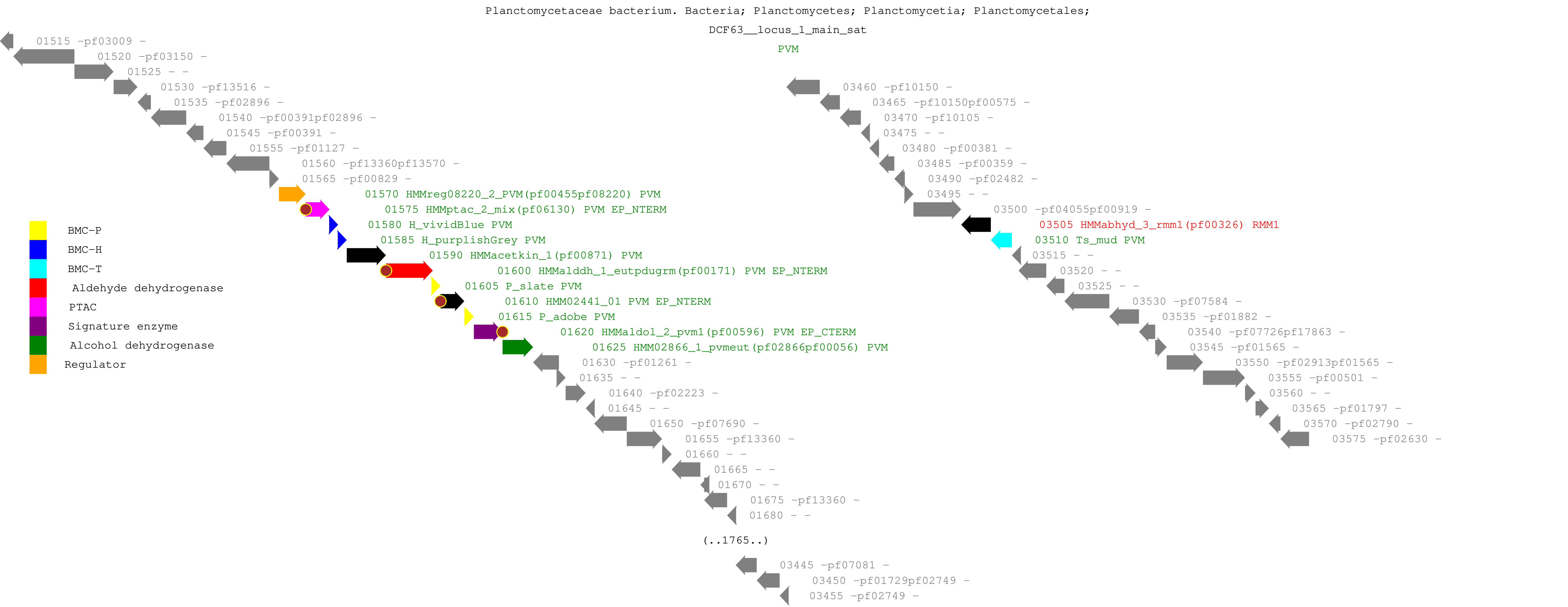
Planctomycetaceae bacterium. Bacteria; Planctomycetes; Planctomycetia; Planctomycetales;
DCF63__locus_1_main_sat
PVM
01515 -pf03009 -
01520 -pf03150 -
01525 - -
01530 -pf13516 -
01535 -pf02896 -
01540 -pf00391pf02896 -
01545 -pf00391 -
01555 -pf01127 -
01560 -pf13360pf13570 -
01565 -pf00829 -
01570 HMMreg08220_2_PVM(pf00455pf08220) PVM
01575 HMMptac_2_mix(pf06130) PVM EP_NTERM
01580 H_vividBlue PVM
01585 H_purplishGrey PVM
01590 HMMacetkin_1(pf00871) PVM
01600 HMMalddh_1_eutpdugrm(pf00171) PVM EP_NTERM
01605 P_slate PVM
01610 HMM02441_01 PVM EP_NTERM
01615 P_adobe PVM
01620 HMMaldol_2_pvm1(pf00596) PVM EP_CTERM
01625 HMM02866_1_pvmeut(pf02866pf00056) PVM
01630 -pf01261 -
01635 - -
01640 -pf02223 -
01645 - -
01650 -pf07690 -
01655 -pf13360 -
01660 - -
01665 - -
01670 - -
01675 -pf13360 -
01680 - -
(..1765..)
03445 -pf07081 -
03450 -pf01729pf02749 -
03455 -pf02749 -
03460 -pf10150 -
03465 -pf10150pf00575 -
03470 -pf10105 -
03475 - -
03480 -pf00381 -
03485 -pf00359 -
03490 -pf02482 -
03495 - -
03500 -pf04055pf00919 -
03505 HMMabhyd_3_rmm1(pf00326) RMM1
03510 Ts_mud PVM
03515 - -
03520 - -
03525 - -
03530 -pf07584 -
03535 -pf01882 -
03540 -pf07726pf17863 -
03545 -pf01565 -
03550 -pf02913pf01565 -
03555 -pf00501 -
03560 - -
03565 -pf01797 -
03570 -pf02790 -
03575 -pf02630 -
BMC-P
BMC-H
BMC-T
Aldehyde dehydrogenase
PTAC
Signature enzyme
Alcohol dehydrogenase
Regulator

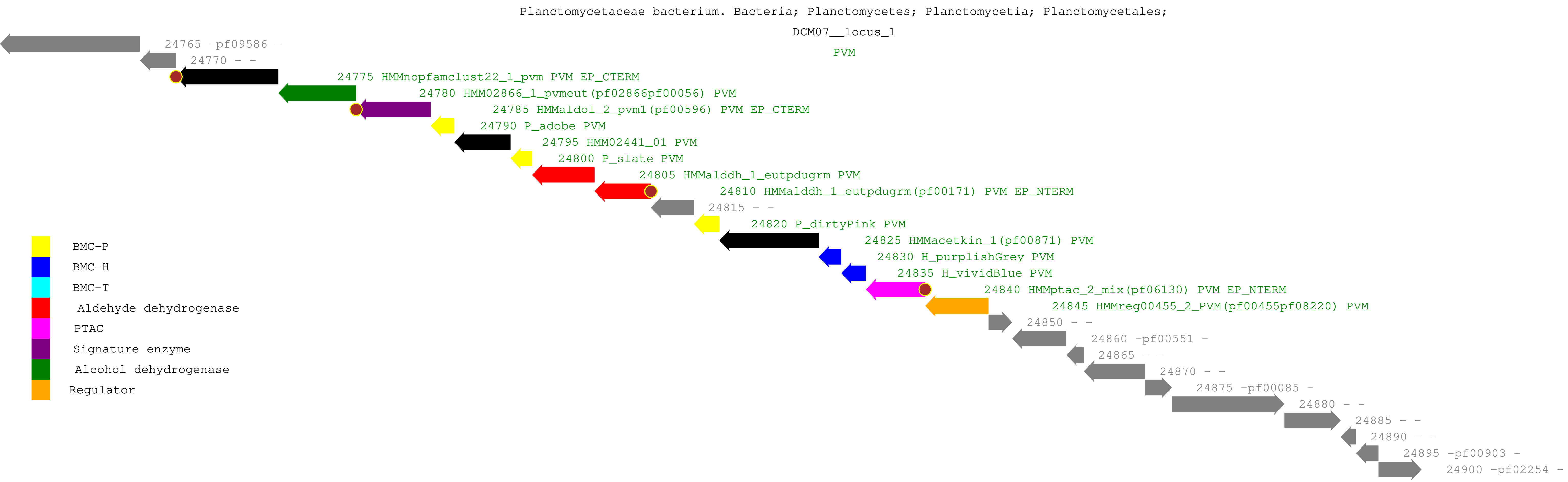

Planctomycetaceae bacterium. Bacteria; Planctomycetes; Planctomycetia; Planctomycetales;
DCM07__locus_1
PVM
24765 -pf09586 -
24770 - -
24775 HMMnopfamclust22_1_pvm PVM EP_CTERM
24780 HMM02866_1_pvmeut(pf02866pf00056) PVM
24785 HMMaldol_2_pvm1(pf00596) PVM EP_CTERM
24790 P_adobe PVM
24795 HMM02441_01 PVM
24800 P_slate PVM
24805 HMMalddh_1_eutpdugrm PVM
24810 HMMalddh_1_eutpdugrm(pf00171) PVM EP_NTERM
24815 - -
24820 P_dirtyPink PVM
24825 HMMacetkin_1(pf00871) PVM
24830 H_purplishGrey PVM
24835 H_vividBlue PVM
24840 HMMptac_2_mix(pf06130) PVM EP_NTERM
24845 HMMreg00455_2_PVM(pf00455pf08220) PVM
24850 - -
24860 -pf00551 -
24865 - -
24870 - -
24875 -pf00085 -
24880 - -
24885 - -
24890 - -
24895 -pf00903 -
24900 -pf02254 -
BMC-P
BMC-H
BMC-T
Aldehyde dehydrogenase
PTAC
Signature enzyme
Alcohol dehydrogenase
Regulator

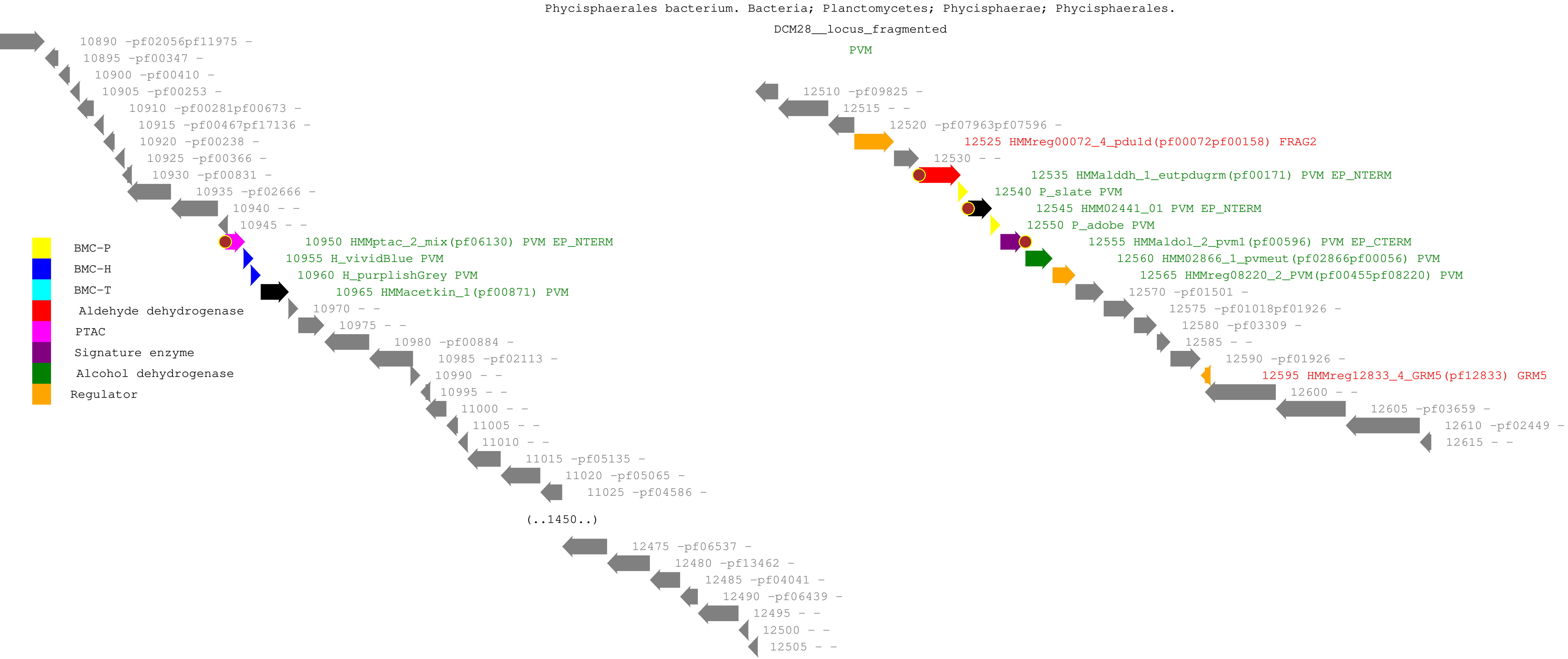
Phycisphaerales bacterium. Bacteria; Planctomycetes; Phycisphaerae; Phycisphaerales.
DCM28__locus_fragmented
PVM
10890 -pf02056pf11975 -
10895 -pf00347 -
10900 -pf00410 -
10905 -pf00253 -
10910 -pf00281pf00673 -
10915 -pf00467pf17136 -
10920 -pf00238 -
10925 -pf00366 -
10930 -pf00831 -
10935 -pf02666 -
10940 - -
10945 - -
10950 HMMptac_2_mix(pf06130) PVM EP_NTERM
10955 H_vividBlue PVM
10960 H_purplishGrey PVM
10965 HMMacetkin_1(pf00871) PVM
10970 - -
10975 - -
10980 -pf00884 -
10985 -pf02113 -
10990 - -
10995 - -
11000 - -
11005 - -
11010 - -
11015 -pf05135 -
11020 -pf05065 -
11025 -pf04586 -
(..1450..)
12475 -pf06537 -
12480 -pf13462 -
12485 -pf04041 -
12490 -pf06439 -
12495 - -
12500 - -
12505 - -
12510 -pf09825 -
12515 - -
12520 -pf07963pf07596 -
12525 HMMreg00072_4_pdu1d(pf00072pf00158) FRAG2
12530 - -
12535 HMMalddh_1_eutpdugrm(pf00171) PVM EP_NTERM
12540 P_slate PVM
12545 HMM02441_01 PVM EP_NTERM
12550 P_adobe PVM
12555 HMMaldol_2_pvm1(pf00596) PVM EP_CTERM
12560 HMM02866_1_pvmeut(pf02866pf00056) PVM
12565 HMMreg08220_2_PVM(pf00455pf08220) PVM
12570 -pf01501 -
12575 -pf01018pf01926 -
12580 -pf03309 -
12585 - -
12590 -pf01926 -
12595 HMMreg12833_4_GRM5(pf12833) GRM5
12600 - -
12605 -pf03659 -
12610 -pf02449 -
12615 - -
BMC-P
BMC-H
BMC-T
Aldehyde dehydrogenase
PTAC
Signature enzyme
Alcohol dehydrogenase
Regulator

Verrucomicrobiales bacterium. Bacteria; Verrucomicrobia; Verrucomicrobiae; Verrucomicrobiales.

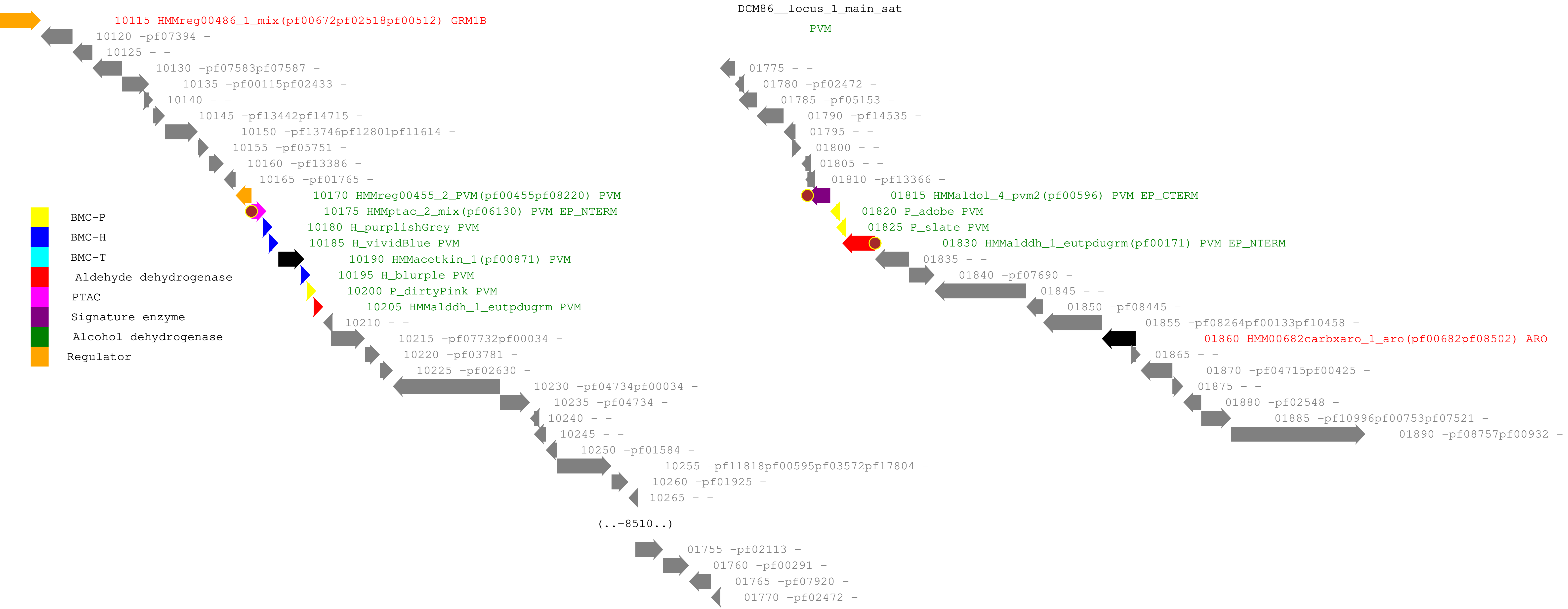

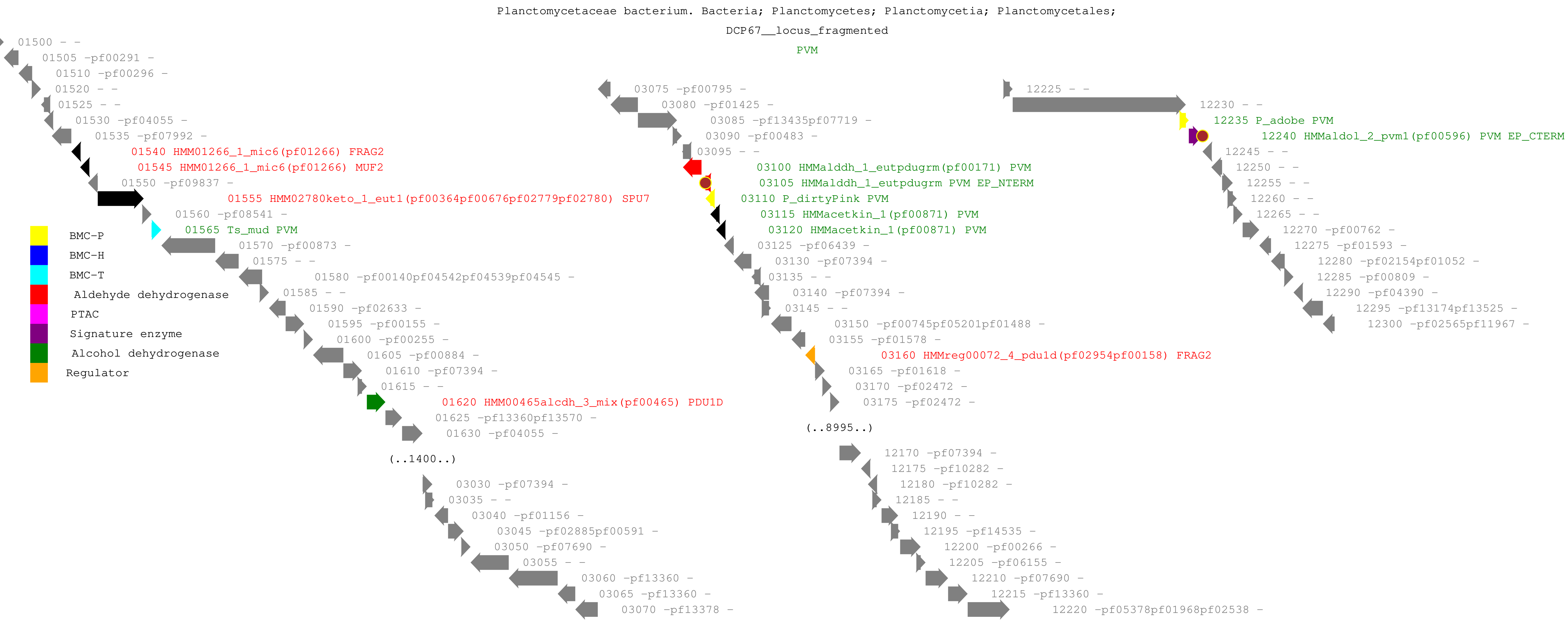
Planctomycetaceae bacterium. Bacteria; Planctomycetes; Planctomycetia; Planctomycetales;
DCP67__locus_fragmented
PVM
01500 - -
01505 -pf00291 -
01510 -pf00296 -
01520 - -
01525 - -
01530 -pf04055 -
01535 -pf07992 -
01540 HMM01266_1_mic6(pf01266) FRAG2
01545 HMM01266_1_mic6(pf01266) MUF2
01550 -pf09837 -
01555 HMM02780keto_1_eut1(pf00364pf00676pf02779pf02780) SPU7
01560 -pf08541 -
01565 Ts_mud PVM
01570 -pf00873 -
01575 - -
01580 -pf00140pf04542pf04539pf04545 -
01585 - -
01590 -pf02633 -
01595 -pf00155 -
01600 -pf00255 -
01605 -pf00884 -
01610 -pf07394 -
01615 - -
01620 HMM00465alcdh_3_mix(pf00465) PDU1D
01625 -pf13360pf13570 -
01630 -pf04055 -
(..1400..)
03030 -pf07394 -
03035 - -
03040 -pf01156 -
03045 -pf02885pf00591 -
03050 -pf07690 -
03055 - -
03060 -pf13360 -
03065 -pf13360 -
03070 -pf13378 -
BMC-P
BMC-H
BMC-T
Aldehyde dehydrogenase
PTAC
Signature enzyme
Alcohol dehydrogenase
Regulator
03075 -pf00795 -
03080 -pf01425 -
03085 -pf13435pf07719 -
03090 -pf00483 -
03095 - -
03100 HMMalddh_1_eutpdugrm(pf00171) PVM
03105 HMMalddh_1_eutpdugrm PVM EP_NTERM
03110 P_dirtyPink PVM
03115 HMMacetkin_1(pf00871) PVM
03120 HMMacetkin_1(pf00871) PVM
03125 -pf06439 -
03130 -pf07394 -
03135 - -
03140 -pf07394 -
03145 - -
03150 -pf00745pf05201pf01488 -
03155 -pf01578 -
03160 HMMreg00072_4_pdu1d(pf02954pf00158) FRAG2
03165 -pf01618 -
03170 -pf02472 -
03175 -pf02472 -
(..8995..)
12170 -pf07394 -
12175 -pf10282 -
12180 -pf10282 -
12185 - -
12190 - -
12195 -pf14535 -
12200 -pf00266 -
12205 -pf06155 -
12210 -pf07690 -
12215 -pf13360 -
12220 -pf05378pf01968pf02538 -
12225 - -
12230 - -
12235 P_adobe PVM
12240 HMMaldol_2_pvm1(pf00596) PVM EP_CTERM
12245 - -
12250 - -
12255 - -
12260 - -
12265 - -
12270 -pf00762 -
12275 -pf01593 -
12280 -pf02154pf01052 -
12285 -pf00809 -
12290 -pf04390 -
12295 -pf13174pf13525 -
12300 -pf02565pf11967 -

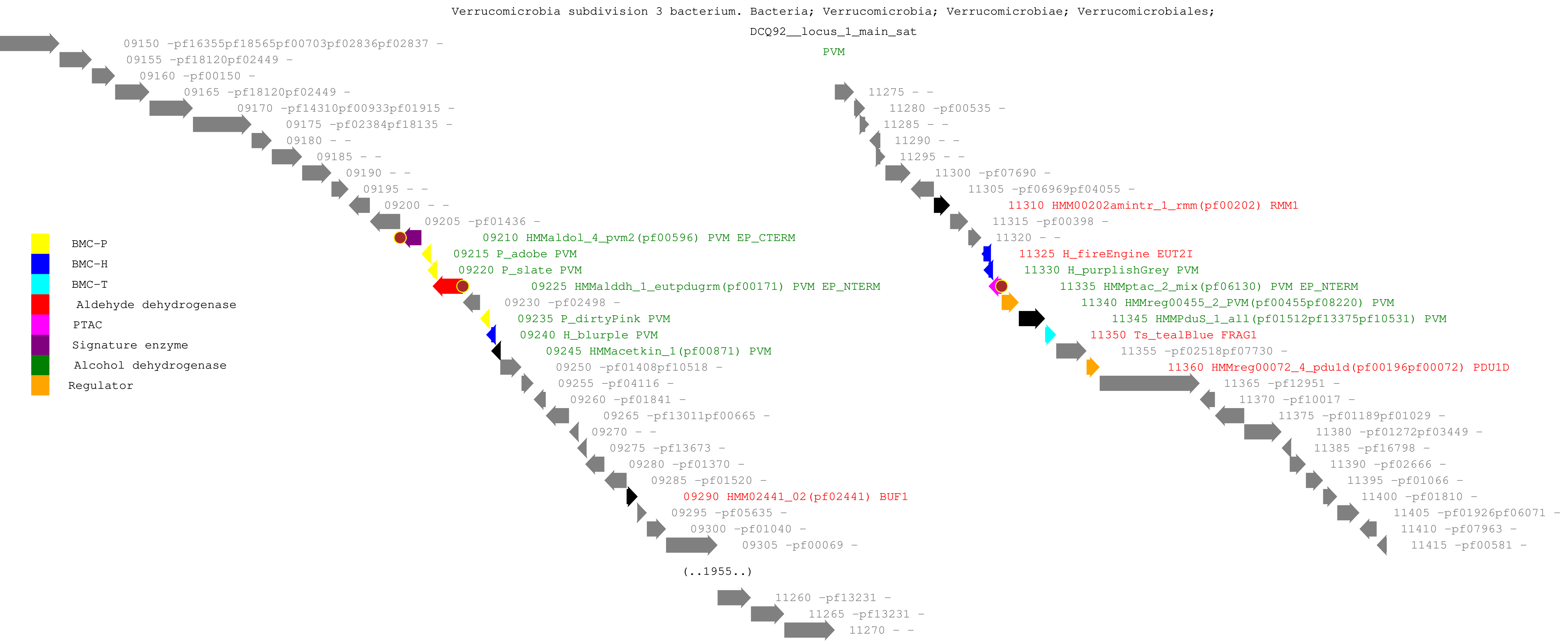

Verrucomicrobia subdivision 3 bacterium. Bacteria; Verrucomicrobia; Verrucomicrobiae; Verrucomicrobiales;
DCQ92__locus_1_main_sat
PVM
09150 -pf16355pf18565pf00703pf02836pf02837 -
09155 -pf18120pf02449 -
09160 -pf00150 -
09165 -pf18120pf02449 -
09170 -pf14310pf00933pf01915 -
09175 -pf02384pf18135 -
09180 - -
09185 - -
09190 - -
09195 - -
09200 - -
09205 -pf01436 -
09210 HMMaldol_4_pvm2(pf00596) PVM EP_CTERM
09215 P_adobe PVM
09220 P_slate PVM
09225 HMMalddh_1_eutpdugrm(pf00171) PVM EP_NTERM
09230 -pf02498 -
09235 P_dirtyPink PVM
09240 H_blurple PVM
09245 HMMacetkin_1(pf00871) PVM
09250 -pf01408pf10518 -
09255 -pf04116 -
09260 -pf01841 -
09265 -pf13011pf00665 -
09270 - -
09275 -pf13673 -
09280 -pf01370 -
09285 -pf01520 -
09290 HMM02441_02(pf02441) BUF1
09295 -pf05635 -
09300 -pf01040 -
09305 -pf00069 -
(..1955..)
11260 -pf13231 -
11265 -pf13231 -
11270 - -
11275 - -
11280 -pf00535 -
11285 - -
11290 - -
11295 - -
11300 -pf07690 -
11305 -pf06969pf04055 -
11310 HMM00202amintr_1_rmm(pf00202) RMM1
11315 -pf00398 -
11320 - -
11325 H_fireEngine EUT2I
11330 H_purplishGrey PVM
11335 HMMptac_2_mix(pf06130) PVM EP_NTERM
11340 HMMreg00455_2_PVM(pf00455pf08220) PVM
11345 HMMPduS_1_all(pf01512pf13375pf10531) PVM
11350 Ts_tealBlue FRAG1
11355 -pf02518pf07730 -
11360 HMMreg00072_4_pdu1d(pf00196pf00072) PDU1D
11365 -pf12951 -
11370 -pf10017 -
11375 -pf01189pf01029 -
11380 -pf01272pf03449 -
11385 -pf16798 -
11390 -pf02666 -
11395 -pf01066 -
11400 -pf01810 -
11405 -pf01926pf06071 -
11410 -pf07963 -
11415 -pf00581 -
BMC-P
BMC-H
BMC-T
Aldehyde dehydrogenase
PTAC
Signature enzyme
Alcohol dehydrogenase
Regulator

Verrucomicrobiales bacterium. Bacteria; Verrucomicrobia; Verrucomicrobiae; Verrucomicrobiales.

DCQ96__locus_1

PVM

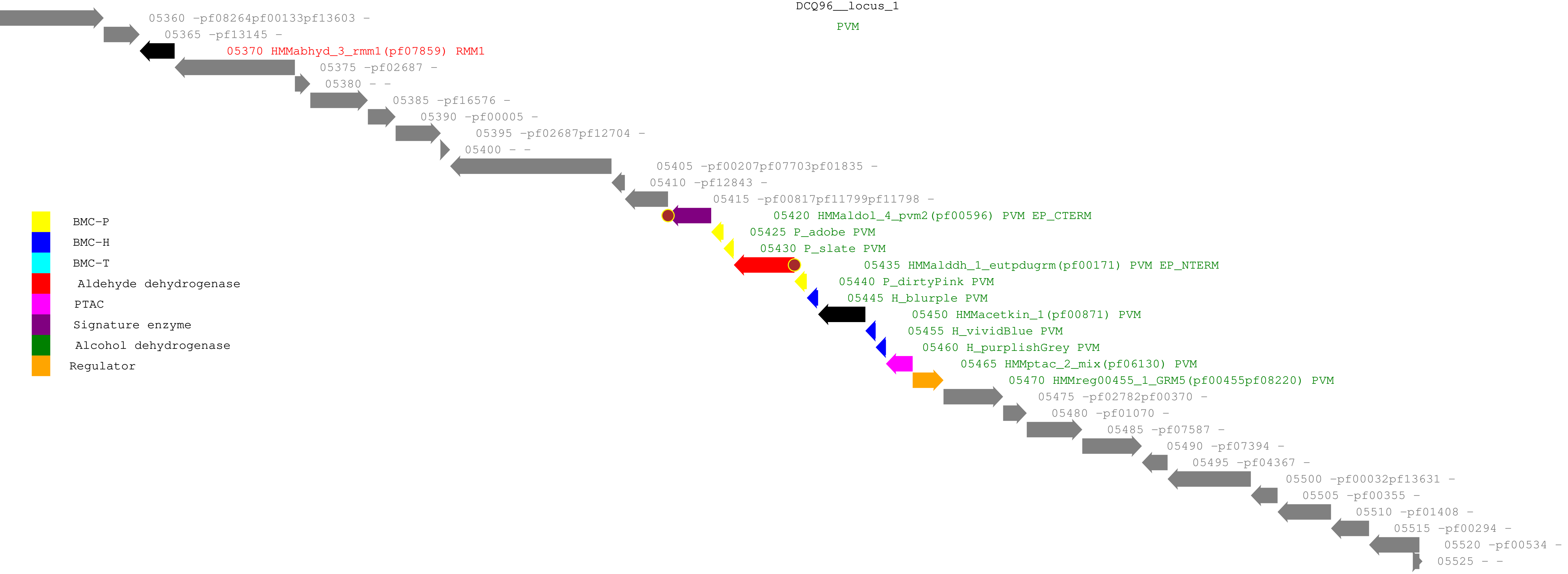

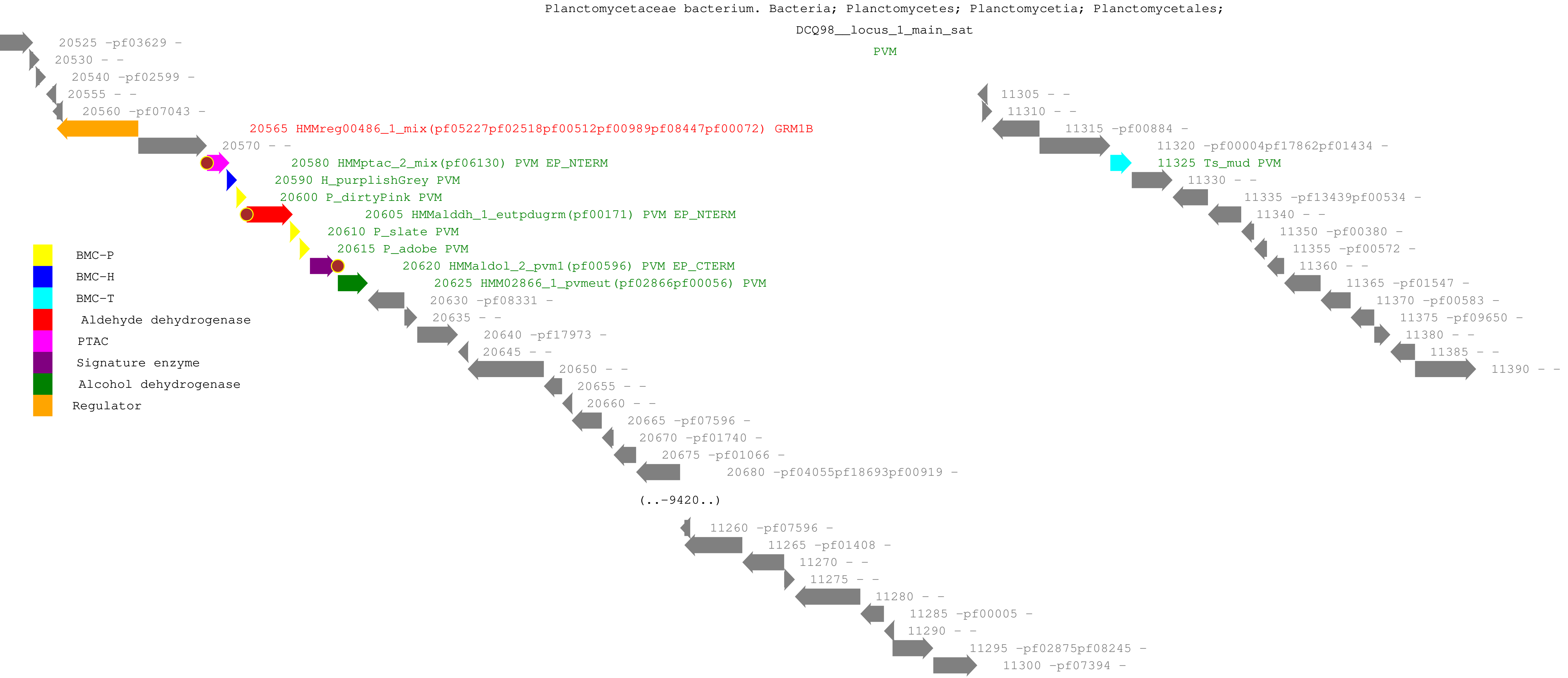

Planctomycetaceae bacterium. Bacteria; Planctomycetes; Planctomycetia; Planctomycetales;
DCQ98__locus_1_main_sat
PVM
20525 -pf03629 -
20530 - -
20540 -pf02599 -
20555 - -
20560 -pf07043 -
20565 HMMreg00486_1_mix(pf05227pf02518pf00512pf00989pf08447pf00072) GRM1B
20570 - -
20580 HMMptac_2_mix(pf06130) PVM EP_NTERM
20590 H_purplishGrey PVM
20600 P_dirtyPink PVM
20605 HMMalddh_1_eutpdugrm(pf00171) PVM EP_NTERM
20610 P_slate PVM
20615 P_adobe PVM
20620 HMMaldol_2_pvm1(pf00596) PVM EP_CTERM
20625 HMM02866_1_pvmeut(pf02866pf00056) PVM
20630 -pf08331 -
20635 - -
20640 -pf17973 -
20645 - -
20650 - -
20655 - -
20660 - -
20665 -pf07596 -
20670 -pf01740 -
20675 -pf01066 -
20680 -pf04055pf18693pf00919 -
(..-9420..)
11260 -pf07596 -
11265 -pf01408 -
11270 - -
11275 - -
11280 - -
11285 -pf00005 -
11290 - -
11295 -pf02875pf08245 -
11300 -pf07394 -
11305 - -
11310 - -
11315 -pf00884 -
11320 -pf00004pf17862pf01434 -
11325 Ts_mud PVM
11330 - -
11335 -pf13439pf00534 -
11340 - -
11350 -pf00380 -
11355 -pf00572 -
11360 - -
11365 -pf01547 -
11370 -pf00583 -
11375 -pf09650 -
11380 - -
11385 - -
11390 - -
BMC-P
BMC-H
BMC-T
Aldehyde dehydrogenase
PTAC
Signature enzyme
Alcohol dehydrogenase
Regulator

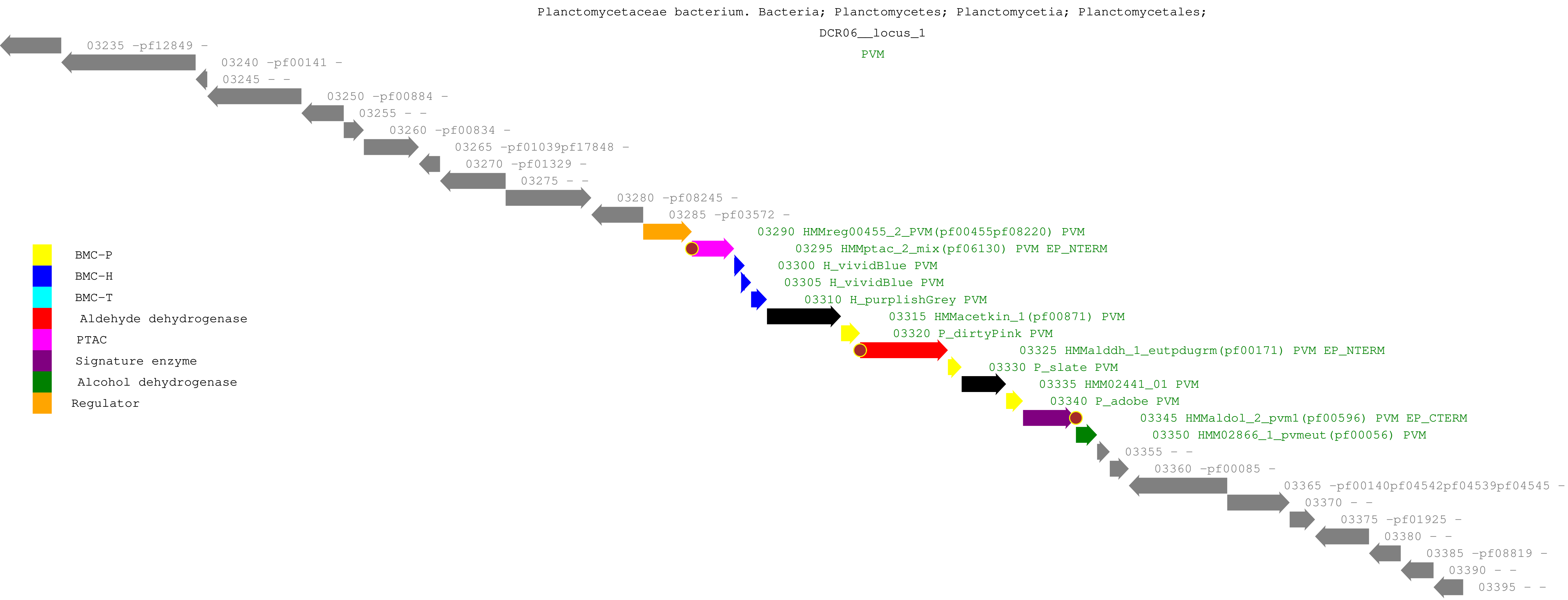
Planctomycetaceae bacterium. Bacteria; Planctomycetes; Planctomycetia; Planctomycetales;
DCR06__locus_1
PVM
03235 -pf12849 -
03240 -pf00141 -
03245 - -
03250 -pf00884 -
03255 - -
03260 -pf00834 -
03265 -pf01039pf17848 -
03270 -pf01329 -
03275 - -
03280 -pf08245 -
03285 -pf03572 -
03290 HMMreg00455_2_PVM(pf00455pf08220) PVM
03295 HMMptac_2_mix(pf06130) PVM EP_NTERM
03300 H_vividBlue PVM
03305 H_vividBlue PVM
03310 H_purplishGrey PVM
03315 HMMacetkin_1(pf00871) PVM
03320 P_dirtyPink PVM
03325 HMMaldh_1_eutpdugrm(pf00171) PVM EP_NTERM
03330 P_slate PVM
03335 HMM02441_01 PVM
03340 P_adobe PVM
03345 HMMaldol_2_pvm1(pf00596) PVM EP_CTERM
03350 HMM02866_1_pvmeut(pf00056) PVM
03355 - -
03360 -pf00085 -
03365 -pf00140pf04542pf04539pf04545 -
03370 - -
03375 -pf01925 -
03380 - -
03385 -pf08819 -
03390 - -
03395 - -
BMC-P
BMC-H
BMC-T
Aldehyde dehydrogenase
PTAC
Signature enzyme
Alcohol dehydrogenase
Regulator

Verrucomicrobiales bacterium. Bacteria; Verrucomicrobia; Verrucomicrobiae; Verrucomicrobiales.

DCR17__locus_1

PVM

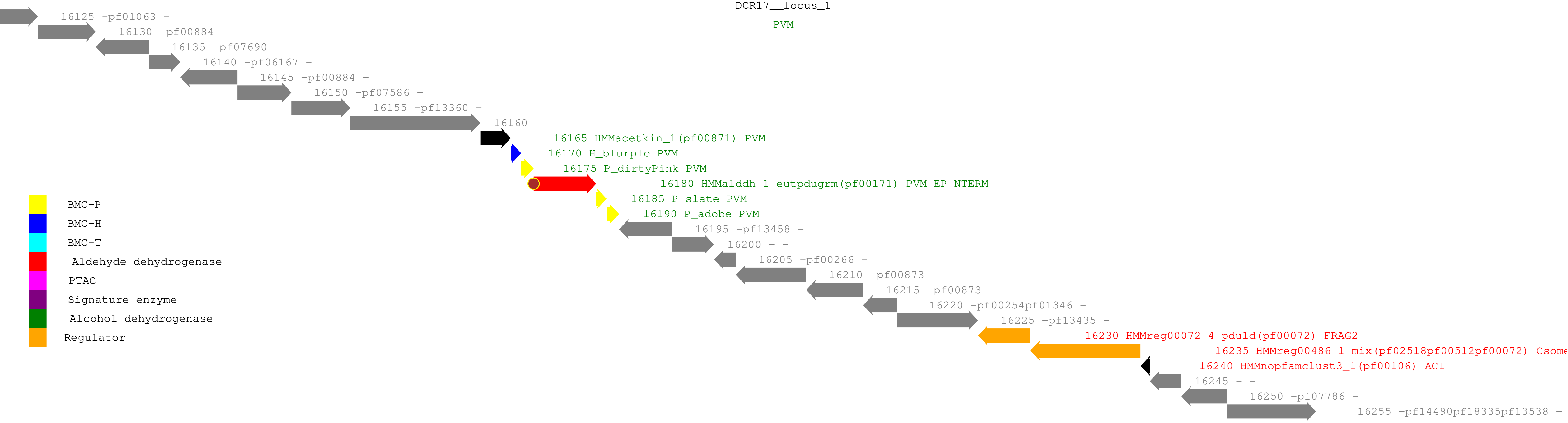

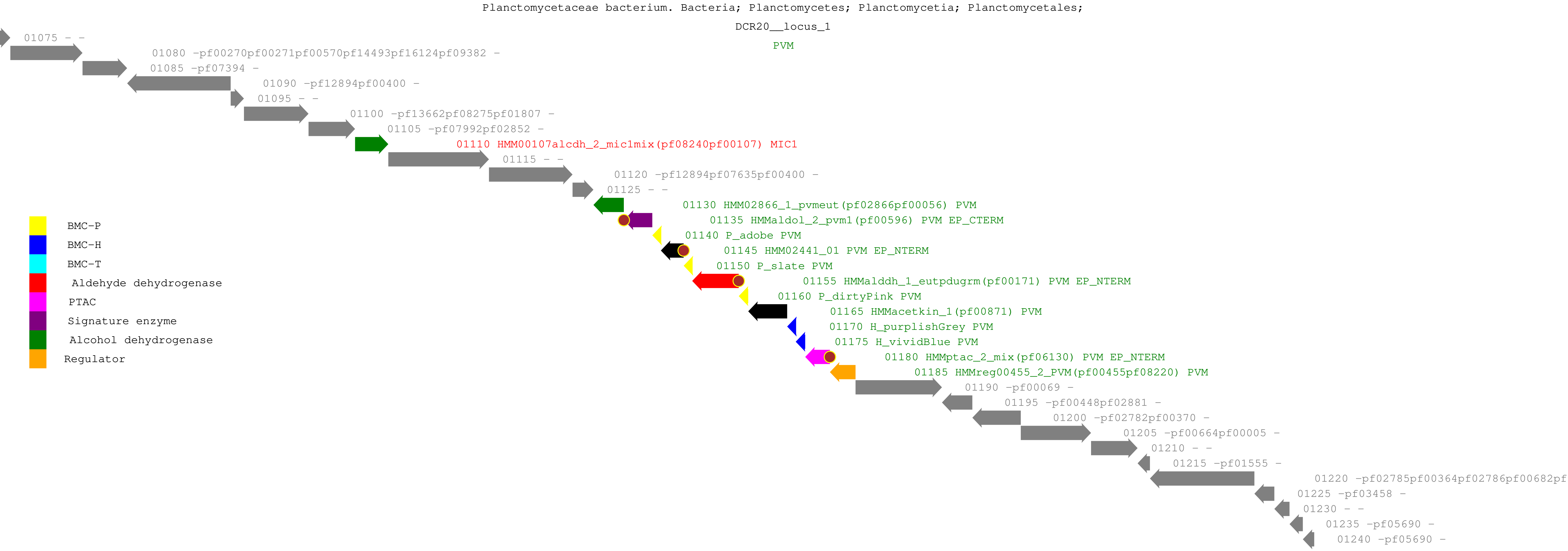
Planctomycetaceae bacterium. Bacteria; Planctomycetes; Planctomycetia; Planctomycetales;
DCR20__locus_1
PVM
01075 - -
01080 -pf00270pf00271pf00570pf14493pf16124pf09382 -
01085 -pf07394 -
01090 -pf12894pf00400 -
01095 - -
01100 -pf13662pf08275pf01807 -
01105 -pf07992pf02852 -
01110 HMM00107alcdh_2_mic1mix(pf08240pf00107) MIC1
01115 - -
01120 -pf12894pf07635pf00400 -
01125 - -
01130 HMM02866_1_pvmeut(pf02866pf00056) PVM
01135 HMMaldol_2_pvm1(pf00596) PVM EP_CTERM
01140 P_adobe PVM
01145 HMM02441_01 PVM EP_NTERM
01150 P_slate PVM
01155 HMMalddh_1_eutpdugrm(pf00171) PVM EP_NTERM
01160 P_dirtyPink PVM
01165 HMMacetkin_1(pf00871) PVM
01170 H_purplishGrey PVM
01175 H_vividBlue PVM
01180 HMMptac_2_mix(pf06130) PVM EP_NTERM
01185 HMMreg00455_2_PVM(pf00455pf08220) PVM
01190 -pf00069 -
01195 -pf00448pf02881 -
01200 -pf02782pf00370 -
01205 -pf00664pf00005 -
01210 - -
01215 -pf01555 -
01220 -pf02785pf00364pf02786pf00682pf
01225 -pf03458 -
01230 - -
01235 -pf05690 -
01240 -pf05690 -
BMC-P
BMC-H
BMC-T
Aldehyde dehydrogenase
PTAC
Signature enzyme
Alcohol dehydrogenase
Regulator

Verrucomicrobiales bacterium. Bacteria; Verrucomicrobia; Verrucomicrobiae; Verrucomicrobiales.

DCS85__locus_1_main_sat

PVM

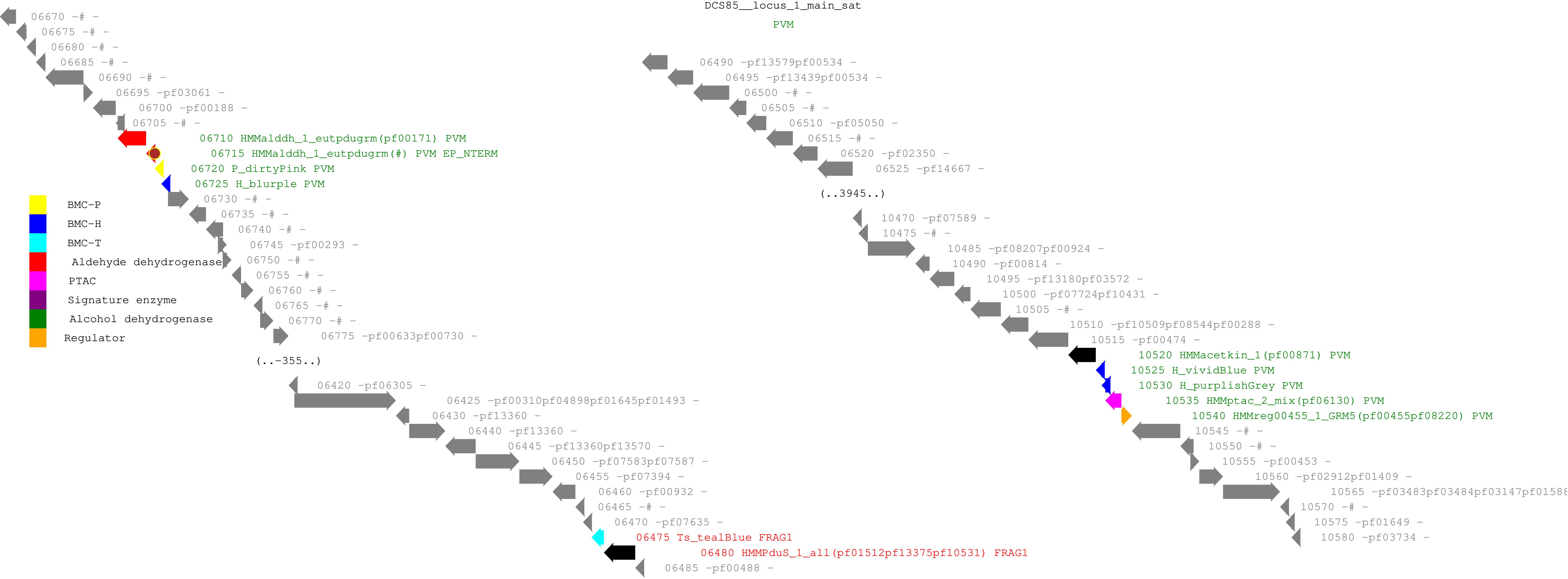

Planctomycetaceae bacterium. Bacteria; Planctomycetes; Planctomycetia; Planctomycetales;

DCW57__locus_1_main_sat

PVM

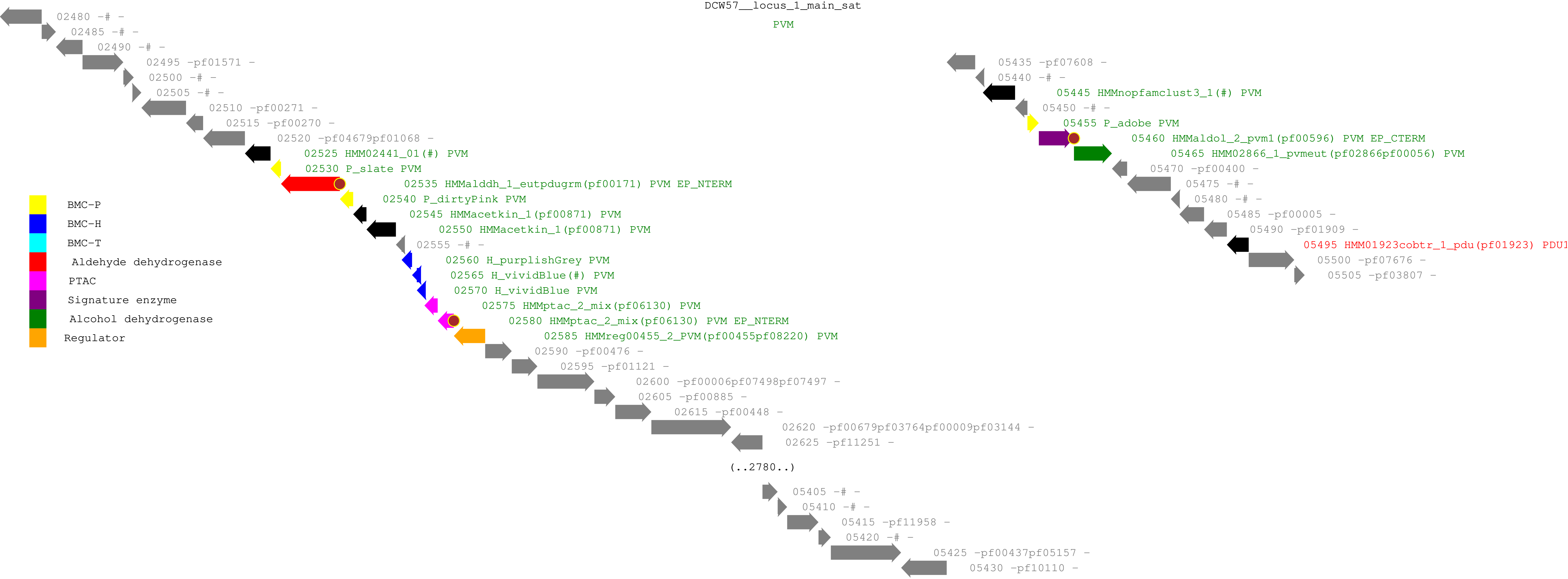

Opitutae bacterium. Bacteria; Verrucomicrobia; Opitutae.

DCX06__locus_1_main_sat

PVM

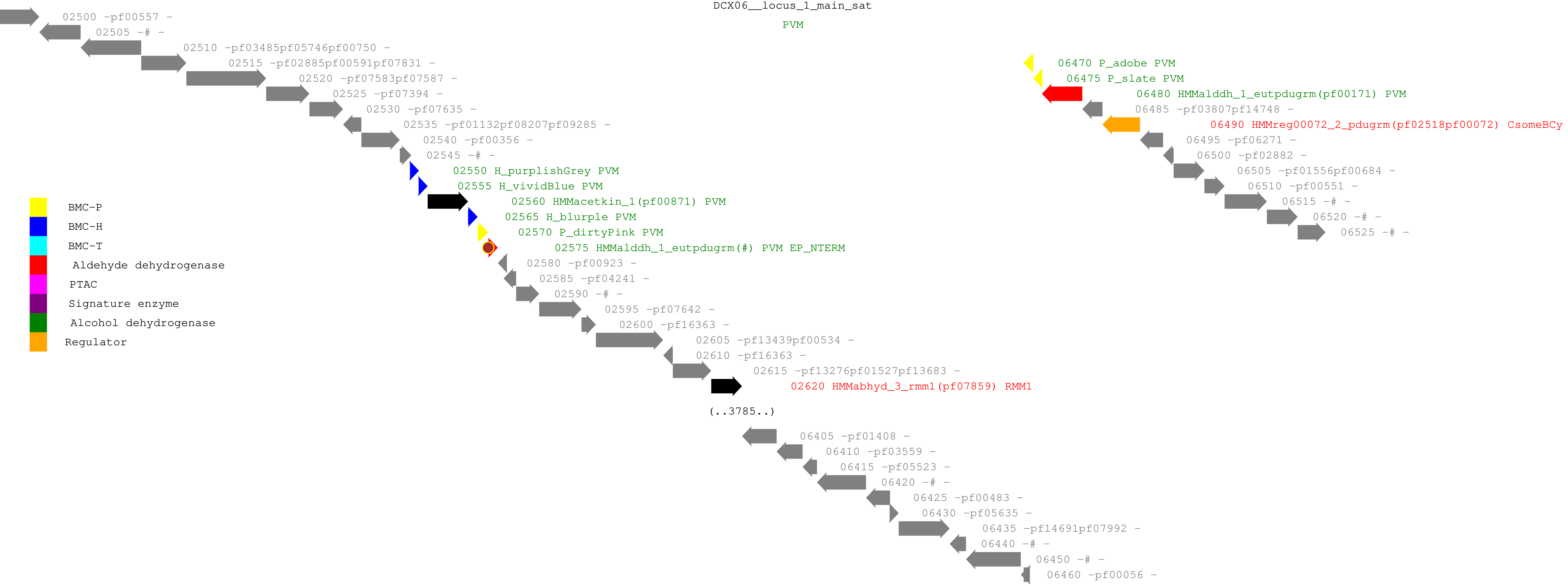

Planctomycetaceae bacterium. Bacteria; Planctomycetes; Planctomycetia; Planctomycetales;

DCX79__locus_1

PVM

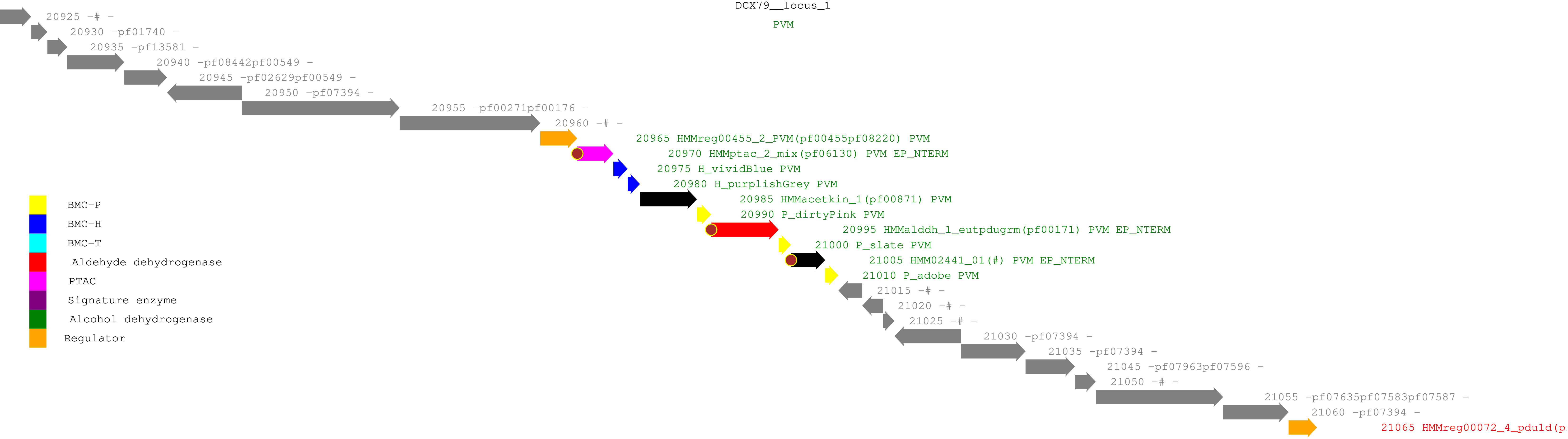

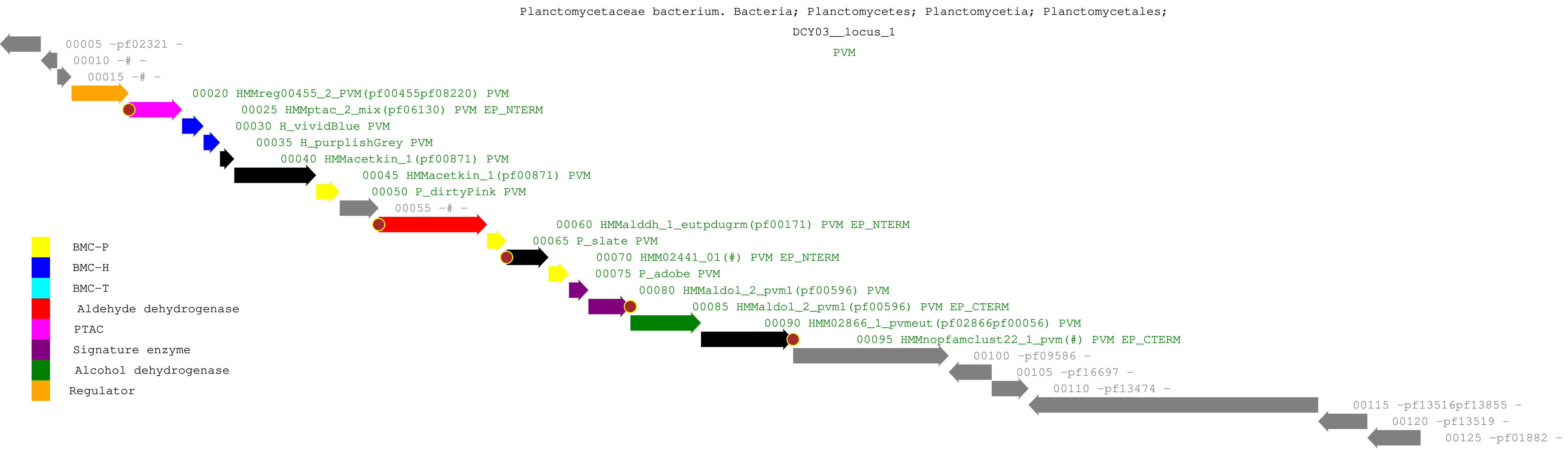
Planctomycetaceae bacterium. Bacteria; Planctomycetes; Planctomycetia; Planctomycetales;
DCY03__locus_1
PVM
00005 -pf02321 -
00010 -# -
00015 -# -
00020 HMMreg00455_2_PVM(pf00455pf08220) PVM
00025 HMMptac_2_mix(pf06130) PVM EP_NTERM
00030 H_vividBlue PVM
00035 H_purplishGrey PVM
00040 HMMacetkin_1(pf00871) PVM
00045 HMMacetkin_1(pf00871) PVM
00050 P_dirtyPink PVM
00055 -# -
00060 HMMaldh_1_eutpdugrm(pf00171) PVM EP_NTERM
00065 P_slate PVM
00070 HMM02441_01(#) PVM EP_NTERM
00075 P_adobe PVM
00080 HMMaldol_2_pvm1(pf00596) PVM
00085 HMMaldol_2_pvm1(pf00596) PVM EP_CTERM
00090 HMM02866_1_pvmeut(pf02866pf00056) PVM
00095 HMMnopfamclust22_1_pvm(#) PVM EP_CTERM
00100 -pf09586 -
00105 -pf16697 -
00110 -pf13474 -
00115 -pf13516pf13855 -
00120 -pf13519 -
00125 -pf01882 -
BMC-P
BMC-H
BMC-T
Aldehyde dehydrogenase
PTAC
Signature enzyme
Alcohol dehydrogenase
Regulator

Verrucomicrobiales bacterium. Bacteria; Verrucomicrobia; Verrucomicrobiae; Verrucomicrobiales.

DCY13__locus_1

PVM

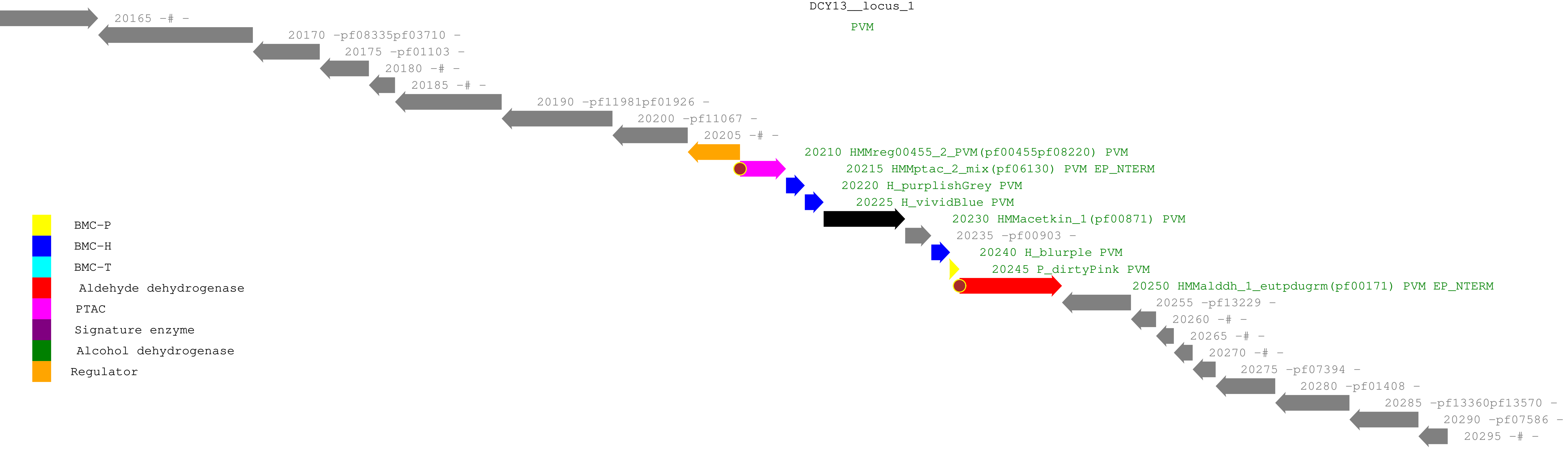

Opitutae bacterium. Bacteria; Verrucomicrobia; Opitutae.

DCY38__locus_1

PVM

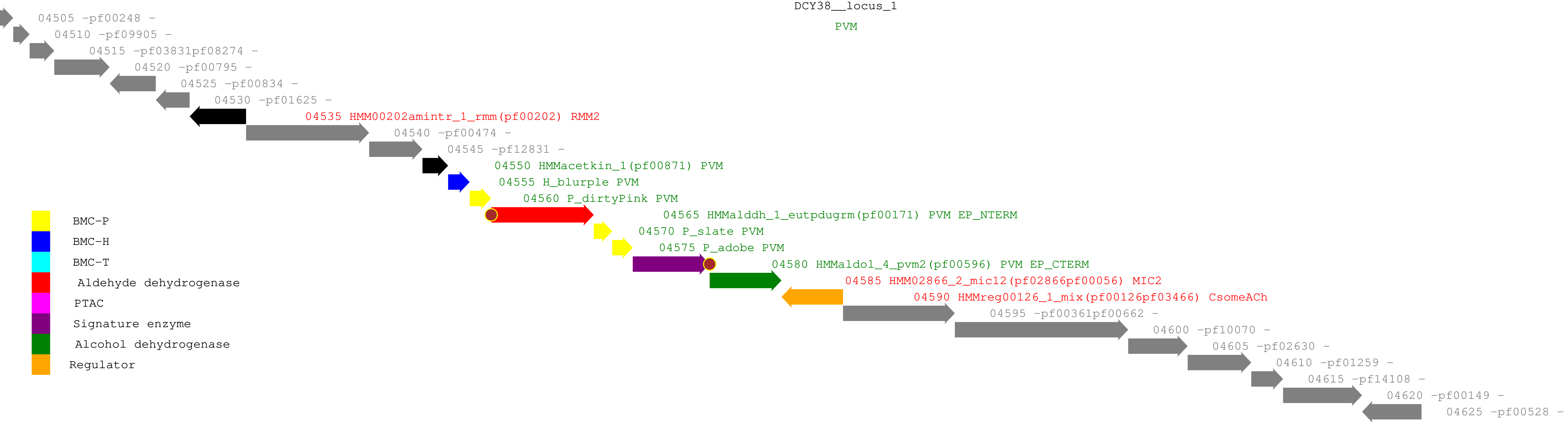

Planctomycetaceae bacterium. Bacteria; Planctomycetes; Planctomycetia; Planctomycetales;

DCY79__locus_1_main_sat

PVM

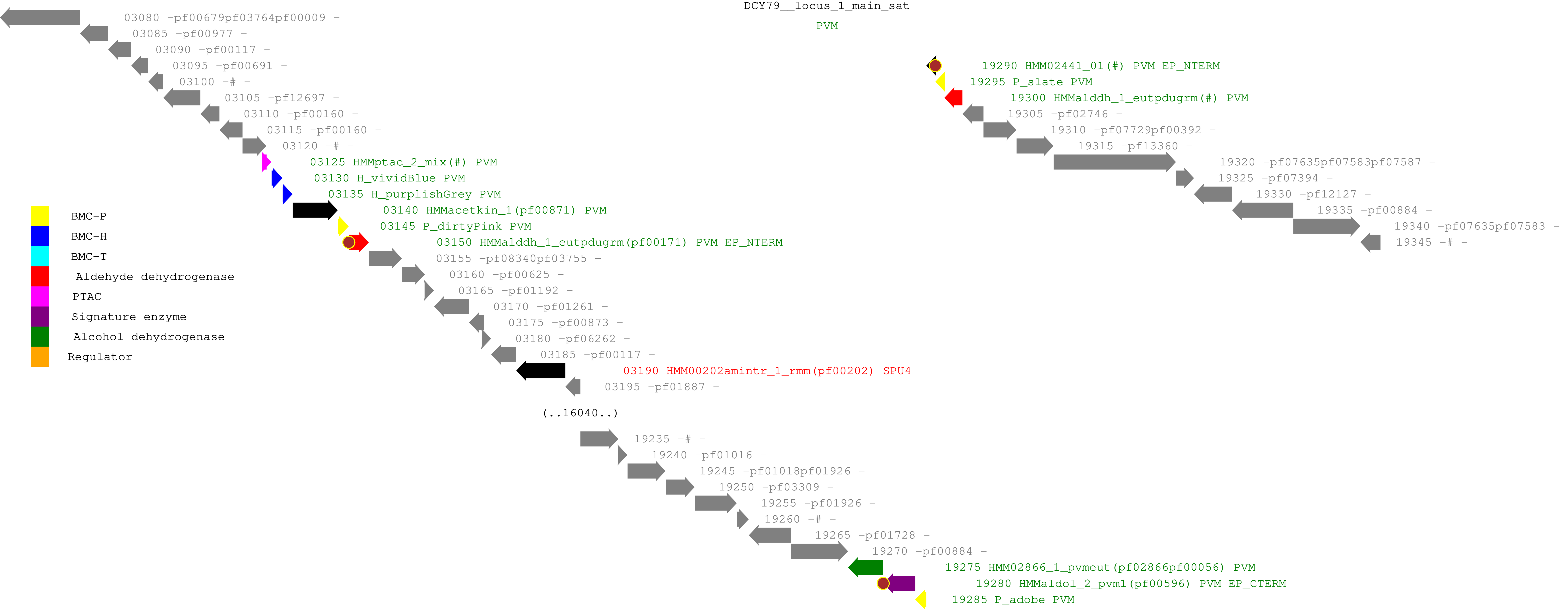

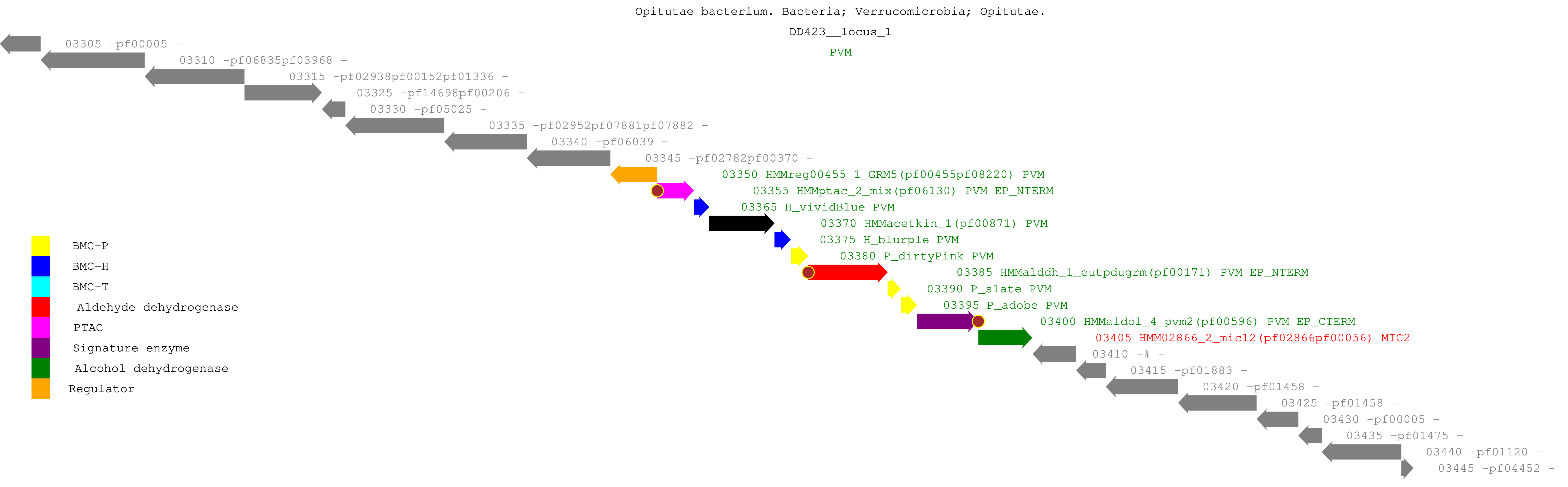

Opitutae bacterium. Bacteria; Verrucomicrobia; Opitutae.
DD423__locus_1
PVM
03305 -pf00005 -
03310 -pf06835pf03968 -
03315 -pf02938pf00152pf01336 -
03325 -pf14698pf00206 -
03330 -pf05025 -
03335 -pf02952pf07881pf07882 -
03340 -pf06039 -
03345 -pf02782pf00370 -
03350 HMMreg00455_1_GRM5(pf00455pf08220) PVM
03355 HMMptac_2_mix(pf06130) PVM EP_NTERM
03365 H_vividBlue PVM
03370 HMMacetkin_1(pf00871) PVM
03375 H_blurple PVM
03380 P_dirtyPink PVM
03385 HMMalddh_1_eutpdugrm(pf00171) PVM EP_NTERM
03390 P_slate PVM
03395 P_adobe PVM
03400 HMMaldol_4_pvm2(pf00596) PVM EP_CTERM
03405 HMM02866_2_mic12(pf02866pf00056) MIC2
03410 -# -
03415 -pf01883 -
03420 -pf01458 -
03425 -pf01458 -
03430 -pf00005 -
03435 -pf01475 -
03440 -pf01120 -
03445 -pf04452 -
BMC-P
BMC-H
BMC-T
Aldehyde dehydrogenase
PTAC
Signature enzyme
Alcohol dehydrogenase
Regulator

Verrucomicrobiales bacterium. Bacteria; Verrucomicrobia; Verrucomicrobiae; Verrucomicrobiales.

DD438__locus_1_main_sat

PVM

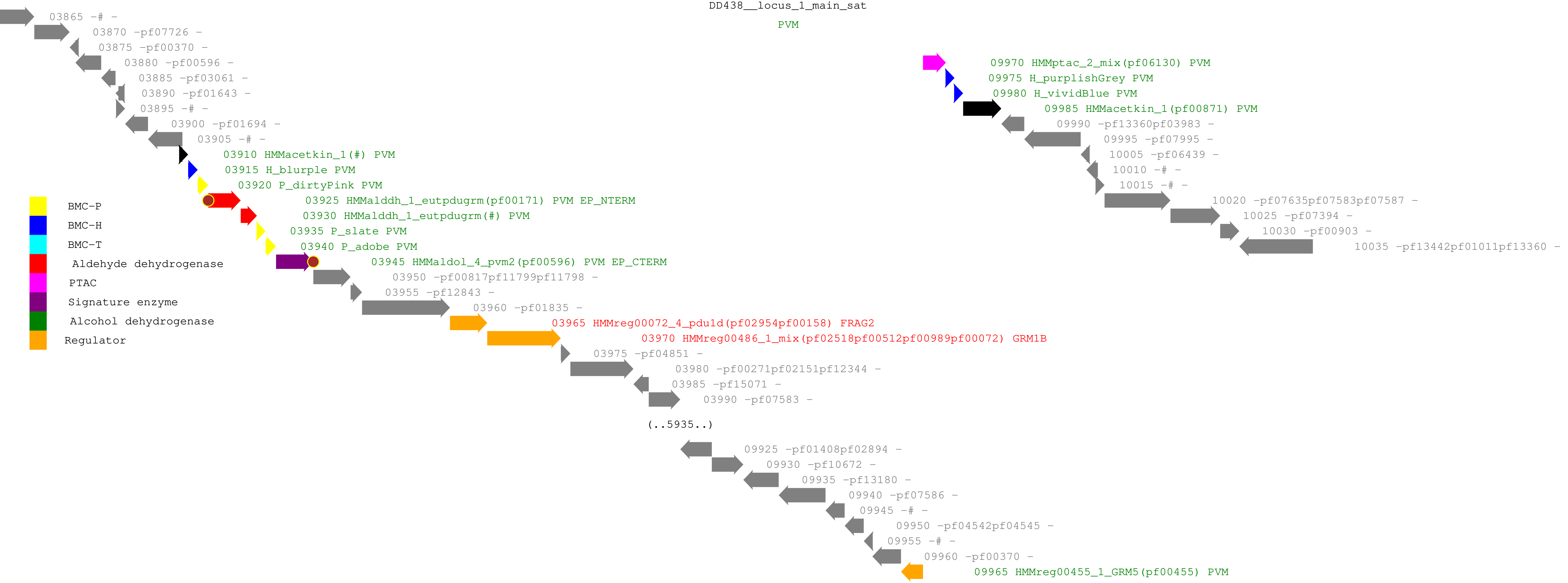

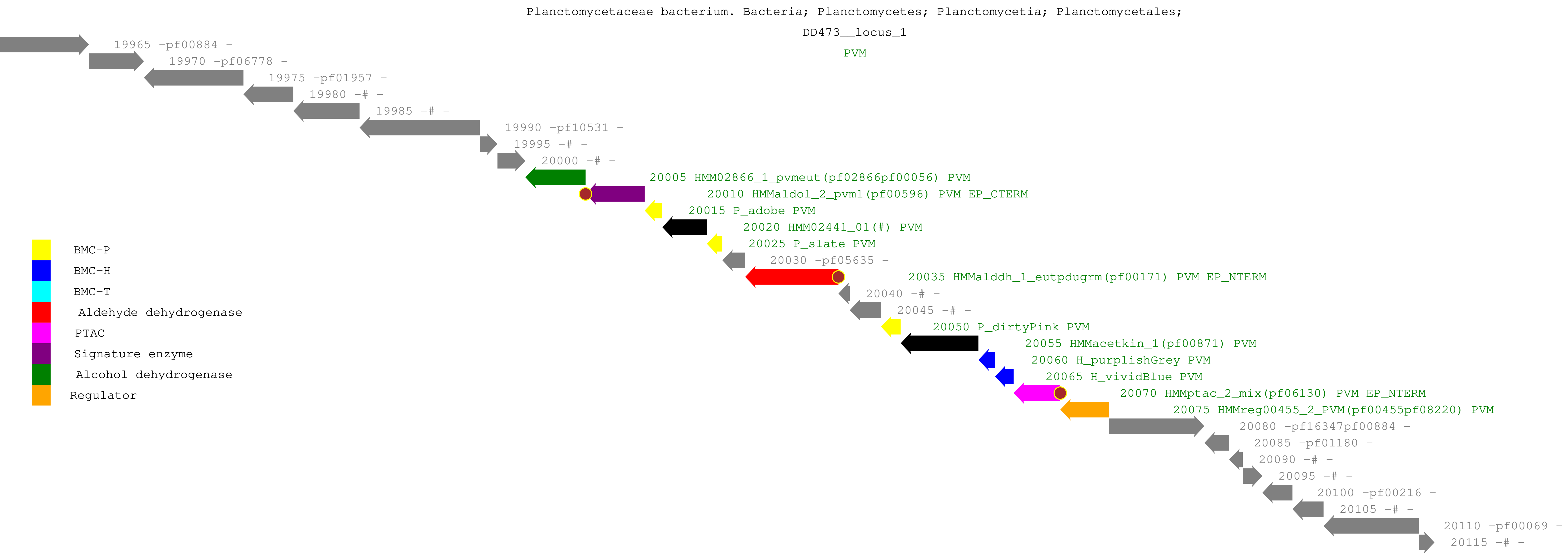

Planctomycetaceae bacterium. Bacteria; Planctomycetes; Planctomycetia; Planctomycetales;
DD473__locus_1
PVM
19965 -pf00884 -
19970 -pf06778 -
19975 -pf01957 -
19980 -# -
19985 -# -
19990 -pf10531 -
19995 -# -
20000 -# -
20005 HMM02866_1_pvmeut(pf02866pf00056) PVM
20010 HMMaldol_2_pvm1(pf00596) PVM EP_CTERM
20015 P_adobe PVM
20020 HMM02441_01(#) PVM
20025 P_slate PVM
20030 -pf05635 -
20035 HMMaldd h_1_eutpdugrm(pf00171) PVM EP_NTERM
20040 -# -
20045 -# -
20050 P_dirtyPink PVM
20055 HMMacetkin_1(pf00871) PVM
20060 H_purplishGrey PVM
20065 H_vividBlue PVM
20070 HMMptac_2_mix(pf06130) PVM EP_NTERM
20075 HMMreg00455_2_PVM(pf00455pf08220) PVM
20080 -pf16347pf00884 -
20085 -pf01180 -
20090 -# -
20095 -# -
20100 -pf00216 -
20105 -# -
20110 -pf00069 -
20115 -# -
BMC-P
BMC-H
BMC-T
Aldehyde dehydrogenase
PTAC
Signature enzyme
Alcohol dehydrogenase
Regulator

Planctomycetaceae bacterium. Bacteria; Planctomycetes; Planctomycetia; Planctomycetales;

DD662__locus_1

PVM

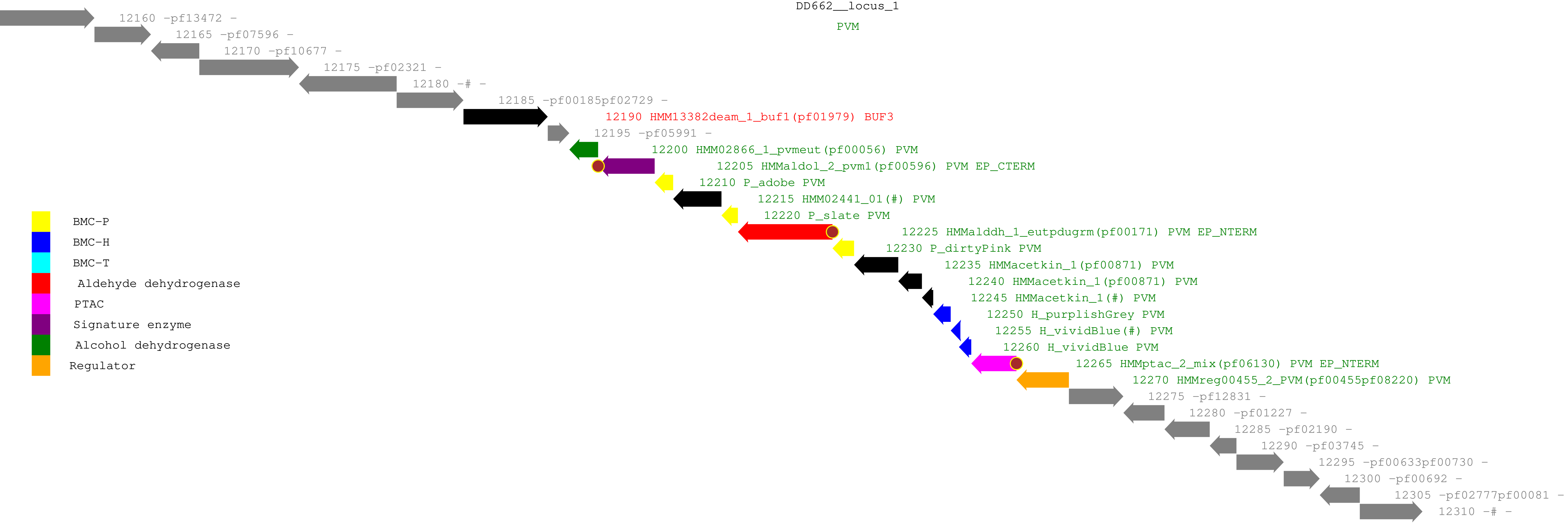

Planctomycetaceae bacterium. Bacteria; Planctomycetes; Planctomycetia; Planctomycetales;

DD670__locus_1_main_sat

PVM

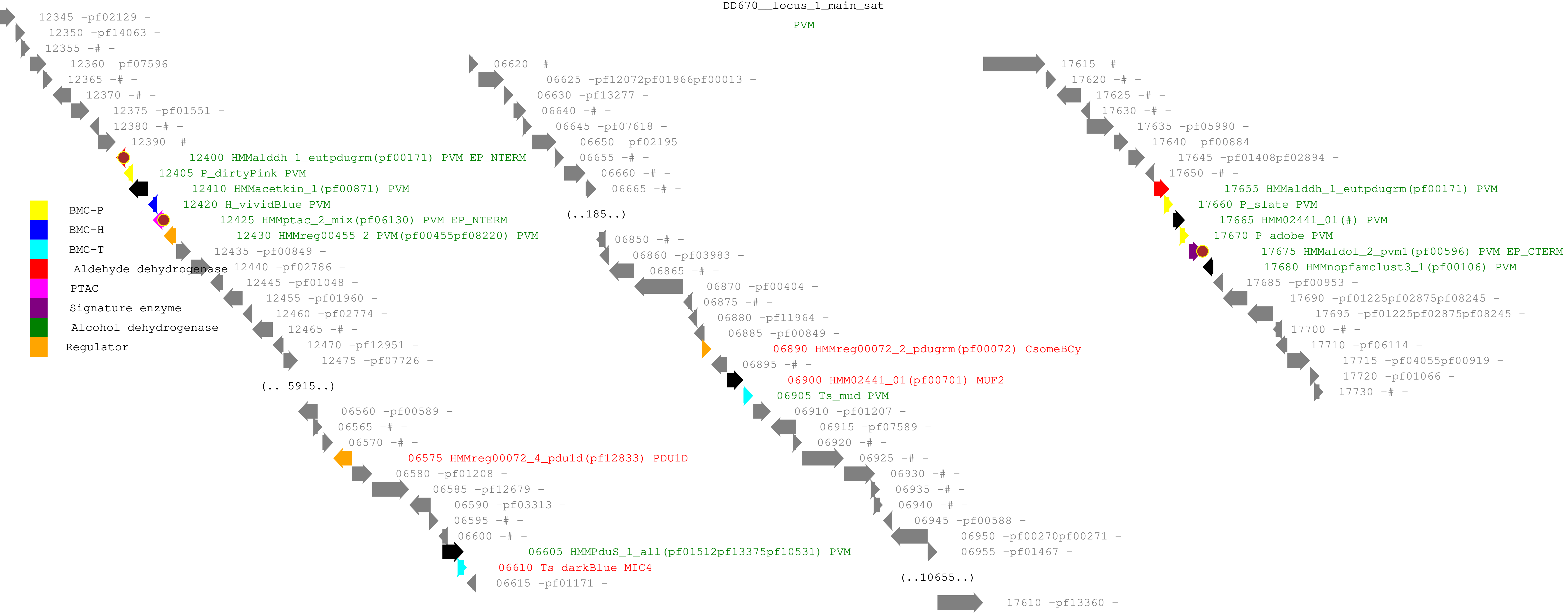

Verrucomicrobiales bacterium. Bacteria; Verrucomicrobia; Verrucomicrobiae; Verrucomicrobiales.

DD687__locus_1_main_sat

PVM

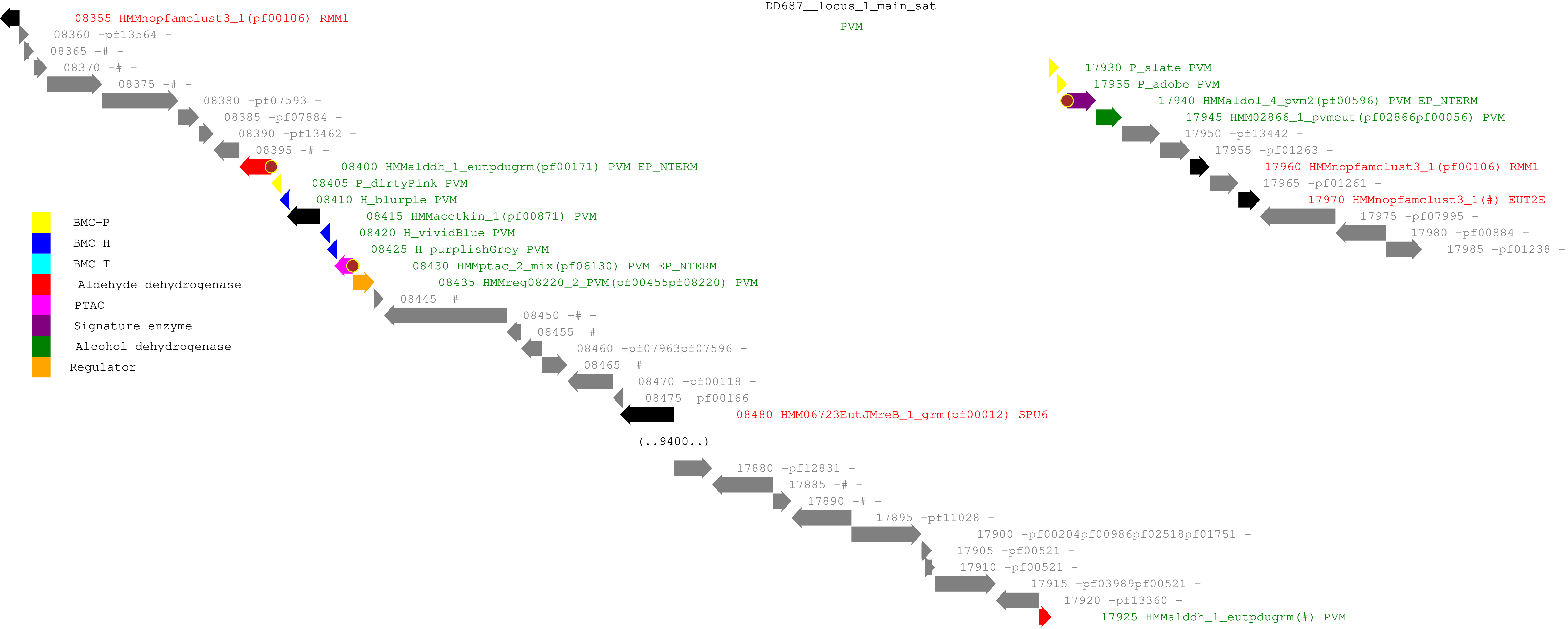

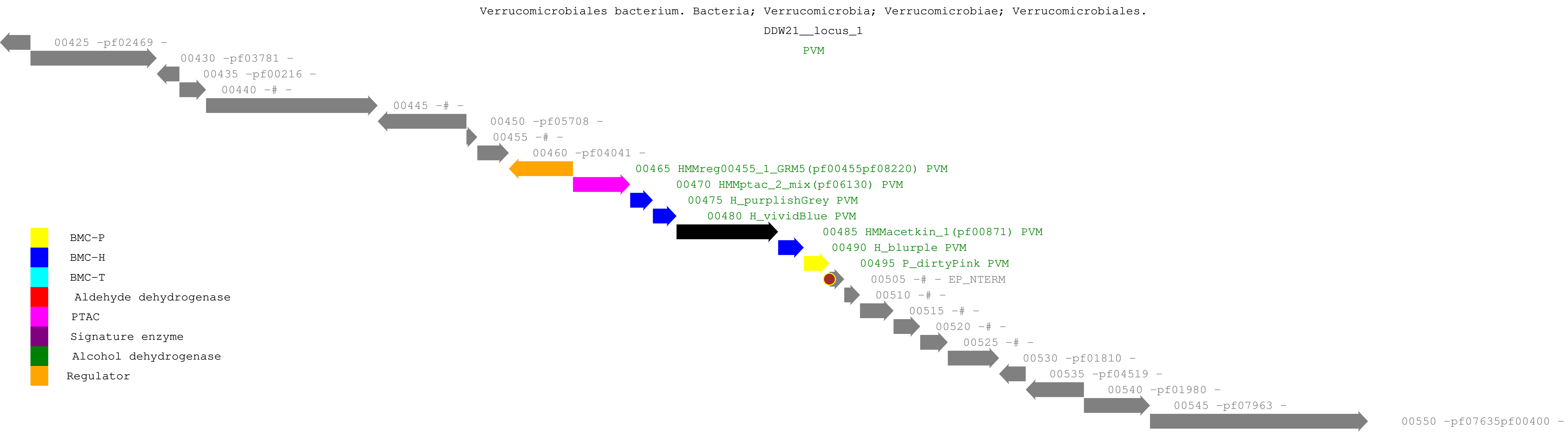
Verrucomicrobiales bacterium. Bacteria; Verrucomicrobia; Verrucomicrobiae; Verrucomicrobiales.
DDW21__locus_1
PVM
00425 -pf02469 -
00430 -pf03781 -
00435 -pf00216 -
00440 -# -
00445 -# -
00450 -pf05708 -
00455 -# -
00460 -pf04041 -
00465 HMMreg00455_1_GRM5(pf00455pf08220) PVM
00470 HMMptac_2_mix(pf06130) PVM
00475 H_purplishGrey PVM
00480 H_vividBlue PVM
00485 HMMacetkin_1(pf00871) PVM
00490 H_blurple PVM
00495 P_dirtyPink PVM
00505 -# - EP_NTERM
00510 -# -
00515 -# -
00520 -# -
00525 -# -
00530 -pf01810 -
00535 -pf04519 -
00540 -pf01980 -
00545 -pf07963 -
00550 -pf07635pf00400 -
BMC-P
BMC-H
BMC-T
Aldehyde dehydrogenase
PTAC
Signature enzyme
Alcohol dehydrogenase
Regulator

Planctomycetaceae bacterium. Bacteria; Planctomycetes; Planctomycetia; Planctomycetales;

DDW52__locus_fragmented

PVM

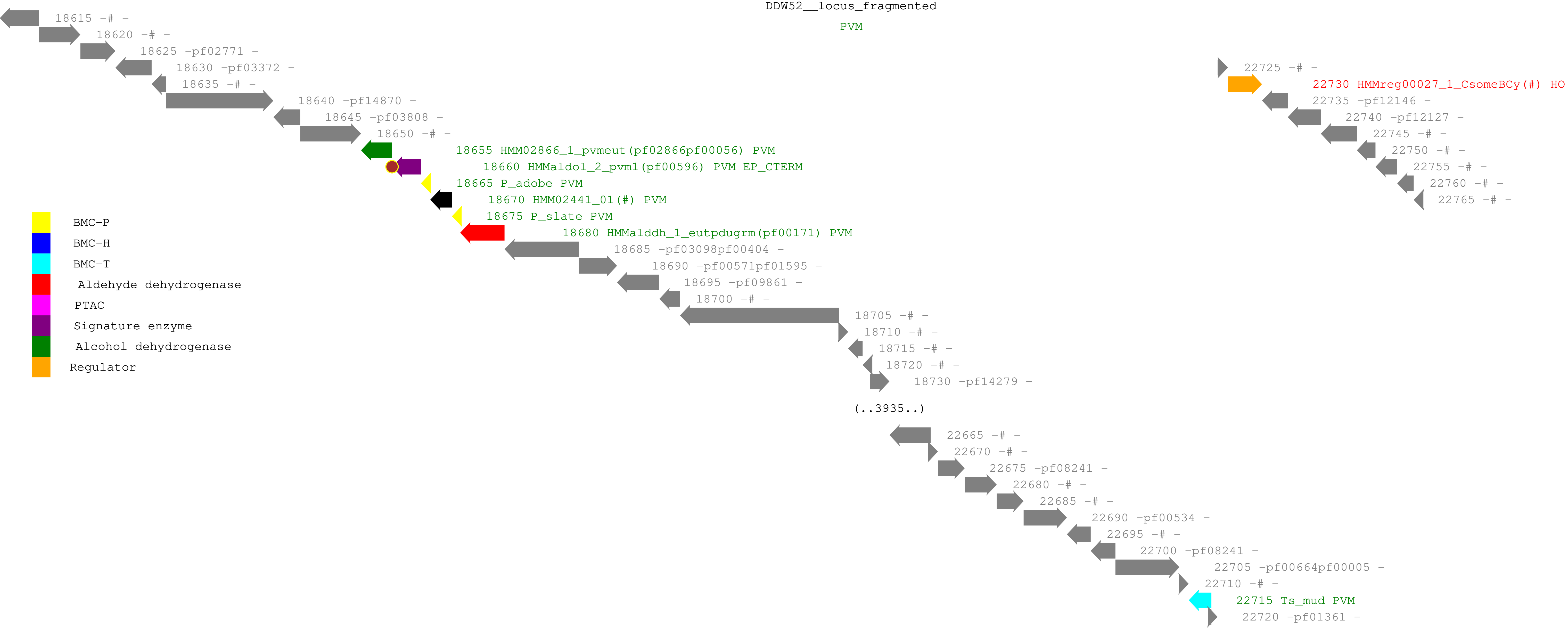

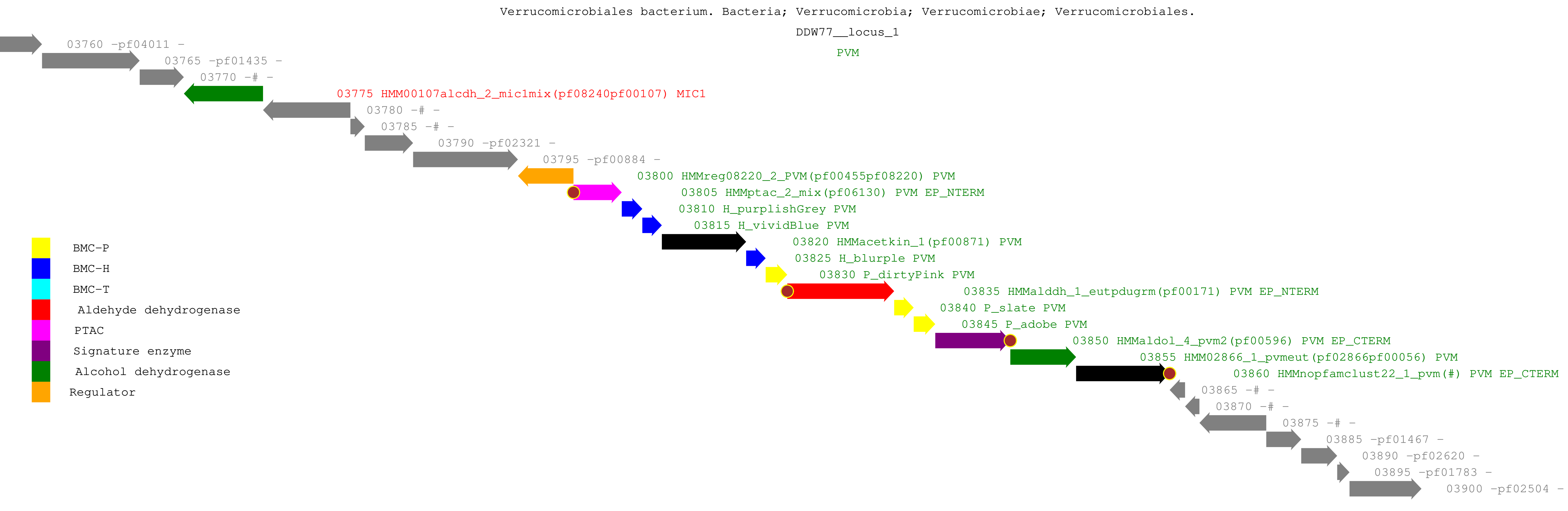
Verrucomicrobiales bacterium. Bacteria; Verrucomicrobia; Verrucomicrobiae; Verrucomicrobiales.
DDW77__locus_1
PVM
03760 -pf04011 -
03765 -pf01435 -
03770 -# -
03775 HMM00107alcdh_2_mic1mix(pf08240pf00107) MIC1
03780 -# -
03785 -# -
03790 -pf02321 -
03795 -pf00884 -
03800 HMMreg08220_2_PVM(pf00455pf08220) PVM
03805 HMMptac_2_mix(pf06130) PVM EP_NTERM
03810 H_purplishGrey PVM
03815 H_vividBlue PVM
03820 HMMacetkin_1(pf00871) PVM
03825 H_blurple PVM
03830 P_dirtyPink PVM
03835 HMMalddh_1_eutpdugrm(pf00171) PVM EP_NTERM
03840 P_slate PVM
03845 P_adobe PVM
03850 HMMaldol_4_pvm2(pf00596) PVM EP_CTERM
03855 HMM02866_1_pvmeut(pf02866pf00056) PVM
03860 HMMnopfamclust22_1_pvm(#) PVM EP_CTERM
03865 -# -
03870 -# -
03875 -# -
03885 -pf01467 -
03890 -pf02620 -
03895 -pf01783 -
03900 -pf02504 -
BMC-P
BMC-H
BMC-T
Aldehyde dehydrogenase
PTAC
Signature enzyme
Alcohol dehydrogenase
Regulator

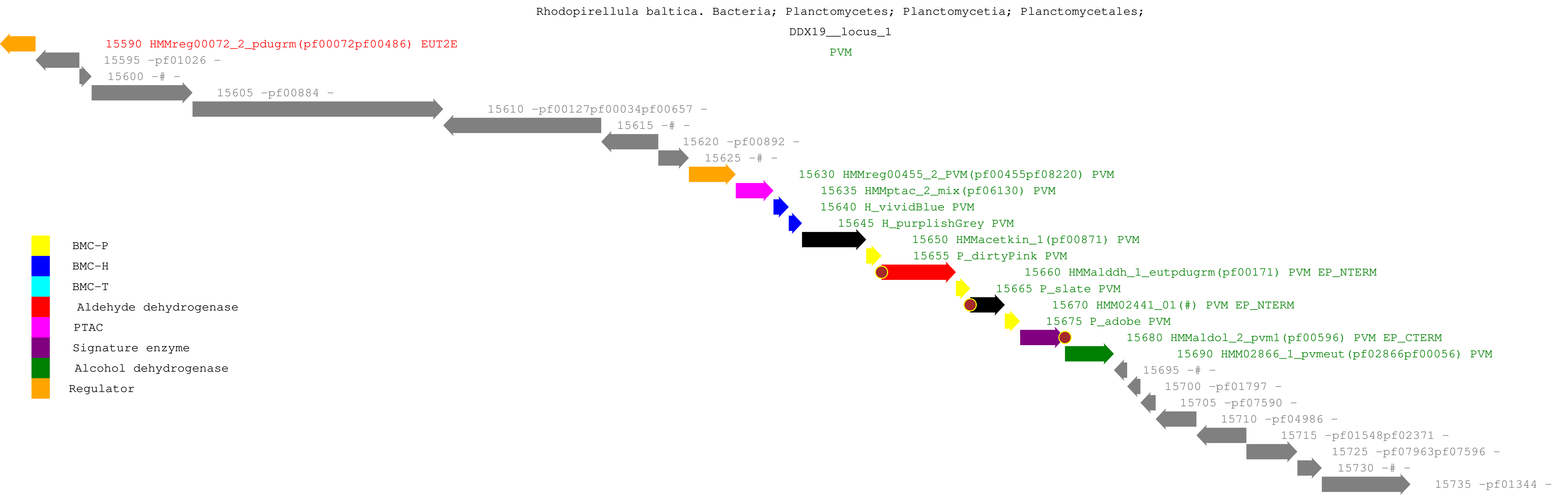
Rhodopirellula baltica. Bacteria; Planctomycetes; Planctomycetia; Planctomycetales;
DDX19__locus_1
PVM
15590 HMMreg00072_2_pdugrm(pf00072pf00486) EUT2E
15595 -pf01026 -
15600 -# -
15605 -pf00884 -
15610 -pf00127pf00034pf00657 -
15615 -# -
15620 -pf00892 -
15625 -# -
15630 HMMreg00455_2_PVM(pf00455pf08220) PVM
15635 HMMptac_2_mix(pf06130) PVM
15640 H_vividBlue PVM
15645 H_purplishGrey PVM
15650 HMMacetkin_1(pf00871) PVM
15655 P_dirtyPink PVM
15660 HMMaldh_1_eutpdugrm(pf00171) PVM EP_NTERM
15665 P_slate PVM
15670 HMM02441_01(#) PVM EP_NTERM
15675 P_adobe PVM
15680 HMMaldol_2_pvm1(pf00596) PVM EP_CTERM
15690 HMM02866_1_pvmeut(pf02866pf00056) PVM
15695 -# -
15700 -pf01797 -
15705 -pf07590 -
15710 -pf04986 -
15715 -pf01548pf02371 -
15725 -pf07963pf07596 -
15730 -# -
15735 -pf01344 -
BMC-P
BMC-H
BMC-T
Aldehyde dehydrogenase
PTAC
Signature enzyme
Alcohol dehydrogenase
Regulator

Planctomycetales bacterium. Bacteria; Planctomycetes; Planctomycetia; Planctomycetales;

DDY78__locus_1

PVM

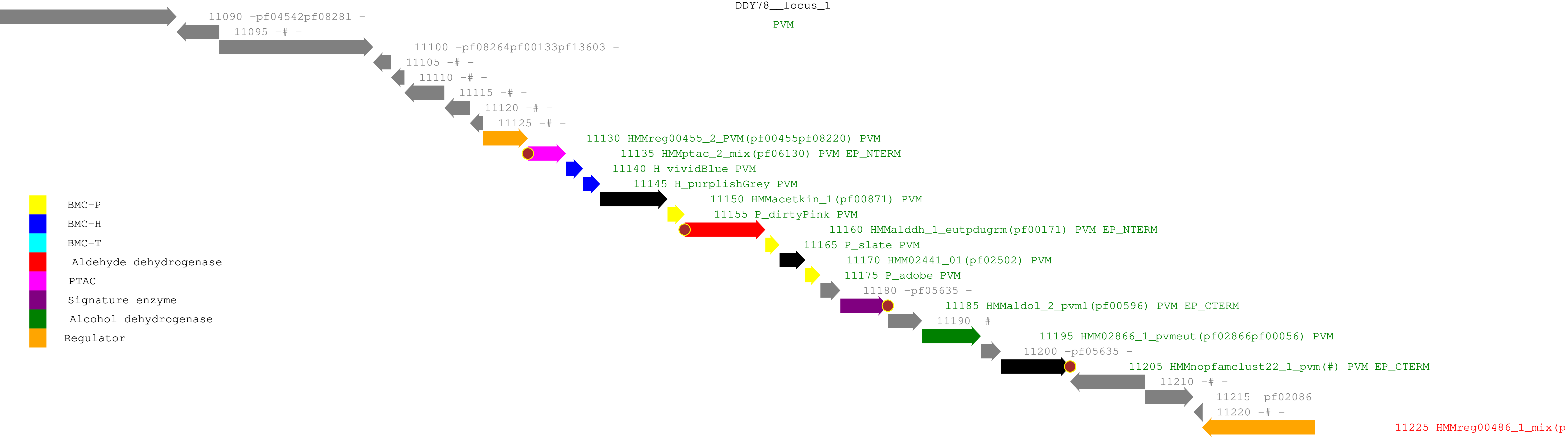

Planctomycetaceae bacterium. Bacteria; Planctomycetes; Planctomycetia; Planctomycetales;

DDY91__locus_1_main_sat

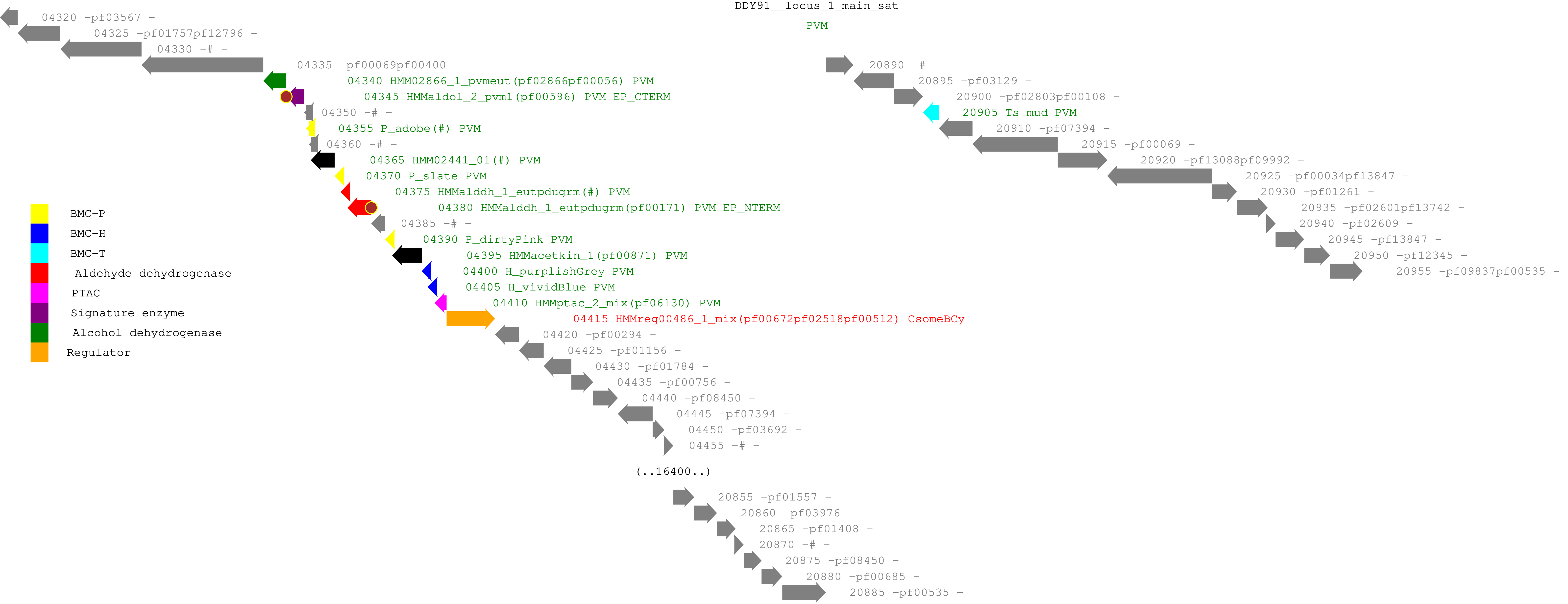

Coraliomargarita sp. WN38. Bacteria; Verrucomicrobia; Opitutae; Puniceicoccales;

DDZ13__locus_1

PVM

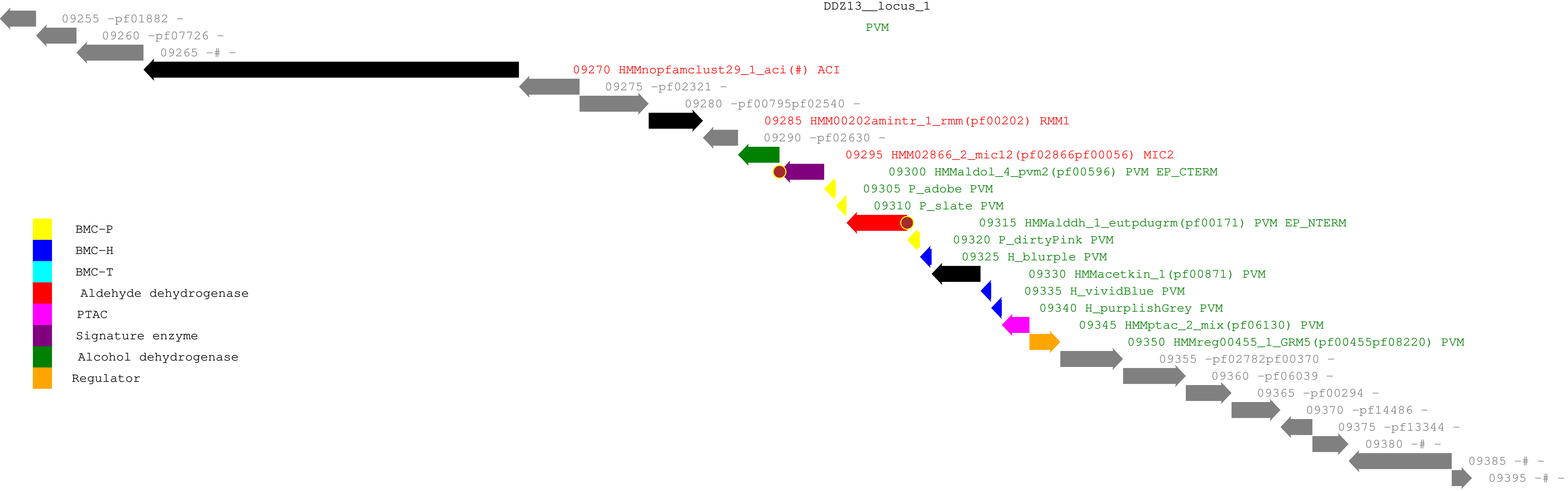

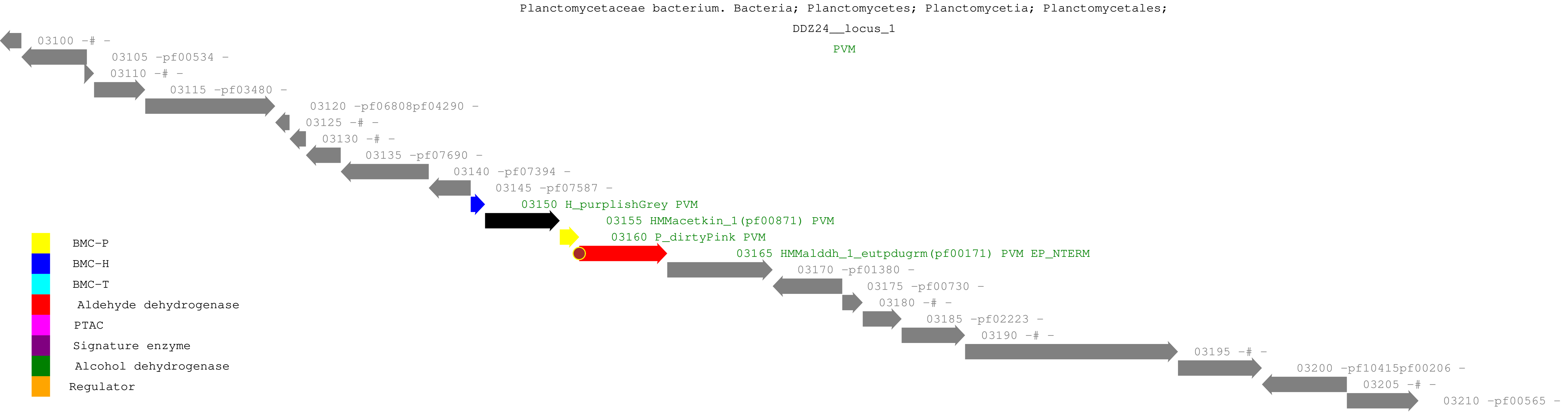
Planctomycetaceae bacterium. Bacteria; Planctomycetes; Planctomycetia; Planctomycetales;
DDZ24__locus_1
PVM
03100 -# -
03105 -pf00534 -
03110 -# -
03115 -pf03480 -
03120 -pf06808pf04290 -
03125 -# -
03130 -# -
03135 -pf07690 -
03140 -pf07394 -
03145 -pf07587 -
03150 H_purplishGrey PVM
03155 HMMacetkin_1(pf00871) PVM
03160 P_dirtyPink PVM
03165 HMMalddh_1_eutpdugrm(pf00171) PVM EP_NTERM
03170 -pf01380 -
03175 -pf00730 -
03180 -# -
03185 -pf02223 -
03190 -# -
03195 -# -
03200 -pf10415pf00206 -
03205 -# -
03210 -pf00565 -
BMC-P
BMC-H
BMC-T
Aldehyde dehydrogenase
PTAC
Signature enzyme
Alcohol dehydrogenase
Regulator

Planctomycetaceae bacterium. Bacteria; Planctomycetes; Planctomycetia; Planctomycetales;

DDZ51__locus_fragmented

PVM

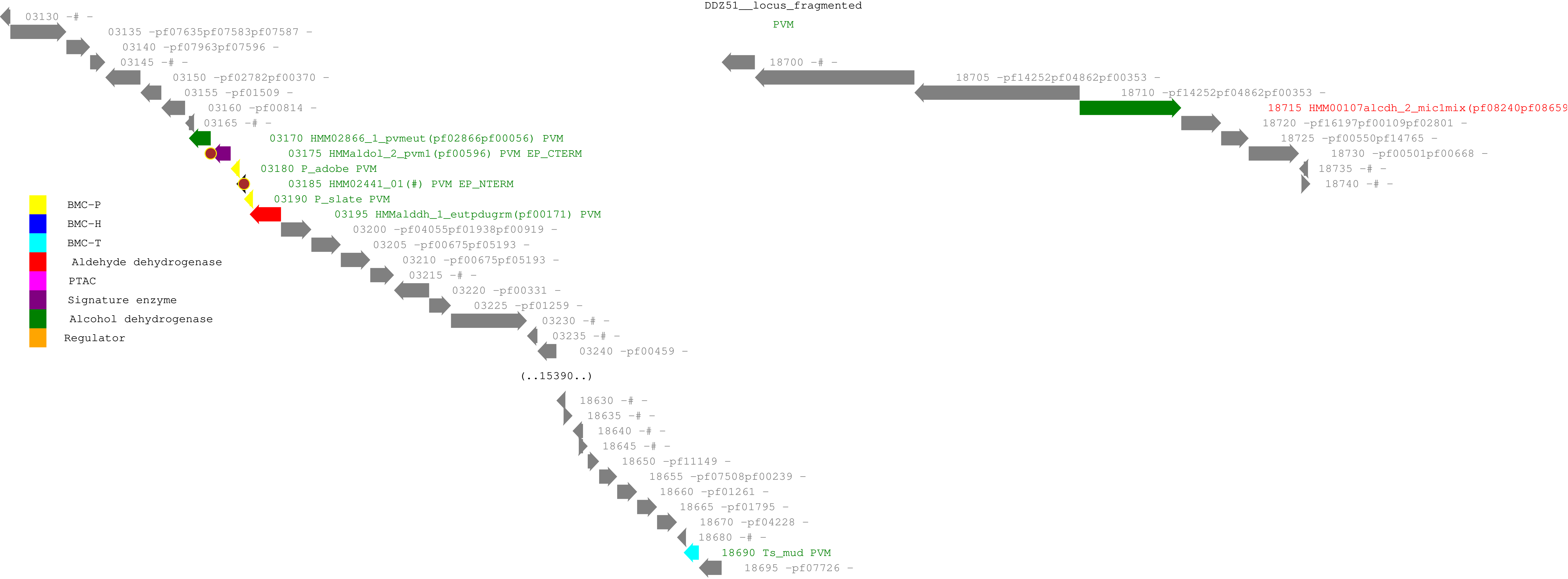

Planctomycetaceae bacterium. Bacteria; Planctomycetes; Planctomycetia; Planctomycetales;

DDZ90__locus_1_main_sat

PVM

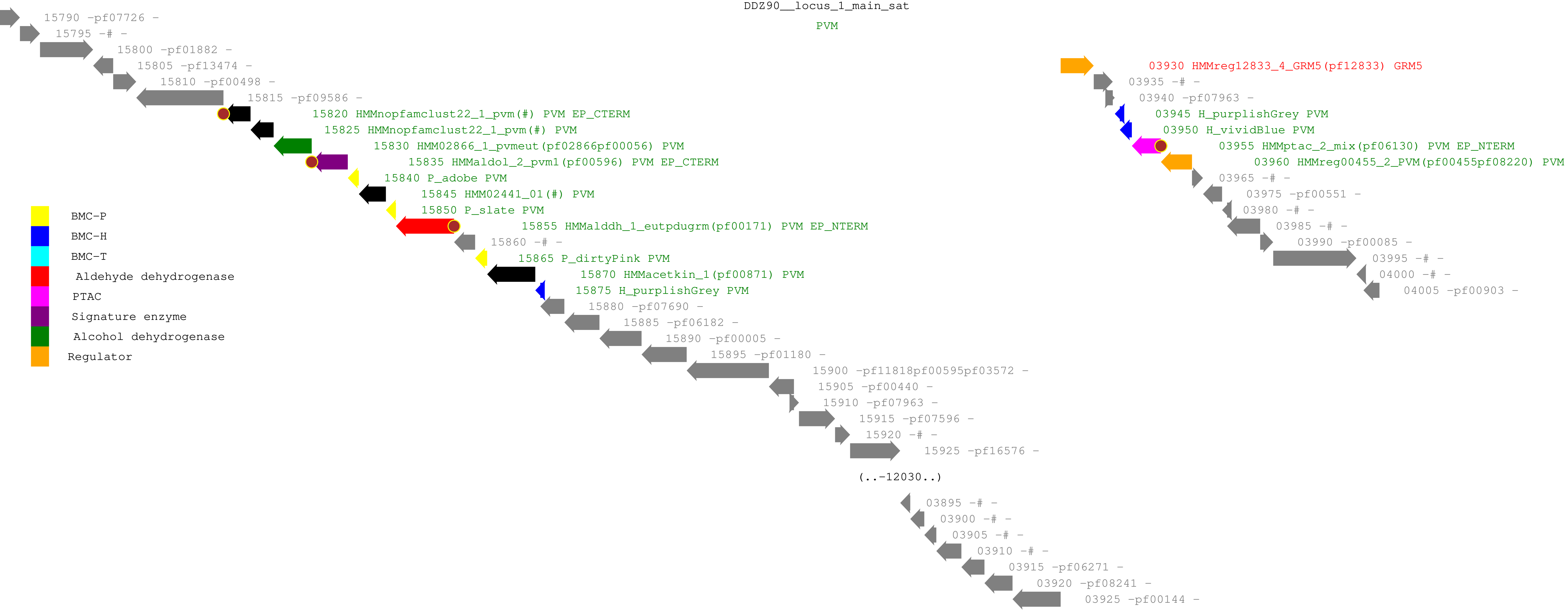

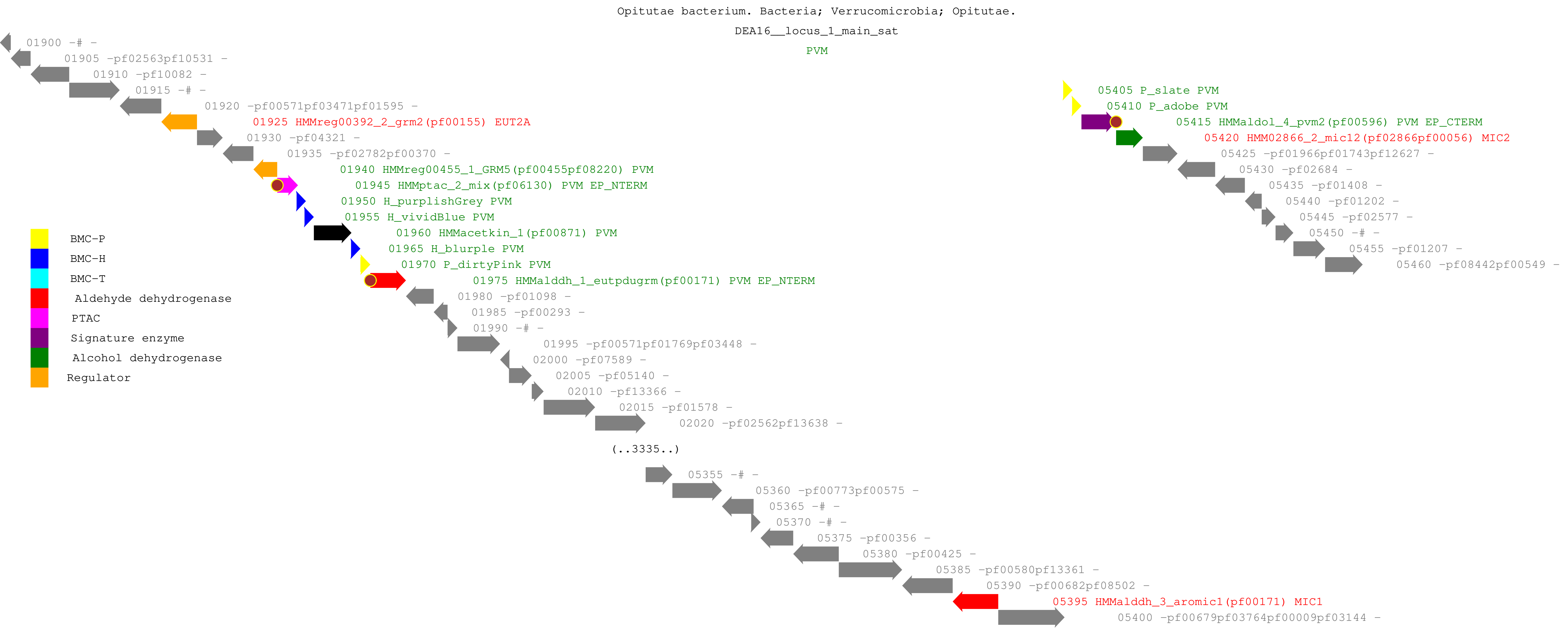
Opitutae bacterium. Bacteria; Verrucomicrobia; Opitutae.
DEA16__locus_1_main_sat
PVM
01900 -# -
01905 -pf02563pf10531 -
01910 -pf10082 -
01915 -# -
01920 -pf00571pf03471pf01595 -
01925 HMMreg00392_2_grm2(pf00155) EUT2A
01930 -pf04321 -
01935 -pf02782pf00370 -
01940 HMMreg00455_1_GRM5(pf00455pf08220) PVM
01945 HMMptac_2_mix(pf06130) PVM EP_NTERM
01950 H_purplishGrey PVM
01955 H_vividBlue PVM
01960 HMMacetkin_1(pf00871) PVM
01965 H_blurple PVM
01970 P_dirtyPink PVM
01975 HMMalddh_1_eutpdugrm(pf00171) PVM EP_NTERM
01980 -pf01098 -
01985 -pf00293 -
01990 -# -
01995 -pf00571pf01769pf03448 -
02000 -pf07589 -
02005 -pf05140 -
02010 -pf13366 -
02015 -pf01578 -
02020 -pf02562pf13638 -
(..3335..)
05355 -# -
05360 -pf00773pf00575 -
05365 -# -
05370 -# -
05375 -pf00356 -
05380 -pf00425 -
05385 -pf00580pf13361 -
05390 -pf00682pf08502 -
05395 HMMalddh_3_aromic1(pf00171) MIC1
05400 -pf00679pf03764pf00009pf03144 -
05405 P_slate PVM
05410 P_adobe PVM
05415 HMMaldol_4_pvm2(pf00596) PVM EP_CTERM
05420 HMM02866_2_mic12(pf02866pf00056) MIC2
05425 -pf01966pf01743pf12627 -
05430 -pf02684 -
05435 -pf01408 -
05440 -pf01202 -
05445 -pf02577 -
05450 -# -
05455 -pf01207 -
05460 -pf08442pf00549 -
BMC-P
BMC-H
BMC-T
Aldehyde dehydrogenase
PTAC
Signature enzyme
Alcohol dehydrogenase
Regulator

Opitutae bacterium. Bacteria; Verrucomicrobia; Opitutae.

DEA90__locus_1_main_sat

PVM

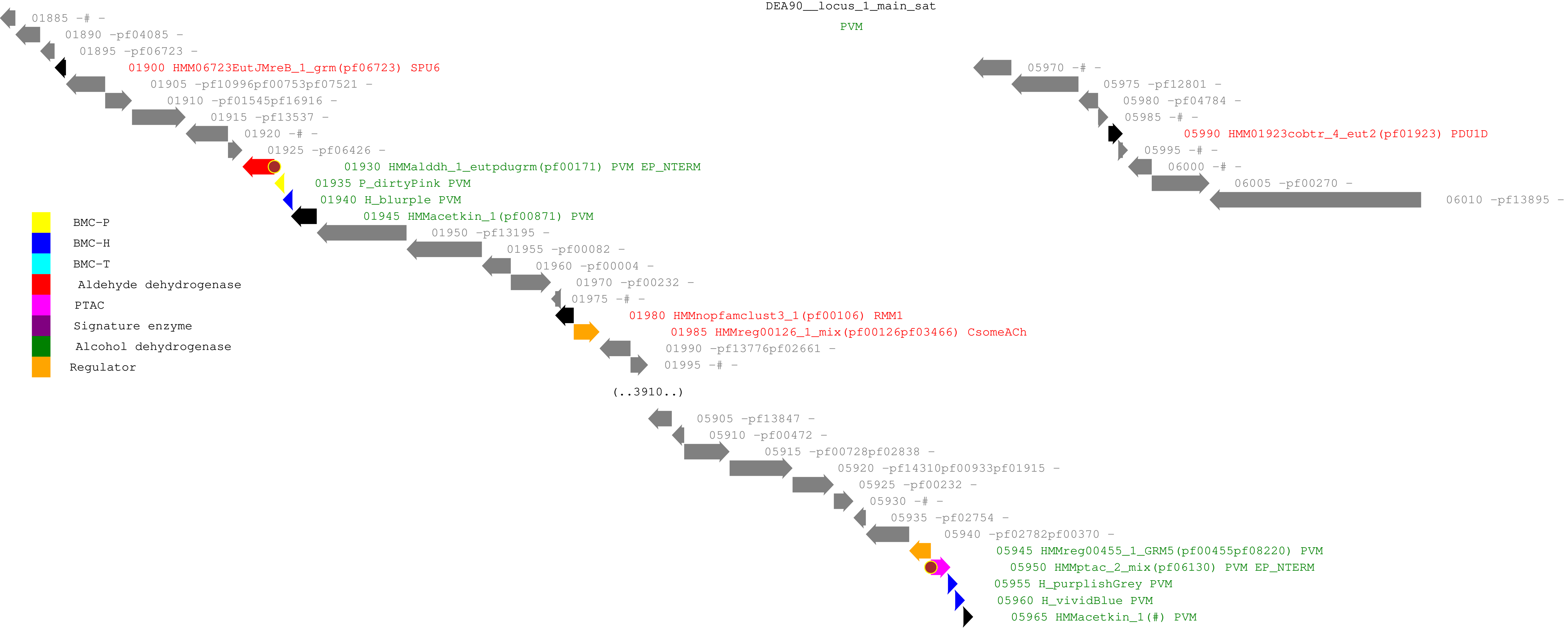

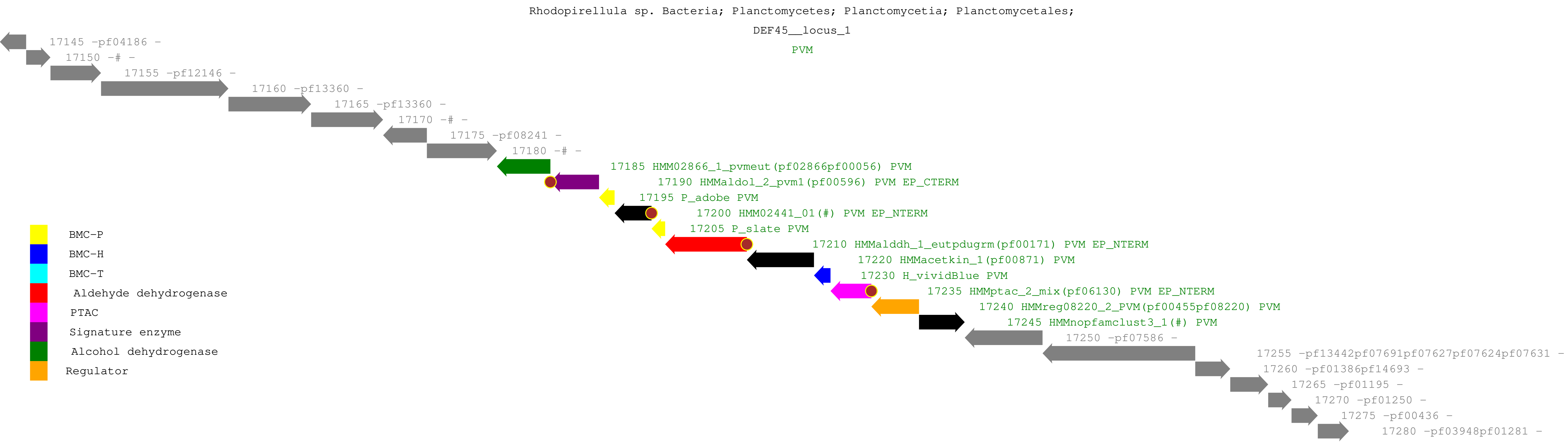

Rhodopirellula sp. Bacteria; Planctomycetes; Planctomycetia; Planctomycetales;
DEF45__locus_1
PVM
17145 -pf04186 -
17150 -# -
17155 -pf12146 -
17160 -pf13360 -
17165 -pf13360 -
17170 -# -
17175 -pf08241 -
17180 -# -
17185 HMM02866_1_pvmeut(pf02866pf00056) PVM
17190 HMMaldol_2_pvm1(pf00596) PVM EP_CTERM
17195 P_adobe PVM
17200 HMM02441_01(#) PVM EP_NTERM
17205 P_slate PVM
17210 HMMalddh_1_eutpdugrm(pf00171) PVM EP_NTERM
17220 HMMacetkin_1(pf00871) PVM
17230 H_vividBlue PVM
17235 HMMptac_2_mix(pf06130) PVM EP_NTERM
17240 HMMreg08220_2_PVM(pf00455pf08220) PVM
17245 HMMnopfamclust3_1(#) PVM
17250 -pf07586 -
17255 -pf13442pf07691pf07627pf07624pf07631 -
17260 -pf01386pf14693 -
17265 -pf01195 -
17270 -pf01250 -
17275 -pf00436 -
17280 -pf03948pf01281 -
BMC-P
BMC-H
BMC-T
Aldehyde dehydrogenase
PTAC
Signature enzyme
Alcohol dehydrogenase
Regulator

Phycisphaerales bacterium. Bacteria; Planctomycetes; Phycisphaerae; Phycisphaerales.

DER01__locus_1

PVM

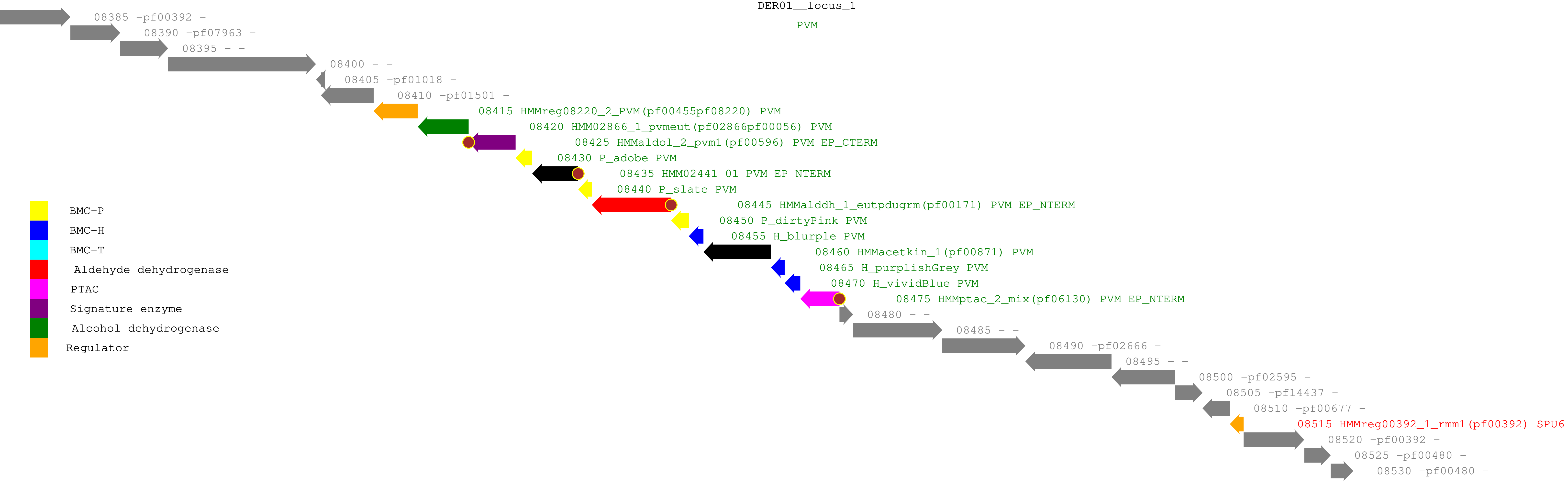

Planctomycetaceae bacterium. Bacteria; Planctomycetes; Planctomycetia; Planctomycetales;

DER64__locus_1

PVM

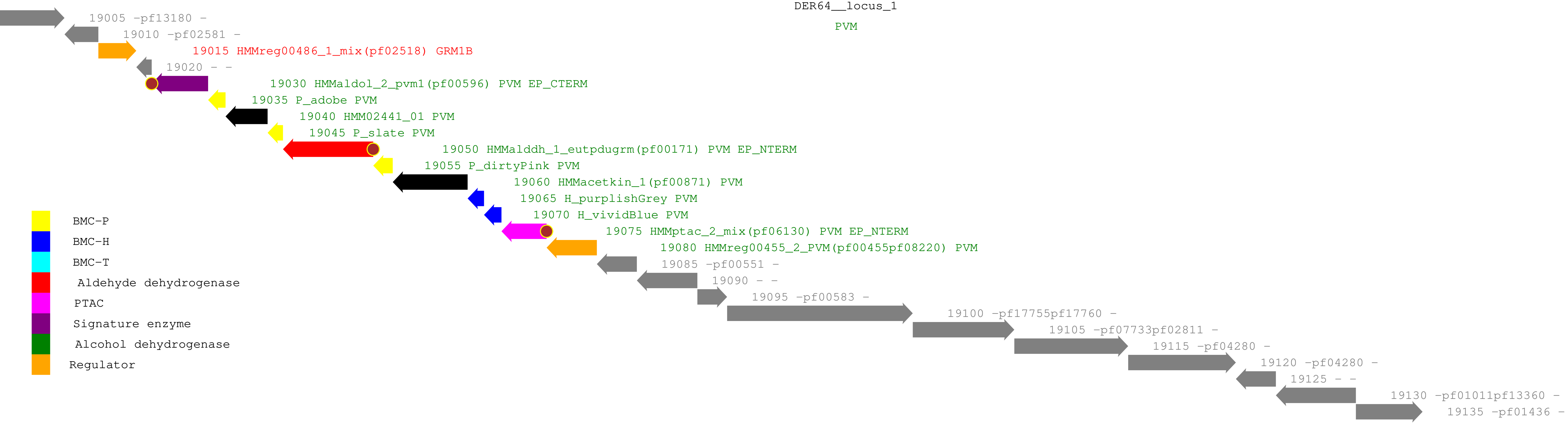

Planctomycetaceae bacterium. Bacteria; Planctomycetes; Planctomycetia; Planctomycetales;

DHW38__locus_1_main_sat

PVM

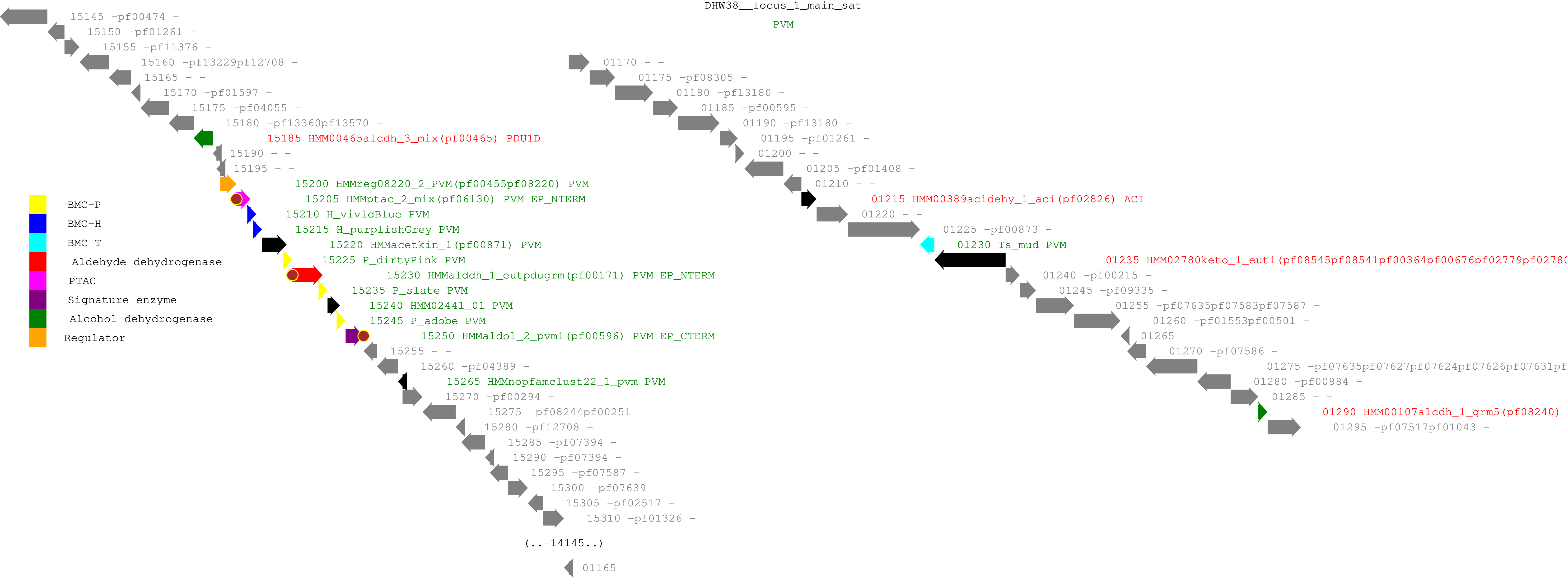

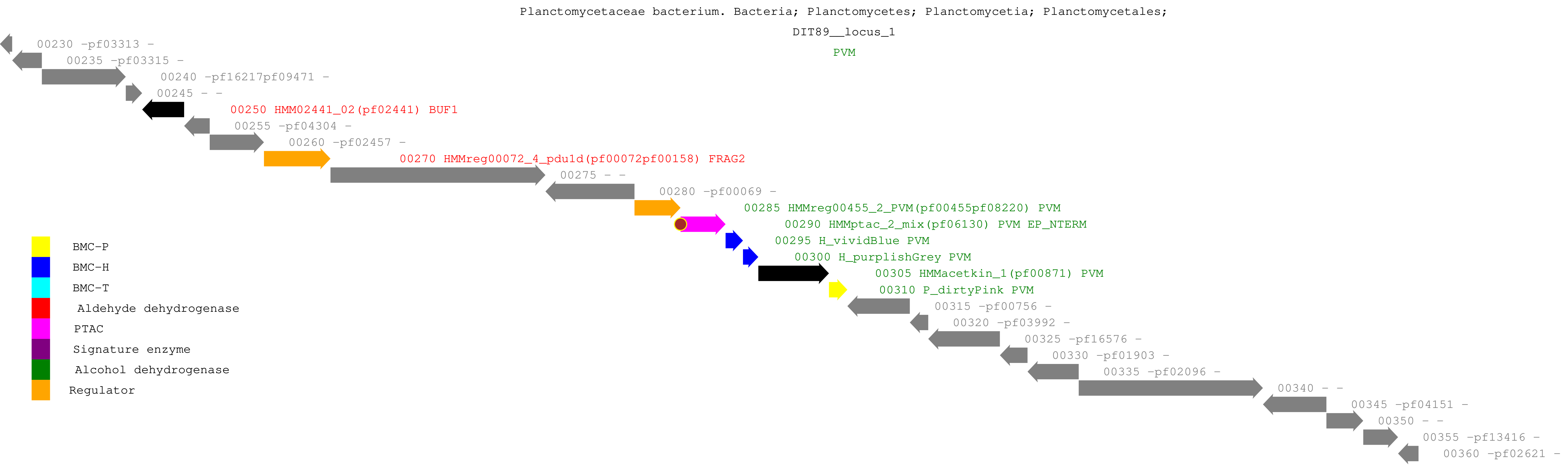
Planctomycetaceae bacterium. Bacteria; Planctomycetes; Planctomycetia; Planctomycetales;
DIT89__locus_1
PVM
00230 -pf03313 -
00235 -pf03315 -
00240 -pf16217pf09471 -
00245 - -
00250 HMM02441_02(pf02441) BUF1
00255 -pf04304 -
00260 -pf02457 -
00270 HMMreg00072_4_pdu1d(pf00072pf00158) FRAG2
00275 - -
00280 -pf00069 -
00285 HMMreg00455_2_PVM(pf00455pf08220) PVM
00290 HMMptac_2_mix(pf06130) PVM EP_NTERM
00295 H_vividBlue PVM
00300 H_purplishGrey PVM
00305 HMMacetkin_1(pf00871) PVM
00310 P_dirtyPink PVM
00315 -pf00756 -
00320 -pf03992 -
00325 -pf16576 -
00330 -pf01903 -
00335 -pf02096 -
00340 - -
00345 -pf04151 -
00350 - -
00355 -pf13416 -
00360 -pf02621 -
BMC-P
BMC-H
BMC-T
Aldehyde dehydrogenase
PTAC
Signature enzyme
Alcohol dehydrogenase
Regulator

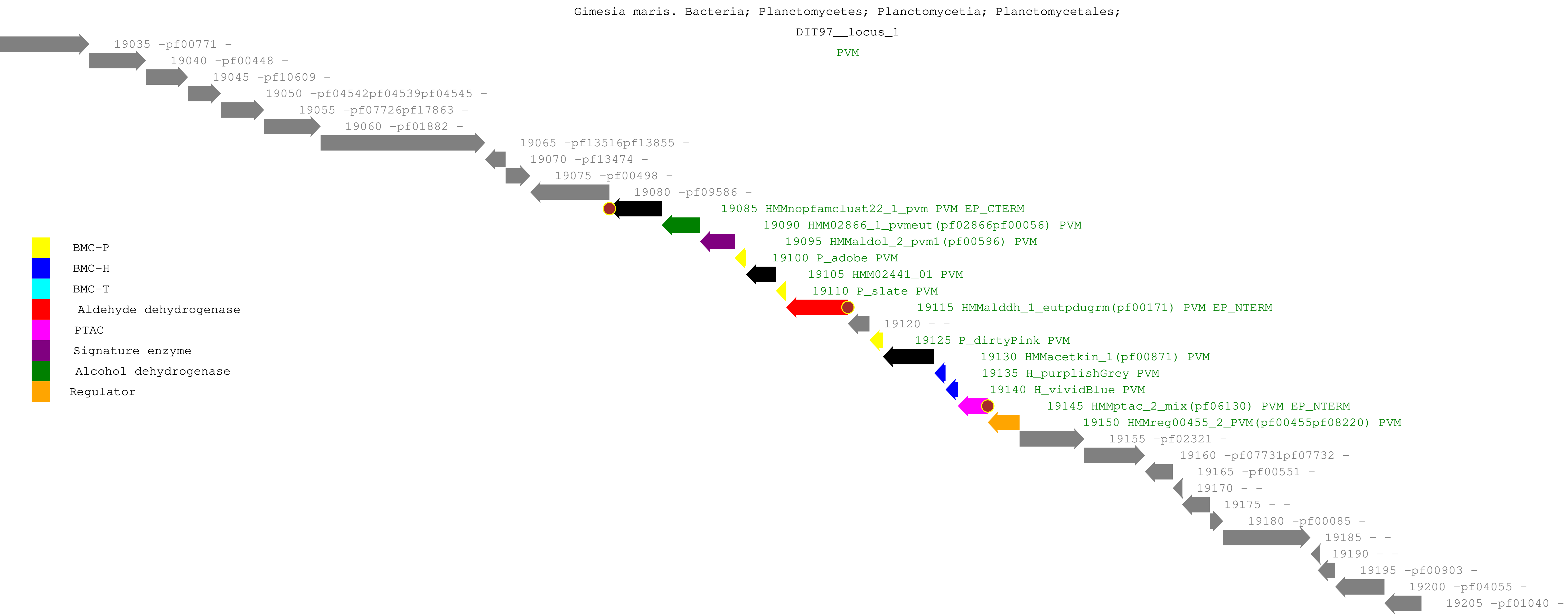
Gimesia maris. Bacteria; Planctomycetes; Planctomycetia; Planctomycetales;
DIT97__locus_1
PVM
19035 -pf00771 -
19040 -pf00448 -
19045 -pf10609 -
19050 -pf04542pf04539pf04545 -
19055 -pf07726pf17863 -
19060 -pf01882 -
19065 -pf13516pf13855 -
19070 -pf13474 -
19075 -pf00498 -
19080 -pf09586 -
19085 HMMnopfamclust22_1_pvm PVM EP_CTERM
19090 HMM02866_1_pvmeut(pf02866pf00056) PVM
19095 HMMaldol_2_pvm1(pf00596) PVM
19100 P_adobe PVM
19105 HMM02441_01 PVM
19110 P_slate PVM
19115 HMMalddh_1_eutpdugrm(pf00171) PVM EP_NTERM
19120 - -
19125 P_dirtyPink PVM
19130 HMMacetkin_1(pf00871) PVM
19135 H_purplishGrey PVM
19140 H_vividBlue PVM
19145 HMMptac_2_mix(pf06130) PVM EP_NTERM
19150 HMMreg00455_2_PVM(pf00455pf08220) PVM
19155 -pf02321 -
19160 -pf07731pf07732 -
19165 -pf00551 -
19170 - -
19175 - -
19180 -pf00085 -
19185 - -
19190 - -
19195 -pf00903 -
19200 -pf04055 -
19205 -pf01040 -
BMC-P
BMC-H
BMC-T
Aldehyde dehydrogenase
PTAC
Signature enzyme
Alcohol dehydrogenase
Regulator

Verrucomicrobiales bacterium. Bacteria; Verrucomicrobia; Verrucomicrobiae; Verrucomicrobiales.

DIV54__locus_1

PVM

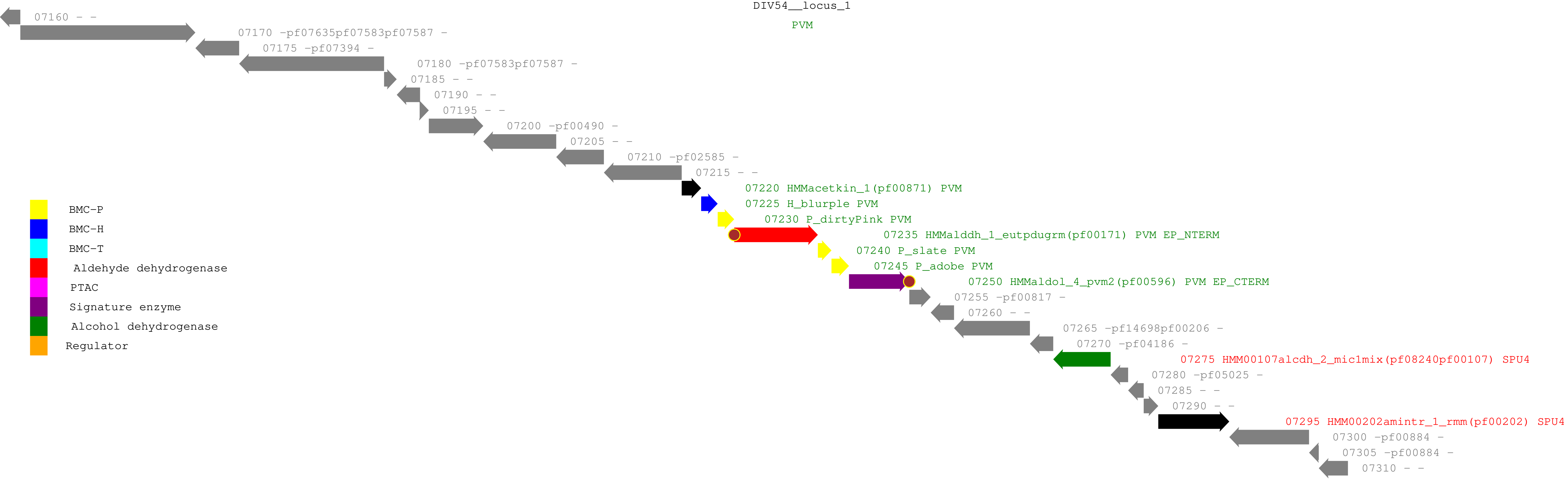

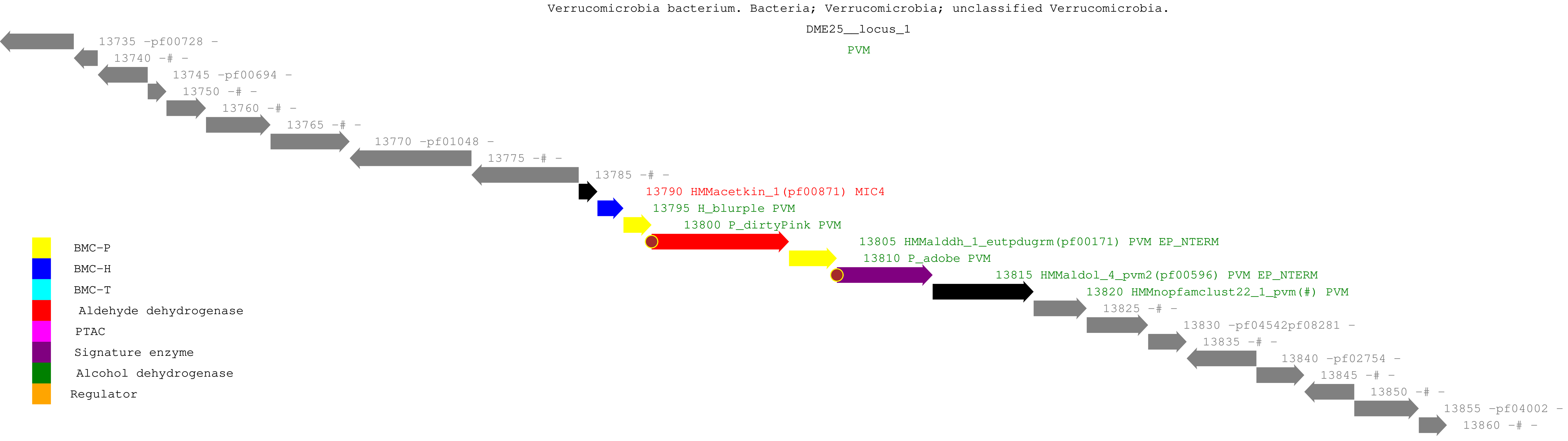
Verrucomicrobia bacterium. Bacteria; Verrucomicrobia; unclassified Verrucomicrobia.
DME25__locus_1
PVM
13735 -pf00728 -
13740 -# -
13745 -pf00694 -
13750 -# -
13760 -# -
13765 -# -
13770 -pf01048 -
13775 -# -
13785 -# -
13790 HMMacetkin_1(pf00871) MIC4
13795 H_blurple PVM
13800 P_dirtyPink PVM
13805 HMMalddh_1_eutpdugrm(pf00171) PVM EP_NTERM
13810 P_adobe PVM
13815 HMMaldol_4_pvm2(pf00596) PVM EP_NTERM
13820 HMMnopfamclust22_1_pvm(#) PVM
13825 -# -
13830 -pf04542pf08281 -
13835 -# -
13840 -pf02754 -
13845 -# -
13850 -# -
13855 -pf04002 -
13860 -# -
BMC-P
BMC-H
BMC-T
Aldehyde dehydrogenase
PTAC
Signature enzyme
Alcohol dehydrogenase
Regulator

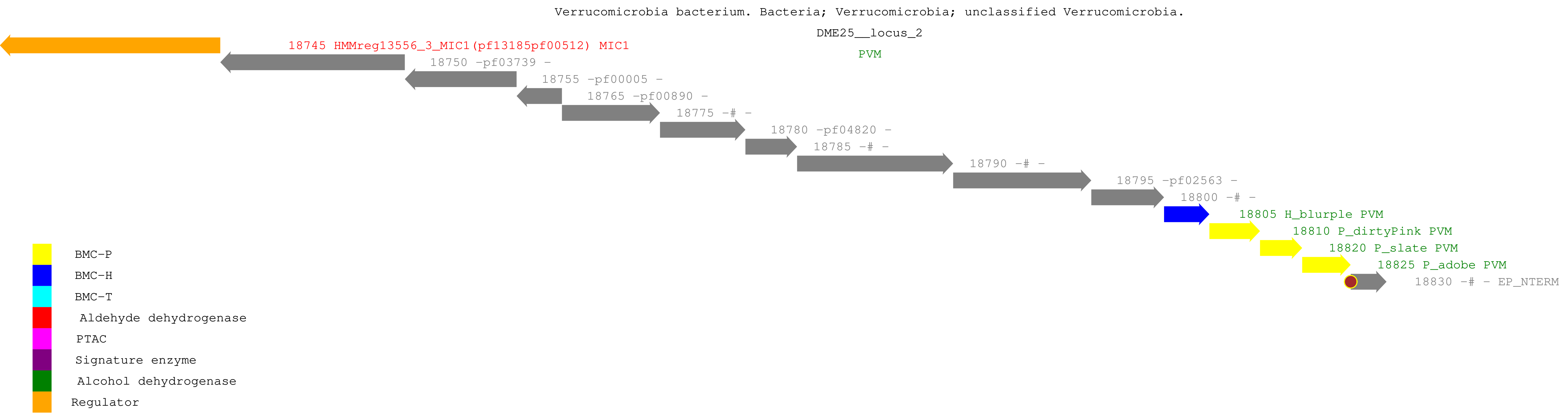
Verrucomicrobia bacterium. Bacteria; Verrucomicrobia; unclassified Verrucomicrobia.
DME25__locus_2
PVM
18745 HMMreg13556_3_MIC1(pf13185pf00512) MIC1
18750 -pf03739 -
18755 -pf00005 -
18765 -pf00890 -
18775 -# -
18780 -pf04820 -
18785 -# -
18790 -# -
18795 -pf02563 -
18800 -# -
18805 H_blurple PVM
18810 P_dirtyPink PVM
18820 P_slate PVM
18825 P_adobe PVM
18830 -# - EP_NTERM
BMC-P
BMC-H
BMC-T
Aldehyde dehydrogenase
PTAC
Signature enzyme
Alcohol dehydrogenase
Regulator

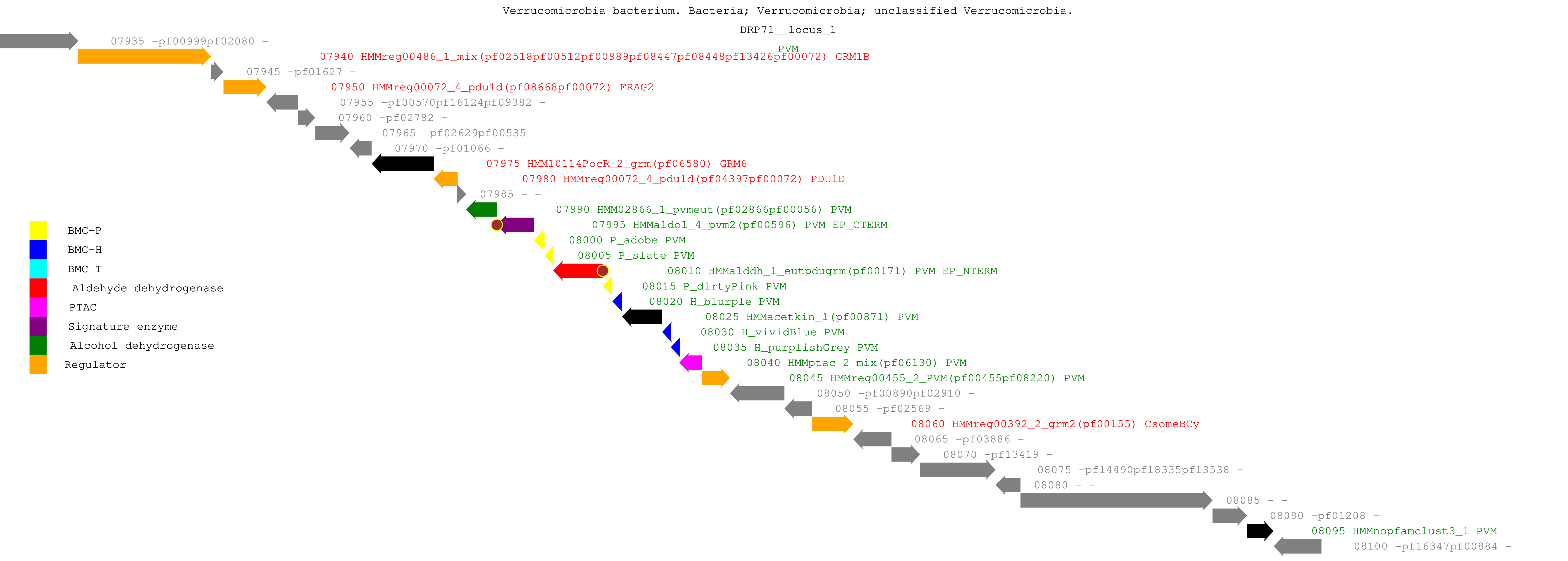

Verrucomicrobia bacterium. Bacteria; Verrucomicrobia; unclassified Verrucomicrobia.
DRP71__locus_1
PVM
07935 -pf00999pf02080 -
07940 HMMreg00486_1_mix(pf02518pf00512pf00989pf08447pf08448pf13426pf00072) GRM1B
07945 -pf01627 -
07950 HMMreg00072_4_pdu1d(pf08668pf00072) FRAG2
07955 -pf00570pf16124pf09382 -
07960 -pf02782 -
07965 -pf02629pf00535 -
07970 -pf01066 -
07975 HMM10114PocR_2_grm(pf06580) GRM6
07980 HMMreg00072_4_pdu1d(pf04397pf00072) PDU1D
07985 - -
07990 HMM02866_1_pvmeut(pf02866pf00056) PVM
07995 HMMaldol_4_pvm2(pf00596) PVM EP_CTERM
08000 P_adobe PVM
08005 P_slate PVM
08010 HMMalddh_1_eutpdugrm(pf00171) PVM EP_NTERM
08015 P_dirtyPink PVM
08020 H_blurple PVM
08025 HMMacetkin_1(pf00871) PVM
08030 H_vividBlue PVM
08035 H_purplishGrey PVM
08040 HMMptac_2_mix(pf06130) PVM
08045 HMMreg00455_2_PVM(pf00455pf08220) PVM
08050 -pf00890pf02910 -
08055 -pf02569 -
08060 HMMreg00392_2_grm2(pf00155) CsomeBCy
08065 -pf03886 -
08070 -pf13419 -
08075 -pf14490pf18335pf13538 -
08080 - -
08085 - -
08090 -pf01208 -
08095 HMMnopfamclust3_1 PVM
08100 -pf16347pf00884 -
BMC-P
BMC-H
BMC-T
Aldehyde dehydrogenase
PTAC
Signature enzyme
Alcohol dehydrogenase
Regulator

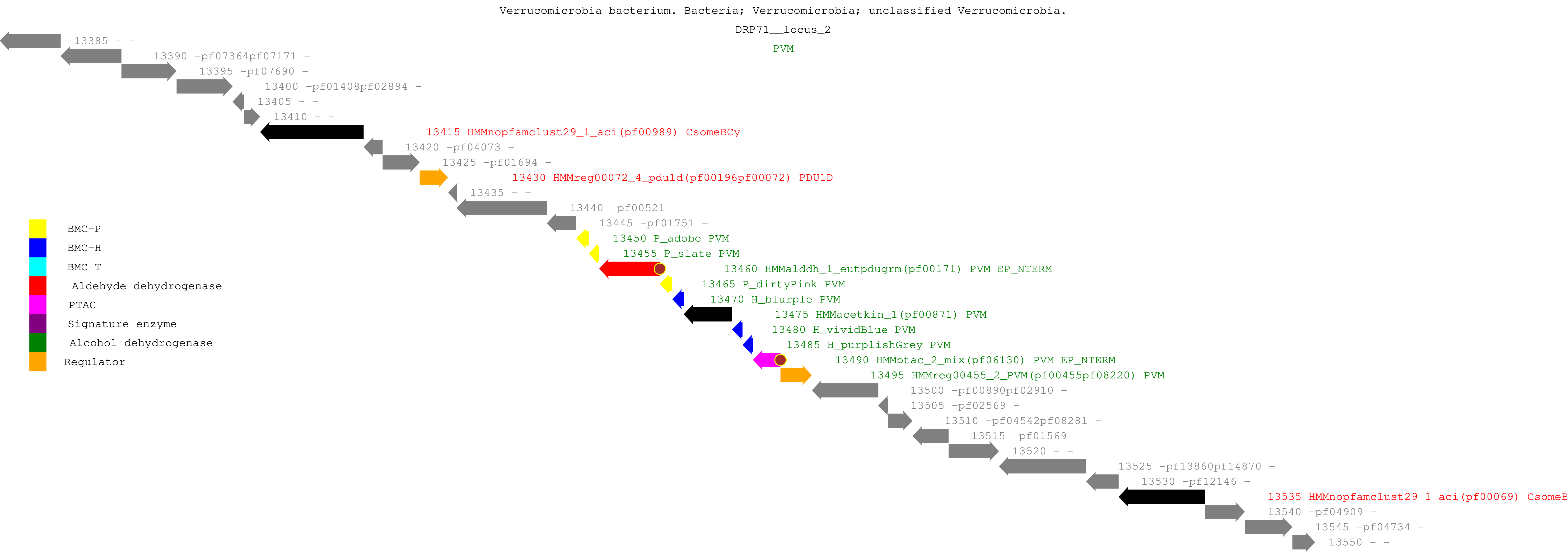

Verrucomicrobia bacterium. Bacteria; Verrucomicrobia; unclassified Verrucomicrobia.
DRP71__locus_2
PVM
13385 - -
13390 -pf07364pf07171 -
13395 -pf07690 -
13400 -pf01408pf02894 -
13405 - -
13410 - -
13415 HMMnopfamclust29_1_aci(pf00989) CsomeBCy
13420 -pf04073 -
13425 -pf01694 -
13430 HMMreg00072_4_pdu1d(pf00196pf00072) PDU1D
13435 - -
13440 -pf00521 -
13445 -pf01751 -
13450 P_adobe PVM
13455 P_slate PVM
13460 HMMaldh_1_eutpdugrm(pf00171) PVM EP_NTERM
13465 P_dirtyPink PVM
13470 H_blurple PVM
13475 HMMacetkin_1(pf00871) PVM
13480 H_vividBlue PVM
13485 H_purplishGrey PVM
13490 HMMptac_2_mix(pf06130) PVM EP_NTERM
13495 HMMreg00455_2_PVM(pf00455pf08220) PVM
13500 -pf00890pf02910 -
13505 -pf02569 -
13510 -pf04542pf08281 -
13515 -pf01569 -
13520 - -
13525 -pf13860pf14870 -
13530 -pf12146 -
13535 HMMnopfamclust29_1_aci(pf00069) CsomeB
13540 -pf04909 -
13545 -pf04734 -
13550 - -
BMC-P
BMC-H
BMC-T
Aldehyde dehydrogenase
PTAC
Signature enzyme
Alcohol dehydrogenase
Regulator

Blastopirellula marina DSM 3645. Bacteria; Planctomycetes; Planctomycetia; Planctomycetales;

DSM3645__locus_1

PVM

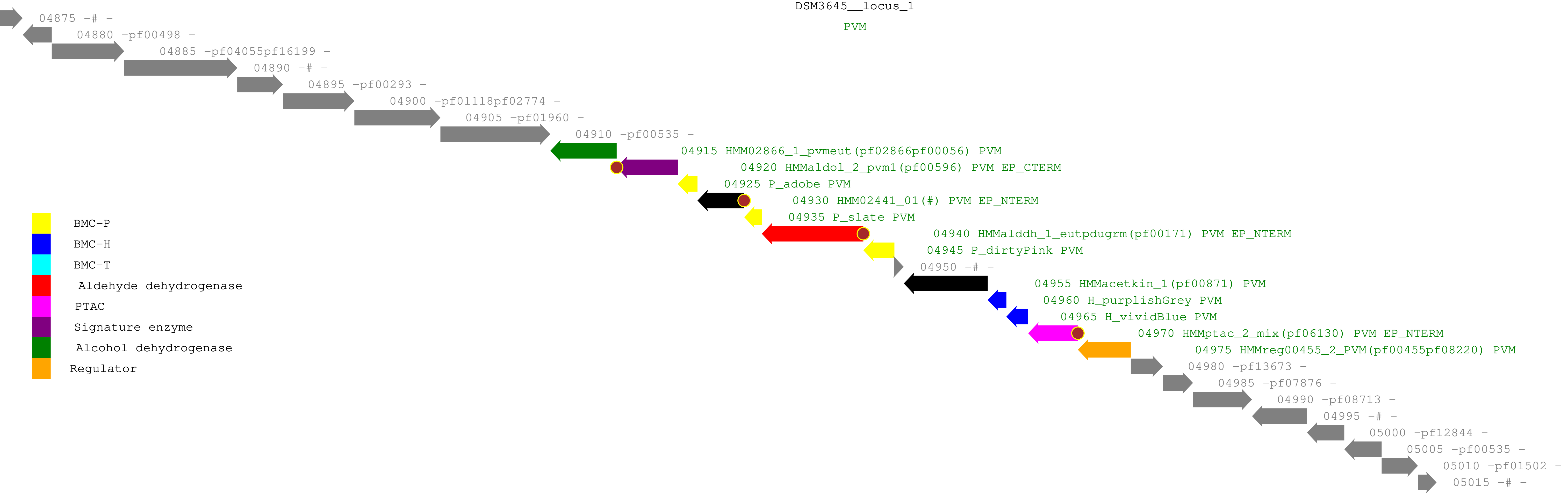

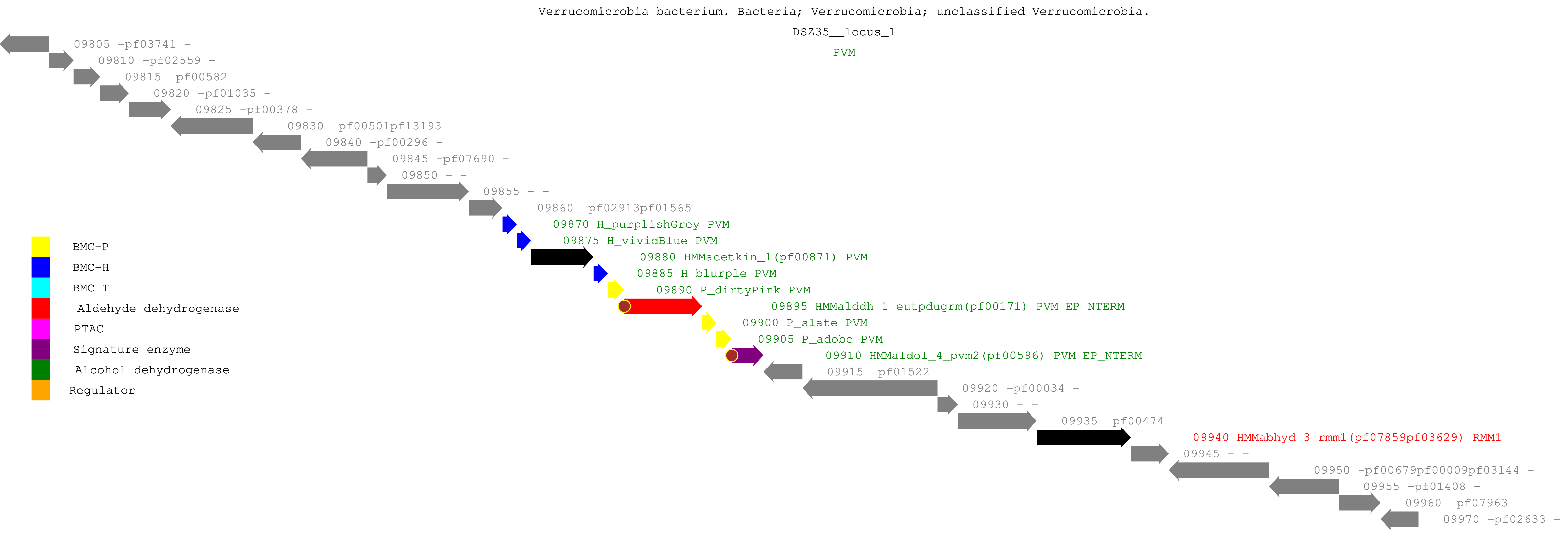
Verrucomicrobia bacterium. Bacteria; Verrucomicrobia; unclassified Verrucomicrobia.
DSZ35__locus_1
PVM
09805 -pf03741 -
09810 -pf02559 -
09815 -pf00582 -
09820 -pf01035 -
09825 -pf00378 -
09830 -pf00501pf13193 -
09840 -pf00296 -
09845 -pf07690 -
09850 - -
09855 - -
09860 -pf02913pf01565 -
09870 H_purplishGrey PVM
09875 H_vividBlue PVM
09880 HMMacetkin_1(pf00871) PVM
09885 H_blurple PVM
09890 P_dirtyPink PVM
09895 HMMalddh_1_eutpdugrm(pf00171) PVM EP_NTERM
09900 P_slate PVM
09905 P_adobe PVM
09910 HMMaldol_4_pvm2(pf00596) PVM EP_NTERM
09915 -pf01522 -
09920 -pf00034 -
09930 - -
09935 -pf00474 -
09940 HMMabhyd_3_rmm1(pf07859pf03629) RMM1
09945 - -
09950 -pf00679pf00009pf03144 -
09955 -pf01408 -
09960 -pf07963 -
09970 -pf02633 -
BMC-P
BMC-H
BMC-T
Aldehyde dehydrogenase
PTAC
Signature enzyme
Alcohol dehydrogenase
Regulator

Blastopirellula cremea. Bacteria; Planctomycetes; Planctomycetia; Planctomycetales;

DTL42__locus_1

PVM

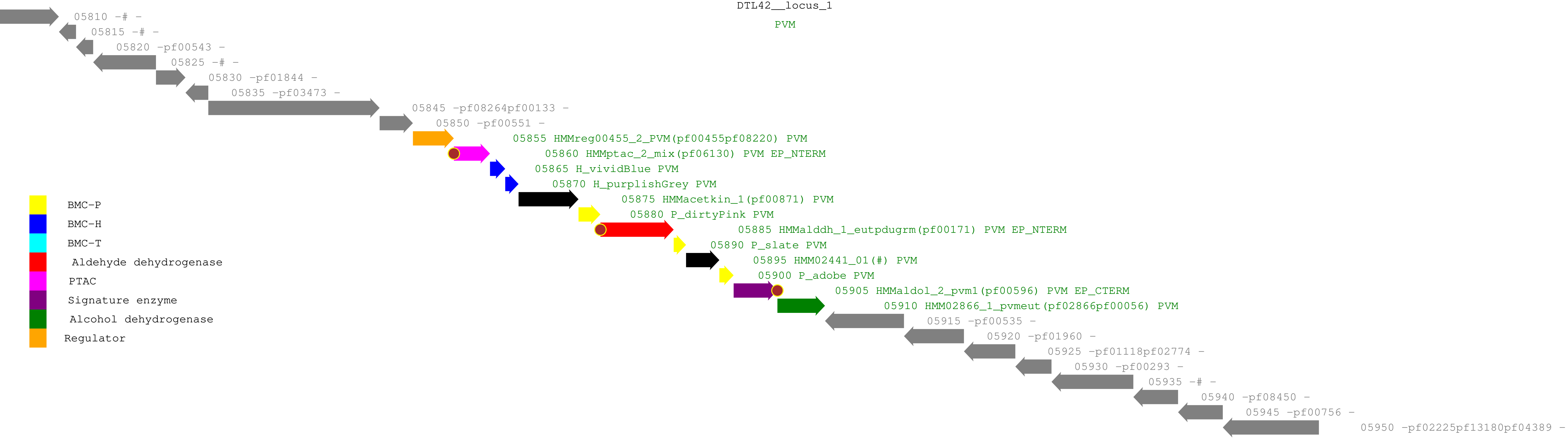

Verrucomicrobia bacterium. Bacteria; Verrucomicrobia; unclassified Verrucomicrobia.

DUW69__locus_1

PVM

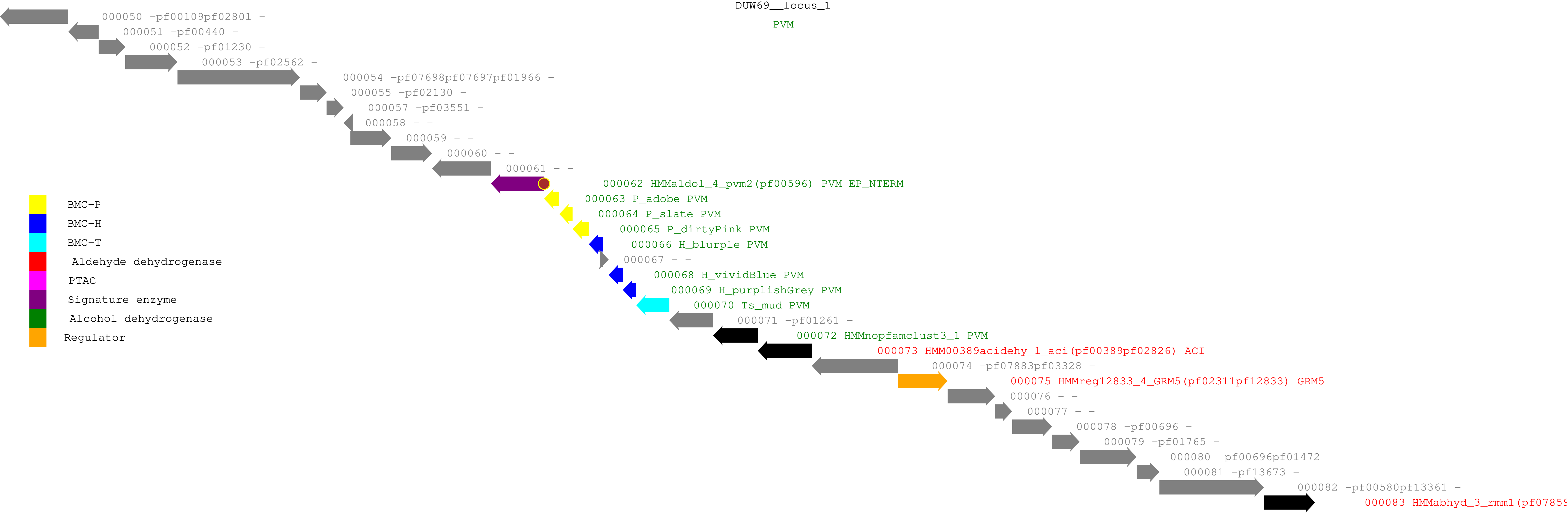

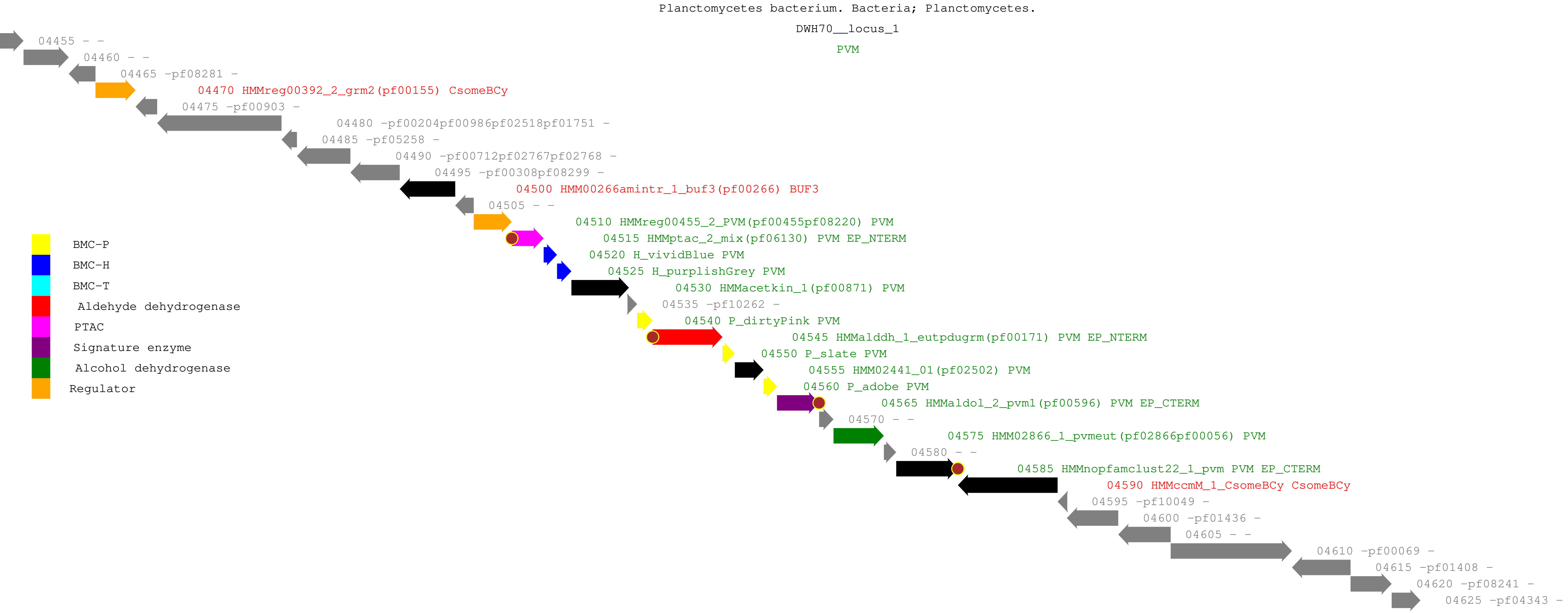
Planctomycetes bacterium. Bacteria; Planctomycetes.
DWH70__locus_1
PVM
04455 - -
04460 - -
04465 -pf08281 -
04470 HMMreg00392_2_grm2(pf00155) CsomeBCy
04475 -pf00903 -
04480 -pf00204pf00986pf02518pf01751 -
04485 -pf05258 -
04490 -pf00712pf02767pf02768 -
04495 -pf00308pf08299 -
04500 HMM00266amintr_1_buf3(pf00266) BUF3
04505 - -
04510 HMMreg00455_2_PVM(pf00455pf08220) PVM
04515 HMMptac_2_mix(pf06130) PVM EP_NTERM
04520 H_vividBlue PVM
04525 H_purplishGrey PVM
04530 HMMacetkin_1(pf00871) PVM
04535 -pf10262 -
04540 P_dirtyPink PVM
04545 HMMalddh_1_eutpdugrm(pf00171) PVM EP_NTERM
04550 P_slate PVM
04555 HMM02441_01(pf02502) PVM
04560 P_adobe PVM
04565 HMMaldol_2_pvm1(pf00596) PVM EP_CTERM
04570 - -
04575 HMM02866_1_pvmeut(pf02866pf00056) PVM
04580 - -
04585 HMMnopfamclust22_1_pvm PVM EP_CTERM
04590 HMMccmM_1_CsomeBCy CsomeBCy
04595 -pf10049 -
04600 -pf01436 -
04605 - -
04610 -pf00069 -
04615 -pf01408 -
04620 -pf08241 -
04625 -pf04343 -
BMC-P
BMC-H
BMC-T
Aldehyde dehydrogenase
PTAC
Signature enzyme
Alcohol dehydrogenase
Regulator

Planctomycetes bacterium. Bacteria; Planctomycetes.

DWH78__locus_1

PVM

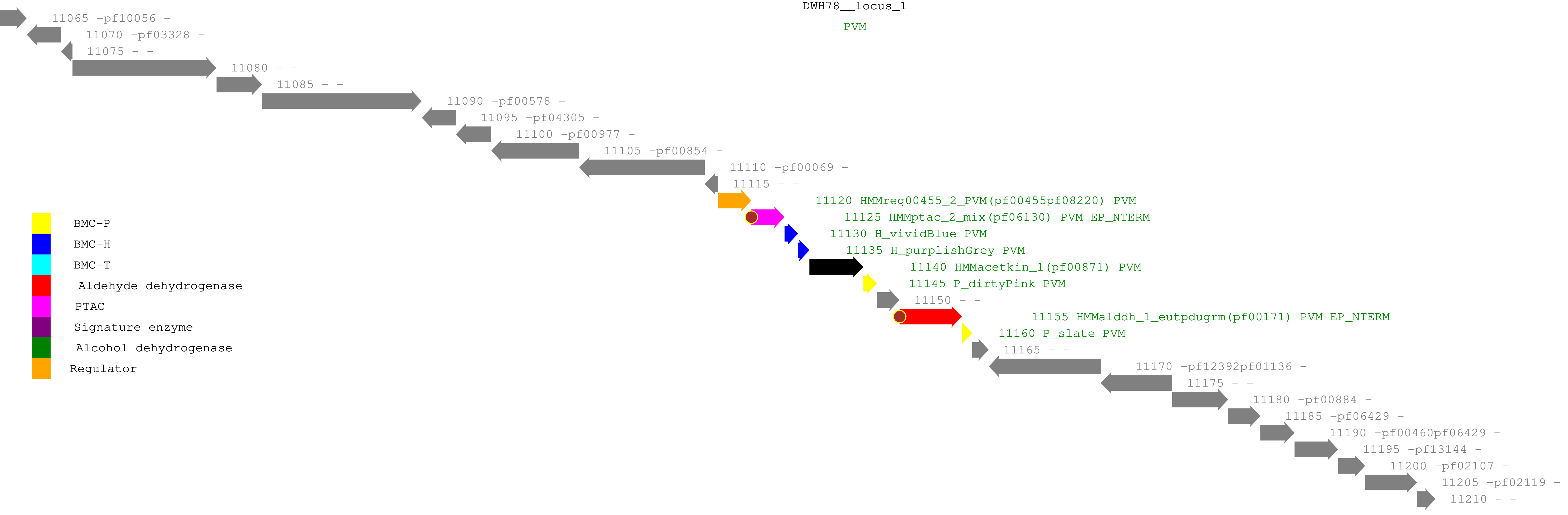

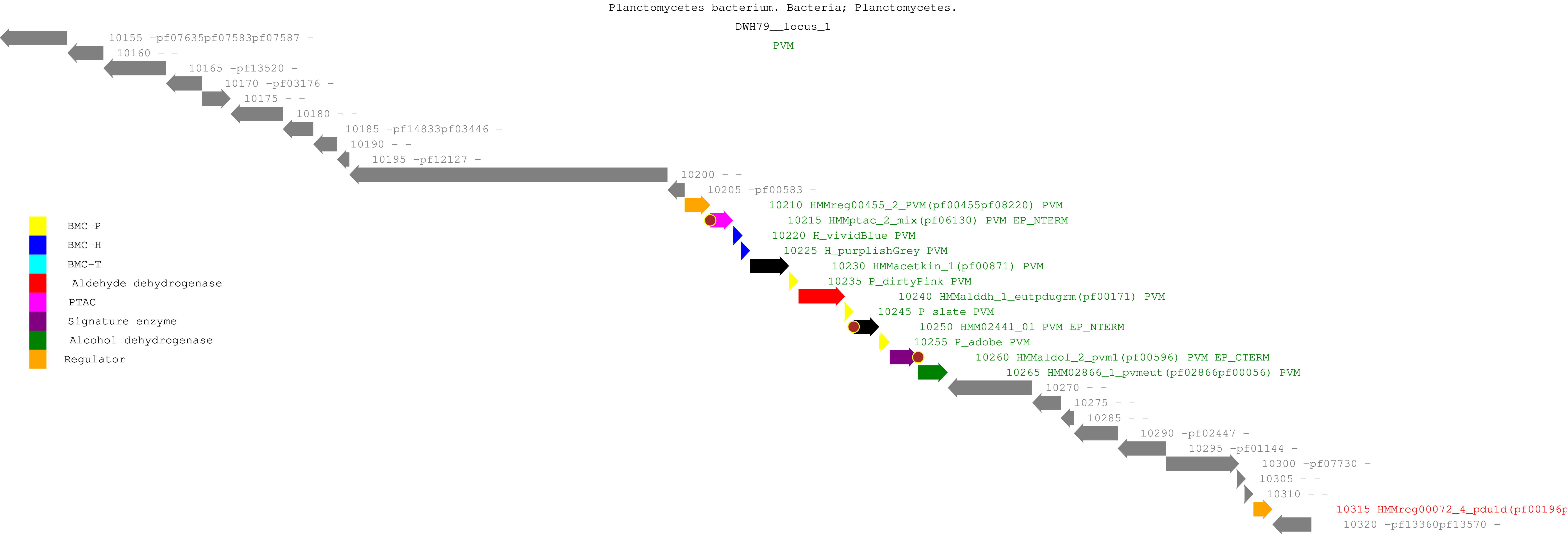
Planctomycetes bacterium. Bacteria; Planctomycetes.
DWH79__locus_1
PVM
10155 -pf07635pf07583pf07587 -
10160 - -
10165 -pf13520 -
10170 -pf03176 -
10175 - -
10180 - -
10185 -pf14833pf03446 -
10190 - -
10195 -pf12127 -
10200 - -
10205 -pf00583 -
10210 HMMreg00455_2_PVM(pf00455pf08220) PVM
10215 HMMptac_2_mix(pf06130) PVM EP_NTERM
10220 H_vividBlue PVM
10225 H_purplishGrey PVM
10230 HMMacetkin_1(pf00871) PVM
10235 P_dirtyPink PVM
10240 HMMaldh_1_eutpdugrm(pf00171) PVM
10245 P_slate PVM
10250 HMM02441_01 PVM EP_NTERM
10255 P_adobe PVM
10260 HMMaldol_2_pvm1(pf00596) PVM EP_CTERM
10265 HMM02866_1_pvmeut(pf02866pf00056) PVM
10270 - -
10275 - -
10285 - -
10290 -pf02447 -
10295 -pf01144 -
10300 -pf07730 -
10305 - -
10310 - -
10315 HMMreg00072_4_pdu1d(pf00196p
10320 -pf13360pf13570 -
BMC-P
BMC-H
BMC-T
Aldehyde dehydrogenase
PTAC
Signature enzyme
Alcohol dehydrogenase
Regulator

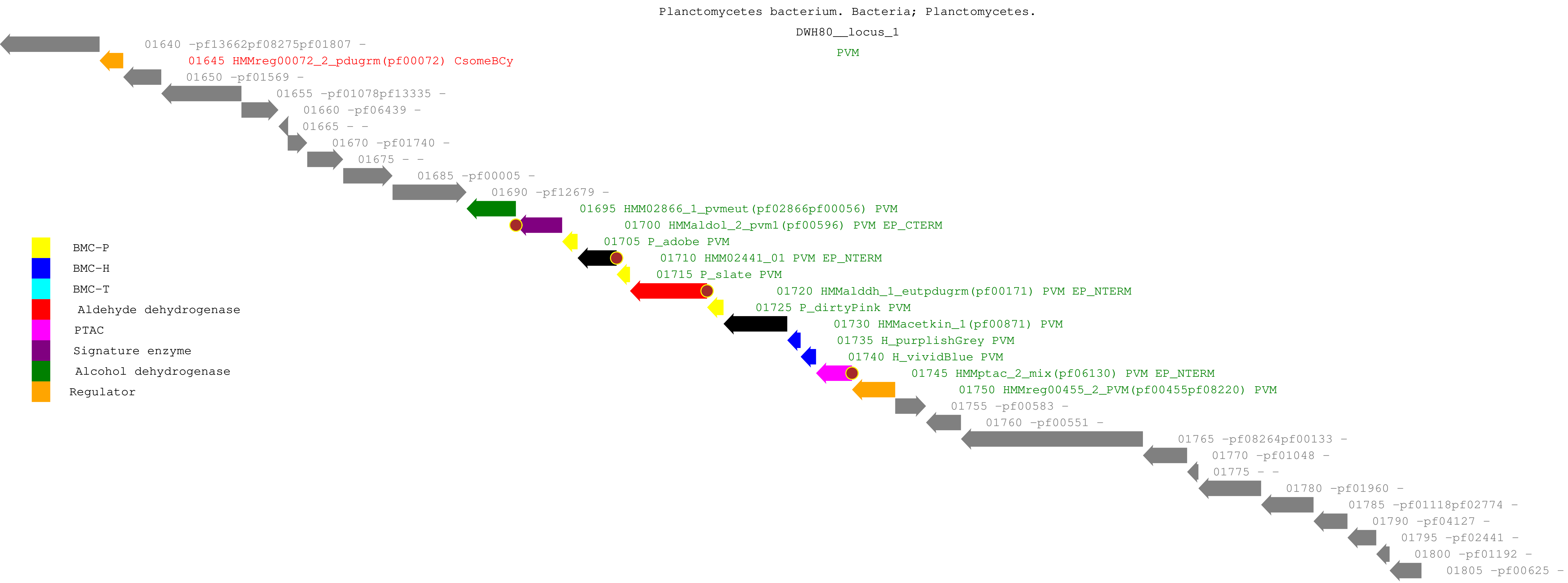
Planctomycetes bacterium. Bacteria; Planctomycetes.
DWH80__locus_1
PVM
01640 -pf13662pf08275pf01807 -
01645 HMMreg00072_2_pdugrm(pf00072) CsomeBCy
01650 -pf01569 -
01655 -pf01078pf13335 -
01660 -pf06439 -
01665 - -
01670 -pf01740 -
01675 - -
01685 -pf00005 -
01690 -pf12679 -
01695 HMM02866_1_pvmeut(pf02866pf00056) PVM
01700 HMMaldol_2_pvm1(pf00596) PVM EP_CTERM
01705 P_adobe PVM
01710 HMM02441_01 PVM EP_NTERM
01715 P_slate PVM
01720 HMMalddh_1_eutpdugrm(pf00171) PVM EP_NTERM
01725 P_dirtyPink PVM
01730 HMMacetkin_1(pf00871) PVM
01735 H_purplishGrey PVM
01740 H_vividBlue PVM
01745 HMMptac_2_mix(pf06130) PVM EP_NTERM
01750 HMMreg00455_2_PVM(pf00455pf08220) PVM
01755 -pf00583 -
01760 -pf00551 -
01765 -pf08264pf00133 -
01770 -pf01048 -
01775 - -
01780 -pf01960 -
01785 -pf01118pf02774 -
01790 -pf04127 -
01795 -pf02441 -
01800 -pf01192 -
01805 -pf00625 -
BMC-P
BMC-H
BMC-T
Aldehyde dehydrogenase
PTAC
Signature enzyme
Alcohol dehydrogenase
Regulator

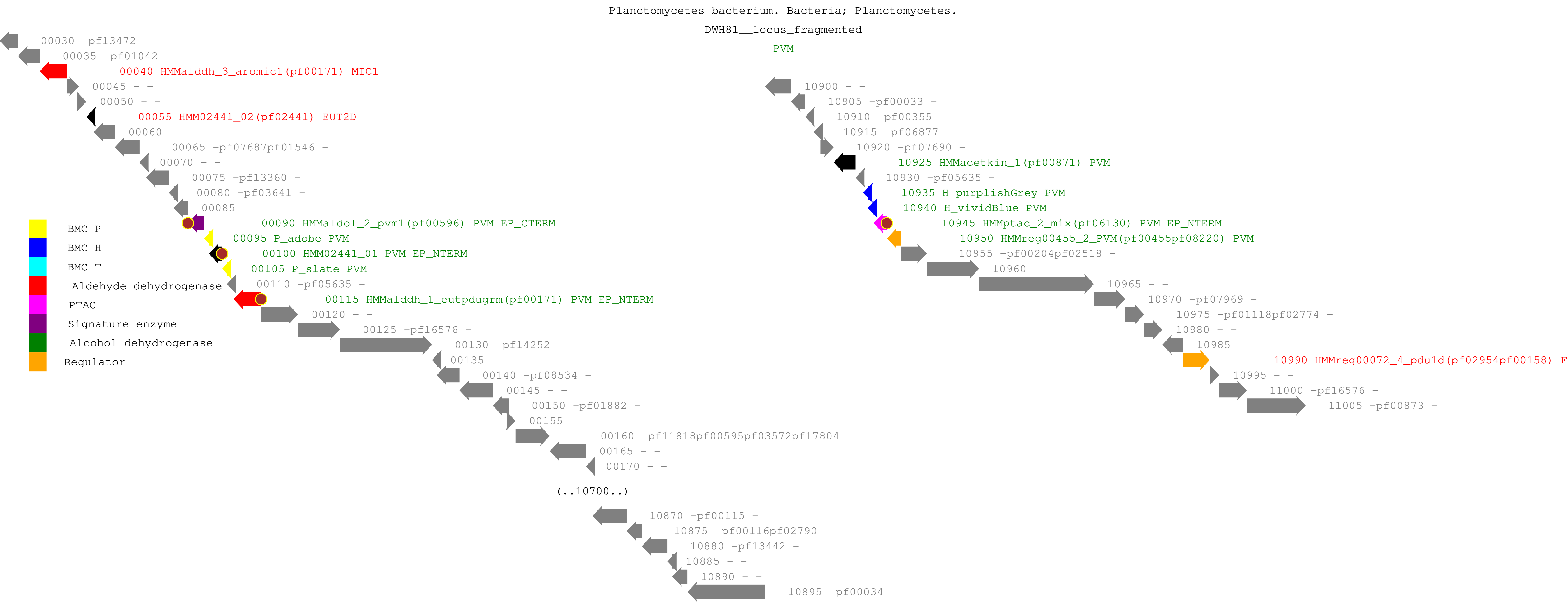

Planctomycetes bacterium. Bacteria; Planctomycetes.
DWH81__locus_fragmented
PVM
00030 -pf13472 -
00035 -pf01042 -
00040 HMMaldh_3_aromic1(pf00171) MIC1
00045 - -
00050 - -
00055 HMM02441_02(pf02441) EUT2D
00060 - -
00065 -pf07687pf01546 -
00070 - -
00075 -pf13360 -
00080 -pf03641 -
00085 - -
00090 HMMaldol_2_pvm1(pf00596) PVM EP_CTERM
00095 P_adobe PVM
00100 HMM02441_01 PVM EP_NTERM
00105 P_slate PVM
00110 -pf05635 -
00115 HMMaldh_1_eutpdugrm(pf00171) PVM EP_NTERM
00120 - -
00125 -pf16576 -
00130 -pf14252 -
00135 - -
00140 -pf08534 -
00145 - -
00150 -pf01882 -
00155 - -
00160 -pf11818pf00595pf03572pf17804 -
00165 - -
00170 - -
(..10700..)
10870 -pf00115 -
10875 -pf00116pf02790 -
10880 -pf13442 -
10885 - -
10890 - -
10895 -pf00034 -
10900 - -
10905 -pf00033 -
10910 -pf00355 -
10915 -pf06877 -
10920 -pf07690 -
10925 HMMacetkin_1(pf00871) PVM
10930 -pf05635 -
10935 H_purplishGrey PVM
10940 H_vividBlue PVM
10945 HMMptac_2_mix(pf06130) PVM EP_NTERM
10950 HMMreg00455_2_PVM(pf00455pf08220) PVM
10955 -pf00204pf02518 -
10960 - -
10965 - -
10970 -pf07969 -
10975 -pf01118pf02774 -
10980 - -
10985 - -
10990 HMMreg00072_4_pdu1d(pf02954pf00158) F
10995 - -
11000 -pf16576 -
11005 -pf00873 -
BMC-P
BMC-H
BMC-T
Aldehyde dehydrogenase
PTAC
Signature enzyme
Alcohol dehydrogenase
Regulator

Planctomycetes bacterium. Bacteria; Planctomycetes.

DWH82__locus_1

PVM

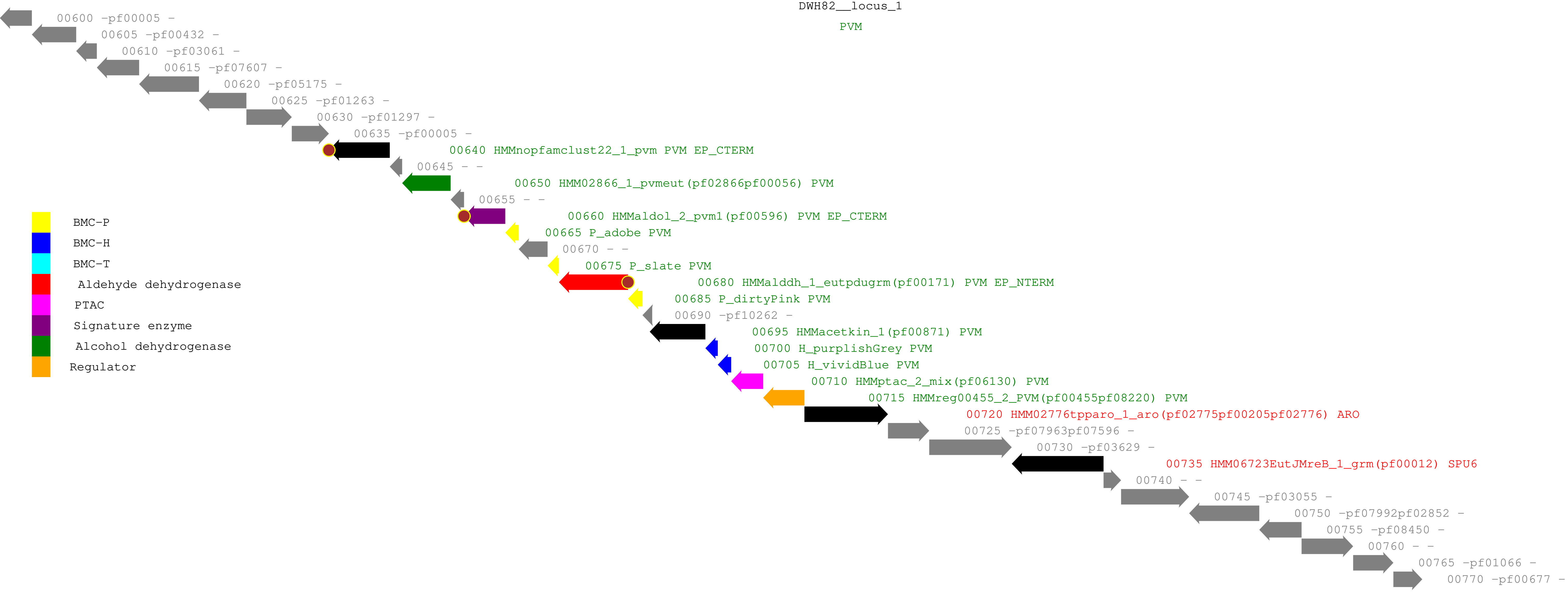

Planctomycetes bacterium. Bacteria; Planctomycetes.

DWH83__locus_1

PVM

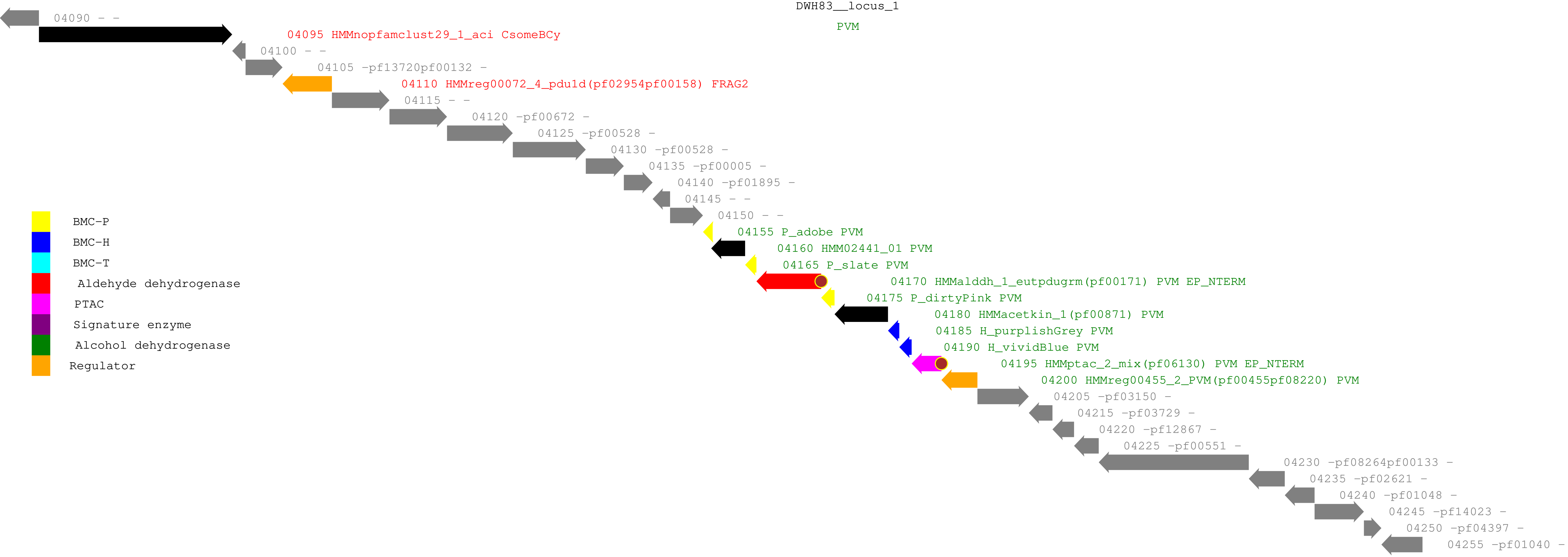

Planctomycetes bacterium. Bacteria; Planctomycetes.

DWH91__locus_1

PVM

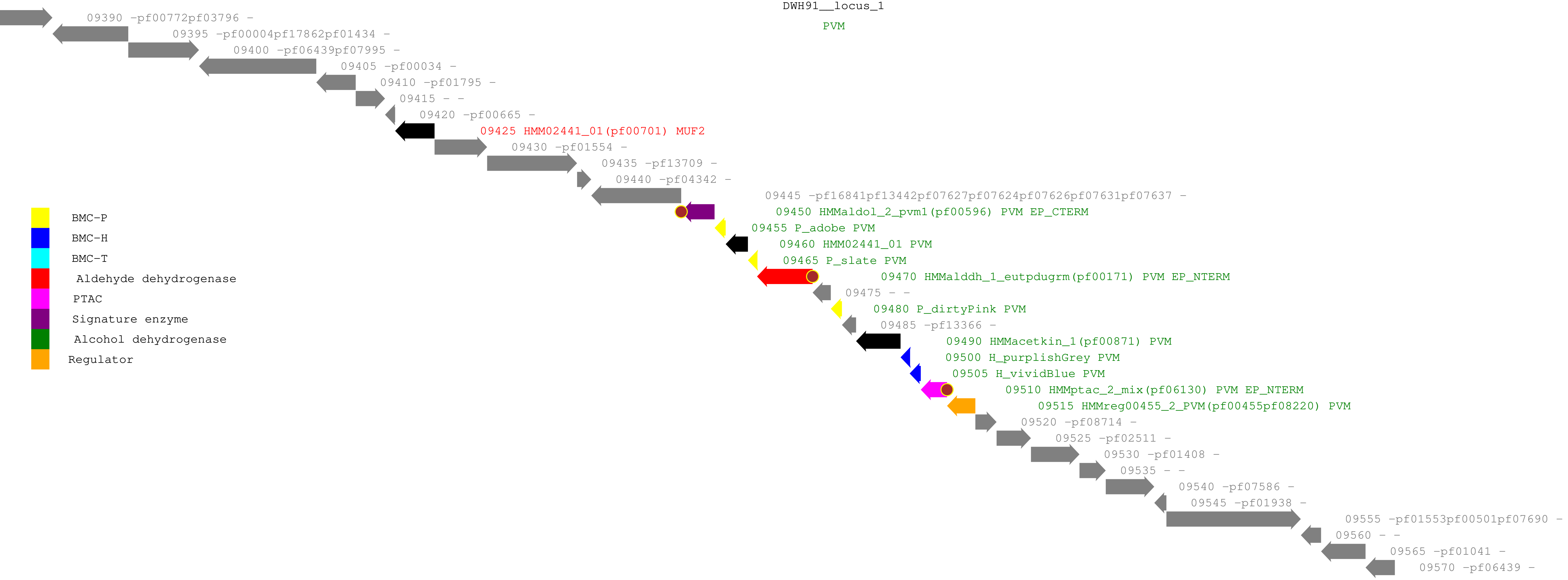

Planctomycetes bacterium. Bacteria; Planctomycetes.

DWH99__locus_1_main_sat

PVM

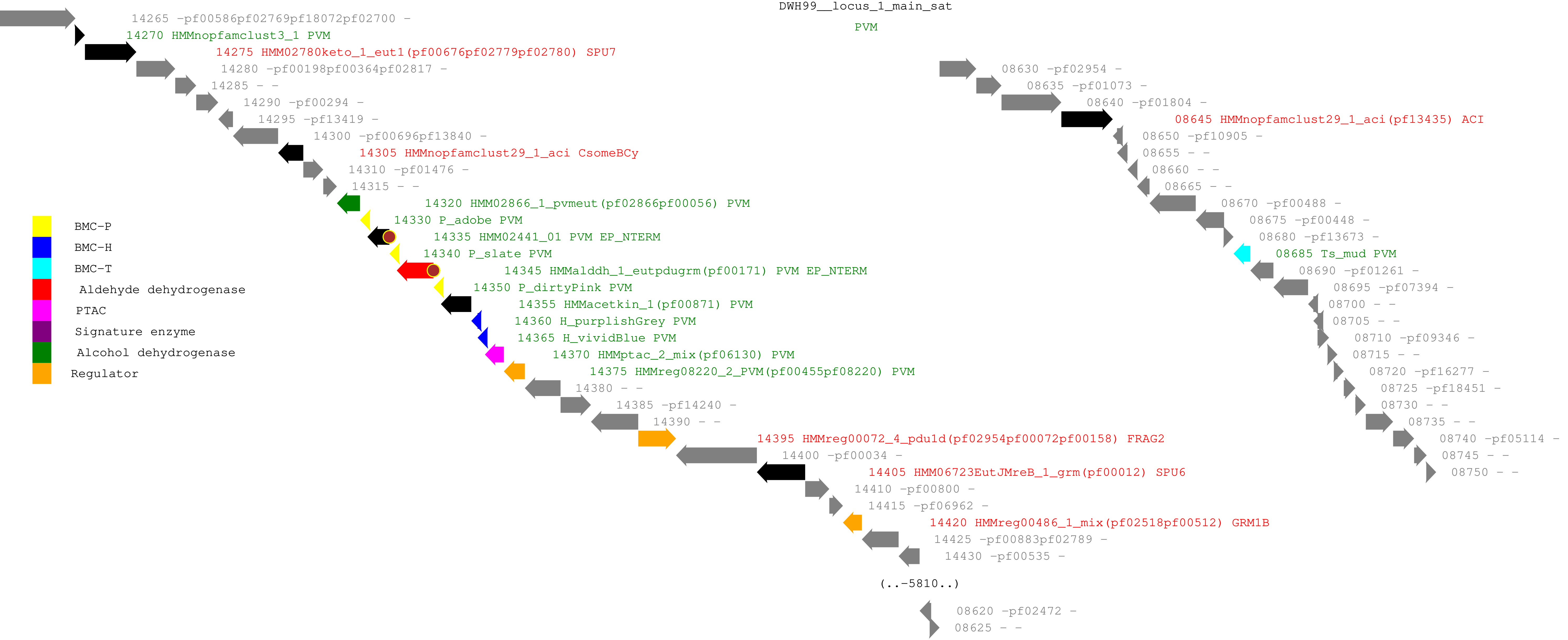

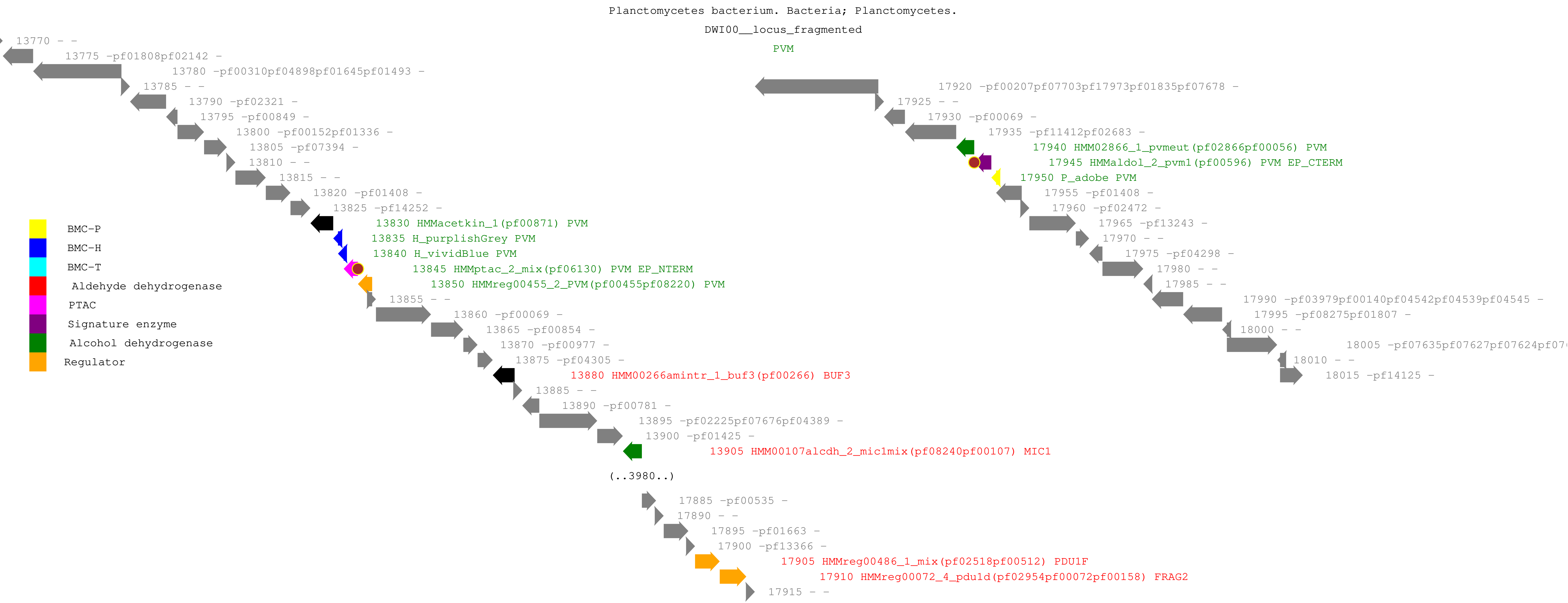

Planctomycetes bacterium. Bacteria; Planctomycetes.
DWI00__locus_fragmented
PVM
13770 - -
13775 -pf01808pf02142 -
13780 -pf00310pf04898pf01645pf01493 -
13785 - -
13790 -pf02321 -
13795 -pf00849 -
13800 -pf00152pf01336 -
13805 -pf07394 -
13810 - -
13815 - -
13820 -pf01408 -
13825 -pf14252 -
13830 HMMacetkin_1(pf00871) PVM
13835 H_purplishGrey PVM
13840 H_vividBlue PVM
13845 HMMptac_2_mix(pf06130) PVM EP_NTERM
13850 HMMreg00455_2_PVM(pf00455pf08220) PVM
13855 - -
13860 -pf00069 -
13865 -pf00854 -
13870 -pf00977 -
13875 -pf04305 -
13880 HMM00266amintr_1_buf3(pf00266) BUF3
13885 - -
13890 -pf00781 -
13895 -pf02225pf07676pf04389 -
13900 -pf01425 -
13905 HMM00107alcdh_2_mic1mix(pf08240pf00107) MIC1
(..3980..)
17885 -pf00535 -
17890 - -
17895 -pf01663 -
17900 -pf13366 -
17905 HMMreg00486_1_mix(pf02518pf00512) PDU1F
17910 HMMreg00072_4_pdu1d(pf02954pf00072pf00158) FRAG2
17915 - -
17920 -pf00207pf07703pf17973pf01835pf07678 -
17925 - -
17930 -pf00069 -
17935 -pf11412pf02683 -
17940 HMM02866_1_pvmeut(pf02866pf00056) PVM
17945 HMMaldol_2_pvm1(pf00596) PVM EP_CTERM
17950 P_adobe PVM
17955 -pf01408 -
17960 -pf02472 -
17965 -pf13243 -
17970 - -
17975 -pf04298 -
17980 - -
17985 - -
17990 -pf03979pf00140pf04542pf04539pf04545 -
17995 -pf08275pf01807 -
18000 - -
18005 -pf07635pf07627pf07624pf07
18010 - -
18015 -pf14125 -
BMC-P
BMC-H
BMC-T
Aldehyde dehydrogenase
PTAC
Signature enzyme
Alcohol dehydrogenase
Regulator

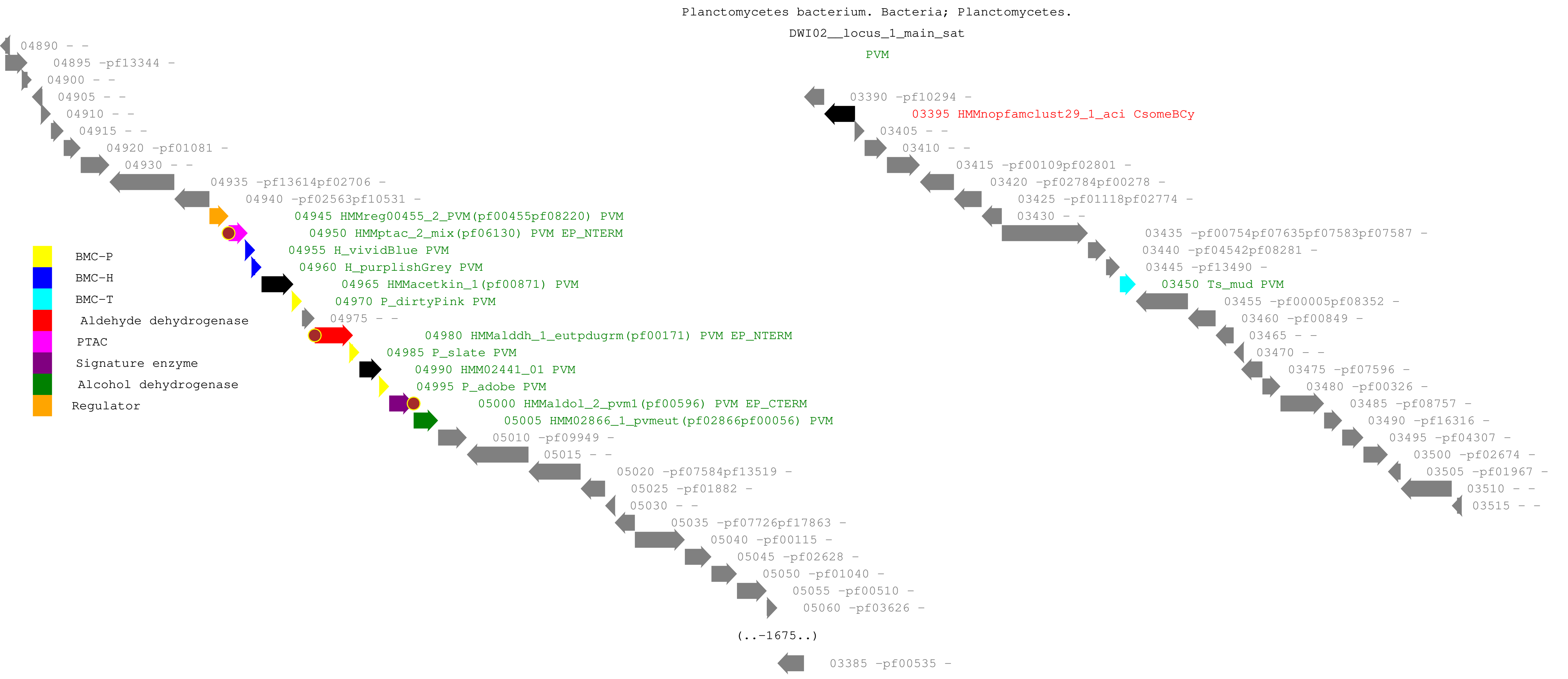
Planctomycetes bacterium. Bacteria; Planctomycetes.
DWI02__locus_1_main_sat
PVM
04890 - -
04895 -pf13344 -
04900 - -
04905 - -
04910 - -
04915 - -
04920 -pf01081 -
04930 - -
04935 -pf13614pf02706 -
04940 -pf02563pf10531 -
04945 HMMreg00455_2_PVM(pf00455pf08220) PVM
04950 HMMptac_2_mix(pf06130) PVM EP_NTERM
04955 H_vividBlue PVM
04960 H_purplishGrey PVM
04965 HMMacetkin_1(pf00871) PVM
04970 P_dirtyPink PVM
04975 - -
04980 HMMaldddh_1_eutpdugrm(pf00171) PVM EP_NTERM
04985 P_slate PVM
04990 HMM02441_01 PVM
04995 P_adobe PVM
05000 HMMaldol_2_pvm1(pf00596) PVM EP_CTERM
05005 HMM02866_1_pvmeut(pf02866pf00056) PVM
05010 -pf09949 -
05015 - -
05020 -pf07584pf13519 -
05025 -pf01882 -
05030 - -
05035 -pf07726pf17863 -
05040 -pf00115 -
05045 -pf02628 -
05050 -pf01040 -
05055 -pf00510 -
05060 -pf03626 -
(..-1675..)
03385 -pf00535 -
03390 -pf10294 -
03395 HMMnopfamclust29_1_aci CsomeBCy
03405 - -
03410 - -
03415 -pf00109pf02801 -
03420 -pf02784pf00278 -
03425 -pf01118pf02774 -
03430 - -
03435 -pf00754pf07635pf07583pf07587 -
03440 -pf04542pf08281 -
03445 -pf13490 -
03450 Ts_mud PVM
03455 -pf00005pf08352 -
03460 -pf00849 -
03465 - -
03470 - -
03475 -pf07596 -
03480 -pf00326 -
03485 -pf08757 -
03490 -pf16316 -
03495 -pf04307 -
03500 -pf02674 -
03505 -pf01967 -
03510 - -
03515 - -
BMC-P
BMC-H
BMC-T
Aldehyde dehydrogenase
PTAC
Signature enzyme
Alcohol dehydrogenase
Regulator

Planctomycetes bacterium. Bacteria; Planctomycetes.

DWI03__locus_1

PVM

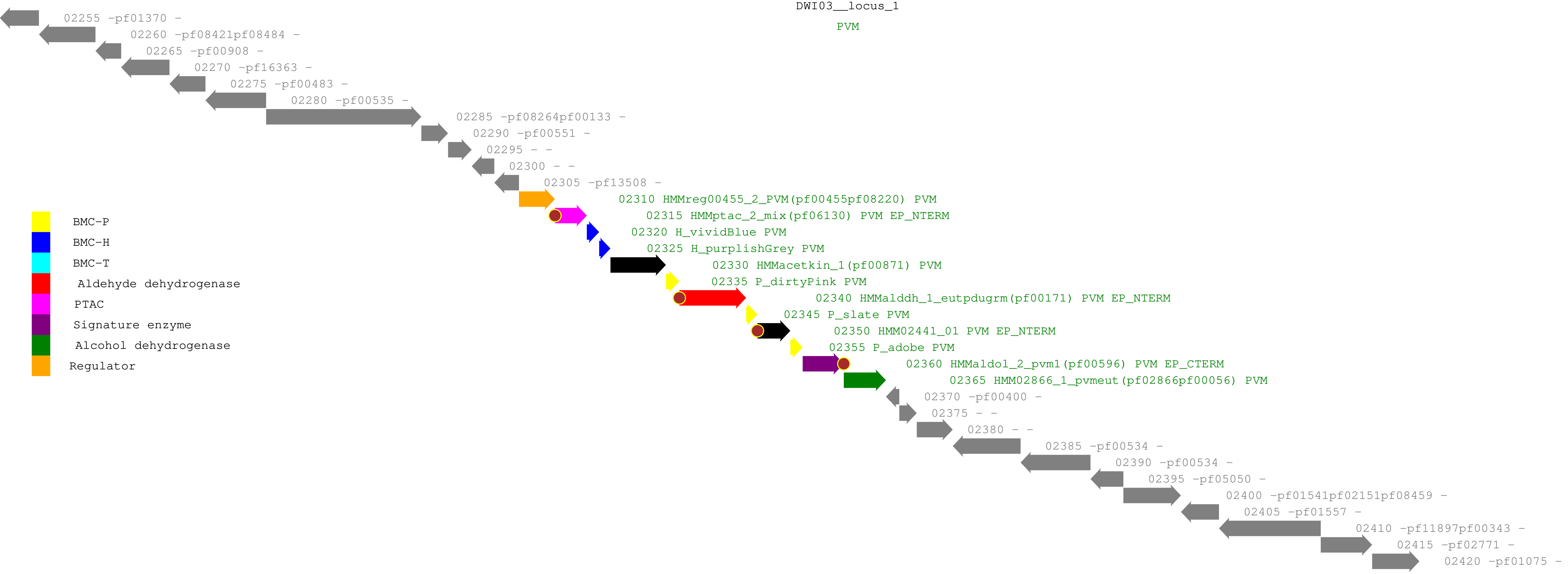

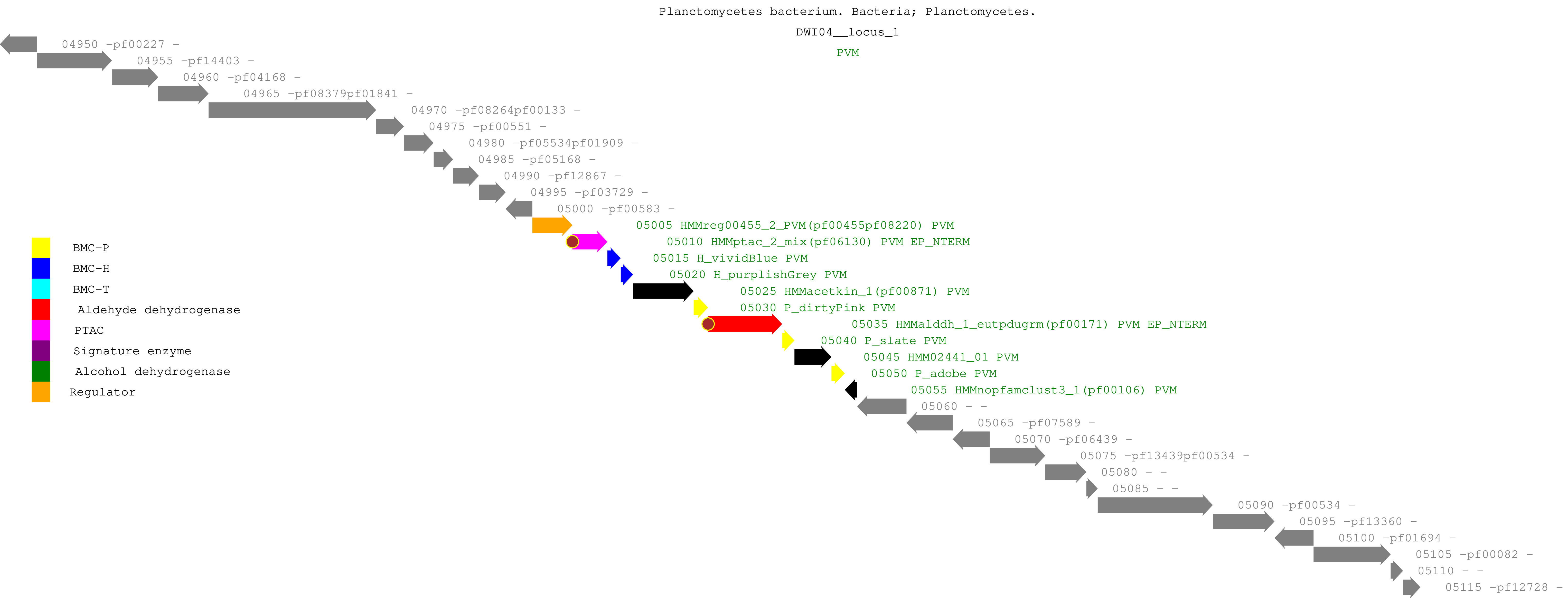
Planctomycetes bacterium. Bacteria; Planctomycetes.
DWI04__locus_1
PVM
04950 -pf00227 -
04955 -pf14403 -
04960 -pf04168 -
04965 -pf08379pf01841 -
04970 -pf08264pf00133 -
04975 -pf00551 -
04980 -pf05534pf01909 -
04985 -pf05168 -
04990 -pf12867 -
04995 -pf03729 -
05000 -pf00583 -
05005 HMMreg00455_2_PVM(pf00455pf08220) PVM
05010 HMMptac_2_mix(pf06130) PVM EP_NTERM
05015 H_vividBlue PVM
05020 H_purplishGrey PVM
05025 HMMacetkin_1(pf00871) PVM
05030 P_dirtyPink PVM
05035 HMMalddh_1_eutpdugrm(pf00171) PVM EP_NTERM
05040 P_slate PVM
05045 HMM02441_01 PVM
05050 P_adobe PVM
05055 HMMnopfamclust3_1(pf00106) PVM
05060 - -
05065 -pf07589 -
05070 -pf06439 -
05075 -pf13439pf00534 -
05080 - -
05085 - -
05090 -pf00534 -
05095 -pf13360 -
05100 -pf01694 -
05105 -pf00082 -
05110 - -
05115 -pf12728 -
BMC-P
BMC-H
BMC-T
Aldehyde dehydrogenase
PTAC
Signature enzyme
Alcohol dehydrogenase
Regulator

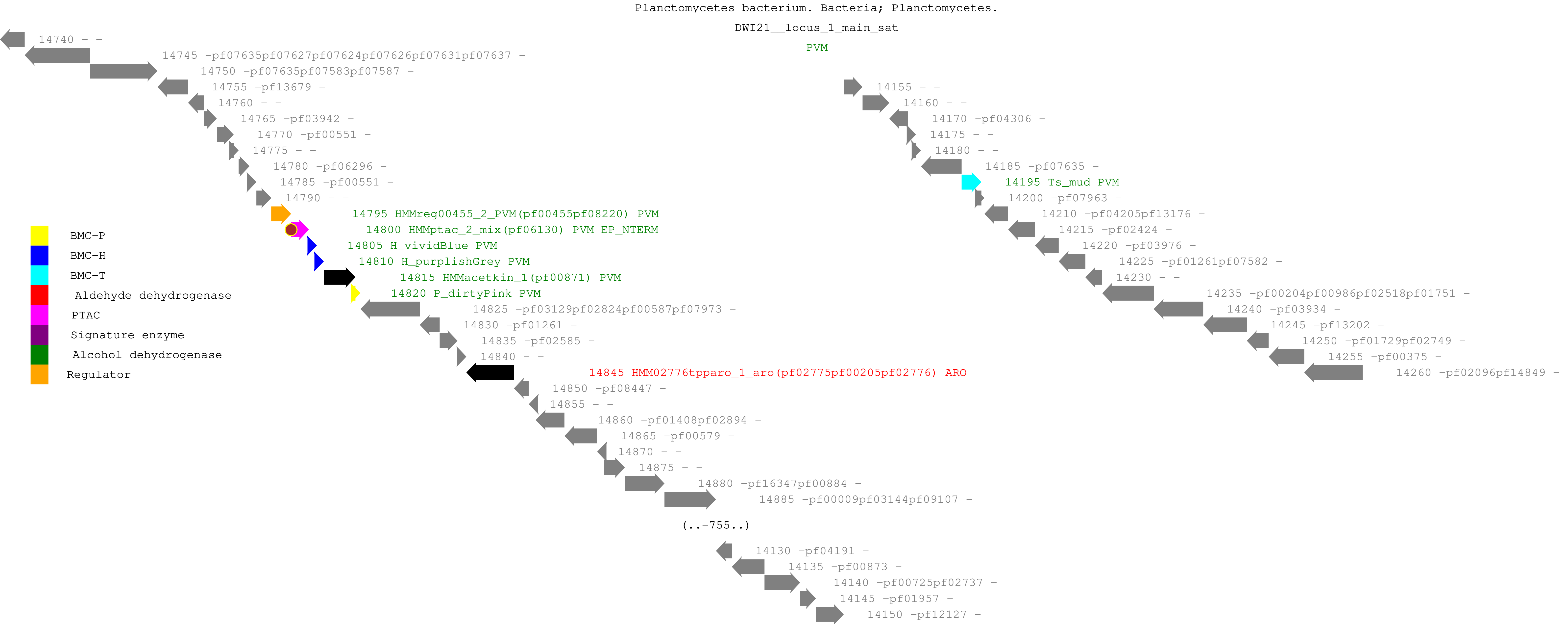
Planctomycetes bacterium. Bacteria; Planctomycetes.
DWI21__locus_1_main_sat
PVM
14740 - -
14745 -pf07635pf07627pf07624pf07626pf07631pf07637 -
14750 -pf07635pf07583pf07587 -
14755 -pf13679 -
14760 - -
14765 -pf03942 -
14770 -pf00551 -
14775 - -
14780 -pf06296 -
14785 -pf00551 -
14790 - -
14795 HMMreg00455_2_PVM(pf00455pf08220) PVM
14800 HMMptac_2_mix(pf06130) PVM EP_NTERM
14805 H_vividBlue PVM
14810 H_purplishGrey PVM
14815 HMMacetkin_1(pf00871) PVM
14820 P_dirtyPink PVM
14825 -pf03129pf02824pf00587pf07973 -
14830 -pf01261 -
14835 -pf02585 -
14840 - -
14845 HMM02776tpparo_1_aro(pf02775pf00205pf02776) ARO
14850 -pf08447 -
14855 - -
14860 -pf01408pf02894 -
14865 -pf00579 -
14870 - -
14875 - -
14880 -pf16347pf00884 -
14885 -pf00009pf03144pf09107 -
(..-755..)
14130 -pf04191 -
14135 -pf00873 -
14140 -pf00725pf02737 -
14145 -pf01957 -
14150 -pf12127 -
14155 - -
14160 - -
14170 -pf04306 -
14175 - -
14180 - -
14185 -pf07635 -
14195 Ts_mud PVM
14200 -pf07963 -
14210 -pf04205pf13176 -
14215 -pf02424 -
14220 -pf03976 -
14225 -pf01261pf07582 -
14230 - -
14235 -pf00204pf00986pf02518pf01751 -
14240 -pf03934 -
14245 -pf13202 -
14250 -pf01729pf02749 -
14255 -pf00375 -
14260 -pf02096pf14849 -
BMC-P
BMC-H
BMC-T
Aldehyde dehydrogenase
PTAC
Signature enzyme
Alcohol dehydrogenase
Regulator

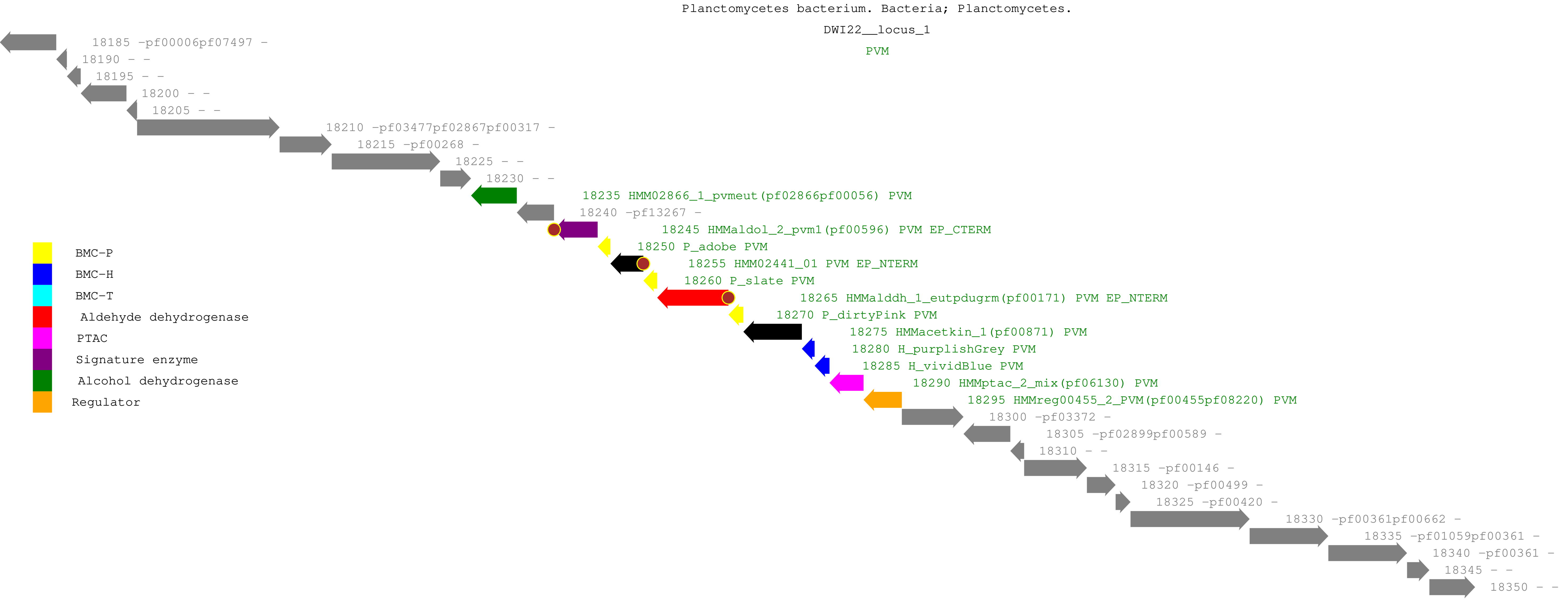
Planctomycetes bacterium. Bacteria; Planctomycetes.
DWI22__locus_1
PVM
18185 -pf00006pf07497 -
18190 - -
18195 - -
18200 - -
18205 - -
18210 -pf03477pf02867pf00317 -
18215 -pf00268 -
18225 - -
18230 - -
18235 HMM02866_1_pvmeut(pf02866pf00056) PVM
18240 -pf13267 -
18245 HMMaldol_2_pvm1(pf00596) PVM EP_CTERM
18250 P_adobe PVM
18255 HMM02441_01 PVM EP_NTERM
18260 P_slate PVM
18265 HMMaldh_1_eutpdugrm(pf00171) PVM EP_NTERM
18270 P_dirtyPink PVM
18275 HMMacetkin_1(pf00871) PVM
18280 H_purplishGrey PVM
18285 H_vividBlue PVM
18290 HMMptac_2_mix(pf06130) PVM
18295 HMMreg00455_2_PVM(pf00455pf08220) PVM
18300 -pf03372 -
18305 -pf02899pf00589 -
18310 - -
18315 -pf00146 -
18320 -pf00499 -
18325 -pf00420 -
18330 -pf00361pf00662 -
18335 -pf01059pf00361 -
18340 -pf00361 -
18345 - -
18350 - -
BMC-P
BMC-H
BMC-T
Aldehyde dehydrogenase
PTAC
Signature enzyme
Alcohol dehydrogenase
Regulator

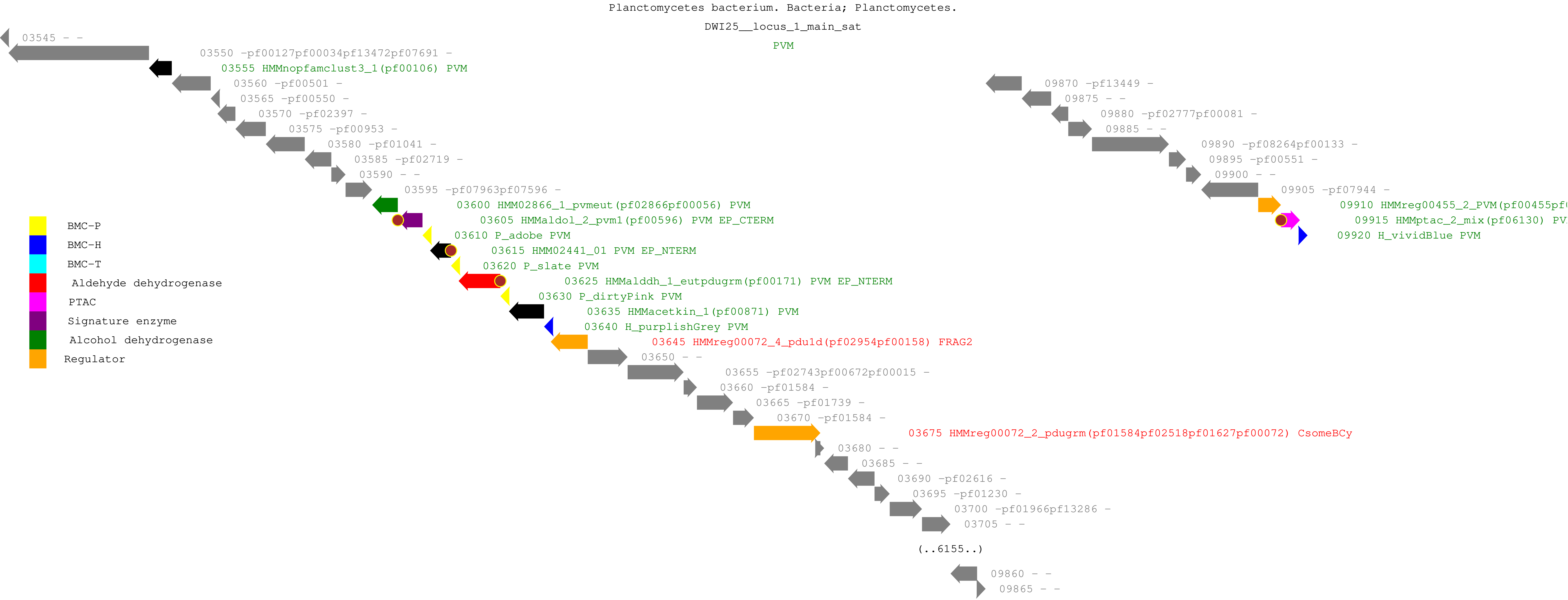
Planctomycetes bacterium. Bacteria; Planctomycetes.
DWI25__locus_1_main_sat
PVM
03545 - -
03550 -pf00127pf00034pf13472pf07691 -
03555 HMMnopfamclust3_1(pf00106) PVM
03560 -pf00501 -
03565 -pf00550 -
03570 -pf02397 -
03575 -pf00953 -
03580 -pf01041 -
03585 -pf02719 -
03590 - -
03595 -pf07963pf07596 -
03600 HMM02866_1_pvmeut(pf02866pf00056) PVM
03605 HMMaldol_2_pvm1(pf00596) PVM EP_CTERM
03610 P_adobe PVM
03615 HMM02441_01 PVM EP_NTERM
03620 P_slate PVM
03625 HMMaldh_1_eutpdugrm(pf00171) PVM EP_NTERM
03630 P_dirtyPink PVM
03635 HMMacetkin_1(pf00871) PVM
03640 H_purplishGrey PVM
03645 HMMreg00072_4_pdu1d(pf02954pf00158) FRAG2
03650 - -
03655 -pf02743pf00672pf00015 -
03660 -pf01584 -
03665 -pf01739 -
03670 -pf01584 -
03675 HMMreg00072_2_pdugrm(pf01584pf02518pf01627pf00072) CsomeBCy
03680 - -
03685 - -
03690 -pf02616 -
03695 -pf01230 -
03700 -pf01966pf13286 -
03705 - -
(..6155..)
09860 - -
09865 - -
09870 -pf13449 -
09875 - -
09880 -pf02777pf00081 -
09885 - -
09890 -pf08264pf00133 -
09895 -pf00551 -
09900 - -
09905 -pf07944 -
09910 HMMreg00455_2_PVM
09915 HMMptac_2_mix(pf06130)
09920 H_vividBlue PVM
BMC-P
BMC-H
BMC-T
Aldehyde dehydrogenase
PTAC
Signature enzyme
Alcohol dehydrogenase
Regulator

Planctomycetes bacterium. Bacteria; Planctomycetes.

DWI26__locus_1

PVM

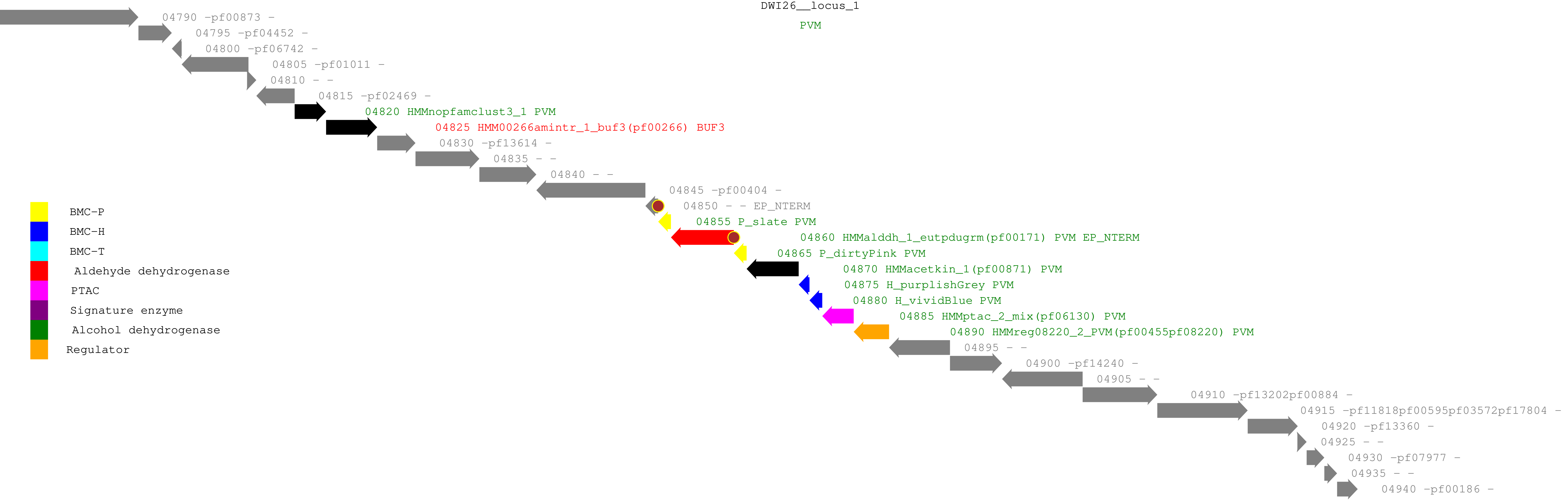

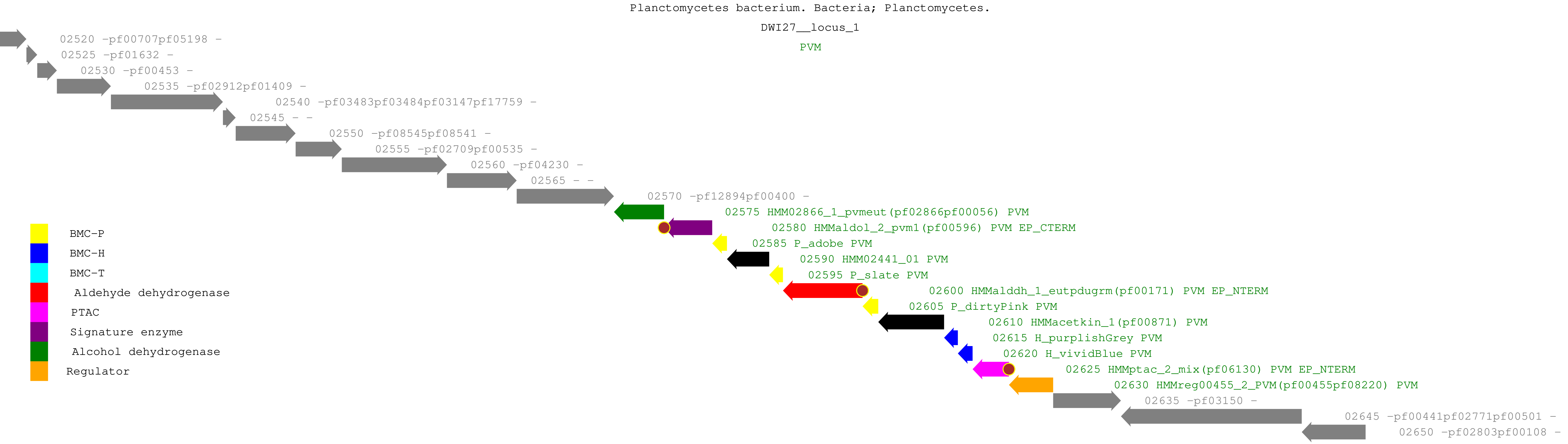
Planctomycetes bacterium. Bacteria; Planctomycetes.
DWI27__locus_1
PVM
02520 -pf00707pf05198 -
02525 -pf01632 -
02530 -pf00453 -
02535 -pf02912pf01409 -
02540 -pf03483pf03484pf03147pf17759 -
02545 - -
02550 -pf08545pf08541 -
02555 -pf02709pf00535 -
02560 -pf04230 -
02565 - -
02570 -pf12894pf00400 -
02575 HMM02866_1_pvmeut(pf02866pf00056) PVM
02580 HMMaldol_2_pvm1(pf00596) PVM EP_CTERM
02585 P_adobe PVM
02590 HMM02441_01 PVM
02595 P_slate PVM
02600 HMMalddh_1_eutpdugrm(pf00171) PVM EP_NTERM
02605 P_dirtyPink PVM
02610 HMMacetkin_1(pf00871) PVM
02615 H_purplishGrey PVM
02620 H_vividBlue PVM
02625 HMMptac_2_mix(pf06130) PVM EP_NTERM
02630 HMMreg00455_2_PVM(pf00455pf08220) PVM
02635 -pf03150 -
02645 -pf00441pf02771pf00501 -
02650 -pf02803pf00108 -
BMC-P
BMC-H
BMC-T
Aldehyde dehydrogenase
PTAC
Signature enzyme
Alcohol dehydrogenase
Regulator

Planctomycetes bacterium. Bacteria; Planctomycetes.

DWQ31__locus_1

PVM

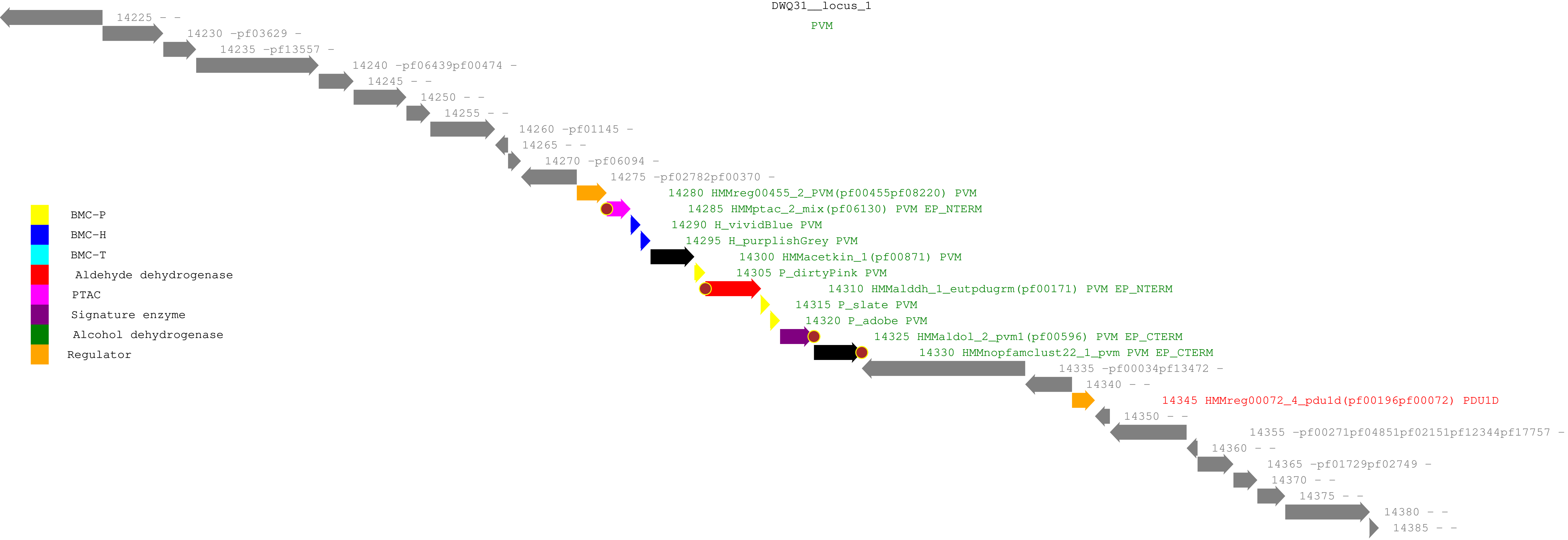

Planctomycetes bacterium. Bacteria; Planctomycetes.

DWQ37__locus_1

PVM

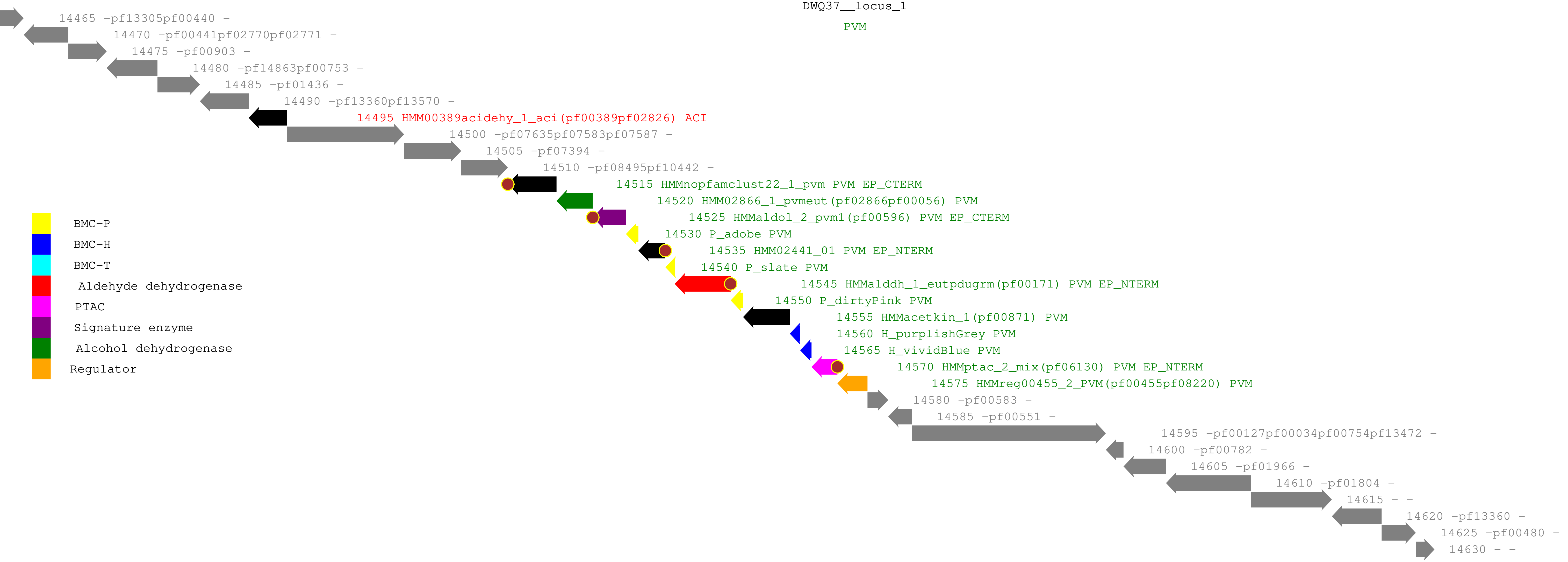

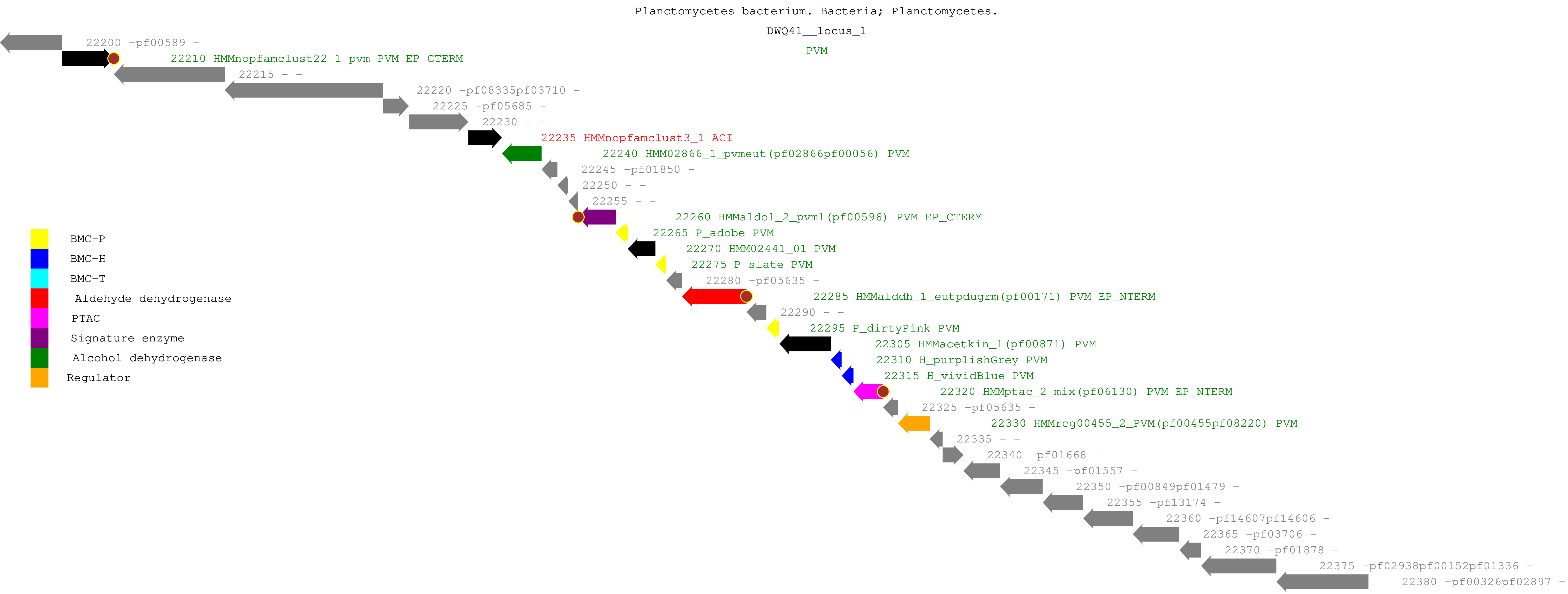

Planctomycetes bacterium. Bacteria; Planctomycetes.
DWQ41__locus_1
PVM
22200 -pf00589 -
22210 HMMnopfamclust22_1_pvm PVM EP_CTERM
22215 - -
22220 -pf08335pf03710 -
22225 -pf05685 -
22230 - -
22235 HMMnopfamclust3_1 ACI
22240 HMM02866_1_pvmeut(pf02866pf00056) PVM
22245 -pf01850 -
22250 - -
22255 - -
22260 HMMaldol_2_pvm1(pf00596) PVM EP_CTERM
22265 P_adobe PVM
22270 HMM02441_01 PVM
22275 P_slate PVM
22280 -pf05635 -
22285 HMMalddh_1_eutpdugrm(pf00171) PVM EP_NTERM
22290 - -
22295 P_dirtyPink PVM
22305 HMMacetkin_1(pf00871) PVM
22310 H_purplishGrey PVM
22315 H_vividBlue PVM
22320 HMMptac_2_mix(pf06130) PVM EP_NTERM
22325 -pf05635 -
22330 HMMreg00455_2_PVM(pf00455pf08220) PVM
22335 - -
22340 -pf01668 -
22345 -pf01557 -
22350 -pf00849pf01479 -
22355 -pf13174 -
22360 -pf14607pf14606 -
22365 -pf03706 -
22370 -pf01878 -
22375 -pf02938pf00152pf01336 -
22380 -pf00326pf02897 -
BMC-P
BMC-H
BMC-T
Aldehyde dehydrogenase
PTAC
Signature enzyme
Alcohol dehydrogenase
Regulator

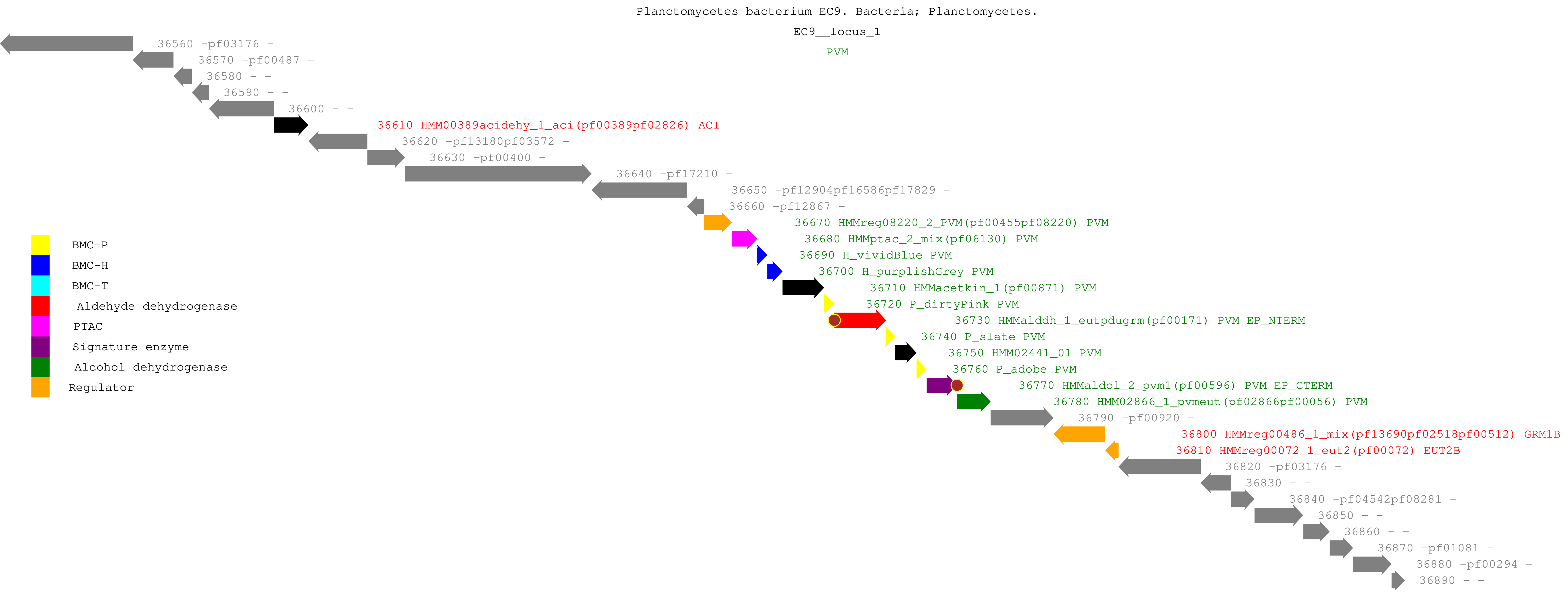

Planctomycetes bacterium EC9. Bacteria; Planctomycetes.
EC9__locus_1
PVM
36560 -pf03176 -
36570 -pf00487 -
36580 - -
36590 - -
36600 - -
36610 HMM00389acidehy_1_aci(pf00389pf02826) ACI
36620 -pf13180pf03572 -
36630 -pf00400 -
36640 -pf17210 -
36650 -pf12904pf16586pf17829 -
36660 -pf12867 -
36670 HMMreg08220_2_PVM(pf00455pf08220) PVM
36680 HMMptac_2_mix(pf06130) PVM
36690 H_vividBlue PVM
36700 H_purplishGrey PVM
36710 HMMacetkin_1(pf00871) PVM
36720 P_dirtyPink PVM
36730 HMMalddh_1_eutpdugrm(pf00171) PVM EP_NTERM
36740 P_slate PVM
36750 HMM02441_01 PVM
36760 P_adobe PVM
36770 HMMaldol_2_pvm1(pf00596) PVM EP_CTERM
36780 HMM02866_1_pvmeut(pf02866pf00056) PVM
36790 -pf00920 -
36800 HMMreg00486_1_mix(pf13690pf02518pf00512) GRM1B
36810 HMMreg00072_1_eut2(pf00072) EUT2B
36820 -pf03176 -
36830 - -
36840 -pf04542pf08281 -
36850 - -
36860 - -
36870 -pf01081 -
36880 -pf00294 -
36890 - -
BMC-P
BMC-H
BMC-T
Aldehyde dehydrogenase
PTAC
Signature enzyme
Alcohol dehydrogenase
Regulator

Verrucomicrobiaceae bacterium. Bacteria; Verrucomicrobia; Verrucomicrobiae; Verrucomicrobiales;

EHM17__locus_1_main_sat

PVM

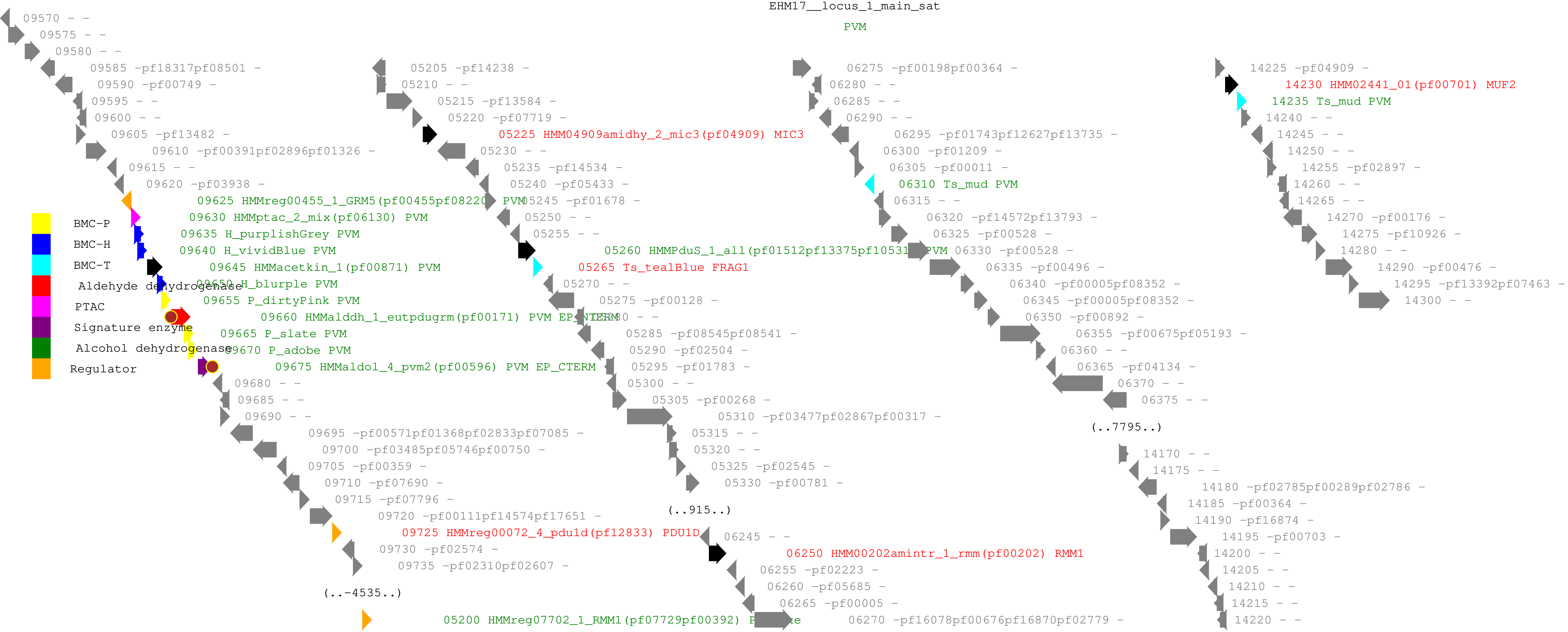

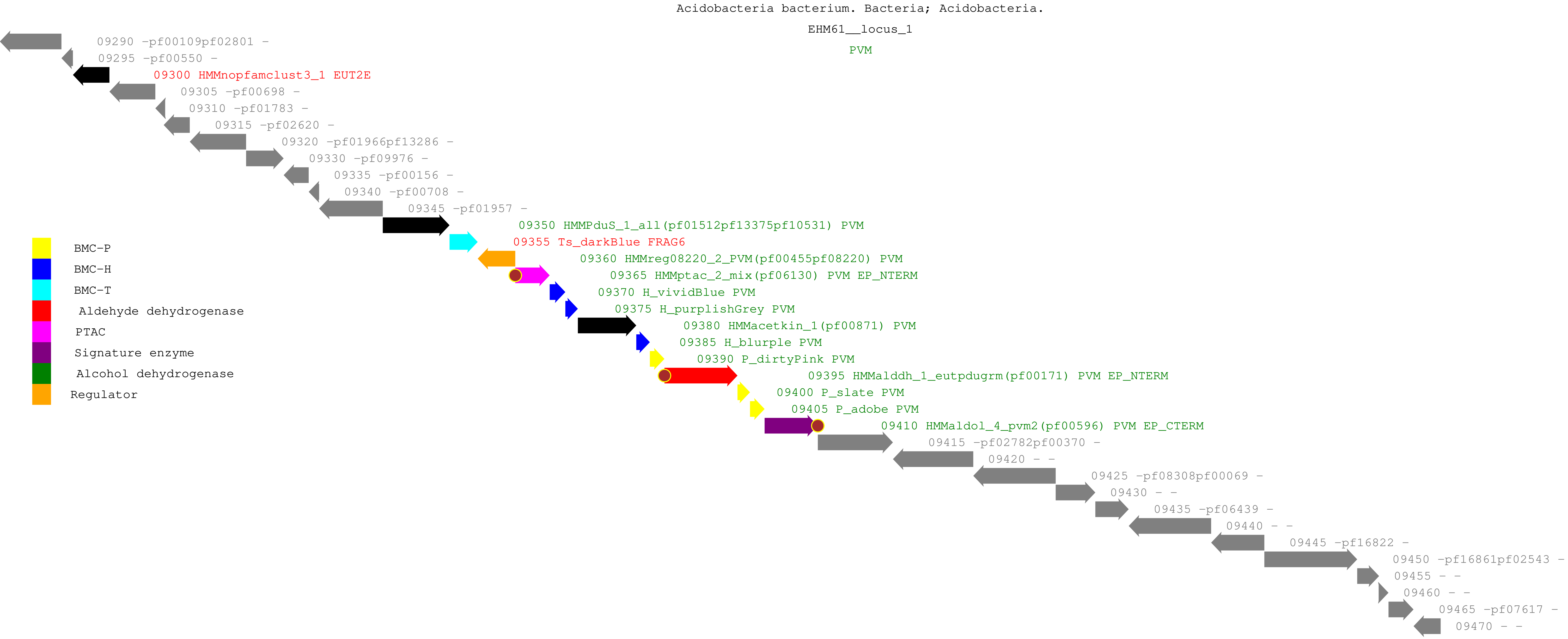

Acidobacteria bacterium. Bacteria; Acidobacteria.
EHM61__locus_1
PVM
09290 -pf00109pf02801 -
09295 -pf00550 -
09300 HMMnopfamclust3_1 EUT2E
09305 -pf00698 -
09310 -pf01783 -
09315 -pf02620 -
09320 -pf01966pf13286 -
09330 -pf09976 -
09335 -pf00156 -
09340 -pf00708 -
09345 -pf01957 -
09350 HMMPduS_1_all(pf01512pf13375pf10531) PVM
09355 Ts_darkBlue FRAG6
09360 HMMreg08220_2_PVM(pf00455pf08220) PVM
09365 HMMptac_2_mix(pf06130) PVM EP_NTERM
09370 H_vividBlue PVM
09375 H_purplishGrey PVM
09380 HMMacetkin_1(pf00871) PVM
09385 H_blurple PVM
09390 P_dirtyPink PVM
09395 HMMalddh_1_eutpdugrm(pf00171) PVM EP_NTERM
09400 P_slate PVM
09405 P_adobe PVM
09410 HMMaldol_4_pvm2(pf00596) PVM EP_CTERM
09415 -pf02782pf00370 -
09420 - -
09425 -pf08308pf00069 -
09430 - -
09435 -pf06439 -
09440 - -
09445 -pf16822 -
09450 -pf16861pf02543 -
09455 - -
09460 - -
09465 -pf07617 -
09470 - -
BMC-P
BMC-H
BMC-T
Aldehyde dehydrogenase
PTAC
Signature enzyme
Alcohol dehydrogenase
Regulator

Planctomycetes bacterium ElP. Bacteria; Planctomycetes.

ElP__locus_1

PVM

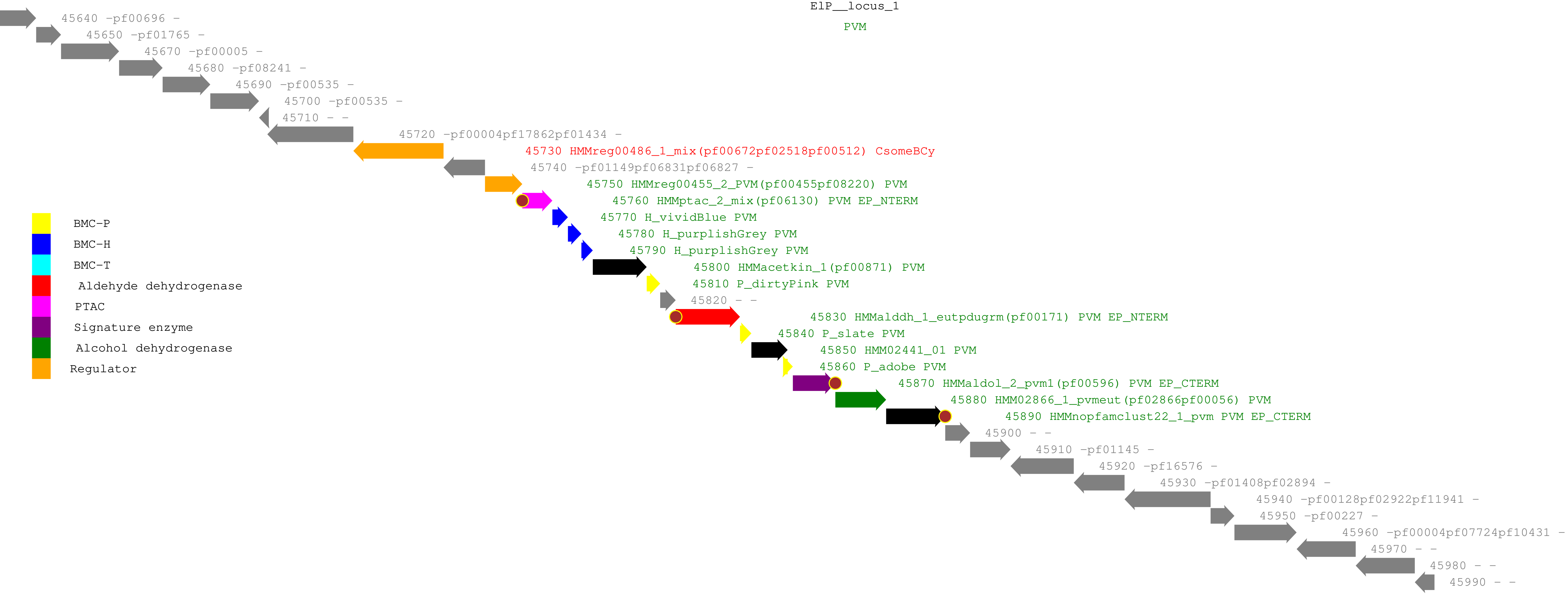

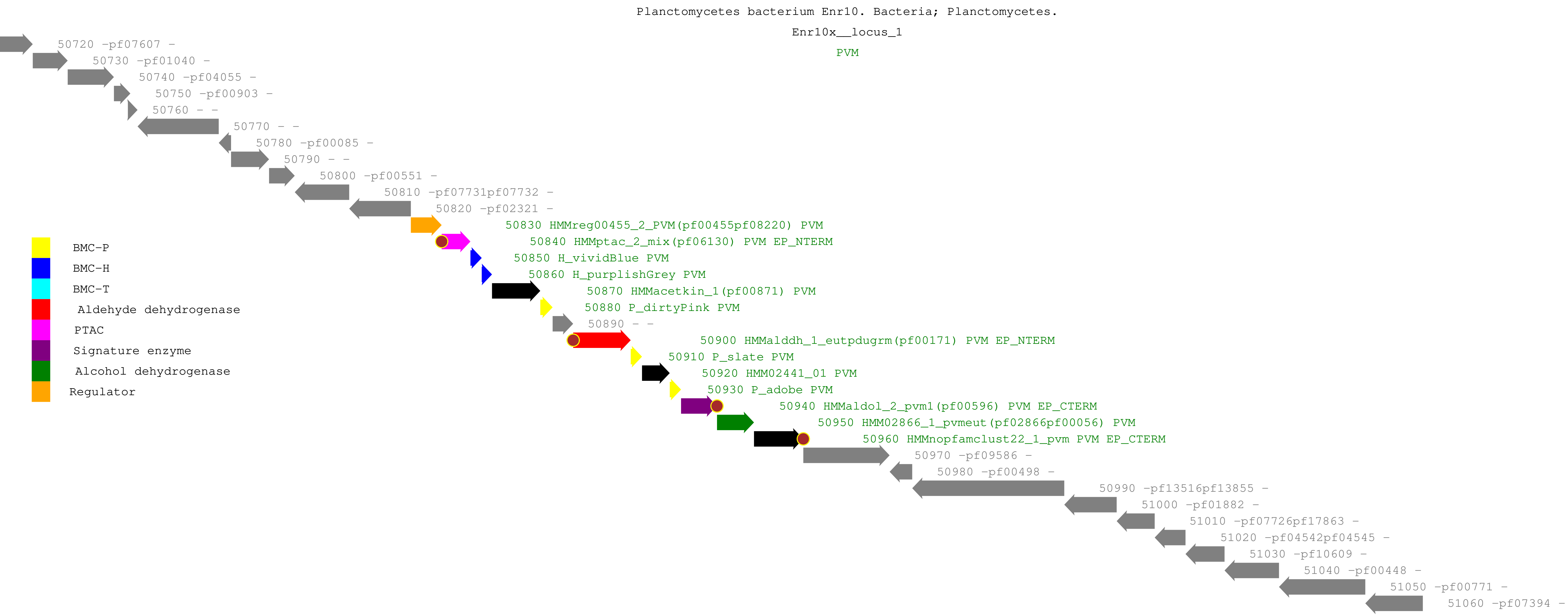

Planctomycetes bacterium Enr10. Bacteria; Planctomycetes.
Enr10x__locus_1
PVM
50720 -pf07607 -
50730 -pf01040 -
50740 -pf04055 -
50750 -pf00903 -
50760 - -
50770 - -
50780 -pf00085 -
50790 - -
50800 -pf00551 -
50810 -pf07731pf07732 -
50820 -pf02321 -
50830 HMMreg00455_2_PVM(pf00455pf08220) PVM
50840 HMMptac_2_mix(pf06130) PVM EP_NTERM
50850 H_vividBlue PVM
50860 H_purplishGrey PVM
50870 HMMacetkin_1(pf00871) PVM
50880 P_dirtyPink PVM
50890 - -
50900 HMMaldh_1_eutpdugrm(pf00171) PVM EP_NTERM
50910 P_slate PVM
50920 HMM02441_01 PVM
50930 P_adobe PVM
50940 HMMaldol_2_pvm1(pf00596) PVM EP_CTERM
50950 HMM02866_1_pvmeut(pf02866pf00056) PVM
50960 HMMnopfamclust22_1_pvm PVM EP_CTERM
50970 -pf09586 -
50980 -pf00498 -
50990 -pf13516pf13855 -
51000 -pf01882 -
51010 -pf07726pf17863 -
51020 -pf04542pf04545 -
51030 -pf10609 -
51040 -pf00448 -
51050 -pf00771 -
51060 -pf07394 -
BMC-P
BMC-H
BMC-T
Aldehyde dehydrogenase
PTAC
Signature enzyme
Alcohol dehydrogenase
Regulator

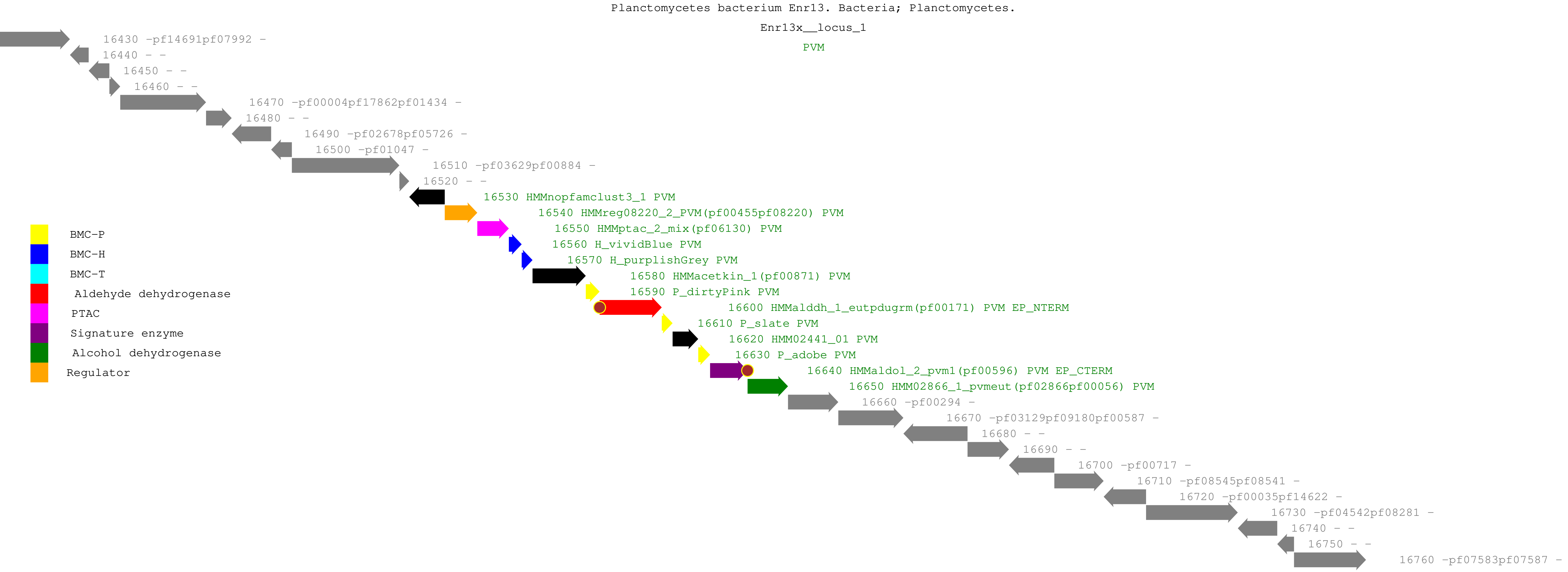

Planctomycetes bacterium Enr13. Bacteria; Planctomycetes.
Enr13x__locus_1
PVM
16430 -pf14691pf07992 -
16440 - -
16450 - -
16460 - -
16470 -pf00004pf17862pf01434 -
16480 - -
16490 -pf02678pf05726 -
16500 -pf01047 -
16510 -pf03629pf00884 -
16520 - -
16530 HMMnopfamclust3_1 PVM
16540 HMMreg08220_2_PVM(pf00455pf08220) PVM
16550 HMMptac_2_mix(pf06130) PVM
16560 H_vividBlue PVM
16570 H_purplishGrey PVM
16580 HMMacetkin_1(pf00871) PVM
16590 P_dirtyPink PVM
16600 HMMalddh_1_eutpdugrm(pf00171) PVM EP_NTERM
16610 P_slate PVM
16620 HMM02441_01 PVM
16630 P_adobe PVM
16640 HMMaldol_2_pvm1(pf00596) PVM EP_CTERM
16650 HMM02866_1_pvmeut(pf02866pf00056) PVM
16660 -pf00294 -
16670 -pf03129pf09180pf00587 -
16680 - -
16690 - -
16700 -pf00717 -
16710 -pf08545pf08541 -
16720 -pf00035pf14622 -
16730 -pf04542pf08281 -
16740 - -
16750 - -
16760 -pf07583pf07587 -
BMC-P
BMC-H
BMC-T
Aldehyde dehydrogenase
PTAC
Signature enzyme
Alcohol dehydrogenase
Regulator

Planctomycetes bacterium Enr17. Bacteria; Planctomycetes.

Enr17x__locus_1

PVM

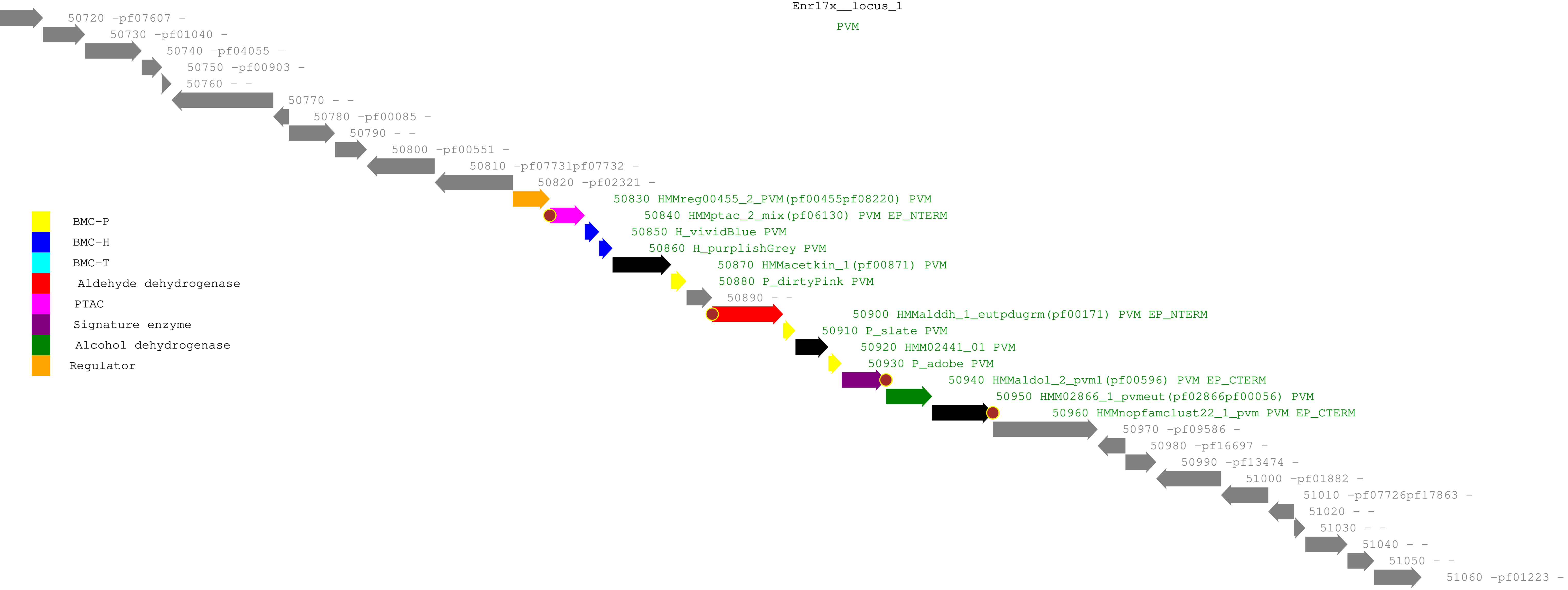

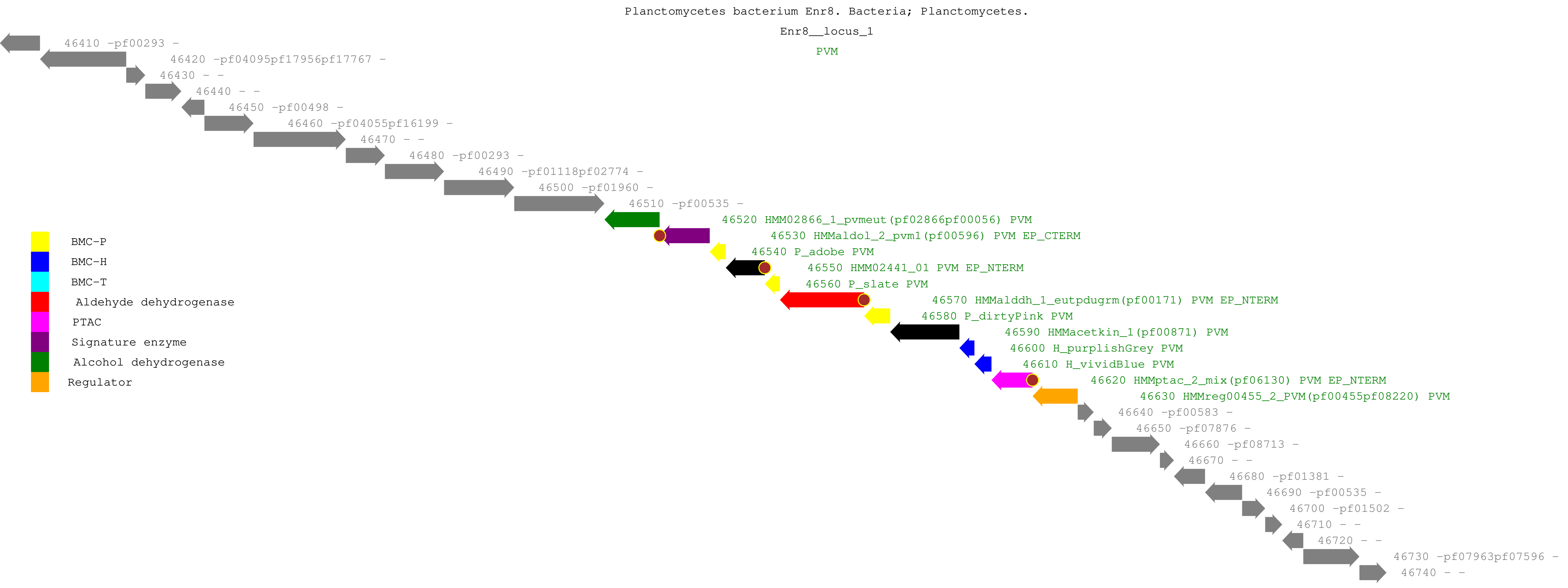
Planctomycetes bacterium Enr8. Bacteria; Planctomycetes.
Enr8__locus_1
PVM
46410 -pf00293 -
46420 -pf04095pf17956pf17767 -
46430 - -
46440 - -
46450 -pf00498 -
46460 -pf04055pf16199 -
46470 - -
46480 -pf00293 -
46490 -pf01118pf02774 -
46500 -pf01960 -
46510 -pf00535 -
46520 HMM02866_1_pvmeut(pf02866pf00056) PVM
46530 HMMaldol_2_pvm1(pf00596) PVM EP_CTERM
46540 P_adobe PVM
46550 HMM02441_01 PVM EP_NTERM
46560 P_slate PVM
46570 HMMalddh_1_eutpdugrm(pf00171) PVM EP_NTERM
46580 P_dirtyPink PVM
46590 HMMacetkin_1(pf00871) PVM
46600 H_purplishGrey PVM
46610 H_vividBlue PVM
46620 HMMptac_2_mix(pf06130) PVM EP_NTERM
46630 HMMreg00455_2_PVM(pf00455pf08220) PVM
46640 -pf00583 -
46650 -pf07876 -
46660 -pf08713 -
46670 - -
46680 -pf01381 -
46690 -pf00535 -
46700 -pf01502 -
46710 - -
46720 - -
46730 -pf07963pf07596 -
46740 - -
BMC-P
BMC-H
BMC-T
Aldehyde dehydrogenase
PTAC
Signature enzyme
Alcohol dehydrogenase
Regulator

Verrucomicrobia bacterium. Bacteria; Verrucomicrobia; unclassified Verrucomicrobia.

EPO07__locus_1_main_sat

PVM

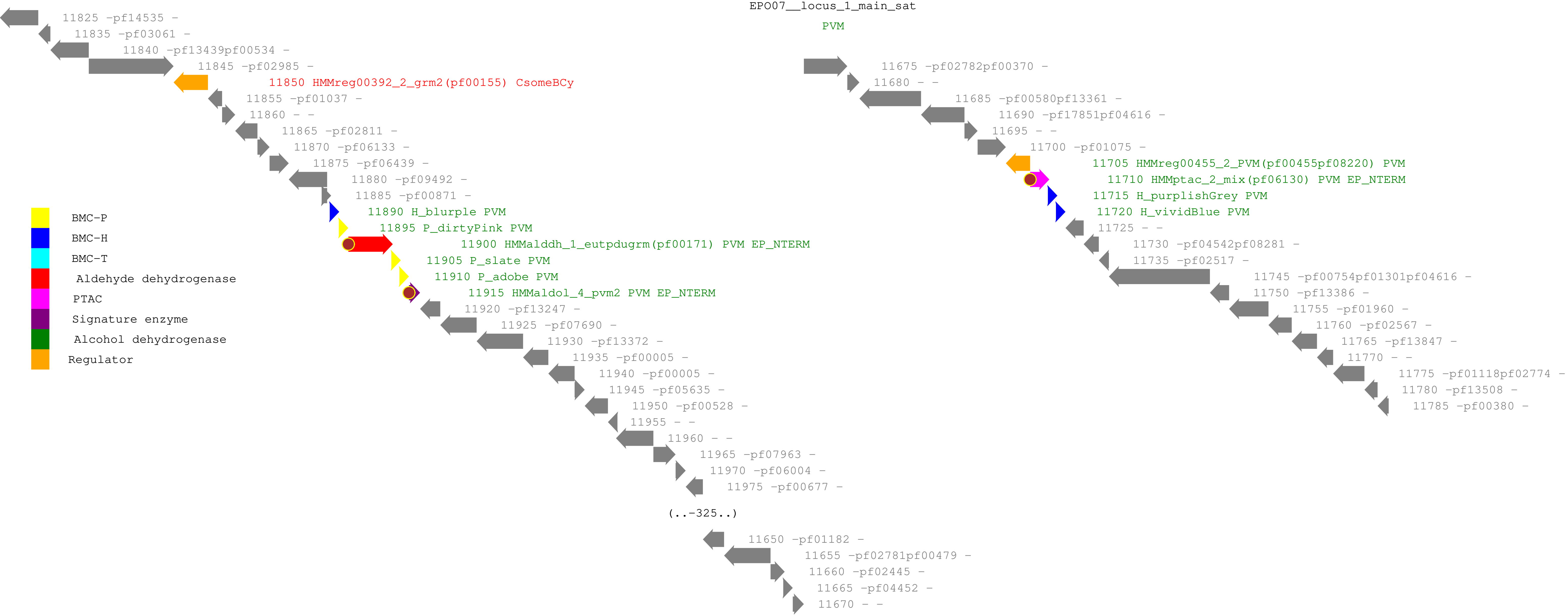

Planctomycetes bacterium ETA_A8. Bacteria; Planctomycetes.

ETAA8__locus_1

PVM

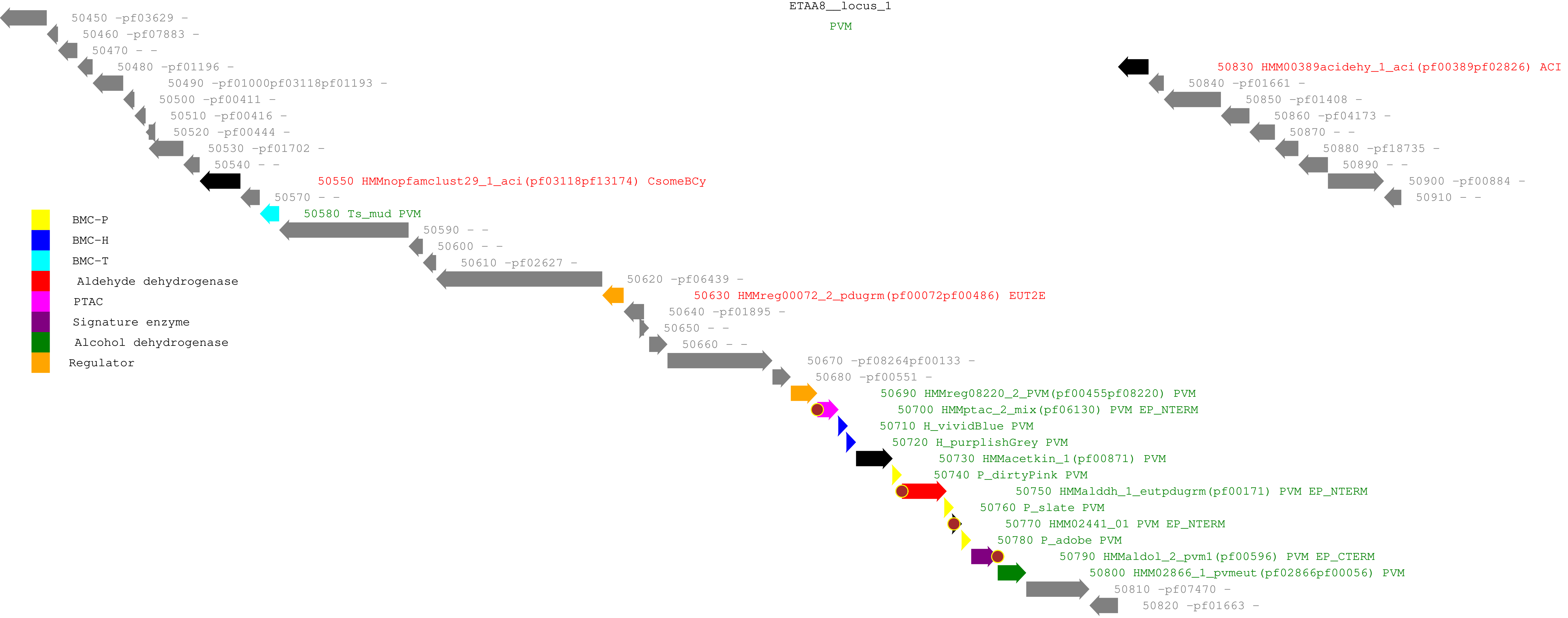

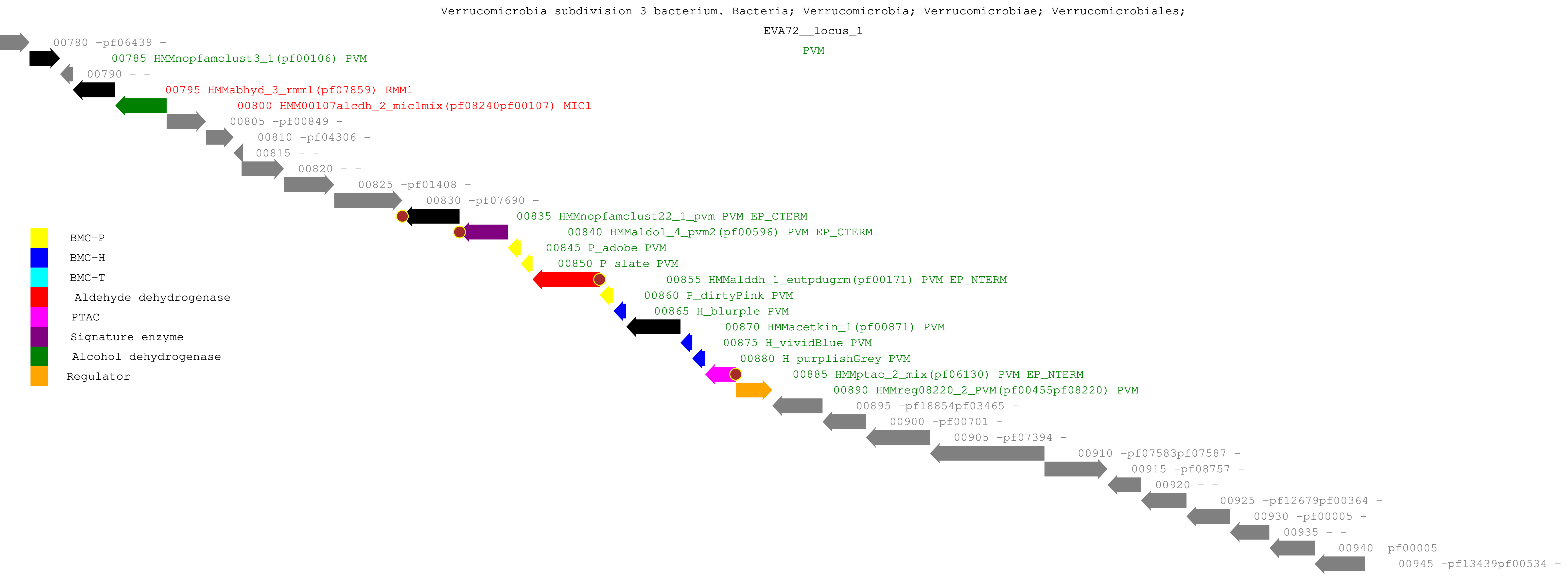
Verrucomicrobia subdivision 3 bacterium. Bacteria; Verrucomicrobia; Verrucomicrobiae; Verrucomicrobiales;
EVA72__locus_1
PVM
00780 -pf06439 -
00785 HMMnopfamclust3_1(pf00106) PVM
00790 - -
00795 HMMabhyd_3_rmm1(pf07859) RMM1
00800 HMM00107alcdh_2_mic1mix(pf08240pf00107) MIC1
00805 -pf00849 -
00810 -pf04306 -
00815 - -
00820 - -
00825 -pf01408 -
00830 -pf07690 -
00835 HMMnopfamclust22_1_pvm PVM EP_CTERM
00840 HMMaldol_4_pvm2(pf00596) PVM EP_CTERM
00845 P_adobe PVM
00850 P_slate PVM
00855 HMMalddh_1_eutpdugrm(pf00171) PVM EP_NTERM
00860 P_dirtyPink PVM
00865 H_blurple PVM
00870 HMMacetkin_1(pf00871) PVM
00875 H_vividBlue PVM
00880 H_purplishGrey PVM
00885 HMMptac_2_mix(pf06130) PVM EP_NTERM
00890 HMMreg08220_2_PVM(pf00455pf08220) PVM
00895 -pf18854pf03465 -
00900 -pf00701 -
00905 -pf07394 -
00910 -pf07583pf07587 -
00915 -pf08757 -
00920 - -
00925 -pf12679pf00364 -
00930 -pf00005 -
00935 - -
00940 -pf00005 -
00945 -pf13439pf00534 -
BMC-P
BMC-H
BMC-T
Aldehyde dehydrogenase
PTAC
Signature enzyme
Alcohol dehydrogenase
Regulator

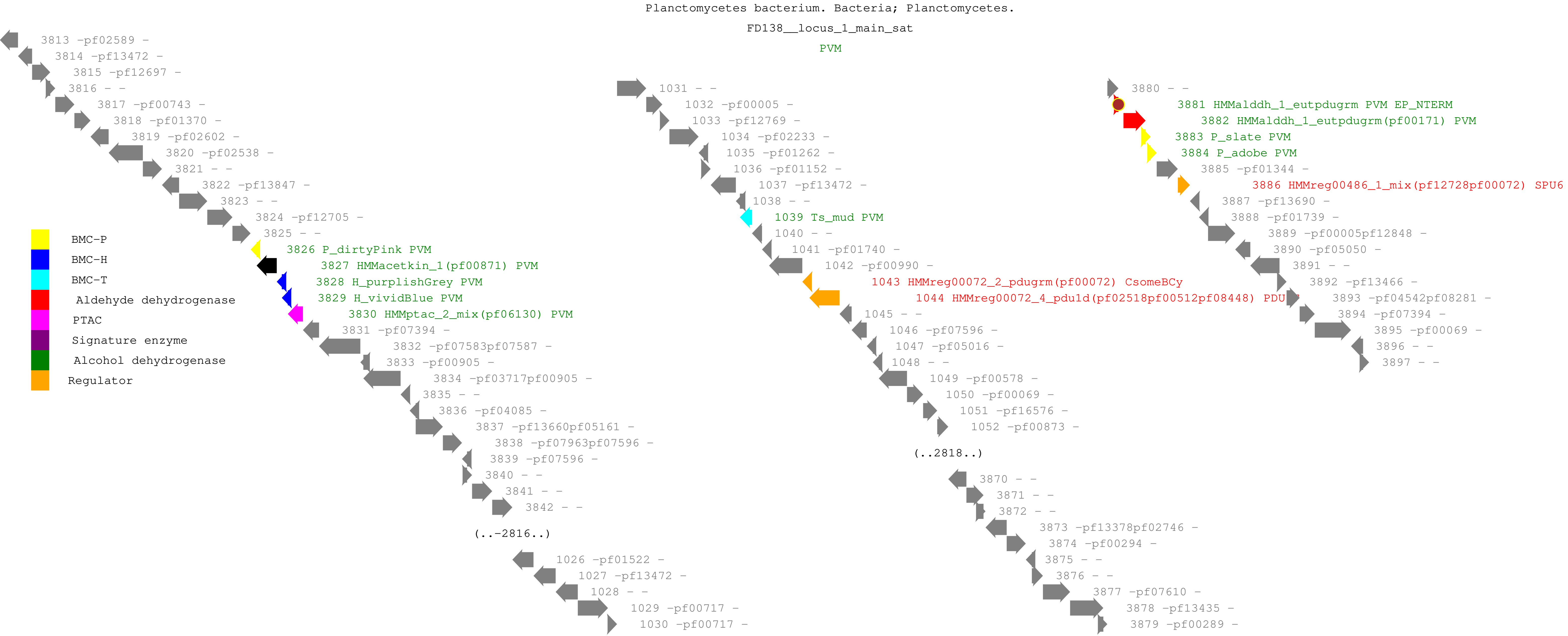
Planctomycetes bacterium. Bacteria; Planctomycetes.
FD138__locus_1_main_sat
PVM
3813 -pf02589 -
3814 -pf13472 -
3815 -pf12697 -
3816 - -
3817 -pf00743 -
3818 -pf01370 -
3819 -pf02602 -
3820 -pf02538 -
3821 - -
3822 -pf13847 -
3823 - -
3824 -pf12705 -
3825 - -
3826 P_dirtyPink PVM
3827 HMMacetkin_1(pf00871) PVM
3828 H_purplishGrey PVM
3829 H_vividBlue PVM
3830 HMMptac_2_mix(pf06130) PVM
3831 -pf07394 -
3832 -pf07583pf07587 -
3833 -pf00905 -
3834 -pf03717pf00905 -
3835 - -
3836 -pf04085 -
3837 -pf13660pf05161 -
3838 -pf07963pf07596 -
3839 -pf07596 -
3840 - -
3841 - -
3842 - -
(..-2816..)
1026 -pf01522 -
1027 -pf13472 -
1028 - -
1029 -pf00717 -
1030 -pf00717 -
1031 - -
1032 -pf00005 -
1033 -pf12769 -
1034 -pf02233 -
1035 -pf01262 -
1036 -pf01152 -
1037 -pf13472 -
1038 - -
1039 Ts_mud PVM
1040 - -
1041 -pf01740 -
1042 -pf00990 -
1043 HMMreg00072_2_pdugrm(pf00072) CsomeBCy
1044 HMMreg00072_4_pdu1d(pf02518pf00512pf08448) PDU
1045 - -
1046 -pf07596 -
1047 -pf05016 -
1048 - -
1049 -pf00578 -
1050 -pf00069 -
1051 -pf16576 -
1052 -pf00873 -
(..2818..)
3870 - -
3871 - -
3872 - -
3873 -pf13378pf02746 -
3874 -pf00294 -
3875 - -
3876 - -
3877 -pf07610 -
3878 -pf13435 -
3879 -pf00289 -
3880 - -
3881 HMMalddh_1_eutpdugrm PVM EP_NTERM
3882 HMMalddh_1_eutpdugrm(pf00171) PVM
3883 P_slate PVM
3884 P_adobe PVM
3885 -pf01344 -
3886 HMMreg00486_1_mix(pf12728pf00072) SPU6
3887 -pf13690 -
3888 -pf01739 -
3889 -pf00005pf12848 -
3890 -pf05050 -
3891 - -
3892 -pf13466 -
3893 -pf04542pf08281 -
3894 -pf07394 -
3895 -pf00069 -
3896 - -
3897 - -
BMC-P
BMC-H
BMC-T
Aldehyde dehydrogenase
PTAC
Signature enzyme
Alcohol dehydrogenase
Regulator

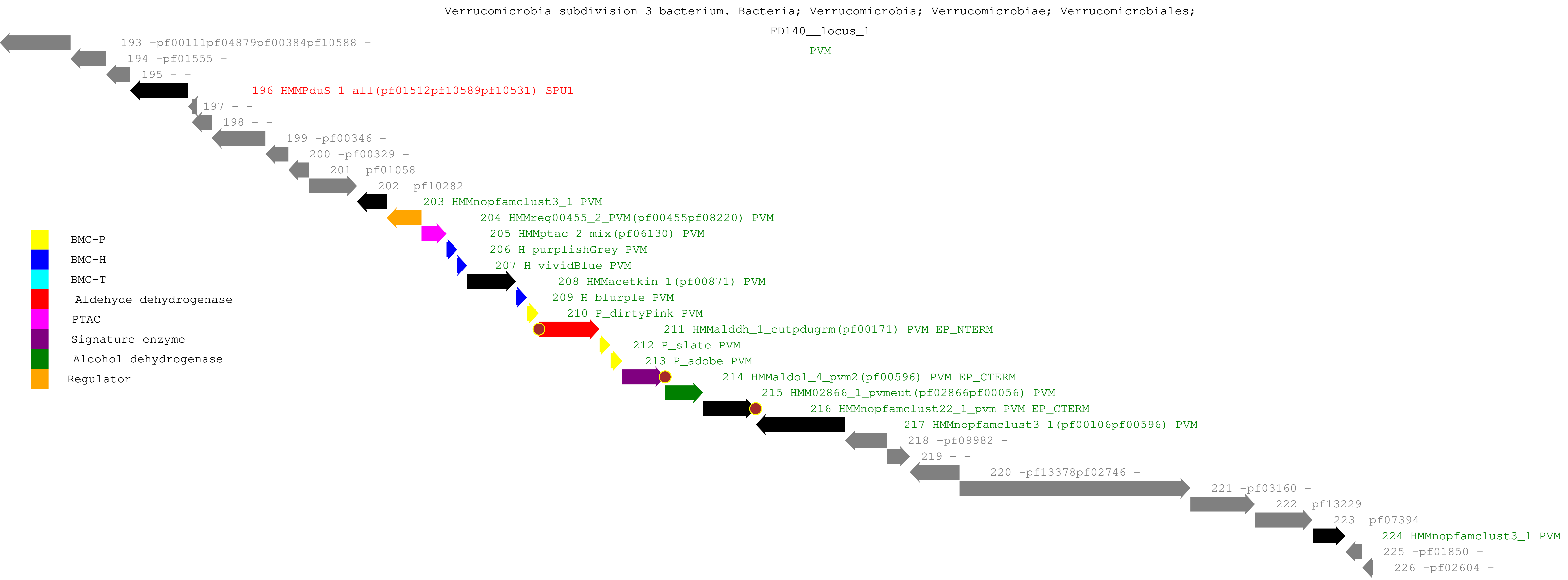

Verrucomicrobia subdivision 3 bacterium. Bacteria; Verrucomicrobia; Verrucomicrobiae; Verrucomicrobiales;
FD140__locus_1
PVM
193 -pf00111pf04879pf00384pf10588 -
194 -pf01555 -
195 - -
196 HMMPduS_1_all(pf01512pf10589pf10531) SPU1
197 - -
198 - -
199 -pf00346 -
200 -pf00329 -
201 -pf01058 -
202 -pf10282 -
203 HMMnopfamclust3_1 PVM
204 HMMreg00455_2_PVM(pf00455pf08220) PVM
205 HMMptac_2_mix(pf06130) PVM
206 H_purplishGrey PVM
207 H_vividBlue PVM
208 HMMacetkin_1(pf00871) PVM
209 H_blurple PVM
210 P_dirtyPink PVM
211 HMMalddh_1_eutpdugrm(pf00171) PVM EP_NTERM
212 P_slate PVM
213 P_adobe PVM
214 HMMaldol_4_pvm2(pf00596) PVM EP_CTERM
215 HMM02866_1_pvmeut(pf02866pf00056) PVM
216 HMMnopfamclust22_1_pvm PVM EP_CTERM
217 HMMnopfamclust3_1(pf00106pf00596) PVM
218 -pf09982 -
219 - -
220 -pf13378pf02746 -
221 -pf03160 -
222 -pf13229 -
223 -pf07394 -
224 HMMnopfamclust3_1 PVM
225 -pf01850 -
226 -pf02604 -
BMC-P
BMC-H
BMC-T
Aldehyde dehydrogenase
PTAC
Signature enzyme
Alcohol dehydrogenase
Regulator

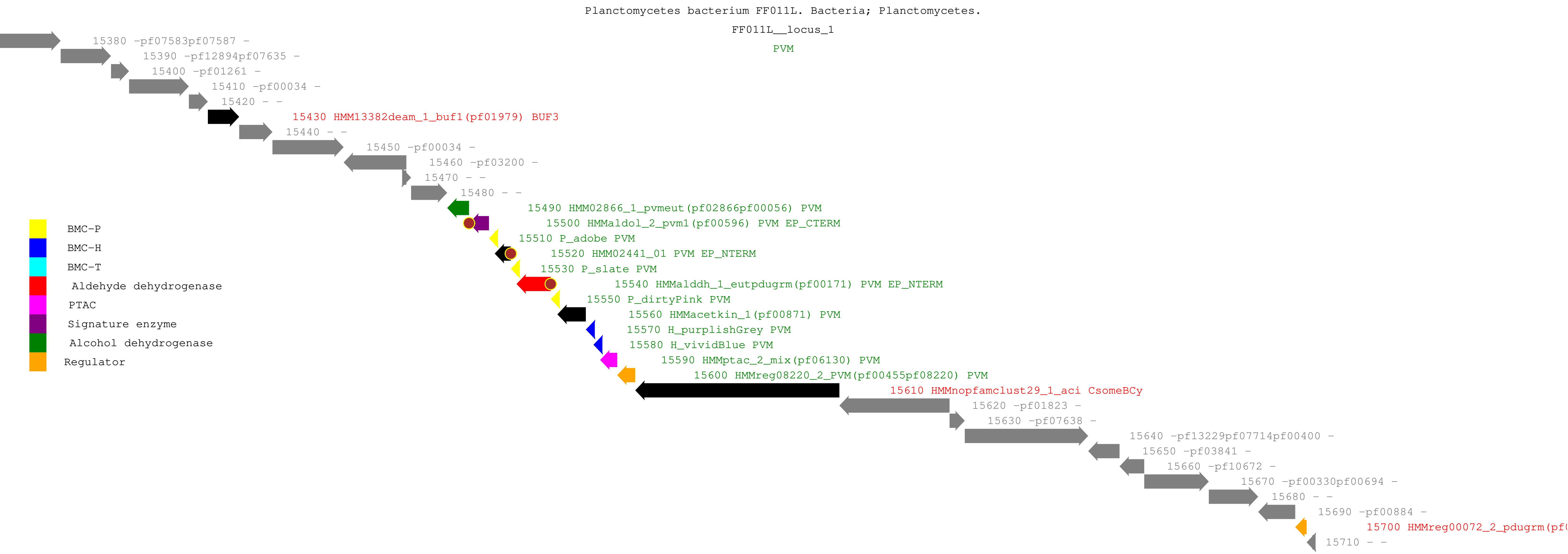

Planctomycetes bacterium FF011L. Bacteria; Planctomycetes.
FF011L__locus_1
PVM
15380 -pf07583pf07587 -
15390 -pf12894pf07635 -
15400 -pf01261 -
15410 -pf00034 -
15420 - -
15430 HMM13382deam_1_buf1(pf01979) BUF3
15440 - -
15450 -pf00034 -
15460 -pf03200 -
15470 - -
15480 - -
15490 HMM02866_1_pvmeut(pf02866pf00056) PVM
15500 HMMaldol_2_pvm1(pf00596) PVM EP_CTERM
15510 P_adobe PVM
15520 HMM02441_01 PVM EP_NTERM
15530 P_slate PVM
15540 HMMalddh_1_eutpdugrm(pf00171) PVM EP_NTERM
15550 P_dirtyPink PVM
15560 HMMacetkin_1(pf00871) PVM
15570 H_purplishGrey PVM
15580 H_vividBlue PVM
15590 HMMptac_2_mix(pf06130) PVM
15600 HMMreg08220_2_PVM(pf00455pf08220) PVM
15610 HMMnopfamclust29_1_aci CsomeBCy
15620 -pf01823 -
15630 -pf07638 -
15640 -pf13229pf07714pf00400 -
15650 -pf03841 -
15660 -pf10672 -
15670 -pf00330pf00694 -
15680 - -
15690 -pf00884 -
15700 HMMreg00072_2_pdugrm(pf
15710 - -
BMC-P
BMC-H
BMC-T
Aldehyde dehydrogenase
PTAC
Signature enzyme
Alcohol dehydrogenase
Regulator

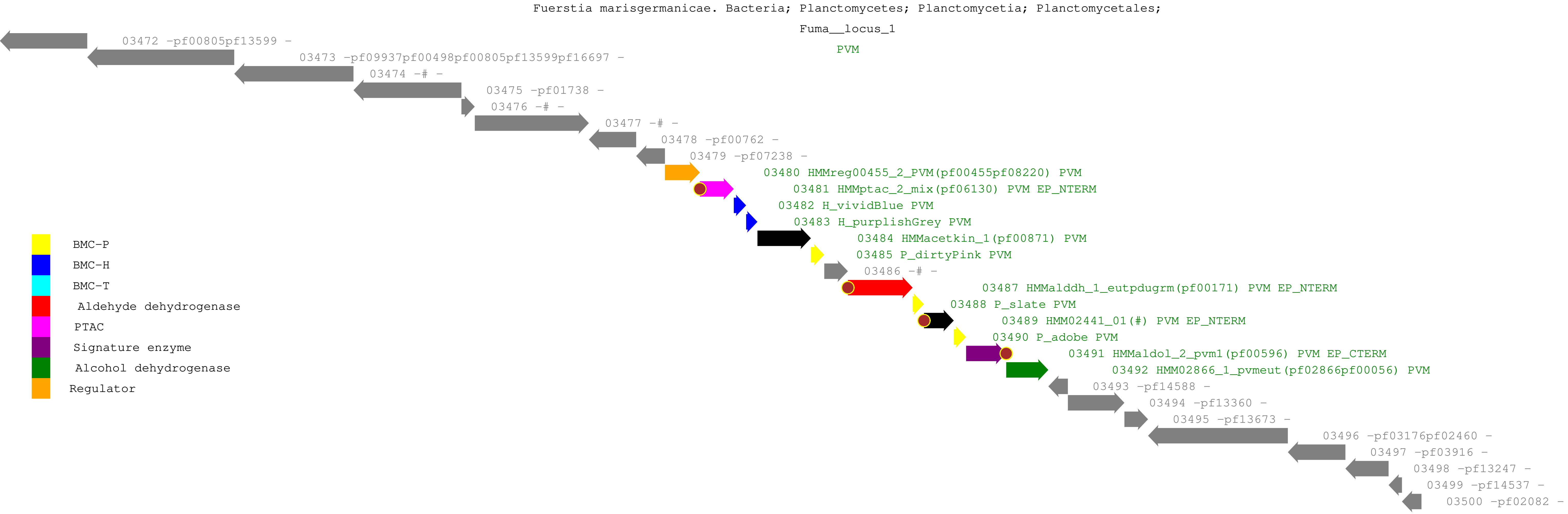

Fuerstia marisgermanicae. Bacteria; Planctomycetes; Planctomycetia; Planctomycetales;
Fuma__locus_1
PVM
03472 -pf00805pf13599 -
03473 -pf09937pf00498pf00805pf13599pf16697 -
03474 -# -
03475 -pf01738 -
03476 -# -
03477 -# -
03478 -pf00762 -
03479 -pf07238 -
03480 HMMreg00455_2_PVM(pf00455pf08220) PVM
03481 HMMptac_2_mix(pf06130) PVM EP_NTERM
03482 H_vividBlue PVM
03483 H_purplishGrey PVM
03484 HMMacetkin_1(pf00871) PVM
03485 P_dirtyPink PVM
03486 -# -
03487 HMMaldh_1_eutpdugrm(pf00171) PVM EP_NTERM
03488 P_slate PVM
03489 HMM02441_01(#) PVM EP_NTERM
03490 P_adobe PVM
03491 HMMaldol_2_pvm1(pf00596) PVM EP_CTERM
03492 HMM02866_1_pvmeut(pf02866pf00056) PVM
03493 -pf14588 -
03494 -pf13360 -
03495 -pf13673 -
03496 -pf03176pf02460 -
03497 -pf03916 -
03498 -pf13247 -
03499 -pf14537 -
03500 -pf02082 -
BMC-P
BMC-H
BMC-T
Aldehyde dehydrogenase
PTAC
Signature enzyme
Alcohol dehydrogenase
Regulator

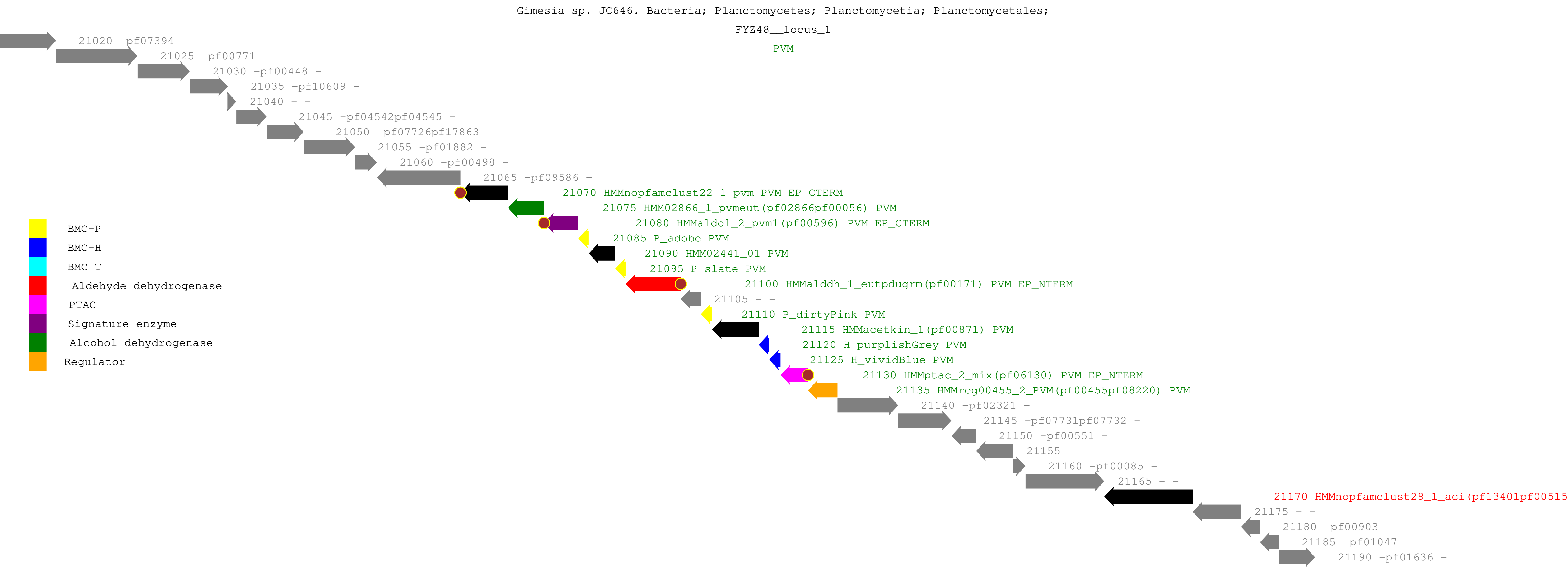

Gimesia sp. JC646. Bacteria; Planctomycetes; Planctomycetia; Planctomycetales;
FYZ48__locus_1
PVM
21020 -pf07394 -
21025 -pf00771 -
21030 -pf00448 -
21035 -pf10609 -
21040 - -
21045 -pf04542pf04545 -
21050 -pf07726pf17863 -
21055 -pf01882 -
21060 -pf00498 -
21065 -pf09586 -
21070 HMMnopfamclust22_1_pvm PVM EP_CTERM
21075 HMM02866_1_pvmeut(pf02866pf00056) PVM
21080 HMMaldol_2_pvm1(pf00596) PVM EP_CTERM
21085 P_adobe PVM
21090 HMM02441_01 PVM
21095 P_slate PVM
21100 HMMalddh_1_eutpdugrm(pf00171) PVM EP_NTERM
21105 - -
21110 P_dirtyPink PVM
21115 HMMacetkin_1(pf00871) PVM
21120 H_purplishGrey PVM
21125 H_vividBlue PVM
21130 HMMptac_2_mix(pf06130) PVM EP_NTERM
21135 HMMreg00455_2_PVM(pf00455pf08220) PVM
21140 -pf02321 -
21145 -pf07731pf07732 -
21150 -pf00551 -
21155 - -
21160 -pf00085 -
21165 - -
21170 HMMnopfamclust29_1_aci(pf13401pf00515
21175 - -
21180 -pf00903 -
21185 -pf01047 -
21190 -pf01636 -
BMC-P
BMC-H
BMC-T
Aldehyde dehydrogenase
PTAC
Signature enzyme
Alcohol dehydrogenase
Regulator

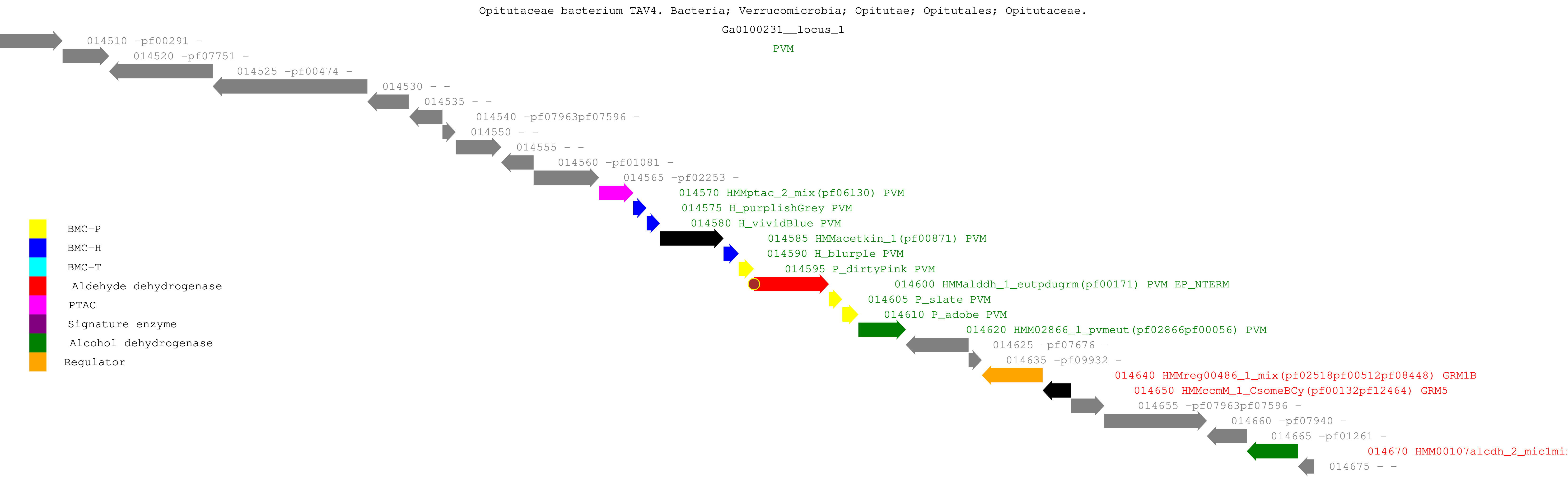
Opitutaceae bacterium TAV4. Bacteria; Verrucomicrobia; Opitutae; Opitutales; Opitutaceae.
Ga0100231__locus_1
PVM
014510 -pf00291 -
014520 -pf07751 -
014525 -pf00474 -
014530 - -
014535 - -
014540 -pf07963pf07596 -
014550 - -
014555 - -
014560 -pf01081 -
014565 -pf02253 -
014570 HMMptac_2_mix(pf06130) PVM
014575 H_purplishGrey PVM
014580 H_vividBlue PVM
014585 HMMacetkin_1(pf00871) PVM
014590 H_blurple PVM
014595 P_dirtyPink PVM
014600 HMMalddh_1_eutpdugrm(pf00171) PVM EP_NTERM
014605 P_slate PVM
014610 P_adobe PVM
014620 HMM02866_1_pvmeut(pf02866pf00056) PVM
014625 -pf07676 -
014635 -pf09932 -
014640 HMMreg00486_1_mix(pf02518pf00512pf08448) GRM1B
014650 HMMccmM_1_CsomeBCy(pf00132pf12464) GRM5
014655 -pf07963pf07596 -
014660 -pf07940 -
014665 -pf01261 -
014670 HMM00107alcdh_2_mic1mi
014675 - -
BMC-P
BMC-H
BMC-T
Aldehyde dehydrogenase
PTAC
Signature enzyme
Alcohol dehydrogenase
Regulator

mine drainage metagenome. unclassified sequences; metagenomes; ecological metagenomes.

GALL__locus_1

PVM

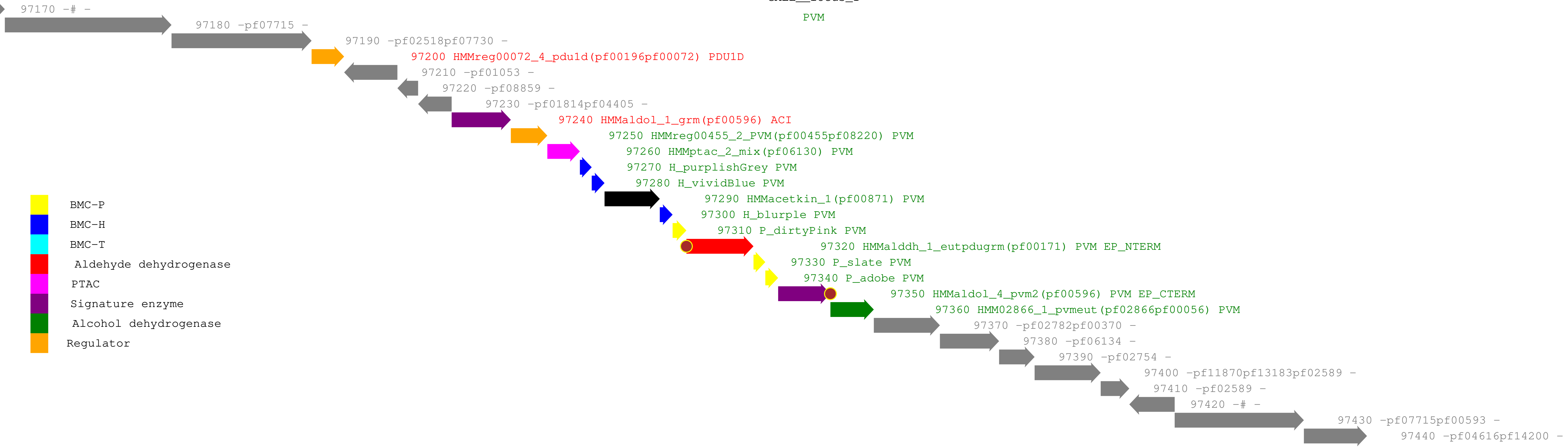

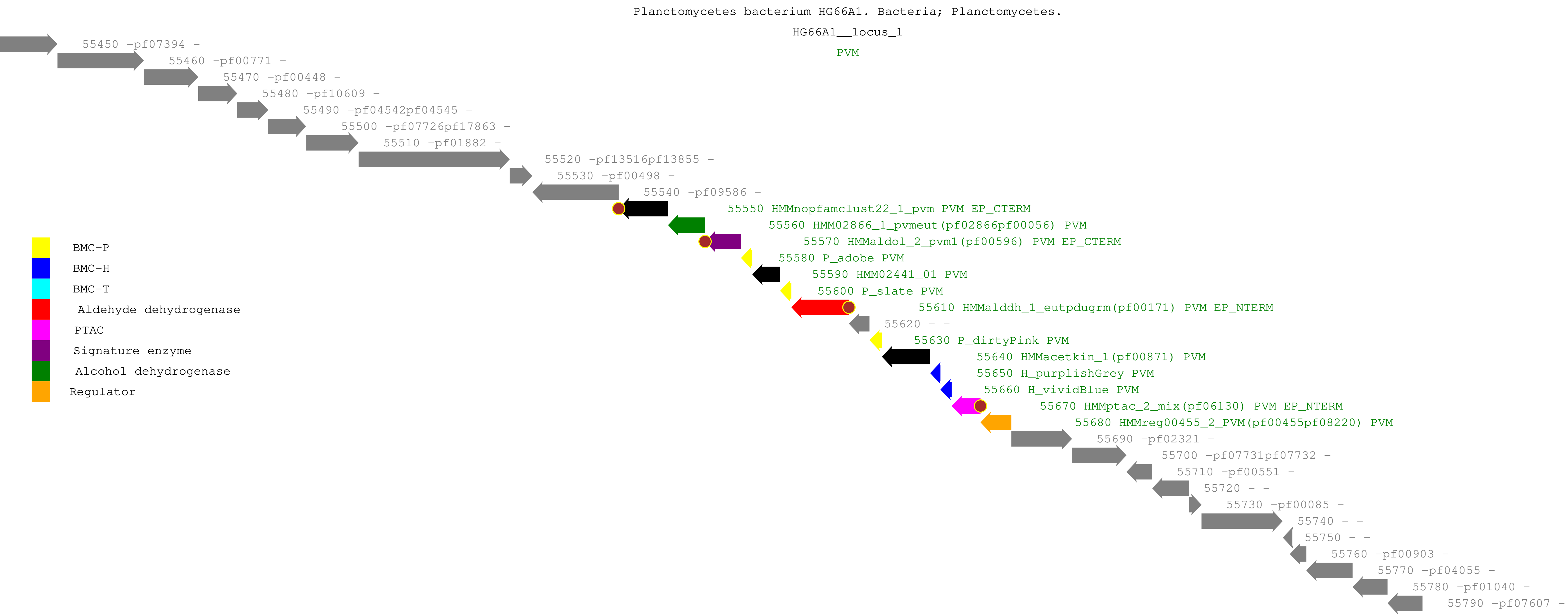
Planctomycetes bacterium HG66A1. Bacteria; Planctomycetes.
HG66A1__locus_1
PVM
55450 -pf07394 -
55460 -pf00771 -
55470 -pf00448 -
55480 -pf10609 -
55490 -pf04542pf04545 -
55500 -pf07726pf17863 -
55510 -pf01882 -
55520 -pf13516pf13855 -
55530 -pf00498 -
55540 -pf09586 -
55550 HMMnopfamclust22_1_pvm PVM EP_CTERM
55560 HMM02866_1_pvmeut(pf02866pf00056) PVM
55570 HMMaldol_2_pvm1(pf00596) PVM EP_CTERM
55580 P_adobe PVM
55590 HMM02441_01 PVM
55600 P_slate PVM
55610 HMMalddh_1_eutpdugrm(pf00171) PVM EP_NTERM
55620 - -
55630 P_dirtyPink PVM
55640 HMMacetkin_1(pf00871) PVM
55650 H_purplishGrey PVM
55660 H_vividBlue PVM
55670 HMMptac_2_mix(pf06130) PVM EP_NTERM
55680 HMMreg00455_2_PVM(pf00455pf08220) PVM
55690 -pf02321 -
55700 -pf07731pf07732 -
55710 -pf00551 -
55720 - -
55730 -pf00085 -
55740 - -
55750 - -
55760 -pf00903 -
55770 -pf04055 -
55780 -pf01040 -
55790 -pf07607 -
BMC-P
BMC-H
BMC-T
Aldehyde dehydrogenase
PTAC
Signature enzyme
Alcohol dehydrogenase
Regulator

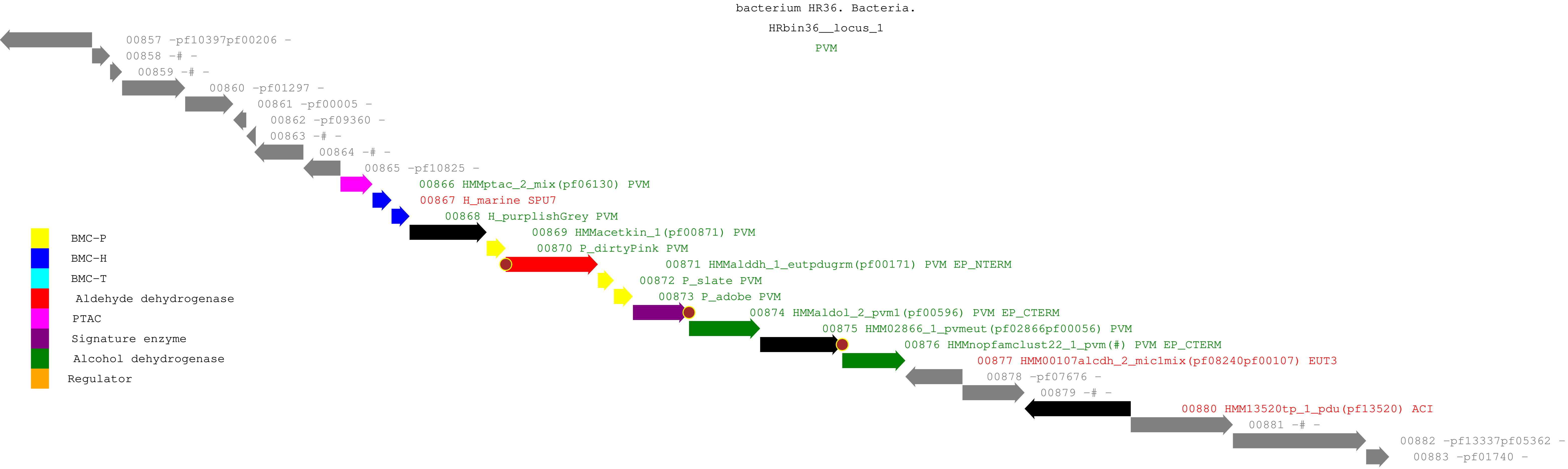
bacterium HR36. Bacteria.
HRbin36__locus_1
PVM
00857 -pf10397pf00206 -
00858 -# -
00859 -# -
00860 -pf01297 -
00861 -pf00005 -
00862 -pf09360 -
00863 -# -
00864 -# -
00865 -pf10825 -
00866 HMMptac_2_mix(pf06130) PVM
00867 H_marine SPU7
00868 H_purplishGrey PVM
00869 HMMacetkin_1(pf00871) PVM
00870 P_dirtyPink PVM
00871 HMMalddh_1_eutpdugrm(pf00171) PVM EP_NTERM
00872 P_slate PVM
00873 P_adobe PVM
00874 HMMaldol_2_pvm1(pf00596) PVM EP_CTERM
00875 HMM02866_1_pvmeut(pf02866pf00056) PVM
00876 HMMnopfamclust22_1_pvm(#) PVM EP_CTERM
00877 HMM00107alcdh_2_mic1mix(pf08240pf00107) EUT3
00878 -pf07676 -
00879 -# -
00880 HMM13520tp_1_pdu(pf13520) ACI
00881 -# -
00882 -pf13337pf05362 -
00883 -pf01740 -
BMC-P
BMC-H
BMC-T
Aldehyde dehydrogenase
PTAC
Signature enzyme
Alcohol dehydrogenase
Regulator

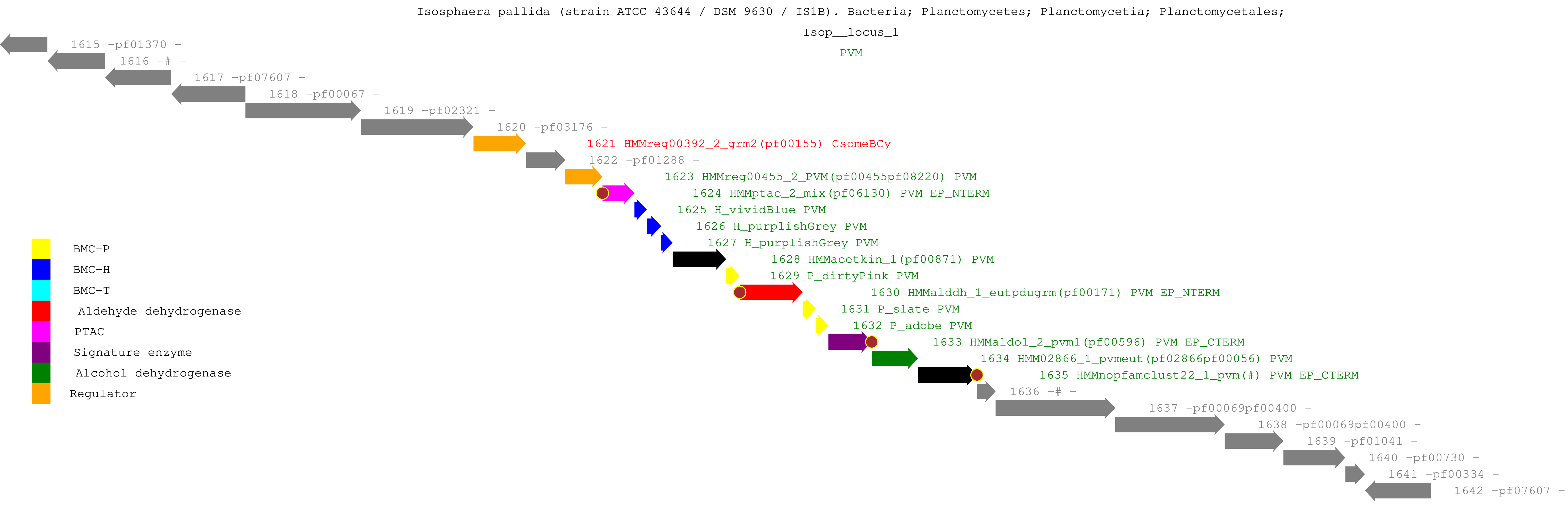

Isosphaera pallida (strain ATCC 43644 / DSM 9630 / IS1B). Bacteria; Planctomycetes; Planctomycetia; Planctomycetales;
Isop__locus_1
PVM
1615 -pf01370 -
1616 -# -
1617 -pf07607 -
1618 -pf00067 -
1619 -pf02321 -
1620 -pf03176 -
1621 HMMreg00392_2_grm2(pf00155) CsomeBCy
1622 -pf01288 -
1623 HMMreg00455_2_PVM(pf00455pf08220) PVM
1624 HMMptac_2_mix(pf06130) PVM EP_NTERM
1625 H_vividBlue PVM
1626 H_purplishGrey PVM
1627 H_purplishGrey PVM
1628 HMMacetkin_1(pf00871) PVM
1629 P_dirtyPink PVM
1630 HMMalddh_1_eutpdugrm(pf00171) PVM EP_NTERM
1631 P_slate PVM
1632 P_adobe PVM
1633 HMMaldol_2_pvm1(pf00596) PVM EP_CTERM
1634 HMM02866_1_pvmeut(pf02866pf00056) PVM
1635 HMMnopfamclust22_1_pvm(#) PVM EP_CTERM
1636 -# -
1637 -pf00069pf00400 -
1638 -pf00069pf00400 -
1639 -pf01041 -
1640 -pf00730 -
1641 -pf00334 -
1642 -pf07607 -
BMC-P
BMC-H
BMC-T
Aldehyde dehydrogenase
PTAC
Signature enzyme
Alcohol dehydrogenase
Regulator

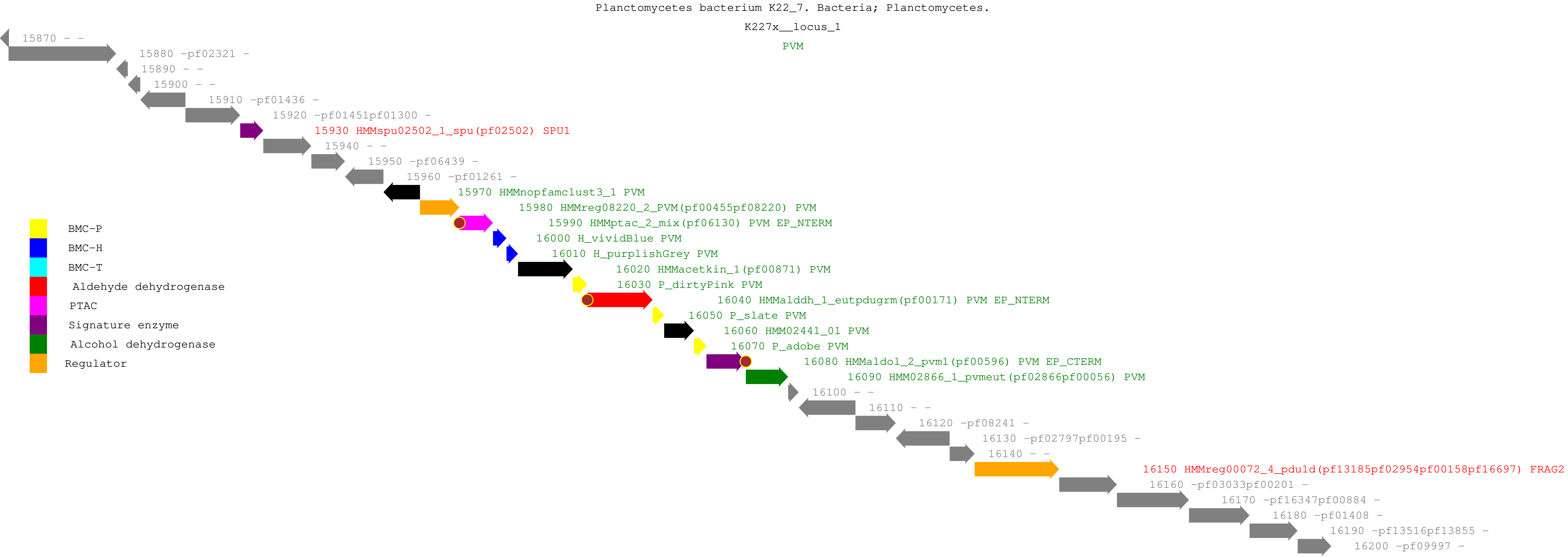
Planctomycetes bacterium K22_7. Bacteria; Planctomycetes.
K227x__locus_1
PVM
15870 - -
15880 -pf02321 -
15890 - -
15900 - -
15910 -pf01436 -
15920 -pf01451pf01300 -
15930 HMMspu02502_1_spu(pf02502) SPU1
15940 - -
15950 -pf06439 -
15960 -pf01261 -
15970 HMMnopfamclust3_1 PVM
15980 HMMreg08220_2_PVM(pf00455pf08220) PVM
15990 HMMptac_2_mix(pf06130) PVM EP_NTERM
16000 H_vividBlue PVM
16010 H_purplishGrey PVM
16020 HMMacetkin_1(pf00871) PVM
16030 P_dirtyPink PVM
16040 HMMalddh_1_eutpdugrm(pf00171) PVM EP_NTERM
16050 P_slate PVM
16060 HMM02441_01 PVM
16070 P_adobe PVM
16080 HMMaldol_2_pvm1(pf00596) PVM EP_CTERM
16090 HMM02866_1_pvmeut(pf02866pf00056) PVM
16100 - -
16110 - -
16120 -pf08241 -
16130 -pf02797pf00195 -
16140 - -
16150 HMMreg00072_4_pdu1d(pf13185pf02954pf00158pf16697) FRAG2
16160 -pf03033pf00201 -
16170 -pf16347pf00884 -
16180 -pf01408 -
16190 -pf13516pf13855 -
16200 -pf09997 -
BMC-P
BMC-H
BMC-T
Aldehyde dehydrogenase
PTAC
Signature enzyme
Alcohol dehydrogenase
Regulator

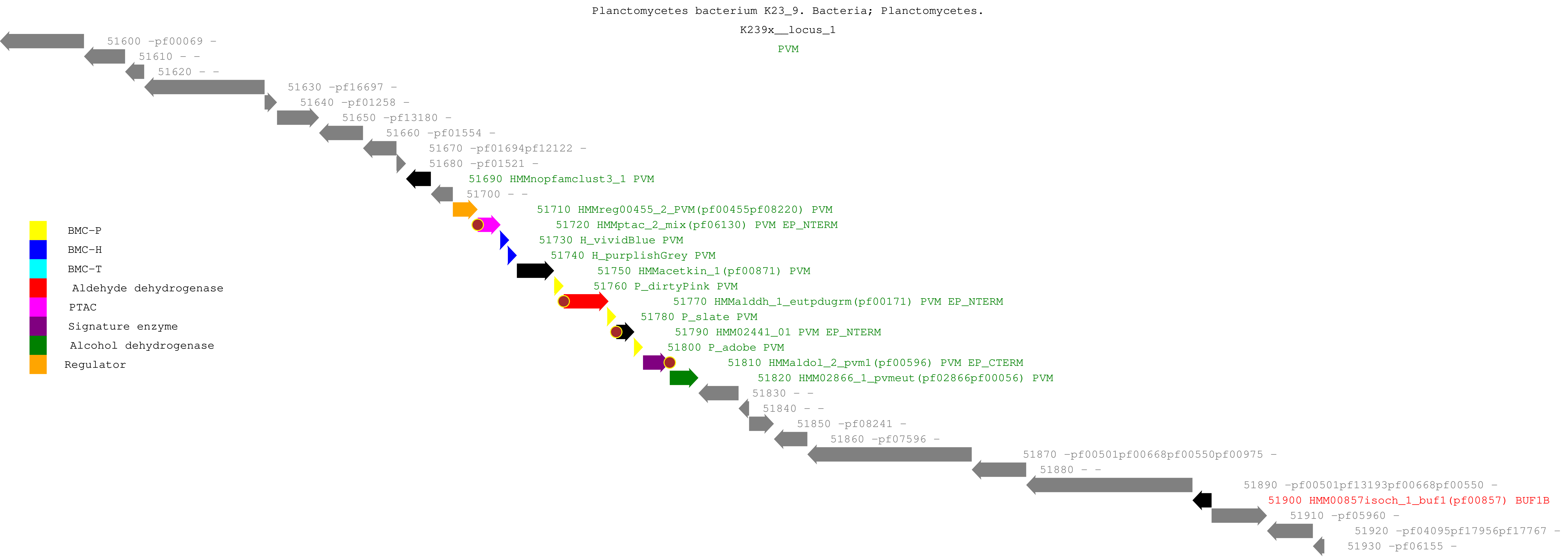
Planctomycetes bacterium K23_9. Bacteria; Planctomycetes.
K239x__locus_1
PVM
51600 -pf00069 -
51610 - -
51620 - -
51630 -pf16697 -
51640 -pf01258 -
51650 -pf13180 -
51660 -pf01554 -
51670 -pf01694pf12122 -
51680 -pf01521 -
51690 HMMnopfamclust3_1 PVM
51700 - -
51710 HMMreg00455_2_PVM(pf00455pf08220) PVM
51720 HMMptac_2_mix(pf06130) PVM EP_NTERM
51730 H_vividBlue PVM
51740 H_purplishGrey PVM
51750 HMMacetkin_1(pf00871) PVM
51760 P_dirtyPink PVM
51770 HMMaldh_1_eutpdugrm(pf00171) PVM EP_NTERM
51780 P_slate PVM
51790 HMM02441_01 PVM EP_NTERM
51800 P_adobe PVM
51810 HMMaldol_2_pvm1(pf00596) PVM EP_CTERM
51820 HMM02866_1_pvmeut(pf02866pf00056) PVM
51830 - -
51840 - -
51850 -pf08241 -
51860 -pf07596 -
51870 -pf00501pf00668pf00550pf00975 -
51880 - -
51890 -pf00501pf13193pf00668pf00550 -
51900 HMM00857isoch_1_buf1(pf00857) BUF1B
51910 -pf05960 -
51920 -pf04095pf17956pf17767 -
51930 -pf06155 -
BMC-P
BMC-H
BMC-T
Aldehyde dehydrogenase
PTAC
Signature enzyme
Alcohol dehydrogenase
Regulator

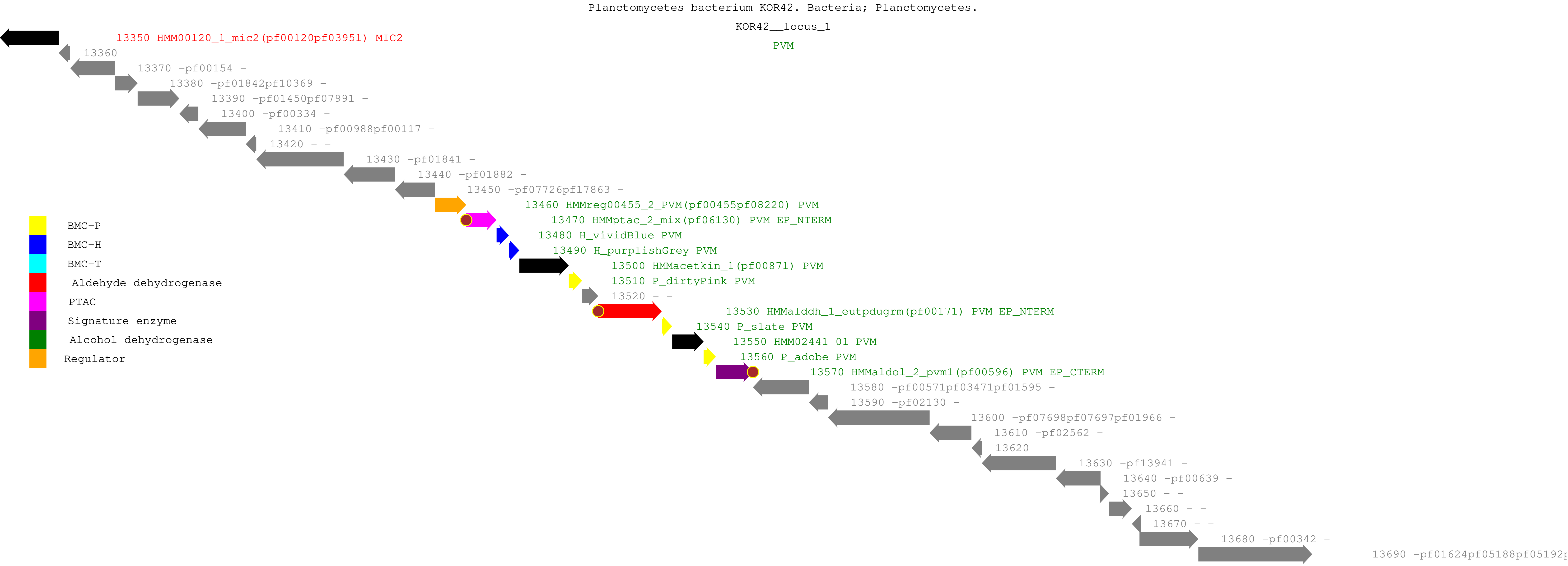
Planctomycetes bacterium KOR42. Bacteria; Planctomycetes.
KOR42__locus_1
PVM
13350 HMM00120_1_mic2(pf00120pf03951) MIC2
13360 - -
13370 -pf00154 -
13380 -pf01842pf10369 -
13390 -pf01450pf07991 -
13400 -pf00334 -
13410 -pf00988pf00117 -
13420 - -
13430 -pf01841 -
13440 -pf01882 -
13450 -pf07726pf17863 -
13460 HMMreg00455_2_PVM(pf00455pf08220) PVM
13470 HMMptac_2_mix(pf06130) PVM EP_NTERM
13480 H_vividBlue PVM
13490 H_purplishGrey PVM
13500 HMMacetkin_1(pf00871) PVM
13510 P_dirtyPink PVM
13520 - -
13530 HMMalddh_1_eutpdugrm(pf00171) PVM EP_NTERM
13540 P_slate PVM
13550 HMM02441_01 PVM
13560 P_adobe PVM
13570 HMMaldol_2_pvm1(pf00596) PVM EP_CTERM
13580 -pf00571pf03471pf01595 -
13590 -pf02130 -
13600 -pf07698pf07697pf01966 -
13610 -pf02562 -
13620 - -
13630 -pf13941 -
13640 -pf00639 -
13650 - -
13660 - -
13670 - -
13680 -pf00342 -
13690 -pf01624pf05188pf05192
BMC-P
BMC-H
BMC-T
Aldehyde dehydrogenase
PTAC
Signature enzyme
Alcohol dehydrogenase
Regulator

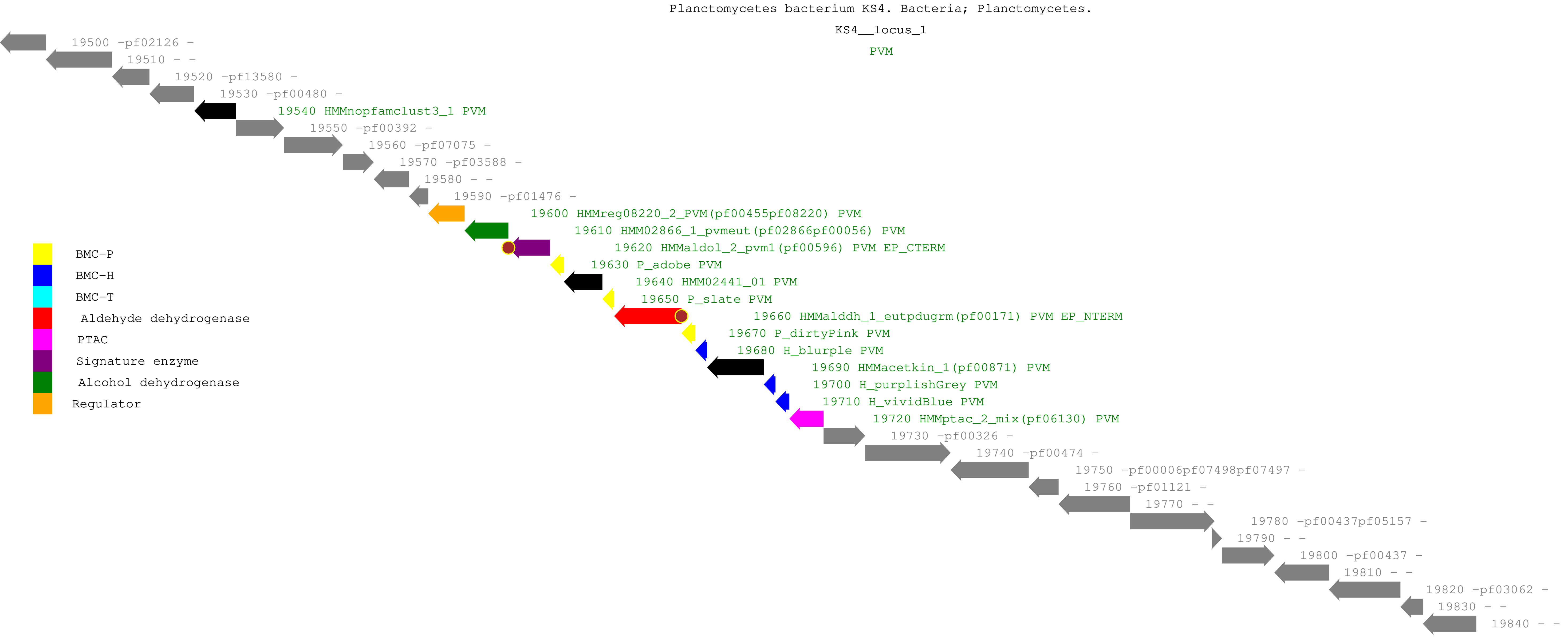

Planctomycetes bacterium KS4. Bacteria; Planctomycetes.
KS4__locus_1
PVM
19500 -pf02126 -
19510 - -
19520 -pf13580 -
19530 -pf00480 -
19540 HMMnopfamclust3_1 PVM
19550 -pf00392 -
19560 -pf07075 -
19570 -pf03588 -
19580 - -
19590 -pf01476 -
19600 HMMreg08220_2_PVM(pf00455pf08220) PVM
19610 HMM02866_1_pvmeut(pf02866pf00056) PVM
19620 HMMaldol_2_pvm1(pf00596) PVM EP_CTERM
19630 P_adobe PVM
19640 HMM02441_01 PVM
19650 P_slate PVM
19660 HMMalddh_1_eutpdugrm(pf00171) PVM EP_NTERM
19670 P_dirtyPink PVM
19680 H_blurple PVM
19690 HMMacetkin_1(pf00871) PVM
19700 H_purplishGrey PVM
19710 H_vividBlue PVM
19720 HMMptac_2_mix(pf06130) PVM
19730 -pf00326 -
19740 -pf00474 -
19750 -pf00006pf07498pf07497 -
19760 -pf01121 -
19770 - -
19780 -pf00437pf05157 -
19790 - -
19800 -pf00437 -
19810 - -
19820 -pf03062 -
19830 - -
19840 - -
BMC-P
BMC-H
BMC-T
Aldehyde dehydrogenase
PTAC
Signature enzyme
Alcohol dehydrogenase
Regulator

Planctomycetia bacterium. Bacteria; Planctomycetes; Planctomycetia; unclassified Planctomycetia.

LBMAG46__locus_1

PVM

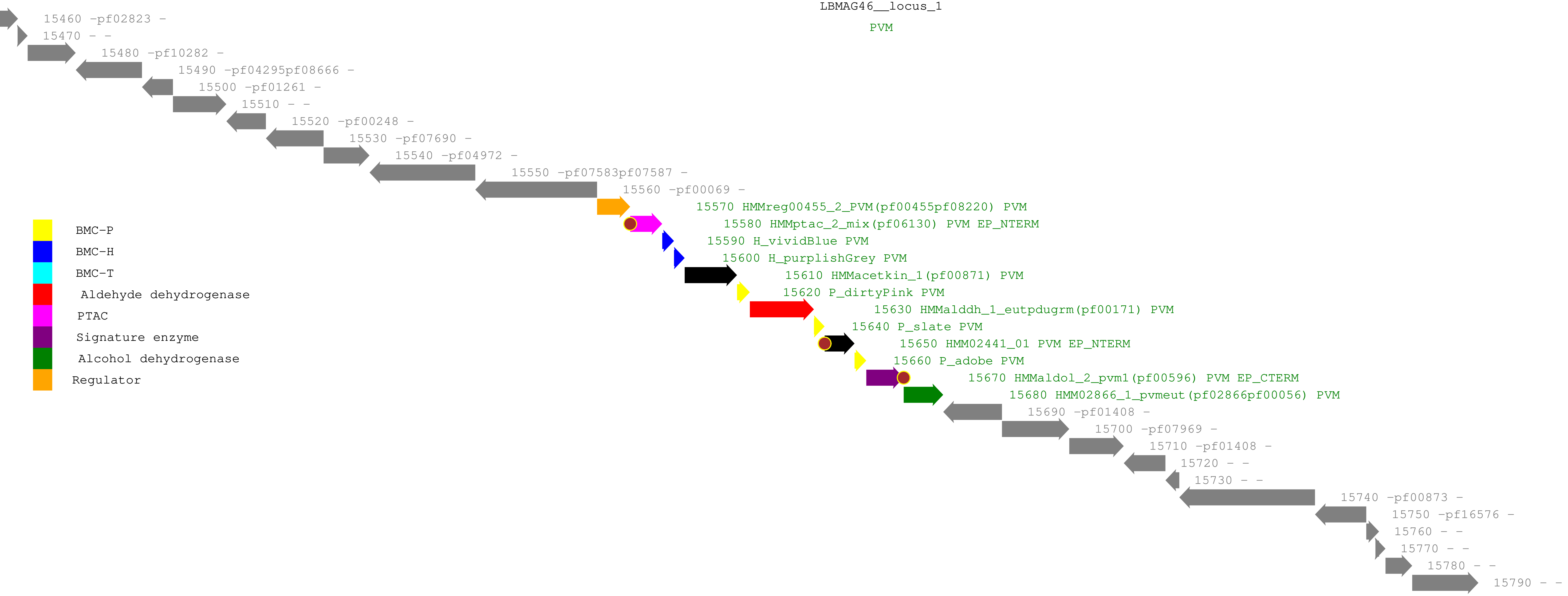

Planctomycetia bacterium. Bacteria; Planctomycetes; Planctomycetia; unclassified Planctomycetia.

LBMAG47__locus_1

PVM

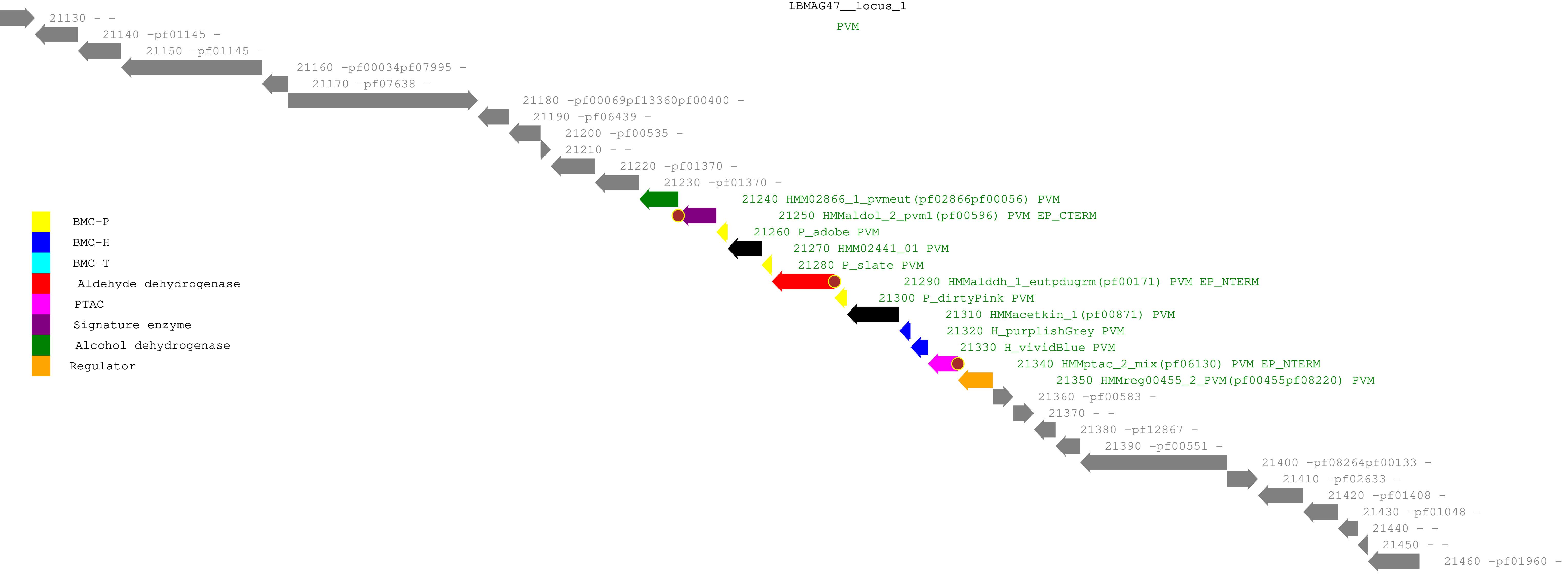

Planctomycetia bacterium. Bacteria; Planctomycetes; Planctomycetia; unclassified Planctomycetia.

LBMAG52__locus_1_main_sat

PVM

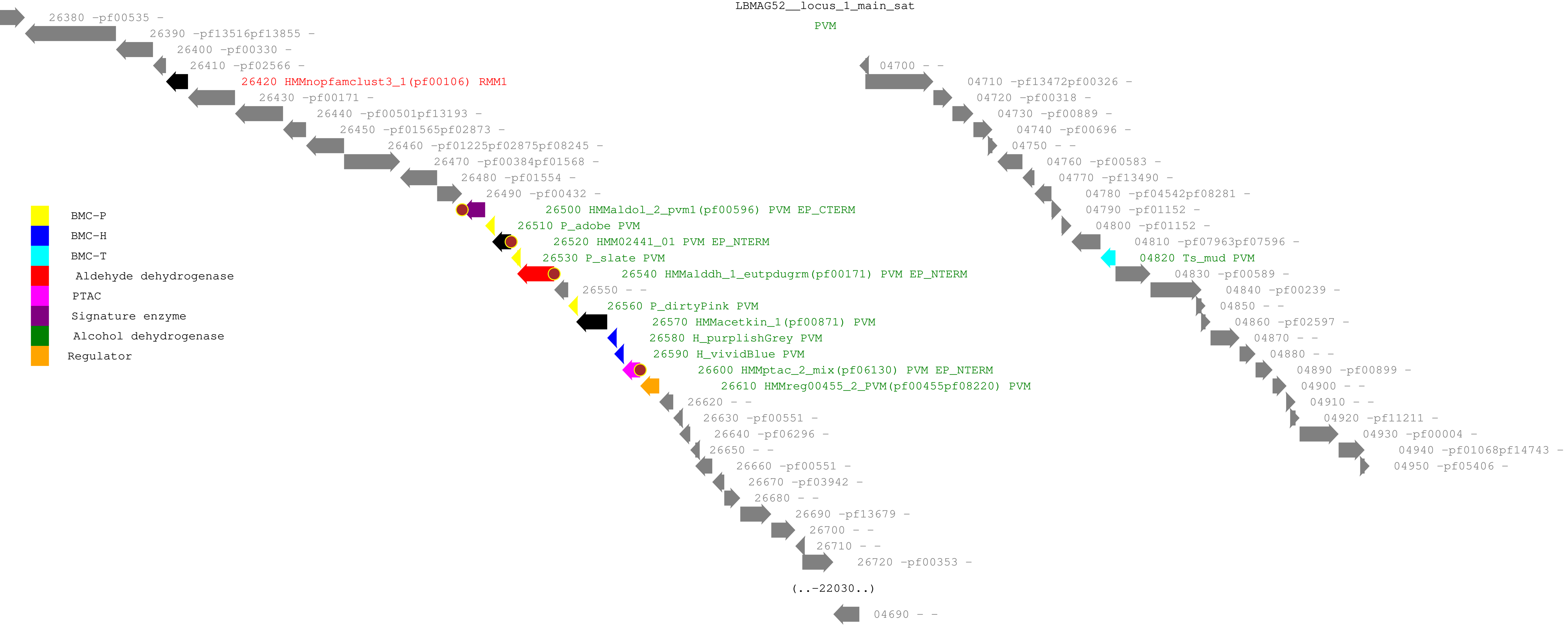

Verrucomicrobia bacterium. Bacteria; Verrucomicrobia; unclassified Verrucomicrobia.

LBMAG56__locus_1

PVM

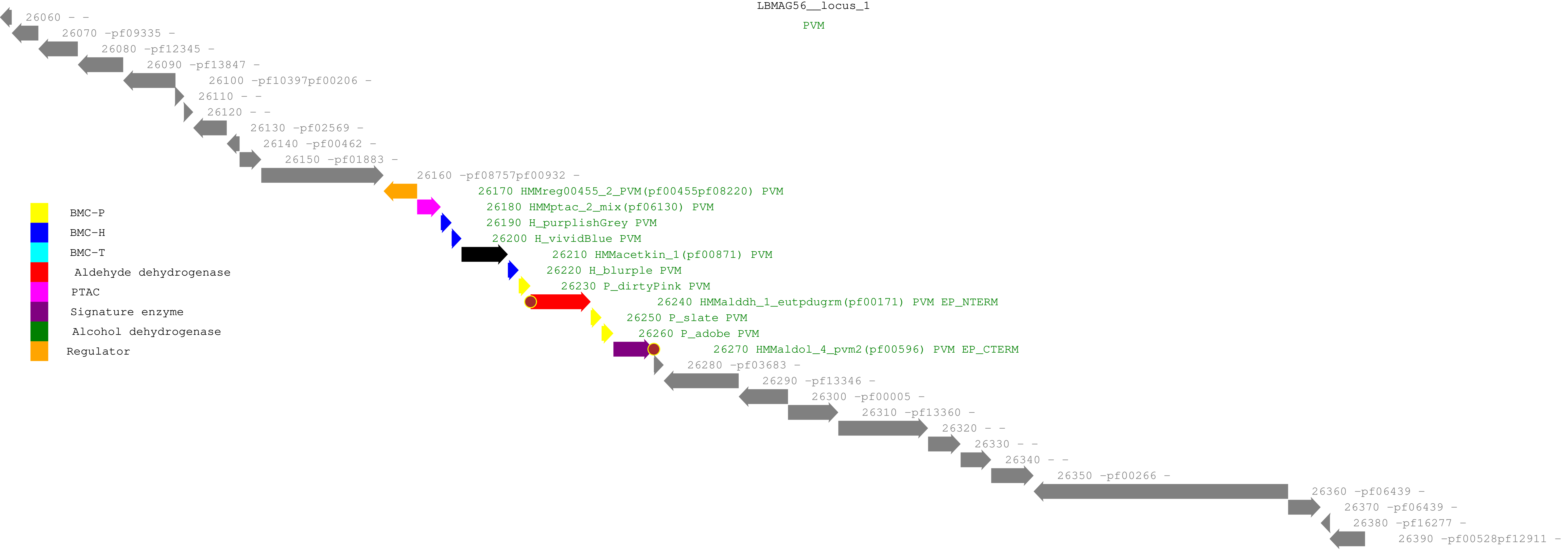

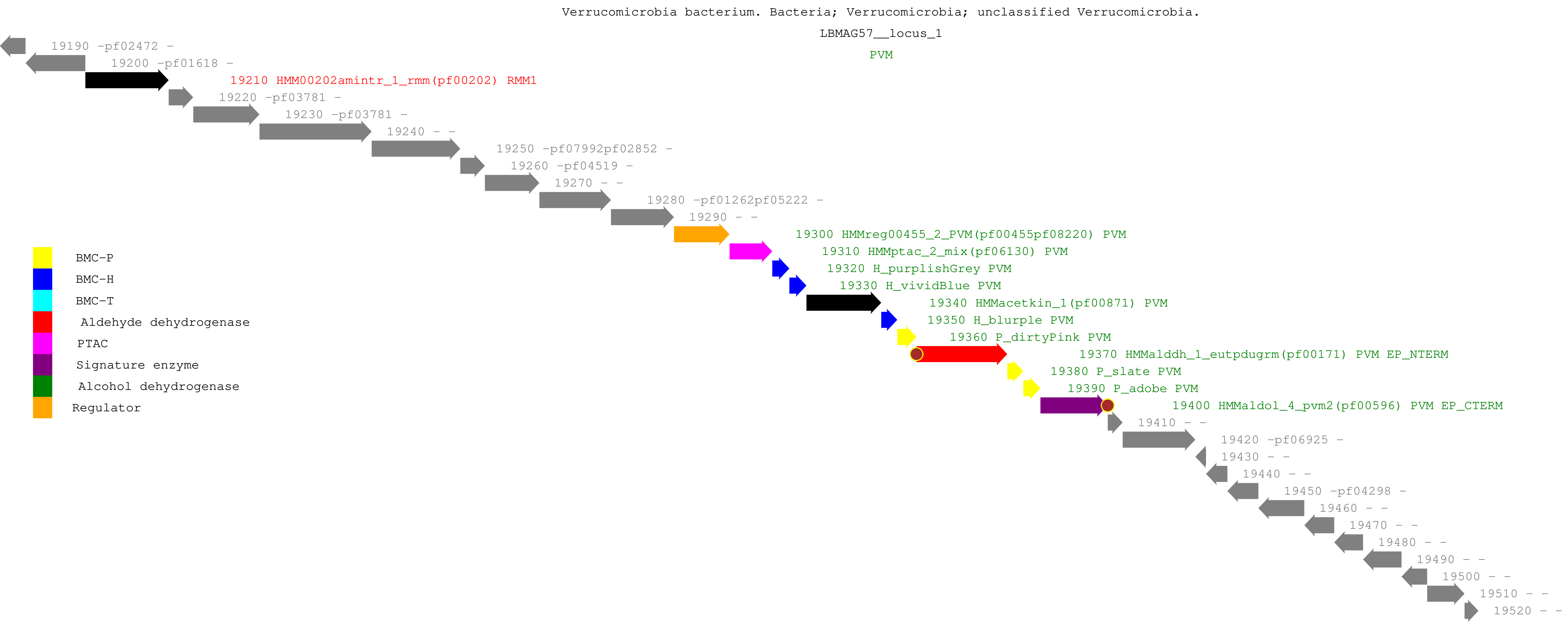
Verrucomicrobia bacterium. Bacteria; Verrucomicrobia; unclassified Verrucomicrobia.
LBMAG57__locus_1
PVM
19190 -pf02472 -
19200 -pf01618 -
19210 HMM00202amintr_1_rmm(pf00202) RMM1
19220 -pf03781 -
19230 -pf03781 -
19240 - -
19250 -pf07992pf02852 -
19260 -pf04519 -
19270 - -
19280 -pf01262pf05222 -
19290 - -
19300 HMMreg00455_2_PVM(pf00455pf08220) PVM
19310 HMMptac_2_mix(pf06130) PVM
19320 H_purplishGrey PVM
19330 H_vividBlue PVM
19340 HMMacetkin_1(pf00871) PVM
19350 H_blurple PVM
19360 P_dirtyPink PVM
19370 HMMalddh_1_eutpdugrm(pf00171) PVM EP_NTERM
19380 P_slate PVM
19390 P_adobe PVM
19400 HMMaldol_4_pvm2(pf00596) PVM EP_CTERM
19410 - -
19420 -pf06925 -
19430 - -
19440 - -
19450 -pf04298 -
19460 - -
19470 - -
19480 - -
19490 - -
19500 - -
19510 - -
19520 - -
BMC-P
BMC-H
BMC-T
Aldehyde dehydrogenase
PTAC
Signature enzyme
Alcohol dehydrogenase
Regulator

marine sediment metagenome. unclassified sequences; metagenomes; ecological metagenomes.

LCGC14__locus_1

PVM

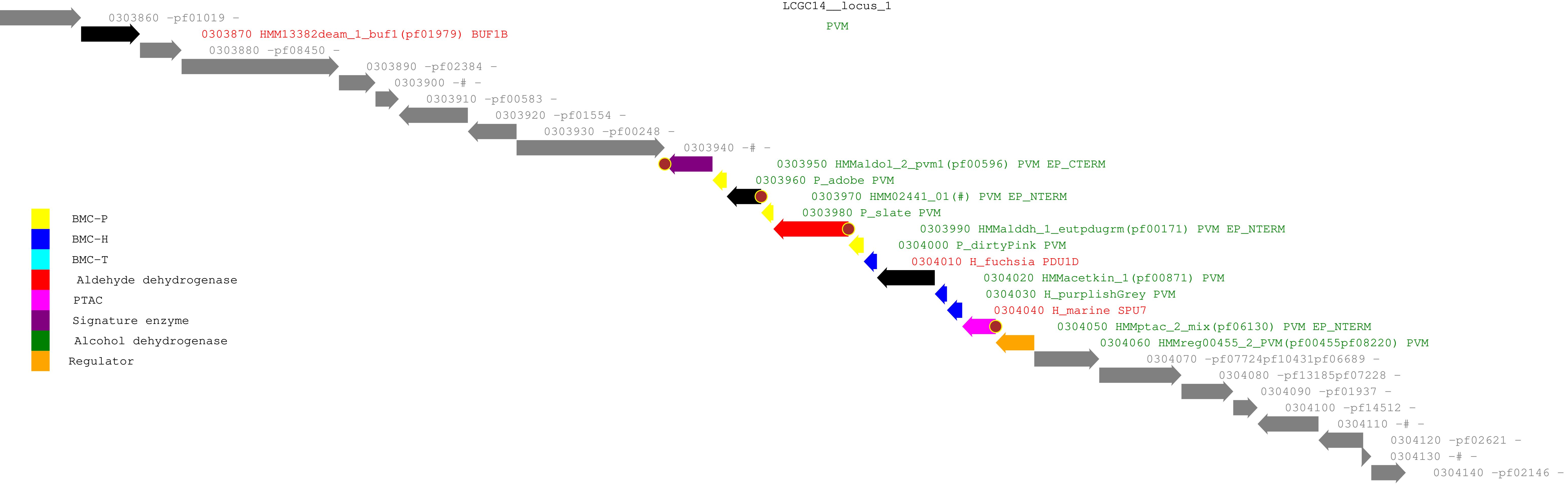

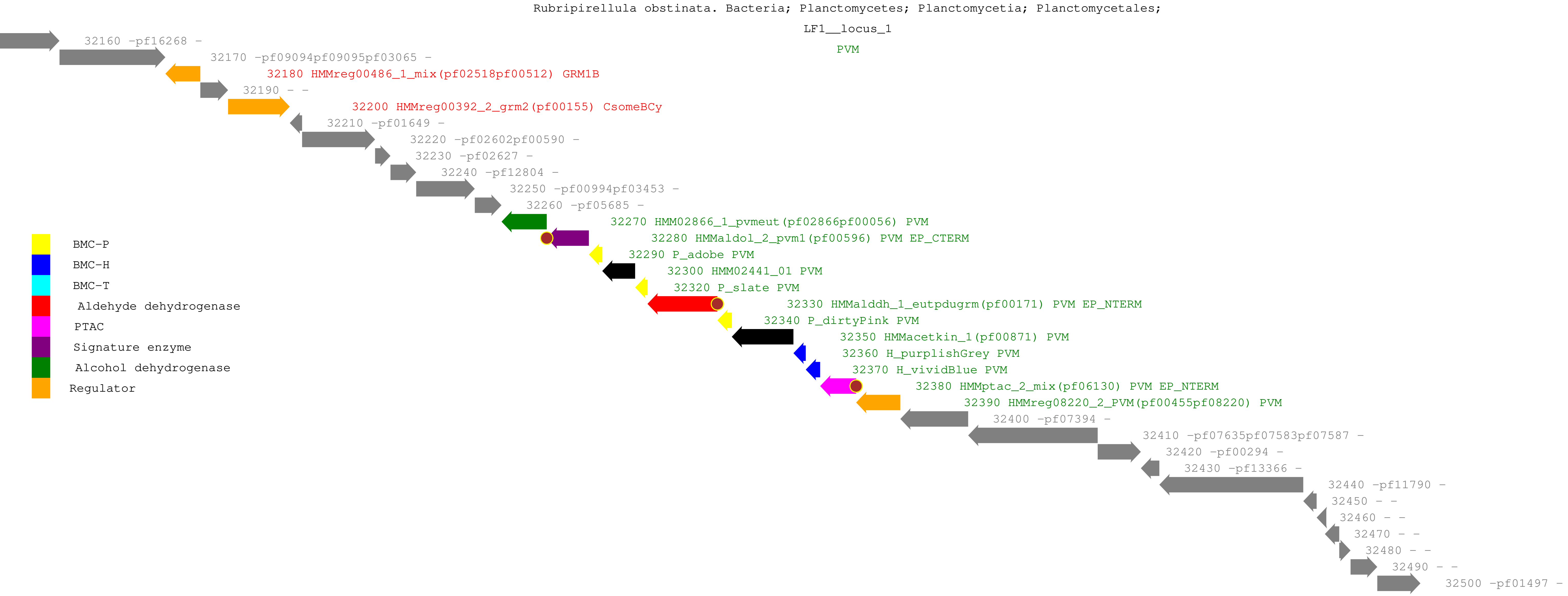
Rubripirellula obstinata. Bacteria; Planctomycetes; Planctomycetia; Planctomycetales;
LF1__locus_1
PVM
32160 -pf16268 -
32170 -pf09094pf09095pf03065 -
32180 HMMreg00486_1_mix(pf02518pf00512) GRM1B
32190 - -
32200 HMMreg00392_2_grm2(pf00155) CsomeBCy
32210 -pf01649 -
32220 -pf02602pf00590 -
32230 -pf02627 -
32240 -pf12804 -
32250 -pf00994pf03453 -
32260 -pf05685 -
32270 HMM02866_1_pvmeut(pf02866pf00056) PVM
32280 HMMaldol_2_pvm1(pf00596) PVM EP_CTERM
32290 P_adobe PVM
32300 HMM02441_01 PVM
32320 P_slate PVM
32330 HMMalddh_1_eutpdugrm(pf00171) PVM EP_NTERM
32340 P_dirtyPink PVM
32350 HMMacetkin_1(pf00871) PVM
32360 H_purplishGrey PVM
32370 H_vividBlue PVM
32380 HMMptac_2_mix(pf06130) PVM EP_NTERM
32390 HMMreg08220_2_PVM(pf00455pf08220) PVM
32400 -pf07394 -
32410 -pf07635pf07583pf07587 -
32420 -pf00294 -
32430 -pf13366 -
32440 -pf11790 -
32450 - -
32460 - -
32470 - -
32480 - -
32490 - -
32500 -pf01497 -
BMC-P
BMC-H
BMC-T
Aldehyde dehydrogenase
PTAC
Signature enzyme
Alcohol dehydrogenase
Regulator

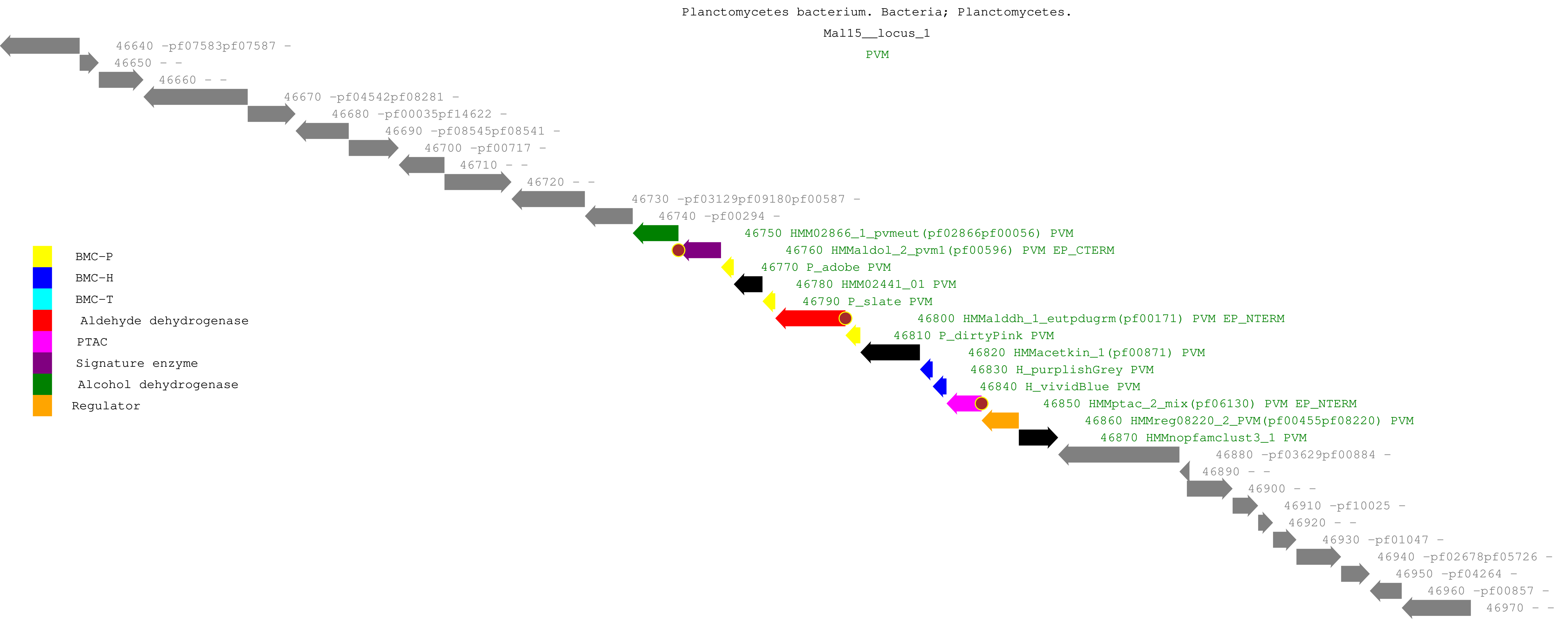
Planctomycetes bacterium. Bacteria; Planctomycetes.
Mal15__locus_1
PVM
46640 -pf07583pf07587 -
46650 - -
46660 - -
46670 -pf04542pf08281 -
46680 -pf00035pf14622 -
46690 -pf08545pf08541 -
46700 -pf00717 -
46710 - -
46720 - -
46730 -pf03129pf09180pf00587 -
46740 -pf00294 -
46750 HMM02866_1_pvmeut(pf02866pf00056) PVM
46760 HMMaldol_2_pvm1(pf00596) PVM EP_CTERM
46770 P_adobe PVM
46780 HMM02441_01 PVM
46790 P_slate PVM
46800 HMMaldh_1_eutpdugrm(pf00171) PVM EP_NTERM
46810 P_dirtyPink PVM
46820 HMMacetkin_1(pf00871) PVM
46830 H_purplishGrey PVM
46840 H_vividBlue PVM
46850 HMMptac_2_mix(pf06130) PVM EP_NTERM
46860 HMMreg08220_2_PVM(pf00455pf08220) PVM
46870 HMMnopfamclust3_1 PVM
46880 -pf03629pf00884 -
46890 - -
46900 - -
46910 -pf10025 -
46920 - -
46930 -pf01047 -
46940 -pf02678pf05726 -
46950 -pf04264 -
46960 -pf00857 -
46970 - -
BMC-P
BMC-H
BMC-T
Aldehyde dehydrogenase
PTAC
Signature enzyme
Alcohol dehydrogenase
Regulator

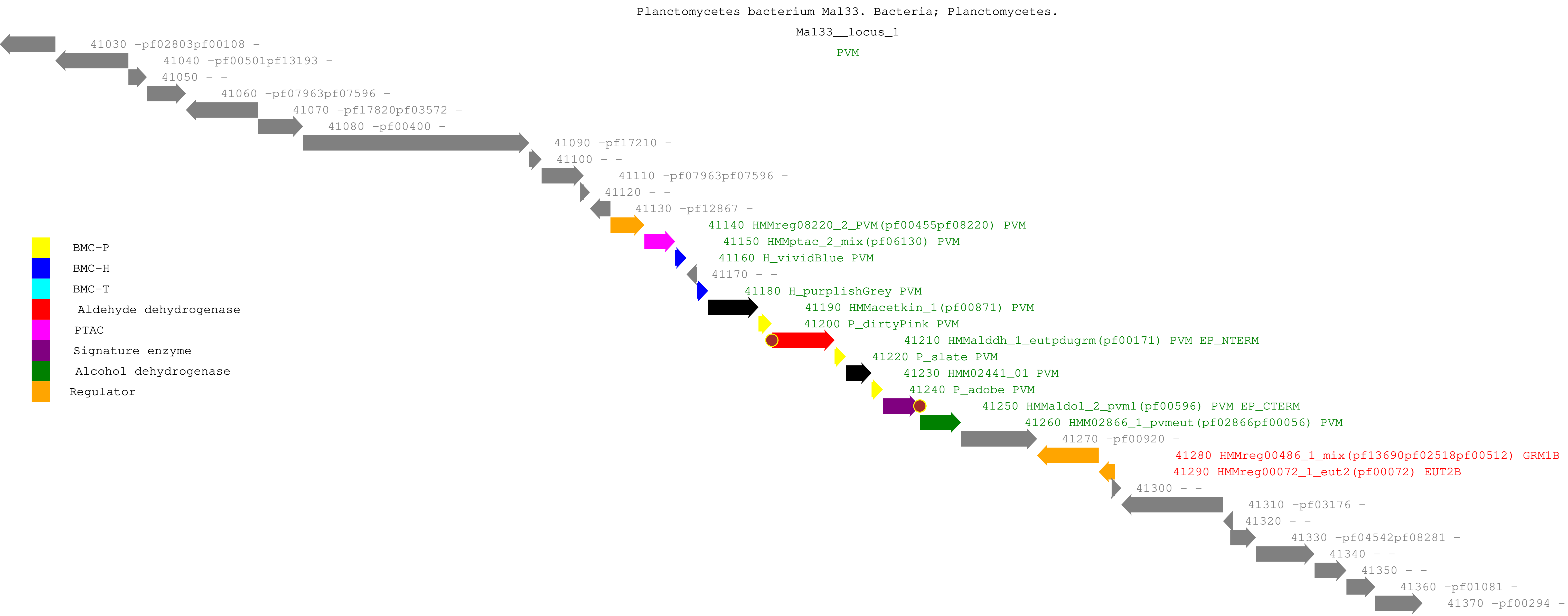

Planctomycetes bacterium Mal33. Bacteria; Planctomycetes.
Mal33__locus_1
PVM
41030 -pf02803pf00108 -
41040 -pf00501pf13193 -
41050 - -
41060 -pf07963pf07596 -
41070 -pf17820pf03572 -
41080 -pf00400 -
41090 -pf17210 -
41100 - -
41110 -pf07963pf07596 -
41120 - -
41130 -pf12867 -
41140 HMMreg08220_2_PVM(pf00455pf08220) PVM
41150 HMMptac_2_mix(pf06130) PVM
41160 H_vividBlue PVM
41170 - -
41180 H_purplishGrey PVM
41190 HMMacetkin_1(pf00871) PVM
41200 P_dirtyPink PVM
41210 HMMalddh_1_eutpdugrm(pf00171) PVM EP_NTERM
41220 P_slate PVM
41230 HMM02441_01 PVM
41240 P_adobe PVM
41250 HMMaldol_2_pvm1(pf00596) PVM EP_CTERM
41260 HMM02866_1_pvmeut(pf02866pf00056) PVM
41270 -pf00920 -
41280 HMMreg00486_1_mix(pf13690pf02518pf00512) GRM1B
41290 HMMreg00072_1_eut2(pf00072) EUT2B
41300 - -
41310 -pf03176 -
41320 - -
41330 -pf04542pf08281 -
41340 - -
41350 - -
41360 -pf01081 -
41370 -pf00294 -
BMC-P
BMC-H
BMC-T
Aldehyde dehydrogenase
PTAC
Signature enzyme
Alcohol dehydrogenase
Regulator

Planctomycetes bacterium Mal48. Bacteria; Planctomycetes.

Mal48__locus_1

PVM

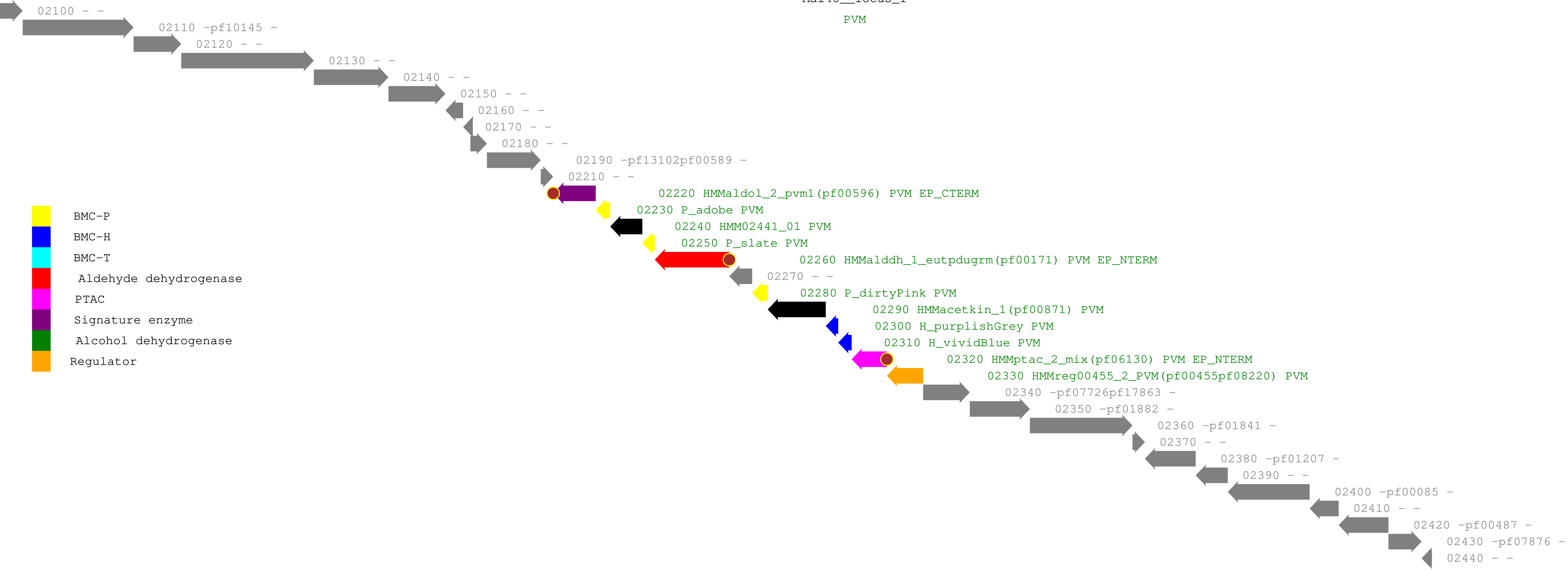

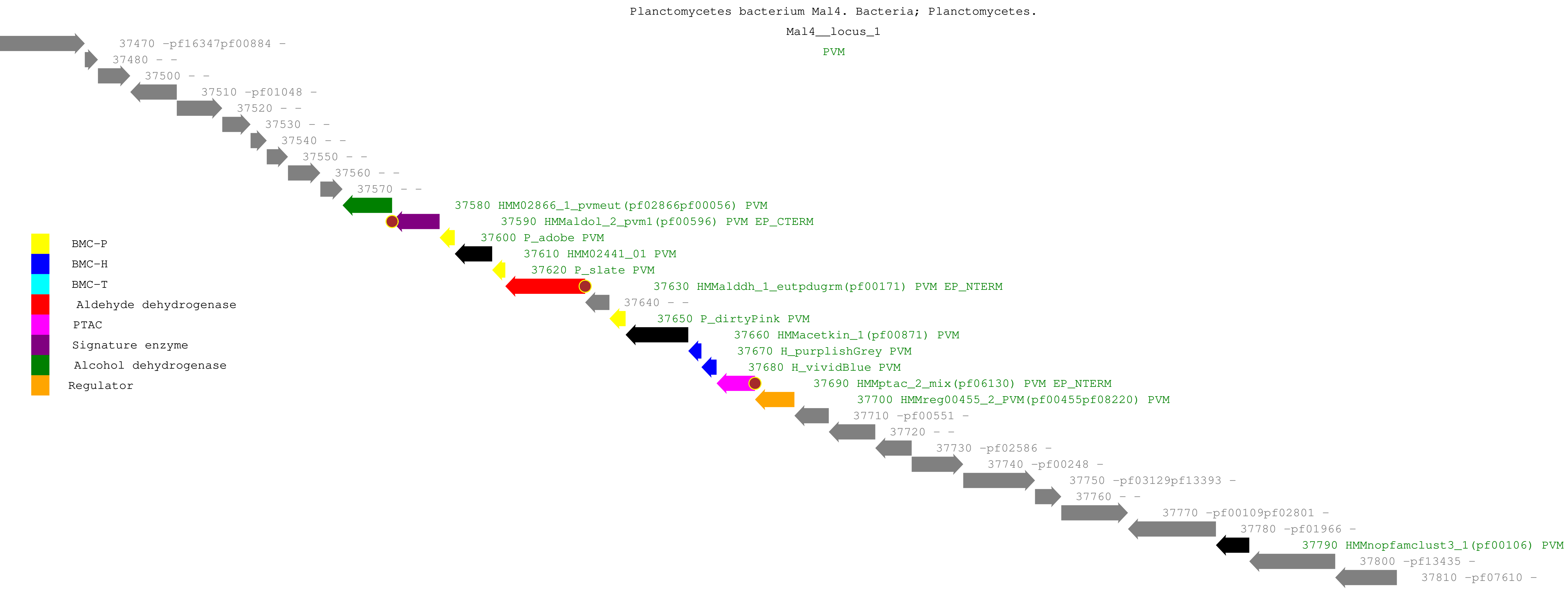
Planctomycetes bacterium Mal4. Bacteria; Planctomycetes.
Mal4__locus_1
PVM
37470 -pf16347pf00884 -
37480 - -
37500 - -
37510 -pf01048 -
37520 - -
37530 - -
37540 - -
37550 - -
37560 - -
37570 - -
37580 HMM02866_1_pvmeut(pf02866pf00056) PVM
37590 HMMaldol_2_pvm1(pf00596) PVM EP_CTERM
37600 P_adobe PVM
37610 HMM02441_01 PVM
37620 P_slate PVM
37630 HMMalddh_1_eutpdugrm(pf00171) PVM EP_NTERM
37640 - -
37650 P_dirtyPink PVM
37660 HMMacetkin_1(pf00871) PVM
37670 H_purplishGrey PVM
37680 H_vividBlue PVM
37690 HMMptac_2_mix(pf06130) PVM EP_NTERM
37700 HMMreg00455_2_PVM(pf00455pf08220) PVM
37710 -pf00551 -
37720 - -
37730 -pf02586 -
37740 -pf00248 -
37750 -pf03129pf13393 -
37760 - -
37770 -pf00109pf02801 -
37780 -pf01966 -
37790 HMMnopfamclust3_1(pf00106) PVM
37800 -pf13435 -
37810 -pf07610 -
BMC-P
BMC-H
BMC-T
Aldehyde dehydrogenase
PTAC
Signature enzyme
Alcohol dehydrogenase
Regulator

Planctomycetes bacterium Mal52. Bacteria; Planctomycetes.

Mal52__locus_1

PVM

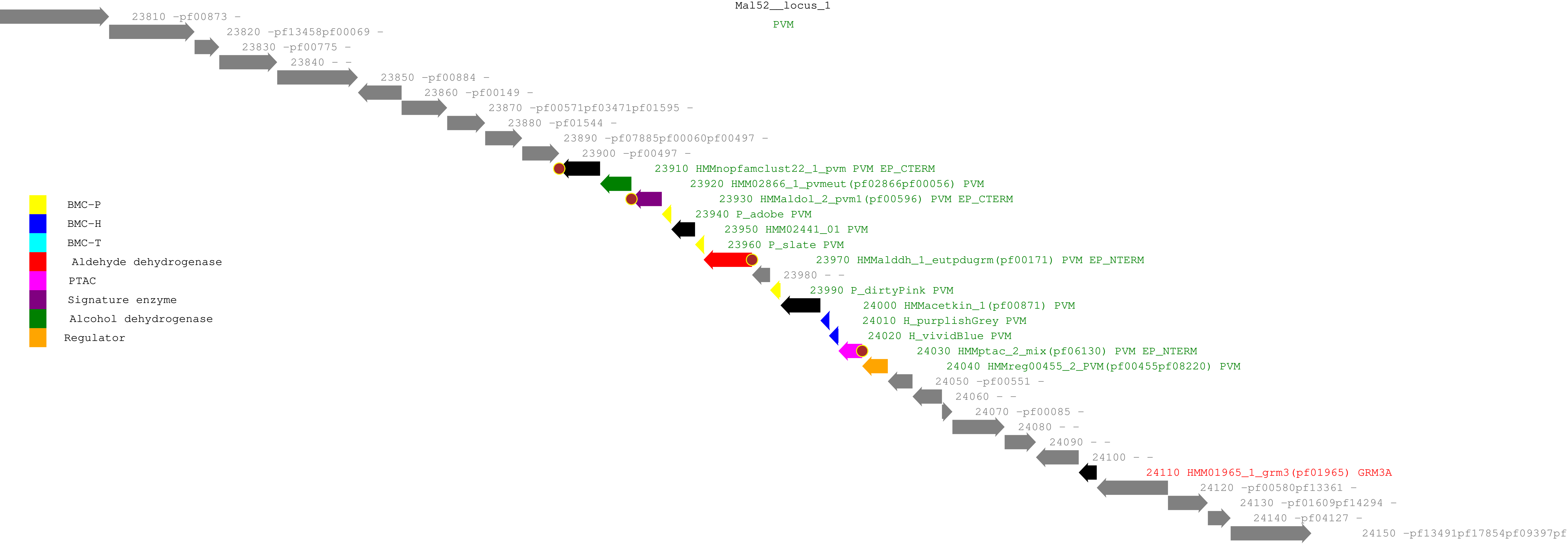

hydrothermal vent metagenome. unclassified sequences; metagenomes; ecological metagenomes.

MNBD_PLANCTO02-_locus_1_main_sat

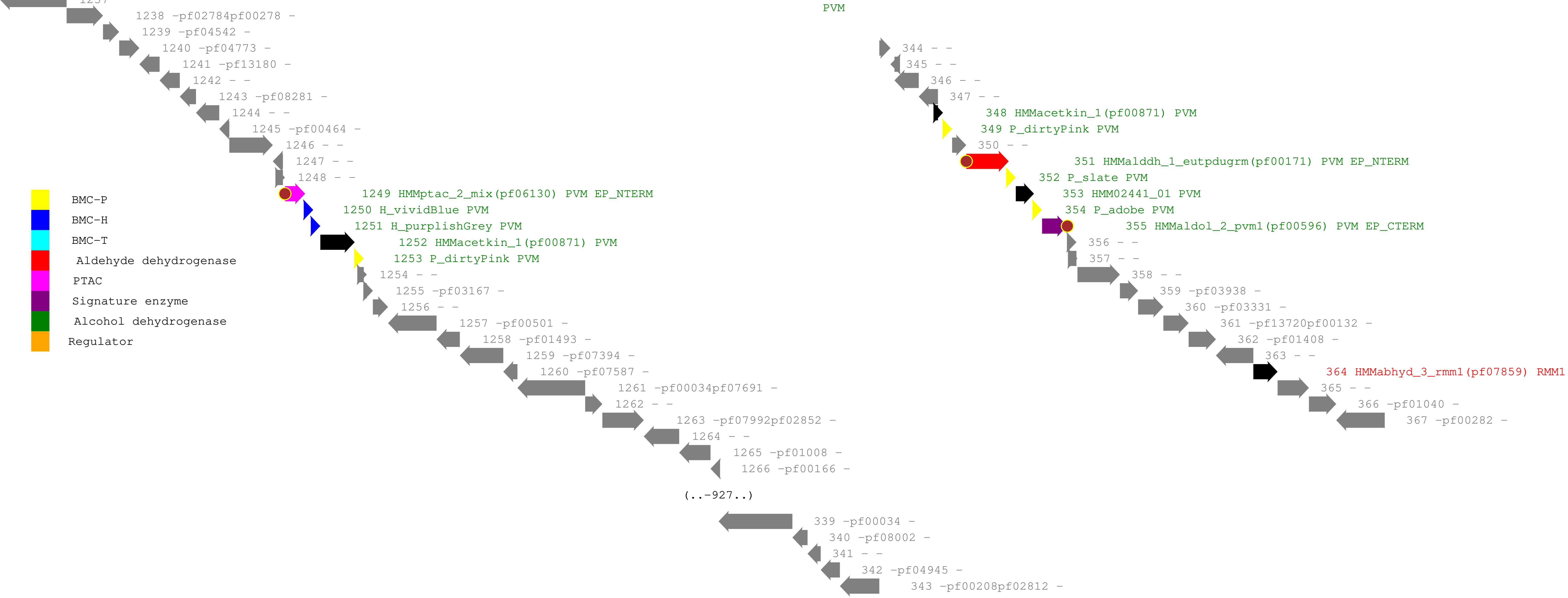

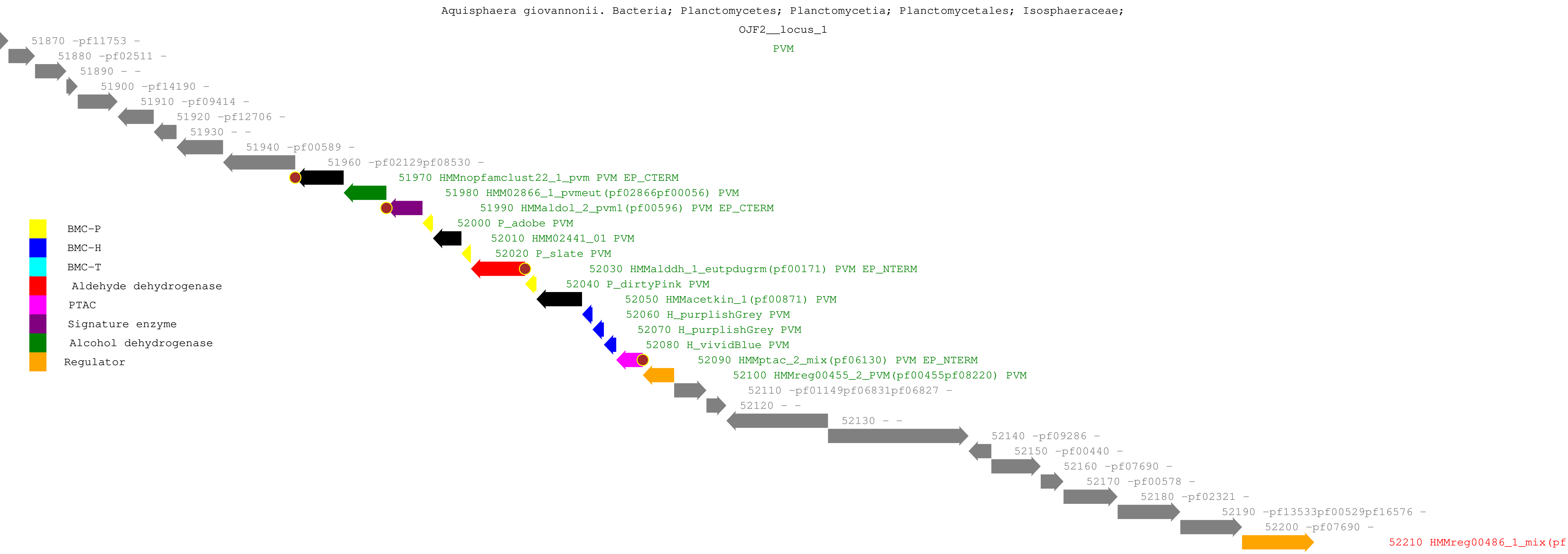
Aquisphaera giovannonii. Bacteria; Planctomycetes; Planctomycetia; Planctomycetales; Isosphaeraceae;
OJF2__locus_1
PVM
51870 -pf11753 -
51880 -pf02511 -
51890 - -
51900 -pf14190 -
51910 -pf09414 -
51920 -pf12706 -
51930 - -
51940 -pf00589 -
51960 -pf02129pf08530 -
51970 HMMnopfamclust22_1_pvm PVM EP_CTERM
51980 HMM02866_1_pvmeut(pf02866pf00056) PVM
51990 HMMaldol_2_pvm1(pf00596) PVM EP_CTERM
52000 P_adobe PVM
52010 HMM02441_01 PVM
52020 P_slate PVM
52030 HMMaldh_1_eutpdugrm(pf00171) PVM EP_NTERM
52040 P_dirtyPink PVM
52050 HMMacetkin_1(pf00871) PVM
52060 H_purplishGrey PVM
52070 H_purplishGrey PVM
52080 H_vividBlue PVM
52090 HMMptac_2_mix(pf06130) PVM EP_NTERM
52100 HMMreg00455_2_PVM(pf00455pf08220) PVM
52110 -pf01149pf06831pf06827 -
52120 - -
52130 - -
52140 -pf09286 -
52150 -pf00440 -
52160 -pf07690 -
52170 -pf00578 -
52180 -pf02321 -
52190 -pf13533pf00529pf16576 -
52200 -pf07690 -
52210 HMMreg00486_1_mix(pf
BMC-P
BMC-H
BMC-T
Aldehyde dehydrogenase
PTAC
Signature enzyme
Alcohol dehydrogenase
Regulator

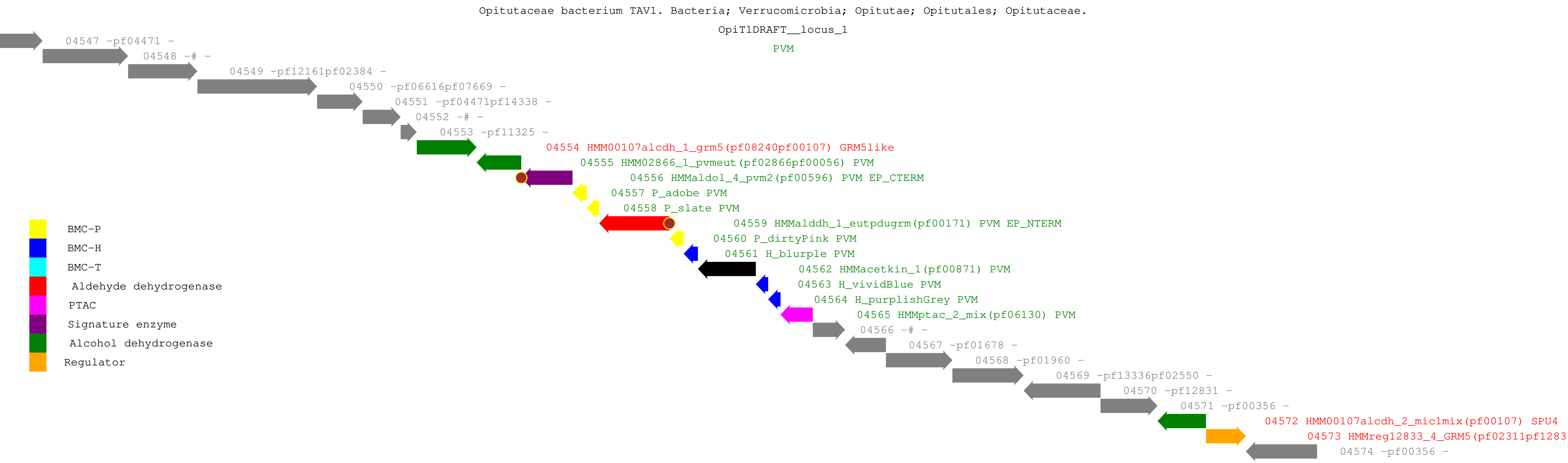
Opitutaceae bacterium TAV1. Bacteria; Verrucomicrobia; Opitutae; Opitutales; Opitutaceae.
OpiT1DRAFT__locus_1
PVM
04547 -pf04471 -
04548 -# -
04549 -pf12161pf02384 -
04550 -pf06616pf07669 -
04551 -pf04471pf14338 -
04552 -# -
04553 -pf11325 -
04554 HMM00107alcdh_1_grm5(pf08240pf00107) GRM5like
04555 HMM02866_1_pvmeut(pf02866pf00056) PVM
04556 HMMaldol_4_pvm2(pf00596) PVM EP_CTERM
04557 P_adobe PVM
04558 P_slate PVM
04559 HMMalddh_1_eutpdugrm(pf00171) PVM EP_NTERM
04560 P_dirtyPink PVM
04561 H_blurple PVM
04562 HMMacetkin_1(pf00871) PVM
04563 H_vividBlue PVM
04564 H_purplishGrey PVM
04565 HMMptac_2_mix(pf06130) PVM
04566 -# -
04567 -pf01678 -
04568 -pf01960 -
04569 -pf13336pf02550 -
04570 -pf12831 -
04571 -pf00356 -
04572 HMM00107alcdh_2_mic1mix(pf00107) SPU4
04573 HMMreg12833_4_GRM5(pf02311pf1283
04574 -pf00356 -
BMC-P
BMC-H
BMC-T
Aldehyde dehydrogenase
PTAC
Signature enzyme
Alcohol dehydrogenase
Regulator

Opitutaceae bacterium TAV5. Bacteria; Verrucomicrobia; Opitutae; Opitutales; Opitutaceae.

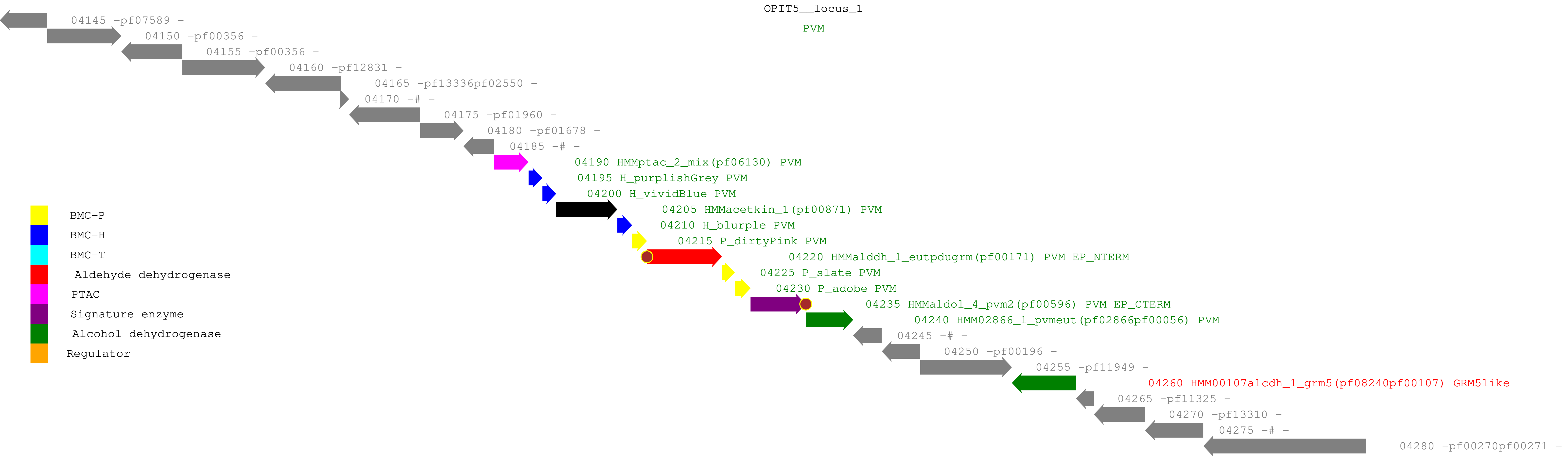

Opitutus terrae (strain DSM 11246 / JCM 15787 / PB90-1). Bacteria; Verrucomicrobia; Opitutae; Opitutales; Opitutaceae;

Oter__locus_1_main_sat

PVM

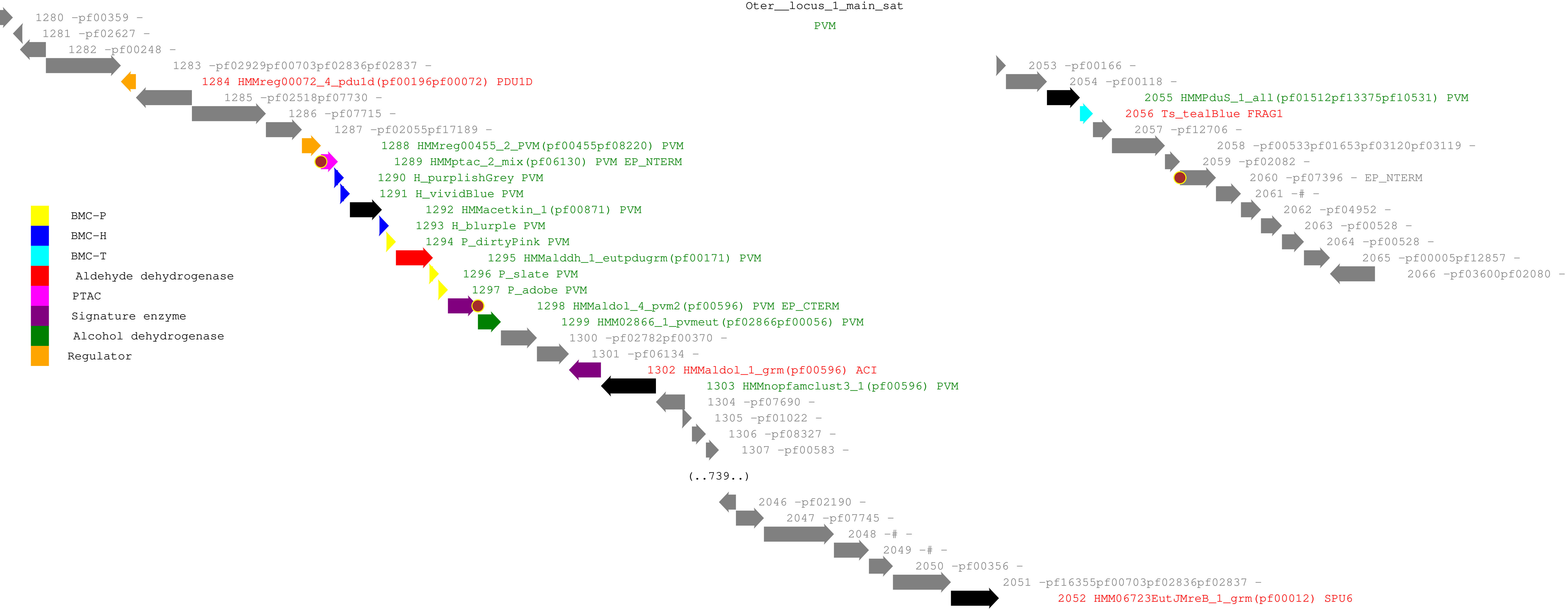

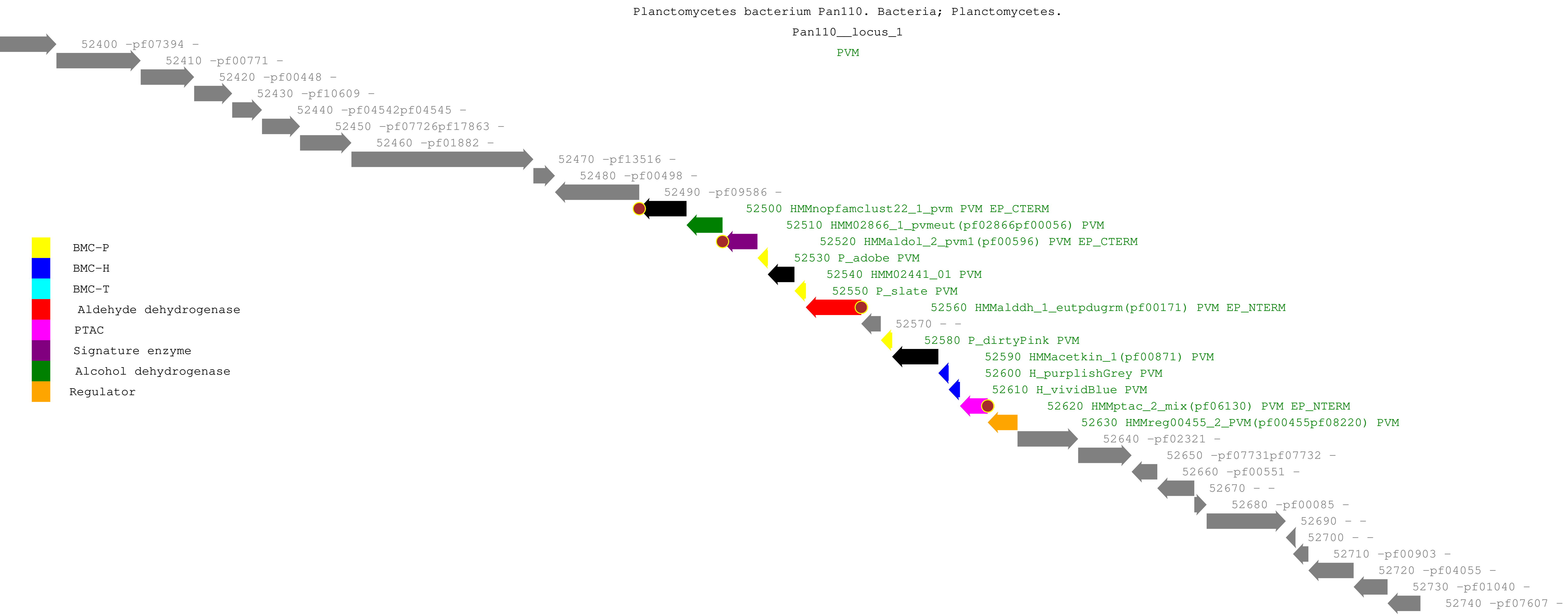
Planctomycetes bacterium Pan110. Bacteria; Planctomycetes.
Pan110__locus_1
PVM
52400 -pf07394 -
52410 -pf00771 -
52420 -pf00448 -
52430 -pf10609 -
52440 -pf04542pf04545 -
52450 -pf07726pf17863 -
52460 -pf01882 -
52470 -pf13516 -
52480 -pf00498 -
52490 -pf09586 -
52500 HMMnopfamclust22_1_pvm PVM EP_CTERM
52510 HMM02866_1_pvmeut(pf02866pf00056) PVM
52520 HMMaldol_2_pvm1(pf00596) PVM EP_CTERM
52530 P_adobe PVM
52540 HMM02441_01 PVM
52550 P_slate PVM
52560 HMMalddh_1_eutpdugrm(pf00171) PVM EP_NTERM
52570 - -
52580 P_dirtyPink PVM
52590 HMMacetkin_1(pf00871) PVM
52600 H_purplishGrey PVM
52610 H_vividBlue PVM
52620 HMMptac_2_mix(pf06130) PVM EP_NTERM
52630 HMMreg00455_2_PVM(pf00455pf08220) PVM
52640 -pf02321 -
52650 -pf07731pf07732 -
52660 -pf00551 -
52670 - -
52680 -pf00085 -
52690 - -
52700 - -
52710 -pf00903 -
52720 -pf04055 -
52730 -pf01040 -
52740 -pf07607 -
BMC-P
BMC-H
BMC-T
Aldehyde dehydrogenase
PTAC
Signature enzyme
Alcohol dehydrogenase
Regulator

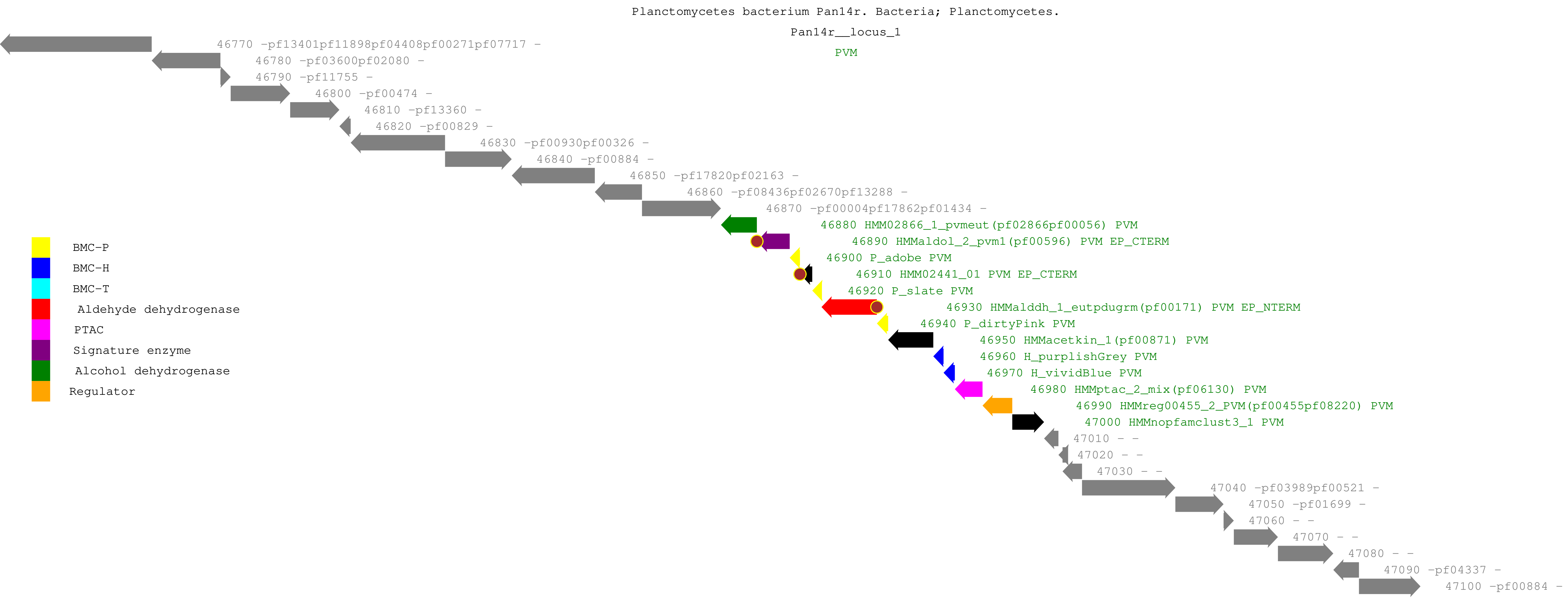

Planctomycetes bacterium Pan14r. Bacteria; Planctomycetes.
Pan14r__locus_1
PVM
46770 -pf13401pf11898pf04408pf00271pf07717 -
46780 -pf03600pf02080 -
46790 -pf11755 -
46800 -pf00474 -
46810 -pf13360 -
46820 -pf00829 -
46830 -pf00930pf00326 -
46840 -pf00884 -
46850 -pf17820pf02163 -
46860 -pf08436pf02670pf13288 -
46870 -pf00004pf17862pf01434 -
46880 HMM02866_1_pvmeut(pf02866pf00056) PVM
46890 HMMaldol_2_pvm1(pf00596) PVM EP_CTERM
46900 P_adobe PVM
46910 HMM02441_01 PVM EP_CTERM
46920 P_slate PVM
46930 HMMalddh_1_eutpdugrm(pf00171) PVM EP_NTERM
46940 P_dirtyPink PVM
46950 HMMacetkin_1(pf00871) PVM
46960 H_purplishGrey PVM
46970 H_vividBlue PVM
46980 HMMptac_2_mix(pf06130) PVM
46990 HMMreg00455_2_PVM(pf00455pf08220) PVM
47000 HMMnopfamclust3_1 PVM
47010 - -
47020 - -
47030 - -
47040 -pf03989pf00521 -
47050 -pf01699 -
47060 - -
47070 - -
47080 - -
47090 -pf04337 -
47100 -pf00884 -
BMC-P
BMC-H
BMC-T
Aldehyde dehydrogenase
PTAC
Signature enzyme
Alcohol dehydrogenase
Regulator

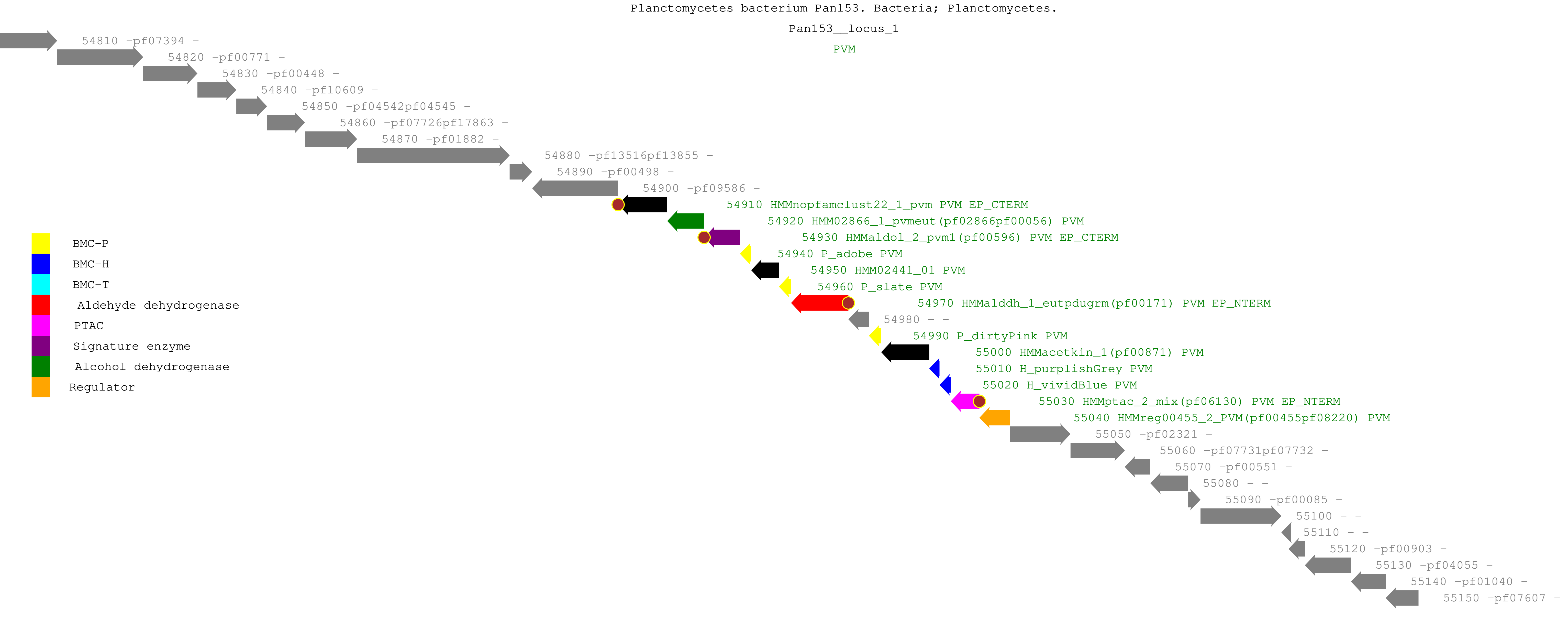
Planctomycetes bacterium Pan153. Bacteria; Planctomycetes.
Pan153__locus_1
PVM
54810 -pf07394 -
54820 -pf00771 -
54830 -pf00448 -
54840 -pf10609 -
54850 -pf04542pf04545 -
54860 -pf07726pf17863 -
54870 -pf01882 -
54880 -pf13516pf13855 -
54890 -pf00498 -
54900 -pf09586 -
54910 HMMnopfamclust22_1_pvm PVM EP_CTERM
54920 HMM02866_1_pvmeut(pf02866pf00056) PVM
54930 HMMaldol_2_pvm1(pf00596) PVM EP_CTERM
54940 P_adobe PVM
54950 HMM02441_01 PVM
54960 P_slate PVM
54970 HMMalddh_1_eutpdugrm(pf00171) PVM EP_NTERM
54980 - -
54990 P_dirtyPink PVM
55000 HMMacetkin_1(pf00871) PVM
55010 H_purplishGrey PVM
55020 H_vividBlue PVM
55030 HMMptac_2_mix(pf06130) PVM EP_NTERM
55040 HMMreg00455_2_PVM(pf00455pf08220) PVM
55050 -pf02321 -
55060 -pf07731pf07732 -
55070 -pf00551 -
55080 - -
55090 -pf00085 -
55100 - -
55110 - -
55120 -pf00903 -
55130 -pf04055 -
55140 -pf01040 -
55150 -pf07607 -
BMC-P
BMC-H
BMC-T
Aldehyde dehydrogenase
PTAC
Signature enzyme
Alcohol dehydrogenase
Regulator

Planctomycetes bacterium Pan161. Bacteria; Planctomycetes.

Pan161__locus_1

PVM

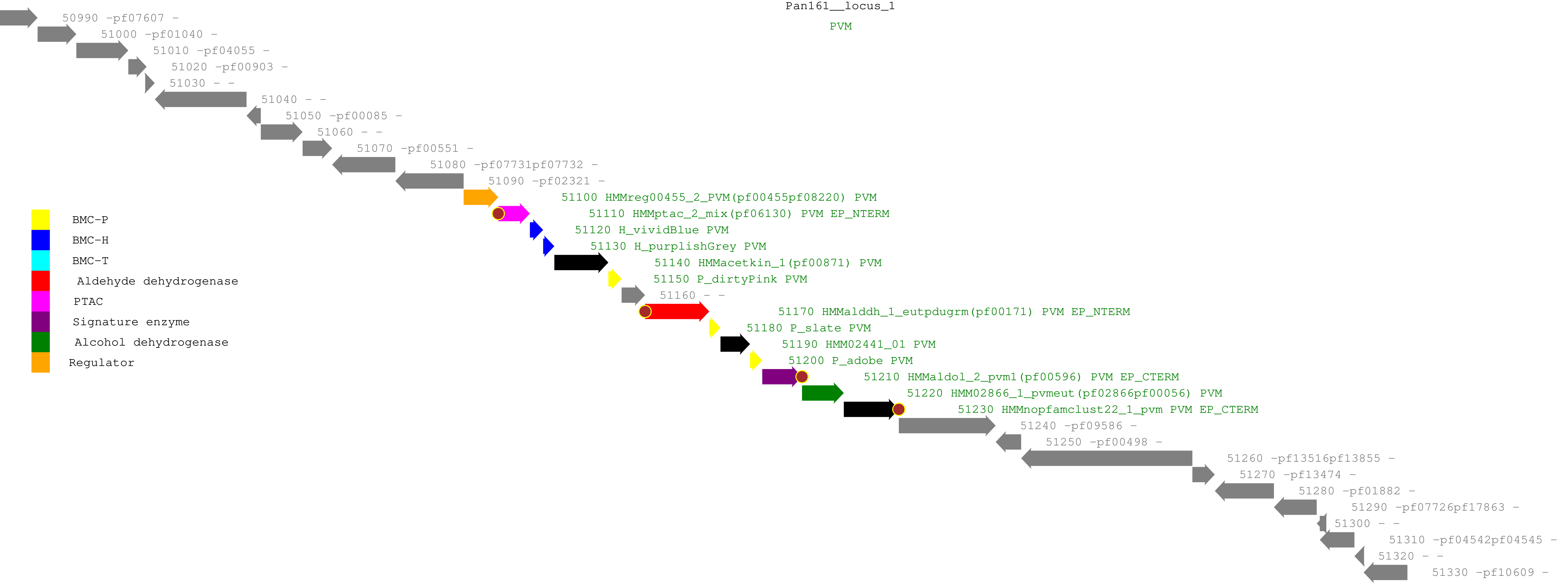

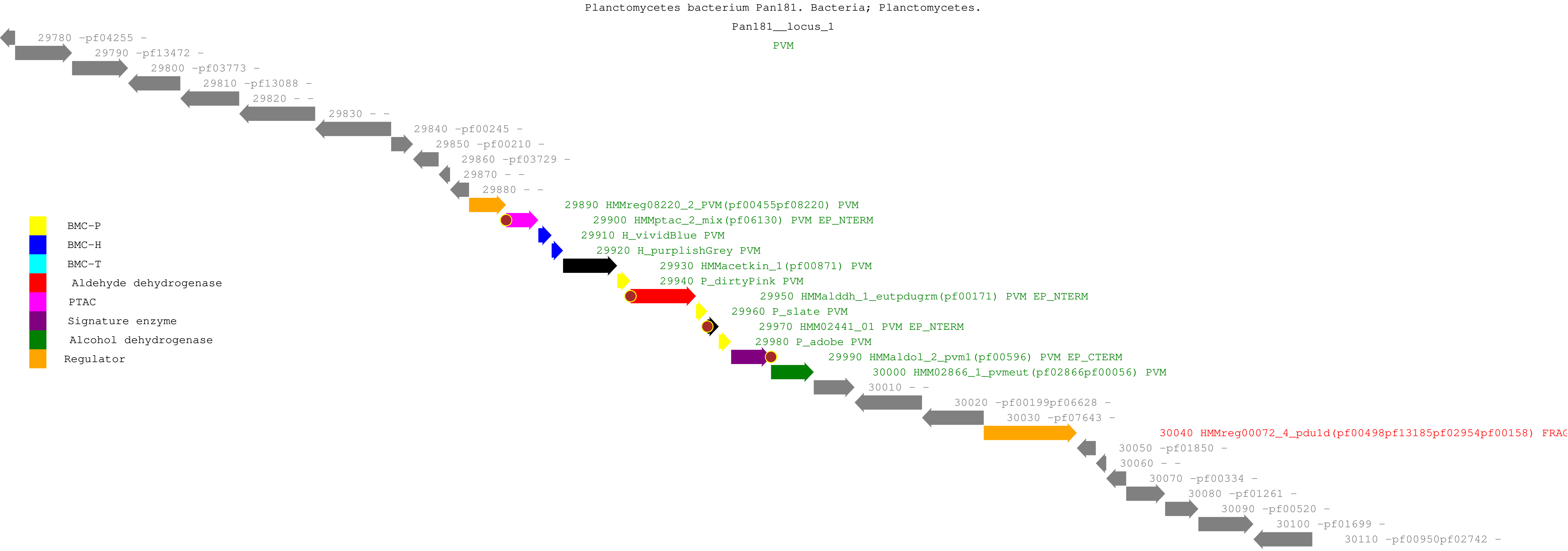
Planctomycetes bacterium Pan181. Bacteria; Planctomycetes.
Pan181__locus_1
PVM
29780 -pf04255 -
29790 -pf13472 -
29800 -pf03773 -
29810 -pf13088 -
29820 - -
29830 - -
29840 -pf00245 -
29850 -pf00210 -
29860 -pf03729 -
29870 - -
29880 - -
29890 HMMreg08220_2_PVM(pf00455pf08220) PVM
29900 HMMptac_2_mix(pf06130) PVM EP_NTERM
29910 H_vividBlue PVM
29920 H_purplishGrey PVM
29930 HMMacetkin_1(pf00871) PVM
29940 P_dirtyPink PVM
29950 HMMalddh_1_eutpdugrm(pf00171) PVM EP_NTERM
29960 P_slate PVM
29970 HMM02441_01 PVM EP_NTERM
29980 P_adobe PVM
29990 HMMaldol_2_pvm1(pf00596) PVM EP_CTERM
30000 HMM02866_1_pvmeut(pf02866pf00056) PVM
30010 - -
30020 -pf00199pf06628 -
30030 -pf07643 -
30040 HMMreg00072_4_pdu1d(pf00498pf13185pf02954pf00158) FRAG
30050 -pf01850 -
30060 - -
30070 -pf00334 -
30080 -pf01261 -
30090 -pf00520 -
30100 -pf01699 -
30110 -pf00950pf02742 -
BMC-P
BMC-H
BMC-T
Aldehyde dehydrogenase
PTAC
Signature enzyme
Alcohol dehydrogenase
Regulator

Planctomycetes bacterium Pan189. Bacteria; Planctomycetes.

Pan189__locus_1

PVM

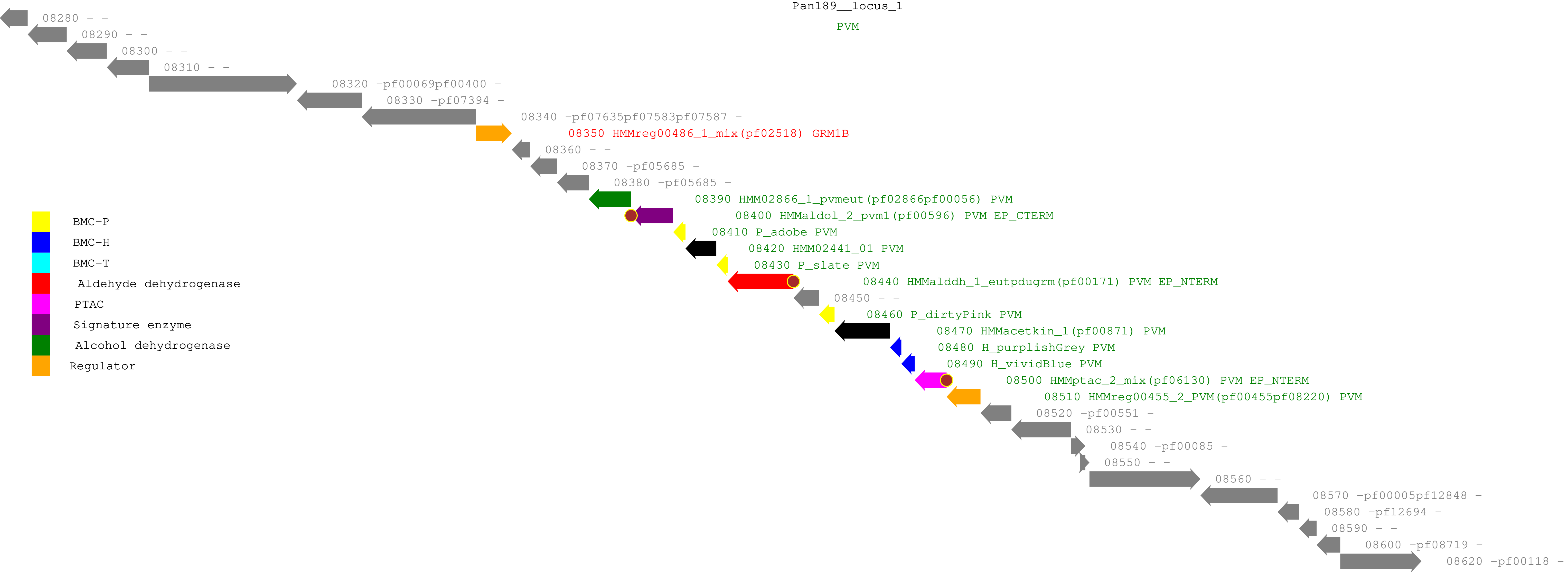

Planctomycetes bacterium Pan216. Bacteria; Planctomycetes.

Pan216__locus_1

PVM

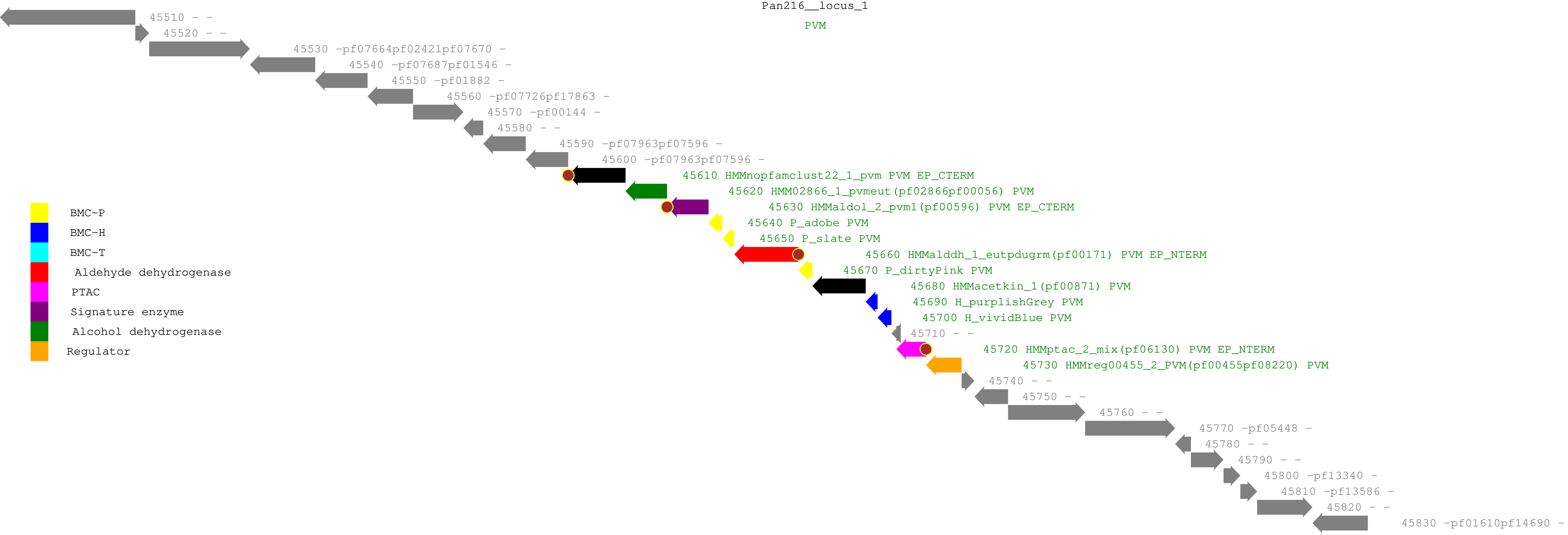

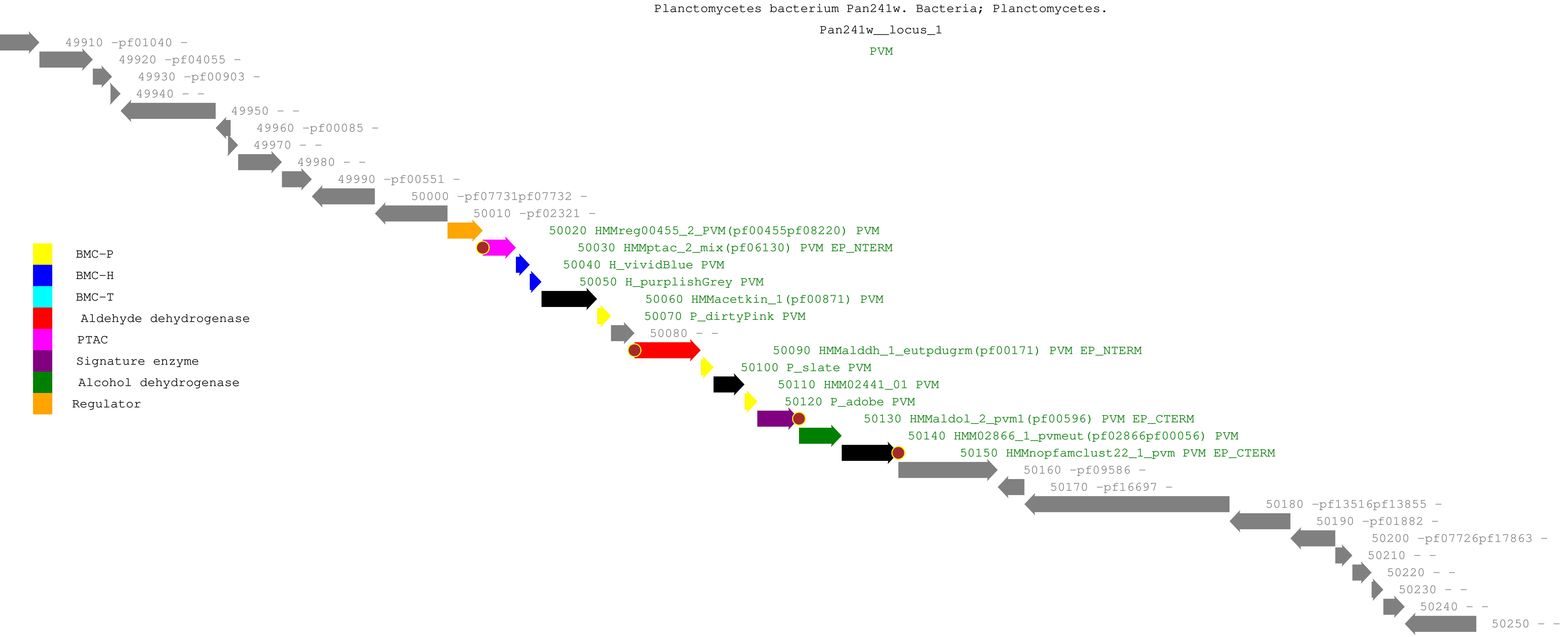

Planctomycetes bacterium Pan241w. Bacteria; Planctomycetes.
Pan241w__locus_1
PVM
49910 -pf01040 -
49920 -pf04055 -
49930 -pf00903 -
49940 - -
49950 - -
49960 -pf00085 -
49970 - -
49980 - -
49990 -pf00551 -
50000 -pf07731pf07732 -
50010 -pf02321 -
50020 HMMreg00455_2_PVM(pf00455pf08220) PVM
50030 HMMptac_2_mix(pf06130) PVM EP_NTERM
50040 H_vividBlue PVM
50050 H_purplishGrey PVM
50060 HMMacetkin_1(pf00871) PVM
50070 P_dirtyPink PVM
50080 - -
50090 HMMaldd h_1_eutpdugrm(pf00171) PVM EP_NTERM
50100 P_slate PVM
50110 HMM02441_01 PVM
50120 P_adobe PVM
50130 HMMaldol_2_pvm1(pf00596) PVM EP_CTERM
50140 HMM02866_1_pvmeut(pf02866pf00056) PVM
50150 HMMnopfamclust22_1_pvm PVM EP_CTERM
50160 -pf09586 -
50170 -pf16697 -
50180 -pf13516pf13855 -
50190 -pf01882 -
50200 -pf07726pf17863 -
50210 - -
50220 - -
50230 - -
50240 - -
50250 - -
BMC-P
BMC-H
BMC-T
Aldehyde dehydrogenase
PTAC
Signature enzyme
Alcohol dehydrogenase
Regulator

Planctomycetes bacterium Pan265. Bacteria; Planctomycetes.

Pan265__locus_1

PVM

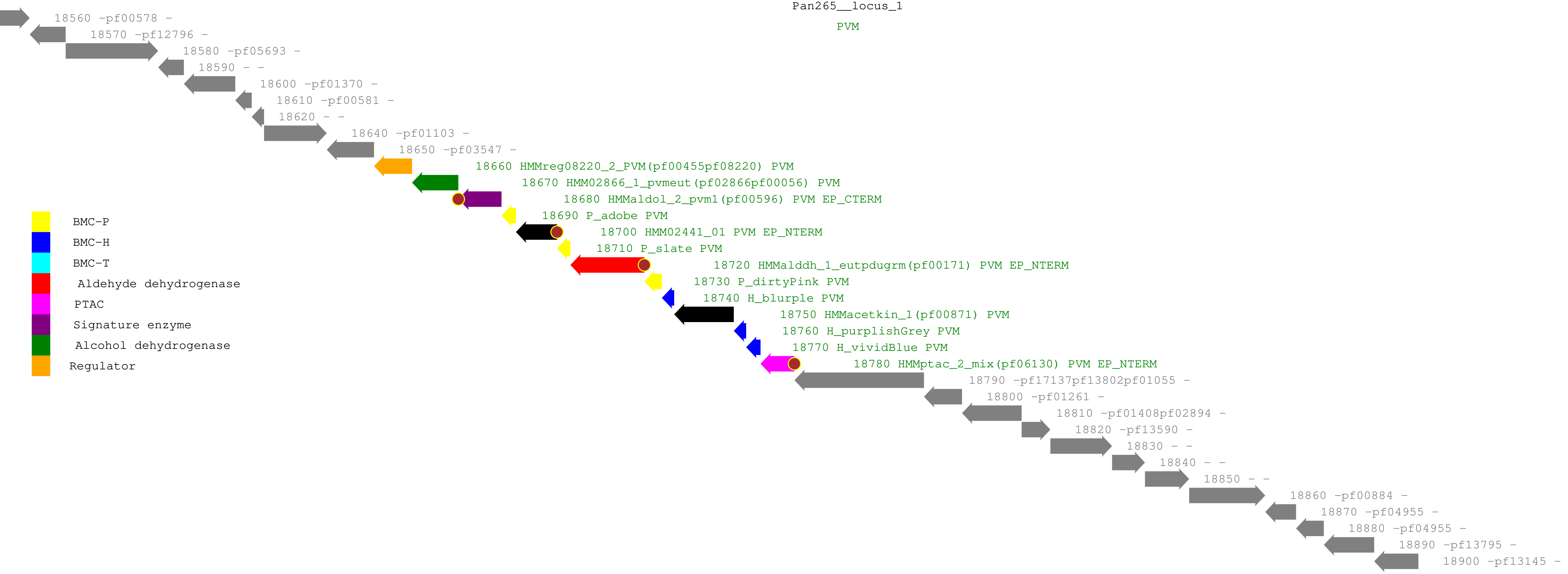

Planctomycetes bacterium Pan44. Bacteria; Planctomycetes.

Pan44__locus_fragmented

PVM

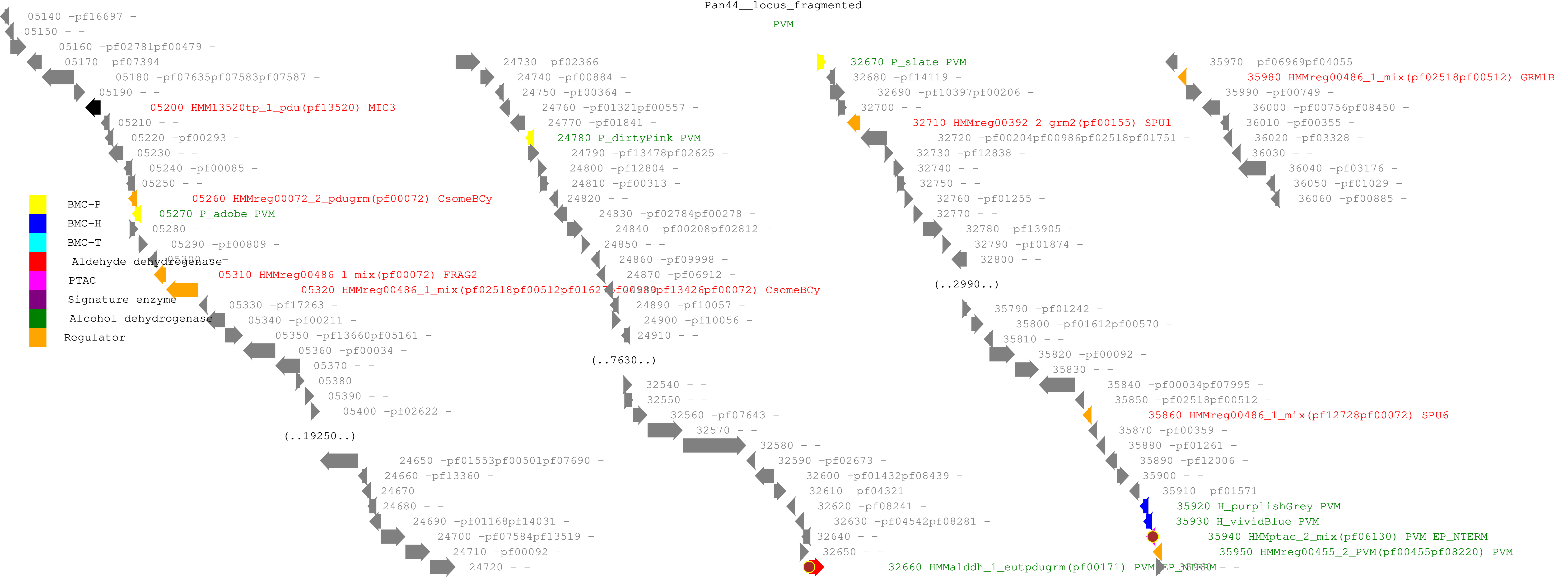

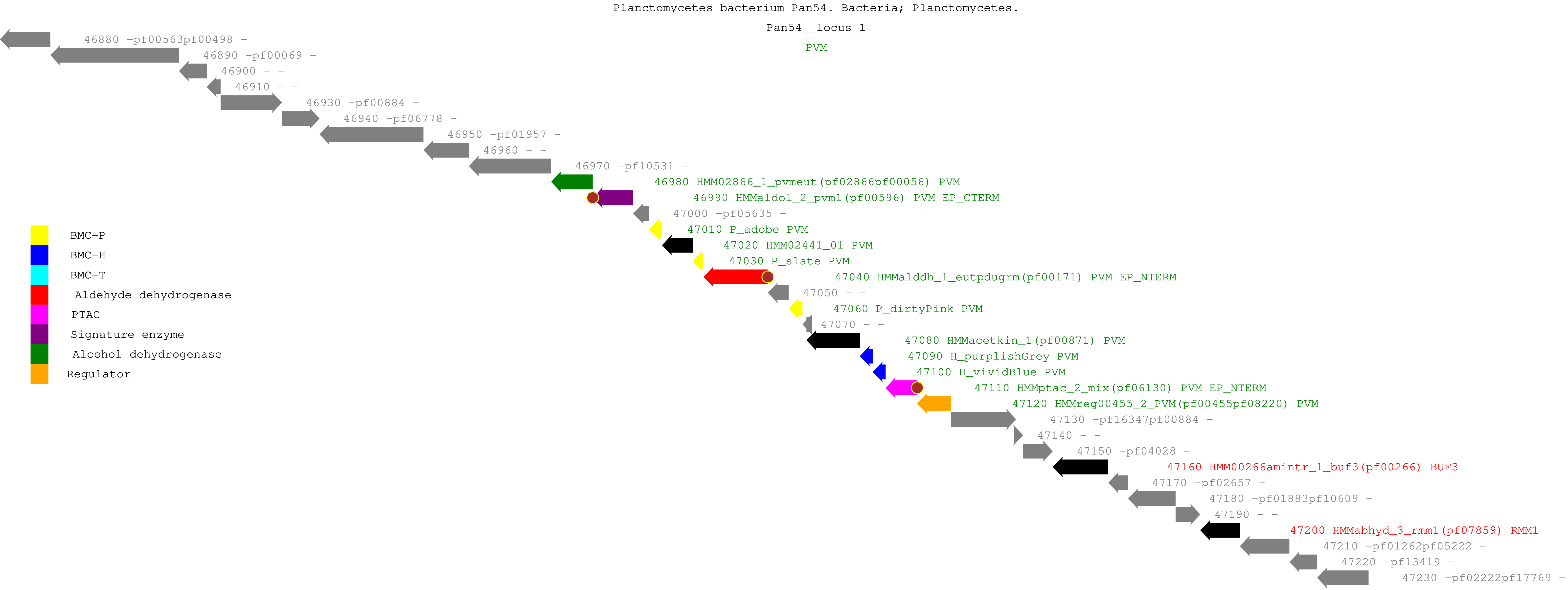

Planctomycetes bacterium Pan54. Bacteria; Planctomycetes.
Pan54__locus_1
PVM
46880 -pf00563pf00498 -
46890 -pf00069 -
46900 - -
46910 - -
46930 -pf00884 -
46940 -pf06778 -
46950 -pf01957 -
46960 - -
46970 -pf10531 -
46980 HMM02866_1_pvmeut(pf02866pf00056) PVM
46990 HMMaldol_2_pvm1(pf00596) PVM EP_CTERM
47000 -pf05635 -
47010 P_adobe PVM
47020 HMM02441_01 PVM
47030 P_slate PVM
47040 HMMalddh_1_eutpdugrm(pf00171) PVM EP_NTERM
47050 - -
47060 P_dirtyPink PVM
47070 - -
47080 HMMacetkin_1(pf00871) PVM
47090 H_purplishGrey PVM
47100 H_vividBlue PVM
47110 HMMptac_2_mix(pf06130) PVM EP_NTERM
47120 HMMreg00455_2_PVM(pf00455pf08220) PVM
47130 -pf16347pf00884 -
47140 - -
47150 -pf04028 -
47160 HMM00266amintr_1_buf3(pf00266) BUF3
47170 -pf02657 -
47180 -pf01883pf10609 -
47190 - -
47200 HMMabhyd_3_rmm1(pf07859) RMM1
47210 -pf01262pf05222 -
47220 -pf13419 -
47230 -pf02222pf17769 -
BMC-P
BMC-H
BMC-T
Aldehyde dehydrogenase
PTAC
Signature enzyme
Alcohol dehydrogenase
Regulator

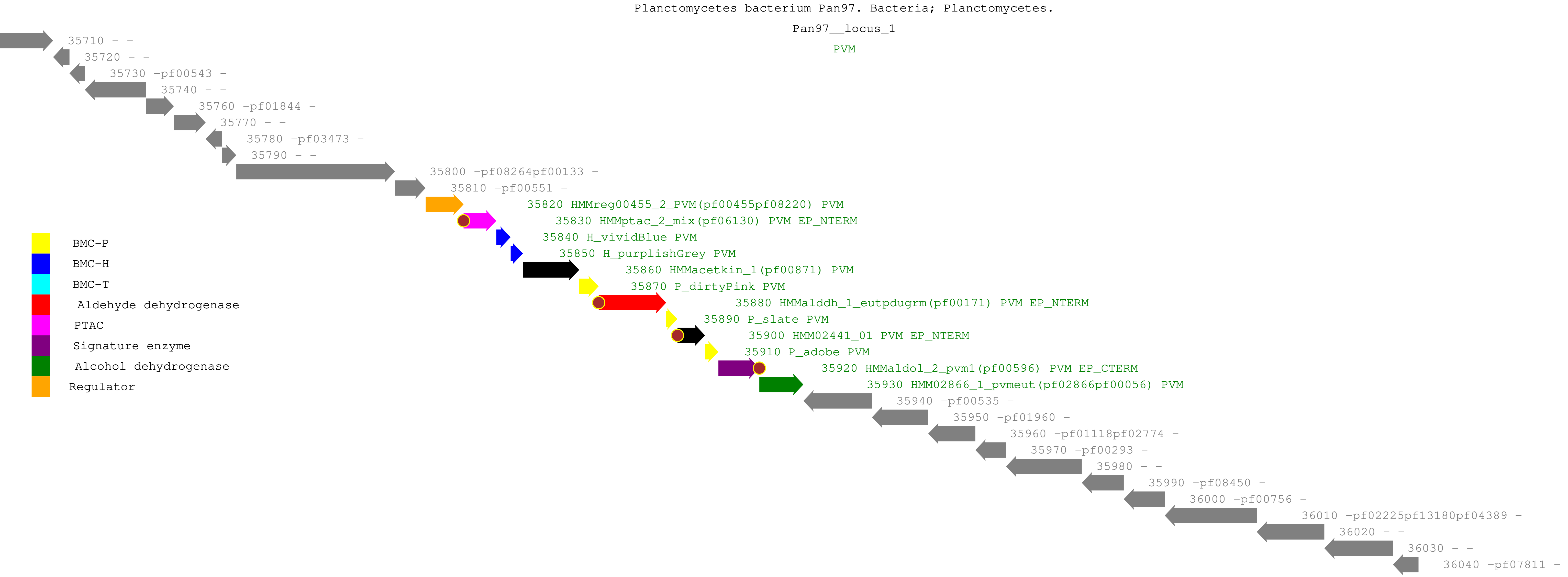
Planctomycetes bacterium Pan97. Bacteria; Planctomycetes.
Pan97__locus_1
PVM
35710 - -
35720 - -
35730 -pf00543 -
35740 - -
35760 -pf01844 -
35770 - -
35780 -pf03473 -
35790 - -
35800 -pf08264pf00133 -
35810 -pf00551 -
35820 HMMreg00455_2_PVM(pf00455pf08220) PVM
35830 HMMptac_2_mix(pf06130) PVM EP_NTERM
35840 H_vividBlue PVM
35850 H_purplishGrey PVM
35860 HMMacetkin_1(pf00871) PVM
35870 P_dirtyPink PVM
35880 HMMaldh_1_eutpdugrm(pf00171) PVM EP_NTERM
35890 P_slate PVM
35900 HMM02441_01 PVM EP_NTERM
35910 P_adobe PVM
35920 HMMaldol_2_pvm1(pf00596) PVM EP_CTERM
35930 HMM02866_1_pvmeut(pf02866pf00056) PVM
35940 -pf00535 -
35950 -pf01960 -
35960 -pf01118pf02774 -
35970 -pf00293 -
35980 - -
35990 -pf08450 -
36000 -pf00756 -
36010 -pf02225pf13180pf04389 -
36020 - -
36030 - -
36040 -pf07811 -
BMC-P
BMC-H
BMC-T
Aldehyde dehydrogenase
PTAC
Signature enzyme
Alcohol dehydrogenase
Regulator

Planctomycetes bacterium Pla100. Bacteria; Planctomycetes.

Pla100__locus_1

PVM

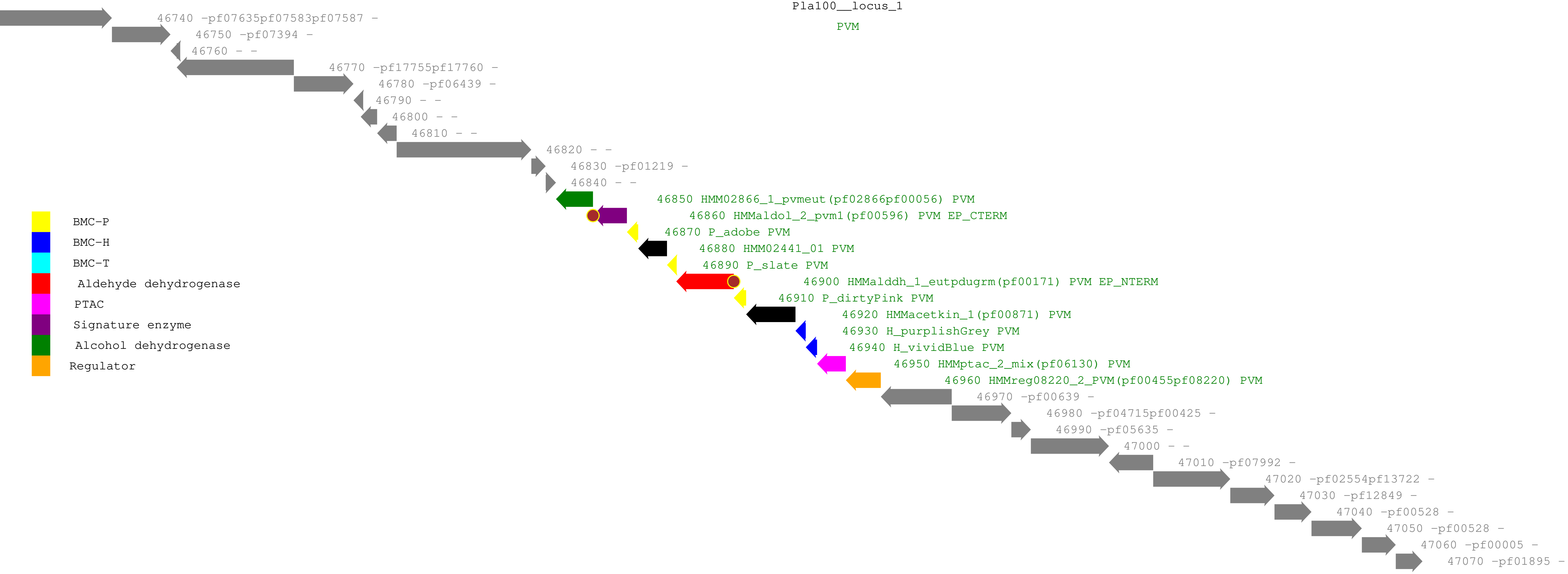

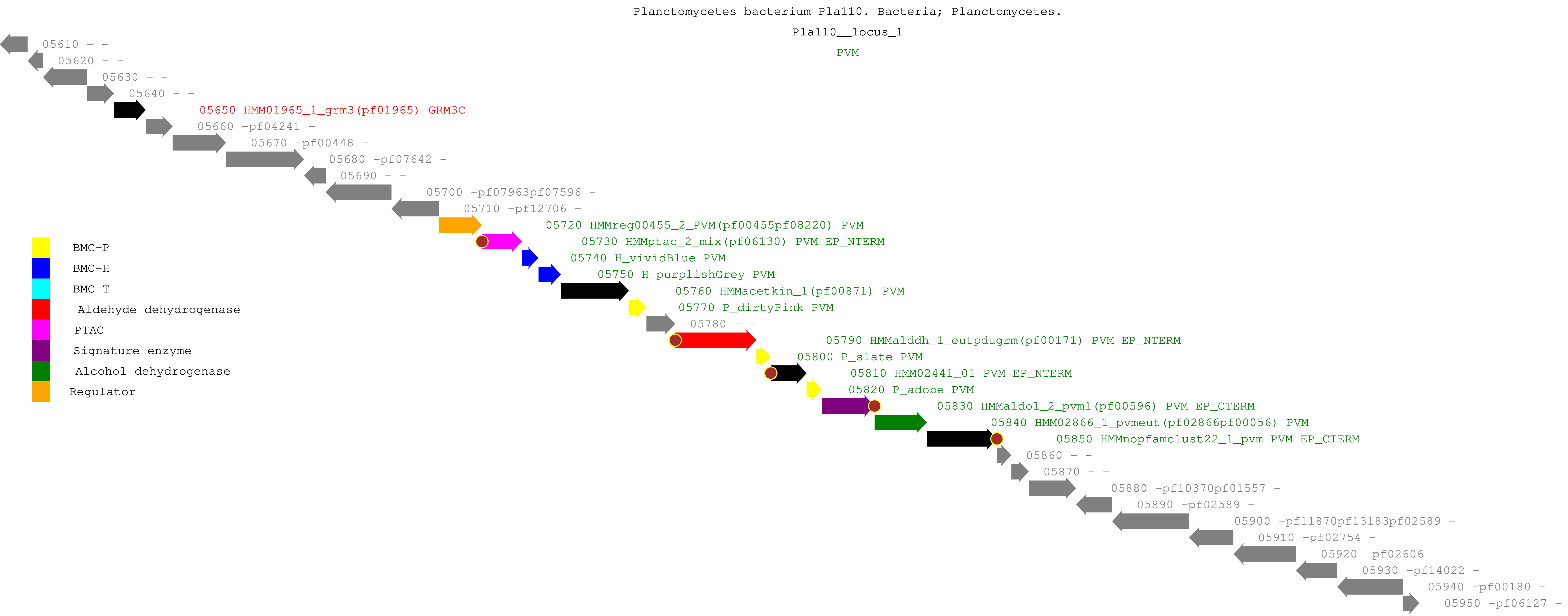

Planctomycetes bacterium Pla110. Bacteria; Planctomycetes.
Pla110__locus_1
PVM
05610 - -
05620 - -
05630 - -
05640 - -
05650 HMM01965_1_grm3(pf01965) GRM3C
05660 -pf04241 -
05670 -pf00448 -
05680 -pf07642 -
05690 - -
05700 -pf07963pf07596 -
05710 -pf12706 -
05720 HMMreg00455_2_PVM(pf00455pf08220) PVM
05730 HMMptac_2_mix(pf06130) PVM EP_NTERM
05740 H_vividBlue PVM
05750 H_purplishGrey PVM
05760 HMMacetkin_1(pf00871) PVM
05770 P_dirtyPink PVM
05780 - -
05790 HMMaldh_1_eutpdugrm(pf00171) PVM EP_NTERM
05800 P_slate PVM
05810 HMM02441_01 PVM EP_NTERM
05820 P_adobe PVM
05830 HMMaldol_2_pvm1(pf00596) PVM EP_CTERM
05840 HMM02866_1_pvmeut(pf02866pf00056) PVM
05850 HMMnopfamclust22_1_pvm PVM EP_CTERM
05860 - -
05870 - -
05880 -pf10370pf01557 -
05890 -pf02589 -
05900 -pf11870pf13183pf02589 -
05910 -pf02754 -
05920 -pf02606 -
05930 -pf14022 -
05940 -pf00180 -
05950 -pf06127 -
BMC-P
BMC-H
BMC-T
Aldehyde dehydrogenase
PTAC
Signature enzyme
Alcohol dehydrogenase
Regulator

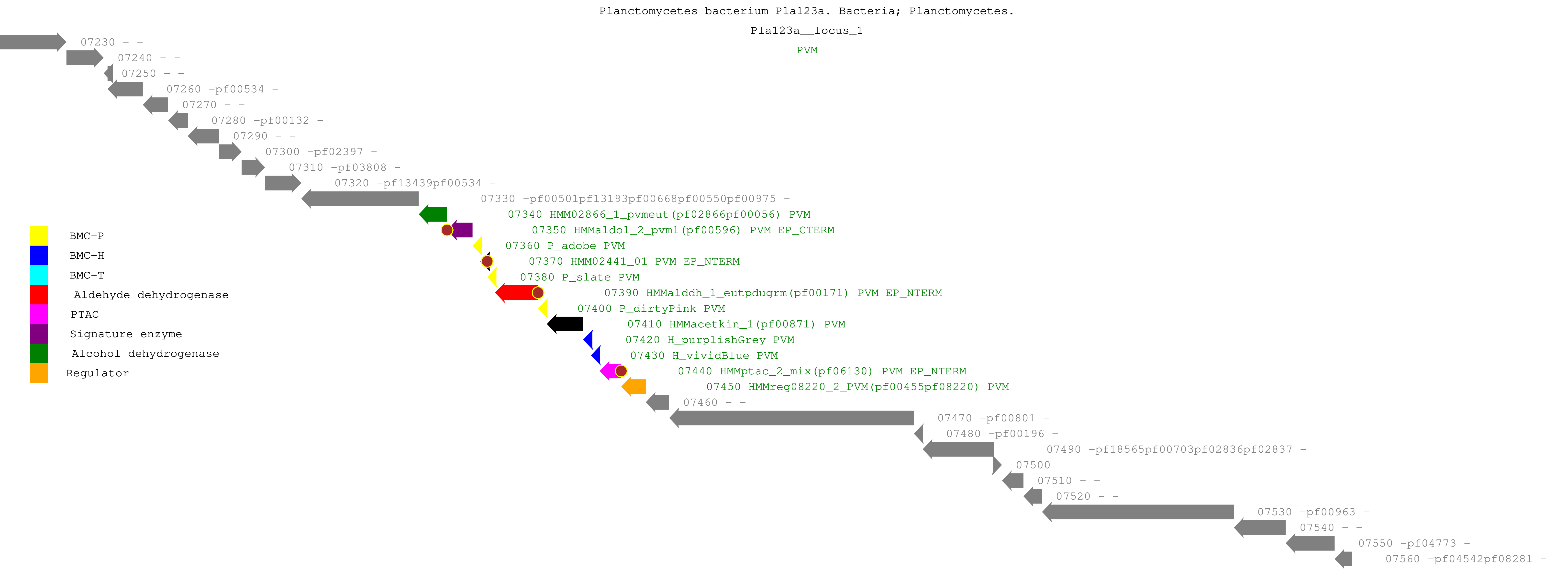
Planctomycetes bacterium Pla123a. Bacteria; Planctomycetes.
Pla123a__locus_1
PVM
07230 - -
07240 - -
07250 - -
07260 -pf00534 -
07270 - -
07280 -pf00132 -
07290 - -
07300 -pf02397 -
07310 -pf03808 -
07320 -pf13439pf00534 -
07330 -pf00501pf13193pf00668pf00550pf00975 -
07340 HMM02866_1_pvmeut(pf02866pf00056) PVM
07350 HMMaldol_2_pvm1(pf00596) PVM EP_CTERM
07360 P_adobe PVM
07370 HMM02441_01 PVM EP_NTERM
07380 P_slate PVM
07390 HMMalddh_1_eutpdugrm(pf00171) PVM EP_NTERM
07400 P_dirtyPink PVM
07410 HMMacetkin_1(pf00871) PVM
07420 H_purplishGrey PVM
07430 H_vividBlue PVM
07440 HMMptac_2_mix(pf06130) PVM EP_NTERM
07450 HMMreg08220_2_PVM(pf00455pf08220) PVM
07460 - -
07470 -pf00801 -
07480 -pf00196 -
07490 -pf18565pf00703pf02836pf02837 -
07500 - -
07510 - -
07520 - -
07530 -pf00963 -
07540 - -
07550 -pf04773 -
07560 -pf04542pf08281 -
BMC-P
BMC-H
BMC-T
Aldehyde dehydrogenase
PTAC
Signature enzyme
Alcohol dehydrogenase
Regulator

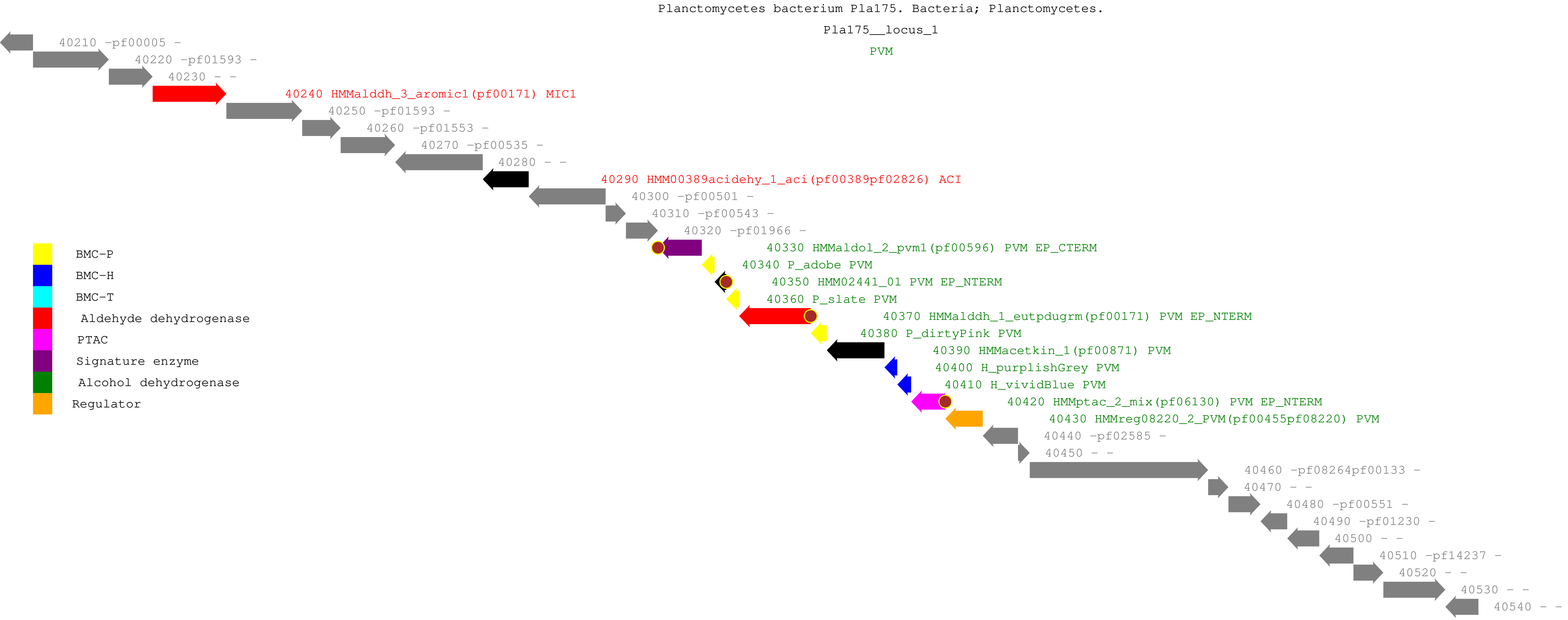
Planctomycetes bacterium Pla175. Bacteria; Planctomycetes.
Pla175__locus_1
PVM
40210 -pf00005 -
40220 -pf01593 -
40230 - -
40240 HMMaldh_3_aromic1(pf00171) MIC1
40250 -pf01593 -
40260 -pf01553 -
40270 -pf00535 -
40280 - -
40290 HMM00389acidehy_1_aci(pf00389pf02826) ACI
40300 -pf00501 -
40310 -pf00543 -
40320 -pf01966 -
40330 HMMaldol_2_pvm1(pf00596) PVM EP_CTERM
40340 P_adobe PVM
40350 HMM02441_01 PVM EP_NTERM
40360 P_slate PVM
40370 HMMaldh_1_eutpdugrm(pf00171) PVM EP_NTERM
40380 P_dirtyPink PVM
40390 HMMacetkin_1(pf00871) PVM
40400 H_purplishGrey PVM
40410 H_vividBlue PVM
40420 HMMptac_2_mix(pf06130) PVM EP_NTERM
40430 HMMreg08220_2_PVM(pf00455pf08220) PVM
40440 -pf02585 -
40450 - -
40460 -pf08264pf00133 -
40470 - -
40480 -pf00551 -
40490 -pf01230 -
40500 - -
40510 -pf14237 -
40520 - -
40530 - -
40540 - -
BMC-P
BMC-H
BMC-T
Aldehyde dehydrogenase
PTAC
Signature enzyme
Alcohol dehydrogenase
Regulator

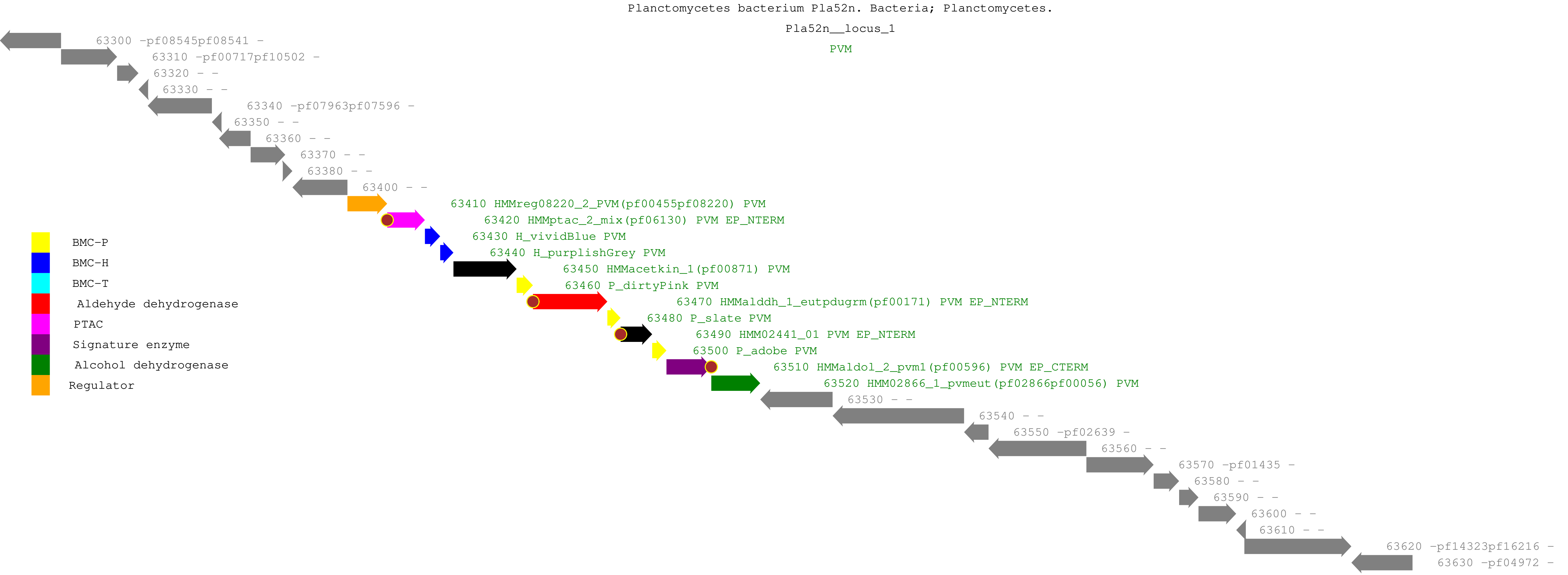
Planctomycetes bacterium Pla52n. Bacteria; Planctomycetes.
Pla52n__locus_1
PVM
63300 -pf08545pf08541 -
63310 -pf00717pf10502 -
63320 - -
63330 - -
63340 -pf07963pf07596 -
63350 - -
63360 - -
63370 - -
63380 - -
63400 - -
63410 HMMreg08220_2_PVM(pf00455pf08220) PVM
63420 HMMptac_2_mix(pf06130) PVM EP_NTERM
63430 H_vividBlue PVM
63440 H_purplishGrey PVM
63450 HMMacetkin_1(pf00871) PVM
63460 P_dirtyPink PVM
63470 HMMaldh_1_eutpdugrm(pf00171) PVM EP_NTERM
63480 P_slate PVM
63490 HMM02441_01 PVM EP_NTERM
63500 P_adobe PVM
63510 HMMaldol_2_pvm1(pf00596) PVM EP_CTERM
63520 HMM02866_1_pvmeut(pf02866pf00056) PVM
63530 - -
63540 - -
63550 -pf02639 -
63560 - -
63570 -pf01435 -
63580 - -
63590 - -
63600 - -
63610 - -
63620 -pf14323pf16216 -
63630 -pf04972 -
BMC-P
BMC-H
BMC-T
Aldehyde dehydrogenase
PTAC
Signature enzyme
Alcohol dehydrogenase
Regulator

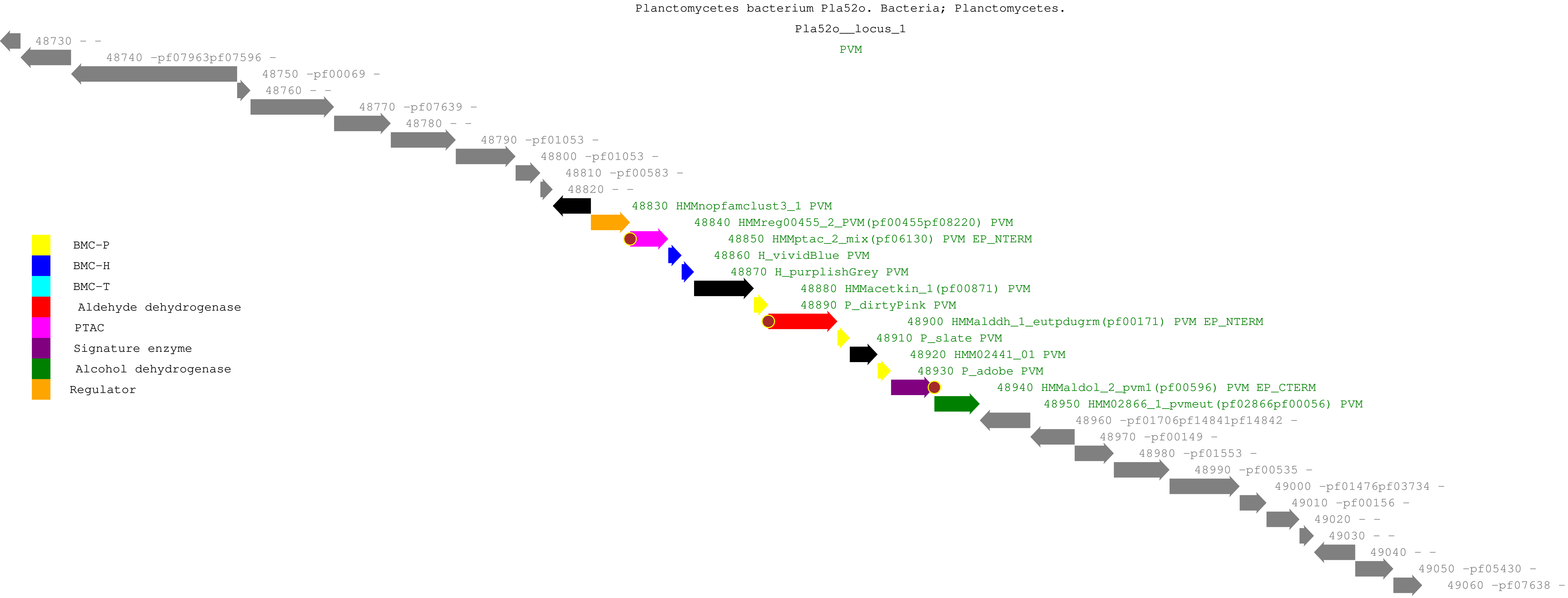
Planctomycetes bacterium Pla52o. Bacteria; Planctomycetes.
Pla52o__locus_1
PVM
48730 - -
48740 -pf07963pf07596 -
48750 -pf00069 -
48760 - -
48770 -pf07639 -
48780 - -
48790 -pf01053 -
48800 -pf01053 -
48810 -pf00583 -
48820 - -
48830 HMMnopfamclust3_1 PVM
48840 HMMreg00455_2_PVM(pf00455pf08220) PVM
48850 HMMptac_2_mix(pf06130) PVM EP_NTERM
48860 H_vividBlue PVM
48870 H_purplishGrey PVM
48880 HMMacetkin_1(pf00871) PVM
48890 P_dirtyPink PVM
48900 HMMalddh_1_eutpdugrm(pf00171) PVM EP_NTERM
48910 P_slate PVM
48920 HMM02441_01 PVM
48930 P_adobe PVM
48940 HMMaldol_2_pvm1(pf00596) PVM EP_CTERM
48950 HMM02866_1_pvmeut(pf02866pf00056) PVM
48960 -pf01706pf14841pf14842 -
48970 -pf00149 -
48980 -pf01553 -
48990 -pf00535 -
49000 -pf01476pf03734 -
49010 -pf00156 -
49020 - -
49030 - -
49040 - -
49050 -pf05430 -
49060 -pf07638 -
BMC-P
BMC-H
BMC-T
Aldehyde dehydrogenase
PTAC
Signature enzyme
Alcohol dehydrogenase
Regulator

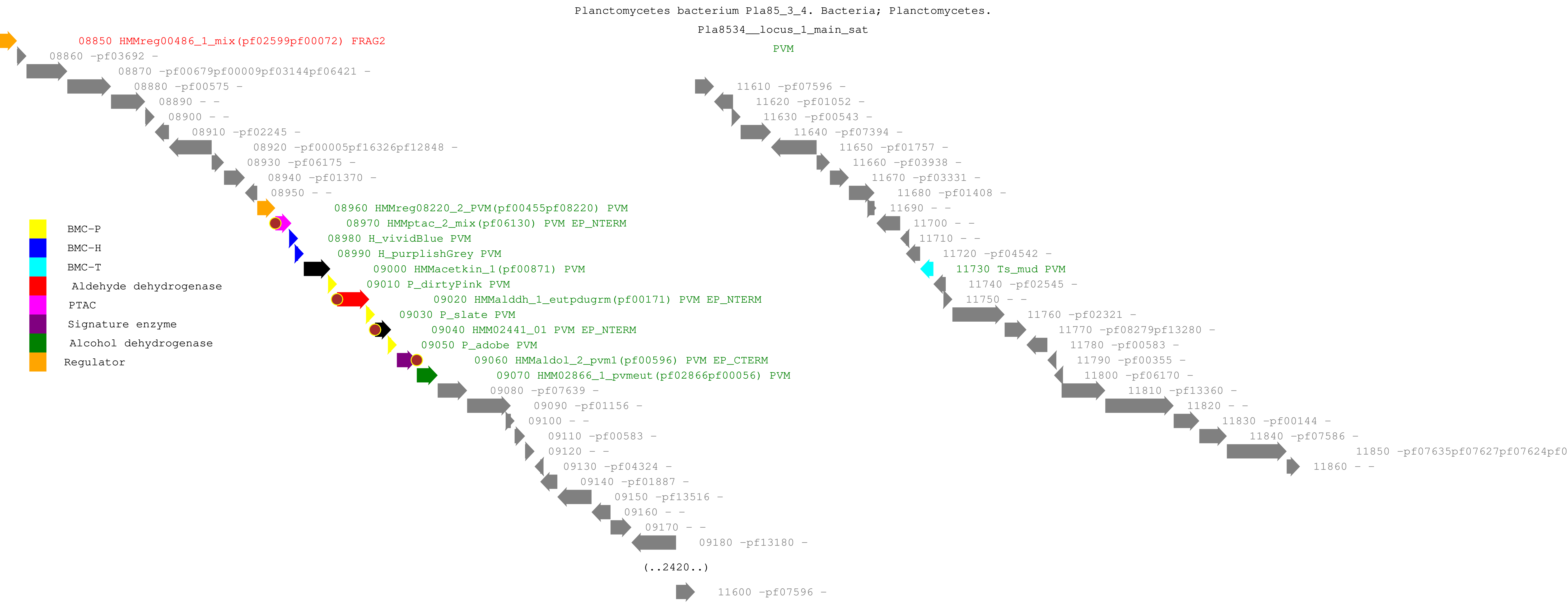
Planctomycetes bacterium Pla85_3_4. Bacteria; Planctomycetes.
Pla8534__locus_1_main_sat
PVM
08850 HMMreg00486_1_mix(pf02599pf00072) FRAG2
08860 -pf03692 -
08870 -pf00679pf00009pf03144pf06421 -
08880 -pf00575 -
08890 - -
08900 - -
08910 -pf02245 -
08920 -pf00005pf16326pf12848 -
08930 -pf06175 -
08940 -pf01370 -
08950 - -
08960 HMMreg08220_2_PVM(pf00455pf08220) PVM
08970 HMMptac_2_mix(pf06130) PVM EP_NTERM
08980 H_vividBlue PVM
08990 H_purplishGrey PVM
09000 HMMacetkin_1(pf00871) PVM
09010 P_dirtyPink PVM
09020 HMMaldh_1_eutpdugrm(pf00171) PVM EP_NTERM
09030 P_slate PVM
09040 HMM02441_01 PVM EP_NTERM
09050 P_adobe PVM
09060 HMMaldol_2_pvm1(pf00596) PVM EP_CTERM
09070 HMM02866_1_pvmeut(pf02866pf00056) PVM
09080 -pf07639 -
09090 -pf01156 -
09100 - -
09110 -pf00583 -
09120 - -
09130 -pf04324 -
09140 -pf01887 -
09150 -pf13516 -
09160 - -
09170 - -
09180 -pf13180 -
(..2420..)
11600 -pf07596 -
11610 -pf07596 -
11620 -pf01052 -
11630 -pf00543 -
11640 -pf07394 -
11650 -pf01757 -
11660 -pf03938 -
11670 -pf03331 -
11680 -pf01408 -
11690 - -
11700 - -
11710 - -
11720 -pf04542 -
11730 Ts_mud PVM
11740 -pf02545 -
11750 - -
11760 -pf02321 -
11770 -pf08279pf13280 -
11780 -pf00583 -
11790 -pf00355 -
11800 -pf06170 -
11810 -pf13360 -
11820 - -
11830 -pf00144 -
11840 -pf07586 -
11850 -pf07635pf07627pf07624pf0
11860 - -
BMC-P
BMC-H
BMC-T
Aldehyde dehydrogenase
PTAC
Signature enzyme
Alcohol dehydrogenase
Regulator

Rubinisphaera brasiliensis (strain ATCC 49424 / DSM 5305 / JCM 21570 / Bacteria; Planctomycetes; Planctomycetia; Planctomycetales;

Plabr__locus_1

PVM

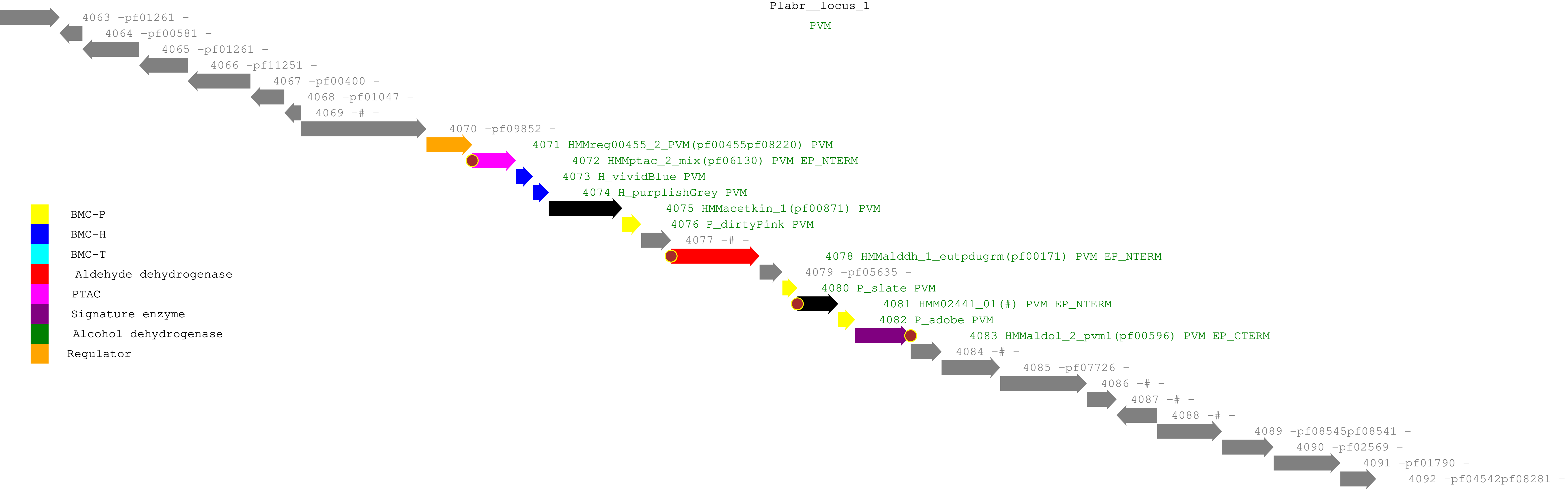

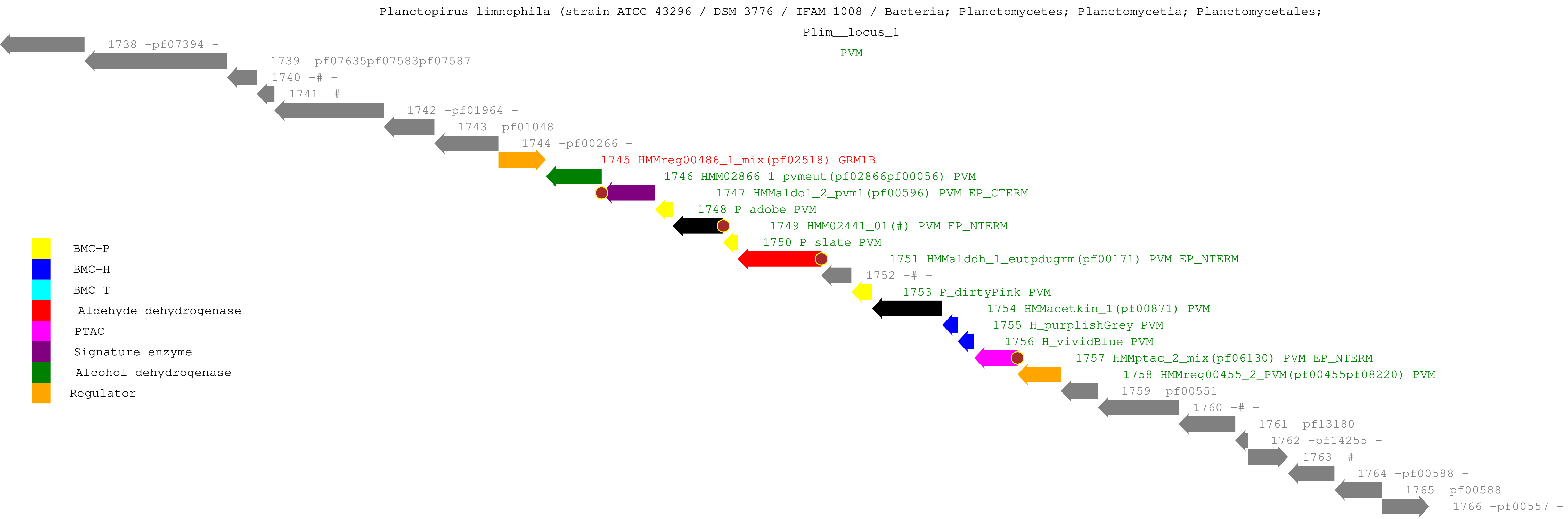

Planctopirus limnophila (strain ATCC 43296 / DSM 3776 / IFAM 1008 / Bacteria; Planctomycetes; Planctomycetia; Planctomycetales;
Plim__locus_1
PVM
1738 -pf07394 -
1739 -pf07635pf07583pf07587 -
1740 -# -
1741 -# -
1742 -pf01964 -
1743 -pf01048 -
1744 -pf00266 -
1745 HMMreg00486_1_mix(pf02518) GRM1B
1746 HMM02866_1_pvmeut(pf02866pf00056) PVM
1747 HMMaldol_2_pvm1(pf00596) PVM EP_CTERM
1748 P_adobe PVM
1749 HMM02441_01(#) PVM EP_NTERM
1750 P_slate PVM
1751 HMMalddh_1_eutpdugrm(pf00171) PVM EP_NTERM
1752 -# -
1753 P_dirtyPink PVM
1754 HMMacetkin_1(pf00871) PVM
1755 H_purplishGrey PVM
1756 H_vividBlue PVM
1757 HMMptac_2_mix(pf06130) PVM EP_NTERM
1758 HMMreg00455_2_PVM(pf00455pf08220) PVM
1759 -pf00551 -
1760 -# -
1761 -pf13180 -
1762 -pf14255 -
1763 -# -
1764 -pf00588 -
1765 -pf00588 -
1766 -pf00557 -
BMC-P
BMC-H
BMC-T
Aldehyde dehydrogenase
PTAC
Signature enzyme
Alcohol dehydrogenase
Regulator

Gimesia maris DSM 8797. Bacteria; Planctomycetes; Planctomycetia; Planctomycetales;

PM8797T__locus_1

PVM

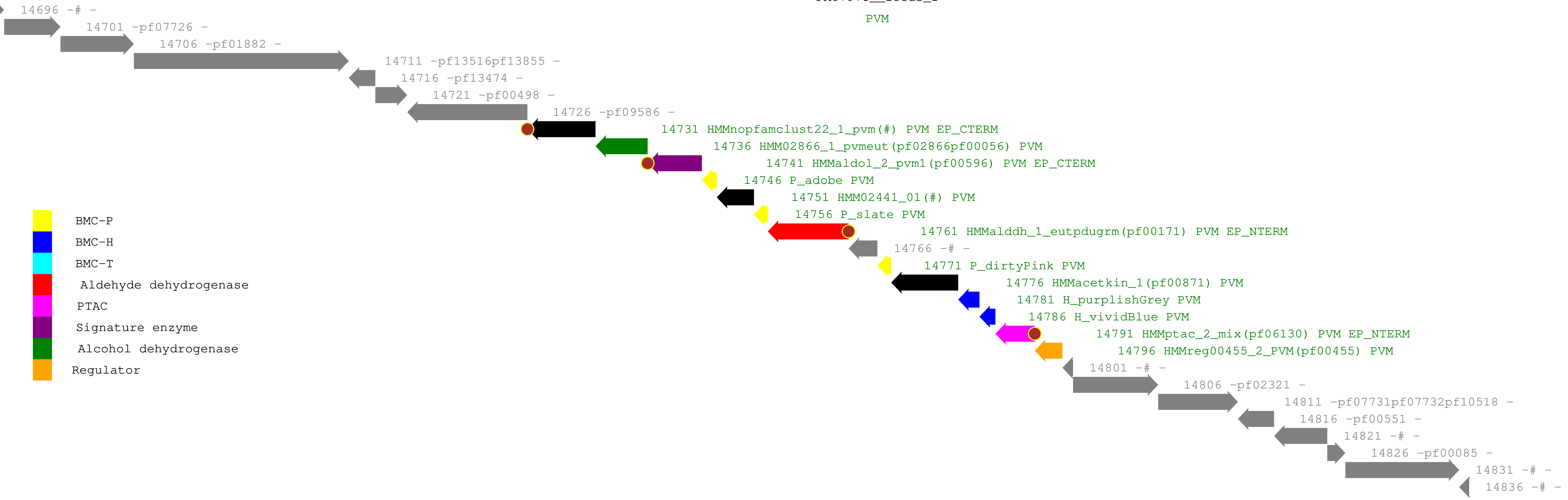

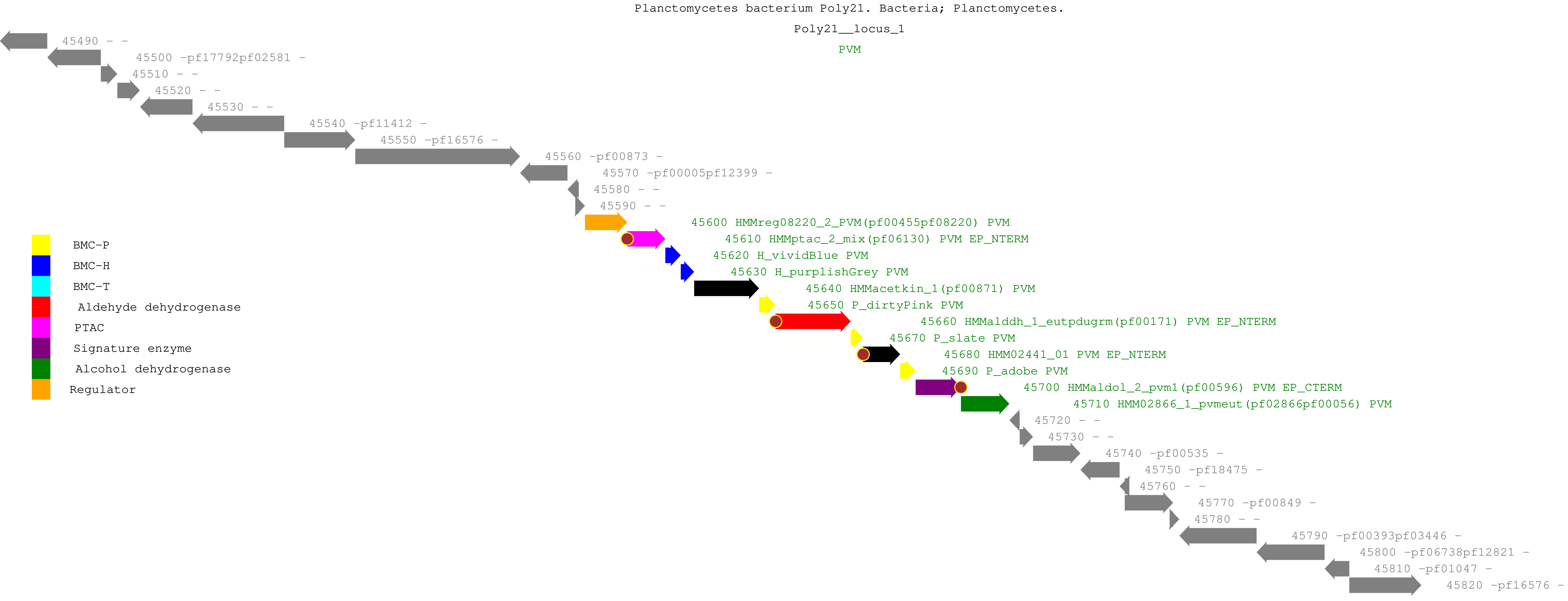
Planctomycetes bacterium Poly21. Bacteria; Planctomycetes.
Poly21__locus_1
PVM
45490 - -
45500 -pf17792pf02581 -
45510 - -
45520 - -
45530 - -
45540 -pf11412 -
45550 -pf16576 -
45560 -pf00873 -
45570 -pf00005pf12399 -
45580 - -
45590 - -
45600 HMMreg08220_2_PVM(pf00455pf08220) PVM
45610 HMMptac_2_mix(pf06130) PVM EP_NTERM
45620 H_vividBlue PVM
45630 H_purplishGrey PVM
45640 HMMacetkin_1(pf00871) PVM
45650 P_dirtyPink PVM
45660 HMMalddh_1_eutpdugrm(pf00171) PVM EP_NTERM
45670 P_slate PVM
45680 HMM02441_01 PVM EP_NTERM
45690 P_adobe PVM
45700 HMMaldol_2_pvm1(pf00596) PVM EP_CTERM
45710 HMM02866_1_pvmeut(pf02866pf00056) PVM
45720 - -
45730 - -
45740 -pf00535 -
45750 -pf18475 -
45760 - -
45770 -pf00849 -
45780 - -
45790 -pf00393pf03446 -
45800 -pf06738pf12821 -
45810 -pf01047 -
45820 -pf16576 -
BMC-P
BMC-H
BMC-T
Aldehyde dehydrogenase
PTAC
Signature enzyme
Alcohol dehydrogenase
Regulator

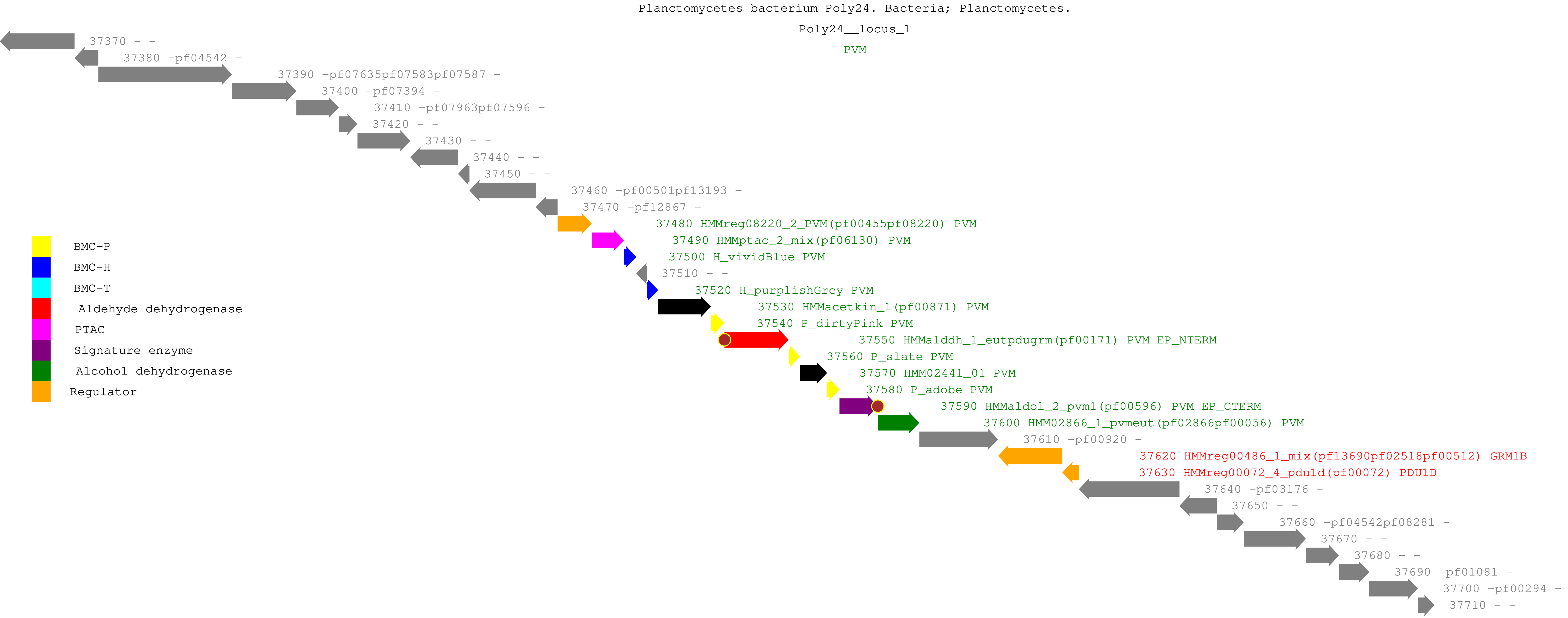
Planctomycetes bacterium Poly24. Bacteria; Planctomycetes.
Poly24__locus_1
PVM
37370 - -
37380 -pf04542 -
37390 -pf07635pf07583pf07587 -
37400 -pf07394 -
37410 -pf07963pf07596 -
37420 - -
37430 - -
37440 - -
37450 - -
37460 -pf00501pf13193 -
37470 -pf12867 -
37480 HMMreg08220_2_PVM(pf00455pf08220) PVM
37490 HMMptac_2_mix(pf06130) PVM
37500 H_vividBlue PVM
37510 - -
37520 H_purplishGrey PVM
37530 HMMacetkin_1(pf00871) PVM
37540 P_dirtyPink PVM
37550 HMMaldh_1_eutpdugrm(pf00171) PVM EP_NTERM
37560 P_slate PVM
37570 HMM02441_01 PVM
37580 P_adobe PVM
37590 HMMaldol_2_pvm1(pf00596) PVM EP_CTERM
37600 HMM02866_1_pvmeut(pf02866pf00056) PVM
37610 -pf00920 -
37620 HMMreg00486_1_mix(pf13690pf02518pf00512) GRM1B
37630 HMMreg00072_4_pdu1d(pf00072) PDU1D
37640 -pf03176 -
37650 - -
37660 -pf04542pf08281 -
37670 - -
37680 - -
37690 -pf01081 -
37700 -pf00294 -
37710 - -
BMC-P
BMC-H
BMC-T
Aldehyde dehydrogenase
PTAC
Signature enzyme
Alcohol dehydrogenase
Regulator

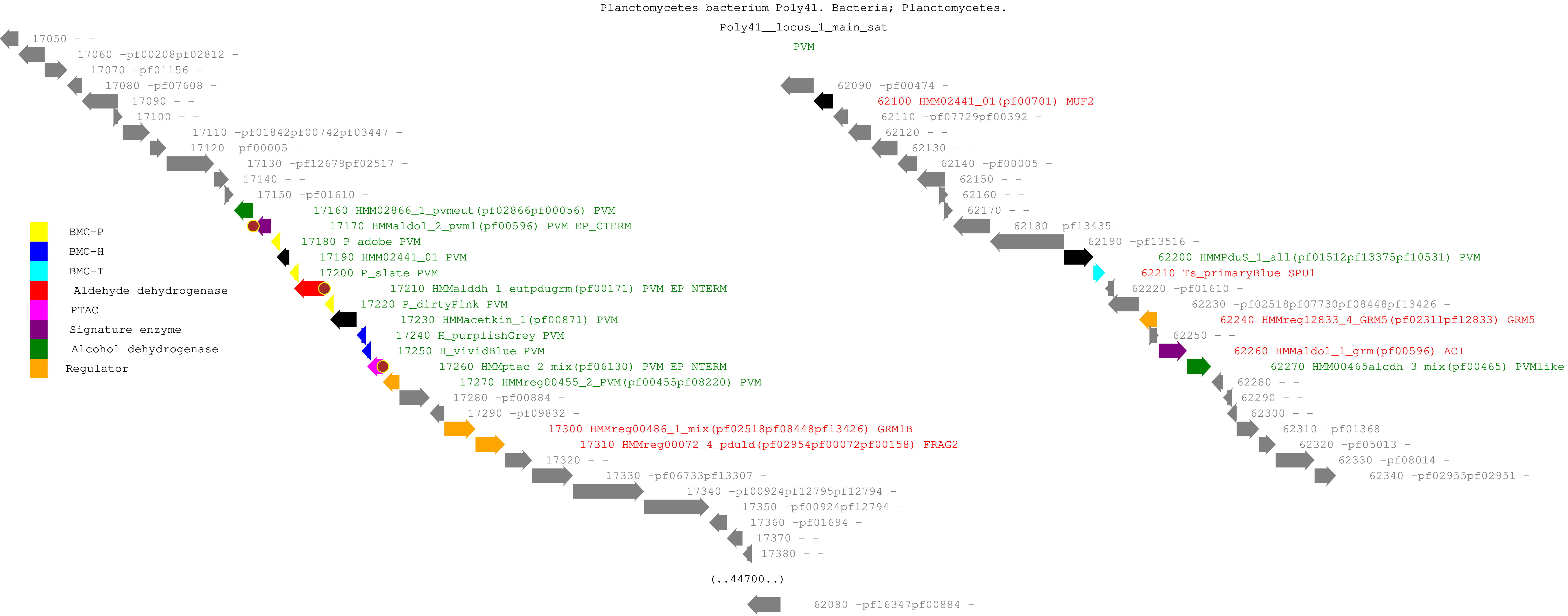
Planctomycetes bacterium Poly41. Bacteria; Planctomycetes.
Poly41__locus_1_main_sat
PVM
17050 - -
17060 -pf00208pf02812 -
17070 -pf01156 -
17080 -pf07608 -
17090 - -
17100 - -
17110 -pf01842pf00742pf03447 -
17120 -pf00005 -
17130 -pf12679pf02517 -
17140 - -
17150 -pf01610 -
17160 HMM02866_1_pvmeut(pf02866pf00056) PVM
17170 HMMaldol_2_pvm1(pf00596) PVM EP_CTERM
17180 P_adobe PVM
17190 HMM02441_01 PVM
17200 P_slate PVM
17210 HMMalddh_1_eutpdugrm(pf00171) PVM EP_NTERM
17220 P_dirtyPink PVM
17230 HMMacetkin_1(pf00871) PVM
17240 H_purplishGrey PVM
17250 H_vividBlue PVM
17260 HMMptac_2_mix(pf06130) PVM EP_NTERM
17270 HMMreg00455_2_PVM(pf00455pf08220) PVM
17280 -pf00884 -
17290 -pf09832 -
17300 HMMreg00486_1_mix(pf02518pf08448pf13426) GRM1B
17310 HMMreg00072_4_pdu1d(pf02954pf00072pf00158) FRAG2
17320 - -
17330 -pf06733pf13307 -
17340 -pf00924pf12795pf12794 -
17350 -pf00924pf12794 -
17360 -pf01694 -
17370 - -
17380 - -
(..44700..)
62080 -pf16347pf00884 -
62090 -pf00474 -
62100 HMM02441_01(pf00701) MUF2
62110 -pf07729pf00392 -
62120 - -
62130 - -
62140 -pf00005 -
62150 - -
62160 - -
62170 - -
62180 -pf13435 -
62190 -pf13516 -
62200 HMMPduS_1_all(pf01512pf13375pf10531) PVM
62210 Ts_primaryBlue SPU1
62220 -pf01610 -
62230 -pf02518pf07730pf08448pf13426 -
62240 HMMreg12833_4_GRM5(pf02311pf12833) GRM5
62250 - -
62260 HMMaldol_1_grm(pf00596) ACI
62270 HMM00465alcdh_3_mix(pf00465) PVMlike
62280 - -
62290 - -
62300 - -
62310 -pf01368 -
62320 -pf05013 -
62330 -pf08014 -
62340 -pf02955pf02951 -
BMC-P
BMC-H
BMC-T
Aldehyde dehydrogenase
PTAC
Signature enzyme
Alcohol dehydrogenase
Regulator

Planctomycetes bacterium Poly51. Bacteria; Planctomycetes.

Poly51__locus_1

PVM

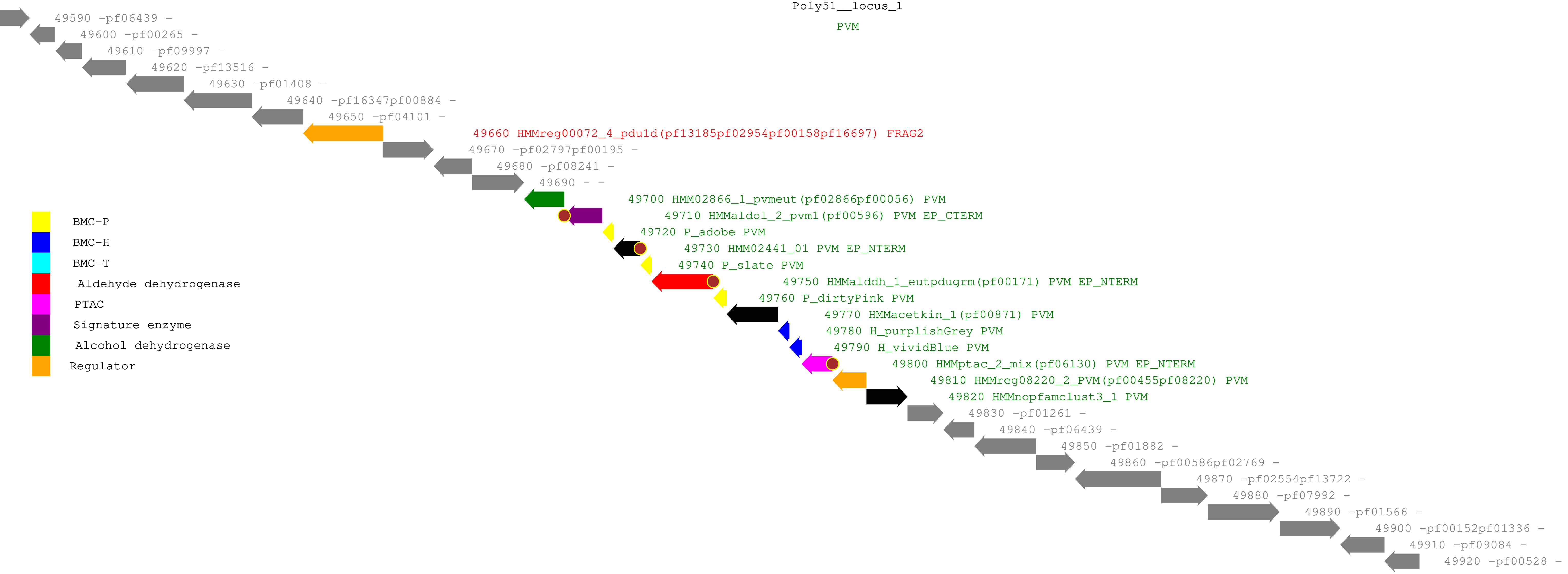

Planctomycetes bacterium Poly59. Bacteria; Planctomycetes.

Poly59__locus_1

PVM

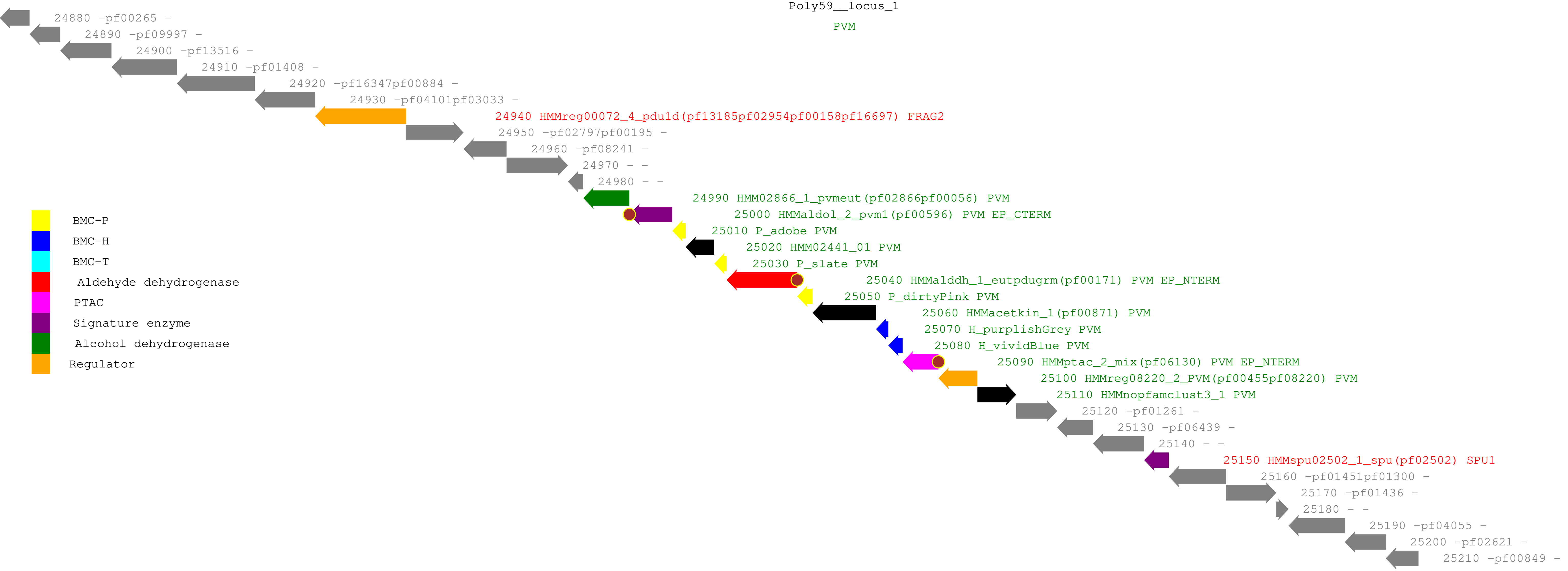

Pirellula staleyi (strain ATCC 27377 / DSM 6068 / ICPB 4128) (Pirella Bacteria; Planctomycetes; Planctomycetia; Planctomycetales;

Psta__locus_1_main_sat

PVM

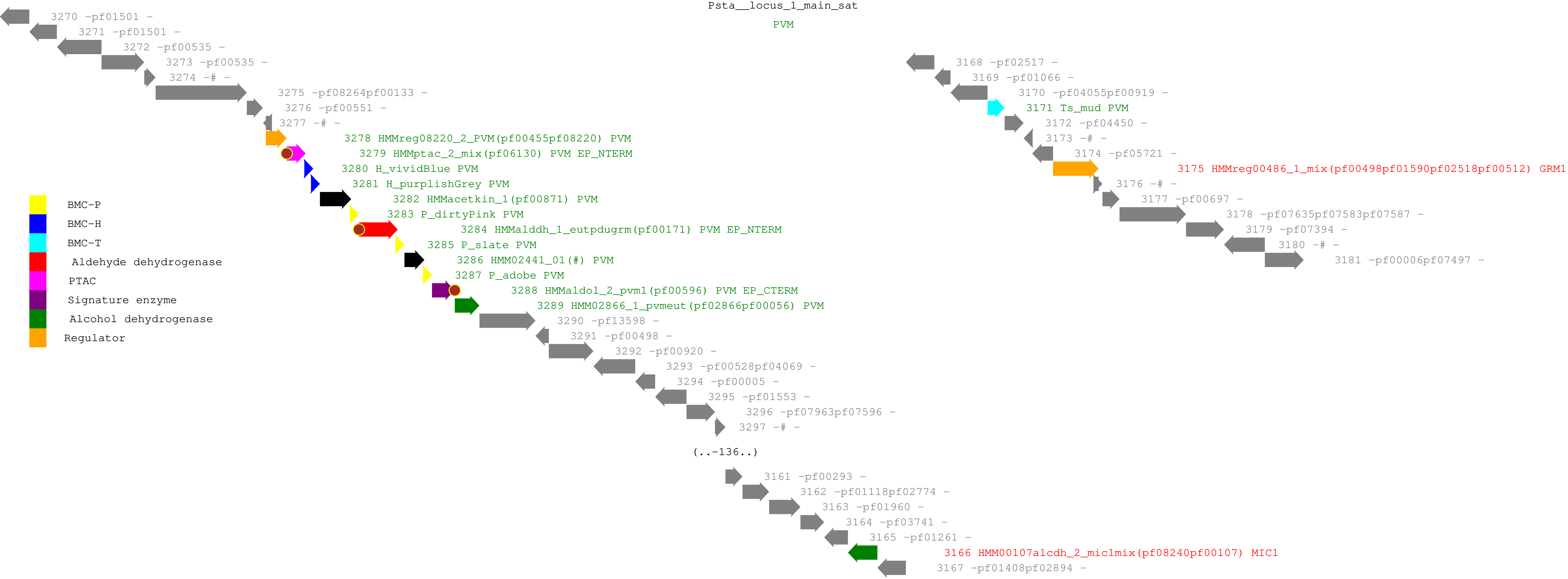

Planctomycetes bacterium Q31a. Bacteria; Planctomycetes.

Q31a__locus_1_main_sat

PVM

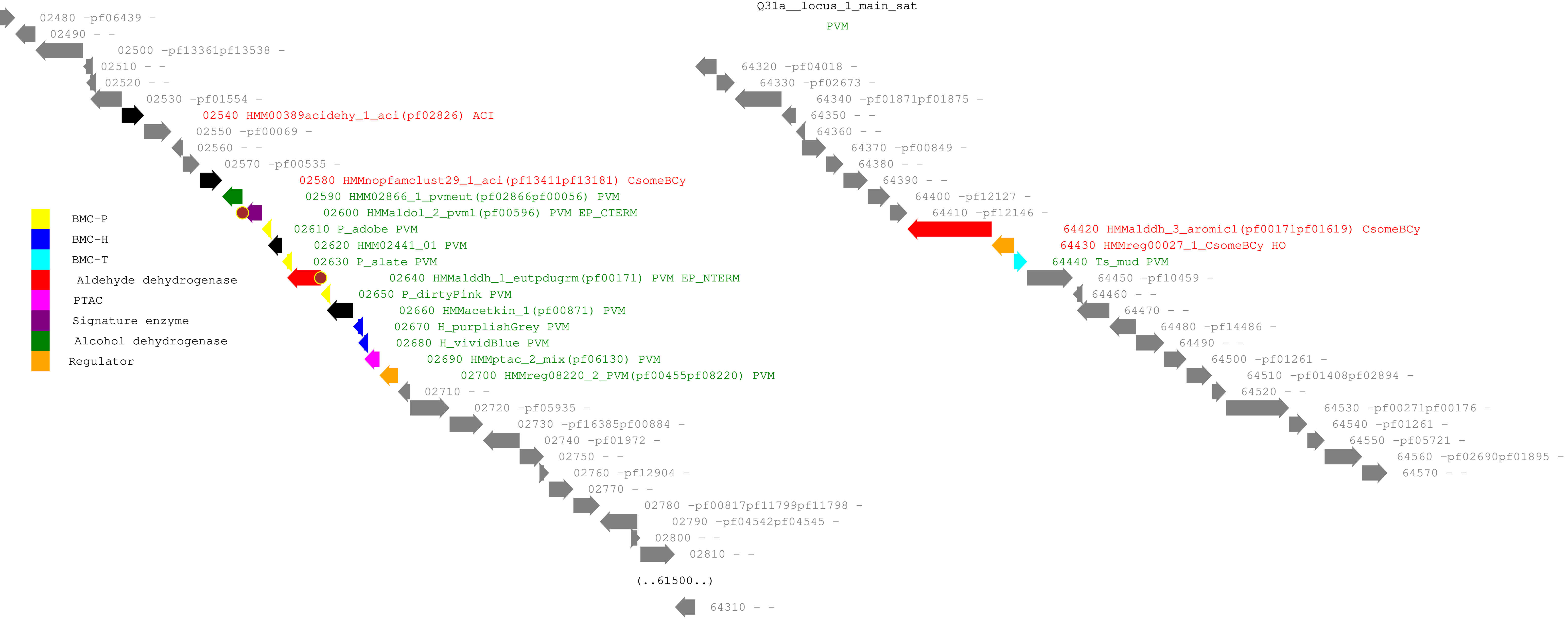

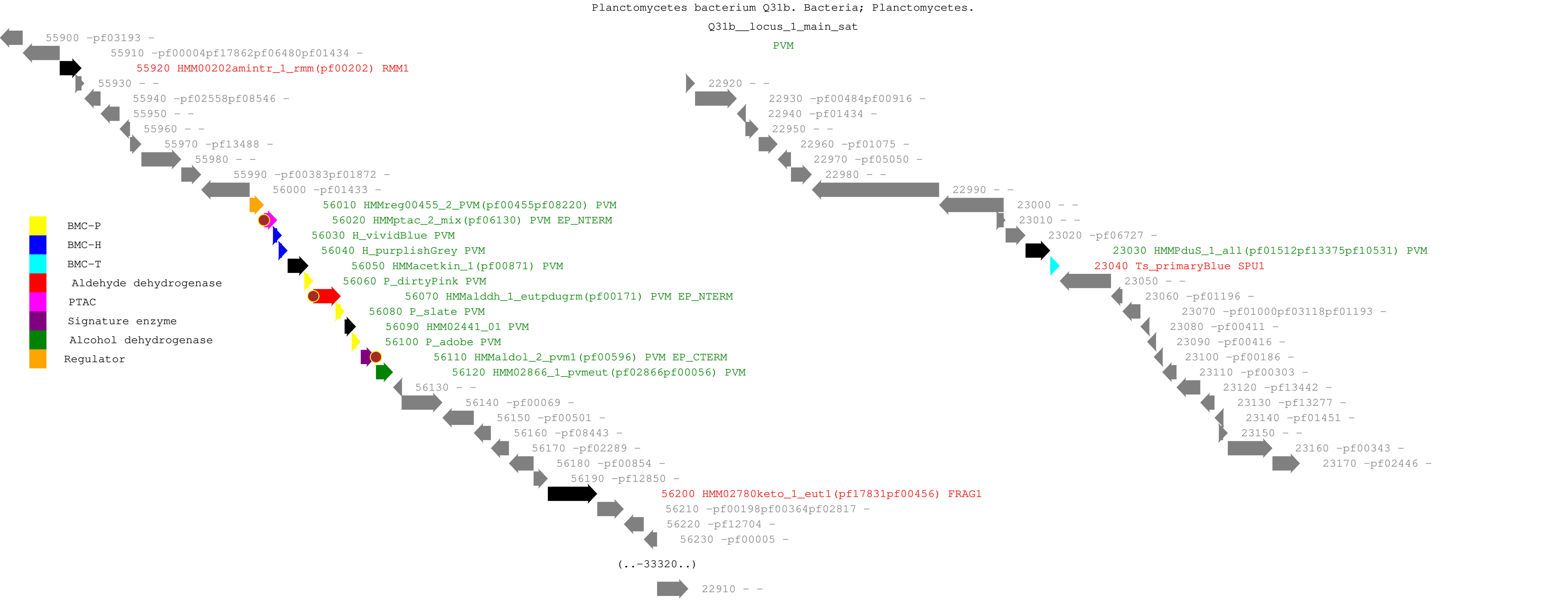

Planctomycetes bacterium Q31b. Bacteria; Planctomycetes.
Q31b__locus_1_main_sat
PVM
55900 -pf03193 -
55910 -pf00004pf17862pf06480pf01434 -
55920 HMM00202amintr_1_rmm(pf00202) RMM1
55930 - -
55940 -pf02558pf08546 -
55950 - -
55960 - -
55970 -pf13488 -
55980 - -
55990 -pf00383pf01872 -
56000 -pf01433 -
56010 HMMreg00455_2_PVM(pf00455pf08220) PVM
56020 HMMptac_2_mix(pf06130) PVM EP_NTERM
56030 H_vividBlue PVM
56040 H_purplishGrey PVM
56050 HMMacetkin_1(pf00871) PVM
56060 P_dirtyPink PVM
56070 HMMalddh_1_eutpdugrm(pf00171) PVM EP_NTERM
56080 P_slate PVM
56090 HMM02441_01 PVM
56100 P_adobe PVM
56110 HMMaldol_2_pvm1(pf00596) PVM EP_CTERM
56120 HMM02866_1_pvmeut(pf02866pf00056) PVM
56130 - -
56140 -pf00069 -
56150 -pf00501 -
56160 -pf08443 -
56170 -pf02289 -
56180 -pf00854 -
56190 -pf12850 -
56200 HMM02780keto_1_eut1(pf17831pf00456) FRAG1
56210 -pf00198pf00364pf02817 -
56220 -pf12704 -
56230 -pf00005 -
(..-33320..)
22910 - -
22920 - -
22930 -pf00484pf00916 -
22940 -pf01434 -
22950 - -
22960 -pf01075 -
22970 -pf05050 -
22980 - -
22990 - -
23000 - -
23010 - -
23020 -pf06727 -
23030 HMMPduS_1_all(pf01512pf13375pf10531) PVM
23040 Ts_primaryBlue SPU1
23050 - -
23060 -pf01196 -
23070 -pf01000pf03118pf01193 -
23080 -pf00411 -
23090 -pf00416 -
23100 -pf00186 -
23110 -pf00303 -
23120 -pf13442 -
23130 -pf13277 -
23140 -pf01451 -
23150 - -
23160 -pf00343 -
23170 -pf02446 -
BMC-P
BMC-H
BMC-T
Aldehyde dehydrogenase
PTAC
Signature enzyme
Alcohol dehydrogenase
Regulator

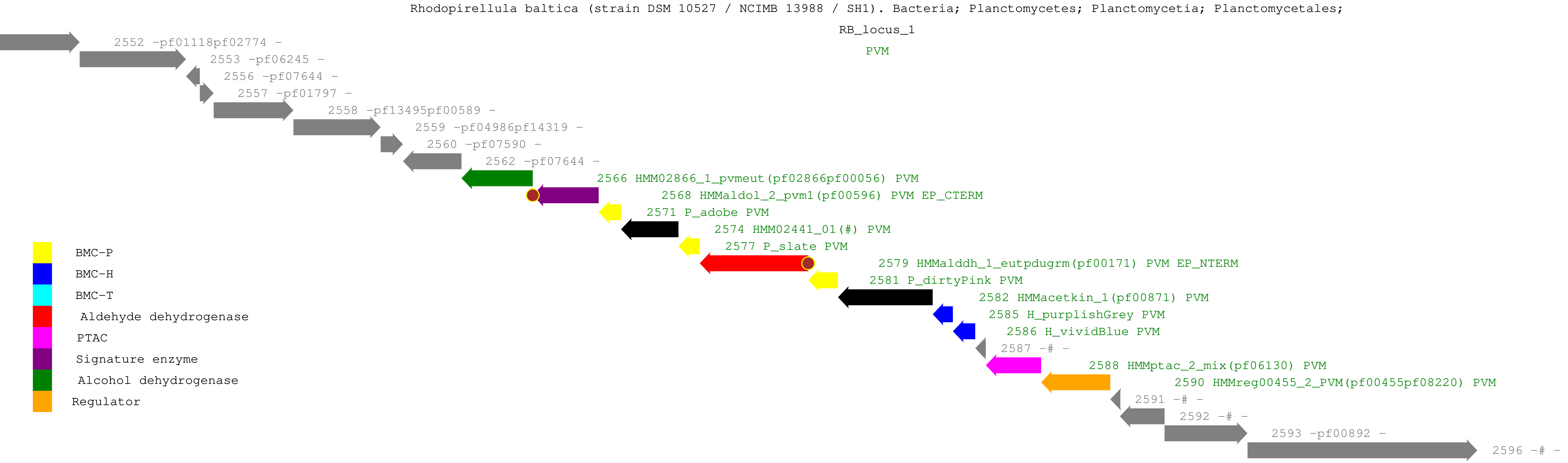
Rhodopirellula baltica (strain DSM 10527 / NCIMB 13988 / SH1). Bacteria; Planctomycetes; Planctomycetia; Planctomycetales;
RB_locus_1
PVM
2552 -pf01118pf02774 -
2553 -pf06245 -
2556 -pf07644 -
2557 -pf01797 -
2558 -pf13495pf00589 -
2559 -pf04986pf14319 -
2560 -pf07590 -
2562 -pf07644 -
2566 HMM02866_1_pvmeut(pf02866pf00056) PVM
2568 HMMaldol_2_pvm1(pf00596) PVM EP_CTERM
2571 P_adobe PVM
2574 HMM02441_01(#) PVM
2577 P_slate PVM
2579 HMMalddh_1_eutpdugrm(pf00171) PVM EP_NTERM
2581 P_dirtyPink PVM
2582 HMMacetkin_1(pf00871) PVM
2585 H_purplishGrey PVM
2586 H_vividBlue PVM
2587 -# -
2588 HMMptac_2_mix(pf06130) PVM
2590 HMMreg00455_2_PVM(pf00455pf08220) PVM
2591 -# -
2592 -# -
2593 -pf00892 -
2596 -# -
BMC-P
BMC-H
BMC-T
Aldehyde dehydrogenase
PTAC
Signature enzyme
Alcohol dehydrogenase
Regulator

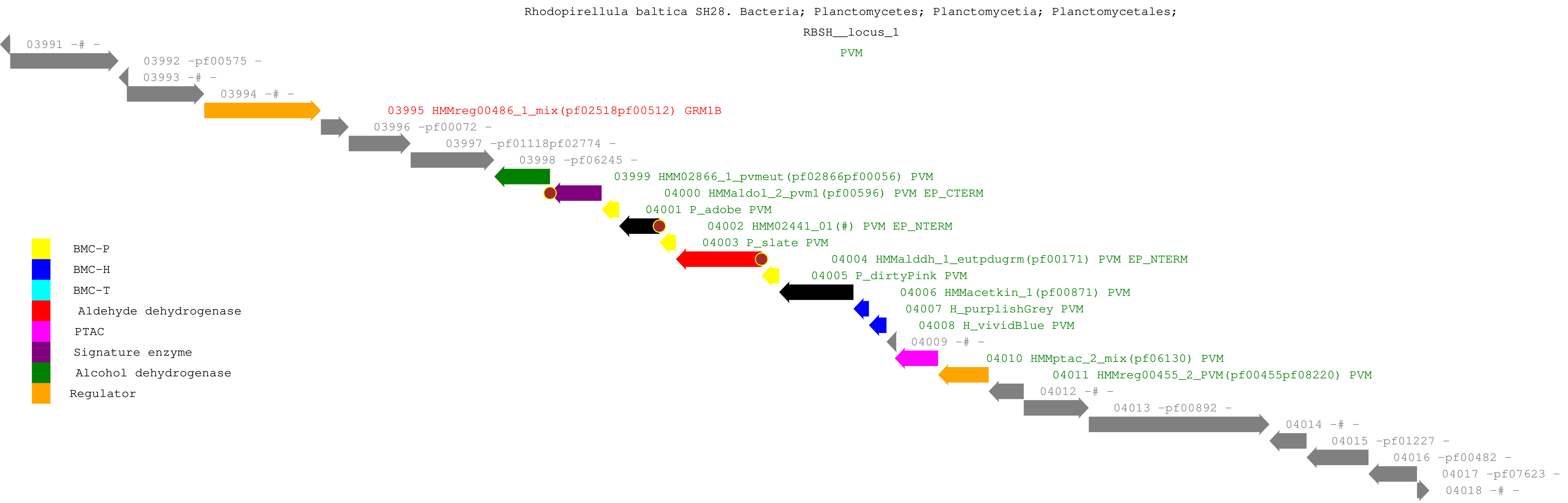
Rhodopirellula baltica SH28. Bacteria; Planctomycetes; Planctomycetia; Planctomycetales;
RBSH__locus_1
PVM
03991 -# -
03992 -pf00575 -
03993 -# -
03994 -# -
03995 HMMreg00486_1_mix(pf02518pf00512) GRM1B
03996 -pf00072 -
03997 -pf01118pf02774 -
03998 -pf06245 -
03999 HMM02866_1_pvmeut(pf02866pf00056) PVM
04000 HMMaldol_2_pvm1(pf00596) PVM EP_CTERM
04001 P_adobe PVM
04002 HMM02441_01(#) PVM EP_NTERM
04003 P_slate PVM
04004 HMMalddh_1_eutpdugrm(pf00171) PVM EP_NTERM
04005 P_dirtyPink PVM
04006 HMMacetkin_1(pf00871) PVM
04007 H_purplishGrey PVM
04008 H_vividBlue PVM
04009 -# -
04010 HMMptac_2_mix(pf06130) PVM
04011 HMMreg00455_2_PVM(pf00455pf08220) PVM
04012 -# -
04013 -pf00892 -
04014 -# -
04015 -pf01227 -
04016 -pf00482 -
04017 -pf07623 -
04018 -# -
BMC-P
BMC-H
BMC-T
Aldehyde dehydrogenase
PTAC
Signature enzyme
Alcohol dehydrogenase
Regulator

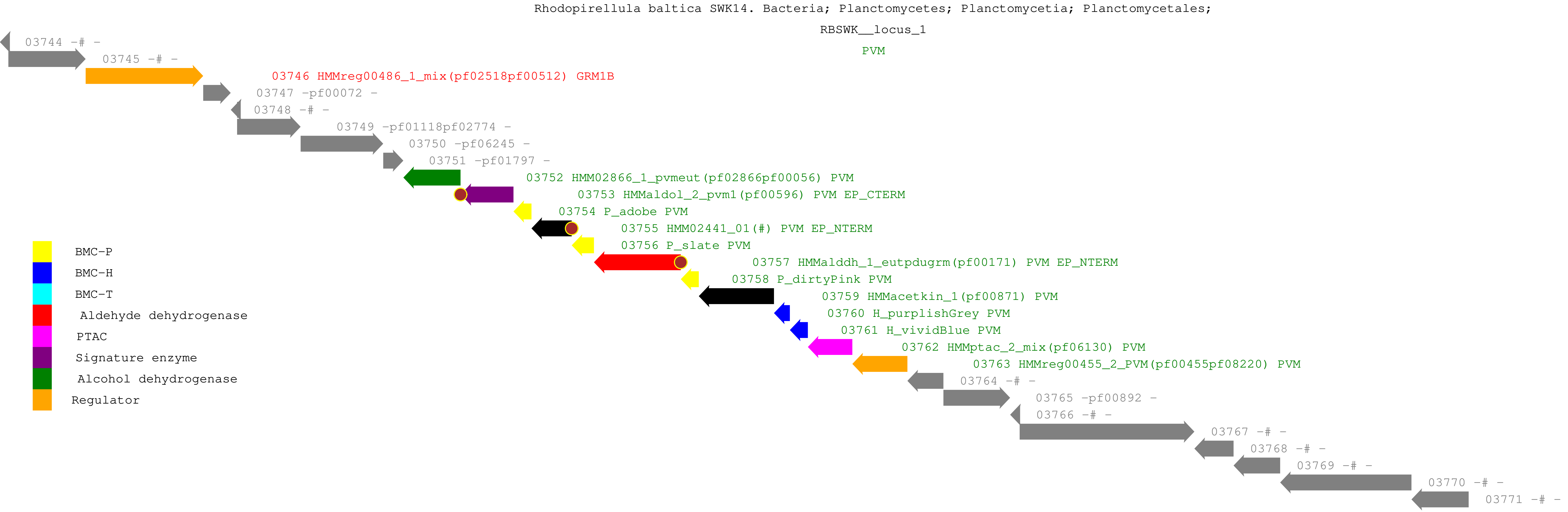
Rhodopirellula baltica SWK14. Bacteria; Planctomycetes; Planctomycetia; Planctomycetales;
RBSWK__locus_1
PVM
03744 -# -
03745 -# -
03746 HMMreg00486_1_mix(pf02518pf00512) GRM1B
03747 -pf00072 -
03748 -# -
03749 -pf01118pf02774 -
03750 -pf06245 -
03751 -pf01797 -
03752 HMM02866_1_pvmeut(pf02866pf00056) PVM
03753 HMMaldol_2_pvm1(pf00596) PVM EP_CTERM
03754 P_adobe PVM
03755 HMM02441_01(#) PVM EP_NTERM
03756 P_slate PVM
03757 HMMalddh_1_eutpdugrm(pf00171) PVM EP_NTERM
03758 P_dirtyPink PVM
03759 HMMacetkin_1(pf00871) PVM
03760 H_purplishGrey PVM
03761 H_vividBlue PVM
03762 HMMptac_2_mix(pf06130) PVM
03763 HMMreg00455_2_PVM(pf00455pf08220) PVM
03764 -# -
03765 -pf00892 -
03766 -# -
03767 -# -
03768 -# -
03769 -# -
03770 -# -
03771 -# -
BMC-P
BMC-H
BMC-T
Aldehyde dehydrogenase
PTAC
Signature enzyme
Alcohol dehydrogenase
Regulator

Rhodopirellula baltica WH47. Bacteria; Planctomycetes; Planctomycetia; Planctomycetales;

RBWH47__locus_1

PVM

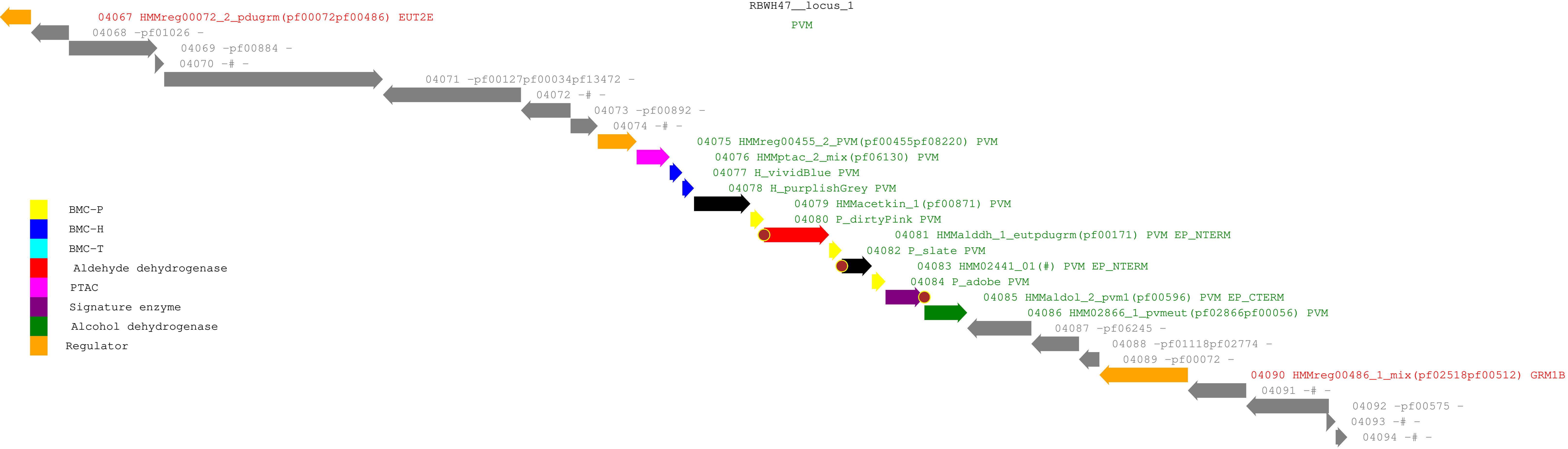

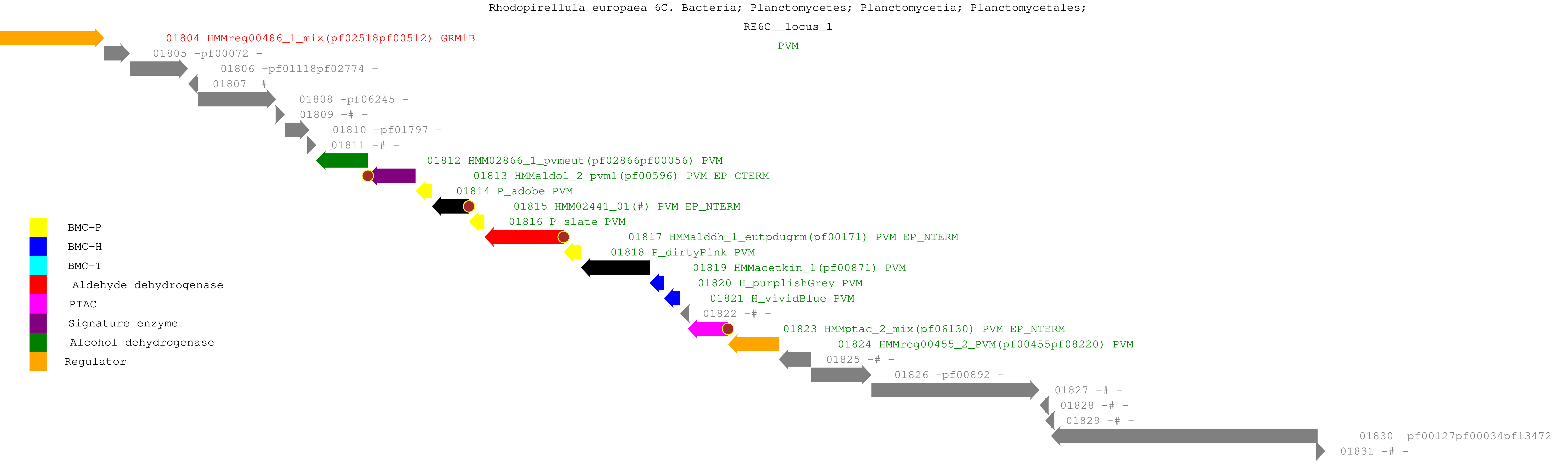
Rhodopirellula europaea 6C. Bacteria; Planctomycetes; Planctomycetia; Planctomycetales;
RE6C__locus_1
PVM
01804 HMMreg00486_1_mix(pf02518pf00512) GRM1B
01805 -pf00072 -
01806 -pf01118pf02774 -
01807 -# -
01808 -pf06245 -
01809 -# -
01810 -pf01797 -
01811 -# -
01812 HMM02866_1_pvmeut(pf02866pf00056) PVM
01813 HMMaldol_2_pvm1(pf00596) PVM EP_CTERM
01814 P_adobe PVM
01815 HMM02441_01(#) PVM EP_NTERM
01816 P_slate PVM
01817 HMMalddh_1_eutpdugrm(pf00171) PVM EP_NTERM
01818 P_dirtyPink PVM
01819 HMMacetkin_1(pf00871) PVM
01820 H_purplishGrey PVM
01821 H_vividBlue PVM
01822 -# -
01823 HMMptac_2_mix(pf06130) PVM EP_NTERM
01824 HMMreg00455_2_PVM(pf00455pf08220) PVM
01825 -# -
01826 -pf00892 -
01827 -# -
01828 -# -
01829 -# -
01830 -pf00127pf00034pf13472 -
01831 -# -
BMC-P
BMC-H
BMC-T
Aldehyde dehydrogenase
PTAC
Signature enzyme
Alcohol dehydrogenase
Regulator

Rhodopirellula europaea SH398. Bacteria; Planctomycetes; Planctomycetia; Planctomycetales;

RESH__locus_1

PVM

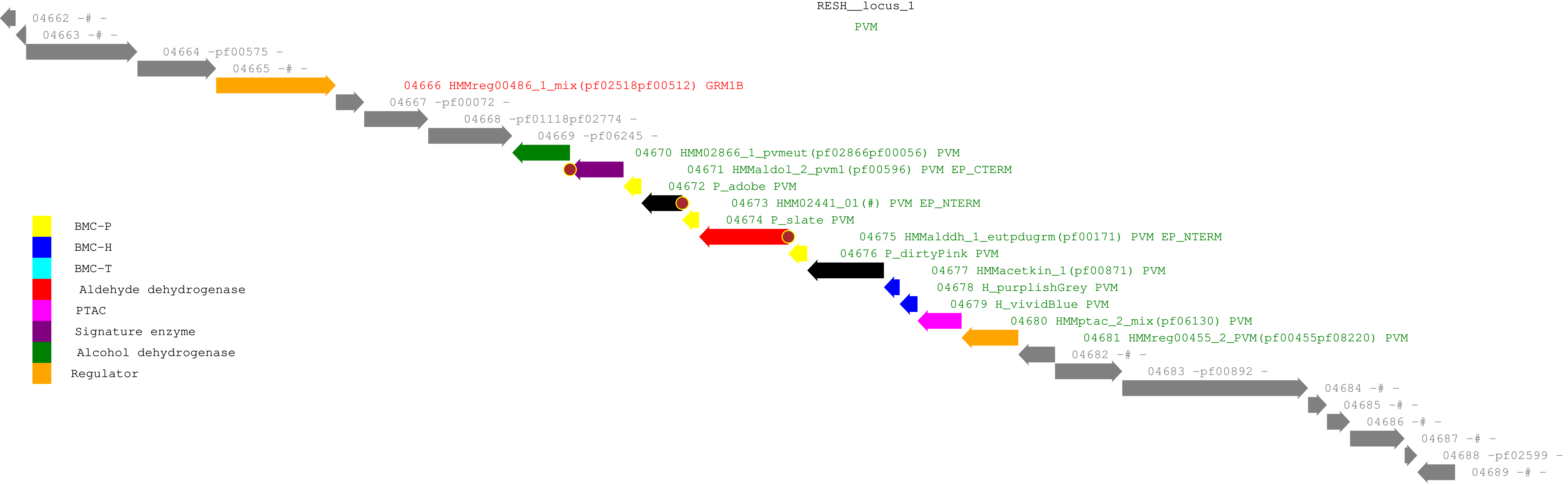

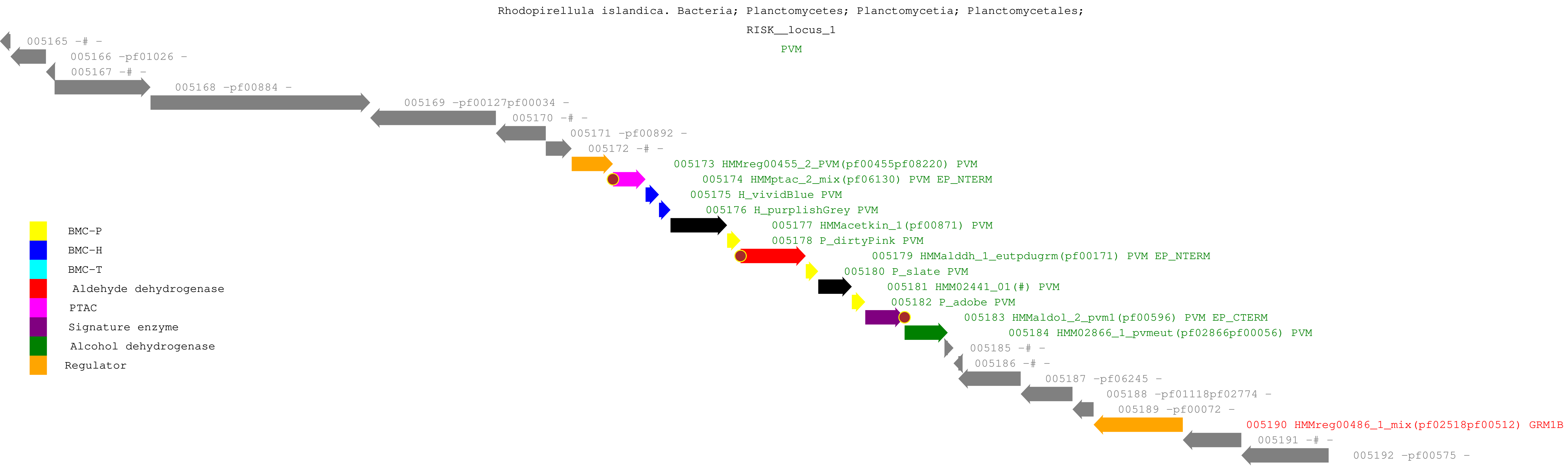
Rhodopirellula islandica. Bacteria; Planctomycetes; Planctomycetia; Planctomycetales;
RISK__locus_1
PVM
005165 -# -
005166 -pf01026 -
005167 -# -
005168 -pf00884 -
005169 -pf00127pf00034 -
005170 -# -
005171 -pf00892 -
005172 -# -
005173 HMMreg00455_2_PVM(pf00455pf08220) PVM
005174 HMMptac_2_mix(pf06130) PVM EP_NTERM
005175 H_vividBlue PVM
005176 H_purplishGrey PVM
005177 HMMacetkin_1(pf00871) PVM
005178 P_dirtyPink PVM
005179 HMMalddh_1_eutpdugrm(pf00171) PVM EP_NTERM
005180 P_slate PVM
005181 HMM02441_01(#) PVM
005182 P_adobe PVM
005183 HMMaldol_2_pvm1(pf00596) PVM EP_CTERM
005184 HMM02866_1_pvmeut(pf02866pf00056) PVM
005185 -# -
005186 -# -
005187 -pf06245 -
005188 -pf01118pf02774 -
005189 -pf00072 -
005190 HMMreg00486_1_mix(pf02518pf00512) GRM1B
005191 -# -
005192 -pf00575 -
BMC-P
BMC-H
BMC-T
Aldehyde dehydrogenase
PTAC
Signature enzyme
Alcohol dehydrogenase
Regulator

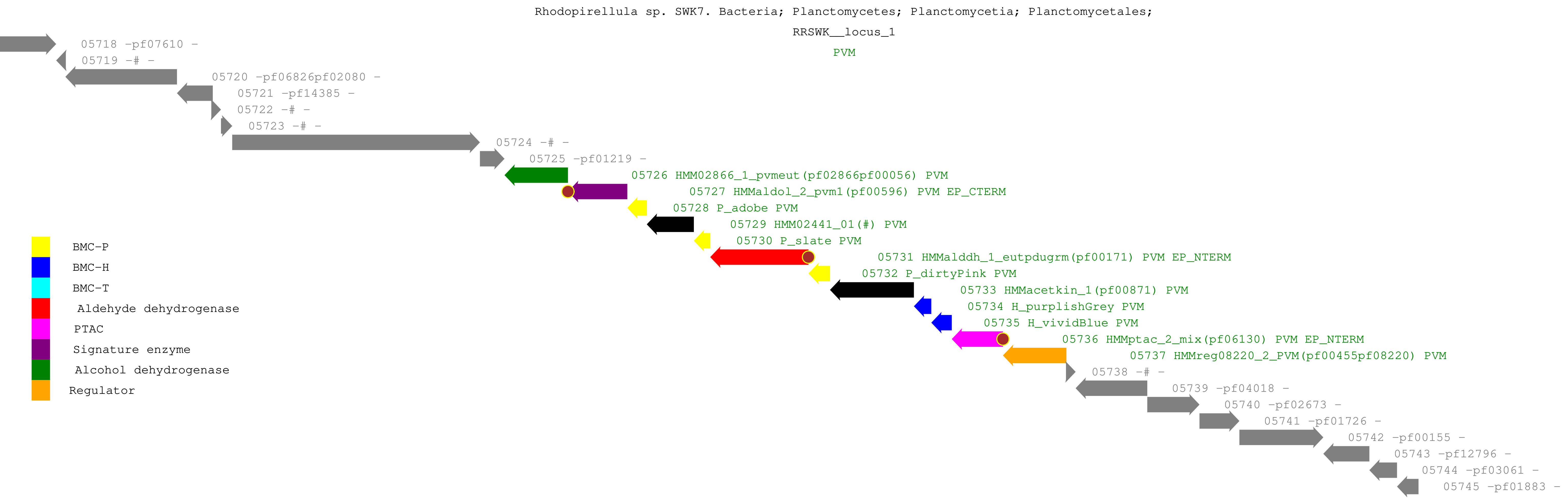

Rhodopirellula sp. SWK7. Bacteria; Planctomycetes; Planctomycetia; Planctomycetales;
RRSWK__locus_1
PVM
05718 -pf07610 -
05719 -# -
05720 -pf06826pf02080 -
05721 -pf14385 -
05722 -# -
05723 -# -
05724 -# -
05725 -pf01219 -
05726 HMM02866_1_pvmeut(pf02866pf00056) PVM
05727 HMMaldol_2_pvm1(pf00596) PVM EP_CTERM
05728 P_adobe PVM
05729 HMM02441_01(#) PVM
05730 P_slate PVM
05731 HMMalddh_1_eutpdugrm(pf00171) PVM EP_NTERM
05732 P_dirtyPink PVM
05733 HMMacetkin_1(pf00871) PVM
05734 H_purplishGrey PVM
05735 H_vividBlue PVM
05736 HMMptac_2_mix(pf06130) PVM EP_NTERM
05737 HMMreg08220_2_PVM(pf00455pf08220) PVM
05738 -# -
05739 -pf04018 -
05740 -pf02673 -
05741 -pf01726 -
05742 -pf00155 -
05743 -pf12796 -
05744 -pf03061 -
05745 -pf01883 -
BMC-P
BMC-H
BMC-T
Aldehyde dehydrogenase
PTAC
Signature enzyme
Alcohol dehydrogenase
Regulator

Rhodopirellula sallentina SM41. Bacteria; Planctomycetes; Planctomycetia; Planctomycetales;

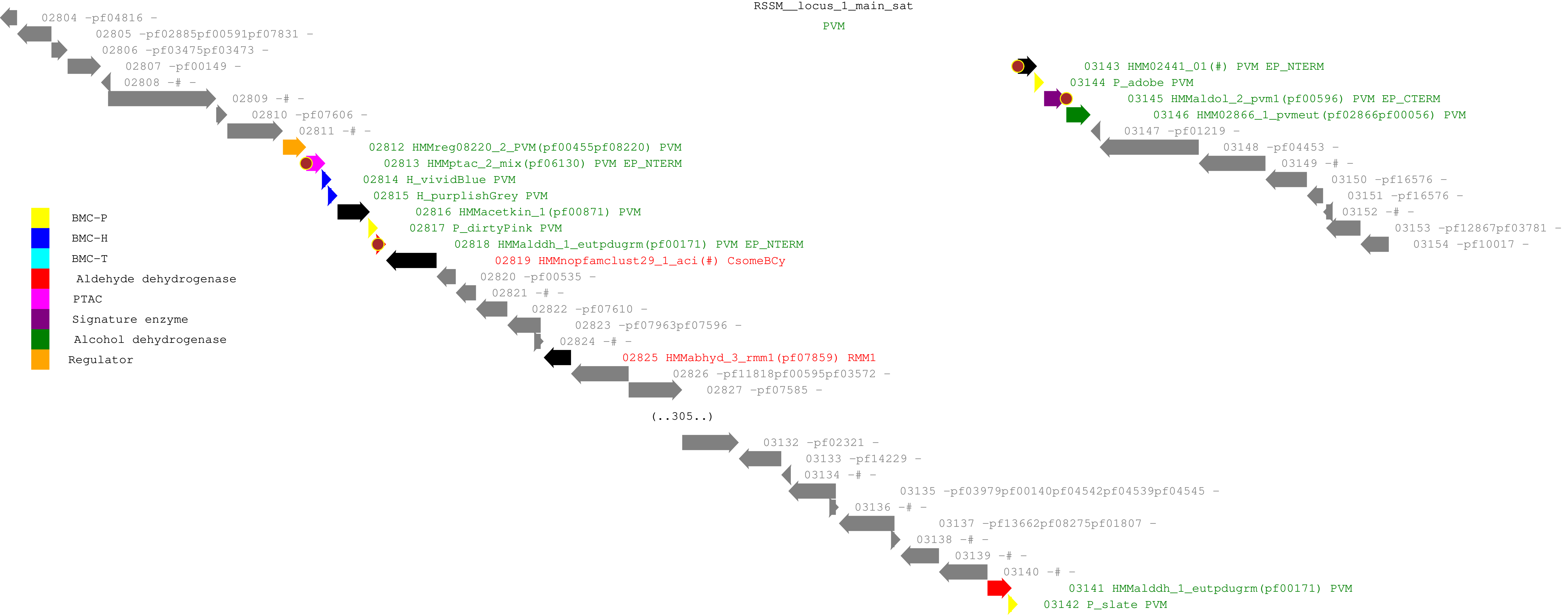

Planctomicrobium piriforme. Bacteria; Planctomycetes; Planctomycetia; Planctomycetales;

SAMN05421753__locus_1

PVM

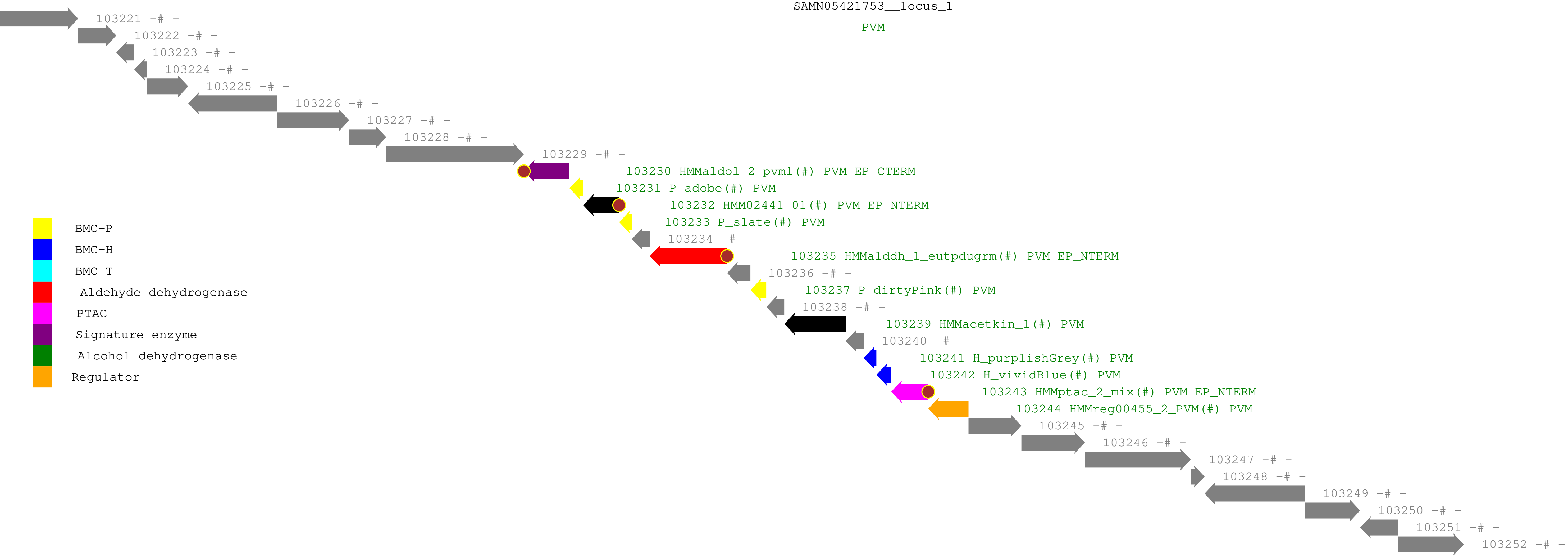

Singulisphaera sp. GP187. Bacteria; Planctomycetes; Planctomycetia; Planctomycetales;

SAMN05444166__locus_1

PVM

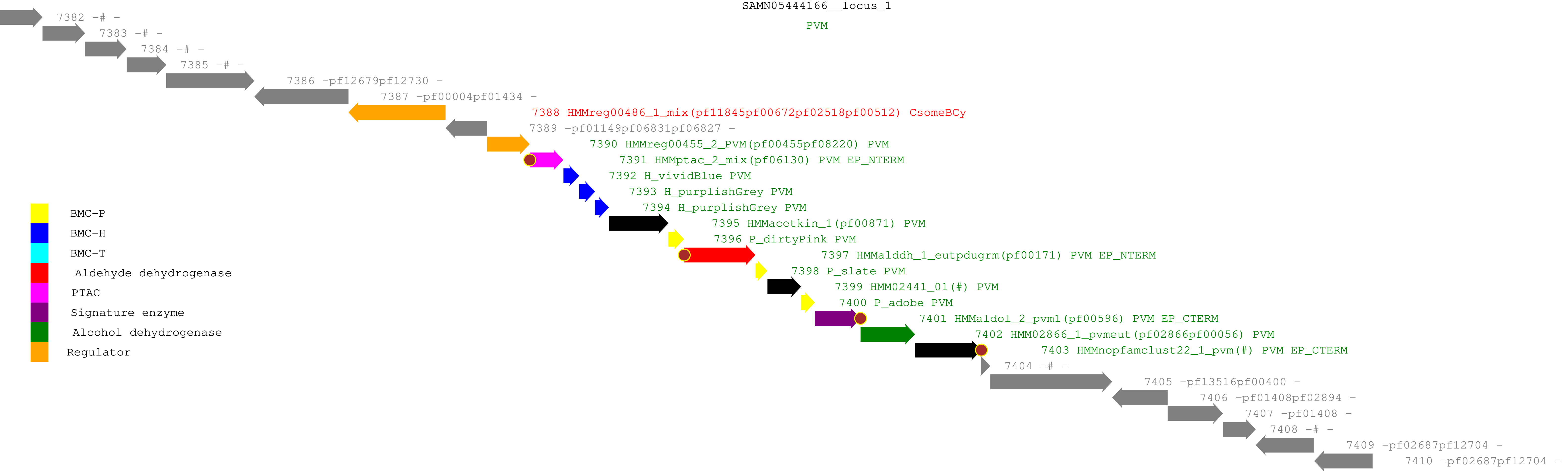

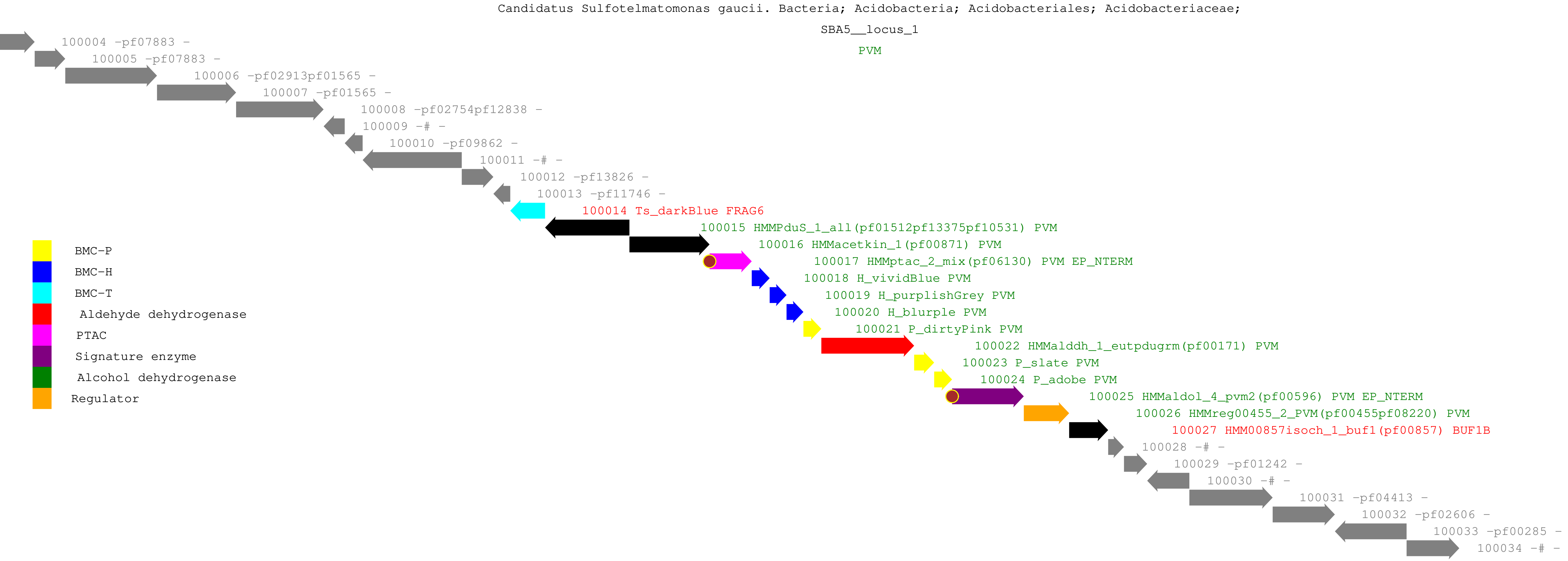
Candidatus Sulfotelmatomonas gaucii. Bacteria; Acidobacteria; Acidobacteriales; Acidobacteriaceae;
SBA5__locus_1
PVM
100004 -pf07883 -
100005 -pf07883 -
100006 -pf02913pf01565 -
100007 -pf01565 -
100008 -pf02754pf12838 -
100009 -# -
100010 -pf09862 -
100011 -# -
100012 -pf13826 -
100013 -pf11746 -
100014 Ts_darkBlue FRAG6
100015 HMMPduS_1_all(pf01512pf13375pf10531) PVM
100016 HMMacetkin_1(pf00871) PVM
100017 HMMptac_2_mix(pf06130) PVM EP_NTERM
100018 H_vividBlue PVM
100019 H_purplishGrey PVM
100020 H_blurple PVM
100021 P_dirtyPink PVM
100022 HMMalddh_1_eutpdugrm(pf00171) PVM
100023 P_slate PVM
100024 P_adobe PVM
100025 HMMaldol_4_pvm2(pf00596) PVM EP_NTERM
100026 HMMreg00455_2_PVM(pf00455pf08220) PVM
100027 HMM00857isoch_1_buf1(pf00857) BUF1B
100028 -# -
100029 -pf01242 -
100030 -# -
100031 -pf04413 -
100032 -pf02606 -
100033 -pf00285 -
100034 -# -
BMC-P
BMC-H
BMC-T
Aldehyde dehydrogenase
PTAC
Signature enzyme
Alcohol dehydrogenase
Regulator

Singulisphaera acidiphila (strain ATCC BAA-1392 / DSM 18658 / VKM Bacteria; Planctomycetes; Planctomycetia; Planctomycetales;

Sinac__locus_1

PVM

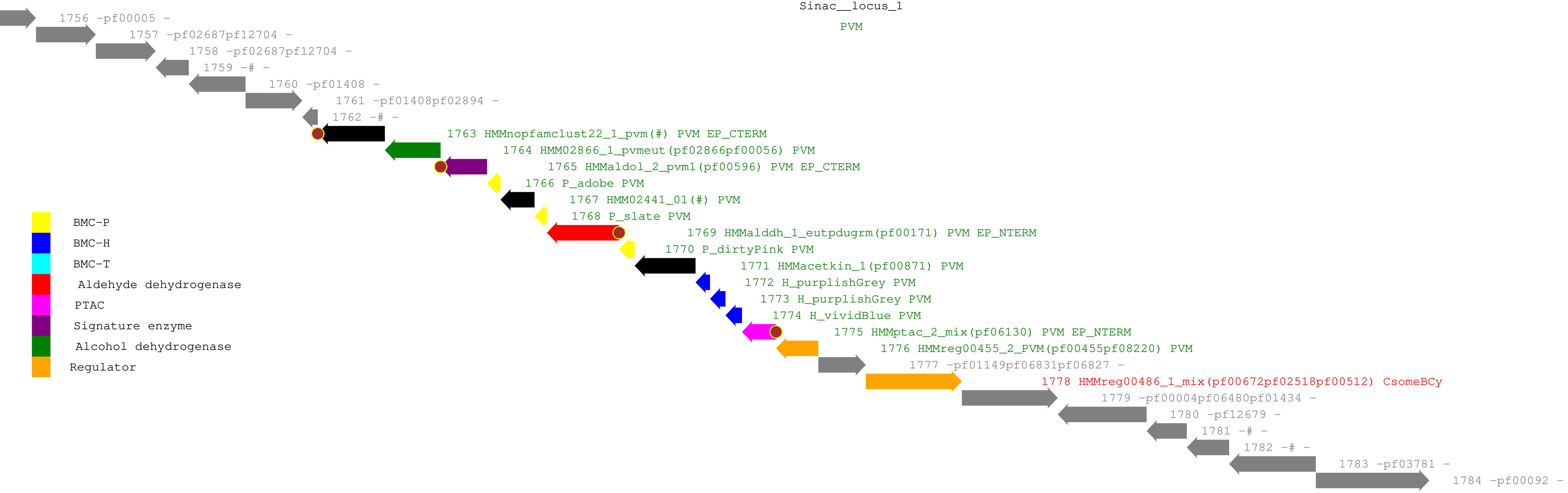

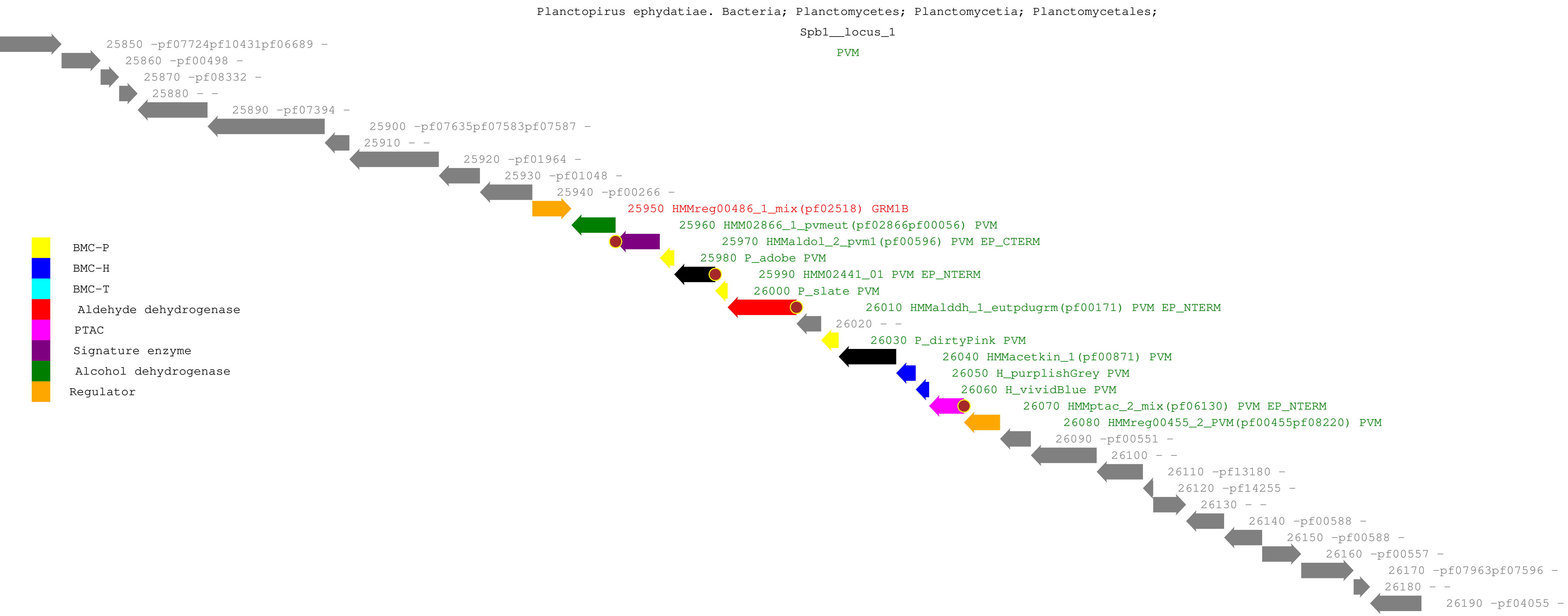
Planctopirus ephydatiae. Bacteria; Planctomycetes; Planctomycetia; Planctomycetales;
Spb1__locus_1
PVM
25850 -pf07724pf10431pf06689 -
25860 -pf00498 -
25870 -pf08332 -
25880 - -
25890 -pf07394 -
25900 -pf07635pf07583pf07587 -
25910 - -
25920 -pf01964 -
25930 -pf01048 -
25940 -pf00266 -
25950 HMMreg00486_1_mix(pf02518) GRM1B
25960 HMM02866_1_pvmeut(pf02866pf00056) PVM
25970 HMMaldol_2_pvm1(pf00596) PVM EP_CTERM
25980 P_adobe PVM
25990 HMM02441_01 PVM EP_NTERM
26000 P_slate PVM
26010 HMMalddh_1_eutpdugrm(pf00171) PVM EP_NTERM
26020 - -
26030 P_dirtyPink PVM
26040 HMMacetkin_1(pf00871) PVM
26050 H_purplishGrey PVM
26060 H_vividBlue PVM
26070 HMMptac_2_mix(pf06130) PVM EP_NTERM
26080 HMMreg00455_2_PVM(pf00455pf08220) PVM
26090 -pf00551 -
26100 - -
26110 -pf13180 -
26120 -pf14255 -
26130 - -
26140 -pf00588 -
26150 -pf00588 -
26160 -pf00557 -
26170 -pf07963pf07596 -
26180 - -
26190 -pf04055 -
BMC-P
BMC-H
BMC-T
Aldehyde dehydrogenase
PTAC
Signature enzyme
Alcohol dehydrogenase
Regulator

Planctomycetes bacterium SV_7m_r. Bacteria; Planctomycetes.

SV7mr__locus_1

PVM

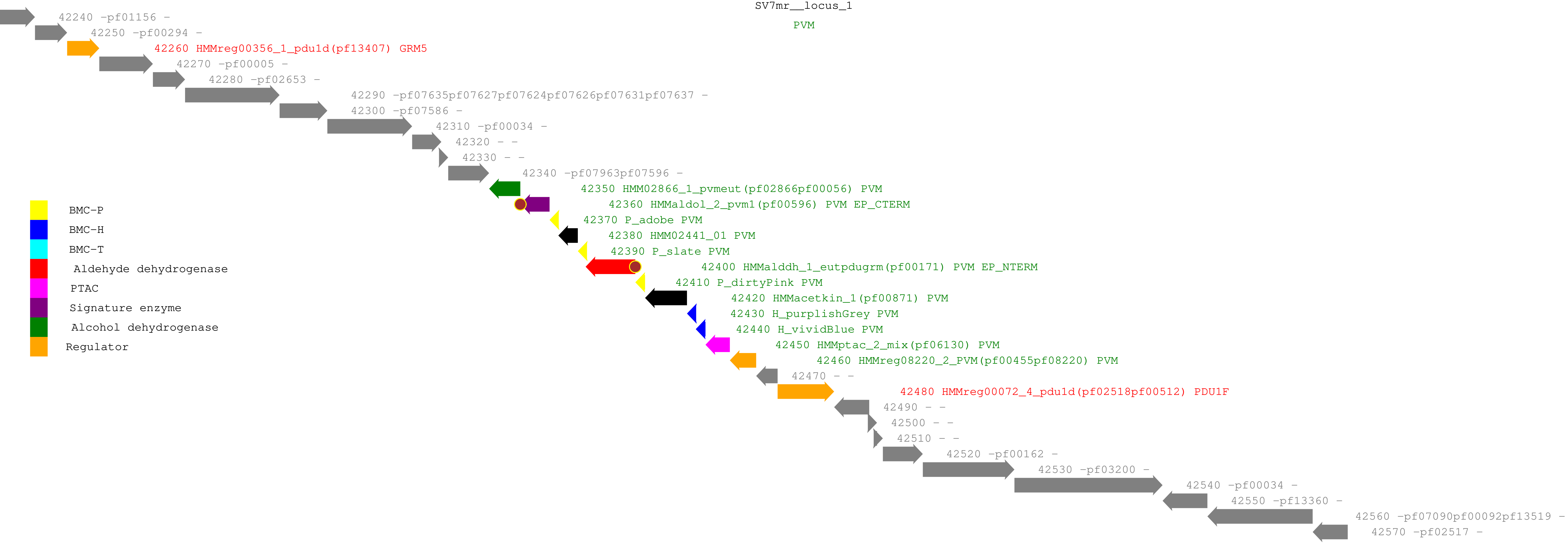

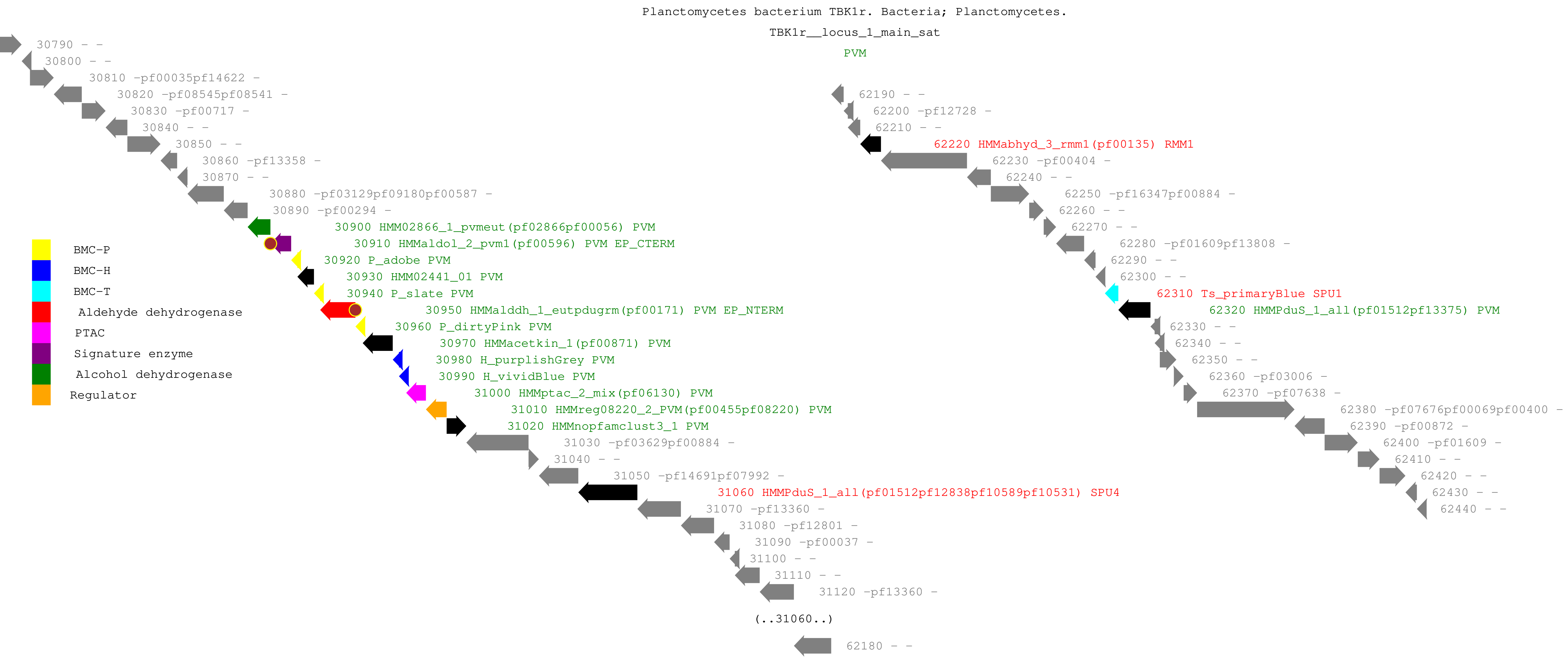

Planctomycetes bacterium TBK1r. Bacteria; Planctomycetes.
TBK1r__locus_1_main_sat
PVM
30790 - -
30800 - -
30810 -pf00035pf14622 -
30820 -pf08545pf08541 -
30830 -pf00717 -
30840 - -
30850 - -
30860 -pf13358 -
30870 - -
30880 -pf03129pf09180pf00587 -
30890 -pf00294 -
30900 HMM02866_1_pvmeut(pf02866pf00056) PVM
30910 HMMaldol_2_pvm1(pf00596) PVM EP_CTERM
30920 P_adobe PVM
30930 HMM02441_01 PVM
30940 P_slate PVM
30950 HMMalddh_1_eutpdugrm(pf00171) PVM EP_NTERM
30960 P_dirtyPink PVM
30970 HMMacetkin_1(pf00871) PVM
30980 H_purplishGrey PVM
30990 H_vividBlue PVM
31000 HMMptac_2_mix(pf06130) PVM
31010 HMMreg08220_2_PVM(pf00455pf08220) PVM
31020 HMMnopfamclust3_1 PVM
31030 -pf03629pf00884 -
31040 - -
31050 -pf14691pf07992 -
31060 HMMPduS_1_all(pf01512pf12838pf10589pf10531) SPU4
31070 -pf13360 -
31080 -pf12801 -
31090 -pf00037 -
31100 - -
31110 - -
31120 -pf13360 -
(..31060..)
62180 - -
62190 - -
62200 -pf12728 -
62210 - -
62220 HMMabhyd_3_rmm1(pf00135) RMM1
62230 -pf00404 -
62240 - -
62250 -pf16347pf00884 -
62260 - -
62270 - -
62280 -pf01609pf13808 -
62290 - -
62300 - -
62310 Ts_primaryBlue SPU1
62320 HMMPduS_1_all(pf01512pf13375) PVM
62330 - -
62340 - -
62350 - -
62360 -pf03006 -
62370 -pf07638 -
62380 -pf07676pf00069pf00400 -
62390 -pf00872 -
62400 -pf01609 -
62410 - -
62420 - -
62430 - -
62440 - -
BMC-P
BMC-H
BMC-T
Aldehyde dehydrogenase
PTAC
Signature enzyme
Alcohol dehydrogenase
Regulator

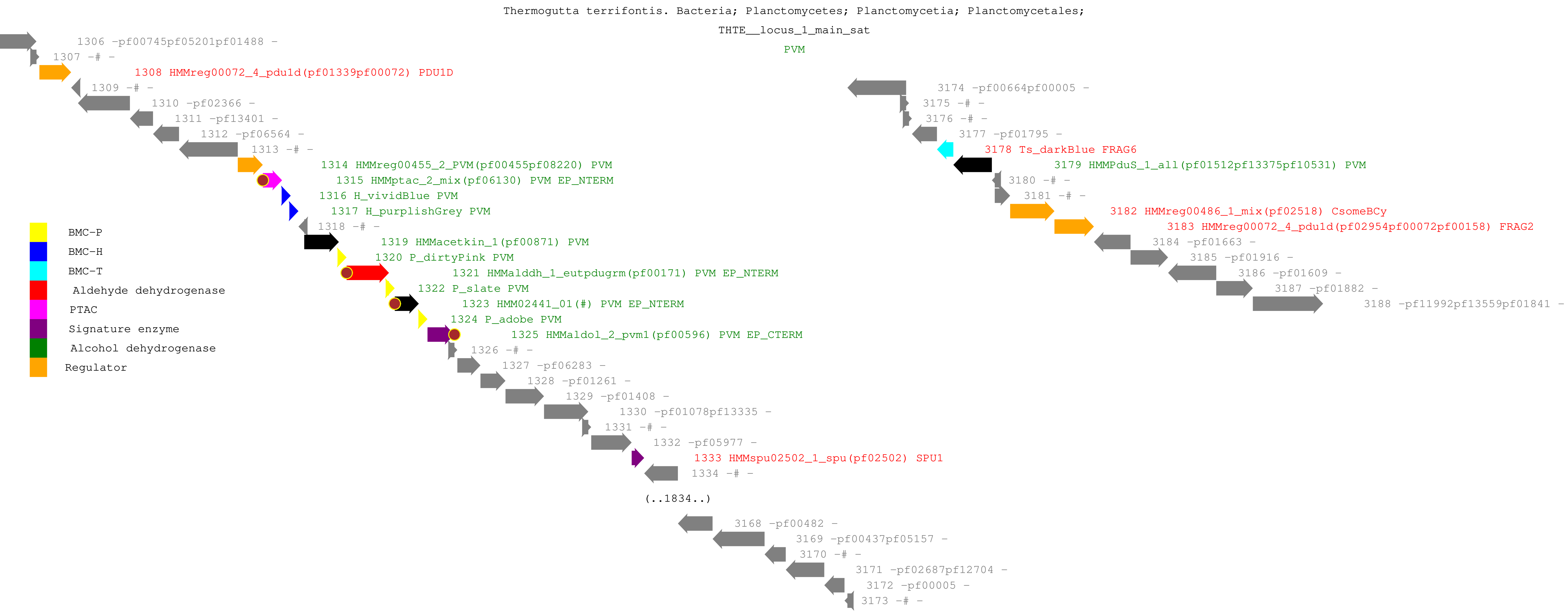
Thermogutta terrifontis. Bacteria; Planctomycetes; Planctomycetia; Planctomycetales;
THTE__locus_1_main_sat
PVM
1306 -pf00745pf05201pf01488 -
1307 -# -
1308 HMMreg00072_4_pdu1d(pf01339pf00072) PDU1D
1309 -# -
1310 -pf02366 -
1311 -pf13401 -
1312 -pf06564 -
1313 -# -
1314 HMMreg00455_2_PVM(pf00455pf08220) PVM
1315 HMMptac_2_mix(pf06130) PVM EP_NTERM
1316 H_vividBlue PVM
1317 H_purplishGrey PVM
1318 -# -
1319 HMMacetkin_1(pf00871) PVM
1320 P_dirtyPink PVM
1321 HMMalddh_1_eutpdugrm(pf00171) PVM EP_NTERM
1322 P_slate PVM
1323 HMM02441_01(#) PVM EP_NTERM
1324 P_adobe PVM
1325 HMMaldol_2_pvm1(pf00596) PVM EP_CTERM
1326 -# -
1327 -pf06283 -
1328 -pf01261 -
1329 -pf01408 -
1330 -pf01078pf13335 -
1331 -# -
1332 -pf05977 -
1333 HMMspu02502_1_spu(pf02502) SPU1
1334 -# -
(..1834..)
3168 -pf00482 -
3169 -pf00437pf05157 -
3170 -# -
3171 -pf02687pf12704 -
3172 -pf00005 -
3173 -# -
3174 -pf00664pf00005 -
3175 -# -
3176 -# -
3177 -pf01795 -
3178 Ts_darkBlue FRAG6
3179 HMMPduS_1_all(pf01512pf13375pf10531) PVM
3180 -# -
3181 -# -
3182 HMMreg00486_1_mix(pf02518) CsomeBCy
3183 HMMreg00072_4_pdu1d(pf02954pf00072pf00158) FRAG2
3184 -pf01663 -
3185 -pf01916 -
3186 -pf01609 -
3187 -pf01882 -
3188 -pf11992pf13559pf01841 -
BMC-P
BMC-H
BMC-T
Aldehyde dehydrogenase
PTAC
Signature enzyme
Alcohol dehydrogenase
Regulator

Tautonia sociabilis. Bacteria; Planctomycetes; Planctomycetia; Planctomycetales; Isosphaeraceae;

TsocGM__locus_1

PVM

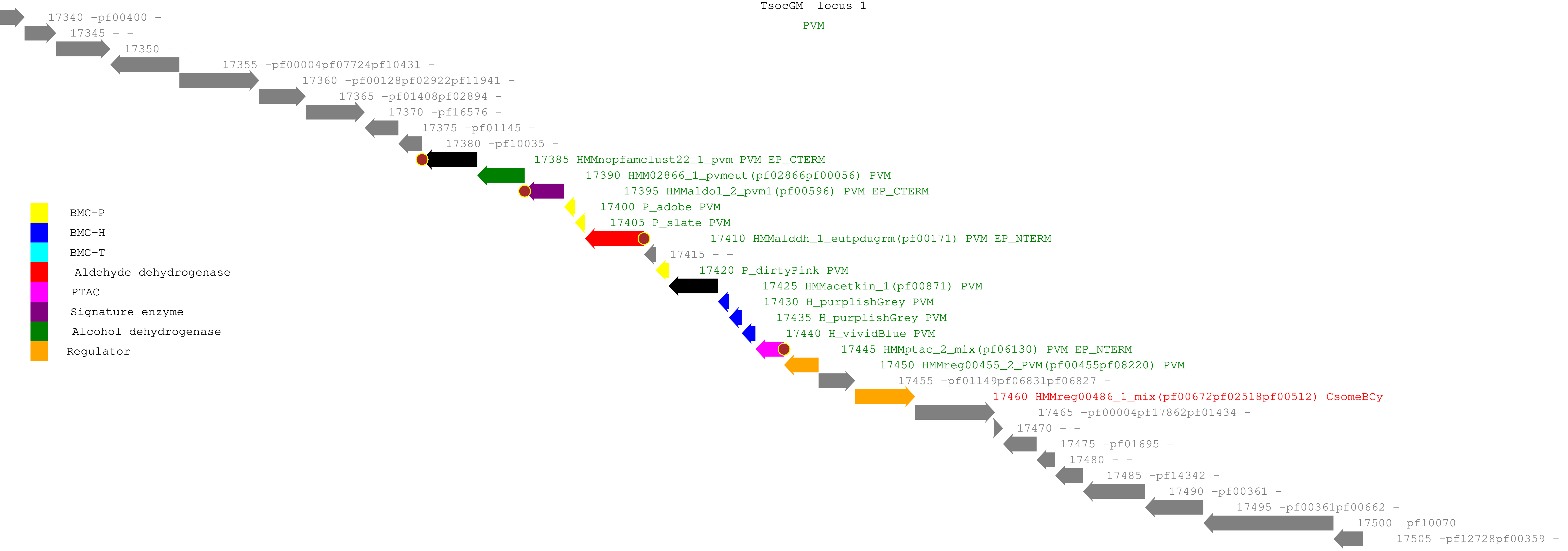

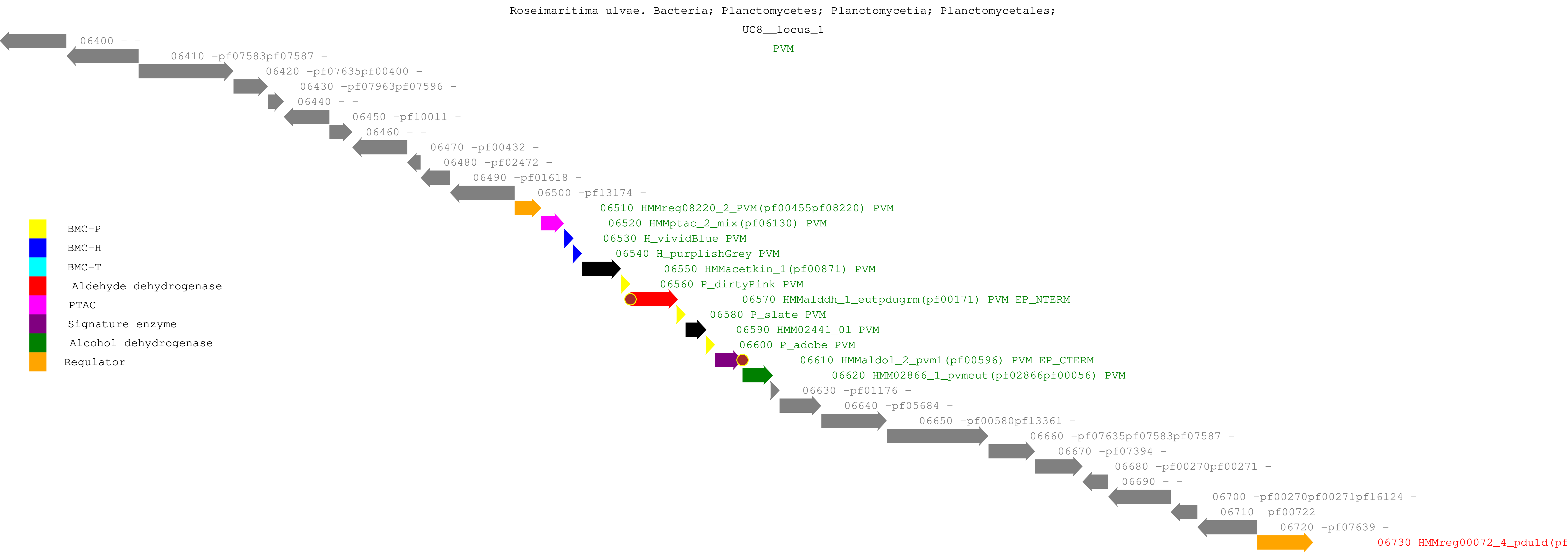
Roseimaritima ulvae. Bacteria; Planctomycetes; Planctomycetia; Planctomycetales;
UC8__locus_1
PVM
06400 - -
06410 -pf07583pf07587 -
06420 -pf07635pf00400 -
06430 -pf07963pf07596 -
06440 - -
06450 -pf10011 -
06460 - -
06470 -pf00432 -
06480 -pf02472 -
06490 -pf01618 -
06500 -pf13174 -
06510 HMMreg08220_2_PVM(pf00455pf08220) PVM
06520 HMMptac_2_mix(pf06130) PVM
06530 H_vividBlue PVM
06540 H_purplishGrey PVM
06550 HMMacetkin_1(pf00871) PVM
06560 P_dirtyPink PVM
06570 HMMalddh_1_eutpdugrm(pf00171) PVM EP_NTERM
06580 P_slate PVM
06590 HMM02441_01 PVM
06600 P_adobe PVM
06610 HMMaldol_2_pvm1(pf00596) PVM EP_CTERM
06620 HMM02866_1_pvmeut(pf02866pf00056) PVM
06630 -pf01176 -
06640 -pf05684 -
06650 -pf00580pf13361 -
06660 -pf07635pf07583pf07587 -
06670 -pf07394 -
06680 -pf00270pf00271 -
06690 - -
06700 -pf00270pf00271pf16124 -
06710 -pf00722 -
06720 -pf07639 -
06730 HMMreg00072_4_pdu1d(pf
BMC-P
BMC-H
BMC-T
Aldehyde dehydrogenase
PTAC
Signature enzyme
Alcohol dehydrogenase
Regulator

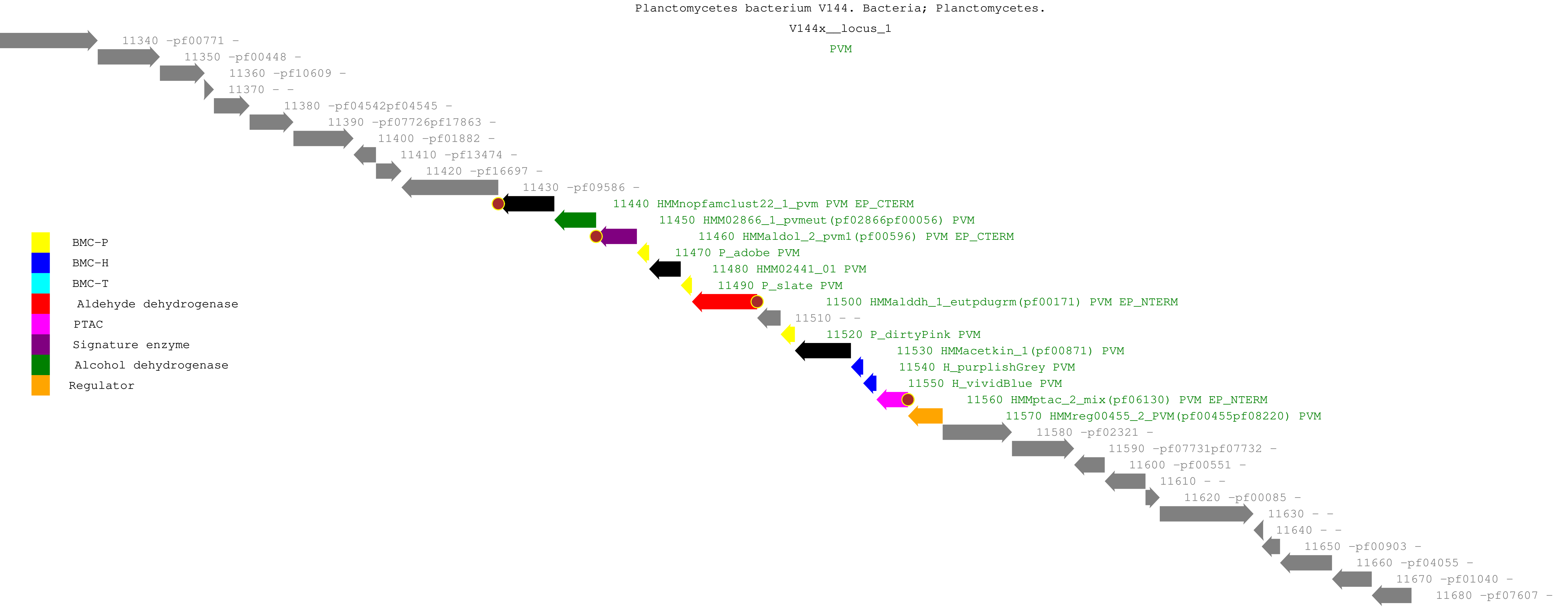

Planctomycetes bacterium V144. Bacteria; Planctomycetes.
V144x__locus_1
PVM
11340 -pf00771 -
11350 -pf00448 -
11360 -pf10609 -
11370 - -
11380 -pf04542pf04545 -
11390 -pf07726pf17863 -
11400 -pf01882 -
11410 -pf13474 -
11420 -pf16697 -
11430 -pf09586 -
11440 HMMnopfamclust22_1_pvm PVM EP_CTERM
11450 HMM02866_1_pvmeut(pf02866pf00056) PVM
11460 HMMaldol_2_pvm1(pf00596) PVM EP_CTERM
11470 P_adobe PVM
11480 HMM02441_01 PVM
11490 P_slate PVM
11500 HMMalddh_1_eutpdugrm(pf00171) PVM EP_NTERM
11510 - -
11520 P_dirtyPink PVM
11530 HMMacetkin_1(pf00871) PVM
11540 H_purplishGrey PVM
11550 H_vividBlue PVM
11560 HMMptac_2_mix(pf06130) PVM EP_NTERM
11570 HMMreg00455_2_PVM(pf00455pf08220) PVM
11580 -pf02321 -
11590 -pf07731pf07732 -
11600 -pf00551 -
11610 - -
11620 -pf00085 -
11630 - -
11640 - -
11650 -pf00903 -
11660 -pf04055 -
11670 -pf01040 -
11680 -pf07607 -
BMC-P
BMC-H
BMC-T
Aldehyde dehydrogenase
PTAC
Signature enzyme
Alcohol dehydrogenase
Regulator

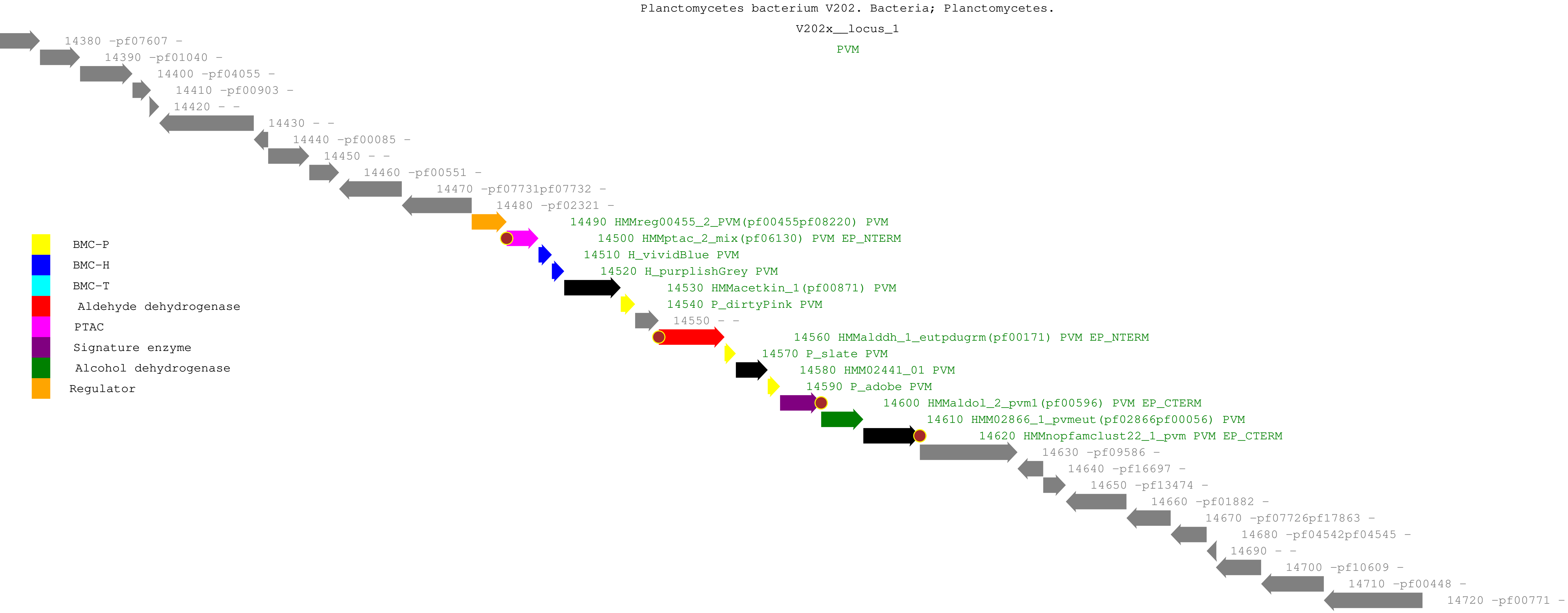
Planctomycetes bacterium V202. Bacteria; Planctomycetes.
V202x__locus_1
PVM
14380 -pf07607 -
14390 -pf01040 -
14400 -pf04055 -
14410 -pf00903 -
14420 - -
14430 - -
14440 -pf00085 -
14450 - -
14460 -pf00551 -
14470 -pf07731pf07732 -
14480 -pf02321 -
14490 HMMreg00455_2_PVM(pf00455pf08220) PVM
14500 HMMptac_2_mix(pf06130) PVM EP_NTERM
14510 H_vividBlue PVM
14520 H_purplishGrey PVM
14530 HMMacetkin_1(pf00871) PVM
14540 P_dirtyPink PVM
14550 - -
14560 HMMalddh_1_eutpdugrm(pf00171) PVM EP_NTERM
14570 P_slate PVM
14580 HMM02441_01 PVM
14590 P_adobe PVM
14600 HMMaldol_2_pvm1(pf00596) PVM EP_CTERM
14610 HMM02866_1_pvmeut(pf02866pf00056) PVM
14620 HMMnopfamclust22_1_pvm PVM EP_CTERM
14630 -pf09586 -
14640 -pf16697 -
14650 -pf13474 -
14660 -pf01882 -
14670 -pf07726pf17863 -
14680 -pf04542pf04545 -
14690 - -
14700 -pf10609 -
14710 -pf00448 -
14720 -pf00771 -
BMC-P
BMC-H
BMC-T
Aldehyde dehydrogenase
PTAC
Signature enzyme
Alcohol dehydrogenase
Regulator

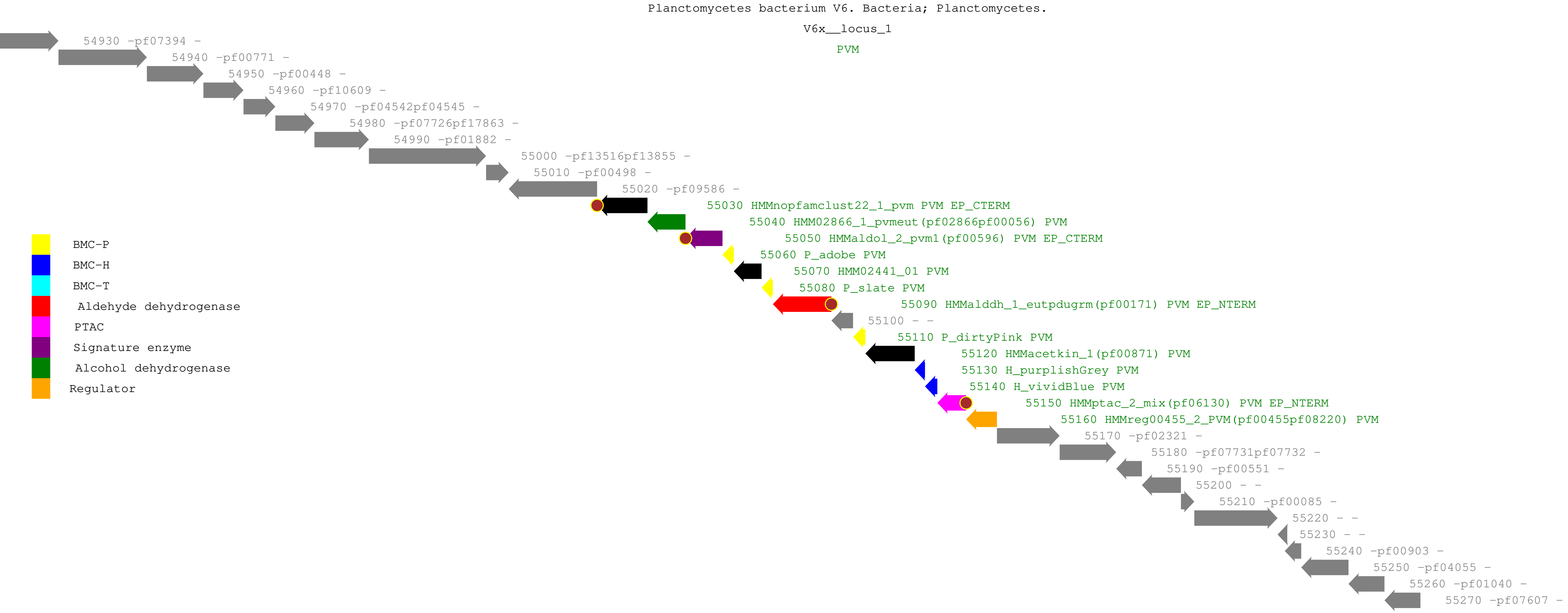
Planctomycetes bacterium V6. Bacteria; Planctomycetes.
V6x__locus_1
PVM
54930 -pf07394 -
54940 -pf00771 -
54950 -pf00448 -
54960 -pf10609 -
54970 -pf04542pf04545 -
54980 -pf07726pf17863 -
54990 -pf01882 -
55000 -pf13516pf13855 -
55010 -pf00498 -
55020 -pf09586 -
55030 HMMnopfamclust22_1_pvm PVM EP_CTERM
55040 HMM02866_1_pvmeut(pf02866pf00056) PVM
55050 HMMaldol_2_pvm1(pf00596) PVM EP_CTERM
55060 P_adobe PVM
55070 HMM02441_01 PVM
55080 P_slate PVM
55090 HMMaldd h_1_eutpdugrm(pf00171) PVM EP_NTERM
55100 - -
55110 P_dirtyPink PVM
55120 HMMacetkin_1(pf00871) PVM
55130 H_purplishGrey PVM
55140 H_vividBlue PVM
55150 HMMptac_2_mix(pf06130) PVM EP_NTERM
55160 HMMreg00455_2_PVM(pf00455pf08220) PVM
55170 -pf02321 -
55180 -pf07731pf07732 -
55190 -pf00551 -
55200 - -
55210 -pf00085 -
55220 - -
55230 - -
55240 -pf00903 -
55250 -pf04055 -
55260 -pf01040 -
55270 -pf07607 -
BMC-P
BMC-H
BMC-T
Aldehyde dehydrogenase
PTAC
Signature enzyme
Alcohol dehydrogenase
Regulator

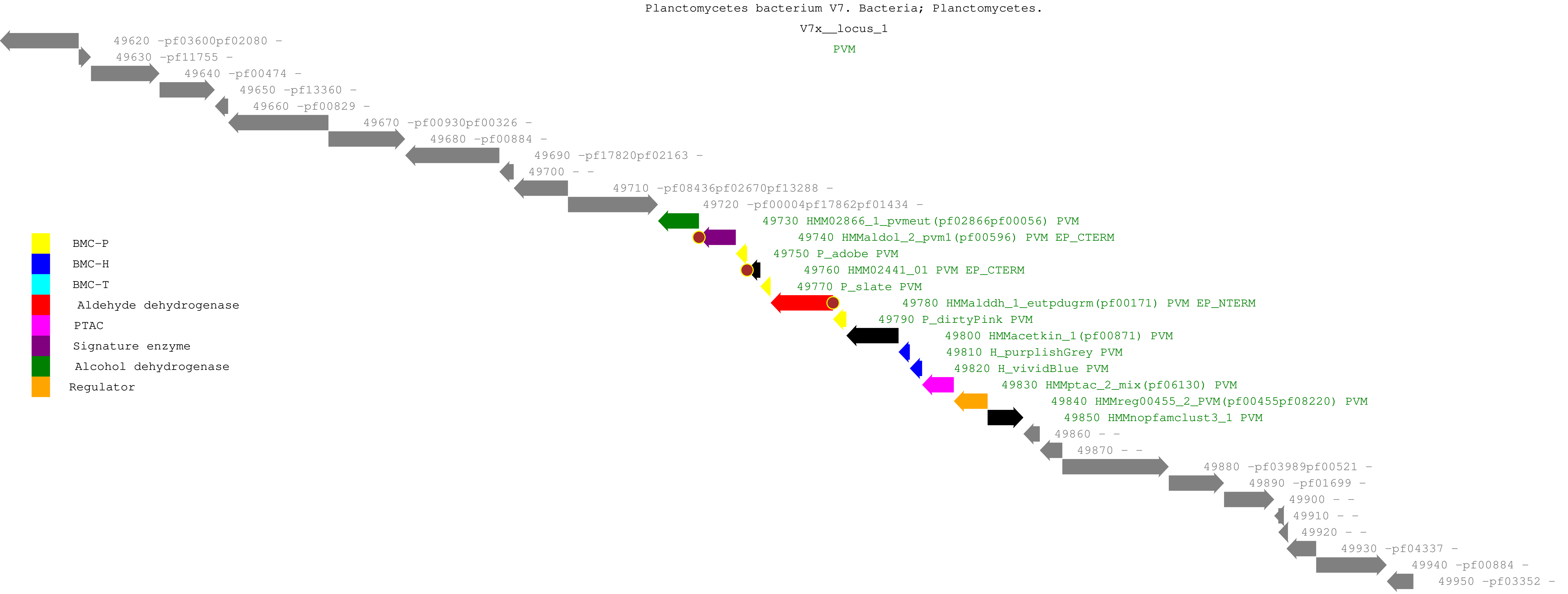
Planctomycetes bacterium V7. Bacteria; Planctomycetes.
V7x__locus_1
PVM
49620 -pf03600pf02080 -
49630 -pf11755 -
49640 -pf00474 -
49650 -pf13360 -
49660 -pf00829 -
49670 -pf00930pf00326 -
49680 -pf00884 -
49690 -pf17820pf02163 -
49700 - -
49710 -pf08436pf02670pf13288 -
49720 -pf00004pf17862pf01434 -
49730 HMM02866_1_pvmeut(pf02866pf00056) PVM
49740 HMMaldol_2_pvm1(pf00596) PVM EP_CTERM
49750 P_adobe PVM
49760 HMM02441_01 PVM EP_CTERM
49770 P_slate PVM
49780 HMMalddh_1_eutpdugrm(pf00171) PVM EP_NTERM
49790 P_dirtyPink PVM
49800 HMMacetkin_1(pf00871) PVM
49810 H_purplishGrey PVM
49820 H_vividBlue PVM
49830 HMMptac_2_mix(pf06130) PVM
49840 HMMreg00455_2_PVM(pf00455pf08220) PVM
49850 HMMnopfamclust3_1 PVM
49860 - -
49870 - -
49880 -pf03989pf00521 -
49890 -pf01699 -
49900 - -
49910 - -
49920 - -
49930 -pf04337 -
49940 -pf00884 -
49950 -pf03352 -
BMC-P
BMC-H
BMC-T
Aldehyde dehydrogenase
PTAC
Signature enzyme
Alcohol dehydrogenase
Regulator

Verrucomicrobiae bacterium DG1235. Bacteria; Verrucomicrobia; Verrucomicrobiae; Verrucomicrobiales.

VDG1235__locus_1_main_sat

PVM

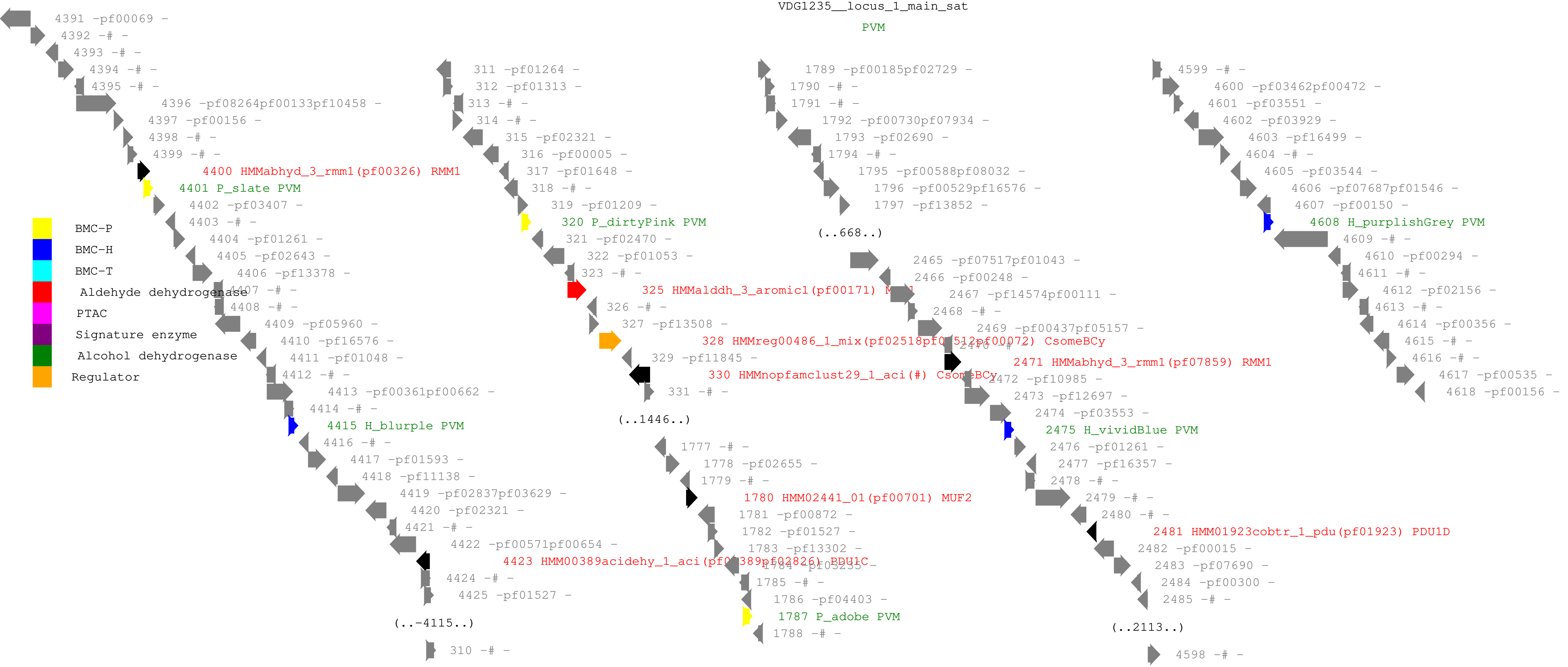

Pirellula sp. SH-Sr6A. Bacteria; Planctomycetes; Planctomycetia; Planctomycetales;

VN12__locus_1_main_sat

PVM

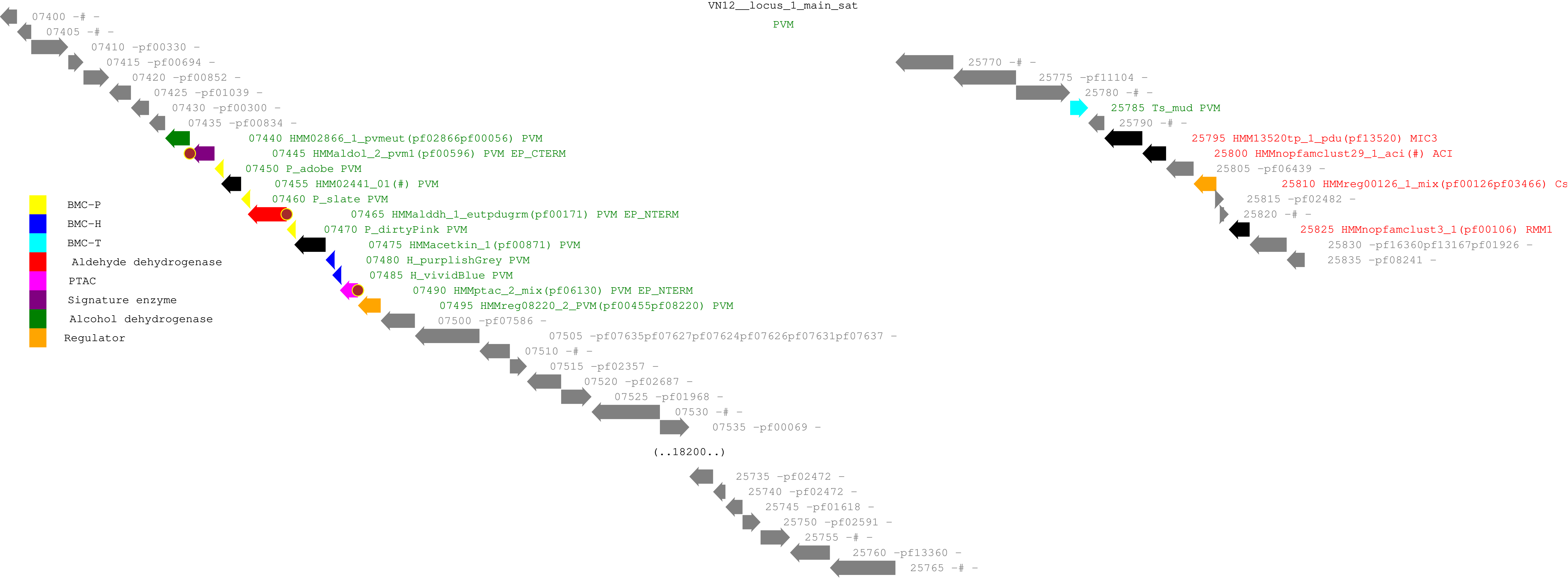

Planctomyces sp. SH-PL14. Bacteria; Planctomycetes; Planctomycetia; Planctomycetales;

VT03__locus_1

PVM

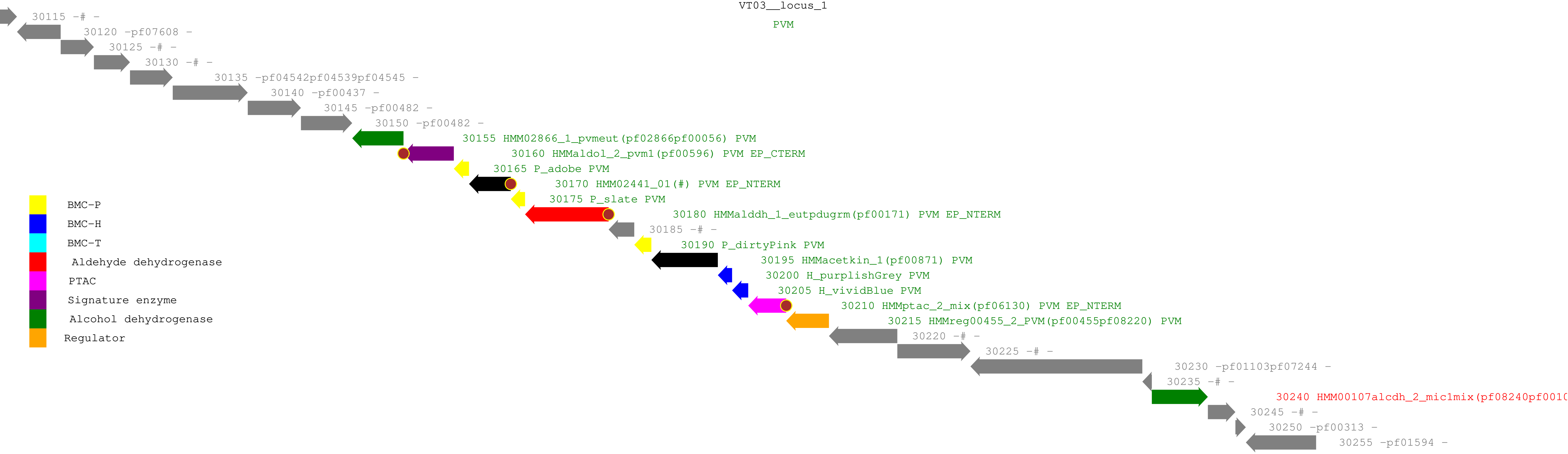

Planctomyces sp. SH-PL62. Bacteria; Planctomycetes; Planctomycetia; Planctomycetales;

VT85__locus_1

PVM

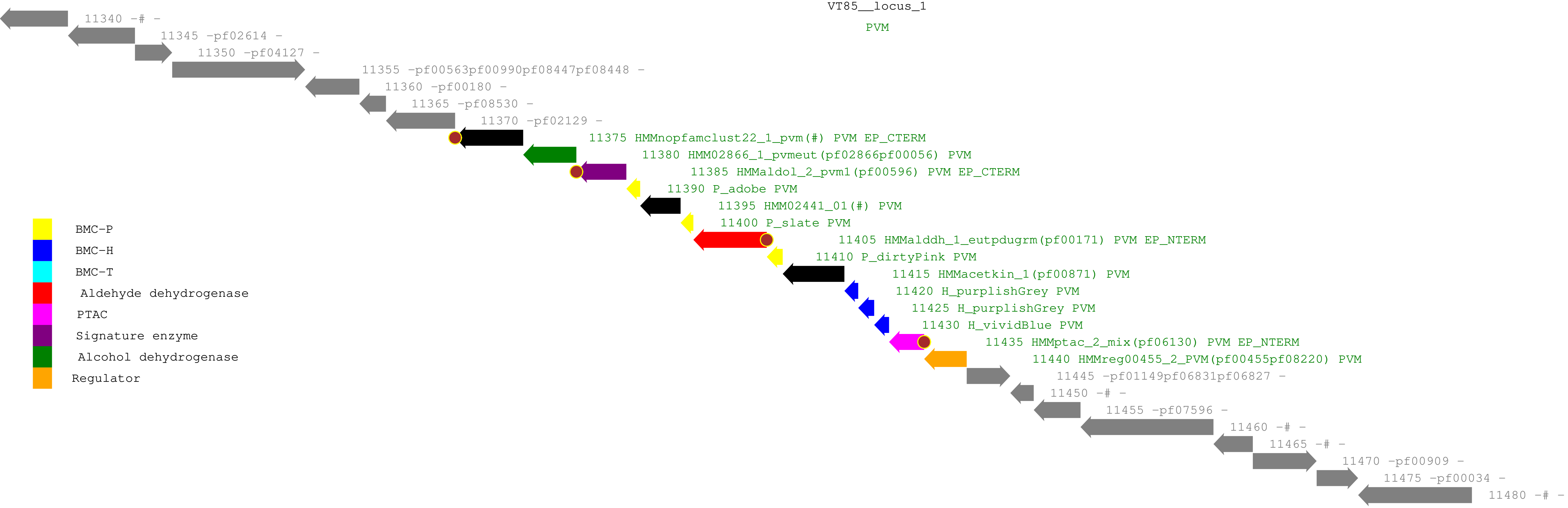

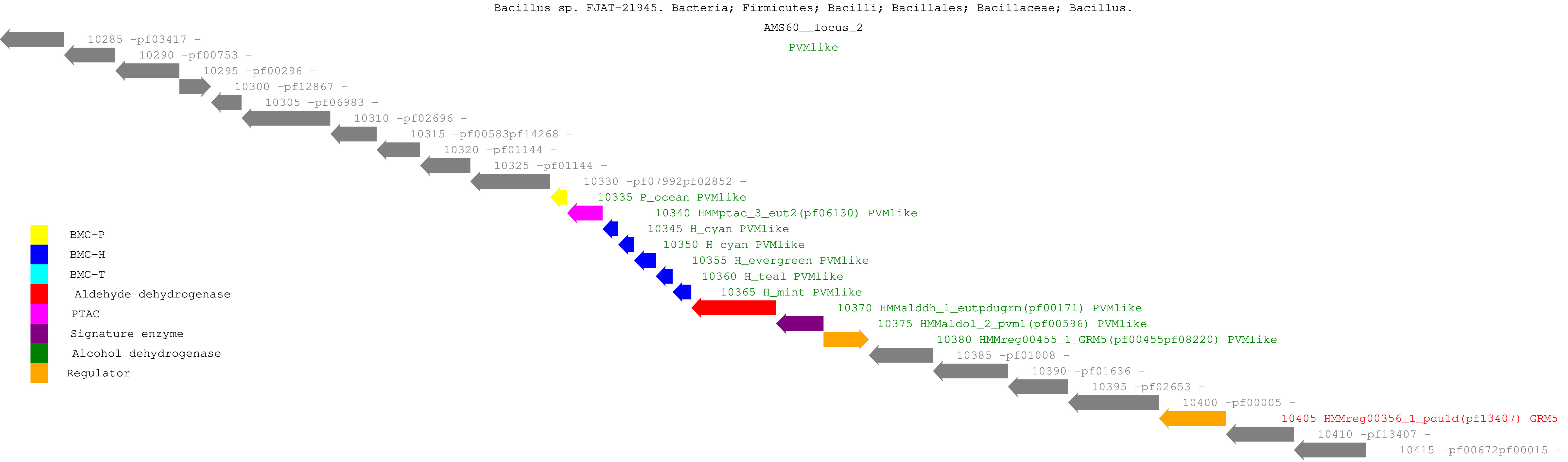
Bacillus sp. FJAT-21945. Bacteria; Firmicutes; Bacilli; Bacillales; Bacillaceae; Bacillus.
AMS60__locus_2
PVMlike
10285 -pf03417 -
10290 -pf00753 -
10295 -pf00296 -
10300 -pf12867 -
10305 -pf06983 -
10310 -pf02696 -
10315 -pf00583pf14268 -
10320 -pf01144 -
10325 -pf01144 -
10330 -pf07992pf02852 -
10335 P_ocean PVMlike
10340 HMMptac_3_eut2(pf06130) PVMlike
10345 H_cyan PVMlike
10350 H_cyan PVMlike
10355 H_evergreen PVMlike
10360 H_teal PVMlike
10365 H_mint PVMlike
10370 HMMalddh_1_eutpdugrm(pf00171) PVMlike
10375 HMMaldol_2_pvm1(pf00596) PVMlike
10380 HMMreg00455_1_GRM5(pf00455pf08220) PVMlike
10385 -pf01008 -
10390 -pf01636 -
10395 -pf02653 -
10400 -pf00005 -
10405 HMMreg00356_1_pdu1d(pf13407) GRM5
10410 -pf13407 -
10415 -pf00672pf00015 -
BMC-P
BMC-H
BMC-T
Aldehyde dehydrogenase
PTAC
Signature enzyme
Alcohol dehydrogenase
Regulator

Bacillus solani. Bacteria; Firmicutes; Bacilli; Bacillales; Bacillaceae; Bacillus.

AN957__locus_2

PVMlike

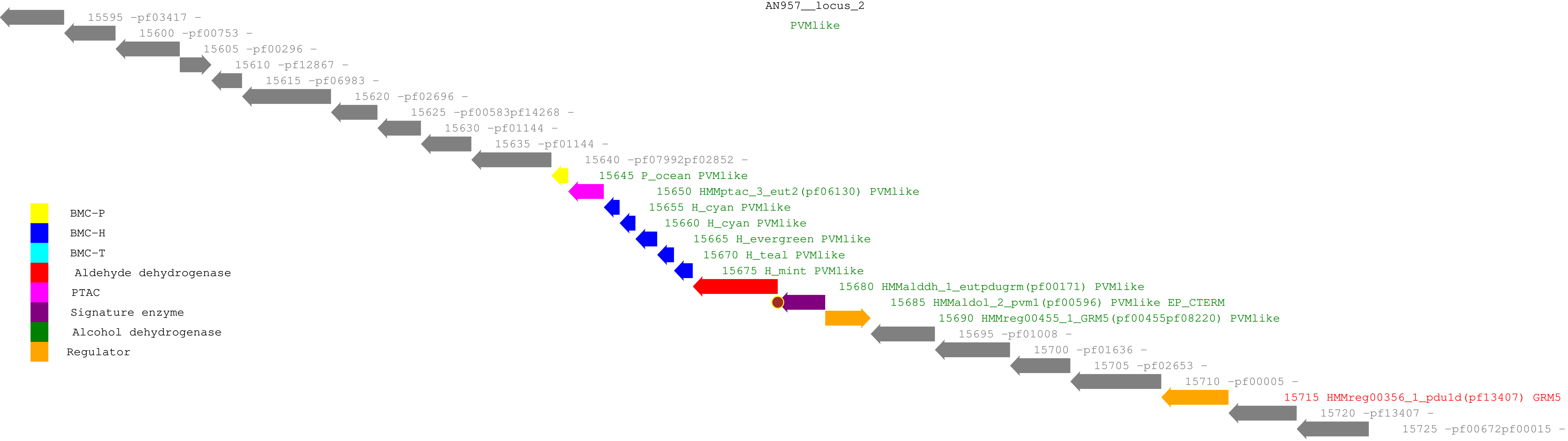

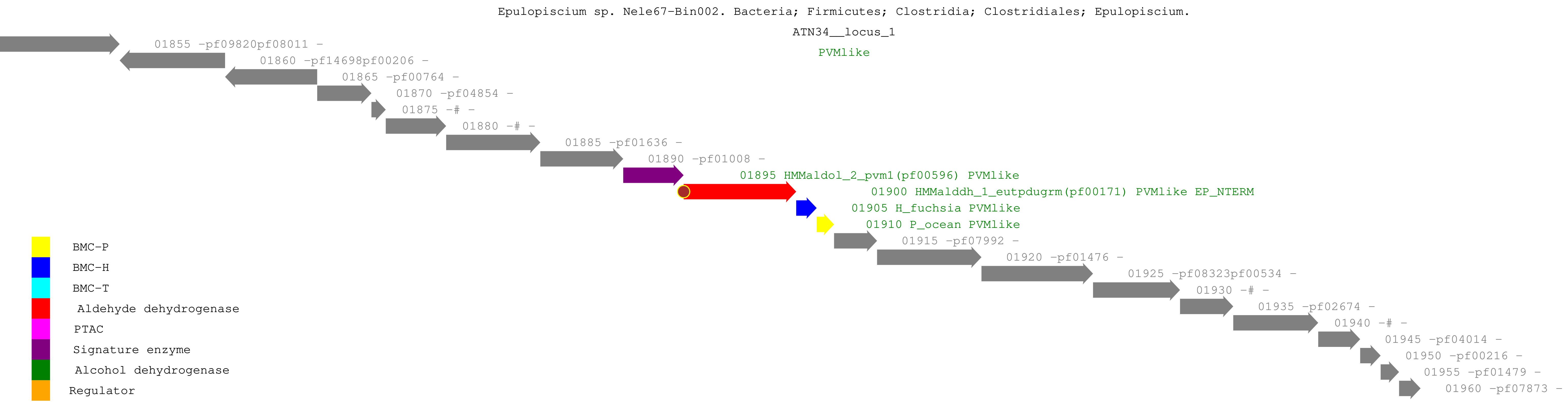
Epulopiscium sp. Nele67-Bin002. Bacteria; Firmicutes; Clostridia; Clostridiales; Epulopiscium.
ATN34__locus_1
PVMlike
01855 -pf09820pf08011 -
01860 -pf14698pf00206 -
01865 -pf00764 -
01870 -pf04854 -
01875 -# -
01880 -# -
01885 -pf01636 -
01890 -pf01008 -
01895 HMMaldol_2_pvm1(pf00596) PVMlike
01900 HMMalddh_1_eutpdugrm(pf00171) PVMlike EP_NTERM
01905 H_fuchsia PVMlike
01910 P_ocean PVMlike
01915 -pf07992 -
01920 -pf01476 -
01925 -pf08323pf00534 -
01930 -# -
01935 -pf02674 -
01940 -# -
01945 -pf04014 -
01950 -pf00216 -
01955 -pf01479 -
01960 -pf07873 -
BMC-P
BMC-H
BMC-T
Aldehyde dehydrogenase
PTAC
Signature enzyme
Alcohol dehydrogenase
Regulator

Paenibacillaceae bacterium ZCTH02-B3. Bacteria; Firmicutes; Bacilli; Bacillales; Paenibacillaceae.

BAA02__locus_1

PVMlike

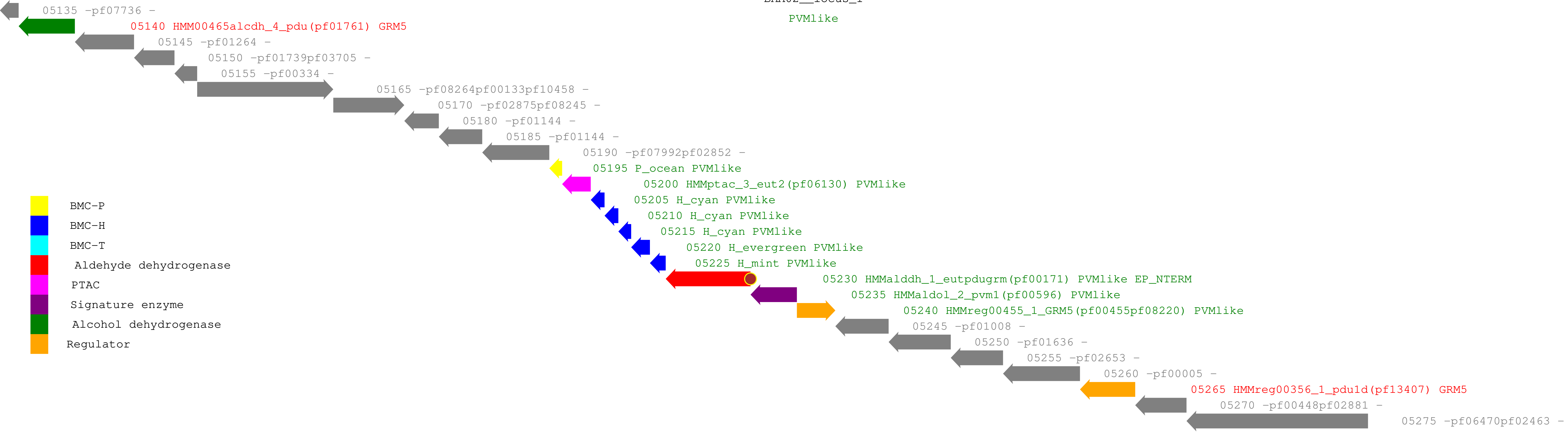

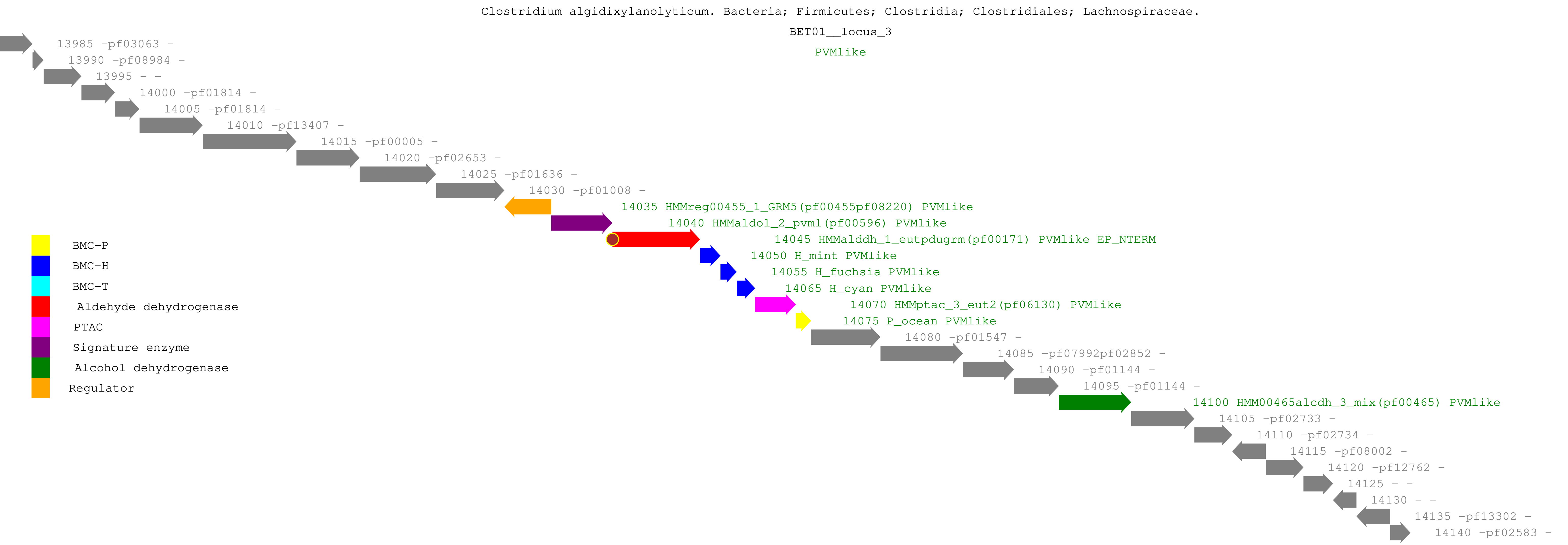
Clostridium algidixylanolyticum. Bacteria; Firmicutes; Clostridia; Clostridiales; Lachnospiraceae.
BET01__locus_3
PVMlike
13985 -pf03063 -
13990 -pf08984 -
13995 - -
14000 -pf01814 -
14005 -pf01814 -
14010 -pf13407 -
14015 -pf00005 -
14020 -pf02653 -
14025 -pf01636 -
14030 -pf01008 -
14035 HMMreg00455_1_GRM5(pf00455pf08220) PVMlike
14040 HMMaldol_2_pvm1(pf00596) PVMlike
14045 HMMalddh_1_eutpdugrm(pf00171) PVMlike EP_NTERM
14050 H_mint PVMlike
14055 H_fuchsia PVMlike
14065 H_cyan PVMlike
14070 HMMptac_3_eut2(pf06130) PVMlike
14075 P_ocean PVMlike
14080 -pf01547 -
14085 -pf07992pf02852 -
14090 -pf01144 -
14095 -pf01144 -
14100 HMM00465alcdh_3_mix(pf00465) PVMlike
14105 -pf02733 -
14110 -pf02734 -
14115 -pf08002 -
14120 -pf12762 -
14125 - -
14130 - -
14135 -pf13302 -
14140 -pf02583 -
BMC-P
BMC-H
BMC-T
Aldehyde dehydrogenase
PTAC
Signature enzyme
Alcohol dehydrogenase
Regulator

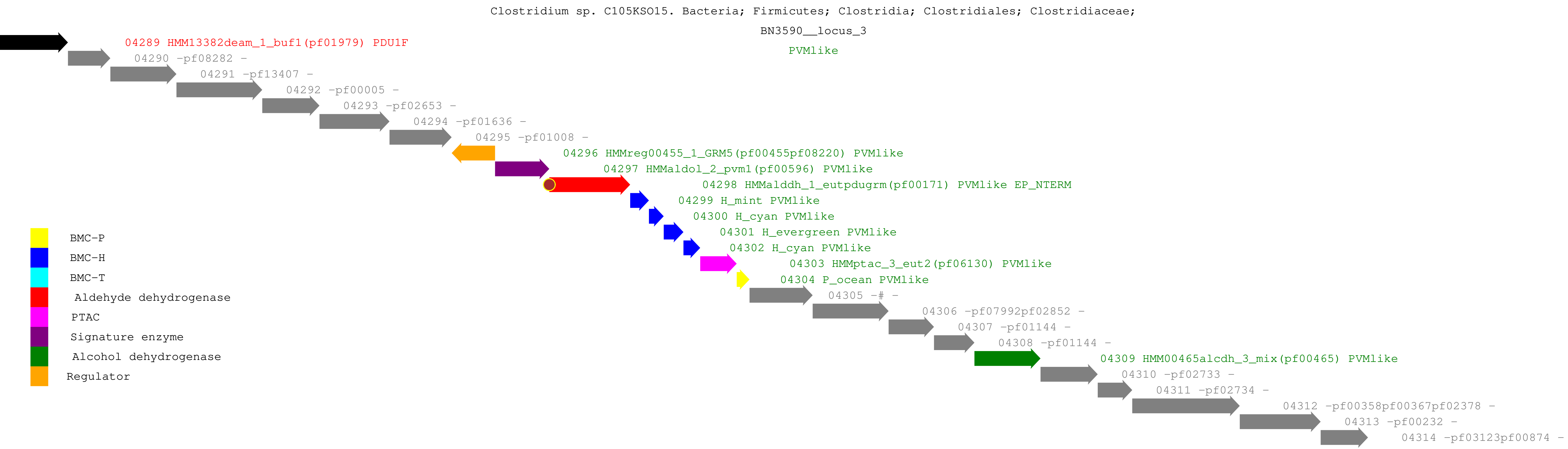

Clostridium sp. C105KSO15. Bacteria; Firmicutes; Clostridia; Clostridiales; Clostridiaceae;
BN3590__locus_3
PVMlike
04289 HMM13382deam_1_buf1(pf01979) PDU1F
04290 -pf08282 -
04291 -pf13407 -
04292 -pf00005 -
04293 -pf02653 -
04294 -pf01636 -
04295 -pf01008 -
04296 HMMreg00455_1_GRM5(pf00455pf08220) PVMlike
04297 HMMaldol_2_pvm1(pf00596) PVMlike
04298 HMMaldh_1_eutpdugrm(pf00171) PVMlike EP_NTERM
04299 H_mint PVMlike
04300 H_cyan PVMlike
04301 H_evergreen PVMlike
04302 H_cyan PVMlike
04303 HMMptac_3_eut2(pf06130) PVMlike
04304 P_ocean PVMlike
04305 -# -
04306 -pf07992pf02852 -
04307 -pf01144 -
04308 -pf01144 -
04309 HMM00465alcdh_3_mix(pf00465) PVMlike
04310 -pf02733 -
04311 -pf02734 -
04312 -pf00358pf00367pf02378 -
04313 -pf00232 -
04314 -pf03123pf00874 -
BMC-P
BMC-H
BMC-T
Aldehyde dehydrogenase
PTAC
Signature enzyme
Alcohol dehydrogenase
Regulator

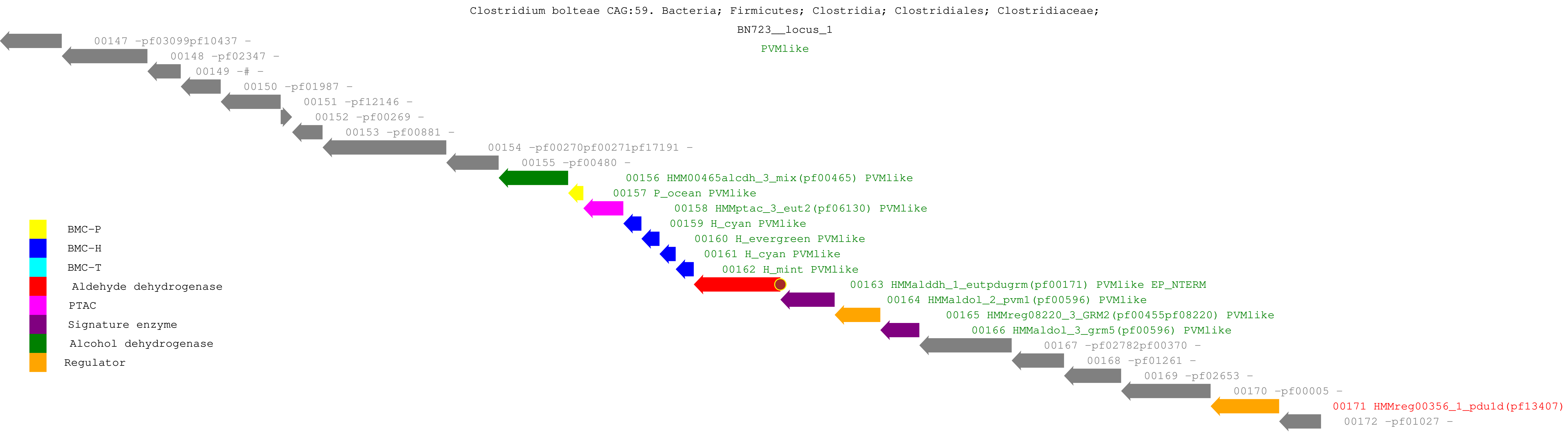
Clostridium bolteae CAG:59. Bacteria; Firmicutes; Clostridia; Clostridiales; Clostridiaceae;
BN723__locus_1
PVMlike
00147 -pf03099pf10437 -
00148 -pf02347 -
00149 -# -
00150 -pf01987 -
00151 -pf12146 -
00152 -pf00269 -
00153 -pf00881 -
00154 -pf00270pf00271pf17191 -
00155 -pf00480 -
00156 HMM00465alcdh_3_mix(pf00465) PVMlike
00157 P_ocean PVMlike
00158 HMMptac_3_eut2(pf06130) PVMlike
00159 H_cyan PVMlike
00160 H_evergreen PVMlike
00161 H_cyan PVMlike
00162 H_mint PVMlike
00163 HMMaldh_1_eutpdugrm(pf00171) PVMlike EP_NTERM
00164 HMMaldol_2_pvm1(pf00596) PVMlike
00165 HMMreg08220_3_GRM2(pf00455pf08220) PVMlike
00166 HMMaldol_3_grm5(pf00596) PVMlike
00167 -pf02782pf00370 -
00168 -pf01261 -
00169 -pf02653 -
00170 -pf00005 -
00171 HMMreg00356_1_pdu1d(pf13407)
00172 -pf01027 -
BMC-P
BMC-H
BMC-T
Aldehyde dehydrogenase
PTAC
Signature enzyme
Alcohol dehydrogenase
Regulator

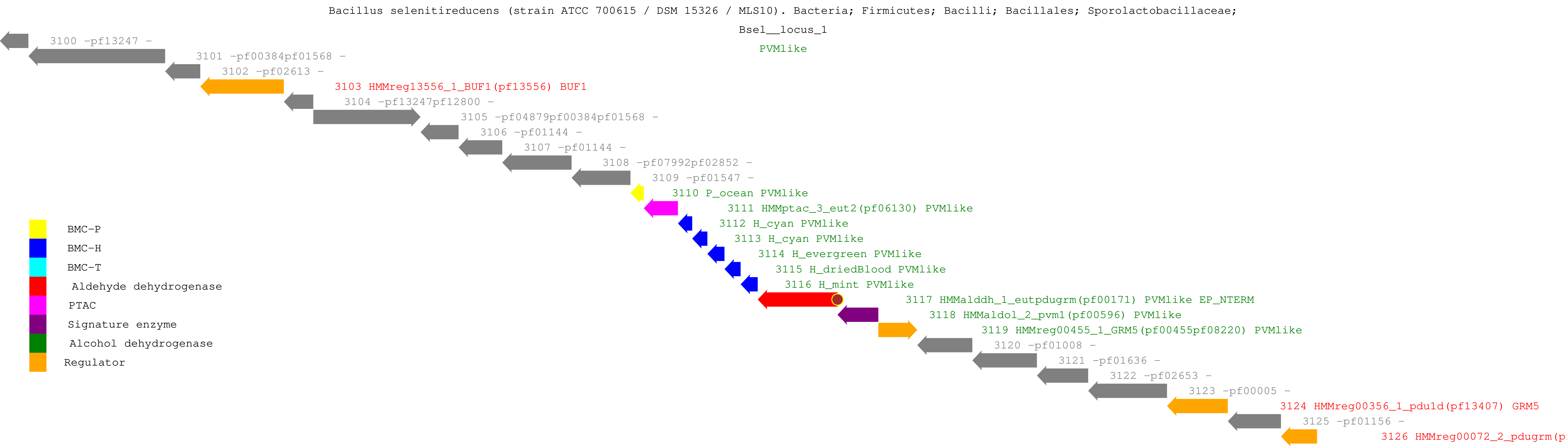
Bacillus selenitireducens (strain ATCC 700615 / DSM 15326 / MLS10). Bacteria; Firmicutes; Bacilli; Bacillales; Sporolactobacillaceae;
Bsel__locus_1
PVMlike
3100 -pf13247 -
3101 -pf00384pf01568 -
3102 -pf02613 -
3103 HMMreg13556_1_BUF1(pf13556) BUF1
3104 -pf13247pf12800 -
3105 -pf04879pf00384pf01568 -
3106 -pf01144 -
3107 -pf01144 -
3108 -pf07992pf02852 -
3109 -pf01547 -
3110 P_ocean PVMlike
3111 HMMptac_3_eut2(pf06130) PVMlike
3112 H_cyan PVMlike
3113 H_cyan PVMlike
3114 H_evergreen PVMlike
3115 H_driedBlood PVMlike
3116 H_mint PVMlike
3117 HMMaldh_1_eutpdugrm(pf00171) PVMlike EP_NTERM
3118 HMMaldol_2_pvm1(pf00596) PVMlike
3119 HMMreg00455_1_GRM5(pf00455pf08220) PVMlike
3120 -pf01008 -
3121 -pf01636 -
3122 -pf02653 -
3123 -pf00005 -
3124 HMMreg00356_1_pdu1d(pf13407) GRM5
3125 -pf01156 -
3126 HMMreg00072_2_pdugrm(p
BMC-P
BMC-H
BMC-T
Aldehyde dehydrogenase
PTAC
Signature enzyme
Alcohol dehydrogenase
Regulator

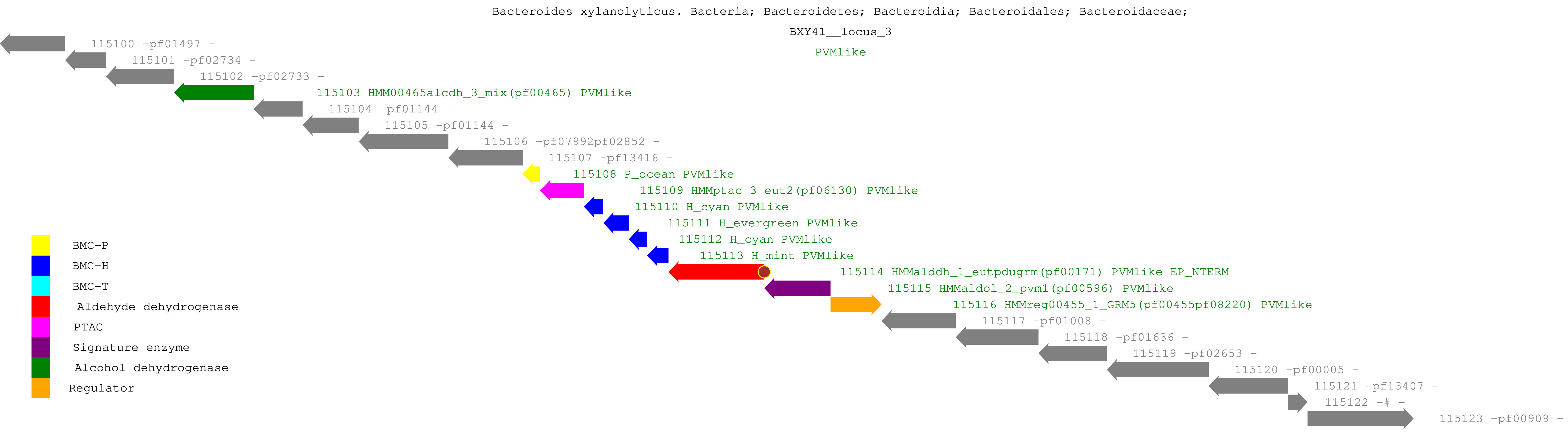
Bacteroides xylanolyticus. Bacteria; Bacteroidetes; Bacteroidia; Bacteroidales; Bacteroidaceae;
BXY41__locus_3
PVMlike
115100 -pf01497 -
115101 -pf02734 -
115102 -pf02733 -
115103 HMM00465alcdh_3_mix(pf00465) PVMlike
115104 -pf01144 -
115105 -pf01144 -
115106 -pf07992pf02852 -
115107 -pf13416 -
115108 P_ocean PVMlike
115109 HMMptac_3_eut2(pf06130) PVMlike
115110 H_cyan PVMlike
115111 H_evergreen PVMlike
115112 H_cyan PVMlike
115113 H_mint PVMlike
115114 HMMalddh_1_eutpdugrm(pf00171) PVMlike EP_NTERM
115115 HMMaldol_2_pvm1(pf00596) PVMlike
115116 HMMreg00455_1_GRM5(pf00455pf08220) PVMlike
115117 -pf01008 -
115118 -pf01636 -
115119 -pf02653 -
115120 -pf00005 -
115121 -pf13407 -
115122 -# -
115123 -pf00909 -
BMC-P
BMC-H
BMC-T
Aldehyde dehydrogenase
PTAC
Signature enzyme
Alcohol dehydrogenase
Regulator

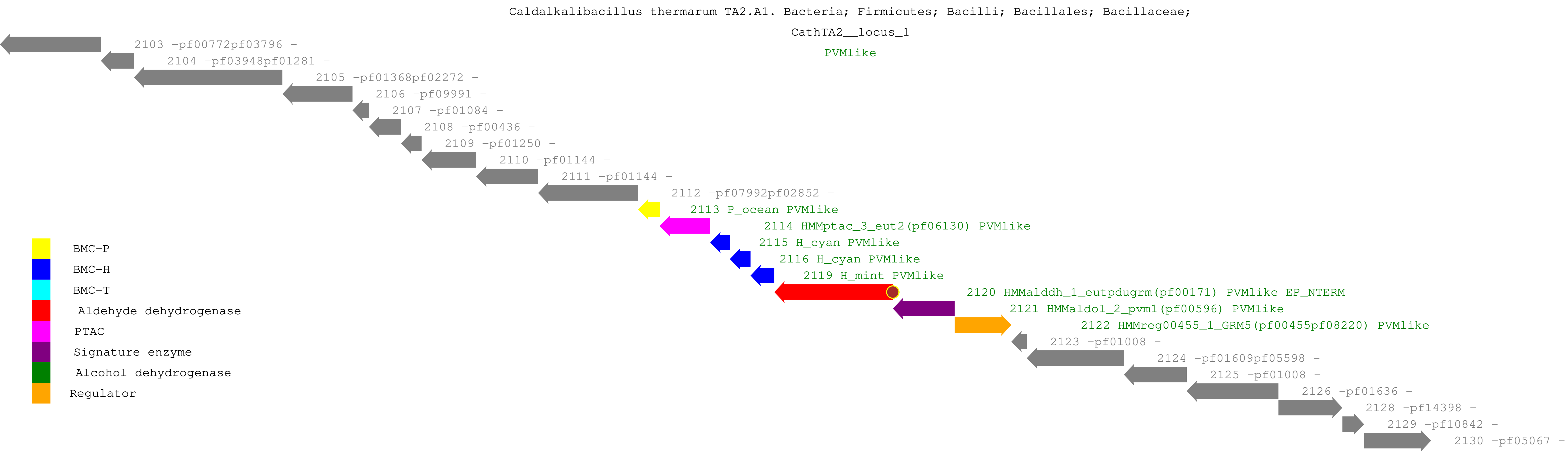
Caldalkalibacillus thermarum TA2.A1. Bacteria; Firmicutes; Bacilli; Bacillales; Bacillaceae;
CathTA2__locus_1
PVMlike
2103 -pf00772pf03796 -
2104 -pf03948pf01281 -
2105 -pf01368pf02272 -
2106 -pf09991 -
2107 -pf01084 -
2108 -pf00436 -
2109 -pf01250 -
2110 -pf01144 -
2111 -pf01144 -
2112 -pf07992pf02852 -
2113 P_ocean PVMlike
2114 HMMptac_3_eut2(pf06130) PVMlike
2115 H_cyan PVMlike
2116 H_cyan PVMlike
2119 H_mint PVMlike
2120 HMMaldh_1_eutpdugrm(pf00171) PVMlike EP_NTERM
2121 HMMaldol_2_pvm1(pf00596) PVMlike
2122 HMMreg00455_1_GRM5(pf00455pf08220) PVMlike
2123 -pf01008 -
2124 -pf01609pf05598 -
2125 -pf01008 -
2126 -pf01636 -
2128 -pf14398 -
2129 -pf10842 -
2130 -pf05067 -
BMC-P
BMC-H
BMC-T
Aldehyde dehydrogenase
PTAC
Signature enzyme
Alcohol dehydrogenase
Regulator

Faecalibacterium prausnitzii. Bacteria; Firmicutes; Clostridia; Clostridiales; Ruminococcaceae;

CGS50__locus_1

PVMlike

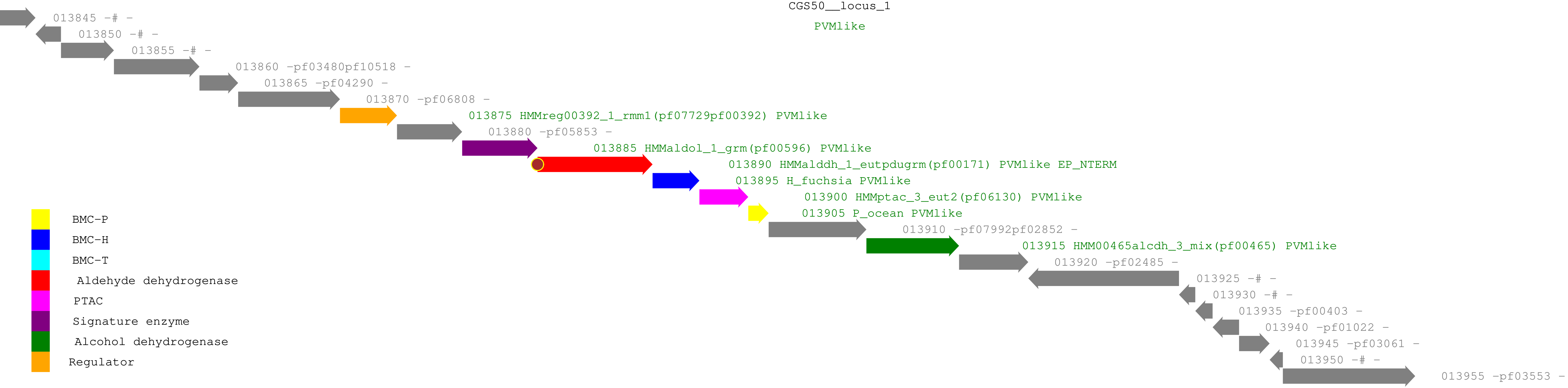

Clostridium homopropionicum DSM 5847. Bacteria; Firmicutes; Clostridia; Clostridiales; Clostridiaceae;

CLHOM__locus_1

PVMlike

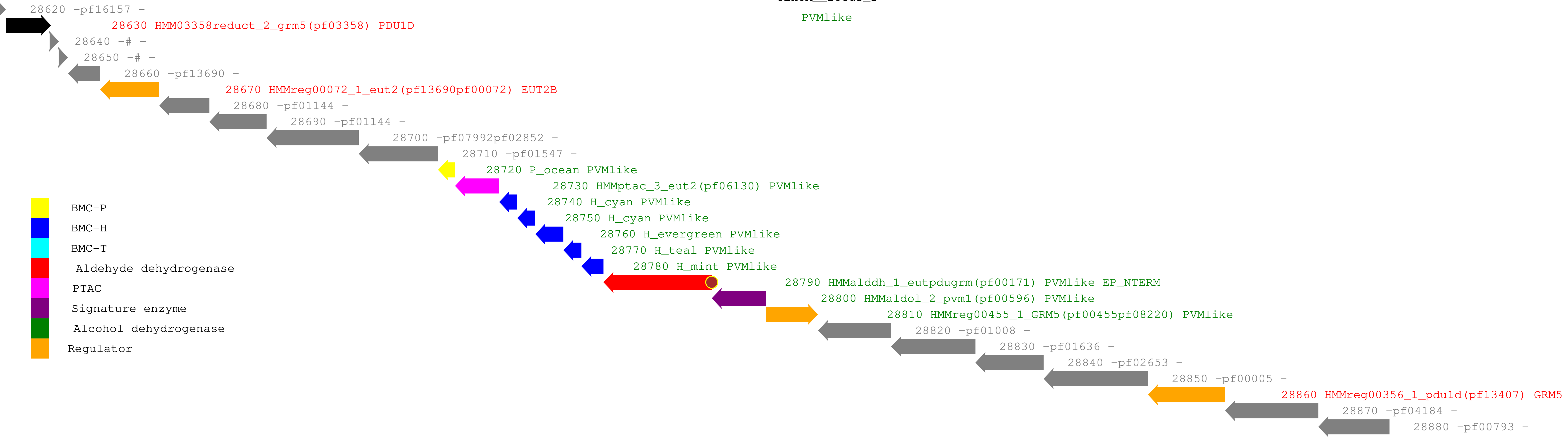

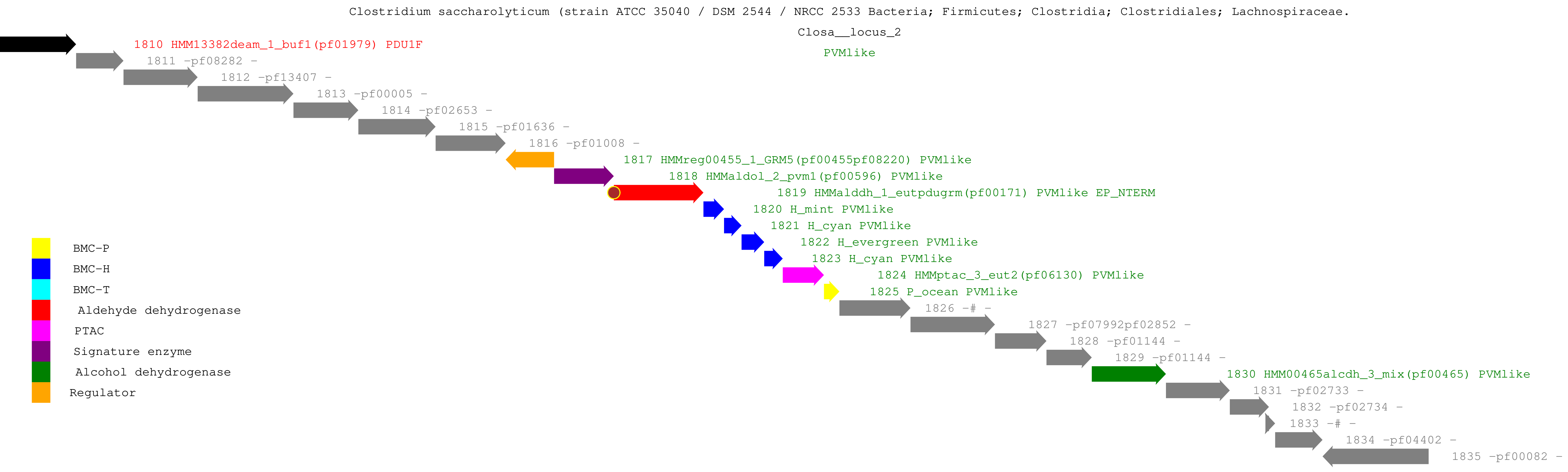
Clostridium saccharolyticum (strain ATCC 35040 / DSM 2544 / NRCC 2533 Bacteria; Firmicutes; Clostridia; Clostridiales; Lachnospiraceae.
Closa__locus_2
PVMlike
1810 HMM13382deam_1_buf1(pf01979) PDU1F
1811 -pf08282 -
1812 -pf13407 -
1813 -pf00005 -
1814 -pf02653 -
1815 -pf01636 -
1816 -pf01008 -
1817 HMMreg00455_1_GRM5(pf00455pf08220) PVMlike
1818 HMMaldol_2_pvm1(pf00596) PVMlike
1819 HMMalddh_1_eutpdugrm(pf00171) PVMlike EP_NTERM
1820 H_mint PVMlike
1821 H_cyan PVMlike
1822 H_evergreen PVMlike
1823 H_cyan PVMlike
1824 HMMptac_3_eut2(pf06130) PVMlike
1825 P_ocean PVMlike
1826 -# -
1827 -pf07992pf02852 -
1828 -pf01144 -
1829 -pf01144 -
1830 HMM00465alcdh_3_mix(pf00465) PVMlike
1831 -pf02733 -
1832 -pf02734 -
1833 -# -
1834 -pf04402 -
1835 -pf00082 -
BMC-P
BMC-H
BMC-T
Aldehyde dehydrogenase
PTAC
Signature enzyme
Alcohol dehydrogenase
Regulator

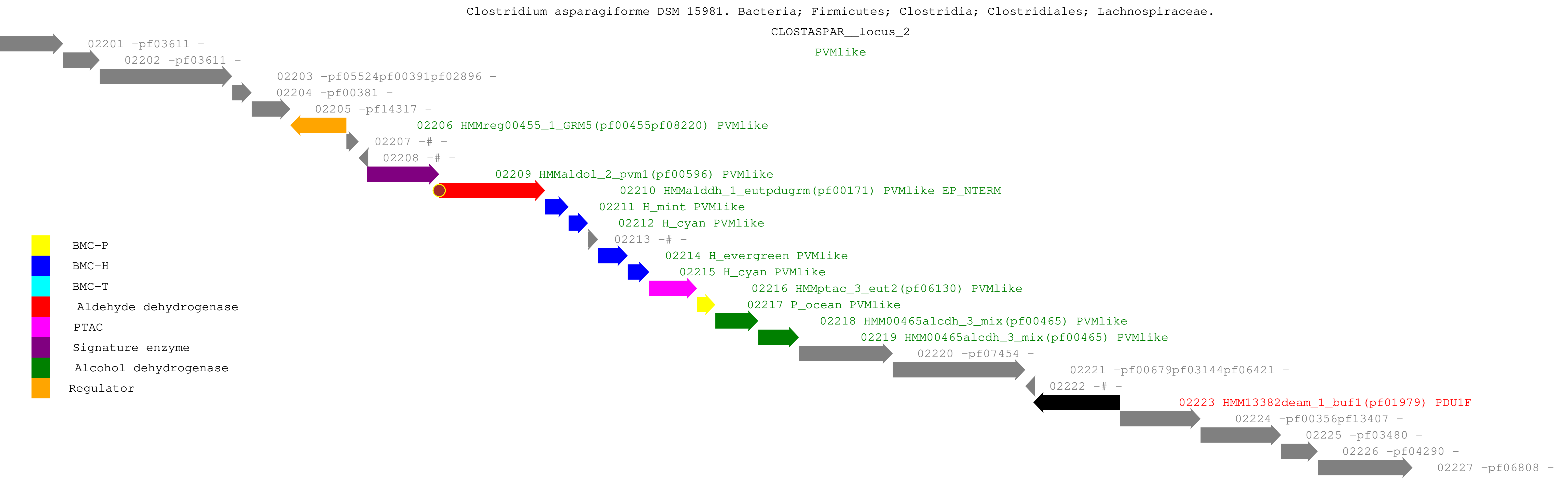
Clostridium asparagiforme DSM 15981. Bacteria; Firmicutes; Clostridia; Clostridiales; Lachnospiraceae.
CLOSTASPAR__locus_2
PVMlike
02201 -pf03611 -
02202 -pf03611 -
02203 -pf05524pf00391pf02896 -
02204 -pf00381 -
02205 -pf14317 -
02206 HMMreg00455_1_GRM5(pf00455pf08220) PVMlike
02207 -# -
02208 -# -
02209 HMMaldol_2_pvm1(pf00596) PVMlike
02210 HMMaldh_1_eutpdugrm(pf00171) PVMlike EP_NTERM
02211 H_mint PVMlike
02212 H_cyan PVMlike
02213 -# -
02214 H_evergreen PVMlike
02215 H_cyan PVMlike
02216 HMMptac_3_eut2(pf06130) PVMlike
02217 P_ocean PVMlike
02218 HMM00465alcdh_3_mix(pf00465) PVMlike
02219 HMM00465alcdh_3_mix(pf00465) PVMlike
02220 -pf07454 -
02221 -pf00679pf03144pf06421 -
02222 -# -
02223 HMM13382deam_1_buf1(pf01979) PDU1F
02224 -pf00356pf13407 -
02225 -pf03480 -
02226 -pf04290 -
02227 -pf06808 -
BMC-P
BMC-H
BMC-T
Aldehyde dehydrogenase
PTAC
Signature enzyme
Alcohol dehydrogenase
Regulator

Acetoanaerobium sticklandii (strain ATCC 12662 / DSM 519 / JCM 1433 / Bacteria; Firmicutes; Clostridia; Clostridiales;

CLOST__locus_2

PVMlike

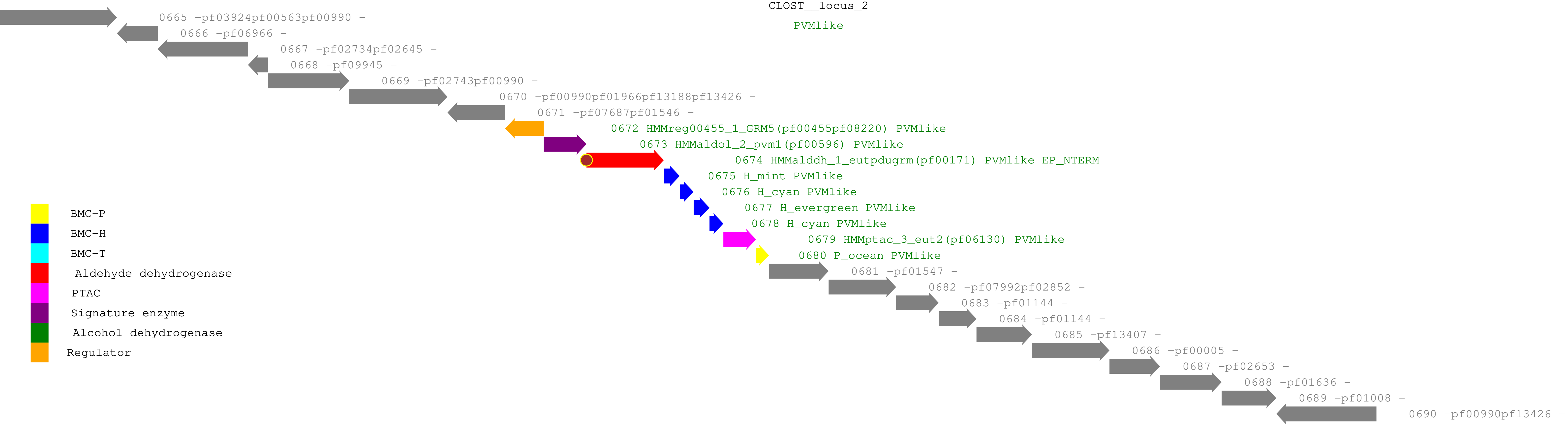

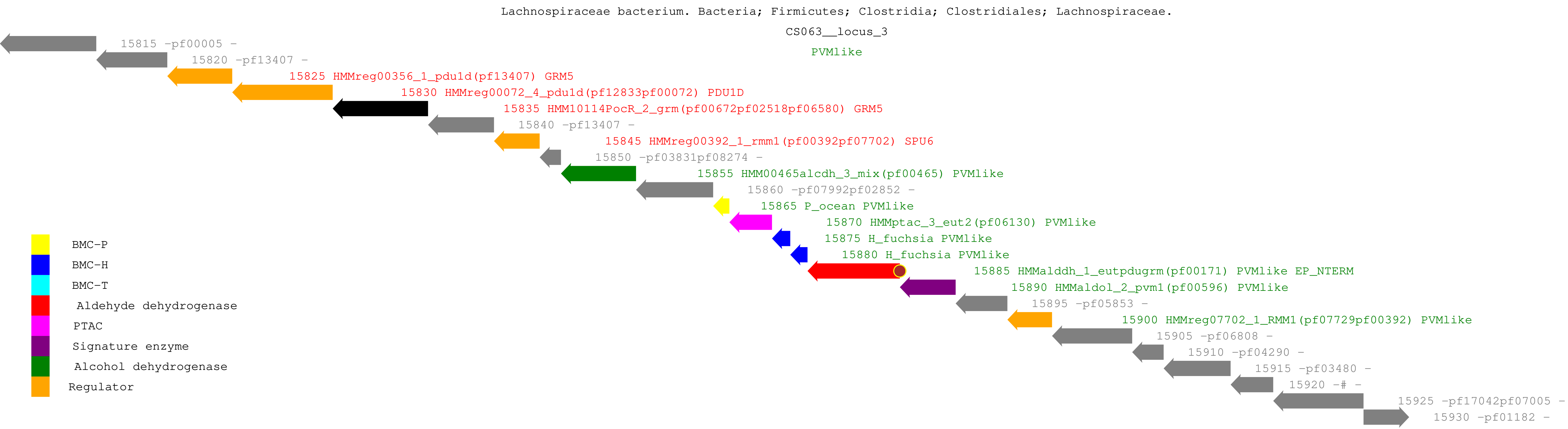
Lachnospiraceae bacterium. Bacteria; Firmicutes; Clostridia; Clostridiales; Lachnospiraceae.
CS063__locus_3
PVMlike
15815 -pf00005 -
15820 -pf13407 -
15825 HMMreg00356_1_pdu1d(pf13407) GRM5
15830 HMMreg00072_4_pdu1d(pf12833pf00072) PDU1D
15835 HMM10114PocR_2_grm(pf00672pf02518pf06580) GRM5
15840 -pf13407 -
15845 HMMreg00392_1_rmm1(pf00392pf07702) SPU6
15850 -pf03831pf08274 -
15855 HMM00465alcdh_3_mix(pf00465) PVMlike
15860 -pf07992pf02852 -
15865 P_ocean PVMlike
15870 HMMptac_3_eut2(pf06130) PVMlike
15875 H_fuchsia PVMlike
15880 H_fuchsia PVMlike
15885 HMMaldh_1_eutpdugrm(pf00171) PVMlike EP_NTERM
15890 HMMaldol_2_pvm1(pf00596) PVMlike
15895 -pf05853 -
15900 HMMreg07702_1_RMM1(pf07729pf00392) PVMlike
15905 -pf06808 -
15910 -pf04290 -
15915 -pf03480 -
15920 -# -
15925 -pf17042pf07005 -
15930 -pf01182 -
BMC-P
BMC-H
BMC-T
Aldehyde dehydrogenase
PTAC
Signature enzyme
Alcohol dehydrogenase
Regulator

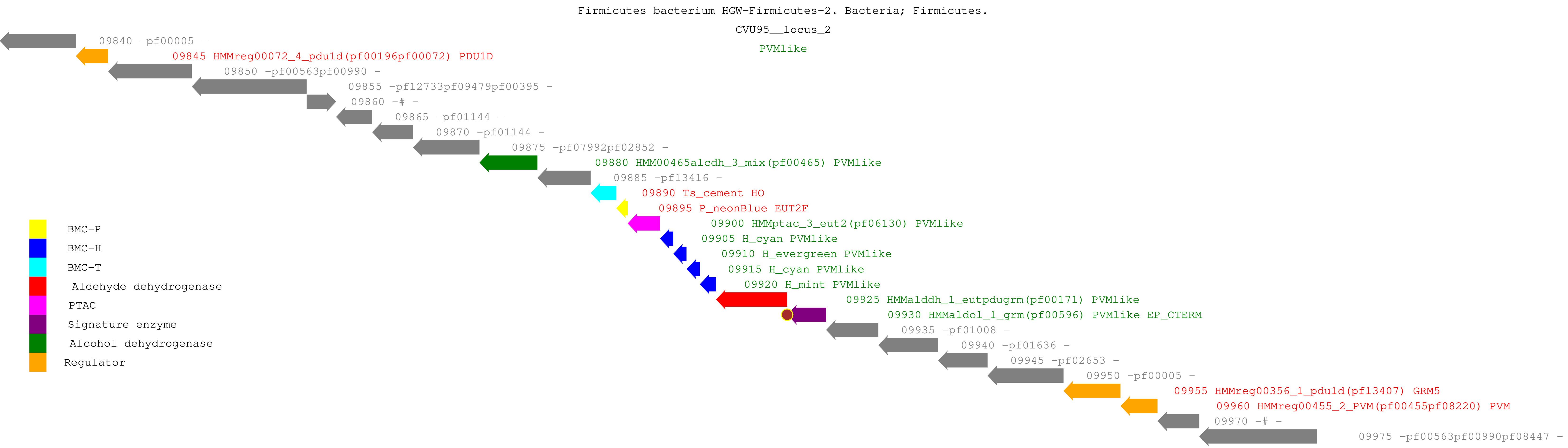
Firmicutes bacterium HGW-Firmicutes-2. Bacteria; Firmicutes.
CVU95__locus_2
PVMlike
09840 -pf00005 -
09845 HMMreg00072_4_pdu1d(pf00196pf00072) PDU1D
09850 -pf00563pf00990 -
09855 -pf12733pf09479pf00395 -
09860 -# -
09865 -pf01144 -
09870 -pf01144 -
09875 -pf07992pf02852 -
09880 HMM00465alcdh_3_mix(pf00465) PVMlike
09885 -pf13416 -
09890 Ts_cement HO
09895 P_neonBlue EUT2F
09900 HMMptac_3_eut2(pf06130) PVMlike
09905 H_cyan PVMlike
09910 H_evergreen PVMlike
09915 H_cyan PVMlike
09920 H_mint PVMlike
09925 HMMalddh_1_eutpdugrm(pf00171) PVMlike
09930 HMMaldol_1_grm(pf00596) PVMlike EP_CTERM
09935 -pf01008 -
09940 -pf01636 -
09945 -pf02653 -
09950 -pf00005 -
09955 HMMreg00356_1_pdu1d(pf13407) GRM5
09960 HMMreg00455_2_PVM(pf00455pf08220) PVM
09970 -# -
09975 -pf00563pf00990pf08447 -
BMC-P
BMC-H
BMC-T
Aldehyde dehydrogenase
PTAC
Signature enzyme
Alcohol dehydrogenase
Regulator

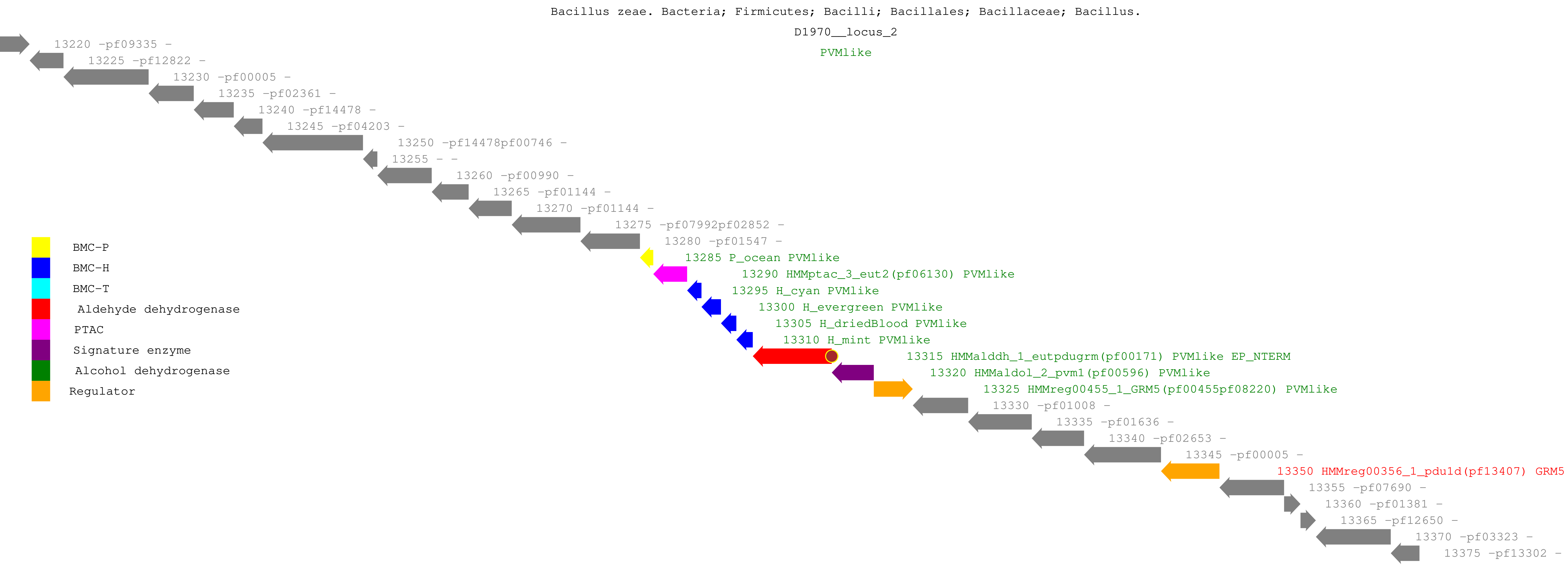
Bacillus zeae. Bacteria; Firmicutes; Bacilli; Bacillales; Bacillaceae; Bacillus.
D1970__locus_2
PVMlike
13220 -pf09335 -
13225 -pf12822 -
13230 -pf00005 -
13235 -pf02361 -
13240 -pf14478 -
13245 -pf04203 -
13250 -pf14478pf00746 -
13255 - -
13260 -pf00990 -
13265 -pf01144 -
13270 -pf01144 -
13275 -pf07992pf02852 -
13280 -pf01547 -
13285 P_ocean PVMlike
13290 HMMptac_3_eut2(pf06130) PVMlike
13295 H_cyan PVMlike
13300 H_evergreen PVMlike
13305 H_driedBlood PVMlike
13310 H_mint PVMlike
13315 HMMalddh_1_eutpdugrm(pf00171) PVMlike EP_NTERM
13320 HMMaldol_2_pvm1(pf00596) PVMlike
13325 HMMreg00455_1_GRM5(pf00455pf08220) PVMlike
13330 -pf01008 -
13335 -pf01636 -
13340 -pf02653 -
13345 -pf00005 -
13350 HMMreg00356_1_pdu1d(pf13407) GRM5
13355 -pf07690 -
13360 -pf01381 -
13365 -pf12650 -
13370 -pf03323 -
13375 -pf13302 -
BMC-P
BMC-H
BMC-T
Aldehyde dehydrogenase
PTAC
Signature enzyme
Alcohol dehydrogenase
Regulator

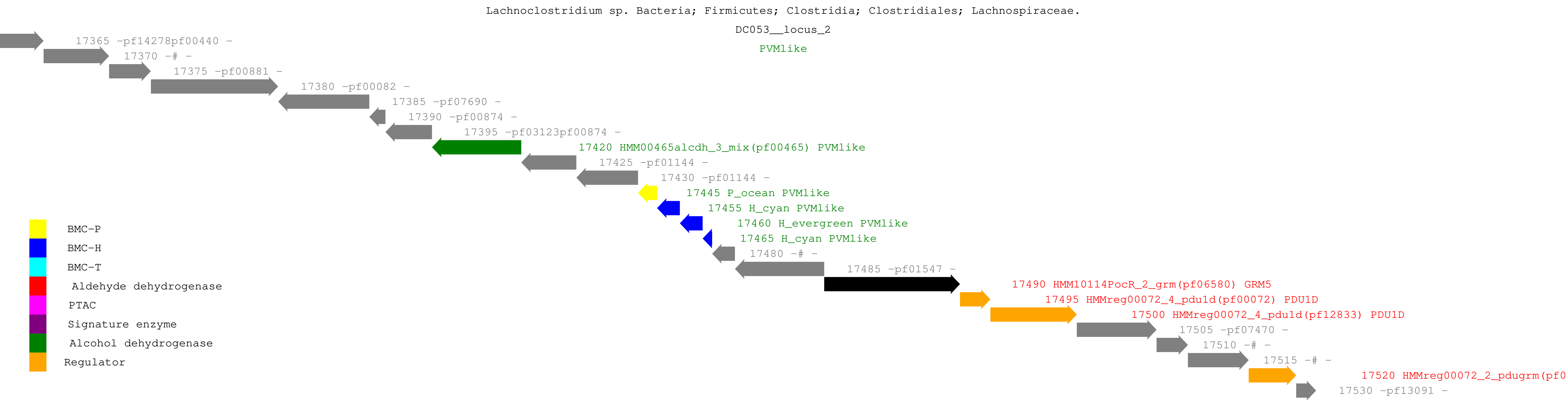
Lachnoclostridium sp. Bacteria; Firmicutes; Clostridia; Clostridiales; Lachnospiraceae.
DC053__locus_2
PVMlike
17365 -pf14278pf00440 -
17370 -# -
17375 -pf00881 -
17380 -pf00082 -
17385 -pf07690 -
17390 -pf00874 -
17395 -pf03123pf00874 -
17420 HMM00465alcdh_3_mix(pf00465) PVMlike
17425 -pf01144 -
17430 -pf01144 -
17445 P_ocean PVMlike
17455 H_cyan PVMlike
17460 H_evergreen PVMlike
17465 H_cyan PVMlike
17480 -# -
17485 -pf01547 -
17490 HMM10114PocR_2_grm(pf06580) GRM5
17495 HMMreg00072_4_pdu1d(pf00072) PDU1D
17500 HMMreg00072_4_pdu1d(pf12833) PDU1D
17505 -pf07470 -
17510 -# -
17515 -# -
17520 HMMreg00072_2_pdugrm(pf0
17530 -pf13091 -
BMC-P
BMC-H
BMC-T
Aldehyde dehydrogenase
PTAC
Signature enzyme
Alcohol dehydrogenase
Regulator

Lachnoclostridium sp. Bacteria; Firmicutes; Clostridia; Clostridiales; Lachnospiraceae.

DDW53__locus_2

PVMlike

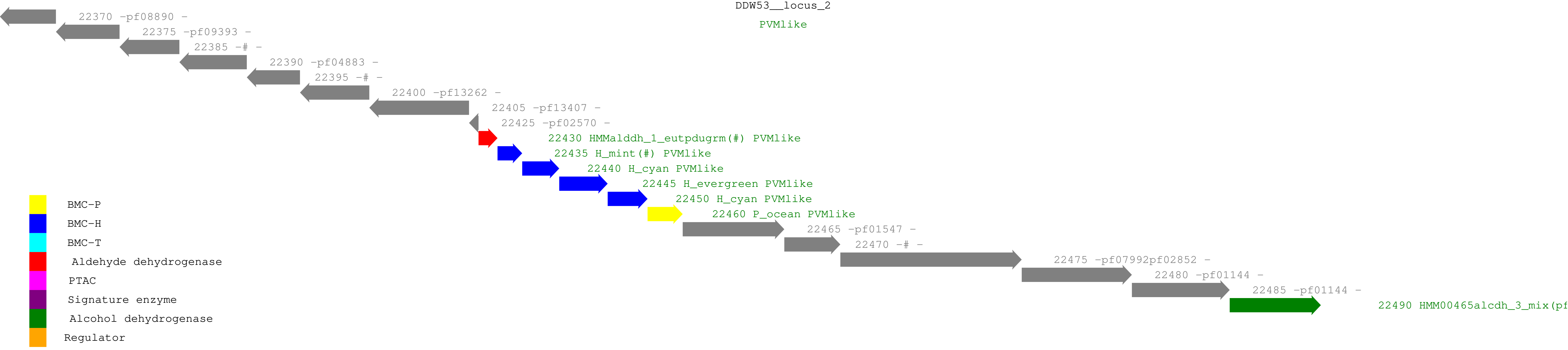

Clostridium sp. Bacteria; Firmicutes; Clostridia; Clostridiales; Clostridiaceae;

DDX68__locus_1

PVMlike

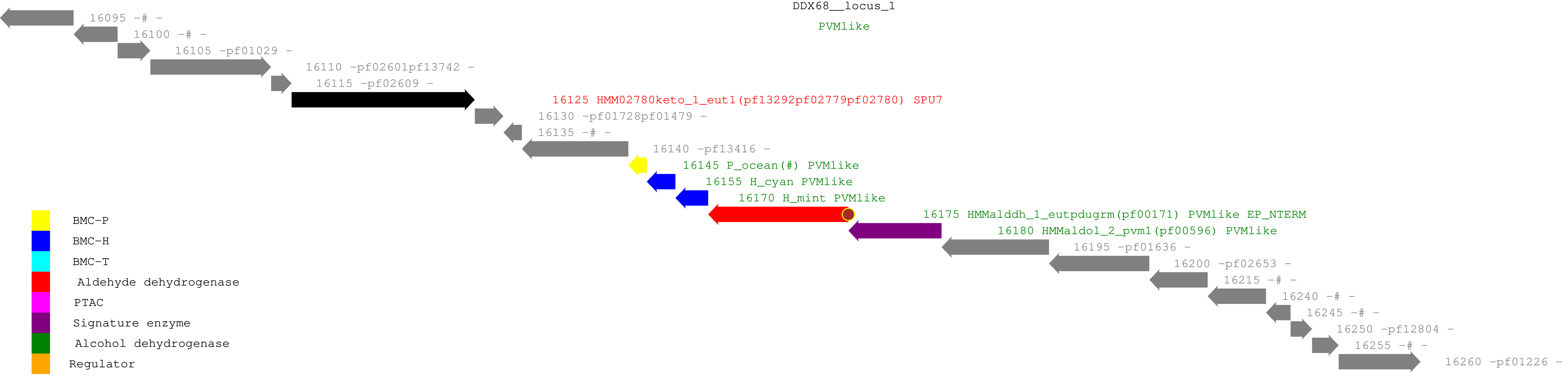

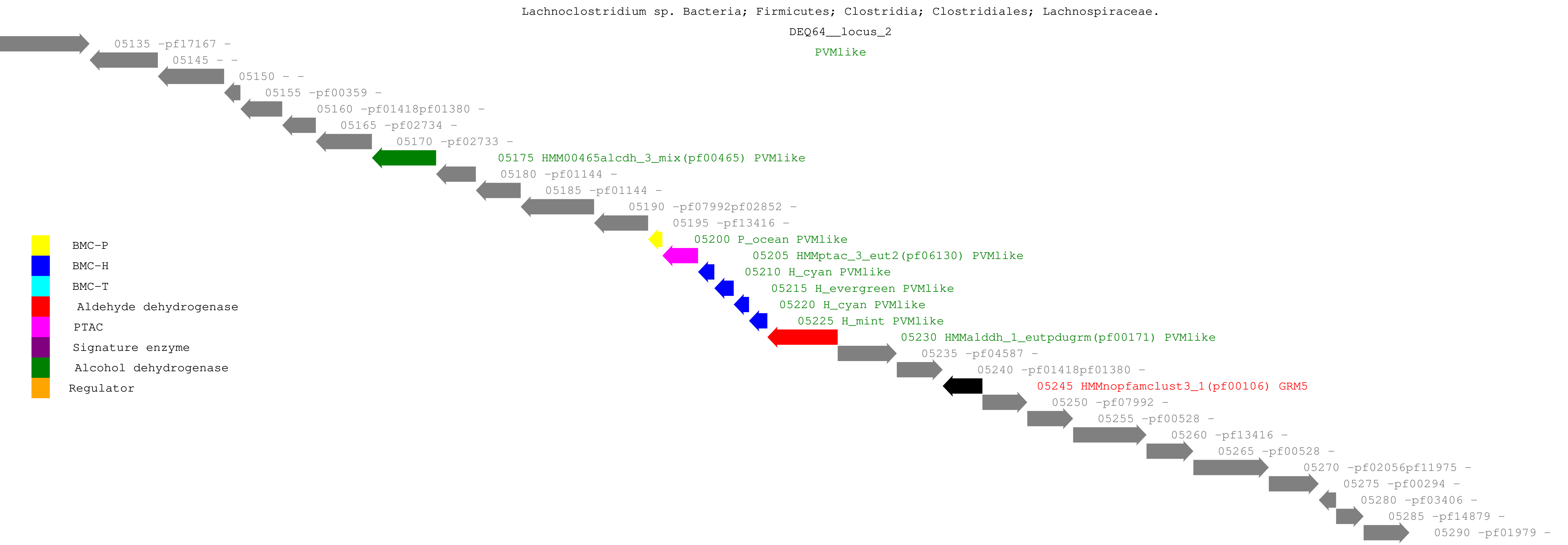
Lachnoclostridium sp. Bacteria; Firmicutes; Clostridia; Clostridiales; Lachnospiraceae.
DEQ64__locus_2
PVMlike
05135 -pf17167 -
05145 - -
05150 - -
05155 -pf00359 -
05160 -pf01418pf01380 -
05165 -pf02734 -
05170 -pf02733 -
05175 HMM00465alcdh_3_mix(pf00465) PVMlike
05180 -pf01144 -
05185 -pf01144 -
05190 -pf07992pf02852 -
05195 -pf13416 -
05200 P_ocean PVMlike
05205 HMMptac_3_eut2(pf06130) PVMlike
05210 H_cyan PVMlike
05215 H_evergreen PVMlike
05220 H_cyan PVMlike
05225 H_mint PVMlike
05230 HMMalddh_1_eutpdugrm(pf00171) PVMlike
05235 -pf04587 -
05240 -pf01418pf01380 -
05245 HMMnopfamclust3_1(pf00106) GRM5
05250 -pf07992 -
05255 -pf00528 -
05260 -pf13416 -
05265 -pf00528 -
05270 -pf02056pf11975 -
05275 -pf00294 -
05280 -pf03406 -
05285 -pf14879 -
05290 -pf01979 -
BMC-P
BMC-H
BMC-T
Aldehyde dehydrogenase
PTAC
Signature enzyme
Alcohol dehydrogenase
Regulator

Saliterribacillus persicus. Bacteria; Firmicutes; Bacilli; Bacillales; Bacillaceae;

DFR57__locus_1

PVMlike

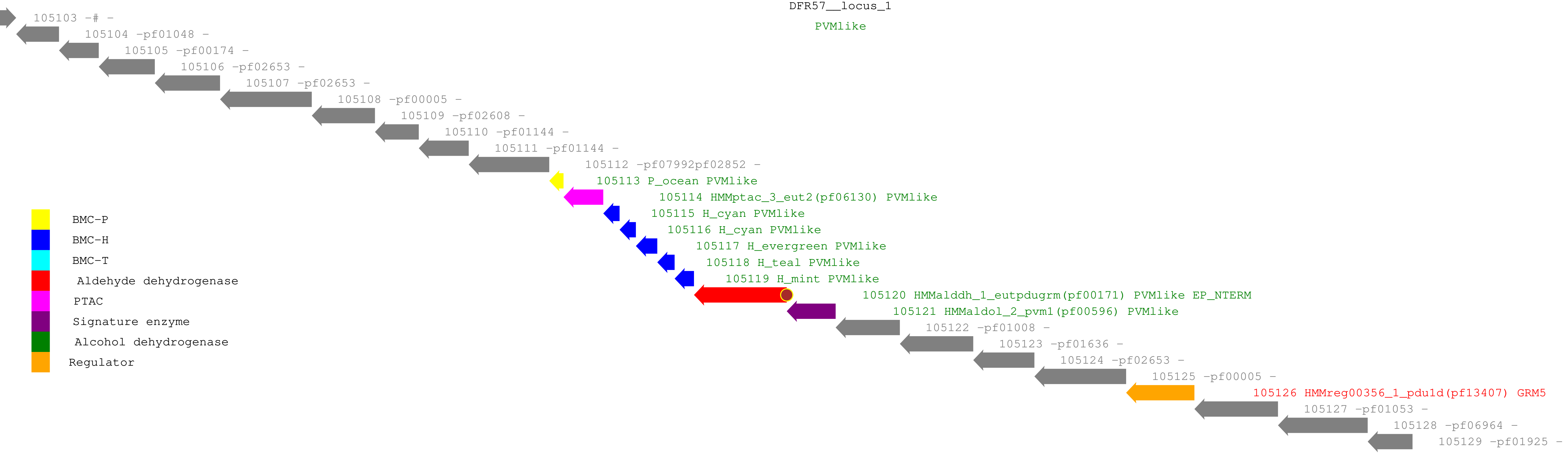

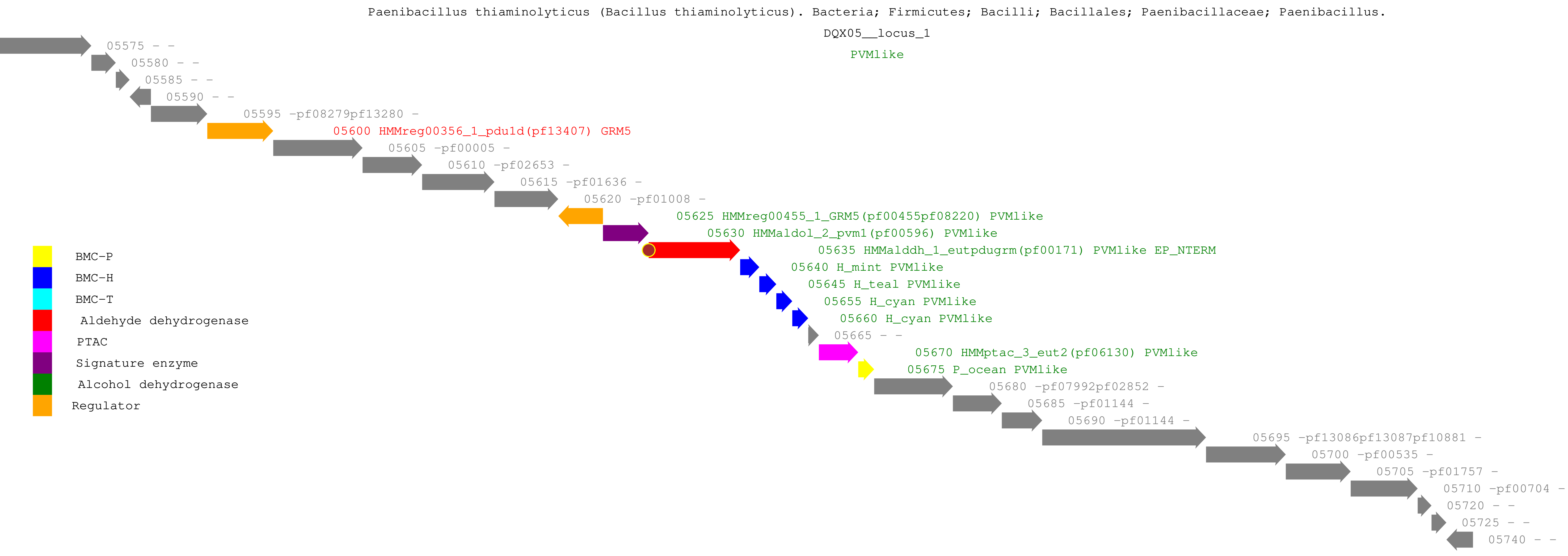
Paenibacillus thiaminolyticus (Bacillus thiaminolyticus). Bacteria; Firmicutes; Bacilli; Bacillales; Paenibacillaceae; Paenibacillus.
DQX05__locus_1
PVMlike
05575 - -
05580 - -
05585 - -
05590 - -
05595 -pf08279pf13280 -
05600 HMMreg00356_1_pdu1d(pf13407) GRM5
05605 -pf00005 -
05610 -pf02653 -
05615 -pf01636 -
05620 -pf01008 -
05625 HMMreg00455_1_GRM5(pf00455pf08220) PVMlike
05630 HMMaldol_2_pvm1(pf00596) PVMlike
05635 HMMalddh_1_eutpdugrm(pf00171) PVMlike EP_NTERM
05640 H_mint PVMlike
05645 H_teal PVMlike
05655 H_cyan PVMlike
05660 H_cyan PVMlike
05665 - -
05670 HMMptac_3_eut2(pf06130) PVMlike
05675 P_ocean PVMlike
05680 -pf07992pf02852 -
05685 -pf01144 -
05690 -pf01144 -
05695 -pf13086pf13087pf10881 -
05700 -pf00535 -
05705 -pf01757 -
05710 -pf00704 -
05720 - -
05725 - -
05740 - -
BMC-P
BMC-H
BMC-T
Aldehyde dehydrogenase
PTAC
Signature enzyme
Alcohol dehydrogenase
Regulator

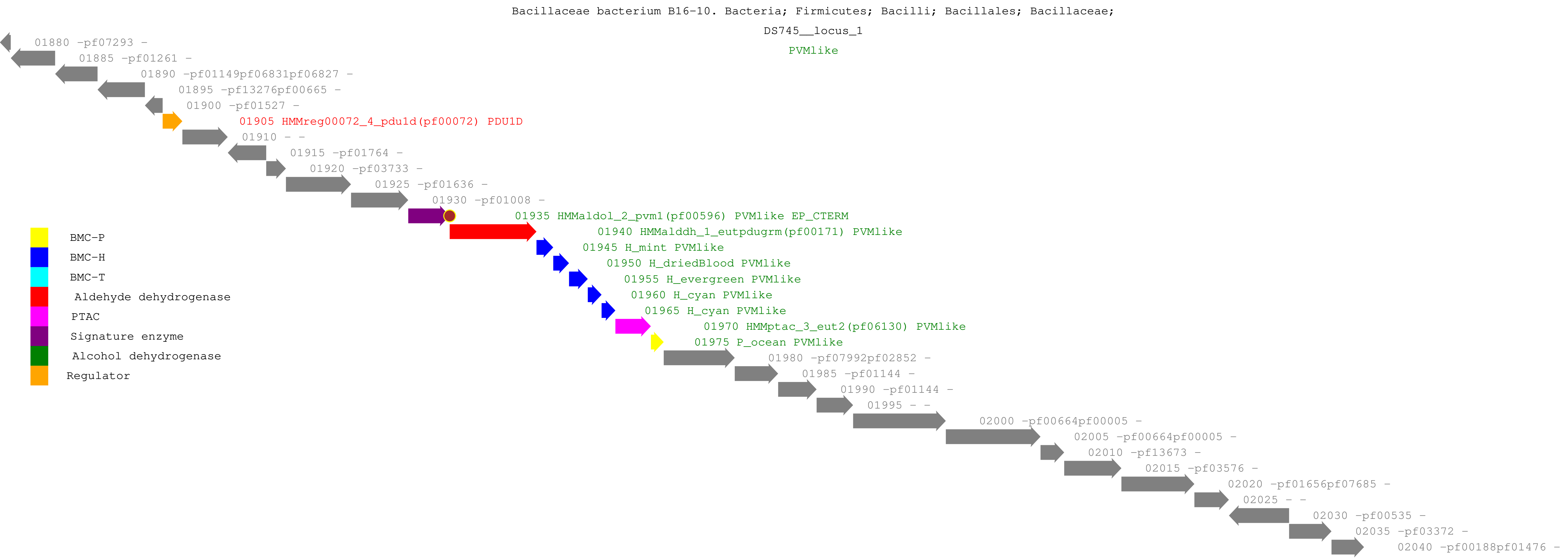

Bacillaceae bacterium B16-10. Bacteria; Firmicutes; Bacilli; Bacillales; Bacillaceae;
DS745__locus_1
PVMlike
01880 -pf07293 -
01885 -pf01261 -
01890 -pf01149pf06831pf06827 -
01895 -pf13276pf00665 -
01900 -pf01527 -
01905 HMMreg00072_4_pdu1d(pf00072) PDU1D
01910 - -
01915 -pf01764 -
01920 -pf03733 -
01925 -pf01636 -
01930 -pf01008 -
01935 HMMaldol_2_pvm1(pf00596) PVMlike EP_CTERM
01940 HMMalddh_1_eutpdugrm(pf00171) PVMlike
01945 H_mint PVMlike
01950 H_driedBlood PVMlike
01955 H_evergreen PVMlike
01960 H_cyan PVMlike
01965 H_cyan PVMlike
01970 HMMptac_3_eut2(pf06130) PVMlike
01975 P_ocean PVMlike
01980 -pf07992pf02852 -
01985 -pf01144 -
01990 -pf01144 -
01995 - -
02000 -pf00664pf00005 -
02005 -pf00664pf00005 -
02010 -pf13673 -
02015 -pf03576 -
02020 -pf01656pf07685 -
02025 - -
02030 -pf00535 -
02035 -pf03372 -
02040 -pf00188pf01476 -
BMC-P
BMC-H
BMC-T
Aldehyde dehydrogenase
PTAC
Signature enzyme
Alcohol dehydrogenase
Regulator

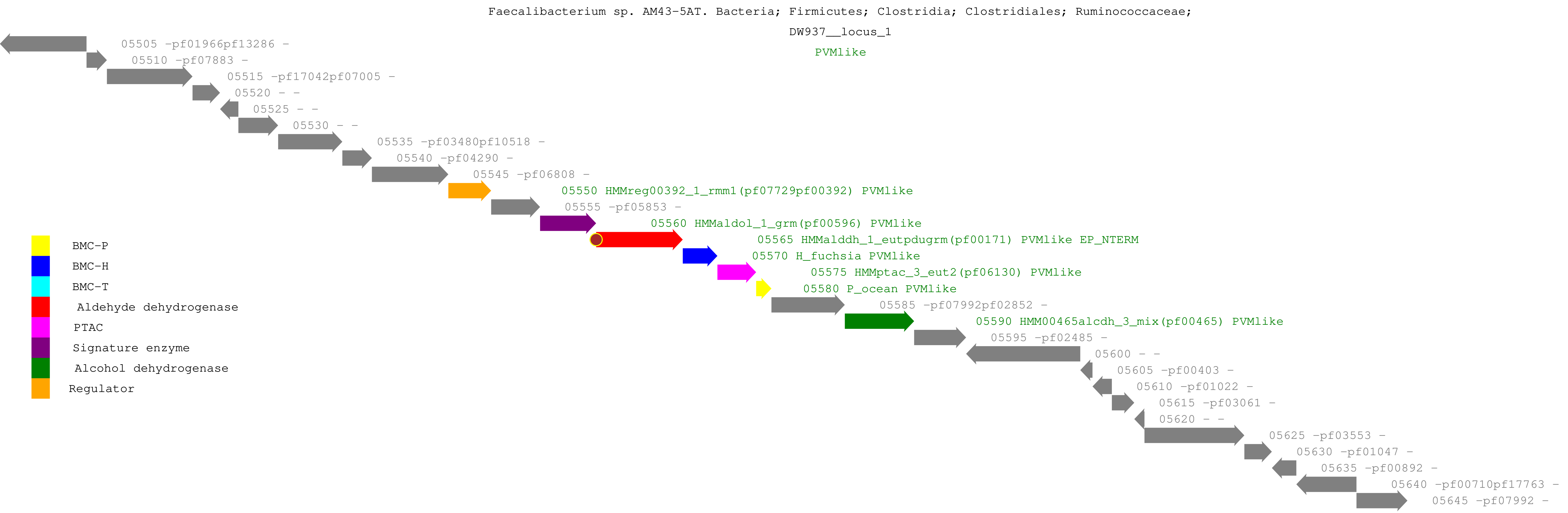

Faecalibacterium sp. AM43-5AT. Bacteria; Firmicutes; Clostridia; Clostridiales; Ruminococcaceae;
DW937__locus_1
PVMlike
05505 -pf01966pf13286 -
05510 -pf07883 -
05515 -pf17042pf07005 -
05520 - -
05525 - -
05530 - -
05535 -pf03480pf10518 -
05540 -pf04290 -
05545 -pf06808 -
05550 HMMreg00392_1_rmm1(pf07729pf00392) PVMlike
05555 -pf05853 -
05560 HMMaldol_1_grm(pf00596) PVMlike
05565 HMMalddh_1_eutpdugrm(pf00171) PVMlike EP_NTERM
05570 H_fuchsia PVMlike
05575 HMMptac_3_eut2(pf06130) PVMlike
05580 P_ocean PVMlike
05585 -pf07992pf02852 -
05590 HMM00465alcdh_3_mix(pf00465) PVMlike
05595 -pf02485 -
05600 - -
05605 -pf00403 -
05610 -pf01022 -
05615 -pf03061 -
05620 - -
05625 -pf03553 -
05630 -pf01047 -
05635 -pf00892 -
05640 -pf00710pf17763 -
05645 -pf07992 -
BMC-P
BMC-H
BMC-T
Aldehyde dehydrogenase
PTAC
Signature enzyme
Alcohol dehydrogenase
Regulator

Clostridium asparagiforme. Bacteria; Firmicutes; Clostridia; Clostridiales; Lachnospiraceae.

DWV29__locus_2

PVMlike

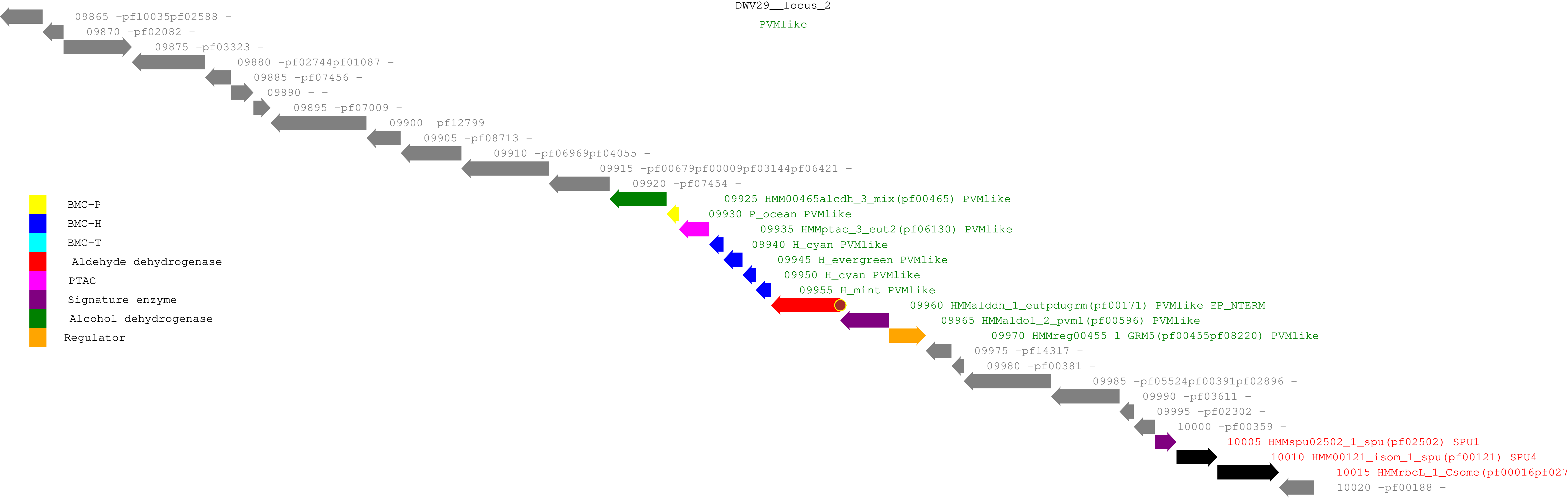

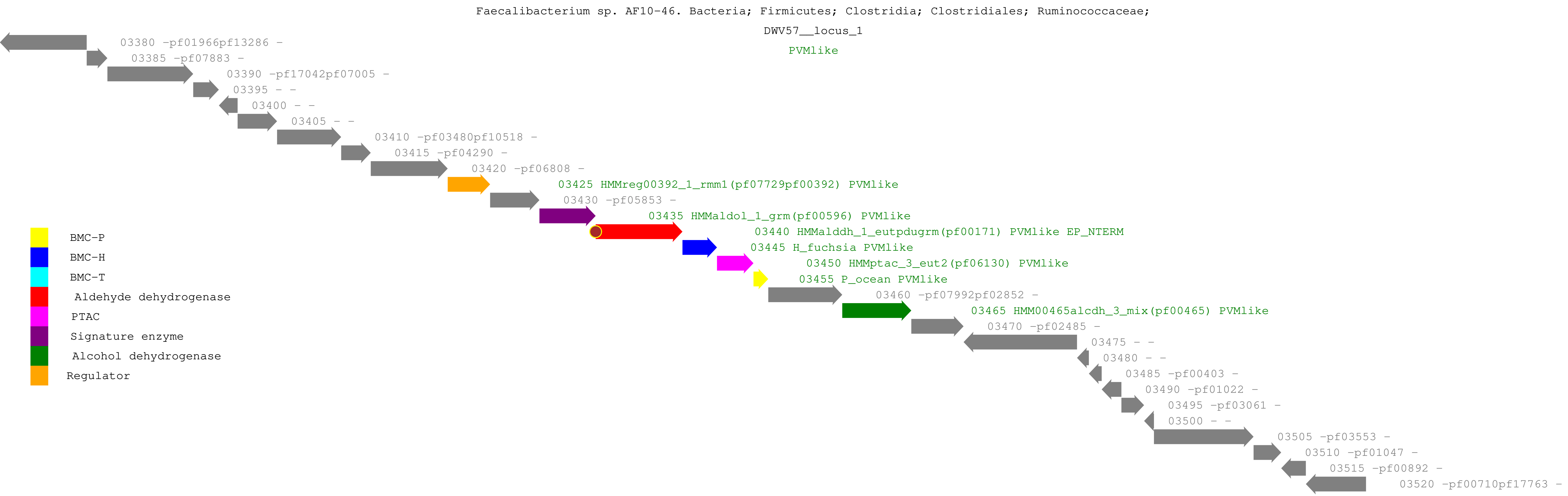
Faecalibacterium sp. AF10-46. Bacteria; Firmicutes; Clostridia; Clostridiales; Ruminococcaceae;
DWV57__locus_1
PVMlike
03380 -pf01966pf13286 -
03385 -pf07883 -
03390 -pf17042pf07005 -
03395 - -
03400 - -
03405 - -
03410 -pf03480pf10518 -
03415 -pf04290 -
03420 -pf06808 -
03425 HMMreg00392_1_rmm1(pf07729pf00392) PVMlike
03430 -pf05853 -
03435 HMMaldol_1_grm(pf00596) PVMlike
03440 HMMalddh_1_eutpdugrm(pf00171) PVMlike EP_NTERM
03445 H_fuchsia PVMlike
03450 HMMptac_3_eut2(pf06130) PVMlike
03455 P_ocean PVMlike
03460 -pf07992pf02852 -
03465 HMM00465alcdh_3_mix(pf00465) PVMlike
03470 -pf02485 -
03475 - -
03480 - -
03485 -pf00403 -
03490 -pf01022 -
03495 -pf03061 -
03500 - -
03505 -pf03553 -
03510 -pf01047 -
03515 -pf00892 -
03520 -pf00710pf17763 -
BMC-P
BMC-H
BMC-T
Aldehyde dehydrogenase
PTAC
Signature enzyme
Alcohol dehydrogenase
Regulator

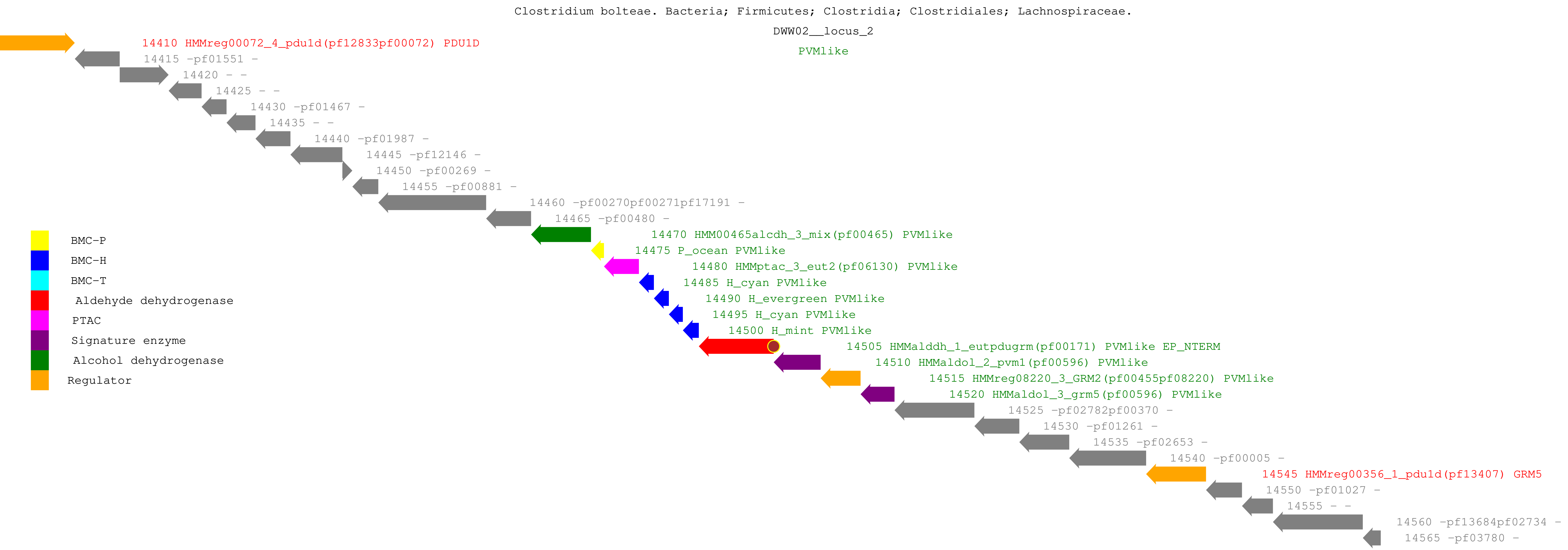
Clostridium bolteae. Bacteria; Firmicutes; Clostridia; Clostridiales; Lachnospiraceae.
DWW02__locus_2
PVMlike
14410 HMMreg00072_4_pdu1d(pf12833pf00072) PDU1D
14415 -pf01551 -
14420 - -
14425 - -
14430 -pf01467 -
14435 - -
14440 -pf01987 -
14445 -pf12146 -
14450 -pf00269 -
14455 -pf00881 -
14460 -pf00270pf00271pf17191 -
14465 -pf00480 -
14470 HMM00465alcdh_3_mix(pf00465) PVMlike
14475 P_ocean PVMlike
14480 HMMptac_3_eut2(pf06130) PVMlike
14485 H_cyan PVMlike
14490 H_evergreen PVMlike
14495 H_cyan PVMlike
14500 H_mint PVMlike
14505 HMMaldh_1_eutpdugrm(pf00171) PVMlike EP_NTERM
14510 HMMaldol_2_pvm1(pf00596) PVMlike
14515 HMMreg08220_3_GRM2(pf00455pf08220) PVMlike
14520 HMMaldol_3_grm5(pf00596) PVMlike
14525 -pf02782pf00370 -
14530 -pf01261 -
14535 -pf02653 -
14540 -pf00005 -
14545 HMMreg00356_1_pdu1d(pf13407) GRM5
14550 -pf01027 -
14555 - -
14560 -pf13684pf02734 -
14565 -pf03780 -
BMC-P
BMC-H
BMC-T
Aldehyde dehydrogenase
PTAC
Signature enzyme
Alcohol dehydrogenase
Regulator

Clostridium sp. AF18-27. Bacteria; Firmicutes; Clostridia; Clostridiales; Clostridiaceae;

DWX10__locus_1

PVMlike

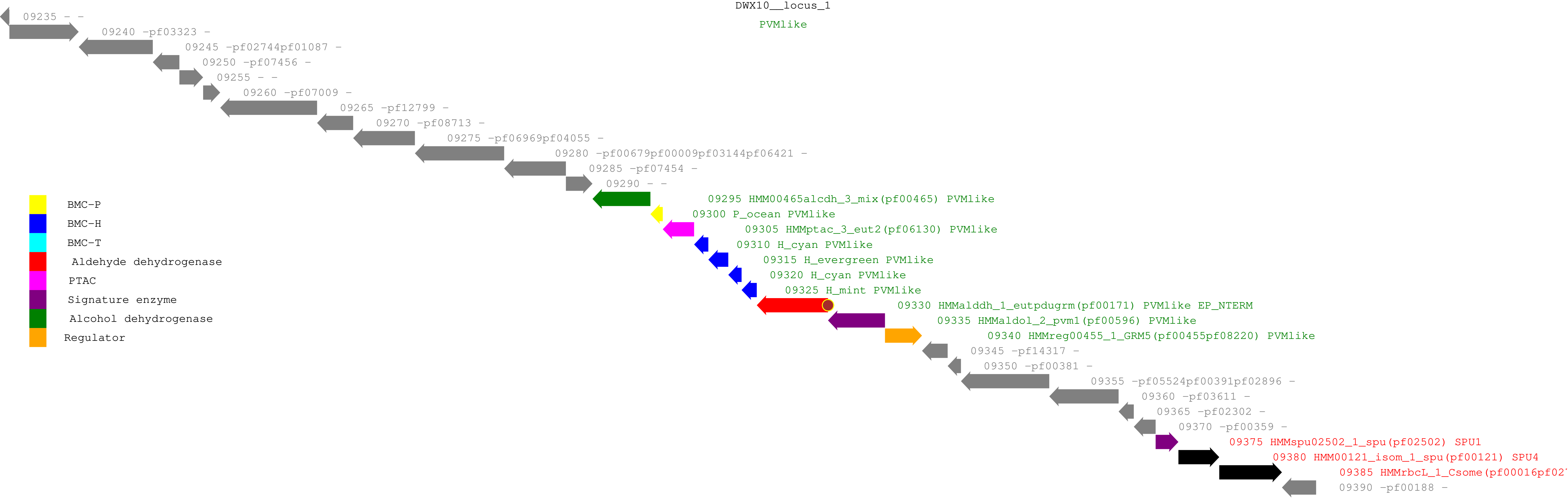

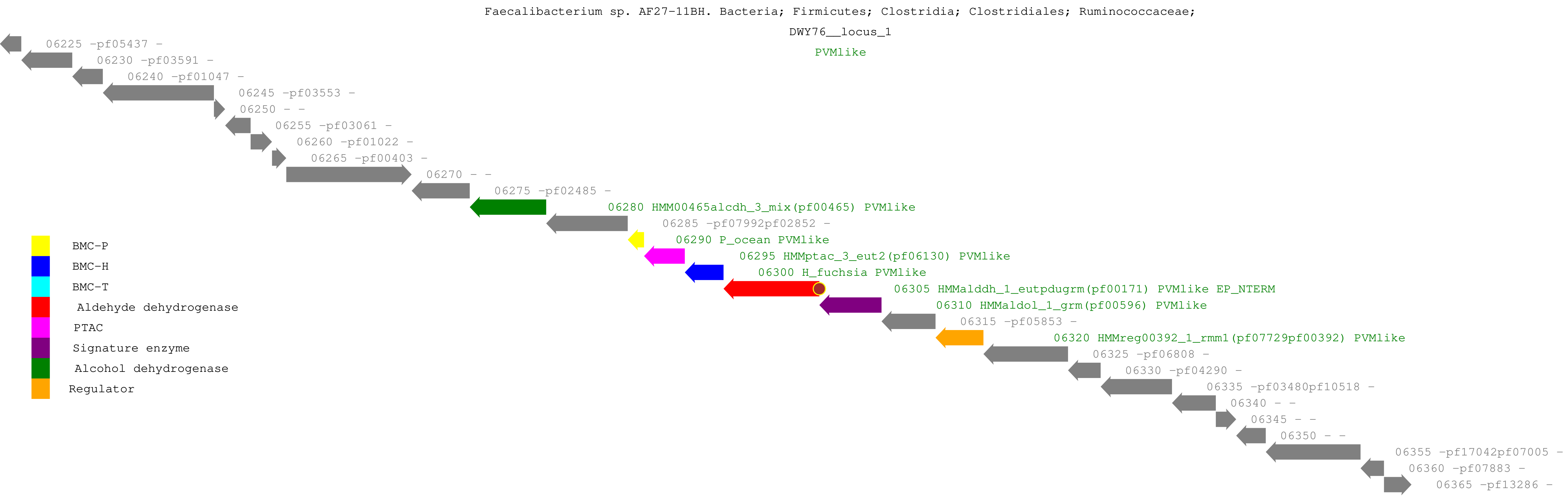
Faecalibacterium sp. AF27-11BH. Bacteria; Firmicutes; Clostridia; Clostridiales; Ruminococcaceae;
DWY76__locus_1
PVMlike
06225 -pf05437 -
06230 -pf03591 -
06240 -pf01047 -
06245 -pf03553 -
06250 - -
06255 -pf03061 -
06260 -pf01022 -
06265 -pf00403 -
06270 - -
06275 -pf02485 -
06280 HMM00465alcdh_3_mix(pf00465) PVMlike
06285 -pf07992pf02852 -
06290 P_ocean PVMlike
06295 HMMptac_3_eut2(pf06130) PVMlike
06300 H_fuchsia PVMlike
06305 HMMalddh_1_eutpdugrm(pf00171) PVMlike EP_NTERM
06310 HMMaldol_1_grm(pf00596) PVMlike
06315 -pf05853 -
06320 HMMreg00392_1_rmm1(pf07729pf00392) PVMlike
06325 -pf06808 -
06330 -pf04290 -
06335 -pf03480pf10518 -
06340 - -
06345 - -
06350 - -
06355 -pf17042pf07005 -
06360 -pf07883 -
06365 -pf13286 -
BMC-P
BMC-H
BMC-T
Aldehyde dehydrogenase
PTAC
Signature enzyme
Alcohol dehydrogenase
Regulator

Faecalibacterium sp. OF03-6AC. Bacteria; Firmicutes; Clostridia; Clostridiales; Ruminococcaceae;

DXA66__locus_1

PVMlike

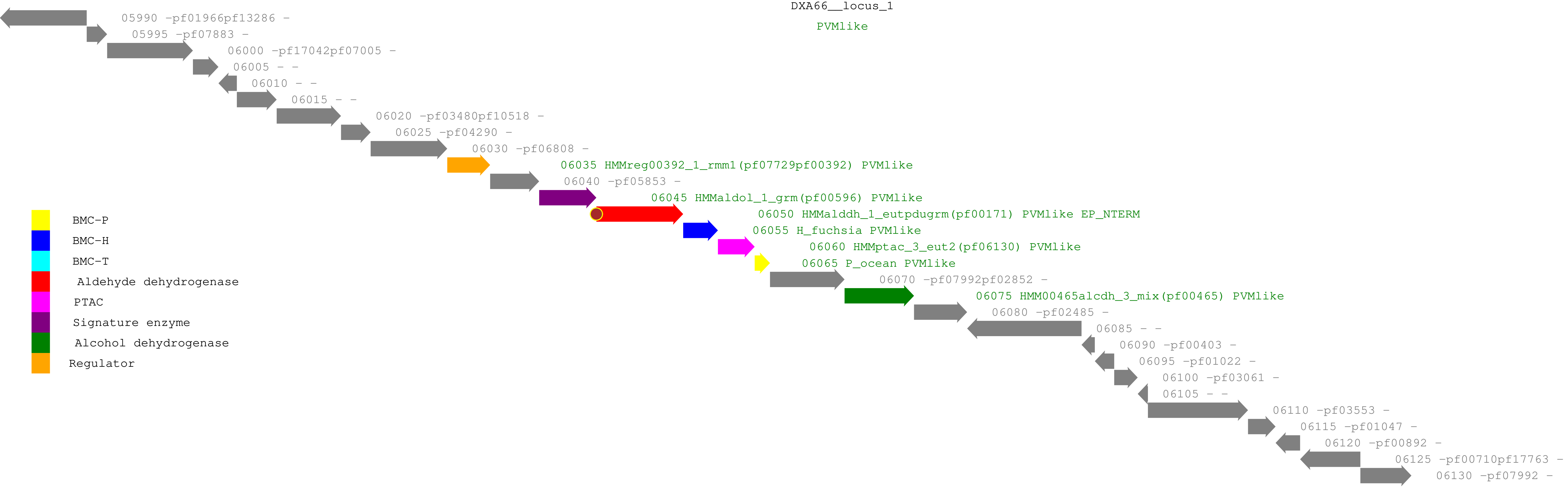

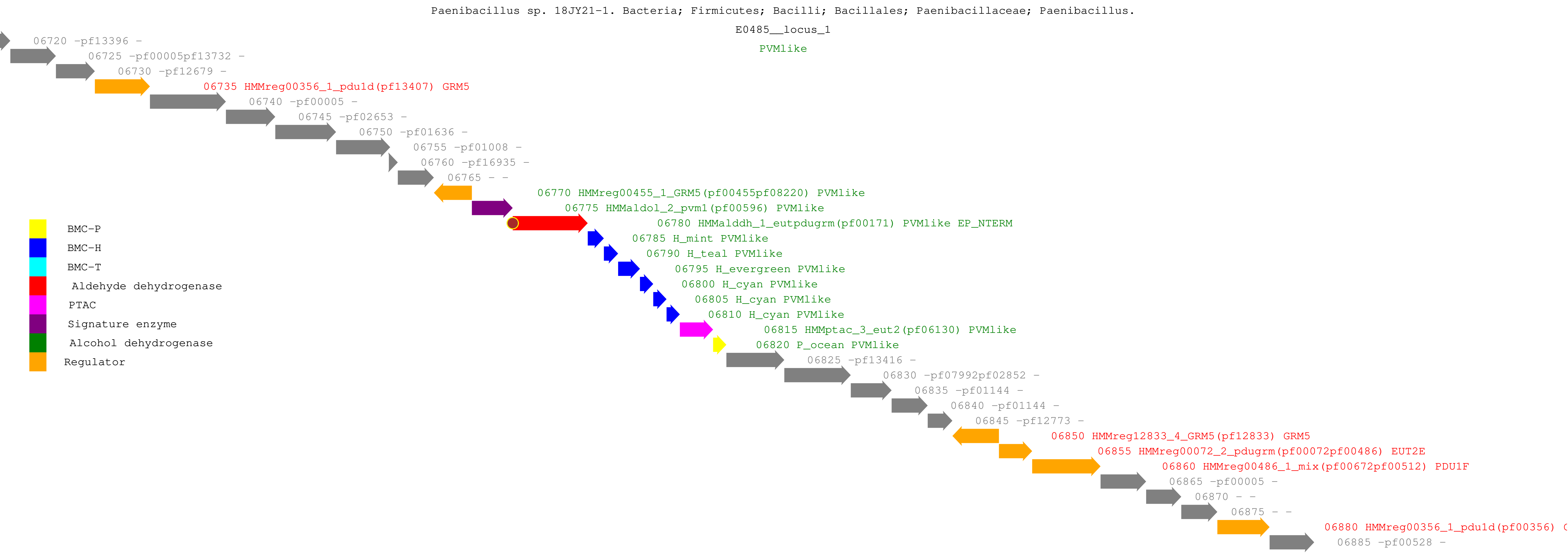
Paenibacillus sp. 18JY21-1. Bacteria; Firmicutes; Bacilli; Bacillales; Paenibacillaceae; Paenibacillus.
E0485__locus_1
PVMlike
06720 -pf13396 -
06725 -pf00005pf13732 -
06730 -pf12679 -
06735 HMMreg00356_1_pdu1d(pf13407) GRM5
06740 -pf00005 -
06745 -pf02653 -
06750 -pf01636 -
06755 -pf01008 -
06760 -pf16935 -
06765 - -
06770 HMMreg00455_1_GRM5(pf00455pf08220) PVMlike
06775 HMMaldol_2_pvm1(pf00596) PVMlike
06780 HMMalddh_1_eutpdugrm(pf00171) PVMlike EP_NTERM
06785 H_mint PVMlike
06790 H_teal PVMlike
06795 H_evergreen PVMlike
06800 H_cyan PVMlike
06805 H_cyan PVMlike
06810 H_cyan PVMlike
06815 HMMptac_3_eut2(pf06130) PVMlike
06820 P_ocean PVMlike
06825 -pf13416 -
06830 -pf07992pf02852 -
06835 -pf01144 -
06840 -pf01144 -
06845 -pf12773 -
06850 HMMreg12833_4_GRM5(pf12833) GRM5
06855 HMMreg00072_2_pdugrm(pf00072pf00486) EUT2E
06860 HMMreg00486_1_mix(pf00672pf00512) PDU1F
06865 -pf00005 -
06870 - -
06875 - -
06880 HMMreg00356_1_pdu1d(pf00356)
06885 -pf00528 -
BMC-P
BMC-H
BMC-T
Aldehyde dehydrogenase
PTAC
Signature enzyme
Alcohol dehydrogenase
Regulator

Bacillus praedii. Bacteria; Firmicutes; Bacilli; Bacillales; Bacillaceae; Bacillus.

E0Y62__locus_1

PVMlike

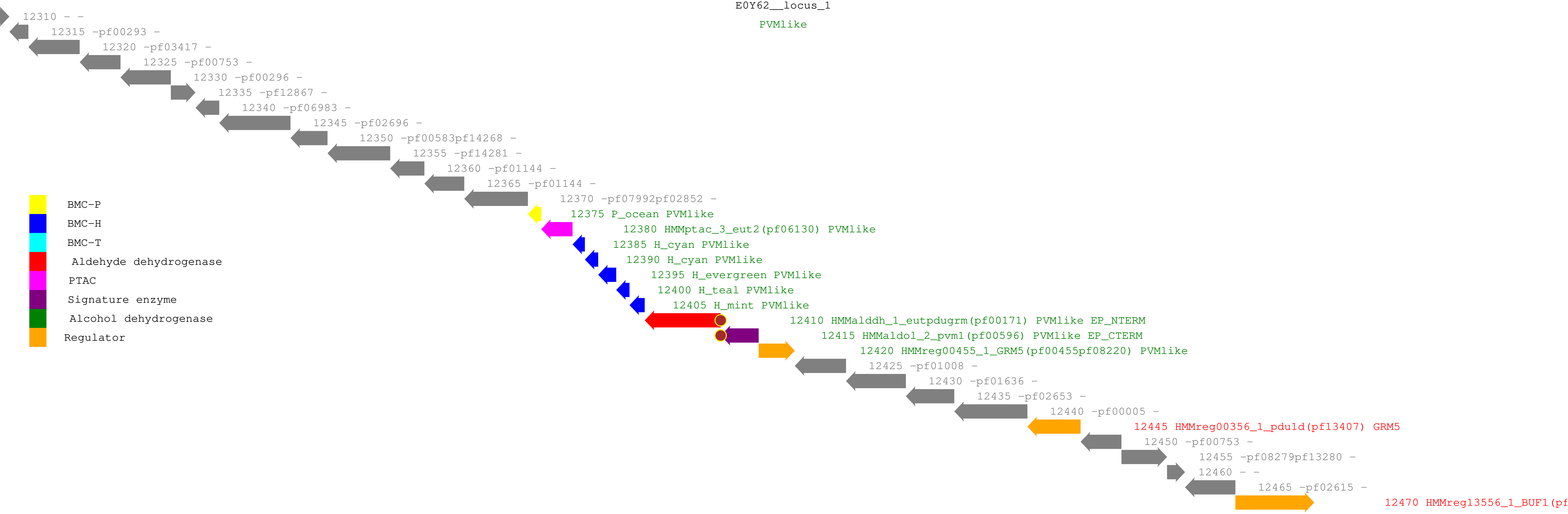

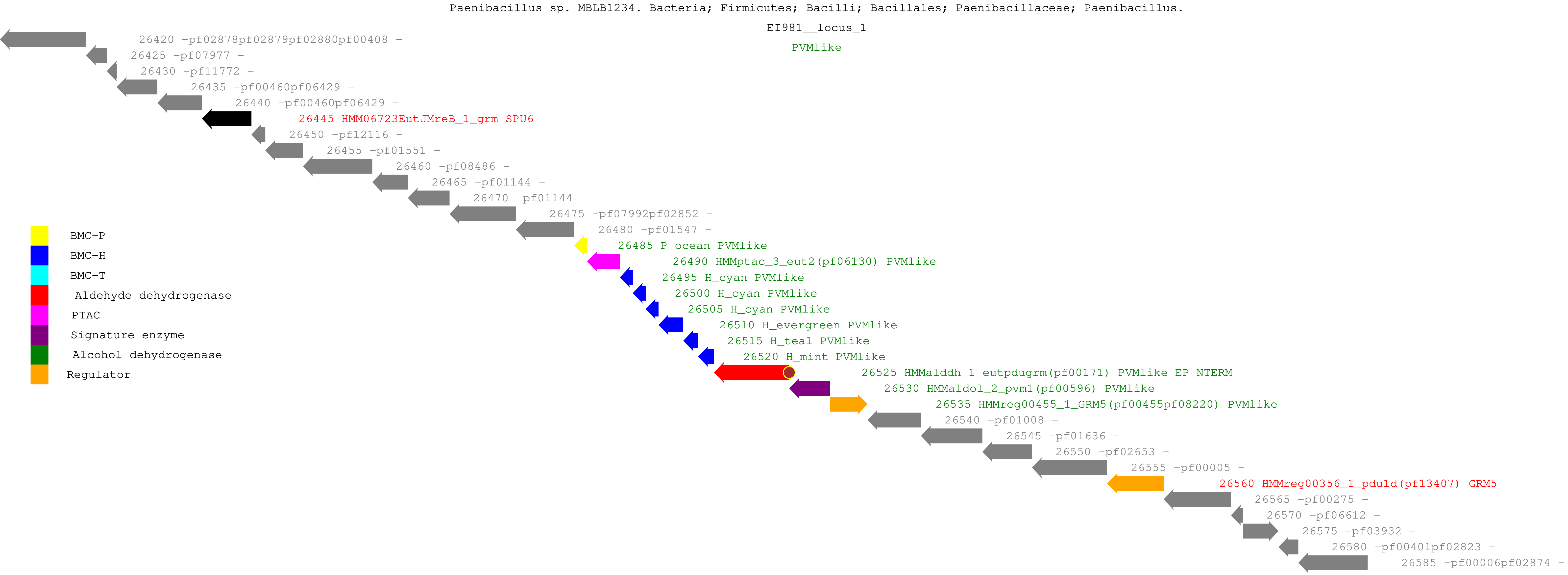
Paenibacillus sp. MBLB1234. Bacteria; Firmicutes; Bacilli; Bacillales; Paenibacillaceae; Paenibacillus.
EI981__locus_1
PVMlike
26420 -pf02878pf02879pf02880pf00408 -
26425 -pf07977 -
26430 -pf11772 -
26435 -pf00460pf06429 -
26440 -pf00460pf06429 -
26445 HMM06723EutJMreB_1_grm SPU6
26450 -pf12116 -
26455 -pf01551 -
26460 -pf08486 -
26465 -pf01144 -
26470 -pf01144 -
26475 -pf07992pf02852 -
26480 -pf01547 -
26485 P_ocean PVMlike
26490 HMMptac_3_eut2(pf06130) PVMlike
26495 H_cyan PVMlike
26500 H_cyan PVMlike
26505 H_cyan PVMlike
26510 H_evergreen PVMlike
26515 H_teal PVMlike
26520 H_mint PVMlike
26525 HMMaldh_1_eutpdugrm(pf00171) PVMlike EP_NTERM
26530 HMMaldol_2_pvm1(pf00596) PVMlike
26535 HMMreg00455_1_GRM5(pf00455pf08220) PVMlike
26540 -pf01008 -
26545 -pf01636 -
26550 -pf02653 -
26555 -pf00005 -
26560 HMMreg00356_1_pdu1d(pf13407) GRM5
26565 -pf00275 -
26570 -pf06612 -
26575 -pf03932 -
26580 -pf00401pf02823 -
26585 -pf00006pf02874 -
BMC-P
BMC-H
BMC-T
Aldehyde dehydrogenase
PTAC
Signature enzyme
Alcohol dehydrogenase
Regulator

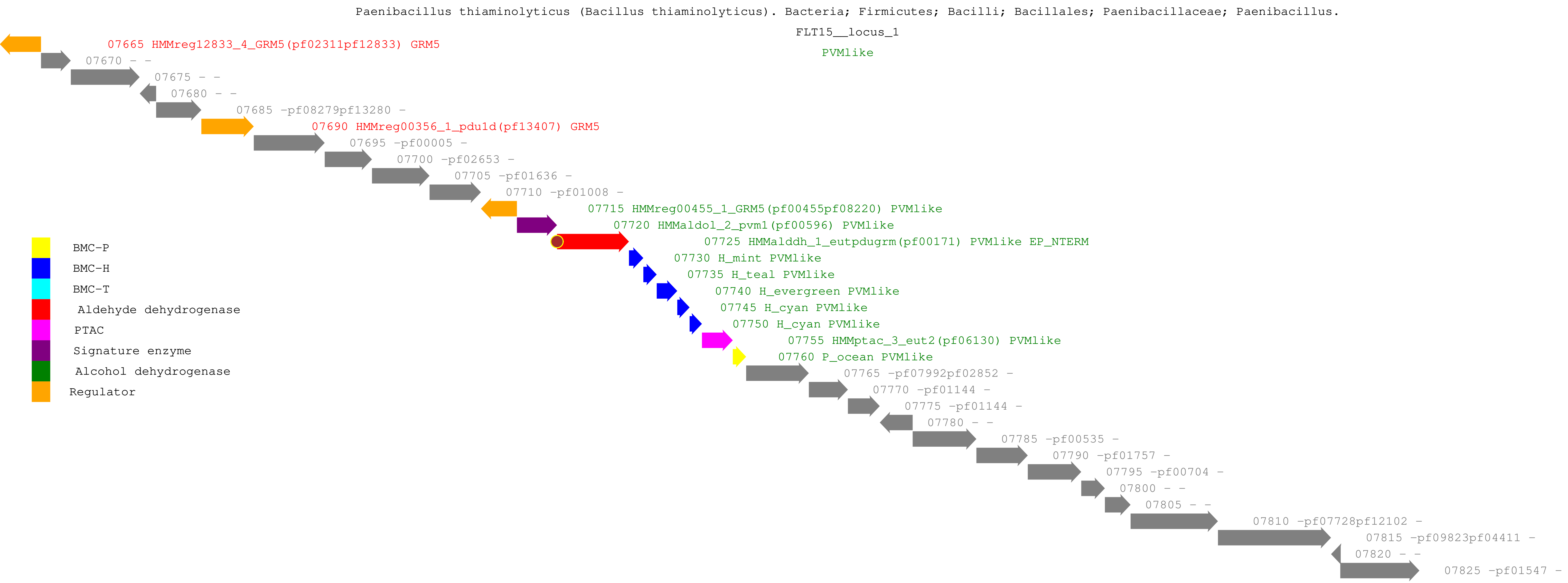
Paenibacillus thiaminolyticus (Bacillus thiaminolyticus). Bacteria; Firmicutes; Bacilli; Bacillales; Paenibacillaceae; Paenibacillus.
FLT15__locus_1
PVMlike
07665 HMMreg12833_4_GRM5(pf02311pf12833) GRM5
07670 - -
07675 - -
07680 - -
07685 -pf08279pf13280 -
07690 HMMreg00356_1_pdu1d(pf13407) GRM5
07695 -pf00005 -
07700 -pf02653 -
07705 -pf01636 -
07710 -pf01008 -
07715 HMMreg00455_1_GRM5(pf00455pf08220) PVMlike
07720 HMMaldol_2_pvm1(pf00596) PVMlike
07725 HMMalddh_1_eutpdugrm(pf00171) PVMlike EP_NTERM
07730 H_mint PVMlike
07735 H_teal PVMlike
07740 H_evergreen PVMlike
07745 H_cyan PVMlike
07750 H_cyan PVMlike
07755 HMMptac_3_eut2(pf06130) PVMlike
07760 P_ocean PVMlike
07765 -pf07992pf02852 -
07770 -pf01144 -
07775 -pf01144 -
07780 - -
07785 -pf00535 -
07790 -pf01757 -
07795 -pf00704 -
07800 - -
07805 - -
07810 -pf07728pf12102 -
07815 -pf09823pf04411 -
07820 - -
07825 -pf01547 -
BMC-P
BMC-H
BMC-T
Aldehyde dehydrogenase
PTAC
Signature enzyme
Alcohol dehydrogenase
Regulator

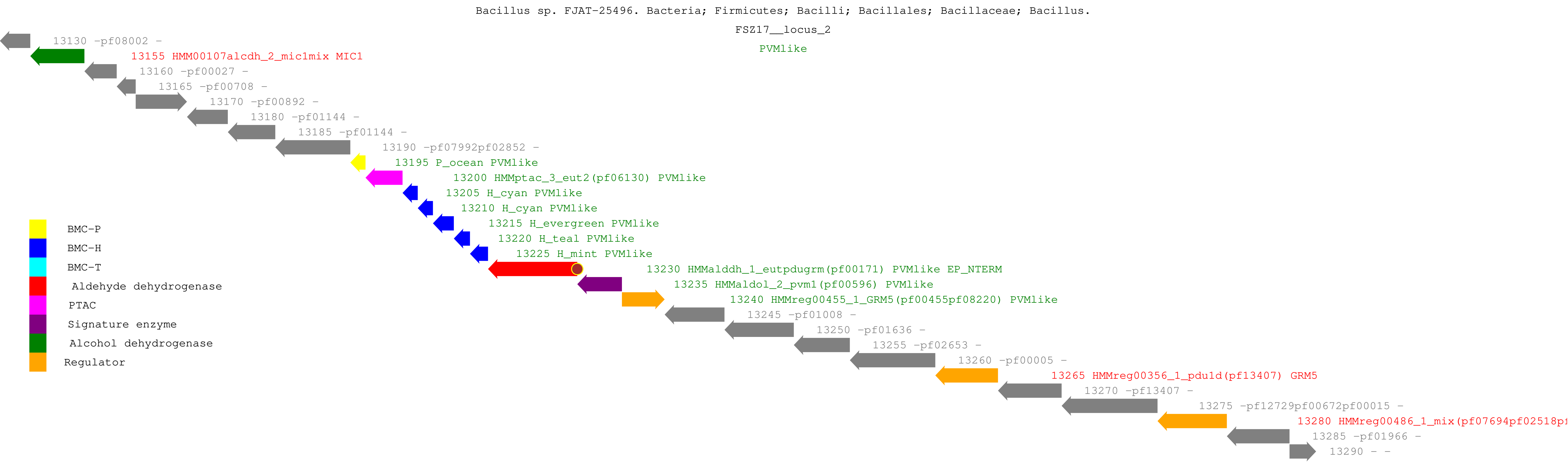
Bacillus sp. FJAT-25496. Bacteria; Firmicutes; Bacilli; Bacillales; Bacillaceae; Bacillus.
FSZ17__locus_2
PVMlike
13130 -pf08002 -
13155 HMM00107alcdh_2_mic1mix MIC1
13160 -pf00027 -
13165 -pf00708 -
13170 -pf00892 -
13180 -pf01144 -
13185 -pf01144 -
13190 -pf07992pf02852 -
13195 P_ocean PVMlike
13200 HMMptac_3_eut2(pf06130) PVMlike
13205 H_cyan PVMlike
13210 H_cyan PVMlike
13215 H_evergreen PVMlike
13220 H_teal PVMlike
13225 H_mint PVMlike
13230 HMMalddh_1_eutpdugrm(pf00171) PVMlike EP_NTERM
13235 HMMaldol_2_pvm1(pf00596) PVMlike
13240 HMMreg00455_1_GRM5(pf00455pf08220) PVMlike
13245 -pf01008 -
13250 -pf01636 -
13255 -pf02653 -
13260 -pf00005 -
13265 HMMreg00356_1_pdu1d(pf13407) GRM5
13270 -pf13407 -
13275 -pf12729pf00672pf00015 -
13280 HMMreg00486_1_mix(pf07694pf02518p
13285 -pf01966 -
13290 - -
BMC-P
BMC-H
BMC-T
Aldehyde dehydrogenase
PTAC
Signature enzyme
Alcohol dehydrogenase
Regulator

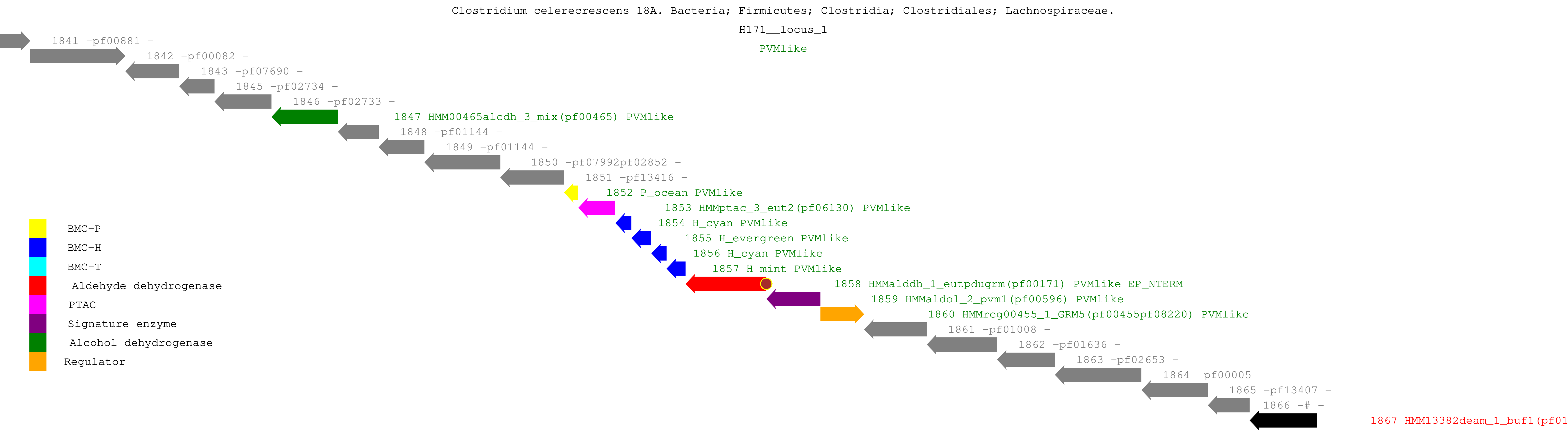
Clostridium celerecrescens 18A. Bacteria; Firmicutes; Clostridia; Clostridiales; Lachnospiraceae.
H171__locus_1
PVMlike
1841 -pf00881 -
1842 -pf00082 -
1843 -pf07690 -
1845 -pf02734 -
1846 -pf02733 -
1847 HMM00465alcdh_3_mix(pf00465) PVMlike
1848 -pf01144 -
1849 -pf01144 -
1850 -pf07992pf02852 -
1851 -pf13416 -
1852 P_ocean PVMlike
1853 HMMptac_3_eut2(pf06130) PVMlike
1854 H_cyan PVMlike
1855 H_evergreen PVMlike
1856 H_cyan PVMlike
1857 H_mint PVMlike
1858 HMMalddh_1_eutpdugrm(pf00171) PVMlike EP_NTERM
1859 HMMaldol_2_pvm1(pf00596) PVMlike
1860 HMMreg00455_1_GRM5(pf00455pf08220) PVMlike
1861 -pf01008 -
1862 -pf01636 -
1863 -pf02653 -
1864 -pf00005 -
1865 -pf13407 -
1866 -# -
1867 HMM13382deam_1_buf1(pf01
BMC-P
BMC-H
BMC-T
Aldehyde dehydrogenase
PTAC
Signature enzyme
Alcohol dehydrogenase
Regulator

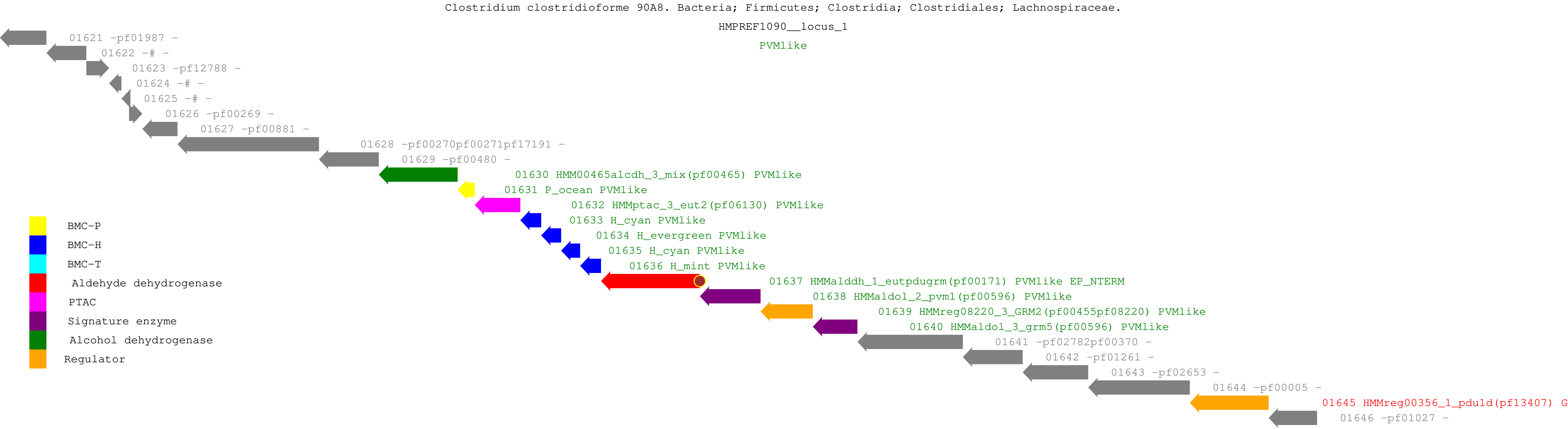

Clostridium clostridioforme 90A8. Bacteria; Firmicutes; Clostridia; Clostridiales; Lachnospiraceae.
HMPREF1090__locus_1
PVMlike
01621 -pf01987 -
01622 -# -
01623 -pf12788 -
01624 -# -
01625 -# -
01626 -pf00269 -
01627 -pf00881 -
01628 -pf00270pf00271pf17191 -
01629 -pf00480 -
01630 HMM00465alcdh_3_mix(pf00465) PVMlike
01631 P_ocean PVMlike
01632 HMMptac_3_eut2(pf06130) PVMlike
01633 H_cyan PVMlike
01634 H_evergreen PVMlike
01635 H_cyan PVMlike
01636 H_mint PVMlike
01637 HMMaldh_1_eutpdugrm(pf00171) PVMlike EP_NTERM
01638 HMMaldol_2_pvm1(pf00596) PVMlike
01639 HMMreg08220_3_GRM2(pf00455pf08220) PVMlike
01640 HMMaldol_3_grm5(pf00596) PVMlike
01641 -pf02782pf00370 -
01642 -pf01261 -
01643 -pf02653 -
01644 -pf00005 -
01645 HMMreg00356_1_pdu1d(pf13407) G
01646 -pf01027 -
BMC-P
BMC-H
BMC-T
Aldehyde dehydrogenase
PTAC
Signature enzyme
Alcohol dehydrogenase
Regulator

Faecalibacterium cf. prausnitzii KLE1255. Bacteria; Firmicutes; Clostridia; Clostridiales; Ruminococcaceae;

HMPREF9436__locus_1

PVMlike

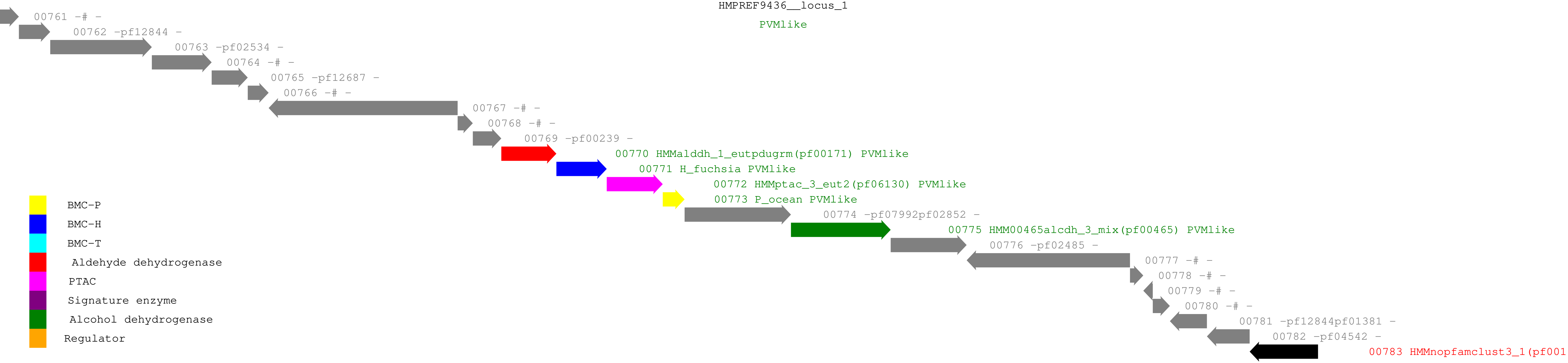

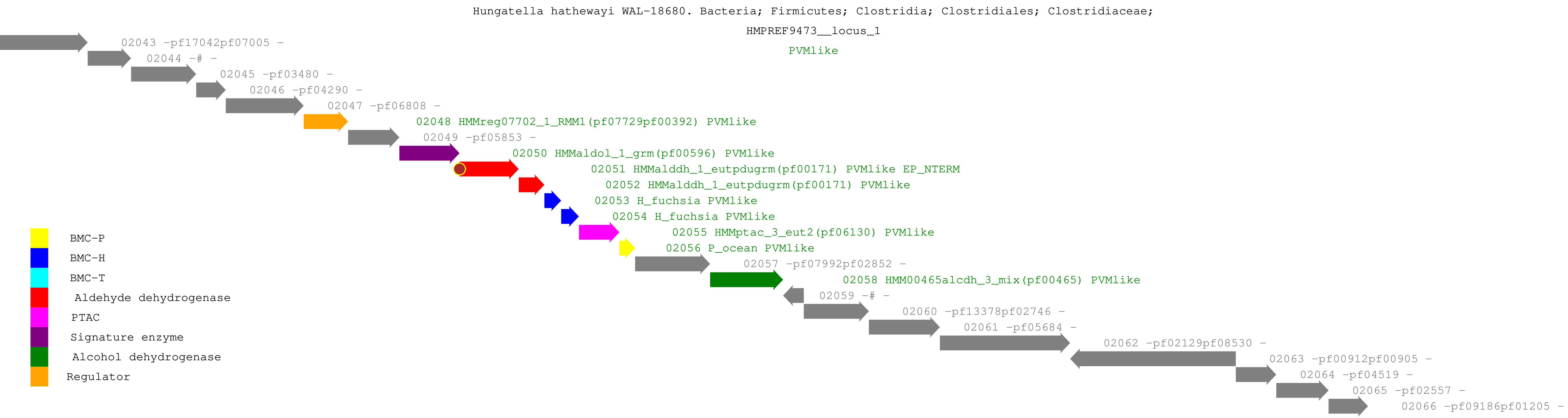

Hungatella hathewayi WAL-18680. Bacteria; Firmicutes; Clostridia; Clostridiales; Clostridiaceae;
HMPREF9473__locus_1
PVMlike
02043 -pf17042pf07005 -
02044 -# -
02045 -pf03480 -
02046 -pf04290 -
02047 -pf06808 -
02048 HMMreg07702_1_RMM1(pf07729pf00392) PVMlike
02049 -pf05853 -
02050 HMMaldol_1_grm(pf00596) PVMlike
02051 HMMaldddh_1_eutpdugrm(pf00171) PVMlike EP_NTERM
02052 HMMaldddh_1_eutpdugrm(pf00171) PVMlike
02053 H_fuchsia PVMlike
02054 H_fuchsia PVMlike
02055 HMMptac_3_eut2(pf06130) PVMlike
02056 P_ocean PVMlike
02057 -pf07992pf02852 -
02058 HMM00465alcdh_3_mix(pf00465) PVMlike
02059 -# -
02060 -pf13378pf02746 -
02061 -pf05684 -
02062 -pf02129pf08530 -
02063 -pf00912pf00905 -
02064 -pf04519 -
02065 -pf02557 -
02066 -pf09186pf01205 -
BMC-P
BMC-H
BMC-T
Aldehyde dehydrogenase
PTAC
Signature enzyme
Alcohol dehydrogenase
Regulator

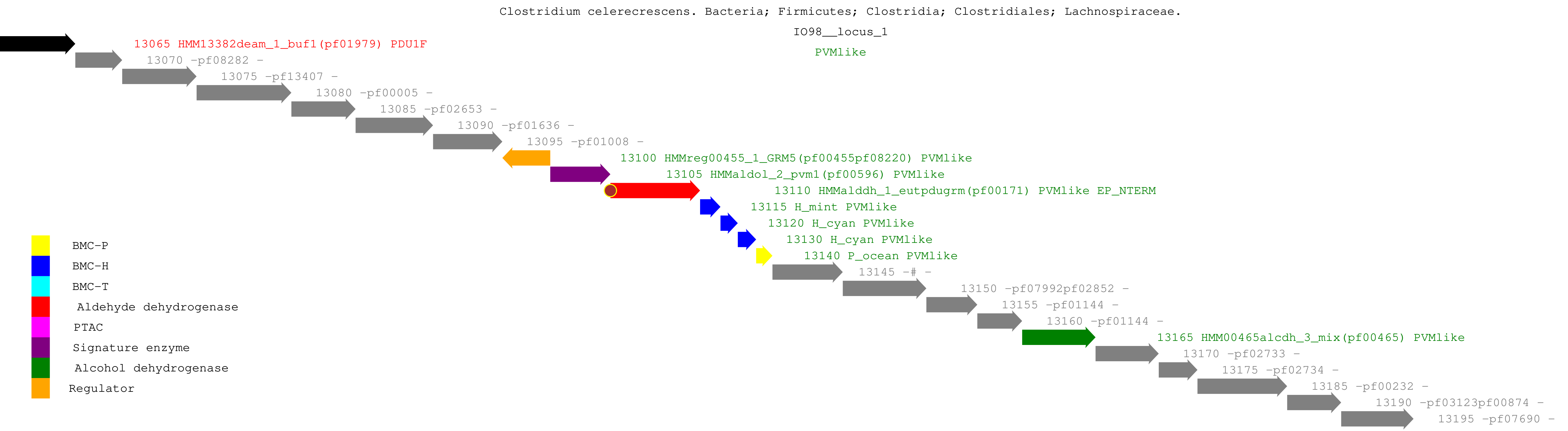
Clostridium celerecrescens. Bacteria; Firmicutes; Clostridia; Clostridiales; Lachnospiraceae.
IO98__locus_1
PVMlike
13065 HMM13382deam_1_buf1(pf01979) PDU1F
13070 -pf08282 -
13075 -pf13407 -
13080 -pf00005 -
13085 -pf02653 -
13090 -pf01636 -
13095 -pf01008 -
13100 HMMreg00455_1_GRM5(pf00455pf08220) PVMlike
13105 HMMaldol_2_pvm1(pf00596) PVMlike
13110 HMMalddh_1_eutpdugrm(pf00171) PVMlike EP_NTERM
13115 H_mint PVMlike
13120 H_cyan PVMlike
13130 H_cyan PVMlike
13140 P_ocean PVMlike
13145 -# -
13150 -pf07992pf02852 -
13155 -pf01144 -
13160 -pf01144 -
13165 HMM00465alcdh_3_mix(pf00465) PVMlike
13170 -pf02733 -
13175 -pf02734 -
13185 -pf00232 -
13190 -pf03123pf00874 -
13195 -pf07690 -
BMC-P
BMC-H
BMC-T
Aldehyde dehydrogenase
PTAC
Signature enzyme
Alcohol dehydrogenase
Regulator

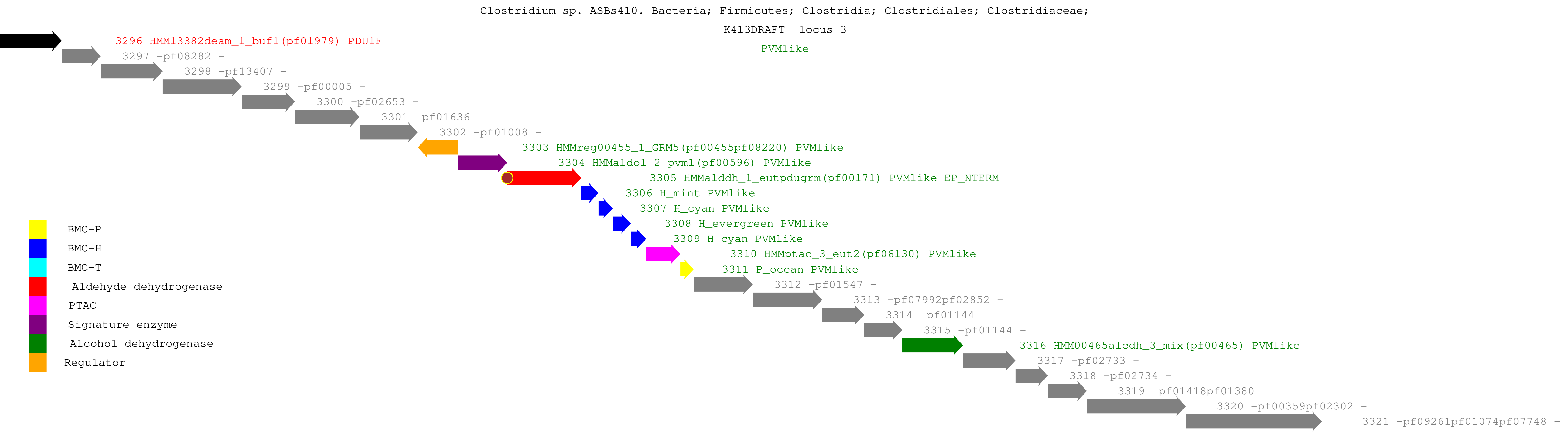

Clostridium sp. ASBs410. Bacteria; Firmicutes; Clostridia; Clostridiales; Clostridiaceae;
K413DRAFT__locus_3
PVMlike
3296 HMM13382deam_1_buf1(pf01979) PDU1F
3297 -pf08282 -
3298 -pf13407 -
3299 -pf00005 -
3300 -pf02653 -
3301 -pf01636 -
3302 -pf01008 -
3303 HMMreg00455_1_GRM5(pf00455pf08220) PVMlike
3304 HMMaldol_2_pvm1(pf00596) PVMlike
3305 HMMalddh_1_eutpdugrm(pf00171) PVMlike EP_NTERM
3306 H_mint PVMlike
3307 H_cyan PVMlike
3308 H_evergreen PVMlike
3309 H_cyan PVMlike
3310 HMMptac_3_eut2(pf06130) PVMlike
3311 P_ocean PVMlike
3312 -pf01547 -
3313 -pf07992pf02852 -
3314 -pf01144 -
3315 -pf01144 -
3316 HMM00465alcdh_3_mix(pf00465) PVMlike
3317 -pf02733 -
3318 -pf02734 -
3319 -pf01418pf01380 -
3320 -pf00359pf02302 -
3321 -pf09261pf01074pf07748 -
BMC-P
BMC-H
BMC-T
Aldehyde dehydrogenase
PTAC
Signature enzyme
Alcohol dehydrogenase
Regulator

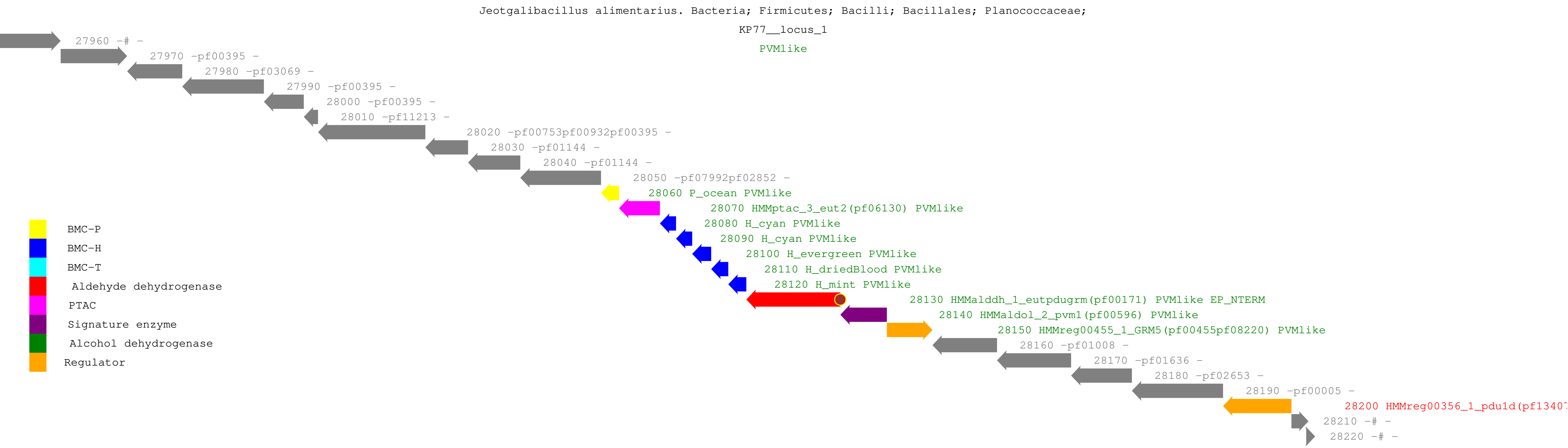
Jeotgalibacillus alimentarius. Bacteria; Firmicutes; Bacilli; Bacillales; Planococcaceae;
KP77__locus_1
PVMlike
27960 -# -
27970 -pf00395 -
27980 -pf03069 -
27990 -pf00395 -
28000 -pf00395 -
28010 -pf11213 -
28020 -pf00753pf00932pf00395 -
28030 -pf01144 -
28040 -pf01144 -
28050 -pf07992pf02852 -
28060 P_ocean PVMlike
28070 HMMptac_3_eut2(pf06130) PVMlike
28080 H_cyan PVMlike
28090 H_cyan PVMlike
28100 H_evergreen PVMlike
28110 H_driedBlood PVMlike
28120 H_mint PVMlike
28130 HMMalddh_1_eutpdugrm(pf00171) PVMlike EP_NTERM
28140 HMMaldol_2_pvm1(pf00596) PVMlike
28150 HMMreg00455_1_GRM5(pf00455pf08220) PVMlike
28160 -pf01008 -
28170 -pf01636 -
28180 -pf02653 -
28190 -pf00005 -
28200 HMMreg00356_1_pdu1d(pf1340
28210 -# -
28220 -# -
BMC-P
BMC-H
BMC-T
Aldehyde dehydrogenase
PTAC
Signature enzyme
Alcohol dehydrogenase
Regulator

Paenibacillus thiaminolyticus (Bacillus thiaminolyticus). Bacteria; Firmicutes; Bacilli; Bacillales; Paenibacillaceae;

NCTC11027__locus_1

PVMlike

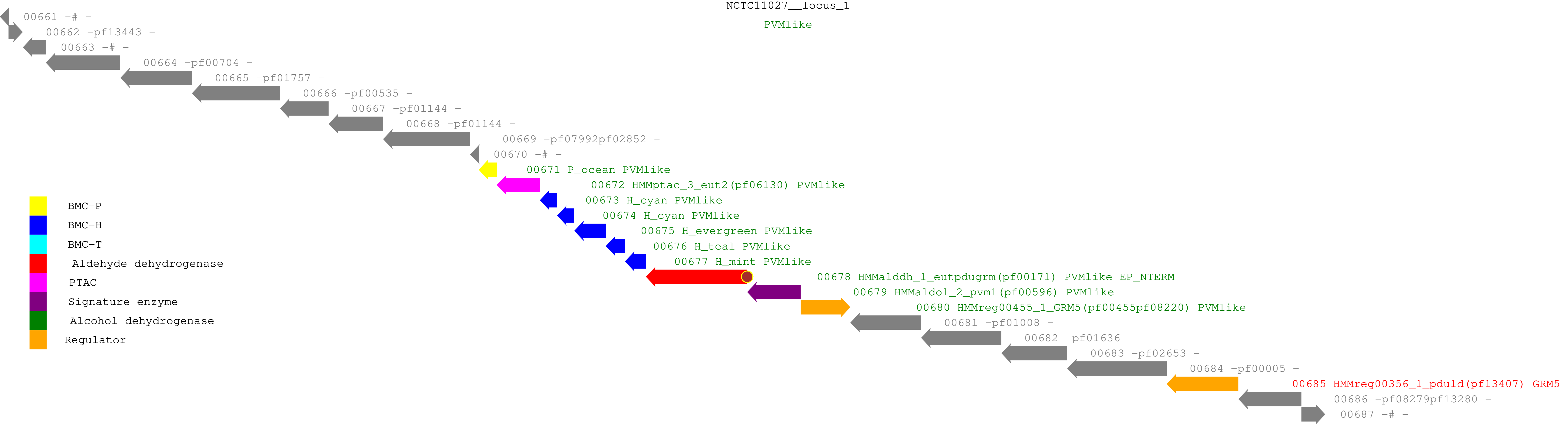

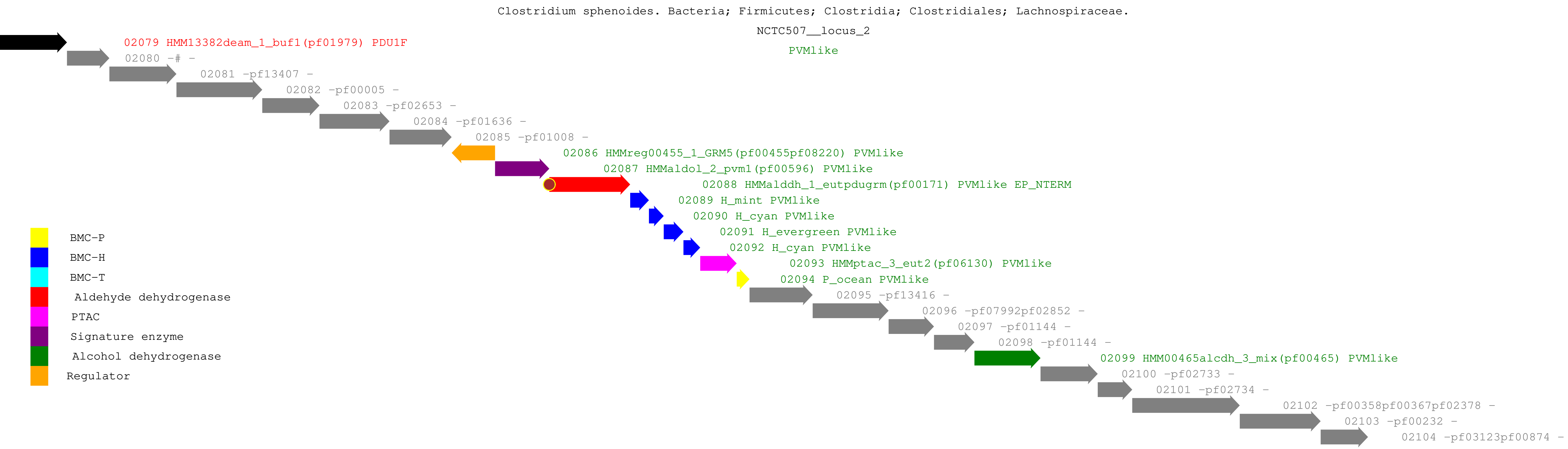
Clostridium sphenoides. Bacteria; Firmicutes; Clostridia; Clostridiales; Lachnospiraceae.
NCTC507__locus_2
PVMlike
02079 HMM13382deam_1_buf1(pf01979) PDU1F
02080 -# -
02081 -pf13407 -
02082 -pf00005 -
02083 -pf02653 -
02084 -pf01636 -
02085 -pf01008 -
02086 HMMreg00455_1_GRM5(pf00455pf08220) PVMlike
02087 HMMaldol_2_pvm1(pf00596) PVMlike
02088 HMMalddh_1_eutpdugrm(pf00171) PVMlike EP_NTERM
02089 H_mint PVMlike
02090 H_cyan PVMlike
02091 H_evergreen PVMlike
02092 H_cyan PVMlike
02093 HMMptac_3_eut2(pf06130) PVMlike
02094 P_ocean PVMlike
02095 -pf13416 -
02096 -pf07992pf02852 -
02097 -pf01144 -
02098 -pf01144 -
02099 HMM00465alcdh_3_mix(pf00465) PVMlike
02100 -pf02733 -
02101 -pf02734 -
02102 -pf00358pf00367pf02378 -
02103 -pf00232 -
02104 -pf03123pf00874 -
BMC-P
BMC-H
BMC-T
Aldehyde dehydrogenase
PTAC
Signature enzyme
Alcohol dehydrogenase
Regulator

Acetoanaerobium noterae. Bacteria; Firmicutes; Clostridia; Clostridiales;

SAMN02745120__locus_2

PVMlike

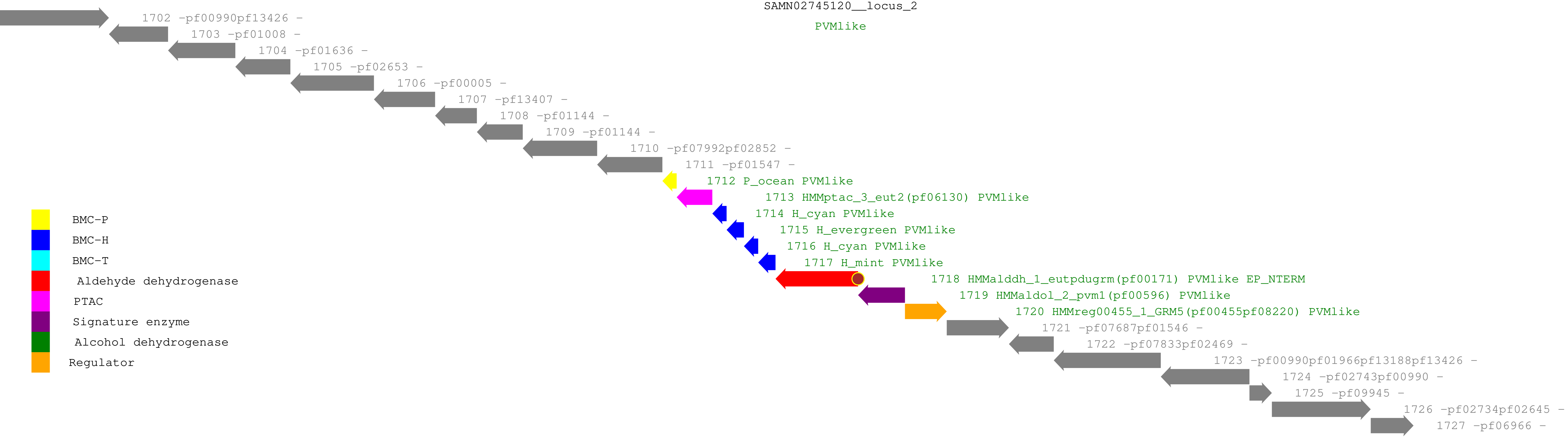

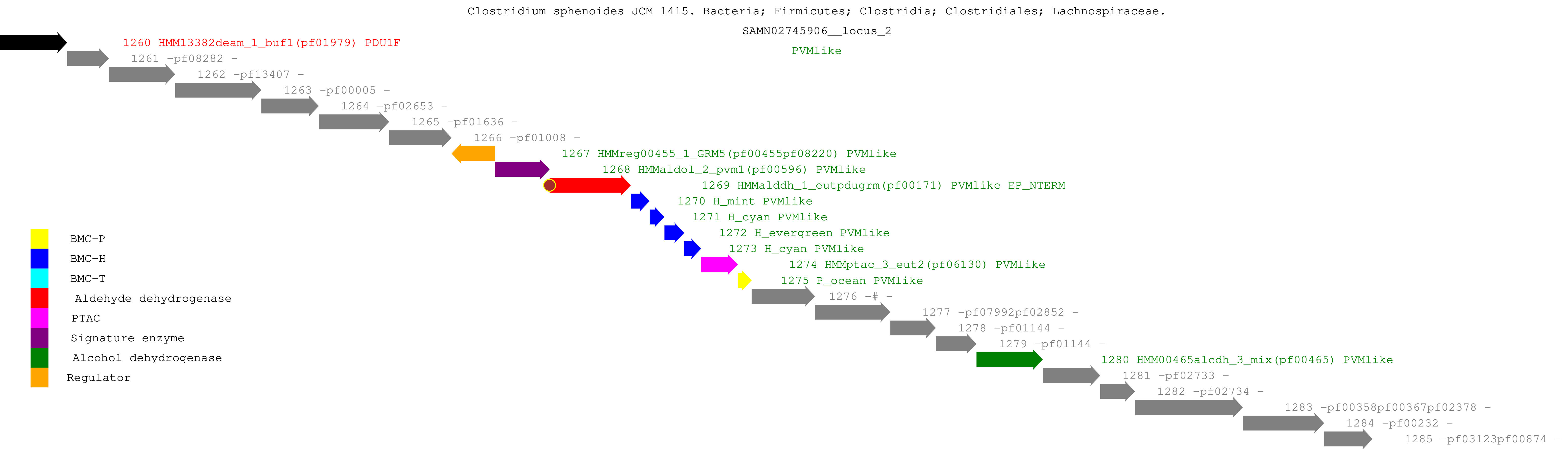
Clostridium sphenoides JCM 1415. Bacteria; Firmicutes; Clostridia; Clostridiales; Lachnospiraceae.
SAMN02745906__locus_2
PVMlike
1260 HMM13382deam_1_buf1(pf01979) PDU1F
1261 -pf08282 -
1262 -pf13407 -
1263 -pf00005 -
1264 -pf02653 -
1265 -pf01636 -
1266 -pf01008 -
1267 HMMreg00455_1_GRM5(pf00455pf08220) PVMlike
1268 HMMaldol_2_pvm1(pf00596) PVMlike
1269 HMMalddh_1_eutpdugrm(pf00171) PVMlike EP_NTERM
1270 H_mint PVMlike
1271 H_cyan PVMlike
1272 H_evergreen PVMlike
1273 H_cyan PVMlike
1274 HMMptac_3_eut2(pf06130) PVMlike
1275 P_ocean PVMlike
1276 -# -
1277 -pf07992pf02852 -
1278 -pf01144 -
1279 -pf01144 -
1280 HMM00465alcdh_3_mix(pf00465) PVMlike
1281 -pf02733 -
1282 -pf02734 -
1283 -pf00358pf00367pf02378 -
1284 -pf00232 -
1285 -pf03123pf00874 -
BMC-P
BMC-H
BMC-T
Aldehyde dehydrogenase
PTAC
Signature enzyme
Alcohol dehydrogenase
Regulator

Clostridium caminithermale DSM 15212. Bacteria; Firmicutes; Clostridia; Clostridiales; Clostridiaceae;

SAMN02745912__locus_1

PVMlike

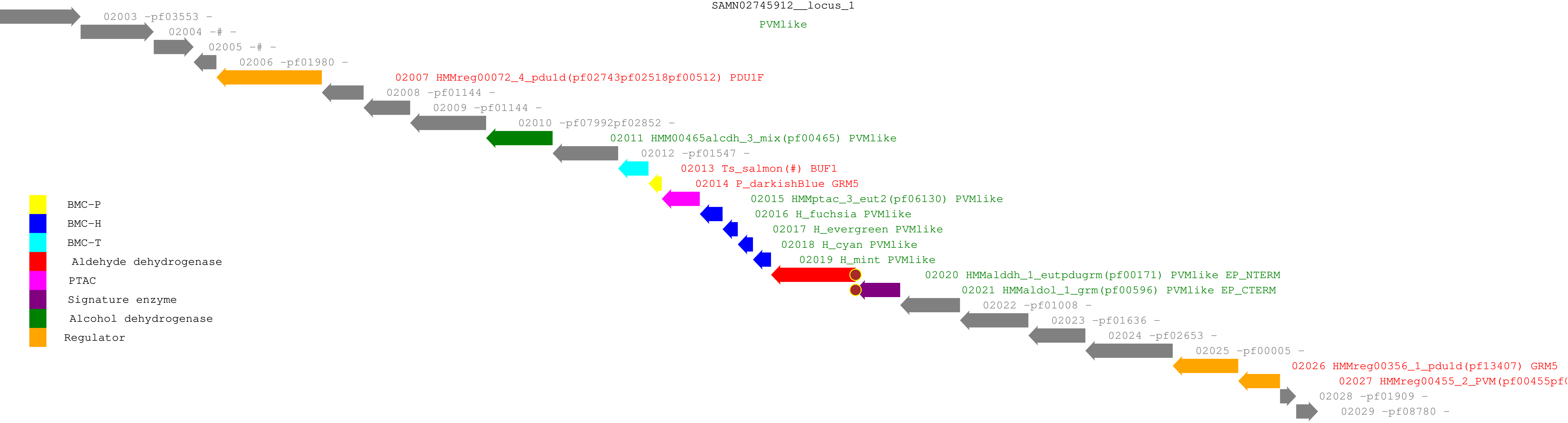

Gracilibacillus ureilyticus. Bacteria; Firmicutes; Bacilli; Bacillales; Bacillaceae;

SAMN04487944__locus_1

PVMlike

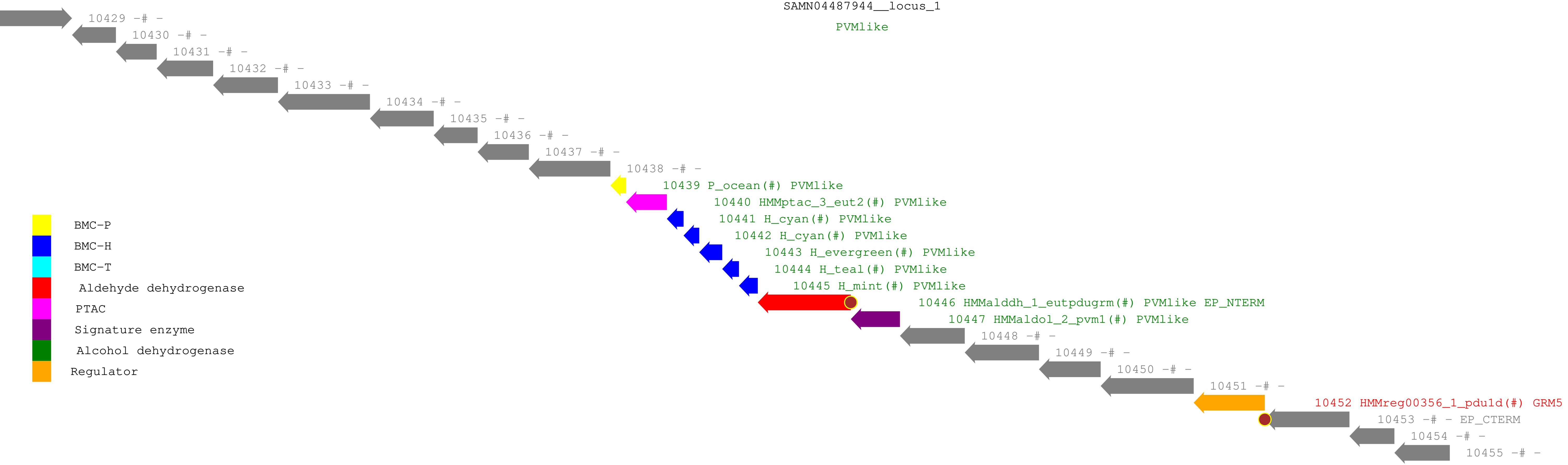

Bacillus daliensis. Bacteria; Firmicutes; Bacilli; Bacillales; Bacillaceae; Bacillus.

SAMN04488053__locus_1

PVMlike

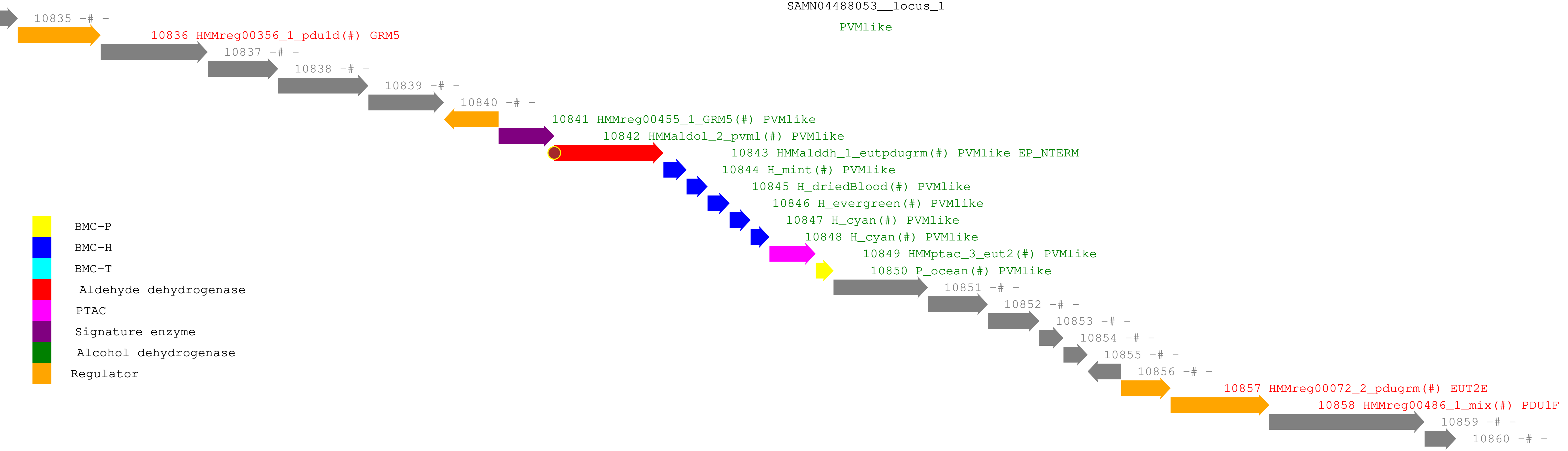

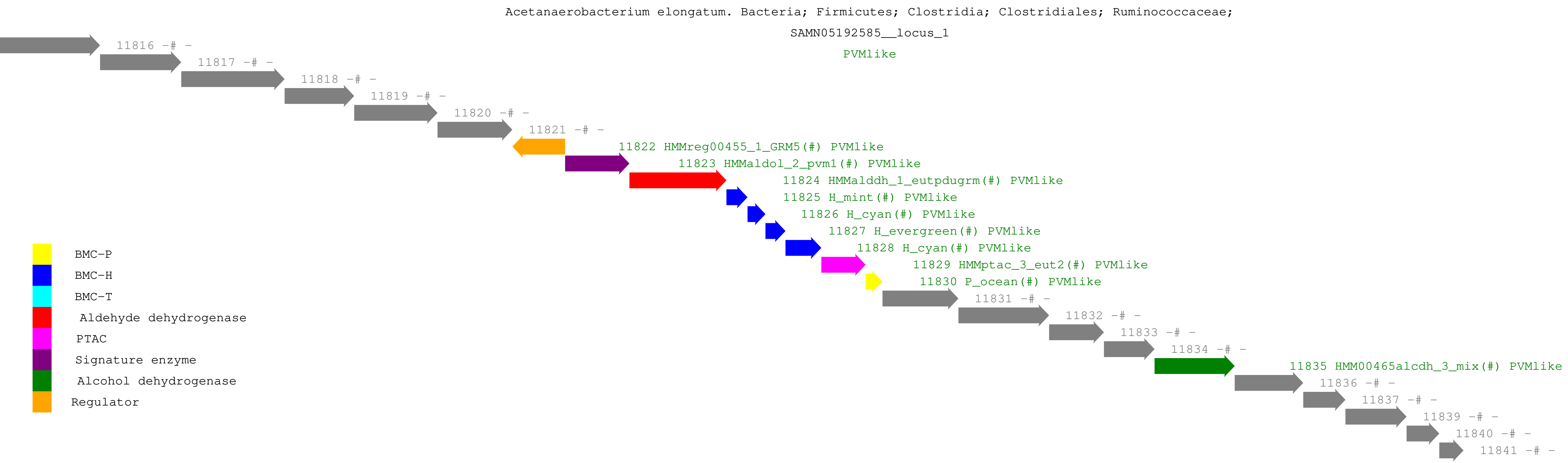
Acetanaerobacterium elongatum. Bacteria; Firmicutes; Clostridia; Clostridiales; Ruminococcaceae;
SAMN05192585__locus_1
PVMlike
11816 -# -
11817 -# -
11818 -# -
11819 -# -
11820 -# -
11821 -# -
11822 HMMreg00455_1_GRM5(#) PVMlike
11823 HMMaldol_2_pvm1(#) PVMlike
11824 HMMaldh_1_eutpdugrm(#) PVMlike
11825 H_mint(#) PVMlike
11826 H_cyan(#) PVMlike
11827 H_evergreen(#) PVMlike
11828 H_cyan(#) PVMlike
11829 HMMptac_3_eut2(#) PVMlike
11830 P_ocean(#) PVMlike
11831 -# -
11832 -# -
11833 -# -
11834 -# -
11835 HMM00465alcdh_3_mix(#) PVMlike
11836 -# -
11837 -# -
11839 -# -
11840 -# -
11841 -# -
BMC-P
BMC-H
BMC-T
Aldehyde dehydrogenase
PTAC
Signature enzyme
Alcohol dehydrogenase
Regulator

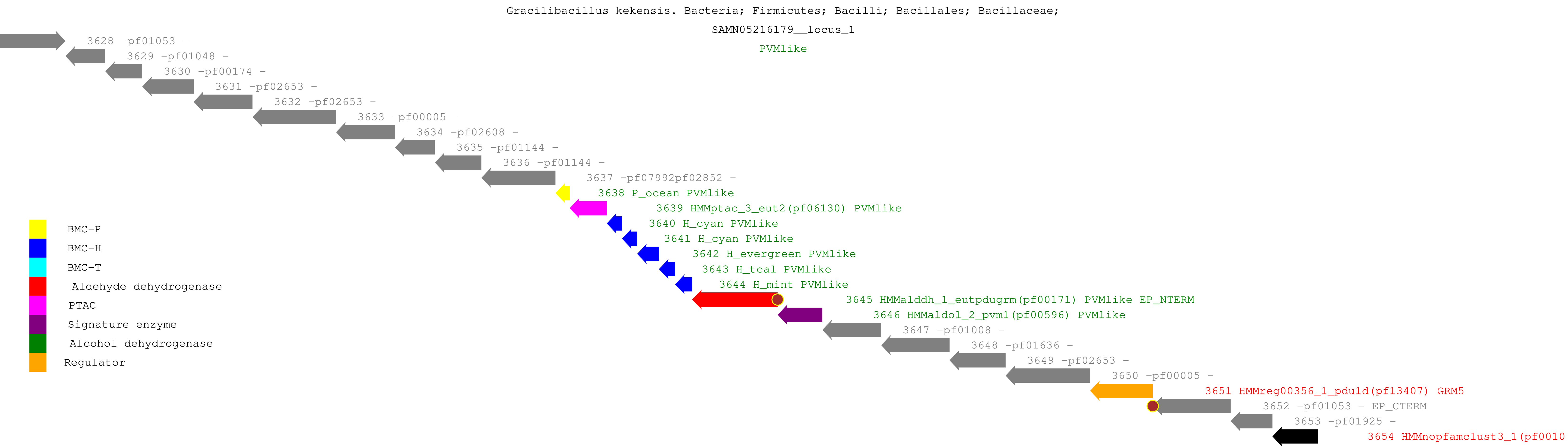
Gracilibacillus kekensis. Bacteria; Firmicutes; Bacilli; Bacillales; Bacillaceae;
SAMN05216179__locus_1
PVMlike
3628 -pf01053 -
3629 -pf01048 -
3630 -pf00174 -
3631 -pf02653 -
3632 -pf02653 -
3633 -pf00005 -
3634 -pf02608 -
3635 -pf01144 -
3636 -pf01144 -
3637 -pf07992pf02852 -
3638 P_ocean PVMlike
3639 HMMptac_3_eut2(pf06130) PVMlike
3640 H_cyan PVMlike
3641 H_cyan PVMlike
3642 H_evergreen PVMlike
3643 H_teal PVMlike
3644 H_mint PVMlike
3645 HMMaldh_1_eutpdugrm(pf00171) PVMlike EP_NTERM
3646 HMMaldol_2_pvm1(pf00596) PVMlike
3647 -pf01008 -
3648 -pf01636 -
3649 -pf02653 -
3650 -pf00005 -
3651 HMMreg00356_1_pdu1d(pf13407) GRM5
3652 -pf01053 - EP_CTERM
3653 -pf01925 -
3654 HMMnopfamclust3_1(pf0010
BMC-P
BMC-H
BMC-T
Aldehyde dehydrogenase
PTAC
Signature enzyme
Alcohol dehydrogenase
Regulator

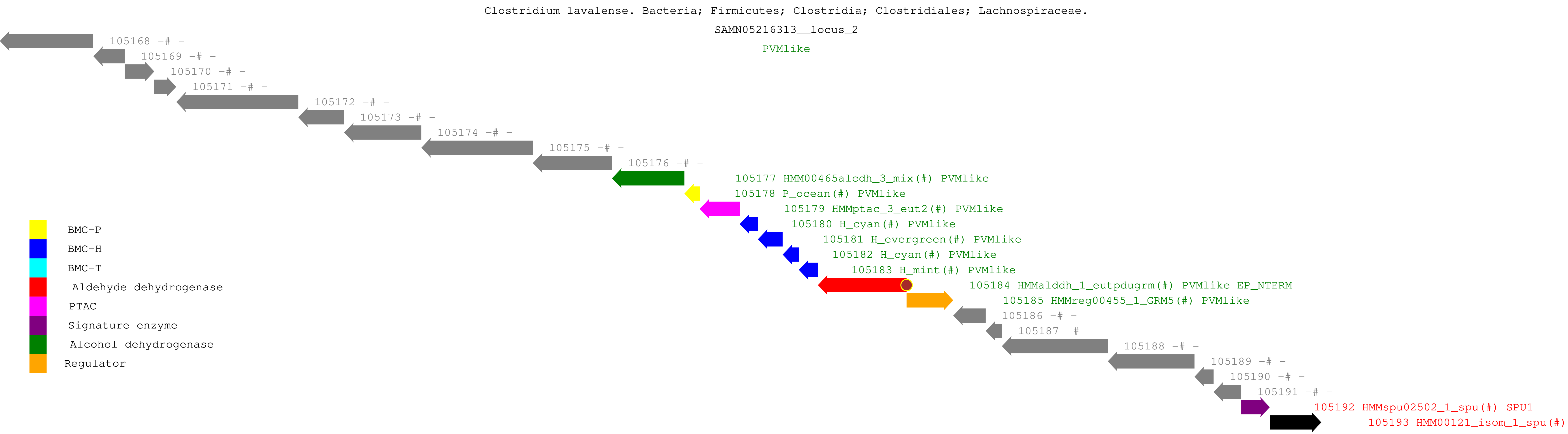
Clostridium lavalense. Bacteria; Firmicutes; Clostridia; Clostridiales; Lachnospiraceae.
SAMN05216313__locus_2
PVMlike
105168 -# -
105169 -# -
105170 -# -
105171 -# -
105172 -# -
105173 -# -
105174 -# -
105175 -# -
105176 -# -
105177 HMM00465alcdh_3_mix(#) PVMlike
105178 P_ocean(#) PVMlike
105179 HMMptac_3_eut2(#) PVMlike
105180 H_cyan(#) PVMlike
105181 H_evergreen(#) PVMlike
105182 H_cyan(#) PVMlike
105183 H_mint(#) PVMlike
105184 HMMalddh_1_eutpdugrm(#) PVMlike EP_NTERM
105185 HMMreg00455_1_GRM5(#) PVMlike
105186 -# -
105187 -# -
105188 -# -
105189 -# -
105190 -# -
105191 -# -
105192 HMMspu02502_1_spu(#) SPU1
105193 HMM00121_isom_1_spu(#)
BMC-P
BMC-H
BMC-T
Aldehyde dehydrogenase
PTAC
Signature enzyme
Alcohol dehydrogenase
Regulator

Desulfotomaculum guttoideum. Bacteria; Firmicutes; Clostridia; Clostridiales; Lachnospiraceae.

SAMN05443270__locus_3

PVMlike

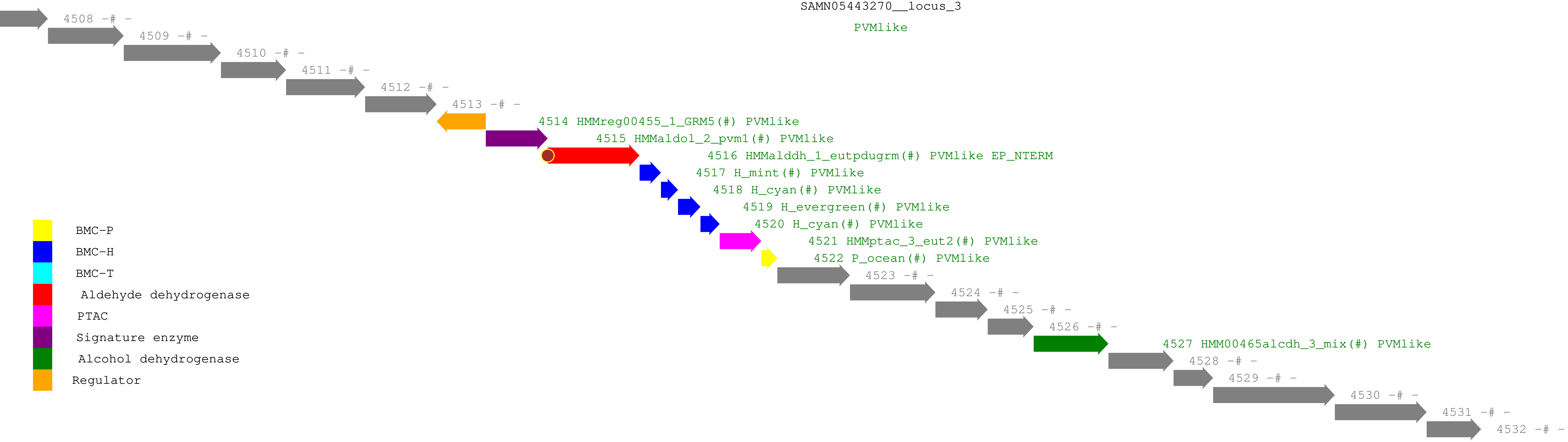

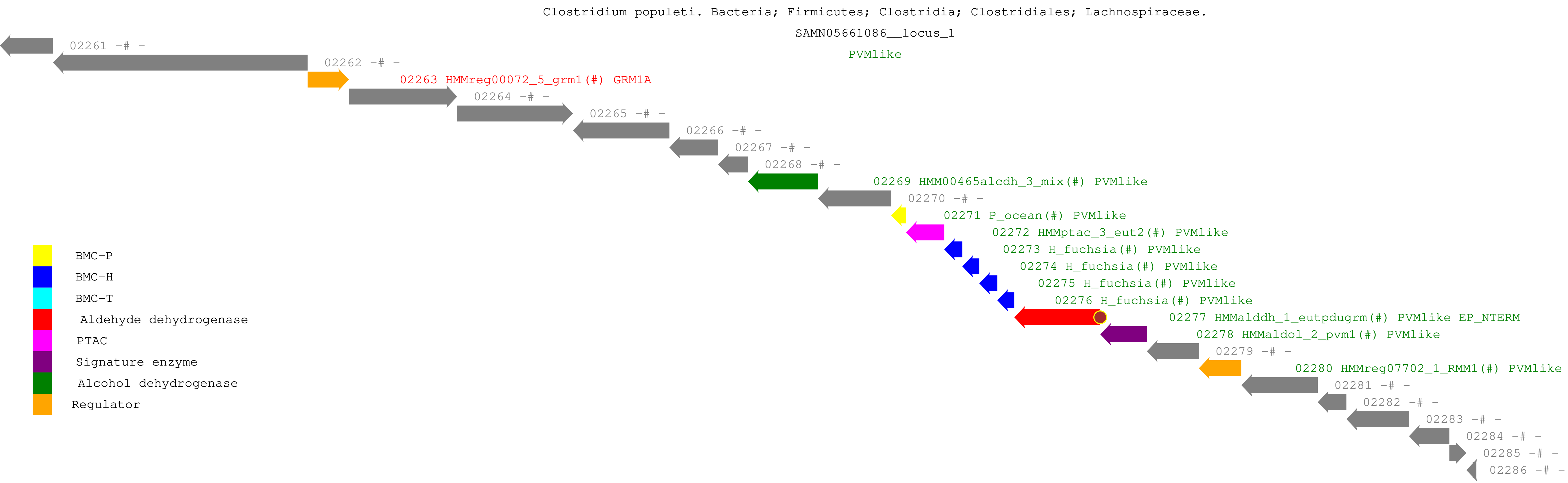
Clostridium populeti. Bacteria; Firmicutes; Clostridia; Clostridiales; Lachnospiraceae.
SAMN05661086__locus_1
PVMlike
02261 -# -
02262 -# -
02263 HMMreg00072_5_grm1(#) GRM1A
02264 -# -
02265 -# -
02266 -# -
02267 -# -
02268 -# -
02269 HMM00465alcdh_3_mix(#) PVMlike
02270 -# -
02271 P_ocean(#) PVMlike
02272 HMMptac_3_eut2(#) PVMlike
02273 H_fuchsia(#) PVMlike
02274 H_fuchsia(#) PVMlike
02275 H_fuchsia(#) PVMlike
02276 H_fuchsia(#) PVMlike
02277 HMMalddh_1_eutpdugrm(#) PVMlike EP_NTERM
02278 HMMaldol_2_pvm1(#) PVMlike
02279 -# -
02280 HMMreg07702_1_RMM1(#) PVMlike
02281 -# -
02282 -# -
02283 -# -
02284 -# -
02285 -# -
02286 -# -
BMC-P
BMC-H
BMC-T
Aldehyde dehydrogenase
PTAC
Signature enzyme
Alcohol dehydrogenase
Regulator

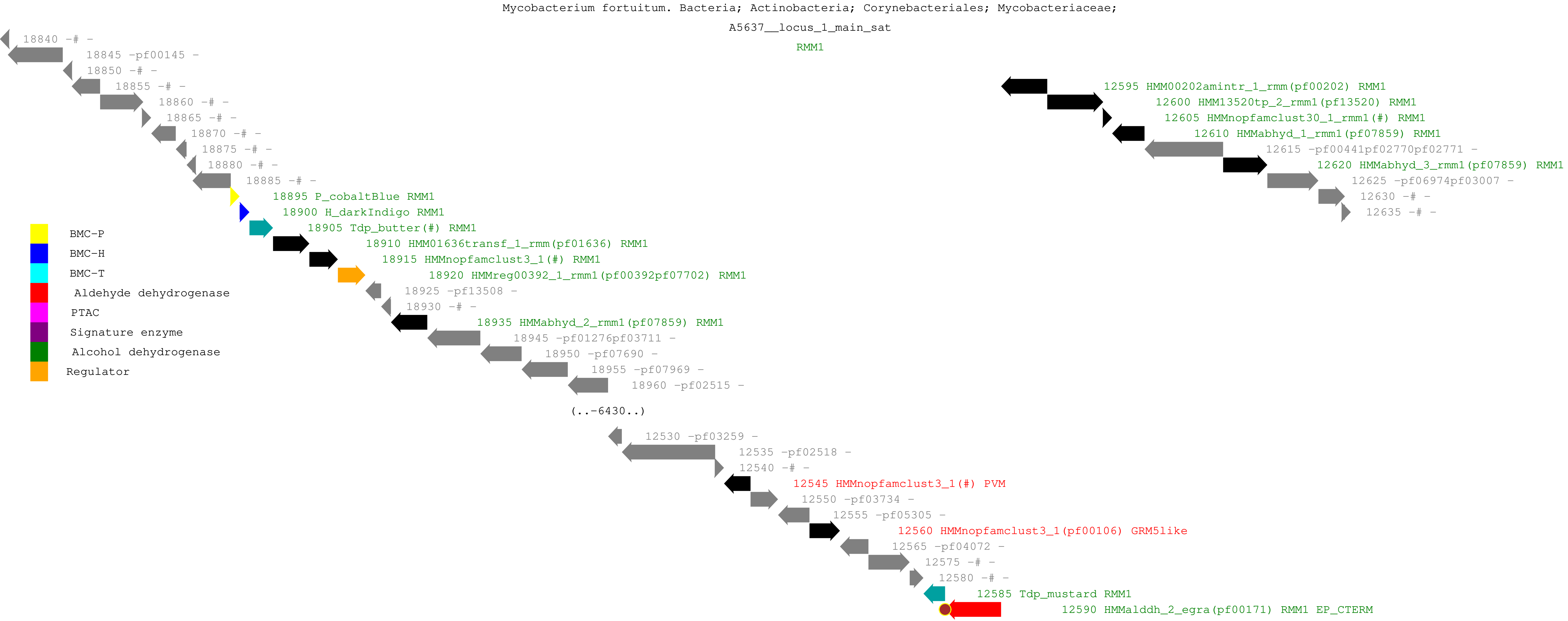
Mycobacterium fortuitum. Bacteria; Actinobacteria; Corynebacteriales; Mycobacteriaceae;
A5637__locus_1_main_sat
RMM1
18840 -# -
18845 -pf00145 -
18850 -# -
18855 -# -
18860 -# -
18865 -# -
18870 -# -
18875 -# -
18880 -# -
18885 -# -
18895 P_cobaltBlue RMM1
18900 H_darkIndigo RMM1
18905 Tdp_butter(#) RMM1
18910 HMM01636transf_1_rmm(pf01636) RMM1
18915 HMMnopfamclust3_1(#) RMM1
18920 HMMreg00392_1_rmm1(pf00392pf07702) RMM1
18925 -pf13508 -
18930 -# -
18935 HMMabhyd_2_rmm1(pf07859) RMM1
18945 -pf01276pf03711 -
18950 -pf07690 -
18955 -pf07969 -
18960 -pf02515 -
(..-6430..)
12530 -pf03259 -
12535 -pf02518 -
12540 -# -
12545 HMMnopfamclust3_1(#) PVM
12550 -pf03734 -
12555 -pf05305 -
12560 HMMnopfamclust3_1(pf00106) GRM5like
12565 -pf04072 -
12575 -# -
12580 -# -
12585 Tdp_mustard RMM1
12590 HMMaldh_2_egra(pf00171) RMM1 EP_CTERM
12595 HMM00202amintr_1_rmm(pf00202) RMM1
12600 HMM13520tp_2_rmm1(pf13520) RMM1
12605 HMMnopfamclust30_1_rmm1(#) RMM1
12610 HMMabhyd_1_rmm1(pf07859) RMM1
12615 -pf00441pf02770pf02771 -
12620 HMMabhyd_3_rmm1(pf07859) RMM1
12625 -pf06974pf03007 -
12630 -# -
12635 -# -
BMC-P
BMC-H
BMC-T
Aldehyde dehydrogenase
PTAC
Signature enzyme
Alcohol dehydrogenase
Regulator

Mycobacterium sp. E802. Bacteria; Actinobacteria; Corynebacteriales; Mycobacteriaceae;

A5699__locus_1

RMM1

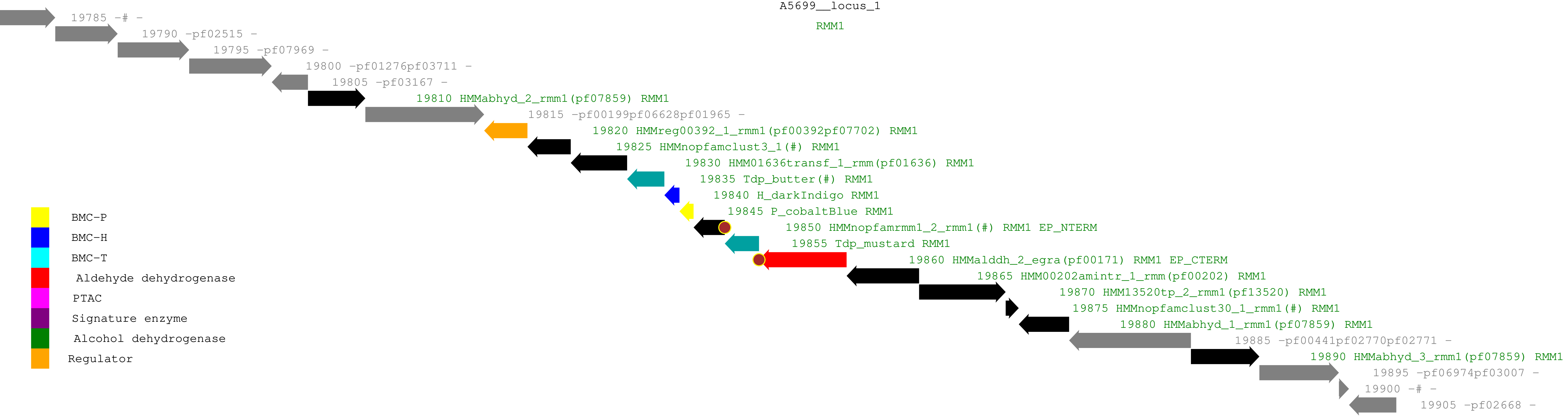

Mycolicibacterium porcinum. Bacteria; Actinobacteria; Corynebacteriales; Mycobacteriaceae;

A5717__locus_1

RMM1

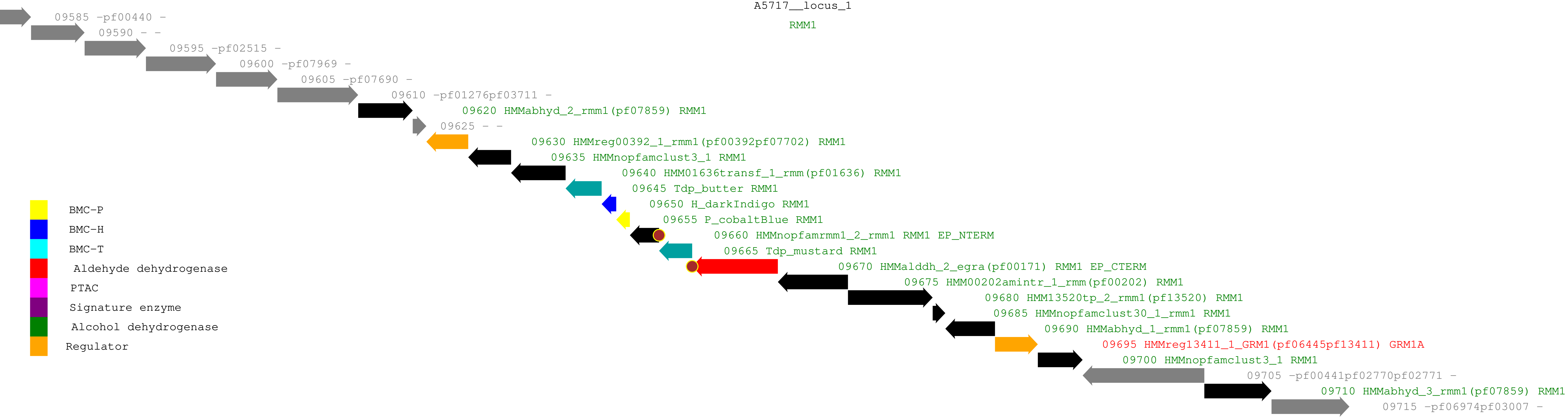

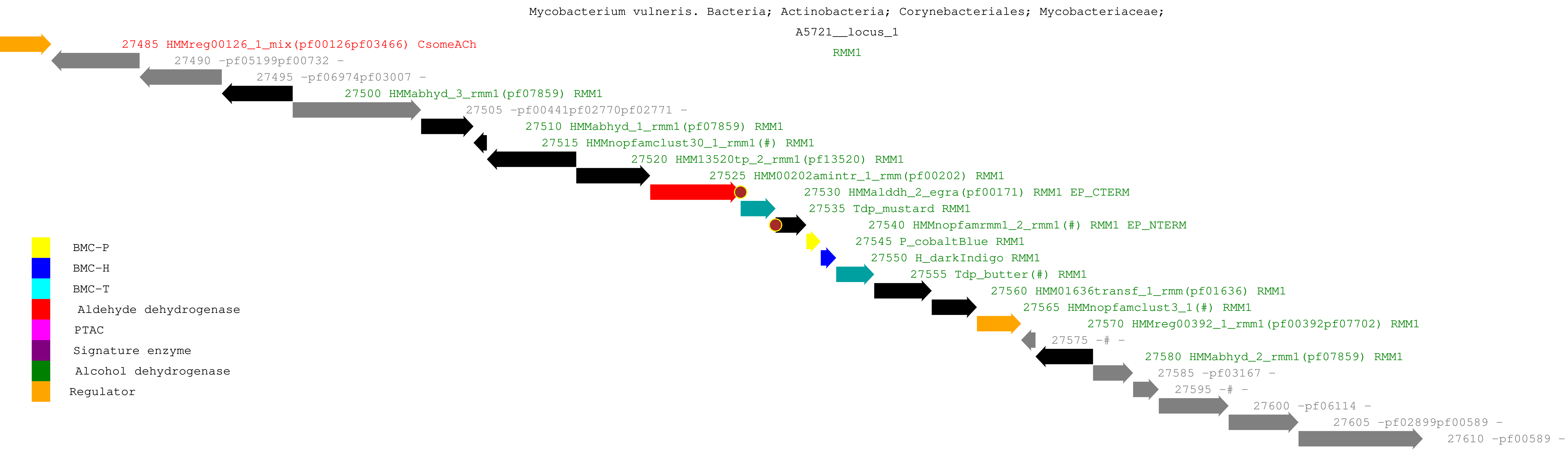
Mycobacterium vulneris. Bacteria; Actinobacteria; Corynebacteriales; Mycobacteriaceae;
A5721__locus_1
RMM1
27485 HMMreg00126_1_mix(pf00126pf03466) CsomeACh
27490 -pf05199pf00732 -
27495 -pf06974pf03007 -
27500 HMMabhyd_3_rmm1(pf07859) RMM1
27505 -pf00441pf02770pf02771 -
27510 HMMabhyd_1_rmm1(pf07859) RMM1
27515 HMMnopfamclust30_1_rmm1(#) RMM1
27520 HMM13520tp_2_rmm1(pf13520) RMM1
27525 HMM00202amintr_1_rmm(pf00202) RMM1
27530 HMMaldh_2_egra(pf00171) RMM1 EP_CTERM
27535 Tdp_mustard RMM1
27540 HMMnopfamrmm1_2_rmm1(#) RMM1 EP_NTERM
27545 P_cobaltBlue RMM1
27550 H_darkIndigo RMM1
27555 Tdp_butter(#) RMM1
27560 HMM01636transf_1_rmm(pf01636) RMM1
27565 HMMnopfamclust3_1(#) RMM1
27570 HMMreg00392_1_rmm1(pf00392pf07702) RMM1
27575 -# -
27580 HMMabhyd_2_rmm1(pf07859) RMM1
27585 -pf03167 -
27595 -# -
27600 -pf06114 -
27605 -pf02899pf00589 -
27610 -pf00589 -
BMC-P
BMC-H
BMC-T
Aldehyde dehydrogenase
PTAC
Signature enzyme
Alcohol dehydrogenase
Regulator

Mycobacterium sp. ACS1612. Bacteria; Actinobacteria; Corynebacteriales; Mycobacteriaceae;

A5724__locus_1

RMM1

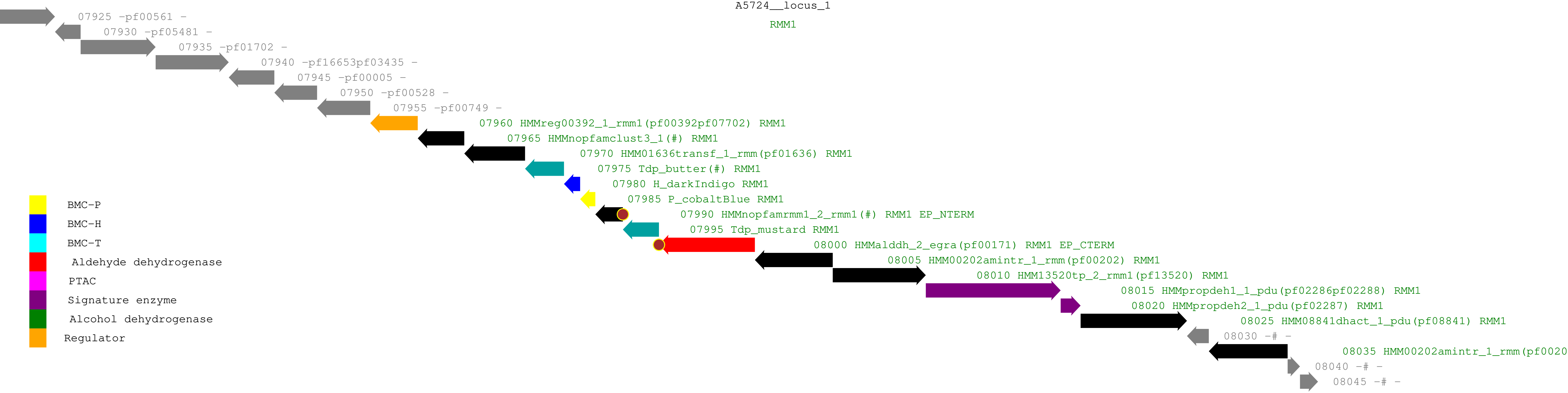

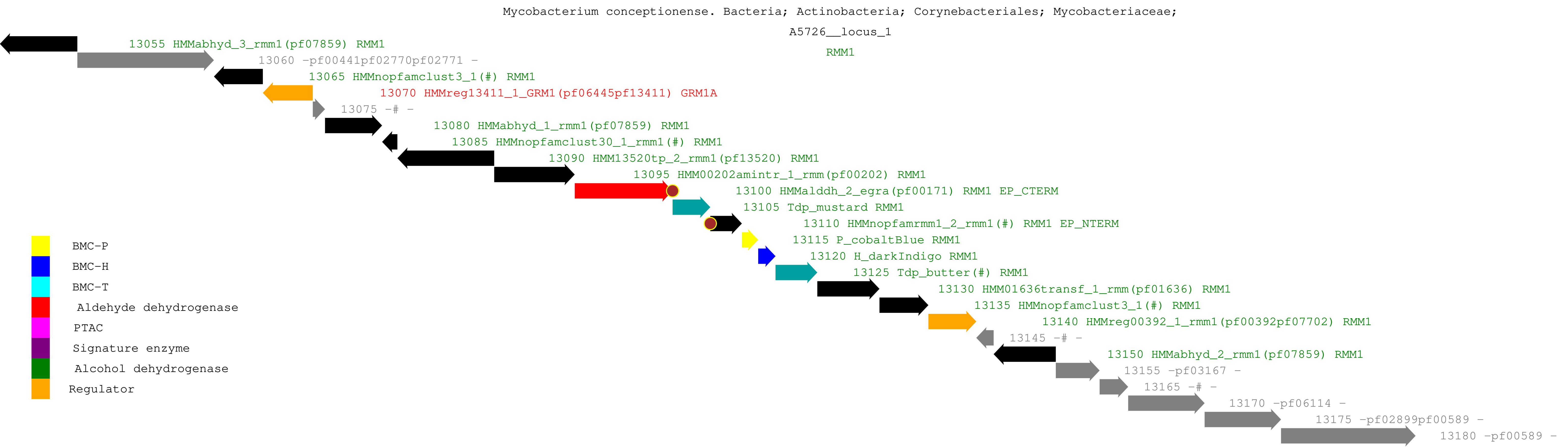

Mycobacterium conceptionense. Bacteria; Actinobacteria; Corynebacteriales; Mycobacteriaceae;
A5726__locus_1
RMM1
13055 HMMabhyd_3_rmm1(pf07859) RMM1
13060 -pf00441pf02770pf02771 -
13065 HMMnopfamclust3_1(#) RMM1
13070 HMMreg13411_1_GRM1(pf06445pf13411) GRM1A
13075 -# -
13080 HMMabhyd_1_rmm1(pf07859) RMM1
13085 HMMnopfamclust30_1_rmm1(#) RMM1
13090 HMM13520tp_2_rmm1(pf13520) RMM1
13095 HMM00202amintr_1_rmm(pf00202) RMM1
13100 HMMaldh_2_egra(pf00171) RMM1 EP_CTERM
13105 Tdp_mustard RMM1
13110 HMMnopfamrmm1_2_rmm1(#) RMM1 EP_NTERM
13115 P_cobaltBlue RMM1
13120 H_darkIndigo RMM1
13125 Tdp_butter(#) RMM1
13130 HMM01636transf_1_rmm(pf01636) RMM1
13135 HMMnopfamclust3_1(#) RMM1
13140 HMMreg00392_1_rmm1(pf00392pf07702) RMM1
13145 -# -
13150 HMMabhyd_2_rmm1(pf07859) RMM1
13155 -pf03167 -
13165 -# -
13170 -pf06114 -
13175 -pf02899pf00589 -
13180 -pf00589 -
BMC-P
BMC-H
BMC-T
Aldehyde dehydrogenase
PTAC
Signature enzyme
Alcohol dehydrogenase
Regulator

Mycobacterium sp. GA-1841. Bacteria; Actinobacteria; Corynebacteriales; Mycobacteriaceae;

A5740__locus_1

RMM1

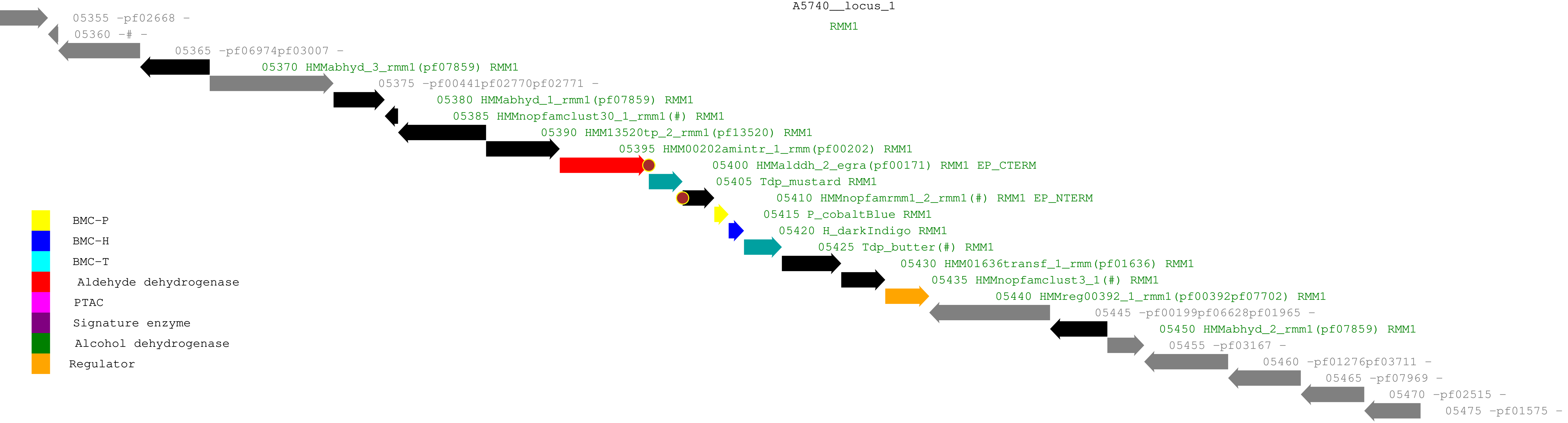

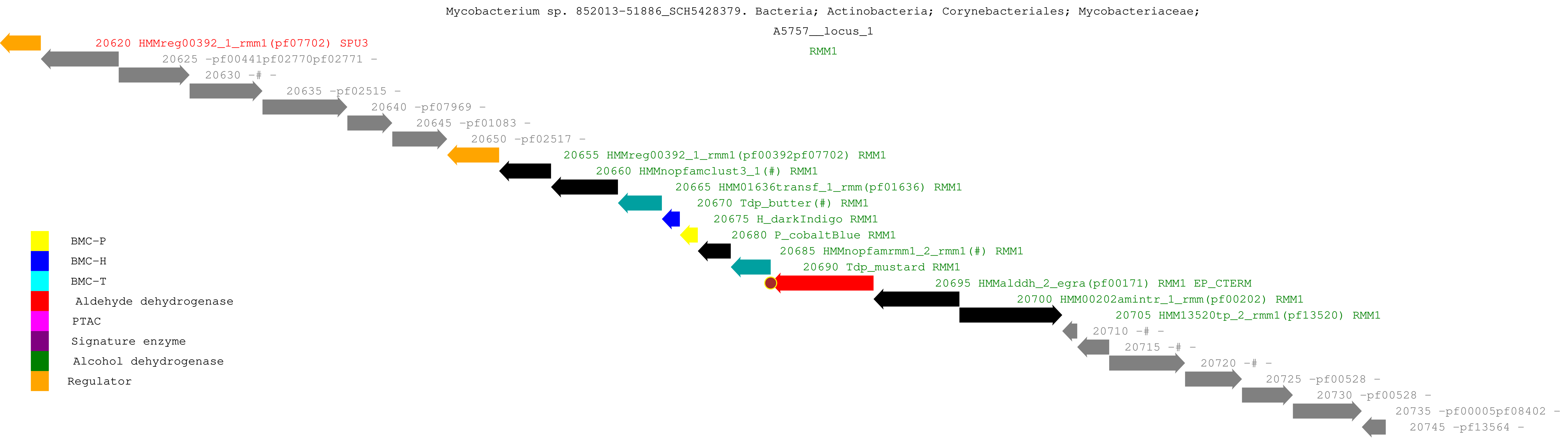
Mycobacterium sp. 852013-51886_SCH5428379. Bacteria; Actinobacteria; Corynebacteriales; Mycobacteriaceae;
A5757__locus_1
RMM1
20620 HMMreg00392_1_rmm1(pf07702) SPU3
20625 -pf00441pf02770pf02771 -
20630 -# -
20635 -pf02515 -
20640 -pf07969 -
20645 -pf01083 -
20650 -pf02517 -
20655 HMMreg00392_1_rmm1(pf00392pf07702) RMM1
20660 HMMnopfamclust3_1(#) RMM1
20665 HMM01636transf_1_rmm(pf01636) RMM1
20670 Tdp_butter(#) RMM1
20675 H_darkIndigo RMM1
20680 P_cobaltBlue RMM1
20685 HMMnopfamrmm1_2_rmm1(#) RMM1
20690 Tdp_mustard RMM1
20695 HMMaldh_2_egra(pf00171) RMM1 EP_CTERM
20700 HMM00202amintr_1_rmm(pf00202) RMM1
20705 HMM13520tp_2_rmm1(pf13520) RMM1
20710 -# -
20715 -# -
20720 -# -
20725 -pf00528 -
20730 -pf00528 -
20735 -pf00005pf08402 -
20745 -pf13564 -
BMC-P
BMC-H
BMC-T
Aldehyde dehydrogenase
PTAC
Signature enzyme
Alcohol dehydrogenase
Regulator

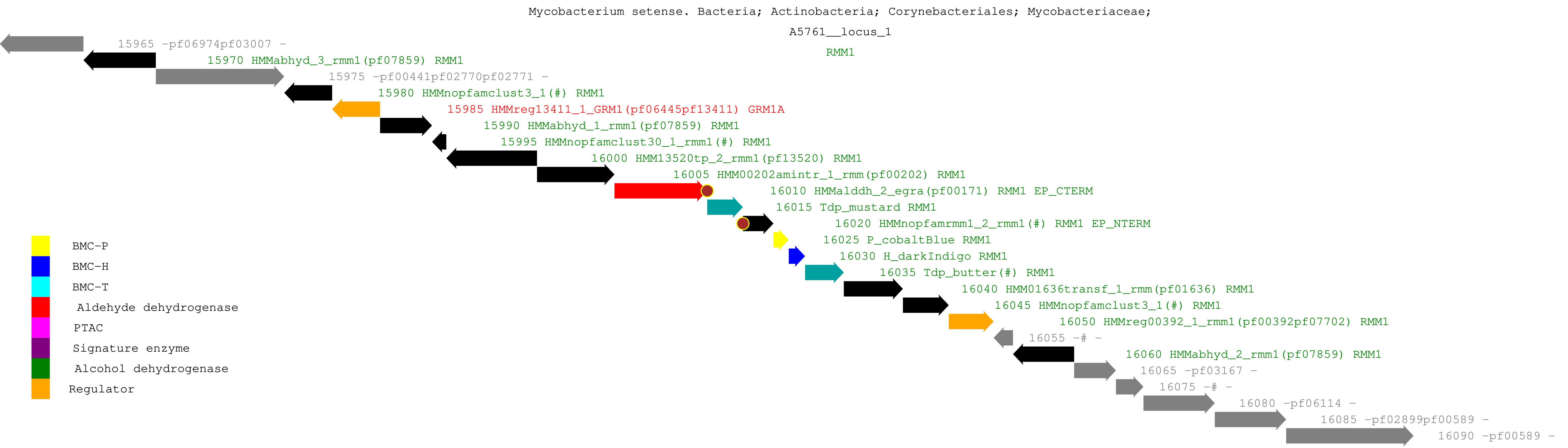
Mycobacterium setense. Bacteria; Actinobacteria; Corynebacteriales; Mycobacteriaceae;
A5761__locus_1
RMM1
15965 -pf06974pf03007 -
15970 HMMabhyd_3_rmm1(pf07859) RMM1
15975 -pf00441pf02770pf02771 -
15980 HMMnopfamclust3_1(#) RMM1
15985 HMMreg13411_1_GRM1(pf06445pf13411) GRM1A
15990 HMMabhyd_1_rmm1(pf07859) RMM1
15995 HMMnopfamclust30_1_rmm1(#) RMM1
16000 HMM13520tp_2_rmm1(pf13520) RMM1
16005 HMM00202amintr_1_rmm(pf00202) RMM1
16010 HMMalddh_2_egra(pf00171) RMM1 EP_CTERM
16015 Tdp_mustard RMM1
16020 HMMnopfamrmm1_2_rmm1(#) RMM1 EP_NTERM
16025 P_cobaltBlue RMM1
16030 H_darkIndigo RMM1
16035 Tdp_butter(#) RMM1
16040 HMM01636transf_1_rmm(pf01636) RMM1
16045 HMMnopfamclust3_1(#) RMM1
16050 HMMreg00392_1_rmm1(pf00392pf07702) RMM1
16055 -# -
16060 HMMabhyd_2_rmm1(pf07859) RMM1
16065 -pf03167 -
16075 -# -
16080 -pf06114 -
16085 -pf02899pf00589 -
16090 -pf00589 -
BMC-P
BMC-H
BMC-T
Aldehyde dehydrogenase
PTAC
Signature enzyme
Alcohol dehydrogenase
Regulator

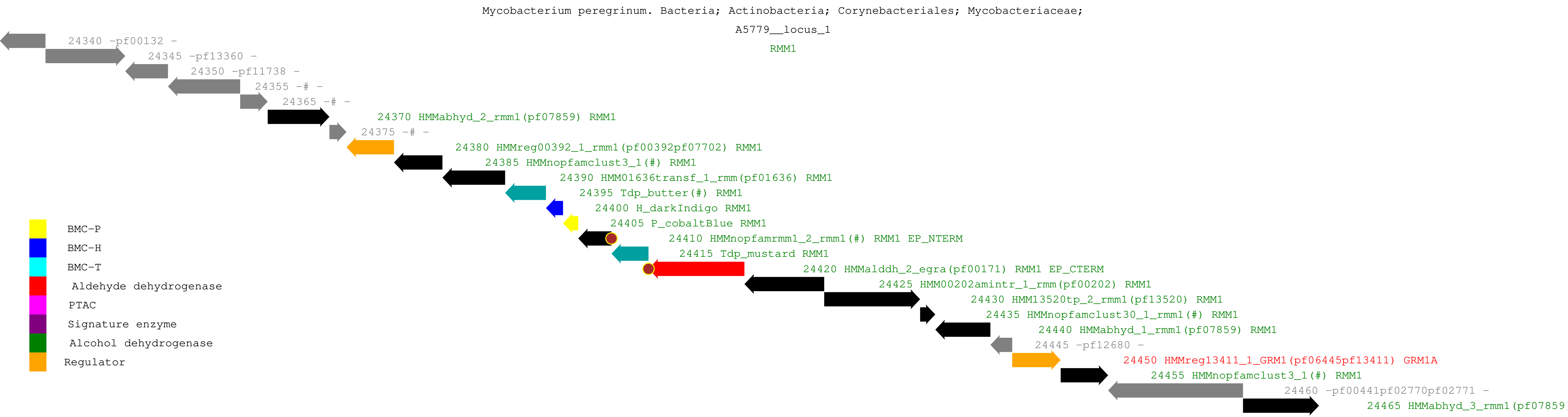
Mycobacterium peregrinum. Bacteria; Actinobacteria; Corynebacteriales; Mycobacteriaceae;
A5779__locus_1
RMM1
24340 -pf00132 -
24345 -pf13360 -
24350 -pf11738 -
24355 -# -
24365 -# -
24370 HMMabhyd_2_rmm1(pf07859) RMM1
24375 -# -
24380 HMMreg00392_1_rmm1(pf00392pf07702) RMM1
24385 HMMnopfamclust3_1(#) RMM1
24390 HMM01636transf_1_rmm(pf01636) RMM1
24395 Tdp_butter(#) RMM1
24400 H_darkIndigo RMM1
24405 P_cobaltBlue RMM1
24410 HMMnopfamrmm1_2_rmm1(#) RMM1 EP_NTERM
24415 Tdp_mustard RMM1
24420 HMMaldd h_2_egra(pf00171) RMM1 EP_CTERM
24425 HMM00202amintr_1_rmm(pf00202) RMM1
24430 HMM13520tp_2_rmm1(pf13520) RMM1
24435 HMMnopfamclust30_1_rmm1(#) RMM1
24440 HMMabhyd_1_rmm1(pf07859) RMM1
24445 -pf12680 -
24450 HMMreg13411_1_GRM1(pf06445pf13411) GRM1A
24455 HMMnopfamclust3_1(#) RMM1
24460 -pf00441pf02770pf02771 -
24465 HMMabhyd_3_rmm1(pf07859)
BMC-P
BMC-H
BMC-T
Aldehyde dehydrogenase
PTAC
Signature enzyme
Alcohol dehydrogenase
Regulator

Mycobacterium sp. 852013-50091_SCH5140682. Bacteria; Actinobacteria; Corynebacteriales; Mycobacteriaceae;

A5784__locus_1

RMM1

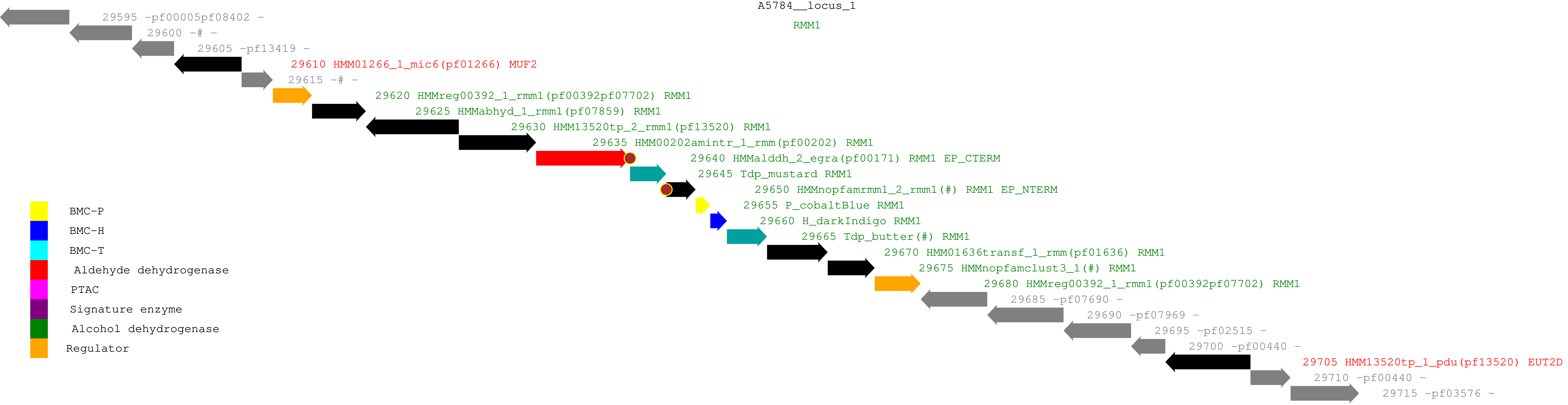

Mycobacterium peregrinum. Bacteria; Actinobacteria; Corynebacteriales; Mycobacteriaceae;

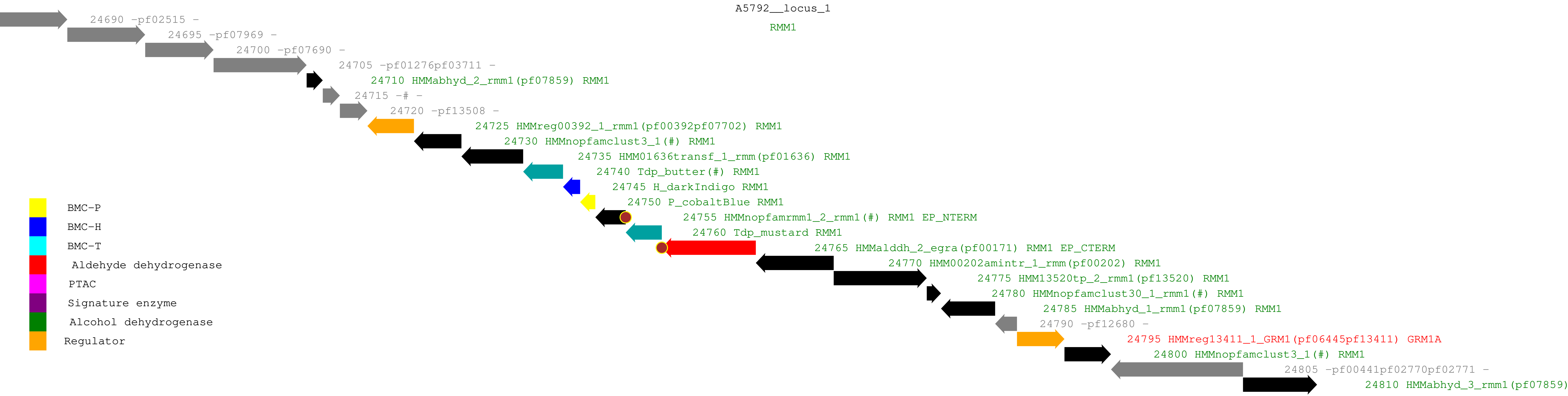

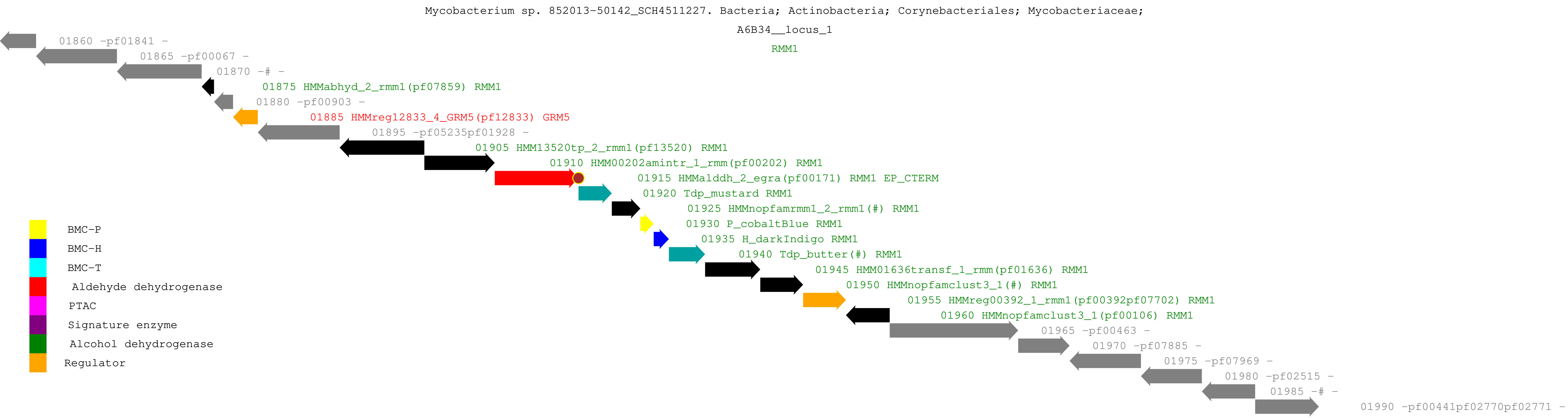
Mycobacterium sp. 852013-50142_SCH4511227. Bacteria; Actinobacteria; Corynebacteriales; Mycobacteriaceae;
A6B34__locus_1
RMM1
01860 -pf01841 -
01865 -pf00067 -
01870 -# -
01875 HMMabhyd_2_rmm1(pf07859) RMM1
01880 -pf00903 -
01885 HMMreg12833_4_GRM5(pf12833) GRM5
01895 -pf05235pf01928 -
01905 HMM13520tp_2_rmm1(pf13520) RMM1
01910 HMM00202amintr_1_rmm(pf00202) RMM1
01915 HMMalddh_2_egra(pf00171) RMM1 EP_CTERM
01920 Tdp_mustard RMM1
01925 HMMnopfamrmm1_2_rmm1(#) RMM1
01930 P_cobaltBlue RMM1
01935 H_darkIndigo RMM1
01940 Tdp_butter(#) RMM1
01945 HMM01636transf_1_rmm(pf01636) RMM1
01950 HMMnopfamclust3_1(#) RMM1
01955 HMMreg00392_1_rmm1(pf00392pf07702) RMM1
01960 HMMnopfamclust3_1(pf00106) RMM1
01965 -pf00463 -
01970 -pf07885 -
01975 -pf07969 -
01980 -pf02515 -
01985 -# -
01990 -pf00441pf02770pf02771 -
BMC-P
BMC-H
BMC-T
Aldehyde dehydrogenase
PTAC
Signature enzyme
Alcohol dehydrogenase
Regulator

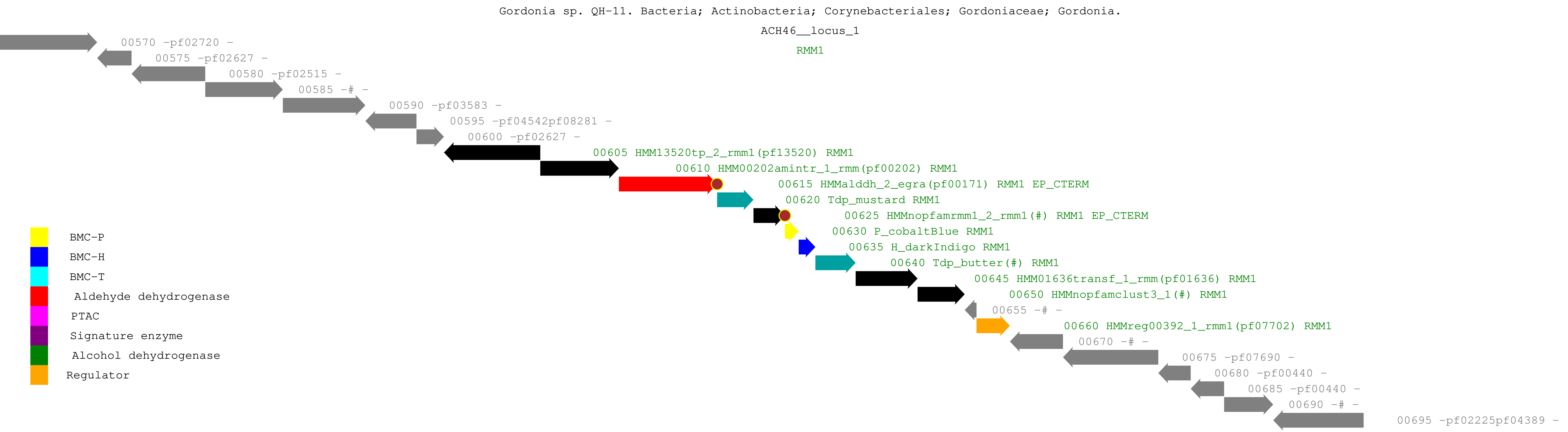
Gordonia sp. QH-11. Bacteria; Actinobacteria; Corynebacteriales; Gordoniaceae; Gordonia.
ACH46__locus_1
RMM1
00570 -pf02720 -
00575 -pf02627 -
00580 -pf02515 -
00585 -# -
00590 -pf03583 -
00595 -pf04542pf08281 -
00600 -pf02627 -
00605 HMM13520tp_2_rmm1(pf13520) RMM1
00610 HMM00202amintr_1_rmm(pf00202) RMM1
00615 HMMalddh_2_egra(pf00171) RMM1 EP_CTERM
00620 Tdp_mustard RMM1
00625 HMMnopfamrmm1_2_rmm1(#) RMM1 EP_CTERM
00630 P_cobaltBlue RMM1
00635 H_darkIndigo RMM1
00640 Tdp_butter(#) RMM1
00645 HMM01636transf_1_rmm(pf01636) RMM1
00650 HMMnopfamclust3_1(#) RMM1
00655 -# -
00660 HMMreg00392_1_rmm1(pf07702) RMM1
00670 -# -
00675 -pf07690 -
00680 -pf00440 -
00685 -pf00440 -
00690 -# -
00695 -pf02225pf04389 -
BMC-P
BMC-H
BMC-T
Aldehyde dehydrogenase
PTAC
Signature enzyme
Alcohol dehydrogenase
Regulator

Mycobacterium conceptionense. Bacteria; Actinobacteria; Corynebacteriales; Mycobacteriaceae;

ACT17__locus_1

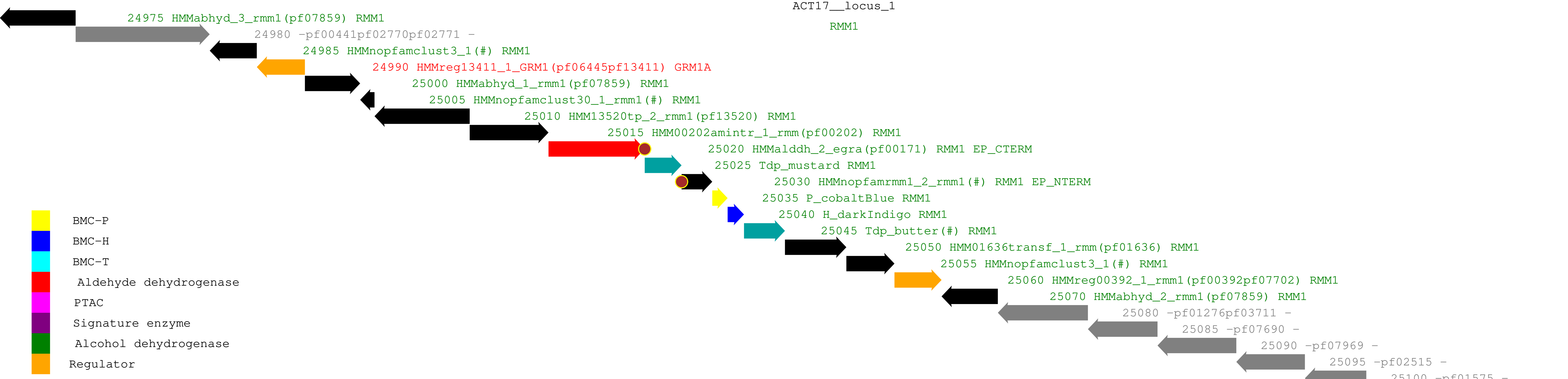

Mycobacterium goodii. Bacteria; Actinobacteria; Corynebacteriales; Mycobacteriaceae;

AFA91__locus_1

RMM1

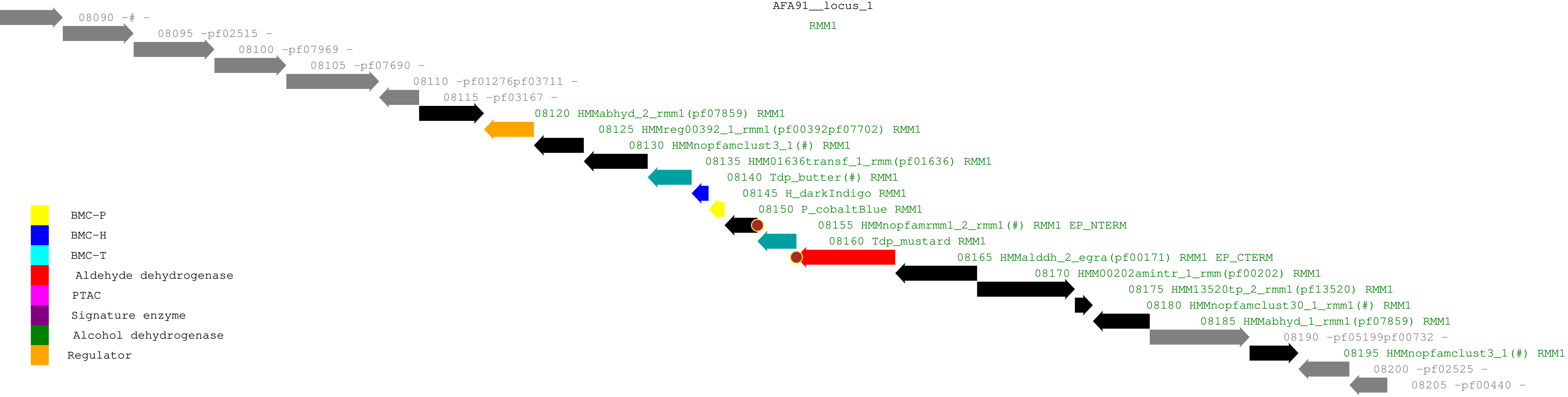

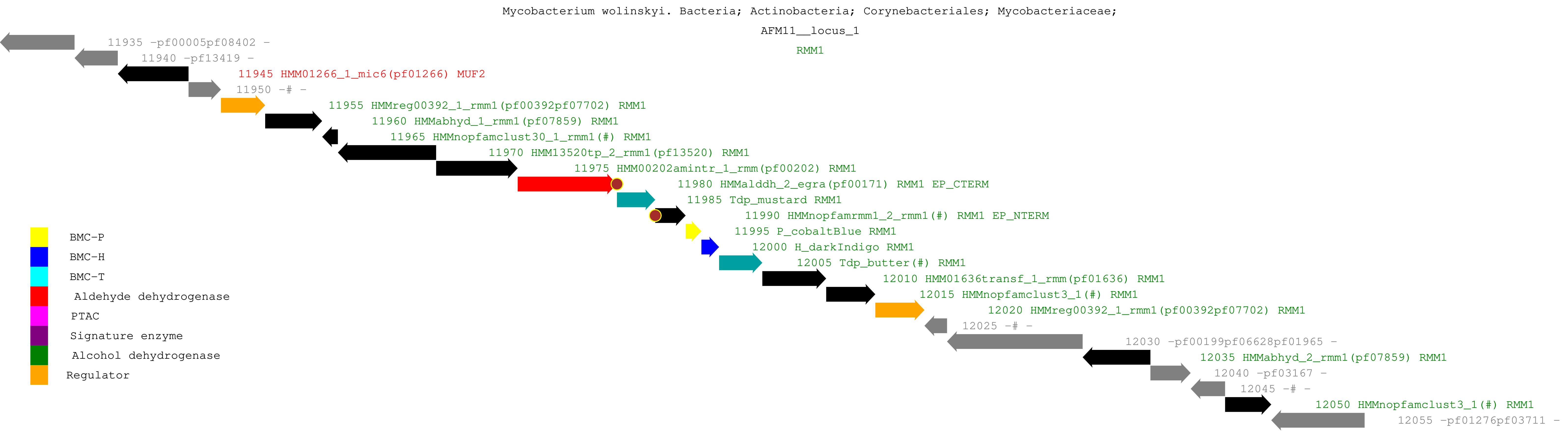
Mycobacterium wolinskyi. Bacteria; Actinobacteria; Corynebacteriales; Mycobacteriaceae;
AFM11__locus_1
RMM1
11935 -pf00005pf08402 -
11940 -pf13419 -
11945 HMM01266_1_mic6(pf01266) MUF2
11950 -# -
11955 HMMreg00392_1_rmm1(pf00392pf07702) RMM1
11960 HMMabhyd_1_rmm1(pf07859) RMM1
11965 HMMnopfamclust30_1_rmm1(#) RMM1
11970 HMM13520tp_2_rmm1(pf13520) RMM1
11975 HMM00202amintr_1_rmm(pf00202) RMM1
11980 HMMalddh_2_egra(pf00171) RMM1 EP_CTERM
11985 Tdp_mustard RMM1
11990 HMMnopfamrmm1_2_rmm1(#) RMM1 EP_NTERM
11995 P_cobaltBlue RMM1
12000 H_darkIndigo RMM1
12005 Tdp_butter(#) RMM1
12010 HMM01636transf_1_rmm(pf01636) RMM1
12015 HMMnopfamclust3_1(#) RMM1
12020 HMMreg00392_1_rmm1(pf00392pf07702) RMM1
12025 -# -
12030 -pf00199pf06628pf01965 -
12035 HMMabhyd_2_rmm1(pf07859) RMM1
12040 -pf03167 -
12045 -# -
12050 HMMnopfamclust3_1(#) RMM1
12055 -pf01276pf03711 -
BMC-P
BMC-H
BMC-T
Aldehyde dehydrogenase
PTAC
Signature enzyme
Alcohol dehydrogenase
Regulator

Nocardioides sp. Root79. Bacteria; Actinobacteria; Propionibacteriales; Nocardioidaceae;

ASE19__locus_1

RMM1

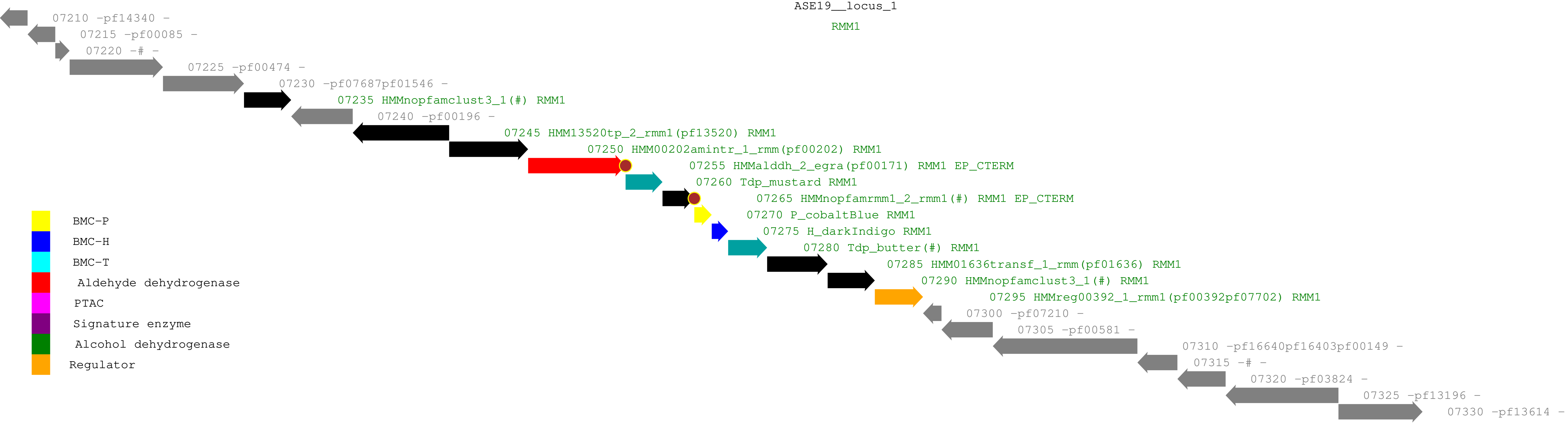

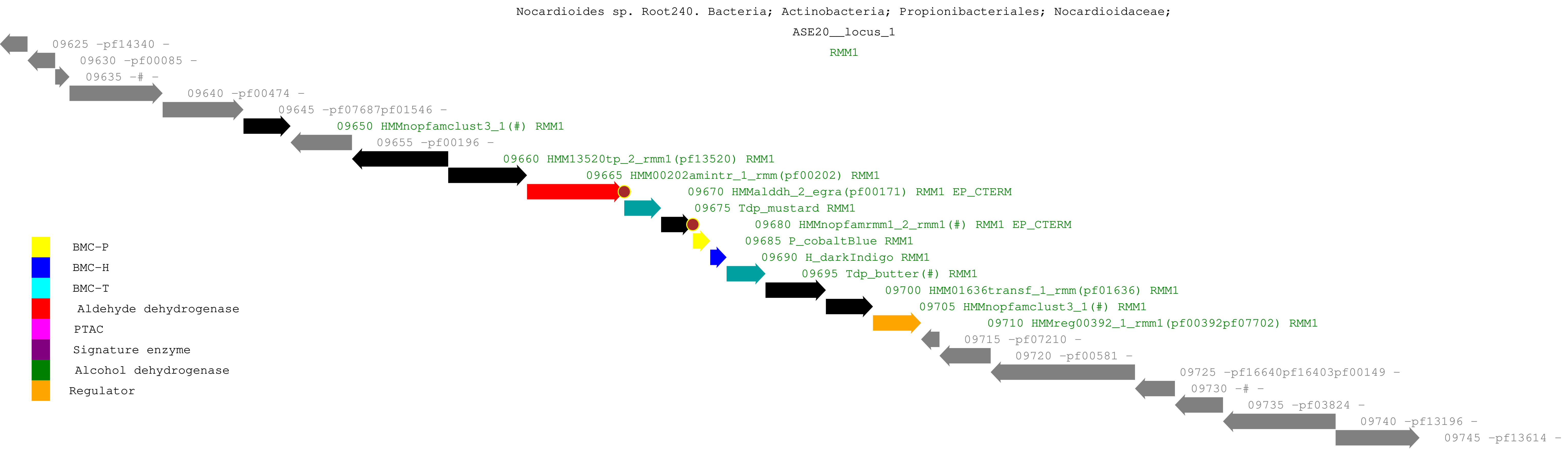
Nocardioides sp. Root240. Bacteria; Actinobacteria; Propionibacteriales; Nocardioidaceae;
ASE20__locus_1
RMM1
09625 -pf14340 -
09630 -pf00085 -
09635 -# -
09640 -pf00474 -
09645 -pf07687pf01546 -
09650 HMMnopfamclust3_1(#) RMM1
09655 -pf00196 -
09660 HMM13520tp_2_rmm1(pf13520) RMM1
09665 HMM00202amintr_1_rmm(pf00202) RMM1
09670 HMMalddh_2_egra(pf00171) RMM1 EP_CTERM
09675 Tdp_mustard RMM1
09680 HMMnopfamrmm1_2_rmm1(#) RMM1 EP_CTERM
09685 P_cobaltBlue RMM1
09690 H_darkIndigo RMM1
09695 Tdp_butter(#) RMM1
09700 HMM01636transf_1_rmm(pf01636) RMM1
09705 HMMnopfamclust3_1(#) RMM1
09710 HMMreg00392_1_rmm1(pf00392pf07702) RMM1
09715 -pf07210 -
09720 -pf00581 -
09725 -pf16640pf16403pf00149 -
09730 -# -
09735 -pf03824 -
09740 -pf13196 -
09745 -pf13614 -
BMC-P
BMC-H
BMC-T
Aldehyde dehydrogenase
PTAC
Signature enzyme
Alcohol dehydrogenase
Regulator

Mycobacterium sp. Soil538. Bacteria; Actinobacteria; Corynebacteriales; Mycobacteriaceae;

ASG82__locus_1

RMM1

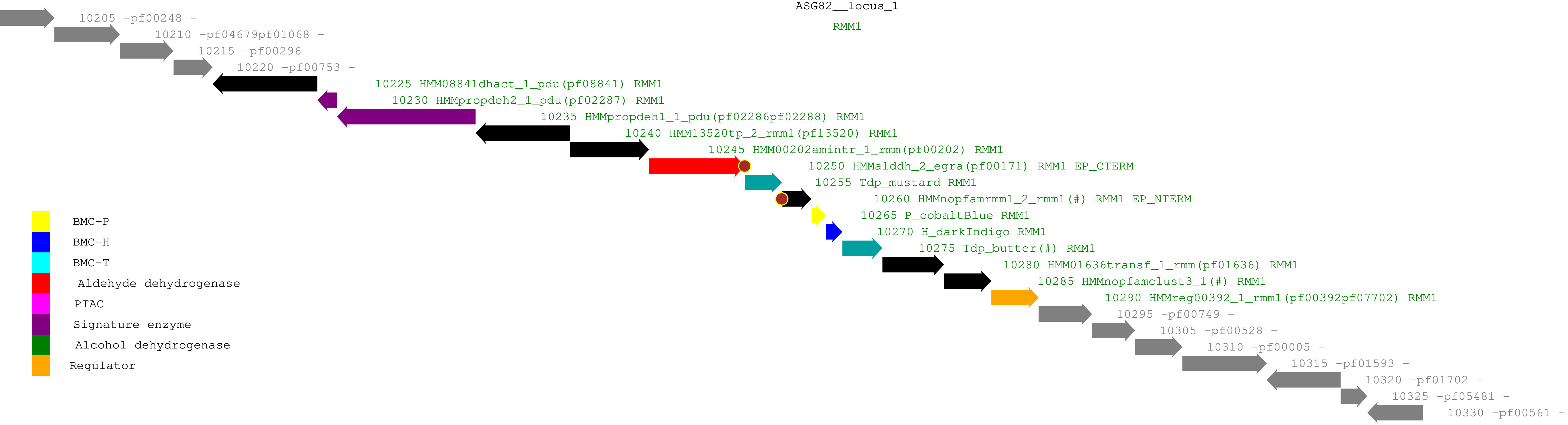

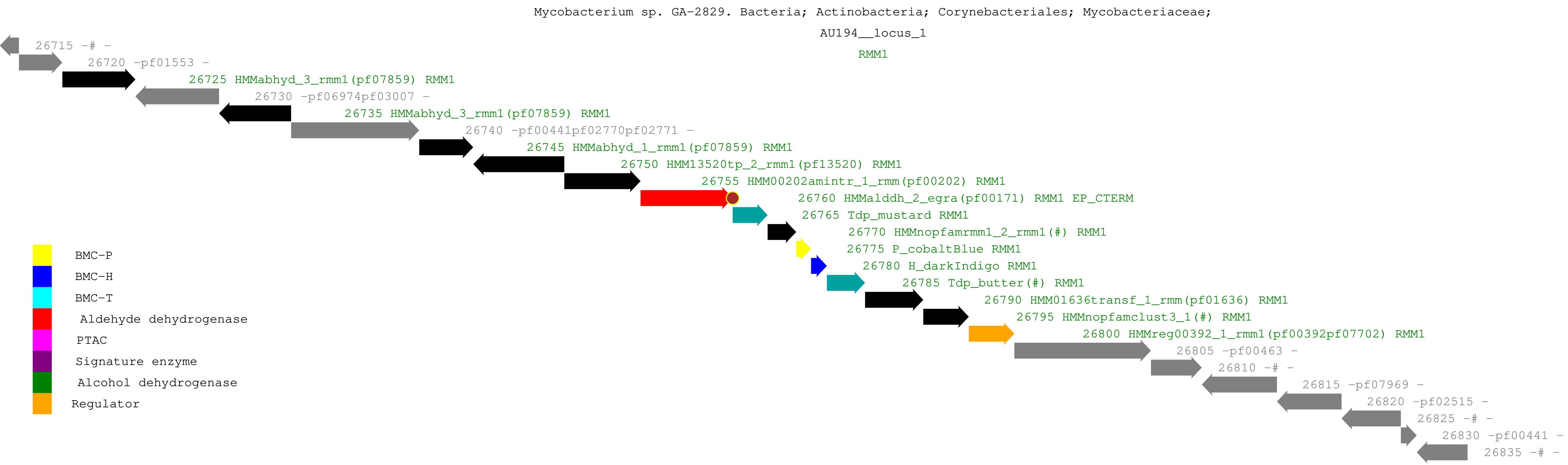
Mycobacterium sp. GA-2829. Bacteria; Actinobacteria; Corynebacteriales; Mycobacteriaceae;
AU194__locus_1
RMM1
26715 -# -
26720 -pf01553 -
26725 HMMabhyd_3_rmm1(pf07859) RMM1
26730 -pf06974pf03007 -
26735 HMMabhyd_3_rmm1(pf07859) RMM1
26740 -pf00441pf02770pf02771 -
26745 HMMabhyd_1_rmm1(pf07859) RMM1
26750 HMM13520tp_2_rmm1(pf13520) RMM1
26755 HMM00202amintr_1_rmm(pf00202) RMM1
26760 HMMalddh_2_egra(pf00171) RMM1 EP_CTERM
26765 Tdp_mustard RMM1
26770 HMMnopfamrmm1_2_rmm1(#) RMM1
26775 P_cobaltBlue RMM1
26780 H_darkIndigo RMM1
26785 Tdp_butter(#) RMM1
26790 HMM01636transf_1_rmm(pf01636) RMM1
26795 HMMnopfamclust3_1(#) RMM1
26800 HMMreg00392_1_rmm1(pf00392pf07702) RMM1
26805 -pf00463 -
26810 -# -
26815 -pf07969 -
26820 -pf02515 -
26825 -# -
26830 -pf00441 -
26835 -# -
BMC-P
BMC-H
BMC-T
Aldehyde dehydrogenase
PTAC
Signature enzyme
Alcohol dehydrogenase
Regulator

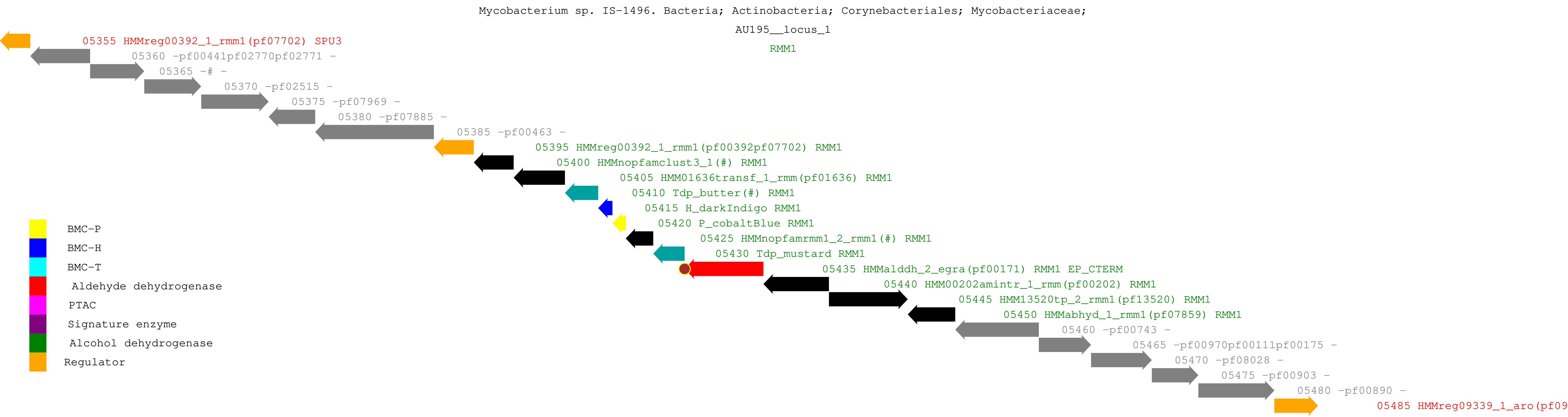
Mycobacterium sp. IS-1496. Bacteria; Actinobacteria; Corynebacteriales; Mycobacteriaceae;
AU195__locus_1
RMM1
05355 HMMreg00392_1_rmm1(pf07702) SPU3
05360 -pf00441pf02770pf02771 -
05365 -# -
05370 -pf02515 -
05375 -pf07969 -
05380 -pf07885 -
05385 -pf00463 -
05395 HMMreg00392_1_rmm1(pf00392pf07702) RMM1
05400 HMMnopfamclust3_1(#) RMM1
05405 HMM01636transf_1_rmm(pf01636) RMM1
05410 Tdp_butter(#) RMM1
05415 H_darkIndigo RMM1
05420 P_cobaltBlue RMM1
05425 HMMnopfamrmm1_2_rmm1(#) RMM1
05430 Tdp_mustard RMM1
05435 HMMalddh_2_egra(pf00171) RMM1 EP_CTERM
05440 HMM00202amintr_1_rmm(pf00202) RMM1
05445 HMM13520tp_2_rmm1(pf13520) RMM1
05450 HMMabhyd_1_rmm1(pf07859) RMM1
05460 -pf00743 -
05465 -pf00970pf00111pf00175 -
05470 -pf08028 -
05475 -pf00903 -
05480 -pf00890 -
05485 HMMreg09339_1_aro(pf09
BMC-P
BMC-H
BMC-T
Aldehyde dehydrogenase
PTAC
Signature enzyme
Alcohol dehydrogenase
Regulator

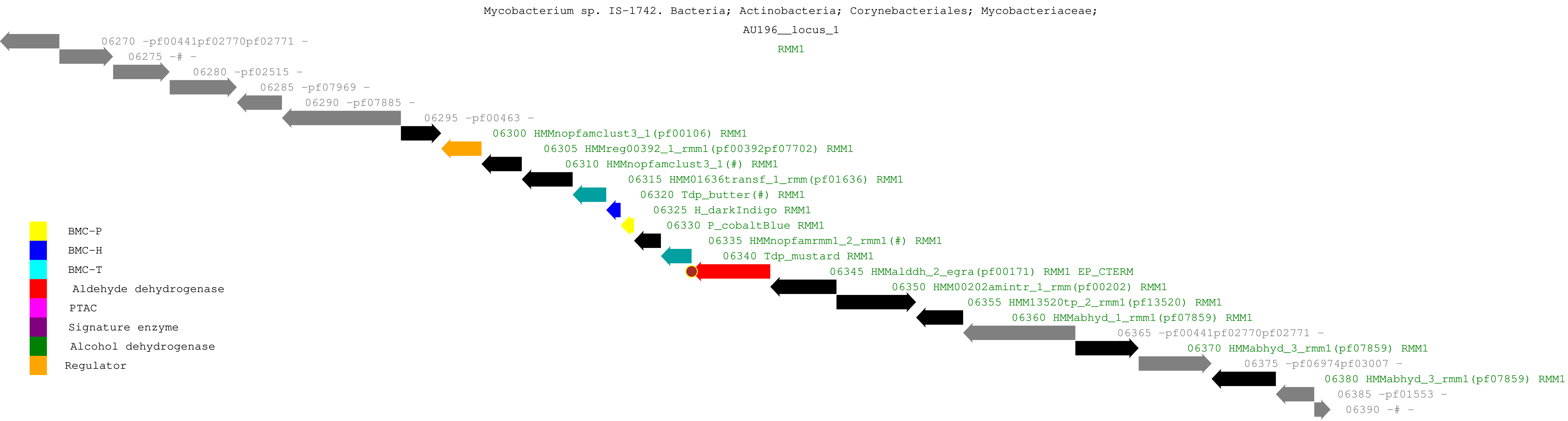
Mycobacterium sp. IS-1742. Bacteria; Actinobacteria; Corynebacteriales; Mycobacteriaceae;
AU196__locus_1
RMM1
06270 -pf00441pf02770pf02771 -
06275 -# -
06280 -pf02515 -
06285 -pf07969 -
06290 -pf07885 -
06295 -pf00463 -
06300 HMMnopfamclust3_1(pf00106) RMM1
06305 HMMreg00392_1_rmm1(pf00392pf07702) RMM1
06310 HMMnopfamclust3_1(#) RMM1
06315 HMM01636transf_1_rmm(pf01636) RMM1
06320 Tdp_butter(#) RMM1
06325 H_darkIndigo RMM1
06330 P_cobaltBlue RMM1
06335 HMMnopfamrmm1_2_rmm1(#) RMM1
06340 Tdp_mustard RMM1
06345 HMMalddh_2_egra(pf00171) RMM1 EP_CTERM
06350 HMM00202amintr_1_rmm(pf00202) RMM1
06355 HMM13520tp_2_rmm1(pf13520) RMM1
06360 HMMabhyd_1_rmm1(pf07859) RMM1
06365 -pf00441pf02770pf02771 -
06370 HMMabhyd_3_rmm1(pf07859) RMM1
06375 -pf06974pf03007 -
06380 HMMabhyd_3_rmm1(pf07859) RMM1
06385 -pf01553 -
06390 -# -
BMC-P
BMC-H
BMC-T
Aldehyde dehydrogenase
PTAC
Signature enzyme
Alcohol dehydrogenase
Regulator

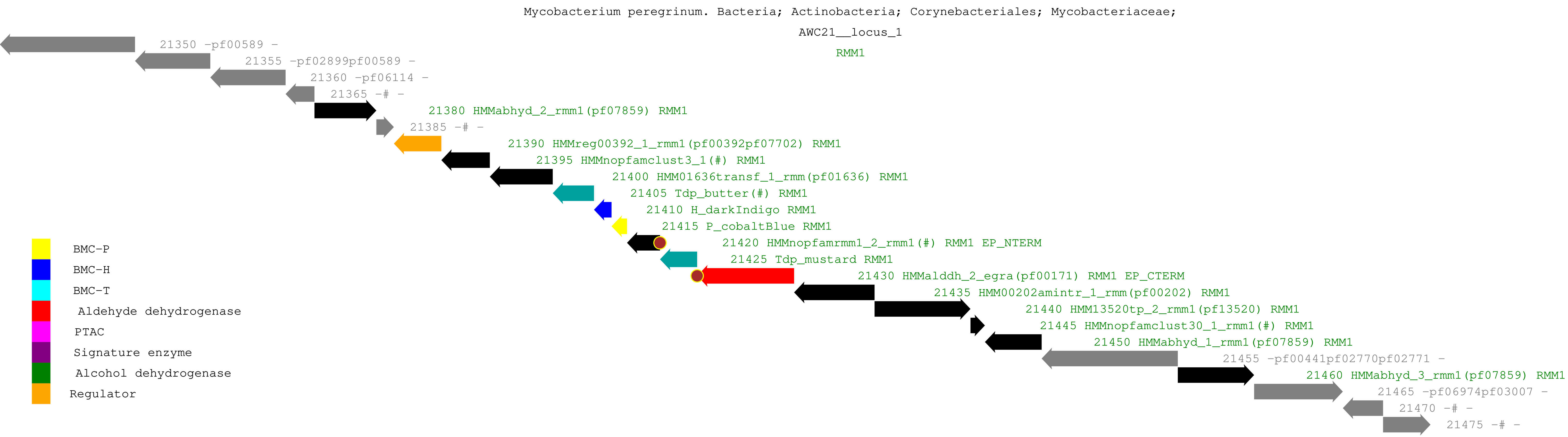
Mycobacterium peregrinum. Bacteria; Actinobacteria; Corynebacteriales; Mycobacteriaceae;
AWC21__locus_1
RMM1
21350 -pf00589 -
21355 -pf02899pf00589 -
21360 -pf06114 -
21365 -# -
21380 HMMabhyd_2_rmm1(pf07859) RMM1
21385 -# -
21390 HMMreg00392_1_rmm1(pf00392pf07702) RMM1
21395 HMMnopfamclust3_1(#) RMM1
21400 HMM01636transf_1_rmm(pf01636) RMM1
21405 Tdp_butter(#) RMM1
21410 H_darkIndigo RMM1
21415 P_cobaltBlue RMM1
21420 HMMnopfamrmm1_2_rmm1(#) RMM1 EP_NTERM
21425 Tdp_mustard RMM1
21430 HMMalddh_2_egra(pf00171) RMM1 EP_CTERM
21435 HMM00202amintr_1_rmm(pf00202) RMM1
21440 HMM13520tp_2_rmm1(pf13520) RMM1
21445 HMMnopfamclust30_1_rmm1(#) RMM1
21450 HMMabhyd_1_rmm1(pf07859) RMM1
21455 -pf00441pf02770pf02771 -
21460 HMMabhyd_3_rmm1(pf07859) RMM1
21465 -pf06974pf03007 -
21470 -# -
21475 -# -
BMC-P
BMC-H
BMC-T
Aldehyde dehydrogenase
PTAC
Signature enzyme
Alcohol dehydrogenase
Regulator

Mycobacterium wolinskyi. Bacteria; Actinobacteria; Corynebacteriales; Mycobacteriaceae;

AWC31__locus_1

RMM1

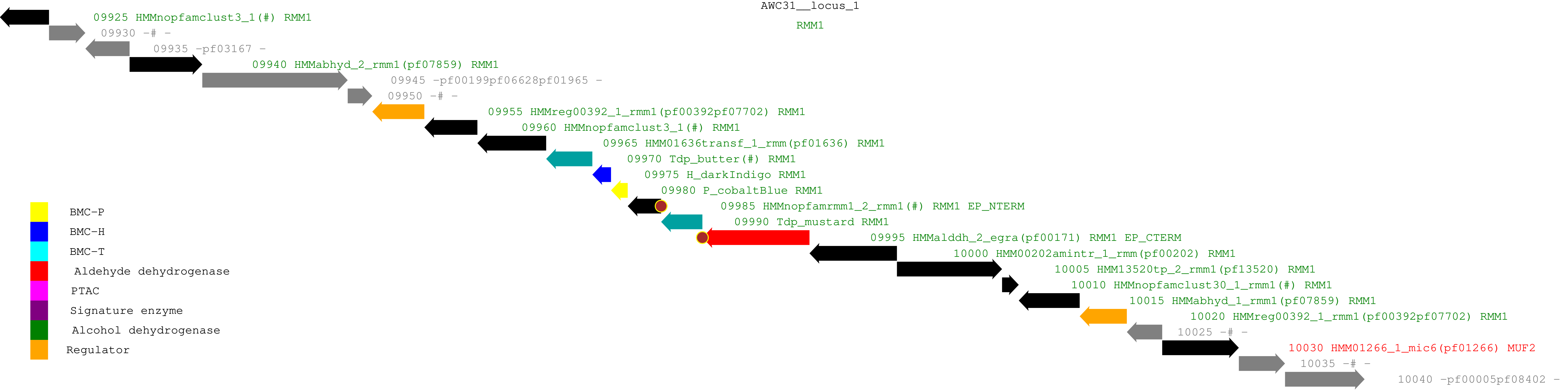

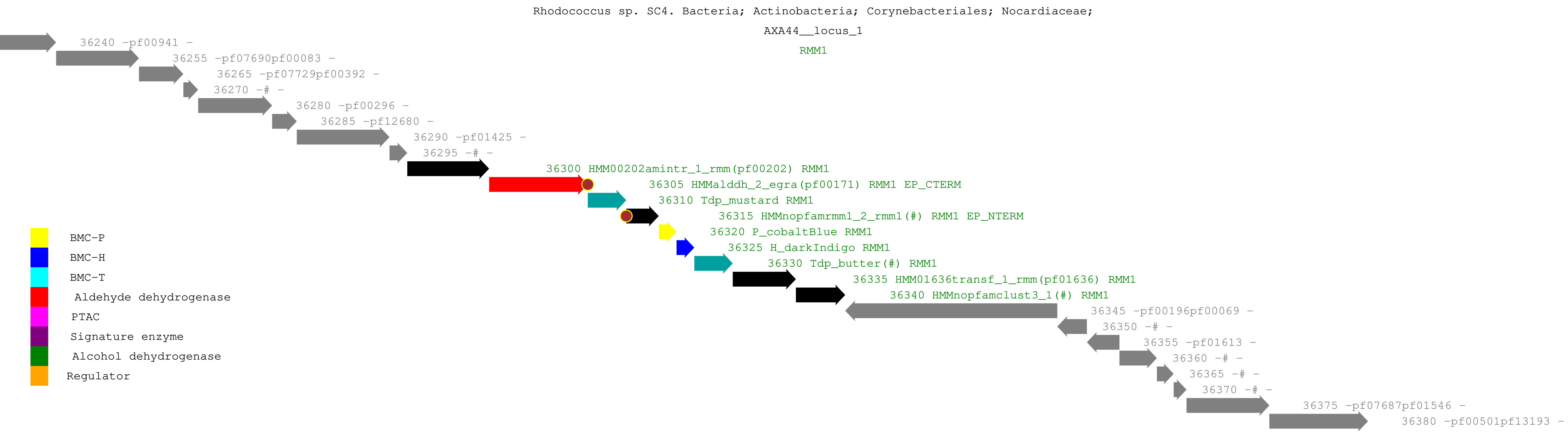
Rhodococcus sp. SC4. Bacteria; Actinobacteria; Corynebacteriales; Nocardiaceae;
AXA44__locus_1
RMM1
36240 -pf00941 -
36255 -pf07690pf00083 -
36265 -pf07729pf00392 -
36270 -# -
36280 -pf00296 -
36285 -pf12680 -
36290 -pf01425 -
36295 -# -
36300 HMM00202amintr_1_rmm(pf00202) RMM1
36305 HMMalddh_2_egra(pf00171) RMM1 EP_CTERM
36310 Tdp_mustard RMM1
36315 HMMnopfamrmm1_2_rmm1(#) RMM1 EP_NTERM
36320 P_cobaltBlue RMM1
36325 H_darkIndigo RMM1
36330 Tdp_butter(#) RMM1
36335 HMM01636transf_1_rmm(pf01636) RMM1
36340 HMMnopfamclust3_1(#) RMM1
36345 -pf00196pf00069 -
36350 -# -
36355 -pf01613 -
36360 -# -
36365 -# -
36370 -# -
36375 -pf07687pf01546 -
36380 -pf00501pf13193 -
BMC-P
BMC-H
BMC-T
Aldehyde dehydrogenase
PTAC
Signature enzyme
Alcohol dehydrogenase
Regulator

Vibrio europaeus. Bacteria; Proteobacteria; Gammaproteobacteria; Vibrionales;

AZ468__locus_1

RMM1

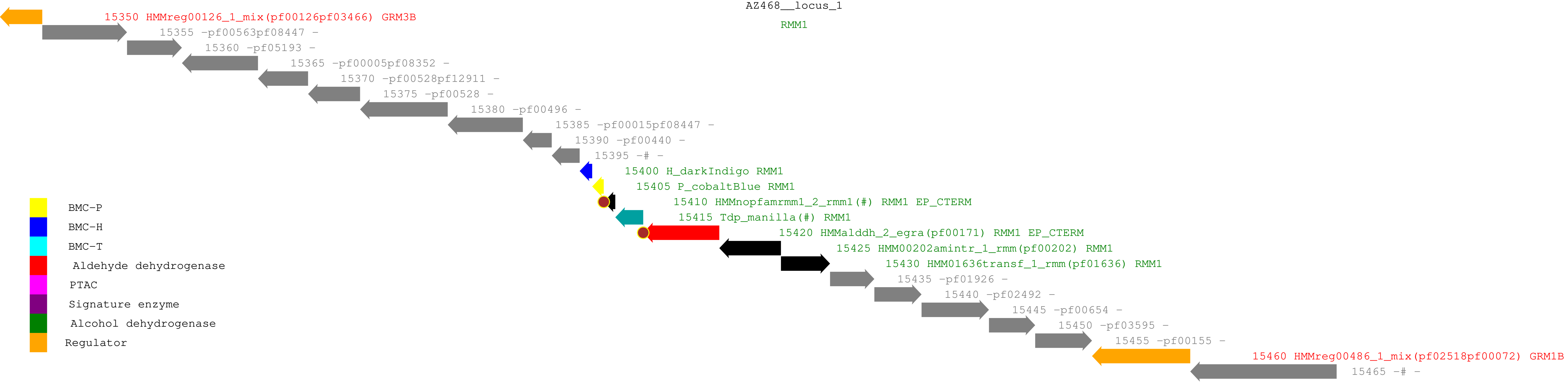

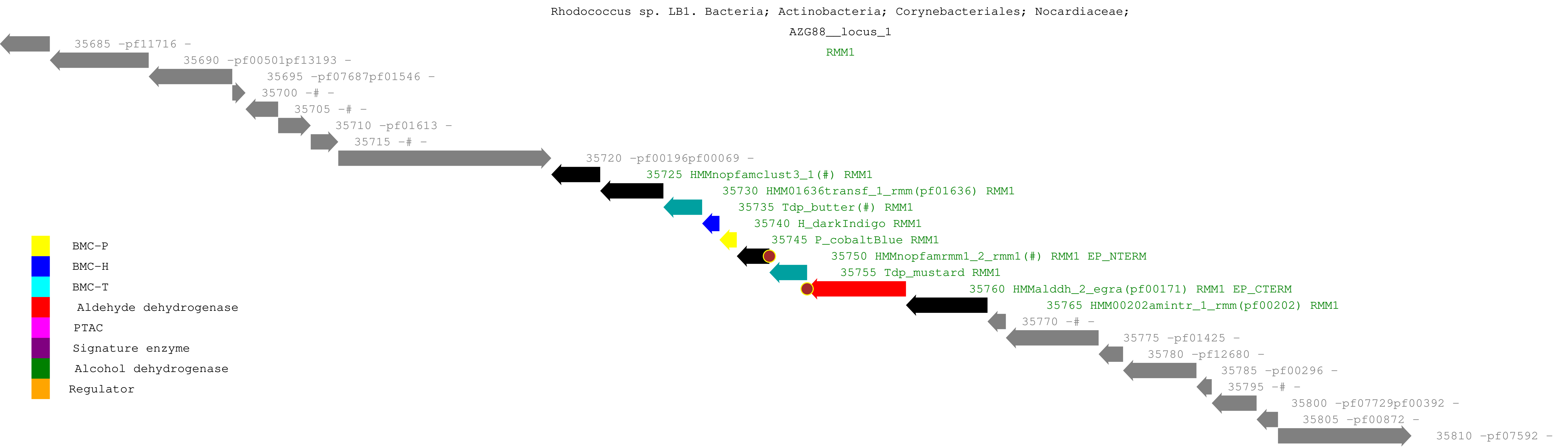
Rhodococcus sp. LB1. Bacteria; Actinobacteria; Corynebacteriales; Nocardiaceae;
AZG88__locus_1
RMM1
35685 -pf11716 -
35690 -pf00501pf13193 -
35695 -pf07687pf01546 -
35700 -# -
35705 -# -
35710 -pf01613 -
35715 -# -
35720 -pf00196pf00069 -
35725 HMMnopfamclust3_1(#) RMM1
35730 HMM01636transf_1_rmm(pf01636) RMM1
35735 Tdp_butter(#) RMM1
35740 H_darkIndigo RMM1
35745 P_cobaltBlue RMM1
35750 HMMnopfamrmm1_2_rmm1(#) RMM1 EP_NTERM
35755 Tdp_mustard RMM1
35760 HMMalddh_2_egra(pf00171) RMM1 EP_CTERM
35765 HMM00202amintr_1_rmm(pf00202) RMM1
35770 -# -
35775 -pf01425 -
35780 -pf12680 -
35785 -pf00296 -
35795 -# -
35800 -pf07729pf00392 -
35805 -pf00872 -
35810 -pf07592 -
BMC-P
BMC-H
BMC-T
Aldehyde dehydrogenase
PTAC
Signature enzyme
Alcohol dehydrogenase
Regulator

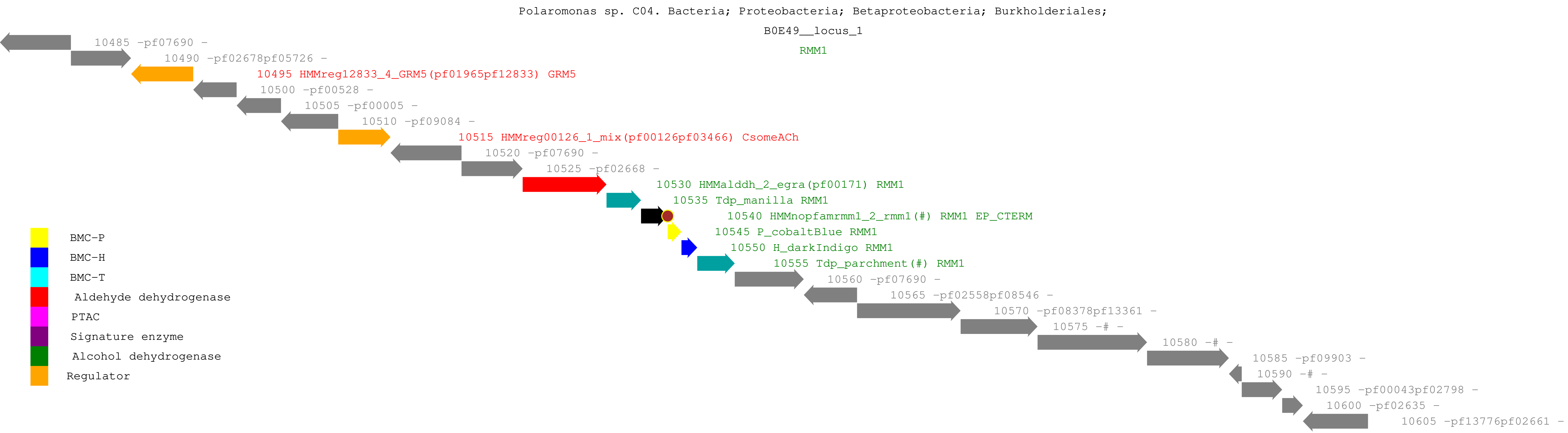
Polaromonas sp. C04. Bacteria; Proteobacteria; Betaproteobacteria; Burkholderiales;
B0E49__locus_1
RMM1
10485 -pf07690 -
10490 -pf02678pf05726 -
10495 HMMreg12833_4_GRM5(pf01965pf12833) GRM5
10500 -pf00528 -
10505 -pf00005 -
10510 -pf09084 -
10515 HMMreg00126_1_mix(pf00126pf03466) CsomeACh
10520 -pf07690 -
10525 -pf02668 -
10530 HMMalddh_2_egra(pf00171) RMM1
10535 Tdp_manilla RMM1
10540 HMMnopfamrmm1_2_rmm1(#) RMM1 EP_CTERM
10545 P_cobaltBlue RMM1
10550 H_darkIndigo RMM1
10555 Tdp_parchment(#) RMM1
10560 -pf07690 -
10565 -pf02558pf08546 -
10570 -pf08378pf13361 -
10575 -# -
10580 -# -
10585 -pf09903 -
10590 -# -
10595 -pf00043pf02798 -
10600 -pf02635 -
10605 -pf13776pf02661 -
BMC-P
BMC-H
BMC-T
Aldehyde dehydrogenase
PTAC
Signature enzyme
Alcohol dehydrogenase
Regulator

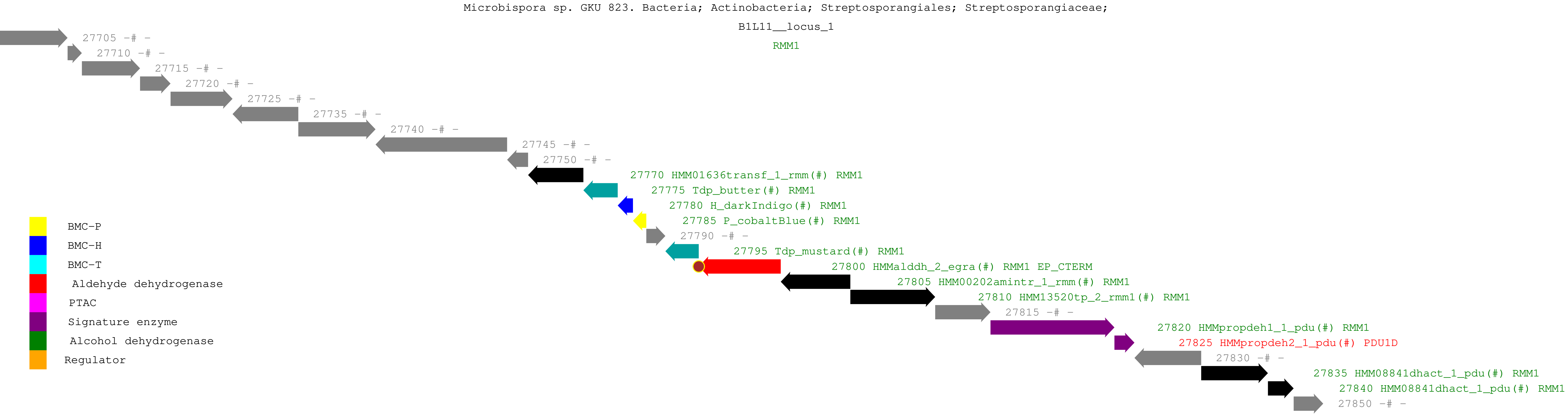

Microbispora sp. GKU 823. Bacteria; Actinobacteria; Streptosporangiales; Streptosporangiaceae;
B1L11__locus_1
RMM1
27705 -# -
27710 -# -
27715 -# -
27720 -# -
27725 -# -
27735 -# -
27740 -# -
27745 -# -
27750 -# -
27770 HMM01636transf_1_rmm(#) RMM1
27775 Tdp_butter(#) RMM1
27780 H_darkIndigo(#) RMM1
27785 P_cobaltBlue(#) RMM1
27790 -# -
27795 Tdp_mustard(#) RMM1
27800 HMMalddh_2_egra(#) RMM1 EP_CTERM
27805 HMM00202amintr_1_rmm(#) RMM1
27810 HMM13520tp_2_rmm1(#) RMM1
27815 -# -
27820 HMMpropdeh1_1_pdu(#) RMM1
27825 HMMpropdeh2_1_pdu(#) PDU1D
27830 -# -
27835 HMM08841dhact_1_pdu(#) RMM1
27840 HMM08841dhact_1_pdu(#) RMM1
27850 -# -
BMC-P
BMC-H
BMC-T
Aldehyde dehydrogenase
PTAC
Signature enzyme
Alcohol dehydrogenase
Regulator

Limnohabitans sp. Rim28. Bacteria; Proteobacteria; Betaproteobacteria; Burkholderiales;

B472__locus_1

RMM1

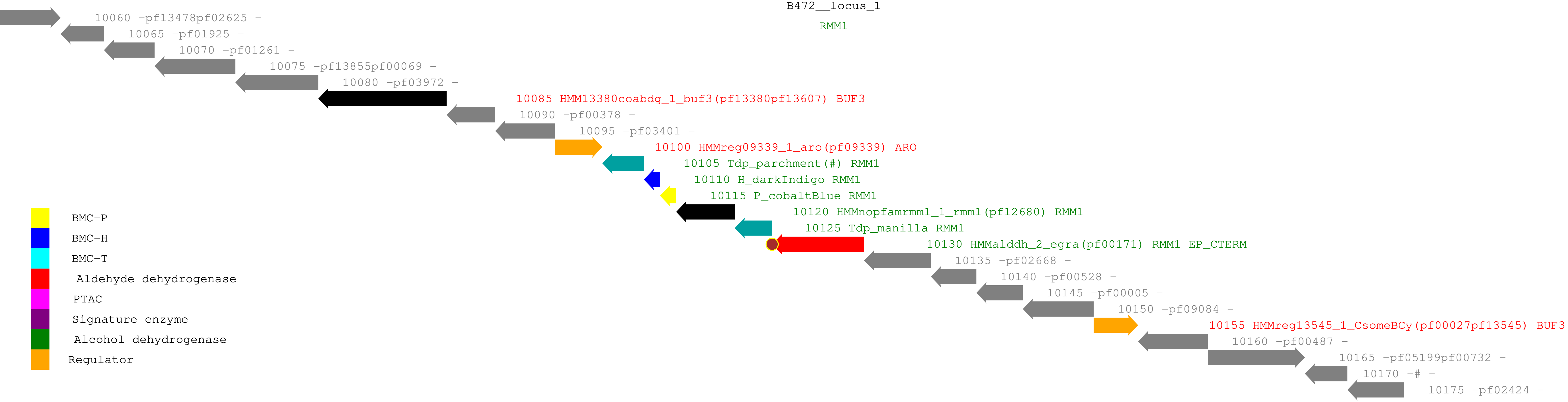

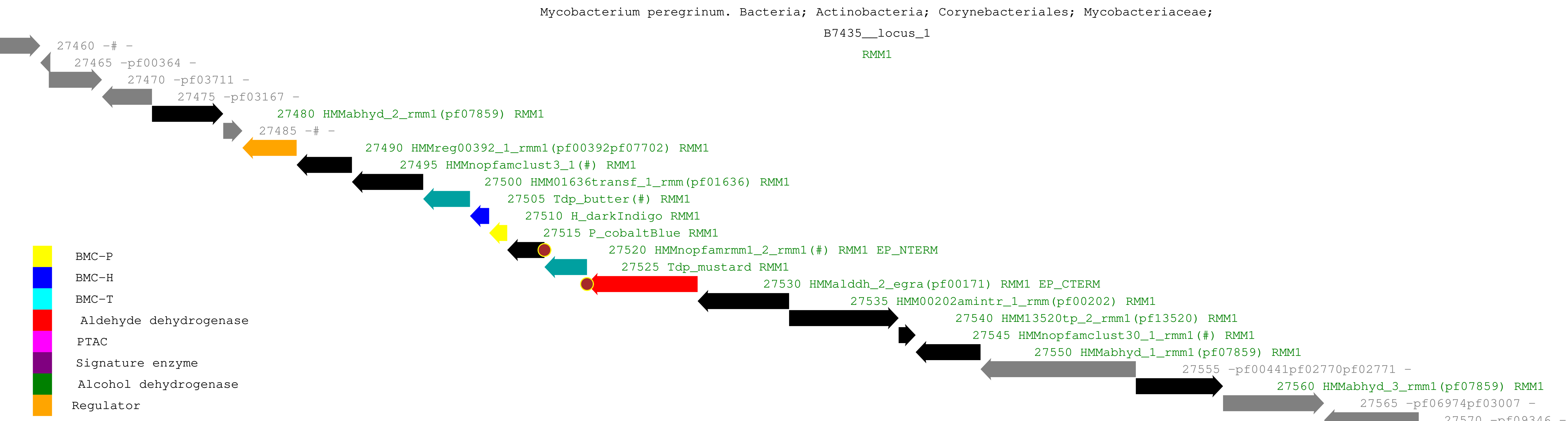
Mycobacterium peregrinum. Bacteria; Actinobacteria; Corynebacteriales; Mycobacteriaceae;
B7435__locus_1
RMM1
27460 -# -
27465 -pf00364 -
27470 -pf03711 -
27475 -pf03167 -
27480 HMMabhyd_2_rmm1(pf07859) RMM1
27485 -# -
27490 HMMreg00392_1_rmm1(pf00392pf07702) RMM1
27495 HMMnopfamclust3_1(#) RMM1
27500 HMM01636transf_1_rmm(pf01636) RMM1
27505 Tdp_butter(#) RMM1
27510 H_darkIndigo RMM1
27515 P_cobaltBlue RMM1
27520 HMMnopfamrmm1_2_rmm1(#) RMM1 EP_NTERM
27525 Tdp_mustard RMM1
27530 HMMalddh_2_egra(pf00171) RMM1 EP_CTERM
27535 HMM00202amintr_1_rmm(pf00202) RMM1
27540 HMM13520tp_2_rmm1(pf13520) RMM1
27545 HMMnopfamclust30_1_rmm1(#) RMM1
27550 HMMabhyd_1_rmm1(pf07859) RMM1
27555 -pf00441pf02770pf02771 -
27560 HMMabhyd_3_rmm1(pf07859) RMM1
27565 -pf06974pf03007 -
27570 -pf09346 -
27575 -# -
BMC-P
BMC-H
BMC-T
Aldehyde dehydrogenase
PTAC
Signature enzyme
Alcohol dehydrogenase
Regulator

Limnohabitans sp. T6-20. Bacteria; Proteobacteria; Betaproteobacteria; Burkholderiales;

B9Z33__locus_1

RMM1

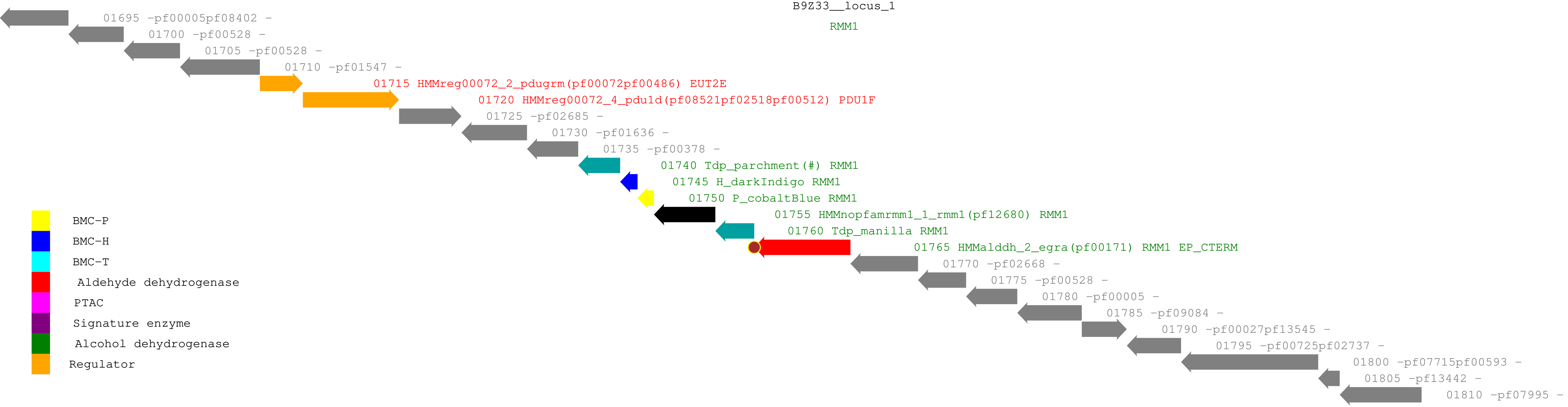

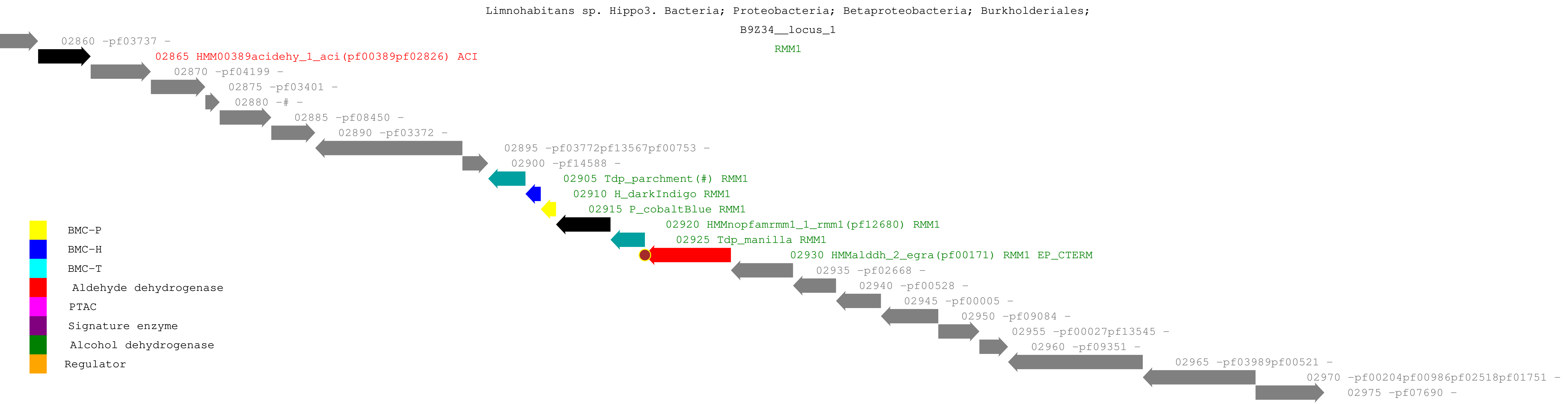
Limnohabitans sp. Hippo3. Bacteria; Proteobacteria; Betaproteobacteria; Burkholderiales;
B9Z34__locus_1
RMM1
02860 -pf03737 -
02865 HMM00389acidehy_1_aci(pf00389pf02826) ACI
02870 -pf04199 -
02875 -pf03401 -
02880 -# -
02885 -pf08450 -
02890 -pf03372 -
02895 -pf03772pf13567pf00753 -
02900 -pf14588 -
02905 Tdp_parchment(#) RMM1
02910 H_darkIndigo RMM1
02915 P_cobaltBlue RMM1
02920 HMMnopfamrmm1_1_rmm1(pf12680) RMM1
02925 Tdp_manilla RMM1
02930 HMMalddh_2_egra(pf00171) RMM1 EP_CTERM
02935 -pf02668 -
02940 -pf00528 -
02945 -pf00005 -
02950 -pf09084 -
02955 -pf00027pf13545 -
02960 -pf09351 -
02965 -pf03989pf00521 -
02970 -pf00204pf00986pf02518pf01751 -
02975 -pf07690 -
BMC-P
BMC-H
BMC-T
Aldehyde dehydrogenase
PTAC
Signature enzyme
Alcohol dehydrogenase
Regulator

Limnohabitans sp. MMS-10A-160. Bacteria; Proteobacteria; Betaproteobacteria; Burkholderiales;

B9Z38__locus_1

RMM1

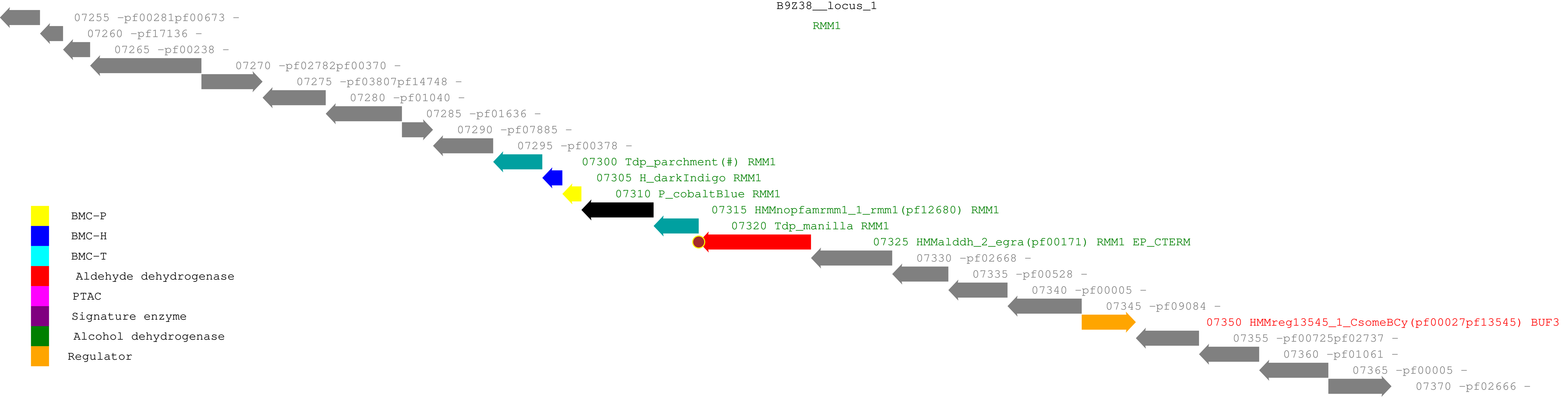

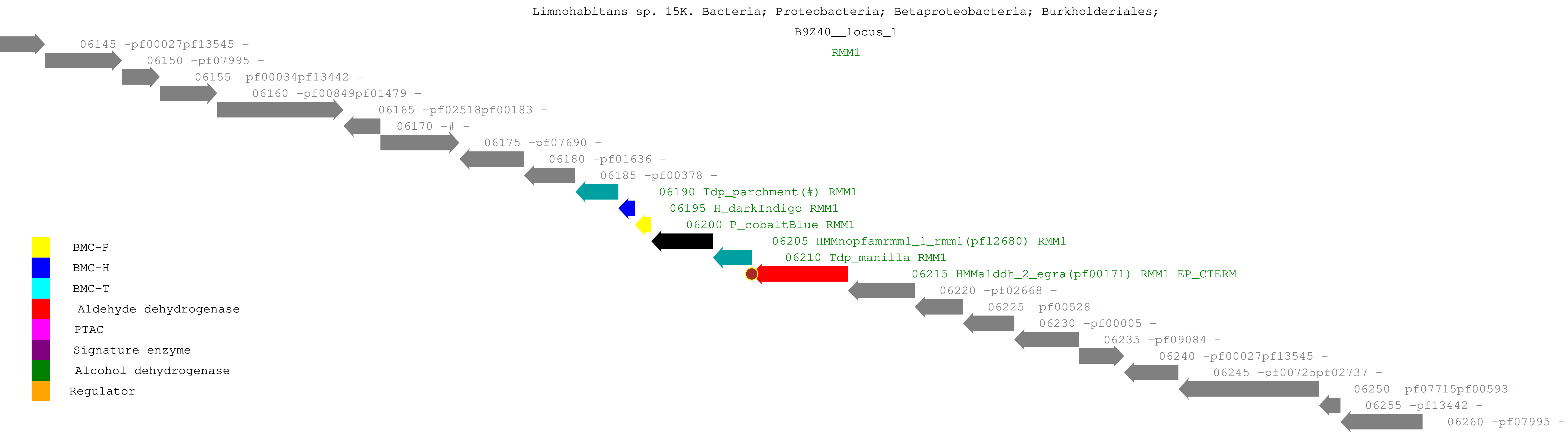

Limnohabitans sp. 15K. Bacteria; Proteobacteria; Betaproteobacteria; Burkholderiales;
B9Z40__locus_1
RMM1
06145 -pf00027pf13545 -
06150 -pf07995 -
06155 -pf00034pf13442 -
06160 -pf00849pf01479 -
06165 -pf02518pf00183 -
06170 -# -
06175 -pf07690 -
06180 -pf01636 -
06185 -pf00378 -
06190 Tdp_parchment(#) RMM1
06195 H_darkIndigo RMM1
06200 P_cobaltBlue RMM1
06205 HMMnopfamrmm1_1_rmm1(pf12680) RMM1
06210 Tdp_manilla RMM1
06215 HMMalddh_2_egra(pf00171) RMM1 EP_CTERM
06220 -pf02668 -
06225 -pf00528 -
06230 -pf00005 -
06235 -pf09084 -
06240 -pf00027pf13545 -
06245 -pf00725pf02737 -
06250 -pf07715pf00593 -
06255 -pf13442 -
06260 -pf07995 -
BMC-P
BMC-H
BMC-T
Aldehyde dehydrogenase
PTAC
Signature enzyme
Alcohol dehydrogenase
Regulator

Limnohabitans sp. MMS-10A-192. Bacteria; Proteobacteria; Betaproteobacteria; Burkholderiales;

B9Z43__locus_1

RMM1

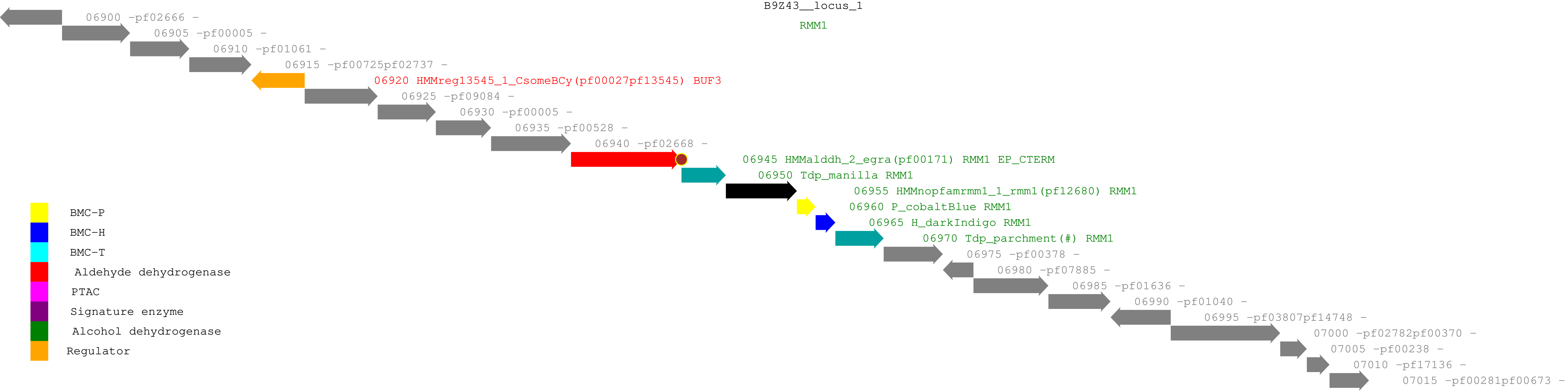

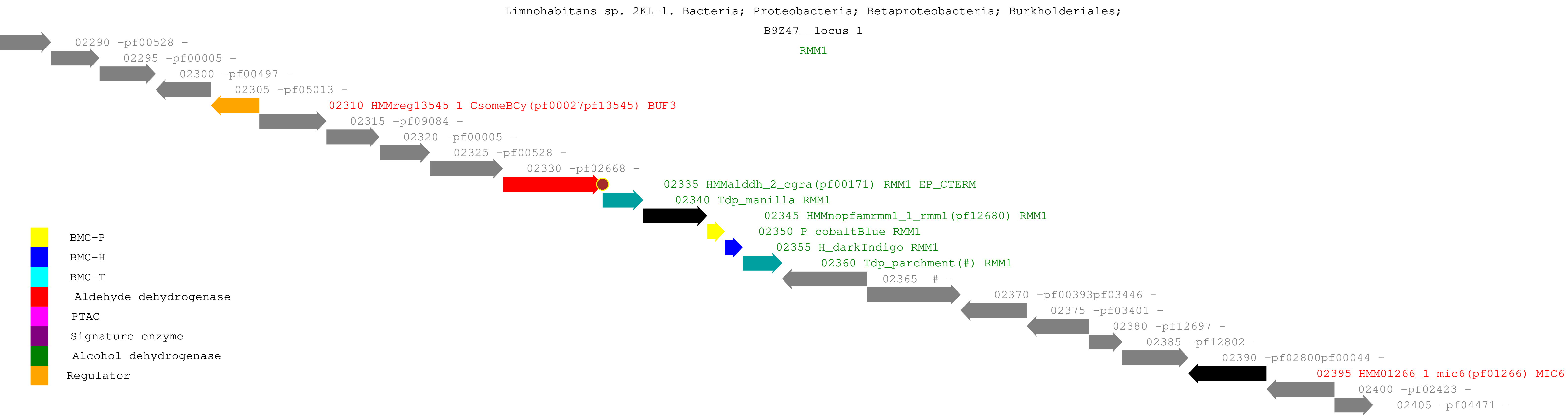

Limnohabitans sp. 2KL-1. Bacteria; Proteobacteria; Betaproteobacteria; Burkholderiales;
B9Z47__locus_1
RMM1
02290 -pf00528 -
02295 -pf00005 -
02300 -pf00497 -
02305 -pf05013 -
02310 HMMreg13545_1_CsomeBCy(pf00027pf13545) BUF3
02315 -pf09084 -
02320 -pf00005 -
02325 -pf00528 -
02330 -pf02668 -
02335 HMMalddh_2_egra(pf00171) RMM1 EP_CTERM
02340 Tdp_manilla RMM1
02345 HMMnopfamrmm1_1_rmm1(pf12680) RMM1
02350 P_cobaltBlue RMM1
02355 H_darkIndigo RMM1
02360 Tdp_parchment(#) RMM1
02365 -# -
02370 -pf00393pf03446 -
02375 -pf03401 -
02380 -pf12697 -
02385 -pf12802 -
02390 -pf02800pf00044 -
02395 HMM01266_1_mic6(pf01266) MIC6
02400 -pf02423 -
02405 -pf04471 -
BMC-P
BMC-H
BMC-T
Aldehyde dehydrogenase
PTAC
Signature enzyme
Alcohol dehydrogenase
Regulator

Limnohabitans sp. WS1. Bacteria; Proteobacteria; Betaproteobacteria; Burkholderiales;

B9Z48__locus_1

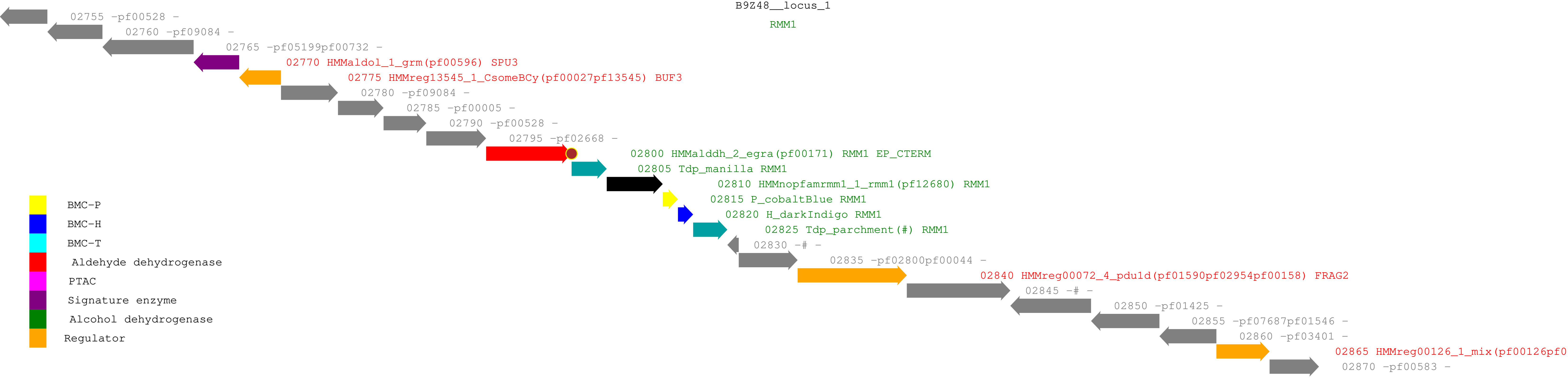

Limnohabitans sp. 2KL-51. Bacteria; Proteobacteria; Betaproteobacteria; Burkholderiales;

B9Z49__locus_1

RMM1

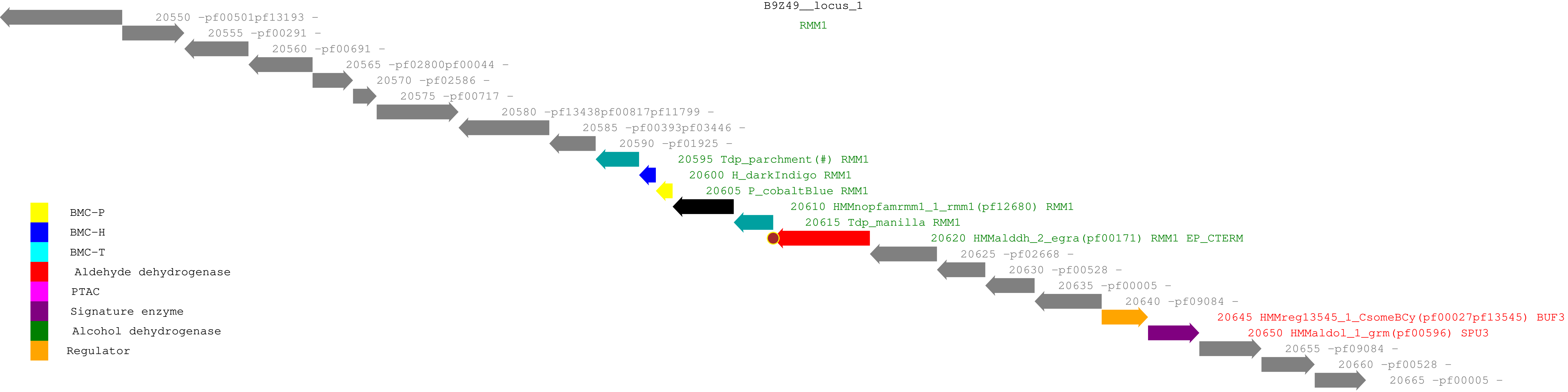

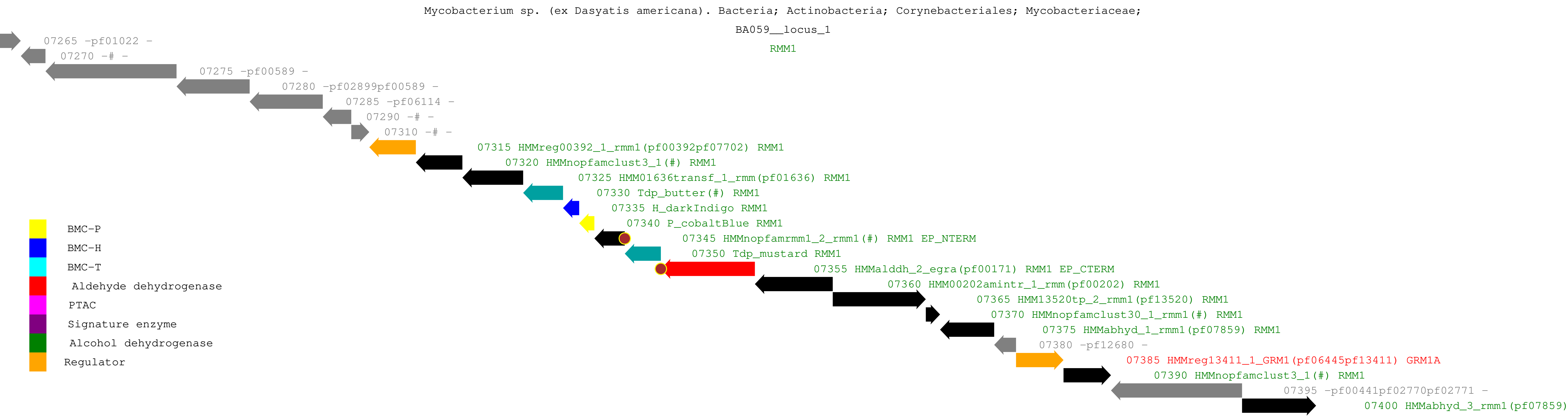

Mycobacterium sp. (ex Dasyatis americana). Bacteria; Actinobacteria; Corynebacteriales; Mycobacteriaceae;
BA059__locus_1
RMM1
07265 -pf01022 -
07270 -# -
07275 -pf00589 -
07280 -pf02899pf00589 -
07285 -pf06114 -
07290 -# -
07310 -# -
07315 HMMreg00392_1_rmm1(pf00392pf07702) RMM1
07320 HMMnopfamclust3_1(#) RMM1
07325 HMM01636transf_1_rmm(pf01636) RMM1
07330 Tdp_butter(#) RMM1
07335 H_darkIndigo RMM1
07340 P_cobaltBlue RMM1
07345 HMMnopfamrmm1_2_rmm1(#) RMM1 EP_NTERM
07350 Tdp_mustard RMM1
07355 HMMalddh_2_egra(pf00171) RMM1 EP_CTERM
07360 HMM00202amintr_1_rmm(pf00202) RMM1
07365 HMM13520tp_2_rmm1(pf13520) RMM1
07370 HMMnopfamclust30_1_rmm1(#) RMM1
07375 HMMabhyd_1_rmm1(pf07859) RMM1
07380 -pf12680 -
07385 HMMreg13411_1_GRM1(pf06445pf13411) GRM1A
07390 HMMnopfamclust3_1(#) RMM1
07395 -pf00441pf02770pf02771 -
07400 HMMabhyd_3_rmm1(pf07859)
BMC-P
BMC-H
BMC-T
Aldehyde dehydrogenase
PTAC
Signature enzyme
Alcohol dehydrogenase
Regulator

Mycobacterium sp. djl-10. Bacteria; Actinobacteria; Corynebacteriales; Mycobacteriaceae;

BCA37__locus_1

RMM1

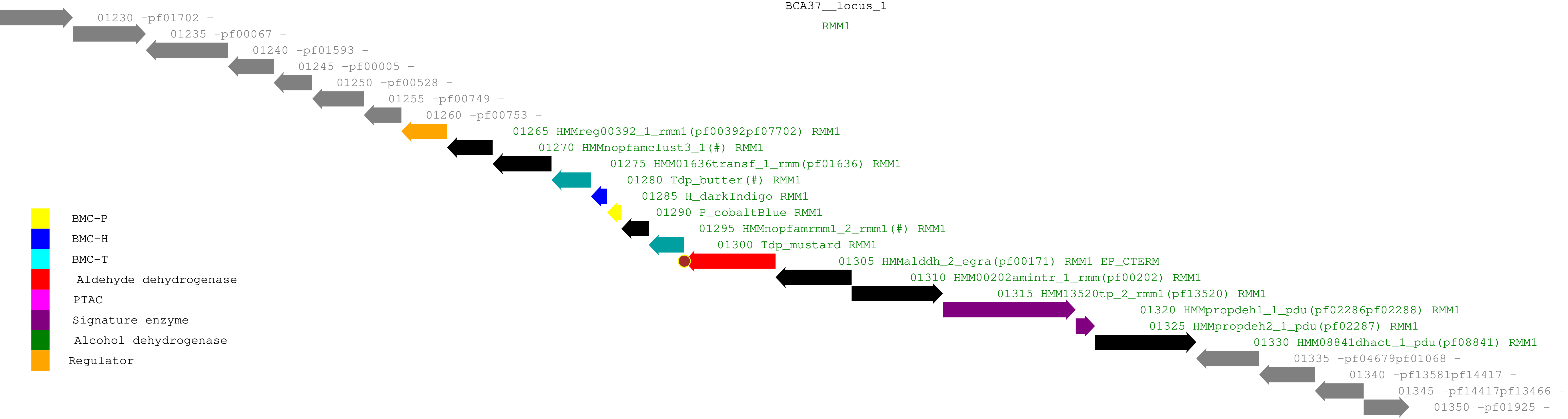

Mycolicibacterium litorale. Bacteria; Actinobacteria; Corynebacteriales; Mycobacteriaceae;

BCL50__locus_1

RMM1

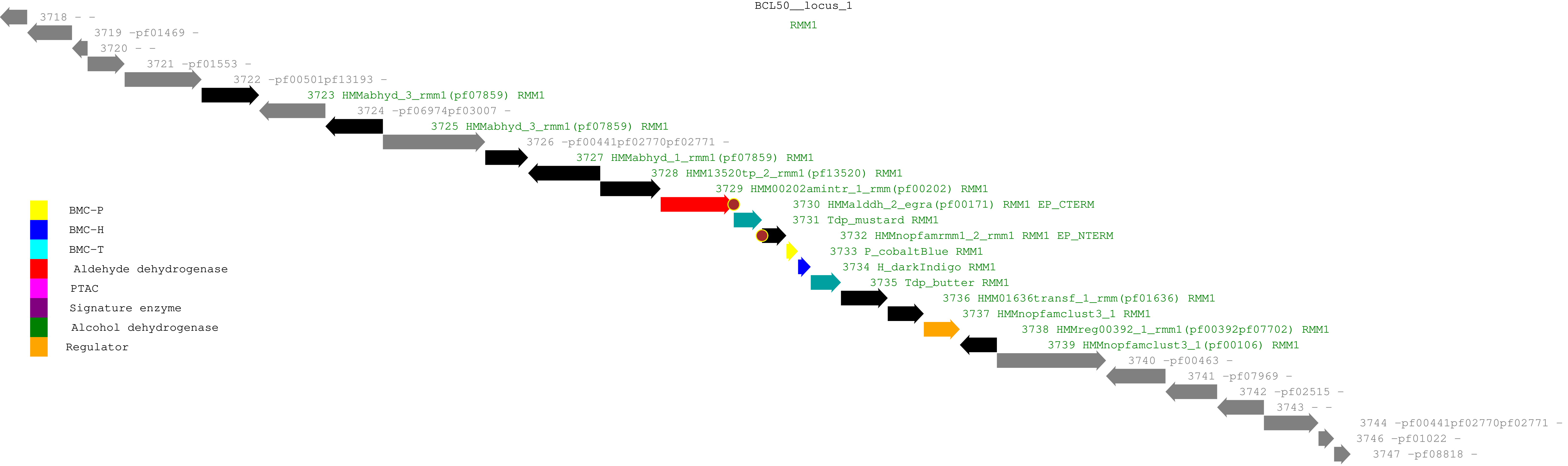

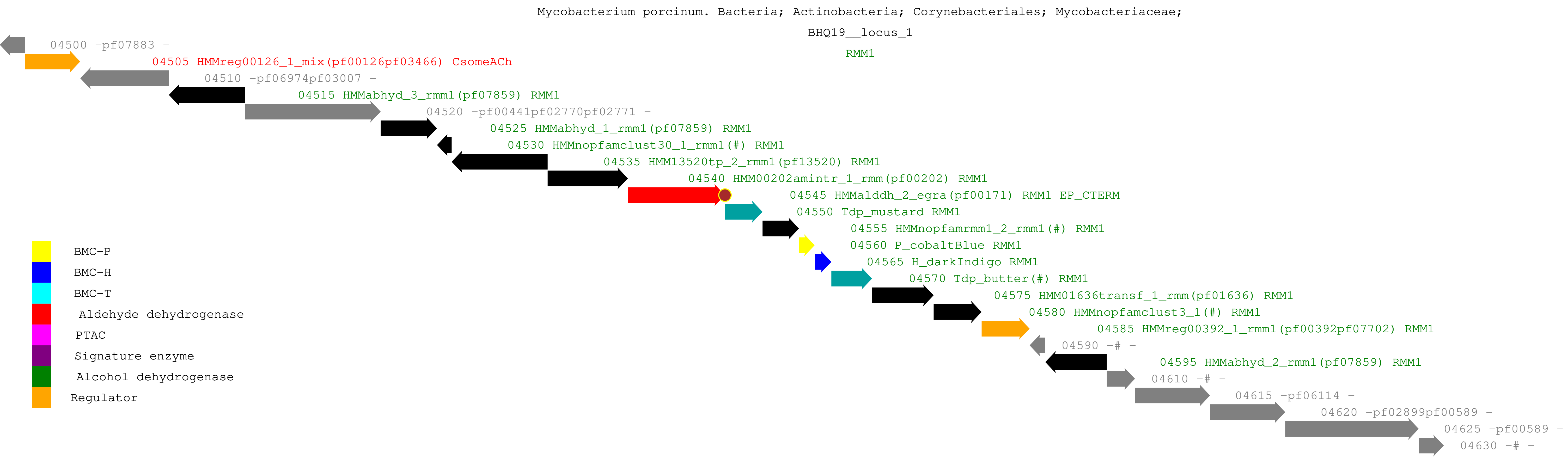
Mycobacterium porcinum. Bacteria; Actinobacteria; Corynebacteriales; Mycobacteriaceae;
BHQ19__locus_1
RMM1
04500 -pf07883 -
04505 HMMreg00126_1_mix(pf00126pf03466) CsomeACh
04510 -pf06974pf03007 -
04515 HMMabhyd_3_rmm1(pf07859) RMM1
04520 -pf00441pf02770pf02771 -
04525 HMMabhyd_1_rmm1(pf07859) RMM1
04530 HMMnopfamclust30_1_rmm1(#) RMM1
04535 HMM13520tp_2_rmm1(pf13520) RMM1
04540 HMM00202amintr_1_rmm(pf00202) RMM1
04545 HMMaldh_2_egra(pf00171) RMM1 EP_CTERM
04550 Tdp_mustard RMM1
04555 HMMnopfamrmm1_2_rmm1(#) RMM1
04560 P_cobaltBlue RMM1
04565 H_darkIndigo RMM1
04570 Tdp_butter(#) RMM1
04575 HMM01636transf_1_rmm(pf01636) RMM1
04580 HMMnopfamclust3_1(#) RMM1
04585 HMMreg00392_1_rmm1(pf00392pf07702) RMM1
04590 -# -
04595 HMMabhyd_2_rmm1(pf07859) RMM1
04610 -# -
04615 -pf06114 -
04620 -pf02899pf00589 -
04625 -pf00589 -
04630 -# -
BMC-P
BMC-H
BMC-T
Aldehyde dehydrogenase
PTAC
Signature enzyme
Alcohol dehydrogenase
Regulator

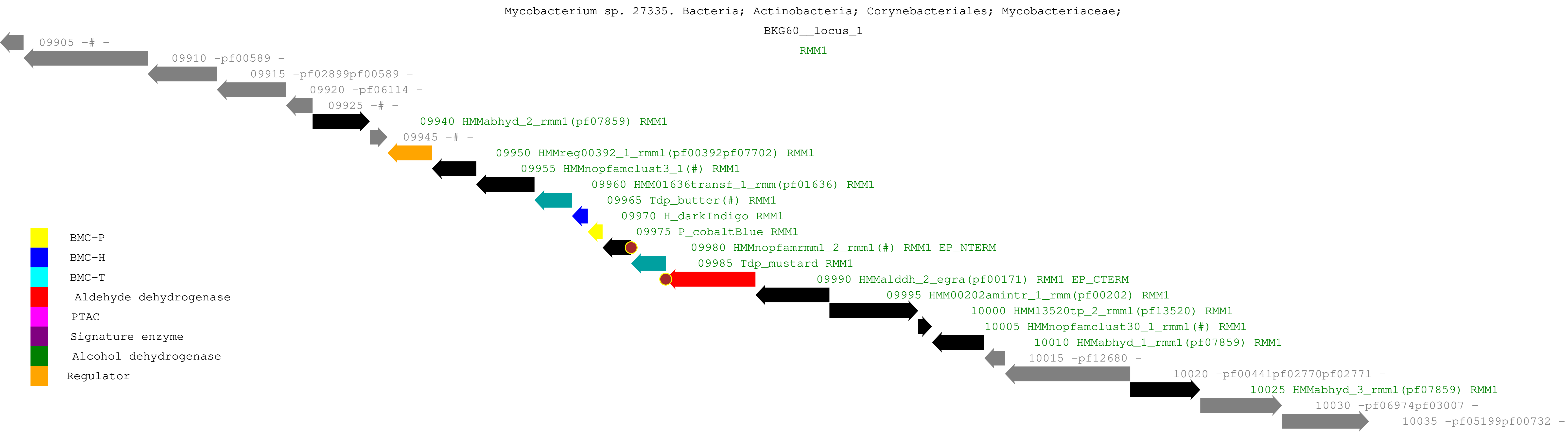
Mycobacterium sp. 27335. Bacteria; Actinobacteria; Corynebacteriales; Mycobacteriaceae;
BKG60__locus_1
RMM1
09905 -# -
09910 -pf00589 -
09915 -pf02899pf00589 -
09920 -pf06114 -
09925 -# -
09940 HMMabhyd_2_rmm1(pf07859) RMM1
09945 -# -
09950 HMMreg00392_1_rmm1(pf00392pf07702) RMM1
09955 HMMnopfamclust3_1(#) RMM1
09960 HMM01636transf_1_rmm(pf01636) RMM1
09965 Tdp_butter(#) RMM1
09970 H_darkIndigo RMM1
09975 P_cobaltBlue RMM1
09980 HMMnopfamrmm1_2_rmm1(#) RMM1 EP_NTERM
09985 Tdp_mustard RMM1
09990 HMMaldh_2_egra(pf00171) RMM1 EP_CTERM
09995 HMM00202amintr_1_rmm(pf00202) RMM1
10000 HMM13520tp_2_rmm1(pf13520) RMM1
10005 HMMnopfamclust30_1_rmm1(#) RMM1
10010 HMMabhyd_1_rmm1(pf07859) RMM1
10015 -pf12680 -
10020 -pf00441pf02770pf02771 -
10025 HMMabhyd_3_rmm1(pf07859) RMM1
10030 -pf06974pf03007 -
10035 -pf05199pf00732 -
BMC-P
BMC-H
BMC-T
Aldehyde dehydrogenase
PTAC
Signature enzyme
Alcohol dehydrogenase
Regulator

Mycobacterium sp. 24999. Bacteria; Actinobacteria; Corynebacteriales; Mycobacteriaceae;

BKG61__locus_1

RMM1

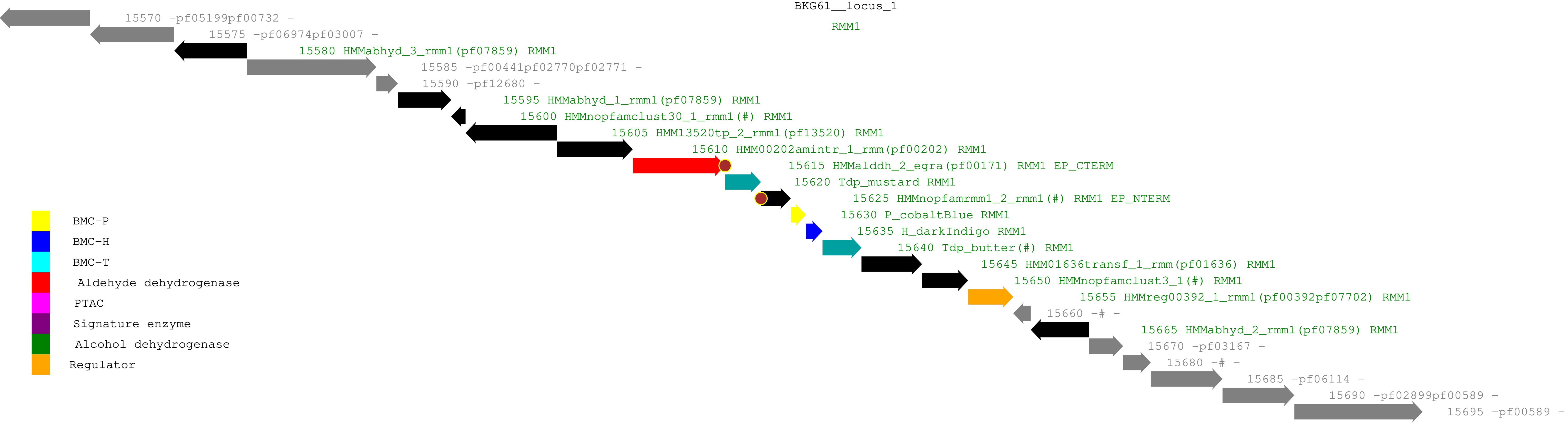

Mycobacterium neoaurum. Bacteria; Actinobacteria; Corynebacteriales; Mycobacteriaceae;

BN1047__locus_1

RMM1

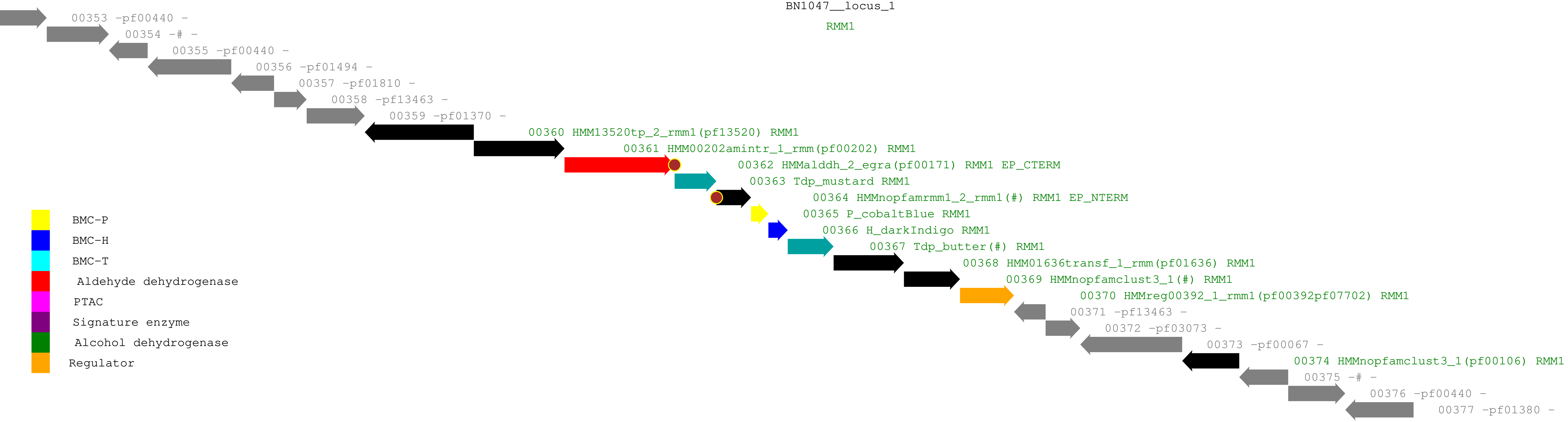

Mycobacterium neworleansense. Bacteria; Actinobacteria; Corynebacteriales; Mycobacteriaceae;

BN2156__locus_1

RMM1

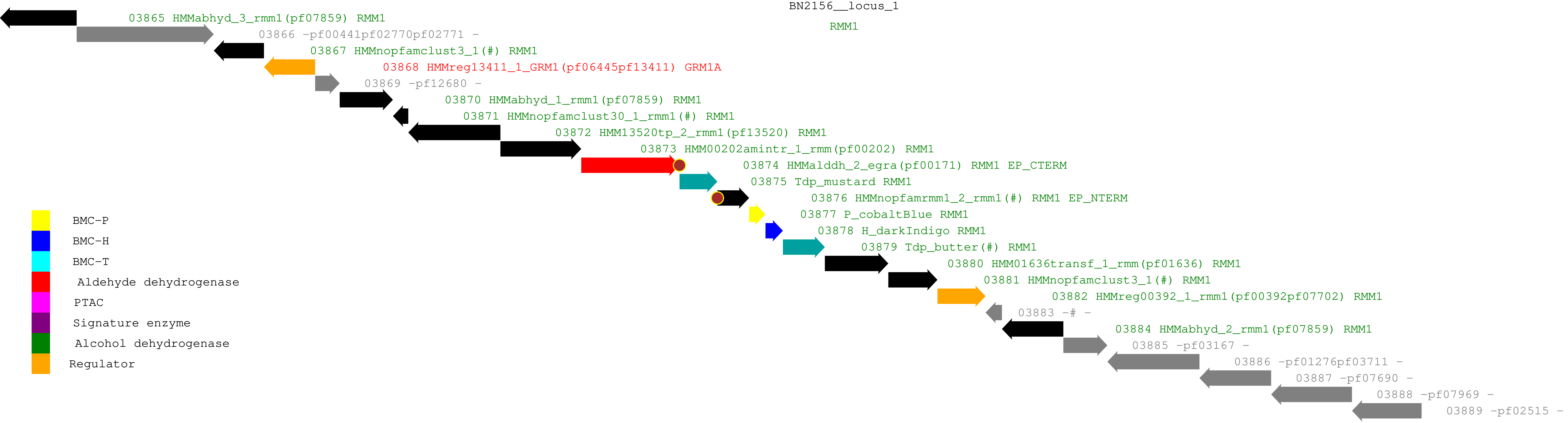

Mycobacterium farcinogenes. Bacteria; Actinobacteria; Corynebacteriales; Mycobacteriaceae;

BN975__locus_1

RMM1

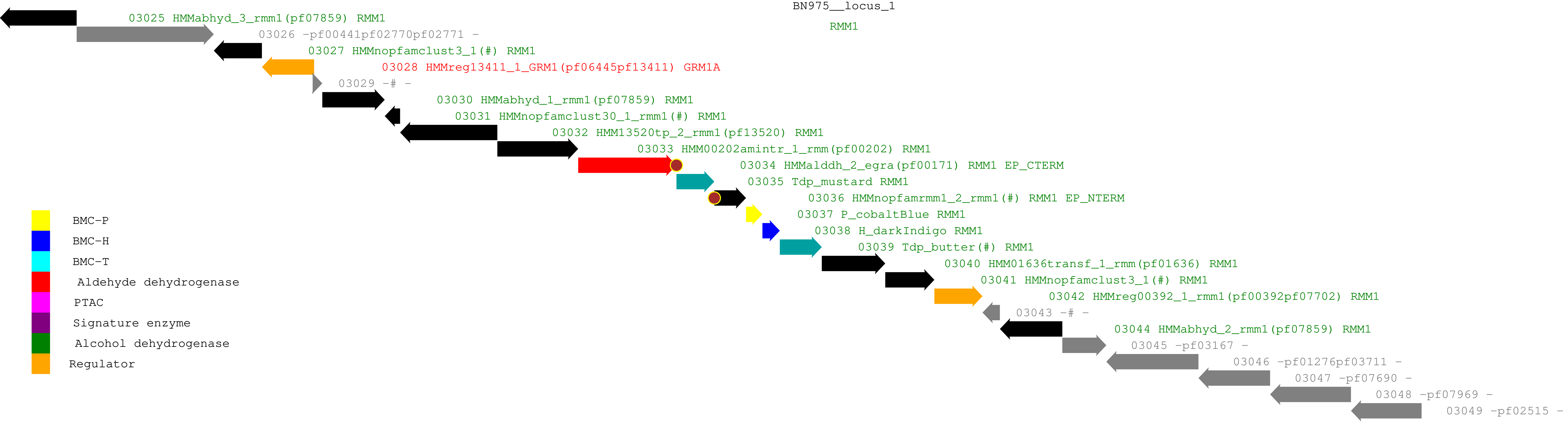

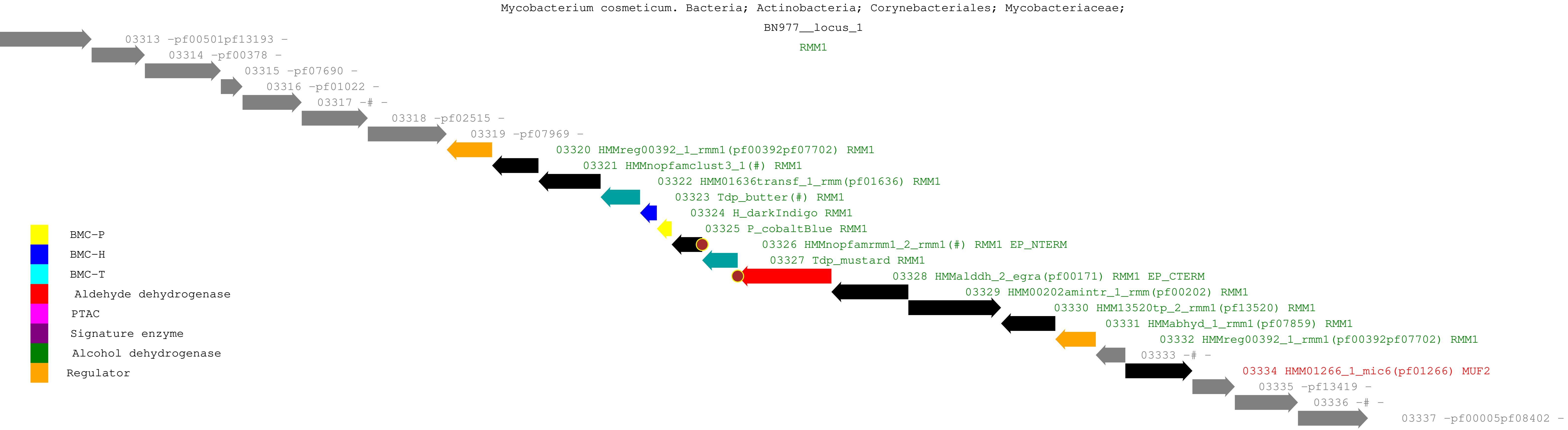
Mycobacterium cosmeticum. Bacteria; Actinobacteria; Corynebacteriales; Mycobacteriaceae;
BN977__locus_1
RMM1
03313 -pf00501pf13193 -
03314 -pf00378 -
03315 -pf07690 -
03316 -pf01022 -
03317 -# -
03318 -pf02515 -
03319 -pf07969 -
03320 HMMreg00392_1_rmm1(pf00392pf07702) RMM1
03321 HMMnopfamclust3_1(#) RMM1
03322 HMM01636transf_1_rmm(pf01636) RMM1
03323 Tdp_butter(#) RMM1
03324 H_darkIndigo RMM1
03325 P_cobaltBlue RMM1
03326 HMMnopfamrmm1_2_rmm1(#) RMM1 EP_NTERM
03327 Tdp_mustard RMM1
03328 HMMalddh_2_egra(pf00171) RMM1 EP_CTERM
03329 HMM00202amintr_1_rmm(pf00202) RMM1
03330 HMM13520tp_2_rmm1(pf13520) RMM1
03331 HMMabhyd_1_rmm1(pf07859) RMM1
03332 HMMreg00392_1_rmm1(pf00392pf07702) RMM1
03333 -# -
03334 HMM01266_1_mic6(pf01266) MUF2
03335 -pf13419 -
03336 -# -
03337 -pf00005pf08402 -
BMC-P
BMC-H
BMC-T
Aldehyde dehydrogenase
PTAC
Signature enzyme
Alcohol dehydrogenase
Regulator

Mycobacterium mageritense DSM 44476 = CIP 104973. Bacteria; Actinobacteria; Corynebacteriales; Mycobacteriaceae;

BN978__locus_1

RMM1

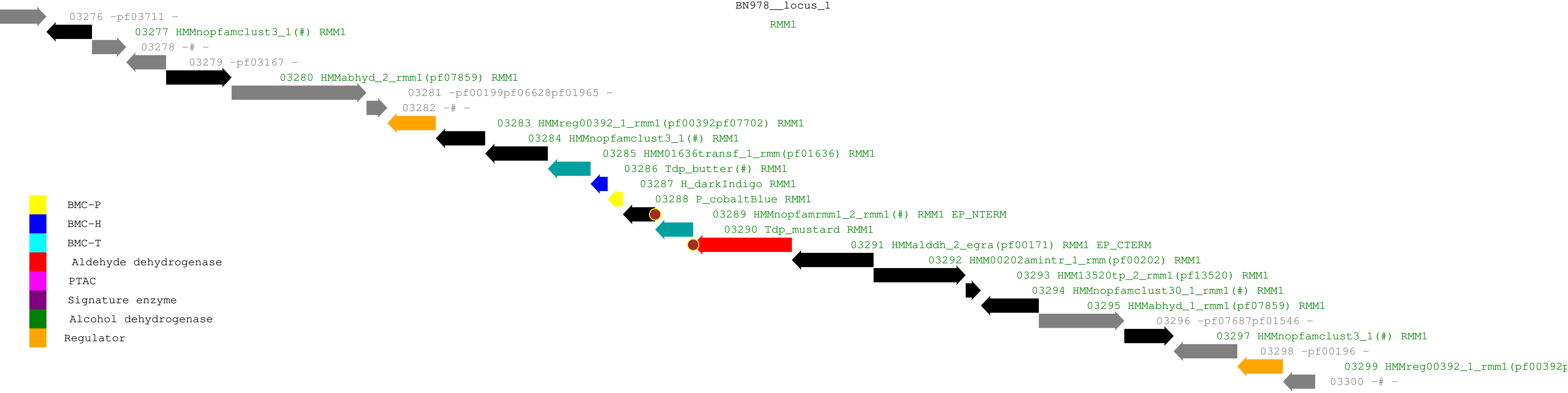

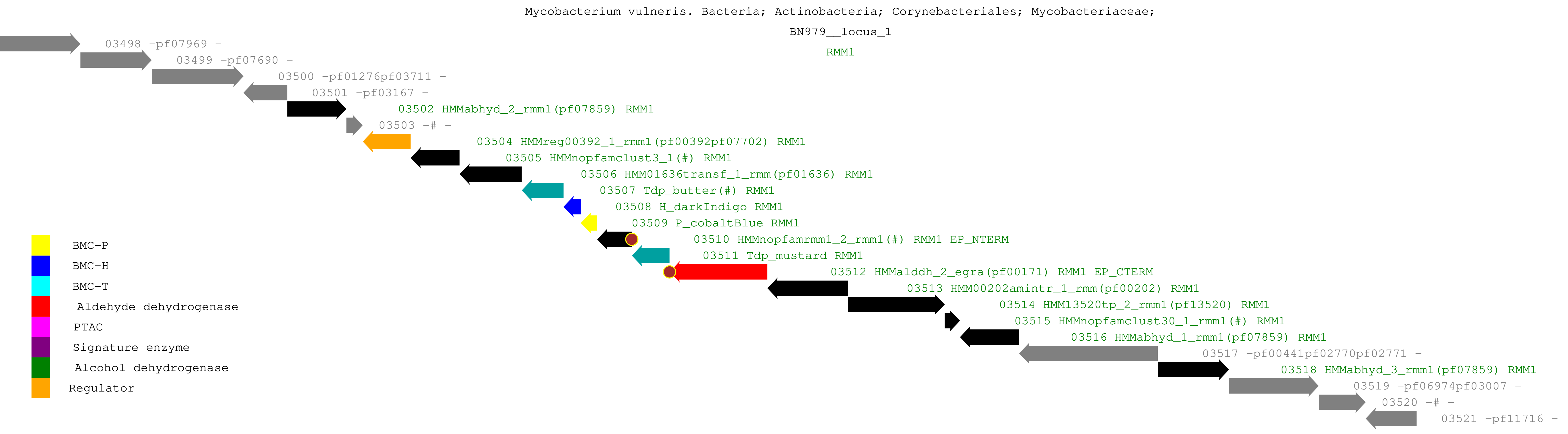
Mycobacterium vulneris. Bacteria; Actinobacteria; Corynebacteriales; Mycobacteriaceae;
BN979__locus_1
RMM1
03498 -pf07969 -
03499 -pf07690 -
03500 -pf01276pf03711 -
03501 -pf03167 -
03502 HMMabhyd_2_rmm1(pf07859) RMM1
03503 -# -
03504 HMMreg00392_1_rmm1(pf00392pf07702) RMM1
03505 HMMnopfamclust3_1(#) RMM1
03506 HMM01636transf_1_rmm(pf01636) RMM1
03507 Tdp_butter(#) RMM1
03508 H_darkIndigo RMM1
03509 P_cobaltBlue RMM1
03510 HMMnopfamrmm1_2_rmm1(#) RMM1 EP_NTERM
03511 Tdp_mustard RMM1
03512 HMMalddh_2_egra(pf00171) RMM1 EP_CTERM
03513 HMM00202amintr_1_rmm(pf00202) RMM1
03514 HMM13520tp_2_rmm1(pf13520) RMM1
03515 HMMnopfamclust30_1_rmm1(#) RMM1
03516 HMMabhyd_1_rmm1(pf07859) RMM1
03517 -pf00441pf02770pf02771 -
03518 HMMabhyd_3_rmm1(pf07859) RMM1
03519 -pf06974pf03007 -
03520 -# -
03521 -pf11716 -
BMC-P
BMC-H
BMC-T
Aldehyde dehydrogenase
PTAC
Signature enzyme
Alcohol dehydrogenase
Regulator

Mycobacterium aquaticum. Bacteria; Actinobacteria; Corynebacteriales; Mycobacteriaceae;

BST13__locus_1

RMM1

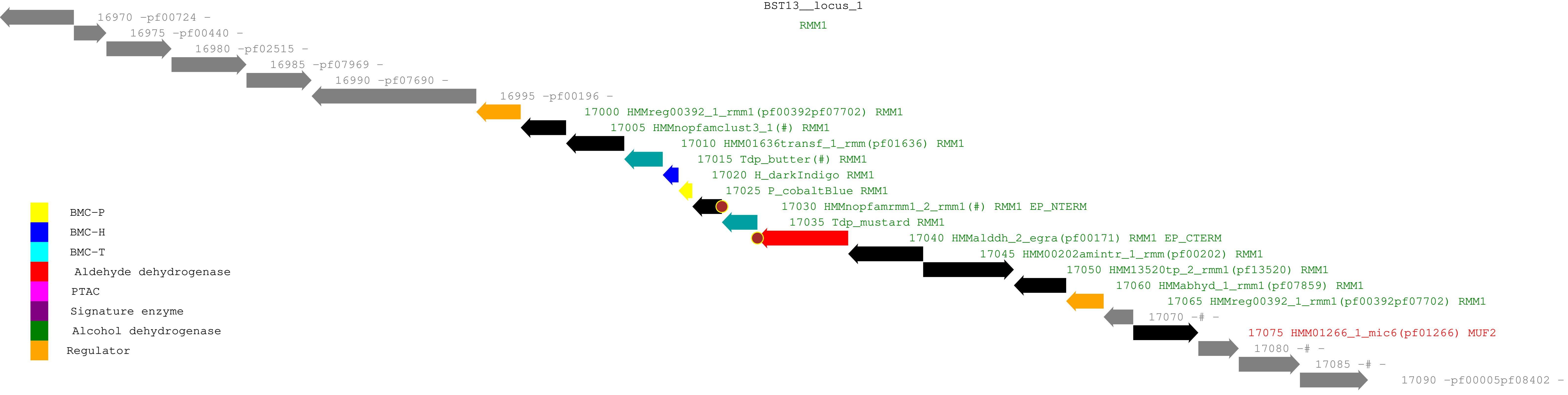

Mycobacterium monacense DSM 44395. Bacteria; Actinobacteria; Corynebacteriales; Mycobacteriaceae;

BST34__locus_1

RMM1

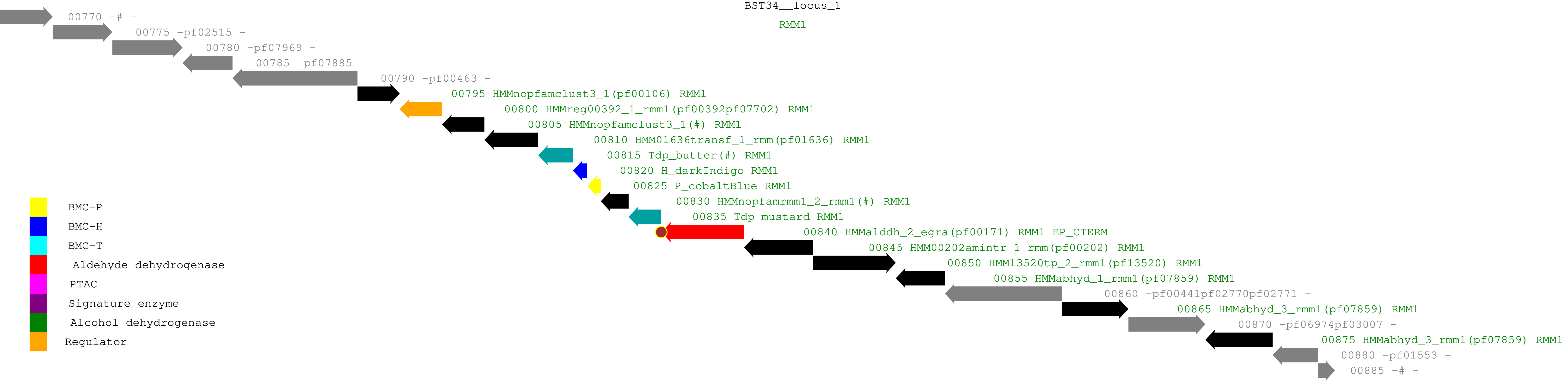

Mycobacterium parafortuitum. Bacteria; Actinobacteria; Corynebacteriales; Mycobacteriaceae;

BST38__locus_1

RMM1

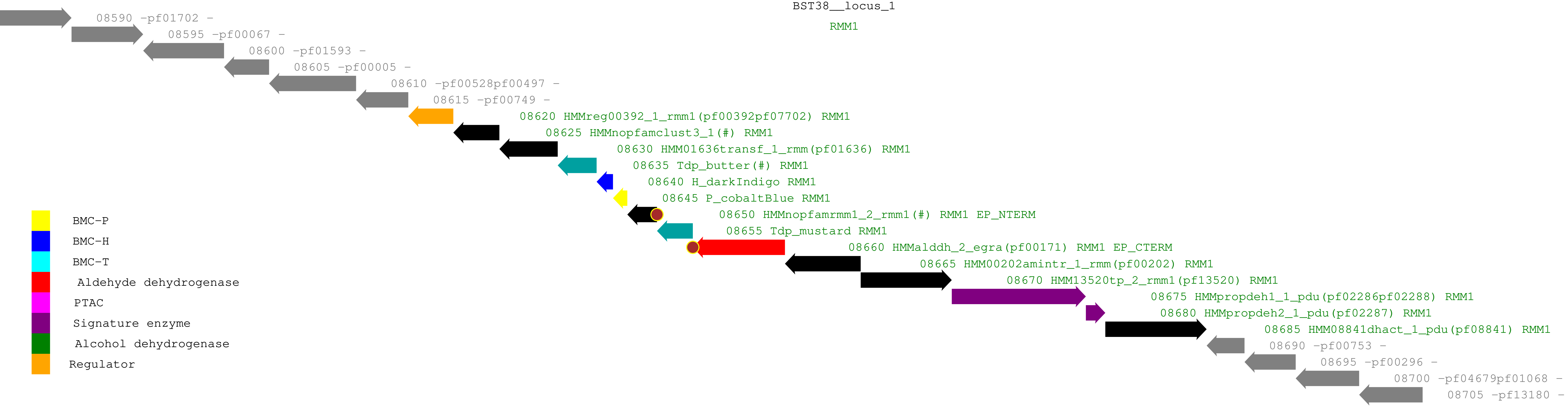

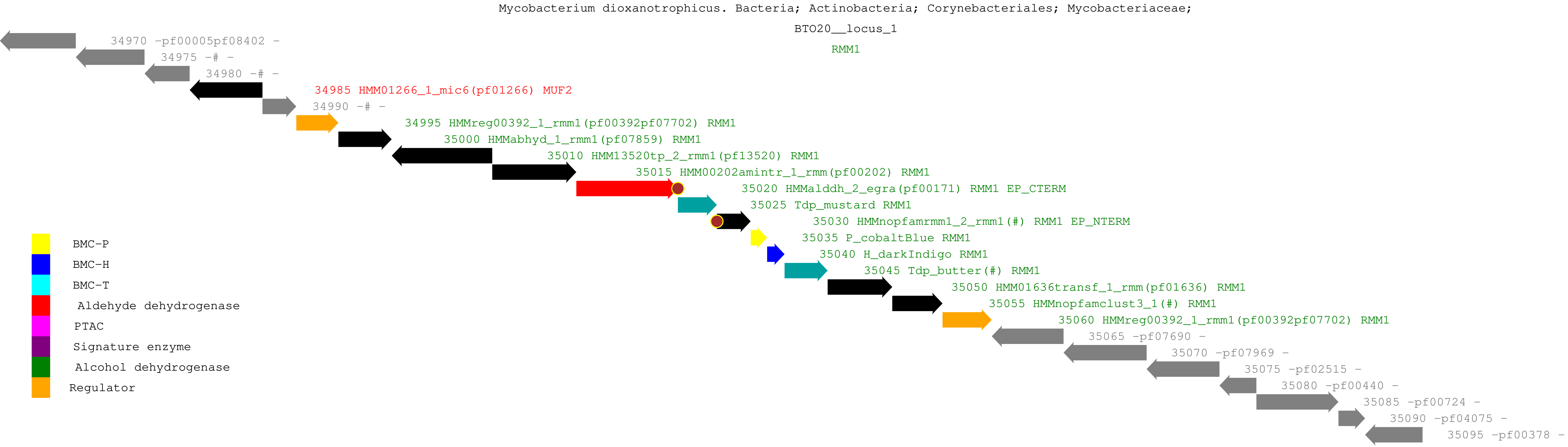

Mycobacterium dioxanotrophicus. Bacteria; Actinobacteria; Corynebacteriales; Mycobacteriaceae;
BTO20__locus_1
RMM1
34970 -pf00005pf08402 -
34975 -# -
34980 -# -
34985 HMM01266_1_mic6(pf01266) MUF2
34990 -# -
34995 HMMreg00392_1_rmm1(pf00392pf07702) RMM1
35000 HMMabhyd_1_rmm1(pf07859) RMM1
35010 HMM13520tp_2_rmm1(pf13520) RMM1
35015 HMM00202amintr_1_rmm(pf00202) RMM1
35020 HMMalddh_2_egra(pf00171) RMM1 EP_CTERM
35025 Tdp_mustard RMM1
35030 HMMnopfamrmm1_2_rmm1(#) RMM1 EP_NTERM
35035 P_cobaltBlue RMM1
35040 H_darkIndigo RMM1
35045 Tdp_butter(#) RMM1
35050 HMM01636transf_1_rmm(pf01636) RMM1
35055 HMMnopfamclust3_1(#) RMM1
35060 HMMreg00392_1_rmm1(pf00392pf07702) RMM1
35065 -pf07690 -
35070 -pf07969 -
35075 -pf02515 -
35080 -pf00440 -
35085 -pf00724 -
35090 -pf04075 -
35095 -pf00378 -
BMC-P
BMC-H
BMC-T
Aldehyde dehydrogenase
PTAC
Signature enzyme
Alcohol dehydrogenase
Regulator

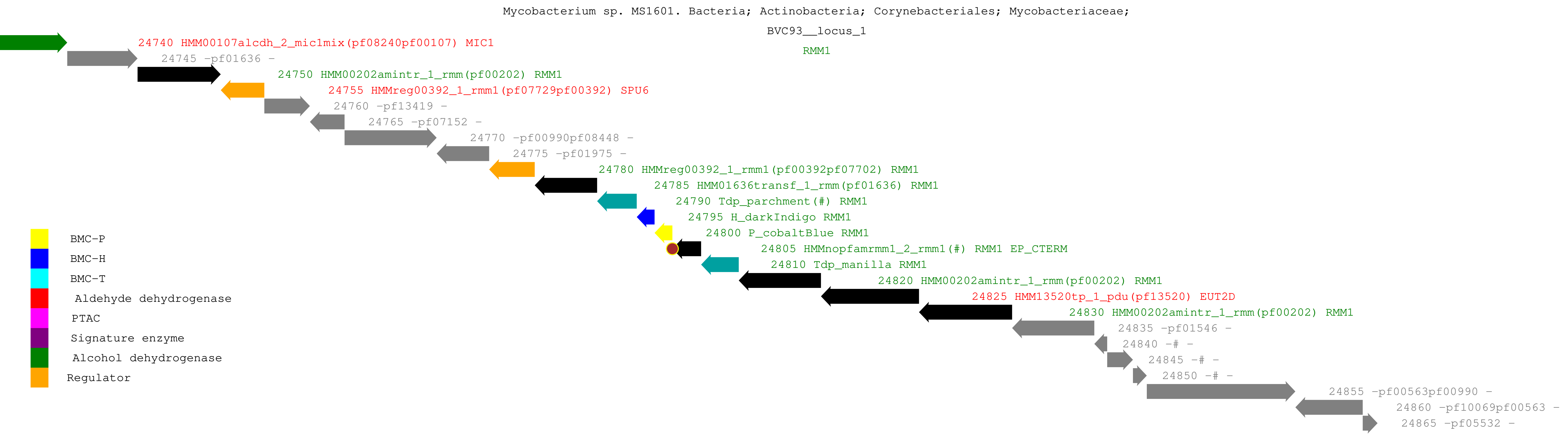
Mycobacterium sp. MS1601. Bacteria; Actinobacteria; Corynebacteriales; Mycobacteriaceae;
BVC93__locus_1
RMM1
24740 HMM00107alcdh_2_mic1mix(pf08240pf00107) MIC1
24745 -pf01636 -
24750 HMM00202amintr_1_rmm(pf00202) RMM1
24755 HMMreg00392_1_rmm1(pf07729pf00392) SPU6
24760 -pf13419 -
24765 -pf07152 -
24770 -pf00990pf08448 -
24775 -pf01975 -
24780 HMMreg00392_1_rmm1(pf00392pf07702) RMM1
24785 HMM01636transf_1_rmm(pf01636) RMM1
24790 Tdp_parchment(#) RMM1
24795 H_darkIndigo RMM1
24800 P_cobaltBlue RMM1
24805 HMMnopfamrmm1_2_rmm1(#) RMM1 EP_CTERM
24810 Tdp_manilla RMM1
24820 HMM00202amintr_1_rmm(pf00202) RMM1
24825 HMM13520tp_1_pdu(pf13520) EUT2D
24830 HMM00202amintr_1_rmm(pf00202) RMM1
24835 -pf01546 -
24840 -# -
24845 -# -
24850 -# -
24855 -pf00563pf00990 -
24860 -pf10069pf00563 -
24865 -pf05532 -
BMC-P
BMC-H
BMC-T
Aldehyde dehydrogenase
PTAC
Signature enzyme
Alcohol dehydrogenase
Regulator

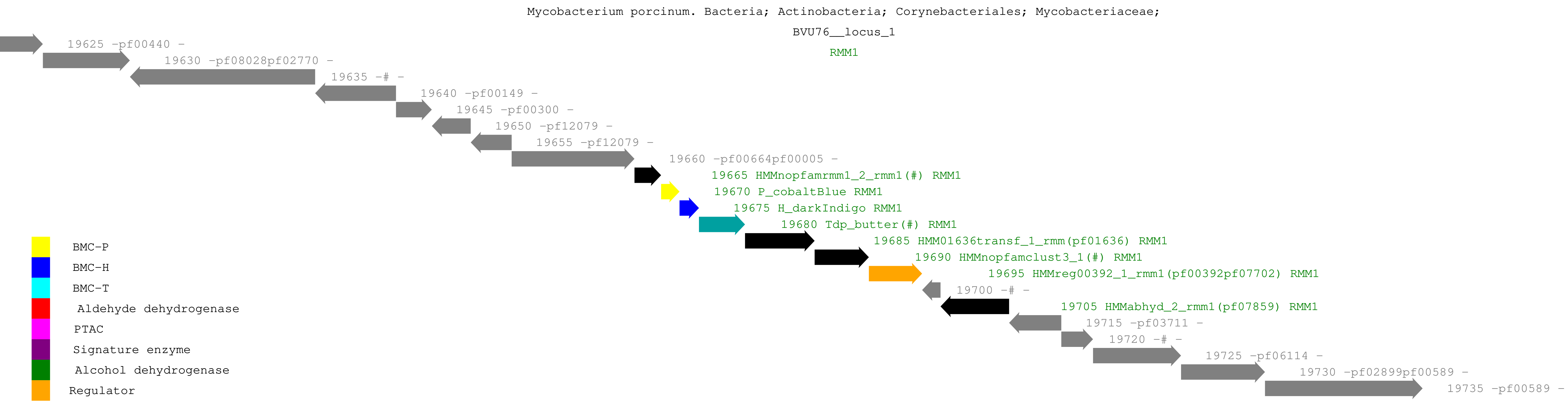

Mycobacterium porcinum. Bacteria; Actinobacteria; Corynebacteriales; Mycobacteriaceae;
BVU76__locus_1
RMM1
19625 -pf00440 -
19630 -pf08028pf02770 -
19635 -# -
19640 -pf00149 -
19645 -pf00300 -
19650 -pf12079 -
19655 -pf12079 -
19660 -pf00664pf00005 -
19665 HMMnopfamrmm1_2_rmm1(#) RMM1
19670 P_cobaltBlue RMM1
19675 H_darkIndigo RMM1
19680 Tdp_butter(#) RMM1
19685 HMM01636transf_1_rmm(pf01636) RMM1
19690 HMMnopfamclust3_1(#) RMM1
19695 HMMreg00392_1_rmm1(pf00392pf07702) RMM1
19700 -# -
19705 HMMabhyd_2_rmm1(pf07859) RMM1
19715 -pf03711 -
19720 -# -
19725 -pf06114 -
19730 -pf02899pf00589 -
19735 -pf00589 -
BMC-P
BMC-H
BMC-T
Aldehyde dehydrogenase
PTAC
Signature enzyme
Alcohol dehydrogenase
Regulator

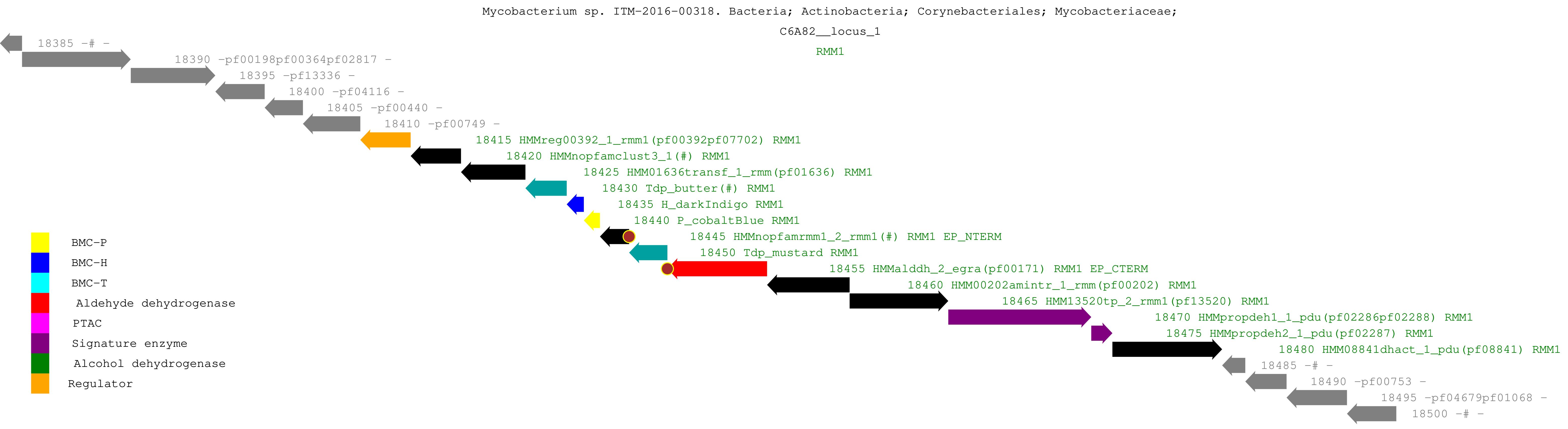
Mycobacterium sp. ITM-2016-00318. Bacteria; Actinobacteria; Corynebacteriales; Mycobacteriaceae;
C6A82__locus_1
RMM1
18385 -# -
18390 -pf00198pf00364pf02817 -
18395 -pf13336 -
18400 -pf04116 -
18405 -pf00440 -
18410 -pf00749 -
18415 HMMreg00392_1_rmm1(pf00392pf07702) RMM1
18420 HMMnopfamclust3_1(#) RMM1
18425 HMM01636transf_1_rmm(pf01636) RMM1
18430 Tdp_butter(#) RMM1
18435 H_darkIndigo RMM1
18440 P_cobaltBlue RMM1
18445 HMMnopfamrmm1_2_rmm1(#) RMM1 EP_NTERM
18450 Tdp_mustard RMM1
18455 HMMaldh_2_egra(pf00171) RMM1 EP_CTERM
18460 HMM00202amintr_1_rmm(pf00202) RMM1
18465 HMM13520tp_2_rmm1(pf13520) RMM1
18470 HMMpropdeh1_1_pdu(pf02286pf02288) RMM1
18475 HMMpropdeh2_1_pdu(pf02287) RMM1
18480 HMM08841dhact_1_pdu(pf08841) RMM1
18485 -# -
18490 -pf00753 -
18495 -pf04679pf01068 -
18500 -# -
BMC-P
BMC-H
BMC-T
Aldehyde dehydrogenase
PTAC
Signature enzyme
Alcohol dehydrogenase
Regulator

Mycobacterium sp. ITM-2016-00317. Bacteria; Actinobacteria; Corynebacteriales; Mycobacteriaceae;

C6A87__locus_1

RMM1

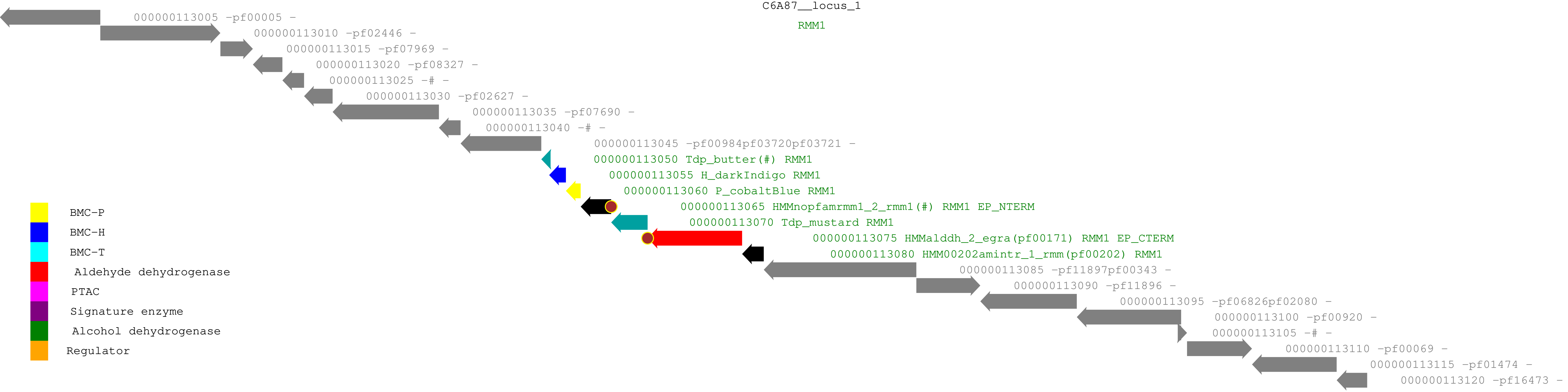

Mycolicibacterium austroafricanum. Bacteria; Actinobacteria; Corynebacteriales; Mycobacteriaceae;

C6A88__locus_1

RMM1

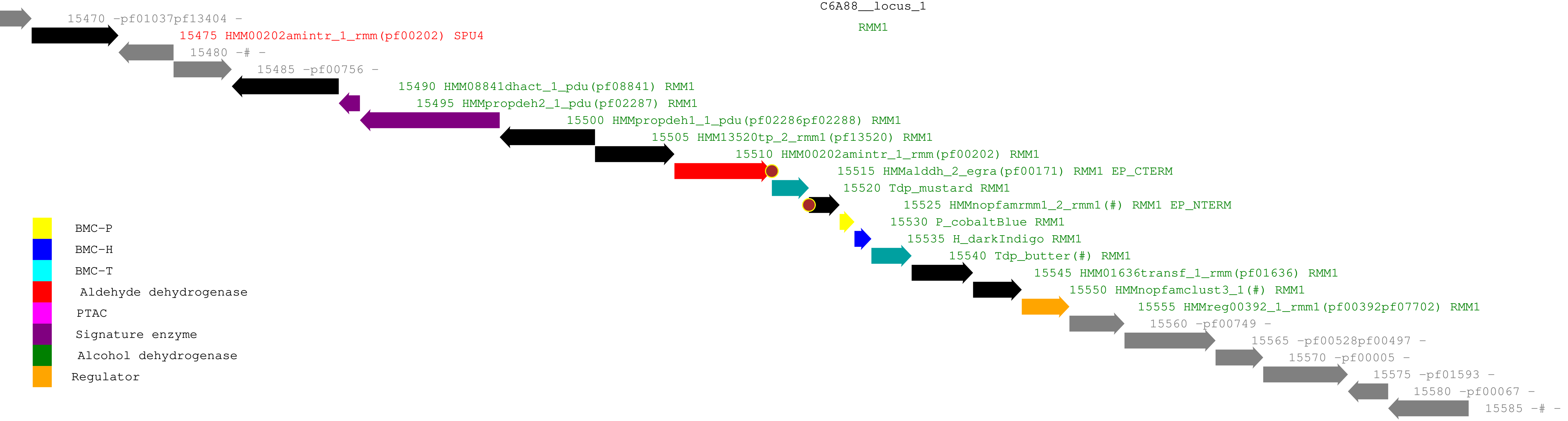

Mycobacterium hassiacum (strain DSM 44199 / CIP 105218 / JCM 12690 / Bacteria; Actinobacteria; Corynebacteriales; Mycobacteriaceae;
C731__locus_1
RMM1

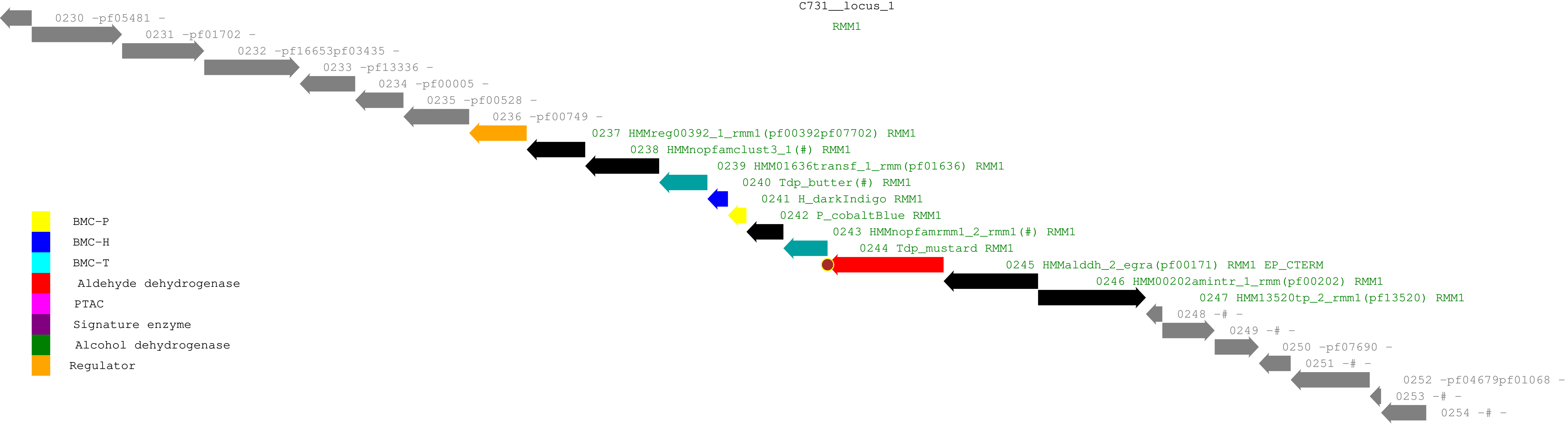

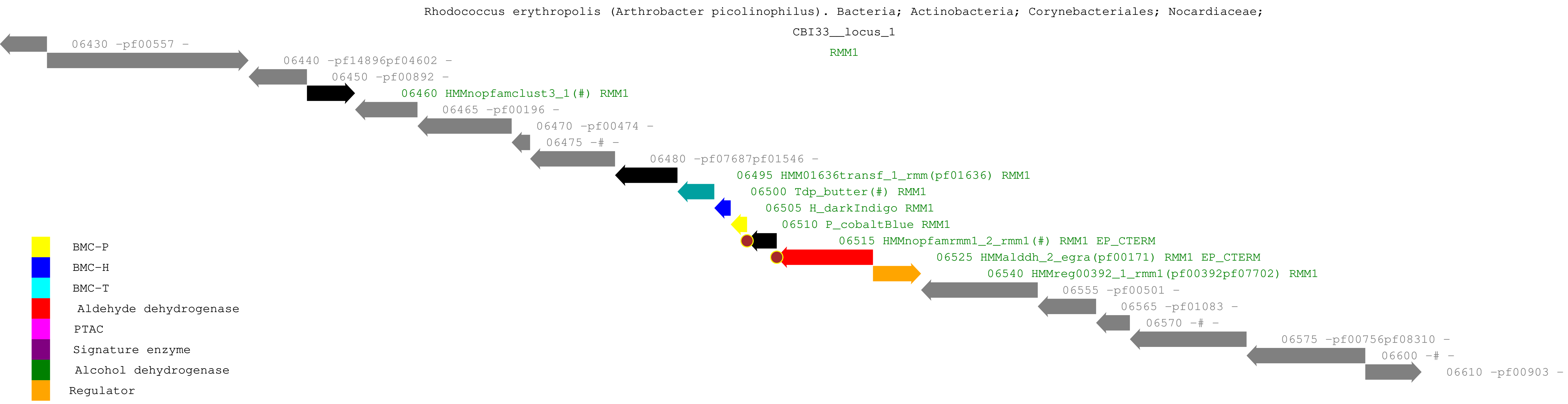
Rhodococcus erythropolis (Arthrobacter picolinophilus). Bacteria; Actinobacteria; Corynebacteriales; Nocardiaceae;
CBI33__locus_1
RMM1
06430 -pf00557 -
06440 -pf14896pf04602 -
06450 -pf00892 -
06460 HMMnopfamclust3_1(#) RMM1
06465 -pf00196 -
06470 -pf00474 -
06475 -# -
06480 -pf07687pf01546 -
06495 HMM01636transf_1_rmm(pf01636) RMM1
06500 Tdp_butter(#) RMM1
06505 H_darkIndigo RMM1
06510 P_cobaltBlue RMM1
06515 HMMnopfamrmm1_2_rmm1(#) RMM1 EP_CTERM
06525 HMMalddh_2_egra(pf00171) RMM1 EP_CTERM
06540 HMMreg00392_1_rmm1(pf00392pf07702) RMM1
06555 -pf00501 -
06565 -pf01083 -
06570 -# -
06575 -pf00756pf08310 -
06600 -# -
06610 -pf00903 -
BMC-P
BMC-H
BMC-T
Aldehyde dehydrogenase
PTAC
Signature enzyme
Alcohol dehydrogenase
Regulator

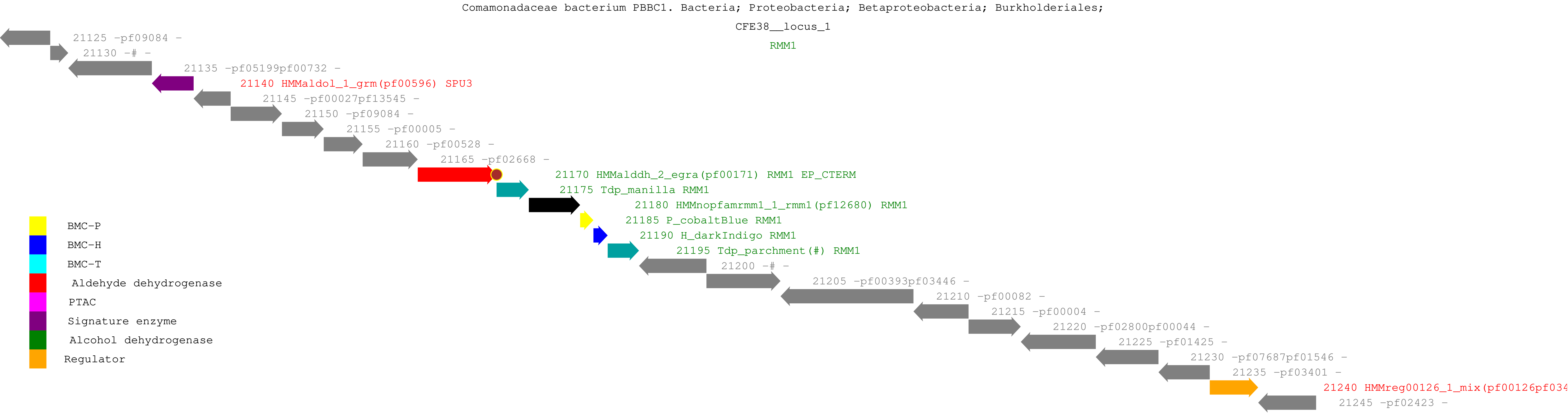
Comamonadaceae bacterium PBBC1. Bacteria; Proteobacteria; Betaproteobacteria; Burkholderiales;
CFE38__locus_1
RMM1
21125 -pf09084 -
21130 -# -
21135 -pf05199pf00732 -
21140 HMMaldol_1_grm(pf00596) SPU3
21145 -pf00027pf13545 -
21150 -pf09084 -
21155 -pf00005 -
21160 -pf00528 -
21165 -pf02668 -
21170 HMMalddh_2_egra(pf00171) RMM1 EP_CTERM
21175 Tdp_manilla RMM1
21180 HMMnopfamrmm1_1_rmm1(pf12680) RMM1
21185 P_cobaltBlue RMM1
21190 H_darkIndigo RMM1
21195 Tdp_parchment(#) RMM1
21200 -# -
21205 -pf00393pf03446 -
21210 -pf00082 -
21215 -pf00004 -
21220 -pf02800pf00044 -
21225 -pf01425 -
21230 -pf07687pf01546 -
21235 -pf03401 -
21240 HMMreg00126_1_mix(pf00126pf034
21245 -pf02423 -
BMC-P
BMC-H
BMC-T
Aldehyde dehydrogenase
PTAC
Signature enzyme
Alcohol dehydrogenase
Regulator

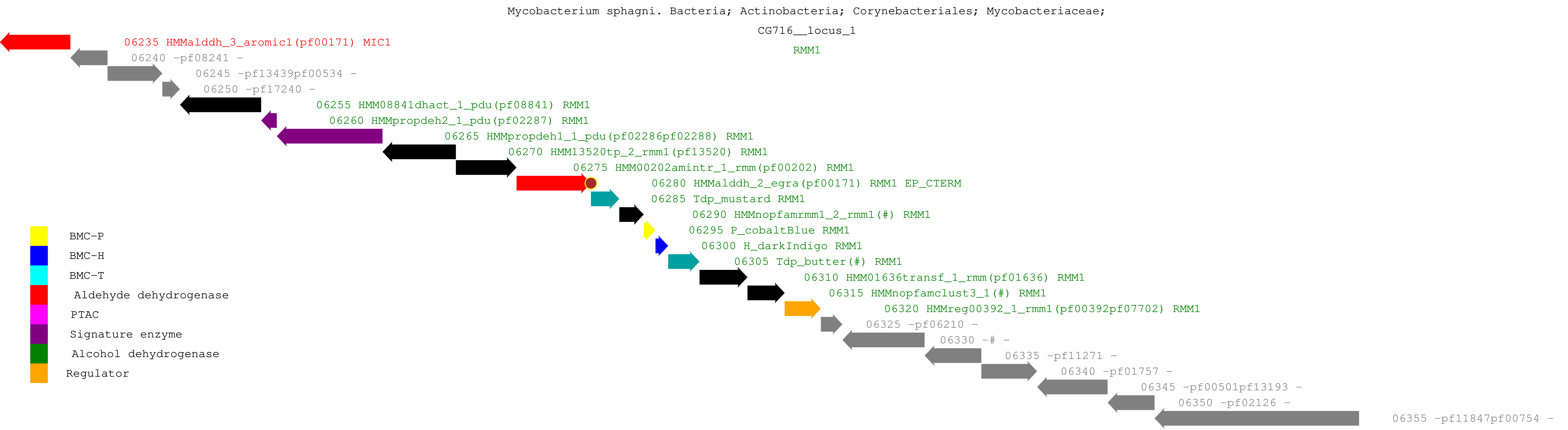
Mycobacterium sphagni. Bacteria; Actinobacteria; Corynebacteriales; Mycobacteriaceae;
CG716__locus_1
RMM1
06235 HMMaldh_3_aromic1(pf00171) MIC1
06240 -pf08241 -
06245 -pf13439pf00534 -
06250 -pf17240 -
06255 HMM08841dhact_1_pdu(pf08841) RMM1
06260 HMMpropdeh2_1_pdu(pf02287) RMM1
06265 HMMpropdeh1_1_pdu(pf02286pf02288) RMM1
06270 HMM13520tp_2_rmm1(pf13520) RMM1
06275 HMM00202amintr_1_rmm(pf00202) RMM1
06280 HMMaldh_2_egra(pf00171) RMM1 EP_CTERM
06285 Tdp_mustard RMM1
06290 HMMnopfamrmm1_2_rmm1(#) RMM1
06295 P_cobaltBlue RMM1
06300 H_darkIndigo RMM1
06305 Tdp_butter(#) RMM1
06310 HMM01636transf_1_rmm(pf01636) RMM1
06315 HMMnopfamclust3_1(#) RMM1
06320 HMMreg00392_1_rmm1(pf00392pf07702) RMM1
06325 -pf06210 -
06330 -# -
06335 -pf11271 -
06340 -pf01757 -
06345 -pf00501pf13193 -
06350 -pf02126 -
06355 -pf11847pf00754 -
BMC-P
BMC-H
BMC-T
Aldehyde dehydrogenase
PTAC
Signature enzyme
Alcohol dehydrogenase
Regulator

Mycobacterium agri. Bacteria; Actinobacteria; Corynebacteriales; Mycobacteriaceae;

CQY20__locus_1

RMM1

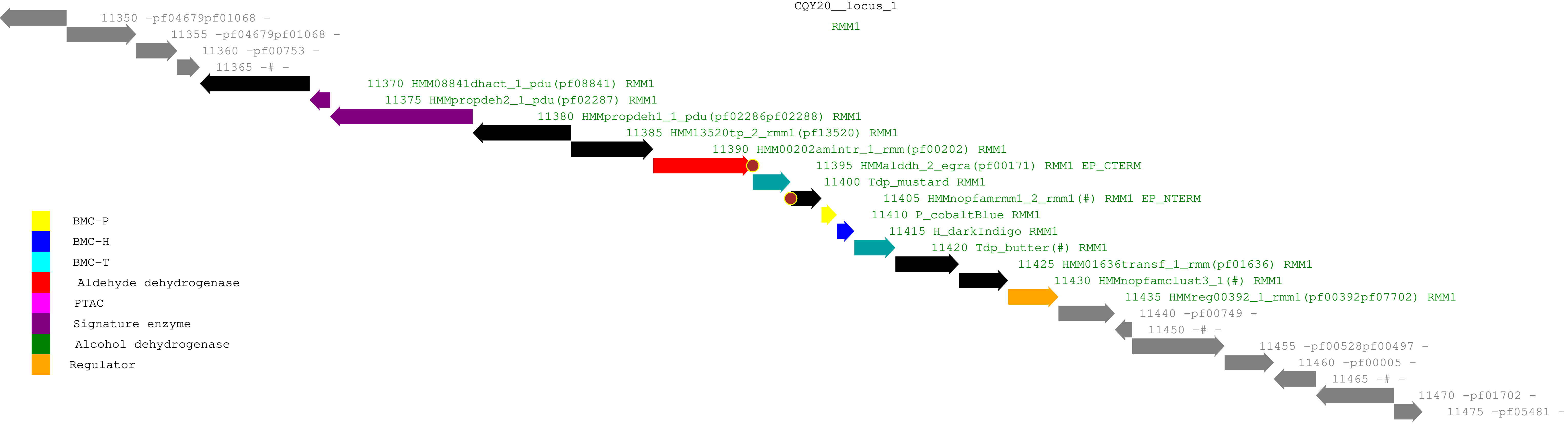

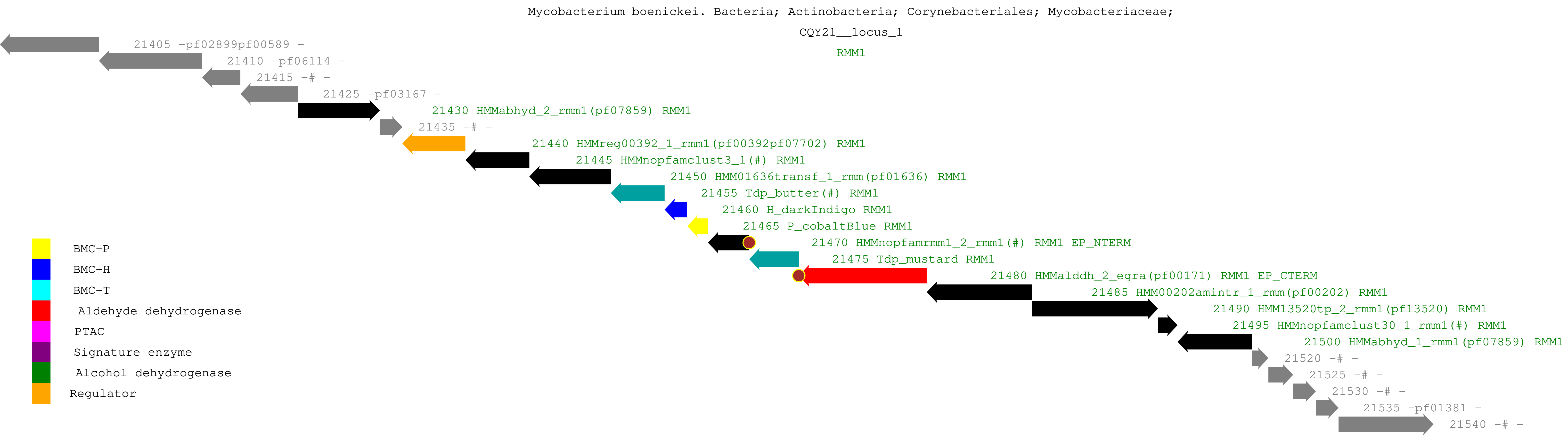
Mycobacterium boenickei. Bacteria; Actinobacteria; Corynebacteriales; Mycobacteriaceae;
CQY21__locus_1
RMM1
21405 -pf02899pf00589 -
21410 -pf06114 -
21415 -# -
21425 -pf03167 -
21430 HMMabhyd_2_rmm1(pf07859) RMM1
21435 -# -
21440 HMMreg00392_1_rmm1(pf00392pf07702) RMM1
21445 HMMnopfamclust3_1(#) RMM1
21450 HMM01636transf_1_rmm(pf01636) RMM1
21455 Tdp_butter(#) RMM1
21460 H_darkIndigo RMM1
21465 P_cobaltBlue RMM1
21470 HMMnopfamrmm1_2_rmm1(#) RMM1 EP_NTERM
21475 Tdp_mustard RMM1
21480 HMMalddh_2_egra(pf00171) RMM1 EP_CTERM
21485 HMM00202amintr_1_rmm(pf00202) RMM1
21490 HMM13520tp_2_rmm1(pf13520) RMM1
21495 HMMnopfamclust30_1_rmm1(#) RMM1
21500 HMMabhyd_1_rmm1(pf07859) RMM1
21520 -# -
21525 -# -
21530 -# -
21535 -pf01381 -
21540 -# -
BMC-P
BMC-H
BMC-T
Aldehyde dehydrogenase
PTAC
Signature enzyme
Alcohol dehydrogenase
Regulator

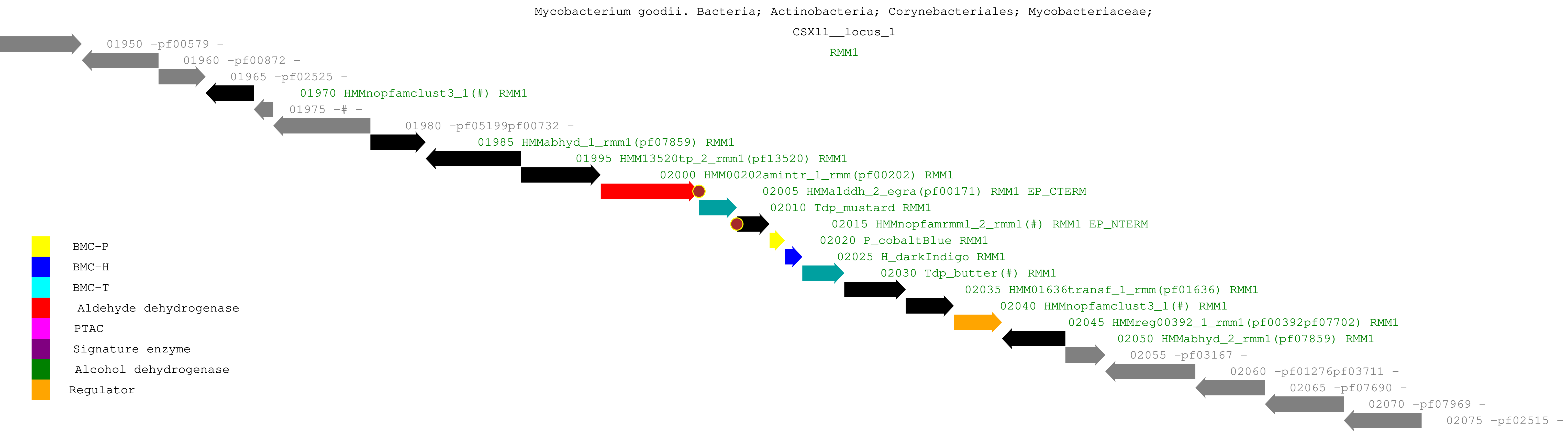

Mycobacterium goodii. Bacteria; Actinobacteria; Corynebacteriales; Mycobacteriaceae;
CSX11__locus_1
RMM1
01950 -pf00579 -
01960 -pf00872 -
01965 -pf02525 -
01970 HMMnopfamclust3_1(#) RMM1
01975 -# -
01980 -pf05199pf00732 -
01985 HMMabhyd_1_rmm1(pf07859) RMM1
01995 HMM13520tp_2_rmm1(pf13520) RMM1
02000 HMM00202amintr_1_rmm(pf00202) RMM1
02005 HMMalddh_2_egra(pf00171) RMM1 EP_CTERM
02010 Tdp_mustard RMM1
02015 HMMnopfamrmm1_2_rmm1(#) RMM1 EP_NTERM
02020 P_cobaltBlue RMM1
02025 H_darkIndigo RMM1
02030 Tdp_butter(#) RMM1
02035 HMM01636transf_1_rmm(pf01636) RMM1
02040 HMMnopfamclust3_1(#) RMM1
02045 HMMreg00392_1_rmm1(pf00392pf07702) RMM1
02050 HMMabhyd_2_rmm1(pf07859) RMM1
02055 -pf03167 -
02060 -pf01276pf03711 -
02065 -pf07690 -
02070 -pf07969 -
02075 -pf02515 -
BMC-P
BMC-H
BMC-T
Aldehyde dehydrogenase
PTAC
Signature enzyme
Alcohol dehydrogenase
Regulator

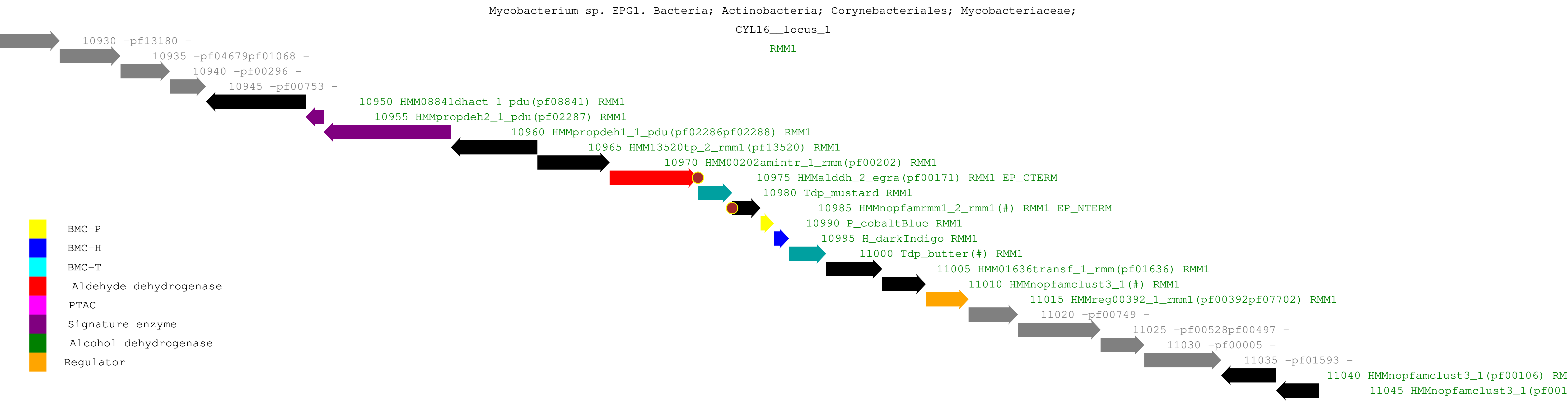
Mycobacterium sp. EPG1. Bacteria; Actinobacteria; Corynebacteriales; Mycobacteriaceae;
CYL16__locus_1
RMM1
10930 -pf13180 -
10935 -pf04679pf01068 -
10940 -pf00296 -
10945 -pf00753 -
10950 HMM08841dhact_1_pdu(pf08841) RMM1
10955 HMMpropdeh2_1_pdu(pf02287) RMM1
10960 HMMpropdeh1_1_pdu(pf02286pf02288) RMM1
10965 HMM13520tp_2_rmm1(pf13520) RMM1
10970 HMM00202amintr_1_rmm(pf00202) RMM1
10975 HMMalddh_2_egra(pf00171) RMM1 EP_CTERM
10980 Tdp_mustard RMM1
10985 HMMnopfamrmm1_2_rmm1(#) RMM1 EP_NTERM
10990 P_cobaltBlue RMM1
10995 H_darkIndigo RMM1
11000 Tdp_butter(#) RMM1
11005 HMM01636transf_1_rmm(pf01636) RMM1
11010 HMMnopfamclust3_1(#) RMM1
11015 HMMreg00392_1_rmm1(pf00392pf07702) RMM1
11020 -pf00749 -
11025 -pf00528pf00497 -
11030 -pf00005 -
11035 -pf01593 -
11040 HMMnopfamclust3_1(pf00106) RM
11045 HMMnopfamclust3_1(pf001
BMC-P
BMC-H
BMC-T
Aldehyde dehydrogenase
PTAC
Signature enzyme
Alcohol dehydrogenase
Regulator

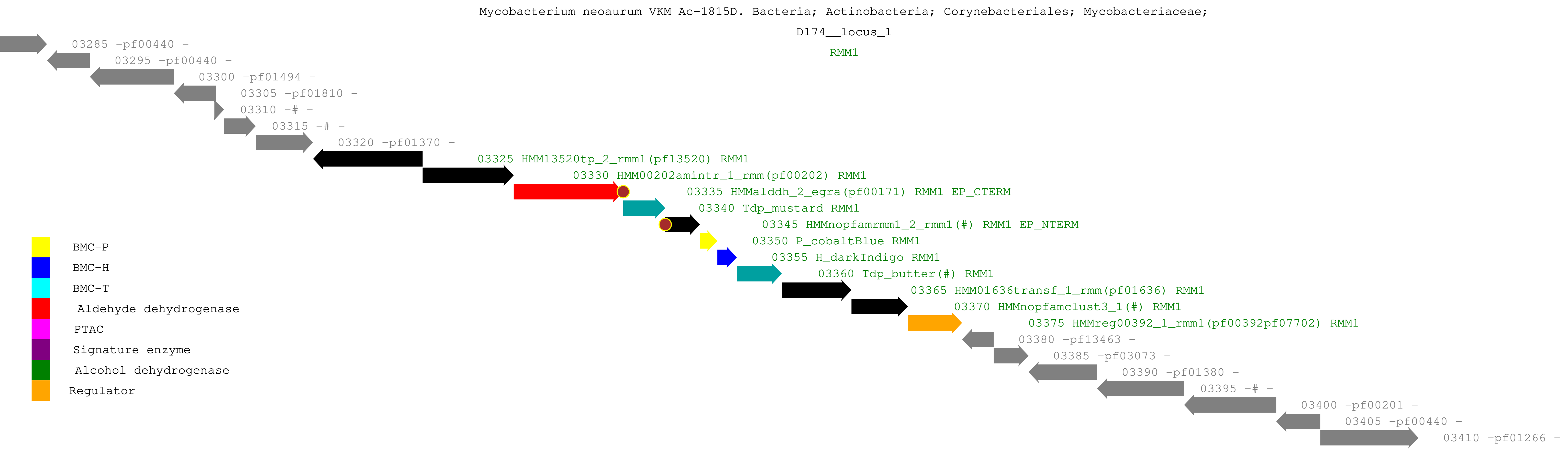

Mycobacterium neoaurum VKM Ac-1815D. Bacteria; Actinobacteria; Corynebacteriales; Mycobacteriaceae;
D174__locus_1
RMM1
03285 -pf00440 -
03295 -pf00440 -
03300 -pf01494 -
03305 -pf01810 -
03310 -# -
03315 -# -
03320 -pf01370 -
03325 HMM13520tp_2_rmm1(pf13520) RMM1
03330 HMM00202amintr_1_rmm(pf00202) RMM1
03335 HMMaldh_2_egra(pf00171) RMM1 EP_CTERM
03340 Tdp_mustard RMM1
03345 HMMnopfamrmm1_2_rmm1(#) RMM1 EP_NTERM
03350 P_cobaltBlue RMM1
03355 H_darkIndigo RMM1
03360 Tdp_butter(#) RMM1
03365 HMM01636transf_1_rmm(pf01636) RMM1
03370 HMMnopfamclust3_1(#) RMM1
03375 HMMreg00392_1_rmm1(pf00392pf07702) RMM1
03380 -pf13463 -
03385 -pf03073 -
03390 -pf01380 -
03395 -# -
03400 -pf00201 -
03405 -pf00440 -
03410 -pf01266 -
BMC-P
BMC-H
BMC-T
Aldehyde dehydrogenase
PTAC
Signature enzyme
Alcohol dehydrogenase
Regulator

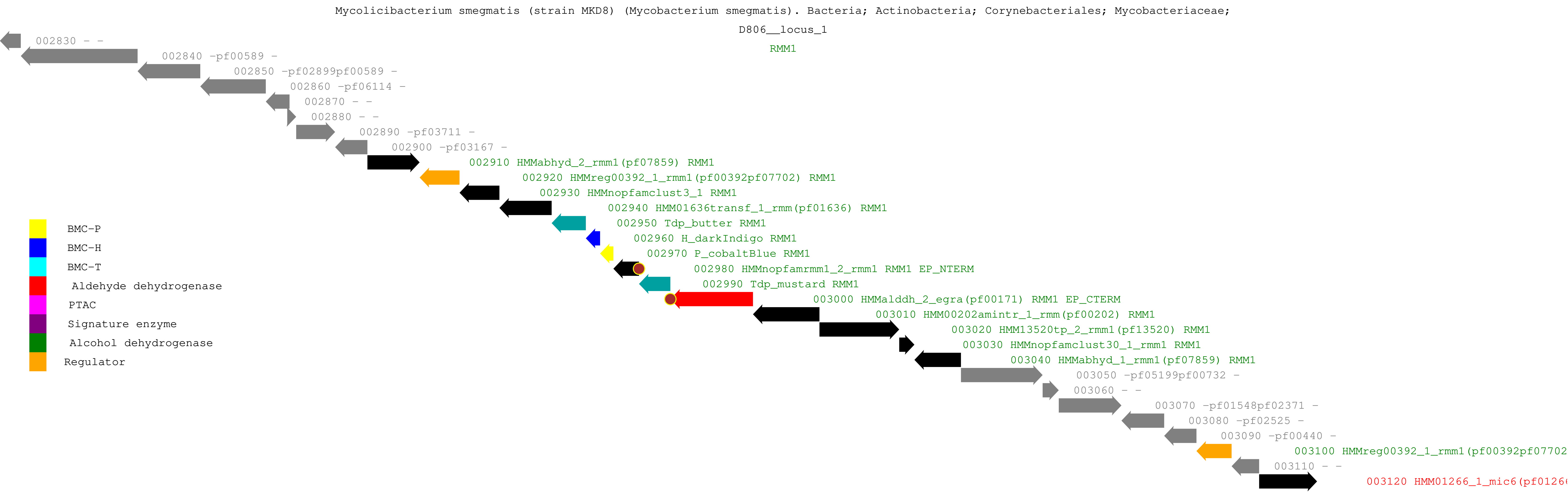
Mycolicibacterium smegmatis (strain MKD8) (Mycobacterium smegmatis). Bacteria; Actinobacteria; Corynebacteriales; Mycobacteriaceae;
D806__locus_1
RMM1
002830 - -
002840 -pf00589 -
002850 -pf02899pf00589 -
002860 -pf06114 -
002870 - -
002880 - -
002890 -pf03711 -
002900 -pf03167 -
002910 HMMabhyd_2_rmm1(pf07859) RMM1
002920 HMMreg00392_1_rmm1(pf00392pf07702) RMM1
002930 HMMnopfamclust3_1 RMM1
002940 HMM01636transf_1_rmm(pf01636) RMM1
002950 Tdp_butter RMM1
002960 H_darkIndigo RMM1
002970 P_cobaltBlue RMM1
002980 HMMnopfamrmm1_2_rmm1 RMM1 EP_NTERM
002990 Tdp_mustard RMM1
003000 HMMaldh_2_egra(pf00171) RMM1 EP_CTERM
003010 HMM00202amintr_1_rmm(pf00202) RMM1
003020 HMM13520tp_2_rmm1(pf13520) RMM1
003030 HMMnopfamclust30_1_rmm1 RMM1
003040 HMMabhyd_1_rmm1(pf07859) RMM1
003050 -pf05199pf00732 -
003060 - -
003070 -pf01548pf02371 -
003080 -pf02525 -
003090 -pf00440 -
003100 HMMreg00392_1_rmm1(pf00392pf07702
003110 - -
003120 HMM01266_1_mic6(pf0126
BMC-P
BMC-H
BMC-T
Aldehyde dehydrogenase
PTAC
Signature enzyme
Alcohol dehydrogenase
Regulator

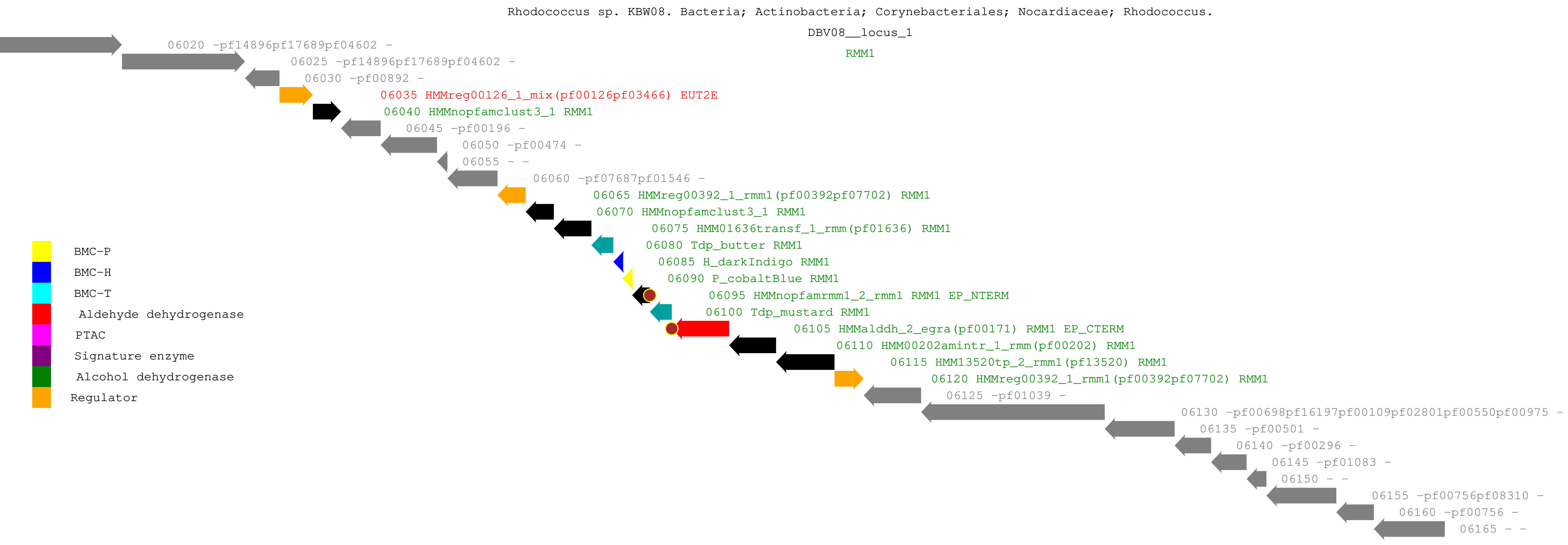
Rhodococcus sp. KBW08. Bacteria; Actinobacteria; Corynebacteriales; Nocardiaceae; Rhodococcus.
DBV08__locus_1
RMM1
06020 -pf14896pf17689pf04602 -
06025 -pf14896pf17689pf04602 -
06030 -pf00892 -
06035 HMMreg00126_1_mix(pf00126pf03466) EUT2E
06040 HMMnopfamclust3_1 RMM1
06045 -pf00196 -
06050 -pf00474 -
06055 - -
06060 -pf07687pf01546 -
06065 HMMreg00392_1_rmm1(pf00392pf07702) RMM1
06070 HMMnopfamclust3_1 RMM1
06075 HMM01636transf_1_rmm(pf01636) RMM1
06080 Tdp_butter RMM1
06085 H_darkIndigo RMM1
06090 P_cobaltBlue RMM1
06095 HMMnopfamrmm1_2_rmm1 RMM1 EP_NTERM
06100 Tdp_mustard RMM1
06105 HMMalddh_2_egra(pf00171) RMM1 EP_CTERM
06110 HMM00202amintr_1_rmm(pf00202) RMM1
06115 HMM13520tp_2_rmm1(pf13520) RMM1
06120 HMMreg00392_1_rmm1(pf00392pf07702) RMM1
06125 -pf01039 -
06130 -pf00698pf16197pf00109pf02801pf00550pf00975 -
06135 -pf00501 -
06140 -pf00296 -
06145 -pf01083 -
06150 - -
06155 -pf00756pf08310 -
06160 -pf00756 -
06165 - -
BMC-P
BMC-H
BMC-T
Aldehyde dehydrogenase
PTAC
Signature enzyme
Alcohol dehydrogenase
Regulator

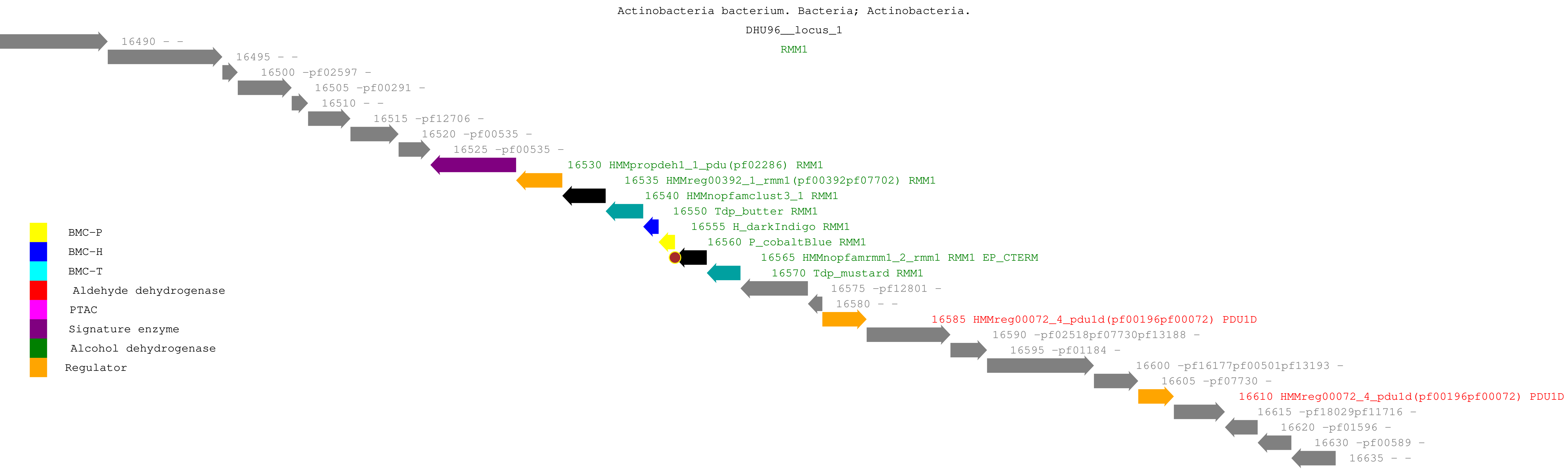
Actinobacteria bacterium. Bacteria; Actinobacteria.
DHU96__locus_1
RMM1
16490 - -
16495 - -
16500 -pf02597 -
16505 -pf00291 -
16510 - -
16515 -pf12706 -
16520 -pf00535 -
16525 -pf00535 -
16530 HMMpropdeh1_1_pdu(pf02286) RMM1
16535 HMMreg00392_1_rmm1(pf00392pf07702) RMM1
16540 HMMnopfamclust3_1 RMM1
16550 Tdp_butter RMM1
16555 H_darkIndigo RMM1
16560 P_cobaltBlue RMM1
16565 HMMnopfamrmm1_2_rmm1 RMM1 EP_CTERM
16570 Tdp_mustard RMM1
16575 -pf12801 -
16580 - -
16585 HMMreg00072_4_pdu1d(pf00196pf00072) PDU1D
16590 -pf02518pf07730pf13188 -
16595 -pf01184 -
16600 -pf16177pf00501pf13193 -
16605 -pf07730 -
16610 HMMreg00072_4_pdu1d(pf00196pf00072) PDU1D
16615 -pf18029pf11716 -
16620 -pf01596 -
16630 -pf00589 -
16635 - -
BMC-P
BMC-H
BMC-T
Aldehyde dehydrogenase
PTAC
Signature enzyme
Alcohol dehydrogenase
Regulator

Mycolicibacterium hassiacum. Bacteria; Actinobacteria; Corynebacteriales; Mycobacteriaceae;

DIU75__locus_1

RMM1

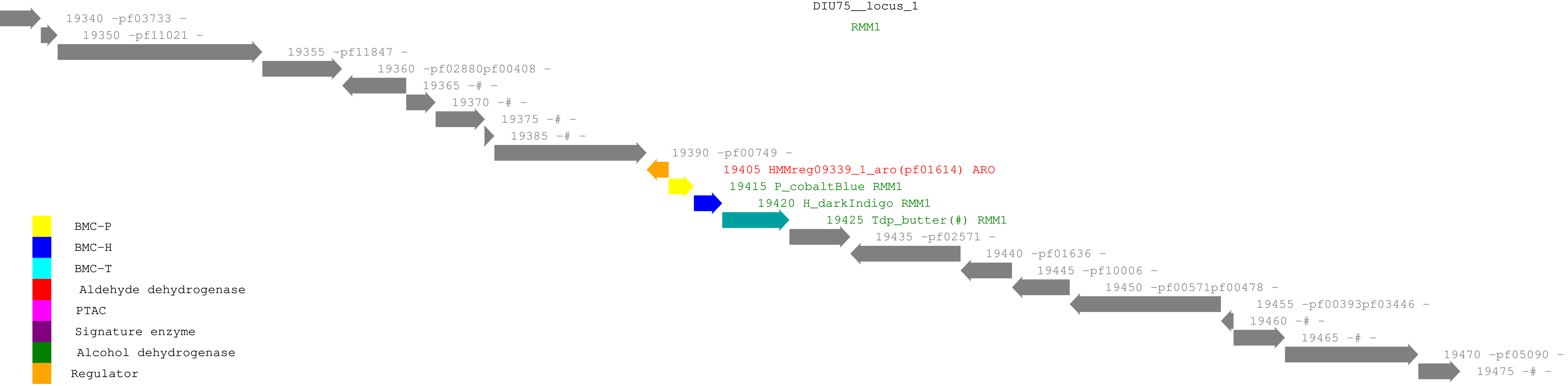

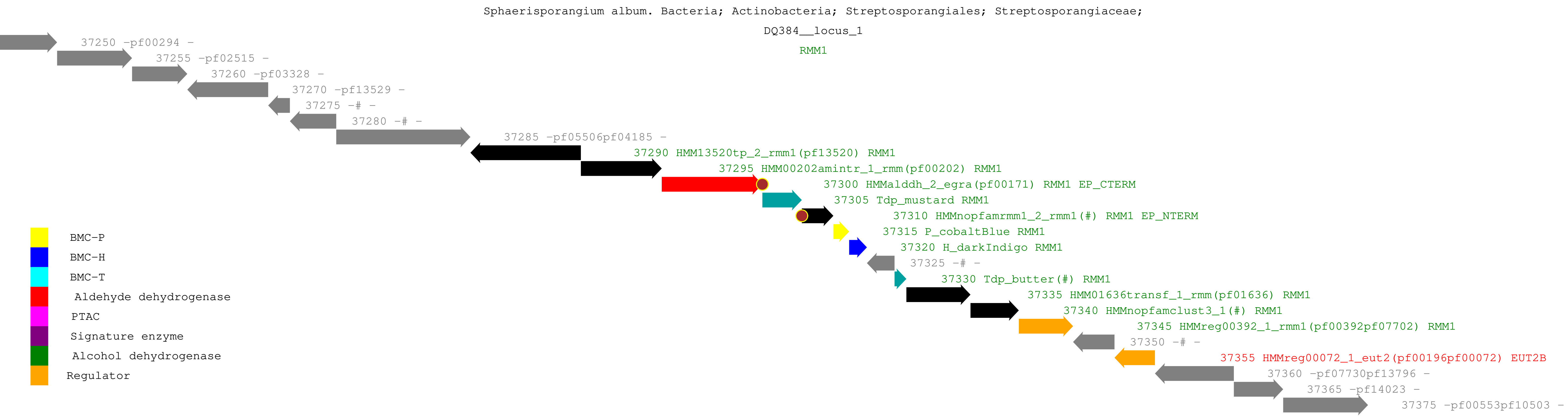
Sphaerisporangium album. Bacteria; Actinobacteria; Streptosporangiales; Streptosporangiaceae;
DQ384__locus_1
RMM1
37250 -pf00294 -
37255 -pf02515 -
37260 -pf03328 -
37270 -pf13529 -
37275 -# -
37280 -# -
37285 -pf05506pf04185 -
37290 HMM13520tp_2_rmm1(pf13520) RMM1
37295 HMM00202amintr_1_rmm(pf00202) RMM1
37300 HMMalddh_2_egra(pf00171) RMM1 EP_CTERM
37305 Tdp_mustard RMM1
37310 HMMnopfamrmm1_2_rmm1(#) RMM1 EP_NTERM
37315 P_cobaltBlue RMM1
37320 H_darkIndigo RMM1
37325 -# -
37330 Tdp_butter(#) RMM1
37335 HMM01636transf_1_rmm(pf01636) RMM1
37340 HMMnopfamclust3_1(#) RMM1
37345 HMMreg00392_1_rmm1(pf00392pf07702) RMM1
37350 -# -
37355 HMMreg00072_1_eut2(pf00196pf00072) EUT2B
37360 -pf07730pf13796 -
37365 -pf14023 -
37375 -pf00553pf10503 -
BMC-P
BMC-H
BMC-T
Aldehyde dehydrogenase
PTAC
Signature enzyme
Alcohol dehydrogenase
Regulator

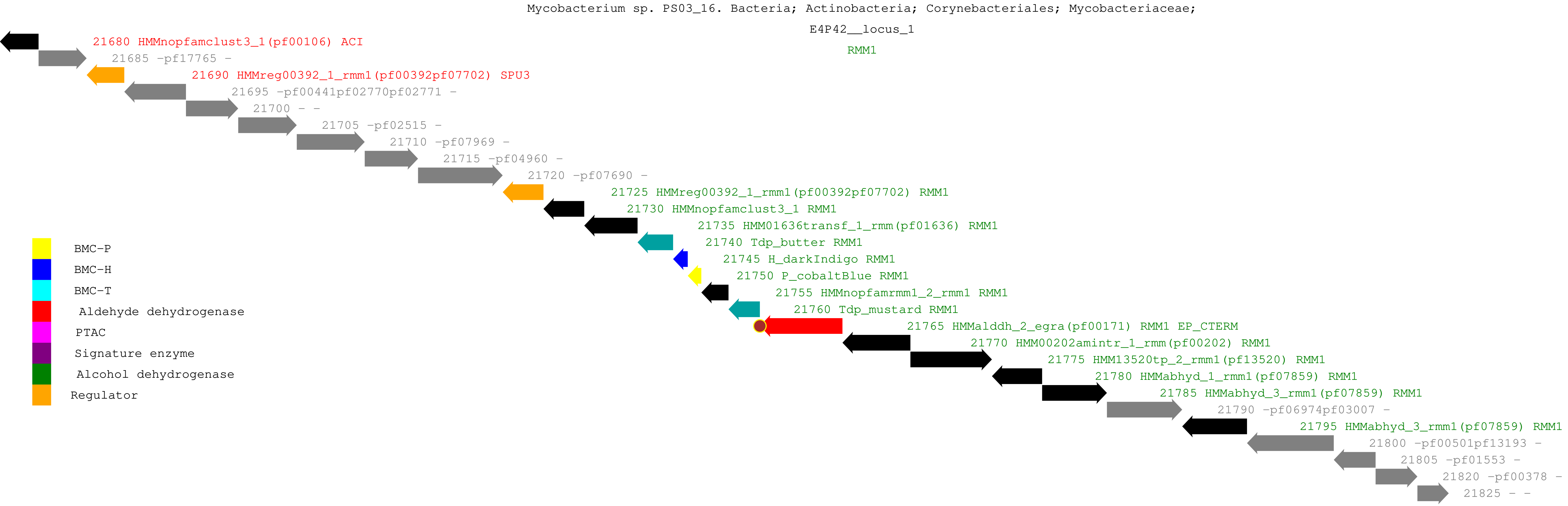
Mycobacterium sp. PS03_16. Bacteria; Actinobacteria; Corynebacteriales; Mycobacteriaceae;
E4P42__locus_1
RMM1
21680 HMMnopfamclust3_1(pf00106) ACI
21685 -pf17765 -
21690 HMMreg00392_1_rmm1(pf00392pf07702) SPU3
21695 -pf00441pf02770pf02771 -
21700 - -
21705 -pf02515 -
21710 -pf07969 -
21715 -pf04960 -
21720 -pf07690 -
21725 HMMreg00392_1_rmm1(pf00392pf07702) RMM1
21730 HMMnopfamclust3_1 RMM1
21735 HMM01636transf_1_rmm(pf01636) RMM1
21740 Tdp_butter RMM1
21745 H_darkIndigo RMM1
21750 P_cobaltBlue RMM1
21755 HMMnopfamrmm1_2_rmm1 RMM1
21760 Tdp_mustard RMM1
21765 HMMalddh_2_egra(pf00171) RMM1 EP_CTERM
21770 HMM00202amintr_1_rmm(pf00202) RMM1
21775 HMM13520tp_2_rmm1(pf13520) RMM1
21780 HMMabhyd_1_rmm1(pf07859) RMM1
21785 HMMabhyd_3_rmm1(pf07859) RMM1
21790 -pf06974pf03007 -
21795 HMMabhyd_3_rmm1(pf07859) RMM1
21800 -pf00501pf13193 -
21805 -pf01553 -
21820 -pf00378 -
21825 - -
BMC-P
BMC-H
BMC-T
Aldehyde dehydrogenase
PTAC
Signature enzyme
Alcohol dehydrogenase
Regulator

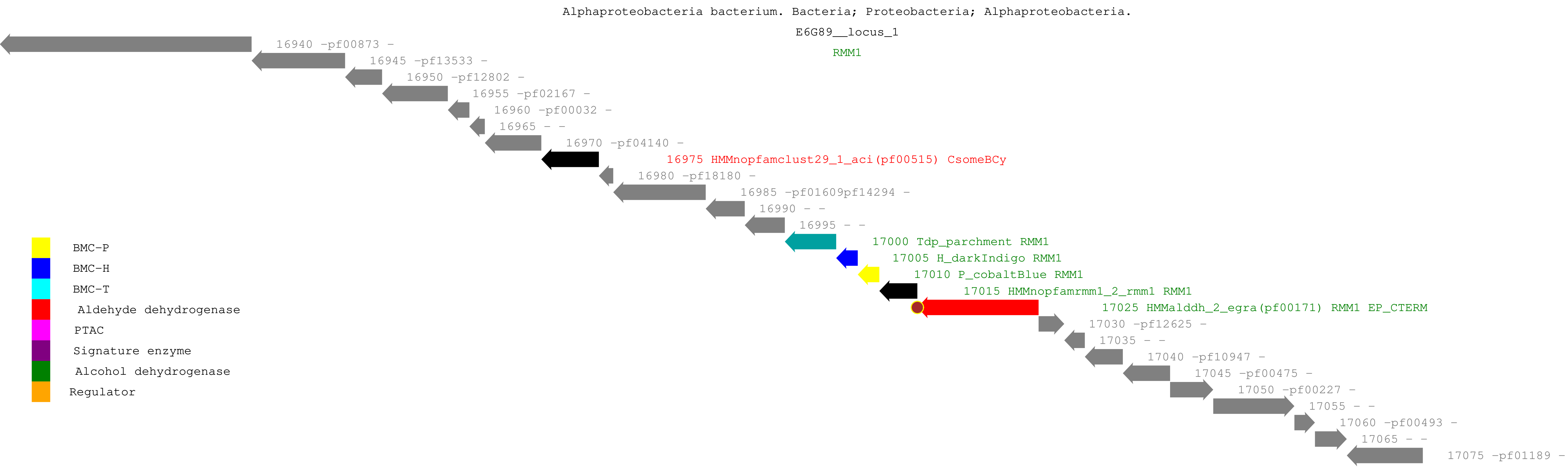

Alphaproteobacteria bacterium. Bacteria; Proteobacteria; Alphaproteobacteria.
E6G89__locus_1
RMM1
16940 -pf00873 -
16945 -pf13533 -
16950 -pf12802 -
16955 -pf02167 -
16960 -pf00032 -
16965 - -
16970 -pf04140 -
16975 HMMnopfamclust29_1_aci(pf00515) CsomeBCy
16980 -pf18180 -
16985 -pf01609pf14294 -
16990 - -
16995 - -
17000 Tdp_parchment RMM1
17005 H_darkIndigo RMM1
17010 P_cobaltBlue RMM1
17015 HMMnopfamrmm1_2_rmm1 RMM1
17025 HMMalddh_2_egra(pf00171) RMM1 EP_CTERM
17030 -pf12625 -
17035 - -
17040 -pf10947 -
17045 -pf00475 -
17050 -pf00227 -
17055 - -
17060 -pf00493 -
17065 - -
17075 -pf01189 -
BMC-P
BMC-H
BMC-T
Aldehyde dehydrogenase
PTAC
Signature enzyme
Alcohol dehydrogenase
Regulator

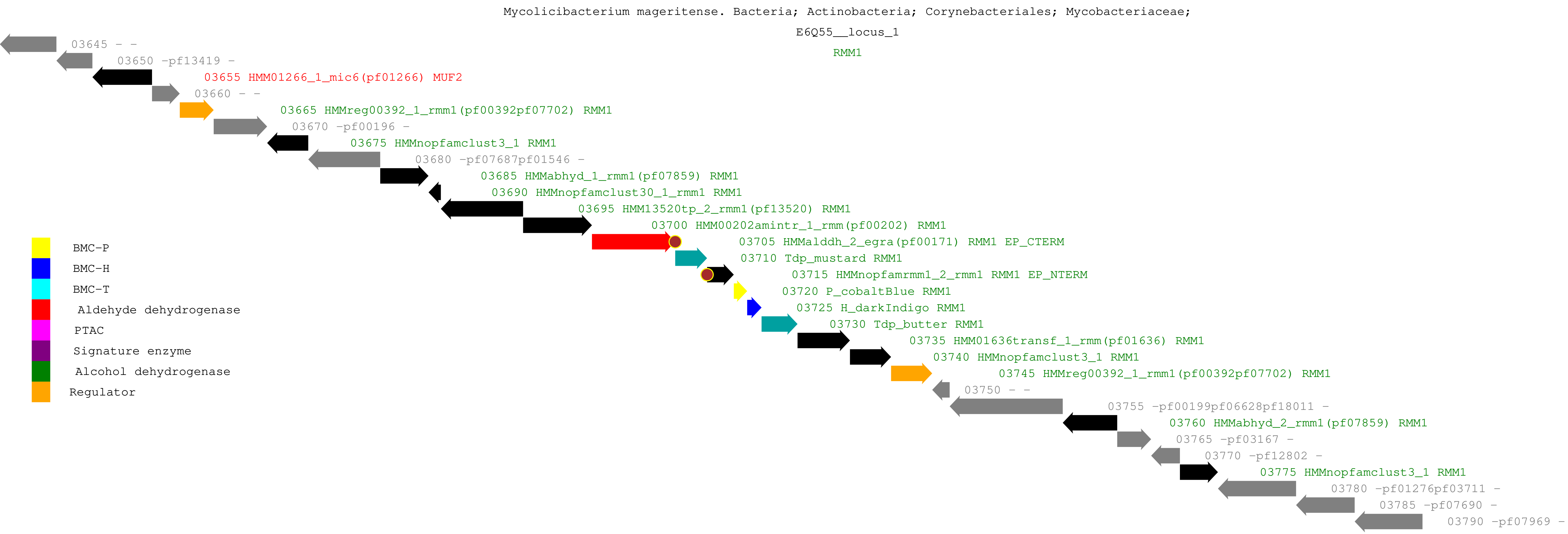
Mycolicibacterium mageritense. Bacteria; Actinobacteria; Corynebacteriales; Mycobacteriaceae;
E6Q55__locus_1
RMM1
03645 - -
03650 -pf13419 -
03655 HMM01266_1_mic6(pf01266) MUF2
03660 - -
03665 HMMreg00392_1_rmm1(pf00392pf07702) RMM1
03670 -pf00196 -
03675 HMMnopfamclust3_1 RMM1
03680 -pf07687pf01546 -
03685 HMMabhyd_1_rmm1(pf07859) RMM1
03690 HMMnopfamclust30_1_rmm1 RMM1
03695 HMM13520tp_2_rmm1(pf13520) RMM1
03700 HMM00202amintr_1_rmm(pf00202) RMM1
03705 HMMalddh_2_egra(pf00171) RMM1 EP_CTERM
03710 Tdp_mustard RMM1
03715 HMMnopfamrmm1_2_rmm1 RMM1 EP_NTERM
03720 P_cobaltBlue RMM1
03725 H_darkIndigo RMM1
03730 Tdp_butter RMM1
03735 HMM01636transf_1_rmm(pf01636) RMM1
03740 HMMnopfamclust3_1 RMM1
03745 HMMreg00392_1_rmm1(pf00392pf07702) RMM1
03750 - -
03755 -pf00199pf06628pf18011 -
03760 HMMabhyd_2_rmm1(pf07859) RMM1
03765 -pf03167 -
03770 -pf12802 -
03775 HMMnopfamclust3_1 RMM1
03780 -pf01276pf03711 -
03785 -pf07690 -
03790 -pf07969 -
BMC-P
BMC-H
BMC-T
Aldehyde dehydrogenase
PTAC
Signature enzyme
Alcohol dehydrogenase
Regulator

Mycobacterium sp. SWH-M1. Bacteria; Actinobacteria; Corynebacteriales; Mycobacteriaceae;

EB72__locus_1

RMM1

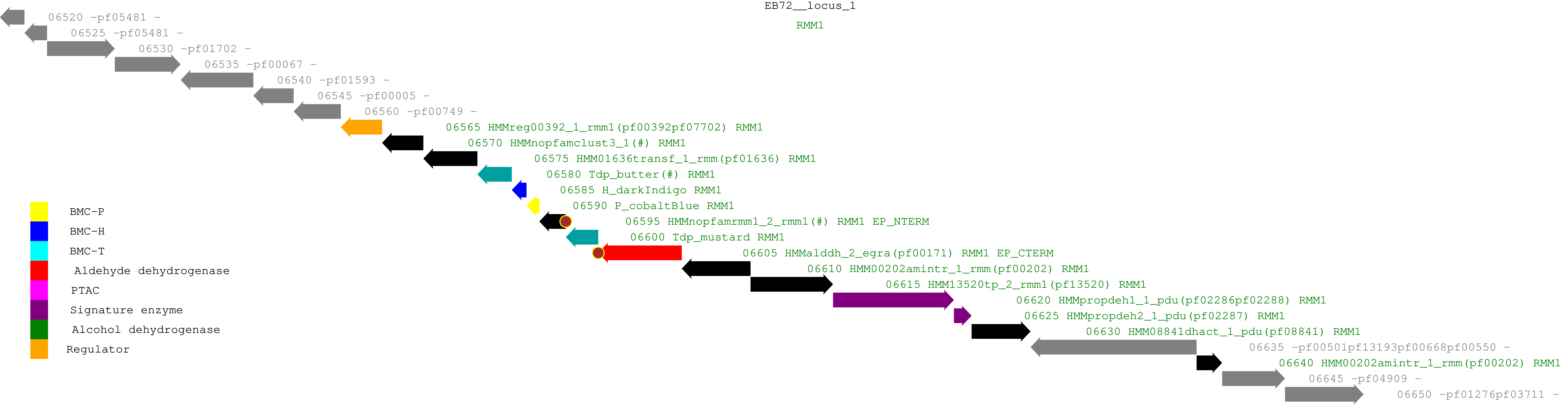

Mycobacterium sp. SWH-M3. Bacteria; Actinobacteria; Corynebacteriales; Mycobacteriaceae;

EB73__locus_1

RMM1

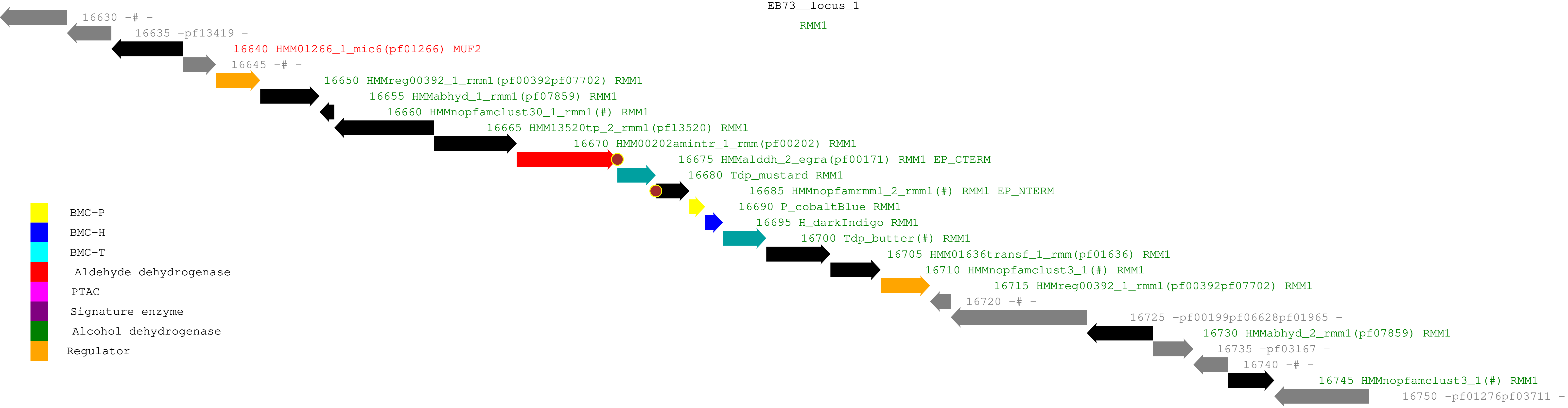

Mycobacterium sp. SWH-M5. Bacteria; Actinobacteria; Corynebacteriales; Mycobacteriaceae;

EB74__locus_1

RMM1

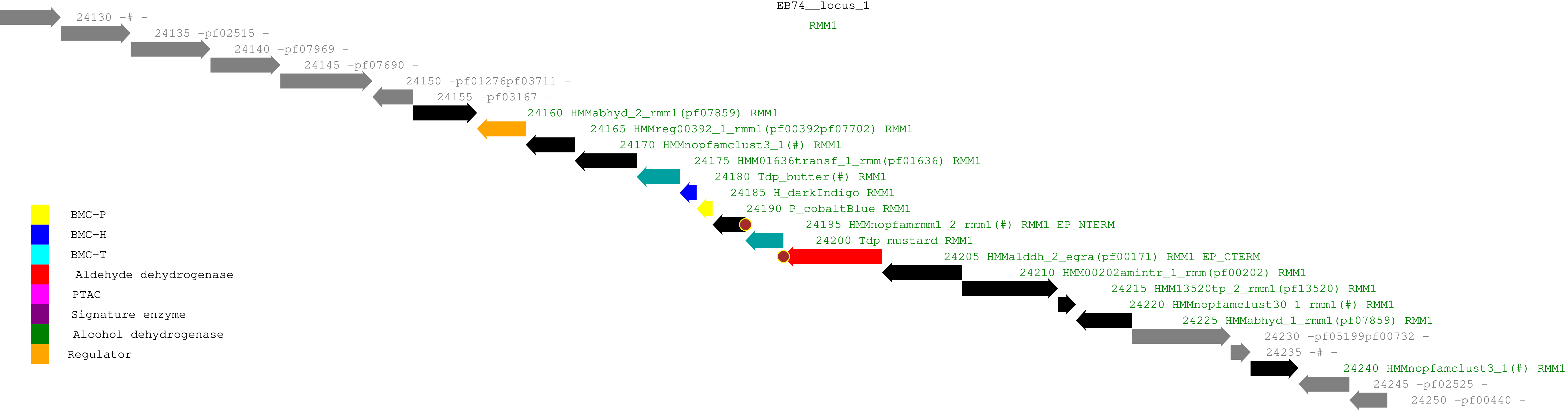

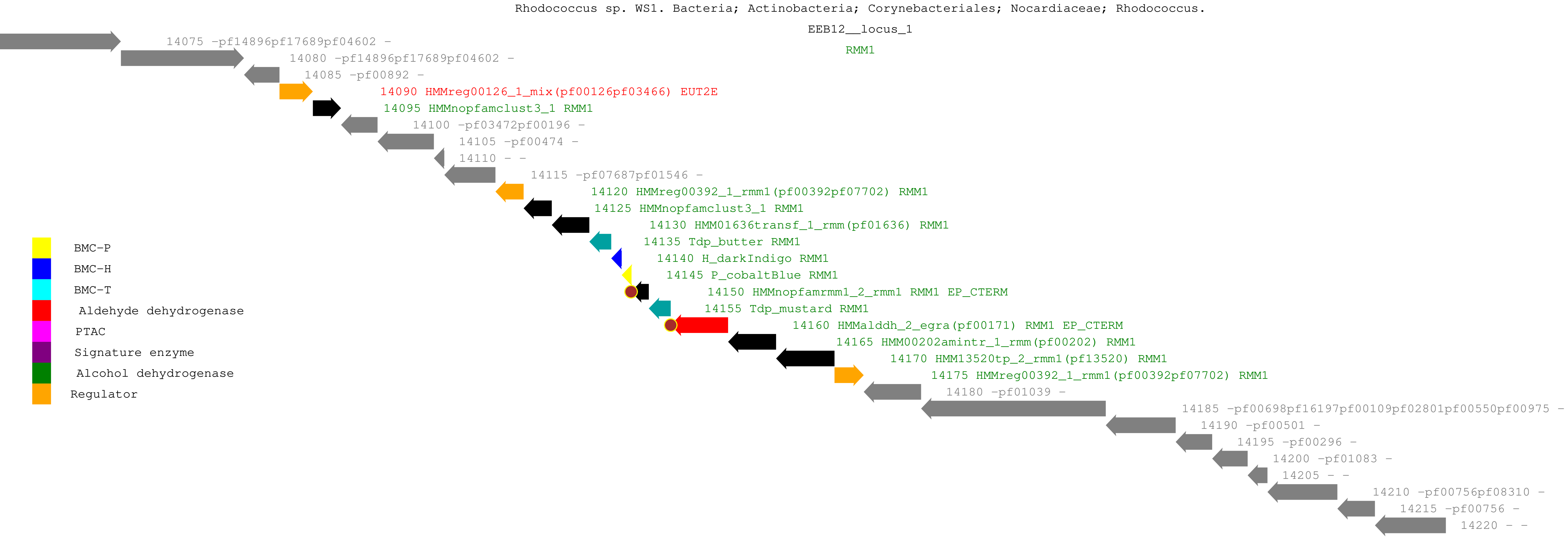
Rhodococcus sp. WS1. Bacteria; Actinobacteria; Corynebacteriales; Nocardiaceae; Rhodococcus.
EEB12__locus_1
RMM1
14075 -pf14896pf17689pf04602 -
14080 -pf14896pf17689pf04602 -
14085 -pf00892 -
14090 HMMreg00126_1_mix(pf00126pf03466) EUT2E
14095 HMMnopfamclust3_1 RMM1
14100 -pf03472pf00196 -
14105 -pf00474 -
14110 - -
14115 -pf07687pf01546 -
14120 HMMreg00392_1_rmm1(pf00392pf07702) RMM1
14125 HMMnopfamclust3_1 RMM1
14130 HMM01636transf_1_rmm(pf01636) RMM1
14135 Tdp_butter RMM1
14140 H_darkIndigo RMM1
14145 P_cobaltBlue RMM1
14150 HMMnopfamrmm1_2_rmm1 RMM1 EP_CTERM
14155 Tdp_mustard RMM1
14160 HMMalddh_2_egra(pf00171) RMM1 EP_CTERM
14165 HMM00202amintr_1_rmm(pf00202) RMM1
14170 HMM13520tp_2_rmm1(pf13520) RMM1
14175 HMMreg00392_1_rmm1(pf00392pf07702) RMM1
14180 -pf01039 -
14185 -pf00698pf16197pf00109pf02801pf00550pf00975 -
14190 -pf00501 -
14195 -pf00296 -
14200 -pf01083 -
14205 - -
14210 -pf00756pf08310 -
14215 -pf00756 -
14220 - -
BMC-P
BMC-H
BMC-T
Aldehyde dehydrogenase
PTAC
Signature enzyme
Alcohol dehydrogenase
Regulator

Rhodococcus sp. WS4. Bacteria; Actinobacteria; Corynebacteriales; Nocardiaceae; Rhodococcus.

EEB14__locus_1

RMM1

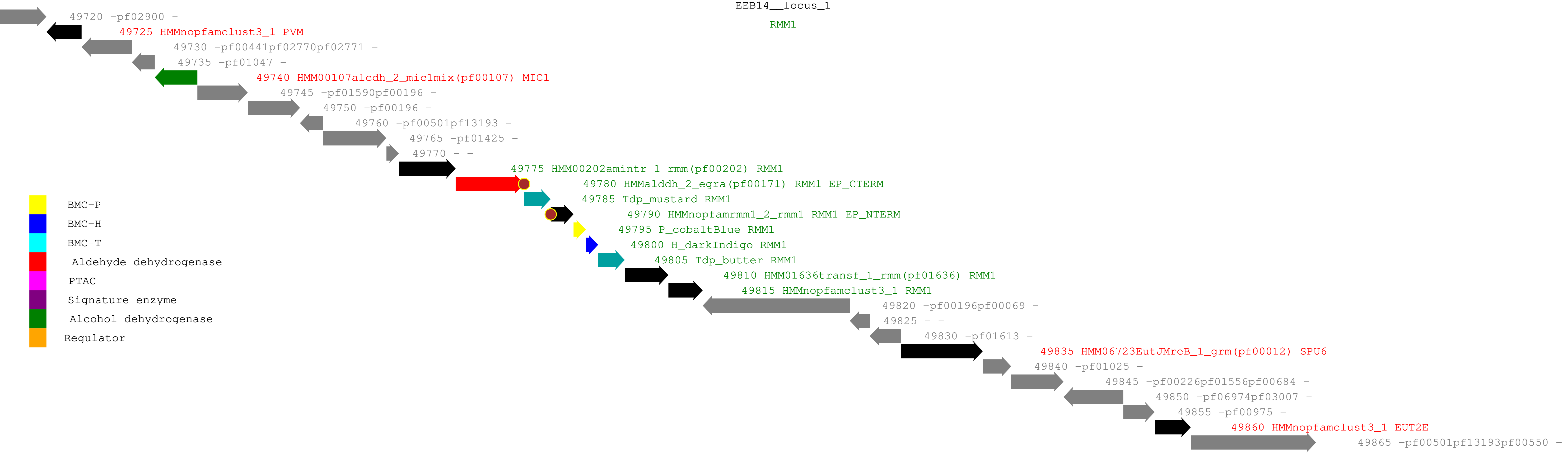

Streptomyces hyalinus. Bacteria; Actinobacteria; Streptomycetales; Streptomycetaceae;

EHYA__locus_1

RMM1

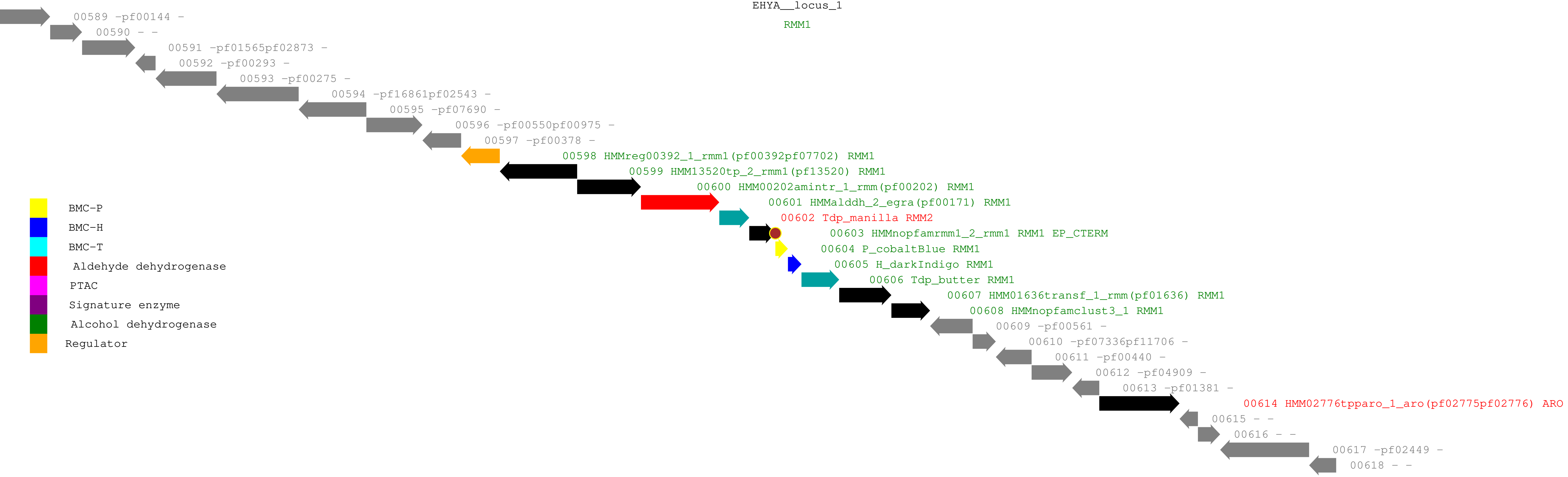

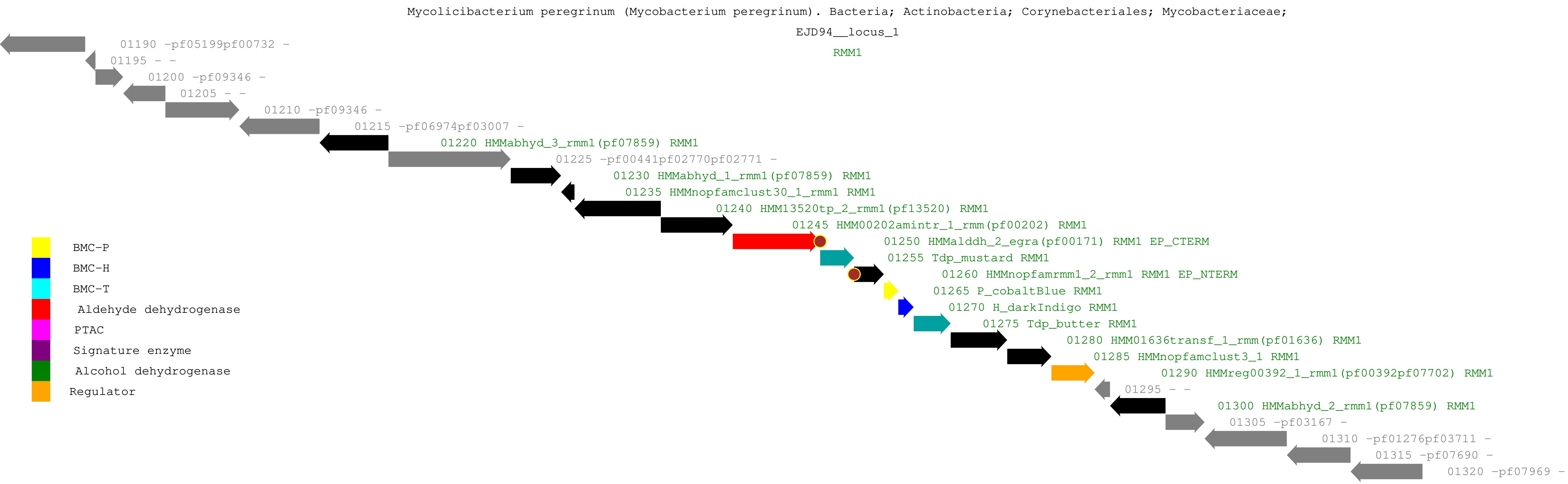
Mycolicibacterium peregrinum (Mycobacterium peregrinum). Bacteria; Actinobacteria; Corynebacteriales; Mycobacteriaceae;
EJD94__locus_1
RMM1
01190 -pf05199pf00732 -
01195 - -
01200 -pf09346 -
01205 - -
01210 -pf09346 -
01215 -pf06974pf03007 -
01220 HMMabhyd_3_rmm1(pf07859) RMM1
01225 -pf00441pf02770pf02771 -
01230 HMMabhyd_1_rmm1(pf07859) RMM1
01235 HMMnopfamclust30_1_rmm1 RMM1
01240 HMM13520tp_2_rmm1(pf13520) RMM1
01245 HMM00202amintr_1_rmm(pf00202) RMM1
01250 HMMalddh_2_egra(pf00171) RMM1 EP_CTERM
01255 Tdp_mustard RMM1
01260 HMMnopfamrmm1_2_rmm1 RMM1 EP_NTERM
01265 P_cobaltBlue RMM1
01270 H_darkIndigo RMM1
01275 Tdp_butter RMM1
01280 HMM01636transf_1_rmm(pf01636) RMM1
01285 HMMnopfamclust3_1 RMM1
01290 HMMreg00392_1_rmm1(pf00392pf07702) RMM1
01295 - -
01300 HMMabhyd_2_rmm1(pf07859) RMM1
01305 -pf03167 -
01310 -pf01276pf03711 -
01315 -pf07690 -
01320 -pf07969 -
BMC-P
BMC-H
BMC-T
Aldehyde dehydrogenase
PTAC
Signature enzyme
Alcohol dehydrogenase
Regulator

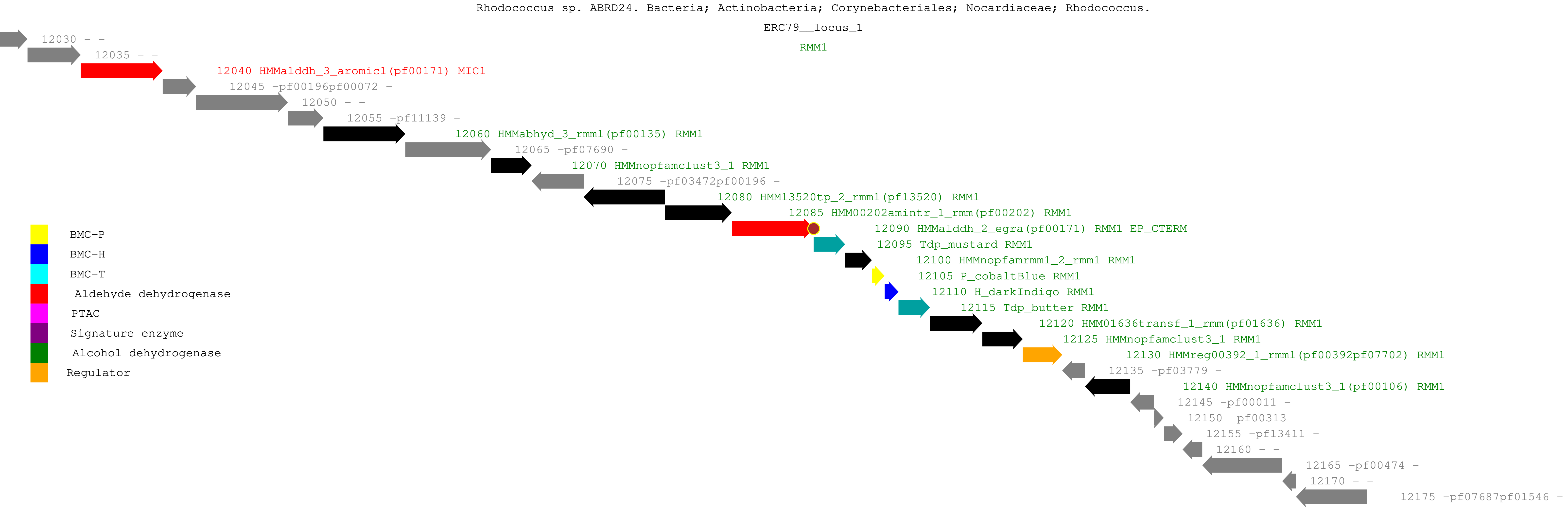

Rhodococcus sp. ABRD24. Bacteria; Actinobacteria; Corynebacteriales; Nocardiaceae; Rhodococcus.
ERC79__locus_1
RMM1
12030 – –
12035 – –
12040 HMMaldh_3_aromic1(pf00171) MIC1
12045 –pf00196pf00072 –
12050 – –
12055 –pf11139 –
12060 HMMabhyd_3_rmm1(pf00135) RMM1
12065 –pf07690 –
12070 HMMnopfamclust3_1 RMM1
12075 –pf03472pf00196 –
12080 HMM13520tp_2_rmm1(pf13520) RMM1
12085 HMM00202amintr_1_rmm(pf00202) RMM1
12090 HMMaldh_2_egra(pf00171) RMM1 EP_CTERM
12095 Tdp_mustard RMM1
12100 HMMnopfamrmm1_2_rmm1 RMM1
12105 P_cobaltBlue RMM1
12110 H_darkIndigo RMM1
12115 Tdp_butter RMM1
12120 HMM01636transf_1_rmm(pf01636) RMM1
12125 HMMnopfamclust3_1 RMM1
12130 HMMreg00392_1_rmm1(pf00392pf07702) RMM1
12135 –pf03779 –
12140 HMMnopfamclust3_1(pf00106) RMM1
12145 –pf00011 –
12150 –pf00313 –
12155 –pf13411 –
12160 – –
12165 –pf00474 –
12170 – –
12175 –pf07687pf01546 –
BMC-P
BMC-H
BMC-T
Aldehyde dehydrogenase
PTAC
Signature enzyme
Alcohol dehydrogenase
Regulator

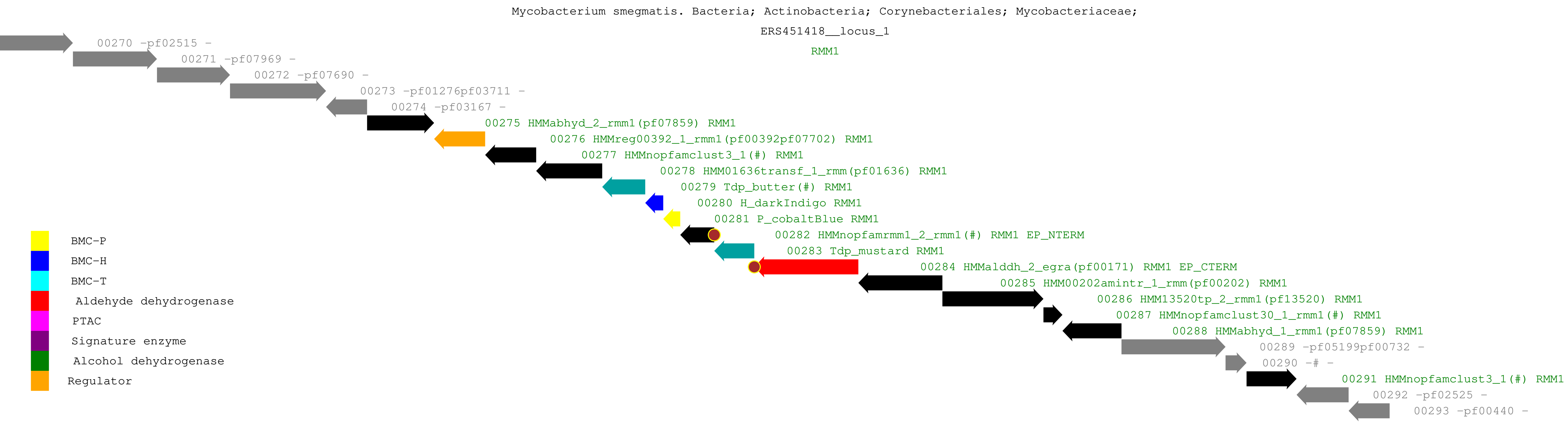
Mycobacterium smegmatis. Bacteria; Actinobacteria; Corynebacteriales; Mycobacteriaceae;
ERS451418__locus_1
RMM1
00270 -pf02515 -
00271 -pf07969 -
00272 -pf07690 -
00273 -pf01276pf03711 -
00274 -pf03167 -
00275 HMMabhyd_2_rmm1(pf07859) RMM1
00276 HMMreg00392_1_rmm1(pf00392pf07702) RMM1
00277 HMMnopfamclust3_1(#) RMM1
00278 HMM01636transf_1_rmm(pf01636) RMM1
00279 Tdp_butter(#) RMM1
00280 H_darkIndigo RMM1
00281 P_cobaltBlue RMM1
00282 HMMnopfamrmm1_2_rmm1(#) RMM1 EP_NTERM
00283 Tdp_mustard RMM1
00284 HMMalddh_2_egra(pf00171) RMM1 EP_CTERM
00285 HMM00202amintr_1_rmm(pf00202) RMM1
00286 HMM13520tp_2_rmm1(pf13520) RMM1
00287 HMMnopfamclust30_1_rmm1(#) RMM1
00288 HMMabhyd_1_rmm1(pf07859) RMM1
00289 -pf05199pf00732 -
00290 -# -
00291 HMMnopfamclust3_1(#) RMM1
00292 -pf02525 -
00293 -pf00440 -
BMC-P
BMC-H
BMC-T
Aldehyde dehydrogenase
PTAC
Signature enzyme
Alcohol dehydrogenase
Regulator

Mycobacterium rufum. Bacteria; Actinobacteria; Corynebacteriales; Mycobacteriaceae;

EU78__locus_1

RMM1

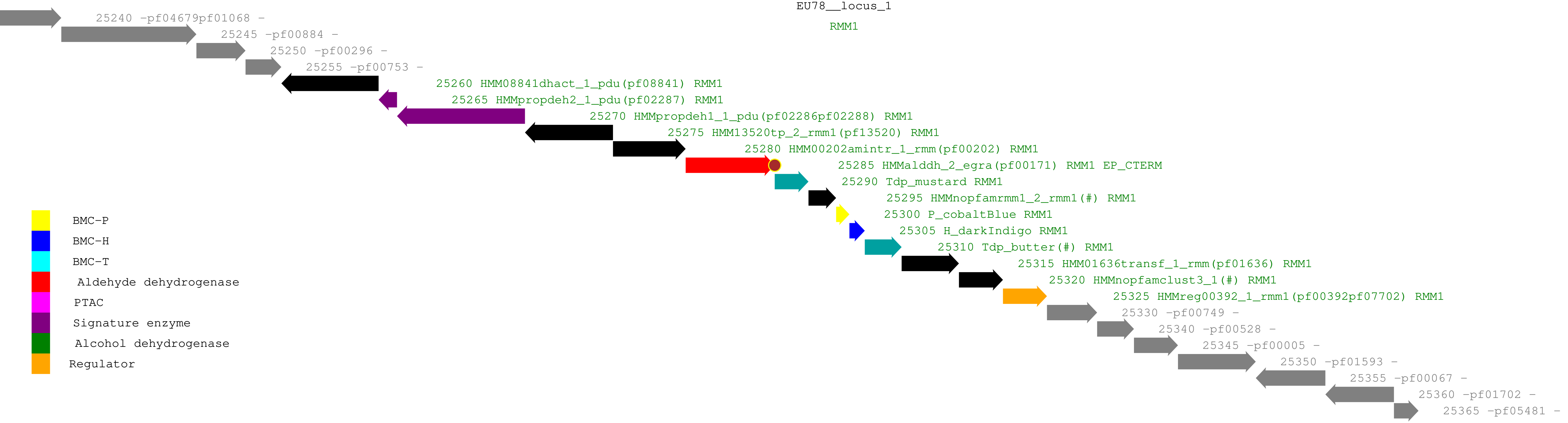

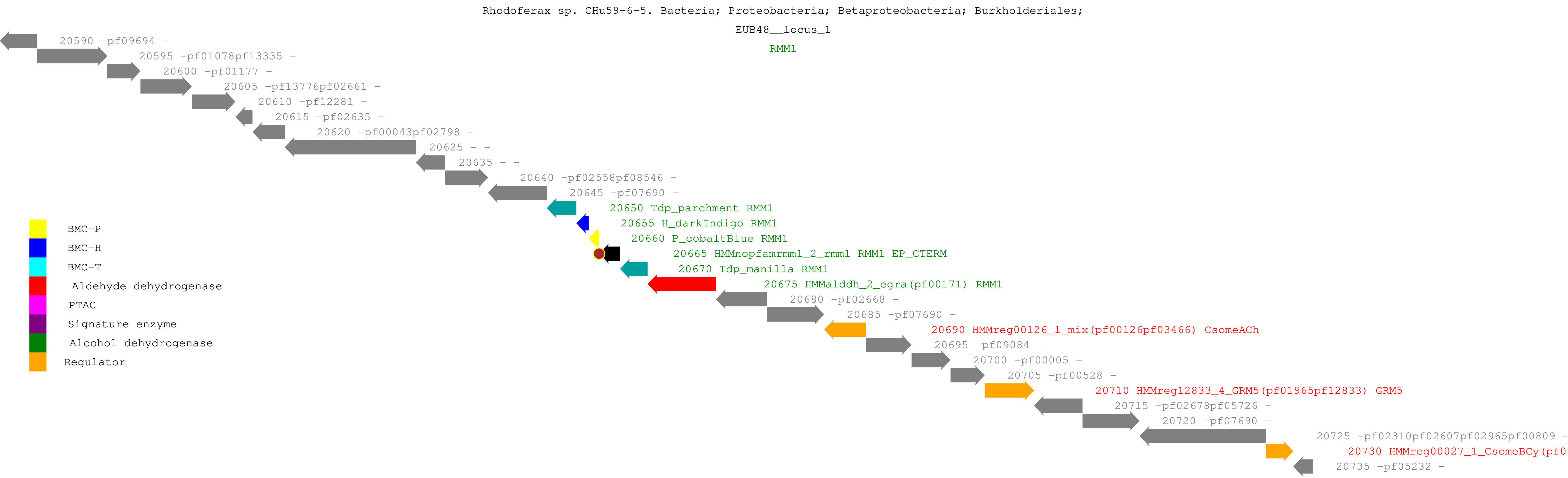
Rhodoferax sp. CHu59-6-5. Bacteria; Proteobacteria; Betaproteobacteria; Burkholderiales;
EUB48__locus_1
RMM1
20590 -pf09694 -
20595 -pf01078pf13335 -
20600 -pf01177 -
20605 -pf13776pf02661 -
20610 -pf12281 -
20615 -pf02635 -
20620 -pf00043pf02798 -
20625 - -
20635 - -
20640 -pf02558pf08546 -
20645 -pf07690 -
20650 Tdp_parchment RMM1
20655 H_darkIndigo RMM1
20660 P_cobaltBlue RMM1
20665 HMMnopfamrmm1_2_rmm1 RMM1 EP_CTERM
20670 Tdp_manilla RMM1
20675 HMMalddh_2_egra(pf00171) RMM1
20680 -pf02668 -
20685 -pf07690 -
20690 HMMreg00126_1_mix(pf00126pf03466) CsomeACh
20695 -pf09084 -
20700 -pf00005 -
20705 -pf00528 -
20710 HMMreg12833_4_GRM5(pf01965pf12833) GRM5
20715 -pf02678pf05726 -
20720 -pf07690 -
20725 -pf02310pf02607pf02965pf00809 -
20730 HMMreg00027_1_CsomeBCy(pf0
20735 -pf05232 -
BMC-P
BMC-H
BMC-T
Aldehyde dehydrogenase
PTAC
Signature enzyme
Alcohol dehydrogenase
Regulator

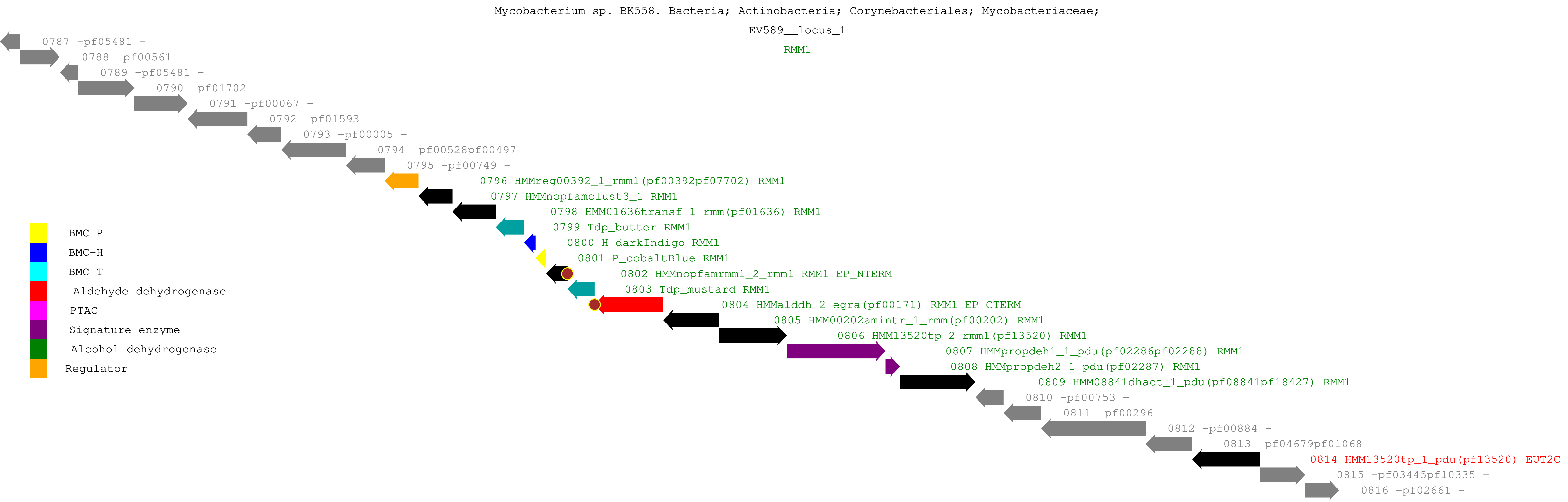

Mycobacterium sp. BK558. Bacteria; Actinobacteria; Corynebacteriales; Mycobacteriaceae;
EV589__locus_1
RMM1
0787 -pf05481 -
0788 -pf00561 -
0789 -pf05481 -
0790 -pf01702 -
0791 -pf00067 -
0792 -pf01593 -
0793 -pf00005 -
0794 -pf00528pf00497 -
0795 -pf00749 -
0796 HMMreg00392_1_rmm1(pf00392pf07702) RMM1
0797 HMMnopfamclust3_1 RMM1
0798 HMM01636transf_1_rmm(pf01636) RMM1
0799 Tdp_butter RMM1
0800 H_darkIndigo RMM1
0801 P_cobaltBlue RMM1
0802 HMMnopfamrmm1_2_rmm1 RMM1 EP_NTERM
0803 Tdp_mustard RMM1
0804 HMMaldh_2_egra(pf00171) RMM1 EP_CTERM
0805 HMM00202amintr_1_rmm(pf00202) RMM1
0806 HMM13520tp_2_rmm1(pf13520) RMM1
0807 HMMpropdeh1_1_pdu(pf02286pf02288) RMM1
0808 HMMpropdeh2_1_pdu(pf02287) RMM1
0809 HMM08841dhact_1_pdu(pf08841pf18427) RMM1
0810 -pf00753 -
0811 -pf00296 -
0812 -pf00884 -
0813 -pf04679pf01068 -
0814 HMM13520tp_1_pdu(pf13520) EUT2C
0815 -pf03445pf10335 -
0816 -pf02661 -
BMC-P
BMC-H
BMC-T
Aldehyde dehydrogenase
PTAC
Signature enzyme
Alcohol dehydrogenase
Regulator

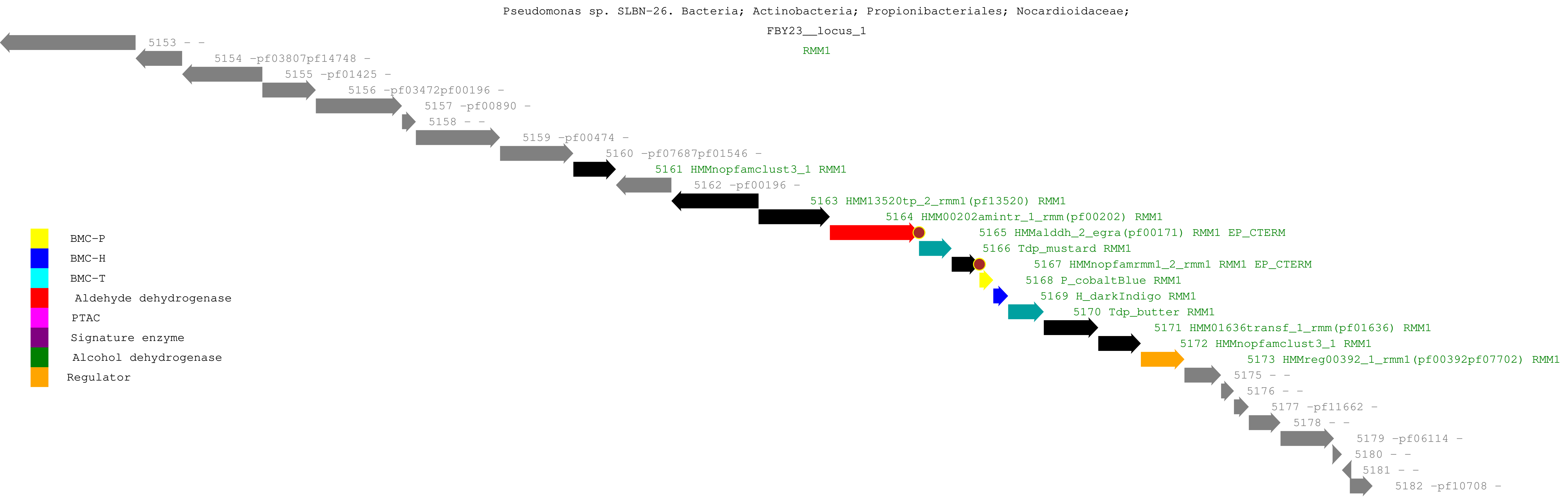
Pseudomonas sp. SLBN-26. Bacteria; Actinobacteria; Propionibacteriales; Nocardioidaceae;
FBY23__locus_1
RMM1
5153 - -
5154 -pf03807pf14748 -
5155 -pf01425 -
5156 -pf03472pf00196 -
5157 -pf00890 -
5158 - -
5159 -pf00474 -
5160 -pf07687pf01546 -
5161 HMMnopfamclust3_1 RMM1
5162 -pf00196 -
5163 HMM13520tp_2_rmm1(pf13520) RMM1
5164 HMM00202amintr_1_rmm(pf00202) RMM1
5165 HMMalddh_2_egra(pf00171) RMM1 EP_CTERM
5166 Tdp_mustard RMM1
5167 HMMnopfamrmm1_2_rmm1 RMM1 EP_CTERM
5168 P_cobaltBlue RMM1
5169 H_darkIndigo RMM1
5170 Tdp_butter RMM1
5171 HMM01636transf_1_rmm(pf01636) RMM1
5172 HMMnopfamclust3_1 RMM1
5173 HMMreg00392_1_rmm1(pf00392pf07702) RMM1
5175 - -
5176 - -
5177 -pf11662 -
5178 - -
5179 -pf06114 -
5180 - -
5181 - -
5182 -pf10708 -
BMC-P
BMC-H
BMC-T
Aldehyde dehydrogenase
PTAC
Signature enzyme
Alcohol dehydrogenase
Regulator

Arthrobacter sp. SLBN-53. Bacteria; Actinobacteria; Corynebacteriales; Mycobacteriaceae;

FBY28__locus_1

RMM1

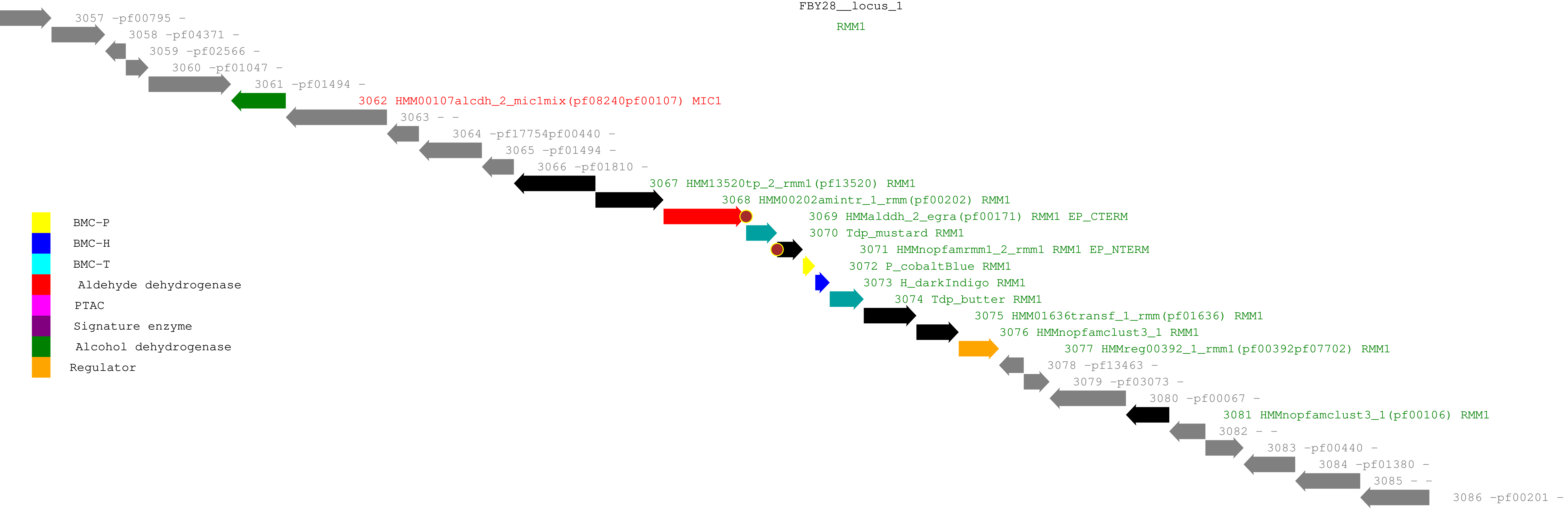

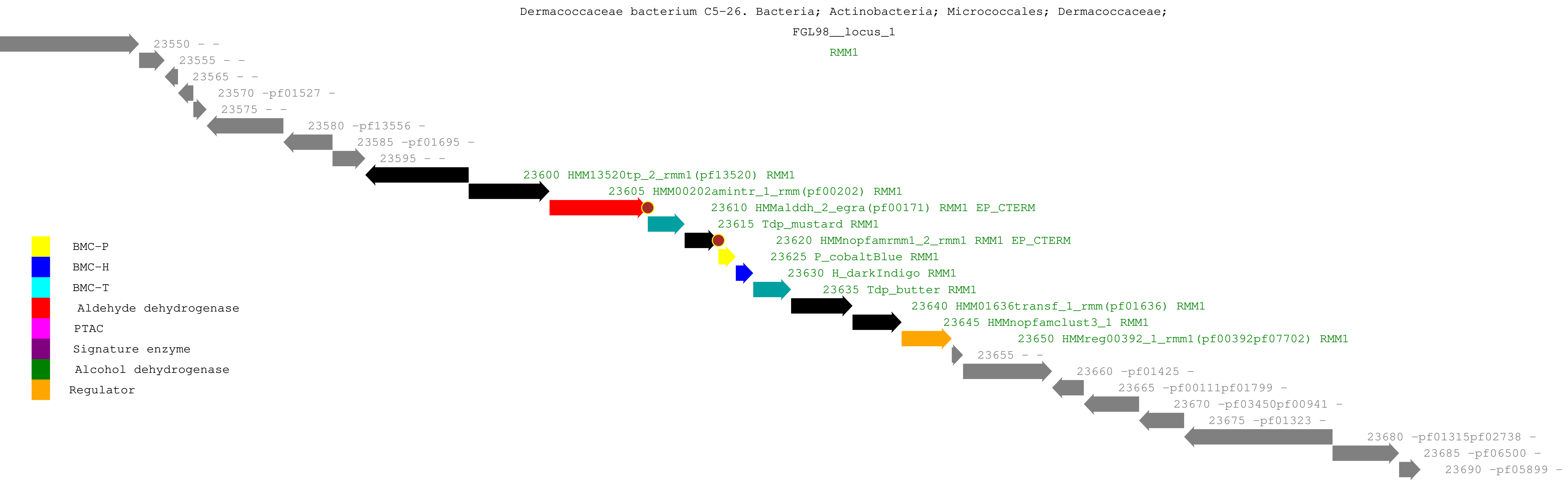
Dermacoccaceae bacterium C5-26. Bacteria; Actinobacteria; Micrococcales; Dermacoccaceae;
FGL98__locus_1
RMM1
23550 - -
23555 - -
23565 - -
23570 -pf01527 -
23575 - -
23580 -pf13556 -
23585 -pf01695 -
23595 - -
23600 HMM13520tp_2_rmm1(pf13520) RMM1
23605 HMM00202amintr_1_rmm(pf00202) RMM1
23610 HMMalddh_2_egra(pf00171) RMM1 EP_CTERM
23615 Tdp_mustard RMM1
23620 HMMnopfamrmm1_2_rmm1 RMM1 EP_CTERM
23625 P_cobaltBlue RMM1
23630 H_darkIndigo RMM1
23635 Tdp_butter RMM1
23640 HMM01636transf_1_rmm(pf01636) RMM1
23645 HMMnopfamclust3_1 RMM1
23650 HMMreg00392_1_rmm1(pf00392pf07702) RMM1
23655 - -
23660 -pf01425 -
23665 -pf00111pf01799 -
23670 -pf03450pf00941 -
23675 -pf01323 -
23680 -pf01315pf02738 -
23685 -pf06500 -
23690 -pf05899 -
BMC-P
BMC-H
BMC-T
Aldehyde dehydrogenase
PTAC
Signature enzyme
Alcohol dehydrogenase
Regulator

Mycolicibacterium fortuitum (Mycobacterium fortuitum). Bacteria; Actinobacteria; Corynebacteriales; Mycobacteriaceae;

FKW78__locus_1

RMM1

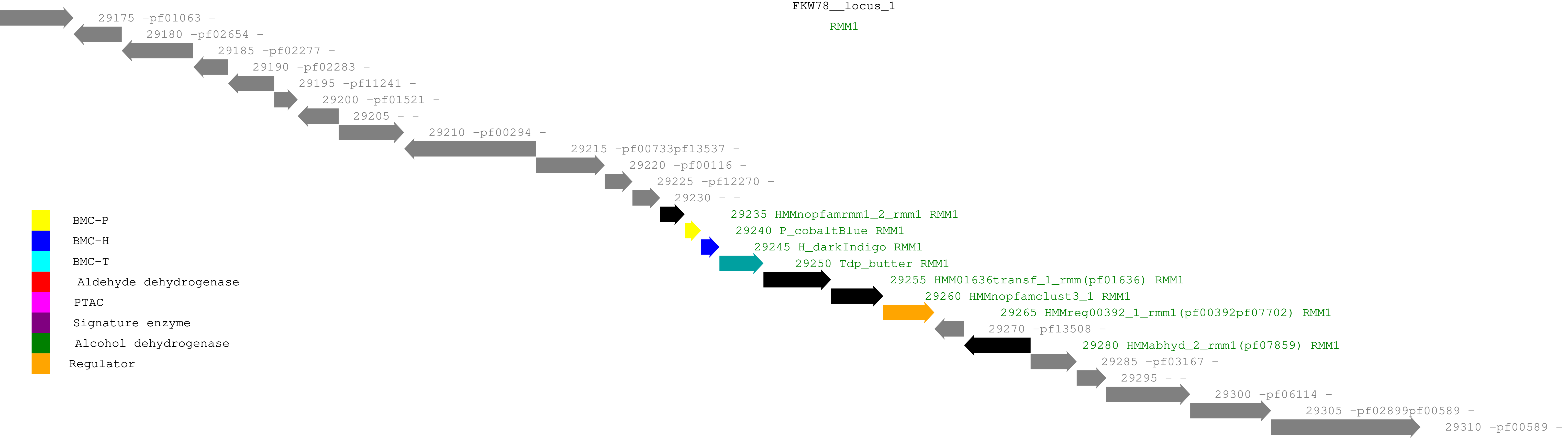

Microbispora sp. KK1-11. Bacteria; Actinobacteria; Streptosporangiales; Streptosporangiaceae;

FLW16__locus_1

RMM1

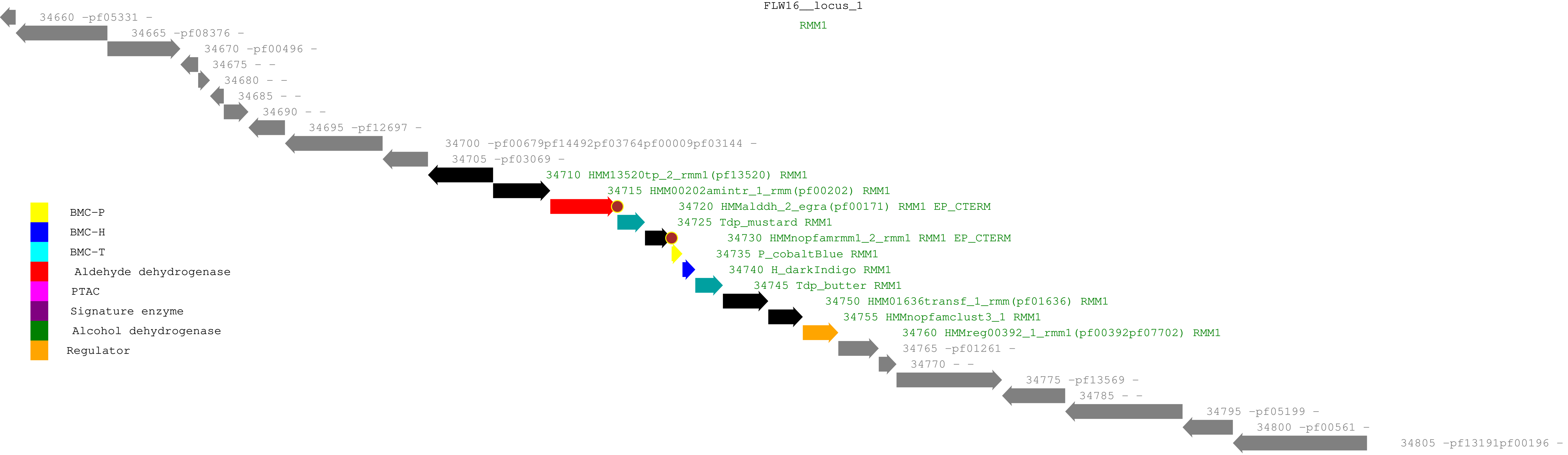

Rhodospirillaceae bacterium R5913. Bacteria; Proteobacteria; Alphaproteobacteria; Rhodospirillales;

FRZ44__locus_1

RMM1

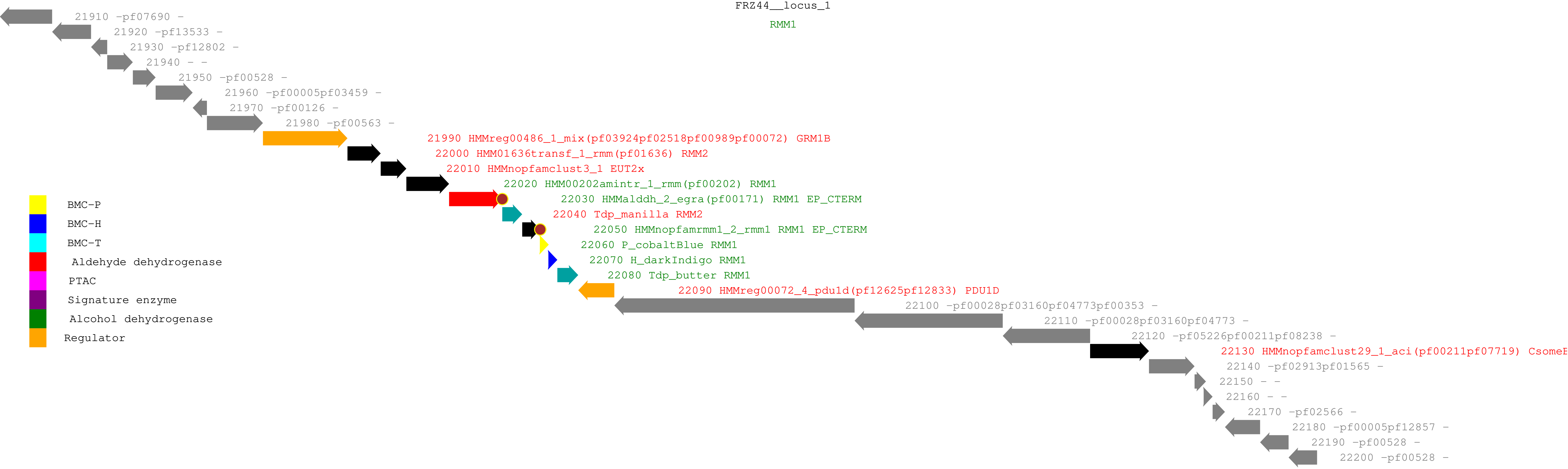

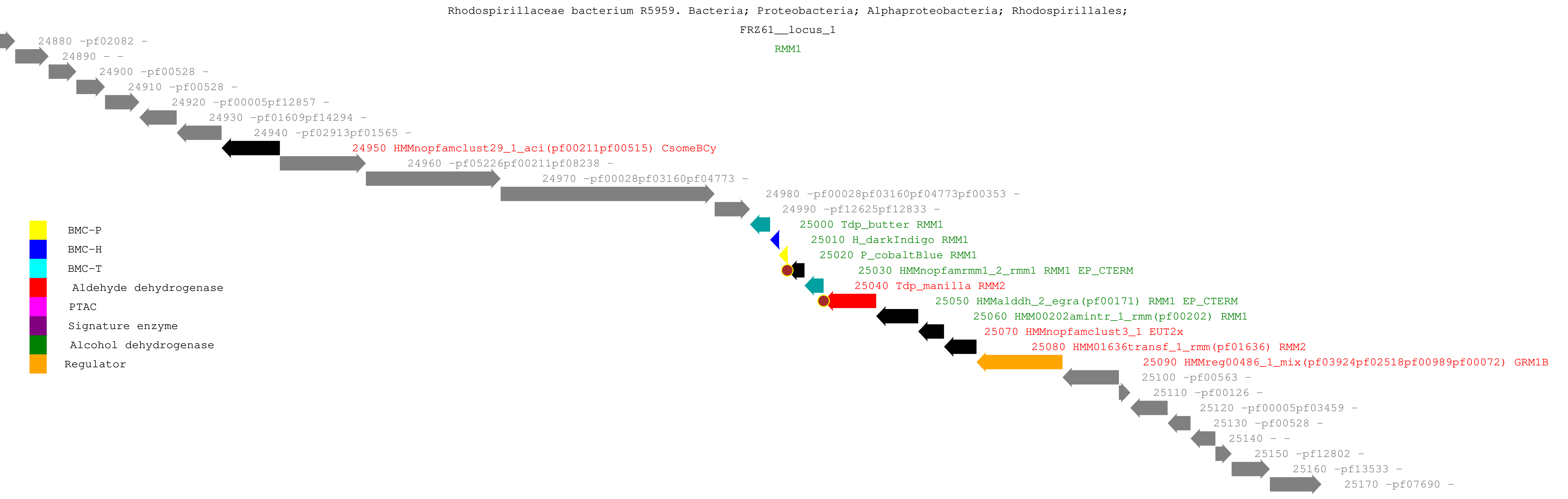
Rhodospirillaceae bacterium R5959. Bacteria; Proteobacteria; Alphaproteobacteria; Rhodospirillales;
FRZ61__locus_1
RMM1
24880 -pf02082 -
24890 - -
24900 -pf00528 -
24910 -pf00528 -
24920 -pf00005pf12857 -
24930 -pf01609pf14294 -
24940 -pf02913pf01565 -
24950 HMMnopfamclust29_1_aci(pf00211pf00515) CsomeBCy
24960 -pf05226pf00211pf08238 -
24970 -pf00028pf03160pf04773 -
24980 -pf00028pf03160pf04773pf00353 -
24990 -pf12625pf12833 -
25000 Tdp_butter RMM1
25010 H_darkIndigo RMM1
25020 P_cobaltBlue RMM1
25030 HMMnopfamrmm1_2_rmm1 RMM1 EP_CTERM
25040 Tdp_manilla RMM2
25050 HMMalddh_2_egra(pf00171) RMM1 EP_CTERM
25060 HMM00202amintr_1_rmm(pf00202) RMM1
25070 HMMnopfamclust3_1 EUT2x
25080 HMM01636transf_1_rmm(pf01636) RMM2
25090 HMMreg00486_1_mix(pf03924pf02518pf00989pf00072) GRM1B
25100 -pf00563 -
25110 -pf00126 -
25120 -pf00005pf03459 -
25130 -pf00528 -
25140 - -
25150 -pf12802 -
25160 -pf13533 -
25170 -pf07690 -
BMC-P
BMC-H
BMC-T
Aldehyde dehydrogenase
PTAC
Signature enzyme
Alcohol dehydrogenase
Regulator

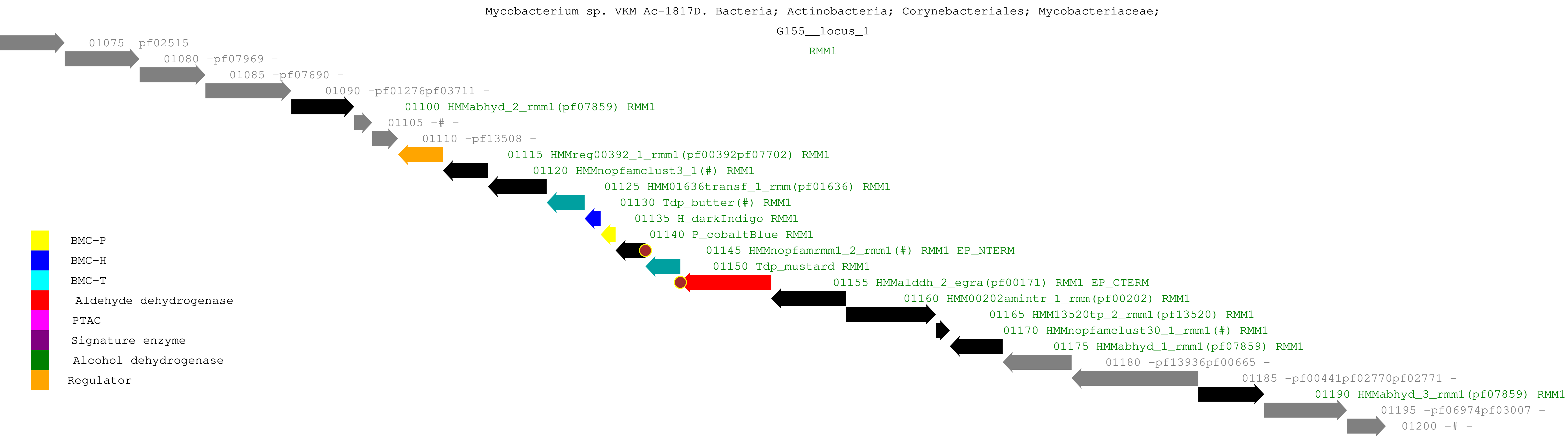
Mycobacterium sp. VKM Ac-1817D. Bacteria; Actinobacteria; Corynebacteriales; Mycobacteriaceae;
G155__locus_1
RMM1
01075 -pf02515 -
01080 -pf07969 -
01085 -pf07690 -
01090 -pf01276pf03711 -
01100 HMMabhyd_2_rmm1(pf07859) RMM1
01105 -# -
01110 -pf13508 -
01115 HMMreg00392_1_rmm1(pf00392pf07702) RMM1
01120 HMMnopfamclust3_1(#) RMM1
01125 HMM01636transf_1_rmm(pf01636) RMM1
01130 Tdp_butter(#) RMM1
01135 H_darkIndigo RMM1
01140 P_cobaltBlue RMM1
01145 HMMnopfamrmm1_2_rmm1(#) RMM1 EP_NTERM
01150 Tdp_mustard RMM1
01155 HMMalddh_2_egra(pf00171) RMM1 EP_CTERM
01160 HMM00202amintr_1_rmm(pf00202) RMM1
01165 HMM13520tp_2_rmm1(pf13520) RMM1
01170 HMMnopfamclust30_1_rmm1(#) RMM1
01175 HMMabhyd_1_rmm1(pf07859) RMM1
01180 -pf13936pf00665 -
01185 -pf00441pf02770pf02771 -
01190 HMMabhyd_3_rmm1(pf07859) RMM1
01195 -pf06974pf03007 -
01200 -# -
BMC-P
BMC-H
BMC-T
Aldehyde dehydrogenase
PTAC
Signature enzyme
Alcohol dehydrogenase
Regulator

Limnohabitans planktonicus II-D5. Bacteria; Proteobacteria; Betaproteobacteria; Burkholderiales;

H663__locus_1

RMM1

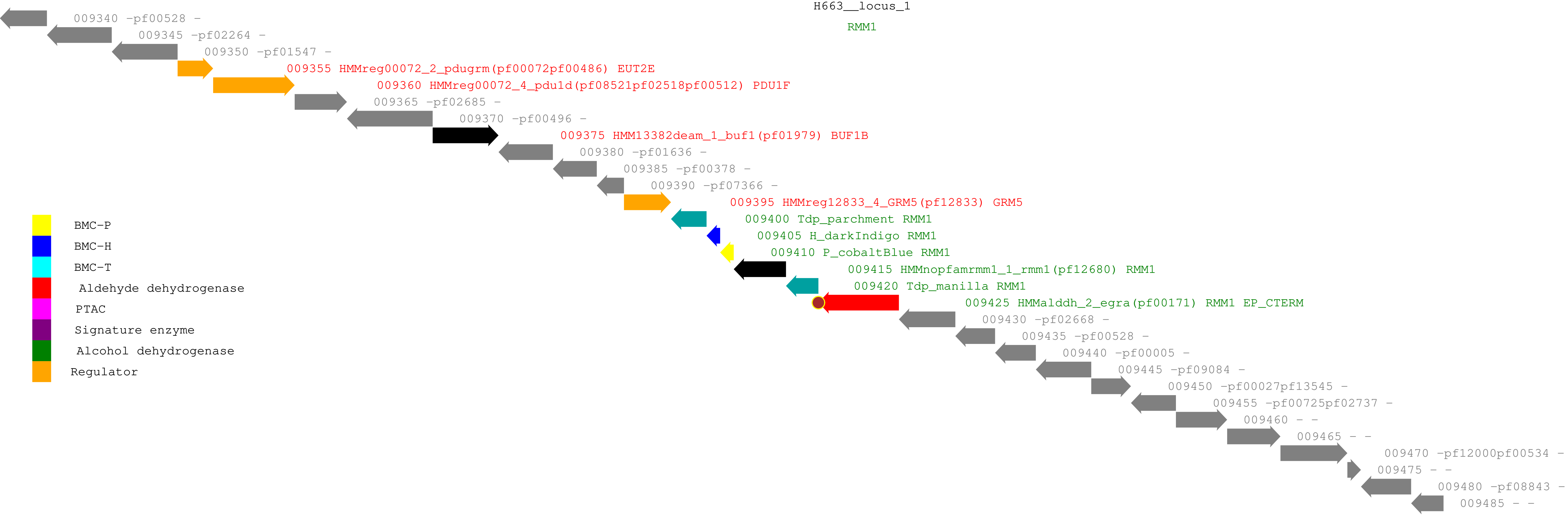

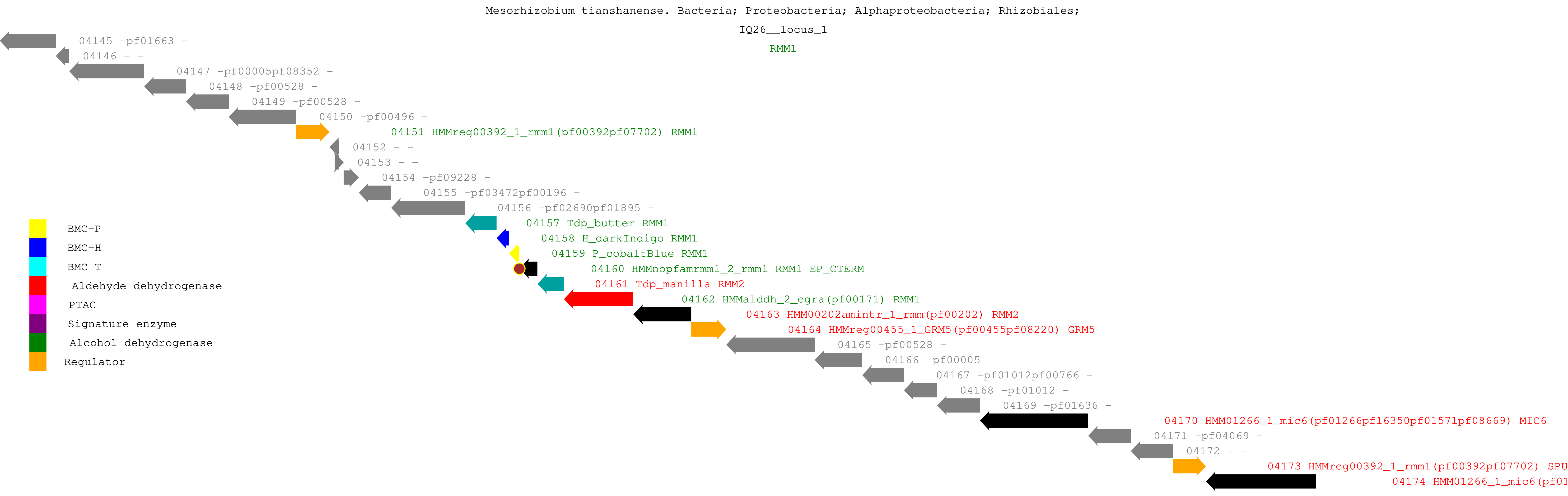

Mesorhizobium tianshanense. Bacteria; Proteobacteria; Alphaproteobacteria; Rhizobiales;
IQ26__locus_1
RMM1
04145 -pf01663 -
04146 - -
04147 -pf00005pf08352 -
04148 -pf00528 -
04149 -pf00528 -
04150 -pf00496 -
04151 HMMreg00392_1_rmm1(pf00392pf07702) RMM1
04152 - -
04153 - -
04154 -pf09228 -
04155 -pf03472pf00196 -
04156 -pf02690pf01895 -
04157 Tdp_butter RMM1
04158 H_darkIndigo RMM1
04159 P_cobaltBlue RMM1
04160 HMMnopfamrmm1_2_rmm1 RMM1 EP_CTERM
04161 Tdp_manilla RMM2
04162 HMMaldh_2_egra(pf00171) RMM1
04163 HMM00202amintr_1_rmm(pf00202) RMM2
04164 HMMreg00455_1_GRM5(pf00455pf08220) GRM5
04165 -pf00528 -
04166 -pf00005 -
04167 -pf01012pf00766 -
04168 -pf01012 -
04169 -pf01636 -
04170 HMM01266_1_mic6(pf01266pf16350pf01571pf08669) MIC6
04171 -pf04069 -
04172 - -
04173 HMMreg00392_1_rmm1(pf00392pf07702) SPU
04174 HMM01266_1_mic6(pf01
BMC-P
BMC-H
BMC-T
Aldehyde dehydrogenase
PTAC
Signature enzyme
Alcohol dehydrogenase
Regulator

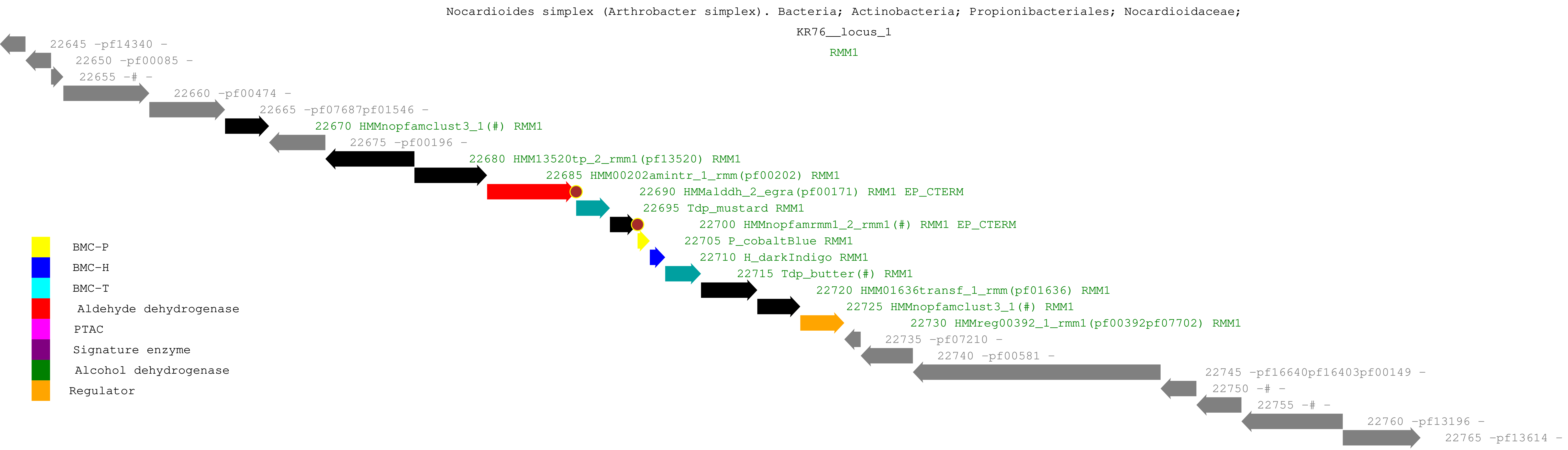

Nocardioides simplex (Arthrobacter simplex). Bacteria; Actinobacteria; Propionibacteriales; Nocardioidaceae;
KR76__locus_1
RMM1
22645 -pf14340 -
22650 -pf00085 -
22655 -# -
22660 -pf00474 -
22665 -pf07687pf01546 -
22670 HMMnopfamclust3_1(#) RMM1
22675 -pf00196 -
22680 HMM13520tp_2_rmm1(pf13520) RMM1
22685 HMM00202amintr_1_rmm(pf00202) RMM1
22690 HMMalddh_2_egra(pf00171) RMM1 EP_CTERM
22695 Tdp_mustard RMM1
22700 HMMnopfamrmm1_2_rmm1(#) RMM1 EP_CTERM
22705 P_cobaltBlue RMM1
22710 H_darkIndigo RMM1
22715 Tdp_butter(#) RMM1
22720 HMM01636transf_1_rmm(pf01636) RMM1
22725 HMMnopfamclust3_1(#) RMM1
22730 HMMreg00392_1_rmm1(pf00392pf07702) RMM1
22735 -pf07210 -
22740 -pf00581 -
22745 -pf16640pf16403pf00149 -
22750 -# -
22755 -# -
22760 -pf13196 -
22765 -pf13614 -
BMC-P
BMC-H
BMC-T
Aldehyde dehydrogenase
PTAC
Signature enzyme
Alcohol dehydrogenase
Regulator

Limnohabitans sp. 63ED37-2. Bacteria; Proteobacteria; Betaproteobacteria; Burkholderiales;

L63ED372__locus_1

RMM1

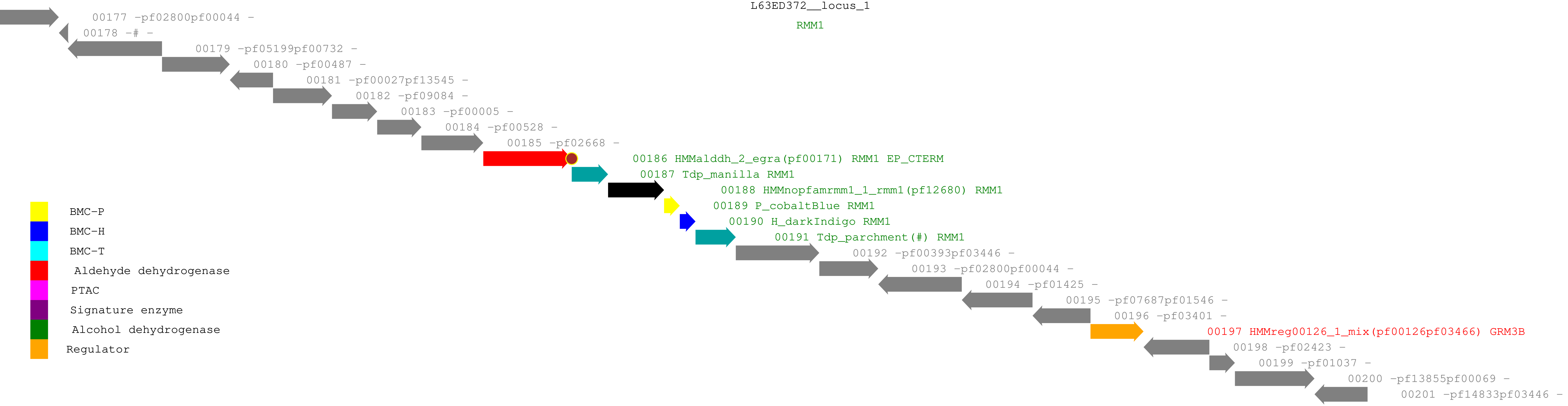

Streptomyces thermoautotrophicus. Bacteria; Actinobacteria; Streptomycetales; Streptomycetaceae;

LI90__locus_1

RMM1

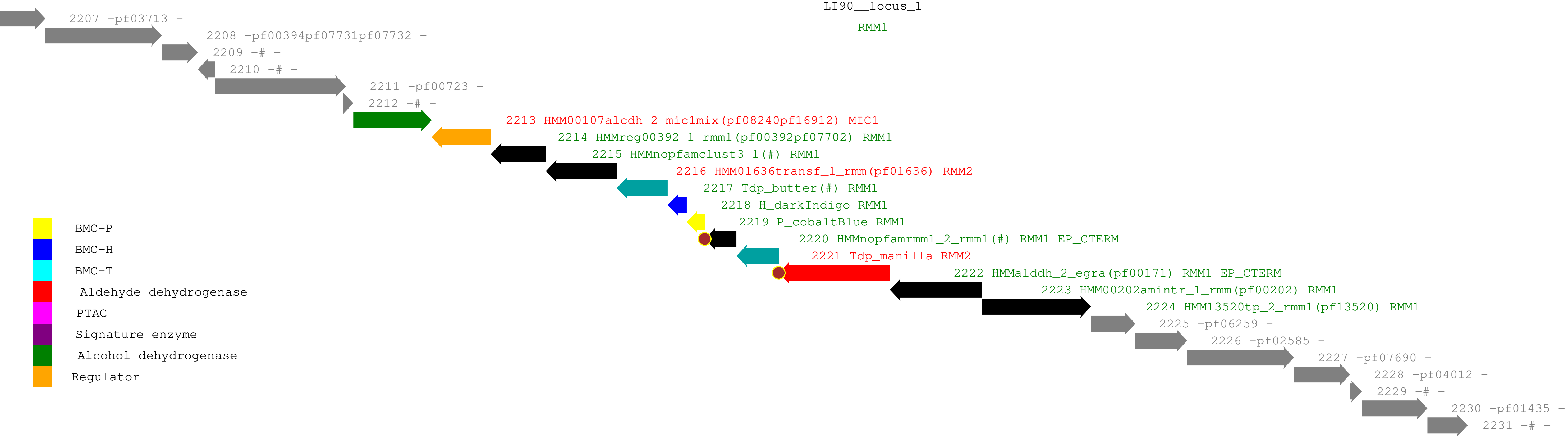

Mycobacterium chlorophenolicum. Bacteria; Actinobacteria; Corynebacteriales; Mycobacteriaceae;

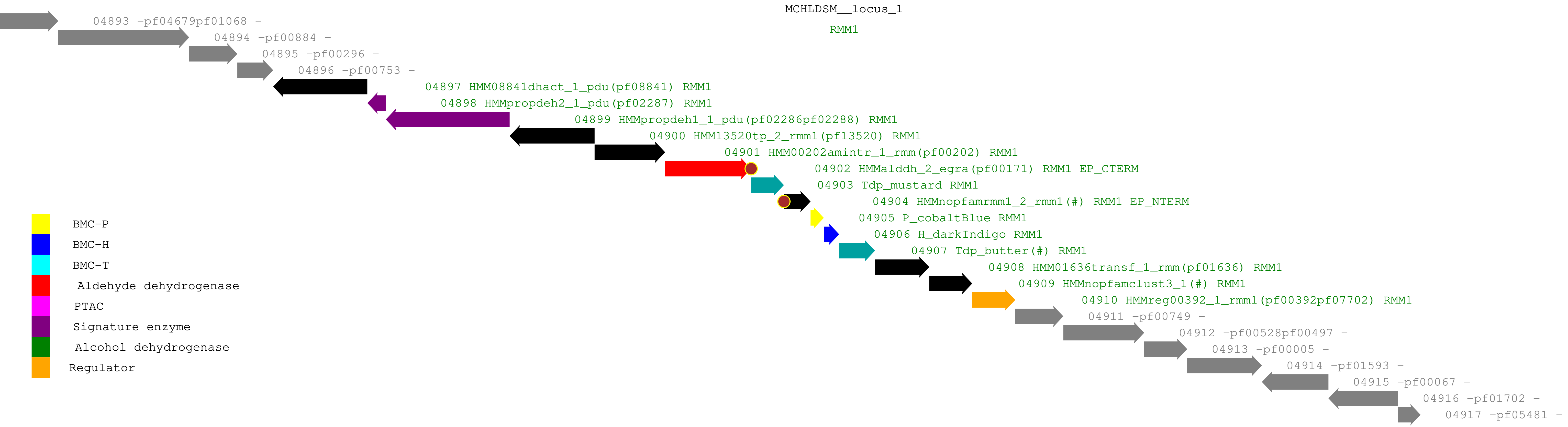

Mycobacterium chubuense. Bacteria; Actinobacteria; Corynebacteriales; Mycobacteriaceae;

MCHUDSM44219__locus_1

RMM1

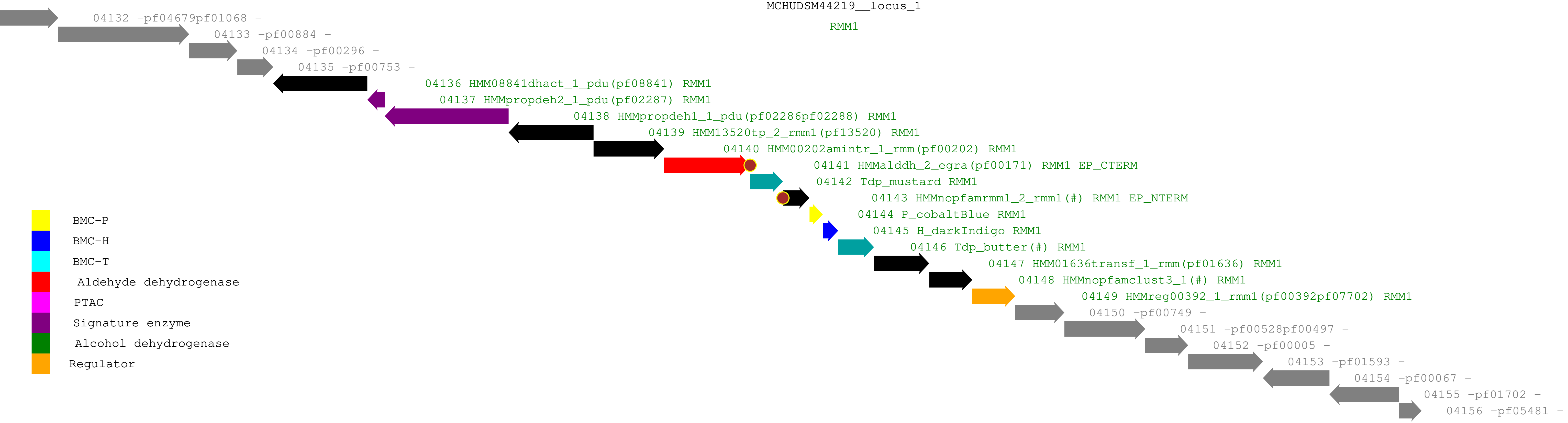

Mycobacterium gilvum (strain PYR-GCK) (Mycobacterium flavescens Bacteria; Actinobacteria; Corynebacteriales; Mycobacteriaceae;

Mflv__locus_1

RMM1

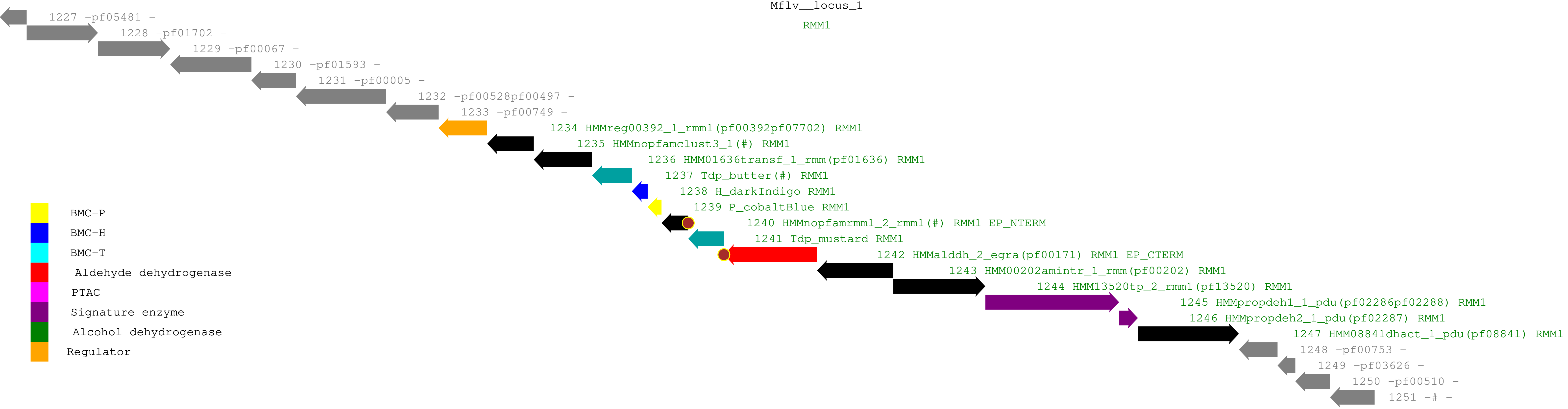

Mycobacterium fortuitum subsp. fortuitum DSM 46621 = ATCC 6841. Bacteria; Actinobacteria; Corynebacteriales; Mycobacteriaceae;

MFORT__locus_1

RMM1

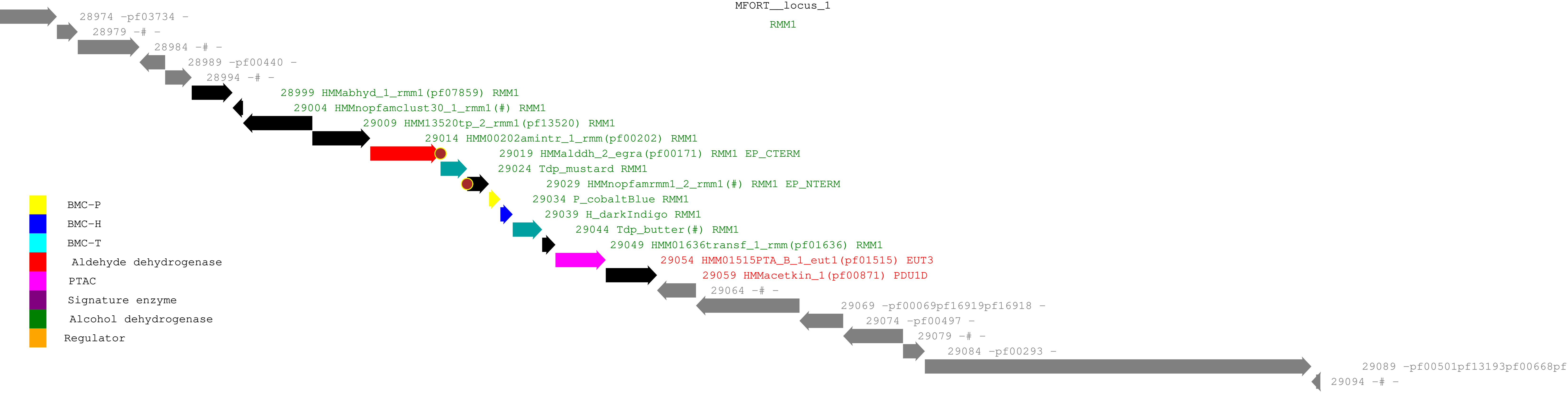

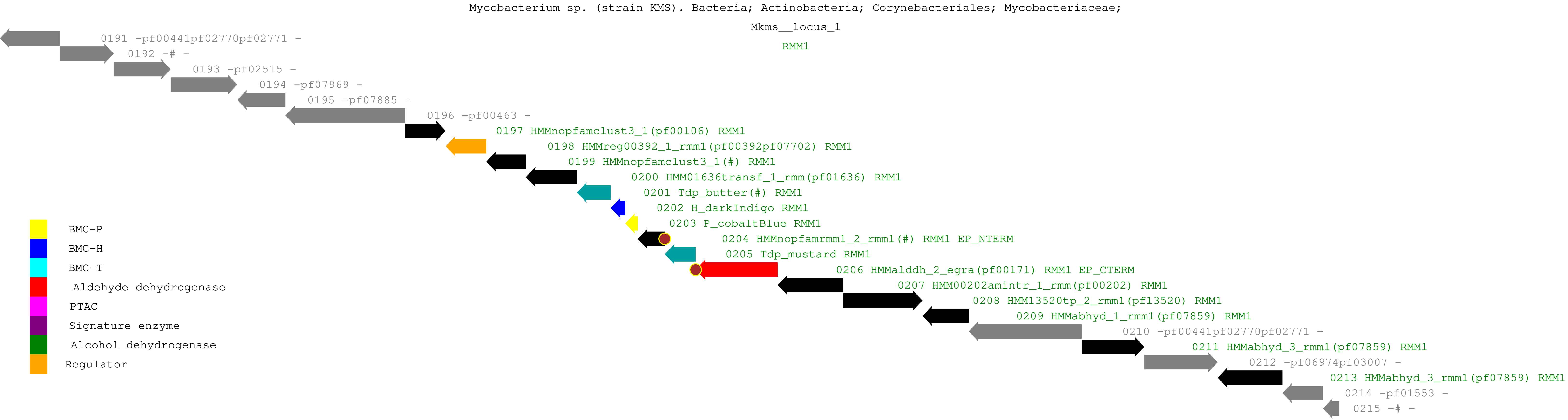
Mycobacterium sp. (strain KMS). Bacteria; Actinobacteria; Corynebacteriales; Mycobacteriaceae;
Mkms__locus_1
RMM1
0191 -pf00441pf02770pf02771 -
0192 -# -
0193 -pf02515 -
0194 -pf07969 -
0195 -pf07885 -
0196 -pf00463 -
0197 HMMnopfamclust3_1(pf00106) RMM1
0198 HMMreg00392_1_rmm1(pf00392pf07702) RMM1
0199 HMMnopfamclust3_1(#) RMM1
0200 HMM01636transf_1_rmm(pf01636) RMM1
0201 Tdp_butter(#) RMM1
0202 H_darkIndigo RMM1
0203 P_cobaltBlue RMM1
0204 HMMnopfamrmm1_2_rmm1(#) RMM1 EP_NTERM
0205 Tdp_mustard RMM1
0206 HMMalddh_2_egra(pf00171) RMM1 EP_CTERM
0207 HMM00202amintr_1_rmm(pf00202) RMM1
0208 HMM13520tp_2_rmm1(pf13520) RMM1
0209 HMMabhyd_1_rmm1(pf07859) RMM1
0210 -pf00441pf02770pf02771 -
0211 HMMabhyd_3_rmm1(pf07859) RMM1
0212 -pf06974pf03007 -
0213 HMMabhyd_3_rmm1(pf07859) RMM1
0214 -pf01553 -
0215 -# -
BMC-P
BMC-H
BMC-T
Aldehyde dehydrogenase
PTAC
Signature enzyme
Alcohol dehydrogenase
Regulator

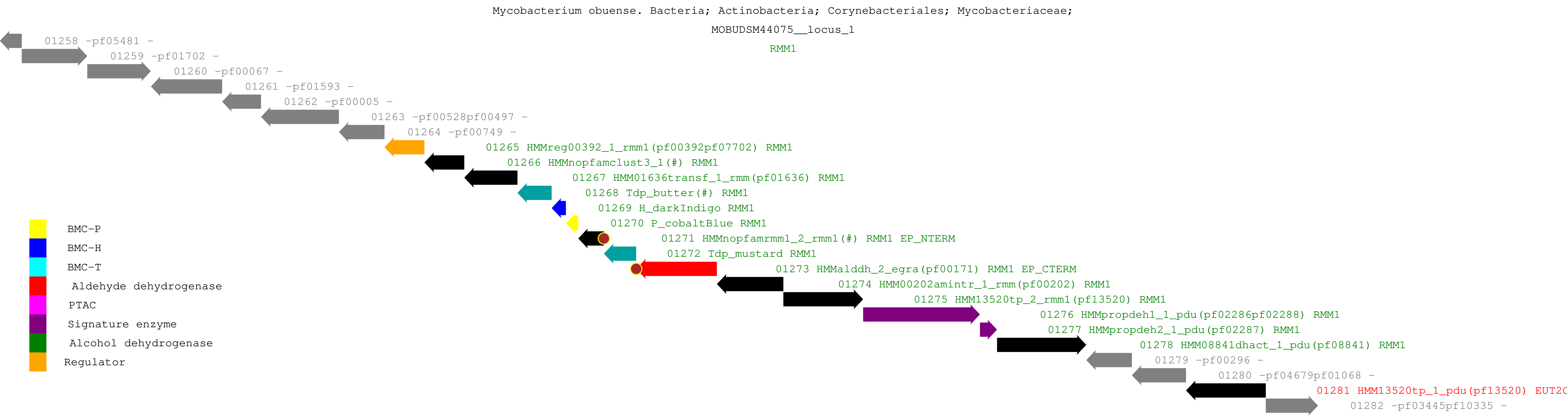
Mycobacterium obuense. Bacteria; Actinobacteria; Corynebacteriales; Mycobacteriaceae;
MOBUDSM44075__locus_1
RMM1
01258 -pf05481 -
01259 -pf01702 -
01260 -pf00067 -
01261 -pf01593 -
01262 -pf00005 -
01263 -pf00528pf00497 -
01264 -pf00749 -
01265 HMMreg00392_1_rmm1(pf00392pf07702) RMM1
01266 HMMnopfamclust3_1(#) RMM1
01267 HMM01636transf_1_rmm(pf01636) RMM1
01268 Tdp_butter(#) RMM1
01269 H_darkIndigo RMM1
01270 P_cobaltBlue RMM1
01271 HMMnopfamrmm1_2_rmm1(#) RMM1 EP_NTERM
01272 Tdp_mustard RMM1
01273 HMMalddh_2_egra(pf00171) RMM1 EP_CTERM
01274 HMM00202amintr_1_rmm(pf00202) RMM1
01275 HMM13520tp_2_rmm1(pf13520) RMM1
01276 HMMpropdeh1_1_pdu(pf02286pf02288) RMM1
01277 HMMpropdeh2_1_pdu(pf02287) RMM1
01278 HMM08841dhact_1_pdu(pf08841) RMM1
01279 -pf00296 -
01280 -pf04679pf01068 -
01281 HMM13520tp_1_pdu(pf13520) EUT2C
01282 -pf03445pf10335 -
BMC-P
BMC-H
BMC-T
Aldehyde dehydrogenase
PTAC
Signature enzyme
Alcohol dehydrogenase
Regulator

Mycobacterium phlei DSM 43239 = CCUG 21000. Bacteria; Actinobacteria; Corynebacteriales; Mycobacteriaceae;

MPHL43239__locus_1

RMM1

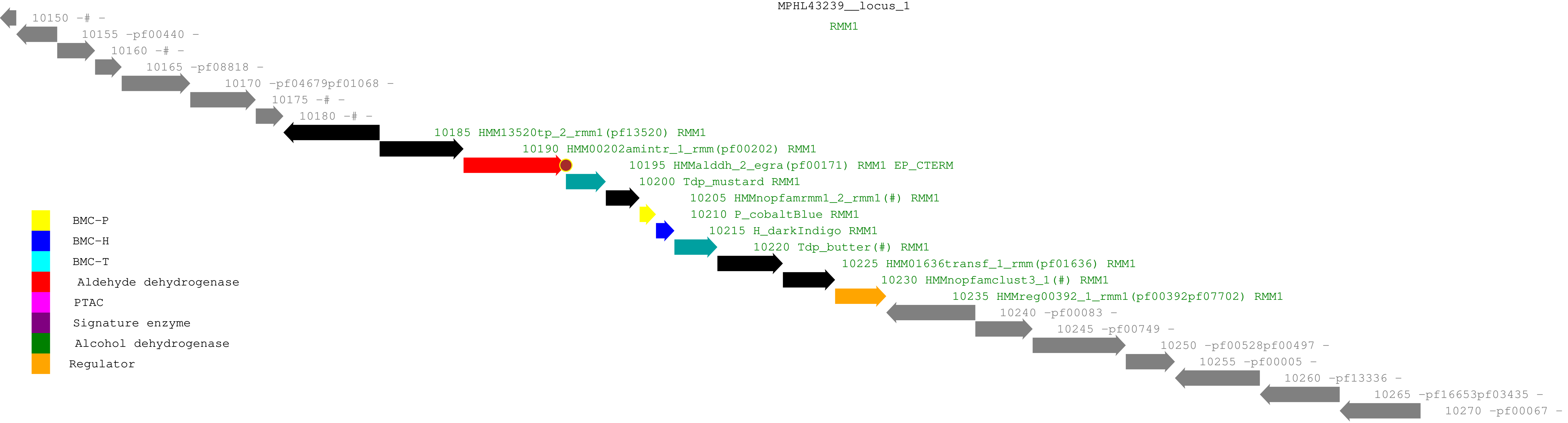

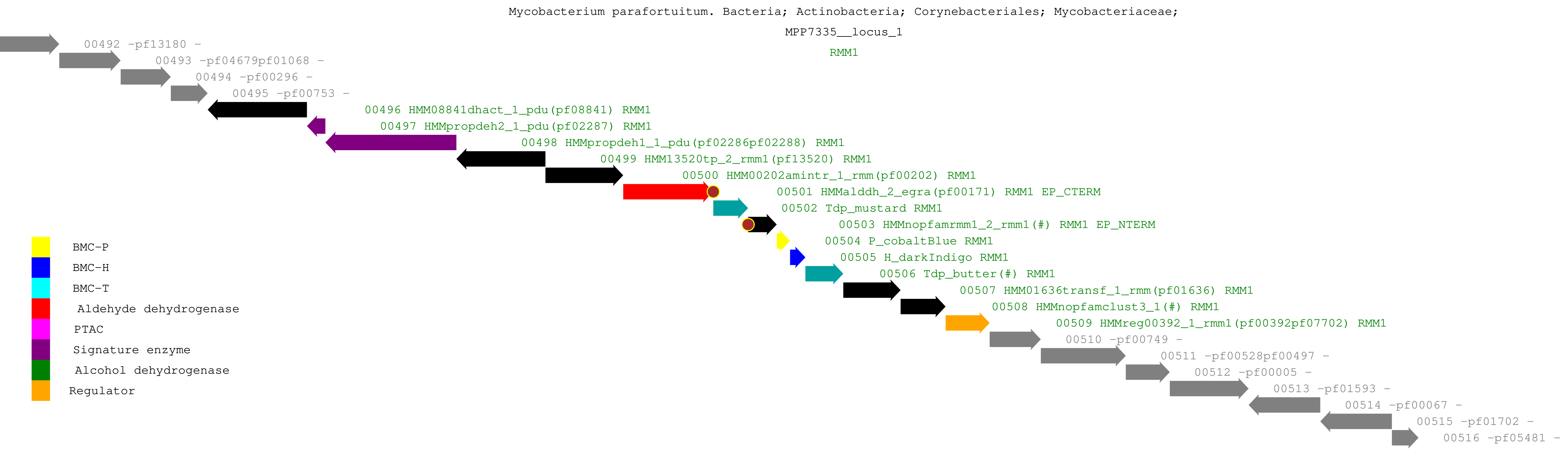

Mycobacterium parafortuitum. Bacteria; Actinobacteria; Corynebacteriales; Mycobacteriaceae;
MPP7335__locus_1
RMM1
00492 -pf13180 -
00493 -pf04679pf01068 -
00494 -pf00296 -
00495 -pf00753 -
00496 HMM08841dhact_1_pdu(pf08841) RMM1
00497 HMMpropdeh2_1_pdu(pf02287) RMM1
00498 HMMpropdeh1_1_pdu(pf02286pf02288) RMM1
00499 HMM13520tp_2_rmm1(pf13520) RMM1
00500 HMM00202amintr_1_rmm(pf00202) RMM1
00501 HMMalddh_2_egra(pf00171) RMM1 EP_CTERM
00502 Tdp_mustard RMM1
00503 HMMnopfamrmm1_2_rmm1(#) RMM1 EP_NTERM
00504 P_cobaltBlue RMM1
00505 H_darkIndigo RMM1
00506 Tdp_butter(#) RMM1
00507 HMM01636transf_1_rmm(pf01636) RMM1
00508 HMMnopfamclust3_1(#) RMM1
00509 HMMreg00392_1_rmm1(pf00392pf07702) RMM1
00510 -pf00749 -
00511 -pf00528pf00497 -
00512 -pf00005 -
00513 -pf01593 -
00514 -pf00067 -
00515 -pf01702 -
00516 -pf05481 -
BMC-P
BMC-H
BMC-T
Aldehyde dehydrogenase
PTAC
Signature enzyme
Alcohol dehydrogenase
Regulator

Mycobacterium smegmatis (strain ATCC 700084 / mc(2)155). Bacteria; Actinobacteria; Corynebacteriales; Mycobacteriaceae;

MSMEG__locus_1

RMM1

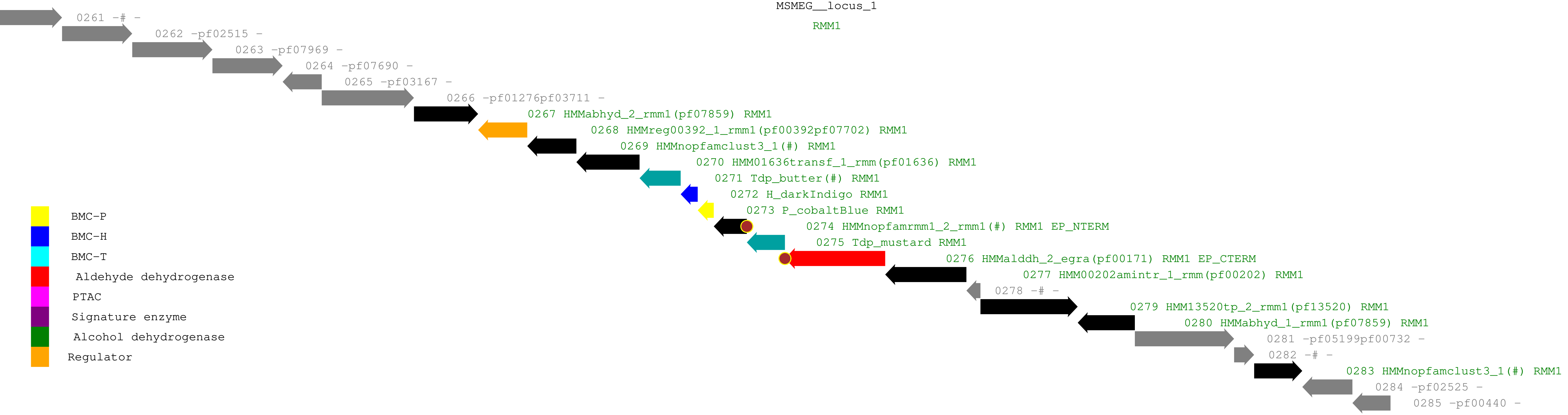

Mycobacterium smegmatis (strain ATCC 700084 / mc(2)155). Bacteria; Actinobacteria; Corynebacteriales; Mycobacteriaceae;

MSMEI__locus_1

RMM1

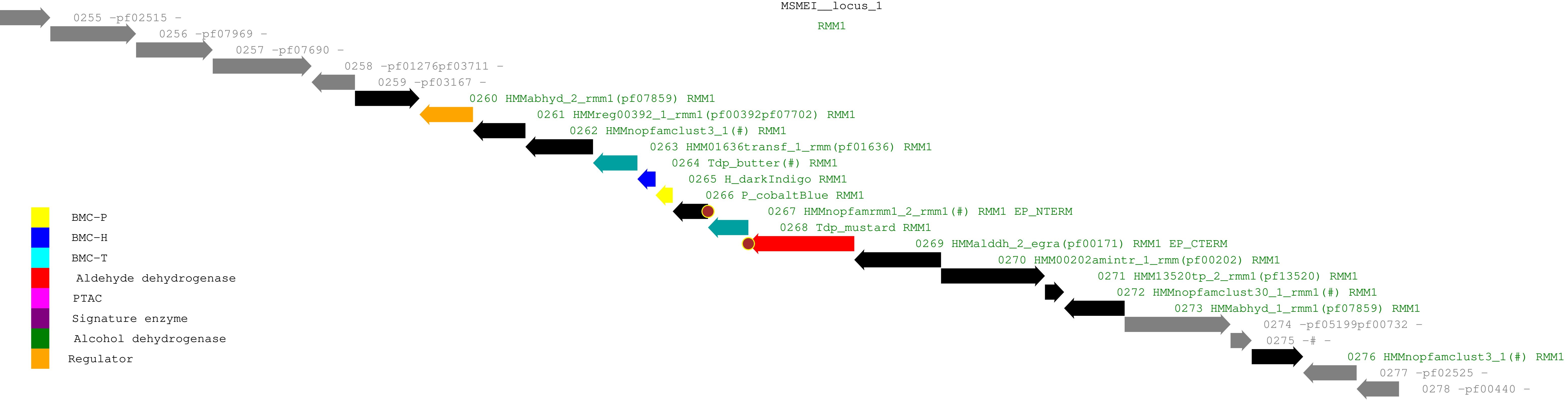

Mycobacterium gilvum (strain DSM 45189 / LMG 24558 / Spyr1). Bacteria; Actinobacteria; Corynebacteriales; Mycobacteriaceae;

Mspyr1__locus_1

RMM1

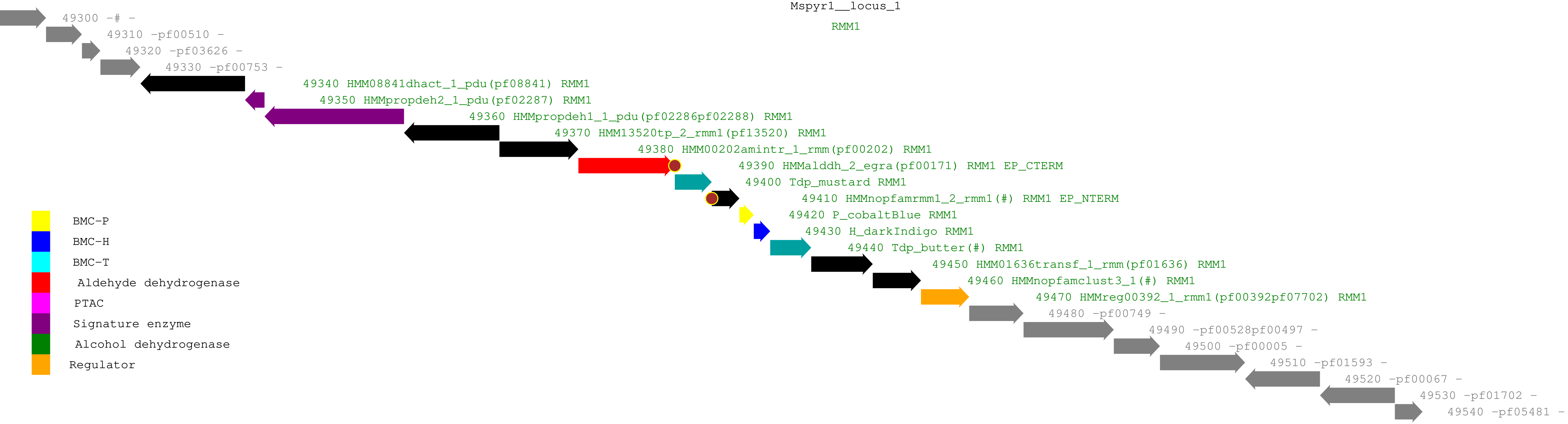

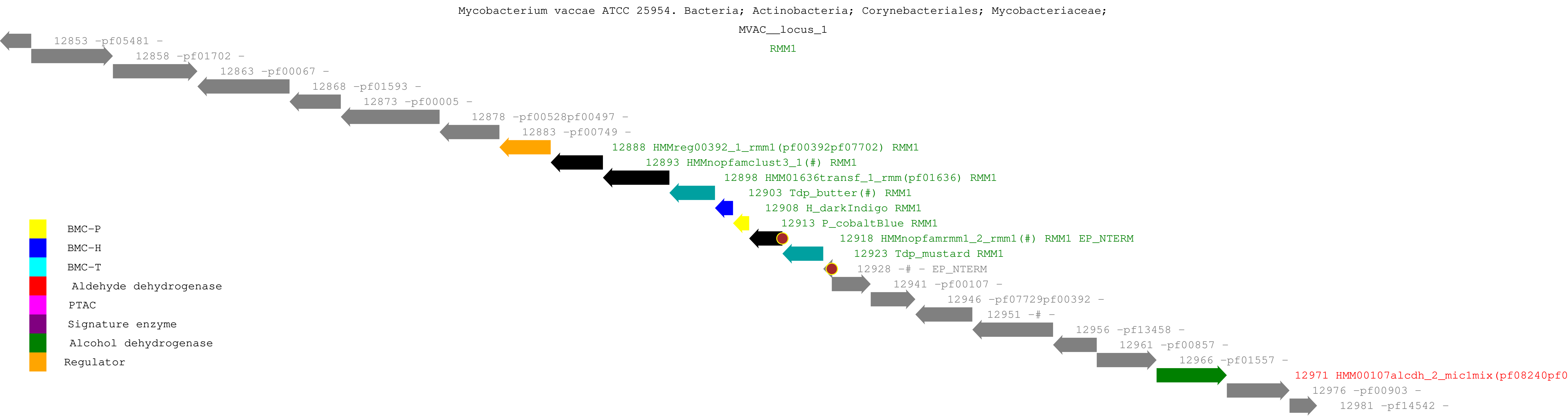

Mycobacterium vaccae ATCC 25954. Bacteria; Actinobacteria; Corynebacteriales; Mycobacteriaceae;
MVAC__locus_1
RMM1
12853 -pf05481 -
12858 -pf01702 -
12863 -pf00067 -
12868 -pf01593 -
12873 -pf00005 -
12878 -pf00528pf00497 -
12883 -pf00749 -
12888 HMMreg00392_1_rmm1(pf00392pf07702) RMM1
12893 HMMnopfamclust3_1(#) RMM1
12898 HMM01636transf_1_rmm(pf01636) RMM1
12903 Tdp_butter(#) RMM1
12908 H_darkIndigo RMM1
12913 P_cobaltBlue RMM1
12918 HMMnopfamrmm1_2_rmm1(#) RMM1 EP_NTERM
12923 Tdp_mustard RMM1
12928 -# - EP_NTERM
12941 -pf00107 -
12946 -pf07729pf00392 -
12951 -# -
12956 -pf13458 -
12961 -pf00857 -
12966 -pf01557 -
12971 HMM00107alcdh_2_mic1mix(pf08240pf0
12976 -pf00903 -
12981 -pf14542 -
BMC-P
BMC-H
BMC-T
Aldehyde dehydrogenase
PTAC
Signature enzyme
Alcohol dehydrogenase
Regulator

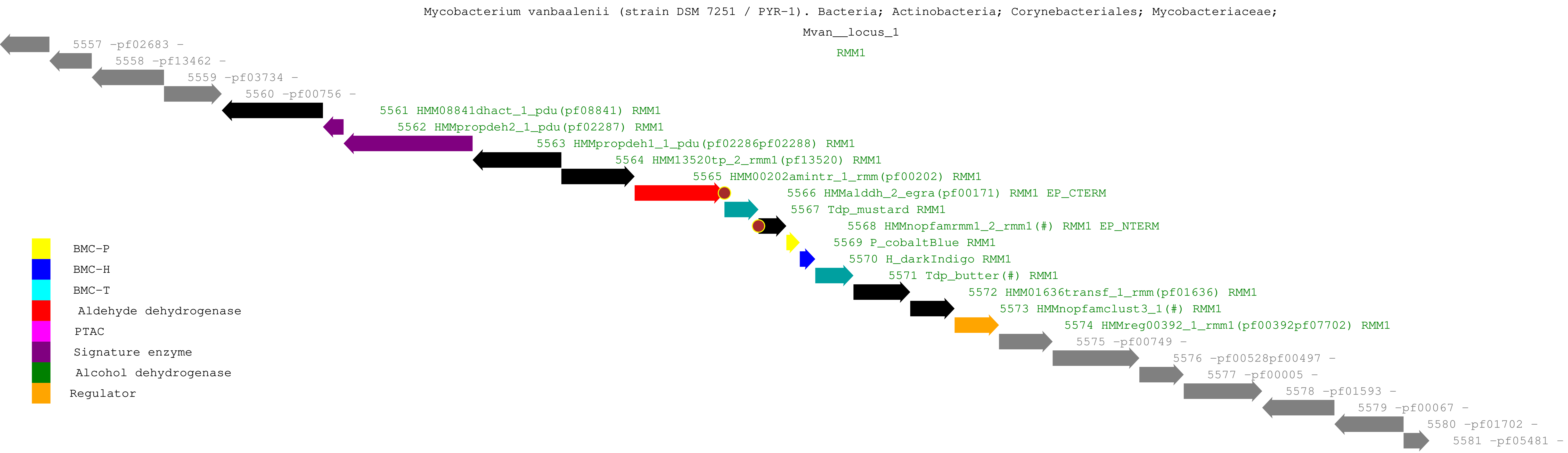

Mycobacterium vanbaalenii (strain DSM 7251 / PYR-1). Bacteria; Actinobacteria; Corynebacteriales; Mycobacteriaceae;
Mvan__locus_1
RMM1
5557 -pf02683 -
5558 -pf13462 -
5559 -pf03734 -
5560 -pf00756 -
5561 HMM08841dhact_1_pdu(pf08841) RMM1
5562 HMMpropdeh2_1_pdu(pf02287) RMM1
5563 HMMpropdeh1_1_pdu(pf02286pf02288) RMM1
5564 HMM13520tp_2_rmm1(pf13520) RMM1
5565 HMM00202amintr_1_rmm(pf00202) RMM1
5566 HMMalddh_2_egra(pf00171) RMM1 EP_CTERM
5567 Tdp_mustard RMM1
5568 HMMnopfamrmm1_2_rmm1(#) RMM1 EP_NTERM
5569 P_cobaltBlue RMM1
5570 H_darkIndigo RMM1
5571 Tdp_butter(#) RMM1
5572 HMM01636transf_1_rmm(pf01636) RMM1
5573 HMMnopfamclust3_1(#) RMM1
5574 HMMreg00392_1_rmm1(pf00392pf07702) RMM1
5575 -pf00749 -
5576 -pf00528pf00497 -
5577 -pf00005 -
5578 -pf01593 -
5579 -pf00067 -
5580 -pf01702 -
5581 -pf05481 -
BMC-P
BMC-H
BMC-T
Aldehyde dehydrogenase
PTAC
Signature enzyme
Alcohol dehydrogenase
Regulator

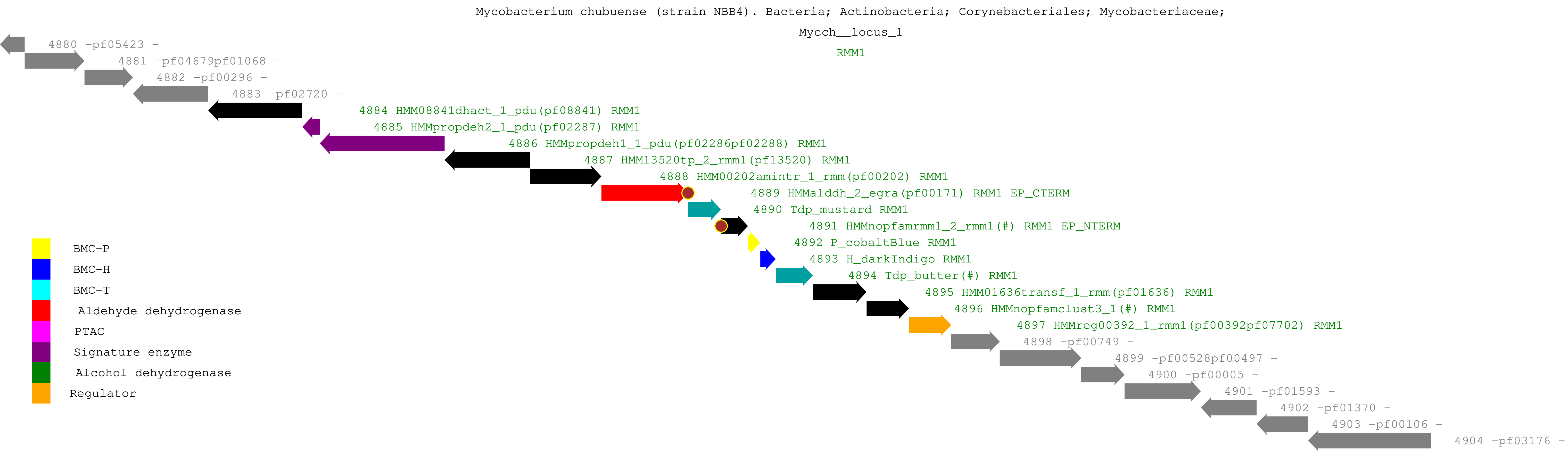

Mycobacterium chubuense (strain NBB4). Bacteria; Actinobacteria; Corynebacteriales; Mycobacteriaceae;
Mycch__locus_1
RMM1
4880 -pf05423 -
4881 -pf04679pf01068 -
4882 -pf00296 -
4883 -pf02720 -
4884 HMM08841dhact_1_pdu(pf08841) RMM1
4885 HMMpropdeh2_1_pdu(pf02287) RMM1
4886 HMMpropdeh1_1_pdu(pf02286pf02288) RMM1
4887 HMM13520tp_2_rmm1(pf13520) RMM1
4888 HMM00202amintr_1_rmm(pf00202) RMM1
4889 HMMalddh_2_egra(pf00171) RMM1 EP_CTERM
4890 Tdp_mustard RMM1
4891 HMMnopfamrmm1_2_rmm1(#) RMM1 EP_NTERM
4892 P_cobaltBlue RMM1
4893 H_darkIndigo RMM1
4894 Tdp_butter(#) RMM1
4895 HMM01636transf_1_rmm(pf01636) RMM1
4896 HMMnopfamclust3_1(#) RMM1
4897 HMMreg00392_1_rmm1(pf00392pf07702) RMM1
4898 -pf00749 -
4899 -pf00528pf00497 -
4900 -pf00005 -
4901 -pf01593 -
4902 -pf01370 -
4903 -pf00106 -
4904 -pf03176 -
BMC-P
BMC-H
BMC-T
Aldehyde dehydrogenase
PTAC
Signature enzyme
Alcohol dehydrogenase
Regulator

Mycolicibacterium aurum (Mycobacterium aurum). Bacteria; Actinobacteria; Corynebacteriales; Mycobacteriaceae;

NCTC10437__locus_1

RMM1

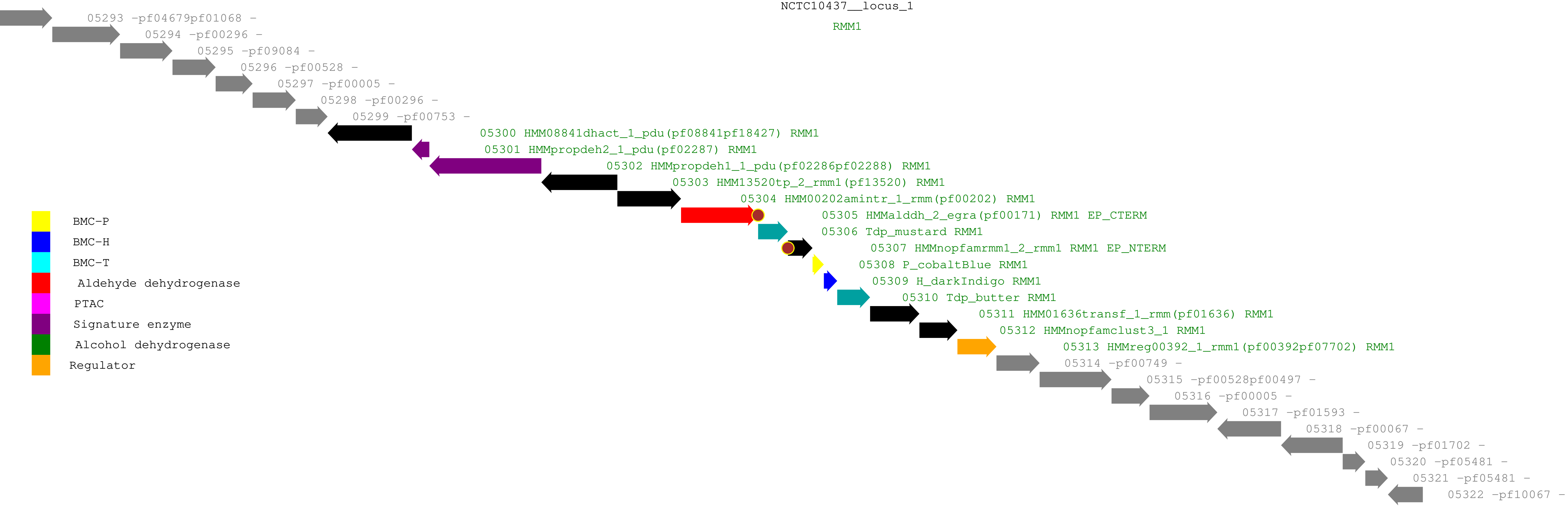

Mycolicibacterium chitae (Mycobacterium chitae). Bacteria; Actinobacteria; Corynebacteriales; Mycobacteriaceae;

NCTC10485__locus_1

RMM1

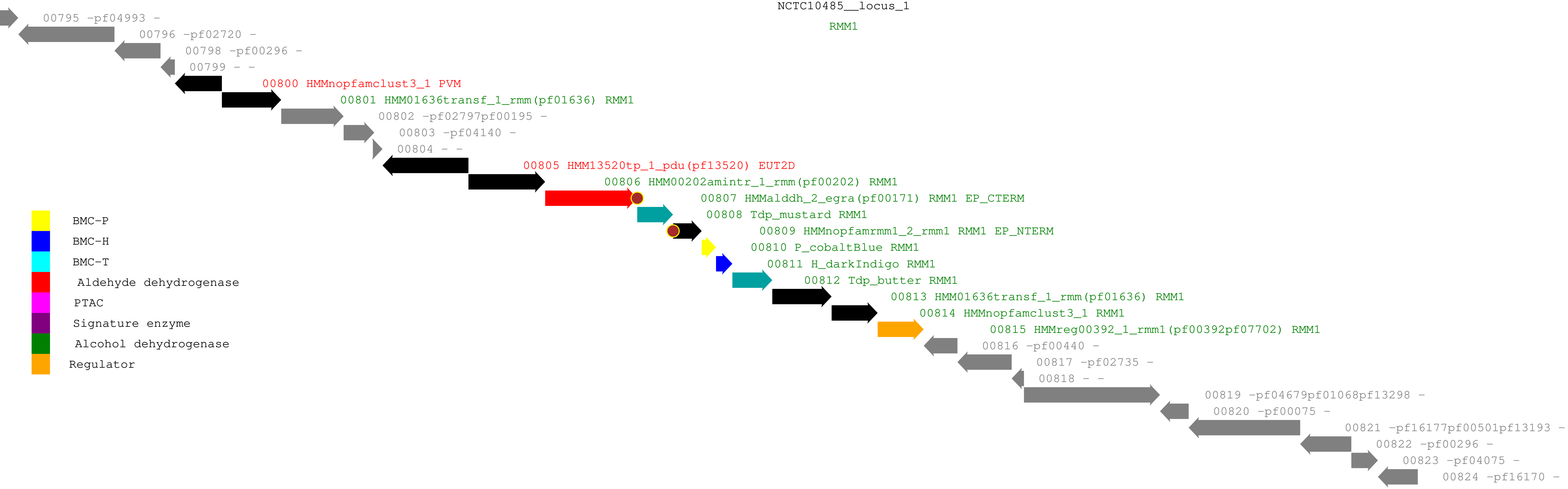

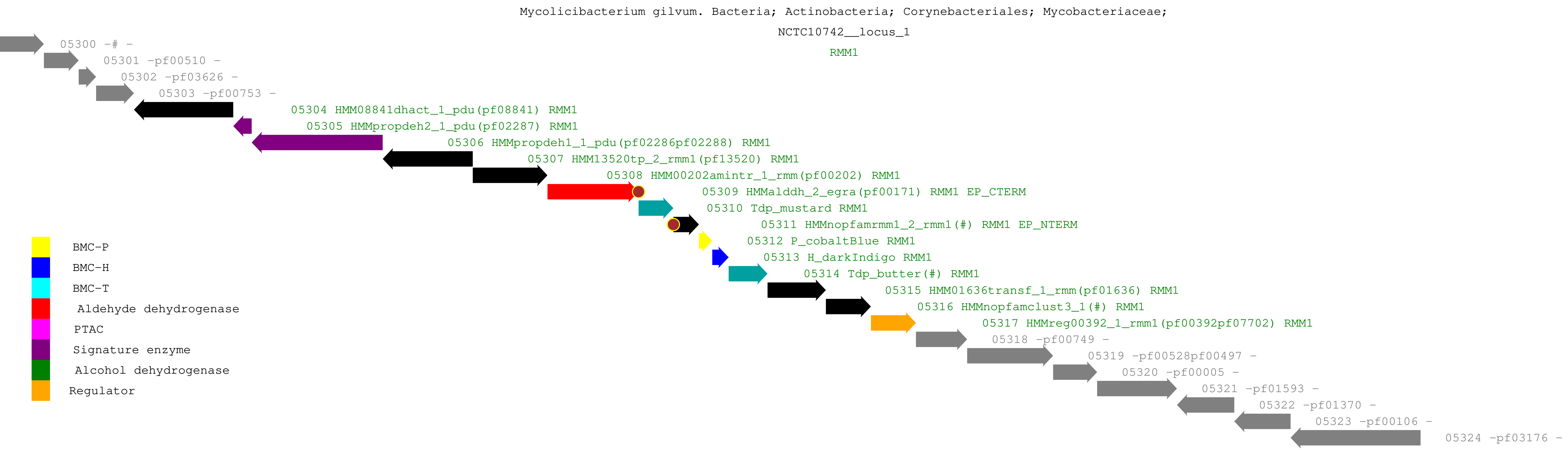
Mycolicibacterium gilvum. Bacteria; Actinobacteria; Corynebacteriales; Mycobacteriaceae;
NCTC10742__locus_1
RMM1
05300 -# -
05301 -pf00510 -
05302 -pf03626 -
05303 -pf00753 -
05304 HMM08841dhact_1_pdu(pf08841) RMM1
05305 HMMpropdeh2_1_pdu(pf02287) RMM1
05306 HMMpropdeh1_1_pdu(pf02286pf02288) RMM1
05307 HMM13520tp_2_rmm1(pf13520) RMM1
05308 HMM00202amintr_1_rmm(pf00202) RMM1
05309 HMMalddh_2_egra(pf00171) RMM1 EP_CTERM
05310 Tdp_mustard RMM1
05311 HMMnopfamrmm1_2_rmm1(#) RMM1 EP_NTERM
05312 P_cobaltBlue RMM1
05313 H_darkIndigo RMM1
05314 Tdp_butter(#) RMM1
05315 HMM01636transf_1_rmm(pf01636) RMM1
05316 HMMnopfamclust3_1(#) RMM1
05317 HMMreg00392_1_rmm1(pf00392pf07702) RMM1
05318 -pf00749 -
05319 -pf00528pf00497 -
05320 -pf00005 -
05321 -pf01593 -
05322 -pf01370 -
05323 -pf00106 -
05324 -pf03176 -
BMC-P
BMC-H
BMC-T
Aldehyde dehydrogenase
PTAC
Signature enzyme
Alcohol dehydrogenase
Regulator

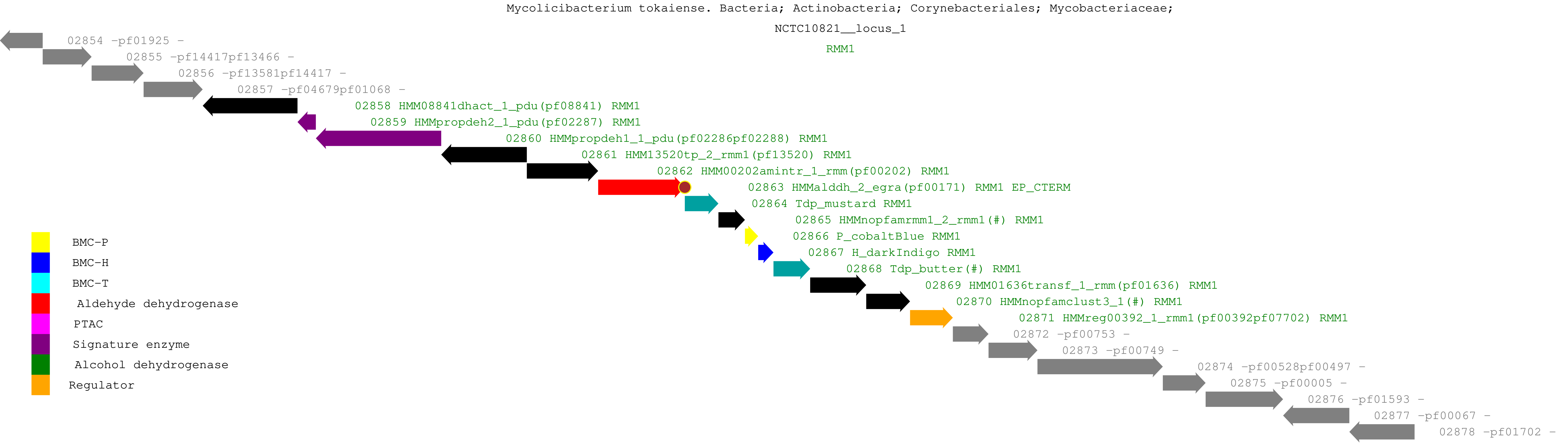
Mycolicibacterium tokaiense. Bacteria; Actinobacteria; Corynebacteriales; Mycobacteriaceae;
NCTC10821__locus_1
RMM1
02854 -pf01925 -
02855 -pf14417pf13466 -
02856 -pf13581pf14417 -
02857 -pf04679pf01068 -
02858 HMM08841dhact_1_pdu(pf08841) RMM1
02859 HMMpropdeh2_1_pdu(pf02287) RMM1
02860 HMMpropdeh1_1_pdu(pf02286pf02288) RMM1
02861 HMM13520tp_2_rmm1(pf13520) RMM1
02862 HMM00202amintr_1_rmm(pf00202) RMM1
02863 HMMalddh_2_egra(pf00171) RMM1 EP_CTERM
02864 Tdp_mustard RMM1
02865 HMMnopfamrmm1_2_rmm1(#) RMM1
02866 P_cobaltBlue RMM1
02867 H_darkIndigo RMM1
02868 Tdp_butter(#) RMM1
02869 HMM01636transf_1_rmm(pf01636) RMM1
02870 HMMnopfamclust3_1(#) RMM1
02871 HMMreg00392_1_rmm1(pf00392pf07702) RMM1
02872 -pf00753 -
02873 -pf00749 -
02874 -pf00528pf00497 -
02875 -pf00005 -
02876 -pf01593 -
02877 -pf00067 -
02878 -pf01702 -
BMC-P
BMC-H
BMC-T
Aldehyde dehydrogenase
PTAC
Signature enzyme
Alcohol dehydrogenase
Regulator

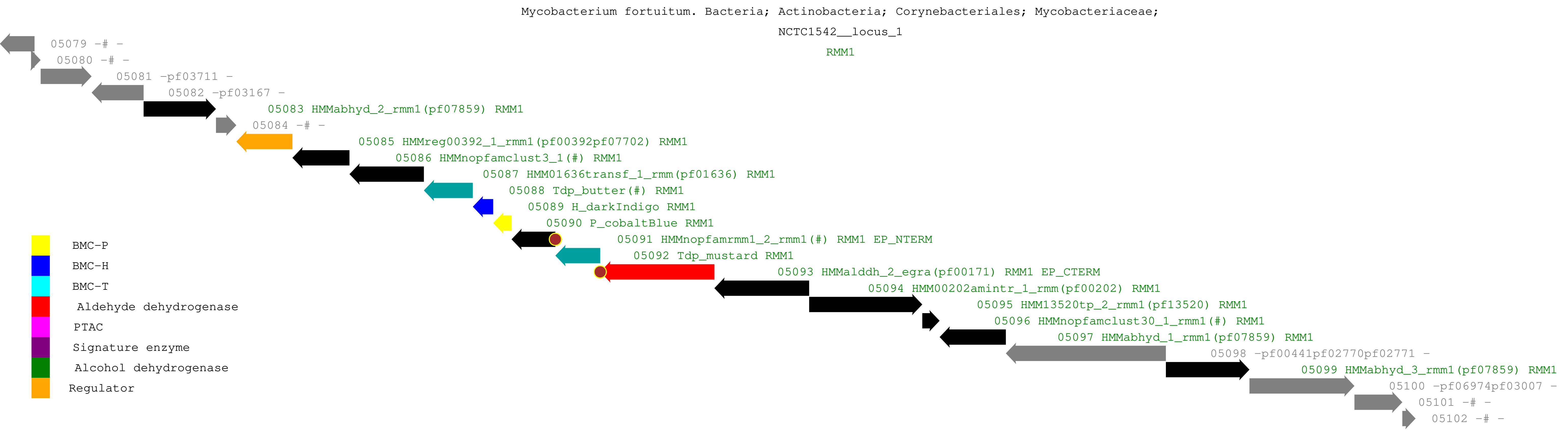
Mycobacterium fortuitum. Bacteria; Actinobacteria; Corynebacteriales; Mycobacteriaceae;
NCTC1542__locus_1
RMM1
05079 -# -
05080 -# -
05081 -pf03711 -
05082 -pf03167 -
05083 HMMabhyd_2_rmm1(pf07859) RMM1
05084 -# -
05085 HMMreg00392_1_rmm1(pf00392pf07702) RMM1
05086 HMMnopfamclust3_1(#) RMM1
05087 HMM01636transf_1_rmm(pf01636) RMM1
05088 Tdp_butter(#) RMM1
05089 H_darkIndigo RMM1
05090 P_cobaltBlue RMM1
05091 HMMnopfamrmm1_2_rmm1(#) RMM1 EP_NTERM
05092 Tdp_mustard RMM1
05093 HMMalddh_2_egra(pf00171) RMM1 EP_CTERM
05094 HMM00202amintr_1_rmm(pf00202) RMM1
05095 HMM13520tp_2_rmm1(pf13520) RMM1
05096 HMMnopfamclust30_1_rmm1(#) RMM1
05097 HMMabhyd_1_rmm1(pf07859) RMM1
05098 -pf00441pf02770pf02771 -
05099 HMMabhyd_3_rmm1(pf07859) RMM1
05100 -pf06974pf03007 -
05101 -# -
05102 -# -
BMC-P
BMC-H
BMC-T
Aldehyde dehydrogenase
PTAC
Signature enzyme
Alcohol dehydrogenase
Regulator

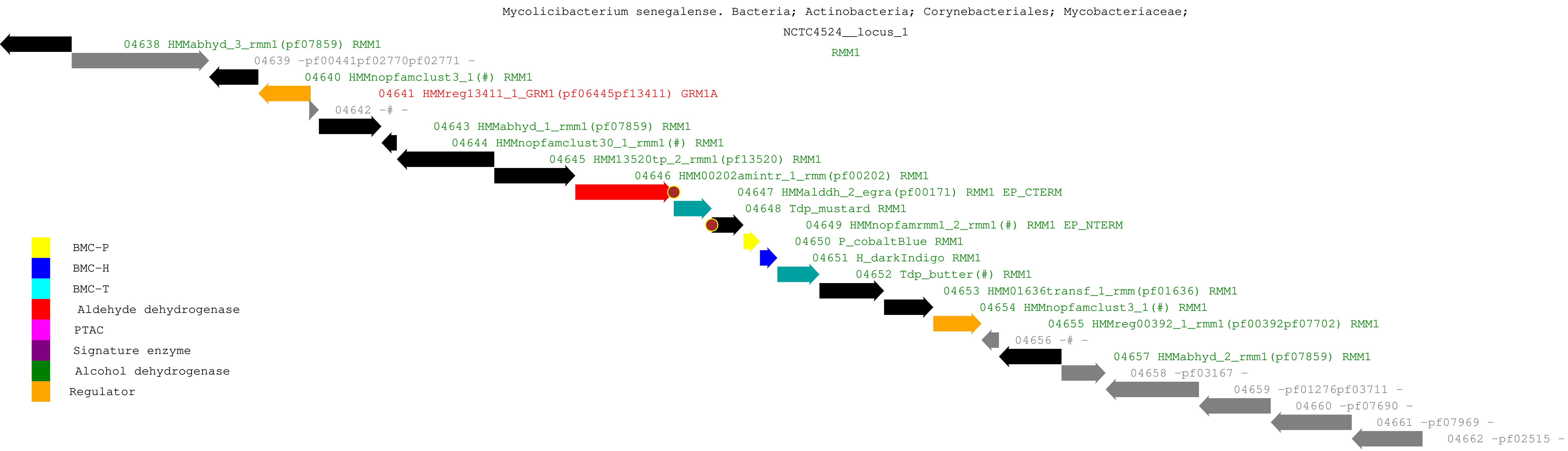

Mycolicibacterium senegalense. Bacteria; Actinobacteria; Corynebacteriales; Mycobacteriaceae;
NCTC4524__locus_1
RMM1
04638 HMMabhyd_3_rmm1(pf07859) RMM1
04639 -pf00441pf02770pf02771 -
04640 HMMnopfamclust3_1(#) RMM1
04641 HMMreg13411_1_GRM1(pf06445pf13411) GRM1A
04642 -# -
04643 HMMabhyd_1_rmm1(pf07859) RMM1
04644 HMMnopfamclust30_1_rmm1(#) RMM1
04645 HMM13520tp_2_rmm1(pf13520) RMM1
04646 HMM00202amintr_1_rmm(pf00202) RMM1
04647 HMMalddh_2_egra(pf00171) RMM1 EP_CTERM
04648 Tdp_mustard RMM1
04649 HMMnopfamrmm1_2_rmm1(#) RMM1 EP_NTERM
04650 P_cobaltBlue RMM1
04651 H_darkIndigo RMM1
04652 Tdp_butter(#) RMM1
04653 HMM01636transf_1_rmm(pf01636) RMM1
04654 HMMnopfamclust3_1(#) RMM1
04655 HMMreg00392_1_rmm1(pf00392pf07702) RMM1
04656 -# -
04657 HMMabhyd_2_rmm1(pf07859) RMM1
04658 -pf03167 -
04659 -pf01276pf03711 -
04660 -pf07690 -
04661 -pf07969 -
04662 -pf02515 -
BMC-P
BMC-H
BMC-T
Aldehyde dehydrogenase
PTAC
Signature enzyme
Alcohol dehydrogenase
Regulator

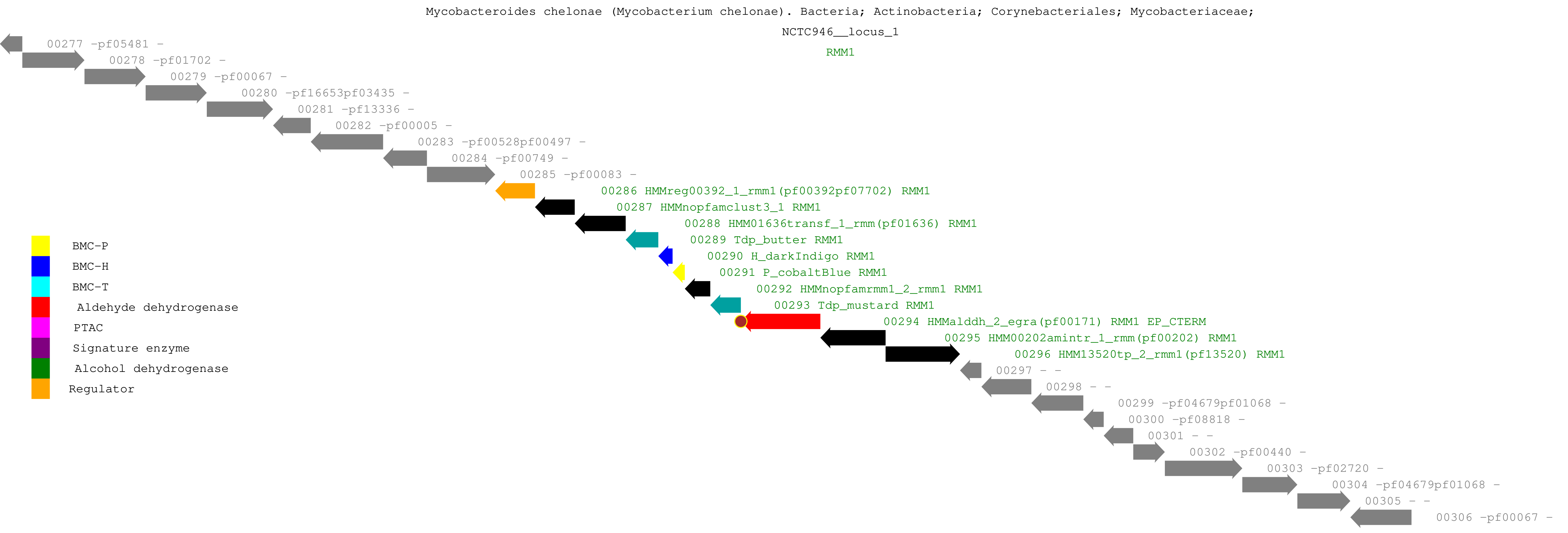
Mycobacteroides chelonae (Mycobacterium chelonae). Bacteria; Actinobacteria; Corynebacteriales; Mycobacteriaceae;
NCTC946__locus_1
RMM1
00277 -pf05481 -
00278 -pf01702 -
00279 -pf00067 -
00280 -pf16653pf03435 -
00281 -pf13336 -
00282 -pf00005 -
00283 -pf00528pf00497 -
00284 -pf00749 -
00285 -pf00083 -
00286 HMMreg00392_1_rmm1(pf00392pf07702) RMM1
00287 HMMnopfamclust3_1 RMM1
00288 HMM01636transf_1_rmm(pf01636) RMM1
00289 Tdp_butter RMM1
00290 H_darkIndigo RMM1
00291 P_cobaltBlue RMM1
00292 HMMnopfamrmm1_2_rmm1 RMM1
00293 Tdp_mustard RMM1
00294 HMMalddh_2_egra(pf00171) RMM1 EP_CTERM
00295 HMM00202amintr_1_rmm(pf00202) RMM1
00296 HMM13520tp_2_rmm1(pf13520) RMM1
00297 - -
00298 - -
00299 -pf04679pf01068 -
00300 -pf08818 -
00301 - -
00302 -pf00440 -
00303 -pf02720 -
00304 -pf04679pf01068 -
00305 - -
00306 -pf00067 -
BMC-P
BMC-H
BMC-T
Aldehyde dehydrogenase
PTAC
Signature enzyme
Alcohol dehydrogenase
Regulator

Nocardioides simplex (Arthrobacter simplex). Bacteria; Actinobacteria; Propionibacteriales; Nocardioidaceae;

NSI01__locus_1

RMM1

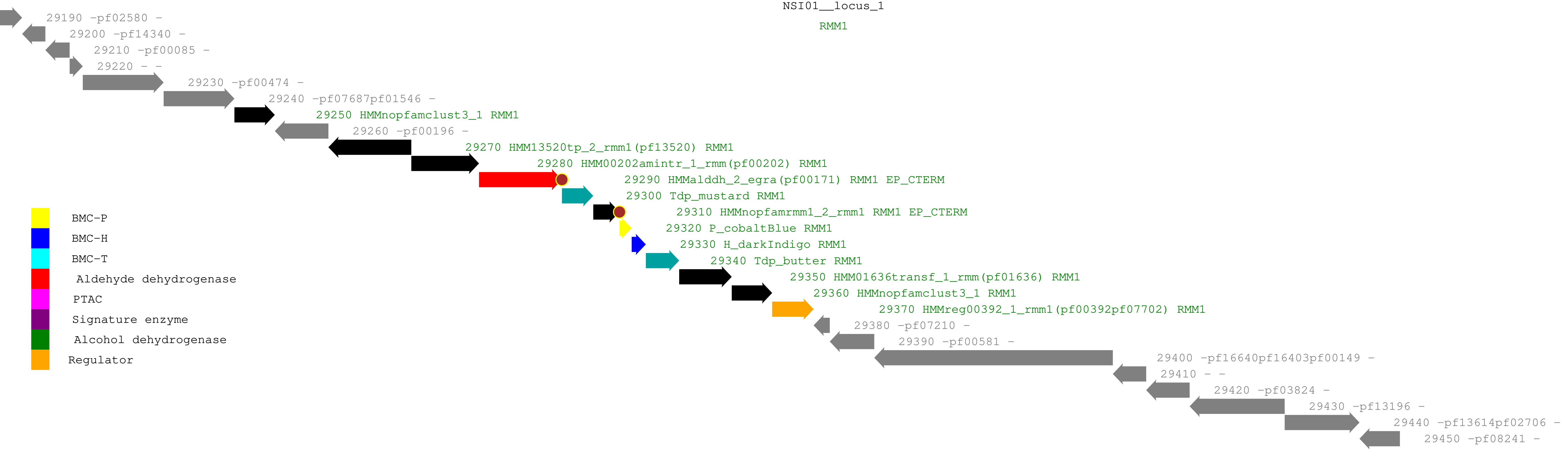

Rhodococcus erythropolis (Arthrobacter picolinophilus). Bacteria; Actinobacteria; Corynebacteriales; Nocardiaceae;

QV65__locus_1

RMM1

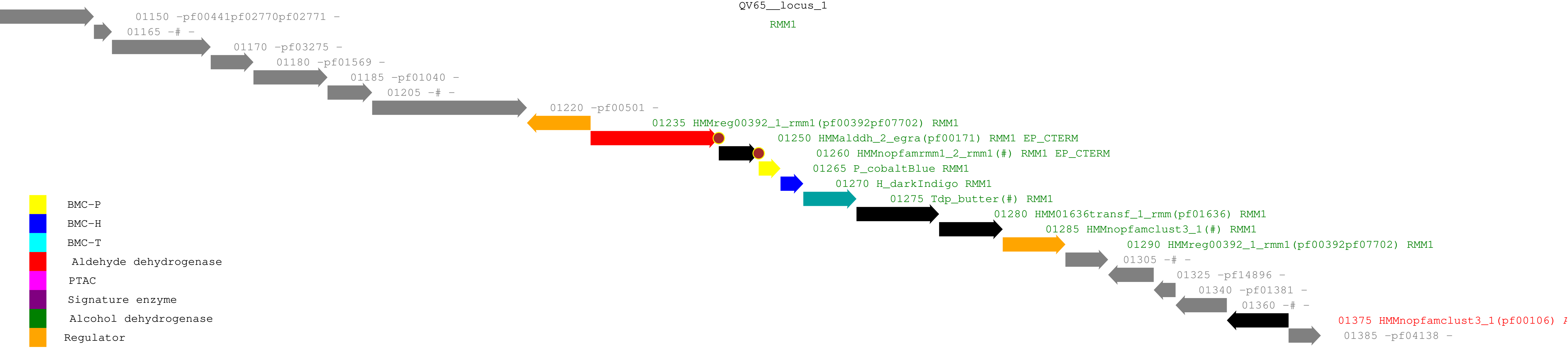

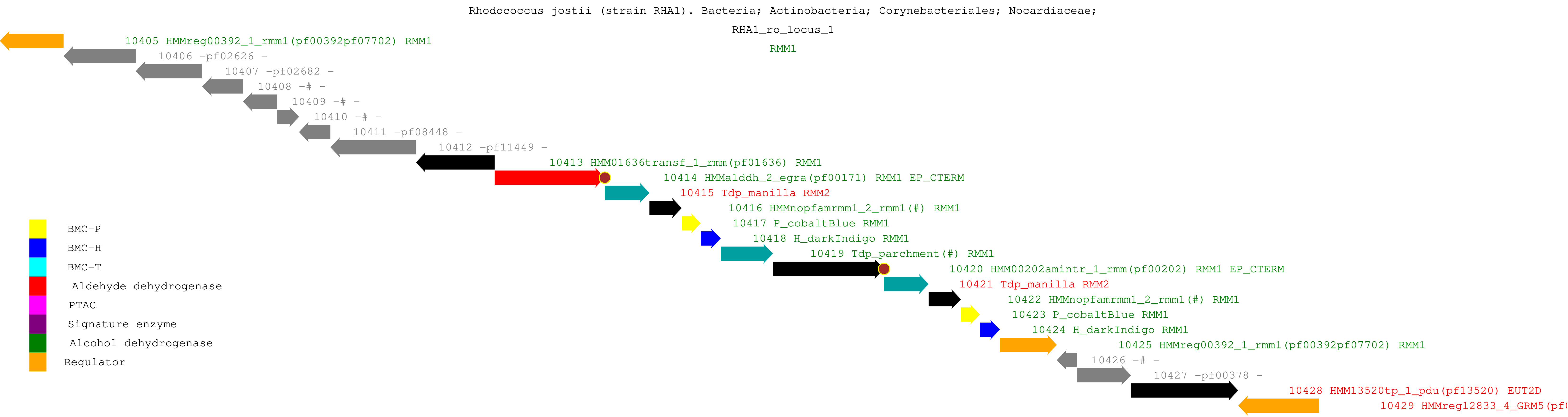
Rhodococcus jostii (strain RHA1). Bacteria; Actinobacteria; Corynebacteriales; Nocardiaceae;
RHA1_ro_locus_1
RMM1
10405 HMMreg00392_1_rmm1(pf00392pf07702) RMM1
10406 -pf02626 -
10407 -pf02682 -
10408 -# -
10409 -# -
10410 -# -
10411 -pf08448 -
10412 -pf11449 -
10413 HMM01636transf_1_rmm(pf01636) RMM1
10414 HMMaldh_2_egra(pf00171) RMM1 EP_CTERM
10415 Tdp_manilla RMM2
10416 HMMnopfamrmm1_2_rmm1(#) RMM1
10417 P_cobaltBlue RMM1
10418 H_darkIndigo RMM1
10419 Tdp_parchment(#) RMM1
10420 HMM00202amintr_1_rmm(pf00202) RMM1 EP_CTERM
10421 Tdp_manilla RMM2
10422 HMMnopfamrmm1_2_rmm1(#) RMM1
10423 P_cobaltBlue RMM1
10424 H_darkIndigo RMM1
10425 HMMreg00392_1_rmm1(pf00392pf07702) RMM1
10426 -# -
10427 -pf00378 -
10428 HMM13520tp_1_pdu(pf13520) EUT2D
10429 HMMreg12833_4_GRM5(pf
BMC-P
BMC-H
BMC-T
Aldehyde dehydrogenase
PTAC
Signature enzyme
Alcohol dehydrogenase
Regulator

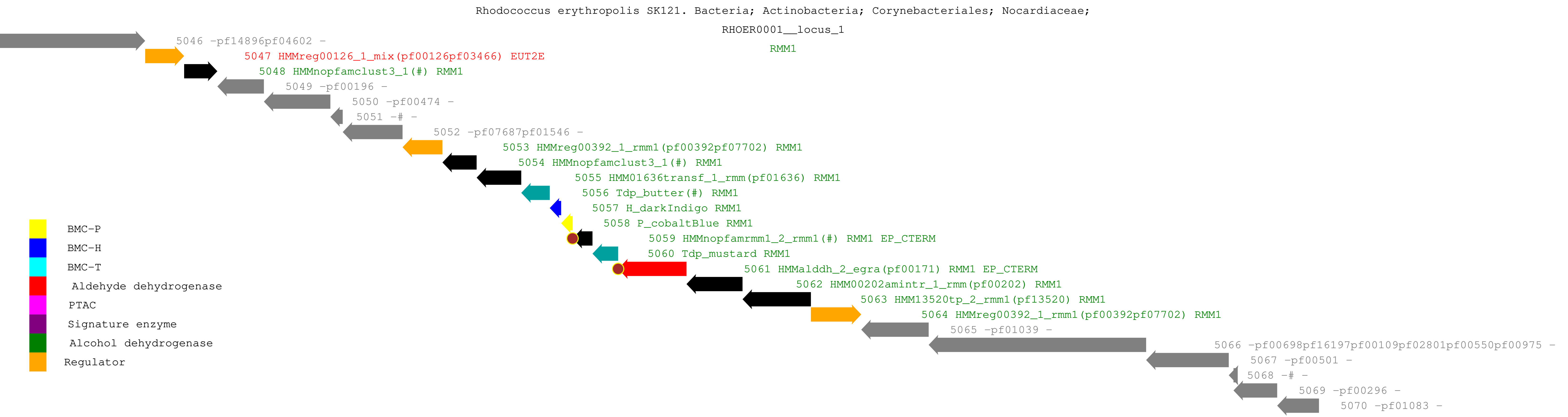

Rhodococcus erythropolis SK121. Bacteria; Actinobacteria; Corynebacteriales; Nocardiaceae;
RHOER0001__locus_1
RMM1
5046 -pf14896pf04602 -
5047 HMMreg00126_1_mix(pf00126pf03466) EUT2E
5048 HMMnopfamclust3_1(#) RMM1
5049 -pf00196 -
5050 -pf00474 -
5051 -# -
5052 -pf07687pf01546 -
5053 HMMreg00392_1_rmm1(pf00392pf07702) RMM1
5054 HMMnopfamclust3_1(#) RMM1
5055 HMM01636transf_1_rmm(pf01636) RMM1
5056 Tdp_butter(#) RMM1
5057 H_darkIndigo RMM1
5058 P_cobaltBlue RMM1
5059 HMMnopfamrmm1_2_rmm1(#) RMM1 EP_CTERM
5060 Tdp_mustard RMM1
5061 HMMaldh_2_egra(pf00171) RMM1 EP_CTERM
5062 HMM00202amintr_1_rmm(pf00202) RMM1
5063 HMM13520tp_2_rmm1(pf13520) RMM1
5064 HMMreg00392_1_rmm1(pf00392pf07702) RMM1
5065 -pf01039 -
5066 -pf00698pf16197pf00109pf02801pf00550pf00975 -
5067 -pf00501 -
5068 -# -
5069 -pf00296 -
5070 -pf01083 -
BMC-P
BMC-H
BMC-T
Aldehyde dehydrogenase
PTAC
Signature enzyme
Alcohol dehydrogenase
Regulator

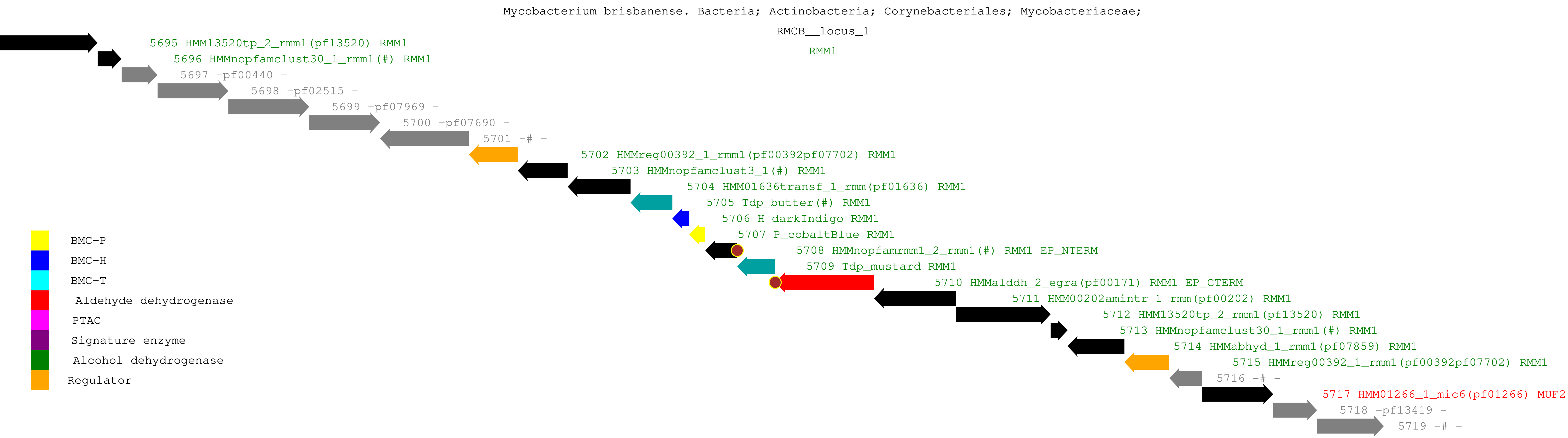
Mycobacterium brisbanense. Bacteria; Actinobacteria; Corynebacteriales; Mycobacteriaceae;
RMCB__locus_1
RMM1
5695 HMM13520tp_2_rmm1(pf13520) RMM1
5696 HMMnopfamclust30_1_rmm1(#) RMM1
5697 -pf00440 -
5698 -pf02515 -
5699 -pf07969 -
5700 -pf07690 -
5701 -# -
5702 HMMreg00392_1_rmm1(pf00392pf07702) RMM1
5703 HMMnopfamclust3_1(#) RMM1
5704 HMM01636transf_1_rmm(pf01636) RMM1
5705 Tdp_butter(#) RMM1
5706 H_darkIndigo RMM1
5707 P_cobaltBlue RMM1
5708 HMMnopfamrmm1_2_rmm1(#) RMM1 EP_NTERM
5709 Tdp_mustard RMM1
5710 HMMalddh_2_egra(pf00171) RMM1 EP_CTERM
5711 HMM00202amintr_1_rmm(pf00202) RMM1
5712 HMM13520tp_2_rmm1(pf13520) RMM1
5713 HMMnopfamclust30_1_rmm1(#) RMM1
5714 HMMabhyd_1_rmm1(pf07859) RMM1
5715 HMMreg00392_1_rmm1(pf00392pf07702) RMM1
5716 -# -
5717 HMM01266_1_mic6(pf01266) MUF2
5718 -pf13419 -
5719 -# -
BMC-P
BMC-H
BMC-T
Aldehyde dehydrogenase
PTAC
Signature enzyme
Alcohol dehydrogenase
Regulator

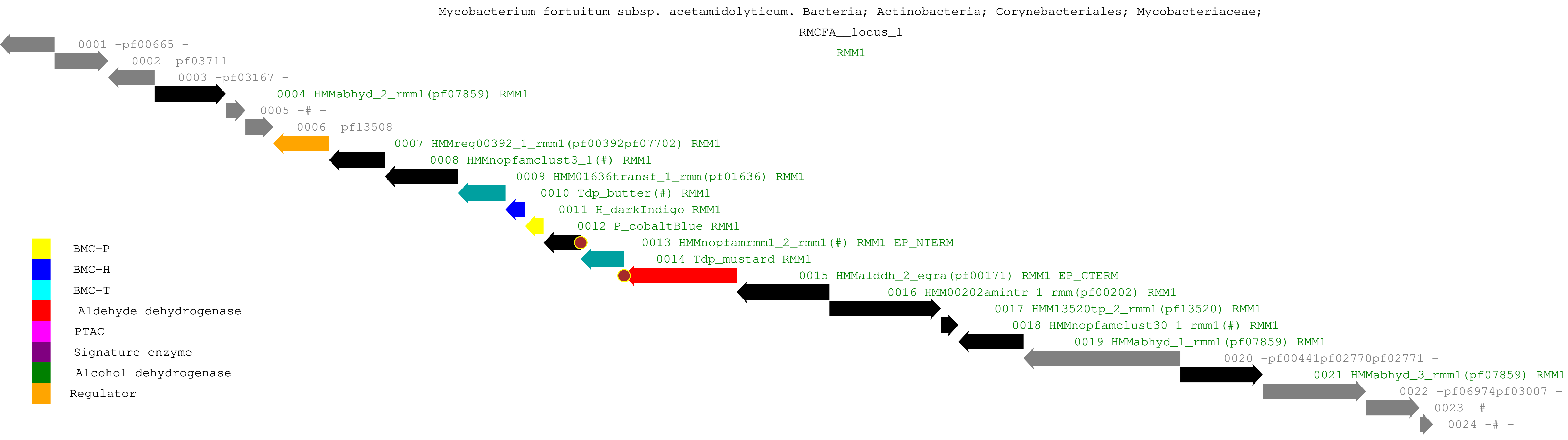
Mycobacterium fortuitum subsp. acetamidolyticum. Bacteria; Actinobacteria; Corynebacteriales; Mycobacteriaceae;
RMCFA__locus_1
RMM1
0001 -pf00665 -
0002 -pf03711 -
0003 -pf03167 -
0004 HMMabhyd_2_rmm1(pf07859) RMM1
0005 -# -
0006 -pf13508 -
0007 HMMreg00392_1_rmm1(pf00392pf07702) RMM1
0008 HMMnopfamclust3_1(#) RMM1
0009 HMM01636transf_1_rmm(pf01636) RMM1
0010 Tdp_butter(#) RMM1
0011 H_darkIndigo RMM1
0012 P_cobaltBlue RMM1
0013 HMMnopfamrmm1_2_rmm1(#) RMM1 EP_NTERM
0014 Tdp_mustard RMM1
0015 HMMalddh_2_egra(pf00171) RMM1 EP_CTERM
0016 HMM00202amintr_1_rmm(pf00202) RMM1
0017 HMM13520tp_2_rmm1(pf13520) RMM1
0018 HMMnopfamclust30_1_rmm1(#) RMM1
0019 HMMabhyd_1_rmm1(pf07859) RMM1
0020 -pf00441pf02770pf02771 -
0021 HMMabhyd_3_rmm1(pf07859) RMM1
0022 -pf06974pf03007 -
0023 -# -
0024 -# -
BMC-P
BMC-H
BMC-T
Aldehyde dehydrogenase
PTAC
Signature enzyme
Alcohol dehydrogenase
Regulator

Mycobacterium sp. UNC267MFSha1.1M11. Bacteria; Actinobacteria; Corynebacteriales; Mycobacteriaceae;

SAMN02799620__locus_1

RMM1

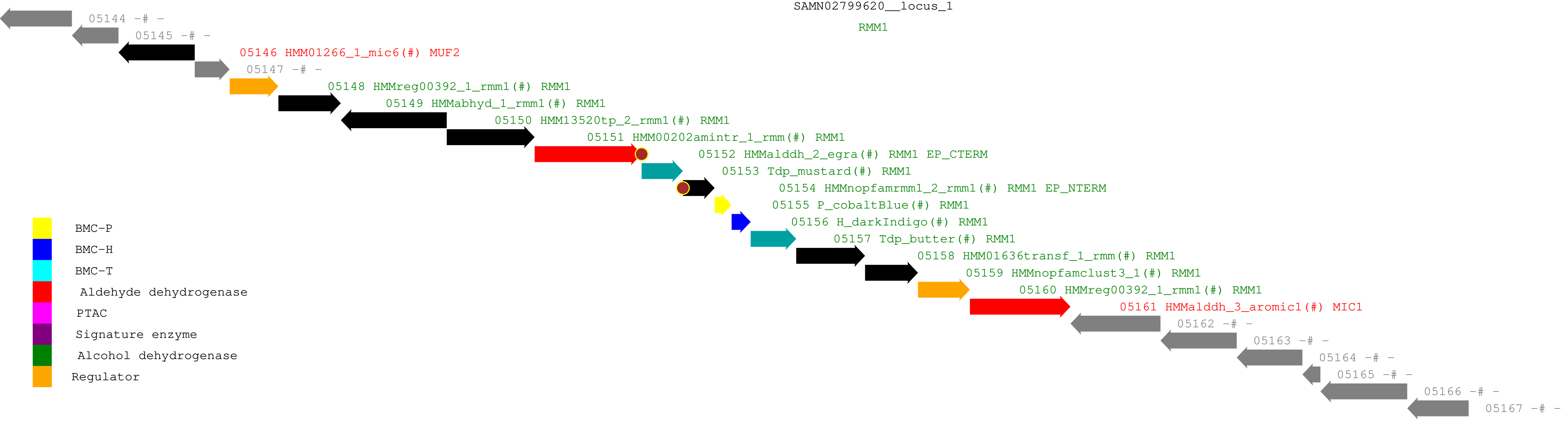

Bauldia litoralis. Bacteria; Proteobacteria; Alphaproteobacteria; Rhizobiales; Bauldia.

SAMN02982931__locus_1

RMM1

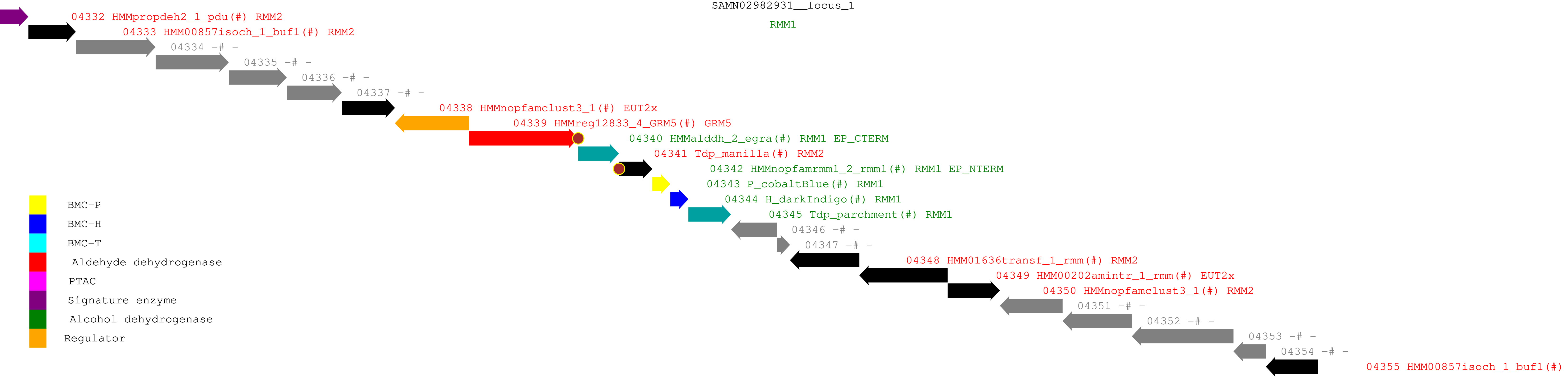

Mycobacterium sp. 455mf. Bacteria; Actinobacteria; Corynebacteriales; Mycobacteriaceae;

SAMN04488582__locus_1

RMM1

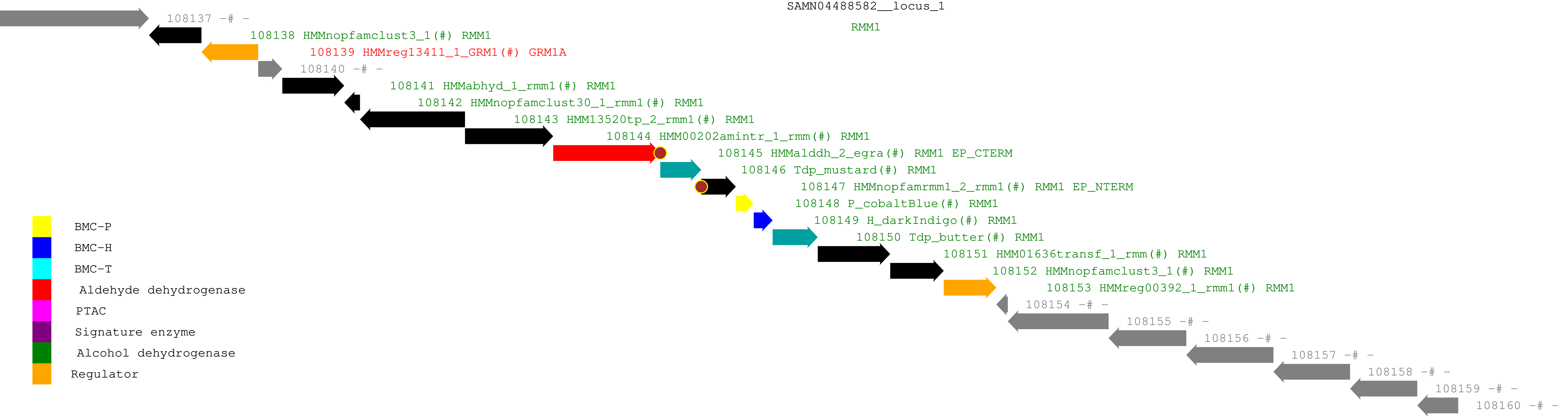

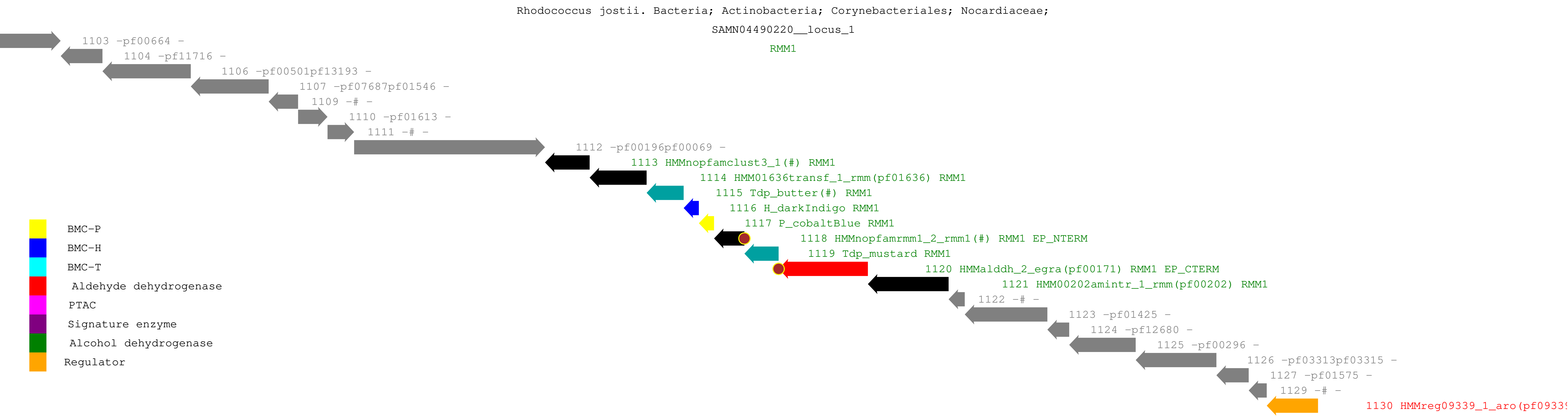

Rhodococcus jostii. Bacteria; Actinobacteria; Corynebacteriales; Nocardiaceae;
SAMN04490220__locus_1
RMM1
1103 -pf00664 -
1104 -pf11716 -
1106 -pf00501pf13193 -
1107 -pf07687pf01546 -
1109 -# -
1110 -pf01613 -
1111 -# -
1112 -pf00196pf00069 -
1113 HMMnopfamclust3_1(#) RMM1
1114 HMM01636transf_1_rmm(pf01636) RMM1
1115 Tdp_butter(#) RMM1
1116 H_darkIndigo RMM1
1117 P_cobaltBlue RMM1
1118 HMMnopfamrmm1_2_rmm1(#) RMM1 EP_NTERM
1119 Tdp_mustard RMM1
1120 HMMalddh_2_egra(pf00171) RMM1 EP_CTERM
1121 HMM00202amintr_1_rmm(pf00202) RMM1
1122 -# -
1123 -pf01425 -
1124 -pf12680 -
1125 -pf00296 -
1126 -pf03313pf03315 -
1127 -pf01575 -
1129 -# -
1130 HMMreg09339_1_aro(pf0933
BMC-P
BMC-H
BMC-T
Aldehyde dehydrogenase
PTAC
Signature enzyme
Alcohol dehydrogenase
Regulator

Nocardioides simplex (Arthrobacter simplex). Bacteria; Actinobacteria; Propionibacteriales; Nocardioidaceae;

SAMN05421671__locus_1

RMM1

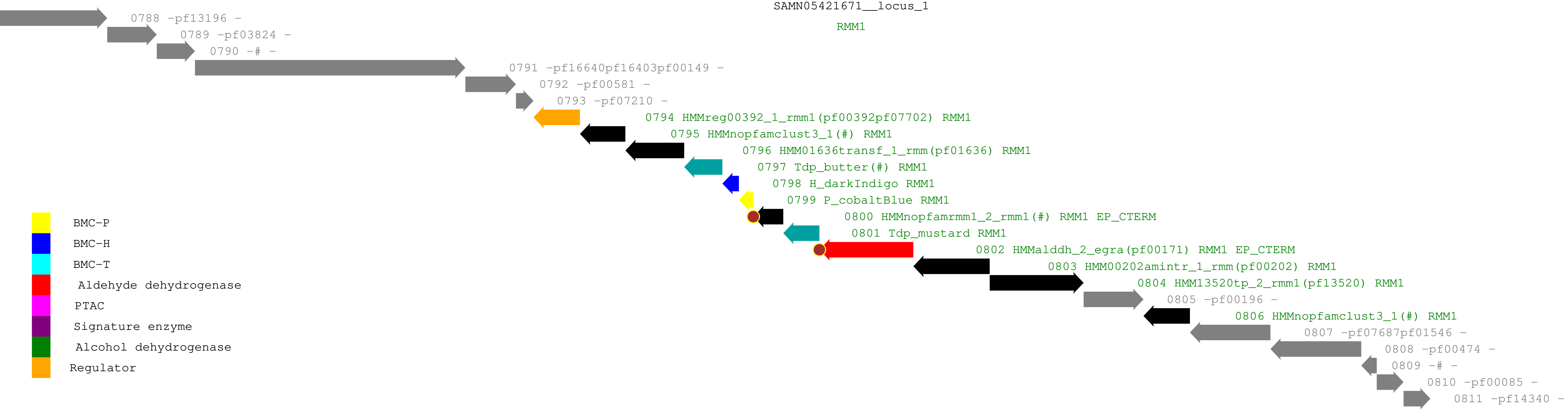

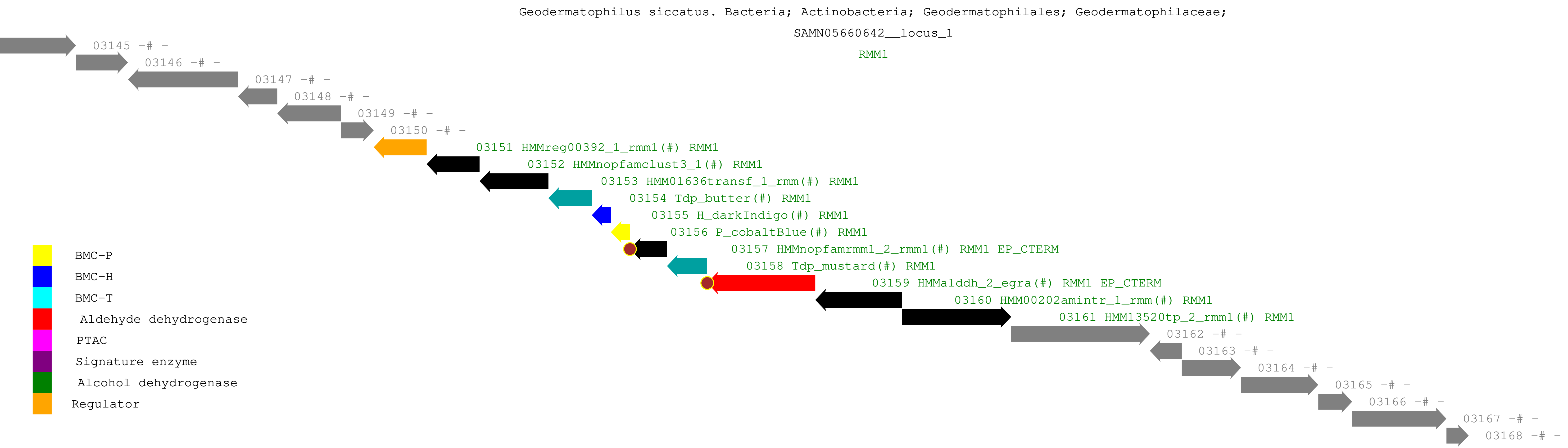
Geodermatophilus siccatus. Bacteria; Actinobacteria; Geodermatophilales; Geodermatophilaceae;
SAMN05660642__locus_1
RMM1
03145 -# -
03146 -# -
03147 -# -
03148 -# -
03149 -# -
03150 -# -
03151 HMMreg00392_1_rmm1(#) RMM1
03152 HMMnopfamclust3_1(#) RMM1
03153 HMM01636transf_1_rmm(#) RMM1
03154 Tdp_butter(#) RMM1
03155 H_darkIndigo(#) RMM1
03156 P_cobaltBlue(#) RMM1
03157 HMMnopfamrmm1_2_rmm1(#) RMM1 EP_CTERM
03158 Tdp_mustard(#) RMM1
03159 HMMalddh_2_egra(#) RMM1 EP_CTERM
03160 HMM00202amintr_1_rmm(#) RMM1
03161 HMM13520tp_2_rmm1(#) RMM1
03162 -# -
03163 -# -
03164 -# -
03165 -# -
03166 -# -
03167 -# -
03168 -# -
BMC-P
BMC-H
BMC-T
Aldehyde dehydrogenase
PTAC
Signature enzyme
Alcohol dehydrogenase
Regulator

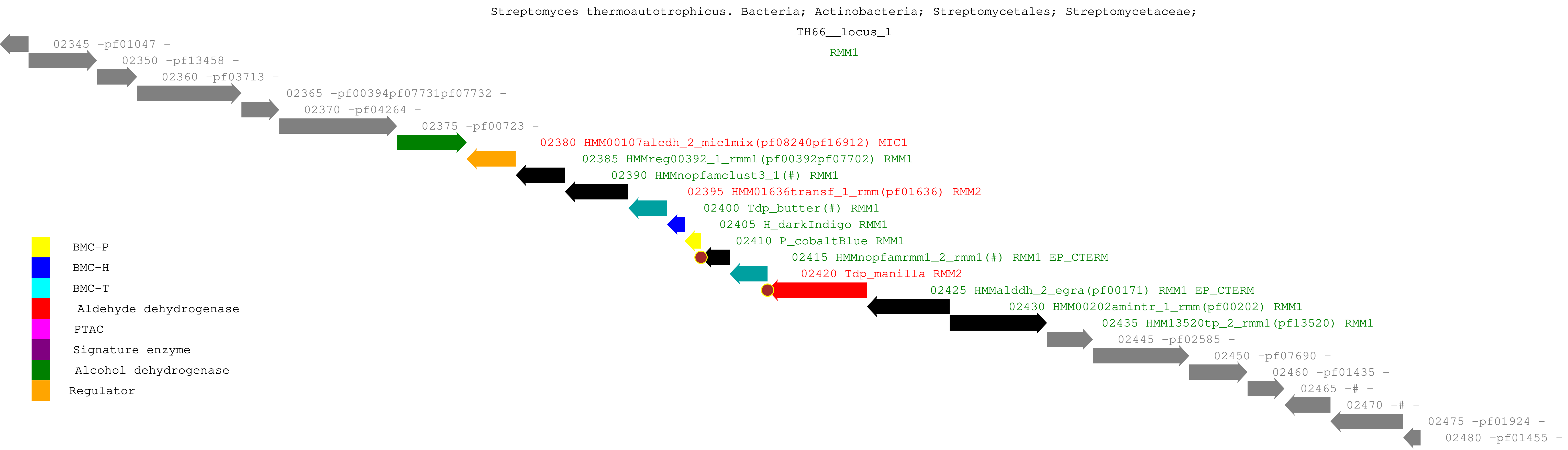
Streptomyces thermoautotrophicus. Bacteria; Actinobacteria; Streptomycetales; Streptomycetaceae;
TH66__locus_1
RMM1
02345 -pf01047 -
02350 -pf13458 -
02360 -pf03713 -
02365 -pf00394pf07731pf07732 -
02370 -pf04264 -
02375 -pf00723 -
02380 HMM00107alcdh_2_mic1mix(pf08240pf16912) MIC1
02385 HMMreg00392_1_rmm1(pf00392pf07702) RMM1
02390 HMMnopfamclust3_1(#) RMM1
02395 HMM01636transf_1_rmm(pf01636) RMM2
02400 Tdp_butter(#) RMM1
02405 H_darkIndigo RMM1
02410 P_cobaltBlue RMM1
02415 HMMnopfamrmm1_2_rmm1(#) RMM1 EP_CTERM
02420 Tdp_manilla RMM2
02425 HMMaldh_2_egra(pf00171) RMM1 EP_CTERM
02430 HMM00202amintr_1_rmm(pf00202) RMM1
02435 HMM13520tp_2_rmm1(pf13520) RMM1
02445 -pf02585 -
02450 -pf07690 -
02460 -pf01435 -
02465 -# -
02470 -# -
02475 -pf01924 -
02480 -pf01455 -
BMC-P
BMC-H
BMC-T
Aldehyde dehydrogenase
PTAC
Signature enzyme
Alcohol dehydrogenase
Regulator

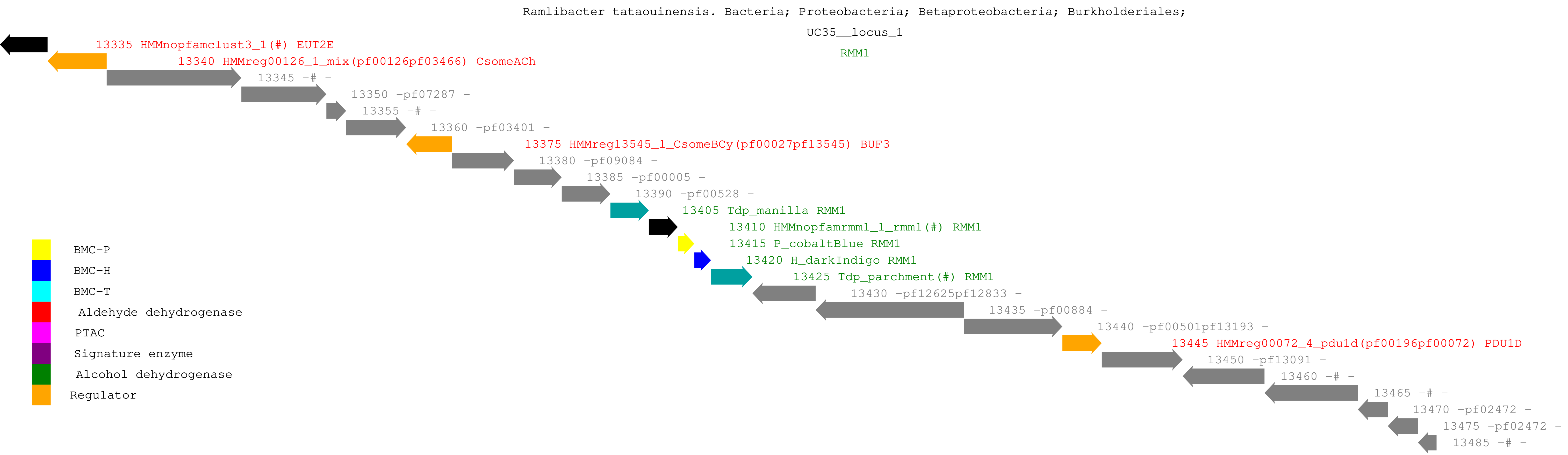
Ramlibacter tataouinensis. Bacteria; Proteobacteria; Betaproteobacteria; Burkholderiales;
UC35__locus_1
RMM1
13335 HMMnopfamclust3_1(#) EUT2E
13340 HMMreg00126_1_mix(pf00126pf03466) CsomeACh
13345 -# -
13350 -pf07287 -
13355 -# -
13360 -pf03401 -
13375 HMMreg13545_1_CsomeBCy(pf00027pf13545) BUF3
13380 -pf09084 -
13385 -pf00005 -
13390 -pf00528 -
13405 Tdp_manilla RMM1
13410 HMMnopfamrmm1_1_rmm1(#) RMM1
13415 P_cobaltBlue RMM1
13420 H_darkIndigo RMM1
13425 Tdp_parchment(#) RMM1
13430 -pf12625pf12833 -
13435 -pf00884 -
13440 -pf00501pf13193 -
13445 HMMreg00072_4_pdu1d(pf00196pf00072) PDU1D
13450 -pf13091 -
13460 -# -
13465 -# -
13470 -pf02472 -
13475 -pf02472 -
13485 -# -
BMC-P
BMC-H
BMC-T
Aldehyde dehydrogenase
PTAC
Signature enzyme
Alcohol dehydrogenase
Regulator

Vibrio tapetis subsp. tapetis. Bacteria; Proteobacteria; Gammaproteobacteria; Vibrionales;

VTAP4600_A_locus_1

RMM1

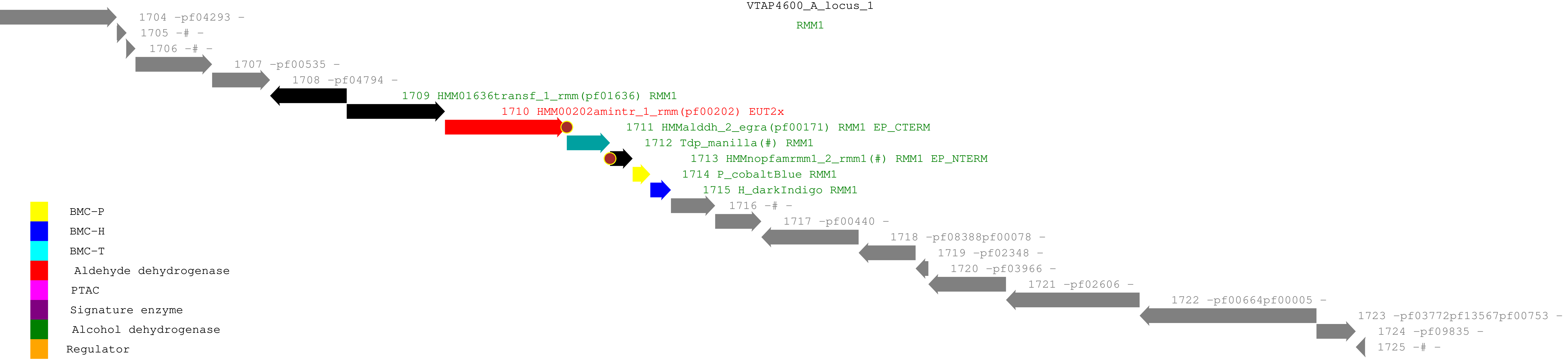

Mycobacterium fortuitum. Bacteria; Actinobacteria; Corynebacteriales; Mycobacteriaceae;

XA26__locus_1

RMM1

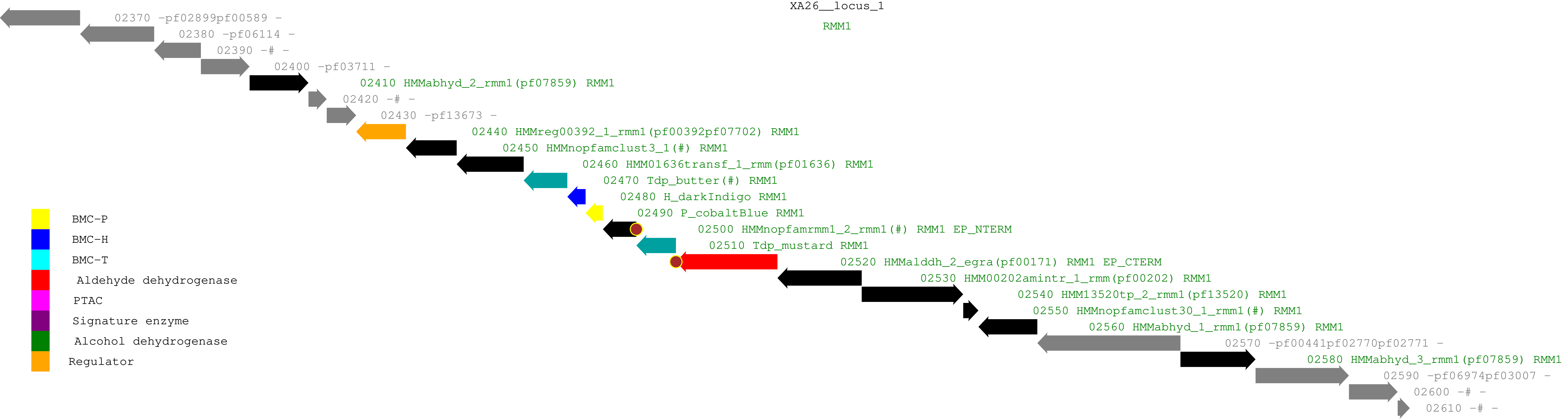

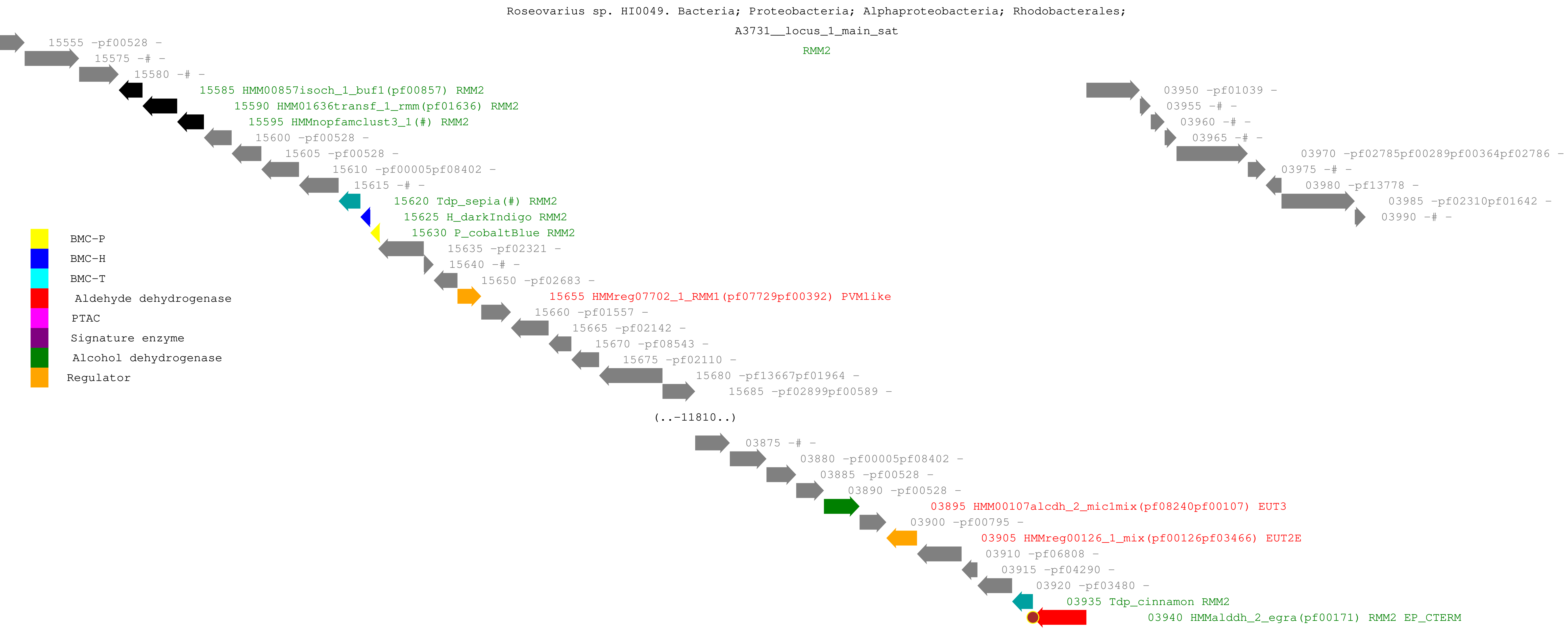
Roseovarius sp. HI0049. Bacteria; Proteobacteria; Alphaproteobacteria; Rhodobacterales;
A3731__locus_1_main_sat
RMM2
15555 -pf00528 -
15575 -# -
15580 -# -
15585 HMM00857isoch_1_buf1(pf00857) RMM2
15590 HMM01636transf_1_rmm(pf01636) RMM2
15595 HMMnopfamclust3_1(#) RMM2
15600 -pf00528 -
15605 -pf00528 -
15610 -pf00005pf08402 -
15615 -# -
15620 Tdp_sepia(#) RMM2
15625 H_darkIndigo RMM2
15630 P_cobaltBlue RMM2
15635 -pf02321 -
15640 -# -
15650 -pf02683 -
15655 HMMreg07702_1_RMM1(pf07729pf00392) PVMlike
15660 -pf01557 -
15665 -pf02142 -
15670 -pf08543 -
15675 -pf02110 -
15680 -pf13667pf01964 -
15685 -pf02899pf00589 -
(..-11810..)
03875 -# -
03880 -pf00005pf08402 -
03885 -pf00528 -
03890 -pf00528 -
03895 HMM00107alcdh_2_mic1mix(pf08240pf00107) EUT3
03900 -pf00795 -
03905 HMMreg00126_1_mix(pf00126pf03466) EUT2E
03910 -pf06808 -
03915 -pf04290 -
03920 -pf03480 -
03935 Tdp_cinnamon RMM2
03940 HMMalddh_2_egra(pf00171) RMM2 EP_CTERM
03950 -pf01039 -
03955 -# -
03960 -# -
03965 -# -
03970 -pf02785pf00289pf00364pf02786 -
03975 -# -
03980 -pf13778 -
03985 -pf02310pf01642 -
03990 -# -
BMC-P
BMC-H
BMC-T
Aldehyde dehydrogenase
PTAC
Signature enzyme
Alcohol dehydrogenase
Regulator

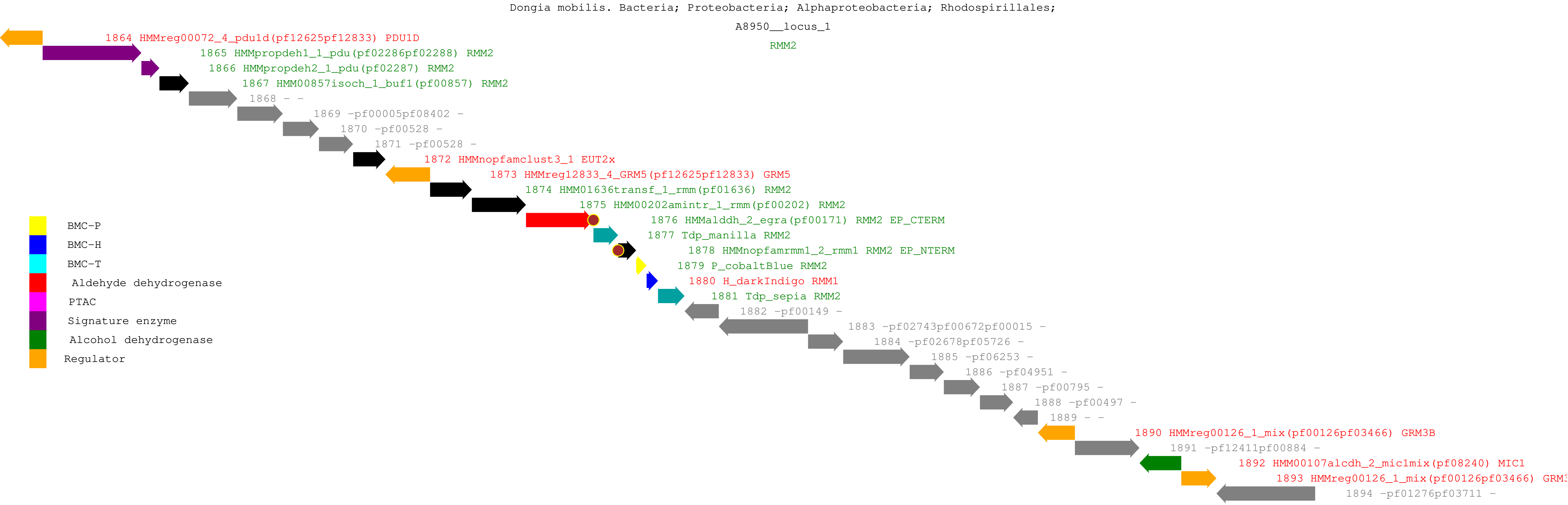

Dongia mobilis. Bacteria; Proteobacteria; Alphaproteobacteria; Rhodospirillales;
A8950__locus_1
RMM2
1864 HMMreg00072_4_pdu1d(pf12625pf12833) PDU1D
1865 HMMpropdeh1_1_pdu(pf02286pf02288) RMM2
1866 HMMpropdeh2_1_pdu(pf02287) RMM2
1867 HMM00857isoch_1_buf1(pf00857) RMM2
1868 - -
1869 -pf00005pf08402 -
1870 -pf00528 -
1871 -pf00528 -
1872 HMMnopfamclust3_1 EUT2x
1873 HMMreg12833_4_GRM5(pf12625pf12833) GRM5
1874 HMM01636transf_1_rmm(pf01636) RMM2
1875 HMM00202amintr_1_rmm(pf00202) RMM2
1876 HMMalddh_2_egra(pf00171) RMM2 EP_CTERM
1877 Tdp_manilla RMM2
1878 HMMnopfamrmm1_2_rmm1 RMM2 EP_NTERM
1879 P_cobaltBlue RMM2
1880 H_darkIndigo RMM1
1881 Tdp_sepia RMM2
1882 -pf00149 -
1883 -pf02743pf00672pf00015 -
1884 -pf02678pf05726 -
1885 -pf06253 -
1886 -pf04951 -
1887 -pf00795 -
1888 -pf00497 -
1889 - -
1890 HMMreg00126_1_mix(pf00126pf03466) GRM3B
1891 -pf12411pf00884 -
1892 HMM00107alcdh_2_mic1mix(pf08240) MIC1
1893 HMMreg00126_1_mix(pf00126pf03466) GRM3
1894 -pf01276pf03711 -
BMC-P
BMC-H
BMC-T
Aldehyde dehydrogenase
PTAC
Signature enzyme
Alcohol dehydrogenase
Regulator

Rhodobacteraceae bacterium (ex Bugula neritina AB1). Bacteria; Proteobacteria; Alphaproteobacteria; Rhodobacterales;

AB838__locus_1

RMM2

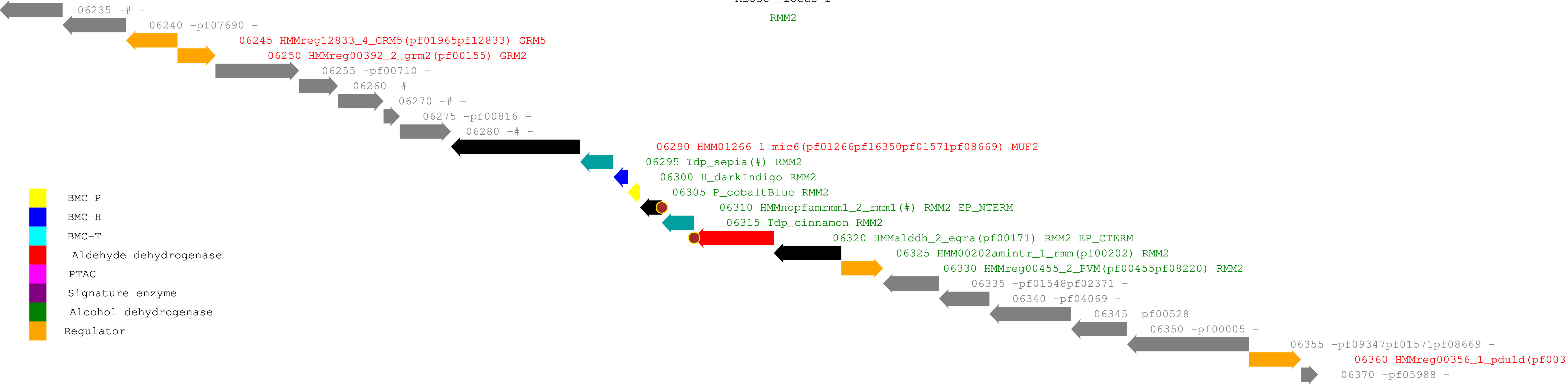

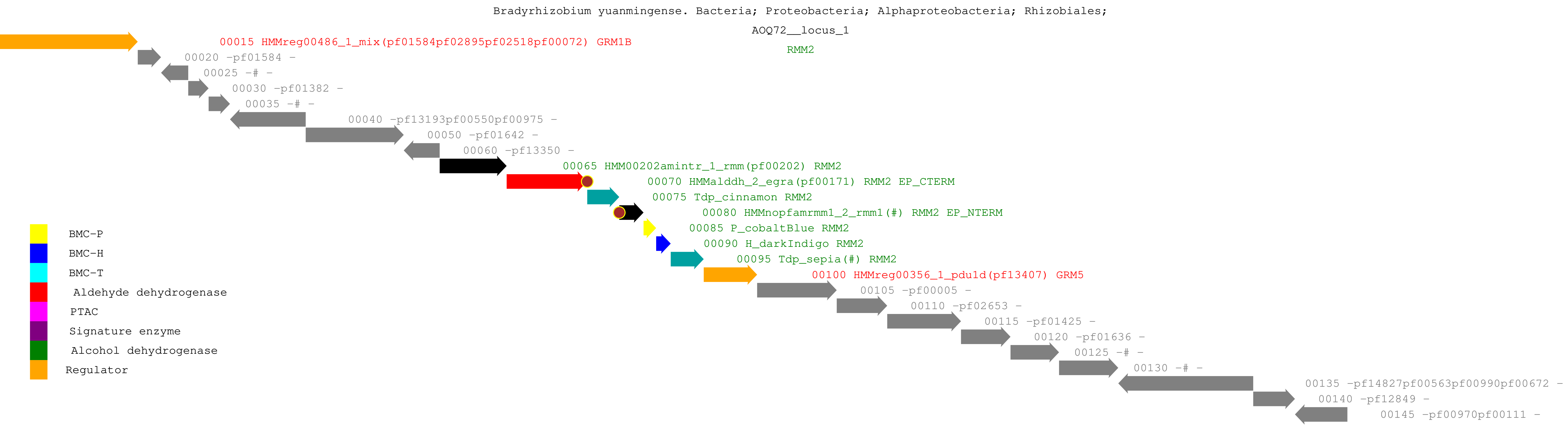
Bradyrhizobium yuanmingense. Bacteria; Proteobacteria; Alphaproteobacteria; Rhizobiales;
AOQ72__locus_1
RMM2
00015 HMMreg00486_1_mix(pf01584pf02895pf02518pf00072) GRM1B
00020 -pf01584 -
00025 -# -
00030 -pf01382 -
00035 -# -
00040 -pf13193pf00550pf00975 -
00050 -pf01642 -
00060 -pf13350 -
00065 HMM00202amintr_1_rmm(pf00202) RMM2
00070 HMMaldd h_2_egra(pf00171) RMM2 EP_CTERM
00075 Tdp_cinnamon RMM2
00080 HMMnopfamrmm1_2_rmm1(#) RMM2 EP_NTERM
00085 P_cobaltBlue RMM2
00090 H_darkIndigo RMM2
00095 Tdp_sepia(#) RMM2
00100 HMMreg00356_1_pdu1d(pf13407) GRM5
00105 -pf00005 -
00110 -pf02653 -
00115 -pf01425 -
00120 -pf01636 -
00125 -# -
00130 -# -
00135 -pf14827pf00563pf00990pf00672 -
00140 -pf12849 -
00145 -pf00970pf00111 -
BMC-P
BMC-H
BMC-T
Aldehyde dehydrogenase
PTAC
Signature enzyme
Alcohol dehydrogenase
Regulator

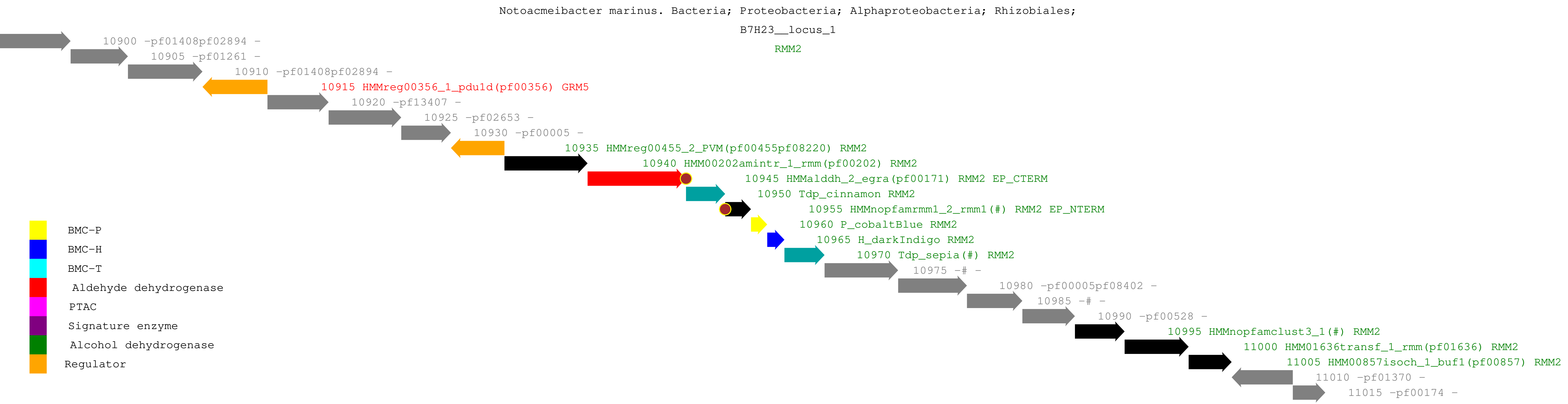
Notoacmeibacter marinus. Bacteria; Proteobacteria; Alphaproteobacteria; Rhizobiales;
B7H23__locus_1
RMM2
10900 -pf01408pf02894 -
10905 -pf01261 -
10910 -pf01408pf02894 -
10915 HMMreg00356_1_pdu1d(pf00356) GRM5
10920 -pf13407 -
10925 -pf02653 -
10930 -pf00005 -
10935 HMMreg00455_2_PVM(pf00455pf08220) RMM2
10940 HMM00202amintr_1_rmm(pf00202) RMM2
10945 HMMalddh_2_egra(pf00171) RMM2 EP_CTERM
10950 Tdp_cinnamon RMM2
10955 HMMnopfamrmm1_2_rmm1(#) RMM2 EP_NTERM
10960 P_cobaltBlue RMM2
10965 H_darkIndigo RMM2
10970 Tdp_sepia(#) RMM2
10975 -# -
10980 -pf00005pf08402 -
10985 -# -
10990 -pf00528 -
10995 HMMnopfamclust3_1(#) RMM2
11000 HMM01636transf_1_rmm(pf01636) RMM2
11005 HMM00857isoch_1_buf1(pf00857) RMM2
11010 -pf01370 -
11015 -pf00174 -
BMC-P
BMC-H
BMC-T
Aldehyde dehydrogenase
PTAC
Signature enzyme
Alcohol dehydrogenase
Regulator

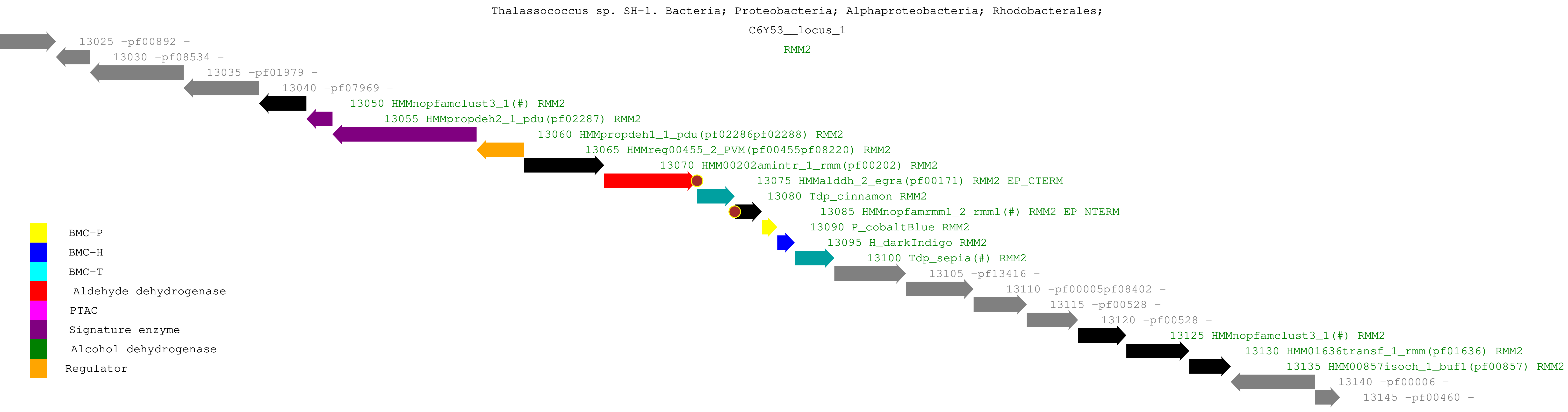
Thalassococcus sp. SH-1. Bacteria; Proteobacteria; Alphaproteobacteria; Rhodobacterales;
C6Y53__locus_1
RMM2
13025 -pf00892 -
13030 -pf08534 -
13035 -pf01979 -
13040 -pf07969 -
13050 HMMnopfamclust3_1(#) RMM2
13055 HMMpropdeh2_1_pdu(pf02287) RMM2
13060 HMMpropdeh1_1_pdu(pf02286pf02288) RMM2
13065 HMMreg00455_2_PVM(pf00455pf08220) RMM2
13070 HMM00202amintr_1_rmm(pf00202) RMM2
13075 HMMalddh_2_egra(pf00171) RMM2 EP_CTERM
13080 Tdp_cinnamon RMM2
13085 HMMnopfamrmm1_2_rmm1(#) RMM2 EP_NTERM
13090 P_cobaltBlue RMM2
13095 H_darkIndigo RMM2
13100 Tdp_sepia(#) RMM2
13105 -pf13416 -
13110 -pf00005pf08402 -
13115 -pf00528 -
13120 -pf00528 -
13125 HMMnopfamclust3_1(#) RMM2
13130 HMM01636transf_1_rmm(pf01636) RMM2
13135 HMM00857isoch_1_buf1(pf00857) RMM2
13140 -pf00006 -
13145 -pf00460 -
BMC-P
BMC-H
BMC-T
Aldehyde dehydrogenase
PTAC
Signature enzyme
Alcohol dehydrogenase
Regulator

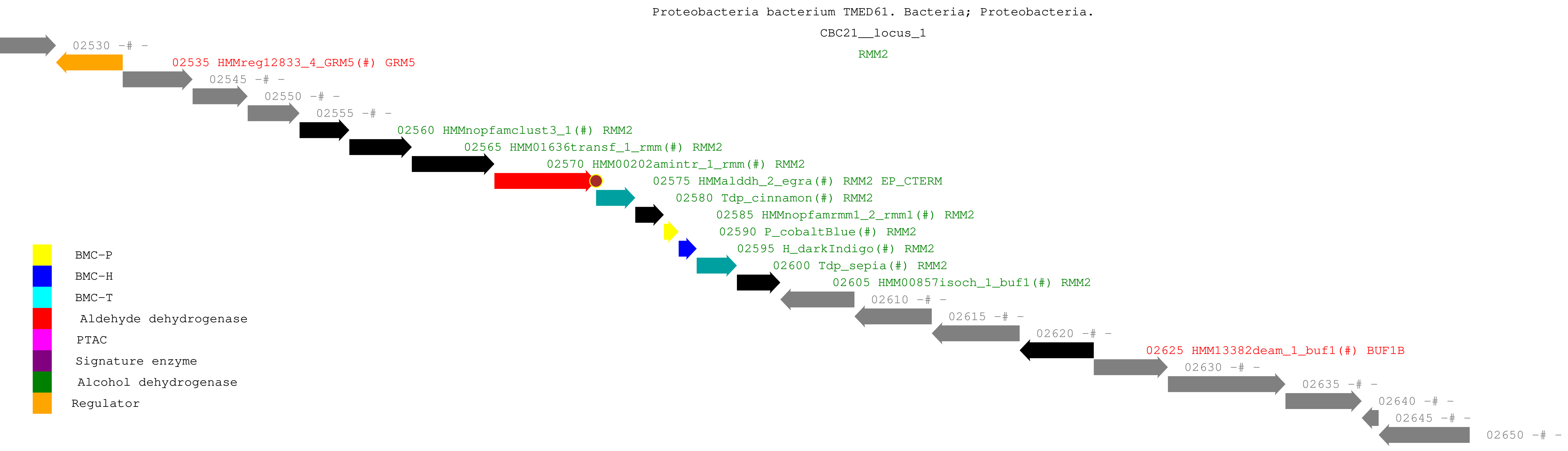
Proteobacteria bacterium TMED61. Bacteria; Proteobacteria.
CBC21__locus_1
RMM2
02530 -# -
02535 HMMreg12833_4_GRM5(#) GRM5
02545 -# -
02550 -# -
02555 -# -
02560 HMMnopfamclust3_1(#) RMM2
02565 HMM01636transf_1_rmm(#) RMM2
02570 HMM00202amintr_1_rmm(#) RMM2
02575 HMMalddh_2_egra(#) RMM2 EP_CTERM
02580 Tdp_cinnamon(#) RMM2
02585 HMMnopfamrmm1_2_rmm1(#) RMM2
02590 P_cobaltBlue(#) RMM2
02595 H_darkIndigo(#) RMM2
02600 Tdp_sepia(#) RMM2
02605 HMM00857isoch_1_buf1(#) RMM2
02610 -# -
02615 -# -
02620 -# -
02625 HMM13382deam_1_buf1(#) BUF1B
02630 -# -
02635 -# -
02640 -# -
02645 -# -
02650 -# -
BMC-P
BMC-H
BMC-T
Aldehyde dehydrogenase
PTAC
Signature enzyme
Alcohol dehydrogenase
Regulator

Acidiferrobacter sp. Bacteria; Proteobacteria; Gammaproteobacteria; Acidiferrobacterales;

CL386__locus_1

RMM2

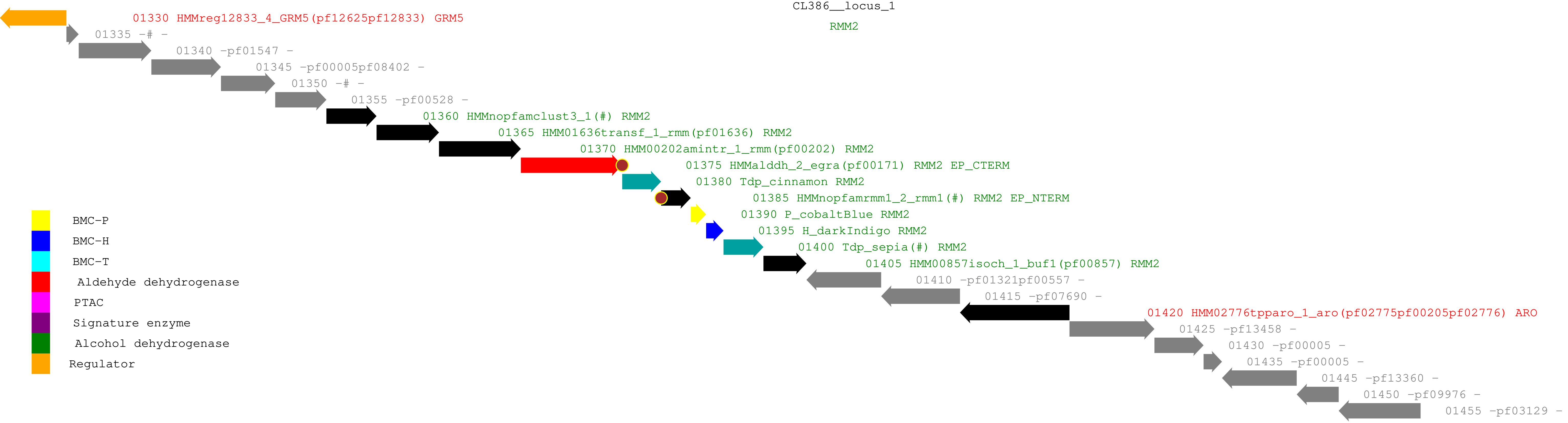

Acidiferrobacteraceae bacterium. Bacteria; Proteobacteria; Gammaproteobacteria; Acidiferrobacterales;

CL389__locus_1

RMM2

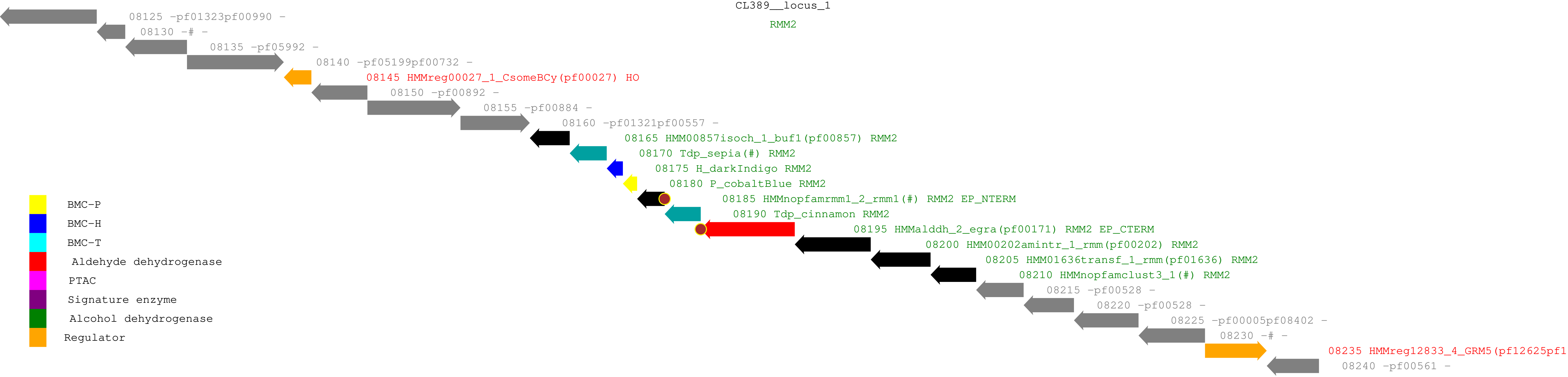

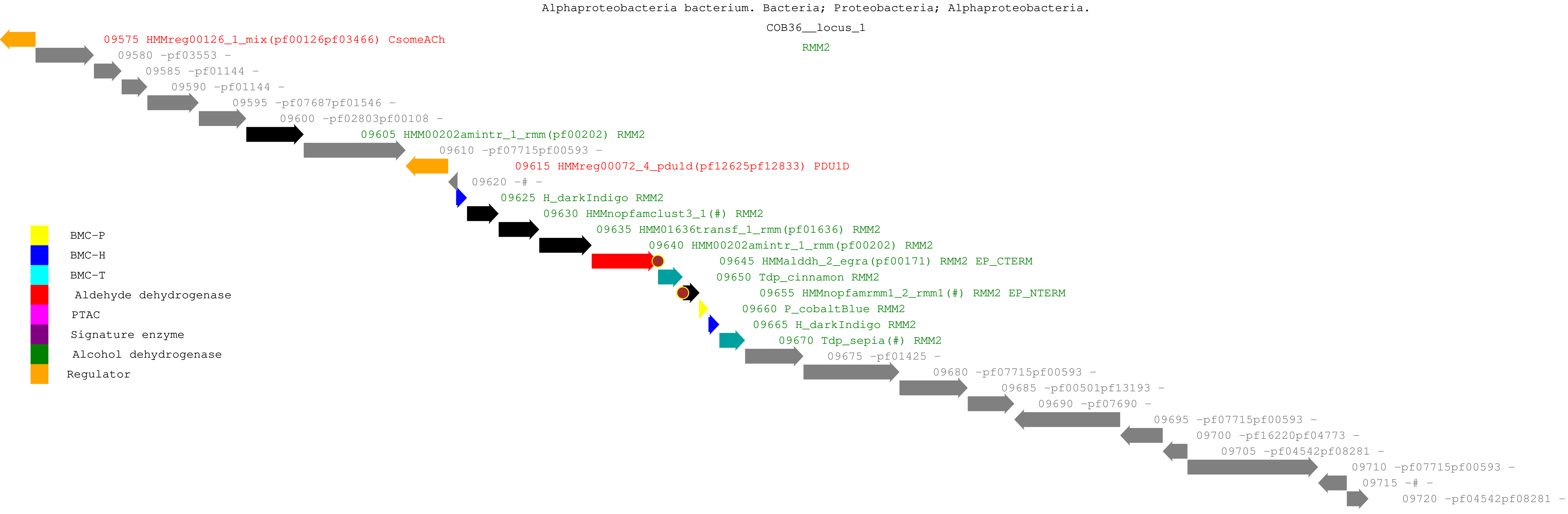

Alphaproteobacteria bacterium. Bacteria; Proteobacteria; Alphaproteobacteria.
COB36__locus_1
RMM2
09575 HMMreg00126_1_mix(pf00126pf03466) CsomeACh
09580 -pf03553 -
09585 -pf01144 -
09590 -pf01144 -
09595 -pf07687pf01546 -
09600 -pf02803pf00108 -
09605 HMM00202amintr_1_rmm(pf00202) RMM2
09610 -pf07715pf00593 -
09615 HMMreg00072_4_pdu1d(pf12625pf12833) PDU1D
09620 -# -
09625 H_darkIndigo RMM2
09630 HMMnopfamclust3_1(#) RMM2
09635 HMM01636transf_1_rmm(pf01636) RMM2
09640 HMM00202amintr_1_rmm(pf00202) RMM2
09645 HMMalddh_2_egra(pf00171) RMM2 EP_CTERM
09650 Tdp_cinnamon RMM2
09655 HMMnopfamrmm1_2_rmm1(#) RMM2 EP_NTERM
09660 P_cobaltBlue RMM2
09665 H_darkIndigo RMM2
09670 Tdp_sepia(#) RMM2
09675 -pf01425 -
09680 -pf07715pf00593 -
09685 -pf00501pf13193 -
09690 -pf07690 -
09695 -pf07715pf00593 -
09700 -pf16220pf04773 -
09705 -pf04542pf08281 -
09710 -pf07715pf00593 -
09715 -# -
09720 -pf04542pf08281 -
BMC-P
BMC-H
BMC-T
Aldehyde dehydrogenase
PTAC
Signature enzyme
Alcohol dehydrogenase
Regulator

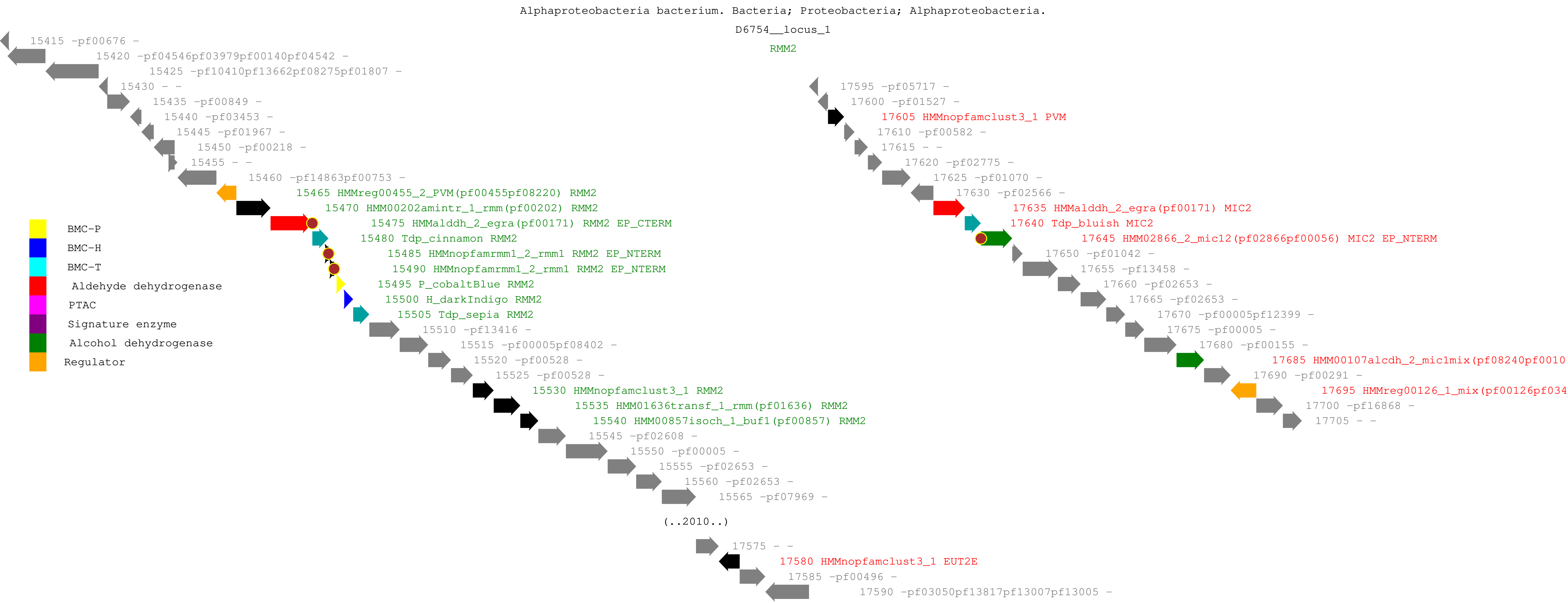

Alphaproteobacteria bacterium. Bacteria; Proteobacteria; Alphaproteobacteria.
D6754__locus_1
RMM2
15415 -pf00676 -
15420 -pf04546pf03979pf00140pf04542 -
15425 -pf10410pf13662pf08275pf01807 -
15430 - -
15435 -pf00849 -
15440 -pf03453 -
15445 -pf01967 -
15450 -pf00218 -
15455 - -
15460 -pf14863pf00753 -
15465 HMMreg00455_2_PVM(pf00455pf08220) RMM2
15470 HMM00202amintr_1_rmm(pf00202) RMM2
15475 HMMalddh_2_egra(pf00171) RMM2 EP_CTERM
15480 Tdp_cinnamon RMM2
15485 HMMnopfamrmm1_2_rmm1 RMM2 EP_NTERM
15490 HMMnopfamrmm1_2_rmm1 RMM2 EP_NTERM
15495 P_cobaltBlue RMM2
15500 H_darkIndigo RMM2
15505 Tdp_sepia RMM2
15510 -pf13416 -
15515 -pf00005pf08402 -
15520 -pf00528 -
15525 -pf00528 -
15530 HMMnopfamclust3_1 RMM2
15535 HMM01636transf_1_rmm(pf01636) RMM2
15540 HMM00857isoch_1_buf1(pf00857) RMM2
15545 -pf02608 -
15550 -pf00005 -
15555 -pf02653 -
15560 -pf02653 -
15565 -pf07969 -
(..2010..)
17575 - -
17580 HMMnopfamclust3_1 EUT2E
17585 -pf00496 -
17590 -pf03050pf13817pf13007pf13005 -
17595 -pf05717 -
17600 -pf01527 -
17605 HMMnopfamclust3_1 PVM
17610 -pf00582 -
17615 - -
17620 -pf02775 -
17625 -pf01070 -
17630 -pf02566 -
17635 HMMalddh_2_egra(pf00171) MIC2
17640 Tdp_bluish MIC2
17645 HMM02866_2_mic12(pf02866pf00056) MIC2 EP_NTERM
17650 -pf01042 -
17655 -pf13458 -
17660 -pf02653 -
17665 -pf02653 -
17670 -pf00005pf12399 -
17675 -pf00005 -
17680 -pf00155 -
17685 HMM00107alcdh_2_mic1mix(pf08240pf0010
17690 -pf00291 -
17695 HMMreg00126_1_mix(pf00126pf034
17700 -pf16868 -
17705 - -
BMC-P
BMC-H
BMC-T
Aldehyde dehydrogenase
PTAC
Signature enzyme
Alcohol dehydrogenase
Regulator

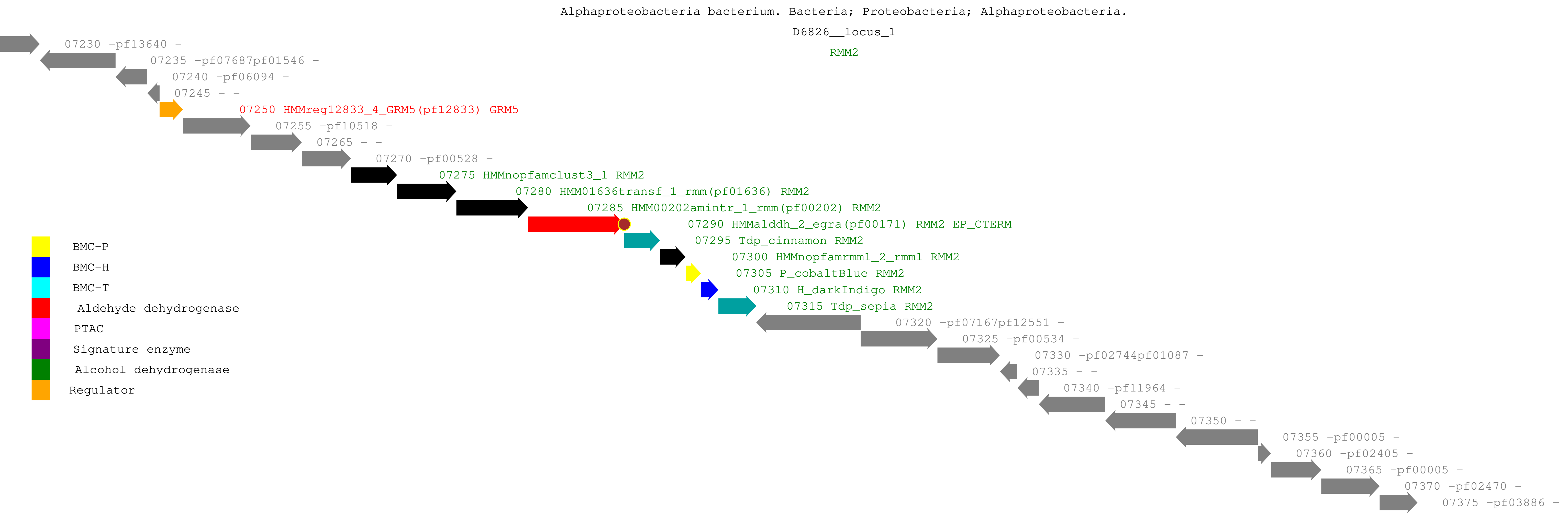
Alphaproteobacteria bacterium. Bacteria; Proteobacteria; Alphaproteobacteria.
D6826__locus_1
RMM2
07230 -pf13640 -
07235 -pf07687pf01546 -
07240 -pf06094 -
07245 - -
07250 HMMreg12833_4_GRM5(pf12833) GRM5
07255 -pf10518 -
07265 - -
07270 -pf00528 -
07275 HMMnopfamclust3_1 RMM2
07280 HMM01636transf_1_rmm(pf01636) RMM2
07285 HMM00202amintr_1_rmm(pf00202) RMM2
07290 HMMalddh_2_egra(pf00171) RMM2 EP_CTERM
07295 Tdp_cinnamon RMM2
07300 HMMnopfamrmm1_2_rmm1 RMM2
07305 P_cobaltBlue RMM2
07310 H_darkIndigo RMM2
07315 Tdp_sepia RMM2
07320 -pf07167pf12551 -
07325 -pf00534 -
07330 -pf02744pf01087 -
07335 - -
07340 -pf11964 -
07345 - -
07350 - -
07355 -pf00005 -
07360 -pf02405 -
07365 -pf00005 -
07370 -pf02470 -
07375 -pf03886 -
BMC-P
BMC-H
BMC-T
Aldehyde dehydrogenase
PTAC
Signature enzyme
Alcohol dehydrogenase
Regulator

Gammaproteobacteria bacterium. Bacteria; Proteobacteria; Gammaproteobacteria.

DCE12__locus_1

RMM2

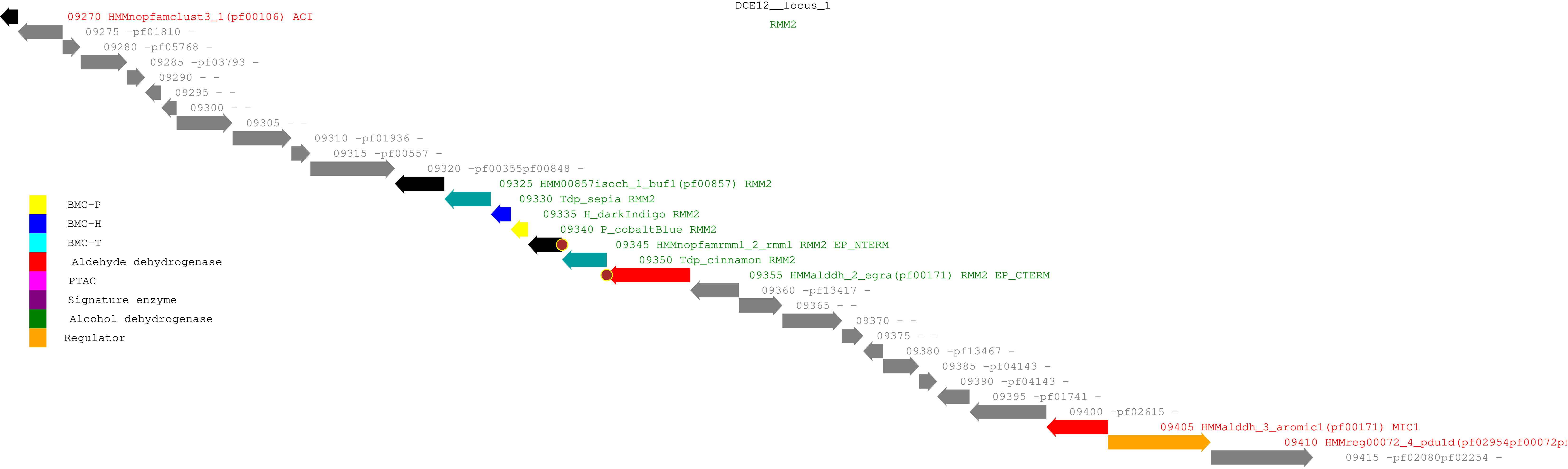

Roseovarius nubinhibens. Bacteria; Proteobacteria; Alphaproteobacteria; Rhodobacterales;

DCS45__locus_1

RMM2

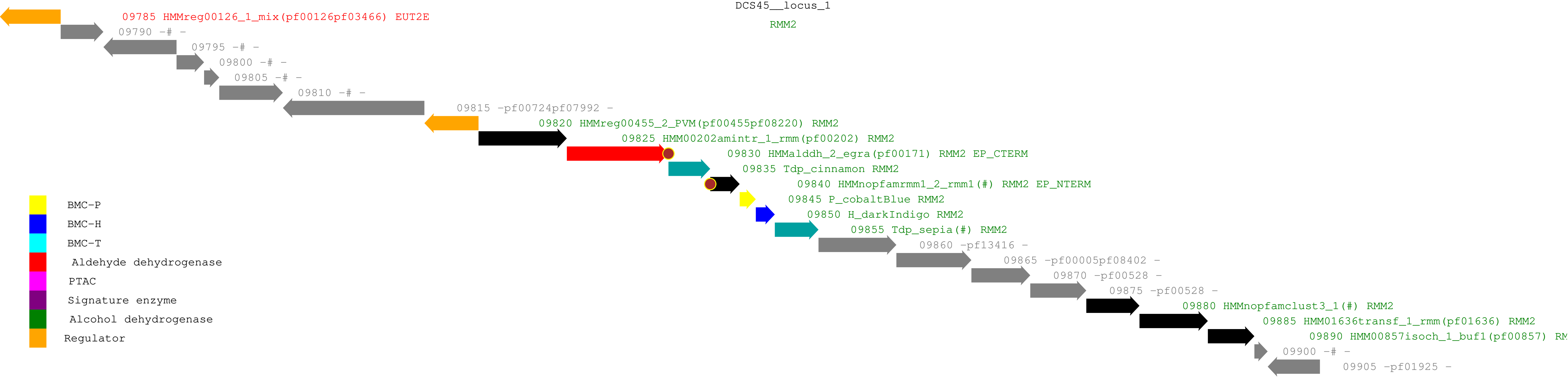

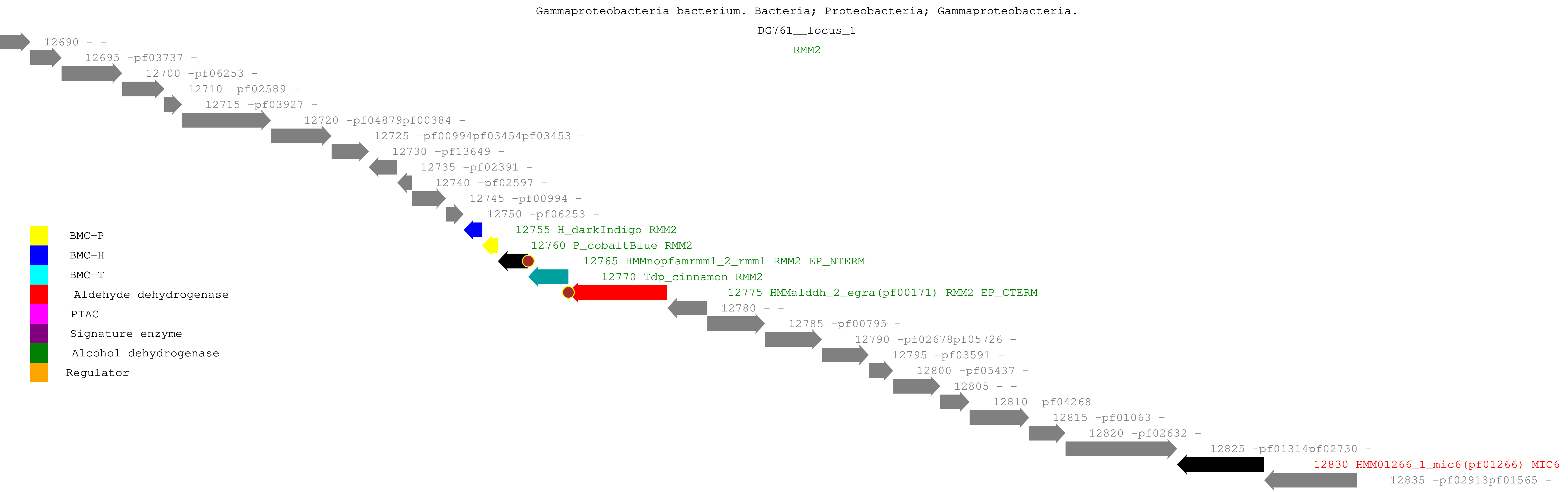

Gammaproteobacteria bacterium. Bacteria; Proteobacteria; Gammaproteobacteria.
DG761__locus_1
RMM2
12690 - -
12695 -pf03737 -
12700 -pf06253 -
12710 -pf02589 -
12715 -pf03927 -
12720 -pf04879pf00384 -
12725 -pf00994pf03454pf03453 -
12730 -pf13649 -
12735 -pf02391 -
12740 -pf02597 -
12745 -pf00994 -
12750 -pf06253 -
12755 H_darkIndigo RMM2
12760 P_cobaltBlue RMM2
12765 HMMnopfamrmm1_2_rmm1 RMM2 EP_NTERM
12770 Tdp_cinnamon RMM2
12775 HMMaldh_2_egra(pf00171) RMM2 EP_CTERM
12780 - -
12785 -pf00795 -
12790 -pf02678pf05726 -
12795 -pf03591 -
12800 -pf05437 -
12805 - -
12810 -pf04268 -
12815 -pf01063 -
12820 -pf02632 -
12825 -pf01314pf02730 -
12830 HMM01266_1_mic6(pf01266) MIC6
12835 -pf02913pf01565 -
BMC-P
BMC-H
BMC-T
Aldehyde dehydrogenase
PTAC
Signature enzyme
Alcohol dehydrogenase
Regulator

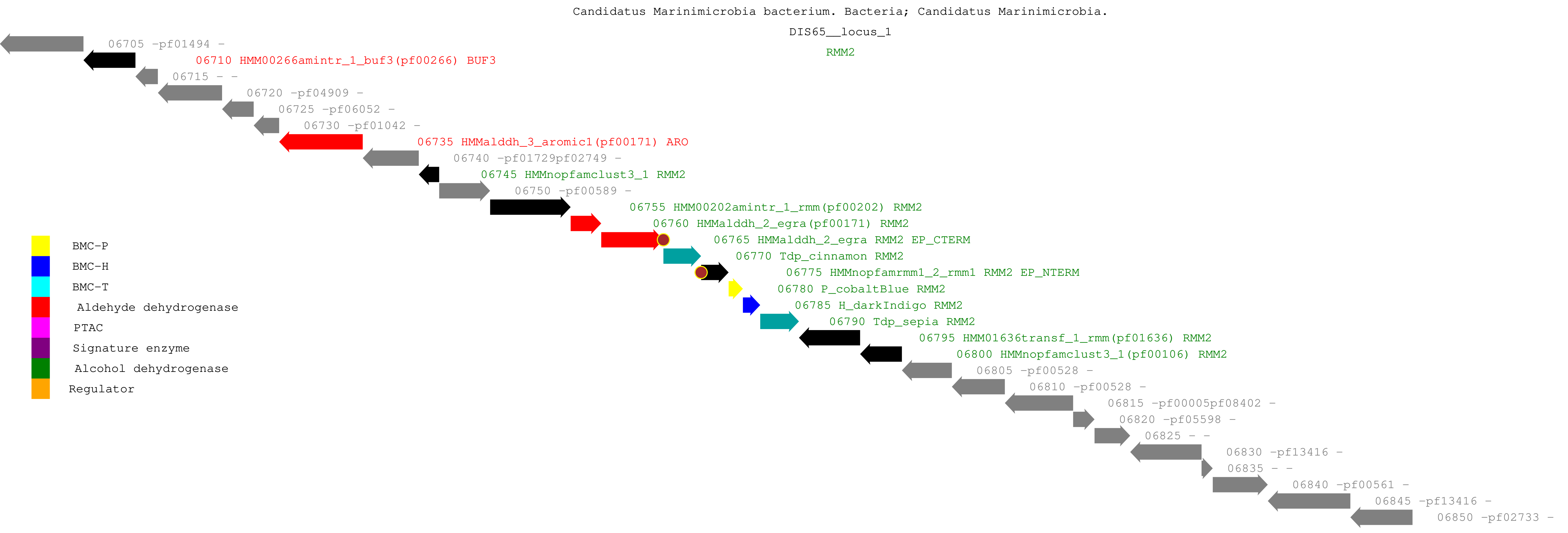
Candidatus Marinimicrobia bacterium. Bacteria; Candidatus Marinimicrobia.
DIS65__locus_1
RMM2
06705 -pf01494 -
06710 HMM00266amintr_1_buf3(pf00266) BUF3
06715 - -
06720 -pf04909 -
06725 -pf06052 -
06730 -pf01042 -
06735 HMMalddh_3_aromic1(pf00171) ARO
06740 -pf01729pf02749 -
06745 HMMnopfamclust3_1 RMM2
06750 -pf00589 -
06755 HMM00202amintr_1_rmm(pf00202) RMM2
06760 HMMalddh_2_egra(pf00171) RMM2
06765 HMMalddh_2_egra RMM2 EP_CTERM
06770 Tdp_cinnamon RMM2
06775 HMMnopfamrmm1_2_rmm1 RMM2 EP_NTERM
06780 P_cobaltBlue RMM2
06785 H_darkIndigo RMM2
06790 Tdp_sepia RMM2
06795 HMM01636transf_1_rmm(pf01636) RMM2
06800 HMMnopfamclust3_1(pf00106) RMM2
06805 -pf00528 -
06810 -pf00528 -
06815 -pf00005pf08402 -
06820 -pf05598 -
06825 - -
06830 -pf13416 -
06835 - -
06840 -pf00561 -
06845 -pf13416 -
06850 -pf02733 -
BMC-P
BMC-H
BMC-T
Aldehyde dehydrogenase
PTAC
Signature enzyme
Alcohol dehydrogenase
Regulator

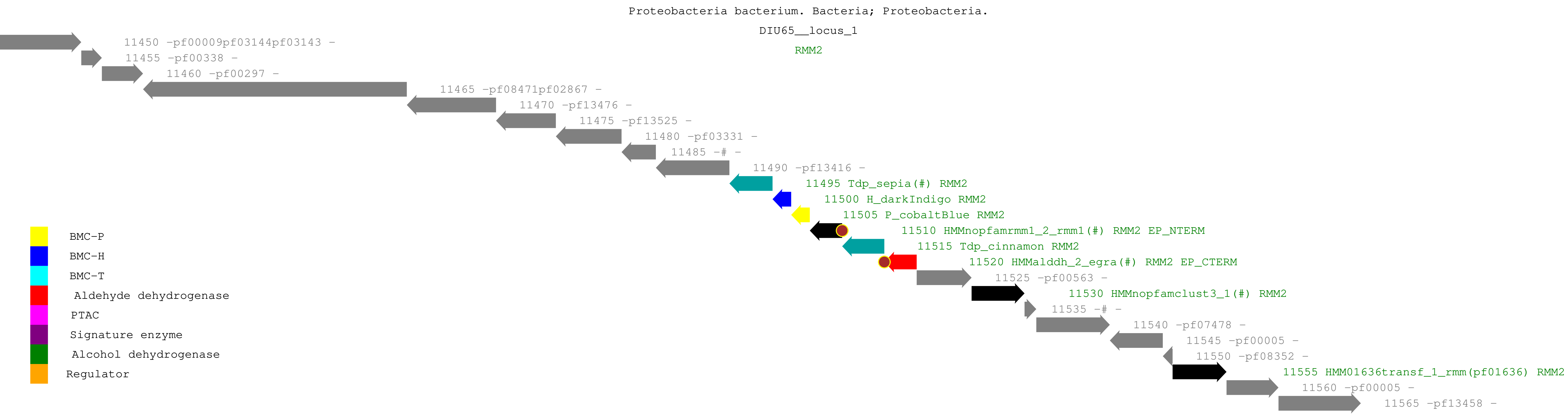
Proteobacteria bacterium. Bacteria; Proteobacteria.
DIU65__locus_1
RMM2
11450 -pf00009pf03144pf03143 -
11455 -pf00338 -
11460 -pf00297 -
11465 -pf08471pf02867 -
11470 -pf13476 -
11475 -pf13525 -
11480 -pf03331 -
11485 -# -
11490 -pf13416 -
11495 Tdp_sepia(#) RMM2
11500 H_darkIndigo RMM2
11505 P_cobaltBlue RMM2
11510 HMMnopfamrmm1_2_rmm1(#) RMM2 EP_NTERM
11515 Tdp_cinnamon RMM2
11520 HMMalddh_2_egra(#) RMM2 EP_CTERM
11525 -pf00563 -
11530 HMMnopfamclust3_1(#) RMM2
11535 -# -
11540 -pf07478 -
11545 -pf00005 -
11550 -pf08352 -
11555 HMM01636transf_1_rmm(pf01636) RMM2
11560 -pf00005 -
11565 -pf13458 -
BMC-P
BMC-H
BMC-T
Aldehyde dehydrogenase
PTAC
Signature enzyme
Alcohol dehydrogenase
Regulator

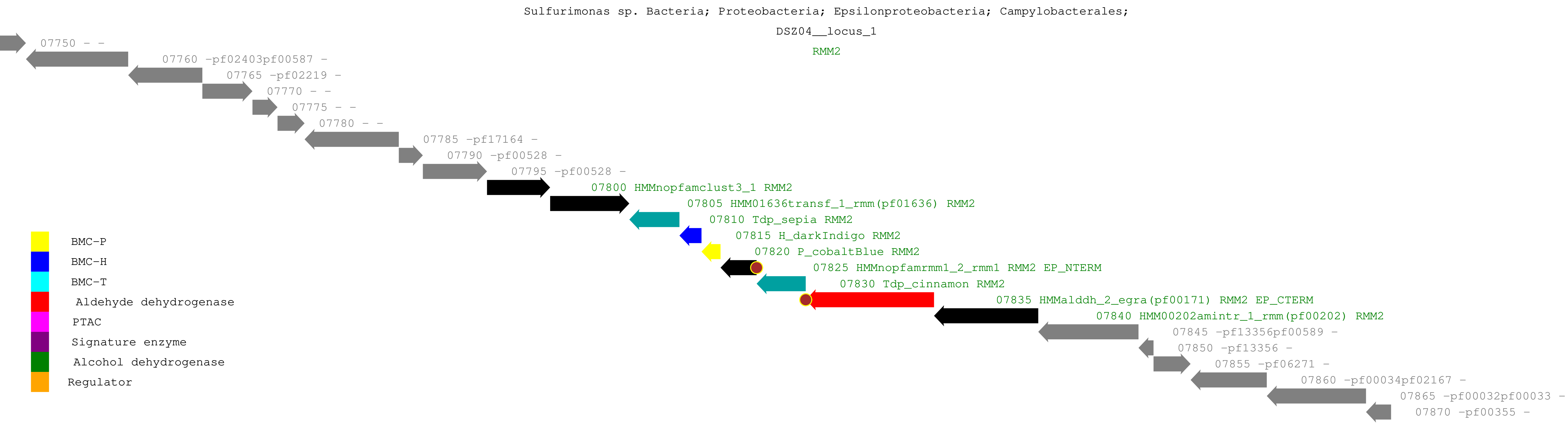
Sulfurimonas sp. Bacteria; Proteobacteria; Epsilonproteobacteria; Campylobacterales;
DSZ04__locus_1
RMM2
07750 – –
07760 -pf02403pf00587 –
07765 -pf02219 –
07770 – –
07775 – –
07780 – –
07785 -pf17164 –
07790 -pf00528 –
07795 -pf00528 –
07800 HMMnopfamclust3_1 RMM2
07805 HMM01636transf_1_rmm(pf01636) RMM2
07810 Tdp_sepia RMM2
07815 H_darkIndigo RMM2
07820 P_cobaltBlue RMM2
07825 HMMnopfamrmm1_2_rmm1 RMM2 EP_NTERM
07830 Tdp_cinnamon RMM2
07835 HMMalddh_2_egra(pf00171) RMM2 EP_CTERM
07840 HMM00202amintr_1_rmm(pf00202) RMM2
07845 -pf13356pf00589 –
07850 -pf13356 –
07855 -pf06271 –
07860 -pf00034pf02167 –
07865 -pf00032pf00033 –
07870 -pf00355 –
BMC-P
BMC-H
BMC-T
Aldehyde dehydrogenase
PTAC
Signature enzyme
Alcohol dehydrogenase
Regulator

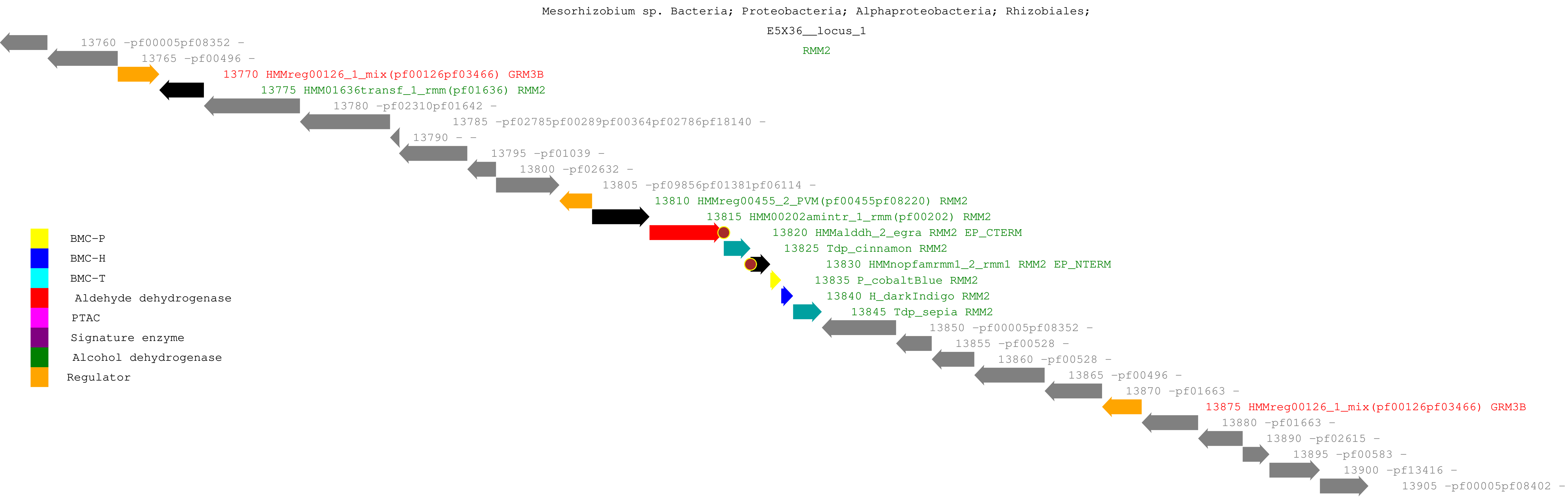
Mesorhizobium sp. Bacteria; Proteobacteria; Alphaproteobacteria; Rhizobiales;
E5X36__locus_1
RMM2
13760 -pf00005pf08352 -
13765 -pf00496 -
13770 HMMreg00126_1_mix(pf00126pf03466) GRM3B
13775 HMM01636transf_1_rmm(pf01636) RMM2
13780 -pf02310pf01642 -
13785 -pf02785pf00289pf00364pf02786pf18140 -
13790 - -
13795 -pf01039 -
13800 -pf02632 -
13805 -pf09856pf01381pf06114 -
13810 HMMreg00455_2_PVM(pf00455pf08220) RMM2
13815 HMM00202amintr_1_rmm(pf00202) RMM2
13820 HMMalddh_2_egra RMM2 EP_CTERM
13825 Tdp_cinnamon RMM2
13830 HMMnopfamrmm1_2_rmm1 RMM2 EP_NTERM
13835 P_cobaltBlue RMM2
13840 H_darkIndigo RMM2
13845 Tdp_sepia RMM2
13850 -pf00005pf08352 -
13855 -pf00528 -
13860 -pf00528 -
13865 -pf00496 -
13870 -pf01663 -
13875 HMMreg00126_1_mix(pf00126pf03466) GRM3B
13880 -pf01663 -
13890 -pf02615 -
13895 -pf00583 -
13900 -pf13416 -
13905 -pf00005pf08402 -
BMC-P
BMC-H
BMC-T
Aldehyde dehydrogenase
PTAC
Signature enzyme
Alcohol dehydrogenase
Regulator

Rhodospirillaceae bacterium. Bacteria; Proteobacteria; Alphaproteobacteria; Rhodospirillales;

E6Q98__locus_1

RMM2

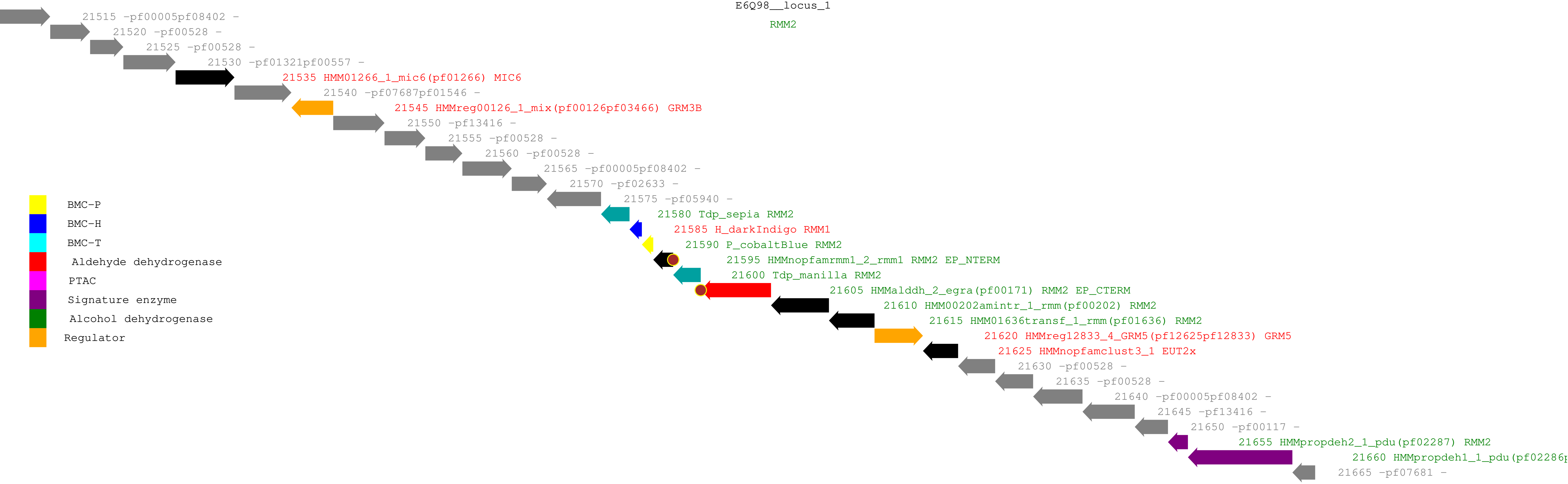

Tepidamorphus gemmatus. Bacteria; Proteobacteria; Alphaproteobacteria; Rhizobiales; Rhodobiaceae;

EDC22__locus_1

RMM2

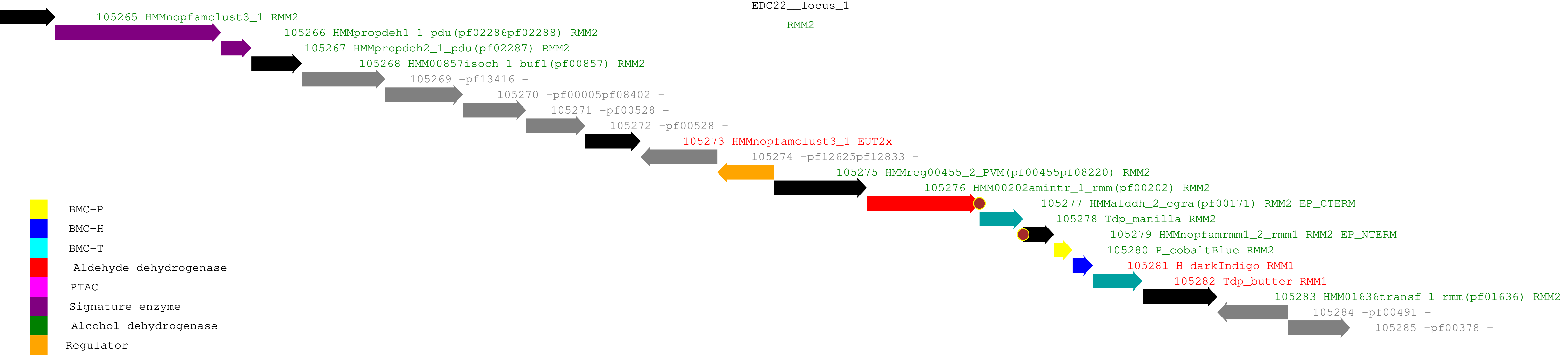

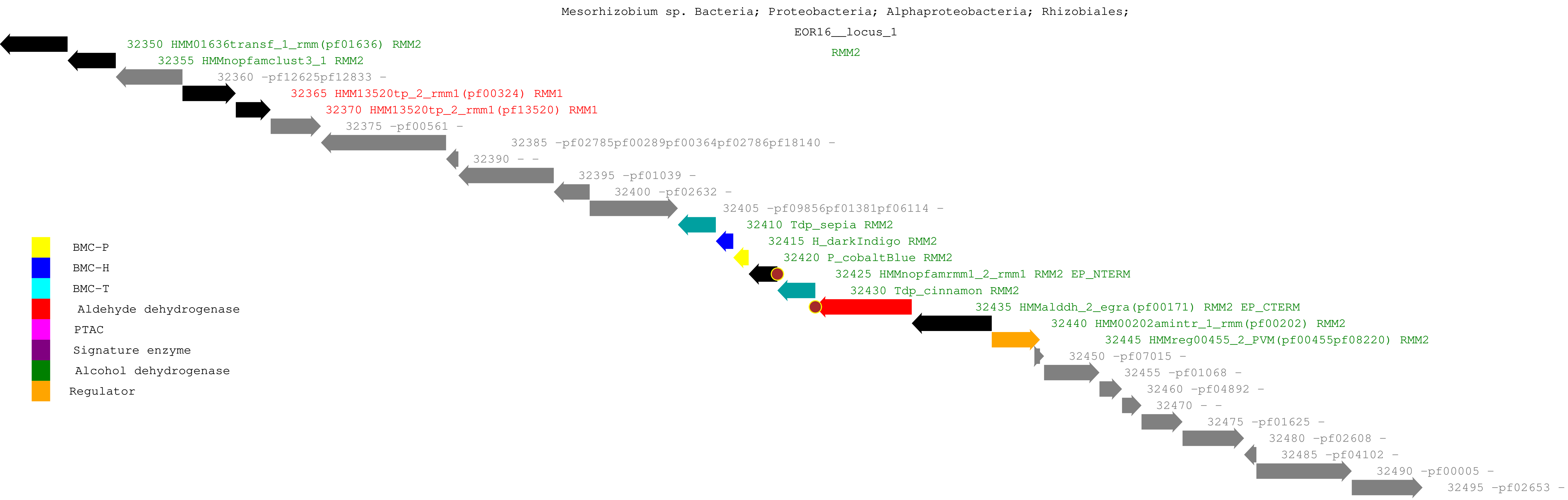
Mesorhizobium sp. Bacteria; Proteobacteria; Alphaproteobacteria; Rhizobiales;
EOR16__locus_1
RMM2
32350 HMM01636transf_1_rmm(pf01636) RMM2
32355 HMMnopfamclust3_1 RMM2
32360 -pf12625pf12833 -
32365 HMM13520tp_2_rmm1(pf00324) RMM1
32370 HMM13520tp_2_rmm1(pf13520) RMM1
32375 -pf00561 -
32385 -pf02785pf00289pf00364pf02786pf18140 -
32390 - -
32395 -pf01039 -
32400 -pf02632 -
32405 -pf09856pf01381pf06114 -
32410 Tdp_sepia RMM2
32415 H_darkIndigo RMM2
32420 P_cobaltBlue RMM2
32425 HMMnopfamrmm1_2_rmm1 RMM2 EP_NTERM
32430 Tdp_cinnamon RMM2
32435 HMMalddh_2_egra(pf00171) RMM2 EP_CTERM
32440 HMM00202amintr_1_rmm(pf00202) RMM2
32445 HMMreg00455_2_PVM(pf00455pf08220) RMM2
32450 -pf07015 -
32455 -pf01068 -
32460 -pf04892 -
32470 - -
32475 -pf01625 -
32480 -pf02608 -
32485 -pf04102 -
32490 -pf00005 -
32495 -pf02653 -
BMC-P
BMC-H
BMC-T
Aldehyde dehydrogenase
PTAC
Signature enzyme
Alcohol dehydrogenase
Regulator

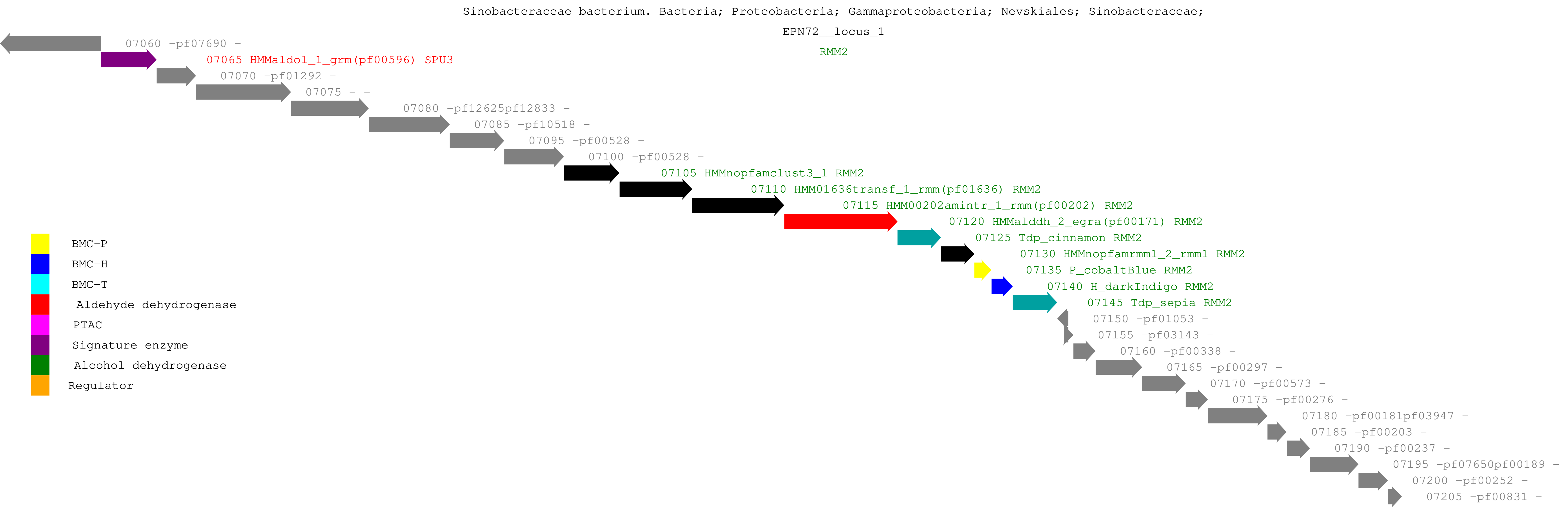
Sinobacteraceae bacterium. Bacteria; Proteobacteria; Gammaproteobacteria; Nevskiales; Sinobacteraceae;
EPN72__locus_1
RMM2
07060 -pf07690 -
07065 HMMaldol_1_grm(pf00596) SPU3
07070 -pf01292 -
07075 - -
07080 -pf12625pf12833 -
07085 -pf10518 -
07095 -pf00528 -
07100 -pf00528 -
07105 HMMnopfamclust3_1 RMM2
07110 HMM01636transf_1_rmm(pf01636) RMM2
07115 HMM00202amintr_1_rmm(pf00202) RMM2
07120 HMMalddh_2_egra(pf00171) RMM2
07125 Tdp_cinnamon RMM2
07130 HMMnopfamrmm1_2_rmm1 RMM2
07135 P_cobaltBlue RMM2
07140 H_darkIndigo RMM2
07145 Tdp_sepia RMM2
07150 -pf01053 -
07155 -pf03143 -
07160 -pf00338 -
07165 -pf00297 -
07170 -pf00573 -
07175 -pf00276 -
07180 -pf00181pf03947 -
07185 -pf00203 -
07190 -pf00237 -
07195 -pf07650pf00189 -
07200 -pf00252 -
07205 -pf00831 -
BMC-P
BMC-H
BMC-T
Aldehyde dehydrogenase
PTAC
Signature enzyme
Alcohol dehydrogenase
Regulator

Verminephrobacter sp. Larva24. Bacteria; Proteobacteria; Betaproteobacteria; Burkholderiales;

ET532__locus_1

RMM2

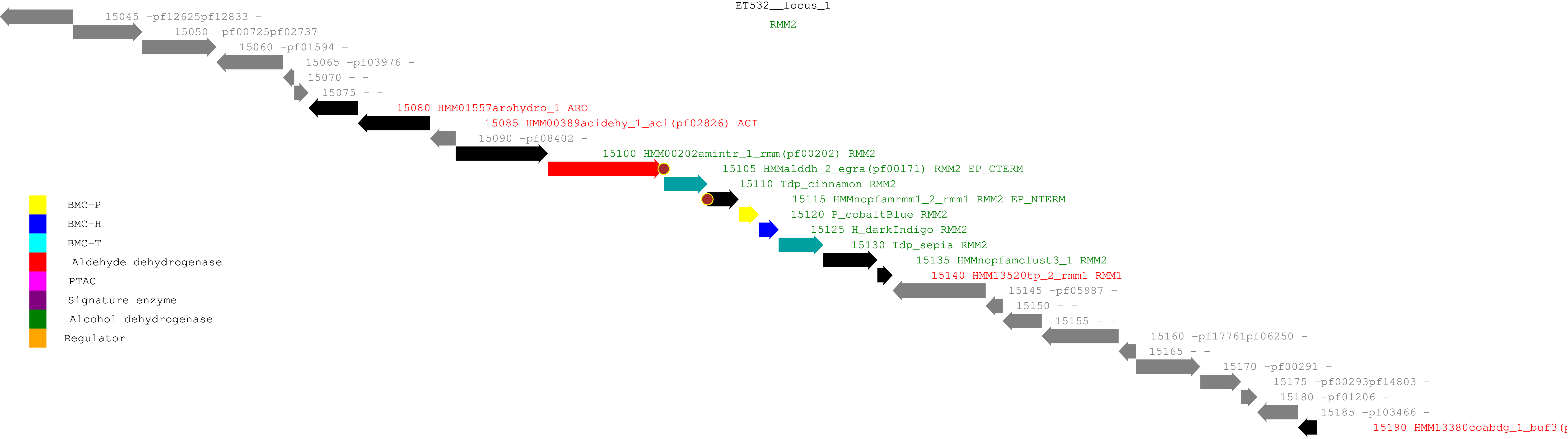

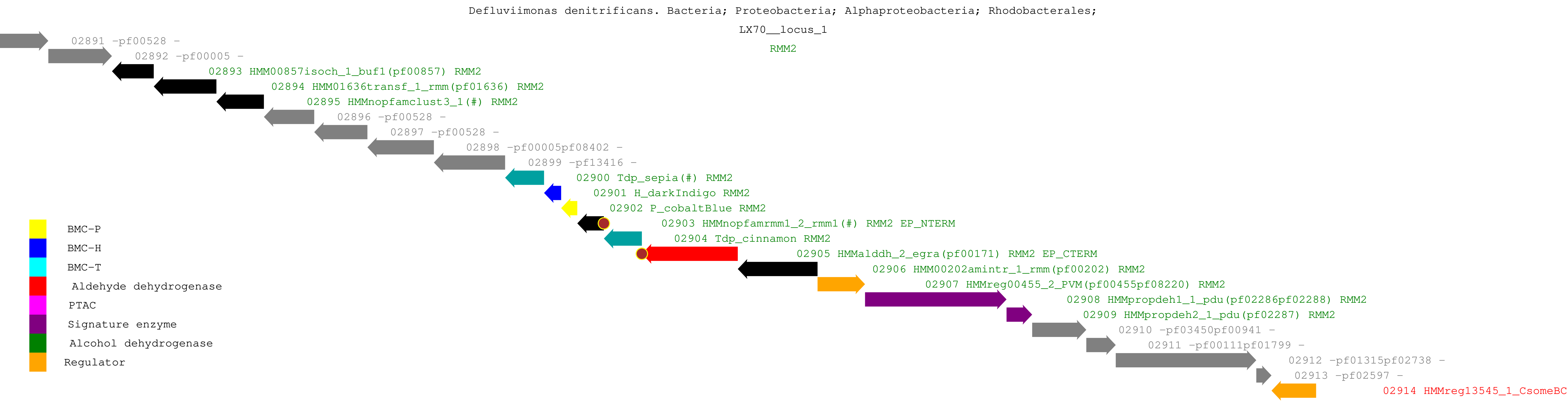
Defluviimonas denitrificans. Bacteria; Proteobacteria; Alphaproteobacteria; Rhodobacterales;
LX70__locus_1
RMM2
02891 -pf00528 -
02892 -pf00005 -
02893 HMM00857isoch_1_buf1(pf00857) RMM2
02894 HMM01636transf_1_rmm(pf01636) RMM2
02895 HMMnopfamclust3_1(#) RMM2
02896 -pf00528 -
02897 -pf00528 -
02898 -pf00005pf08402 -
02899 -pf13416 -
02900 Tdp_sepia(#) RMM2
02901 H_darkIndigo RMM2
02902 P_cobaltBlue RMM2
02903 HMMnopfamrmm1_2_rmm1(#) RMM2 EP_NTERM
02904 Tdp_cinnamon RMM2
02905 HMMaldh_2_egra(pf00171) RMM2 EP_CTERM
02906 HMM00202amintr_1_rmm(pf00202) RMM2
02907 HMMreg00455_2_PVM(pf00455pf08220) RMM2
02908 HMMpropdeh1_1_pdu(pf02286pf02288) RMM2
02909 HMMpropdeh2_1_pdu(pf02287) RMM2
02910 -pf03450pf00941 -
02911 -pf00111pf01799 -
02912 -pf01315pf02738 -
02913 -pf02597 -
02914 HMMreg13545_1_CsomeBC
BMC-P
BMC-H
BMC-T
Aldehyde dehydrogenase
PTAC
Signature enzyme
Alcohol dehydrogenase
Regulator

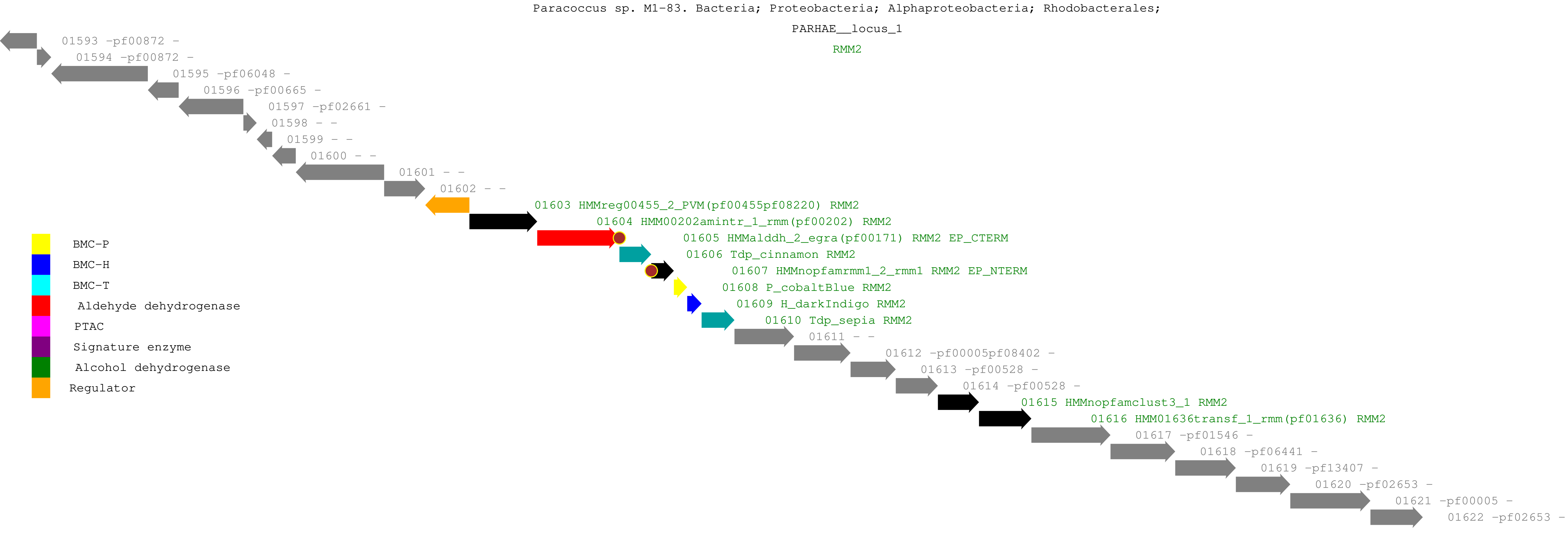

Paracoccus sp. M1-83. Bacteria; Proteobacteria; Alphaproteobacteria; Rhodobacterales;
PARHAE__locus_1
RMM2
01593 -pf00872 -
01594 -pf00872 -
01595 -pf06048 -
01596 -pf00665 -
01597 -pf02661 -
01598 - -
01599 - -
01600 - -
01601 - -
01602 - -
01603 HMMreg00455_2_PVM(pf00455pf08220) RMM2
01604 HMM00202amintr_1_rmm(pf00202) RMM2
01605 HMMalddh_2_egra(pf00171) RMM2 EP_CTERM
01606 Tdp_cinnamon RMM2
01607 HMMnopfamrmm1_2_rmm1 RMM2 EP_NTERM
01608 P_cobaltBlue RMM2
01609 H_darkIndigo RMM2
01610 Tdp_sepia RMM2
01611 - -
01612 -pf00005pf08402 -
01613 -pf00528 -
01614 -pf00528 -
01615 HMMnopfamclust3_1 RMM2
01616 HMM01636transf_1_rmm(pf01636) RMM2
01617 -pf01546 -
01618 -pf06441 -
01619 -pf13407 -
01620 -pf02653 -
01621 -pf00005 -
01622 -pf02653 -
BMC-P
BMC-H
BMC-T
Aldehyde dehydrogenase
PTAC
Signature enzyme
Alcohol dehydrogenase
Regulator

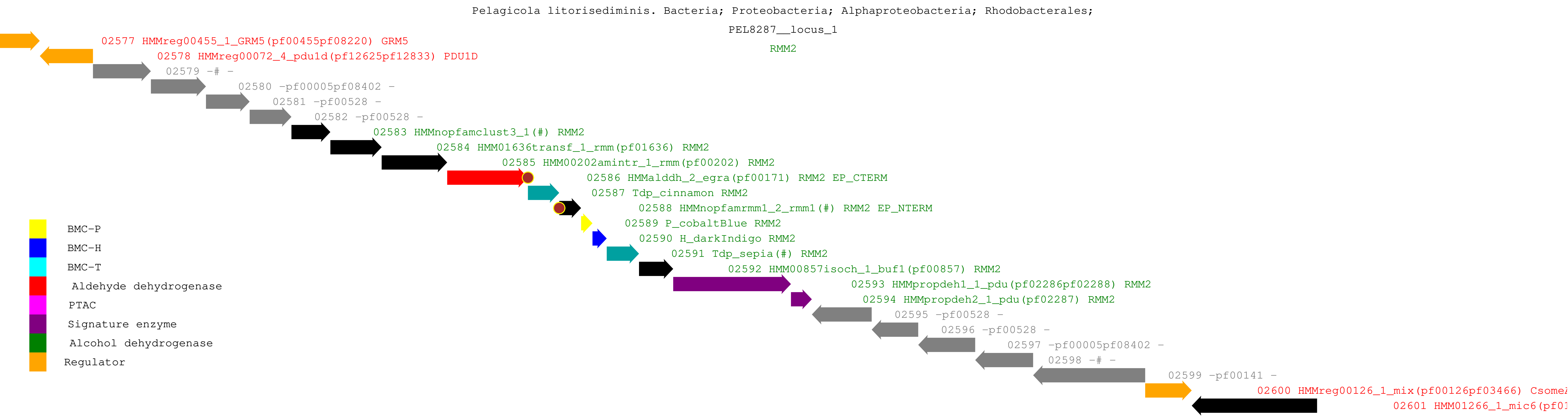

Pelagicola litorisediminis. Bacteria; Proteobacteria; Alphaproteobacteria; Rhodobacterales;
PEL8287__locus_1
RMM2
02577 HMMreg00455_1_GRM5(pf00455pf08220) GRM5
02578 HMMreg00072_4_pdu1d(pf12625pf12833) PDU1D
02579 -# -
02580 -pf00005pf08402 -
02581 -pf00528 -
02582 -pf00528 -
02583 HMMnopfamclust3_1(#) RMM2
02584 HMM01636transf_1_rmm(pf01636) RMM2
02585 HMM00202amintr_1_rmm(pf00202) RMM2
02586 HMMalddh_2_egra(pf00171) RMM2 EP_CTERM
02587 Tdp_cinnamon RMM2
02588 HMMnopfamrmm1_2_rmm1(#) RMM2 EP_NTERM
02589 P_cobaltBlue RMM2
02590 H_darkIndigo RMM2
02591 Tdp_sepia(#) RMM2
02592 HMM00857isoch_1_buf1(pf00857) RMM2
02593 HMMpropdeh1_1_pdu(pf02286pf02288) RMM2
02594 HMMpropdeh2_1_pdu(pf02287) RMM2
02595 -pf00528 -
02596 -pf00528 -
02597 -pf00005pf08402 -
02598 -# -
02599 -pf00141 -
02600 HMMreg00126_1_mix(pf00126pf03466) Csome
02601 HMM01266_1_mic6(pf01
BMC-P
BMC-H
BMC-T
Aldehyde dehydrogenase
PTAC
Signature enzyme
Alcohol dehydrogenase
Regulator

Paracoccus yeei. Bacteria; Proteobacteria; Alphaproteobacteria; Rhodobacterales;

PYTT13__locus_1

RMM2

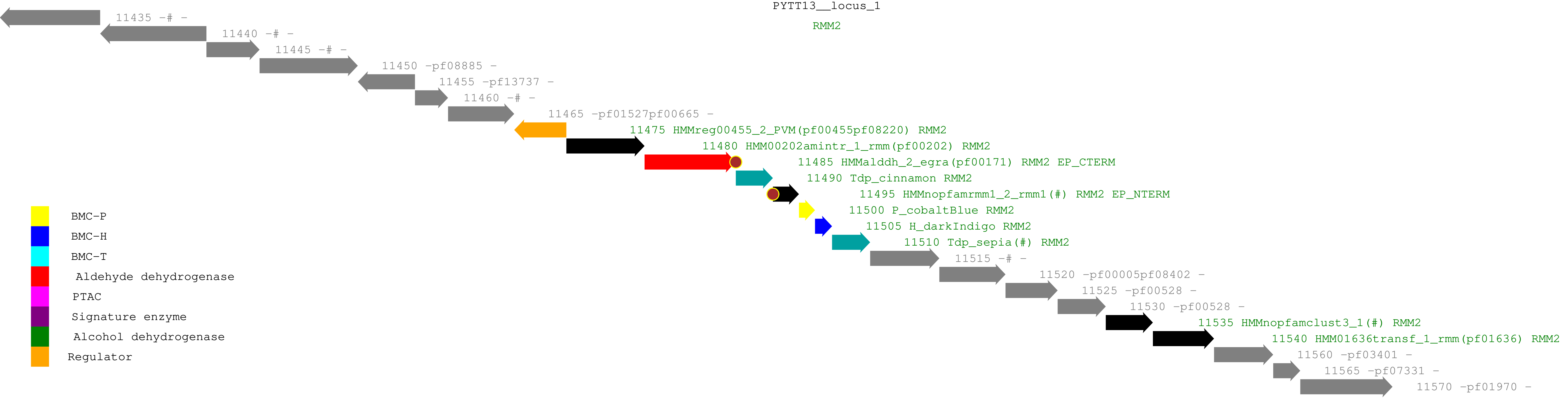

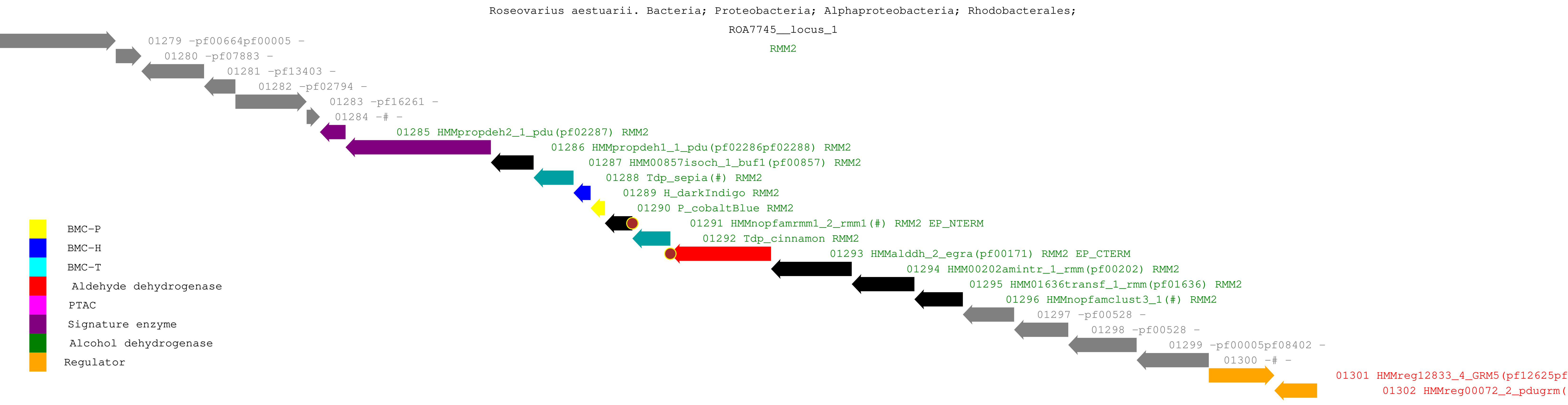

Roseovarius aestuarii. Bacteria; Proteobacteria; Alphaproteobacteria; Rhodobacterales;
ROA7745__locus_1
RMM2
01279 -pf00664pf00005 -
01280 -pf07883 -
01281 -pf13403 -
01282 -pf02794 -
01283 -pf16261 -
01284 -# -
01285 HMMpropdeh2_1_pdu(pf02287) RMM2
01286 HMMpropdeh1_1_pdu(pf02286pf02288) RMM2
01287 HMM00857isoch_1_buf1(pf00857) RMM2
01288 Tdp_sepia(#) RMM2
01289 H_darkIndigo RMM2
01290 P_cobaltBlue RMM2
01291 HMMnopfamrmm1_2_rmm1(#) RMM2 EP_NTERM
01292 Tdp_cinnamon RMM2
01293 HMMalddh_2_egra(pf00171) RMM2 EP_CTERM
01294 HMM00202amintr_1_rmm(pf00202) RMM2
01295 HMM01636transf_1_rmm(pf01636) RMM2
01296 HMMnopfamclust3_1(#) RMM2
01297 -pf00528 -
01298 -pf00528 -
01299 -pf00005pf08402 -
01300 -# -
01301 HMMreg12833_4_GRM5(pf12625pf
01302 HMMreg00072_2_pdugrm(
BMC-P
BMC-H
BMC-T
Aldehyde dehydrogenase
PTAC
Signature enzyme
Alcohol dehydrogenase
Regulator

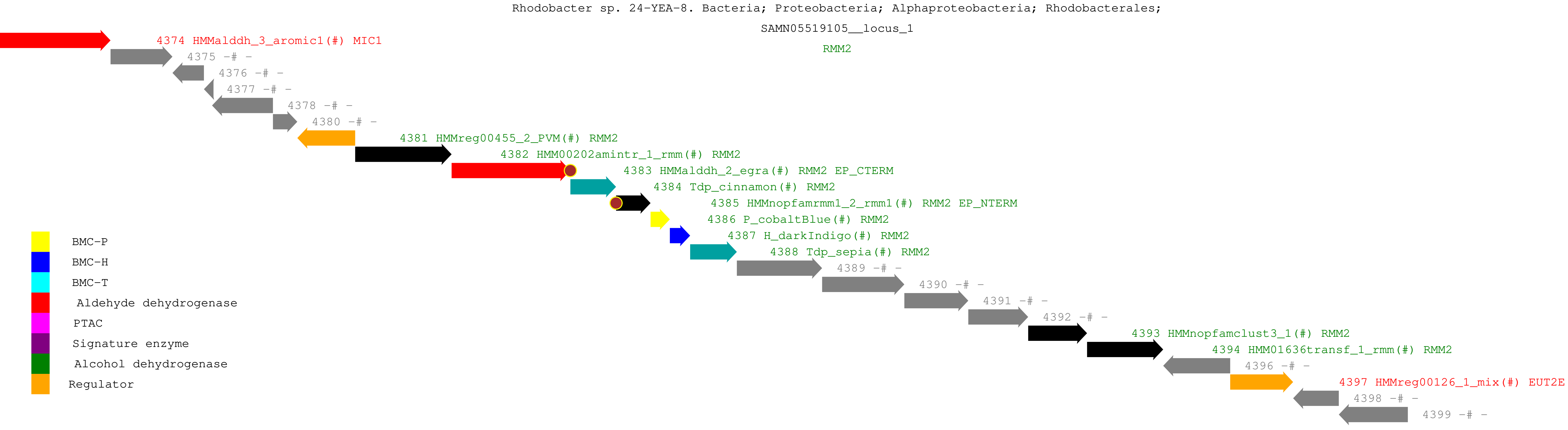

Rhodobacter sp. 24-YEA-8. Bacteria; Proteobacteria; Alphaproteobacteria; Rhodobacterales;
SAMN05519105__locus_1
RMM2
4374 HMMalddh_3_aromic1(#) MIC1
4375 -# -
4376 -# -
4377 -# -
4378 -# -
4380 -# -
4381 HMMreg00455_2_PVM(#) RMM2
4382 HMM00202amintr_1_rmm(#) RMM2
4383 HMMalddh_2_egra(#) RMM2 EP_CTERM
4384 Tdp_cinnamon(#) RMM2
4385 HMMnopfamrmm1_2_rmm1(#) RMM2 EP_NTERM
4386 P_cobaltBlue(#) RMM2
4387 H_darkIndigo(#) RMM2
4388 Tdp_sepia(#) RMM2
4389 -# -
4390 -# -
4391 -# -
4392 -# -
4393 HMMnopfamclust3_1(#) RMM2
4394 HMM01636transf_1_rmm(#) RMM2
4396 -# -
4397 HMMreg00126_1_mix(#) EUT2E
4398 -# -
4399 -# -
BMC-P
BMC-H
BMC-T
Aldehyde dehydrogenase
PTAC
Signature enzyme
Alcohol dehydrogenase
Regulator

Verminephrobacter eiseniae (strain EF01-2). Bacteria; Proteobacteria; Betaproteobacteria; Burkholderiales;

Veis__locus_1

RMM2

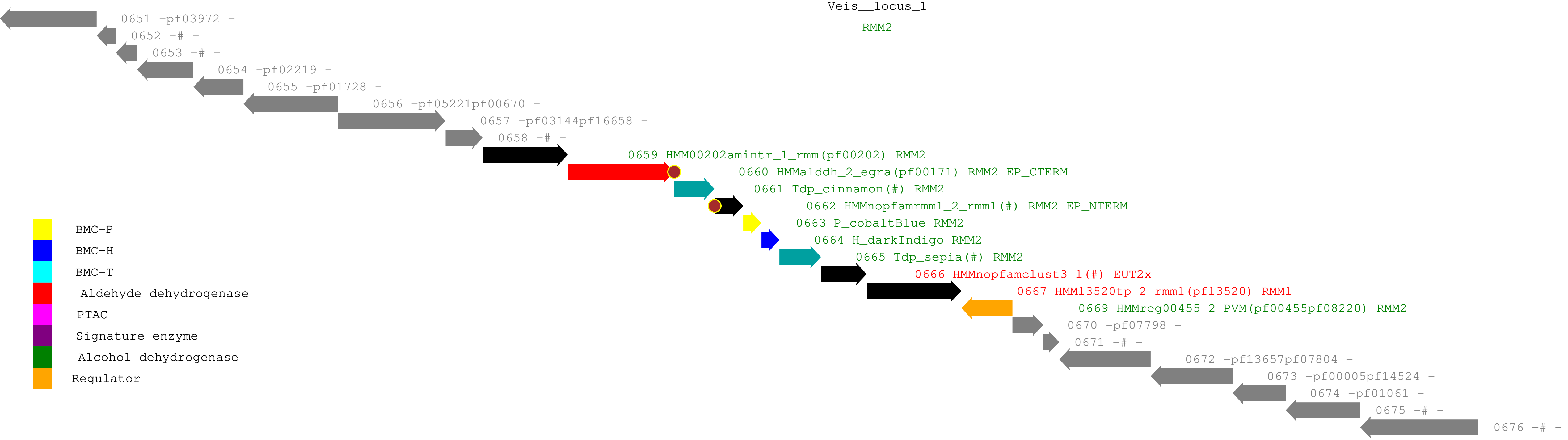

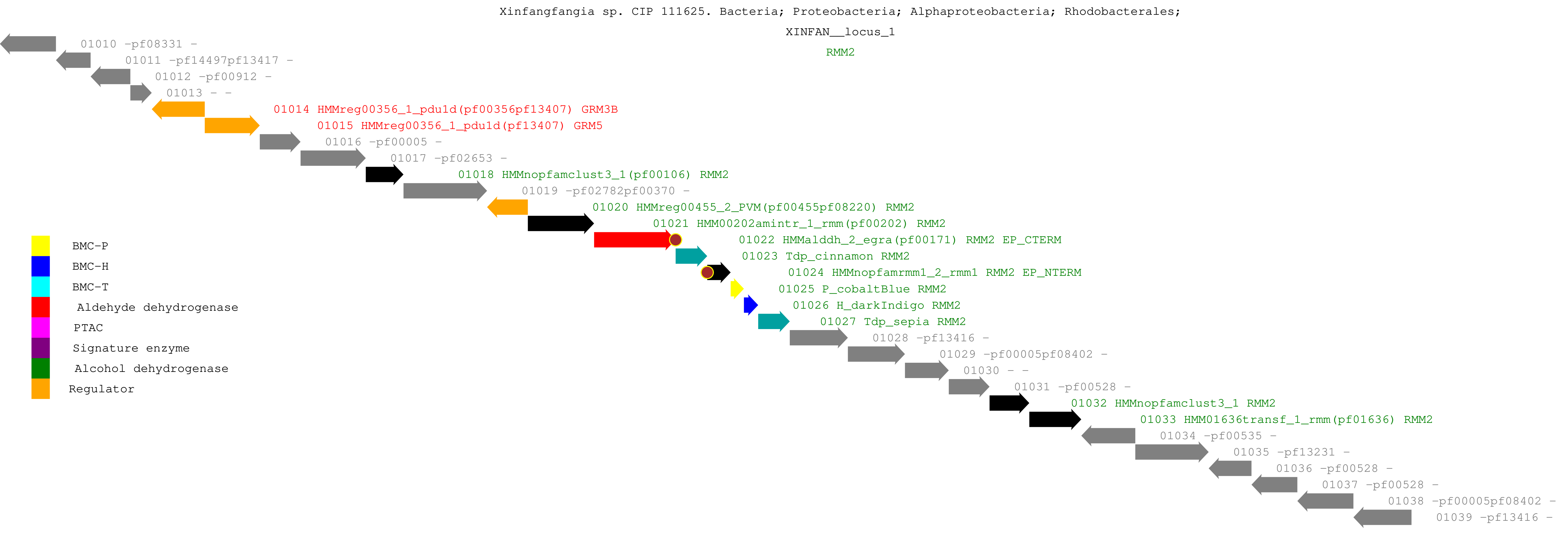
Xinfangfangia sp. CIP 111625. Bacteria; Proteobacteria; Alphaproteobacteria; Rhodobacterales;
XINFAN__locus_1
RMM2
01010 -pf08331 -
01011 -pf14497pf13417 -
01012 -pf00912 -
01013 - -
01014 HMMreg00356_1_pdu1d(pf00356pf13407) GRM3B
01015 HMMreg00356_1_pdu1d(pf13407) GRM5
01016 -pf00005 -
01017 -pf02653 -
01018 HMMnopfamclust3_1(pf00106) RMM2
01019 -pf02782pf00370 -
01020 HMMreg00455_2_PVM(pf00455pf08220) RMM2
01021 HMM00202amintr_1_rmm(pf00202) RMM2
01022 HMMalddh_2_egra(pf00171) RMM2 EP_CTERM
01023 Tdp_cinnamon RMM2
01024 HMMnopfamrmm1_2_rmm1 RMM2 EP_NTERM
01025 P_cobaltBlue RMM2
01026 H_darkIndigo RMM2
01027 Tdp_sepia RMM2
01028 -pf13416 -
01029 -pf00005pf08402 -
01030 - -
01031 -pf00528 -
01032 HMMnopfamclust3_1 RMM2
01033 HMM01636transf_1_rmm(pf01636) RMM2
01034 -pf00535 -
01035 -pf13231 -
01036 -pf00528 -
01037 -pf00528 -
01038 -pf00005pf08402 -
01039 -pf13416 -
BMC-P
BMC-H
BMC-T
Aldehyde dehydrogenase
PTAC
Signature enzyme
Alcohol dehydrogenase
Regulator

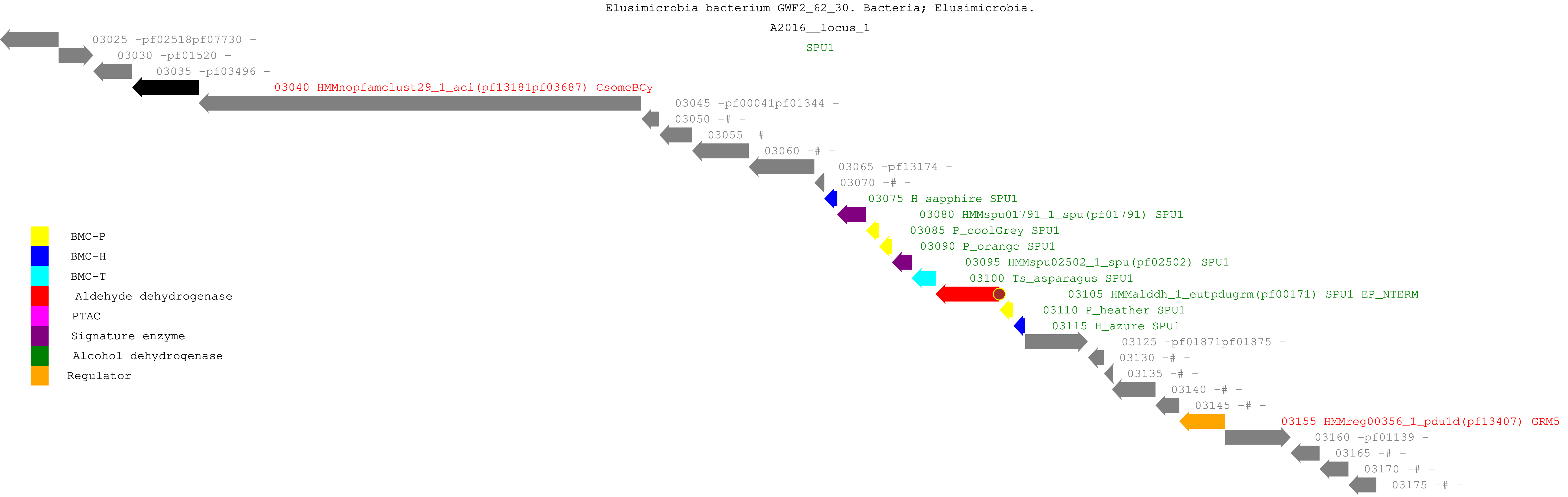

Elusimicrobia bacterium GWF2_62_30. Bacteria; Elusimicrobia.
A2016__locus_1
SPU1
03025 -pf02518pf07730 -
03030 -pf01520 -
03035 -pf03496 -
03040 HMMnopfamclust29_1_aci(pf13181pf03687) CsomeBCy
03045 -pf00041pf01344 -
03050 -# -
03055 -# -
03060 -# -
03065 -pf13174 -
03070 -# -
03075 H_sapphire SPU1
03080 HMMspu01791_1_spu(pf01791) SPU1
03085 P_coolGrey SPU1
03090 P_orange SPU1
03095 HMMspu02502_1_spu(pf02502) SPU1
03100 Ts_asparagus SPU1
03105 HMMalddh_1_eutpdugrm(pf00171) SPU1 EP_NTERM
03110 P_heather SPU1
03115 H_azure SPU1
03125 -pf01871pf01875 -
03130 -# -
03135 -# -
03140 -# -
03145 -# -
03155 HMMreg00356_1_pdu1d(pf13407) GRM5
03160 -pf01139 -
03165 -# -
03170 -# -
03175 -# -
BMC-P
BMC-H
BMC-T
Aldehyde dehydrogenase
PTAC
Signature enzyme
Alcohol dehydrogenase
Regulator

Elusimicrobia bacterium GWA2_62_23. Bacteria; Elusimicrobia.

A2049__locus_1

SPU1

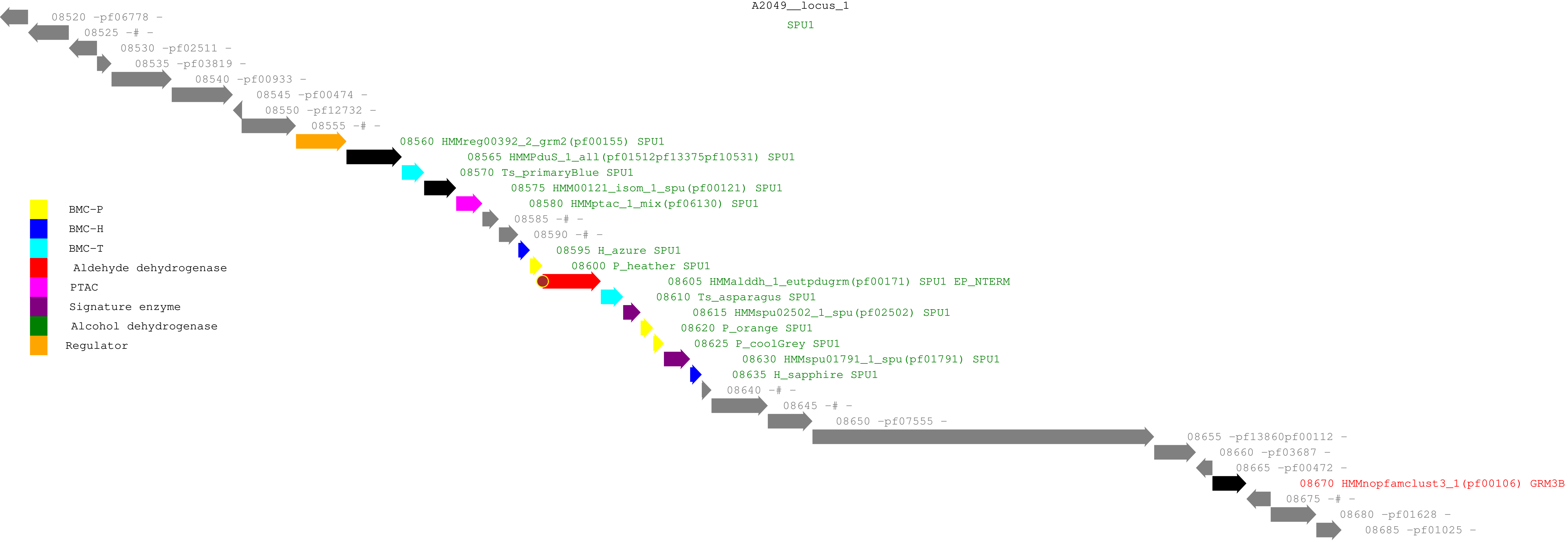

Elusimicrobia bacterium GWC2_61_19. Bacteria; Elusimicrobia.

A2081__locus_1

SPU1

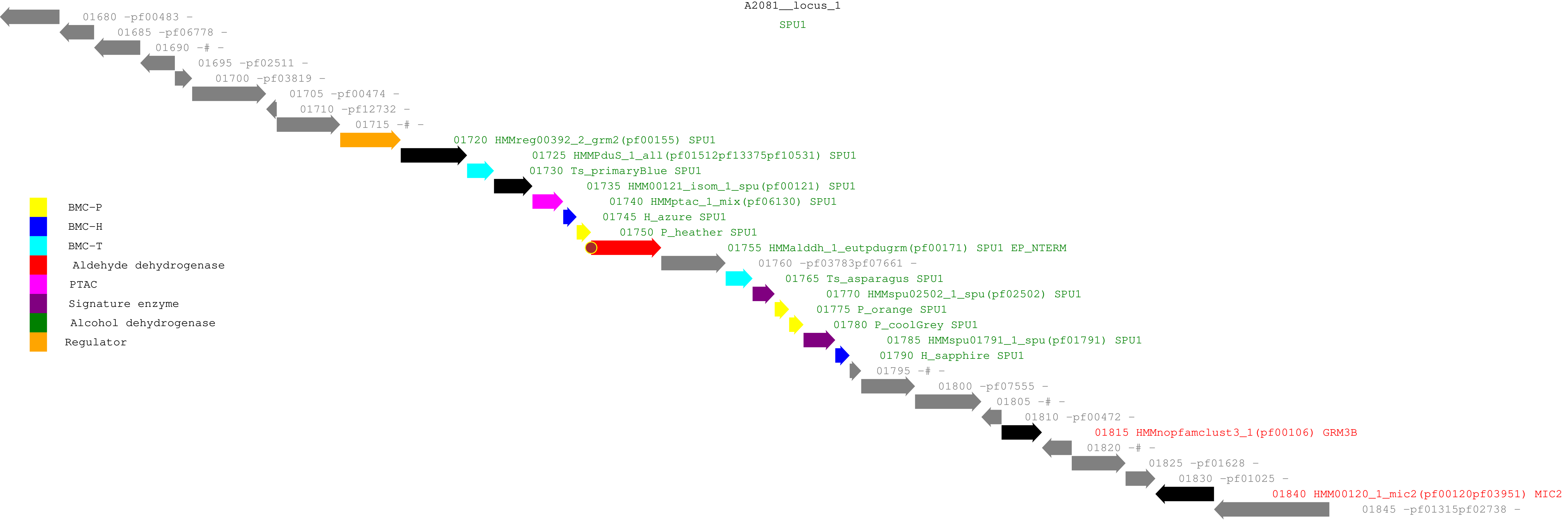

Elusimicrobia bacterium GWD2_63_28. Bacteria; Elusimicrobia.

A2089__locus_1_main_sat

SPU1

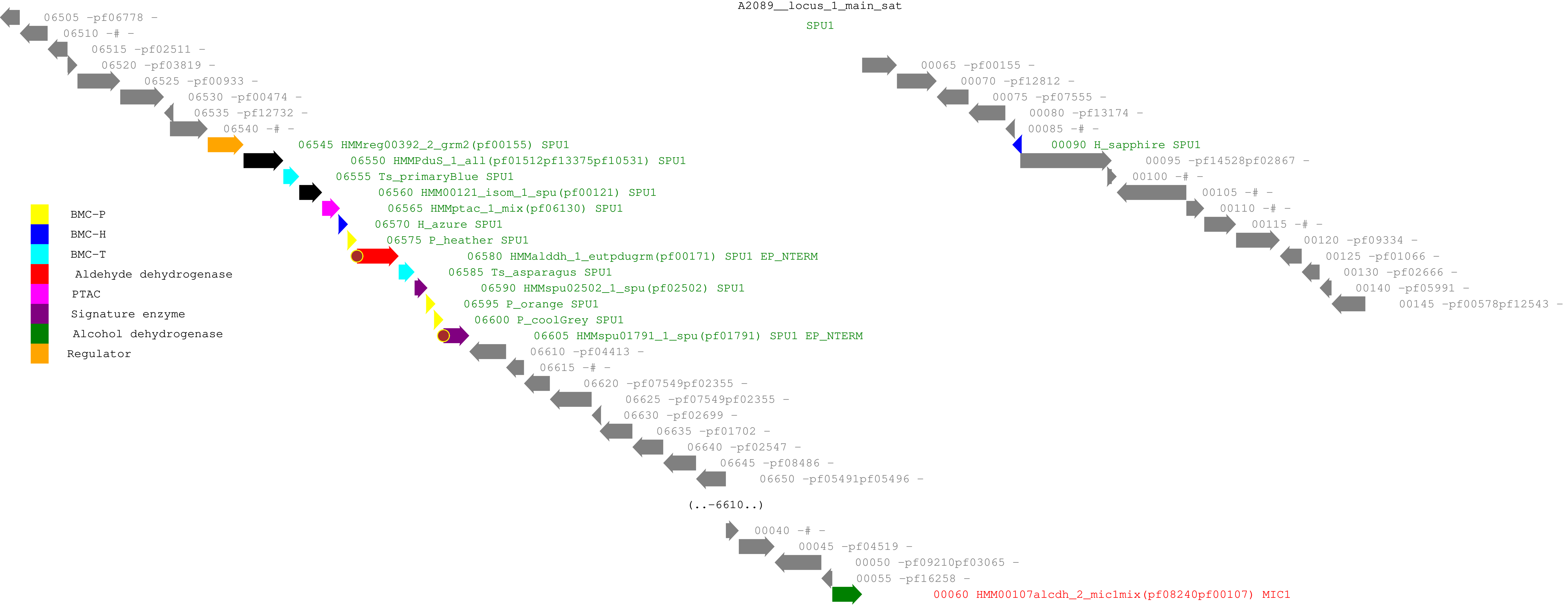

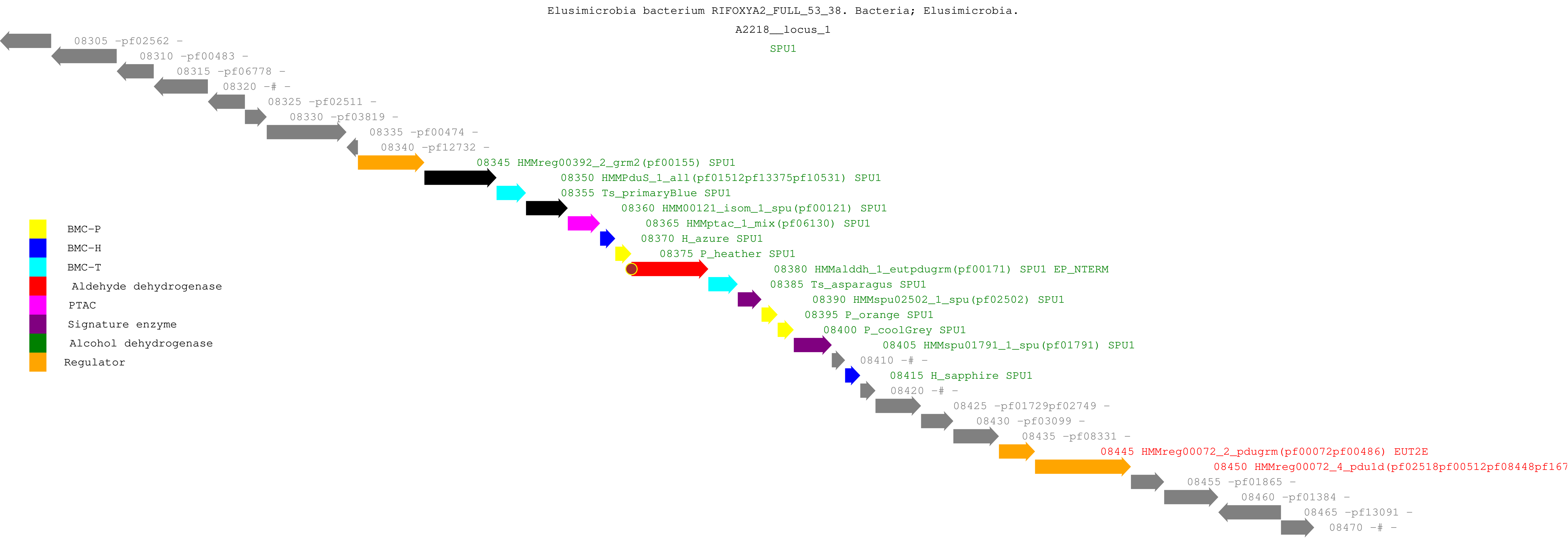
Elusimicrobia bacterium RIFOXYA2_FULL_53_38. Bacteria; Elusimicrobia.
A2218__locus_1
SPU1
08305 -pf02562 -
08310 -pf00483 -
08315 -pf06778 -
08320 -# -
08325 -pf02511 -
08330 -pf03819 -
08335 -pf00474 -
08340 -pf12732 -
08345 HMMreg00392_2_grm2(pf00155) SPU1
08350 HMMPduS_1_all(pf01512pf13375pf10531) SPU1
08355 Ts_primaryBlue SPU1
08360 HMM00121_isom_1_spu(pf00121) SPU1
08365 HMMptac_1_mix(pf06130) SPU1
08370 H_azure SPU1
08375 P_heather SPU1
08380 HMMaldd h_1_eutpdugrm(pf00171) SPU1 EP_NTERM
08385 Ts_asparagus SPU1
08390 HMMspu02502_1_spu(pf02502) SPU1
08395 P_orange SPU1
08400 P_coolGrey SPU1
08405 HMMspu01791_1_spu(pf01791) SPU1
08410 -# -
08415 H_sapphire SPU1
08420 -# -
08425 -pf01729pf02749 -
08430 -pf03099 -
08435 -pf08331 -
08445 HMMreg00072_2_pdugrm(pf00072pf00486) EUT2E
08450 HMMreg00072_4_pdu1d(pf02518pf00512pf08448pf167
08455 -pf01865 -
08460 -pf01384 -
08465 -pf13091 -
08470 -# -
BMC-P
BMC-H
BMC-T
Aldehyde dehydrogenase
PTAC
Signature enzyme
Alcohol dehydrogenase
Regulator

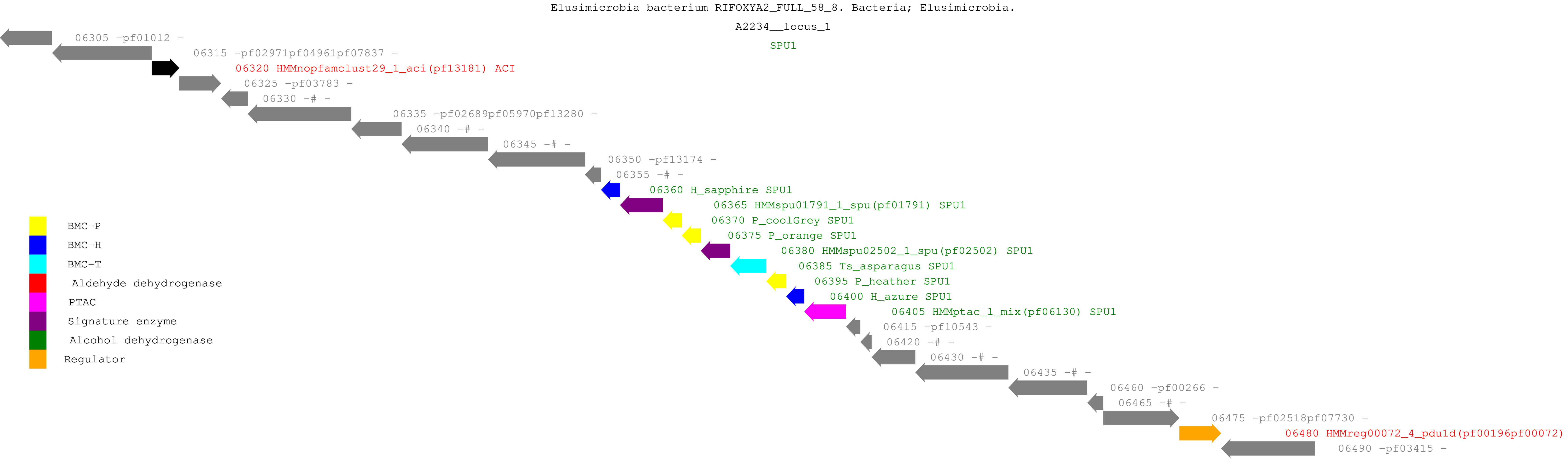
Elusimicrobia bacterium RIFOXYA2_FULL_58_8. Bacteria; Elusimicrobia.
A2234__locus_1
SPU1
06305 -pf01012 -
06315 -pf02971pf04961pf07837 -
06320 HMMnopfamclust29_1_aci(pf13181) ACI
06325 -pf03783 -
06330 -# -
06335 -pf02689pf05970pf13280 -
06340 -# -
06345 -# -
06350 -pf13174 -
06355 -# -
06360 H_sapphire SPU1
06365 HMMspu01791_1_spu(pf01791) SPU1
06370 P_coolGrey SPU1
06375 P_orange SPU1
06380 HMMspu02502_1_spu(pf02502) SPU1
06385 Ts_asparagus SPU1
06395 P_heather SPU1
06400 H_azure SPU1
06405 HMMptac_1_mix(pf06130) SPU1
06415 -pf10543 -
06420 -# -
06430 -# -
06435 -# -
06460 -pf00266 -
06465 -# -
06475 -pf02518pf07730 -
06480 HMMreg00072_4_pdu1d(pf00196pf00072)
06490 -pf03415 -
BMC-P
BMC-H
BMC-T
Aldehyde dehydrogenase
PTAC
Signature enzyme
Alcohol dehydrogenase
Regulator

Elusimicrobia bacterium RIFOXYA12_FULL_57_11. Bacteria; Elusimicrobia.

A2285__locus_1_main_sat

SPU1

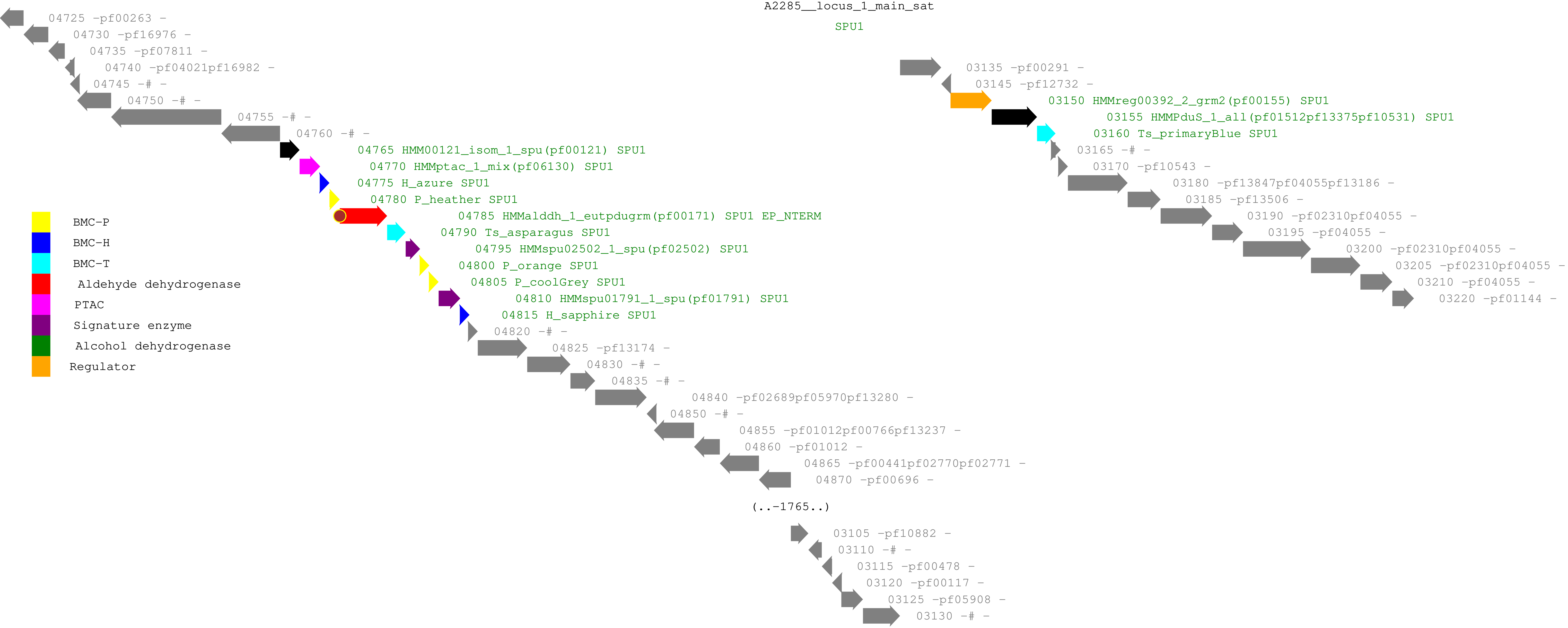

Elusimicrobia bacterium RIFOXYD12_FULL_66_9. Bacteria; Elusimicrobia.

A2506__locus_1

SPU1

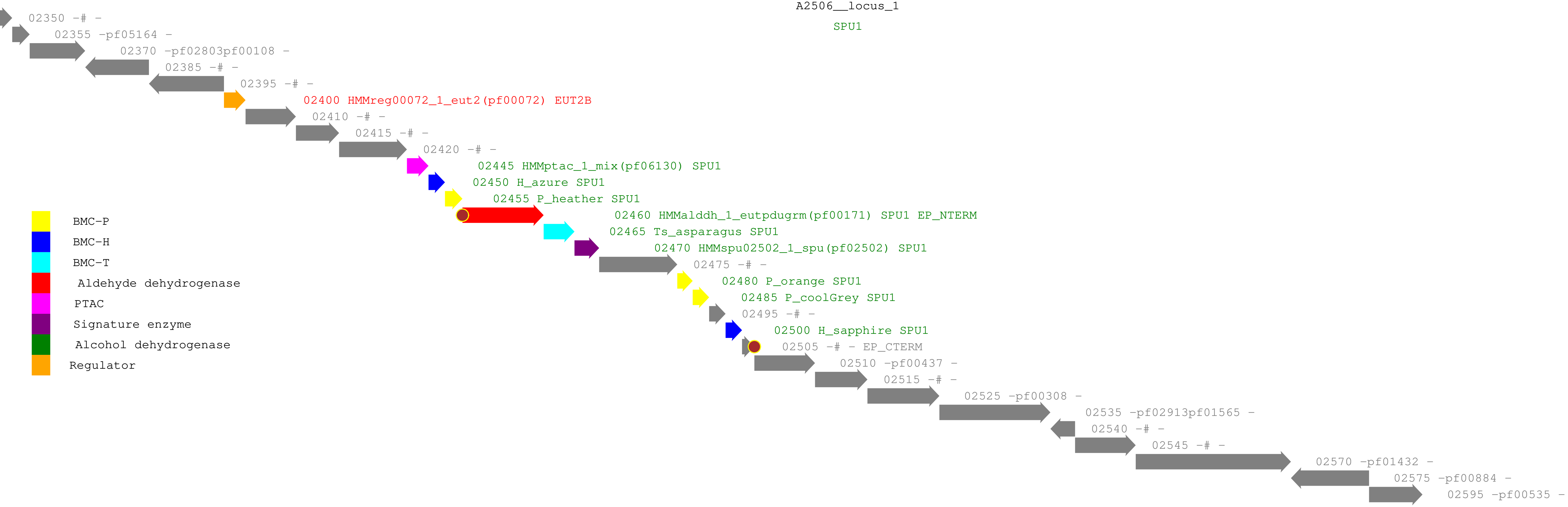

Elusimicrobia bacterium RIFCSPHIGHO2_01_FULL_64_10. Bacteria; Elusimicrobia.

A2636__locus_1

SPU1

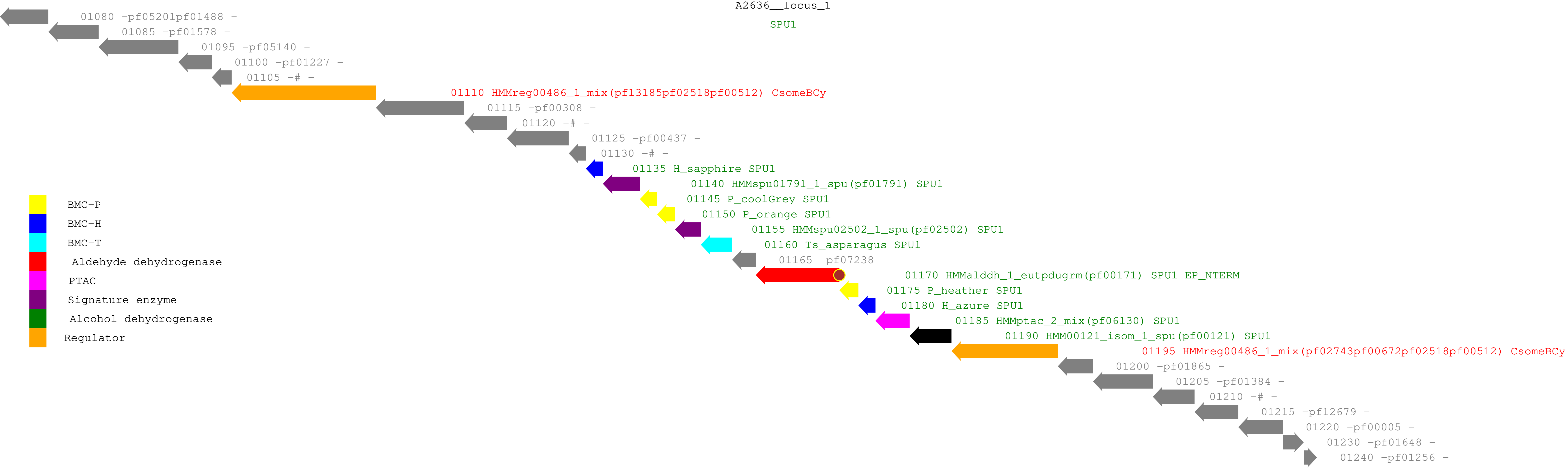

Elusimicrobia bacterium RIFCSPLOWO2_01_FULL_64_13. Bacteria; Elusimicrobia.

A2902__locus_1

SPU1

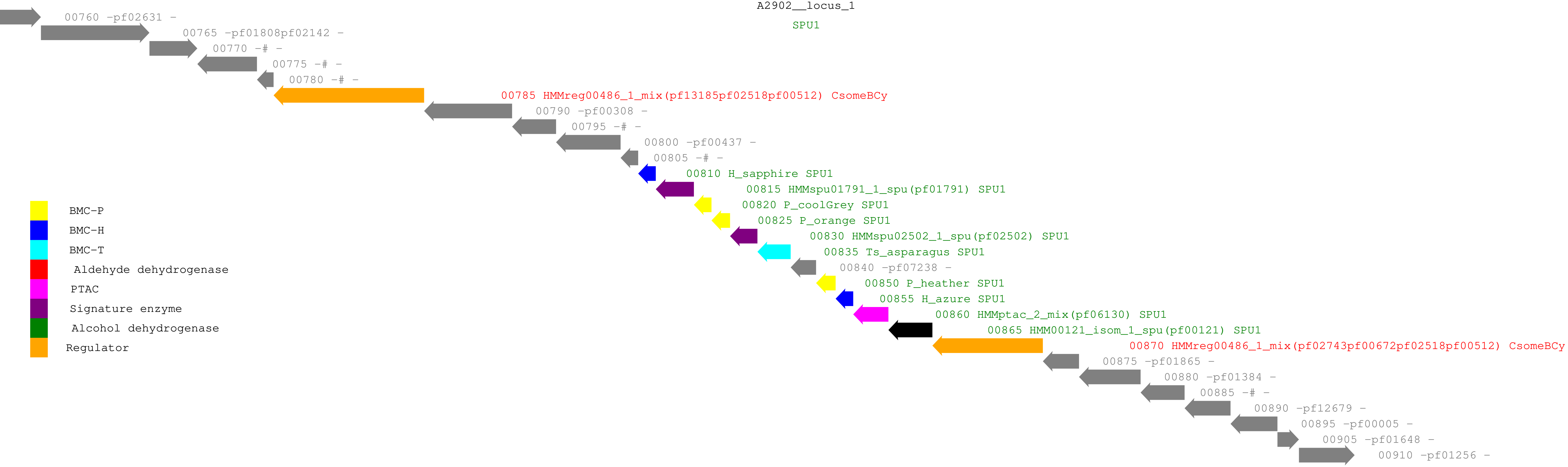

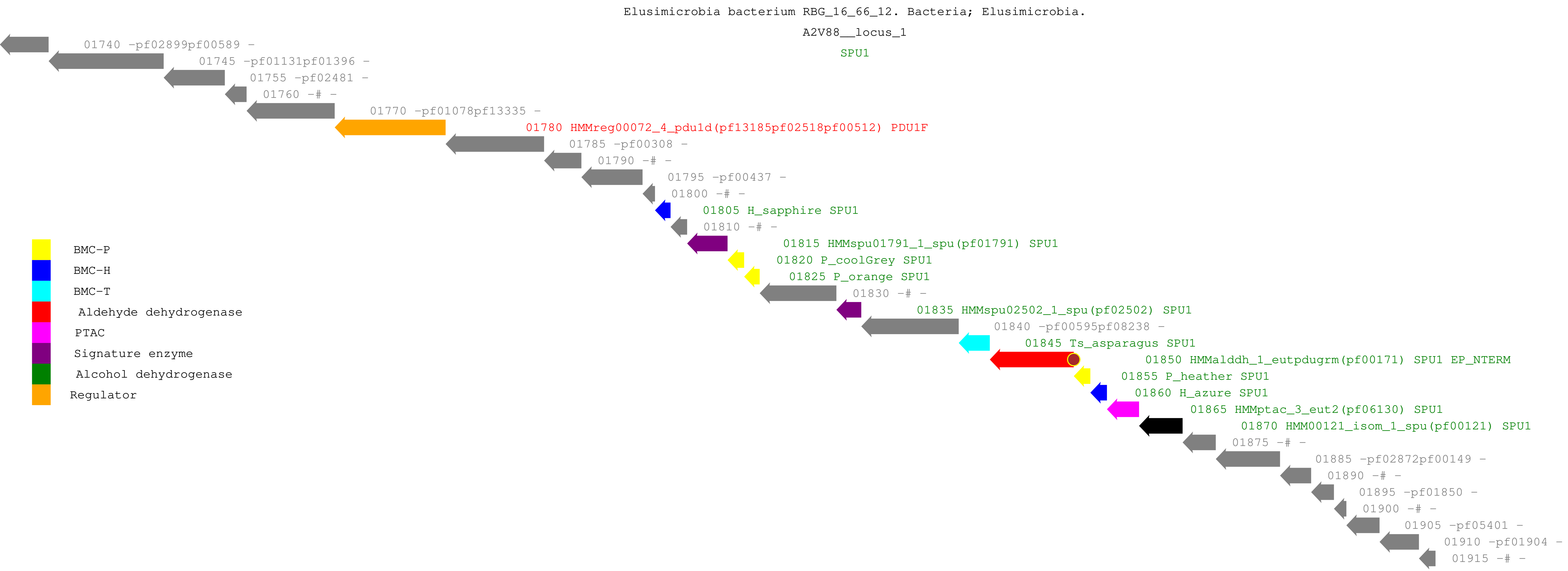
Elusimicrobia bacterium RBG_16_66_12. Bacteria; Elusimicrobia.
A2V88__locus_1
SPU1
01740 -pf02899pf00589 -
01745 -pf01131pf01396 -
01755 -pf02481 -
01760 -# -
01770 -pf01078pf13335 -
01780 HMMreg00072_4_pdu1d(pf13185pf02518pf00512) PDU1F
01785 -pf00308 -
01790 -# -
01795 -pf00437 -
01800 -# -
01805 H_sapphire SPU1
01810 -# -
01815 HMMspu01791_1_spu(pf01791) SPU1
01820 P_coolGrey SPU1
01825 P_orange SPU1
01830 -# -
01835 HMMspu02502_1_spu(pf02502) SPU1
01840 -pf00595pf08238 -
01845 Ts_asparagus SPU1
01850 HMMaldh_1_eutpdugrm(pf00171) SPU1 EP_NTERM
01855 P_heather SPU1
01860 H_azure SPU1
01865 HMMptac_3_eut2(pf06130) SPU1
01870 HMM00121_isom_1_spu(pf00121) SPU1
01875 -# -
01885 -pf02872pf00149 -
01890 -# -
01895 -pf01850 -
01900 -# -
01905 -pf05401 -
01910 -pf01904 -
01915 -# -
BMC-P
BMC-H
BMC-T
Aldehyde dehydrogenase
PTAC
Signature enzyme
Alcohol dehydrogenase
Regulator

Elusimicrobia bacterium GWA2_56_46. Bacteria; Elusimicrobia.

A2X28__locus_1

SPU1

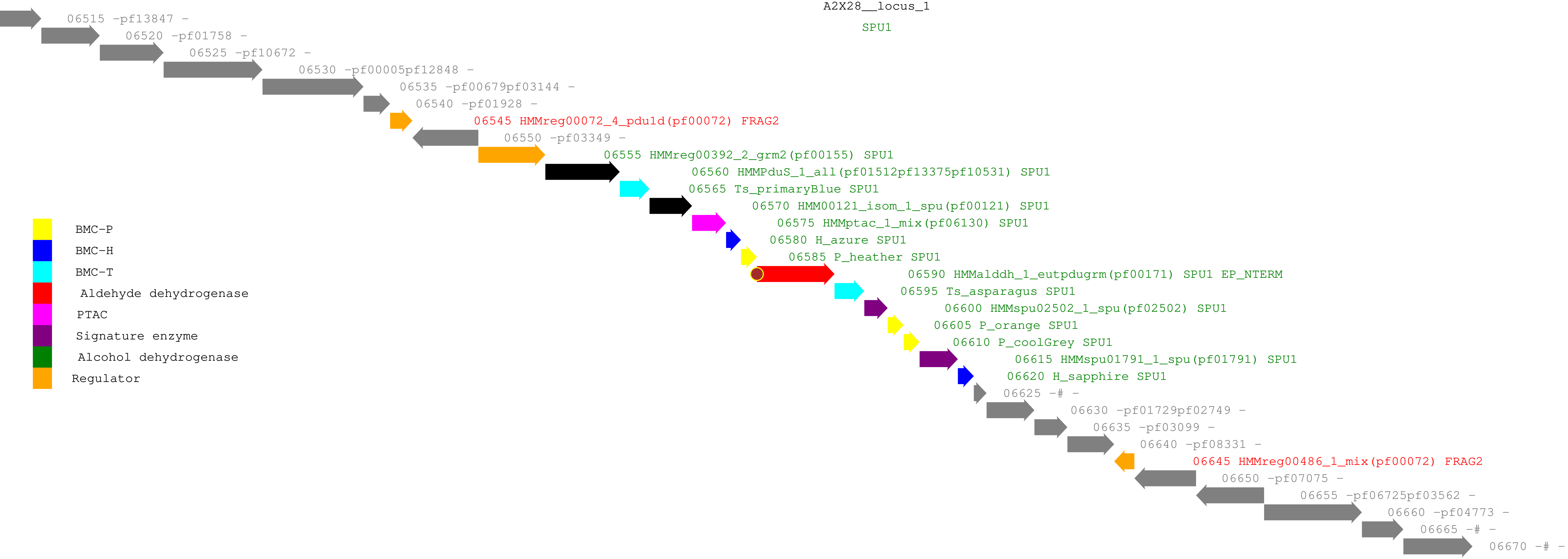

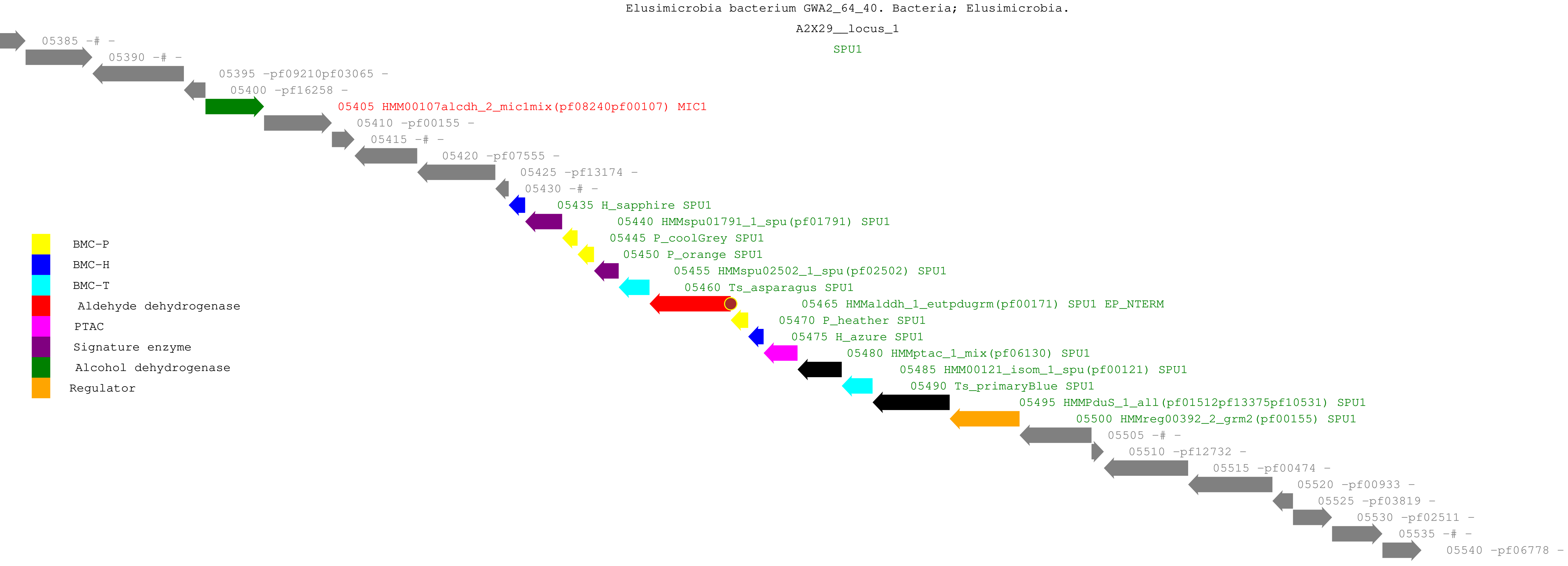
Elusimicrobia bacterium GWA2_64_40. Bacteria; Elusimicrobia.
A2X29__locus_1
SPU1
05385 -# -
05390 -# -
05395 -pf09210pf03065 -
05400 -pf16258 -
05405 HMM00107alcdh_2_mic1mix(pf08240pf00107) MIC1
05410 -pf00155 -
05415 -# -
05420 -pf07555 -
05425 -pf13174 -
05430 -# -
05435 H_sapphire SPU1
05440 HMMspu01791_1_spu(pf01791) SPU1
05445 P_coolGrey SPU1
05450 P_orange SPU1
05455 HMMspu02502_1_spu(pf02502) SPU1
05460 Ts_asparagus SPU1
05465 HMMalddh_1_eutpdugrm(pf00171) SPU1 EP_NTERM
05470 P_heather SPU1
05475 H_azure SPU1
05480 HMMptac_1_mix(pf06130) SPU1
05485 HMM00121_isom_1_spu(pf00121) SPU1
05490 Ts_primaryBlue SPU1
05495 HMMPduS_1_all(pf01512pf13375pf10531) SPU1
05500 HMMreg00392_2_grm2(pf00155) SPU1
05505 -# -
05510 -pf12732 -
05515 -pf00474 -
05520 -pf00933 -
05525 -pf03819 -
05530 -pf02511 -
05535 -# -
05540 -pf06778 -
BMC-P
BMC-H
BMC-T
Aldehyde dehydrogenase
PTAC
Signature enzyme
Alcohol dehydrogenase
Regulator

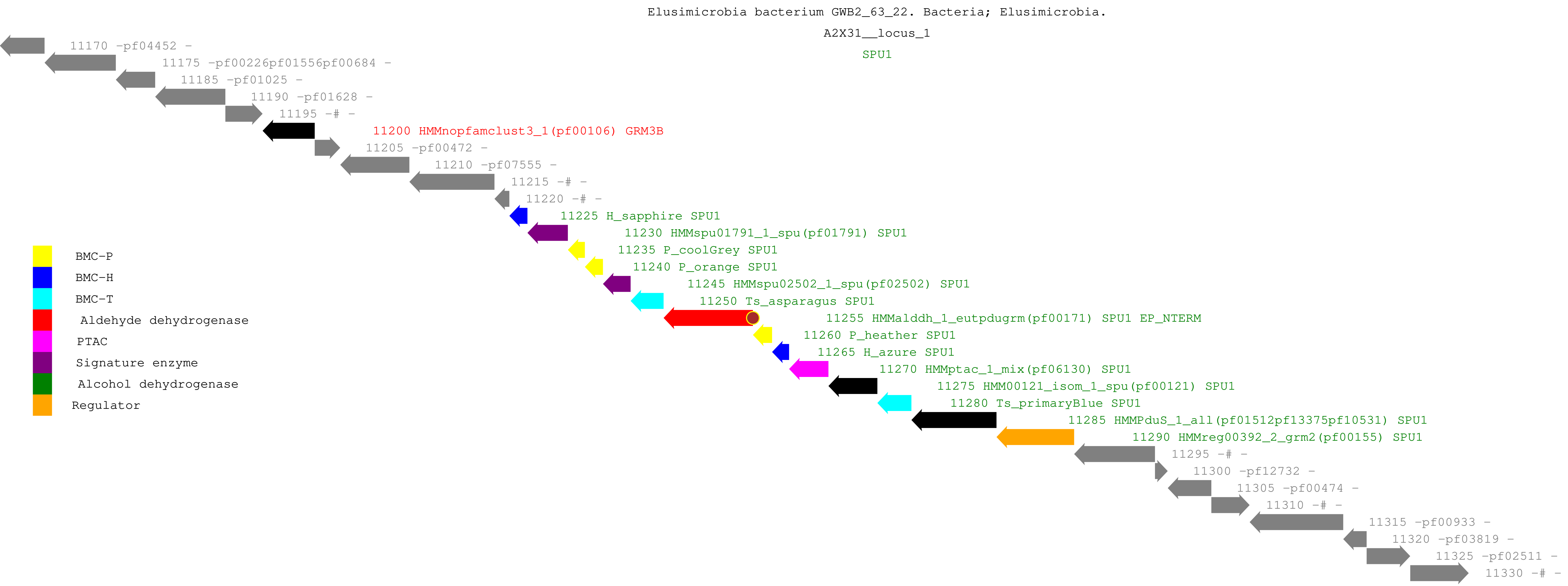
Elusimicrobia bacterium GWB2_63_22. Bacteria; Elusimicrobia.
A2X31__locus_1
SPU1
11170 -pf04452 -
11175 -pf00226pf01556pf00684 -
11185 -pf01025 -
11190 -pf01628 -
11195 -# -
11200 HMMnopfamclust3_1(pf00106) GRM3B
11205 -pf00472 -
11210 -pf07555 -
11215 -# -
11220 -# -
11225 H_sapphire SPU1
11230 HMMspu01791_1_spu(pf01791) SPU1
11235 P_coolGrey SPU1
11240 P_orange SPU1
11245 HMMspu02502_1_spu(pf02502) SPU1
11250 Ts_asparagus SPU1
11255 HMMaldh_1_eutpdugrm(pf00171) SPU1 EP_NTERM
11260 P_heather SPU1
11265 H_azure SPU1
11270 HMMptac_1_mix(pf06130) SPU1
11275 HMM00121_isom_1_spu(pf00121) SPU1
11280 Ts_primaryBlue SPU1
11285 HMMPduS_1_all(pf01512pf13375pf10531) SPU1
11290 HMMreg00392_2_grm2(pf00155) SPU1
11295 -# -
11300 -pf12732 -
11305 -pf00474 -
11310 -# -
11315 -pf00933 -
11320 -pf03819 -
11325 -pf02511 -
11330 -# -
BMC-P
BMC-H
BMC-T
Aldehyde dehydrogenase
PTAC
Signature enzyme
Alcohol dehydrogenase
Regulator

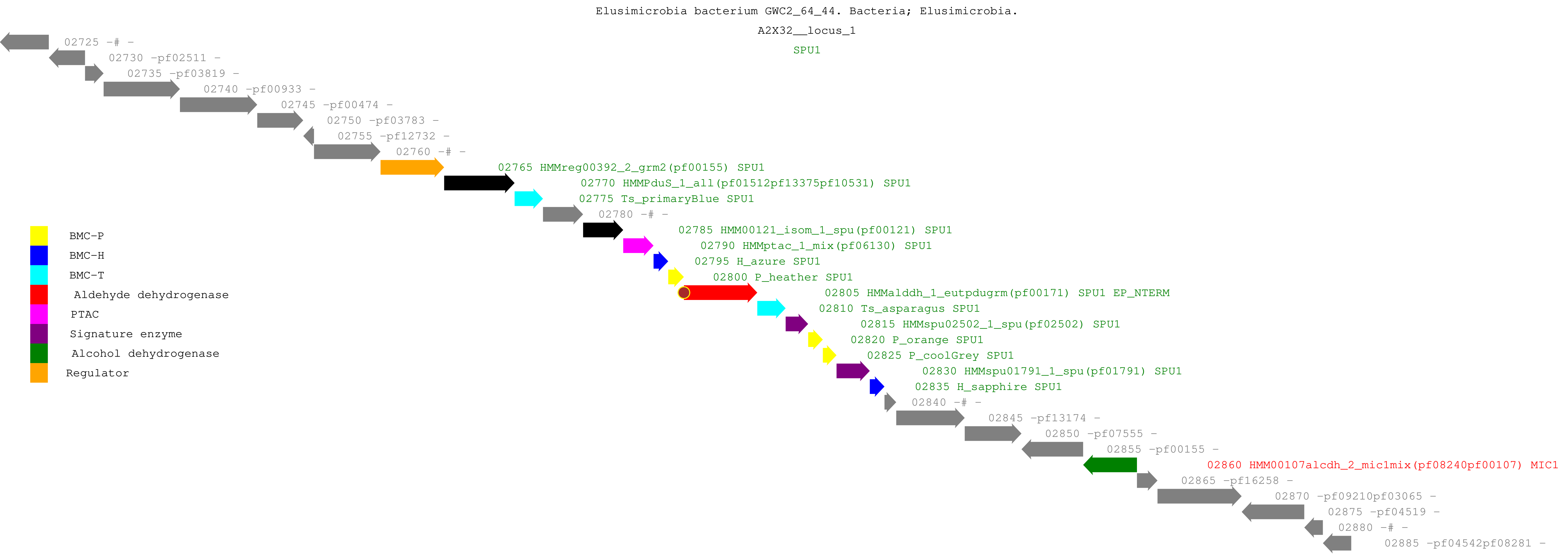
Elusimicrobia bacterium GWC2_64_44. Bacteria; Elusimicrobia.
A2X32__locus_1
SPU1
02725 -# -
02730 -pf02511 -
02735 -pf03819 -
02740 -pf00933 -
02745 -pf00474 -
02750 -pf03783 -
02755 -pf12732 -
02760 -# -
02765 HMMreg00392_2_grm2(pf00155) SPU1
02770 HMMPduS_1_all(pf01512pf13375pf10531) SPU1
02775 Ts_primaryBlue SPU1
02780 -# -
02785 HMM00121_isom_1_spu(pf00121) SPU1
02790 HMMptac_1_mix(pf06130) SPU1
02795 H_azure SPU1
02800 P_heather SPU1
02805 HMMalddh_1_eutpdugrm(pf00171) SPU1 EP_NTERM
02810 Ts_asparagus SPU1
02815 HMMspu02502_1_spu(pf02502) SPU1
02820 P_orange SPU1
02825 P_coolGrey SPU1
02830 HMMspu01791_1_spu(pf01791) SPU1
02835 H_sapphire SPU1
02840 -# -
02845 -pf13174 -
02850 -pf07555 -
02855 -pf00155 -
02860 HMM00107alcdh_2_mic1mix(pf08240pf00107) MIC1
02865 -pf16258 -
02870 -pf09210pf03065 -
02875 -pf04519 -
02880 -# -
02885 -pf04542pf08281 -
BMC-P
BMC-H
BMC-T
Aldehyde dehydrogenase
PTAC
Signature enzyme
Alcohol dehydrogenase
Regulator

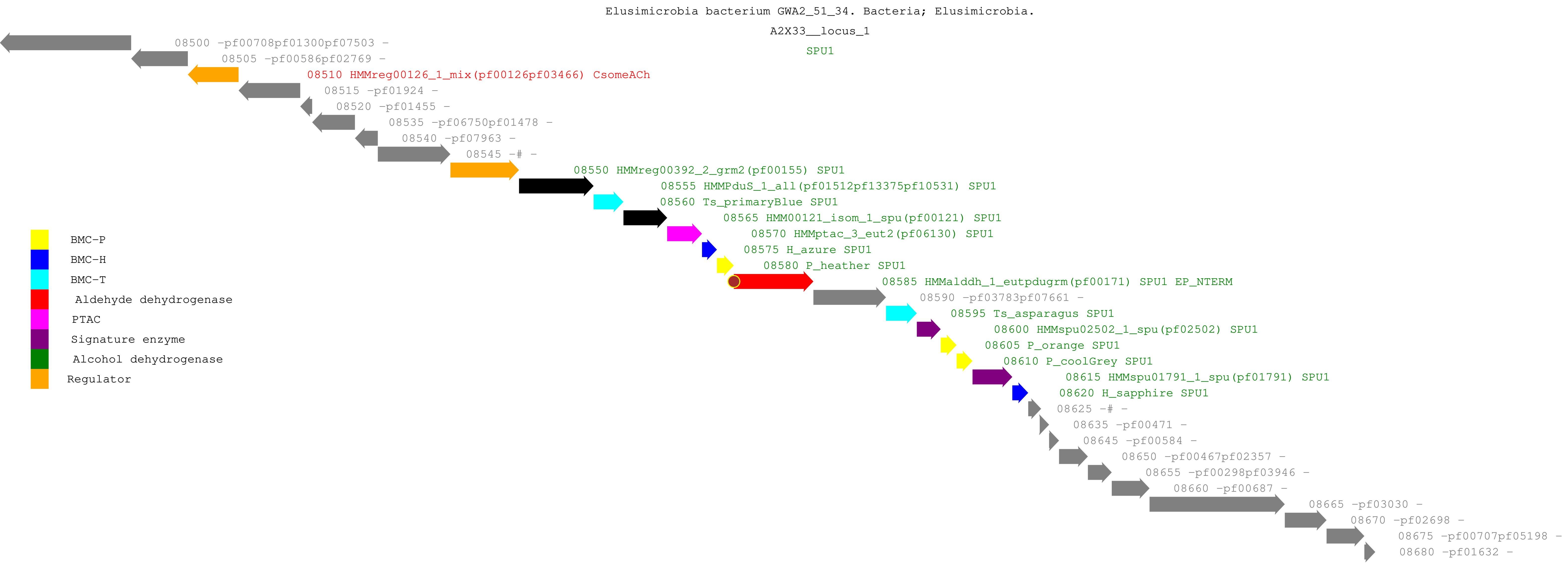
Elusimicrobia bacterium GWA2_51_34. Bacteria; Elusimicrobia.
A2X33__locus_1
SPU1
08500 -pf00708pf01300pf07503 -
08505 -pf00586pf02769 -
08510 HMMreg00126_1_mix(pf00126pf03466) CsomeACh
08515 -pf01924 -
08520 -pf01455 -
08535 -pf06750pf01478 -
08540 -pf07963 -
08545 -# -
08550 HMMreg00392_2_grm2(pf00155) SPU1
08555 HMMPduS_1_all(pf01512pf13375pf10531) SPU1
08560 Ts_primaryBlue SPU1
08565 HMM00121_isom_1_spu(pf00121) SPU1
08570 HMMptac_3_eut2(pf06130) SPU1
08575 H_azure SPU1
08580 P_heather SPU1
08585 HMMalddh_1_eutpdugrm(pf00171) SPU1 EP_NTERM
08590 -pf03783pf07661 -
08595 Ts_asparagus SPU1
08600 HMMspu02502_1_spu(pf02502) SPU1
08605 P_orange SPU1
08610 P_coolGrey SPU1
08615 HMMspu01791_1_spu(pf01791) SPU1
08620 H_sapphire SPU1
08625 -# -
08635 -pf00471 -
08645 -pf00584 -
08650 -pf00467pf02357 -
08655 -pf00298pf03946 -
08660 -pf00687 -
08665 -pf03030 -
08670 -pf02698 -
08675 -pf00707pf05198 -
08680 -pf01632 -
BMC-P
BMC-H
BMC-T
Aldehyde dehydrogenase
PTAC
Signature enzyme
Alcohol dehydrogenase
Regulator

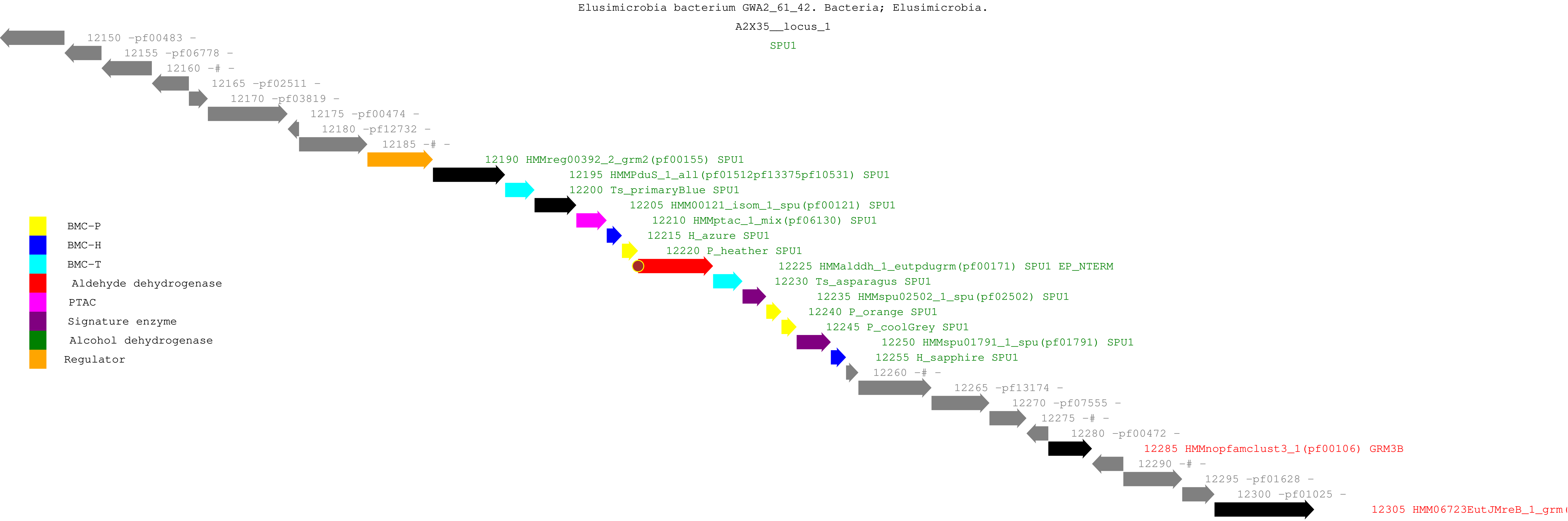

Elusimicrobia bacterium GWA2_61_42. Bacteria; Elusimicrobia.
A2X35__locus_1
SPU1
12150 -pf00483 -
12155 -pf06778 -
12160 -# -
12165 -pf02511 -
12170 -pf03819 -
12175 -pf00474 -
12180 -pf12732 -
12185 -# -
12190 HMMreg00392_2_grm2(pf00155) SPU1
12195 HMMPduS_1_all(pf01512pf13375pf10531) SPU1
12200 Ts_primaryBlue SPU1
12205 HMM00121_isom_1_spu(pf00121) SPU1
12210 HMMptac_1_mix(pf06130) SPU1
12215 H_azure SPU1
12220 P_heather SPU1
12225 HMMalddh_1_eutpdugrm(pf00171) SPU1 EP_NTERM
12230 Ts_asparagus SPU1
12235 HMMspu02502_1_spu(pf02502) SPU1
12240 P_orange SPU1
12245 P_coolGrey SPU1
12250 HMMspu01791_1_spu(pf01791) SPU1
12255 H_sapphire SPU1
12260 -# -
12265 -pf13174 -
12270 -pf07555 -
12275 -# -
12280 -pf00472 -
12285 HMMnopfamclust3_1(pf00106) GRM3B
12290 -# -
12295 -pf01628 -
12300 -pf01025 -
12305 HMM06723EutJMreB_1_grm
BMC-P
BMC-H
BMC-T
Aldehyde dehydrogenase
PTAC
Signature enzyme
Alcohol dehydrogenase
Regulator

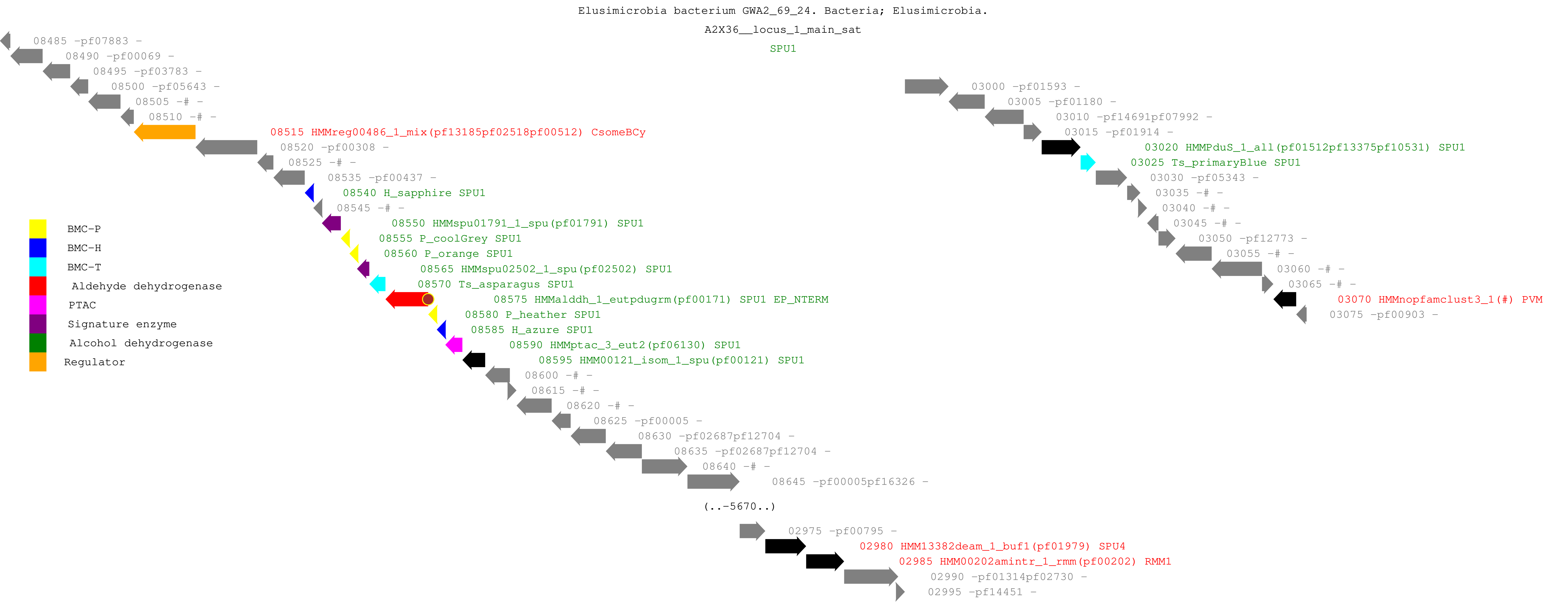
Elusimicrobia bacterium GWA2_69_24. Bacteria; Elusimicrobia.
A2X36__locus_1_main_sat
SPU1
08485 -pf07883 -
08490 -pf00069 -
08495 -pf03783 -
08500 -pf05643 -
08505 -# -
08510 -# -
08515 HMMreg00486_1_mix(pf13185pf02518pf00512) CsomeBCy
08520 -pf00308 -
08525 -# -
08535 -pf00437 -
08540 H_sapphire SPU1
08545 -# -
08550 HMMspu01791_1_spu(pf01791) SPU1
08555 P_coolGrey SPU1
08560 P_orange SPU1
08565 HMMspu02502_1_spu(pf02502) SPU1
08570 Ts_asparagus SPU1
08575 HMMalddh_1_eutpdugrm(pf00171) SPU1 EP_NTERM
08580 P_heather SPU1
08585 H_azure SPU1
08590 HMMptac_3_eut2(pf06130) SPU1
08595 HMM00121_isom_1_spu(pf00121) SPU1
08600 -# -
08615 -# -
08620 -# -
08625 -pf00005 -
08630 -pf02687pf12704 -
08635 -pf02687pf12704 -
08640 -# -
08645 -pf00005pf16326 -
(..-5670..)
02975 -pf00795 -
02980 HMM13382deam_1_buf1(pf01979) SPU4
02985 HMM00202amintr_1_rmm(pf00202) RMM1
02990 -pf01314pf02730 -
02995 -pf14451 -
03000 -pf01593 -
03005 -pf01180 -
03010 -pf14691pf07992 -
03015 -pf01914 -
03020 HMMPduS_1_all(pf01512pf13375pf10531) SPU1
03025 Ts_primaryBlue SPU1
03030 -pf05343 -
03035 -# -
03040 -# -
03045 -# -
03050 -pf12773 -
03055 -# -
03060 -# -
03065 -# -
03070 HMMnopfamclust3_1(#) PVM
03075 -pf00903 -
BMC-P
BMC-H
BMC-T
Aldehyde dehydrogenase
PTAC
Signature enzyme
Alcohol dehydrogenase
Regulator

Elusimicrobia bacterium GWA2_66_18. Bacteria; Elusimicrobia.

A2X37__locus_1

SPU1

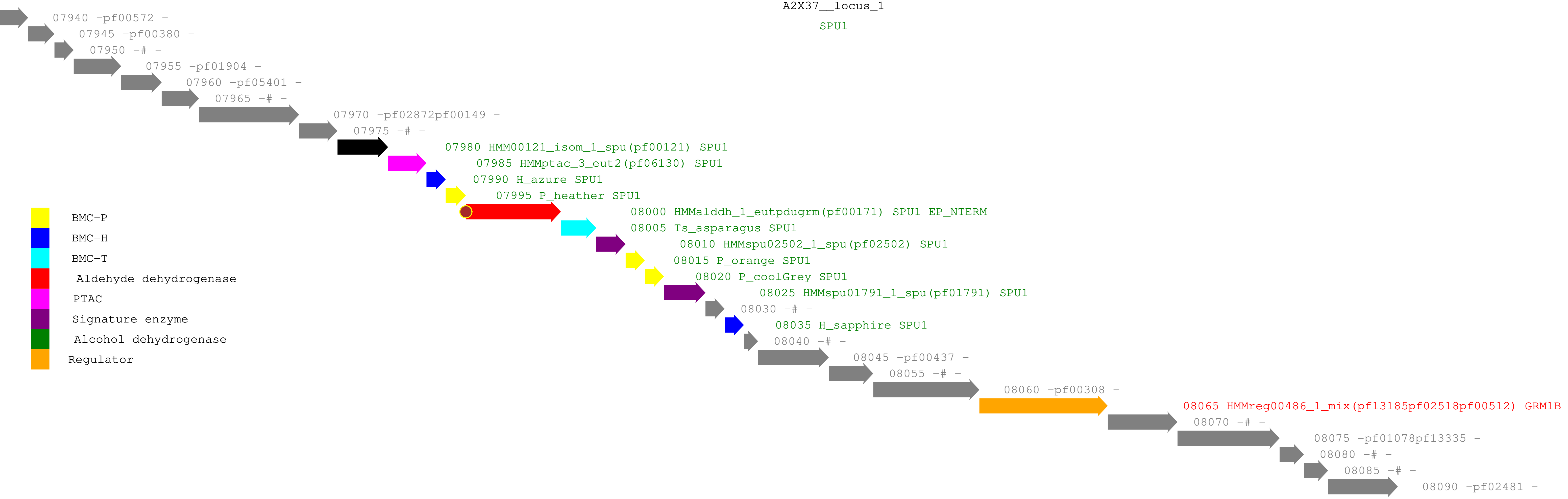

Elusimicrobia bacterium RIFCSPLOWO2_01_FULL_60_11. Bacteria; Elusimicrobia.

A3A86__locus_1

SPU1

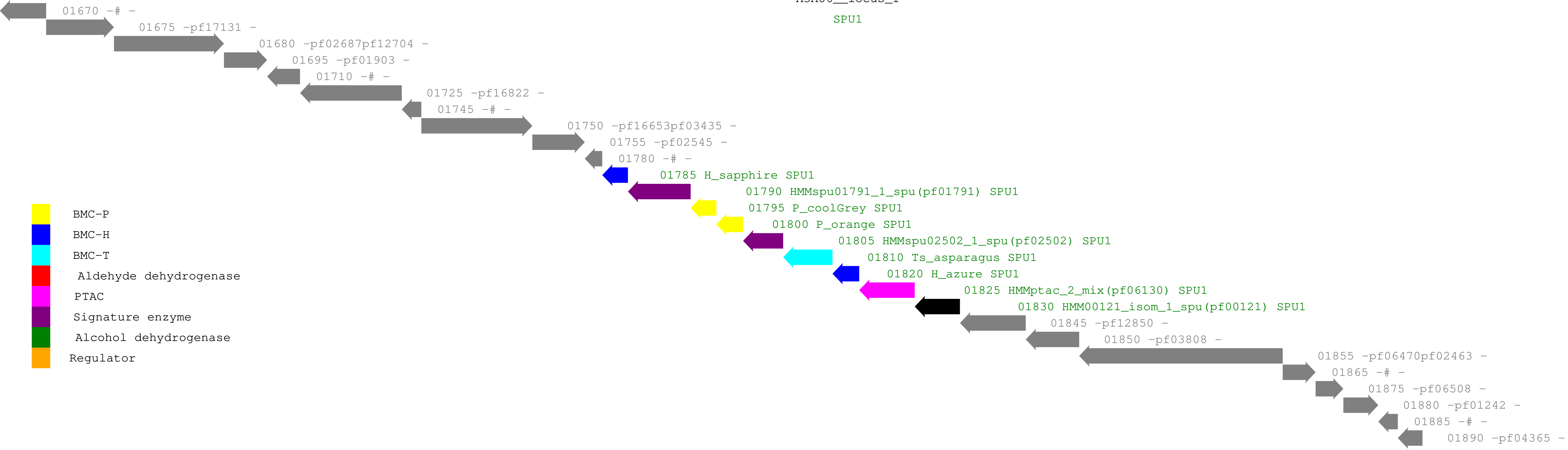

Elusimicrobia bacterium RIFCSPLOWO2_12_FULL_59_9. Bacteria; Elusimicrobia.

A3G41__locus_1

SPU1

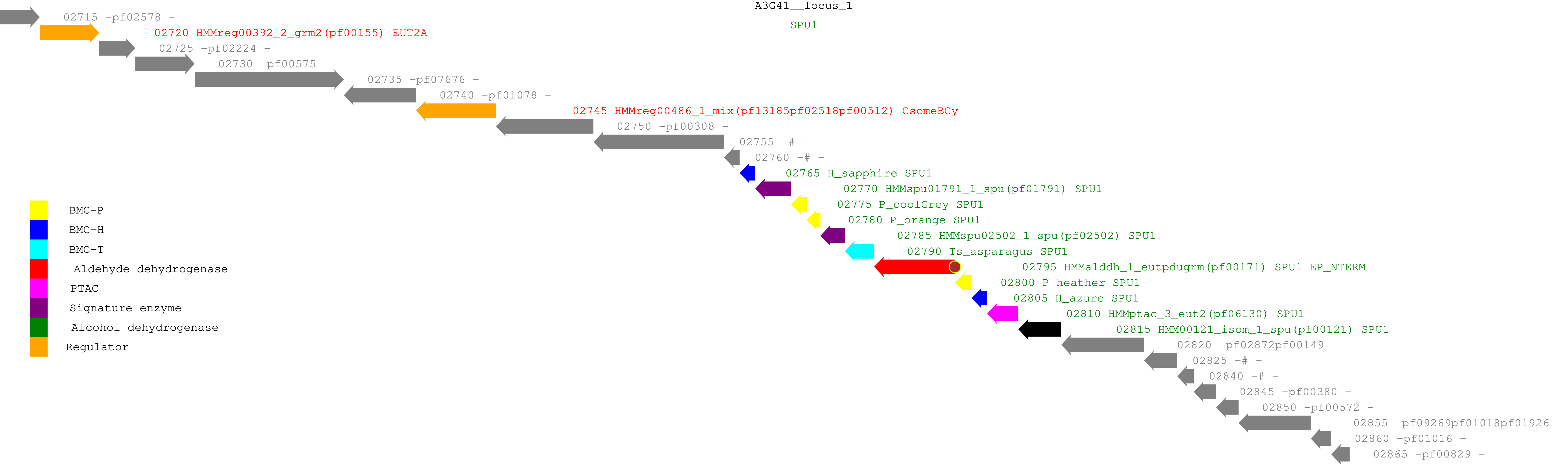

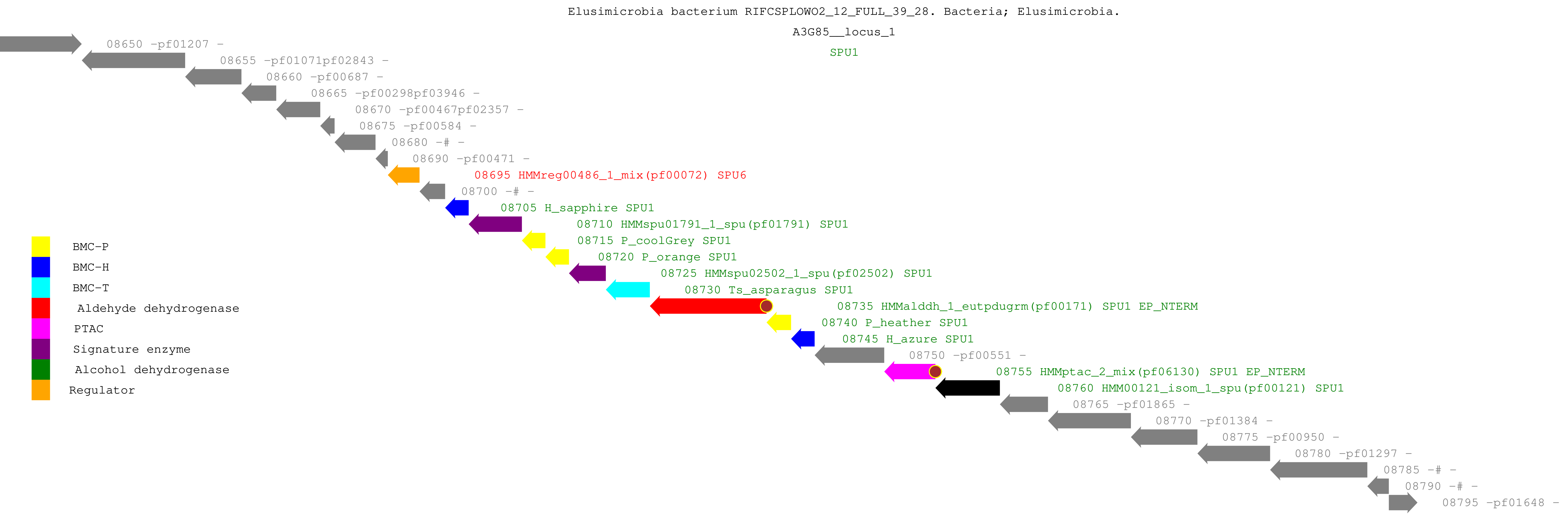
Elusimicrobia bacterium RIFCSPLOWO2_12_FULL_39_28. Bacteria; Elusimicrobia.
A3G85__locus_1
SPU1
08650 -pf01207 -
08655 -pf01071pf02843 -
08660 -pf00687 -
08665 -pf00298pf03946 -
08670 -pf00467pf02357 -
08675 -pf00584 -
08680 -# -
08690 -pf00471 -
08695 HMMreg00486_1_mix(pf00072) SPU6
08700 -# -
08705 H_sapphire SPU1
08710 HMMspu01791_1_spu(pf01791) SPU1
08715 P_coolGrey SPU1
08720 P_orange SPU1
08725 HMMspu02502_1_spu(pf02502) SPU1
08730 Ts_asparagus SPU1
08735 HMMalddh_1_eutpdugrm(pf00171) SPU1 EP_NTERM
08740 P_heather SPU1
08745 H_azure SPU1
08750 -pf00551 -
08755 HMMptac_2_mix(pf06130) SPU1 EP_NTERM
08760 HMM00121_isom_1_spu(pf00121) SPU1
08765 -pf01865 -
08770 -pf01384 -
08775 -pf00950 -
08780 -pf01297 -
08785 -# -
08790 -# -
08795 -pf01648 -
BMC-P
BMC-H
BMC-T
Aldehyde dehydrogenase
PTAC
Signature enzyme
Alcohol dehydrogenase
Regulator

Elusimicrobia bacterium RIFCSPLOWO2_02_FULL_61_11. Bacteria; Elusimicrobia.

A3I76__locus_1

SPU1

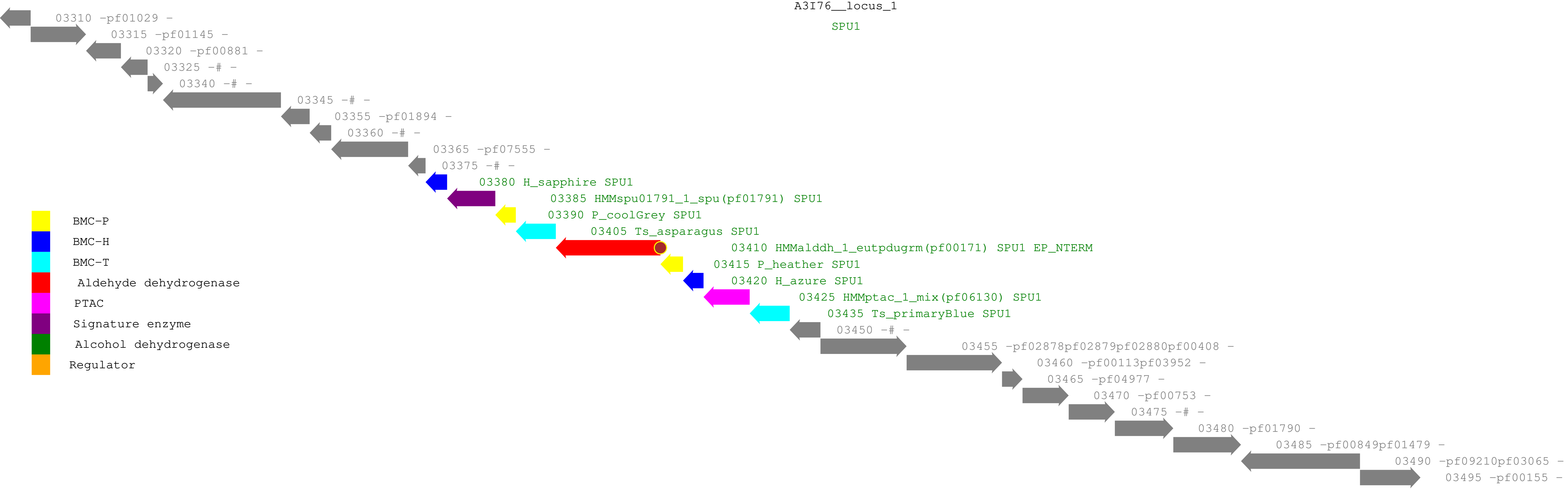

Elusimicrobia bacterium RIFCSPHIGHO2_02_FULL_61_10. Bacteria; Elusimicrobia.

A3J70__locus_1_main_sat

SPU1

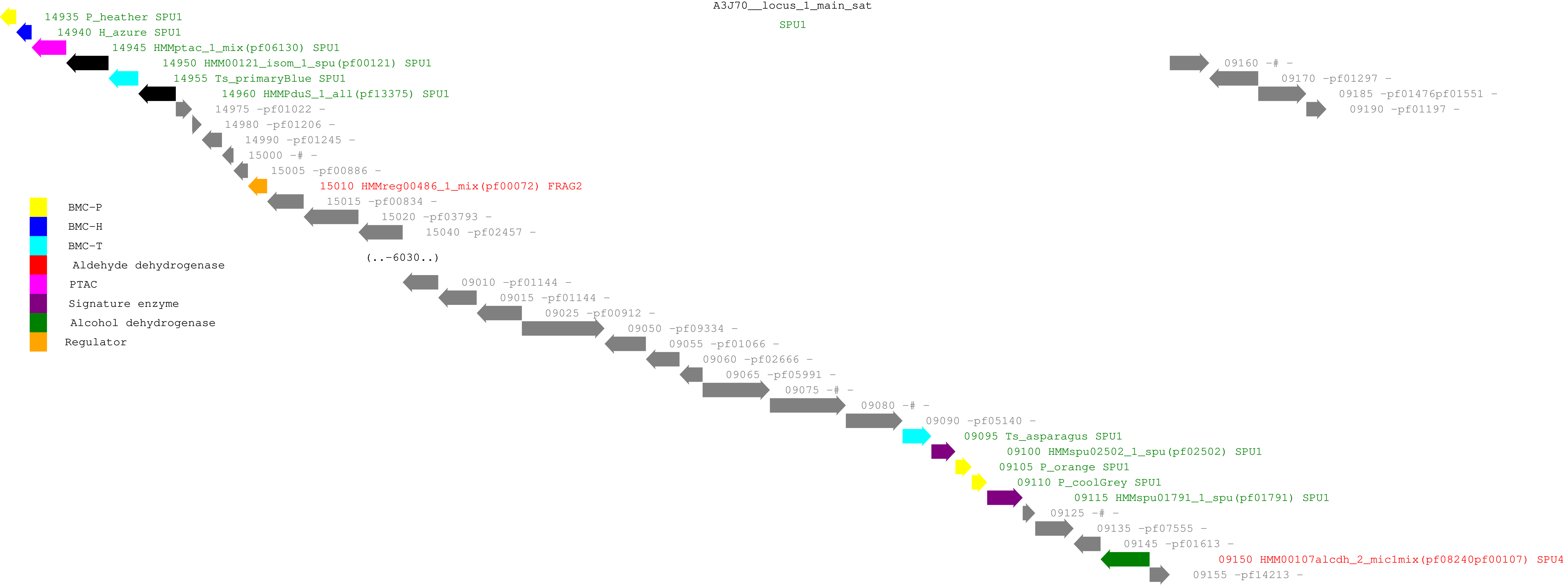

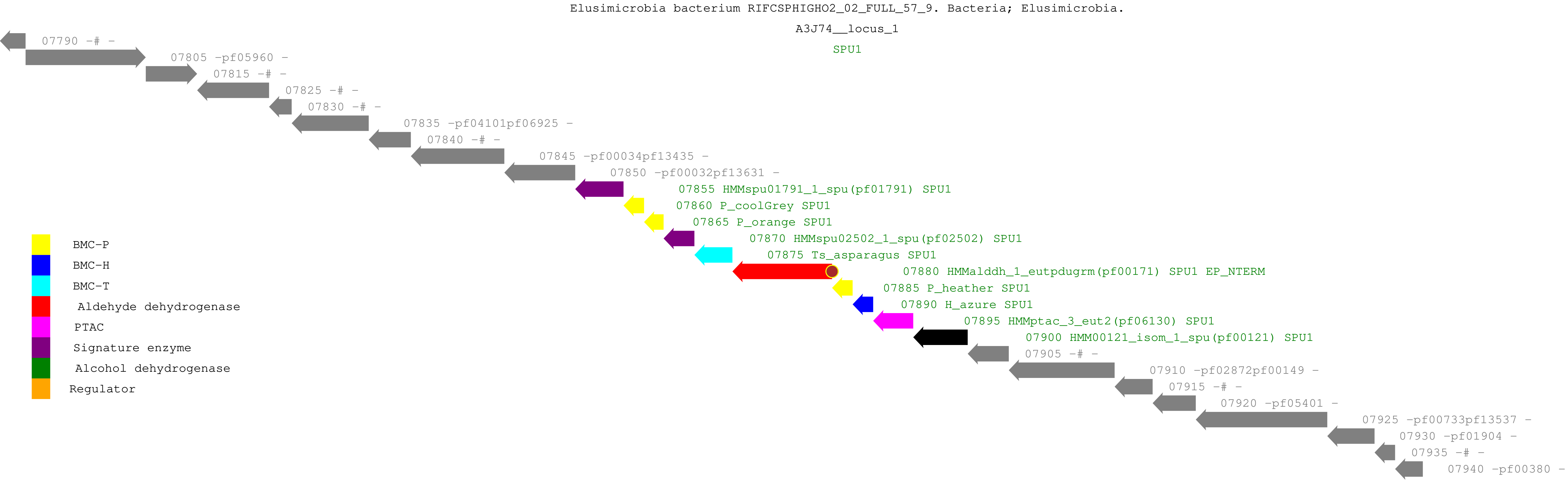
Elusimicrobia bacterium RIFCSPHIGHO2_02_FULL_57_9. Bacteria; Elusimicrobia.
A3J74__locus_1
SPU1
07790 -# -
07805 -pf05960 -
07815 -# -
07825 -# -
07830 -# -
07835 -pf04101pf06925 -
07840 -# -
07845 -pf00034pf13435 -
07850 -pf00032pf13631 -
07855 HMMspu01791_1_spu(pf01791) SPU1
07860 P_coolGrey SPU1
07865 P_orange SPU1
07870 HMMspu02502_1_spu(pf02502) SPU1
07875 Ts_asparagus SPU1
07880 HMMaldde_1_eutpdugrm(pf00171) SPU1 EP_NTERM
07885 P_heather SPU1
07890 H_azure SPU1
07895 HMMptac_3_eut2(pf06130) SPU1
07900 HMM00121_isom_1_spu(pf00121) SPU1
07905 -# -
07910 -pf02872pf00149 -
07915 -# -
07920 -pf05401 -
07925 -pf00733pf13537 -
07930 -pf01904 -
07935 -# -
07940 -pf00380 -
BMC-P
BMC-H
BMC-T
Aldehyde dehydrogenase
PTAC
Signature enzyme
Alcohol dehydrogenase
Regulator

Elusimicrobia bacterium RIFOXYB2_FULL_62_6. Bacteria; Elusimicrobia.

A3J79__locus_1_main_sat

SPU1

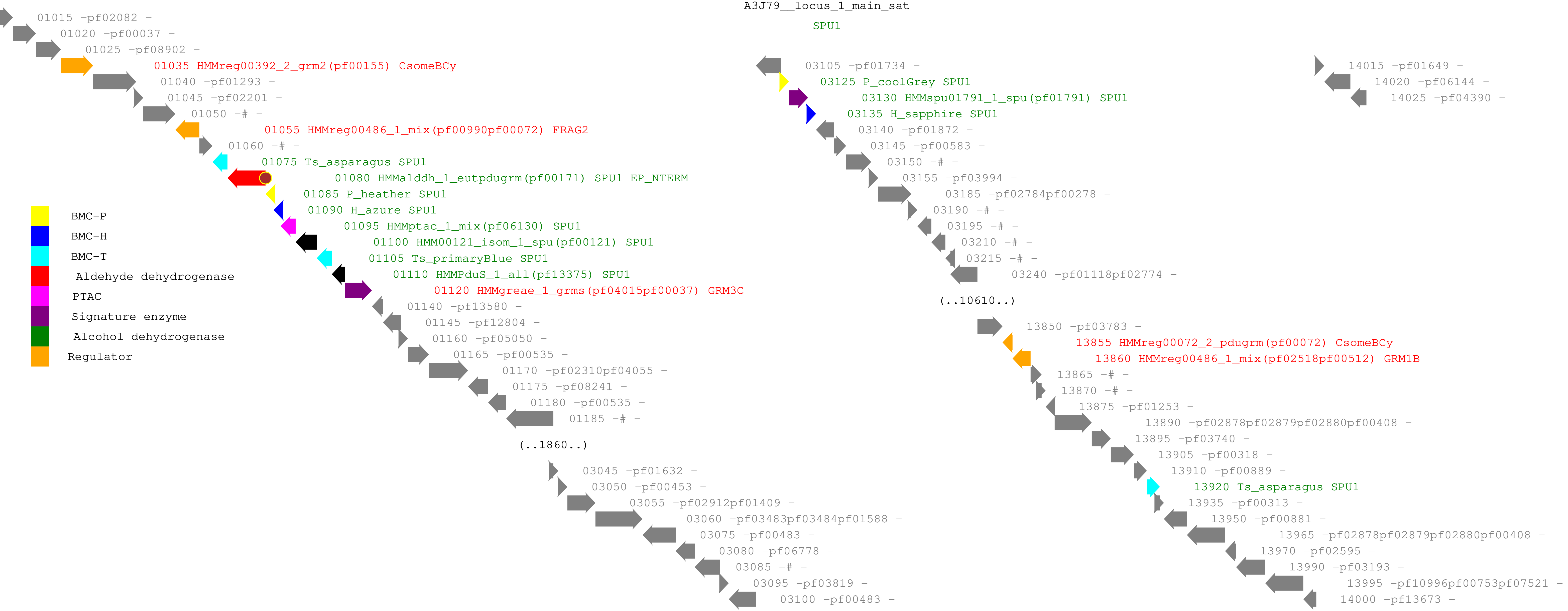

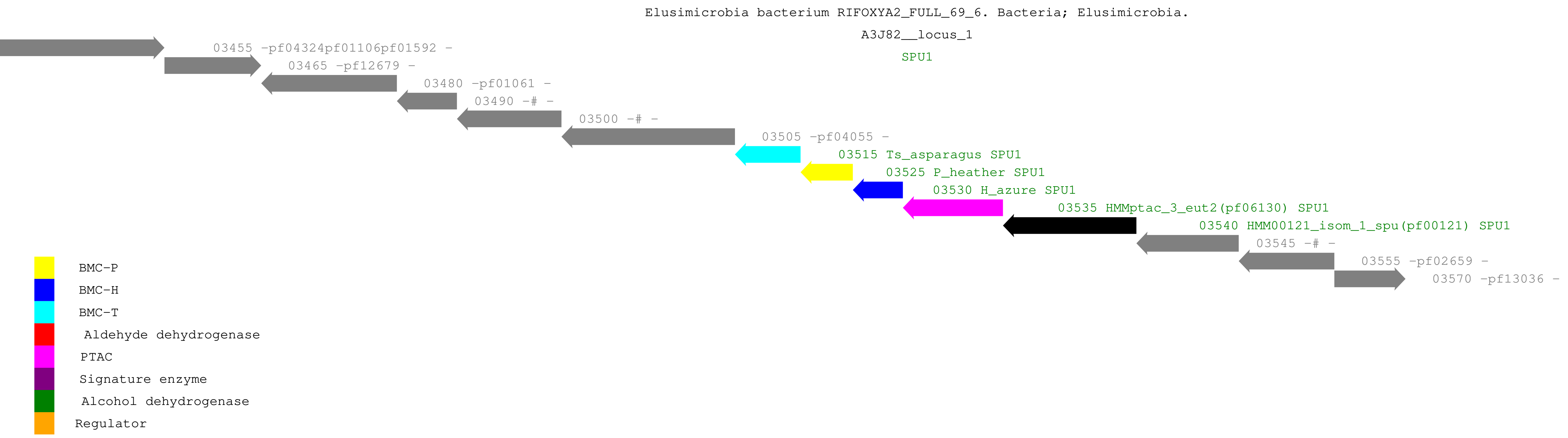
Elusimicrobia bacterium RIFOXYA2_FULL_69_6. Bacteria; Elusimicrobia.
A3J82__locus_1
SPU1
03455 -pf04324pf01106pf01592 -
03465 -pf12679 -
03480 -pf01061 -
03490 -# -
03500 -# -
03505 -pf04055 -
03515 Ts_asparagus SPU1
03525 P_heather SPU1
03530 H_azure SPU1
03535 HMMptac_3_eut2(pf06130) SPU1
03540 HMM00121_isom_1_spu(pf00121) SPU1
03545 -# -
03555 -pf02659 -
03570 -pf13036 -
BMC-P
BMC-H
BMC-T
Aldehyde dehydrogenase
PTAC
Signature enzyme
Alcohol dehydrogenase
Regulator

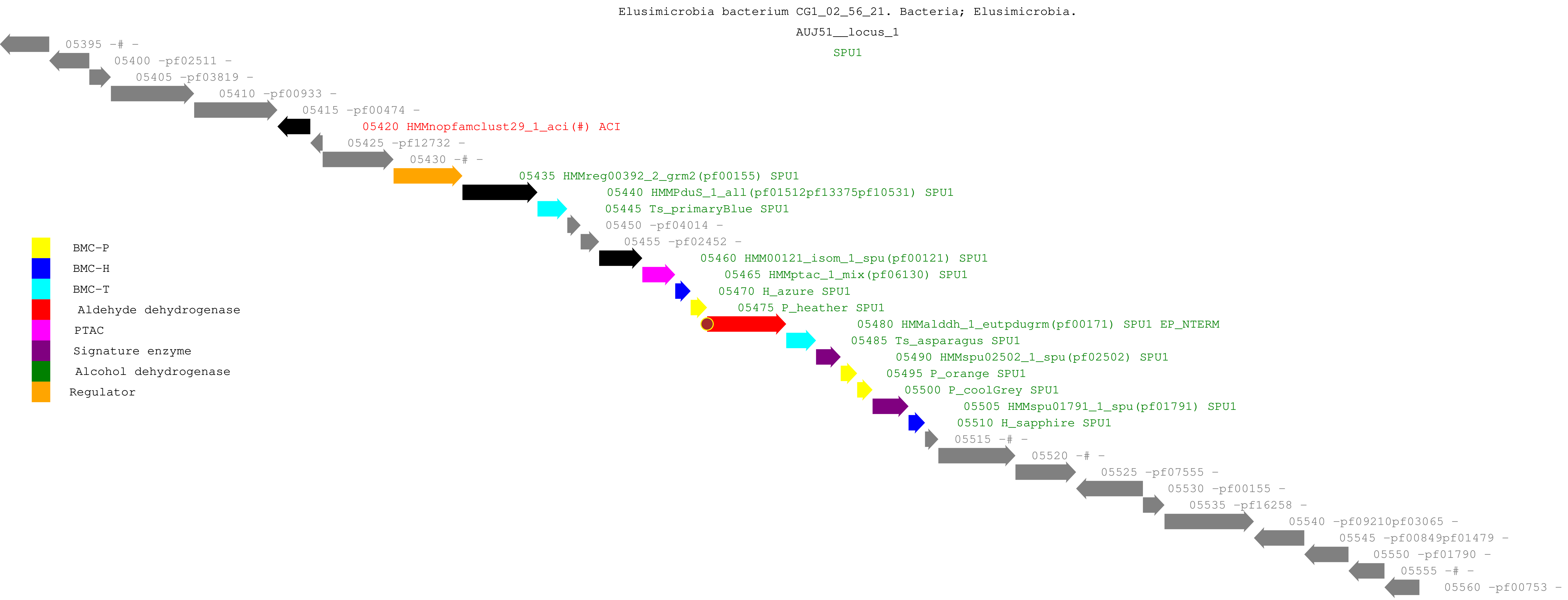
Elusimicrobia bacterium CG1_02_56_21. Bacteria; Elusimicrobia.
AUJ51__locus_1
SPU1
05395 -# -
05400 -pf02511 -
05405 -pf03819 -
05410 -pf00933 -
05415 -pf00474 -
05420 HMMnopfamclust29_1_aci(#) ACI
05425 -pf12732 -
05430 -# -
05435 HMMreg00392_2_grm2(pf00155) SPU1
05440 HMMPduS_1_all(pf01512pf13375pf10531) SPU1
05445 Ts_primaryBlue SPU1
05450 -pf04014 -
05455 -pf02452 -
05460 HMM00121_isom_1_spu(pf00121) SPU1
05465 HMMptac_1_mix(pf06130) SPU1
05470 H_azure SPU1
05475 P_heather SPU1
05480 HMMalddh_1_eutpdugrm(pf00171) SPU1 EP_NTERM
05485 Ts_asparagus SPU1
05490 HMMspu02502_1_spu(pf02502) SPU1
05495 P_orange SPU1
05500 P_coolGrey SPU1
05505 HMMspu01791_1_spu(pf01791) SPU1
05510 H_sapphire SPU1
05515 -# -
05520 -# -
05525 -pf07555 -
05530 -pf00155 -
05535 -pf16258 -
05540 -pf09210pf03065 -
05545 -pf00849pf01479 -
05550 -pf01790 -
05555 -# -
05560 -pf00753 -
BMC-P
BMC-H
BMC-T
Aldehyde dehydrogenase
PTAC
Signature enzyme
Alcohol dehydrogenase
Regulator

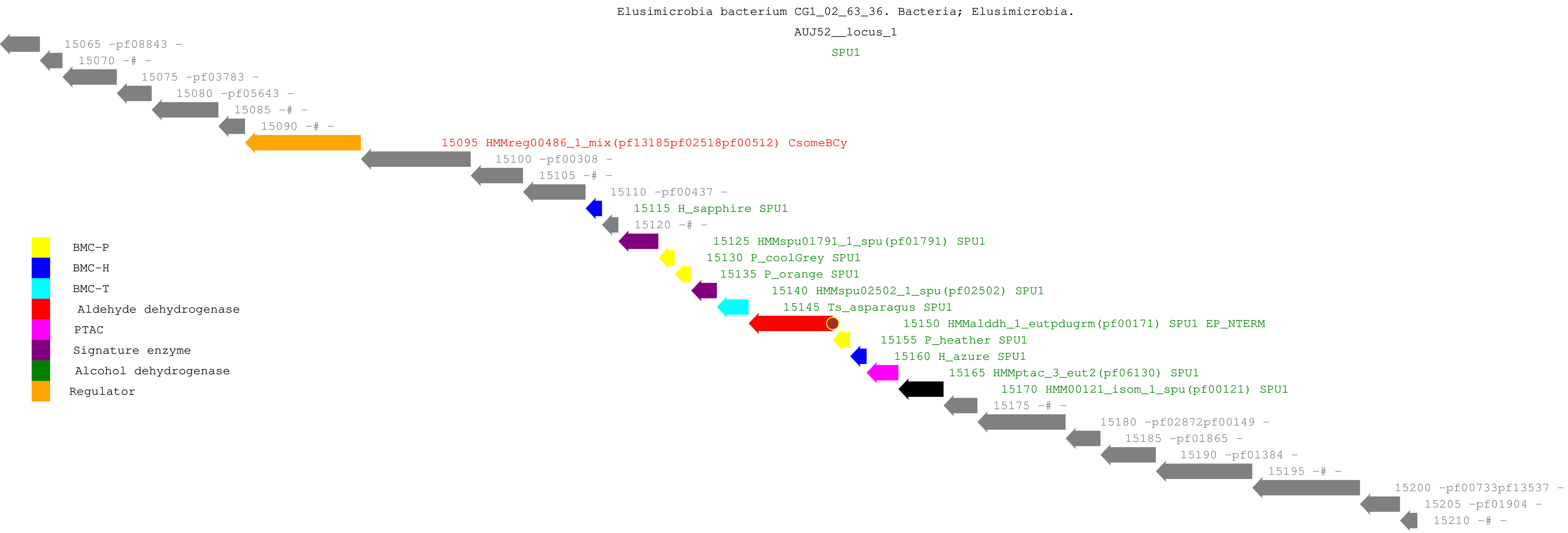

Elusimicrobia bacterium CG1_02_63_36. Bacteria; Elusimicrobia.
AUJ52__locus_1
SPU1
15065 -pf08843 -
15070 -# -
15075 -pf03783 -
15080 -pf05643 -
15085 -# -
15090 -# -
15095 HMMreg00486_1_mix(pf13185pf02518pf00512) CsomeBCy
15100 -pf00308 -
15105 -# -
15110 -pf00437 -
15115 H_sapphire SPU1
15120 -# -
15125 HMMspu01791_1_spu(pf01791) SPU1
15130 P_coolGrey SPU1
15135 P_orange SPU1
15140 HMMspu02502_1_spu(pf02502) SPU1
15145 Ts_asparagus SPU1
15150 HMMalddh_1_eutpdugrm(pf00171) SPU1 EP_NTERM
15155 P_heather SPU1
15160 H_azure SPU1
15165 HMMptac_3_eut2(pf06130) SPU1
15170 HMM00121_isom_1_spu(pf00121) SPU1
15175 -# -
15180 -pf02872pf00149 -
15185 -pf01865 -
15190 -pf01384 -
15195 -# -
15200 -pf00733pf13537 -
15205 -pf01904 -
15210 -# -
BMC-P
BMC-H
BMC-T
Aldehyde dehydrogenase
PTAC
Signature enzyme
Alcohol dehydrogenase
Regulator

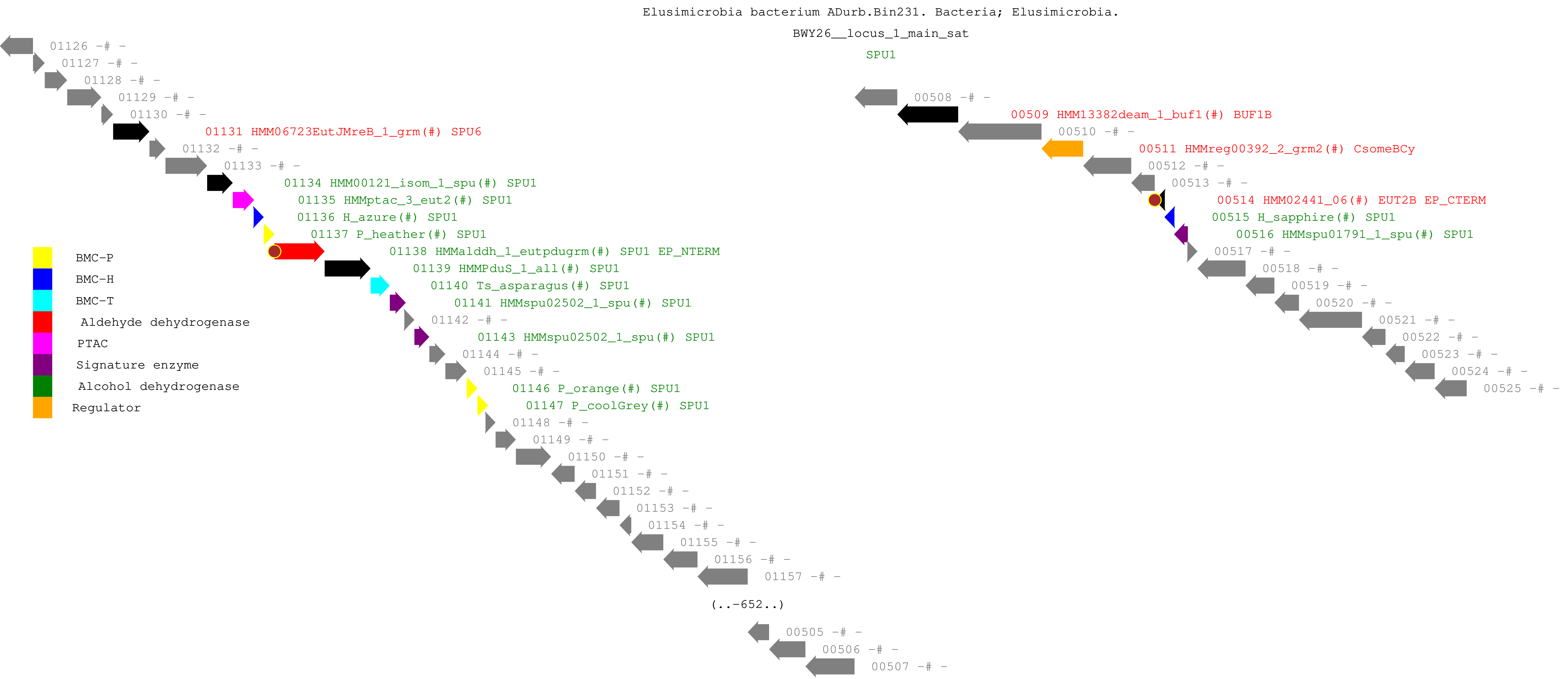
Elusimicrobia bacterium ADurb.Bin231. Bacteria; Elusimicrobia.
BWY26__locus_1_main_sat
SPU1
01126 -# -
01127 -# -
01128 -# -
01129 -# -
01130 -# -
01131 HMM06723EutJMreB_1_grm(#) SPU6
01132 -# -
01133 -# -
01134 HMM00121_isom_1_spu(#) SPU1
01135 HMMptac_3_eut2(#) SPU1
01136 H_azure(#) SPU1
01137 P_heather(#) SPU1
01138 HMMalddh_1_eutpdugrm(#) SPU1 EP_NTERM
01139 HMMPduS_1_all(#) SPU1
01140 Ts_asparagus(#) SPU1
01141 HMMspu02502_1_spu(#) SPU1
01142 -# -
01143 HMMspu02502_1_spu(#) SPU1
01144 -# -
01145 -# -
01146 P_orange(#) SPU1
01147 P_coolGrey(#) SPU1
01148 -# -
01149 -# -
01150 -# -
01151 -# -
01152 -# -
01153 -# -
01154 -# -
01155 -# -
01156 -# -
01157 -# -
(..-652..)
00505 -# -
00506 -# -
00507 -# -
00508 -# -
00509 HMM13382deam_1_buf1(#) BUF1B
00510 -# -
00511 HMMreg00392_2_grm2(#) CsomeBCy
00512 -# -
00513 -# -
00514 HMM02441_06(#) EUT2B EP_CTERM
00515 H_sapphire(#) SPU1
00516 HMMspu01791_1_spu(#) SPU1
00517 -# -
00518 -# -
00519 -# -
00520 -# -
00521 -# -
00522 -# -
00523 -# -
00524 -# -
00525 -# -
BMC-P
BMC-H
BMC-T
Aldehyde dehydrogenase
PTAC
Signature enzyme
Alcohol dehydrogenase
Regulator

bacterium BM706. Bacteria.
C0601__locus_1
SPU1

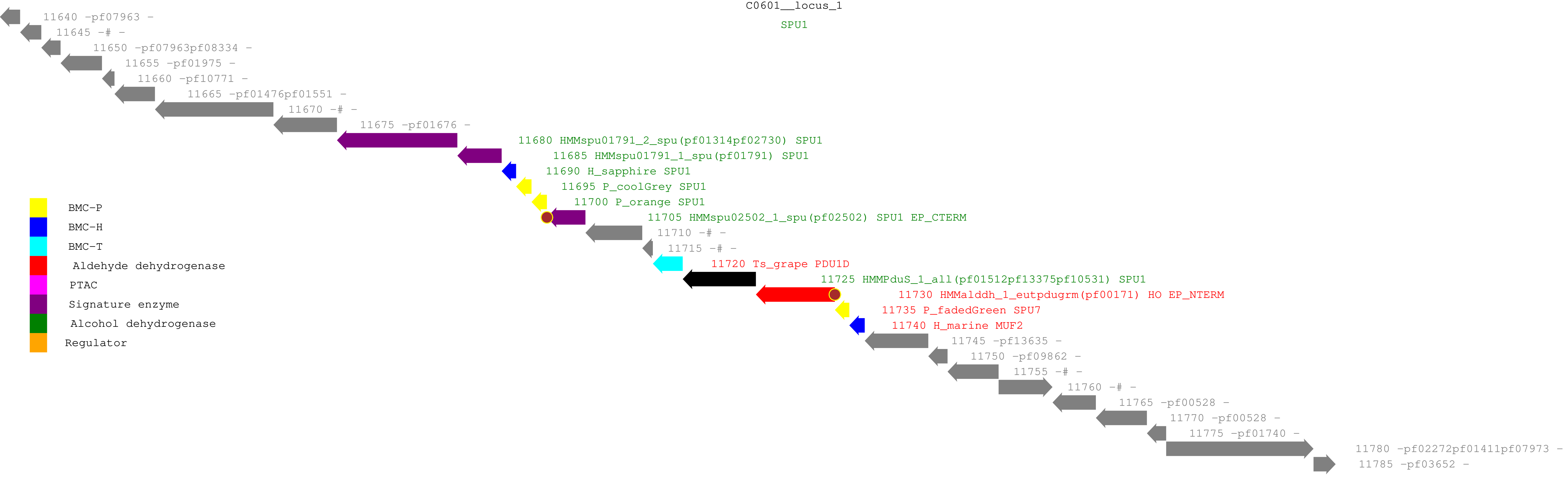

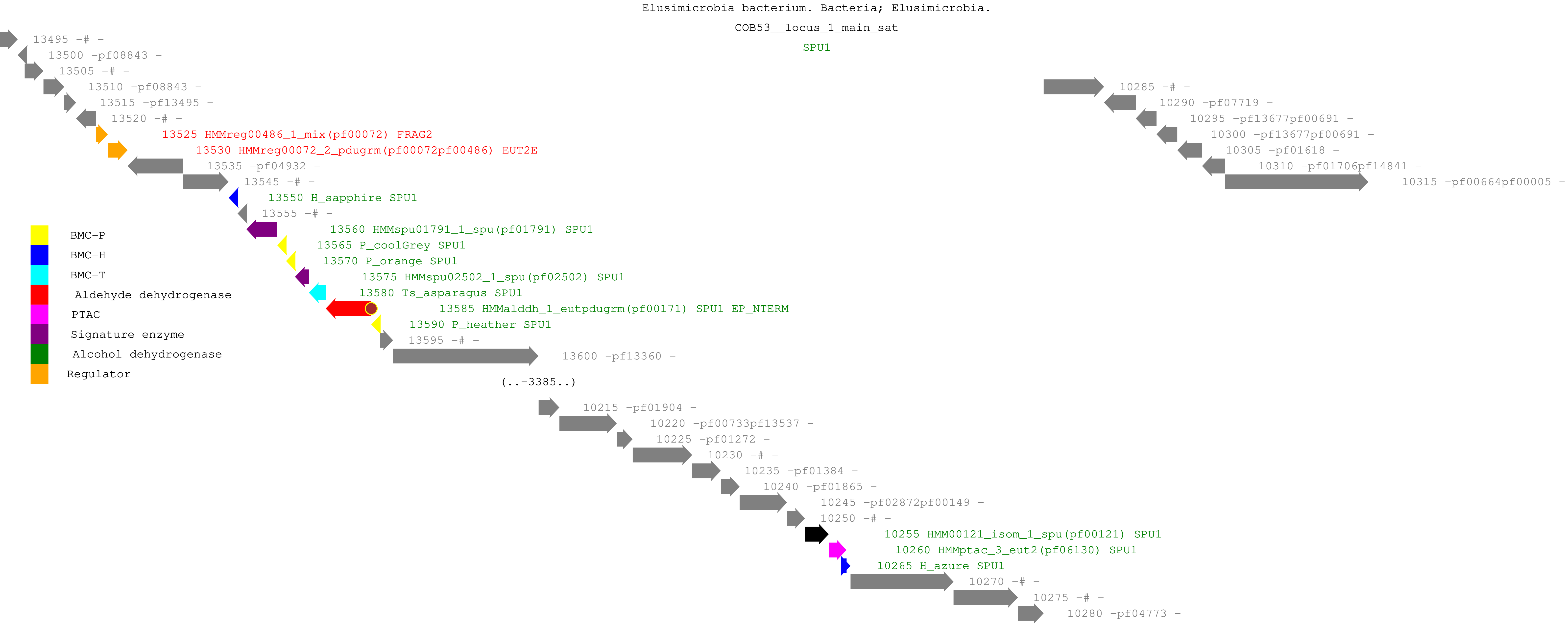
Elusimicrobia bacterium. Bacteria; Elusimicrobia.
COB53__locus_1_main_sat
SPU1
13495 -# -
13500 -pf08843 -
13505 -# -
13510 -pf08843 -
13515 -pf13495 -
13520 -# -
13525 HMMreg00486_1_mix(pf00072) FRAG2
13530 HMMreg00072_2_pdugrm(pf00072pf00486) EUT2E
13535 -pf04932 -
13545 -# -
13550 H_sapphire SPU1
13555 -# -
13560 HMMspu01791_1_spu(pf01791) SPU1
13565 P_coolGrey SPU1
13570 P_orange SPU1
13575 HMMspu02502_1_spu(pf02502) SPU1
13580 Ts_asparagus SPU1
13585 HMMaldh_1_eutpdugrm(pf00171) SPU1 EP_NTERM
13590 P_heather SPU1
13595 -# -
13600 -pf13360 -
(..-3385..)
10215 -pf01904 -
10220 -pf00733pf13537 -
10225 -pf01272 -
10230 -# -
10235 -pf01384 -
10240 -pf01865 -
10245 -pf02872pf00149 -
10250 -# -
10255 HMM00121_isom_1_spu(pf00121) SPU1
10260 HMMptac_3_eut2(pf06130) SPU1
10265 H_azure SPU1
10270 -# -
10275 -# -
10280 -pf04773 -
10285 -# -
10290 -pf07719 -
10295 -pf13677pf00691 -
10300 -pf13677pf00691 -
10305 -pf01618 -
10310 -pf01706pf14841 -
10315 -pf00664pf00005 -
BMC-P
BMC-H
BMC-T
Aldehyde dehydrogenase
PTAC
Signature enzyme
Alcohol dehydrogenase
Regulator

Elusimicrobia bacterium CG22_combo_CG10-13_8_21_14_all_63_91. Bacteria; Elusimicrobia.

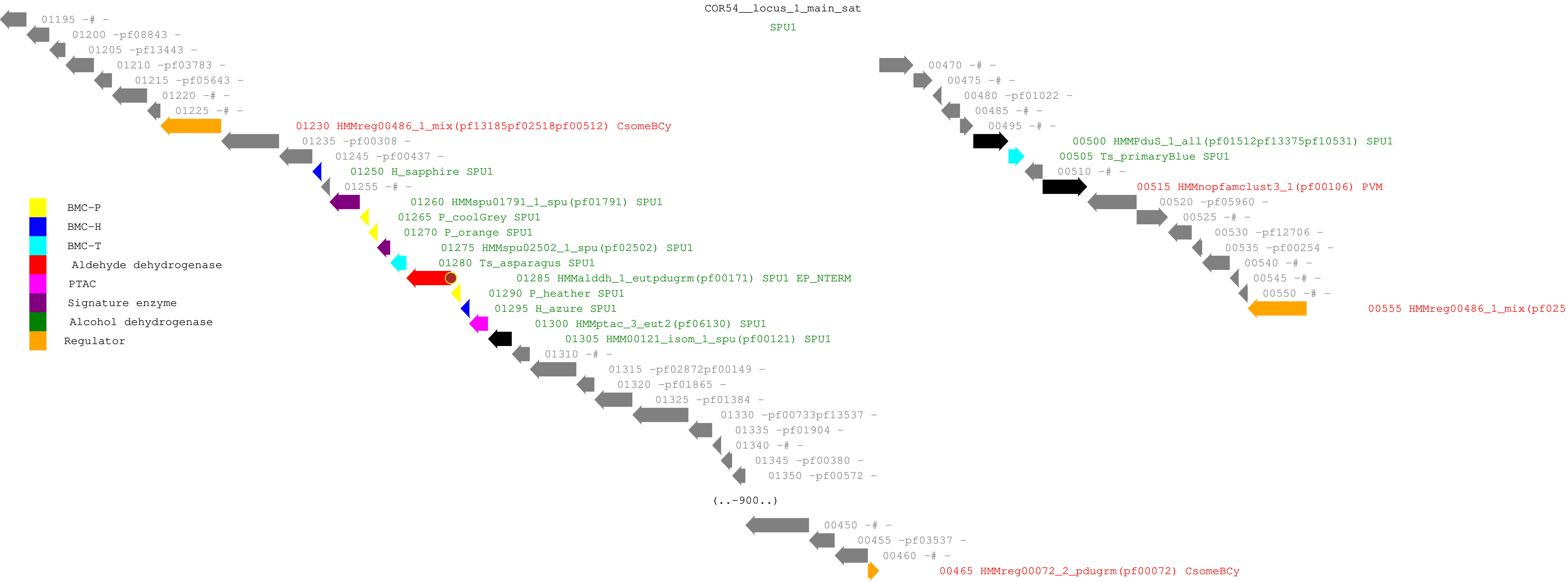

Elusimicrobia bacterium CG03_land_8_20_14_0_80_50_18. Bacteria; Elusimicrobia.

COS41__locus_1

SPU1

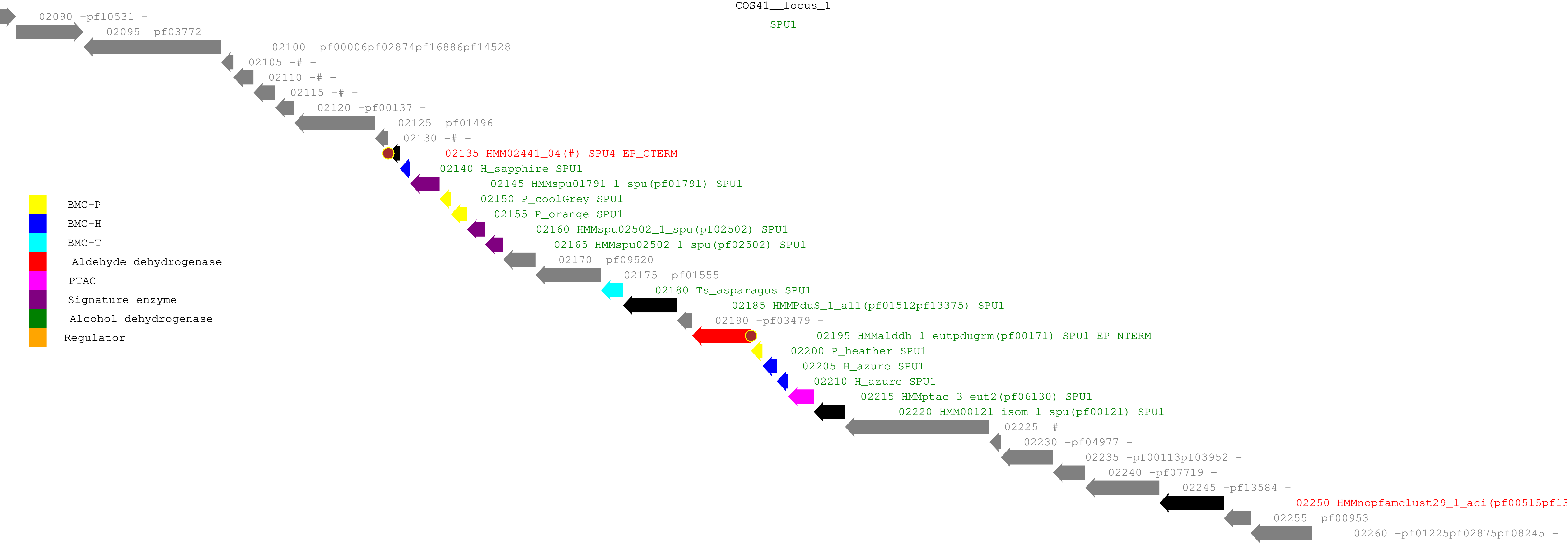

Elusimicrobia bacterium CG06_land_8_20_14_3_00_38_11. Bacteria; Elusimicrobia.

COS68__locus_1_main_sat

SPU1

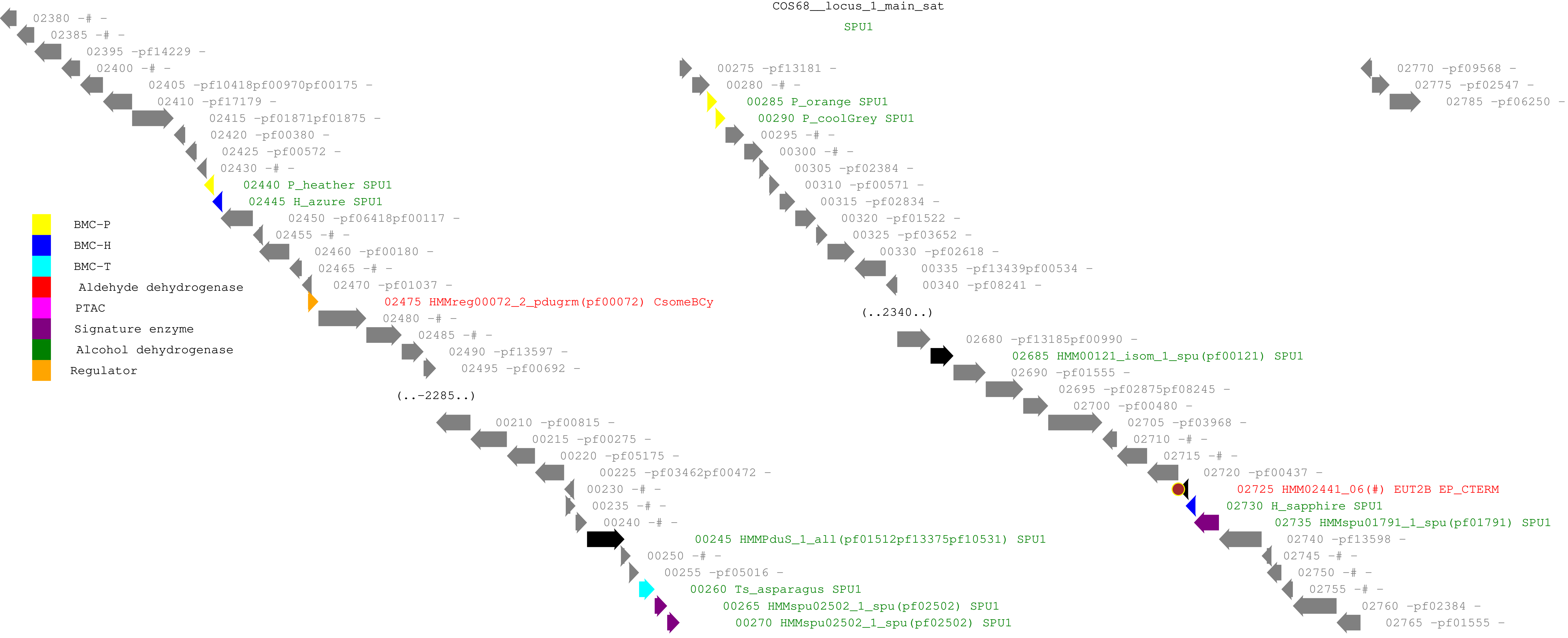

Elusimicrobia bacterium CG08_land_8_20_14_0_20_44_26. Bacteria; Elusimicrobia.

COT16__locus_1

SPU1

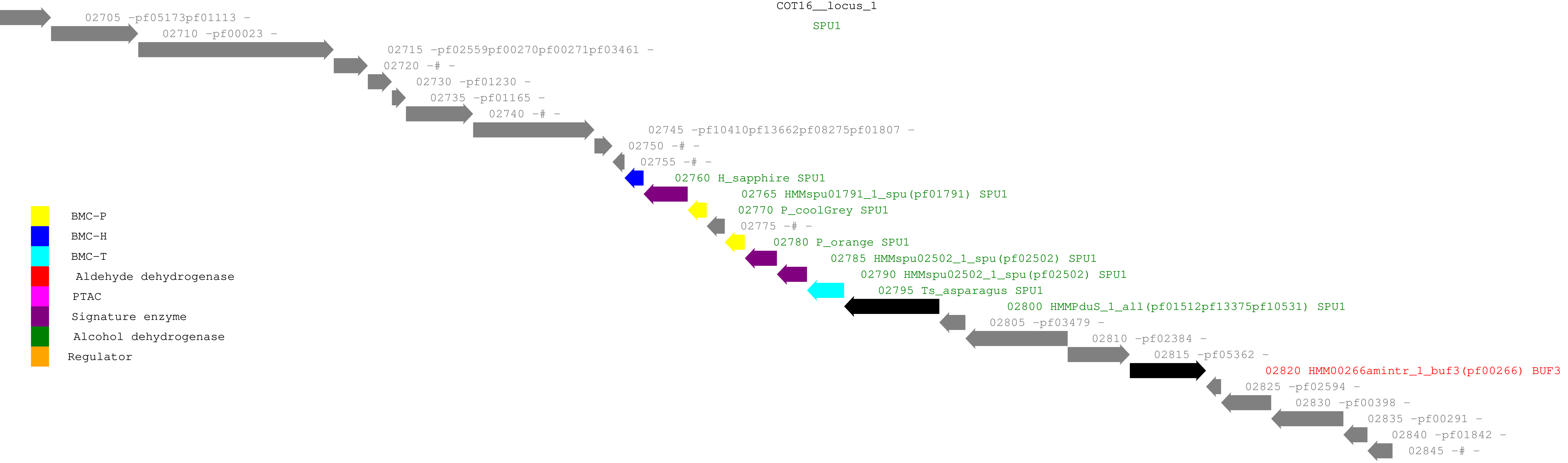

Elusimicrobia bacterium CG08_land_8_20_14_0_20_51_18. Bacteria; Elusimicrobia.

COT17__locus_fragmented

SPU1

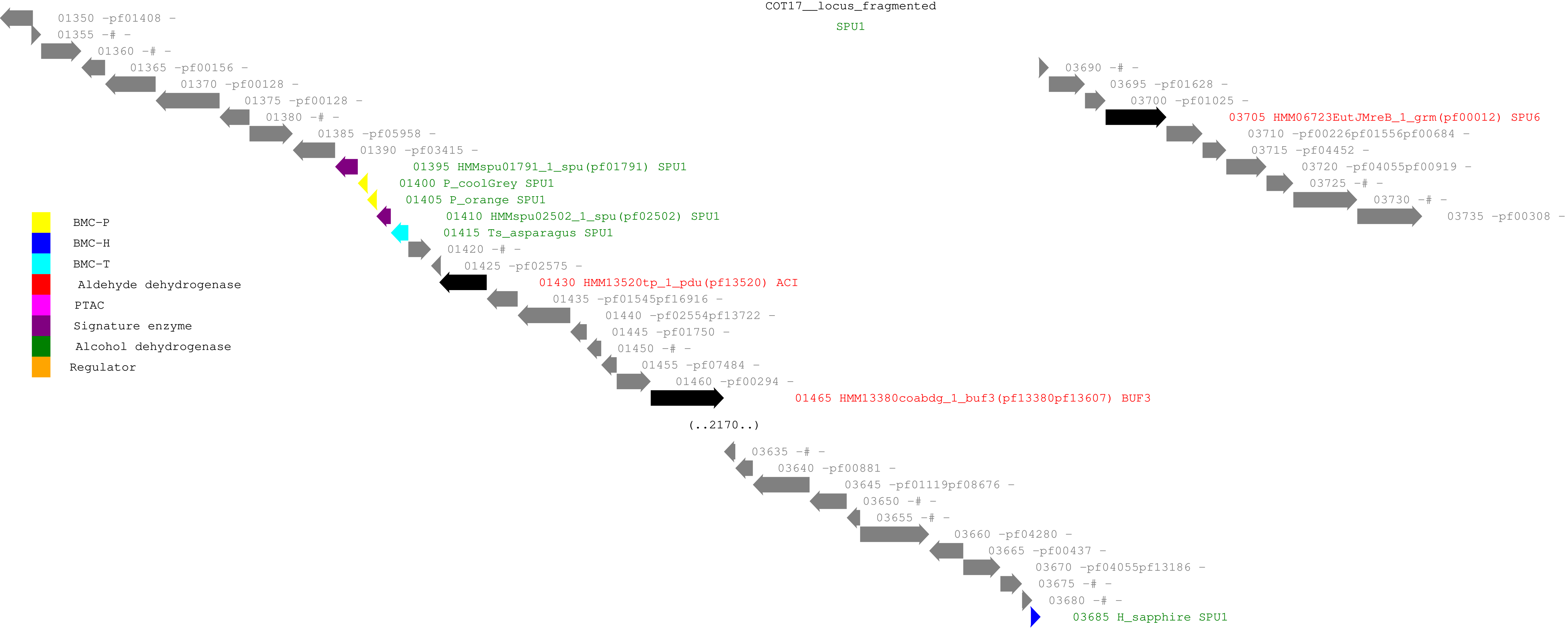

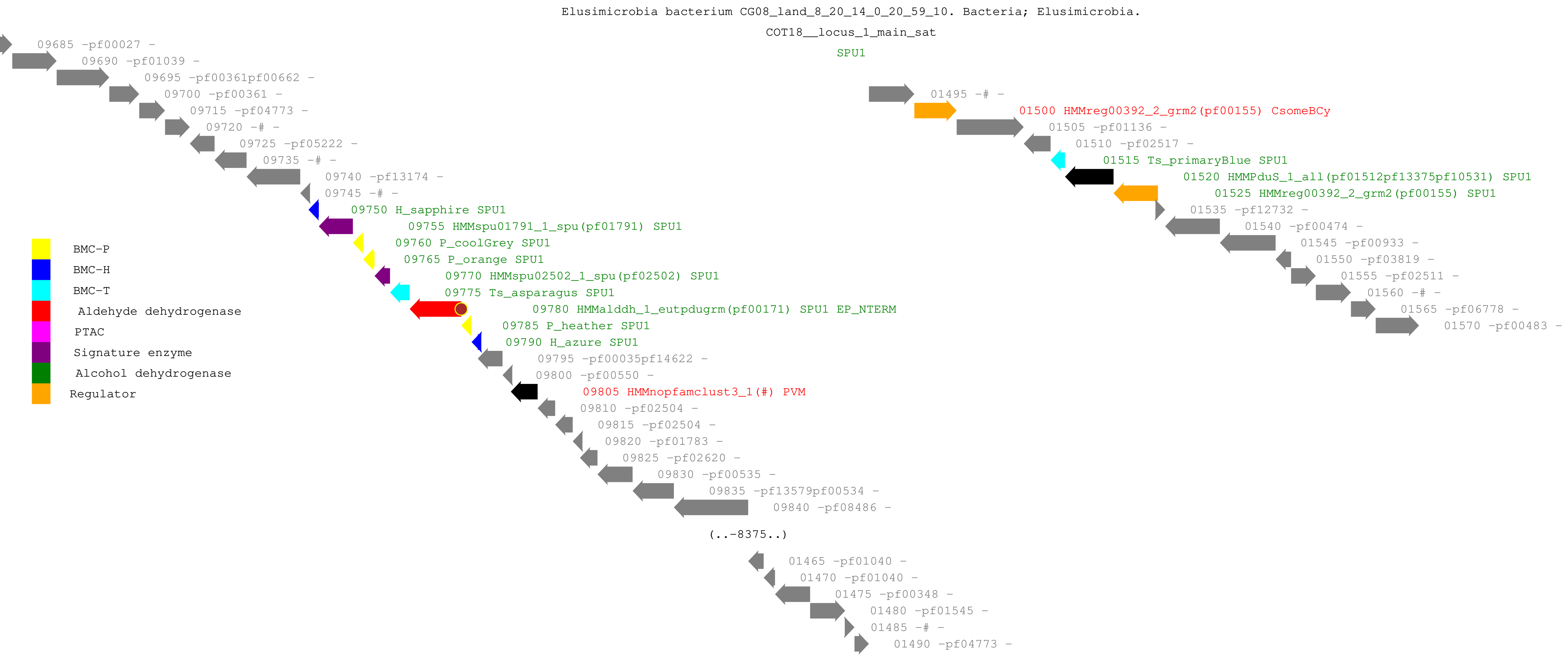
Elusimicrobia bacterium CG08_land_8_20_14_0_20_59_10. Bacteria; Elusimicrobia.
COT18__locus_1_main_sat
SPU1
09685 -pf00027 -
09690 -pf01039 -
09695 -pf00361pf00662 -
09700 -pf00361 -
09715 -pf04773 -
09720 -# -
09725 -pf05222 -
09735 -# -
09740 -pf13174 -
09745 -# -
09750 H_sapphire SPU1
09755 HMMspu01791_1_spu(pf01791) SPU1
09760 P_coolGrey SPU1
09765 P_orange SPU1
09770 HMMspu02502_1_spu(pf02502) SPU1
09775 Ts_asparagus SPU1
09780 HMMaldh_1_eutpdugrm(pf00171) SPU1 EP_NTERM
09785 P_heather SPU1
09790 H_azure SPU1
09795 -pf00035pf14622 -
09800 -pf00550 -
09805 HMMnopfamclust3_1(#) PVM
09810 -pf02504 -
09815 -pf02504 -
09820 -pf01783 -
09825 -pf02620 -
09830 -pf00535 -
09835 -pf13579pf00534 -
09840 -pf08486 -
(..-8375..)
01465 -pf01040 -
01470 -pf01040 -
01475 -pf00348 -
01480 -pf01545 -
01485 -# -
01490 -pf04773 -
01495 -# -
01500 HMMreg00392_2_grm2(pf00155) CsomeBCy
01505 -pf01136 -
01510 -pf02517 -
01515 Ts_primaryBlue SPU1
01520 HMMPduS_1_all(pf01512pf13375pf10531) SPU1
01525 HMMreg00392_2_grm2(pf00155) SPU1
01535 -pf12732 -
01540 -pf00474 -
01545 -pf00933 -
01550 -pf03819 -
01555 -pf02511 -
01560 -# -
01565 -pf06778 -
01570 -pf00483 -
BMC-P
BMC-H
BMC-T
Aldehyde dehydrogenase
PTAC
Signature enzyme
Alcohol dehydrogenase
Regulator

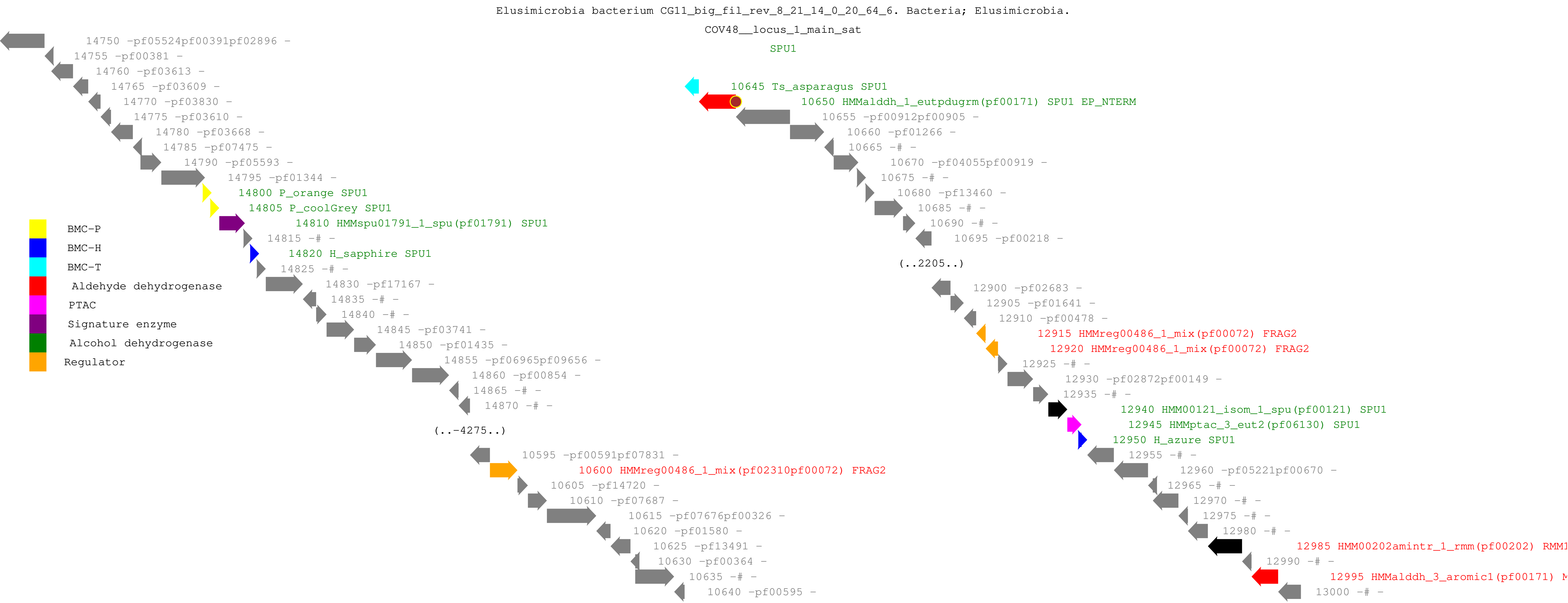

Elusimicrobia bacterium CG11_big_fil_rev_8_21_14_0_20_64_6. Bacteria; Elusimicrobia.
COV48__locus_1_main_sat
SPU1
14750 -pf05524pf00391pf02896 -
14755 -pf00381 -
14760 -pf03613 -
14765 -pf03609 -
14770 -pf03830 -
14775 -pf03610 -
14780 -pf03668 -
14785 -pf07475 -
14790 -pf05593 -
14795 -pf01344 -
14800 P_orange SPU1
14805 P_coolGrey SPU1
14810 HMMspu01791_1_spu(pf01791) SPU1
14815 -# -
14820 H_sapphire SPU1
14825 -# -
14830 -pf17167 -
14835 -# -
14840 -# -
14845 -pf03741 -
14850 -pf01435 -
14855 -pf06965pf09656 -
14860 -pf00854 -
14865 -# -
14870 -# -
(..-4275..)
10595 -pf00591pf07831 -
10600 HMMreg00486_1_mix(pf02310pf00072) FRAG2
10605 -pf14720 -
10610 -pf07687 -
10615 -pf07676pf00326 -
10620 -pf01580 -
10625 -pf13491 -
10630 -pf00364 -
10635 -# -
10640 -pf00595 -
BMC-P
BMC-H
BMC-T
Aldehyde dehydrogenase
PTAC
Signature enzyme
Alcohol dehydrogenase
Regulator
10645 Ts_asparagus SPU1
10650 HMMaldddh_1_eutpdugrm(pf00171) SPU1 EP_NTERM
10655 -pf00912pf00905 -
10660 -pf01266 -
10665 -# -
10670 -pf04055pf00919 -
10675 -# -
10680 -pf13460 -
10685 -# -
10690 -# -
10695 -pf00218 -
(..2205..)
12900 -pf02683 -
12905 -pf01641 -
12910 -pf00478 -
12915 HMMreg00486_1_mix(pf00072) FRAG2
12920 HMMreg00486_1_mix(pf00072) FRAG2
12925 -# -
12930 -pf02872pf00149 -
12935 -# -
12940 HMM00121_isom_1_spu(pf00121) SPU1
12945 HMMptac_3_eut2(pf06130) SPU1
12950 H_azure SPU1
12955 -# -
12960 -pf05221pf00670 -
12965 -# -
12970 -# -
12975 -# -
12980 -# -
12985 HMM00202amintr_1_rmm(pf00202) RMM1
12990 -# -
12995 HMMaldddh_3_aromic1(pf00171)
13000 -# -

Elusimicrobia bacterium CG_4_10_14_3_um_filter_49_12_50_7. Bacteria; Elusimicrobia.

COZ15__locus_1_main_sat

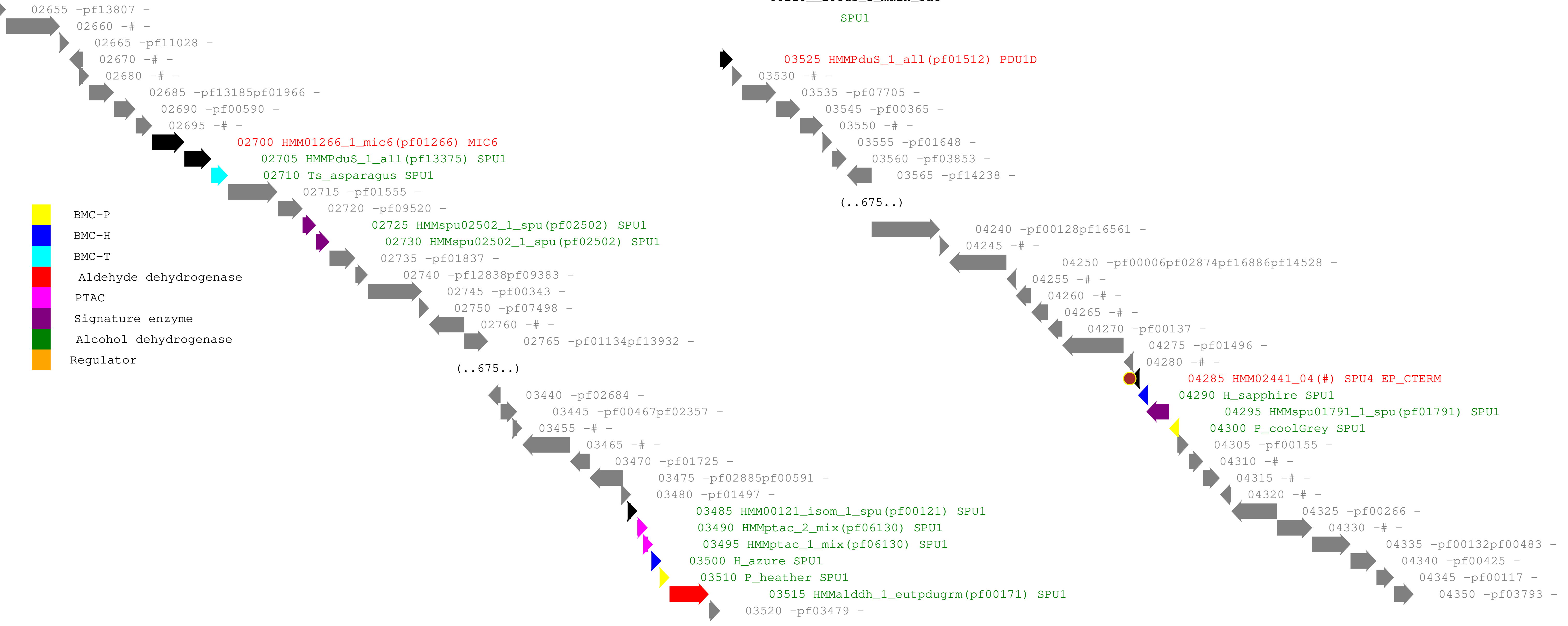

Elusimicrobia bacterium CG_4_8_14_3_um_filter_50_9. Bacteria; Elusimicrobia.

COZ72__locus_1

SPU1

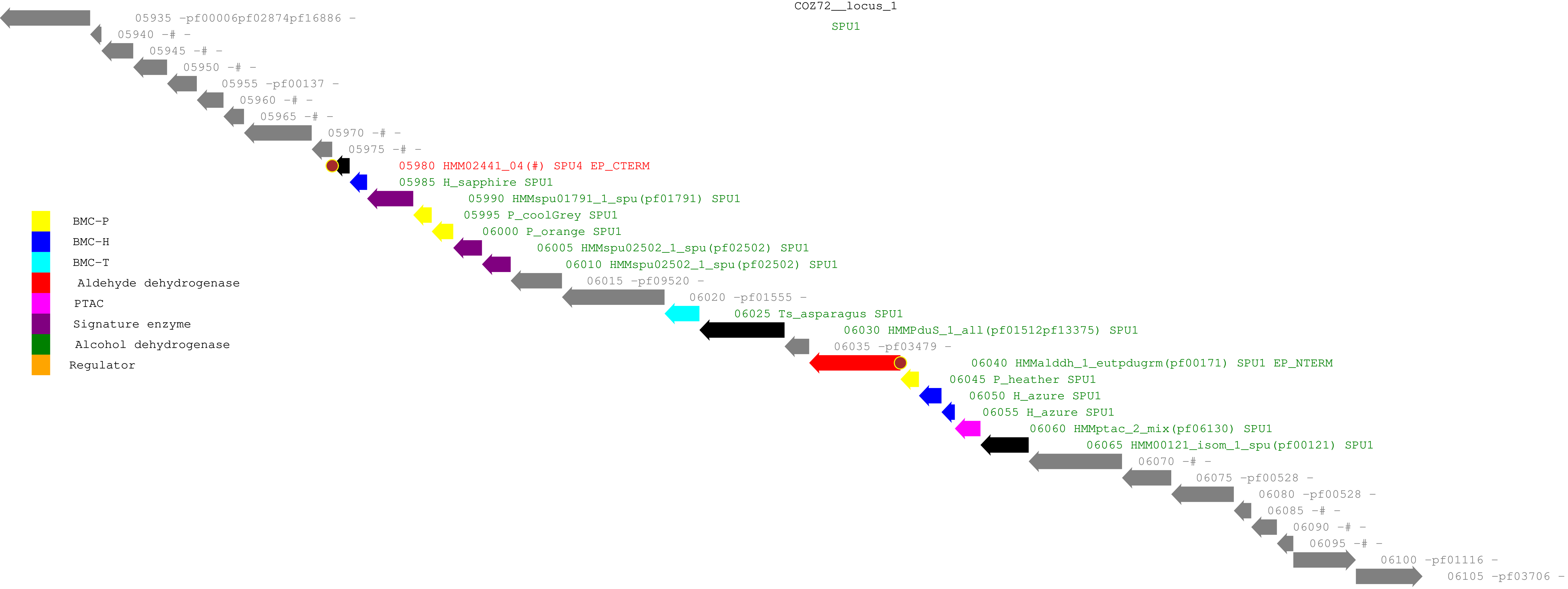

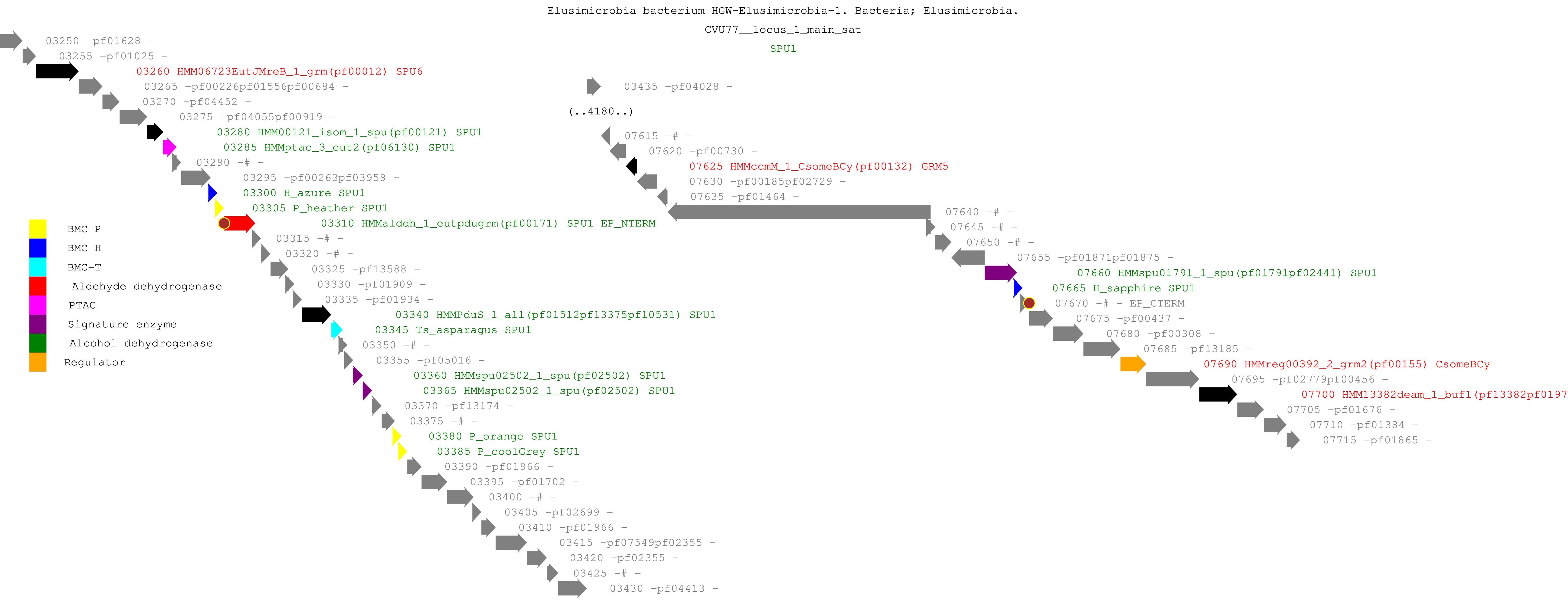

Elusimicrobia bacterium HGW-Elusimicrobia-1. Bacteria; Elusimicrobia.
CVU77__locus_1_main_sat
SPU1
03250 -pf01628 -
03255 -pf01025 -
03260 HMM06723EutJMreB_1_grm(pf00012) SPU6
03265 -pf00226pf01556pf00684 -
03270 -pf04452 -
03275 -pf04055pf00919 -
03280 HMM00121_isom_1_spu(pf00121) SPU1
03285 HMMptac_3_eut2(pf06130) SPU1
03290 -# -
03295 -pf00263pf03958 -
03300 H_azure SPU1
03305 P_heather SPU1
03310 HMMalddh_1_eutpdugrm(pf00171) SPU1 EP_NTERM
03315 -# -
03320 -# -
03325 -pf13588 -
03330 -pf01909 -
03335 -pf01934 -
03340 HMMPduS_1_all(pf01512pf13375pf10531) SPU1
03345 Ts_asparagus SPU1
03350 -# -
03355 -pf05016 -
03360 HMMspu02502_1_spu(pf02502) SPU1
03365 HMMspu02502_1_spu(pf02502) SPU1
03370 -pf13174 -
03375 -# -
03380 P_orange SPU1
03385 P_coolGrey SPU1
03390 -pf01966 -
03395 -pf01702 -
03400 -# -
03405 -pf02699 -
03410 -pf01966 -
03415 -pf07549pf02355 -
03420 -pf02355 -
03425 -# -
03430 -pf04413 -
03435 -pf04028 -
(..4180..)
07615 -# -
07620 -pf00730 -
07625 HMMccmM_1_CsomeBCy(pf00132) GRM5
07630 -pf00185pf02729 -
07635 -pf01464 -
07640 -# -
07645 -# -
07650 -# -
07655 -pf01871pf01875 -
07660 HMMspu01791_1_spu(pf01791pf02441) SPU1
07665 H_sapphire SPU1
07670 -# - EP_CTERM
07675 -pf00437 -
07680 -pf00308 -
07685 -pf13185 -
07690 HMMreg00392_2_grm2(pf00155) CsomeBCy
07695 -pf02779pf00456 -
07700 HMM13382deam_1_buf1(pf13382pf0197
07705 -pf01676 -
07710 -pf01384 -
07715 -pf01865 -
BMC-P
BMC-H
BMC-T
Aldehyde dehydrogenase
PTAC
Signature enzyme
Alcohol dehydrogenase
Regulator

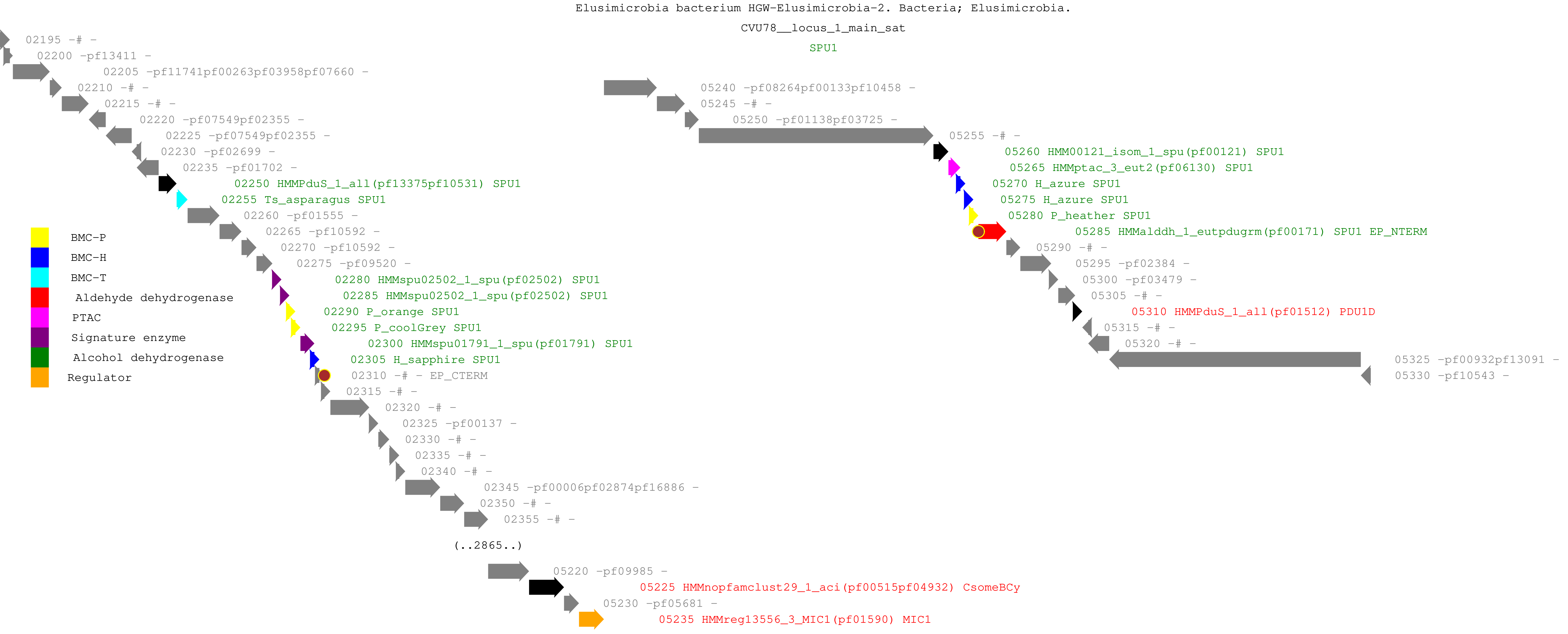
Elusimicrobia bacterium HGW-Elusimicrobia-2. Bacteria; Elusimicrobia.
CVU78__locus_1_main_sat
SPU1
02195 -# -
02200 -pf13411 -
02205 -pf11741pf00263pf03958pf07660 -
02210 -# -
02215 -# -
02220 -pf07549pf02355 -
02225 -pf07549pf02355 -
02230 -pf02699 -
02235 -pf01702 -
02250 HMMPduS_1_all(pf13375pf10531) SPU1
02255 Ts_asparagus SPU1
02260 -pf01555 -
02265 -pf10592 -
02270 -pf10592 -
02275 -pf09520 -
02280 HMMspu02502_1_spu(pf02502) SPU1
02285 HMMspu02502_1_spu(pf02502) SPU1
02290 P_orange SPU1
02295 P_coolGrey SPU1
02300 HMMspu01791_1_spu(pf01791) SPU1
02305 H_sapphire SPU1
02310 -# - EP_CTERM
02315 -# -
02320 -# -
02325 -pf00137 -
02330 -# -
02335 -# -
02340 -# -
02345 -pf00006pf02874pf16886 -
02350 -# -
02355 -# -
(..2865..)
05220 -pf09985 -
05225 HMMnopfamclust29_1_aci(pf00515pf04932) CsomeBCy
05230 -pf05681 -
05235 HMMreg13556_3_MIC1(pf01590) MIC1
05240 -pf08264pf00133pf10458 -
05245 -# -
05250 -pf01138pf03725 -
05255 -# -
05260 HMM00121_isom_1_spu(pf00121) SPU1
05265 HMMptac_3_eut2(pf06130) SPU1
05270 H_azure SPU1
05275 H_azure SPU1
05280 P_heather SPU1
05285 HMMalddh_1_eutpdugrm(pf00171) SPU1 EP_NTERM
05290 -# -
05295 -pf02384 -
05300 -pf03479 -
05305 -# -
05310 HMMPduS_1_all(pf01512) PDU1D
05315 -# -
05320 -# -
05325 -pf00932pf13091 -
05330 -pf10543 -
BMC-P
BMC-H
BMC-T
Aldehyde dehydrogenase
PTAC
Signature enzyme
Alcohol dehydrogenase
Regulator

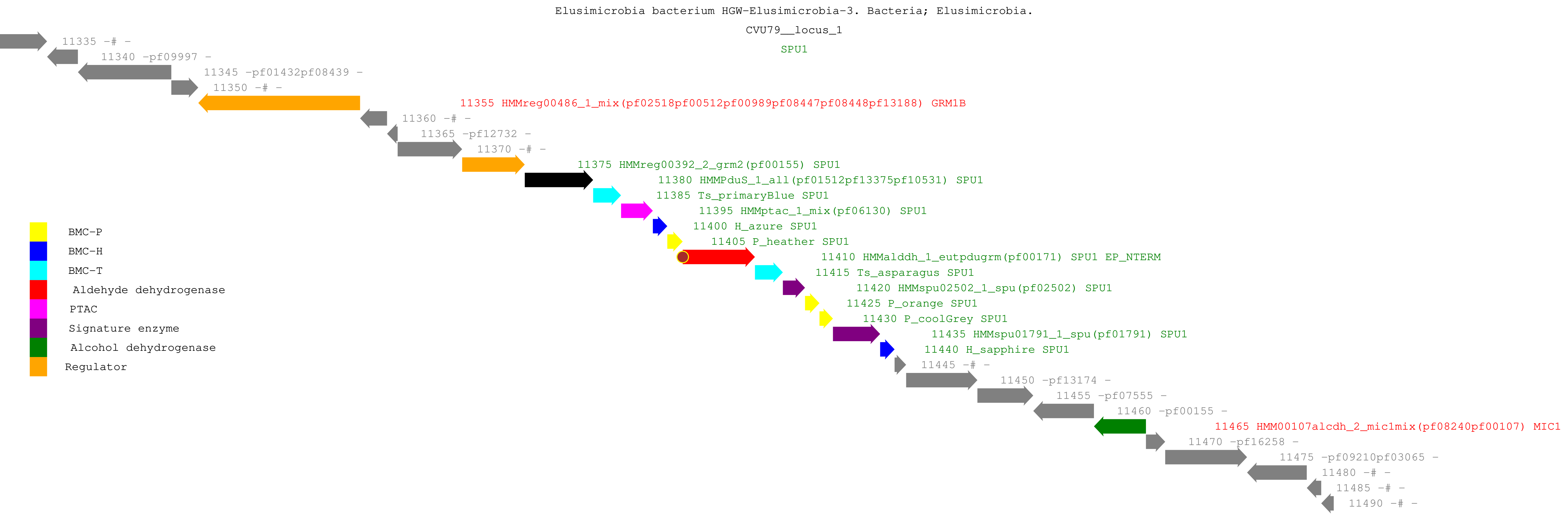

Elusimicrobia bacterium HGW-Elusimicrobia-3. Bacteria; Elusimicrobia.
CVU79__locus_1
SPU1
11335 -# -
11340 -pf09997 -
11345 -pf01432pf08439 -
11350 -# -
11355 HMMreg00486_1_mix(pf02518pf00512pf00989pf08447pf08448pf13188) GRM1B
11360 -# -
11365 -pf12732 -
11370 -# -
11375 HMMreg00392_2_grm2(pf00155) SPU1
11380 HMMPduS_1_all(pf01512pf13375pf10531) SPU1
11385 Ts_primaryBlue SPU1
11395 HMMptac_1_mix(pf06130) SPU1
11400 H_azure SPU1
11405 P_heather SPU1
11410 HMMaldh_1_eutpdugrm(pf00171) SPU1 EP_NTERM
11415 Ts_asparagus SPU1
11420 HMMspu02502_1_spu(pf02502) SPU1
11425 P_orange SPU1
11430 P_coolGrey SPU1
11435 HMMspu01791_1_spu(pf01791) SPU1
11440 H_sapphire SPU1
11445 -# -
11450 -pf13174 -
11455 -pf07555 -
11460 -pf00155 -
11465 HMM00107alcdh_2_mic1mix(pf08240pf00107) MIC1
11470 -pf16258 -
11475 -pf09210pf03065 -
11480 -# -
11485 -# -
11490 -# -
BMC-P
BMC-H
BMC-T
Aldehyde dehydrogenase
PTAC
Signature enzyme
Alcohol dehydrogenase
Regulator

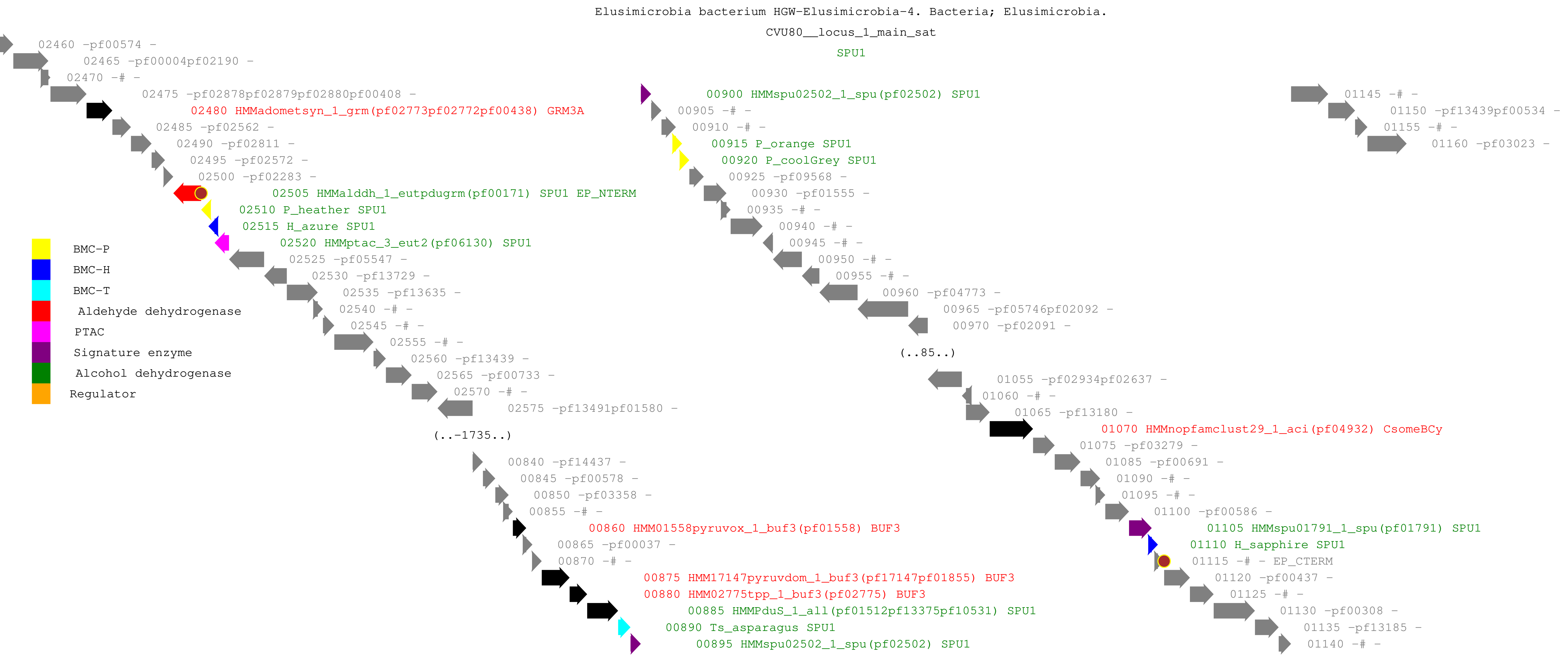
Elusimicrobia bacterium HGW-Elusimicrobia-4. Bacteria; Elusimicrobia.
CVU80__locus_1_main_sat
SPU1
02460 -pf00574 -
02465 -pf00004pf02190 -
02470 -# -
02475 -pf02878pf02879pf02880pf00408 -
02480 HMMadometsyn_1_grm(pf02773pf02772pf00438) GRM3A
02485 -pf02562 -
02490 -pf02811 -
02495 -pf02572 -
02500 -pf02283 -
02505 HMMalddh_1_eutpdugrm(pf00171) SPU1 EP_NTERM
02510 P_heather SPU1
02515 H_azure SPU1
02520 HMMptac_3_eut2(pf06130) SPU1
02525 -pf05547 -
02530 -pf13729 -
02535 -pf13635 -
02540 -# -
02545 -# -
02555 -# -
02560 -pf13439 -
02565 -pf00733 -
02570 -# -
02575 -pf13491pf01580 -
(..-1735..)
00840 -pf14437 -
00845 -pf00578 -
00850 -pf03358 -
00855 -# -
00860 HMM01558pyruvox_1_buf3(pf01558) BUF3
00865 -pf00037 -
00870 -# -
00875 HMM17147pyruvdom_1_buf3(pf17147pf01855) BUF3
00880 HMM02775tpp_1_buf3(pf02775) BUF3
00885 HMMPduS_1_all(pf01512pf13375pf10531) SPU1
00890 Ts_asparagus SPU1
00895 HMMspu02502_1_spu(pf02502) SPU1
00900 HMMspu02502_1_spu(pf02502) SPU1
00905 -# -
00910 -# -
00915 P_orange SPU1
00920 P_coolGrey SPU1
00925 -pf09568 -
00930 -pf01555 -
00935 -# -
00940 -# -
00945 -# -
00950 -# -
00955 -# -
00960 -pf04773 -
00965 -pf05746pf02092 -
00970 -pf02091 -
(..85..)
01055 -pf02934pf02637 -
01060 -# -
01065 -pf13180 -
01070 HMMnopfamclust29_1_aci(pf04932) CsomeBCy
01075 -pf03279 -
01085 -pf00691 -
01090 -# -
01095 -# -
01100 -pf00586 -
01105 HMMspu01791_1_spu(pf01791) SPU1
01110 H_sapphire SPU1
01115 -# - EP_CTERM
01120 -pf00437 -
01125 -# -
01130 -pf00308 -
01135 -pf13185 -
01140 -# -
01145 -# -
01150 -pf13439pf00534 -
01155 -# -
01160 -pf03023 -
BMC-P
BMC-H
BMC-T
Aldehyde dehydrogenase
PTAC
Signature enzyme
Alcohol dehydrogenase
Regulator

Elusimicrobia bacterium. Bacteria; Elusimicrobia.

DCL44__locus_1

SPU1

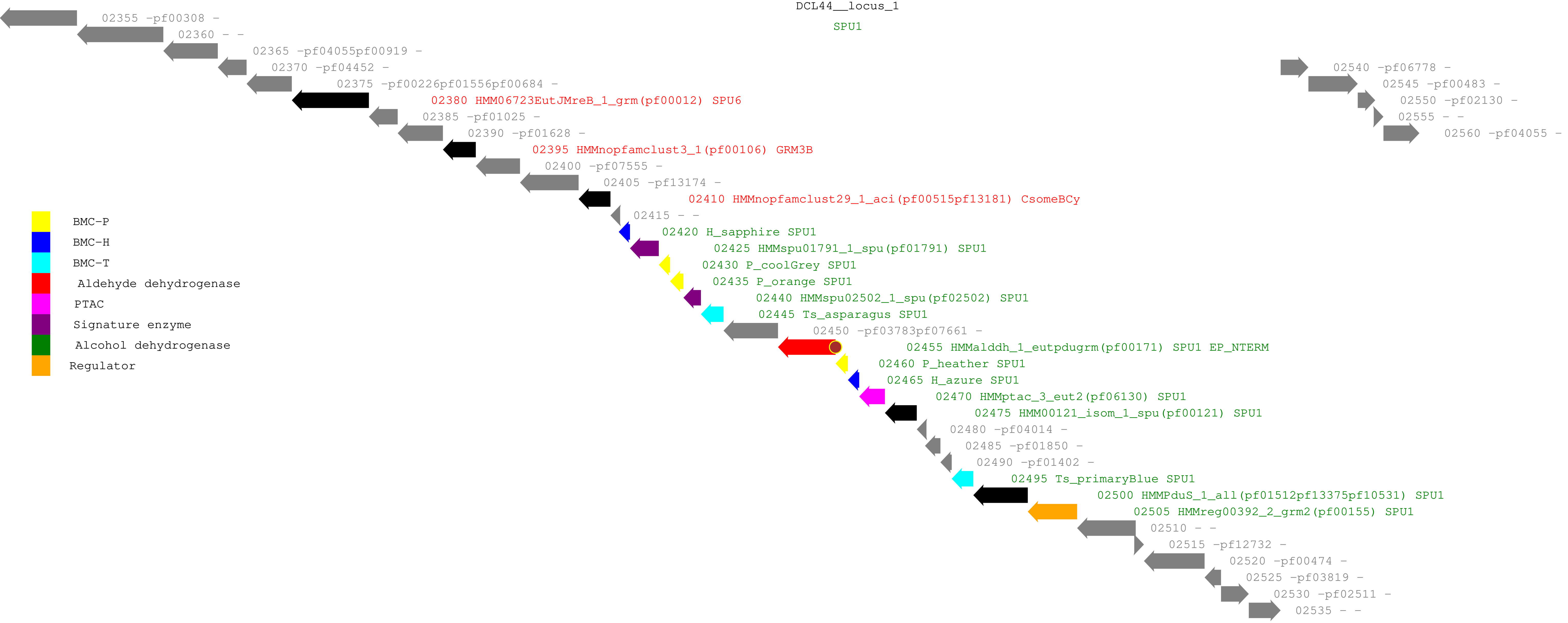

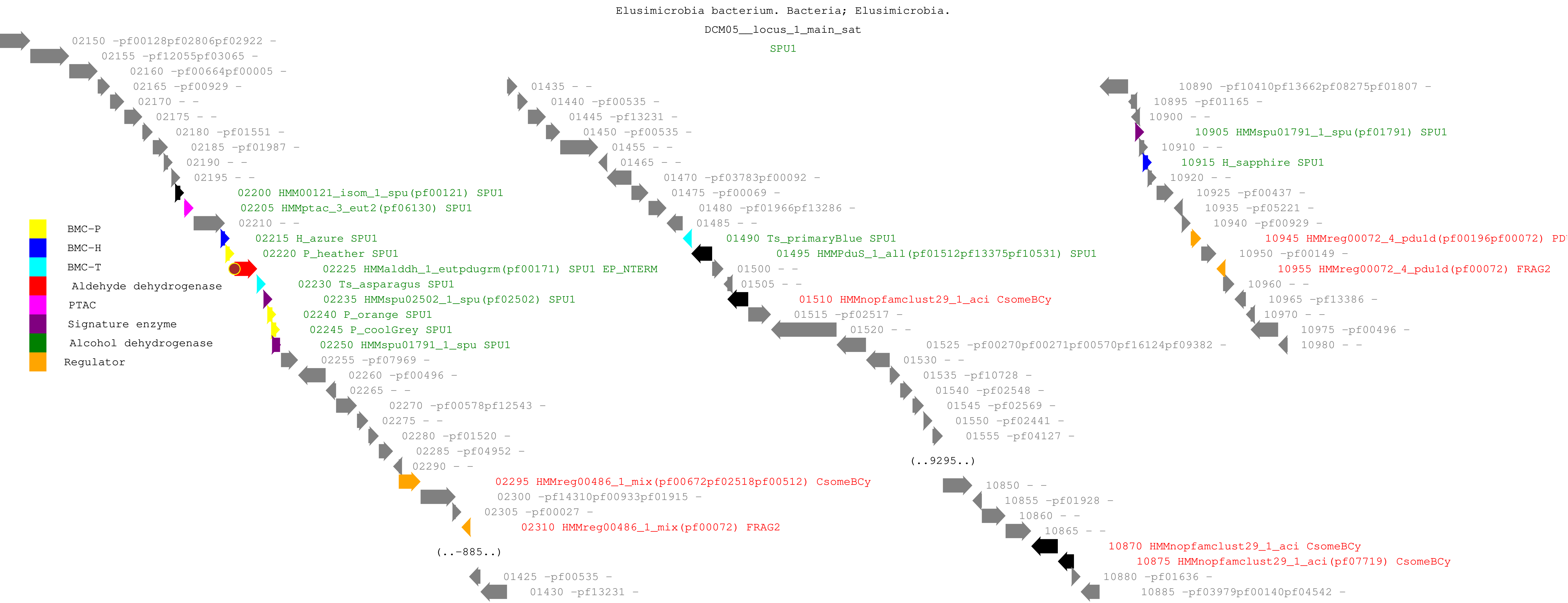

Elusimicrobia bacterium. Bacteria; Elusimicrobia.
DCM05__locus_1_main_sat
SPU1
02150 -pf00128pf02806pf02922 -
02155 -pf12055pf03065 -
02160 -pf00664pf00005 -
02165 -pf00929 -
02170 - -
02175 - -
02180 -pf01551 -
02185 -pf01987 -
02190 - -
02195 - -
02200 HMM00121_isom_1_spu(pf00121) SPU1
02205 HMMptac_3_eut2(pf06130) SPU1
02210 - -
02215 H_azure SPU1
02220 P_heather SPU1
02225 HMMalddh_1_eutpdugrm(pf00171) SPU1 EP_NTERM
02230 Ts_asparagus SPU1
02235 HMMspu02502_1_spu(pf02502) SPU1
02240 P_orange SPU1
02245 P_coolGrey SPU1
02250 HMMspu01791_1_spu SPU1
02255 -pf07969 -
02260 -pf00496 -
02265 - -
02270 -pf00578pf12543 -
02275 - -
02280 -pf01520 -
02285 -pf04952 -
02290 - -
02295 HMMreg00486_1_mix(pf00672pf02518pf00512) CsomeBCy
02300 -pf14310pf00933pf01915 -
02305 -pf00027 -
02310 HMMreg00486_1_mix(pf00072) FRAG2
(..-885..)
01425 -pf00535 -
01430 -pf13231 -
BMC-P
BMC-H
BMC-T
Aldehyde dehydrogenase
PTAC
Signature enzyme
Alcohol dehydrogenase
Regulator
01435 - -
01440 -pf00535 -
01445 -pf13231 -
01450 -pf00535 -
01455 - -
01465 - -
01470 -pf03783pf00092 -
01475 -pf00069 -
01480 -pf01966pf13286 -
01485 - -
01490 Ts_primaryBlue SPU1
01495 HMMPduS_1_all(pf01512pf13375pf10531) SPU1
01500 - -
01505 - -
01510 HMMnopfamclust29_1_aci CsomeBCy
01515 -pf02517 -
01520 - -
01525 -pf00270pf00271pf00570pf16124pf09382 -
01530 - -
01535 -pf10728 -
01540 -pf02548 -
01545 -pf02569 -
01550 -pf02441 -
01555 -pf04127 -
(..9295..)
10850 - -
10855 -pf01928 -
10860 - -
10865 - -
10870 HMMnopfamclust29_1_aci CsomeBCy
10875 HMMnopfamclust29_1_aci(pf07719) CsomeBCy
10880 -pf01636 -
10885 -pf03979pf00140pf04542 -
10890 -pf10410pf13662pf08275pf01807 -
10895 -pf01165 -
10900 - -
10905 HMMspu01791_1_spu(pf01791) SPU1
10910 - -
10915 H_sapphire SPU1
10920 - -
10925 -pf00437 -
10935 -pf05221 -
10940 -pf00929 -
10945 HMMreg00072_4_pdu1d(pf00196pf00072) PD
10950 -pf00149 -
10955 HMMreg00072_4_pdu1d(pf00072) FRAG2
10960 - -
10965 -pf13386 -
10970 - -
10975 -pf00496 -
10980 - -

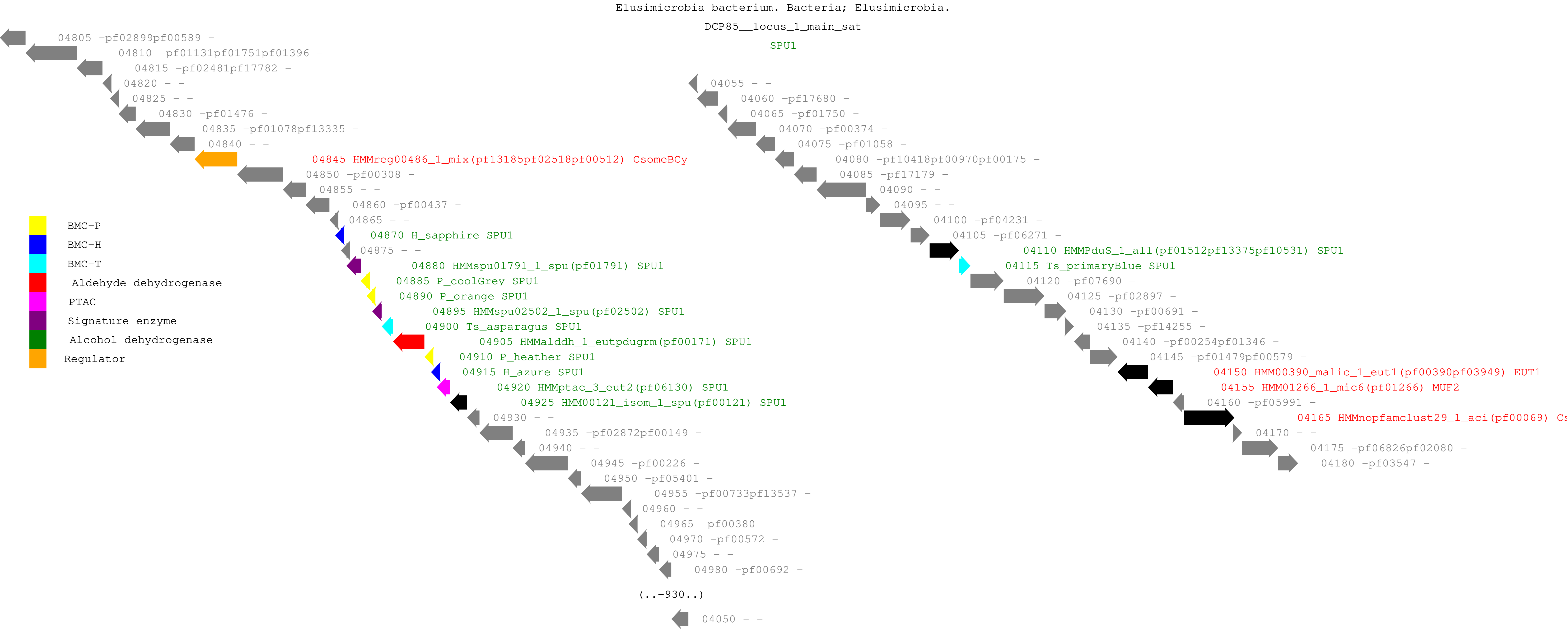
Elusimicrobia bacterium. Bacteria; Elusimicrobia.
DCP85__locus_1_main_sat
SPU1
04805 -pf02899pf00589 -
04810 -pf01131pf01751pf01396 -
04815 -pf02481pf17782 -
04820 - -
04825 - -
04830 -pf01476 -
04835 -pf01078pf13335 -
04840 - -
04845 HMMreg00486_1_mix(pf13185pf02518pf00512) CsomeBCy
04850 -pf00308 -
04855 - -
04860 -pf00437 -
04865 - -
04870 H_sapphire SPU1
04875 - -
04880 HMMspu01791_1_spu(pf01791) SPU1
04885 P_coolGrey SPU1
04890 P_orange SPU1
04895 HMMspu02502_1_spu(pf02502) SPU1
04900 Ts_asparagus SPU1
04905 HMMalddh_1_eutpdugrm(pf00171) SPU1
04910 P_heather SPU1
04915 H_azure SPU1
04920 HMMptac_3_eut2(pf06130) SPU1
04925 HMM00121_isom_1_spu(pf00121) SPU1
04930 - -
04935 -pf02872pf00149 -
04940 - -
04945 -pf00226 -
04950 -pf05401 -
04955 -pf00733pf13537 -
04960 - -
04965 -pf00380 -
04970 -pf00572 -
04975 - -
04980 -pf00692 -
(..-930..)
04050 - -
04055 - -
04060 -pf17680 -
04065 -pf01750 -
04070 -pf00374 -
04075 -pf01058 -
04080 -pf10418pf00970pf00175 -
04085 -pf17179 -
04090 - -
04095 - -
04100 -pf04231 -
04105 -pf06271 -
04110 HMMPduS_1_all(pf01512pf13375pf10531) SPU1
04115 Ts_primaryBlue SPU1
04120 -pf07690 -
04125 -pf02897 -
04130 -pf00691 -
04135 -pf14255 -
04140 -pf00254pf01346 -
04145 -pf01479pf00579 -
04150 HMM00390_malic_1_eut1(pf00390pf03949) EUT1
04155 HMM01266_1_mic6(pf01266) MUF2
04160 -pf05991 -
04165 HMMnopfamclust29_1_aci(pf00069) Cs
04170 - -
04175 -pf06826pf02080 -
04180 -pf03547 -
BMC-P
BMC-H
BMC-T
Aldehyde dehydrogenase
PTAC
Signature enzyme
Alcohol dehydrogenase
Regulator

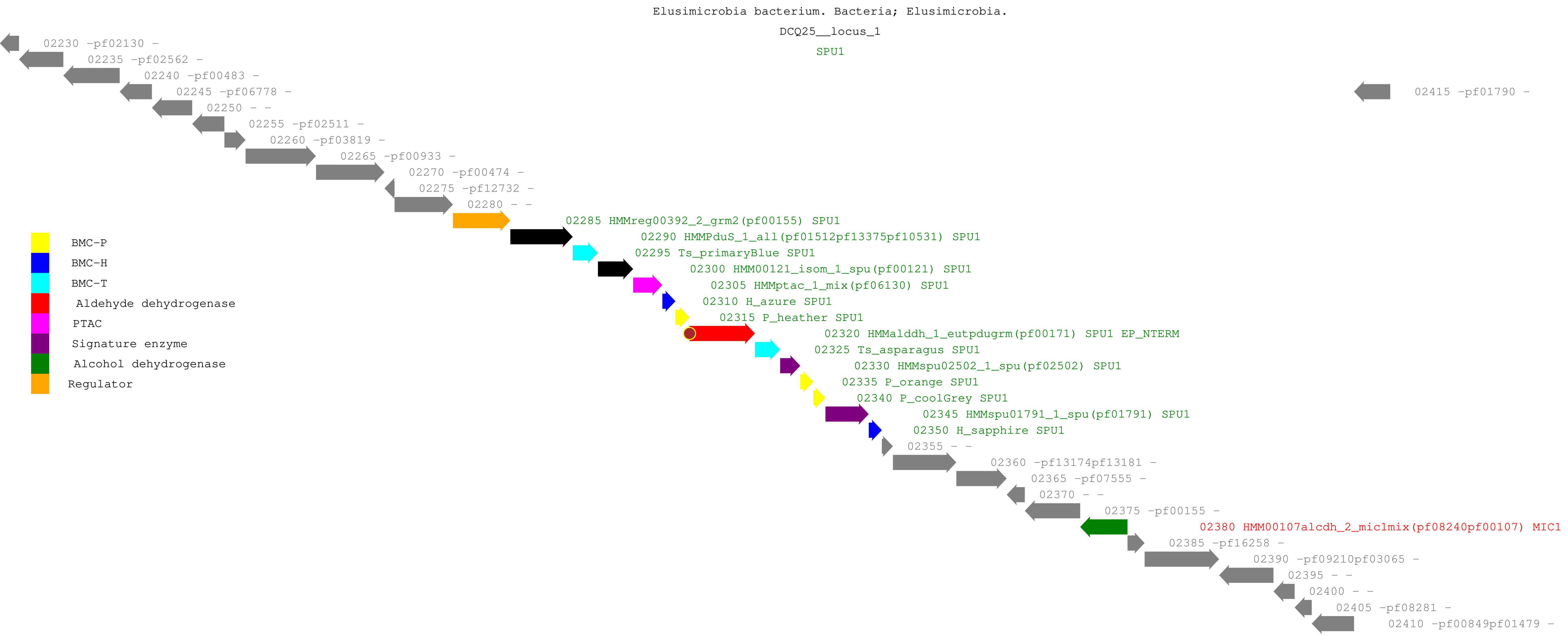
Elusimicrobia bacterium. Bacteria; Elusimicrobia.
DCQ25__locus_1
SPU1
02230 -pf02130 -
02235 -pf02562 -
02240 -pf00483 -
02245 -pf06778 -
02250 - -
02255 -pf02511 -
02260 -pf03819 -
02265 -pf00933 -
02270 -pf00474 -
02275 -pf12732 -
02280 - -
02285 HMMreg00392_2_grm2(pf00155) SPU1
02290 HMMPduS_1_all(pf01512pf13375pf10531) SPU1
02295 Ts_primaryBlue SPU1
02300 HMM00121_isom_1_spu(pf00121) SPU1
02305 HMMptac_1_mix(pf06130) SPU1
02310 H_azure SPU1
02315 P_heather SPU1
02320 HMMalddh_1_eutpdugrm(pf00171) SPU1 EP_NTERM
02325 Ts_asparagus SPU1
02330 HMMspu02502_1_spu(pf02502) SPU1
02335 P_orange SPU1
02340 P_coolGrey SPU1
02345 HMMspu01791_1_spu(pf01791) SPU1
02350 H_sapphire SPU1
02355 - -
02360 -pf13174pf13181 -
02365 -pf07555 -
02370 - -
02375 -pf00155 -
02380 HMM00107alcdh_2_mic1mix(pf08240pf00107) MIC1
02385 -pf16258 -
02390 -pf09210pf03065 -
02395 - -
02400 - -
02405 -pf08281 -
02410 -pf00849pf01479 -
02415 -pf01790 -
BMC-P
BMC-H
BMC-T
Aldehyde dehydrogenase
PTAC
Signature enzyme
Alcohol dehydrogenase
Regulator

Elusimicrobia bacterium. Bacteria; Elusimicrobia.

DCS63__locus_1

SPU1

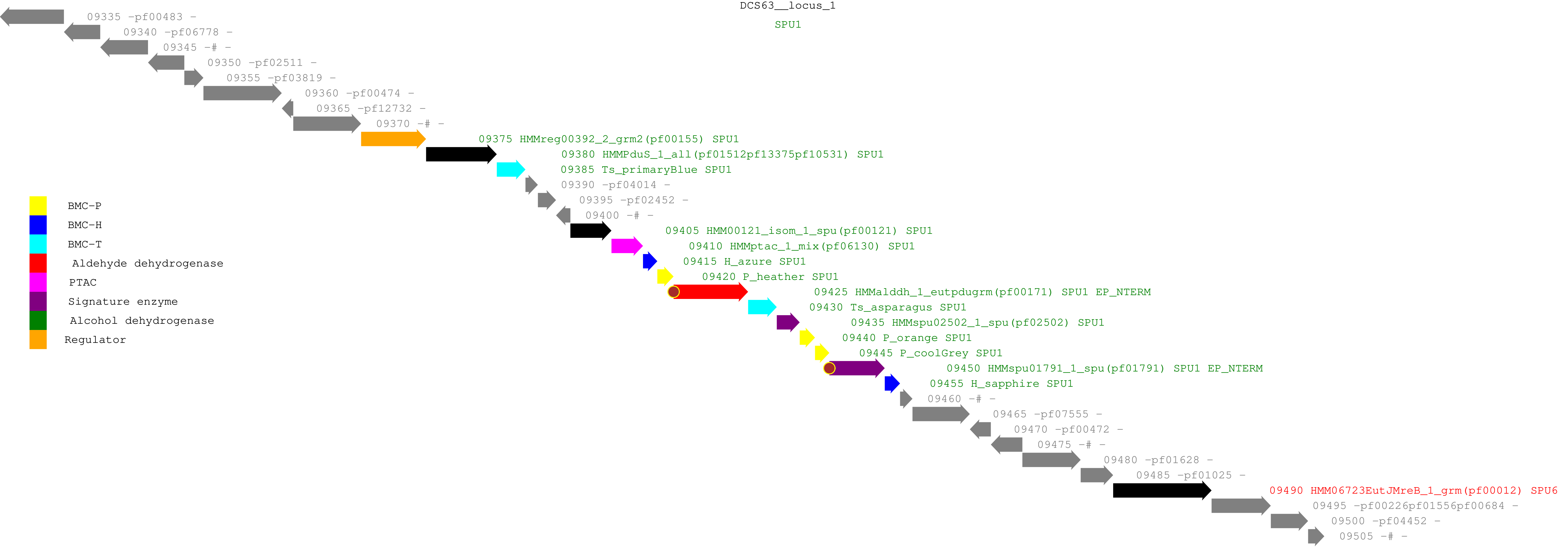

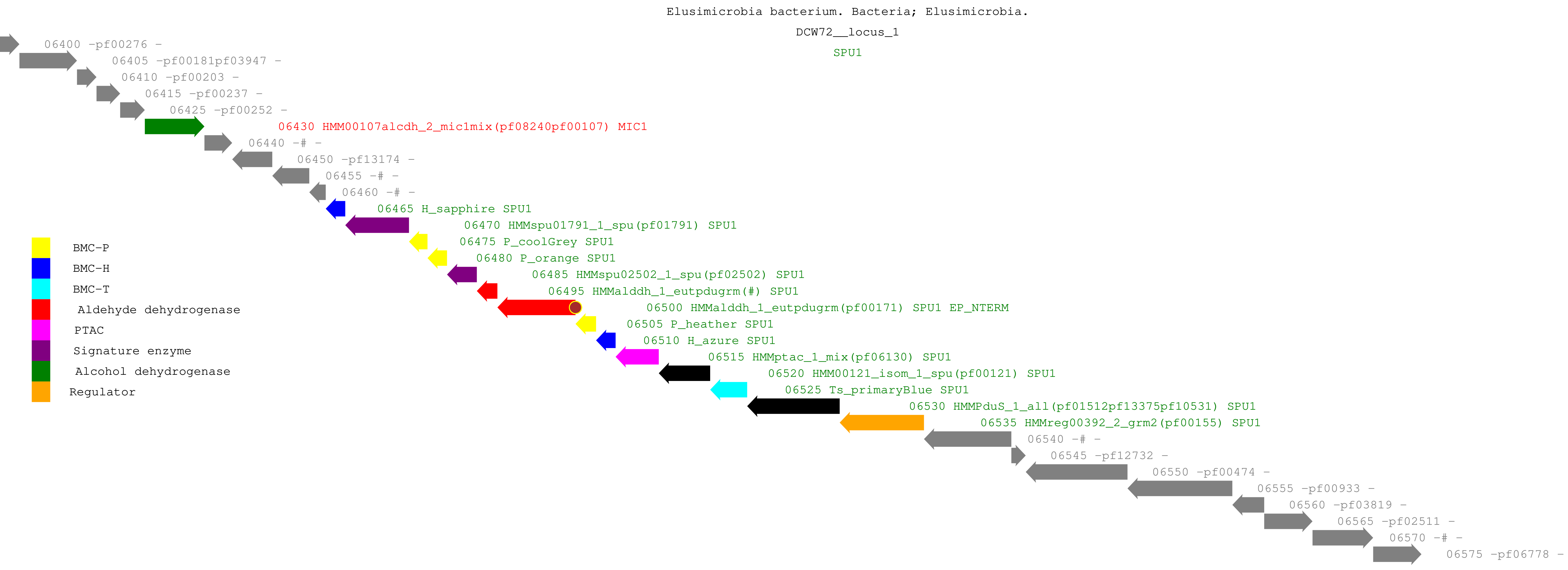
Elusimicrobia bacterium. Bacteria; Elusimicrobia.
DCW72__locus_1
SPU1
06400 -pf00276 -
06405 -pf00181pf03947 -
06410 -pf00203 -
06415 -pf00237 -
06425 -pf00252 -
06430 HMM00107alcdh_2_mic1mix(pf08240pf00107) MIC1
06440 -# -
06450 -pf13174 -
06455 -# -
06460 -# -
06465 H_sapphire SPU1
06470 HMMspu01791_1_spu(pf01791) SPU1
06475 P_coolGrey SPU1
06480 P_orange SPU1
06485 HMMspu02502_1_spu(pf02502) SPU1
06495 HMMalddh_1_eutpdugrm(#) SPU1
06500 HMMalddh_1_eutpdugrm(pf00171) SPU1 EP_NTERM
06505 P_heather SPU1
06510 H_azure SPU1
06515 HMMptac_1_mix(pf06130) SPU1
06520 HMM00121_isom_1_spu(pf00121) SPU1
06525 Ts_primaryBlue SPU1
06530 HMMPduS_1_all(pf01512pf13375pf10531) SPU1
06535 HMMreg00392_2_grm2(pf00155) SPU1
06540 -# -
06545 -pf12732 -
06550 -pf00474 -
06555 -pf00933 -
06560 -pf03819 -
06565 -pf02511 -
06570 -# -
06575 -pf06778 -
BMC-P
BMC-H
BMC-T
Aldehyde dehydrogenase
PTAC
Signature enzyme
Alcohol dehydrogenase
Regulator

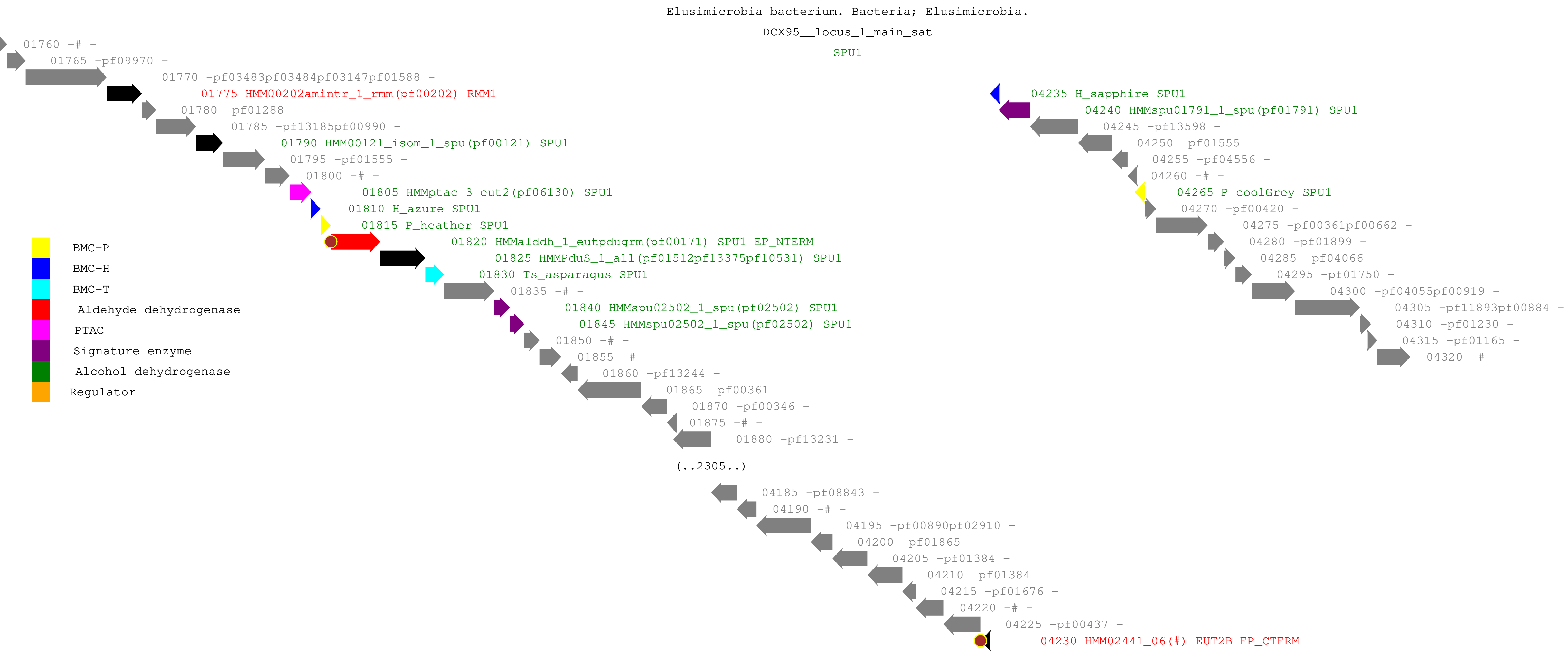
Elusimicrobia bacterium. Bacteria; Elusimicrobia.
DCX95__locus_1_main_sat
SPU1
01760 -# -
01765 -pf09970 -
01770 -pf03483pf03484pf03147pf01588 -
01775 HMM00202amintr_1_rmm(pf00202) RMM1
01780 -pf01288 -
01785 -pf13185pf00990 -
01790 HMM00121_isom_1_spu(pf00121) SPU1
01795 -pf01555 -
01800 -# -
01805 HMMptac_3_eut2(pf06130) SPU1
01810 H_azure SPU1
01815 P_heather SPU1
01820 HMMalddh_1_eutpdugrm(pf00171) SPU1 EP_NTERM
01825 HMMPduS_1_all(pf01512pf13375pf10531) SPU1
01830 Ts_asparagus SPU1
01835 -# -
01840 HMMspu02502_1_spu(pf02502) SPU1
01845 HMMspu02502_1_spu(pf02502) SPU1
01850 -# -
01855 -# -
01860 -pf13244 -
01865 -pf00361 -
01870 -pf00346 -
01875 -# -
01880 -pf13231 -
(..2305..)
04185 -pf08843 -
04190 -# -
04195 -pf00890pf02910 -
04200 -pf01865 -
04205 -pf01384 -
04210 -pf01384 -
04215 -pf01676 -
04220 -# -
04225 -pf00437 -
04230 HMM02441_06(#) EUT2B EP_CTERM
04235 H_sapphire SPU1
04240 HMMspu01791_1_spu(pf01791) SPU1
04245 -pf13598 -
04250 -pf01555 -
04255 -pf04556 -
04260 -# -
04265 P_coolGrey SPU1
04270 -pf00420 -
04275 -pf00361pf00662 -
04280 -pf01899 -
04285 -pf04066 -
04295 -pf01750 -
04300 -pf04055pf00919 -
04305 -pf11893pf00884 -
04310 -pf01230 -
04315 -pf01165 -
04320 -# -
BMC-P
BMC-H
BMC-T
Aldehyde dehydrogenase
PTAC
Signature enzyme
Alcohol dehydrogenase
Regulator

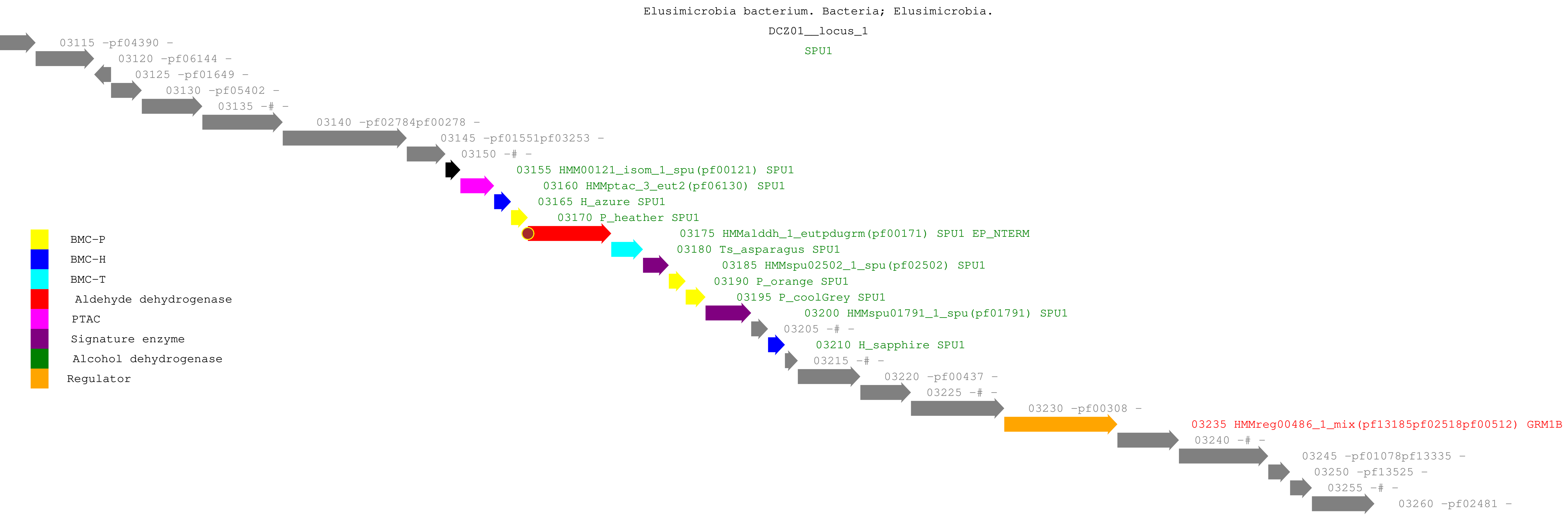
Elusimicrobia bacterium. Bacteria; Elusimicrobia.
DCZ01__locus_1
SPU1
03115 -pf04390 -
03120 -pf06144 -
03125 -pf01649 -
03130 -pf05402 -
03135 -# -
03140 -pf02784pf00278 -
03145 -pf01551pf03253 -
03150 -# -
03155 HMM00121_isom_1_spu(pf00121) SPU1
03160 HMMptac_3_eut2(pf06130) SPU1
03165 H_azure SPU1
03170 P_heather SPU1
03175 HMMalddh_1_eutpdugrm(pf00171) SPU1 EP_NTERM
03180 Ts_asparagus SPU1
03185 HMMspu02502_1_spu(pf02502) SPU1
03190 P_orange SPU1
03195 P_coolGrey SPU1
03200 HMMspu01791_1_spu(pf01791) SPU1
03205 -# -
03210 H_sapphire SPU1
03215 -# -
03220 -pf00437 -
03225 -# -
03230 -pf00308 -
03235 HMMreg00486_1_mix(pf13185pf02518pf00512) GRM1B
03240 -# -
03245 -pf01078pf13335 -
03250 -pf13525 -
03255 -# -
03260 -pf02481 -
BMC-P
BMC-H
BMC-T
Aldehyde dehydrogenase
PTAC
Signature enzyme
Alcohol dehydrogenase
Regulator

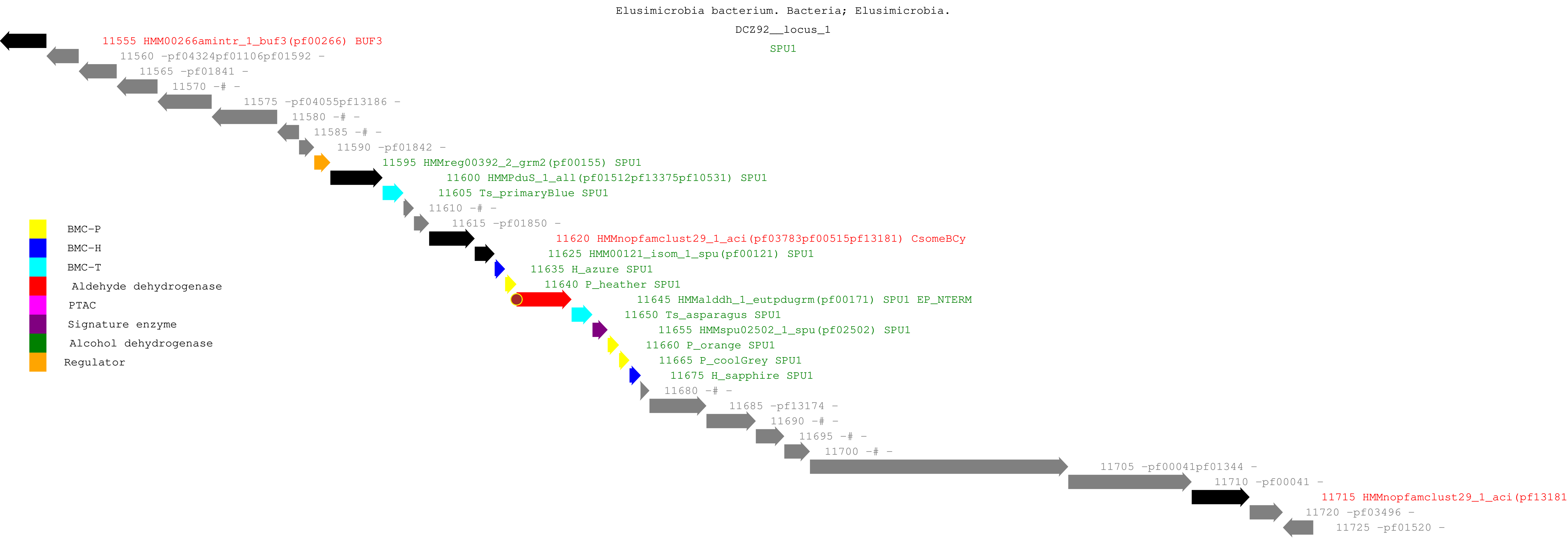
Elusimicrobia bacterium. Bacteria; Elusimicrobia.
DCZ92__locus_1
SPU1
11555 HMM00266amintr_1_buf3(pf00266) BUF3
11560 -pf04324pf01106pf01592 -
11565 -pf01841 -
11570 -# -
11575 -pf04055pf13186 -
11580 -# -
11585 -# -
11590 -pf01842 -
11595 HMMreg00392_2_grm2(pf00155) SPU1
11600 HMMPduS_1_all(pf01512pf13375pf10531) SPU1
11605 Ts_primaryBlue SPU1
11610 -# -
11615 -pf01850 -
11620 HMMnopfamclust29_1_aci(pf03783pf00515pf13181) CsomeBCy
11625 HMM00121_isom_1_spu(pf00121) SPU1
11635 H_azure SPU1
11640 P_heather SPU1
11645 HMMalddh_1_eutpdugrm(pf00171) SPU1 EP_NTERM
11650 Ts_asparagus SPU1
11655 HMMspu02502_1_spu(pf02502) SPU1
11660 P_orange SPU1
11665 P_coolGrey SPU1
11675 H_sapphire SPU1
11680 -# -
11685 -pf13174 -
11690 -# -
11695 -# -
11700 -# -
11705 -pf00041pf01344 -
11710 -pf00041 -
11715 HMMnopfamclust29_1_aci(pf13181
11720 -pf03496 -
11725 -pf01520 -
BMC-P
BMC-H
BMC-T
Aldehyde dehydrogenase
PTAC
Signature enzyme
Alcohol dehydrogenase
Regulator

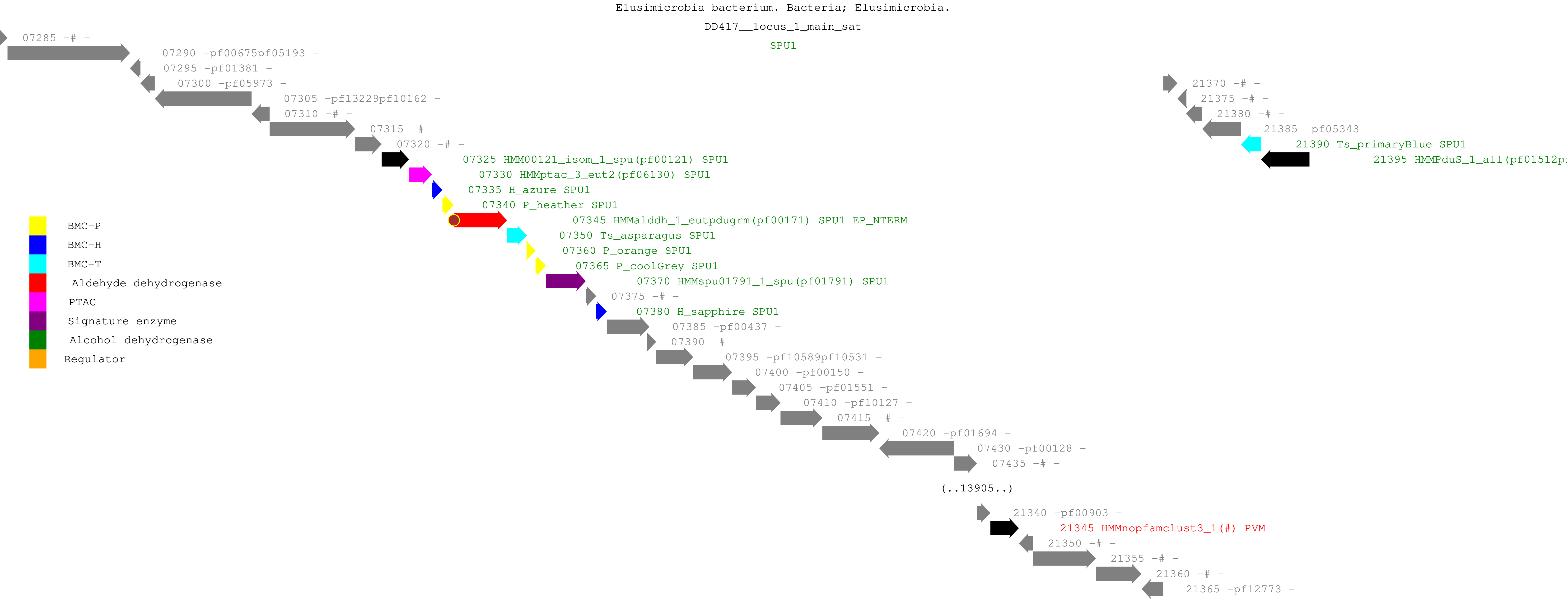
Elusimicrobia bacterium. Bacteria; Elusimicrobia.
DD417__locus_1_main_sat
SPU1
07285 -# -
07290 -pf00675pf05193 -
07295 -pf01381 -
07300 -pf05973 -
07305 -pf13229pf10162 -
07310 -# -
07315 -# -
07320 -# -
07325 HMM00121_isom_1_spu(pf00121) SPU1
07330 HMMptac_3_eut2(pf06130) SPU1
07335 H_azure SPU1
07340 P_heather SPU1
07345 HMMalddh_1_eutpdugrm(pf00171) SPU1 EP_NTERM
07350 Ts_asparagus SPU1
07360 P_orange SPU1
07365 P_coolGrey SPU1
07370 HMMspu01791_1_spu(pf01791) SPU1
07375 -# -
07380 H_sapphire SPU1
07385 -pf00437 -
07390 -# -
07395 -pf10589pf10531 -
07400 -pf00150 -
07405 -pf01551 -
07410 -pf10127 -
07415 -# -
07420 -pf01694 -
07430 -pf00128 -
07435 -# -
(..13905..)
21340 -pf00903 -
21345 HMMnopfamclust3_1(#) PVM
21350 -# -
21355 -# -
21360 -# -
21365 -pf12773 -
21370 -# -
21375 -# -
21380 -# -
21385 -pf05343 -
21390 Ts_primaryBlue SPU1
21395 HMMPduS_1_all(pf01512p
BMC-P
BMC-H
BMC-T
Aldehyde dehydrogenase
PTAC
Signature enzyme
Alcohol dehydrogenase
Regulator

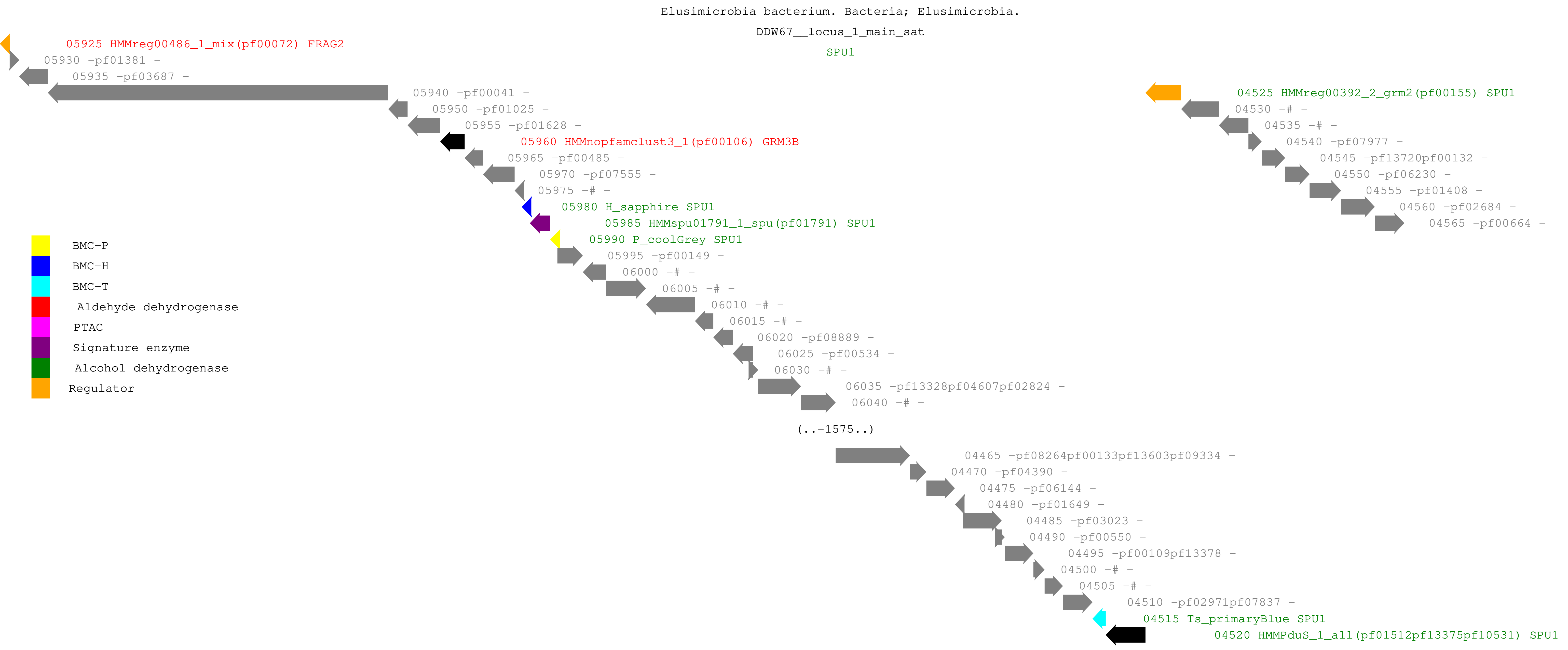
Elusimicrobia bacterium. Bacteria; Elusimicrobia.
DDW67__locus_1_main_sat
SPU1
05925 HMMreg00486_1_mix(pf00072) FRAG2
05930 -pf01381 -
05935 -pf03687 -
05940 -pf00041 -
05950 -pf01025 -
05955 -pf01628 -
05960 HMMnopfamclust3_1(pf00106) GRM3B
05965 -pf00485 -
05970 -pf07555 -
05975 -# -
05980 H_sapphire SPU1
05985 HMMspu01791_1_spu(pf01791) SPU1
05990 P_coolGrey SPU1
05995 -pf00149 -
06000 -# -
06005 -# -
06010 -# -
06015 -# -
06020 -pf08889 -
06025 -pf00534 -
06030 -# -
06035 -pf13328pf04607pf02824 -
06040 -# -
(..-1575..)
04465 -pf08264pf00133pf13603pf09334 -
04470 -pf04390 -
04475 -pf06144 -
04480 -pf01649 -
04485 -pf03023 -
04490 -pf00550 -
04495 -pf00109pf13378 -
04500 -# -
04505 -# -
04510 -pf02971pf07837 -
04515 Ts_primaryBlue SPU1
04520 HMMPduS_1_all(pf01512pf13375pf10531) SPU1
04525 HMMreg00392_2_grm2(pf00155) SPU1
04530 -# -
04535 -# -
04540 -pf07977 -
04545 -pf13720pf00132 -
04550 -pf06230 -
04555 -pf01408 -
04560 -pf02684 -
04565 -pf00664 -
BMC-P
BMC-H
BMC-T
Aldehyde dehydrogenase
PTAC
Signature enzyme
Alcohol dehydrogenase
Regulator

Elusimicrobia bacterium. Bacteria; Elusimicrobia.

DEB40__locus_1_main_sat

SPU1

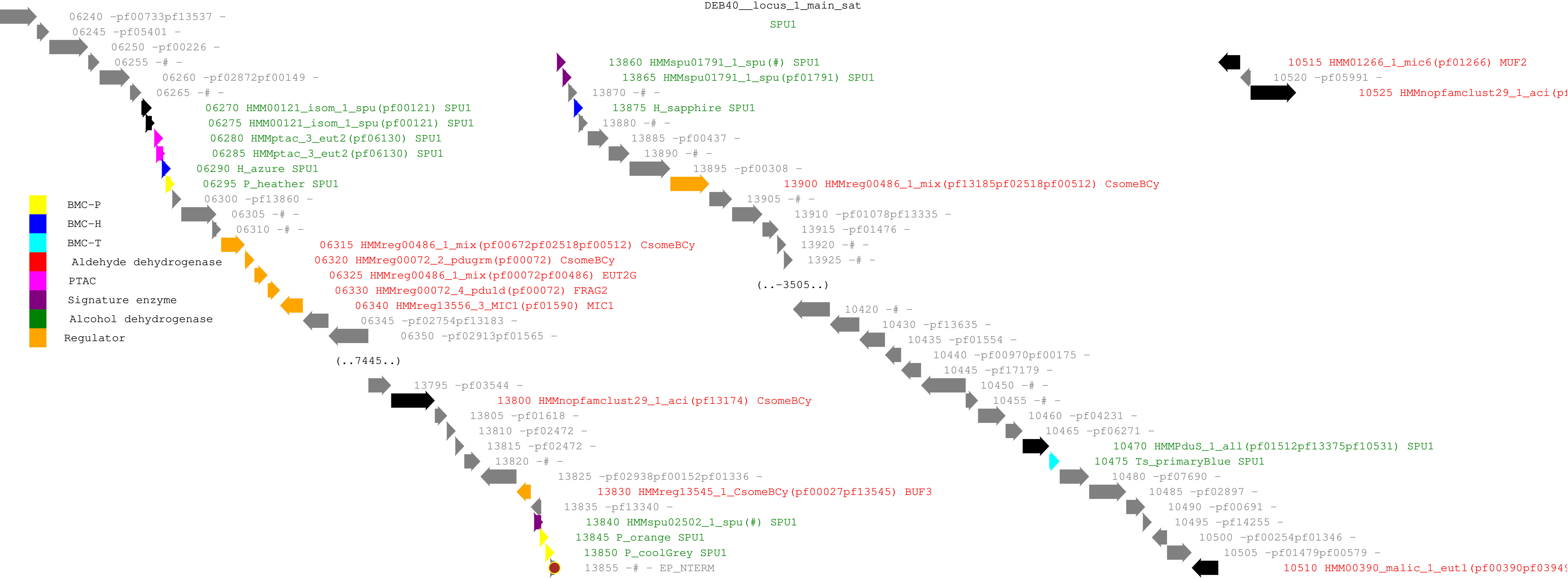

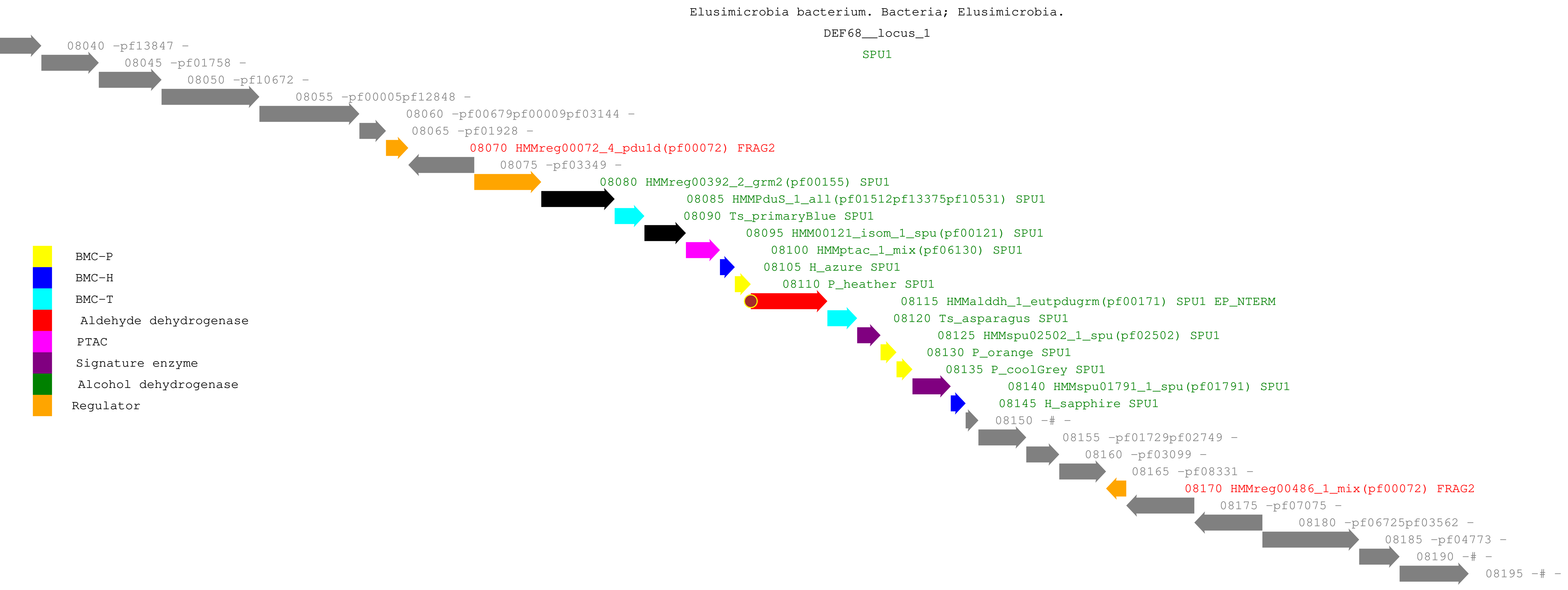
Elusimicrobia bacterium. Bacteria; Elusimicrobia.
DEF68__locus_1
SPU1
08040 -pf13847 -
08045 -pf01758 -
08050 -pf10672 -
08055 -pf00005pf12848 -
08060 -pf00679pf00009pf03144 -
08065 -pf01928 -
08070 HMMreg00072_4_pdu1d(pf00072) FRAG2
08075 -pf03349 -
08080 HMMreg00392_2_grm2(pf00155) SPU1
08085 HMMPduS_1_all(pf01512pf13375pf10531) SPU1
08090 Ts_primaryBlue SPU1
08095 HMM00121_isom_1_spu(pf00121) SPU1
08100 HMMptac_1_mix(pf06130) SPU1
08105 H_azure SPU1
08110 P_heather SPU1
08115 HMMaldh_1_eutpdugrm(pf00171) SPU1 EP_NTERM
08120 Ts_asparagus SPU1
08125 HMMspu02502_1_spu(pf02502) SPU1
08130 P_orange SPU1
08135 P_coolGrey SPU1
08140 HMMspu01791_1_spu(pf01791) SPU1
08145 H_sapphire SPU1
08150 -# -
08155 -pf01729pf02749 -
08160 -pf03099 -
08165 -pf08331 -
08170 HMMreg00486_1_mix(pf00072) FRAG2
08175 -pf07075 -
08180 -pf06725pf03562 -
08185 -pf04773 -
08190 -# -
08195 -# -
BMC-P
BMC-H
BMC-T
Aldehyde dehydrogenase
PTAC
Signature enzyme
Alcohol dehydrogenase
Regulator

Elusimicrobia bacterium. Bacteria; Elusimicrobia.

DEQ38__locus_1_main_sat

SPU1

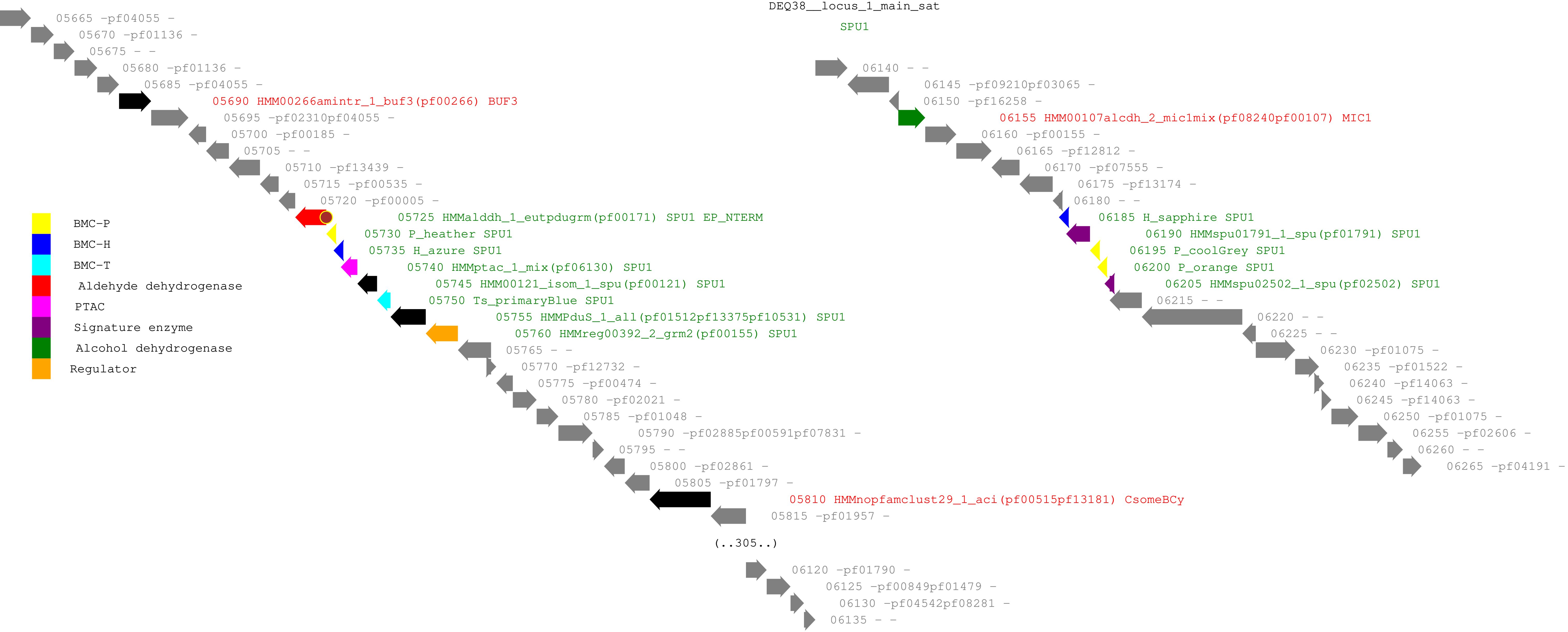

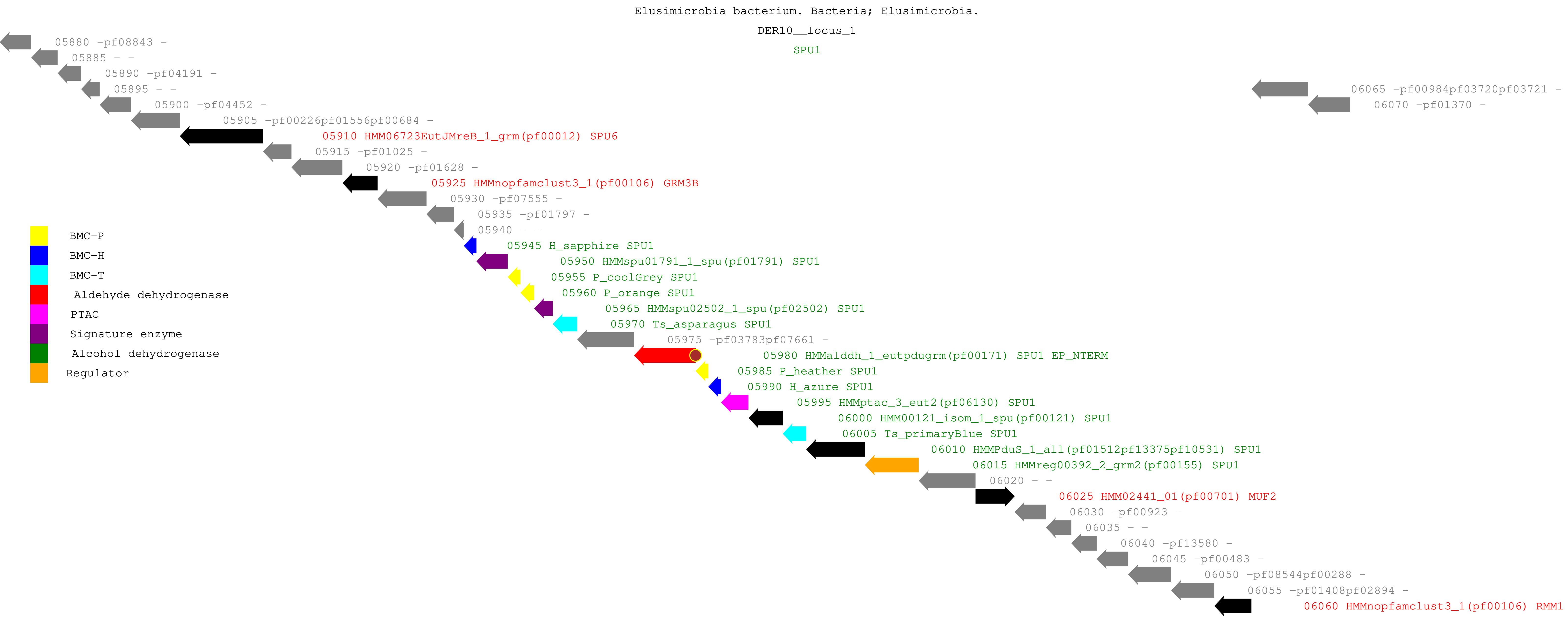
Elusimicrobia bacterium. Bacteria; Elusimicrobia.
DER10__locus_1
SPU1
05880 -pf08843 -
05885 - -
05890 -pf04191 -
05895 - -
05900 -pf04452 -
05905 -pf00226pf01556pf00684 -
05910 HMM06723EutJMreB_1_grm(pf00012) SPU6
05915 -pf01025 -
05920 -pf01628 -
05925 HMMnopfamclust3_1(pf00106) GRM3B
05930 -pf07555 -
05935 -pf01797 -
05940 - -
05945 H_sapphire SPU1
05950 HMMspu01791_1_spu(pf01791) SPU1
05955 P_coolGrey SPU1
05960 P_orange SPU1
05965 HMMspu02502_1_spu(pf02502) SPU1
05970 Ts_asparagus SPU1
05975 -pf03783pf07661 -
05980 HMMalddh_1_eutpdugrm(pf00171) SPU1 EP_NTERM
05985 P_heather SPU1
05990 H_azure SPU1
05995 HMMptac_3_eut2(pf06130) SPU1
06000 HMM00121_isom_1_spu(pf00121) SPU1
06005 Ts_primaryBlue SPU1
06010 HMMPduS_1_all(pf01512pf13375pf10531) SPU1
06015 HMMreg00392_2_grm2(pf00155) SPU1
06020 - -
06025 HMM02441_01(pf00701) MUF2
06030 -pf00923 -
06035 - -
06040 -pf13580 -
06045 -pf00483 -
06050 -pf08544pf00288 -
06055 -pf01408pf02894 -
06060 HMMnopfamclust3_1(pf00106) RMM1
06065 -pf00984pf03720pf03721 -
06070 -pf01370 -
BMC-P
BMC-H
BMC-T
Aldehyde dehydrogenase
PTAC
Signature enzyme
Alcohol dehydrogenase
Regulator

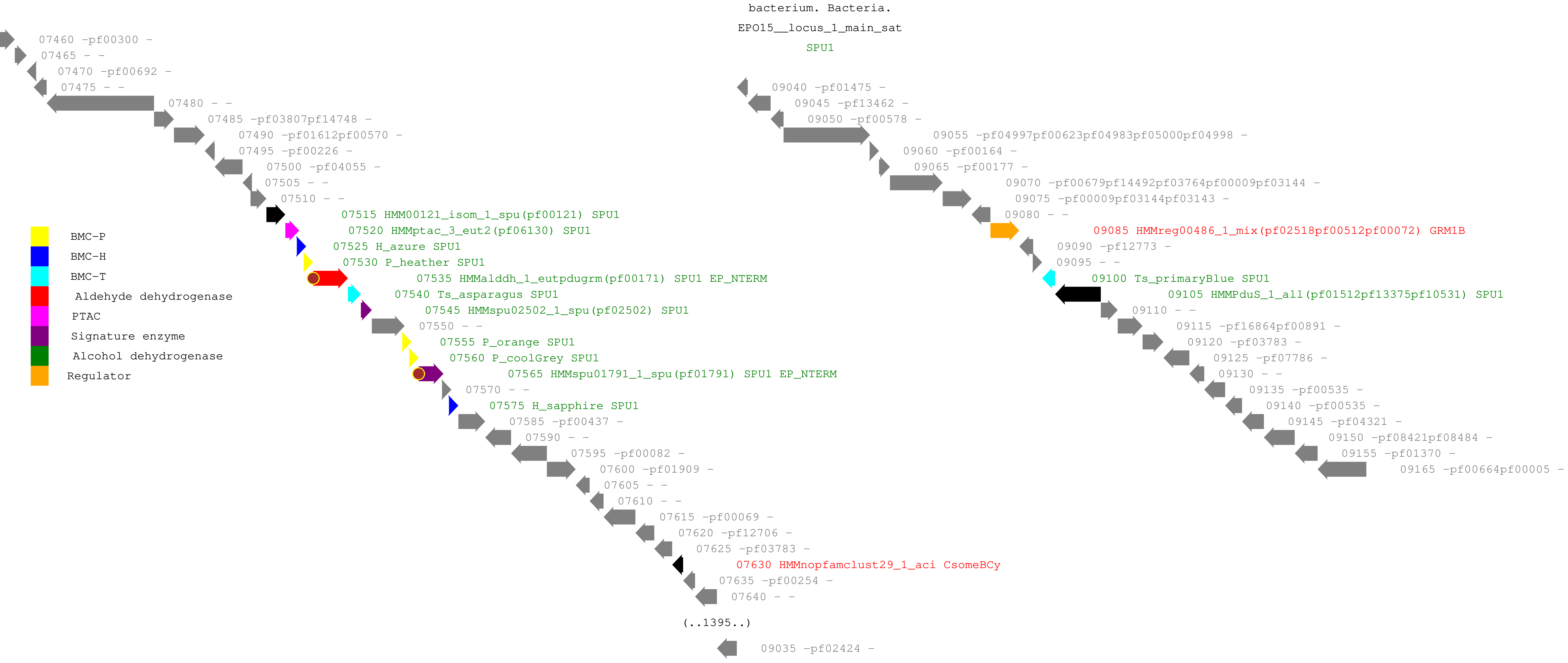

bacterium. Bacteria.
EPO15__locus_1_main_sat
SPU1
07460 -pf00300 -
07465 - -
07470 -pf00692 -
07475 - -
07480 - -
07485 -pf03807pf14748 -
07490 -pf01612pf00570 -
07495 -pf00226 -
07500 -pf04055 -
07505 - -
07510 - -
07515 HMM00121_isom_1_spu(pf00121) SPU1
07520 HMMptac_3_eut2(pf06130) SPU1
07525 H_azure SPU1
07530 P_heather SPU1
07535 HMMalddh_1_eutpdugrm(pf00171) SPU1 EP_NTERM
07540 Ts_asparagus SPU1
07545 HMMspu02502_1_spu(pf02502) SPU1
07550 - -
07555 P_orange SPU1
07560 P_coolGrey SPU1
07565 HMMspu01791_1_spu(pf01791) SPU1 EP_NTERM
07570 - -
07575 H_sapphire SPU1
07585 -pf00437 -
07590 - -
07595 -pf00082 -
07600 -pf01909 -
07605 - -
07610 - -
07615 -pf00069 -
07620 -pf12706 -
07625 -pf03783 -
07630 HMMnopfamclust29_1_aci CsomeBCy
07635 -pf00254 -
07640 - -
(..1395..)
09035 -pf02424 -
09040 -pf01475 -
09045 -pf13462 -
09050 -pf00578 -
09055 -pf04997pf00623pf04983pf05000pf04998 -
09060 -pf00164 -
09065 -pf00177 -
09070 -pf00679pf14492pf03764pf00009pf03144 -
09075 -pf00009pf03144pf03143 -
09080 - -
09085 HMMreg00486_1_mix(pf02518pf00512pf00072) GRM1B
09090 -pf12773 -
09095 - -
09100 Ts_primaryBlue SPU1
09105 HMMPduS_1_all(pf01512pf13375pf10531) SPU1
09110 - -
09115 -pf16864pf00891 -
09120 -pf03783 -
09125 -pf07786 -
09130 - -
09135 -pf00535 -
09140 -pf00535 -
09145 -pf04321 -
09150 -pf08421pf08484 -
09155 -pf01370 -
09165 -pf00664pf00005 -
BMC-P
BMC-H
BMC-T
Aldehyde dehydrogenase
PTAC
Signature enzyme
Alcohol dehydrogenase
Regulator

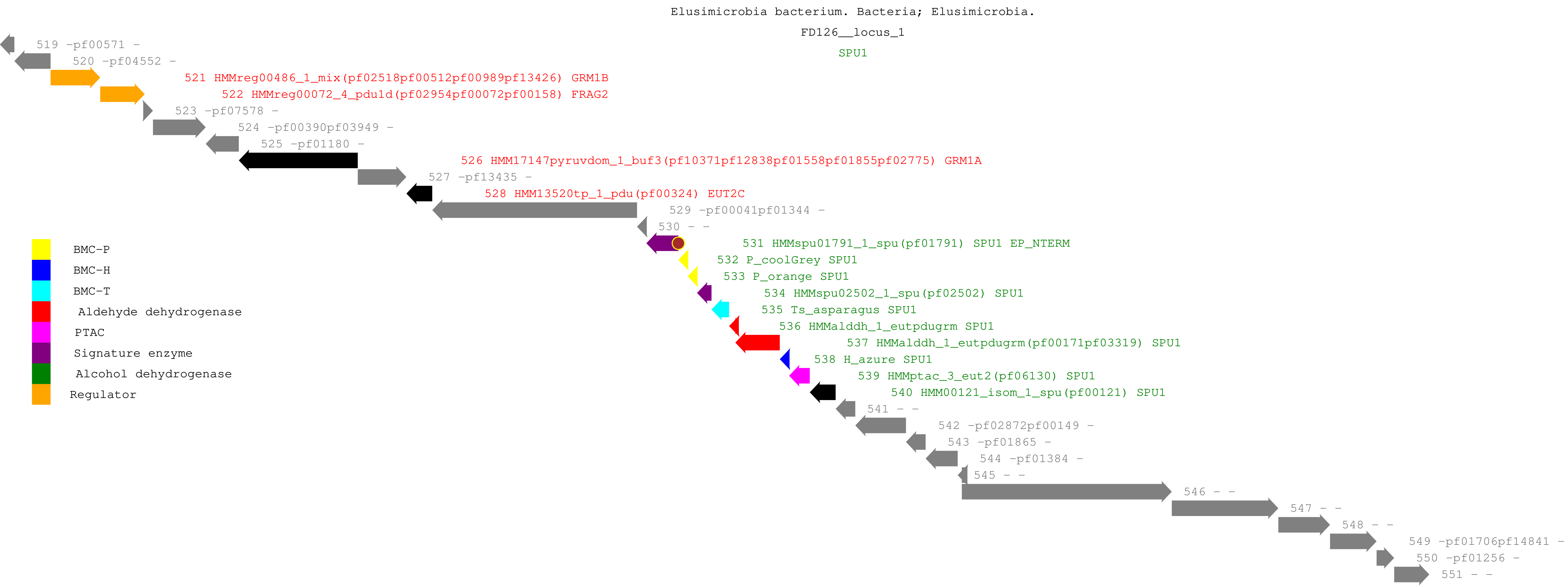
Elusimicrobia bacterium. Bacteria; Elusimicrobia.
FD126__locus_1
SPU1
519 -pf00571 -
520 -pf04552 -
521 HMMreg00486_1_mix(pf02518pf00512pf00989pf13426) GRM1B
522 HMMreg00072_4_pdu1d(pf02954pf00072pf00158) FRAG2
523 -pf07578 -
524 -pf00390pf03949 -
525 -pf01180 -
526 HMM17147pyruvdom_1_buf3(pf10371pf12838pf01558pf01855pf02775) GRM1A
527 -pf13435 -
528 HMM13520tp_1_pdu(pf00324) EUT2C
529 -pf00041pf01344 -
530 - -
531 HMMspu01791_1_spu(pf01791) SPU1 EP_NTERM
532 P_coolGrey SPU1
533 P_orange SPU1
534 HMMspu02502_1_spu(pf02502) SPU1
535 Ts_asparagus SPU1
536 HMMalddh_1_eutpdugrm SPU1
537 HMMalddh_1_eutpdugrm(pf00171pf03319) SPU1
538 H_azure SPU1
539 HMMptac_3_eut2(pf06130) SPU1
540 HMM00121_isom_1_spu(pf00121) SPU1
541 - -
542 -pf02872pf00149 -
543 -pf01865 -
544 -pf01384 -
545 - -
546 - -
547 - -
548 - -
549 -pf01706pf14841 -
550 -pf01256 -
551 - -
BMC-P
BMC-H
BMC-T
Aldehyde dehydrogenase
PTAC
Signature enzyme
Alcohol dehydrogenase
Regulator

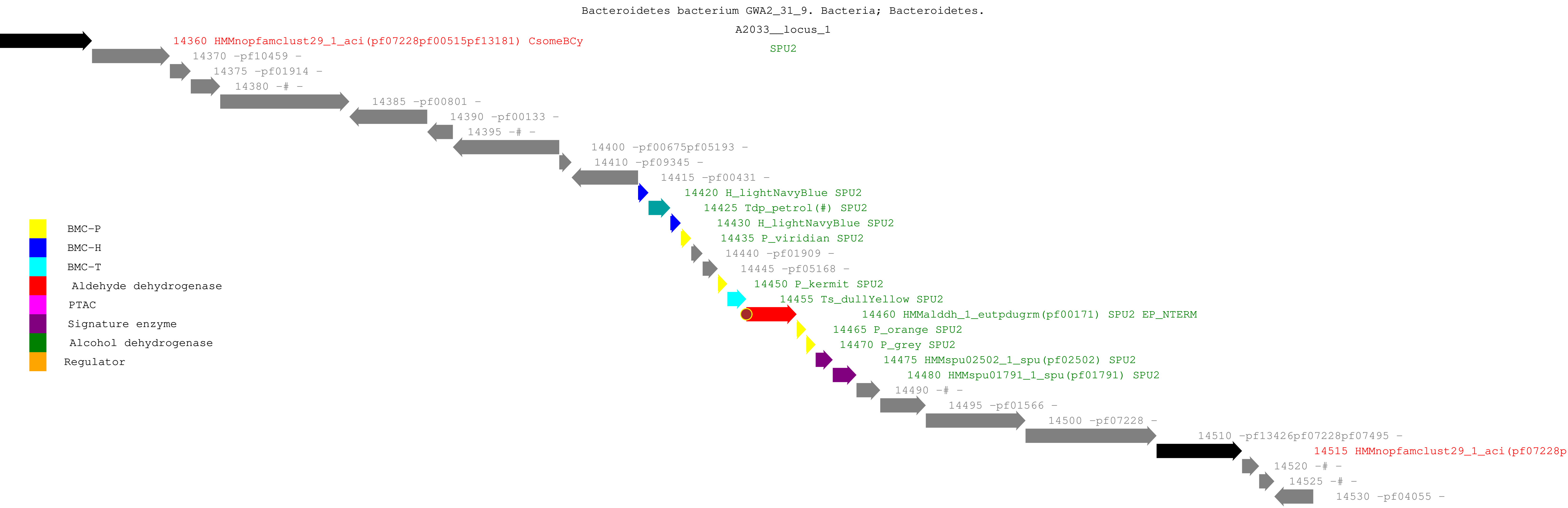
Bacteroidetes bacterium GWA2_31_9. Bacteria; Bacteroidetes.
A2033__locus_1
SPU2
14360 HMMnopfamclust29_1_aci(pf07228pf00515pf13181) CsomeBCy
14370 -pf10459 -
14375 -pf01914 -
14380 -# -
14385 -pf00801 -
14390 -pf00133 -
14395 -# -
14400 -pf00675pf05193 -
14410 -pf09345 -
14415 -pf00431 -
14420 H_lightNavyBlue SPU2
14425 Tdp_petrol(#) SPU2
14430 H_lightNavyBlue SPU2
14435 P_viridian SPU2
14440 -pf01909 -
14445 -pf05168 -
14450 P_kermit SPU2
14455 Ts_dullYellow SPU2
14460 HMMalddh_1_eutpdugrm(pf00171) SPU2 EP_NTERM
14465 P_orange SPU2
14470 P_grey SPU2
14475 HMMspu02502_1_spu(pf02502) SPU2
14480 HMMspu01791_1_spu(pf01791) SPU2
14490 -# -
14495 -pf01566 -
14500 -pf07228 -
14510 -pf13426pf07228pf07495 -
14515 HMMnopfamclust29_1_aci(pf07228p
14520 -# -
14525 -# -
14530 -pf04055 -
BMC-P
BMC-H
BMC-T
Aldehyde dehydrogenase
PTAC
Signature enzyme
Alcohol dehydrogenase
Regulator

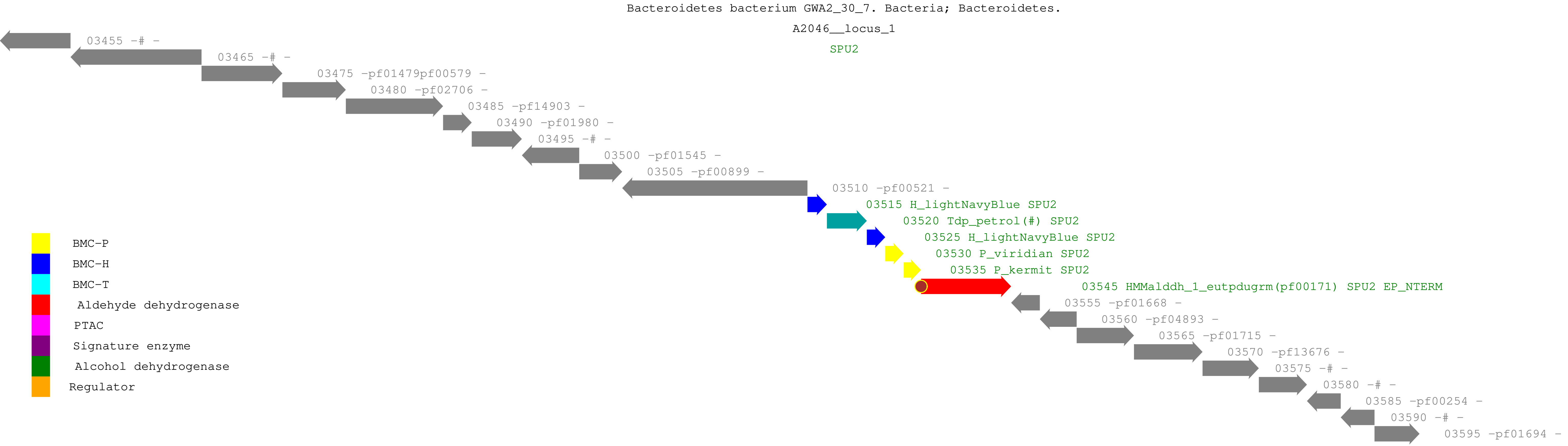
Bacteroidetes bacterium GWA2_30_7. Bacteria; Bacteroidetes.
A2046__locus_1
SPU2
03455 -# -
03465 -# -
03475 -pf01479pf00579 -
03480 -pf02706 -
03485 -pf14903 -
03490 -pf01980 -
03495 -# -
03500 -pf01545 -
03505 -pf00899 -
03510 -pf00521 -
03515 H_lightNavyBlue SPU2
03520 Tdp_petrol(#) SPU2
03525 H_lightNavyBlue SPU2
03530 P_viridian SPU2
03535 P_kermit SPU2
03545 HMMalddh_1_eutpdugrm(pf00171) SPU2 EP_NTERM
03555 -pf01668 -
03560 -pf04893 -
03565 -pf01715 -
03570 -pf13676 -
03575 -# -
03580 -# -
03585 -pf00254 -
03590 -# -
03595 -pf01694 -
BMC-P
BMC-H
BMC-T
Aldehyde dehydrogenase
PTAC
Signature enzyme
Alcohol dehydrogenase
Regulator

Candidatus Atribacteria bacterium RBG_19FT_COMBO_35_14. Bacteria; Candidatus Atribacteria.

A2V47__locus_1

SPU2

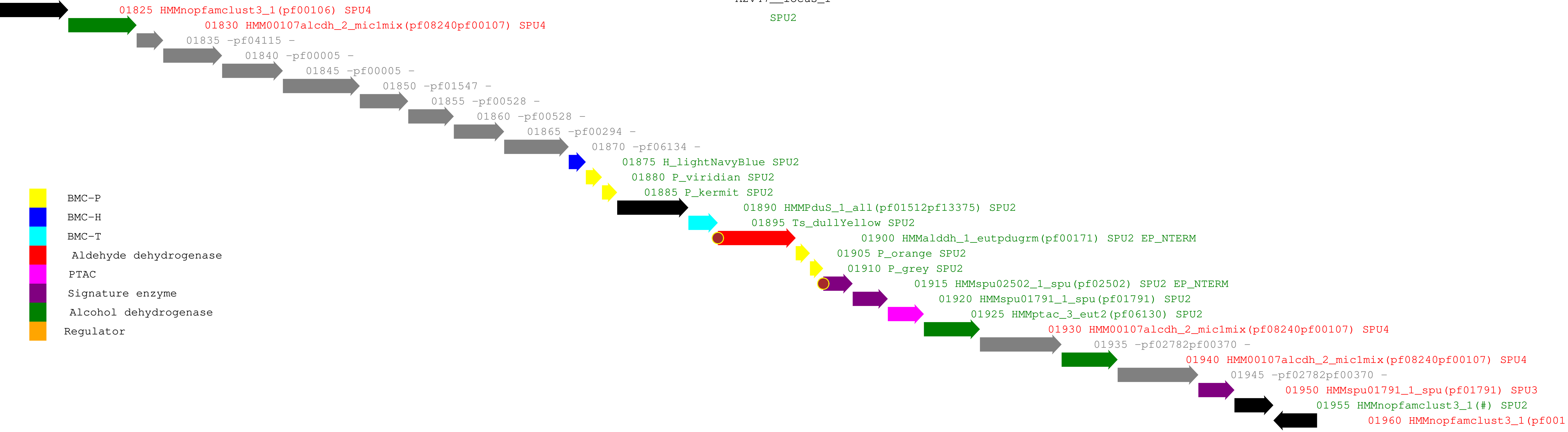

Bacteroidetes bacterium GWF2_41_31. Bacteria; Bacteroidetes.

A2W85__locus_1

SPU2

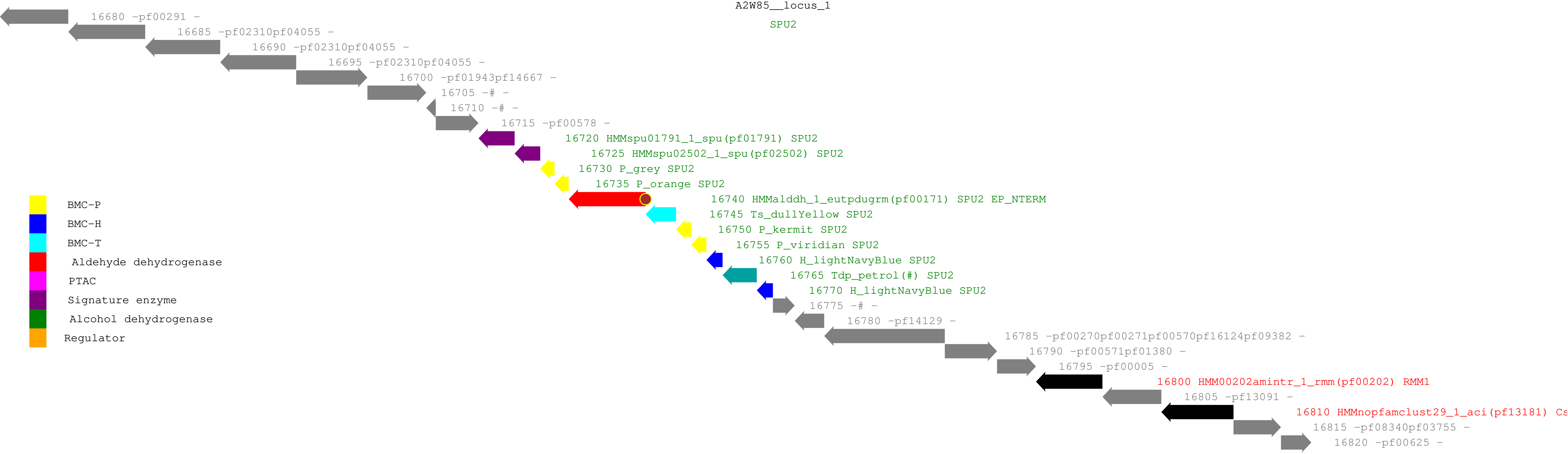

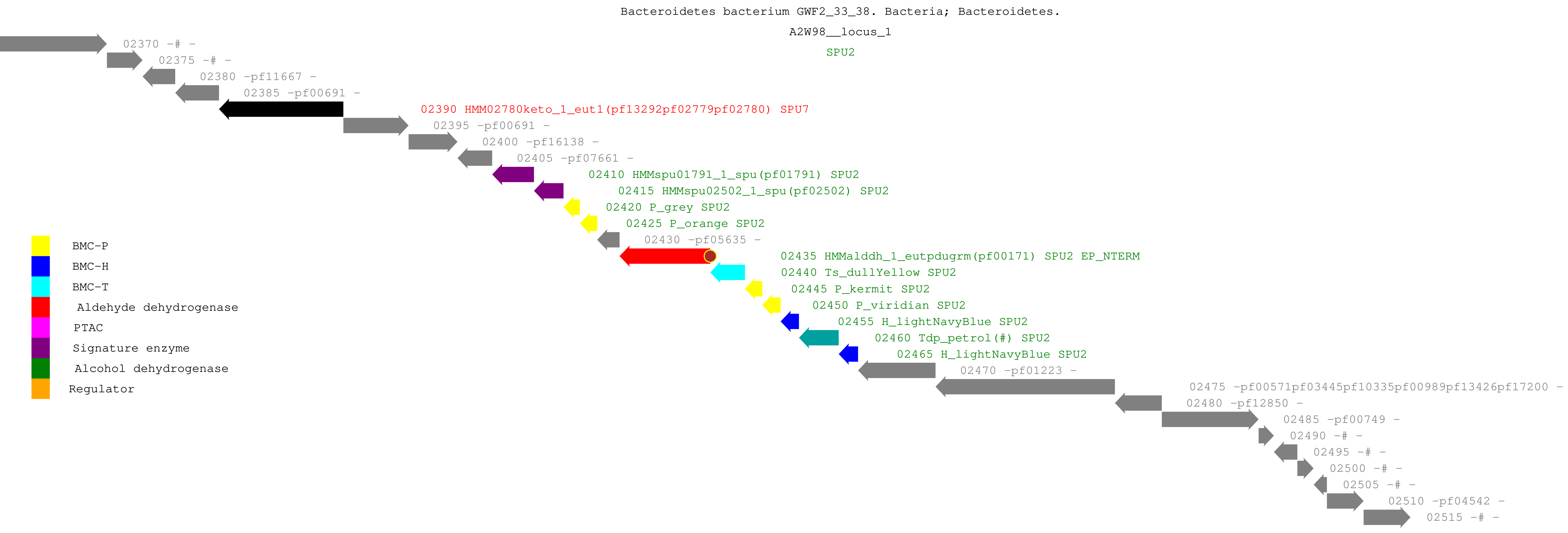

Bacteroidetes bacterium GWF2_33_38. Bacteria; Bacteroidetes.
A2W98__locus_1
SPU2
02370 -# -
02375 -# -
02380 -pf11667 -
02385 -pf00691 -
02390 HMM02780keto_1_eut1(pf13292pf02779pf02780) SPU7
02395 -pf00691 -
02400 -pf16138 -
02405 -pf07661 -
02410 HMMspu01791_1_spu(pf01791) SPU2
02415 HMMspu02502_1_spu(pf02502) SPU2
02420 P_grey SPU2
02425 P_orange SPU2
02430 -pf05635 -
02435 HMMalddh_1_eutpdugrm(pf00171) SPU2 EP_NTERM
02440 Ts_dullYellow SPU2
02445 P_kermit SPU2
02450 P_viridian SPU2
02455 H_lightNavyBlue SPU2
02460 Tdp_petrol(#) SPU2
02465 H_lightNavyBlue SPU2
02470 -pf01223 -
02475 -pf00571pf03445pf10335pf00989pf13426pf17200 -
02480 -pf12850 -
02485 -pf00749 -
02490 -# -
02495 -# -
02500 -# -
02505 -# -
02510 -pf04542 -
02515 -# -
BMC-P
BMC-H
BMC-T
Aldehyde dehydrogenase
PTAC
Signature enzyme
Alcohol dehydrogenase
Regulator

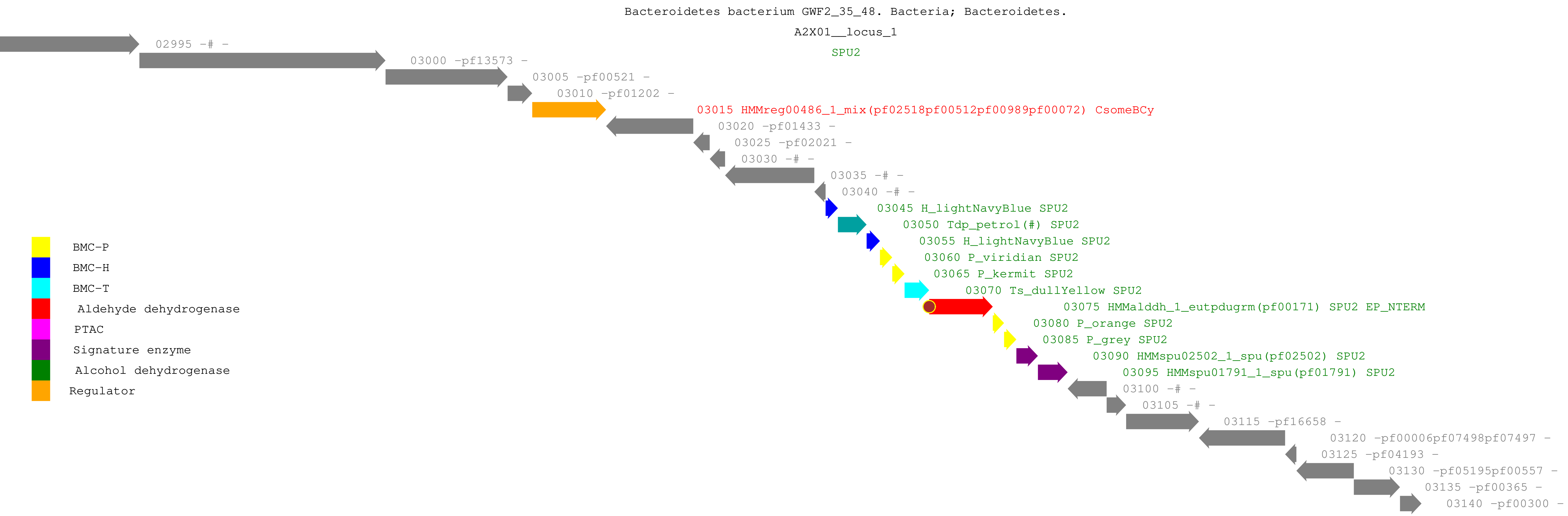
Bacteroidetes bacterium GWF2_35_48. Bacteria; Bacteroidetes.
A2X01__locus_1
SPU2
02995 -# -
03000 -pf13573 -
03005 -pf00521 -
03010 -pf01202 -
03015 HMMreg00486_1_mix(pf02518pf00512pf00989pf00072) CsomeBCy
03020 -pf01433 -
03025 -pf02021 -
03030 -# -
03035 -# -
03040 -# -
03045 H_lightNavyBlue SPU2
03050 Tdp_petrol(#) SPU2
03055 H_lightNavyBlue SPU2
03060 P_viridian SPU2
03065 P_kermit SPU2
03070 Ts_dullYellow SPU2
03075 HMMalddh_1_eutpdugrm(pf00171) SPU2 EP_NTERM
03080 P_orange SPU2
03085 P_grey SPU2
03090 HMMspu02502_1_spu(pf02502) SPU2
03095 HMMspu01791_1_spu(pf01791) SPU2
03100 -# -
03105 -# -
03115 -pf16658 -
03120 -pf00006pf07498pf07497 -
03125 -pf04193 -
03130 -pf05195pf00557 -
03135 -pf00365 -
03140 -pf00300 -
BMC-P
BMC-H
BMC-T
Aldehyde dehydrogenase
PTAC
Signature enzyme
Alcohol dehydrogenase
Regulator

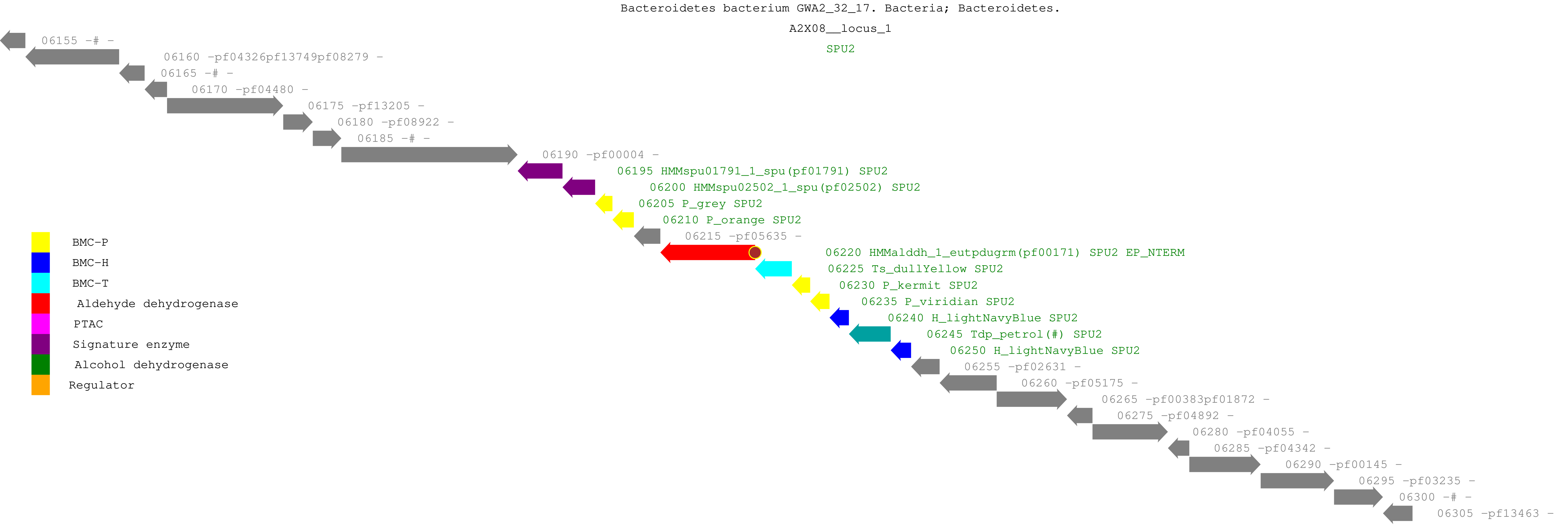
Bacteroidetes bacterium GWA2_32_17. Bacteria; Bacteroidetes.
A2X08__locus_1
SPU2
06155 -# -
06160 -pf04326pf13749pf08279 -
06165 -# -
06170 -pf04480 -
06175 -pf13205 -
06180 -pf08922 -
06185 -# -
06190 -pf00004 -
06195 HMMspu01791_1_spu(pf01791) SPU2
06200 HMMspu02502_1_spu(pf02502) SPU2
06205 P_grey SPU2
06210 P_orange SPU2
06215 -pf05635 -
06220 HMMaldh_1_eutpdugrm(pf00171) SPU2 EP_NTERM
06225 Ts_dullYellow SPU2
06230 P_kermit SPU2
06235 P_viridian SPU2
06240 H_lightNavyBlue SPU2
06245 Tdp_petrol(#) SPU2
06250 H_lightNavyBlue SPU2
06255 -pf02631 -
06260 -pf05175 -
06265 -pf00383pf01872 -
06275 -pf04892 -
06280 -pf04055 -
06285 -pf04342 -
06290 -pf00145 -
06295 -pf03235 -
06300 -# -
06305 -pf13463 -
BMC-P
BMC-H
BMC-T
Aldehyde dehydrogenase
PTAC
Signature enzyme
Alcohol dehydrogenase
Regulator

Spirochaetes bacterium GWC2_52_13. Bacteria; Spirochaetes.

A2Y31__locus_1

SPU2

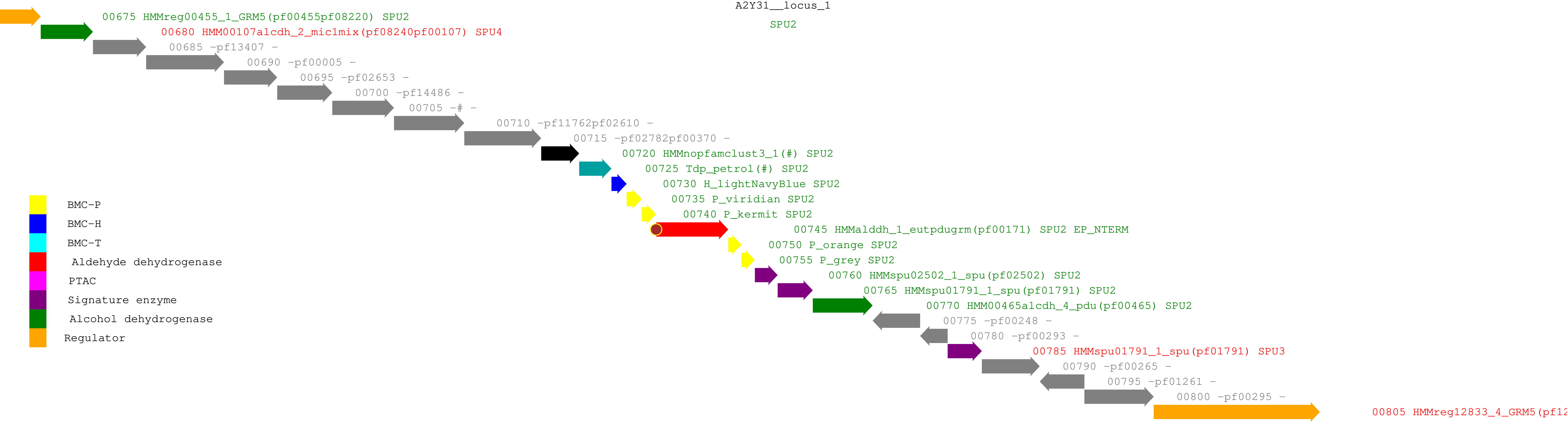

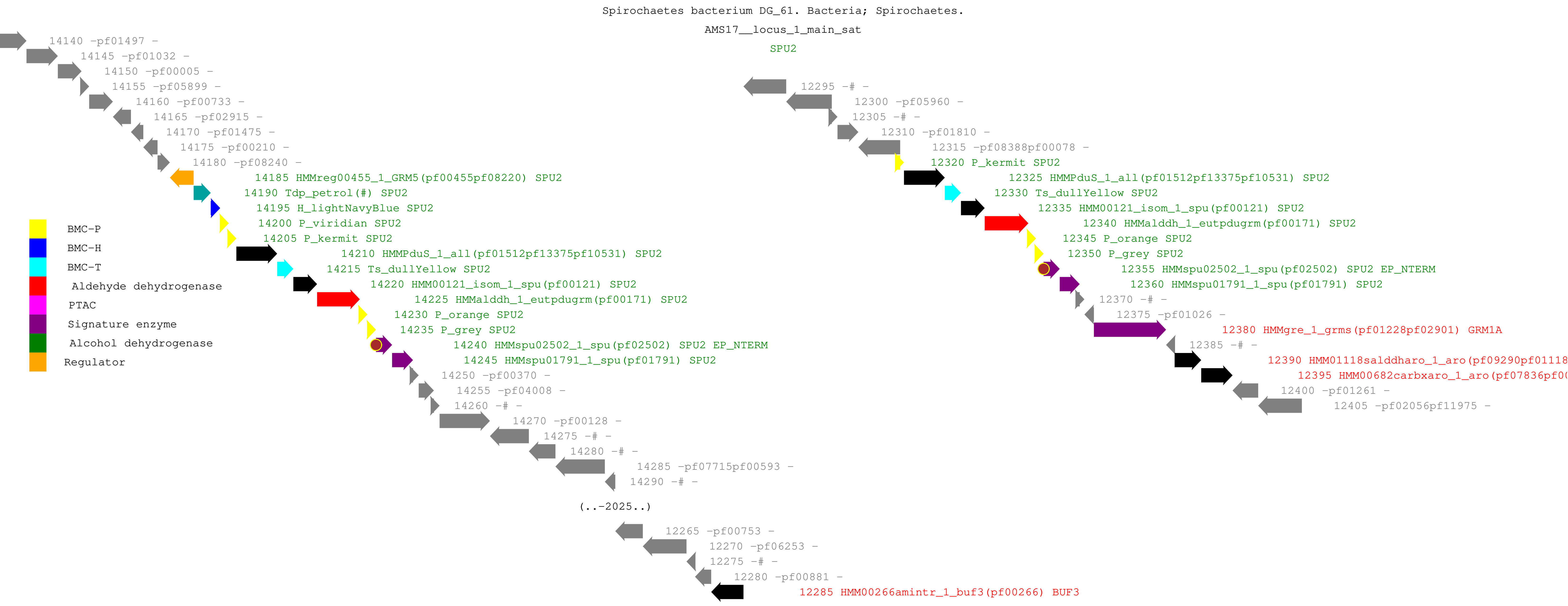

Spirochaetes bacterium DG_61. Bacteria; Spirochaetes.
AMS17__locus_1_main_sat
SPU2
14140 -pf01497 -
14145 -pf01032 -
14150 -pf00005 -
14155 -pf05899 -
14160 -pf00733 -
14165 -pf02915 -
14170 -pf01475 -
14175 -pf00210 -
14180 -pf08240 -
14185 HMMreg00455_1_GRM5(pf00455pf08220) SPU2
14190 Tdp_petrol(#) SPU2
14195 H_lightNavyBlue SPU2
14200 P_viridian SPU2
14205 P_kermit SPU2
14210 HMMPduS_1_all(pf01512pf13375pf10531) SPU2
14215 Ts_dullYellow SPU2
14220 HMM00121_isom_1_spu(pf00121) SPU2
14225 HMMalddh_1_eutpdugrm(pf00171) SPU2
14230 P_orange SPU2
14235 P_grey SPU2
14240 HMMspu02502_1_spu(pf02502) SPU2 EP_NTERM
14245 HMMspu01791_1_spu(pf01791) SPU2
14250 -pf00370 -
14255 -pf04008 -
14260 -# -
14270 -pf00128 -
14275 -# -
14280 -# -
14285 -pf07715pf00593 -
14290 -# -
(..-2025..)
12265 -pf00753 -
12270 -pf06253 -
12275 -# -
12280 -pf00881 -
12285 HMM00266amintr_1_buf3(pf00266) BUF3
12295 -# -
12300 -pf05960 -
12305 -# -
12310 -pf01810 -
12315 -pf08388pf00078 -
12320 P_kermit SPU2
12325 HMMPduS_1_all(pf01512pf13375pf10531) SPU2
12330 Ts_dullYellow SPU2
12335 HMM00121_isom_1_spu(pf00121) SPU2
12340 HMMalddh_1_eutpdugrm(pf00171) SPU2
12345 P_orange SPU2
12350 P_grey SPU2
12355 HMMspu02502_1_spu(pf02502) SPU2 EP_NTERM
12360 HMMspu01791_1_spu(pf01791) SPU2
12370 -# -
12375 -pf01026 -
12380 HMMgre_1_grms(pf01228pf02901) GRM1A
12385 -# -
12390 HMM01118salddharo_1_aro(pf09290pf01118
12395 HMM00682carbxaro_1_aro(pf07836pf0
12400 -pf01261 -
12405 -pf02056pf11975 -
BMC-P
BMC-H
BMC-T
Aldehyde dehydrogenase
PTAC
Signature enzyme
Alcohol dehydrogenase
Regulator

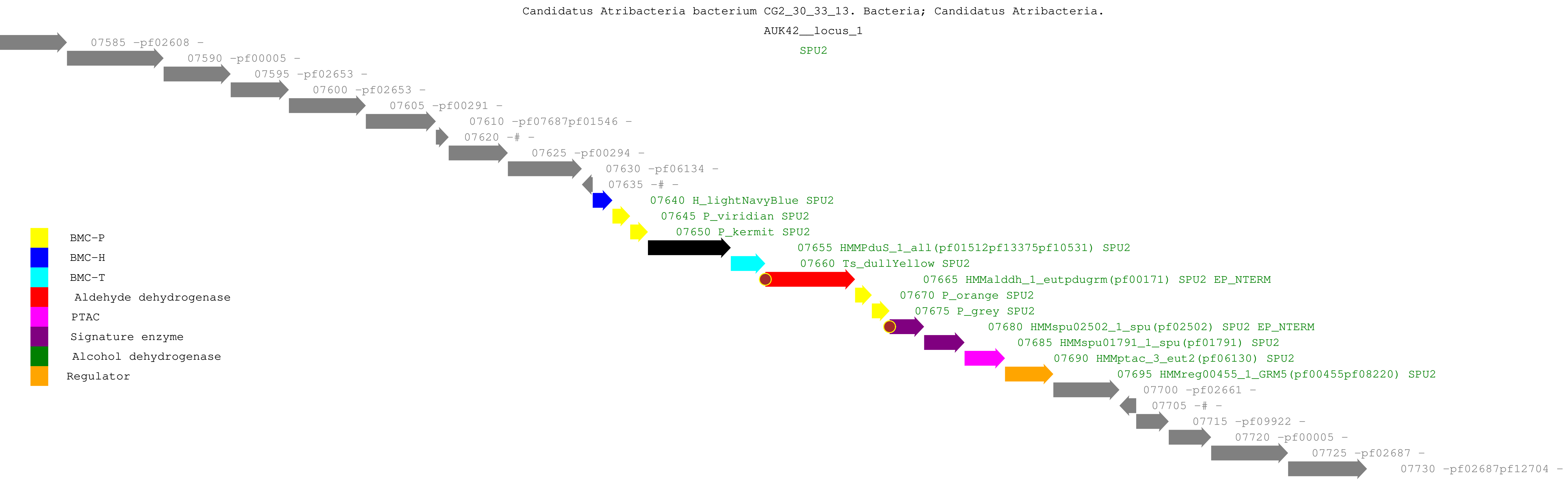
Candidatus Atribacteria bacterium CG2_30_33_13. Bacteria; Candidatus Atribacteria.
AUK42__locus_1
SPU2
07585 -pf02608 -
07590 -pf00005 -
07595 -pf02653 -
07600 -pf02653 -
07605 -pf00291 -
07610 -pf07687pf01546 -
07620 -# -
07625 -pf00294 -
07630 -pf06134 -
07635 -# -
07640 H_lightNavyBlue SPU2
07645 P_viridian SPU2
07650 P_kermit SPU2
07655 HMMPduS_1_all(pf01512pf13375pf10531) SPU2
07660 Ts_dullYellow SPU2
07665 HMMalddh_1_eutpdugrm(pf00171) SPU2 EP_NTERM
07670 P_orange SPU2
07675 P_grey SPU2
07680 HMMspu02502_1_spu(pf02502) SPU2 EP_NTERM
07685 HMMspu01791_1_spu(pf01791) SPU2
07690 HMMptac_3_eut2(pf06130) SPU2
07695 HMMreg00455_1_GRM5(pf00455pf08220) SPU2
07700 -pf02661 -
07705 -# -
07715 -pf09922 -
07720 -pf00005 -
07725 -pf02687 -
07730 -pf02687pf12704 -
BMC-P
BMC-H
BMC-T
Aldehyde dehydrogenase
PTAC
Signature enzyme
Alcohol dehydrogenase
Regulator

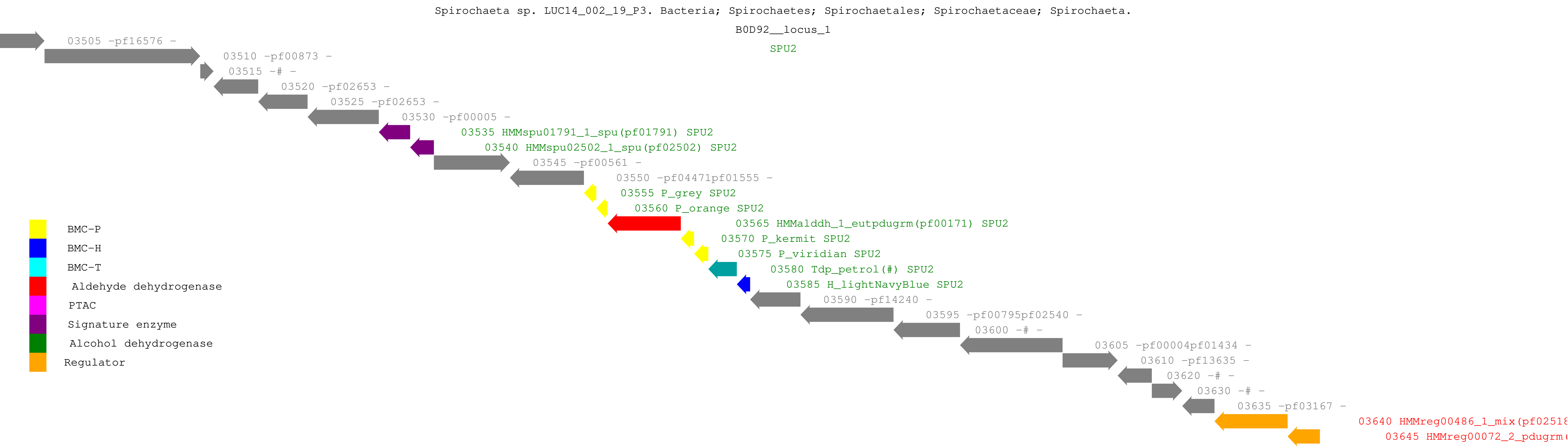
Spirochaeta sp. LUC14_002_19_P3. Bacteria; Spirochaetes; Spirochaetales; Spirochaetaceae; Spirochaeta.
B0D92__locus_1
SPU2
03505 -pf16576 -
03510 -pf00873 -
03515 -# -
03520 -pf02653 -
03525 -pf02653 -
03530 -pf00005 -
03535 HMMspu01791_1_spu(pf01791) SPU2
03540 HMMspu02502_1_spu(pf02502) SPU2
03545 -pf00561 -
03550 -pf04471pf01555 -
03555 P_grey SPU2
03560 P_orange SPU2
03565 HMMaldh_1_eutpdugrm(pf00171) SPU2
03570 P_kermit SPU2
03575 P_viridian SPU2
03580 Tdp_petrol(#) SPU2
03585 H_lightNavyBlue SPU2
03590 -pf14240 -
03595 -pf00795pf02540 -
03600 -# -
03605 -pf00004pf01434 -
03610 -pf13635 -
03620 -# -
03630 -# -
03635 -pf03167 -
BMC-P
BMC-H
BMC-T
Aldehyde dehydrogenase
PTAC
Signature enzyme
Alcohol dehydrogenase
Regulator

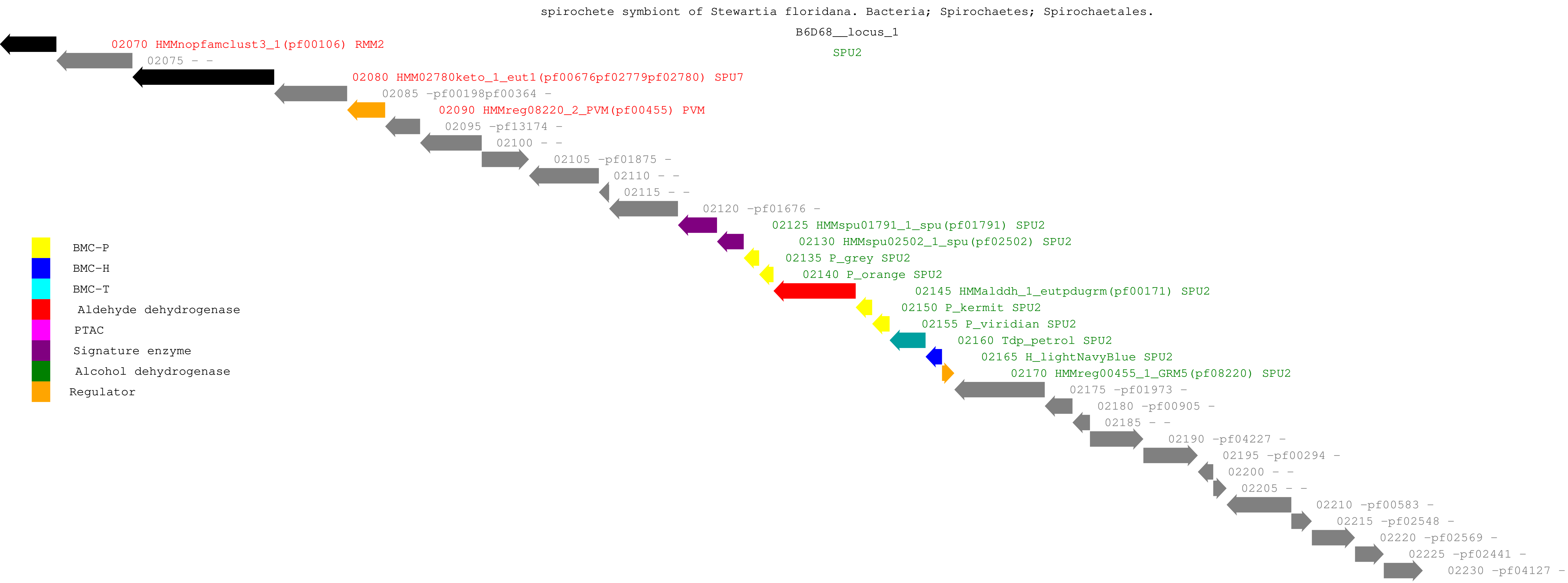
spirochete symbiont of Stewartia floridana. Bacteria; Spirochaetes; Spirochaetales.
B6D68__locus_1
SPU2
02070 HMMnopfamclust3_1(pf00106) RMM2
02075 - -
02080 HMM02780keto_1_eut1(pf00676pf02779pf02780) SPU7
02085 -pf00198pf00364 -
02090 HMMreg08220_2_PVM(pf00455) PVM
02095 -pf13174 -
02100 - -
02105 -pf01875 -
02110 - -
02115 - -
02120 -pf01676 -
02125 HMMspu01791_1_spu(pf01791) SPU2
02130 HMMspu02502_1_spu(pf02502) SPU2
02135 P_grey SPU2
02140 P_orange SPU2
02145 HMMalddh_1_eutpdugrm(pf00171) SPU2
02150 P_kermit SPU2
02155 P_viridian SPU2
02160 Tdp_petrol SPU2
02165 H_lightNavyBlue SPU2
02170 HMMreg00455_1_GRM5(pf08220) SPU2
02175 -pf01973 -
02180 -pf00905 -
02185 - -
02190 -pf04227 -
02195 -pf00294 -
02200 - -
02205 - -
02210 -pf00583 -
02215 -pf02548 -
02220 -pf02569 -
02225 -pf02441 -
02230 -pf04127 -
BMC-P
BMC-H
BMC-T
Aldehyde dehydrogenase
PTAC
Signature enzyme
Alcohol dehydrogenase
Regulator

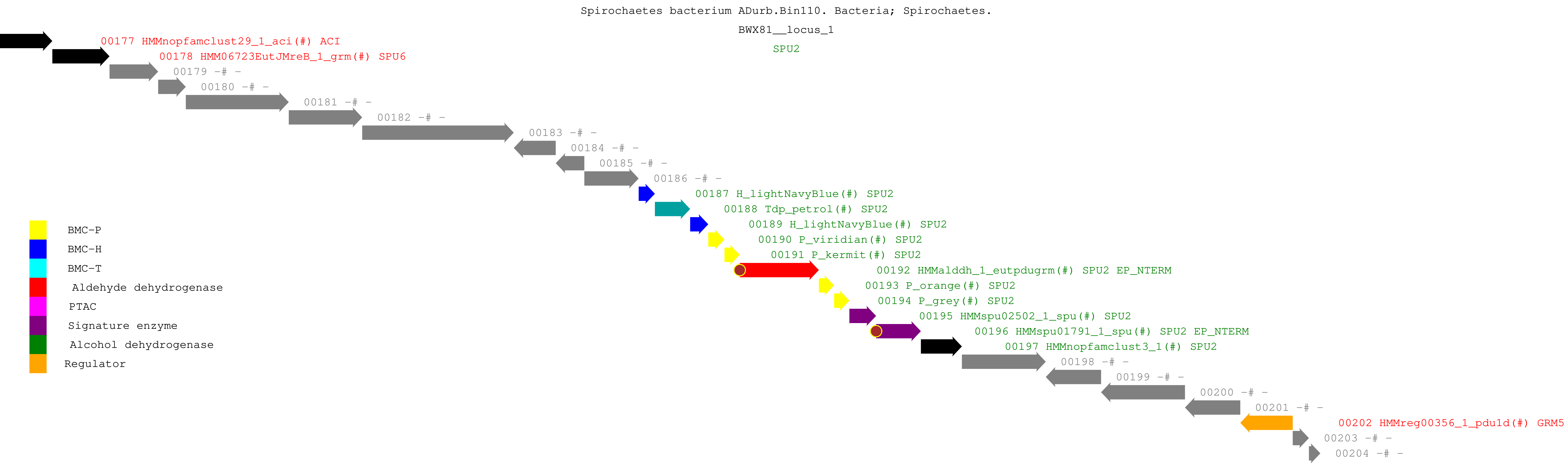
Spirochaetes bacterium ADurb.Bin110. Bacteria; Spirochaetes.
BWX81__locus_1
SPU2
00177 HMMnopfamclust29_1_aci(#) ACI
00178 HMM06723EutJMreB_1_grm(#) SPU6
00179 -# -
00180 -# -
00181 -# -
00182 -# -
00183 -# -
00184 -# -
00185 -# -
00186 -# -
00187 H_lightNavyBlue(#) SPU2
00188 Tdp_petrol(#) SPU2
00189 H_lightNavyBlue(#) SPU2
00190 P_viridian(#) SPU2
00191 P_kermit(#) SPU2
00192 HMMalddh_1_eutpdugrm(#) SPU2 EP_NTERM
00193 P_orange(#) SPU2
00194 P_grey(#) SPU2
00195 HMMspu02502_1_spu(#) SPU2
00196 HMMspu01791_1_spu(#) SPU2 EP_NTERM
00197 HMMnopfamclust3_1(#) SPU2
00198 -# -
00199 -# -
00200 -# -
00201 -# -
00202 HMMreg00356_1_pdu1d(#) GRM5
00203 -# -
00204 -# -
BMC-P
BMC-H
BMC-T
Aldehyde dehydrogenase
PTAC
Signature enzyme
Alcohol dehydrogenase
Regulator

Candidatus Atribacteria bacterium ADurb.Bin276. Bacteria; Candidatus Atribacteria.

BWY41__locus_1

SPU2

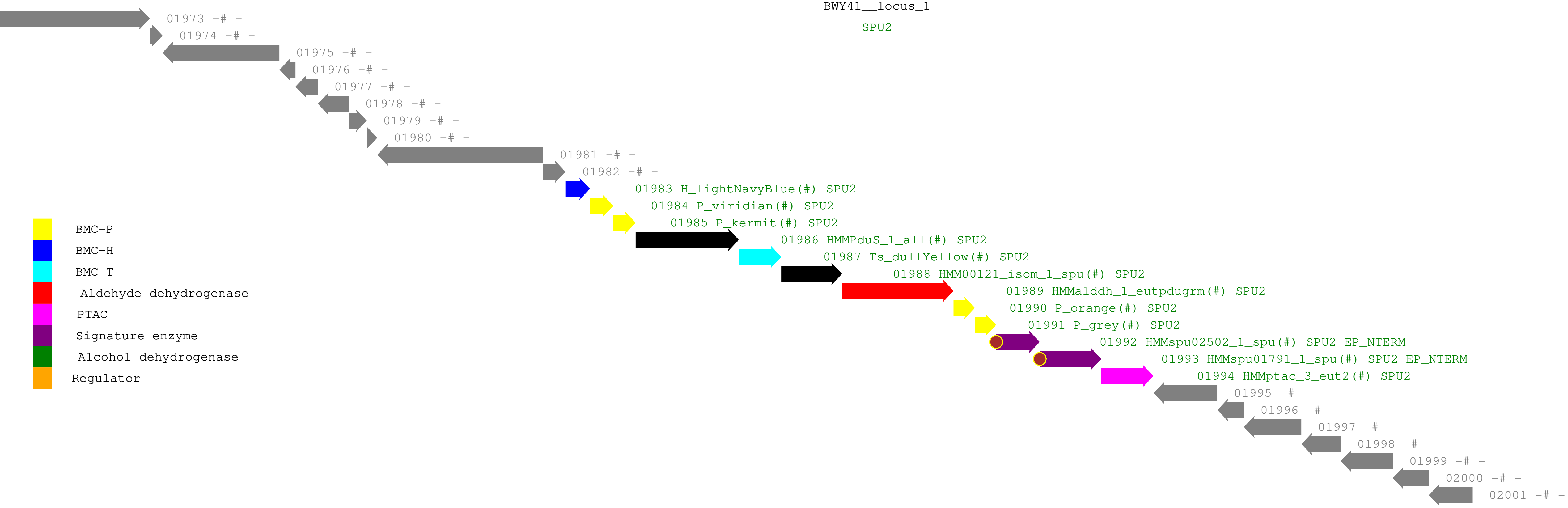

Marinilabiliales bacterium. Bacteria; Bacteroidetes; Bacteroidia; Marinilabiliales;

C0596__locus_1

SPU2

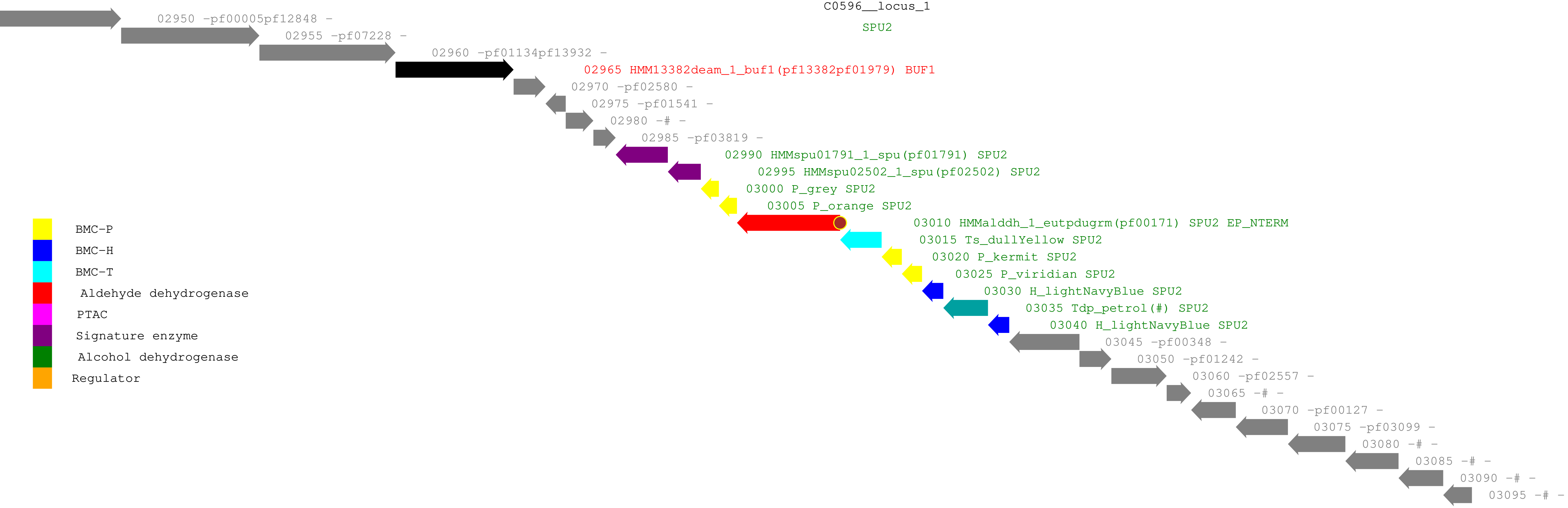

Candidatus Atribacteria bacterium CG_4_9_14_3_um_filter_33_16. Bacteria; Candidatus Atribacteria.

CO097__locus_1

SPU2

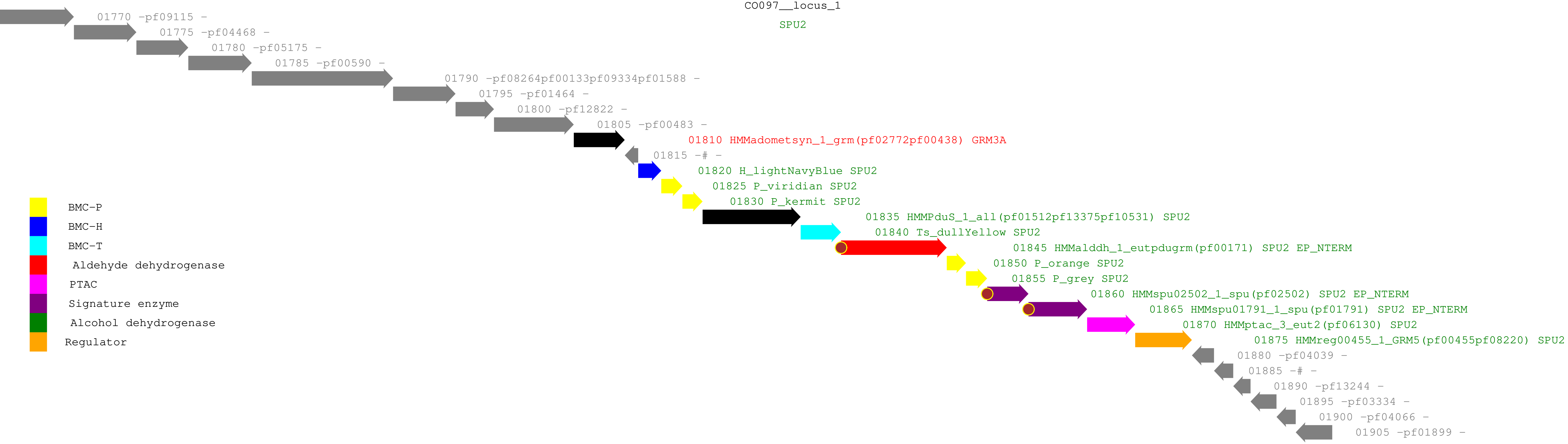

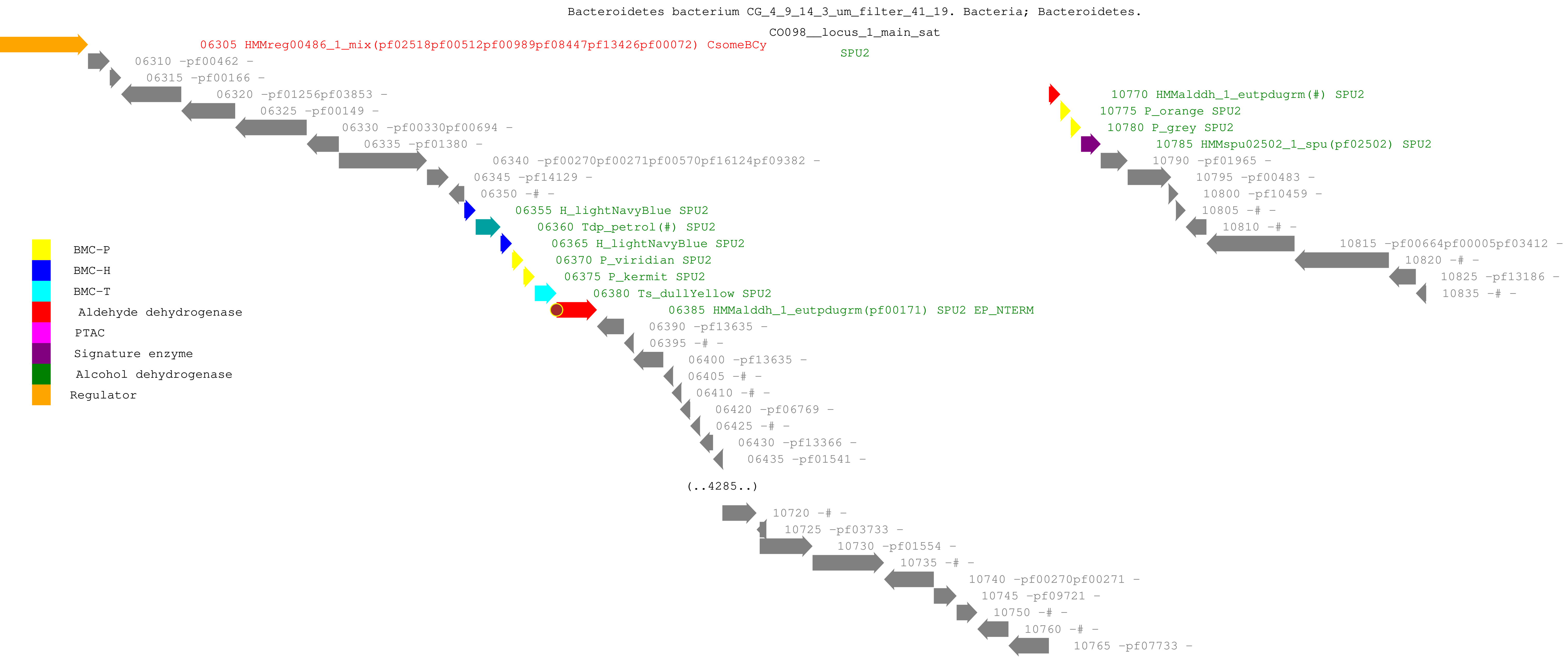
Bacteroidetes bacterium CG_4_9_14_3_um_filter_41_19. Bacteria; Bacteroidetes.
CO098__locus_1_main_sat
SPU2
06305 HMMreg00486_1_mix(pf02518pf00512pf00989pf08447pf13426pf00072) CsomeBCy
06310 -pf00462 -
06315 -pf00166 -
06320 -pf01256pf03853 -
06325 -pf00149 -
06330 -pf00330pf00694 -
06335 -pf01380 -
06340 -pf00270pf00271pf00570pf16124pf09382 -
06345 -pf14129 -
06350 -# -
06355 H_lightNavyBlue SPU2
06360 Tdp_petrol(#) SPU2
06365 H_lightNavyBlue SPU2
06370 P_viridian SPU2
06375 P_kermit SPU2
06380 Ts_dullYellow SPU2
06385 HMMaldddh_1_eutpdugrm(pf00171) SPU2 EP_NTERM
06390 -pf13635 -
06395 -# -
06400 -pf13635 -
06405 -# -
06410 -# -
06420 -pf06769 -
06425 -# -
06430 -pf13366 -
06435 -pf01541 -
(..4285..)
10720 -# -
10725 -pf03733 -
10730 -pf01554 -
10735 -# -
10740 -pf00270pf00271 -
10745 -pf09721 -
10750 -# -
10760 -# -
10765 -pf07733 -
10770 HMMaldddh_1_eutpdugrm(#) SPU2
10775 P_orange SPU2
10780 P_grey SPU2
10785 HMMspu02502_1_spu(pf02502) SPU2
10790 -pf01965 -
10795 -pf00483 -
10800 -pf10459 -
10805 -# -
10810 -# -
10815 -pf00664pf00005pf03412 -
10820 -# -
10825 -pf13186 -
10835 -# -
BMC-P
BMC-H
BMC-T
Aldehyde dehydrogenase
PTAC
Signature enzyme
Alcohol dehydrogenase
Regulator

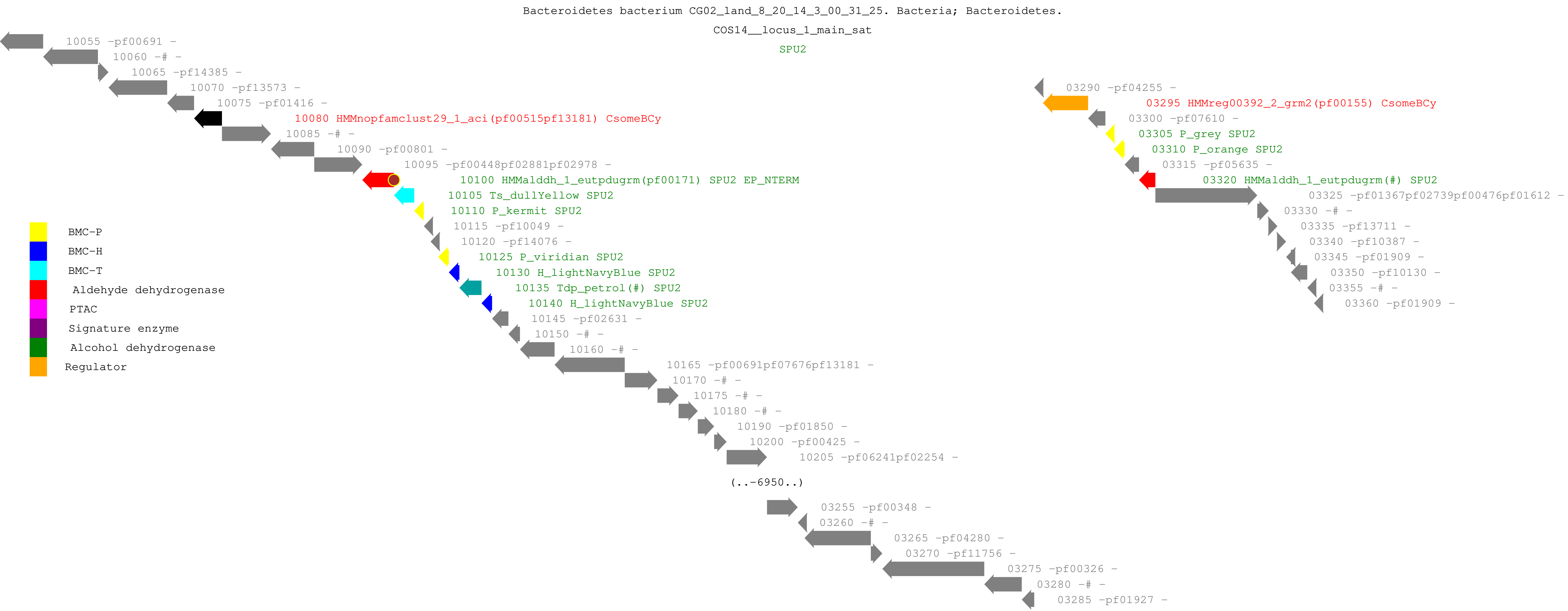
Bacteroidetes bacterium CG02_land_8_20_14_3_00_31_25. Bacteria; Bacteroidetes.
COS14__locus_1_main_sat
SPU2
10055 -pf00691 -
10060 -# -
10065 -pf14385 -
10070 -pf13573 -
10075 -pf01416 -
10080 HMMnopfamclust29_1_aci(pf00515pf13181) CsomeBCy
10085 -# -
10090 -pf00801 -
10095 -pf00448pf02881pf02978 -
10100 HMMalddh_1_eutpdugrm(pf00171) SPU2 EP_NTERM
10105 Ts_dullYellow SPU2
10110 P_kermit SPU2
10115 -pf10049 -
10120 -pf14076 -
10125 P_viridian SPU2
10130 H_lightNavyBlue SPU2
10135 Tdp_petrol(#) SPU2
10140 H_lightNavyBlue SPU2
10145 -pf02631 -
10150 -# -
10160 -# -
10165 -pf00691pf07676pf13181 -
10170 -# -
10175 -# -
10180 -# -
10190 -pf01850 -
10200 -pf00425 -
10205 -pf06241pf02254 -
(..-6950..)
03255 -pf00348 -
03260 -# -
03265 -pf04280 -
03270 -pf11756 -
03275 -pf00326 -
03280 -# -
03285 -pf01927 -
03290 -pf04255 -
03295 HMMreg00392_2_grm2(pf00155) CsomeBCy
03300 -pf07610 -
03305 P_grey SPU2
03310 P_orange SPU2
03315 -pf05635 -
03320 HMMaldh_1_eutpdugrm(#) SPU2
03325 -pf01367pf02739pf00476pf01612 -
03330 -# -
03335 -pf13711 -
03340 -pf10387 -
03345 -pf01909 -
03350 -pf10130 -
03355 -# -
03360 -pf01909 -
BMC-P
BMC-H
BMC-T
Aldehyde dehydrogenase
PTAC
Signature enzyme
Alcohol dehydrogenase
Regulator

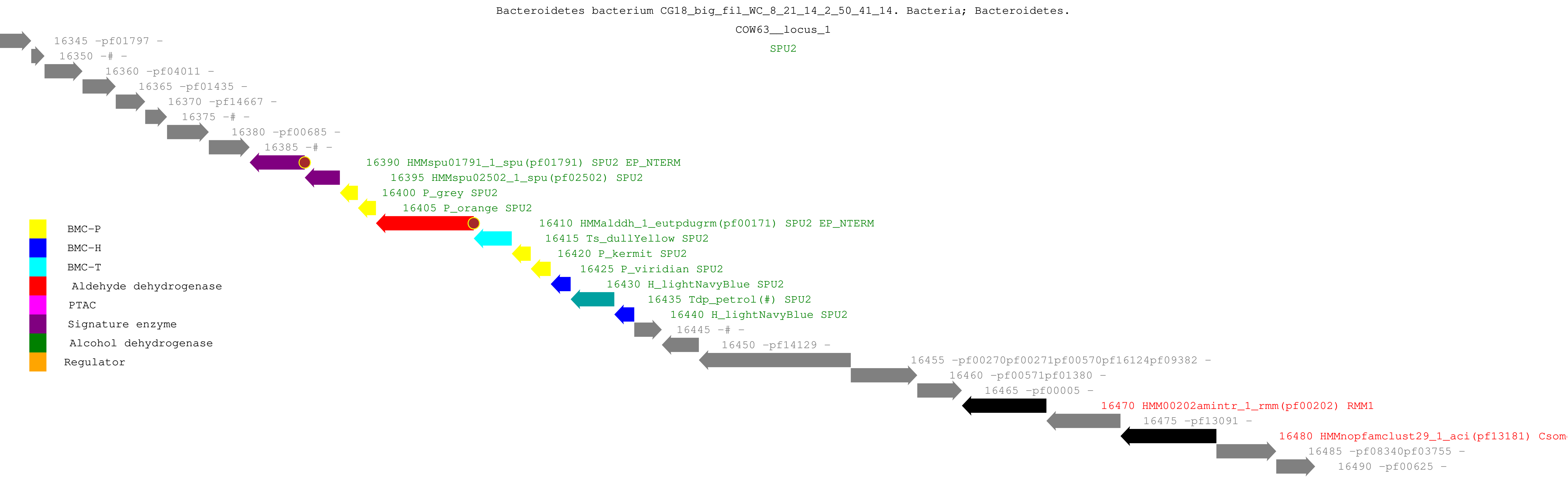
Bacteroidetes bacterium CG18_big_fil_WC_8_21_14_2_50_41_14. Bacteria; Bacteroidetes.
COW63__locus_1
SPU2
16345 -pf01797 -
16350 -# -
16360 -pf04011 -
16365 -pf01435 -
16370 -pf14667 -
16375 -# -
16380 -pf00685 -
16385 -# -
16390 HMMspu01791_1_spu(pf01791) SPU2 EP_NTERM
16395 HMMspu02502_1_spu(pf02502) SPU2
16400 P_grey SPU2
16405 P_orange SPU2
16410 HMMaldh_1_eutpdugrm(pf00171) SPU2 EP_NTERM
16415 Ts_dullYellow SPU2
16420 P_kermit SPU2
16425 P_viridian SPU2
16430 H_lightNavyBlue SPU2
16435 Tdp_petrol(#) SPU2
16440 H_lightNavyBlue SPU2
16445 -# -
16450 -pf14129 -
16455 -pf00270pf00271pf00570pf16124pf09382 -
16460 -pf00571pf01380 -
16465 -pf00005 -
16470 HMM00202amintr_1_rmm(pf00202) RMM1
16475 -pf13091 -
16480 HMMnopfamclust29_1_aci(pf13181) Csom
16485 -pf08340pf03755 -
16490 -pf00625 -
BMC-P
BMC-H
BMC-T
Aldehyde dehydrogenase
PTAC
Signature enzyme
Alcohol dehydrogenase
Regulator

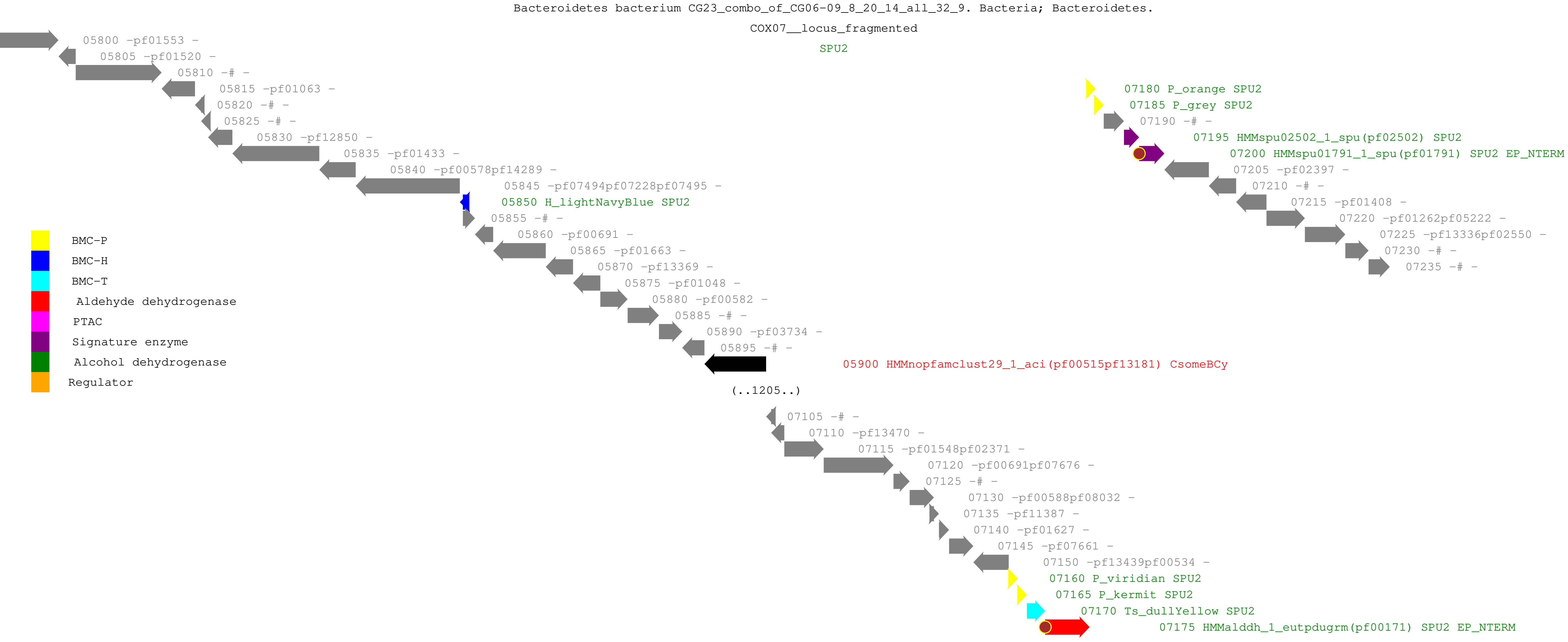

Bacteroidetes bacterium CG23_combo_of_CG06-09_8_20_14_all_32_9. Bacteria; Bacteroidetes.
COX07__locus_fragmented
SPU2
05800 -pf01553 -
05805 -pf01520 -
05810 -# -
05815 -pf01063 -
05820 -# -
05825 -# -
05830 -pf12850 -
05835 -pf01433 -
05840 -pf00578pf14289 -
05845 -pf07494pf07228pf07495 -
05850 H_lightNavyBlue SPU2
05855 -# -
05860 -pf00691 -
05865 -pf01663 -
05870 -pf13369 -
05875 -pf01048 -
05880 -pf00582 -
05885 -# -
05890 -pf03734 -
05895 -# -
05900 HMMnopfamclust29_1_aci(pf00515pf13181) CsomeBCy
(..1205..)
07105 -# -
07110 -pf13470 -
07115 -pf01548pf02371 -
07120 -pf00691pf07676 -
07125 -# -
07130 -pf00588pf08032 -
07135 -pf11387 -
07140 -pf01627 -
07145 -pf07661 -
07150 -pf13439pf00534 -
07160 P_viridian SPU2
07165 P_kermit SPU2
07170 Ts_dullYellow SPU2
07175 HMMalddh_1_eutpdugrm(pf00171) SPU2 EP_NTERM
07180 P_orange SPU2
07185 P_grey SPU2
07190 -# -
07195 HMMspu02502_1_spu(pf02502) SPU2
07200 HMMspu01791_1_spu(pf01791) SPU2 EP_NTERM
07205 -pf02397 -
07210 -# -
07215 -pf01408 -
07220 -pf01262pf05222 -
07225 -pf13336pf02550 -
07230 -# -
07235 -# -
BMC-P
BMC-H
BMC-T
Aldehyde dehydrogenase
PTAC
Signature enzyme
Alcohol dehydrogenase
Regulator

Candidatus Atribacteria bacterium CG_4_10_14_3_um_filter_34_13. Bacteria; Candidatus Atribacteria.

COZ07__locus_1

SPU2

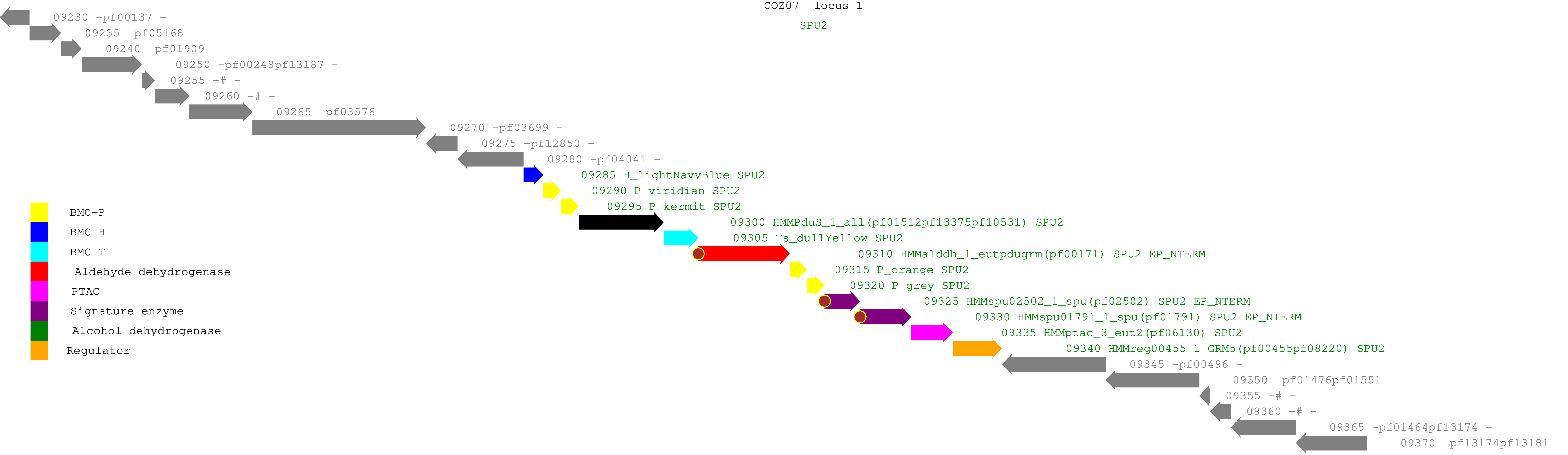

Bacteroidetes bacterium CG_4_10_14_3_um_filter_42_6. Bacteria; Bacteroidetes.

COZ08__locus_fragmented

SPU2

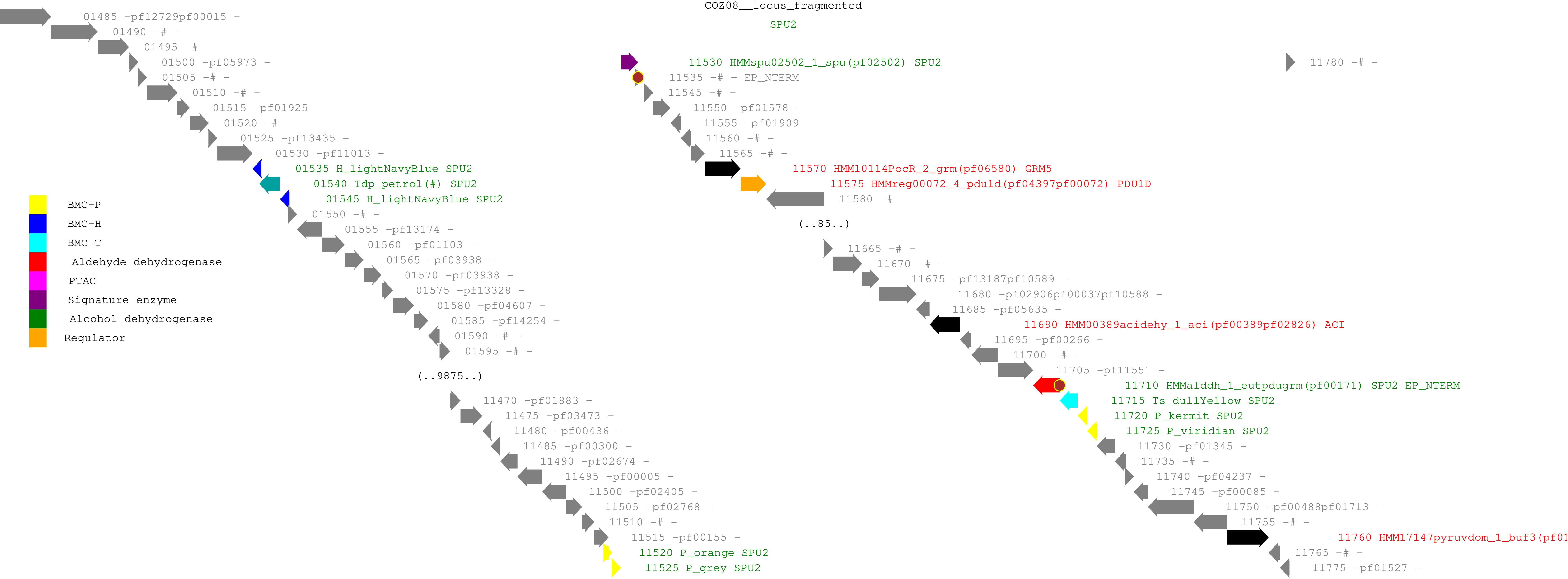

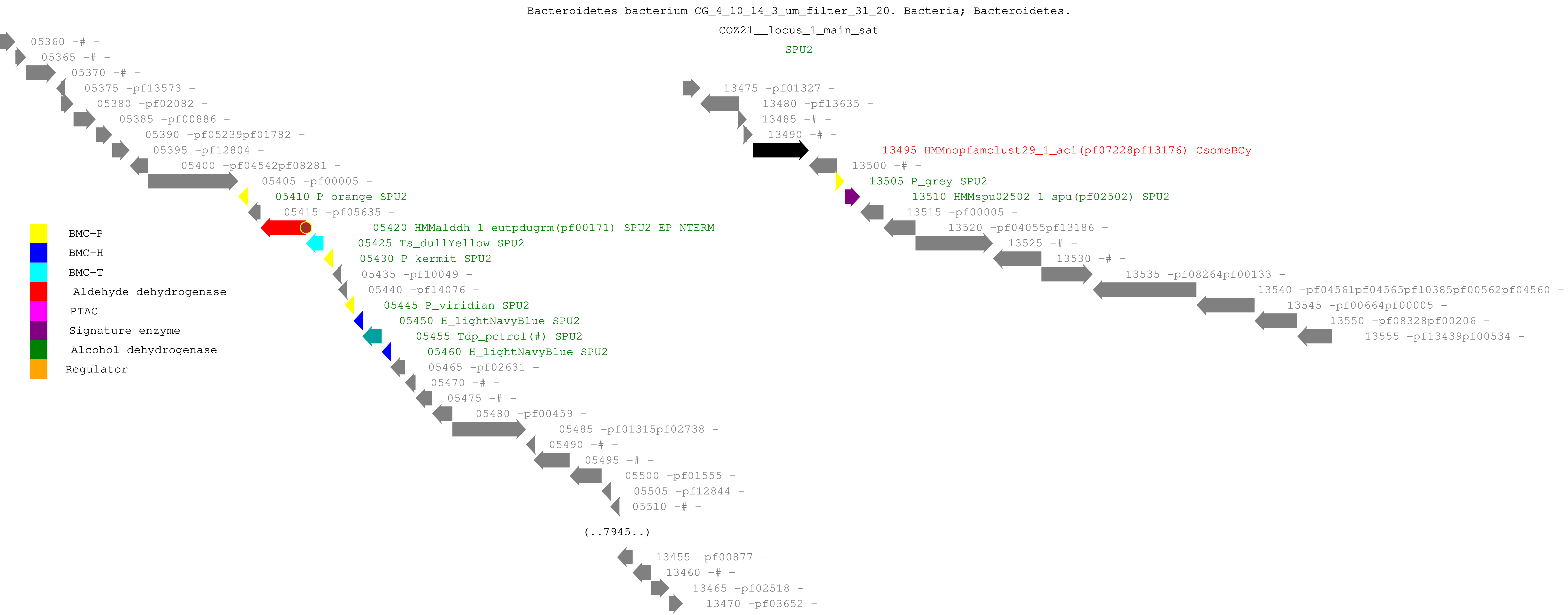
Bacteroidetes bacterium CG_4_10_14_3_um_filter_31_20. Bacteria; Bacteroidetes.
COZ21__locus_1_main_sat
SPU2
05360 -# -
05365 -# -
05370 -# -
05375 -pf13573 -
05380 -pf02082 -
05385 -pf00886 -
05390 -pf05239pf01782 -
05395 -pf12804 -
05400 -pf04542pf08281 -
05405 -pf00005 -
05410 P_orange SPU2
05415 -pf05635 -
05420 HMMalddh_1_eutpdugrm(pf00171) SPU2 EP_NTERM
05425 Ts_dullYellow SPU2
05430 P_kermit SPU2
05435 -pf10049 -
05440 -pf14076 -
05445 P_viridian SPU2
05450 H_lightNavyBlue SPU2
05455 Tdp_petrol(#) SPU2
05460 H_lightNavyBlue SPU2
05465 -pf02631 -
05470 -# -
05475 -# -
05480 -pf00459 -
05485 -pf01315pf02738 -
05490 -# -
05495 -# -
05500 -pf01555 -
05505 -pf12844 -
05510 -# -
(..7945..)
13455 -pf00877 -
13460 -# -
13465 -pf02518 -
13470 -pf03652 -
13475 -pf01327 -
13480 -pf13635 -
13485 -# -
13490 -# -
13495 HMMnopfamclust29_1_aci(pf07228pf13176) CsomeBCy
13500 -# -
13505 P_grey SPU2
13510 HMMspu02502_1_spu(pf02502) SPU2
13515 -pf00005 -
13520 -pf04055pf13186 -
13525 -# -
13530 -# -
13535 -pf08264pf00133 -
13540 -pf04561pf04565pf10385pf00562pf04560 -
13545 -pf00664pf00005 -
13550 -pf08328pf00206 -
13555 -pf13439pf00534 -
BMC-P
BMC-H
BMC-T
Aldehyde dehydrogenase
PTAC
Signature enzyme
Alcohol dehydrogenase
Regulator

Candidatus Atribacteria bacterium CG_4_8_14_3_um_filter_34_18. Bacteria; Candidatus Atribacteria.

COZ58__locus_1

SPU2

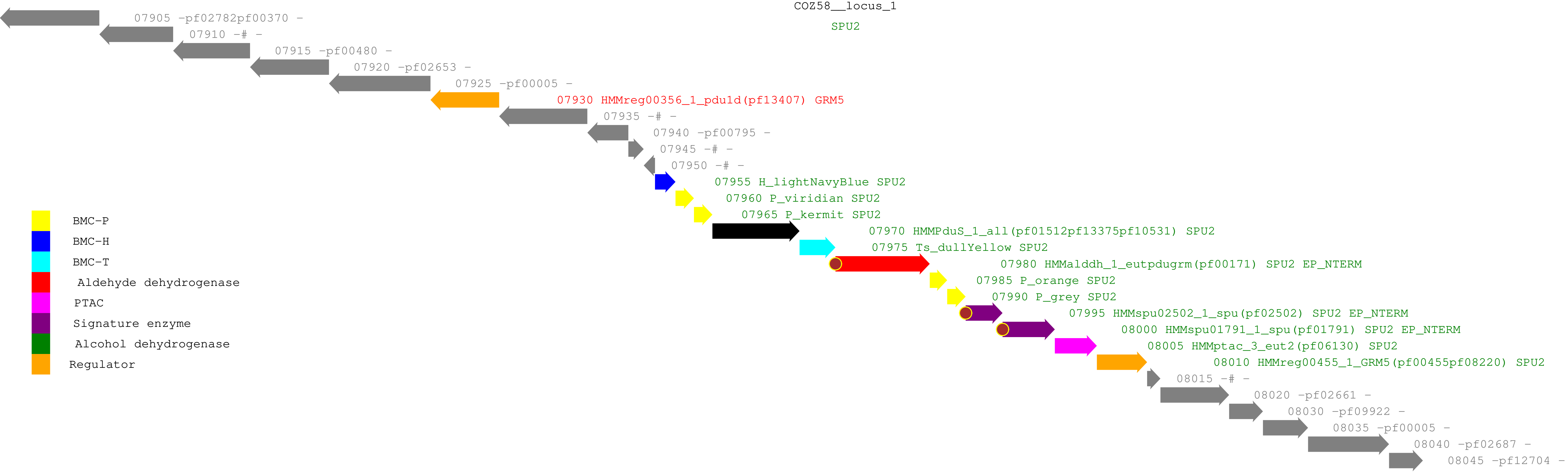

Bacteroidetes bacterium CG_4_8_14_3_um_filter_31_14. Bacteria; Bacteroidetes.

COZ59__locus_1_main_sat

SPU2

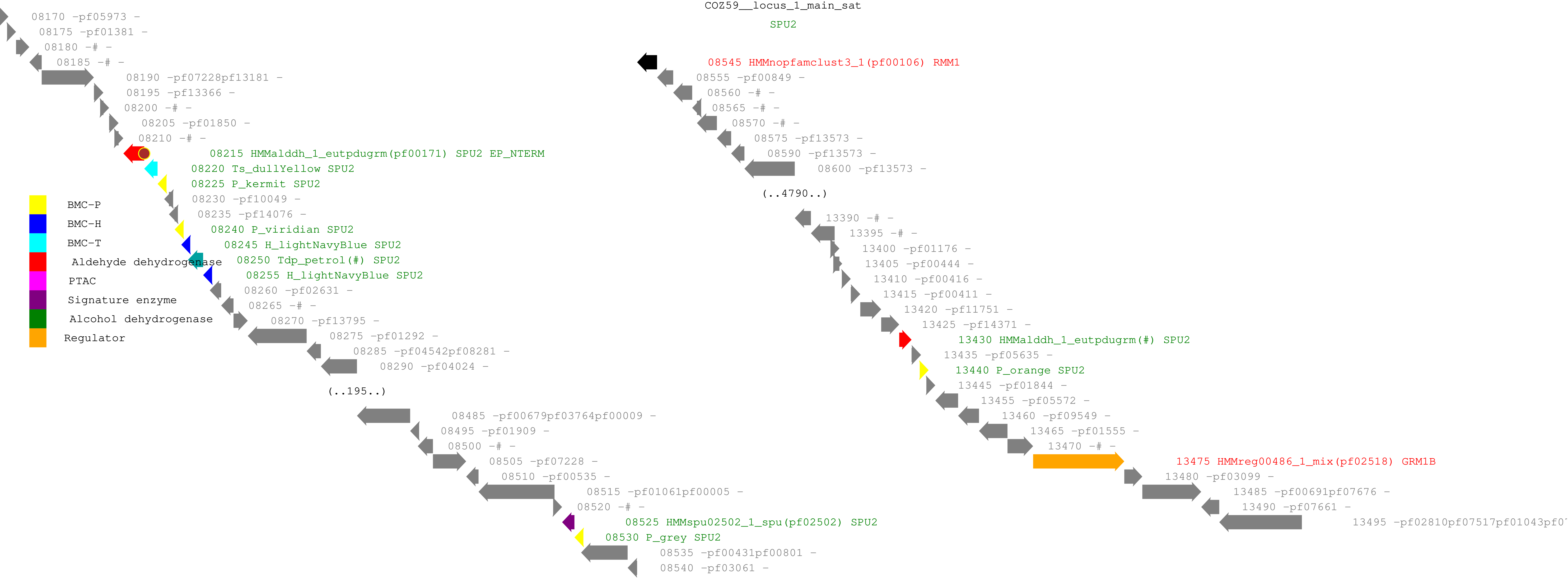

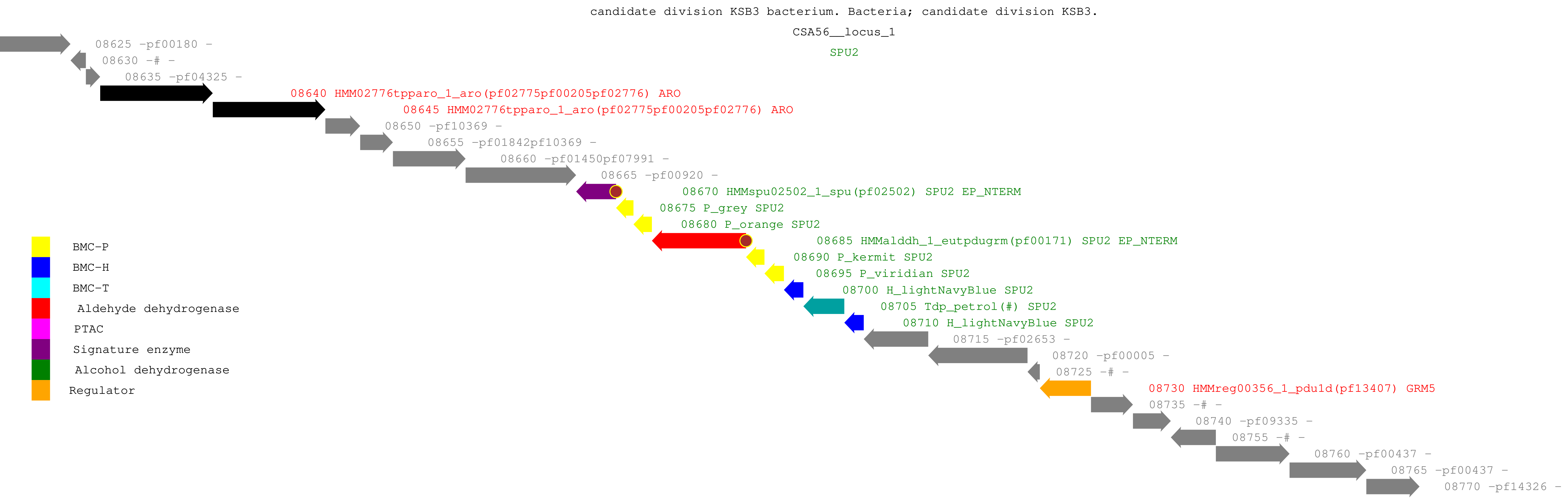
candidate division KSB3 bacterium. Bacteria; candidate division KSB3.
CSA56__locus_1
SPU2
08625 -pf00180 -
08630 -# -
08635 -pf04325 -
08640 HMM02776tpparo_1_aro(pf02775pf00205pf02776) ARO
08645 HMM02776tpparo_1_aro(pf02775pf00205pf02776) ARO
08650 -pf10369 -
08655 -pf01842pf10369 -
08660 -pf01450pf07991 -
08665 -pf00920 -
08670 HMMspu02502_1_spu(pf02502) SPU2 EP_NTERM
08675 P_grey SPU2
08680 P_orange SPU2
08685 HMMaldh_1_eutpdugrm(pf00171) SPU2 EP_NTERM
08690 P_kermit SPU2
08695 P_viridian SPU2
08700 H_lightNavyBlue SPU2
08705 Tdp_petrol(#) SPU2
08710 H_lightNavyBlue SPU2
08715 -pf02653 -
08720 -pf00005 -
08725 -# -
08730 HMMreg00356_1_pdu1d(pf13407) GRM5
08735 -# -
08740 -pf09335 -
08755 -# -
08760 -pf00437 -
08765 -pf00437 -
08770 -pf14326 -
BMC-P
BMC-H
BMC-T
Aldehyde dehydrogenase
PTAC
Signature enzyme
Alcohol dehydrogenase
Regulator

candidate division KSB3 bacterium. Bacteria; candidate division KSB3.

CSB45__locus_1

SPU2

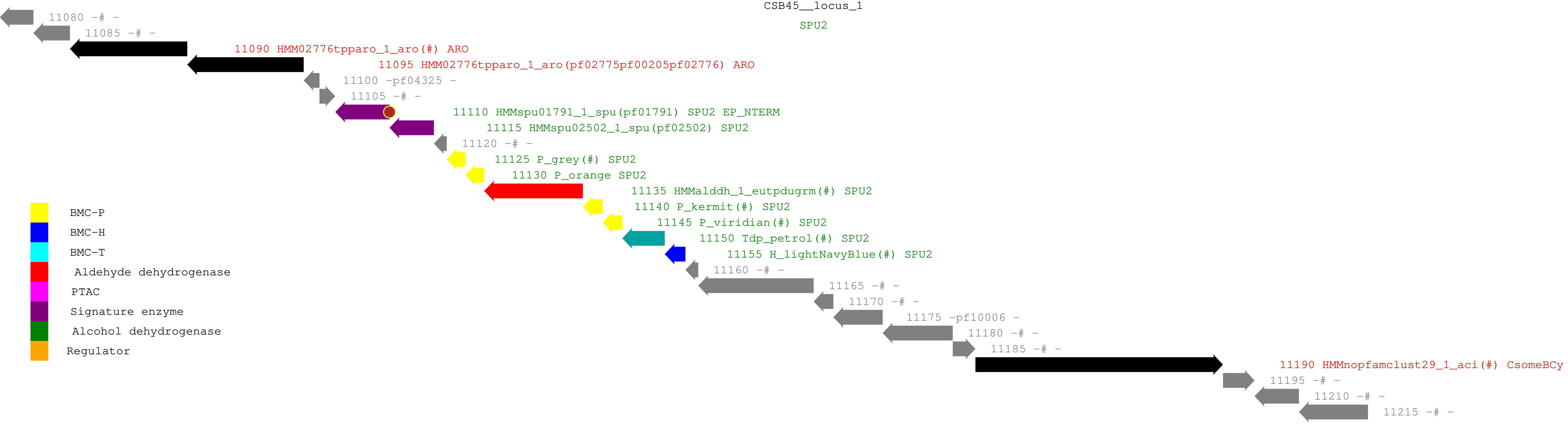

Candidatus Atribacteria bacterium HGW-Atribacteria-1. Bacteria; Candidatus Atribacteria.

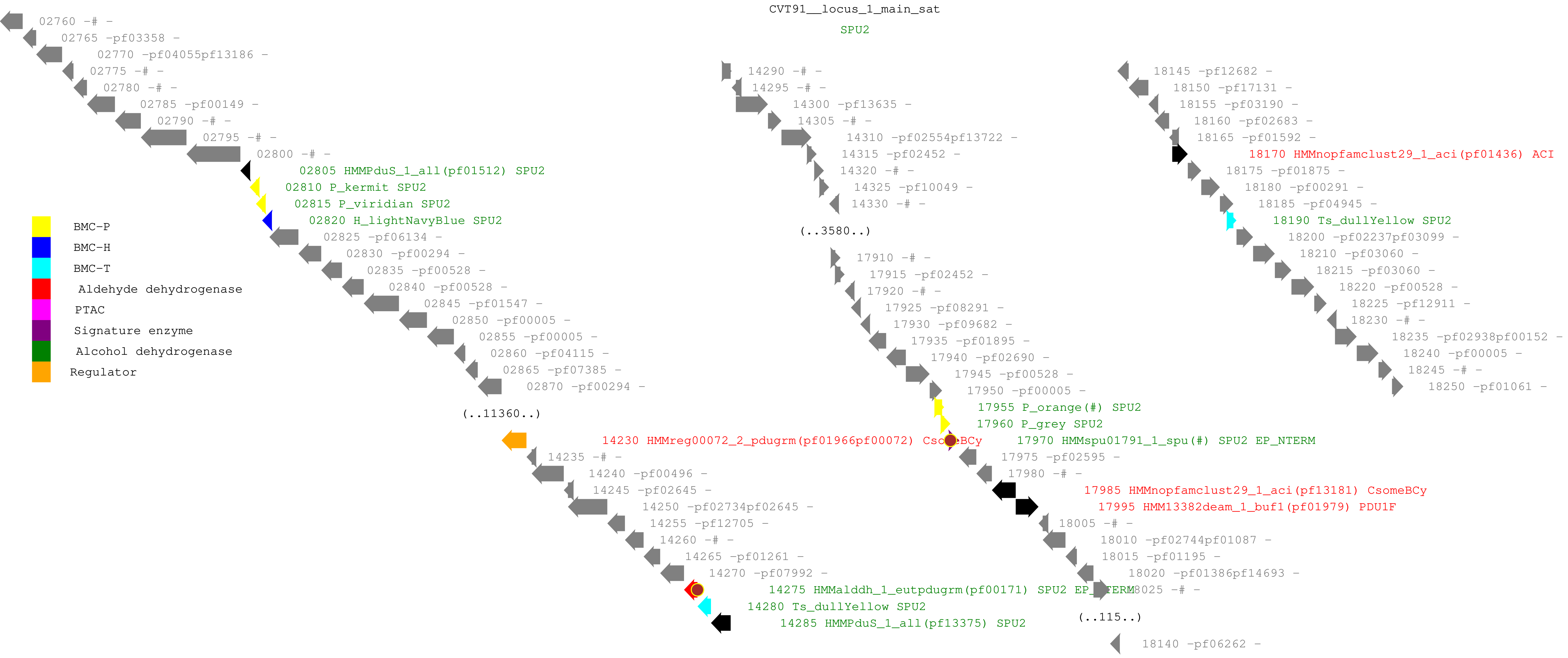

Bacteroidetes bacterium HGW-Bacteroidetes-15. Bacteria; Bacteroidetes.

CVT98__locus_1

SPU2

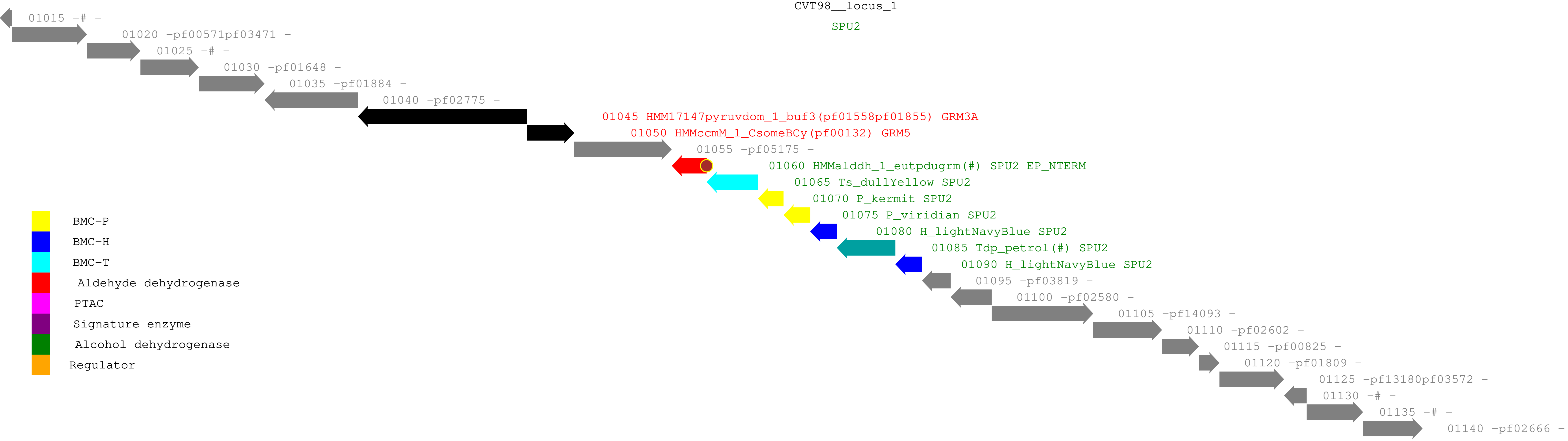

Bacteroidetes bacterium HGW-Bacteroidetes-16. Bacteria; Bacteroidetes.

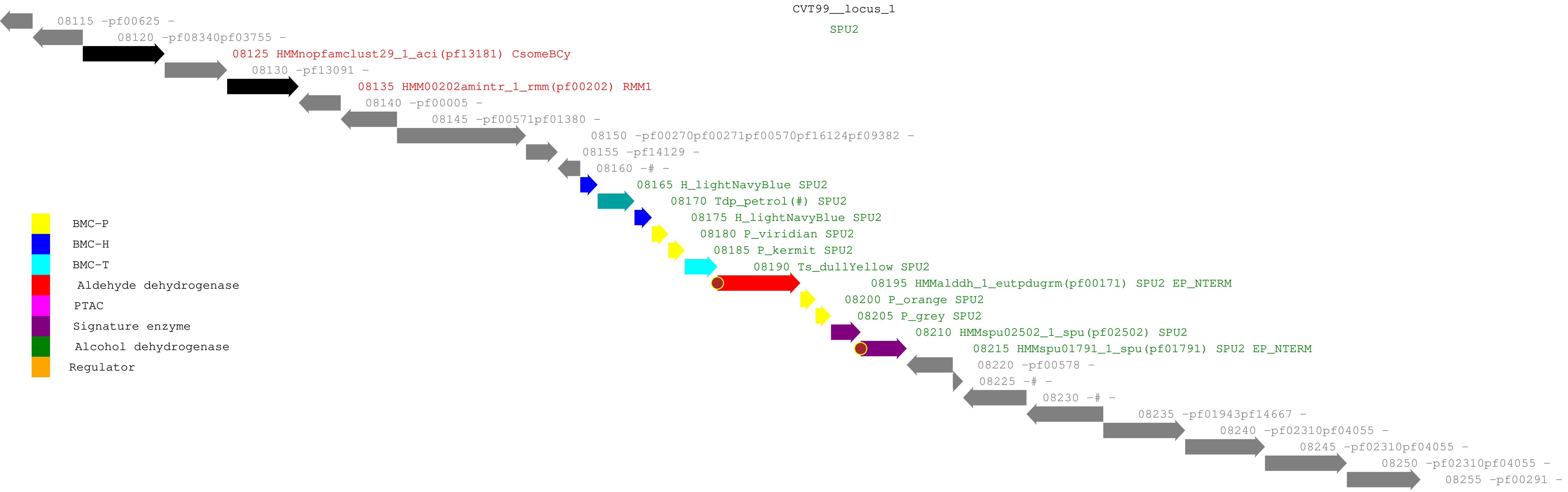

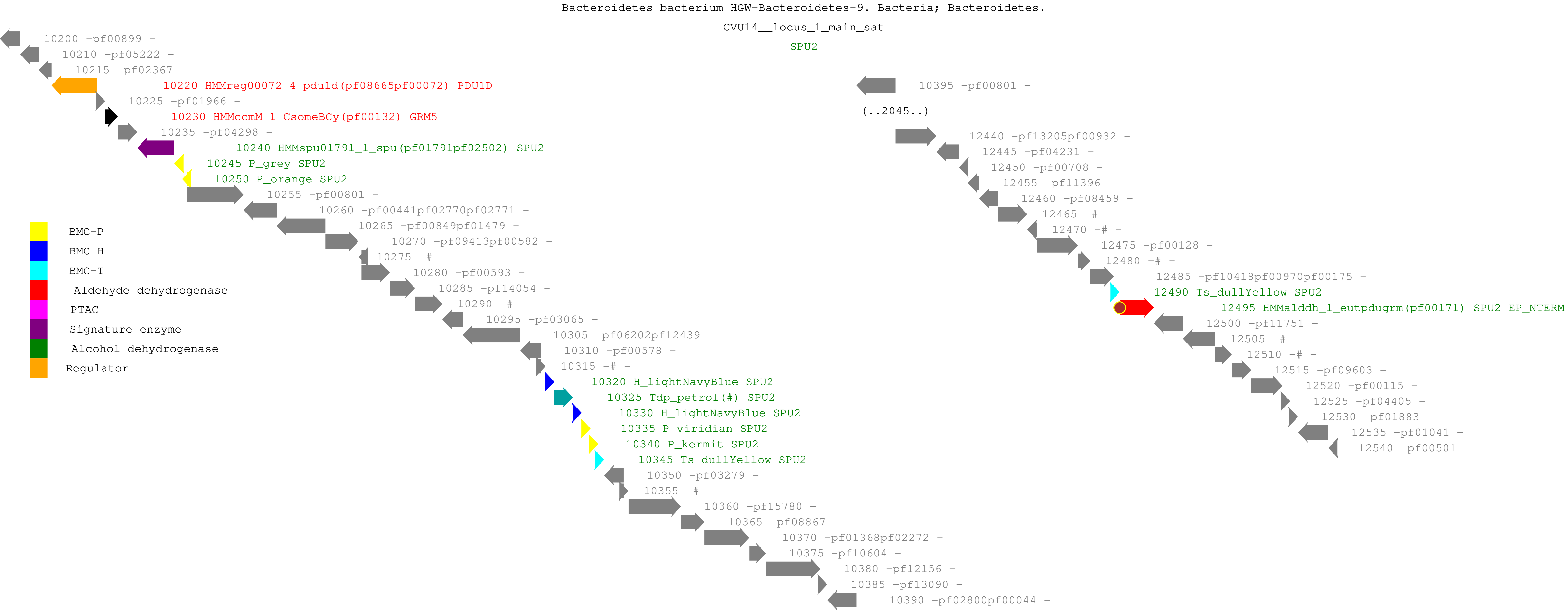

Bacteroidetes bacterium HGW-Bacteroidetes-9. Bacteria; Bacteroidetes.
CVU14__locus_1_main_sat
SPU2
10200 -pf00899 -
10210 -pf05222 -
10215 -pf02367 -
10220 HMMreg00072_4_pdu1d(pf08665pf00072) PDU1D
10225 -pf01966 -
10230 HMMccmM_1_CsomeBCy(pf00132) GRM5
10235 -pf04298 -
10240 HMMspu01791_1_spu(pf01791pf02502) SPU2
10245 P_grey SPU2
10250 P_orange SPU2
10255 -pf00801 -
10260 -pf00441pf02770pf02771 -
10265 -pf00849pf01479 -
10270 -pf09413pf00582 -
10275 -# -
10280 -pf00593 -
10285 -pf14054 -
10290 -# -
10295 -pf03065 -
10305 -pf06202pf12439 -
10310 -pf00578 -
10315 -# -
10320 H_lightNavyBlue SPU2
10325 Tdp_petrol(#) SPU2
10330 H_lightNavyBlue SPU2
10335 P_viridian SPU2
10340 P_kermit SPU2
10345 Ts_dullYellow SPU2
10350 -pf03279 -
10355 -# -
10360 -pf15780 -
10365 -pf08867 -
10370 -pf01368pf02272 -
10375 -pf10604 -
10380 -pf12156 -
10385 -pf13090 -
10390 -pf02800pf00044 -
10395 -pf00801 -
(..2045..)
12440 -pf13205pf00932 -
12445 -pf04231 -
12450 -pf00708 -
12455 -pf11396 -
12460 -pf08459 -
12465 -# -
12470 -# -
12475 -pf00128 -
12480 -# -
12485 -pf10418pf00970pf00175 -
12490 Ts_dullYellow SPU2
12495 HMMalddh_1_eutpdugrm(pf00171) SPU2 EP_NTERM
12500 -pf11751 -
12505 -# -
12510 -# -
12515 -pf09603 -
12520 -pf00115 -
12525 -pf04405 -
12530 -pf01883 -
12535 -pf01041 -
12540 -pf00501 -
BMC-P
BMC-H
BMC-T
Aldehyde dehydrogenase
PTAC
Signature enzyme
Alcohol dehydrogenase
Regulator

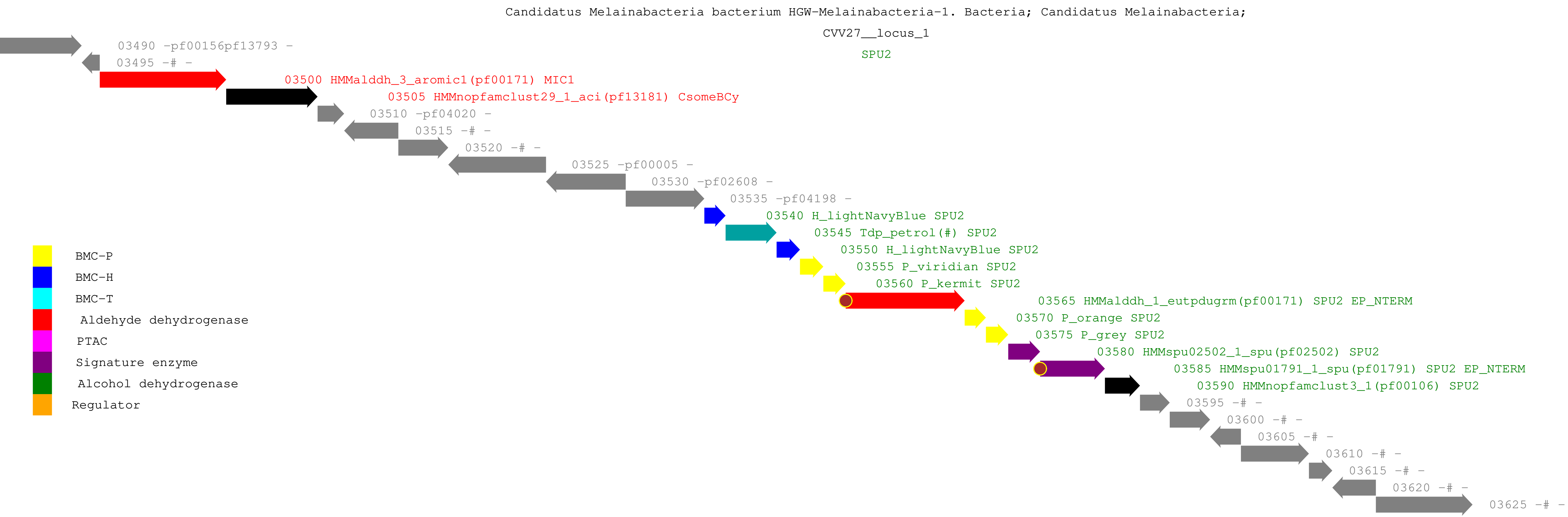
Candidatus Melainabacteria bacterium HGW-Melainabacteria-1. Bacteria; Candidatus Melainabacteria;
CVV27__locus_1
SPU2
03490 -pf00156pf13793 -
03495 -# -
03500 HMMaldDh_3_aromic1(pf00171) MIC1
03505 HMMnopfamclust29_1_aci(pf13181) CsomeBCy
03510 -pf04020 -
03515 -# -
03520 -# -
03525 -pf00005 -
03530 -pf02608 -
03535 -pf04198 -
03540 H_lightNavyBlue SPU2
03545 Tdp_petrol(#) SPU2
03550 H_lightNavyBlue SPU2
03555 P_viridian SPU2
03560 P_kermit SPU2
03565 HMMaldDh_1_eutpdugrm(pf00171) SPU2 EP_NTERM
03570 P_orange SPU2
03575 P_grey SPU2
03580 HMMspu02502_1_spu(pf02502) SPU2
03585 HMMspu01791_1_spu(pf01791) SPU2 EP_NTERM
03590 HMMnopfamclust3_1(pf00106) SPU2
03595 -# -
03600 -# -
03605 -# -
03610 -# -
03615 -# -
03620 -# -
03625 -# -
BMC-P
BMC-H
BMC-T
Aldehyde dehydrogenase
PTAC
Signature enzyme
Alcohol dehydrogenase
Regulator

Spirochaetae bacterium HGW-Spirochaetae-2. Bacteria; Spirochaetes.

CVV46__locus_1

SPU2

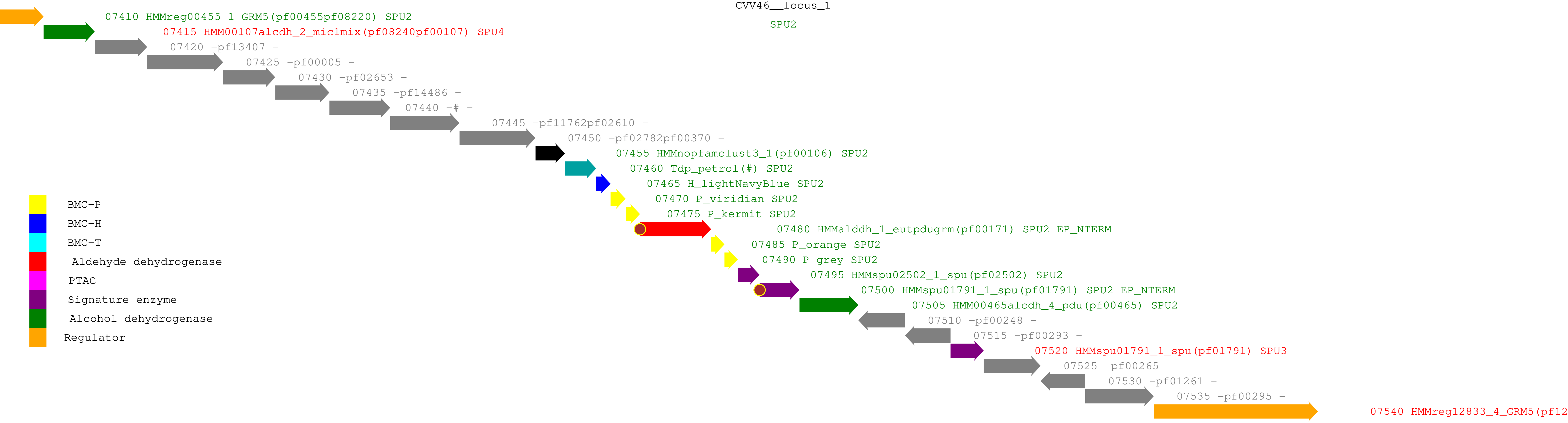

Spirochaetae bacterium HGW-Spirochaetae-4. Bacteria; Spirochaetes.

CVV48__locus_1

SPU2

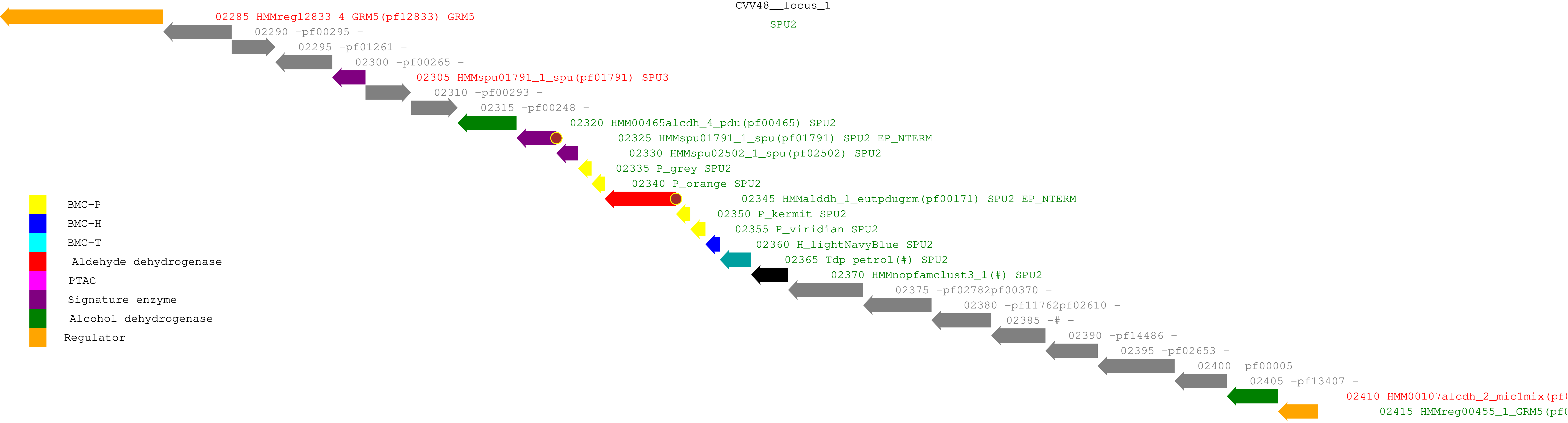

Candidatus Wallbacteria bacterium HGW-Wallbacteria-1. Bacteria; Candidatus Wallbacteria.

CVV64__locus_2

SPU2

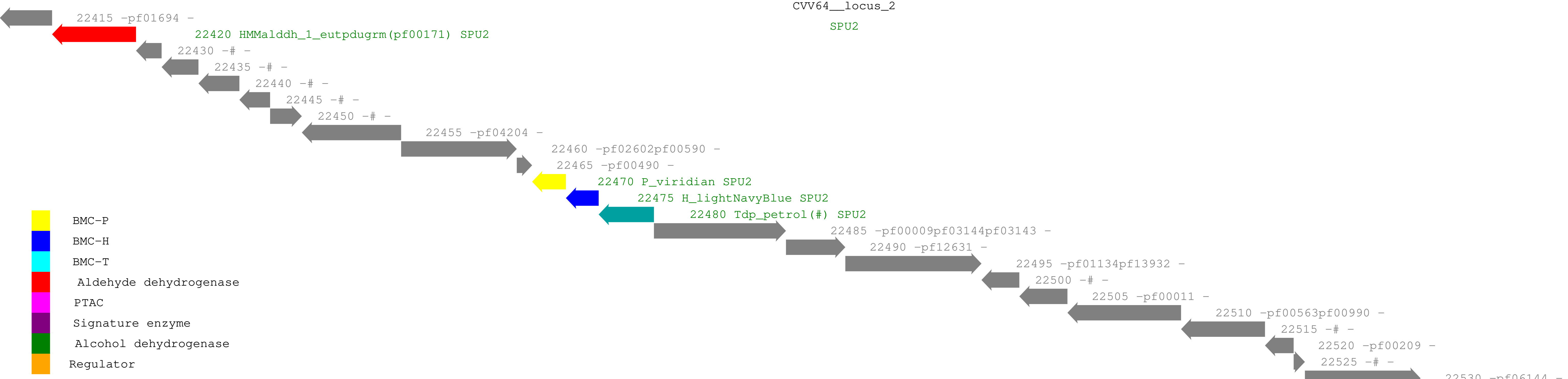

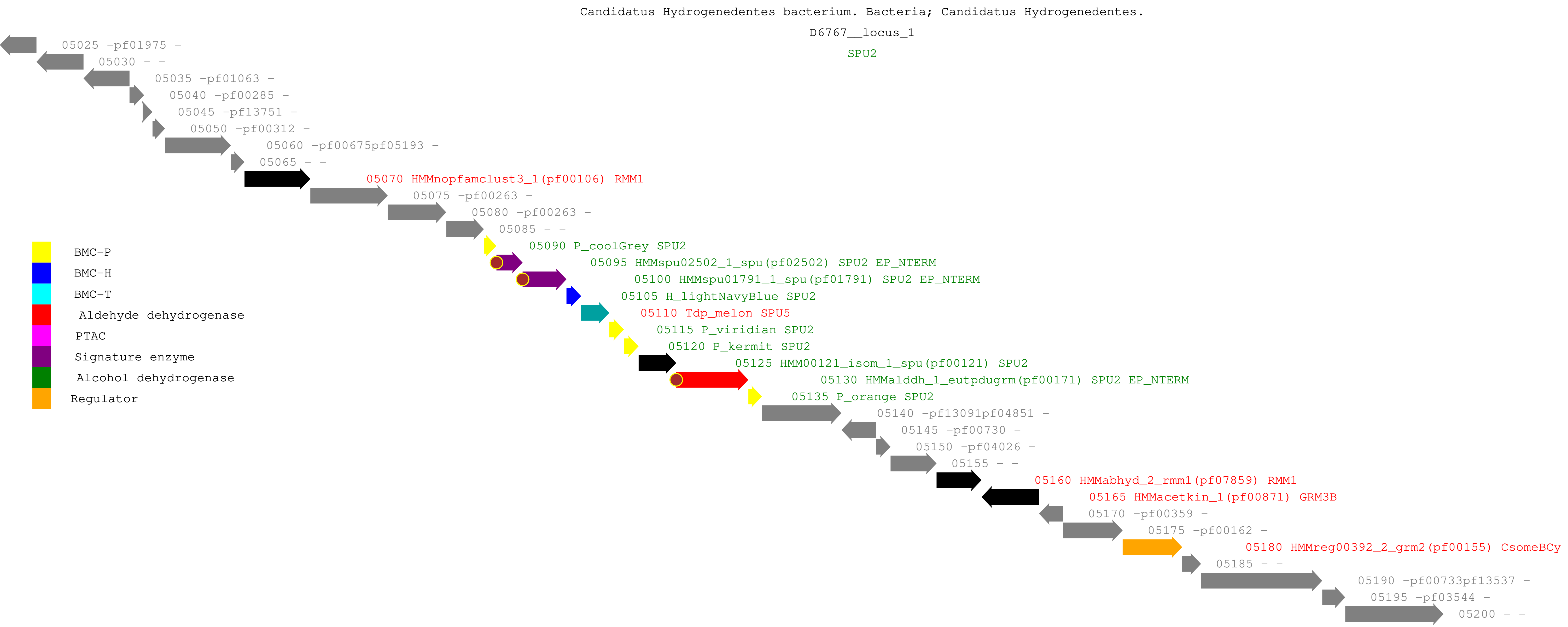
Candidatus Hydrogenedentes bacterium. Bacteria; Candidatus Hydrogenedentes.
D6767__locus_1
SPU2
05025 -pf01975 -
05030 - -
05035 -pf01063 -
05040 -pf00285 -
05045 -pf13751 -
05050 -pf00312 -
05060 -pf00675pf05193 -
05065 - -
05070 HMMnopfamclust3_1(pf00106) RMM1
05075 -pf00263 -
05080 -pf00263 -
05085 - -
05090 P_coolGrey SPU2
05095 HMMspu02502_1_spu(pf02502) SPU2 EP_NTERM
05100 HMMspu01791_1_spu(pf01791) SPU2 EP_NTERM
05105 H_lightNavyBlue SPU2
05110 Tdp_melon SPU5
05115 P_viridian SPU2
05120 P_kermit SPU2
05125 HMM00121_isom_1_spu(pf00121) SPU2
05130 HMMalddh_1_eutpdugrm(pf00171) SPU2 EP_NTERM
05135 P_orange SPU2
05140 -pf13091pf04851 -
05145 -pf00730 -
05150 -pf04026 -
05155 - -
05160 HMMabhyd_2_rmm1(pf07859) RMM1
05165 HMMacetkin_1(pf00871) GRM3B
05170 -pf00359 -
05175 -pf00162 -
05180 HMMreg00392_2_grm2(pf00155) CsomeBCy
05185 - -
05190 -pf00733pf13537 -
05195 -pf03544 -
05200 - -
BMC-P
BMC-H
BMC-T
Aldehyde dehydrogenase
PTAC
Signature enzyme
Alcohol dehydrogenase
Regulator

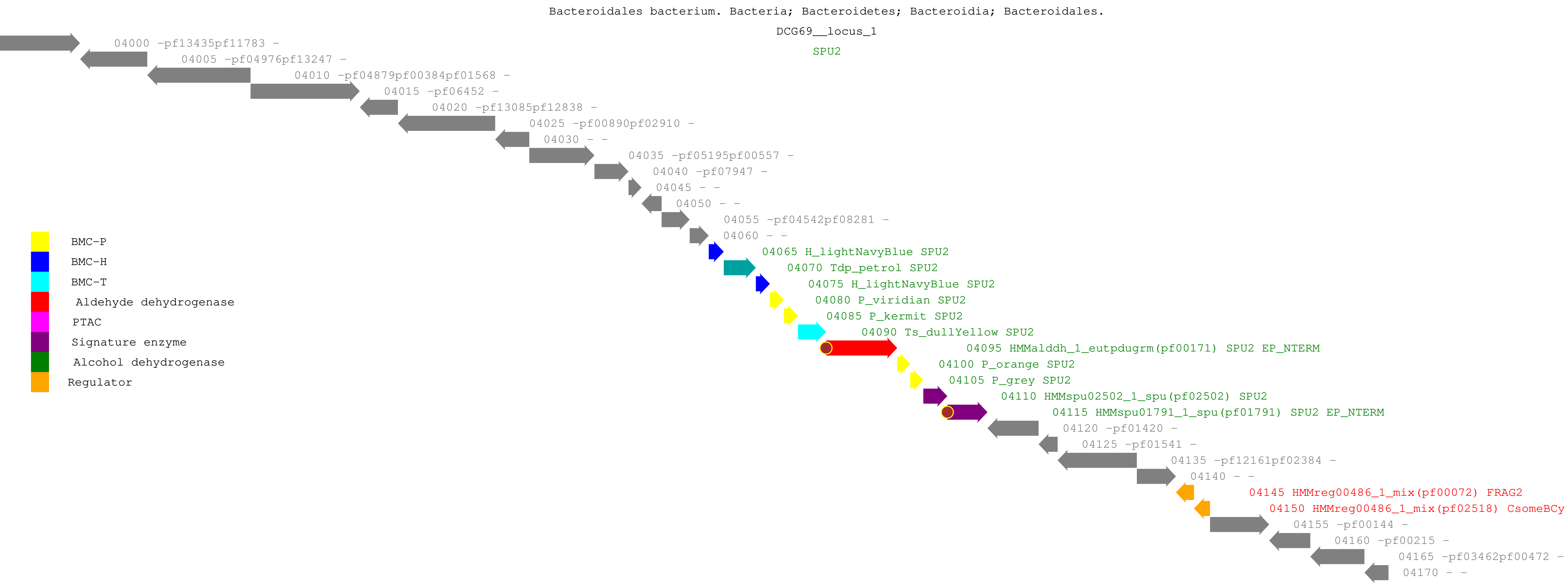

Bacteroidales bacterium. Bacteria; Bacteroidetes; Bacteroidia; Bacteroidales.
DCG69__locus_1
SPU2
04000 -pf13435pf11783 -
04005 -pf04976pf13247 -
04010 -pf04879pf00384pf01568 -
04015 -pf06452 -
04020 -pf13085pf12838 -
04025 -pf00890pf02910 -
04030 - -
04035 -pf05195pf00557 -
04040 -pf07947 -
04045 - -
04050 - -
04055 -pf04542pf08281 -
04060 - -
04065 H_lightNavyBlue SPU2
04070 Tdp_petrol SPU2
04075 H_lightNavyBlue SPU2
04080 P_viridian SPU2
04085 P_kermit SPU2
04090 Ts_dullYellow SPU2
04095 HMMalddh_1_eutpdugrm(pf00171) SPU2 EP_NTERM
04100 P_orange SPU2
04105 P_grey SPU2
04110 HMMspu02502_1_spu(pf02502) SPU2
04115 HMMspu01791_1_spu(pf01791) SPU2 EP_NTERM
04120 -pf01420 -
04125 -pf01541 -
04135 -pf12161pf02384 -
04140 - -
04145 HMMreg00486_1_mix(pf00072) FRAG2
04150 HMMreg00486_1_mix(pf02518) CsomeBCy
04155 -pf00144 -
04160 -pf00215 -
04165 -pf03462pf00472 -
04170 - -
BMC-P
BMC-H
BMC-T
Aldehyde dehydrogenase
PTAC
Signature enzyme
Alcohol dehydrogenase
Regulator

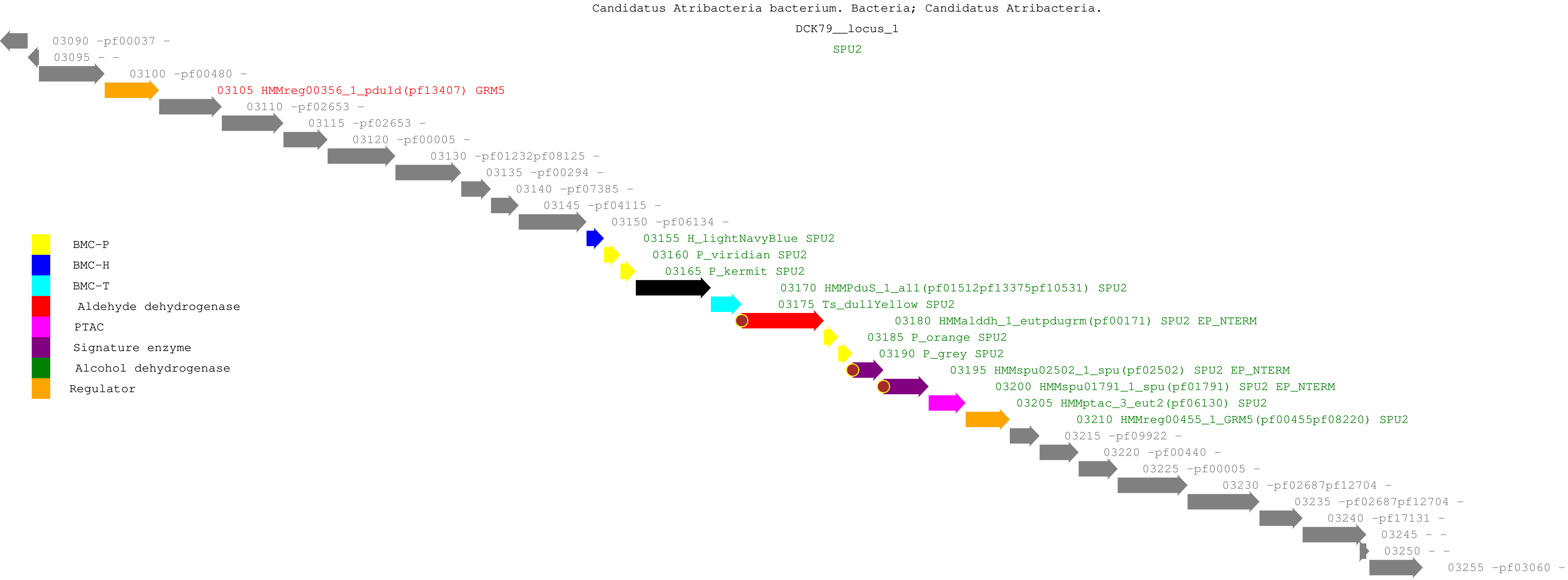
Candidatus Atribacteria bacterium. Bacteria; Candidatus Atribacteria.
DCK79__locus_1
SPU2
03090 -pf00037 -
03095 - -
03100 -pf00480 -
03105 HMMreg00356_1_pdu1d(pf13407) GRM5
03110 -pf02653 -
03115 -pf02653 -
03120 -pf00005 -
03130 -pf01232pf08125 -
03135 -pf00294 -
03140 -pf07385 -
03145 -pf04115 -
03150 -pf06134 -
03155 H_lightNavyBlue SPU2
03160 P_viridian SPU2
03165 P_kermit SPU2
03170 HMMPduS_1_all(pf01512pf13375pf10531) SPU2
03175 Ts_dullYellow SPU2
03180 HMMalddh_1_eutpdugrm(pf00171) SPU2 EP_NTERM
03185 P_orange SPU2
03190 P_grey SPU2
03195 HMMspu02502_1_spu(pf02502) SPU2 EP_NTERM
03200 HMMspu01791_1_spu(pf01791) SPU2 EP_NTERM
03205 HMMptac_3_eut2(pf06130) SPU2
03210 HMMreg00455_1_GRM5(pf00455pf08220) SPU2
03215 -pf09922 -
03220 -pf00440 -
03225 -pf00005 -
03230 -pf02687pf12704 -
03235 -pf02687pf12704 -
03240 -pf17131 -
03245 - -
03250 - -
03255 -pf03060 -
BMC-P
BMC-H
BMC-T
Aldehyde dehydrogenase
PTAC
Signature enzyme
Alcohol dehydrogenase
Regulator

Bacteroidales bacterium. Bacteria; Bacteroidetes; Bacteroidia; Bacteroidales.

DCM62__locus_1

SPU2

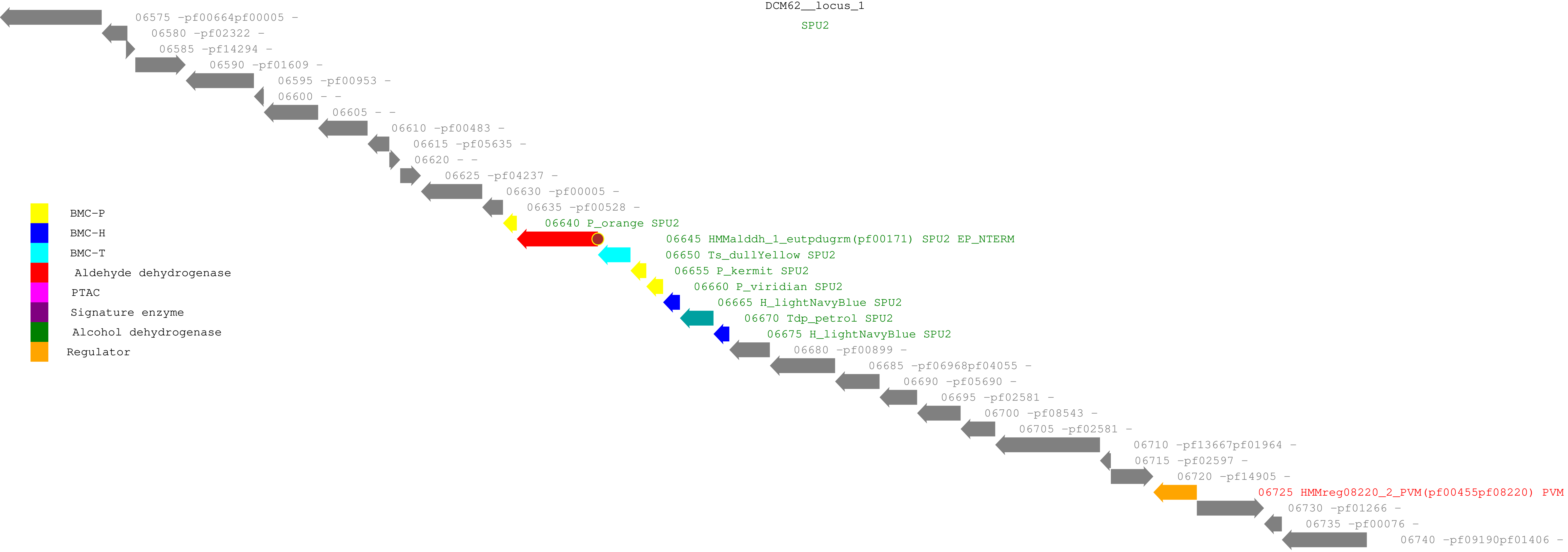

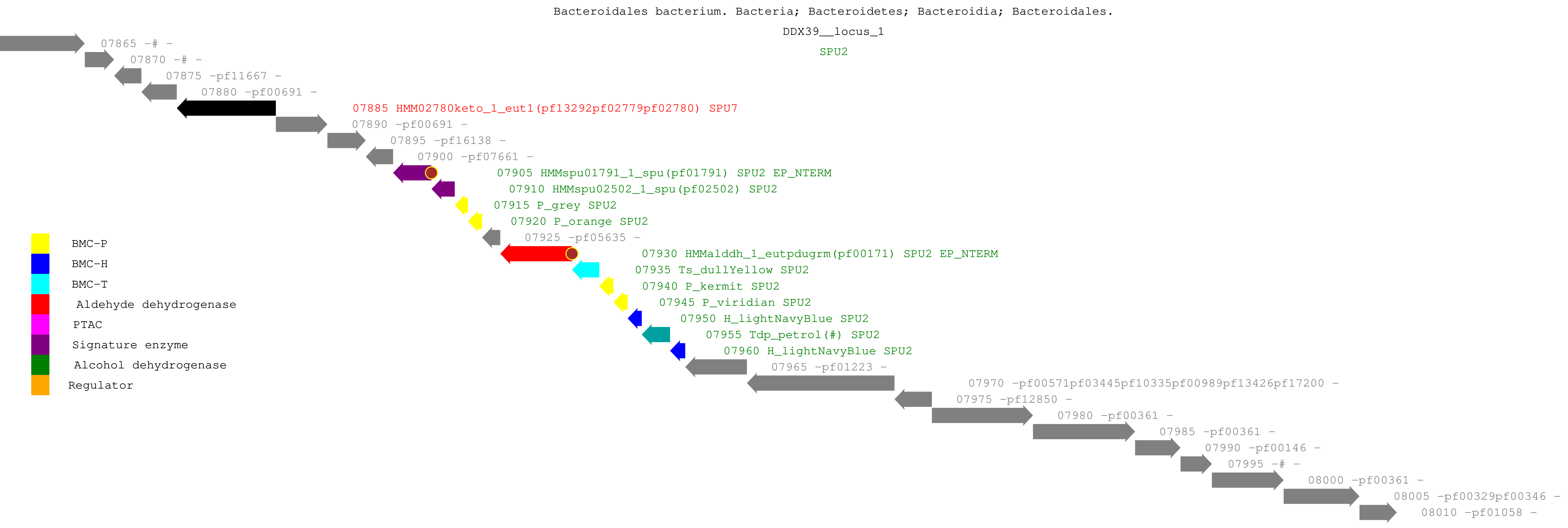
Bacteroidales bacterium. Bacteria; Bacteroidetes; Bacteroidia; Bacteroidales.
DDX39__locus_1
SPU2
07865 -# -
07870 -# -
07875 -pf11667 -
07880 -pf00691 -
07885 HMM02780keto_1_eut1(pf13292pf02779pf02780) SPU7
07890 -pf00691 -
07895 -pf16138 -
07900 -pf07661 -
07905 HMMspu01791_1_spu(pf01791) SPU2 EP_NTERM
07910 HMMspu02502_1_spu(pf02502) SPU2
07915 P_grey SPU2
07920 P_orange SPU2
07925 -pf05635 -
07930 HMMalddh_1_eutpdugrm(pf00171) SPU2 EP_NTERM
07935 Ts_dullYellow SPU2
07940 P_kermit SPU2
07945 P_viridian SPU2
07950 H_lightNavyBlue SPU2
07955 Tdp_petrol(#) SPU2
07960 H_lightNavyBlue SPU2
07965 -pf01223 -
07970 -pf00571pf03445pf10335pf00989pf13426pf17200 -
07975 -pf12850 -
07980 -pf00361 -
07985 -pf00361 -
07990 -pf00146 -
07995 -# -
08000 -pf00361 -
08005 -pf00329pf00346 -
08010 -pf01058 -
BMC-P
BMC-H
BMC-T
Aldehyde dehydrogenase
PTAC
Signature enzyme
Alcohol dehydrogenase
Regulator

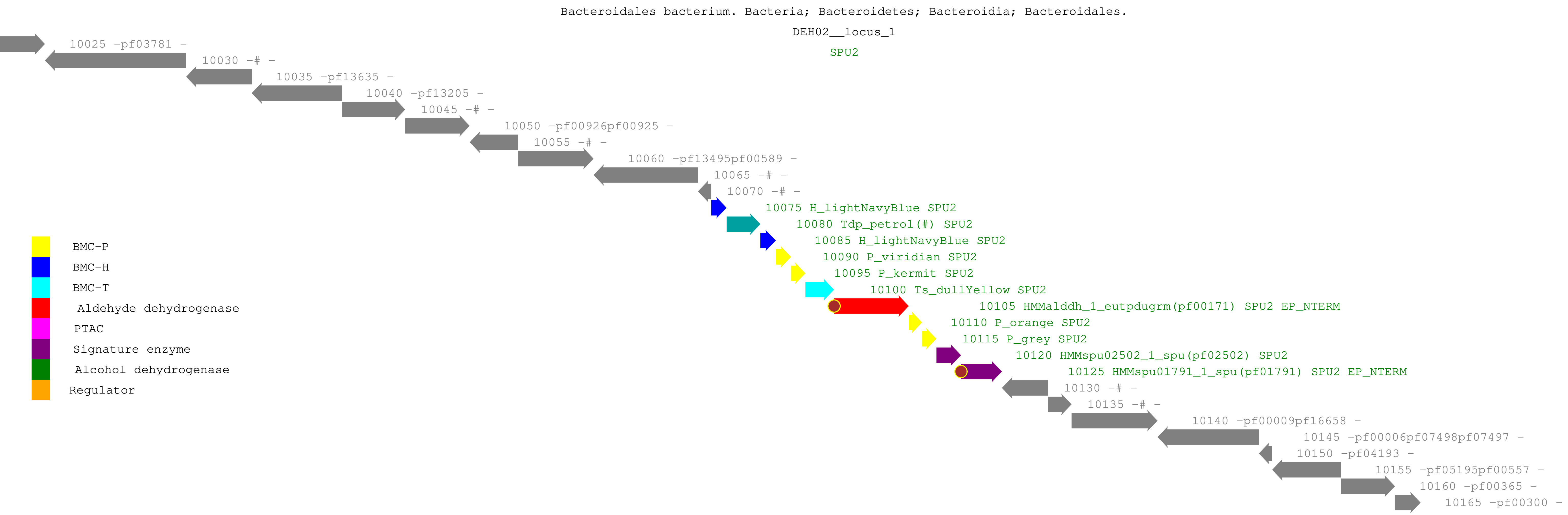
Bacteroidales bacterium. Bacteria; Bacteroidetes; Bacteroidia; Bacteroidales.
DEH02__locus_1
SPU2
10025 -pf03781 -
10030 -# -
10035 -pf13635 -
10040 -pf13205 -
10045 -# -
10050 -pf00926pf00925 -
10055 -# -
10060 -pf13495pf00589 -
10065 -# -
10070 -# -
10075 H_lightNavyBlue SPU2
10080 Tdp_petrol(#) SPU2
10085 H_lightNavyBlue SPU2
10090 P_viridian SPU2
10095 P_kermit SPU2
10100 Ts_dullYellow SPU2
10105 HMMalddh_1_eutpdugrm(pf00171) SPU2 EP_NTERM
10110 P_orange SPU2
10115 P_grey SPU2
10120 HMMspu02502_1_spu(pf02502) SPU2
10125 HMMspu01791_1_spu(pf01791) SPU2 EP_NTERM
10130 -# -
10135 -# -
10140 -pf00009pf16658 -
10145 -pf00006pf07498pf07497 -
10150 -pf04193 -
10155 -pf05195pf00557 -
10160 -pf00365 -
10165 -pf00300 -
BMC-P
BMC-H
BMC-T
Aldehyde dehydrogenase
PTAC
Signature enzyme
Alcohol dehydrogenase
Regulator

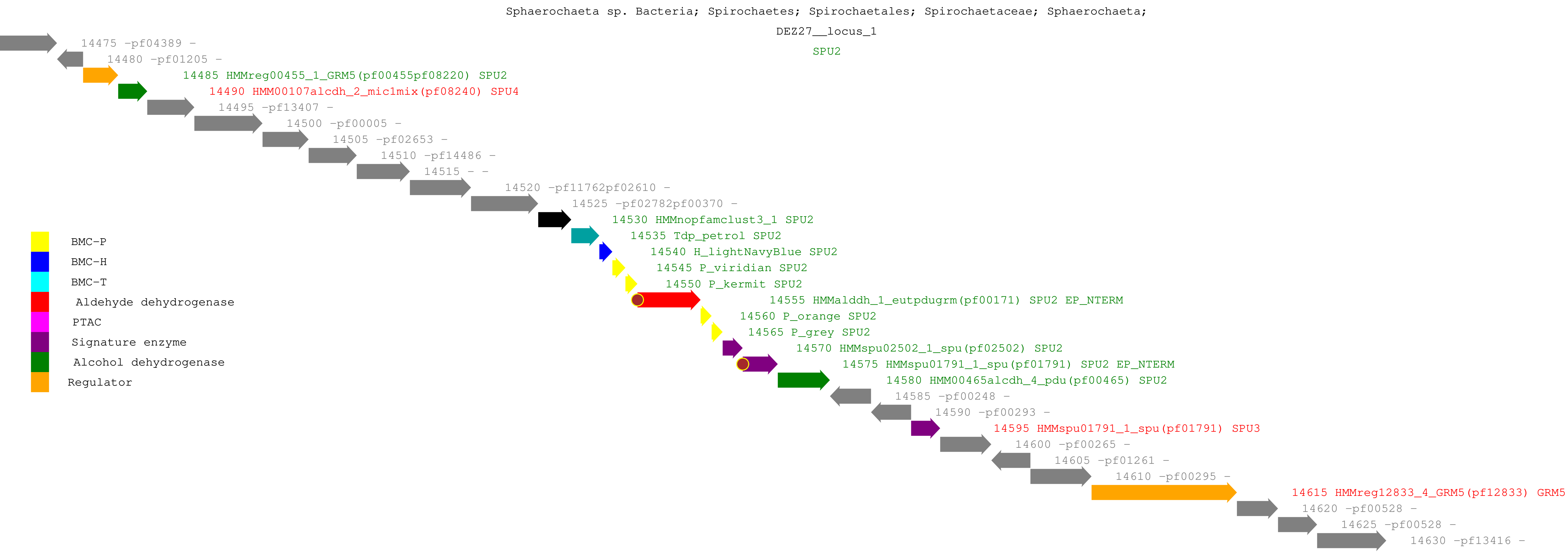

Sphaerochaeta sp. Bacteria; Spirochaetes; Spirochaetales; Spirochaetaceae; Sphaerochaeta;
DEZ27__locus_1
SPU2
14475 -pf04389 -
14480 -pf01205 -
14485 HMMreg00455_1_GRM5(pf00455pf08220) SPU2
14490 HMM00107alcdh_2_mic1mix(pf08240) SPU4
14495 -pf13407 -
14500 -pf00005 -
14505 -pf02653 -
14510 -pf14486 -
14515 - -
14520 -pf11762pf02610 -
14525 -pf02782pf00370 -
14530 HMMnopfamclust3_1 SPU2
14535 Tdp_petrol SPU2
14540 H_lightNavyBlue SPU2
14545 P_viridian SPU2
14550 P_kermit SPU2
14555 HMMaldh_1_eutpdugrm(pf00171) SPU2 EP_NTERM
14560 P_orange SPU2
14565 P_grey SPU2
14570 HMMspu02502_1_spu(pf02502) SPU2
14575 HMMspu01791_1_spu(pf01791) SPU2 EP_NTERM
14580 HMM00465alcdh_4_pdu(pf00465) SPU2
14585 -pf00248 -
14590 -pf00293 -
14595 HMMspu01791_1_spu(pf01791) SPU3
14600 -pf00265 -
14605 -pf01261 -
14610 -pf00295 -
14615 HMMreg12833_4_GRM5(pf12833) GRM5
14620 -pf00528 -
14625 -pf00528 -
14630 -pf13416 -
BMC-P
BMC-H
BMC-T
Aldehyde dehydrogenase
PTAC
Signature enzyme
Alcohol dehydrogenase
Regulator

Candidatus Atribacteria bacterium. Bacteria; Candidatus Atribacteria.

DF698__locus_1

SPU2

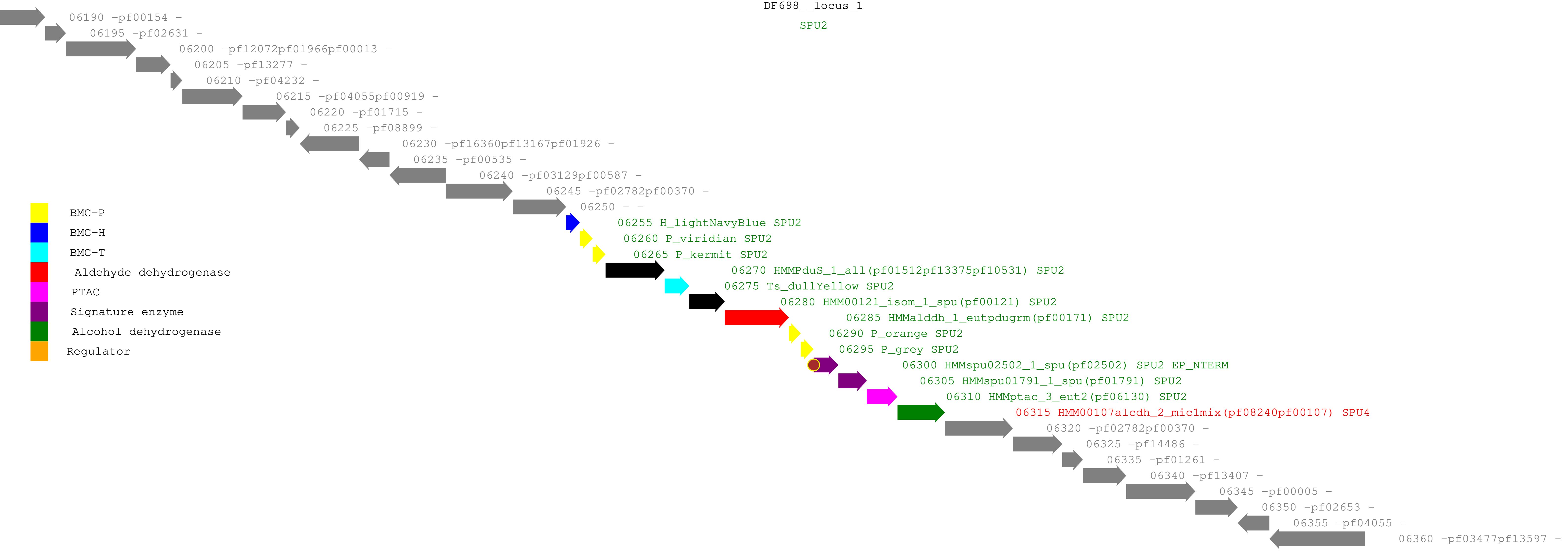

Sphaerochaeta sp. Bacteria; Spirochaetes; Spirochaetales; Spirochaetaceae; Sphaerochaeta;

DHV69__locus_1

SPU2

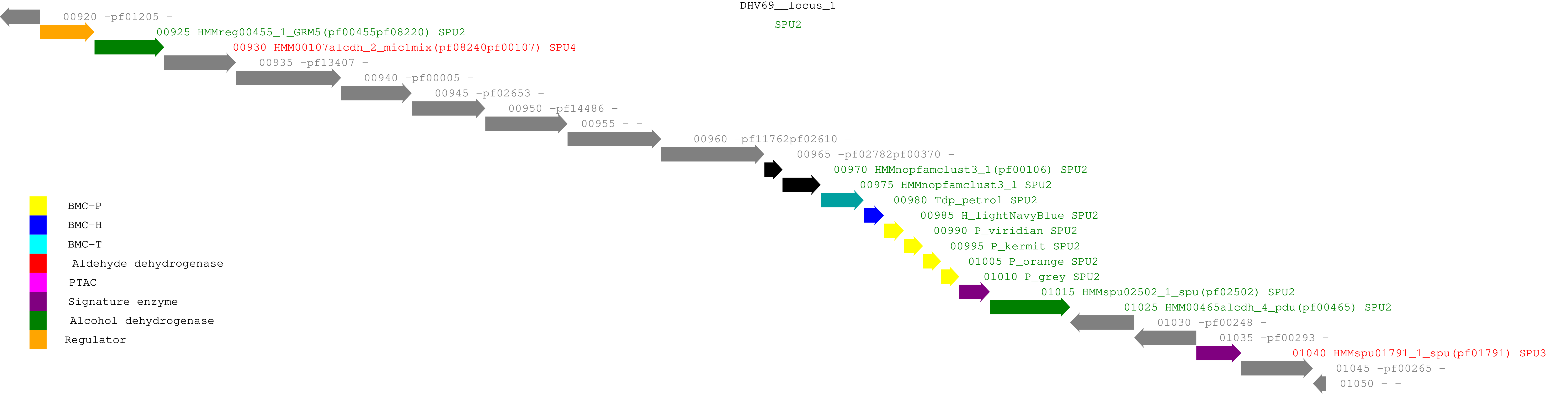

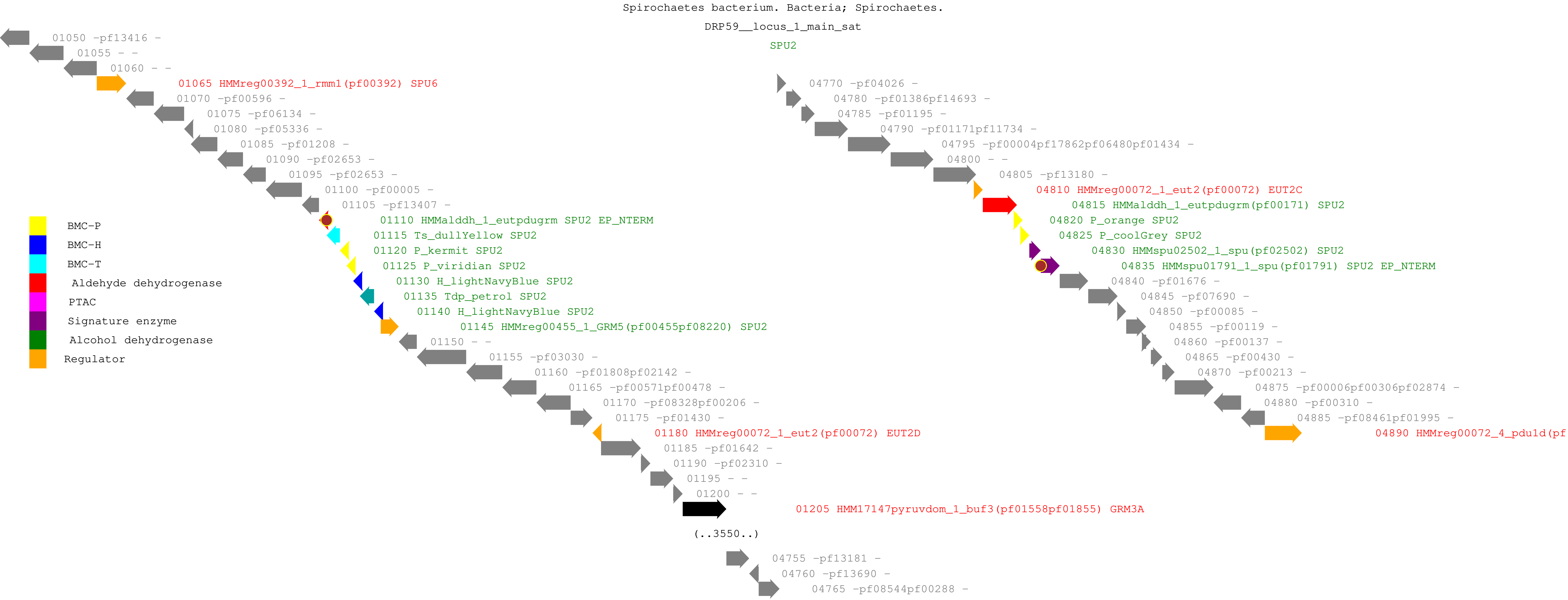
Spirochaetes bacterium. Bacteria; Spirochaetes.
DRP59__locus_1_main_sat
SPU2
01050 -pf13416 -
01055 - -
01060 - -
01065 HMMreg00392_1_rmm1(pf00392) SPU6
01070 -pf00596 -
01075 -pf06134 -
01080 -pf05336 -
01085 -pf01208 -
01090 -pf02653 -
01095 -pf02653 -
01100 -pf00005 -
01105 -pf13407 -
01110 HMMalddh_1_eutpdugrm SPU2 EP_NTERM
01115 Ts_dullYellow SPU2
01120 P_kermit SPU2
01125 P_viridian SPU2
01130 H_lightNavyBlue SPU2
01135 Tdp_petrol SPU2
01140 H_lightNavyBlue SPU2
01145 HMMreg00455_1_GRM5(pf00455pf08220) SPU2
01150 - -
01155 -pf03030 -
01160 -pf01808pf02142 -
01165 -pf00571pf00478 -
01170 -pf08328pf00206 -
01175 -pf01430 -
01180 HMMreg00072_1_eut2(pf00072) EUT2D
01185 -pf01642 -
01190 -pf02310 -
01195 - -
01200 - -
01205 HMM17147pyruvdom_1_buf3(pf01558pf01855) GRM3A
(..3550..)
04755 -pf13181 -
04760 -pf13690 -
04765 -pf08544pf00288 -
04770 -pf04026 -
04780 -pf01386pf14693 -
04785 -pf01195 -
04790 -pf01171pf11734 -
04795 -pf00004pf17862pf06480pf01434 -
04800 - -
04805 -pf13180 -
04810 HMMreg00072_1_eut2(pf00072) EUT2C
04815 HMMalddh_1_eutpdugrm(pf00171) SPU2
04820 P_orange SPU2
04825 P_coolGrey SPU2
04830 HMMspu02502_1_spu(pf02502) SPU2
04835 HMMspu01791_1_spu(pf01791) SPU2 EP_NTERM
04840 -pf01676 -
04845 -pf07690 -
04850 -pf00085 -
04855 -pf00119 -
04860 -pf00137 -
04865 -pf00430 -
04870 -pf00213 -
04875 -pf00006pf00306pf02874 -
04880 -pf00310 -
04885 -pf08461pf01995 -
04890 HMMreg00072_4_pdu1d(pf
BMC-P
BMC-H
BMC-T
Aldehyde dehydrogenase
PTAC
Signature enzyme
Alcohol dehydrogenase
Regulator

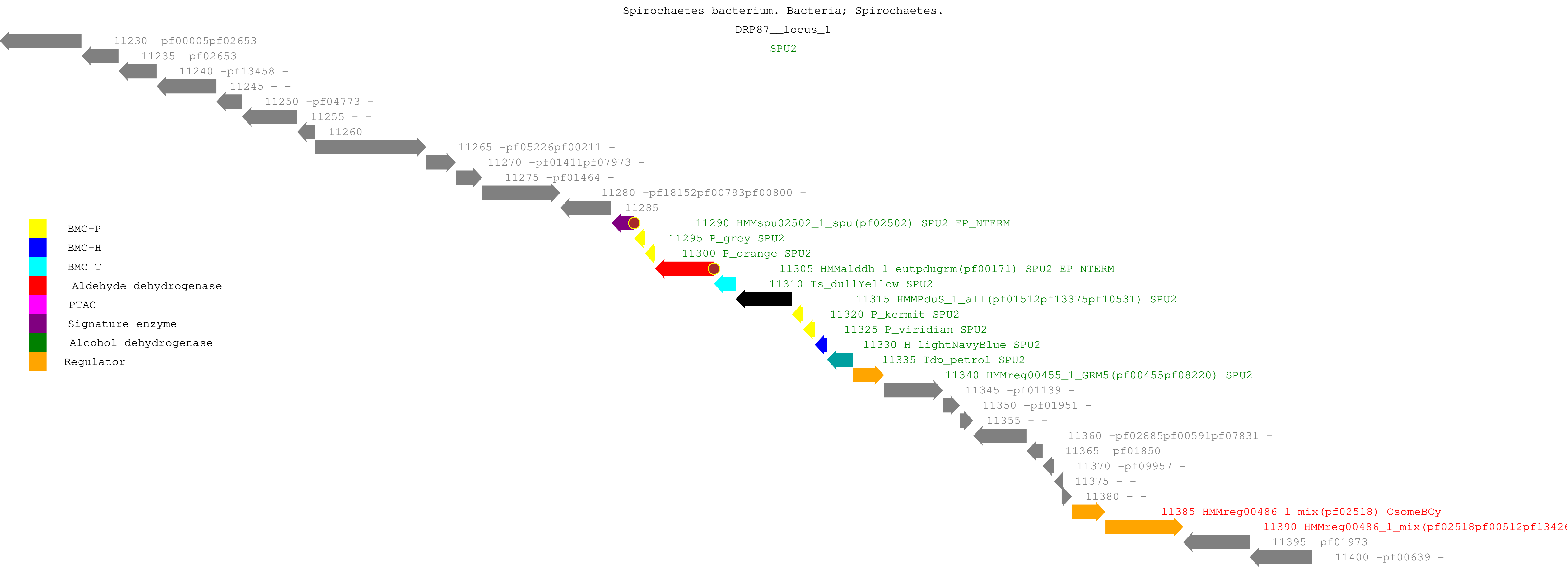

Spirochaetes bacterium. Bacteria; Spirochaetes.
DRP87__locus_1
SPU2
11230 -pf00005pf02653 -
11235 -pf02653 -
11240 -pf13458 -
11245 - -
11250 -pf04773 -
11255 - -
11260 - -
11265 -pf05226pf00211 -
11270 -pf01411pf07973 -
11275 -pf01464 -
11280 -pf18152pf00793pf00800 -
11285 - -
11290 HMMspu02502_1_spu(pf02502) SPU2 EP_NTERM
11295 P_grey SPU2
11300 P_orange SPU2
11305 HMMaldh_1_eutpdugrm(pf00171) SPU2 EP_NTERM
11310 Ts_dullYellow SPU2
11315 HMMPduS_1_all(pf01512pf13375pf10531) SPU2
11320 P_kermit SPU2
11325 P_viridian SPU2
11330 H_lightNavyBlue SPU2
11335 Tdp_petrol SPU2
11340 HMMreg00455_1_GRM5(pf00455pf08220) SPU2
11345 -pf01139 -
11350 -pf01951 -
11355 - -
11360 -pf02885pf00591pf07831 -
11365 -pf01850 -
11370 -pf09957 -
11375 - -
11380 - -
11385 HMMreg00486_1_mix(pf02518) CsomeBCy
11390 HMMreg00486_1_mix(pf02518pf00512pf1342
11395 -pf01973 -
11400 -pf00639 -
BMC-P
BMC-H
BMC-T
Aldehyde dehydrogenase
PTAC
Signature enzyme
Alcohol dehydrogenase
Regulator

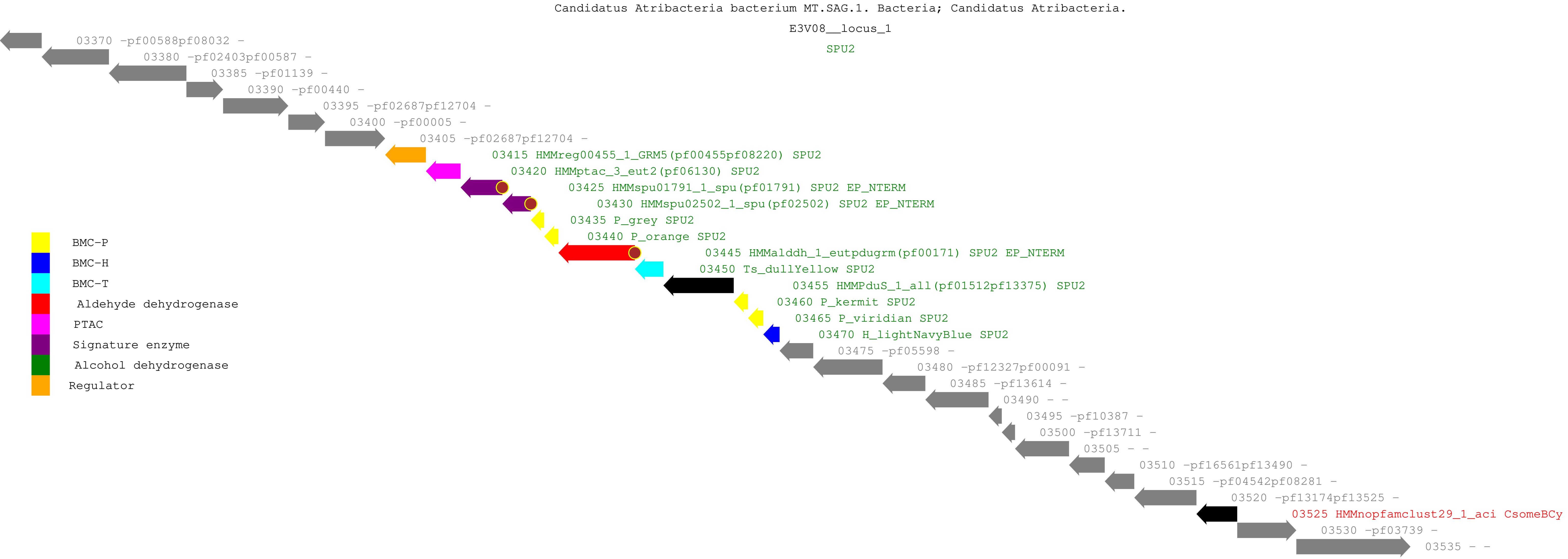
Candidatus Atribacteria bacterium MT.SAG.1. Bacteria; Candidatus Atribacteria.
E3V08__locus_1
SPU2
03370 -pf00588pf08032 -
03380 -pf02403pf00587 -
03385 -pf01139 -
03390 -pf00440 -
03395 -pf02687pf12704 -
03400 -pf00005 -
03405 -pf02687pf12704 -
03415 HMMreg00455_1_GRM5(pf00455pf08220) SPU2
03420 HMMptac_3_eut2(pf06130) SPU2
03425 HMMspu01791_1_spu(pf01791) SPU2 EP_NTERM
03430 HMMspu02502_1_spu(pf02502) SPU2 EP_NTERM
03435 P_grey SPU2
03440 P_orange SPU2
03445 HMMaldh_1_eutpdugrm(pf00171) SPU2 EP_NTERM
03450 Ts_dullYellow SPU2
03455 HMMPduS_1_all(pf01512pf13375) SPU2
03460 P_kermit SPU2
03465 P_viridian SPU2
03470 H_lightNavyBlue SPU2
03475 -pf05598 -
03480 -pf12327pf00091 -
03485 -pf13614 -
03490 - -
03495 -pf10387 -
03500 -pf13711 -
03505 - -
03510 -pf16561pf13490 -
03515 -pf04542pf08281 -
03520 -pf13174pf13525 -
03525 HMMnopfamclust29_1_aci CsomeBCy
03530 -pf03739 -
03535 - -
BMC-P
BMC-H
BMC-T
Aldehyde dehydrogenase
PTAC
Signature enzyme
Alcohol dehydrogenase
Regulator

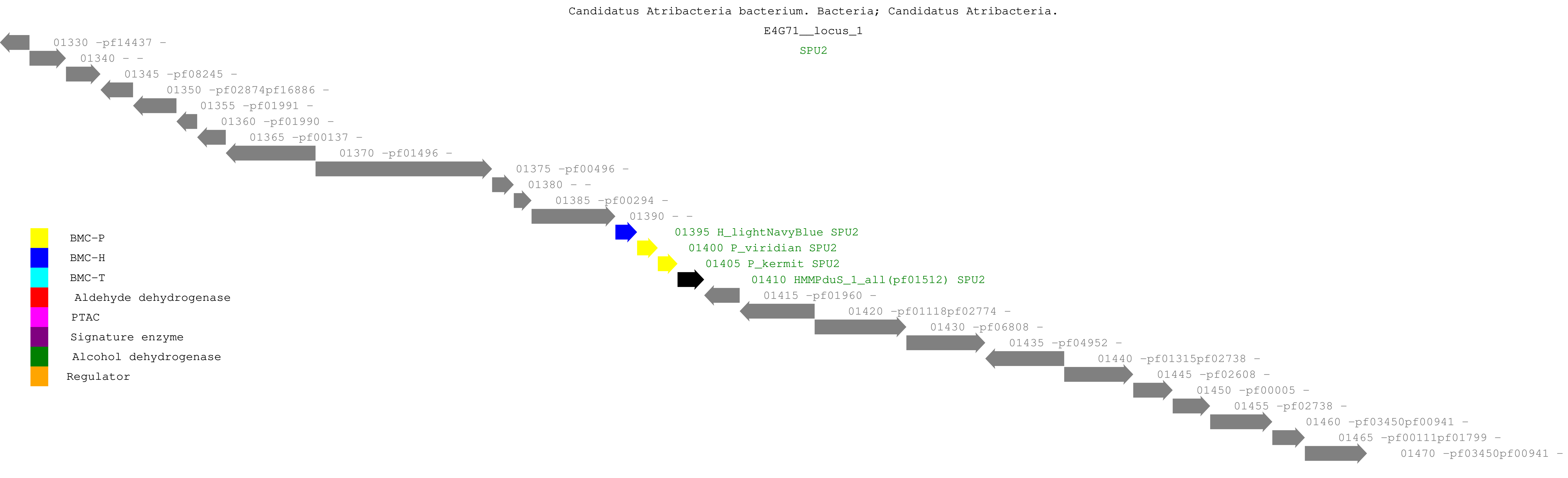
Candidatus Atribacteria bacterium. Bacteria; Candidatus Atribacteria.
E4G71__locus_1
SPU2
01330 -pf14437 -
01340 - -
01345 -pf08245 -
01350 -pf02874pf16886 -
01355 -pf01991 -
01360 -pf01990 -
01365 -pf00137 -
01370 -pf01496 -
01375 -pf00496 -
01380 - -
01385 -pf00294 -
01390 - -
01395 H_lightNavyBlue SPU2
01400 P_viridian SPU2
01405 P_kermit SPU2
01410 HMMPduS_1_all(pf01512) SPU2
01415 -pf01960 -
01420 -pf01118pf02774 -
01430 -pf06808 -
01435 -pf04952 -
01440 -pf01315pf02738 -
01445 -pf02608 -
01450 -pf00005 -
01455 -pf02738 -
01460 -pf03450pf00941 -
01465 -pf00111pf01799 -
01470 -pf03450pf00941 -
BMC-P
BMC-H
BMC-T
Aldehyde dehydrogenase
PTAC
Signature enzyme
Alcohol dehydrogenase
Regulator

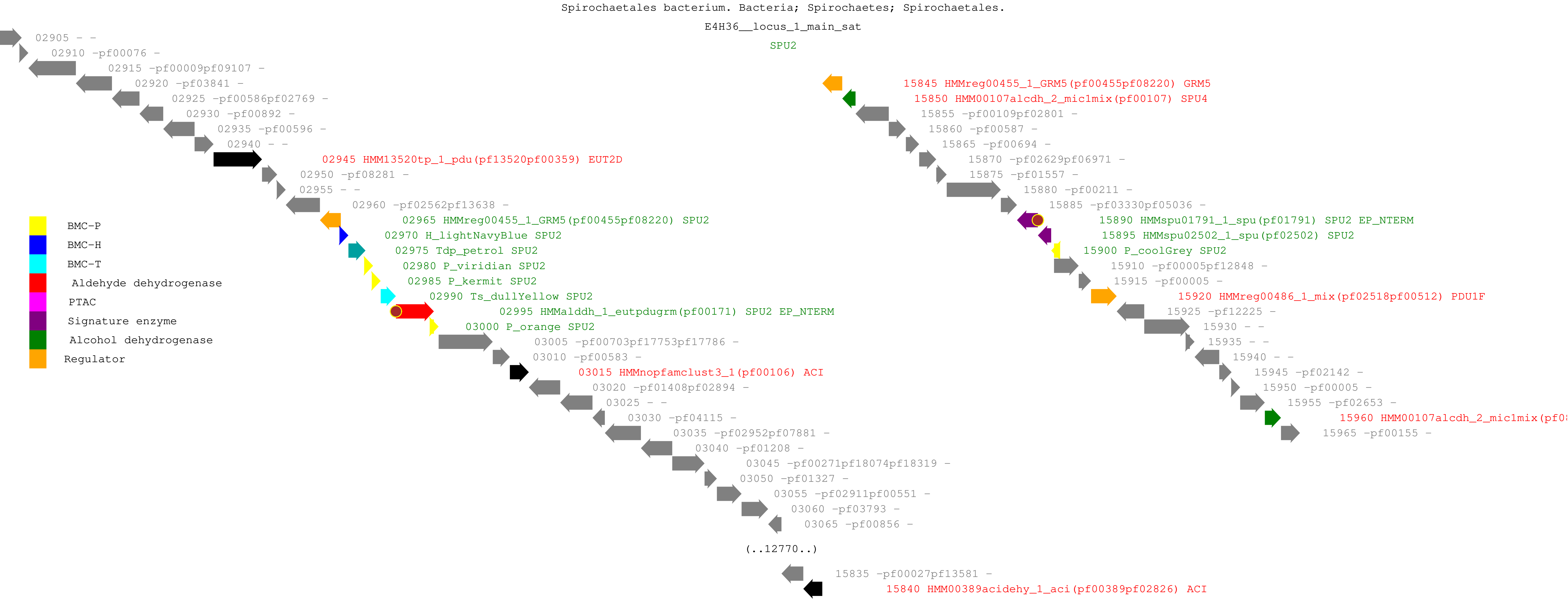
Spirochaetales bacterium. Bacteria; Spirochaetes; Spirochaetales.
E4H36__locus_1_main_sat
SPU2
02905 - -
02910 -pf00076 -
02915 -pf00009pf09107 -
02920 -pf03841 -
02925 -pf00586pf02769 -
02930 -pf00892 -
02935 -pf00596 -
02940 - -
02945 HMM13520tp_1_pdu(pf13520pf00359) EUT2D
02950 -pf08281 -
02955 - -
02960 -pf02562pf13638 -
02965 HMMreg00455_1_GRM5(pf00455pf08220) SPU2
02970 H_lightNavyBlue SPU2
02975 Tdp_petrol SPU2
02980 P_viridian SPU2
02985 P_kermit SPU2
02990 Ts_dullYellow SPU2
02995 HMMalddh_1_eutpdugrm(pf00171) SPU2 EP_NTERM
03000 P_orange SPU2
03005 -pf00703pf17753pf17786 -
03010 -pf00583 -
03015 HMMnopfamclust3_1(pf00106) ACI
03020 -pf01408pf02894 -
03025 - -
03030 -pf04115 -
03035 -pf02952pf07881 -
03040 -pf01208 -
03045 -pf00271pf18074pf18319 -
03050 -pf01327 -
03055 -pf02911pf00551 -
03060 -pf03793 -
03065 -pf00856 -
(..12770..)
15835 -pf00027pf13581 -
15840 HMM00389acidehy_1_aci(pf00389pf02826) ACI
15845 HMMreg00455_1_GRM5(pf00455pf08220) GRM5
15850 HMM00107alcdh_2_mic1mix(pf00107) SPU4
15855 -pf00109pf02801 -
15860 -pf00587 -
15865 -pf00694 -
15870 -pf02629pf06971 -
15875 -pf01557 -
15880 -pf00211 -
15885 -pf03330pf05036 -
15890 HMMspu01791_1_spu(pf01791) SPU2 EP_NTERM
15895 HMMspu02502_1_spu(pf02502) SPU2
15900 P_coolGrey SPU2
15910 -pf00005pf12848 -
15915 -pf00005 -
15920 HMMreg00486_1_mix(pf02518pf00512) PDU1F
15925 -pf12225 -
15930 - -
15935 - -
15940 - -
15945 -pf02142 -
15950 -pf00005 -
15955 -pf02653 -
15960 HMM00107alcdh_2_mic1mix(pf0
15965 -pf00155 -
BMC-P
BMC-H
BMC-T
Aldehyde dehydrogenase
PTAC
Signature enzyme
Alcohol dehydrogenase
Regulator

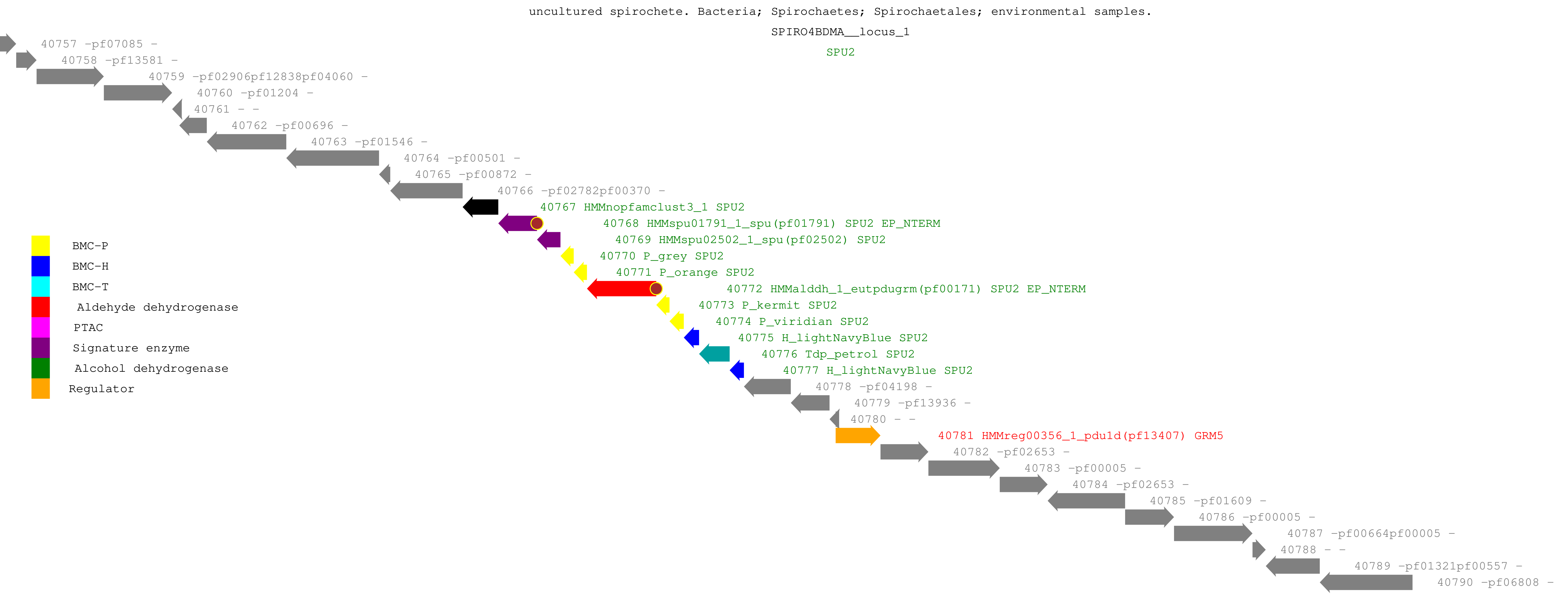

uncultured spirochete. Bacteria; Spirochaetes; Spirochaetales; environmental samples.
SPIRO4BDMA__locus_1
SPU2
40757 -pf07085 -
40758 -pf13581 -
40759 -pf02906pf12838pf04060 -
40760 -pf01204 -
40761 - -
40762 -pf00696 -
40763 -pf01546 -
40764 -pf00501 -
40765 -pf00872 -
40766 -pf02782pf00370 -
40767 HMMnopfamclust3_1 SPU2
40768 HMMspu01791_1_spu(pf01791) SPU2 EP_NTERM
40769 HMMspu02502_1_spu(pf02502) SPU2
40770 P_grey SPU2
40771 P_orange SPU2
40772 HMMaldh_1_eutpdugrm(pf00171) SPU2 EP_NTERM
40773 P_kermit SPU2
40774 P_viridian SPU2
40775 H_lightNavyBlue SPU2
40776 Tdp_petrol SPU2
40777 H_lightNavyBlue SPU2
40778 -pf04198 -
40779 -pf13936 -
40780 - -
40781 HMMreg00356_1_pdu1d(pf13407) GRM5
40782 -pf02653 -
40783 -pf00005 -
40784 -pf02653 -
40785 -pf01609 -
40786 -pf00005 -
40787 -pf00664pf00005 -
40788 - -
40789 -pf01321pf00557 -
40790 -pf06808 -
BMC-P
BMC-H
BMC-T
Aldehyde dehydrogenase
PTAC
Signature enzyme
Alcohol dehydrogenase
Regulator

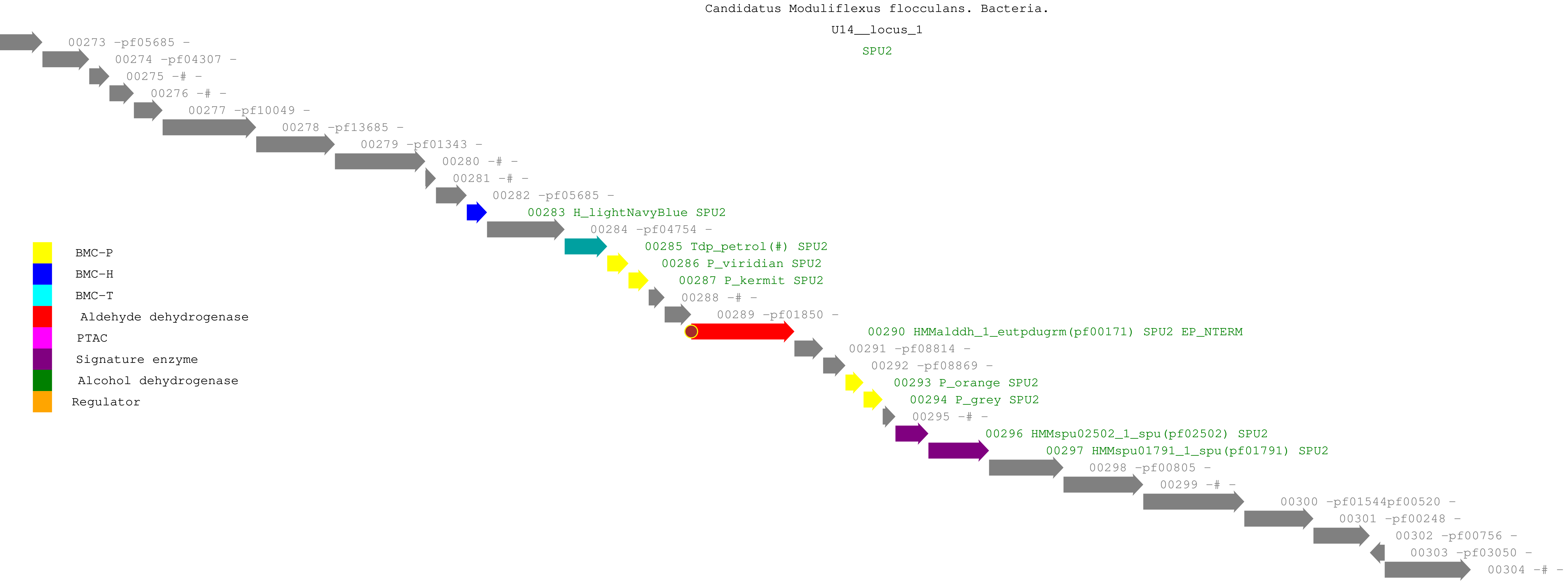
Candidatus Moduliflexus flocculans. Bacteria.
U14__locus_1
SPU2
00273 -pf05685 -
00274 -pf04307 -
00275 -# -
00276 -# -
00277 -pf10049 -
00278 -pf13685 -
00279 -pf01343 -
00280 -# -
00281 -# -
00282 -pf05685 -
00283 H_lightNavyBlue SPU2
00284 -pf04754 -
00285 Tdp_petrol(#) SPU2
00286 P_viridian SPU2
00287 P_kermit SPU2
00288 -# -
00289 -pf01850 -
00290 HMMalddh_1_eutpdugrm(pf00171) SPU2 EP_NTERM
00291 -pf08814 -
00292 -pf08869 -
00293 P_orange SPU2
00294 P_grey SPU2
00295 -# -
00296 HMMspu02502_1_spu(pf02502) SPU2
00297 HMMspu01791_1_spu(pf01791) SPU2
00298 -pf00805 -
00299 -# -
00300 -pf01544pf00520 -
00301 -pf00248 -
00302 -pf00756 -
00303 -pf03050 -
00304 -# -
BMC-P
BMC-H
BMC-T
Aldehyde dehydrogenase
PTAC
Signature enzyme
Alcohol dehydrogenase
Regulator

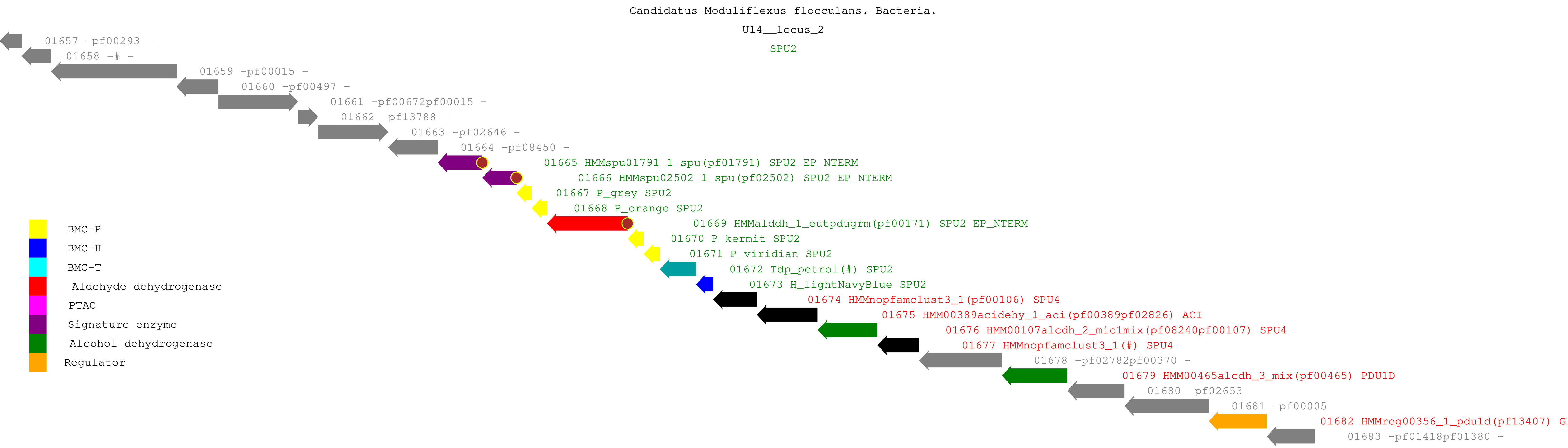
Candidatus Moduliflexus flocculans. Bacteria.
U14__locus_2
SPU2
01657 -pf00293 -
01658 -# -
01659 -pf00015 -
01660 -pf00497 -
01661 -pf00672pf00015 -
01662 -pf13788 -
01663 -pf02646 -
01664 -pf08450 -
01665 HMMspu01791_1_spu(pf01791) SPU2 EP_NTERM
01666 HMMspu02502_1_spu(pf02502) SPU2 EP_NTERM
01667 P_grey SPU2
01668 P_orange SPU2
01669 HMMaldh_1_eutpdugrm(pf00171) SPU2 EP_NTERM
01670 P_kermit SPU2
01671 P_viridian SPU2
01672 Tdp_petrol(#) SPU2
01673 H_lightNavyBlue SPU2
01674 HMMnopfamclust3_1(pf00106) SPU4
01675 HMM00389acidehy_1_aci(pf00389pf02826) ACI
01676 HMM00107alcdh_2_mic1mix(pf08240pf00107) SPU4
01677 HMMnopfamclust3_1(#) SPU4
01678 -pf02782pf00370 -
01679 HMM00465alcdh_3_mix(pf00465) PDU1D
01680 -pf02653 -
01681 -pf00005 -
01682 HMMreg00356_1_pdu1d(pf13407)
01683 -pf01418pf01380 -
BMC-P
BMC-H
BMC-T
Aldehyde dehydrogenase
PTAC
Signature enzyme
Alcohol dehydrogenase
Regulator

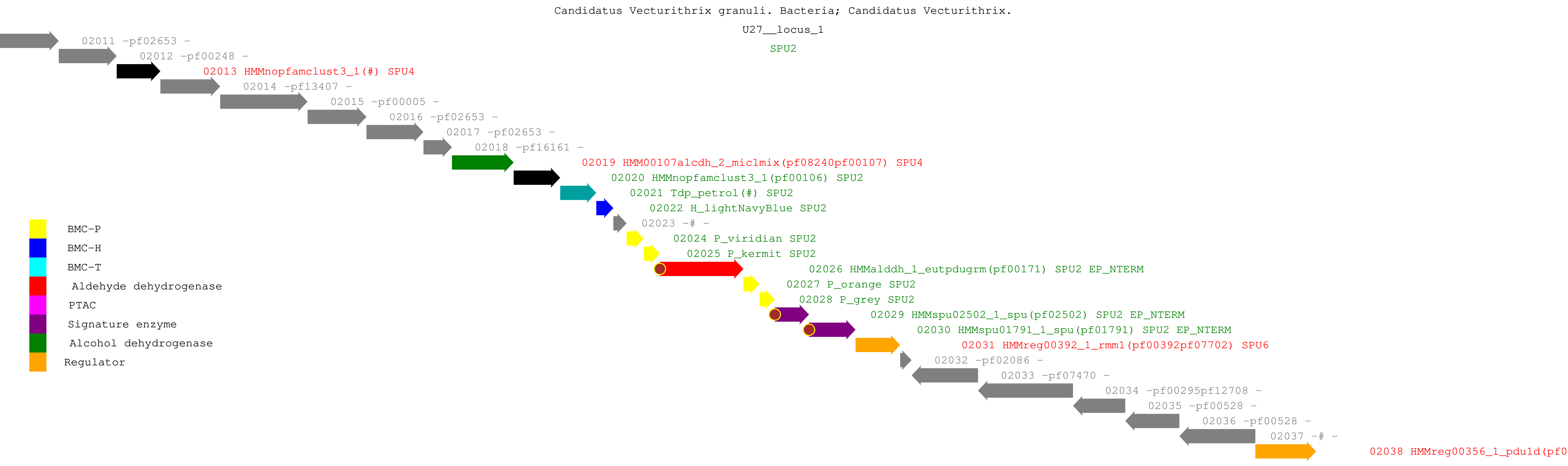
Candidatus Vecturithrix granuli. Bacteria; Candidatus Vecturithrix.
U27__locus_1
SPU2
02011 -pf02653 -
02012 -pf00248 -
02013 HMMnopfamclust3_1(#) SPU4
02014 -pf13407 -
02015 -pf00005 -
02016 -pf02653 -
02017 -pf02653 -
02018 -pf16161 -
02019 HMM00107alcdh_2_mic1mix(pf08240pf00107) SPU4
02020 HMMnopfamclust3_1(pf00106) SPU2
02021 Tdp_petrol(#) SPU2
02022 H_lightNavyBlue SPU2
02023 -# -
02024 P_viridian SPU2
02025 P_kermit SPU2
02026 HMMaldh_1_eutpdugrm(pf00171) SPU2 EP_NTERM
02027 P_orange SPU2
02028 P_grey SPU2
02029 HMMspu02502_1_spu(pf02502) SPU2 EP_NTERM
02030 HMMspu01791_1_spu(pf01791) SPU2 EP_NTERM
02031 HMMreg00392_1_rmm1(pf00392pf07702) SPU6
02032 -pf02086 -
02033 -pf07470 -
02034 -pf00295pf12708 -
02035 -pf00528 -
02036 -pf00528 -
02037 -# -
02038 HMMreg00356_1_pdu1d(pf0
BMC-P
BMC-H
BMC-T
Aldehyde dehydrogenase
PTAC
Signature enzyme
Alcohol dehydrogenase
Regulator

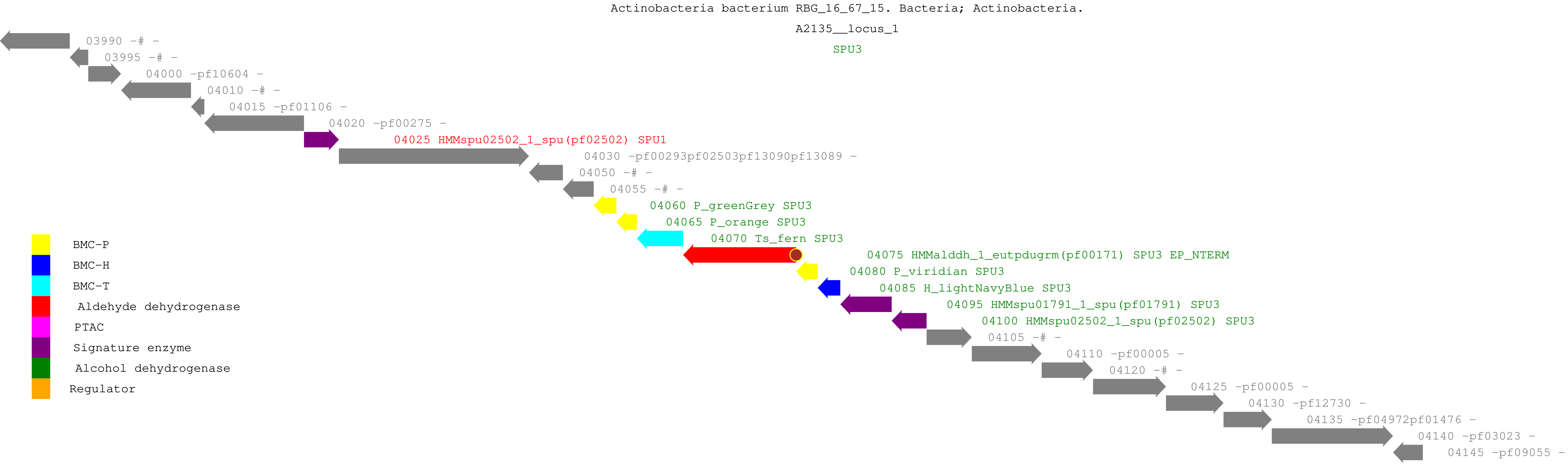

Actinobacteria bacterium RBG_16_67_15. Bacteria; Actinobacteria.
A2135__locus_1
SPU3
03990 -# -
03995 -# -
04000 -pf10604 -
04010 -# -
04015 -pf01106 -
04020 -pf00275 -
04025 HMMspu02502_1_spu(pf02502) SPU1
04030 -pf00293pf02503pf13090pf13089 -
04050 -# -
04055 -# -
04060 P_greenGrey SPU3
04065 P_orange SPU3
04070 Ts_fern SPU3
04075 HMMalddh_1_eutpdugrm(pf00171) SPU3 EP_NTERM
04080 P_viridian SPU3
04085 H_lightNavyBlue SPU3
04095 HMMspu01791_1_spu(pf01791) SPU3
04100 HMMspu02502_1_spu(pf02502) SPU3
04105 -# -
04110 -pf00005 -
04120 -# -
04125 -pf00005 -
04130 -pf12730 -
04135 -pf04972pf01476 -
04140 -pf03023 -
04145 -pf09055 -
BMC-P
BMC-H
BMC-T
Aldehyde dehydrogenase
PTAC
Signature enzyme
Alcohol dehydrogenase
Regulator

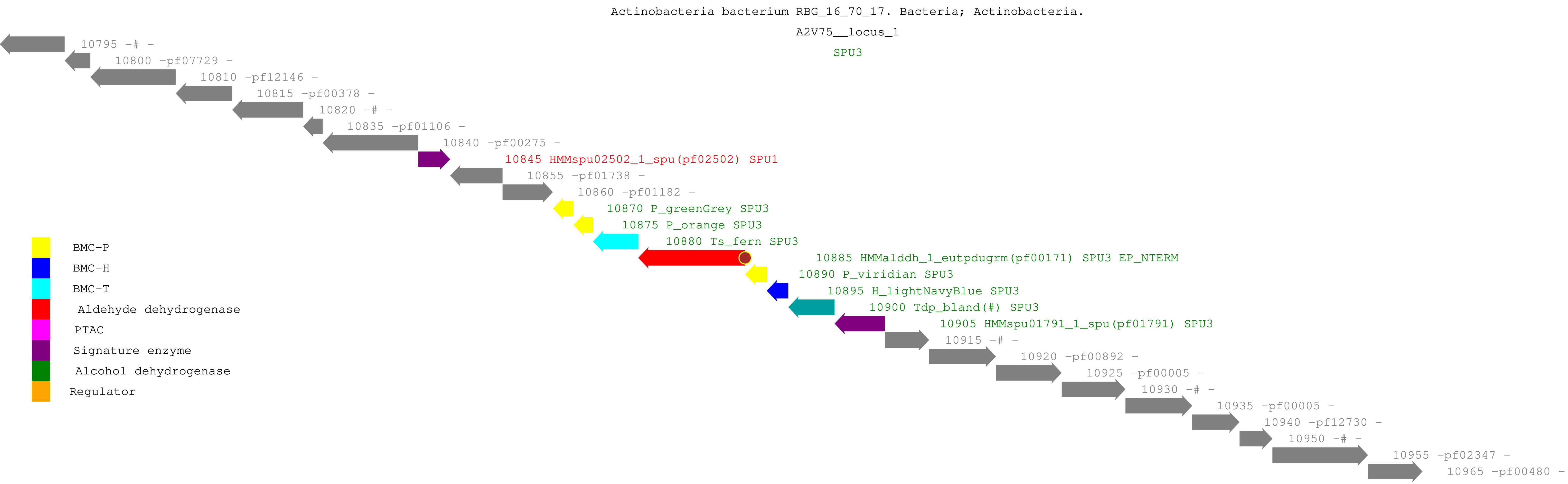
Actinobacteria bacterium RBG_16_70_17. Bacteria; Actinobacteria.
A2V75__locus_1
SPU3
10795 -# -
10800 -pf07729 -
10810 -pf12146 -
10815 -pf00378 -
10820 -# -
10835 -pf01106 -
10840 -pf00275 -
10845 HMMspu02502_1_spu(pf02502) SPU1
10855 -pf01738 -
10860 -pf01182 -
10870 P_greenGrey SPU3
10875 P_orange SPU3
10880 Ts_fern SPU3
10885 HMMalddh_1_eutpdugrm(pf00171) SPU3 EP_NTERM
10890 P_viridian SPU3
10895 H_lightNavyBlue SPU3
10900 Tdp_bland(#) SPU3
10905 HMMspu01791_1_spu(pf01791) SPU3
10915 -# -
10920 -pf00892 -
10925 -pf00005 -
10930 -# -
10935 -pf00005 -
10940 -pf12730 -
10950 -# -
10955 -pf02347 -
10965 -pf00480 -
BMC-P
BMC-H
BMC-T
Aldehyde dehydrogenase
PTAC
Signature enzyme
Alcohol dehydrogenase
Regulator

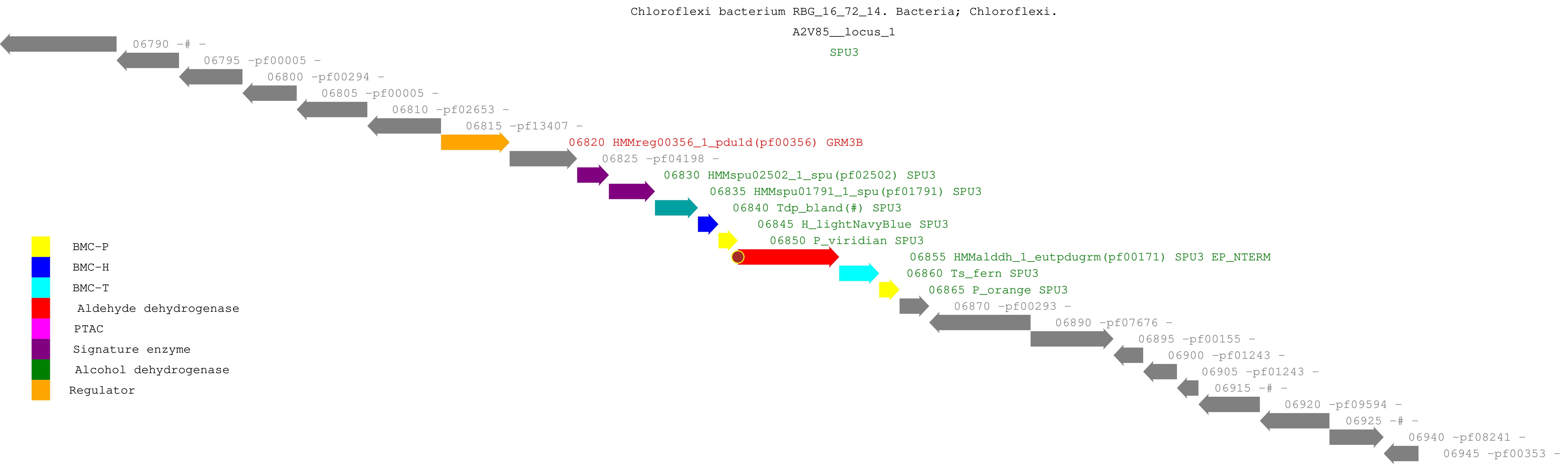
Chloroflexi bacterium RBG_16_72_14. Bacteria; Chloroflexi.
A2V85__locus_1
SPU3
06790 -# -
06795 -pf00005 -
06800 -pf00294 -
06805 -pf00005 -
06810 -pf02653 -
06815 -pf13407 -
06820 HMMreg00356_1_pdu1d(pf00356) GRM3B
06825 -pf04198 -
06830 HMMspu02502_1_spu(pf02502) SPU3
06835 HMMspu01791_1_spu(pf01791) SPU3
06840 Tdp_bland(#) SPU3
06845 H_lightNavyBlue SPU3
06850 P_viridian SPU3
06855 HMMalddh_1_eutpdugrm(pf00171) SPU3 EP_NTERM
06860 Ts_fern SPU3
06865 P_orange SPU3
06870 -pf00293 -
06890 -pf07676 -
06895 -pf00155 -
06900 -pf01243 -
06905 -pf01243 -
06915 -# -
06920 -pf09594 -
06925 -# -
06940 -pf08241 -
06945 -pf00353 -
BMC-P
BMC-H
BMC-T
Aldehyde dehydrogenase
PTAC
Signature enzyme
Alcohol dehydrogenase
Regulator

Lentisphaerae bacterium GWF2_57_35. Bacteria; Lentisphaerae; unclassified Lentisphaerae (miscellaneous).

A2X46__locus_1

SPU3

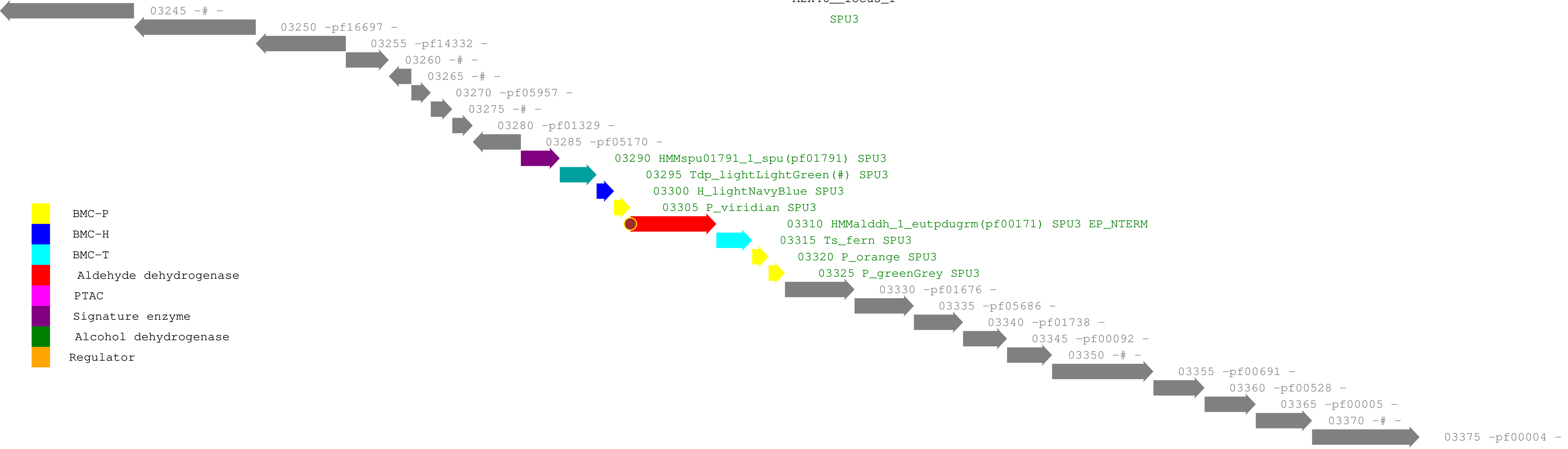

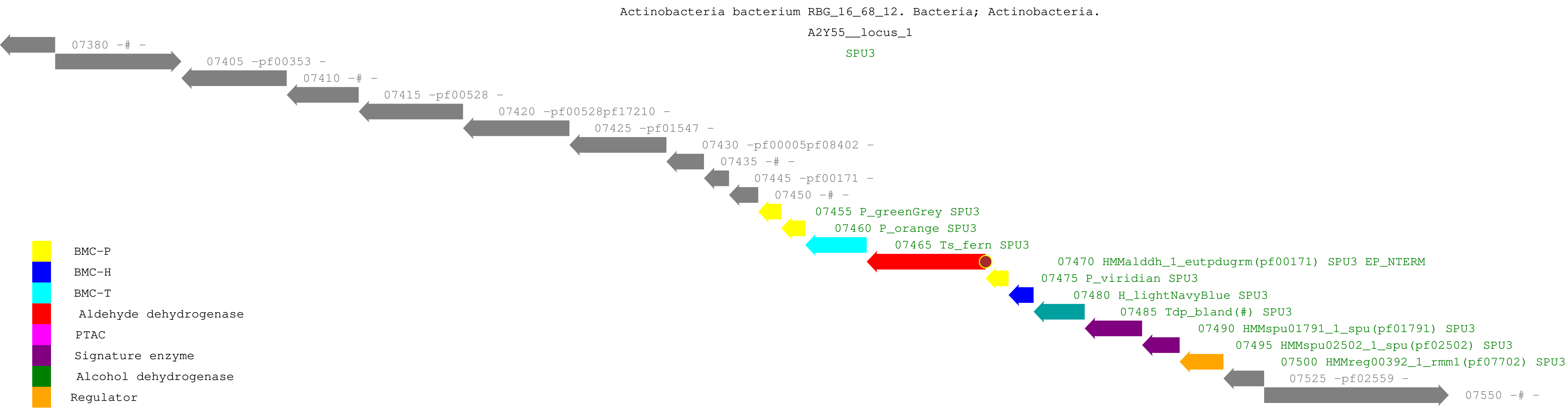

Actinobacteria bacterium RBG_16_68_12. Bacteria; Actinobacteria.
A2Y55__locus_1
SPU3
07380 -# -
07405 -pf00353 -
07410 -# -
07415 -pf00528 -
07420 -pf00528pf17210 -
07425 -pf01547 -
07430 -pf00005pf08402 -
07435 -# -
07445 -pf00171 -
07450 -# -
07455 P_greenGrey SPU3
07460 P_orange SPU3
07465 Ts_fern SPU3
07470 HMMalddh_1_eutpdugrm(pf00171) SPU3 EP_NTERM
07475 P_viridian SPU3
07480 H_lightNavyBlue SPU3
07485 Tdp_bland(#) SPU3
07490 HMMspu01791_1_spu(pf01791) SPU3
07495 HMMspu02502_1_spu(pf02502) SPU3
07500 HMMreg00392_1_rmm1(pf07702) SPU3
07525 -pf02559 -
07550 -# -
BMC-P
BMC-H
BMC-T
Aldehyde dehydrogenase
PTAC
Signature enzyme
Alcohol dehydrogenase
Regulator

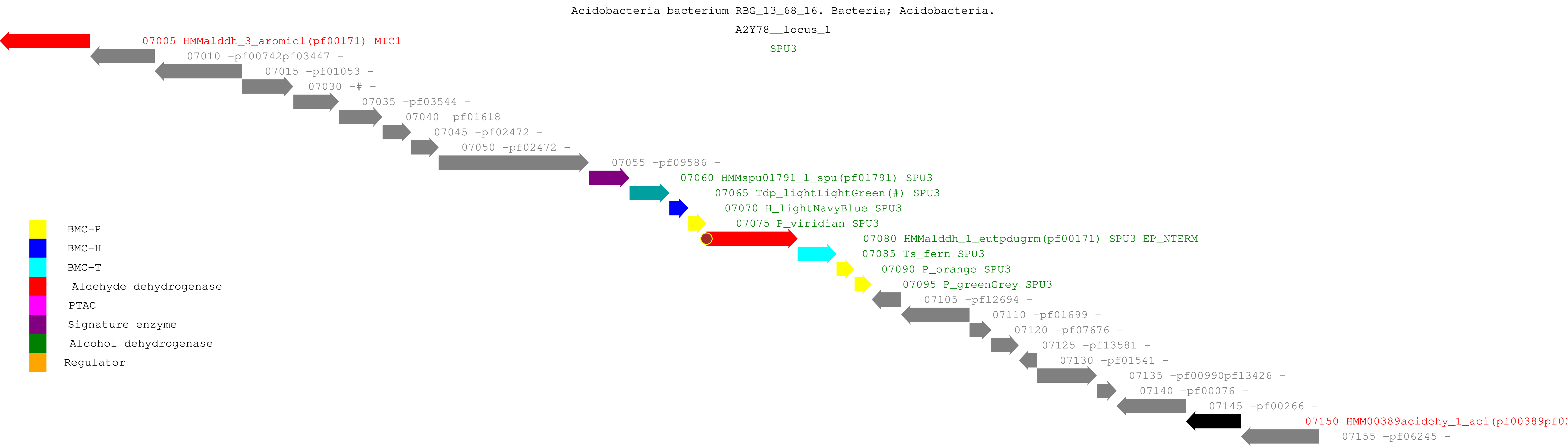

Acidobacteria bacterium RBG_13_68_16. Bacteria; Acidobacteria.
A2Y78__locus_1
SPU3
07005 HMMalddh_3_aromic1(pf00171) MIC1
07010 -pf00742pf03447 -
07015 -pf01053 -
07030 -# -
07035 -pf03544 -
07040 -pf01618 -
07045 -pf02472 -
07050 -pf02472 -
07055 -pf09586 -
07060 HMMspu01791_1_spu(pf01791) SPU3
07065 Tdp_lightLightGreen(#) SPU3
07070 H_lightNavyBlue SPU3
07075 P_viridian SPU3
07080 HMMalddh_1_eutpdugrm(pf00171) SPU3 EP_NTERM
07085 Ts_fern SPU3
07090 P_orange SPU3
07095 P_greenGrey SPU3
07105 -pf12694 -
07110 -pf01699 -
07120 -pf07676 -
07125 -pf13581 -
07130 -pf01541 -
07135 -pf00990pf13426 -
07140 -pf00076 -
07145 -pf00266 -
07150 HMM00389acidehy_1_aci(pf00389pf0
07155 -pf06245 -
BMC-P
BMC-H
BMC-T
Aldehyde dehydrogenase
PTAC
Signature enzyme
Alcohol dehydrogenase
Regulator

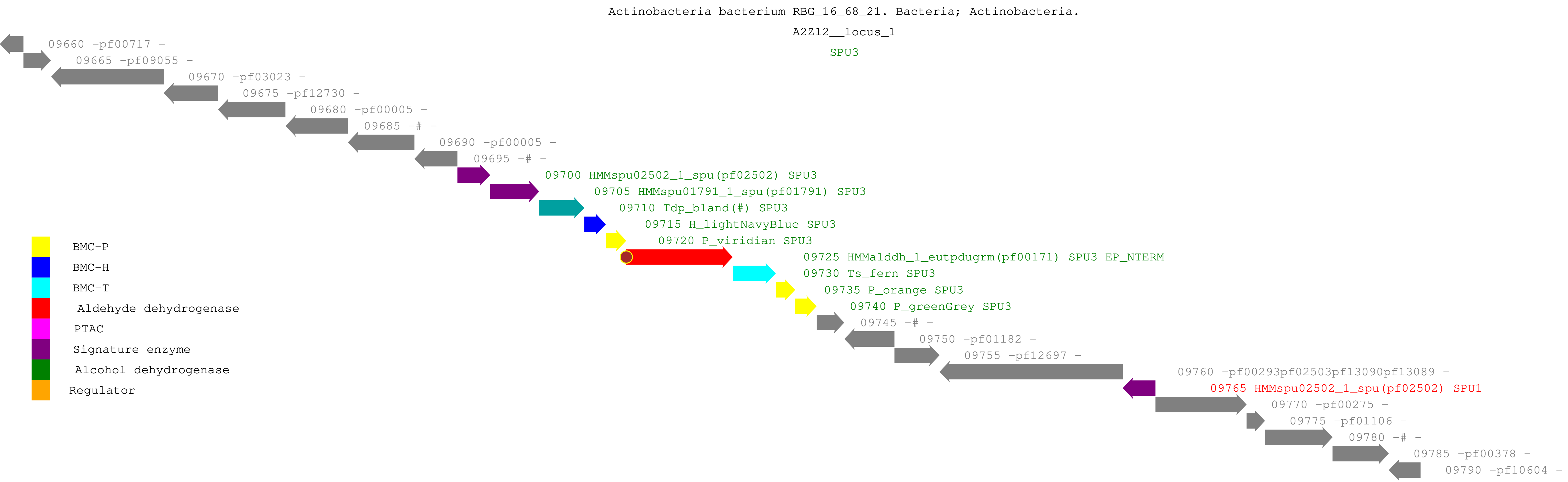

Actinobacteria bacterium RBG_16_68_21. Bacteria; Actinobacteria.
A2Z12__locus_1
SPU3
09660 -pf00717 -
09665 -pf09055 -
09670 -pf03023 -
09675 -pf12730 -
09680 -pf00005 -
09685 -# -
09690 -pf00005 -
09695 -# -
09700 HMMspu02502_1_spu(pf02502) SPU3
09705 HMMspu01791_1_spu(pf01791) SPU3
09710 Tdp_bland(#) SPU3
09715 H_lightNavyBlue SPU3
09720 P_viridian SPU3
09725 HMMalddh_1_eutpdugrm(pf00171) SPU3 EP_NTERM
09730 Ts_fern SPU3
09735 P_orange SPU3
09740 P_greenGrey SPU3
09745 -# -
09750 -pf01182 -
09755 -pf12697 -
09760 -pf00293pf02503pf13090pf13089 -
09765 HMMspu02502_1_spu(pf02502) SPU1
09770 -pf00275 -
09775 -pf01106 -
09780 -# -
09785 -pf00378 -
09790 -pf10604 -
BMC-P
BMC-H
BMC-T
Aldehyde dehydrogenase
PTAC
Signature enzyme
Alcohol dehydrogenase
Regulator

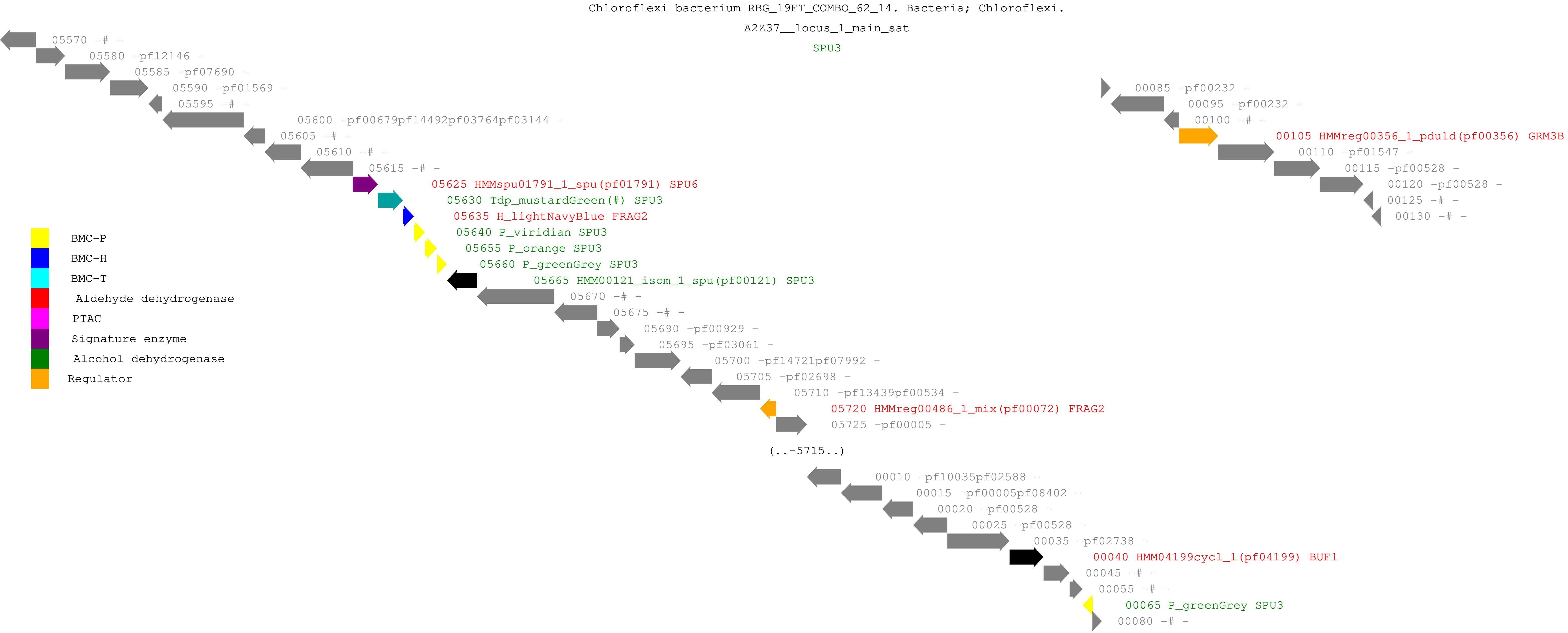
Chloroflexi bacterium RBG_19FT_COMBO_62_14. Bacteria; Chloroflexi.
A2Z37__locus_1_main_sat
SPU3
05570 -# -
05580 -pf12146 -
05585 -pf07690 -
05590 -pf01569 -
05595 -# -
05600 -pf00679pf14492pf03764pf03144 -
05605 -# -
05610 -# -
05615 -# -
05625 HMMspu01791_1_spu(pf01791) SPU6
05630 Tdp_mustardGreen(#) SPU3
05635 H_lightNavyBlue FRAG2
05640 P_viridian SPU3
05655 P_orange SPU3
05660 P_greenGrey SPU3
05665 HMM00121_isom_1_spu(pf00121) SPU3
05670 -# -
05675 -# -
05690 -pf00929 -
05695 -pf03061 -
05700 -pf14721pf07992 -
05705 -pf02698 -
05710 -pf13439pf00534 -
05720 HMMreg00486_1_mix(pf00072) FRAG2
05725 -pf00005 -
(..-5715..)
00010 -pf10035pf02588 -
00015 -pf00005pf08402 -
00020 -pf00528 -
00025 -pf00528 -
00035 -pf02738 -
00040 HMM04199cycl_1(pf04199) BUF1
00045 -# -
00055 -# -
00065 P_greenGrey SPU3
00080 -# -
00085 -pf00232 -
00095 -pf00232 -
00100 -# -
00105 HMMreg00356_1_pdu1d(pf00356) GRM3B
00110 -pf01547 -
00115 -pf00528 -
00120 -pf00528 -
00125 -# -
00130 -# -
BMC-P
BMC-H
BMC-T
Aldehyde dehydrogenase
PTAC
Signature enzyme
Alcohol dehydrogenase
Regulator

Minicystis rosea. Bacteria; Proteobacteria; Deltaproteobacteria; Myxococcales;

A7982__locus_1_main_sat

SPU3

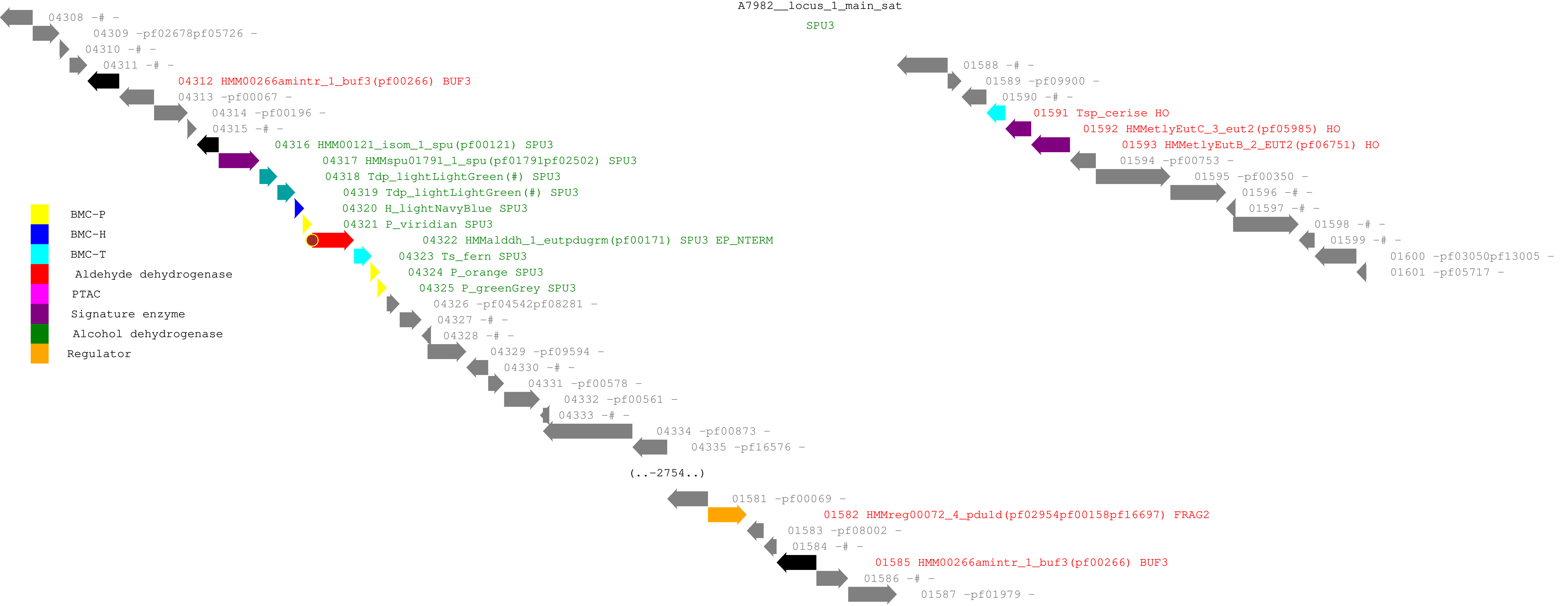

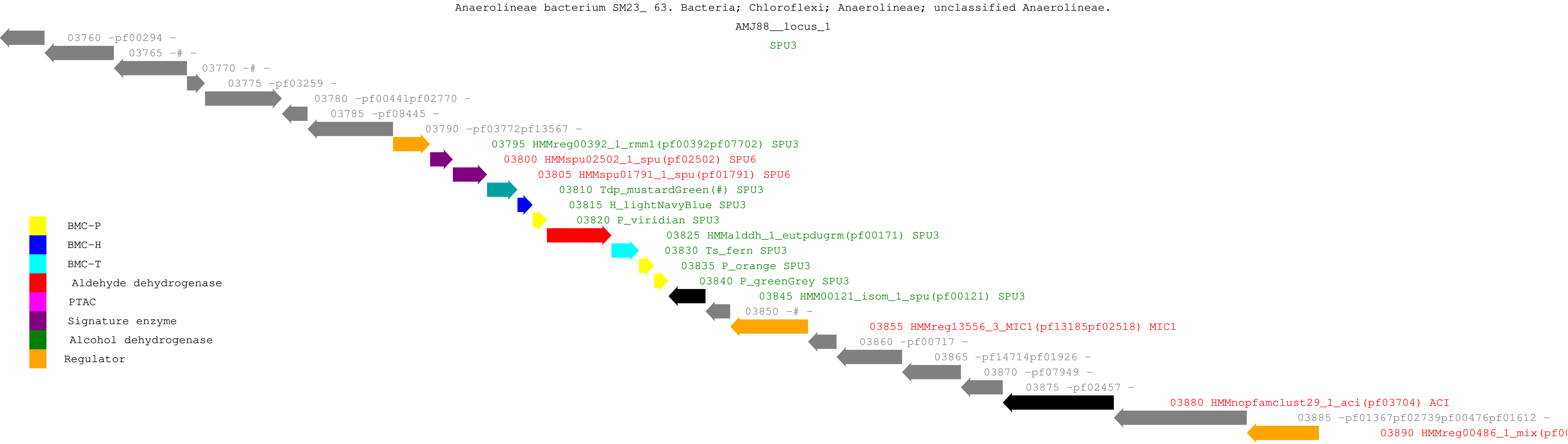
Anaerolineae bacterium SM23_ 63. Bacteria; Chloroflexi; Anaerolineae; unclassified Anaerolineae.
AMJ88__locus_1
SPU3
03760 -pf00294 -
03765 -# -
03770 -# -
03775 -pf03259 -
03780 -pf00441pf02770 -
03785 -pf08445 -
03790 -pf03772pf13567 -
03795 HMMreg00392_1_rmm1(pf00392pf07702) SPU3
03800 HMMspu02502_1_spu(pf02502) SPU6
03805 HMMspu01791_1_spu(pf01791) SPU6
03810 Tdp_mustardGreen(#) SPU3
03815 H_lightNavyBlue SPU3
03820 P_viridian SPU3
03825 HMMalddh_1_eutpdugrm(pf00171) SPU3
03830 Ts_fern SPU3
03835 P_orange SPU3
03840 P_greenGrey SPU3
03845 HMM00121_isom_1_spu(pf00121) SPU3
03850 -# -
03855 HMMreg13556_3_MIC1(pf13185pf02518) MIC1
03860 -pf00717 -
03865 -pf14714pf01926 -
03870 -pf07949 -
03875 -pf02457 -
03880 HMMnopfamclust29_1_aci(pf03704) ACI
03885 -pf01367pf02739pf00476pf01612 -
03890 HMMreg00486_1_mix(pf0
BMC-P
BMC-H
BMC-T
Aldehyde dehydrogenase
PTAC
Signature enzyme
Alcohol dehydrogenase
Regulator

Anaerolineae bacterium SM23_84. Bacteria; Chloroflexi; Anaerolineae; unclassified Anaerolineae.

AMJ93__locus_1

SPU3

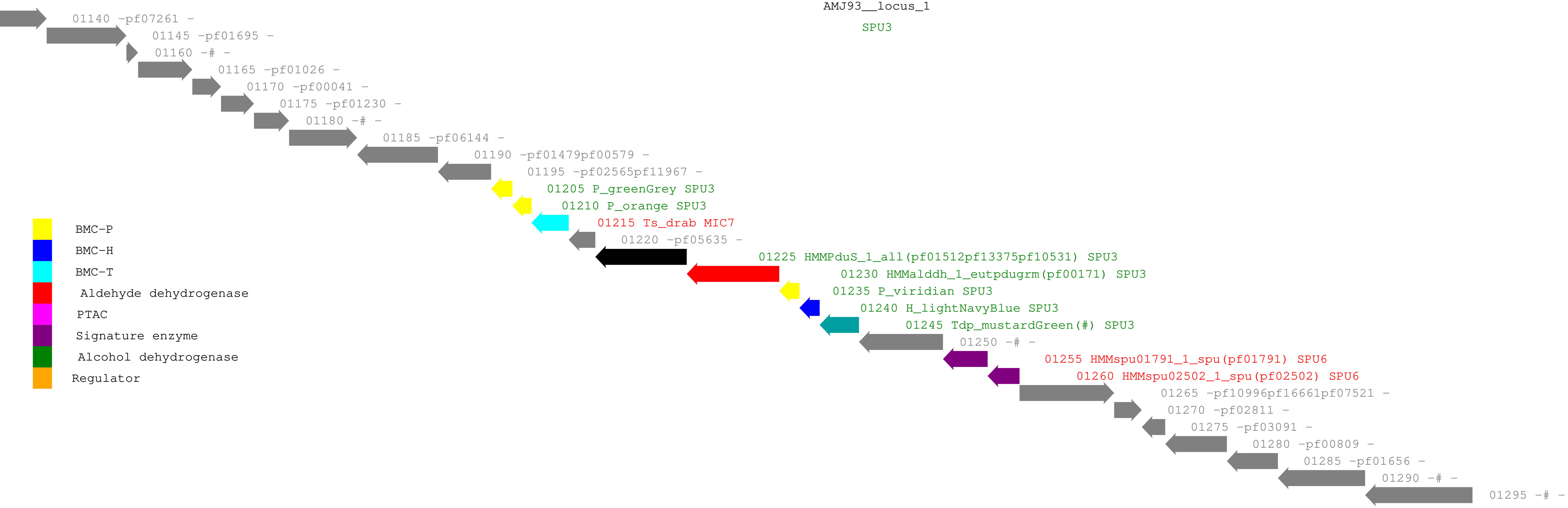

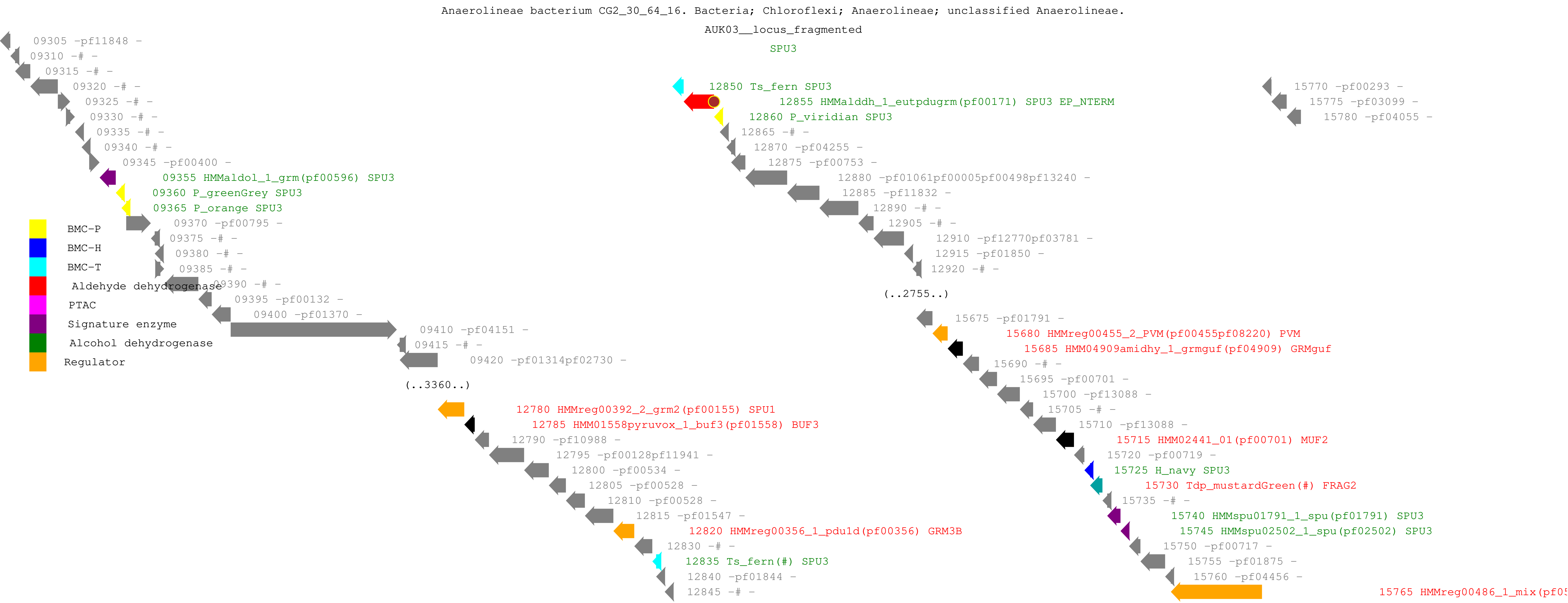
Anaerolineae bacterium CG2_30_64_16. Bacteria; Chloroflexi; Anaerolineae; unclassified Anaerolineae.
AUK03__locus_fragmented
SPU3
09305 -pf11848 -
09310 -# -
09315 -# -
09320 -# -
09325 -# -
09330 -# -
09335 -# -
09340 -# -
09345 -pf00400 -
09355 HMMaldol_1_grm(pf00596) SPU3
09360 P_greenGrey SPU3
09365 P_orange SPU3
09370 -pf00795 -
09375 -# -
09380 -# -
09385 -# -
09390 -# -
09395 -pf00132 -
09400 -pf01370 -
09410 -pf04151 -
09415 -# -
09420 -pf01314pf02730 -
(..3360..)
12780 HMMreg00392_2_grm2(pf00155) SPU1
12785 HMM01558pyruvox_1_buf3(pf01558) BUF3
12790 -pf10988 -
12795 -pf00128pf11941 -
12800 -pf00534 -
12805 -pf00528 -
12810 -pf00528 -
12815 -pf01547 -
12820 HMMreg00356_1_pdu1d(pf00356) GRM3B
12830 -# -
12835 Ts_fern(#) SPU3
12840 -pf01844 -
12845 -# -
12850 Ts_fern SPU3
12855 HMMalddh_1_eutpdugrm(pf00171) SPU3 EP_NTERM
12860 P_viridian SPU3
12865 -# -
12870 -pf04255 -
12875 -pf00753 -
12880 -pf01061pf00005pf00498pf13240 -
12885 -pf11832 -
12890 -# -
12905 -# -
12910 -pf12770pf03781 -
12915 -pf01850 -
12920 -# -
(..2755..)
15675 -pf01791 -
15680 HMMreg00455_2_PVM(pf00455pf08220) PVM
15685 HMM04909amidhy_1_grmguf(pf04909) GRMguf
15690 -# -
15695 -pf00701 -
15700 -pf13088 -
15705 -# -
15710 -pf13088 -
15715 HMM02441_01(pf00701) MUF2
15720 -pf00719 -
15725 H_navy SPU3
15730 Tdp_mustardGreen(#) FRAG2
15735 -# -
15740 HMMspu01791_1_spu(pf01791) SPU3
15745 HMMspu02502_1_spu(pf02502) SPU3
15750 -pf00717 -
15755 -pf01875 -
15760 -pf04456 -
15765 HMMreg00486_1_mix(pf0
15770 -pf00293 -
15775 -pf03099 -
15780 -pf04055 -
BMC-P
BMC-H
BMC-T
Aldehyde dehydrogenase
PTAC
Signature enzyme
Alcohol dehydrogenase
Regulator

candidate division KSB1 4572_119. Bacteria; candidate division KSB1.

B6I21__locus_fragmented

SPU3

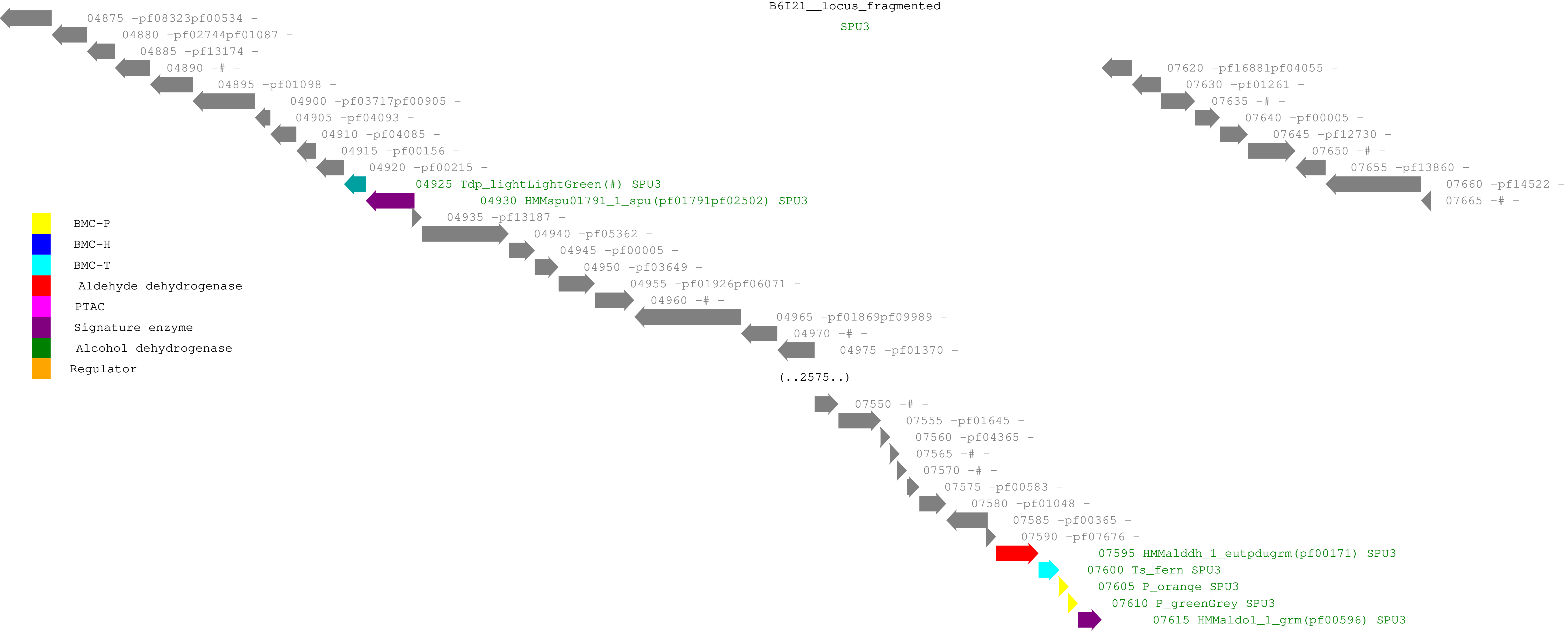

Anaerolineaceae bacterium 4572_32.1. Bacteria; Chloroflexi; Anaerolineae; Anaerolineales; Anaerolineaceae.

B6I34__locus_1_main_sat

SPU3

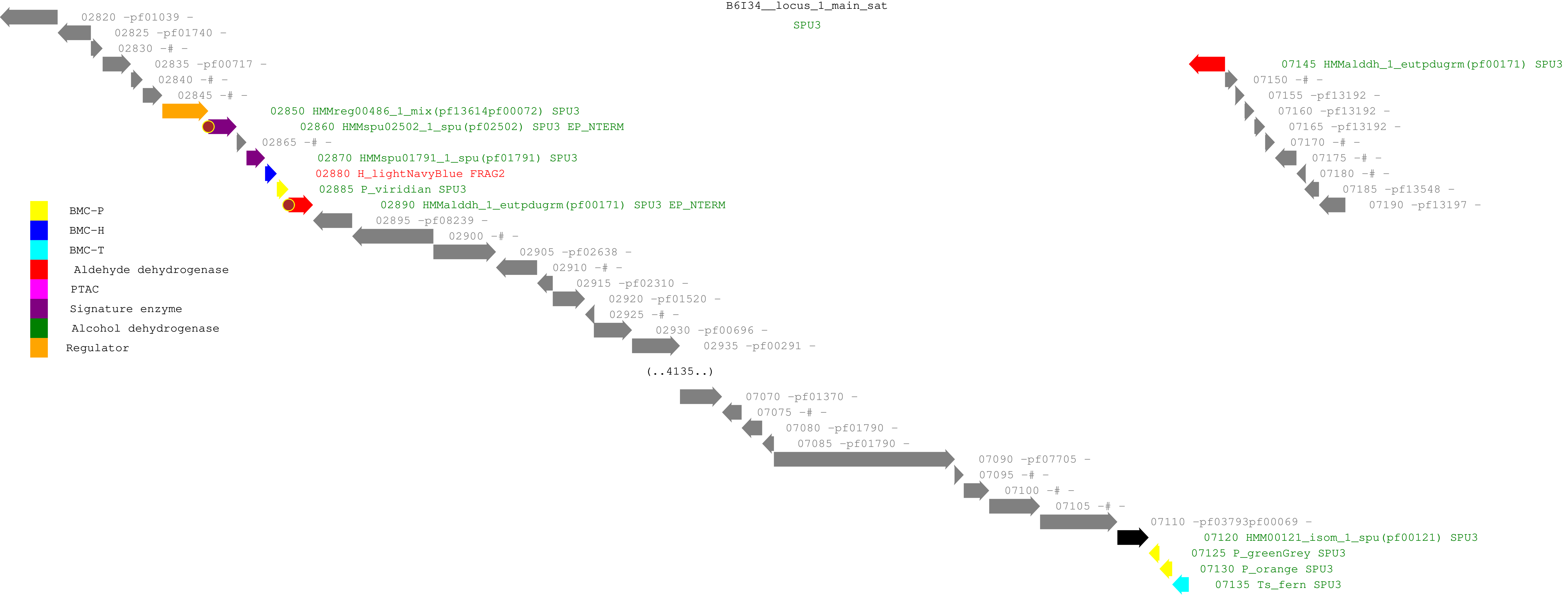

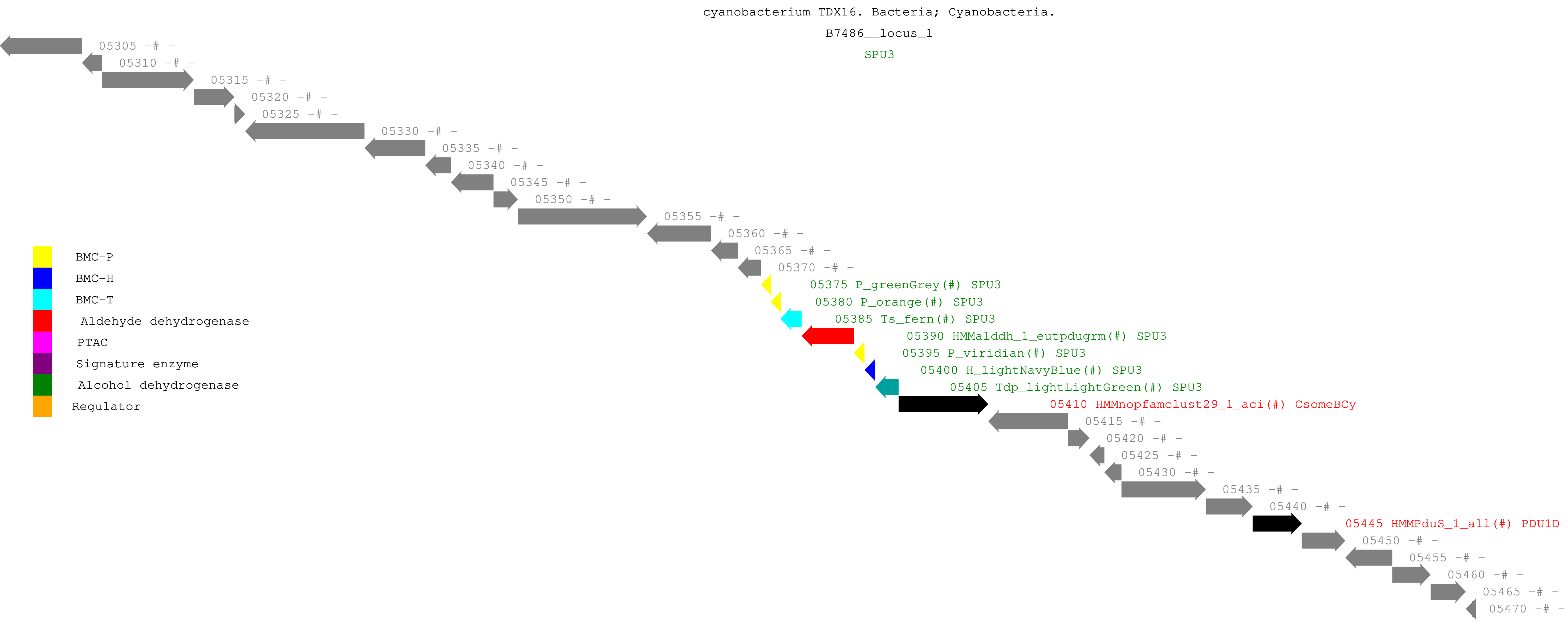
cyanobacterium TDX16. Bacteria; Cyanobacteria.
B7486__locus_1
SPU3
05305 -# -
05310 -# -
05315 -# -
05320 -# -
05325 -# -
05330 -# -
05335 -# -
05340 -# -
05345 -# -
05350 -# -
05355 -# -
05360 -# -
05365 -# -
05370 -# -
05375 P_greenGrey(#) SPU3
05380 P_orange(#) SPU3
05385 Ts_fern(#) SPU3
05390 HMMalddh_1_eutpdugrm(#) SPU3
05395 P_viridian(#) SPU3
05400 H_lightNavyBlue(#) SPU3
05405 Tdp_lightLightGreen(#) SPU3
05410 HMMnopfamclust29_1_aci(#) CsomeBCy
05415 -# -
05420 -# -
05425 -# -
05430 -# -
05435 -# -
05440 -# -
05445 HMMPduS_1_all(#) PDU1D
05450 -# -
05455 -# -
05460 -# -
05465 -# -
05470 -# -
BMC-P
BMC-H
BMC-T
Aldehyde dehydrogenase
PTAC
Signature enzyme
Alcohol dehydrogenase
Regulator

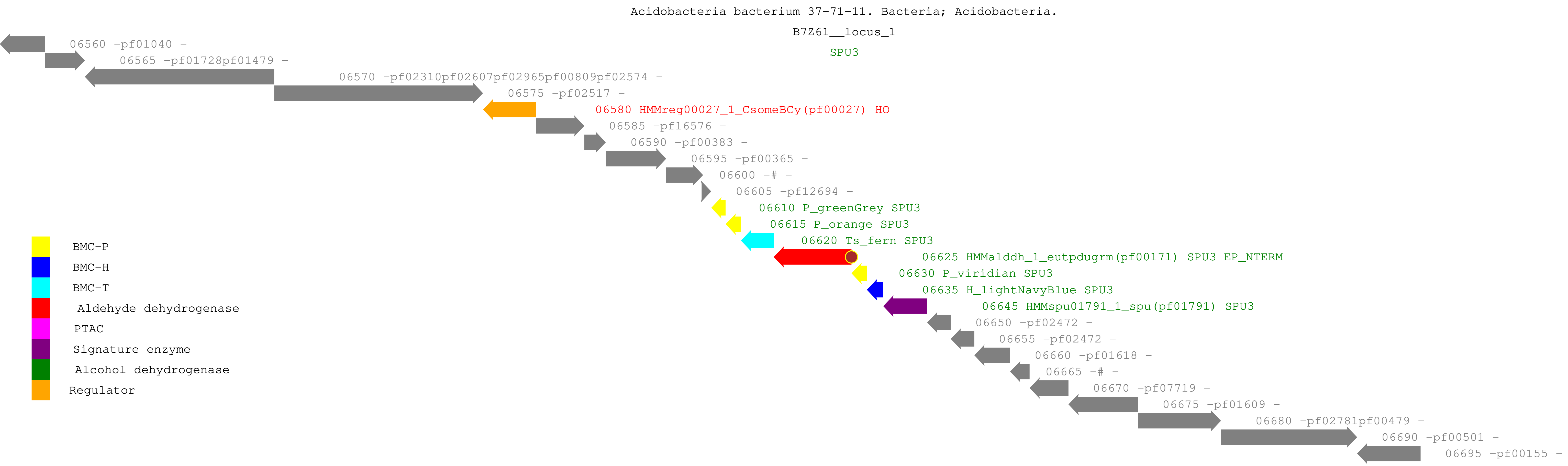
Acidobacteria bacterium 37-71-11. Bacteria; Acidobacteria.
B7Z61__locus_1
SPU3
06560 -pf01040 -
06565 -pf01728pf01479 -
06570 -pf02310pf02607pf02965pf00809pf02574 -
06575 -pf02517 -
06580 HMMreg00027_1_CsomeBCy(pf00027) HO
06585 -pf16576 -
06590 -pf00383 -
06595 -pf00365 -
06600 -# -
06605 -pf12694 -
06610 P_greenGrey SPU3
06615 P_orange SPU3
06620 Ts_fern SPU3
06625 HMMalddh_1_eutpdugrm(pf00171) SPU3 EP_NTERM
06630 P_viridian SPU3
06635 H_lightNavyBlue SPU3
06645 HMMspu01791_1_spu(pf01791) SPU3
06650 -pf02472 -
06655 -pf02472 -
06660 -pf01618 -
06665 -# -
06670 -pf07719 -
06675 -pf01609 -
06680 -pf02781pf00479 -
06690 -pf00501 -
06695 -pf00155 -
BMC-P
BMC-H
BMC-T
Aldehyde dehydrogenase
PTAC
Signature enzyme
Alcohol dehydrogenase
Regulator

Acidobacteria bacterium 21-70-11. Bacteria; Acidobacteria.

B7Z68__locus_1

SPU3

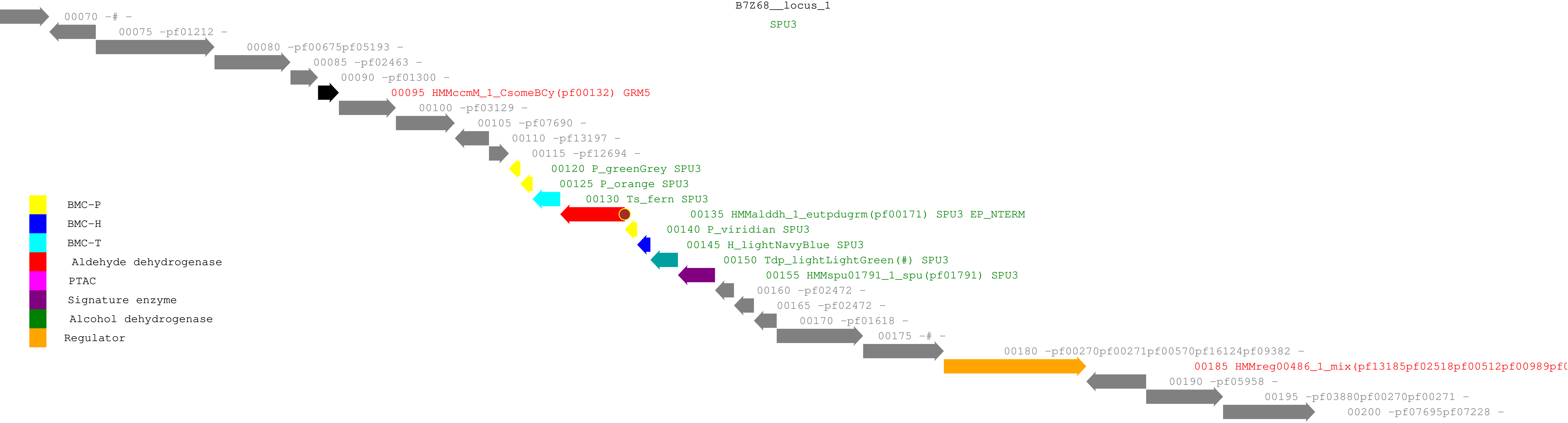

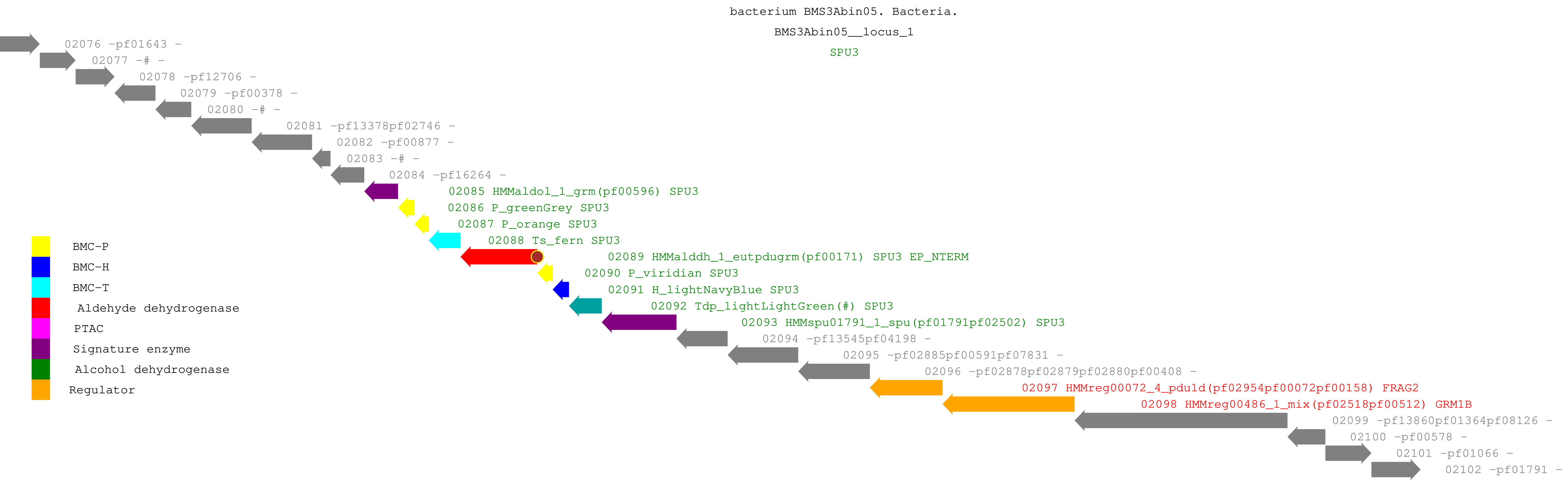
bacterium BMS3Abin05. Bacteria.
BMS3Abin05__locus_1
SPU3
02076 -pf01643 -
02077 -# -
02078 -pf12706 -
02079 -pf00378 -
02080 -# -
02081 -pf13378pf02746 -
02082 -pf00877 -
02083 -# -
02084 -pf16264 -
02085 HMMaldol_1_grm(pf00596) SPU3
02086 P_greenGrey SPU3
02087 P_orange SPU3
02088 Ts_fern SPU3
02089 HMMalddh_1_eutpdugrm(pf00171) SPU3 EP_NTERM
02090 P_viridian SPU3
02091 H_lightNavyBlue SPU3
02092 Tdp_lightLightGreen(#) SPU3
02093 HMMspu01791_1_spu(pf01791pf02502) SPU3
02094 -pf13545pf04198 -
02095 -pf02885pf00591pf07831 -
02096 -pf02878pf02879pf02880pf00408 -
02097 HMMreg00072_4_pdu1d(pf02954pf00072pf00158) FRAG2
02098 HMMreg00486_1_mix(pf02518pf00512) GRM1B
02099 -pf13860pf01364pf08126 -
02100 -pf00578 -
02101 -pf01066 -
02102 -pf01791 -
BMC-P
BMC-H
BMC-T
Aldehyde dehydrogenase
PTAC
Signature enzyme
Alcohol dehydrogenase
Regulator

bacterium BMS3Bbin01. Bacteria.

BMS3Bbin01__locus_1_main_sat

SPU3

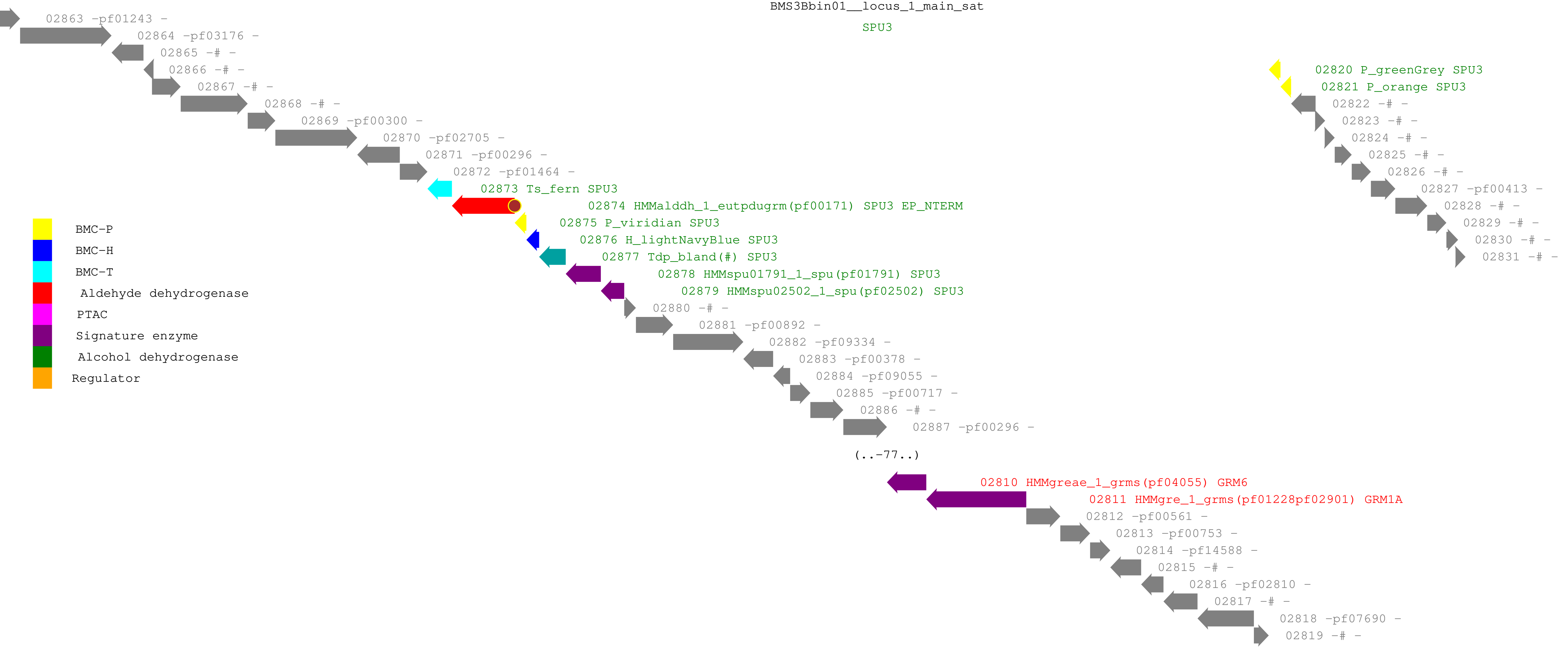

Verrucomicrobia bacterium ADurb.Bin018. Bacteria; Verrucomicrobia; unclassified Verrucomicrobia.

BWX54__locus_1

SPU3

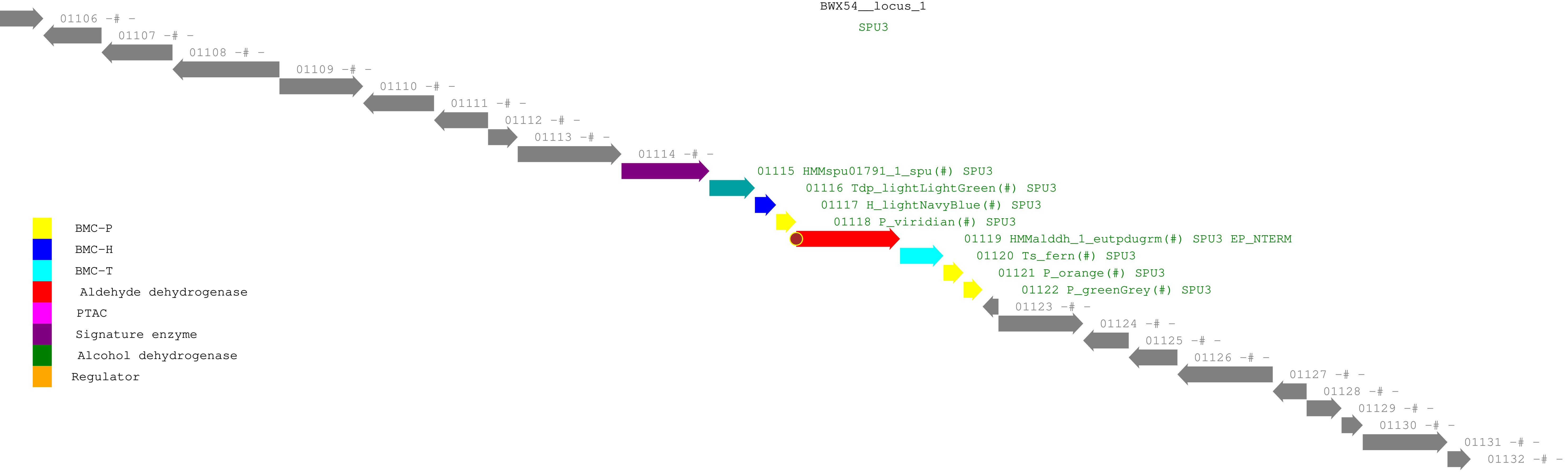

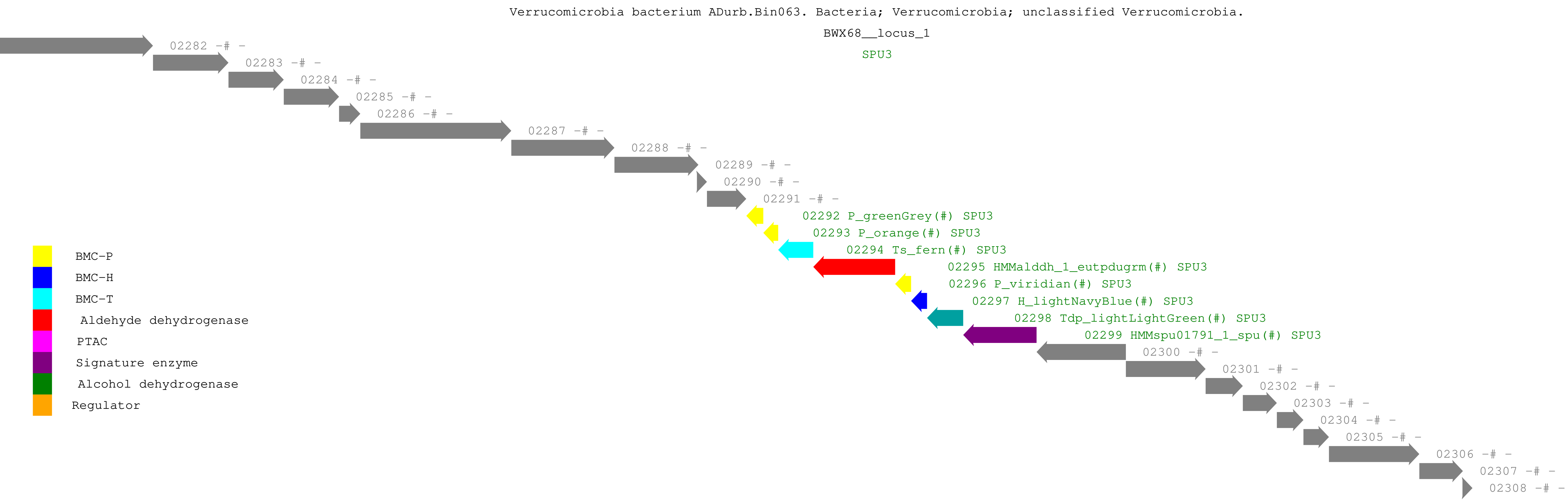
Verrucomicrobia bacterium ADurb.Bin063. Bacteria; Verrucomicrobia; unclassified Verrucomicrobia.
BWX68__locus_1
SPU3
02282 -# -
02283 -# -
02284 -# -
02285 -# -
02286 -# -
02287 -# -
02288 -# -
02289 -# -
02290 -# -
02291 -# -
02292 P_greenGrey(#) SPU3
02293 P_orange(#) SPU3
02294 Ts_fern(#) SPU3
02295 HMMalddh_1_eutpdugrm(#) SPU3
02296 P_viridian(#) SPU3
02297 H_lightNavyBlue(#) SPU3
02298 Tdp_lightLightGreen(#) SPU3
02299 HMMspu01791_1_spu(#) SPU3
02300 -# -
02301 -# -
02302 -# -
02303 -# -
02304 -# -
02305 -# -
02306 -# -
02307 -# -
02308 -# -
BMC-P
BMC-H
BMC-T
Aldehyde dehydrogenase
PTAC
Signature enzyme
Alcohol dehydrogenase
Regulator

Deltaproteobacteria bacterium ADurb.Bin207. Bacteria; Proteobacteria; Deltaproteobacteria.

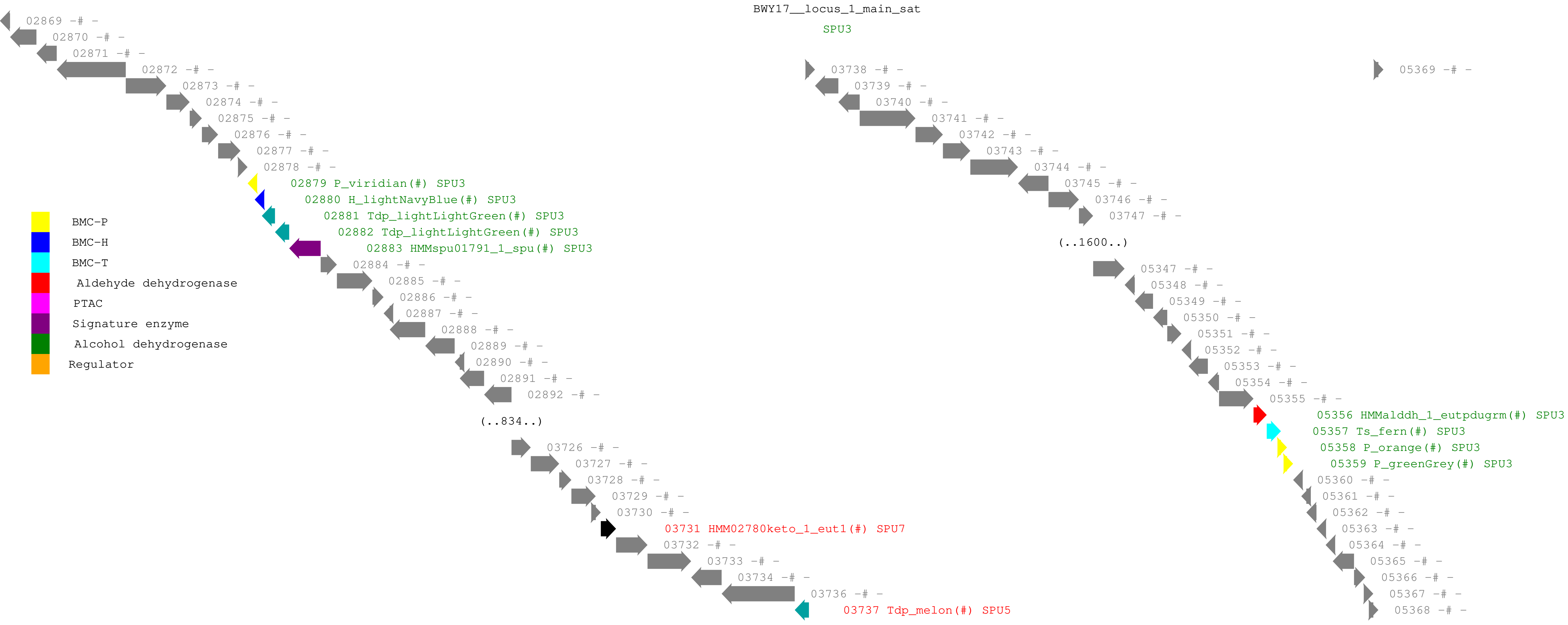

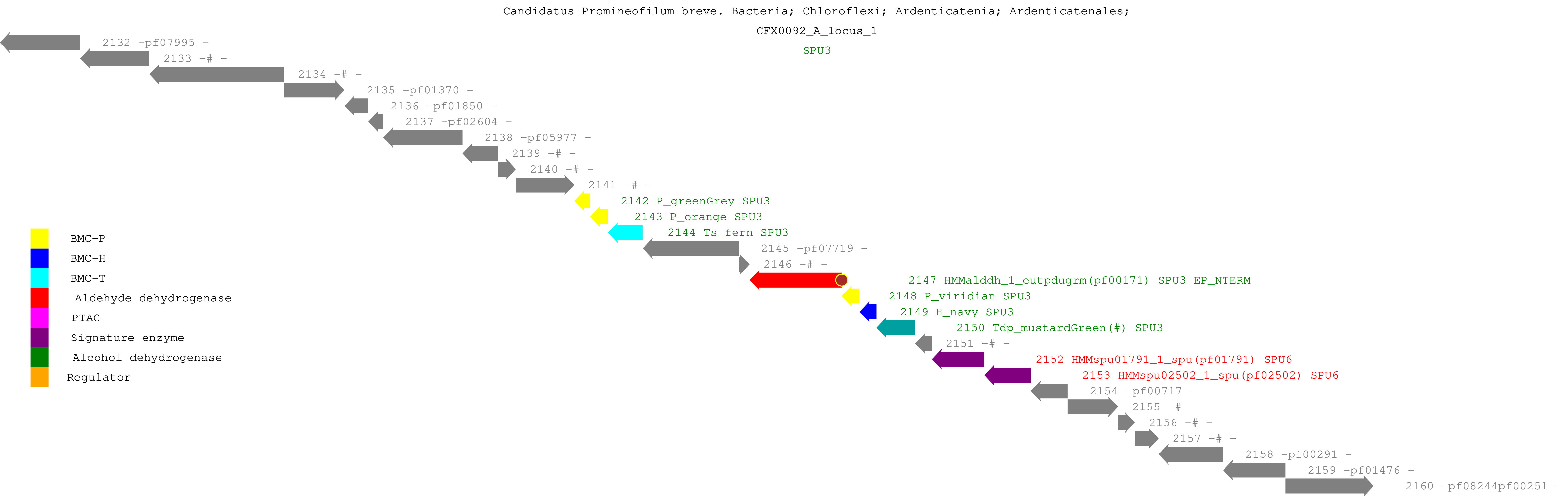
Candidatus Promineofilum breve. Bacteria; Chloroflexi; Ardenticatenia; Ardenticatenales;
CFX0092_A_locus_1
SPU3
2132 -pf07995 -
2133 -# -
2134 -# -
2135 -pf01370 -
2136 -pf01850 -
2137 -pf02604 -
2138 -pf05977 -
2139 -# -
2140 -# -
2141 -# -
2142 P_greenGrey SPU3
2143 P_orange SPU3
2144 Ts_fern SPU3
2145 -pf07719 -
2146 -# -
2147 HMMaldh_1_eutpdugrm(pf00171) SPU3 EP_NTERM
2148 P_viridian SPU3
2149 H_navy SPU3
2150 Tdp_mustardGreen(#) SPU3
2151 -# -
2152 HMMspu01791_1_spu(pf01791) SPU6
2153 HMMspu02502_1_spu(pf02502) SPU6
2154 -pf00717 -
2155 -# -
2156 -# -
2157 -# -
2158 -pf00291 -
2159 -pf01476 -
2160 -pf08244pf00251 -
BMC-P
BMC-H
BMC-T
Aldehyde dehydrogenase
PTAC
Signature enzyme
Alcohol dehydrogenase
Regulator

Anaerolineaceae bacterium. Bacteria; Chloroflexi; Anaerolineae; Anaerolineales; Anaerolineaceae.

CL608__locus_1

SPU3

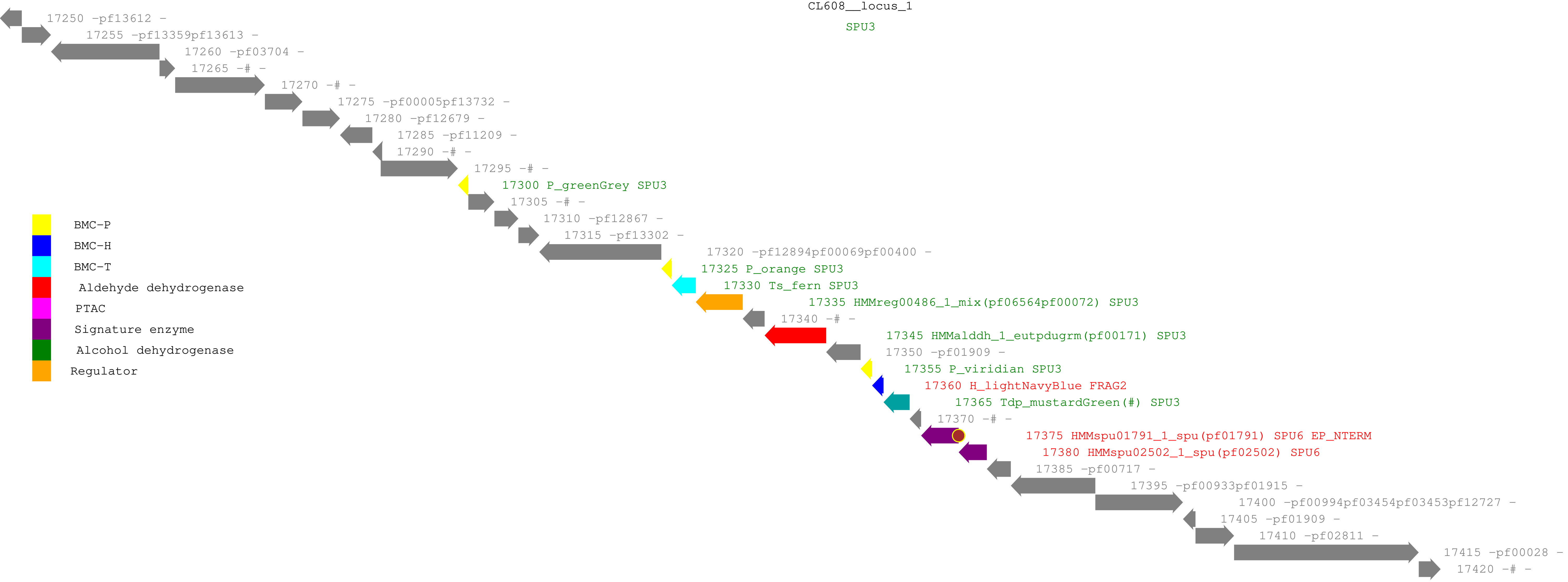

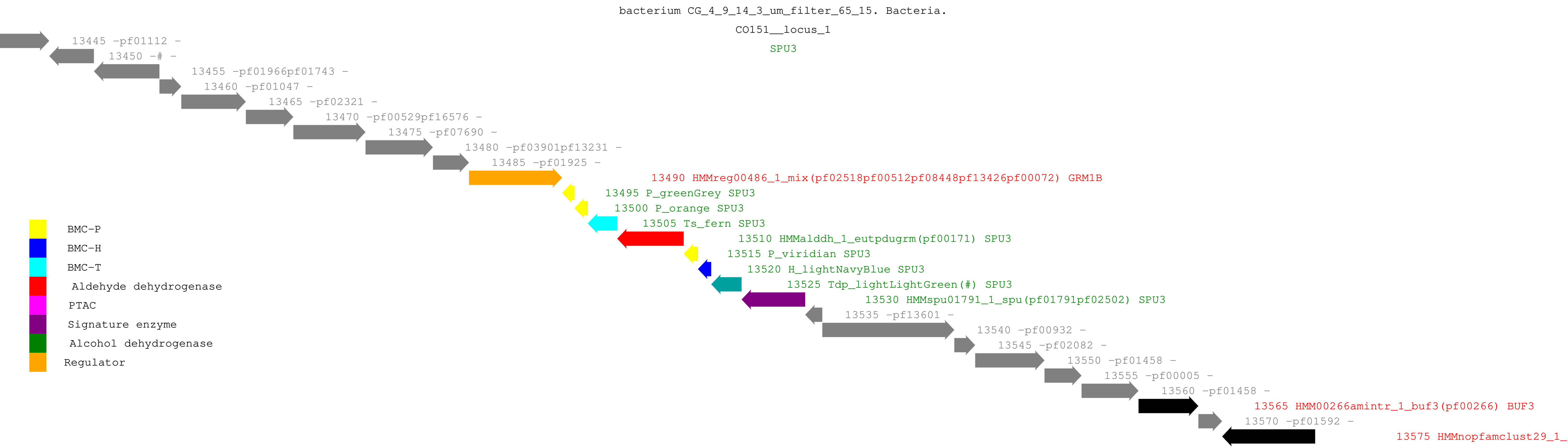
bacterium CG_4_9_14_3_um_filter_65_15. Bacteria.
CO151__locus_1
SPU3
13445 -pf01112 -
13450 -# -
13455 -pf01966pf01743 -
13460 -pf01047 -
13465 -pf02321 -
13470 -pf00529pf16576 -
13475 -pf07690 -
13480 -pf03901pf13231 -
13485 -pf01925 -
13490 HMMreg00486_1_mix(pf02518pf00512pf08448pf13426pf00072) GRM1B
13495 P_greenGrey SPU3
13500 P_orange SPU3
13505 Ts_fern SPU3
13510 HMMaldh_1_eutpdugrm(pf00171) SPU3
13515 P_viridian SPU3
13520 H_lightNavyBlue SPU3
13525 Tdp_lightLightGreen(#) SPU3
13530 HMMspu01791_1_spu(pf01791pf02502) SPU3
13535 -pf13601 -
13540 -pf00932 -
13545 -pf02082 -
13550 -pf01458 -
13555 -pf00005 -
13560 -pf01458 -
13565 HMM00266amintr_1_buf3(pf00266) BUF3
13570 -pf01592 -
13575 HMMnopfamclust29_1_
BMC-P
BMC-H
BMC-T
Aldehyde dehydrogenase
PTAC
Signature enzyme
Alcohol dehydrogenase
Regulator

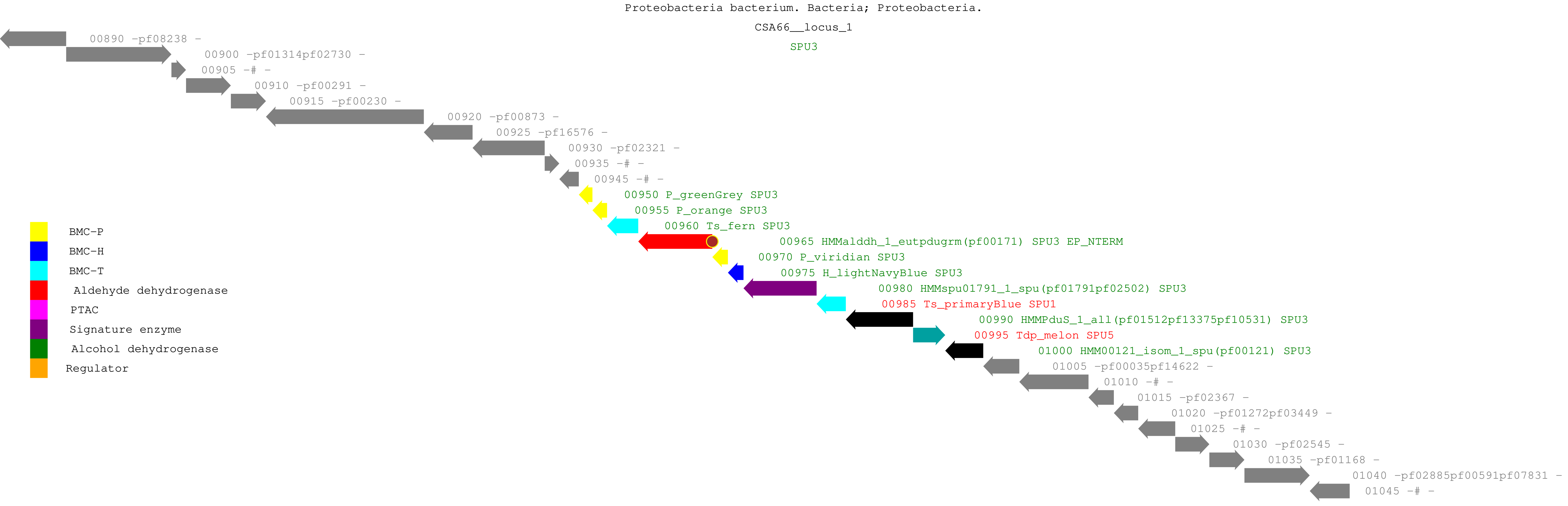
Proteobacteria bacterium. Bacteria; Proteobacteria.
CSA66__locus_1
SPU3
00890 -pf08238 -
00900 -pf01314pf02730 -
00905 -# -
00910 -pf00291 -
00915 -pf00230 -
00920 -pf00873 -
00925 -pf16576 -
00930 -pf02321 -
00935 -# -
00945 -# -
00950 P_greenGrey SPU3
00955 P_orange SPU3
00960 Ts_fern SPU3
00965 HMMalddh_1_eutpdugrm(pf00171) SPU3 EP_NTERM
00970 P_viridian SPU3
00975 H_lightNavyBlue SPU3
00980 HMMspu01791_1_spu(pf01791pf02502) SPU3
00985 Ts_primaryBlue SPU1
00990 HMMPduS_1_all(pf01512pf13375pf10531) SPU3
00995 Tdp_melon SPU5
01000 HMM00121_isom_1_spu(pf00121) SPU3
01005 -pf00035pf14622 -
01010 -# -
01015 -pf02367 -
01020 -pf01272pf03449 -
01025 -# -
01030 -pf02545 -
01035 -pf01168 -
01040 -pf02885pf00591pf07831 -
01045 -# -
BMC-P
BMC-H
BMC-T
Aldehyde dehydrogenase
PTAC
Signature enzyme
Alcohol dehydrogenase
Regulator

Deltaproteobacteria bacterium HGW-Deltaproteobacteria-14. Bacteria; Proteobacteria; Deltaproteobacteria.

CVU56__locus_1_main_sat

SPU3

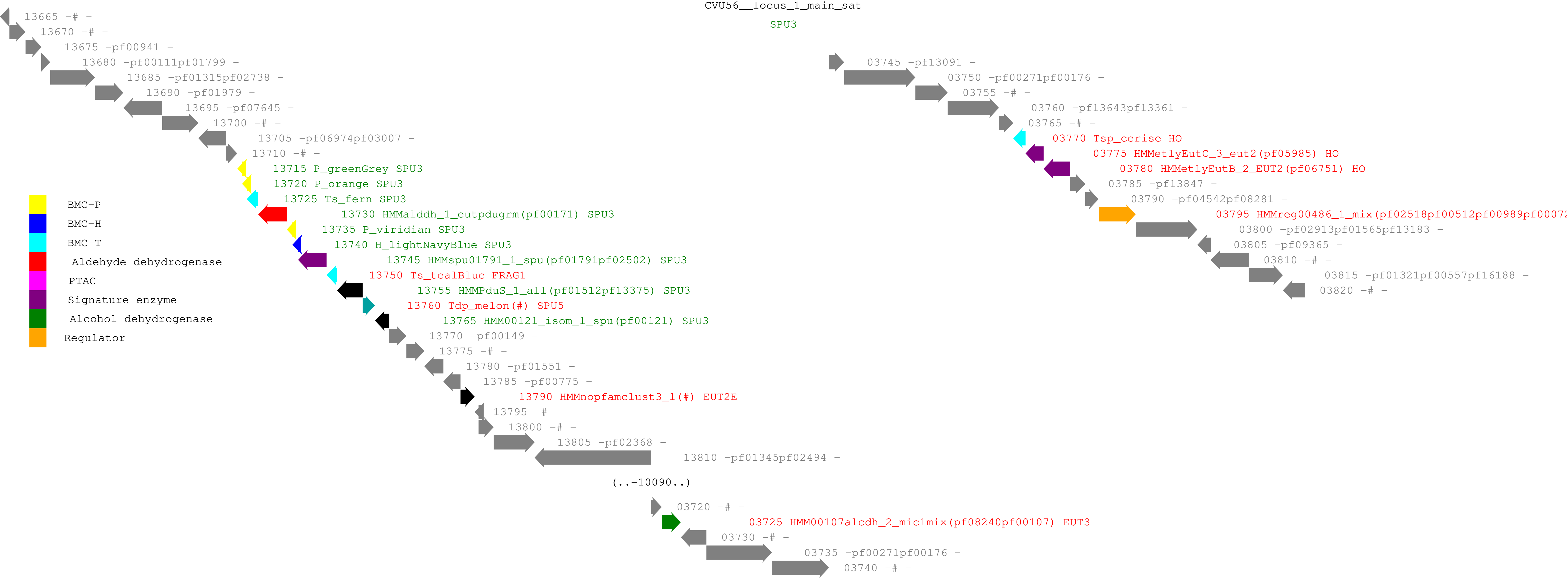

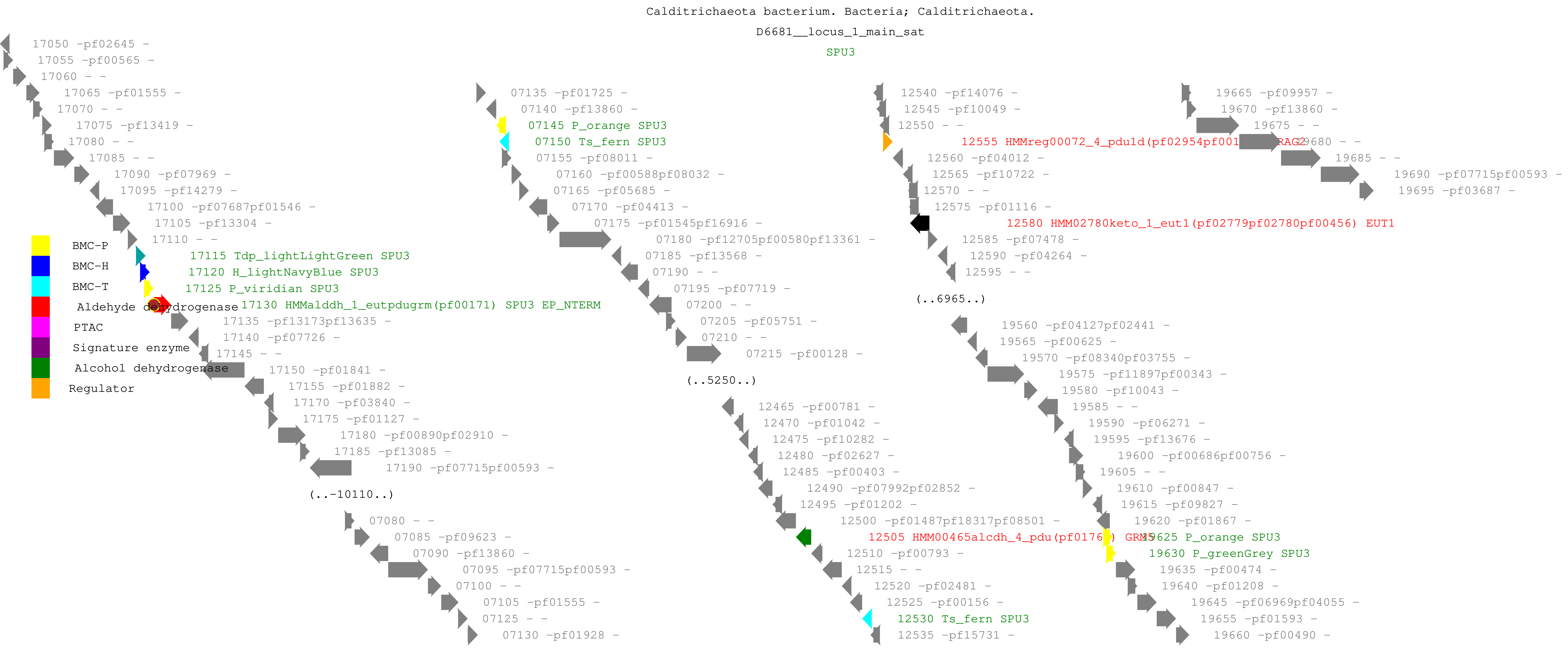
Calditrichaeota bacterium. Bacteria; Calditrichaeota.
D6681__locus_1_main_sat
SPU3
17050 -pf02645 -
17055 -pf00565 -
17060 - -
17065 -pf01555 -
17070 - -
17075 -pf13419 -
17080 - -
17085 - -
17090 -pf07969 -
17095 -pf14279 -
17100 -pf07687pf01546 -
17105 -pf13304 -
17110 - -
17115 Tdp_lightLightGreen SPU3
17120 H_lightNavyBlue SPU3
17125 P_viridian SPU3
17130 HMMalddh_1_eutpdugrm(pf00171) SPU3 EP_NTERM
17135 -pf13173pf13635 -
17140 -pf07726 -
17145 - -
17150 -pf01841 -
17155 -pf01882 -
17170 -pf03840 -
17175 -pf01127 -
17180 -pf00890pf02910 -
17185 -pf13085 -
17190 -pf07715pf00593 -
(..-10110..)
07080 - -
07085 -pf09623 -
07090 -pf13860 -
07095 -pf07715pf00593 -
07100 - -
07105 -pf01555 -
07125 - -
07130 -pf01928 -
BMC-P
BMC-H
BMC-T
Aldehyde dehydrogenase
PTAC
Signature enzyme
Alcohol dehydrogenase
Regulator
07135 -pf01725 -
07140 -pf13860 -
07145 P_orange SPU3
07150 Ts_fern SPU3
07155 -pf08011 -
07160 -pf00588pf08032 -
07165 -pf05685 -
07170 -pf04413 -
07175 -pf01545pf16916 -
07180 -pf12705pf00580pf13361 -
07185 -pf13568 -
07190 - -
07195 -pf07719 -
07200 - -
07205 -pf05751 -
07210 - -
07215 -pf00128 -
(..5250..)
12465 -pf00781 -
12470 -pf01042 -
12475 -pf10282 -
12480 -pf02627 -
12485 -pf00403 -
12490 -pf07992pf02852 -
12495 -pf01202 -
12500 -pf01487pf18317pf08501 -
12505 HMM00465alcdh_4_pdu(pf0176 ) GRM
12510 -pf00793 -
12515 - -
12520 -pf02481 -
12525 -pf00156 -
12530 Ts_fern SPU3
12535 -pf15731 -
12540 -pf14076 -
12545 -pf10049 -
12550 - -
12555 HMMreg00072_4_pdu1d(pf02954pf001 RAG2
12560 -pf04012 -
12565 -pf10722 -
12570 - -
12575 -pf01116 -
12580 HMM02780keto_1_eut1(pf02779pf02780pf00456) EUT1
12585 -pf07478 -
12590 -pf04264 -
12595 - -
(..6965..)
19560 -pf04127pf02441 -
19565 -pf00625 -
19570 -pf08340pf03755 -
19575 -pf11897pf00343 -
19580 -pf10043 -
19585 - -
19590 -pf06271 -
19595 -pf13676 -
19600 -pf00686pf00756 -
19605 - -
19610 -pf00847 -
19615 -pf09827 -
19620 -pf01867 -
19625 P_orange SPU3
19630 P_greenGrey SPU3
19635 -pf00474 -
19640 -pf01208 -
19645 -pf06969pf04055 -
19655 -pf01593 -
19660 -pf00490 -
19665 -pf09957 -
19670 -pf13860 -
19675 - -
19680 - -
19685 - -
19690 -pf07715pf00593 -
19695 -pf03687 -

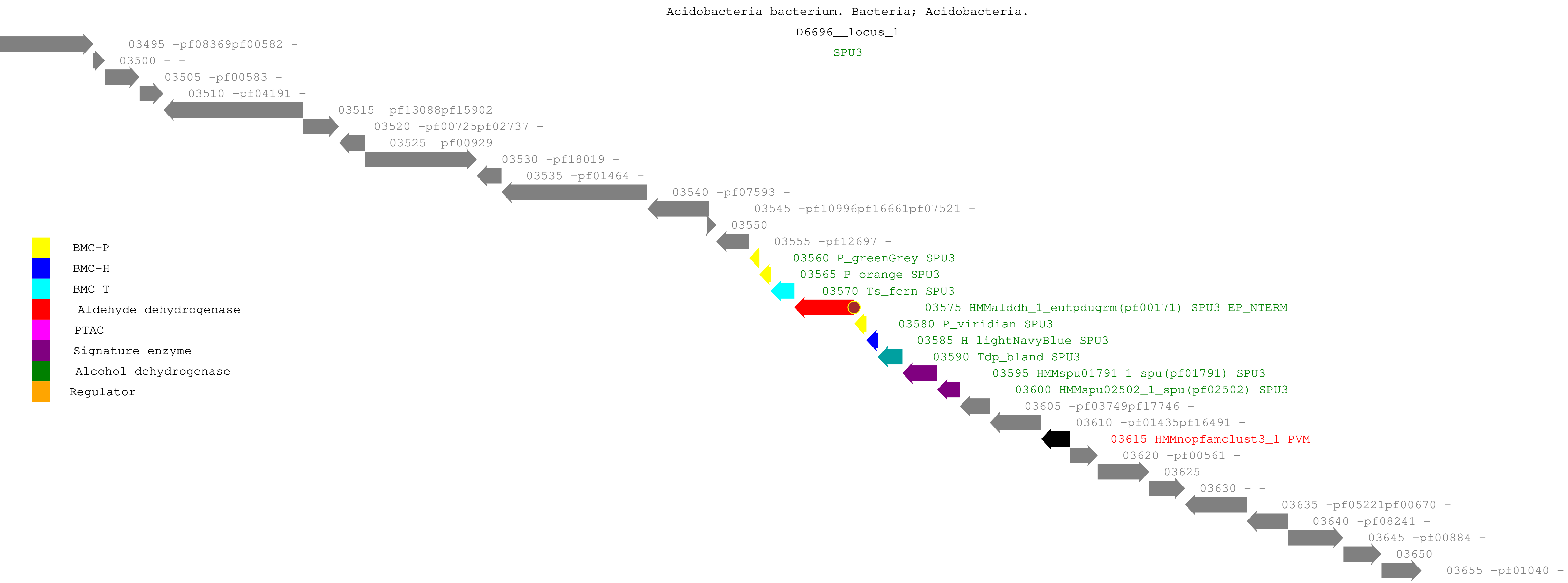
Acidobacteria bacterium. Bacteria; Acidobacteria.
D6696__locus_1
SPU3
03495 -pf08369pf00582 -
03500 - -
03505 -pf00583 -
03510 -pf04191 -
03515 -pf13088pf15902 -
03520 -pf00725pf02737 -
03525 -pf00929 -
03530 -pf18019 -
03535 -pf01464 -
03540 -pf07593 -
03545 -pf10996pf16661pf07521 -
03550 - -
03555 -pf12697 -
03560 P_greenGrey SPU3
03565 P_orange SPU3
03570 Ts_fern SPU3
03575 HMMaldh_1_eutpdugrm(pf00171) SPU3 EP_NTERM
03580 P_viridian SPU3
03585 H_lightNavyBlue SPU3
03590 Tdp_bland SPU3
03595 HMMspu01791_1_spu(pf01791) SPU3
03600 HMMspu02502_1_spu(pf02502) SPU3
03605 -pf03749pf17746 -
03610 -pf01435pf16491 -
03615 HMMnopfamclust3_1 PVM
03620 -pf00561 -
03625 - -
03630 - -
03635 -pf05221pf00670 -
03640 -pf08241 -
03645 -pf00884 -
03650 - -
03655 -pf01040 -
BMC-P
BMC-H
BMC-T
Aldehyde dehydrogenase
PTAC
Signature enzyme
Alcohol dehydrogenase
Regulator

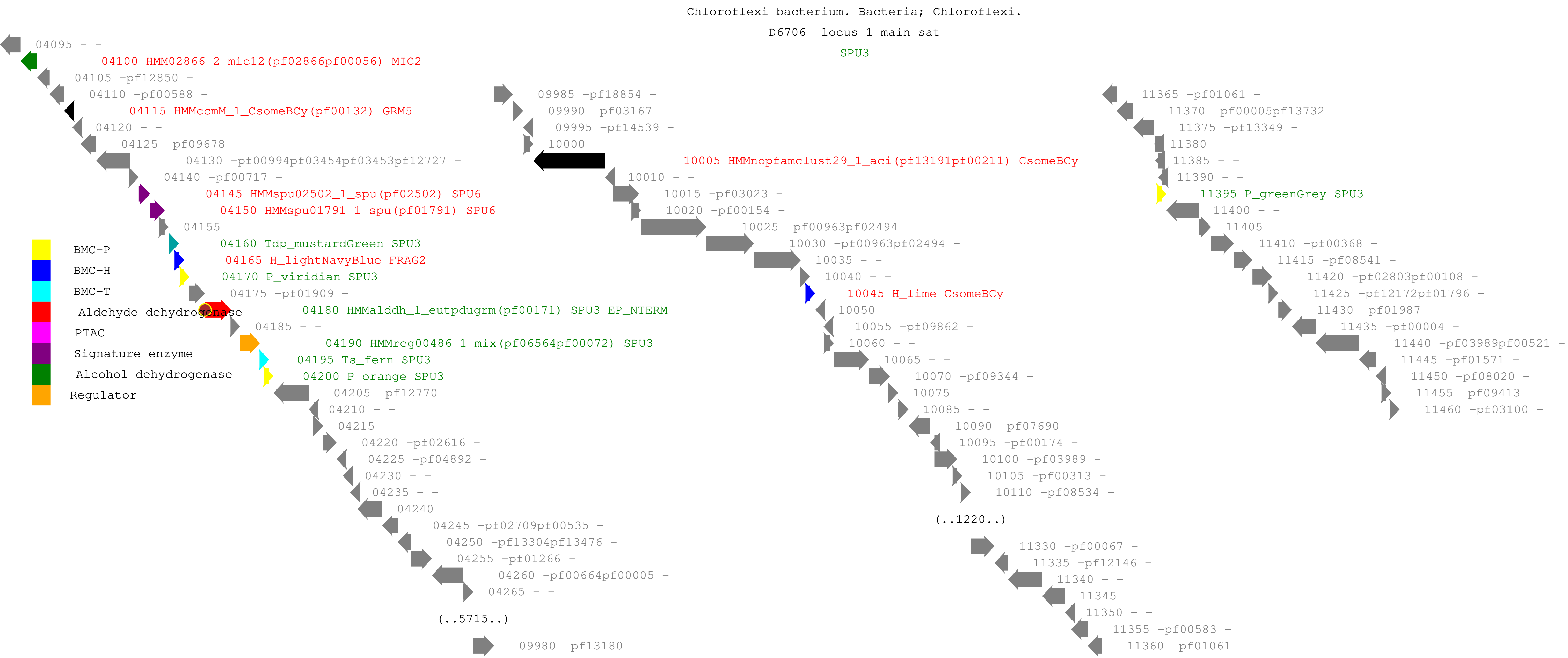

Chloroflexi bacterium. Bacteria; Chloroflexi.
D6706__locus_1_main_sat
SPU3
04095 - -
04100 HMM02866_2_mic12(pf02866pf00056) MIC2
04105 -pf12850 -
04110 -pf00588 -
04115 HMMccmM_1_CsomeBCy(pf00132) GRM5
04120 - -
04125 -pf09678 -
04130 -pf00994pf03454pf03453pf12727 -
04140 -pf00717 -
04145 HMMspu02502_1_spu(pf02502) SPU6
04150 HMMspu01791_1_spu(pf01791) SPU6
04155 - -
04160 Tdp_mustardGreen SPU3
04165 H_lightNavyBlue FRAG2
04170 P_viridian SPU3
04175 -pf01909 -
04180 HMMalddh_1_eutpdugrm(pf00171) SPU3 EP_NTERM
04185 - -
04190 HMMreg00486_1_mix(pf06564pf00072) SPU3
04195 Ts_fern SPU3
04200 P_orange SPU3
04205 -pf12770 -
04210 - -
04215 - -
04220 -pf02616 -
04225 -pf04892 -
04230 - -
04235 - -
04240 - -
04245 -pf02709pf00535 -
04250 -pf13304pf13476 -
04255 -pf01266 -
04260 -pf00664pf00005 -
04265 - -
(..5715..)
09980 -pf13180 -
BMC-P
BMC-H
BMC-T
Aldehyde dehydrogenase
PTAC
Signature enzyme
Alcohol dehydrogenase
Regulator
09985 -pf18854 -
09990 -pf03167 -
09995 -pf14539 -
10000 - -
10005 HMMnopfamclust29_1_aci(pf13191pf00211) CsomeBCy
10010 - -
10015 -pf03023 -
10020 -pf00154 -
10025 -pf00963pf02494 -
10030 -pf00963pf02494 -
10035 - -
10040 - -
10045 H_lime CsomeBCy
10050 - -
10055 -pf09862 -
10060 - -
10065 - -
10070 -pf09344 -
10075 - -
10085 - -
10090 -pf07690 -
10095 -pf00174 -
10100 -pf03989 -
10105 -pf00313 -
10110 -pf08534 -
(..1220..)
11330 -pf00067 -
11335 -pf12146 -
11340 - -
11345 - -
11350 - -
11355 -pf00583 -
11360 -pf01061 -
11365 -pf01061 -
11370 -pf00005pf13732 -
11375 -pf13349 -
11380 - -
11385 - -
11390 - -
11395 P_greenGrey SPU3
11400 - -
11405 - -
11410 -pf00368 -
11415 -pf08541 -
11420 -pf02803pf00108 -
11425 -pf12172pf01796 -
11430 -pf01987 -
11435 -pf00004 -
11440 -pf03989pf00521 -
11445 -pf01571 -
11450 -pf08020 -
11455 -pf09413 -
11460 -pf03100 -

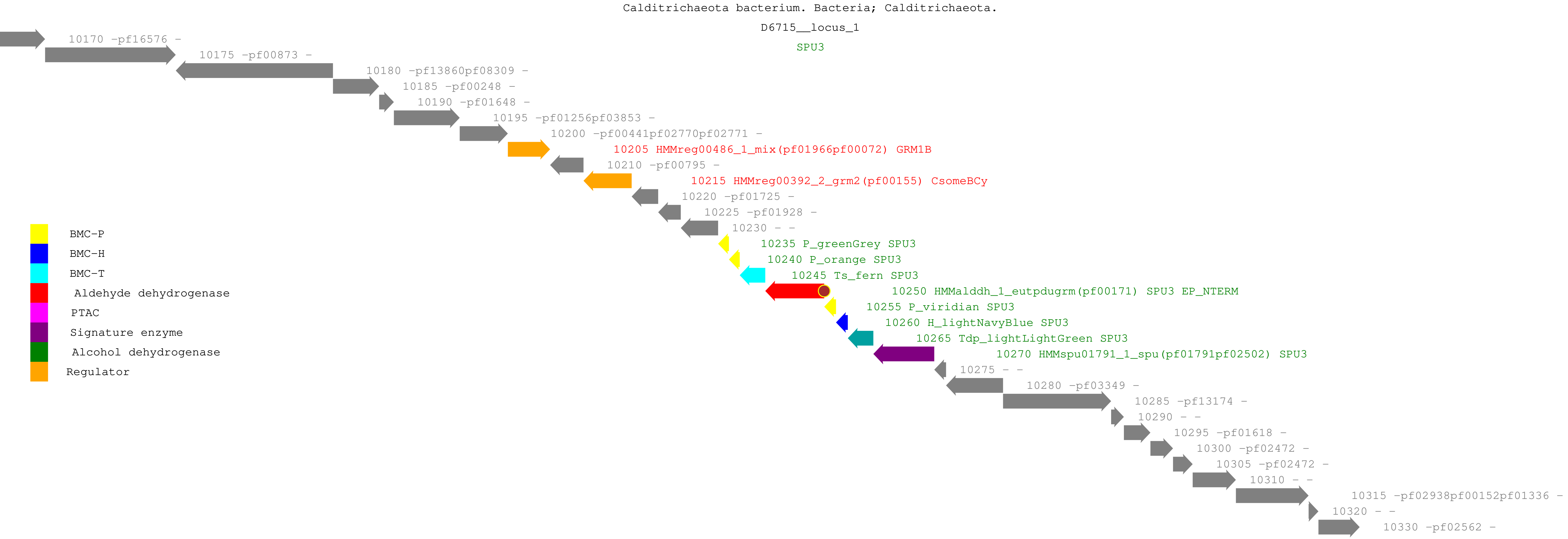
Calditrichaeota bacterium. Bacteria; Calditrichaeota.
D6715__locus_1
SPU3
10170 -pf16576 -
10175 -pf00873 -
10180 -pf13860pf08309 -
10185 -pf00248 -
10190 -pf01648 -
10195 -pf01256pf03853 -
10200 -pf00441pf02770pf02771 -
10205 HMMreg00486_1_mix(pf01966pf00072) GRM1B
10210 -pf00795 -
10215 HMMreg00392_2_grm2(pf00155) CsomeBCy
10220 -pf01725 -
10225 -pf01928 -
10230 - -
10235 P_greenGrey SPU3
10240 P_orange SPU3
10245 Ts_fern SPU3
10250 HMMaldh_1_eutpdugrm(pf00171) SPU3 EP_NTERM
10255 P_viridian SPU3
10260 H_lightNavyBlue SPU3
10265 Tdp_lightLightGreen SPU3
10270 HMMspu01791_1_spu(pf01791pf02502) SPU3
10275 - -
10280 -pf03349 -
10285 -pf13174 -
10290 - -
10295 -pf01618 -
10300 -pf02472 -
10305 -pf02472 -
10310 - -
10315 -pf02938pf00152pf01336 -
10320 - -
10330 -pf02562 -
BMC-P
BMC-H
BMC-T
Aldehyde dehydrogenase
PTAC
Signature enzyme
Alcohol dehydrogenase
Regulator

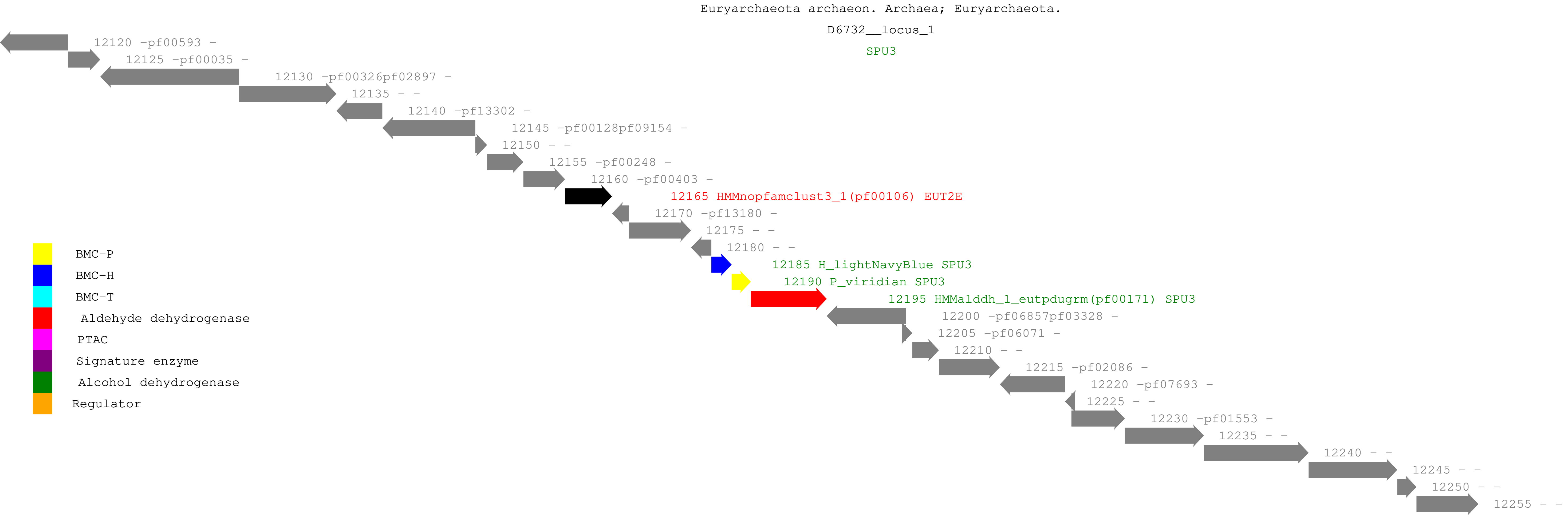

Euryarchaeota archaeon. Archaea; Euryarchaeota.
D6732__locus_1
SPU3
12120 -pf00593 -
12125 -pf00035 -
12130 -pf00326pf02897 -
12135 - -
12140 -pf13302 -
12145 -pf00128pf09154 -
12150 - -
12155 -pf00248 -
12160 -pf00403 -
12165 HMMnopfamclust3_1(pf00106) EUT2E
12170 -pf13180 -
12175 - -
12180 - -
12185 H_lightNavyBlue SPU3
12190 P_viridian SPU3
12195 HMMalddh_1_eutpdugrm(pf00171) SPU3
12200 -pf06857pf03328 -
12205 -pf06071 -
12210 - -
12215 -pf02086 -
12220 -pf07693 -
12225 - -
12230 -pf01553 -
12235 - -
12240 - -
12245 - -
12250 - -
12255 - -
BMC-P
BMC-H
BMC-T
Aldehyde dehydrogenase
PTAC
Signature enzyme
Alcohol dehydrogenase
Regulator

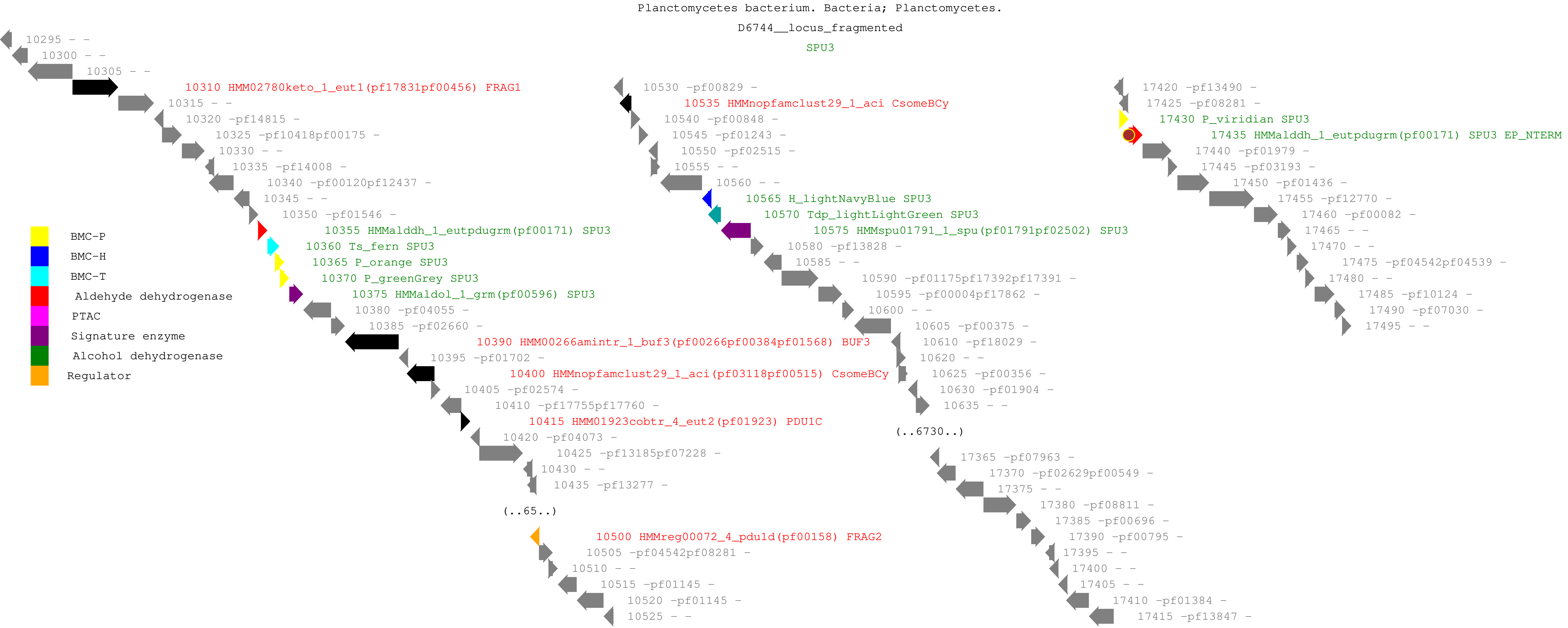

Planctomycetes bacterium. Bacteria; Planctomycetes.
D6744__locus_fragmented
SPU3
10295 - -
10300 - -
10305 - -
10310 HMM02780keto_1_eut1(pf17831pf00456) FRAG1
10315 - -
10320 -pf14815 -
10325 -pf10418pf00175 -
10330 - -
10335 -pf14008 -
10340 -pf00120pf12437 -
10345 - -
10350 -pf01546 -
10355 HMMaldh_1_eutpdugrm(pf00171) SPU3
10360 Ts_fern SPU3
10365 P_orange SPU3
10370 P_greenGrey SPU3
10375 HMMaldol_1_grm(pf00596) SPU3
10380 -pf04055 -
10385 -pf02660 -
10390 HMM00266amintr_1_buf3(pf00266pf00384pf01568) BUF3
10395 -pf01702 -
10400 HMMnopfamclust29_1_aci(pf03118pf00515) CsomeBCy
10405 -pf02574 -
10410 -pf17755pf17760 -
10415 HMM01923cobtr_4_eut2(pf01923) PDU1C
10420 -pf04073 -
10425 -pf13185pf07228 -
10430 - -
10435 -pf13277 -
(..65..)
10500 HMMreg00072_4_pdu1d(pf00158) FRAG2
10505 -pf04542pf08281 -
10510 - -
10515 -pf01145 -
10520 -pf01145 -
10525 - -
10530 -pf00829 -
10535 HMMnopfamclust29_1_aci CsomeBCy
10540 -pf00848 -
10545 -pf01243 -
10550 -pf02515 -
10555 - -
10560 - -
10565 H_lightNavyBlue SPU3
10570 Tdp_lightLightGreen SPU3
10575 HMMspu01791_1_spu(pf01791pf02502) SPU3
10580 -pf13828 -
10585 - -
10590 -pf01175pf17392pf17391 -
10595 -pf00004pf17862 -
10600 - -
10605 -pf00375 -
10610 -pf18029 -
10620 - -
10625 -pf00356 -
10630 -pf01904 -
10635 - -
(..6730..)
17365 -pf07963 -
17370 -pf02629pf00549 -
17375 - -
17380 -pf08811 -
17385 -pf00696 -
17390 -pf00795 -
17395 - -
17400 - -
17405 - -
17410 -pf01384 -
17415 -pf13847 -
17420 -pf13490 -
17425 -pf08281 -
17430 P_viridian SPU3
17435 HMMaldh_1_eutpdugrm(pf00171) SPU3 EP_NTERM
17440 -pf01979 -
17445 -pf03193 -
17450 -pf01436 -
17455 -pf12770 -
17460 -pf00082 -
17465 - -
17470 - -
17475 -pf04542pf04539 -
17480 - -
17485 -pf10124 -
17490 -pf07030 -
17495 - -
BMC-P
BMC-H
BMC-T
Aldehyde dehydrogenase
PTAC
Signature enzyme
Alcohol dehydrogenase
Regulator

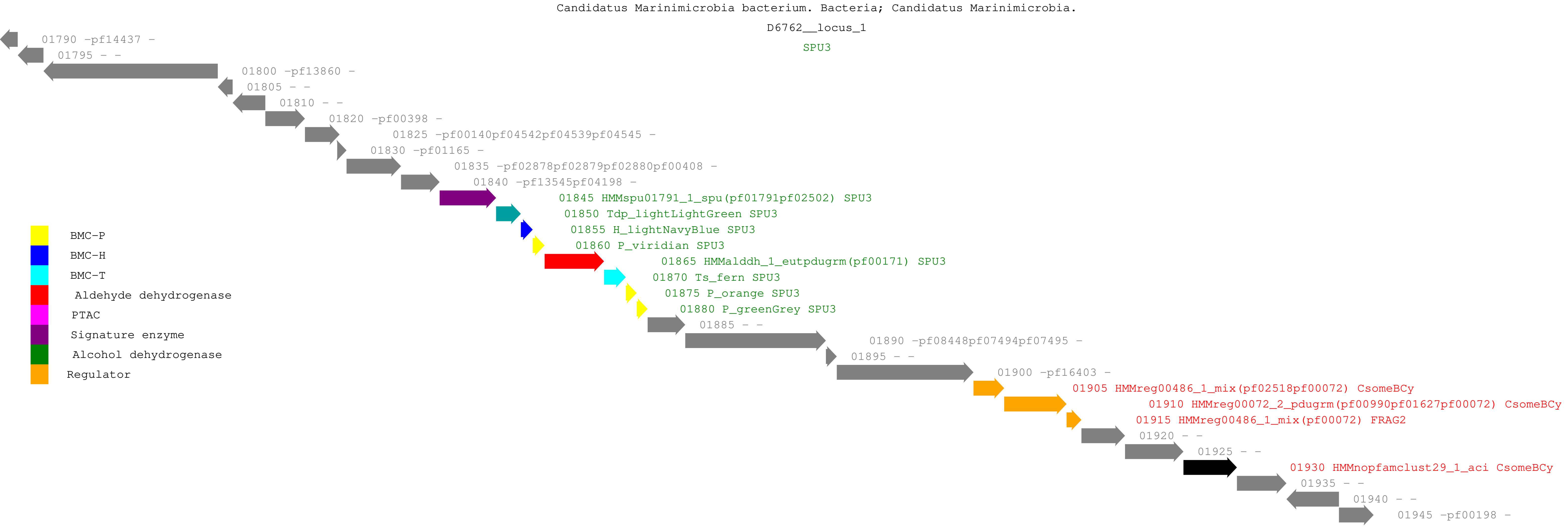

Candidatus Marinimicrobia bacterium. Bacteria; Candidatus Marinimicrobia.
D6762__locus_1
SPU3
01790 -pf14437 -
01795 - -
01800 -pf13860 -
01805 - -
01810 - -
01820 -pf00398 -
01825 -pf00140pf04542pf04539pf04545 -
01830 -pf01165 -
01835 -pf02878pf02879pf02880pf00408 -
01840 -pf13545pf04198 -
01845 HMMspu01791_1_spu(pf01791pf02502) SPU3
01850 Tdp_lightLightGreen SPU3
01855 H_lightNavyBlue SPU3
01860 P_viridian SPU3
01865 HMMalddh_1_eutpdugrm(pf00171) SPU3
01870 Ts_fern SPU3
01875 P_orange SPU3
01880 P_greenGrey SPU3
01885 - -
01890 -pf08448pf07494pf07495 -
01895 - -
01900 -pf16403 -
01905 HMMreg00486_1_mix(pf02518pf00072) CsomeBCy
01910 HMMreg00072_2_pdugrm(pf00990pf01627pf00072) CsomeBCy
01915 HMMreg00486_1_mix(pf00072) FRAG2
01920 - -
01925 - -
01930 HMMnopfamclust29_1_aci CsomeBCy
01935 - -
01940 - -
01945 -pf00198 -
BMC-P
BMC-H
BMC-T
Aldehyde dehydrogenase
PTAC
Signature enzyme
Alcohol dehydrogenase
Regulator

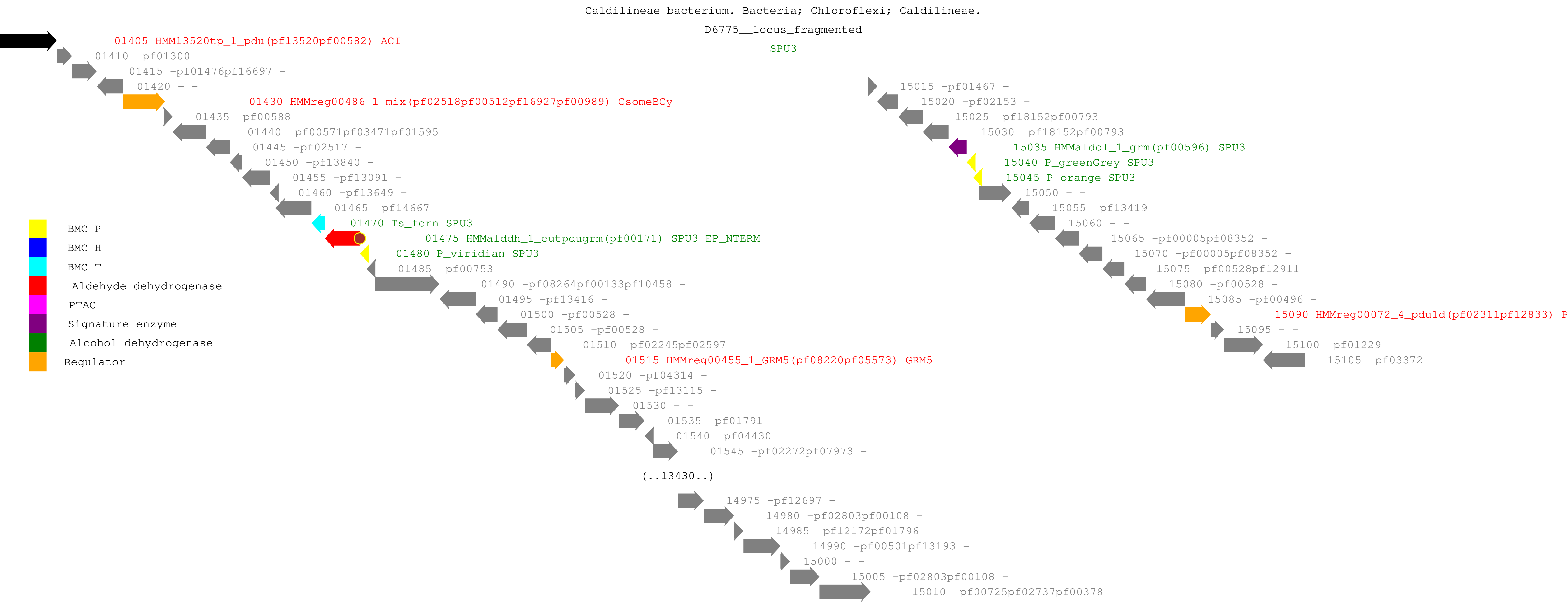

Caldilineae bacterium. Bacteria; Chloroflexi; Caldilineae.
D6775__locus_fragmented
SPU3
01405 HMM13520tp_1_pdu(pf13520pf00582) ACI
01410 -pf01300 -
01415 -pf01476pf16697 -
01420 - -
01430 HMMreg00486_1_mix(pf02518pf00512pf16927pf00989) CsomeBCy
01435 -pf00588 -
01440 -pf00571pf03471pf01595 -
01445 -pf02517 -
01450 -pf13840 -
01455 -pf13091 -
01460 -pf13649 -
01465 -pf14667 -
01470 Ts_fern SPU3
01475 HMMaldh_1_eutpdugrm(pf00171) SPU3 EP_NTERM
01480 P_viridian SPU3
01485 -pf00753 -
01490 -pf08264pf00133pf10458 -
01495 -pf13416 -
01500 -pf00528 -
01505 -pf00528 -
01510 -pf02245pf02597 -
01515 HMMreg00455_1_GRM5(pf08220pf05573) GRM5
01520 -pf04314 -
01525 -pf13115 -
01530 - -
01535 -pf01791 -
01540 -pf04430 -
01545 -pf02272pf07973 -
(..13430..)
14975 -pf12697 -
14980 -pf02803pf00108 -
14985 -pf12172pf01796 -
14990 -pf00501pf13193 -
15000 - -
15005 -pf02803pf00108 -
15010 -pf00725pf02737pf00378 -
15015 -pf01467 -
15020 -pf02153 -
15025 -pf18152pf00793 -
15030 -pf18152pf00793 -
15035 HMMaldol_1_grm(pf00596) SPU3
15040 P_greenGrey SPU3
15045 P_orange SPU3
15050 - -
15055 -pf13419 -
15060 - -
15065 -pf00005pf08352 -
15070 -pf00005pf08352 -
15075 -pf00528pf12911 -
15080 -pf00528 -
15085 -pf00496 -
15090 HMMreg00072_4_pdu1d(pf02311pf12833) P
15095 - -
15100 -pf01229 -
15105 -pf03372 -
BMC-P
BMC-H
BMC-T
Aldehyde dehydrogenase
PTAC
Signature enzyme
Alcohol dehydrogenase
Regulator

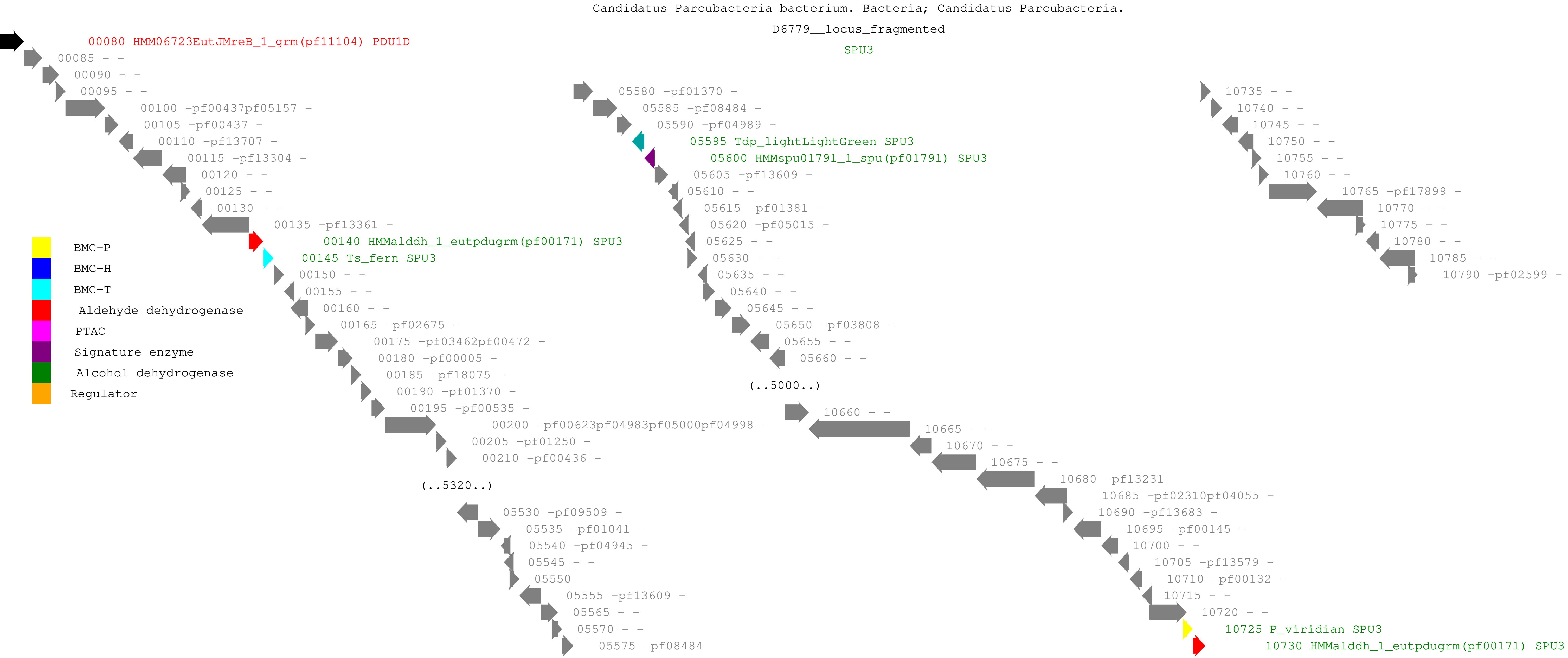

Candidatus Parcubacteria bacterium. Bacteria; Candidatus Parcubacteria.
D6779__locus_fragmented
SPU3
00080 HMM06723EutJMreB_1_grm(pf11104) PDU1D
00085 - -
00090 - -
00095 - -
00100 -pf00437pf05157 -
00105 -pf00437 -
00110 -pf13707 -
00115 -pf13304 -
00120 - -
00125 - -
00130 - -
00135 -pf13361 -
00140 HMMalddh_1_eutpdugrm(pf00171) SPU3
00145 Ts_fern SPU3
00150 - -
00155 - -
00160 - -
00165 -pf02675 -
00175 -pf03462pf00472 -
00180 -pf00005 -
00185 -pf18075 -
00190 -pf01370 -
00195 -pf00535 -
00200 -pf00623pf04983pf05000pf04998 -
00205 -pf01250 -
00210 -pf00436 -
(..5320..)
05530 -pf09509 -
05535 -pf01041 -
05540 -pf04945 -
05545 - -
05550 - -
05555 -pf13609 -
05565 - -
05570 - -
05575 -pf08484 -
05580 -pf01370 -
05585 -pf08484 -
05590 -pf04989 -
05595 Tdp_lightLightGreen SPU3
05600 HMMspu01791_1_spu(pf01791) SPU3
05605 -pf13609 -
05610 - -
05615 -pf01381 -
05620 -pf05015 -
05625 - -
05630 - -
05635 - -
05640 - -
05645 - -
05650 -pf03808 -
05655 - -
05660 - -
(..5000..)
10660 - -
10665 - -
10670 - -
10675 - -
10680 -pf13231 -
10685 -pf02310pf04055 -
10690 -pf13683 -
10695 -pf00145 -
10700 - -
10705 -pf13579 -
10710 -pf00132 -
10715 - -
10720 - -
10725 P_viridian SPU3
10730 HMMalddh_1_eutpdugrm(pf00171) SPU3
10735 - -
10740 - -
10745 - -
10750 - -
10755 - -
10760 - -
10765 -pf17899 -
10770 - -
10775 - -
10780 - -
10785 - -
10790 -pf02599 -
BMC-P
BMC-H
BMC-T
Aldehyde dehydrogenase
PTAC
Signature enzyme
Alcohol dehydrogenase
Regulator

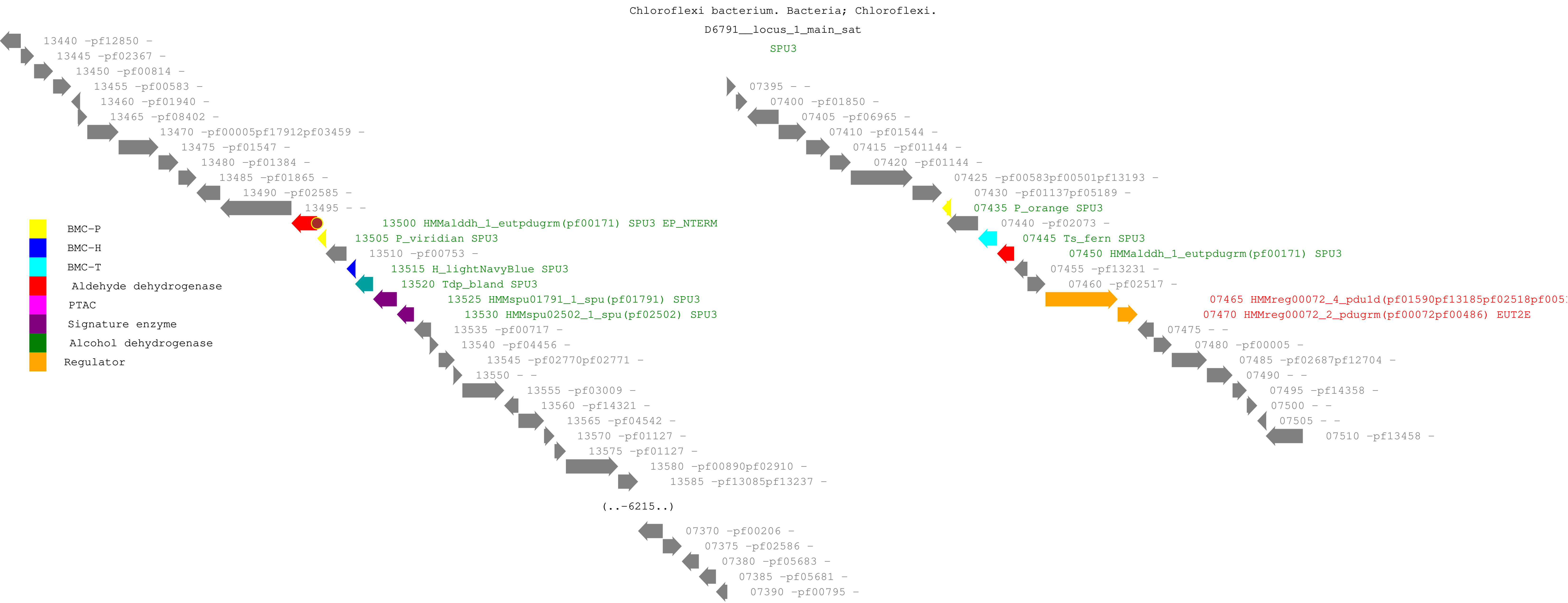
Chloroflexi bacterium. Bacteria; Chloroflexi.
D6791__locus_1_main_sat
SPU3
13440 -pf12850 -
13445 -pf02367 -
13450 -pf00814 -
13455 -pf00583 -
13460 -pf01940 -
13465 -pf08402 -
13470 -pf00005pf17912pf03459 -
13475 -pf01547 -
13480 -pf01384 -
13485 -pf01865 -
13490 -pf02585 -
13495 - -
13500 HMMaldh_1_eutpdugrm(pf00171) SPU3 EP_NTERM
13505 P_viridian SPU3
13510 -pf00753 -
13515 H_lightNavyBlue SPU3
13520 Tdp_bland SPU3
13525 HMMspu01791_1_spu(pf01791) SPU3
13530 HMMspu02502_1_spu(pf02502) SPU3
13535 -pf00717 -
13540 -pf04456 -
13545 -pf02770pf02771 -
13550 - -
13555 -pf03009 -
13560 -pf14321 -
13565 -pf04542 -
13570 -pf01127 -
13575 -pf01127 -
13580 -pf00890pf02910 -
13585 -pf13085pf13237 -
(..-6215..)
07370 -pf00206 -
07375 -pf02586 -
07380 -pf05683 -
07385 -pf05681 -
07390 -pf00795 -
07395 - -
07400 -pf01850 -
07405 -pf06965 -
07410 -pf01544 -
07415 -pf01144 -
07420 -pf01144 -
07425 -pf00583pf00501pf13193 -
07430 -pf01137pf05189 -
07435 P_orange SPU3
07440 -pf02073 -
07445 Ts_fern SPU3
07450 HMMalddh_1_eutpdugrm(pf00171) SPU3
07455 -pf13231 -
07460 -pf02517 -
07465 HMMreg00072_4_pdu1d(pf01590pf13185pf02518pf005
07470 HMMreg00072_2_pdugrm(pf00072pf00486) EUT2E
07475 - -
07480 -pf00005 -
07485 -pf02687pf12704 -
07490 - -
07495 -pf14358 -
07500 - -
07505 - -
07510 -pf13458 -
BMC-P
BMC-H
BMC-T
Aldehyde dehydrogenase
PTAC
Signature enzyme
Alcohol dehydrogenase
Regulator

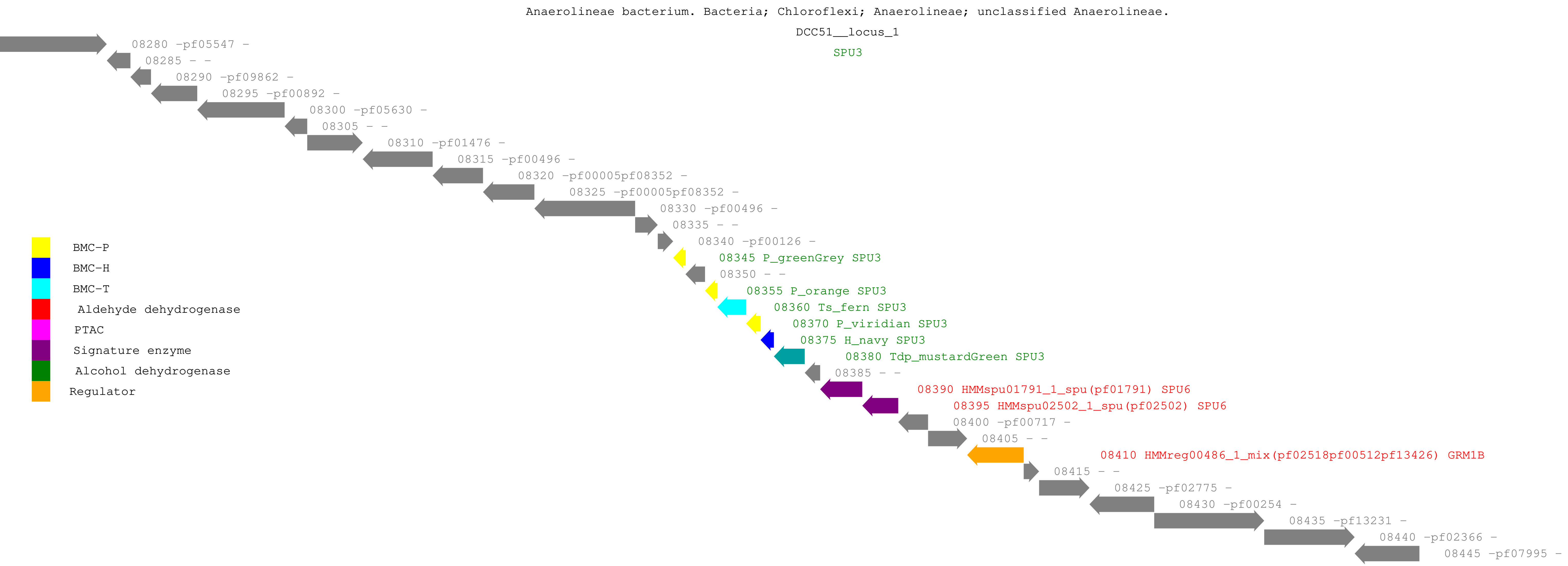
Anaerolineae bacterium. Bacteria; Chloroflexi; Anaerolineae; unclassified Anaerolineae.
DCC51__locus_1
SPU3
08280 -pf05547 -
08285 - -
08290 -pf09862 -
08295 -pf00892 -
08300 -pf05630 -
08305 - -
08310 -pf01476 -
08315 -pf00496 -
08320 -pf00005pf08352 -
08325 -pf00005pf08352 -
08330 -pf00496 -
08335 - -
08340 -pf00126 -
08345 P_greenGrey SPU3
08350 - -
08355 P_orange SPU3
08360 Ts_fern SPU3
08370 P_viridian SPU3
08375 H_navy SPU3
08380 Tdp_mustardGreen SPU3
08385 - -
08390 HMMspu01791_1_spu(pf01791) SPU6
08395 HMMspu02502_1_spu(pf02502) SPU6
08400 -pf00717 -
08405 - -
08410 HMMreg00486_1_mix(pf02518pf00512pf13426) GRM1B
08415 - -
08425 -pf02775 -
08430 -pf00254 -
08435 -pf13231 -
08440 -pf02366 -
08445 -pf07995 -
BMC-P
BMC-H
BMC-T
Aldehyde dehydrogenase
PTAC
Signature enzyme
Alcohol dehydrogenase
Regulator

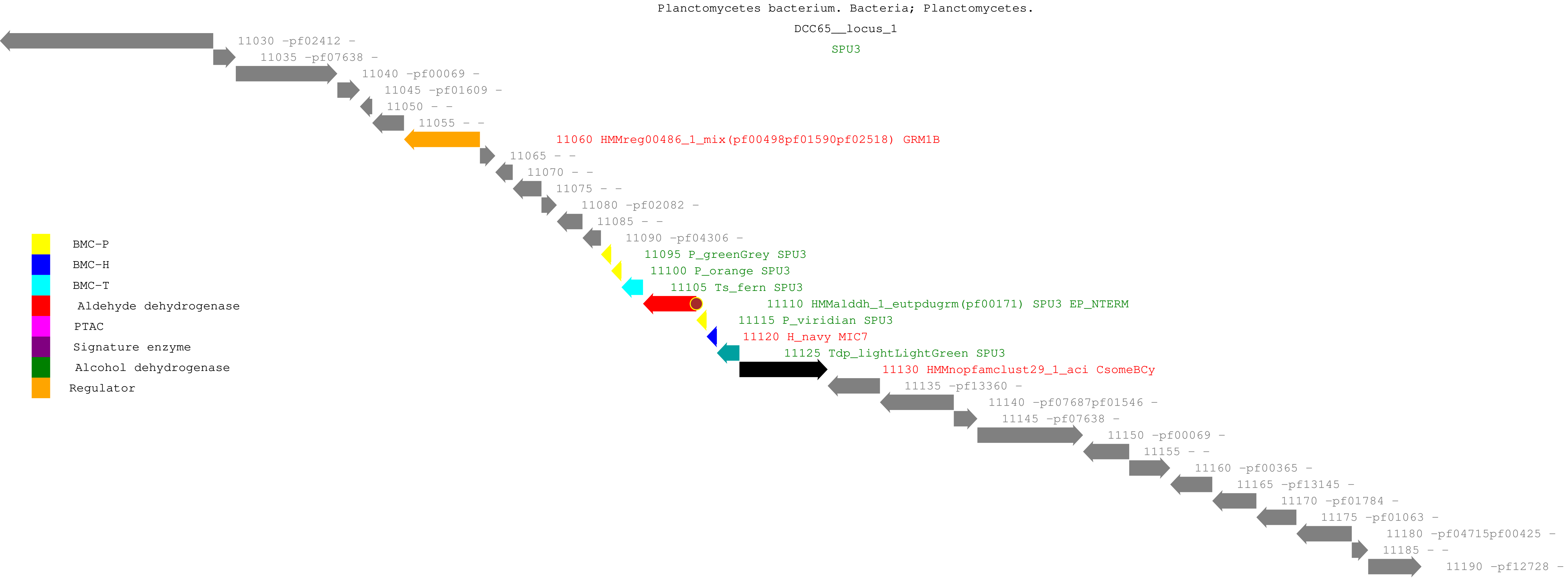
Planctomycetes bacterium. Bacteria; Planctomycetes.
DCC65__locus_1
SPU3
11030 -pf02412 -
11035 -pf07638 -
11040 -pf00069 -
11045 -pf01609 -
11050 - -
11055 - -
11060 HMMreg00486_1_mix(pf00498pf01590pf02518) GRM1B
11065 - -
11070 - -
11075 - -
11080 -pf02082 -
11085 - -
11090 -pf04306 -
11095 P_greenGrey SPU3
11100 P_orange SPU3
11105 Ts_fern SPU3
11110 HMMaldh_1_eutpdugrm(pf00171) SPU3 EP_NTERM
11115 P_viridian SPU3
11120 H_navy MIC7
11125 Tdp_lightLightGreen SPU3
11130 HMMnopfamclust29_1_aci CsomeBCy
11135 -pf13360 -
11140 -pf07687pf01546 -
11145 -pf07638 -
11150 -pf00069 -
11155 - -
11160 -pf00365 -
11165 -pf13145 -
11170 -pf01784 -
11175 -pf01063 -
11180 -pf04715pf00425 -
11185 - -
11190 -pf12728 -
BMC-P
BMC-H
BMC-T
Aldehyde dehydrogenase
PTAC
Signature enzyme
Alcohol dehydrogenase
Regulator

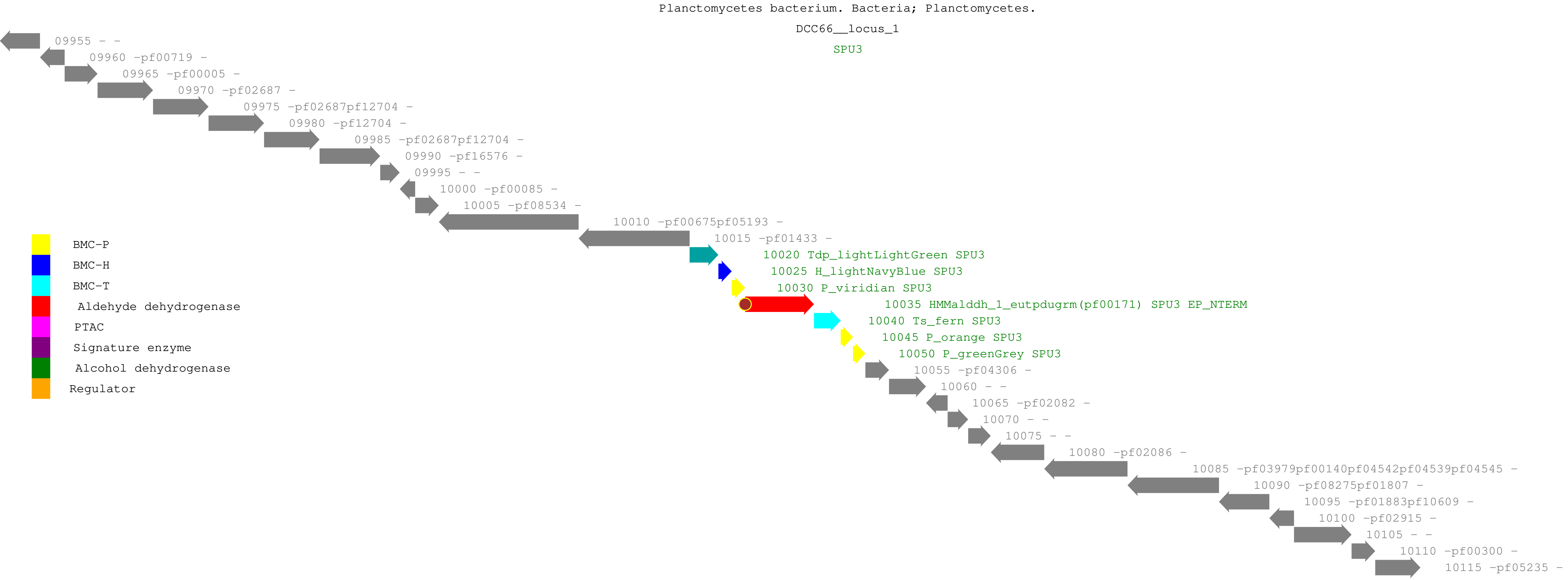
Planctomycetes bacterium. Bacteria; Planctomycetes.
DCC66__locus_1
SPU3
09955 - -
09960 -pf00719 -
09965 -pf00005 -
09970 -pf02687 -
09975 -pf02687pf12704 -
09980 -pf12704 -
09985 -pf02687pf12704 -
09990 -pf16576 -
09995 - -
10000 -pf00085 -
10005 -pf08534 -
10010 -pf00675pf05193 -
10015 -pf01433 -
10020 Tdp_lightLightGreen SPU3
10025 H_lightNavyBlue SPU3
10030 P_viridian SPU3
10035 HMMaldh_1_eutpdugrm(pf00171) SPU3 EP_NTERM
10040 Ts_fern SPU3
10045 P_orange SPU3
10050 P_greenGrey SPU3
10055 -pf04306 -
10060 - -
10065 -pf02082 -
10070 - -
10075 - -
10080 -pf02086 -
10085 -pf03979pf00140pf04542pf04539pf04545 -
10090 -pf08275pf01807 -
10095 -pf01883pf10609 -
10100 -pf02915 -
10105 - -
10110 -pf00300 -
10115 -pf05235 -
BMC-P
BMC-H
BMC-T
Aldehyde dehydrogenase
PTAC
Signature enzyme
Alcohol dehydrogenase
Regulator

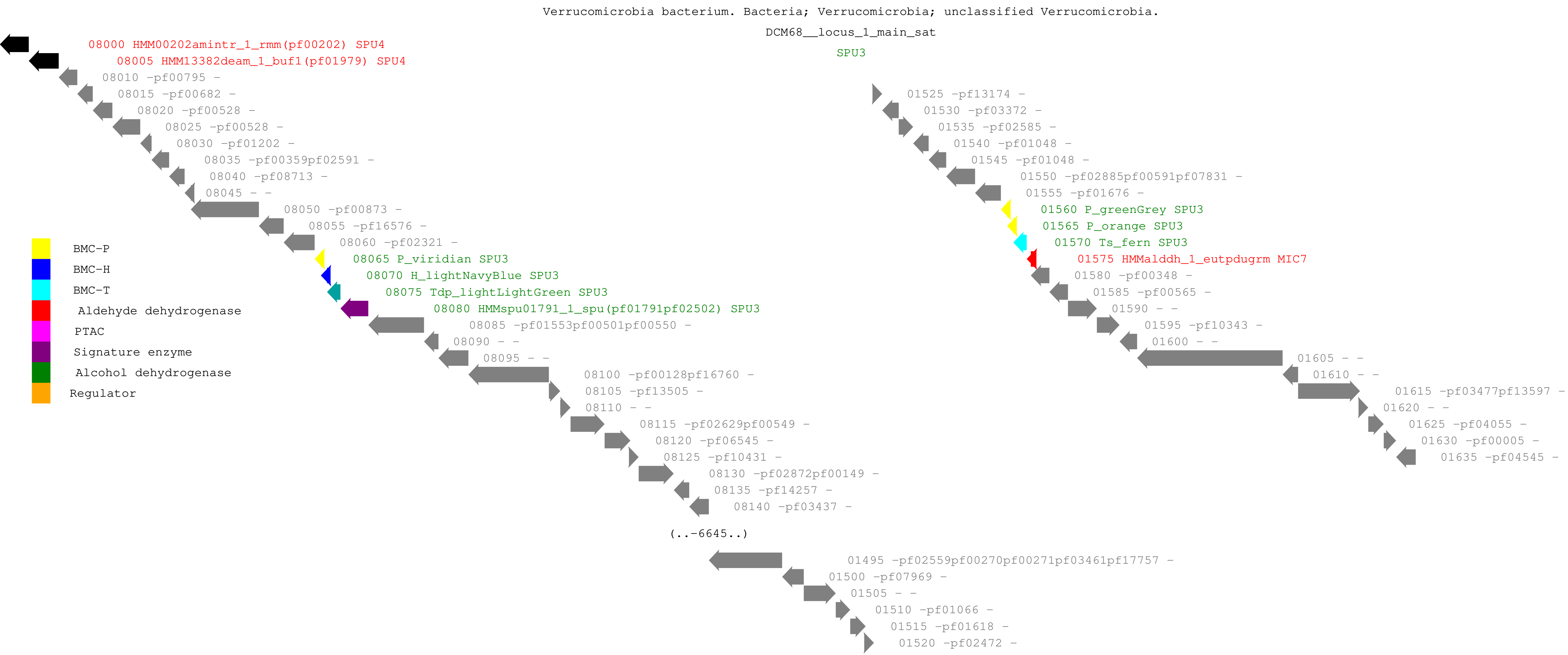
Verrucomicrobia bacterium. Bacteria; Verrucomicrobia; unclassified Verrucomicrobia.
DCM68__locus_1_main_sat
SPU3
08000 HMM00202amintr_1_rmm(pf00202) SPU4
08005 HMM13382deam_1_buf1(pf01979) SPU4
08010 -pf00795 -
08015 -pf00682 -
08020 -pf00528 -
08025 -pf00528 -
08030 -pf01202 -
08035 -pf00359pf02591 -
08040 -pf08713 -
08045 - -
08050 -pf00873 -
08055 -pf16576 -
08060 -pf02321 -
08065 P_viridian SPU3
08070 H_lightNavyBlue SPU3
08075 Tdp_lightLightGreen SPU3
08080 HMMspu01791_1_spu(pf01791pf02502) SPU3
08085 -pf01553pf00501pf00550 -
08090 - -
08095 - -
08100 -pf00128pf16760 -
08105 -pf13505 -
08110 - -
08115 -pf02629pf00549 -
08120 -pf06545 -
08125 -pf10431 -
08130 -pf02872pf00149 -
08135 -pf14257 -
08140 -pf03437 -
(..-6645..)
01495 -pf02559pf00270pf00271pf03461pf17757 -
01500 -pf07969 -
01505 - -
01510 -pf01066 -
01515 -pf01618 -
01520 -pf02472 -
01525 -pf13174 -
01530 -pf03372 -
01535 -pf02585 -
01540 -pf01048 -
01545 -pf01048 -
01550 -pf02885pf00591pf07831 -
01555 -pf01676 -
01560 P_greenGrey SPU3
01565 P_orange SPU3
01570 Ts_fern SPU3
01575 HMMalddh_1_eutpdugrm MIC7
01580 -pf00348 -
01585 -pf00565 -
01590 - -
01595 -pf10343 -
01600 - -
01605 - -
01610 - -
01615 -pf03477pf13597 -
01620 - -
01625 -pf04055 -
01630 -pf00005 -
01635 -pf04545 -
BMC-P
BMC-H
BMC-T
Aldehyde dehydrogenase
PTAC
Signature enzyme
Alcohol dehydrogenase
Regulator

Actinobacteria bacterium. Bacteria; Actinobacteria.

DCY40__locus_1

SPU3

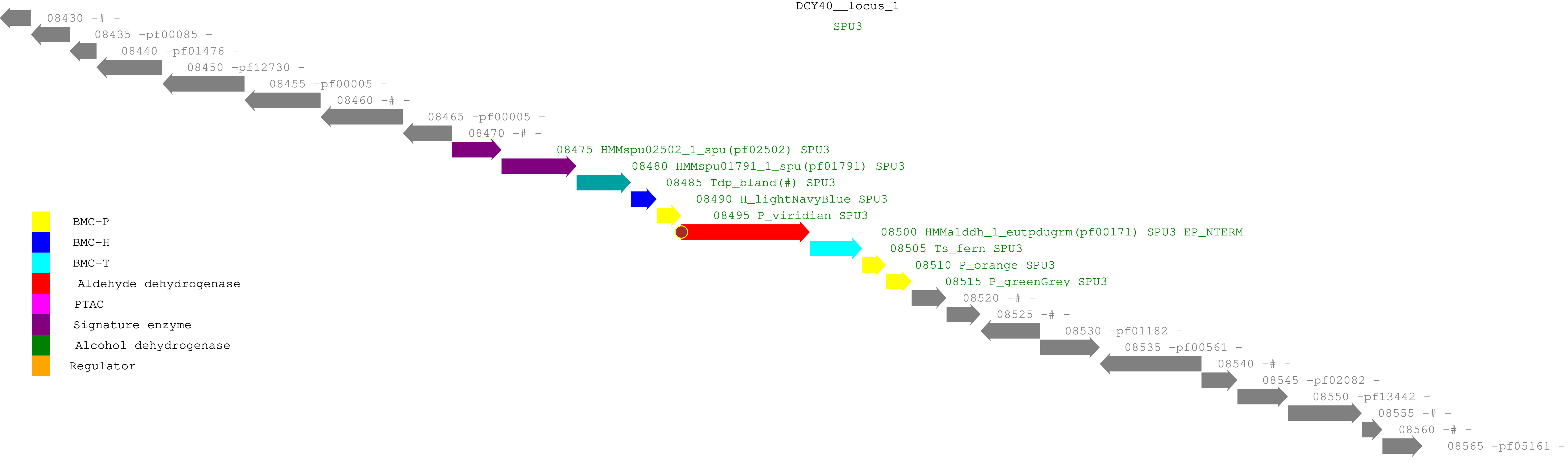

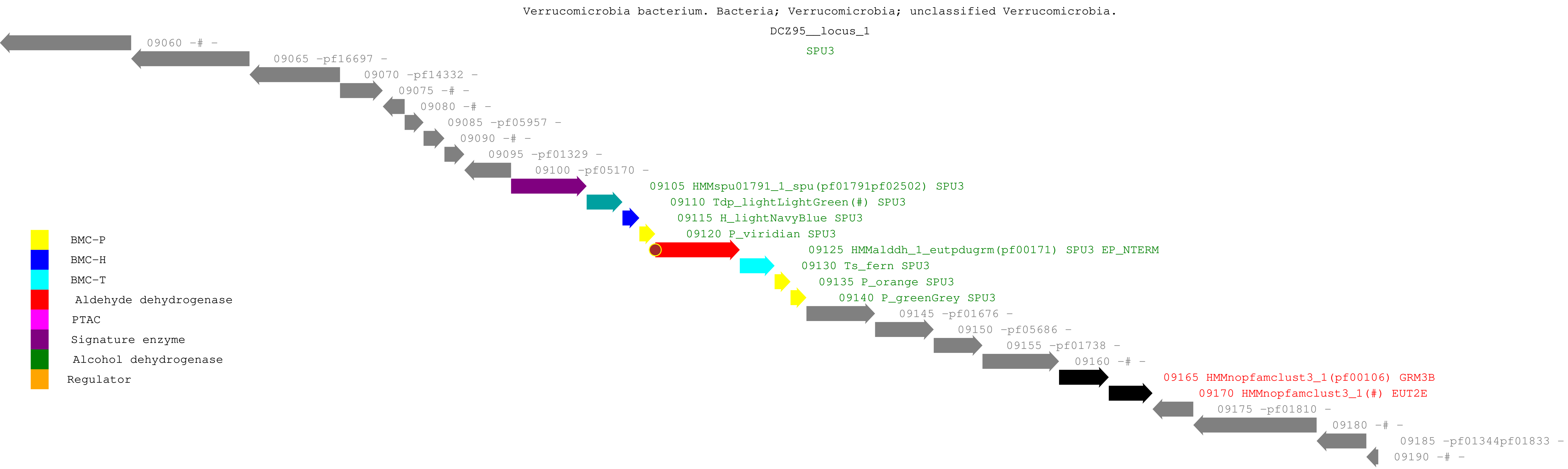

Verrucomicrobia bacterium. Bacteria; Verrucomicrobia; unclassified Verrucomicrobia.
DCZ95__locus_1
SPU3
09060 -# -
09065 -pf16697 -
09070 -pf14332 -
09075 -# -
09080 -# -
09085 -pf05957 -
09090 -# -
09095 -pf01329 -
09100 -pf05170 -
09105 HMMspu01791_1_spu(pf01791pf02502) SPU3
09110 Tdp_lightLightGreen(#) SPU3
09115 H_lightNavyBlue SPU3
09120 P_viridian SPU3
09125 HMMalddh_1_eutpdugrm(pf00171) SPU3 EP_NTERM
09130 Ts_fern SPU3
09135 P_orange SPU3
09140 P_greenGrey SPU3
09145 -pf01676 -
09150 -pf05686 -
09155 -pf01738 -
09160 -# -
09165 HMMnopfamclust3_1(pf00106) GRM3B
09170 HMMnopfamclust3_1(#) EUT2E
09175 -pf01810 -
09180 -# -
09185 -pf01344pf01833 -
09190 -# -
BMC-P
BMC-H
BMC-T
Aldehyde dehydrogenase
PTAC
Signature enzyme
Alcohol dehydrogenase
Regulator

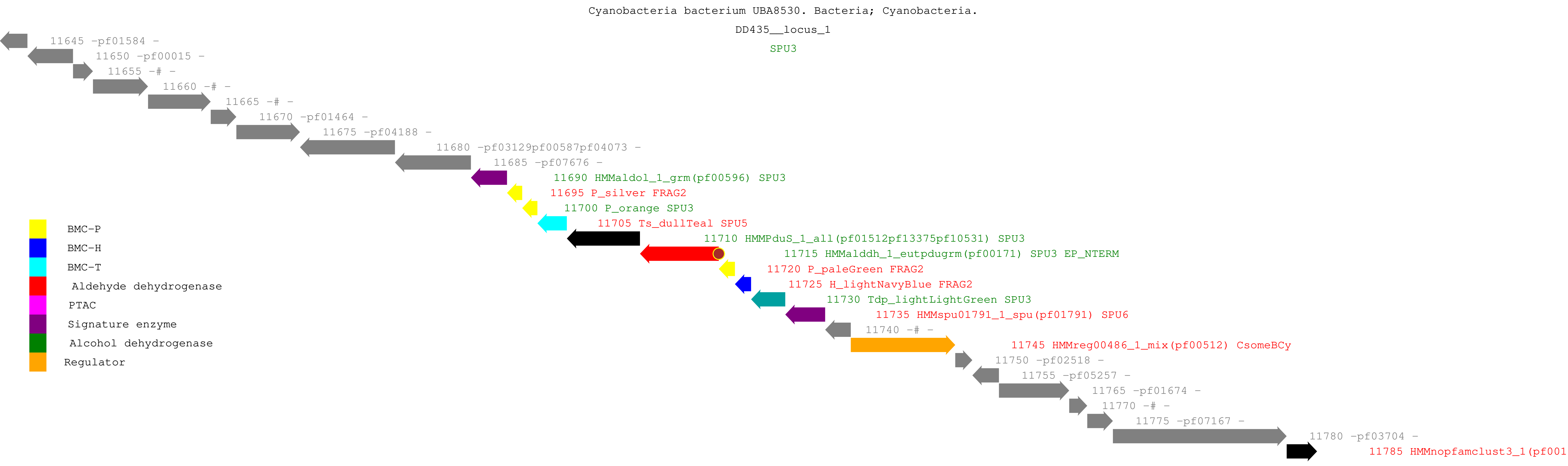

Cyanobacteria bacterium UBA8530. Bacteria; Cyanobacteria.
DD435__locus_1
SPU3
11645 -pf01584 -
11650 -pf00015 -
11655 -# -
11660 -# -
11665 -# -
11670 -pf01464 -
11675 -pf04188 -
11680 -pf03129pf00587pf04073 -
11685 -pf07676 -
11690 HMMaldol_1_grm(pf00596) SPU3
11695 P_silver FRAG2
11700 P_orange SPU3
11705 Ts_dullTeal SPU5
11710 HMMPduS_1_all(pf01512pf13375pf10531) SPU3
11715 HMMalddh_1_eutpdugrm(pf00171) SPU3 EP_NTERM
11720 P_paleGreen FRAG2
11725 H_lightNavyBlue FRAG2
11730 Tdp_lightLightGreen SPU3
11735 HMMspu01791_1_spu(pf01791) SPU6
11740 -# -
11745 HMMreg00486_1_mix(pf00512) CsomeBCy
11750 -pf02518 -
11755 -pf05257 -
11765 -pf01674 -
11770 -# -
11775 -pf07167 -
11780 -pf03704 -
11785 HMMnopfamclust3_1(pf001
BMC-P
BMC-H
BMC-T
Aldehyde dehydrogenase
PTAC
Signature enzyme
Alcohol dehydrogenase
Regulator

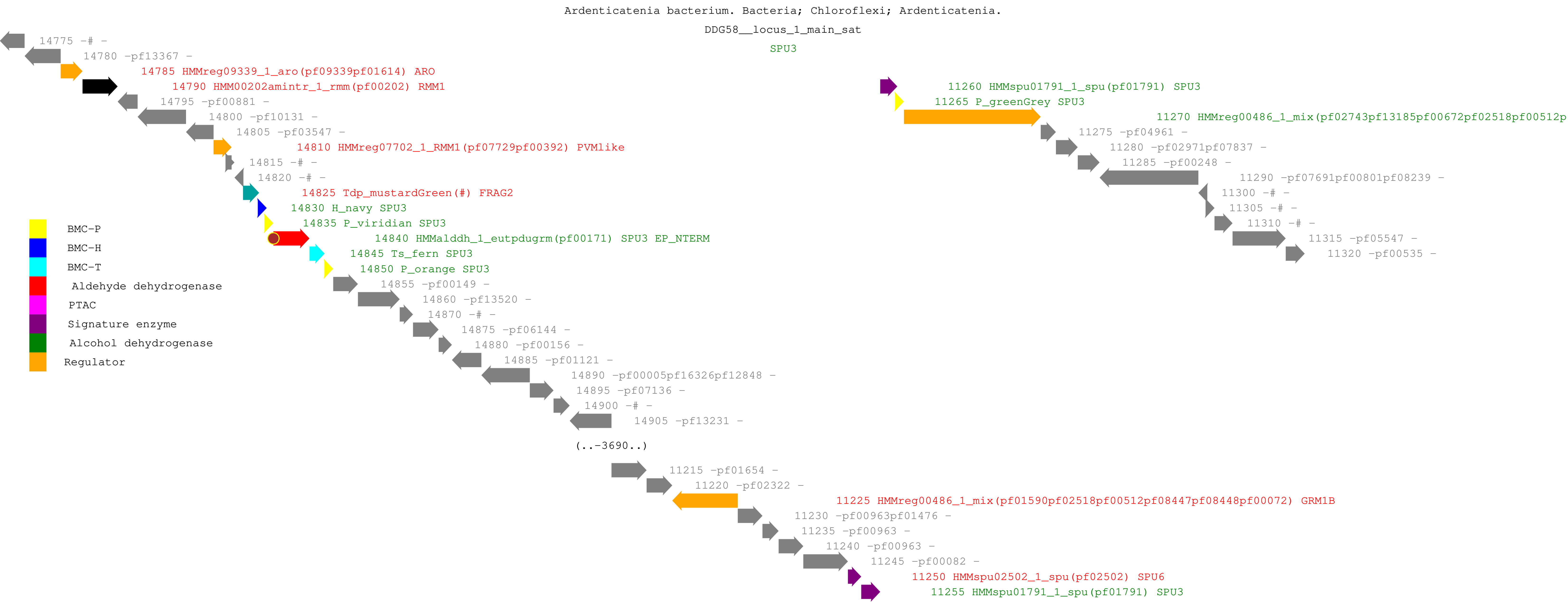
Ardenticatenia bacterium. Bacteria; Chloroflexi; Ardenticatenia.
DDG58__locus_1_main_sat
SPU3
14775 -# -
14780 -pf13367 -
14785 HMMreg09339_1_aro(pf09339pf01614) ARO
14790 HMM00202amintr_1_rmm(pf00202) RMM1
14795 -pf00881 -
14800 -pf10131 -
14805 -pf03547 -
14810 HMMreg07702_1_RMM1(pf07729pf00392) PVMlike
14815 -# -
14820 -# -
14825 Tdp_mustardGreen(#) FRAG2
14830 H_navy SPU3
14835 P_viridian SPU3
14840 HMMalddh_1_eutpdugrm(pf00171) SPU3 EP_NTERM
14845 Ts_fern SPU3
14850 P_orange SPU3
14855 -pf00149 -
14860 -pf13520 -
14870 -# -
14875 -pf06144 -
14880 -pf00156 -
14885 -pf01121 -
14890 -pf00005pf16326pf12848 -
14895 -pf07136 -
14900 -# -
14905 -pf13231 -
(..-3690..)
11215 -pf01654 -
11220 -pf02322 -
11225 HMMreg00486_1_mix(pf01590pf02518pf00512pf08447pf08448pf00072) GRM1B
11230 -pf00963pf01476 -
11235 -pf00963 -
11240 -pf00963 -
11245 -pf00082 -
11250 HMMspu02502_1_spu(pf02502) SPU6
11255 HMMspu01791_1_spu(pf01791) SPU3
11260 HMMspu01791_1_spu(pf01791) SPU3
11265 P_greenGrey SPU3
11270 HMMreg00486_1_mix(pf02743pf13185pf00672pf02518pf00512p
11275 -pf04961 -
11280 -pf02971pf07837 -
11285 -pf00248 -
11290 -pf07691pf00801pf08239 -
11300 -# -
11305 -# -
11310 -# -
11315 -pf05547 -
11320 -pf00535 -
BMC-P
BMC-H
BMC-T
Aldehyde dehydrogenase
PTAC
Signature enzyme
Alcohol dehydrogenase
Regulator

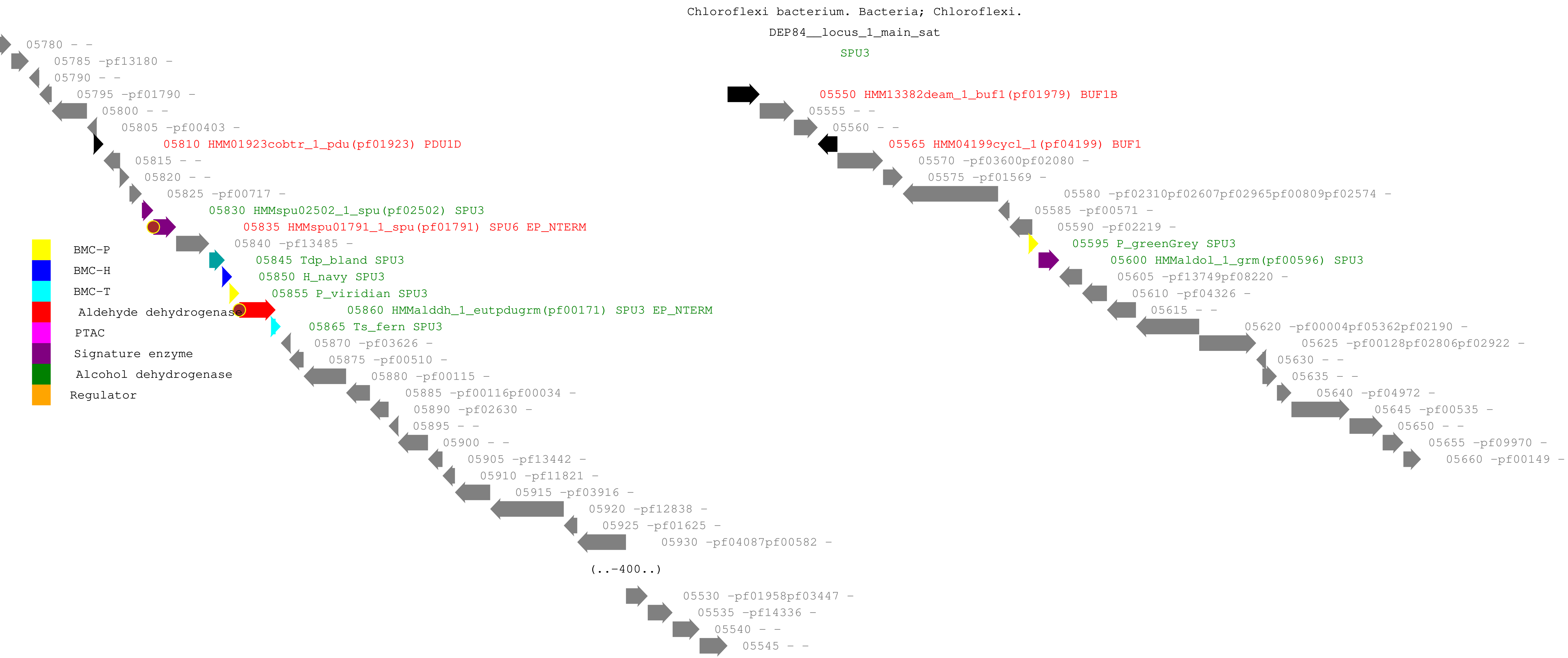
Chloroflexi bacterium. Bacteria; Chloroflexi.
DEP84__locus_1_main_sat
SPU3
05780 - -
05785 -pf13180 -
05790 - -
05795 -pf01790 -
05800 - -
05805 -pf00403 -
05810 HMM01923cobtr_1_pdu(pf01923) PDU1D
05815 - -
05820 - -
05825 -pf00717 -
05830 HMMspu02502_1_spu(pf02502) SPU3
05835 HMMspu01791_1_spu(pf01791) SPU6 EP_NTERM
05840 -pf13485 -
05845 Tdp_bland SPU3
05850 H_navy SPU3
05855 P_viridian SPU3
05860 HMMalddh_1_eutpdugrm(pf00171) SPU3 EP_NTERM
05865 Ts_fern SPU3
05870 -pf03626 -
05875 -pf00510 -
05880 -pf00115 -
05885 -pf00116pf00034 -
05890 -pf02630 -
05895 - -
05900 - -
05905 -pf13442 -
05910 -pf11821 -
05915 -pf03916 -
05920 -pf12838 -
05925 -pf01625 -
05930 -pf04087pf00582 -
(..-400..)
05530 -pf01958pf03447 -
05535 -pf14336 -
05540 - -
05545 - -
05550 HMM13382deam_1_buf1(pf01979) BUF1B
05555 - -
05560 - -
05565 HMM04199cycl_1(pf04199) BUF1
05570 -pf03600pf02080 -
05575 -pf01569 -
05580 -pf02310pf02607pf02965pf00809pf02574 -
05585 -pf00571 -
05590 -pf02219 -
05595 P_greenGrey SPU3
05600 HMMaldol_1_grm(pf00596) SPU3
05605 -pf13749pf08220 -
05610 -pf04326 -
05615 - -
05620 -pf00004pf05362pf02190 -
05625 -pf00128pf02806pf02922 -
05630 - -
05635 - -
05640 -pf04972 -
05645 -pf00535 -
05650 - -
05655 -pf09970 -
05660 -pf00149 -
BMC-P
BMC-H
BMC-T
Aldehyde dehydrogenase
PTAC
Signature enzyme
Alcohol dehydrogenase
Regulator

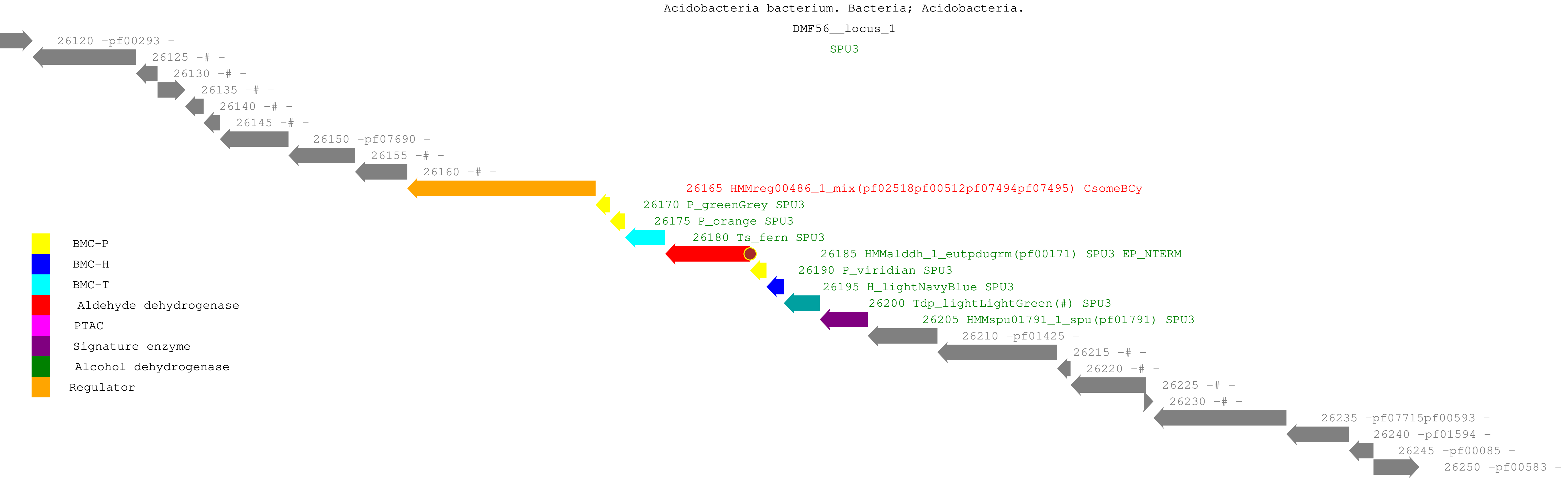

Acidobacteria bacterium. Bacteria; Acidobacteria.
DMF56__locus_1
SPU3
26120 -pf00293 -
26125 -# -
26130 -# -
26135 -# -
26140 -# -
26145 -# -
26150 -pf07690 -
26155 -# -
26160 -# -
26165 HMMreg00486_1_mix(pf02518pf00512pf07494pf07495) CsomeBCy
26170 P_greenGrey SPU3
26175 P_orange SPU3
26180 Ts_fern SPU3
26185 HMMaldh_1_eutpdugrm(pf00171) SPU3 EP_NTERM
26190 P_viridian SPU3
26195 H_lightNavyBlue SPU3
26200 Tdp_lightLightGreen(#) SPU3
26205 HMMspu01791_1_spu(pf01791) SPU3
26210 -pf01425 -
26215 -# -
26220 -# -
26225 -# -
26230 -# -
26235 -pf07715pf00593 -
26240 -pf01594 -
26245 -pf00085 -
26250 -pf00583 -
BMC-P
BMC-H
BMC-T
Aldehyde dehydrogenase
PTAC
Signature enzyme
Alcohol dehydrogenase
Regulator

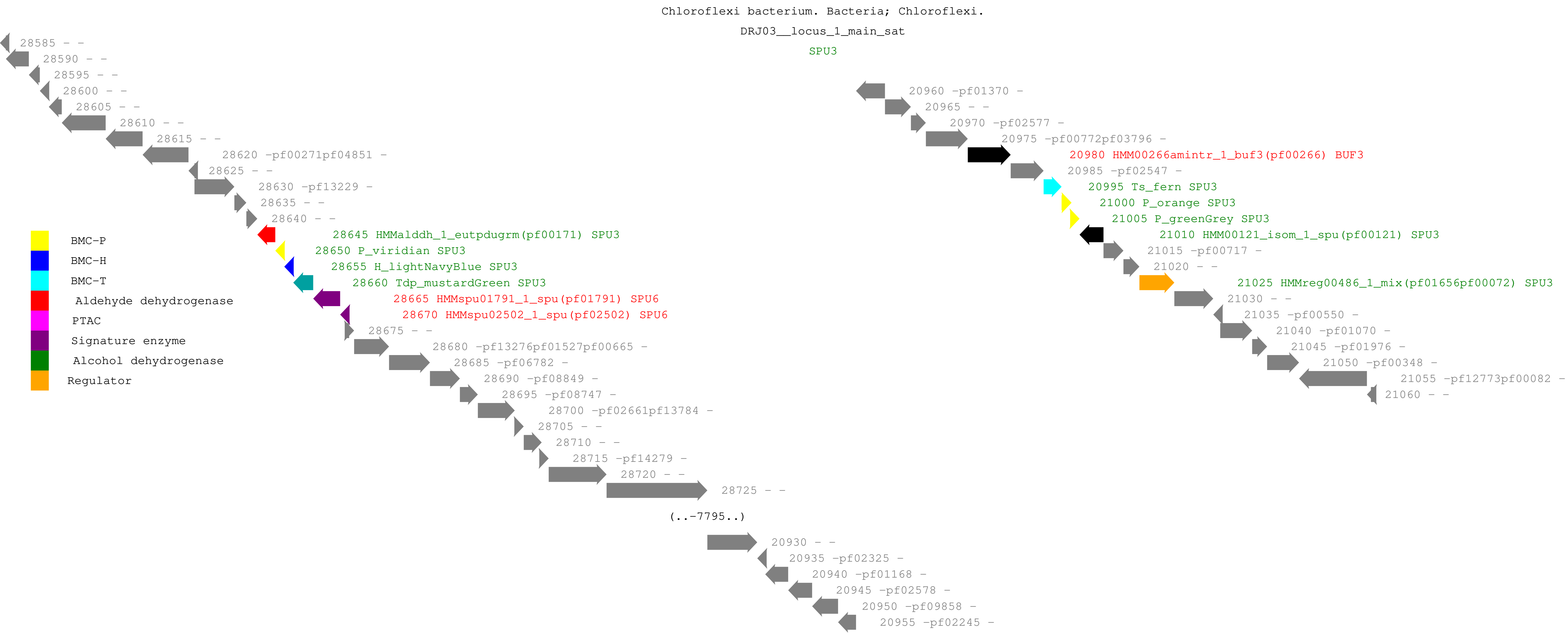
Chloroflexi bacterium. Bacteria; Chloroflexi.
DRJ03__locus_1_main_sat
SPU3
28585 - -
28590 - -
28595 - -
28600 - -
28605 - -
28610 - -
28615 - -
28620 -pf00271pf04851 -
28625 - -
28630 -pf13229 -
28635 - -
28640 - -
28645 HMMalddh_1_eutpdugrm(pf00171) SPU3
28650 P_viridian SPU3
28655 H_lightNavyBlue SPU3
28660 Tdp_mustardGreen SPU3
28665 HMMspu01791_1_spu(pf01791) SPU6
28670 HMMspu02502_1_spu(pf02502) SPU6
28675 - -
28680 -pf13276pf01527pf00665 -
28685 -pf06782 -
28690 -pf08849 -
28695 -pf08747 -
28700 -pf02661pf13784 -
28705 - -
28710 - -
28715 -pf14279 -
28720 - -
28725 - -
(..-7795..)
20930 - -
20935 -pf02325 -
20940 -pf01168 -
20945 -pf02578 -
20950 -pf09858 -
20955 -pf02245 -
20960 -pf01370 -
20965 - -
20970 -pf02577 -
20975 -pf00772pf03796 -
20980 HMM00266amintr_1_buf3(pf00266) BUF3
20985 -pf02547 -
20995 Ts_fern SPU3
21000 P_orange SPU3
21005 P_greenGrey SPU3
21010 HMM00121_isom_1_spu(pf00121) SPU3
21015 -pf00717 -
21020 - -
21025 HMMreg00486_1_mix(pf01656pf00072) SPU3
21030 - -
21035 -pf00550 -
21040 -pf01070 -
21045 -pf01976 -
21050 -pf00348 -
21055 -pf12773pf00082 -
21060 - -
BMC-P
BMC-H
BMC-T
Aldehyde dehydrogenase
PTAC
Signature enzyme
Alcohol dehydrogenase
Regulator

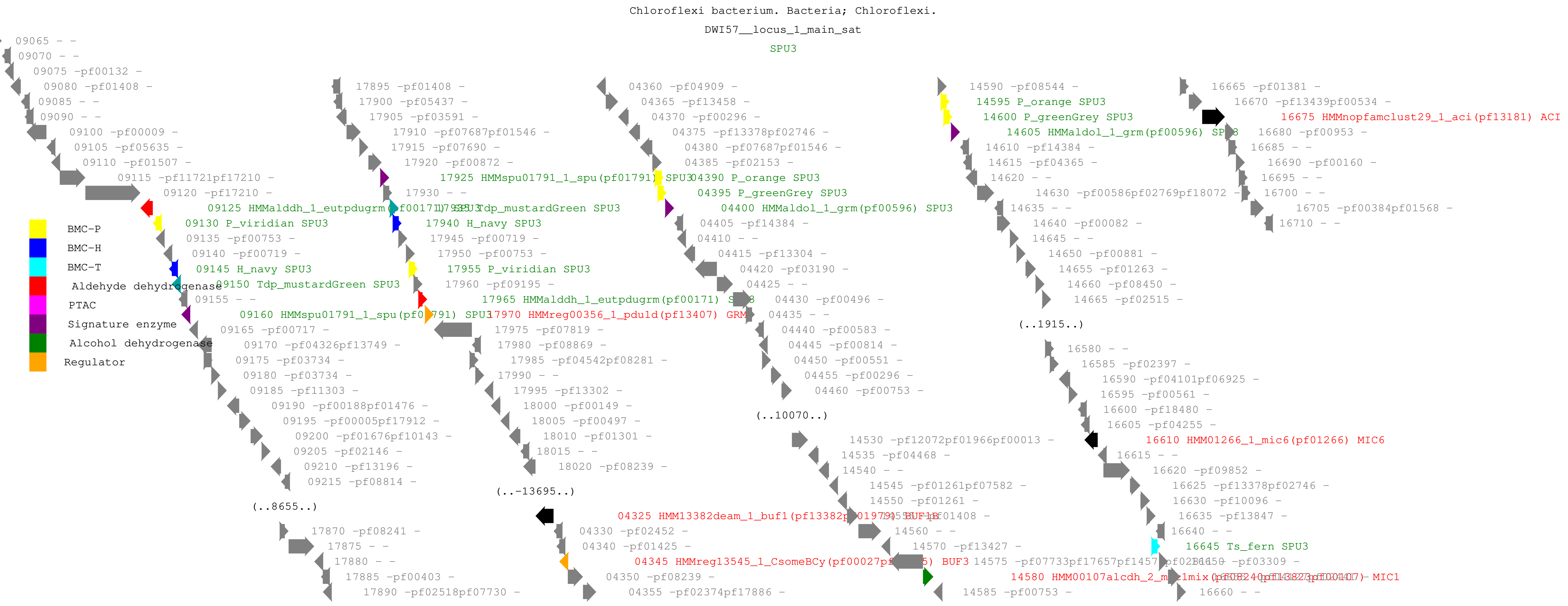
Chloroflexi bacterium. Bacteria; Chloroflexi.
DWI57__locus_1_main_sat
SPU3
09065 - -
09070 - -
09075 -pf00132 -
09080 -pf01408 -
09085 - -
09090 - -
09100 -pf00009 -
09105 -pf05635 -
09110 -pf01507 -
09115 -pf11721pf17210 -
09120 -pf17210 -
09125 HMMaldh_1_eutpdugrm(pf00171) SPU3
09130 P_viridian SPU3
09135 -pf00753 -
09140 -pf00719 -
09145 H_navy SPU3
09150 Tdp_mustardGreen SPU3
09155 - -
09160 HMMspu01791_1_spu(pf01791) SPU3
09165 -pf00717 -
09170 -pf04326pf13749 -
09175 -pf03734 -
09180 -pf03734 -
09185 -pf11303 -
09190 -pf00188pf01476 -
09195 -pf00005pf17912 -
09200 -pf01676pf10143 -
09205 -pf02146 -
09210 -pf13196 -
09215 -pf08814 -
(..8655..)
17870 -pf08241 -
17875 - -
17880 - -
17885 -pf00403 -
17890 -pf02518pf07730 -
BMC-P
BMC-H
BMC-T
Aldehyde dehydrogenase
PTAC
Signature enzyme
Alcohol dehydrogenase
Regulator
17895 -pf01408 -
17900 -pf05437 -
17905 -pf03591 -
17910 -pf07687pf01546 -
17915 -pf07690 -
17920 -pf00872 -
17925 HMMspu01791_1_spu(pf01791) SPU3
17930 - -
17935 Tdp_mustardGreen SPU3
17940 H_navy SPU3
17945 -pf00719 -
17950 -pf00753 -
17955 P_viridian SPU3
17960 -pf09195 -
17965 HMMaldh_1_eutpdugrm(pf00171) SPU3
17970 HMMreg00356_1_pdu1d(pf13407) GRM
17975 -pf07819 -
17980 -pf08869 -
17985 -pf04542pf08281 -
17990 - -
17995 -pf13302 -
18000 -pf00149 -
18005 -pf00497 -
18010 -pf01301 -
18015 - -
18020 -pf08239 -
(..-13695..)
04325 HMM13382deam_1_buf1(pf13382pf01979) BUF1
04330 -pf02452 -
04340 -pf01425 -
04345 HMMreg13545_1_CsomeBCy(pf00027pf [illegible]) BUF3
04350 -pf08239 -
04355 -pf02374pf17886 -
04360 -pf04909 -
04365 -pf13458 -
04370 -pf00296 -
04375 -pf13378pf02746 -
04380 -pf07687pf01546 -
04385 -pf02153 -
04390 P_orange SPU3
04395 P_greenGrey SPU3
04400 HMMaldol_1_grm(pf00596) SPU3
04405 -pf14384 -
04410 - -
04415 -pf13304 -
04420 -pf03190 -
04425 - -
04430 -pf00496 -
04435 - -
04440 -pf00583 -
04445 -pf00814 -
04450 -pf00551 -
04455 -pf00296 -
04460 -pf00753 -
(..10070..)
14530 -pf12072pf01966pf00013 -
14535 -pf04468 -
14540 - -
14545 -pf01261pf07582 -
14550 -pf01261 -
14555 -pf01408 -
14560 - -
14570 -pf13427 -
14575 -pf07733pf17657pf1457 [illegible]
14580 HMM00107alcdh_2_m[illegible]1mix(pf08240pf13823pf00107) MIC1
14585 -pf00753 -
14590 -pf08544 -
14595 P_orange SPU3
14600 P_greenGrey SPU3
14605 HMMaldol_1_grm(pf00596) SPU3
14610 -pf14384 -
14615 -pf04365 -
14620 - -
14630 -pf00586pf02769pf18072 -
14635 - -
14640 -pf00082 -
14645 - -
14650 -pf00881 -
14655 -pf01263 -
14660 -pf08450 -
14665 -pf02515 -
(..1915..)
16580 - -
16585 -pf02397 -
16590 -pf04101pf06925 -
16595 -pf00561 -
16600 -pf18480 -
16605 -pf04255 -
16610 HMM01266_1_mic6(pf01266) MIC6
16615 - -
16620 -pf09852 -
16625 -pf13378pf02746 -
16630 -pf10096 -
16635 -pf13847 -
16640 - -
16645 Ts_fern SPU3
16650 -pf03309 -
16660 - -
16665 -pf01381 -
16670 -pf13439pf00534 -
16675 HMMnopfamclust29_1_aci(pf13181) ACI
16680 -pf00953 -
16685 - -
16690 -pf00160 -
16695 - -
16700 - -
16705 -pf00384pf01568 -
16710 - -

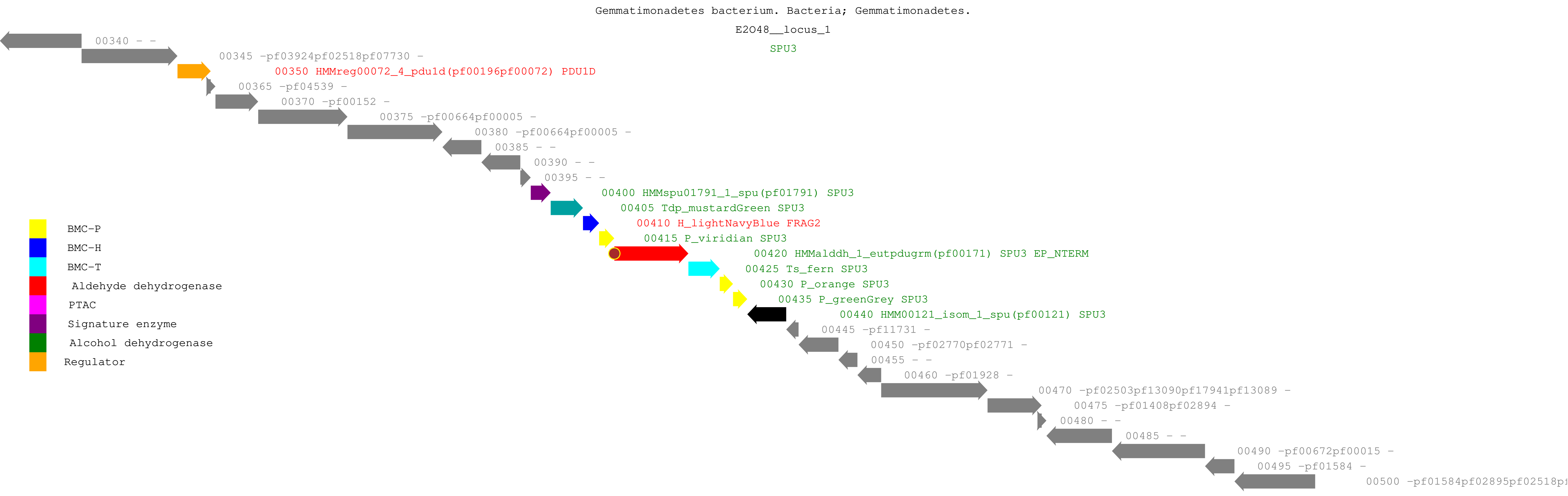
Gemmatimonadetes bacterium. Bacteria; Gemmatimonadetes.
E2O48__locus_1
SPU3
00340 - -
00345 -pf03924pf02518pf07730 -
00350 HMMreg00072_4_pdu1d(pf00196pf00072) PDU1D
00365 -pf04539 -
00370 -pf00152 -
00375 -pf00664pf00005 -
00380 -pf00664pf00005 -
00385 - -
00390 - -
00395 - -
00400 HMMspu01791_1_spu(pf01791) SPU3
00405 Tdp_mustardGreen SPU3
00410 H_lightNavyBlue FRAG2
00415 P_viridian SPU3
00420 HMMalddh_1_eutpdugrm(pf00171) SPU3 EP_NTERM
00425 Ts_fern SPU3
00430 P_orange SPU3
00435 P_greenGrey SPU3
00440 HMM00121_isom_1_spu(pf00121) SPU3
00445 -pf11731 -
00450 -pf02770pf02771 -
00455 - -
00460 -pf01928 -
00470 -pf02503pf13090pf17941pf13089 -
00475 -pf01408pf02894 -
00480 - -
00485 - -
00490 -pf00672pf00015 -
00495 -pf01584 -
00500 -pf01584pf02895pf02518pf
BMC-P
BMC-H
BMC-T
Aldehyde dehydrogenase
PTAC
Signature enzyme
Alcohol dehydrogenase
Regulator

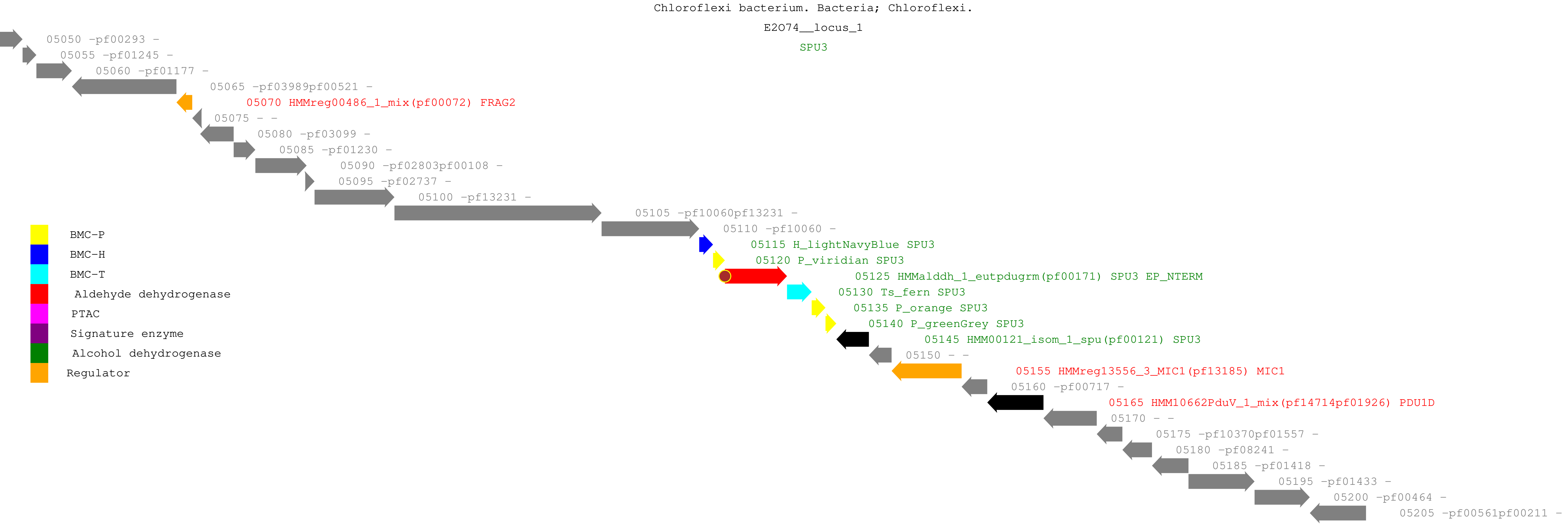
Chloroflexi bacterium. Bacteria; Chloroflexi.
E2074__locus_1
SPU3
05050 -pf00293 -
05055 -pf01245 -
05060 -pf01177 -
05065 -pf03989pf00521 -
05070 HMMreg00486_1_mix(pf00072) FRAG2
05075 - -
05080 -pf03099 -
05085 -pf01230 -
05090 -pf02803pf00108 -
05095 -pf02737 -
05100 -pf13231 -
05105 -pf10060pf13231 -
05110 -pf10060 -
05115 H_lightNavyBlue SPU3
05120 P_viridian SPU3
05125 HMMalddh_1_eutpdugrm(pf00171) SPU3 EP_NTERM
05130 Ts_fern SPU3
05135 P_orange SPU3
05140 P_greenGrey SPU3
05145 HMM00121_isom_1_spu(pf00121) SPU3
05150 - -
05155 HMMreg13556_3_MIC1(pf13185) MIC1
05160 -pf00717 -
05165 HMM10662PduV_1_mix(pf14714pf01926) PDU1D
05170 - -
05175 -pf10370pf01557 -
05180 -pf08241 -
05185 -pf01418 -
05195 -pf01433 -
05200 -pf00464 -
05205 -pf00561pf00211 -
BMC-P
BMC-H
BMC-T
Aldehyde dehydrogenase
PTAC
Signature enzyme
Alcohol dehydrogenase
Regulator

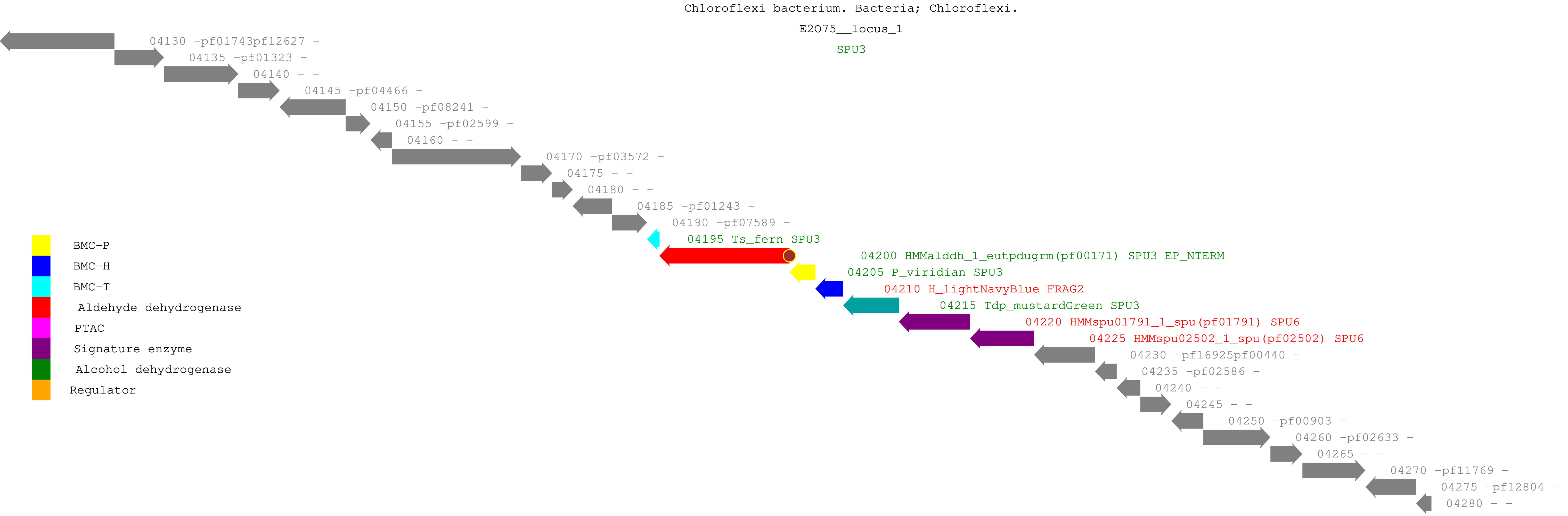

Chloroflexi bacterium. Bacteria; Chloroflexi.
E2075__locus_1
SPU3
04130 -pf01743pf12627 -
04135 -pf01323 -
04140 - -
04145 -pf04466 -
04150 -pf08241 -
04155 -pf02599 -
04160 - -
04170 -pf03572 -
04175 - -
04180 - -
04185 -pf01243 -
04190 -pf07589 -
04195 Ts_fern SPU3
04200 HMMalddh_1_eutpdugrm(pf00171) SPU3 EP_NTERM
04205 P_viridian SPU3
04210 H_lightNavyBlue FRAG2
04215 Tdp_mustardGreen SPU3
04220 HMMspu01791_1_spu(pf01791) SPU6
04225 HMMspu02502_1_spu(pf02502) SPU6
04230 -pf16925pf00440 -
04235 -pf02586 -
04240 - -
04245 - -
04250 -pf00903 -
04260 -pf02633 -
04265 - -
04270 -pf11769 -
04275 -pf12804 -
04280 - -
BMC-P
BMC-H
BMC-T
Aldehyde dehydrogenase
PTAC
Signature enzyme
Alcohol dehydrogenase
Regulator

Acidobacteria bacterium. Bacteria; Acidobacteria.

E2O95__locus_1

SPU3

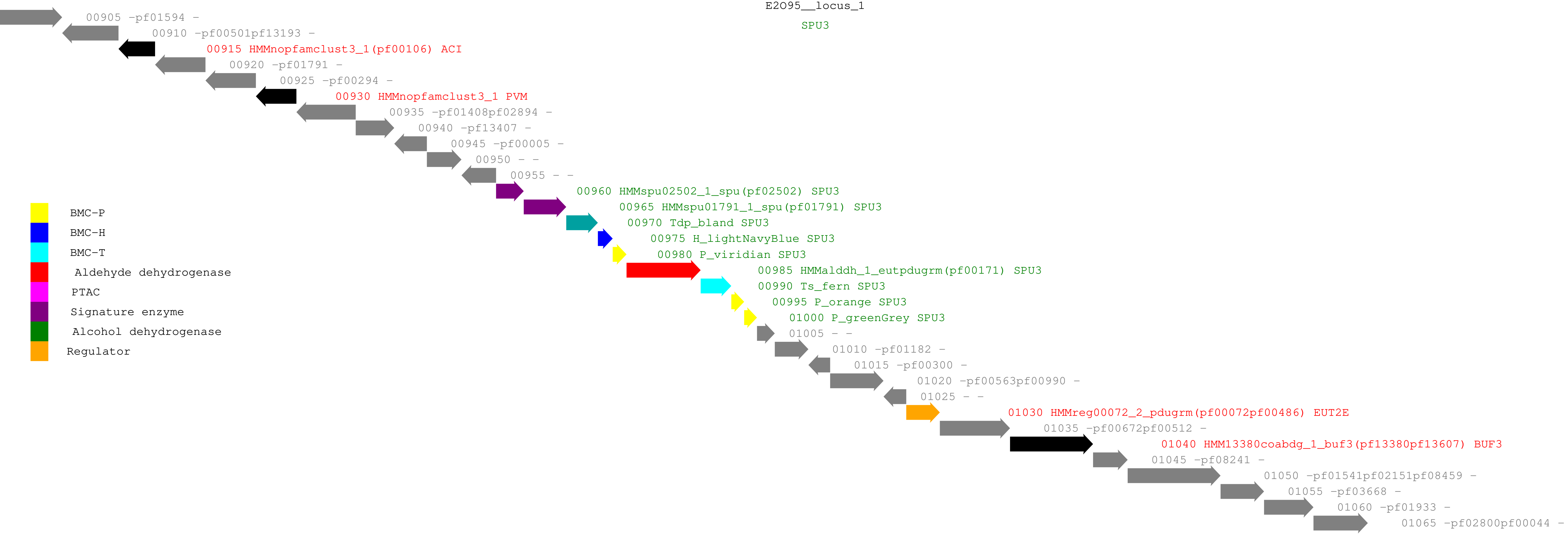

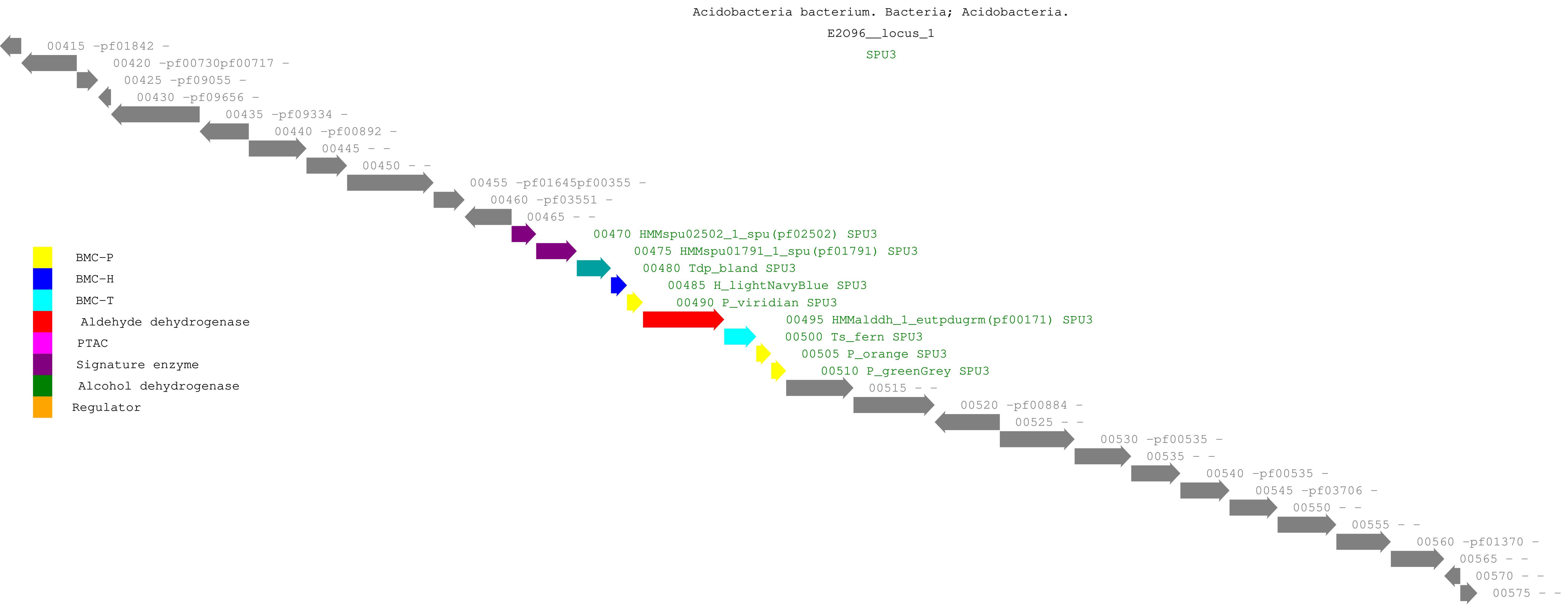
Acidobacteria bacterium. Bacteria; Acidobacteria.
E2O96__locus_1
SPU3
00415 -pf01842 -
00420 -pf00730pf00717 -
00425 -pf09055 -
00430 -pf09656 -
00435 -pf09334 -
00440 -pf00892 -
00445 - -
00450 - -
00455 -pf01645pf00355 -
00460 -pf03551 -
00465 - -
00470 HMMspu02502_1_spu(pf02502) SPU3
00475 HMMspu01791_1_spu(pf01791) SPU3
00480 Tdp_bland SPU3
00485 H_lightNavyBlue SPU3
00490 P_viridian SPU3
00495 HMMalddh_1_eutpdugrm(pf00171) SPU3
00500 Ts_fern SPU3
00505 P_orange SPU3
00510 P_greenGrey SPU3
00515 - -
00520 -pf00884 -
00525 - -
00530 -pf00535 -
00535 - -
00540 -pf00535 -
00545 -pf03706 -
00550 - -
00555 - -
00560 -pf01370 -
00565 - -
00570 - -
00575 - -
BMC-P
BMC-H
BMC-T
Aldehyde dehydrogenase
PTAC
Signature enzyme
Alcohol dehydrogenase
Regulator

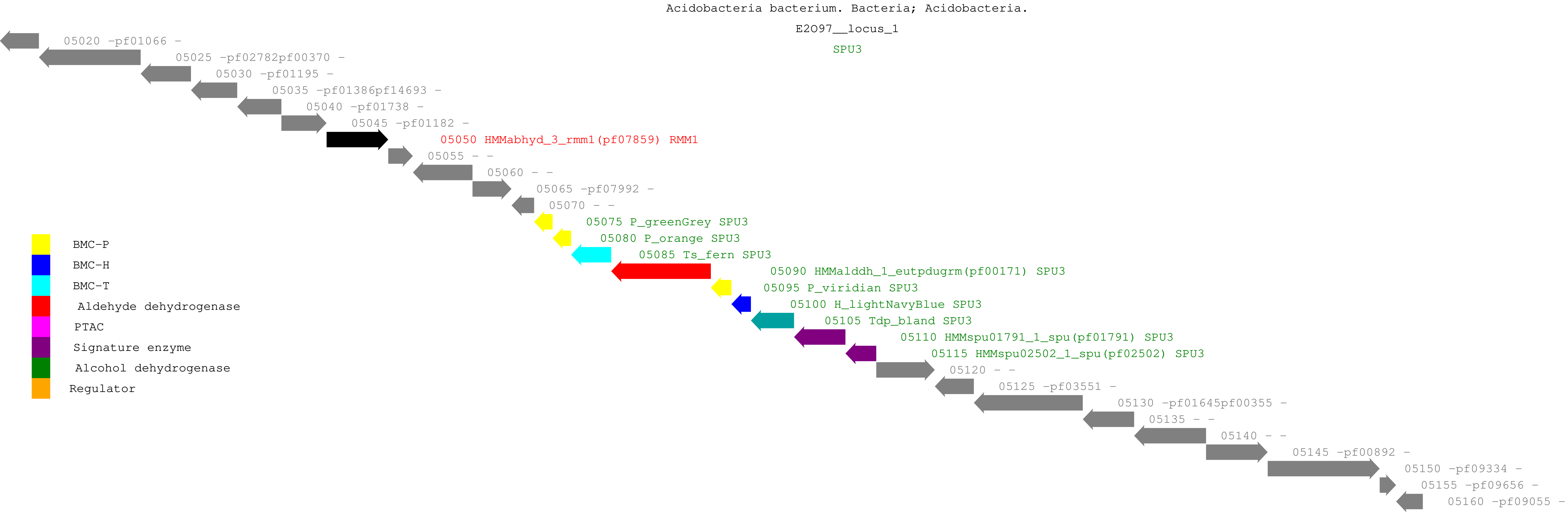

Acidobacteria bacterium. Bacteria; Acidobacteria.
E2O97__locus_1
SPU3
05020 -pf01066 -
05025 -pf02782pf00370 -
05030 -pf01195 -
05035 -pf01386pf14693 -
05040 -pf01738 -
05045 -pf01182 -
05050 HMMabhyd_3_rmm1(pf07859) RMM1
05055 - -
05060 - -
05065 -pf07992 -
05070 - -
05075 P_greenGrey SPU3
05080 P_orange SPU3
05085 Ts_fern SPU3
05090 HMMaldh_1_eutpdugrm(pf00171) SPU3
05095 P_viridian SPU3
05100 H_lightNavyBlue SPU3
05105 Tdp_bland SPU3
05110 HMMspu01791_1_spu(pf01791) SPU3
05115 HMMspu02502_1_spu(pf02502) SPU3
05120 - -
05125 -pf03551 -
05130 -pf01645pf00355 -
05135 - -
05140 - -
05145 -pf00892 -
05150 -pf09334 -
05155 -pf09656 -
05160 -pf09055 -
BMC-P
BMC-H
BMC-T
Aldehyde dehydrogenase
PTAC
Signature enzyme
Alcohol dehydrogenase
Regulator

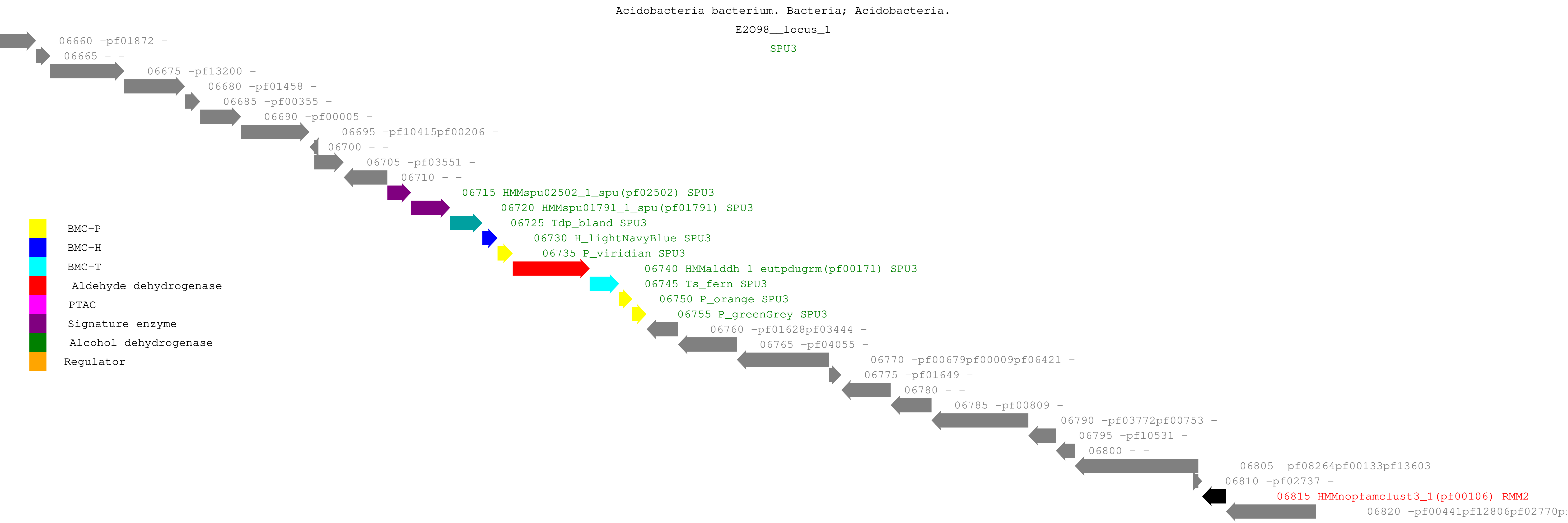

Acidobacteria bacterium. Bacteria; Acidobacteria.
E2098__locus_1
SPU3
06660 -pf01872 -
06665 - -
06675 -pf13200 -
06680 -pf01458 -
06685 -pf00355 -
06690 -pf00005 -
06695 -pf10415pf00206 -
06700 - -
06705 -pf03551 -
06710 - -
06715 HMMspu02502_1_spu(pf02502) SPU3
06720 HMMspu01791_1_spu(pf01791) SPU3
06725 Tdp_bland SPU3
06730 H_lightNavyBlue SPU3
06735 P_viridian SPU3
06740 HMMaldh_1_eutpdugrm(pf00171) SPU3
06745 Ts_fern SPU3
06750 P_orange SPU3
06755 P_greenGrey SPU3
06760 -pf01628pf03444 -
06765 -pf04055 -
06770 -pf00679pf00009pf06421 -
06775 -pf01649 -
06780 - -
06785 -pf00809 -
06790 -pf03772pf00753 -
06795 -pf10531 -
06800 - -
06805 -pf08264pf00133pf13603 -
06810 -pf02737 -
06815 HMMnopfamclust3_1(pf00106) RMM2
06820 -pf00441pf12806pf02770p
BMC-P
BMC-H
BMC-T
Aldehyde dehydrogenase
PTAC
Signature enzyme
Alcohol dehydrogenase
Regulator

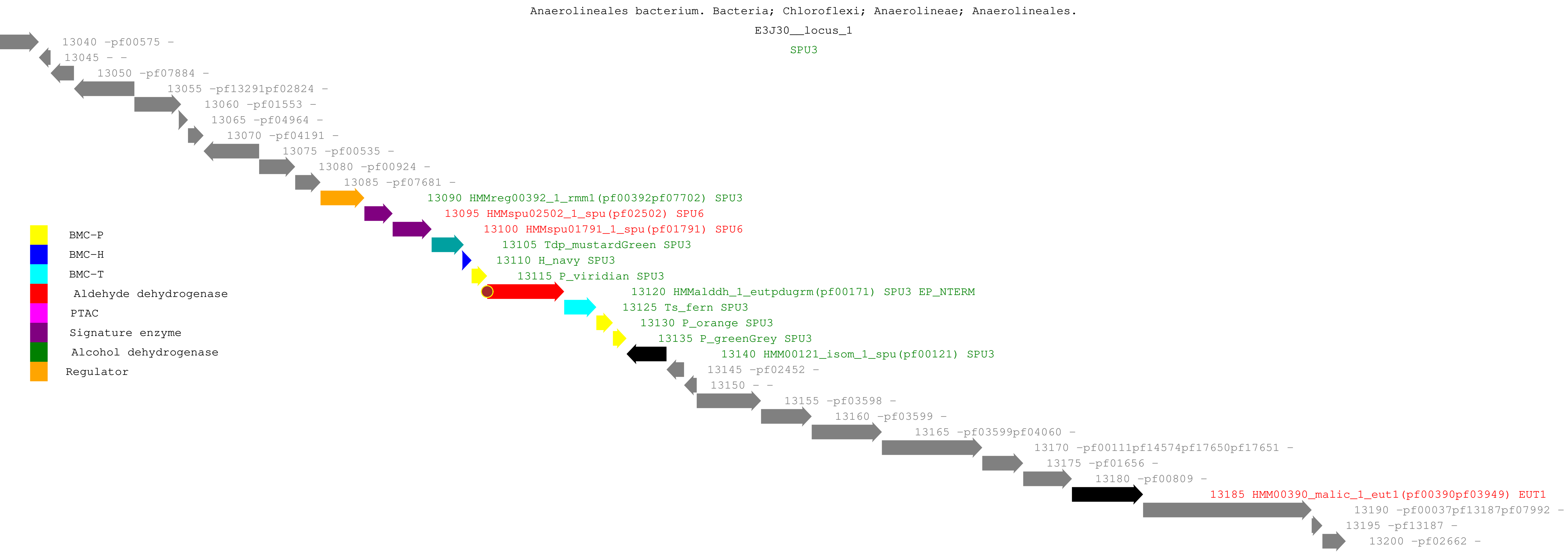
Anaerolineales bacterium. Bacteria; Chloroflexi; Anaerolineae; Anaerolineales.
E3J30__locus_1
SPU3
13040 -pf00575 -
13045 - -
13050 -pf07884 -
13055 -pf13291pf02824 -
13060 -pf01553 -
13065 -pf04964 -
13070 -pf04191 -
13075 -pf00535 -
13080 -pf00924 -
13085 -pf07681 -
13090 HMMreg00392_1_rmm1(pf00392pf07702) SPU3
13095 HMMspu02502_1_spu(pf02502) SPU6
13100 HMMspu01791_1_spu(pf01791) SPU6
13105 Tdp_mustardGreen SPU3
13110 H_navy SPU3
13115 P_viridian SPU3
13120 HMMalddh_1_eutpdugrm(pf00171) SPU3 EP_NTERM
13125 Ts_fern SPU3
13130 P_orange SPU3
13135 P_greenGrey SPU3
13140 HMM00121_isom_1_spu(pf00121) SPU3
13145 -pf02452 -
13150 - -
13155 -pf03598 -
13160 -pf03599 -
13165 -pf03599pf04060 -
13170 -pf00111pf14574pf17650pf17651 -
13175 -pf01656 -
13180 -pf00809 -
13185 HMM00390_malic_1_eut1(pf00390pf03949) EUT1
13190 -pf00037pf13187pf07992 -
13195 -pf13187 -
13200 -pf02662 -
BMC-P
BMC-H
BMC-T
Aldehyde dehydrogenase
PTAC
Signature enzyme
Alcohol dehydrogenase
Regulator

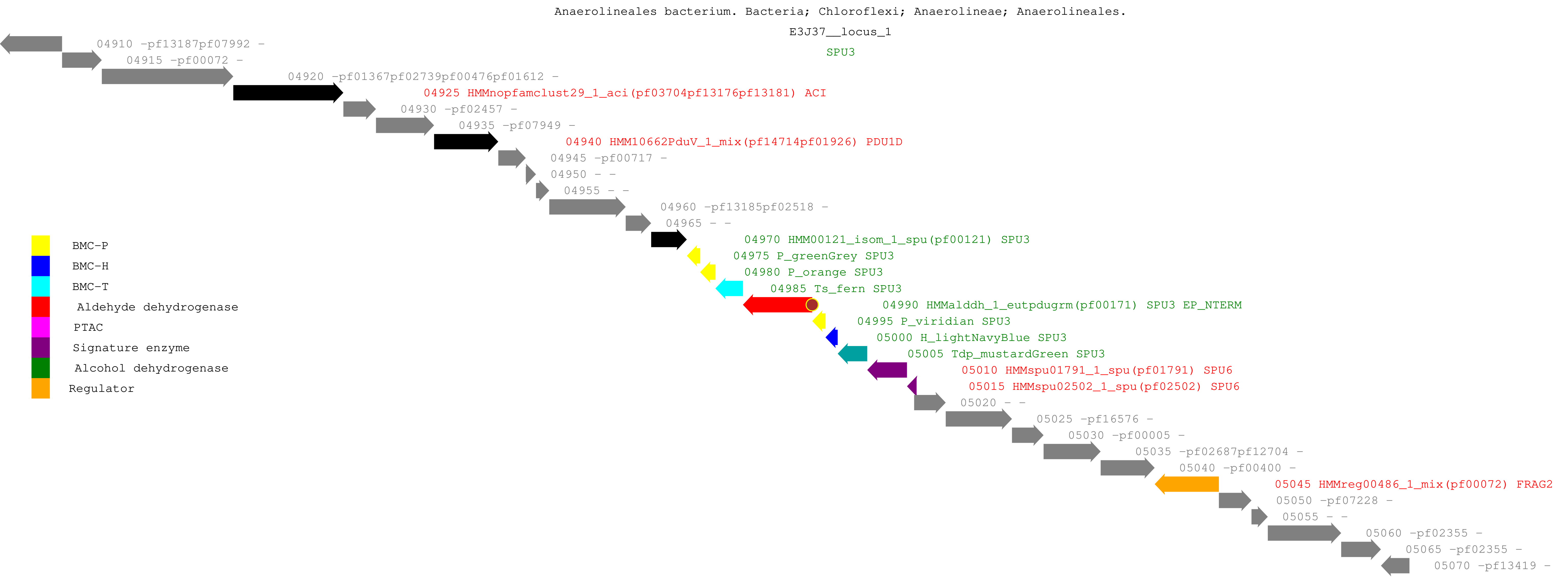
Anaerolineales bacterium. Bacteria; Chloroflexi; Anaerolineae; Anaerolineales.
E3J37__locus_1
SPU3
04910 -pf13187pf07992 -
04915 -pf00072 -
04920 -pf01367pf02739pf00476pf01612 -
04925 HMMnopfamclust29_1_aci(pf03704pf13176pf13181) ACI
04930 -pf02457 -
04935 -pf07949 -
04940 HMM10662PduV_1_mix(pf14714pf01926) PDU1D
04945 -pf00717 -
04950 - -
04955 - -
04960 -pf13185pf02518 -
04965 - -
04970 HMM00121_isom_1_spu(pf00121) SPU3
04975 P_greenGrey SPU3
04980 P_orange SPU3
04985 Ts_fern SPU3
04990 HMMalddh_1_eutpdugrm(pf00171) SPU3 EP_NTERM
04995 P_viridian SPU3
05000 H_lightNavyBlue SPU3
05005 Tdp_mustardGreen SPU3
05010 HMMspu01791_1_spu(pf01791) SPU6
05015 HMMspu02502_1_spu(pf02502) SPU6
05020 - -
05025 -pf16576 -
05030 -pf00005 -
05035 -pf02687pf12704 -
05040 -pf00400 -
05045 HMMreg00486_1_mix(pf00072) FRAG2
05050 -pf07228 -
05055 - -
05060 -pf02355 -
05065 -pf02355 -
05070 -pf13419 -
BMC-P
BMC-H
BMC-T
Aldehyde dehydrogenase
PTAC
Signature enzyme
Alcohol dehydrogenase
Regulator

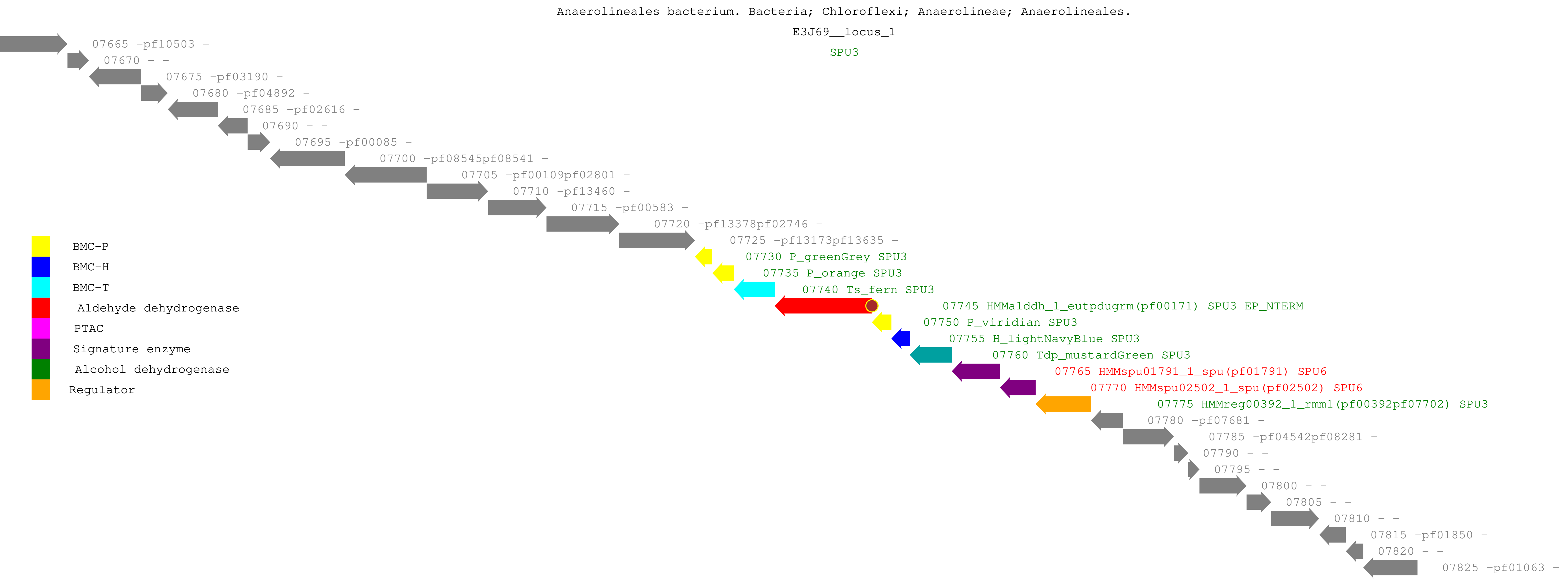
Anaerolineales bacterium. Bacteria; Chloroflexi; Anaerolineae; Anaerolineales.
E3J69__locus_1
SPU3
07665 -pf10503 -
07670 - -
07675 -pf03190 -
07680 -pf04892 -
07685 -pf02616 -
07690 - -
07695 -pf00085 -
07700 -pf08545pf08541 -
07705 -pf00109pf02801 -
07710 -pf13460 -
07715 -pf00583 -
07720 -pf13378pf02746 -
07725 -pf13173pf13635 -
07730 P_greenGrey SPU3
07735 P_orange SPU3
07740 Ts_fern SPU3
07745 HMMalddh_1_eutpdugrm(pf00171) SPU3 EP_NTERM
07750 P_viridian SPU3
07755 H_lightNavyBlue SPU3
07760 Tdp_mustardGreen SPU3
07765 HMMspu01791_1_spu(pf01791) SPU6
07770 HMMspu02502_1_spu(pf02502) SPU6
07775 HMMreg00392_1_rmm1(pf00392pf07702) SPU3
07780 -pf07681 -
07785 -pf04542pf08281 -
07790 - -
07795 - -
07800 - -
07805 - -
07810 - -
07815 -pf01850 -
07820 - -
07825 -pf01063 -
BMC-P
BMC-H
BMC-T
Aldehyde dehydrogenase
PTAC
Signature enzyme
Alcohol dehydrogenase
Regulator

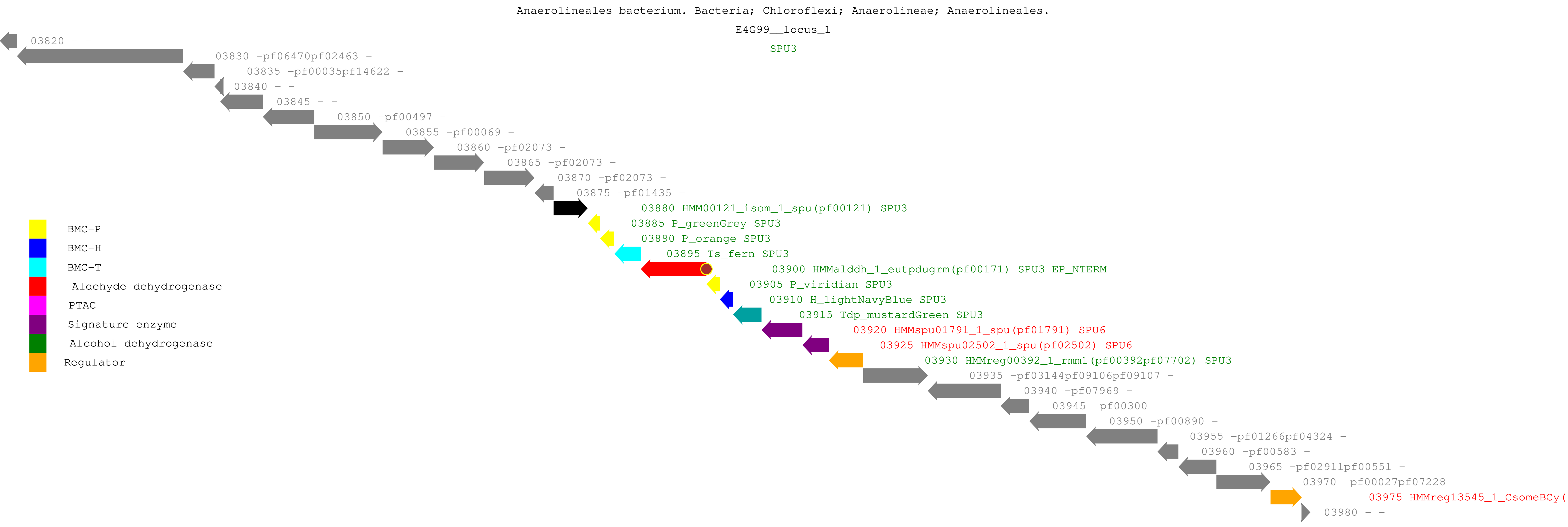
Anaerolineales bacterium. Bacteria; Chloroflexi; Anaerolineae; Anaerolineales.
E4G99__locus_1
SPU3
03820 - -
03830 -pf06470pf02463 -
03835 -pf00035pf14622 -
03840 - -
03845 - -
03850 -pf00497 -
03855 -pf00069 -
03860 -pf02073 -
03865 -pf02073 -
03870 -pf02073 -
03875 -pf01435 -
03880 HMM00121_isom_1_spu(pf00121) SPU3
03885 P_greenGrey SPU3
03890 P_orange SPU3
03895 Ts_fern SPU3
03900 HMMaldh_1_eutpdugrm(pf00171) SPU3 EP_NTERM
03905 P_viridian SPU3
03910 H_lightNavyBlue SPU3
03915 Tdp_mustardGreen SPU3
03920 HMMspu01791_1_spu(pf01791) SPU6
03925 HMMspu02502_1_spu(pf02502) SPU6
03930 HMMreg00392_1_rmm1(pf00392pf07702) SPU3
03935 -pf03144pf09106pf09107 -
03940 -pf07969 -
03945 -pf00300 -
03950 -pf00890 -
03955 -pf01266pf04324 -
03960 -pf00583 -
03965 -pf02911pf00551 -
03970 -pf00027pf07228 -
03975 HMMreg13545_1_CsomeBCy(
03980 - -
BMC-P
BMC-H
BMC-T
Aldehyde dehydrogenase
PTAC
Signature enzyme
Alcohol dehydrogenase
Regulator

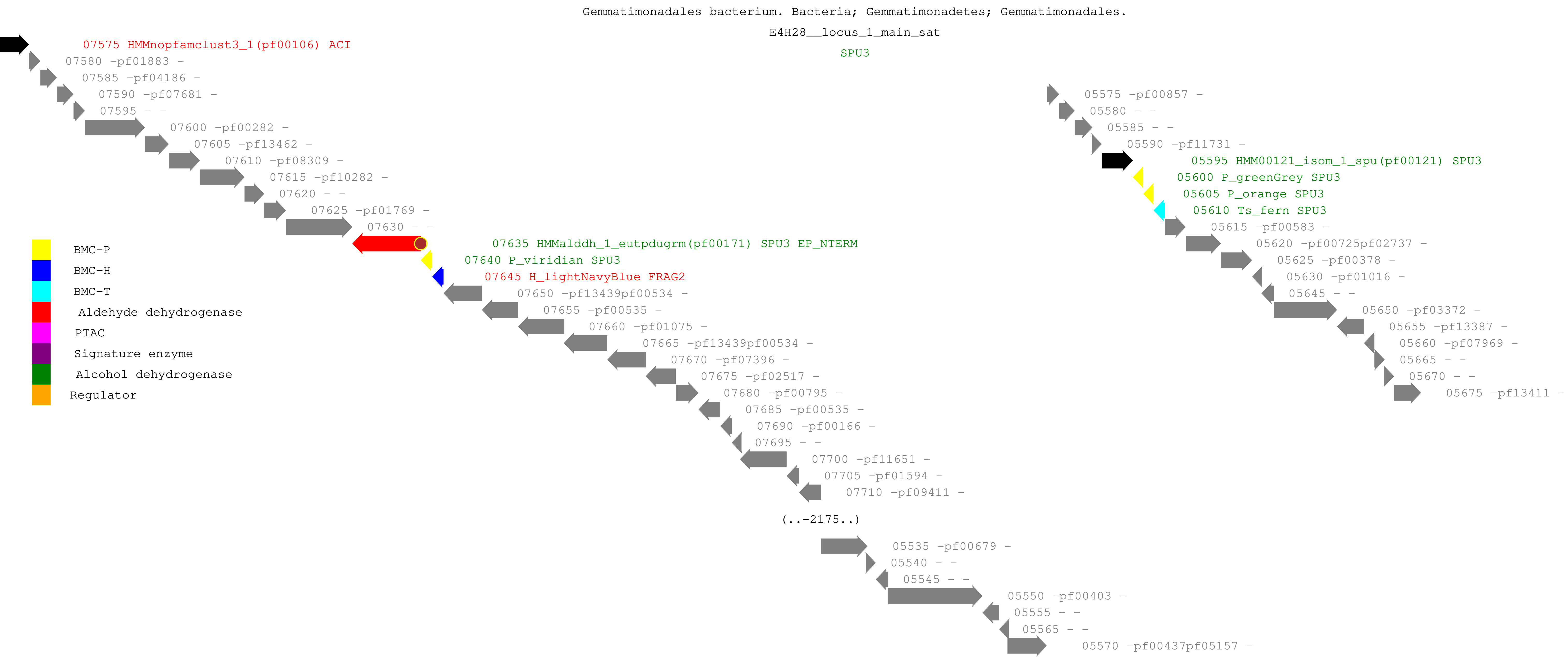
Gemmatimonadales bacterium. Bacteria; Gemmatimonadetes; Gemmatimonadales.
E4H28__locus_1_main_sat
SPU3
07575 HMMnopfamclust3_1(pf00106) ACI
07580 -pf01883 -
07585 -pf04186 -
07590 -pf07681 -
07595 - -
07600 -pf00282 -
07605 -pf13462 -
07610 -pf08309 -
07615 -pf10282 -
07620 - -
07625 -pf01769 -
07630 - -
07635 HMMalddh_1_eutpdugrm(pf00171) SPU3 EP_NTERM
07640 P_viridian SPU3
07645 H_lightNavyBlue FRAG2
07650 -pf13439pf00534 -
07655 -pf00535 -
07660 -pf01075 -
07665 -pf13439pf00534 -
07670 -pf07396 -
07675 -pf02517 -
07680 -pf00795 -
07685 -pf00535 -
07690 -pf00166 -
07695 - -
07700 -pf11651 -
07705 -pf01594 -
07710 -pf09411 -
(..-2175..)
05535 -pf00679 -
05540 - -
05545 - -
05550 -pf00403 -
05555 - -
05565 - -
05570 -pf00437pf05157 -
05575 -pf00857 -
05580 - -
05585 - -
05590 -pf11731 -
05595 HMM00121_isom_1_spu(pf00121) SPU3
05600 P_greenGrey SPU3
05605 P_orange SPU3
05610 Ts_fern SPU3
05615 -pf00583 -
05620 -pf00725pf02737 -
05625 -pf00378 -
05630 -pf01016 -
05645 - -
05650 -pf03372 -
05655 -pf13387 -
05660 -pf07969 -
05665 - -
05670 - -
05675 -pf13411 -
BMC-P
BMC-H
BMC-T
Aldehyde dehydrogenase
PTAC
Signature enzyme
Alcohol dehydrogenase
Regulator

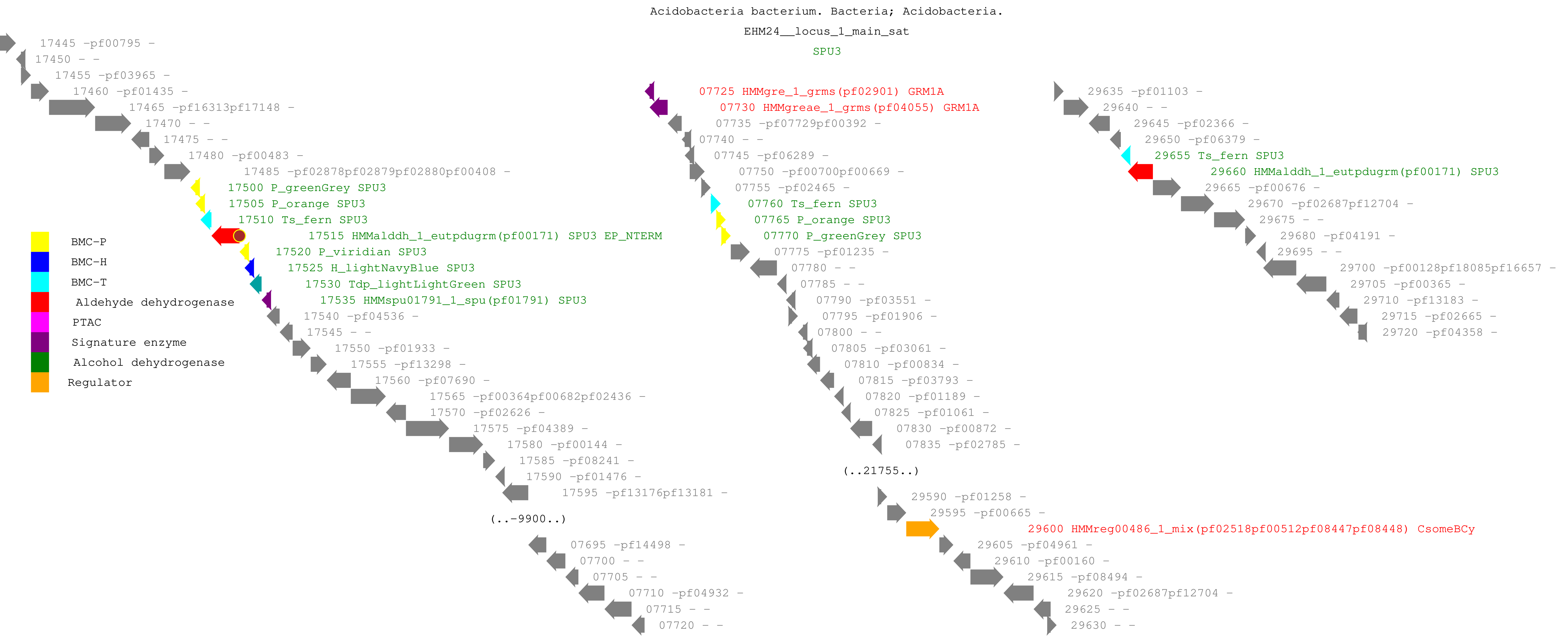

Acidobacteria bacterium. Bacteria; Acidobacteria.
EHM24__locus_1_main_sat
SPU3
17445 -pf00795 -
17450 - -
17455 -pf03965 -
17460 -pf01435 -
17465 -pf16313pf17148 -
17470 - -
17475 - -
17480 -pf00483 -
17485 -pf02878pf02879pf02880pf00408 -
17500 P_greenGrey SPU3
17505 P_orange SPU3
17510 Ts_fern SPU3
17515 HMMalddh_1_eutpdugrm(pf00171) SPU3 EP_NTERM
17520 P_viridian SPU3
17525 H_lightNavyBlue SPU3
17530 Tdp_lightLightGreen SPU3
17535 HMMspu01791_1_spu(pf01791) SPU3
17540 -pf04536 -
17545 - -
17550 -pf01933 -
17555 -pf13298 -
17560 -pf07690 -
17565 -pf00364pf00682pf02436 -
17570 -pf02626 -
17575 -pf04389 -
17580 -pf00144 -
17585 -pf08241 -
17590 -pf01476 -
17595 -pf13176pf13181 -
(..-9900..)
07695 -pf14498 -
07700 - -
07705 - -
07710 -pf04932 -
07715 - -
07720 - -
BMC-P
BMC-H
BMC-T
Aldehyde dehydrogenase
PTAC
Signature enzyme
Alcohol dehydrogenase
Regulator
07725 HMMgre_1_grms(pf02901) GRM1A
07730 HMMgreae_1_grms(pf04055) GRM1A
07735 -pf07729pf00392 -
07740 - -
07745 -pf06289 -
07750 -pf00700pf00669 -
07755 -pf02465 -
07760 Ts_fern SPU3
07765 P_orange SPU3
07770 P_greenGrey SPU3
07775 -pf01235 -
07780 - -
07785 - -
07790 -pf03551 -
07795 -pf01906 -
07800 - -
07805 -pf03061 -
07810 -pf00834 -
07815 -pf03793 -
07820 -pf01189 -
07825 -pf01061 -
07830 -pf00872 -
07835 -pf02785 -
(..21755..)
29590 -pf01258 -
29595 -pf00665 -
29600 HMMreg00486_1_mix(pf02518pf00512pf08447pf08448) CsomeBCy
29605 -pf04961 -
29610 -pf00160 -
29615 -pf08494 -
29620 -pf02687pf12704 -
29625 - -
29630 - -
29635 -pf01103 -
29640 - -
29645 -pf02366 -
29650 -pf06379 -
29655 Ts_fern SPU3
29660 HMMaldDH_1_eutpdugrm(pf00171) SPU3
29665 -pf00676 -
29670 -pf02687pf12704 -
29675 - -
29680 -pf04191 -
29695 - -
29700 -pf00128pf18085pf16657 -
29705 -pf00365 -
29710 -pf13183 -
29715 -pf02665 -
29720 -pf04358 -

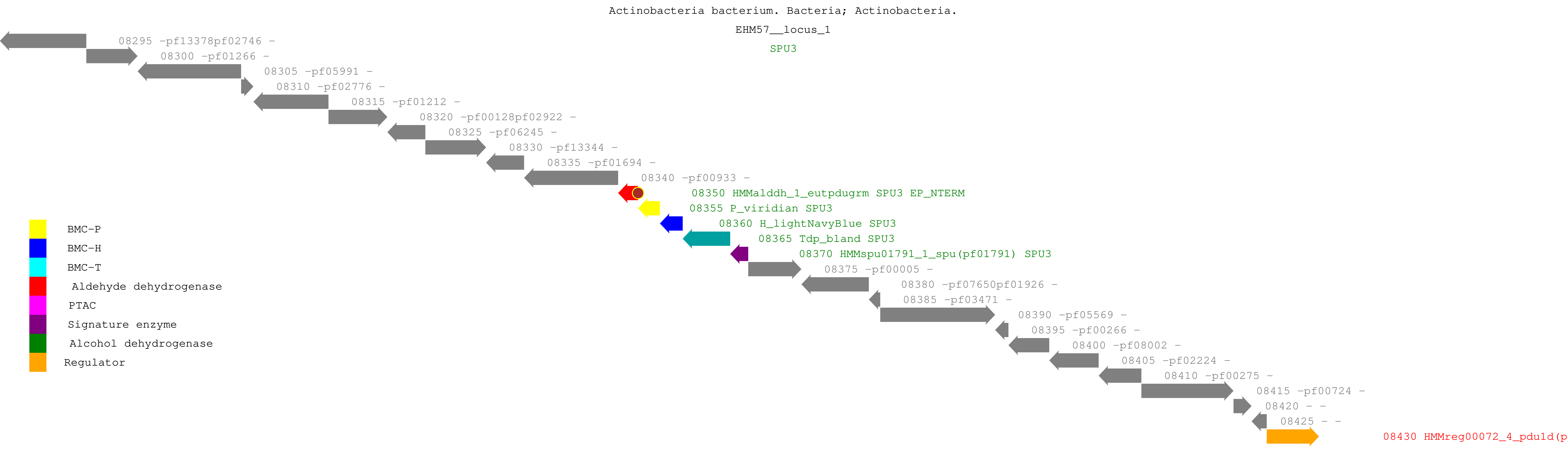
Actinobacteria bacterium. Bacteria; Actinobacteria.
EHM57__locus_1
SPU3
08295 -pf13378pf02746 -
08300 -pf01266 -
08305 -pf05991 -
08310 -pf02776 -
08315 -pf01212 -
08320 -pf00128pf02922 -
08325 -pf06245 -
08330 -pf13344 -
08335 -pf01694 -
08340 -pf00933 -
08350 HMMalddh_1_eutpdugrm SPU3 EP_NTERM
08355 P_viridian SPU3
08360 H_lightNavyBlue SPU3
08365 Tdp_bland SPU3
08370 HMMspu01791_1_spu(pf01791) SPU3
08375 -pf00005 -
08380 -pf07650pf01926 -
08385 -pf03471 -
08390 -pf05569 -
08395 -pf00266 -
08400 -pf08002 -
08405 -pf02224 -
08410 -pf00275 -
08415 -pf00724 -
08420 - -
08425 - -
08430 HMMreg00072_4_pdu1d(p
BMC-P
BMC-H
BMC-T
Aldehyde dehydrogenase
PTAC
Signature enzyme
Alcohol dehydrogenase
Regulator

Flavobacterium sp. Bacteria; Bacteroidetes; Flavobacteriia; Flavobacteriales;

EO043__locus_1

SPU3

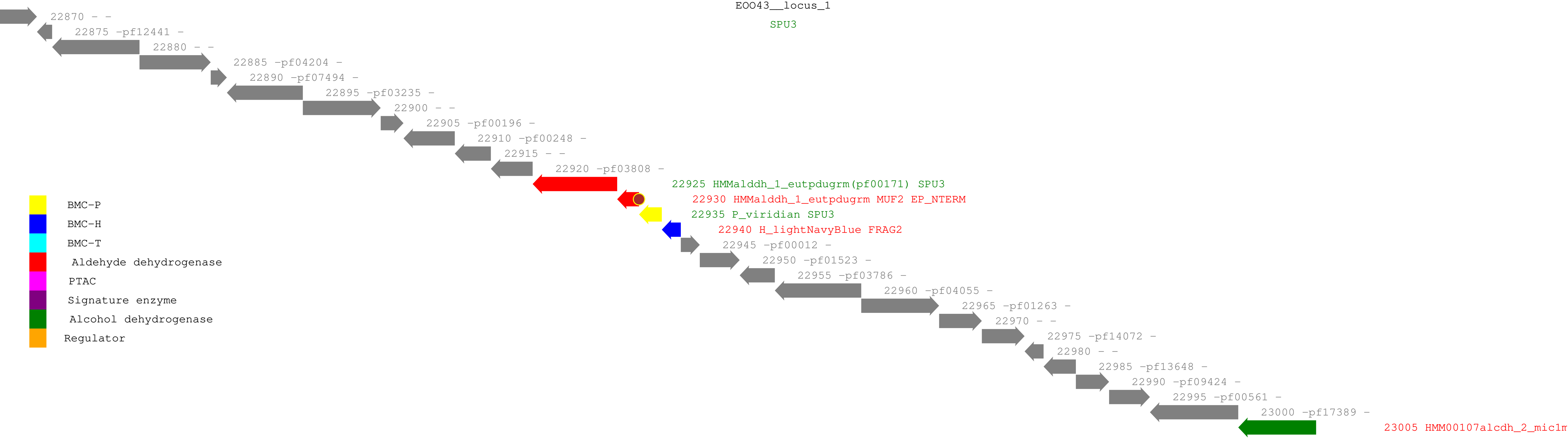

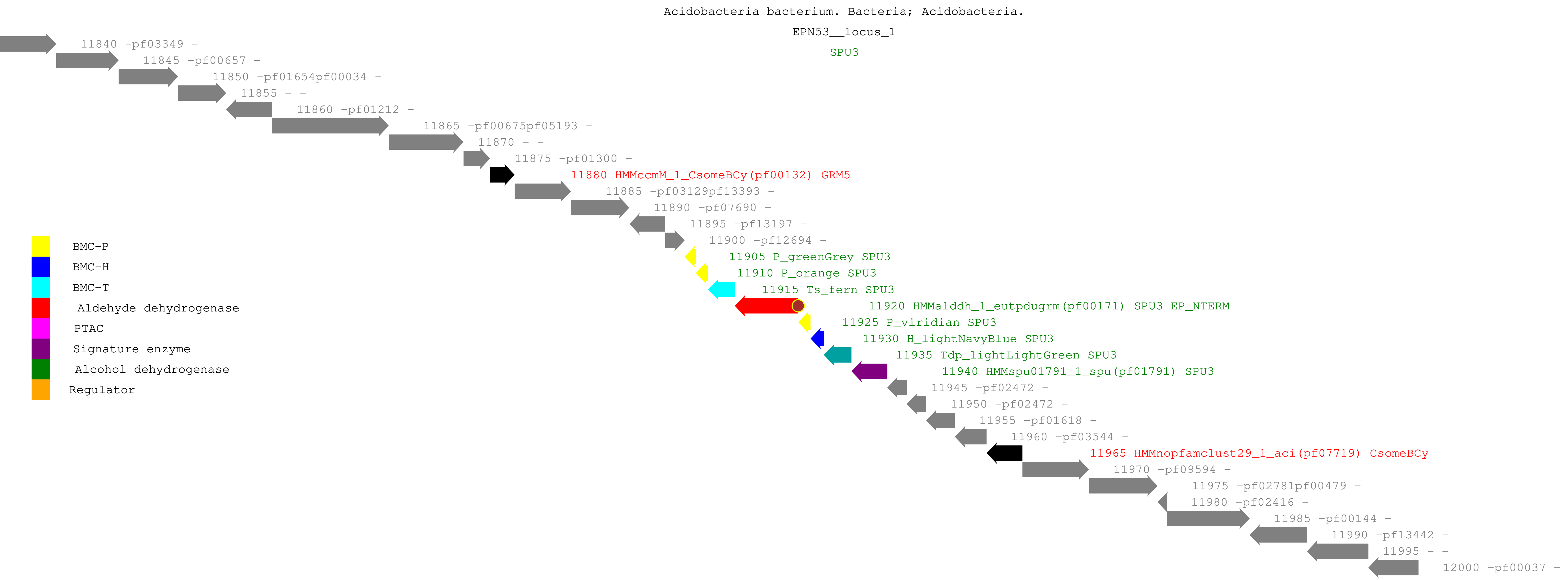
Acidobacteria bacterium. Bacteria; Acidobacteria.
EPN53__locus_1
SPU3
11840 -pf03349 -
11845 -pf00657 -
11850 -pf01654pf00034 -
11855 - -
11860 -pf01212 -
11865 -pf00675pf05193 -
11870 - -
11875 -pf01300 -
11880 HMMccmM_1_CsomeBCy(pf00132) GRM5
11885 -pf03129pf13393 -
11890 -pf07690 -
11895 -pf13197 -
11900 -pf12694 -
11905 P_greenGrey SPU3
11910 P_orange SPU3
11915 Ts_fern SPU3
11920 HMMalddh_1_eutpdugrm(pf00171) SPU3 EP_NTERM
11925 P_viridian SPU3
11930 H_lightNavyBlue SPU3
11935 Tdp_lightLightGreen SPU3
11940 HMMspu01791_1_spu(pf01791) SPU3
11945 -pf02472 -
11950 -pf02472 -
11955 -pf01618 -
11960 -pf03544 -
11965 HMMnopfamclust29_1_aci(pf07719) CsomeBCy
11970 -pf09594 -
11975 -pf02781pf00479 -
11980 -pf02416 -
11985 -pf00144 -
11990 -pf13442 -
11995 - -
12000 -pf00037 -
BMC-P
BMC-H
BMC-T
Aldehyde dehydrogenase
PTAC
Signature enzyme
Alcohol dehydrogenase
Regulator

Egibacter rhizosphaerae. Bacteria; Actinobacteria; Nitriliruptoria; Egibacterales; Egibacteraceae;

ER308__locus_1

SPU3

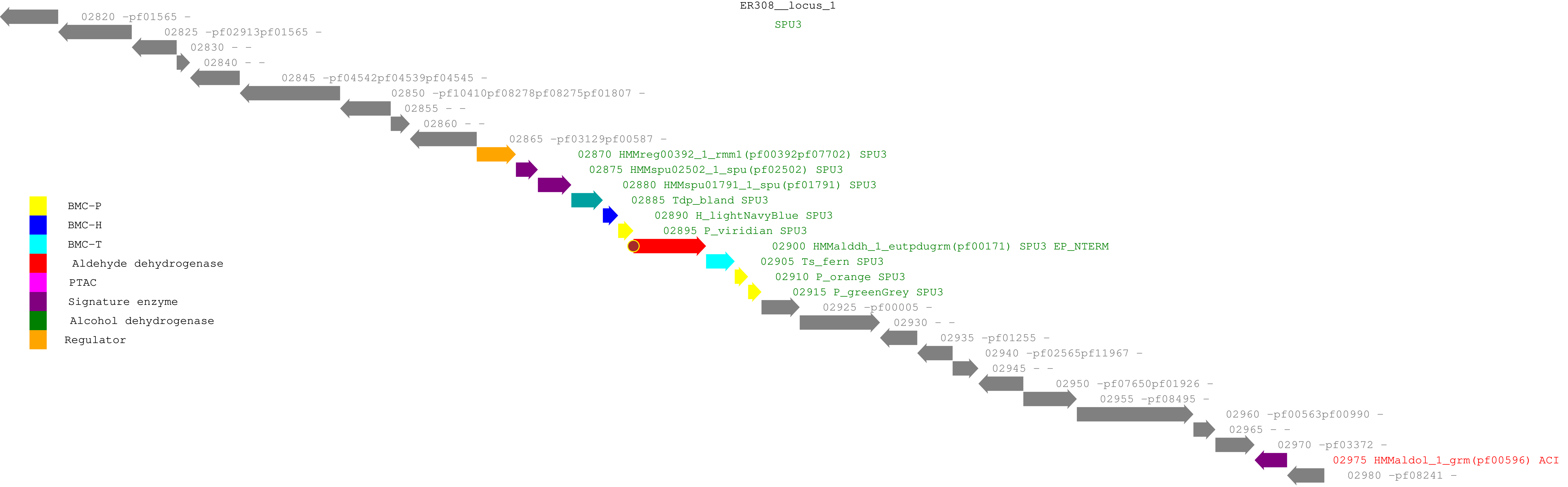

Acidimicrobiia bacterium. Bacteria; Actinobacteria; Acidimicrobiia.

EX267__locus_1

SPU3

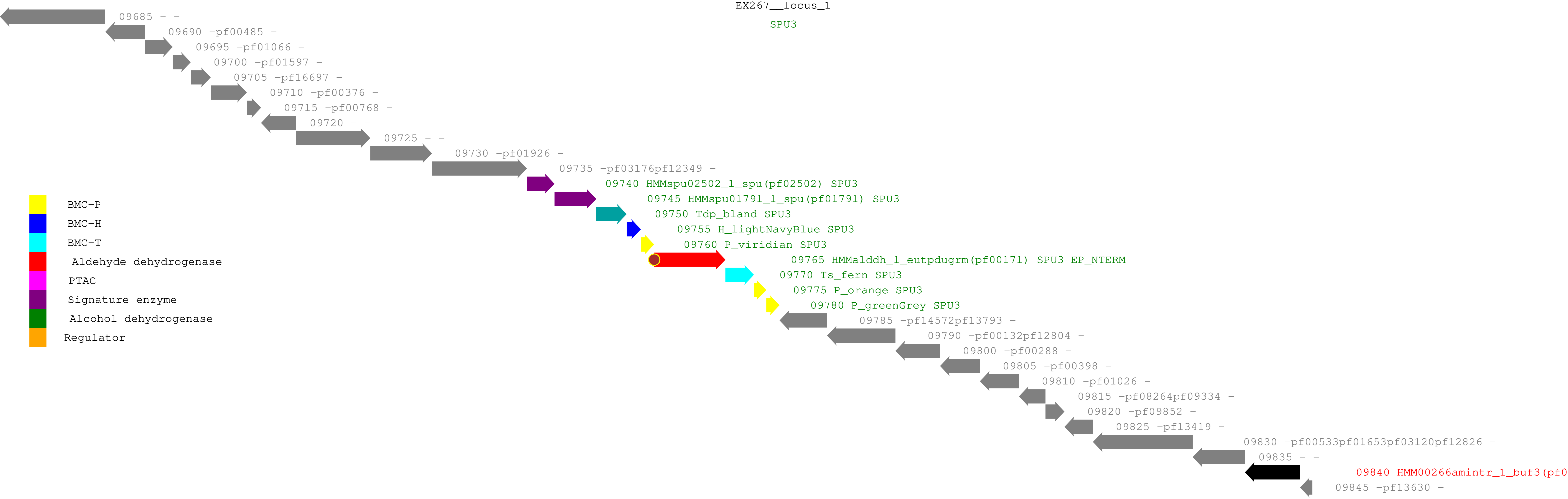

Chromatiales bacterium. Bacteria; Proteobacteria; Gammaproteobacteria; Chromatiales;

EX272__locus_1

SPU3

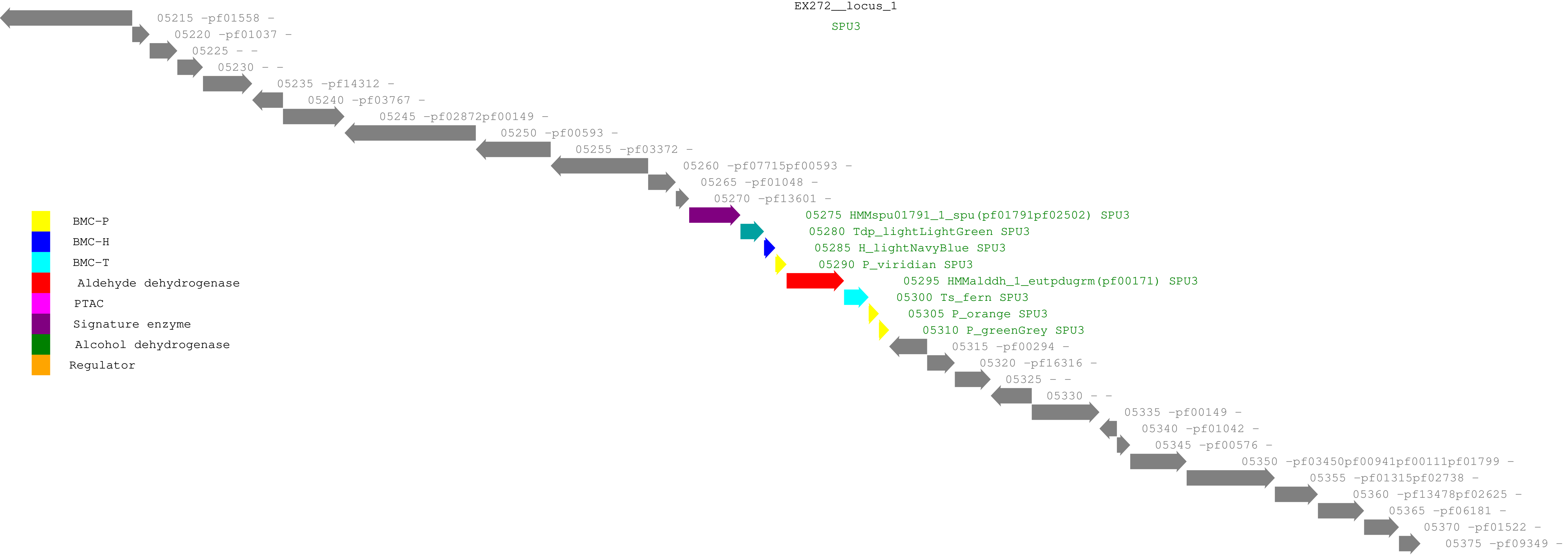

hydrothermal vent metagenome. unclassified sequences; metagenomes; ecological metagenomes.

MNBD_ACTINO01-_locus_fragmented

SPU3

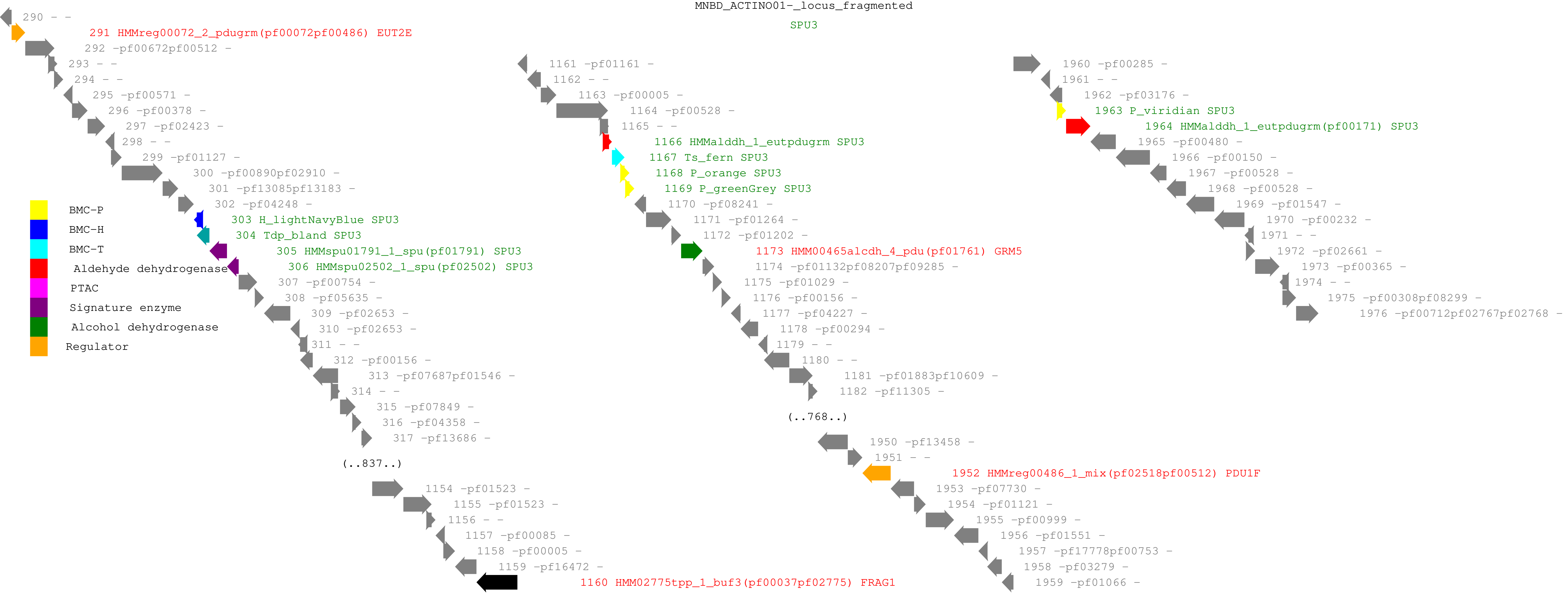

Phycisphaerae bacterium RAS2. Bacteria; Planctomycetes; Phycisphaerae.

RAS2__locus_1

SPU3

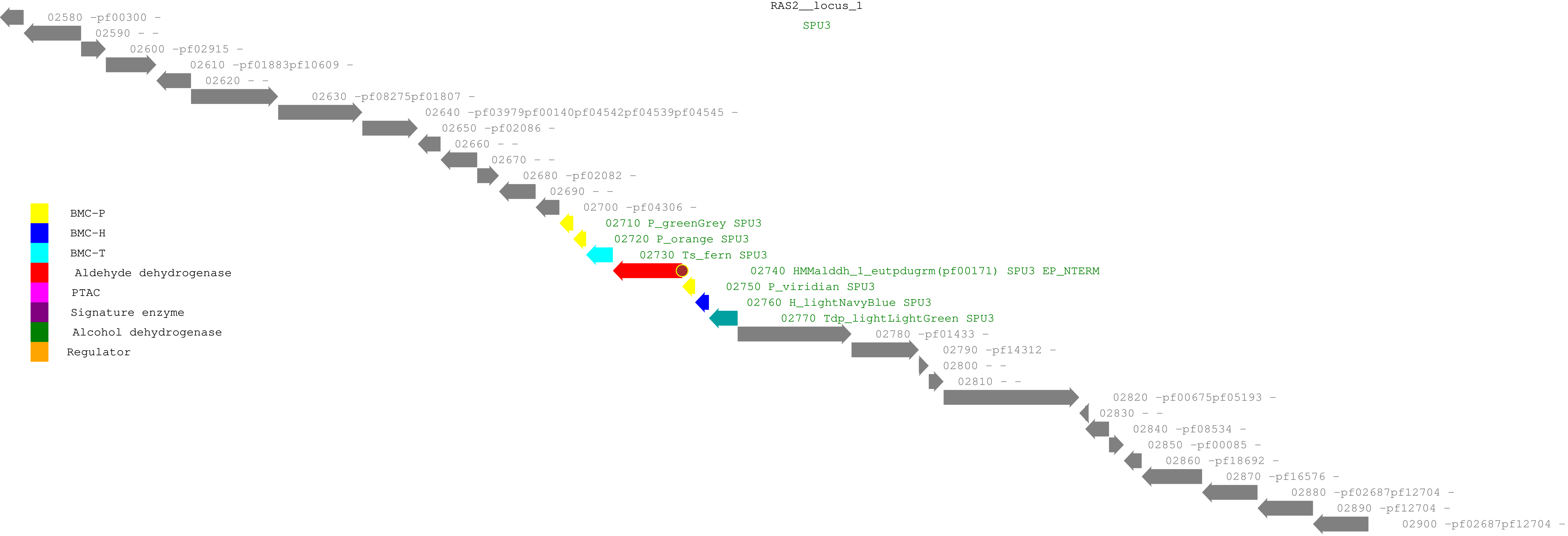

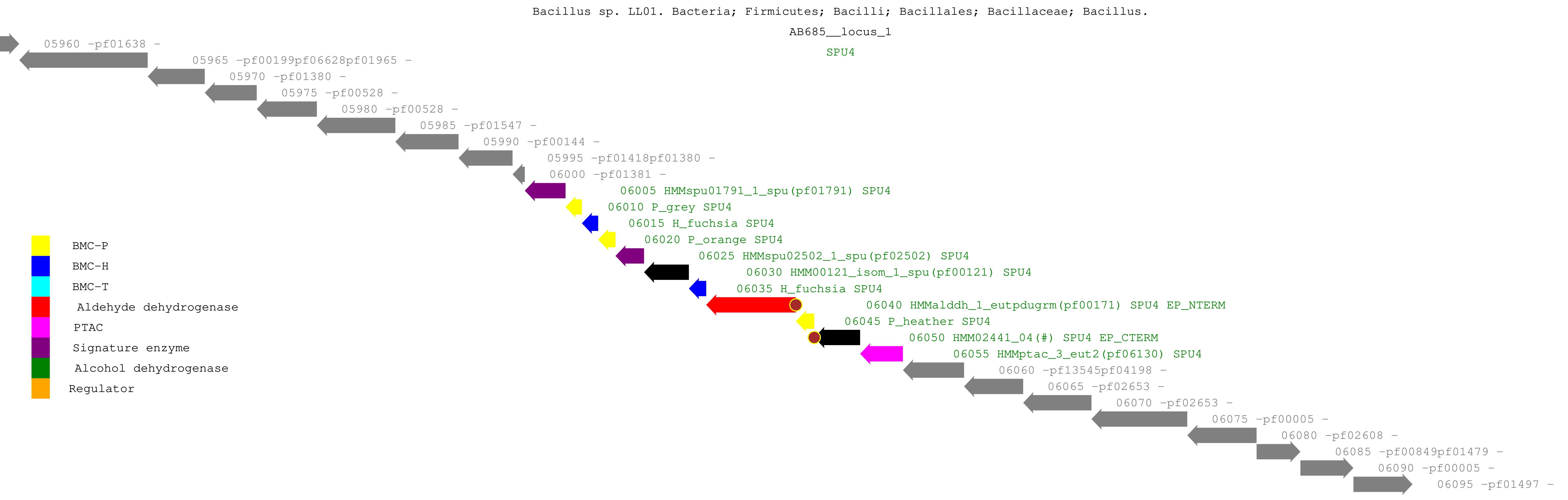
Bacillus sp. LL01. Bacteria; Firmicutes; Bacilli; Bacillales; Bacillaceae; Bacillus.
AB685__locus_1
SPU4
05960 -pf01638 -
05965 -pf00199pf06628pf01965 -
05970 -pf01380 -
05975 -pf00528 -
05980 -pf00528 -
05985 -pf01547 -
05990 -pf00144 -
05995 -pf01418pf01380 -
06000 -pf01381 -
06005 HMMspu01791_1_spu(pf01791) SPU4
06010 P_grey SPU4
06015 H_fuchsia SPU4
06020 P_orange SPU4
06025 HMMspu02502_1_spu(pf02502) SPU4
06030 HMM00121_isom_1_spu(pf00121) SPU4
06035 H_fuchsia SPU4
06040 HMMalddh_1_eutpdugrm(pf00171) SPU4 EP_NTERM
06045 P_heather SPU4
06050 HMM02441_04(#) SPU4 EP_CTERM
06055 HMMptac_3_eut2(pf06130) SPU4
06060 -pf13545pf04198 -
06065 -pf02653 -
06070 -pf02653 -
06075 -pf00005 -
06080 -pf02608 -
06085 -pf00849pf01479 -
06090 -pf00005 -
06095 -pf01497 -
BMC-P
BMC-H
BMC-T
Aldehyde dehydrogenase
PTAC
Signature enzyme
Alcohol dehydrogenase
Regulator

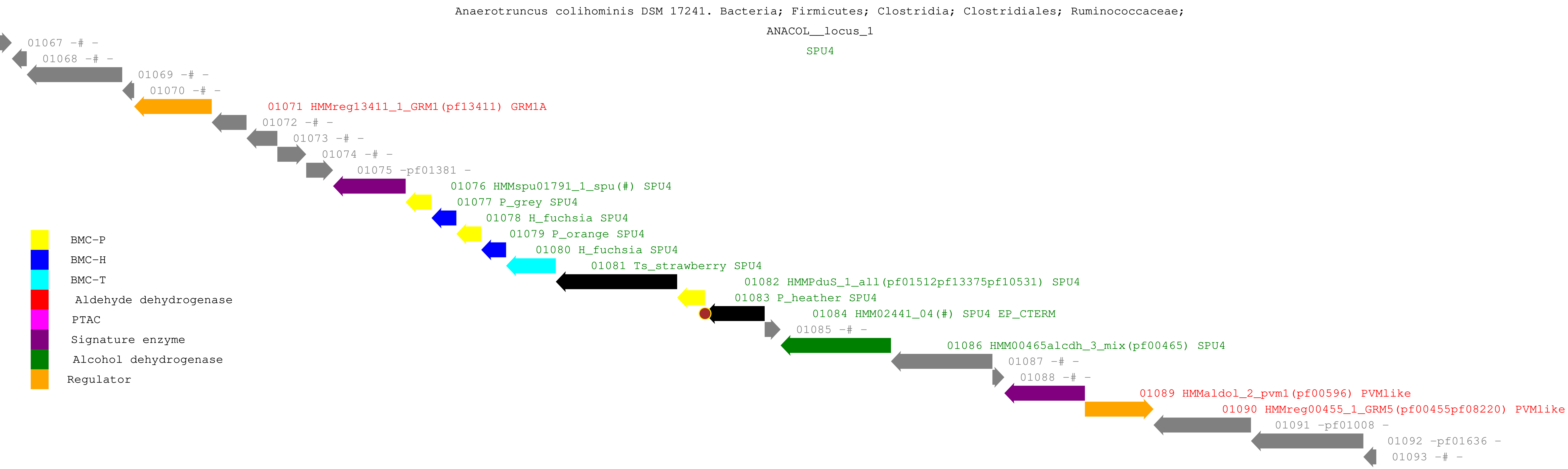

Anaerotruncus colihominis DSM 17241. Bacteria; Firmicutes; Clostridia; Clostridiales; Ruminococcaceae;
ANACOL__locus_1
SPU4
01067 -# -
01068 -# -
01069 -# -
01070 -# -
01071 HMMreg13411_1_GRM1(pf13411) GRM1A
01072 -# -
01073 -# -
01074 -# -
01075 -pf01381 -
01076 HMMspu01791_1_spu(#) SPU4
01077 P_grey SPU4
01078 H_fuchsia SPU4
01079 P_orange SPU4
01080 H_fuchsia SPU4
01081 Ts_strawberry SPU4
01082 HMMPduS_1_all(pf01512pf13375pf10531) SPU4
01083 P_heather SPU4
01084 HMM02441_04(#) SPU4 EP_CTERM
01085 -# -
01086 HMM00465alcdh_3_mix(pf00465) SPU4
01087 -# -
01088 -# -
01089 HMMaldol_2_pvm1(pf00596) PVMlike
01090 HMMreg00455_1_GRM5(pf00455pf08220) PVMlike
01091 -pf01008 -
01092 -pf01636 -
01093 -# -
BMC-P
BMC-H
BMC-T
Aldehyde dehydrogenase
PTAC
Signature enzyme
Alcohol dehydrogenase
Regulator

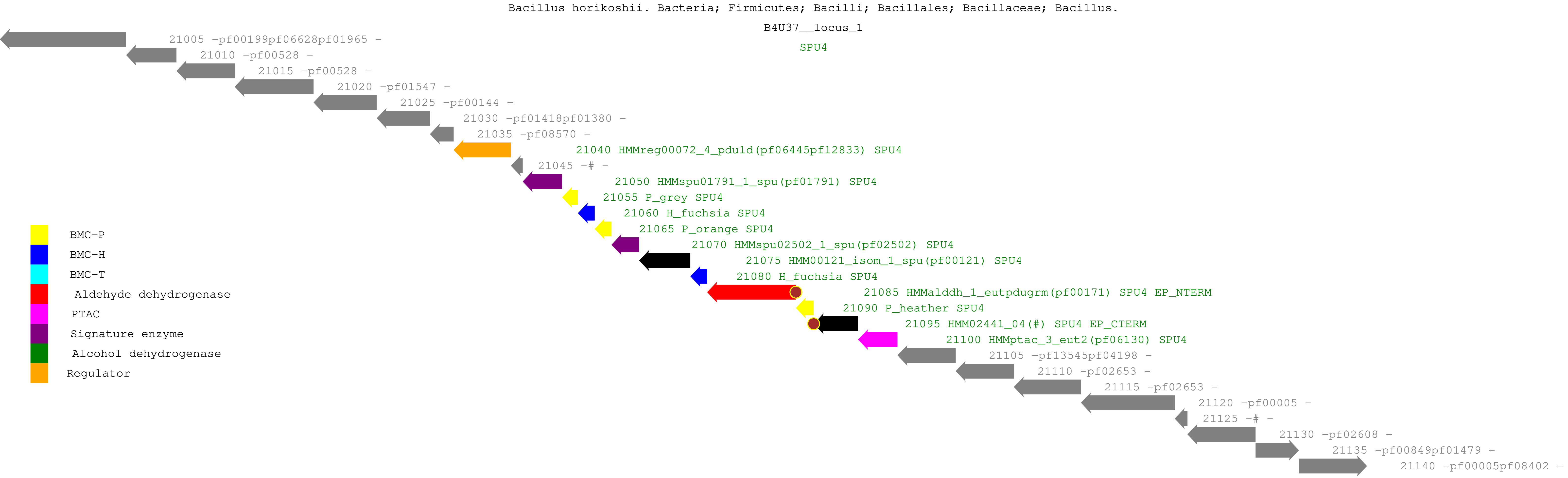
Bacillus horikoshii. Bacteria; Firmicutes; Bacilli; Bacillales; Bacillaceae; Bacillus.
B4U37__locus_1
SPU4
21005 -pf00199pf06628pf01965 -
21010 -pf00528 -
21015 -pf00528 -
21020 -pf01547 -
21025 -pf00144 -
21030 -pf01418pf01380 -
21035 -pf08570 -
21040 HMMreg00072_4_pdu1d(pf06445pf12833) SPU4
21045 -# -
21050 HMMspu01791_1_spu(pf01791) SPU4
21055 P_grey SPU4
21060 H_fuchsia SPU4
21065 P_orange SPU4
21070 HMMspu02502_1_spu(pf02502) SPU4
21075 HMM00121_isom_1_spu(pf00121) SPU4
21080 H_fuchsia SPU4
21085 HMMalddh_1_eutpdugrm(pf00171) SPU4 EP_NTERM
21090 P_heather SPU4
21095 HMM02441_04(#) SPU4 EP_CTERM
21100 HMMptac_3_eut2(pf06130) SPU4
21105 -pf13545pf04198 -
21110 -pf02653 -
21115 -pf02653 -
21120 -pf00005 -
21125 -# -
21130 -pf02608 -
21135 -pf00849pf01479 -
21140 -pf00005pf08402 -
BMC-P
BMC-H
BMC-T
Aldehyde dehydrogenase
PTAC
Signature enzyme
Alcohol dehydrogenase
Regulator

Paenibacillus sp. MEC069. Bacteria; Firmicutes; Bacilli; Bacillales; Paenibacillaceae; Paenibacillus.

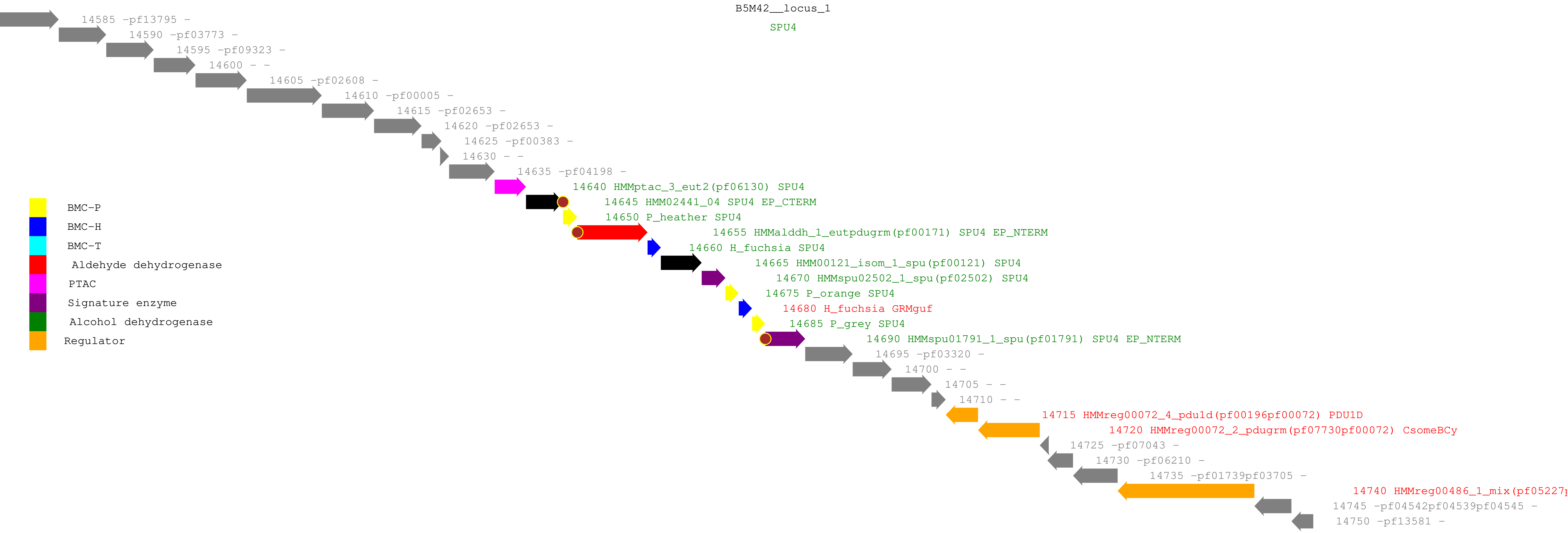

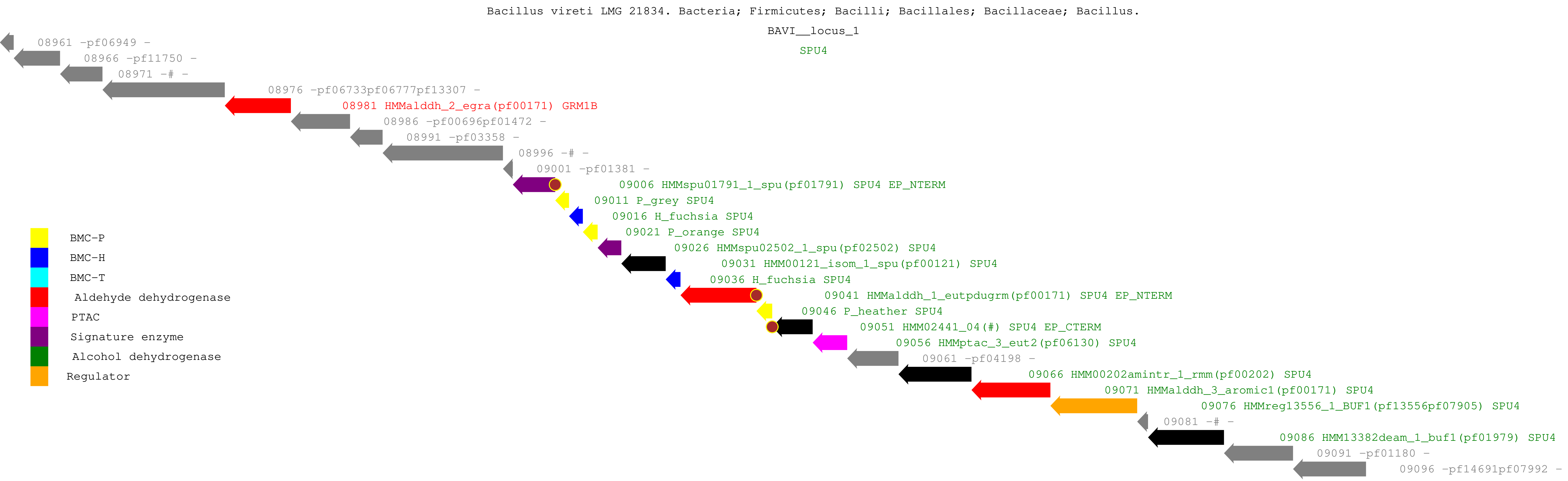
Bacillus vireti LMG 21834. Bacteria; Firmicutes; Bacilli; Bacillales; Bacillaceae; Bacillus.
BAVI__locus_1
SPU4
08961 -pf06949 -
08966 -pf11750 -
08971 -# -
08976 -pf06733pf06777pf13307 -
08981 HMMaldh_2_egra(pf00171) GRM1B
08986 -pf00696pf01472 -
08991 -pf03358 -
08996 -# -
09001 -pf01381 -
09006 HMMspu01791_1_spu(pf01791) SPU4 EP_NTERM
09011 P_grey SPU4
09016 H_fuchsia SPU4
09021 P_orange SPU4
09026 HMMspu02502_1_spu(pf02502) SPU4
09031 HMM00121_isom_1_spu(pf00121) SPU4
09036 H_fuchsia SPU4
09041 HMMaldh_1_eutpdugrm(pf00171) SPU4 EP_NTERM
09046 P_heather SPU4
09051 HMM02441_04(#) SPU4 EP_CTERM
09056 HMMptac_3_eut2(pf06130) SPU4
09061 -pf04198 -
09066 HMM00202amintr_1_rmm(pf00202) SPU4
09071 HMMaldh_3_aromic1(pf00171) SPU4
09076 HMMreg13556_1_BUF1(pf13556pf07905) SPU4
09081 -# -
09086 HMM13382deam_1_buf1(pf01979) SPU4
09091 -pf01180 -
09096 -pf14691pf07992 -
BMC-P
BMC-H
BMC-T
Aldehyde dehydrogenase
PTAC
Signature enzyme
Alcohol dehydrogenase
Regulator

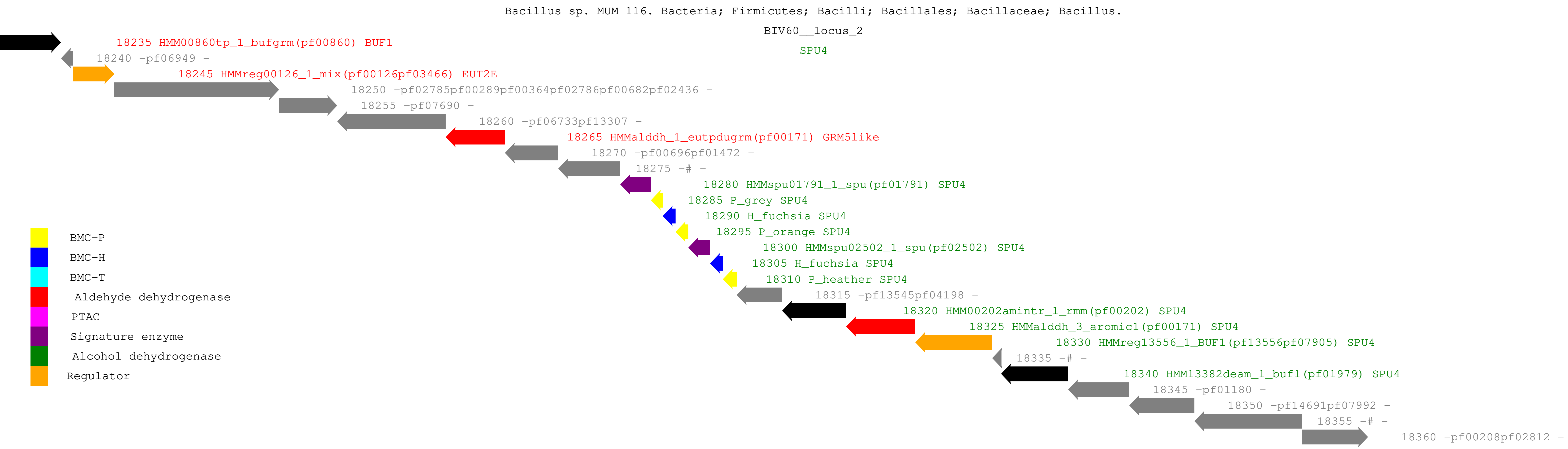

Bacillus sp. MUM 116. Bacteria; Firmicutes; Bacilli; Bacillales; Bacillaceae; Bacillus.
BIV60__locus_2
SPU4
18235 HMM00860tp_1_bufgrm(pf00860) BUF1
18240 -pf06949 -
18245 HMMreg00126_1_mix(pf00126pf03466) EUT2E
18250 -pf02785pf00289pf00364pf02786pf00682pf02436 -
18255 -pf07690 -
18260 -pf06733pf13307 -
18265 HMMalddh_1_eutpdugrm(pf00171) GRM5like
18270 -pf00696pf01472 -
18275 -# -
18280 HMMspu01791_1_spu(pf01791) SPU4
18285 P_grey SPU4
18290 H_fuchsia SPU4
18295 P_orange SPU4
18300 HMMspu02502_1_spu(pf02502) SPU4
18305 H_fuchsia SPU4
18310 P_heather SPU4
18315 -pf13545pf04198 -
18320 HMM00202amintr_1_rmm(pf00202) SPU4
18325 HMMalddh_3_aromic1(pf00171) SPU4
18330 HMMreg13556_1_BUF1(pf13556pf07905) SPU4
18335 -# -
18340 HMM13382deam_1_buf1(pf01979) SPU4
18345 -pf01180 -
18350 -pf14691pf07992 -
18355 -# -
18360 -pf00208pf02812 -
BMC-P
BMC-H
BMC-T
Aldehyde dehydrogenase
PTAC
Signature enzyme
Alcohol dehydrogenase
Regulator

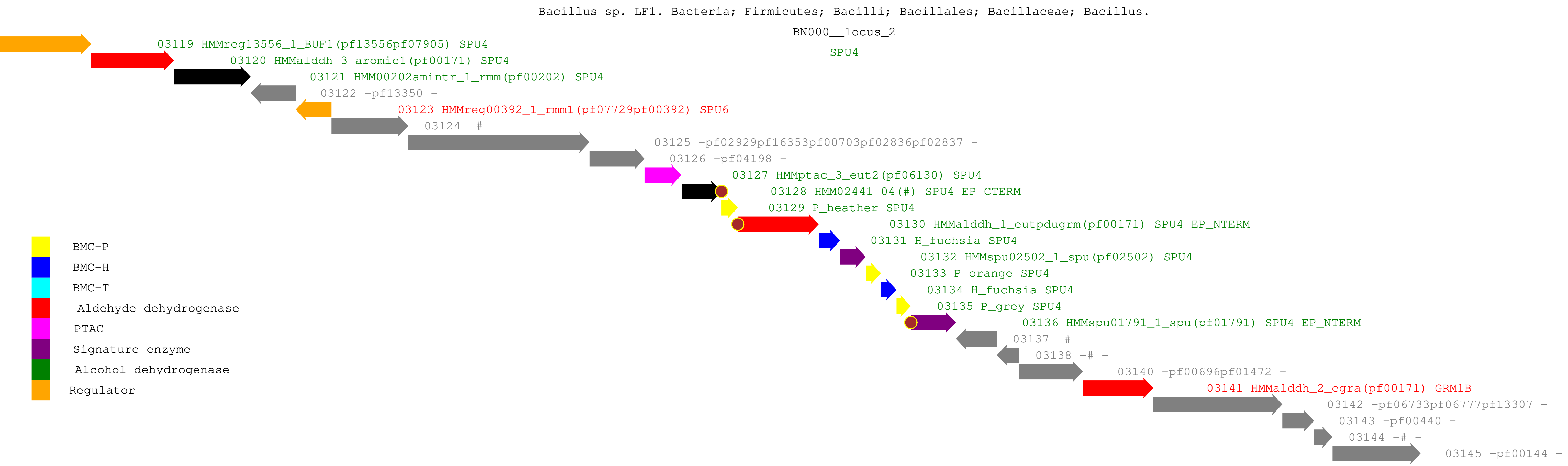
Bacillus sp. LF1. Bacteria; Firmicutes; Bacilli; Bacillales; Bacillaceae; Bacillus.
BN000__locus_2
SPU4
03119 HMMreg13556_1_BUF1(pf13556pf07905) SPU4
03120 HMMalddh_3_aromic1(pf00171) SPU4
03121 HMM00202amintr_1_rmm(pf00202) SPU4
03122 -pf13350 -
03123 HMMreg00392_1_rmm1(pf07729pf00392) SPU6
03124 -# -
03125 -pf02929pf16353pf00703pf02836pf02837 -
03126 -pf04198 -
03127 HMMptac_3_eut2(pf06130) SPU4
03128 HMM02441_04(#) SPU4 EP_CTERM
03129 P_heather SPU4
03130 HMMalddh_1_eutpdugrm(pf00171) SPU4 EP_NTERM
03131 H_fuchsia SPU4
03132 HMMspu02502_1_spu(pf02502) SPU4
03133 P_orange SPU4
03134 H_fuchsia SPU4
03135 P_grey SPU4
03136 HMMspu01791_1_spu(pf01791) SPU4 EP_NTERM
03137 -# -
03138 -# -
03140 -pf00696pf01472 -
03141 HMMalddh_2_egra(pf00171) GRM1B
03142 -pf06733pf06777pf13307 -
03143 -pf00440 -
03144 -# -
03145 -pf00144 -
BMC-P
BMC-H
BMC-T
Aldehyde dehydrogenase
PTAC
Signature enzyme
Alcohol dehydrogenase
Regulator

Bacillus sp. MRMR6. Bacteria; Firmicutes; Bacilli; Bacillales; Bacillaceae; Bacillus.

BTR25__locus_1

SPU4

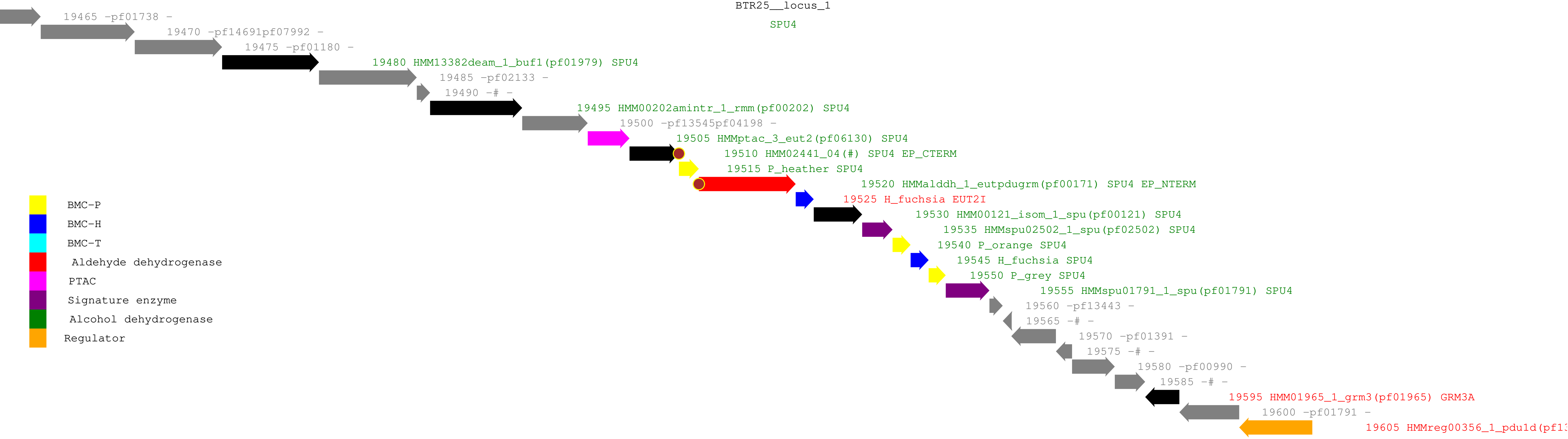

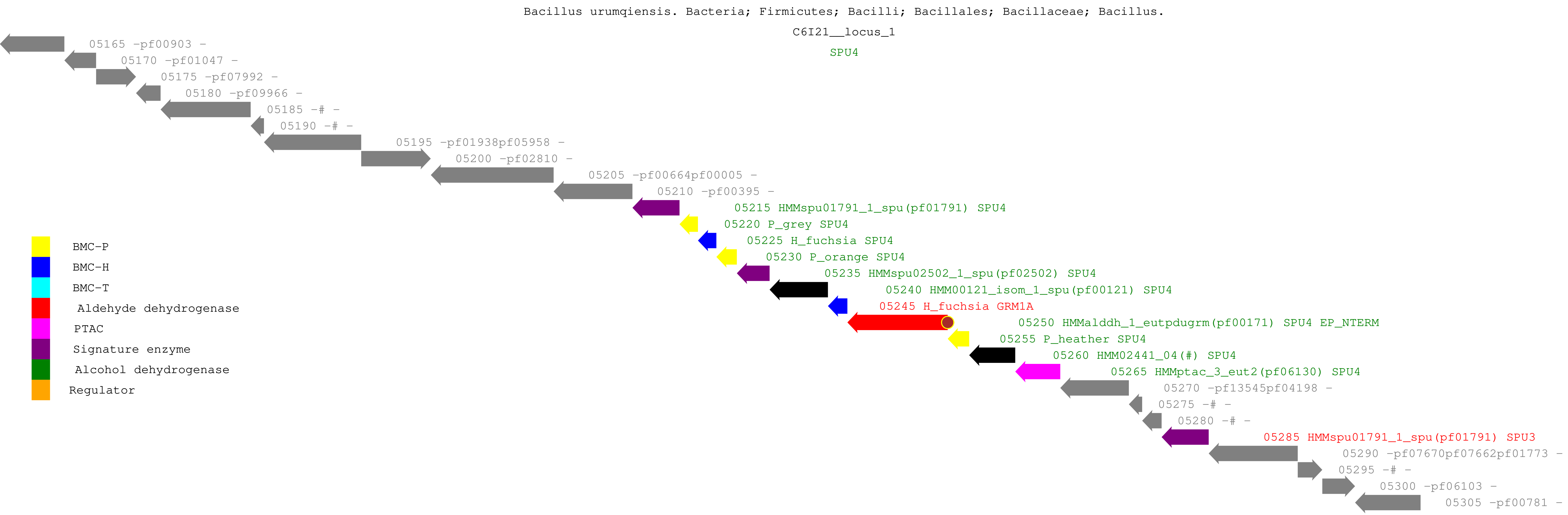

Bacillus urumqiensis. Bacteria; Firmicutes; Bacilli; Bacillales; Bacillaceae; Bacillus.
C6I21__locus_1
SPU4
05165 -pf00903 -
05170 -pf01047 -
05175 -pf07992 -
05180 -pf09966 -
05185 -# -
05190 -# -
05195 -pf01938pf05958 -
05200 -pf02810 -
05205 -pf00664pf00005 -
05210 -pf00395 -
05215 HMMspu01791_1_spu(pf01791) SPU4
05220 P_grey SPU4
05225 H_fuchsia SPU4
05230 P_orange SPU4
05235 HMMspu02502_1_spu(pf02502) SPU4
05240 HMM00121_isom_1_spu(pf00121) SPU4
05245 H_fuchsia GRM1A
05250 HMMalddh_1_eutpdugrm(pf00171) SPU4 EP_NTERM
05255 P_heather SPU4
05260 HMM02441_04(#) SPU4
05265 HMMptac_3_eut2(pf06130) SPU4
05270 -pf13545pf04198 -
05275 -# -
05280 -# -
05285 HMMspu01791_1_spu(pf01791) SPU3
05290 -pf07670pf07662pf01773 -
05295 -# -
05300 -pf06103 -
05305 -pf00781 -
BMC-P
BMC-H
BMC-T
Aldehyde dehydrogenase
PTAC
Signature enzyme
Alcohol dehydrogenase
Regulator

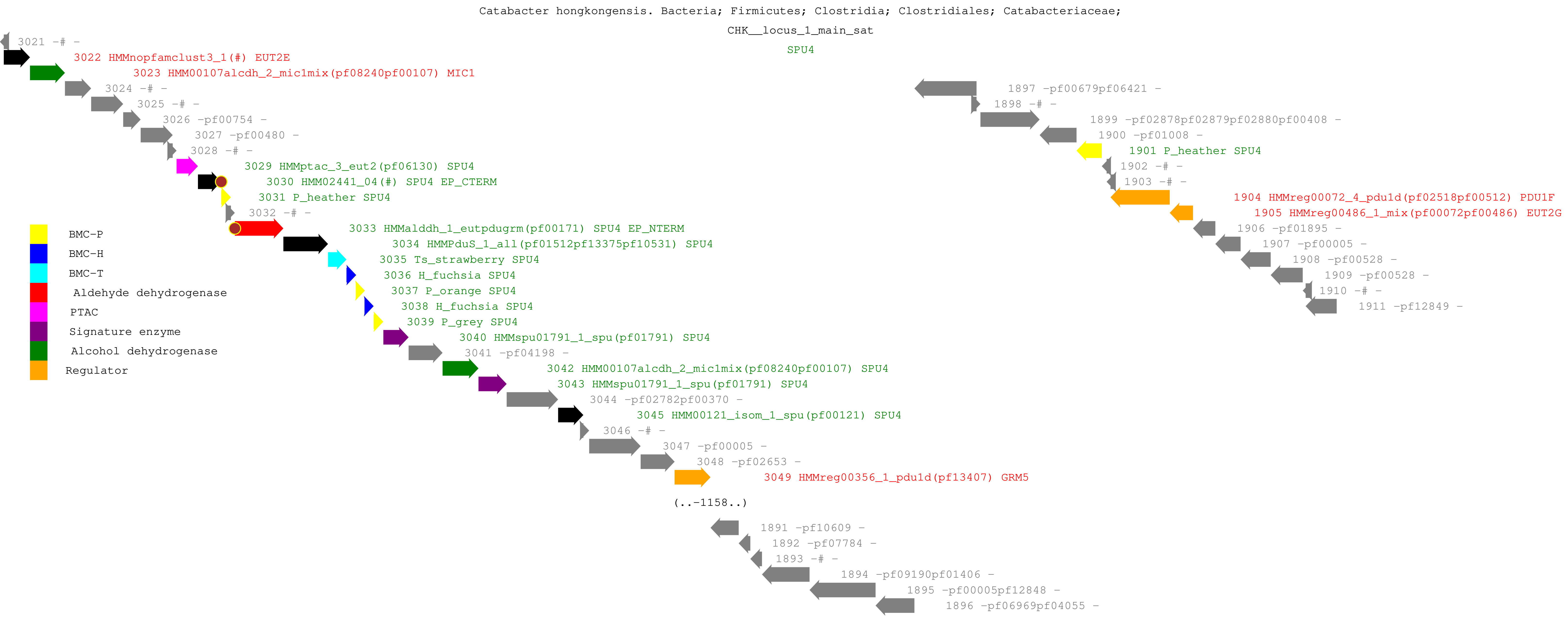

Catabacter hongkongensis. Bacteria; Firmicutes; Clostridia; Clostridiales; Catabacteriaceae;
CHK__locus_1_main_sat
SPU4
3021 -# -
3022 HMMnopfamclust3_1(#) EUT2E
3023 HMM00107alcdh_2_mic1mix(pf08240pf00107) MIC1
3024 -# -
3025 -# -
3026 -pf00754 -
3027 -pf00480 -
3028 -# -
3029 HMMptac_3_eut2(pf06130) SPU4
3030 HMM02441_04(#) SPU4 EP_CTERM
3031 P_heather SPU4
3032 -# -
3033 HMMaldh_1_eutpdugrm(pf00171) SPU4 EP_NTERM
3034 HMMPduS_1_all(pf01512pf13375pf10531) SPU4
3035 Ts_strawberry SPU4
3036 H_fuchsia SPU4
3037 P_orange SPU4
3038 H_fuchsia SPU4
3039 P_grey SPU4
3040 HMMspu01791_1_spu(pf01791) SPU4
3041 -pf04198 -
3042 HMM00107alcdh_2_mic1mix(pf08240pf00107) SPU4
3043 HMMspu01791_1_spu(pf01791) SPU4
3044 -pf02782pf00370 -
3045 HMM00121_isom_1_spu(pf00121) SPU4
3046 -# -
3047 -pf00005 -
3048 -pf02653 -
3049 HMMreg00356_1_pdu1d(pf13407) GRM5
(..-1158..)
1891 -pf10609 -
1892 -pf07784 -
1893 -# -
1894 -pf09190pf01406 -
1895 -pf00005pf12848 -
1896 -pf06969pf04055 -
1897 -pf00679pf06421 -
1898 -# -
1899 -pf02878pf02879pf02880pf00408 -
1900 -pf01008 -
1901 P_heather SPU4
1902 -# -
1903 -# -
1904 HMMreg00072_4_pdu1d(pf02518pf00512) PDU1F
1905 HMMreg00486_1_mix(pf00072pf00486) EUT2G
1906 -pf01895 -
1907 -pf00005 -
1908 -pf00528 -
1909 -pf00528 -
1910 -# -
1911 -pf12849 -
BMC-P
BMC-H
BMC-T
Aldehyde dehydrogenase
PTAC
Signature enzyme
Alcohol dehydrogenase
Regulator

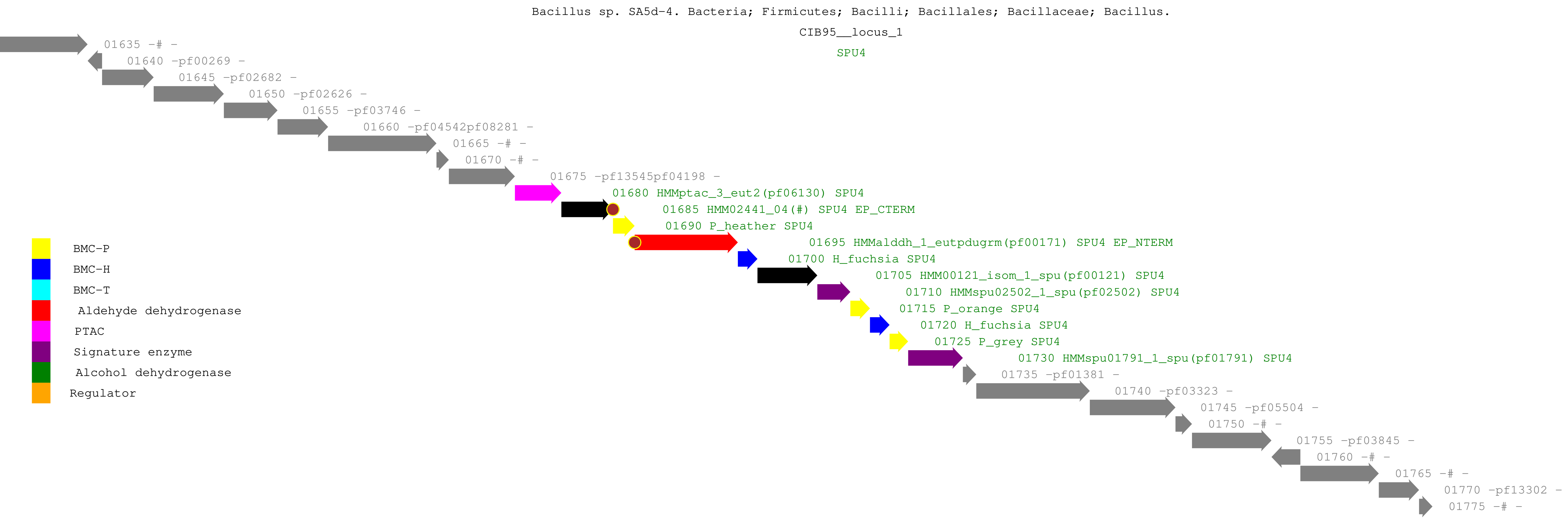
Bacillus sp. SA5d-4. Bacteria; Firmicutes; Bacilli; Bacillales; Bacillaceae; Bacillus.
CIB95__locus_1
SPU4
01635 -# -
01640 -pf00269 -
01645 -pf02682 -
01650 -pf02626 -
01655 -pf03746 -
01660 -pf04542pf08281 -
01665 -# -
01670 -# -
01675 -pf13545pf04198 -
01680 HMMptac_3_eut2(pf06130) SPU4
01685 HMM02441_04(#) SPU4 EP_CTERM
01690 P_heather SPU4
01695 HMMalddh_1_eutpdugrm(pf00171) SPU4 EP_NTERM
01700 H_fuchsia SPU4
01705 HMM00121_isom_1_spu(pf00121) SPU4
01710 HMMspu02502_1_spu(pf02502) SPU4
01715 P_orange SPU4
01720 H_fuchsia SPU4
01725 P_grey SPU4
01730 HMMspu01791_1_spu(pf01791) SPU4
01735 -pf01381 -
01740 -pf03323 -
01745 -pf05504 -
01750 -# -
01755 -pf03845 -
01760 -# -
01765 -# -
01770 -pf13302 -
01775 -# -
BMC-P
BMC-H
BMC-T
Aldehyde dehydrogenase
PTAC
Signature enzyme
Alcohol dehydrogenase
Regulator

Bacillus sp. AFS006103. Bacteria; Firmicutes; Bacilli; Bacillales; Bacillaceae; Bacillus.

CN481__locus_1

SPU4

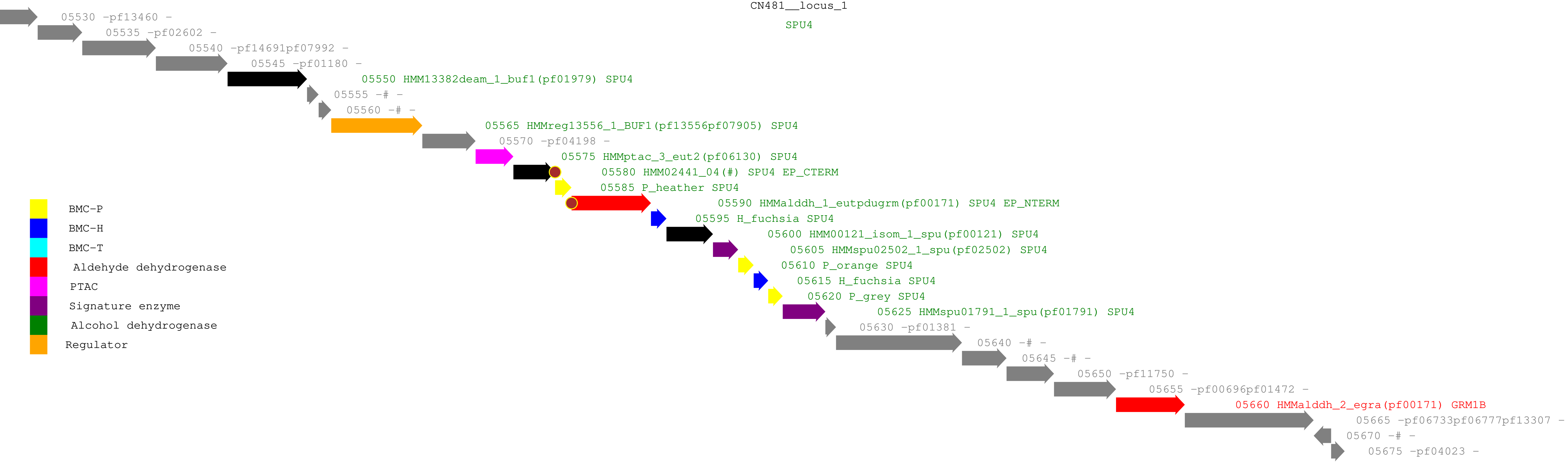

Bacillus sp. AFS073361. Bacteria; Firmicutes; Bacilli; Bacillales; Bacillaceae; Bacillus.

COJ96__locus_1

SPU4

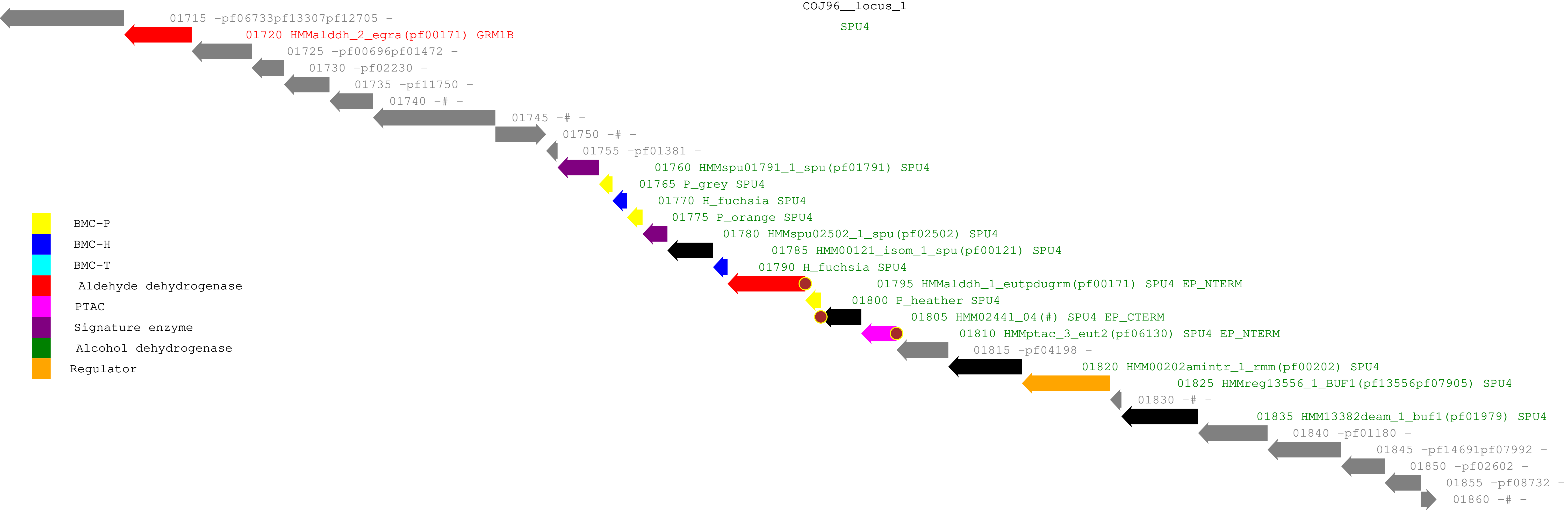

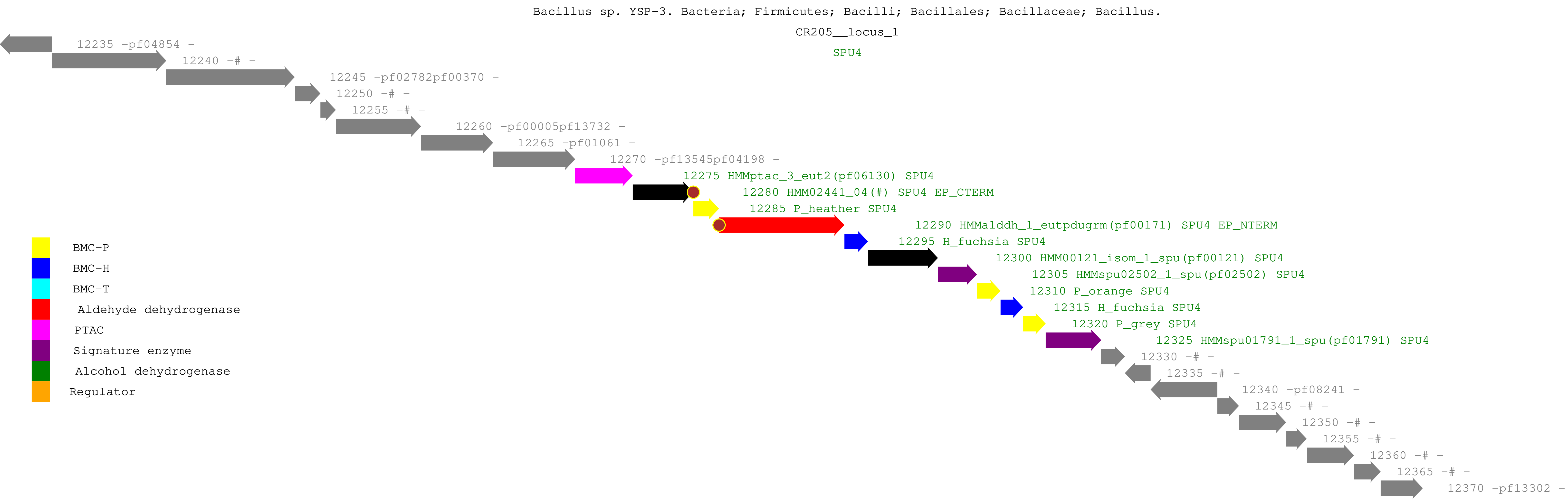

Bacillus sp. YSP-3. Bacteria; Firmicutes; Bacilli; Bacillales; Bacillaceae; Bacillus.
CR205__locus_1
SPU4
12235 -pf04854 -
12240 -# -
12245 -pf02782pf00370 -
12250 -# -
12255 -# -
12260 -pf00005pf13732 -
12265 -pf01061 -
12270 -pf13545pf04198 -
12275 HMMptac_3_eut2(pf06130) SPU4
12280 HMM02441_04(#) SPU4 EP_CTERM
12285 P_heather SPU4
12290 HMMaldh_1_eutpdugrm(pf00171) SPU4 EP_NTERM
12295 H_fuchsia SPU4
12300 HMM00121_isom_1_spu(pf00121) SPU4
12305 HMMspu02502_1_spu(pf02502) SPU4
12310 P_orange SPU4
12315 H_fuchsia SPU4
12320 P_grey SPU4
12325 HMMspu01791_1_spu(pf01791) SPU4
12330 -# -
12335 -# -
12340 -pf08241 -
12345 -# -
12350 -# -
12355 -# -
12360 -# -
12365 -# -
12370 -pf13302 -
BMC-P
BMC-H
BMC-T
Aldehyde dehydrogenase
PTAC
Signature enzyme
Alcohol dehydrogenase
Regulator

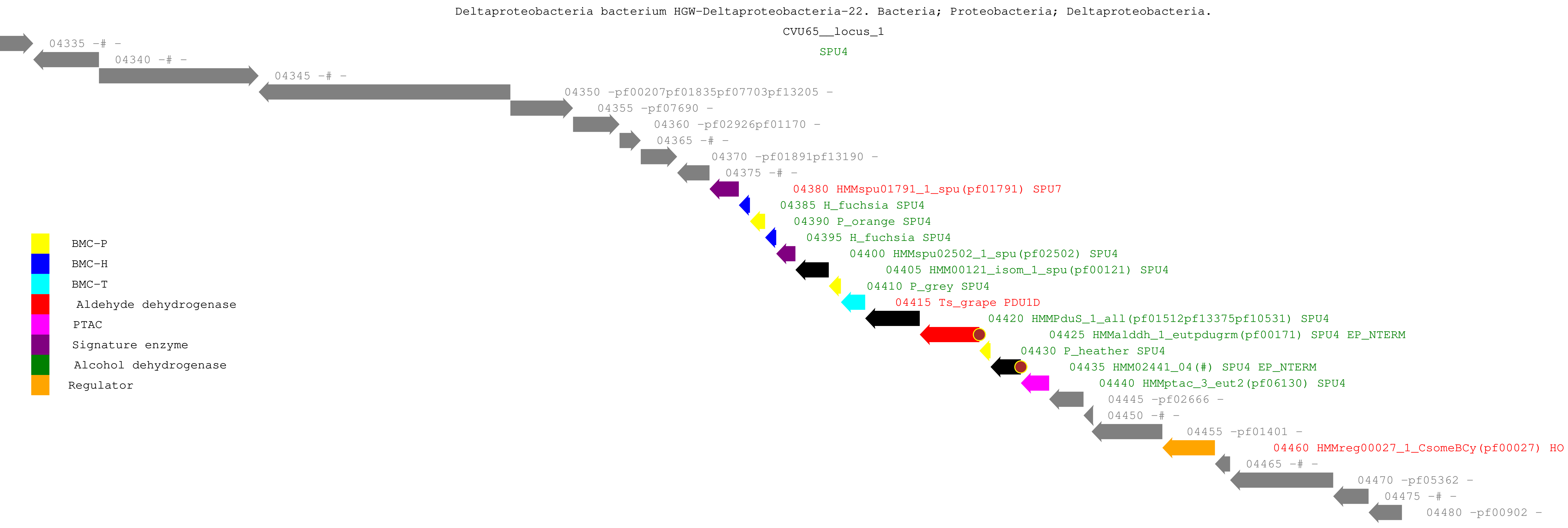
Deltaproteobacteria bacterium HGW-Deltaproteobacteria-22. Bacteria; Proteobacteria; Deltaproteobacteria.
CVU65__locus_1
SPU4
04335 -# -
04340 -# -
04345 -# -
04350 -pf00207pf01835pf07703pf13205 -
04355 -pf07690 -
04360 -pf02926pf01170 -
04365 -# -
04370 -pf01891pf13190 -
04375 -# -
04380 HMMspu01791_1_spu(pf01791) SPU7
04385 H_fuchsia SPU4
04390 P_orange SPU4
04395 H_fuchsia SPU4
04400 HMMspu02502_1_spu(pf02502) SPU4
04405 HMM00121_isom_1_spu(pf00121) SPU4
04410 P_grey SPU4
04415 Ts_grape PDU1D
04420 HMMPduS_1_all(pf01512pf13375pf10531) SPU4
04425 HMMalddh_1_eutpdugrm(pf00171) SPU4 EP_NTERM
04430 P_heather SPU4
04435 HMM02441_04(#) SPU4 EP_NTERM
04440 HMMptac_3_eut2(pf06130) SPU4
04445 -pf02666 -
04450 -# -
04455 -pf01401 -
04460 HMMreg00027_1_CsomeBCy(pf00027) HO
04465 -# -
04470 -pf05362 -
04475 -# -
04480 -pf00902 -
BMC-P
BMC-H
BMC-T
Aldehyde dehydrogenase
PTAC
Signature enzyme
Alcohol dehydrogenase
Regulator

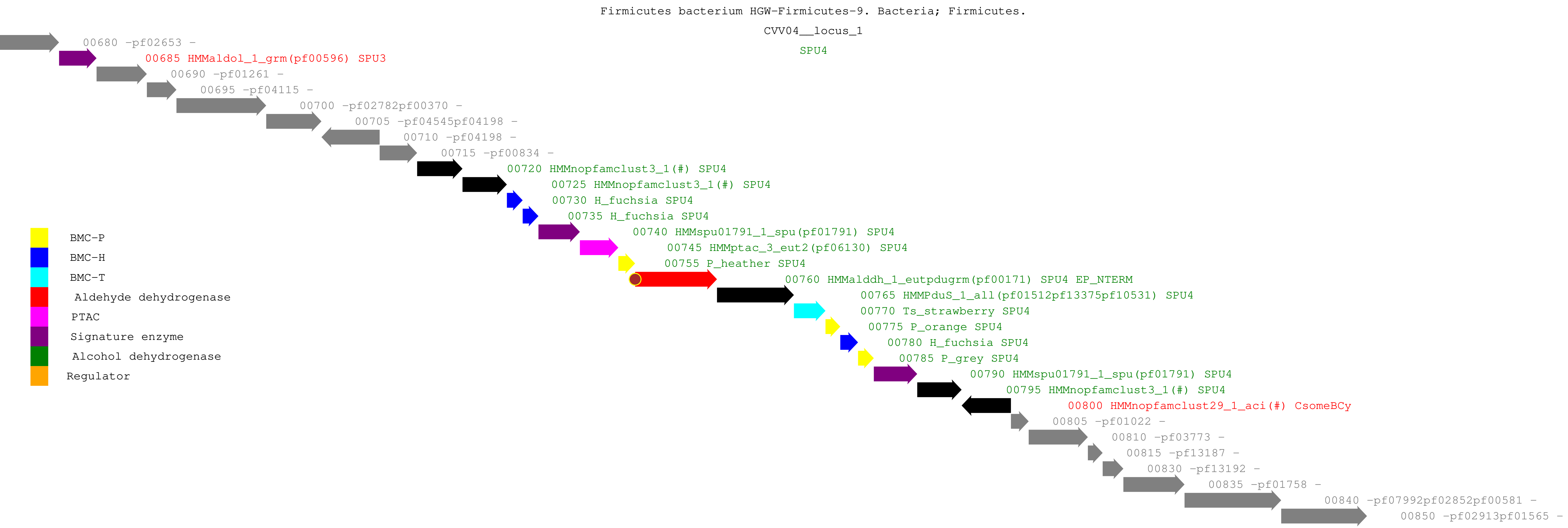
Firmicutes bacterium HGW-Firmicutes-9. Bacteria; Firmicutes.
CVV04__locus_1
SPU4
00680 -pf02653 -
00685 HMMaldol_1_grm(pf00596) SPU3
00690 -pf01261 -
00695 -pf04115 -
00700 -pf02782pf00370 -
00705 -pf04545pf04198 -
00710 -pf04198 -
00715 -pf00834 -
00720 HMMnopfamclust3_1(#) SPU4
00725 HMMnopfamclust3_1(#) SPU4
00730 H_fuchsia SPU4
00735 H_fuchsia SPU4
00740 HMMspu01791_1_spu(pf01791) SPU4
00745 HMMptac_3_eut2(pf06130) SPU4
00755 P_heather SPU4
00760 HMMaldh_1_eutpdugrm(pf00171) SPU4 EP_NTERM
00765 HMMPduS_1_all(pf01512pf13375pf10531) SPU4
00770 Ts_strawberry SPU4
00775 P_orange SPU4
00780 H_fuchsia SPU4
00785 P_grey SPU4
00790 HMMspu01791_1_spu(pf01791) SPU4
00795 HMMnopfamclust3_1(#) SPU4
00800 HMMnopfamclust29_1_aci(#) CsomeBCy
00805 -pf01022 -
00810 -pf03773 -
00815 -pf13187 -
00830 -pf13192 -
00835 -pf01758 -
00840 -pf07992pf02852pf00581 -
00850 -pf02913pf01565 -
BMC-P
BMC-H
BMC-T
Aldehyde dehydrogenase
PTAC
Signature enzyme
Alcohol dehydrogenase
Regulator

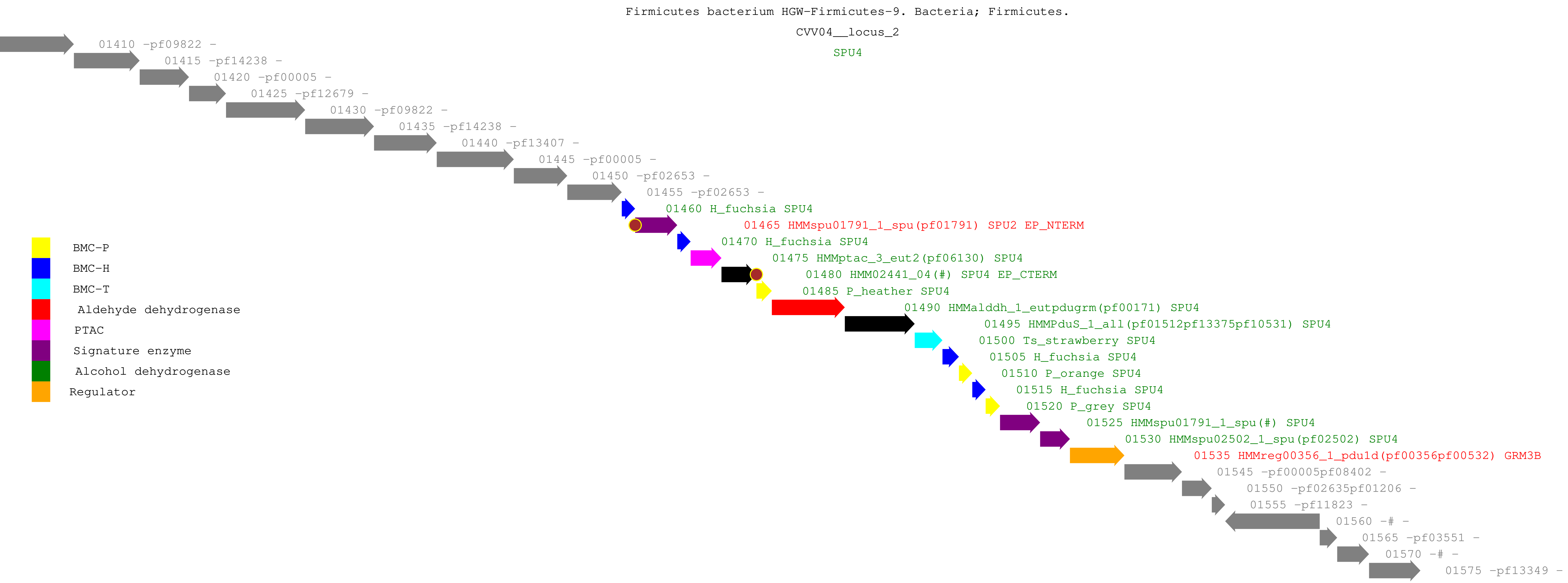

Firmicutes bacterium HGW-Firmicutes-9. Bacteria; Firmicutes.
CVV04__locus_2
SPU4
01410 -pf09822 -
01415 -pf14238 -
01420 -pf00005 -
01425 -pf12679 -
01430 -pf09822 -
01435 -pf14238 -
01440 -pf13407 -
01445 -pf00005 -
01450 -pf02653 -
01455 -pf02653 -
01460 H_fuchsia SPU4
01465 HMMspu01791_1_spu(pf01791) SPU2 EP_NTERM
01470 H_fuchsia SPU4
01475 HMMptac_3_eut2(pf06130) SPU4
01480 HMM02441_04(#) SPU4 EP_CTERM
01485 P_heather SPU4
01490 HMMaldh_1_eutpdugrm(pf00171) SPU4
01495 HMMPduS_1_all(pf01512pf13375pf10531) SPU4
01500 Ts_strawberry SPU4
01505 H_fuchsia SPU4
01510 P_orange SPU4
01515 H_fuchsia SPU4
01520 P_grey SPU4
01525 HMMspu01791_1_spu(#) SPU4
01530 HMMspu02502_1_spu(pf02502) SPU4
01535 HMMreg00356_1_pdu1d(pf00356pf00532) GRM3B
01545 -pf00005pf08402 -
01550 -pf02635pf01206 -
01555 -pf11823 -
01560 -# -
01565 -pf03551 -
01570 -# -
01575 -pf13349 -
BMC-P
BMC-H
BMC-T
Aldehyde dehydrogenase
PTAC
Signature enzyme
Alcohol dehydrogenase
Regulator

Clostridiales bacterium. Bacteria; Firmicutes; Clostridia; Clostridiales.

DBX46__locus_1_main_sat

SPU4

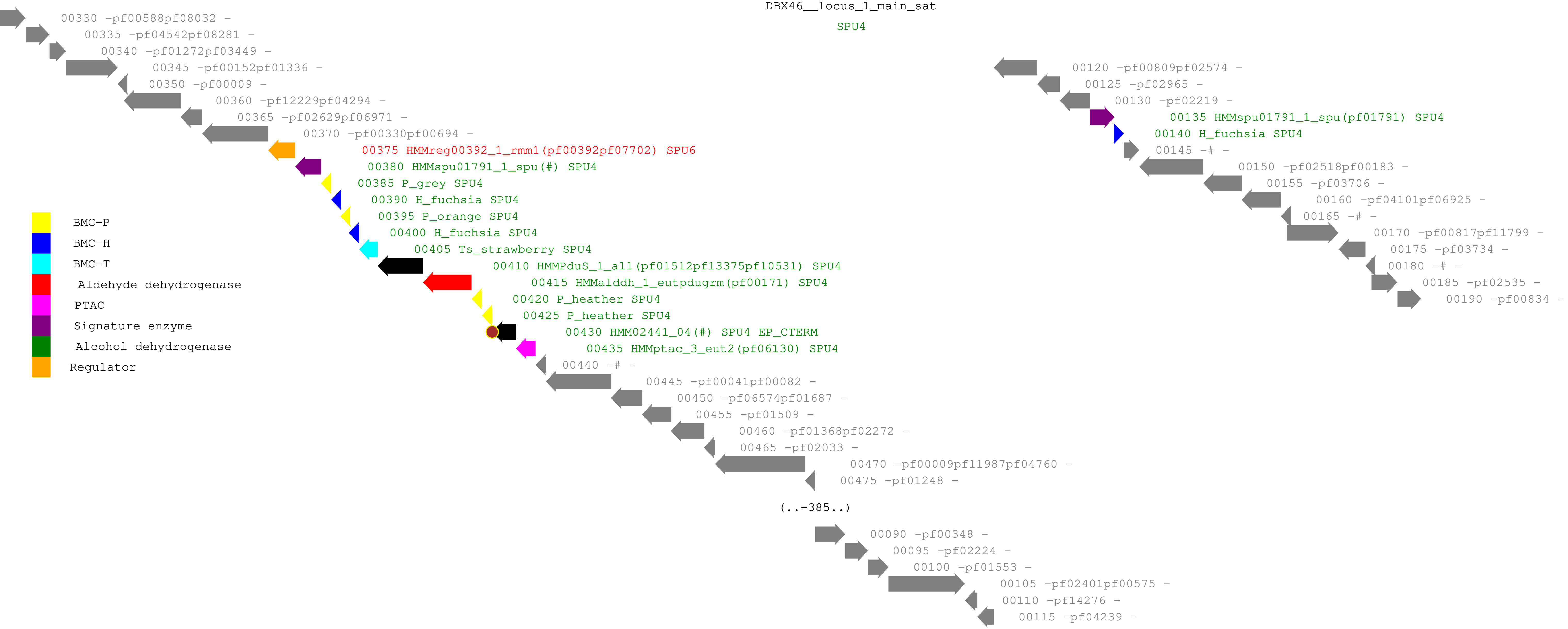

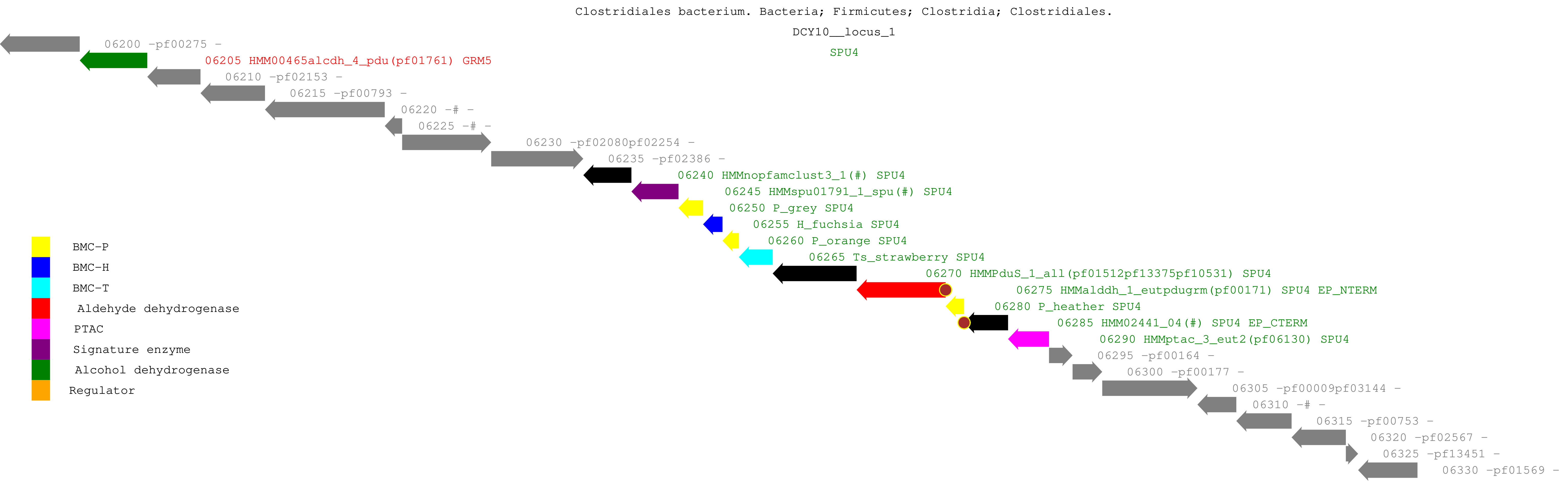

Clostridiales bacterium. Bacteria; Firmicutes; Clostridia; Clostridiales.
DCY10__locus_1
SPU4
06200 -pf00275 -
06205 HMM00465alcdh_4_pdu(pf01761) GRM5
06210 -pf02153 -
06215 -pf00793 -
06220 -# -
06225 -# -
06230 -pf02080pf02254 -
06235 -pf02386 -
06240 HMMnopfamclust3_1(#) SPU4
06245 HMMspu01791_1_spu(#) SPU4
06250 P_grey SPU4
06255 H_fuchsia SPU4
06260 P_orange SPU4
06265 Ts_strawberry SPU4
06270 HMMPduS_1_all(pf01512pf13375pf10531) SPU4
06275 HMMalddh_1_eutpdugrm(pf00171) SPU4 EP_NTERM
06280 P_heather SPU4
06285 HMM02441_04(#) SPU4 EP_CTERM
06290 HMMptac_3_eut2(pf06130) SPU4
06295 -pf00164 -
06300 -pf00177 -
06305 -pf00009pf03144 -
06310 -# -
06315 -pf00753 -
06320 -pf02567 -
06325 -pf13451 -
06330 -pf01569 -
BMC-P
BMC-H
BMC-T
Aldehyde dehydrogenase
PTAC
Signature enzyme
Alcohol dehydrogenase
Regulator

Clostridiales bacterium. Bacteria; Firmicutes; Clostridia; Clostridiales.

DEB31__locus_1

SPU4

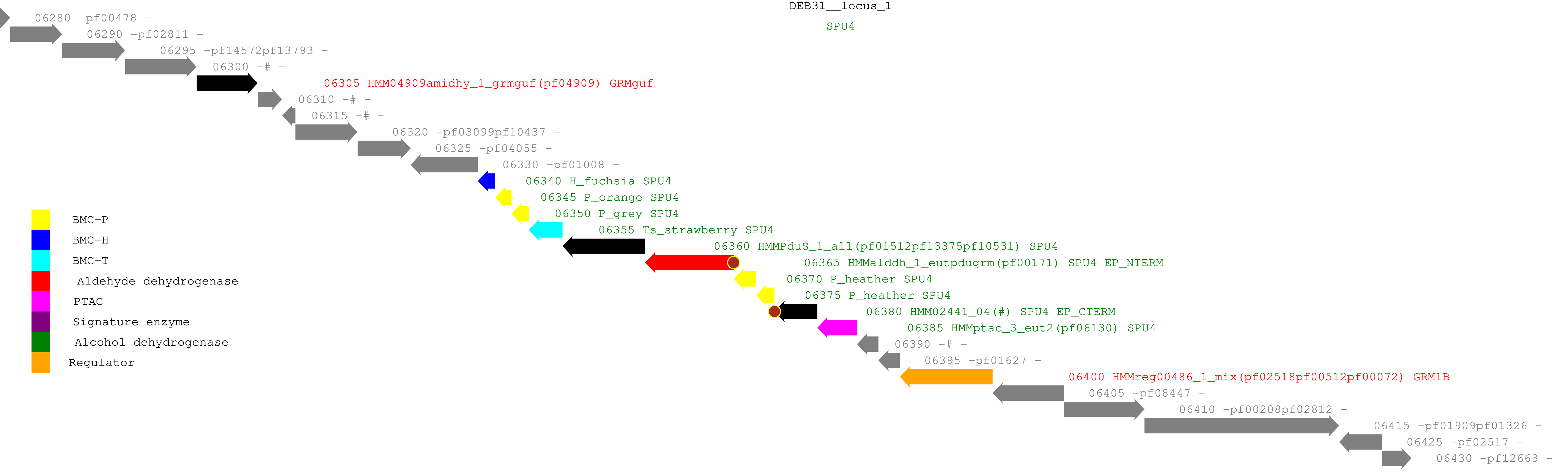

Pseudogracilibacillus auburnensis. Bacteria; Firmicutes; Bacilli; Bacillales; Bacillaceae;

DFR56__locus_1

SPU4

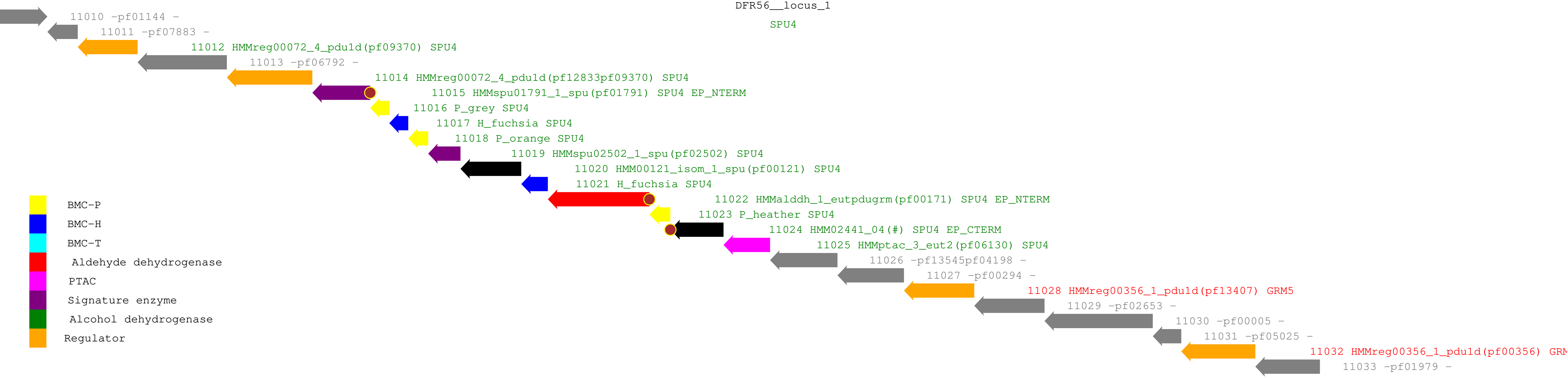

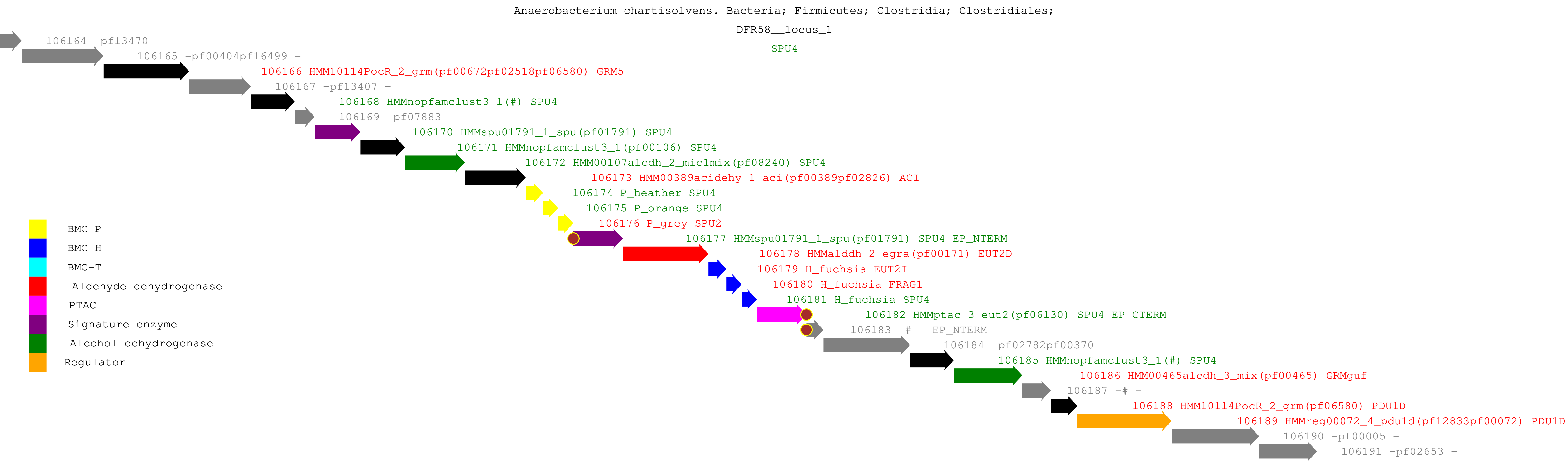
Anaerobacterium chartisolvens. Bacteria; Firmicutes; Clostridia; Clostridiales;
DFR58__locus_1
SPU4
106164 -pf13470 -
106165 -pf00404pf16499 -
106166 HMM10114PocR_2_grm(pf00672pf02518pf06580) GRM5
106167 -pf13407 -
106168 HMMnopfamclust3_1(#) SPU4
106169 -pf07883 -
106170 HMMspu01791_1_spu(pf01791) SPU4
106171 HMMnopfamclust3_1(pf00106) SPU4
106172 HMM00107alcdh_2_mic1mix(pf08240) SPU4
106173 HMM00389acidehy_1_aci(pf00389pf02826) ACI
106174 P_heather SPU4
106175 P_orange SPU4
106176 P_grey SPU2
106177 HMMspu01791_1_spu(pf01791) SPU4 EP_NTERM
106178 HMMalddh_2_egra(pf00171) EUT2D
106179 H_fuchsia EUT2I
106180 H_fuchsia FRAG1
106181 H_fuchsia SPU4
106182 HMMptac_3_eut2(pf06130) SPU4 EP_CTERM
106183 -# - EP_NTERM
106184 -pf02782pf00370 -
106185 HMMnopfamclust3_1(#) SPU4
106186 HMM00465alcdh_3_mix(pf00465) GRMguf
106187 -# -
106188 HMM10114PocR_2_grm(pf06580) PDU1D
106189 HMMreg00072_4_pdu1d(pf12833pf00072) PDU1D
106190 -pf00005 -
106191 -pf02653 -
BMC-P
BMC-H
BMC-T
Aldehyde dehydrogenase
PTAC
Signature enzyme
Alcohol dehydrogenase
Regulator

Clostridiales bacterium. Bacteria; Firmicutes; Clostridia; Clostridiales.

DIW49__locus_1

SPU4

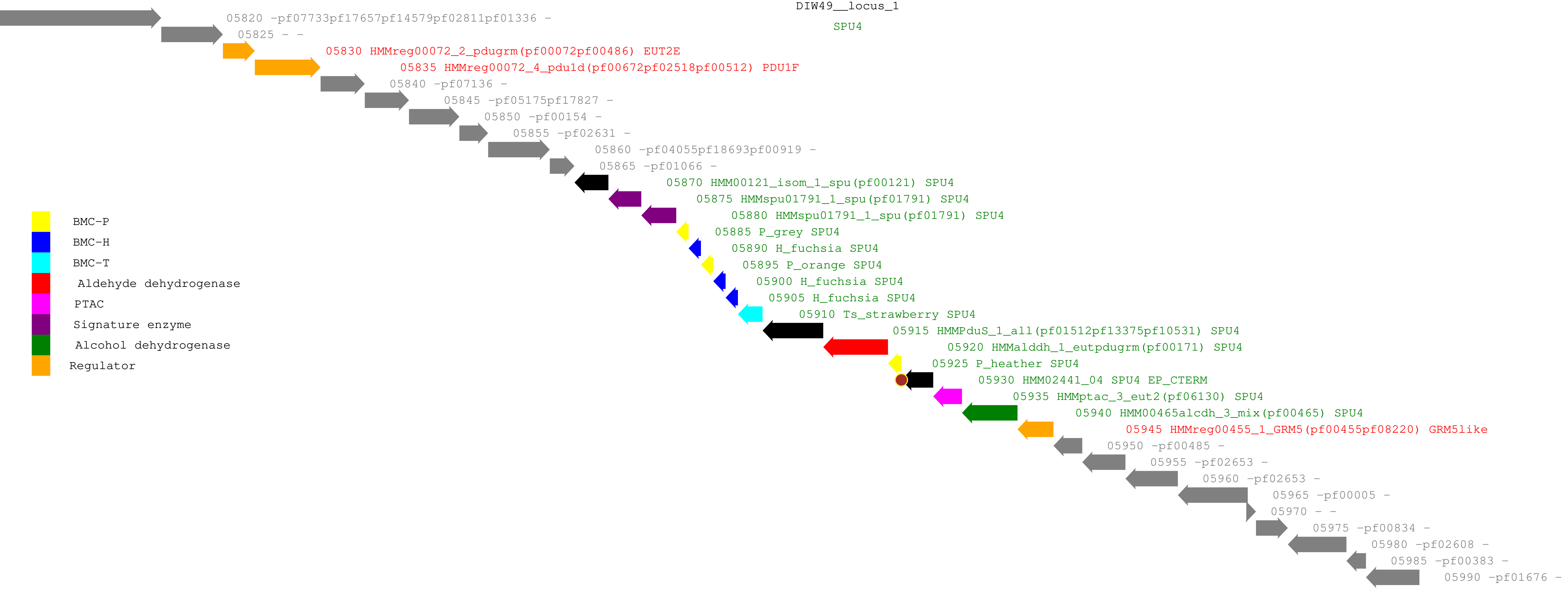

Clostridium sp. AF15-17LB. Bacteria; Firmicutes; Clostridia; Clostridiales; Clostridiaceae;

DWW31__locus_1

SPU4

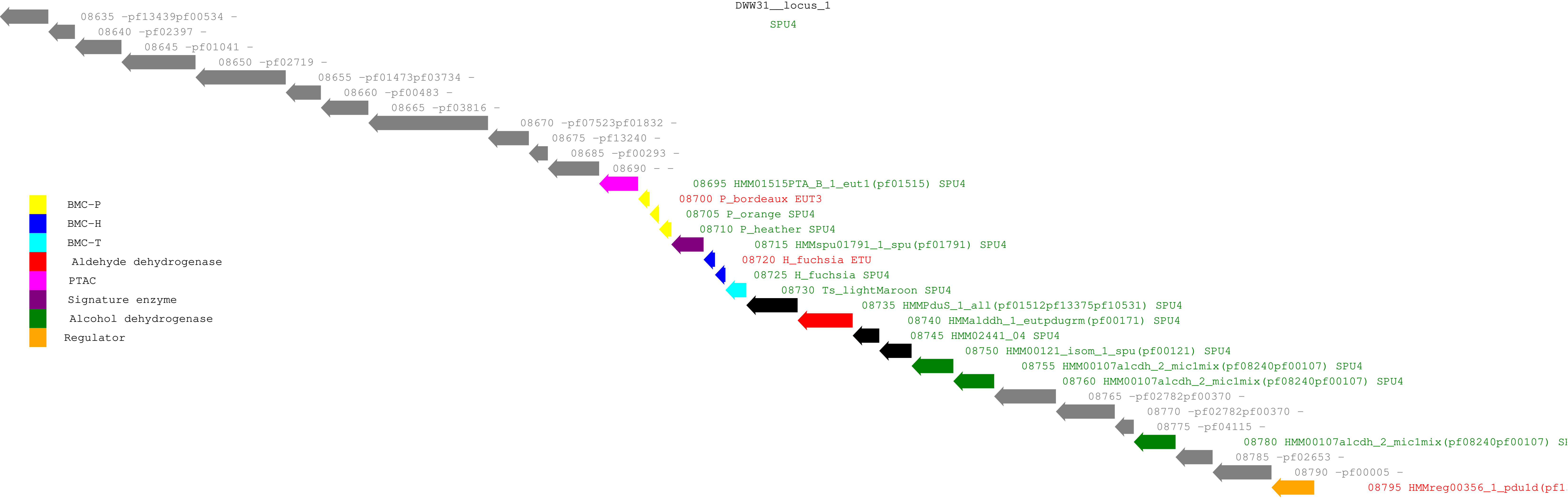

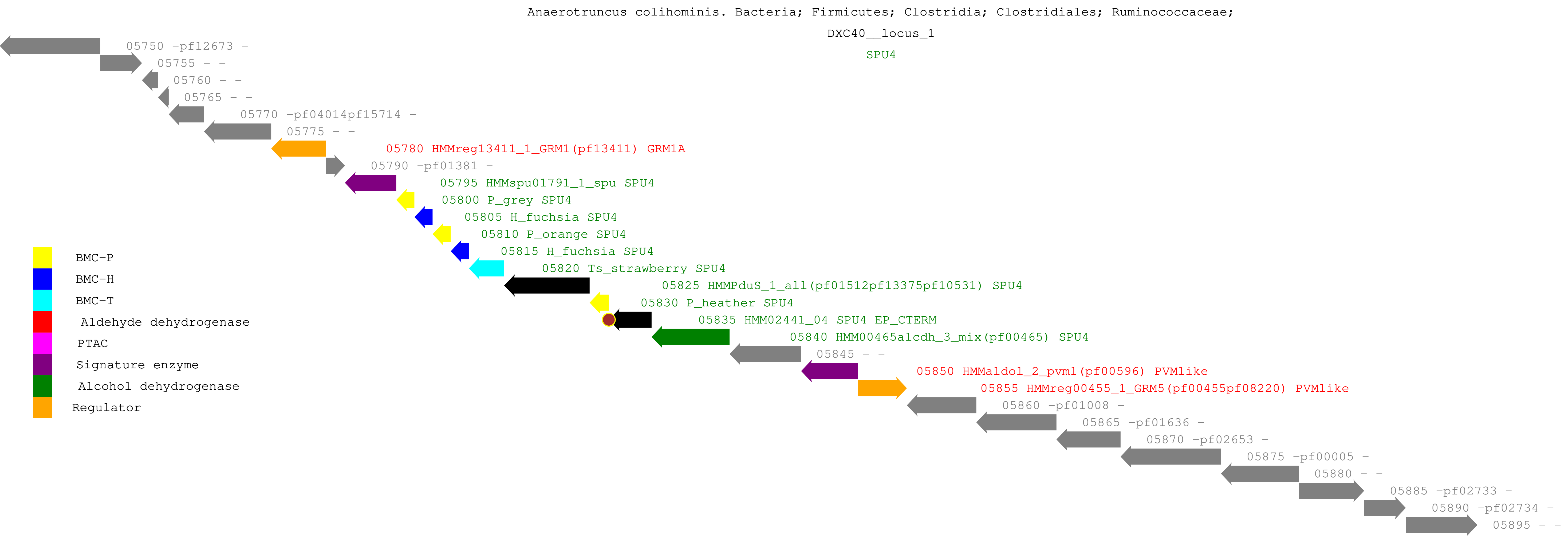
Anaerotruncus colihominis. Bacteria; Firmicutes; Clostridia; Clostridiales; Ruminococcaceae;
DXC40__locus_1
SPU4
05750 -pf12673 -
05755 - -
05760 - -
05765 - -
05770 -pf04014pf15714 -
05775 - -
05780 HMMreg13411_1_GRM1(pf13411) GRM1A
05790 -pf01381 -
05795 HMMspu01791_1_spu SPU4
05800 P_grey SPU4
05805 H_fuchsia SPU4
05810 P_orange SPU4
05815 H_fuchsia SPU4
05820 Ts_strawberry SPU4
05825 HMMPduS_1_all(pf01512pf13375pf10531) SPU4
05830 P_heather SPU4
05835 HMM02441_04 SPU4 EP_CTERM
05840 HMM00465alcdh_3_mix(pf00465) SPU4
05845 - -
05850 HMMaldol_2_pvm1(pf00596) PVMlike
05855 HMMreg00455_1_GRM5(pf00455pf08220) PVMlike
05860 -pf01008 -
05865 -pf01636 -
05870 -pf02653 -
05875 -pf00005 -
05880 - -
05885 -pf02733 -
05890 -pf02734 -
05895 - -
BMC-P
BMC-H
BMC-T
Aldehyde dehydrogenase
PTAC
Signature enzyme
Alcohol dehydrogenase
Regulator

Bacillus sp. WN066. Bacteria; Firmicutes; Bacilli; Bacillales; Bacillaceae; Bacillus.

E2K98__locus_1

SPU4

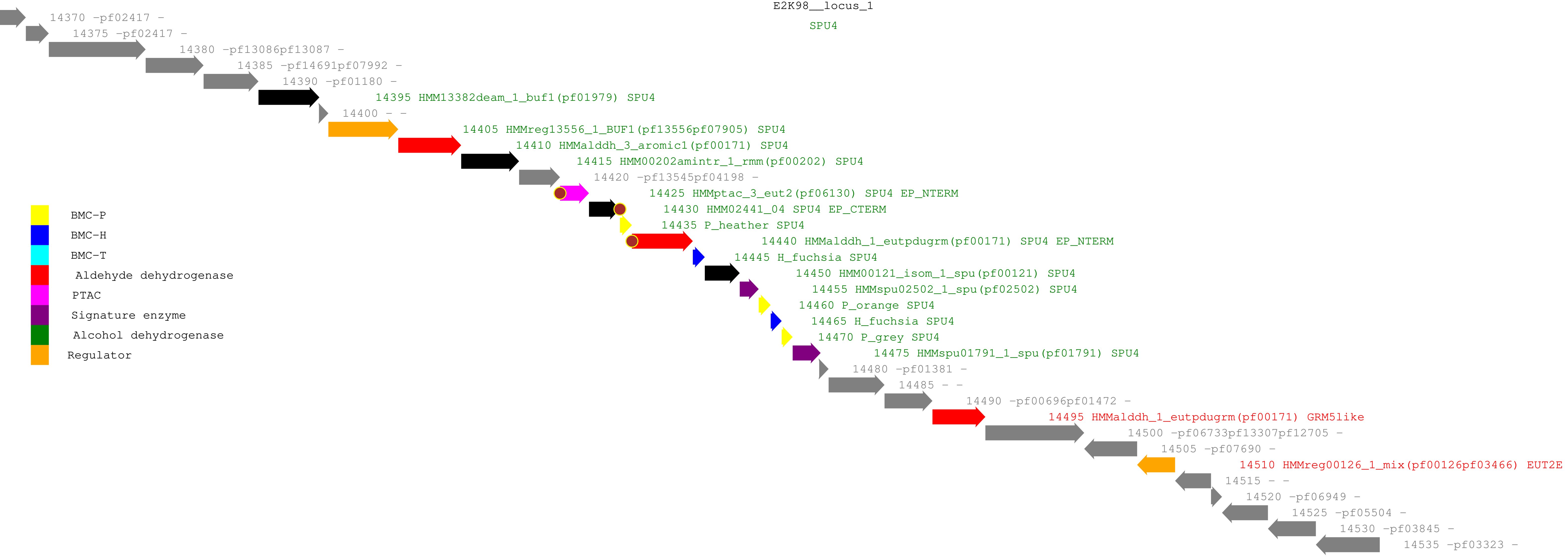

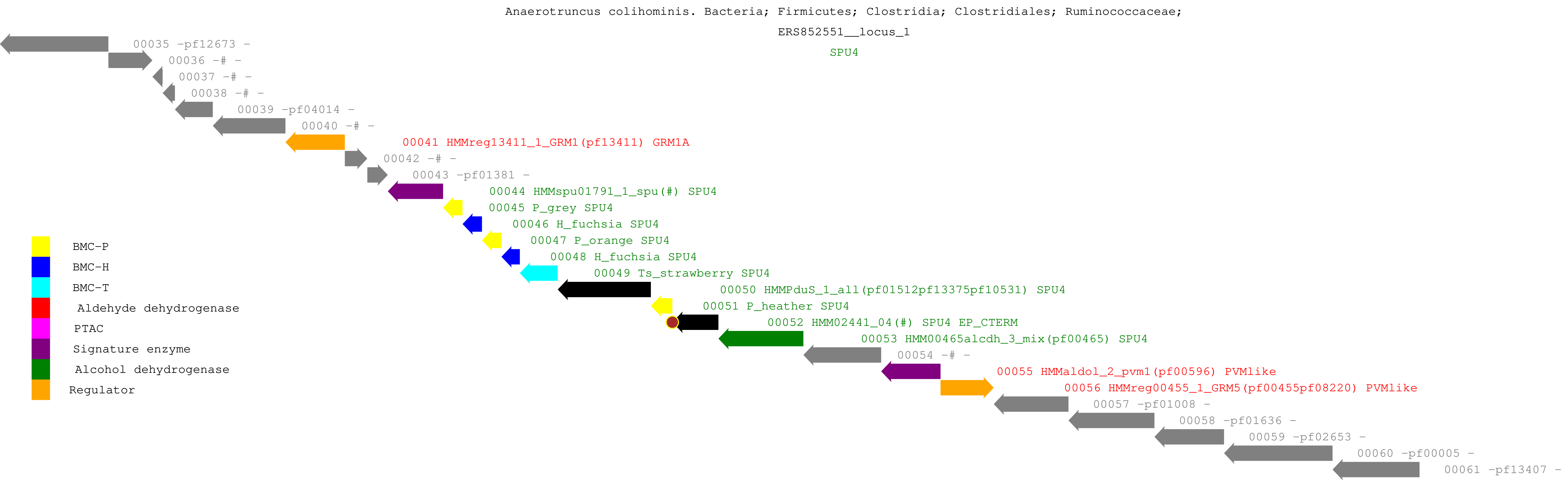
Anaerotruncus colihominis. Bacteria; Firmicutes; Clostridia; Clostridiales; Ruminococcaceae;
ERS852551__locus_1
SPU4
00035 -pf12673 -
00036 -# -
00037 -# -
00038 -# -
00039 -pf04014 -
00040 -# -
00041 HMMreg13411_1_GRM1(pf13411) GRM1A
00042 -# -
00043 -pf01381 -
00044 HMMspu01791_1_spu(#) SPU4
00045 P_grey SPU4
00046 H_fuchsia SPU4
00047 P_orange SPU4
00048 H_fuchsia SPU4
00049 Ts_strawberry SPU4
00050 HMMPduS_1_all(pf01512pf13375pf10531) SPU4
00051 P_heather SPU4
00052 HMM02441_04(#) SPU4 EP_CTERM
00053 HMM00465alcdh_3_mix(pf00465) SPU4
00054 -# -
00055 HMMaldol_2_pvm1(pf00596) PVMlike
00056 HMMreg00455_1_GRM5(pf00455pf08220) PVMlike
00057 -pf01008 -
00058 -pf01636 -
00059 -pf02653 -
00060 -pf00005 -
00061 -pf13407 -
BMC-P
BMC-H
BMC-T
Aldehyde dehydrogenase
PTAC
Signature enzyme
Alcohol dehydrogenase
Regulator

Bacillus sp. S3. Bacteria; Firmicutes; Bacilli; Bacillales; Bacillaceae; Bacillus.

FAY30__locus_1

SPU4

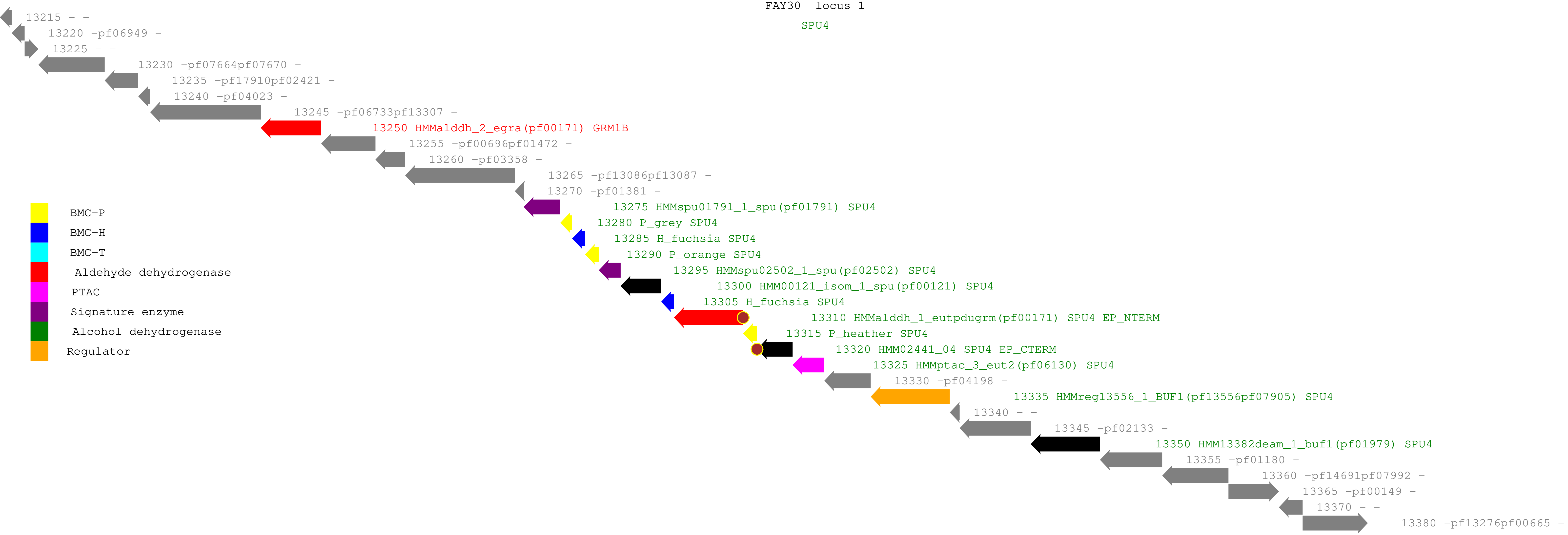

Bacillus horikoshii. Bacteria; Firmicutes; Bacilli; Bacillales; Bacillaceae; Bacillus.

FZC75__locus_1

SPU4

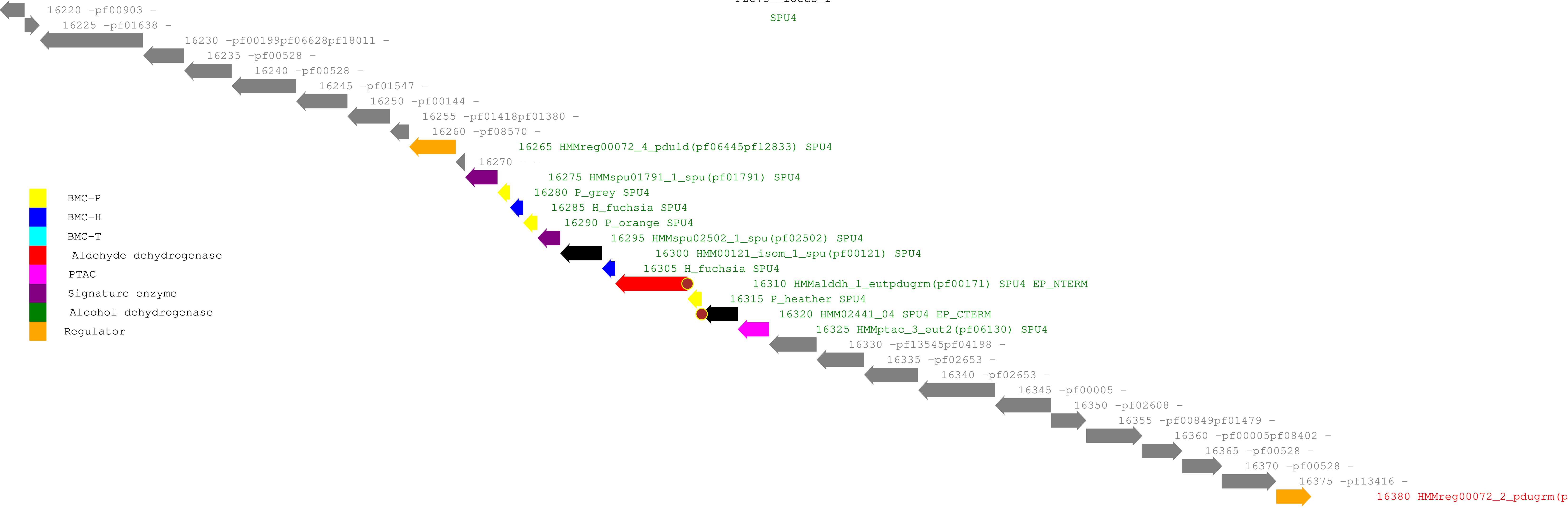

Bacillus horikoshii. Bacteria; Firmicutes; Bacilli; Bacillales; Bacillaceae; Bacillus.

FZC76__locus_1

SPU4

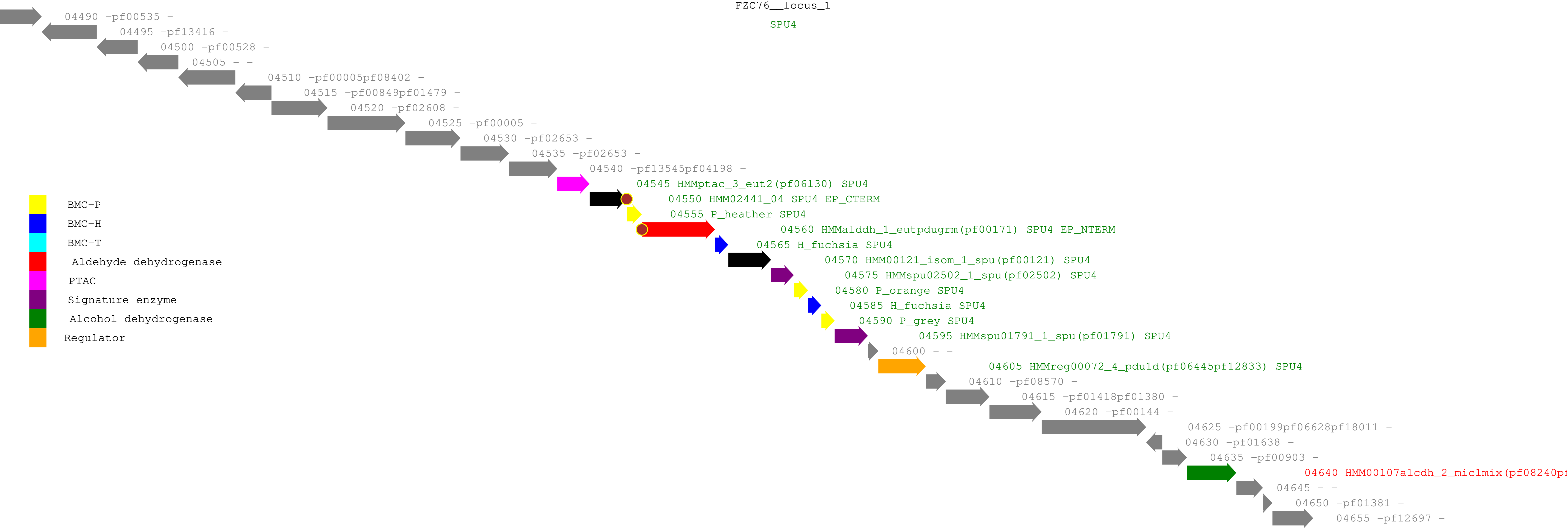

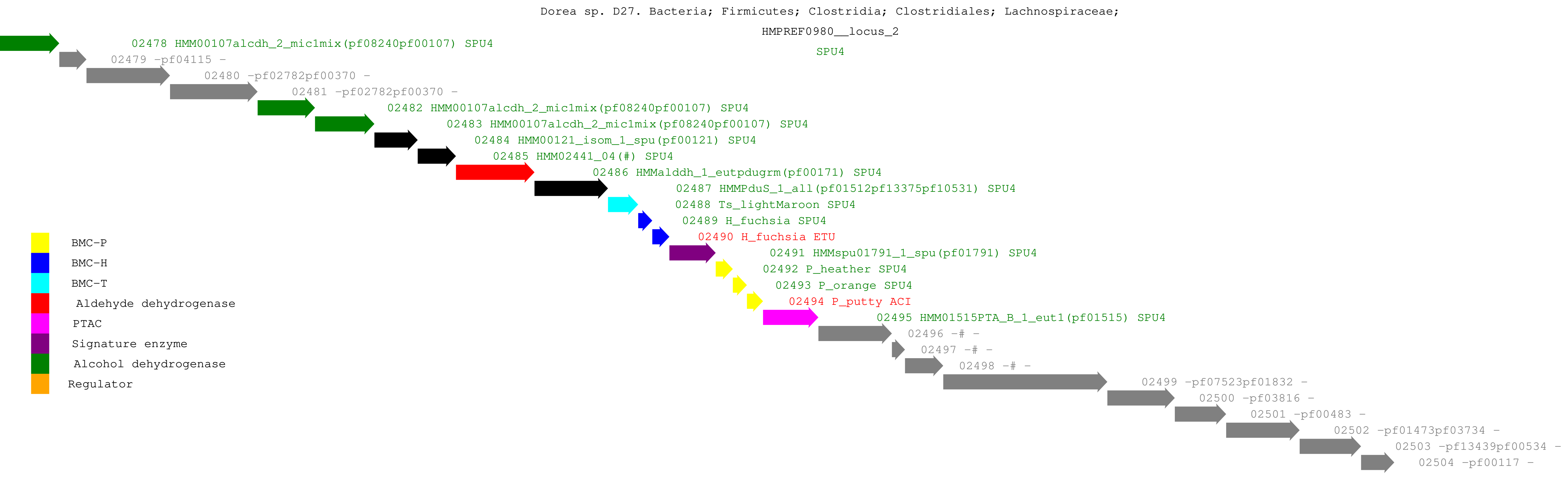
Dorea sp. D27. Bacteria; Firmicutes; Clostridia; Clostridiales; Lachnospiraceae;
HMPREF0980__locus_2
SPU4
02478 HMM00107alcdh_2_mic1mix(pf08240pf00107) SPU4
02479 -pf04115 -
02480 -pf02782pf00370 -
02481 -pf02782pf00370 -
02482 HMM00107alcdh_2_mic1mix(pf08240pf00107) SPU4
02483 HMM00107alcdh_2_mic1mix(pf08240pf00107) SPU4
02484 HMM00121_isom_1_spu(pf00121) SPU4
02485 HMM02441_04(#) SPU4
02486 HMMaldh_1_eutpdugrm(pf00171) SPU4
02487 HMMPduS_1_all(pf01512pf13375pf10531) SPU4
02488 Ts_lightMaroon SPU4
02489 H_fuchsia SPU4
02490 H_fuchsia ETU
02491 HMMspu01791_1_spu(pf01791) SPU4
02492 P_heather SPU4
02493 P_orange SPU4
02494 P_putty ACI
02495 HMM01515PTA_B_1_eut1(pf01515) SPU4
02496 -# -
02497 -# -
02498 -# -
02499 -pf07523pf01832 -
02500 -pf03816 -
02501 -pf00483 -
02502 -pf01473pf03734 -
02503 -pf13439pf00534 -
02504 -pf00117 -
BMC-P
BMC-H
BMC-T
Aldehyde dehydrogenase
PTAC
Signature enzyme
Alcohol dehydrogenase
Regulator

Candidatus Izimaplasma sp. HR2. Bacteria; Tenericutes; Candidatus Izimaplasma.

KQ78__locus_1

SPU4

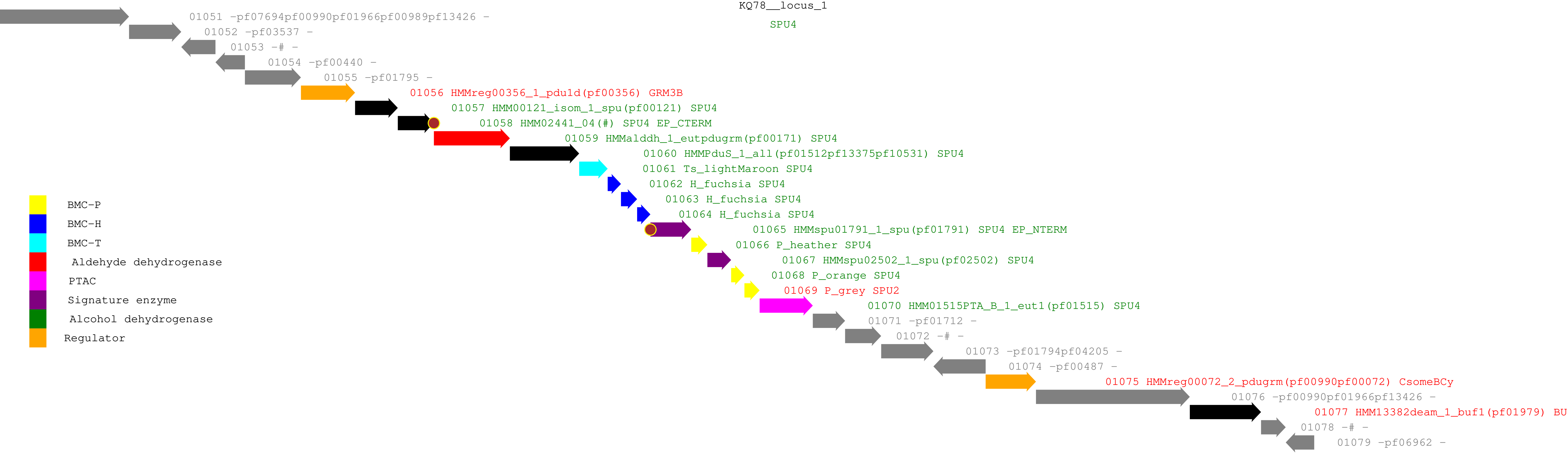

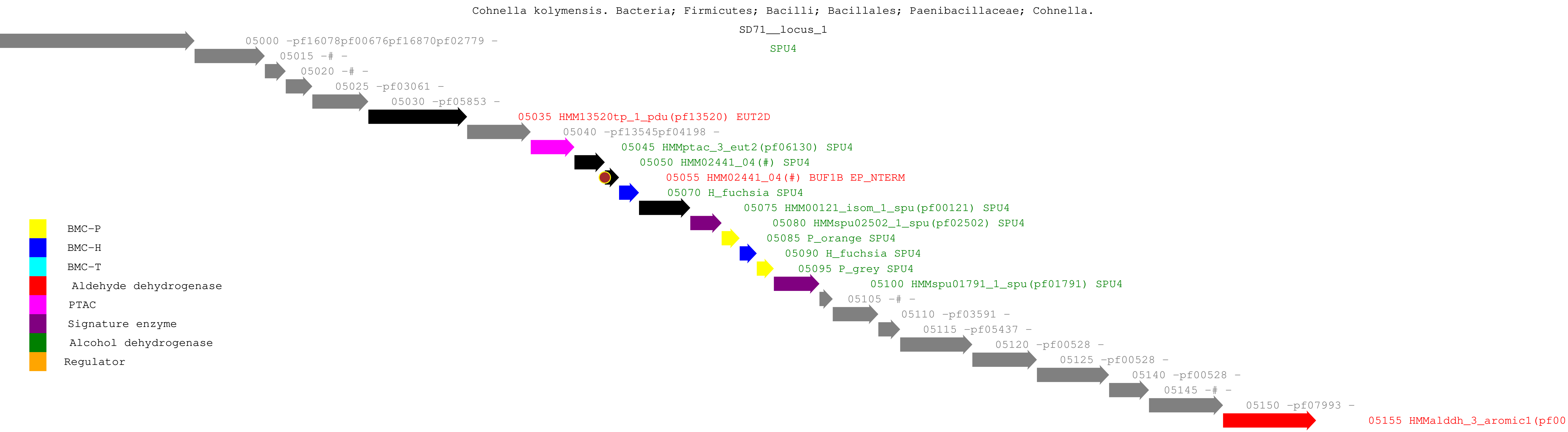
Cohnella kolymensis. Bacteria; Firmicutes; Bacilli; Bacillales; Paenibacillaceae; Cohnella.
SD71__locus_1
SPU4
05000 -pf16078pf00676pf16870pf02779 -
05015 -# -
05020 -# -
05025 -pf03061 -
05030 -pf05853 -
05035 HMM13520tp_1_pdu(pf13520) EUT2D
05040 -pf13545pf04198 -
05045 HMMptac_3_eut2(pf06130) SPU4
05050 HMM02441_04(#) SPU4
05055 HMM02441_04(#) BUF1B EP_NTERM
05070 H_fuchsia SPU4
05075 HMM00121_isom_1_spu(pf00121) SPU4
05080 HMMspu02502_1_spu(pf02502) SPU4
05085 P_orange SPU4
05090 H_fuchsia SPU4
05095 P_grey SPU4
05100 HMMspu01791_1_spu(pf01791) SPU4
05105 -# -
05110 -pf03591 -
05115 -pf05437 -
05120 -pf00528 -
05125 -pf00528 -
05140 -pf00528 -
05145 -# -
05150 -pf07993 -
05155 HMMaldd h_3_aromic1(pf00
BMC-P
BMC-H
BMC-T
Aldehyde dehydrogenase
PTAC
Signature enzyme
Alcohol dehydrogenase
Regulator

Gammaproteobacteria bacterium GWE2_42_36. Bacteria; Proteobacteria; Gammaproteobacteria.

A2X77__locus_1

SPU5

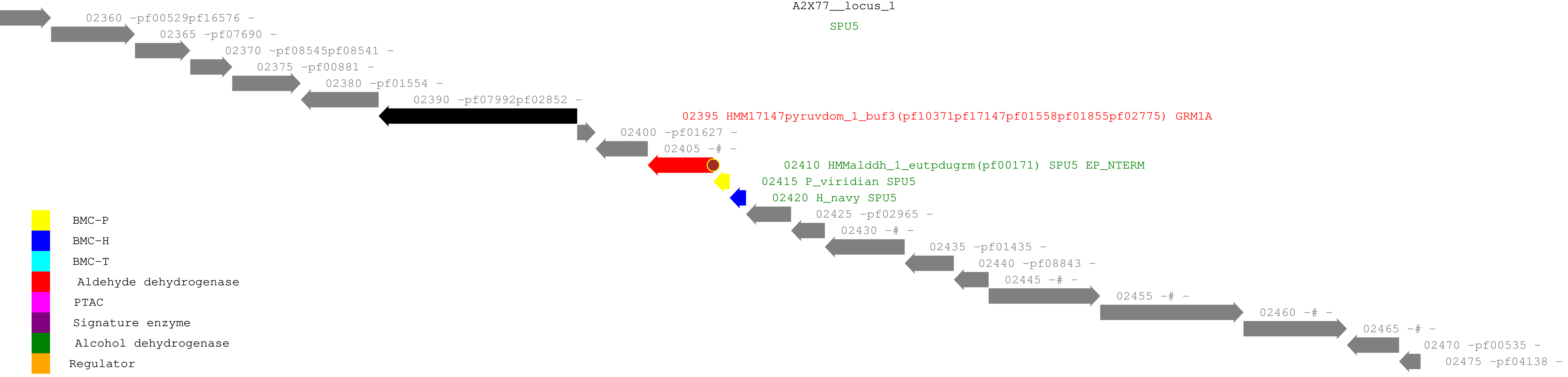

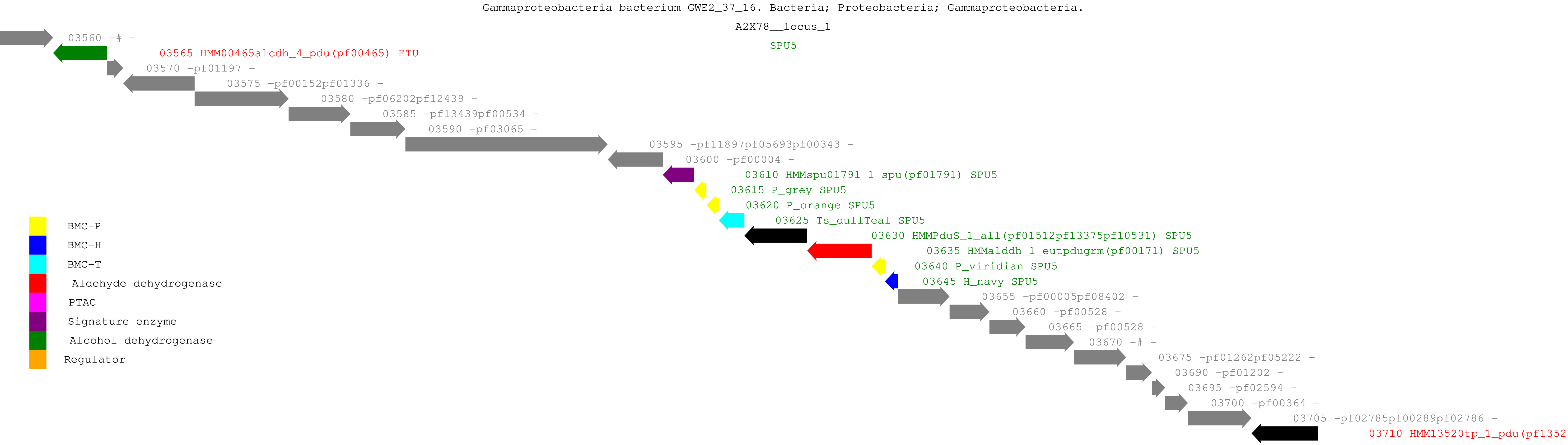
Gammaproteobacteria bacterium GWE2_37_16. Bacteria; Proteobacteria; Gammaproteobacteria.
A2X78__locus_1
SPU5
03560 -# -
03565 HMM00465alcdh_4_pdu(pf00465) ETU
03570 -pf01197 -
03575 -pf00152pf01336 -
03580 -pf06202pf12439 -
03585 -pf13439pf00534 -
03590 -pf03065 -
03595 -pf11897pf05693pf00343 -
03600 -pf00004 -
03610 HMMspu01791_1_spu(pf01791) SPU5
03615 P_grey SPU5
03620 P_orange SPU5
03625 Ts_dullTeal SPU5
03630 HMMPduS_1_all(pf01512pf13375pf10531) SPU5
03635 HMMalddh_1_eutpdugrm(pf00171) SPU5
03640 P_viridian SPU5
03645 H_navy SPU5
03655 -pf00005pf08402 -
03660 -pf00528 -
03665 -pf00528 -
03670 -# -
03675 -pf01262pf05222 -
03690 -pf01202 -
03695 -pf02594 -
03700 -pf00364 -
03705 -pf02785pf00289pf02786 -
03710 HMM13520tp_1_pdu(pf1352
BMC-P
BMC-H
BMC-T
Aldehyde dehydrogenase
PTAC
Signature enzyme
Alcohol dehydrogenase
Regulator

Bdellovibrionales bacterium GWA2_49_15. Bacteria; Proteobacteria; Deltaproteobacteria; Bdellovibrionales.

A2X86__locus_1

SPU5

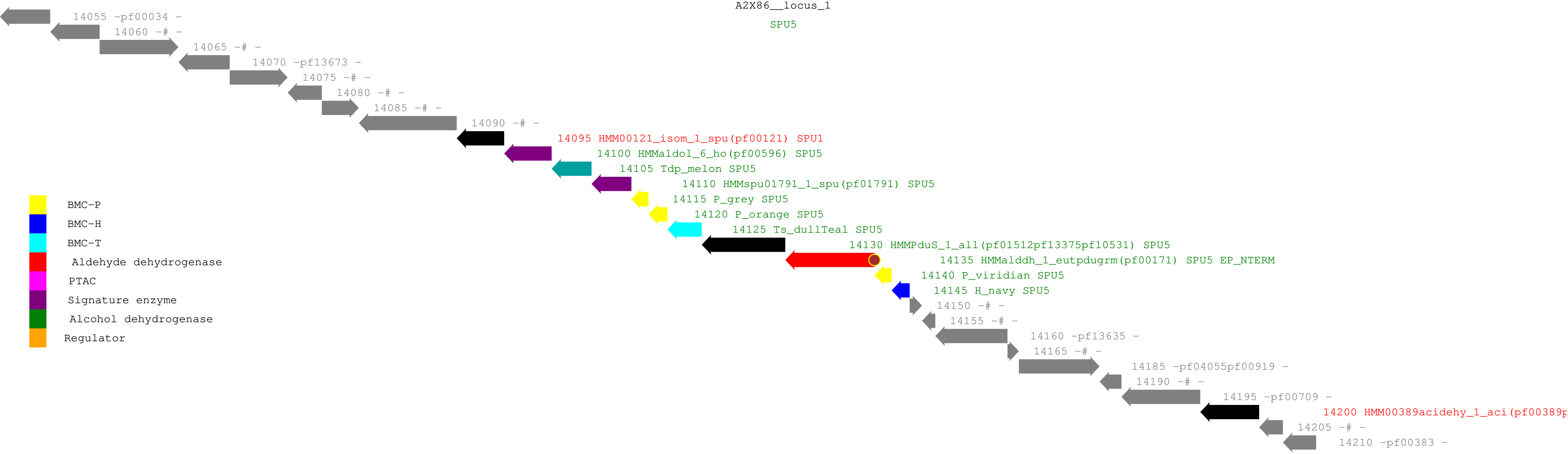

Gammaproteobacteria bacterium RIFCSPHIGHO2_02_FULL_42_13. Bacteria; Proteobacteria; Gammaproteobacteria.

A3C55__locus_1

SPU5

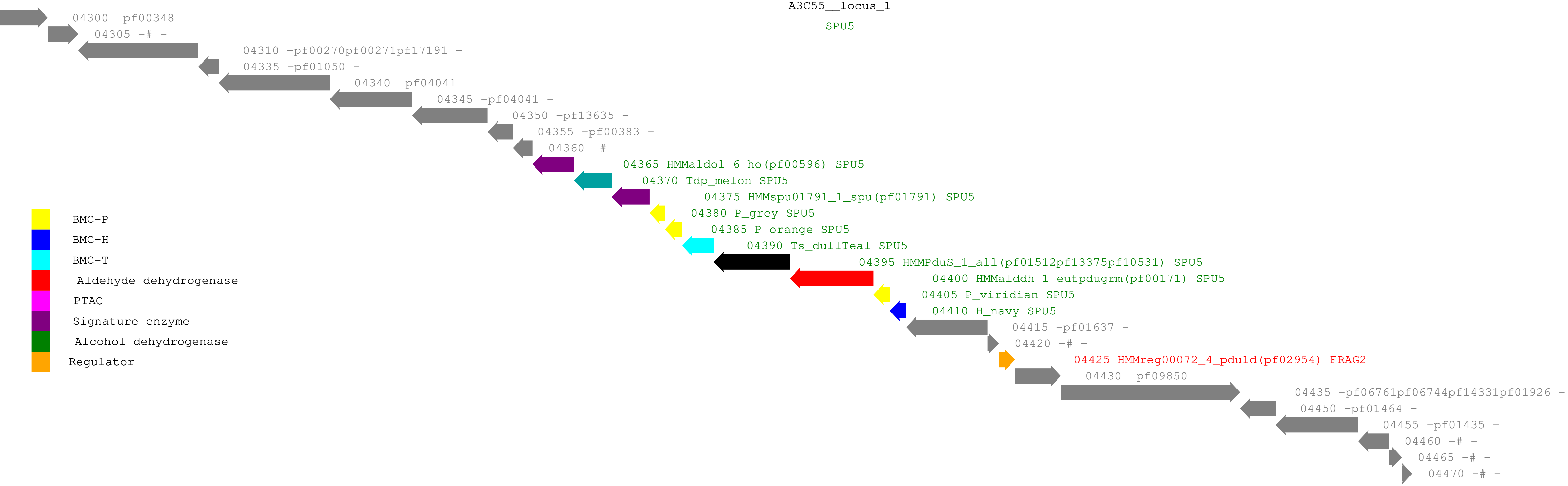

Deltaproteobacteria bacterium RIFCSPLOWO2_02_FULL_47_10. Bacteria; Proteobacteria; Deltaproteobacteria.

A3I09__locus_1

SPU5

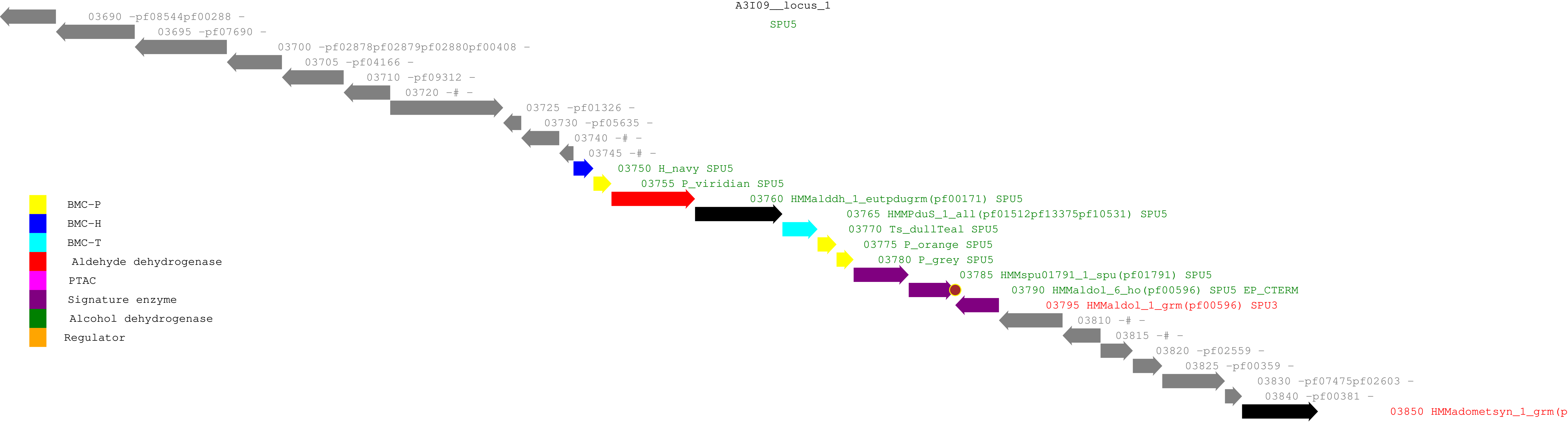

Dehalococcoidia bacterium. Bacteria; Chloroflexi; Dehalococcoidia.

CL889__locus_1

SPU5

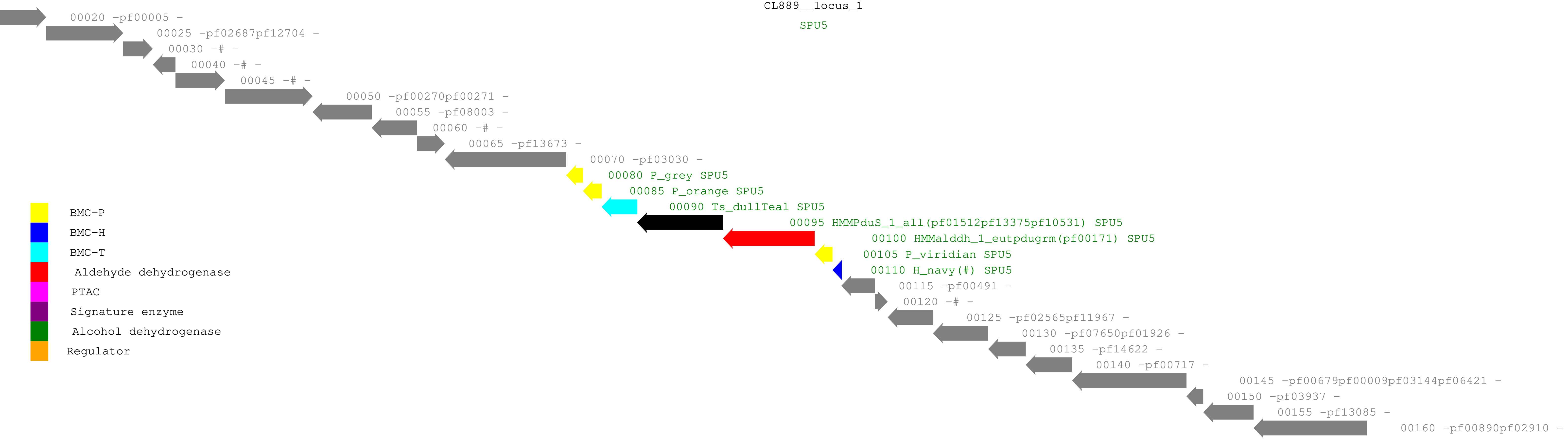

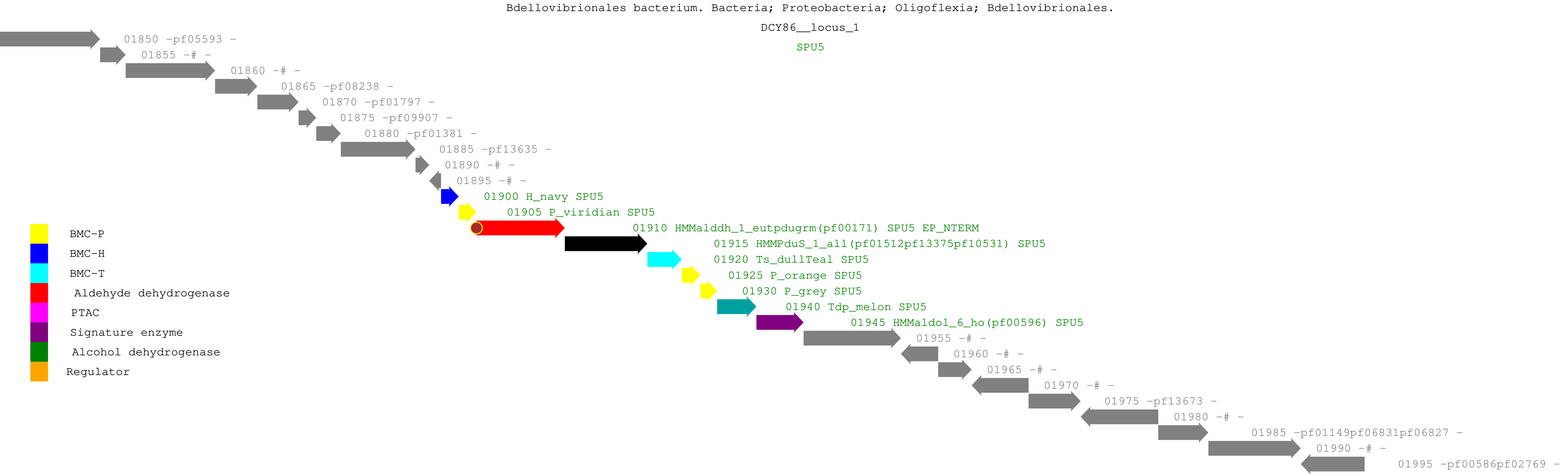
Bdellovibrionales bacterium. Bacteria; Proteobacteria; Oligoflexia; Bdellovibrionales.
DCY86__locus_1
SPU5
01850 -pf05593 -
01855 -# -
01860 -# -
01865 -pf08238 -
01870 -pf01797 -
01875 -pf09907 -
01880 -pf01381 -
01885 -pf13635 -
01890 -# -
01895 -# -
01900 H_navy SPU5
01905 P_viridian SPU5
01910 HMMalddh_1_eutpdugrm(pf00171) SPU5 EP_NTERM
01915 HMMPduS_1_all(pf01512pf13375pf10531) SPU5
01920 Ts_dullTeal SPU5
01925 P_orange SPU5
01930 P_grey SPU5
01940 Tdp_melon SPU5
01945 HMMaldol_6_ho(pf00596) SPU5
01955 -# -
01960 -# -
01965 -# -
01970 -# -
01975 -pf13673 -
01980 -# -
01985 -pf01149pf06831pf06827 -
01990 -# -
01995 -pf00586pf02769 -
BMC-P
BMC-H
BMC-T
Aldehyde dehydrogenase
PTAC
Signature enzyme
Alcohol dehydrogenase
Regulator

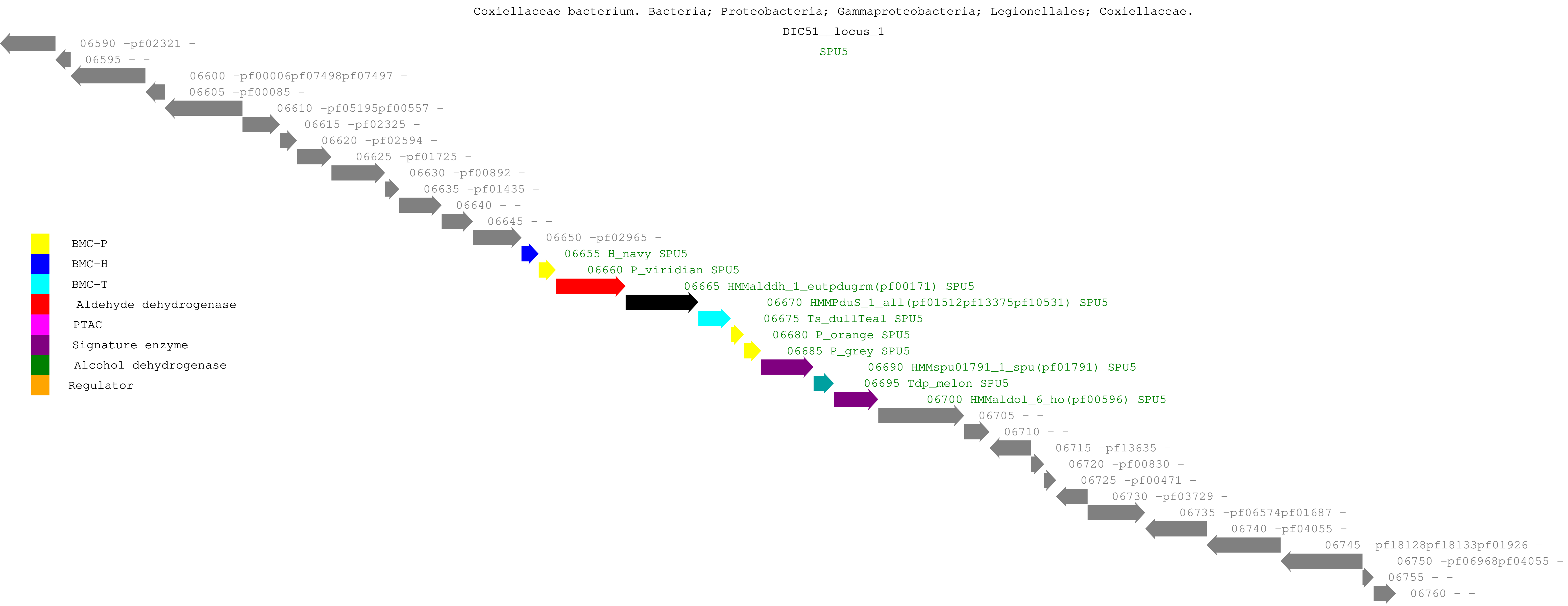
Coxiellaceae bacterium. Bacteria; Proteobacteria; Gammaproteobacteria; Legionellales; Coxiellaceae.
DIC51__locus_1
SPU5
06590 -pf02321 -
06595 - -
06600 -pf00006pf07498pf07497 -
06605 -pf00085 -
06610 -pf05195pf00557 -
06615 -pf02325 -
06620 -pf02594 -
06625 -pf01725 -
06630 -pf00892 -
06635 -pf01435 -
06640 - -
06645 - -
06650 -pf02965 -
06655 H_navy SPU5
06660 P_viridian SPU5
06665 HMMaldh_1_eutpdugrm(pf00171) SPU5
06670 HMMPduS_1_all(pf01512pf13375pf10531) SPU5
06675 Ts_dullTeal SPU5
06680 P_orange SPU5
06685 P_grey SPU5
06690 HMMspu01791_1_spu(pf01791) SPU5
06695 Tdp_melon SPU5
06700 HMMaldol_6_ho(pf00596) SPU5
06705 - -
06710 - -
06715 -pf13635 -
06720 -pf00830 -
06725 -pf00471 -
06730 -pf03729 -
06735 -pf06574pf01687 -
06740 -pf04055 -
06745 -pf18128pf18133pf01926 -
06750 -pf06968pf04055 -
06755 - -
06760 - -
BMC-P
BMC-H
BMC-T
Aldehyde dehydrogenase
PTAC
Signature enzyme
Alcohol dehydrogenase
Regulator

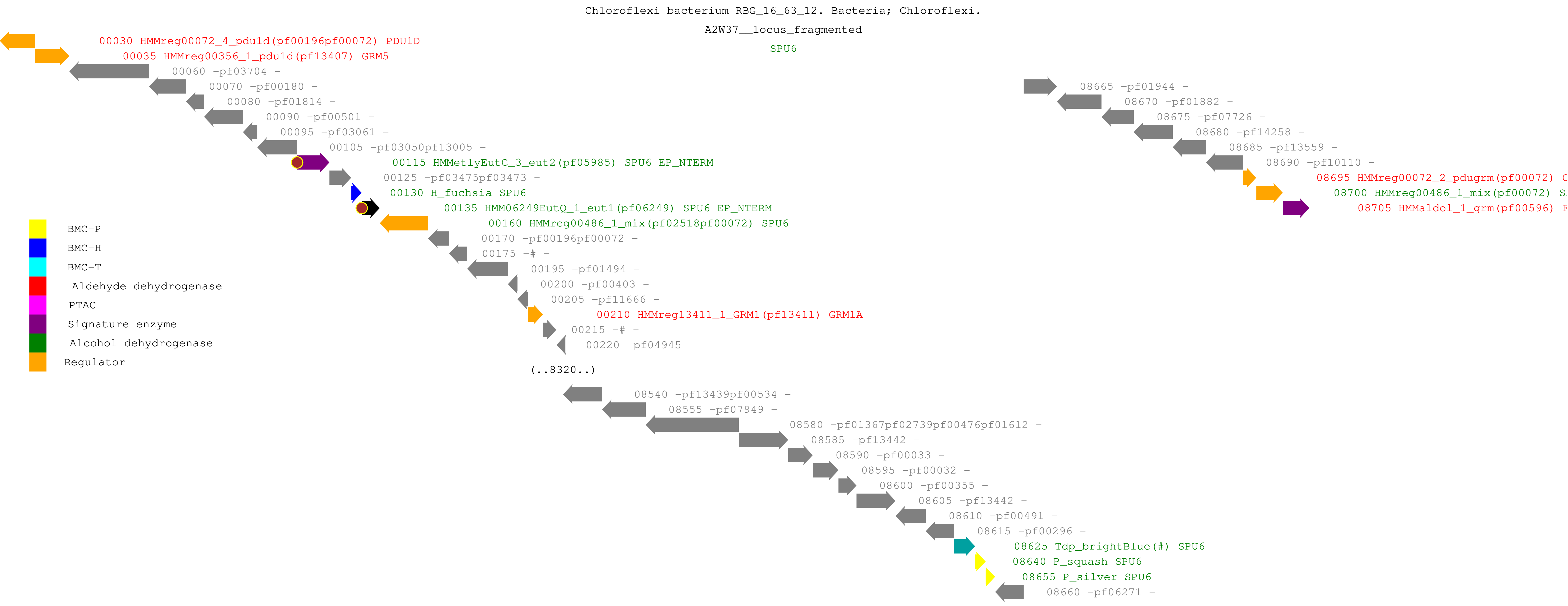
Chloroflexi bacterium RBG_16_63_12. Bacteria; Chloroflexi.
A2W37__locus_fragmented
SPU6
00030 HMMreg00072_4_pdu1d(pf00196pf00072) PDU1D
00035 HMMreg00356_1_pdu1d(pf13407) GRM5
00060 -pf03704 -
00070 -pf00180 -
00080 -pf01814 -
00090 -pf00501 -
00095 -pf03061 -
00105 -pf03050pf13005 -
00115 HMMetlyEutC_3_eut2(pf05985) SPU6 EP_NTERM
00125 -pf03475pf03473 -
00130 H_fuchsia SPU6
00135 HMM06249EutQ_1_eut1(pf06249) SPU6 EP_NTERM
00160 HMMreg00486_1_mix(pf02518pf00072) SPU6
00170 -pf00196pf00072 -
00175 -# -
00195 -pf01494 -
00200 -pf00403 -
00205 -pf11666 -
00210 HMMreg13411_1_GRM1(pf13411) GRM1A
00215 -# -
00220 -pf04945 -
(..8320..)
08540 -pf13439pf00534 -
08555 -pf07949 -
08580 -pf01367pf02739pf00476pf01612 -
08585 -pf13442 -
08590 -pf00033 -
08595 -pf00032 -
08600 -pf00355 -
08605 -pf13442 -
08610 -pf00491 -
08615 -pf00296 -
08625 Tdp_brightBlue(#) SPU6
08640 P_squash SPU6
08655 P_silver SPU6
08660 -pf06271 -
08665 -pf01944 -
08670 -pf01882 -
08675 -pf07726 -
08680 -pf14258 -
08685 -pf13559 -
08690 -pf10110 -
08695 HMMreg00072_2_pdugrm(pf00072)
08700 HMMreg00486_1_mix(pf00072)
08705 HMMaldol_1_grm(pf00596)
BMC-P
BMC-H
BMC-T
Aldehyde dehydrogenase
PTAC
Signature enzyme
Alcohol dehydrogenase
Regulator

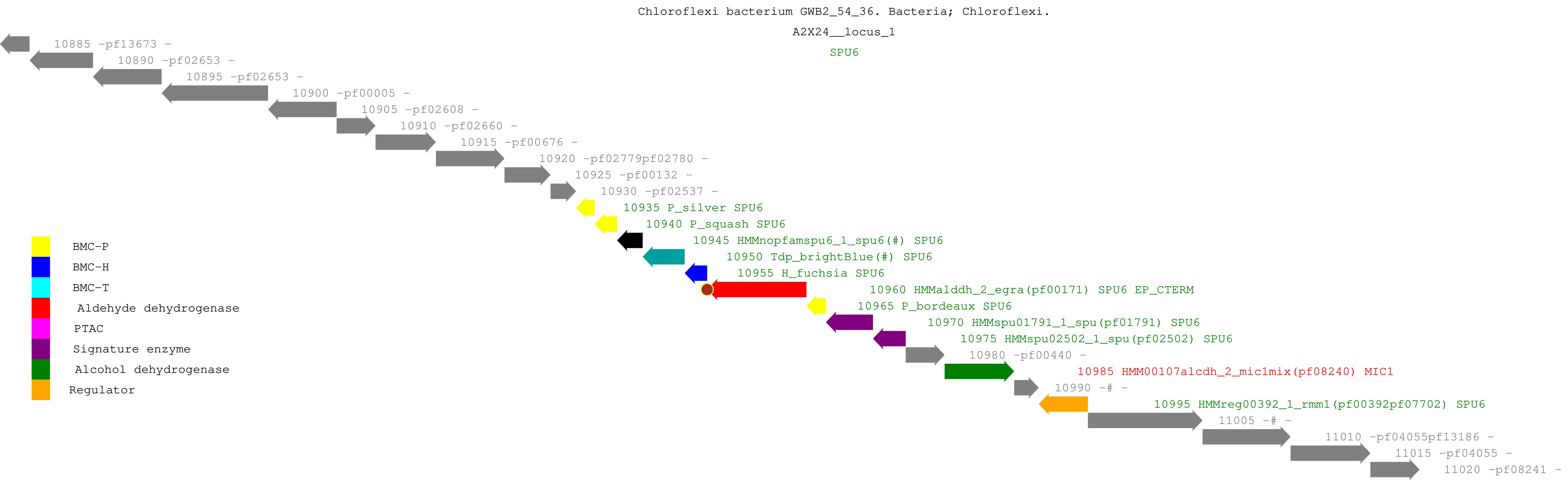
Chloroflexi bacterium GWB2_54_36. Bacteria; Chloroflexi.
A2X24__locus_1
SPU6
10885 -pf13673 -
10890 -pf02653 -
10895 -pf02653 -
10900 -pf00005 -
10905 -pf02608 -
10910 -pf02660 -
10915 -pf00676 -
10920 -pf02779pf02780 -
10925 -pf00132 -
10930 -pf02537 -
10935 P_silver SPU6
10940 P_squash SPU6
10945 HMMnopfamspu6_1_spu6(#) SPU6
10950 Tdp_brightBlue(#) SPU6
10955 H_fuchsia SPU6
10960 HMMalddh_2_egra(pf00171) SPU6 EP_CTERM
10965 P_bordeaux SPU6
10970 HMMspu01791_1_spu(pf01791) SPU6
10975 HMMspu02502_1_spu(pf02502) SPU6
10980 -pf00440 -
10985 HMM00107alcdh_2_mic1mix(pf08240) MIC1
10990 -# -
10995 HMMreg00392_1_rmm1(pf00392pf07702) SPU6
11005 -# -
11010 -pf04055pf13186 -
11015 -pf04055 -
11020 -pf08241 -
BMC-P
BMC-H
BMC-T
Aldehyde dehydrogenase
PTAC
Signature enzyme
Alcohol dehydrogenase
Regulator

Chloroflexi bacterium RBG_16_51_16. Bacteria; Chloroflexi.

A2Y54__locus_1

SPU6

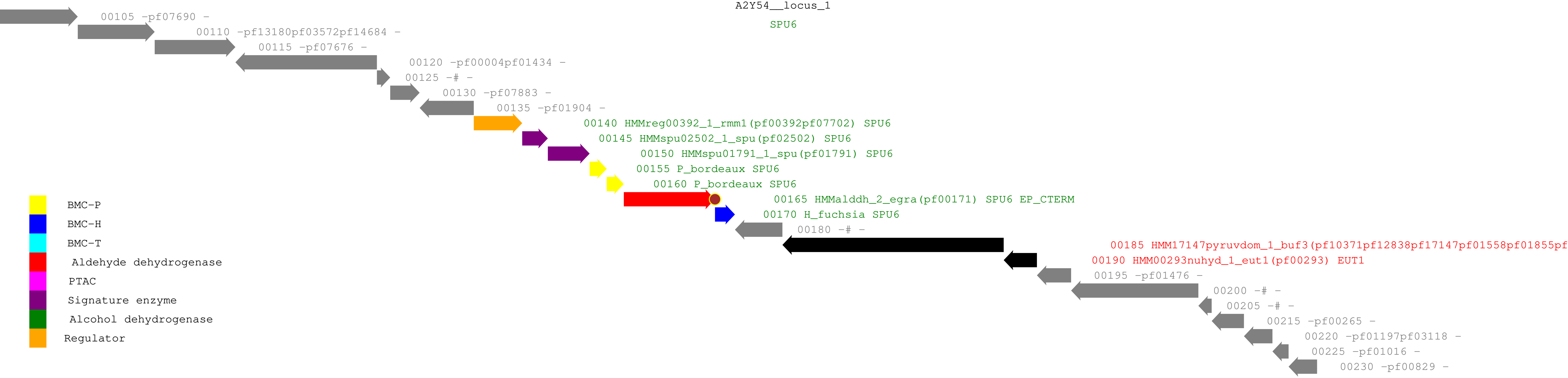

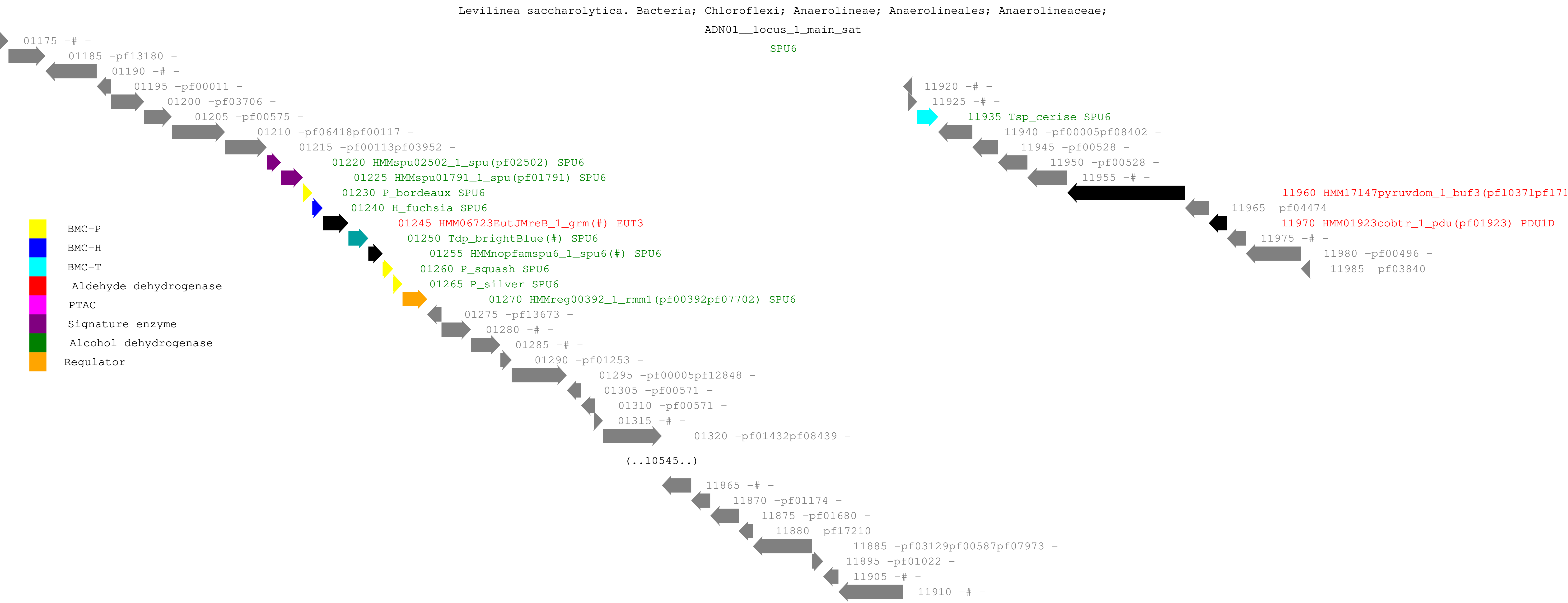
Levilinea saccharolytica. Bacteria; Chloroflexi; Anaerolineae; Anaerolineales; Anaerolineaceae;
ADN01__locus_1_main_sat
SPU6
01175 -# -
01185 -pf13180 -
01190 -# -
01195 -pf00011 -
01200 -pf03706 -
01205 -pf00575 -
01210 -pf06418pf00117 -
01215 -pf00113pf03952 -
01220 HMMspu02502_1_spu(pf02502) SPU6
01225 HMMspu01791_1_spu(pf01791) SPU6
01230 P_bordeaux SPU6
01240 H_fuchsia SPU6
01245 HMM06723EutJMreB_1_grm(#) EUT3
01250 Tdp_brightBlue(#) SPU6
01255 HMMnopfamspu6_1_spu6(#) SPU6
01260 P_squash SPU6
01265 P_silver SPU6
01270 HMMreg00392_1_rmm1(pf00392pf07702) SPU6
01275 -pf13673 -
01280 -# -
01285 -# -
01290 -pf01253 -
01295 -pf00005pf12848 -
01305 -pf00571 -
01310 -pf00571 -
01315 -# -
01320 -pf01432pf08439 -
(..10545..)
11865 -# -
11870 -pf01174 -
11875 -pf01680 -
11880 -pf17210 -
11885 -pf03129pf00587pf07973 -
11895 -pf01022 -
11905 -# -
11910 -# -
11920 -# -
11925 -# -
11935 Tsp_cerise SPU6
11940 -pf00005pf08402 -
11945 -pf00528 -
11950 -pf00528 -
11955 -# -
11960 HMM17147pyruvdom_1_buf3(pf10371pf171
11965 -pf04474 -
11970 HMM01923cobtr_1_pdu(pf01923) PDU1D
11975 -# -
11980 -pf00496 -
11985 -pf03840 -
BMC-P
BMC-H
BMC-T
Aldehyde dehydrogenase
PTAC
Signature enzyme
Alcohol dehydrogenase
Regulator

Anaerolineae bacterium CG1_02_58_13. Bacteria; Chloroflexi; Anaerolineae; unclassified Anaerolineae.

AUJ21__locus_1_main_sat

SPU6

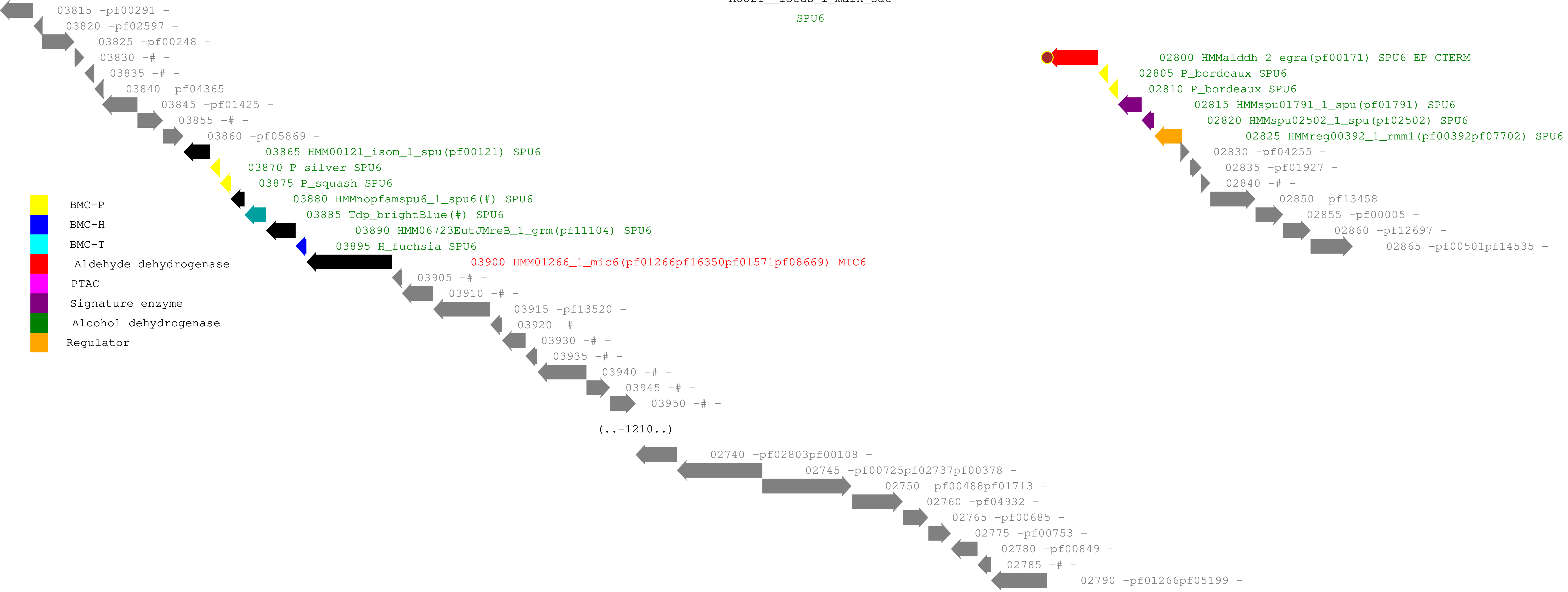

Anaerolineaceae bacterium 4572_5.2. Bacteria; Chloroflexi; Anaerolineae; Anaerolineales; Anaerolineaceae.

B6243__locus_1

SPU6

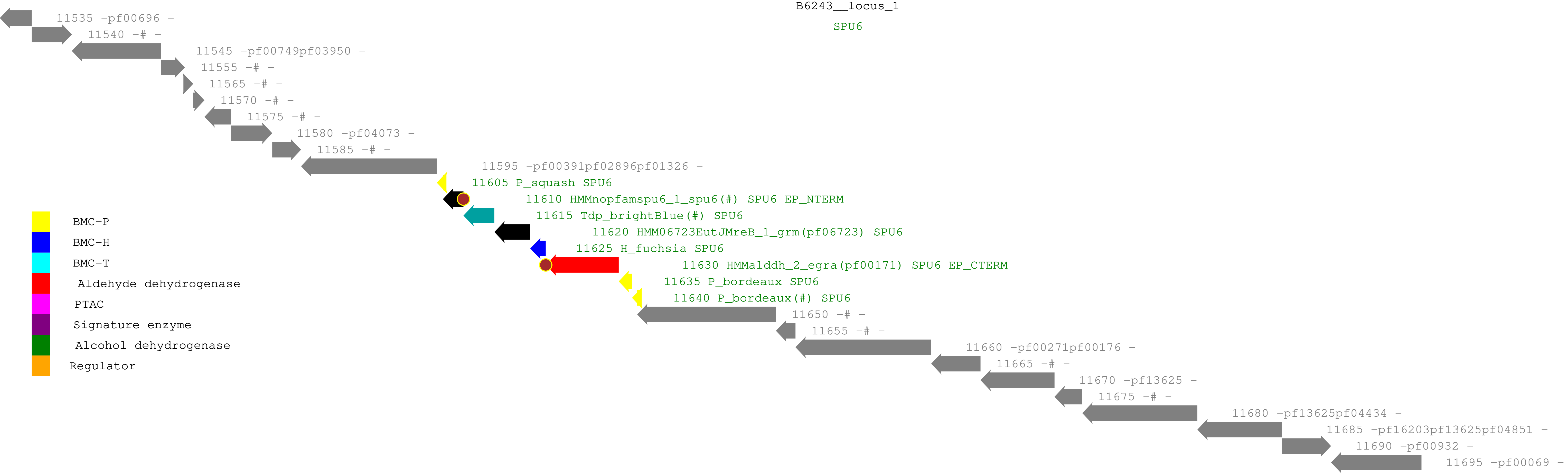

Chloroflexi bacterium 44-23. Bacteria; Chloroflexi.

BGO78__locus_1_main_sat

SPU6

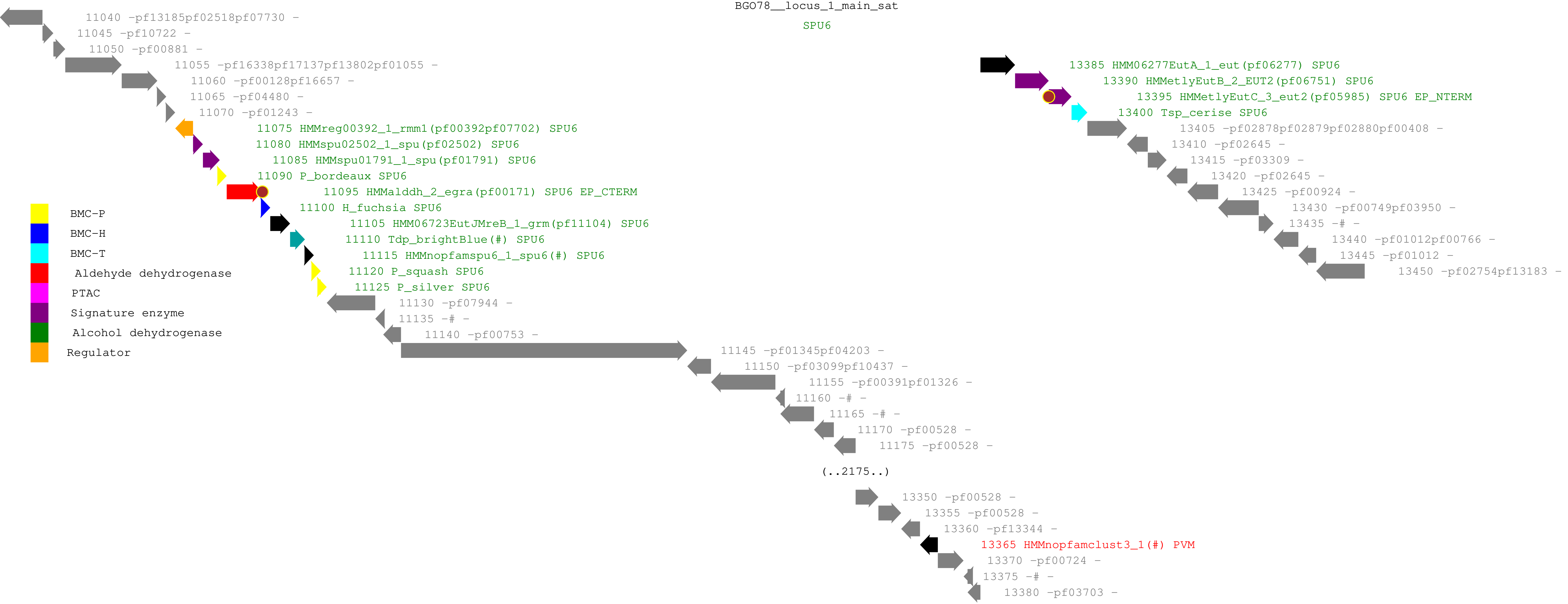

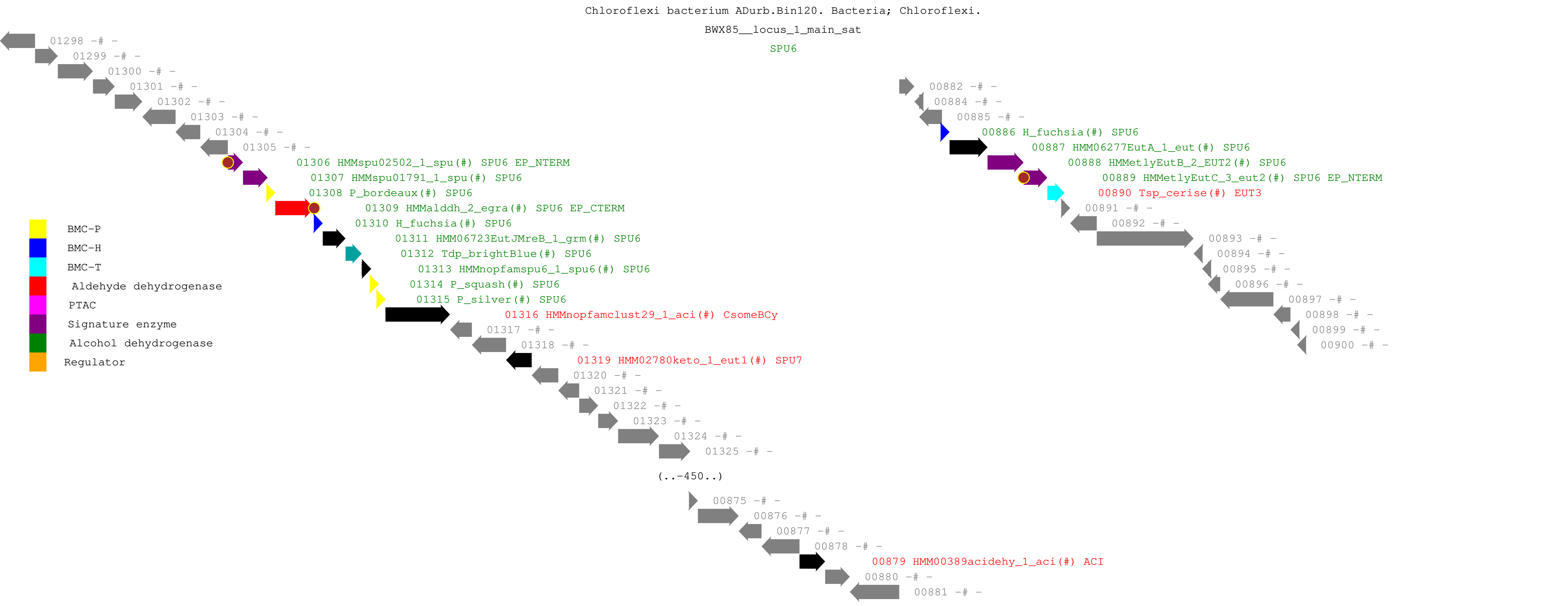
Chloroflexi bacterium ADurb.Bin120. Bacteria; Chloroflexi.
BWX85__locus_1_main_sat
SPU6
01298 -# -
01299 -# -
01300 -# -
01301 -# -
01302 -# -
01303 -# -
01304 -# -
01305 -# -
01306 HMMspu02502_1_spu(#) SPU6 EP_NTERM
01307 HMMspu01791_1_spu(#) SPU6
01308 P_bordeaux(#) SPU6
01309 HMMalddh_2_egra(#) SPU6 EP_CTERM
01310 H_fuchsia(#) SPU6
01311 HMM06723EutJMreB_1_grm(#) SPU6
01312 Tdp_brightBlue(#) SPU6
01313 HMMnopfamspu6_1_spu6(#) SPU6
01314 P_squash(#) SPU6
01315 P_silver(#) SPU6
01316 HMMnopfamclust29_1_aci(#) CsomeBCy
01317 -# -
01318 -# -
01319 HMM02780keto_1_eut1(#) SPU7
01320 -# -
01321 -# -
01322 -# -
01323 -# -
01324 -# -
01325 -# -
(..-450..)
00875 -# -
00876 -# -
00877 -# -
00878 -# -
00879 HMM00389acidehy_1_aci(#) ACI
00880 -# -
00881 -# -
00882 -# -
00884 -# -
00885 -# -
00886 H_fuchsia(#) SPU6
00887 HMM06277EutA_1_eut(#) SPU6
00888 HMMetlyEutB_2_EUT2(#) SPU6
00889 HMMetlyEutC_3_eut2(#) SPU6 EP_NTERM
00890 Tsp_cerise(#) EUT3
00891 -# -
00892 -# -
00893 -# -
00894 -# -
00895 -# -
00896 -# -
00897 -# -
00898 -# -
00899 -# -
00900 -# -
BMC-P
BMC-H
BMC-T
Aldehyde dehydrogenase
PTAC
Signature enzyme
Alcohol dehydrogenase
Regulator

Anaerolineaceae bacterium. Bacteria; Chloroflexi; Anaerolineae; Anaerolineales; Anaerolineaceae.

C4545__locus_1

SPU6

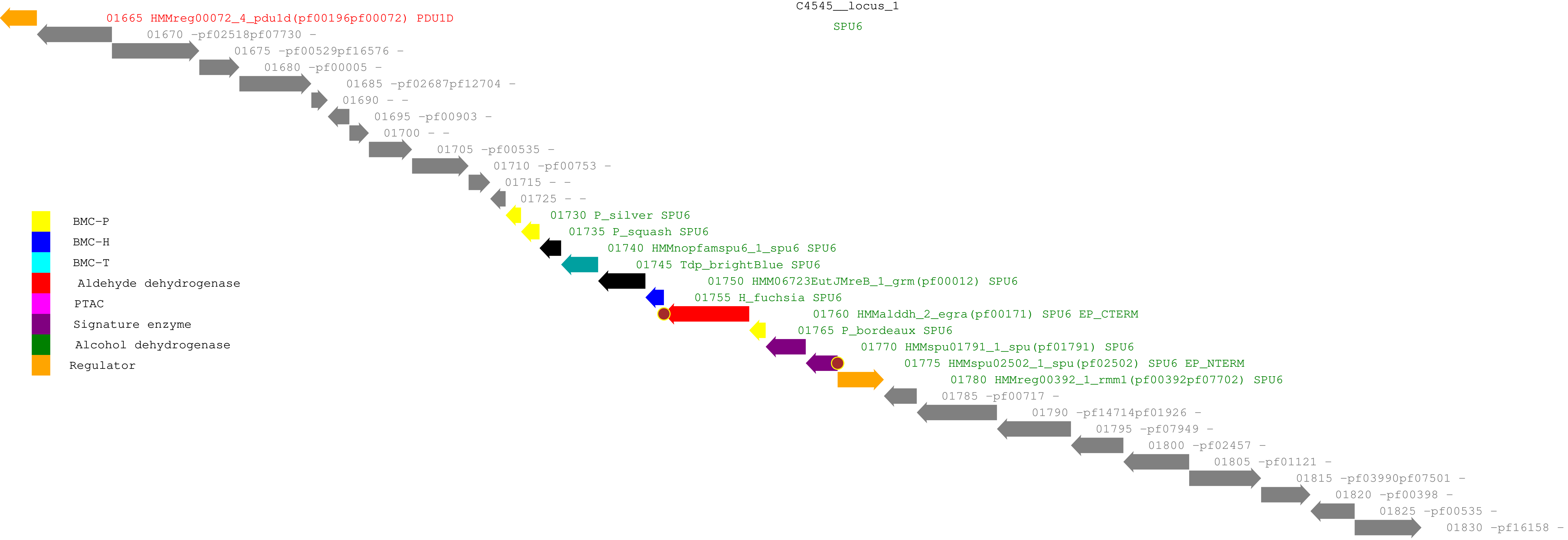

Brevefilum fermentans. Bacteria; Chloroflexi; Anaerolineae; Anaerolineales; Anaerolineaceae;

CFX1CAM__locus_1_main_sat

SPU6

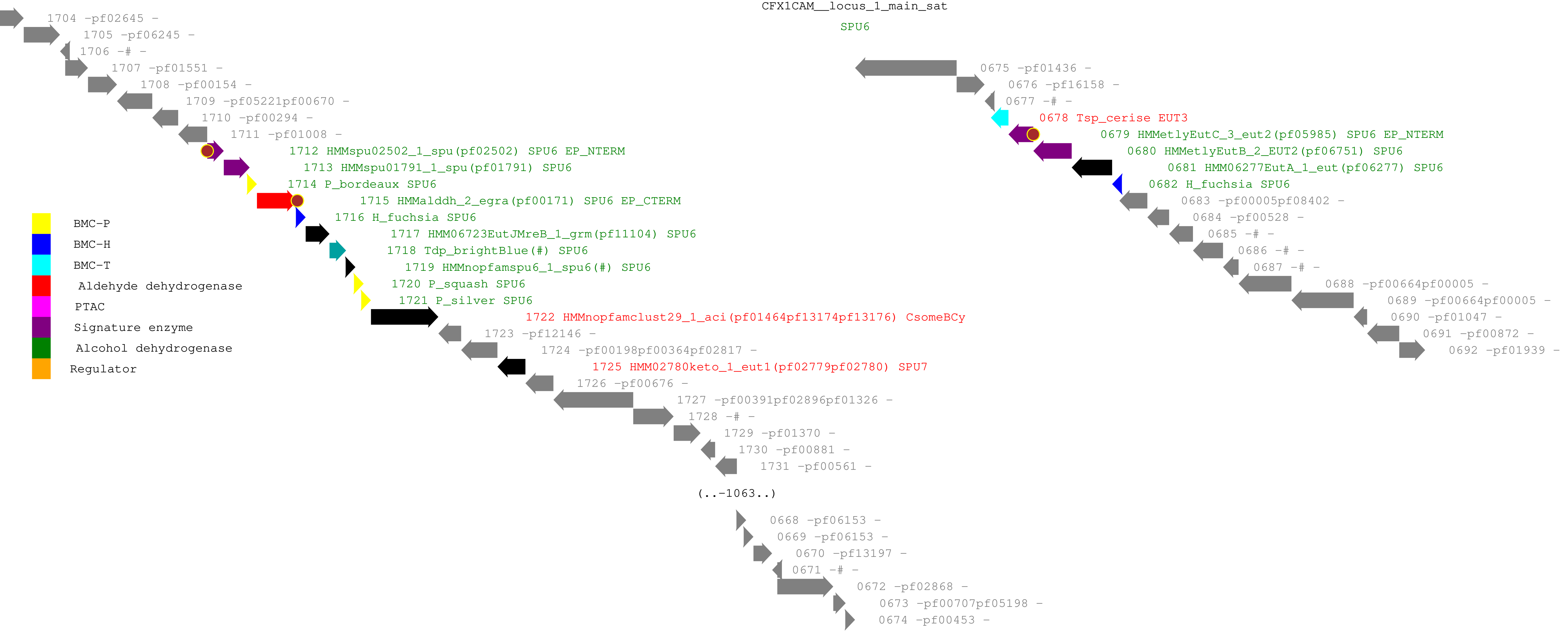

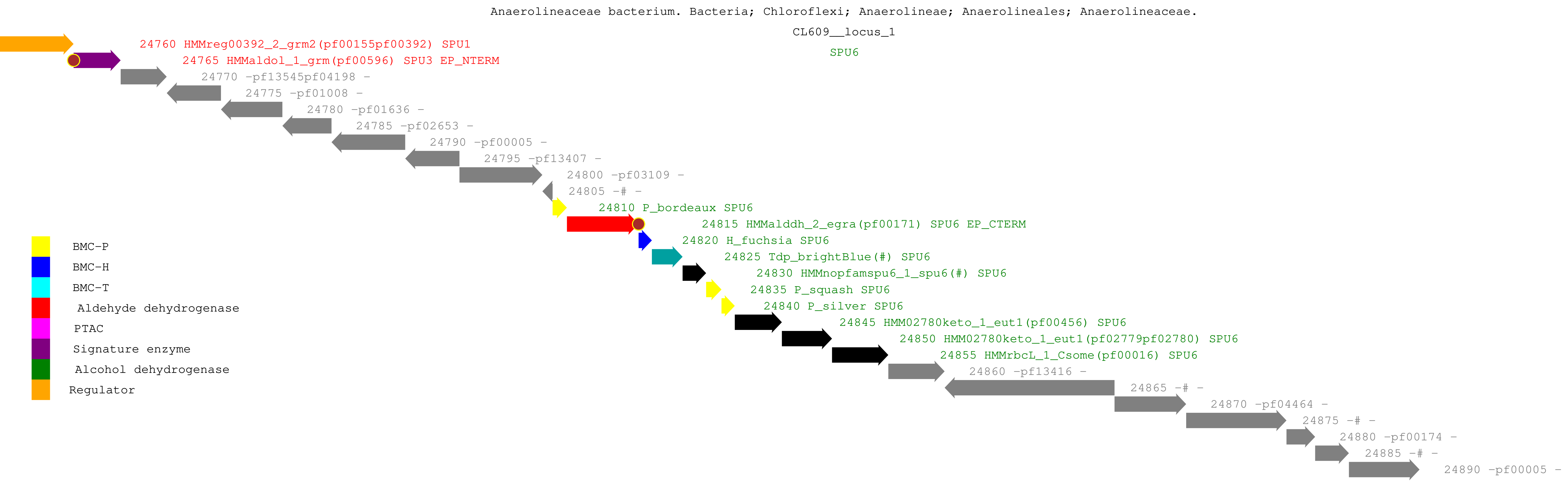
Anaerolineaceae bacterium. Bacteria; Chloroflexi; Anaerolineae; Anaerolineales; Anaerolineaceae.
CL609__locus_1
SPU6
24760 HMMreg00392_2_grm2(pf00155pf00392) SPU1
24765 HMMaldol_1_grm(pf00596) SPU3 EP_NTERM
24770 -pf13545pf04198 -
24775 -pf01008 -
24780 -pf01636 -
24785 -pf02653 -
24790 -pf00005 -
24795 -pf13407 -
24800 -pf03109 -
24805 -# -
24810 P_bordeaux SPU6
24815 HMMaldh_2_egra(pf00171) SPU6 EP_CTERM
24820 H_fuchsia SPU6
24825 Tdp_brightBlue(#) SPU6
24830 HMMnopfamspu6_1_spu6(#) SPU6
24835 P_squash SPU6
24840 P_silver SPU6
24845 HMM02780keto_1_eut1(pf00456) SPU6
24850 HMM02780keto_1_eut1(pf02779pf02780) SPU6
24855 HMMrbcL_1_Csome(pf00016) SPU6
24860 -pf13416 -
24865 -# -
24870 -pf04464 -
24875 -# -
24880 -pf00174 -
24885 -# -
24890 -pf00005 -
BMC-P
BMC-H
BMC-T
Aldehyde dehydrogenase
PTAC
Signature enzyme
Alcohol dehydrogenase
Regulator

Anaerolineae bacterium CG_4_9_14_0_8_um_filter_58_9. Bacteria; Chloroflexi; Anaerolineae; unclassified Anaerolineae.

CO064__locus_fragmented

SPU6

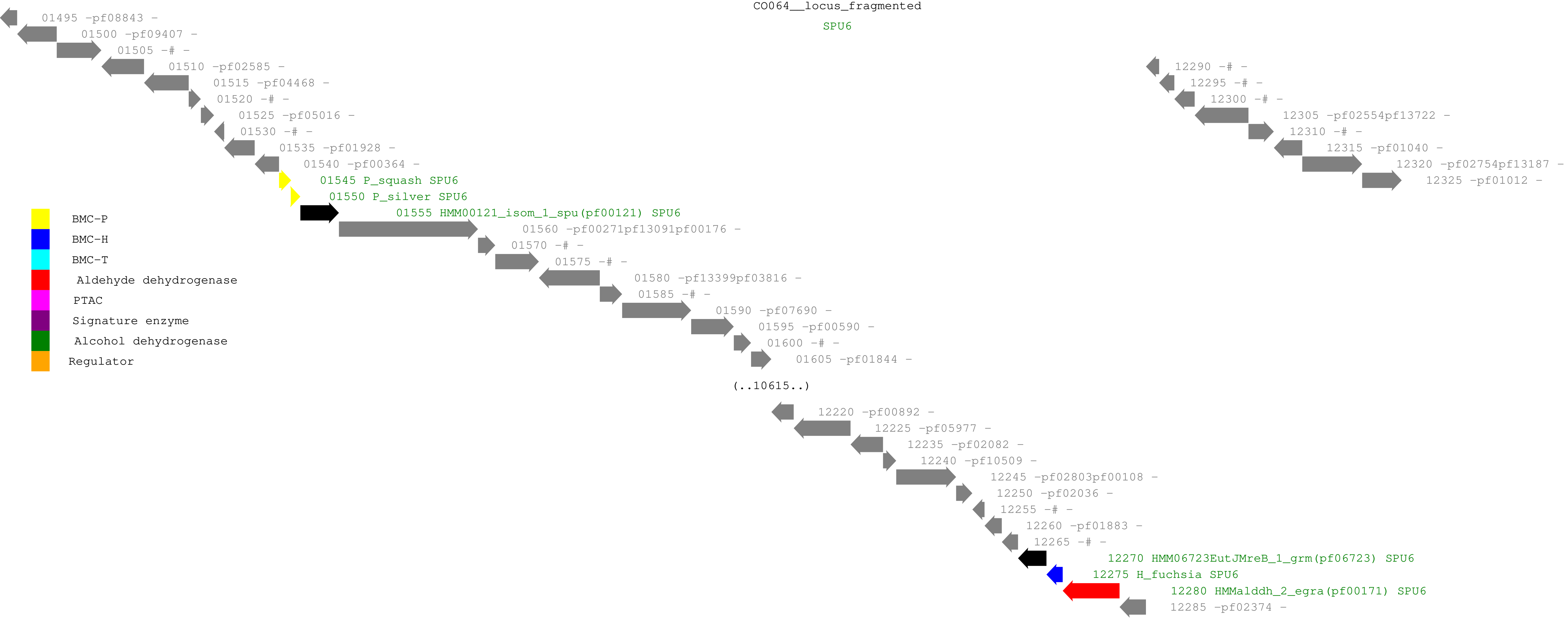

Anaerolineae bacterium CG03_land_8_20_14_0_80_58_20. Bacteria; Chloroflexi; Anaerolineae; unclassified Anaerolineae.

COS37__locus_fragmented

SPU6

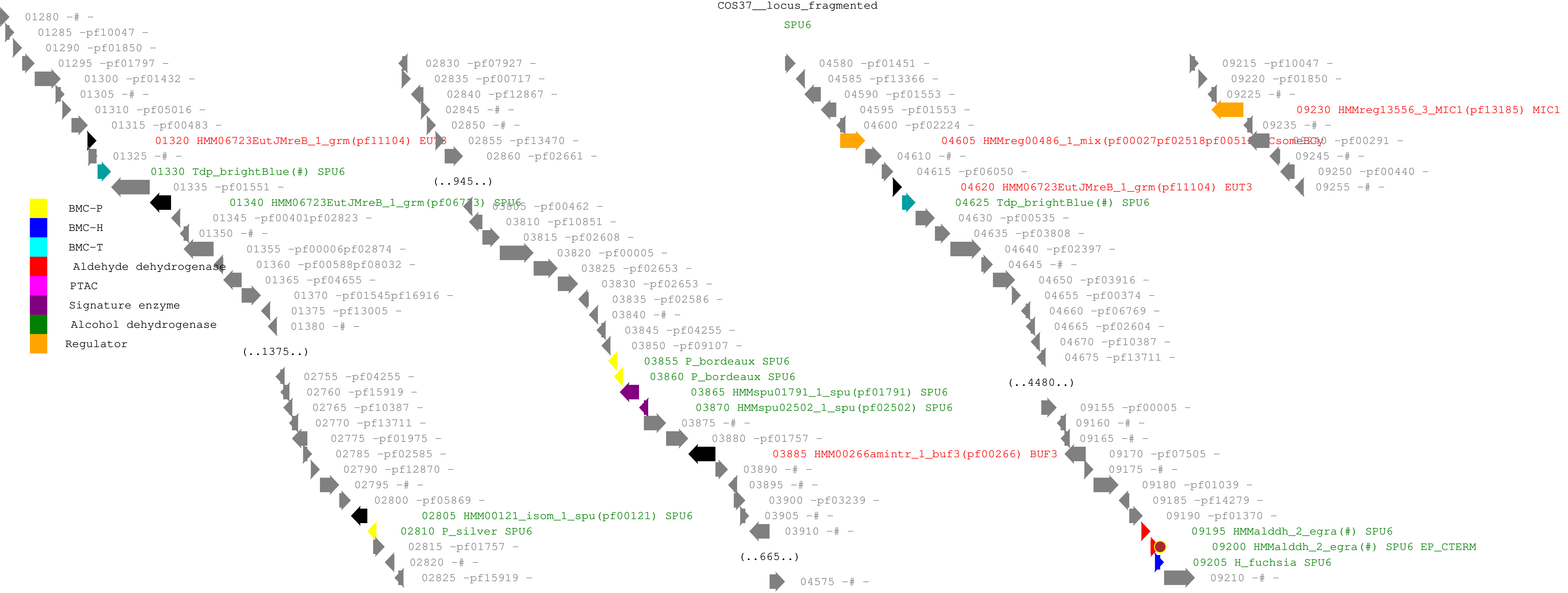

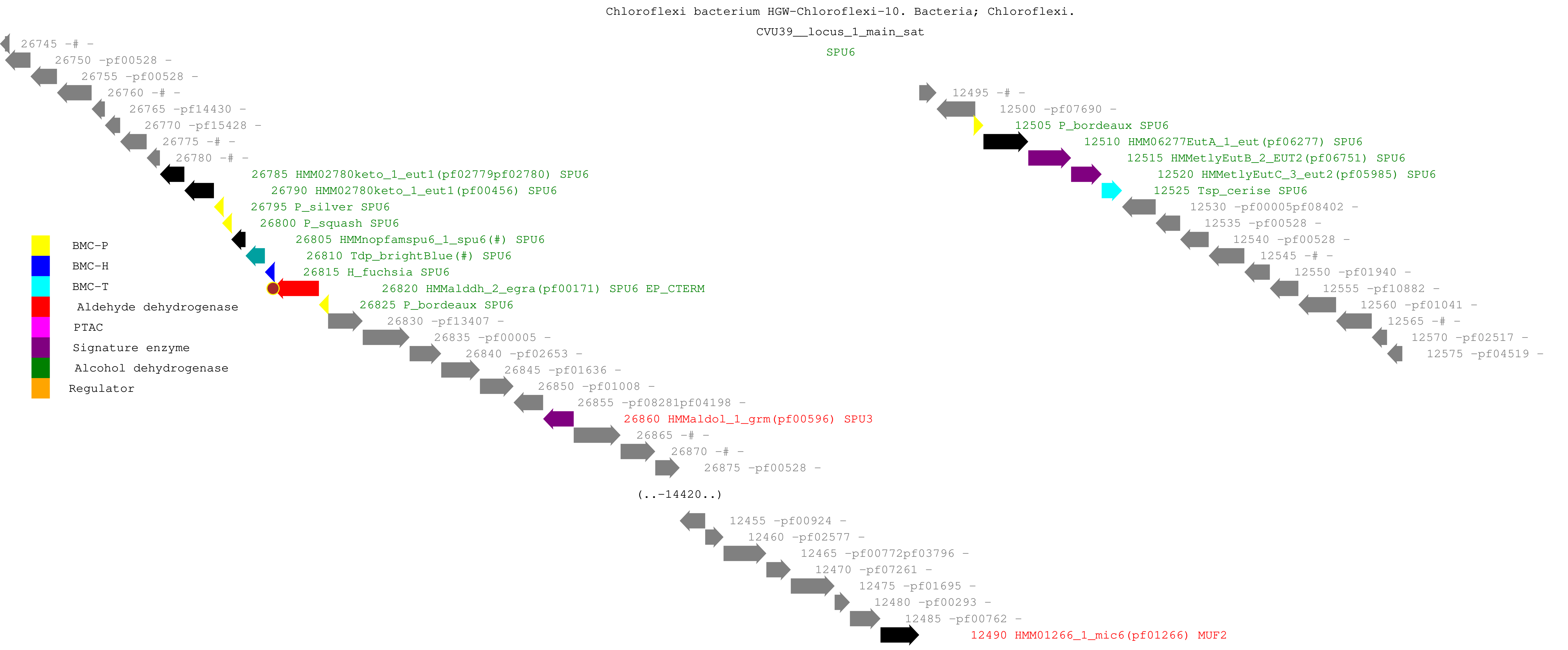
Chloroflexi bacterium HGW-Chloroflexi-10. Bacteria; Chloroflexi.
CVU39__locus_1_main_sat
SPU6
26745 -# -
26750 -pf00528 -
26755 -pf00528 -
26760 -# -
26765 -pf14430 -
26770 -pf15428 -
26775 -# -
26780 -# -
26785 HMM02780keto_1_eut1(pf02779pf02780) SPU6
26790 HMM02780keto_1_eut1(pf00456) SPU6
26795 P_silver SPU6
26800 P_squash SPU6
26805 HMMnopfamspu6_1_spu6(#) SPU6
26810 Tdp_brightBlue(#) SPU6
26815 H_fuchsia SPU6
26820 HMMalddh_2_egra(pf00171) SPU6 EP_CTERM
26825 P_bordeaux SPU6
26830 -pf13407 -
26835 -pf00005 -
26840 -pf02653 -
26845 -pf01636 -
26850 -pf01008 -
26855 -pf08281pf04198 -
26860 HMMaldol_1_grm(pf00596) SPU3
26865 -# -
26870 -# -
26875 -pf00528 -
(..-14420..)
12455 -pf00924 -
12460 -pf02577 -
12465 -pf00772pf03796 -
12470 -pf07261 -
12475 -pf01695 -
12480 -pf00293 -
12485 -pf00762 -
12490 HMM01266_1_mic6(pf01266) MUF2
12495 -# -
12500 -pf07690 -
12505 P_bordeaux SPU6
12510 HMM06277EutA_1_eut(pf06277) SPU6
12515 HMMetlyEutB_2_EUT2(pf06751) SPU6
12520 HMMetlyEutC_3_eut2(pf05985) SPU6
12525 Tsp_cerise SPU6
12530 -pf00005pf08402 -
12535 -pf00528 -
12540 -pf00528 -
12545 -# -
12550 -pf01940 -
12555 -pf10882 -
12560 -pf01041 -
12565 -# -
12570 -pf02517 -
12575 -pf04519 -
BMC-P
BMC-H
BMC-T
Aldehyde dehydrogenase
PTAC
Signature enzyme
Alcohol dehydrogenase
Regulator

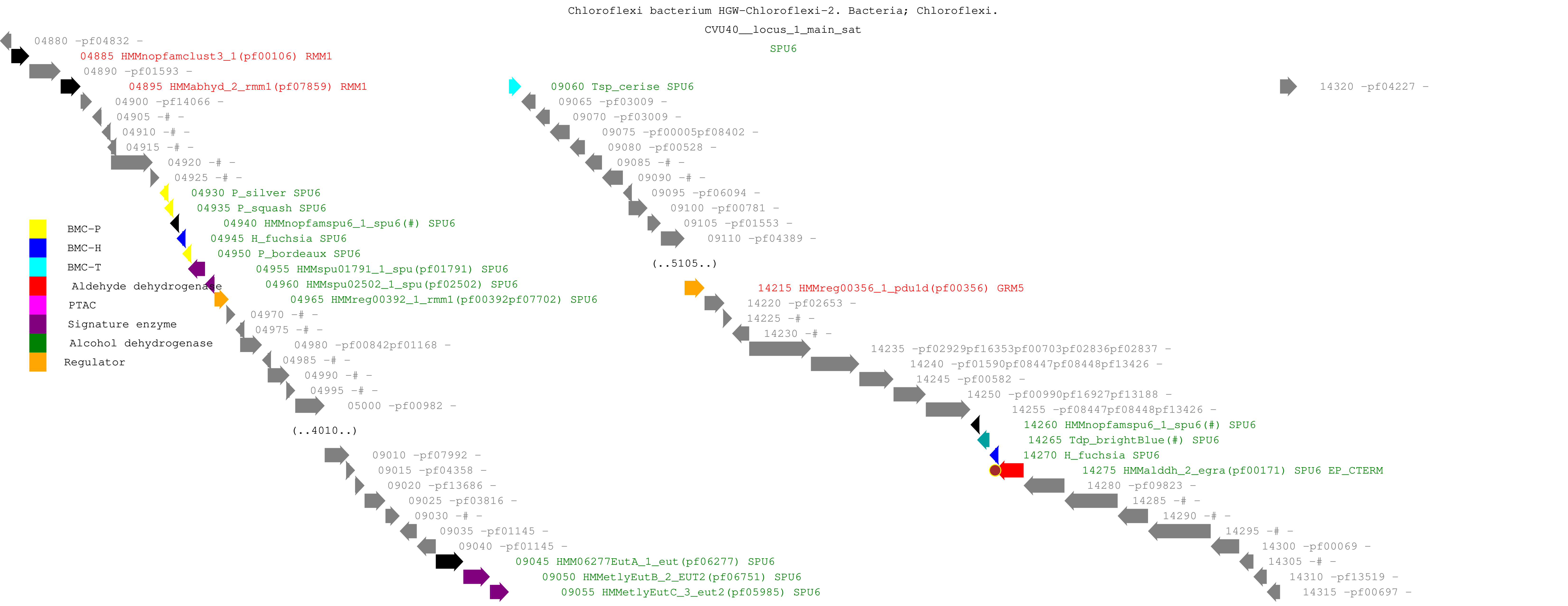
Chloroflexi bacterium HGW-Chloroflexi-2. Bacteria; Chloroflexi.
CVU40__locus_1_main_sat
SPU6
04880 -pf04832 -
04885 HMMnopfamclust3_1(pf00106) RMM1
04890 -pf01593 -
04895 HMMabhyd_2_rmm1(pf07859) RMM1
04900 -pf14066 -
04905 -# -
04910 -# -
04915 -# -
04920 -# -
04925 -# -
04930 P_silver SPU6
04935 P_squash SPU6
04940 HMMnopfamspu6_1_spu6(#) SPU6
04945 H_fuchsia SPU6
04950 P_bordeaux SPU6
04955 HMMspu01791_1_spu(pf01791) SPU6
04960 HMMspu02502_1_spu(pf02502) SPU6
04965 HMMreg00392_1_rmm1(pf00392pf07702) SPU6
04970 -# -
04975 -# -
04980 -pf00842pf01168 -
04985 -# -
04990 -# -
04995 -# -
05000 -pf00982 -
(..4010..)
09010 -pf07992 -
09015 -pf04358 -
09020 -pf13686 -
09025 -pf03816 -
09030 -# -
09035 -pf01145 -
09040 -pf01145 -
09045 HMM06277EutA_1_eut(pf06277) SPU6
09050 HMMetlyEutB_2_EUT2(pf06751) SPU6
09055 HMMetlyEutC_3_eut2(pf05985) SPU6
09060 Tsp_cerise SPU6
09065 -pf03009 -
09070 -pf03009 -
09075 -pf00005pf08402 -
09080 -pf00528 -
09085 -# -
09090 -# -
09095 -pf06094 -
09100 -pf00781 -
09105 -pf01553 -
09110 -pf04389 -
(..5105..)
14215 HMMreg00356_1_pdu1d(pf00356) GRM5
14220 -pf02653 -
14225 -# -
14230 -# -
14235 -pf02929pf16353pf00703pf02836pf02837 -
14240 -pf01590pf08447pf08448pf13426 -
14245 -pf00582 -
14250 -pf00990pf16927pf13188 -
14255 -pf08447pf08448pf13426 -
14260 HMMnopfamspu6_1_spu6(#) SPU6
14265 Tdp_brightBlue(#) SPU6
14270 H_fuchsia SPU6
14275 HMMalddh_2_egra(pf00171) SPU6 EP_CTERM
14280 -pf09823 -
14285 -# -
14290 -# -
14295 -# -
14300 -pf00069 -
14305 -# -
14310 -pf13519 -
14315 -pf00697 -
14320 -pf04227 -
BMC-P
BMC-H
BMC-T
Aldehyde dehydrogenase
PTAC
Signature enzyme
Alcohol dehydrogenase
Regulator

Chloroflexi bacterium HGW-Chloroflexi-3. Bacteria; Chloroflexi.

CVU41__locus_1_main_sat

SPU6

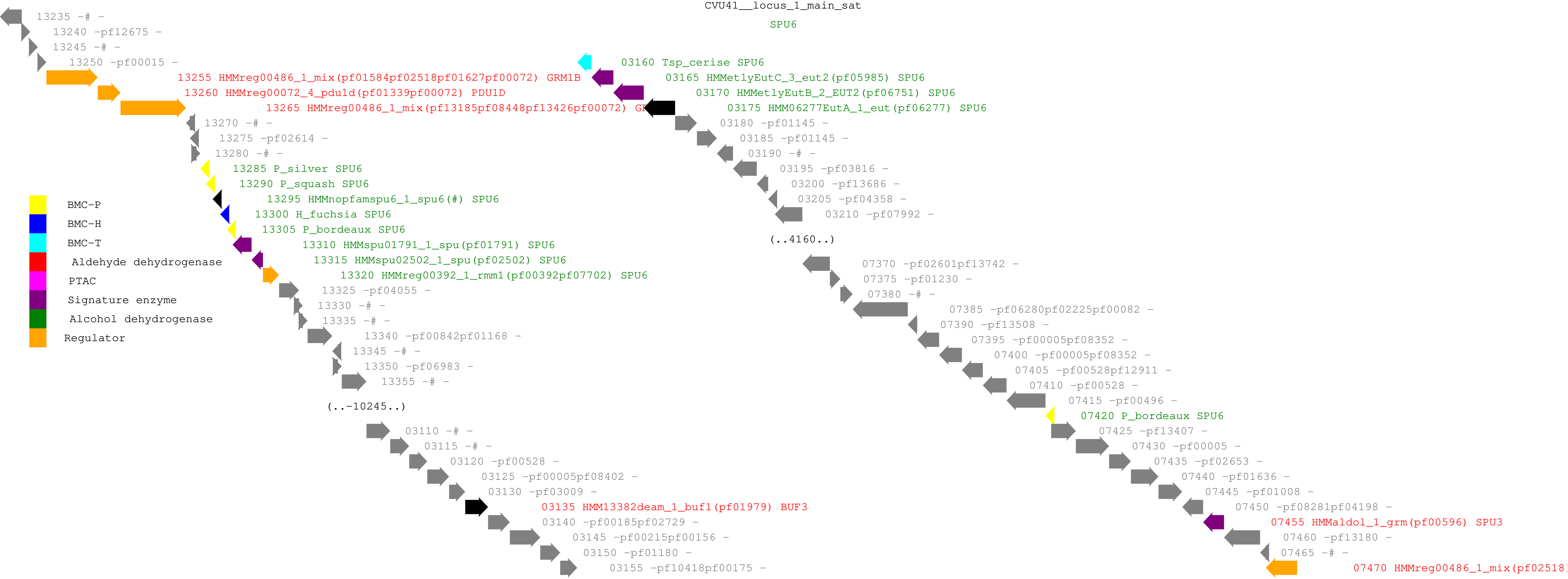

Chloroflexi bacterium HGW-Chloroflexi-4. Bacteria; Chloroflexi.

CVU42__locus_1

SPU6

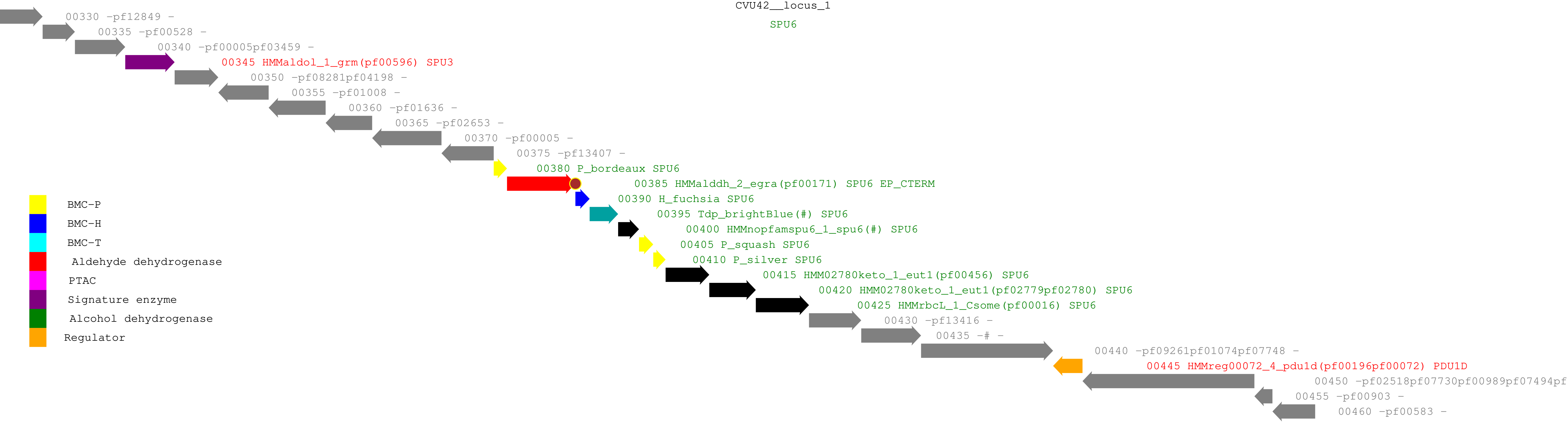

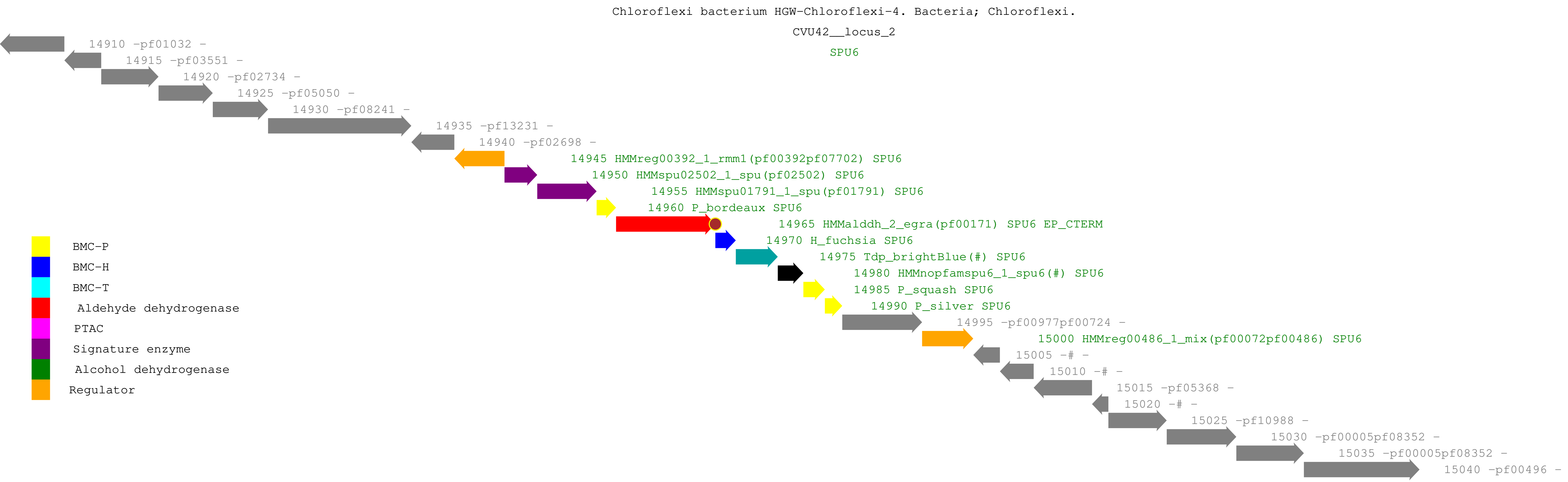
Chloroflexi bacterium HGW-Chloroflexi-4. Bacteria; Chloroflexi.
CVU42__locus_2
SPU6
14910 -pf01032 -
14915 -pf03551 -
14920 -pf02734 -
14925 -pf05050 -
14930 -pf08241 -
14935 -pf13231 -
14940 -pf02698 -
14945 HMMreg00392_1_rmm1(pf00392pf07702) SPU6
14950 HMMspu02502_1_spu(pf02502) SPU6
14955 HMMspu01791_1_spu(pf01791) SPU6
14960 P_bordeaux SPU6
14965 HMMaldh_2_egra(pf00171) SPU6 EP_CTERM
14970 H_fuchsia SPU6
14975 Tdp_brightBlue(#) SPU6
14980 HMMnopfamspu6_1_spu6(#) SPU6
14985 P_squash SPU6
14990 P_silver SPU6
14995 -pf00977pf00724 -
15000 HMMreg00486_1_mix(pf00072pf00486) SPU6
15005 -# -
15010 -# -
15015 -pf05368 -
15020 -# -
15025 -pf10988 -
15030 -pf00005pf08352 -
15035 -pf00005pf08352 -
15040 -pf00496 -
BMC-P
BMC-H
BMC-T
Aldehyde dehydrogenase
PTAC
Signature enzyme
Alcohol dehydrogenase
Regulator

Chloroflexi bacterium HGW-Chloroflexi-5. Bacteria; Chloroflexi.

CVU43__locus_1

SPU6

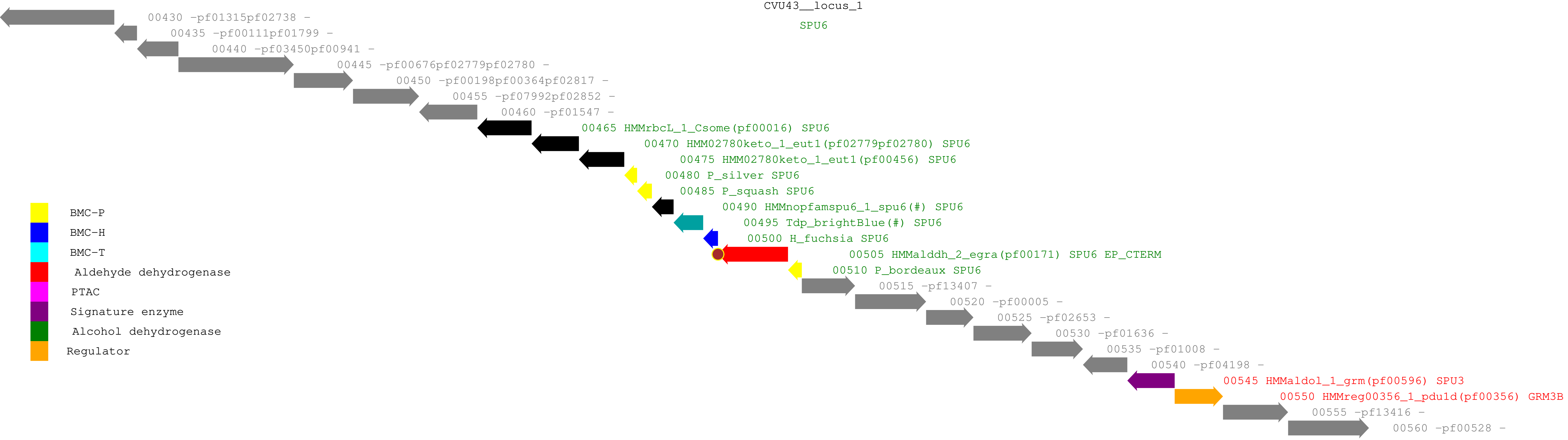

Chloroflexi bacterium HGW-Chloroflexi-5. Bacteria; Chloroflexi.

CVU43__locus_2

SPU6

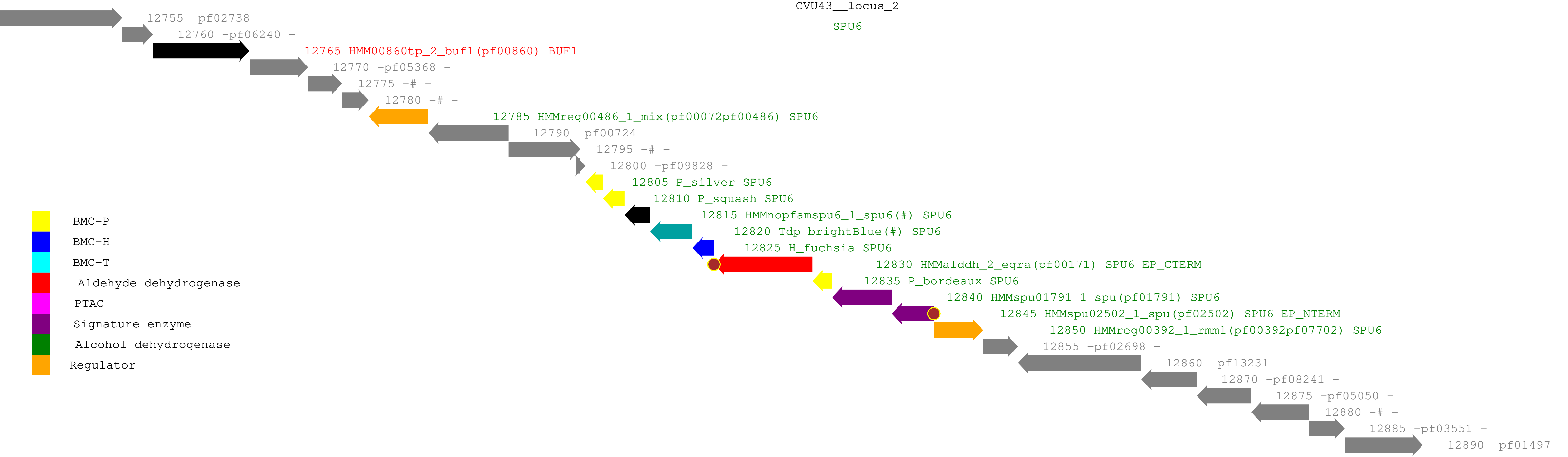

Chloroflexi bacterium HGW-Chloroflexi-7. Bacteria; Chloroflexi.

CVU45__locus_1_main_sat

SPU6

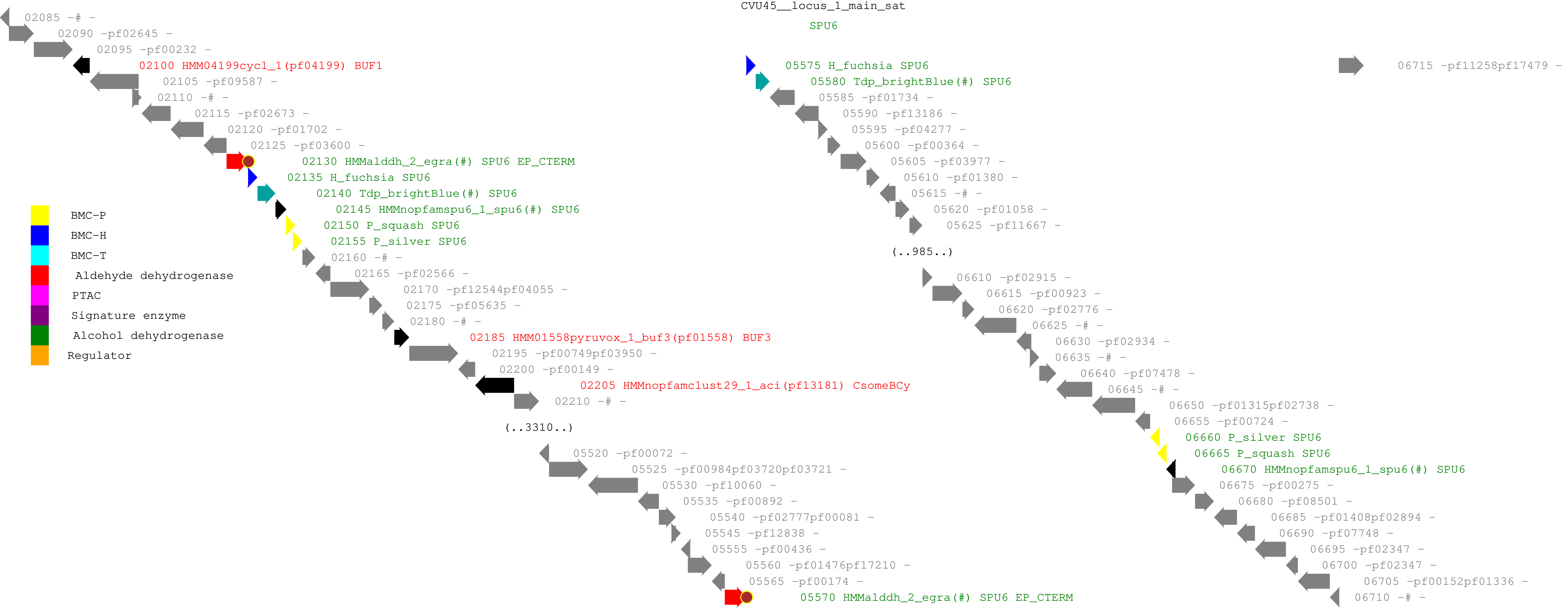

Chloroflexi bacterium HGW-Chloroflexi-8. Bacteria; Chloroflexi.

CVU46__locus_1_main_sat

SPU6

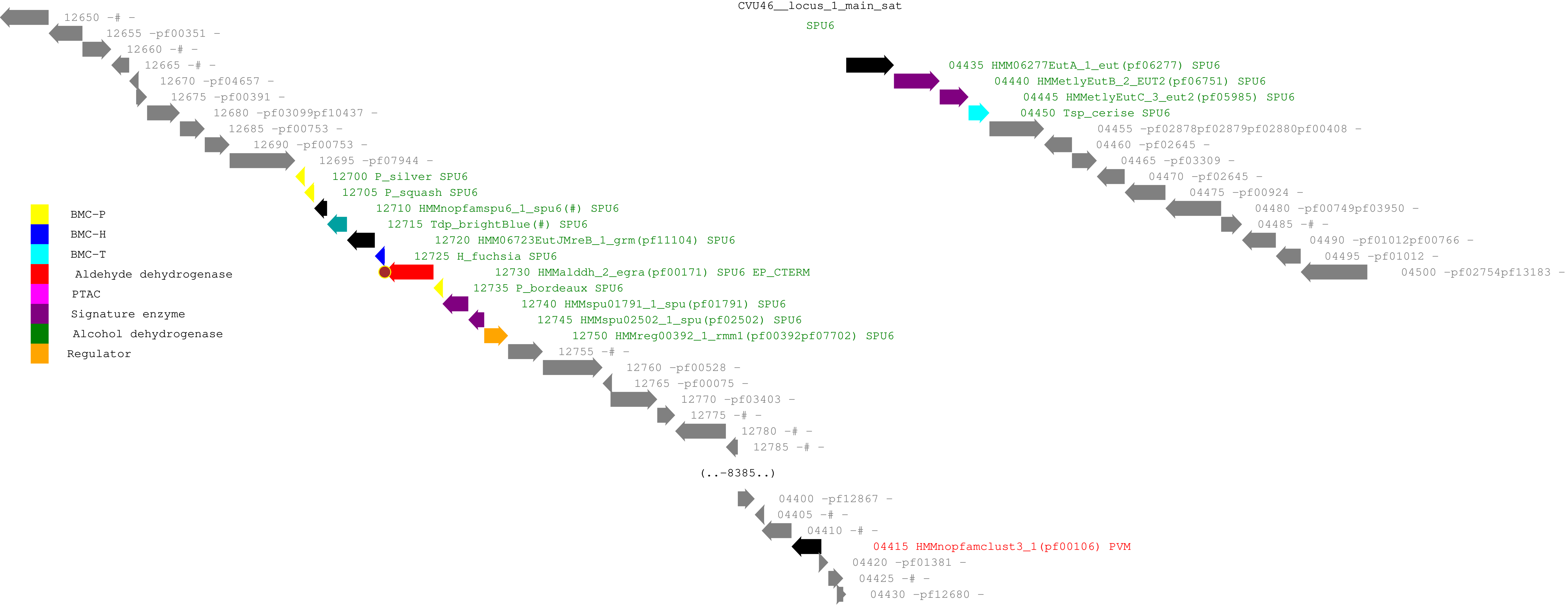

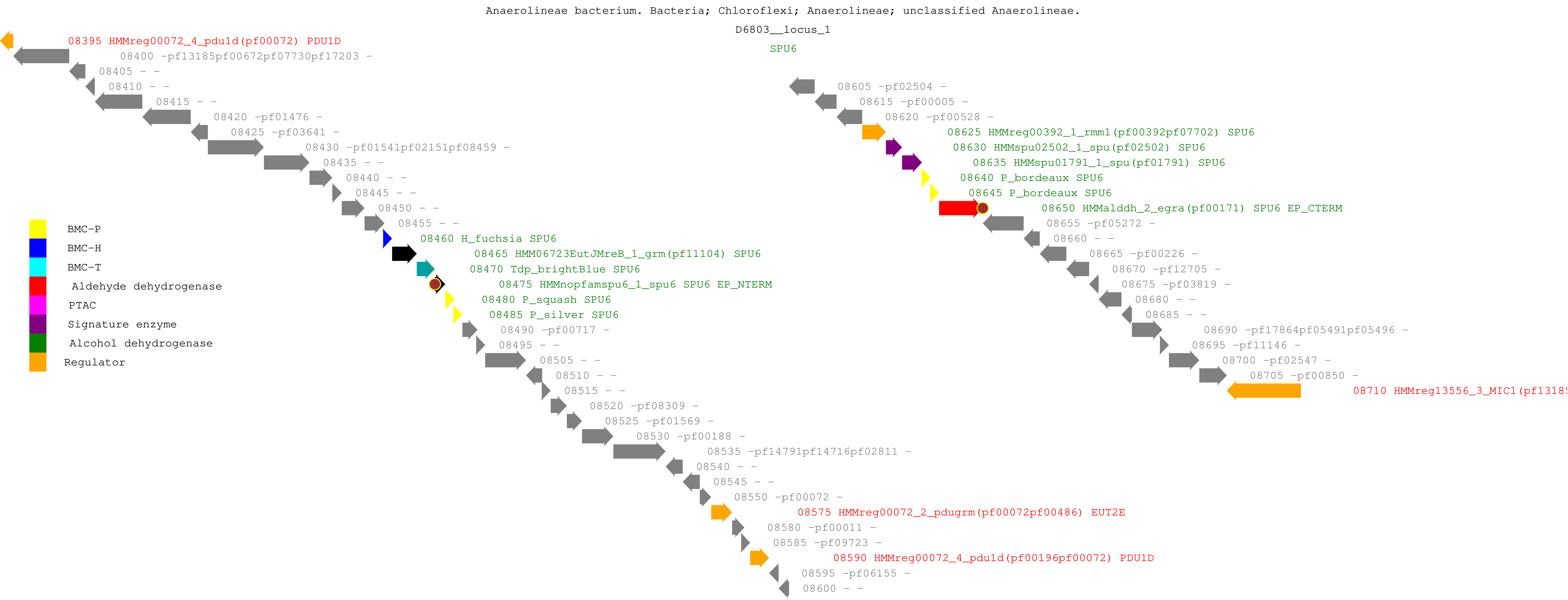
Anaerolineae bacterium. Bacteria; Chloroflexi; Anaerolineae; unclassified Anaerolineae.
D6803__locus_1
SPU6
08395 HMMreg00072_4_pdu1d(pf00072) PDU1D
08400 -pf13185pf00672pf07730pf17203 -
08405 - -
08410 - -
08415 - -
08420 -pf01476 -
08425 -pf03641 -
08430 -pf01541pf02151pf08459 -
08435 - -
08440 - -
08445 - -
08450 - -
08455 - -
08460 H_fuchsia SPU6
08465 HMM06723EutJMreB_1_grm(pf11104) SPU6
08470 Tdp_brightBlue SPU6
08475 HMMnopfamspu6_1_spu6 SPU6 EP_NTERM
08480 P_squash SPU6
08485 P_silver SPU6
08490 -pf00717 -
08495 - -
08505 - -
08510 - -
08515 - -
08520 -pf08309 -
08525 -pf01569 -
08530 -pf00188 -
08535 -pf14791pf14716pf02811 -
08540 - -
08545 - -
08550 -pf00072 -
08575 HMMreg00072_2_pdugrm(pf00072pf00486) EUT2E
08580 -pf00011 -
08585 -pf09723 -
08590 HMMreg00072_4_pdu1d(pf00196pf00072) PDU1D
08595 -pf06155 -
08600 - -
08605 -pf02504 -
08615 -pf00005 -
08620 -pf00528 -
08625 HMMreg00392_1_rmm1(pf00392pf07702) SPU6
08630 HMMspu02502_1_spu(pf02502) SPU6
08635 HMMspu01791_1_spu(pf01791) SPU6
08640 P_bordeaux SPU6
08645 P_bordeaux SPU6
08650 HMMalddh_2_egra(pf00171) SPU6 EP_CTERM
08655 -pf05272 -
08660 - -
08665 -pf00226 -
08670 -pf12705 -
08675 -pf03819 -
08680 - -
08685 - -
08690 -pf17864pf05491pf05496 -
08695 -pf11146 -
08700 -pf02547 -
08705 -pf00850 -
08710 HMMreg13556_3_MIC1(pf1318
BMC-P
BMC-H
BMC-T
Aldehyde dehydrogenase
PTAC
Signature enzyme
Alcohol dehydrogenase
Regulator

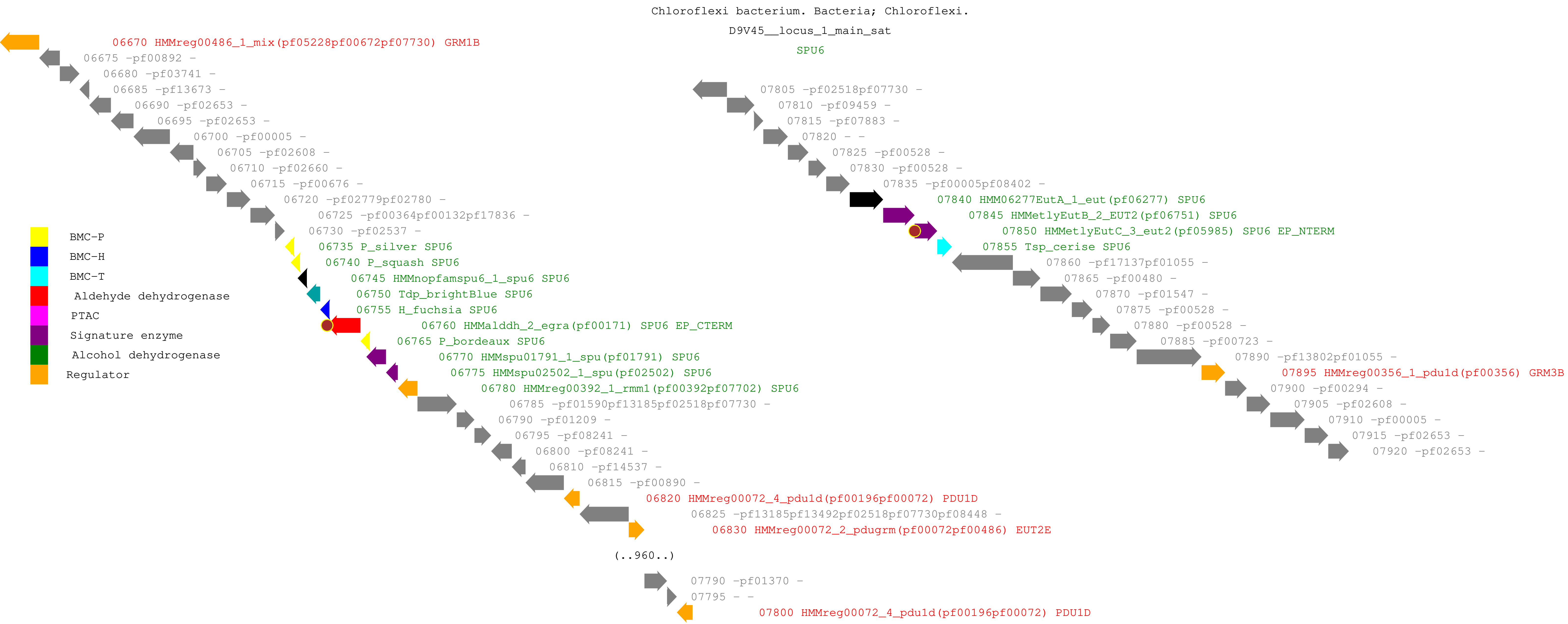
Chloroflexi bacterium. Bacteria; Chloroflexi.
D9V45__locus_1_main_sat
SPU6
06670 HMMreg00486_1_mix(pf05228pf00672pf07730) GRM1B
06675 -pf00892 -
06680 -pf03741 -
06685 -pf13673 -
06690 -pf02653 -
06695 -pf02653 -
06700 -pf00005 -
06705 -pf02608 -
06710 -pf02660 -
06715 -pf00676 -
06720 -pf02779pf02780 -
06725 -pf00364pf00132pf17836 -
06730 -pf02537 -
06735 P_silver SPU6
06740 P_squash SPU6
06745 HMMnopfamspu6_1_spu6 SPU6
06750 Tdp_brightBlue SPU6
06755 H_fuchsia SPU6
06760 HMMalddh_2_egra(pf00171) SPU6 EP_CTERM
06765 P_bordeaux SPU6
06770 HMMspu01791_1_spu(pf01791) SPU6
06775 HMMspu02502_1_spu(pf02502) SPU6
06780 HMMreg00392_1_rmm1(pf00392pf07702) SPU6
06785 -pf01590pf13185pf02518pf07730 -
06790 -pf01209 -
06795 -pf08241 -
06800 -pf08241 -
06810 -pf14537 -
06815 -pf00890 -
06820 HMMreg00072_4_pdu1d(pf00196pf00072) PDU1D
06825 -pf13185pf13492pf02518pf07730pf08448 -
06830 HMMreg00072_2_pdugrm(pf00072pf00486) EUT2E
(..960..)
07790 -pf01370 -
07795 - -
07800 HMMreg00072_4_pdu1d(pf00196pf00072) PDU1D
07805 -pf02518pf07730 -
07810 -pf09459 -
07815 -pf07883 -
07820 - -
07825 -pf00528 -
07830 -pf00528 -
07835 -pf00005pf08402 -
07840 HMM06277EutA_1_eut(pf06277) SPU6
07845 HMMetlyEutB_2_EUT2(pf06751) SPU6
07850 HMMetlyEutC_3_eut2(pf05985) SPU6 EP_NTERM
07855 Tsp_cerise SPU6
07860 -pf17137pf01055 -
07865 -pf00480 -
07870 -pf01547 -
07875 -pf00528 -
07880 -pf00528 -
07885 -pf00723 -
07890 -pf13802pf01055 -
07895 HMMreg00356_1_pdu1d(pf00356) GRM3B
07900 -pf00294 -
07905 -pf02608 -
07910 -pf00005 -
07915 -pf02653 -
07920 -pf02653 -
BMC-P
BMC-H
BMC-T
Aldehyde dehydrogenase
PTAC
Signature enzyme
Alcohol dehydrogenase
Regulator

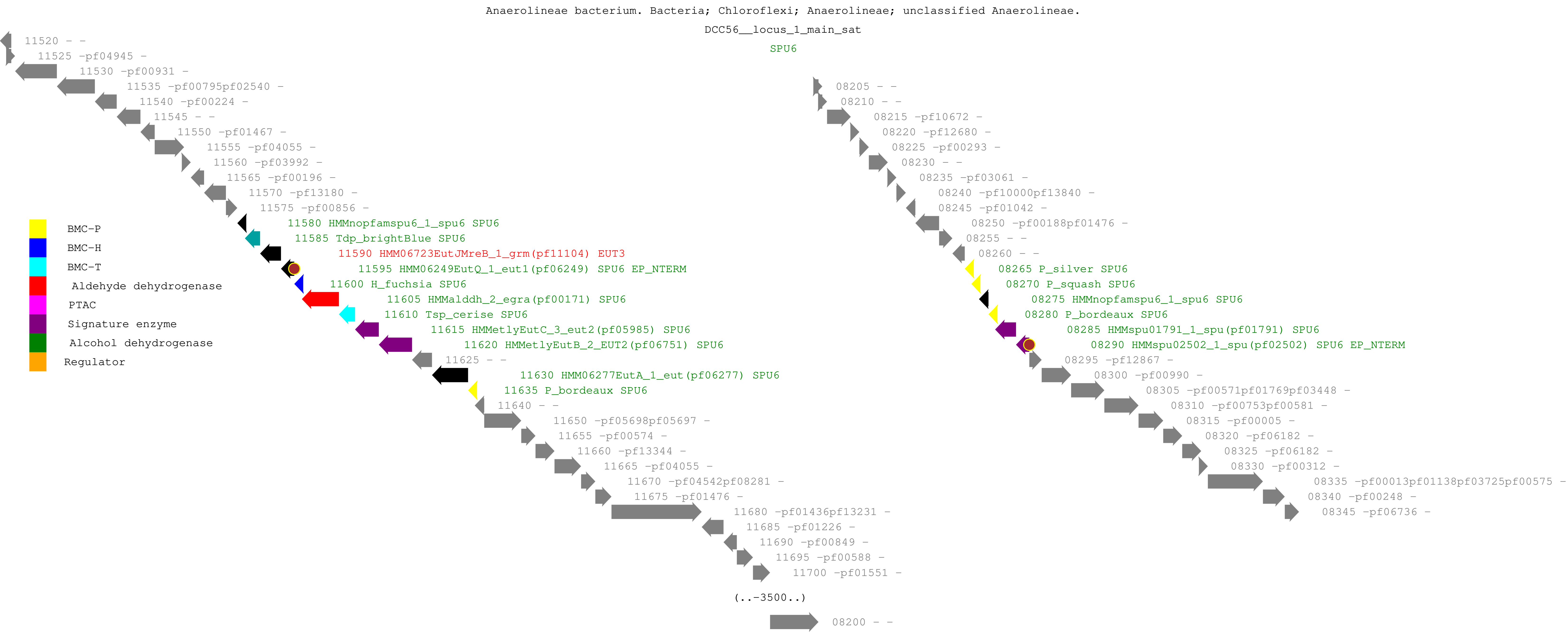
Anaerolineae bacterium. Bacteria; Chloroflexi; Anaerolineae; unclassified Anaerolineae.
DCC56__locus_1_main_sat
SPU6
11520 - -
11525 -pf04945 -
11530 -pf00931 -
11535 -pf00795pf02540 -
11540 -pf00224 -
11545 - -
11550 -pf01467 -
11555 -pf04055 -
11560 -pf03992 -
11565 -pf00196 -
11570 -pf13180 -
11575 -pf00856 -
11580 HMMnopfamspu6_1_spu6 SPU6
11585 Tdp_brightBlue SPU6
11590 HMM06723EutJMreB_1_grm(pf11104) EUT3
11595 HMM06249EutQ_1_eut1(pf06249) SPU6 EP_NTERM
11600 H_fuchsia SPU6
11605 HMMalddh_2_egra(pf00171) SPU6
11610 Tsp_cerise SPU6
11615 HMMetlyEutC_3_eut2(pf05985) SPU6
11620 HMMetlyEutB_2_EUT2(pf06751) SPU6
11625 - -
11630 HMM06277EutA_1_eut(pf06277) SPU6
11635 P_bordeaux SPU6
11640 - -
11650 -pf05698pf05697 -
11655 -pf00574 -
11660 -pf13344 -
11665 -pf04055 -
11670 -pf04542pf08281 -
11675 -pf01476 -
11680 -pf01436pf13231 -
11685 -pf01226 -
11690 -pf00849 -
11695 -pf00588 -
11700 -pf01551 -
(..-3500..)
08200 - -
08205 - -
08210 - -
08215 -pf10672 -
08220 -pf12680 -
08225 -pf00293 -
08230 - -
08235 -pf03061 -
08240 -pf10000pf13840 -
08245 -pf01042 -
08250 -pf00188pf01476 -
08255 - -
08260 - -
08265 P_silver SPU6
08270 P_squash SPU6
08275 HMMnopfamspu6_1_spu6 SPU6
08280 P_bordeaux SPU6
08285 HMMspu01791_1_spu(pf01791) SPU6
08290 HMMspu02502_1_spu(pf02502) SPU6 EP_NTERM
08295 -pf12867 -
08300 -pf00990 -
08305 -pf00571pf01769pf03448 -
08310 -pf00753pf00581 -
08315 -pf00005 -
08320 -pf06182 -
08325 -pf06182 -
08330 -pf00312 -
08335 -pf00013pf01138pf03725pf00575 -
08340 -pf00248 -
08345 -pf06736 -
BMC-P
BMC-H
BMC-T
Aldehyde dehydrogenase
PTAC
Signature enzyme
Alcohol dehydrogenase
Regulator

Anaerolineaceae bacterium. Bacteria; Chloroflexi; Anaerolineae; Anaerolineales; Anaerolineaceae.

DCK95__locus_1_main_sat

SPU6

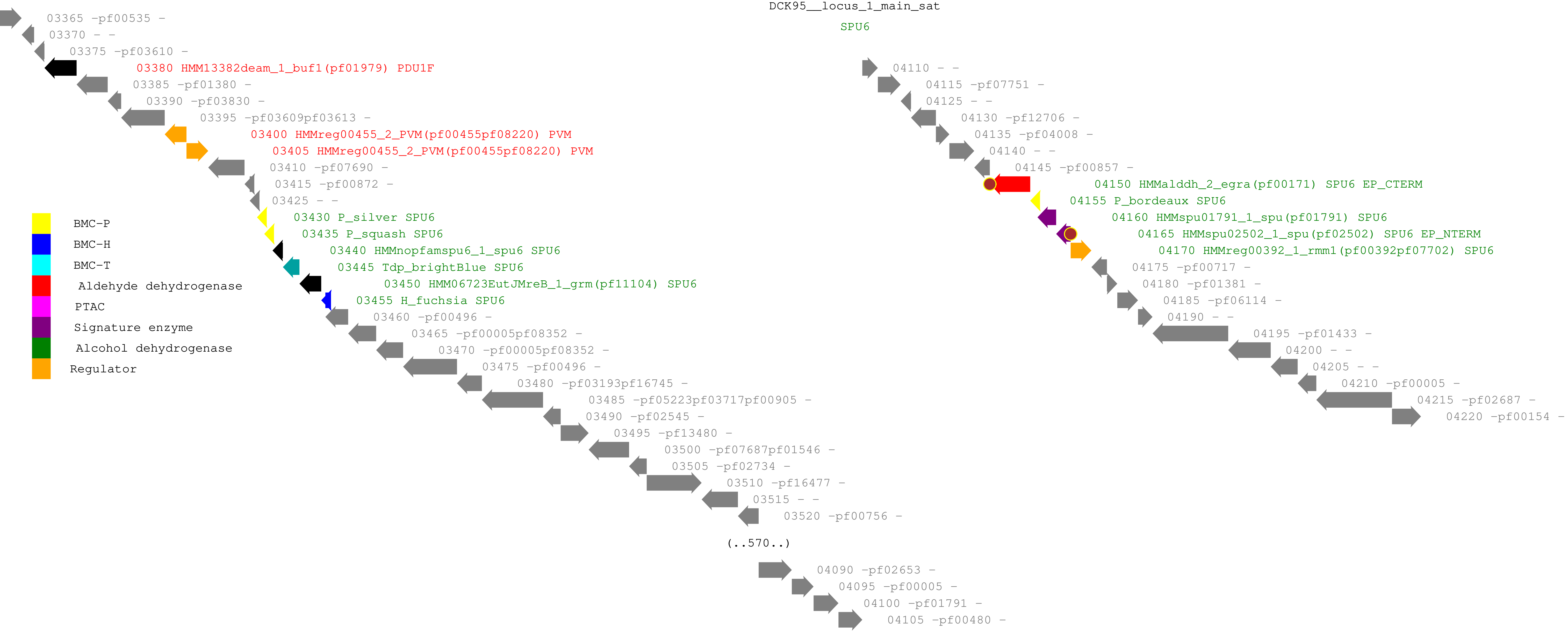

Anaerolineaceae bacterium. Bacteria; Chloroflexi; Anaerolineae; Anaerolineales; Anaerolineaceae.

DCL08__locus_1

SPU6

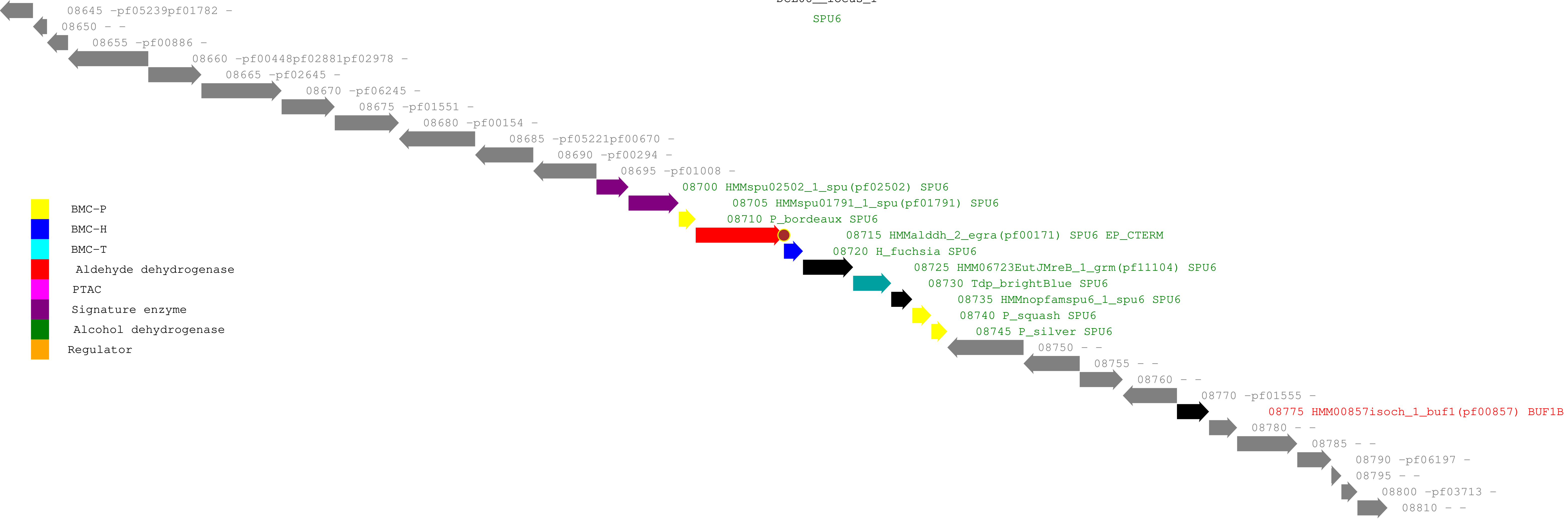

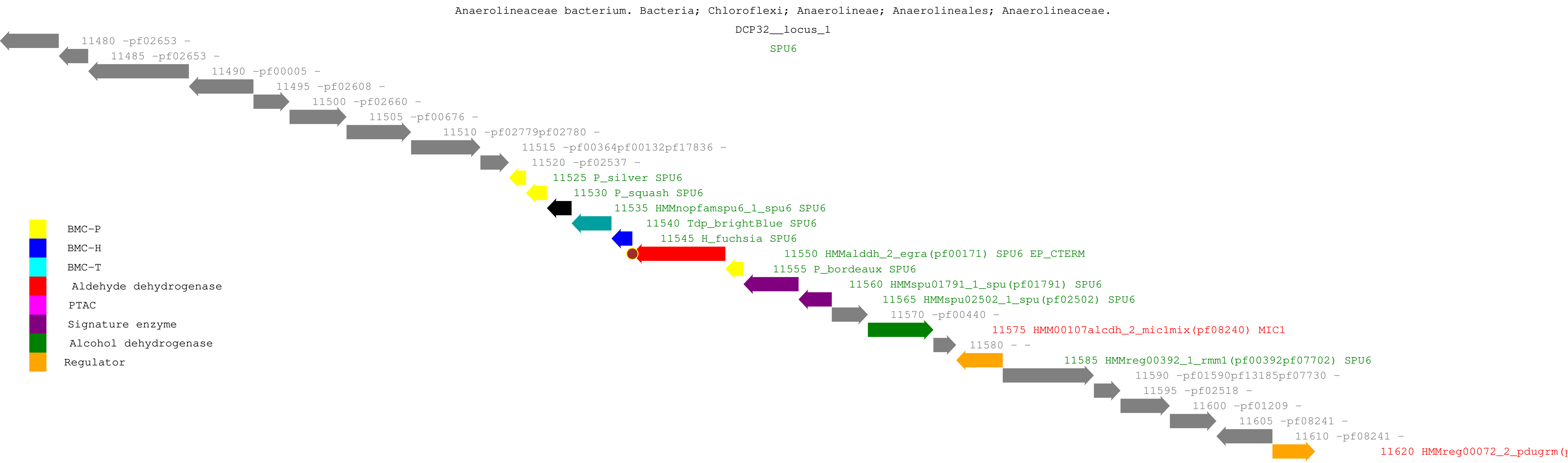

Anaerolineaceae bacterium. Bacteria; Chloroflexi; Anaerolineae; Anaerolineales; Anaerolineaceae.
DCP32__locus_1
SPU6
11480 -pf02653 -
11485 -pf02653 -
11490 -pf00005 -
11495 -pf02608 -
11500 -pf02660 -
11505 -pf00676 -
11510 -pf02779pf02780 -
11515 -pf00364pf00132pf17836 -
11520 -pf02537 -
11525 P_silver SPU6
11530 P_squash SPU6
11535 HMMnopfamspu6_1_spu6 SPU6
11540 Tdp_brightBlue SPU6
11545 H_fuchsia SPU6
11550 HMMaldh_2_egra(pf00171) SPU6 EP_CTERM
11555 P_bordeaux SPU6
11560 HMMspu01791_1_spu(pf01791) SPU6
11565 HMMspu02502_1_spu(pf02502) SPU6
11570 -pf00440 -
11575 HMM00107alcdh_2_mic1mix(pf08240) MIC1
11580 - -
11585 HMMreg00392_1_rmm1(pf00392pf07702) SPU6
11590 -pf01590pf13185pf07730 -
11595 -pf02518 -
11600 -pf01209 -
11605 -pf08241 -
11610 -pf08241 -
11620 HMMreg00072_2_pdugrm(
BMC-P
BMC-H
BMC-T
Aldehyde dehydrogenase
PTAC
Signature enzyme
Alcohol dehydrogenase
Regulator

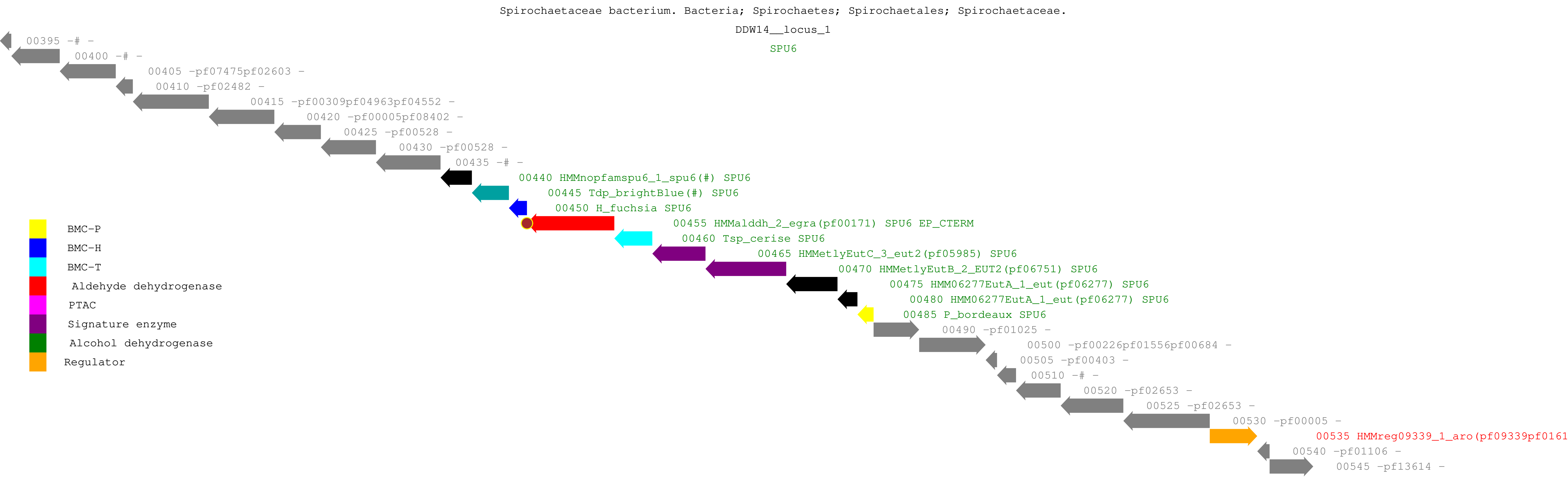
Spirochaetaceae bacterium. Bacteria; Spirochaetes; Spirochaetales; Spirochaetaceae.
DDW14__locus_1
SPU6
00395 -# -
00400 -# -
00405 -pf07475pf02603 -
00410 -pf02482 -
00415 -pf00309pf04963pf04552 -
00420 -pf00005pf08402 -
00425 -pf00528 -
00430 -pf00528 -
00435 -# -
00440 HMMnopfamspu6_1_spu6(#) SPU6
00445 Tdp_brightBlue(#) SPU6
00450 H_fuchsia SPU6
00455 HMMaldh_2_egra(pf00171) SPU6 EP_CTERM
00460 Tsp_cerise SPU6
00465 HMMetlyEutC_3_eut2(pf05985) SPU6
00470 HMMetlyEutB_2_EUT2(pf06751) SPU6
00475 HMM06277EutA_1_eut(pf06277) SPU6
00480 HMM06277EutA_1_eut(pf06277) SPU6
00485 P_bordeaux SPU6
00490 -pf01025 -
00500 -pf00226pf01556pf00684 -
00505 -pf00403 -
00510 -# -
00520 -pf02653 -
00525 -pf02653 -
00530 -pf00005 -
00535 HMMreg09339_1_aro(pf09339pf0161
00540 -pf01106 -
00545 -pf13614 -
BMC-P
BMC-H
BMC-T
Aldehyde dehydrogenase
PTAC
Signature enzyme
Alcohol dehydrogenase
Regulator

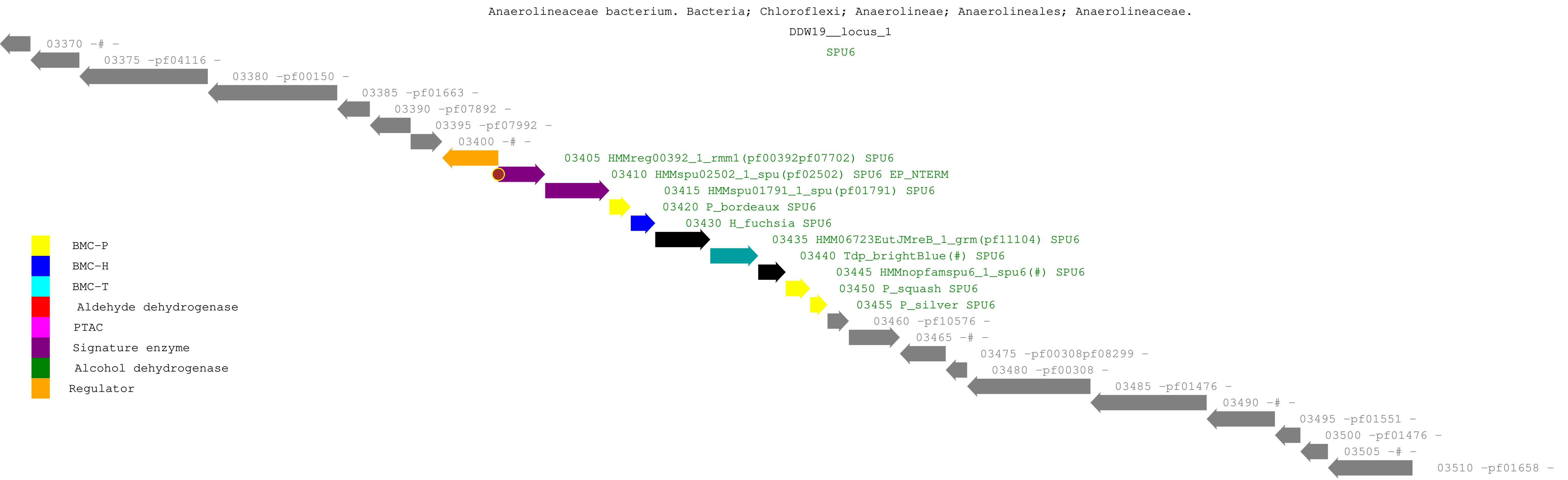
Anaerolineaceae bacterium. Bacteria; Chloroflexi; Anaerolineae; Anaerolineales; Anaerolineaceae.
DDW19__locus_1
SPU6
03370 -# -
03375 -pf04116 -
03380 -pf00150 -
03385 -pf01663 -
03390 -pf07892 -
03395 -pf07992 -
03400 -# -
03405 HMMreg00392_1_rmm1(pf00392pf07702) SPU6
03410 HMMspu02502_1_spu(pf02502) SPU6 EP_NTERM
03415 HMMspu01791_1_spu(pf01791) SPU6
03420 P_bordeaux SPU6
03430 H_fuchsia SPU6
03435 HMM06723EutJMreB_1_grm(pf11104) SPU6
03440 Tdp_brightBlue(#) SPU6
03445 HMMnopfamspu6_1_spu6(#) SPU6
03450 P_squash SPU6
03455 P_silver SPU6
03460 -pf10576 -
03465 -# -
03475 -pf00308pf08299 -
03480 -pf00308 -
03485 -pf01476 -
03490 -# -
03495 -pf01551 -
03500 -pf01476 -
03505 -# -
03510 -pf01658 -
BMC-P
BMC-H
BMC-T
Aldehyde dehydrogenase
PTAC
Signature enzyme
Alcohol dehydrogenase
Regulator

Anaerolineaceae bacterium. Bacteria; Chloroflexi; Anaerolineae; Anaerolineales; Anaerolineaceae.

DF984__locus_1_main_sat

SPU6

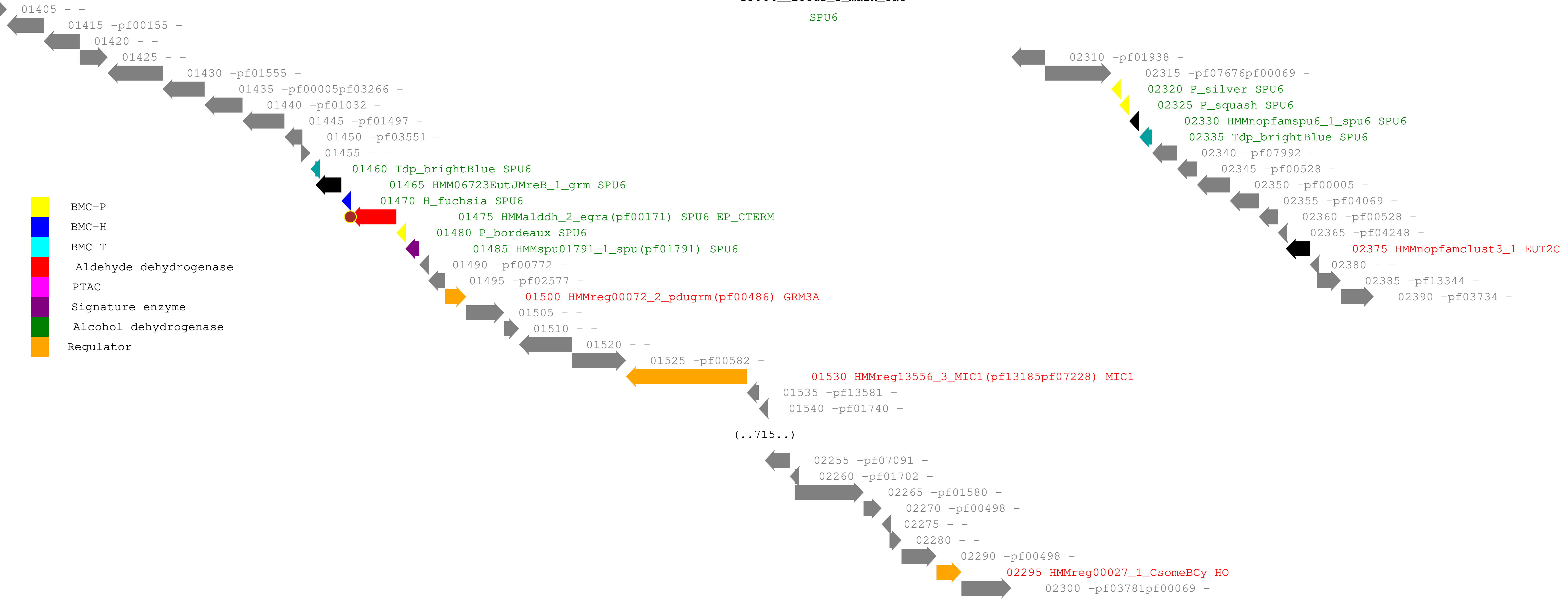

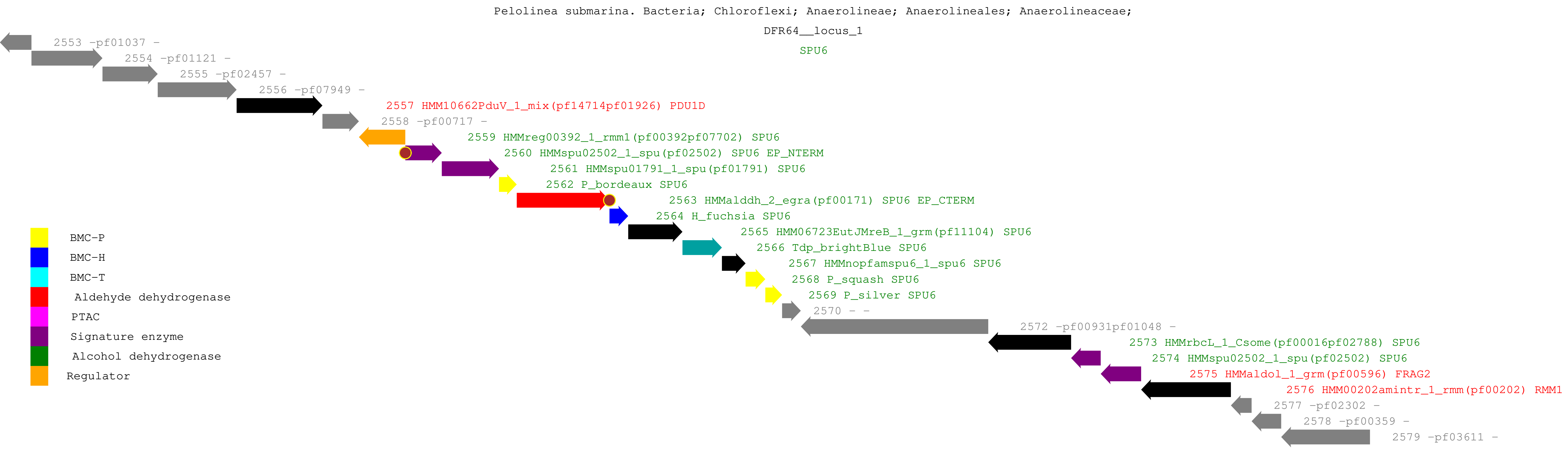
Pelolinea submarina. Bacteria; Chloroflexi; Anaerolineae; Anaerolineales; Anaerolineaceae;
DFR64__locus_1
SPU6
2553 -pf01037 -
2554 -pf01121 -
2555 -pf02457 -
2556 -pf07949 -
2557 HMM10662PduV_1_mix(pf14714pf01926) PDU1D
2558 -pf00717 -
2559 HMMreg00392_1_rmm1(pf00392pf07702) SPU6
2560 HMMspu02502_1_spu(pf02502) SPU6 EP_NTERM
2561 HMMspu01791_1_spu(pf01791) SPU6
2562 P_bordeaux SPU6
2563 HMMalddh_2_egra(pf00171) SPU6 EP_CTERM
2564 H_fuchsia SPU6
2565 HMM06723EutJMreB_1_grm(pf11104) SPU6
2566 Tdp_brightBlue SPU6
2567 HMMnopfamspu6_1_spu6 SPU6
2568 P_squash SPU6
2569 P_silver SPU6
2570 - -
2572 -pf00931pf01048 -
2573 HMMrbcL_1_Csome(pf00016pf02788) SPU6
2574 HMMspu02502_1_spu(pf02502) SPU6
2575 HMMaldol_1_grm(pf00596) FRAG2
2576 HMM00202amintr_1_rmm(pf00202) RMM1
2577 -pf02302 -
2578 -pf00359 -
2579 -pf03611 -
BMC-P
BMC-H
BMC-T
Aldehyde dehydrogenase
PTAC
Signature enzyme
Alcohol dehydrogenase
Regulator

Anaerolineaceae bacterium. Bacteria; Chloroflexi; Anaerolineae; Anaerolineales; Anaerolineaceae.

DIW44__locus_1

SPU6

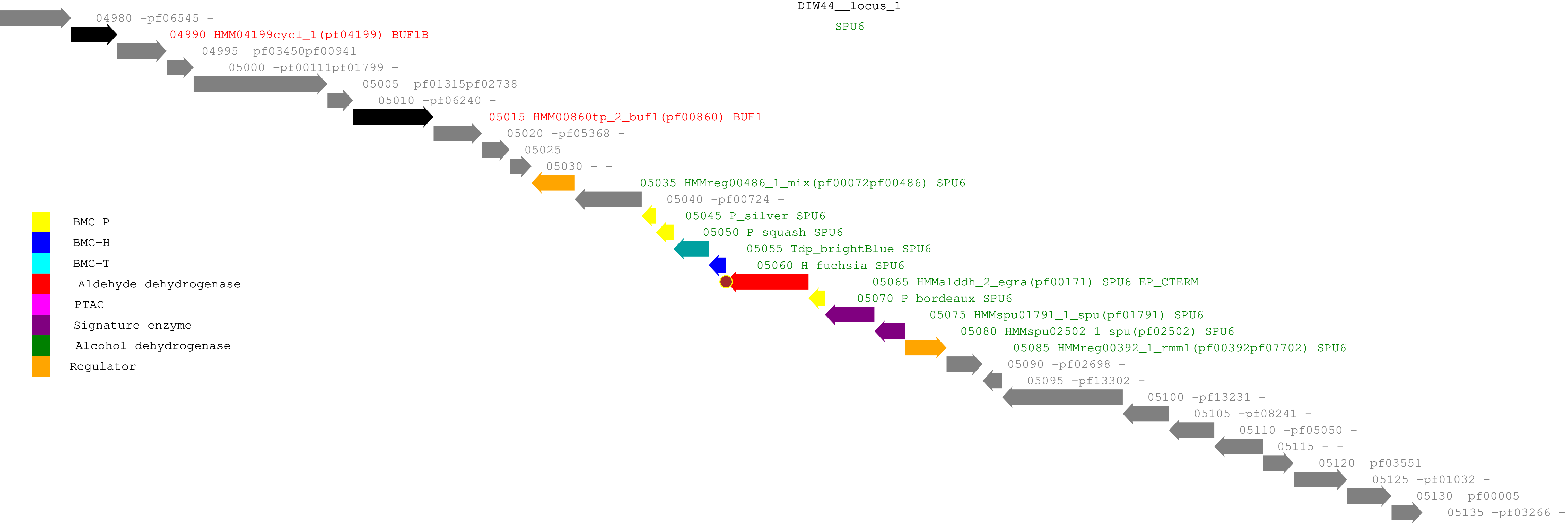

Chloroflexi bacterium. Bacteria; Chloroflexi.

DRI46__locus_1

SPU6

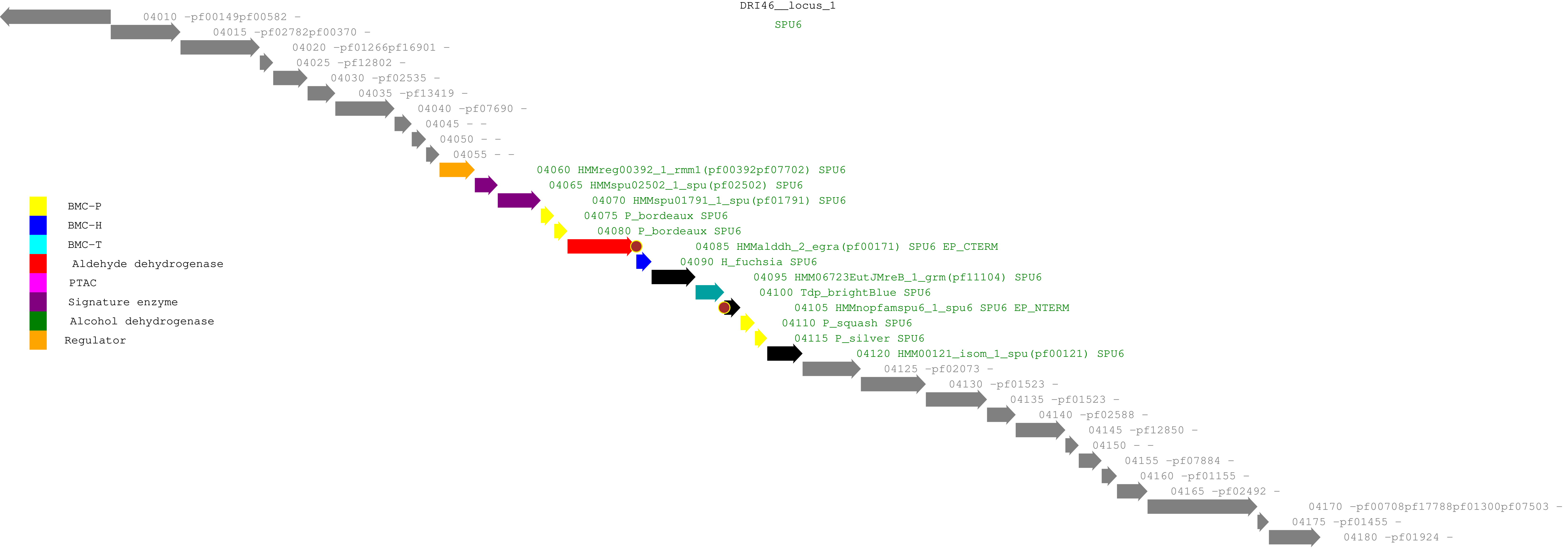

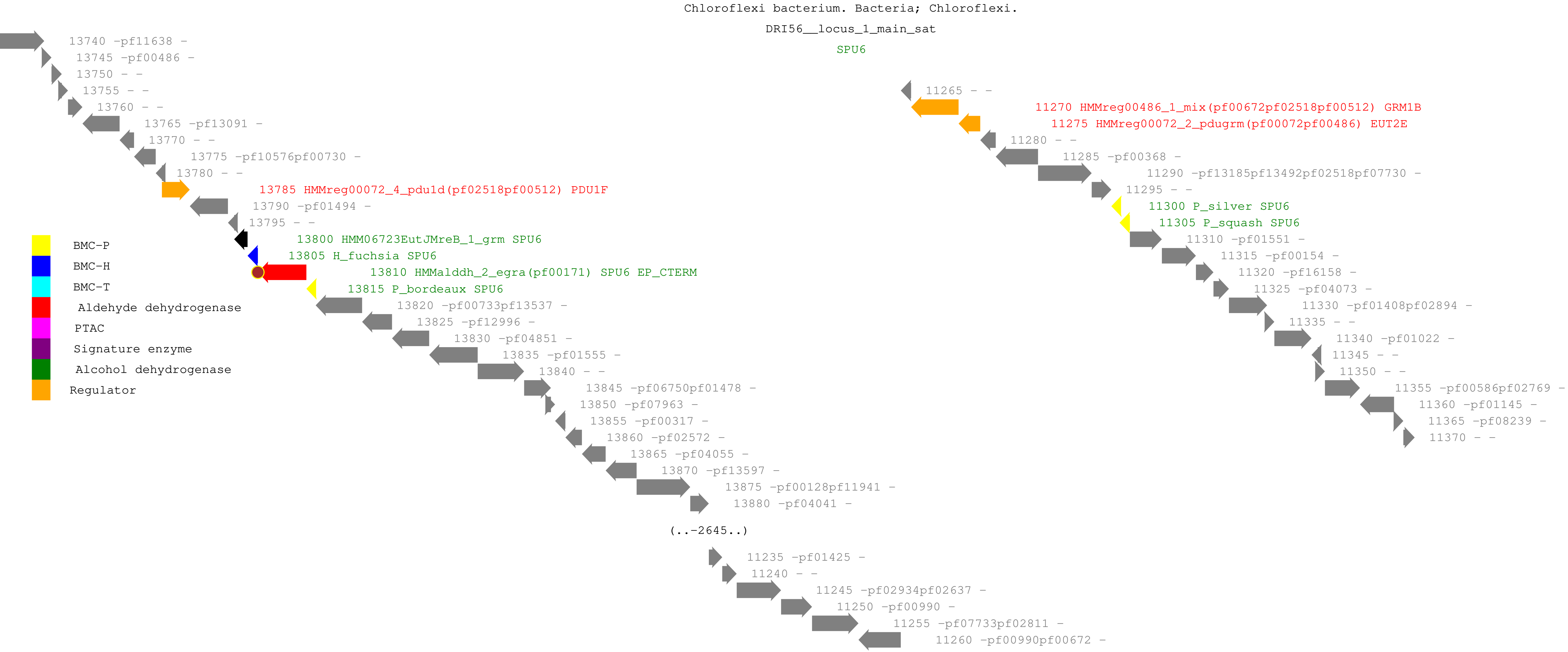

Chloroflexi bacterium. Bacteria; Chloroflexi.
DRI56__locus_1_main_sat
SPU6
13740 -pf11638 -
13745 -pf00486 -
13750 - -
13755 - -
13760 - -
13765 -pf13091 -
13770 - -
13775 -pf10576pf00730 -
13780 - -
13785 HMMreg00072_4_pdu1d(pf02518pf00512) PDU1F
13790 -pf01494 -
13795 - -
13800 HMM06723EutJMreB_1_grm SPU6
13805 H_fuchsia SPU6
13810 HMMalddh_2_egra(pf00171) SPU6 EP_CTERM
13815 P_bordeaux SPU6
13820 -pf00733pf13537 -
13825 -pf12996 -
13830 -pf04851 -
13835 -pf01555 -
13840 - -
13845 -pf06750pf01478 -
13850 -pf07963 -
13855 -pf00317 -
13860 -pf02572 -
13865 -pf04055 -
13870 -pf13597 -
13875 -pf00128pf11941 -
13880 -pf04041 -
(..-2645..)
11235 -pf01425 -
11240 - -
11245 -pf02934pf02637 -
11250 -pf00990 -
11255 -pf07733pf02811 -
11260 -pf00990pf00672 -
11265 - -
11270 HMMreg00486_1_mix(pf00672pf02518pf00512) GRM1B
11275 HMMreg00072_2_pdugrm(pf00072pf00486) EUT2E
11280 - -
11285 -pf00368 -
11290 -pf13185pf13492pf02518pf07730 -
11295 - -
11300 P_silver SPU6
11305 P_squash SPU6
11310 -pf01551 -
11315 -pf00154 -
11320 -pf16158 -
11325 -pf04073 -
11330 -pf01408pf02894 -
11335 - -
11340 -pf01022 -
11345 - -
11350 - -
11355 -pf00586pf02769 -
11360 -pf01145 -
11365 -pf08239 -
11370 - -
BMC-P
BMC-H
BMC-T
Aldehyde dehydrogenase
PTAC
Signature enzyme
Alcohol dehydrogenase
Regulator

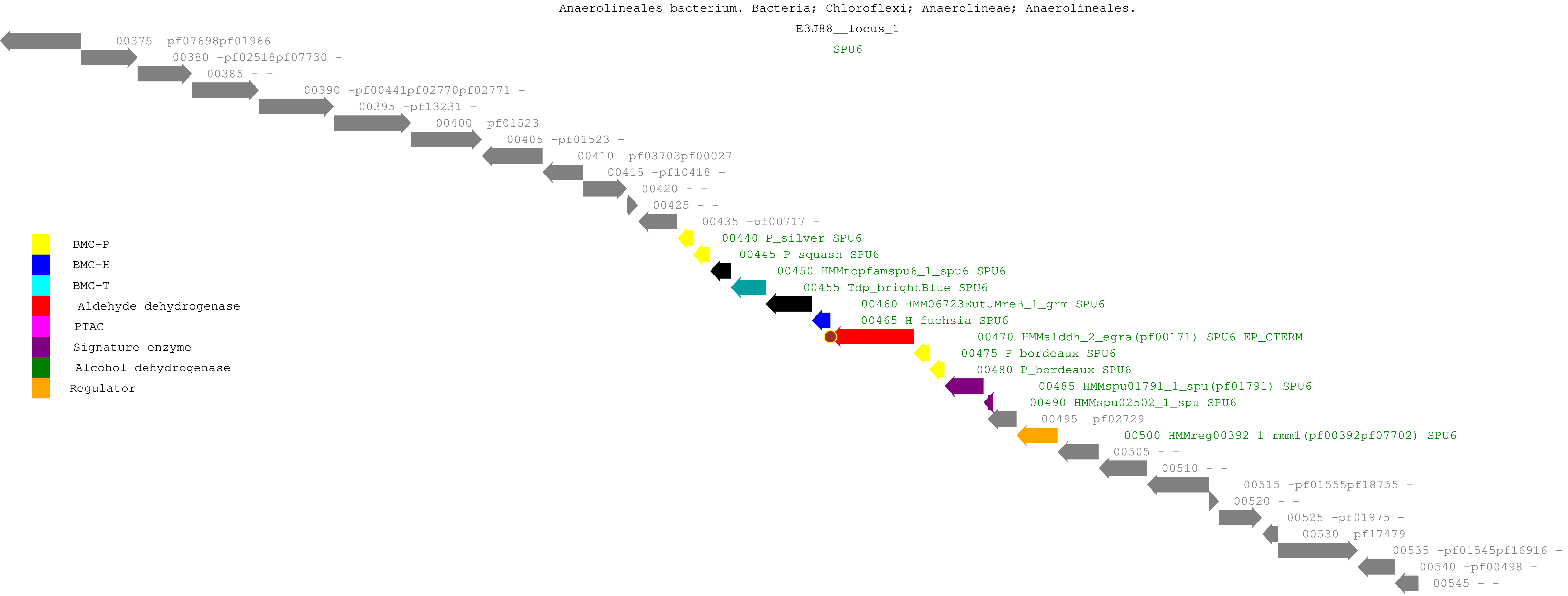
Anaerolineales bacterium. Bacteria; Chloroflexi; Anaerolineae; Anaerolineales.
E3J88__locus_1
SPU6
00375 -pf07698pf01966 -
00380 -pf02518pf07730 -
00385 - -
00390 -pf00441pf02770pf02771 -
00395 -pf13231 -
00400 -pf01523 -
00405 -pf01523 -
00410 -pf03703pf00027 -
00415 -pf10418 -
00420 - -
00425 - -
00435 -pf00717 -
00440 P_silver SPU6
00445 P_squash SPU6
00450 HMMnopfamspu6_1_spu6 SPU6
00455 Tdp_brightBlue SPU6
00460 HMM06723EutJMreB_1_grm SPU6
00465 H_fuchsia SPU6
00470 HMMaldh_2_egra(pf00171) SPU6 EP_CTERM
00475 P_bordeaux SPU6
00480 P_bordeaux SPU6
00485 HMMspu01791_1_spu(pf01791) SPU6
00490 HMMspu02502_1_spu SPU6
00495 -pf02729 -
00500 HMMreg00392_1_rmm1(pf00392pf07702) SPU6
00505 - -
00510 - -
00515 -pf01555pf18755 -
00520 - -
00525 -pf01975 -
00530 -pf17479 -
00535 -pf01545pf16916 -
00540 -pf00498 -
00545 - -
BMC-P
BMC-H
BMC-T
Aldehyde dehydrogenase
PTAC
Signature enzyme
Alcohol dehydrogenase
Regulator

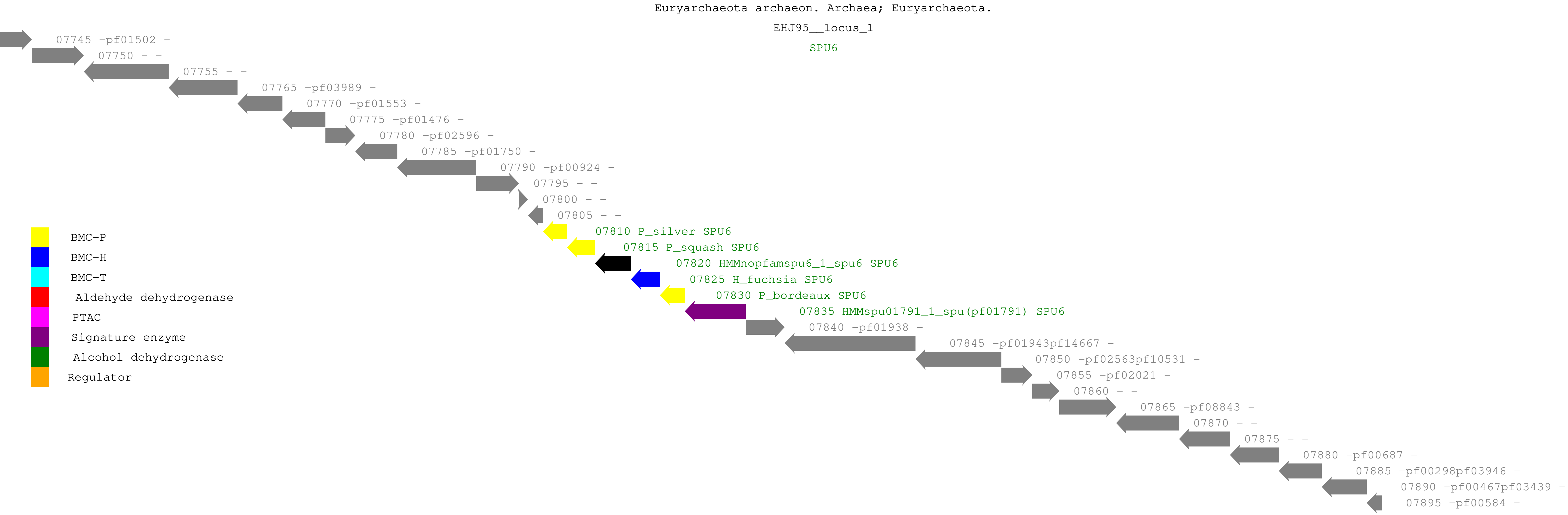
Euryarchaeota archaeon. Archaea; Euryarchaeota.
EHJ95__locus_1
SPU6
07745 -pf01502 -
07750 - -
07755 - -
07765 -pf03989 -
07770 -pf01553 -
07775 -pf01476 -
07780 -pf02596 -
07785 -pf01750 -
07790 -pf00924 -
07795 - -
07800 - -
07805 - -
07810 P_silver SPU6
07815 P_squash SPU6
07820 HMMnopfamspu6_1_spu6 SPU6
07825 H_fuchsia SPU6
07830 P_bordeaux SPU6
07835 HMMspu01791_1_spu(pf01791) SPU6
07840 -pf01938 -
07845 -pf01943pf14667 -
07850 -pf02563pf10531 -
07855 -pf02021 -
07860 - -
07865 -pf08843 -
07870 - -
07875 - -
07880 -pf00687 -
07885 -pf00298pf03946 -
07890 -pf00467pf03439 -
07895 -pf00584 -
BMC-P
BMC-H
BMC-T
Aldehyde dehydrogenase
PTAC
Signature enzyme
Alcohol dehydrogenase
Regulator

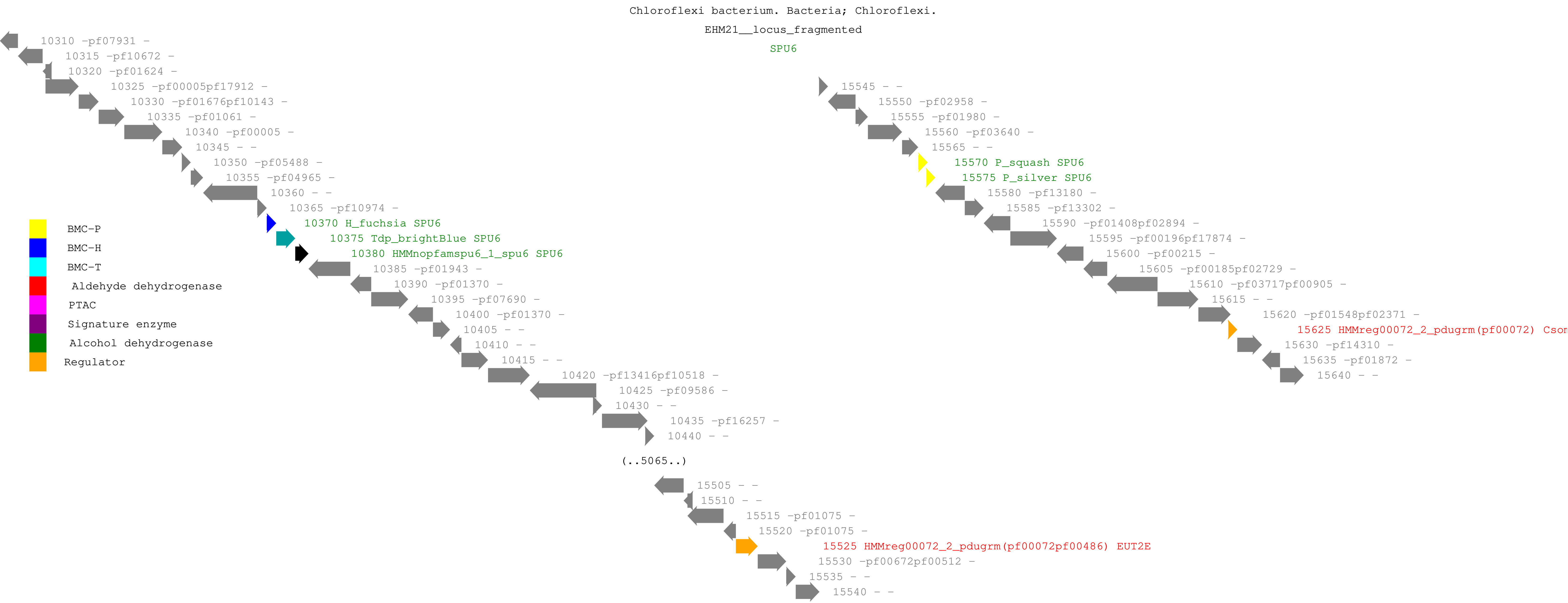
Chloroflexi bacterium. Bacteria; Chloroflexi.
EHM21__locus_fragmented
SPU6
10310 -pf07931 -
10315 -pf10672 -
10320 -pf01624 -
10325 -pf00005pf17912 -
10330 -pf01676pf10143 -
10335 -pf01061 -
10340 -pf00005 -
10345 - -
10350 -pf05488 -
10355 -pf04965 -
10360 - -
10365 -pf10974 -
10370 H_fuchsia SPU6
10375 Tdp_brightBlue SPU6
10380 HMMnopfamspu6_1_spu6 SPU6
10385 -pf01943 -
10390 -pf01370 -
10395 -pf07690 -
10400 -pf01370 -
10405 - -
10410 - -
10415 - -
10420 -pf13416pf10518 -
10425 -pf09586 -
10430 - -
10435 -pf16257 -
10440 - -
(..5065..)
15505 - -
15510 - -
15515 -pf01075 -
15520 -pf01075 -
15525 HMMreg00072_2_pdugrm(pf00072pf00486) EUT2E
15530 -pf00672pf00512 -
15535 - -
15540 - -
15545 - -
15550 -pf02958 -
15555 -pf01980 -
15560 -pf03640 -
15565 - -
15570 P_squash SPU6
15575 P_silver SPU6
15580 -pf13180 -
15585 -pf13302 -
15590 -pf01408pf02894 -
15595 -pf00196pf17874 -
15600 -pf00215 -
15605 -pf00185pf02729 -
15610 -pf03717pf00905 -
15615 - -
15620 -pf01548pf02371 -
15625 HMMreg00072_2_pdugrm(pf00072) Cso
15630 -pf14310 -
15635 -pf01872 -
15640 - -
BMC-P
BMC-H
BMC-T
Aldehyde dehydrogenase
PTAC
Signature enzyme
Alcohol dehydrogenase
Regulator

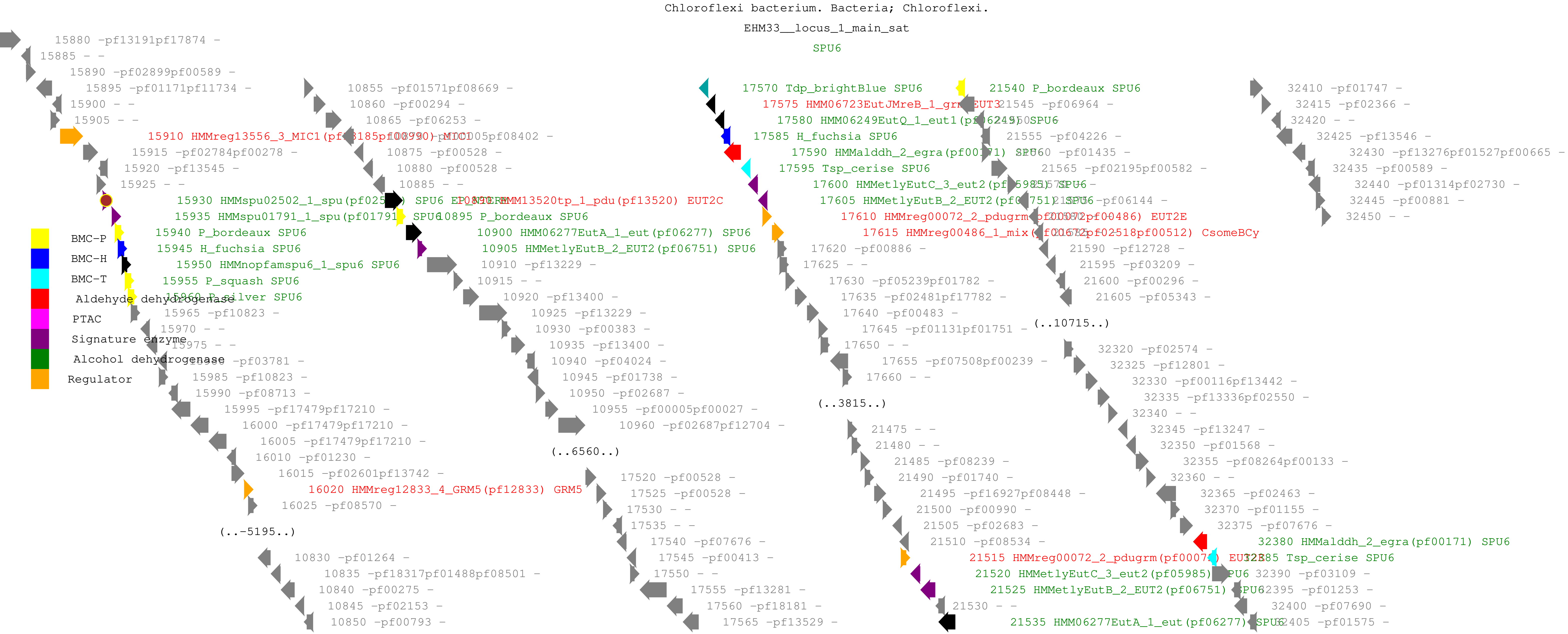
Chloroflexi bacterium. Bacteria; Chloroflexi.
EHM33__locus_1_main_sat
SPU6
BMC-P
BMC-H
BMC-T
Aldehyde dehydrogenase
PTAC
Signature enzyme
Alcohol dehydrogenase
Regulator
15880 -pf13191pf17874 -
15885 - -
15890 -pf02899pf00589 -
15895 -pf01171pf11734 -
15900 - -
15905 - -
15910 HMMreg13556_3_MIC1
15915 -pf02784pf00278 -
15920 -pf13545 -
15925 - -
15930 HMMspu02502_1_spu
15935 HMMspu01791_1_spu
15940 P_bordeaux SPU6
15945 H_fuchsia SPU6
15950 HMMnopfamspu6_1_spu6 SPU6
15955 P_squash SPU6
15960 P_silver SPU6
15965 -pf10823 -
15970 - -
15975 - -
-pf03781 -
15985 -pf10823 -
15990 -pf08713 -
15995 -pf17479pf17210 -
16000 -pf17479pf17210 -
16005 -pf17479pf17210 -
16010 -pf01230 -
16015 -pf02601pf13742 -
16020 HMMreg12833_4_GRM5(pf12833) GRM5
16025 -pf08570 -
(..-5195..)
10830 -pf01264 -
10835 -pf18317pf01488pf08501 -
10840 -pf00275 -
10845 -pf02153 -
10850 -pf00793 -
10855 -pf01571pf08669 -
10860 -pf00294 -
10865 -pf06253 -
10875 -pf00528 -
10880 -pf00528 -
10885 - -
HMM13520tp_1_pdu(pf13520) EUT2C
10895 P_bordeaux SPU6
10900 HMM06277EutA_1_eut(pf06277) SPU6
10905 HMMetlyEutB_2_EUT2(pf06751) SPU6
10910 -pf13229 -
10915 - -
10920 -pf13400 -
10925 -pf13229 -
10930 -pf00383 -
10935 -pf13400 -
10940 -pf04024 -
10945 -pf01738 -
10950 -pf02687 -
10955 -pf00005pf00027 -
10960 -pf02687pf12704 -
(..6560..)
17520 -pf00528 -
17525 -pf00528 -
17530 - -
17535 - -
17540 -pf07676 -
17545 -pf00413 -
17550 - -
17555 -pf13281 -
17560 -pf18181 -
17565 -pf13529 -
17570 Tdp_brightBlue SPU6
17575 HMM06723EutJMreB_1_gr
17580 HMM06249EutQ_1_eut1
17585 H_fuchsia SPU6
17590 HMMalddh_2_egra(pf00171) SPU6
17595 Tsp_cerise SPU6
17600 HMMetlyEutC_3_eut2
17605 HMMetlyEutB_2_EUT2
17610 HMMreg00072_2_pdugrm
17615 HMMreg00486_1_mix
17620 -pf00886 -
17625 - -
17630 -pf05239pf01782 -
17635 -pf02481pf17782 -
17640 -pf00483 -
17645 -pf01131pf01751 -
17650 - -
17655 -pf07508pf00239 -
17660 - -
(..3815..)
21475 - -
21480 - -
21485 -pf08239 -
21490 -pf01740 -
21495 -pf16927pf08448 -
21500 -pf00990 -
21505 -pf02683 -
21510 -pf08534 -
21515 HMMreg00072_2_pdugrm EUT2E
21520 HMMetlyEutC_3_eut2(pf05985)
21525 HMMetlyEutB_2_EUT2(pf06751)
21530 - -
21535 HMM06277EutA_1_eut(pf06277) SPU6
21540 P_bordeaux SPU6
21545 -pf06964 -
21555 -pf04226 -
21560 -pf01435 -
21565 -pf02195pf00582 -
21590 -pf12728 -
21595 -pf03209 -
21600 -pf00296 -
21605 -pf05343 -
(..10715..)
32320 -pf02574 -
32325 -pf12801 -
32330 -pf00116pf13442 -
32335 -pf13336pf02550 -
32340 - -
32345 -pf13247 -
32350 -pf01568 -
32355 -pf08264pf00133 -
32360 - -
32365 -pf02463 -
32370 -pf01155 -
32375 -pf07676 -
32380 HMMalddh_2_egra(pf00171) SPU6
Tsp_cerise SPU6
32390 -pf03109 -
-pf01253 -
32400 -pf07690 -
-pf01575 -
32410 -pf01747 -
32415 -pf02366 -
32420 - -
32425 -pf13546 -
32430 -pf13276pf01527pf00665 -
32435 -pf00589 -
32440 -pf01314pf02730 -
32445 -pf00881 -
32450 - -

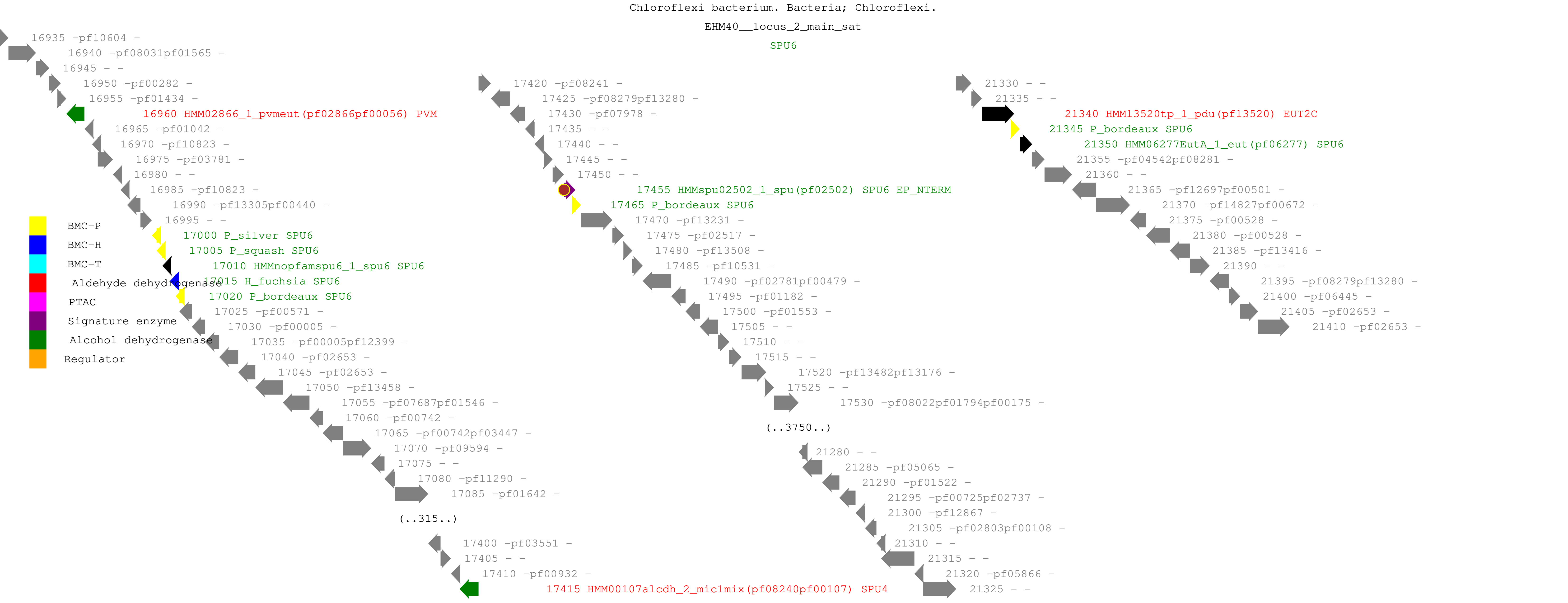

Chloroflexi bacterium. Bacteria; Chloroflexi.
EHM40__locus_2_main_sat
SPU6
16935 -pf10604 -
16940 -pf08031pf01565 -
16945 - -
16950 -pf00282 -
16955 -pf01434 -
16960 HMM02866_1_pvmeut(pf02866pf00056) PVM
16965 -pf01042 -
16970 -pf10823 -
16975 -pf03781 -
16980 - -
16985 -pf10823 -
16990 -pf13305pf00440 -
16995 - -
17000 P_silver SPU6
17005 P_squash SPU6
17010 HMMnopfamspu6_1_spu6 SPU6
17015 H_fuchsia SPU6
17020 P_bordeaux SPU6
17025 -pf00571 -
17030 -pf00005 -
17035 -pf00005pf12399 -
17040 -pf02653 -
17045 -pf02653 -
17050 -pf13458 -
17055 -pf07687pf01546 -
17060 -pf00742 -
17065 -pf00742pf03447 -
17070 -pf09594 -
17075 - -
17080 -pf11290 -
17085 -pf01642 -
(..315..)
17400 -pf03551 -
17405 - -
17410 -pf00932 -
17415 HMM00107alcdh_2_mic1mix(pf08240pf00107) SPU4
17420 -pf08241 -
17425 -pf08279pf13280 -
17430 -pf07978 -
17435 - -
17440 - -
17445 - -
17450 - -
17455 HMMspu02502_1_spu(pf02502) SPU6 EP_NTERM
17465 P_bordeaux SPU6
17470 -pf13231 -
17475 -pf02517 -
17480 -pf13508 -
17485 -pf10531 -
17490 -pf02781pf00479 -
17495 -pf01182 -
17500 -pf01553 -
17505 - -
17510 - -
17515 - -
17520 -pf13482pf13176 -
17525 - -
17530 -pf08022pf01794pf00175 -
(..3750..)
21280 - -
21285 -pf05065 -
21290 -pf01522 -
21295 -pf00725pf02737 -
21300 -pf12867 -
21305 -pf02803pf00108 -
21310 - -
21315 - -
21320 -pf05866 -
21325 - -
21330 - -
21335 - -
21340 HMM13520tp_1_pdu(pf13520) EUT2C
21345 P_bordeaux SPU6
21350 HMM06277EutA_1_eut(pf06277) SPU6
21355 -pf04542pf08281 -
21360 - -
21365 -pf12697pf00501 -
21370 -pf14827pf00672 -
21375 -pf00528 -
21380 -pf00528 -
21385 -pf13416 -
21390 - -
21395 -pf08279pf13280 -
21400 -pf06445 -
21405 -pf02653 -
21410 -pf02653 -
BMC-P
BMC-H
BMC-T
Aldehyde dehydrogenase
PTAC
Signature enzyme
Alcohol dehydrogenase
Regulator

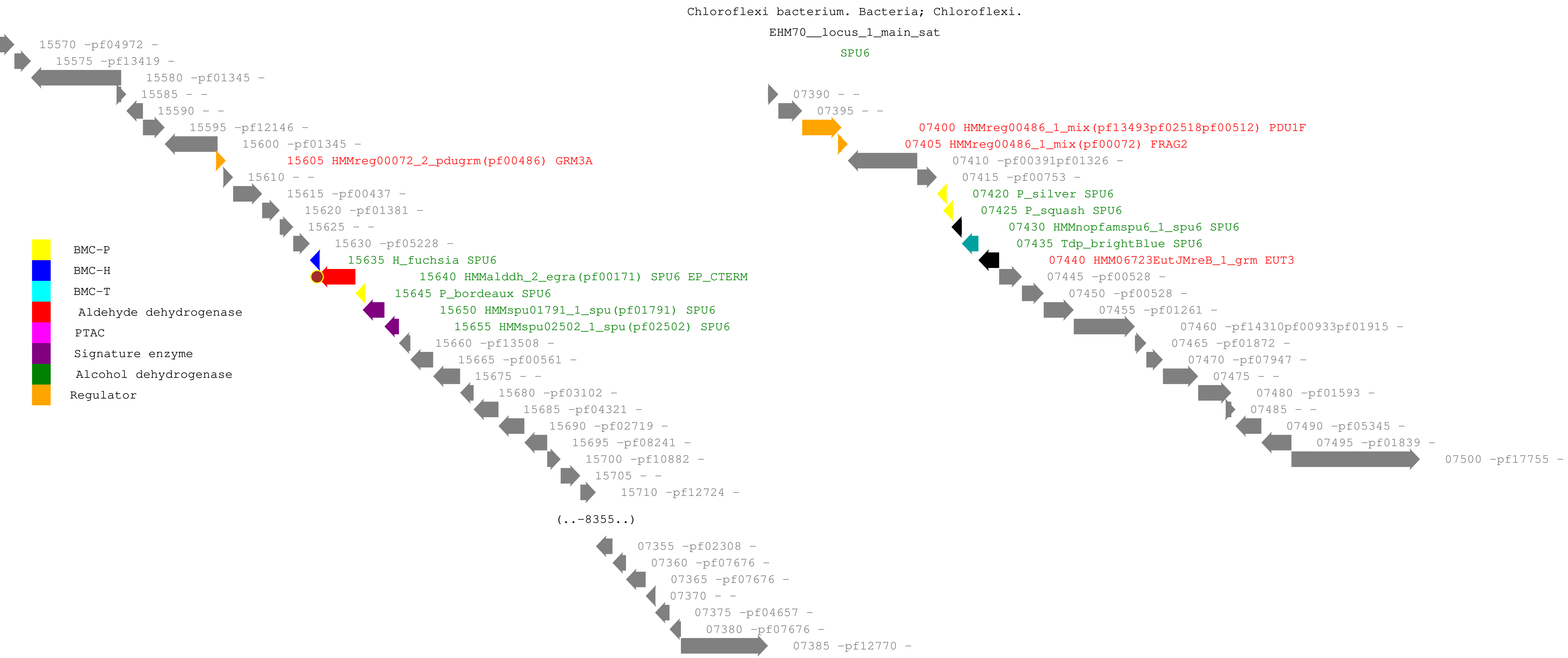
Chloroflexi bacterium. Bacteria; Chloroflexi.
EHM70__locus_1_main_sat
SPU6
15570 -pf04972 -
15575 -pf13419 -
15580 -pf01345 -
15585 - -
15590 - -
15595 -pf12146 -
15600 -pf01345 -
15605 HMMreg00072_2_pdugrm(pf00486) GRM3A
15610 - -
15615 -pf00437 -
15620 -pf01381 -
15625 - -
15630 -pf05228 -
15635 H_fuchsia SPU6
15640 HMMalddh_2_egra(pf00171) SPU6 EP_CTERM
15645 P_bordeaux SPU6
15650 HMMspu01791_1_spu(pf01791) SPU6
15655 HMMspu02502_1_spu(pf02502) SPU6
15660 -pf13508 -
15665 -pf00561 -
15675 - -
15680 -pf03102 -
15685 -pf04321 -
15690 -pf02719 -
15695 -pf08241 -
15700 -pf10882 -
15705 - -
15710 -pf12724 -
(..-8355..)
07355 -pf02308 -
07360 -pf07676 -
07365 -pf07676 -
07370 - -
07375 -pf04657 -
07380 -pf07676 -
07385 -pf12770 -
07390 - -
07395 - -
07400 HMMreg00486_1_mix(pf13493pf02518pf00512) PDU1F
07405 HMMreg00486_1_mix(pf00072) FRAG2
07410 -pf00391pf01326 -
07415 -pf00753 -
07420 P_silver SPU6
07425 P_squash SPU6
07430 HMMnopfamspu6_1_spu6 SPU6
07435 Tdp_brightBlue SPU6
07440 HMM06723EutJMreB_1_grm EUT3
07445 -pf00528 -
07450 -pf00528 -
07455 -pf01261 -
07460 -pf14310pf00933pf01915 -
07465 -pf01872 -
07470 -pf07947 -
07475 - -
07480 -pf01593 -
07485 - -
07490 -pf05345 -
07495 -pf01839 -
07500 -pf17755 -
BMC-P
BMC-H
BMC-T
Aldehyde dehydrogenase
PTAC
Signature enzyme
Alcohol dehydrogenase
Regulator

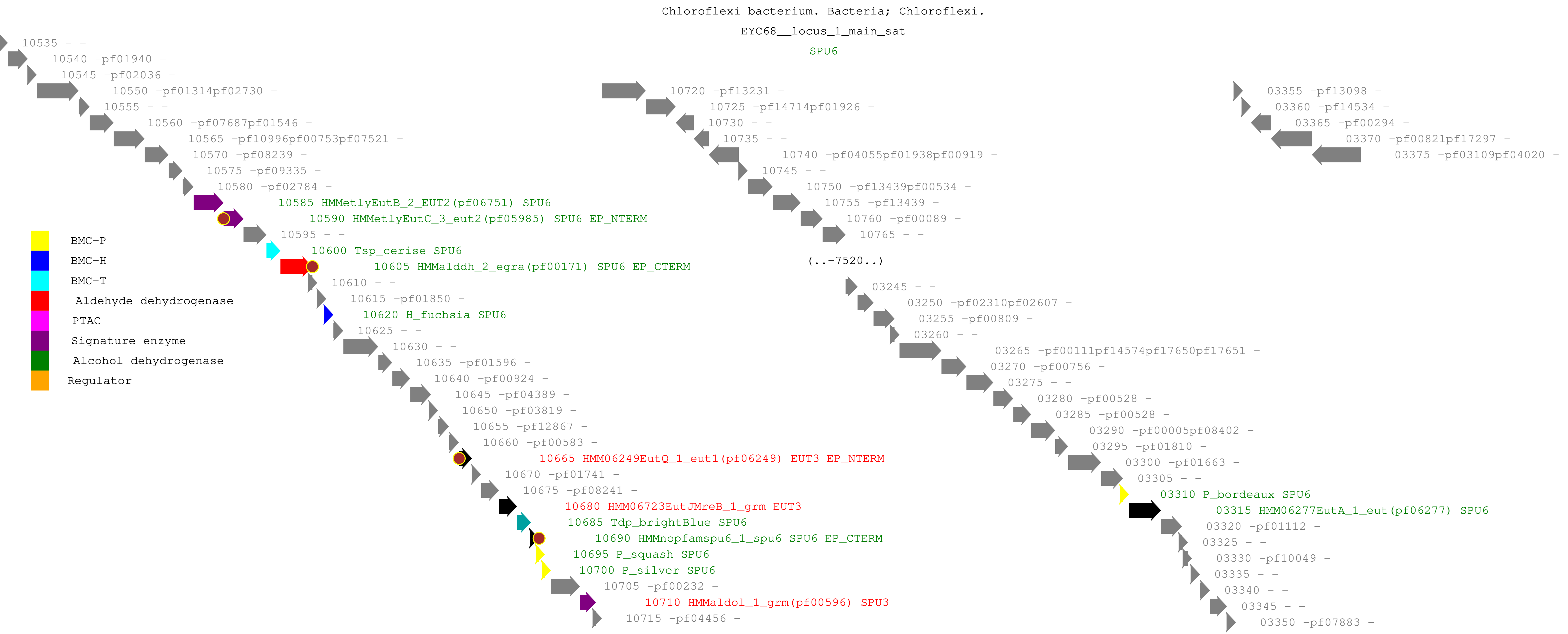
Chloroflexi bacterium. Bacteria; Chloroflexi.
EYC68__locus_1_main_sat
SPU6
10535 - -
10540 -pf01940 -
10545 -pf02036 -
10550 -pf01314pf02730 -
10555 - -
10560 -pf07687pf01546 -
10565 -pf10996pf00753pf07521 -
10570 -pf08239 -
10575 -pf09335 -
10580 -pf02784 -
10585 HMMetlyEutB_2_EUT2(pf06751) SPU6
10590 HMMetlyEutC_3_eut2(pf05985) SPU6 EP_NTERM
10595 - -
10600 Tsp_cerise SPU6
10605 HMMalddh_2_egra(pf00171) SPU6 EP_CTERM
10610 - -
10615 -pf01850 -
10620 H_fuchsia SPU6
10625 - -
10630 - -
10635 -pf01596 -
10640 -pf00924 -
10645 -pf04389 -
10650 -pf03819 -
10655 -pf12867 -
10660 -pf00583 -
10665 HMM06249EutQ_1_eut1(pf06249) EUT3 EP_NTERM
10670 -pf01741 -
10675 -pf08241 -
10680 HMM06723EutJMreB_1_grm EUT3
10685 Tdp_brightBlue SPU6
10690 HMMnopfamspu6_1_spu6 SPU6 EP_CTERM
10695 P_squash SPU6
10700 P_silver SPU6
10705 -pf00232 -
10710 HMMaldol_1_grm(pf00596) SPU3
10715 -pf04456 -
BMC-P
BMC-H
BMC-T
Aldehyde dehydrogenase
PTAC
Signature enzyme
Alcohol dehydrogenase
Regulator
10720 -pf13231 -
10725 -pf14714pf01926 -
10730 - -
10735 - -
10740 -pf04055pf01938pf00919 -
10745 - -
10750 -pf13439pf00534 -
10755 -pf13439 -
10760 -pf00089 -
10765 - -
(..-7520..)
03245 - -
03250 -pf02310pf02607 -
03255 -pf00809 -
03260 - -
03265 -pf00111pf14574pf17650pf17651 -
03270 -pf00756 -
03275 - -
03280 -pf00528 -
03285 -pf00528 -
03290 -pf00005pf08402 -
03295 -pf01810 -
03300 -pf01663 -
03305 - -
03310 P_bordeaux SPU6
03315 HMM06277EutA_1_eut(pf06277) SPU6
03320 -pf01112 -
03325 - -
03330 -pf10049 -
03335 - -
03340 - -
03345 - -
03350 -pf07883 -
03355 -pf13098 -
03360 -pf14534 -
03365 -pf00294 -
03370 -pf00821pf17297 -
03375 -pf03109pf04020 -

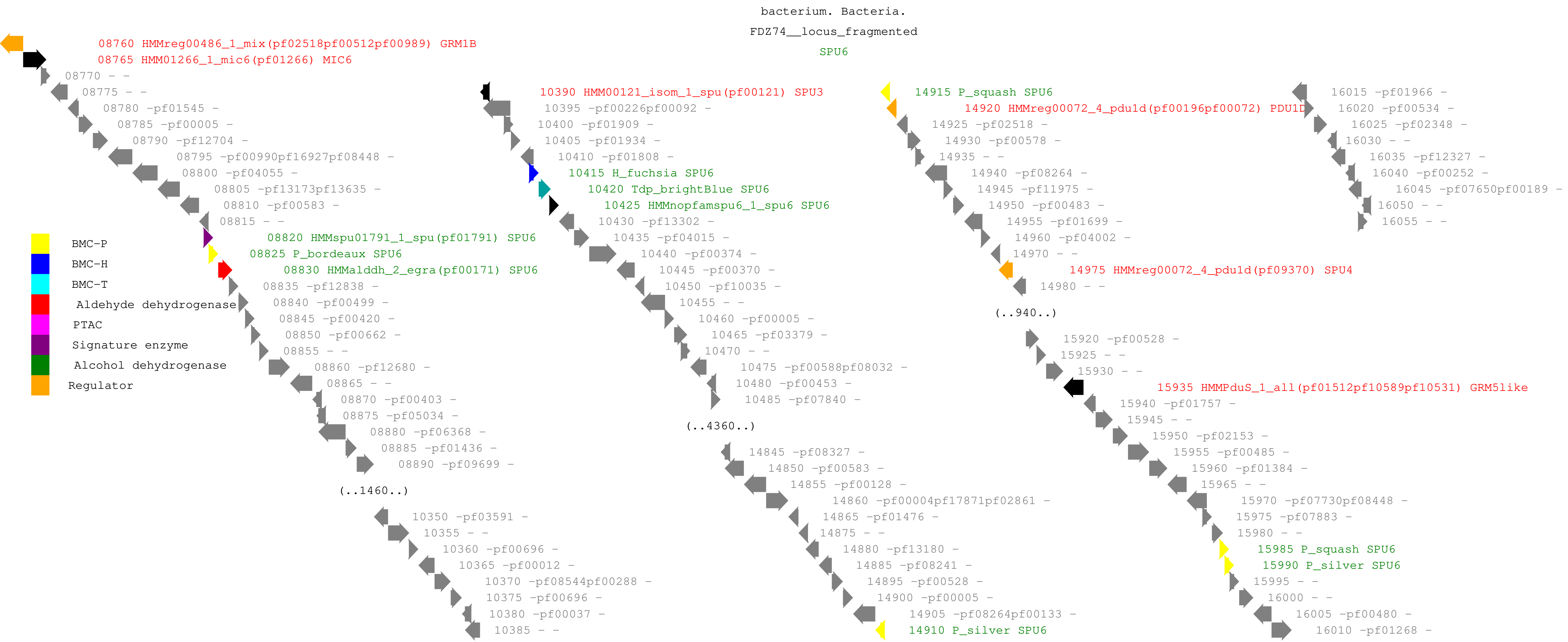
bacterium. Bacteria.
FDZ74__locus_fragmented
SPU6
08760 HMMreg00486_1_mix(pf02518pf00512pf00989) GRM1B
08765 HMM01266_1_mic6(pf01266) MIC6
08770 - -
08775 - -
08780 -pf01545 -
08785 -pf00005 -
08790 -pf12704 -
08795 -pf00990pf16927pf08448 -
08800 -pf04055 -
08805 -pf13173pf13635 -
08810 -pf00583 -
08815 - -
08820 HMMspu01791_1_spu(pf01791) SPU6
08825 P_bordeaux SPU6
08830 HMMaldh_2_egra(pf00171) SPU6
08835 -pf12838 -
08840 -pf00499 -
08845 -pf00420 -
08850 -pf00662 -
08855 - -
08860 -pf12680 -
08865 - -
08870 -pf00403 -
08875 -pf05034 -
08880 -pf06368 -
08885 -pf01436 -
08890 -pf09699 -
(..1460..)
10350 -pf03591 -
10355 - -
10360 -pf00696 -
10365 -pf00012 -
10370 -pf08544pf00288 -
10375 -pf00696 -
10380 -pf00037 -
10385 - -
BMC-P
BMC-H
BMC-T
Aldehyde dehydrogenase
PTAC
Signature enzyme
Alcohol dehydrogenase
Regulator
10390 HMM00121_isom_1_spu(pf00121) SPU3
10395 -pf00226pf00092 -
10400 -pf01909 -
10405 -pf01934 -
10410 -pf01808 -
10415 H_fuchsia SPU6
10420 Tdp_brightBlue SPU6
10425 HMMnopfamspu6_1_spu6 SPU6
10430 -pf13302 -
10435 -pf04015 -
10440 -pf00374 -
10445 -pf00370 -
10450 -pf10035 -
10455 - -
10460 -pf00005 -
10465 -pf03379 -
10470 - -
10475 -pf00588pf08032 -
10480 -pf00453 -
10485 -pf07840 -
(..4360..)
14845 -pf08327 -
14850 -pf00583 -
14855 -pf00128 -
14860 -pf00004pf17871pf02861 -
14865 -pf01476 -
14875 - -
14880 -pf13180 -
14885 -pf08241 -
14895 -pf00528 -
14900 -pf00005 -
14905 -pf08264pf00133 -
14910 P_silver SPU6
14915 P_squash SPU6
14920 HMMreg00072_4_pdu1d(pf00196pf00072) PDU1D
14925 -pf02518 -
14930 -pf00578 -
14935 - -
14940 -pf08264 -
14945 -pf11975 -
14950 -pf00483 -
14955 -pf01699 -
14960 -pf04002 -
14970 - -
14975 HMMreg00072_4_pdu1d(pf09370) SPU4
14980 - -
(..940..)
15920 -pf00528 -
15925 - -
15930 - -
15935 HMMPduS_1_all(pf01512pf10589pf10531) GRM5like
15940 -pf01757 -
15945 - -
15950 -pf02153 -
15955 -pf00485 -
15960 -pf01384 -
15965 - -
15970 -pf07730pf08448 -
15975 -pf07883 -
15980 - -
15985 P_squash SPU6
15990 P_silver SPU6
15995 - -
16000 - -
16005 -pf00480 -
16010 -pf01268 -
16015 -pf01966 -
16020 -pf00534 -
16025 -pf02348 -
16030 - -
16035 -pf12327 -
16040 -pf00252 -
16045 -pf07650pf00189 -
16050 - -
16055 - -

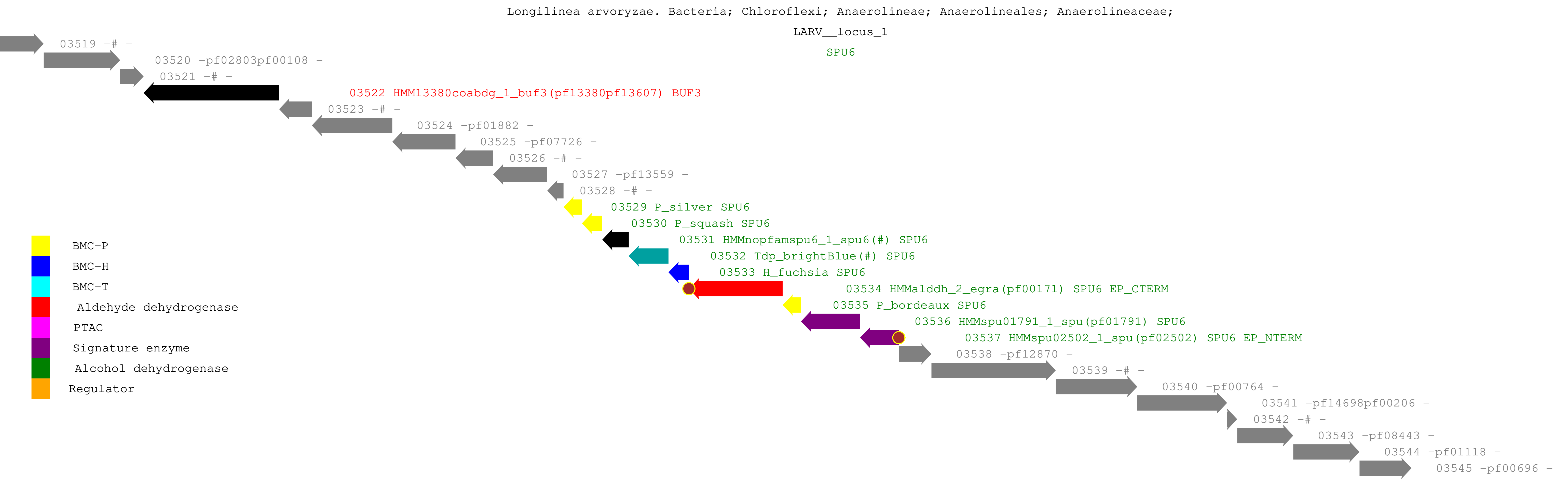

Longilinea arvoryzae. Bacteria; Chloroflexi; Anaerolineae; Anaerolineales; Anaerolineaceae;
LARV__locus_1
SPU6
03519 -# -
03520 -pf02803pf00108 -
03521 -# -
03522 HMM13380coabdg_1_buf3(pf13380pf13607) BUF3
03523 -# -
03524 -pf01882 -
03525 -pf07726 -
03526 -# -
03527 -pf13559 -
03528 -# -
03529 P_silver SPU6
03530 P_squash SPU6
03531 HMMnopfamspu6_1_spu6(#) SPU6
03532 Tdp_brightBlue(#) SPU6
03533 H_fuchsia SPU6
03534 HMMalddh_2_egra(pf00171) SPU6 EP_CTERM
03535 P_bordeaux SPU6
03536 HMMspu01791_1_spu(pf01791) SPU6
03537 HMMspu02502_1_spu(pf02502) SPU6 EP_NTERM
03538 -pf12870 -
03539 -# -
03540 -pf00764 -
03541 -pf14698pf00206 -
03542 -# -
03543 -pf08443 -
03544 -pf01118 -
03545 -pf00696 -
BMC-P
BMC-H
BMC-T
Aldehyde dehydrogenase
PTAC
Signature enzyme
Alcohol dehydrogenase
Regulator

Pelolinea submarina. Bacteria; Chloroflexi; Anaerolineae; Anaerolineales; Anaerolineaceae;

Pelsub_P_locus_1

SPU6

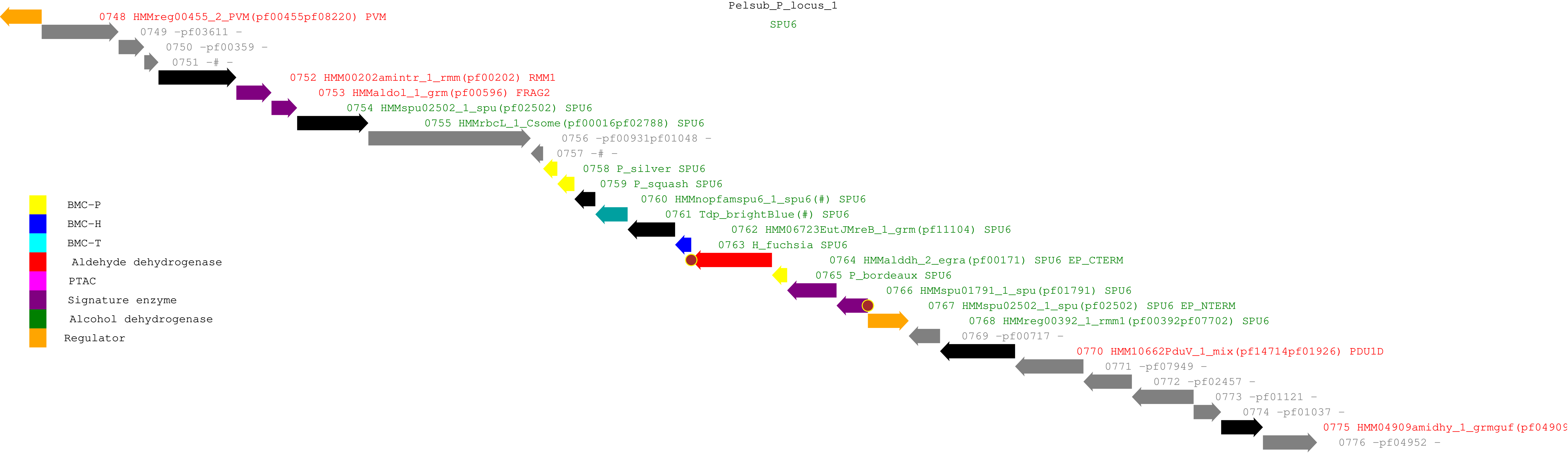

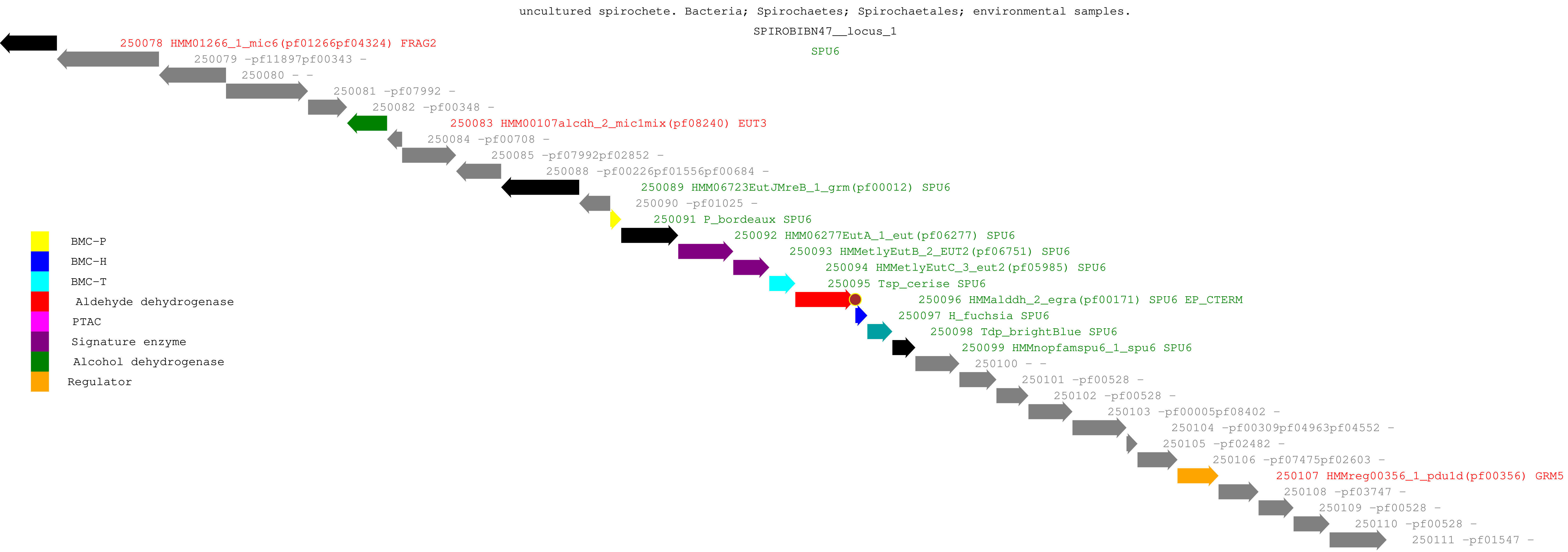
uncultured spirochete. Bacteria; Spirochaetes; Spirochaetales; environmental samples.
SPIROBIBN47__locus_1
SPU6
250078 HMM01266_1_mic6(pf01266pf04324) FRAG2
250079 -pf11897pf00343 -
250080 - -
250081 -pf07992 -
250082 -pf00348 -
250083 HMM00107alcdh_2_mic1mix(pf08240) EUT3
250084 -pf00708 -
250085 -pf07992pf02852 -
250088 -pf00226pf01556pf00684 -
250089 HMM06723EutJMreB_1_grm(pf00012) SPU6
250090 -pf01025 -
250091 P_bordeaux SPU6
250092 HMM06277EutA_1_eut(pf06277) SPU6
250093 HMMetlyEutB_2_EUT2(pf06751) SPU6
250094 HMMetlyEutC_3_eut2(pf05985) SPU6
250095 Tsp_cerise SPU6
250096 HMMalddh_2_egra(pf00171) SPU6 EP_CTERM
250097 H_fuchsia SPU6
250098 Tdp_brightBlue SPU6
250099 HMMnopfamspu6_1_spu6 SPU6
250100 - -
250101 -pf00528 -
250102 -pf00528 -
250103 -pf00005pf08402 -
250104 -pf00309pf04963pf04552 -
250105 -pf02482 -
250106 -pf07475pf02603 -
250107 HMMreg00356_1_pdu1d(pf00356) GRM5
250108 -pf03747 -
250109 -pf00528 -
250110 -pf00528 -
250111 -pf01547 -
BMC-P
BMC-H
BMC-T
Aldehyde dehydrogenase
PTAC
Signature enzyme
Alcohol dehydrogenase
Regulator

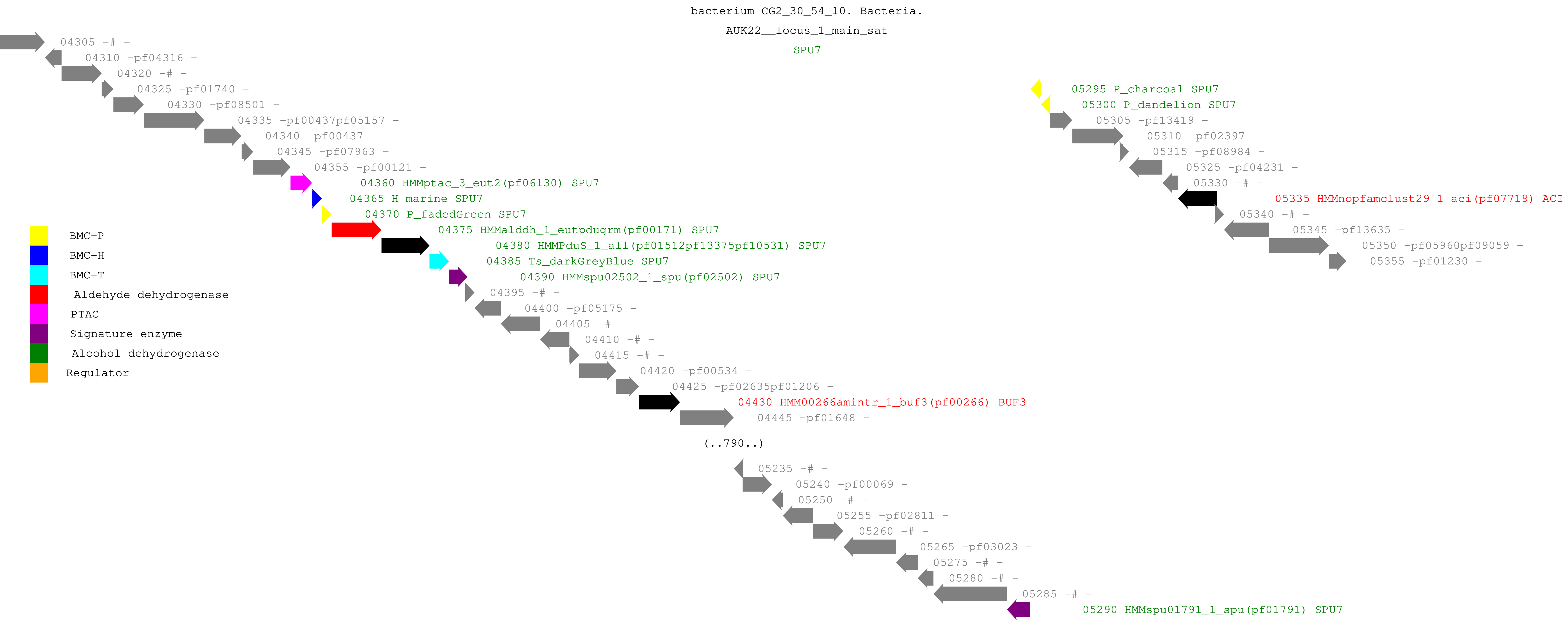
bacterium CG2_30_54_10. Bacteria.
AUK22__locus_1_main_sat
SPU7
04305 -# -
04310 -pf04316 -
04320 -# -
04325 -pf01740 -
04330 -pf08501 -
04335 -pf00437pf05157 -
04340 -pf00437 -
04345 -pf07963 -
04355 -pf00121 -
04360 HMMptac_3_eut2(pf06130) SPU7
04365 H_marine SPU7
04370 P_fadedGreen SPU7
04375 HMMalddh_1_eutpdugrm(pf00171) SPU7
04380 HMMPduS_1_all(pf01512pf13375pf10531) SPU7
04385 Ts_darkGreyBlue SPU7
04390 HMMspu02502_1_spu(pf02502) SPU7
04395 -# -
04400 -pf05175 -
04405 -# -
04410 -# -
04415 -# -
04420 -pf00534 -
04425 -pf02635pf01206 -
04430 HMM00266amintr_1_buf3(pf00266) BUF3
04445 -pf01648 -
(..790..)
05235 -# -
05240 -pf00069 -
05250 -# -
05255 -pf02811 -
05260 -# -
05265 -pf03023 -
05275 -# -
05280 -# -
05285 -# -
05290 HMMspu01791_1_spu(pf01791) SPU7
05295 P_charcoal SPU7
05300 P_dandelion SPU7
05305 -pf13419 -
05310 -pf02397 -
05315 -pf08984 -
05325 -pf04231 -
05330 -# -
05335 HMMnopfamclust29_1_aci(pf07719) ACI
05340 -# -
05345 -pf13635 -
05350 -pf05960pf09059 -
05355 -pf01230 -
BMC-P
BMC-H
BMC-T
Aldehyde dehydrogenase
PTAC
Signature enzyme
Alcohol dehydrogenase
Regulator

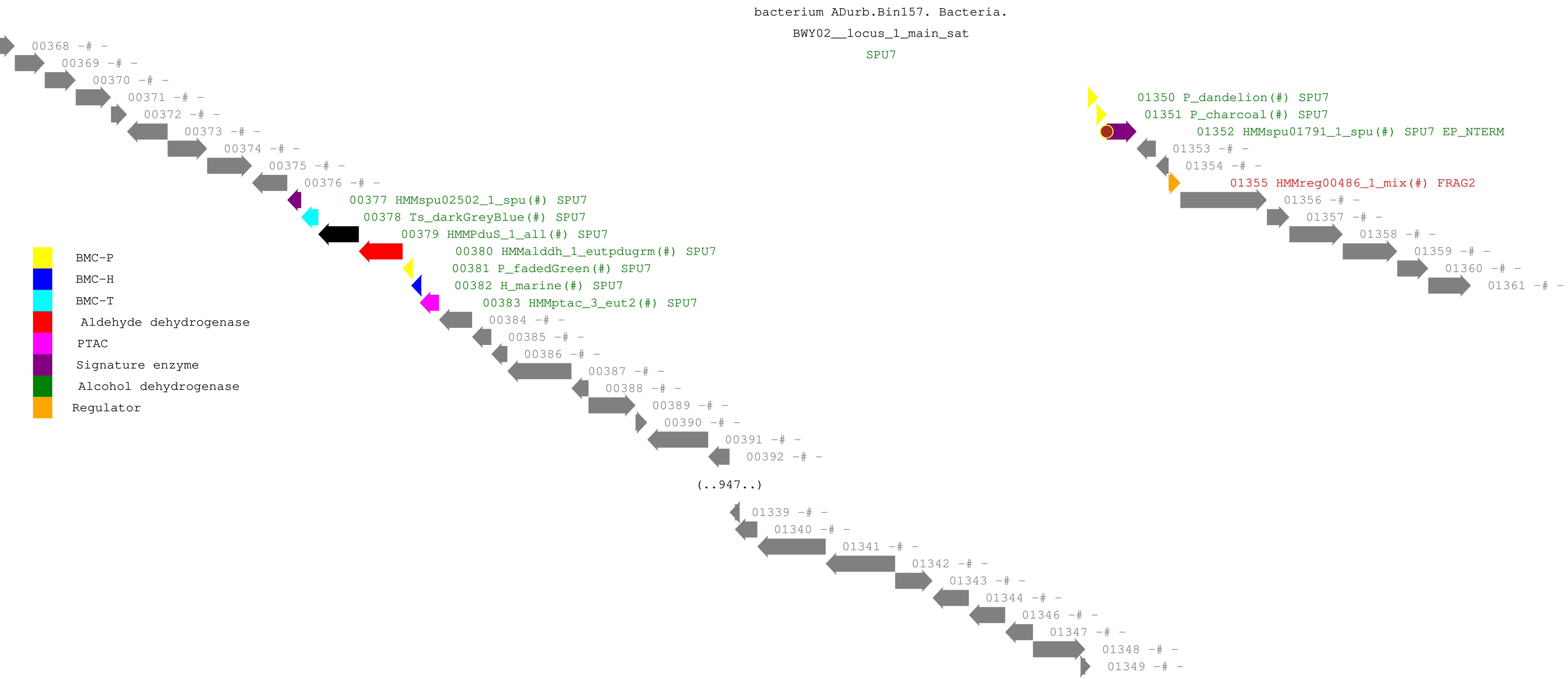
bacterium ADurb.Bin157. Bacteria.
BWY02__locus_1_main_sat
SPU7
00368 -# -
00369 -# -
00370 -# -
00371 -# -
00372 -# -
00373 -# -
00374 -# -
00375 -# -
00376 -# -
00377 HMMspu02502_1_spu(#) SPU7
00378 Ts_darkGreyBlue(#) SPU7
00379 HMMPduS_1_all(#) SPU7
00380 HMMaldh_1_eutpdugrm(#) SPU7
00381 P_fadedGreen(#) SPU7
00382 H_marine(#) SPU7
00383 HMMptac_3_eut2(#) SPU7
00384 -# -
00385 -# -
00386 -# -
00387 -# -
00388 -# -
00389 -# -
00390 -# -
00391 -# -
00392 -# -
(..947..)
01339 -# -
01340 -# -
01341 -# -
01342 -# -
01343 -# -
01344 -# -
01346 -# -
01347 -# -
01348 -# -
01349 -# -
01350 P_dandelion(#) SPU7
01351 P_charcoal(#) SPU7
01352 HMMspu01791_1_spu(#) SPU7 EP_NTERM
01353 -# -
01354 -# -
01355 HMMreg00486_1_mix(#) FRAG2
01356 -# -
01357 -# -
01358 -# -
01359 -# -
01360 -# -
01361 -# -
BMC-P
BMC-H
BMC-T
Aldehyde dehydrogenase
PTAC
Signature enzyme
Alcohol dehydrogenase
Regulator

Candidatus Cloacimonetes bacterium. Bacteria; Candidatus Cloacimonetes.

COB02__locus_1_main_sat

SPU7

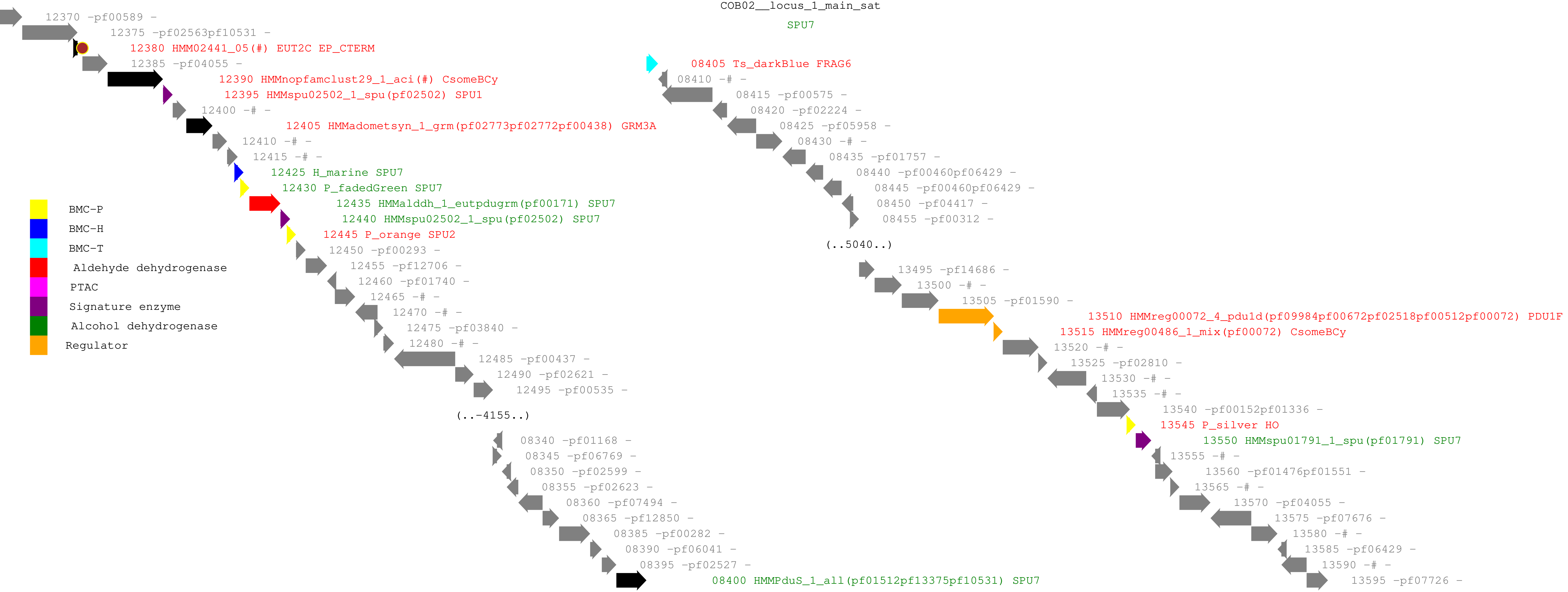

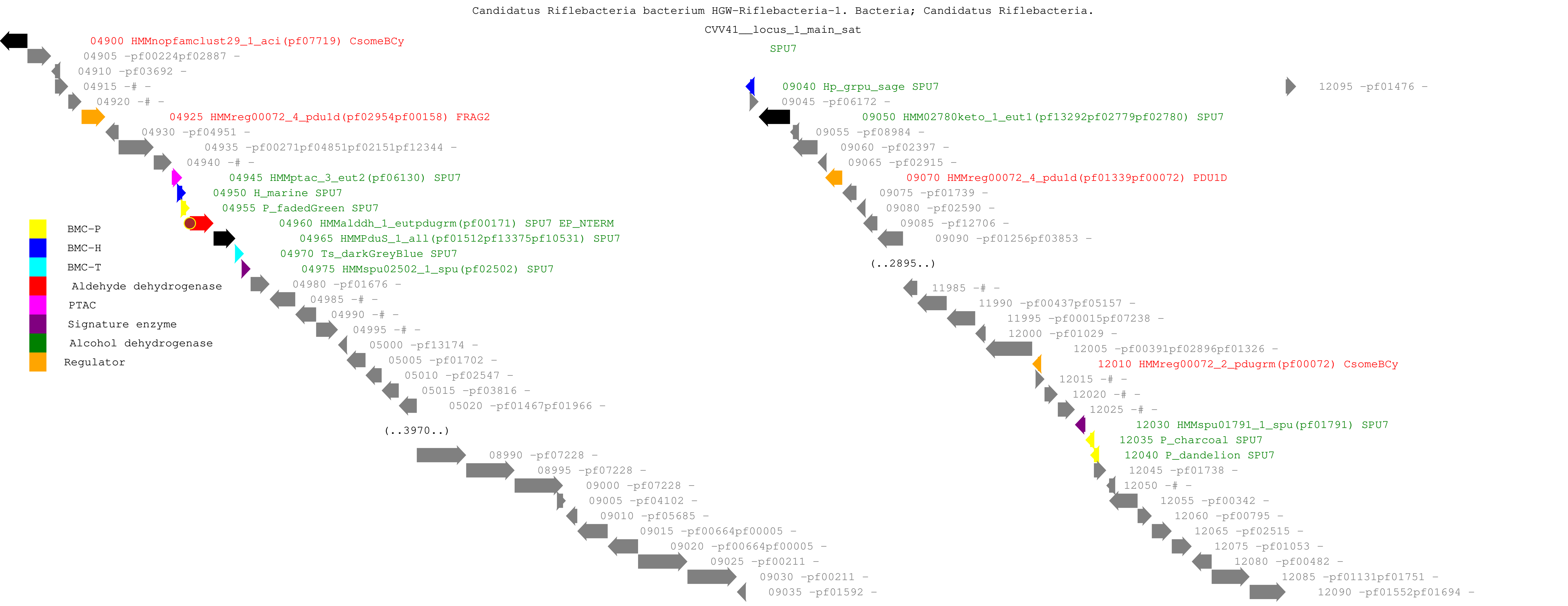
Candidatus Riflebacteria bacterium HGW-Riflebacteria-1. Bacteria; Candidatus Riflebacteria.
CVV41__locus_1_main_sat
SPU7
04900 HMMnopfamclust29_1_aci(pf07719) CsomeBCy
04905 -pf00224pf02887 -
04910 -pf03692 -
04915 -# -
04920 -# -
04925 HMMreg00072_4_pdu1d(pf02954pf00158) FRAG2
04930 -pf04951 -
04935 -pf00271pf04851pf02151pf12344 -
04940 -# -
04945 HMMptac_3_eut2(pf06130) SPU7
04950 H_marine SPU7
04955 P_fadedGreen SPU7
04960 HMMalddh_1_eutpdugrm(pf00171) SPU7 EP_NTERM
04965 HMMPduS_1_all(pf01512pf13375pf10531) SPU7
04970 Ts_darkGreyBlue SPU7
04975 HMMspu02502_1_spu(pf02502) SPU7
04980 -pf01676 -
04985 -# -
04990 -# -
04995 -# -
05000 -pf13174 -
05005 -pf01702 -
05010 -pf02547 -
05015 -pf03816 -
05020 -pf01467pf01966 -
(..3970..)
08990 -pf07228 -
08995 -pf07228 -
09000 -pf07228 -
09005 -pf04102 -
09010 -pf05685 -
09015 -pf00664pf00005 -
09020 -pf00664pf00005 -
09025 -pf00211 -
09030 -pf00211 -
09035 -pf01592 -
BMC-P
BMC-H
BMC-T
Aldehyde dehydrogenase
PTAC
Signature enzyme
Alcohol dehydrogenase
Regulator
09040 Hp_grpu_sage SPU7
09045 -pf06172 -
09050 HMM02780keto_1_eut1(pf13292pf02779pf02780) SPU7
09055 -pf08984 -
09060 -pf02397 -
09065 -pf02915 -
09070 HMMreg00072_4_pdu1d(pf01339pf00072) PDU1D
09075 -pf01739 -
09080 -pf02590 -
09085 -pf12706 -
09090 -pf01256pf03853 -
(..2895..)
11985 -# -
11990 -pf00437pf05157 -
11995 -pf00015pf07238 -
12000 -pf01029 -
12005 -pf00391pf02896pf01326 -
12010 HMMreg00072_2_pdugrm(pf00072) CsomeBCy
12015 -# -
12020 -# -
12025 -# -
12030 HMMspu01791_1_spu(pf01791) SPU7
12035 P_charcoal SPU7
12040 P_dandelion SPU7
12045 -pf01738 -
12050 -# -
12055 -pf00342 -
12060 -pf00795 -
12065 -pf02515 -
12075 -pf01053 -
12080 -pf00482 -
12085 -pf01131pf01751 -
12090 -pf01552pf01694 -
12095 -pf01476 -

Candidatus Riflebacteria bacterium HGW-Riflebacteria-2. Bacteria; Candidatus Riflebacteria.

CVV42__locus_1_main_sat

SPU7

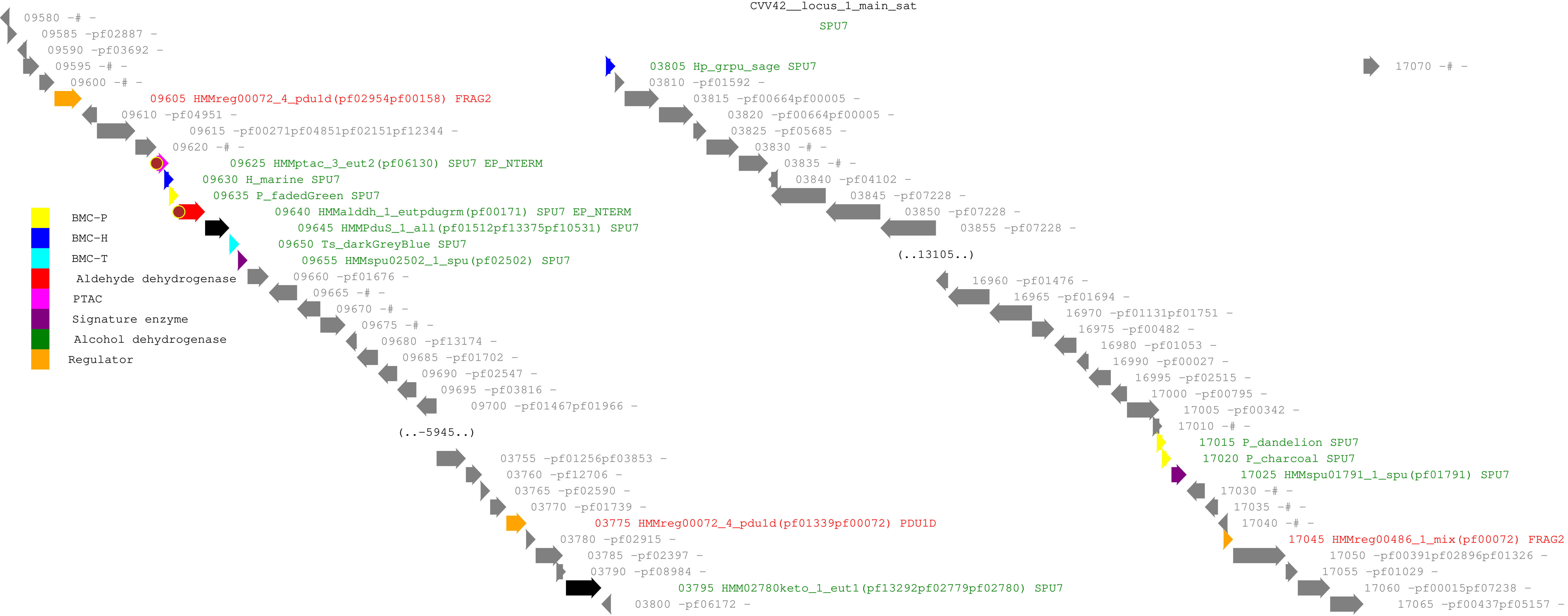

Candidatus Wallbacteria bacterium HGW-Wallbacteria-1. Bacteria; Candidatus Wallbacteria.

CVV64__locus_1

SPU7

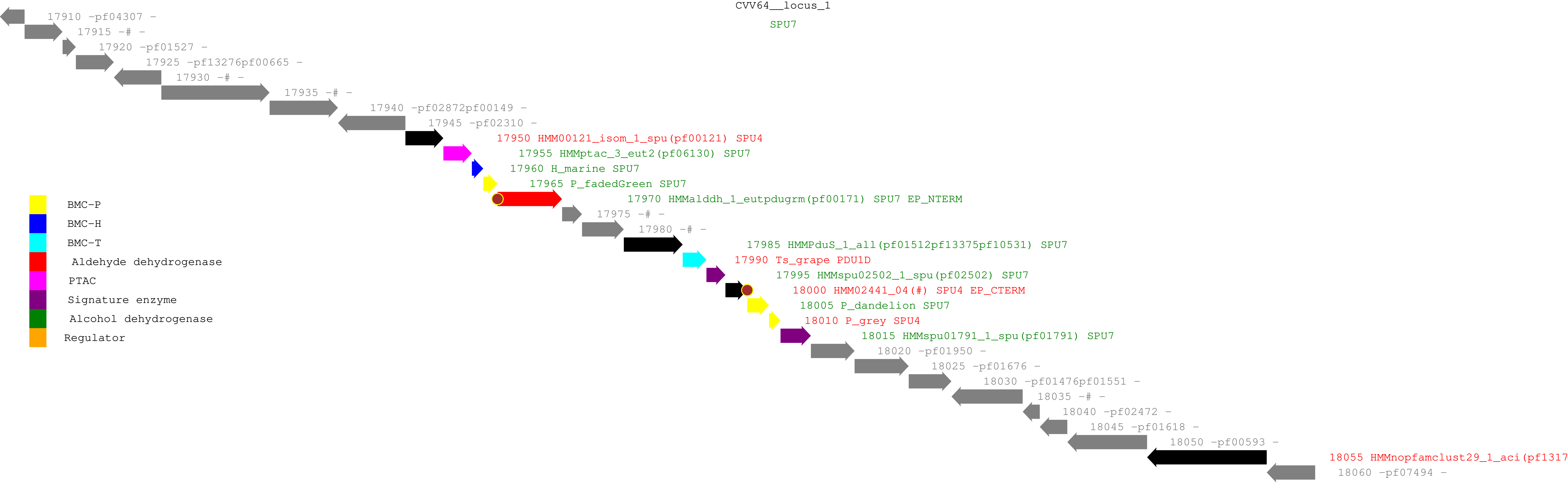

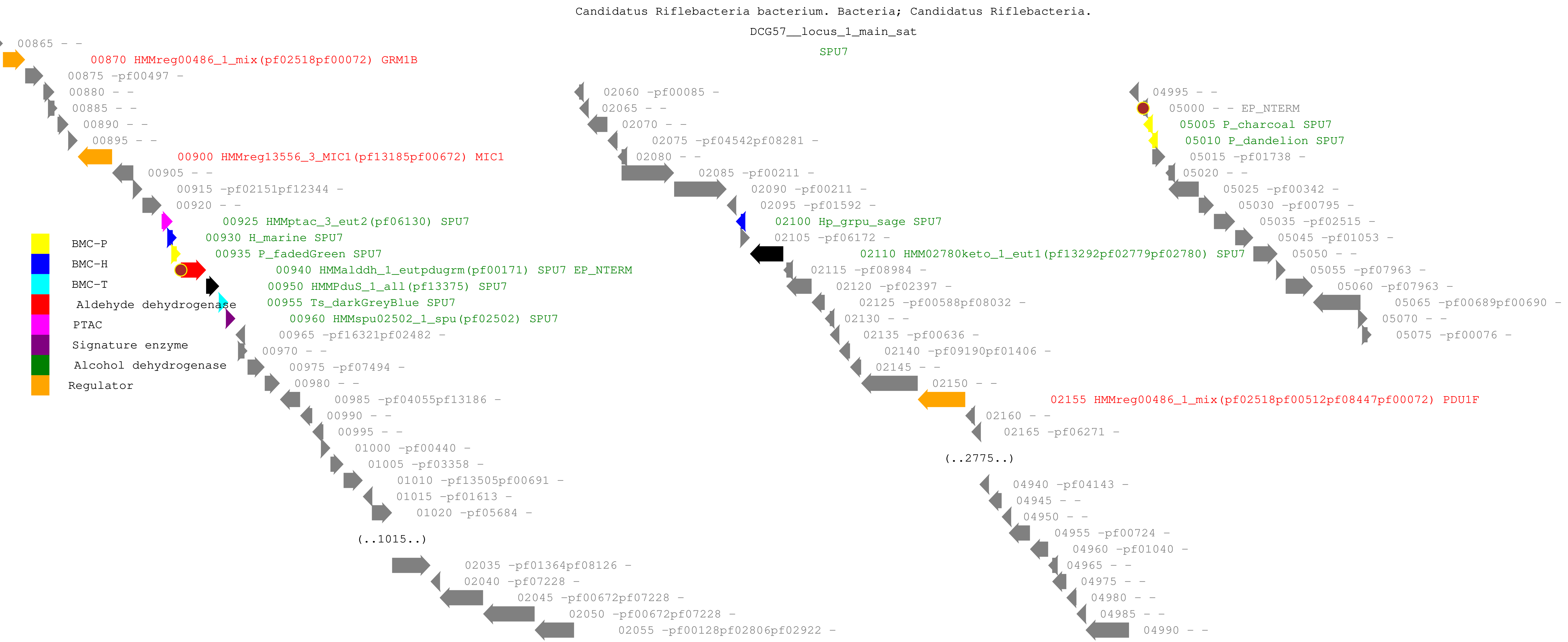
Candidatus Riflebacteria bacterium. Bacteria; Candidatus Riflebacteria.
DCG57__locus_1_main_sat
SPU7
00865 - -
00870 HMMreg00486_1_mix(pf02518pf00072) GRM1B
00875 -pf00497 -
00880 - -
00885 - -
00890 - -
00895 - -
00900 HMMreg13556_3_MIC1(pf13185pf00672) MIC1
00905 - -
00915 -pf02151pf12344 -
00920 - -
00925 HMMptac_3_eut2(pf06130) SPU7
00930 H_marine SPU7
00935 P_fadedGreen SPU7
00940 HMMaldh_1_eutpdugrm(pf00171) SPU7 EP_NTERM
00950 HMMPduS_1_all(pf13375) SPU7
00955 Ts_darkGreyBlue SPU7
00960 HMMspu02502_1_spu(pf02502) SPU7
00965 -pf16321pf02482 -
00970 - -
00975 -pf07494 -
00980 - -
00985 -pf04055pf13186 -
00990 - -
00995 - -
01000 -pf00440 -
01005 -pf03358 -
01010 -pf13505pf00691 -
01015 -pf01613 -
01020 -pf05684 -
(..1015..)
02035 -pf01364pf08126 -
02040 -pf07228 -
02045 -pf00672pf07228 -
02050 -pf00672pf07228 -
02055 -pf00128pf02806pf02922 -
BMC-P
BMC-H
BMC-T
Aldehyde dehydrogenase
PTAC
Signature enzyme
Alcohol dehydrogenase
Regulator
02060 -pf00085 -
02065 - -
02070 - -
02075 -pf04542pf08281 -
02080 - -
02085 -pf00211 -
02090 -pf00211 -
02095 -pf01592 -
02100 Hp_grpu_sage SPU7
02105 -pf06172 -
02110 HMM02780keto_1_eut1(pf13292pf02779pf02780) SPU7
02115 -pf08984 -
02120 -pf02397 -
02125 -pf00588pf08032 -
02130 - -
02135 -pf00636 -
02140 -pf09190pf01406 -
02145 - -
02150 - -
02155 HMMreg00486_1_mix(pf02518pf00512pf08447pf00072) PDU1F
02160 - -
02165 -pf06271 -
(..2775..)
04940 -pf04143 -
04945 - -
04950 - -
04955 -pf00724 -
04960 -pf01040 -
04965 - -
04975 - -
04980 - -
04985 - -
04990 - -
04995 - -
05000 - - EP_NTERM
05005 P_charcoal SPU7
05010 P_dandelion SPU7
05015 -pf01738 -
05020 - -
05025 -pf00342 -
05030 -pf00795 -
05035 -pf02515 -
05045 -pf01053 -
05050 - -
05055 -pf07963 -
05060 -pf07963 -
05065 -pf00689pf00690 -
05070 - -
05075 -pf00076 -

Candidatus Ozemobacter sibiricus. Bacteria; Candidatus Riflebacteria; Candidatus Ozemobacteria;

OZSIB__locus_1_main_sat

SPU7

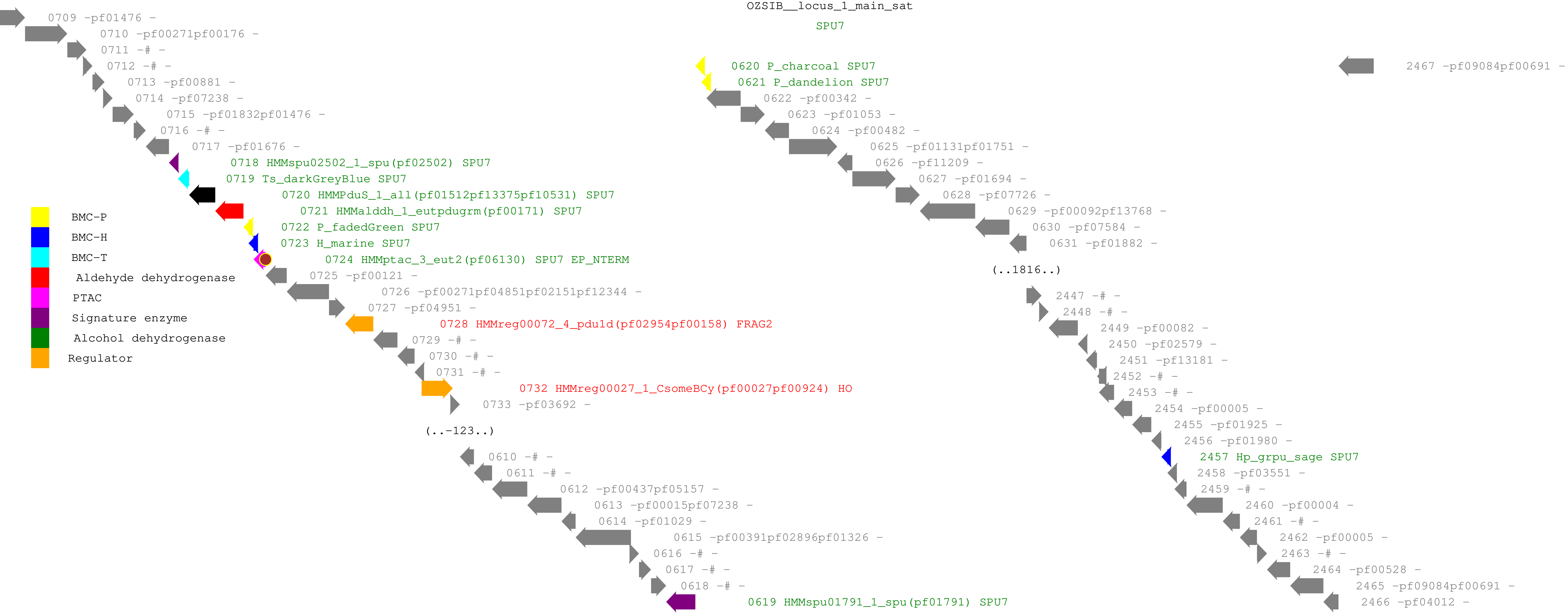
