## Supplementary Data: BMC types for "A Catalog of the Diversity and Ubiquity of Metabolic Organelles in Bacteria"

Descriptions of the names of HMMs can be found in Supplementary Data: HMM names table.

Color key for locus diagrams:

- BMC-P
- BMC-H, BMC-H$^{p}$
- BMC-T$^{s}$, BMC-T$^{sp}$
- BMC-T$^{dp}$
- Aldehyde dehydrogenase / AldDh
- Phosphotransacylase / PTAC
- Signature enzyme
- Alcohol dehydrogenase / AlcDh
- Regulator
- Gene for protein with assigned HMM
- Gene for protein without HMM (pfam shown if available)

Abbreviations:

EP: Encapsulation peptide

AldDh: aldehyde dehydrogenase (pfam00171)

PTAC: phosphotransacylase (pfam06130)

### New loci

#### *ACI – 125 loci*

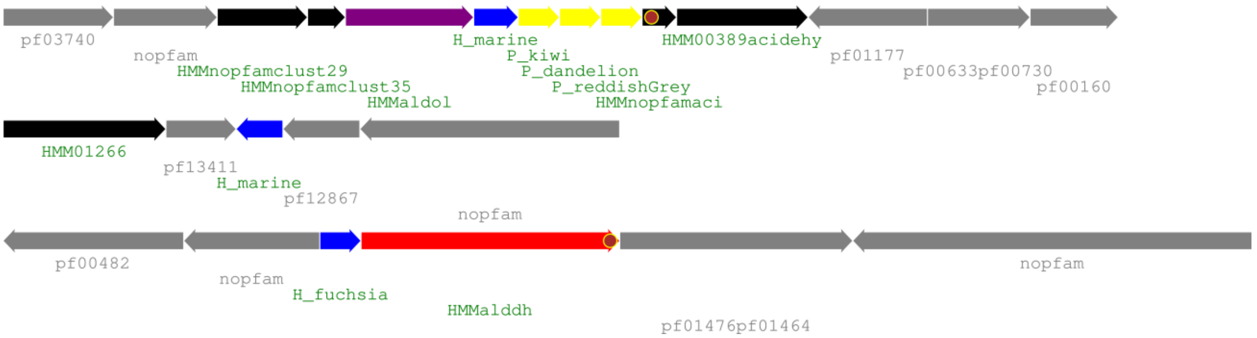


The ACI locus is found exclusively in Acidobacteria. The main locus contains a pfam00596 aldolase with a C-terminal EP, a hydroxyacid dehydrogenase (pfam00389/02826), and three proteins that have no known homologs; one of them is a very short protein ~80aa with an N-terminal EP and three conserved cysteines in its C-terminal domain. Interestingly, another one of the unknown domain proteins contains 3 conserved histidines, indicating that metal binding could be an important aspect of this locus type. A satellite locus found in most genomes contains an aldehyde dehydrogenase which has an EP close to the C-terminus; however there is a small ~50aa domain of unknown function following the EP in most of the members. Its aldehyde dehydrogenase is unique and phylogenetically associated with that of MIC3 (**Error! Reference source not found.**), while its aldolase is quite distinct and most similar to that of HO/MIC3. Homologs of PduS and the BMC-T$^{s}$ type PduT are occasionally found as pairs in a satellite location. The locus is always on its own in the genome.

The shell of ACI BMCs consists of a BMC-H with typical interface motifs and a BMC-P triplet in the main locus. The triplet BMC-P contains a BMC-P each from the green, grey and orange major clades and they are almost always encoded next to each other chromosomally. Occasionally there is a fourth BMC-P (from the blue major clade) on a satellite locus together with the aldehyde dehydrogenase. Two other BMC-H with mostly standard interaction motifs are occasionally found either by itself (H_marine) or with an AldDh (H_fuchsia or sometimes H_royalpurple with an unusual PRPF motif).

#### *ARO – 17 loci*

type I

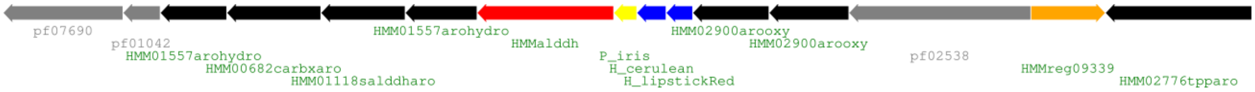


type II

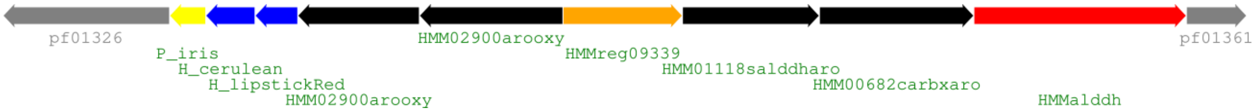


ARO type loci are found in two orders of Actinobacteria, Micromonosporales (type I) and Pseudonocardiales (type II). Core enzymes are two pfam02900 ring-opening oxygenases and a set of enzymes related to the degradation of aromatic aldehyde compounds, possibly starting with 2-aminophenol based on the assignment of related pfam00171 AldDh as aminomuconate-semialdehyde dehydrogenase. There is also a pfam02538 family protein in the type I operon that is annotated as Hydantoinase B/oxoprolinase that could be the first transcribed protein if the protein containing a pfam09339 helix-turn-helix DNA binding motif at the 5' end is responsible for the operon regulation. No encapsulation peptides are found on any proteins and there are no satellite loci except one Pseudonocardia member (ABS81__locus_1, this one also does not contain all genes so it is either a remnant or a related subtype). There is a small N-terminal extension on one of the ring-opening oxygenases but it is much shorter than typical EPs (only ~20aa) and also predicted to be a beta-strand rather than alpha-helix.

The shell proteins of the ARO consist of one BMC-P and two distinct BMC-H. Their sequences are very unique and on their respective trees they are found on their own on branches with very long stems. This indicates divergent evolution from other shell proteins, and supports them acquiring a novel function which has undergone evolutionary constraint. While the H_cerulean BMC-H contains typical residue motifs for interfaces, the H_lipstickRed one seems more specialized; it lacks the PRPH motif and has at least one insertion compared to regular BMC-H proteins.

#### *BUF2 – 8 loci*

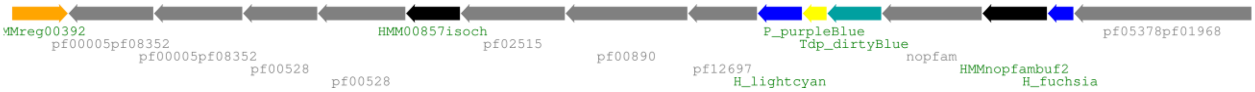


The BUF2 locus occurs in eight members of Actinobacteria/Micromonosporales. There are two conserved proteins located between the shell proteins, one of them resembles a methyltransferase and the other is a protein of about 260 aa length that has no known homologs; a sequence alignment reveals a conserved CDxxDxCSCGC motif that most likely binds a metal ion. There are no detectable encapsulation peptides and no satellite loci for this BMC type. The loci are on their own in the genomes except for two co-occurrences with ARO type BMCs.

The shell of this type consists of two BMC-H, one BMC-P and a BMC-$T^{dp}$ protein. The H_fuchsia BMC-H contains the standard interface residue motifs and the H_lightcyan one has a C-terminal extension of about 80aa that contains many prolines and charged residues. The Tdp_dirtyblue type BMC-$T^{dp}$ are unique to BUF2 and are located on a long stem in the blue major clade; its closest relatives are found in BUF3 and SPU6 type loci.

#### *BUF3 – 10 loci*

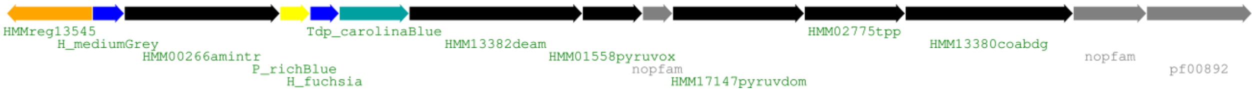


BUF3 loci are mostly found in the Clostridiales order of the Clostridia class but one instance of this type is found in an organism of the Spirochaetes phylum. The enzyme set consist of a pfam00266 aminotransferase, a pfam13382 deamidase, pfam01558 and pfam17147 domains of a pyruvate ferredoxin/flavodoxin oxidoreductase, a pfam02775 thiamine pyrophosphate enzyme and a pfam13380 CoA binding domain. There are no detectable typical encapsulation peptides, but there are N-terminal extensions of about 70-80 aa found on two enzymes (pfam00266 and pfam13382) that are only present in BMC-loci members. Their predicted secondary structure contains helical segments (and the pfam00266 one is separated from the main protein by a proline rich, likely flexible linker) so it is possible that they represent a targeting signal or have a role in BMC assembly.

One member of this locus type has been noted previously [1] as a potentially new type and our finding of several new occurrences confirms this prediction. BUF3 loci are found mostly on their own with no satellite loci, except one is found in a Maledivibacter species that contains six loci and two that co-occur with a SPU4-type locus.

Most BUF3 loci contain two BMC-H, one BMC-P and one BMC-T$^{dp}$ protein. The two BMC-H proteins have the standard motifs for interface interaction and belong to a basal type of BMC-H class. The BMC-P protein has an unusual amount of cysteine residues, two absolutely conserved ones and another two to three in more variable positions. Homology models indicate that all conserved ones and most of the others are oriented towards the protein core except for one close to the C-terminus. The BMC-T$^{dp}$ protein is unique to BUF3 loci and found in nine out of ten loci (replaced by three BMC-H that are different from the BUF3 typical ones in one locus).

#### *EUT3 - 17 loci*

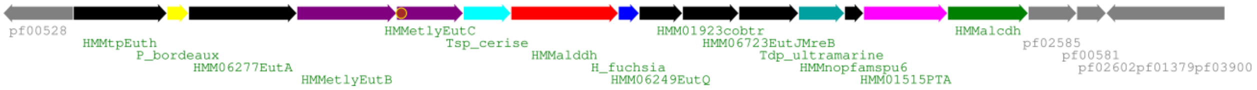


EUT3 type loci are found in diverse species, most are found in the Micrococcales and Propionibacteriales orders of Actinobacteria but several are from the Chloroflexi and Spirochaetes phyla. EUT3 loci are mostly on their own in the genome except for two co-occurrences with a PDU1D type locus. Protein sequences of the EUT3 type enzymes and shell proteins are distinct from EUT1 and EUT2, e.g. on a phylogenetic tree of EutB / the ethanolamine ammonia-lyase heavy chain is distinct and the only other EutB members in the same clade are found in satellite loci of SPU6 and HO. The AldDh is most closely related to homologs in SPU6.

The shell of EUT3 contains one protein each of a BMC-H, BMC-P, BMC-T$^{sp}$ and BMC-T$^{dp}$. The presence of a BMC-T$^{dp}$ is very unusual for a EUT type BMC. The BMC-T$^{dp}$ protein sequence is another aspect of the parallels of this type with SPU6 type BMCs since their BMC-T$^{dp}$ are neighbors on the BMC-T$^{dp}$ tree. The BMC-T$^{sp}$ is of the EutL type, consistent with a ethanolamine utilizing type and its closest relatives are found in SPU3, SPU6 and HO type BMCs.

#### *Fragmented loci*

This group of loci is characterized by a lack of locus organization. The components are found in up to six different genomic locations and some of the loci lack any identifiable enzymes. Searches with encapsulation peptide HMMs in those genomes have not yielded positive hits either. Many of those types are found in a wide variety of phyla and candidate phyla are common among those. An interesting common feature is the presence of BMC-P triplets proteins from the green/grey/orange major clades.

#### *FRAG1 – 57 loci*

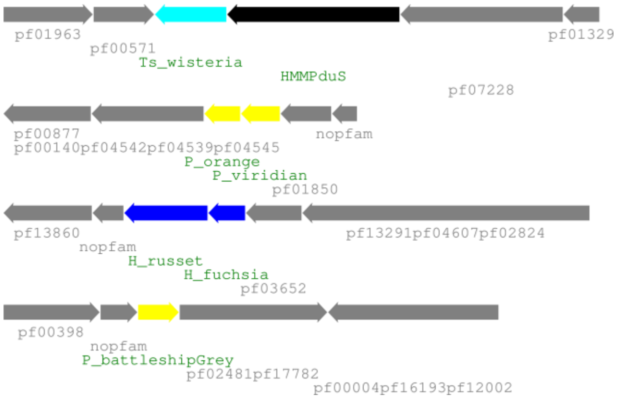


This shell protein only locus type is most prominent in Ignavibacteria but also found in several Candidatus Kryptonia and Bacteroidetes, a few each in Chlorobi, Rhodothermaeota and one in a Proteobacterium. A PduS/T homolog pair is found in 24 of those loci. Besides a BMC-P triplet from the green/grey/orange clades there are two BMC-H, one with standard interface residue motifs and another with a large C-terminal extension that seems to contain an ordered domain analogous to EutK.

#### *FRAG2 – 33 loci*

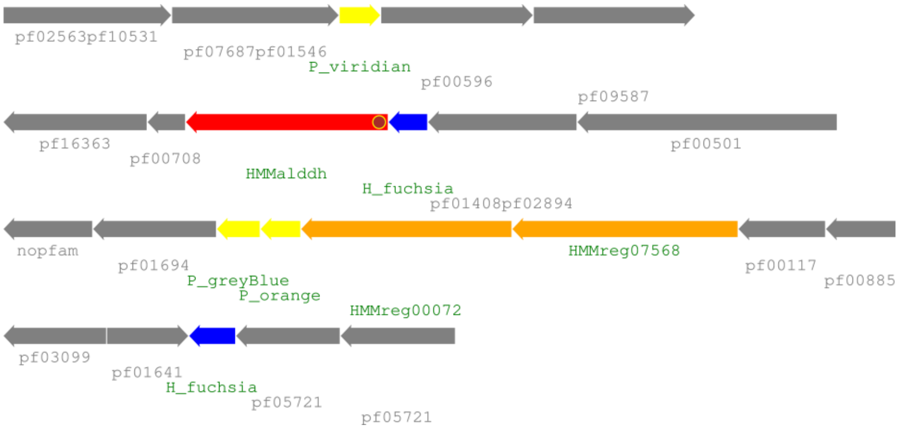


The most common phyla in this locus type is Candidatus Poribaceria with 21 loci; some of the other represented phyla which include Calditrichaeota, Candidatus Omnitrophica, Chlodoflexi, Planctomycetes and Spirochaetes have some type HMM mismatches so more sequenced genomes they could with their own type similar to FRAG2. The shell proteins consist of a green/grey/orange BMC-P triplet as well as two BMC-H with standard interface residues.

#### *FRAG3 – 8 loci*

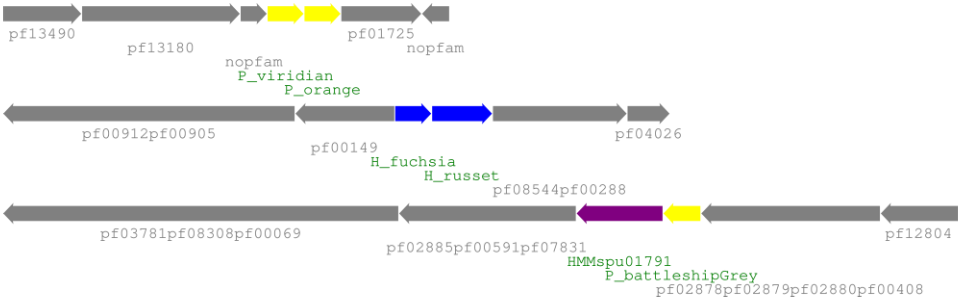


FRAG3 loci are most common in the Calditrichaeota phylum; one member each is found in candidate division KB1 and LCP-89. A DeoC/LacD family pfam01791 aldolase (a signature enzyme in SPU loci) is found next to a BMC-P protein and could give a clue to this type's function. The FRAG3 shell contains a green/grey/orange BMC-P triplet and two BMC-H, one with standard interface residues and another with a C-terminal extension of about 50 residues.

#### *FRAG4 – 4 loci*

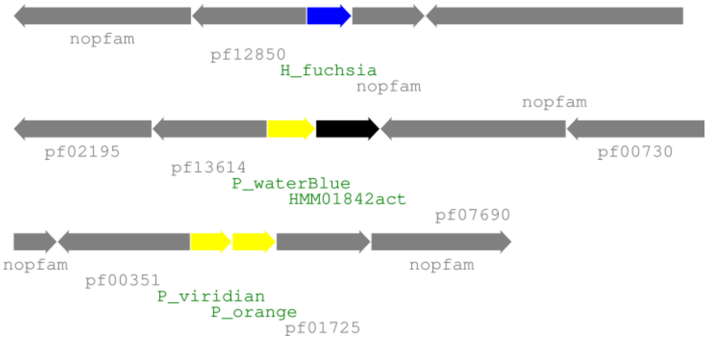


FRAG4 loci are only found in Candidatus Marinimicrobia. The only BMC typical protein besides shell proteins is with a pfam01842 ACT domain of about 90 aa at its C-terminus. ACT domains are commonly found as regulators that bind amino acids [2]. The N-terminal domain of that protein seems specific to this BMC and has no significant homology to any other currently available protein sequence. The shell protein set is not consistent across the loci and some BMC-P might be missing ; besides a standard BMC-H there are BMC-P from the blue, green, orange and grey major clades.

#### *FRAG5 – 3 loci*

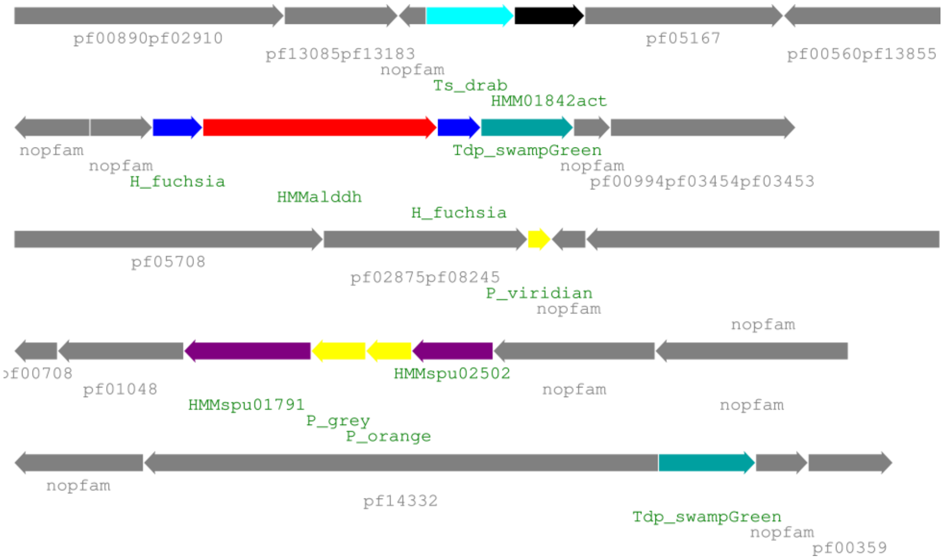


All of the FRAG5 loci are found in Planctomycetes; the presence of the SPU signature enzymes (pfam01791/pfam02502) indicates a similar type of reaction. The aldehyde dehydrogenases has a C-terminal EP and the closest relatives are found in FRAG8 and ACI types. The shell proteins consist of a BMC-P triplet from green/grey/orange clades, two standard BMC-H, two BMC-T$^{dp}$ from the major clade containing CcmP as well as a BMC-T$^{s}$ without pore cysteine motif and no co-occurring PduS homolog.

#### *FRAG6 – 5 loci*

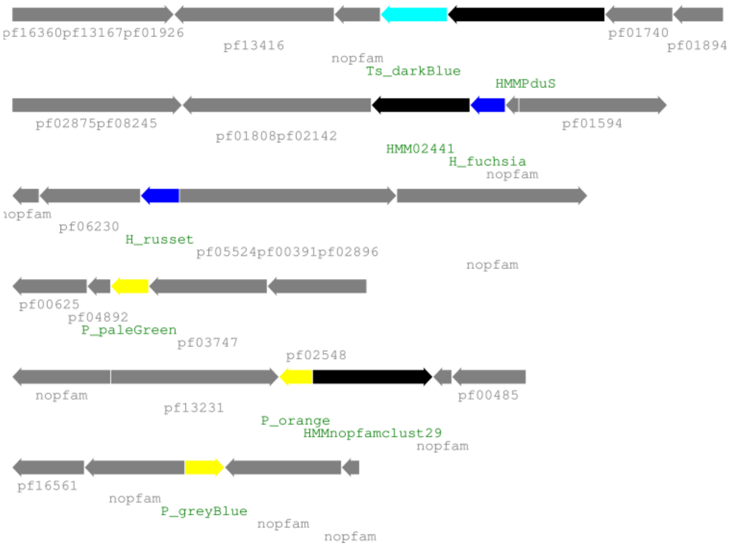


FRAG6 loci are found in three candidate phyla: Candidatus Hydrogenedentes, Candidatus Sumerlaeota and candidate division BRC1. The BMC-P triplet from the green/grey/orange clades are co-localized in two loci but in separate locations in all other genomes. A BMC-H with standard interface residues is accompanied by a pfam02441 flavoprotein but no other typical BMC enzymes can be found in the proximity of shell proteins. Another BMC-H with mostly standard interface residues is also found by itself and a PduS-T homolog pair is found in two genomes.

### *FRAG7 – 2 loci*

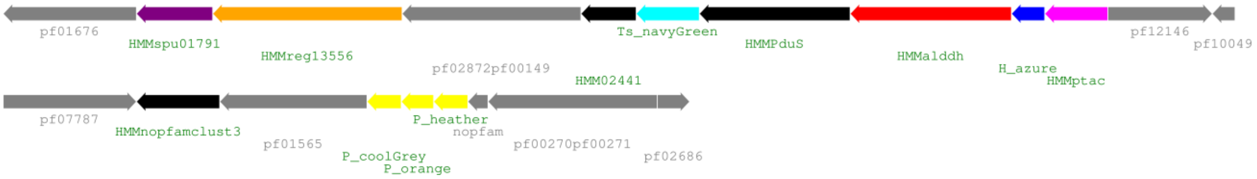


The FRAG7 loci are found in two Candidatus Hydrogenedentes strains. The locus is split in two locations, one containing a BMC-P triplet from green/grey/orange clades and the other a standard BMC-H, a PduS-T homolog pair, an AldDh and PTAC as well as a pfam02441 flavoprotein. The closest relatives of the AldDh with an N-terminal EP are found in the HO and SPU3 types.

### *FRAG8 – 7 loci*

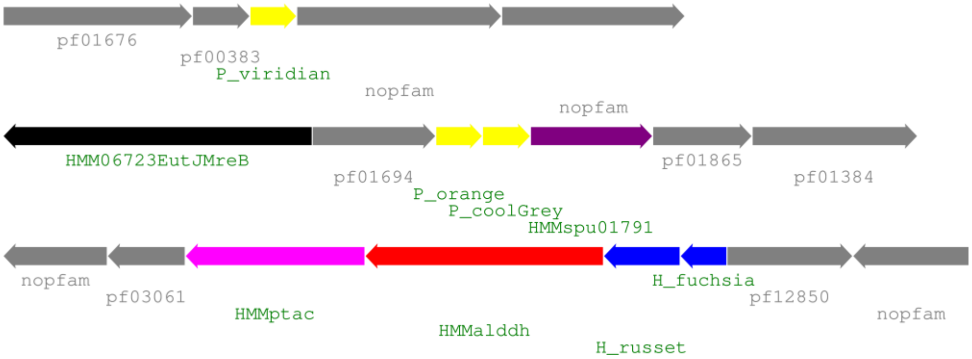


FRAG8 BMCs are mostly found in Gemmatimonadetes but two taxonomic outliers are from a Planctomycete and a Deltaproteobacterium. The AldDh with a C-terminal EP is closest related to the ACI one but it is only found in three loci; while the SPU-type signature enzyme pfam01791 is found in almost all loci, the pfam02502 enzyme is only found in two loci. Shell proteins consists of a BMC-P triplet from green/grey/orange clades as well as a BMC-H with standard interface residues; two loci also additionally contain a BMC-H protein with a 50 amino acid C-terminal extension .

### *GRM loci*

GRM loci are characterized by the presence of glycyl radical enzymes (GREs); there are six different previously characterized members (GRM1-5, GRMguf; see [3] and below). GRM1 and GRM2 use choline as an initial substrate and are sometimes referred to as Cut (choline utilizing). GRM3 and GRM4 utilize 1,2-propanediol and GRM5 fuculose phosphate as initial substrate; no function has yet been proposed for the GRMguf. We introduce here several subtypes for those already established types (most additional ones are also described in [4]) and add a new GRM6 type.

### *GRM6 – 19 loci*

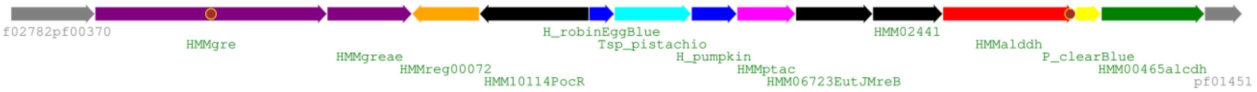

GRM6 loci occur mostly in various Firmicutes and a single member is from the Bacteroidetes phylum. The loci contain a complete enzyme set that matches the degradation pathway of 1,2-propanediol using a glycyl radical enzyme analogous to GRM3 and GRM4. Like its GRM3/4/5 counterparts the GRE contains an internal encapsulation peptide and on a GRE phylogenetic tree it is closest related to GRM4. The AldDh however has a C-terminal EP, which is usually found in enzymes related to ethanolamine degradation and their closest homologs are enzymes found in EUT2 loci.

The shell of GRM6 loci generally contains a shell protein set of two BMC-H, one BMC-P and a BMC-$T^{sp}$ protein. The H_robineggblue BMC-H contains the standard interface residues and the more specialized H_pumpkin BMC-H has a C-terminal extension that contains a conserved 4xCys motif likely to bind a metal ion. The BMC-$T^{sp}$ shell protein is of the PduB type and its closest related members are from GRM1A and PDU1D BMC types.

#### *HO – 41 loci*

type I – 30 loci

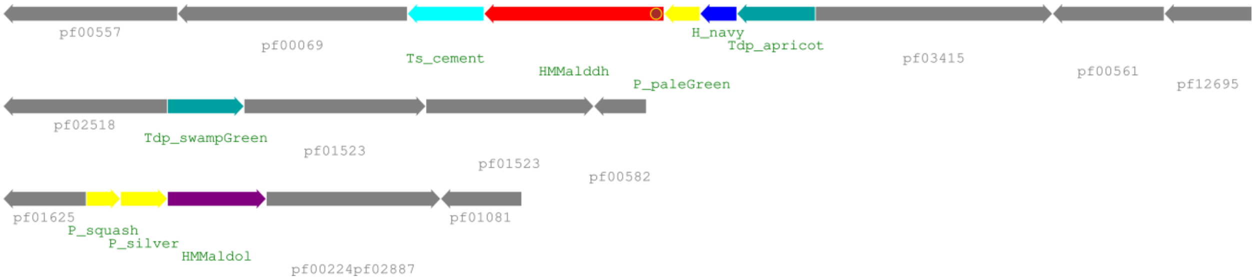


type I – with PduS/T homologs and EutBC

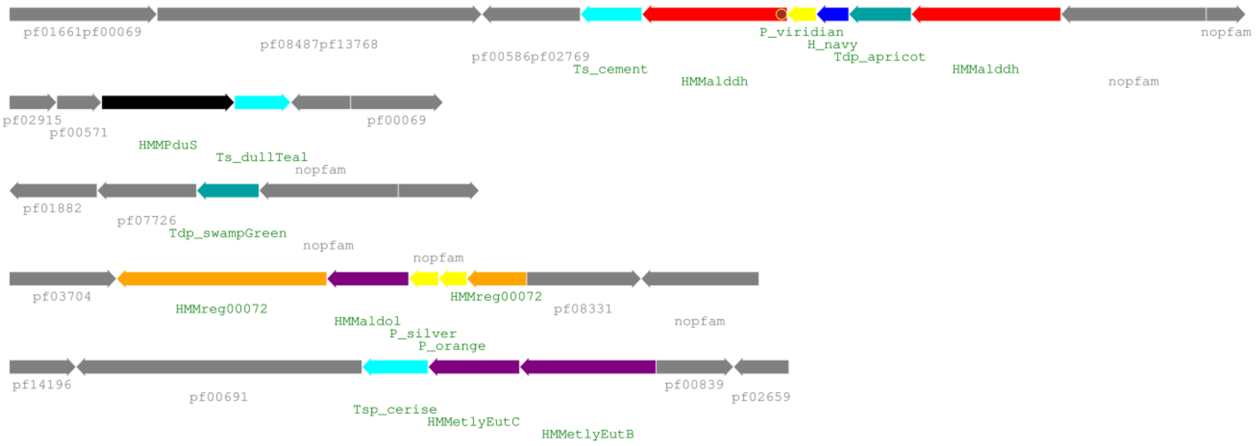


type II – 11 loci

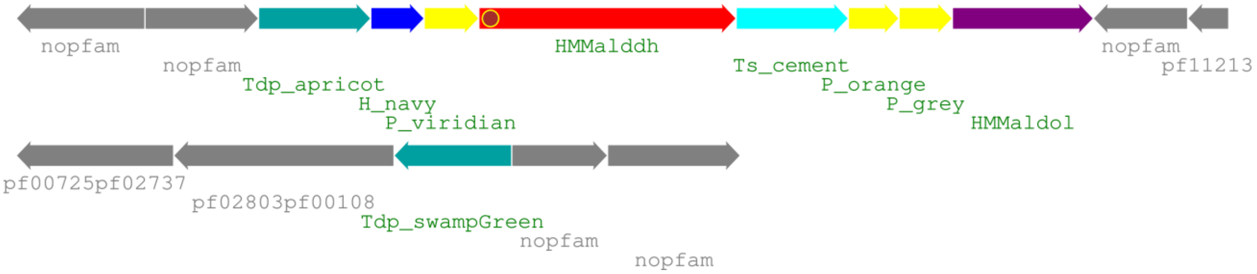


The HO type BMC is found mostly in Deltaproteobacteria, with one member also from Alphaproteobacteria (Rickettsiales bacterium) and another found in a Candidatus Latescibacteria bacterium. The typical main locus of type I (that includes the *Haliangium ochraceum* model organism) contains only shell proteins along with an AldDh. In a second type, almost all components are found in the main locus besides a satellite BMC-T$^{dp}$ protein. A pfam00596 aldolase is found on a satellite locus of many type I loci (24) and in the main locus of about half type II loci (5). The AldDh always has an N-terminal EP but only about a third of the pfam00596 aldolases have a C-terminal extension that is possibly an EP. Based on the AldDh tree, the closest related BMC types are SPU5 and SPU7. Interestingly, in four loci there is a BMC-T$^{sp}$ (EutL type) together with the EUT signature enzyme ethanolamine lyase EutB/C; this is possibly a EUT module that could process ethanolamine. PduS/T homolog copies are found in three genomes in a satellite location.

The combined shell components of both HO subtypes consist of a single BMC-H, three BMC-P, two BMC-T$^{dp}$ and a BMC-T$^{s}$. The BMC-P are part of a typical BMC-P triplet; they are found split into main and satellite loci in type I and in the same main operon in type II. The two BMC-T$^{dp}$ are found in two distinct clades on the tree, the main locus one is in the clade that also contains the alpha-carboxysome CsoS1D and the satellite BMC-T$^{dp}$ is found in the clade that contains the beta-carboxysomal CcmP member.

#### *MIC2 – 22 loci*

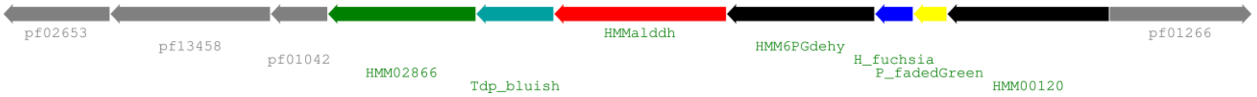


MIC2 type BMCs are found in different classes of Proteobacteria (Acidithiobacillia, Alpha- and Gammaproteobacteria) as well as two members from Actinobacteria. The enzymes in the locus consist of an aldehyde dehydrogenase, a pfam02866 lactate/malate dehydrogenase, a protein that belongs to a 6-phosphogluconate dehydrogenase-like superfamily (http://www.ebi.ac.uk/interpro/entry/IPR008927) and a pfam00120 glutamine synthetase. There is an N-terminal extension of about 70 amino acids on the pfam02866 protein that is only present in the context of a BMC locus and it has detectable homology with the EutQ/cupin-like superfamily, possibly interacting with a ligand involved in this BMC's pathway. A similar domain is also found in the MIC1 members of the same protein. There is a short N-terminal extension on the 6-phosphogluconate dehydrogenase-like protein that could be a ~10aa long predicted helical EP but there is no EP on the aldehyde dehydrogenase. There are no satellite loci and no other BMC types in the genomes that have a MIC2 locus. The closest relative of this type is MIC1 and the next closest characterized member is RMM1 based on AldDh phylogeny.

There is one shell protein each of the BMC-H, BMC-P and BMC-$T^{dp}$ type in the locus, representing a minimal set of shell proteins. Both the BMC-H and BMC-P have standard sequence motifs and the BMC-$T^{dp}$ is from the blue major clade that contains other uncharacterized BMC types.

### *MIC3 – 11 loci*

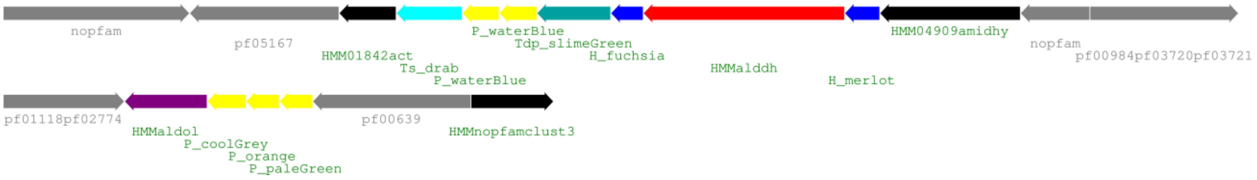


All MIC3 loci are found in Planctomycetes with the exception of one from a candidate phylum archaeon, Candidatus Woesearchaeota archaeon (CMO84__locus_1); this however seems to be an annotation mistake as it is only found in one Woesearchaeota archaeon (ARS1170), and the phylogenetic distribution of the proteins in this species analyzed with IMG shows many BLAST matches against Planctomycetes and Proteobacteria. The enzymes in the main locus consist of an AldDh as well as a pfam04909 amidohydrolase. In a satellite locus that is present in all members there is a BMC-P triplet with a pfam00596 aldolase. There are no detectable encapsulation peptides on any of the proteins. The AldDh however has a unique C-terminal extension that consists of a flexible linker of up to about 120 amino acids followed by a domain of about 40 aa with four conserved cysteines. The AldDh is phylogenetically closest to EUT3 members.

The shell proteins of the main locus consist of two BMC-P that are in the same P_waterblue clade yet they show distinct sequence differences. Similarly, the BMC-H have standard interface residue motifs yet the two copies are distinct. A BMC-$T^{s}$ protein is also found in the main locus along with a BMC-$T^{dp}$ that is in the same major clade as the beta-carboxysomal CcmP. In a satellite locus there is an additional BMC-P triplet from the green, grey and orange major clades. This adds up to an unusual total number of five pentamers for the consolidated locus.

### *MIC4 – 15 loci*

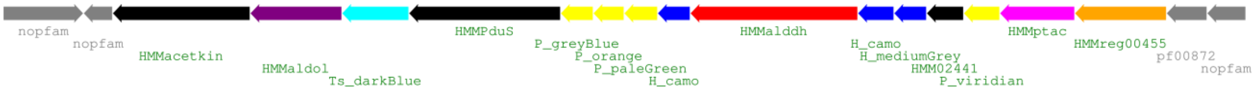


MIC4 type BMCs are found in a diverse group of organisms, there are members from Actinobacteria, Bacteroidetes, Calditrichaeota, candidate division KSB1, Firmicutes and Ignavibacteria. The enzymes of the main locus consist of an AldDh, a PTAC, a pfam00596 aldolase as well as a pfam00871 acetate kinase. The AldDh with an N-terminal EP is closest related to MUF2 and MUF3 (it is found in two places on the AldDh tree).

The BMC-H shell protein set is not consistent across the loci, some contain a single BMC-H with standard interface residues(H_fuchsia), others contain two mostly standard H_camo type BMC-H as well as an H_mediumGrey with unusual interface residues. All of the loci have a BMC-P triplet of the green/grey/orange major clades with sometimes an additional BMC-P from the green major clade. A PduS homolog associated with a PduT-type BMC-T$^{s}$ is also found in the operon with a conserved cysteine in the pore.

#### *MIC6 – 3 loci*

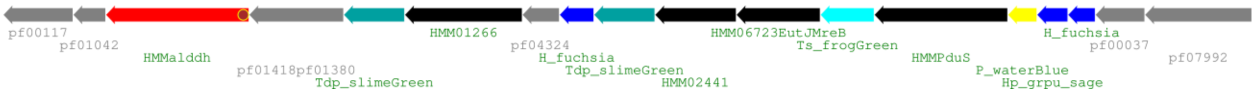


MIC6 is present in anaerobic Deltaproteobacteria and contains a variety of common BMC proteins of unknown function and an aldehyde dehydrogenase with an N-terminal EP. The locus contains a predicted [2Fe-2S] binding protein of pfam04324 and a FAD dependent oxidoreductase of pfam01266 with a BMC-specific 30 amino acid C-terminal extension that contains a conserved 4xCys motif. The AldDh is closest related to the SPU2 homolog.

The shell protein set of this type contains two BMC-H and a BMC-P protein with standard sequence motifs as well as two distinct copies of a BMC-T$^{dp}$ from the green major clade that contains the beta-carboxysomal CcmP. Additionally there are two shell proteins that are expected to bind an iron-sulfur cluster at its pore, a PduT-type BMC-T$^{s}$ and a GrpU-type BMC-H$^{p}$.

#### *MIC7 – 6 loci*

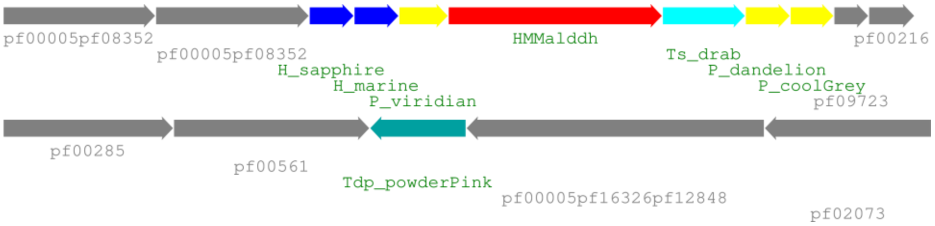


The MIC7 locus is found in *Candidatus Latescibacteria* and *Candidatus Eisenbacteria* and contains only an AldDh besides shell proteins. The AldDh with an N-terminal EP is closest related to the SPU3 one.

The shell proteins of the MIC7 locus contain a BMC-H with standard interface motifs (H_marine) as well as a more specialized BMC-H and a BMC-P triplet from the green/grey/orange major clades. There are two types of BMC-T$^{s}$; in two loci there is a Ts_dullteal with a predicted pore cysteine located next to a PduS homolog and in four loci there is a Ts_drab that lacks the pore cysteine (and does not co-occur with a PduS homolog). A BMC-T$^{dp}$ protein is also found in a satellite location; its sequence places it at the base of the major clade containing the alpha-carboxysomal CsoS1D homolog.

#### *MUF2 – 13 loci*

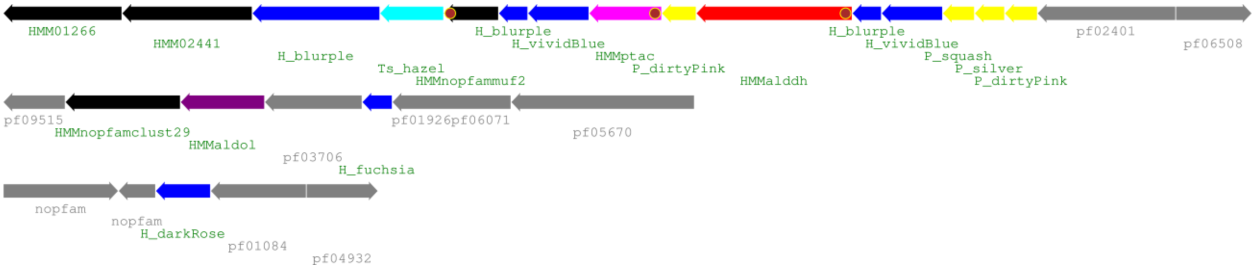


MUF2 type BMC loci are primarily found in Gemmatimonadetes and additionally in a Candidatus Handelsmanbacteria bacterium and an Armatimonadetes bacterium. There is an AldDh and a PTAC in the main locus; a pfam00596 aldolase in a satellite locus (sometimes also in the main locus) together with a BMC-H. Five of the loci have an uncharacterized protein located between the genes for the BMC-H and BMC-T$^s$. This protein has a predicted C-terminal EP and its middle domain (~60aa long) contains six conserved cysteines; the N-terminal domain is also found in a nearby pfam00701 class I aldolase. EPs are also found on the N-terminus of all the AldDh, on the N-terminus of some of the PTAC and on the C-terminus of all pfam00596 aldolases. The MUF2 loci occur mostly on their own, only one genome has an additional PVM locus. The AldDh is most closely related to the PVM one; considering also that the PVM contain the same set of enzymes, it seems likely that they have similar substrates.

There is a large variety of different shell proteins in this BMC type. All loci contain a BMC-P triplet from the purple, grey and orange major clades and most have another copy of the more basic purple clade member. There are at least two normal length BMC-H in the main locus and several organisms have double (Ts_hazel) or very unusual quadruple linear fusions of the BMC-H domains (large H_blurple). Another BMC-H with an unknown C-terminal extension domain is also found by itself in a satellite locus (H_darkrose).

#### *MUF3 – 8 loci*

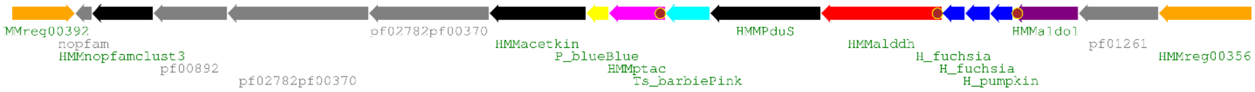


All MUF3 BMC loci are found in the Clostridiales and Thermoanaerobacterales orders of the Clostridia class. An AldDh, a PTAC, a pfam00871 acetate kinase and a pfam00596 aldolase form the enzymatic core. EPs are found one the N-terminus of the AldDh and the PTAC as well as on the C-terminus of the aldolase. There are no satellite loci besides a BMC-T$^s$ by itself in one genome and no other BMCs are found in the organisms that contain MUF3 loci. The MUF2 AldDh is its own branch on a phylogenetic tree, located between the major clades that contain most SPU types and PVM.

The main locus contains one BMC-P, three different BMC-H proteins, two of them with standard interface sequence motifs and a third, more unusual BMC-H (H_pumpkin). A PduT-type BMC-T$^s$ with a conserved cysteine pore residue is found next to a PduS homolog in the center of the locus.

### *SPU – general information*

While the SPU type has been described in Axen et al, at that time there were only eight members of this type. The more recent genome sequencing has increased that number to more than 150, which enabled us to make a more thorough analysis of the type in general as well as determine seven subtypes. They generally contain two enzymes related to sugar phosphate reactions, pfam01791 and pfam02502 and additionally they all contain at least three distinct BMC-P proteins, one each from the grey and orange major clades that are always next to each other as well as a third BMC-P from the green, blue or purple clades.

### *SPU1 – 62 loci*

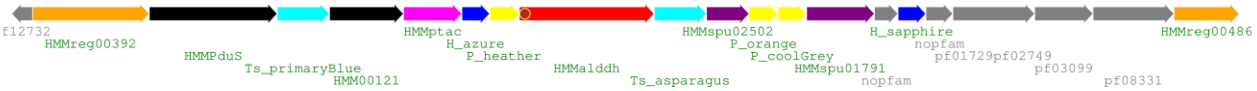


The SPU1 type BMC loci are found exclusively in Elusimicrobia species. The SPU signature enzymes pfam01791 and pfam02502 are adjacent to two BMC-P and there is also an AldDh, a PTAC and pfam00121 isomerase. EPs are found on all AldDh enzymes and on about half of the PTAC; the pfam01791 proteins sometimes have N-terminal extensions but those do not resemble EPs. A few organisms have the locus split into two locations and PduS/T homologs are found in satellite loci of five genomes but there are no other BMC loci in those genomes.

The shell proteins of the main locus consist of a BMC-P triplet and two BMC-H paralogs that have charge complementary interface residues (KAA**K**/P**R**PH vs KAA(**N/D**)/P**Q**PH). There are two BMC-$T^{s}$, a PduT-type with a pore cysteine associated with a PduS homolog and a second BMC-$T^{s}$ related to PduT but that lacks the pore cysteine and seems to fulfill a purely structural function.

### *SPU2 – 53 loci*

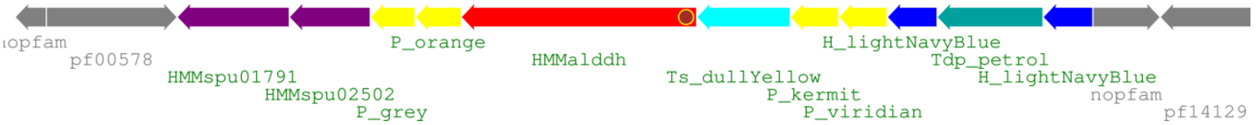


SPU2 type BMC loci are found in a variety of bacterial species, most members are in the Bacteroidetes and Spirochaetes phyla but many are also from the various candidate phyla: Atribacteria, Melainabacteria, Moduliflexus flocculans, division KSB3, Wallbacteria and Vecturithrix. The enzyme set in this locus type is more limited and consists of only an AldDh and the pfam01791 and pfam02502 signature enzymes. Only few loci also contain a pfam00121 triosephosphate isomerase, a PTAC or an alcohol dehydrogenase. There are no satellite loci but a few loci are split into 2-3 genomic locations. EPs are found on the N-terminus of all AldDh and on about half of the pfam01791 and a third of the pfam02502 enzymes. There are no other BMC types co-occurring with SPU2 in these genomes.

The shell proteins of this locus consist of one or two similar BMC-H with standard interface motifs, a BMC-P triplet with an additional copy of a green clade member, a BMC-$T^{dp}$ from the CcmP-like major clade as well as a BMC-T[s]. The BMC-T[s] co-occurs with a PduS homolog in about a quarter of the loci; however the cysteine in the pore is still found in 80% of the BMC-T[s], possibly indicating that many of them have lost PduS more recently but still retained the BMC-T[s] for structural reasons.

#### *SPU3 – 31 loci*

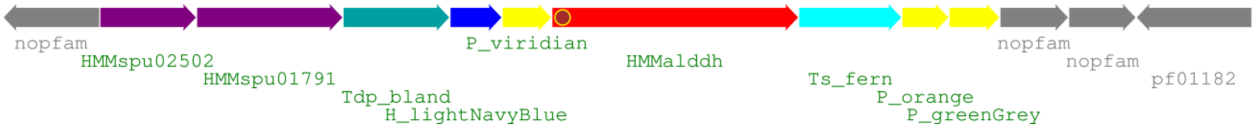


SPU3 loci are found in very diverse species, there are loci in the phyla Acidobacteria, Actinobacteria, Calditrichaeota, Chloroflexi, Gemmatimonadetes, Lentisphaerae, Planctomycetes, Proteobacteria and Verrucomicrobia as well as the candidate phyla Marinimicrobia, Parcubacteria and candidate division KSB1. The enzymes of this locus consists of an aldehyde dehydrogenase and the pfam01791 and pfam02502 enzymes, the latter two are found fused on one polypeptide chain in 12 observed loci (pfam02502 is the N-terminal domain, which matches the order in the non-fused loci). EPs are found on the N-termini of the AldDh as well as the pfam01791; for the fused pfam02502-pfam01791 a sequence that could be an EP is found internally between the two domains. Satellite loci are not common in this BMC type; there are two cases where the EUT signature enzymes are found together with a EutL type BMC-T[sp] ("EUT module") and sometimes the locus is split into two locations.

The shell components of this locus consists of a BMC-H with standard interface residues, a BMC-P triplet from the green, grey and orange major clades, a BMC-T[dp] from the CcmP-like major clade as well as a BMC-T[s] without co-occurring PduS homolog and no cysteine in the pore.

#### *SPU4 – 33 loci*

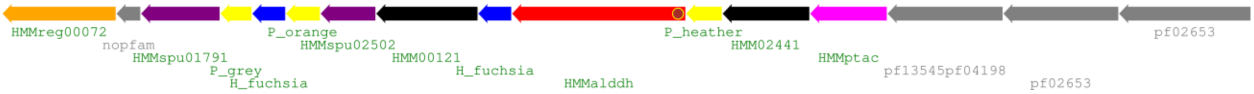


SPU4 type BMCs occur mostly in the Firmicute orders Bacillales and Clostridiales, one member is found in a Deltaproteobacterium (Deltaproteobacteria bacterium HGW-Deltaproteobacteria-22) and one in a Tenericute (Candidatus Izimaplasma sp. HR2). There is an aldehyde dehydrogenase, the two SPU signature enzymes pfam02502/pfam01791, a PTAC as well as a pfam00121 triosephosphate isomerase in the locus. In almost all loci there is also a protein that belongs to the eut_hyp type (https://www.ncbi.nlm.nih.gov/Structure/cdd/TIGR02536) that is related to the pfam02441 flavoprotein family. The presence of a 30-40 amino acid N-terminal extension with a predicted amphipathic helix followed by a region of low conservation indicates an EP and suggests a role for this protein in the BMC function. N-terminal EPs are also found the AldDh and most of the pfam01791 proteins, PTACs and about half of the pfam00121 isomerases. There are generally no satellite loci for SPU4, only in two cases there are 1-2 locus components in a different location. SPU4 loci occasionally co-occur with other BMCs, with EUT2-type BMCs (7x), GRM1 (2x) and BUF3 (2x).

The shell proteins in the locus consist of a BMC-P triplet and two similar BMC-H with standard interface motifs. Unusually for SPU loci there is no BMC-T of any type consistently in the main locus, there is only a PduS homolog-associated BMC-T$^{s}$ in 14 loci.

### *SPU5 – 8 loci*

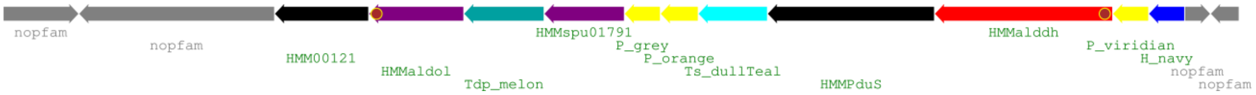


SPU5 type BMCs are found in eight Proteobacteria and one Chloroflexi class bacterium. There is always an AldDh but no pfam02502 and only four have the pfam1791 SPU signature enzyme; in five loci there is a pfam00596 aldolase. Despite lacking some SPU type signature enzymes it is likely still closely related to other SPU due to their partial presence and the shell protein composition. EPs are found on the N-termini of the AldDh on the C-terminus of the pfam00596 aldolase. The AldDh is closest related to the HO type and the next closest SPU relative is SPU3. No satellite loci are observed in those genomes and also no other BMC types.

The shell proteins of this locus consist of a BMC-P triplet from the green/grey/orange clades, a BMC-H with standard interface motifs and a PduS homolog-associated BMC-T$^{s}$ with a cysteine pore motif. Four loci additionally contain a BMC-T$^{dp}$ from the major clade that contains the alpha-carboxysomal CsoS1D.

### *SPU6 – 44 loci*

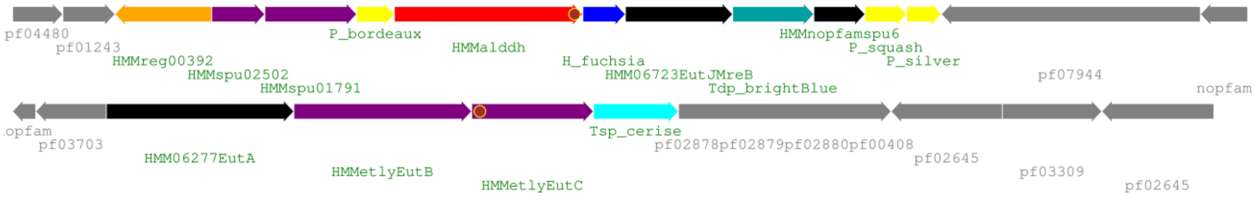


related Spirochaetes – 2 loci

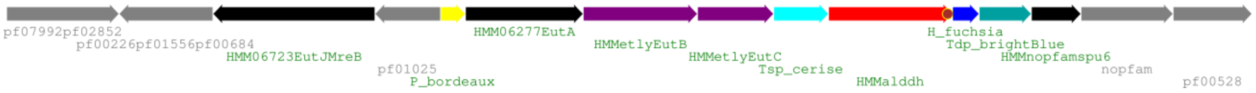


SPU6 BMCs occur only in Chloroflexi class bacteria. The core enzymes are an aldehyde dehydrogenase, the SPU signature enzymes pfam01791/02502 and a short ~130 amino acid protein with no pfam that has some homology with the EutQ type protein. A EutJ type protein occurs in eight loci. The locus is occasionally split into two locations or contains a duplicate set that lacks some enzymes. A “EUT module” satellite locus that consists of a BMC-T$^{sp}$/EutLPduB and EUT signature enzymes EutA/B/C is found in seven loci but there are no other full BMC loci found in the genomes. A SPU6 related BMC type is found in two Spirochaetes genomes where the EUT signature enzymes are in the locus and replace the SPU signature enzymes. This could represent an intermediate form of a transition between SPU6 and EUT3 that exchanged the signature enzymes.

EPs are found on the C-terminus of all AldDh, on the N-terminus of six pfam01791 enzymes and on six of the seven EutC on the satellite locus. The AldDh of SPU6 is the only one of any SPU type that is found on the EUT23/GRM12 major branch of the AldDh tree; the closest related ones are EUT3 and MIC1/2.

The shell proteins consist of a BMC-P triplet, one BMC-H with standard interface residues and a BMC-$T^{dp}$ of the blue major clade. Interestingly, the BMC-$T^{dp}$ of SPU6 and EUT3 are closely related (Tdp_brightBlue / Tdp_ultramarine) and the Spirochaetes members mentioned above are found basal to the SPU6 branch, consistent with a potential intermediate form in between EUT3 and SPU6.

### *SPU7 – 8 loci*

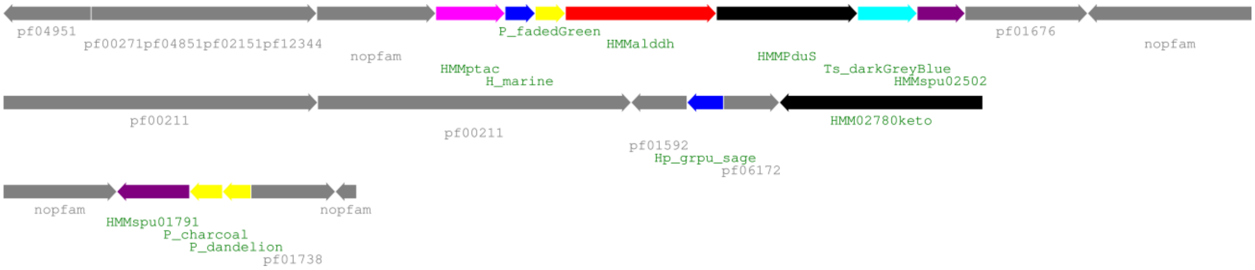


SPU7 type BMC loci are only found in unclassified and candidate bacterial phyla (candidate phyla Cloacimonetes, Riflebacteria, Wallbacteria and Ozemobacter sibiricus). The organisms where this BMC is found seem to be mainly from anaerobic environments (Candidatus Cloacimonetes [5], Candidatus Riflebacteria/Ozemobacter sibiricus [6]. The enzyme set of the locus consists of an AldDh, a PTAC and a pfam02502 SPU enzyme in the main locus as well as a pfam01791 SPU signature enzyme in a satellite location. There is an EP on the N-terminus of all aldehyde dehydrogenases and PTACs. The SPU7 generally occurs by itself, only one MIC5 locus is in one of the genomes. The closest relatives of the AldDh are from the SPU5/HO-type.

The locus shell proteins consists of one BMC-H with standard interface motifs, a BMC-P triplet with grey/orange clade members in a satellite location and a green clade BMC-P in the main locus. A PduS/T homolog pair is found in the main locus and another shell protein with a cysteine in the central pore, a GrpU type BMC-$H^p$ is found by itself in a satellite location of half of the loci.

### Previously characterized BMC types and additional subtypes

#### *BUF1 – 46 loci*

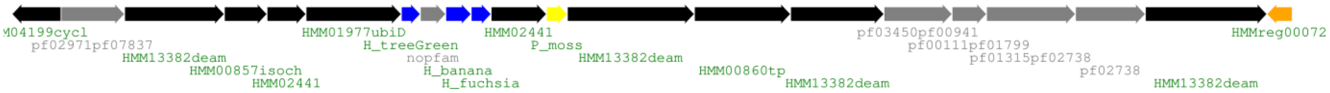


BUF1 is one of a few BMCs that do not have an aldehyde dehydrogenase; recently a role in the degradation of xanthine was proposed [1]. This BMC is found most commonly in Firmicutes but single members are found in the phyla Bacteroidetes, Proteobacteria and Synergistetes, however those do not have the full set of enzyme seen in Firmicutes.

#### *BUF1B – 55 loci*

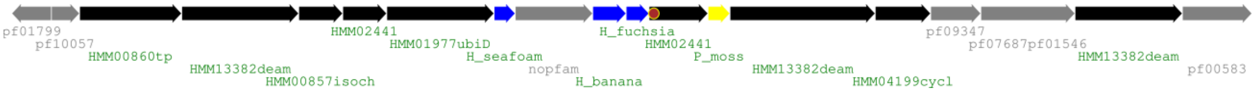


BUF1B loci are mostly identical to BUF1 but a few genes are shuffled (e.g. the membrane transporter HMM0860 is at the beginning of the locus). One of the BMC-H shell proteins (H_treeGreen vs H_seafoam) is similar but distinct. The BUF1B locus is almost exclusively found in the Bacilli order of Firmicutes.

#### *CsomeACh – 214 loci*

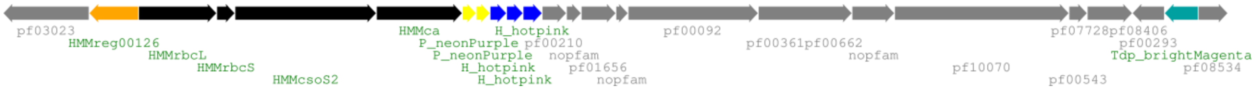


Chemoautotrophic carboxysome loci have been extensively characterized; the main locus consists of two BMC-P, 2-3 BMC-H, the catalytic enzymes RbcL/S and carbonic anhydrase and the interior organizing protein CsoS2. A BMC-$T^{dp}$ (CsoS1D) is frequently found in proximity to the locus or in a satellite location. These carboxysome loci are found mostly in Proteobacteria but also in five genome of the Actinobacteria phylum.

#### *CsomeACy – 125 loci*

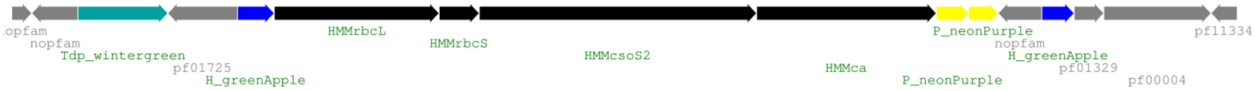


Cyanobacterial alpha-carboxysomes (alpha because of the type of RuBisCO) are very similar to their chemoautotrophic counterparts with a slightly different gene order, e.g. the two BMC-H are typically located before and after the rest of the main locus. BMC-$T^{dp}$ (CsoS1D) are found next to the main locus or in a satellite location; infrequently one of the BMC-H shell proteins are also in a satellite location. The locus is also found in three genomes of the eukaryotic Paulinella, an amoeba that has taken up a symbiont containing an alpha-carboxysome.

#### *CsomeBCy – 394 loci*

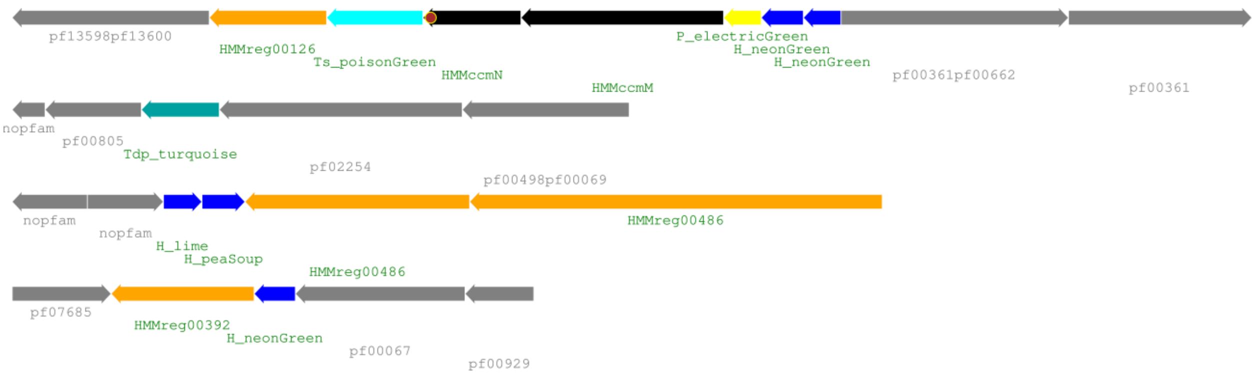


Beta-cyanobacterial carboxysomes are also highly studied and literature is available on all of its components. The main locus consists of genes coding for two BMC-H (named CcmK2 and CcmK1), a BMC-P (CcmL), the interior proteins CcmM and CcmN as well as a BMC-$T^{s}$ (CcmO; sometimes in a satellite location). On separate satellite loci are a BMC-$T^{dp}$ (CcmP), a pair of BMC-H (CcmK3 and CcmK4) and sometimes a single BMC-H (CcmK5 or CcmK6). All Beta-carboxysomes occur only in Cyanobacteria (one exception found in searches is an engineered *Bacillus subtilis* strain).

#### *ETU – 3 loci*

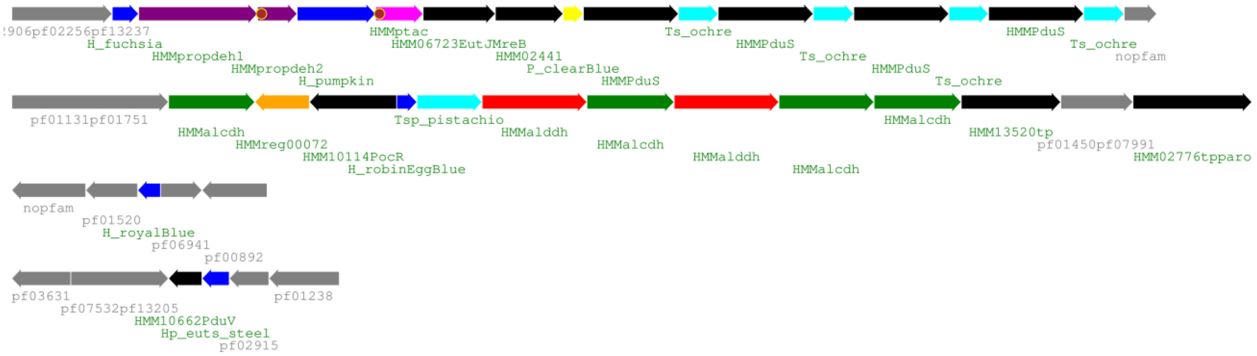


This locus type is found in three strains of *Clostridium kluivery* that has been shown to be able to process ethanol [7]. The described locus contains three alcohol dehydrogenases and two aldehyde dehydrogenases as well as a BMC-H (EtuA) and a structurally characterized BMC-$T^{sp}$ (EtuB). However in all strains there is another potential locus that contains a PDU signature enzymes, a PTAC and several PduS/T homolog pairs as well as two BMC-H and a BMC-P that are likely also involved in BMC formation since the other locus does not contain a BMC-P which might be necessary to form a closed shell. This locus was referred to as PDU-like/PDU3 in Axen et al but since it always co-occurs with the ETU type we conclude that they are likely to function together and are not a separate type. There are two satellite loci, one with a single BMC-H protein and another with a BMC-$H^{p}$ (EutS type) and a PduV homolog.

#### *EUT1 – 1378 loci*

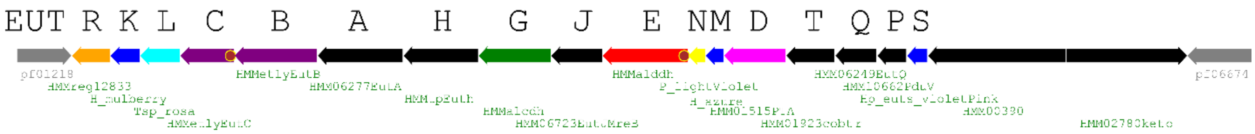


EUT1-split locus

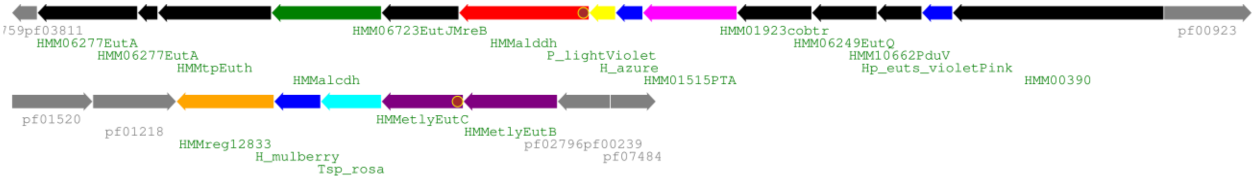


EUT1 type loci occur exclusively in Beta- and Gammaproteobacteria, most members are from very commonly sequenced Enterobacterial strains like *Escherichia coli* and various Klebsiella, Salmonella, Shigella. There is a wealth of biochemical characterization available on the EUT1 locus from *Salmonella enterica*. Most loci are organized in one operon but in 57 cases the locus is split into two genomic locations (typically like shown in the second locus diagram).

#### *EUT2*

EUT2 loci are much more diverse than EUT1 loci and there several subtypes with distinct gene composition and order. The main difference to EUT1 is the presence of the BMC specific pfam06130 PTAC instead of the "housekeeping" pfam01515 PTAC usually found outside of BMC loci. We added five other EUT2 subtypes based on distinct groups of the EUT clustering. We also designate an incomplete locus named EUT2x, which lacks the EUT signature enzymes but seems related to the EUT2 type. Some of those however co-occur with another EUT2 type so it could be a functional extension of that locus. The previously named EUT3 in Axen et al seems to be a fragment of a EUT2D type; almost all protein sequences are most closely related to EUT2D homologs; we renamed it to EUT2K since a EUT3 type should be reserved for a more distinct type.

#### *EUT2A – 169 loci*

EUT2A type loci are found mostly in Firmicutes but two members are also found in Actinobacteria, one in a Bacteroidetes and one hit in a Chlamydiae genome (which might be a mis-annotation of a shotgun sequencing gene fragment). EUT2A loci are mostly found on their own in genomes but about 5-10% of them co-occur with GRM1A, PDU1C, PDU1E or PVMlike BMC loci.

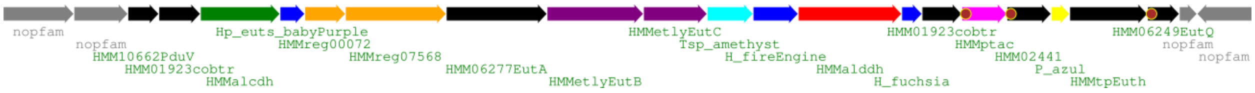


#### *EUT2B – 147 loci*

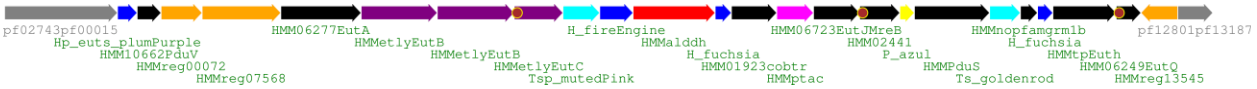


EUT2B type BMCs are found widely distributed among the phyla Firmicutes, Bacteroidetes, Fusobacteria, Proteobacteria and Synergistetes. The most interesting difference to EUT2A is the presence of a PduS/T homolog pair in most loci and absence of an alcohol dehydrogenase. EUT2B type loci frequently co-occur with other BMC types, about 20% with PDU1D and about 5-15% each with BUF1, EUT2B, EUT2G, GRM1A, GRM5/5like and PVMlike.

#### *EUT2C – 88 loci*

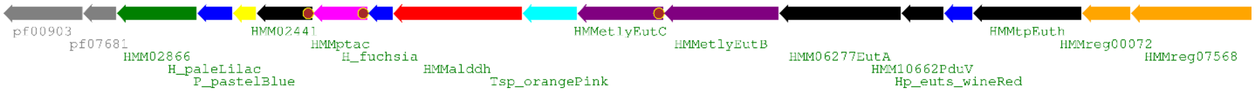


The loci of EUT2C are well conserved in order and composition and only found in the Bacillales order of Firmicutes. They do not contain satellite loci and about 10% co-occur with BUF1B type loci and four of them with SPU4, which is quite unusual as the two EUT and SPU locus types only co-occur in three other cases. This locus has an alcohol dehydrogenase of the pf02866 type.

#### *EUT2D – 28 loci*

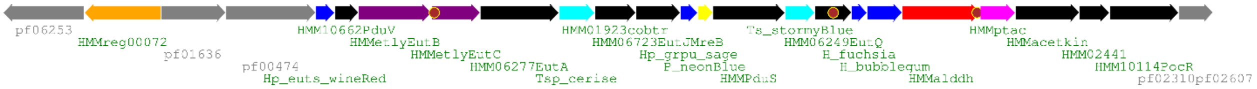


EUT2D loci are found only in the Firmicute orders Clostridia and Negativicutes. Like EUT2B, it contains a PduS/T homolog pair but lacks an alcohol dehydrogenase.

#### *EUT2E – 75 loci*

EUT2E type loci are only found in Bacillales. It is very similar to EUT2C in components but with a different gene order.

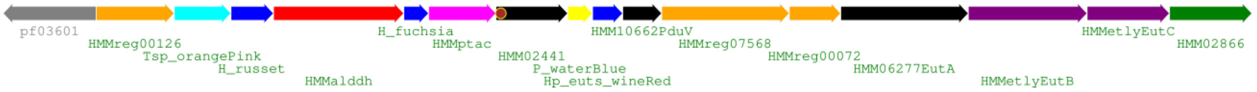


#### *EUT2F - 59 loci*

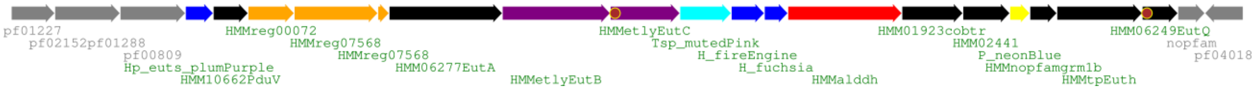


EUT2F loci are found exclusively in Fusobacteria. Fusobacteria are common in periodontal diseases and they are also found in the gut and have been implicated in colorectal cancer [8]. More than 60% of the loci co-occur with PDU1F type loci. The locus is similar to EUT2E but with a different order and additional EutQ and EutH type  proteins as well as a flavoprotein homolog of the HMM02441 family.

#### *EUT2G – 29 loci*

EUT2G type loci are found exclusively in Clostridia and are a locus type with minimal components. Unusual for a EUT type locus is the presence of only one BMC-H type protein along a single BMC-P and BMC-T$^{sp}$.

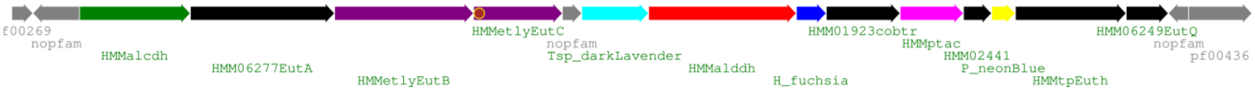


#### *EUT2H – 28 loci*

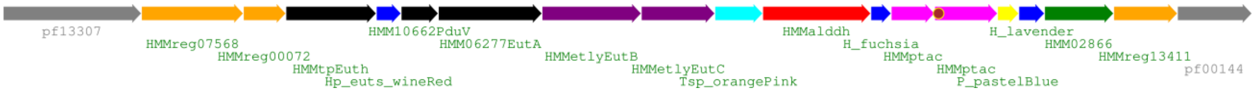


EUT2H type loci are only found in the Bacillales order of Firmicutes. A distinctive feature of this subtype is the presence of two pfam061030 PTAC enzymes, only one of them with an EP. This locus has an alcohol dehydrogenase of the pf02866 type.

#### *EUT2I – 162 loci*

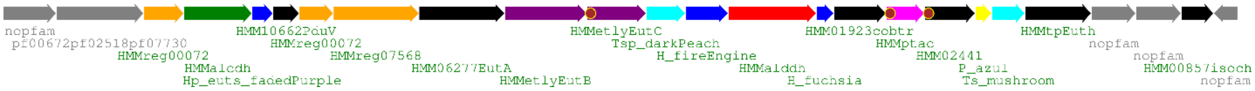


EUT2I loci are found mostly in Firmicutes but single members are also found in a Proteobacterium and one, very surprisingly, in a Thermotogae. This could represent a HGT event and this locus is the only one found in any Thermotogae species. The BMC-T$^{s}$ is related to PduT but there is no PduS homolog in this locus type.

#### *EUT2K – 4 members; previously EUT3*

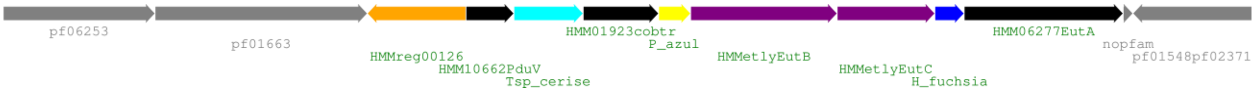


This locus was named EUT3 in Axen et al, however we have renamed it to EUT2K since it seems to be only a fragment occurring in only four closely related strains of Desulfitobacterium and does not represent a significantly different type than other EUT2 BMCs. The enzyme set is very reduced and only contains a cobalamin adenosyltransferase besides the EUT signature enzymes EutABC. They all co-occur with GRM1A and GRMguf so this could be just an extra module to process ethanolamine, integrated in the GRM1A BMC shell that has a matching pathway starting from acetaldehyde. In two of the loci even the BMC-H is missing which would support such a function as a supplementary module dependent on a main BMC locus.

#### *EUT2x – 24 loci*

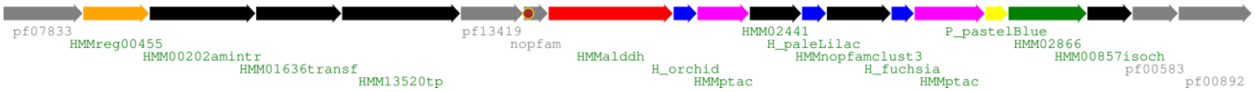

This BMC type lacks the EUT signature enzymes but most of the proteins are closest related to the ones from EUT2C. All of them occur in Bacillales and all except five of them co-occur with another EUT2 type so it could be an extension of those. Interestingly, in 60% of the loci there is a short-chain dehydrogenase usually found in RMM1; this could indicate a different initial substrate for this BMC type. Five of the loci contain an additional, unusual BMC-H (H_bloodorange) that nests within beta-carboxysomal BMC-H on the phylogenetic tree.

#### *GRM1A – 294 loci*

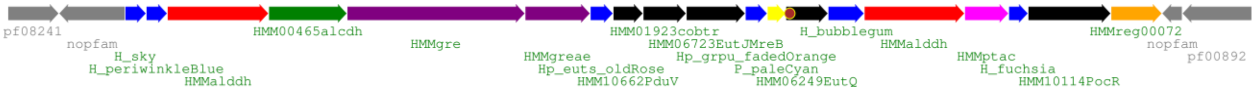


GRM1A type loci are found predominantly in various Firmicutes but a few loci are found in the commensal Coriobacteriales order from the phylum Actinobacteria and 1-2 members each in Bacteroidetes, Chlamydiae, Fusobacteria and Proteobacteria. Only one of the two AldDh is likely to be active since the other lacks an active site cysteine [9]. GRM1A type loci sometimes occur with other types; EUT2, PDU1 and PVMlike BMC types are most common.

#### GRM1B – 25 loci

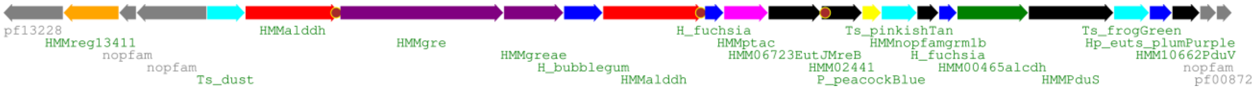


GRM1B loci are all found in Delta- or Gammaproteobacteria with one exception of a member from the Bacteroidetes phylum. The overall content is similar to GRM1A but with a different gene order. In addition to the PduS/T homolog pair there are two extra BMC-$T^s$ in this locus type that do not seem related to PduS. There is also a GRM1B specific protein that has not been assigned a protein family.

#### *GRM2 – 253 loci*

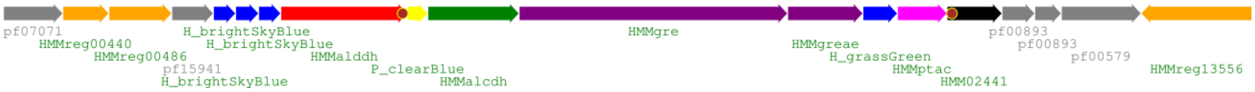


The choline degrading GRM2 BMCs are found exclusively in Gammaproteobacterial orders Aeromonadales, Enterobacterales and Vibrionales. The shell proteins consist of four BMC-H and a BMC-P; no BMC-T of any type are found in GRM2 loci. GRM2 commonly co-occur with EUT1 and PDU1AB.

#### *GRM3A – 84 loci*

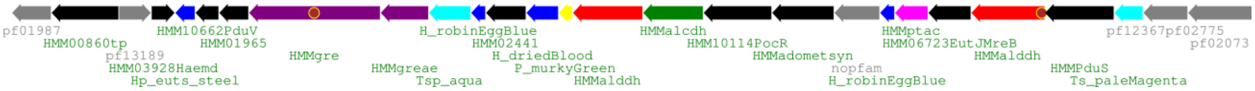


GRM3A loci are found exclusively in the Firmicute classes Bacilli, Clostridia, Negativicutes and Tissierellia. The shell protein set consists of one BMC-$H^p$, three BMC-H, one BMC-P, a BMC-$T^s$ and a BMC-$T^{sp}$. One of the BMC-H (H_driedblood) has a long C-terminal extension with a large number of positive and negative charges. Similar to GRM1 loci there are two aldehyde dehydrogenases; only one of them has an N-terminal EP but the active site cysteine seems mostly conserved in the one lacking the EP. Nevertheless, there could be other mutations that render it inactive and maybe fulfill a similar role as the inactive AldDh in GRM1 type loci.

#### *GRM3B – 92 loci*

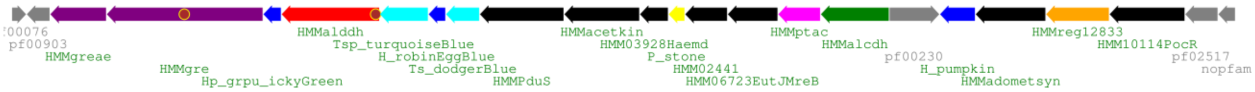


GRM3B type BMCs are all from Beta-, Gamma- or Epsilonproteobacteria. Most components are similar to GRM3A but in a different order; however there is no second aldehyde dehydrogenase in GRM3B. The BMC-H$^{p}$ in GRM3B however is a GrpU type with a conserved cysteine at the pore as opposed to a EutS type BMC-H$^{p}$ in GRM3A. GRM3B loci sometimes co-occur with EUT1 and GRM2.

#### *GRM3C – 24 loci*

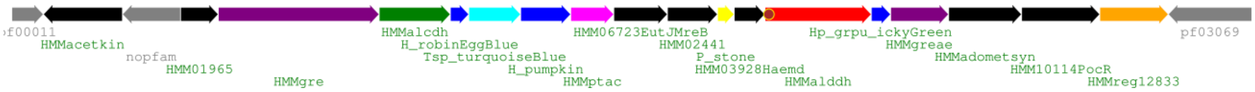


This GRM3 subtype is found exclusively in Alphaproteobacteria. In this locus type there is an unusual separation of GRE and GRE-activating enzyme (that are usually adjacent) and it lacks the PduS/T homolog pair observed in other GRM3 types.

#### *GRM4 – 8 loci*

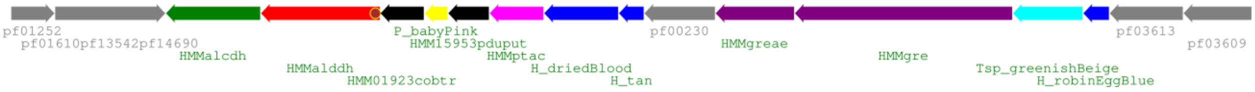


GRM4 loci were previously only observed in Shewanella strains. We have now found this BMC type in another seven types of Gammaproteobacteria, among them strains of *E. coli* and *Salmonella*, establishing it as a specific GRM BMC type. Half of the GRM4 loci co-occur with EUT1 and PDU1AB.

#### *GRM5 – 202 loci*

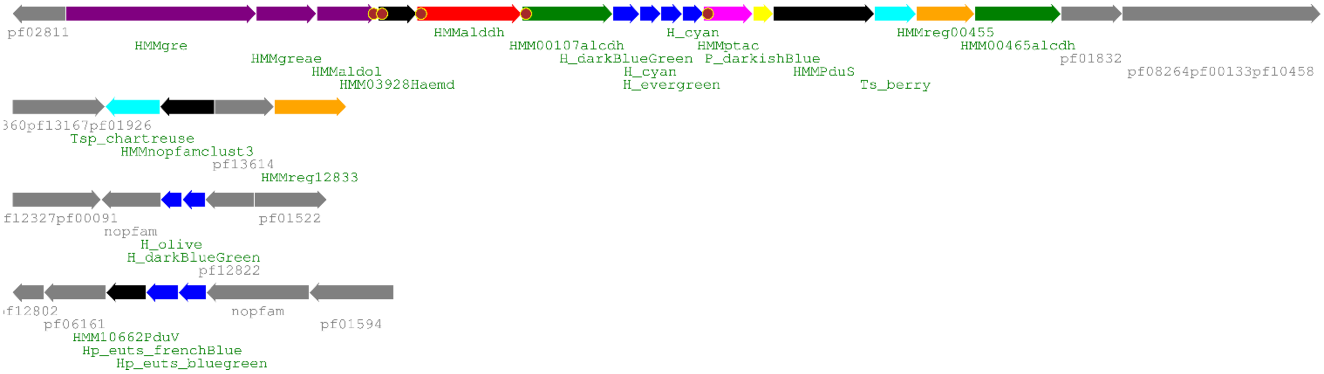

Occurrence of GRM5 type BMCs is limited to Clostridia class of Firmicutes. The main locus shell proteins consist of four BMC-H, one BMC-P and a BMC-$T^{s}$ of the PduT type next to a PduS homolog. While this locus is mostly on its own in a genome, more than 75% of the loci have one or more satellite loci. The three types of satellite loci are a PduB-like BMC-$T^{sp}$ next to a short-chain dehydrogenase, two BMC-H and two PduU related BMC-$H^{p}$ next to a PduV homolog.

#### *GRM5like – 44 loci*

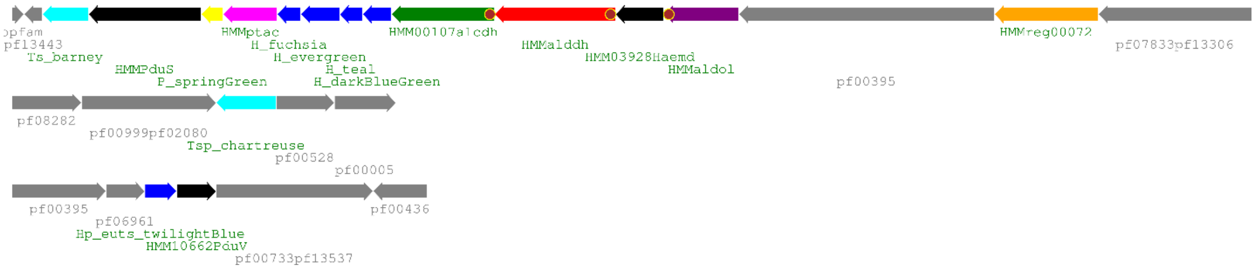


The GRM5like locus type is highly similar to the GRM5 type but with the notable absence of the GRE and GRE activating enzyme. Like its counterpart, it is found exclusively in Clostridia with one exception of a member in an Alphaproteobacterium. It is mostly on its own in the genome, indicating that it is likely functional. Indeed the GRE and activating enzyme can be found in a different location and the protein sequences are highly homologous to the GRM5 ones and contain the BMC-specific EP. While it is unusual that the central enzyme of the pathway is not in the main locus it is not unprecedented; for example RuBisCO in beta-carboxysomes, which makes up the bulk of the BMC interior, is only sometimes found in or close to the main locus.

#### *GRMguf – 18 loci*

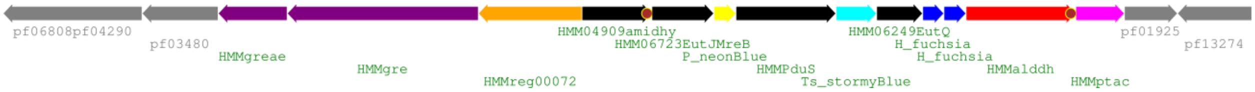


The GRMguf BMC type, named for GRE of unknown function because it does not cluster with other GRM GREs and also it does not have an encapsulation peptide sequence [3]. The genes for the GRE and its activating enzyme are also in the opposite direction than the rest of the locus. It is found in Clostridia as well as in Deltaproteobacteria. The shell proteins of the locus consist of two BMC-H, one BMC-P and a PduT-like BMC-$T^{s}$. 11 of the loci co-occur with GRM1A loci and in six of those there is also a EUT type BMC in the genome. This could mean that this locus fulfills some ancillary function instead of being a standalone BMC.

#### *MIC1 = MIC – 45 loci*

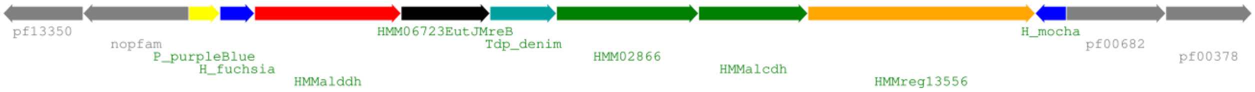


MIC1 loci are found in various types of Actinobacteria. The main locus shell proteins consist of two BMC-H, one BMC-P and a BMC-$T^{dp}$. There are no satellite loci observed for this type but there are three separate co-occurrences each with an ARO, RMM1 or CsomeACh BMC type. The closest related AldDh is found in the MIC2 type and both share the absence of an EP.

#### *MUF1 - 7 loci*

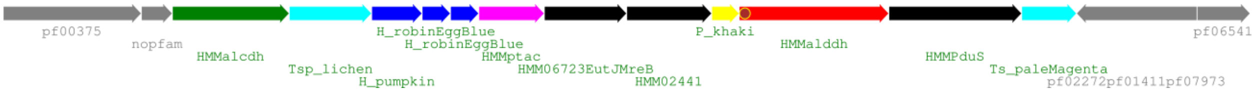


MUF1 type BMCs occur only in strains of Clostridium. The locus contains genes for three BMC-H, one BMC-P, one PduT-like BMC-$T^{s}$ and one BMC-$T^{sp}$. Both the aldehyde dehydrogenase and the PduB-like BMC-$T^{sp}$ are closest related to the ones found in PDU1D, indicating possible similarities regarding substrates or intermediates. Based on this 1,2-propanediol as a substrate would seem likely, however it lacks a signature enzyme. Enzymes for the both the B12-dependent and independent degradation of propanediol are found for example in the genome of *Clostridium botulinum E1 str. 'BoNT E Beluga'* but they do not contain EP motifs. Five of the seven loci co- occur with GRM1A.

#### *PDU1AB – 1293 loci*

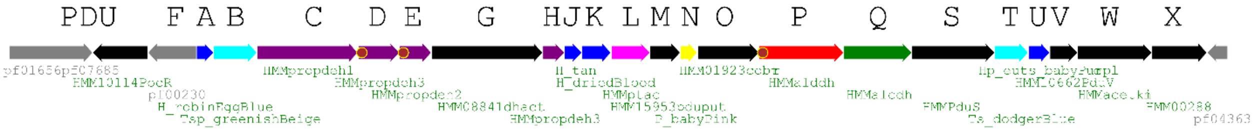


PDU1AB loci are found exclusively in Gammaproteobacteria, most of them of the frequently sequenced Enterobacterales order. The locus from *Salmonella enterica* Serovar Typhimurium LT2 has been extensively investigated and there is a wealth of biochemical studies available in the literature. This locus was also the basis of naming the PDU type proteins (see locus diagram). The loci are very rarely split into two locations and in more 60% of the genomes they co-occur with EUT1; rarely (~5%) they co-occur with GRM2 (and in 2/3 of those EUT1 is also present).

#### *PDU1C – 89 loci*

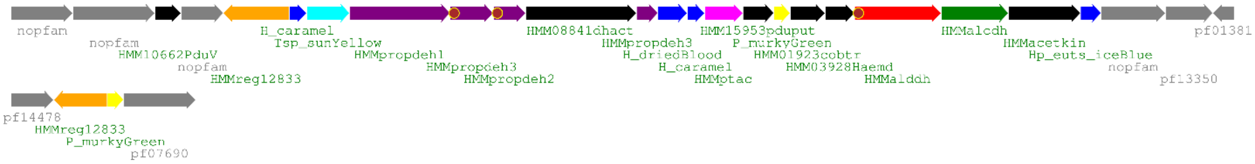


PDU1C type BMCs are specific to Lactobacillales. In comparison to PDU1AB they lack the PduS/T pair. In about 10% of the loci there is an extra BMC-P in a satellite location. 10 of the PDU1C loci co-occur with EUT2A in the genome.

#### *PDU1D – 182 loci*

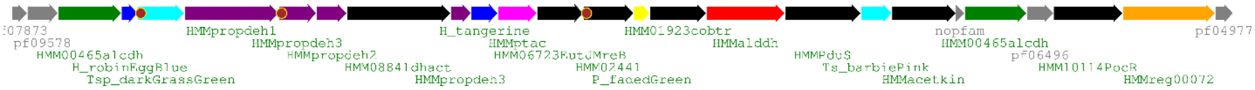

PDU1D loci are primarily found in Firmicutes but they are also present in the phyla Acidobacteria (one single member), Actinobacteria, Chloroflexi, Proteobacteria, Spirochaetes and Synergistetes. The widespread use indicates that this is likely a very efficient metabolic module that works independently across different phyla. In about 15% of the loci there is an extra BMC-$T^{sp}$ (sometimes duplicated) in a satellite location. Interestingly that one is homologous to EutL, unlike the one in the main locus which is homologous to PduB. The locus co-occurs with a EUT2 type in about 30% of the genomes and with GRM1A in less than 10%.

#### *PDU1E – 44 loci*

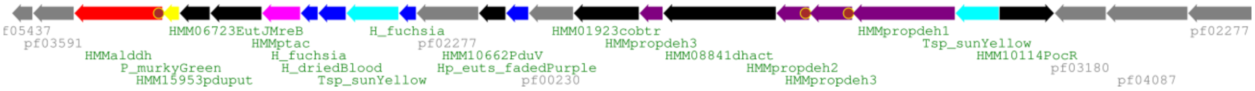


The loci for PDU1E are only found in the Firmicute classes Bacilli and Negativicutes. There are no satellite loci for this type except for a single PduS/T pair and it co-occurs with a EUT2 in about 30%, and with GRM1A in about 15% of the genomes. About half of the loci contain two BMC-$T^{sp}$, which are usually only found as one copy per locus.

#### *PDU1F – 44 loci*

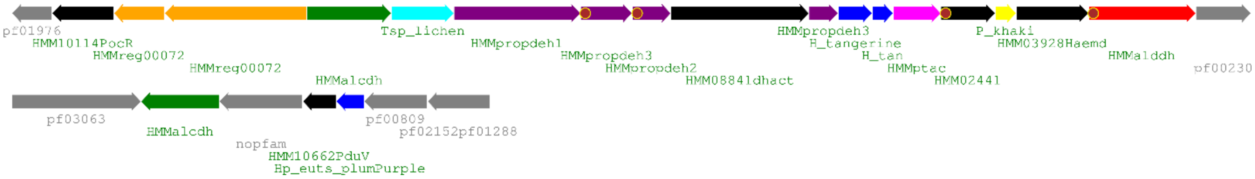


The PDU1F type locus is only found in different strains of the anaerobic Fusobacterium genus. More than 80% of the loci co-occur with a EUT2F type BMC in the genome, indicating a possible co-dependence.

#### *PVM – 285 loci*

*Planctomycetes limnophilus* characterized locus

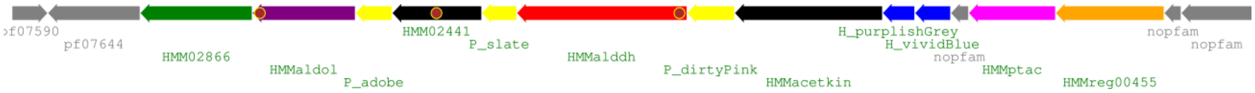


organization with satellite loci

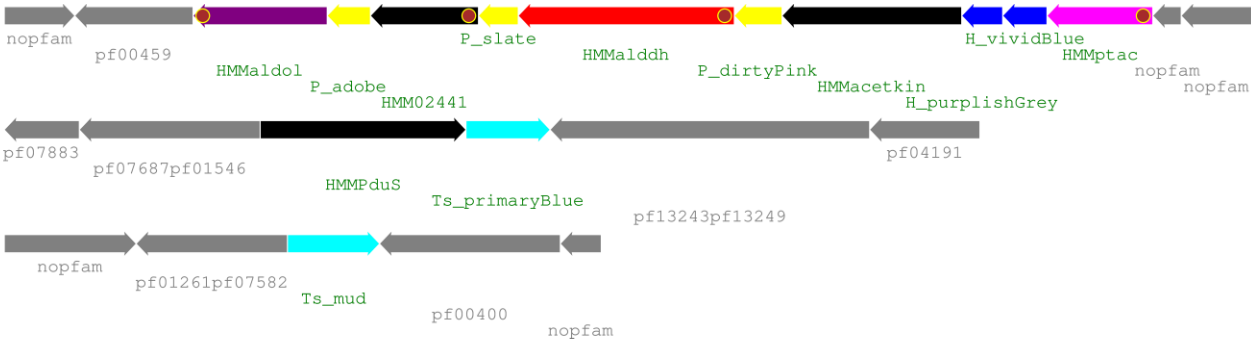


PVM (Planctomycetes and Verrucomicrobia microcompartment, [10]) type BMCs are only found in Planctomycetes and Verrucomicrobia where they degrade plant saccharides. The shell proteins of this locus consists of two BMC-H and three distinct BMC-P. This is the first characterized locus that features a BMC-P triplet, with members from the purple, grey and orange major clades.

#### *PVMlike –36 loci*

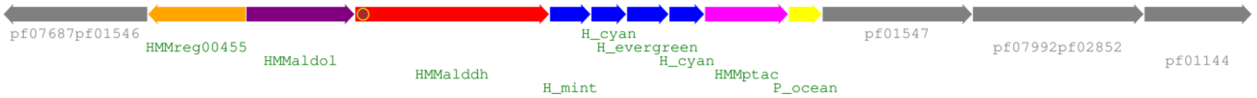


PVMlike are similar to PVM with regards to the potential signature aldolase enzyme, however other components like the aldehyde dehydrogenase, the PTAC and BMC-P are closest related to the GRM5/GRM5like homologs. PVMlike BMCs are found mostly in the Firmicute classes Clostridia and Bacilli and one member is found in *Bacteroides xylanolyticus*. The presence in that single Bacteroidetes strain [11] possibly indicates a more recent HGT event and the PVMlike BMC locus might assist with processing some type of sugar derivative based on the homology of the aldolase with the one from PVM loci. PVM type loci do not have satellite loci and in more than half of the genomes they co-occur with EUT2 and/or GRM1A BMC loci.

#### *RMM1 / AAUM1 – 140 loci*

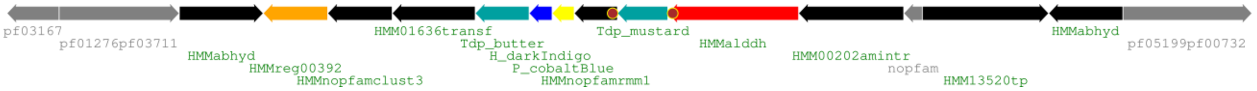


RMM1 (Rhodococcus and Mycobacterium microcompartment), or AAUM (aminoacetone utilization microcompartment) type BMCs are found in various Actinobacteria and Proteobacteria. The best studied locus is from *Mycobacterium smegmatis* MC2 155 where the crystal structures of all shell proteins were determined and enzymatic studies of the short-chain dehydrogenase/reductase were performed [12,13]. RMM1 loci contain only one BMC-H and one BMC-P but two different BMC-$T^{dp}$. There are no satellite loci and besides three instances of a co-occurrence with MIC1 the loci are the only type of BMC in the genomes.

#### *RMM2 / AAUM2– 32 loci*

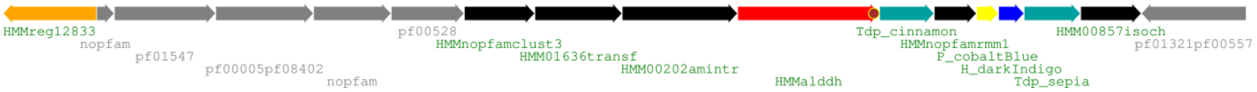


These loci were named RMM2/PDU2 in Axen because PDU signature enzymes were observed in proximity. However, despite the increased number of genomes (from 4 to 32) we only observe this in 4 genomes so it is unlikely to be an important part of the locus. The locus is very similar to RMM1 but with a different gene order. It is found in a variety of Proteobacteria and in *Candidatus Marinimicrobia*. The RMM2 type locus does not have satellite loci and always occurs by itself.

### References


1. Ravcheev DA, Moussu L, Smajic S, Thiele I: **Comparative Genomic Analysis Reveals Novel Microcompartment-Associated Metabolic Pathways in the Human Gut Microbiome**. *Front Genet* 2019, **10**:636.
2. Lang EJ, Cross PJ, Mittelstadt G, Jameson GB, Parker EJ: **Allosteric ACTion: the varied ACT domains regulating enzymes of amino-acid metabolism**. *Curr Opin Struct Biol* 2014, **29**:102-111.
3. Zarzycki J, Erbilgin O, Kerfeld CA: **Bioinformatic characterization of glycyl radical enzyme-associated bacterial microcompartments**. *Appl Environ Microbiol* 2015, **81**:8315-8329.
4. Ferlez B, Sutter M, Kerfeld CA: **Glycyl Radical Enzyme-Associated Microcompartments: Redox-Replete Bacterial Organelles**. *mBio* 2019, **10**.
5. Solli L, Havelsrud OE, Horn SJ, Rike AG: **A metagenomic study of the microbial communities in four parallel biogas reactors**. *Biotechnology for Biofuels* 2014, **14**.
6. Kadnikov VV, Mardanov AV, Beletsky AV, Karnachuk OV, Ravin NV: **Genome Analysis of a Member of the Uncultured Phylum Riflebacteria Revealed Pathways of Organotrophic Metabolism and Dissimilatory Iron Reduction**. *Microbiology* 2020, **89**:328-336.
7. Heldt D, Frank S, Seyedarabi A, Ladikis D, Parsons JB, Warren MJ, Pickersgill RW: **Structure of a trimeric bacterial microcompartment shell protein, EtuB, associated with ethanol utilization in *Clostridium kluyveri***. *The Biochemical journal* 2009, **423**:199-207.
8. Zhou Z, Chen J, Yao H, Hu H: **Fusobacterium and Colorectal Cancer**. *Front Oncol* 2018, **8**:371.
9. Axen SD, Erbilgin O, Kerfeld CA: **A taxonomy of bacterial microcompartment loci constructed by a novel scoring method**. *PLoS Comput Biol* 2014, **10**:e1003898.
10. Erbilgin O, McDonald KL, Kerfeld CA: **Characterization of a planctomycetal organelle: a novel bacterial microcompartment for the aerobic degradation of plant saccharides**. *Appl Environ Microbiol* 2014, **80**:2193-2205.
11. Scholten-Koerselman I, Houwaard F, Janssen P, Zehnder AJ: **Bacteroides xylanolyticus sp. nov., a xylanolytic bacterium from methane producing cattle manure**. *Antonie Van Leeuwenhoek* 1986, **52**:543-554.
12. Mallette E, Kimber MS: **A Complete Structural Inventory of the Mycobacterial Microcompartment Shell Proteins Constrains Models of Global Architecture and Transport**. *J Biol Chem* 2017, **292**:1197-1210.
13. Mallette E, Kimber MS: **Structural and kinetic characterization of (S)-1-amino-2-propanol kinase from the aminoacetone utilization microcompartment of Mycobacterium smegmatis**. *J Biol Chem* 2018, **293**:19909-19918.
