## Supplementary Figures and Tables for "A Catalog of the Diversity and Ubiquity of Metabolic Organelles in Bacteria"

| | total # | ARO | BUF1 | BUF1B | BUF2 | BUF3 | EUT1 | EUT2A | EUT2B | EUT2C | EUT2D | EUT2E | EUT2F | EUT2G | EUT2H | EUT2I | EUT2x | EUT3 | GRM1A | GRM2 | GRM3A | GRM3B | GRM4 | GRM5 | GRM6 | GRMguf | MIC1 | MUF1 | PDU1AB | PDU1C | PDU1D | PDU1E | PDU1F | PVM | PVMlike | SPU1 | SPU4 |
|---|---|---|---|---|---|---|---|---|---|---|---|---|---|---|---|---|---|---|---|---|---|---|---|---|---|---|---|---|---|---|---|---|---|---|---|---|---|
| ARO | 17 | | | | 2 | | | | | | | | | | | | | | | | | | | | | | 3 | | | | | | | | | | |
| BUF1 | 46 | | | | | 2 | | 2 | 14 | | | | | | | 7 | | | 4 | | | | | | | | | | | | 4 | | | | | | |
| BUF1B | 55 | | | | | | | | | 11 | | 1 | | | | | | | | | | | | | 1 | | | | | | | | | | 4 | | |
| BUF2 | 8 | 2 | | | | | | | | | | | | | | | | | | | | | | | | | | | | | | | | | | | |
| BUF3 | 10 | | 2 | | | | | | 2 | | | | | | | | | | 1 | | | | | | | | | | | | 1 | | | | | | 2 |
| EUT1 | 1408 | | | | | | 2 | | | | | | | | | | | | | 68 | | 28 | 4 | | | | | | 817 | | | 1 | | | | | |
| EUT2A | 177 | | 2 | | | | | | | | | | | | | | | | 19 | | | | | | | | | | | 10 | | 13 | | | 9 | | |
| EUT2B | 147 | | 14 | | | 2 | | | 6 | | | | | 14 | | | | | 17 | | | | | 4 | 1 | | | | | | 27 | | | | 8 | | |
| EUT2C | 88 | | | 11 | | | | | | | | 2 | | | | | 8 | | | | | | | | | | | | | | 1 | | | | | | 4 |
| EUT2D | 28 | | | | | | | | | | 18 | | | | | | | | 14 | | | | | | | 3 | | | | | 15 | | | | | | |
| EUT2E | 75 | | | 1 | | | | | | 2 | | 10 | | | | 1 | 3 | | | | | | | | | | | | | | | | | | | | 1 |
| EUT2F | 59 | | | | | | | | | | | | | | | | | | 1 | | | | | | | | | | | | | | 37 | | | | |
| EUT2G | 29 | | | | | | | | 14 | | | | | 2 | | | | | | | | | | 1 | | | | | | | 10 | | | | | | |
| EUT2H | 38 | | | | | | | | | | | | | | | | 8 | | | | | | | | | | | | | | | | | | | | |
| EUT2I | 162 | | 7 | | | | | | | | | 1 | | | | 24 | | | 36 | | 2 | | | 1 | | | | | | | 3 | | | | 4 | | |
| EUT2x | 24 | | | | | | | | | 8 | | 3 | | | 8 | | | | | | | | | | | | | | | | 1 | | | | | | 2 |
| EUT3 | 21 | | | | | | | | | | | | | | | | | | 4 | | | | | | | 4 | | | | | 2 | | | | | | |
| GRM1A | 297 | | 4 | | | 1 | | 19 | 17 | | 14 | | 1 | | | 36 | | 4 | 4 | | 9 | | | 1 | 3 | 11 | | 5 | | 1 | 14 | 8 | | | 19 | | 2 |
| GRM2 | 261 | | | | | | 68 | | | | | | | | | | | | | 2 | | 17 | 1 | | | | | | 71 | | | | | | | | |
| GRM3A | 84 | | | | | | | | | | | | | | | 2 | | | 9 | | 2 | | | | | | | | | | | 2 | | | 3 | | |
| GRM3B | 94 | | | | | | 28 | | | | | | | | | | | | | 17 | | | | | | | | | 2 | | | | | | | | |
| GRM4 | 8 | | | | | | 4 | | | | | | | | | | | | | 1 | | | | | | | | | 4 | | | | | | | | |
| GRM5 | 216 | | | | | | | | 4 | | | | | 1 | | 1 | | | 1 | | | | | 2 | | | | | | | 1 | | | | | | |
| GRM6 | 19 | | | 1 | | | | | 1 | | | | | | | | | | 3 | | | | | | | | | | | | 2 | 1 | | | 1 | | |
| GRMguf | 18 | | | | | | | | | | 3 | | | | | | | 4 | 11 | | | | | | | | | | | | 1 | | | | | | |
| MIC1 | 46 | 3 | | | | | | | | | | | | | | | | | | | | | | | | | 4 | | | | | | | | | | |
| MUF1 | 7 | | | | | | | | | | | | | | | | | | 5 | | | | | | | | | | | | | | | | | | |
| PDU1AB | 1315 | | | | | | 817 | | | | | | | | | | | | | 71 | | 2 | 4 | | | | | | 18 | | | 1 | | | | | |
| PDU1C | 89 | | | | | | | 10 | | | | | | | | | | | 1 | | | | | | | | | | | | | 1 | | | | | |
| PDU1D | 183 | | 4 | | | 1 | | | 27 | 1 | 15 | | | 10 | | 3 | 1 | 2 | 14 | | | | | 1 | 2 | 1 | | | | | 8 | | | | | | |
| PDU1E | 46 | | | | | | 1 | 13 | | | | | | | | | | | 8 | | 2 | | | | 1 | | | | 1 | 1 | | | | | | | |
| PDU1F | 44 | | | | | | | | | | | | 37 | | | | | | | | | | | | | | | | | | | | | | | | |
| PVM | 305 | | | | | | | | | | | | | | | | | | | | | | | | | | | | | | | | | 10 | | | |
| PVMlike | 57 | | | 4 | | | | 9 | 8 | | | | | | | 4 | | | 19 | | 3 | | | | 1 | | | | | | | | | | | | |
| SPU1 | 65 | | | | | | | | | | | | | | | | | | | | | | | | | | | | | | | | | | | 6 | |
| SPU4 | 34 | | | | | 2 | | | | 4 | | 1 | | | | | 2 | | 2 | | | | | | | | | | | | | | | | | | 2 |

Supplementary Table 1. Co-occurrence of BMC types (BMC types with less than 3 total co-occurrences are not shown). Green and dark green colors indicate 50% or higher co-occurrence of the BMC type on the left with the corresponding second type in the rows.

| # genomes | | | | | |
|---|---|---|---|---|---|
| 1 | BUF1 | BUF3 | EUT2 | GRM1 | PDU1 |
| 3 | BUF1 | EUT2 | GRM1 | | |
| 1 | BUF1 | EUT2 | PDU1 | | |
| 1 | EUT1 | GRM2 | GRM4 | | |
| 43 | EUT1 | GRM2 | PDU1 | | |
| 3 | EUT1 | GRM4 | PDU1 | | |
| 1 | EUT2 | GRM1 | GRM5 | | |
| 1 | EUT2 | GRM1 | GRM6 | | |
| 6 | EUT2 | GRM1 | GRMguf | | |
| 7 | EUT2 | GRM1 | PDU1 | | |
| 16 | EUT2 | GRM1 | PVMlike | | |
| 1 | GRM1 | GRM3 | PVMlike | | |
| 1 | GRM1 | GRM6 | PDU1 | | |
| 1 | GRM1 | GRM6 | PVMlike | | |
| 1 | GRM1 | GRMguf | PDU1 | | |

Supplementary Table 2. Co-occurrence loci for distinct BMC functional types in genomes that contain three or more BMC types (not counting duplicates).

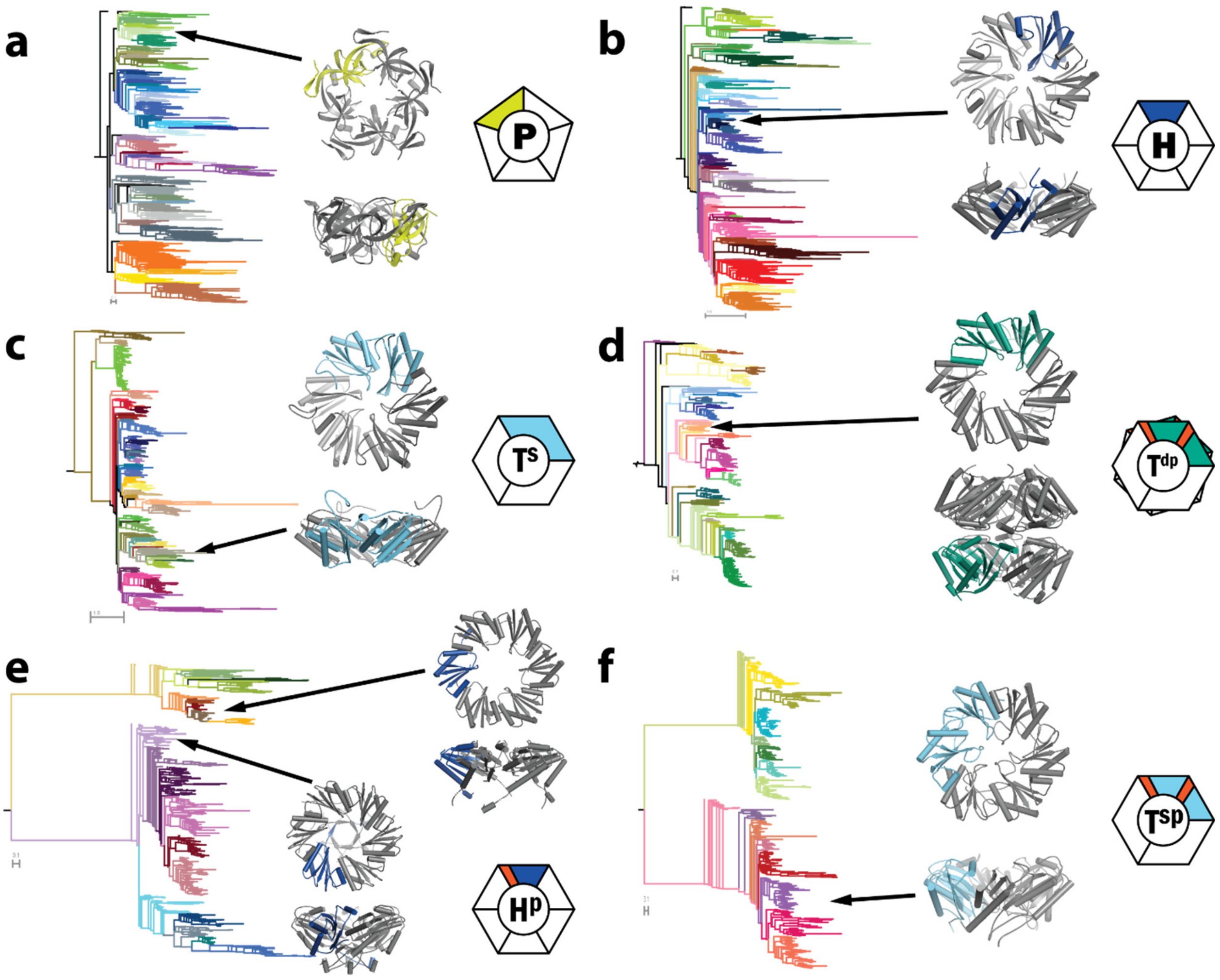


Supplementary Fig. 1. BMC shell protein phylogenetic trees with representative protein structures. **a**, BMC-P: Hoch_5814 PDB ID 5V74 (HO complete shell). **b**, BMC-H: Hoch_5815, PDB ID 5V74 (HO complete shell). **c**, BMC-$T^s$: Hoch_5812, PDB ID 5DIH. **d**, BMC-$T^{dp}$ : Hoch_5816, PDB ID 5V75. **e**, BMC-$H^p$ : (top): GrpU, PDB ID 4OLO; (bottom): EutS, PDB ID 4AXI. **f**, BMC-Tsp: EutL, PDB ID 3GFH.

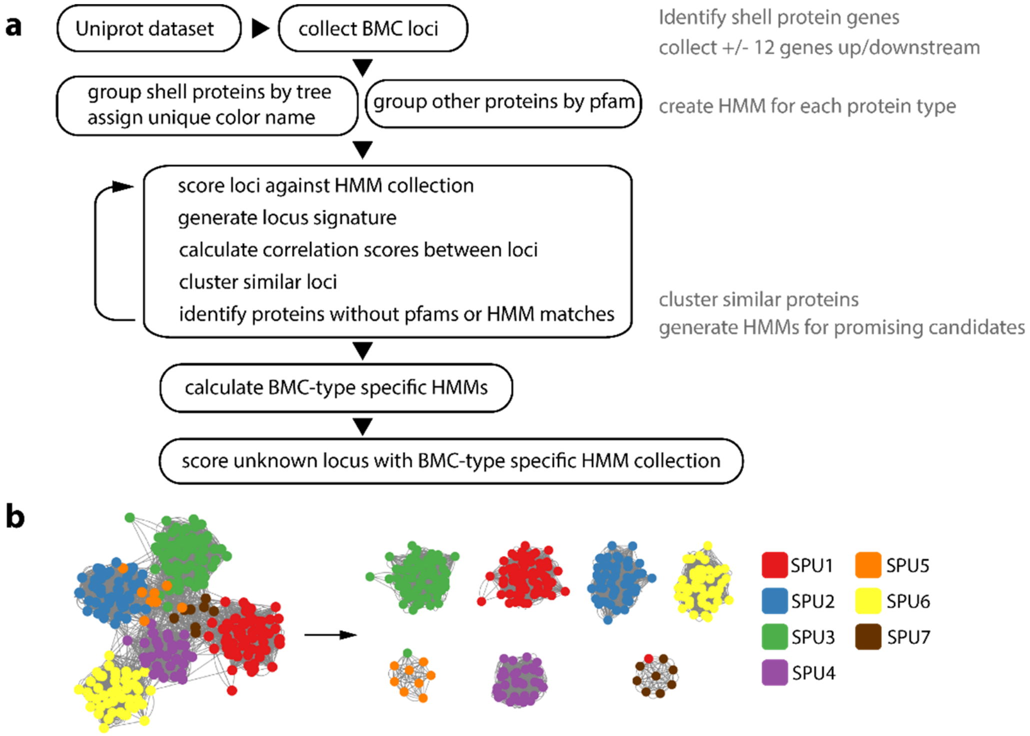


Supplementary Fig. 2. BMC classification methodology and clustering example. **a**, Flowchart of the methodology of BMC locus classification. **b**, Example of clustering by locus scoring, shown for all SPU (sugar phosphate utilization) BMC types: Force directed layout (left) and MCL clustering result (right) shows consistency of the BMC type clusters.

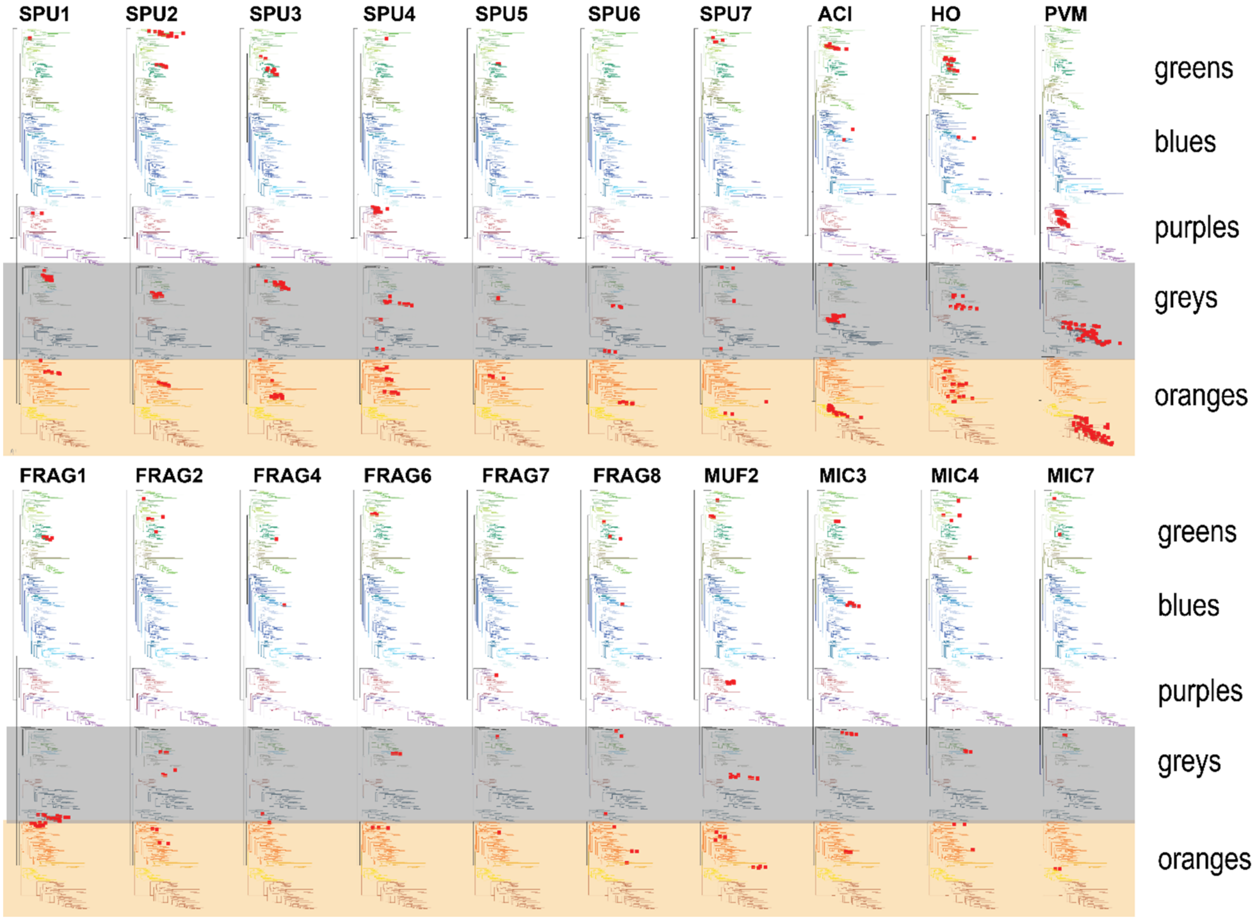


Supplementary Fig. 3. Distribution of BMC-P proteins in BMC types that have characteristic BMC-P triplets, with one member each from the grey (shaded) and orange (shaded) major clades and a third from the green/blue/purple clades. Positions of the BMC-P on the tree for each type are marked with red squares.

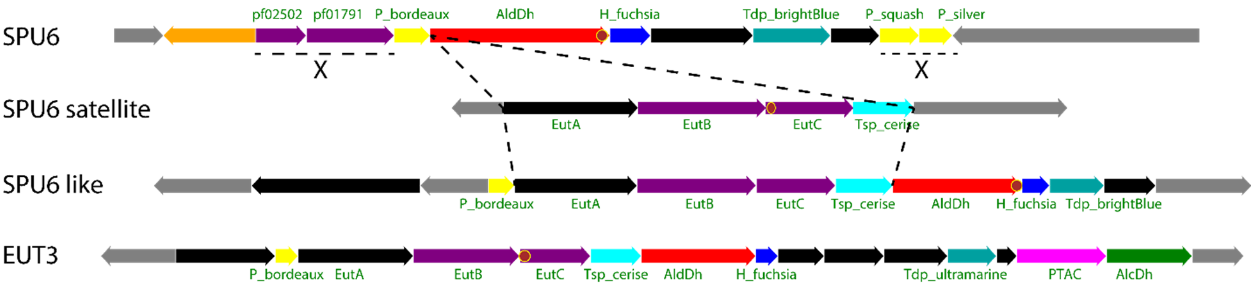


Supplementary Fig. 4. Locus diagrams of an SPU6 locus, its satellite, a SPU6 like locus and a EUT3 locus. The satellite of SPU6 inserting into the main locus would result in a SPU6 like locus. The genes for SPU signature enzymes pf02502 and pf01791 as well as two BMC-P would be deleted. Unlabeled genes in black are conserved among loci, in grey non-conserved proteins.

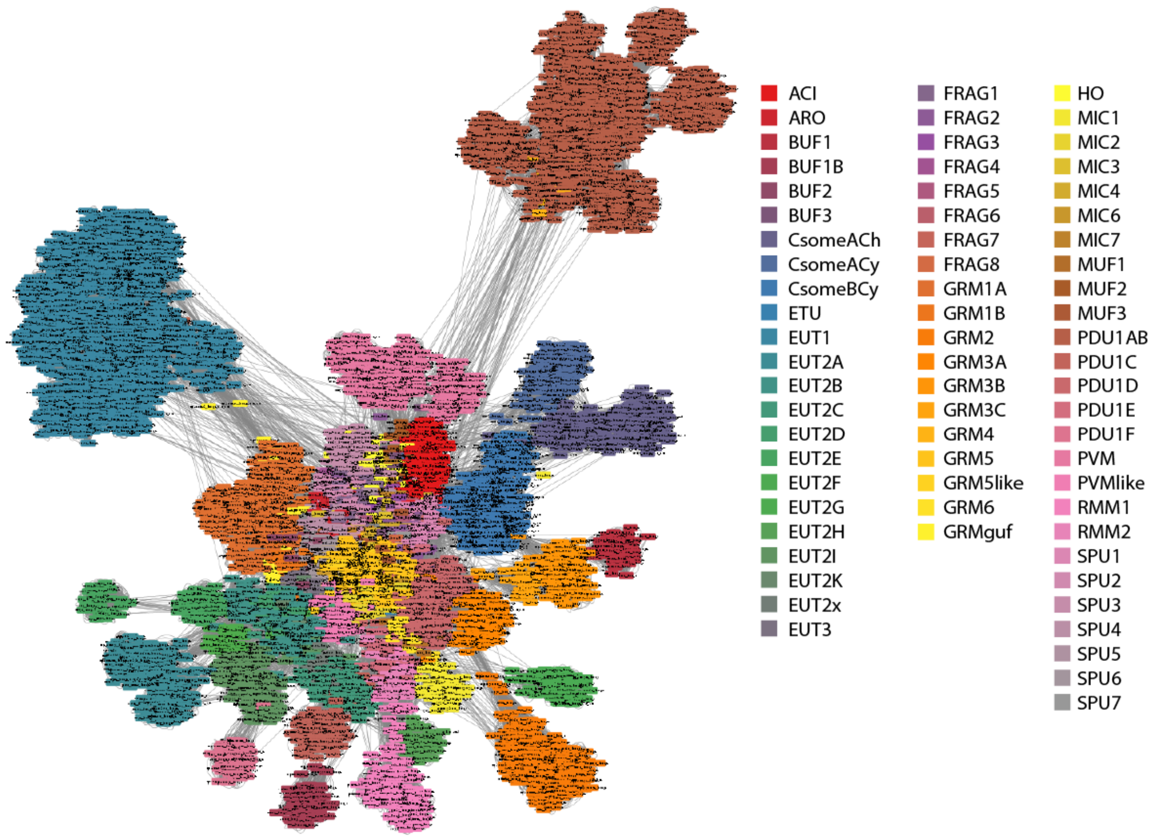


Supplementary Fig. 5. Visualization of all vs all BMC type scoring in a force-directed layout.

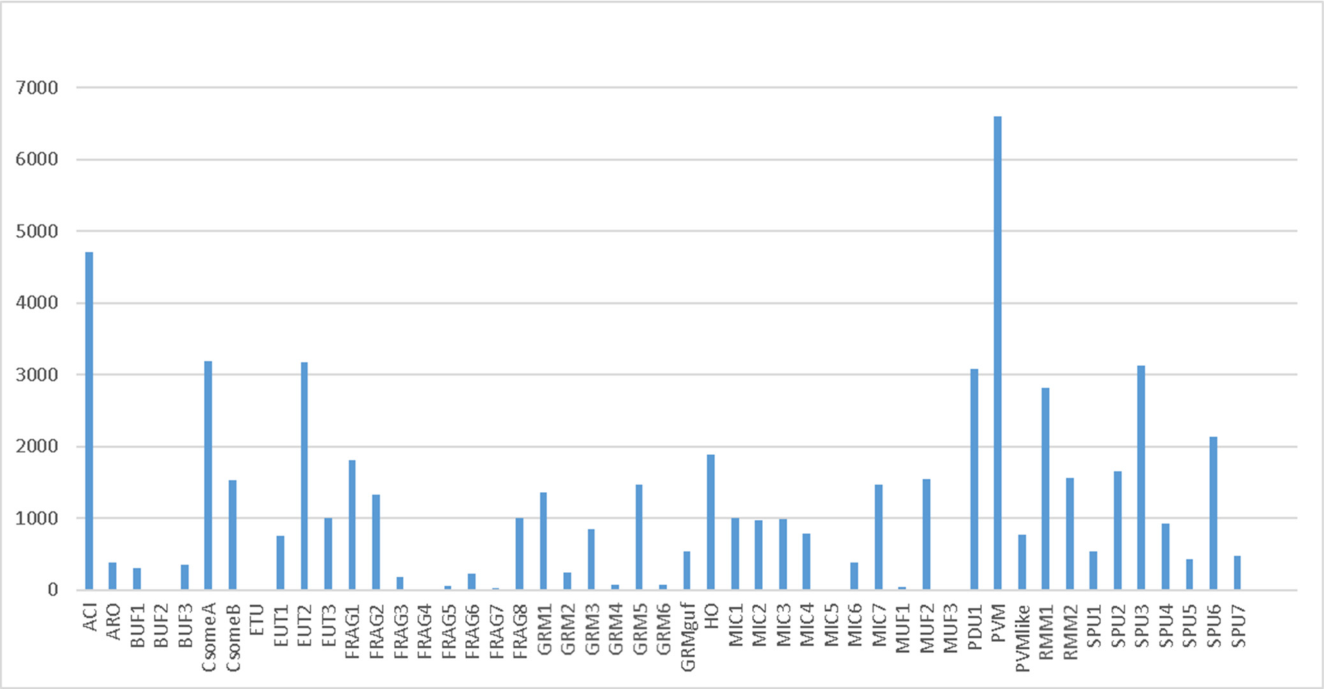


Supplementary Fig. 6. Presence of BMC types in metagenomes. Y-axis represents the number of metagenomes where both the BMC-H and BMC-P of a certain type are found as top hits of an HMM search.
